## Supplementary material for "Single-nucleotide Differences and Cell Type Decide the Subcellular Localization of miRNA Isoforms (isomiRs), tRNA-derived Fragments (tRFs) and rRNA-derived Fragments (rRFs)": Supp. Table S8 (2 of 5)

### Supp Table S8.2.Columns_AM_to_BW

**Supplemental Table S8:** The abundances

#### 111 Short RNA (Figure 5)

| | >MI0000542\|hsa-mir-320a&WithFlank&8\|-\|22244960\|22245043@48.69.22(+2U)&[MIMAT0000510&hsa-miR-320a-3p&offsets\|0\|0(+2U);m-60&8\|-\|22244975\|22244996&offsets\|0\|0(+2U)]\|AAAAGCTGGGTTGAGAGGGCGATT | >hg38_21_+_8433172_8446622_RNA45SN1_45S_50ntFlanks@10738.10755.18\|AAATCTCGCGCCGGGCCG | >MI0000289\|hsa-mir-181a-1&WithFlank&1\|-\|198859038\|198859159@30.53.24;MI0000269\|hsa-mir-181a-2&WithFlank&9\|+\|124692436\|124692557@45.68.24&[MIMAT0000256&hsa-miR-181a-5p&offsets\|0\|+1;m-32&1\|-\|198859108\|198859130&offsets\|0\|+1];[MIMAT0000256_1& | >hg38_21_+_8433172_8446622_RNA45SN1_45S_50ntFlanks@11718.11740.23\|AACGGCGGGAGTAACTATGACTC | >hg38_21_+_8433172_8446622_RNA45SN1_45S_50ntFlanks@11718.11742.25\|AACGGCGGGAGTAACTATGACTCTC | >hg38_21_+_8433172_8446622_RNA45SN1_45S_50ntFlanks@6650.6683.34\|ACGACTCTTAGCGGTGGATCACTCGGCTCGTGCG | >trna10_AspGTC_12_-_125424193_125424264@37.73.37,trna12_AspGTC_12_-_125411891_125411962@37.73.37,trna144_AspGTC_6_-_27551236_27551307@37.73.37,trna38_AspGTC_17_-_8125556_8125627@37.73.37,trna45_AspGTC_6_+_27447453_27447524@37.73.37,trna48_AspGTC_6_+_27471523_2 | >hg38_21_+_8433172_8446622_RNA45SN1_45S_50ntFlanks@5472.5516.45\|ACGGCCCTGGCGGAGCGCTGAGAAGACGGTCGAACTTGACTATCT | >hg38_21_+_8433172_8446622_RNA45SN1_45S_50ntFlanks@6653.6683.31\|ACTCTTAGCGGTGGATCACTCGGCTCGTGCG | >hg38_21_+_8433172_8446622_RNA45SN1_45S_50ntFlanks@7989.8006.18\|AGACGTGGCGACCCGCTG | >hg38_21_+_8433172_8446622_RNA45SN1_45S_50ntFlanks@7989.8007.19\|AGACGTGGCGACCCGCTGA | >MI0000298\|hsa-mir-221&WithFlank&X\|-\|45746151\|45746272@71.93.23(+1U)&[MIMAT0000278&hsa-miR-221-3p&offsets\|0\|0(+1U);m-91&X\|-\|45746181\|45746202&offsets\|0\|+1(+1U)]\|AGCTACATTGTCTGCTGGGTTTCT | >multi-hg38_MT_+_598_1651_RNR1_12S_50ntFlanks@52.77.26\|ATAGGTTTGGTCCTAGCCTTTCTATT | >hg38_21_+_8433172_8446622_RNA45SN1_45S_50ntFlanks@11784.11803.20\|ATGAATGGATGAACGAGATT | >trnaMT_PheGAA_MT_+_577_647@39.56.18\|ATGTTTAGACGGGCTCAC | >hg38_21_+_8433172_8446622_RNA45SN1_45S_50ntFlanks@6722.6744.23\|ATTGCAGGACACATTGATCATCG | >MI0000095\|hsa-mir-93&WithFlank&7\|-\|100093762\|100093853@17.38.22(+1A)&[MIMAT0000093&hsa-miR-93-5p&offsets\|0\|-1(+1A);m-29&7\|-\|100093815\|100093837&offsets\|0\|-1(+1A)]\|CAAAGTGCTGTTCGTGCAGGTAA | >multi-am_trnaMT_ValTAC_MT_+_1602_1670@48.67.20\|CAACTTAACTTGACCGCTCT | >trna10_AspGTC_12_-_125424193_125424264@36.74.39,trna12_AspGTC_12_-_125411891_125411962@36.74.39,trna144_AspGTC_6_-_27551236_27551307@36.74.39,trna38_AspGTC_17_-_8125556_8125627@36.74.39,trna45_AspGTC_6_+_27447453_27447524@36.74.39,trna48_AspGTC_6_+_27471523_2 | >trna10_AspGTC_12_-_125424193_125424264@36.75.40,trna12_AspGTC_12_-_125411891_125411962@36.75.40,trna144_AspGTC_6_-_27551236_27551307@36.75.40,trna38_AspGTC_17_-_8125556_8125627@36.75.40,trna45_AspGTC_6_+_27447453_27447524@36.75.40,trna48_AspGTC_6_+_27471523_2 | >trnaMT_GluTTC_MT_-_14674_14742@38.72.35,trnalookalike8_GluTTC_5_-_93905172_93905240@38.72.35\|CATTGGTCGTGGTTGTAGTCCGTGCGAGAATACCA | >hg38_21_+_8433172_8446622_RNA45SN1_45S_50ntFlanks@11670.11701.32\|CCCAGTGCTCTGAATGTCAAAGTGAAGAAATT | >hg38_21_+_8433172_8446622_RNA45SN1_45S_50ntFlanks@12072.12095.24\|CCCGAGGGGCTCTCGCTTCTGGCG | >hg38_21_+_8433172_8446622_RNA45SN1_45S_50ntFlanks@12993.13039.47\|CCCTCGCTGCGATCTATTGAAAGTCAGCCCTCGACACAAGGGTTTGT | >hg38_21_+_8433172_8446622_RNA45SN1_45S_50ntFlanks@6651.6670.20\|CGACTCTTAGCGGTGGATCA | >hg38_21_+_8433172_8446622_RNA45SN1_45S_50ntFlanks@6651.6683.33\|CGACTCTTAGCGGTGGATCACTCGGCTCGTGCG | >hg38_21_+_8433172_8446622_RNA45SN1_45S_50ntFlanks@6651.6688.38\|CGACTCTTAGCGGTGGATCACTCGGCTCGTGCGTCGAT | >hg38_21_+_8433172_8446622_RNA45SN1_45S_50ntFlanks@11781.11803.23\|CGCATGAATGGATGAACGAGATT | >hg38_21_+_8433172_8446622_RNA45SN1_45S_50ntFlanks@8115.8143.29\|CGCGGGACATGTGGCGTACGGAAGACCCG | >hg38_21_+_8433172_8446622_RNA45SN1_45S_50ntFlanks@5473.5516.44\|CGGCCCTGGCGGAGCGCTGAGAAGACGGTCGAACTTGACTATCT | >hg38_21_+_8433172_8446622_RNA45SN1_45S_50ntFlanks@6661.6683.23\|CGGTGGATCACTCGGCTCGTGCG | >hg38_21_+_8433172_8446622_RNA45SN1_45S_50ntFlanks@12991.13016.26\|CTCCCTCGCTGCGATCTATTGAAAGT | >hg38_1_-_228634819_228635039_RNA5S12_5S_50ntFlanks@84.101.18\|CTCGTCTGATCTCGGAAG | >hg38_21_+_8433172_8446622_RNA45SN1_45S_50ntFlanks@5489.5516.28\|CTGAGAAGACGGTCGAACTTGACTATCT | >hg38_21_+_8433172_8446622_RNA45SN1_45S_50ntFlanks@475.503.29\|CTTCGTGATCGATGTGGTGACGTCGTGCT | >hg38_21_+_8433172_8446622_RNA45SN1_45S_50ntFlanks@475.504.30\|CTTCGTGATCGATGTGGTGACGTCGTGCTC | >hg38_21_+_8433172_8446622_RNA45SN1_45S_50ntFlanks@475.505.31\|CTTCGTGATCGATGTGGTGACGTCGTGCTCT |
|---|---|---|---|---|---|---|---|---|---|---|---|---|---|---|---|---|---|---|---|---|---|---|---|---|---|---|---|---|---|---|---|---|---|---|---|---|---|
| SRR10024150 | 40 | 0 | 0 | 0 | 15 | 271 | 23 | 51 | 194 | 0 | 31 | 20 | 0 | 0 | 70 | 0 | 0 | 0 | 23 | 116 | 116 | 0 | 0 | 56 | 82 | 1293 | 210 | 0 | 0 | 112 | 97 | 0 | 15 | 1150 | 465 | 82 | 976 |
| SRR10024151 | 125 | 0 | 0 | 0 | 0 | 16 | 29 | 28 | 14 | 0 | 0 | 97 | 19 | 0 | 39 | 0 | 24 | 0 | 58 | 204 | 717 | 0 | 0 | 22 | 0 | 74 | 133 | 0 | 0 | 75 | 99 | 0 | 0 | 39 | 34 | 199 | 52 |
| SRR10024152 | 22 | 0 | 0 | 0 | 0 | 298 | 12 | 21 | 159 | 21 | 77 | 16 | 10 | 15 | 140 | 0 | 0 | 0 | 12 | 81 | 291 | 0 | 0 | 36 | 87 | 1070 | 67 | 15 | 0 | 62 | 46 | 21 | 36 | 982 | 180 | 51 | 812 |
| SRR10024153 | 52 | 0 | 0 | 0 | 0 | 433 | 0 | 0 | 344 | 0 | 20 | 25 | 40 | 0 | 200 | 0 | 15 | 0 | 0 | 31 | 278 | 0 | 0 | 139 | 70 | 2111 | 80 | 30 | 0 | 55 | 75 | 15 | 0 | 1150 | 1284 | 80 | 1733 |
| SRR10024154 | 25 | 0 | 0 | 0 | 0 | 145 | 0 | 19 | 130 | 22 | 136 | 28 | 19 | 32 | 144 | 0 | 10 | 0 | 16 | 112 | 514 | 0 | 0 | 22 | 35 | 14 | 54 | 0 | 0 | 57 | 95 | 13 | 32 | 11 | 136 | 25 | 383 |
| SRR10024155 | 36 | 0 | 0 | 0 | 0 | 219 | 0 | 16 | 110 | 26 | 439 | 27 | 0 | 0 | 153 | 27 | 15 | 0 | 26 | 77 | 574 | 0 | 0 | 27 | 628 | 1024 | 56 | 40 | 0 | 45 | 67 | 24 | 0 | 259 | 279 | 38 | 556 |
| SRR10024156 | 108 | 0 | 0 | 0 | 0 | 11 | 47 | 46 | 216 | 0 | 38 | 40 | 15 | 0 | 109 | 0 | 30 | 0 | 0 | 62 | 1355 | 0 | 0 | 91 | 23 | 47 | 99 | 0 | 0 | 69 | 56 | 0 | 0 | 39 | 19 | 112 | 35 |
| SRR10024158 | 46 | 0 | 0 | 0 | 0 | 140 | 0 | 0 | 65 | 17 | 34 | 64 | 39 | 39 | 215 | 0 | 35 | 0 | 0 | 43 | 230 | 0 | 0 | 24 | 36 | 580 | 39 | 0 | 12 | 12 | 85 | 22 | 0 | 1357 | 307 | 34 | 338 |
| SRR10024159 | 99 | 11 | 0 | 0 | 0 | 185 | 0 | 0 | 246 | 0 | 53 | 54 | 0 | 0 | 14 | 20 | 24 | 0 | 0 | 0 | 226 | 0 | 0 | 0 | 53 | 1368 | 20 | 0 | 14 | 0 | 73 | 42 | 14 | 571 | 654 | 70 | 1127 |
| SRR10024161 | 23 | 0 | 0 | 0 | 0 | 206 | 0 | 0 | 130 | 0 | 46 | 32 | 27 | 0 | 77 | 61 | 16 | 0 | 17 | 43 | 77 | 0 | 0 | 19 | 466 | 13 | 73 | 19 | 0 | 23 | 57 | 0 | 19 | 511 | 584 | 57 | 527 |
| SRR10024162 | 74 | 0 | 0 | 0 | 0 | 14 | 11 | 25 | 11 | 0 | 22 | 43 | 25 | 0 | 179 | 0 | 26 | 0 | 0 | 21 | 527 | 0 | 0 | 25 | 40 | 62 | 165 | 0 | 0 | 75 | 118 | 0 | 0 | 17 | 30 | 190 | 51 |
| SRR10024163 | 40 | 0 | 0 | 0 | 0 | 10 | 0 | 0 | 592 | 0 | 23 | 19 | 14 | 0 | 63 | 0 | 0 | 0 | 14 | 49 | 216 | 0 | 0 | 33 | 56 | 47 | 117 | 0 | 0 | 23 | 159 | 14 | 23 | 760 | 760 | 131 | 22 |
| SRR10024164 | 116 | 0 | 0 | 0 | 0 | 303 | 0 | 0 | 139 | 0 | 43 | 32 | 0 | 43 | 0 | 0 | 15 | 0 | 57 | 86 | 570 | 0 | 0 | 17 | 35 | 1585 | 78 | 26 | 0 | 0 | 87 | 0 | 0 | 728 | 139 | 121 | 1282 |
| SRR10024166 | 100 | 0 | 0 | 0 | 0 | 528 | 0 | 33 | 392 | 0 | 87 | 18 | 33 | 0 | 42 | 0 | 0 | 0 | 0 | 0 | 92 | 0 | 0 | 87 | 37 | 56 | 17 | 0 | 0 | 74 | 178 | 12 | 21 | 26 | 14 | 235 | 41 |
| SRR10024167 | 29 | 10 | 0 | 0 | 0 | 65 | 16 | 130 | 82 | 17 | 82 | 39 | 31 | 14 | 304 | 0 | 18 | 0 | 0 | 0 | 48 | 0 | 0 | 68 | 58 | 287 | 41 | 0 | 0 | 44 | 27 | 17 | 0 | 546 | 277 | 51 | 294 |
| SRR10024168 | 25 | 0 | 0 | 0 | 0 | 136 | 0 | 0 | 174 | 0 | 24 | 53 | 0 | 0 | 113 | 0 | 56 | 0 | 0 | 129 | 804 | 0 | 0 | 58 | 44 | 547 | 73 | 15 | 0 | 0 | 53 | 24 | 19 | 310 | 121 | 29 | 305 |
| SRR10024169 | 138 | 0 | 0 | 0 | 0 | 12 | 0 | 0 | 299 | 0 | 31 | 87 | 24 | 0 | 69 | 0 | 23 | 0 | 23 | 0 | 240 | 0 | 0 | 13 | 29 | 63 | 24 | 27 | 0 | 0 | 73 | 16 | 0 | 36 | 45 | 109 | 45 |
| SRR10024170 | 89 | 13 | 0 | 0 | 0 | 334 | 11 | 0 | 230 | 0 | 35 | 37 | 13 | 0 | 171 | 0 | 18 | 0 | 0 | 57 | 205 | 0 | 0 | 30 | 43 | 33 | 82 | 0 | 0 | 22 | 104 | 17 | 0 | 12 | 273 | 121 | 21 |
| SRR10024171 | 36 | 17 | 0 | 0 | 0 | 215 | 0 | 62 | 197 | 0 | 47 | 55 | 25 | 0 | 74 | 0 | 21 | 0 | 37 | 49 | 246 | 0 | 0 | 99 | 49 | 21 | 74 | 0 | 0 | 106 | 69 | 0 | 0 | 23 | 291 | 89 | 11 |
| SRR10024173 | 114 | 0 | 0 | 0 | 0 | 144 | 17 | 0 | 122 | 0 | 32 | 74 | 18 | 27 | 68 | 0 | 54 | 0 | 0 | 154 | 1061 | 0 | 0 | 45 | 63 | 13 | 81 | 0 | 0 | 0 | 32 | 0 | 0 | 554 | 108 | 90 | 653 |
| SRR10024176 | 43 | 0 | 0 | 0 | 0 | 70 | 0 | 0 | 70 | 0 | 0 | 17 | 14 | 0 | 22 | 0 | 21 | 0 | 0 | 0 | 22 | 0 | 0 | 0 | 77 | 325 | 38 | 0 | 10 | 10 | 27 | 0 | 10 | 1804 | 218 | 24 | 147 |
| SRR10024178 | 56 | 14 | 0 | 0 | 0 | 359 | 0 | 14 | 345 | 0 | 31 | 32 | 0 | 0 | 28 | 0 | 11 | 0 | 0 | 35 | 206 | 0 | 0 | 14 | 61 | 31 | 78 | 0 | 0 | 33 | 78 | 0 | 33 | 27 | 228 | 64 | 412 |
| SRR10024179 | 25 | 0 | 0 | 11 | 0 | 188 | 17 | 0 | 116 | 17 | 108 | 46 | 0 | 0 | 150 | 34 | 13 | 0 | 0 | 17 | 316 | 0 | 0 | 0 | 410 | 832 | 13 | 61 | 0 | 0 | 70 | 27 | 0 | 142 | 169 | 17 | 215 |
| SRR10024180 | 39 | 11 | 0 | 0 | 0 | 158 | 0 | 26 | 173 | 0 | 44 | 23 | 33 | 18 | 181 | 0 | 0 | 0 | 0 | 0 | 251 | 0 | 0 | 74 | 48 | 10 | 44 | 0 | 11 | 18 | 63 | 0 | 15 | 13 | 409 | 70 | 11 |
| SRR10024181 | 35 | 0 | 0 | 0 | 0 | 483 | 26 | 151 | 270 | 0 | 0 | 49 | 22 | 0 | 26 | 0 | 16 | 0 | 26 | 65 | 496 | 0 | 0 | 126 | 22 | 2025 | 72 | 0 | 16 | 399 | 107 | 0 | 13 | 1001 | 640 | 91 | 1033 |
| SRR10024182 | 77 | 0 | 0 | 0 | 0 | 705 | 0 | 24 | 473 | 0 | 34 | 18 | 58 | 0 | 97 | 0 | 10 | 0 | 64 | 97 | 661 | 0 | 0 | 29 | 97 | 48 | 159 | 0 | 0 | 53 | 97 | 43 | 0 | 22 | 11 | 101 | 31 |
| SRR10024183 | 73 | 0 | 0 | 0 | 0 | 489 | 0 | 12 | 199 | 0 | 37 | 61 | 12 | 0 | 51 | 18 | 19 | 0 | 0 | 51 | 243 | 0 | 0 | 46 | 104 | 2336 | 119 | 0 | 0 | 15 | 128 | 21 | 34 | 516 | 205 | 86 | 931 |
| SRR10024185 | 68 | 0 | 0 | 0 | 0 | 313 | 0 | 18 | 145 | 0 | 12 | 29 | 15 | 0 | 138 | 0 | 14 | 0 | 13 | 25 | 1151 | 0 | 0 | 45 | 33 | 1591 | 57 | 0 | 0 | 12 | 63 | 12 | 0 | 3633 | 464 | 111 | 1380 |
| SRR10024186 | 30 | 0 | 0 | 0 | 0 | 52 | 0 | 30 | 96 | 0 | 11 | 25 | 39 | 0 | 26 | 0 | 12 | 0 | 0 | 0 | 77 | 0 | 0 | 41 | 77 | 311 | 14 | 0 | 0 | 19 | 17 | 0 | 17 | 638 | 344 | 39 | 352 |
| SRR10024187 | 405 | 0 | 11 | 0 | 0 | 296 | 0 | 0 | 101 | 0 | 13 | 33 | 13 | 31 | 18 | 0 | 21 | 0 | 88 | 263 | 263 | 0 | 0 | 31 | 107 | 28 | 164 | 25 | 0 | 0 | 69 | 0 | 0 | 208 | 82 | 183 | 19 |
| SRR10024188 | 31 | 16 | 0 | 0 | 0 | 258 | 0 | 0 | 101 | 12 | 69 | 0 | 24 | 16 | 102 | 0 | 0 | 0 | 0 | 18 | 166 | 0 | 0 | 0 | 24 | 1214 | 20 | 0 | 0 | 16 | 77 | 0 | 28 | 1335 | 210 | 165 | 1915 |
| SRR10024189 | 48 | 0 | 0 | 0 | 0 | 534 | 0 | 13 | 175 | 0 | 42 | 23 | 21 | 13 | 35 | 0 | 15 | 0 | 24 | 12 | 188 | 0 | 0 | 13 | 38 | 3215 | 46 | 0 | 17 | 0 | 200 | 17 | 25 | 842 | 700 | 125 | 1901 |
| SRR10024190 | 29 | 12 | 0 | 0 | 0 | 149 | 0 | 32 | 165 | 0 | 52 | 0 | 20 | 12 | 218 | 0 | 0 | 0 | 0 | 0 | 1487 | 24 | 0 | 0 | 16 | 15 | 0 | 16 | 0 | 40 | 84 | 20 | 16 | 25 | 125 | 277 | 74 |
| SRR10024191 | 32 | 0 | 0 | 0 | 0 | 315 | 52 | 0 | 184 | 0 | 0 | 32 | 0 | 11 | 79 | 0 | 29 | 0 | 0 | 209 | 288 | 11 | 11 | 195 | 43 | 1746 | 108 | 0 | 0 | 0 | 54 | 11 | 11 | 206 | 206 | 98 | 662 |
| SRR10024192 | 37 | 11 | 0 | 0 | 21 | 1110 | 0 | 0 | 406 | 0 | 0 | 16 | 21 | 0 | 106 | 0 | 13 | 0 | 35 | 70 | 633 | 11 | 0 | 21 | 85 | 5627 | 192 | 0 | 0 | 11 | 117 | 64 | 0 | 192 | 438 | 171 | 3374 |
| SRR10024193 | 54 | 0 | 0 | 0 | 0 | 507 | 0 | 11 | 428 | 0 | 15 | 40 | 51 | 0 | 0 | 0 | 24 | 0 | 18 | 55 | 273 | 0 | 0 | 51 | 54 | 41 | 210 | 0 | 0 | 40 | 91 | 0 | 0 | 13 | 22 | 120 | 22 |
| SRR10024194 | 28 | 0 | 0 | 0 | 0 | 111 | 10 | 79 | 76 | 0 | 38 | 41 | 0 | 18 | 50 | 0 | 12 | 0 | 10 | 20 | 230 | 0 | 0 | 0 | 67 | 428 | 56 | 0 | 0 | 26 | 35 | 0 | 12 | 10 | 138 | 41 | 352 |
| SRR10024195 | 32 | 0 | 0 | 0 | 0 | 452 | 12 | 130 | 314 | 0 | 35 | 22 | 0 | 0 | 185 | 0 | 15 | 0 | 58 | 35 | 381 | 0 | 0 | 114 | 0 | 1910 | 51 | 0 | 0 | 200 | 157 | 0 | 24 | 1387 | 534 | 55 | 994 |
| SRR10024196 | 52 | 0 | 0 | 0 | 16 | 336 | 0 | 86 | 219 | 0 | 23 | 15 | 70 | 16 | 53 | 0 | 13 | 0 | 0 | 18 | 177 | 23 | 0 | 235 | 23 | 2402 | 16 | 0 | 0 | 47 | 125 | 23 | 23 | 1487 | 509 | 180 | 1432 |
| SRR10024197 | 12 | 13 | 0 | 0 | 0 | 74 | 0 | 26 | 121 | 0 | 26 | 24 | 0 | 13 | 309 | 0 | 10 | 0 | 0 | 0 | 185 | 0 | 0 | 52 | 65 | 355 | 56 | 0 | 0 | 95 | 56 | 0 | 0 | 553 | 216 | 52 | 311 |
| SRR10024198 | 26 | 0 | 0 | 0 | 11 | 178 | 13 | 24 | 178 | 0 | 21 | 36 | 0 | 0 | 13 | 13 | 22 | 0 | 13 | 105 | 262 | 0 | 0 | 66 | 58 | 15 | 72 | 0 | 11 | 24 | 85 | 11 | 0 | 284 | 202 | 48 | 326 |
| SRR10024199 | 70 | 0 | 0 | 0 | 0 | 439 | 53 | 0 | 161 | 0 | 71 | 14 | 45 | 19 | 53 | 0 | 14 | 0 | 18 | 71 | 463 | 26 | 0 | 110 | 26 | 2389 | 39 | 0 | 0 | 0 | 123 | 39 | 0 | 2383 | 155 | 65 | 988 |
| SRR10024201 | 27 | 23 | 0 | 0 | 0 | 370 | 0 | 36 | 370 | 0 | 32 | 27 | 41 | 14 | 123 | 0 | 0 | 0 | 35 | 0 | 228 | 0 | 0 | 68 | 41 | 1795 | 45 | 0 | 0 | 72 | 149 | 14 | 14 | 613 | 1037 | 99 | 1222 |
| SRR10024202 | 77 | 0 | 0 | 0 | 0 | 375 | 0 | 16 | 159 | 0 | 16 | 13 | 24 | 0 | 102 | 0 | 0 | 0 | 17 | 102 | 546 | 0 | 0 | 40 | 32 | 1626 | 56 | 0 | 0 | 64 | 64 | 24 | 48 | 1523 | 151 | 191 | 1674 |
| SRR10024203 | 94 | 0 | 0 | 0 | 0 | 11 | 60 | 14 | 570 | 0 | 14 | 70 | 14 | 0 | 119 | 0 | 23 | 0 | 0 | 60 | 775 | 0 | 0 | 74 | 83 | 39 | 147 | 0 | 0 | 55 | 184 | 0 | 0 | 566 | 21 | 143 | 25 |
| SRR10024204 | 28 | 0 | 0 | 0 | 0 | 192 | 14 | 0 | 241 | 20 | 65 | 35 | 0 | 0 | 349 | 0 | 11 | 0 | 14 | 0 | 223 | 0 | 0 | 16 | 39 | 17 | 36 | 0 | 0 | 29 | 137 | 0 | 26 | 11 | 17 | 78 | 30 |
| SRR10024205 | 28 | 0 | 0 | 0 | 0 | 1069 | 38 | 0 | 418 | 12 | 70 | 15 | 0 | 12 | 840 | 0 | 0 | 0 | 0 | 0 | 382 | 0 | 0 | 23 | 58 | 4995 | 58 | 0 | 0 | 0 | 256 | 0 | 58 | 441 | 709 | 244 | 4437 |
| SRR10024206 | 50 | 10 | 0 | 0 | 0 | 14 | 0 | 10 | 10 | 15 | 50 | 58 | 31 | 27 | 11 | 0 | 21 | 0 | 32 | 21 | 275 | 0 | 0 | 0 | 42 | 69 | 96 | 10 | 13 | 19 | 103 | 0 | 13 | 28 | 23 | 84 | 25 |
| SRR10024207 | 69 | 0 | 0 | 0 | 0 | 138 | 0 | 0 | 122 | 0 | 15 | 38 | 36 | 10 | 16 | 0 | 11 | 0 | 0 | 16 | 79 | 0 | 0 | 10 | 25 | 800 | 25 | 0 | 10 | 0 | 46 | 0 | 10 | 16 | 260 | 46 | 566 |
| SRR10024208 | 21 | 0 | 0 | 0 | 0 | 219 | 0 | 28 | 191 | 12 | 48 | 13 | 0 | 12 | 79 | 32 | 13 | 0 | 16 | 47 | 189 | 0 | 0 | 20 | 88 | 11 | 56 | 12 | 0 | 16 | 104 | 0 | 28 | 626 | 207 | 36 | 682 |
| SRR1003025 | 30 | 31 | 8001 | 0 | 0 | 57 | 87 | 44 | 29 | 0 | 0 | 0 | 0 | 0 | 73 | 0 | 43 | 0 | 0 | 1364 | 499 | 0 | 0 | 0 | 0 | 66 | 136 | 0 | 0 | 0 | 64 | 0 | 0 | 14 | 77 | 22 | 50 |
| SRR1005875 | 279 | 0 | 1558 | 0 | 19 | 0 | 0 | 29 | 0 | 0 | 0 | 1555 | 0 | 0 | 0 | 0 | 13 | 0 | 0 | 729 | 364 | 0 | 0 | 0 | 0 | 39 | 0 | 0 | 0 | 0 | 0 | 0 | 0 | 0 | 0 | 0 | 0 |
| SRR10072377 | 44 | 0 | 69 | 0 | 0 | 116 | 457 | 42 | 0 | 19 | 51 | 12 | 130 | 16 | 37818 | 0 | 43 | 73 | 67 | 287 | 7444 | 0 | 0 | 576 | 408 | 666 | 2035 | 0 | 0 | 22 | 40 | 0 | 0 | 147 | 34 | 0 | 42 |
| SRR10072378 | 44 | 0 | 65 | 0 | 0 | 18 | 373 | 32 | 16 | 14 | 42 | 11 | 17 | 16 | 35144 | 12 | 48 | 24 | 76 | 309 | 7796 | 0 | 0 | 80 | 56 | 90 | 275 | 0 | 0 | 16 | 51 | 0 | 0 | 21 | 31 | 15 | 38 |
| SRR10072379 | 49 | 0 | 78 | 0 | 0 | 100 | 67 | 0 | 16 | 15 | 62 | 17 | 329 | 18 | 49177 | 0 | 58 | 344 | 13 | 253 | 7711 | 0 | 0 | 651 | 400 | 478 | 1596 | 0 | 0 | 0 | 33 | 0 | 0 | 23 | 46 | 13 | 63 |
| SRR10072380 | 45 | 0 | 68 | 0 | 0 | 94 | 33 | 0 | 10 | 25 | 52 | 20 | 329 | 17 | 48456 | 0 | 55 | 363 | 26 | 150 | 7547 | 0 | 0 | 596 | 386 | 452 | 1594 | 0 | 0 | 0 | 30 | 0 | 0 | 30 | 43 | 13 | 67 |
| SRR10072381 | 63 | 0 | 61 | 0 | 0 | 15 | 156 | 39 | 21 | 18 | 43 | 0 | 34 | 14 | 13448 | 0 | 33 | 21 | 78 | 524 | 3862 | 0 | 0 | 170 | 51 | 84 | 254 | 0 | 0 | 18 | 44 | 0 | 0 | 15 | 38 | 16 | 61 |
| SRR10072382 | 49 | 0 | 63 | 0 | 0 | 16 | 194 | 31 | 12 | 19 | 41 | 13 | 31 | 12 | 13337 | 0 | 34 | 40 | 56 | 500 | 3525 | 0 | 0 | 170 | 50 | 85 | 230 | 0 | 0 | 19 | 39 | 0 | 0 | 13 | 24 | 10 | 52 |
| SRR10072383 | 37 | 0 | 152 | 0 | 0 | 75 | 856 | 26 | 0 | 16 | 28 | 12 | 43 | 0 | 39959 | 0 | 22 | 32 | 43 | 271 | 5993 | 0 | 0 | 117 | 30 | 30 | 95 | 0 | 0 | 14 | 23 | 0 | 0 | 12 | 99 | 33 | 124 |
| SRR10072384 | 38 | 0 | 129 | 0 | 0 | 81 | 771 | 18 | 0 | 0 | 28 | 12 | 49 | 0 | 39691 | 0 | 31 | 38 | 53 | 348 | 5869 | 0 | 0 | 118 | 30 | 32 | 100 | 0 | 0 | 12 | 18 | 0 | 0 | 14 | 97 | 28 | 12 |
| SRR10072385 | 44 | 0 | 78 | 0 | 0 | 11 | 1181 | 12 | 18 | 15 | 41 | 13 | 18 | 19 | 8201 | 0 | 37 | 71 | 82 | 239 | 3443 | 0 | 0 | 191 | 32 | 65 | 168 | 0 | 0 | 45 | 19 | 0 | 0 | 47 | 55 | 30 | 13 |
| SRR10072386 | 51 | 0 | 90 | 0 | 0 | 112 | 997 | 137 | 23 | 0 | 44 | 14 | 153 | 0 | 7965 | 0 | 36 | 64 | 123 | 275 | 3321 | 0 | 0 | 1927 | 287 | 582 | 1472 | 0 | 0 | 29 | 37 | 0 | 0 | 42 | 67 | 29 | 90 |
| SRR10072387 | 57 | 0 | 73 | 0 | 0 | 84 | 787 | 153 | 14 | 17 | 63 | 13 | 232 | 34 | 6225 | 0 | 39 | 66 | 59 | 659 | 1467 | 0 | 0 | 1328 | 299 | 462 | 1324 | 0 | 0 | 27 | 24 | 0 | 0 | 108 | 21 | 12 | 44 |
| SRR10072388 | 54 | 0 | 66 | 0 | 0 | 14 | 777 | 20 | 12 | 13 | 53 | 0 | 27 | 28 | 5746 | 0 | 34 | 65 | 62 | 692 | 1509 | 0 | 0 | 185 | 40 | 66 | 172 | 0 | 0 | 21 | 35 | 0 | 0 | 12 | 27 | 15 | 39 |
| SRR10072389 | 22 | 0 | 181 | 0 | 0 | 64 | 259 | 0 | 0 | 0 | 20 | 0 | 18 | 0 | 75067 | 0 | 11 | 22 | 26 | 272 | 22281 | 0 | 0 | 12 | 22 | 26 | 46 | 0 | 0 | 0 | 32 | 0 | 0 | 34 | 38 | 19 | 65 |
| SRR10072390 | 24 | 0 | 176 | 0 | 0 | 51 | 112 | 0 | 0 | 0 | 0 | 0 | 175 | 0 | 73190 | 0 | 0 | 11 | 74 | 198 | 21362 | 0 | 0 | 185 | 304 | 362 | 523 | 0 | 0 | 0 | 39 | 0 | 24 | 36 | 372 | 175 | 821 |
| SRR10072391 | 39 | 0 | 114 | 0 | 0 | 10 | 107 | 17 | 0 | 0 | 19 | 18 | 70 | 10 | 34138 | 0 | 36 | 31 | 33 | 296 | 6066 | 0 | 0 | 262 | 41 | 48 | 147 | 0 | 0 | 0 | 23 | 0 | 0 | 16 | 59 | 36 | 12 |
| SRR10072392 | 44 | 0 | 117 | 0 | 0 | 64 | 161 | 11 | 0 | 0 | 16 | 18 | 511 | 0 | 33675 | 0 | 40 | 59 | 0 | 194 | 6346 | 0 | 0 | 2003 | 330 | 395 | 1266 | 0 | 0 | 0 | 16 | 0 | 0 | 18 | 68 | 30 | 77 |
| SRR10072393 | 28 | 0 | 86 | 0 | 0 | 133 | 450 | 48 | 25 | 11 | 28 | 16 | 169 | 12 | 12231 | 12 | 52 | 184 | 56 | 177 | 5906 | 0 | 0 | 1296 | 394 | 708 | 1582 | 11 | 0 | 32 | 50 | 0 | 0 | 69 | 44 | 16 | 50 |
| SRR10072394 | 31 | 0 | 77 | 0 | 0 | 135 | 482 | 47 | 16 | 0 | 39 | 12 | 161 | 18 | 11687 | 0 | 49 | 197 | 63 | 146 | 5924 | 0 | 0 | 1269 | 372 | 686 | 1576 | 0 | 0 | 35 | 39 | 0 | 0 | 60 | 51 | 28 | 72 |
| SRR10072395 | 24 | 0 | 125 | 0 | 0 | 129 | 659 | 0 | 14 | 20 | 48 | 23 | 381 | 0 | 328866 | 14 | 16 | 43 | 51 | 203 | 10451 | 0 | 0 | 395 | 231 | 347 | 14 | 0 | 0 | 0 | 34 | 0 | 0 | 0 | 225 | 109 | 12 |
| SRR10072396 | 23 | 0 | 129 | 0 | 0 | 94 | 587 | 0 | 13 | 34 | 20 | 22 | 302 | 13 | 324424 | 13 | 17 | 35 | 0 | 200 | 10621 | 0 | 0 | 315 | 194 | 476 | 617 | 0 | 0 | 0 | 20 | 0 | 27 | 0 | 235 | 141 | 456 |
| SRR10072397 | 44 | 0 | 115 | 0 | 0 | 125 | 807 | 44 | 15 | 0 | 30 | 26 | 206 | 15 | 15545 | 0 | 50 | 14 | 32 | 144 | 6111 | 0 | 0 | 555 | 334 | 896 | 1407 | 0 | 0 | 17 | 37 | 0 | 15 | 98 | 74 | 51 | 204 |
| SRR10072398 | 43 | 0 | 132 | 0 | 0 | 12 | 660 | 55 | 20 | 13 | 32 | 23 | 20 | 17 | 15464 | 0 | 59 | 23 | 36 | 223 | 5878 | 0 | 0 | 71 | 32 | 106 | 179 | 0 | 0 | 12 | 37 | 0 | 13 | 12 | 12 | 32 | 26 |
| SRR10072399 | 53 | 0 | 52 | 0 | 0 | 136 | 1443 | 30 | 21 | 13 | 40 | 17 | 286 | 16 | 10122 | 0 | 28 | 45 | 51 | 314 | 3611 | 0 | 0 | 1142 | 300 | 635 | 1581 | 0 | 0 | 18 | 48 | 0 | 0 | 70 | 32 | 0 | 39 |
| SRR10072400 | 52 | 0 | 49 | 0 | 0 | 121 | 1236 | 28 | 21 | 0 | 63 | 17 | 273 | 17 | 10066 | 0 | 26 | 28 | 22 | 337 | 3649 | 0 | 0 | 1183 | 297 | 628 | 1401 | 0 | 0 | 22 | 45 | 0 | 0 | 115 | 36 | 12 | 36 |
| SRR10077412 | 29 | 0 | 4053 | 0 | 0 | 0 | 0 | 0 | 19 | 0 | 0 | 712 | 0 | 0 | 19 | 0 | 12 | 0 | 0 | 76 | 191 | 0 | 0 | 14 | 0 | 222 | 188 | 0 | 0 | 0 | 68 | 0 | 0 | 0 | 608 | 53 | 319 |
| SRR10077413 | 30 | 0 | 4039 | 0 | 0 | 0 | 0 | 0 | 35 | 0 | 0 | 718 | 0 | 0 | 39 | 0 | 12 | 0 | 0 | 156 | 234 | 0 | 0 | 15 | 0 | 192 | 183 | 0 | 0 | 0 | 113 | 0 | 0 | 0 | 30 | 79 | 17 |
| SRR10077414 | 37 | 0 | 3977 | 0 | 0 | 0 | 0 | 0 | 32 | 0 | 0 | 696 | 0 | 0 | 81 | 0 | 15 | 0 | 0 | 121 | 121 | 0 | 0 | 11 | 0 | 248 | 174 | 0 | 0 | 0 | 90 | 0 | 0 | 0 | 706 | 90 | 395 |
| SRR10077415 | 31 | 0 | 4114 | 0 | 0 | 20 | 0 | 0 | 25 | 0 | 0 | 716 | 0 | 0 | 20 | 0 | 15 | 0 | 0 | 137 | 118 | 0 | 0 | 0 | 0 | 229 | 189 | 0 | 0 | 0 | 104 | 0 | 0 | 0 | 776 | 60 | 303 |
| SRR10077416 | 26 | 0 | 3045 | 0 | 0 | 0 | 0 | 0 | 0 | 0 | 0 | 591 | 0 | 0 | 109 | 0 | 14 | 0 | 0 | 22 | 88 | 0 | 0 | 0 | 0 | 39 | 43 | 0 | 0 | 0 | 58 | 0 | 0 | 0 | 207 | 14 | 106 |
| SRR10077417 | 18 | 0 | 3067 | 0 | 0 | 0 | 0 | 0 | 0 | 0 | 0 | 577 | 0 | 0 | 110 | 0 | 13 | 0 | 0 | 66 | 0 | 0 | 0 | 0 | 0 | 20 | 39 | 0 | 0 | 0 | 68 | 0 | 0 | 0 | 205 | 0 | 88 |
| SRR10077418 | 21 | 0 | 3012 | 0 | 0 | 0 | 0 | 0 | 0 | 0 | 0 | 580 | 0 | 0 | 96 | 0 | 15 | 0 | 0 | 72 | 48 | 0 | 0 | 0 | 0 | 58 | 79 | 0 | 0 | 0 | 63 | 0 | 0 | 0 | 189 | 21 | 58 |
| SRR10077419 | 22 | 0 | 3028 | 0 | 0 | 0 | 0 | 0 | 0 | 0 | 0 | 580 | 0 | 0 | 111 | 0 | 13 | 0 | 0 | 22 | 67 | 0 | 0 | 0 | 0 | 25 | 74 | 15 | 0 | 0 | 79 | 0 | 0 | 0 | 168 | 25 | 49 |
| SRR10077420 | 35 | 0 | 3393 | 0 | 0 | 0 | 0 | 0 | 0 | 0 | 0 | 830 | 0 | 0 | 101 | 0 | 13 | 0 | 0 | 34 | 84 | 0 | 0 | 0 | 0 | 52 | 82 | 0 | 0 | 0 | 67 | 0 | 0 | 0 | 248 | 17 | 99 |
| SRR10077421 | 32 | 0 | 3369 | 0 | 0 | 0 | 0 | 0 | 0 | 0 | 0 | 838 | 0 | 0 | 86 | 0 | 14 | 0 | 0 | 121 | 138 | 0 | 0 | 0 | 0 | 39 | 72 | 0 | 0 | 0 | 60 | 0 | 0 | 0 | 14 | 21 | 54 |
| SRR10077422 | 32 | 0 | 3296 | 0 | 0 | 0 | 0 | 0 | 0 | 0 | 0 | 825 | 0 | 0 | 18 | 0 | 16 | 0 | 0 | 110 | 110 | 0 | 0 | 0 | 0 | 48 | 70 | 0 | 0 | 0 | 32 | 0 | 0 | 0 | 166 | 19 | 61 |
| SRR10077423 | 36 | 0 | 3427 | 0 | 0 | 0 | 0 | 0 | 0 | 0 | 0 | 837 | 0 | 0 | 70 | 0 | 17 | 0 | 0 | 122 | 52 | 0 | 0 | 0 | 0 | 60 | 57 | 0 | 0 | 0 | 78 | 0 | 0 | 0 | 11 | 24 | 78 |
| SRR10077424 | 28 | 0 | 2862 | 0 | 0 | 0 | 0 | 0 | 0 | 0 | 0 | 696 | 0 | 0 | 56 | 0 | 15 | 0 | 0 | 40 | 24 | 0 | 0 | 0 | 11 | 90 | 195 | 32 | 0 | 0 | 162 | 0 | 0 | 0 | 350 | 51 | 191 |
| SRR10077425 | 24 | 0 | 2896 | 0 | 0 | 0 | 0 | 0 | 0 | 0 | 0 | 709 | 0 | 0 | 65 | 0 | 17 | 16 | 0 | 65 | 41 | 0 | 0 | 0 | 0 | 92 | 125 | 0 | 0 | 0 | 158 | 0 | 11 | 0 | 468 | 44 | 180 |
| SRR10077426 | 28 | 0 | 2880 | 0 | 0 | 0 | 0 | 0 | 0 | 0 | 0 | 697 | 0 | 0 | 26 | 0 | 19 | 0 | 0 | 61 | 44 | 0 | 0 | 0 | 0 | 98 | 169 | 20 | 0 | 0 | 106 | 0 | 0 | 0 | 448 | 59 | 161 |
| SRR10077427 | 30 | 0 | 2864 | 0 | 0 | 0 | 0 | 0 | 0 | 0 | 0 | 686 | 0 | 15 | 75 | 0 | 16 | 0 | 0 | 100 | 33 | 0 | 0 | 0 | 0 | 122 | 188 | 0 | 0 | 0 | 181 | 0 | 0 | 0 | 14 | 48 | 144 |

### Supp Table S8.2.Columns_AM_to_BW

| | | | | | | | | | | | | | | | | | | | | | | | | | | | | | | | | | | | | | |
|---|---|---|---|---|---|---|---|---|---|---|---|---|---|---|---|---|---|---|---|---|---|---|---|---|---|---|---|---|---|---|---|---|---|---|---|---|---|
| SRR10077428 | 24 | 0 | 3141 | 0 | 0 | 28 | 0 | 0 | 21 | 0 | 0 | 693 | 0 | 0 | 12 | 0 | 16 | 0 | 0 | 128 | 360 | 0 | 0 | 0 | 0 | 26 | 20 | 28 | 0 | 0 | 92 | 0 | 0 | 0 | 33 | 177 | 21 |
| SRR10077429 | 29 | 0 | 3105 | 0 | 0 | 29 | 0 | 0 | 36 | 0 | 0 | 644 | 15 | 0 | 24 | 0 | 16 | 0 | 0 | 95 | 343 | 0 | 0 | 0 | 0 | 847 | 598 | 0 | 0 | 0 | 139 | 0 | 0 | 0 | 825 | 153 | 803 |
| SRR10077430 | 30 | 0 | 3125 | 0 | 0 | 39 | 0 | 0 | 31 | 0 | 0 | 682 | 0 | 0 | 25 | 0 | 15 | 0 | 0 | 137 | 287 | 0 | 0 | 0 | 0 | 774 | 555 | 0 | 0 | 0 | 141 | 0 | 0 | 16 | 828 | 141 | 774 |
| SRR10077431 | 24 | 0 | 3133 | 0 | 0 | 22 | 0 | 0 | 29 | 0 | 0 | 699 | 0 | 0 | 12 | 0 | 16 | 0 | 0 | 118 | 294 | 0 | 0 | 0 | 0 | 25 | 23 | 0 | 0 | 0 | 124 | 0 | 0 | 0 | 27 | 87 | 21 |
| SRR10077432 | 24 | 0 | 3204 | 0 | 14 | 0 | 0 | 0 | 0 | 0 | 0 | 696 | 0 | 14 | 43 | 0 | 16 | 0 | 0 | 93 | 93 | 0 | 0 | 0 | 14 | 218 | 253 | 0 | 0 | 0 | 189 | 0 | 0 | 0 | 561 | 81 | 274 |
| SRR10077433 | 27 | 0 | 3224 | 0 | 0 | 15 | 0 | 0 | 0 | 0 | 0 | 714 | 0 | 11 | 26 | 0 | 18 | 0 | 0 | 96 | 77 | 0 | 0 | 0 | 0 | 196 | 236 | 0 | 0 | 0 | 174 | 0 | 0 | 0 | 21 | 105 | 294 |
| SRR10077434 | 25 | 0 | 3162 | 0 | 0 | 23 | 0 | 0 | 19 | 0 | 0 | 684 | 12 | 0 | 21 | 0 | 15 | 0 | 0 | 84 | 91 | 0 | 0 | 0 | 12 | 281 | 200 | 19 | 0 | 0 | 173 | 0 | 0 | 0 | 21 | 104 | 270 |
| SRR10077435 | 26 | 0 | 3212 | 0 | 0 | 15 | 0 | 0 | 0 | 0 | 0 | 697 | 0 | 11 | 79 | 0 | 16 | 18 | 0 | 131 | 59 | 0 | 0 | 0 | 15 | 204 | 219 | 18 | 0 | 0 | 157 | 0 | 0 | 0 | 712 | 91 | 318 |
| SRR10116271 | 30 | 0 | 662 | 21 | 0 | 18 | 236 | 0 | 26 | 26 | 134 | 76 | 0 | 34 | 19 | 0 | 0 | 37 | 12 | 31 | 50 | 0 | 0 | 0 | 42 | 192 | 29 | 0 | 15 | 0 | 208 | 0 | 0 | 13 | 260 | 0 | 37 |
| SRR10116272 | 55 | 0 | 222 | 16 | 0 | 0 | 13 | 0 | 24 | 37 | 73 | 105 | 0 | 65 | 175 | 0 | 0 | 451 | 0 | 22 | 175 | 0 | 0 | 0 | 113 | 99 | 28 | 0 | 97 | 0 | 171 | 0 | 11 | 184 | 83 | 0 | 40 |
| SRR10116273 | 26 | 0 | 388 | 29 | 0 | 14 | 44 | 0 | 0 | 79 | 42 | 403 | 0 | 44 | 31 | 10 | 0 | 157 | 0 | 13 | 149 | 0 | 32 | 0 | 23 | 13 | 20 | 0 | 50 | 0 | 15 | 0 | 0 | 39 | 48 | 10 | 72 |
| SRR10116274 | 32 | 0 | 157 | 11 | 0 | 11 | 20 | 0 | 17 | 11 | 20 | 143 | 0 | 264 | 259 | 0 | 0 | 268 | 0 | 60 | 129 | 0 | 0 | 0 | 71 | 139 | 40 | 20 | 0 | 0 | 173 | 0 | 0 | 51 | 201 | 26 | 60 |
| SRR10116275 | 85 | 0 | 1799 | 35 | 14 | 11 | 33 | 0 | 42 | 0 | 68 | 164 | 0 | 73 | 45 | 15 | 0 | 34 | 0 | 20 | 106 | 0 | 0 | 0 | 69 | 180 | 33 | 40 | 133 | 0 | 321 | 0 | 0 | 25 | 204 | 17 | 73 |
| SRR10119011 | 259 | 0 | 419 | 0 | 0 | 0 | 19 | 18 | 0 | 0 | 0 | 26 | 0 | 26 | 214 | 0 | 0 | 0 | 0 | 466 | 9245 | 0 | 0 | 39 | 0 | 23 | 33 | 0 | 0 | 0 | 0 | 0 | 0 | 0 | 0 | 0 | 13 |
| SRR10119012 | 44 | 52 | 17 | 18 | 158 | 0 | 0 | 0 | 0 | 0 | 0 | 0 | 0 | 0 | 0 | 55 | 0 | 0 | 0 | 0 | 8611 | 30 | 0 | 0 | 0 | 0 | 143 | 0 | 0 | 0 | 0 | 0 | 0 | 0 | 0 | 0 | 0 |
| SRR10119014 | 0 | 0 | 0 | 0 | 0 | 0 | 0 | 16 | 0 | 0 | 0 | 0 | 0 | 0 | 0 | 0 | 0 | 0 | 0 | 0 | 0 | 0 | 0 | 0 | 0 | 0 | 35 | 0 | 0 | 0 | 0 | 0 | 0 | 0 | 0 | 0 | 0 |
| SRR10119022 | 316 | 0 | 625 | 0 | 0 | 0 | 86 | 132 | 0 | 0 | 0 | 0 | 0 | 0 | 84 | 0 | 0 | 0 | 0 | 304 | 1048 | 0 | 0 | 11 | 0 | 52 | 155 | 0 | 0 | 0 | 0 | 0 | 10 | 0 | 0 | 0 | 0 |
| SRR10119033 | 166 | 0 | 62 | 764 | 195 | 0 | 0 | 0 | 0 | 0 | 0 | 28 | 0 | 0 | 0 | 0 | 0 | 0 | 0 | 0 | 2337 | 0 | 0 | 0 | 0 | 0 | 0 | 0 | 0 | 0 | 0 | 0 | 0 | 0 | 0 | 0 | 0 |
| SRR10119034 | 12 | 0 | 22 | 2254 | 376 | 0 | 0 | 0 | 0 | 0 | 0 | 0 | 0 | 0 | 0 | 0 | 0 | 0 | 0 | 0 | 3540 | 0 | 0 | 0 | 0 | 0 | 139 | 0 | 0 | 0 | 0 | 0 | 0 | 0 | 0 | 0 | 0 |
| SRR10119035 | 269 | 0 | 771 | 0 | 0 | 0 | 388 | 53 | 0 | 0 | 0 | 23 | 0 | 0 | 0 | 0 | 0 | 0 | 51 | 1045 | 1744 | 0 | 0 | 12 | 0 | 67 | 89 | 0 | 0 | 0 | 0 | 0 | 0 | 0 | 0 | 0 | 0 |
| SRR10119037 | 55 | 0 | 37 | 1913 | 638 | 0 | 0 | 0 | 0 | 0 | 0 | 15 | 0 | 0 | 0 | 0 | 0 | 0 | 0 | 0 | 0 | 98 | 0 | 0 | 0 | 0 | 0 | 0 | 0 | 0 | 0 | 65 | 0 | 0 | 0 | 0 | 0 |
| SRR10119040 | 31 | 0 | 27 | 25 | 535 | 0 | 0 | 52 | 0 | 0 | 0 | 24 | 0 | 0 | 0 | 0 | 0 | 0 | 0 | 0 | 9543 | 120 | 0 | 0 | 0 | 0 | 92 | 114 | 123 | 0 | 0 | 0 | 0 | 0 | 0 | 0 | 0 |
| SRR10119044 | 27 | 0 | 13 | 15 | 58 | 0 | 0 | 0 | 0 | 0 | 0 | 15 | 0 | 0 | 0 | 0 | 0 | 0 | 0 | 1575 | 15308 | 0 | 0 | 0 | 0 | 0 | 0 | 0 | 0 | 27 | 0 | 0 | 0 | 0 | 0 | 0 | 0 |
| SRR10119046 | 293 | 13 | 722 | 0 | 0 | 0 | 293 | 18 | 0 | 0 | 0 | 15 | 0 | 0 | 35 | 0 | 0 | 0 | 18 | 928 | 1892 | 0 | 0 | 20 | 0 | 16 | 29 | 0 | 0 | 0 | 0 | 0 | 0 | 0 | 0 | 0 | 0 |
| SRR10119048 | 151 | 0 | 33 | 999 | 49 | 0 | 0 | 80 | 0 | 0 | 0 | 46 | 0 | 0 | 0 | 0 | 0 | 0 | 0 | 377 | 3093 | 0 | 0 | 0 | 0 | 0 | 217 | 0 | 0 | 0 | 0 | 0 | 0 | 0 | 0 | 0 | 0 |
| SRR10119051 | 64 | 55 | 36 | 22 | 455 | 0 | 0 | 0 | 0 | 0 | 0 | 0 | 0 | 0 | 0 | 0 | 0 | 0 | 0 | 0 | 1511 | 18 | 0 | 0 | 0 | 0 | 111 | 0 | 0 | 0 | 0 | 0 | 0 | 0 | 0 | 0 | 0 |
| SRR10119052 | 26 | 0 | 20 | 19 | 276 | 0 | 0 | 0 | 0 | 0 | 0 | 14 | 0 | 0 | 0 | 0 | 0 | 0 | 0 | 876 | 3004 | 11 | 0 | 0 | 0 | 0 | 322 | 0 | 0 | 0 | 0 | 0 | 0 | 0 | 0 | 0 | 0 |
| SRR10119056 | 44 | 90 | 0 | 35 | 415 | 0 | 0 | 0 | 0 | 0 | 0 | 0 | 0 | 50 | 0 | 0 | 0 | 0 | 0 | 0 | 2582 | 0 | 0 | 0 | 0 | 0 | 61 | 0 | 0 | 0 | 0 | 0 | 0 | 0 | 0 | 0 | 0 |
| SRR10119057 | 390 | 0 | 643 | 0 | 0 | 0 | 0 | 12 | 0 | 0 | 0 | 0 | 0 | 0 | 12 | 0 | 0 | 0 | 0 | 1389 | 32111 | 0 | 0 | 39 | 0 | 21 | 13 | 0 | 0 | 0 | 0 | 0 | 0 | 0 | 0 | 0 | 0 |
| SRR10119058 | 383 | 0 | 929 | 0 | 0 | 0 | 28 | 10 | 0 | 0 | 0 | 0 | 0 | 0 | 0 | 0 | 0 | 0 | 14 | 1514 | 31543 | 0 | 0 | 119 | 0 | 52 | 40 | 0 | 0 | 0 | 0 | 0 | 0 | 0 | 0 | 0 | 0 |
| SRR10119059 | 314 | 15 | 429 | 0 | 11 | 18 | 64 | 22 | 0 | 0 | 0 | 15 | 0 | 0 | 0 | 0 | 0 | 0 | 0 | 180 | 865 | 0 | 0 | 0 | 18 | 29 | 43 | 11 | 0 | 0 | 0 | 0 | 30 | 0 | 0 | 11 | 22 |
| SRR10119063 | 16 | 0 | 0 | 0 | 0 | 0 | 392 | 514 | 0 | 0 | 0 | 0 | 0 | 25 | 0 | 0 | 0 | 0 | 0 | 0 | 0 | 0 | 0 | 0 | 0 | 0 | 0 | 0 | 0 | 20 | 0 | 0 | 0 | 28 | 0 | 0 | 0 |
| SRR10119066 | 307 | 0 | 1551 | 0 | 0 | 0 | 41 | 62 | 0 | 0 | 0 | 0 | 0 | 0 | 43 | 0 | 0 | 0 | 0 | 566 | 14036 | 0 | 0 | 105 | 0 | 80 | 40 | 0 | 0 | 0 | 0 | 0 | 0 | 0 | 0 | 0 | 0 |
| SRR10119067 | 280 | 0 | 1383 | 0 | 0 | 0 | 48 | 27 | 0 | 0 | 0 | 0 | 0 | 0 | 49 | 0 | 0 | 0 | 16 | 681 | 17251 | 0 | 0 | 48 | 0 | 23 | 12 | 0 | 0 | 0 | 0 | 0 | 0 | 0 | 0 | 0 | 0 |
| SRR10119068 | 192 | 0 | 789 | 0 | 0 | 0 | 150 | 12 | 0 | 0 | 0 | 0 | 0 | 0 | 0 | 0 | 0 | 0 | 0 | 561 | 3584 | 0 | 0 | 20 | 0 | 19 | 65 | 0 | 0 | 0 | 0 | 0 | 0 | 0 | 0 | 0 | 0 |
| SRR10119069 | 259 | 0 | 915 | 0 | 0 | 0 | 82 | 18 | 0 | 0 | 0 | 0 | 0 | 0 | 0 | 0 | 0 | 0 | 0 | 612 | 3034 | 0 | 0 | 13 | 0 | 15 | 50 | 0 | 0 | 0 | 0 | 0 | 0 | 0 | 0 | 0 | 0 |
| SRR10119070 | 275 | 0 | 329 | 0 | 0 | 0 | 0 | 17 | 0 | 0 | 0 | 39 | 0 | 0 | 83 | 0 | 0 | 0 | 54 | 540 | 16462 | 0 | 0 | 20 | 0 | 19 | 20 | 0 | 0 | 0 | 0 | 0 | 0 | 0 | 0 | 0 | 0 |
| SRR10119071 | 159 | 0 | 1357 | 0 | 0 | 0 | 0 | 0 | 0 | 0 | 0 | 0 | 0 | 0 | 0 | 0 | 0 | 0 | 0 | 920 | 29939 | 0 | 0 | 292 | 0 | 35 | 19 | 0 | 0 | 0 | 0 | 0 | 0 | 0 | 0 | 0 | 0 |
| SRR10119072 | 350 | 0 | 1085 | 0 | 0 | 0 | 0 | 0 | 0 | 0 | 0 | 0 | 0 | 0 | 0 | 0 | 0 | 0 | 0 | 669 | 26033 | 0 | 0 | 140 | 0 | 0 | 17 | 0 | 0 | 0 | 0 | 0 | 0 | 0 | 0 | 0 | 0 |
| SRR10119073 | 215 | 11 | 451 | 0 | 0 | 0 | 0 | 0 | 0 | 0 | 0 | 0 | 0 | 0 | 54 | 0 | 0 | 0 | 0 | 611 | 22401 | 0 | 0 | 170 | 0 | 0 | 11 | 0 | 0 | 0 | 0 | 0 | 0 | 0 | 0 | 0 | 0 |
| SRR10119074 | 349 | 0 | 331 | 0 | 0 | 0 | 15 | 0 | 0 | 0 | 0 | 0 | 0 | 0 | 0 | 0 | 0 | 0 | 0 | 635 | 25267 | 0 | 0 | 111 | 0 | 0 | 11 | 0 | 0 | 0 | 0 | 0 | 0 | 0 | 0 | 0 | 0 |
| SRR10119075 | 213 | 0 | 578 | 0 | 0 | 0 | 34 | 13 | 0 | 0 | 0 | 19 | 0 | 0 | 15 | 0 | 0 | 15 | 0 | 619 | 4909 | 0 | 0 | 17 | 0 | 13 | 19 | 0 | 0 | 0 | 0 | 0 | 0 | 0 | 10 | 0 | 0 |
| SRR10119076 | 288 | 10 | 470 | 0 | 0 | 0 | 32 | 20 | 0 | 0 | 0 | 15 | 0 | 0 | 123 | 0 | 0 | 0 | 36 | 624 | 4352 | 0 | 0 | 20 | 0 | 10 | 23 | 0 | 0 | 0 | 0 | 0 | 0 | 0 | 0 | 0 | 0 |
| SRR10119077 | 116 | 0 | 712 | 0 | 0 | 0 | 67 | 16 | 0 | 0 | 0 | 11 | 0 | 0 | 449 | 0 | 0 | 0 | 31 | 1316 | 13358 | 0 | 0 | 110 | 0 | 76 | 96 | 0 | 0 | 0 | 0 | 0 | 0 | 0 | 0 | 0 | 0 |
| SRR10119078 | 135 | 0 | 830 | 0 | 0 | 0 | 93 | 21 | 0 | 0 | 0 | 0 | 0 | 0 | 529 | 0 | 0 | 0 | 25 | 1448 | 18911 | 0 | 0 | 29 | 0 | 32 | 30 | 0 | 0 | 0 | 0 | 0 | 0 | 0 | 0 | 0 | 0 |
| SRR10119079 | 153 | 17 | 416 | 0 | 0 | 0 | 23 | 109 | 0 | 0 | 0 | 0 | 0 | 0 | 39 | 0 | 0 | 0 | 0 | 273 | 2151 | 0 | 0 | 11 | 0 | 22 | 63 | 0 | 0 | 0 | 0 | 0 | 0 | 0 | 16 | 0 | 0 |
| SRR10119080 | 160 | 0 | 469 | 12 | 0 | 0 | 27 | 30 | 0 | 0 | 0 | 0 | 0 | 0 | 54 | 0 | 0 | 0 | 0 | 284 | 2715 | 0 | 0 | 0 | 0 | 34 | 22 | 0 | 0 | 0 | 0 | 0 | 0 | 0 | 0 | 0 | 12 |
| SRR10119081 | 132 | 0 | 564 | 0 | 0 | 0 | 0 | 0 | 0 | 0 | 0 | 14 | 0 | 0 | 25 | 0 | 0 | 0 | 0 | 970 | 23347 | 0 | 0 | 340 | 0 | 11 | 19 | 0 | 0 | 0 | 0 | 0 | 0 | 0 | 0 | 0 | 0 |
| SRR10119082 | 206 | 0 | 384 | 0 | 0 | 0 | 17 | 0 | 0 | 0 | 0 | 28 | 0 | 0 | 62 | 0 | 0 | 0 | 15 | 658 | 19366 | 0 | 0 | 106 | 0 | 0 | 17 | 0 | 0 | 0 | 0 | 0 | 0 | 0 | 0 | 0 | 0 |
| SRR1014666 | 59 | 0 | 88 | 0 | 0 | 0 | 0 | 0 | 0 | 0 | 0 | 40 | 0 | 34 | 44 | 0 | 19 | 29 | 0 | 0 | 0 | 0 | 0 | 0 | 20 | 0 | 0 | 31 | 36 | 0 | 22 | 0 | 0 | 0 | 20 | 17 | 0 |
| SRR1014667 | 46 | 17 | 62 | 0 | 0 | 0 | 12 | 0 | 0 | 0 | 0 | 26 | 0 | 14 | 18 | 0 | 21 | 134 | 0 | 0 | 29 | 0 | 0 | 0 | 14 | 0 | 0 | 19 | 10 | 0 | 0 | 0 | 0 | 13 | 44 | 10 | 11 |
| SRR1014669 | 84 | 0 | 111 | 0 | 0 | 0 | 11 | 0 | 0 | 0 | 0 | 37 | 0 | 37 | 0 | 0 | 22 | 0 | 11 | 11 | 0 | 0 | 0 | 0 | 34 | 0 | 0 | 19 | 38 | 0 | 0 | 0 | 0 | 110 | 12 | 0 | 0 |
| SRR10197184 | 14 | 0 | 157 | 0 | 0 | 0 | 0 | 10 | 12 | 0 | 45 | 832 | 467 | 20 | 1081 | 11 | 81 | 194 | 0 | 0 | 48 | 0 | 0 | 0 | 721 | 17 | 0 | 0 | 0 | 0 | 183 | 0 | 76 | 1581 | 52 | 62 | 97 |
| SRR10197185 | 33 | 0 | 0 | 0 | 0 | 0 | 0 | 91 | 0 | 0 | 26 | 108 | 92 | 0 | 6697 | 0 | 0 | 0 | 0 | 0 | 148 | 0 | 0 | 0 | 0 | 0 | 0 | 0 | 0 | 64 | 47 | 0 | 24 | 787 | 0 | 15 | 94 |
| SRR10197186 | 17 | 0 | 33 | 0 | 0 | 0 | 51 | 13 | 0 | 0 | 19 | 526 | 60 | 20 | 1742 | 22 | 10 | 38 | 0 | 0 | 174 | 0 | 0 | 0 | 114 | 0 | 0 | 0 | 0 | 0 | 35 | 0 | 14 | 314 | 19 | 17 | 41 |
| SRR10197187 | 16 | 0 | 42 | 0 | 0 | 0 | 19 | 13 | 0 | 14 | 138 | 900 | 80 | 347 | 1417 | 0 | 14 | 16 | 0 | 0 | 34 | 0 | 0 | 0 | 222 | 0 | 0 | 30 | 0 | 0 | 63 | 0 | 42 | 735 | 0 | 16 | 37 |
| SRR10197188 | 16 | 0 | 117 | 0 | 0 | 0 | 0 | 12 | 10 | 23 | 225 | 1050 | 150 | 289 | 797 | 0 | 46 | 55 | 0 | 0 | 19 | 0 | 0 | 0 | 290 | 0 | 0 | 39 | 0 | 0 | 73 | 0 | 78 | 939 | 10 | 13 | 28 |
| SRR10197189 | 10 | 0 | 17 | 0 | 0 | 0 | 27 | 20 | 0 | 28 | 282 | 547 | 82 | 204 | 2345 | 14 | 0 | 10 | 0 | 0 | 96 | 0 | 0 | 0 | 45 | 0 | 0 | 19 | 0 | 13 | 50 | 0 | 47 | 692 | 0 | 17 | 51 |
| SRR10197190 | 16 | 0 | 131 | 0 | 0 | 0 | 0 | 0 | 0 | 25 | 133 | 519 | 19 | 105 | 573 | 0 | 34 | 36 | 0 | 0 | 47 | 0 | 0 | 0 | 60 | 0 | 0 | 13 | 12 | 0 | 14 | 0 | 44 | 281 | 0 | 0 | 11 |
| SRR10197191 | 21 | 0 | 107 | 0 | 0 | 0 | 33 | 0 | 0 | 15 | 149 | 684 | 24 | 152 | 511 | 0 | 32 | 18 | 0 | 0 | 66 | 0 | 0 | 0 | 117 | 0 | 0 | 18 | 15 | 0 | 14 | 0 | 29 | 390 | 0 | 0 | 14 |
| SRR10197192 | 22 | 0 | 107 | 0 | 0 | 0 | 0 | 0 | 0 | 16 | 147 | 634 | 49 | 117 | 792 | 0 | 28 | 18 | 0 | 0 | 183 | 0 | 0 | 0 | 48 | 0 | 0 | 14 | 21 | 0 | 11 | 0 | 23 | 583 | 0 | 0 | 15 |
| SRR10225092 | 525 | 0 | 4597 | 0 | 0 | 88 | 499 | 263 | 0 | 0 | 0 | 263 | 0 | 0 | 0 | 0 | 22 | 0 | 166 | 12146 | 19800 | 0 | 0 | 0 | 88 | 1314 | 1664 | 0 | 0 | 0 | 526 | 0 | 0 | 0 | 876 | 263 | 1139 |
| SRR10225093 | 21 | 0 | 6492 | 0 | 0 | 0 | 24 | 3863 | 14 | 0 | 0 | 681 | 0 | 0 | 0 | 0 | 42 | 0 | 12 | 59 | 83 | 0 | 0 | 0 | 14 | 64 | 423 | 0 | 37 | 294 | 0 | 0 | 0 | 165 | 0 | 0 | 0 |
| SRR10225094 | 326 | 0 | 16230 | 0 | 0 | 22 | 303 | 143 | 11 | 0 | 0 | 199 | 0 | 0 | 0 | 0 | 0 | 0 | 38 | 2876 | 10143 | 0 | 0 | 0 | 0 | 179 | 255 | 0 | 0 | 11 | 61 | 22 | 0 | 0 | 40 | 11 | 40 |
| SRR10225096 | 563 | 0 | 9006 | 0 | 0 | 10 | 138 | 20 | 0 | 0 | 0 | 281 | 0 | 0 | 0 | 0 | 141 | 0 | 138 | 2492 | 646 | 0 | 0 | 0 | 0 | 354 | 269 | 0 | 15 | 0 | 48 | 0 | 0 | 0 | 12 | 0 | 10 |
| SRR10282881 | 148 | 25 | 1955 | 0 | 0 | 0 | 0 | 18 | 16 | 0 | 0 | 490 | 0 | 17 | 0 | 0 | 0 | 0 | 0 | 0 | 0 | 0 | 0 | 0 | 13 | 87 | 10 | 14 | 0 | 0 | 13 | 0 | 0 | 14 | 0 | 0 | 0 |
| SRR10282882 | 165 | 25 | 2219 | 0 | 0 | 0 | 0 | 12 | 13 | 0 | 0 | 517 | 0 | 20 | 0 | 0 | 0 | 0 | 0 | 0 | 0 | 12 | 0 | 0 | 24 | 74 | 0 | 162 | 0 | 0 | 13 | 0 | 0 | 12 | 0 | 0 | 0 |
| SRR10282883 | 235 | 24 | 2402 | 0 | 0 | 0 | 16 | 0 | 0 | 0 | 0 | 588 | 0 | 12 | 0 | 0 | 0 | 0 | 0 | 0 | 51 | 0 | 0 | 0 | 0 | 82 | 48 | 79 | 0 | 0 | 0 | 0 | 0 | 0 | 0 | 0 | 0 |
| SRR10282884 | 228 | 15 | 2657 | 0 | 0 | 0 | 17 | 0 | 15 | 0 | 0 | 623 | 0 | 0 | 0 | 0 | 0 | 0 | 0 | 0 | 66 | 0 | 0 | 0 | 0 | 114 | 61 | 101 | 0 | 0 | 0 | 0 | 0 | 0 | 0 | 0 | 0 |
| SRR10282885 | 94 | 13 | 2274 | 11 | 0 | 0 | 12 | 0 | 33 | 0 | 0 | 623 | 0 | 19 | 0 | 0 | 0 | 0 | 0 | 0 | 46 | 0 | 0 | 0 | 19 | 15 | 20 | 15 | 0 | 0 | 17 | 0 | 0 | 23 | 0 | 0 | 0 |
| SRR10282886 | 137 | 0 | 2578 | 0 | 0 | 0 | 14 | 0 | 33 | 0 | 0 | 608 | 0 | 17 | 0 | 0 | 0 | 0 | 0 | 0 | 53 | 0 | 0 | 0 | 18 | 223 | 19 | 202 | 0 | 0 | 15 | 0 | 0 | 15 | 0 | 0 | 0 |
| SRR10282890 | 144 | 713 | 1320 | 0 | 0 | 0 | 60 | 30 | 19 | 0 | 0 | 556 | 0 | 287 | 32 | 0 | 0 | 0 | 0 | 0 | 126 | 0 | 0 | 0 | 0 | 139 | 11 | 124 | 0 | 0 | 0 | 0 | 0 | 0 | 15 | 0 | 15 |
| SRR10282891 | 135 | 679 | 1552 | 0 | 0 | 0 | 53 | 34 | 22 | 0 | 0 | 599 | 0 | 287 | 29 | 0 | 0 | 0 | 0 | 0 | 126 | 0 | 0 | 0 | 0 | 131 | 0 | 136 | 0 | 0 | 0 | 0 | 0 | 0 | 15 | 0 | 0 |
| SRR10282892 | 77 | 151 | 1393 | 0 | 0 | 0 | 0 | 51 | 18 | 0 | 0 | 473 | 0 | 12 | 10 | 0 | 0 | 0 | 0 | 0 | 0 | 0 | 0 | 0 | 0 | 12 | 0 | 20 | 0 | 0 | 0 | 0 | 0 | 11 | 28 | 0 | 14 |
| SRR10282893 | 98 | 1078 | 1576 | 0 | 0 | 0 | 0 | 35 | 14 | 0 | 0 | 469 | 0 | 111 | 20 | 0 | 0 | 0 | 0 | 0 | 0 | 0 | 0 | 0 | 0 | 112 | 14 | 165 | 0 | 0 | 0 | 0 | 10 | 13 | 28 | 0 | 12 |
| SRR10282894 | 16 | 121 | 411 | 0 | 0 | 0 | 0 | 0 | 0 | 0 | 0 | 169 | 0 | 25 | 0 | 0 | 0 | 0 | 0 | 0 | 28 | 0 | 0 | 0 | 0 | 33 | 0 | 59 | 0 | 0 | 0 | 0 | 0 | 0 | 11 | 0 | 14 |
| SRR10282895 | 19 | 111 | 463 | 0 | 0 | 0 | 0 | 0 | 0 | 0 | 0 | 200 | 0 | 27 | 0 | 0 | 0 | 0 | 0 | 0 | 26 | 0 | 0 | 0 | 0 | 34 | 0 | 71 | 0 | 0 | 0 | 0 | 0 | 0 | 0 | 0 | 12 |
| SRR10323931 | 74 | 0 | 106 | 0 | 0 | 61 | 0 | 0 | 63 | 66 | 12 | 838 | 155 | 0 | 114 | 0 | 477 | 10 | 0 | 0 | 20 | 0 | 0 | 15 | 168 | 42 | 13 | 13 | 0 | 0 | 194 | 15 | 0 | 15 | 18 | 83 | 198 |
| SRR10323932 | 53 | 10 | 24 | 0 | 0 | 67 | 16 | 0 | 142 | 79 | 553 | 696 | 108 | 17 | 67 | 12 | 693 | 0 | 0 | 0 | 27 | 0 | 0 | 0 | 116 | 792 | 14 | 0 | 43 | 0 | 78 | 20 | 0 | 332 | 442 | 1605 | 4219 |
| SRR10323933 | 46 | 0 | 35 | 0 | 12 | 38 | 19 | 0 | 23 | 14 | 75 | 866 | 125 | 0 | 0 | 0 | 757 | 0 | 0 | 11 | 102 | 0 | 0 | 10 | 78 | 452 | 28 | 0 | 18 | 0 | 69 | 17 | 0 | 19 | 26 | 135 | 290 |
| SRR10323939 | 78 | 0 | 220 | 0 | 0 | 34 | 0 | 0 | 89 | 74 | 458 | 2130 | 155 | 24 | 100 | 0 | 391 | 37 | 10 | 10 | 25 | 0 | 0 | 0 | 254 | 398 | 0 | 0 | 0 | 0 | 219 | 15 | 13 | 192 | 383 | 841 | 1683 |
| SRR10323940 | 83 | 14 | 168 | 0 | 0 | 186 | 19 | 0 | 369 | 194 | 1373 | 1365 | 106 | 30 | 50 | 0 | 692 | 12 | 0 | 0 | 131 | 0 | 0 | 17 | 161 | 1954 | 100 | 13 | 41 | 0 | 142 | 45 | 0 | 207 | 596 | 1215 | 3837 |
| SRR10323941 | 85 | 61 | 72 | 0 | 0 | 213 | 0 | 0 | 726 | 644 | 1411 | 859 | 101 | 25 | 0 | 0 | 459 | 0 | 14 | 0 | 64 | 0 | 0 | 0 | 111 | 1731 | 49 | 15 | 124 | 0 | 237 | 16 | 0 | 98 | 480 | 703 | 2063 |
| SRR10385125 | 13 | 0 | 643 | 0 | 0 | 94 | 14 | 0 | 60 | 56 | 106 | 1401 | 29 | 25 | 287 | 0 | 422 | 0 | 22 | 0 | 216 | 0 | 0 | 16 | 26 | 557 | 113 | 23 | 23 | 0 | 334 | 0 | 16 | 66 | 67 | 33 | 75 |
| SRR10385126 | 32 | 0 | 771 | 0 | 0 | 1868 | 0 | 0 | 885 | 24 | 75 | 1524 | 63 | 17 | 120 | 11 | 582 | 23 | 0 | 20 | 691 | 0 | 0 | 19 | 251 | 10580 | 319 | 16 | 0 | 0 | 2012 | 13 | 22 | 90 | 1565 | 647 | 1747 |
| SRR10416439 | 287 | 0 | 291 | 0 | 0 | 317 | 0 | 0 | 69 | 0 | 19 | 0 | 31 | 11 | 126 | 0 | 0 | 0 | 0 | 13 | 419 | 0 | 0 | 0 | 25 | 1951 | 39 | 0 | 0 | 0 | 88 | 0 | 0 | 58 | 995 | 56 | 798 |
| SRR10416440 | 218 | 0 | 406 | 0 | 0 | 39 | 0 | 10 | 17 | 0 | 0 | 0 | 16 | 10 | 31 | 0 | 0 | 0 | 0 | 0 | 211 | 0 | 0 | 0 | 16 | 265 | 28 | 0 | 0 | 25 | 34 | 0 | 0 | 16 | 142 | 17 | 113 |
| SRR10416441 | 199 | 0 | 139 | 0 | 0 | 67 | 0 | 0 | 16 | 0 | 14 | 0 | 33 | 0 | 114 | 0 | 0 | 23 | 0 | 11 | 228 | 0 | 0 | 14 | 11 | 397 | 14 | 0 | 0 | 0 | 18 | 0 | 0 | 24 | 170 | 30 | 114 |
| SRR10416442 | 143 | 0 | 658 | 0 | 0 | 888 | 0 | 0 | 201 | 0 | 23 | 0 | 57 | 0 | 83 | 0 | 39 | 20 | 0 | 34 | 179 | 0 | 0 | 14 | 131 | 6056 | 205 | 0 | 0 | 0 | 242 | 0 | 0 | 54 | 1051 | 311 | 1121 |
| SRR10416496 | 0 | 0 | 0 | 0 | 0 | 0 | 0 | 10 | 0 | 0 | 0 | 0 | 0 | 0 | 0 | 21 | 0 | 0 | 0 | 265 | 0 | 0 | 0 | 10 | 10 | 31 | 164 | 10 | 0 | 10 | 0 | 0 | 0 | 0 | 0 | 0 | 0 |
| SRR10416497 | 0 | 0 | 0 | 18 | 0 | 0 | 359 | 36 | 0 | 0 | 0 | 0 | 0 | 0 | 0 | 18 | 0 | 0 | 0 | 717 | 0 | 0 | 0 | 0 | 0 | 0 | 181 | 18 | 0 | 0 | 0 | 0 | 18 | 0 | 0 | 0 | 18 |
| SRR10416498 | 0 | 0 | 0 | 0 | 0 | 0 | 0 | 0 | 0 | 0 | 0 | 0 | 0 | 0 | 0 | 0 | 0 | 0 | 0 | 502 | 0 | 0 | 0 | 18 | 0 | 0 | 12 | 0 | 0 | 0 | 0 | 0 | 0 | 0 | 0 | 0 | 0 |
| SRR10416499 | 0 | 0 | 0 | 0 | 0 | 0 | 0 | 0 | 0 | 0 | 0 | 0 | 0 | 0 | 0 | 0 | 0 | 0 | 0 | 792 | 0 | 0 | 0 | 23 | 0 | 45 | 17 | 0 | 0 | 23 | 0 | 0 | 0 | 0 | 0 | 0 | 0 |
| SRR10416500 | 0 | 0 | 0 | 0 | 0 | 0 | 0 | 0 | 0 | 0 | 0 | 0 | 0 | 0 | 0 | 0 | 0 | 0 | 0 | 0 | 287 | 0 | 0 | 0 | 0 | 45 | 208 | 0 | 0 | 0 | 0 | 0 | 0 | 0 | 0 | 0 | 0 |
| SRR10416501 | 0 | 0 | 0 | 0 | 0 | 0 | 0 | 74 | 0 | 0 | 0 | 0 | 0 | 0 | 0 | 0 | 0 | 0 | 0 | 1250 | 313 | 0 | 0 | 45 | 0 | 45 | 23 | 0 | 0 | 0 | 0 | 0 | 0 | 0 | 0 | 0 | 15 |
| SRR10416502 | 0 | 0 | 0 | 0 | 0 | 20 | 0 | 0 | 0 | 0 | 0 | 0 | 0 | 0 | 0 | 0 | 0 | 0 | 0 | 0 | 0 | 0 | 0 | 60 | 0 | 20 | 516 | 0 | 0 | 0 | 0 | 0 | 20 | 0 | 0 | 0 | 0 |
| SRR10416503 | 0 | 0 | 0 | 0 | 0 | 0 | 0 | 0 | 0 | 0 | 0 | 0 | 0 | 0 | 0 | 0 | 0 | 0 | 0 | 832 | 0 | 0 | 0 | 44 | 0 | 22 | 15 | 0 | 0 | 0 | 0 | 0 | 0 | 22 | 22 | 0 | 0 |
| SRR10416504 | 0 | 0 | 0 | 0 | 0 | 0 | 0 | 0 | 0 | 0 | 0 | 0 | 0 | 0 | 0 | 0 | 0 | 0 | 0 | 0 | 0 | 0 | 0 | 0 | 21 | 42 | 148 | 0 | 0 | 21 | 0 | 0 | 0 | 0 | 0 | 0 | 0 |
| SRR10416505 | 0 | 0 | 0 | 0 | 0 | 0 | 157 | 26 | 0 | 0 | 0 | 0 | 0 | 0 | 0 | 0 | 0 | 0 | 0 | 1732 | 0 | 0 | 0 | 31 | 0 | 20 | 158 | 0 | 0 | 0 | 0 | 0 | 10 | 0 | 0 | 0 | 0 |
| SRR10416506 | 0 | 0 | 10 | 0 | 0 | 0 | 0 | 18 | 0 | 0 | 0 | 0 | 0 | 0 | 0 | 18 | 0 | 0 | 0 | 772 | 386 | 0 | 0 | 0 | 0 | 88 | 194 | 0 | 0 | 0 | 0 | 0 | 0 | 0 | 0 | 0 | 0 |
| SRR10416507 | 0 | 0 | 0 | 0 | 0 | 0 | 0 | 15 | 0 | 0 | 0 | 0 | 0 | 0 | 0 | 15 | 0 | 0 | 0 | 461 | 461 | 0 | 0 | 45 | 30 | 74 | 12 | 0 | 0 | 15 | 0 | 0 | 15 | 0 | 0 | 0 | 0 |
| SRR10416508 | 0 | 0 | 0 | 0 | 0 | 0 | 0 | 68 | 0 | 0 | 0 | 0 | 0 | 0 | 0 | 0 | 0 | 0 | 0 | 379 | 0 | 0 | 0 | 0 | 0 | 23 | 158 | 23 | 0 | 0 | 0 | 0 | 0 | 0 | 0 | 0 | 0 |
| SRR10416509 | 0 | 0 | 0 | 0 | 0 | 0 | 0 | 0 | 0 | 0 | 0 | 0 | 0 | 0 | 0 | 0 | 0 | 0 | 0 | 863 | 216 | 0 | 0 | 30 | 0 | 0 | 119 | 0 | 0 | 0 | 0 | 0 | 0 | 0 | 0 | 0 | 0 |
| SRR10416510 | 15 | 0 | 0 | 0 | 0 | 0 | 0 | 0 | 0 | 0 | 0 | 0 | 0 | 0 | 0 | 0 | 0 | 0 | 0 | 1178 | 0 | 0 | 0 | 13 | 0 | 63 | 12 | 0 | 0 | 0 | 0 | 0 | 31 | 31 | 0 | 0 | 0 |
| SRR10416513 | 13 | 0 | 0 | 0 | 0 | 0 | 0 | 71 | 0 | 0 | 0 | 0 | 0 | 0 | 0 | 0 | 0 | 0 | 0 | 133 | 0 | 0 | 0 | 39 | 0 | 0 | 175 | 0 | 0 | 18 | 0 | 0 | 0 | 0 | 0 | 0 | 0 |
| SRR10416514 | 17 | 0 | 0 | 0 | 0 | 0 | 0 | 70 | 0 | 0 | 0 | 0 | 0 | 0 | 513 | 0 | 0 | 0 | 0 | 513 | 0 | 0 | 0 | 47 | 0 | 94 | 258 | 0 | 0 | 0 | 0 | 0 | 0 | 0 | 23 | 0 | 0 |
| SRR10416515 | 0 | 0 | 0 | 0 | 0 | 0 | 0 | 0 | 0 | 0 | 0 | 0 | 0 | 0 | 0 | 0 | 0 | 0 | 0 | 2155 | 431 | 0 | 0 | 23 | 0 | 113 | 473 | 0 | 0 | 0 | 0 | 0 | 0 | 23 | 0 | 0 | 0 |
| SRR10416516 | 13 | 0 | 0 | 0 | 0 | 0 | 0 | 0 | 0 | 0 | 0 | 0 | 0 | 0 | 400 | 0 | 0 | 0 | 0 | 800 | 400 | 0 | 0 | 22 | 22 | 0 | 16 | 0 | 0 | 0 | 0 | 0 | 0 | 0 | 0 | 0 | 0 |
| SRR10416517 | 0 | 0 | 0 | 0 | 0 | 0 | 0 | 0 | 0 | 0 | 0 | 0 | 0 | 0 | 0 | 0 | 0 | 0 | 0 | 1433 | 0 | 15 | 0 | 15 | 0 | 15 | 24 | 0 | 0 | 0 | 0 | 0 | 0 | 0 | 0 | 0 | 0 |
| SRR10416518 | 19 | 0 | 0 | 0 | 0 | 0 | 0 | 28 | 0 | 0 | 0 | 0 | 0 | 0 | 0 | 0 | 0 | 0 | 0 | 0 | 645 | 0 | 0 | 111 | 0 | 83 | 18 | 0 | 0 | 0 | 0 | 0 | 0 | 28 | 0 | 0 | 0 |
| SRR10416519 | 14 | 16 | 0 | 0 | 16 | 0 | 0 | 0 | 0 | 0 | 0 | 0 | 0 | 0 | 0 | 16 | 0 | 0 | 0 | 1563 | 1042 | 0 | 0 | 290 | 0 | 306 | 709 | 0 | 0 | 16 | 0 | 16 | 0 | 16 | 0 | 0 | 16 |
| SRR10416520 | 12 | 0 | 0 | 0 | 0 | 39 | 0 | 0 | 0 | 0 | 0 | 0 | 0 | 0 | 0 | 0 | 0 | 0 | 0 | 3245 | 649 | 0 | 0 | 116 | 0 | 387 | 10 | 0 | 0 | 0 | 0 | 0 | 0 | 39 | 0 | 0 | 0 |
| SRR10416521 | 20 | 0 | 0 | 0 | 74 | 0 | 0 | 74 | 0 | 0 | 0 | 0 | 0 | 0 | 0 | 0 | 0 | 0 | 0 | 485 | 0 | 0 | 0 | 148 | 0 | 148 | 813 | 0 | 74 | 0 | 0 | 0 | 74 | 0 | 0 | 0 | 0 |
| SRR10416522 | 0 | 0 | 0 | 0 | 0 | 0 | 0 | 0 | 0 | 0 | 0 | 0 | 0 | 0 | 0 | 0 | 0 | 0 | 0 | 1581 | 0 | 0 | 0 | 0 | 0 | 0 | 135 | 0 | 0 | 0 | 0 | 0 | 0 | 0 | 0 | 0 | 0 |
| SRR10416523 | 21 | 0 | 0 | 0 | 0 | 0 | 0 | 0 | 0 | 0 | 0 | 0 | 0 | 0 | 0 | 0 | 0 | 0 | 0 | 2044 | 1635 | 0 | 0 | 63 | 0 | 42 | 230 | 0 | 0 | 0 | 0 | 0 | 21 | 0 | 0 | 0 | 21 |
| SRR10416524 | 0 | 0 | 0 | 0 | 0 | 30 | 0 | 0 | 0 | 0 | 0 | 0 | 0 | 0 | 0 | 0 | 0 | 0 | 0 | 1988 | 0 | 0 | 0 | 178 | 0 | 89 | 148 | 0 | 0 | 0 | 0 | 0 | 0 | 0 | 0 | 0 | 0 |
| SRR10416525 | 0 | 0 | 0 | 0 | 0 | 43 | 0 | 0 | 0 | 0 | 0 | 0 | 0 | 0 | 0 | 0 | 0 | 0 | 0 | 1453 | 3634 | 0 | 0 | 43 | 0 | 129 | 172 | 0 | 0 | 0 | 0 | 0 | 0 | 43 | 0 | 0 | 0 |
| SRR10416526 | 0 | 0 | 0 | 0 | 0 | 39 | 0 | 0 | 0 | 0 | 0 | 0 | 0 | 0 | 0 | 0 | 0 | 0 | 0 | 2705 | 2028 | 0 | 0 | 347 | 0 | 193 | 425 | 0 | 0 | 0 | 0 | 0 | 39 | 0 | 0 | 0 | 0 |
| SRR10416527 | 14 | 0 | 0 | 0 | 0 | 0 | 0 | 47 | 0 | 0 | 0 | 0 | 0 | 0 | 0 | 0 | 0 | 0 | 0 | 5155 | 0 | 0 | 0 | 95 | 0 | 142 | 95 | 0 | 0 | 0 | 0 | 0 | 0 | 0 | 0 | 0 | 0 |
| SRR10416528 | 20 | 0 | 0 | 0 | 0 | 0 | 0 | 41 | 81 | 0 | 0 | 0 | 0 | 0 | 0 | 0 | 0 | 0 | 0 | 1518 | 1012 | 0 | 0 | 81 | 0 | 81 | 41 | 0 | 0 | 0 | 0 | 0 | 0 | 0 | 0 | 0 | 0 |
| SRR10416529 | 0 | 0 | 0 | 0 | 0 | 0 | 0 | 0 | 0 | 0 | 0 | 0 | 0 | 0 | 0 | 0 | 0 | 0 | 903 | 0 | 903 | 0 | 0 | 55 | 0 | 218 | 0 | 0 | 0 | 0 | 0 | 0 | 0 | 0 | 0 | 0 | 0 |
| SRR10416530 | 0 | 0 | 0 | 0 | 0 | 0 | 0 | 60 | 0 | 0 | 0 | 0 | 0 | 0 | 0 | 0 | 0 | 0 | 0 | 2994 | 749 | 0 | 0 | 60 | 0 | 0 | 179 | 0 | 0 | 0 | 0 | 0 | 0 | 0 | 0 | 0 | 0 |
| SRR10416531 | 0 | 0 | 0 | 0 | 0 | 0 | 0 | 99 | 0 | 0 | 0 | 0 | 0 | 0 | 0 | 0 | 0 | 0 | 0 | 1105 | 0 | 0 | 0 | 0 | 0 | 397 | 199 | 0 | 0 | 0 | 0 | 0 | 0 | 0 | 0 | 0 | 0 |
| SRR10416532 | 0 | 0 | 0 | 0 | 0 | 0 | 0 | 0 | 0 | 0 | 0 | 0 | 0 | 0 | 0 | 0 | 0 | 0 | 0 | 0 | 0 | 0 | 0 | 0 | 0 | 0 | 69 | 0 | 0 | 0 | 0 | 0 | 0 | 0 | 0 | 0 | 0 |

### Supp Table S8.2.Columns_AM_to_BW

| | | | | | | | | | | | | | | | | | | | | | | | | | | | | | | | | | | | | |
|---|---|---|---|---|---|---|---|---|---|---|---|---|---|---|---|---|---|---|---|---|---|---|---|---|---|---|---|---|---|---|---|---|---|---|---|---|---|
| SRR10416533 | 28 | 44 | 0 | 0 | 0 | 0 | 0 | 0 | 0 | 0 | 0 | 0 | 0 | 0 | 0 | 0 | 0 | 0 | 0 | 2010 | 1005 | 0 | 0 | 175 | 0 | 87 | 306 | 0 | 0 | 0 | 0 | 0 | 0 | 0 | 0 | 0 | 0 |
| SRR10416534 | 10 | 0 | 0 | 0 | 0 | 0 | 0 | 85 | 0 | 0 | 0 | 0 | 0 | 0 | 0 | 0 | 0 | 0 | 0 | 3468 | 1156 | 0 | 0 | 0 | 0 | 0 | 426 | 0 | 0 | 0 | 0 | 0 | 0 | 0 | 0 | 0 | 0 |
| SRR10416535 | 0 | 0 | 0 | 0 | 0 | 0 | 0 | 46 | 0 | 0 | 0 | 0 | 0 | 0 | 0 | 0 | 0 | 0 | 0 | 840 | 1679 | 0 | 0 | 91 | 46 | 0 | 274 | 0 | 0 | 0 | 46 | 0 | 0 | 0 | 0 | 0 | 0 |
| SRR10416536 | 0 | 0 | 0 | 0 | 0 | 0 | 0 | 0 | 0 | 0 | 0 | 0 | 0 | 0 | 0 | 0 | 0 | 0 | 0 | 4326 | 2884 | 0 | 0 | 89 | 0 | 133 | 177 | 0 | 0 | 0 | 0 | 0 | 0 | 0 | 0 | 0 | 0 |
| SRR10416537 | 0 | 0 | 0 | 0 | 0 | 0 | 0 | 45 | 0 | 0 | 0 | 0 | 0 | 0 | 0 | 0 | 0 | 0 | 0 | 3836 | 0 | 0 | 0 | 134 | 0 | 90 | 90 | 0 | 0 | 0 | 45 | 0 | 0 | 0 | 0 | 0 | 0 |
| SRR10416538 | 0 | 0 | 0 | 0 | 0 | 0 | 0 | 34 | 0 | 0 | 0 | 0 | 0 | 0 | 0 | 0 | 0 | 0 | 661 | 1982 | 661 | 0 | 0 | 34 | 0 | 205 | 273 | 0 | 0 | 0 | 0 | 0 | 0 | 0 | 0 | 0 | 0 |
| SRR10416539 | 0 | 0 | 0 | 0 | 0 | 0 | 0 | 67 | 0 | 0 | 0 | 0 | 0 | 67 | 0 | 0 | 0 | 0 | 0 | 5630 | 704 | 0 | 0 | 0 | 0 | 0 | 67 | 0 | 0 | 0 | 0 | 0 | 0 | 0 | 0 | 0 | 0 |
| SRR10416540 | 0 | 0 | 0 | 0 | 0 | 0 | 0 | 0 | 0 | 0 | 0 | 0 | 0 | 0 | 0 | 0 | 0 | 0 | 0 | 806 | 0 | 0 | 0 | 50 | 50 | 50 | 151 | 0 | 0 | 0 | 0 | 50 | 0 | 0 | 0 | 0 | 0 |
| SRR10416541 | 0 | 0 | 0 | 0 | 0 | 0 | 0 | 114 | 0 | 0 | 0 | 0 | 0 | 0 | 0 | 0 | 0 | 0 | 0 | 0 | 0 | 0 | 0 | 114 | 0 | 0 | 456 | 0 | 0 | 0 | 0 | 0 | 0 | 0 | 228 | 0 | 0 |
| SRR10416542 | 0 | 0 | 0 | 90 | 0 | 0 | 0 | 0 | 0 | 0 | 0 | 0 | 0 | 0 | 0 | 0 | 0 | 0 | 0 | 6033 | 1508 | 0 | 0 | 0 | 0 | 0 | 271 | 0 | 0 | 0 | 0 | 0 | 0 | 0 | 0 | 0 | 0 |
| SRR10416543 | 0 | 0 | 0 | 0 | 0 | 10 | 0 | 42 | 0 | 0 | 0 | 0 | 0 | 0 | 0 | 10 | 0 | 0 | 0 | 526 | 0 | 0 | 0 | 21 | 10 | 21 | 426 | 0 | 0 | 10 | 0 | 0 | 0 | 0 | 0 | 0 | 0 |
| SRR10416544 | 0 | 0 | 0 | 0 | 0 | 0 | 0 | 0 | 0 | 0 | 0 | 0 | 0 | 0 | 0 | 0 | 0 | 0 | 0 | 4854 | 0 | 0 | 0 | 234 | 0 | 78 | 156 | 0 | 0 | 0 | 0 | 0 | 0 | 0 | 0 | 0 | 0 |
| SRR10416545 | 0 | 19 | 0 | 38 | 0 | 0 | 0 | 19 | 0 | 0 | 0 | 0 | 0 | 0 | 0 | 0 | 0 | 0 | 0 | 834 | 0 | 0 | 0 | 94 | 19 | 19 | 357 | 0 | 0 | 0 | 0 | 0 | 0 | 0 | 0 | 0 | 0 |
| SRR10416546 | 0 | 33 | 0 | 0 | 0 | 0 | 0 | 66 | 0 | 0 | 0 | 0 | 0 | 0 | 0 | 0 | 0 | 0 | 0 | 2448 | 612 | 0 | 0 | 66 | 0 | 33 | 99 | 0 | 0 | 0 | 0 | 0 | 0 | 0 | 0 | 0 | 0 |
| SRR10416547 | 0 | 0 | 0 | 0 | 0 | 0 | 0 | 0 | 0 | 0 | 0 | 0 | 0 | 0 | 0 | 0 | 0 | 0 | 0 | 1577 | 0 | 0 | 0 | 145 | 0 | 48 | 193 | 0 | 0 | 0 | 0 | 0 | 0 | 0 | 0 | 0 | 0 |
| SRR10416549 | 15 | 0 | 0 | 23 | 0 | 0 | 0 | 0 | 0 | 0 | 0 | 0 | 0 | 0 | 0 | 0 | 0 | 0 | 0 | 4487 | 280 | 0 | 0 | 68 | 0 | 68 | 295 | 0 | 0 | 0 | 0 | 0 | 0 | 0 | 0 | 0 | 0 |
| SRR10416550 | 0 | 21 | 0 | 0 | 0 | 0 | 0 | 0 | 0 | 0 | 0 | 0 | 0 | 0 | 0 | 0 | 0 | 0 | 0 | 3943 | 493 | 21 | 0 | 104 | 0 | 104 | 208 | 21 | 0 | 0 | 0 | 0 | 21 | 0 | 21 | 0 | 0 |
| SRR10416551 | 0 | 0 | 0 | 0 | 0 | 0 | 0 | 27 | 0 | 0 | 0 | 0 | 0 | 0 | 0 | 0 | 0 | 0 | 0 | 3294 | 941 | 0 | 0 | 27 | 0 | 27 | 318 | 0 | 0 | 0 | 0 | 0 | 0 | 0 | 0 | 0 | 0 |
| SRR10416553 | 0 | 0 | 0 | 0 | 0 | 0 | 0 | 65 | 0 | 0 | 0 | 0 | 0 | 0 | 0 | 0 | 0 | 0 | 0 | 0 | 0 | 0 | 0 | 0 | 0 | 324 | 259 | 0 | 0 | 0 | 0 | 0 | 0 | 0 | 0 | 0 | 0 |
| SRR10416554 | 31 | 0 | 0 | 0 | 0 | 71 | 0 | 0 | 0 | 0 | 0 | 0 | 0 | 0 | 0 | 0 | 0 | 0 | 0 | 1397 | 698 | 0 | 0 | 0 | 71 | 71 | 11 | 0 | 0 | 0 | 71 | 71 | 0 | 0 | 0 | 0 | 0 |
| SRR10416555 | 0 | 0 | 0 | 0 | 0 | 0 | 0 | 0 | 0 | 0 | 0 | 0 | 0 | 0 | 0 | 0 | 0 | 0 | 0 | 2336 | 0 | 0 | 0 | 0 | 0 | 0 | 343 | 0 | 0 | 0 | 0 | 0 | 0 | 0 | 0 | 0 | 0 |
| SRR10416557 | 0 | 0 | 0 | 0 | 0 | 0 | 0 | 0 | 0 | 0 | 0 | 0 | 0 | 0 | 0 | 0 | 0 | 0 | 0 | 1862 | 0 | 0 | 0 | 0 | 0 | 115 | 0 | 0 | 0 | 0 | 0 | 0 | 0 | 0 | 0 | 0 | 0 |
| SRR10416558 | 16 | 0 | 0 | 0 | 0 | 0 | 0 | 51 | 0 | 0 | 0 | 0 | 0 | 0 | 0 | 0 | 0 | 0 | 0 | 4714 | 673 | 0 | 0 | 51 | 0 | 101 | 152 | 0 | 0 | 0 | 0 | 0 | 0 | 0 | 0 | 0 | 0 |
| SRR10416559 | 0 | 0 | 0 | 0 | 0 | 0 | 0 | 0 | 0 | 0 | 0 | 0 | 0 | 0 | 0 | 0 | 0 | 0 | 0 | 2181 | 545 | 0 | 0 | 39 | 0 | 78 | 234 | 0 | 0 | 0 | 0 | 0 | 0 | 0 | 0 | 0 | 0 |
| SRR10416560 | 20 | 0 | 0 | 0 | 0 | 0 | 0 | 23 | 0 | 0 | 0 | 0 | 0 | 0 | 0 | 0 | 0 | 0 | 0 | 6076 | 434 | 0 | 0 | 161 | 23 | 0 | 115 | 0 | 0 | 0 | 0 | 0 | 0 | 0 | 0 | 0 | 0 |
| SRR10416562 | 25 | 35 | 0 | 0 | 0 | 0 | 0 | 0 | 0 | 0 | 0 | 0 | 0 | 0 | 0 | 0 | 0 | 0 | 0 | 722 | 0 | 0 | 0 | 173 | 0 | 0 | 243 | 0 | 0 | 0 | 0 | 0 | 0 | 0 | 0 | 0 | 0 |
| SRR10416563 | 24 | 0 | 0 | 15 | 0 | 15 | 0 | 60 | 0 | 0 | 0 | 0 | 0 | 0 | 0 | 0 | 0 | 0 | 0 | 4379 | 0 | 0 | 0 | 135 | 0 | 75 | 465 | 0 | 0 | 0 | 0 | 0 | 0 | 15 | 0 | 0 | 0 |
| SRR10416564 | 0 | 0 | 0 | 0 | 0 | 0 | 0 | 0 | 0 | 0 | 0 | 0 | 0 | 0 | 0 | 0 | 0 | 0 | 0 | 3494 | 0 | 0 | 0 | 129 | 0 | 129 | 172 | 0 | 0 | 0 | 0 | 0 | 0 | 0 | 0 | 0 | 0 |
| SRR10416565 | 0 | 31 | 0 | 0 | 0 | 0 | 0 | 0 | 0 | 0 | 0 | 0 | 0 | 31 | 0 | 0 | 0 | 0 | 0 | 3161 | 0 | 0 | 0 | 0 | 0 | 31 | 187 | 0 | 0 | 0 | 0 | 0 | 0 | 0 | 0 | 0 | 0 |
| SRR10416566 | 0 | 0 | 0 | 0 | 0 | 0 | 0 | 0 | 0 | 0 | 0 | 0 | 0 | 0 | 0 | 0 | 0 | 0 | 0 | 0 | 0 | 0 | 0 | 0 | 0 | 72 | 0 | 0 | 0 | 0 | 0 | 0 | 0 | 0 | 0 | 0 | 0 |
| SRR10416567 | 0 | 0 | 10 | 0 | 0 | 0 | 0 | 107 | 0 | 0 | 0 | 0 | 0 | 0 | 0 | 0 | 0 | 0 | 0 | 528 | 0 | 0 | 0 | 107 | 0 | 107 | 214 | 0 | 0 | 0 | 0 | 0 | 0 | 0 | 0 | 0 | 0 |
| SRR10416568 | 0 | 0 | 0 | 0 | 0 | 0 | 0 | 158 | 0 | 0 | 0 | 0 | 0 | 0 | 0 | 0 | 0 | 0 | 0 | 9267 | 0 | 0 | 0 | 0 | 0 | 236 | 158 | 0 | 0 | 0 | 0 | 0 | 0 | 0 | 0 | 0 | 0 |
| SRR10416569 | 15 | 0 | 0 | 0 | 0 | 0 | 0 | 0 | 0 | 0 | 0 | 0 | 0 | 0 | 0 | 0 | 0 | 0 | 0 | 1274 | 0 | 0 | 0 | 0 | 0 | 143 | 143 | 0 | 0 | 0 | 0 | 0 | 0 | 0 | 0 | 0 | 0 |
| SRR10416570 | 0 | 0 | 0 | 0 | 0 | 53 | 0 | 0 | 0 | 0 | 0 | 0 | 0 | 0 | 644 | 0 | 0 | 0 | 0 | 0 | 644 | 0 | 0 | 0 | 0 | 106 | 17 | 0 | 0 | 0 | 0 | 0 | 0 | 0 | 0 | 0 | 53 |
| SRR10416571 | 0 | 0 | 0 | 0 | 0 | 0 | 0 | 183 | 0 | 0 | 0 | 0 | 0 | 0 | 0 | 0 | 0 | 0 | 0 | 4623 | 770 | 0 | 0 | 0 | 0 | 306 | 978 | 0 | 0 | 0 | 0 | 0 | 0 | 0 | 0 | 0 | 0 |
| SRR10416572 | 16 | 0 | 0 | 0 | 0 | 0 | 0 | 44 | 0 | 0 | 0 | 0 | 0 | 0 | 0 | 0 | 0 | 0 | 0 | 1092 | 546 | 0 | 0 | 87 | 0 | 218 | 741 | 0 | 0 | 0 | 0 | 0 | 0 | 0 | 0 | 0 | 0 |
| SRR10416573 | 0 | 0 | 0 | 0 | 0 | 0 | 0 | 0 | 0 | 0 | 0 | 0 | 0 | 0 | 0 | 0 | 0 | 0 | 0 | 5102 | 0 | 0 | 0 | 204 | 0 | 102 | 409 | 0 | 0 | 0 | 0 | 0 | 0 | 0 | 0 | 0 | 0 |
| SRR10416574 | 0 | 0 | 0 | 0 | 0 | 0 | 0 | 0 | 0 | 0 | 0 | 0 | 0 | 0 | 0 | 0 | 0 | 0 | 0 | 1505 | 0 | 0 | 0 | 0 | 0 | 0 | 191 | 0 | 0 | 0 | 0 | 0 | 0 | 0 | 0 | 0 | 0 |
| SRR10416575 | 41 | 0 | 0 | 0 | 0 | 0 | 0 | 42 | 0 | 0 | 0 | 0 | 0 | 0 | 0 | 0 | 0 | 0 | 0 | 0 | 1297 | 0 | 0 | 170 | 42 | 42 | 425 | 0 | 0 | 0 | 0 | 0 | 0 | 0 | 0 | 0 | 0 |
| SRR10416576 | 48 | 0 | 0 | 0 | 0 | 0 | 0 | 0 | 0 | 0 | 0 | 0 | 0 | 0 | 0 | 0 | 0 | 0 | 0 | 2210 | 0 | 0 | 0 | 169 | 0 | 56 | 169 | 0 | 0 | 28 | 0 | 0 | 0 | 0 | 0 | 0 | 0 |
| SRR10416577 | 10 | 71 | 0 | 0 | 0 | 0 | 0 | 71 | 0 | 0 | 0 | 0 | 0 | 0 | 0 | 71 | 0 | 0 | 0 | 3408 | 1363 | 0 | 0 | 0 | 0 | 214 | 427 | 0 | 0 | 0 | 0 | 0 | 0 | 0 | 0 | 0 | 0 |
| SRR10416578 | 0 | 0 | 0 | 0 | 292 | 0 | 0 | 146 | 0 | 0 | 0 | 0 | 0 | 0 | 0 | 146 | 0 | 0 | 0 | 0 | 0 | 0 | 0 | 146 | 0 | 146 | 1023 | 0 | 0 | 0 | 0 | 0 | 0 | 0 | 0 | 0 | 0 |
| SRR10416579 | 0 | 0 | 0 | 0 | 0 | 0 | 0 | 0 | 0 | 0 | 0 | 0 | 0 | 0 | 1124 | 0 | 0 | 0 | 0 | 2247 | 1124 | 0 | 0 | 0 | 0 | 137 | 137 | 0 | 0 | 0 | 0 | 0 | 0 | 0 | 0 | 0 | 0 |
| SRR10416580 | 0 | 0 | 0 | 0 | 0 | 0 | 0 | 0 | 0 | 0 | 0 | 0 | 0 | 0 | 0 | 0 | 0 | 0 | 0 | 2315 | 0 | 0 | 0 | 156 | 0 | 52 | 363 | 52 | 0 | 0 | 0 | 0 | 0 | 0 | 0 | 0 | 0 |
| SRR10416581 | 19 | 0 | 0 | 0 | 0 | 0 | 0 | 0 | 0 | 0 | 0 | 0 | 0 | 0 | 0 | 0 | 0 | 0 | 0 | 3731 | 1493 | 0 | 0 | 61 | 0 | 263 | 444 | 0 | 0 | 0 | 0 | 0 | 20 | 20 | 0 | 0 | 0 |
| SRR10416582 | 0 | 0 | 0 | 0 | 0 | 0 | 0 | 0 | 0 | 0 | 0 | 0 | 0 | 0 | 0 | 0 | 0 | 0 | 0 | 1869 | 935 | 0 | 0 | 92 | 0 | 92 | 276 | 0 | 0 | 0 | 0 | 0 | 0 | 0 | 0 | 0 | 0 |
| SRR10416583 | 29 | 0 | 0 | 0 | 0 | 0 | 0 | 167 | 0 | 0 | 0 | 0 | 0 | 0 | 0 | 0 | 0 | 0 | 0 | 1905 | 0 | 0 | 0 | 84 | 0 | 0 | 84 | 0 | 0 | 0 | 0 | 0 | 0 | 0 | 0 | 0 | 0 |
| SRR10416584 | 0 | 44 | 0 | 0 | 0 | 0 | 0 | 0 | 0 | 0 | 0 | 0 | 0 | 0 | 0 | 0 | 0 | 0 | 0 | 1672 | 0 | 0 | 0 | 89 | 0 | 89 | 178 | 0 | 0 | 0 | 0 | 0 | 44 | 0 | 0 | 0 | 0 |
| SRR10416585 | 0 | 0 | 0 | 165 | 0 | 0 | 0 | 0 | 0 | 0 | 0 | 0 | 0 | 0 | 0 | 0 | 0 | 0 | 0 | 2028 | 0 | 0 | 0 | 0 | 0 | 0 | 0 | 0 | 0 | 0 | 0 | 0 | 0 | 0 | 0 | 0 | 0 |
| SRR10416586 | 0 | 0 | 0 | 0 | 0 | 0 | 0 | 0 | 0 | 0 | 0 | 0 | 0 | 0 | 0 | 0 | 0 | 0 | 0 | 1131 | 1131 | 0 | 0 | 94 | 0 | 94 | 282 | 0 | 0 | 0 | 0 | 0 | 0 | 0 | 0 | 0 | 0 |
| SRR10416587 | 0 | 0 | 0 | 0 | 0 | 0 | 0 | 0 | 0 | 0 | 0 | 0 | 0 | 0 | 0 | 0 | 0 | 0 | 0 | 6012 | 1002 | 0 | 0 | 129 | 0 | 129 | 258 | 65 | 0 | 0 | 0 | 0 | 0 | 0 | 0 | 0 | 0 |
| SRR10416588 | 0 | 0 | 0 | 0 | 0 | 0 | 0 | 78 | 0 | 0 | 0 | 0 | 0 | 0 | 0 | 78 | 0 | 0 | 0 | 1982 | 991 | 0 | 0 | 0 | 0 | 0 | 155 | 0 | 0 | 0 | 0 | 0 | 0 | 0 | 0 | 0 | 0 |
| SRR10416589 | 0 | 0 | 0 | 0 | 0 | 0 | 0 | 49 | 0 | 0 | 0 | 0 | 0 | 0 | 0 | 0 | 0 | 0 | 0 | 794 | 7143 | 0 | 0 | 49 | 0 | 123 | 197 | 0 | 0 | 0 | 0 | 0 | 0 | 0 | 0 | 0 | 0 |
| SRR10416590 | 0 | 0 | 0 | 0 | 0 | 0 | 0 | 0 | 0 | 0 | 0 | 0 | 0 | 0 | 0 | 0 | 0 | 0 | 0 | 4762 | 794 | 0 | 0 | 67 | 0 | 0 | 67 | 0 | 0 | 0 | 0 | 0 | 0 | 0 | 0 | 0 | 0 |
| SRR10416591 | 0 | 0 | 0 | 0 | 0 | 0 | 0 | 90 | 0 | 0 | 0 | 0 | 0 | 0 | 0 | 0 | 0 | 0 | 0 | 9063 | 0 | 0 | 0 | 90 | 0 | 179 | 359 | 0 | 0 | 0 | 0 | 0 | 0 | 0 | 0 | 0 | 0 |
| SRR10416592 | 22 | 0 | 0 | 0 | 0 | 0 | 0 | 0 | 0 | 0 | 0 | 0 | 0 | 0 | 0 | 0 | 0 | 0 | 0 | 743 | 371 | 0 | 0 | 227 | 0 | 227 | 16 | 0 | 0 | 0 | 0 | 0 | 0 | 0 | 0 | 0 | 0 |
| SRR10416593 | 0 | 0 | 0 | 0 | 0 | 0 | 0 | 0 | 0 | 0 | 0 | 0 | 0 | 0 | 0 | 0 | 0 | 0 | 0 | 7484 | 0 | 0 | 0 | 47 | 0 | 0 | 140 | 0 | 0 | 0 | 47 | 0 | 0 | 0 | 0 | 0 | 0 |
| SRR10416594 | 0 | 0 | 0 | 0 | 0 | 0 | 0 | 0 | 0 | 0 | 0 | 0 | 0 | 0 | 0 | 0 | 0 | 0 | 0 | 1862 | 0 | 0 | 0 | 0 | 0 | 75 | 0 | 0 | 0 | 0 | 0 | 0 | 0 | 0 | 0 | 0 | 0 |
| SRR10416595 | 31 | 0 | 0 | 42 | 42 | 0 | 0 | 0 | 0 | 0 | 0 | 0 | 0 | 42 | 0 | 0 | 0 | 0 | 0 | 2462 | 1477 | 42 | 0 | 0 | 0 | 42 | 502 | 0 | 0 | 0 | 0 | 0 | 0 | 0 | 0 | 0 | 0 |
| SRR10416596 | 22 | 0 | 0 | 0 | 0 | 19 | 0 | 0 | 0 | 0 | 0 | 0 | 0 | 0 | 0 | 19 | 0 | 0 | 0 | 1048 | 2096 | 0 | 0 | 289 | 39 | 173 | 694 | 0 | 0 | 0 | 0 | 0 | 0 | 39 | 0 | 0 | 0 |
| SRR10416597 | 27 | 0 | 0 | 0 | 0 | 0 | 0 | 27 | 0 | 0 | 0 | 0 | 0 | 0 | 0 | 0 | 0 | 0 | 0 | 6027 | 0 | 0 | 0 | 82 | 0 | 82 | 329 | 0 | 0 | 0 | 0 | 0 | 0 | 0 | 0 | 0 | 0 |
| SRR10416598 | 18 | 0 | 0 | 0 | 0 | 0 | 0 | 0 | 0 | 0 | 0 | 0 | 0 | 0 | 0 | 0 | 0 | 0 | 0 | 0 | 1182 | 0 | 0 | 165 | 0 | 110 | 523 | 0 | 0 | 0 | 0 | 0 | 0 | 28 | 0 | 0 | 0 |
| SRR10416599 | 0 | 0 | 0 | 0 | 0 | 0 | 0 | 0 | 0 | 0 | 0 | 0 | 0 | 0 | 0 | 0 | 0 | 0 | 0 | 0 | 0 | 0 | 0 | 63 | 0 | 189 | 946 | 0 | 0 | 0 | 0 | 0 | 63 | 63 | 0 | 0 | 0 |
| SRR10416600 | 25 | 0 | 0 | 0 | 0 | 0 | 0 | 0 | 0 | 0 | 0 | 0 | 0 | 0 | 0 | 0 | 0 | 0 | 0 | 3769 | 838 | 0 | 0 | 100 | 0 | 100 | 18 | 0 | 0 | 0 | 0 | 0 | 0 | 20 | 0 | 0 | 0 |
| SRR10428553 | 63 | 20 | 2597 | 15 | 0 | 0 | 11 | 16 | 10 | 0 | 0 | 36 | 0 | 23 | 30 | 0 | 0 | 0 | 0 | 0 | 14 | 0 | 0 | 0 | 18 | 188 | 226 | 34 | 0 | 21 | 19 | 0 | 0 | 33 | 0 | 0 | 0 |
| SRR10428554 | 20 | 23 | 2479 | 0 | 0 | 16 | 62 | 37 | 21 | 0 | 0 | 14 | 0 | 39 | 1037 | 0 | 0 | 0 | 0 | 71 | 481 | 0 | 0 | 0 | 17 | 232 | 127 | 26 | 0 | 15 | 0 | 0 | 0 | 16 | 53 | 0 | 25 |
| SRR10428555 | 38 | 36 | 1781 | 20 | 0 | 0 | 20 | 0 | 0 | 0 | 0 | 15 | 0 | 39 | 456 | 0 | 10 | 0 | 0 | 16 | 93 | 0 | 0 | 0 | 21 | 12 | 59 | 56 | 0 | 0 | 0 | 0 | 14 | 20 | 13 | 0 | 15 |
| SRR10428556 | 36 | 13 | 2647 | 11 | 0 | 0 | 0 | 0 | 0 | 0 | 0 | 18 | 0 | 29 | 4393 | 0 | 0 | 0 | 0 | 218 | 325 | 0 | 0 | 0 | 0 | 145 | 479 | 31 | 0 | 0 | 14 | 0 | 0 | 14 | 54 | 0 | 29 |
| SRR10428557 | 31 | 79 | 906 | 36 | 12 | 0 | 25 | 0 | 0 | 0 | 0 | 70 | 0 | 53 | 224 | 0 | 0 | 13 | 12 | 54 | 318 | 18 | 0 | 0 | 12 | 29 | 63 | 61 | 0 | 0 | 0 | 0 | 13 | 19 | 0 | 0 | 10 |
| SRR10428558 | 38 | 24 | 1247 | 15 | 12 | 0 | 16 | 0 | 0 | 0 | 0 | 21 | 0 | 41 | 520 | 0 | 0 | 0 | 0 | 51 | 152 | 10 | 0 | 0 | 0 | 105 | 85 | 38 | 0 | 0 | 11 | 0 | 0 | 17 | 0 | 0 | 0 |
| SRR10428559 | 24 | 23 | 3145 | 14 | 0 | 0 | 29 | 0 | 0 | 0 | 0 | 45 | 0 | 37 | 883 | 0 | 0 | 0 | 0 | 31 | 185 | 12 | 0 | 0 | 11 | 142 | 148 | 43 | 0 | 0 | 13 | 0 | 0 | 26 | 37 | 0 | 31 |
| SRR10428560 | 18 | 0 | 1490 | 0 | 0 | 0 | 12 | 26 | 0 | 0 | 0 | 55 | 0 | 16 | 266 | 0 | 17 | 0 | 0 | 0 | 21 | 0 | 0 | 0 | 13 | 92 | 214 | 0 | 0 | 0 | 0 | 0 | 0 | 23 | 46 | 0 | 29 |
| SRR10446207 | 183 | 0 | 832 | 0 | 0 | 148 | 2556 | 0 | 111 | 0 | 0 | 28 | 20 | 0 | 54 | 0 | 39 | 0 | 125 | 510 | 1913 | 0 | 0 | 295 | 1182 | 2236 | 1528 | 0 | 0 | 0 | 199 | 11 | 0 | 0 | 136 | 11 | 43 |
| SRR10446208 | 153 | 0 | 780 | 0 | 0 | 115 | 1273 | 0 | 61 | 0 | 0 | 85 | 16 | 16 | 147 | 0 | 58 | 0 | 316 | 5319 | 11 | 0 | 0 | 48 | 92 | 133 | 186 | 0 | 0 | 0 | 115 | 0 | 0 | 0 | 83 | 26 | 42 |
| SRR10446209 | 322 | 0 | 514 | 0 | 0 | 290 | 4329 | 25 | 226 | 0 | 0 | 73 | 15 | 29 | 2213 | 0 | 34 | 14 | 316 | 1654 | 1727 | 0 | 0 | 447 | 658 | 4761 | 1140 | 0 | 0 | 0 | 865 | 0 | 0 | 0 | 275 | 54 | 167 |
| SRR10446210 | 78 | 0 | 236 | 0 | 0 | 19 | 450 | 0 | 256 | 0 | 0 | 96 | 72 | 0 | 536 | 0 | 42 | 54 | 279 | 150 | 1136 | 0 | 0 | 11 | 118 | 327 | 159 | 20 | 26 | 0 | 19 | 0 | 0 | 0 | 282 | 52 | 138 |
| SRR10446211 | 460 | 16 | 1390 | 0 | 0 | 101 | 65 | 23 | 71 | 0 | 0 | 113 | 0 | 23 | 74 | 0 | 37 | 0 | 37 | 18 | 1763 | 0 | 0 | 96 | 101 | 1499 | 172 | 13 | 13 | 0 | 87 | 17 | 0 | 0 | 29 | 15 | 27 |
| SRR10446212 | 769 | 0 | 378 | 0 | 0 | 382 | 159 | 41 | 245 | 0 | 0 | 233 | 0 | 13 | 569 | 0 | 17 | 0 | 13 | 251 | 6044 | 0 | 0 | 525 | 436 | 6778 | 897 | 0 | 0 | 0 | 465 | 0 | 0 | 0 | 156 | 64 | 108 |
| SRR10531363 | 12 | 0 | 0 | 170 | 0 | 0 | 0 | 457 | 0 | 0 | 0 | 0 | 95 | 0 | 0 | 0 | 0 | 0 | 0 | 0 | 2180 | 0 | 0 | 0 | 0 | 0 | 0 | 0 | 0 | 1810 | 0 | 0 | 0 | 208 | 0 | 0 | 21 |
| SRR10531364 | 13 | 0 | 0 | 0 | 0 | 0 | 0 | 272 | 0 | 0 | 0 | 18 | 0 | 62 | 829 | 0 | 0 | 0 | 0 | 581 | 16 | 15 | 0 | 43 | 0 | 0 | 0 | 67 | 0 | 77 | 0 | 0 | 0 | 0 | 0 | 0 | 0 |
| SRR10531365 | 0 | 139 | 0 | 0 | 0 | 0 | 0 | 0 | 0 | 0 | 0 | 0 | 0 | 0 | 0 | 0 | 0 | 0 | 0 | 0 | 1704 | 0 | 0 | 261 | 0 | 0 | 0 | 177 | 0 | 13 | 0 | 0 | 0 | 337 | 0 | 0 | 0 |
| SRR10531367 | 14 | 0 | 0 | 0 | 0 | 0 | 0 | 215 | 0 | 0 | 0 | 33 | 0 | 0 | 1181 | 0 | 0 | 0 | 0 | 0 | 12500 | 0 | 0 | 109 | 0 | 0 | 0 | 0 | 0 | 47 | 0 | 0 | 0 | 129 | 0 | 0 | 0 |
| SRR10531368 | 16 | 0 | 0 | 131 | 0 | 0 | 0 | 40 | 0 | 0 | 0 | 21 | 0 | 0 | 0 | 0 | 0 | 0 | 0 | 862 | 4689 | 0 | 0 | 0 | 0 | 0 | 0 | 0 | 0 | 113 | 0 | 0 | 0 | 0 | 0 | 0 | 0 |
| SRR10531369 | 26 | 0 | 0 | 0 | 0 | 0 | 11 | 0 | 0 | 0 | 0 | 19 | 0 | 0 | 11 | 0 | 0 | 0 | 0 | 0 | 8488 | 0 | 0 | 0 | 0 | 0 | 0 | 0 | 0 | 512 | 0 | 0 | 0 | 0 | 0 | 0 | 0 |
| SRR10531370 | 38 | 0 | 0 | 0 | 0 | 0 | 0 | 2977 | 0 | 0 | 0 | 82 | 0 | 0 | 642 | 0 | 0 | 0 | 0 | 0 | 6331 | 0 | 0 | 0 | 0 | 0 | 0 | 0 | 0 | 6035 | 0 | 0 | 0 | 0 | 0 | 0 | 0 |
| SRR10544939 | 937 | 160 | 2204 | 0 | 0 | 24 | 100075 | 462 | 0 | 0 | 0 | 27 | 0 | 25 | 0 | 0 | 105 | 0 | 1000 | 6726 | 0 | 0 | 0 | 1255 | 0 | 46 | 11 | 0 | 0 | 0 | 0 | 0 | 0 | 0 | 0 | 0 | 0 |
| SRR10544940 | 1085 | 76 | 2076 | 0 | 0 | 24 | 64706 | 416 | 13 | 0 | 13 | 28 | 17 | 28 | 0 | 0 | 85 | 0 | 435 | 2695 | 0 | 0 | 0 | 166 | 21 | 27 | 16 | 0 | 0 | 0 | 0 | 0 | 0 | 0 | 0 | 0 | 0 |
| SRR10544941 | 739 | 170 | 944 | 0 | 0 | 66 | 65221 | 859 | 37 | 0 | 18 | 18 | 11 | 41 | 0 | 0 | 144 | 0 | 1089 | 8024 | 0 | 0 | 0 | 932 | 22 | 166 | 47 | 0 | 0 | 0 | 11 | 0 | 0 | 0 | 0 | 15 | 19 |
| SRR10544942 | 1513 | 197 | 3517 | 0 | 0 | 25 | 49331 | 800 | 12 | 10 | 24 | 27 | 20 | 61 | 0 | 0 | 88 | 0 | 336 | 3132 | 0 | 0 | 0 | 552 | 0 | 27 | 0 | 0 | 0 | 0 | 0 | 0 | 0 | 0 | 0 | 0 | 0 |
| SRR10544943 | 1434 | 65 | 1623 | 0 | 0 | 13 | 94941 | 44 | 22 | 0 | 13 | 42 | 0 | 12 | 0 | 0 | 110 | 0 | 611 | 4599 | 0 | 0 | 0 | 178 | 11 | 17 | 13 | 0 | 0 | 0 | 0 | 0 | 0 | 0 | 0 | 0 | 0 |
| SRR10544944 | 1187 | 49 | 2391 | 0 | 0 | 12 | 58936 | 271 | 0 | 0 | 11 | 16 | 10 | 16 | 0 | 0 | 282 | 0 | 1886 | 9657 | 0 | 0 | 0 | 333 | 18 | 12 | 0 | 0 | 0 | 0 | 0 | 0 | 0 | 0 | 0 | 0 | 0 |
| SRR10544945 | 928 | 85 | 8328 | 0 | 0 | 13 | 76076 | 37 | 0 | 0 | 13 | 8677 | 0 | 0 | 28 | 0 | 56 | 0 | 337 | 1146 | 51 | 0 | 0 | 110 | 0 | 0 | 0 | 0 | 0 | 0 | 0 | 0 | 0 | 0 | 0 | 0 | 0 |
| SRR10544946 | 809 | 148 | 4666 | 0 | 0 | 43 | 80074 | 37 | 12 | 0 | 51 | 10693 | 0 | 0 | 16 | 0 | 30 | 0 | 157 | 1032 | 37 | 0 | 0 | 242 | 0 | 0 | 0 | 0 | 0 | 0 | 0 | 0 | 0 | 0 | 0 | 0 | 0 |
| SRR10544947 | 560 | 17 | 6633 | 0 | 0 | 17 | 75048 | 10 | 0 | 0 | 22 | 9270 | 0 | 0 | 20 | 0 | 39 | 0 | 678 | 2389 | 60 | 0 | 0 | 57 | 0 | 0 | 0 | 0 | 0 | 0 | 0 | 0 | 0 | 0 | 0 | 0 | 0 |
| SRR10544948 | 519 | 305 | 9300 | 0 | 0 | 26 | 116388 | 181 | 0 | 33 | 169 | 9878 | 0 | 0 | 31 | 0 | 36 | 0 | 170 | 466 | 20 | 0 | 0 | 337 | 15 | 0 | 0 | 0 | 0 | 0 | 0 | 0 | 0 | 0 | 0 | 0 | 0 |
| SRR10544949 | 570 | 40 | 4272 | 0 | 0 | 21 | 116096 | 23 | 11 | 0 | 29 | 10850 | 0 | 0 | 0 | 0 | 48 | 0 | 223 | 1937 | 83 | 0 | 0 | 65 | 29 | 0 | 0 | 0 | 0 | 0 | 0 | 0 | 0 | 0 | 0 | 0 | 0 |
| SRR10544950 | 504 | 27 | 6605 | 0 | 0 | 23 | 113649 | 25 | 0 | 0 | 15 | 8748 | 0 | 0 | 52 | 0 | 140 | 0 | 274 | 1336 | 98 | 0 | 0 | 102 | 26 | 0 | 0 | 0 | 0 | 0 | 0 | 0 | 0 | 0 | 0 | 0 | 0 |
| SRR10756575 | 221 | 14 | 101 | 0 | 0 | 98 | 20 | 20 | 141 | 52 | 22 | 1008 | 10 | 44 | 436 | 0 | 28 | 0 | 0 | 23 | 651 | 17 | 0 | 17 | 27 | 390 | 37 | 0 | 14 | 95 | 62 | 17 | 37 | 99 | 118 | 87 | 173 |
| SRR10756576 | 161 | 0 | 141 | 15 | 24 | 69 | 21 | 49 | 100 | 26 | 15 | 651 | 109 | 49 | 1031 | 0 | 33 | 93 | 12 | 0 | 1134 | 60 | 0 | 31 | 27 | 339 | 24 | 18 | 0 | 29 | 90 | 15 | 18 | 17 | 193 | 182 | 222 |
| SRR10756577 | 135 | 33 | 153 | 10 | 0 | 810 | 16 | 534 | 1523 | 60 | 1097 | 1012 | 98 | 76 | 1576 | 10 | 26 | 57 | 0 | 0 | 233 | 0 | 0 | 55 | 58 | 3836 | 0 | 13 | 10 | 88 | 1634 | 18 | 28 | 571 | 3781 | 2207 | 3552 |
| SRR10756578 | 156 | 18 | 207 | 0 | 13 | 662 | 102 | 123 | 793 | 37 | 349 | 1406 | 134 | 58 | 308 | 0 | 46 | 12 | 0 | 28 | 428 | 30 | 0 | 21 | 75 | 3061 | 39 | 0 | 14 | 28 | 639 | 0 | 10 | 482 | 3271 | 1784 | 2668 |
| SRR10756579 | 231 | 0 | 130 | 0 | 0 | 42 | 77 | 28 | 46 | 32 | 17 | 1566 | 19 | 41 | 537 | 0 | 38 | 12 | 0 | 14 | 387 | 17 | 0 | 32 | 69 | 200 | 32 | 23 | 20 | 122 | 35 | 0 | 29 | 93 | 151 | 112 | 161 |
| SRR10756581 | 157 | 0 | 161 | 0 | 17 | 52 | 599 | 10 | 38 | 0 | 66 | 886 | 12 | 12 | 2019 | 0 | 60 | 28 | 17 | 19 | 1236 | 0 | 0 | 28 | 69 | 260 | 145 | 0 | 0 | 57 | 29 | 0 | 76 | 23 | 256 | 264 | 435 |
| SRR10756582 | 300 | 22 | 67 | 0 | 22 | 2588 | 42 | 725 | 3578 | 0 | 148 | 1964 | 148 | 0 | 577 | 0 | 37 | 0 | 0 | 0 | 512 | 22 | 0 | 133 | 0 | 12029 | 37 | 0 | 0 | 244 | 1967 | 0 | 22 | 710 | 5634 | 7401 | 12421 |
| SRR10756583 | 203 | 25 | 79 | 17 | 34 | 1449 | 24 | 0 | 1592 | 0 | 184 | 1325 | 226 | 0 | 382 | 0 | 39 | 0 | 0 | 24 | 470 | 0 | 0 | 17 | 134 | 8151 | 50 | 0 | 0 | 17 | 2480 | 0 | 84 | 117 | 5939 | 3728 | 6149 |
| SRR10875785 | 444 | 0 | 201 | 0 | 50 | 204 | 163 | 0 | 84 | 285 | 790 | 139 | 0 | 203 | 12376 | 28 | 84 | 14 | 33 | 0 | 763 | 21 | 18 | 0 | 67 | 1311 | 36 | 26 | 57 | 0 | 259 | 1406 | 355 | 137 | 1683 | 5201 | 9410 |
| SRR10875786 | 246 | 19 | 113 | 0 | 13 | 16 | 37 | 0 | 0 | 352 | 781 | 87 | 0 | 92 | 2603 | 94 | 37 | 17 | 0 | 37 | 289 | 24 | 15 | 0 | 46 | 53 | 12 | 42 | 85 | 0 | 14 | 113 | 863 | 16 | 29 | 107 | 227 |
| SRR10953234 | 59 | 0 | 168 | 0 | 0 | 290 | 142 | 172 | 136 | 0 | 0 | 16 | 0 | 0 | 20 | 0 | 103 | 0 | 212 | 2838 | 471 | 0 | 0 | 5152 | 10 | 1129 | 1063 | 0 | 0 | 90 | 37 | 0 | 0 | 16 | 253 | 259 | 654 |
| SRR10953235 | 48 | 0 | 156 | 0 | 0 | 262 | 120 | 180 | 133 | 0 | 0 | 38 | 0 | 0 | 0 | 0 | 111 | 0 | 185 | 2283 | 315 | 0 | 0 | 6389 | 10 | 959 | 955 | 0 | 0 | 116 | 45 | 0 | 0 | 13 | 240 | 329 | 640 |
| SRR10953236 | 57 | 0 | 172 | 0 | 0 | 268 | 79 | 257 | 144 | 0 | 0 | 17 | 0 | 0 | 14 | 0 | 83 | 0 | 157 | 2258 | 292 | 0 | 0 | 6433 | 12 | 970 | 888 | 0 | 0 | 127 | 48 | 0 | 0 | 22 | 284 | 250 | 597 |
| SRR10953237 | 49 | 0 | 215 | 0 | 0 | 105 | 273 | 71 | 27 | 0 | 0 | 27 | 0 | 0 | 21 | 0 | 113 | 0 | 373 | 4733 | 461 | 0 | 0 | 1854 | 0 | 488 | 472 | 0 | 0 | 27 | 12 | 0 | 0 | 12 | 111 | 147 | 274 |
| SRR10953238 | 85 | 0 | 66 | 0 | 0 | 64 | 144 | 76 | 33 | 0 | 0 | 44 | 0 | 0 | 20 | 0 | 75 | 0 | 123 | 1292 | 281 | 0 | 0 | 1933 | 12 | 259 | 313 | 0 | 0 | 39 | 27 | 0 | 0 | 26 | 90 | 91 | 207 |
| SRR10953239 | 64 | 0 | 45 | 0 | 0 | 190 | 130 | 215 | 103 | 0 | 0 | 95 | 11 | 0 | 30 | 0 | 55 | 0 | 110 | 1195 | 339 | 0 | 0 | 3977 | 12 | 742 | 1034 | 0 | 40 | 143 | 39 | 0 | 0 | 30 | 305 | 414 | 752 |
| SRR1107492 | 34 | 0 | 400 | 0 | 0 | 0 | 0 | 0 | 0 | 0 | 0 | 45 | 0 | 25 | 19 | 0 | 33 | 0 | 0 | 0 | 446 | 33 | 0 | 0 | 0 | 0 | 0 | 12 | 0 | 0 | 0 | 0 | 0 | 19 | 993 | 60 | 635 |
| SRR1107493 | 39 | 0 | 1405 | 0 | 0 | 0 | 0 | 53 | 0 | 0 | 0 | 53 | 0 | 20 | 39 | 0 | 19 | 0 | 0 | 135 | 771 | 22 | 0 | 0 | 0 | 41 | 14 | 0 | 13 | 30 | 0 | 0 | 0 | 35 | 21 | 0 | 17 |
| SRR1107494 | 38 | 17 | 1997 | 0 | 0 | 51 | 176 | 81 | 37 | 0 | 0 | 35 | 0 | 0 | 213 | 0 | 24 | 0 | 38 | 991 | 703 | 11 | 0 | 0 | 15 | 100 | 72 | 0 | 13 | 27 | 11 | 0 | 0 | 15 | 57 | 24 | 29 |
| SRR1107495 | 37 | 21 | 1966 | 0 | 0 | 35 | 150 | 85 | 37 | 0 | 0 | 40 | 0 | 0 | 55 | 0 | 28 | 0 | 0 | 846 | 641 | 17 | 0 | 0 | 0 | 90 | 63 | 0 | 14 | 25 | 29 | 0 | 0 | 21 | 72 | 34 | 26 |
| SRR1107496 | 44 | 21 | 1872 | 0 | 0 | 53 | 349 | 853 | 32 | 0 | 0 | 34 | 0 | 0 | 48 | 0 | 39 | 0 | 48 | 1499 | 650 | 19 | 0 | 0 | 0 | 813 | 529 | 0 | 148 | 345 | 11 | 0 | 0 | 14 | 396 | 29 | 160 |
| SRR1107497 | 46 | 0 | 1794 | 0 | 0 | 47 | 180 | 68 | 34 | 0 | 0 | 34 | 0 | 17 | 45 | 0 | 25 | 0 | 0 | 1167 | 763 | 25 | 0 | 0 | 0 | 75 | 47 | 0 | 13 | 26 | 13 | 0 | 0 | 0 | 45 | 21 | 17 |
| SRR1159512 | 45 | 0 | 1716 | 0 | 0 | 64 | 24 | 0 | 29 | 0 | 0 | 108 | 0 | 10 | 37 | 0 | 15 | 0 | 10 | 2297 | 652 | 0 | 0 | 0 | 48 | 116 | 491 | 0 | 0 | 0 | 11 | 0 | 0 | 39 | 187 | 29 | 111 |
| SRR1207050 | 284 | 0 | 386 | 0 | 0 | 55 | 0 | 0 | 15 | 0 | 0 | 25 | 0 | 0 | 11 | 86 | 18 | 28 | 0 | 24 | 44 | 11 | 45 | 0 | 15 | 447 | 34 | 0 | 616 | 0 | 11 | 0 | 45 | 0 | 0 | 0 | 0 |
| SRR1207051 | 381 | 0 | 710 | 0 | 0 | 11 | 0 | 0 | 178 | 0 | 0 | 697 | 0 | 0 | 127 | 11 | 0 | 0 | 28 | 443 | 174 | 25 | 0 | 0 | 22 | 81 | 58 | 15 | 98 | 0 | 255 | 0 | 23 | 0 | 34 | 22 | 0 |
| SRR1207053 | 455 | 0 | 428 | 0 | 0 | 247 | 0 | 0 | 51 | 0 | 0 | 479 | 0 | 0 | 11 | 261 | 0 | 0 | 0 | 0 | 22 | 14 | 11 | 0 | 0 | 591 | 0 | 23 | 253 | 0 | 207 | 0 | 205 | 0 | 20 | 0 | 0 |
| SRR1207054 | 416 | 0 | 671 | 0 | 0 | 358 | 0 | 0 | 69 | 0 | 0 | 205 | 0 | 0 | 0 | 34 | 0 | 0 | 0 | 0 | 26 | 0 | 32 | 0 | 0 | 2266 | 0 | 0 | 18463 | 0 | 59 | 0 | 89 | 0 | 0 | 0 | 0 |

### Supp Table S8.2.Columns_AM_to_BW

| | | | | | | | | | | | | | | | | | | | | | | | | | | | | | | | | | | | | | |
|---|---|---|---|---|---|---|---|---|---|---|---|---|---|---|---|---|---|---|---|---|---|---|---|---|---|---|---|---|---|---|---|---|---|---|---|---|---|
| SRR1207055 | 391 | 0 | 546 | 0 | 0 | 154 | 0 | 0 | 57 | 0 | 39 | 656 | 0 | 0 | 0 | 129 | 0 | 0 | 0 | 0 | 0 | 0 | 14 | 0 | 10 | 61 | 0 | 11 | 16 | 0 | 34 | 0 | 106 | 0 | 0 | 0 | 0 |
| SRR1232072 | 36 | 0 | 472 | 0 | 0 | 0 | 157 | 467 | 0 | 0 | 0 | 18 | 0 | 22 | 0 | 0 | 22 | 0 | 24 | 2412 | 100 | 0 | 0 | 0 | 0 | 22 | 94 | 0 | 24 | 21 | 0 | 0 | 0 | 17 | 0 | 0 | 0 |
| SRR1232073 | 66 | 15 | 2836 | 0 | 0 | 0 | 429 | 1584 | 0 | 0 | 0 | 89 | 0 | 32 | 14 | 0 | 17 | 0 | 26 | 2220 | 82 | 0 | 0 | 0 | 0 | 99 | 334 | 15 | 51 | 12 | 0 | 0 | 0 | 0 | 54 | 0 | 17 |
| SRR1240812 | 28 | 0 | 3500 | 0 | 0 | 12 | 41 | 13 | 76 | 0 | 0 | 141 | 15 | 0 | 23 | 0 | 47 | 0 | 44 | 16688 | 711 | 0 | 0 | 72 | 0 | 176 | 1030 | 0 | 0 | 0 | 96 | 0 | 0 | 0 | 277 | 25 | 135 |
| SRR1240813 | 25 | 0 | 3155 | 0 | 0 | 54 | 34 | 0 | 16 | 0 | 0 | 103 | 12 | 0 | 17 | 0 | 35 | 0 | 68 | 15708 | 799 | 0 | 0 | 38 | 0 | 160 | 914 | 0 | 0 | 0 | 46 | 0 | 0 | 0 | 197 | 22 | 111 |
| SRR1240814 | 69 | 0 | 2522 | 0 | 0 | 49 | 20 | 11 | 18 | 0 | 0 | 247 | 0 | 0 | 13 | 0 | 16 | 0 | 26 | 16339 | 3863 | 0 | 0 | 48 | 0 | 1079 | 10868 | 0 | 0 | 0 | 66 | 0 | 0 | 0 | 874 | 129 | 808 |
| SRR1240815 | 66 | 0 | 2416 | 0 | 0 | 49 | 0 | 0 | 17 | 0 | 0 | 236 | 0 | 0 | 15 | 0 | 19 | 0 | 23 | 15621 | 4070 | 0 | 0 | 46 | 0 | 1127 | 11061 | 0 | 0 | 0 | 75 | 0 | 0 | 0 | 966 | 156 | 834 |
| SRR1240816 | 39 | 0 | 402 | 0 | 0 | 60 | 0 | 0 | 11 | 0 | 0 | 525 | 23 | 0 | 0 | 0 | 39 | 0 | 60 | 4166 | 587 | 0 | 0 | 91 | 0 | 107 | 866 | 0 | 0 | 0 | 61 | 0 | 0 | 0 | 257 | 57 | 168 |
| SRR1240817 | 34 | 0 | 324 | 0 | 0 | 61 | 14 | 0 | 17 | 0 | 0 | 501 | 15 | 0 | 0 | 0 | 29 | 0 | 60 | 3240 | 559 | 0 | 0 | 83 | 0 | 1076 | 7799 | 10 | 0 | 0 | 84 | 0 | 0 | 0 | 2437 | 567 | 1529 |
| SRR1240818 | 78 | 76 | 5944 | 0 | 0 | 0 | 1652 | 612 | 0 | 0 | 0 | 176 | 0 | 11 | 544 | 0 | 0 | 0 | 282 | 11255 | 2843 | 0 | 0 | 19 | 0 | 82 | 1466 | 0 | 0 | 16 | 0 | 12 | 0 | 14 | 193 | 22 | 143 |
| SRR1241600 | 142 | 0 | 348 | 0 | 0 | 14 | 10 | 0 | 11 | 0 | 0 | 516 | 0 | 11 | 294 | 0 | 123 | 0 | 15 | 308 | 369 | 0 | 0 | 0 | 27 | 112 | 41 | 0 | 11 | 0 | 22 | 0 | 0 | 0 | 17 | 39 | 16 |
| SRR1241601 | 101 | 14 | 1572 | 0 | 0 | 95 | 72 | 0 | 27 | 14 | 0 | 0 | 0 | 0 | 340 | 0 | 14 | 0 | 51 | 385 | 1104 | 0 | 0 | 0 | 35 | 1180 | 163 | 0 | 0 | 0 | 111 | 0 | 31 | 0 | 17 | 39 | 13 |
| SRR1241602 | 187 | 0 | 2412 | 0 | 0 | 38 | 42 | 0 | 21 | 11 | 0 | 13 | 0 | 12 | 110 | 0 | 160 | 0 | 34 | 225 | 286 | 0 | 0 | 0 | 37 | 396 | 25 | 10 | 0 | 0 | 164 | 0 | 0 | 0 | 0 | 14 | 0 |
| SRR1241603 | 39 | 18 | 1268 | 0 | 0 | 125 | 0 | 171 | 16 | 0 | 0 | 36 | 0 | 0 | 36 | 0 | 0 | 0 | 0 | 239 | 1586 | 0 | 0 | 0 | 0 | 1241 | 1524 | 47 | 0 | 0 | 152 | 0 | 0 | 0 | 8458 | 798 | 3314 |
| SRR1241604 | 26 | 12 | 1284 | 0 | 0 | 113 | 0 | 141 | 31 | 0 | 0 | 55 | 0 | 0 | 45 | 0 | 10 | 0 | 0 | 371 | 1445 | 0 | 0 | 0 | 0 | 1627 | 1264 | 41 | 0 | 0 | 132 | 0 | 0 | 0 | 4986 | 452 | 1934 |
| SRR1241605 | 12 | 10 | 1163 | 0 | 0 | 123 | 0 | 217 | 22 | 0 | 0 | 26 | 0 | 0 | 20 | 0 | 0 | 0 | 10 | 583 | 4669 | 0 | 0 | 0 | 0 | 1448 | 1558 | 40 | 0 | 0 | 196 | 0 | 0 | 0 | 5528 | 675 | 2292 |
| SRR1264353 | 80 | 0 | 538 | 0 | 0 | 0 | 481 | 0 | 0 | 0 | 0 | 844 | 0 | 0 | 64 | 0 | 0 | 0 | 192 | 4198 | 1186 | 0 | 0 | 0 | 14 | 23 | 15 | 0 | 0 | 0 | 0 | 0 | 0 | 0 | 17 | 16 | 31 |
| SRR1264354 | 190 | 0 | 221 | 0 | 0 | 0 | 282 | 0 | 0 | 0 | 12 | 3316 | 0 | 0 | 0 | 0 | 12 | 0 | 141 | 2115 | 740 | 0 | 0 | 0 | 13 | 17 | 15 | 0 | 0 | 0 | 0 | 0 | 0 | 0 | 51 | 22 | 44 |
| SRR1270153 | 23 | 0 | 46 | 0 | 0 | 17 | 627 | 19 | 0 | 0 | 0 | 30 | 0 | 0 | 0 | 0 | 0 | 0 | 206 | 81 | 0 | 0 | 0 | 0 | 12 | 578 | 0 | 0 | 0 | 0 | 427 | 0 | 12 | 0 | 0 | 0 | 0 |
| SRR1290784 | 27 | 0 | 36433 | 0 | 0 | 10 | 24 | 0 | 40 | 0 | 0 | 72 | 0 | 0 | 32 | 0 | 66 | 0 | 0 | 2398 | 1151 | 0 | 0 | 0 | 0 | 183 | 570 | 0 | 0 | 0 | 14 | 0 | 0 | 0 | 102 | 56 | 353 |
| SRR1290785 | 24 | 0 | 37335 | 0 | 0 | 75 | 30 | 34 | 20 | 0 | 0 | 59 | 0 | 0 | 24 | 0 | 83 | 18 | 24 | 3123 | 997 | 0 | 0 | 26 | 0 | 146 | 567 | 0 | 0 | 0 | 13 | 0 | 0 | 0 | 74 | 38 | 274 |
| SRR1290786 | 24 | 0 | 42741 | 0 | 0 | 74 | 0 | 11 | 38 | 0 | 0 | 71 | 0 | 0 | 35 | 0 | 75 | 0 | 0 | 2587 | 605 | 0 | 0 | 31 | 0 | 1639 | 4922 | 0 | 0 | 0 | 130 | 0 | 0 | 0 | 816 | 371 | 3119 |
| SRR1290787 | 21 | 0 | 42502 | 0 | 0 | 99 | 33 | 0 | 27 | 0 | 0 | 78 | 0 | 0 | 191 | 0 | 71 | 17 | 0 | 2646 | 3525 | 24 | 0 | 0 | 0 | 2176 | 5958 | 0 | 0 | 0 | 104 | 0 | 0 | 0 | 611 | 330 | 2951 |
| SRR1290788 | 23 | 0 | 41461 | 0 | 0 | 86 | 23 | 0 | 23 | 0 | 0 | 75 | 0 | 0 | 256 | 0 | 72 | 34 | 0 | 2263 | 2733 | 18 | 0 | 0 | 0 | 1606 | 4389 | 0 | 0 | 0 | 133 | 0 | 0 | 0 | 666 | 325 | 2653 |
| SRR1290789 | 26 | 0 | 37670 | 0 | 0 | 43 | 11 | 0 | 25 | 0 | 0 | 63 | 0 | 0 | 128 | 0 | 66 | 0 | 0 | 399 | 415 | 0 | 0 | 0 | 0 | 1590 | 1844 | 0 | 0 | 0 | 200 | 0 | 0 | 0 | 1297 | 547 | 4477 |
| SRR1290790 | 15 | 0 | 39123 | 0 | 0 | 93 | 42 | 14 | 11 | 0 | 0 | 53 | 0 | 0 | 30 | 0 | 70 | 0 | 0 | 2377 | 942 | 11 | 0 | 0 | 0 | 1514 | 5812 | 0 | 0 | 0 | 149 | 0 | 0 | 0 | 678 | 330 | 2329 |
| SRR1290791 | 18 | 0 | 31234 | 0 | 0 | 12 | 35 | 0 | 36 | 0 | 0 | 53 | 0 | 0 | 31 | 0 | 80 | 0 | 0 | 1435 | 1183 | 0 | 0 | 0 | 0 | 182 | 364 | 0 | 0 | 0 | 14 | 0 | 0 | 0 | 104 | 70 | 457 |
| SRR1290792 | 27 | 0 | 39759 | 0 | 0 | 108 | 24 | 31 | 27 | 0 | 0 | 88 | 0 | 0 | 42 | 0 | 75 | 0 | 0 | 2801 | 872 | 0 | 0 | 0 | 16 | 176 | 455 | 0 | 0 | 0 | 12 | 0 | 0 | 0 | 93 | 50 | 345 |
| SRR1290793 | 27 | 0 | 41470 | 0 | 0 | 88 | 0 | 0 | 32 | 0 | 0 | 61 | 0 | 0 | 23 | 0 | 82 | 0 | 0 | 977 | 575 | 0 | 0 | 0 | 0 | 1970 | 4777 | 0 | 0 | 0 | 171 | 0 | 0 | 0 | 1665 | 606 | 4432 |
| SRR1290794 | 33 | 0 | 62602 | 0 | 0 | 64 | 54 | 16 | 18 | 0 | 0 | 65 | 0 | 0 | 27 | 0 | 46 | 0 | 0 | 876 | 424 | 0 | 0 | 0 | 0 | 1091 | 1051 | 0 | 0 | 0 | 87 | 0 | 0 | 0 | 543 | 337 | 1916 |
| SRR1290795 | 36 | 0 | 66537 | 0 | 0 | 53 | 0 | 0 | 22 | 0 | 0 | 58 | 0 | 0 | 19 | 0 | 46 | 0 | 0 | 867 | 647 | 0 | 0 | 0 | 0 | 130 | 174 | 0 | 0 | 0 | 71 | 0 | 0 | 0 | 80 | 35 | 247 |
| SRR1290796 | 27 | 0 | 54573 | 0 | 0 | 47 | 13 | 30 | 18 | 0 | 0 | 57 | 0 | 0 | 99 | 0 | 48 | 0 | 0 | 693 | 648 | 0 | 0 | 61 | 0 | 1337 | 1562 | 0 | 0 | 0 | 212 | 0 | 0 | 0 | 794 | 221 | 1739 |
| SRR1290797 | 21 | 0 | 49576 | 0 | 0 | 87 | 17 | 27 | 31 | 0 | 0 | 44 | 0 | 0 | 133 | 0 | 53 | 0 | 0 | 2414 | 2034 | 17 | 0 | 0 | 0 | 228 | 568 | 0 | 0 | 0 | 28 | 0 | 0 | 0 | 132 | 32 | 263 |
| SRR1290798 | 21 | 0 | 41116 | 0 | 0 | 122 | 34 | 106 | 30 | 0 | 0 | 34 | 0 | 0 | 71 | 0 | 47 | 0 | 0 | 2060 | 1952 | 21 | 0 | 32 | 0 | 2929 | 5814 | 0 | 0 | 0 | 242 | 0 | 0 | 0 | 3603 | 870 | 6672 |
| SRR1290799 | 25 | 13 | 48931 | 0 | 0 | 42 | 26 | 17 | 45 | 0 | 0 | 19 | 0 | 0 | 55 | 0 | 27 | 0 | 0 | 494 | 1213 | 24 | 0 | 0 | 0 | 275 | 518 | 0 | 0 | 0 | 63 | 0 | 0 | 0 | 62 | 18 | 164 |
| SRR1290800 | 16 | 10 | 49709 | 0 | 0 | 46 | 50 | 553 | 19 | 0 | 0 | 24 | 0 | 0 | 57 | 0 | 28 | 0 | 0 | 1044 | 1280 | 122 | 0 | 20 | 0 | 1165 | 2846 | 0 | 0 | 0 | 45 | 0 | 0 | 0 | 225 | 64 | 585 |
| SRR1290801 | 27 | 12 | 53346 | 0 | 0 | 14 | 131 | 134 | 29 | 0 | 0 | 82 | 0 | 0 | 20 | 0 | 46 | 0 | 0 | 2044 | 669 | 24 | 0 | 29 | 0 | 235 | 393 | 0 | 0 | 0 | 47 | 0 | 0 | 0 | 35 | 52 | 112 |
| SRR1290802 | 29 | 0 | 41702 | 0 | 0 | 97 | 0 | 0 | 12 | 0 | 0 | 82 | 0 | 0 | 42 | 0 | 78 | 0 | 0 | 4177 | 1271 | 13 | 0 | 30 | 16 | 132 | 691 | 0 | 0 | 0 | 105 | 0 | 0 | 0 | 143 | 39 | 210 |
| SRR1290803 | 27 | 0 | 45564 | 0 | 0 | 77 | 0 | 0 | 12 | 0 | 0 | 91 | 0 | 0 | 19 | 0 | 66 | 0 | 0 | 4878 | 769 | 0 | 0 | 82 | 0 | 1297 | 8976 | 0 | 0 | 0 | 70 | 0 | 0 | 0 | 1266 | 332 | 2061 |
| SRR1290804 | 24 | 0 | 32747 | 0 | 0 | 63 | 13 | 0 | 23 | 0 | 0 | 68 | 19 | 0 | 218 | 0 | 95 | 13 | 0 | 2775 | 8155 | 0 | 0 | 34 | 0 | 100 | 492 | 0 | 0 | 0 | 11 | 0 | 0 | 0 | 165 | 62 | 314 |
| SRR1290805 | 37 | 0 | 38194 | 0 | 0 | 74 | 0 | 0 | 20 | 0 | 0 | 70 | 0 | 0 | 163 | 0 | 86 | 10 | 0 | 3251 | 9112 | 11 | 0 | 17 | 0 | 157 | 784 | 0 | 0 | 0 | 76 | 0 | 0 | 0 | 196 | 62 | 479 |
| SRR1290806 | 33 | 0 | 34963 | 0 | 0 | 61 | 0 | 0 | 12 | 0 | 0 | 109 | 0 | 0 | 138 | 0 | 75 | 12 | 0 | 1759 | 3404 | 12 | 0 | 0 | 0 | 143 | 480 | 0 | 0 | 0 | 10 | 0 | 0 | 0 | 219 | 68 | 349 |
| SRR1290807 | 16 | 0 | 39898 | 0 | 0 | 128 | 0 | 0 | 21 | 0 | 0 | 73 | 0 | 0 | 21 | 0 | 66 | 0 | 0 | 2306 | 939 | 0 | 0 | 0 | 0 | 2023 | 7061 | 0 | 0 | 0 | 118 | 0 | 0 | 0 | 2497 | 824 | 4559 |
| SRR1290808 | 24 | 0 | 31363 | 0 | 0 | 131 | 95 | 0 | 39 | 0 | 0 | 60 | 0 | 0 | 0 | 0 | 92 | 0 | 0 | 1814 | 1218 | 20 | 0 | 0 | 0 | 2157 | 7442 | 0 | 0 | 0 | 98 | 0 | 0 | 0 | 2060 | 758 | 4142 |
| SRR1290809 | 28 | 0 | 35478 | 0 | 0 | 116 | 28 | 12 | 33 | 0 | 0 | 73 | 0 | 0 | 36 | 0 | 72 | 0 | 10 | 3980 | 1519 | 20 | 0 | 0 | 0 | 131 | 723 | 0 | 0 | 0 | 114 | 0 | 0 | 0 | 150 | 51 | 285 |
| SRR1292461 | 624 | 0 | 20987 | 0 | 0 | 14 | 0 | 0 | 322 | 37 | 22 | 342 | 0 | 0 | 180 | 48 | 32 | 0 | 0 | 0 | 27 | 18 | 0 | 0 | 384 | 82 | 22 | 0 | 238 | 0 | 191 | 0 | 15 | 0 | 523 | 48 | 0 |
| SRR1304304 | 20 | 60 | 624 | 0 | 28 | 0 | 0 | 0 | 0 | 0 | 0 | 104 | 0 | 27 | 150 | 0 | 0 | 0 | 0 | 0 | 0 | 0 | 0 | 0 | 0 | 0 | 48 | 13 | 0 | 0 | 25 | 0 | 0 | 0 | 24 | 276 | 17 |
| SRR1304307 | 36 | 14 | 1542 | 0 | 14 | 0 | 0 | 0 | 0 | 0 | 0 | 329 | 0 | 47 | 185 | 0 | 21 | 0 | 0 | 0 | 12 | 0 | 0 | 0 | 0 | 0 | 0 | 30 | 0 | 0 | 83 | 0 | 0 | 15 | 511 | 17 | 24 |
| SRR1304309 | 102 | 28 | 1022 | 45 | 40 | 0 | 0 | 0 | 0 | 10 | 0 | 69 | 0 | 10 | 424 | 0 | 0 | 0 | 0 | 0 | 64 | 0 | 0 | 0 | 15 | 23 | 13 | 63 | 0 | 0 | 277 | 0 | 0 | 0 | 1906 | 169 | 176 |
| SRR1304313 | 21 | 0 | 8419 | 0 | 0 | 40 | 46 | 393 | 18 | 0 | 0 | 0 | 0 | 0 | 67 | 0 | 33 | 0 | 11 | 3573 | 1327 | 0 | 0 | 22 | 0 | 839 | 5688 | 14 | 35 | 30 | 165 | 0 | 0 | 35 | 1356 | 139 | 814 |
| SRR1393733 | 25 | 0 | 643 | 100 | 127 | 0 | 0 | 0 | 57 | 0 | 0 | 2101 | 0 | 0 | 118 | 0 | 41 | 15 | 0 | 0 | 51 | 49 | 0 | 0 | 290 | 285 | 0 | 0 | 11 | 0 | 803 | 0 | 0 | 0 | 20041 | 1873 | 5891 |
| SRR1510077 | 20 | 0 | 3559 | 0 | 0 | 0 | 66 | 116 | 0 | 0 | 0 | 289 | 0 | 0 | 609 | 0 | 0 | 0 | 101 | 301 | 2207 | 0 | 0 | 59 | 13 | 148 | 125 | 13 | 0 | 13 | 0 | 0 | 0 | 44 | 0 | 0 | 11 |
| SRR1510078 | 17 | 0 | 3593 | 0 | 0 | 0 | 390 | 48 | 0 | 0 | 0 | 262 | 0 | 20 | 2259 | 0 | 0 | 0 | 116 | 107 | 1201 | 0 | 0 | 32 | 45 | 74 | 69 | 33 | 0 | 11 | 0 | 0 | 25 | 52 | 13 | 0 | 0 |
| SRR1510079 | 13 | 0 | 3727 | 0 | 0 | 36 | 70 | 41 | 23 | 0 | 0 | 161 | 0 | 0 | 347 | 0 | 0 | 0 | 88 | 28 | 755 | 0 | 0 | 14 | 0 | 757 | 182 | 18 | 0 | 0 | 0 | 0 | 14 | 32 | 14 | 0 | 0 |
| SRR1510080 | 14 | 0 | 4201 | 0 | 0 | 0 | 840 | 111 | 0 | 0 | 0 | 105 | 0 | 0 | 3834 | 0 | 0 | 0 | 241 | 90 | 1737 | 0 | 0 | 0 | 0 | 128 | 57 | 74 | 0 | 25 | 0 | 0 | 0 | 48 | 14 | 0 | 12 |
| SRR1510081 | 0 | 0 | 1114 | 0 | 0 | 11 | 358 | 68 | 0 | 0 | 0 | 51 | 0 | 0 | 332 | 0 | 0 | 0 | 130 | 79 | 732 | 0 | 0 | 28 | 0 | 15 | 53 | 11 | 0 | 0 | 15 | 0 | 13 | 19 | 17 | 0 | 0 |
| SRR1510082 | 44 | 0 | 2711 | 0 | 0 | 0 | 1040 | 154 | 0 | 0 | 0 | 198 | 0 | 0 | 417 | 0 | 0 | 0 | 279 | 227 | 2300 | 0 | 0 | 91 | 0 | 197 | 81 | 0 | 0 | 15 | 0 | 0 | 0 | 0 | 12 | 0 | 23 |
| SRR1510083 | 38 | 0 | 4638 | 0 | 0 | 0 | 370 | 26 | 0 | 0 | 0 | 155 | 0 | 0 | 1638 | 0 | 0 | 0 | 727 | 316 | 3658 | 0 | 0 | 70 | 0 | 96 | 94 | 0 | 0 | 0 | 0 | 0 | 0 | 0 | 10 | 0 | 12 |
| SRR1510084 | 19 | 11 | 2739 | 0 | 0 | 23 | 476 | 42 | 0 | 0 | 0 | 363 | 0 | 11 | 118 | 0 | 0 | 0 | 176 | 55 | 352 | 0 | 0 | 31 | 0 | 119 | 54 | 25 | 11 | 23 | 0 | 0 | 11 | 37 | 45 | 0 | 37 |
| SRR1510085 | 19 | 10 | 2396 | 0 | 0 | 10 | 274 | 0 | 0 | 0 | 0 | 208 | 0 | 0 | 1019 | 0 | 0 | 0 | 251 | 145 | 1058 | 0 | 0 | 12 | 24 | 81 | 110 | 24 | 0 | 0 | 0 | 0 | 0 | 24 | 16 | 0 | 0 |
| SRR1510086 | 11 | 13 | 2755 | 13 | 0 | 10 | 1695 | 11 | 0 | 0 | 0 | 109 | 0 | 13 | 95 | 0 | 0 | 0 | 95 | 33 | 172 | 0 | 0 | 0 | 0 | 180 | 58 | 0 | 0 | 41 | 0 | 0 | 13 | 48 | 20 | 0 | 10 |
| SRR1510087 | 39 | 11 | 2732 | 0 | 0 | 24 | 148 | 17 | 0 | 0 | 0 | 270 | 0 | 0 | 1001 | 0 | 0 | 0 | 234 | 673 | 8714 | 0 | 0 | 50 | 0 | 32 | 19 | 0 | 0 | 0 | 0 | 0 | 0 | 0 | 21 | 0 | 25 |
| SRR1510088 | 17 | 0 | 3653 | 0 | 0 | 0 | 1285 | 89 | 0 | 0 | 0 | 216 | 0 | 0 | 2469 | 0 | 0 | 0 | 143 | 34 | 469 | 0 | 0 | 0 | 13 | 52 | 66 | 0 | 0 | 20 | 0 | 0 | 0 | 21 | 67 | 0 | 27 |
| SRR1510089 | 18 | 0 | 4715 | 0 | 0 | 11 | 47 | 14 | 0 | 0 | 0 | 289 | 0 | 0 | 950 | 0 | 0 | 0 | 120 | 137 | 3144 | 0 | 0 | 21 | 0 | 16 | 14 | 0 | 0 | 17 | 11 | 0 | 0 | 19 | 25 | 0 | 11 |
| SRR1510090 | 26 | 0 | 2274 | 0 | 0 | 26 | 98 | 10 | 0 | 0 | 0 | 171 | 0 | 0 | 60 | 0 | 0 | 0 | 82 | 67 | 259 | 0 | 0 | 83 | 16 | 268 | 153 | 18 | 0 | 0 | 10 | 10 | 0 | 18 | 42 | 0 | 44 |
| SRR1510091 | 13 | 0 | 2362 | 0 | 0 | 19 | 474 | 99 | 0 | 0 | 0 | 117 | 0 | 0 | 654 | 0 | 0 | 0 | 236 | 72 | 540 | 0 | 0 | 22 | 16 | 189 | 25 | 0 | 0 | 25 | 0 | 0 | 0 | 0 | 12 | 0 | 19 |
| SRR1510092 | 71 | 0 | 2107 | 0 | 0 | 21 | 255 | 31 | 0 | 0 | 0 | 103 | 0 | 0 | 81 | 0 | 0 | 0 | 168 | 814 | 798 | 0 | 0 | 131 | 0 | 139 | 86 | 0 | 0 | 0 | 0 | 0 | 0 | 0 | 0 | 0 | 0 |
| SRR1510093 | 29 | 0 | 2798 | 0 | 0 | 18 | 150 | 39 | 0 | 0 | 0 | 114 | 0 | 0 | 953 | 0 | 0 | 0 | 77 | 154 | 44 | 0 | 0 | 67 | 22 | 150 | 75 | 20 | 0 | 0 | 0 | 0 | 0 | 18 | 10 | 0 | 20 |
| SRR1510094 | 19 | 0 | 3754 | 0 | 0 | 24 | 207 | 72 | 0 | 0 | 0 | 58 | 0 | 0 | 774 | 0 | 0 | 0 | 467 | 172 | 1441 | 0 | 0 | 17 | 14 | 245 | 95 | 0 | 0 | 10 | 10 | 0 | 14 | 0 | 14 | 0 | 27 |
| SRR1510095 | 0 | 11 | 2761 | 0 | 0 | 22 | 251 | 108 | 0 | 0 | 0 | 122 | 0 | 0 | 318 | 0 | 0 | 0 | 99 | 23 | 160 | 0 | 0 | 11 | 11 | 264 | 141 | 0 | 0 | 0 | 15 | 0 | 0 | 15 | 56 | 0 | 26 |
| SRR1510096 | 11 | 15 | 1863 | 0 | 0 | 0 | 85 | 0 | 12 | 0 | 0 | 199 | 0 | 24 | 1049 | 0 | 0 | 0 | 52 | 340 | 1998 | 0 | 0 | 85 | 18 | 124 | 121 | 12 | 0 | 0 | 0 | 0 | 18 | 18 | 21 | 0 | 18 |
| SRR1510097 | 80 | 0 | 10972 | 0 | 0 | 32 | 143 | 23 | 0 | 0 | 0 | 98 | 0 | 0 | 104 | 0 | 21 | 0 | 68 | 610 | 698 | 14 | 0 | 23 | 50 | 372 | 113 | 36 | 0 | 0 | 14 | 0 | 23 | 14 | 18 | 0 | 50 |
| SRR1510098 | 81 | 10 | 8857 | 0 | 13 | 10 | 127 | 33 | 0 | 0 | 0 | 222 | 0 | 0 | 69 | 0 | 0 | 0 | 816 | 583 | 1277 | 0 | 0 | 28 | 25 | 25 | 17 | 15 | 0 | 0 | 0 | 0 | 0 | 30 | 43 | 0 | 45 |
| SRR1510099 | 356 | 0 | 4273 | 14 | 11 | 11 | 181 | 75 | 0 | 0 | 0 | 78 | 0 | 0 | 32 | 0 | 0 | 0 | 781 | 646 | 980 | 18 | 0 | 61 | 0 | 14 | 12 | 20 | 0 | 25 | 0 | 18 | 0 | 16 | 11 | 0 | 32 |
| SRR1510100 | 74 | 27 | 10417 | 0 | 0 | 11 | 613 | 14 | 0 | 0 | 0 | 273 | 0 | 17 | 124 | 0 | 0 | 40 | 1963 | 810 | 387 | 0 | 0 | 12 | 22 | 17 | 31 | 22 | 0 | 0 | 0 | 10 | 0 | 0 | 10 | 0 | 14 |
| SRR1510101 | 47 | 0 | 5436 | 0 | 0 | 0 | 351 | 0 | 0 | 11 | 0 | 313 | 0 | 0 | 177 | 0 | 11 | 0 | 471 | 104 | 665 | 0 | 0 | 19 | 49 | 163 | 166 | 0 | 0 | 0 | 0 | 0 | 0 | 0 | 24 | 0 | 24 |
| SRR1510102 | 17 | 16 | 5201 | 0 | 0 | 12 | 146 | 0 | 0 | 0 | 0 | 312 | 0 | 10 | 536 | 0 | 0 | 0 | 411 | 216 | 913 | 0 | 0 | 24 | 32 | 107 | 109 | 11 | 0 | 0 | 0 | 0 | 27 | 24 | 35 | 0 | 22 |
| SRR1510103 | 18 | 0 | 4679 | 0 | 0 | 0 | 658 | 69 | 0 | 0 | 0 | 360 | 0 | 0 | 93 | 0 | 0 | 0 | 204 | 44 | 150 | 0 | 0 | 13 | 12 | 172 | 153 | 0 | 0 | 0 | 0 | 0 | 0 | 0 | 77 | 0 | 41 |
| SRR1510104 | 46 | 0 | 4952 | 0 | 0 | 0 | 768 | 119 | 0 | 0 | 0 | 280 | 0 | 0 | 187 | 0 | 0 | 0 | 306 | 131 | 742 | 0 | 0 | 35 | 0 | 133 | 131 | 0 | 0 | 12 | 10 | 0 | 0 | 0 | 0 | 0 | 0 |
| SRR1510105 | 19 | 0 | 4897 | 0 | 0 | 20 | 65 | 0 | 0 | 0 | 0 | 300 | 0 | 0 | 5978 | 0 | 11 | 0 | 260 | 242 | 2065 | 0 | 0 | 72 | 16 | 165 | 13 | 0 | 0 | 0 | 0 | 0 | 0 | 0 | 32 | 0 | 12 |
| SRR1510106 | 24 | 0 | 4867 | 0 | 0 | 10 | 659 | 11 | 0 | 0 | 0 | 284 | 0 | 0 | 525 | 0 | 0 | 0 | 335 | 106 | 1334 | 0 | 0 | 24 | 0 | 108 | 85 | 0 | 0 | 28 | 0 | 0 | 0 | 16 | 26 | 0 | 21 |
| SRR1510107 | 0 | 0 | 4203 | 0 | 0 | 14 | 121 | 0 | 14 | 0 | 0 | 242 | 0 | 0 | 380 | 0 | 0 | 0 | 175 | 103 | 559 | 0 | 0 | 34 | 17 | 221 | 146 | 29 | 0 | 0 | 0 | 0 | 0 | 0 | 12 | 0 | 22 |
| SRR1510108 | 24 | 0 | 4088 | 0 | 0 | 11 | 317 | 207 | 0 | 0 | 0 | 276 | 0 | 0 | 29 | 0 | 0 | 0 | 548 | 173 | 839 | 0 | 0 | 86 | 49 | 233 | 83 | 0 | 0 | 40 | 0 | 0 | 0 | 0 | 14 | 0 | 11 |
| SRR1510109 | 26 | 0 | 4234 | 0 | 0 | 26 | 236 | 119 | 0 | 0 | 0 | 306 | 0 | 0 | 145 | 0 | 0 | 11 | 165 | 82 | 888 | 0 | 0 | 23 | 38 | 17 | 12 | 0 | 0 | 11 | 0 | 0 | 0 | 0 | 25 | 0 | 26 |
| SRR1510110 | 78 | 0 | 2535 | 0 | 0 | 15 | 1113 | 19 | 0 | 0 | 0 | 366 | 0 | 0 | 312 | 0 | 0 | 0 | 253 | 167 | 1677 | 0 | 0 | 39 | 0 | 17 | 14 | 0 | 0 | 15 | 0 | 0 | 0 | 0 | 0 | 0 | 15 |
| SRR1510111 | 0 | 0 | 1777 | 0 | 0 | 21 | 186 | 85 | 0 | 0 | 0 | 170 | 0 | 0 | 41 | 0 | 0 | 0 | 369 | 94 | 311 | 0 | 0 | 13 | 55 | 226 | 128 | 21 | 0 | 0 | 0 | 0 | 0 | 0 | 64 | 0 | 38 |
| SRR1510112 | 19 | 0 | 2545 | 0 | 0 | 24 | 910 | 110 | 0 | 0 | 0 | 91 | 0 | 0 | 5418 | 0 | 0 | 0 | 158 | 74 | 3322 | 0 | 0 | 11 | 0 | 11 | 81 | 0 | 0 | 0 | 0 | 0 | 0 | 48 | 48 | 0 | 35 |
| SRR1510113 | 22 | 0 | 3162 | 0 | 0 | 17 | 164 | 15 | 0 | 0 | 0 | 125 | 0 | 0 | 345 | 0 | 0 | 0 | 224 | 411 | 786 | 0 | 0 | 19 | 0 | 11 | 113 | 0 | 0 | 0 | 19 | 0 | 0 | 0 | 72 | 0 | 41 |
| SRR1510114 | 11 | 0 | 1863 | 0 | 0 | 11 | 901 | 107 | 0 | 0 | 0 | 160 | 0 | 15 | 1056 | 0 | 0 | 0 | 150 | 54 | 948 | 0 | 0 | 33 | 15 | 13 | 80 | 18 | 0 | 41 | 0 | 0 | 14 | 39 | 20 | 0 | 27 |
| SRR1510115 | 12 | 0 | 2508 | 0 | 0 | 0 | 1117 | 12 | 0 | 0 | 0 | 210 | 0 | 0 | 1026 | 0 | 0 | 0 | 174 | 23 | 579 | 0 | 0 | 32 | 0 | 93 | 68 | 0 | 0 | 24 | 0 | 0 | 0 | 24 | 19 | 0 | 29 |
| SRR1510116 | 38 | 0 | 1909 | 0 | 0 | 16 | 345 | 51 | 0 | 0 | 0 | 156 | 0 | 0 | 226 | 0 | 0 | 11 | 283 | 190 | 1489 | 0 | 0 | 0 | 56 | 337 | 111 | 14 | 0 | 12 | 14 | 0 | 0 | 0 | 14 | 0 | 23 |
| SRR1510117 | 115 | 0 | 1261 | 0 | 0 | 19 | 289 | 12 | 0 | 0 | 0 | 152 | 0 | 0 | 85 | 0 | 0 | 46 | 567 | 310 | 3293 | 0 | 0 | 11 | 33 | 28 | 15 | 0 | 0 | 14 | 0 | 0 | 0 | 11 | 0 | 0 | 13 |
| SRR1510118 | 40 | 0 | 2174 | 0 | 0 | 0 | 1058 | 135 | 0 | 0 | 0 | 165 | 0 | 14 | 2565 | 0 | 0 | 0 | 281 | 119 | 2721 | 0 | 0 | 33 | 0 | 202 | 100 | 0 | 0 | 21 | 0 | 0 | 0 | 13 | 0 | 0 | 38 |
| SRR1510119 | 18 | 0 | 2924 | 0 | 0 | 23 | 212 | 31 | 0 | 0 | 0 | 73 | 0 | 0 | 3206 | 0 | 0 | 0 | 244 | 173 | 1396 | 0 | 0 | 31 | 23 | 15 | 94 | 10 | 0 | 0 | 0 | 0 | 0 | 14 | 31 | 0 | 30 |
| SRR1510120 | 23 | 0 | 3122 | 0 | 0 | 12 | 537 | 89 | 0 | 0 | 0 | 137 | 0 | 0 | 4982 | 0 | 0 | 0 | 233 | 85 | 1545 | 0 | 0 | 18 | 11 | 14 | 11 | 0 | 0 | 0 | 0 | 0 | 0 | 12 | 66 | 0 | 37 |
| SRR1510121 | 10 | 0 | 953 | 0 | 0 | 0 | 423 | 118 | 0 | 0 | 0 | 65 | 0 | 0 | 89 | 0 | 0 | 0 | 200 | 142 | 370 | 0 | 0 | 19 | 60 | 23 | 153 | 0 | 0 | 16 | 28 | 0 | 0 | 12 | 58 | 14 | 51 |
| SRR1519906 | 53 | 130 | 25 | 108 | 29 | 0 | 0 | 0 | 0 | 0 | 0 | 19 | 0 | 79 | 392 | 0 | 0 | 0 | 0 | 131 | 0 | 0 | 0 | 0 | 0 | 0 | 14 | 0 | 0 | 0 | 0 | 0 | 0 | 14 | 0 | 0 | 0 |
| SRR1519919 | 182 | 21 | 254 | 117 | 128 | 0 | 70 | 480 | 0 | 0 | 0 | 187 | 0 | 21 | 0 | 0 | 0 | 0 | 0 | 0 | 246 | 0 | 0 | 0 | 11 | 11 | 149 | 0 | 0 | 0 | 0 | 0 | 0 | 0 | 0 | 0 | 0 |
| SRR1525198 | 18 | 11 | 335 | 0 | 0 | 39 | 0 | 869 | 14 | 0 | 0 | 236 | 0 | 35 | 96 | 0 | 0 | 0 | 0 | 230 | 80 | 0 | 0 | 0 | 0 | 918 | 1995 | 49 | 0 | 11 | 35 | 0 | 0 | 0 | 17100 | 1384 | 4262 |
| SRR1525199 | 13 | 14 | 331 | 0 | 0 | 59 | 0 | 410 | 11 | 0 | 0 | 451 | 0 | 28 | 97 | 0 | 13 | 0 | 0 | 301 | 38 | 0 | 0 | 0 | 0 | 1236 | 1903 | 42 | 0 | 17 | 59 | 0 | 0 | 0 | 18489 | 1325 | 3974 |
| SRR1525200 | 14 | 20 | 573 | 0 | 0 | 20 | 0 | 1038 | 14 | 0 | 0 | 60 | 0 | 14 | 117 | 0 | 11 | 0 | 0 | 325 | 44 | 0 | 0 | 0 | 0 | 410 | 1906 | 75 | 0 | 0 | 0 | 0 | 0 | 0 | 17513 | 1940 | 6011 |
| SRR1525201 | 14 | 15 | 619 | 0 | 0 | 78 | 0 | 346 | 20 | 0 | 0 | 100 | 0 | 20 | 272 | 0 | 26 | 0 | 0 | 377 | 109 | 0 | 0 | 0 | 0 | 1250 | 2187 | 43 | 0 | 10 | 33 | 0 | 0 | 10 | 14516 | 1453 | 3968 |
| SRR1525202 | 25 | 0 | 433 | 0 | 0 | 0 | 0 | 27 | 0 | 0 | 0 | 138 | 0 | 0 | 423 | 0 | 14 | 0 | 0 | 297 | 214 | 0 | 0 | 0 | 15 | 17 | 82 | 0 | 0 | 0 | 0 | 0 | 0 | 0 | 445 | 51 | 156 |
| SRR1525203 | 19 | 0 | 305 | 0 | 0 | 48 | 0 | 16 | 0 | 0 | 0 | 244 | 0 | 21 | 294 | 0 | 13 | 0 | 0 | 443 | 181 | 0 | 0 | 0 | 0 | 35 | 84 | 54 | 0 | 0 | 51 | 0 | 0 | 0 | 555 | 53 | 165 |
| SRR1563015 | 21 | 0 | 6094 | 0 | 0 | 42 | 27 | 39 | 23 | 0 | 0 | 0 | 0 | 12 | 63 | 0 | 25 | 0 | 0 | 270 | 654 | 0 | 0 | 0 | 0 | 293 | 443 | 0 | 252 | 0 | 13 | 0 | 0 | 27 | 2694 | 229 | 1637 |
| SRR1563018 | 58 | 30 | 6280 | 0 | 0 | 13 | 0 | 72 | 26 | 0 | 0 | 0 | 0 | 0 | 71 | 0 | 20 | 0 | 0 | 527 | 737 | 0 | 0 | 20 | 0 | 11 | 91 | 39 | 125 | 16 | 13 | 0 | 0 | 13 | 498 | 58 | 473 |
| SRR1563057 | 11 | 31 | 2591 | 0 | 0 | 0 | 25 | 36 | 31 | 0 | 0 | 0 | 0 | 52 | 83 | 0 | 33 | 0 | 0 | 1537 | 1006 | 10 | 0 | 0 | 0 | 15 | 256 | 52 | 21 | 0 | 0 | 0 | 10 | 21 | 530 | 84 | 459 |
| SRR1563058 | 53 | 0 | 4339 | 18 | 13 | 0 | 155 | 372 | 0 | 0 | 0 | 115 | 0 | 27 | 0 | 0 | 29 | 0 | 379 | 1802 | 38 | 0 | 0 | 11 | 0 | 79 | 683 | 17 | 0 | 36 | 0 | 0 | 0 | 27 | 32 | 0 | 21 |
| SRR1563059 | 42 | 24 | 16238 | 10 | 0 | 219 | 10 | 199 | 192 | 0 | 0 | 0 | 0 | 13 | 80 | 0 | 17 | 0 | 0 | 1059 | 924 | 0 | 0 | 17 | 0 | 6000 | 11002 | 13 | 0 | 0 | 297 | 0 | 0 | 0 | 11487 | 563 | 4230 |
| SRR1563060 | 33 | 12 | 14958 | 0 | 0 | 0 | 185 | 76 | 0 | 0 | 0 | 15 | 0 | 10 | 15 | 0 | 0 | 0 | 139 | 703 | 108 | 0 | 0 | 0 | 17 | 22 | 433 | 14 | 0 | 31 | 0 | 0 | 0 | 16 | 38 | 0 | 20 |
| SRR1563061 | 87 | 0 | 9242 | 22 | 0 | 17 | 0 | 54 | 0 | 0 | 0 | 0 | 0 | 0 | 38 | 0 | 15 | 0 | 0 | 206 | 701 | 0 | 0 | 0 | 0 | 42 | 33 | 0 | 122 | 14 | 0 | 0 | 0 | 0 | 205 | 10 | 82 |
| SRR1563062 | 0 | 0 | 11237 | 14 | 22 | 0 | 312 | 345 | 0 | 0 | 0 | 0 | 0 | 0 | 15 | 0 | 30 | 0 | 372 | 1279 | 30 | 0 | 0 | 0 | 0 | 12 | 166 | 11 | 0 | 27 | 0 | 0 | 0 | 17 | 14 | 0 | 0 |
| SRR1563063 | 56 | 26 | 9497 | 0 | 0 | 231 | 19 | 26 | 242 | 0 | 0 | 19 | 0 | 0 | 66 | 0 | 29 | 0 | 0 | 2102 | 599 | 0 | 0 | 0 | 0 | 5125 | 3508 | 18 | 24 | 0 | 402 | 0 | 0 | 13 | 6241 | 1092 | 3812 |
| SRR1563064 | 50 | 0 | 12195 | 0 | 0 | 0 | 338 | 806 | 0 | 0 | 0 | 74 | 0 | 11 | 0 | 0 | 28 | 0 | 547 | 2500 | 74 | 0 | 0 | 0 | 0 | 62 | 1599 | 0 | 0 | 77 | 0 | 0 | 0 | 56 | 46 | 0 | 26 |
| SRR1575598 | 11 | 20 | 481 | 0 | 0 | 0 | 0 | 0 | 0 | 0 | 0 | 102 | 0 | 34 | 85 | 18 | 11 | 0 | 0 | 0 | 0 | 0 | 0 | 0 | 12 | 0 | 0 | 46 | 0 | 0 | 0 | 0 | 1278 | 0 | 0 | 0 | 0 |
| SRR1575599 | 70 | 23 | 748 | 0 | 0 | 0 | 0 | 0 | 0 | 0 | 0 | 174 | 0 | 13 | 0 | 0 | 16 | 0 | 0 | 0 | 0 | 0 | 0 | 0 | 0 | 0 | 0 | 16 | 10 | 0 | 0 | 0 | 48 | 0 | 181 | 19 | 74 |
| SRR1575601 | 0 | 24 | 95 | 0 | 0 | 0 | 0 | 0 | 0 | 0 | 0 | 0 | 0 | 13 | 13 | 27 | 12 | 0 | 0 | 0 | 0 | 0 | 0 | 0 | 0 | 0 | 0 | 19 | 0 | 0 | 0 | 0 | 678 | 0 | 0 | 0 | 0 |
| SRR1575602 | 52 | 22 | 747 | 13 | 0 | 0 | 0 | 0 | 0 | 0 | 0 | 125 | 0 | 21 | 27 | 11 | 0 | 0 | 0 | 0 | 0 | 0 | 0 | 0 | 0 | 19 | 0 | 32 | 0 | 0 | 0 | 0 | 211 | 14 | 283 | 26 | 178 |
| SRR1592244 | 937 | 0 | 957 | 0 | 0 | 0 | 55 | 24 | 11 | 0 | 0 | 929 | 0 | 0 | 11 | 0 | 0 | 15 | 62 | 733 | 12515 | 0 | 0 | 0 | 0 | 20 | 136 | 0 | 0 | 0 | 0 | 0 | 26 | 0 | 29 | 0 | 24 |
| SRR1592245 | 1237 | 0 | 734 | 0 | 0 | 0 | 27 | 12 | 0 | 0 | 0 | 1256 | 0 | 0 | 12 | 0 | 0 | 18 | 30 | 106 | 8424 | 0 | 0 | 0 | 14 | 38 | 14 | 0 | 0 | 16 | 0 | 0 | 30 | 12 | 0 | 0 | 0 |
| SRR1592246 | 1657 | 0 | 854 | 0 | 0 | 0 | 32 | 238 | 0 | 0 | 0 | 1178 | 0 | 0 | 0 | 0 | 0 | 12 | 43 | 53 | 3099 | 0 | 0 | 0 | 0 | 58 | 182 | 0 | 0 | 61 | 0 | 0 | 10 | 28 | 0 | 0 | 0 |
| SRR1592247 | 2570 | 0 | 896 | 0 | 0 | 0 | 36 | 345 | 0 | 0 | 0 | 797 | 0 | 0 | 0 | 0 | 0 | 15 | 23 | 40 | 6353 | 0 | 0 | 0 | 16 | 39 | 11 | 0 | 0 | 64 | 0 | 0 | 0 | 17 | 0 | 0 | 0 |
| SRR1592248 | 1851 | 0 | 1061 | 0 | 0 | 0 | 132 | 1523 | 0 | 0 | 0 | 927 | 0 | 0 | 0 | 0 | 0 | 15 | 12 | 28 | 3849 | 0 | 0 | 0 | 0 | 86 | 109 | 0 | 0 | 266 | 0 | 0 | 0 | 29 | 0 | 0 | 0 |
| SRR1596226 | 150 | 0 | 2217 | 0 | 0 | 11 | 0 | 0 | 0 | 0 | 0 | 69 | 0 | 0 | 0 | 0 | 0 | 0 | 17 | 390 | 1161 | 0 | 0 | 0 | 88 | 147 | 500 | 0 | 0 | 0 | 11 | 0 | 0 | 0 | 18 | 0 | 18 |

### Supp Table S8.2.Columns_AM_to_BW

| | | | | | | | | | | | | | | | | | | | | | | | | | | | | | | | | | | | | | |
|---|---|---|---|---|---|---|---|---|---|---|---|---|---|---|---|---|---|---|---|---|---|---|---|---|---|---|---|---|---|---|---|---|---|---|---|---|---|
| SRR1596227 | 137 | 0 | 2227 | 0 | 0 | 12 | 0 | 0 | 11 | 0 | 0 | 102 | 0 | 0 | 13 | 0 | 0 | 0 | 13 | 804 | 1331 | 0 | 0 | 0 | 78 | 132 | 510 | 0 | 0 | 0 | 15 | 0 | 0 | 0 | 22 | 0 | 12 |
| SRR1596228 | 53 | 0 | 1205 | 0 | 0 | 13 | 0 | 0 | 0 | 0 | 0 | 67 | 13 | 0 | 0 | 0 | 22 | 0 | 21 | 600 | 1062 | 0 | 0 | 13 | 73 | 427 | 1169 | 0 | 0 | 0 | 0 | 0 | 10 | 10 | 17 | 0 | 0 |
| SRR1596229 | 59 | 0 | 1129 | 0 | 0 | 21 | 0 | 0 | 0 | 0 | 0 | 69 | 0 | 0 | 0 | 0 | 20 | 0 | 44 | 435 | 1546 | 0 | 0 | 0 | 29 | 132 | 246 | 0 | 0 | 0 | 11 | 0 | 0 | 14 | 0 | 0 | 0 |
| SRR1596230 | 31 | 0 | 685 | 0 | 0 | 81 | 90 | 0 | 10 | 0 | 0 | 14 | 0 | 0 | 0 | 0 | 0 | 11 | 54 | 189 | 1410 | 0 | 0 | 0 | 108 | 1041 | 1200 | 0 | 0 | 0 | 47 | 0 | 0 | 10 | 47 | 10 | 14 |
| SRR1596231 | 35 | 0 | 747 | 0 | 0 | 81 | 127 | 0 | 20 | 0 | 0 | 14 | 12 | 0 | 0 | 0 | 11 | 0 | 53 | 137 | 1417 | 0 | 0 | 0 | 94 | 1112 | 1222 | 0 | 0 | 0 | 49 | 0 | 0 | 12 | 65 | 24 | 41 |
| SRR1608128 | 0 | 0 | 0 | 0 | 0 | 0 | 0 | 0 | 0 | 0 | 0 | 0 | 0 | 0 | 0 | 0 | 0 | 504 | 0 | 0 | 232 | 0 | 0 | 0 | 45 | 0 | 0 | 0 | 10 | 0 | 20 | 0 | 34 | 0 | 0 | 12 | 10 |
| SRR1608138 | 0 | 0 | 30 | 0 | 0 | 0 | 0 | 0 | 0 | 0 | 0 | 0 | 0 | 0 | 37 | 0 | 0 | 83 | 0 | 0 | 186 | 0 | 0 | 0 | 13 | 23 | 0 | 0 | 0 | 0 | 15 | 0 | 23 | 0 | 0 | 10 | 0 |
| SRR1608143 | 0 | 0 | 0 | 0 | 0 | 0 | 0 | 0 | 0 | 0 | 0 | 0 | 0 | 0 | 0 | 0 | 0 | 210 | 0 | 0 | 170 | 0 | 0 | 0 | 148 | 0 | 0 | 0 | 0 | 0 | 0 | 0 | 72 | 0 | 0 | 0 | 0 |
| SRR1608150 | 0 | 0 | 10 | 0 | 0 | 0 | 0 | 0 | 0 | 0 | 0 | 0 | 0 | 0 | 0 | 0 | 0 | 209 | 0 | 0 | 68 | 0 | 0 | 0 | 87 | 0 | 0 | 0 | 0 | 0 | 0 | 0 | 84 | 0 | 0 | 0 | 0 |
| SRR1608158 | 0 | 0 | 11 | 0 | 0 | 0 | 0 | 0 | 0 | 0 | 0 | 0 | 0 | 0 | 30 | 0 | 0 | 81 | 0 | 0 | 193 | 0 | 0 | 0 | 39 | 0 | 0 | 0 | 0 | 0 | 14 | 0 | 0 | 0 | 0 | 0 | 0 |
| SRR1608160 | 0 | 0 | 0 | 0 | 0 | 0 | 17 | 0 | 0 | 0 | 0 | 0 | 0 | 0 | 17 | 0 | 0 | 192 | 0 | 0 | 555 | 0 | 0 | 0 | 39 | 0 | 0 | 0 | 0 | 0 | 42 | 0 | 19 | 0 | 0 | 0 | 0 |
| SRR1608163 | 0 | 0 | 16 | 0 | 0 | 0 | 0 | 0 | 0 | 0 | 0 | 0 | 0 | 0 | 0 | 0 | 0 | 109 | 0 | 0 | 37 | 0 | 0 | 0 | 74 | 0 | 0 | 0 | 0 | 0 | 33 | 0 | 0 | 0 | 0 | 0 | 0 |
| SRR1608165 | 0 | 0 | 0 | 0 | 0 | 0 | 0 | 0 | 0 | 0 | 0 | 0 | 0 | 0 | 0 | 0 | 0 | 130 | 0 | 0 | 58 | 0 | 0 | 0 | 45 | 0 | 0 | 0 | 0 | 0 | 31 | 0 | 20 | 0 | 0 | 0 | 0 |
| SRR1608166 | 0 | 0 | 0 | 0 | 0 | 0 | 0 | 0 | 0 | 0 | 0 | 0 | 0 | 0 | 25 | 0 | 0 | 277 | 0 | 0 | 325 | 0 | 0 | 0 | 59 | 0 | 0 | 0 | 0 | 0 | 16 | 0 | 69 | 0 | 0 | 0 | 0 |
| SRR1608167 | 0 | 0 | 0 | 0 | 0 | 0 | 0 | 0 | 0 | 0 | 0 | 0 | 0 | 0 | 30 | 0 | 0 | 101 | 0 | 0 | 273 | 0 | 0 | 0 | 52 | 0 | 0 | 0 | 0 | 0 | 35 | 0 | 22 | 0 | 0 | 0 | 0 |
| SRR1608168 | 0 | 0 | 0 | 0 | 0 | 0 | 0 | 0 | 0 | 0 | 0 | 0 | 0 | 0 | 0 | 0 | 0 | 179 | 0 | 0 | 38 | 0 | 0 | 0 | 102 | 12 | 0 | 0 | 0 | 0 | 38 | 0 | 34 | 0 | 0 | 0 | 0 |
| SRR1608169 | 0 | 0 | 0 | 0 | 0 | 0 | 0 | 0 | 0 | 0 | 0 | 0 | 0 | 0 | 51 | 0 | 0 | 210 | 10 | 0 | 1997 | 0 | 0 | 0 | 75 | 11 | 0 | 0 | 0 | 0 | 11 | 0 | 45 | 0 | 0 | 0 | 0 |
| SRR1608170 | 0 | 0 | 20 | 0 | 0 | 0 | 0 | 0 | 0 | 0 | 0 | 0 | 0 | 0 | 29 | 0 | 0 | 209 | 0 | 0 | 72 | 0 | 0 | 0 | 52 | 0 | 0 | 0 | 0 | 0 | 0 | 0 | 18 | 0 | 0 | 0 | 0 |
| SRR1608171 | 0 | 0 | 12 | 0 | 0 | 0 | 0 | 0 | 0 | 0 | 0 | 0 | 0 | 0 | 17 | 0 | 0 | 129 | 0 | 0 | 188 | 0 | 0 | 0 | 88 | 0 | 0 | 0 | 0 | 0 | 25 | 0 | 27 | 0 | 0 | 0 | 0 |
| SRR1608172 | 0 | 0 | 27 | 0 | 0 | 0 | 0 | 0 | 0 | 0 | 0 | 0 | 0 | 0 | 18 | 0 | 0 | 217 | 0 | 0 | 587 | 0 | 0 | 0 | 210 | 0 | 0 | 0 | 0 | 0 | 0 | 0 | 65 | 0 | 0 | 0 | 0 |
| SRR1608174 | 0 | 0 | 11 | 0 | 0 | 0 | 0 | 0 | 0 | 0 | 0 | 0 | 0 | 0 | 0 | 0 | 0 | 215 | 0 | 0 | 807 | 0 | 0 | 0 | 35 | 0 | 0 | 0 | 0 | 0 | 23 | 0 | 62 | 0 | 18 | 12 | 0 |
| SRR1608175 | 0 | 0 | 0 | 0 | 0 | 0 | 0 | 0 | 0 | 0 | 0 | 0 | 0 | 0 | 0 | 0 | 0 | 138 | 0 | 0 | 52 | 0 | 0 | 0 | 112 | 19 | 0 | 0 | 0 | 0 | 27 | 0 | 0 | 0 | 0 | 0 | 0 |
| SRR1608177 | 0 | 0 | 0 | 0 | 0 | 0 | 0 | 0 | 0 | 0 | 0 | 0 | 0 | 0 | 0 | 0 | 0 | 636 | 0 | 0 | 125 | 0 | 0 | 0 | 57 | 0 | 0 | 0 | 0 | 0 | 0 | 0 | 15 | 0 | 0 | 0 | 0 |
| SRR1608178 | 0 | 0 | 18 | 0 | 0 | 0 | 0 | 0 | 0 | 0 | 0 | 0 | 0 | 0 | 36 | 0 | 0 | 100 | 0 | 0 | 417 | 0 | 0 | 0 | 70 | 12 | 0 | 0 | 0 | 0 | 50 | 0 | 31 | 0 | 19 | 35 | 12 |
| SRR1608179 | 0 | 0 | 16 | 0 | 0 | 0 | 0 | 0 | 0 | 0 | 0 | 0 | 0 | 0 | 14 | 0 | 0 | 177 | 0 | 0 | 557 | 0 | 0 | 0 | 115 | 19 | 11 | 0 | 0 | 0 | 18 | 0 | 27 | 0 | 0 | 19 | 0 |
| SRR1608180 | 0 | 0 | 22 | 0 | 0 | 0 | 0 | 0 | 0 | 0 | 0 | 0 | 0 | 0 | 15 | 0 | 0 | 246 | 0 | 0 | 131 | 0 | 0 | 0 | 75 | 0 | 0 | 0 | 0 | 0 | 89 | 0 | 18 | 0 | 0 | 14 | 14 |
| SRR1608181 | 0 | 0 | 21 | 0 | 0 | 0 | 0 | 0 | 0 | 0 | 0 | 0 | 0 | 0 | 0 | 0 | 0 | 174 | 0 | 0 | 447 | 0 | 0 | 0 | 230 | 0 | 0 | 0 | 0 | 0 | 15 | 0 | 25 | 0 | 0 | 0 | 0 |
| SRR1608182 | 0 | 0 | 0 | 0 | 0 | 0 | 0 | 0 | 0 | 0 | 0 | 0 | 0 | 0 | 0 | 0 | 0 | 165 | 0 | 0 | 67 | 0 | 0 | 0 | 47 | 0 | 0 | 0 | 0 | 0 | 14 | 0 | 35 | 0 | 0 | 0 | 0 |
| SRR1608184 | 0 | 0 | 20 | 0 | 0 | 0 | 0 | 0 | 0 | 0 | 0 | 0 | 14 | 0 | 0 | 0 | 0 | 437 | 0 | 0 | 259 | 0 | 0 | 0 | 14 | 0 | 0 | 0 | 0 | 0 | 101 | 0 | 66 | 0 | 0 | 14 | 0 |
| SRR1608185 | 0 | 0 | 26 | 0 | 0 | 0 | 0 | 0 | 0 | 0 | 0 | 0 | 0 | 0 | 13 | 0 | 0 | 208 | 0 | 0 | 238 | 0 | 0 | 0 | 69 | 0 | 0 | 0 | 0 | 0 | 94 | 0 | 25 | 0 | 0 | 0 | 0 |
| SRR1608186 | 0 | 0 | 0 | 0 | 0 | 0 | 0 | 0 | 0 | 0 | 0 | 0 | 0 | 0 | 59 | 0 | 0 | 199 | 0 | 0 | 119 | 0 | 0 | 0 | 105 | 0 | 0 | 0 | 0 | 0 | 44 | 0 | 27 | 0 | 0 | 14 | 0 |
| SRR1608189 | 0 | 0 | 17 | 0 | 0 | 0 | 0 | 0 | 0 | 0 | 0 | 0 | 0 | 0 | 67 | 0 | 0 | 203 | 0 | 0 | 118 | 0 | 0 | 0 | 10 | 0 | 0 | 0 | 0 | 0 | 15 | 0 | 32 | 0 | 0 | 0 | 0 |
| SRR1608190 | 0 | 0 | 13 | 0 | 0 | 0 | 0 | 0 | 0 | 0 | 0 | 0 | 0 | 0 | 43 | 0 | 0 | 160 | 0 | 0 | 0 | 0 | 0 | 0 | 87 | 0 | 0 | 0 | 0 | 0 | 67 | 0 | 38 | 0 | 0 | 12 | 0 |
| SRR1608191 | 0 | 0 | 15 | 0 | 0 | 0 | 0 | 0 | 0 | 0 | 0 | 0 | 0 | 0 | 122 | 0 | 0 | 212 | 0 | 0 | 465 | 0 | 0 | 0 | 55 | 0 | 0 | 0 | 0 | 0 | 49 | 0 | 18 | 0 | 0 | 12 | 0 |
| SRR1608192 | 0 | 0 | 14 | 0 | 0 | 0 | 0 | 0 | 0 | 0 | 0 | 0 | 0 | 0 | 0 | 0 | 0 | 275 | 0 | 0 | 192 | 0 | 0 | 0 | 56 | 0 | 0 | 0 | 0 | 0 | 33 | 0 | 33 | 0 | 0 | 0 | 0 |
| SRR1608193 | 0 | 0 | 37 | 0 | 0 | 0 | 0 | 0 | 0 | 0 | 0 | 0 | 0 | 0 | 0 | 0 | 0 | 237 | 0 | 0 | 47 | 0 | 0 | 0 | 60 | 0 | 0 | 0 | 0 | 0 | 39 | 0 | 26 | 0 | 0 | 0 | 0 |
| SRR1608194 | 0 | 0 | 17 | 0 | 0 | 0 | 0 | 0 | 0 | 0 | 0 | 0 | 0 | 0 | 15 | 0 | 0 | 228 | 0 | 0 | 119 | 0 | 0 | 0 | 89 | 11 | 0 | 0 | 0 | 0 | 167 | 0 | 11 | 0 | 0 | 26 | 0 |
| SRR1608196 | 0 | 0 | 0 | 0 | 0 | 0 | 0 | 0 | 0 | 0 | 0 | 0 | 0 | 0 | 53 | 0 | 0 | 118 | 0 | 0 | 127 | 0 | 0 | 0 | 80 | 0 | 0 | 0 | 0 | 0 | 13 | 0 | 14 | 0 | 0 | 0 | 0 |
| SRR1608201 | 0 | 0 | 12 | 0 | 0 | 0 | 0 | 0 | 0 | 0 | 0 | 0 | 0 | 0 | 0 | 0 | 0 | 104 | 0 | 0 | 35 | 0 | 0 | 0 | 77 | 0 | 0 | 0 | 0 | 0 | 54 | 0 | 15 | 0 | 0 | 23 | 0 |
| SRR1608208 | 0 | 0 | 0 | 0 | 0 | 0 | 0 | 0 | 0 | 0 | 0 | 0 | 0 | 0 | 23 | 0 | 0 | 99 | 0 | 0 | 70 | 0 | 0 | 0 | 115 | 0 | 0 | 0 | 11 | 0 | 17 | 0 | 17 | 0 | 0 | 0 | 0 |
| SRR1608213 | 0 | 0 | 0 | 0 | 0 | 0 | 0 | 0 | 0 | 0 | 0 | 0 | 0 | 0 | 19 | 0 | 0 | 158 | 0 | 0 | 229 | 0 | 0 | 0 | 41 | 0 | 0 | 0 | 0 | 0 | 132 | 0 | 17 | 0 | 19 | 14 | 0 |
| SRR1608214 | 0 | 0 | 12 | 0 | 0 | 0 | 0 | 0 | 0 | 0 | 0 | 0 | 0 | 0 | 14 | 0 | 0 | 32 | 0 | 0 | 28 | 0 | 0 | 0 | 10 | 0 | 0 | 0 | 0 | 0 | 12 | 0 | 0 | 0 | 0 | 0 | 0 |
| SRR1608222 | 0 | 0 | 0 | 0 | 0 | 0 | 0 | 0 | 0 | 0 | 0 | 0 | 0 | 0 | 32 | 0 | 0 | 86 | 0 | 0 | 256 | 0 | 0 | 0 | 37 | 0 | 0 | 0 | 0 | 0 | 16 | 0 | 0 | 0 | 0 | 0 | 0 |
| SRR1608232 | 0 | 0 | 0 | 0 | 0 | 0 | 0 | 0 | 0 | 0 | 0 | 0 | 0 | 0 | 0 | 0 | 0 | 89 | 0 | 0 | 325 | 0 | 0 | 0 | 90 | 0 | 0 | 0 | 0 | 0 | 56 | 0 | 21 | 0 | 10 | 31 | 0 |
| SRR1608237 | 0 | 0 | 0 | 0 | 0 | 0 | 0 | 0 | 0 | 0 | 0 | 0 | 0 | 0 | 0 | 0 | 0 | 31 | 0 | 0 | 33 | 0 | 0 | 0 | 81 | 0 | 0 | 0 | 0 | 0 | 23 | 0 | 15 | 0 | 0 | 0 | 0 |
| SRR1608254 | 0 | 0 | 0 | 0 | 0 | 0 | 0 | 0 | 0 | 0 | 0 | 0 | 0 | 0 | 0 | 0 | 0 | 118 | 0 | 0 | 497 | 0 | 0 | 0 | 104 | 0 | 0 | 0 | 0 | 0 | 27 | 0 | 89 | 0 | 0 | 0 | 0 |
| SRR1608255 | 0 | 0 | 0 | 0 | 0 | 0 | 0 | 0 | 0 | 0 | 0 | 0 | 0 | 0 | 20 | 0 | 0 | 155 | 0 | 0 | 416 | 0 | 0 | 0 | 68 | 0 | 0 | 0 | 0 | 0 | 26 | 0 | 66 | 0 | 0 | 0 | 0 |
| SRR1608256 | 0 | 0 | 0 | 0 | 0 | 0 | 0 | 0 | 0 | 0 | 0 | 0 | 0 | 0 | 29 | 0 | 0 | 148 | 0 | 0 | 336 | 0 | 0 | 0 | 231 | 0 | 0 | 0 | 0 | 0 | 14 | 0 | 94 | 0 | 0 | 0 | 0 |
| SRR1608257 | 0 | 0 | 15 | 0 | 0 | 0 | 0 | 0 | 0 | 0 | 0 | 0 | 0 | 0 | 0 | 0 | 0 | 133 | 0 | 0 | 232 | 0 | 0 | 0 | 74 | 0 | 0 | 0 | 0 | 0 | 14 | 0 | 66 | 0 | 0 | 0 | 0 |
| SRR1608260 | 0 | 10 | 10 | 0 | 0 | 0 | 0 | 0 | 0 | 0 | 0 | 0 | 0 | 0 | 19 | 0 | 0 | 122 | 0 | 0 | 305 | 0 | 0 | 0 | 146 | 0 | 0 | 0 | 0 | 0 | 10 | 0 | 74 | 0 | 0 | 0 | 0 |
| SRR1608263 | 0 | 0 | 0 | 0 | 0 | 0 | 0 | 0 | 0 | 0 | 0 | 0 | 0 | 0 | 0 | 0 | 0 | 78 | 0 | 0 | 196 | 0 | 0 | 0 | 146 | 0 | 0 | 0 | 0 | 0 | 27 | 0 | 11 | 0 | 0 | 0 | 0 |
| SRR1608264 | 0 | 0 | 0 | 0 | 0 | 0 | 0 | 0 | 0 | 0 | 0 | 0 | 0 | 0 | 26 | 0 | 0 | 109 | 0 | 0 | 207 | 0 | 0 | 0 | 36 | 0 | 0 | 0 | 0 | 0 | 29 | 0 | 16 | 0 | 11 | 11 | 11 |
| SRR1608266 | 0 | 0 | 0 | 0 | 0 | 0 | 0 | 0 | 0 | 0 | 0 | 0 | 0 | 0 | 0 | 0 | 0 | 60 | 0 | 0 | 412 | 0 | 0 | 0 | 131 | 0 | 0 | 0 | 0 | 0 | 35 | 0 | 11 | 0 | 0 | 14 | 0 |
| SRR1608267 | 0 | 0 | 0 | 0 | 0 | 0 | 0 | 0 | 0 | 0 | 0 | 0 | 0 | 0 | 0 | 0 | 0 | 117 | 0 | 0 | 54 | 0 | 0 | 0 | 81 | 0 | 0 | 0 | 0 | 0 | 30 | 0 | 0 | 0 | 0 | 0 | 12 |
| SRR1608269 | 0 | 0 | 0 | 0 | 0 | 0 | 0 | 0 | 0 | 0 | 0 | 0 | 0 | 0 | 0 | 0 | 0 | 0 | 0 | 0 | 640 | 0 | 0 | 0 | 84 | 0 | 0 | 0 | 0 | 0 | 57 | 0 | 13 | 0 | 0 | 22 | 0 |
| SRR1608270 | 0 | 0 | 0 | 0 | 0 | 0 | 0 | 0 | 0 | 0 | 0 | 0 | 0 | 0 | 0 | 0 | 0 | 46 | 0 | 0 | 97 | 0 | 0 | 0 | 249 | 0 | 0 | 0 | 0 | 0 | 54 | 0 | 33 | 0 | 0 | 37 | 33 |
| SRR1608271 | 0 | 16 | 0 | 0 | 0 | 0 | 0 | 0 | 0 | 0 | 0 | 0 | 0 | 0 | 0 | 0 | 0 | 85 | 0 | 0 | 40 | 0 | 0 | 0 | 59 | 0 | 0 | 0 | 0 | 0 | 0 | 0 | 0 | 0 | 0 | 0 | 0 |
| SRR1608274 | 0 | 0 | 0 | 0 | 0 | 0 | 0 | 0 | 0 | 0 | 0 | 0 | 0 | 0 | 0 | 0 | 0 | 228 | 0 | 0 | 63 | 0 | 0 | 0 | 79 | 0 | 0 | 0 | 17 | 0 | 53 | 0 | 0 | 0 | 0 | 0 | 13 |
| SRR1608276 | 0 | 0 | 0 | 0 | 0 | 0 | 0 | 0 | 0 | 0 | 0 | 0 | 0 | 0 | 29 | 0 | 0 | 39 | 0 | 0 | 705 | 0 | 0 | 0 | 127 | 0 | 0 | 0 | 0 | 0 | 42 | 0 | 0 | 0 | 0 | 19 | 28 |
| SRR1608277 | 0 | 0 | 0 | 0 | 0 | 0 | 0 | 0 | 0 | 0 | 0 | 0 | 0 | 0 | 0 | 0 | 0 | 106 | 0 | 0 | 0 | 0 | 0 | 0 | 101 | 0 | 0 | 0 | 0 | 0 | 22 | 0 | 0 | 0 | 0 | 16 | 14 |
| SRR1608278 | 0 | 0 | 0 | 0 | 0 | 0 | 0 | 0 | 0 | 0 | 0 | 0 | 0 | 0 | 33 | 0 | 0 | 73 | 0 | 0 | 0 | 0 | 0 | 0 | 39 | 0 | 0 | 0 | 0 | 0 | 20 | 0 | 0 | 0 | 0 | 0 | 0 |
| SRR1608281 | 0 | 0 | 0 | 0 | 0 | 0 | 0 | 0 | 0 | 0 | 0 | 0 | 0 | 0 | 0 | 0 | 0 | 141 | 0 | 0 | 525 | 0 | 0 | 0 | 98 | 0 | 0 | 0 | 0 | 0 | 20 | 0 | 0 | 0 | 0 | 15 | 0 |
| SRR1608282 | 0 | 0 | 0 | 0 | 0 | 0 | 0 | 0 | 0 | 0 | 0 | 0 | 0 | 0 | 13 | 0 | 0 | 342 | 0 | 0 | 229 | 0 | 0 | 0 | 58 | 0 | 0 | 0 | 0 | 0 | 42 | 0 | 15 | 0 | 0 | 11 | 0 |
| SRR1608283 | 0 | 0 | 0 | 0 | 0 | 0 | 0 | 0 | 0 | 0 | 0 | 0 | 0 | 0 | 0 | 0 | 0 | 93 | 0 | 0 | 153 | 0 | 0 | 0 | 71 | 0 | 0 | 0 | 0 | 0 | 18 | 0 | 0 | 0 | 0 | 12 | 0 |
| SRR1608286 | 0 | 0 | 0 | 0 | 0 | 0 | 0 | 0 | 0 | 0 | 0 | 0 | 0 | 0 | 39 | 0 | 0 | 194 | 0 | 0 | 473 | 0 | 0 | 0 | 28 | 0 | 0 | 0 | 0 | 0 | 28 | 0 | 18 | 0 | 0 | 0 | 0 |
| SRR1608287 | 0 | 0 | 0 | 0 | 0 | 0 | 0 | 0 | 0 | 0 | 0 | 0 | 0 | 0 | 0 | 0 | 0 | 241 | 0 | 0 | 250 | 0 | 0 | 0 | 159 | 0 | 0 | 0 | 0 | 0 | 23 | 0 | 11 | 0 | 0 | 0 | 0 |
| SRR1608289 | 0 | 0 | 0 | 0 | 0 | 0 | 0 | 0 | 0 | 0 | 0 | 0 | 0 | 0 | 0 | 0 | 0 | 105 | 0 | 0 | 233 | 0 | 0 | 0 | 199 | 0 | 0 | 0 | 0 | 0 | 50 | 0 | 17 | 0 | 0 | 17 | 0 |
| SRR1608290 | 0 | 0 | 0 | 0 | 0 | 0 | 0 | 0 | 0 | 0 | 0 | 0 | 0 | 0 | 0 | 0 | 0 | 224 | 0 | 0 | 134 | 0 | 0 | 0 | 143 | 0 | 0 | 0 | 0 | 0 | 60 | 0 | 11 | 0 | 0 | 22 | 0 |
| SRR1608302 | 0 | 0 | 0 | 0 | 0 | 0 | 0 | 0 | 0 | 0 | 0 | 0 | 0 | 0 | 59 | 0 | 0 | 131 | 0 | 0 | 118 | 0 | 0 | 0 | 78 | 0 | 0 | 0 | 0 | 0 | 31 | 0 | 16 | 0 | 0 | 0 | 0 |
| SRR1608307 | 0 | 0 | 0 | 0 | 0 | 0 | 0 | 0 | 0 | 0 | 0 | 0 | 0 | 0 | 0 | 0 | 0 | 191 | 0 | 0 | 50 | 0 | 0 | 0 | 78 | 0 | 0 | 0 | 0 | 0 | 27 | 0 | 15 | 0 | 0 | 0 | 0 |
| SRR1608308 | 0 | 0 | 0 | 0 | 0 | 0 | 0 | 0 | 0 | 0 | 0 | 0 | 0 | 0 | 37 | 0 | 0 | 102 | 0 | 0 | 187 | 0 | 0 | 0 | 166 | 0 | 0 | 0 | 0 | 0 | 85 | 0 | 0 | 0 | 0 | 11 | 0 |
| SRR1634379 | 172 | 0 | 429 | 0 | 0 | 33 | 37 | 26 | 0 | 0 | 0 | 86 | 0 | 0 | 14 | 0 | 0 | 0 | 0 | 188 | 1169 | 0 | 0 | 0 | 225 | 585 | 1029 | 0 | 0 | 25 | 15 | 0 | 0 | 38 | 13 | 0 | 11 |
| SRR1634381 | 87 | 0 | 381 | 0 | 0 | 46 | 0 | 15 | 0 | 0 | 0 | 64 | 0 | 0 | 0 | 0 | 19 | 0 | 0 | 455 | 967 | 0 | 0 | 0 | 419 | 1154 | 1761 | 0 | 0 | 0 | 36 | 0 | 0 | 14 | 23 | 15 | 23 |
| SRR1634384 | 188 | 0 | 405 | 0 | 0 | 48 | 27 | 12 | 0 | 0 | 0 | 64 | 0 | 0 | 12 | 0 | 0 | 0 | 0 | 1491 | 2016 | 0 | 0 | 21 | 86 | 909 | 904 | 0 | 0 | 0 | 17 | 0 | 0 | 0 | 0 | 0 | 0 |
| SRR1634388 | 210 | 0 | 255 | 0 | 0 | 27 | 58 | 12 | 0 | 0 | 0 | 12 | 0 | 0 | 0 | 0 | 21 | 14 | 0 | 1455 | 828 | 0 | 0 | 15 | 68 | 518 | 550 | 0 | 0 | 0 | 15 | 0 | 0 | 0 | 0 | 0 | 0 |
| SRR1634390 | 90 | 0 | 302 | 0 | 0 | 34 | 0 | 0 | 0 | 0 | 0 | 140 | 0 | 11 | 0 | 0 | 0 | 0 | 0 | 218 | 508 | 0 | 0 | 0 | 548 | 664 | 992 | 0 | 0 | 0 | 40 | 0 | 0 | 23 | 42 | 11 | 33 |
| SRR1634392 | 168 | 0 | 241 | 0 | 0 | 45 | 17 | 10 | 0 | 0 | 0 | 56 | 0 | 0 | 0 | 0 | 17 | 0 | 0 | 478 | 539 | 0 | 0 | 0 | 303 | 830 | 1171 | 0 | 0 | 0 | 40 | 0 | 0 | 10 | 18 | 0 | 18 |
| SRR1634394 | 248 | 0 | 243 | 0 | 0 | 33 | 0 | 18 | 0 | 0 | 0 | 40 | 0 | 0 | 0 | 0 | 23 | 0 | 0 | 359 | 416 | 0 | 0 | 0 | 300 | 739 | 1265 | 0 | 0 | 0 | 33 | 0 | 0 | 16 | 0 | 0 | 13 |
| SRR1634395 | 76 | 0 | 965 | 0 | 0 | 15 | 0 | 0 | 0 | 0 | 0 | 221 | 0 | 0 | 0 | 0 | 21 | 0 | 0 | 336 | 604 | 0 | 0 | 0 | 226 | 273 | 435 | 0 | 0 | 0 | 10 | 0 | 0 | 22 | 33 | 13 | 16 |
| SRR1634398 | 38 | 0 | 556 | 0 | 0 | 47 | 16 | 10 | 0 | 0 | 0 | 30 | 0 | 0 | 0 | 0 | 26 | 0 | 0 | 515 | 265 | 0 | 0 | 0 | 259 | 730 | 716 | 0 | 0 | 0 | 35 | 0 | 0 | 24 | 56 | 12 | 28 |
| SRR1634399 | 67 | 0 | 862 | 0 | 0 | 10 | 23 | 0 | 0 | 0 | 0 | 115 | 0 | 0 | 0 | 0 | 18 | 0 | 0 | 455 | 320 | 0 | 0 | 0 | 184 | 235 | 379 | 0 | 0 | 0 | 11 | 0 | 0 | 13 | 14 | 10 | 11 |
| SRR1634400 | 68 | 0 | 838 | 0 | 0 | 27 | 22 | 0 | 0 | 0 | 0 | 98 | 0 | 0 | 0 | 0 | 20 | 0 | 0 | 446 | 349 | 0 | 0 | 0 | 521 | 674 | 1075 | 0 | 0 | 0 | 38 | 0 | 0 | 15 | 45 | 18 | 30 |
| SRR1634406 | 90 | 0 | 342 | 0 | 0 | 46 | 13 | 0 | 0 | 0 | 0 | 128 | 0 | 0 | 0 | 0 | 19 | 0 | 0 | 464 | 1322 | 0 | 0 | 0 | 259 | 722 | 992 | 0 | 0 | 0 | 78 | 0 | 0 | 21 | 23 | 0 | 24 |
| SRR1634409 | 21 | 0 | 811 | 0 | 0 | 18 | 0 | 0 | 0 | 0 | 0 | 29 | 0 | 13 | 0 | 0 | 18 | 0 | 0 | 241 | 398 | 0 | 0 | 0 | 320 | 277 | 479 | 10 | 0 | 0 | 12 | 0 | 0 | 17 | 27 | 14 | 15 |
| SRR1634411 | 67 | 0 | 614 | 0 | 0 | 27 | 30 | 0 | 0 | 0 | 0 | 39 | 0 | 0 | 0 | 0 | 0 | 0 | 0 | 746 | 2250 | 0 | 0 | 0 | 132 | 187 | 261 | 0 | 0 | 0 | 23 | 0 | 0 | 21 | 19 | 16 | 13 |
| SRR1634412 | 69 | 0 | 608 | 0 | 0 | 11 | 23 | 0 | 0 | 0 | 0 | 31 | 0 | 0 | 0 | 0 | 12 | 0 | 0 | 697 | 2017 | 0 | 0 | 11 | 137 | 175 | 258 | 0 | 0 | 0 | 26 | 0 | 0 | 20 | 20 | 16 | 11 |
| SRR1634414 | 52 | 0 | 492 | 0 | 0 | 17 | 0 | 0 | 0 | 0 | 0 | 149 | 0 | 0 | 0 | 0 | 13 | 0 | 0 | 310 | 350 | 0 | 0 | 0 | 198 | 355 | 486 | 0 | 0 | 0 | 18 | 0 | 0 | 23 | 19 | 13 | 13 |
| SRR1634415 | 62 | 0 | 1287 | 0 | 0 | 16 | 13 | 0 | 0 | 0 | 0 | 135 | 0 | 0 | 0 | 0 | 0 | 0 | 0 | 267 | 555 | 0 | 0 | 0 | 257 | 424 | 616 | 0 | 0 | 0 | 19 | 0 | 0 | 21 | 20 | 14 | 13 |
| SRR1634417 | 55 | 0 | 524 | 0 | 0 | 29 | 0 | 11 | 0 | 0 | 0 | 46 | 0 | 11 | 11 | 0 | 14 | 0 | 0 | 178 | 766 | 0 | 0 | 0 | 682 | 620 | 1188 | 0 | 0 | 0 | 20 | 0 | 0 | 32 | 58 | 15 | 27 |
| SRR1636959 | 19 | 0 | 912 | 0 | 0 | 28 | 27 | 545 | 0 | 0 | 0 | 28 | 0 | 0 | 576 | 0 | 0 | 56 | 48 | 1114 | 5303 | 0 | 0 | 30 | 28 | 417 | 373 | 0 | 0 | 0 | 0 | 0 | 0 | 0 | 146 | 22 | 121 |
| SRR1636960 | 0 | 0 | 448 | 0 | 0 | 16 | 15 | 243 | 0 | 0 | 0 | 15 | 0 | 0 | 109 | 0 | 0 | 243 | 58 | 884 | 1257 | 0 | 0 | 19 | 12 | 200 | 177 | 0 | 0 | 0 | 0 | 0 | 0 | 16 | 290 | 35 | 179 |
| SRR1636961 | 0 | 0 | 479 | 0 | 11 | 0 | 0 | 310 | 0 | 0 | 0 | 17 | 0 | 0 | 106 | 0 | 0 | 251 | 17 | 611 | 1150 | 0 | 0 | 39 | 0 | 194 | 246 | 0 | 0 | 0 | 0 | 0 | 0 | 28 | 285 | 11 | 141 |
| SRR1636962 | 12 | 0 | 301 | 0 | 0 | 62 | 18 | 149 | 0 | 0 | 0 | 0 | 0 | 0 | 259 | 0 | 0 | 21 | 32 | 2710 | 1150 | 0 | 0 | 56 | 12 | 1081 | 724 | 0 | 0 | 12 | 16 | 0 | 0 | 0 | 301 | 40 | 149 |
| SRR1636963 | 13 | 0 | 338 | 0 | 0 | 26 | 21 | 58 | 26 | 0 | 0 | 0 | 0 | 0 | 230 | 0 | 0 | 0 | 51 | 3670 | 2208 | 0 | 0 | 35 | 0 | 799 | 814 | 0 | 0 | 0 | 19 | 0 | 0 | 0 | 271 | 27 | 159 |
| SRR1636964 | 17 | 0 | 276 | 0 | 0 | 42 | 19 | 24 | 0 | 0 | 0 | 0 | 0 | 0 | 162 | 0 | 0 | 0 | 23 | 3734 | 1262 | 0 | 0 | 33 | 39 | 40 | 37 | 0 | 0 | 0 | 27 | 0 | 0 | 0 | 215 | 21 | 147 |
| SRR1636965 | 45 | 0 | 691 | 0 | 22 | 0 | 200 | 660 | 0 | 0 | 0 | 0 | 0 | 105 | 12 | 14 | 0 | 0 | 73 | 9031 | 260 | 13 | 0 | 0 | 18 | 51 | 188 | 13 | 0 | 16 | 0 | 0 | 0 | 183 | 78 | 0 | 35 |
| SRR1636966 | 22 | 0 | 698 | 0 | 10 | 0 | 250 | 102 | 0 | 0 | 0 | 0 | 0 | 42 | 0 | 0 | 0 | 0 | 50 | 2948 | 67 | 13 | 0 | 0 | 15 | 26 | 55 | 0 | 0 | 18 | 0 | 0 | 0 | 22 | 12 | 0 | 0 |
| SRR1636967 | 14 | 0 | 569 | 0 | 0 | 14 | 215 | 1577 | 14 | 0 | 0 | 14 | 0 | 55 | 22 | 0 | 14 | 85 | 43 | 4093 | 2284 | 0 | 0 | 0 | 0 | 295 | 161 | 0 | 0 | 0 | 0 | 0 | 0 | 0 | 46 | 14 | 14 |
| SRR1636968 | 15 | 0 | 596 | 0 | 0 | 54 | 69 | 3738 | 15 | 0 | 0 | 28 | 0 | 13 | 12 | 0 | 10 | 16 | 30 | 2330 | 393 | 0 | 0 | 30 | 12 | 567 | 173 | 0 | 15 | 24 | 12 | 0 | 0 | 128 | 1051 | 81 | 699 |
| SRR1636969 | 20 | 0 | 623 | 0 | 0 | 27 | 109 | 4249 | 14 | 0 | 0 | 42 | 0 | 18 | 72 | 0 | 0 | 36 | 24 | 1418 | 438 | 0 | 0 | 23 | 10 | 306 | 162 | 0 | 0 | 11 | 19 | 0 | 0 | 87 | 1320 | 108 | 830 |
| SRR1636970 | 20 | 28 | 496 | 0 | 0 | 31 | 93 | 3924 | 0 | 0 | 0 | 78 | 0 | 17 | 47 | 0 | 0 | 0 | 34 | 1602 | 392 | 0 | 0 | 16 | 14 | 301 | 241 | 0 | 0 | 17 | 18 | 0 | 0 | 59 | 1360 | 130 | 968 |
| SRR1642941 | 105 | 0 | 126 | 0 | 38 | 29 | 95 | 0 | 16 | 47 | 201 | 44 | 0 | 14 | 102 | 0 | 14 | 33 | 0 | 548 | 278 | 0 | 0 | 949 | 125 | 165 | 480 | 0 | 0 | 14 | 27 | 0 | 0 | 63 | 0 | 0 | 0 |
| SRR1642942 | 209 | 0 | 66 | 18 | 70 | 40 | 0 | 0 | 0 | 352 | 841 | 504 | 12 | 18 | 201 | 0 | 46 | 72 | 0 | 452 | 1219 | 0 | 0 | 700 | 250 | 302 | 1072 | 0 | 0 | 14 | 45 | 0 | 0 | 85 | 16 | 13 | 29 |
| SRR1642943 | 22 | 0 | 66 | 12 | 17 | 12 | 140 | 11 | 0 | 45 | 120 | 14 | 18 | 10 | 32 | 0 | 25 | 20 | 0 | 156 | 252 | 0 | 0 | 247 | 82 | 67 | 370 | 0 | 0 | 0 | 16 | 0 | 0 | 32 | 0 | 0 | 0 |
| SRR1642944 | 29 | 0 | 160 | 0 | 40 | 41 | 46 | 12 | 17 | 56 | 264 | 68 | 21 | 16 | 3874 | 0 | 10 | 85 | 0 | 205 | 296 | 0 | 0 | 1123 | 126 | 199 | 574 | 0 | 0 | 19 | 22 | 0 | 0 | 33 | 12 | 10 | 24 |
| SRR1642945 | 39 | 0 | 46 | 0 | 29 | 32 | 158 | 0 | 16 | 39 | 178 | 19 | 0 | 18 | 64 | 0 | 19 | 0 | 10 | 117 | 71 | 0 | 0 | 831 | 170 | 162 | 1083 | 0 | 0 | 0 | 25 | 0 | 0 | 129 | 0 | 0 | 0 |
| SRR1642946 | 36 | 0 | 70 | 0 | 26 | 14 | 96 | 0 | 19 | 16 | 61 | 21 | 18 | 0 | 265 | 0 | 22 | 30 | 12 | 187 | 127 | 0 | 0 | 460 | 61 | 56 | 404 | 0 | 0 | 12 | 13 | 0 | 0 | 23 | 0 | 0 | 0 |
| SRR1642947 | 48 | 0 | 33 | 0 | 11 | 12 | 85 | 0 | 14 | 20 | 109 | 30 | 14 | 15 | 148 | 0 | 26 | 42 | 15 | 359 | 144 | 0 | 0 | 387 | 69 | 63 | 301 | 0 | 0 | 0 | 15 | 0 | 0 | 20 | 0 | 0 | 0 |
| SRR1642948 | 47 | 0 | 87 | 0 | 19 | 16 | 73 | 0 | 20 | 18 | 86 | 82 | 11 | 15 | 161 | 0 | 14 | 26 | 15 | 197 | 186 | 0 | 0 | 422 | 55 | 90 | 217 | 0 | 0 | 0 | 22 | 0 | 0 | 15 | 13 | 0 | 15 |
| SRR1642949 | 77 | 0 | 80 | 0 | 29 | 22 | 34 | 0 | 12 | 62 | 301 | 60 | 0 | 16 | 3962 | 0 | 23 | 27 | 20 | 365 | 284 | 0 | 0 | 1025 | 152 | 112 | 701 | 0 | 0 | 0 | 31 | 0 | 0 | 23 | 0 | 0 | 0 |
| SRR1642950 | 103 | 0 | 74 | 0 | 28 | 25 | 77 | 0 | 13 | 51 | 251 | 576 | 25 | 13 | 1173 | 0 | 27 | 43 | 14 | 349 | 119 | 0 | 0 | 1051 | 174 | 160 | 932 | 0 | 0 | 0 | 16 | 0 | 0 | 11 | 0 | 0 | 0 |
| SRR1642951 | 51 | 0 | 118 | 0 | 12 | 11 | 107 | 0 | 11 | 18 | 86 | 19 | 17 | 13 | 2172 | 0 | 22 | 51 | 12 | 438 | 290 | 0 | 0 | 341 | 80 | 82 | 328 | 0 | 0 | 0 | 11 | 0 | 0 | 11 | 0 | 0 | 10 |
| SRR1642952 | 71 | 0 | 61 | 0 | 17 | 12 | 279 | 0 | 10 | 58 | 200 | 60 | 14 | 17 | 387 | 0 | 31 | 42 | 18 | 481 | 256 | 0 | 0 | 297 | 65 | 70 | 358 | 0 | 0 | 0 | 11 | 0 | 0 | 13 | 0 | 0 | 10 |
| SRR1642953 | 61 | 0 | 100 | 11 | 15 | 19 | 133 | 0 | 24 | 39 | 110 | 19 | 29 | 14 | 454 | 0 | 15 | 36 | 13 | 87 | 187 | 0 | 0 | 15 | 77 | 143 | 110 | 0 | 0 | 0 | 14 | 0 | 0 | 136 | 0 | 0 | 14 |
| SRR1642954 | 42 | 0 | 311 | 0 | 30 | 39 | 31 | 0 | 22 | 14 | 42 | 45 | 26 | 0 | 997 | 0 | 32 | 11 | 15 | 57 | 416 | 0 | 0 | 16 | 51 | 328 | 90 | 0 | 0 | 0 | 23 | 0 | 0 | 99 | 18 | 13 | 36 |
| SRR1642955 | 30 | 0 | 59 | 0 | 40 | 39 | 64 | 21 | 12 | 50 | 149 | 31 | 19 | 31 | 41 | 0 | 21 | 0 | 0 | 0 | 131 | 0 | 0 | 15 | 145 | 233 | 371 | 0 | 0 | 0 | 41 | 0 | 0 | 2126 | 0 | 0 | 0 |
| SRR1642956 | 109 | 0 | 455 | 26 | 43 | 60 | 157 | 0 | 20 | 163 | 511 | 152 | 24 | 19 | 202 | 0 | 208 | 0 | 0 | 90 | 175 | 0 | 0 | 28 | 203 | 500 | 335 | 13 | 0 | 0 | 56 | 0 | 0 | 412 | 23 | 21 | 55 |
| SRR1642957 | 104 | 0 | 131 | 0 | 42 | 258 | 56 | 0 | 73 | 71 | 241 | 188 | 54 | 0 | 208 | 0 | 57 | 29 | 13 | 69 | 1081 | 0 | 0 | 59 | 311 | 1290 | 322 | 0 | 0 | 0 | 124 | 0 | 0 | 161 | 24 | 19 | 34 |
| SRR1642958 | 90 | 0 | 113 | 14 | 12 | 18 | 50 | 0 | 20 | 34 | 98 | 30 | 11 | 22 | 5578 | 0 | 17 | 49 | 29 | 36 | 465 | 0 | 0 | 15 | 71 | 134 | 113 | 11 | 0 | 0 | 16 | 0 | 0 | 138 | 14 | 11 | 15 |
| SRR1642959 | 58 | 0 | 109 | 0 | 59 | 39 | 45 | 0 | 13 | 58 | 235 | 36 | 16 | 17 | 722 | 0 | 18 | 0 | 0 | 54 | 262 | 0 | 0 | 48 | 203 | 266 | 191 | 0 | 0 | 0 | 39 | 0 | 0 | 146 | 0 | 0 | 11 |
| SRR1642960 | 62 | 0 | 65 | 11 | 22 | 18 | 84 | 0 | 14 | 24 | 101 | 33 | 11 | 10 | 911 | 0 | 22 | 76 | 23 | 61 | 283 | 0 | 0 | 20 | 78 | 130 | 84 | 13 | 0 | 0 | 14 | 0 | 0 | 55 | 0 | 0 | 14 |
| SRR1642961 | 32 | 0 | 57 | 0 | 11 | 12 | 89 | 0 | 17 | 18 | 72 | 22 | 25 | 17 | 96 | 0 | 0 | 0 | 0 | 16 | 75 | 0 | 0 | 12 | 35 | 82 | 67 | 0 | 0 | 0 | 10 | 0 | 0 | 210 | 0 | 0 | 13 |
| SRR1642962 | 81 | 0 | 70 | 0 | 17 | 15 | 84 | 0 | 0 | 12 | 57 | 230 | 14 | 0 | 1493 | 0 | 34 | 19 | 10 | 94 | 188 | 0 | 0 | 23 | 91 | 123 | 110 | 0 | 0 | 0 | 25 | 0 | 0 | 29 | 0 | 0 | 17 |
| SRR1642963 | 44 | 0 | 53 | 0 | 30 | 58 | 87 | 0 | 24 | 113 | 403 | 55 | 58 | 13 | 91 | 0 | 37 | 17 | 0 | 65 | 239 | 0 | 0 | 72 | 273 | 305 | 263 | 0 | 0 | 0 | 28 | 0 | 0 | 317 | 10 | 0 | 12 |

### Supp Table S8.2.Columns_AM_to_BW

| | | | | | | | | | | | | | | | | | | | | | | | | | | | | | | | | | | | | | |
|---|---|---|---|---|---|---|---|---|---|---|---|---|---|---|---|---|---|---|---|---|---|---|---|---|---|---|---|---|---|---|---|---|---|---|---|---|---|
| SRR1642964 | 74 | 0 | 12 | 12 | 11 | 43 | 146 | 0 | 24 | 28 | 92 | 694 | 19 | 11 | 369 | 0 | 44 | 94 | 39 | 97 | 457 | 0 | 0 | 15 | 80 | 229 | 76 | 0 | 0 | 0 | 34 | 0 | 0 | 80 | 15 | 11 | 13 |
| SRR1642965 | 55 | 0 | 187 | 17 | 19 | 0 | 0 | 0 | 0 | 54 | 170 | 36 | 15 | 26 | 397 | 0 | 24 | 102 | 0 | 0 | 0 | 0 | 0 | 27 | 79 | 33 | 0 | 13 | 0 | 0 | 19 | 0 | 0 | 74 | 0 | 0 | 0 |
| SRR1642966 | 72 | 0 | 171 | 14 | 56 | 0 | 0 | 0 | 0 | 112 | 368 | 20 | 33 | 21 | 307 | 0 | 77 | 60 | 0 | 19 | 42 | 0 | 0 | 35 | 248 | 112 | 13 | 14 | 0 | 0 | 21 | 0 | 0 | 101 | 0 | 0 | 14 |
| SRR1642967 | 46 | 0 | 149 | 12 | 19 | 0 | 60 | 0 | 0 | 32 | 112 | 17 | 10 | 25 | 589 | 0 | 21 | 48 | 0 | 0 | 32 | 0 | 0 | 19 | 76 | 34 | 13 | 13 | 0 | 0 | 15 | 0 | 0 | 88 | 0 | 0 | 0 |
| SRR1642968 | 88 | 0 | 140 | 12 | 54 | 10 | 16 | 0 | 10 | 121 | 398 | 40 | 26 | 24 | 956 | 0 | 26 | 47 | 0 | 16 | 0 | 0 | 0 | 36 | 253 | 109 | 18 | 0 | 0 | 0 | 61 | 0 | 0 | 183 | 0 | 0 | 0 |
| SRR1642969 | 48 | 0 | 97 | 13 | 17 | 0 | 0 | 0 | 0 | 44 | 131 | 28 | 15 | 22 | 309 | 0 | 22 | 0 | 0 | 0 | 71 | 0 | 0 | 25 | 80 | 26 | 17 | 0 | 0 | 0 | 12 | 0 | 0 | 157 | 0 | 0 | 0 |
| SRR1642970 | 19 | 0 | 179 | 14 | 11 | 21 | 0 | 0 | 27 | 37 | 39 | 55 | 28 | 28 | 523 | 0 | 37 | 86 | 0 | 0 | 49 | 0 | 0 | 33 | 65 | 63 | 14 | 12 | 0 | 0 | 12 | 0 | 0 | 75 | 18 | 13 | 24 |
| SRR1642971 | 41 | 0 | 115 | 0 | 16 | 18 | 21 | 0 | 12 | 20 | 78 | 39 | 31 | 22 | 276 | 0 | 33 | 0 | 0 | 0 | 0 | 0 | 0 | 31 | 95 | 44 | 22 | 0 | 0 | 0 | 23 | 0 | 0 | 106 | 0 | 0 | 11 |
| SRR1642972 | 38 | 0 | 216 | 14 | 23 | 0 | 0 | 0 | 0 | 30 | 89 | 196 | 11 | 28 | 1824 | 0 | 60 | 53 | 0 | 10 | 21 | 0 | 0 | 26 | 71 | 27 | 11 | 12 | 0 | 0 | 28 | 0 | 0 | 72 | 14 | 11 | 21 |
| SRR1642973 | 61 | 0 | 182 | 13 | 22 | 12 | 15 | 0 | 14 | 20 | 84 | 22 | 14 | 23 | 4364 | 0 | 19 | 88 | 0 | 0 | 45 | 0 | 0 | 23 | 101 | 46 | 22 | 0 | 0 | 0 | 16 | 0 | 0 | 96 | 0 | 0 | 10 |
| SRR1642974 | 120 | 0 | 193 | 11 | 40 | 12 | 14 | 0 | 16 | 78 | 225 | 74 | 42 | 24 | 9076 | 0 | 15 | 37 | 14 | 0 | 99 | 0 | 0 | 23 | 292 | 144 | 18 | 0 | 0 | 0 | 46 | 0 | 0 | 368 | 13 | 0 | 20 |
| SRR1642975 | 54 | 0 | 91 | 0 | 35 | 18 | 0 | 0 | 23 | 35 | 41 | 38 | 22 | 12 | 1169 | 0 | 35 | 35 | 0 | 20 | 73 | 0 | 0 | 11 | 157 | 43 | 27 | 0 | 0 | 0 | 24 | 0 | 0 | 64 | 14 | 0 | 12 |
| SRR1642976 | 36 | 0 | 139 | 15 | 22 | 30 | 0 | 0 | 11 | 13 | 56 | 39 | 14 | 21 | 402 | 0 | 24 | 54 | 0 | 0 | 24 | 0 | 0 | 18 | 93 | 143 | 22 | 0 | 0 | 0 | 62 | 0 | 0 | 88 | 12 | 0 | 12 |
| SRR1642977 | 81 | 0 | 140 | 0 | 21 | 13 | 91 | 0 | 14 | 36 | 125 | 48 | 14 | 23 | 44 | 0 | 23 | 13 | 0 | 0 | 63 | 0 | 0 | 11 | 85 | 85 | 83 | 11 | 0 | 0 | 12 | 0 | 0 | 120 | 12 | 13 | 11 |
| SRR1642978 | 32 | 0 | 62 | 0 | 38 | 41 | 35 | 0 | 26 | 61 | 209 | 393 | 46 | 11 | 135 | 0 | 29 | 81 | 0 | 35 | 262 | 0 | 0 | 22 | 205 | 281 | 135 | 0 | 0 | 0 | 51 | 0 | 0 | 253 | 18 | 15 | 24 |
| SRR1642979 | 78 | 0 | 133 | 0 | 29 | 35 | 27 | 0 | 25 | 54 | 150 | 100 | 34 | 13 | 81 | 0 | 25 | 38 | 0 | 18 | 322 | 0 | 0 | 24 | 204 | 300 | 158 | 11 | 0 | 0 | 71 | 0 | 0 | 192 | 17 | 19 | 32 |
| SRR1642980 | 73 | 0 | 40 | 17 | 67 | 36 | 14 | 0 | 25 | 65 | 306 | 499 | 45 | 23 | 289 | 0 | 23 | 20 | 14 | 14 | 524 | 0 | 0 | 15 | 185 | 272 | 124 | 11 | 0 | 0 | 33 | 0 | 0 | 452 | 39 | 30 | 64 |
| SRR1642981 | 69 | 0 | 112 | 0 | 74 | 28 | 62 | 0 | 19 | 68 | 292 | 55 | 28 | 14 | 36 | 0 | 14 | 52 | 0 | 15 | 73 | 0 | 0 | 19 | 264 | 232 | 141 | 0 | 0 | 0 | 41 | 0 | 0 | 259 | 0 | 0 | 0 |
| SRR1642982 | 36 | 0 | 117 | 11 | 53 | 50 | 68 | 0 | 34 | 45 | 197 | 172 | 75 | 17 | 121 | 0 | 28 | 78 | 0 | 0 | 47 | 0 | 0 | 15 | 240 | 539 | 160 | 0 | 0 | 0 | 39 | 0 | 0 | 644 | 24 | 13 | 42 |
| SRR1642983 | 47 | 0 | 89 | 15 | 29 | 42 | 76 | 0 | 15 | 192 | 476 | 48 | 41 | 34 | 55 | 0 | 15 | 19 | 0 | 34 | 122 | 0 | 0 | 0 | 287 | 375 | 264 | 13 | 0 | 0 | 54 | 0 | 0 | 397 | 0 | 0 | 12 |
| SRR1642984 | 19 | 0 | 21 | 17 | 11 | 27 | 62 | 0 | 33 | 64 | 143 | 137 | 20 | 12 | 467 | 0 | 21 | 38 | 0 | 0 | 249 | 0 | 0 | 13 | 89 | 196 | 74 | 11 | 0 | 0 | 19 | 0 | 0 | 115 | 14 | 0 | 10 |
| SRR1642985 | 81 | 0 | 142 | 20 | 47 | 18 | 247 | 0 | 16 | 39 | 165 | 19 | 14 | 18 | 1655 | 0 | 18 | 58 | 0 | 31 | 145 | 0 | 0 | 15 | 104 | 143 | 47 | 0 | 0 | 0 | 15 | 0 | 0 | 143 | 11 | 12 | 15 |
| SRR1642986 | 89 | 0 | 163 | 12 | 29 | 15 | 121 | 0 | 12 | 70 | 221 | 24 | 20 | 18 | 472 | 0 | 26 | 78 | 0 | 33 | 159 | 0 | 0 | 16 | 123 | 148 | 72 | 11 | 0 | 0 | 13 | 0 | 0 | 160 | 12 | 0 | 19 |
| SRR1642987 | 54 | 0 | 89 | 14 | 33 | 15 | 50 | 0 | 19 | 27 | 117 | 36 | 21 | 10 | 175 | 0 | 16 | 27 | 33 | 192 | 83 | 0 | 0 | 16 | 83 | 116 | 44 | 12 | 0 | 0 | 20 | 0 | 0 | 74 | 0 | 0 | 13 |
| SRR1642988 | 53 | 0 | 450 | 16 | 56 | 34 | 0 | 0 | 15 | 32 | 120 | 40 | 57 | 21 | 1250 | 0 | 54 | 23 | 13 | 19 | 119 | 0 | 0 | 22 | 269 | 240 | 121 | 10 | 0 | 0 | 149 | 0 | 0 | 488 | 18 | 18 | 34 |
| SRR1657194 | 60 | 0 | 1419 | 0 | 0 | 102 | 15 | 18 | 76 | 0 | 0 | 513 | 18 | 0 | 58 | 0 | 34 | 34 | 10 | 1117 | 119 | 0 | 0 | 0 | 0 | 103 | 148 | 10 | 0 | 0 | 11 | 0 | 0 | 0 | 148 | 129 | 71 |
| SRR1657195 | 70 | 0 | 1305 | 0 | 0 | 0 | 0 | 30 | 0 | 0 | 0 | 313 | 0 | 0 | 110 | 0 | 44 | 126 | 0 | 822 | 303 | 0 | 0 | 17 | 0 | 15 | 51 | 22 | 0 | 0 | 23 | 0 | 0 | 0 | 940 | 44 | 364 |
| SRR1657196 | 72 | 0 | 1278 | 0 | 0 | 65 | 0 | 18 | 17 | 0 | 0 | 313 | 0 | 0 | 35 | 0 | 37 | 44 | 0 | 733 | 166 | 0 | 0 | 11 | 13 | 948 | 1267 | 0 | 0 | 0 | 77 | 0 | 0 | 0 | 3952 | 211 | 1483 |
| SRR1657197 | 65 | 0 | 1603 | 0 | 0 | 111 | 29 | 345 | 14 | 0 | 0 | 597 | 0 | 14 | 59 | 0 | 61 | 26 | 15 | 4877 | 221 | 0 | 0 | 180 | 0 | 705 | 3883 | 0 | 0 | 0 | 124 | 0 | 0 | 0 | 1382 | 138 | 746 |
| SRR1657198 | 66 | 23 | 2320 | 0 | 0 | 0 | 0 | 0 | 0 | 0 | 0 | 542 | 0 | 0 | 46 | 0 | 78 | 26 | 0 | 260 | 31 | 0 | 0 | 0 | 0 | 29 | 26 | 0 | 0 | 0 | 12 | 0 | 0 | 0 | 154 | 48 | 53 |
| SRR1686806 | 40 | 0 | 912 | 0 | 0 | 0 | 293 | 0 | 0 | 0 | 0 | 125 | 0 | 50 | 0 | 0 | 0 | 19 | 3104 | 5413 | 5303 | 0 | 0 | 0 | 0 | 0 | 146 | 0 | 0 | 0 | 0 | 0 | 0 | 18 | 46 | 73 | 45 |
| SRR1707574 | 28 | 0 | 0 | 0 | 0 | 0 | 0 | 82 | 0 | 0 | 0 | 459 | 0 | 40 | 0 | 0 | 0 | 0 | 48 | 3681 | 297 | 0 | 0 | 0 | 15 | 0 | 211 | 81 | 0 | 0 | 0 | 0 | 0 | 74 | 36 | 0 | 24 |
| SRR1707575 | 0 | 26 | 807 | 17 | 12 | 19 | 11 | 0 | 44 | 0 | 0 | 11 | 0 | 0 | 26 | 0 | 0 | 0 | 12 | 2142 | 339 | 0 | 0 | 0 | 0 | 586 | 0 | 0 | 29 | 0 | 17 | 0 | 41 | 0 | 0 | 13 | 0 |
| SRR1708906 | 83 | 0 | 676 | 0 | 0 | 16 | 0 | 0 | 10 | 0 | 0 | 758 | 0 | 26 | 18 | 0 | 17 | 0 | 36 | 3770 | 1278 | 0 | 0 | 0 | 0 | 784 | 1885 | 0 | 0 | 0 | 55 | 0 | 0 | 16 | 2194 | 693 | 3564 |
| SRR1708907 | 141 | 0 | 725 | 0 | 0 | 24 | 0 | 0 | 12 | 0 | 0 | 811 | 0 | 0 | 32 | 0 | 17 | 12 | 0 | 140 | 295 | 0 | 0 | 0 | 0 | 31 | 11 | 17 | 0 | 0 | 70 | 0 | 0 | 0 | 92 | 20 | 135 |
| SRR1708908 | 116 | 0 | 532 | 0 | 0 | 0 | 0 | 0 | 0 | 0 | 0 | 422 | 0 | 0 | 22 | 0 | 20 | 0 | 0 | 72 | 79 | 0 | 0 | 0 | 0 | 173 | 77 | 0 | 0 | 0 | 74 | 0 | 0 | 17 | 44 | 11 | 58 |
| SRR1708909 | 283 | 0 | 938 | 0 | 0 | 26 | 0 | 0 | 0 | 0 | 0 | 455 | 0 | 0 | 10 | 0 | 20 | 0 | 0 | 1693 | 2040 | 0 | 0 | 0 | 0 | 56 | 116 | 0 | 0 | 0 | 53 | 0 | 24 | 0 | 77 | 90 | 78 |
| SRR1708910 | 146 | 0 | 551 | 0 | 0 | 20 | 0 | 0 | 13 | 0 | 0 | 968 | 0 | 0 | 0 | 0 | 20 | 0 | 0 | 59 | 129 | 0 | 0 | 0 | 13 | 17 | 109 | 13 | 0 | 0 | 52 | 0 | 0 | 0 | 31 | 13 | 48 |
| SRR1708911 | 97 | 0 | 609 | 0 | 0 | 0 | 0 | 0 | 0 | 0 | 0 | 588 | 0 | 0 | 34 | 0 | 29 | 0 | 0 | 91 | 57 | 0 | 0 | 0 | 0 | 12 | 94 | 24 | 0 | 0 | 126 | 0 | 0 | 16 | 27 | 175 | 30 |
| SRR1708912 | 66 | 0 | 474 | 0 | 0 | 12 | 0 | 0 | 0 | 0 | 0 | 766 | 0 | 0 | 30 | 0 | 14 | 33 | 0 | 242 | 545 | 0 | 0 | 0 | 0 | 419 | 554 | 14 | 0 | 0 | 61 | 0 | 0 | 0 | 727 | 323 | 1097 |
| SRR1708913 | 112 | 0 | 496 | 0 | 0 | 0 | 0 | 0 | 0 | 0 | 0 | 591 | 0 | 12 | 44 | 0 | 16 | 0 | 0 | 68 | 83 | 0 | 0 | 0 | 0 | 145 | 171 | 0 | 0 | 0 | 75 | 0 | 0 | 16 | 17 | 206 | 31 |
| SRR1708914 | 59 | 0 | 362 | 20 | 0 | 45 | 0 | 0 | 35 | 0 | 0 | 268 | 0 | 0 | 87 | 0 | 21 | 58 | 0 | 246 | 825 | 0 | 0 | 0 | 25 | 20 | 37 | 0 | 0 | 0 | 60 | 0 | 0 | 15 | 32 | 243 | 32 |
| SRR1708915 | 94 | 0 | 355 | 0 | 0 | 15 | 14 | 0 | 0 | 0 | 0 | 249 | 0 | 13 | 35 | 0 | 18 | 0 | 0 | 1585 | 1655 | 0 | 0 | 0 | 0 | 522 | 1373 | 13 | 0 | 0 | 67 | 0 | 0 | 0 | 503 | 165 | 784 |
| SRR1708916 | 106 | 11 | 400 | 0 | 0 | 19 | 0 | 0 | 14 | 0 | 0 | 651 | 0 | 14 | 72 | 0 | 18 | 0 | 0 | 126 | 260 | 0 | 0 | 0 | 0 | 415 | 206 | 16 | 0 | 0 | 79 | 0 | 0 | 0 | 848 | 293 | 992 |
| SRR1708917 | 75 | 0 | 576 | 0 | 0 | 57 | 0 | 0 | 12 | 0 | 0 | 511 | 0 | 17 | 71 | 0 | 20 | 0 | 0 | 478 | 1514 | 0 | 0 | 0 | 0 | 780 | 1216 | 0 | 0 | 0 | 80 | 0 | 0 | 0 | 608 | 231 | 931 |
| SRR1708918 | 83 | 0 | 465 | 0 | 0 | 13 | 0 | 0 | 0 | 0 | 0 | 521 | 0 | 23 | 88 | 0 | 11 | 0 | 0 | 77 | 387 | 0 | 0 | 0 | 0 | 258 | 127 | 13 | 0 | 0 | 124 | 0 | 0 | 0 | 418 | 163 | 725 |
| SRR1708919 | 73 | 0 | 508 | 0 | 0 | 53 | 0 | 0 | 30 | 0 | 0 | 330 | 0 | 11 | 47 | 0 | 17 | 0 | 0 | 410 | 392 | 14 | 0 | 0 | 11 | 42 | 27 | 0 | 0 | 0 | 111 | 0 | 0 | 0 | 28 | 158 | 37 |
| SRR1708920 | 80 | 0 | 402 | 0 | 0 | 31 | 0 | 0 | 30 | 0 | 0 | 249 | 0 | 10 | 42 | 0 | 17 | 0 | 0 | 358 | 473 | 10 | 0 | 0 | 0 | 648 | 421 | 18 | 0 | 0 | 71 | 0 | 0 | 0 | 362 | 100 | 435 |
| SRR1708921 | 79 | 0 | 350 | 0 | 0 | 51 | 0 | 0 | 28 | 0 | 0 | 162 | 0 | 12 | 78 | 0 | 19 | 0 | 0 | 729 | 1259 | 0 | 0 | 0 | 26 | 847 | 736 | 12 | 0 | 0 | 109 | 0 | 0 | 0 | 475 | 107 | 576 |
| SRR1708922 | 78 | 0 | 415 | 0 | 0 | 39 | 0 | 0 | 23 | 0 | 0 | 484 | 0 | 0 | 29 | 0 | 13 | 0 | 0 | 818 | 995 | 0 | 0 | 0 | 0 | 27 | 35 | 23 | 0 | 0 | 84 | 0 | 0 | 0 | 15 | 195 | 34 |
| SRR1708923 | 72 | 15 | 475 | 0 | 0 | 49 | 12 | 0 | 33 | 0 | 0 | 405 | 0 | 0 | 24 | 0 | 23 | 0 | 0 | 358 | 2384 | 0 | 0 | 0 | 0 | 38 | 26 | 27 | 0 | 0 | 97 | 0 | 0 | 0 | 17 | 122 | 32 |
| SRR1708924 | 121 | 0 | 913 | 0 | 0 | 19 | 0 | 0 | 0 | 0 | 0 | 938 | 0 | 0 | 27 | 0 | 17 | 0 | 15 | 385 | 296 | 0 | 0 | 0 | 0 | 33 | 34 | 15 | 0 | 0 | 32 | 0 | 0 | 0 | 111 | 31 | 147 |
| SRR1708925 | 105 | 0 | 830 | 0 | 0 | 20 | 23 | 0 | 18 | 0 | 0 | 592 | 0 | 12 | 27 | 0 | 15 | 0 | 27 | 871 | 1036 | 18 | 0 | 0 | 0 | 32 | 37 | 14 | 0 | 0 | 64 | 0 | 0 | 0 | 124 | 22 | 117 |
| SRR1708926 | 141 | 0 | 568 | 0 | 0 | 36 | 0 | 21 | 24 | 0 | 0 | 496 | 0 | 0 | 17 | 0 | 16 | 0 | 28 | 3011 | 626 | 19 | 0 | 0 | 13 | 39 | 140 | 21 | 0 | 0 | 56 | 0 | 0 | 0 | 46 | 21 | 100 |
| SRR1708927 | 92 | 0 | 413 | 0 | 0 | 45 | 0 | 0 | 16 | 0 | 0 | 310 | 0 | 0 | 38 | 0 | 24 | 0 | 0 | 1395 | 917 | 0 | 0 | 0 | 0 | 789 | 1085 | 14 | 11 | 0 | 104 | 0 | 0 | 20 | 1974 | 342 | 1666 |
| SRR1708928 | 78 | 0 | 483 | 14 | 0 | 29 | 0 | 0 | 14 | 0 | 0 | 666 | 0 | 17 | 38 | 0 | 22 | 0 | 0 | 324 | 166 | 24 | 0 | 0 | 12 | 727 | 277 | 0 | 0 | 0 | 96 | 0 | 0 | 38 | 2236 | 524 | 3057 |
| SRR1708929 | 99 | 0 | 800 | 0 | 13 | 13 | 0 | 0 | 11 | 0 | 0 | 646 | 0 | 0 | 50 | 0 | 29 | 0 | 0 | 132 | 149 | 20 | 0 | 0 | 0 | 27 | 19 | 13 | 0 | 0 | 64 | 0 | 0 | 0 | 46 | 13 | 58 |
| SRR1708930 | 61 | 15 | 907 | 0 | 0 | 27 | 22 | 10 | 0 | 0 | 0 | 389 | 0 | 13 | 44 | 0 | 12 | 0 | 0 | 1730 | 535 | 0 | 0 | 0 | 13 | 24 | 78 | 19 | 0 | 0 | 39 | 0 | 0 | 19 | 63 | 10 | 46 |
| SRR1708931 | 69 | 0 | 757 | 0 | 0 | 33 | 17 | 0 | 0 | 0 | 0 | 358 | 0 | 0 | 55 | 0 | 13 | 0 | 0 | 1916 | 1226 | 0 | 0 | 0 | 0 | 454 | 2092 | 0 | 0 | 0 | 36 | 0 | 0 | 0 | 1082 | 187 | 1346 |
| SRR1708932 | 57 | 0 | 499 | 0 | 0 | 0 | 0 | 0 | 0 | 0 | 0 | 204 | 0 | 23 | 73 | 0 | 17 | 114 | 0 | 67 | 152 | 0 | 0 | 0 | 17 | 158 | 122 | 0 | 11 | 0 | 85 | 0 | 0 | 0 | 56 | 133 | 26 |
| SRR1708933 | 85 | 0 | 432 | 0 | 0 | 24 | 0 | 0 | 0 | 0 | 0 | 119 | 0 | 24 | 108 | 0 | 24 | 0 | 0 | 177 | 279 | 0 | 0 | 0 | 0 | 341 | 402 | 15 | 0 | 0 | 136 | 0 | 0 | 18 | 2161 | 278 | 1212 |
| SRR1708934 | 88 | 0 | 620 | 0 | 0 | 44 | 0 | 0 | 17 | 0 | 0 | 537 | 0 | 0 | 22 | 0 | 19 | 0 | 0 | 2358 | 581 | 0 | 0 | 0 | 0 | 914 | 4320 | 13 | 0 | 0 | 70 | 0 | 0 | 0 | 2870 | 404 | 2163 |
| SRR1708935 | 60 | 0 | 749 | 10 | 0 | 0 | 0 | 0 | 0 | 0 | 0 | 297 | 0 | 0 | 145 | 0 | 23 | 0 | 0 | 41 | 139 | 0 | 0 | 0 | 0 | 210 | 96 | 0 | 0 | 0 | 73 | 0 | 0 | 0 | 758 | 129 | 698 |
| SRR1708936 | 41 | 11 | 827 | 0 | 0 | 58 | 40 | 15 | 34 | 0 | 0 | 48 | 0 | 0 | 76 | 0 | 18 | 0 | 22 | 4451 | 1163 | 0 | 0 | 0 | 0 | 1018 | 1657 | 0 | 0 | 0 | 69 | 0 | 0 | 0 | 718 | 122 | 707 |
| SRR1708937 | 38 | 24 | 604 | 0 | 0 | 0 | 0 | 0 | 0 | 0 | 0 | 41 | 0 | 0 | 100 | 0 | 16 | 0 | 0 | 29 | 47 | 0 | 0 | 0 | 0 | 96 | 40 | 12 | 0 | 0 | 76 | 0 | 0 | 0 | 21 | 36 | 133 |
| SRR1708938 | 44 | 0 | 334 | 0 | 0 | 12 | 0 | 0 | 15 | 0 | 0 | 37 | 0 | 28 | 132 | 0 | 18 | 0 | 0 | 66 | 234 | 0 | 0 | 0 | 0 | 350 | 108 | 28 | 0 | 0 | 183 | 0 | 0 | 12 | 718 | 84 | 412 |
| SRR1708939 | 75 | 0 | 386 | 0 | 0 | 33 | 0 | 0 | 23 | 0 | 0 | 26 | 0 | 20 | 110 | 0 | 32 | 0 | 0 | 191 | 294 | 0 | 0 | 0 | 13 | 29 | 198 | 13 | 36 | 0 | 176 | 0 | 0 | 0 | 30 | 120 | 22 |
| SRR1708940 | 62 | 0 | 562 | 0 | 0 | 39 | 0 | 0 | 44 | 0 | 0 | 158 | 0 | 29 | 58 | 0 | 22 | 0 | 0 | 246 | 472 | 0 | 0 | 0 | 0 | 829 | 255 | 25 | 19 | 0 | 75 | 0 | 0 | 12 | 1111 | 193 | 904 |
| SRR1708941 | 57 | 52 | 571 | 0 | 0 | 30 | 0 | 0 | 39 | 0 | 0 | 144 | 0 | 30 | 54 | 0 | 28 | 0 | 0 | 132 | 278 | 0 | 0 | 0 | 0 | 36 | 153 | 13 | 0 | 0 | 133 | 0 | 0 | 15 | 23 | 153 | 27 |
| SRR1708942 | 91 | 0 | 742 | 0 | 0 | 11 | 0 | 0 | 17 | 0 | 0 | 986 | 0 | 0 | 90 | 0 | 18 | 0 | 0 | 214 | 255 | 0 | 0 | 0 | 0 | 28 | 11 | 0 | 0 | 0 | 63 | 0 | 0 | 20 | 156 | 36 | 158 |
| SRR1708943 | 193 | 0 | 817 | 0 | 0 | 0 | 0 | 0 | 0 | 0 | 0 | 1010 | 0 | 0 | 30 | 0 | 13 | 0 | 0 | 34 | 188 | 0 | 0 | 0 | 0 | 104 | 30 | 0 | 0 | 0 | 38 | 0 | 0 | 0 | 61 | 13 | 38 |
| SRR1708944 | 143 | 0 | 816 | 11 | 0 | 27 | 0 | 0 | 16 | 0 | 0 | 547 | 0 | 0 | 23 | 0 | 25 | 0 | 11 | 110 | 141 | 0 | 0 | 0 | 19 | 529 | 177 | 0 | 0 | 0 | 56 | 0 | 11 | 16 | 744 | 204 | 1325 |
| SRR1708945 | 175 | 0 | 589 | 0 | 0 | 16 | 10 | 0 | 21 | 0 | 0 | 410 | 0 | 0 | 30 | 0 | 23 | 0 | 36 | 1452 | 584 | 24 | 0 | 0 | 11 | 52 | 110 | 0 | 11 | 0 | 72 | 0 | 0 | 0 | 90 | 22 | 81 |
| SRR1708946 | 80 | 0 | 433 | 0 | 0 | 11 | 0 | 0 | 0 | 0 | 0 | 769 | 0 | 0 | 47 | 0 | 20 | 0 | 0 | 126 | 226 | 20 | 0 | 0 | 11 | 22 | 103 | 0 | 0 | 0 | 60 | 0 | 0 | 26 | 43 | 12 | 58 |
| SRR1708947 | 149 | 0 | 682 | 0 | 0 | 41 | 15 | 0 | 12 | 0 | 0 | 609 | 0 | 18 | 15 | 0 | 22 | 0 | 28 | 801 | 513 | 24 | 0 | 0 | 12 | 846 | 931 | 0 | 0 | 0 | 98 | 0 | 0 | 0 | 573 | 167 | 1052 |
| SRR1708948 | 69 | 13 | 884 | 0 | 0 | 0 | 0 | 0 | 11 | 0 | 0 | 562 | 0 | 11 | 74 | 0 | 16 | 15 | 0 | 84 | 99 | 0 | 0 | 0 | 0 | 304 | 146 | 21 | 0 | 0 | 38 | 0 | 0 | 11 | 1004 | 146 | 800 |
| SRR1708949 | 82 | 0 | 746 | 0 | 0 | 34 | 0 | 0 | 11 | 0 | 0 | 402 | 11 | 14 | 53 | 0 | 17 | 0 | 0 | 386 | 386 | 0 | 0 | 0 | 0 | 20 | 36 | 0 | 11 | 0 | 34 | 0 | 0 | 0 | 45 | 230 | 54 |
| SRR1708950 | 69 | 0 | 586 | 0 | 0 | 0 | 0 | 0 | 0 | 0 | 0 | 300 | 0 | 11 | 62 | 0 | 18 | 0 | 0 | 93 | 251 | 0 | 0 | 0 | 0 | 12 | 11 | 11 | 0 | 0 | 17 | 0 | 0 | 0 | 54 | 12 | 41 |
| SRR1708951 | 85 | 0 | 469 | 0 | 0 | 14 | 0 | 0 | 0 | 0 | 0 | 109 | 0 | 23 | 270 | 0 | 22 | 16 | 14 | 1745 | 1181 | 0 | 0 | 0 | 0 | 346 | 945 | 0 | 0 | 0 | 41 | 0 | 0 | 0 | 1261 | 172 | 800 |
| SRR1708952 | 70 | 0 | 628 | 0 | 17 | 22 | 0 | 0 | 0 | 0 | 0 | 372 | 0 | 20 | 165 | 0 | 16 | 0 | 0 | 177 | 300 | 0 | 0 | 0 | 0 | 18 | 13 | 25 | 0 | 0 | 39 | 0 | 0 | 0 | 78 | 10 | 44 |
| SRR1708953 | 54 | 0 | 652 | 0 | 0 | 18 | 0 | 0 | 18 | 0 | 0 | 284 | 0 | 18 | 212 | 0 | 22 | 0 | 0 | 154 | 503 | 0 | 0 | 0 | 20 | 314 | 271 | 18 | 0 | 0 | 47 | 0 | 0 | 18 | 1204 | 147 | 951 |
| SRR1708954 | 43 | 0 | 938 | 0 | 0 | 26 | 0 | 0 | 18 | 0 | 0 | 68 | 0 | 26 | 262 | 0 | 22 | 0 | 0 | 152 | 290 | 0 | 0 | 0 | 0 | 550 | 181 | 0 | 0 | 0 | 51 | 0 | 0 | 0 | 772 | 124 | 611 |
| SRR1708955 | 41 | 0 | 859 | 0 | 0 | 36 | 0 | 0 | 24 | 0 | 0 | 39 | 0 | 14 | 174 | 0 | 19 | 0 | 0 | 212 | 514 | 0 | 0 | 0 | 0 | 688 | 227 | 0 | 0 | 0 | 62 | 0 | 0 | 0 | 676 | 71 | 450 |
| SRR1708956 | 34 | 17 | 541 | 0 | 0 | 67 | 14 | 0 | 25 | 0 | 0 | 19 | 0 | 19 | 451 | 0 | 23 | 0 | 0 | 845 | 1222 | 0 | 0 | 0 | 10 | 1053 | 620 | 15 | 0 | 0 | 118 | 0 | 0 | 0 | 898 | 64 | 425 |
| SRR1708957 | 36 | 0 | 288 | 0 | 0 | 87 | 31 | 0 | 32 | 0 | 0 | 0 | 0 | 16 | 94 | 0 | 28 | 0 | 27 | 3635 | 1419 | 0 | 0 | 0 | 0 | 78 | 58 | 16 | 0 | 0 | 145 | 0 | 0 | 14 | 38 | 106 | 20 |
| SRR1708958 | 39 | 22 | 555 | 0 | 0 | 29 | 0 | 0 | 17 | 0 | 0 | 23 | 0 | 27 | 231 | 0 | 23 | 0 | 0 | 196 | 479 | 0 | 0 | 0 | 11 | 617 | 109 | 0 | 0 | 0 | 54 | 0 | 0 | 0 | 988 | 54 | 449 |
| SRR1708959 | 36 | 0 | 605 | 0 | 0 | 55 | 0 | 0 | 26 | 0 | 0 | 16 | 0 | 0 | 100 | 0 | 31 | 0 | 0 | 228 | 548 | 0 | 0 | 0 | 0 | 1143 | 403 | 0 | 0 | 0 | 68 | 0 | 0 | 0 | 645 | 90 | 563 |
| SRR1708960 | 67 | 0 | 747 | 0 | 0 | 55 | 0 | 14 | 0 | 0 | 0 | 615 | 0 | 10 | 83 | 0 | 13 | 0 | 41 | 1943 | 847 | 10 | 0 | 0 | 12 | 579 | 960 | 16 | 0 | 0 | 39 | 0 | 0 | 18 | 1804 | 536 | 2073 |
| SRR1708961 | 93 | 0 | 772 | 0 | 0 | 24 | 35 | 0 | 0 | 0 | 0 | 598 | 0 | 0 | 35 | 0 | 19 | 20 | 32 | 1382 | 425 | 18 | 0 | 0 | 12 | 609 | 1434 | 0 | 0 | 0 | 44 | 0 | 0 | 30 | 2216 | 454 | 2558 |
| SRR1708962 | 74 | 0 | 430 | 0 | 0 | 14 | 0 | 0 | 0 | 0 | 0 | 372 | 0 | 10 | 71 | 0 | 17 | 0 | 18 | 165 | 254 | 29 | 0 | 0 | 0 | 16 | 147 | 0 | 0 | 0 | 55 | 0 | 0 | 10 | 45 | 15 | 72 |
| SRR1708963 | 135 | 0 | 448 | 0 | 0 | 12 | 0 | 0 | 0 | 0 | 0 | 340 | 0 | 0 | 43 | 0 | 20 | 0 | 0 | 259 | 187 | 23 | 0 | 0 | 14 | 18 | 142 | 0 | 0 | 0 | 85 | 0 | 0 | 0 | 94 | 16 | 77 |
| SRR1708964 | 59 | 15 | 351 | 11 | 15 | 56 | 0 | 0 | 0 | 0 | 0 | 543 | 0 | 0 | 15 | 0 | 23 | 0 | 22 | 601 | 400 | 23 | 0 | 0 | 15 | 31 | 18 | 19 | 11 | 0 | 72 | 0 | 0 | 11 | 43 | 16 | 56 |
| SRR1708965 | 59 | 0 | 667 | 0 | 0 | 12 | 0 | 0 | 14 | 0 | 0 | 480 | 0 | 0 | 88 | 0 | 24 | 0 | 0 | 51 | 95 | 16 | 0 | 0 | 0 | 331 | 67 | 0 | 0 | 0 | 83 | 0 | 0 | 24 | 904 | 189 | 849 |
| SRR1708966 | 49 | 13 | 823 | 0 | 0 | 19 | 57 | 46 | 0 | 0 | 0 | 477 | 0 | 38 | 76 | 0 | 19 | 0 | 0 | 2232 | 739 | 0 | 0 | 0 | 0 | 423 | 1552 | 17 | 0 | 0 | 11 | 0 | 0 | 0 | 994 | 163 | 860 |
| SRR1708967 | 59 | 19 | 679 | 0 | 0 | 0 | 0 | 0 | 13 | 0 | 0 | 392 | 0 | 10 | 98 | 0 | 18 | 0 | 0 | 74 | 61 | 0 | 0 | 0 | 15 | 61 | 69 | 10 | 0 | 0 | 29 | 0 | 0 | 0 | 48 | 106 | 27 |
| SRR1708968 | 50 | 28 | 514 | 0 | 16 | 0 | 0 | 0 | 0 | 0 | 0 | 165 | 0 | 44 | 320 | 0 | 22 | 38 | 0 | 100 | 119 | 0 | 0 | 0 | 0 | 128 | 53 | 34 | 0 | 0 | 31 | 0 | 0 | 19 | 1853 | 141 | 616 |
| SRR1708969 | 69 | 19 | 567 | 19 | 0 | 15 | 0 | 0 | 0 | 0 | 0 | 107 | 0 | 31 | 359 | 0 | 26 | 0 | 0 | 366 | 400 | 0 | 0 | 0 | 19 | 380 | 312 | 40 | 12 | 0 | 49 | 0 | 0 | 0 | 1709 | 158 | 1014 |
| SRR1708970 | 86 | 0 | 626 | 0 | 0 | 0 | 0 | 0 | 0 | 0 | 0 | 397 | 0 | 27 | 256 | 0 | 19 | 0 | 0 | 38 | 218 | 11 | 0 | 0 | 13 | 160 | 51 | 16 | 0 | 0 | 27 | 0 | 0 | 0 | 73 | 10 | 40 |
| SRR1708971 | 57 | 14 | 793 | 0 | 0 | 0 | 0 | 0 | 11 | 0 | 0 | 260 | 0 | 14 | 308 | 0 | 29 | 0 | 0 | 50 | 208 | 0 | 0 | 0 | 17 | 216 | 146 | 17 | 0 | 0 | 77 | 0 | 0 | 0 | 1335 | 149 | 752 |
| SRR1708972 | 47 | 31 | 913 | 0 | 0 | 16 | 0 | 0 | 0 | 0 | 0 | 42 | 13 | 13 | 287 | 0 | 20 | 11 | 0 | 55 | 82 | 0 | 0 | 0 | 0 | 22 | 43 | 11 | 0 | 0 | 52 | 0 | 0 | 16 | 60 | 52 | 30 |
| SRR1708973 | 30 | 46 | 780 | 0 | 0 | 0 | 0 | 0 | 0 | 0 | 0 | 24 | 0 | 35 | 407 | 0 | 18 | 24 | 0 | 21 | 45 | 0 | 0 | 0 | 23 | 67 | 0 | 23 | 0 | 0 | 67 | 0 | 0 | 0 | 322 | 15 | 78 |
| SRR1708974 | 27 | 18 | 518 | 0 | 0 | 22 | 0 | 0 | 17 | 0 | 0 | 0 | 0 | 15 | 392 | 0 | 25 | 10 | 0 | 116 | 345 | 0 | 0 | 0 | 0 | 32 | 141 | 0 | 0 | 0 | 62 | 0 | 0 | 0 | 42 | 36 | 13 |
| SRR1708975 | 36 | 23 | 334 | 0 | 0 | 42 | 0 | 0 | 37 | 0 | 0 | 0 | 0 | 11 | 124 | 0 | 29 | 0 | 0 | 170 | 408 | 0 | 0 | 0 | 0 | 49 | 126 | 0 | 0 | 0 | 104 | 0 | 0 | 12 | 45 | 76 | 19 |
| SRR1708976 | 41 | 36 | 542 | 0 | 0 | 21 | 0 | 0 | 14 | 0 | 0 | 40 | 0 | 55 | 197 | 0 | 23 | 0 | 0 | 235 | 368 | 0 | 0 | 0 | 15 | 575 | 179 | 30 | 0 | 0 | 57 | 0 | 0 | 0 | 1167 | 74 | 616 |
| SRR1708977 | 26 | 38 | 519 | 0 | 0 | 67 | 25 | 11 | 29 | 0 | 0 | 21 | 0 | 15 | 115 | 0 | 29 | 0 | 0 | 1928 | 903 | 0 | 0 | 0 | 0 | 76 | 54 | 11 | 0 | 0 | 96 | 0 | 0 | 15 | 31 | 88 | 29 |
| SRR1708978 | 141 | 0 | 852 | 0 | 0 | 23 | 0 | 0 | 19 | 0 | 0 | 1061 | 0 | 24 | 30 | 0 | 19 | 0 | 0 | 156 | 260 | 0 | 0 | 0 | 13 | 34 | 12 | 11 | 0 | 0 | 77 | 0 | 13 | 11 | 122 | 31 | 118 |
| SRR1708979 | 199 | 0 | 777 | 0 | 0 | 17 | 0 | 0 | 24 | 0 | 0 | 785 | 0 | 0 | 17 | 0 | 15 | 0 | 0 | 403 | 483 | 13 | 0 | 0 | 13 | 510 | 556 | 0 | 0 | 0 | 65 | 0 | 0 | 11 | 1529 | 330 | 1613 |
| SRR1708980 | 63 | 0 | 504 | 0 | 0 | 17 | 0 | 0 | 0 | 0 | 0 | 343 | 0 | 0 | 37 | 0 | 15 | 0 | 0 | 157 | 315 | 0 | 0 | 0 | 0 | 345 | 205 | 0 | 0 | 0 | 46 | 0 | 12 | 0 | 653 | 217 | 777 |
| SRR1708981 | 197 | 0 | 577 | 0 | 0 | 15 | 0 | 0 | 12 | 0 | 0 | 530 | 0 | 0 | 19 | 0 | 25 | 0 | 0 | 164 | 228 | 0 | 0 | 0 | 22 | 27 | 15 | 0 | 0 | 0 | 82 | 0 | 0 | 0 | 56 | 15 | 69 |
| SRR1708982 | 71 | 0 | 445 | 24 | 15 | 24 | 0 | 0 | 24 | 0 | 0 | 651 | 0 | 0 | 0 | 0 | 17 | 0 | 0 | 395 | 353 | 0 | 0 | 0 | 12 | 745 | 337 | 20 | 0 | 0 | 98 | 0 | 0 | 22 | 1068 | 323 | 1845 |
| SRR1708983 | 153 | 0 | 731 | 0 | 0 | 0 | 0 | 0 | 0 | 0 | 0 | 791 | 0 | 0 | 0 | 0 | 23 | 0 | 0 | 250 | 289 | 0 | 0 | 0 | 11 | 545 | 541 | 17 | 0 | 0 | 60 | 0 | 0 | 0 | 426 | 188 | 879 |
| SRR1708984 | 64 | 0 | 652 | 0 | 0 | 0 | 0 | 0 | 0 | 0 | 0 | 526 | 0 | 14 | 30 | 0 | 14 | 0 | 0 | 311 | 288 | 0 | 0 | 0 | 14 | 20 | 33 | 0 | 0 | 0 | 63 | 0 | 0 | 0 | 28 | 10 | 60 |
| SRR1708985 | 76 | 0 | 889 | 0 | 0 | 28 | 16 | 0 | 0 | 0 | 0 | 221 | 0 | 0 | 66 | 0 | 16 | 0 | 12 | 3191 | 1005 | 0 | 0 | 0 | 0 | 460 | 2514 | 0 | 0 | 0 | 38 | 0 | 0 | 0 | 656 | 220 | 993 |
| SRR1708986 | 73 | 0 | 736 | 0 | 0 | 24 | 0 | 0 | 10 | 0 | 0 | 173 | 10 | 0 | 54 | 0 | 22 | 41 | 0 | 702 | 702 | 0 | 0 | 0 | 10 | 26 | 118 | 0 | 0 | 0 | 40 | 0 | 0 | 0 | 54 | 14 | 69 |
| SRR1708987 | 77 | 0 | 545 | 0 | 11 | 15 | 0 | 0 | 0 | 0 | 0 | 54 | 0 | 11 | 44 | 0 | 21 | 0 | 0 | 356 | 737 | 0 | 0 | 0 | 0 | 16 | 32 | 0 | 0 | 0 | 94 | 0 | 0 | 11 | 54 | 195 | 42 |
| SRR1708988 | 109 | 0 | 812 | 0 | 0 | 14 | 0 | 0 | 10 | 0 | 0 | 181 | 0 | 19 | 74 | 0 | 22 | 10 | 0 | 203 | 285 | 0 | 0 | 0 | 42 | 381 | 304 | 11 | 12 | 0 | 39 | 0 | 0 | 14 | 1576 | 85 | 832 |
| SRR1708989 | 49 | 11 | 893 | 0 | 0 | 13 | 0 | 0 | 11 | 0 | 0 | 36 | 0 | 25 | 73 | 0 | 26 | 0 | 0 | 56 | 266 | 0 | 0 | 0 | 0 | 11 | 152 | 34 | 17 | 0 | 63 | 0 | 0 | 0 | 86 | 173 | 51 |
| SRR1708990 | 80 | 0 | 510 | 0 | 0 | 51 | 14 | 0 | 25 | 0 | 0 | 325 | 0 | 27 | 56 | 0 | 16 | 0 | 14 | 1885 | 562 | 60 | 0 | 0 | 12 | 40 | 87 | 23 | 0 | 0 | 49 | 0 | 0 | 0 | 29 | 11 | 50 |
| SRR1708991 | 49 | 0 | 686 | 0 | 0 | 26 | 0 | 0 | 22 | 0 | 0 | 52 | 0 | 39 | 78 | 0 | 15 | 0 | 0 | 558 | 558 | 16 | 0 | 0 | 0 | 42 | 24 | 12 | 0 | 0 | 88 | 0 | 0 | 14 | 28 | 118 | 36 |
| SRR1708992 | 61 | 0 | 365 | 0 | 0 | 68 | 52 | 0 | 26 | 0 | 0 | 59 | 0 | 36 | 162 | 0 | 19 | 0 | 13 | 2974 | 1348 | 32 | 0 | 0 | 0 | 53 | 74 | 19 | 13 | 0 | 152 | 0 | 0 | 0 | 32 | 133 | 28 |
| SRR1708993 | 53 | 0 | 331 | 0 | 0 | 12 | 0 | 0 | 34 | 0 | 0 | 0 | 0 | 31 | 88 | 0 | 34 | 0 | 0 | 163 | 203 | 0 | 0 | 0 | 0 | 29 | 168 | 22 | 19 | 0 | 144 | 0 | 0 | 16 | 31 | 47 | 22 |
| SRR1708994 | 50 | 0 | 663 | 10 | 0 | 17 | 0 | 0 | 0 | 0 | 0 | 44 | 0 | 29 | 70 | 0 | 20 | 0 | 0 | 154 | 299 | 0 | 0 | 0 | 11 | 25 | 90 | 50 | 42 | 0 | 38 | 0 | 20 | 11 | 55 | 67 | 36 |
| SRR1708995 | 19 | 0 | 382 | 0 | 0 | 49 | 0 | 0 | 28 | 0 | 0 | 0 | 0 | 16 | 35 | 0 | 42 | 0 | 0 | 361 | 721 | 0 | 0 | 0 | 12 | 852 | 584 | 24 | 0 | 0 | 70 | 0 | 0 | 10 | 758 | 86 | 738 |
| SRR1708996 | 74 | 0 | 718 | 0 | 0 | 33 | 0 | 0 | 12 | 0 | 0 | 643 | 0 | 0 | 33 | 0 | 15 | 0 | 0 | 450 | 800 | 0 | 0 | 0 | 15 | 511 | 341 | 12 | 0 | 0 | 56 | 0 | 0 | 27 | 1777 | 409 | 1668 |

### Supp Table S8.2.Columns_AM_to_BW

| | | | | | | | | | | | | | | | | | | | | | | | | | | | | | | | | | | | | | |
|---|---|---|---|---|---|---|---|---|---|---|---|---|---|---|---|---|---|---|---|---|---|---|---|---|---|---|---|---|---|---|---|---|---|---|---|---|---|
| SRR1708997 | 82 | 0 | 518 | 0 | 21 | 17 | 69 | 23 | 0 | 0 | 0 | 522 | 0 | 23 | 116 | 0 | 13 | 0 | 39 | 4176 | 894 | 13 | 0 | 0 | 0 | 422 | 881 | 0 | 0 | 0 | 29 | 0 | 0 | 0 | 884 | 271 | 1110 |
| SRR1708998 | 73 | 0 | 482 | 0 | 0 | 28 | 0 | 0 | 28 | 0 | 0 | 365 | 0 | 0 | 60 | 0 | 20 | 0 | 0 | 371 | 821 | 13 | 0 | 0 | 10 | 777 | 529 | 15 | 0 | 0 | 65 | 0 | 0 | 0 | 1211 | 405 | 1748 |
| SRR1708999 | 185 | 0 | 528 | 0 | 0 | 0 | 0 | 0 | 0 | 0 | 0 | 416 | 0 | 0 | 0 | 0 | 21 | 0 | 0 | 79 | 120 | 0 | 0 | 0 | 25 | 161 | 66 | 0 | 0 | 0 | 120 | 0 | 0 | 0 | 1072 | 234 | 945 |
| SRR1709000 | 69 | 0 | 830 | 0 | 0 | 43 | 0 | 0 | 18 | 0 | 0 | 701 | 0 | 25 | 65 | 0 | 16 | 23 | 0 | 850 | 413 | 0 | 0 | 0 | 0 | 731 | 2953 | 0 | 0 | 0 | 59 | 0 | 0 | 13 | 997 | 331 | 1577 |
| SRR1709001 | 75 | 15 | 522 | 0 | 0 | 0 | 0 | 0 | 0 | 0 | 0 | 443 | 0 | 23 | 83 | 0 | 11 | 39 | 0 | 201 | 153 | 0 | 0 | 0 | 0 | 218 | 218 | 15 | 0 | 0 | 44 | 0 | 0 | 0 | 883 | 241 | 1097 |
| SRR1709002 | 63 | 0 | 600 | 0 | 0 | 31 | 0 | 0 | 0 | 0 | 0 | 259 | 0 | 0 | 249 | 0 | 19 | 528 | 0 | 2268 | 1239 | 0 | 0 | 19 | 0 | 21 | 176 | 0 | 0 | 0 | 37 | 0 | 0 | 0 | 63 | 174 | 59 |
| SRR1709003 | 75 | 0 | 514 | 0 | 0 | 12 | 0 | 0 | 0 | 0 | 0 | 109 | 0 | 12 | 134 | 0 | 29 | 24 | 0 | 120 | 374 | 0 | 0 | 0 | 0 | 371 | 392 | 0 | 0 | 0 | 77 | 0 | 0 | 15 | 1427 | 166 | 762 |
| SRR1709004 | 150 | 0 | 975 | 0 | 0 | 33 | 0 | 0 | 16 | 0 | 0 | 918 | 0 | 0 | 12 | 0 | 15 | 0 | 0 | 112 | 186 | 0 | 0 | 0 | 0 | 27 | 215 | 0 | 0 | 0 | 107 | 0 | 0 | 0 | 104 | 28 | 135 |
| SRR1709005 | 113 | 0 | 768 | 0 | 0 | 30 | 0 | 0 | 12 | 0 | 0 | 721 | 0 | 0 | 27 | 0 | 12 | 0 | 0 | 1259 | 1506 | 0 | 0 | 0 | 0 | 35 | 64 | 0 | 0 | 0 | 33 | 0 | 0 | 0 | 113 | 32 | 128 |
| SRR1709006 | 87 | 0 | 553 | 0 | 0 | 61 | 0 | 0 | 0 | 0 | 0 | 360 | 0 | 0 | 0 | 0 | 19 | 0 | 13 | 3109 | 1201 | 15 | 0 | 0 | 13 | 46 | 97 | 0 | 0 | 0 | 69 | 0 | 0 | 17 | 51 | 19 | 69 |
| SRR1709007 | 113 | 0 | 516 | 0 | 0 | 32 | 0 | 0 | 19 | 0 | 0 | 346 | 0 | 0 | 48 | 0 | 19 | 0 | 0 | 3143 | 1644 | 16 | 0 | 0 | 0 | 783 | 1472 | 11 | 0 | 0 | 67 | 0 | 0 | 11 | 1709 | 507 | 2094 |
| SRR1709008 | 104 | 0 | 503 | 0 | 0 | 24 | 0 | 0 | 13 | 0 | 0 | 861 | 0 | 0 | 11 | 0 | 20 | 0 | 0 | 109 | 153 | 0 | 0 | 0 | 11 | 24 | 118 | 0 | 0 | 0 | 57 | 0 | 0 | 13 | 47 | 15 | 71 |
| SRR1709009 | 88 | 0 | 615 | 0 | 0 | 22 | 0 | 0 | 15 | 0 | 0 | 452 | 0 | 0 | 61 | 0 | 22 | 0 | 0 | 227 | 681 | 15 | 0 | 0 | 0 | 655 | 272 | 0 | 13 | 0 | 99 | 0 | 0 | 11 | 1019 | 327 | 1535 |
| SRR1709010 | 63 | 13 | 1401 | 0 | 0 | 28 | 0 | 0 | 13 | 0 | 0 | 144 | 0 | 10 | 57 | 0 | 15 | 0 | 0 | 184 | 259 | 0 | 0 | 0 | 23 | 24 | 26 | 0 | 0 | 0 | 71 | 0 | 0 | 10 | 73 | 186 | 57 |
| SRR1709011 | 82 | 0 | 1446 | 0 | 0 | 31 | 0 | 0 | 10 | 0 | 0 | 82 | 0 | 0 | 20 | 0 | 23 | 0 | 0 | 824 | 780 | 0 | 0 | 0 | 0 | 541 | 1787 | 0 | 0 | 0 | 54 | 0 | 0 | 0 | 1886 | 177 | 1618 |
| SRR1709012 | 57 | 0 | 795 | 0 | 0 | 34 | 0 | 0 | 0 | 0 | 0 | 33 | 0 | 0 | 79 | 0 | 19 | 44 | 0 | 899 | 1117 | 0 | 0 | 0 | 0 | 19 | 63 | 0 | 0 | 0 | 47 | 0 | 0 | 0 | 69 | 162 | 42 |
| SRR1709013 | 70 | 0 | 781 | 0 | 0 | 0 | 0 | 0 | 0 | 0 | 0 | 28 | 0 | 0 | 220 | 0 | 21 | 0 | 0 | 45 | 189 | 0 | 0 | 0 | 0 | 14 | 172 | 15 | 0 | 0 | 82 | 0 | 0 | 0 | 96 | 97 | 42 |
| SRR1709014 | 83 | 0 | 851 | 0 | 15 | 27 | 0 | 0 | 0 | 0 | 0 | 140 | 0 | 15 | 11 | 0 | 20 | 0 | 0 | 350 | 530 | 15 | 0 | 0 | 0 | 638 | 703 | 15 | 0 | 0 | 80 | 0 | 0 | 0 | 4926 | 396 | 2428 |
| SRR1709015 | 55 | 0 | 945 | 0 | 0 | 16 | 0 | 0 | 13 | 0 | 0 | 70 | 0 | 0 | 110 | 0 | 29 | 0 | 0 | 80 | 314 | 0 | 0 | 0 | 0 | 17 | 16 | 0 | 0 | 0 | 53 | 0 | 0 | 0 | 64 | 145 | 59 |
| SRR1709016 | 36 | 0 | 719 | 0 | 0 | 79 | 19 | 0 | 16 | 0 | 0 | 0 | 0 | 14 | 113 | 0 | 19 | 0 | 0 | 2082 | 1069 | 0 | 0 | 0 | 0 | 1220 | 1179 | 12 | 0 | 0 | 79 | 0 | 0 | 0 | 1244 | 187 | 806 |
| SRR1709017 | 21 | 0 | 570 | 0 | 0 | 100 | 0 | 0 | 21 | 0 | 0 | 0 | 0 | 11 | 44 | 0 | 23 | 0 | 0 | 1691 | 1399 | 0 | 0 | 0 | 0 | 92 | 94 | 0 | 0 | 0 | 109 | 0 | 0 | 0 | 78 | 102 | 43 |
| SRR1709018 | 20 | 0 | 269 | 0 | 0 | 41 | 0 | 0 | 16 | 0 | 0 | 0 | 0 | 0 | 132 | 0 | 31 | 0 | 0 | 158 | 506 | 0 | 0 | 0 | 0 | 955 | 126 | 0 | 18 | 0 | 102 | 0 | 0 | 0 | 1234 | 51 | 314 |
| SRR1709019 | 16 | 0 | 231 | 0 | 0 | 44 | 0 | 0 | 16 | 0 | 0 | 0 | 0 | 16 | 13 | 0 | 36 | 0 | 0 | 261 | 189 | 0 | 0 | 0 | 0 | 39 | 11 | 16 | 0 | 0 | 180 | 0 | 0 | 0 | 51 | 78 | 28 |
| SRR1709020 | 39 | 0 | 650 | 0 | 0 | 52 | 0 | 0 | 27 | 0 | 0 | 19 | 0 | 0 | 37 | 0 | 29 | 0 | 11 | 2743 | 716 | 0 | 0 | 0 | 0 | 63 | 111 | 0 | 0 | 0 | 44 | 0 | 0 | 0 | 65 | 99 | 50 |
| SRR1709021 | 20 | 0 | 473 | 0 | 0 | 67 | 0 | 0 | 34 | 0 | 0 | 0 | 0 | 0 | 42 | 0 | 30 | 0 | 0 | 438 | 700 | 0 | 0 | 0 | 0 | 75 | 33 | 0 | 0 | 0 | 104 | 0 | 0 | 0 | 50 | 148 | 39 |
| SRR1732015 | 71 | 13 | 369 | 0 | 0 | 0 | 0 | 0 | 0 | 0 | 0 | 73 | 0 | 32 | 250 | 16 | 22 | 0 | 17 | 4329 | 979 | 14 | 12 | 0 | 21 | 26 | 20 | 39 | 20 | 0 | 0 | 17 | 10 | 11 | 24 | 15 | 18 |
| SRR1732016 | 43 | 51 | 428 | 14 | 0 | 0 | 39 | 0 | 0 | 0 | 0 | 37 | 0 | 74 | 76 | 47 | 32 | 0 | 13 | 317 | 248 | 0 | 55 | 0 | 67 | 56 | 46 | 77 | 863 | 0 | 0 | 0 | 160 | 46 | 0 | 0 | 0 |
| SRR1748132 | 187 | 15 | 816 | 25 | 15 | 0 | 88 | 12 | 0 | 0 | 0 | 30 | 0 | 15 | 18 | 0 | 0 | 0 | 0 | 50 | 397 | 0 | 0 | 0 | 88 | 259 | 77 | 25 | 0 | 0 | 14 | 0 | 0 | 92 | 247 | 16 | 59 |
| SRR1748133 | 95 | 0 | 1985 | 11 | 18 | 11 | 91 | 0 | 0 | 0 | 0 | 133 | 0 | 22 | 184 | 0 | 22 | 0 | 0 | 213 | 1795 | 0 | 0 | 0 | 26 | 88 | 45 | 16 | 0 | 0 | 0 | 0 | 0 | 70 | 14 | 0 | 22 |
| SRR1759212 | 12 | 0 | 2365 | 0 | 0 | 33 | 63 | 14 | 16 | 0 | 0 | 263 | 13 | 0 | 329 | 0 | 0 | 0 | 62 | 322 | 14 | 0 | 0 | 100 | 14 | 322 | 229 | 0 | 0 | 0 | 11 | 0 | 0 | 0 | 25 | 0 | 16 |
| SRR1759213 | 33 | 0 | 2404 | 0 | 0 | 0 | 24 | 0 | 0 | 0 | 0 | 145 | 0 | 0 | 705 | 0 | 11 | 0 | 115 | 1238 | 1745 | 0 | 0 | 59 | 23 | 17 | 24 | 0 | 0 | 0 | 0 | 0 | 0 | 0 | 0 | 0 | 0 |
| SRR1759214 | 24 | 0 | 2528 | 0 | 0 | 18 | 46 | 22 | 0 | 0 | 0 | 157 | 0 | 0 | 1200 | 0 | 0 | 0 | 130 | 609 | 2663 | 0 | 0 | 62 | 0 | 18 | 19 | 0 | 0 | 0 | 0 | 0 | 0 | 0 | 26 | 0 | 21 |
| SRR1759215 | 30 | 0 | 2299 | 0 | 0 | 14 | 184 | 13 | 10 | 0 | 0 | 151 | 0 | 0 | 0 | 0 | 0 | 0 | 183 | 330 | 2746 | 0 | 0 | 50 | 24 | 197 | 246 | 0 | 0 | 0 | 0 | 0 | 0 | 0 | 24 | 0 | 0 |
| SRR1759216 | 23 | 0 | 2483 | 0 | 0 | 13 | 60 | 0 | 0 | 0 | 0 | 139 | 0 | 0 | 880 | 0 | 0 | 0 | 140 | 426 | 1116 | 0 | 0 | 54 | 24 | 216 | 250 | 0 | 0 | 0 | 0 | 0 | 0 | 0 | 28 | 0 | 17 |
| SRR1759217 | 26 | 0 | 2388 | 0 | 0 | 14 | 112 | 0 | 0 | 0 | 0 | 214 | 0 | 0 | 1227 | 0 | 0 | 0 | 142 | 536 | 2182 | 0 | 0 | 140 | 76 | 145 | 163 | 0 | 0 | 0 | 0 | 0 | 0 | 0 | 18 | 0 | 11 |
| SRR1759218 | 34 | 0 | 3646 | 0 | 0 | 20 | 103 | 33 | 0 | 0 | 0 | 154 | 0 | 0 | 2868 | 0 | 10 | 39 | 245 | 816 | 2534 | 0 | 0 | 22 | 33 | 281 | 277 | 0 | 0 | 0 | 11 | 0 | 0 | 0 | 31 | 0 | 31 |
| SRR1759219 | 20 | 0 | 2372 | 0 | 0 | 21 | 82 | 19 | 0 | 0 | 0 | 165 | 0 | 0 | 2992 | 0 | 0 | 15 | 53 | 365 | 3471 | 0 | 0 | 64 | 29 | 28 | 25 | 0 | 0 | 0 | 12 | 0 | 0 | 0 | 25 | 0 | 19 |
| SRR1759220 | 22 | 0 | 2806 | 0 | 0 | 16 | 71 | 0 | 0 | 0 | 0 | 170 | 0 | 0 | 6234 | 0 | 0 | 11 | 93 | 529 | 5066 | 0 | 0 | 12 | 32 | 174 | 266 | 0 | 0 | 0 | 0 | 0 | 0 | 0 | 35 | 0 | 19 |
| SRR1759221 | 12 | 0 | 3026 | 0 | 0 | 12 | 162 | 57 | 0 | 0 | 0 | 153 | 0 | 0 | 715 | 0 | 0 | 13 | 57 | 30 | 872 | 0 | 0 | 0 | 0 | 11 | 178 | 0 | 0 | 0 | 13 | 0 | 0 | 0 | 90 | 0 | 50 |
| SRR1759222 | 26 | 65 | 3421 | 0 | 0 | 20 | 36 | 0 | 17 | 0 | 0 | 67 | 0 | 0 | 13580 | 0 | 0 | 0 | 111 | 81 | 2205 | 0 | 0 | 0 | 0 | 362 | 51 | 0 | 0 | 0 | 48 | 0 | 0 | 0 | 80 | 0 | 37 |
| SRR1759223 | 24 | 0 | 2876 | 0 | 0 | 26 | 497 | 133 | 0 | 0 | 0 | 156 | 0 | 0 | 526 | 0 | 0 | 23 | 146 | 258 | 3321 | 0 | 0 | 77 | 0 | 15 | 17 | 0 | 0 | 13 | 13 | 0 | 0 | 0 | 71 | 0 | 46 |
| SRR1759224 | 18 | 0 | 3186 | 0 | 0 | 14 | 124 | 0 | 0 | 0 | 0 | 82 | 0 | 0 | 2321 | 0 | 0 | 0 | 33 | 44 | 4426 | 0 | 0 | 0 | 0 | 184 | 77 | 0 | 0 | 0 | 0 | 0 | 0 | 0 | 141 | 0 | 70 |
| SRR1759225 | 22 | 0 | 2440 | 0 | 0 | 0 | 98 | 12 | 0 | 0 | 0 | 132 | 0 | 0 | 532 | 0 | 0 | 0 | 54 | 34 | 552 | 0 | 0 | 0 | 0 | 134 | 70 | 0 | 0 | 0 | 16 | 0 | 0 | 12 | 100 | 0 | 43 |
| SRR1759226 | 13 | 0 | 4135 | 0 | 0 | 0 | 78 | 0 | 0 | 0 | 0 | 50 | 0 | 0 | 1837 | 0 | 0 | 0 | 32 | 61 | 4861 | 0 | 0 | 0 | 0 | 160 | 62 | 0 | 0 | 0 | 0 | 0 | 0 | 0 | 72 | 0 | 33 |
| SRR1759227 | 15 | 0 | 3211 | 0 | 0 | 0 | 74 | 25 | 0 | 0 | 0 | 126 | 0 | 10 | 1910 | 0 | 0 | 0 | 47 | 45 | 1725 | 0 | 0 | 0 | 0 | 221 | 212 | 0 | 0 | 0 | 0 | 0 | 0 | 20 | 86 | 0 | 43 |
| SRR1759228 | 15 | 0 | 3272 | 0 | 0 | 43 | 234 | 12 | 0 | 0 | 0 | 81 | 0 | 0 | 698 | 0 | 0 | 0 | 165 | 64 | 277 | 0 | 0 | 0 | 0 | 10 | 97 | 0 | 0 | 0 | 14 | 0 | 0 | 25 | 88 | 0 | 60 |
| SRR1759229 | 19 | 0 | 3063 | 0 | 0 | 13 | 450 | 103 | 0 | 0 | 0 | 148 | 0 | 0 | 995 | 0 | 0 | 18 | 63 | 138 | 1092 | 0 | 0 | 25 | 0 | 261 | 216 | 0 | 0 | 11 | 13 | 0 | 0 | 10 | 55 | 10 | 29 |
| SRR1759230 | 11 | 0 | 2694 | 0 | 0 | 0 | 673 | 16 | 0 | 0 | 0 | 112 | 0 | 0 | 1480 | 0 | 0 | 11 | 56 | 14 | 635 | 0 | 0 | 0 | 0 | 43 | 36 | 0 | 0 | 0 | 0 | 0 | 0 | 17 | 77 | 0 | 31 |
| SRR1759231 | 18 | 14 | 3587 | 0 | 0 | 41 | 35 | 0 | 0 | 0 | 0 | 84 | 0 | 0 | 148 | 0 | 0 | 0 | 60 | 152 | 263 | 0 | 0 | 0 | 0 | 19 | 143 | 0 | 0 | 0 | 25 | 0 | 0 | 0 | 100 | 0 | 43 |
| SRR1759232 | 18 | 0 | 2196 | 0 | 0 | 22 | 475 | 0 | 11 | 0 | 0 | 126 | 0 | 0 | 2362 | 0 | 0 | 16 | 58 | 71 | 2688 | 0 | 0 | 0 | 0 | 12 | 193 | 0 | 0 | 0 | 0 | 0 | 0 | 0 | 213 | 0 | 113 |
| SRR1759233 | 11 | 0 | 3106 | 0 | 0 | 0 | 77 | 0 | 0 | 0 | 0 | 110 | 0 | 0 | 817 | 0 | 0 | 0 | 24 | 0 | 263 | 0 | 0 | 0 | 0 | 13 | 38 | 0 | 0 | 0 | 16 | 0 | 0 | 0 | 106 | 0 | 57 |
| SRR1759234 | 26 | 0 | 3054 | 0 | 0 | 33 | 200 | 52 | 12 | 0 | 0 | 159 | 0 | 0 | 527 | 0 | 0 | 0 | 114 | 184 | 829 | 0 | 0 | 19 | 0 | 24 | 18 | 0 | 0 | 0 | 0 | 0 | 0 | 0 | 60 | 0 | 37 |
| SRR1759235 | 19 | 0 | 4204 | 0 | 0 | 35 | 258 | 52 | 0 | 0 | 0 | 209 | 0 | 0 | 2277 | 0 | 0 | 0 | 272 | 186 | 594 | 0 | 0 | 25 | 0 | 27 | 20 | 0 | 0 | 0 | 12 | 0 | 0 | 0 | 44 | 0 | 19 |
| SRR1759236 | 25 | 0 | 3351 | 0 | 0 | 22 | 130 | 71 | 0 | 0 | 0 | 179 | 0 | 0 | 1512 | 0 | 0 | 0 | 158 | 213 | 1764 | 0 | 0 | 73 | 0 | 346 | 284 | 0 | 0 | 0 | 0 | 0 | 0 | 26 | 72 | 0 | 42 |
| SRR1759237 | 16 | 0 | 4674 | 0 | 0 | 22 | 146 | 106 | 0 | 0 | 0 | 104 | 0 | 0 | 2577 | 0 | 0 | 0 | 422 | 392 | 2093 | 0 | 0 | 86 | 0 | 236 | 269 | 0 | 0 | 0 | 0 | 0 | 0 | 17 | 73 | 0 | 52 |
| SRR1759238 | 11 | 0 | 3130 | 0 | 0 | 30 | 108 | 34 | 11 | 0 | 0 | 103 | 0 | 0 | 995 | 0 | 0 | 0 | 145 | 262 | 1655 | 0 | 0 | 55 | 0 | 26 | 15 | 0 | 0 | 0 | 17 | 0 | 0 | 11 | 94 | 0 | 72 |
| SRR1759239 | 26 | 0 | 1096 | 0 | 0 | 0 | 3254 | 11 | 0 | 0 | 0 | 201 | 0 | 11 | 4957 | 0 | 0 | 44 | 409 | 78 | 261 | 0 | 0 | 0 | 0 | 10 | 41 | 0 | 12 | 0 | 0 | 0 | 0 | 19 | 67 | 21 | 34 |
| SRR1759240 | 30 | 0 | 4621 | 0 | 0 | 24 | 84 | 61 | 0 | 0 | 0 | 193 | 0 | 0 | 1982 | 0 | 0 | 0 | 187 | 387 | 2440 | 0 | 0 | 76 | 0 | 21 | 19 | 0 | 0 | 0 | 18 | 0 | 0 | 0 | 66 | 0 | 52 |
| SRR1759241 | 27 | 0 | 3932 | 0 | 0 | 23 | 271 | 74 | 0 | 0 | 0 | 196 | 0 | 0 | 1593 | 0 | 0 | 0 | 169 | 95 | 3092 | 0 | 0 | 29 | 0 | 20 | 15 | 0 | 0 | 0 | 13 | 0 | 0 | 14 | 110 | 0 | 76 |
| SRR1759242 | 18 | 0 | 4979 | 0 | 0 | 54 | 214 | 196 | 19 | 0 | 0 | 105 | 0 | 0 | 101 | 0 | 0 | 0 | 273 | 150 | 330 | 0 | 0 | 24 | 0 | 916 | 442 | 0 | 0 | 33 | 22 | 0 | 0 | 19 | 104 | 16 | 101 |
| SRR1759243 | 22 | 0 | 4019 | 0 | 0 | 15 | 58 | 25 | 0 | 0 | 0 | 102 | 0 | 0 | 497 | 0 | 0 | 0 | 66 | 102 | 356 | 0 | 0 | 0 | 0 | 248 | 223 | 0 | 0 | 0 | 27 | 0 | 0 | 10 | 117 | 13 | 30 |
| SRR1759244 | 24 | 0 | 2503 | 0 | 0 | 15 | 140 | 0 | 0 | 0 | 0 | 219 | 11 | 0 | 279 | 0 | 0 | 11 | 94 | 1261 | 629 | 0 | 0 | 67 | 47 | 227 | 320 | 0 | 0 | 0 | 0 | 0 | 0 | 0 | 20 | 0 | 12 |
| SRR1759245 | 18 | 0 | 2281 | 0 | 0 | 25 | 172 | 39 | 0 | 0 | 0 | 145 | 16 | 0 | 362 | 0 | 0 | 0 | 125 | 167 | 411 | 0 | 0 | 88 | 0 | 235 | 162 | 0 | 0 | 0 | 16 | 0 | 0 | 0 | 25 | 0 | 15 |
| SRR1759246 | 16 | 0 | 1188 | 0 | 0 | 37 | 63 | 0 | 13 | 0 | 0 | 107 | 0 | 0 | 856 | 0 | 0 | 66 | 44 | 120 | 911 | 0 | 0 | 16 | 0 | 12 | 13 | 0 | 0 | 0 | 19 | 0 | 0 | 0 | 55 | 0 | 23 |
| SRR1759247 | 14 | 0 | 2610 | 0 | 0 | 11 | 167 | 35 | 0 | 0 | 0 | 118 | 0 | 0 | 354 | 0 | 0 | 0 | 79 | 46 | 301 | 0 | 0 | 0 | 0 | 145 | 71 | 0 | 0 | 0 | 0 | 0 | 0 | 11 | 96 | 0 | 54 |
| SRR1759248 | 24 | 0 | 2470 | 0 | 0 | 36 | 380 | 174 | 18 | 0 | 0 | 143 | 0 | 0 | 1954 | 0 | 0 | 12 | 148 | 251 | 1676 | 0 | 0 | 61 | 0 | 20 | 13 | 0 | 0 | 16 | 21 | 0 | 0 | 0 | 74 | 0 | 32 |
| SRR1759249 | 35 | 0 | 2594 | 0 | 0 | 22 | 82 | 25 | 14 | 0 | 0 | 171 | 0 | 0 | 307 | 0 | 0 | 0 | 16 | 354 | 935 | 0 | 0 | 45 | 0 | 11 | 15 | 0 | 0 | 0 | 0 | 0 | 0 | 14 | 32 | 0 | 30 |
| SRR1759250 | 40 | 0 | 3693 | 0 | 0 | 31 | 40 | 0 | 11 | 0 | 0 | 123 | 0 | 0 | 1914 | 0 | 12 | 16 | 69 | 1434 | 4041 | 0 | 0 | 0 | 13 | 15 | 15 | 0 | 0 | 0 | 0 | 0 | 0 | 0 | 29 | 0 | 18 |
| SRR1759251 | 24 | 0 | 3685 | 0 | 0 | 18 | 153 | 20 | 0 | 0 | 0 | 122 | 0 | 0 | 254 | 0 | 0 | 0 | 54 | 268 | 975 | 0 | 0 | 112 | 0 | 196 | 282 | 0 | 0 | 0 | 0 | 0 | 0 | 0 | 26 | 0 | 11 |
| SRR1759252 | 30 | 0 | 1588 | 0 | 0 | 30 | 562 | 35 | 16 | 0 | 0 | 135 | 0 | 0 | 719 | 0 | 0 | 0 | 128 | 302 | 1380 | 0 | 0 | 64 | 0 | 516 | 488 | 0 | 0 | 0 | 0 | 0 | 0 | 0 | 90 | 0 | 39 |
| SRR1759253 | 40 | 0 | 3380 | 0 | 0 | 22 | 103 | 24 | 19 | 0 | 0 | 159 | 0 | 0 | 2929 | 0 | 0 | 0 | 189 | 482 | 3785 | 0 | 0 | 38 | 0 | 193 | 282 | 0 | 0 | 0 | 0 | 0 | 0 | 0 | 51 | 0 | 30 |
| SRR1759254 | 29 | 0 | 2720 | 0 | 0 | 18 | 161 | 0 | 0 | 0 | 0 | 129 | 0 | 0 | 224 | 0 | 0 | 0 | 157 | 1168 | 2120 | 0 | 0 | 80 | 0 | 196 | 292 | 0 | 0 | 0 | 0 | 0 | 0 | 0 | 24 | 0 | 15 |
| SRR1759255 | 40 | 0 | 2961 | 0 | 0 | 11 | 40 | 12 | 0 | 0 | 0 | 204 | 0 | 0 | 417 | 0 | 0 | 0 | 35 | 652 | 1123 | 0 | 0 | 64 | 0 | 108 | 14 | 0 | 0 | 0 | 0 | 0 | 0 | 0 | 22 | 0 | 0 |
| SRR1759256 | 26 | 0 | 2973 | 0 | 0 | 21 | 82 | 32 | 18 | 0 | 0 | 157 | 0 | 0 | 4330 | 0 | 0 | 0 | 156 | 287 | 3736 | 0 | 0 | 26 | 12 | 325 | 220 | 0 | 0 | 0 | 36 | 0 | 0 | 13 | 127 | 12 | 88 |
| SRR1759257 | 19 | 0 | 3565 | 0 | 0 | 25 | 39 | 0 | 0 | 0 | 0 | 103 | 0 | 0 | 66 | 0 | 0 | 0 | 43 | 108 | 143 | 0 | 0 | 0 | 11 | 158 | 239 | 0 | 0 | 0 | 0 | 0 | 0 | 25 | 37 | 0 | 42 |
| SRR1759258 | 13 | 0 | 1679 | 0 | 0 | 206 | 1458 | 39 | 25 | 0 | 0 | 229 | 0 | 35 | 581 | 0 | 0 | 0 | 240 | 154 | 406 | 0 | 0 | 21 | 0 | 2213 | 953 | 13 | 0 | 0 | 62 | 0 | 0 | 11 | 414 | 40 | 228 |
| SRR1759259 | 19 | 0 | 3171 | 0 | 0 | 106 | 280 | 0 | 26 | 0 | 0 | 142 | 0 | 0 | 2756 | 0 | 0 | 0 | 311 | 150 | 2008 | 0 | 0 | 0 | 0 | 67 | 33 | 0 | 0 | 0 | 117 | 0 | 0 | 0 | 12 | 22 | 202 |
| SRR1759260 | 18 | 0 | 2786 | 0 | 0 | 0 | 192 | 177 | 0 | 0 | 0 | 164 | 0 | 0 | 531 | 0 | 0 | 11 | 97 | 71 | 246 | 0 | 0 | 23 | 0 | 101 | 91 | 0 | 0 | 0 | 0 | 0 | 0 | 0 | 61 | 0 | 14 |
| SRR1759261 | 14 | 0 | 2589 | 0 | 0 | 14 | 767 | 98 | 13 | 0 | 0 | 157 | 0 | 10 | 1410 | 0 | 0 | 0 | 165 | 114 | 747 | 0 | 0 | 21 | 0 | 190 | 197 | 0 | 0 | 10 | 0 | 0 | 0 | 12 | 116 | 0 | 49 |
| SRR1759262 | 16 | 0 | 2722 | 0 | 0 | 15 | 597 | 13 | 0 | 0 | 0 | 175 | 0 | 0 | 336 | 0 | 0 | 23 | 91 | 125 | 237 | 0 | 0 | 26 | 0 | 11 | 12 | 0 | 0 | 0 | 0 | 0 | 0 | 0 | 42 | 0 | 13 |
| SRR1759263 | 26 | 0 | 2050 | 0 | 0 | 18 | 33 | 0 | 0 | 0 | 0 | 144 | 0 | 0 | 1632 | 0 | 0 | 16 | 94 | 238 | 618 | 0 | 0 | 0 | 0 | 191 | 113 | 0 | 0 | 0 | 23 | 0 | 0 | 0 | 85 | 0 | 58 |
| SRR1759264 | 17 | 0 | 2667 | 0 | 0 | 0 | 40 | 0 | 0 | 0 | 0 | 107 | 0 | 0 | 1024 | 0 | 0 | 0 | 49 | 261 | 1278 | 0 | 0 | 0 | 0 | 167 | 179 | 0 | 0 | 0 | 0 | 0 | 0 | 0 | 87 | 0 | 37 |
| SRR1759265 | 16 | 12 | 3020 | 0 | 0 | 12 | 29 | 31 | 27 | 0 | 0 | 56 | 0 | 0 | 67 | 0 | 0 | 0 | 38 | 152 | 169 | 0 | 0 | 0 | 0 | 251 | 157 | 0 | 0 | 12 | 0 | 0 | 0 | 0 | 39 | 0 | 24 |
| SRR1759266 | 22 | 0 | 2122 | 0 | 0 | 18 | 66 | 0 | 0 | 0 | 0 | 115 | 11 | 0 | 36 | 0 | 17 | 0 | 31 | 103 | 215 | 0 | 0 | 66 | 22 | 217 | 10 | 0 | 0 | 0 | 15 | 0 | 0 | 0 | 0 | 0 | 13 |
| SRR1759267 | 19 | 0 | 1200 | 0 | 0 | 20 | 98 | 0 | 25 | 0 | 0 | 86 | 11 | 0 | 238 | 0 | 0 | 29 | 21 | 26 | 337 | 0 | 0 | 0 | 16 | 11 | 72 | 0 | 0 | 0 | 16 | 0 | 0 | 0 | 110 | 11 | 52 |
| SRR1759268 | 18 | 0 | 1483 | 0 | 0 | 34 | 298 | 0 | 37 | 0 | 0 | 84 | 0 | 0 | 3381 | 0 | 0 | 0 | 71 | 141 | 1744 | 0 | 0 | 0 | 0 | 438 | 187 | 0 | 0 | 0 | 24 | 0 | 0 | 12 | 90 | 0 | 51 |
| SRR1759269 | 26 | 0 | 2819 | 0 | 0 | 0 | 36 | 38 | 0 | 0 | 0 | 156 | 0 | 0 | 245 | 0 | 0 | 0 | 118 | 526 | 824 | 0 | 0 | 0 | 0 | 139 | 130 | 0 | 0 | 0 | 0 | 0 | 0 | 0 | 43 | 0 | 16 |
| SRR1759270 | 17 | 0 | 2231 | 0 | 0 | 45 | 53 | 12 | 0 | 0 | 0 | 168 | 12 | 0 | 67 | 0 | 0 | 0 | 90 | 313 | 270 | 0 | 0 | 100 | 0 | 236 | 228 | 0 | 0 | 0 | 15 | 0 | 0 | 0 | 37 | 0 | 32 |
| SRR1759271 | 16 | 0 | 2531 | 0 | 0 | 0 | 44 | 0 | 0 | 0 | 0 | 113 | 0 | 0 | 768 | 0 | 0 | 0 | 78 | 161 | 723 | 0 | 0 | 0 | 0 | 133 | 101 | 0 | 0 | 0 | 0 | 0 | 0 | 10 | 79 | 18 | 40 |
| SRR1759272 | 18 | 0 | 2795 | 0 | 0 | 0 | 70 | 0 | 11 | 0 | 0 | 85 | 0 | 0 | 529 | 0 | 0 | 0 | 28 | 131 | 1029 | 0 | 0 | 0 | 19 | 136 | 57 | 0 | 0 | 0 | 0 | 0 | 0 | 0 | 114 | 11 | 89 |
| SRR1759273 | 56 | 0 | 1449 | 0 | 0 | 24 | 700 | 25 | 0 | 0 | 0 | 74 | 0 | 0 | 747 | 0 | 0 | 0 | 331 | 295 | 960 | 0 | 0 | 0 | 23 | 15 | 11 | 0 | 0 | 0 | 16 | 0 | 0 | 0 | 59 | 0 | 37 |
| SRR1759274 | 12 | 0 | 1952 | 0 | 0 | 0 | 919 | 208 | 0 | 0 | 0 | 127 | 0 | 0 | 559 | 0 | 0 | 0 | 157 | 63 | 325 | 0 | 0 | 38 | 10 | 67 | 80 | 0 | 15 | 10 | 0 | 0 | 0 | 16 | 80 | 0 | 18 |
| SRR1759275 | 28 | 0 | 2755 | 0 | 0 | 13 | 403 | 130 | 0 | 0 | 0 | 162 | 0 | 0 | 444 | 0 | 0 | 14 | 353 | 292 | 268 | 0 | 0 | 67 | 0 | 154 | 160 | 0 | 0 | 0 | 12 | 0 | 0 | 0 | 59 | 0 | 27 |
| SRR1777167 | 96 | 32 | 174 | 0 | 0 | 28 | 535 | 165 | 25 | 24 | 27 | 38 | 32 | 17 | 970 | 0 | 0 | 0 | 17 | 67 | 218 | 0 | 0 | 18 | 19 | 139 | 63 | 0 | 0 | 115 | 25 | 0 | 39 | 51 | 37 | 51 | 24 |
| SRR1820679 | 28 | 0 | 93 | 0 | 45 | 78 | 140 | 21 | 21 | 12 | 108 | 36 | 25 | 13 | 260 | 0 | 0 | 34 | 520 | 6246 | 520 | 0 | 0 | 4403 | 54 | 264 | 2020 | 0 | 0 | 0 | 30 | 0 | 0 | 27 | 14 | 12 | 23 |
| SRR1820680 | 13 | 0 | 19 | 11 | 44 | 18 | 0 | 0 | 16 | 48 | 187 | 0 | 68 | 18 | 1219 | 0 | 0 | 353 | 0 | 21 | 86 | 0 | 0 | 86 | 520 | 90 | 20 | 0 | 0 | 0 | 49 | 0 | 0 | 53 | 0 | 0 | 14 |
| SRR1820681 | 0 | 0 | 32 | 0 | 17 | 94 | 166 | 32 | 17 | 0 | 54 | 0 | 68 | 15 | 83 | 0 | 0 | 0 | 182 | 5158 | 862 | 0 | 0 | 4509 | 87 | 302 | 2209 | 0 | 0 | 18 | 30 | 0 | 0 | 123 | 0 | 0 | 13 |
| SRR1820682 | 10 | 11 | 21 | 0 | 62 | 21 | 0 | 0 | 16 | 36 | 172 | 0 | 117 | 26 | 1268 | 0 | 0 | 649 | 20 | 82 | 184 | 0 | 0 | 124 | 561 | 120 | 47 | 10 | 0 | 0 | 87 | 0 | 0 | 109 | 14 | 10 | 21 |
| SRR1820683 | 21 | 0 | 35 | 0 | 32 | 83 | 116 | 27 | 39 | 15 | 83 | 0 | 0 | 0 | 290 | 0 | 0 | 62 | 503 | 6072 | 928 | 0 | 0 | 4697 | 68 | 283 | 2022 | 0 | 0 | 0 | 40 | 0 | 0 | 47 | 14 | 0 | 17 |
| SRR1820684 | 21 | 0 | 18 | 0 | 52 | 47 | 26 | 0 | 47 | 36 | 19 | 65 | 0 | 49 | 5054 | 0 | 0 | 153 | 0 | 26 | 306 | 0 | 0 | 54 | 43 | 23 | 33 | 18 | 0 | 0 | 13 | 0 | 15 | 16 | 33 | 21 | 23 |
| SRR1820685 | 14 | 0 | 45 | 0 | 12 | 29 | 128 | 21 | 12 | 0 | 18 | 0 | 14 | 0 | 224 | 0 | 0 | 79 | 481 | 7275 | 705 | 0 | 0 | 1490 | 21 | 102 | 814 | 0 | 0 | 0 | 32 | 0 | 0 | 14 | 12 | 0 | 19 |
| SRR1820686 | 0 | 0 | 21 | 0 | 51 | 18 | 0 | 0 | 29 | 34 | 23 | 0 | 14 | 38 | 540 | 0 | 0 | 69 | 0 | 57 | 28 | 0 | 0 | 70 | 64 | 14 | 45 | 15 | 0 | 0 | 11 | 0 | 0 | 63 | 18 | 0 | 18 |
| SRR1820687 | 19 | 12 | 15 | 0 | 22 | 39 | 0 | 0 | 13 | 18 | 23 | 40 | 22 | 17 | 11383 | 0 | 0 | 55 | 0 | 22 | 649 | 0 | 0 | 11 | 31 | 38 | 30 | 20 | 0 | 0 | 29 | 0 | 0 | 44 | 12 | 12 | 21 |
| SRR1820688 | 0 | 0 | 11 | 0 | 21 | 34 | 0 | 0 | 36 | 26 | 23 | 0 | 25 | 28 | 2054 | 0 | 0 | 73 | 0 | 24 | 325 | 0 | 0 | 42 | 34 | 29 | 29 | 0 | 0 | 0 | 19 | 0 | 0 | 51 | 0 | 0 | 19 |
| SRR1820689 | 25 | 0 | 25 | 0 | 24 | 0 | 0 | 0 | 19 | 32 | 155 | 13 | 60 | 49 | 2943 | 0 | 0 | 12 | 0 | 28 | 87 | 0 | 0 | 37 | 107 | 36 | 0 | 13 | 0 | 0 | 99 | 0 | 0 | 498 | 0 | 11 | 15 |
| SRR1820690 | 0 | 13 | 12 | 0 | 26 | 0 | 0 | 0 | 12 | 36 | 173 | 0 | 130 | 49 | 3197 | 0 | 0 | 28 | 0 | 138 | 120 | 0 | 0 | 93 | 112 | 32 | 13 | 19 | 0 | 0 | 60 | 0 | 0 | 186 | 0 | 0 | 0 |
| SRR1853804 | 71 | 0 | 893 | 0 | 0 | 0 | 0 | 0 | 0 | 0 | 0 | 0 | 0 | 12 | 0 | 0 | 0 | 0 | 0 | 1471 | 787 | 0 | 0 | 0 | 0 | 107 | 31 | 12 | 0 | 0 | 12 | 0 | 0 | 0 | 41 | 42 | 21 |
| SRR1853805 | 80 | 0 | 573 | 0 | 0 | 0 | 0 | 18 | 31 | 0 | 0 | 0 | 0 | 0 | 0 | 0 | 0 | 0 | 0 | 1863 | 518 | 0 | 0 | 13 | 0 | 408 | 75 | 13 | 0 | 0 | 0 | 0 | 0 | 0 | 197 | 31 | 88 |
| SRR1853806 | 63 | 0 | 998 | 0 | 0 | 25 | 416 | 21 | 0 | 0 | 0 | 46 | 0 | 0 | 33 | 0 | 0 | 0 | 17 | 2531 | 982 | 0 | 0 | 0 | 0 | 80 | 13 | 0 | 0 | 0 | 20 | 0 | 0 | 0 | 55 | 19 | 26 |
| SRR1853807 | 56 | 0 | 1804 | 0 | 0 | 15 | 295 | 11 | 11 | 0 | 0 | 0 | 0 | 13 | 33 | 0 | 0 | 0 | 0 | 426 | 1113 | 0 | 0 | 0 | 0 | 50 | 42 | 15 | 0 | 0 | 29 | 0 | 0 | 0 | 30 | 14 | 30 |
| SRR1853808 | 64 | 0 | 2236 | 0 | 0 | 99 | 977 | 0 | 22 | 0 | 0 | 0 | 0 | 0 | 0 | 0 | 14 | 0 | 0 | 2104 | 2631 | 16 | 0 | 0 | 0 | 1594 | 221 | 0 | 0 | 0 | 45 | 0 | 0 | 0 | 221 | 109 | 388 |
| SRR1853809 | 48 | 0 | 2743 | 0 | 0 | 32 | 987 | 17 | 0 | 0 | 0 | 10 | 0 | 0 | 0 | 0 | 11 | 0 | 0 | 1073 | 1717 | 0 | 0 | 0 | 0 | 496 | 117 | 0 | 0 | 0 | 0 | 0 | 0 | 0 | 279 | 95 | 346 |
| SRR1917323 | 86 | 12 | 272 | 0 | 0 | 88 | 42 | 11 | 29 | 13 | 39 | 609 | 28 | 0 | 978 | 0 | 134 | 19 | 18 | 189 | 1114 | 0 | 0 | 84 | 136 | 689 | 62 | 0 | 0 | 147 | 44 | 0 | 18 | 41 | 24 | 207 | 360 |
| SRR1917324 | 53 | 20 | 145 | 0 | 0 | 1749 | 36 | 62 | 523 | 11 | 53 | 326 | 123 | 0 | 2924 | 0 | 50 | 11 | 21 | 153 | 2043 | 14 | 0 | 49 | 128 | 11216 | 713 | 0 | 0 | 37 | 1064 | 0 | 36 | 318 | 455 | 3167 | 6227 |
| SRR1917325 | 99 | 0 | 247 | 0 | 0 | 99 | 45 | 31 | 41 | 0 | 34 | 908 | 26 | 11 | 924 | 0 | 202 | 0 | 24 | 184 | 1742 | 0 | 0 | 36 | 13 | 731 | 57 | 0 | 0 | 50 | 38 | 0 | 0 | 46 | 32 | 220 | 509 |
| SRR1917326 | 238 | 0 | 393 | 0 | 0 | 67 | 363 | 0 | 296 | 42 | 139 | 590 | 0 | 0 | 0 | 0 | 106 | 0 | 51 | 46 | 0 | 0 | 0 | 0 | 29 | 489 | 63 | 0 | 329 | 0 | 49 | 0 | 0 | 74 | 0 | 0 | 0 |
| SRR1917327 | 164 | 0 | 280 | 0 | 0 | 14 | 506 | 0 | 320 | 33 | 83 | 595 | 0 | 0 | 0 | 0 | 114 | 0 | 39 | 48 | 0 | 0 | 0 | 0 | 12 | 96 | 19 | 0 | 61 | 0 | 20 | 0 | 0 | 19 | 0 | 0 | 0 |
| SRR1917328 | 160 | 0 | 15 | 0 | 0 | 10 | 360 | 0 | 16 | 0 | 18 | 56 | 0 | 0 | 0 | 0 | 30 | 0 | 18 | 0 | 0 | 0 | 0 | 0 | 0 | 20 | 0 | 0 | 0 | 0 | 12 | 0 | 0 | 0 | 0 | 0 | 0 |
| SRR1917329 | 102 | 0 | 126 | 0 | 0 | 112 | 112 | 19 | 38 | 13 | 36 | 451 | 25 | 0 | 904 | 0 | 126 | 21 | 53 | 461 | 1300 | 0 | 0 | 42 | 22 | 948 | 57 | 0 | 0 | 24 | 34 | 0 | 15 | 25 | 24 | 108 | 212 |
| SRR1917330 | 90 | 0 | 128 | 0 | 0 | 101 | 93 | 46 | 33 | 17 | 56 | 361 | 25 | 0 | 1129 | 0 | 125 | 17 | 71 | 405 | 1753 | 13 | 0 | 36 | 15 | 830 | 50 | 0 | 0 | 71 | 29 | 0 | 22 | 40 | 25 | 140 | 318 |
| SRR1917331 | 126 | 0 | 188 | 0 | 0 | 1015 | 72 | 367 | 299 | 0 | 43 | 482 | 728 | 0 | 1126 | 0 | 206 | 26 | 107 | 800 | 1496 | 0 | 0 | 308 | 201 | 10158 | 996 | 0 | 0 | 215 | 276 | 0 | 0 | 934 | 275 | 976 | 2671 |
| SRR1917332 | 50 | 0 | 21 | 0 | 0 | 18 | 1135 | 24 | 30 | 18 | 77 | 171 | 0 | 0 | 24 | 0 | 14 | 0 | 284 | 284 | 47 | 0 | 0 | 0 | 0 | 154 | 17 | 0 | 33 | 20 | 50 | 0 | 0 | 20 | 0 | 0 | 0 |
| SRR1917333 | 63 | 0 | 57 | 0 | 0 | 13 | 582 | 29 | 57 | 23 | 59 | 297 | 0 | 0 | 0 | 0 | 17 | 0 | 241 | 100 | 0 | 0 | 0 | 0 | 0 | 75 | 18 | 0 | 21 | 11 | 51 | 0 | 0 | 15 | 0 | 0 | 0 |
| SRR1917334 | 105 | 0 | 60 | 0 | 0 | 34 | 1248 | 378 | 112 | 25 | 113 | 311 | 0 | 0 | 0 | 0 | 25 | 0 | 401 | 446 | 0 | 0 | 0 | 21 | 0 | 206 | 62 | 0 | 132 | 70 | 49 | 0 | 0 | 45 | 0 | 0 | 0 |

### Supp Table S8.2.Columns_AM_to_BW

| | | | | | | | | | | | | | | | | | | | | | | | | | | | | | | | | | | | | | |
|---|---|---|---|---|---|---|---|---|---|---|---|---|---|---|---|---|---|---|---|---|---|---|---|---|---|---|---|---|---|---|---|---|---|---|---|---|---|
| SRR1917335 | 97 | 0 | 225 | 0 | 0 | 101 | 59 | 90 | 37 | 14 | 38 | 602 | 35 | 0 | 2254 | 0 | 134 | 0 | 33 | 189 | 1621 | 0 | 0 | 46 | 13 | 697 | 67 | 0 | 0 | 157 | 36 | 0 | 27 | 88 | 24 | 166 | 316 |
| SRR1917336 | 87 | 0 | 285 | 0 | 0 | 1685 | 43 | 62 | 613 | 15 | 76 | 493 | 362 | 12 | 2816 | 0 | 138 | 26 | 14 | 111 | 1691 | 11 | 0 | 35 | 108 | 14035 | 800 | 0 | 0 | 47 | 698 | 0 | 30 | 640 | 810 | 4170 | 8356 |
| SRR1917337 | 121 | 0 | 404 | 0 | 0 | 79 | 44 | 59 | 31 | 21 | 47 | 689 | 30 | 0 | 1862 | 0 | 246 | 66 | 39 | 219 | 1539 | 0 | 0 | 27 | 16 | 666 | 70 | 0 | 0 | 65 | 28 | 0 | 19 | 91 | 48 | 147 | 316 |
| SRR1917338 | 81 | 0 | 0 | 0 | 0 | 21 | 275 | 212 | 112 | 11 | 61 | 641 | 0 | 0 | 0 | 0 | 16 | 0 | 81 | 152 | 0 | 0 | 0 | 22 | 0 | 36 | 12 | 0 | 36 | 23 | 23 | 0 | 0 | 15 | 0 | 0 | 0 |
| SRR1917339 | 62 | 0 | 0 | 0 | 0 | 11 | 329 | 135 | 45 | 14 | 88 | 746 | 0 | 0 | 0 | 0 | 14 | 0 | 82 | 93 | 0 | 0 | 0 | 0 | 0 | 28 | 15 | 0 | 18 | 10 | 19 | 0 | 0 | 10 | 0 | 0 | 22 |
| SRR1917340 | 29 | 0 | 21 | 0 | 0 | 11 | 409 | 461 | 115 | 26 | 195 | 566 | 0 | 0 | 0 | 0 | 0 | 0 | 162 | 263 | 11 | 0 | 0 | 0 | 0 | 86 | 28 | 0 | 71 | 30 | 68 | 0 | 0 | 56 | 0 | 0 | 21 |
| SRR1922516 | 196 | 0 | 4383 | 0 | 0 | 0 | 26 | 0 | 0 | 0 | 0 | 258 | 0 | 0 | 435 | 0 | 23 | 0 | 30 | 379 | 1231 | 0 | 0 | 0 | 0 | 24 | 40 | 0 | 0 | 0 | 0 | 0 | 0 | 0 | 17 | 0 | 14 |
| SRR1922517 | 78 | 0 | 2744 | 0 | 0 | 0 | 22 | 0 | 0 | 0 | 0 | 214 | 0 | 0 | 885 | 0 | 14 | 0 | 11 | 291 | 1169 | 0 | 0 | 0 | 0 | 34 | 32 | 0 | 0 | 0 | 0 | 0 | 0 | 0 | 23 | 0 | 17 |
| SRR1922518 | 259 | 0 | 4453 | 0 | 0 | 0 | 0 | 0 | 0 | 0 | 0 | 256 | 0 | 0 | 484 | 0 | 29 | 0 | 20 | 327 | 1369 | 0 | 0 | 0 | 0 | 35 | 37 | 0 | 0 | 0 | 0 | 0 | 0 | 0 | 33 | 0 | 25 |
| SRR1922519 | 100 | 0 | 2581 | 0 | 0 | 0 | 0 | 0 | 0 | 0 | 0 | 213 | 0 | 0 | 895 | 0 | 0 | 0 | 13 | 339 | 716 | 0 | 0 | 0 | 0 | 14 | 15 | 0 | 0 | 0 | 0 | 0 | 0 | 0 | 21 | 0 | 15 |
| SRR1922520 | 87 | 0 | 2481 | 0 | 0 | 0 | 28 | 0 | 0 | 0 | 0 | 126 | 0 | 0 | 479 | 0 | 18 | 0 | 0 | 232 | 1083 | 0 | 0 | 0 | 0 | 15 | 16 | 0 | 0 | 0 | 0 | 0 | 0 | 0 | 29 | 0 | 27 |
| SRR1922521 | 132 | 0 | 4246 | 0 | 0 | 0 | 22 | 0 | 0 | 0 | 0 | 225 | 0 | 0 | 472 | 0 | 72 | 0 | 29 | 580 | 890 | 0 | 0 | 0 | 0 | 28 | 38 | 0 | 0 | 0 | 0 | 0 | 0 | 0 | 14 | 0 | 12 |
| SRR1922522 | 112 | 0 | 3915 | 0 | 0 | 0 | 0 | 0 | 0 | 0 | 0 | 221 | 0 | 0 | 450 | 0 | 27 | 11 | 0 | 330 | 703 | 0 | 0 | 0 | 0 | 26 | 27 | 0 | 0 | 0 | 0 | 0 | 0 | 0 | 41 | 0 | 35 |
| SRR1922523 | 102 | 0 | 4494 | 0 | 0 | 0 | 10 | 0 | 0 | 0 | 0 | 196 | 0 | 0 | 436 | 0 | 64 | 0 | 10 | 190 | 512 | 0 | 0 | 0 | 13 | 23 | 16 | 0 | 0 | 0 | 0 | 0 | 0 | 0 | 16 | 0 | 15 |
| SRR1922524 | 179 | 0 | 4010 | 0 | 0 | 0 | 13 | 0 | 0 | 0 | 0 | 319 | 0 | 0 | 612 | 0 | 30 | 0 | 16 | 431 | 825 | 0 | 0 | 0 | 11 | 40 | 50 | 0 | 0 | 0 | 0 | 0 | 0 | 0 | 50 | 0 | 37 |
| SRR1922525 | 100 | 0 | 5300 | 0 | 0 | 0 | 0 | 0 | 0 | 0 | 0 | 263 | 0 | 0 | 382 | 0 | 28 | 10 | 17 | 488 | 870 | 0 | 0 | 0 | 0 | 13 | 10 | 0 | 0 | 0 | 0 | 0 | 0 | 0 | 20 | 0 | 14 |
| SRR1922526 | 227 | 0 | 4240 | 0 | 0 | 0 | 22 | 0 | 0 | 0 | 0 | 307 | 0 | 0 | 580 | 0 | 27 | 0 | 14 | 420 | 638 | 0 | 0 | 0 | 20 | 47 | 43 | 0 | 0 | 0 | 0 | 0 | 0 | 0 | 66 | 0 | 47 |
| SRR1922527 | 163 | 0 | 7330 | 0 | 0 | 0 | 0 | 0 | 0 | 0 | 0 | 270 | 0 | 0 | 420 | 0 | 35 | 0 | 0 | 316 | 691 | 0 | 0 | 0 | 0 | 14 | 14 | 0 | 0 | 0 | 0 | 0 | 0 | 0 | 30 | 0 | 19 |
| SRR1973407 | 41 | 0 | 3693 | 0 | 0 | 85 | 407 | 127 | 53 | 0 | 0 | 104 | 0 | 11 | 0 | 0 | 0 | 0 | 136 | 4207 | 543 | 0 | 0 | 0 | 32 | 1081 | 954 | 0 | 11 | 21 | 32 | 0 | 0 | 21 | 201 | 21 | 106 |
| SRR1973408 | 49 | 0 | 5213 | 0 | 12 | 71 | 542 | 166 | 83 | 0 | 0 | 80 | 0 | 0 | 0 | 0 | 0 | 0 | 0 | 2867 | 852 | 0 | 0 | 0 | 12 | 1159 | 1064 | 12 | 71 | 47 | 59 | 12 | 0 | 0 | 154 | 12 | 35 |
| SRR1973409 | 50 | 0 | 4318 | 0 | 0 | 158 | 378 | 197 | 89 | 0 | 0 | 130 | 0 | 0 | 0 | 0 | 11 | 0 | 47 | 2694 | 567 | 0 | 0 | 0 | 0 | 1467 | 1319 | 0 | 20 | 59 | 30 | 0 | 0 | 0 | 256 | 39 | 148 |
| SRR1973410 | 50 | 0 | 4503 | 0 | 0 | 179 | 862 | 248 | 70 | 0 | 0 | 106 | 0 | 0 | 0 | 0 | 0 | 0 | 172 | 5403 | 575 | 0 | 0 | 0 | 0 | 1917 | 2344 | 0 | 0 | 60 | 50 | 0 | 0 | 0 | 238 | 30 | 109 |
| SRR1973411 | 37 | 0 | 3890 | 0 | 0 | 62 | 724 | 16 | 93 | 0 | 0 | 94 | 0 | 31 | 0 | 0 | 30 | 0 | 241 | 6517 | 483 | 0 | 0 | 0 | 0 | 114 | 130 | 0 | 62 | 93 | 31 | 0 | 0 | 31 | 13 | 0 | 93 |
| SRR1973412 | 57 | 0 | 5350 | 15 | 0 | 11 | 184 | 16 | 135 | 0 | 0 | 184 | 0 | 0 | 0 | 0 | 17 | 0 | 276 | 5622 | 645 | 0 | 0 | 0 | 45 | 111 | 118 | 0 | 15 | 30 | 30 | 0 | 0 | 0 | 15 | 0 | 150 |
| SRR1973413 | 36 | 0 | 4826 | 0 | 0 | 247 | 387 | 450 | 101 | 0 | 0 | 85 | 0 | 0 | 0 | 0 | 14 | 0 | 387 | 4571 | 1007 | 0 | 0 | 0 | 22 | 1979 | 2215 | 0 | 34 | 56 | 0 | 11 | 0 | 22 | 382 | 0 | 90 |
| SRR1973414 | 42 | 0 | 5399 | 0 | 0 | 132 | 377 | 557 | 44 | 0 | 0 | 75 | 0 | 0 | 0 | 0 | 21 | 0 | 188 | 3768 | 1130 | 0 | 0 | 0 | 0 | 1920 | 2608 | 0 | 29 | 44 | 88 | 0 | 0 | 59 | 396 | 44 | 103 |
| SRR1973415 | 38 | 0 | 6989 | 0 | 0 | 177 | 484 | 645 | 81 | 0 | 0 | 64 | 0 | 0 | 0 | 0 | 22 | 0 | 121 | 3510 | 1937 | 0 | 0 | 0 | 32 | 1756 | 2288 | 0 | 48 | 64 | 64 | 0 | 0 | 16 | 548 | 16 | 129 |
| SRR1973952 | 223 | 0 | 474 | 0 | 0 | 90 | 195 | 26 | 26 | 13 | 64 | 0 | 39 | 13 | 292 | 13 | 83 | 0 | 0 | 341 | 292 | 0 | 0 | 52 | 193 | 696 | 787 | 0 | 0 | 13 | 400 | 0 | 0 | 64 | 1444 | 11117 | 29443 |
| SRR1973953 | 37 | 0 | 439 | 0 | 10 | 23 | 149 | 120 | 10 | 60 | 182 | 17 | 28 | 0 | 30 | 0 | 39 | 0 | 0 | 199 | 507 | 0 | 0 | 205 | 166 | 371 | 385 | 0 | 0 | 0 | 163 | 0 | 24 | 89 | 131 | 138 | 201 |
| SRR1973956 | 193 | 0 | 0 | 18 | 25 | 0 | 0 | 24 | 0 | 13 | 21 | 0 | 0 | 13 | 0 | 34 | 0 | 0 | 0 | 0 | 0 | 0 | 0 | 41 | 11 | 27 | 25 | 13 | 0 | 14 | 14 | 0 | 14 | 57 | 0 | 0 | 0 |
| SRR1973958 | 88 | 0 | 0 | 0 | 0 | 0 | 13 | 18 | 0 | 0 | 0 | 876 | 0 | 13 | 0 | 0 | 0 | 0 | 0 | 0 | 0 | 0 | 0 | 0 | 0 | 0 | 0 | 0 | 0 | 10 | 0 | 0 | 13 | 26 | 0 | 0 | 0 |
| SRR1973964 | 0 | 0 | 0 | 0 | 0 | 0 | 0 | 0 | 0 | 0 | 0 | 0 | 0 | 0 | 0 | 0 | 0 | 0 | 0 | 0 | 0 | 0 | 0 | 0 | 0 | 0 | 0 | 0 | 0 | 0 | 0 | 0 | 153 | 30 | 0 | 0 | 0 |
| SRR1973966 | 96 | 0 | 1138 | 11 | 17 | 87 | 25 | 33 | 32 | 157 | 312 | 1472 | 59 | 0 | 25 | 0 | 0 | 19 | 25 | 507 | 717 | 0 | 0 | 1357 | 408 | 625 | 732 | 0 | 0 | 87 | 50 | 0 | 24 | 324 | 37 | 33 | 53 |
| SRR1973967 | 402 | 0 | 4905 | 13 | 16 | 42 | 516 | 12 | 0 | 53 | 160 | 5325 | 0 | 12 | 32 | 0 | 0 | 0 | 64 | 483 | 145 | 0 | 0 | 12 | 213 | 527 | 550 | 16 | 0 | 0 | 51 | 0 | 31 | 79 | 72 | 54 | 86 |
| SRR1979367 | 66 | 0 | 16 | 0 | 0 | 0 | 0 | 0 | 0 | 0 | 0 | 33 | 91 | 75 | 601 | 0 | 274 | 49 | 0 | 0 | 17 | 0 | 0 | 0 | 17 | 108 | 124 | 33 | 0 | 0 | 1077 | 0 | 0 | 0 | 0 | 17 | 0 |
| SRR1988280 | 52 | 27 | 418 | 0 | 0 | 12 | 119 | 0 | 0 | 0 | 0 | 134 | 0 | 0 | 79 | 0 | 60 | 12 | 49 | 1948 | 979 | 0 | 0 | 0 | 20 | 52 | 38 | 0 | 0 | 0 | 0 | 0 | 0 | 0 | 185 | 37 | 121 |
| SRR1988281 | 63 | 21 | 515 | 0 | 0 | 12 | 141 | 0 | 12 | 0 | 0 | 163 | 0 | 0 | 94 | 0 | 68 | 0 | 78 | 1494 | 1076 | 0 | 0 | 0 | 26 | 235 | 176 | 0 | 0 | 0 | 0 | 0 | 0 | 0 | 762 | 176 | 574 |
| SRR1988282 | 55 | 0 | 325 | 0 | 0 | 14 | 103 | 0 | 0 | 0 | 0 | 157 | 0 | 0 | 144 | 0 | 76 | 16 | 78 | 1741 | 1045 | 0 | 0 | 0 | 19 | 60 | 35 | 0 | 0 | 0 | 11 | 0 | 0 | 0 | 218 | 47 | 122 |
| SRR1988283 | 27 | 15 | 1420 | 0 | 0 | 0 | 159 | 16 | 21 | 0 | 0 | 597 | 0 | 0 | 2468 | 0 | 0 | 16 | 0 | 2011 | 3115 | 0 | 0 | 0 | 0 | 18 | 106 | 0 | 0 | 0 | 0 | 0 | 0 | 0 | 97 | 14 | 74 |
| SRR1988284 | 26 | 59 | 1672 | 0 | 0 | 0 | 147 | 83 | 27 | 0 | 0 | 690 | 0 | 0 | 3010 | 0 | 0 | 0 | 10 | 1759 | 2884 | 0 | 0 | 0 | 0 | 114 | 560 | 0 | 0 | 0 | 0 | 0 | 0 | 0 | 513 | 55 | 404 |
| SRR1988285 | 24 | 73 | 1501 | 0 | 0 | 0 | 176 | 69 | 19 | 0 | 0 | 756 | 0 | 0 | 4978 | 0 | 0 | 27 | 0 | 1755 | 3703 | 0 | 0 | 0 | 0 | 106 | 526 | 0 | 0 | 0 | 0 | 0 | 0 | 0 | 440 | 53 | 340 |
| SRR1988286 | 20 | 13 | 1484 | 0 | 0 | 0 | 212 | 12 | 26 | 0 | 0 | 650 | 0 | 0 | 2452 | 0 | 0 | 26 | 0 | 2476 | 2814 | 0 | 0 | 0 | 0 | 19 | 99 | 0 | 0 | 0 | 0 | 0 | 0 | 0 | 93 | 11 | 70 |
| SRR1988287 | 0 | 24 | 302 | 0 | 0 | 22 | 69 | 0 | 11 | 0 | 0 | 161 | 0 | 0 | 511 | 0 | 40 | 0 | 30 | 1078 | 297 | 0 | 0 | 0 | 15 | 156 | 114 | 0 | 0 | 0 | 0 | 0 | 0 | 0 | 113 | 15 | 91 |
| SRR1988288 | 0 | 25 | 320 | 0 | 0 | 14 | 46 | 0 | 0 | 0 | 0 | 110 | 0 | 0 | 598 | 0 | 44 | 0 | 33 | 987 | 383 | 0 | 0 | 0 | 0 | 505 | 390 | 0 | 0 | 0 | 0 | 0 | 0 | 0 | 360 | 63 | 285 |
| SRR1988289 | 0 | 69 | 573 | 0 | 0 | 12 | 97 | 0 | 18 | 0 | 0 | 281 | 0 | 0 | 317 | 0 | 35 | 108 | 17 | 2299 | 470 | 0 | 0 | 0 | 15 | 318 | 242 | 0 | 0 | 0 | 0 | 0 | 0 | 0 | 272 | 36 | 258 |
| SRR1988290 | 11 | 29 | 332 | 0 | 0 | 14 | 91 | 0 | 20 | 0 | 0 | 128 | 0 | 0 | 145 | 0 | 43 | 84 | 15 | 2794 | 278 | 0 | 0 | 0 | 12 | 128 | 83 | 0 | 0 | 0 | 0 | 0 | 0 | 0 | 86 | 13 | 80 |
| SRR1988291 | 68 | 115 | 983 | 0 | 0 | 22 | 920 | 0 | 63 | 0 | 0 | 130 | 0 | 0 | 1064 | 0 | 33 | 27 | 96 | 1703 | 3180 | 0 | 0 | 0 | 11 | 497 | 207 | 0 | 0 | 0 | 18 | 0 | 0 | 0 | 1254 | 148 | 574 |
| SRR1988292 | 76 | 105 | 1101 | 0 | 0 | 24 | 1134 | 15 | 56 | 0 | 0 | 134 | 0 | 0 | 1525 | 0 | 39 | 63 | 98 | 1641 | 4105 | 0 | 0 | 0 | 20 | 524 | 252 | 0 | 0 | 0 | 28 | 0 | 0 | 0 | 1530 | 169 | 651 |
| SRR1988293 | 71 | 129 | 962 | 0 | 0 | 24 | 1289 | 12 | 40 | 0 | 0 | 129 | 0 | 0 | 1359 | 0 | 40 | 26 | 97 | 1810 | 4822 | 0 | 0 | 0 | 0 | 487 | 267 | 0 | 0 | 0 | 28 | 0 | 0 | 0 | 1322 | 191 | 596 |
| SRR1988294 | 71 | 22 | 997 | 0 | 0 | 26 | 1326 | 0 | 50 | 0 | 0 | 140 | 0 | 0 | 797 | 0 | 30 | 27 | 95 | 1908 | 3555 | 0 | 0 | 0 | 0 | 86 | 45 | 0 | 0 | 0 | 24 | 0 | 0 | 0 | 226 | 29 | 111 |
| SRR1988295 | 30 | 455 | 1262 | 0 | 0 | 0 | 209 | 52 | 25 | 0 | 0 | 332 | 0 | 15 | 1386 | 0 | 24 | 0 | 35 | 1028 | 1167 | 0 | 0 | 0 | 0 | 299 | 413 | 0 | 0 | 0 | 25 | 0 | 0 | 0 | 305 | 32 | 400 |
| SRR1988296 | 30 | 420 | 1504 | 0 | 0 | 10 | 225 | 37 | 33 | 0 | 0 | 289 | 0 | 17 | 768 | 0 | 30 | 11 | 28 | 1218 | 956 | 0 | 0 | 0 | 0 | 281 | 391 | 0 | 0 | 0 | 28 | 0 | 0 | 0 | 262 | 26 | 347 |
| SRR1988297 | 26 | 415 | 1495 | 0 | 0 | 0 | 157 | 71 | 33 | 0 | 0 | 377 | 0 | 15 | 854 | 0 | 26 | 0 | 0 | 1038 | 1066 | 0 | 0 | 0 | 0 | 267 | 437 | 0 | 0 | 0 | 11 | 18 | 0 | 0 | 295 | 35 | 371 |
| SRR1988298 | 30 | 84 | 1275 | 0 | 0 | 0 | 289 | 13 | 31 | 0 | 0 | 292 | 0 | 27 | 507 | 0 | 41 | 14 | 20 | 789 | 661 | 0 | 0 | 0 | 0 | 56 | 66 | 0 | 0 | 0 | 17 | 0 | 0 | 0 | 49 | 16 | 52 |
| SRR1988299 | 61 | 130 | 1147 | 0 | 0 | 31 | 839 | 253 | 67 | 0 | 0 | 154 | 0 | 0 | 4313 | 0 | 48 | 0 | 120 | 2067 | 9075 | 0 | 0 | 0 | 12 | 732 | 268 | 0 | 0 | 0 | 34 | 0 | 0 | 0 | 1305 | 222 | 713 |
| SRR1988300 | 58 | 18 | 1418 | 0 | 0 | 27 | 519 | 30 | 74 | 0 | 0 | 195 | 0 | 0 | 3047 | 0 | 59 | 15 | 33 | 2961 | 6960 | 0 | 0 | 0 | 16 | 90 | 39 | 0 | 0 | 0 | 25 | 0 | 0 | 0 | 213 | 26 | 99 |
| SRR1988301 | 71 | 16 | 1473 | 0 | 0 | 25 | 762 | 30 | 72 | 0 | 0 | 170 | 0 | 0 | 2617 | 0 | 56 | 0 | 99 | 3365 | 8162 | 0 | 0 | 0 | 0 | 90 | 37 | 0 | 0 | 0 | 41 | 0 | 0 | 0 | 217 | 36 | 88 |
| SRR1988302 | 65 | 137 | 1372 | 0 | 0 | 31 | 781 | 221 | 57 | 0 | 0 | 203 | 0 | 0 | 3647 | 0 | 52 | 21 | 74 | 2918 | 9169 | 0 | 0 | 0 | 0 | 783 | 285 | 0 | 0 | 0 | 29 | 0 | 0 | 0 | 1666 | 194 | 754 |
| SRR1988303 | 38 | 143 | 1380 | 0 | 0 | 19 | 842 | 252 | 64 | 0 | 0 | 76 | 0 | 0 | 2737 | 0 | 34 | 30 | 86 | 3797 | 6779 | 0 | 0 | 0 | 11 | 640 | 247 | 0 | 0 | 0 | 24 | 0 | 0 | 0 | 1288 | 161 | 603 |
| SRR1988304 | 30 | 149 | 1457 | 0 | 0 | 20 | 734 | 363 | 60 | 0 | 0 | 55 | 0 | 0 | 5970 | 0 | 25 | 11 | 79 | 1625 | 7392 | 0 | 0 | 0 | 16 | 636 | 275 | 0 | 0 | 0 | 23 | 0 | 0 | 0 | 1616 | 190 | 764 |
| SRR1988305 | 33 | 166 | 1272 | 0 | 0 | 16 | 814 | 479 | 60 | 0 | 0 | 63 | 0 | 0 | 3149 | 0 | 28 | 0 | 87 | 2348 | 6909 | 0 | 0 | 0 | 0 | 651 | 249 | 0 | 0 | 0 | 25 | 0 | 0 | 0 | 1373 | 168 | 648 |
| SRR1988306 | 50 | 16 | 1307 | 0 | 0 | 22 | 319 | 66 | 62 | 0 | 0 | 91 | 0 | 13 | 3861 | 0 | 23 | 21 | 44 | 1694 | 5411 | 0 | 0 | 0 | 0 | 64 | 32 | 0 | 0 | 0 | 33 | 0 | 0 | 11 | 168 | 19 | 89 |
| SRR1988307 | 65 | 22 | 5791 | 0 | 0 | 28 | 194 | 623 | 64 | 0 | 0 | 96 | 0 | 0 | 4857 | 0 | 32 | 22 | 23 | 1721 | 7410 | 0 | 0 | 0 | 31 | 96 | 51 | 0 | 0 | 12 | 48 | 0 | 0 | 48 | 213 | 33 | 155 |
| SRR1988308 | 101 | 11 | 4404 | 0 | 0 | 28 | 268 | 22 | 69 | 0 | 0 | 138 | 0 | 16 | 1858 | 0 | 45 | 22 | 66 | 1975 | 2912 | 0 | 0 | 0 | 25 | 98 | 39 | 0 | 0 | 0 | 34 | 0 | 0 | 0 | 129 | 20 | 100 |
| SRR1988309 | 74 | 30 | 6317 | 0 | 0 | 22 | 229 | 240 | 33 | 0 | 0 | 165 | 0 | 0 | 8792 | 0 | 27 | 0 | 28 | 1294 | 7838 | 0 | 0 | 0 | 33 | 76 | 73 | 0 | 0 | 74 | 31 | 0 | 0 | 37 | 209 | 30 | 141 |
| SRR1988310 | 95 | 104 | 2986 | 0 | 0 | 27 | 654 | 288 | 38 | 0 | 0 | 141 | 0 | 0 | 1592 | 0 | 45 | 0 | 61 | 2050 | 2948 | 0 | 0 | 0 | 51 | 655 | 356 | 0 | 0 | 0 | 30 | 0 | 0 | 0 | 2438 | 282 | 998 |
| SRR1988311 | 88 | 39 | 1601 | 0 | 0 | 45 | 104 | 293 | 125 | 0 | 0 | 101 | 0 | 16 | 624 | 0 | 31 | 18 | 40 | 2215 | 2655 | 0 | 0 | 0 | 27 | 898 | 293 | 0 | 0 | 0 | 25 | 10 | 0 | 10 | 2152 | 231 | 1054 |
| SRR1988312 | 65 | 52 | 1819 | 0 | 0 | 33 | 128 | 18 | 96 | 0 | 0 | 104 | 0 | 0 | 618 | 0 | 25 | 0 | 96 | 1855 | 2578 | 0 | 0 | 0 | 27 | 56 | 19 | 0 | 0 | 0 | 25 | 0 | 0 | 0 | 149 | 17 | 71 |
| SRR2052873 | 111 | 0 | 3761 | 26 | 0 | 213 | 41 | 62 | 53 | 0 | 0 | 0 | 0 | 17 | 1126 | 0 | 25 | 642 | 0 | 125 | 5286 | 0 | 0 | 0 | 21 | 4478 | 796 | 0 | 0 | 0 | 320 | 0 | 0 | 0 | 7931 | 368 | 3247 |
| SRR2052874 | 128 | 0 | 3769 | 21 | 0 | 268 | 46 | 69 | 66 | 0 | 0 | 0 | 0 | 21 | 1247 | 0 | 24 | 707 | 0 | 153 | 4487 | 0 | 0 | 0 | 19 | 5083 | 749 | 0 | 0 | 0 | 454 | 12 | 0 | 0 | 7597 | 293 | 2852 |
| SRR2052875 | 198 | 0 | 4538 | 24 | 0 | 336 | 12 | 62 | 128 | 0 | 0 | 0 | 0 | 36 | 1546 | 0 | 19 | 582 | 0 | 91 | 6277 | 0 | 0 | 0 | 0 | 5896 | 1109 | 11 | 27 | 0 | 565 | 0 | 0 | 14 | 7665 | 360 | 2465 |
| SRR2052877 | 507 | 0 | 3567 | 20 | 0 | 141 | 28 | 105 | 54 | 0 | 0 | 0 | 0 | 23 | 1344 | 0 | 22 | 889 | 0 | 115 | 6106 | 0 | 0 | 0 | 12 | 2924 | 646 | 0 | 0 | 0 | 154 | 11 | 0 | 0 | 4630 | 260 | 1967 |
| SRR2052878 | 173 | 0 | 4901 | 13 | 0 | 27 | 23 | 14 | 68 | 0 | 0 | 0 | 0 | 29 | 1658 | 0 | 22 | 738 | 0 | 116 | 6211 | 0 | 0 | 0 | 13 | 481 | 108 | 0 | 22 | 0 | 24 | 0 | 0 | 0 | 935 | 50 | 419 |
| SRR2053027 | 302 | 0 | 5319 | 0 | 0 | 283 | 0 | 113 | 100 | 0 | 0 | 0 | 0 | 11 | 1621 | 0 | 30 | 720 | 0 | 278 | 8321 | 0 | 0 | 0 | 14 | 4415 | 814 | 0 | 0 | 0 | 313 | 19 | 0 | 0 | 4324 | 251 | 1900 |
| SRR2053028 | 238 | 0 | 5926 | 0 | 10 | 220 | 23 | 141 | 92 | 0 | 0 | 0 | 0 | 17 | 1736 | 0 | 21 | 895 | 0 | 135 | 8087 | 0 | 0 | 0 | 30 | 3984 | 786 | 0 | 0 | 0 | 251 | 14 | 0 | 0 | 5973 | 277 | 2220 |
| SRR2053029 | 227 | 0 | 5343 | 13 | 0 | 392 | 10 | 111 | 164 | 0 | 0 | 0 | 0 | 19 | 2740 | 0 | 23 | 1177 | 0 | 103 | 12846 | 0 | 0 | 0 | 18 | 7475 | 962 | 0 | 0 | 0 | 610 | 0 | 0 | 0 | 5055 | 283 | 2300 |
| SRR2061931 | 61 | 25 | 2777 | 0 | 0 | 29 | 84 | 1087 | 0 | 0 | 0 | 14 | 0 | 0 | 48 | 0 | 10 | 0 | 36 | 1819 | 723 | 13 | 0 | 21 | 0 | 755 | 1918 | 0 | 0 | 46 | 0 | 0 | 13 | 25 | 63 | 0 | 25 |
| SRR2061932 | 81 | 14 | 1617 | 0 | 0 | 27 | 0 | 1242 | 0 | 0 | 0 | 0 | 0 | 14 | 0 | 14 | 10 | 0 | 45 | 2003 | 668 | 0 | 0 | 54 | 14 | 716 | 918 | 0 | 0 | 270 | 27 | 0 | 27 | 0 | 27 | 0 | 0 |
| SRR2061933 | 119 | 13 | 1732 | 0 | 0 | 35 | 0 | 99 | 13 | 0 | 0 | 11 | 0 | 26 | 0 | 0 | 15 | 0 | 0 | 1321 | 1421 | 0 | 0 | 0 | 0 | 185 | 356 | 0 | 0 | 13 | 0 | 0 | 39 | 0 | 0 | 0 | 0 |
| SRR2061934 | 56 | 0 | 1108 | 0 | 0 | 23 | 374 | 543 | 0 | 0 | 0 | 14 | 0 | 10 | 0 | 0 | 0 | 0 | 0 | 2710 | 1308 | 0 | 0 | 17 | 23 | 115 | 125 | 0 | 0 | 34 | 0 | 0 | 23 | 0 | 0 | 0 | 0 |
| SRR2061935 | 106 | 0 | 1057 | 0 | 0 | 51 | 212 | 164 | 25 | 0 | 0 | 0 | 0 | 0 | 0 | 0 | 0 | 0 | 0 | 1380 | 743 | 0 | 0 | 76 | 25 | 60 | 101 | 25 | 25 | 11 | 0 | 0 | 25 | 25 | 25 | 0 | 25 |
| SRR2061936 | 218 | 16 | 514 | 0 | 0 | 80 | 34 | 102 | 43 | 0 | 0 | 11 | 0 | 11 | 238 | 0 | 0 | 0 | 0 | 1224 | 1054 | 16 | 0 | 16 | 0 | 983 | 1736 | 0 | 0 | 0 | 0 | 0 | 59 | 0 | 32 | 0 | 11 |
| SRR2061937 | 367 | 22 | 723 | 0 | 0 | 123 | 80 | 101 | 0 | 0 | 0 | 49 | 0 | 22 | 121 | 0 | 0 | 0 | 0 | 804 | 884 | 0 | 0 | 0 | 0 | 1291 | 1500 | 0 | 0 | 22 | 14 | 0 | 58 | 0 | 36 | 0 | 0 |
| SRR2061938 | 227 | 0 | 675 | 0 | 0 | 59 | 0 | 142 | 34 | 0 | 0 | 45 | 0 | 20 | 55 | 0 | 11 | 0 | 27 | 2684 | 1397 | 0 | 0 | 29 | 0 | 661 | 2487 | 0 | 0 | 0 | 15 | 0 | 29 | 15 | 24 | 0 | 0 |
| SRR2061940 | 89 | 0 | 311 | 0 | 0 | 24 | 0 | 142 | 0 | 0 | 0 | 11 | 0 | 24 | 146 | 0 | 0 | 0 | 0 | 1166 | 1603 | 0 | 0 | 24 | 0 | 897 | 1180 | 0 | 0 | 24 | 24 | 24 | 0 | 0 | 24 | 0 | 0 |
| SRR2061941 | 22 | 0 | 8988 | 0 | 0 | 741 | 0 | 838 | 182 | 0 | 0 | 67 | 0 | 0 | 13 | 0 | 0 | 0 | 0 | 999 | 89 | 0 | 0 | 73 | 0 | 6351 | 5549 | 0 | 0 | 73 | 73 | 0 | 12 | 36 | 522 | 49 | 364 |
| SRR2061942 | 21 | 0 | 9257 | 0 | 11 | 36 | 0 | 112 | 11 | 0 | 0 | 54 | 0 | 0 | 0 | 0 | 0 | 0 | 0 | 1095 | 240 | 0 | 0 | 41 | 11 | 394 | 622 | 0 | 0 | 11 | 12 | 0 | 11 | 33 | 66 | 44 | 41 |
| SRR2061943 | 19 | 0 | 5057 | 0 | 0 | 20 | 0 | 94 | 15 | 0 | 0 | 41 | 0 | 0 | 0 | 0 | 0 | 0 | 0 | 511 | 267 | 0 | 0 | 0 | 0 | 246 | 504 | 10 | 0 | 28 | 23 | 0 | 13 | 13 | 63 | 46 | 40 |
| SRR2061944 | 19 | 0 | 14193 | 0 | 0 | 158 | 24 | 583 | 45 | 0 | 0 | 65 | 0 | 0 | 0 | 0 | 0 | 0 | 12 | 838 | 413 | 0 | 0 | 35 | 0 | 2309 | 3402 | 0 | 0 | 59 | 54 | 0 | 0 | 15 | 183 | 0 | 173 |
| SRR2061945 | 36 | 0 | 8729 | 0 | 0 | 246 | 18 | 1152 | 76 | 0 | 0 | 34 | 0 | 0 | 0 | 0 | 0 | 0 | 0 | 970 | 242 | 0 | 0 | 55 | 10 | 2815 | 4472 | 0 | 0 | 73 | 69 | 0 | 21 | 10 | 304 | 24 | 277 |
| SRR2061946 | 20 | 0 | 6365 | 0 | 0 | 119 | 0 | 1325 | 42 | 0 | 0 | 28 | 0 | 12 | 0 | 0 | 0 | 0 | 0 | 310 | 127 | 0 | 0 | 0 | 0 | 1752 | 2529 | 0 | 0 | 125 | 22 | 0 | 14 | 104 | 156 | 0 | 82 |
| SRR2061947 | 29 | 0 | 9069 | 0 | 0 | 261 | 14 | 759 | 84 | 0 | 0 | 43 | 0 | 0 | 28 | 0 | 0 | 0 | 14 | 1056 | 248 | 0 | 0 | 79 | 0 | 3168 | 4672 | 0 | 0 | 61 | 56 | 0 | 0 | 23 | 312 | 33 | 275 |
| SRR2061948 | 16 | 0 | 7636 | 0 | 0 | 157 | 21 | 851 | 31 | 0 | 0 | 53 | 0 | 0 | 14 | 0 | 0 | 0 | 0 | 849 | 392 | 0 | 0 | 55 | 0 | 2157 | 3875 | 0 | 0 | 43 | 59 | 0 | 27 | 27 | 271 | 20 | 161 |
| SRR2061949 | 14 | 0 | 4661 | 0 | 0 | 19 | 0 | 171 | 31 | 0 | 0 | 33 | 0 | 0 | 0 | 0 | 0 | 0 | 0 | 337 | 242 | 0 | 0 | 11 | 17 | 253 | 414 | 0 | 0 | 15 | 33 | 0 | 20 | 22 | 25 | 0 | 56 |
| SRR2061950 | 66 | 0 | 2718 | 0 | 0 | 131 | 0 | 71 | 29 | 0 | 0 | 186 | 0 | 0 | 0 | 0 | 12 | 0 | 0 | 923 | 519 | 0 | 0 | 41 | 14 | 1468 | 241 | 0 | 0 | 14 | 84 | 0 | 0 | 0 | 137 | 23 | 100 |
| SRR2061951 | 66 | 0 | 3264 | 0 | 0 | 1284 | 24 | 515 | 218 | 0 | 0 | 211 | 0 | 0 | 12 | 0 | 23 | 0 | 0 | 925 | 231 | 0 | 0 | 66 | 0 | 12984 | 2422 | 0 | 0 | 0 | 605 | 0 | 14 | 0 | 1021 | 163 | 685 |
| SRR2061952 | 40 | 0 | 1787 | 0 | 0 | 108 | 10 | 58 | 21 | 0 | 0 | 119 | 0 | 0 | 14 | 0 | 13 | 0 | 14 | 623 | 292 | 0 | 0 | 15 | 0 | 1234 | 249 | 0 | 0 | 18 | 71 | 0 | 0 | 20 | 128 | 11 | 54 |
| SRR2061953 | 77 | 0 | 3348 | 0 | 0 | 302 | 19 | 78 | 42 | 0 | 0 | 213 | 0 | 0 | 26 | 0 | 20 | 0 | 0 | 1476 | 226 | 0 | 0 | 19 | 0 | 2375 | 323 | 0 | 14 | 51 | 107 | 0 | 19 | 14 | 196 | 16 | 115 |
| SRR2061954 | 57 | 0 | 2647 | 0 | 0 | 152 | 0 | 111 | 32 | 0 | 0 | 144 | 0 | 0 | 0 | 0 | 16 | 0 | 0 | 855 | 223 | 0 | 0 | 36 | 0 | 1747 | 275 | 0 | 0 | 40 | 71 | 0 | 0 | 0 | 151 | 28 | 116 |
| SRR2061955 | 42 | 0 | 1557 | 0 | 0 | 106 | 11 | 72 | 18 | 0 | 0 | 99 | 0 | 0 | 17 | 0 | 12 | 0 | 0 | 505 | 210 | 0 | 0 | 18 | 0 | 989 | 178 | 14 | 0 | 21 | 58 | 0 | 0 | 32 | 101 | 49 | 45 |
| SRR2061956 | 76 | 0 | 3394 | 0 | 0 | 2444 | 0 | 709 | 333 | 0 | 0 | 240 | 0 | 0 | 16 | 0 | 22 | 0 | 0 | 1099 | 419 | 0 | 0 | 46 | 0 | 18541 | 2307 | 0 | 0 | 101 | 908 | 0 | 0 | 20 | 1640 | 144 | 925 |
| SRR2061957 | 60 | 0 | 2727 | 0 | 0 | 154 | 0 | 102 | 27 | 0 | 0 | 144 | 0 | 0 | 13 | 0 | 15 | 0 | 0 | 804 | 340 | 0 | 0 | 60 | 0 | 1415 | 234 | 0 | 0 | 13 | 71 | 0 | 0 | 22 | 160 | 17 | 86 |
| SRR2061958 | 48 | 0 | 1891 | 0 | 0 | 1093 | 0 | 888 | 208 | 0 | 0 | 120 | 0 | 0 | 20 | 0 | 0 | 0 | 0 | 435 | 355 | 0 | 0 | 20 | 0 | 11279 | 1718 | 0 | 0 | 52 | 517 | 0 | 0 | 33 | 817 | 46 | 384 |
| SRR2069629 | 352 | 0 | 277 | 0 | 0 | 66 | 20 | 0 | 19 | 0 | 0 | 4789 | 0 | 0 | 17 | 0 | 1138 | 10 | 11 | 64 | 156 | 0 | 0 | 0 | 136 | 244 | 154 | 0 | 0 | 0 | 88 | 0 | 10 | 19 | 17 | 10 | 17 |
| SRR2069630 | 582 | 0 | 404 | 0 | 0 | 45 | 16 | 0 | 0 | 17 | 0 | 2411 | 0 | 0 | 11 | 0 | 862 | 0 | 0 | 21 | 16 | 0 | 0 | 106 | 251 | 190 | 100 | 0 | 0 | 0 | 86 | 0 | 1221 | 15 | 0 | 0 | 15 |
| SRR2084358 | 0 | 0 | 70 | 0 | 0 | 0 | 34 | 0 | 196 | 0 | 0 | 15 | 0 | 49 | 34 | 98 | 0 | 0 | 0 | 4990 | 547 | 0 | 0 | 49 | 0 | 11 | 343 | 0 | 49 | 0 | 22 | 0 | 49 | 0 | 55 | 17 | 82 |
| SRR2084359 | 0 | 0 | 98 | 0 | 0 | 247 | 72 | 70 | 106 | 0 | 0 | 23 | 0 | 176 | 72 | 35 | 0 | 91 | 0 | 6443 | 1134 | 0 | 0 | 0 | 35 | 21 | 37 | 35 | 0 | 0 | 26 | 0 | 35 | 0 | 108 | 29 | 140 |
| SRR2085173 | 41 | 0 | 246 | 0 | 0 | 0 | 0 | 0 | 0 | 0 | 0 | 16 | 0 | 0 | 12 | 66 | 0 | 0 | 0 | 0 | 16 | 0 | 0 | 0 | 0 | 0 | 0 | 0 | 68 | 0 | 69 | 0 | 116 | 0 | 105 | 40 | 0 |
| SRR2103379 | 26 | 213 | 38 | 0 | 0 | 74 | 935 | 1008 | 26 | 10 | 24 | 0 | 0 | 143 | 36 | 0 | 30 | 35 | 33 | 2156 | 0 | 0 | 0 | 0 | 0 | 919 | 772 | 0 | 183 | 18 | 39 | 0 | 15 | 23 | 1112 | 142 | 684 |
| SRR2103380 | 24 | 230 | 33 | 0 | 0 | 70 | 956 | 1021 | 23 | 12 | 29 | 0 | 0 | 124 | 66 | 0 | 31 | 36 | 38 | 2311 | 0 | 0 | 0 | 0 | 11 | 989 | 831 | 0 | 184 | 20 | 42 | 0 | 11 | 14 | 987 | 117 | 646 |
| SRR2103381 | 31 | 39 | 33 | 0 | 0 | 15 | 919 | 210 | 21 | 0 | 22 | 0 | 0 | 22 | 54 | 0 | 33 | 34 | 30 | 2445 | 0 | 0 | 0 | 0 | 12 | 226 | 181 | 0 | 43 | 22 | 44 | 0 | 15 | 15 | 227 | 28 | 149 |
| SRR2103382 | 29 | 117 | 206 | 14 | 21 | 68 | 257 | 561 | 30 | 0 | 0 | 0 | 0 | 52 | 439 | 0 | 34 | 237 | 27 | 5434 | 0 | 0 | 0 | 0 | 35 | 1136 | 746 | 0 | 0 | 0 | 40 | 0 | 0 | 0 | 2475 | 277 | 797 |
| SRR2103383 | 30 | 120 | 178 | 0 | 11 | 72 | 218 | 617 | 21 | 0 | 0 | 0 | 0 | 39 | 350 | 0 | 39 | 274 | 23 | 4475 | 0 | 0 | 0 | 0 | 40 | 1139 | 756 | 0 | 0 | 0 | 31 | 0 | 0 | 0 | 2377 | 273 | 752 |
| SRR2103384 | 25 | 133 | 159 | 0 | 0 | 142 | 282 | 1149 | 47 | 0 | 0 | 0 | 0 | 50 | 145 | 0 | 32 | 127 | 27 | 4837 | 0 | 0 | 0 | 0 | 25 | 2168 | 1182 | 0 | 0 | 0 | 99 | 0 | 0 | 11 | 4234 | 359 | 1168 |
| SRR2105940 | 249 | 0 | 870 | 14 | 0 | 0 | 0 | 0 | 0 | 0 | 0 | 21 | 0 | 0 | 0 | 0 | 15 | 0 | 0 | 0 | 133 | 0 | 0 | 0 | 0 | 0 | 0 | 0 | 0 | 0 | 0 | 0 | 0 | 0 | 55 | 0 | 0 |
| SRR2105943 | 311 | 0 | 1325 | 13 | 0 | 0 | 0 | 0 | 0 | 0 | 0 | 0 | 0 | 0 | 50 | 0 | 0 | 0 | 0 | 0 | 56 | 0 | 0 | 0 | 0 | 0 | 0 | 0 | 0 | 0 | 13 | 0 | 0 | 0 | 40 | 0 | 0 |
| SRR2105944 | 114 | 0 | 399 | 0 | 0 | 0 | 0 | 0 | 0 | 0 | 0 | 85 | 0 | 0 | 70 | 0 | 0 | 0 | 0 | 0 | 18 | 0 | 0 | 0 | 0 | 0 | 0 | 0 | 0 | 0 | 0 | 0 | 0 | 0 | 13 | 0 | 0 |
| SRR2105949 | 147 | 0 | 486 | 19 | 0 | 0 | 0 | 0 | 0 | 0 | 0 | 28 | 0 | 0 | 7222 | 0 | 0 | 500 | 0 | 0 | 0 | 0 | 0 | 0 | 0 | 0 | 0 | 11 | 0 | 0 | 0 | 0 | 0 | 0 | 0 | 0 | 0 |
| SRR2105951 | 82 | 0 | 478 | 35 | 0 | 0 | 0 | 0 | 0 | 0 | 0 | 23 | 0 | 0 | 2868 | 0 | 0 | 0 | 0 | 0 | 0 | 0 | 0 | 0 | 0 | 0 | 0 | 13 | 0 | 0 | 0 | 0 | 0 | 0 | 38 | 0 | 0 |
| SRR2106873 | 87 | 0 | 2271 | 0 | 0 | 18 | 0 | 0 | 14 | 0 | 0 | 151 | 0 | 0 | 0 | 0 | 0 | 0 | 0 | 192 | 105 | 0 | 0 | 0 | 0 | 394 | 920 | 0 | 0 | 0 | 482 | 0 | 0 | 0 | 1522 | 217 | 1600 |
| SRR2106874 | 63 | 0 | 1703 | 0 | 0 | 40 | 0 | 0 | 31 | 0 | 0 | 150 | 0 | 0 | 54 | 0 | 0 | 0 | 0 | 194 | 475 | 0 | 0 | 0 | 0 | 59 | 54 | 0 | 0 | 0 | 42 | 0 | 0 | 0 | 142 | 18 | 98 |
| SRR2106875 | 58 | 0 | 1661 | 0 | 0 | 73 | 0 | 0 | 23 | 0 | 0 | 180 | 0 | 0 | 14 | 0 | 12 | 0 | 14 | 1133 | 1190 | 0 | 0 | 0 | 0 | 108 | 435 | 0 | 0 | 0 | 30 | 0 | 0 | 0 | 113 | 10 | 91 |
| SRR2106876 | 71 | 0 | 2989 | 0 | 0 | 136 | 0 | 16 | 39 | 0 | 0 | 137 | 0 | 0 | 21 | 0 | 11 | 0 | 84 | 4096 | 1710 | 0 | 0 | 70 | 0 | 1348 | 2437 | 0 | 0 | 0 | 275 | 0 | 0 | 0 | 337 | 50 | 186 |
| SRR2106877 | 65 | 0 | 894 | 0 | 0 | 15 | 18 | 12 | 73 | 0 | 0 | 168 | 0 | 0 | 0 | 0 | 13 | 0 | 36 | 5717 | 1894 | 0 | 0 | 110 | 0 | 134 | 595 | 0 | 0 | 49 | 50 | 0 | 0 | 14 | 56 | 14 | 80 |

### Supp Table S8.2.Columns_AM_to_BW

| | | | | | | | | | | | | | | | | | | | | | | | | | | | | | | | | | | | | | |
|---|---|---|---|---|---|---|---|---|---|---|---|---|---|---|---|---|---|---|---|---|---|---|---|---|---|---|---|---|---|---|---|---|---|---|---|---|---|
| SRR2106878 | 89 | 0 | 1690 | 0 | 0 | 172 | 21 | 55 | 37 | 0 | 0 | 252 | 0 | 12 | 21 | 0 | 11 | 0 | 21 | 7038 | 2023 | 0 | 0 | 215 | 0 | 2144 | 9430 | 0 | 0 | 31 | 448 | 0 | 0 | 0 | 1481 | 141 | 1259 |
| SRR2106879 | 79 | 0 | 531 | 0 | 0 | 10 | 0 | 0 | 59 | 0 | 0 | 180 | 0 | 0 | 86 | 0 | 15 | 0 | 0 | 830 | 1336 | 0 | 0 | 0 | 0 | 154 | 128 | 12 | 0 | 0 | 58 | 0 | 0 | 15 | 28 | 44 | 24 |
| SRR2106880 | 124 | 0 | 2980 | 0 | 0 | 25 | 25 | 0 | 16 | 0 | 0 | 120 | 0 | 0 | 0 | 0 | 12 | 0 | 0 | 562 | 830 | 0 | 0 | 0 | 0 | 94 | 106 | 0 | 0 | 0 | 60 | 0 | 0 | 0 | 138 | 16 | 80 |
| SRR2106881 | 85 | 0 | 1072 | 0 | 0 | 70 | 0 | 0 | 15 | 0 | 0 | 237 | 0 | 0 | 58 | 0 | 14 | 0 | 14 | 434 | 666 | 0 | 0 | 0 | 0 | 81 | 89 | 0 | 0 | 0 | 50 | 0 | 0 | 15 | 47 | 48 | 39 |
| SRR2106882 | 57 | 11 | 5688 | 0 | 0 | 0 | 0 | 0 | 0 | 0 | 0 | 462 | 0 | 0 | 111 | 0 | 17 | 22 | 0 | 400 | 271 | 0 | 0 | 0 | 0 | 268 | 347 | 0 | 0 | 0 | 160 | 0 | 0 | 0 | 609 | 72 | 577 |
| SRR2106883 | 48 | 0 | 4433 | 0 | 0 | 23 | 28 | 0 | 25 | 0 | 0 | 359 | 0 | 0 | 21 | 0 | 20 | 0 | 35 | 4502 | 595 | 0 | 0 | 28 | 0 | 418 | 4453 | 16 | 0 | 0 | 259 | 0 | 0 | 0 | 616 | 72 | 476 |
| SRR2106884 | 55 | 0 | 4734 | 0 | 0 | 21 | 0 | 0 | 13 | 0 | 0 | 384 | 0 | 0 | 66 | 0 | 18 | 0 | 0 | 2255 | 436 | 0 | 0 | 16 | 0 | 359 | 2147 | 0 | 0 | 0 | 203 | 0 | 0 | 0 | 583 | 69 | 544 |
| SRR2124760 | 1232 | 0 | 969 | 0 | 0 | 10 | 24 | 15 | 0 | 0 | 0 | 0 | 0 | 0 | 38 | 0 | 12 | 0 | 14 | 2763 | 3600 | 0 | 0 | 16 | 0 | 45 | 30 | 0 | 0 | 0 | 0 | 0 | 0 | 0 | 0 | 0 | 0 |
| SRR2124761 | 672 | 22 | 857 | 0 | 0 | 21 | 113 | 22 | 0 | 0 | 0 | 20 | 0 | 0 | 13 | 0 | 0 | 0 | 15 | 1507 | 1034 | 0 | 0 | 17 | 0 | 34 | 22 | 0 | 0 | 0 | 0 | 0 | 0 | 0 | 14 | 0 | 0 |
| SRR2124762 | 597 | 25 | 1160 | 0 | 0 | 26 | 30 | 131 | 0 | 0 | 0 | 26 | 0 | 0 | 0 | 0 | 0 | 0 | 0 | 1202 | 521 | 11 | 0 | 78 | 0 | 255 | 162 | 0 | 0 | 0 | 0 | 0 | 0 | 0 | 0 | 0 | 0 |
| SRR2124763 | 433 | 15 | 1206 | 12 | 0 | 37 | 111 | 20 | 13 | 0 | 0 | 0 | 0 | 11 | 0 | 0 | 0 | 0 | 13 | 692 | 555 | 13 | 0 | 54 | 0 | 59 | 25 | 0 | 0 | 0 | 14 | 0 | 0 | 0 | 14 | 0 | 18 |
| SRR2124764 | 864 | 18 | 716 | 0 | 0 | 18 | 21 | 16 | 0 | 0 | 0 | 16 | 0 | 0 | 34 | 0 | 0 | 0 | 10 | 1833 | 2452 | 0 | 0 | 17 | 0 | 84 | 67 | 0 | 0 | 0 | 0 | 0 | 0 | 0 | 10 | 0 | 0 |
| SRR2124765 | 897 | 20 | 537 | 14 | 0 | 14 | 14 | 24 | 0 | 0 | 0 | 12 | 0 | 16 | 60 | 0 | 0 | 0 | 43 | 2319 | 4499 | 17 | 0 | 83 | 0 | 147 | 164 | 16 | 0 | 0 | 0 | 0 | 12 | 0 | 0 | 0 | 0 |
| SRR2124766 | 662 | 20 | 550 | 0 | 0 | 0 | 0 | 55 | 0 | 0 | 0 | 12 | 0 | 0 | 26 | 0 | 0 | 0 | 13 | 2145 | 2021 | 14 | 0 | 42 | 0 | 125 | 113 | 0 | 0 | 0 | 0 | 0 | 0 | 0 | 0 | 0 | 0 |
| SRR2124767 | 654 | 26 | 562 | 20 | 0 | 11 | 53 | 19 | 0 | 0 | 0 | 12 | 0 | 0 | 13 | 0 | 0 | 0 | 15 | 2730 | 1108 | 22 | 0 | 15 | 0 | 25 | 30 | 11 | 0 | 0 | 0 | 0 | 11 | 0 | 11 | 0 | 10 |
| SRR2124768 | 803 | 25 | 834 | 20 | 0 | 16 | 0 | 60 | 0 | 0 | 0 | 13 | 0 | 12 | 0 | 0 | 0 | 0 | 0 | 1915 | 502 | 11 | 0 | 106 | 0 | 162 | 157 | 11 | 0 | 0 | 0 | 0 | 0 | 0 | 10 | 0 | 0 |
| SRR2124769 | 808 | 10 | 673 | 21 | 0 | 14 | 17 | 0 | 0 | 0 | 0 | 0 | 0 | 14 | 93 | 0 | 0 | 0 | 47 | 2376 | 2598 | 20 | 0 | 18 | 0 | 49 | 52 | 0 | 0 | 0 | 0 | 0 | 0 | 0 | 0 | 0 | 12 |
| SRR2124770 | 1006 | 37 | 1096 | 15 | 0 | 0 | 11 | 44 | 0 | 0 | 0 | 0 | 0 | 0 | 11 | 0 | 13 | 0 | 0 | 1847 | 721 | 15 | 0 | 41 | 0 | 69 | 110 | 0 | 0 | 0 | 0 | 0 | 0 | 0 | 0 | 0 | 0 |
| SRR2124771 | 498 | 21 | 556 | 28 | 0 | 10 | 60 | 33 | 0 | 0 | 0 | 22 | 0 | 13 | 33 | 0 | 0 | 12 | 50 | 2162 | 2476 | 10 | 0 | 51 | 0 | 107 | 122 | 0 | 0 | 0 | 0 | 0 | 0 | 0 | 10 | 0 | 0 |
| SRR2124772 | 746 | 22 | 490 | 19 | 0 | 12 | 0 | 39 | 0 | 0 | 0 | 0 | 0 | 0 | 37 | 0 | 0 | 0 | 0 | 3130 | 1260 | 18 | 0 | 52 | 0 | 160 | 126 | 0 | 0 | 0 | 12 | 0 | 0 | 0 | 0 | 0 | 0 |
| SRR2124773 | 549 | 15 | 546 | 12 | 0 | 13 | 40 | 15 | 0 | 0 | 0 | 12 | 0 | 0 | 104 | 0 | 0 | 44 | 48 | 5265 | 8249 | 18 | 0 | 29 | 0 | 58 | 60 | 0 | 0 | 0 | 22 | 0 | 16 | 0 | 0 | 0 | 0 |
| SRR2124774 | 484 | 19 | 772 | 28 | 0 | 16 | 0 | 40 | 0 | 0 | 0 | 0 | 0 | 14 | 0 | 0 | 0 | 0 | 0 | 2375 | 877 | 12 | 0 | 14 | 0 | 49 | 30 | 14 | 0 | 0 | 0 | 0 | 0 | 0 | 12 | 0 | 0 |
| SRR2124775 | 704 | 25 | 653 | 18 | 0 | 16 | 13 | 14 | 0 | 0 | 0 | 14 | 0 | 23 | 0 | 0 | 0 | 0 | 26 | 2511 | 1551 | 21 | 0 | 85 | 0 | 263 | 153 | 12 | 0 | 0 | 11 | 0 | 0 | 0 | 28 | 0 | 25 |
| SRR2124776 | 879 | 23 | 675 | 17 | 0 | 23 | 26 | 39 | 0 | 0 | 0 | 18 | 0 | 0 | 21 | 0 | 0 | 0 | 12 | 3344 | 1648 | 14 | 0 | 92 | 0 | 234 | 129 | 0 | 0 | 0 | 0 | 0 | 0 | 0 | 0 | 0 | 0 |
| SRR2124777 | 706 | 23 | 713 | 18 | 0 | 31 | 252 | 183 | 16 | 0 | 0 | 15 | 0 | 0 | 20 | 0 | 0 | 0 | 23 | 2789 | 1137 | 15 | 0 | 59 | 0 | 357 | 131 | 0 | 0 | 0 | 10 | 0 | 0 | 0 | 31 | 0 | 14 |
| SRR2124778 | 1397 | 28 | 966 | 19 | 0 | 11 | 0 | 29 | 0 | 0 | 0 | 0 | 0 | 0 | 26 | 0 | 0 | 0 | 0 | 3943 | 993 | 19 | 0 | 27 | 0 | 44 | 42 | 0 | 0 | 0 | 0 | 0 | 0 | 0 | 0 | 0 | 0 |
| SRR2124779 | 624 | 20 | 824 | 0 | 0 | 12 | 31 | 99 | 0 | 0 | 0 | 0 | 0 | 0 | 0 | 0 | 0 | 0 | 0 | 1585 | 871 | 16 | 0 | 63 | 0 | 203 | 131 | 0 | 0 | 0 | 0 | 0 | 0 | 0 | 0 | 0 | 11 |
| SRR2126527 | 60 | 0 | 83 | 0 | 0 | 14 | 127 | 31 | 0 | 0 | 0 | 24 | 0 | 0 | 0 | 0 | 17 | 0 | 44 | 8279 | 4162 | 0 | 0 | 50 | 14 | 137 | 1707 | 0 | 0 | 0 | 0 | 0 | 0 | 0 | 497 | 45 | 298 |
| SRR2126528 | 67 | 0 | 64 | 0 | 0 | 82 | 130 | 61 | 0 | 0 | 0 | 26 | 0 | 0 | 0 | 0 | 16 | 0 | 52 | 9822 | 4492 | 0 | 0 | 49 | 14 | 888 | 10996 | 0 | 0 | 0 | 0 | 0 | 0 | 0 | 3035 | 328 | 1899 |
| SRR2126529 | 64 | 0 | 80 | 0 | 0 | 88 | 68 | 41 | 13 | 0 | 0 | 25 | 0 | 0 | 0 | 0 | 18 | 0 | 37 | 7342 | 4209 | 0 | 0 | 43 | 17 | 875 | 9795 | 0 | 0 | 0 | 0 | 0 | 0 | 0 | 3218 | 351 | 2131 |
| SRR2126530 | 76 | 0 | 71 | 0 | 0 | 17 | 153 | 14 | 11 | 0 | 0 | 24 | 0 | 0 | 0 | 0 | 15 | 0 | 61 | 7495 | 3584 | 0 | 0 | 27 | 17 | 158 | 1956 | 0 | 0 | 0 | 0 | 0 | 0 | 0 | 346 | 28 | 224 |
| SRR2126531 | 60 | 0 | 64 | 0 | 0 | 81 | 97 | 64 | 10 | 0 | 0 | 17 | 0 | 0 | 0 | 0 | 16 | 0 | 58 | 7453 | 3718 | 0 | 0 | 35 | 18 | 729 | 8781 | 0 | 0 | 0 | 0 | 0 | 0 | 0 | 1719 | 169 | 1101 |
| SRR2126532 | 69 | 0 | 70 | 0 | 0 | 20 | 78 | 42 | 0 | 0 | 0 | 22 | 0 | 0 | 0 | 0 | 14 | 0 | 62 | 8987 | 3745 | 0 | 0 | 27 | 16 | 193 | 2426 | 0 | 0 | 0 | 0 | 0 | 0 | 0 | 579 | 59 | 407 |
| SRR2135530 | 42 | 15 | 379 | 0 | 14 | 36 | 344 | 109 | 19 | 11 | 16 | 1041 | 29 | 13 | 1036 | 0 | 189 | 13 | 269 | 1860 | 242 | 0 | 0 | 0 | 27 | 165 | 32 | 0 | 0 | 0 | 35 | 0 | 0 | 128 | 59 | 34 | 137 |
| SRR2135531 | 53 | 55 | 729 | 0 | 12 | 337 | 569 | 9250 | 97 | 12 | 72 | 1381 | 78 | 0 | 522 | 0 | 371 | 39 | 1110 | 17227 | 309 | 0 | 0 | 302 | 214 | 2110 | 1238 | 0 | 0 | 77 | 153 | 0 | 0 | 312 | 169 | 210 | 452 |
| SRR2135532 | 58 | 12 | 722 | 0 | 0 | 46 | 272 | 72 | 17 | 10 | 63 | 1556 | 18 | 0 | 616 | 0 | 722 | 48 | 193 | 2131 | 240 | 0 | 0 | 0 | 67 | 298 | 111 | 0 | 0 | 0 | 35 | 0 | 0 | 64 | 29 | 34 | 68 |
| SRR2229584 | 176 | 0 | 35 | 0 | 0 | 0 | 0 | 0 | 0 | 0 | 0 | 0 | 0 | 0 | 70 | 0 | 0 | 0 | 0 | 351 | 18 | 0 | 0 | 75 | 0 | 70 | 15 | 0 | 0 | 0 | 15 | 0 | 0 | 0 | 34 | 16 | 48 |
| SRR2229585 | 170 | 0 | 56 | 0 | 0 | 0 | 0 | 0 | 0 | 0 | 0 | 0 | 0 | 10 | 43 | 0 | 0 | 0 | 0 | 2005 | 0 | 0 | 0 | 122 | 0 | 44 | 45 | 0 | 0 | 0 | 0 | 0 | 0 | 0 | 20 | 0 | 16 |
| SRR2229586 | 0 | 0 | 0 | 0 | 0 | 43 | 17 | 0 | 32 | 0 | 0 | 0 | 0 | 0 | 0 | 0 | 0 | 30 | 34 | 18853 | 0 | 0 | 0 | 111 | 0 | 43 | 22 | 0 | 0 | 1295 | 0 | 0 | 0 | 18 | 0 | 0 | 22 |
| SRR2229587 | 0 | 0 | 0 | 0 | 0 | 26 | 27 | 0 | 0 | 0 | 0 | 0 | 0 | 0 | 0 | 0 | 0 | 0 | 27 | 13645 | 0 | 0 | 0 | 153 | 0 | 17 | 31 | 0 | 0 | 529 | 0 | 0 | 0 | 0 | 0 | 0 | 0 |
| SRR2229588 | 34 | 0 | 100 | 0 | 0 | 0 | 130 | 131 | 0 | 0 | 0 | 0 | 0 | 15 | 0 | 0 | 0 | 0 | 104 | 8490 | 0 | 0 | 0 | 13 | 0 | 24 | 33 | 0 | 0 | 15 | 0 | 0 | 0 | 0 | 34 | 24 | 49 |
| SRR2229589 | 26 | 0 | 54 | 0 | 0 | 0 | 72 | 75 | 0 | 0 | 0 | 0 | 0 | 0 | 96 | 0 | 0 | 0 | 48 | 9325 | 0 | 0 | 0 | 10 | 0 | 69 | 15 | 0 | 0 | 20 | 0 | 0 | 0 | 12 | 16 | 16 | 69 |
| SRR2287868 | 16 | 0 | 1301 | 0 | 0 | 24 | 211 | 0 | 0 | 0 | 0 | 145 | 0 | 0 | 300 | 0 | 0 | 38 | 58 | 37 | 2321 | 0 | 0 | 21 | 0 | 402 | 96 | 0 | 0 | 0 | 16 | 0 | 0 | 11 | 73 | 21 | 39 |
| SRR2289624 | 0 | 0 | 1678 | 0 | 0 | 0 | 446 | 178 | 0 | 0 | 0 | 86 | 0 | 0 | 972 | 0 | 0 | 0 | 172 | 241 | 2511 | 0 | 0 | 0 | 31 | 95 | 224 | 0 | 52 | 72 | 0 | 0 | 0 | 22 | 0 | 0 | 0 |
| SRR2289625 | 12 | 0 | 2053 | 0 | 0 | 30 | 474 | 0 | 18 | 0 | 0 | 111 | 0 | 0 | 8876 | 0 | 0 | 0 | 71 | 65 | 5415 | 0 | 0 | 0 | 0 | 11 | 96 | 0 | 0 | 0 | 32 | 0 | 0 | 16 | 50 | 0 | 50 |
| SRR2289626 | 0 | 0 | 1210 | 0 | 0 | 0 | 33 | 0 | 0 | 0 | 0 | 44 | 0 | 0 | 61 | 0 | 0 | 0 | 25 | 201 | 304 | 0 | 0 | 0 | 0 | 68 | 39 | 0 | 0 | 0 | 0 | 0 | 0 | 0 | 0 | 0 | 0 |
| SRR2294377 | 0 | 0 | 1682 | 0 | 0 | 34 | 131 | 0 | 0 | 0 | 0 | 60 | 0 | 0 | 643 | 0 | 0 | 0 | 315 | 1082 | 951 | 0 | 0 | 0 | 0 | 461 | 140 | 0 | 0 | 0 | 23 | 0 | 0 | 0 | 31 | 0 | 41 |
| SRR2294380 | 11 | 0 | 2760 | 0 | 0 | 45 | 455 | 10 | 12 | 0 | 0 | 46 | 0 | 0 | 39 | 0 | 16 | 0 | 517 | 281 | 291 | 0 | 0 | 0 | 10 | 602 | 166 | 0 | 109 | 0 | 27 | 0 | 0 | 25 | 100 | 23 | 79 |
| SRR2294739 | 11 | 0 | 2467 | 0 | 0 | 100 | 107 | 33 | 0 | 0 | 0 | 20 | 0 | 0 | 44 | 0 | 11 | 29 | 157 | 110 | 427 | 0 | 0 | 12 | 0 | 1232 | 198 | 0 | 0 | 0 | 37 | 0 | 0 | 14 | 74 | 0 | 44 |
| SRR2294742 | 11 | 14 | 2814 | 0 | 0 | 95 | 168 | 16 | 11 | 0 | 0 | 38 | 0 | 0 | 260 | 0 | 12 | 0 | 746 | 679 | 530 | 0 | 0 | 0 | 0 | 19 | 237 | 0 | 0 | 0 | 53 | 0 | 0 | 22 | 92 | 24 | 72 |
| SRR2294746 | 14 | 0 | 3186 | 0 | 0 | 33 | 227 | 22 | 44 | 0 | 0 | 24 | 0 | 0 | 175 | 0 | 0 | 0 | 462 | 315 | 388 | 0 | 0 | 0 | 15 | 977 | 144 | 0 | 11 | 0 | 33 | 0 | 0 | 0 | 74 | 0 | 33 |
| SRR2294747 | 0 | 10 | 2501 | 0 | 0 | 0 | 21 | 30 | 0 | 0 | 0 | 35 | 0 | 0 | 11 | 0 | 0 | 0 | 220 | 821 | 122 | 0 | 0 | 17 | 0 | 83 | 75 | 0 | 0 | 0 | 0 | 0 | 0 | 12 | 16 | 0 | 51 |
| SRR2294748 | 10 | 21 | 1775 | 0 | 0 | 49 | 304 | 41 | 14 | 0 | 0 | 29 | 0 | 14 | 315 | 0 | 0 | 0 | 1734 | 1989 | 798 | 0 | 0 | 0 | 0 | 637 | 113 | 0 | 0 | 0 | 18 | 0 | 0 | 0 | 33 | 0 | 34 |
| SRR2294820 | 14 | 0 | 3882 | 0 | 0 | 59 | 721 | 48 | 10 | 0 | 0 | 36 | 0 | 0 | 189 | 0 | 12 | 0 | 1030 | 1533 | 479 | 0 | 0 | 0 | 10 | 14 | 117 | 0 | 14 | 0 | 32 | 0 | 0 | 0 | 110 | 27 | 103 |
| SRR2296788 | 46 | 0 | 880 | 0 | 0 | 54 | 152 | 193 | 12 | 0 | 0 | 1297 | 0 | 0 | 0 | 0 | 36 | 0 | 76 | 6464 | 1800 | 0 | 0 | 0 | 59 | 87 | 579 | 0 | 0 | 24 | 24 | 0 | 0 | 0 | 72 | 36 | 34 |
| SRR2296789 | 44 | 0 | 1098 | 0 | 0 | 61 | 0 | 42 | 30 | 0 | 0 | 1160 | 15 | 0 | 0 | 0 | 29 | 0 | 38 | 5780 | 1436 | 0 | 0 | 0 | 61 | 89 | 1092 | 0 | 0 | 0 | 46 | 0 | 0 | 0 | 110 | 16 | 65 |
| SRR2296790 | 53 | 0 | 1058 | 0 | 0 | 0 | 0 | 3137 | 0 | 0 | 0 | 1058 | 0 | 0 | 0 | 0 | 36 | 0 | 0 | 4082 | 1521 | 0 | 0 | 0 | 83 | 432 | 3120 | 0 | 0 | 0 | 33 | 0 | 0 | 0 | 730 | 66 | 465 |
| SRR2296791 | 46 | 0 | 1105 | 0 | 13 | 11 | 42 | 229 | 13 | 0 | 0 | 1188 | 0 | 0 | 42 | 0 | 35 | 0 | 0 | 7513 | 918 | 0 | 0 | 0 | 26 | 111 | 685 | 0 | 0 | 13 | 13 | 0 | 0 | 13 | 100 | 40 | 42 |
| SRR2296792 | 72 | 12 | 1131 | 0 | 0 | 11 | 0 | 27 | 12 | 0 | 0 | 1612 | 0 | 0 | 0 | 0 | 32 | 0 | 0 | 7576 | 2841 | 0 | 0 | 0 | 25 | 106 | 1448 | 0 | 0 | 0 | 37 | 0 | 0 | 12 | 114 | 74 | 71 |
| SRR2296793 | 37 | 0 | 929 | 0 | 0 | 27 | 122 | 122 | 0 | 0 | 0 | 1064 | 0 | 0 | 0 | 0 | 38 | 0 | 41 | 6269 | 2890 | 0 | 0 | 0 | 76 | 55 | 479 | 0 | 0 | 0 | 14 | 0 | 0 | 0 | 62 | 48 | 30 |
| SRR2296794 | 44 | 0 | 1200 | 0 | 0 | 46 | 30 | 431 | 13 | 0 | 0 | 1570 | 0 | 0 | 30 | 0 | 49 | 0 | 0 | 8255 | 2498 | 0 | 0 | 0 | 99 | 795 | 4697 | 0 | 0 | 13 | 13 | 0 | 0 | 0 | 629 | 40 | 219 |
| SRR2296795 | 72 | 0 | 817 | 0 | 0 | 27 | 0 | 10 | 0 | 0 | 0 | 2158 | 0 | 0 | 71 | 0 | 22 | 0 | 0 | 4608 | 1679 | 0 | 0 | 0 | 27 | 60 | 318 | 0 | 0 | 0 | 18 | 0 | 0 | 0 | 60 | 36 | 33 |
| SRR2296797 | 39 | 0 | 1265 | 0 | 0 | 47 | 16 | 122 | 0 | 0 | 0 | 1647 | 0 | 0 | 16 | 14 | 41 | 0 | 62 | 4930 | 1326 | 0 | 0 | 0 | 11 | 92 | 400 | 28 | 0 | 19 | 33 | 0 | 0 | 0 | 63 | 52 | 35 |
| SRR2296798 | 41 | 0 | 1228 | 0 | 0 | 38 | 132 | 1897 | 13 | 0 | 0 | 1216 | 0 | 0 | 0 | 0 | 31 | 0 | 0 | 5231 | 1390 | 0 | 0 | 0 | 63 | 341 | 2339 | 0 | 0 | 0 | 0 | 0 | 25 | 0 | 645 | 51 | 316 |
| SRR2296799 | 52 | 10 | 844 | 0 | 0 | 73 | 51 | 354 | 0 | 0 | 0 | 1774 | 10 | 0 | 0 | 0 | 35 | 0 | 17 | 6859 | 1920 | 0 | 0 | 0 | 52 | 1093 | 10292 | 0 | 0 | 0 | 16 | 0 | 10 | 0 | 755 | 89 | 380 |
| SRR2296800 | 75 | 0 | 1086 | 0 | 0 | 14 | 90 | 26 | 0 | 0 | 0 | 1763 | 0 | 0 | 0 | 0 | 31 | 0 | 45 | 8443 | 3296 | 0 | 0 | 0 | 28 | 134 | 1524 | 0 | 0 | 0 | 0 | 0 | 0 | 0 | 122 | 84 | 64 |
| SRR2296801 | 92 | 0 | 815 | 0 | 0 | 43 | 97 | 19 | 0 | 0 | 0 | 2638 | 0 | 0 | 39 | 0 | 26 | 0 | 58 | 5127 | 1930 | 0 | 0 | 0 | 62 | 616 | 6516 | 0 | 0 | 0 | 28 | 0 | 0 | 0 | 337 | 0 | 133 |
| SRR2296802 | 61 | 0 | 1019 | 0 | 0 | 48 | 0 | 48 | 0 | 0 | 0 | 1727 | 0 | 0 | 0 | 0 | 0 | 0 | 0 | 3832 | 1299 | 0 | 0 | 0 | 97 | 1135 | 7847 | 24 | 0 | 0 | 0 | 0 | 24 | 0 | 507 | 48 | 217 |
| SRR2296805 | 66 | 0 | 977 | 0 | 0 | 42 | 71 | 32 | 0 | 0 | 0 | 1380 | 0 | 0 | 27 | 0 | 38 | 0 | 27 | 5956 | 1453 | 0 | 0 | 0 | 84 | 79 | 861 | 0 | 0 | 12 | 35 | 0 | 0 | 12 | 67 | 52 | 38 |
| SRR2296806 | 40 | 0 | 1220 | 13 | 0 | 40 | 43 | 422 | 0 | 0 | 0 | 905 | 0 | 0 | 0 | 0 | 39 | 0 | 86 | 5917 | 1887 | 26 | 0 | 0 | 40 | 646 | 5746 | 0 | 0 | 0 | 40 | 0 | 13 | 0 | 1291 | 79 | 738 |
| SRR2296807 | 73 | 0 | 979 | 0 | 0 | 52 | 29 | 121 | 12 | 0 | 0 | 1652 | 0 | 0 | 14 | 0 | 35 | 0 | 58 | 7583 | 2123 | 0 | 0 | 0 | 52 | 98 | 1019 | 0 | 0 | 12 | 0 | 0 | 0 | 0 | 55 | 32 | 29 |
| SRR2296808 | 60 | 0 | 1170 | 0 | 0 | 69 | 0 | 11 | 35 | 0 | 0 | 1465 | 17 | 0 | 0 | 0 | 29 | 0 | 0 | 5224 | 1979 | 0 | 0 | 0 | 13 | 82 | 772 | 0 | 0 | 0 | 17 | 0 | 0 | 0 | 57 | 17 | 43 |
| SRR2296809 | 70 | 0 | 973 | 0 | 0 | 20 | 32 | 20 | 0 | 0 | 0 | 1949 | 0 | 0 | 0 | 0 | 28 | 0 | 0 | 4070 | 1454 | 0 | 0 | 0 | 46 | 61 | 670 | 0 | 0 | 0 | 18 | 0 | 0 | 10 | 39 | 23 | 24 |
| SRR2296810 | 75 | 0 | 1096 | 0 | 0 | 52 | 68 | 12 | 0 | 0 | 0 | 1253 | 0 | 0 | 23 | 0 | 29 | 0 | 0 | 4898 | 1111 | 0 | 0 | 0 | 44 | 135 | 708 | 0 | 0 | 0 | 52 | 0 | 0 | 0 | 83 | 87 | 39 |
| SRR2353417 | 24 | 0 | 1826 | 0 | 0 | 23 | 89 | 0 | 0 | 0 | 0 | 198 | 0 | 0 | 2510 | 0 | 0 | 0 | 64 | 313 | 5364 | 0 | 0 | 57 | 14 | 21 | 17 | 0 | 0 | 0 | 11 | 0 | 0 | 0 | 17 | 0 | 0 |
| SRR2353418 | 19 | 0 | 2340 | 0 | 0 | 0 | 71 | 0 | 0 | 0 | 0 | 189 | 0 | 0 | 389 | 0 | 0 | 0 | 185 | 715 | 0 | 0 | 0 | 82 | 0 | 93 | 172 | 0 | 0 | 0 | 0 | 0 | 0 | 0 | 14 | 0 | 0 |
| SRR2353419 | 23 | 0 | 2683 | 0 | 0 | 26 | 111 | 15 | 0 | 0 | 0 | 171 | 0 | 0 | 3345 | 0 | 0 | 0 | 64 | 479 | 2893 | 0 | 0 | 62 | 0 | 20 | 22 | 0 | 0 | 0 | 17 | 0 | 0 | 0 | 35 | 0 | 22 |
| SRR2353420 | 23 | 0 | 2328 | 0 | 0 | 27 | 99 | 41 | 0 | 0 | 0 | 185 | 0 | 0 | 4594 | 0 | 0 | 47 | 115 | 454 | 3371 | 0 | 0 | 72 | 0 | 260 | 267 | 0 | 0 | 0 | 11 | 0 | 0 | 0 | 16 | 0 | 18 |
| SRR2353421 | 19 | 0 | 2940 | 0 | 0 | 22 | 92 | 52 | 0 | 0 | 0 | 185 | 0 | 0 | 1891 | 0 | 0 | 0 | 84 | 595 | 1838 | 0 | 0 | 76 | 0 | 21 | 16 | 0 | 0 | 0 | 17 | 0 | 0 | 0 | 43 | 0 | 21 |
| SRR2353422 | 25 | 0 | 2305 | 0 | 0 | 13 | 56 | 12 | 0 | 0 | 0 | 217 | 0 | 0 | 874 | 0 | 11 | 10 | 64 | 557 | 1211 | 0 | 0 | 68 | 14 | 14 | 14 | 0 | 0 | 0 | 0 | 0 | 0 | 12 | 14 | 0 | 0 |
| SRR2353423 | 27 | 0 | 2774 | 0 | 0 | 0 | 47 | 0 | 0 | 0 | 0 | 138 | 0 | 0 | 3150 | 0 | 0 | 0 | 87 | 510 | 2765 | 0 | 0 | 37 | 0 | 221 | 251 | 0 | 0 | 0 | 13 | 0 | 0 | 0 | 20 | 0 | 23 |
| SRR2353424 | 33 | 0 | 3179 | 0 | 0 | 21 | 76 | 0 | 0 | 0 | 0 | 166 | 0 | 0 | 406 | 0 | 11 | 0 | 98 | 1064 | 1163 | 0 | 0 | 61 | 12 | 18 | 12 | 0 | 0 | 0 | 0 | 0 | 0 | 0 | 20 | 0 | 12 |
| SRR2353425 | 19 | 0 | 2724 | 0 | 0 | 11 | 126 | 30 | 0 | 0 | 0 | 175 | 0 | 0 | 1644 | 0 | 0 | 0 | 101 | 104 | 1174 | 0 | 0 | 0 | 0 | 15 | 17 | 0 | 0 | 0 | 0 | 0 | 0 | 0 | 45 | 0 | 48 |
| SRR2353426 | 24 | 0 | 2668 | 0 | 0 | 16 | 43 | 20 | 0 | 0 | 0 | 184 | 0 | 0 | 2251 | 0 | 0 | 17 | 79 | 346 | 1719 | 0 | 0 | 52 | 0 | 20 | 19 | 0 | 0 | 0 | 0 | 0 | 0 | 0 | 35 | 0 | 33 |
| SRR2353427 | 18 | 0 | 2220 | 0 | 0 | 11 | 77 | 22 | 0 | 0 | 0 | 195 | 0 | 0 | 1018 | 0 | 0 | 0 | 98 | 175 | 833 | 0 | 0 | 0 | 0 | 14 | 12 | 0 | 0 | 0 | 0 | 0 | 0 | 0 | 31 | 0 | 11 |
| SRR2353428 | 25 | 0 | 2741 | 0 | 0 | 15 | 69 | 21 | 0 | 0 | 0 | 159 | 0 | 0 | 906 | 0 | 0 | 12 | 82 | 206 | 788 | 0 | 0 | 0 | 16 | 13 | 118 | 0 | 0 | 0 | 0 | 0 | 0 | 0 | 45 | 0 | 0 |
| SRR2353429 | 25 | 0 | 2551 | 0 | 0 | 12 | 389 | 13 | 0 | 0 | 0 | 176 | 0 | 0 | 1740 | 0 | 0 | 25 | 110 | 108 | 628 | 0 | 0 | 13 | 0 | 17 | 13 | 0 | 0 | 0 | 0 | 0 | 0 | 0 | 35 | 0 | 15 |
| SRR2353430 | 31 | 0 | 2429 | 0 | 0 | 11 | 233 | 18 | 0 | 0 | 0 | 127 | 0 | 0 | 463 | 0 | 10 | 11 | 52 | 231 | 393 | 0 | 0 | 92 | 23 | 142 | 13 | 0 | 0 | 0 | 0 | 0 | 0 | 0 | 20 | 0 | 0 |
| SRR2353431 | 24 | 0 | 2064 | 0 | 0 | 26 | 193 | 67 | 0 | 0 | 0 | 138 | 0 | 0 | 3395 | 0 | 0 | 15 | 188 | 111 | 1069 | 0 | 0 | 51 | 31 | 28 | 20 | 0 | 0 | 0 | 16 | 0 | 0 | 0 | 45 | 0 | 24 |
| SRR2353432 | 27 | 0 | 2377 | 0 | 0 | 11 | 93 | 27 | 0 | 0 | 0 | 176 | 0 | 0 | 1170 | 0 | 0 | 0 | 62 | 166 | 566 | 0 | 0 | 27 | 0 | 218 | 185 | 0 | 0 | 0 | 16 | 0 | 0 | 0 | 69 | 11 | 14 |
| SRR2353433 | 21 | 0 | 2571 | 0 | 0 | 0 | 76 | 0 | 0 | 0 | 0 | 164 | 0 | 0 | 189 | 0 | 10 | 0 | 62 | 148 | 347 | 0 | 0 | 65 | 0 | 118 | 180 | 0 | 0 | 0 | 0 | 0 | 0 | 0 | 40 | 0 | 0 |
| SRR2353434 | 25 | 0 | 2738 | 0 | 0 | 34 | 147 | 67 | 0 | 0 | 0 | 152 | 0 | 0 | 1810 | 0 | 10 | 15 | 114 | 477 | 895 | 0 | 0 | 66 | 0 | 27 | 19 | 0 | 0 | 0 | 11 | 0 | 0 | 0 | 36 | 0 | 22 |
| SRR2353435 | 21 | 0 | 2769 | 0 | 0 | 23 | 147 | 28 | 17 | 0 | 0 | 165 | 0 | 0 | 1413 | 0 | 0 | 0 | 300 | 445 | 1061 | 0 | 0 | 98 | 0 | 28 | 20 | 0 | 0 | 0 | 14 | 0 | 0 | 0 | 50 | 0 | 21 |
| SRR2353436 | 23 | 0 | 2710 | 0 | 0 | 10 | 76 | 19 | 0 | 0 | 0 | 112 | 0 | 0 | 52 | 0 | 10 | 0 | 175 | 333 | 190 | 0 | 0 | 95 | 22 | 64 | 178 | 0 | 0 | 0 | 0 | 0 | 0 | 0 | 17 | 0 | 0 |
| SRR2353437 | 33 | 0 | 2319 | 0 | 0 | 0 | 84 | 0 | 0 | 0 | 0 | 170 | 0 | 0 | 1362 | 0 | 11 | 12 | 37 | 802 | 1188 | 0 | 0 | 40 | 0 | 234 | 173 | 0 | 0 | 0 | 13 | 0 | 0 | 0 | 16 | 0 | 13 |
| SRR2353438 | 18 | 0 | 3262 | 0 | 0 | 27 | 30 | 0 | 12 | 0 | 0 | 175 | 12 | 0 | 477 | 0 | 0 | 0 | 99 | 857 | 1296 | 0 | 0 | 24 | 0 | 24 | 13 | 0 | 0 | 0 | 13 | 0 | 0 | 0 | 28 | 0 | 0 |
| SRR2353439 | 27 | 13 | 3042 | 0 | 0 | 0 | 33 | 15 | 0 | 0 | 0 | 116 | 0 | 0 | 59 | 0 | 13 | 0 | 122 | 619 | 220 | 0 | 0 | 78 | 0 | 123 | 150 | 0 | 0 | 0 | 0 | 0 | 0 | 0 | 0 | 0 | 0 |
| SRR2353440 | 20 | 0 | 3180 | 0 | 0 | 27 | 88 | 0 | 11 | 0 | 0 | 144 | 0 | 13 | 429 | 0 | 14 | 17 | 64 | 327 | 464 | 0 | 0 | 40 | 0 | 18 | 12 | 0 | 0 | 0 | 0 | 0 | 0 | 0 | 16 | 0 | 16 |
| SRR2353441 | 22 | 0 | 3144 | 0 | 0 | 27 | 59 | 13 | 0 | 0 | 0 | 118 | 0 | 0 | 364 | 0 | 13 | 0 | 73 | 308 | 342 | 0 | 0 | 80 | 0 | 13 | 12 | 0 | 0 | 0 | 20 | 0 | 0 | 0 | 32 | 0 | 22 |
| SRR2353442 | 23 | 0 | 2431 | 0 | 0 | 10 | 181 | 10 | 0 | 0 | 0 | 142 | 0 | 0 | 181 | 0 | 0 | 12 | 85 | 268 | 302 | 0 | 0 | 35 | 12 | 107 | 195 | 0 | 0 | 0 | 0 | 0 | 0 | 0 | 14 | 0 | 0 |
| SRR2353443 | 15 | 0 | 2741 | 0 | 0 | 11 | 183 | 46 | 0 | 0 | 0 | 154 | 0 | 0 | 388 | 0 | 0 | 0 | 90 | 106 | 242 | 0 | 0 | 32 | 0 | 203 | 245 | 0 | 0 | 0 | 0 | 0 | 0 | 0 | 43 | 0 | 25 |
| SRR2353444 | 35 | 0 | 2538 | 0 | 0 | 14 | 154 | 10 | 14 | 0 | 0 | 133 | 0 | 0 | 599 | 0 | 13 | 0 | 48 | 544 | 607 | 0 | 0 | 51 | 22 | 102 | 192 | 0 | 0 | 0 | 0 | 0 | 0 | 0 | 31 | 0 | 12 |
| SRR2353445 | 31 | 0 | 2401 | 0 | 0 | 17 | 46 | 0 | 0 | 0 | 0 | 155 | 0 | 0 | 172 | 0 | 0 | 19 | 64 | 351 | 323 | 0 | 0 | 55 | 0 | 19 | 11 | 0 | 0 | 0 | 17 | 0 | 0 | 0 | 20 | 0 | 17 |
| SRR2422969 | 48 | 64 | 626 | 0 | 0 | 32 | 657 | 860 | 127 | 0 | 0 | 66 | 0 | 32 | 1387 | 0 | 0 | 0 | 146 | 14969 | 1387 | 0 | 0 | 478 | 64 | 828 | 4011 | 0 | 0 | 159 | 32 | 0 | 0 | 0 | 287 | 223 | 159 |
| SRR2422970 | 64 | 33 | 767 | 0 | 0 | 67 | 552 | 899 | 33 | 0 | 0 | 55 | 0 | 0 | 1263 | 0 | 14 | 0 | 395 | 14837 | 1657 | 0 | 0 | 533 | 33 | 1065 | 3363 | 0 | 0 | 67 | 100 | 0 | 0 | 0 | 333 | 67 | 133 |
| SRR2422971 | 34 | 30 | 844 | 0 | 0 | 91 | 627 | 1331 | 91 | 30 | 0 | 101 | 0 | 0 | 1532 | 0 | 0 | 91 | 139 | 16153 | 1323 | 0 | 0 | 484 | 91 | 1089 | 3478 | 0 | 0 | 121 | 272 | 0 | 0 | 30 | 484 | 181 | 242 |
| SRR2422972 | 56 | 0 | 699 | 0 | 0 | 62 | 436 | 57 | 62 | 0 | 0 | 73 | 31 | 62 | 1742 | 0 | 17 | 93 | 290 | 15101 | 1307 | 0 | 0 | 14 | 0 | 39 | 140 | 0 | 0 | 124 | 93 | 0 | 0 | 31 | 11 | 10 | 16 |
| SRR2454328 | 34 | 170 | 1430 | 0 | 0 | 51 | 12 | 126 | 33 | 0 | 0 | 96 | 0 | 14 | 24 | 0 | 26 | 0 | 0 | 65 | 81 | 0 | 0 | 0 | 12 | 1386 | 501 | 0 | 33 | 0 | 706 | 0 | 0 | 134 | 876 | 101 | 368 |
| SRR2454329 | 39 | 20 | 1441 | 0 | 0 | 44 | 24 | 13 | 39 | 0 | 0 | 94 | 0 | 0 | 21 | 0 | 26 | 0 | 0 | 45 | 60 | 0 | 0 | 0 | 0 | 142 | 45 | 11 | 23 | 0 | 77 | 0 | 0 | 16 | 96 | 11 | 38 |
| SRR2454330 | 39 | 173 | 1679 | 0 | 0 | 60 | 0 | 169 | 34 | 0 | 0 | 95 | 0 | 12 | 19 | 0 | 38 | 0 | 0 | 65 | 52 | 0 | 0 | 0 | 16 | 1399 | 452 | 19 | 31 | 0 | 535 | 0 | 0 | 137 | 947 | 94 | 407 |
| SRR2454331 | 48 | 167 | 1656 | 0 | 0 | 62 | 22 | 177 | 31 | 0 | 0 | 104 | 0 | 18 | 11 | 0 | 34 | 0 | 0 | 71 | 69 | 0 | 0 | 0 | 14 | 1549 | 462 | 12 | 20 | 0 | 512 | 0 | 0 | 131 | 900 | 87 | 369 |
| SRR2454332 | 63 | 0 | 1208 | 0 | 0 | 11 | 0 | 24 | 75 | 0 | 0 | 183 | 0 | 19 | 0 | 0 | 24 | 0 | 0 | 143 | 75 | 0 | 0 | 0 | 0 | 236 | 91 | 0 | 0 | 0 | 30 | 0 | 0 | 189 | 81 | 225 | 53 |
| SRR2454333 | 61 | 16 | 1135 | 0 | 0 | 195 | 0 | 28 | 94 | 0 | 0 | 168 | 0 | 0 | 0 | 0 | 24 | 0 | 0 | 129 | 86 | 0 | 0 | 0 | 10 | 252 | 112 | 10 | 14 | 0 | 36 | 0 | 0 | 134 | 105 | 13 | 54 |
| SRR2454334 | 71 | 14 | 1533 | 0 | 0 | 186 | 16 | 0 | 63 | 0 | 0 | 164 | 0 | 30 | 14 | 0 | 15 | 0 | 0 | 140 | 91 | 0 | 0 | 0 | 0 | 5505 | 2841 | 27 | 36 | 0 | 612 | 0 | 0 | 164 | 2333 | 216 | 1186 |
| SRR2454335 | 53 | 11 | 1621 | 0 | 0 | 52 | 0 | 19 | 34 | 0 | 0 | 140 | 15 | 34 | 0 | 0 | 20 | 0 | 0 | 64 | 64 | 0 | 0 | 0 | 0 | 1468 | 609 | 34 | 15 | 0 | 934 | 0 | 0 | 108 | 1920 | 164 | 613 |
| SRR2454336 | 45 | 14 | 1513 | 0 | 0 | 45 | 0 | 17 | 28 | 0 | 0 | 140 | 0 | 14 | 11 | 0 | 19 | 0 | 0 | 86 | 37 | 0 | 0 | 0 | 0 | 1291 | 812 | 14 | 14 | 0 | 833 | 0 | 0 | 115 | 1599 | 154 | 595 |
| SRR2454337 | 48 | 28 | 1640 | 0 | 0 | 49 | 0 | 31 | 28 | 0 | 0 | 134 | 17 | 14 | 15 | 0 | 23 | 0 | 0 | 51 | 46 | 0 | 0 | 0 | 14 | 1012 | 596 | 28 | 21 | 0 | 776 | 0 | 0 | 94 | 1566 | 187 | 544 |
| SRR2487412 | 127 | 0 | 960 | 0 | 0 | 37 | 0 | 0 | 16 | 0 | 0 | 21 | 0 | 0 | 13898 | 0 | 11 | 0 | 0 | 82 | 0 | 0 | 0 | 0 | 16 | 40 | 79 | 0 | 0 | 0 | 53 | 0 | 0 | 21 | 14 | 11 | 64 |
| SRR2487413 | 101 | 0 | 919 | 0 | 0 | 30 | 0 | 0 | 24 | 0 | 0 | 24 | 0 | 18 | 12632 | 0 | 11 | 0 | 0 | 153 | 0 | 0 | 0 | 0 | 0 | 29 | 49 | 0 | 0 | 0 | 42 | 0 | 0 | 0 | 11 | 12 | 79 |
| SRR2487414 | 115 | 0 | 919 | 10 | 0 | 26 | 0 | 0 | 10 | 0 | 0 | 22 | 0 | 10 | 13375 | 0 | 11 | 0 | 0 | 55 | 0 | 0 | 0 | 0 | 0 | 31 | 88 | 0 | 0 | 0 | 36 | 0 | 0 | 0 | 12 | 21 | 62 |
| SRR2487415 | 61 | 0 | 606 | 0 | 28 | 111 | 0 | 0 | 69 | 14 | 0 | 50 | 0 | 28 | 9310 | 0 | 0 | 0 | 35 | 105 | 0 | 0 | 0 | 0 | 14 | 32 | 194 | 0 | 0 | 0 | 69 | 14 | 0 | 0 | 18 | 28 | 347 |
| SRR2487416 | 57 | 0 | 595 | 0 | 16 | 112 | 0 | 0 | 96 | 0 | 0 | 44 | 0 | 0 | 10847 | 0 | 0 | 0 | 40 | 80 | 0 | 0 | 0 | 0 | 0 | 31 | 257 | 0 | 0 | 0 | 177 | 0 | 0 | 32 | 21 | 32 | 594 |
| SRR2487417 | 53 | 0 | 585 | 14 | 14 | 82 | 0 | 0 | 109 | 0 | 0 | 44 | 0 | 55 | 10894 | 0 | 0 | 0 | 0 | 105 | 0 | 0 | 0 | 0 | 14 | 34 | 274 | 14 | 0 | 0 | 123 | 0 | 0 | 0 | 20 | 55 | 438 |
| SRR2487418 | 135 | 0 | 1313 | 20 | 20 | 60 | 0 | 0 | 20 | 0 | 0 | 16 | 0 | 20 | 3021 | 0 | 16 | 30 | 0 | 166 | 786 | 0 | 0 | 0 | 0 | 28 | 201 | 20 | 0 | 0 | 121 | 0 | 0 | 0 | 12 | 0 | 121 |

### Supp Table S8.2.Columns_AM_to_BW

| | | | | | | | | | | | | | | | | | | | | | | | | | | | | | | | | | | | | | |
|---|---|---|---|---|---|---|---|---|---|---|---|---|---|---|---|---|---|---|---|---|---|---|---|---|---|---|---|---|---|---|---|---|---|---|---|---|---|
| SRR2487419 | 129 | 0 | 1315 | 0 | 46 | 92 | 0 | 0 | 0 | 0 | 0 | 15 | 0 | 23 | 2975 | 0 | 0 | 0 | 0 | 186 | 325 | 0 | 0 | 0 | 0 | 26 | 230 | 0 | 23 | 0 | 115 | 46 | 0 | 23 | 13 | 0 | 92 |
| SRR2487420 | 117 | 0 | 1316 | 0 | 40 | 119 | 0 | 0 | 40 | 0 | 0 | 21 | 0 | 20 | 2800 | 0 | 10 | 0 | 0 | 41 | 412 | 0 | 0 | 0 | 60 | 28 | 199 | 40 | 20 | 0 | 119 | 0 | 0 | 20 | 10 | 0 | 99 |
| SRR2487422 | 73 | 0 | 745 | 25 | 74 | 49 | 0 | 0 | 98 | 0 | 0 | 74 | 0 | 0 | 1234 | 0 | 0 | 0 | 0 | 99 | 2862 | 25 | 0 | 0 | 49 | 15 | 319 | 0 | 0 | 0 | 49 | 0 | 0 | 25 | 10 | 0 | 196 |
| SRR2487423 | 76 | 0 | 713 | 0 | 55 | 28 | 0 | 0 | 28 | 0 | 0 | 66 | 0 | 0 | 846 | 0 | 12 | 0 | 0 | 169 | 1862 | 28 | 0 | 55 | 0 | 24 | 524 | 55 | 0 | 0 | 28 | 0 | 0 | 0 | 469 | 0 | 55 |
| SRR2487424 | 75 | 0 | 723 | 0 | 167 | 24 | 0 | 0 | 48 | 0 | 0 | 68 | 0 | 0 | 1807 | 24 | 0 | 0 | 0 | 301 | 2811 | 0 | 0 | 0 | 72 | 22 | 239 | 24 | 24 | 0 | 120 | 0 | 0 | 24 | 13 | 24 | 215 |
| SRR2487426 | 112 | 12 | 1257 | 12 | 24 | 177 | 42 | 0 | 106 | 0 | 0 | 29 | 0 | 24 | 5994 | 12 | 12 | 11 | 0 | 42 | 3621 | 0 | 0 | 0 | 12 | 72 | 284 | 0 | 12 | 12 | 154 | 0 | 0 | 12 | 17 | 0 | 236 |
| SRR2487427 | 106 | 27 | 1288 | 27 | 54 | 121 | 95 | 0 | 54 | 0 | 0 | 36 | 0 | 54 | 4917 | 0 | 13 | 0 | 0 | 167 | 3031 | 27 | 0 | 0 | 27 | 62 | 282 | 0 | 27 | 0 | 148 | 0 | 0 | 0 | 350 | 13 | 296 |
| SRR2487428 | 99 | 12 | 1280 | 23 | 23 | 211 | 84 | 12 | 129 | 0 | 0 | 29 | 0 | 35 | 4987 | 0 | 0 | 11 | 0 | 84 | 3367 | 0 | 0 | 0 | 23 | 2479 | 386 | 12 | 35 | 0 | 152 | 0 | 0 | 0 | 491 | 12 | 222 |
| SRR2487430 | 79 | 0 | 686 | 16 | 33 | 66 | 22 | 16 | 66 | 0 | 0 | 72 | 16 | 82 | 1904 | 0 | 0 | 0 | 0 | 44 | 359 | 0 | 33 | 0 | 16 | 2140 | 181 | 16 | 0 | 0 | 230 | 0 | 0 | 0 | 675 | 16 | 263 |
| SRR2487431 | 71 | 0 | 690 | 0 | 19 | 56 | 74 | 0 | 113 | 0 | 0 | 63 | 0 | 38 | 2035 | 0 | 0 | 0 | 0 | 49 | 518 | 0 | 0 | 0 | 0 | 2269 | 188 | 19 | 0 | 0 | 319 | 0 | 0 | 0 | 825 | 19 | 244 |
| SRR2487432 | 81 | 0 | 712 | 32 | 16 | 130 | 0 | 0 | 49 | 0 | 0 | 80 | 0 | 81 | 1954 | 0 | 0 | 0 | 0 | 22 | 521 | 0 | 16 | 16 | 0 | 28 | 309 | 0 | 0 | 0 | 244 | 0 | 0 | 0 | 11 | 0 | 292 |
| SRR2487434 | 39 | 0 | 604 | 0 | 27 | 95 | 19 | 0 | 14 | 0 | 0 | 14 | 0 | 34 | 869 | 0 | 24 | 0 | 0 | 48 | 76 | 0 | 0 | 0 | 0 | 992 | 162 | 0 | 0 | 0 | 68 | 0 | 0 | 14 | 223 | 0 | 41 |
| SRR2487435 | 40 | 0 | 623 | 23 | 39 | 54 | 22 | 0 | 0 | 0 | 0 | 13 | 0 | 0 | 705 | 0 | 24 | 0 | 0 | 33 | 158 | 0 | 0 | 0 | 0 | 20 | 171 | 0 | 0 | 0 | 31 | 0 | 0 | 23 | 311 | 16 | 62 |
| SRR2487436 | 43 | 0 | 573 | 27 | 0 | 46 | 33 | 0 | 0 | 0 | 0 | 14 | 0 | 33 | 977 | 0 | 27 | 0 | 0 | 33 | 138 | 0 | 0 | 0 | 27 | 25 | 80 | 0 | 0 | 0 | 86 | 0 | 0 | 0 | 219 | 0 | 60 |
| SRR2487438 | 31 | 11 | 413 | 0 | 22 | 100 | 0 | 0 | 33 | 0 | 0 | 48 | 22 | 45 | 315 | 0 | 0 | 0 | 0 | 10 | 519 | 0 | 0 | 11 | 22 | 28 | 245 | 11 | 0 | 0 | 112 | 0 | 0 | 11 | 391 | 0 | 100 |
| SRR2487439 | 32 | 0 | 415 | 0 | 25 | 64 | 0 | 0 | 64 | 0 | 0 | 46 | 13 | 51 | 295 | 0 | 0 | 0 | 0 | 23 | 699 | 0 | 0 | 0 | 76 | 1169 | 229 | 13 | 0 | 0 | 114 | 0 | 0 | 0 | 178 | 13 | 140 |
| SRR2487441 | 114 | 0 | 1644 | 13 | 0 | 26 | 0 | 0 | 0 | 0 | 0 | 34 | 0 | 0 | 4162 | 0 | 26 | 0 | 41 | 775 | 4856 | 13 | 0 | 0 | 0 | 1280 | 758 | 39 | 0 | 0 | 118 | 0 | 13 | 13 | 222 | 0 | 157 |
| SRR2487443 | 105 | 0 | 1602 | 0 | 0 | 40 | 42 | 13 | 0 | 0 | 0 | 33 | 13 | 0 | 3956 | 0 | 22 | 0 | 0 | 250 | 4581 | 0 | 0 | 0 | 0 | 1222 | 940 | 27 | 0 | 0 | 27 | 0 | 0 | 13 | 403 | 0 | 161 |
| SRR2487445 | 108 | 0 | 1533 | 0 | 0 | 51 | 0 | 0 | 67 | 0 | 0 | 32 | 17 | 17 | 4180 | 0 | 19 | 0 | 0 | 265 | 5080 | 17 | 0 | 0 | 0 | 1634 | 943 | 0 | 0 | 0 | 34 | 0 | 0 | 17 | 303 | 0 | 168 |
| SRR2487447 | 137 | 0 | 1591 | 0 | 0 | 83 | 51 | 0 | 17 | 0 | 0 | 30 | 0 | 17 | 4789 | 0 | 26 | 0 | 0 | 257 | 4583 | 17 | 0 | 17 | 0 | 1330 | 1031 | 0 | 0 | 0 | 67 | 0 | 0 | 0 | 283 | 17 | 133 |
| SRR2487449 | 53 | 0 | 558 | 0 | 11 | 157 | 30 | 0 | 123 | 0 | 0 | 27 | 0 | 0 | 443 | 0 | 0 | 0 | 12 | 12 | 2234 | 22 | 0 | 0 | 34 | 4951 | 1467 | 11 | 0 | 0 | 78 | 0 | 0 | 0 | 381 | 11 | 358 |
| SRR2487451 | 52 | 0 | 553 | 0 | 0 | 172 | 12 | 0 | 138 | 0 | 0 | 37 | 23 | 11 | 335 | 0 | 0 | 0 | 0 | 0 | 2135 | 0 | 0 | 0 | 0 | 136 | 33 | 0 | 0 | 0 | 92 | 0 | 0 | 11 | 12 | 11 | 332 |
| SRR2487453 | 42 | 0 | 486 | 0 | 0 | 72 | 23 | 0 | 115 | 0 | 0 | 34 | 0 | 14 | 398 | 0 | 0 | 0 | 39 | 0 | 2037 | 0 | 0 | 0 | 0 | 132 | 27 | 0 | 0 | 0 | 86 | 0 | 0 | 0 | 11 | 14 | 158 |
| SRR2487455 | 45 | 0 | 552 | 0 | 0 | 184 | 15 | 14 | 28 | 0 | 0 | 37 | 0 | 14 | 477 | 0 | 0 | 0 | 15 | 15 | 1999 | 0 | 0 | 0 | 0 | 118 | 34 | 0 | 0 | 0 | 113 | 0 | 14 | 0 | 383 | 0 | 255 |
| SRR2487457 | 41 | 0 | 1452 | 0 | 0 | 122 | 62 | 14 | 27 | 0 | 0 | 18 | 0 | 0 | 3662 | 0 | 25 | 25 | 0 | 228 | 4304 | 14 | 0 | 0 | 0 | 3699 | 1003 | 14 | 0 | 14 | 108 | 0 | 0 | 54 | 203 | 14 | 122 |
| SRR2487459 | 43 | 0 | 1509 | 14 | 0 | 128 | 22 | 28 | 0 | 0 | 0 | 13 | 0 | 14 | 3638 | 0 | 24 | 0 | 0 | 307 | 4668 | 14 | 0 | 0 | 0 | 3732 | 1040 | 14 | 0 | 28 | 100 | 0 | 0 | 0 | 185 | 0 | 157 |
| SRR2487461 | 60 | 0 | 1318 | 0 | 0 | 180 | 29 | 0 | 54 | 0 | 0 | 23 | 0 | 0 | 3981 | 0 | 32 | 0 | 0 | 205 | 4507 | 18 | 0 | 0 | 0 | 4634 | 1118 | 18 | 0 | 36 | 126 | 0 | 18 | 18 | 162 | 0 | 90 |
| SRR2487463 | 57 | 0 | 1372 | 0 | 0 | 106 | 28 | 0 | 18 | 0 | 0 | 20 | 0 | 0 | 3878 | 18 | 46 | 0 | 0 | 138 | 4648 | 35 | 0 | 0 | 0 | 3995 | 866 | 18 | 0 | 18 | 88 | 0 | 18 | 18 | 141 | 18 | 141 |
| SRR2487465 | 51 | 0 | 481 | 0 | 0 | 100 | 0 | 17 | 0 | 0 | 0 | 50 | 0 | 33 | 979 | 0 | 0 | 0 | 15 | 37 | 4918 | 0 | 0 | 17 | 0 | 2707 | 1611 | 0 | 0 | 0 | 166 | 0 | 0 | 0 | 581 | 0 | 249 |
| SRR2487467 | 49 | 0 | 450 | 0 | 17 | 69 | 0 | 17 | 0 | 0 | 0 | 40 | 0 | 0 | 995 | 0 | 0 | 0 | 15 | 30 | 5286 | 0 | 0 | 0 | 0 | 51 | 23 | 0 | 0 | 0 | 103 | 0 | 0 | 0 | 16 | 51 | 172 |
| SRR2487469 | 50 | 0 | 452 | 0 | 0 | 150 | 29 | 0 | 64 | 0 | 0 | 41 | 0 | 21 | 988 | 0 | 0 | 0 | 19 | 0 | 5303 | 0 | 0 | 0 | 43 | 3675 | 1419 | 21 | 0 | 0 | 150 | 0 | 0 | 0 | 301 | 0 | 301 |
| SRR2487471 | 53 | 21 | 428 | 0 | 0 | 85 | 0 | 0 | 0 | 0 | 0 | 47 | 0 | 21 | 909 | 0 | 0 | 0 | 0 | 0 | 5245 | 0 | 0 | 0 | 21 | 46 | 20 | 0 | 0 | 0 | 128 | 0 | 0 | 0 | 449 | 21 | 256 |
| SRR2487473 | 73 | 0 | 469 | 0 | 0 | 220 | 0 | 0 | 34 | 0 | 0 | 0 | 0 | 0 | 7205 | 0 | 0 | 0 | 0 | 2070 | 7950 | 0 | 0 | 0 | 0 | 106 | 42 | 17 | 0 | 0 | 288 | 0 | 0 | 0 | 13 | 17 | 475 |
| SRR2487475 | 96 | 0 | 423 | 0 | 0 | 244 | 42 | 0 | 70 | 0 | 0 | 0 | 0 | 0 | 6953 | 0 | 11 | 0 | 42 | 1550 | 8335 | 0 | 0 | 0 | 0 | 6652 | 2908 | 0 | 17 | 0 | 331 | 0 | 0 | 0 | 818 | 17 | 383 |
| SRR2487477 | 89 | 22 | 412 | 0 | 0 | 219 | 0 | 0 | 0 | 0 | 0 | 0 | 0 | 0 | 7130 | 0 | 16 | 0 | 0 | 1662 | 7237 | 0 | 0 | 0 | 0 | 6916 | 3217 | 0 | 0 | 0 | 241 | 0 | 0 | 0 | 1094 | 22 | 350 |
| SRR2487479 | 81 | 0 | 409 | 22 | 0 | 196 | 0 | 0 | 22 | 0 | 0 | 0 | 0 | 0 | 7192 | 0 | 0 | 0 | 52 | 1407 | 6723 | 22 | 0 | 0 | 0 | 6628 | 2781 | 22 | 22 | 0 | 348 | 0 | 0 | 0 | 1043 | 22 | 239 |
| SRR2487481 | 58 | 0 | 661 | 14 | 0 | 266 | 0 | 0 | 126 | 0 | 0 | 34 | 0 | 0 | 1130 | 0 | 0 | 0 | 39 | 26 | 5549 | 0 | 0 | 0 | 0 | 215 | 46 | 0 | 0 | 0 | 196 | 0 | 0 | 0 | 20 | 0 | 266 |
| SRR2487483 | 53 | 0 | 686 | 14 | 0 | 144 | 39 | 0 | 43 | 0 | 0 | 38 | 0 | 0 | 1352 | 0 | 11 | 0 | 13 | 51 | 4918 | 0 | 0 | 0 | 14 | 213 | 52 | 0 | 0 | 0 | 215 | 0 | 0 | 0 | 21 | 29 | 244 |
| SRR2487485 | 48 | 0 | 566 | 0 | 0 | 198 | 17 | 0 | 36 | 0 | 0 | 44 | 0 | 36 | 1161 | 0 | 0 | 0 | 17 | 34 | 5634 | 18 | 0 | 0 | 0 | 8094 | 1767 | 18 | 0 | 0 | 252 | 0 | 0 | 0 | 523 | 0 | 288 |
| SRR2487487 | 71 | 0 | 643 | 0 | 0 | 179 | 0 | 0 | 36 | 0 | 0 | 51 | 0 | 0 | 1273 | 0 | 0 | 0 | 33 | 17 | 5044 | 0 | 0 | 0 | 0 | 7079 | 1918 | 0 | 0 | 0 | 251 | 0 | 0 | 0 | 412 | 18 | 251 |
| SRR2487489 | 153 | 0 | 2573 | 0 | 0 | 22 | 41 | 62 | 0 | 0 | 0 | 41 | 0 | 0 | 151 | 0 | 27 | 0 | 0 | 76 | 232 | 0 | 0 | 0 | 0 | 818 | 581 | 0 | 34 | 37 | 41 | 0 | 0 | 31 | 91 | 0 | 62 |
| SRR2487491 | 119 | 0 | 2750 | 0 | 0 | 35 | 0 | 29 | 0 | 0 | 0 | 28 | 0 | 0 | 138 | 0 | 11 | 0 | 0 | 42 | 306 | 0 | 0 | 0 | 0 | 128 | 102 | 0 | 55 | 13 | 42 | 0 | 0 | 32 | 58 | 0 | 10 |
| SRR2487493 | 165 | 0 | 2250 | 0 | 0 | 43 | 22 | 59 | 0 | 0 | 0 | 27 | 0 | 0 | 177 | 0 | 21 | 0 | 0 | 66 | 250 | 0 | 0 | 0 | 0 | 123 | 83 | 0 | 39 | 12 | 31 | 0 | 0 | 31 | 11 | 0 | 43 |
| SRR2487495 | 150 | 0 | 2507 | 0 | 0 | 36 | 0 | 44 | 20 | 0 | 0 | 30 | 0 | 0 | 157 | 0 | 17 | 0 | 0 | 50 | 286 | 0 | 0 | 0 | 0 | 1020 | 660 | 0 | 40 | 20 | 20 | 0 | 0 | 16 | 68 | 0 | 52 |
| SRR2487497 | 40 | 0 | 550 | 0 | 0 | 198 | 30 | 0 | 52 | 0 | 0 | 57 | 0 | 0 | 510 | 0 | 11 | 0 | 12 | 71 | 1530 | 0 | 0 | 0 | 43 | 245 | 70 | 34 | 0 | 0 | 155 | 0 | 0 | 0 | 207 | 17 | 86 |
| SRR2487499 | 44 | 0 | 557 | 0 | 0 | 231 | 37 | 0 | 35 | 0 | 0 | 63 | 0 | 0 | 399 | 0 | 0 | 0 | 0 | 49 | 1499 | 0 | 0 | 0 | 71 | 238 | 73 | 18 | 18 | 0 | 266 | 0 | 0 | 0 | 142 | 0 | 106 |
| SRR2487501 | 31 | 0 | 471 | 33 | 0 | 188 | 0 | 0 | 55 | 0 | 0 | 49 | 11 | 0 | 558 | 0 | 0 | 0 | 0 | 38 | 1444 | 0 | 0 | 11 | 33 | 6960 | 1782 | 44 | 0 | 0 | 321 | 0 | 0 | 11 | 144 | 22 | 122 |
| SRR2487503 | 46 | 0 | 500 | 0 | 0 | 121 | 22 | 11 | 22 | 0 | 0 | 51 | 0 | 0 | 641 | 0 | 0 | 0 | 0 | 15 | 1603 | 22 | 0 | 11 | 66 | 6396 | 2114 | 33 | 0 | 0 | 231 | 0 | 0 | 11 | 154 | 11 | 154 |
| SRR2487504 | 55 | 0 | 1455 | 0 | 0 | 480 | 0 | 0 | 60 | 0 | 0 | 19 | 0 | 0 | 476 | 0 | 19 | 0 | 0 | 272 | 816 | 0 | 0 | 0 | 0 | 6957 | 2219 | 0 | 0 | 0 | 120 | 0 | 0 | 0 | 360 | 0 | 120 |
| SRR2487505 | 67 | 0 | 1180 | 0 | 0 | 248 | 0 | 0 | 0 | 0 | 0 | 12 | 0 | 0 | 476 | 0 | 42 | 26 | 0 | 106 | 1958 | 0 | 0 | 0 | 0 | 165 | 62 | 0 | 0 | 0 | 0 | 0 | 0 | 41 | 289 | 0 | 124 |
| SRR2487507 | 58 | 0 | 1377 | 0 | 0 | 165 | 0 | 0 | 0 | 0 | 0 | 11 | 0 | 0 | 255 | 0 | 24 | 0 | 0 | 102 | 1480 | 0 | 0 | 0 | 0 | 6612 | 1777 | 0 | 0 | 0 | 248 | 0 | 0 | 41 | 248 | 41 | 83 |
| SRR2487508 | 72 | 0 | 1431 | 63 | 0 | 379 | 71 | 0 | 0 | 0 | 0 | 16 | 0 | 0 | 1283 | 0 | 16 | 0 | 0 | 143 | 927 | 0 | 0 | 0 | 0 | 6259 | 1833 | 0 | 0 | 0 | 126 | 0 | 0 | 0 | 443 | 63 | 253 |
| SRR2487509 | 76 | 0 | 1310 | 0 | 66 | 11 | 75 | 0 | 0 | 0 | 0 | 24 | 0 | 0 | 601 | 0 | 24 | 0 | 0 | 225 | 751 | 0 | 0 | 0 | 0 | 142 | 41 | 66 | 0 | 0 | 264 | 0 | 0 | 0 | 330 | 0 | 132 |
| SRR2487510 | 80 | 0 | 1484 | 68 | 0 | 405 | 149 | 0 | 0 | 0 | 0 | 13 | 0 | 0 | 1268 | 0 | 30 | 28 | 0 | 149 | 1044 | 0 | 0 | 0 | 0 | 124 | 33 | 0 | 0 | 0 | 0 | 0 | 0 | 0 | 11 | 0 | 135 |
| SRR2487512 | 64 | 0 | 1423 | 0 | 0 | 256 | 78 | 0 | 32 | 0 | 0 | 18 | 0 | 32 | 626 | 0 | 20 | 0 | 0 | 235 | 1486 | 0 | 0 | 0 | 0 | 188 | 61 | 0 | 0 | 0 | 64 | 0 | 0 | 0 | 288 | 0 | 96 |
| SRR2487514 | 57 | 0 | 1370 | 33 | 0 | 233 | 82 | 0 | 33 | 0 | 0 | 0 | 0 | 0 | 368 | 0 | 35 | 0 | 0 | 245 | 1552 | 0 | 0 | 0 | 33 | 150 | 56 | 0 | 0 | 0 | 166 | 0 | 0 | 0 | 299 | 33 | 166 |
| SRR2487516 | 38 | 0 | 550 | 0 | 0 | 101 | 0 | 0 | 0 | 0 | 0 | 64 | 0 | 50 | 479 | 0 | 0 | 0 | 44 | 0 | 2132 | 0 | 0 | 0 | 0 | 1859 | 653 | 0 | 0 | 0 | 352 | 0 | 0 | 0 | 452 | 0 | 151 |
| SRR2487517 | 23 | 0 | 547 | 0 | 0 | 33 | 0 | 0 | 0 | 0 | 0 | 44 | 0 | 0 | 379 | 0 | 0 | 0 | 0 | 29 | 3115 | 0 | 0 | 0 | 0 | 62 | 21 | 98 | 0 | 0 | 163 | 0 | 0 | 0 | 457 | 0 | 392 |
| SRR2487519 | 39 | 0 | 551 | 0 | 0 | 32 | 0 | 0 | 0 | 0 | 0 | 32 | 0 | 0 | 546 | 0 | 0 | 0 | 0 | 29 | 3448 | 0 | 0 | 0 | 0 | 1554 | 1036 | 0 | 0 | 32 | 129 | 0 | 0 | 0 | 259 | 0 | 227 |
| SRR2487520 | 45 | 0 | 517 | 0 | 0 | 103 | 0 | 0 | 51 | 0 | 0 | 58 | 0 | 0 | 678 | 0 | 0 | 0 | 0 | 0 | 2714 | 0 | 0 | 0 | 0 | 2105 | 411 | 0 | 0 | 0 | 205 | 0 | 0 | 0 | 411 | 0 | 51 |
| SRR2487521 | 50 | 0 | 612 | 0 | 0 | 56 | 0 | 0 | 0 | 0 | 0 | 50 | 0 | 0 | 803 | 0 | 0 | 73 | 50 | 50 | 2360 | 0 | 0 | 0 | 0 | 39 | 28 | 56 | 0 | 0 | 0 | 0 | 0 | 0 | 16 | 0 | 223 |
| SRR2487522 | 32 | 0 | 643 | 0 | 0 | 0 | 0 | 0 | 0 | 0 | 0 | 51 | 0 | 0 | 727 | 0 | 14 | 72 | 0 | 48 | 1744 | 0 | 0 | 0 | 0 | 2417 | 899 | 0 | 0 | 0 | 112 | 0 | 0 | 0 | 450 | 0 | 281 |
| SRR2487524 | 40 | 0 | 517 | 0 | 0 | 127 | 0 | 0 | 0 | 0 | 0 | 62 | 0 | 0 | 542 | 0 | 0 | 0 | 22 | 0 | 3449 | 0 | 0 | 0 | 0 | 2233 | 964 | 0 | 0 | 0 | 127 | 25 | 0 | 0 | 508 | 0 | 127 |
| SRR2487526 | 49 | 0 | 530 | 0 | 0 | 182 | 0 | 0 | 52 | 0 | 0 | 49 | 0 | 26 | 536 | 0 | 0 | 42 | 22 | 45 | 3438 | 26 | 0 | 26 | 52 | 2084 | 1120 | 0 | 0 | 0 | 156 | 0 | 0 | 0 | 313 | 0 | 260 |
| SRR2487529 | 47 | 0 | 951 | 0 | 0 | 35 | 81 | 0 | 18 | 0 | 0 | 12 | 0 | 0 | 3525 | 0 | 21 | 0 | 27 | 203 | 2102 | 0 | 0 | 0 | 0 | 34 | 19 | 0 | 0 | 0 | 89 | 0 | 0 | 44 | 11 | 53 | 231 |
| SRR2487531 | 43 | 0 | 970 | 0 | 0 | 55 | 14 | 0 | 0 | 0 | 0 | 0 | 0 | 0 | 3633 | 0 | 19 | 11 | 14 | 224 | 2222 | 0 | 0 | 0 | 0 | 40 | 18 | 0 | 0 | 0 | 92 | 0 | 0 | 0 | 15 | 0 | 293 |
| SRR2487533 | 43 | 0 | 890 | 0 | 0 | 125 | 71 | 0 | 11 | 0 | 0 | 10 | 0 | 0 | 3428 | 0 | 18 | 0 | 0 | 300 | 2085 | 11 | 0 | 0 | 0 | 1826 | 850 | 0 | 0 | 0 | 91 | 0 | 0 | 0 | 590 | 11 | 249 |
| SRR2487535 | 42 | 0 | 898 | 0 | 0 | 67 | 69 | 0 | 11 | 0 | 0 | 13 | 0 | 0 | 4265 | 0 | 20 | 0 | 0 | 225 | 2341 | 0 | 0 | 0 | 0 | 34 | 19 | 11 | 0 | 0 | 34 | 0 | 0 | 22 | 11 | 56 | 336 |
| SRR2487537 | 38 | 0 | 582 | 0 | 0 | 91 | 39 | 15 | 76 | 0 | 0 | 50 | 15 | 0 | 232 | 0 | 14 | 0 | 0 | 23 | 2176 | 0 | 0 | 0 | 0 | 3005 | 850 | 15 | 0 | 0 | 243 | 0 | 0 | 0 | 334 | 30 | 258 |
| SRR2487539 | 35 | 0 | 573 | 0 | 0 | 47 | 32 | 0 | 32 | 0 | 0 | 38 | 0 | 47 | 268 | 0 | 0 | 0 | 0 | 55 | 1986 | 0 | 0 | 0 | 0 | 2821 | 1008 | 16 | 0 | 0 | 126 | 0 | 0 | 0 | 299 | 47 | 221 |
| SRR2487541 | 32 | 0 | 562 | 0 | 0 | 98 | 20 | 0 | 20 | 0 | 0 | 36 | 0 | 20 | 278 | 0 | 16 | 0 | 30 | 20 | 1997 | 0 | 0 | 0 | 0 | 2952 | 997 | 20 | 0 | 0 | 98 | 0 | 0 | 0 | 352 | 20 | 176 |
| SRR2487543 | 41 | 0 | 551 | 0 | 0 | 116 | 19 | 0 | 39 | 0 | 0 | 35 | 0 | 0 | 360 | 0 | 0 | 0 | 0 | 19 | 2148 | 19 | 0 | 19 | 0 | 61 | 20 | 19 | 0 | 0 | 116 | 0 | 0 | 0 | 446 | 58 | 136 |
| SRR2487545 | 104 | 0 | 1426 | 0 | 0 | 183 | 61 | 0 | 0 | 0 | 0 | 14 | 0 | 0 | 101 | 0 | 29 | 0 | 0 | 37 | 57 | 31 | 0 | 0 | 0 | 25 | 18 | 0 | 0 | 0 | 61 | 0 | 0 | 31 | 764 | 31 | 397 |
| SRR2487547 | 91 | 0 | 1493 | 31 | 0 | 156 | 42 | 93 | 0 | 0 | 0 | 23 | 0 | 0 | 121 | 0 | 29 | 0 | 0 | 45 | 21 | 0 | 0 | 0 | 0 | 23 | 17 | 0 | 0 | 0 | 31 | 0 | 0 | 31 | 748 | 31 | 280 |
| SRR2487549 | 114 | 0 | 1238 | 0 | 0 | 0 | 17 | 118 | 0 | 0 | 0 | 31 | 0 | 0 | 111 | 0 | 18 | 0 | 0 | 43 | 51 | 39 | 0 | 0 | 39 | 2511 | 1334 | 0 | 0 | 0 | 196 | 0 | 0 | 0 | 824 | 0 | 392 |
| SRR2487551 | 88 | 0 | 1356 | 0 | 0 | 117 | 38 | 0 | 0 | 0 | 0 | 17 | 0 | 39 | 113 | 0 | 19 | 0 | 0 | 33 | 50 | 0 | 0 | 0 | 0 | 23 | 17 | 0 | 0 | 0 | 195 | 0 | 0 | 0 | 10 | 39 | 428 |
| SRR2487553 | 58 | 0 | 403 | 0 | 0 | 101 | 203 | 404 | 0 | 0 | 0 | 115 | 0 | 0 | 0 | 0 | 0 | 0 | 11 | 133 | 339 | 0 | 0 | 81 | 0 | 42 | 49 | 0 | 0 | 0 | 40 | 0 | 0 | 0 | 445 | 0 | 465 |
| SRR2487555 | 78 | 0 | 566 | 0 | 0 | 103 | 169 | 538 | 62 | 0 | 0 | 72 | 0 | 0 | 11 | 0 | 13 | 0 | 11 | 180 | 347 | 0 | 0 | 0 | 0 | 41 | 46 | 0 | 0 | 0 | 0 | 0 | 0 | 0 | 269 | 62 | 10 |
| SRR2487560 | 112 | 0 | 1406 | 0 | 0 | 208 | 36 | 69 | 69 | 0 | 0 | 60 | 0 | 0 | 516 | 0 | 37 | 0 | 0 | 107 | 142 | 0 | 0 | 0 | 0 | 8854 | 1937 | 0 | 0 | 0 | 138 | 69 | 0 | 0 | 277 | 0 | 138 |
| SRR2487561 | 212 | 0 | 1562 | 0 | 0 | 12 | 57 | 151 | 114 | 0 | 0 | 59 | 0 | 0 | 26 | 0 | 25 | 0 | 0 | 307 | 400 | 0 | 0 | 38 | 0 | 290 | 116 | 0 | 0 | 0 | 19 | 0 | 0 | 0 | 151 | 19 | 189 |
| SRR2487563 | 190 | 0 | 1849 | 0 | 0 | 15 | 31 | 233 | 78 | 0 | 0 | 74 | 0 | 0 | 41 | 0 | 50 | 0 | 10 | 232 | 392 | 0 | 0 | 39 | 0 | 278 | 115 | 0 | 0 | 0 | 19 | 0 | 0 | 19 | 117 | 19 | 155 |
| SRR2487564 | 106 | 0 | 1520 | 0 | 0 | 447 | 19 | 0 | 74 | 0 | 0 | 39 | 0 | 74 | 412 | 0 | 12 | 0 | 0 | 131 | 206 | 0 | 0 | 0 | 0 | 8484 | 1637 | 0 | 0 | 0 | 223 | 0 | 0 | 0 | 744 | 0 | 223 |
| SRR2487565 | 101 | 0 | 1357 | 0 | 0 | 844 | 20 | 0 | 77 | 0 | 0 | 73 | 0 | 0 | 451 | 0 | 28 | 0 | 0 | 98 | 196 | 0 | 0 | 0 | 0 | 8748 | 2225 | 77 | 0 | 0 | 307 | 0 | 0 | 77 | 384 | 0 | 460 |
| SRR2487566 | 94 | 0 | 1397 | 0 | 0 | 10 | 0 | 0 | 157 | 0 | 0 | 59 | 0 | 79 | 473 | 0 | 31 | 0 | 0 | 118 | 256 | 0 | 0 | 0 | 0 | 105 | 32 | 0 | 0 | 0 | 393 | 0 | 0 | 0 | 10 | 79 | 79 |
| SRR2487568 | 240 | 0 | 1988 | 0 | 0 | 412 | 70 | 191 | 44 | 0 | 0 | 48 | 0 | 0 | 17 | 0 | 21 | 0 | 12 | 331 | 384 | 0 | 0 | 15 | 0 | 7952 | 3131 | 0 | 0 | 0 | 15 | 0 | 0 | 0 | 220 | 29 | 250 |
| SRR2487570 | 141 | 0 | 2151 | 0 | 0 | 14 | 38 | 213 | 46 | 0 | 0 | 78 | 0 | 0 | 21 | 0 | 14 | 0 | 0 | 316 | 469 | 0 | 0 | 15 | 0 | 302 | 149 | 0 | 0 | 0 | 0 | 0 | 0 | 15 | 213 | 15 | 137 |
| SRR2487572 | 38 | 0 | 536 | 0 | 0 | 260 | 41 | 0 | 32 | 0 | 0 | 35 | 0 | 0 | 368 | 0 | 0 | 0 | 0 | 65 | 327 | 0 | 0 | 0 | 0 | 6590 | 1266 | 32 | 0 | 0 | 162 | 0 | 0 | 0 | 941 | 32 | 260 |
| SRR2487573 | 94 | 0 | 594 | 0 | 0 | 298 | 31 | 239 | 0 | 0 | 0 | 47 | 0 | 0 | 15 | 0 | 0 | 0 | 15 | 199 | 566 | 0 | 0 | 0 | 0 | 6800 | 2476 | 0 | 0 | 30 | 30 | 0 | 0 | 0 | 298 | 0 | 447 |
| SRR2487575 | 128 | 0 | 722 | 0 | 0 | 180 | 15 | 300 | 120 | 0 | 0 | 82 | 0 | 0 | 30 | 0 | 0 | 0 | 23 | 106 | 551 | 0 | 0 | 60 | 0 | 6539 | 2880 | 0 | 0 | 0 | 0 | 0 | 0 | 0 | 570 | 60 | 360 |
| SRR2487576 | 40 | 0 | 614 | 0 | 0 | 201 | 17 | 34 | 34 | 0 | 0 | 42 | 0 | 0 | 313 | 0 | 15 | 0 | 0 | 59 | 491 | 0 | 0 | 0 | 0 | 7370 | 1474 | 0 | 0 | 0 | 168 | 0 | 0 | 0 | 804 | 34 | 436 |
| SRR2487577 | 54 | 0 | 551 | 36 | 0 | 362 | 0 | 0 | 72 | 0 | 0 | 37 | 0 | 0 | 293 | 0 | 13 | 0 | 0 | 37 | 311 | 0 | 0 | 0 | 72 | 6741 | 1305 | 0 | 0 | 0 | 36 | 0 | 0 | 0 | 761 | 36 | 399 |
| SRR2487578 | 40 | 0 | 661 | 0 | 0 | 255 | 0 | 36 | 36 | 0 | 0 | 40 | 0 | 0 | 321 | 0 | 0 | 0 | 0 | 73 | 476 | 0 | 0 | 0 | 0 | 6493 | 1094 | 0 | 0 | 0 | 182 | 0 | 0 | 0 | 766 | 0 | 292 |
| SRR2487580 | 44 | 0 | 819 | 0 | 0 | 159 | 24 | 68 | 0 | 0 | 0 | 44 | 0 | 0 | 18 | 0 | 0 | 0 | 12 | 130 | 549 | 0 | 0 | 0 | 0 | 227 | 70 | 0 | 0 | 0 | 23 | 0 | 0 | 23 | 295 | 23 | 136 |
| SRR2487582 | 76 | 0 | 884 | 0 | 0 | 324 | 18 | 139 | 116 | 0 | 0 | 61 | 0 | 0 | 36 | 0 | 23 | 0 | 12 | 151 | 627 | 0 | 0 | 0 | 0 | 7162 | 2526 | 0 | 0 | 0 | 0 | 0 | 0 | 0 | 301 | 23 | 93 |
| SRR2487584 | 152 | 0 | 1024 | 0 | 0 | 25 | 0 | 0 | 0 | 0 | 0 | 29 | 0 | 0 | 3671 | 0 | 0 | 0 | 0 | 432 | 1242 | 0 | 0 | 25 | 0 | 148 | 104 | 0 | 0 | 0 | 22 | 0 | 0 | 0 | 47 | 0 | 33 |
| SRR2487585 | 246 | 0 | 1320 | 0 | 0 | 23 | 0 | 0 | 0 | 0 | 0 | 45 | 0 | 0 | 77 | 0 | 0 | 0 | 0 | 922 | 1767 | 0 | 0 | 0 | 0 | 179 | 284 | 0 | 0 | 0 | 0 | 0 | 0 | 0 | 10 | 0 | 10 |
| SRR2487587 | 320 | 0 | 1258 | 0 | 0 | 14 | 0 | 0 | 0 | 0 | 0 | 21 | 0 | 0 | 151 | 0 | 43 | 0 | 76 | 1211 | 2271 | 0 | 0 | 0 | 0 | 560 | 870 | 0 | 0 | 0 | 0 | 0 | 0 | 0 | 37 | 0 | 28 |
| SRR2487588 | 167 | 0 | 1026 | 0 | 0 | 23 | 0 | 0 | 0 | 0 | 0 | 28 | 0 | 19 | 2525 | 0 | 0 | 0 | 0 | 673 | 1010 | 0 | 0 | 0 | 19 | 605 | 434 | 0 | 0 | 0 | 16 | 0 | 0 | 0 | 66 | 0 | 43 |
| SRR2487589 | 148 | 0 | 1042 | 0 | 12 | 45 | 0 | 0 | 0 | 0 | 0 | 27 | 0 | 0 | 2521 | 0 | 0 | 0 | 0 | 300 | 1321 | 0 | 0 | 0 | 24 | 600 | 438 | 0 | 0 | 0 | 37 | 0 | 0 | 0 | 57 | 0 | 37 |
| SRR2487590 | 148 | 0 | 1071 | 0 | 0 | 29 | 0 | 0 | 12 | 0 | 0 | 14 | 0 | 12 | 3948 | 0 | 19 | 0 | 0 | 648 | 1473 | 0 | 0 | 0 | 33 | 538 | 447 | 0 | 0 | 0 | 0 | 0 | 0 | 0 | 70 | 0 | 12 |
| SRR2487592 | 285 | 0 | 1389 | 0 | 0 | 14 | 60 | 0 | 0 | 0 | 0 | 36 | 0 | 0 | 241 | 0 | 36 | 0 | 0 | 1267 | 2414 | 0 | 0 | 11 | 0 | 539 | 803 | 0 | 0 | 0 | 0 | 0 | 0 | 0 | 32 | 0 | 25 |
| SRR2487594 | 148 | 0 | 1294 | 0 | 0 | 26 | 0 | 0 | 0 | 0 | 0 | 37 | 0 | 0 | 184 | 0 | 18 | 0 | 0 | 1041 | 2449 | 0 | 0 | 0 | 0 | 503 | 813 | 0 | 0 | 0 | 0 | 0 | 0 | 0 | 29 | 0 | 29 |
| SRR2487597 | 73 | 0 | 629 | 21 | 21 | 169 | 13 | 0 | 63 | 0 | 0 | 46 | 0 | 11 | 1776 | 0 | 18 | 0 | 64 | 51 | 4491 | 0 | 0 | 0 | 11 | 129 | 30 | 0 | 0 | 0 | 148 | 0 | 0 | 11 | 15 | 21 | 254 |
| SRR2487598 | 64 | 0 | 643 | 0 | 0 | 226 | 0 | 0 | 102 | 0 | 0 | 48 | 11 | 11 | 1569 | 0 | 0 | 0 | 42 | 28 | 4298 | 0 | 0 | 0 | 23 | 116 | 26 | 0 | 0 | 0 | 215 | 0 | 0 | 11 | 11 | 0 | 215 |
| SRR2487599 | 71 | 0 | 606 | 0 | 0 | 182 | 0 | 0 | 102 | 0 | 0 | 40 | 0 | 34 | 1797 | 11 | 0 | 0 | 28 | 56 | 4457 | 0 | 0 | 0 | 11 | 3768 | 820 | 11 | 0 | 0 | 114 | 0 | 0 | 0 | 421 | 11 | 205 |
| SRR2487600 | 94 | 0 | 622 | 0 | 0 | 16 | 0 | 0 | 79 | 0 | 0 | 65 | 16 | 0 | 108 | 0 | 29 | 0 | 43 | 151 | 7794 | 16 | 0 | 79 | 0 | 365 | 198 | 0 | 0 | 0 | 32 | 0 | 0 | 0 | 27 | 32 | 20 |
| SRR2487602 | 112 | 0 | 786 | 0 | 0 | 24 | 0 | 0 | 81 | 0 | 0 | 120 | 0 | 0 | 154 | 0 | 30 | 0 | 0 | 309 | 7537 | 0 | 0 | 0 | 0 | 425 | 193 | 0 | 0 | 0 | 16 | 0 | 0 | 0 | 21 | 0 | 19 |
| SRR2487604 | 73 | 0 | 684 | 0 | 0 | 13 | 0 | 0 | 20 | 0 | 0 | 64 | 0 | 20 | 175 | 0 | 0 | 0 | 58 | 292 | 8759 | 0 | 0 | 0 | 0 | 400 | 186 | 0 | 0 | 0 | 20 | 0 | 0 | 0 | 15 | 0 | 16 |
| SRR2487606 | 124 | 0 | 807 | 0 | 0 | 21 | 0 | 0 | 41 | 0 | 0 | 98 | 0 | 0 | 114 | 0 | 27 | 0 | 86 | 257 | 8292 | 0 | 0 | 21 | 0 | 401 | 181 | 0 | 0 | 0 | 0 | 0 | 0 | 0 | 29 | 0 | 16 |
| SRR2487607 | 62 | 0 | 635 | 0 | 0 | 151 | 13 | 0 | 131 | 0 | 0 | 31 | 10 | 0 | 1801 | 0 | 14 | 0 | 25 | 75 | 4402 | 0 | 0 | 0 | 10 | 4415 | 855 | 0 | 0 | 0 | 181 | 0 | 0 | 10 | 503 | 50 | 201 |
| SRR2487609 | 69 | 0 | 1015 | 35 | 0 | 69 | 151 | 12 | 23 | 0 | 0 | 0 | 0 | 0 | 534 | 0 | 14 | 0 | 0 | 375 | 191 | 23 | 0 | 0 | 0 | 61 | 28 | 12 | 0 | 0 | 69 | 0 | 0 | 0 | 15 | 12 | 207 |
| SRR2487610 | 59 | 12 | 998 | 23 | 0 | 69 | 191 | 12 | 35 | 0 | 0 | 0 | 0 | 0 | 415 | 0 | 21 | 0 | 0 | 290 | 141 | 12 | 0 | 0 | 12 | 72 | 24 | 12 | 0 | 0 | 104 | 0 | 0 | 0 | 13 | 0 | 93 |
| SRR2487611 | 74 | 0 | 1064 | 23 | 12 | 94 | 217 | 0 | 70 | 0 | 0 | 0 | 0 | 12 | 409 | 0 | 19 | 0 | 0 | 359 | 175 | 23 | 0 | 0 | 0 | 2324 | 669 | 0 | 0 | 0 | 70 | 0 | 0 | 0 | 528 | 23 | 188 |
| SRR2487612 | 39 | 0 | 1278 | 0 | 0 | 67 | 265 | 0 | 0 | 0 | 0 | 20 | 0 | 0 | 22 | 0 | 59 | 0 | 0 | 530 | 221 | 34 | 0 | 0 | 0 | 2685 | 1712 | 0 | 0 | 0 | 0 | 0 | 0 | 0 | 302 | 0 | 134 |
| SRR2487614 | 62 | 0 | 1662 | 0 | 0 | 35 | 228 | 0 | 0 | 0 | 0 | 0 | 0 | 0 | 23 | 0 | 21 | 0 | 0 | 433 | 456 | 0 | 0 | 0 | 0 | 148 | 124 | 0 | 0 | 0 | 0 | 0 | 0 | 0 | 27 | 0 | 15 |
| SRR2487616 | 75 | 0 | 1218 | 0 | 0 | 0 | 112 | 45 | 0 | 0 | 0 | 0 | 0 | 0 | 0 | 0 | 25 | 0 | 0 | 815 | 84 | 45 | 0 | 0 | 0 | 138 | 132 | 0 | 0 | 0 | 45 | 0 | 0 | 0 | 19 | 45 | 91 |
| SRR2487618 | 73 | 0 | 1041 | 0 | 0 | 0 | 141 | 0 | 0 | 0 | 0 | 0 | 0 | 0 | 28 | 0 | 24 | 0 | 0 | 506 | 281 | 0 | 0 | 46 | 0 | 170 | 113 | 0 | 0 | 0 | 46 | 0 | 0 | 0 | 16 | 0 | 46 |
| SRR2487619 | 63 | 0 | 950 | 21 | 0 | 42 | 181 | 0 | 11 | 0 | 0 | 0 | 0 | 11 | 415 | 0 | 15 | 0 | 0 | 340 | 106 | 11 | 0 | 0 | 11 | 66 | 23 | 21 | 0 | 0 | 117 | 0 | 0 | 0 | 17 | 42 | 159 |
| SRR2487621 | 68 | 0 | 745 | 0 | 0 | 26 | 157 | 0 | 0 | 0 | 0 | 34 | 0 | 51 | 157 | 0 | 13 | 0 | 0 | 183 | 730 | 0 | 0 | 0 | 0 | 1357 | 768 | 26 | 0 | 0 | 179 | 0 | 0 | 0 | 307 | 0 | 77 |
| SRR2487622 | 65 | 0 | 724 | 27 | 0 | 55 | 56 | 0 | 27 | 0 | 0 | 26 | 0 | 27 | 112 | 0 | 0 | 0 | 0 | 223 | 558 | 0 | 0 | 0 | 0 | 1717 | 845 | 27 | 0 | 0 | 109 | 0 | 0 | 0 | 327 | 0 | 191 |
| SRR2487623 | 63 | 0 | 755 | 0 | 0 | 55 | 0 | 0 | 55 | 0 | 0 | 39 | 0 | 0 | 421 | 0 | 0 | 0 | 0 | 168 | 869 | 0 | 0 | 0 | 0 | 19 | 20 | 0 | 0 | 0 | 83 | 0 | 0 | 0 | 415 | 28 | 249 |
| SRR2487624 | 104 | 0 | 894 | 0 | 0 | 46 | 51 | 31 | 31 | 0 | 0 | 52 | 0 | 0 | 51 | 0 | 0 | 0 | 0 | 445 | 1473 | 0 | 0 | 15 | 0 | 72 | 97 | 0 | 0 | 0 | 15 | 0 | 0 | 0 | 107 | 0 | 77 |
| SRR2487626 | 100 | 0 | 967 | 0 | 0 | 31 | 18 | 16 | 16 | 0 | 0 | 46 | 0 | 0 | 18 | 0 | 0 | 0 | 0 | 370 | 1497 | 0 | 0 | 31 | 0 | 77 | 108 | 0 | 0 | 0 | 0 | 0 | 0 | 0 | 110 | 0 | 126 |
| SRR2487628 | 89 | 0 | 799 | 0 | 0 | 79 | 152 | 40 | 40 | 0 | 0 | 79 | 0 | 0 | 0 | 0 | 14 | 0 | 0 | 564 | 1280 | 0 | 0 | 20 | 0 | 75 | 94 | 0 | 20 | 0 | 20 | 0 | 0 | 0 | 11 | 40 | 79 |

### Supp Table S8.2.Columns_AM_to_BW

| | | | | | | | | | | | | | | | | | | | | | | | | | | | | | | | | | | | | | |
|---|---|---|---|---|---|---|---|---|---|---|---|---|---|---|---|---|---|---|---|---|---|---|---|---|---|---|---|---|---|---|---|---|---|---|---|---|---|
| SRR2487630 | 119 | 0 | 863 | 0 | 0 | 20 | 65 | 60 | 60 | 0 | 0 | 82 | 0 | 0 | 44 | 0 | 0 | 0 | 0 | 502 | 1288 | 0 | 0 | 20 | 0 | 1033 | 1768 | 0 | 0 | 20 | 20 | 0 | 0 | 0 | 199 | 0 | 60 |
| SRR2487631 | 65 | 0 | 729 | 0 | 0 | 24 | 0 | 0 | 48 | 0 | 0 | 28 | 0 | 24 | 199 | 0 | 12 | 0 | 0 | 124 | 893 | 0 | 0 | 0 | 0 | 28 | 18 | 0 | 0 | 0 | 217 | 0 | 0 | 0 | 338 | 0 | 121 |
| SRR2487633 | 97 | 0 | 1811 | 0 | 0 | 74 | 169 | 111 | 0 | 0 | 0 | 32 | 0 | 0 | 22 | 0 | 26 | 0 | 0 | 331 | 188 | 0 | 0 | 55 | 0 | 1218 | 1365 | 18 | 0 | 37 | 37 | 0 | 0 | 0 | 369 | 18 | 221 |
| SRR2487635 | 133 | 0 | 1727 | 0 | 0 | 19 | 204 | 154 | 0 | 0 | 0 | 20 | 0 | 0 | 13 | 0 | 13 | 0 | 0 | 250 | 191 | 0 | 0 | 39 | 0 | 28 | 41 | 0 | 0 | 39 | 19 | 0 | 0 | 0 | 11 | 19 | 367 |
| SRR2487637 | 163 | 0 | 1433 | 0 | 0 | 48 | 187 | 167 | 0 | 0 | 0 | 65 | 0 | 0 | 28 | 0 | 41 | 0 | 0 | 303 | 183 | 0 | 0 | 71 | 0 | 1547 | 1762 | 0 | 0 | 0 | 0 | 0 | 0 | 24 | 238 | 0 | 167 |
| SRR2487639 | 188 | 0 | 1538 | 0 | 0 | 48 | 135 | 169 | 0 | 0 | 0 | 63 | 0 | 0 | 0 | 0 | 24 | 0 | 0 | 227 | 155 | 0 | 0 | 48 | 0 | 23 | 34 | 0 | 0 | 24 | 0 | 0 | 0 | 0 | 290 | 0 | 338 |
| SRR2487641 | 98 | 0 | 496 | 0 | 0 | 91 | 35 | 23 | 23 | 0 | 0 | 52 | 0 | 0 | 17 | 0 | 12 | 0 | 30 | 196 | 1061 | 0 | 0 | 69 | 0 | 78 | 50 | 0 | 0 | 23 | 0 | 0 | 0 | 0 | 10 | 46 | 206 |
| SRR2487643 | 72 | 0 | 596 | 0 | 0 | 119 | 40 | 47 | 0 | 0 | 0 | 72 | 24 | 0 | 22 | 0 | 0 | 0 | 18 | 152 | 1179 | 0 | 0 | 71 | 0 | 3230 | 2517 | 0 | 0 | 24 | 24 | 0 | 0 | 0 | 427 | 24 | 285 |
| SRR2487645 | 29 | 0 | 417 | 0 | 0 | 151 | 55 | 91 | 121 | 0 | 0 | 51 | 0 | 0 | 0 | 0 | 0 | 0 | 33 | 183 | 1112 | 0 | 0 | 0 | 0 | 69 | 38 | 0 | 0 | 0 | 0 | 0 | 0 | 0 | 303 | 0 | 454 |
| SRR2487647 | 50 | 0 | 647 | 0 | 0 | 61 | 72 | 61 | 123 | 0 | 0 | 50 | 0 | 0 | 22 | 0 | 14 | 0 | 0 | 165 | 1064 | 0 | 0 | 61 | 0 | 75 | 40 | 0 | 0 | 0 | 61 | 0 | 0 | 0 | 461 | 92 | 369 |
| SRR2487648 | 55 | 0 | 1512 | 0 | 21 | 126 | 13 | 0 | 21 | 0 | 0 | 23 | 0 | 0 | 0 | 0 | 25 | 0 | 0 | 253 | 114 | 0 | 0 | 0 | 21 | 3036 | 1235 | 0 | 0 | 0 | 503 | 0 | 0 | 0 | 670 | 0 | 314 |
| SRR2487649 | 136 | 0 | 2052 | 0 | 0 | 143 | 73 | 96 | 24 | 0 | 0 | 0 | 0 | 0 | 0 | 0 | 34 | 0 | 0 | 508 | 203 | 0 | 0 | 0 | 0 | 200 | 138 | 0 | 0 | 0 | 48 | 0 | 0 | 0 | 17 | 48 | 119 |
| SRR2487651 | 74 | 0 | 2275 | 0 | 0 | 24 | 71 | 71 | 0 | 0 | 0 | 33 | 0 | 0 | 0 | 0 | 16 | 0 | 14 | 455 | 156 | 24 | 0 | 48 | 0 | 2875 | 2756 | 0 | 0 | 0 | 48 | 0 | 0 | 0 | 309 | 0 | 143 |
| SRR2487652 | 51 | 0 | 1621 | 21 | 0 | 84 | 26 | 21 | 63 | 0 | 0 | 27 | 0 | 42 | 0 | 0 | 19 | 0 | 0 | 261 | 91 | 0 | 0 | 0 | 21 | 2824 | 1159 | 21 | 0 | 0 | 485 | 0 | 0 | 0 | 695 | 0 | 337 |
| SRR2487653 | 56 | 0 | 1543 | 46 | 0 | 114 | 55 | 0 | 0 | 0 | 0 | 20 | 0 | 0 | 0 | 0 | 21 | 0 | 14 | 152 | 69 | 0 | 0 | 0 | 23 | 59 | 24 | 0 | 23 | 0 | 366 | 0 | 0 | 23 | 11 | 0 | 229 |
| SRR2487654 | 54 | 0 | 1651 | 0 | 0 | 115 | 42 | 0 | 23 | 0 | 0 | 25 | 0 | 46 | 0 | 0 | 23 | 0 | 0 | 168 | 126 | 0 | 0 | 0 | 0 | 61 | 23 | 0 | 0 | 0 | 344 | 0 | 0 | 0 | 13 | 23 | 275 |
| SRR2487656 | 109 | 0 | 2401 | 0 | 0 | 111 | 69 | 111 | 56 | 0 | 0 | 62 | 0 | 0 | 0 | 0 | 14 | 0 | 0 | 821 | 197 | 0 | 0 | 0 | 0 | 177 | 124 | 0 | 0 | 0 | 0 | 0 | 0 | 0 | 13 | 0 | 11 |
| SRR2487658 | 127 | 0 | 2262 | 0 | 0 | 115 | 106 | 38 | 19 | 0 | 0 | 71 | 0 | 0 | 0 | 0 | 14 | 0 | 12 | 730 | 153 | 0 | 0 | 19 | 0 | 160 | 142 | 0 | 0 | 0 | 38 | 19 | 0 | 0 | 28 | 0 | 134 |
| SRR2487661 | 97 | 0 | 655 | 0 | 0 | 113 | 55 | 254 | 0 | 0 | 0 | 38 | 0 | 0 | 30 | 0 | 0 | 0 | 0 | 178 | 984 | 0 | 0 | 28 | 0 | 1159 | 1554 | 0 | 0 | 0 | 28 | 0 | 0 | 28 | 424 | 28 | 254 |
| SRR2487663 | 69 | 0 | 608 | 0 | 0 | 146 | 49 | 117 | 0 | 0 | 0 | 61 | 0 | 0 | 34 | 0 | 13 | 0 | 11 | 187 | 906 | 0 | 0 | 0 | 0 | 962 | 1691 | 0 | 0 | 0 | 29 | 0 | 0 | 29 | 204 | 0 | 233 |
| SRR2487665 | 79 | 0 | 518 | 0 | 0 | 37 | 38 | 111 | 37 | 0 | 0 | 48 | 0 | 0 | 14 | 0 | 11 | 0 | 0 | 132 | 1048 | 0 | 0 | 74 | 0 | 20 | 28 | 0 | 0 | 0 | 0 | 0 | 0 | 74 | 185 | 0 | 334 |
| SRR2487667 | 107 | 0 | 577 | 0 | 0 | 37 | 42 | 112 | 0 | 0 | 0 | 46 | 0 | 0 | 51 | 0 | 0 | 0 | 0 | 149 | 943 | 0 | 0 | 37 | 0 | 1084 | 1869 | 0 | 0 | 0 | 0 | 0 | 0 | 37 | 262 | 0 | 262 |
| SRR2487668 | 58 | 0 | 1213 | 0 | 0 | 182 | 14 | 36 | 73 | 0 | 0 | 23 | 0 | 0 | 1305 | 0 | 14 | 0 | 0 | 139 | 1027 | 0 | 0 | 0 | 0 | 106 | 24 | 36 | 0 | 0 | 219 | 0 | 0 | 146 | 182 | 109 | 291 |
| SRR2487669 | 32 | 0 | 1236 | 0 | 0 | 232 | 72 | 66 | 66 | 0 | 0 | 0 | 0 | 0 | 166 | 0 | 32 | 0 | 0 | 280 | 1625 | 0 | 0 | 0 | 0 | 6388 | 2813 | 0 | 0 | 33 | 33 | 0 | 0 | 33 | 66 | 0 | 232 |
| SRR2487671 | 54 | 0 | 1210 | 0 | 0 | 100 | 10 | 367 | 67 | 0 | 0 | 39 | 0 | 0 | 154 | 0 | 23 | 0 | 0 | 257 | 1285 | 0 | 0 | 0 | 0 | 6705 | 3336 | 0 | 0 | 67 | 67 | 0 | 0 | 67 | 167 | 0 | 167 |
| SRR2487672 | 80 | 0 | 1180 | 0 | 0 | 263 | 57 | 0 | 75 | 0 | 0 | 29 | 0 | 0 | 1003 | 0 | 20 | 0 | 14 | 72 | 1074 | 0 | 0 | 0 | 0 | 93 | 28 | 0 | 0 | 0 | 188 | 0 | 0 | 113 | 338 | 0 | 263 |
| SRR2487673 | 60 | 0 | 1154 | 0 | 0 | 197 | 0 | 0 | 0 | 0 | 0 | 15 | 0 | 0 | 1163 | 0 | 13 | 0 | 0 | 107 | 980 | 0 | 0 | 0 | 0 | 4621 | 1738 | 40 | 0 | 0 | 237 | 0 | 0 | 0 | 79 | 79 | 316 |
| SRR2487674 | 54 | 0 | 1178 | 0 | 0 | 319 | 46 | 0 | 200 | 0 | 0 | 17 | 0 | 0 | 1171 | 0 | 15 | 0 | 0 | 123 | 878 | 0 | 0 | 0 | 0 | 91 | 22 | 80 | 0 | 0 | 160 | 0 | 0 | 0 | 399 | 0 | 120 |
| SRR2487676 | 77 | 0 | 1609 | 0 | 0 | 235 | 66 | 104 | 130 | 0 | 0 | 38 | 0 | 0 | 166 | 0 | 0 | 0 | 0 | 365 | 1677 | 0 | 0 | 0 | 0 | 245 | 124 | 26 | 0 | 52 | 26 | 0 | 0 | 0 | 13 | 0 | 235 |
| SRR2487678 | 118 | 0 | 1420 | 0 | 0 | 108 | 25 | 215 | 81 | 0 | 0 | 0 | 0 | 0 | 110 | 0 | 26 | 0 | 0 | 339 | 1677 | 0 | 0 | 27 | 0 | 5838 | 3524 | 0 | 0 | 27 | 27 | 0 | 0 | 0 | 161 | 27 | 296 |
| SRR2487688 | 38 | 0 | 488 | 0 | 0 | 521 | 0 | 0 | 136 | 0 | 0 | 35 | 0 | 23 | 5552 | 0 | 0 | 0 | 44 | 87 | 2098 | 0 | 0 | 0 | 0 | 11826 | 1359 | 0 | 0 | 0 | 1019 | 0 | 0 | 23 | 725 | 68 | 272 |
| SRR2487689 | 43 | 0 | 650 | 0 | 0 | 589 | 0 | 0 | 59 | 0 | 0 | 32 | 0 | 0 | 894 | 0 | 0 | 0 | 0 | 223 | 3183 | 0 | 0 | 0 | 0 | 15898 | 2296 | 0 | 0 | 0 | 353 | 0 | 0 | 0 | 530 | 0 | 294 |
| SRR2487691 | 63 | 0 | 439 | 0 | 0 | 537 | 0 | 0 | 0 | 0 | 0 | 52 | 0 | 0 | 667 | 0 | 0 | 0 | 0 | 111 | 4000 | 0 | 0 | 0 | 0 | 15766 | 3225 | 0 | 0 | 0 | 299 | 0 | 0 | 0 | 478 | 0 | 418 |
| SRR2487692 | 56 | 0 | 492 | 46 | 0 | 11 | 23 | 0 | 92 | 0 | 0 | 28 | 0 | 23 | 5142 | 0 | 12 | 0 | 0 | 46 | 2275 | 0 | 0 | 0 | 0 | 210 | 27 | 23 | 0 | 0 | 15 | 0 | 0 | 0 | 14 | 23 | 369 |
| SRR2487693 | 42 | 0 | 512 | 0 | 0 | 462 | 0 | 0 | 128 | 0 | 0 | 38 | 0 | 0 | 6324 | 0 | 0 | 0 | 0 | 50 | 2083 | 0 | 0 | 0 | 26 | 12789 | 1079 | 0 | 0 | 0 | 1284 | 0 | 0 | 26 | 642 | 26 | 282 |
| SRR2487694 | 64 | 0 | 456 | 0 | 0 | 459 | 0 | 0 | 153 | 0 | 0 | 37 | 0 | 0 | 5705 | 0 | 0 | 0 | 0 | 25 | 1893 | 0 | 0 | 0 | 0 | 11877 | 1351 | 25 | 0 | 0 | 1045 | 0 | 0 | 0 | 663 | 25 | 459 |
| SRR2487696 | 34 | 0 | 544 | 0 | 0 | 638 | 43 | 0 | 182 | 0 | 0 | 94 | 0 | 0 | 851 | 0 | 17 | 0 | 43 | 85 | 3320 | 0 | 0 | 0 | 46 | 14396 | 2506 | 46 | 0 | 0 | 228 | 0 | 0 | 46 | 683 | 0 | 364 |
| SRR2487698 | 44 | 0 | 612 | 0 | 0 | 26 | 0 | 0 | 138 | 0 | 0 | 26 | 0 | 0 | 957 | 0 | 0 | 0 | 0 | 131 | 3872 | 0 | 0 | 0 | 0 | 726 | 190 | 0 | 0 | 46 | 184 | 0 | 0 | 0 | 26 | 0 | 19 |
| SRR2487700 | 40 | 0 | 1852 | 0 | 0 | 156 | 18 | 13 | 26 | 0 | 0 | 48 | 0 | 0 | 1896 | 0 | 20 | 0 | 0 | 128 | 456 | 0 | 0 | 0 | 0 | 3714 | 1316 | 13 | 0 | 13 | 156 | 0 | 0 | 0 | 626 | 13 | 130 |
| SRR2487701 | 126 | 0 | 2315 | 0 | 0 | 22 | 99 | 24 | 24 | 0 | 0 | 52 | 0 | 0 | 99 | 0 | 31 | 0 | 0 | 395 | 888 | 0 | 0 | 0 | 0 | 315 | 202 | 0 | 0 | 24 | 49 | 0 | 0 | 0 | 17 | 0 | 11 |
| SRR2487703 | 91 | 0 | 2288 | 0 | 0 | 316 | 32 | 24 | 24 | 0 | 0 | 81 | 0 | 0 | 64 | 0 | 0 | 0 | 0 | 447 | 766 | 0 | 0 | 24 | 0 | 3939 | 2820 | 0 | 0 | 73 | 24 | 0 | 0 | 0 | 219 | 0 | 49 |
| SRR2487704 | 45 | 0 | 1817 | 0 | 0 | 249 | 0 | 0 | 66 | 0 | 0 | 42 | 0 | 0 | 1874 | 0 | 18 | 0 | 0 | 94 | 525 | 0 | 0 | 0 | 0 | 3972 | 1219 | 0 | 0 | 0 | 92 | 0 | 0 | 0 | 406 | 0 | 210 |
| SRR2487705 | 53 | 14 | 1834 | 0 | 0 | 286 | 20 | 0 | 57 | 0 | 0 | 46 | 0 | 0 | 1499 | 0 | 20 | 0 | 0 | 100 | 400 | 0 | 0 | 0 | 0 | 109 | 33 | 0 | 0 | 14 | 186 | 0 | 0 | 14 | 13 | 0 | 200 |
| SRR2487706 | 57 | 0 | 1830 | 0 | 0 | 229 | 61 | 14 | 57 | 0 | 0 | 49 | 0 | 0 | 1516 | 0 | 22 | 0 | 0 | 202 | 404 | 0 | 0 | 0 | 0 | 3683 | 1104 | 14 | 0 | 0 | 158 | 0 | 0 | 0 | 545 | 14 | 215 |
| SRR2487708 | 75 | 0 | 2590 | 0 | 0 | 15 | 0 | 94 | 19 | 0 | 0 | 84 | 0 | 0 | 156 | 0 | 25 | 0 | 0 | 521 | 756 | 0 | 0 | 19 | 0 | 365 | 258 | 0 | 0 | 57 | 0 | 0 | 0 | 0 | 26 | 19 | 12 |
| SRR2487710 | 85 | 0 | 2795 | 19 | 0 | 194 | 0 | 78 | 0 | 0 | 0 | 68 | 0 | 0 | 79 | 0 | 26 | 0 | 26 | 526 | 763 | 0 | 0 | 0 | 0 | 3837 | 2442 | 0 | 0 | 58 | 39 | 0 | 0 | 0 | 291 | 0 | 116 |
| SRR2529523 | 81 | 0 | 4824 | 0 | 0 | 24 | 0 | 0 | 23 | 0 | 0 | 347 | 0 | 0 | 45 | 0 | 19 | 0 | 15 | 4297 | 706 | 0 | 0 | 0 | 29 | 46 | 144 | 0 | 0 | 0 | 15 | 0 | 0 | 0 | 44 | 0 | 24 |
| SRR2529524 | 83 | 0 | 4763 | 0 | 0 | 40 | 0 | 0 | 23 | 0 | 0 | 333 | 0 | 0 | 15 | 0 | 22 | 0 | 31 | 4107 | 540 | 0 | 0 | 0 | 22 | 50 | 129 | 0 | 0 | 0 | 0 | 0 | 0 | 0 | 45 | 0 | 32 |
| SRR2529525 | 42 | 0 | 3195 | 0 | 0 | 22 | 0 | 0 | 0 | 0 | 0 | 519 | 0 | 0 | 0 | 0 | 0 | 0 | 0 | 815 | 1238 | 0 | 0 | 0 | 0 | 37 | 81 | 0 | 0 | 0 | 25 | 0 | 0 | 0 | 34 | 49 | 31 |
| SRR2529526 | 36 | 0 | 3178 | 0 | 0 | 21 | 0 | 0 | 0 | 0 | 0 | 498 | 0 | 0 | 0 | 0 | 10 | 0 | 0 | 783 | 1476 | 0 | 0 | 0 | 12 | 216 | 490 | 0 | 0 | 0 | 17 | 0 | 0 | 0 | 203 | 40 | 134 |
| SRR2529527 | 18 | 0 | 3952 | 0 | 0 | 13 | 0 | 0 | 0 | 0 | 0 | 252 | 0 | 37 | 0 | 0 | 22 | 0 | 0 | 792 | 651 | 0 | 0 | 0 | 0 | 19 | 92 | 0 | 0 | 0 | 19 | 0 | 0 | 0 | 45 | 58 | 31 |
| SRR2529528 | 17 | 0 | 3888 | 0 | 0 | 0 | 0 | 0 | 0 | 0 | 0 | 256 | 0 | 30 | 0 | 0 | 18 | 0 | 0 | 704 | 725 | 0 | 0 | 0 | 0 | 135 | 588 | 0 | 0 | 0 | 19 | 0 | 0 | 0 | 291 | 60 | 225 |
| SRR2529529 | 53 | 0 | 5464 | 0 | 0 | 62 | 0 | 0 | 21 | 0 | 0 | 519 | 0 | 0 | 0 | 0 | 23 | 0 | 20 | 2766 | 626 | 0 | 0 | 0 | 23 | 632 | 1566 | 0 | 0 | 0 | 30 | 0 | 0 | 0 | 46 | 0 | 32 |
| SRR2529530 | 69 | 0 | 5503 | 0 | 0 | 52 | 21 | 0 | 21 | 0 | 0 | 530 | 0 | 0 | 62 | 0 | 21 | 0 | 41 | 3480 | 352 | 0 | 0 | 0 | 35 | 63 | 172 | 0 | 0 | 0 | 40 | 0 | 0 | 0 | 80 | 0 | 56 |
| SRR2529531 | 29 | 0 | 4654 | 0 | 0 | 29 | 0 | 12 | 11 | 0 | 0 | 390 | 0 | 0 | 12 | 0 | 25 | 0 | 12 | 2798 | 403 | 0 | 0 | 0 | 34 | 504 | 1770 | 0 | 0 | 0 | 21 | 0 | 0 | 0 | 228 | 51 | 130 |
| SRR2529532 | 37 | 0 | 4592 | 0 | 0 | 24 | 0 | 16 | 0 | 0 | 0 | 415 | 0 | 0 | 12 | 0 | 21 | 0 | 0 | 3136 | 366 | 0 | 0 | 0 | 38 | 454 | 1745 | 0 | 0 | 0 | 16 | 0 | 0 | 0 | 256 | 60 | 149 |
| SRR2529533 | 67 | 0 | 4428 | 0 | 0 | 50 | 0 | 0 | 23 | 0 | 0 | 308 | 0 | 0 | 0 | 0 | 23 | 77 | 0 | 3394 | 516 | 0 | 0 | 0 | 51 | 82 | 187 | 0 | 0 | 0 | 27 | 0 | 0 | 0 | 15 | 20 | 54 |
| SRR2529534 | 76 | 0 | 4429 | 0 | 0 | 52 | 0 | 0 | 22 | 0 | 0 | 311 | 0 | 0 | 15 | 0 | 23 | 0 | 15 | 3761 | 468 | 0 | 0 | 0 | 44 | 610 | 1419 | 0 | 0 | 0 | 28 | 0 | 0 | 0 | 109 | 11 | 61 |
| SRR2529535 | 31 | 0 | 5218 | 28 | 12 | 0 | 103 | 15 | 0 | 0 | 0 | 170 | 0 | 198 | 47 | 0 | 13 | 0 | 0 | 260 | 106 | 0 | 0 | 0 | 0 | 153 | 1288 | 172 | 0 | 0 | 0 | 0 | 0 | 0 | 38 | 0 | 11 |
| SRR2529536 | 32 | 0 | 5290 | 20 | 0 | 15 | 101 | 0 | 11 | 0 | 0 | 166 | 0 | 21 | 61 | 0 | 0 | 0 | 0 | 279 | 113 | 11 | 0 | 0 | 0 | 13 | 112 | 15 | 0 | 0 | 0 | 0 | 0 | 0 | 35 | 0 | 13 |
| SRR2529537 | 16 | 0 | 3598 | 0 | 0 | 20 | 530 | 19 | 0 | 0 | 0 | 37 | 0 | 16 | 0 | 0 | 0 | 0 | 0 | 252 | 512 | 0 | 0 | 0 | 0 | 26 | 57 | 15 | 0 | 0 | 13 | 0 | 0 | 0 | 37 | 18 | 54 |
| SRR2529538 | 22 | 0 | 3546 | 0 | 0 | 12 | 516 | 0 | 0 | 0 | 0 | 30 | 0 | 18 | 0 | 0 | 0 | 0 | 0 | 272 | 556 | 0 | 0 | 0 | 0 | 137 | 331 | 19 | 0 | 0 | 0 | 0 | 0 | 0 | 224 | 22 | 89 |
| SRR2529539 | 0 | 0 | 4394 | 0 | 0 | 22 | 0 | 0 | 0 | 0 | 0 | 15 | 0 | 15 | 0 | 0 | 12 | 0 | 0 | 394 | 477 | 0 | 0 | 0 | 12 | 181 | 402 | 16 | 0 | 0 | 11 | 0 | 0 | 0 | 255 | 22 | 87 |
| SRR2529540 | 0 | 0 | 4427 | 0 | 0 | 16 | 14 | 0 | 0 | 0 | 0 | 18 | 0 | 18 | 23 | 0 | 14 | 0 | 0 | 373 | 566 | 0 | 0 | 0 | 0 | 169 | 399 | 16 | 0 | 0 | 23 | 0 | 0 | 0 | 262 | 16 | 123 |
| SRR2529541 | 57 | 0 | 5527 | 0 | 0 | 19 | 57 | 0 | 17 | 0 | 0 | 271 | 0 | 38 | 32 | 0 | 12 | 0 | 0 | 719 | 208 | 0 | 0 | 0 | 0 | 349 | 1603 | 38 | 0 | 0 | 0 | 0 | 0 | 0 | 60 | 0 | 30 |
| SRR2529542 | 48 | 0 | 5510 | 0 | 0 | 16 | 77 | 0 | 14 | 0 | 0 | 270 | 0 | 27 | 32 | 0 | 16 | 0 | 0 | 779 | 255 | 0 | 0 | 12 | 16 | 297 | 1666 | 37 | 0 | 0 | 0 | 0 | 0 | 0 | 47 | 0 | 21 |
| SRR2529543 | 33 | 0 | 6693 | 15 | 0 | 11 | 54 | 0 | 15 | 0 | 0 | 62 | 0 | 196 | 27 | 0 | 18 | 0 | 0 | 299 | 156 | 0 | 0 | 0 | 46 | 234 | 712 | 179 | 0 | 0 | 0 | 0 | 0 | 0 | 114 | 0 | 15 |
| SRR2529544 | 26 | 0 | 6789 | 0 | 0 | 12 | 63 | 0 | 0 | 0 | 0 | 77 | 0 | 34 | 14 | 0 | 13 | 0 | 14 | 300 | 174 | 0 | 0 | 0 | 33 | 35 | 95 | 25 | 0 | 0 | 14 | 0 | 0 | 0 | 14 | 0 | 20 |
| SRR2529545 | 82 | 0 | 4592 | 0 | 0 | 18 | 87 | 0 | 11 | 0 | 0 | 272 | 0 | 16 | 0 | 0 | 14 | 0 | 0 | 1112 | 425 | 0 | 0 | 0 | 16 | 359 | 1456 | 0 | 0 | 0 | 18 | 0 | 0 | 0 | 65 | 0 | 29 |
| SRR2529546 | 70 | 0 | 4651 | 0 | 0 | 44 | 102 | 0 | 0 | 0 | 0 | 252 | 0 | 0 | 25 | 0 | 17 | 0 | 0 | 1031 | 458 | 0 | 0 | 0 | 16 | 374 | 1605 | 11 | 0 | 0 | 16 | 0 | 0 | 0 | 71 | 0 | 44 |
| SRR2529547 | 36 | 0 | 4384 | 0 | 0 | 32 | 0 | 0 | 23 | 0 | 0 | 360 | 0 | 0 | 27 | 0 | 18 | 0 | 0 | 2971 | 382 | 0 | 0 | 0 | 0 | 443 | 1226 | 0 | 0 | 0 | 0 | 0 | 0 | 0 | 107 | 20 | 95 |
| SRR2529548 | 42 | 0 | 4373 | 0 | 0 | 36 | 0 | 0 | 15 | 0 | 0 | 356 | 0 | 0 | 56 | 0 | 22 | 0 | 0 | 2369 | 334 | 0 | 0 | 0 | 0 | 420 | 1143 | 0 | 0 | 0 | 12 | 0 | 0 | 0 | 92 | 15 | 89 |
| SRR2529549 | 41 | 0 | 3414 | 0 | 0 | 26 | 0 | 13 | 11 | 0 | 0 | 514 | 0 | 0 | 0 | 0 | 0 | 0 | 0 | 799 | 1149 | 0 | 0 | 0 | 13 | 276 | 572 | 0 | 0 | 0 | 11 | 0 | 0 | 0 | 215 | 35 | 118 |
| SRR2529550 | 52 | 0 | 3308 | 0 | 0 | 16 | 0 | 0 | 0 | 0 | 0 | 491 | 0 | 0 | 51 | 0 | 13 | 0 | 0 | 678 | 1318 | 0 | 0 | 0 | 0 | 39 | 109 | 0 | 0 | 0 | 16 | 0 | 0 | 0 | 38 | 34 | 32 |
| SRR2529551 | 20 | 0 | 3892 | 0 | 0 | 12 | 18 | 0 | 0 | 0 | 0 | 286 | 0 | 0 | 73 | 0 | 16 | 0 | 0 | 1123 | 1232 | 0 | 0 | 0 | 21 | 22 | 124 | 0 | 0 | 0 | 18 | 0 | 0 | 0 | 44 | 44 | 37 |
| SRR2529552 | 18 | 0 | 4007 | 0 | 0 | 13 | 19 | 0 | 0 | 0 | 0 | 270 | 0 | 0 | 0 | 0 | 22 | 0 | 0 | 1249 | 1173 | 0 | 0 | 0 | 13 | 154 | 698 | 0 | 0 | 0 | 20 | 0 | 0 | 0 | 240 | 38 | 238 |
| SRR2529553 | 86 | 0 | 5292 | 13 | 0 | 52 | 64 | 0 | 16 | 0 | 0 | 503 | 0 | 21 | 32 | 0 | 20 | 0 | 64 | 3494 | 635 | 0 | 0 | 0 | 41 | 73 | 125 | 0 | 0 | 0 | 34 | 0 | 0 | 0 | 57 | 13 | 49 |
| SRR2529554 | 91 | 0 | 5333 | 0 | 0 | 67 | 0 | 0 | 27 | 0 | 0 | 475 | 0 | 0 | 32 | 0 | 15 | 0 | 0 | 3516 | 452 | 0 | 0 | 0 | 27 | 599 | 934 | 0 | 0 | 0 | 11 | 0 | 0 | 0 | 61 | 0 | 37 |
| SRR2529555 | 46 | 0 | 5752 | 0 | 0 | 34 | 0 | 0 | 18 | 0 | 0 | 417 | 0 | 0 | 39 | 0 | 22 | 0 | 20 | 3666 | 431 | 0 | 0 | 0 | 38 | 504 | 972 | 0 | 0 | 0 | 31 | 0 | 0 | 0 | 246 | 36 | 148 |
| SRR2529556 | 38 | 0 | 5705 | 0 | 0 | 40 | 0 | 0 | 13 | 0 | 0 | 418 | 0 | 0 | 19 | 0 | 21 | 0 | 37 | 3408 | 559 | 0 | 0 | 0 | 33 | 544 | 1017 | 0 | 0 | 0 | 35 | 0 | 10 | 0 | 211 | 33 | 142 |
| SRR2529557 | 80 | 0 | 4571 | 0 | 0 | 58 | 0 | 0 | 35 | 0 | 0 | 372 | 0 | 0 | 26 | 0 | 29 | 40 | 0 | 3191 | 818 | 0 | 0 | 14 | 16 | 811 | 938 | 0 | 0 | 0 | 30 | 0 | 0 | 0 | 174 | 12 | 107 |
| SRR2529558 | 86 | 0 | 4630 | 0 | 0 | 57 | 0 | 0 | 29 | 0 | 0 | 336 | 0 | 0 | 54 | 0 | 31 | 0 | 0 | 3029 | 757 | 0 | 0 | 0 | 24 | 91 | 131 | 0 | 0 | 0 | 38 | 0 | 0 | 0 | 24 | 17 | 17 |
| SRR2529559 | 44 | 0 | 4568 | 0 | 0 | 25 | 0 | 0 | 28 | 0 | 0 | 288 | 0 | 0 | 61 | 0 | 16 | 20 | 0 | 3620 | 389 | 0 | 0 | 0 | 13 | 257 | 579 | 0 | 0 | 0 | 15 | 0 | 0 | 0 | 59 | 0 | 25 |
| SRR2529560 | 39 | 0 | 4506 | 0 | 0 | 28 | 0 | 0 | 30 | 0 | 0 | 310 | 0 | 0 | 41 | 0 | 15 | 0 | 41 | 2949 | 639 | 0 | 0 | 0 | 13 | 311 | 589 | 0 | 0 | 0 | 15 | 0 | 0 | 0 | 65 | 0 | 30 |
| SRR2529561 | 23 | 0 | 2684 | 0 | 0 | 21 | 11 | 0 | 0 | 0 | 0 | 396 | 0 | 0 | 11 | 0 | 0 | 0 | 0 | 1239 | 1464 | 0 | 0 | 0 | 0 | 240 | 428 | 0 | 0 | 0 | 13 | 0 | 0 | 0 | 93 | 17 | 100 |
| SRR2529562 | 24 | 0 | 2685 | 0 | 0 | 19 | 0 | 0 | 0 | 0 | 0 | 411 | 0 | 0 | 23 | 0 | 0 | 0 | 0 | 982 | 1536 | 0 | 0 | 0 | 0 | 206 | 377 | 0 | 0 | 0 | 13 | 0 | 0 | 0 | 110 | 16 | 94 |
| SRR2529563 | 18 | 0 | 3770 | 0 | 0 | 14 | 0 | 0 | 0 | 0 | 0 | 271 | 0 | 0 | 0 | 0 | 14 | 0 | 0 | 899 | 920 | 0 | 0 | 0 | 12 | 188 | 379 | 0 | 0 | 0 | 16 | 0 | 0 | 0 | 97 | 36 | 145 |
| SRR2529564 | 18 | 0 | 3713 | 0 | 0 | 0 | 0 | 0 | 0 | 0 | 0 | 276 | 0 | 0 | 0 | 0 | 16 | 0 | 0 | 978 | 1066 | 0 | 0 | 0 | 0 | 183 | 430 | 0 | 0 | 0 | 21 | 0 | 0 | 0 | 97 | 30 | 122 |
| SRR2529565 | 96 | 0 | 5041 | 12 | 0 | 31 | 0 | 0 | 50 | 0 | 0 | 485 | 0 | 0 | 40 | 0 | 23 | 0 | 40 | 4542 | 1474 | 0 | 0 | 0 | 37 | 418 | 640 | 0 | 0 | 0 | 25 | 0 | 0 | 0 | 31 | 0 | 19 |
| SRR2529566 | 92 | 0 | 10 | 0 | 0 | 22 | 0 | 0 | 22 | 0 | 0 | 483 | 0 | 0 | 124 | 0 | 16 | 0 | 83 | 3937 | 912 | 0 | 0 | 13 | 32 | 450 | 703 | 0 | 0 | 0 | 42 | 0 | 0 | 0 | 35 | 0 | 19 |
| SRR2529567 | 80 | 0 | 5104 | 0 | 0 | 28 | 0 | 0 | 40 | 0 | 0 | 415 | 0 | 0 | 0 | 0 | 15 | 0 | 0 | 4452 | 728 | 0 | 0 | 0 | 38 | 510 | 569 | 0 | 0 | 0 | 30 | 0 | 0 | 0 | 93 | 0 | 55 |
| SRR2529568 | 63 | 0 | 5105 | 0 | 0 | 28 | 57 | 0 | 35 | 0 | 0 | 446 | 0 | 0 | 0 | 0 | 15 | 0 | 28 | 4191 | 651 | 0 | 0 | 0 | 46 | 120 | 92 | 0 | 0 | 0 | 33 | 0 | 0 | 0 | 18 | 0 | 28 |
| SRR2529569 | 95 | 0 | 4452 | 0 | 0 | 28 | 0 | 0 | 28 | 0 | 0 | 394 | 0 | 0 | 47 | 0 | 18 | 62 | 0 | 2506 | 851 | 0 | 0 | 0 | 38 | 76 | 81 | 0 | 0 | 0 | 0 | 0 | 0 | 0 | 0 | 0 | 0 |
| SRR2529570 | 92 | 0 | 4431 | 0 | 0 | 26 | 0 | 0 | 49 | 0 | 0 | 402 | 0 | 0 | 0 | 0 | 17 | 0 | 0 | 3001 | 350 | 0 | 0 | 0 | 45 | 433 | 566 | 0 | 0 | 0 | 13 | 0 | 0 | 0 | 19 | 0 | 0 |
| SRR2529571 | 60 | 0 | 4589 | 0 | 0 | 31 | 59 | 0 | 12 | 0 | 0 | 304 | 0 | 0 | 20 | 0 | 12 | 0 | 20 | 3710 | 707 | 0 | 0 | 0 | 24 | 51 | 109 | 0 | 0 | 0 | 14 | 0 | 0 | 0 | 11 | 0 | 37 |
| SRR2529572 | 64 | 0 | 4561 | 0 | 0 | 22 | 20 | 0 | 28 | 0 | 0 | 293 | 0 | 0 | 40 | 0 | 18 | 0 | 0 | 3946 | 341 | 0 | 0 | 0 | 34 | 54 | 105 | 0 | 0 | 0 | 14 | 0 | 0 | 0 | 71 | 0 | 56 |
| SRR2529573 | 28 | 0 | 3235 | 0 | 0 | 0 | 0 | 0 | 0 | 0 | 0 | 473 | 0 | 0 | 35 | 0 | 14 | 0 | 0 | 600 | 1130 | 0 | 0 | 0 | 0 | 217 | 603 | 0 | 0 | 0 | 0 | 0 | 0 | 0 | 228 | 47 | 199 |
| SRR2529574 | 41 | 0 | 3227 | 0 | 0 | 24 | 0 | 0 | 0 | 0 | 0 | 491 | 0 | 0 | 0 | 0 | 10 | 0 | 0 | 603 | 1046 | 0 | 0 | 0 | 11 | 37 | 101 | 0 | 0 | 0 | 0 | 0 | 0 | 0 | 42 | 33 | 25 |
| SRR2529575 | 17 | 0 | 3343 | 0 | 0 | 0 | 0 | 0 | 0 | 0 | 0 | 277 | 0 | 0 | 27 | 0 | 12 | 0 | 18 | 935 | 1153 | 0 | 0 | 0 | 13 | 172 | 608 | 0 | 0 | 0 | 16 | 0 | 0 | 0 | 271 | 59 | 200 |
| SRR2529576 | 20 | 0 | 3251 | 0 | 0 | 17 | 0 | 0 | 0 | 0 | 0 | 295 | 0 | 0 | 29 | 0 | 20 | 0 | 0 | 951 | 1095 | 0 | 0 | 0 | 12 | 140 | 614 | 0 | 0 | 0 | 15 | 0 | 0 | 0 | 295 | 38 | 225 |
| SRR2529577 | 54 | 0 | 5472 | 0 | 0 | 31 | 0 | 0 | 0 | 0 | 0 | 446 | 0 | 0 | 130 | 0 | 17 | 0 | 26 | 3229 | 807 | 0 | 0 | 0 | 24 | 60 | 120 | 0 | 0 | 0 | 28 | 0 | 0 | 0 | 55 | 11 | 66 |
| SRR2529578 | 68 | 0 | 5454 | 0 | 0 | 38 | 0 | 0 | 17 | 0 | 0 | 445 | 0 | 0 | 74 | 0 | 21 | 0 | 0 | 2414 | 788 | 0 | 0 | 0 | 26 | 549 | 1112 | 0 | 0 | 0 | 26 | 0 | 0 | 0 | 91 | 13 | 40 |
| SRR2529579 | 41 | 0 | 5093 | 0 | 0 | 32 | 16 | 0 | 0 | 0 | 0 | 298 | 0 | 0 | 32 | 0 | 17 | 0 | 32 | 3121 | 512 | 0 | 0 | 0 | 34 | 44 | 120 | 0 | 0 | 0 | 21 | 0 | 0 | 0 | 38 | 18 | 16 |
| SRR2529580 | 38 | 0 | 5034 | 0 | 0 | 21 | 0 | 0 | 0 | 0 | 0 | 282 | 0 | 0 | 0 | 0 | 20 | 0 | 0 | 3615 | 554 | 0 | 0 | 0 | 32 | 384 | 965 | 0 | 0 | 0 | 21 | 0 | 0 | 0 | 330 | 40 | 111 |
| SRR2529581 | 74 | 0 | 4659 | 0 | 0 | 43 | 0 | 0 | 16 | 0 | 0 | 351 | 0 | 0 | 0 | 0 | 25 | 0 | 0 | 2154 | 792 | 0 | 0 | 0 | 48 | 70 | 129 | 0 | 0 | 0 | 21 | 0 | 0 | 0 | 22 | 18 | 10 |
| SRR2529582 | 69 | 0 | 4659 | 0 | 0 | 46 | 0 | 0 | 19 | 0 | 0 | 332 | 0 | 0 | 21 | 0 | 24 | 0 | 0 | 2678 | 851 | 0 | 0 | 0 | 57 | 578 | 1039 | 0 | 0 | 0 | 15 | 0 | 0 | 0 | 159 | 19 | 71 |
| SRR2529583 | 33 | 0 | 5894 | 0 | 0 | 18 | 0 | 0 | 18 | 0 | 0 | 77 | 0 | 0 | 0 | 0 | 12 | 0 | 0 | 1411 | 245 | 0 | 0 | 0 | 0 | 32 | 73 | 11 | 0 | 0 | 0 | 0 | 0 | 0 | 74 | 0 | 14 |
| SRR2529584 | 38 | 0 | 5858 | 0 | 0 | 29 | 0 | 0 | 0 | 0 | 0 | 70 | 0 | 11 | 12 | 0 | 14 | 0 | 0 | 1276 | 385 | 0 | 0 | 0 | 11 | 369 | 628 | 0 | 0 | 0 | 0 | 0 | 0 | 0 | 51 | 0 | 15 |
| SRR2529585 | 0 | 0 | 2248 | 12 | 0 | 24 | 0 | 0 | 0 | 0 | 0 | 0 | 0 | 21 | 0 | 0 | 10 | 0 | 0 | 333 | 977 | 0 | 0 | 0 | 0 | 33 | 49 | 0 | 0 | 0 | 0 | 0 | 0 | 0 | 44 | 12 | 12 |
| SRR2529586 | 11 | 0 | 2273 | 0 | 0 | 19 | 0 | 0 | 0 | 0 | 0 | 0 | 0 | 0 | 23 | 0 | 10 | 0 | 0 | 315 | 1054 | 0 | 0 | 0 | 14 | 188 | 335 | 0 | 0 | 0 | 14 | 0 | 0 | 0 | 297 | 14 | 83 |
| SRR2529587 | 0 | 0 | 2000 | 0 | 0 | 16 | 0 | 0 | 0 | 0 | 0 | 0 | 0 | 13 | 0 | 0 | 11 | 0 | 0 | 322 | 952 | 0 | 0 | 0 | 11 | 183 | 426 | 16 | 0 | 0 | 0 | 0 | 0 | 0 | 341 | 14 | 108 |
| SRR2529588 | 0 | 0 | 1972 | 0 | 0 | 0 | 0 | 0 | 0 | 0 | 0 | 0 | 0 | 13 | 14 | 0 | 11 | 0 | 0 | 357 | 826 | 0 | 0 | 0 | 13 | 160 | 386 | 0 | 0 | 0 | 0 | 0 | 0 | 0 | 324 | 18 | 111 |
| SRR2529589 | 43 | 0 | 6275 | 0 | 0 | 33 | 0 | 0 | 0 | 0 | 0 | 79 | 0 | 0 | 58 | 0 | 11 | 0 | 19 | 2180 | 502 | 0 | 0 | 0 | 0 | 50 | 102 | 0 | 0 | 0 | 0 | 0 | 0 | 0 | 80 | 0 | 27 |
| SRR2529590 | 45 | 0 | 6266 | 0 | 0 | 41 | 39 | 0 | 0 | 0 | 0 | 66 | 0 | 0 | 0 | 0 | 14 | 0 | 20 | 2442 | 985 | 0 | 0 | 0 | 25 | 51 | 103 | 0 | 0 | 0 | 18 | 0 | 0 | 0 | 73 | 0 | 23 |
| SRR2529591 | 27 | 0 | 6395 | 0 | 0 | 19 | 0 | 0 | 0 | 0 | 0 | 20 | 0 | 0 | 0 | 0 | 0 | 0 | 58 | 1337 | 1250 | 0 | 0 | 0 | 27 | 477 | 544 | 0 | 0 | 0 | 0 | 0 | 0 | 0 | 128 | 11 | 53 |
| SRR2529592 | 27 | 0 | 6492 | 0 | 0 | 27 | 0 | 0 | 0 | 0 | 0 | 16 | 0 | 0 | 0 | 0 | 12 | 0 | 28 | 1563 | 995 | 0 | 0 | 0 | 27 | 527 | 502 | 14 | 0 | 0 | 0 | 0 | 0 | 0 | 145 | 0 | 49 |
| SRR2529593 | 55 | 0 | 5751 | 0 | 0 | 38 | 0 | 0 | 21 | 0 | 0 | 90 | 0 | 0 | 58 | 0 | 16 | 0 | 0 | 3004 | 635 | 0 | 0 | 12 | 18 | 494 | 778 | 0 | 0 | 0 | 12 | 0 | 0 | 0 | 77 | 0 | 21 |
| SRR2529594 | 74 | 0 | 5683 | 0 | 0 | 24 | 0 | 0 | 15 | 0 | 0 | 90 | 0 | 0 | 0 | 0 | 16 | 0 | 0 | 2418 | 735 | 0 | 0 | 0 | 49 | 530 | 834 | 0 | 0 | 0 | 21 | 0 | 0 | 0 | 67 | 0 | 33 |
| SRR2648184 | 106 | 11 | 4377 | 0 | 0 | 15 | 0 | 0 | 14 | 0 | 0 | 60 | 0 | 0 | 277 | 0 | 18 | 140 | 0 | 53 | 138 | 0 | 0 | 0 | 10 | 166 | 99 | 0 | 72 | 0 | 119 | 0 | 0 | 14 | 216 | 15 | 78 |
| SRR2648185 | 97 | 15 | 4181 | 0 | 0 | 12 | 0 | 0 | 15 | 0 | 0 | 47 | 0 | 0 | 246 | 0 | 21 | 95 | 0 | 67 | 106 | 0 | 0 | 0 | 54 | 21 | 11 | 0 | 10 | 0 | 12 | 0 | 0 | 14 | 20 | 14 | 51 |
| SRR2648186 | 128 | 10 | 4994 | 0 | 0 | 19 | 0 | 0 | 15 | 0 | 0 | 37 | 0 | 0 | 238 | 0 | 22 | 93 | 0 | 63 | 159 | 0 | 0 | 0 | 11 | 144 | 78 | 0 | 68 | 0 | 82 | 0 | 0 | 18 | 118 | 12 | 51 |
| SRR2648187 | 94 | 0 | 3211 | 0 | 0 | 97 | 0 | 0 | 85 | 0 | 0 | 50 | 0 | 0 | 40 | 0 | 25 | 0 | 0 | 65 | 11 | 0 | 0 | 0 | 11 | 43 | 258 | 0 | 12 | 0 | 19 | 0 | 0 | 0 | 68 | 63 | 19 |
| SRR2648188 | 89 | 0 | 3431 | 0 | 0 | 126 | 0 | 0 | 91 | 0 | 0 | 53 | 0 | 0 | 44 | 0 | 26 | 0 | 0 | 91 | 27 | 0 | 0 | 0 | 0 | 1212 | 282 | 13 | 14 | 0 | 513 | 0 | 0 | 0 | 1944 | 99 | 579 |
| SRR2648189 | 98 | 0 | 3409 | 0 | 0 | 112 | 0 | 0 | 68 | 0 | 0 | 48 | 0 | 0 | 53 | 0 | 21 | 0 | 0 | 122 | 47 | 0 | 0 | 0 | 11 | 1092 | 256 | 0 | 14 | 0 | 464 | 0 | 0 | 0 | 1239 | 57 | 415 |

### Supp Table S8.2.Columns_AM_to_BW

| | | | | | | | | | | | | | | | | | | | | | | | | | | | | | | | | | | | | | |
|---|---|---|---|---|---|---|---|---|---|---|---|---|---|---|---|---|---|---|---|---|---|---|---|---|---|---|---|---|---|---|---|---|---|---|---|---|---|
| SRR2728221 | 50 | 13 | 18 | 0 | 0 | 0 | 0 | 0 | 0 | 0 | 0 | 0 | 0 | 0 | 0 | 0 | 0 | 0 | 0 | 0 | 0 | 0 | 0 | 0 | 61 | 0 | 0 | 0 | 0 | 0 | 0 | 0 | 0 | 0 | 0 | 0 | 0 |
| SRR2731278 | 96 | 0 | 192 | 0 | 0 | 206 | 121 | 0 | 16 | 0 | 0 | 69 | 0 | 0 | 0 | 0 | 192 | 0 | 55 | 69 | 431 | 0 | 0 | 184 | 36 | 2314 | 5165 | 0 | 20 | 0 | 0 | 0 | 0 | 101 | 0 | 0 | 17 |
| SRR2731279 | 370 | 0 | 851 | 0 | 0 | 72 | 618 | 0 | 22 | 0 | 0 | 253 | 0 | 0 | 0 | 0 | 137 | 0 | 36 | 394 | 446 | 0 | 0 | 88 | 15 | 621 | 1084 | 0 | 0 | 0 | 11 | 0 | 0 | 30 | 20 | 23 | 15 |
| SRR2731280 | 65 | 0 | 363 | 0 | 0 | 134 | 0 | 0 | 12 | 0 | 17 | 68 | 11 | 0 | 36 | 0 | 47 | 21 | 30 | 286 | 1541 | 0 | 0 | 0 | 50 | 1017 | 147 | 0 | 0 | 0 | 50 | 0 | 0 | 162 | 21 | 17 | 36 |
| SRR2731281 | 77 | 0 | 507 | 0 | 20 | 575 | 53 | 0 | 67 | 0 | 0 | 29 | 20 | 0 | 0 | 0 | 124 | 0 | 46 | 261 | 549 | 0 | 0 | 272 | 150 | 3838 | 4559 | 0 | 0 | 0 | 23 | 0 | 0 | 81 | 41 | 26 | 57 |
| SRR2731282 | 170 | 0 | 1345 | 0 | 0 | 294 | 40 | 0 | 26 | 0 | 0 | 123 | 13 | 0 | 0 | 0 | 379 | 0 | 78 | 310 | 1298 | 0 | 0 | 903 | 38 | 2327 | 5668 | 0 | 0 | 0 | 0 | 0 | 0 | 53 | 44 | 28 | 83 |
| SRR2731283 | 212 | 0 | 604 | 0 | 0 | 115 | 93 | 0 | 23 | 0 | 0 | 111 | 12 | 0 | 0 | 0 | 161 | 0 | 71 | 739 | 1236 | 0 | 0 | 141 | 12 | 922 | 1661 | 0 | 0 | 12 | 0 | 0 | 0 | 44 | 15 | 10 | 27 |
| SRR2731284 | 211 | 0 | 879 | 0 | 14 | 78 | 87 | 0 | 17 | 0 | 0 | 114 | 20 | 0 | 0 | 0 | 274 | 0 | 121 | 752 | 938 | 0 | 0 | 144 | 23 | 699 | 2206 | 0 | 0 | 0 | 0 | 0 | 0 | 23 | 13 | 12 | 10 |
| SRR2731285 | 85 | 0 | 818 | 0 | 18 | 79 | 37 | 0 | 17 | 0 | 0 | 91 | 17 | 0 | 0 | 0 | 301 | 0 | 55 | 424 | 1670 | 0 | 0 | 61 | 39 | 642 | 1479 | 0 | 0 | 0 | 22 | 0 | 0 | 37 | 19 | 10 | 29 |
| SRR2731286 | 459 | 0 | 1207 | 0 | 22 | 86 | 230 | 0 | 15 | 0 | 14 | 189 | 12 | 0 | 0 | 0 | 99 | 0 | 54 | 189 | 246 | 0 | 0 | 14 | 52 | 919 | 2589 | 0 | 0 | 0 | 15 | 0 | 0 | 59 | 22 | 14 | 17 |
| SRR2731287 | 142 | 0 | 457 | 0 | 13 | 79 | 74 | 0 | 14 | 0 | 0 | 122 | 21 | 0 | 0 | 0 | 133 | 0 | 81 | 190 | 997 | 0 | 0 | 163 | 14 | 514 | 1527 | 0 | 0 | 24 | 0 | 0 | 0 | 80 | 24 | 18 | 26 |
| SRR2731288 | 107 | 0 | 775 | 0 | 0 | 275 | 101 | 10 | 29 | 0 | 0 | 137 | 100 | 0 | 0 | 0 | 128 | 0 | 57 | 111 | 435 | 0 | 0 | 119 | 98 | 2455 | 6406 | 0 | 13 | 0 | 93 | 0 | 0 | 80 | 54 | 33 | 94 |
| SRR2731289 | 65 | 0 | 354 | 0 | 0 | 89 | 33 | 11 | 46 | 0 | 0 | 68 | 0 | 0 | 0 | 0 | 14 | 0 | 11 | 83 | 1236 | 0 | 0 | 151 | 0 | 497 | 465 | 0 | 0 | 17 | 10 | 0 | 0 | 64 | 20 | 22 | 107 |
| SRR2731290 | 219 | 0 | 380 | 0 | 0 | 146 | 42 | 22 | 20 | 0 | 0 | 61 | 0 | 0 | 0 | 0 | 35 | 0 | 21 | 258 | 1840 | 0 | 0 | 92 | 0 | 756 | 633 | 0 | 0 | 11 | 0 | 0 | 0 | 60 | 13 | 17 | 56 |
| SRR2731291 | 122 | 0 | 472 | 0 | 20 | 175 | 43 | 12 | 160 | 0 | 0 | 61 | 0 | 0 | 0 | 0 | 29 | 0 | 15 | 48 | 293 | 0 | 0 | 447 | 0 | 975 | 1658 | 0 | 0 | 20 | 11 | 0 | 0 | 101 | 69 | 48 | 256 |
| SRR2731292 | 149 | 0 | 261 | 0 | 0 | 225 | 32 | 30 | 154 | 0 | 0 | 59 | 0 | 0 | 0 | 0 | 11 | 0 | 22 | 114 | 6134 | 27 | 0 | 499 | 0 | 1235 | 571 | 0 | 0 | 35 | 10 | 0 | 0 | 189 | 59 | 80 | 514 |
| SRR2731293 | 177 | 0 | 767 | 0 | 14 | 81 | 20 | 0 | 12 | 0 | 0 | 118 | 13 | 0 | 0 | 0 | 487 | 0 | 77 | 427 | 950 | 0 | 0 | 177 | 17 | 654 | 2603 | 0 | 0 | 0 | 0 | 0 | 0 | 55 | 12 | 0 | 11 |
| SRR2731294 | 147 | 0 | 1384 | 0 | 17 | 430 | 55 | 0 | 33 | 0 | 0 | 124 | 30 | 0 | 0 | 0 | 156 | 0 | 50 | 891 | 2097 | 0 | 0 | 110 | 131 | 3245 | 4285 | 0 | 0 | 0 | 28 | 0 | 0 | 72 | 29 | 28 | 49 |
| SRR2731295 | 105 | 0 | 1003 | 0 | 0 | 82 | 47 | 0 | 33 | 0 | 0 | 392 | 13 | 0 | 0 | 0 | 343 | 0 | 90 | 128 | 2544 | 0 | 0 | 35 | 29 | 648 | 988 | 0 | 0 | 0 | 24 | 0 | 0 | 36 | 15 | 15 | 29 |
| SRR2731296 | 250 | 0 | 1933 | 0 | 0 | 312 | 31 | 0 | 36 | 0 | 0 | 125 | 12 | 0 | 0 | 0 | 587 | 0 | 84 | 453 | 776 | 0 | 0 | 641 | 74 | 2317 | 8501 | 0 | 0 | 0 | 11 | 0 | 0 | 14 | 26 | 14 | 47 |
| SRR2731297 | 183 | 0 | 1112 | 0 | 14 | 171 | 63 | 0 | 19 | 0 | 0 | 134 | 19 | 0 | 0 | 0 | 452 | 11 | 93 | 640 | 948 | 0 | 0 | 366 | 44 | 1423 | 5322 | 0 | 0 | 0 | 0 | 0 | 0 | 71 | 0 | 0 | 16 |
| SRR2731298 | 26 | 0 | 394 | 0 | 0 | 12 | 131 | 0 | 16 | 0 | 55 | 45 | 0 | 0 | 71 | 0 | 28 | 47 | 21 | 0 | 330 | 0 | 0 | 23 | 42 | 114 | 19 | 0 | 32 | 0 | 74 | 0 | 45 | 0 | 78 | 11 | 21 |
| SRR2731299 | 307 | 0 | 859 | 0 | 0 | 108 | 73 | 0 | 21 | 0 | 0 | 105 | 24 | 0 | 0 | 0 | 210 | 0 | 93 | 448 | 1541 | 0 | 0 | 273 | 28 | 658 | 1435 | 0 | 0 | 0 | 0 | 0 | 0 | 18 | 18 | 0 | 24 |
| SRR2731300 | 126 | 0 | 1181 | 0 | 22 | 77 | 35 | 0 | 21 | 0 | 0 | 101 | 13 | 0 | 0 | 0 | 590 | 0 | 71 | 833 | 3452 | 0 | 0 | 75 | 45 | 639 | 1432 | 0 | 0 | 0 | 18 | 0 | 0 | 66 | 19 | 14 | 30 |
| SRR2731301 | 165 | 0 | 1000 | 0 | 18 | 411 | 83 | 0 | 35 | 0 | 0 | 91 | 46 | 0 | 0 | 0 | 190 | 0 | 101 | 398 | 1023 | 0 | 0 | 150 | 172 | 3312 | 5168 | 0 | 0 | 0 | 48 | 0 | 0 | 156 | 22 | 15 | 34 |
| SRR2731302 | 269 | 0 | 1683 | 0 | 12 | 399 | 27 | 19 | 36 | 0 | 0 | 73 | 73 | 0 | 0 | 0 | 245 | 0 | 37 | 194 | 470 | 0 | 0 | 902 | 55 | 2524 | 6022 | 0 | 0 | 35 | 10 | 0 | 0 | 225 | 41 | 29 | 79 |
| SRR2731303 | 41 | 0 | 310 | 0 | 0 | 299 | 86 | 68 | 26 | 0 | 62 | 53 | 23 | 0 | 39 | 0 | 51 | 0 | 75 | 454 | 885 | 0 | 0 | 623 | 383 | 2188 | 686 | 0 | 0 | 75 | 113 | 0 | 0 | 371 | 15 | 19 | 41 |
| SRR2731304 | 113 | 0 | 611 | 0 | 14 | 482 | 65 | 0 | 45 | 0 | 0 | 316 | 15 | 0 | 0 | 0 | 118 | 0 | 116 | 894 | 1017 | 0 | 0 | 179 | 152 | 3813 | 4534 | 0 | 0 | 0 | 37 | 0 | 0 | 21 | 35 | 33 | 48 |
| SRR2731305 | 300 | 0 | 1424 | 0 | 0 | 308 | 260 | 0 | 26 | 0 | 0 | 52 | 38 | 0 | 0 | 0 | 519 | 0 | 51 | 525 | 1101 | 0 | 0 | 511 | 47 | 2293 | 5532 | 0 | 0 | 0 | 12 | 0 | 0 | 105 | 13 | 0 | 29 |
| SRR2731306 | 186 | 0 | 829 | 0 | 12 | 320 | 67 | 13 | 26 | 0 | 0 | 62 | 25 | 0 | 0 | 0 | 220 | 0 | 151 | 1021 | 2986 | 0 | 0 | 992 | 27 | 1930 | 3279 | 0 | 0 | 25 | 0 | 0 | 0 | 183 | 19 | 14 | 42 |
| SRR2731307 | 168 | 0 | 956 | 0 | 0 | 76 | 48 | 0 | 31 | 0 | 0 | 2379 | 15 | 0 | 0 | 0 | 276 | 0 | 21 | 244 | 2359 | 0 | 0 | 138 | 36 | 639 | 1477 | 0 | 0 | 0 | 0 | 0 | 0 | 18 | 29 | 24 | 15 |
| SRR2731308 | 153 | 0 | 1123 | 0 | 0 | 138 | 55 | 0 | 12 | 0 | 0 | 143 | 25 | 0 | 0 | 0 | 232 | 0 | 50 | 422 | 986 | 0 | 0 | 301 | 13 | 1033 | 2182 | 0 | 0 | 16 | 10 | 0 | 0 | 30 | 29 | 14 | 14 |
| SRR2731309 | 182 | 0 | 1200 | 0 | 0 | 242 | 35 | 0 | 19 | 0 | 0 | 79 | 27 | 0 | 0 | 0 | 269 | 0 | 116 | 998 | 2405 | 0 | 0 | 755 | 42 | 1772 | 6254 | 0 | 0 | 10 | 0 | 0 | 0 | 116 | 18 | 11 | 34 |
| SRR2731310 | 160 | 0 | 1358 | 0 | 0 | 718 | 42 | 0 | 82 | 0 | 0 | 113 | 31 | 0 | 0 | 0 | 185 | 0 | 45 | 345 | 1368 | 0 | 0 | 629 | 52 | 4985 | 6657 | 0 | 0 | 0 | 15 | 0 | 0 | 27 | 45 | 29 | 92 |
| SRR2731311 | 272 | 0 | 953 | 0 | 14 | 79 | 93 | 0 | 16 | 0 | 12 | 81 | 13 | 0 | 0 | 0 | 162 | 0 | 93 | 223 | 422 | 0 | 0 | 15 | 16 | 791 | 1772 | 0 | 0 | 0 | 0 | 0 | 0 | 52 | 19 | 12 | 11 |
| SRR2731312 | 192 | 0 | 859 | 0 | 0 | 125 | 86 | 0 | 29 | 0 | 0 | 55 | 18 | 0 | 0 | 0 | 220 | 0 | 99 | 421 | 816 | 0 | 0 | 15 | 15 | 923 | 1912 | 0 | 0 | 0 | 11 | 0 | 0 | 28 | 22 | 14 | 14 |
| SRR2731313 | 127 | 0 | 894 | 0 | 0 | 337 | 226 | 0 | 27 | 0 | 0 | 109 | 23 | 0 | 0 | 0 | 137 | 0 | 163 | 430 | 1254 | 0 | 0 | 237 | 81 | 2775 | 4857 | 0 | 0 | 17 | 27 | 0 | 0 | 236 | 19 | 12 | 36 |
| SRR2731314 | 127 | 0 | 780 | 0 | 23 | 54 | 41 | 0 | 15 | 0 | 11 | 127 | 26 | 0 | 0 | 0 | 473 | 0 | 40 | 570 | 1048 | 0 | 0 | 28 | 48 | 453 | 1346 | 0 | 0 | 0 | 18 | 0 | 0 | 16 | 12 | 0 | 24 |
| SRR2731315 | 202 | 0 | 929 | 0 | 0 | 162 | 89 | 0 | 17 | 0 | 0 | 121 | 24 | 0 | 0 | 0 | 111 | 0 | 75 | 332 | 804 | 0 | 0 | 32 | 35 | 1392 | 1810 | 0 | 0 | 0 | 17 | 0 | 0 | 18 | 35 | 28 | 18 |
| SRR2731316 | 409 | 0 | 1104 | 0 | 0 | 428 | 47 | 0 | 26 | 0 | 0 | 49 | 30 | 0 | 0 | 0 | 346 | 0 | 47 | 711 | 1823 | 0 | 0 | 40 | 92 | 3526 | 5713 | 0 | 0 | 0 | 19 | 0 | 0 | 54 | 16 | 14 | 40 |
| SRR2731317 | 272 | 0 | 1321 | 0 | 0 | 474 | 29 | 0 | 30 | 0 | 0 | 21 | 21 | 0 | 0 | 0 | 221 | 0 | 32 | 291 | 1409 | 0 | 0 | 64 | 99 | 3506 | 7282 | 0 | 0 | 0 | 11 | 0 | 0 | 85 | 35 | 17 | 59 |
| SRR2731318 | 100 | 0 | 1248 | 0 | 0 | 65 | 53 | 0 | 18 | 0 | 0 | 193 | 0 | 0 | 0 | 0 | 125 | 0 | 25 | 98 | 224 | 14 | 0 | 18 | 23 | 617 | 543 | 0 | 15 | 0 | 18 | 19 | 0 | 12 | 19 | 30 | 42 |
| SRR2731319 | 275 | 0 | 641 | 0 | 0 | 277 | 68 | 0 | 21 | 0 | 0 | 76 | 24 | 0 | 0 | 0 | 190 | 0 | 127 | 920 | 2406 | 0 | 0 | 37 | 33 | 1953 | 2099 | 0 | 0 | 0 | 24 | 0 | 0 | 33 | 23 | 17 | 12 |
| SRR2731320 | 81 | 0 | 987 | 0 | 0 | 364 | 13 | 0 | 64 | 0 | 0 | 141 | 0 | 0 | 0 | 0 | 214 | 0 | 11 | 91 | 775 | 14 | 0 | 37 | 42 | 2786 | 3049 | 0 | 0 | 0 | 33 | 0 | 0 | 22 | 65 | 48 | 154 |
| SRR2731321 | 64 | 0 | 1103 | 0 | 14 | 29 | 0 | 0 | 17 | 0 | 0 | 41 | 17 | 0 | 0 | 0 | 209 | 0 | 0 | 63 | 640 | 11 | 0 | 14 | 29 | 315 | 238 | 0 | 14 | 0 | 12 | 25 | 0 | 15 | 16 | 20 | 33 |
| SRR2731322 | 268 | 0 | 860 | 0 | 13 | 645 | 12 | 0 | 58 | 14 | 18 | 61 | 15 | 0 | 15 | 0 | 356 | 0 | 35 | 385 | 3023 | 17 | 0 | 14 | 177 | 5583 | 3735 | 0 | 19 | 0 | 43 | 0 | 0 | 68 | 16 | 29 | 43 |
| SRR2731323 | 733 | 0 | 420 | 0 | 16 | 236 | 205 | 0 | 16 | 11 | 20 | 120 | 10 | 0 | 0 | 0 | 96 | 0 | 119 | 1240 | 694 | 0 | 0 | 12 | 57 | 2713 | 5719 | 0 | 0 | 0 | 0 | 0 | 0 | 76 | 41 | 27 | 66 |
| SRR2731324 | 105 | 0 | 767 | 0 | 14 | 226 | 129 | 0 | 21 | 0 | 0 | 67 | 21 | 0 | 0 | 0 | 300 | 0 | 167 | 802 | 1249 | 0 | 0 | 221 | 105 | 1651 | 3743 | 0 | 0 | 0 | 17 | 0 | 0 | 60 | 16 | 0 | 23 |
| SRR2731325 | 121 | 0 | 826 | 0 | 14 | 78 | 45 | 0 | 29 | 0 | 0 | 86 | 14 | 0 | 0 | 0 | 583 | 10 | 74 | 749 | 1176 | 0 | 0 | 62 | 39 | 621 | 1789 | 0 | 0 | 0 | 15 | 0 | 0 | 17 | 19 | 16 | 11 |
| SRR2731326 | 236 | 0 | 831 | 0 | 0 | 340 | 78 | 0 | 17 | 0 | 0 | 91 | 10 | 0 | 0 | 0 | 177 | 0 | 82 | 620 | 2339 | 0 | 0 | 90 | 96 | 2929 | 7002 | 0 | 0 | 0 | 15 | 0 | 0 | 91 | 23 | 13 | 40 |
| SRR2731327 | 158 | 0 | 897 | 0 | 26 | 96 | 112 | 0 | 19 | 0 | 0 | 101 | 14 | 0 | 0 | 0 | 309 | 0 | 149 | 681 | 1102 | 0 | 0 | 36 | 43 | 1008 | 1931 | 0 | 0 | 0 | 25 | 0 | 0 | 45 | 19 | 11 | 24 |
| SRR2731328 | 198 | 0 | 801 | 0 | 0 | 92 | 78 | 0 | 28 | 0 | 13 | 164 | 0 | 0 | 0 | 0 | 191 | 0 | 117 | 281 | 629 | 0 | 0 | 22 | 33 | 882 | 2858 | 0 | 0 | 0 | 15 | 0 | 0 | 13 | 18 | 0 | 30 |
| SRR2731329 | 187 | 0 | 1791 | 0 | 12 | 392 | 541 | 109 | 23 | 0 | 0 | 649 | 46 | 0 | 0 | 0 | 125 | 0 | 232 | 190 | 354 | 0 | 0 | 183 | 86 | 2515 | 7064 | 0 | 0 | 120 | 0 | 0 | 0 | 142 | 111 | 51 | 186 |
| SRR2731330 | 4006 | 0 | 953 | 0 | 0 | 218 | 453 | 0 | 0 | 0 | 0 | 379 | 0 | 0 | 0 | 0 | 58 | 0 | 144 | 123 | 477 | 16 | 0 | 11 | 49 | 2514 | 5808 | 0 | 0 | 0 | 24 | 0 | 0 | 47 | 271 | 157 | 492 |
| SRR2731331 | 112 | 0 | 1010 | 0 | 0 | 83 | 0 | 0 | 21 | 0 | 0 | 257 | 24 | 0 | 0 | 13 | 65 | 0 | 15 | 144 | 2407 | 10 | 0 | 28 | 29 | 623 | 538 | 0 | 13 | 0 | 12 | 13 | 0 | 12 | 60 | 27 | 72 |
| SRR2731332 | 377 | 0 | 697 | 0 | 0 | 562 | 82 | 0 | 54 | 0 | 0 | 102 | 20 | 0 | 0 | 0 | 76 | 0 | 88 | 1213 | 4266 | 0 | 0 | 126 | 59 | 3858 | 4842 | 0 | 0 | 0 | 12 | 0 | 0 | 36 | 53 | 30 | 84 |
| SRR2731333 | 242 | 0 | 2285 | 0 | 0 | 259 | 99 | 0 | 19 | 0 | 0 | 96 | 21 | 0 | 0 | 0 | 377 | 0 | 53 | 161 | 479 | 0 | 0 | 22 | 60 | 3252 | 4143 | 0 | 0 | 0 | 25 | 0 | 0 | 40 | 165 | 104 | 279 |
| SRR2731335 | 57 | 0 | 701 | 0 | 0 | 78 | 56 | 0 | 24 | 0 | 0 | 54 | 23 | 0 | 0 | 0 | 257 | 0 | 63 | 306 | 921 | 0 | 0 | 114 | 14 | 587 | 1514 | 0 | 0 | 0 | 0 | 0 | 0 | 10 | 16 | 13 | 22 |
| SRR2731336 | 907 | 0 | 487 | 0 | 0 | 208 | 351 | 0 | 18 | 0 | 0 | 322 | 10 | 0 | 0 | 0 | 33 | 0 | 141 | 709 | 3346 | 10 | 0 | 26 | 47 | 2038 | 2871 | 0 | 12 | 0 | 13 | 0 | 0 | 17 | 65 | 22 | 71 |
| SRR2731337 | 189 | 0 | 931 | 0 | 0 | 141 | 39 | 0 | 32 | 0 | 0 | 166 | 11 | 0 | 0 | 0 | 96 | 0 | 46 | 502 | 2150 | 0 | 0 | 83 | 49 | 932 | 1670 | 0 | 0 | 0 | 34 | 0 | 0 | 0 | 32 | 30 | 23 |
| SRR2731338 | 47 | 0 | 494 | 0 | 0 | 52 | 0 | 0 | 16 | 0 | 0 | 215 | 12 | 0 | 0 | 0 | 108 | 0 | 0 | 14 | 1115 | 28 | 0 | 31 | 19 | 427 | 258 | 0 | 21 | 0 | 10 | 12 | 0 | 13 | 45 | 42 | 89 |
| SRR2731339 | 247 | 0 | 1485 | 25 | 40 | 42 | 24 | 0 | 10 | 0 | 0 | 157 | 0 | 0 | 24 | 32 | 45 | 0 | 12 | 60 | 191 | 92 | 0 | 0 | 68 | 559 | 317 | 39 | 88 | 0 | 56 | 48 | 0 | 124 | 26 | 22 | 44 |
| SRR2731340 | 197 | 0 | 934 | 0 | 0 | 514 | 87 | 0 | 47 | 0 | 0 | 87 | 12 | 0 | 0 | 0 | 119 | 0 | 21 | 190 | 1217 | 0 | 0 | 94 | 72 | 4802 | 5839 | 0 | 11 | 0 | 38 | 0 | 0 | 13 | 149 | 90 | 237 |
| SRR2731341 | 172 | 0 | 834 | 0 | 0 | 92 | 39 | 0 | 20 | 0 | 13 | 104 | 16 | 0 | 0 | 0 | 287 | 0 | 29 | 548 | 1570 | 0 | 0 | 16 | 29 | 758 | 1971 | 0 | 0 | 0 | 13 | 0 | 0 | 28 | 16 | 11 | 31 |
| SRR2731342 | 199 | 0 | 889 | 0 | 44 | 307 | 72 | 0 | 26 | 0 | 12 | 57 | 17 | 0 | 0 | 0 | 291 | 0 | 90 | 726 | 1009 | 0 | 0 | 136 | 94 | 2144 | 3305 | 0 | 0 | 0 | 33 | 0 | 0 | 84 | 0 | 0 | 18 |
| SRR2731343 | 107 | 0 | 1002 | 0 | 26 | 63 | 153 | 0 | 18 | 0 | 0 | 307 | 11 | 0 | 0 | 0 | 290 | 0 | 99 | 1398 | 923 | 0 | 0 | 40 | 34 | 603 | 1236 | 0 | 0 | 0 | 26 | 0 | 0 | 21 | 0 | 0 | 13 |
| SRR2731344 | 135 | 0 | 1175 | 0 | 16 | 141 | 50 | 0 | 15 | 0 | 0 | 73 | 14 | 0 | 0 | 0 | 464 | 0 | 50 | 619 | 731 | 0 | 0 | 30 | 76 | 1116 | 1812 | 0 | 0 | 0 | 30 | 0 | 0 | 19 | 17 | 31 | 23 |
| SRR2731345 | 72 | 0 | 207 | 0 | 0 | 288 | 13 | 0 | 42 | 0 | 35 | 39 | 58 | 0 | 60 | 0 | 97 | 0 | 13 | 304 | 748 | 0 | 0 | 35 | 403 | 1722 | 1152 | 0 | 0 | 0 | 163 | 0 | 19 | 129 | 17 | 19 | 61 |
| SRR2731346 | 128 | 0 | 1434 | 0 | 0 | 56 | 36 | 0 | 23 | 0 | 0 | 131 | 16 | 0 | 0 | 0 | 321 | 0 | 41 | 137 | 444 | 0 | 0 | 48 | 28 | 515 | 1139 | 0 | 0 | 0 | 25 | 0 | 0 | 19 | 29 | 22 | 11 |
| SRR2731347 | 107 | 0 | 467 | 0 | 14 | 223 | 74 | 0 | 24 | 0 | 0 | 164 | 52 | 0 | 0 | 0 | 345 | 24 | 77 | 745 | 2692 | 0 | 0 | 283 | 76 | 1665 | 2945 | 0 | 0 | 0 | 18 | 0 | 0 | 40 | 28 | 18 | 55 |
| SRR2731348 | 86 | 0 | 686 | 0 | 0 | 218 | 48 | 0 | 26 | 0 | 15 | 115 | 21 | 0 | 0 | 0 | 112 | 0 | 72 | 273 | 755 | 0 | 0 | 41 | 68 | 1574 | 1788 | 0 | 0 | 0 | 16 | 0 | 0 | 23 | 22 | 14 | 10 |
| SRR2731349 | 36 | 0 | 219 | 0 | 0 | 32 | 18 | 0 | 35 | 16 | 67 | 168 | 31 | 0 | 56 | 0 | 37 | 13 | 15 | 146 | 553 | 0 | 0 | 15 | 114 | 271 | 281 | 0 | 0 | 0 | 23 | 0 | 11 | 25 | 25 | 16 | 38 |
| SRR2731350 | 141 | 0 | 716 | 0 | 0 | 546 | 23 | 0 | 63 | 0 | 0 | 75 | 67 | 0 | 0 | 0 | 252 | 0 | 39 | 305 | 706 | 0 | 0 | 389 | 154 | 3653 | 5431 | 0 | 0 | 0 | 18 | 0 | 0 | 57 | 29 | 17 | 59 |
| SRR2910706 | 37 | 17 | 1997 | 0 | 0 | 45 | 171 | 547 | 27 | 0 | 0 | 53 | 0 | 12 | 69 | 0 | 31 | 0 | 20 | 1033 | 525 | 22 | 0 | 0 | 10 | 771 | 560 | 0 | 118 | 162 | 17 | 0 | 0 | 11 | 531 | 40 | 196 |
| SRR2910737 | 33 | 25 | 1964 | 0 | 0 | 42 | 128 | 349 | 36 | 0 | 0 | 43 | 0 | 21 | 202 | 0 | 21 | 0 | 0 | 440 | 348 | 15 | 0 | 0 | 0 | 748 | 291 | 0 | 130 | 108 | 25 | 0 | 0 | 13 | 509 | 28 | 213 |
| SRR2910738 | 37 | 18 | 2084 | 0 | 0 | 45 | 146 | 120 | 32 | 0 | 0 | 50 | 0 | 10 | 106 | 0 | 24 | 0 | 10 | 848 | 475 | 21 | 0 | 0 | 0 | 143 | 77 | 10 | 15 | 39 | 22 | 0 | 0 | 16 | 65 | 21 | 32 |
| SRR2912507 | 0 | 248 | 10 | 0 | 17 | 0 | 14946 | 32 | 0 | 0 | 0 | 0 | 0 | 0 | 0 | 0 | 51 | 0 | 92879 | 12 | 0 | 0 | 0 | 0 | 0 | 32 | 149 | 0 | 0 | 0 | 0 | 0 | 0 | 0 | 750 | 338 | 285 |
| SRR2912508 | 0 | 170 | 48 | 0 | 92 | 16 | 6140 | 0 | 12 | 0 | 0 | 21 | 0 | 0 | 0 | 0 | 96 | 0 | 121510 | 10 | 0 | 0 | 0 | 0 | 24 | 95 | 667 | 0 | 0 | 0 | 0 | 0 | 0 | 0 | 2038 | 862 | 624 |
| SRR2912510 | 0 | 136 | 275 | 0 | 0 | 16 | 6832 | 16 | 0 | 0 | 0 | 0 | 0 | 0 | 0 | 0 | 107 | 0 | 26634 | 0 | 0 | 0 | 0 | 0 | 0 | 108 | 489 | 0 | 0 | 0 | 0 | 0 | 0 | 0 | 18 | 0 | 0 |
| SRR2912511 | 0 | 129 | 116 | 0 | 0 | 33 | 5347 | 16 | 0 | 0 | 0 | 0 | 0 | 0 | 0 | 0 | 64 | 0 | 20858 | 0 | 0 | 0 | 0 | 0 | 0 | 152 | 460 | 0 | 0 | 0 | 0 | 0 | 0 | 0 | 29 | 0 | 11 |
| SRR2912513 | 0 | 86 | 258 | 0 | 0 | 16 | 5760 | 0 | 0 | 0 | 0 | 39 | 0 | 0 | 0 | 0 | 93 | 0 | 17251 | 0 | 0 | 0 | 0 | 0 | 0 | 17 | 29 | 0 | 0 | 0 | 0 | 0 | 0 | 0 | 11 | 0 | 0 |
| SRR2912514 | 0 | 231 | 13 | 0 | 32 | 0 | 4521 | 27 | 0 | 0 | 0 | 11 | 0 | 11 | 0 | 0 | 59 | 0 | 151600 | 14 | 0 | 0 | 0 | 0 | 0 | 33 | 287 | 0 | 0 | 0 | 0 | 0 | 0 | 0 | 577 | 330 | 206 |
| SRR2912515 | 0 | 172 | 189 | 0 | 0 | 24 | 6239 | 14 | 0 | 0 | 0 | 21 | 0 | 0 | 0 | 0 | 58 | 0 | 29079 | 0 | 0 | 0 | 0 | 0 | 0 | 44 | 198 | 0 | 0 | 0 | 0 | 0 | 0 | 0 | 13 | 0 | 0 |
| SRR2913202 | 1187 | 0 | 826 | 0 | 0 | 19 | 0 | 16 | 0 | 0 | 0 | 13 | 0 | 0 | 0 | 0 | 0 | 0 | 0 | 2674 | 3093 | 0 | 0 | 43 | 0 | 160 | 97 | 0 | 0 | 0 | 0 | 0 | 10 | 0 | 0 | 0 | 0 |
| SRR2913203 | 367 | 0 | 345 | 0 | 0 | 41 | 63 | 21 | 22 | 0 | 0 | 0 | 0 | 0 | 11 | 0 | 0 | 0 | 11 | 2235 | 1170 | 0 | 0 | 16 | 0 | 98 | 45 | 0 | 0 | 0 | 0 | 0 | 0 | 0 | 0 | 0 | 11 |
| SRR2913204 | 846 | 19 | 957 | 12 | 0 | 36 | 59 | 37 | 19 | 0 | 0 | 26 | 0 | 0 | 19 | 0 | 0 | 0 | 0 | 1162 | 993 | 17 | 0 | 17 | 0 | 89 | 31 | 0 | 0 | 19 | 11 | 0 | 0 | 0 | 0 | 0 | 0 |
| SRR2913205 | 1138 | 26 | 642 | 0 | 0 | 19 | 41 | 13 | 0 | 0 | 0 | 15 | 0 | 0 | 11 | 0 | 15 | 14 | 16 | 1349 | 1462 | 14 | 0 | 13 | 0 | 73 | 25 | 0 | 0 | 0 | 11 | 0 | 0 | 0 | 0 | 0 | 0 |
| SRR2913206 | 1246 | 13 | 721 | 11 | 0 | 0 | 0 | 18 | 0 | 0 | 0 | 0 | 0 | 0 | 42 | 0 | 0 | 15 | 0 | 2177 | 2620 | 14 | 0 | 19 | 0 | 41 | 41 | 0 | 0 | 0 | 0 | 0 | 0 | 0 | 0 | 0 | 0 |
| SRR2913207 | 485 | 27 | 725 | 11 | 0 | 16 | 0 | 485 | 0 | 0 | 0 | 15 | 0 | 0 | 0 | 0 | 0 | 0 | 0 | 3724 | 82 | 12 | 0 | 59 | 0 | 262 | 249 | 0 | 0 | 23 | 0 | 0 | 0 | 0 | 28 | 0 | 15 |
| SRR2913208 | 862 | 21 | 644 | 12 | 0 | 0 | 39 | 13 | 0 | 0 | 0 | 0 | 0 | 0 | 83 | 0 | 0 | 74 | 0 | 1676 | 3418 | 18 | 0 | 34 | 0 | 73 | 58 | 0 | 0 | 0 | 0 | 0 | 0 | 0 | 0 | 0 | 0 |
| SRR2913209 | 457 | 10 | 533 | 0 | 0 | 0 | 37 | 15 | 0 | 0 | 0 | 19 | 0 | 0 | 33 | 0 | 0 | 0 | 0 | 704 | 3108 | 0 | 0 | 30 | 0 | 40 | 18 | 0 | 0 | 0 | 0 | 0 | 0 | 0 | 0 | 0 | 0 |
| SRR2913210 | 697 | 17 | 820 | 0 | 0 | 0 | 24 | 37 | 0 | 0 | 0 | 18 | 0 | 0 | 34 | 0 | 0 | 20 | 0 | 1129 | 2089 | 0 | 0 | 46 | 0 | 123 | 60 | 0 | 0 | 0 | 0 | 0 | 0 | 0 | 0 | 0 | 0 |
| SRR2913211 | 1394 | 13 | 1385 | 14 | 0 | 14 | 11 | 14 | 10 | 0 | 0 | 0 | 0 | 10 | 77 | 0 | 0 | 0 | 11 | 2452 | 2507 | 0 | 0 | 25 | 0 | 54 | 35 | 11 | 0 | 0 | 0 | 0 | 0 | 0 | 0 | 0 | 0 |
| SRR2913212 | 404 | 25 | 658 | 0 | 0 | 0 | 132 | 19 | 0 | 0 | 0 | 45 | 0 | 0 | 39 | 0 | 0 | 0 | 28 | 931 | 25 | 0 | 0 | 15 | 0 | 21 | 20 | 0 | 0 | 0 | 0 | 0 | 0 | 0 | 0 | 0 | 0 |
| SRR2913213 | 714 | 0 | 571 | 0 | 0 | 0 | 0 | 28 | 0 | 0 | 0 | 25 | 0 | 0 | 36 | 0 | 0 | 17 | 0 | 1383 | 0 | 0 | 0 | 18 | 0 | 82 | 45 | 0 | 0 | 40 | 0 | 0 | 0 | 0 | 0 | 0 | 0 |
| SRR2913214 | 1254 | 38 | 589 | 14 | 0 | 12 | 23 | 0 | 0 | 0 | 0 | 0 | 0 | 0 | 16 | 0 | 0 | 24 | 23 | 3487 | 5605 | 16 | 0 | 69 | 0 | 113 | 86 | 10 | 0 | 0 | 0 | 12 | 0 | 0 | 0 | 0 | 0 |
| SRR2913215 | 913 | 37 | 820 | 21 | 0 | 10 | 50 | 18 | 0 | 0 | 0 | 12 | 0 | 12 | 32 | 0 | 0 | 0 | 0 | 2732 | 1391 | 30 | 0 | 88 | 0 | 137 | 139 | 11 | 0 | 16 | 0 | 0 | 0 | 0 | 0 | 0 | 0 |
| SRR2913216 | 984 | 15 | 802 | 11 | 0 | 0 | 28 | 202 | 0 | 0 | 0 | 21 | 0 | 0 | 0 | 0 | 0 | 0 | 14 | 584 | 812 | 15 | 0 | 49 | 0 | 115 | 63 | 0 | 0 | 18 | 0 | 0 | 0 | 0 | 0 | 0 | 0 |
| SRR2913217 | 737 | 18 | 670 | 20 | 0 | 0 | 49 | 16 | 0 | 0 | 0 | 0 | 0 | 0 | 35 | 0 | 0 | 16 | 21 | 3226 | 2425 | 21 | 0 | 28 | 0 | 53 | 45 | 0 | 0 | 0 | 0 | 0 | 0 | 0 | 0 | 0 | 0 |
| SRR2913218 | 1420 | 39 | 962 | 17 | 0 | 0 | 19 | 33 | 0 | 0 | 0 | 19 | 0 | 0 | 0 | 0 | 0 | 0 | 0 | 5679 | 713 | 19 | 0 | 27 | 0 | 18 | 18 | 0 | 0 | 0 | 0 | 0 | 0 | 0 | 0 | 0 | 0 |
| SRR2913219 | 1599 | 29 | 948 | 24 | 0 | 0 | 0 | 0 | 0 | 0 | 0 | 16 | 0 | 10 | 55 | 0 | 0 | 0 | 34 | 4118 | 2667 | 15 | 0 | 63 | 0 | 79 | 106 | 0 | 0 | 0 | 0 | 0 | 0 | 0 | 0 | 0 | 0 |
| SRR2913220 | 988 | 11 | 1036 | 23 | 0 | 23 | 77 | 30 | 0 | 0 | 0 | 0 | 0 | 0 | 20 | 0 | 0 | 0 | 0 | 2790 | 707 | 12 | 0 | 19 | 0 | 37 | 41 | 0 | 0 | 17 | 0 | 0 | 0 | 0 | 0 | 0 | 0 |
| SRR2913221 | 594 | 30 | 1006 | 17 | 0 | 14 | 45 | 46 | 0 | 0 | 0 | 0 | 0 | 14 | 13 | 0 | 0 | 0 | 0 | 2852 | 224 | 25 | 0 | 100 | 0 | 135 | 127 | 16 | 0 | 10 | 0 | 0 | 0 | 0 | 0 | 0 | 0 |
| SRR2913222 | 1182 | 19 | 788 | 24 | 0 | 0 | 58 | 17 | 0 | 0 | 0 | 15 | 0 | 0 | 16 | 0 | 0 | 0 | 0 | 1559 | 421 | 19 | 0 | 18 | 0 | 50 | 33 | 0 | 0 | 0 | 0 | 0 | 0 | 0 | 0 | 0 | 0 |
| SRR2913223 | 342 | 25 | 283 | 0 | 0 | 18 | 0 | 23 | 0 | 0 | 0 | 0 | 0 | 0 | 11 | 0 | 0 | 0 | 0 | 1015 | 261 | 14 | 0 | 31 | 0 | 23 | 12 | 0 | 0 | 0 | 0 | 0 | 0 | 0 | 12 | 0 | 0 |
| SRR2913224 | 719 | 37 | 744 | 21 | 0 | 10 | 15 | 19 | 0 | 0 | 0 | 17 | 0 | 0 | 13 | 0 | 11 | 0 | 0 | 2118 | 593 | 19 | 0 | 68 | 0 | 126 | 138 | 0 | 0 | 0 | 0 | 0 | 0 | 0 | 0 | 0 | 0 |
| SRR2913225 | 678 | 11 | 1006 | 16 | 0 | 11 | 123 | 22 | 0 | 0 | 0 | 13 | 0 | 13 | 14 | 0 | 10 | 0 | 36 | 2801 | 734 | 20 | 0 | 27 | 0 | 49 | 47 | 11 | 0 | 0 | 0 | 0 | 0 | 0 | 10 | 0 | 0 |
| SRR2915342 | 21 | 0 | 404 | 13 | 17 | 27 | 47 | 39 | 14 | 0 | 11 | 1121 | 26 | 20 | 48 | 0 | 0 | 0 | 34 | 435 | 1060 | 0 | 0 | 94 | 28 | 136 | 152 | 21 | 0 | 32 | 32 | 0 | 0 | 101 | 97 | 19 | 57 |
| SRR2915343 | 26 | 0 | 493 | 13 | 27 | 32 | 45 | 25 | 14 | 0 | 24 | 1173 | 17 | 18 | 77 | 0 | 0 | 15 | 43 | 449 | 709 | 0 | 0 | 121 | 39 | 175 | 221 | 20 | 0 | 21 | 34 | 0 | 0 | 153 | 85 | 27 | 64 |
| SRR2915344 | 21 | 0 | 540 | 10 | 52 | 79 | 42 | 55 | 38 | 0 | 28 | 1812 | 15 | 29 | 62 | 0 | 0 | 15 | 64 | 689 | 1970 | 0 | 0 | 383 | 103 | 415 | 499 | 11 | 0 | 58 | 73 | 0 | 0 | 232 | 336 | 64 | 209 |
| SRR2915345 | 73 | 0 | 390 | 14 | 117 | 278 | 14 | 15 | 109 | 0 | 36 | 439 | 26 | 24 | 65 | 0 | 0 | 34 | 31 | 379 | 357 | 0 | 0 | 104 | 163 | 1543 | 538 | 0 | 0 | 11 | 159 | 0 | 0 | 184 | 831 | 882 | 680 |
| SRR2915346 | 74 | 0 | 393 | 15 | 48 | 78 | 29 | 17 | 36 | 16 | 23 | 505 | 10 | 19 | 42 | 0 | 0 | 19 | 43 | 456 | 515 | 0 | 0 | 116 | 81 | 432 | 305 | 0 | 0 | 15 | 58 | 0 | 0 | 102 | 156 | 144 | 126 |
| SRR2915347 | 84 | 0 | 526 | 0 | 46 | 97 | 22 | 16 | 27 | 0 | 14 | 484 | 11 | 14 | 70 | 0 | 0 | 71 | 51 | 579 | 455 | 0 | 0 | 192 | 93 | 582 | 495 | 0 | 0 | 11 | 51 | 0 | 0 | 82 | 218 | 194 | 148 |
| SRR2915348 | 28 | 0 | 233 | 10 | 11 | 68 | 0 | 43 | 26 | 0 | 22 | 1166 | 11 | 18 | 161 | 0 | 0 | 29 | 45 | 721 | 1571 | 0 | 0 | 78 | 25 | 364 | 144 | 0 | 0 | 12 | 119 | 0 | 0 | 40 | 60 | 56 | 52 |
| SRR2915349 | 35 | 0 | 303 | 0 | 14 | 67 | 13 | 0 | 32 | 0 | 19 | 1391 | 43 | 19 | 94 | 0 | 0 | 36 | 31 | 396 | 1125 | 0 | 0 | 20 | 47 | 394 | 145 | 0 | 0 | 11 | 130 | 0 | 0 | 34 | 61 | 38 | 36 |
| SRR2915350 | 36 | 0 | 305 | 0 | 42 | 74 | 16 | 12 | 33 | 0 | 18 | 1767 | 30 | 34 | 168 | 0 | 0 | 67 | 31 | 540 | 1347 | 0 | 0 | 45 | 48 | 424 | 275 | 13 | 0 | 10 | 135 | 0 | 0 | 17 | 78 | 71 | 61 |
| SRR2915905 | 0 | 0 | 173 | 0 | 0 | 55 | 88 | 11 | 31 | 0 | 0 | 11 | 0 | 15 | 0 | 0 | 0 | 0 | 63 | 53 | 137 | 0 | 0 | 0 | 13 | 18 | 114 | 0 | 20 | 0 | 62 | 0 | 0 | 26 | 142 | 0 | 106 |
| SRR2915915 | 19 | 0 | 75 | 0 | 0 | 27 | 37 | 0 | 15 | 0 | 0 | 0 | 0 | 12 | 11 | 0 | 0 | 0 | 22 | 23 | 105 | 0 | 0 | 0 | 23 | 286 | 19 | 0 | 0 | 0 | 15 | 0 | 0 | 25 | 271 | 17 | 172 |
| SRR2915918 | 39 | 0 | 1255 | 0 | 0 | 38 | 220 | 33 | 14 | 0 | 0 | 87 | 0 | 0 | 231 | 0 | 16 | 0 | 46 | 577 | 226 | 0 | 0 | 16 | 11 | 11 | 18 | 0 | 59 | 187 | 20 | 0 | 0 | 44 | 191 | 12 | 129 |
| SRR2915921 | 38 | 13 | 1670 | 0 | 0 | 13 | 632 | 108 | 12 | 0 | 0 | 158 | 0 | 16 | 316 | 0 | 21 | 0 | 0 | 427 | 196 | 0 | 0 | 0 | 19 | 109 | 398 | 0 | 311 | 41 | 0 | 0 | 0 | 69 | 22 | 0 | 14 |
| SRR2915923 | 21 | 11 | 610 | 0 | 0 | 14 | 475 | 222 | 11 | 0 | 0 | 73 | 0 | 16 | 94 | 0 | 0 | 0 | 27 | 138 | 90 | 0 | 0 | 0 | 11 | 320 | 355 | 12 | 53 | 0 | 33 | 0 | 0 | 44 | 34 | 0 | 23 |
| SRR2915926 | 51 | 0 | 3822 | 0 | 0 | 29 | 320 | 14 | 11 | 0 | 0 | 79 | 0 | 0 | 15 | 0 | 12 | 0 | 126 | 310 | 69 | 0 | 0 | 0 | 27 | 244 | 262 | 0 | 88 | 74 | 0 | 0 | 0 | 68 | 67 | 0 | 40 |
| SRR2915928 | 43 | 27 | 1712 | 0 | 0 | 60 | 101 | 38 | 36 | 0 | 0 | 64 | 0 | 38 | 25 | 0 | 20 | 0 | 11 | 742 | 253 | 0 | 0 | 0 | 35 | 71 | 60 | 32 | 13 | 0 | 59 | 0 | 16 | 82 | 90 | 0 | 35 |
| SRR2960039 | 595 | 0 | 866 | 0 | 45 | 0 | 0 | 0 | 0 | 22 | 0 | 129 | 0 | 34 | 350 | 0 | 72 | 217 | 0 | 0 | 0 | 0 | 11 | 0 | 0 | 0 | 0 | 17 | 34 | 0 | 151 | 0 | 0 | 0 | 308 | 101 | 0 |

### Supp Table S8.2.Columns_AM_to_BW

| | | | | | | | | | | | | | | | | | | | | | | | | | | | | | | | | | | | | | |
|---|---|---|---|---|---|---|---|---|---|---|---|---|---|---|---|---|---|---|---|---|---|---|---|---|---|---|---|---|---|---|---|---|---|---|---|---|---|
| SRR2976280 | 77 | 11 | 1501 | 0 | 0 | 0 | 0 | 0 | 0 | 0 | 0 | 0 | 0 | 115 | 0 | 0 | 0 | 0 | 0 | 0 | 0 | 0 | 0 | 0 | 0 | 0 | 0 | 0 | 0 | 0 | 0 | 0 | 0 | 0 | 0 | 0 | 0 |
| SRR2976305 | 205 | 0 | 1565 | 0 | 0 | 0 | 0 | 0 | 0 | 0 | 0 | 0 | 0 | 0 | 113 | 0 | 0 | 0 | 0 | 0 | 0 | 0 | 0 | 0 | 0 | 0 | 0 | 20 | 0 | 0 | 0 | 14 | 0 | 0 | 0 | 0 | 0 |
| SRR3092114 | 76 | 0 | 1182 | 0 | 0 | 0 | 0 | 0 | 0 | 0 | 0 | 578 | 0 | 0 | 1997 | 0 | 30 | 0 | 0 | 37 | 256 | 0 | 0 | 0 | 12 | 63 | 0 | 0 | 0 | 0 | 55 | 0 | 0 | 0 | 611 | 925 | 875 |
| SRR3092115 | 109 | 0 | 1746 | 0 | 0 | 0 | 0 | 0 | 0 | 0 | 0 | 381 | 0 | 0 | 4011 | 0 | 23 | 0 | 0 | 40 | 359 | 0 | 0 | 0 | 0 | 16 | 0 | 0 | 0 | 0 | 20 | 0 | 0 | 0 | 198 | 315 | 314 |
| SRR3103222 | 27 | 0 | 486 | 0 | 0 | 21 | 30 | 0 | 23 | 0 | 0 | 16 | 0 | 26 | 5000 | 0 | 0 | 0 | 0 | 34 | 109 | 0 | 0 | 0 | 26 | 12 | 86 | 10 | 0 | 0 | 110 | 0 | 0 | 0 | 193 | 31 | 76 |
| SRR3103223 | 47 | 28 | 751 | 0 | 0 | 28 | 836 | 99 | 14 | 0 | 0 | 0 | 0 | 28 | 251 | 14 | 0 | 0 | 59 | 964 | 339 | 0 | 0 | 0 | 0 | 57 | 0 | 0 | 0 | 0 | 0 | 0 | 0 | 57 | 142 | 0 | 71 |
| SRR3103224 | 45 | 0 | 552 | 0 | 0 | 45 | 229 | 13 | 45 | 0 | 0 | 0 | 0 | 32 | 458 | 0 | 0 | 10 | 164 | 2127 | 851 | 0 | 0 | 0 | 0 | 84 | 0 | 0 | 0 | 0 | 13 | 0 | 0 | 0 | 90 | 13 | 77 |
| SRR3103225 | 31 | 14 | 379 | 0 | 0 | 76 | 503 | 303 | 43 | 0 | 0 | 22 | 0 | 19 | 0 | 0 | 0 | 0 | 16 | 193 | 89 | 0 | 0 | 0 | 0 | 960 | 709 | 0 | 0 | 0 | 57 | 0 | 0 | 0 | 128 | 0 | 57 |
| SRR3103226 | 82 | 0 | 931 | 0 | 0 | 42 | 625 | 42 | 0 | 0 | 0 | 0 | 0 | 21 | 2647 | 0 | 0 | 0 | 37 | 1176 | 2500 | 0 | 0 | 0 | 0 | 42 | 21 | 0 | 21 | 21 | 0 | 0 | 0 | 42 | 106 | 0 | 42 |
| SRR3103227 | 21 | 0 | 254 | 0 | 0 | 30 | 90 | 51 | 91 | 0 | 0 | 12 | 0 | 0 | 294 | 0 | 0 | 0 | 34 | 296 | 378 | 0 | 0 | 0 | 10 | 1586 | 606 | 20 | 0 | 0 | 91 | 0 | 10 | 0 | 192 | 10 | 151 |
| SRR3103228 | 18 | 20 | 162 | 0 | 0 | 34 | 40 | 30 | 44 | 0 | 0 | 11 | 0 | 13 | 176 | 0 | 0 | 0 | 18 | 892 | 426 | 0 | 0 | 0 | 20 | 772 | 329 | 0 | 0 | 0 | 34 | 0 | 0 | 0 | 54 | 17 | 44 |
| SRR3103229 | 33 | 11 | 375 | 0 | 0 | 35 | 383 | 0 | 0 | 0 | 0 | 0 | 0 | 35 | 409 | 0 | 0 | 0 | 368 | 1318 | 626 | 0 | 0 | 0 | 0 | 11 | 91 | 0 | 0 | 0 | 67 | 0 | 0 | 14 | 151 | 11 | 105 |
| SRR3103230 | 13 | 17 | 165 | 0 | 0 | 17 | 369 | 345 | 67 | 0 | 0 | 0 | 0 | 34 | 0 | 0 | 0 | 0 | 0 | 265 | 92 | 0 | 0 | 0 | 17 | 20 | 514 | 0 | 0 | 0 | 34 | 0 | 0 | 25 | 109 | 0 | 67 |
| SRR3103231 | 19 | 0 | 224 | 0 | 0 | 18 | 334 | 0 | 0 | 0 | 0 | 0 | 0 | 0 | 132 | 0 | 0 | 0 | 65 | 1345 | 216 | 0 | 0 | 0 | 14 | 501 | 127 | 0 | 0 | 0 | 55 | 0 | 0 | 0 | 77 | 0 | 59 |
| SRR3103232 | 11 | 18 | 196 | 0 | 0 | 0 | 168 | 54 | 54 | 0 | 0 | 0 | 0 | 0 | 672 | 0 | 0 | 0 | 24 | 648 | 1319 | 0 | 0 | 0 | 18 | 947 | 518 | 0 | 0 | 0 | 125 | 0 | 0 | 0 | 125 | 0 | 54 |
| SRR3103233 | 19 | 0 | 195 | 0 | 0 | 36 | 36 | 0 | 24 | 0 | 0 | 11 | 0 | 0 | 72 | 0 | 0 | 0 | 0 | 347 | 215 | 0 | 0 | 0 | 12 | 14 | 379 | 12 | 12 | 0 | 83 | 0 | 0 | 0 | 112 | 12 | 59 |
| SRR3103234 | 47 | 33 | 220 | 0 | 0 | 33 | 772 | 312 | 37 | 0 | 0 | 0 | 0 | 41 | 0 | 0 | 0 | 0 | 26 | 267 | 118 | 0 | 0 | 0 | 0 | 21 | 304 | 0 | 0 | 0 | 30 | 0 | 0 | 0 | 152 | 19 | 45 |
| SRR3103235 | 36 | 0 | 295 | 0 | 0 | 52 | 215 | 77 | 29 | 0 | 0 | 0 | 0 | 39 | 25 | 0 | 0 | 0 | 24 | 512 | 290 | 0 | 0 | 0 | 0 | 580 | 309 | 0 | 0 | 0 | 26 | 0 | 0 | 0 | 97 | 13 | 55 |
| SRR3103236 | 38 | 0 | 317 | 0 | 0 | 13 | 61 | 10 | 24 | 0 | 0 | 0 | 0 | 10 | 37 | 0 | 0 | 0 | 0 | 166 | 131 | 0 | 0 | 0 | 0 | 13 | 14 | 10 | 0 | 0 | 37 | 0 | 0 | 0 | 114 | 0 | 40 |
| SRR3103237 | 32 | 0 | 256 | 0 | 0 | 119 | 173 | 52 | 38 | 0 | 0 | 0 | 0 | 0 | 0 | 0 | 0 | 0 | 0 | 94 | 66 | 0 | 0 | 0 | 19 | 38 | 466 | 0 | 0 | 0 | 105 | 0 | 0 | 24 | 133 | 29 | 109 |
| SRR3103238 | 22 | 44 | 211 | 0 | 0 | 35 | 66 | 96 | 15 | 0 | 0 | 15 | 0 | 29 | 18 | 0 | 0 | 0 | 0 | 57 | 117 | 0 | 0 | 0 | 0 | 27 | 36 | 0 | 0 | 15 | 53 | 0 | 0 | 11 | 140 | 26 | 90 |
| SRR3103239 | 32 | 0 | 256 | 0 | 0 | 0 | 118 | 0 | 0 | 0 | 0 | 0 | 0 | 0 | 604 | 0 | 0 | 0 | 162 | 20085 | 1488 | 0 | 0 | 0 | 0 | 221 | 109 | 0 | 0 | 0 | 16 | 0 | 0 | 0 | 100 | 0 | 56 |
| SRR3103240 | 41 | 0 | 627 | 0 | 0 | 0 | 83 | 0 | 0 | 0 | 0 | 34 | 0 | 18 | 279 | 0 | 0 | 0 | 50 | 1190 | 405 | 0 | 0 | 0 | 0 | 102 | 46 | 0 | 0 | 0 | 30 | 0 | 0 | 0 | 125 | 12 | 72 |
| SRR3103241 | 38 | 0 | 791 | 0 | 0 | 0 | 56 | 0 | 0 | 0 | 0 | 61 | 0 | 13 | 1392 | 0 | 0 | 0 | 33 | 5012 | 1615 | 0 | 0 | 0 | 0 | 139 | 69 | 0 | 13 | 0 | 76 | 0 | 0 | 0 | 151 | 0 | 177 |
| SRR3103242 | 31 | 0 | 325 | 0 | 0 | 46 | 386 | 186 | 37 | 0 | 0 | 14 | 0 | 23 | 1316 | 14 | 0 | 0 | 24 | 1457 | 1536 | 0 | 0 | 0 | 19 | 29 | 18 | 0 | 19 | 23 | 14 | 0 | 0 | 37 | 16 | 37 | 167 |
| SRR3103243 | 34 | 37 | 295 | 0 | 0 | 12 | 89 | 0 | 15 | 0 | 0 | 11 | 0 | 108 | 250 | 25 | 0 | 15 | 53 | 303 | 143 | 0 | 0 | 0 | 0 | 35 | 0 | 0 | 37 | 0 | 0 | 0 | 0 | 133 | 277 | 0 | 87 |
| SRR3103244 | 42 | 202 | 952 | 0 | 0 | 24 | 1032 | 499 | 42 | 0 | 0 | 29 | 0 | 154 | 31 | 0 | 12 | 11 | 37 | 1168 | 644 | 0 | 0 | 0 | 0 | 59 | 12 | 0 | 0 | 12 | 0 | 0 | 0 | 95 | 344 | 42 | 214 |
| SRR3103245 | 35 | 15 | 624 | 0 | 0 | 83 | 299 | 12 | 253 | 0 | 0 | 20 | 0 | 0 | 4661 | 18 | 18 | 0 | 0 | 668 | 1315 | 0 | 0 | 0 | 0 | 429 | 149 | 0 | 0 | 0 | 60 | 0 | 0 | 12 | 324 | 15 | 185 |
| SRR3103246 | 74 | 19 | 1022 | 0 | 0 | 0 | 1158 | 170 | 12 | 0 | 0 | 47 | 0 | 23 | 270 | 0 | 0 | 0 | 71 | 1675 | 1192 | 0 | 0 | 0 | 0 | 21 | 23 | 0 | 30 | 46 | 15 | 0 | 0 | 97 | 226 | 11 | 138 |
| SRR3103247 | 24 | 20 | 332 | 0 | 0 | 39 | 262 | 44 | 24 | 0 | 0 | 61 | 0 | 0 | 15704 | 0 | 0 | 0 | 0 | 1687 | 1786 | 0 | 0 | 0 | 0 | 886 | 431 | 0 | 0 | 0 | 59 | 0 | 0 | 0 | 377 | 44 | 161 |
| SRR3103248 | 13 | 0 | 264 | 0 | 0 | 78 | 33 | 42 | 44 | 0 | 0 | 0 | 0 | 0 | 31 | 0 | 0 | 0 | 10 | 275 | 126 | 0 | 0 | 0 | 0 | 100 | 23 | 0 | 145 | 23 | 38 | 0 | 0 | 0 | 105 | 0 | 55 |
| SRR3103249 | 12 | 14 | 754 | 0 | 0 | 16 | 156 | 14 | 0 | 0 | 0 | 88 | 0 | 19 | 17 | 0 | 14 | 0 | 66 | 540 | 118 | 0 | 14 | 0 | 0 | 352 | 55 | 0 | 144 | 0 | 33 | 0 | 0 | 68 | 81 | 0 | 53 |
| SRR3103250 | 0 | 0 | 674 | 0 | 0 | 38 | 113 | 22 | 19 | 0 | 0 | 34 | 0 | 0 | 255 | 0 | 18 | 0 | 0 | 496 | 673 | 0 | 0 | 0 | 16 | 40 | 29 | 0 | 25 | 0 | 19 | 0 | 0 | 0 | 224 | 0 | 107 |
| SRR3103251 | 29 | 0 | 168 | 0 | 0 | 97 | 72 | 0 | 47 | 0 | 0 | 15 | 0 | 14 | 556 | 0 | 0 | 0 | 48 | 4215 | 918 | 0 | 0 | 0 | 18 | 63 | 108 | 0 | 0 | 0 | 168 | 11 | 0 | 0 | 369 | 25 | 158 |
| SRR3103252 | 11 | 77 | 323 | 0 | 0 | 15 | 65 | 15 | 15 | 0 | 0 | 35 | 0 | 0 | 406 | 15 | 14 | 0 | 32 | 244 | 747 | 0 | 0 | 0 | 46 | 385 | 77 | 15 | 0 | 0 | 31 | 0 | 0 | 0 | 139 | 0 | 92 |
| SRR3103253 | 0 | 0 | 316 | 0 | 0 | 17 | 20 | 0 | 17 | 0 | 0 | 24 | 0 | 0 | 145 | 0 | 0 | 0 | 0 | 95 | 277 | 0 | 0 | 0 | 30 | 619 | 164 | 0 | 17 | 0 | 69 | 0 | 0 | 0 | 78 | 17 | 39 |
| SRR3103254 | 31 | 13 | 375 | 0 | 0 | 23 | 111 | 32 | 13 | 0 | 0 | 0 | 0 | 0 | 361 | 0 | 0 | 0 | 17 | 722 | 355 | 0 | 0 | 0 | 0 | 326 | 310 | 0 | 13 | 0 | 26 | 0 | 0 | 0 | 377 | 23 | 126 |
| SRR3103255 | 0 | 0 | 615 | 0 | 0 | 16 | 41 | 0 | 18 | 0 | 0 | 12 | 0 | 0 | 2154 | 0 | 0 | 0 | 22 | 1010 | 3628 | 0 | 0 | 0 | 13 | 29 | 27 | 0 | 0 | 0 | 23 | 0 | 0 | 0 | 37 | 24 | 16 |
| SRR3103256 | 13 | 0 | 231 | 0 | 0 | 25 | 113 | 14 | 22 | 0 | 0 | 14 | 0 | 14 | 55 | 0 | 0 | 0 | 0 | 53 | 156 | 0 | 0 | 0 | 30 | 13 | 13 | 0 | 0 | 0 | 33 | 0 | 0 | 27 | 235 | 44 | 145 |
| SRR3103257 | 33 | 14 | 1106 | 0 | 0 | 58 | 70 | 119 | 31 | 0 | 0 | 0 | 0 | 0 | 525 | 0 | 11 | 0 | 0 | 345 | 644 | 0 | 0 | 0 | 17 | 1119 | 379 | 10 | 26 | 29 | 41 | 0 | 0 | 0 | 116 | 12 | 72 |
| SRR3103258 | 36 | 0 | 452 | 0 | 0 | 25 | 274 | 25 | 15 | 0 | 0 | 74 | 0 | 10 | 222 | 0 | 23 | 0 | 0 | 616 | 343 | 0 | 0 | 0 | 50 | 38 | 34 | 0 | 30 | 0 | 35 | 0 | 0 | 15 | 84 | 181 | 45 |
| SRR3103259 | 30 | 21 | 191 | 0 | 0 | 10 | 366 | 36 | 10 | 0 | 0 | 0 | 0 | 18 | 34 | 0 | 0 | 0 | 145 | 2319 | 172 | 0 | 0 | 0 | 0 | 293 | 108 | 0 | 0 | 0 | 10 | 0 | 0 | 64 | 67 | 0 | 39 |
| SRR3103260 | 31 | 11 | 837 | 0 | 0 | 0 | 134 | 0 | 0 | 0 | 0 | 22 | 0 | 11 | 79 | 0 | 0 | 0 | 84 | 3299 | 484 | 0 | 12 | 0 | 0 | 145 | 66 | 0 | 131 | 0 | 0 | 0 | 0 | 14 | 216 | 0 | 136 |
| SRR3103261 | 13 | 0 | 468 | 0 | 0 | 12 | 79 | 52 | 0 | 0 | 0 | 17 | 0 | 13 | 217 | 0 | 0 | 0 | 104 | 3420 | 814 | 0 | 13 | 0 | 0 | 209 | 100 | 11 | 366 | 13 | 0 | 0 | 0 | 34 | 289 | 28 | 258 |
| SRR3103262 | 50 | 46 | 159 | 0 | 0 | 31 | 322 | 31 | 19 | 0 | 0 | 27 | 0 | 100 | 379 | 12 | 0 | 0 | 0 | 497 | 333 | 0 | 0 | 0 | 23 | 28 | 13 | 0 | 69 | 120 | 73 | 0 | 0 | 46 | 11 | 23 | 135 |
| SRR3103263 | 34 | 42 | 768 | 0 | 0 | 19 | 96 | 23 | 42 | 0 | 0 | 0 | 0 | 79 | 3977 | 14 | 11 | 0 | 0 | 105 | 332 | 0 | 0 | 0 | 0 | 19 | 0 | 0 | 56 | 0 | 0 | 0 | 0 | 213 | 16 | 19 | 185 |
| SRR3103264 | 50 | 20 | 293 | 0 | 0 | 18 | 414 | 3220 | 38 | 0 | 0 | 65 | 0 | 16 | 0 | 0 | 0 | 0 | 13 | 407 | 376 | 0 | 0 | 0 | 0 | 71 | 18 | 0 | 101 | 476 | 0 | 0 | 0 | 128 | 33 | 0 | 18 |
| SRR3103265 | 23 | 34 | 135 | 0 | 0 | 166 | 92 | 163 | 221 | 0 | 0 | 15 | 0 | 29 | 863 | 0 | 0 | 0 | 0 | 111 | 121 | 0 | 0 | 0 | 20 | 2307 | 863 | 0 | 0 | 32 | 49 | 0 | 0 | 40 | 138 | 0 | 103 |
| SRR3103266 | 78 | 0 | 857 | 0 | 0 | 38 | 314 | 21 | 38 | 0 | 0 | 65 | 0 | 38 | 0 | 0 | 0 | 0 | 0 | 175 | 309 | 0 | 0 | 0 | 0 | 96 | 38 | 0 | 48 | 48 | 67 | 0 | 0 | 105 | 191 | 29 | 239 |
| SRR3103267 | 25 | 18 | 227 | 0 | 0 | 36 | 57 | 152 | 0 | 0 | 0 | 62 | 0 | 0 | 500 | 0 | 0 | 0 | 0 | 206 | 135 | 0 | 0 | 0 | 12 | 24 | 17 | 15 | 45 | 36 | 161 | 0 | 0 | 12 | 15 | 33 | 209 |
| SRR3103268 | 13 | 0 | 222 | 0 | 0 | 98 | 29 | 75 | 82 | 0 | 0 | 19 | 0 | 26 | 0 | 0 | 0 | 0 | 12 | 126 | 44 | 0 | 0 | 0 | 0 | 68 | 16 | 13 | 127 | 36 | 65 | 0 | 0 | 13 | 189 | 20 | 108 |
| SRR3103269 | 15 | 14 | 183 | 0 | 0 | 112 | 112 | 14 | 46 | 0 | 0 | 19 | 0 | 14 | 33 | 0 | 0 | 0 | 23 | 566 | 87 | 0 | 0 | 0 | 0 | 22 | 112 | 0 | 42 | 0 | 163 | 0 | 0 | 0 | 17 | 65 | 446 |
| SRR3103270 | 13 | 24 | 152 | 0 | 0 | 96 | 244 | 132 | 72 | 0 | 0 | 22 | 0 | 24 | 0 | 0 | 0 | 0 | 0 | 129 | 100 | 0 | 0 | 0 | 24 | 1793 | 490 | 0 | 0 | 0 | 24 | 0 | 0 | 12 | 335 | 0 | 108 |
| SRR3103271 | 21 | 19 | 240 | 0 | 0 | 136 | 224 | 0 | 45 | 0 | 0 | 38 | 0 | 0 | 54 | 0 | 0 | 0 | 22 | 824 | 114 | 0 | 0 | 0 | 19 | 2457 | 220 | 0 | 0 | 0 | 116 | 0 | 0 | 0 | 362 | 52 | 317 |
| SRR3103272 | 15 | 19 | 258 | 0 | 0 | 19 | 129 | 114 | 38 | 0 | 0 | 41 | 0 | 57 | 11 | 0 | 0 | 0 | 0 | 57 | 55 | 0 | 0 | 0 | 38 | 534 | 362 | 38 | 0 | 0 | 0 | 0 | 0 | 0 | 57 | 0 | 38 |
| SRR3103273 | 23 | 17 | 361 | 0 | 0 | 26 | 177 | 104 | 0 | 0 | 0 | 64 | 0 | 17 | 11 | 0 | 0 | 0 | 25 | 288 | 79 | 0 | 0 | 0 | 17 | 883 | 1333 | 0 | 0 | 0 | 0 | 0 | 0 | 0 | 190 | 17 | 95 |
| SRR3103274 | 37 | 0 | 452 | 0 | 0 | 14 | 32 | 11 | 0 | 0 | 0 | 0 | 0 | 11 | 184 | 0 | 0 | 0 | 0 | 424 | 216 | 0 | 0 | 0 | 0 | 185 | 304 | 0 | 27 | 0 | 19 | 0 | 0 | 0 | 11 | 27 | 190 |
| SRR3103275 | 0 | 0 | 401 | 0 | 0 | 20 | 42 | 144 | 0 | 0 | 0 | 19 | 0 | 14 | 14 | 0 | 0 | 0 | 0 | 94 | 187 | 0 | 0 | 0 | 15 | 190 | 152 | 0 | 38 | 38 | 24 | 0 | 0 | 124 | 302 | 20 | 121 |
| SRR3103276 | 15 | 16 | 240 | 0 | 0 | 19 | 169 | 33 | 19 | 0 | 0 | 0 | 0 | 14 | 182 | 0 | 0 | 0 | 0 | 87 | 144 | 0 | 0 | 0 | 28 | 14 | 227 | 0 | 12 | 0 | 44 | 0 | 0 | 40 | 246 | 28 | 94 |
| SRR3103277 | 27 | 18 | 897 | 0 | 0 | 66 | 13 | 24 | 18 | 0 | 0 | 28 | 0 | 0 | 28 | 0 | 0 | 0 | 0 | 90 | 79 | 0 | 0 | 0 | 0 | 1072 | 667 | 12 | 24 | 0 | 66 | 0 | 0 | 24 | 137 | 0 | 125 |
| SRR3103278 | 28 | 0 | 472 | 0 | 0 | 103 | 139 | 86 | 17 | 0 | 0 | 79 | 0 | 26 | 25 | 0 | 22 | 0 | 0 | 94 | 45 | 0 | 0 | 0 | 17 | 20 | 379 | 0 | 17 | 26 | 34 | 0 | 0 | 17 | 29 | 155 | 11 |
| SRR3103279 | 22 | 13 | 271 | 0 | 0 | 15 | 127 | 0 | 10 | 0 | 0 | 0 | 0 | 13 | 61 | 0 | 0 | 0 | 172 | 1383 | 108 | 0 | 0 | 0 | 0 | 280 | 62 | 0 | 0 | 0 | 37 | 0 | 0 | 0 | 106 | 0 | 49 |
| SRR3103280 | 16 | 0 | 333 | 0 | 0 | 20 | 151 | 0 | 48 | 0 | 0 | 0 | 0 | 0 | 35 | 0 | 0 | 0 | 0 | 1501 | 287 | 0 | 0 | 0 | 0 | 323 | 137 | 0 | 0 | 0 | 18 | 0 | 0 | 0 | 77 | 81 | 30 |
| SRR3103281 | 20 | 0 | 574 | 0 | 0 | 372 | 151 | 0 | 71 | 0 | 0 | 14 | 0 | 0 | 44 | 0 | 0 | 0 | 0 | 1337 | 286 | 0 | 0 | 0 | 0 | 6560 | 196 | 0 | 0 | 0 | 267 | 0 | 0 | 0 | 1803 | 67 | 577 |
| SRR3103386 | 70 | 0 | 648 | 0 | 0 | 65 | 196 | 0 | 0 | 0 | 0 | 227 | 0 | 0 | 0 | 0 | 22 | 0 | 0 | 392 | 441 | 0 | 0 | 0 | 0 | 1295 | 130 | 0 | 0 | 0 | 0 | 0 | 0 | 0 | 389 | 32 | 97 |
| SRR3103387 | 51 | 0 | 471 | 0 | 0 | 32 | 456 | 0 | 0 | 0 | 0 | 145 | 0 | 0 | 0 | 0 | 0 | 0 | 91 | 821 | 1186 | 0 | 0 | 0 | 0 | 631 | 126 | 0 | 0 | 0 | 32 | 0 | 0 | 0 | 284 | 32 | 126 |
| SRR3103388 | 46 | 0 | 518 | 0 | 0 | 44 | 2240 | 0 | 0 | 0 | 0 | 319 | 0 | 0 | 0 | 0 | 0 | 0 | 102 | 68 | 1086 | 0 | 0 | 0 | 0 | 438 | 44 | 0 | 0 | 0 | 0 | 0 | 0 | 0 | 131 | 0 | 88 |
| SRR3103389 | 103 | 0 | 1620 | 0 | 0 | 0 | 205 | 0 | 0 | 0 | 0 | 778 | 0 | 0 | 0 | 0 | 28 | 0 | 41 | 328 | 410 | 0 | 0 | 0 | 0 | 16 | 11 | 0 | 0 | 0 | 0 | 0 | 0 | 0 | 222 | 0 | 0 |
| SRR3103390 | 95 | 0 | 1207 | 0 | 0 | 56 | 0 | 0 | 0 | 0 | 0 | 390 | 0 | 0 | 0 | 0 | 12 | 0 | 0 | 517 | 1552 | 0 | 0 | 0 | 0 | 474 | 139 | 0 | 0 | 0 | 0 | 0 | 0 | 0 | 195 | 28 | 111 |
| SRR3103391 | 102 | 0 | 657 | 0 | 0 | 42 | 328 | 0 | 0 | 0 | 0 | 380 | 0 | 0 | 0 | 0 | 15 | 0 | 0 | 562 | 1826 | 0 | 0 | 0 | 0 | 20 | 166 | 0 | 0 | 0 | 42 | 0 | 0 | 0 | 125 | 42 | 166 |
| SRR3103392 | 101 | 0 | 1600 | 0 | 0 | 58 | 171 | 0 | 39 | 0 | 0 | 664 | 19 | 0 | 0 | 0 | 33 | 0 | 13 | 39 | 341 | 0 | 0 | 0 | 19 | 35 | 12 | 0 | 0 | 0 | 77 | 0 | 0 | 0 | 16 | 97 | 252 |
| SRR3103393 | 39 | 0 | 152 | 0 | 0 | 0 | 0 | 0 | 0 | 0 | 0 | 397 | 0 | 0 | 0 | 0 | 24 | 0 | 0 | 0 | 1576 | 0 | 0 | 0 | 0 | 343 | 0 | 0 | 0 | 0 | 0 | 0 | 0 | 0 | 86 | 0 | 86 |
| SRR3103394 | 40 | 0 | 983 | 0 | 0 | 109 | 369 | 0 | 67 | 0 | 0 | 183 | 0 | 0 | 0 | 0 | 37 | 15 | 0 | 427 | 812 | 0 | 0 | 0 | 0 | 1089 | 553 | 0 | 0 | 0 | 17 | 0 | 0 | 0 | 394 | 42 | 159 |
| SRR3103395 | 46 | 0 | 441 | 0 | 0 | 0 | 422 | 0 | 21 | 0 | 0 | 520 | 0 | 0 | 0 | 0 | 23 | 0 | 0 | 543 | 1629 | 0 | 0 | 0 | 21 | 15 | 127 | 0 | 0 | 0 | 0 | 0 | 0 | 0 | 42 | 21 | 63 |
| SRR3103396 | 45 | 0 | 846 | 0 | 0 | 24 | 626 | 0 | 24 | 0 | 0 | 317 | 0 | 0 | 0 | 0 | 17 | 44 | 37 | 589 | 1435 | 0 | 0 | 0 | 0 | 10 | 71 | 0 | 0 | 0 | 0 | 0 | 0 | 24 | 119 | 48 | 95 |
| SRR3103397 | 88 | 0 | 509 | 0 | 0 | 23 | 1012 | 0 | 23 | 0 | 0 | 468 | 0 | 0 | 0 | 0 | 16 | 0 | 35 | 244 | 663 | 0 | 0 | 0 | 0 | 116 | 23 | 0 | 0 | 0 | 23 | 0 | 0 | 0 | 70 | 0 | 70 |
| SRR3103400 | 62 | 0 | 1165 | 0 | 0 | 290 | 370 | 0 | 41 | 0 | 0 | 974 | 0 | 0 | 0 | 0 | 41 | 0 | 0 | 103 | 103 | 0 | 0 | 0 | 0 | 49 | 22 | 0 | 0 | 0 | 83 | 0 | 0 | 0 | 15 | 41 | 166 |
| SRR3103401 | 37 | 0 | 1348 | 0 | 0 | 109 | 202 | 0 | 219 | 0 | 0 | 353 | 0 | 0 | 0 | 0 | 64 | 0 | 0 | 168 | 135 | 0 | 0 | 0 | 0 | 2187 | 1640 | 0 | 0 | 0 | 0 | 0 | 0 | 0 | 1093 | 0 | 875 |
| SRR3103402 | 99 | 0 | 2012 | 0 | 0 | 81 | 84 | 0 | 0 | 0 | 0 | 758 | 0 | 0 | 0 | 0 | 32 | 0 | 0 | 84 | 84 | 0 | 0 | 0 | 0 | 43 | 21 | 0 | 0 | 0 | 0 | 0 | 0 | 0 | 17 | 0 | 649 |
| SRR3103403 | 103 | 0 | 1381 | 0 | 0 | 323 | 280 | 0 | 108 | 0 | 0 | 1290 | 54 | 0 | 0 | 0 | 62 | 0 | 0 | 380 | 120 | 0 | 0 | 0 | 0 | 41 | 22 | 0 | 0 | 0 | 0 | 0 | 0 | 0 | 12 | 0 | 431 |
| SRR3103404 | 66 | 0 | 1311 | 0 | 0 | 228 | 401 | 0 | 0 | 0 | 0 | 1200 | 0 | 0 | 0 | 0 | 40 | 0 | 12 | 122 | 365 | 0 | 0 | 0 | 0 | 30 | 16 | 0 | 57 | 0 | 0 | 0 | 0 | 0 | 742 | 57 | 114 |
| SRR3103405 | 48 | 0 | 726 | 0 | 0 | 214 | 376 | 0 | 54 | 0 | 0 | 805 | 0 | 0 | 0 | 0 | 48 | 0 | 14 | 56 | 279 | 0 | 0 | 0 | 0 | 46 | 14 | 0 | 0 | 0 | 54 | 0 | 0 | 0 | 13 | 321 | 375 |
| SRR3103406 | 191 | 0 | 1052 | 0 | 0 | 33 | 175 | 0 | 0 | 0 | 0 | 1403 | 0 | 0 | 0 | 0 | 24 | 14 | 0 | 263 | 731 | 0 | 0 | 0 | 0 | 501 | 12 | 0 | 0 | 0 | 0 | 0 | 0 | 0 | 534 | 67 | 200 |
| SRR3103407 | 67 | 0 | 819 | 0 | 0 | 175 | 613 | 0 | 0 | 0 | 0 | 1343 | 0 | 0 | 0 | 0 | 31 | 0 | 20 | 20 | 121 | 0 | 0 | 0 | 0 | 23 | 700 | 0 | 0 | 0 | 0 | 0 | 0 | 0 | 700 | 0 | 58 |
| SRR3103408 | 140 | 0 | 920 | 0 | 0 | 114 | 360 | 0 | 0 | 0 | 0 | 1291 | 0 | 114 | 0 | 0 | 17 | 0 | 60 | 240 | 360 | 0 | 0 | 0 | 114 | 2056 | 571 | 0 | 0 | 0 | 0 | 0 | 0 | 0 | 685 | 0 | 228 |
| SRR3103409 | 176 | 0 | 528 | 0 | 0 | 124 | 417 | 0 | 0 | 0 | 0 | 858 | 0 | 0 | 0 | 0 | 18 | 0 | 0 | 417 | 536 | 0 | 0 | 0 | 0 | 19 | 186 | 0 | 0 | 0 | 0 | 0 | 0 | 0 | 249 | 62 | 124 |
| SRR3138807 | 0 | 82 | 65 | 0 | 0 | 16 | 59 | 17 | 56 | 27 | 31 | 0 | 0 | 18 | 0 | 0 | 0 | 0 | 0 | 0 | 0 | 0 | 0 | 0 | 0 | 237 | 369 | 0 | 0 | 0 | 22 | 0 | 0 | 12 | 11 | 49 | 20 |
| SRR3138808 | 18 | 18 | 24 | 0 | 0 | 0 | 37 | 0 | 30 | 20 | 13 | 0 | 0 | 17 | 0 | 0 | 0 | 0 | 0 | 0 | 0 | 0 | 0 | 0 | 0 | 36 | 26 | 0 | 0 | 0 | 13 | 0 | 23 | 0 | 0 | 22 | 16 |
| SRR3138809 | 0 | 0 | 32 | 0 | 0 | 0 | 54 | 0 | 32 | 31 | 17 | 0 | 0 | 17 | 0 | 0 | 0 | 0 | 0 | 0 | 0 | 0 | 0 | 0 | 0 | 34 | 25 | 0 | 0 | 0 | 0 | 0 | 12 | 0 | 0 | 24 | 17 |
| SRR3138810 | 0 | 315 | 72 | 0 | 0 | 0 | 56 | 0 | 35 | 32 | 53 | 0 | 0 | 20 | 0 | 0 | 0 | 0 | 0 | 0 | 0 | 0 | 0 | 0 | 0 | 115 | 140 | 0 | 0 | 0 | 12 | 0 | 107 | 0 | 0 | 20 | 16 |
| SRR3199254 | 44 | 0 | 160 | 0 | 0 | 66 | 44 | 0 | 43 | 0 | 70 | 413 | 0 | 14 | 14 | 11 | 125 | 0 | 90 | 698 | 929 | 0 | 0 | 0 | 657 | 913 | 204 | 0 | 20 | 0 | 63 | 0 | 0 | 0 | 342 | 354 | 471 |
| SRR3199255 | 73 | 13 | 70 | 0 | 0 | 20 | 66 | 11 | 16 | 16 | 12 | 211 | 0 | 0 | 10 | 11 | 59 | 0 | 86 | 264 | 84 | 0 | 0 | 0 | 30 | 11 | 10 | 0 | 22 | 0 | 16 | 0 | 0 | 62 | 44 | 40 | 57 |
| SRR3199256 | 57 | 12 | 192 | 0 | 0 | 70 | 259 | 66 | 21 | 54 | 270 | 511 | 0 | 0 | 0 | 0 | 111 | 0 | 115 | 397 | 259 | 0 | 0 | 17 | 303 | 348 | 539 | 0 | 79 | 21 | 66 | 0 | 0 | 12 | 261 | 328 | 323 |
| SRR3199257 | 0 | 10 | 44 | 0 | 0 | 30 | 841 | 20 | 80 | 0 | 110 | 343 | 0 | 20 | 56 | 10 | 21 | 0 | 33 | 156 | 17 | 0 | 0 | 0 | 80 | 451 | 30 | 0 | 70 | 0 | 191 | 0 | 0 | 50 | 4553 | 1083 | 742 |
| SRR3199258 | 28 | 11 | 180 | 0 | 0 | 25 | 836 | 0 | 28 | 0 | 112 | 153 | 0 | 22 | 12 | 0 | 68 | 0 | 44 | 67 | 0 | 0 | 0 | 0 | 67 | 280 | 151 | 0 | 87 | 0 | 28 | 0 | 11 | 34 | 84 | 115 | 67 |
| SRR3199259 | 23 | 13 | 132 | 0 | 0 | 21 | 1038 | 29 | 32 | 0 | 106 | 121 | 0 | 0 | 0 | 0 | 41 | 0 | 14 | 67 | 36 | 0 | 0 | 0 | 124 | 154 | 75 | 0 | 75 | 18 | 14 | 0 | 0 | 108 | 79 | 64 | 85 |
| SRR3199260 | 25 | 0 | 249 | 0 | 0 | 23 | 245 | 0 | 18 | 0 | 133 | 81 | 0 | 11 | 0 | 0 | 126 | 0 | 0 | 32 | 0 | 0 | 11 | 0 | 155 | 191 | 133 | 0 | 81 | 0 | 15 | 0 | 11 | 55 | 156 | 73 | 240 |
| SRR3199261 | 29 | 0 | 331 | 0 | 0 | 15 | 1858 | 0 | 0 | 0 | 73 | 668 | 0 | 0 | 41 | 0 | 21 | 0 | 41 | 41 | 0 | 0 | 0 | 0 | 73 | 104 | 137 | 0 | 64 | 24 | 0 | 0 | 0 | 61 | 46 | 43 | 34 |
| SRR3199262 | 20 | 12 | 205 | 20 | 0 | 23 | 1678 | 12 | 20 | 12 | 140 | 727 | 0 | 0 | 15 | 20 | 24 | 0 | 25 | 25 | 0 | 0 | 0 | 0 | 179 | 62 | 58 | 0 | 47 | 20 | 16 | 0 | 0 | 39 | 148 | 86 | 129 |
| SRR3199263 | 0 | 0 | 223 | 0 | 0 | 89 | 0 | 0 | 0 | 0 | 0 | 1497 | 0 | 89 | 0 | 0 | 82 | 0 | 0 | 411 | 43 | 0 | 0 | 0 | 178 | 979 | 1068 | 0 | 0 | 0 | 267 | 0 | 0 | 0 | 20 | 178 | 934 |
| SRR3235777 | 183 | 15 | 260 | 0 | 0 | 382 | 170 | 1533 | 614 | 31 | 35 | 521 | 12 | 0 | 3663 | 0 | 122 | 0 | 110 | 399 | 1377 | 0 | 0 | 3120 | 0 | 1722 | 815 | 19 | 0 | 62 | 487 | 27 | 0 | 143 | 174 | 131 | 85 |
| SRR3235778 | 73 | 0 | 325 | 0 | 0 | 1556 | 110 | 575 | 709 | 0 | 0 | 299 | 0 | 0 | 13227 | 0 | 84 | 0 | 55 | 344 | 2324 | 0 | 0 | 3668 | 0 | 6584 | 1238 | 0 | 15 | 27 | 943 | 0 | 0 | 61 | 253 | 195 | 165 |
| SRR3235779 | 116 | 17 | 220 | 0 | 0 | 542 | 337 | 687 | 206 | 0 | 0 | 646 | 0 | 0 | 7201 | 0 | 439 | 0 | 70 | 915 | 9750 | 0 | 0 | 2245 | 0 | 3256 | 2619 | 0 | 0 | 177 | 310 | 12 | 0 | 123 | 605 | 423 | 397 |
| SRR3235780 | 62 | 0 | 548 | 0 | 0 | 744 | 177 | 345 | 393 | 13 | 85 | 192 | 11 | 27 | 18246 | 37 | 119 | 0 | 165 | 530 | 4818 | 48 | 0 | 1135 | 0 | 3295 | 1020 | 0 | 0 | 66 | 311 | 11 | 11 | 199 | 234 | 106 | 114 |
| SRR3235781 | 127 | 0 | 283 | 0 | 0 | 32 | 142 | 16 | 11 | 0 | 21 | 178 | 0 | 0 | 6686 | 0 | 217 | 0 | 133 | 645 | 19185 | 0 | 0 | 130 | 0 | 152 | 105 | 0 | 0 | 177 | 15 | 0 | 0 | 80 | 13 | 22 | 24 |
| SRR3279553 | 0 | 0 | 473 | 0 | 0 | 0 | 0 | 0 | 0 | 0 | 0 | 0 | 0 | 0 | 0 | 0 | 0 | 0 | 0 | 0 | 0 | 0 | 0 | 0 | 54 | 18 | 36 | 0 | 0 | 0 | 18 | 18 | 0 | 0 | 0 | 0 | 0 |
| SRR3279554 | 0 | 0 | 88 | 0 | 0 | 0 | 0 | 0 | 0 | 0 | 0 | 29 | 0 | 0 | 0 | 0 | 0 | 0 | 0 | 0 | 195 | 0 | 0 | 0 | 25 | 0 | 0 | 0 | 0 | 0 | 0 | 0 | 0 | 0 | 59 | 0 | 0 |
| SRR3279555 | 84 | 0 | 0 | 0 | 0 | 0 | 0 | 0 | 0 | 0 | 0 | 0 | 0 | 0 | 0 | 0 | 0 | 0 | 0 | 0 | 232 | 0 | 0 | 0 | 0 | 0 | 0 | 0 | 0 | 0 | 0 | 0 | 0 | 0 | 0 | 0 | 0 |
| SRR3279556 | 0 | 0 | 52 | 0 | 0 | 0 | 0 | 0 | 0 | 0 | 0 | 13 | 0 | 14 | 0 | 0 | 0 | 0 | 0 | 0 | 117 | 0 | 0 | 0 | 95 | 0 | 10 | 10 | 0 | 0 | 0 | 0 | 0 | 20 | 27 | 27 | 0 |
| SRR3279557 | 0 | 0 | 132 | 0 | 0 | 0 | 0 | 0 | 0 | 0 | 0 | 25 | 0 | 15 | 0 | 0 | 0 | 0 | 0 | 11 | 304 | 0 | 0 | 0 | 168 | 0 | 0 | 0 | 0 | 0 | 0 | 0 | 0 | 41 | 159 | 39 | 57 |
| SRR3279558 | 0 | 0 | 38 | 0 | 0 | 0 | 91 | 0 | 0 | 0 | 0 | 38 | 0 | 67 | 0 | 0 | 0 | 0 | 0 | 0 | 91 | 0 | 0 | 0 | 19 | 0 | 0 | 0 | 0 | 0 | 0 | 0 | 0 | 29 | 48 | 29 | 39 |
| SRR3279559 | 0 | 0 | 95 | 0 | 0 | 0 | 0 | 0 | 0 | 0 | 0 | 79 | 0 | 36 | 14 | 0 | 0 | 0 | 0 | 0 | 0 | 0 | 0 | 0 | 42 | 0 | 0 | 0 | 0 | 0 | 0 | 0 | 0 | 12 | 13 | 18 | 48 |
| SRR3279560 | 0 | 0 | 121 | 0 | 0 | 0 | 0 | 0 | 0 | 0 | 0 | 28 | 0 | 12 | 0 | 0 | 0 | 0 | 0 | 19 | 210 | 0 | 0 | 0 | 0 | 0 | 0 | 0 | 0 | 0 | 0 | 0 | 0 | 0 | 0 | 12 | 12 |
| SRR3279561 | 0 | 0 | 79 | 0 | 0 | 0 | 0 | 0 | 0 | 0 | 0 | 64 | 0 | 24 | 0 | 0 | 0 | 0 | 0 | 15 | 133 | 0 | 0 | 0 | 36 | 0 | 0 | 0 | 0 | 0 | 0 | 0 | 0 | 0 | 88 | 16 | 40 |
| SRR3279562 | 0 | 0 | 41 | 0 | 0 | 0 | 0 | 0 | 0 | 0 | 0 | 102 | 0 | 12 | 0 | 12 | 0 | 0 | 0 | 0 | 268 | 0 | 0 | 0 | 10 | 0 | 0 | 0 | 0 | 0 | 0 | 0 | 0 | 0 | 43 | 0 | 0 |
| SRR3279563 | 62 | 0 | 92 | 0 | 0 | 0 | 0 | 0 | 0 | 0 | 0 | 92 | 0 | 23 | 0 | 0 | 31 | 0 | 0 | 0 | 460 | 0 | 0 | 0 | 12 | 0 | 0 | 0 | 0 | 0 | 0 | 0 | 0 | 0 | 22 | 0 | 0 |
| SRR3279564 | 0 | 0 | 254 | 0 | 0 | 0 | 0 | 0 | 0 | 0 | 0 | 55 | 0 | 17 | 0 | 0 | 0 | 0 | 0 | 0 | 99 | 0 | 0 | 0 | 37 | 0 | 0 | 10 | 0 | 0 | 0 | 0 | 0 | 20 | 41 | 0 | 48 |
| SRR3279565 | 0 | 0 | 108 | 0 | 0 | 0 | 12 | 0 | 0 | 0 | 0 | 36 | 0 | 0 | 0 | 0 | 0 | 0 | 0 | 35 | 1221 | 0 | 0 | 0 | 91 | 0 | 14 | 0 | 0 | 0 | 11 | 0 | 0 | 23 | 14 | 11 | 17 |
| SRR3279566 | 0 | 0 | 123 | 0 | 0 | 0 | 0 | 0 | 0 | 0 | 0 | 0 | 0 | 0 | 0 | 0 | 0 | 0 | 0 | 0 | 167 | 0 | 0 | 0 | 11 | 11 | 11 | 0 | 0 | 0 | 0 | 0 | 0 | 0 | 0 | 0 | 0 |
| SRR3279567 | 0 | 0 | 38 | 0 | 0 | 0 | 0 | 0 | 0 | 0 | 0 | 24 | 0 | 45 | 0 | 0 | 0 | 0 | 0 | 0 | 63 | 0 | 0 | 0 | 11 | 0 | 0 | 0 | 0 | 0 | 0 | 0 | 12 | 36 | 68 | 27 | 37 |
| SRR3279568 | 34 | 0 | 119 | 0 | 0 | 0 | 0 | 0 | 0 | 14 | 0 | 102 | 0 | 0 | 0 | 0 | 0 | 0 | 0 | 25 | 369 | 0 | 0 | 0 | 28 | 42 | 14 | 0 | 0 | 0 | 0 | 0 | 0 | 28 | 56 | 0 | 0 |
| SRR3279569 | 77 | 0 | 115 | 0 | 0 | 0 | 0 | 0 | 0 | 0 | 0 | 192 | 0 | 32 | 0 | 0 | 0 | 0 | 0 | 18 | 183 | 0 | 0 | 0 | 16 | 0 | 0 | 16 | 0 | 0 | 0 | 0 | 0 | 0 | 64 | 0 | 32 |
| SRR3279570 | 0 | 0 | 103 | 0 | 0 | 0 | 0 | 0 | 0 | 0 | 0 | 0 | 0 | 0 | 0 | 0 | 19 | 0 | 0 | 0 | 1006 | 0 | 0 | 0 | 0 | 0 | 0 | 0 | 0 | 0 | 0 | 0 | 0 | 0 | 0 | 0 | 0 |
| SRR3279571 | 12 | 0 | 139 | 14 | 0 | 0 | 17 | 0 | 0 | 0 | 0 | 36 | 0 | 27 | 0 | 0 | 0 | 0 | 0 | 33 | 117 | 0 | 0 | 0 | 82 | 0 | 0 | 41 | 0 | 0 | 0 | 0 | 14 | 14 | 41 | 0 | 41 |

### Supp Table S8.2.Columns_AM_to_BW

| | | | | | | | | | | | | | | | | | | | | | | | | | | | | | | | | | | | | | |
|---|---|---|---|---|---|---|---|---|---|---|---|---|---|---|---|---|---|---|---|---|---|---|---|---|---|---|---|---|---|---|---|---|---|---|---|---|---|
| SRR3279572 | 0 | 0 | 96 | 0 | 0 | 0 | 0 | 0 | 0 | 0 | 0 | 16 | 0 | 67 | 0 | 0 | 13 | 0 | 0 | 0 | 157 | 0 | 0 | 0 | 22 | 0 | 0 | 17 | 0 | 0 | 0 | 0 | 0 | 39 | 12 | 22 | 51 |
| SRR3279573 | 31 | 0 | 63 | 0 | 0 | 0 | 0 | 0 | 0 | 0 | 0 | 24 | 0 | 0 | 0 | 0 | 24 | 0 | 0 | 0 | 808 | 0 | 0 | 0 | 20 | 0 | 0 | 0 | 0 | 0 | 0 | 0 | 0 | 0 | 16 | 0 | 0 |
| SRR3279574 | 0 | 0 | 47 | 0 | 0 | 0 | 0 | 0 | 0 | 0 | 0 | 23 | 0 | 0 | 0 | 0 | 0 | 0 | 0 | 47 | 681 | 0 | 0 | 0 | 15 | 0 | 0 | 0 | 0 | 0 | 0 | 0 | 0 | 0 | 0 | 15 | 15 |
| SRR3279575 | 0 | 0 | 145 | 0 | 0 | 0 | 0 | 0 | 0 | 0 | 0 | 18 | 0 | 16 | 0 | 0 | 0 | 0 | 0 | 0 | 402 | 0 | 0 | 0 | 29 | 0 | 13 | 0 | 0 | 0 | 0 | 0 | 0 | 26 | 34 | 11 | 18 |
| SRR3279576 | 0 | 0 | 191 | 0 | 0 | 0 | 0 | 0 | 0 | 0 | 0 | 17 | 0 | 15 | 0 | 0 | 0 | 0 | 0 | 0 | 396 | 0 | 0 | 0 | 130 | 0 | 0 | 0 | 0 | 0 | 0 | 0 | 0 | 12 | 15 | 0 | 15 |
| SRR3279577 | 0 | 0 | 358 | 0 | 0 | 0 | 0 | 39 | 0 | 0 | 0 | 53 | 0 | 28 | 0 | 0 | 11 | 0 | 0 | 16 | 478 | 0 | 0 | 0 | 20 | 16 | 0 | 0 | 12 | 0 | 0 | 0 | 0 | 16 | 24 | 0 | 16 |
| SRR3279578 | 0 | 0 | 50 | 0 | 0 | 0 | 0 | 0 | 0 | 0 | 0 | 114 | 0 | 10 | 14 | 0 | 0 | 0 | 0 | 14 | 246 | 0 | 0 | 0 | 18 | 0 | 0 | 0 | 0 | 0 | 0 | 0 | 0 | 0 | 52 | 18 | 24 |
| SRR3279579 | 0 | 0 | 157 | 0 | 0 | 0 | 0 | 0 | 0 | 0 | 0 | 0 | 0 | 0 | 0 | 0 | 0 | 0 | 0 | 41 | 534 | 0 | 0 | 0 | 0 | 23 | 0 | 0 | 0 | 0 | 0 | 0 | 0 | 0 | 0 | 0 | 0 |
| SRR3279580 | 0 | 0 | 21 | 0 | 0 | 0 | 0 | 0 | 0 | 0 | 0 | 26 | 0 | 27 | 0 | 0 | 0 | 0 | 0 | 0 | 110 | 0 | 0 | 0 | 22 | 0 | 0 | 0 | 0 | 0 | 0 | 0 | 12 | 39 | 45 | 12 | 18 |
| SRR3279581 | 0 | 0 | 117 | 0 | 0 | 0 | 0 | 0 | 0 | 0 | 0 | 91 | 0 | 0 | 0 | 0 | 0 | 0 | 0 | 0 | 251 | 0 | 0 | 0 | 69 | 0 | 0 | 0 | 0 | 0 | 0 | 0 | 0 | 26 | 77 | 13 | 21 |
| SRR3279582 | 0 | 0 | 270 | 0 | 0 | 0 | 12 | 0 | 0 | 0 | 0 | 68 | 0 | 0 | 0 | 0 | 0 | 0 | 0 | 12 | 62 | 0 | 0 | 0 | 39 | 0 | 0 | 0 | 0 | 0 | 0 | 0 | 0 | 11 | 0 | 0 | 11 |
| SRR3279583 | 0 | 0 | 55 | 0 | 0 | 0 | 16 | 0 | 0 | 0 | 0 | 62 | 0 | 0 | 0 | 0 | 0 | 0 | 0 | 48 | 386 | 0 | 0 | 0 | 69 | 0 | 0 | 0 | 0 | 0 | 0 | 0 | 0 | 0 | 51 | 0 | 23 |
| SRR3279584 | 11 | 0 | 876 | 0 | 0 | 0 | 0 | 11 | 0 | 0 | 0 | 25 | 0 | 0 | 0 | 0 | 0 | 0 | 0 | 210 | 1077 | 0 | 0 | 31 | 19 | 29 | 30 | 0 | 0 | 0 | 0 | 0 | 0 | 22 | 30 | 11 | 19 |
| SRR3279585 | 0 | 0 | 224 | 0 | 0 | 0 | 0 | 0 | 0 | 0 | 0 | 0 | 0 | 0 | 0 | 0 | 0 | 0 | 0 | 32 | 517 | 0 | 0 | 0 | 15 | 0 | 11 | 0 | 0 | 0 | 0 | 0 | 0 | 0 | 0 | 0 | 0 |
| SRR3279586 | 32 | 0 | 32 | 0 | 0 | 0 | 0 | 0 | 0 | 0 | 0 | 32 | 0 | 0 | 0 | 0 | 32 | 0 | 0 | 81 | 610 | 0 | 0 | 0 | 14 | 0 | 0 | 0 | 0 | 0 | 0 | 0 | 0 | 0 | 0 | 0 | 0 |
| SRR3317221 | 0 | 0 | 34 | 36 | 0 | 0 | 0 | 0 | 36 | 36 | 72 | 12 | 72 | 0 | 2852 | 0 | 49 | 0 | 0 | 0 | 0 | 0 | 0 | 0 | 501 | 143 | 72 | 0 | 0 | 0 | 15 | 0 | 0 | 215 | 501 | 251 | 286 |
| SRR3477269 | 131 | 12 | 131 | 0 | 0 | 0 | 627 | 104856 | 0 | 0 | 0 | 0 | 0 | 14 | 0 | 0 | 0 | 0 | 139 | 106 | 0 | 0 | 0 | 0 | 0 | 0 | 11 | 0 | 0 | 79 | 0 | 0 | 0 | 0 | 0 | 0 | 0 |
| SRR3477271 | 0 | 10 | 140 | 0 | 0 | 0 | 622 | 111958 | 0 | 0 | 0 | 140 | 0 | 12 | 0 | 0 | 0 | 0 | 156 | 97 | 0 | 0 | 0 | 0 | 0 | 0 | 0 | 0 | 0 | 89 | 0 | 0 | 0 | 0 | 0 | 0 | 0 |
| SRR3490374 | 0 | 0 | 635 | 0 | 0 | 0 | 0 | 0 | 189 | 0 | 0 | 0 | 0 | 0 | 0 | 0 | 0 | 0 | 0 | 296 | 6807 | 0 | 0 | 0 | 0 | 39 | 126 | 0 | 0 | 0 | 63 | 0 | 0 | 0 | 0 | 0 | 63 |
| SRR3490375 | 0 | 0 | 91 | 0 | 0 | 290 | 0 | 0 | 0 | 0 | 0 | 0 | 0 | 0 | 0 | 0 | 0 | 0 | 0 | 0 | 1778 | 0 | 0 | 0 | 0 | 33 | 290 | 0 | 0 | 0 | 580 | 0 | 0 | 0 | 1160 | 0 | 1160 |
| SRR3490376 | 18 | 0 | 278 | 0 | 0 | 0 | 0 | 0 | 30 | 0 | 0 | 10 | 0 | 0 | 0 | 0 | 0 | 0 | 0 | 104 | 9107 | 0 | 0 | 0 | 0 | 164 | 89 | 15 | 0 | 0 | 59 | 0 | 0 | 0 | 44 | 0 | 44 |
| SRR3490377 | 12 | 0 | 203 | 0 | 0 | 0 | 0 | 18 | 0 | 0 | 0 | 12 | 0 | 0 | 0 | 0 | 0 | 0 | 0 | 50 | 18017 | 0 | 0 | 0 | 0 | 18 | 18 | 0 | 0 | 0 | 0 | 0 | 0 | 0 | 0 | 0 | 0 |
| SRR3490378 | 23 | 0 | 434 | 0 | 0 | 0 | 0 | 0 | 0 | 0 | 0 | 14 | 0 | 0 | 0 | 0 | 0 | 0 | 0 | 0 | 10406 | 0 | 0 | 0 | 24 | 95 | 48 | 0 | 24 | 0 | 0 | 0 | 0 | 0 | 24 | 0 | 24 |
| SRR3490379 | 10 | 0 | 337 | 0 | 0 | 0 | 0 | 0 | 0 | 0 | 0 | 0 | 0 | 92 | 0 | 0 | 0 | 0 | 0 | 0 | 4312 | 0 | 0 | 0 | 0 | 10 | 0 | 0 | 0 | 0 | 275 | 0 | 0 | 92 | 183 | 0 | 92 |
| SRR3490380 | 40 | 0 | 427 | 0 | 0 | 0 | 0 | 0 | 0 | 0 | 0 | 0 | 0 | 0 | 0 | 0 | 0 | 0 | 0 | 467 | 6695 | 0 | 0 | 0 | 0 | 45 | 86 | 0 | 0 | 0 | 43 | 0 | 0 | 0 | 0 | 0 | 43 |
| SRR3490382 | 12 | 0 | 140 | 0 | 0 | 0 | 0 | 0 | 0 | 0 | 0 | 0 | 0 | 0 | 0 | 0 | 23 | 0 | 0 | 285 | 4556 | 0 | 0 | 0 | 0 | 29 | 11 | 0 | 0 | 0 | 11 | 0 | 0 | 202 | 0 | 202 | 0 |
| SRR3490383 | 99 | 0 | 539 | 0 | 0 | 0 | 0 | 13 | 0 | 0 | 0 | 51 | 0 | 25 | 0 | 25 | 0 | 0 | 0 | 430 | 11820 | 0 | 0 | 50 | 17 | 41 | 50 | 38 | 0 | 0 | 13 | 0 | 0 | 0 | 63 | 0 | 13 |
| SRR3490384 | 21 | 0 | 606 | 0 | 0 | 0 | 91 | 0 | 47 | 0 | 0 | 57 | 0 | 0 | 0 | 0 | 0 | 0 | 0 | 272 | 2902 | 0 | 0 | 0 | 0 | 3164 | 558 | 47 | 0 | 0 | 233 | 0 | 0 | 0 | 93 | 0 | 47 |
| SRR3490385 | 140 | 0 | 683 | 0 | 0 | 51 | 0 | 0 | 0 | 0 | 0 | 36 | 0 | 51 | 0 | 0 | 0 | 0 | 0 | 987 | 10201 | 0 | 0 | 51 | 51 | 18 | 101 | 0 | 0 | 0 | 51 | 0 | 0 | 51 | 16 | 51 | 16 |
| SRR3490386 | 131 | 0 | 739 | 0 | 0 | 0 | 0 | 100 | 0 | 0 | 0 | 20 | 0 | 0 | 0 | 0 | 15 | 71 | 0 | 924 | 3565 | 50 | 0 | 150 | 50 | 499 | 200 | 0 | 0 | 0 | 0 | 0 | 0 | 0 | 300 | 0 | 50 |
| SRR3490387 | 59 | 0 | 778 | 0 | 0 | 0 | 0 | 36 | 0 | 0 | 0 | 22 | 0 | 36 | 0 | 0 | 0 | 0 | 0 | 151 | 18017 | 0 | 0 | 109 | 0 | 1562 | 254 | 0 | 0 | 0 | 36 | 0 | 0 | 0 | 218 | 0 | 145 |
| SRR3490388 | 15 | 0 | 161 | 0 | 0 | 200 | 0 | 0 | 0 | 0 | 0 | 0 | 0 | 0 | 0 | 0 | 0 | 0 | 0 | 0 | 2725 | 0 | 0 | 0 | 0 | 113 | 400 | 0 | 0 | 0 | 0 | 0 | 0 | 0 | 14 | 0 | 200 |
| SRR3490389 | 18 | 0 | 203 | 0 | 0 | 0 | 0 | 77 | 77 | 0 | 0 | 0 | 0 | 0 | 0 | 0 | 0 | 0 | 0 | 0 | 8070 | 0 | 0 | 0 | 77 | 3294 | 153 | 0 | 0 | 0 | 383 | 0 | 0 | 0 | 230 | 153 | 230 |
| SRR3490390 | 87 | 0 | 633 | 0 | 0 | 65 | 0 | 0 | 0 | 0 | 0 | 0 | 0 | 0 | 0 | 0 | 0 | 0 | 0 | 0 | 7670 | 0 | 0 | 0 | 0 | 400 | 12 | 0 | 0 | 0 | 24 | 0 | 0 | 0 | 12 | 0 | 65 |
| SRR3490391 | 0 | 0 | 95 | 0 | 0 | 0 | 0 | 0 | 0 | 0 | 0 | 0 | 0 | 0 | 0 | 0 | 0 | 0 | 0 | 0 | 3176 | 0 | 0 | 0 | 0 | 86 | 0 | 0 | 0 | 0 | 0 | 0 | 0 | 0 | 0 | 0 | 242 |
| SRR3490392 | 33 | 0 | 287 | 0 | 0 | 34 | 0 | 17 | 51 | 0 | 0 | 0 | 0 | 17 | 0 | 0 | 0 | 0 | 0 | 105 | 3616 | 0 | 0 | 34 | 17 | 2334 | 34 | 0 | 0 | 0 | 69 | 0 | 0 | 34 | 120 | 17 | 69 |
| SRR3490393 | 47 | 0 | 348 | 0 | 0 | 23 | 0 | 23 | 23 | 0 | 0 | 0 | 0 | 0 | 0 | 0 | 0 | 25 | 0 | 58 | 1332 | 0 | 0 | 0 | 0 | 3170 | 282 | 0 | 23 | 0 | 117 | 0 | 0 | 23 | 94 | 0 | 141 |
| SRR3490394 | 25 | 0 | 222 | 0 | 0 | 199 | 0 | 0 | 33 | 0 | 0 | 0 | 0 | 0 | 0 | 0 | 0 | 0 | 0 | 41 | 2198 | 0 | 0 | 0 | 0 | 5380 | 266 | 0 | 0 | 0 | 166 | 0 | 0 | 33 | 531 | 0 | 299 |
| SRR3493372 | 33 | 26 | 139 | 0 | 0 | 0 | 19 | 469 | 0 | 12 | 57 | 436 | 0 | 18 | 14 | 0 | 69 | 0 | 0 | 0 | 0 | 0 | 0 | 0 | 12 | 44 | 21 | 0 | 339 | 16 | 0 | 0 | 22 | 455 | 0 | 0 | 0 |
| SRR3493373 | 42 | 16 | 138 | 0 | 0 | 0 | 33 | 33 | 0 | 21 | 18 | 625 | 0 | 18 | 18 | 0 | 39 | 0 | 0 | 0 | 0 | 0 | 0 | 0 | 12 | 21 | 0 | 0 | 34 | 0 | 0 | 0 | 106 | 248 | 0 | 0 | 0 |
| SRR3534933 | 10 | 24 | 5120 | 14 | 25 | 10 | 83 | 60 | 0 | 0 | 0 | 77 | 0 | 33 | 1425 | 23 | 20 | 0 | 0 | 793 | 1453 | 17 | 0 | 0 | 41 | 152 | 239 | 14 | 0 | 0 | 20 | 12 | 32 | 28 | 125 | 179 | 52 |
| SRR3534934 | 11 | 23 | 4888 | 0 | 0 | 14 | 422 | 158 | 0 | 0 | 0 | 66 | 0 | 41 | 1507 | 0 | 23 | 0 | 0 | 752 | 1608 | 16 | 0 | 0 | 20 | 16 | 285 | 0 | 0 | 0 | 33 | 0 | 0 | 0 | 127 | 136 | 58 |
| SRR3534935 | 20 | 23 | 4072 | 0 | 0 | 31 | 113 | 465 | 16 | 0 | 0 | 477 | 0 | 32 | 2697 | 0 | 13 | 0 | 0 | 743 | 3855 | 0 | 0 | 0 | 0 | 442 | 5471 | 0 | 0 | 0 | 76 | 0 | 0 | 0 | 1285 | 125 | 652 |
| SRR3534936 | 16 | 17 | 3892 | 0 | 0 | 46 | 28 | 73 | 21 | 0 | 0 | 543 | 0 | 0 | 4267 | 0 | 14 | 0 | 0 | 826 | 5545 | 0 | 0 | 16 | 18 | 34 | 454 | 0 | 0 | 0 | 109 | 0 | 0 | 0 | 67 | 188 | 47 |
| SRR3534937 | 28 | 0 | 4474 | 0 | 0 | 56 | 38 | 38 | 27 | 0 | 0 | 509 | 0 | 14 | 999 | 11 | 20 | 0 | 63 | 1670 | 2902 | 14 | 0 | 10 | 26 | 80 | 300 | 0 | 0 | 0 | 95 | 11 | 0 | 11 | 51 | 58 | 33 |
| SRR3534938 | 25 | 0 | 4327 | 0 | 21 | 68 | 27 | 0 | 25 | 0 | 0 | 506 | 0 | 13 | 2223 | 29 | 15 | 0 | 12 | 1277 | 6984 | 32 | 0 | 0 | 71 | 1210 | 4871 | 24 | 11 | 0 | 126 | 14 | 0 | 0 | 898 | 132 | 785 |
| SRR3566024 | 142 | 0 | 58 | 0 | 0 | 0 | 1194 | 0 | 0 | 0 | 42 | 2137 | 0 | 14 | 0 | 0 | 0 | 0 | 334 | 7402 | 1337 | 0 | 0 | 0 | 13 | 34 | 42 | 0 | 0 | 0 | 0 | 0 | 0 | 0 | 43 | 14 | 51 |
| SRR3566026 | 165 | 0 | 31 | 0 | 0 | 0 | 874 | 47 | 19 | 0 | 16 | 2300 | 0 | 23 | 0 | 0 | 0 | 0 | 1061 | 6833 | 1248 | 0 | 0 | 19 | 28 | 19 | 11 | 0 | 0 | 0 | 28 | 0 | 0 | 0 | 64 | 19 | 48 |
| SRR3566150 | 150 | 0 | 24 | 0 | 0 | 26 | 607 | 0 | 0 | 0 | 131 | 1739 | 0 | 225 | 20 | 0 | 0 | 0 | 324 | 6130 | 1598 | 0 | 0 | 21 | 16 | 31 | 58 | 0 | 0 | 0 | 16 | 0 | 0 | 0 | 125 | 42 | 131 |
| SRR3566151 | 174 | 0 | 49 | 0 | 0 | 11 | 617 | 0 | 44 | 27 | 77 | 1386 | 0 | 99 | 0 | 0 | 0 | 0 | 23 | 5626 | 915 | 0 | 0 | 33 | 38 | 16 | 38 | 0 | 0 | 0 | 11 | 0 | 0 | 0 | 88 | 33 | 164 |
| SRR3566152 | 167 | 0 | 0 | 0 | 0 | 0 | 223 | 0 | 0 | 31 | 140 | 1084 | 0 | 342 | 0 | 0 | 0 | 0 | 574 | 6666 | 1116 | 0 | 0 | 39 | 39 | 39 | 39 | 0 | 0 | 0 | 0 | 0 | 0 | 0 | 70 | 23 | 16 |
| SRR3566153 | 203 | 0 | 0 | 0 | 0 | 0 | 278 | 0 | 0 | 0 | 0 | 849 | 0 | 32 | 0 | 0 | 0 | 0 | 56 | 3086 | 445 | 0 | 0 | 27 | 13 | 0 | 16 | 15 | 0 | 0 | 0 | 0 | 0 | 0 | 19 | 16 | 15 |
| SRR3566154 | 0 | 0 | 0 | 0 | 0 | 0 | 301 | 0 | 0 | 0 | 0 | 156 | 0 | 30 | 0 | 74 | 0 | 0 | 0 | 4216 | 0 | 0 | 0 | 0 | 15 | 0 | 15 | 0 | 0 | 0 | 0 | 0 | 0 | 0 | 0 | 0 | 0 |
| SRR3566155 | 0 | 0 | 0 | 0 | 0 | 0 | 1169 | 0 | 0 | 0 | 53 | 268 | 0 | 212 | 0 | 159 | 0 | 0 | 0 | 3272 | 1402 | 0 | 0 | 0 | 53 | 0 | 0 | 212 | 0 | 0 | 0 | 0 | 0 | 0 | 0 | 0 | 0 |
| SRR3582392 | 108 | 0 | 74 | 0 | 0 | 0 | 914 | 0 | 18 | 0 | 18 | 3478 | 0 | 88 | 0 | 0 | 17 | 0 | 492 | 8580 | 1196 | 0 | 0 | 35 | 18 | 71 | 71 | 18 | 0 | 0 | 35 | 0 | 0 | 0 | 672 | 265 | 548 |
| SRR3582393 | 136 | 0 | 63 | 0 | 0 | 0 | 1582 | 0 | 19 | 0 | 39 | 3106 | 0 | 13 | 0 | 0 | 19 | 0 | 870 | 8782 | 1741 | 0 | 0 | 0 | 13 | 0 | 16 | 0 | 0 | 0 | 58 | 0 | 0 | 0 | 108 | 63 | 97 |
| SRR3623374 | 18 | 0 | 76 | 0 | 0 | 0 | 0 | 0 | 0 | 0 | 21 | 128 | 0 | 0 | 0 | 21 | 38 | 183 | 0 | 17 | 164 | 0 | 0 | 0 | 12 | 31 | 21 | 0 | 0 | 0 | 62 | 0 | 21 | 0 | 0 | 0 | 0 |
| SRR3624069 | 132 | 0 | 470 | 11 | 17 | 29 | 24 | 47 | 17 | 37 | 98 | 93 | 25 | 207 | 925 | 0 | 168 | 0 | 12 | 47 | 854 | 18 | 0 | 324 | 130 | 164 | 1447 | 17 | 0 | 0 | 10 | 0 | 47 | 147 | 55 | 71 | 370 |
| SRR3624070 | 144 | 0 | 499 | 0 | 12 | 17 | 50 | 24 | 36 | 15 | 16 | 88 | 14 | 11 | 174 | 0 | 108 | 0 | 0 | 46 | 569 | 0 | 0 | 118 | 33 | 87 | 533 | 17 | 0 | 33 | 22 | 0 | 11 | 53 | 34 | 76 | 530 |
| SRR3624071 | 141 | 0 | 446 | 0 | 12 | 22 | 62 | 20 | 28 | 24 | 26 | 94 | 13 | 24 | 812 | 0 | 128 | 41 | 0 | 54 | 1017 | 0 | 0 | 19 | 58 | 104 | 694 | 13 | 0 | 0 | 18 | 0 | 0 | 26 | 39 | 78 | 496 |
| SRR3624072 | 136 | 0 | 274 | 0 | 17 | 18 | 71 | 85 | 11 | 20 | 17 | 207 | 30 | 13 | 178 | 0 | 61 | 0 | 0 | 34 | 990 | 0 | 0 | 136 | 13 | 56 | 449 | 0 | 0 | 13 | 16 | 0 | 0 | 35 | 56 | 170 | 893 |
| SRR3624073 | 142 | 0 | 417 | 0 | 17 | 19 | 77 | 26 | 36 | 17 | 15 | 222 | 13 | 18 | 216 | 0 | 109 | 0 | 0 | 11 | 414 | 0 | 0 | 28 | 57 | 84 | 535 | 26 | 0 | 0 | 21 | 0 | 0 | 37 | 36 | 46 | 278 |
| SRR3624074 | 149 | 0 | 464 | 0 | 29 | 77 | 43 | 15 | 34 | 27 | 97 | 284 | 67 | 38 | 317 | 0 | 156 | 0 | 0 | 12 | 689 | 0 | 0 | 51 | 257 | 401 | 2255 | 17 | 0 | 0 | 18 | 0 | 0 | 163 | 242 | 678 | 3206 |
| SRR3675003 | 0 | 71 | 849 | 32 | 78 | 0 | 2061 | 2835 | 56 | 0 | 0 | 0 | 0 | 0 | 28 | 0 | 0 | 170 | 67 | 342 | 393 | 0 | 0 | 0 | 11 | 125 | 63 | 0 | 0 | 30 | 0 | 0 | 0 | 39 | 112 | 12 | 51 |
| SRR3675004 | 0 | 20 | 866 | 22 | 15 | 0 | 1409 | 834 | 20 | 0 | 0 | 0 | 0 | 18 | 33 | 0 | 0 | 138 | 56 | 113 | 226 | 0 | 0 | 0 | 0 | 38 | 15 | 0 | 0 | 37 | 0 | 0 | 0 | 33 | 29 | 19 | 15 |
| SRR3675005 | 77 | 27 | 1338 | 0 | 0 | 82 | 16 | 1149 | 28 | 0 | 0 | 0 | 0 | 13 | 19 | 0 | 0 | 0 | 19 | 1573 | 286 | 0 | 0 | 15 | 22 | 2109 | 909 | 24 | 0 | 18 | 60 | 0 | 0 | 12 | 4398 | 349 | 2112 |
| SRR3675006 | 101 | 68 | 304 | 27 | 75 | 0 | 1617 | 3812 | 57 | 0 | 0 | 0 | 0 | 15 | 33 | 0 | 0 | 105 | 88 | 306 | 264 | 0 | 0 | 0 | 0 | 135 | 59 | 0 | 0 | 40 | 0 | 0 | 0 | 43 | 124 | 27 | 66 |
| SRR3675007 | 96 | 24 | 957 | 19 | 16 | 0 | 4587 | 1065 | 31 | 0 | 0 | 0 | 0 | 0 | 41 | 0 | 0 | 100 | 29 | 0 | 283 | 0 | 0 | 0 | 0 | 61 | 24 | 0 | 0 | 20 | 0 | 0 | 0 | 20 | 48 | 11 | 26 |
| SRR3675008 | 58 | 22 | 693 | 17 | 16 | 0 | 1341 | 1068 | 22 | 0 | 0 | 0 | 0 | 12 | 30 | 0 | 0 | 124 | 51 | 153 | 353 | 0 | 0 | 0 | 0 | 54 | 19 | 0 | 0 | 28 | 0 | 0 | 0 | 19 | 20 | 12 | 10 |
| SRR3713941 | 73 | 21 | 245 | 0 | 21 | 17 | 256 | 0 | 21 | 51 | 32 | 1007 | 13 | 58 | 35 | 58 | 139 | 0 | 0 | 12 | 12 | 0 | 28 | 15 | 98 | 100 | 69 | 41 | 62 | 0 | 18 | 13 | 443 | 24 | 18 | 30 | 31 |
| SRR3713942 | 281 | 0 | 22 | 0 | 0 | 14 | 498 | 0 | 16 | 81 | 62 | 1622 | 0 | 0 | 149 | 0 | 87 | 0 | 0 | 0 | 349 | 0 | 21 | 0 | 30 | 93 | 56 | 0 | 393 | 0 | 0 | 0 | 0 | 122 | 0 | 0 | 0 |
| SRR3713943 | 0 | 44 | 80 | 18 | 72 | 0 | 17 | 0 | 0 | 255 | 135 | 0 | 0 | 478 | 83 | 271 | 91 | 0 | 0 | 33 | 0 | 59 | 66 | 0 | 490 | 96 | 79 | 186 | 268 | 0 | 52 | 0 | 264 | 20 | 45 | 52 | 37 |
| SRR3713945 | 81 | 14 | 337 | 0 | 0 | 126 | 72 | 40 | 83 | 30 | 48 | 1897 | 0 | 81 | 215 | 148 | 70 | 0 | 27 | 986 | 502 | 31 | 10 | 278 | 210 | 441 | 428 | 51 | 194 | 17 | 121 | 11 | 362 | 54 | 17 | 37 | 44 |
| SRR3713946 | 146 | 0 | 1390 | 0 | 0 | 45 | 533 | 22 | 30 | 18 | 23 | 1417 | 0 | 26 | 96 | 21 | 39 | 0 | 178 | 1504 | 1107 | 18 | 28 | 19 | 78 | 236 | 150 | 17 | 97 | 0 | 13 | 0 | 125 | 28 | 21 | 15 | 15 |
| SRR3879751 | 132 | 14 | 791 | 0 | 12 | 13 | 94 | 21 | 21 | 14 | 50 | 1438 | 0 | 10 | 145 | 0 | 25 | 0 | 0 | 0 | 13 | 0 | 0 | 0 | 11 | 35 | 0 | 0 | 17 | 0 | 13 | 0 | 25 | 11 | 0 | 0 | 0 |
| SRR3879752 | 198 | 11 | 744 | 0 | 0 | 11 | 65 | 91 | 73 | 48 | 194 | 1922 | 40 | 0 | 74 | 0 | 24 | 0 | 0 | 0 | 13 | 0 | 0 | 0 | 24 | 92 | 12 | 0 | 12 | 19 | 62 | 0 | 26 | 71 | 0 | 0 | 0 |
| SRR3879754 | 54 | 0 | 973 | 0 | 0 | 21 | 21 | 38 | 42 | 241 | 1378 | 242 | 82 | 0 | 0 | 0 | 86 | 0 | 14 | 49 | 55 | 0 | 0 | 0 | 103 | 112 | 21 | 0 | 21 | 0 | 57 | 0 | 12 | 78 | 0 | 0 | 0 |
| SRR3879755 | 63 | 0 | 1055 | 0 | 0 | 14 | 45 | 19 | 13 | 77 | 453 | 303 | 45 | 0 | 45 | 0 | 37 | 0 | 36 | 27 | 72 | 0 | 0 | 0 | 41 | 33 | 12 | 0 | 22 | 0 | 21 | 0 | 19 | 50 | 0 | 0 | 0 |
| SRR3879756 | 65 | 0 | 927 | 0 | 16 | 25 | 102 | 21 | 41 | 138 | 1233 | 401 | 37 | 0 | 0 | 0 | 33 | 0 | 45 | 40 | 0 | 0 | 0 | 0 | 117 | 131 | 18 | 0 | 34 | 0 | 80 | 0 | 39 | 42 | 0 | 0 | 0 |
| SRR3879757 | 73 | 0 | 171 | 0 | 0 | 16 | 0 | 11 | 18 | 219 | 431 | 220 | 280 | 0 | 0 | 0 | 24 | 0 | 0 | 0 | 265 | 0 | 0 | 0 | 172 | 103 | 32 | 0 | 35 | 0 | 562 | 0 | 21 | 156 | 0 | 0 | 0 |
| SRR3879758 | 124 | 15 | 274 | 0 | 0 | 0 | 37 | 13 | 24 | 41 | 105 | 149 | 106 | 0 | 75 | 0 | 25 | 100 | 0 | 37 | 224 | 0 | 0 | 0 | 34 | 27 | 15 | 0 | 31 | 0 | 136 | 0 | 11 | 48 | 0 | 0 | 0 |
| SRR3879759 | 94 | 12 | 141 | 0 | 0 | 39 | 84 | 0 | 63 | 158 | 509 | 282 | 191 | 0 | 135 | 0 | 28 | 0 | 17 | 0 | 506 | 0 | 0 | 0 | 225 | 164 | 41 | 0 | 69 | 0 | 787 | 0 | 32 | 83 | 0 | 0 | 0 |
| SRR3879760 | 56 | 0 | 307 | 0 | 0 | 0 | 65 | 50 | 21 | 143 | 368 | 908 | 110 | 0 | 0 | 0 | 0 | 0 | 0 | 129 | 0 | 0 | 0 | 0 | 194 | 126 | 17 | 0 | 17 | 0 | 178 | 0 | 29 | 143 | 0 | 0 | 0 |
| SRR3879761 | 65 | 0 | 282 | 0 | 0 | 12 | 0 | 36 | 28 | 177 | 482 | 912 | 63 | 0 | 105 | 0 | 11 | 0 | 158 | 0 | 53 | 0 | 0 | 0 | 98 | 85 | 15 | 0 | 19 | 0 | 145 | 0 | 32 | 111 | 0 | 0 | 0 |
| SRR3879766 | 57 | 0 | 679 | 0 | 0 | 208 | 0 | 18 | 52 | 43 | 101 | 765 | 282 | 27 | 439 | 0 | 18 | 10 | 0 | 20 | 0 | 0 | 0 | 0 | 336 | 2005 | 16 | 0 | 0 | 0 | 1125 | 0 | 47 | 37 | 306 | 172 | 438 |
| SRR3879767 | 45 | 0 | 30 | 0 | 0 | 0 | 0 | 33 | 0 | 10 | 124 | 30 | 0 | 26 | 20300 | 0 | 0 | 0 | 0 | 0 | 57 | 16 | 0 | 0 | 0 | 0 | 0 | 0 | 0 | 0 | 58 | 0 | 0 | 251 | 0 | 16 | 297 |
| SRR3879768 | 25 | 10 | 14 | 0 | 0 | 18 | 0 | 10 | 0 | 14 | 44 | 45 | 87 | 11 | 605 | 0 | 0 | 0 | 0 | 11 | 0 | 0 | 0 | 0 | 34 | 28 | 0 | 0 | 0 | 0 | 137 | 0 | 10 | 19 | 22 | 13 | 47 |
| SRR3879769 | 14 | 0 | 21 | 0 | 0 | 26 | 13 | 0 | 0 | 14 | 78 | 124 | 45 | 34 | 858 | 0 | 0 | 0 | 0 | 0 | 40 | 0 | 0 | 0 | 0 | 55 | 0 | 0 | 0 | 0 | 159 | 0 | 11 | 24 | 27 | 11 | 60 |
| SRR3879770 | 43 | 0 | 27 | 0 | 0 | 0 | 20 | 95 | 14 | 32 | 186 | 35 | 0 | 52 | 3429 | 0 | 0 | 0 | 0 | 0 | 61 | 11 | 0 | 0 | 0 | 29 | 0 | 0 | 0 | 0 | 64 | 0 | 0 | 176 | 0 | 139 | 1143 |
| SRR3923813 | 457 | 14 | 1573 | 16 | 0 | 0 | 0 | 0 | 0 | 0 | 0 | 43 | 0 | 0 | 0 | 0 | 0 | 0 | 0 | 0 | 0 | 0 | 0 | 0 | 0 | 0 | 0 | 0 | 0 | 0 | 12 | 0 | 0 | 0 | 11 | 0 | 0 |
| SRR3931357 | 28 | 0 | 1635 | 0 | 0 | 102 | 0 | 0 | 62 | 0 | 0 | 0 | 12 | 0 | 107 | 0 | 22 | 0 | 0 | 459 | 760 | 0 | 0 | 0 | 0 | 1499 | 1764 | 0 | 0 | 0 | 175 | 0 | 0 | 0 | 1996 | 265 | 1602 |
| SRR3931358 | 17 | 0 | 1037 | 0 | 0 | 97 | 0 | 0 | 57 | 0 | 0 | 0 | 0 | 0 | 121 | 0 | 20 | 0 | 0 | 380 | 864 | 0 | 0 | 0 | 0 | 1345 | 1471 | 20 | 0 | 0 | 176 | 0 | 0 | 0 | 2034 | 369 | 1663 |
| SRR3931359 | 16 | 0 | 1617 | 0 | 0 | 12 | 23 | 11 | 20 | 0 | 0 | 0 | 0 | 11 | 653 | 11 | 12 | 72 | 28 | 83 | 8280 | 0 | 0 | 0 | 13 | 200 | 347 | 0 | 0 | 18 | 13 | 0 | 0 | 20 | 304 | 60 | 471 |
| SRR3931360 | 19 | 0 | 1587 | 0 | 0 | 38 | 26 | 29 | 21 | 0 | 0 | 0 | 0 | 25 | 609 | 13 | 0 | 17 | 0 | 53 | 10501 | 0 | 0 | 0 | 39 | 624 | 1065 | 0 | 0 | 12 | 47 | 0 | 0 | 66 | 1167 | 220 | 1642 |
| SRR3931984 | 130 | 0 | 3219 | 27 | 19 | 0 | 0 | 0 | 0 | 0 | 0 | 16 | 0 | 32 | 44 | 17 | 67 | 173 | 0 | 11 | 11 | 0 | 27 | 0 | 56 | 24 | 0 | 14 | 0 | 0 | 485 | 0 | 14 | 42 | 1055 | 69 | 218 |
| SRR3931987 | 178 | 15 | 2574 | 29 | 24 | 0 | 0 | 0 | 0 | 0 | 0 | 39 | 0 | 55 | 45 | 20 | 41 | 27 | 0 | 52 | 0 | 0 | 0 | 0 | 50 | 27 | 0 | 17 | 0 | 0 | 248 | 0 | 14 | 20 | 182 | 0 | 21 |
| SRR3931988 | 233 | 0 | 5073 | 15 | 15 | 0 | 0 | 0 | 11 | 0 | 0 | 11 | 0 | 0 | 12 | 0 | 85 | 0 | 0 | 12 | 0 | 0 | 28 | 0 | 35 | 57 | 0 | 22 | 11 | 0 | 48 | 0 | 0 | 38 | 90 | 71 | 17 |
| SRR3992213 | 237 | 0 | 1621 | 0 | 0 | 203 | 100 | 0 | 74 | 18 | 68 | 3094 | 71 | 15 | 400 | 0 | 0 | 0 | 0 | 549 | 499 | 0 | 0 | 0 | 27 | 2661 | 97 | 0 | 0 | 0 | 50 | 0 | 0 | 162 | 109 | 88 | 725 |
| SRR3992215 | 148 | 0 | 4581 | 0 | 12 | 70 | 0 | 0 | 37 | 0 | 23 | 2160 | 12 | 0 | 283 | 0 | 10 | 0 | 0 | 57 | 113 | 0 | 0 | 0 | 85 | 438 | 21 | 0 | 0 | 0 | 21 | 0 | 0 | 40 | 23 | 13 | 41 |
| SRR3992474 | 21 | 0 | 81 | 0 | 0 | 104 | 14 | 11 | 36 | 0 | 0 | 70 | 0 | 22 | 0 | 0 | 0 | 0 | 0 | 564 | 0 | 0 | 0 | 32 | 0 | 2003 | 1844 | 0 | 0 | 0 | 196 | 0 | 81 | 0 | 0 | 0 | 0 |
| SRR3992475 | 24 | 0 | 78 | 0 | 0 | 80 | 17 | 0 | 37 | 0 | 0 | 70 | 0 | 20 | 0 | 0 | 14 | 0 | 0 | 15462 | 0 | 0 | 0 | 2064 | 0 | 1345 | 1281 | 0 | 0 | 0 | 103 | 0 | 112 | 24 | 0 | 0 | 0 |
| SRR3992476 | 20 | 0 | 721 | 13 | 0 | 22 | 13 | 50 | 11 | 0 | 0 | 334 | 0 | 15 | 110 | 24 | 14 | 52 | 0 | 1279 | 1531 | 13 | 0 | 38 | 41 | 60 | 116 | 13 | 0 | 0 | 25 | 0 | 30 | 29 | 29 | 38 | 19 |
| SRR3992477 | 19 | 0 | 757 | 0 | 0 | 33 | 62 | 59 | 10 | 0 | 0 | 276 | 0 | 0 | 106 | 0 | 15 | 0 | 44 | 26687 | 2610 | 0 | 0 | 924 | 0 | 603 | 1559 | 0 | 0 | 0 | 397 | 0 | 22 | 0 | 470 | 103 | 306 |
| SRR3996364 | 0 | 0 | 2629 | 0 | 0 | 0 | 0 | 67 | 0 | 0 | 0 | 360 | 0 | 20 | 0 | 0 | 0 | 0 | 20 | 1953 | 769 | 0 | 0 | 0 | 0 | 80 | 1468 | 40 | 0 | 87 | 0 | 0 | 0 | 0 | 657 | 27 | 322 |
| SRR3996365 | 0 | 0 | 3813 | 0 | 0 | 0 | 17 | 193 | 0 | 0 | 0 | 394 | 0 | 0 | 0 | 0 | 0 | 0 | 0 | 1456 | 458 | 0 | 0 | 23 | 0 | 139 | 1516 | 15 | 0 | 124 | 0 | 0 | 0 | 0 | 913 | 70 | 209 |
| SRR3996366 | 0 | 0 | 3715 | 0 | 0 | 0 | 0 | 93 | 0 | 0 | 0 | 425 | 0 | 0 | 0 | 0 | 0 | 0 | 20 | 1951 | 985 | 0 | 0 | 25 | 0 | 173 | 84 | 17 | 0 | 13 | 17 | 0 | 0 | 0 | 52 | 110 | 21 |
| SRR3996367 | 10 | 0 | 4189 | 0 | 0 | 0 | 0 | 62 | 0 | 0 | 0 | 406 | 0 | 0 | 0 | 0 | 0 | 0 | 14 | 3305 | 1197 | 0 | 0 | 0 | 0 | 12 | 91 | 0 | 0 | 35 | 14 | 0 | 0 | 0 | 64 | 124 | 20 |
| SRR3996368 | 0 | 0 | 6603 | 0 | 0 | 0 | 39 | 17 | 0 | 0 | 0 | 274 | 0 | 0 | 0 | 0 | 0 | 0 | 156 | 6480 | 455 | 0 | 0 | 0 | 0 | 50 | 175 | 0 | 0 | 12 | 0 | 0 | 0 | 0 | 150 | 17 | 75 |
| SRR3996369 | 0 | 0 | 9675 | 0 | 0 | 0 | 19 | 73 | 0 | 0 | 0 | 255 | 0 | 0 | 0 | 0 | 0 | 0 | 19 | 1044 | 674 | 0 | 0 | 0 | 0 | 51 | 160 | 0 | 0 | 40 | 0 | 0 | 0 | 0 | 258 | 25 | 94 |
| SRR4000625 | 266 | 0 | 627 | 0 | 0 | 18 | 10 | 0 | 585 | 22 | 132 | 540 | 40 | 27 | 40779 | 0 | 0 | 17 | 0 | 31 | 710 | 0 | 0 | 16 | 55 | 191 | 0 | 11 | 0 | 0 | 139 | 0 | 0 | 61 | 0 | 0 | 58 |
| SRR4000626 | 273 | 11 | 634 | 0 | 0 | 21 | 10 | 0 | 603 | 23 | 121 | 603 | 37 | 36 | 38494 | 0 | 0 | 0 | 0 | 21 | 881 | 0 | 0 | 16 | 48 | 206 | 0 | 10 | 0 | 0 | 140 | 0 | 0 | 50 | 0 | 0 | 60 |
| SRR4000627 | 230 | 22 | 383 | 0 | 0 | 0 | 0 | 0 | 11 | 17 | 106 | 798 | 26 | 31 | 108190 | 0 | 0 | 0 | 0 | 19 | 367 | 0 | 0 | 14 | 46 | 14 | 0 | 11 | 0 | 0 | 70 | 0 | 0 | 15 | 0 | 0 | 0 |
| SRR4000628 | 251 | 23 | 346 | 0 | 0 | 0 | 39 | 0 | 11 | 18 | 104 | 853 | 27 | 30 | 109689 | 0 | 0 | 0 | 0 | 19 | 387 | 0 | 0 | 13 | 49 | 15 | 0 | 11 | 0 | 0 | 62 | 0 | 0 | 12 | 0 | 0 | 0 |
| SRR4000629 | 164 | 44 | 222 | 0 | 0 | 0 | 69 | 0 | 176 | 22 | 108 | 936 | 31 | 45 | 32189 | 0 | 0 | 0 | 0 | 69 | 1860 | 0 | 0 | 15 | 45 | 96 | 0 | 12 | 0 | 0 | 97 | 0 | 0 | 39 | 0 | 35 | 126 |
| SRR4000630 | 158 | 31 | 219 | 0 | 0 | 0 | 93 | 0 | 109 | 11 | 64 | 971 | 26 | 27 | 32309 | 0 | 0 | 0 | 23 | 12 | 1513 | 0 | 0 | 15 | 30 | 67 | 0 | 16 | 0 | 0 | 50 | 0 | 0 | 28 | 0 | 22 | 85 |
| SRR4000631 | 242 | 12 | 283 | 0 | 0 | 19 | 24 | 0 | 119 | 24 | 118 | 929 | 49 | 50 | 44219 | 0 | 13 | 0 | 0 | 12 | 3813 | 0 | 0 | 0 | 37 | 202 | 0 | 16 | 0 | 0 | 72 | 0 | 0 | 48 | 0 | 12 | 68 |
| SRR4000632 | 289 | 13 | 289 | 0 | 0 | 11 | 37 | 0 | 77 | 19 | 69 | 848 | 29 | 26 | 46975 | 0 | 23 | 0 | 12 | 25 | 4257 | 0 | 0 | 0 | 27 | 113 | 0 | 16 | 0 | 0 | 54 | 0 | 0 | 32 | 0 | 12 | 45 |
| SRR4000633 | 222 | 44 | 165 | 0 | 0 | 11 | 47 | 0 | 194 | 13 | 77 | 976 | 18 | 21 | 22716 | 0 | 0 | 0 | 12 | 47 | 0 | 0 | 0 | 18 | 24 | 43 | 0 | 13 | 0 | 0 | 46 | 0 | 0 | 19 | 0 | 15 | 54 |
| SRR4000634 | 198 | 70 | 164 | 0 | 0 | 0 | 24 | 0 | 312 | 23 | 134 | 1069 | 35 | 27 | 23418 | 0 | 0 | 0 | 12 | 72 | 0 | 0 | 0 | 33 | 45 | 80 | 0 | 13 | 0 | 0 | 75 | 0 | 0 | 25 | 0 | 17 | 94 |
| SRR4000635 | 330 | 13 | 405 | 0 | 0 | 0 | 10 | 0 | 206 | 26 | 98 | 1222 | 19 | 25 | 34032 | 0 | 0 | 0 | 0 | 0 | 958 | 0 | 0 | 0 | 45 | 31 | 0 | 15 | 0 | 0 | 109 | 0 | 0 | 23 | 0 | 0 | 11 |
| SRR4000636 | 343 | 14 | 395 | 0 | 0 | 0 | 30 | 0 | 213 | 21 | 106 | 1200 | 25 | 22 | 31205 | 0 | 10 | 20 | 0 | 30 | 945 | 0 | 0 | 0 | 44 | 32 | 0 | 15 | 0 | 0 | 107 | 0 | 0 | 22 | 0 | 0 | 15 |
| SRR4000638 | 245 | 16 | 192 | 0 | 0 | 0 | 23 | 0 | 12 | 17 | 75 | 618 | 11 | 35 | 44715 | 0 | 67 | 47 | 0 | 0 | 1418 | 0 | 0 | 0 | 58 | 10 | 0 | 16 | 0 | 0 | 50 | 0 | 0 | 0 | 0 | 0 | 0 |
| SRR4000639 | 272 | 35 | 120 | 0 | 0 | 0 | 47 | 0 | 11 | 28 | 189 | 521 | 11 | 67 | 23130 | 0 | 52 | 0 | 24 | 71 | 801 | 0 | 0 | 129 | 98 | 12 | 0 | 26 | 0 | 0 | 74 | 0 | 0 | 0 | 0 | 0 | 0 |
| SRR4000640 | 227 | 31 | 155 | 0 | 0 | 0 | 119 | 0 | 13 | 40 | 208 | 566 | 12 | 79 | 23323 | 0 | 44 | 54 | 0 | 95 | 788 | 0 | 0 | 137 | 90 | 0 | 0 | 29 | 0 | 0 | 68 | 0 | 0 | 0 | 0 | 0 | 0 |

### Supp Table S8.2.Columns_AM_to_BW

| | | | | | | | | | | | | | | | | | | | | | | | | | | | | | | | | | | | | | |
|---|---|---|---|---|---|---|---|---|---|---|---|---|---|---|---|---|---|---|---|---|---|---|---|---|---|---|---|---|---|---|---|---|---|---|---|---|---|
| SRR4000641 | 271 | 42 | 193 | 0 | 0 | 0 | 120 | 20 | 32 | 24 | 134 | 1000 | 31 | 70 | 17709 | 0 | 54 | 0 | 0 | 201 | 642 | 0 | 0 | 358 | 54 | 0 | 0 | 29 | 0 | 0 | 90 | 0 | 0 | 0 | 0 | 0 | 0 |
| SRR4000642 | 310 | 34 | 181 | 0 | 0 | 0 | 244 | 21 | 29 | 31 | 149 | 997 | 31 | 77 | 18112 | 0 | 42 | 0 | 67 | 177 | 998 | 0 | 0 | 370 | 57 | 0 | 0 | 32 | 0 | 0 | 72 | 0 | 0 | 12 | 0 | 0 | 0 |
| SRR4000643 | 404 | 0 | 151 | 0 | 0 | 0 | 146 | 14 | 0 | 26 | 108 | 678 | 24 | 33 | 15076 | 0 | 79 | 169 | 0 | 379 | 1895 | 0 | 0 | 246 | 46 | 13 | 12 | 13 | 0 | 0 | 20 | 0 | 0 | 33 | 0 | 0 | 0 |
| SRR4000644 | 519 | 0 | 238 | 0 | 0 | 0 | 132 | 21 | 0 | 52 | 207 | 519 | 44 | 59 | 15957 | 0 | 22 | 0 | 0 | 66 | 1550 | 0 | 0 | 428 | 83 | 29 | 23 | 25 | 0 | 0 | 36 | 0 | 0 | 52 | 0 | 0 | 0 |
| SRR4000645 | 358 | 14 | 278 | 0 | 0 | 0 | 0 | 0 | 36 | 13 | 63 | 891 | 22 | 31 | 25201 | 0 | 37 | 0 | 0 | 0 | 460 | 0 | 0 | 0 | 32 | 13 | 0 | 13 | 0 | 0 | 58 | 0 | 0 | 12 | 0 | 0 | 0 |
| SRR4000649 | 138 | 33 | 275 | 0 | 0 | 0 | 14 | 0 | 682 | 31 | 132 | 421 | 18 | 51 | 22019 | 0 | 47 | 15 | 0 | 23 | 656 | 0 | 0 | 0 | 58 | 48 | 0 | 19 | 0 | 0 | 246 | 0 | 0 | 31 | 0 | 0 | 0 |
| SRR4000650 | 131 | 29 | 269 | 0 | 0 | 0 | 14 | 0 | 667 | 31 | 120 | 441 | 21 | 55 | 22612 | 0 | 55 | 0 | 0 | 27 | 696 | 0 | 0 | 0 | 56 | 52 | 0 | 22 | 0 | 0 | 259 | 0 | 0 | 34 | 0 | 0 | 0 |
| SRR4011144 | 1161 | 0 | 787 | 0 | 0 | 15 | 0 | 0 | 18 | 0 | 0 | 797 | 0 | 0 | 0 | 0 | 15 | 0 | 0 | 0 | 1665 | 0 | 0 | 0 | 16 | 89 | 0 | 0 | 0 | 0 | 29 | 0 | 0 | 0 | 11 | 0 | 0 |
| SRR4044175 | 69 | 44 | 257 | 26 | 13 | 0 | 13 | 0 | 0 | 15 | 0 | 280 | 0 | 65 | 0 | 35 | 10 | 0 | 0 | 45 | 36 | 20 | 20 | 0 | 13 | 13 | 115 | 42 | 18 | 0 | 13 | 0 | 177 | 23 | 44 | 0 | 21 |
| SRR4044176 | 60 | 117 | 312 | 129 | 48 | 0 | 20 | 0 | 0 | 0 | 0 | 336 | 0 | 178 | 0 | 113 | 15 | 0 | 0 | 53 | 41 | 24 | 69 | 0 | 25 | 27 | 185 | 113 | 34 | 0 | 10 | 13 | 561 | 43 | 87 | 0 | 39 |
| SRR4044177 | 62 | 35 | 359 | 42 | 19 | 0 | 69 | 0 | 0 | 0 | 0 | 239 | 0 | 31 | 0 | 27 | 18 | 0 | 0 | 94 | 123 | 11 | 21 | 0 | 0 | 18 | 77 | 20 | 15 | 0 | 13 | 11 | 126 | 24 | 49 | 0 | 31 |
| SRR4044178 | 76 | 38 | 257 | 22 | 12 | 0 | 10 | 0 | 0 | 13 | 0 | 393 | 0 | 60 | 0 | 39 | 20 | 0 | 0 | 37 | 85 | 21 | 22 | 0 | 13 | 19 | 83 | 63 | 16 | 0 | 16 | 0 | 164 | 33 | 62 | 15 | 29 |
| SRR4044179 | 56 | 61 | 376 | 79 | 38 | 0 | 24 | 0 | 0 | 0 | 0 | 435 | 0 | 119 | 0 | 75 | 19 | 0 | 0 | 87 | 70 | 34 | 31 | 0 | 15 | 31 | 233 | 92 | 23 | 0 | 11 | 10 | 356 | 47 | 107 | 0 | 51 |
| SRR4044180 | 96 | 86 | 325 | 64 | 12 | 0 | 68 | 0 | 0 | 0 | 0 | 225 | 0 | 92 | 0 | 46 | 14 | 0 | 0 | 42 | 69 | 12 | 58 | 0 | 10 | 104 | 100 | 44 | 180 | 0 | 39 | 0 | 521 | 17 | 77 | 0 | 47 |
| SRR4044206 | 269 | 0 | 0 | 0 | 0 | 18 | 1070 | 10 | 0 | 11 | 22 | 0 | 0 | 10 | 113 | 0 | 35 | 0 | 155 | 101 | 83 | 0 | 0 | 18 | 37 | 46 | 56 | 0 | 0 | 24 | 39 | 0 | 0 | 36 | 0 | 0 | 0 |
| SRR4044210 | 92 | 0 | 0 | 0 | 0 | 22 | 456 | 6144 | 16 | 0 | 48 | 591 | 0 | 0 | 41 | 0 | 25 | 0 | 0 | 83 | 996 | 0 | 0 | 203 | 38 | 58 | 275 | 0 | 0 | 736 | 14 | 0 | 0 | 2322 | 0 | 0 | 0 |
| SRR4044211 | 78 | 0 | 94 | 0 | 0 | 0 | 111 | 159 | 33 | 0 | 26 | 157 | 48 | 0 | 48 | 0 | 0 | 0 | 0 | 32 | 1858 | 0 | 0 | 704 | 11 | 70 | 216 | 0 | 17 | 187 | 35 | 0 | 0 | 288 | 0 | 0 | 31 |
| SRR4270461 | 60 | 0 | 99 | 0 | 36 | 325 | 35 | 461 | 103 | 22 | 41 | 829 | 151 | 43 | 25 | 13 | 520 | 0 | 152 | 1335 | 182 | 0 | 0 | 2513 | 144 | 2063 | 1652 | 29 | 11 | 111 | 258 | 0 | 0 | 316 | 322 | 253 | 447 |
| SRR4270462 | 55 | 0 | 98 | 0 | 54 | 427 | 45 | 86 | 124 | 18 | 34 | 730 | 107 | 60 | 27 | 0 | 252 | 0 | 113 | 839 | 149 | 0 | 0 | 1528 | 152 | 2786 | 1066 | 33 | 15 | 44 | 418 | 0 | 0 | 114 | 371 | 389 | 603 |
| SRR4292754 | 0 | 0 | 1822 | 0 | 0 | 0 | 0 | 0 | 0 | 0 | 0 | 609 | 0 | 0 | 106 | 0 | 0 | 0 | 128 | 6793 | 384 | 0 | 0 | 10 | 15 | 33 | 108 | 0 | 0 | 0 | 27 | 0 | 50 | 0 | 17 | 24 | 66 |
| SRR4292755 | 0 | 0 | 2716 | 0 | 0 | 0 | 0 | 0 | 0 | 0 | 0 | 546 | 0 | 0 | 168 | 0 | 0 | 0 | 60 | 4543 | 322 | 0 | 0 | 65 | 17 | 31 | 766 | 0 | 0 | 0 | 34 | 0 | 67 | 0 | 141 | 19 | 72 |
| SRR4292756 | 21 | 0 | 1810 | 0 | 0 | 0 | 0 | 0 | 0 | 0 | 0 | 624 | 0 | 0 | 43 | 0 | 0 | 0 | 63 | 3331 | 379 | 0 | 0 | 88 | 13 | 45 | 68 | 0 | 0 | 0 | 48 | 0 | 0 | 0 | 11 | 19 | 57 |
| SRR4292757 | 24 | 0 | 1638 | 0 | 0 | 0 | 0 | 0 | 0 | 0 | 0 | 617 | 11 | 0 | 51 | 0 | 0 | 0 | 56 | 2719 | 385 | 0 | 0 | 51 | 15 | 45 | 75 | 0 | 0 | 0 | 34 | 0 | 0 | 0 | 16 | 27 | 67 |
| SRR4296311 | 68 | 0 | 317 | 0 | 0 | 517 | 54 | 61 | 228 | 53 | 105 | 210 | 26 | 0 | 378 | 0 | 107 | 92 | 324 | 1564 | 3829 | 0 | 0 | 626 | 105 | 8101 | 1962 | 26 | 31 | 83 | 1160 | 0 | 0 | 201 | 1730 | 1773 | 5543 |
| SRR4296312 | 72 | 0 | 325 | 23 | 0 | 552 | 140 | 79 | 203 | 113 | 56 | 187 | 45 | 0 | 560 | 0 | 137 | 0 | 560 | 1120 | 5041 | 11 | 0 | 710 | 79 | 7705 | 2061 | 0 | 23 | 90 | 1048 | 0 | 23 | 293 | 1667 | 1633 | 5666 |
| SRR4296313 | 75 | 0 | 670 | 0 | 0 | 48 | 113 | 61 | 23 | 65 | 105 | 219 | 32 | 0 | 567 | 0 | 175 | 0 | 283 | 1927 | 2380 | 0 | 0 | 54 | 12 | 925 | 283 | 20 | 57 | 77 | 110 | 0 | 16 | 85 | 156 | 97 | 281 |
| SRR4296314 | 68 | 0 | 598 | 0 | 11 | 535 | 75 | 43 | 209 | 27 | 59 | 199 | 37 | 32 | 522 | 0 | 192 | 0 | 298 | 2088 | 2982 | 0 | 0 | 514 | 118 | 9566 | 2950 | 0 | 54 | 75 | 1194 | 0 | 16 | 91 | 1638 | 974 | 2714 |
| SRR4296315 | 42 | 11 | 650 | 0 | 0 | 412 | 195 | 39 | 184 | 67 | 84 | 115 | 67 | 11 | 130 | 0 | 111 | 0 | 455 | 2208 | 844 | 0 | 0 | 200 | 178 | 8094 | 3346 | 11 | 67 | 67 | 718 | 0 | 16 | 395 | 1659 | 1592 | 5684 |
| SRR4296316 | 38 | 0 | 635 | 0 | 0 | 443 | 0 | 22 | 188 | 89 | 122 | 118 | 66 | 0 | 478 | 0 | 75 | 0 | 358 | 1792 | 1553 | 11 | 0 | 299 | 144 | 7780 | 3469 | 0 | 33 | 22 | 543 | 0 | 0 | 465 | 1795 | 1651 | 5397 |
| SRR4296317 | 54 | 0 | 511 | 0 | 0 | 282 | 386 | 94 | 237 | 61 | 204 | 133 | 44 | 0 | 110 | 0 | 85 | 0 | 166 | 1766 | 1214 | 0 | 0 | 651 | 144 | 6072 | 3235 | 11 | 44 | 127 | 552 | 0 | 0 | 613 | 1971 | 2401 | 8148 |
| SRR4296318 | 52 | 0 | 456 | 0 | 0 | 16 | 0 | 103 | 174 | 103 | 11 | 133 | 79 | 16 | 159 | 0 | 95 | 0 | 476 | 2064 | 1508 | 24 | 0 | 30 | 11 | 339 | 154 | 16 | 40 | 166 | 42 | 0 | 0 | 25 | 93 | 120 | 421 |
| SRR4296319 | 33 | 0 | 367 | 0 | 0 | 15 | 280 | 14 | 100 | 14 | 18 | 94 | 14 | 0 | 210 | 0 | 60 | 0 | 70 | 1331 | 1261 | 0 | 0 | 15 | 53 | 155 | 65 | 0 | 37 | 53 | 18 | 0 | 0 | 37 | 45 | 49 | 212 |
| SRR4296320 | 29 | 14 | 346 | 0 | 0 | 125 | 51 | 15 | 117 | 143 | 225 | 87 | 43 | 10 | 227 | 0 | 62 | 0 | 76 | 1112 | 1339 | 0 | 0 | 103 | 76 | 1870 | 890 | 0 | 321 | 52 | 218 | 0 | 0 | 526 | 569 | 639 | 2413 |
| SRR4296321 | 29 | 0 | 519 | 0 | 0 | 158 | 156 | 26 | 150 | 106 | 162 | 74 | 26 | 0 | 249 | 0 | 57 | 0 | 62 | 1310 | 1996 | 0 | 0 | 114 | 76 | 2556 | 1152 | 0 | 435 | 62 | 220 | 0 | 0 | 230 | 477 | 483 | 1795 |
| SRR4296322 | 29 | 0 | 515 | 0 | 0 | 146 | 304 | 34 | 87 | 107 | 146 | 86 | 29 | 0 | 304 | 0 | 57 | 0 | 228 | 911 | 1518 | 0 | 0 | 117 | 53 | 2347 | 1132 | 0 | 369 | 15 | 253 | 0 | 0 | 292 | 340 | 505 | 2152 |
| SRR4296979 | 576 | 225 | 13 | 12 | 0 | 0 | 280 | 2006 | 0 | 0 | 0 | 328 | 0 | 12 | 2546 | 0 | 0 | 0 | 13 | 733 | 2600 | 0 | 0 | 124 | 0 | 78 | 1282 | 0 | 0 | 0 | 0 | 0 | 0 | 0 | 225 | 52 | 309 |
| SRR4296985 | 38 | 0 | 285 | 0 | 0 | 0 | 120 | 0 | 0 | 0 | 0 | 98 | 0 | 0 | 0 | 0 | 38 | 0 | 180 | 2583 | 60 | 0 | 0 | 0 | 25 | 102 | 767 | 0 | 112 | 0 | 0 | 0 | 0 | 0 | 0 | 0 | 0 |
| SRR4296986 | 26 | 0 | 419 | 0 | 19 | 0 | 151 | 0 | 0 | 0 | 0 | 233 | 0 | 0 | 0 | 0 | 17 | 46 | 363 | 1663 | 60 | 0 | 0 | 0 | 24 | 46 | 531 | 0 | 84 | 0 | 0 | 0 | 10 | 0 | 0 | 0 | 0 |
| SRR4296987 | 379 | 0 | 98 | 15 | 0 | 0 | 42 | 70 | 0 | 0 | 0 | 336 | 0 | 0 | 0 | 0 | 13 | 0 | 43 | 40 | 48 | 0 | 0 | 0 | 0 | 38 | 364 | 0 | 0 | 22 | 0 | 0 | 0 | 0 | 0 | 0 | 0 |
| SRR4296988 | 334 | 0 | 150 | 12 | 11 | 0 | 46 | 85 | 0 | 0 | 0 | 362 | 0 | 0 | 13 | 0 | 0 | 0 | 51 | 35 | 53 | 0 | 0 | 0 | 0 | 33 | 441 | 0 | 0 | 23 | 0 | 0 | 0 | 11 | 0 | 0 | 0 |
| SRR4328127 | 287 | 0 | 148 | 0 | 0 | 0 | 2911 | 0 | 0 | 0 | 35 | 573 | 0 | 34 | 1338 | 0 | 0 | 0 | 33 | 284 | 284 | 0 | 0 | 0 | 0 | 0 | 0 | 0 | 0 | 0 | 0 | 0 | 0 | 0 | 0 | 0 | 0 |
| SRR4328130 | 868 | 0 | 0 | 0 | 0 | 0 | 3464 | 0 | 0 | 0 | 27 | 2186 | 0 | 13 | 2474 | 0 | 32 | 0 | 1237 | 2227 | 1484 | 0 | 0 | 0 | 0 | 0 | 0 | 0 | 0 | 0 | 0 | 0 | 0 | 12 | 0 | 0 | 0 |
| SRR4328135 | 172 | 0 | 29 | 0 | 0 | 0 | 4624 | 0 | 0 | 0 | 54 | 574 | 0 | 17 | 3391 | 0 | 0 | 0 | 0 | 231 | 540 | 0 | 0 | 0 | 0 | 0 | 0 | 0 | 0 | 0 | 0 | 0 | 0 | 0 | 0 | 0 | 0 |
| SRR4328136 | 1166 | 0 | 38 | 0 | 0 | 0 | 1120 | 0 | 0 | 0 | 24 | 879 | 0 | 35 | 920 | 0 | 38 | 0 | 80 | 380 | 660 | 0 | 0 | 0 | 0 | 0 | 11 | 0 | 0 | 0 | 0 | 0 | 0 | 0 | 0 | 0 | 0 |
| SRR4328137 | 873 | 0 | 998 | 0 | 0 | 0 | 5400 | 0 | 0 | 0 | 12 | 1996 | 0 | 13 | 277 | 0 | 62 | 0 | 254 | 300 | 2031 | 0 | 0 | 0 | 0 | 0 | 11 | 0 | 0 | 0 | 0 | 0 | 0 | 0 | 0 | 0 | 0 |
| SRR4409201 | 38 | 0 | 185 | 0 | 0 | 24 | 18 | 0 | 17 | 0 | 0 | 0 | 0 | 0 | 246 | 0 | 0 | 0 | 0 | 28 | 0 | 0 | 0 | 0 | 0 | 58 | 68 | 0 | 0 | 0 | 19 | 0 | 0 | 0 | 28 | 0 | 35 |
| SRR4409202 | 599 | 0 | 0 | 0 | 0 | 53 | 601 | 16 | 11 | 0 | 0 | 47 | 0 | 0 | 80 | 0 | 52 | 0 | 646 | 38531 | 10 | 0 | 0 | 11 | 0 | 867 | 472 | 0 | 0 | 12 | 0 | 0 | 0 | 0 | 18 | 0 | 22 |
| SRR4409203 | 29 | 0 | 159 | 0 | 0 | 21 | 47 | 0 | 10 | 0 | 0 | 0 | 0 | 0 | 1001 | 0 | 0 | 0 | 0 | 102 | 770 | 0 | 0 | 0 | 0 | 41 | 68 | 0 | 0 | 0 | 12 | 0 | 10 | 0 | 21 | 0 | 24 |
| SRR4409204 | 558 | 0 | 3437 | 0 | 0 | 0 | 576 | 22 | 0 | 0 | 0 | 186 | 0 | 0 | 35 | 0 | 50 | 0 | 882 | 47364 | 2934 | 0 | 0 | 0 | 0 | 43 | 225 | 0 | 0 | 0 | 0 | 0 | 0 | 0 | 0 | 0 | 0 |
| SRR4417245 | 1036 | 0 | 784 | 0 | 0 | 0 | 0 | 0 | 0 | 0 | 120 | 2612 | 1012 | 13 | 13 | 0 | 61 | 282 | 0 | 0 | 57 | 0 | 0 | 0 | 331 | 34 | 13 | 0 | 0 | 0 | 23 | 0 | 0 | 0 | 0 | 0 | 0 |
| SRR4417246 | 772 | 0 | 471 | 0 | 0 | 0 | 0 | 0 | 0 | 0 | 191 | 1558 | 18 | 0 | 73 | 0 | 56 | 298 | 0 | 0 | 29 | 0 | 0 | 0 | 10 | 0 | 0 | 0 | 0 | 0 | 0 | 0 | 0 | 0 | 0 | 0 | 0 |
| SRR4417247 | 3595 | 0 | 614 | 0 | 0 | 0 | 0 | 0 | 0 | 0 | 42 | 3433 | 147 | 0 | 0 | 12 | 56 | 86 | 0 | 0 | 152 | 0 | 0 | 0 | 105 | 29 | 0 | 0 | 0 | 0 | 0 | 0 | 0 | 21 | 0 | 0 | 0 |
| SRR4417248 | 1321 | 0 | 564 | 0 | 0 | 0 | 0 | 0 | 0 | 0 | 101 | 2969 | 68 | 0 | 0 | 0 | 41 | 49 | 0 | 0 | 82 | 0 | 0 | 0 | 181 | 0 | 0 | 0 | 0 | 0 | 0 | 0 | 0 | 21 | 0 | 0 | 0 |
| SRR4417249 | 1156 | 0 | 359 | 0 | 0 | 0 | 0 | 0 | 0 | 0 | 121 | 1238 | 995 | 0 | 10 | 0 | 39 | 347 | 0 | 0 | 0 | 0 | 0 | 0 | 230 | 11 | 0 | 0 | 0 | 0 | 0 | 0 | 0 | 0 | 0 | 0 | 0 |
| SRR4417250 | 2608 | 0 | 779 | 0 | 24 | 0 | 0 | 0 | 0 | 0 | 77 | 3878 | 0 | 0 | 0 | 0 | 46 | 135 | 0 | 0 | 338 | 0 | 0 | 0 | 173 | 17 | 11 | 0 | 0 | 0 | 0 | 0 | 0 | 34 | 0 | 0 | 0 |
| SRR4423247 | 15 | 28 | 1057 | 0 | 0 | 78 | 92 | 345 | 0 | 0 | 0 | 63 | 0 | 94 | 953 | 0 | 38 | 0 | 14 | 2123 | 2137 | 0 | 0 | 0 | 0 | 1044 | 2112 | 0 | 0 | 0 | 333 | 0 | 0 | 0 | 3429 | 272 | 880 |
| SRR4423248 | 27 | 0 | 2282 | 0 | 0 | 17 | 37 | 96 | 0 | 0 | 0 | 181 | 0 | 76 | 776 | 0 | 24 | 0 | 16 | 1004 | 1594 | 0 | 0 | 0 | 19 | 309 | 910 | 0 | 0 | 0 | 43 | 0 | 0 | 25 | 556 | 56 | 430 |
| SRR4423250 | 73 | 13 | 251 | 0 | 0 | 100 | 51 | 432 | 26 | 0 | 0 | 709 | 0 | 0 | 905 | 0 | 27 | 58 | 46 | 8710 | 5397 | 0 | 0 | 54 | 11 | 1421 | 2414 | 0 | 0 | 13 | 30 | 0 | 0 | 10 | 11865 | 2465 | 4896 |
| SRR4423251 | 10 | 0 | 771 | 0 | 0 | 55 | 200 | 156 | 23 | 0 | 0 | 211 | 0 | 0 | 4374 | 0 | 0 | 16 | 135 | 10370 | 30607 | 0 | 0 | 74 | 36 | 914 | 3923 | 0 | 0 | 0 | 89 | 0 | 0 | 17 | 5745 | 1932 | 7143 |
| SRR4473670 | 140 | 32 | 10506 | 0 | 0 | 0 | 255 | 121 | 19 | 0 | 12 | 37 | 0 | 16 | 81 | 0 | 17 | 127 | 56 | 1920 | 2245 | 0 | 0 | 0 | 17 | 24 | 43 | 0 | 14 | 31 | 13 | 0 | 0 | 0 | 13 | 35 | 26 |
| SRR4734676 | 93 | 0 | 1874 | 0 | 0 | 32 | 0 | 0 | 12 | 0 | 0 | 297 | 0 | 0 | 92 | 148 | 43 | 0 | 0 | 267 | 49 | 0 | 0 | 0 | 33 | 454 | 67 | 0 | 0 | 0 | 63 | 0 | 0 | 0 | 1242 | 66 | 228 |
| SRR4734677 | 82 | 0 | 2569 | 0 | 0 | 17 | 0 | 0 | 17 | 0 | 0 | 262 | 0 | 0 | 193 | 34 | 40 | 0 | 0 | 83 | 111 | 0 | 0 | 0 | 0 | 1157 | 184 | 17 | 0 | 0 | 101 | 0 | 0 | 17 | 3235 | 134 | 520 |
| SRR4734678 | 142 | 0 | 836 | 0 | 13 | 27 | 0 | 0 | 13 | 0 | 0 | 147 | 0 | 0 | 43 | 0 | 12 | 0 | 0 | 303 | 563 | 0 | 0 | 0 | 40 | 16 | 80 | 0 | 0 | 0 | 172 | 0 | 0 | 13 | 32 | 40 | 106 |
| SRR4734679 | 211 | 14 | 1253 | 0 | 0 | 82 | 0 | 0 | 41 | 0 | 0 | 268 | 0 | 0 | 47 | 0 | 17 | 0 | 0 | 664 | 190 | 0 | 0 | 0 | 0 | 684 | 96 | 0 | 0 | 0 | 41 | 0 | 0 | 0 | 561 | 82 | 178 |
| SRR4734680 | 181 | 34 | 1405 | 0 | 0 | 67 | 0 | 0 | 51 | 0 | 0 | 185 | 0 | 0 | 0 | 0 | 18 | 0 | 0 | 676 | 406 | 0 | 0 | 0 | 34 | 36 | 17 | 17 | 17 | 0 | 51 | 0 | 0 | 17 | 19 | 0 | 84 |
| SRR4734681 | 127 | 0 | 1948 | 0 | 0 | 61 | 0 | 0 | 31 | 0 | 0 | 247 | 0 | 0 | 24 | 0 | 27 | 0 | 0 | 283 | 378 | 0 | 0 | 0 | 23 | 1158 | 192 | 0 | 0 | 0 | 169 | 0 | 0 | 0 | 2155 | 153 | 437 |
| SRR4734682 | 177 | 0 | 1369 | 0 | 24 | 48 | 0 | 0 | 0 | 0 | 0 | 206 | 0 | 0 | 85 | 16 | 14 | 0 | 0 | 721 | 212 | 0 | 0 | 0 | 48 | 646 | 97 | 0 | 0 | 0 | 48 | 0 | 0 | 0 | 646 | 48 | 154 |
| SRR4734683 | 180 | 0 | 1105 | 0 | 0 | 11 | 0 | 0 | 20 | 0 | 0 | 218 | 0 | 0 | 168 | 11 | 0 | 0 | 0 | 343 | 222 | 0 | 0 | 0 | 15 | 23 | 55 | 0 | 0 | 0 | 29 | 0 | 0 | 21 | 24 | 38 | 122 |
| SRR4734684 | 110 | 13 | 1407 | 0 | 0 | 48 | 0 | 0 | 16 | 0 | 0 | 325 | 0 | 0 | 187 | 13 | 26 | 0 | 0 | 241 | 232 | 0 | 0 | 0 | 0 | 41 | 109 | 0 | 0 | 0 | 100 | 0 | 0 | 10 | 60 | 157 | 14 |
| SRR4734685 | 127 | 0 | 720 | 14 | 0 | 31 | 0 | 0 | 10 | 0 | 0 | 193 | 0 | 17 | 59 | 0 | 17 | 0 | 0 | 259 | 761 | 0 | 0 | 0 | 12 | 30 | 43 | 16 | 0 | 0 | 53 | 0 | 0 | 28 | 32 | 47 | 11 |
| SRR4734686 | 65 | 0 | 1443 | 0 | 0 | 38 | 0 | 0 | 34 | 0 | 0 | 166 | 0 | 0 | 108 | 0 | 39 | 0 | 0 | 173 | 94 | 0 | 0 | 0 | 0 | 19 | 164 | 13 | 0 | 0 | 38 | 0 | 0 | 0 | 67 | 202 | 14 |
| SRR4787084 | 35 | 0 | 3131 | 0 | 0 | 69 | 123 | 0 | 0 | 0 | 0 | 101 | 0 | 0 | 1976 | 0 | 30 | 0 | 741 | 494 | 2593 | 0 | 0 | 345 | 2487 | 138 | 898 | 0 | 0 | 0 | 69 | 0 | 0 | 0 | 0 | 0 | 0 |
| SRR4787085 | 26 | 0 | 3002 | 0 | 36 | 0 | 0 | 0 | 0 | 0 | 0 | 32 | 0 | 0 | 2676 | 0 | 0 | 0 | 357 | 803 | 2052 | 0 | 0 | 0 | 71 | 107 | 17 | 0 | 0 | 0 | 0 | 0 | 0 | 36 | 0 | 0 | 0 |
| SRR4787086 | 50 | 0 | 3470 | 0 | 0 | 64 | 0 | 0 | 64 | 0 | 0 | 114 | 64 | 64 | 2663 | 0 | 30 | 0 | 666 | 533 | 3596 | 0 | 0 | 129 | 119 | 22 | 31 | 0 | 0 | 0 | 0 | 0 | 0 | 0 | 0 | 0 | 0 |
| SRR4787087 | 24 | 0 | 2574 | 0 | 0 | 45 | 0 | 0 | 0 | 0 | 0 | 89 | 224 | 0 | 2833 | 90 | 27 | 0 | 531 | 620 | 3099 | 0 | 0 | 537 | 3000 | 313 | 672 | 0 | 0 | 0 | 45 | 0 | 0 | 0 | 0 | 45 | 134 |
| SRR4787088 | 37 | 0 | 2744 | 0 | 0 | 66 | 126 | 0 | 66 | 0 | 0 | 110 | 0 | 0 | 3391 | 0 | 12 | 0 | 879 | 1130 | 1507 | 0 | 0 | 199 | 861 | 265 | 927 | 0 | 0 | 0 | 0 | 0 | 0 | 0 | 0 | 0 | 0 |
| SRR4787089 | 32 | 0 | 2463 | 0 | 0 | 0 | 0 | 0 | 89 | 0 | 0 | 97 | 267 | 0 | 2010 | 0 | 24 | 0 | 914 | 548 | 1827 | 0 | 0 | 267 | 43 | 178 | 53 | 0 | 0 | 0 | 0 | 0 | 0 | 0 | 0 | 0 | 0 |
| SRR4787090 | 24 | 0 | 341 | 0 | 13 | 94 | 22 | 107 | 18 | 0 | 201 | 107 | 72 | 0 | 9594 | 0 | 0 | 0 | 216 | 108 | 1902 | 0 | 0 | 886 | 188 | 421 | 1003 | 0 | 0 | 81 | 0 | 0 | 0 | 1168 | 18 | 13 | 27 |
| SRR4787091 | 21 | 0 | 2956 | 0 | 0 | 0 | 0 | 0 | 0 | 0 | 157 | 63 | 157 | 0 | 3059 | 0 | 10 | 0 | 834 | 278 | 834 | 0 | 0 | 0 | 23 | 314 | 32 | 0 | 0 | 0 | 0 | 0 | 0 | 0 | 0 | 0 | 0 |
| SRR4787092 | 29 | 0 | 408 | 0 | 14 | 115 | 0 | 101 | 0 | 24 | 245 | 86 | 67 | 0 | 8898 | 0 | 15 | 34 | 137 | 412 | 1830 | 0 | 0 | 923 | 207 | 380 | 1534 | 0 | 0 | 125 | 24 | 0 | 0 | 976 | 0 | 14 | 34 |
| SRR4787093 | 33 | 0 | 358 | 0 | 18 | 170 | 44 | 115 | 0 | 14 | 147 | 96 | 92 | 0 | 9405 | 14 | 0 | 33 | 241 | 329 | 2105 | 0 | 0 | 1053 | 211 | 349 | 1274 | 0 | 0 | 124 | 14 | 0 | 0 | 1191 | 23 | 14 | 18 |
| SRR4787095 | 71 | 0 | 12965 | 0 | 0 | 77 | 94 | 17 | 52 | 0 | 0 | 368 | 0 | 0 | 1318 | 0 | 19 | 0 | 241 | 143 | 2291 | 0 | 0 | 288 | 305 | 1534 | 808 | 0 | 0 | 13 | 137 | 0 | 0 | 0 | 112 | 39 | 99 |
| SRR4787096 | 75 | 0 | 13211 | 0 | 0 | 77 | 97 | 36 | 46 | 0 | 0 | 416 | 10 | 10 | 1314 | 0 | 27 | 0 | 268 | 108 | 2155 | 0 | 0 | 415 | 328 | 1132 | 507 | 0 | 0 | 0 | 133 | 0 | 0 | 0 | 97 | 82 | 108 |
| SRR4787097 | 671 | 0 | 334 | 0 | 0 | 11 | 172 | 46 | 29 | 0 | 0 | 259 | 0 | 0 | 22 | 0 | 22 | 0 | 559 | 387 | 456 | 0 | 0 | 126 | 506 | 297 | 108 | 18 | 59 | 0 | 0 | 0 | 0 | 0 | 0 | 0 | 13 |
| SRR4787098 | 668 | 0 | 401 | 0 | 0 | 15 | 214 | 84 | 16 | 0 | 0 | 290 | 0 | 16 | 26 | 0 | 21 | 21 | 597 | 222 | 504 | 0 | 0 | 180 | 300 | 300 | 64 | 0 | 58 | 0 | 49 | 0 | 0 | 0 | 0 | 10 | 16 |
| SRR4787099 | 580 | 0 | 394 | 0 | 0 | 25 | 236 | 51 | 17 | 0 | 0 | 275 | 0 | 0 | 0 | 0 | 34 | 18 | 535 | 340 | 590 | 0 | 0 | 222 | 326 | 287 | 59 | 0 | 14 | 0 | 43 | 0 | 0 | 0 | 0 | 0 | 0 |
| SRR4787100 | 462 | 0 | 819 | 0 | 0 | 61 | 222 | 0 | 83 | 0 | 0 | 788 | 0 | 12 | 94 | 0 | 125 | 118 | 963 | 882 | 1219 | 0 | 0 | 415 | 2866 | 768 | 104 | 0 | 104 | 0 | 114 | 0 | 0 | 0 | 25 | 18 | 55 |
| SRR4787101 | 468 | 0 | 753 | 0 | 0 | 40 | 273 | 0 | 91 | 0 | 0 | 786 | 0 | 0 | 45 | 0 | 109 | 89 | 991 | 807 | 1113 | 0 | 0 | 21 | 98 | 49 | 10 | 0 | 125 | 16 | 88 | 0 | 0 | 0 | 24 | 0 | 91 |
| SRR4787102 | 481 | 0 | 886 | 0 | 0 | 51 | 206 | 12 | 65 | 0 | 0 | 837 | 0 | 0 | 178 | 0 | 133 | 89 | 785 | 682 | 1042 | 0 | 0 | 22 | 198 | 57 | 15 | 0 | 80 | 12 | 97 | 0 | 0 | 0 | 22 | 17 | 51 |
| SRR4787103 | 173 | 0 | 1640 | 0 | 0 | 46 | 162 | 0 | 40 | 15 | 0 | 1097 | 10 | 0 | 97 | 25 | 280 | 221 | 340 | 340 | 1035 | 0 | 0 | 23 | 105 | 30 | 11 | 0 | 91 | 0 | 91 | 0 | 0 | 20 | 106 | 61 | 131 |
| SRR4787104 | 222 | 0 | 1748 | 0 | 0 | 42 | 90 | 0 | 46 | 0 | 0 | 1020 | 28 | 0 | 90 | 0 | 247 | 187 | 265 | 253 | 922 | 0 | 0 | 21 | 94 | 50 | 21 | 0 | 92 | 0 | 74 | 0 | 0 | 0 | 56 | 42 | 106 |
| SRR4787105 | 322 | 21 | 2391 | 0 | 0 | 12 | 15 | 12 | 0 | 0 | 0 | 242 | 0 | 0 | 109 | 0 | 16 | 28 | 25 | 59 | 545 | 0 | 0 | 0 | 0 | 16 | 79 | 0 | 0 | 0 | 17 | 0 | 0 | 29 | 83 | 0 | 75 |
| SRR4787106 | 314 | 0 | 2482 | 0 | 0 | 19 | 0 | 38 | 0 | 0 | 0 | 255 | 0 | 13 | 92 | 0 | 23 | 14 | 0 | 46 | 382 | 0 | 0 | 0 | 0 | 18 | 108 | 0 | 25 | 0 | 0 | 0 | 0 | 38 | 121 | 0 | 102 |
| SRR4787107 | 125 | 21 | 2498 | 0 | 0 | 24 | 12 | 18 | 15 | 0 | 0 | 372 | 0 | 12 | 36 | 0 | 21 | 14 | 48 | 159 | 469 | 0 | 0 | 0 | 0 | 279 | 112 | 0 | 30 | 0 | 0 | 0 | 0 | 21 | 70 | 0 | 61 |
| SRR4787108 | 111 | 0 | 3055 | 0 | 0 | 22 | 17 | 13 | 0 | 0 | 0 | 246 | 0 | 13 | 85 | 0 | 12 | 0 | 45 | 119 | 741 | 0 | 0 | 0 | 0 | 27 | 28 | 0 | 19 | 0 | 0 | 0 | 19 | 0 | 44 | 0 | 53 |
| SRR4787109 | 327 | 0 | 2368 | 0 | 0 | 22 | 0 | 0 | 0 | 0 | 0 | 278 | 0 | 25 | 43 | 0 | 15 | 0 | 17 | 78 | 338 | 0 | 0 | 0 | 0 | 17 | 115 | 0 | 29 | 0 | 0 | 0 | 14 | 11 | 97 | 0 | 75 |
| SRR4787110 | 127 | 0 | 2441 | 0 | 0 | 27 | 32 | 18 | 0 | 0 | 0 | 357 | 0 | 15 | 32 | 0 | 31 | 15 | 32 | 103 | 343 | 0 | 0 | 0 | 0 | 260 | 128 | 0 | 57 | 0 | 0 | 0 | 0 | 12 | 116 | 0 | 51 |
| SRR4787118 | 0 | 0 | 17038 | 0 | 0 | 0 | 0 | 0 | 0 | 0 | 0 | 0 | 0 | 0 | 0 | 0 | 0 | 0 | 0 | 0 | 0 | 0 | 0 | 0 | 240 | 1002 | 0 | 0 | 0 | 0 | 0 | 0 | 0 | 0 | 0 | 0 | 0 |
| SRR4787119 | 3180 | 0 | 7696 | 0 | 0 | 0 | 0 | 0 | 0 | 0 | 0 | 0 | 0 | 0 | 0 | 0 | 0 | 0 | 0 | 0 | 1662 | 0 | 0 | 0 | 126 | 0 | 0 | 0 | 0 | 0 | 0 | 0 | 0 | 0 | 0 | 0 | 0 |
| SRR4787120 | 51 | 0 | 9949 | 0 | 0 | 64 | 133 | 0 | 14 | 0 | 0 | 501 | 46 | 0 | 1874 | 0 | 38 | 0 | 189 | 124 | 2629 | 0 | 0 | 13 | 32 | 31 | 44 | 0 | 0 | 0 | 147 | 0 | 0 | 0 | 129 | 74 | 101 |
| SRR4787121 | 251 | 0 | 2519 | 0 | 0 | 41 | 12 | 61 | 34 | 0 | 0 | 102 | 0 | 0 | 576 | 0 | 0 | 0 | 93 | 74 | 1555 | 0 | 0 | 82 | 949 | 618 | 232 | 0 | 0 | 32 | 44 | 0 | 0 | 0 | 0 | 0 | 15 |
| SRR4787122 | 262 | 0 | 2065 | 0 | 0 | 35 | 35 | 11 | 35 | 0 | 0 | 96 | 0 | 0 | 851 | 0 | 0 | 11 | 53 | 33 | 2037 | 0 | 0 | 84 | 1442 | 699 | 196 | 0 | 0 | 13 | 44 | 0 | 0 | 0 | 0 | 13 | 0 |
| SRR4787123 | 215 | 0 | 1955 | 0 | 0 | 41 | 12 | 34 | 20 | 0 | 0 | 88 | 0 | 0 | 621 | 0 | 0 | 0 | 92 | 62 | 1494 | 0 | 0 | 73 | 86 | 52 | 12 | 0 | 0 | 0 | 96 | 0 | 0 | 0 | 16 | 0 | 0 |
| SRR4787124 | 438 | 0 | 4250 | 0 | 0 | 303 | 0 | 112 | 146 | 0 | 0 | 147 | 0 | 0 | 137 | 0 | 0 | 0 | 0 | 175 | 1348 | 0 | 0 | 234 | 0 | 2179 | 156 | 0 | 0 | 0 | 0 | 0 | 0 | 0 | 0 | 0 | 0 |
| SRR4787125 | 212 | 0 | 4283 | 0 | 0 | 0 | 0 | 293 | 0 | 0 | 0 | 0 | 0 | 0 | 43 | 0 | 0 | 0 | 218 | 0 | 820 | 0 | 0 | 226 | 538 | 1782 | 773 | 0 | 0 | 0 | 0 | 0 | 0 | 0 | 0 | 0 | 0 |
| SRR4787126 | 236 | 0 | 3164 | 0 | 0 | 0 | 90 | 0 | 0 | 0 | 0 | 0 | 0 | 0 | 0 | 0 | 0 | 0 | 94 | 0 | 724 | 0 | 0 | 0 | 20 | 191 | 23 | 0 | 0 | 0 | 0 | 0 | 0 | 0 | 0 | 0 | 0 |
| SRR4787127 | 0 | 0 | 31 | 0 | 0 | 0 | 0 | 0 | 0 | 0 | 0 | 0 | 0 | 0 | 0 | 0 | 0 | 0 | 0 | 0 | 4163 | 0 | 0 | 0 | 0 | 711 | 0 | 0 | 0 | 0 | 0 | 0 | 0 | 0 | 0 | 0 | 0 |
| SRR4787132 | 25 | 0 | 2913 | 0 | 24 | 24 | 56 | 0 | 0 | 0 | 0 | 70 | 24 | 0 | 1746 | 0 | 12 | 0 | 338 | 1183 | 2366 | 0 | 0 | 71 | 165 | 94 | 18 | 0 | 0 | 0 | 0 | 0 | 0 | 0 | 24 | 0 | 0 |
| SRR4787133 | 17 | 0 | 2769 | 0 | 0 | 0 | 0 | 0 | 0 | 0 | 0 | 28 | 0 | 0 | 1428 | 0 | 0 | 0 | 79 | 873 | 3253 | 0 | 0 | 0 | 100 | 166 | 16 | 0 | 0 | 0 | 0 | 0 | 0 | 0 | 0 | 0 | 0 |
| SRR4787134 | 460 | 0 | 661 | 0 | 0 | 13 | 102 | 35 | 22 | 0 | 0 | 68 | 0 | 0 | 19 | 0 | 0 | 0 | 359 | 105 | 492 | 0 | 0 | 0 | 0 | 39 | 28 | 0 | 13 | 0 | 0 | 0 | 0 | 0 | 35 | 0 | 0 |
| SRR4787135 | 179 | 0 | 1377 | 0 | 0 | 60 | 86 | 0 | 28 | 0 | 0 | 1029 | 18 | 0 | 143 | 0 | 307 | 234 | 278 | 378 | 870 | 0 | 0 | 458 | 1955 | 518 | 222 | 14 | 88 | 0 | 69 | 0 | 0 | 0 | 69 | 23 | 111 |
| SRR4787136 | 458 | 0 | 1371 | 0 | 0 | 69 | 136 | 11 | 12 | 0 | 0 | 492 | 11 | 0 | 16 | 0 | 28 | 51 | 465 | 222 | 662 | 0 | 0 | 19 | 34 | 96 | 862 | 0 | 13 | 0 | 88 | 0 | 0 | 29 | 13 | 50 | 37 |
| SRR4787137 | 674 | 0 | 991 | 0 | 0 | 26 | 52 | 52 | 90 | 0 | 0 | 121 | 17 | 0 | 12 | 0 | 0 | 0 | 141 | 56 | 362 | 0 | 0 | 0 | 202 | 434 | 611 | 0 | 172 | 0 | 69 | 0 | 0 | 39 | 13 | 0 | 0 |
| SRR4787138 | 554 | 0 | 1345 | 0 | 0 | 49 | 255 | 40 | 60 | 0 | 0 | 418 | 11 | 14 | 20 | 0 | 0 | 14 | 850 | 208 | 358 | 0 | 0 | 12 | 88 | 82 | 75 | 0 | 98 | 0 | 63 | 0 | 0 | 17 | 34 | 11 | 25 |
| SRR4787139 | 568 | 0 | 1409 | 0 | 0 | 68 | 257 | 13 | 75 | 0 | 0 | 618 | 20 | 12 | 30 | 0 | 19 | 33 | 745 | 223 | 426 | 0 | 0 | 13 | 363 | 1016 | 942 | 0 | 93 | 0 | 59 | 0 | 0 | 0 | 50 | 33 | 46 |
| SRR4787140 | 356 | 0 | 2382 | 0 | 0 | 85 | 264 | 11 | 107 | 0 | 0 | 1054 | 38 | 0 | 34 | 0 | 52 | 62 | 723 | 203 | 551 | 0 | 0 | 32 | 605 | 1472 | 1904 | 13 | 92 | 11 | 92 | 0 | 0 | 26 | 60 | 36 | 90 |
| SRR4787141 | 604 | 0 | 1284 | 0 | 0 | 48 | 222 | 24 | 55 | 0 | 0 | 625 | 14 | 11 | 0 | 0 | 19 | 38 | 706 | 213 | 422 | 0 | 0 | 11 | 64 | 122 | 151 | 0 | 10 | 0 | 55 | 0 | 0 | 0 | 42 | 27 | 46 |
| SRR4787142 | 350 | 0 | 2405 | 0 | 0 | 91 | 212 | 0 | 107 | 0 | 0 | 1192 | 28 | 0 | 60 | 0 | 66 | 84 | 649 | 204 | 601 | 0 | 0 | 23 | 766 | 1484 | 1707 | 0 | 125 | 0 | 100 | 0 | 0 | 0 | 65 | 67 | 128 |
| SRR4787143 | 413 | 0 | 2303 | 0 | 0 | 103 | 277 | 24 | 90 | 0 | 0 | 1011 | 35 | 0 | 45 | 0 | 37 | 64 | 858 | 221 | 521 | 0 | 0 | 20 | 37 | 145 | 162 | 0 | 10 | 0 | 105 | 0 | 11 | 0 | 66 | 11 | 88 |
| SRR4836291 | 22 | 29 | 2012 | 0 | 0 | 10 | 134 | 488 | 10 | 0 | 0 | 205 | 0 | 13 | 3662 | 0 | 19 | 68 | 13 | 2562 | 2562 | 0 | 0 | 0 | 0 | 41 | 35 | 0 | 0 | 12 | 18 | 0 | 0 | 0 | 64 | 10 | 43 |
| SRR4836293 | 15 | 76 | 2196 | 0 | 13 | 16 | 132 | 1467 | 63 | 0 | 0 | 212 | 0 | 19 | 4643 | 0 | 0 | 55 | 0 | 2157 | 2272 | 0 | 0 | 0 | 0 | 114 | 139 | 0 | 0 | 19 | 28 | 0 | 0 | 0 | 177 | 16 | 123 |
| SRR4836294 | 36 | 25 | 2135 | 0 | 0 | 0 | 187 | 463 | 23 | 0 | 0 | 216 | 0 | 21 | 3974 | 0 | 25 | 33 | 30 | 2426 | 2998 | 0 | 0 | 0 | 0 | 33 | 43 | 0 | 0 | 15 | 15 | 0 | 0 | 0 | 70 | 23 | 52 |
| SRR4836295 | 26 | 36 | 727 | 0 | 0 | 10 | 418 | 1076 | 10 | 0 | 0 | 97 | 0 | 41 | 786 | 0 | 0 | 54 | 0 | 2162 | 2432 | 0 | 0 | 0 | 88 | 129 | 139 | 52 | 0 | 10 | 10 | 10 | 0 | 26 | 206 | 175 | 350 |
| SRR4836297 | 31 | 50 | 739 | 0 | 0 | 0 | 427 | 970 | 13 | 0 | 0 | 99 | 0 | 25 | 549 | 0 | 0 | 64 | 0 | 1617 | 2532 | 0 | 0 | 19 | 94 | 176 | 157 | 38 | 0 | 13 | 13 | 0 | 0 | 31 | 189 | 126 | 372 |

### Supp Table S8.2.Columns_AM_to_BW

| | | | | | | | | | | | | | | | | | | | | | | | | | | | | | | | | | | | | | |
|---|---|---|---|---|---|---|---|---|---|---|---|---|---|---|---|---|---|---|---|---|---|---|---|---|---|---|---|---|---|---|---|---|---|---|---|---|---|
| SRR4836299 | 47 | 41 | 672 | 0 | 0 | 11 | 392 | 195 | 0 | 0 | 0 | 91 | 0 | 26 | 962 | 0 | 0 | 157 | 0 | 1711 | 1889 | 0 | 0 | 11 | 18 | 33 | 27 | 49 | 0 | 0 | 26 | 11 | 0 | 45 | 55 | 33 | 77 |
| SRR4836301 | 35 | 20 | 2577 | 0 | 0 | 20 | 409 | 538 | 14 | 0 | 0 | 257 | 0 | 23 | 444 | 0 | 16 | 618 | 0 | 922 | 2774 | 0 | 0 | 11 | 0 | 66 | 41 | 0 | 0 | 11 | 27 | 11 | 0 | 12 | 106 | 23 | 81 |
| SRR4836302 | 36 | 24 | 2582 | 0 | 0 | 0 | 483 | 1403 | 0 | 0 | 0 | 244 | 0 | 48 | 494 | 0 | 0 | 713 | 0 | 914 | 2626 | 0 | 0 | 11 | 0 | 135 | 122 | 0 | 0 | 28 | 15 | 0 | 0 | 28 | 306 | 39 | 219 |
| SRR4836303 | 29 | 44 | 2415 | 0 | 0 | 14 | 371 | 1502 | 0 | 0 | 0 | 253 | 0 | 28 | 563 | 0 | 25 | 552 | 26 | 755 | 2892 | 0 | 0 | 10 | 0 | 185 | 106 | 0 | 0 | 35 | 21 | 0 | 0 | 30 | 314 | 54 | 217 |
| SRR4836305 | 35 | 21 | 4073 | 0 | 0 | 20 | 0 | 54 | 15 | 0 | 0 | 289 | 0 | 0 | 723 | 0 | 15 | 591 | 0 | 124 | 1239 | 0 | 0 | 0 | 0 | 63 | 80 | 0 | 0 | 15 | 25 | 0 | 0 | 15 | 53 | 11 | 38 |
| SRR4836307 | 51 | 66 | 4282 | 0 | 0 | 12 | 0 | 344 | 12 | 0 | 0 | 288 | 0 | 36 | 819 | 0 | 19 | 949 | 0 | 125 | 1491 | 0 | 0 | 0 | 0 | 271 | 325 | 0 | 0 | 18 | 0 | 0 | 0 | 24 | 325 | 36 | 368 |
| SRR4836308 | 50 | 15 | 3815 | 0 | 0 | 22 | 0 | 50 | 0 | 0 | 0 | 314 | 0 | 25 | 879 | 0 | 13 | 778 | 0 | 177 | 1463 | 0 | 0 | 18 | 0 | 56 | 71 | 0 | 0 | 25 | 22 | 0 | 0 | 15 | 65 | 47 | 50 |
| SRR4836310 | 15 | 105 | 875 | 0 | 0 | 23 | 47 | 1122 | 12 | 0 | 0 | 132 | 0 | 12 | 899 | 0 | 20 | 432 | 0 | 8605 | 1065 | 0 | 0 | 19 | 16 | 413 | 288 | 0 | 0 | 16 | 62 | 0 | 0 | 16 | 810 | 47 | 405 |
| SRR4836311 | 0 | 21 | 918 | 0 | 0 | 33 | 71 | 189 | 0 | 0 | 0 | 141 | 0 | 0 | 958 | 0 | 18 | 306 | 0 | 9269 | 1099 | 0 | 0 | 24 | 33 | 79 | 53 | 0 | 0 | 0 | 15 | 0 | 0 | 0 | 149 | 52 | 79 |
| SRR4836312 | 14 | 18 | 851 | 0 | 0 | 26 | 65 | 182 | 0 | 0 | 0 | 154 | 0 | 0 | 992 | 0 | 19 | 479 | 0 | 8796 | 992 | 0 | 0 | 23 | 11 | 69 | 45 | 0 | 0 | 0 | 20 | 0 | 0 | 11 | 151 | 11 | 64 |
| SRR4836313 | 10 | 64 | 2438 | 0 | 16 | 10 | 52 | 86 | 27 | 0 | 0 | 465 | 0 | 0 | 850 | 0 | 18 | 334 | 10 | 664 | 5049 | 0 | 0 | 0 | 0 | 284 | 324 | 0 | 0 | 0 | 24 | 0 | 0 | 13 | 2046 | 145 | 878 |
| SRR4836314 | 17 | 17 | 2416 | 0 | 15 | 21 | 20 | 26 | 12 | 0 | 0 | 391 | 0 | 16 | 861 | 0 | 19 | 336 | 0 | 851 | 4484 | 0 | 0 | 0 | 0 | 88 | 86 | 0 | 0 | 0 | 21 | 0 | 0 | 21 | 574 | 43 | 284 |
| SRR4836315 | 18 | 80 | 2246 | 0 | 0 | 13 | 50 | 75 | 22 | 0 | 0 | 429 | 0 | 0 | 1072 | 0 | 20 | 331 | 0 | 924 | 4636 | 0 | 0 | 0 | 0 | 267 | 264 | 0 | 0 | 0 | 24 | 0 | 0 | 15 | 2026 | 126 | 937 |
| SRR4836316 | 39 | 15 | 2323 | 0 | 0 | 0 | 94 | 101 | 14 | 0 | 0 | 109 | 0 | 45 | 704 | 0 | 20 | 46 | 0 | 17922 | 857 | 0 | 0 | 0 | 0 | 15 | 44 | 0 | 0 | 0 | 0 | 0 | 0 | 0 | 124 | 45 | 83 |
| SRR4836317 | 41 | 83 | 2264 | 0 | 0 | 0 | 130 | 517 | 0 | 0 | 0 | 95 | 0 | 43 | 810 | 0 | 28 | 73 | 0 | 17389 | 987 | 0 | 0 | 0 | 0 | 72 | 195 | 0 | 0 | 0 | 18 | 0 | 0 | 0 | 741 | 36 | 445 |
| SRR4836318 | 41 | 15 | 2021 | 0 | 0 | 0 | 101 | 90 | 19 | 0 | 0 | 106 | 0 | 29 | 802 | 0 | 19 | 113 | 11 | 18526 | 1059 | 0 | 0 | 0 | 0 | 16 | 43 | 13 | 0 | 0 | 11 | 0 | 0 | 0 | 129 | 46 | 79 |
| SRR4836320 | 10 | 66 | 765 | 0 | 0 | 33 | 19 | 13 | 12 | 0 | 0 | 0 | 0 | 17 | 63 | 0 | 21 | 277 | 27 | 36480 | 170 | 0 | 0 | 0 | 0 | 70 | 56 | 0 | 0 | 0 | 21 | 0 | 0 | 0 | 192 | 87 | 73 |
| SRR4836321 | 16 | 14 | 670 | 0 | 0 | 40 | 18 | 16 | 24 | 0 | 0 | 10 | 0 | 0 | 70 | 0 | 24 | 195 | 17 | 36468 | 199 | 0 | 0 | 0 | 0 | 92 | 55 | 0 | 0 | 0 | 56 | 0 | 0 | 0 | 185 | 72 | 57 |
| SRR4836323 | 0 | 12 | 594 | 0 | 0 | 10 | 19 | 11 | 20 | 0 | 0 | 12 | 0 | 0 | 78 | 0 | 31 | 261 | 19 | 39811 | 182 | 0 | 0 | 0 | 0 | 68 | 39 | 0 | 0 | 0 | 30 | 0 | 0 | 0 | 165 | 74 | 59 |
| SRR4836325 | 13 | 24 | 557 | 0 | 0 | 0 | 0 | 16 | 16 | 0 | 0 | 291 | 0 | 0 | 1152 | 0 | 0 | 114 | 0 | 696 | 3684 | 0 | 0 | 0 | 0 | 17 | 68 | 0 | 0 | 0 | 19 | 0 | 0 | 11 | 361 | 33 | 295 |
| SRR4836326 | 16 | 26 | 539 | 0 | 0 | 0 | 11 | 15 | 13 | 0 | 0 | 297 | 0 | 10 | 839 | 0 | 0 | 48 | 0 | 816 | 3666 | 0 | 0 | 0 | 0 | 18 | 61 | 0 | 0 | 0 | 0 | 0 | 0 | 13 | 385 | 35 | 278 |
| SRR4836327 | 13 | 24 | 481 | 0 | 0 | 0 | 0 | 73 | 0 | 0 | 0 | 322 | 0 | 13 | 1102 | 0 | 0 | 102 | 0 | 924 | 3874 | 0 | 0 | 0 | 0 | 71 | 408 | 0 | 0 | 0 | 31 | 0 | 0 | 0 | 2218 | 209 | 1612 |
| SRR4836328 | 44 | 99 | 747 | 0 | 0 | 0 | 134 | 79 | 16 | 0 | 0 | 150 | 0 | 13 | 2973 | 0 | 22 | 116 | 34 | 2000 | 3967 | 0 | 0 | 0 | 0 | 61 | 69 | 0 | 0 | 0 | 19 | 0 | 0 | 13 | 60 | 23 | 40 |
| SRR4836330 | 33 | 354 | 745 | 0 | 0 | 20 | 139 | 262 | 0 | 0 | 0 | 163 | 0 | 13 | 2937 | 0 | 24 | 63 | 0 | 1794 | 3898 | 0 | 0 | 0 | 0 | 222 | 276 | 0 | 0 | 0 | 14 | 0 | 0 | 18 | 203 | 20 | 107 |
| SRR4836332 | 41 | 369 | 624 | 0 | 0 | 21 | 156 | 218 | 11 | 0 | 0 | 156 | 0 | 17 | 3055 | 0 | 14 | 228 | 0 | 1712 | 3663 | 0 | 0 | 0 | 0 | 212 | 269 | 0 | 0 | 0 | 15 | 0 | 0 | 17 | 170 | 16 | 121 |
| SRR4836334 | 24 | 30 | 2209 | 0 | 0 | 11 | 102 | 45 | 13 | 0 | 0 | 33 | 0 | 13 | 1212 | 0 | 17 | 247 | 0 | 1434 | 4421 | 0 | 0 | 0 | 0 | 103 | 114 | 0 | 0 | 0 | 14 | 0 | 0 | 13 | 516 | 20 | 211 |
| SRR4836335 | 21 | 37 | 2356 | 0 | 0 | 26 | 99 | 45 | 12 | 0 | 0 | 46 | 0 | 14 | 1174 | 0 | 18 | 237 | 0 | 1522 | 4284 | 0 | 0 | 0 | 0 | 104 | 114 | 0 | 0 | 0 | 12 | 0 | 0 | 0 | 513 | 25 | 207 |
| SRR4836336 | 29 | 95 | 2086 | 0 | 10 | 21 | 69 | 126 | 10 | 0 | 0 | 41 | 0 | 12 | 1391 | 0 | 11 | 332 | 14 | 1515 | 4311 | 0 | 0 | 0 | 0 | 269 | 311 | 0 | 0 | 0 | 38 | 0 | 0 | 14 | 1751 | 88 | 678 |
| SRR4836337 | 22 | 17 | 2492 | 0 | 12 | 22 | 64 | 263 | 0 | 0 | 0 | 289 | 0 | 40 | 2520 | 0 | 11 | 292 | 0 | 591 | 3395 | 0 | 0 | 13 | 0 | 63 | 93 | 10 | 0 | 11 | 32 | 10 | 0 | 38 | 49 | 28 | 34 |
| SRR4836339 | 22 | 19 | 2221 | 0 | 0 | 0 | 102 | 957 | 0 | 0 | 0 | 317 | 0 | 46 | 2871 | 0 | 15 | 340 | 0 | 718 | 3811 | 0 | 0 | 0 | 0 | 233 | 357 | 0 | 0 | 63 | 39 | 0 | 0 | 35 | 203 | 23 | 131 |
| SRR4836341 | 24 | 16 | 2479 | 0 | 0 | 14 | 79 | 1220 | 13 | 0 | 0 | 291 | 0 | 41 | 2398 | 0 | 22 | 322 | 0 | 578 | 3078 | 0 | 0 | 16 | 0 | 298 | 442 | 0 | 0 | 43 | 33 | 12 | 0 | 30 | 259 | 0 | 179 |
| SRR4836343 | 26 | 19 | 2121 | 0 | 0 | 0 | 80 | 206 | 0 | 0 | 0 | 313 | 0 | 46 | 2666 | 0 | 0 | 321 | 0 | 702 | 3440 | 0 | 0 | 11 | 0 | 59 | 80 | 0 | 0 | 13 | 26 | 12 | 0 | 11 | 59 | 22 | 34 |
| SRR4836344 | 14 | 34 | 1369 | 0 | 0 | 19 | 65 | 484 | 24 | 0 | 0 | 336 | 0 | 11 | 1780 | 0 | 0 | 87 | 0 | 5082 | 3317 | 0 | 0 | 34 | 0 | 26 | 92 | 0 | 0 | 0 | 29 | 0 | 0 | 15 | 204 | 16 | 112 |
| SRR4836345 | 15 | 20 | 1211 | 10 | 0 | 0 | 75 | 3246 | 0 | 0 | 0 | 310 | 0 | 36 | 1819 | 0 | 0 | 194 | 0 | 5819 | 3728 | 0 | 0 | 31 | 0 | 203 | 610 | 0 | 0 | 20 | 41 | 0 | 0 | 0 | 1501 | 97 | 773 |
| SRR4836346 | 20 | 23 | 1423 | 0 | 0 | 11 | 66 | 3887 | 15 | 0 | 0 | 342 | 0 | 84 | 1519 | 0 | 15 | 75 | 0 | 4934 | 3170 | 0 | 0 | 54 | 0 | 161 | 760 | 0 | 0 | 42 | 73 | 0 | 0 | 0 | 1827 | 122 | 910 |
| SRR4836347 | 17 | 15 | 1276 | 0 | 0 | 15 | 56 | 3388 | 19 | 0 | 0 | 306 | 0 | 81 | 1619 | 0 | 11 | 93 | 0 | 5357 | 3456 | 0 | 0 | 39 | 0 | 139 | 580 | 12 | 0 | 27 | 46 | 0 | 0 | 0 | 1690 | 104 | 816 |
| SRR4836348 | 34 | 45 | 1850 | 0 | 0 | 20 | 87 | 229 | 12 | 0 | 0 | 119 | 0 | 0 | 548 | 0 | 28 | 1231 | 0 | 692 | 4585 | 0 | 0 | 12 | 0 | 78 | 201 | 0 | 0 | 0 | 12 | 0 | 0 | 0 | 94 | 41 | 87 |
| SRR4836349 | 18 | 56 | 1689 | 0 | 0 | 26 | 78 | 195 | 26 | 0 | 0 | 110 | 0 | 0 | 838 | 0 | 20 | 1122 | 0 | 978 | 4842 | 0 | 0 | 17 | 0 | 59 | 160 | 0 | 0 | 30 | 11 | 0 | 0 | 0 | 83 | 34 | 87 |
| SRR4836350 | 22 | 10 | 1875 | 0 | 0 | 35 | 43 | 231 | 23 | 0 | 0 | 105 | 0 | 0 | 649 | 0 | 25 | 979 | 0 | 984 | 4250 | 0 | 0 | 19 | 0 | 70 | 217 | 0 | 0 | 0 | 45 | 0 | 0 | 0 | 96 | 26 | 93 |
| SRR4836351 | 31 | 13 | 1614 | 0 | 0 | 13 | 11 | 194 | 20 | 0 | 0 | 125 | 0 | 0 | 540 | 0 | 20 | 1222 | 11 | 771 | 4254 | 0 | 0 | 13 | 0 | 70 | 178 | 0 | 0 | 17 | 13 | 0 | 0 | 0 | 88 | 10 | 100 |
| SRR4836352 | 19 | 16 | 2111 | 0 | 0 | 18 | 24 | 208 | 0 | 0 | 0 | 80 | 0 | 0 | 1487 | 0 | 10 | 49 | 0 | 775 | 4049 | 0 | 0 | 18 | 0 | 96 | 159 | 0 | 0 | 26 | 10 | 0 | 0 | 0 | 248 | 13 | 117 |
| SRR4836353 | 28 | 16 | 1802 | 0 | 0 | 12 | 17 | 163 | 21 | 0 | 0 | 78 | 0 | 0 | 1630 | 0 | 10 | 76 | 17 | 849 | 3981 | 0 | 0 | 12 | 0 | 91 | 124 | 0 | 0 | 20 | 34 | 0 | 0 | 10 | 223 | 19 | 86 |
| SRR4836354 | 21 | 15 | 2166 | 0 | 0 | 14 | 12 | 231 | 0 | 0 | 0 | 88 | 0 | 0 | 1179 | 0 | 15 | 86 | 12 | 841 | 3844 | 0 | 0 | 14 | 0 | 102 | 157 | 0 | 0 | 28 | 14 | 0 | 0 | 16 | 259 | 16 | 112 |
| SRR4836355 | 29 | 50 | 1919 | 0 | 0 | 16 | 18 | 631 | 12 | 0 | 0 | 79 | 0 | 0 | 1386 | 0 | 0 | 68 | 0 | 849 | 3792 | 0 | 0 | 11 | 0 | 350 | 477 | 0 | 0 | 105 | 41 | 0 | 0 | 0 | 846 | 33 | 355 |
| SRR4836356 | 22 | 25 | 1550 | 0 | 0 | 20 | 586 | 2995 | 0 | 0 | 0 | 81 | 0 | 57 | 2989 | 0 | 24 | 1618 | 0 | 912 | 4480 | 0 | 0 | 12 | 0 | 234 | 216 | 0 | 0 | 42 | 32 | 32 | 0 | 13 | 321 | 40 | 211 |
| SRR4836357 | 20 | 10 | 1350 | 0 | 0 | 0 | 594 | 826 | 15 | 0 | 0 | 78 | 0 | 21 | 3297 | 0 | 21 | 1611 | 16 | 766 | 4735 | 0 | 0 | 0 | 0 | 66 | 50 | 0 | 0 | 12 | 11 | 26 | 0 | 0 | 99 | 13 | 70 |
| SRR4836359 | 23 | 20 | 1565 | 0 | 0 | 15 | 512 | 3042 | 12 | 0 | 0 | 88 | 0 | 76 | 2597 | 0 | 23 | 1581 | 17 | 840 | 4471 | 0 | 0 | 0 | 0 | 251 | 216 | 0 | 0 | 35 | 28 | 24 | 0 | 0 | 320 | 43 | 216 |
| SRR4836361 | 24 | 28 | 1365 | 0 | 0 | 12 | 494 | 2670 | 0 | 0 | 0 | 71 | 0 | 58 | 3285 | 0 | 16 | 1721 | 0 | 916 | 4768 | 0 | 0 | 0 | 0 | 217 | 178 | 0 | 0 | 40 | 31 | 29 | 0 | 16 | 297 | 42 | 230 |
| SRR4836362 | 22 | 241 | 2811 | 0 | 0 | 52 | 40 | 4070 | 16 | 0 | 0 | 173 | 0 | 23 | 968 | 0 | 15 | 208 | 13 | 2009 | 2674 | 0 | 0 | 12 | 0 | 754 | 450 | 0 | 0 | 102 | 115 | 0 | 0 | 41 | 397 | 21 | 205 |
| SRR4836364 | 21 | 64 | 2398 | 0 | 0 | 10 | 14 | 759 | 10 | 0 | 0 | 199 | 0 | 12 | 1049 | 0 | 12 | 272 | 0 | 2055 | 2858 | 0 | 0 | 14 | 0 | 156 | 93 | 0 | 0 | 16 | 26 | 0 | 0 | 40 | 72 | 27 | 38 |
| SRR4836366 | 16 | 251 | 2697 | 0 | 0 | 78 | 42 | 4138 | 17 | 0 | 0 | 202 | 0 | 44 | 916 | 0 | 0 | 272 | 26 | 1985 | 2461 | 0 | 0 | 11 | 0 | 804 | 492 | 0 | 0 | 90 | 101 | 0 | 0 | 30 | 433 | 24 | 198 |
| SRR4836368 | 23 | 246 | 2482 | 0 | 0 | 57 | 43 | 3701 | 19 | 0 | 0 | 183 | 0 | 34 | 845 | 0 | 10 | 365 | 11 | 2091 | 2700 | 0 | 0 | 0 | 0 | 756 | 395 | 0 | 0 | 72 | 121 | 0 | 0 | 33 | 326 | 23 | 171 |
| SRR4836369 | 32 | 65 | 2062 | 0 | 0 | 0 | 104 | 1645 | 14 | 0 | 0 | 123 | 0 | 0 | 868 | 0 | 24 | 60 | 19 | 689 | 2245 | 0 | 0 | 0 | 0 | 284 | 432 | 0 | 0 | 25 | 44 | 0 | 0 | 26 | 412 | 25 | 510 |
| SRR4836370 | 35 | 60 | 1875 | 0 | 0 | 22 | 161 | 1490 | 19 | 0 | 0 | 137 | 0 | 12 | 1079 | 0 | 28 | 130 | 0 | 769 | 2169 | 0 | 0 | 0 | 0 | 294 | 455 | 0 | 0 | 29 | 56 | 0 | 0 | 25 | 446 | 36 | 562 |
| SRR4836371 | 35 | 64 | 2033 | 0 | 0 | 21 | 66 | 1737 | 15 | 0 | 0 | 127 | 0 | 10 | 1064 | 0 | 36 | 170 | 11 | 866 | 2215 | 0 | 0 | 0 | 0 | 277 | 473 | 0 | 0 | 36 | 58 | 0 | 0 | 12 | 423 | 28 | 504 |
| SRR4836373 | 39 | 67 | 1955 | 0 | 0 | 13 | 131 | 1607 | 0 | 0 | 0 | 125 | 0 | 0 | 971 | 0 | 34 | 165 | 0 | 822 | 2161 | 0 | 0 | 15 | 0 | 304 | 473 | 0 | 0 | 73 | 59 | 0 | 0 | 25 | 479 | 38 | 548 |
| SRR4836375 | 55 | 13 | 1311 | 0 | 0 | 0 | 59 | 235 | 18 | 0 | 0 | 37 | 0 | 47 | 753 | 0 | 61 | 1158 | 0 | 979 | 3808 | 0 | 0 | 13 | 0 | 28 | 70 | 0 | 0 | 27 | 10 | 0 | 0 | 13 | 138 | 32 | 112 |
| SRR4836376 | 34 | 52 | 1262 | 0 | 10 | 0 | 50 | 1022 | 15 | 0 | 0 | 51 | 0 | 42 | 666 | 0 | 71 | 989 | 10 | 1114 | 4123 | 0 | 0 | 0 | 0 | 134 | 330 | 0 | 0 | 25 | 15 | 0 | 0 | 0 | 642 | 48 | 487 |
| SRR4836378 | 58 | 54 | 1378 | 0 | 0 | 0 | 39 | 1109 | 12 | 0 | 0 | 53 | 0 | 48 | 746 | 0 | 54 | 1009 | 0 | 1040 | 4091 | 0 | 0 | 28 | 0 | 145 | 298 | 0 | 0 | 28 | 10 | 0 | 0 | 14 | 640 | 28 | 485 |
| SRR4836380 | 45 | 12 | 1369 | 0 | 0 | 0 | 60 | 221 | 0 | 0 | 0 | 60 | 0 | 42 | 699 | 0 | 74 | 1195 | 11 | 899 | 3837 | 0 | 0 | 0 | 0 | 30 | 66 | 0 | 0 | 21 | 16 | 0 | 0 | 16 | 138 | 12 | 112 |
| SRR4836382 | 50 | 29 | 462 | 0 | 0 | 10 | 21 | 452 | 13 | 0 | 0 | 27 | 0 | 0 | 654 | 0 | 13 | 327 | 11 | 11988 | 1732 | 0 | 0 | 0 | 0 | 210 | 54 | 0 | 0 | 0 | 25 | 0 | 0 | 0 | 242 | 18 | 125 |
| SRR4836383 | 24 | 38 | 410 | 0 | 0 | 12 | 13 | 404 | 14 | 0 | 0 | 31 | 0 | 13 | 708 | 0 | 0 | 300 | 13 | 12789 | 1897 | 0 | 0 | 0 | 0 | 217 | 58 | 0 | 0 | 0 | 21 | 0 | 0 | 0 | 239 | 0 | 115 |
| SRR4836384 | 41 | 40 | 529 | 0 | 0 | 24 | 0 | 456 | 22 | 0 | 0 | 39 | 0 | 0 | 738 | 0 | 13 | 286 | 13 | 12690 | 1722 | 0 | 0 | 0 | 0 | 198 | 42 | 0 | 0 | 0 | 19 | 0 | 0 | 0 | 220 | 23 | 119 |
| SRR4836386 | 40 | 41 | 435 | 0 | 0 | 12 | 16 | 480 | 21 | 0 | 0 | 29 | 0 | 0 | 649 | 0 | 14 | 265 | 0 | 12163 | 1807 | 0 | 0 | 0 | 0 | 225 | 53 | 0 | 0 | 0 | 14 | 0 | 0 | 0 | 263 | 17 | 140 |
| SRR4836388 | 41 | 16 | 2861 | 0 | 13 | 0 | 41 | 39 | 19 | 0 | 0 | 34 | 0 | 13 | 3120 | 0 | 32 | 1128 | 0 | 1580 | 3762 | 0 | 0 | 19 | 0 | 52 | 94 | 0 | 0 | 11 | 29 | 0 | 0 | 11 | 180 | 12 | 137 |
| SRR4836390 | 35 | 22 | 2714 | 0 | 12 | 16 | 17 | 159 | 28 | 0 | 0 | 38 | 0 | 12 | 3332 | 0 | 45 | 1190 | 17 | 1679 | 4440 | 0 | 0 | 0 | 0 | 254 | 414 | 0 | 0 | 0 | 24 | 0 | 0 | 16 | 846 | 26 | 626 |
| SRR4836391 | 47 | 23 | 3016 | 0 | 0 | 12 | 0 | 154 | 46 | 0 | 0 | 51 | 0 | 0 | 2889 | 0 | 45 | 870 | 0 | 1534 | 3779 | 0 | 0 | 0 | 0 | 241 | 378 | 0 | 0 | 12 | 19 | 0 | 0 | 14 | 866 | 62 | 586 |
| SRR4836392 | 36 | 20 | 2771 | 0 | 11 | 13 | 0 | 32 | 20 | 0 | 0 | 35 | 0 | 0 | 2955 | 0 | 37 | 1176 | 13 | 1826 | 4103 | 0 | 0 | 22 | 0 | 59 | 93 | 0 | 0 | 11 | 17 | 0 | 0 | 14 | 212 | 14 | 149 |
| SRR4836394 | 94 | 144 | 1086 | 0 | 0 | 23 | 220 | 897 | 12 | 0 | 0 | 751 | 0 | 13 | 5147 | 0 | 45 | 657 | 12 | 887 | 3142 | 0 | 0 | 0 | 0 | 365 | 260 | 0 | 0 | 43 | 34 | 0 | 0 | 18 | 204 | 20 | 123 |
| SRR4836396 | 92 | 41 | 1043 | 0 | 0 | 21 | 90 | 235 | 21 | 0 | 0 | 770 | 0 | 14 | 5199 | 0 | 53 | 655 | 30 | 850 | 3183 | 0 | 0 | 0 | 0 | 109 | 65 | 0 | 0 | 13 | 11 | 0 | 0 | 18 | 57 | 29 | 38 |
| SRR4836398 | 100 | 181 | 1118 | 0 | 0 | 38 | 115 | 807 | 11 | 0 | 0 | 802 | 0 | 21 | 4876 | 0 | 52 | 673 | 43 | 762 | 3100 | 0 | 0 | 0 | 0 | 401 | 229 | 0 | 0 | 55 | 41 | 0 | 0 | 15 | 208 | 28 | 136 |
| SRR4836399 | 104 | 46 | 1043 | 0 | 0 | 23 | 154 | 242 | 16 | 0 | 0 | 757 | 0 | 0 | 5173 | 0 | 54 | 649 | 27 | 797 | 2771 | 0 | 0 | 0 | 0 | 114 | 65 | 0 | 0 | 11 | 31 | 0 | 0 | 20 | 55 | 28 | 46 |
| SRR4836400 | 25 | 38 | 1375 | 0 | 0 | 63 | 71 | 265 | 17 | 0 | 0 | 157 | 0 | 0 | 204 | 0 | 25 | 64 | 0 | 1481 | 1567 | 0 | 0 | 49 | 0 | 1099 | 779 | 0 | 0 | 17 | 124 | 0 | 0 | 0 | 2080 | 144 | 978 |
| SRR4836402 | 30 | 54 | 1280 | 0 | 11 | 15 | 100 | 46 | 36 | 0 | 0 | 129 | 0 | 14 | 201 | 0 | 31 | 80 | 20 | 1555 | 1535 | 0 | 0 | 43 | 0 | 191 | 143 | 0 | 0 | 14 | 26 | 0 | 0 | 11 | 413 | 25 | 174 |
| SRR4836404 | 24 | 28 | 1413 | 0 | 0 | 66 | 67 | 294 | 21 | 0 | 0 | 168 | 0 | 10 | 163 | 0 | 26 | 0 | 38 | 1606 | 1404 | 0 | 0 | 41 | 0 | 1078 | 774 | 0 | 0 | 17 | 149 | 0 | 0 | 10 | 2159 | 93 | 898 |
| SRR4836406 | 19 | 14 | 1213 | 0 | 0 | 36 | 84 | 47 | 25 | 0 | 0 | 158 | 0 | 17 | 214 | 0 | 27 | 37 | 23 | 1400 | 1423 | 0 | 0 | 33 | 0 | 189 | 144 | 0 | 0 | 17 | 30 | 0 | 0 | 0 | 404 | 27 | 188 |
| SRR4836407 | 32 | 81 | 1663 | 0 | 0 | 19 | 293 | 1755 | 14 | 0 | 0 | 171 | 0 | 16 | 548 | 0 | 15 | 43 | 0 | 587 | 2670 | 0 | 0 | 0 | 0 | 68 | 73 | 0 | 0 | 24 | 15 | 0 | 0 | 10 | 94 | 11 | 158 |
| SRR4836408 | 27 | 79 | 1703 | 0 | 0 | 16 | 333 | 2073 | 14 | 0 | 0 | 166 | 0 | 18 | 475 | 0 | 20 | 63 | 0 | 855 | 2566 | 0 | 0 | 0 | 0 | 77 | 90 | 0 | 0 | 28 | 16 | 0 | 0 | 0 | 114 | 30 | 156 |
| SRR4836410 | 41 | 291 | 1554 | 0 | 0 | 13 | 308 | 6497 | 0 | 0 | 0 | 179 | 0 | 69 | 521 | 0 | 23 | 103 | 0 | 741 | 2713 | 0 | 0 | 0 | 0 | 280 | 274 | 0 | 0 | 31 | 62 | 0 | 0 | 0 | 376 | 47 | 586 |
| SRR4836412 | 41 | 224 | 1813 | 0 | 0 | 23 | 31 | 46 | 0 | 0 | 0 | 102 | 0 | 0 | 138 | 0 | 16 | 491 | 0 | 2169 | 352 | 0 | 0 | 0 | 0 | 283 | 333 | 0 | 0 | 0 | 19 | 0 | 0 | 0 | 297 | 44 | 153 |
| SRR4836414 | 67 | 167 | 1916 | 0 | 0 | 14 | 21 | 54 | 0 | 0 | 0 | 118 | 0 | 12 | 142 | 0 | 13 | 426 | 0 | 2030 | 412 | 0 | 0 | 0 | 0 | 313 | 386 | 0 | 0 | 0 | 19 | 0 | 0 | 12 | 311 | 41 | 146 |
| SRR4836415 | 33 | 56 | 1633 | 0 | 0 | 0 | 16 | 14 | 0 | 0 | 0 | 85 | 0 | 0 | 167 | 0 | 10 | 473 | 0 | 2025 | 337 | 0 | 0 | 0 | 0 | 68 | 80 | 0 | 0 | 0 | 0 | 0 | 0 | 10 | 92 | 22 | 37 |
| SRR4836416 | 33 | 35 | 1627 | 0 | 0 | 12 | 218 | 472 | 12 | 0 | 0 | 152 | 0 | 0 | 733 | 0 | 25 | 221 | 52 | 4117 | 2163 | 0 | 0 | 0 | 0 | 73 | 126 | 0 | 0 | 21 | 14 | 0 | 0 | 0 | 228 | 17 | 134 |
| SRR4836417 | 28 | 38 | 1600 | 0 | 0 | 11 | 353 | 565 | 0 | 0 | 0 | 135 | 0 | 0 | 706 | 0 | 32 | 146 | 34 | 3846 | 2160 | 0 | 0 | 0 | 0 | 87 | 139 | 0 | 0 | 15 | 17 | 0 | 0 | 0 | 288 | 26 | 146 |
| SRR4836419 | 41 | 80 | 1596 | 0 | 0 | 10 | 234 | 1027 | 0 | 0 | 0 | 153 | 0 | 0 | 681 | 0 | 41 | 173 | 21 | 3823 | 2263 | 0 | 0 | 0 | 0 | 180 | 287 | 0 | 0 | 31 | 15 | 0 | 0 | 0 | 593 | 42 | 291 |
| SRR4836421 | 13 | 25 | 1136 | 0 | 0 | 0 | 15 | 65 | 29 | 0 | 0 | 329 | 0 | 41 | 600 | 0 | 17 | 756 | 11 | 446 | 4912 | 0 | 0 | 18 | 0 | 47 | 38 | 0 | 0 | 12 | 0 | 0 | 0 | 15 | 76 | 25 | 41 |
| SRR4836422 | 0 | 15 | 1190 | 0 | 11 | 25 | 11 | 373 | 17 | 0 | 0 | 357 | 0 | 63 | 553 | 0 | 20 | 835 | 0 | 452 | 4633 | 0 | 0 | 17 | 13 | 312 | 207 | 0 | 0 | 61 | 19 | 0 | 0 | 70 | 476 | 30 | 238 |
| SRR4836423 | 25 | 17 | 1059 | 0 | 16 | 20 | 0 | 56 | 24 | 0 | 0 | 388 | 0 | 49 | 620 | 0 | 18 | 883 | 12 | 482 | 4549 | 0 | 0 | 13 | 0 | 48 | 34 | 0 | 0 | 14 | 25 | 0 | 0 | 15 | 77 | 41 | 48 |
| SRR4836425 | 48 | 16 | 3692 | 0 | 0 | 0 | 35 | 618 | 12 | 0 | 0 | 1222 | 0 | 23 | 246 | 0 | 11 | 329 | 0 | 2128 | 3245 | 0 | 0 | 0 | 0 | 145 | 1287 | 0 | 0 | 78 | 39 | 0 | 0 | 121 | 794 | 63 | 332 |
| SRR4836427 | 55 | 0 | 3697 | 0 | 15 | 19 | 0 | 642 | 0 | 0 | 0 | 1138 | 0 | 42 | 228 | 0 | 12 | 342 | 0 | 1983 | 3062 | 0 | 0 | 42 | 0 | 306 | 1193 | 0 | 0 | 45 | 23 | 0 | 0 | 42 | 1034 | 49 | 340 |
| SRR4836428 | 58 | 22 | 3449 | 0 | 0 | 13 | 14 | 616 | 0 | 0 | 0 | 1110 | 0 | 16 | 154 | 0 | 15 | 506 | 14 | 2009 | 3107 | 0 | 0 | 0 | 0 | 159 | 1141 | 0 | 0 | 59 | 13 | 0 | 0 | 69 | 1004 | 56 | 381 |
| SRR4836429 | 35 | 58 | 379 | 0 | 0 | 32 | 12 | 533 | 17 | 0 | 0 | 243 | 0 | 0 | 1086 | 0 | 25 | 370 | 0 | 1850 | 2880 | 0 | 0 | 0 | 12 | 392 | 767 | 0 | 0 | 0 | 14 | 0 | 0 | 0 | 334 | 29 | 265 |
| SRR4836431 | 39 | 37 | 407 | 0 | 0 | 48 | 23 | 86 | 23 | 0 | 0 | 224 | 0 | 0 | 1060 | 0 | 16 | 280 | 0 | 1807 | 2592 | 0 | 0 | 0 | 0 | 71 | 125 | 0 | 0 | 14 | 23 | 0 | 0 | 11 | 61 | 37 | 48 |
| SRR4836433 | 34 | 64 | 381 | 0 | 0 | 21 | 16 | 73 | 24 | 0 | 0 | 201 | 0 | 0 | 1135 | 0 | 20 | 201 | 0 | 2093 | 2727 | 0 | 0 | 0 | 0 | 48 | 106 | 0 | 0 | 0 | 26 | 0 | 0 | 12 | 50 | 50 | 44 |
| SRR4836434 | 16 | 56 | 1355 | 0 | 13 | 11 | 0 | 35 | 11 | 0 | 0 | 519 | 0 | 24 | 3017 | 0 | 16 | 1955 | 0 | 483 | 2586 | 0 | 0 | 0 | 0 | 141 | 273 | 0 | 0 | 0 | 35 | 0 | 0 | 13 | 175 | 16 | 106 |
| SRR4836435 | 21 | 21 | 1377 | 0 | 0 | 10 | 33 | 23 | 18 | 0 | 0 | 469 | 0 | 18 | 3005 | 0 | 0 | 1701 | 17 | 631 | 3271 | 0 | 0 | 0 | 0 | 63 | 94 | 0 | 0 | 0 | 16 | 0 | 0 | 10 | 74 | 24 | 29 |
| SRR4836437 | 27 | 24 | 1343 | 0 | 13 | 0 | 13 | 22 | 13 | 0 | 0 | 481 | 0 | 0 | 2753 | 0 | 11 | 1864 | 27 | 671 | 2591 | 0 | 0 | 0 | 0 | 62 | 105 | 0 | 0 | 0 | 15 | 0 | 0 | 25 | 86 | 15 | 26 |
| SRR4836439 | 0 | 18 | 1371 | 0 | 0 | 0 | 13 | 18 | 23 | 0 | 0 | 516 | 0 | 17 | 3129 | 0 | 12 | 1796 | 13 | 639 | 2823 | 0 | 0 | 0 | 0 | 54 | 102 | 0 | 0 | 0 | 12 | 0 | 0 | 11 | 73 | 23 | 32 |
| SRR4836440 | 14 | 106 | 1025 | 19 | 0 | 19 | 0 | 86 | 29 | 0 | 0 | 212 | 0 | 38 | 753 | 0 | 0 | 101 | 0 | 502 | 1351 | 0 | 0 | 0 | 0 | 69 | 93 | 0 | 0 | 0 | 16 | 0 | 0 | 0 | 147 | 13 | 125 |
| SRR4836441 | 20 | 159 | 938 | 0 | 0 | 19 | 19 | 141 | 47 | 0 | 0 | 220 | 0 | 19 | 790 | 0 | 0 | 143 | 0 | 433 | 1373 | 0 | 0 | 0 | 0 | 591 | 928 | 0 | 0 | 0 | 169 | 0 | 0 | 19 | 1556 | 159 | 1612 |
| SRR4836443 | 26 | 14 | 1054 | 0 | 0 | 31 | 0 | 13 | 15 | 0 | 0 | 228 | 0 | 23 | 892 | 0 | 0 | 72 | 15 | 535 | 1368 | 0 | 0 | 0 | 0 | 64 | 90 | 23 | 0 | 0 | 22 | 0 | 0 | 15 | 152 | 19 | 132 |
| SRR4836445 | 11 | 214 | 993 | 0 | 0 | 84 | 30 | 191 | 61 | 0 | 0 | 185 | 0 | 31 | 1101 | 23 | 11 | 81 | 0 | 483 | 1222 | 0 | 0 | 0 | 0 | 596 | 909 | 0 | 0 | 0 | 214 | 0 | 0 | 15 | 1735 | 176 | 1773 |
| SRR4836446 | 0 | 361 | 444 | 0 | 0 | 13 | 11 | 30 | 16 | 0 | 0 | 0 | 0 | 24 | 451 | 0 | 35 | 181 | 22 | 539 | 5208 | 0 | 0 | 0 | 0 | 509 | 719 | 11 | 0 | 0 | 164 | 0 | 0 | 16 | 854 | 46 | 582 |
| SRR4836447 | 17 | 299 | 457 | 0 | 13 | 21 | 11 | 26 | 0 | 0 | 0 | 0 | 0 | 16 | 430 | 0 | 35 | 280 | 11 | 752 | 4911 | 0 | 0 | 0 | 11 | 418 | 718 | 16 | 0 | 0 | 130 | 0 | 0 | 11 | 927 | 50 | 538 |
| SRR4836448 | 14 | 268 | 536 | 0 | 11 | 19 | 44 | 17 | 0 | 0 | 0 | 0 | 0 | 11 | 559 | 0 | 37 | 367 | 0 | 586 | 5386 | 0 | 0 | 0 | 0 | 469 | 610 | 0 | 15 | 0 | 137 | 0 | 0 | 0 | 837 | 36 | 537 |
| SRR4836450 | 17 | 91 | 505 | 0 | 0 | 20 | 26 | 10 | 15 | 0 | 0 | 0 | 0 | 20 | 439 | 0 | 40 | 180 | 18 | 606 | 5332 | 0 | 0 | 0 | 15 | 149 | 262 | 0 | 0 | 0 | 39 | 0 | 0 | 0 | 298 | 17 | 190 |
| SRR4836452 | 44 | 45 | 482 | 0 | 0 | 0 | 184 | 112 | 36 | 0 | 0 | 713 | 0 | 0 | 7165 | 0 | 0 | 439 | 0 | 659 | 3898 | 0 | 0 | 0 | 0 | 21 | 108 | 0 | 0 | 0 | 20 | 0 | 0 | 0 | 60 | 15 | 91 |
| SRR4836453 | 48 | 27 | 458 | 16 | 0 | 0 | 283 | 99 | 20 | 0 | 0 | 774 | 0 | 12 | 7644 | 0 | 0 | 535 | 0 | 849 | 3912 | 0 | 0 | 0 | 0 | 31 | 99 | 0 | 0 | 0 | 0 | 0 | 0 | 0 | 62 | 27 | 83 |
| SRR4836454 | 43 | 140 | 494 | 0 | 0 | 0 | 351 | 327 | 16 | 0 | 0 | 751 | 0 | 13 | 7396 | 0 | 0 | 281 | 0 | 723 | 3430 | 0 | 0 | 0 | 0 | 76 | 407 | 0 | 0 | 0 | 13 | 0 | 0 | 22 | 286 | 35 | 289 |
| SRR4836456 | 46 | 38 | 460 | 0 | 0 | 0 | 167 | 84 | 13 | 0 | 0 | 809 | 0 | 19 | 8001 | 0 | 0 | 253 | 0 | 1063 | 4271 | 0 | 0 | 0 | 0 | 37 | 111 | 0 | 0 | 0 | 0 | 0 | 0 | 0 | 70 | 13 | 82 |
| SRR4836458 | 62 | 16 | 3803 | 0 | 0 | 11 | 54 | 45 | 17 | 0 | 0 | 322 | 0 | 17 | 1360 | 0 | 32 | 19 | 0 | 163 | 1796 | 0 | 0 | 0 | 0 | 25 | 96 | 0 | 0 | 11 | 0 | 0 | 0 | 0 | 203 | 17 | 149 |
| SRR4836459 | 48 | 11 | 3626 | 0 | 0 | 0 | 26 | 45 | 17 | 0 | 0 | 340 | 0 | 14 | 1506 | 0 | 26 | 166 | 0 | 132 | 2272 | 0 | 0 | 0 | 0 | 39 | 107 | 0 | 0 | 0 | 0 | 0 | 0 | 0 | 188 | 15 | 147 |
| SRR4836461 | 54 | 32 | 3569 | 0 | 0 | 0 | 41 | 185 | 0 | 0 | 0 | 377 | 0 | 11 | 1388 | 0 | 18 | 88 | 21 | 124 | 2113 | 0 | 0 | 0 | 0 | 87 | 363 | 0 | 0 | 0 | 0 | 0 | 0 | 0 | 651 | 80 | 445 |
| SRR4836463 | 42 | 37 | 3618 | 0 | 0 | 0 | 0 | 183 | 12 | 0 | 0 | 322 | 0 | 0 | 1965 | 0 | 16 | 45 | 0 | 143 | 1863 | 0 | 0 | 0 | 0 | 85 | 374 | 0 | 0 | 0 | 0 | 0 | 0 | 12 | 709 | 55 | 580 |
| SRR4836464 | 11 | 21 | 1265 | 0 | 0 | 237 | 72 | 0 | 56 | 0 | 0 | 11 | 0 | 0 | 374 | 0 | 24 | 1229 | 13 | 992 | 2837 | 0 | 0 | 0 | 0 | 1260 | 772 | 0 | 0 | 0 | 174 | 0 | 0 | 0 | 607 | 45 | 242 |
| SRR4836466 | 21 | 30 | 1252 | 0 | 0 | 234 | 96 | 0 | 58 | 0 | 0 | 11 | 0 | 0 | 365 | 0 | 29 | 1003 | 19 | 987 | 2725 | 0 | 0 | 0 | 0 | 1317 | 832 | 0 | 0 | 0 | 208 | 0 | 0 | 0 | 622 | 27 | 251 |
| SRR4836468 | 15 | 29 | 1219 | 0 | 0 | 62 | 125 | 0 | 14 | 0 | 0 | 11 | 0 | 0 | 360 | 0 | 22 | 1066 | 16 | 980 | 2680 | 0 | 0 | 0 | 0 | 316 | 211 | 0 | 0 | 0 | 47 | 0 | 0 | 0 | 144 | 35 | 60 |
| SRR4836469 | 17 | 42 | 1238 | 0 | 0 | 210 | 93 | 0 | 54 | 0 | 0 | 0 | 0 | 0 | 422 | 0 | 17 | 1023 | 15 | 792 | 2535 | 0 | 0 | 0 | 0 | 1413 | 782 | 0 | 0 | 0 | 153 | 0 | 0 | 0 | 597 | 28 | 265 |
| SRR4836471 | 71 | 37 | 5319 | 0 | 0 | 13 | 0 | 60 | 33 | 0 | 0 | 56 | 0 | 16 | 532 | 0 | 39 | 365 | 0 | 376 | 925 | 0 | 0 | 0 | 0 | 245 | 78 | 0 | 0 | 0 | 54 | 0 | 0 | 0 | 156 | 57 | 97 |
| SRR4836473 | 61 | 227 | 6185 | 0 | 0 | 93 | 23 | 66 | 36 | 0 | 0 | 57 | 0 | 20 | 555 | 0 | 41 | 234 | 0 | 330 | 775 | 0 | 0 | 0 | 0 | 1621 | 551 | 0 | 0 | 0 | 111 | 0 | 0 | 14 | 1134 | 86 | 640 |
| SRR4836474 | 68 | 28 | 6134 | 0 | 10 | 16 | 17 | 11 | 27 | 0 | 0 | 68 | 0 | 23 | 512 | 0 | 39 | 325 | 12 | 285 | 954 | 0 | 0 | 0 | 0 | 261 | 86 | 0 | 0 | 0 | 11 | 0 | 0 | 12 | 190 | 11 | 125 |
| SRR4836475 | 23 | 522 | 3788 | 0 | 13 | 38 | 12 | 79 | 17 | 0 | 0 | 232 | 0 | 13 | 7779 | 0 | 19 | 717 | 46 | 542 | 1754 | 0 | 0 | 0 | 0 | 409 | 484 | 0 | 0 | 0 | 63 | 0 | 0 | 0 | 580 | 84 | 447 |
| SRR4836477 | 44 | 91 | 4276 | 0 | 15 | 40 | 25 | 15 | 0 | 0 | 0 | 173 | 0 | 12 | 7308 | 0 | 13 | 627 | 0 | 434 | 1219 | 0 | 0 | 0 | 0 | 106 | 90 | 0 | 0 | 0 | 13 | 0 | 0 | 12 | 137 | 14 | 107 |
| SRR4836479 | 40 | 447 | 4289 | 0 | 0 | 13 | 0 | 88 | 22 | 0 | 0 | 204 | 0 | 19 | 7906 | 0 | 10 | 620 | 0 | 424 | 1438 | 0 | 0 | 0 | 0 | 569 | 513 | 0 | 0 | 0 | 66 | 0 | 0 | 0 | 710 | 138 | 713 |

### Supp Table S8.2.Columns_AM_to_BW

| | | | | | | | | | | | | | | | | | | | | | | | | | | | | | | | | | | | | | |
|---|---|---|---|---|---|---|---|---|---|---|---|---|---|---|---|---|---|---|---|---|---|---|---|---|---|---|---|---|---|---|---|---|---|---|---|---|---|
| SRR4836481 | 43 | 24 | 362 | 0 | 0 | 21 | 39 | 0 | 0 | 0 | 0 | 29 | 0 | 0 | 429 | 0 | 11 | 178 | 19 | 1831 | 351 | 0 | 0 | 0 | 0 | 114 | 21 | 0 | 0 | 0 | 27 | 0 | 0 | 0 | 230 | 21 | 114 |
| SRR4836482 | 32 | 49 | 471 | 0 | 0 | 28 | 28 | 0 | 0 | 0 | 0 | 17 | 0 | 0 | 460 | 0 | 15 | 259 | 0 | 1518 | 446 | 0 | 0 | 0 | 0 | 471 | 110 | 0 | 0 | 0 | 67 | 0 | 0 | 0 | 1002 | 131 | 428 |
| SRR4836483 | 26 | 23 | 470 | 0 | 0 | 25 | 14 | 0 | 13 | 0 | 0 | 21 | 0 | 0 | 446 | 0 | 0 | 225 | 0 | 1185 | 418 | 0 | 0 | 0 | 0 | 153 | 19 | 0 | 0 | 0 | 24 | 0 | 0 | 0 | 279 | 27 | 138 |
| SRR4836484 | 22 | 206 | 634 | 11 | 0 | 34 | 24 | 0 | 11 | 0 | 0 | 19 | 0 | 11 | 76262 | 0 | 0 | 80 | 0 | 358 | 12754 | 0 | 0 | 0 | 0 | 800 | 1337 | 11 | 11 | 0 | 126 | 0 | 0 | 11 | 3862 | 171 | 1497 |
| SRR4836485 | 22 | 13 | 737 | 0 | 0 | 69 | 0 | 0 | 0 | 0 | 0 | 13 | 0 | 26 | 72068 | 0 | 11 | 20 | 0 | 427 | 11517 | 0 | 0 | 0 | 0 | 88 | 139 | 34 | 0 | 0 | 16 | 0 | 0 | 0 | 334 | 18 | 154 |
| SRR4836487 | 31 | 87 | 722 | 0 | 0 | 43 | 0 | 0 | 35 | 0 | 0 | 19 | 0 | 0 | 69487 | 0 | 15 | 118 | 0 | 416 | 12173 | 0 | 0 | 0 | 0 | 71 | 130 | 35 | 35 | 0 | 95 | 0 | 0 | 0 | 334 | 16 | 174 |
| SRR4836489 | 47 | 29 | 329 | 0 | 0 | 0 | 40 | 27 | 12 | 0 | 0 | 581 | 0 | 16 | 767 | 0 | 11 | 191 | 0 | 862 | 1171 | 0 | 0 | 0 | 0 | 16 | 39 | 0 | 0 | 0 | 24 | 0 | 0 | 19 | 71 | 30 | 60 |
| SRR4836491 | 42 | 25 | 380 | 0 | 0 | 0 | 38 | 11 | 10 | 0 | 0 | 537 | 0 | 12 | 724 | 0 | 18 | 204 | 0 | 866 | 993 | 0 | 0 | 0 | 0 | 25 | 39 | 0 | 0 | 0 | 17 | 0 | 0 | 10 | 83 | 35 | 66 |
| SRR4836492 | 35 | 11 | 390 | 0 | 0 | 0 | 57 | 13 | 16 | 0 | 0 | 576 | 0 | 14 | 745 | 0 | 12 | 193 | 0 | 826 | 1039 | 0 | 0 | 0 | 0 | 20 | 40 | 0 | 0 | 0 | 12 | 0 | 0 | 21 | 90 | 21 | 81 |
| SRR4836493 | 66 | 98 | 2187 | 0 | 0 | 27 | 53 | 163 | 31 | 0 | 0 | 255 | 0 | 0 | 1030 | 0 | 13 | 31 | 0 | 350 | 1168 | 0 | 0 | 0 | 0 | 599 | 182 | 0 | 0 | 0 | 70 | 0 | 0 | 0 | 572 | 74 | 499 |
| SRR4836494 | 61 | 81 | 2530 | 0 | 0 | 42 | 52 | 171 | 38 | 0 | 0 | 244 | 0 | 0 | 865 | 0 | 0 | 114 | 0 | 246 | 977 | 0 | 0 | 0 | 0 | 730 | 225 | 0 | 0 | 0 | 76 | 0 | 0 | 0 | 678 | 89 | 610 |
| SRR4836495 | 53 | 71 | 2465 | 0 | 0 | 39 | 45 | 184 | 32 | 0 | 0 | 258 | 0 | 0 | 1118 | 0 | 14 | 76 | 0 | 345 | 1223 | 0 | 0 | 0 | 0 | 705 | 239 | 0 | 0 | 0 | 67 | 0 | 0 | 0 | 710 | 101 | 609 |
| SRR4836496 | 44 | 29 | 2664 | 0 | 0 | 0 | 69 | 1234 | 18 | 0 | 0 | 782 | 0 | 22 | 618 | 0 | 0 | 94 | 11 | 1408 | 1786 | 0 | 0 | 0 | 0 | 36 | 68 | 0 | 0 | 25 | 14 | 0 | 0 | 13 | 177 | 17 | 101 |
| SRR4836498 | 38 | 83 | 2443 | 0 | 0 | 17 | 46 | 4393 | 21 | 0 | 0 | 815 | 0 | 17 | 657 | 0 | 12 | 121 | 0 | 1765 | 1971 | 0 | 0 | 0 | 0 | 139 | 326 | 0 | 0 | 33 | 0 | 0 | 0 | 14 | 724 | 98 | 402 |
| SRR4836500 | 36 | 24 | 2576 | 0 | 0 | 12 | 86 | 1182 | 12 | 0 | 0 | 830 | 0 | 25 | 582 | 0 | 0 | 132 | 0 | 1575 | 1798 | 0 | 0 | 0 | 0 | 33 | 80 | 0 | 0 | 25 | 14 | 0 | 0 | 14 | 183 | 21 | 121 |
| SRR4836502 | 44 | 23 | 2812 | 0 | 0 | 10 | 74 | 1319 | 18 | 0 | 0 | 792 | 0 | 19 | 619 | 0 | 0 | 79 | 0 | 1571 | 1581 | 0 | 0 | 0 | 0 | 36 | 89 | 0 | 0 | 10 | 10 | 0 | 0 | 0 | 213 | 22 | 130 |
| SRR4836503 | 19 | 114 | 1788 | 0 | 0 | 0 | 25 | 1268 | 0 | 0 | 0 | 205 | 0 | 11 | 296 | 0 | 18 | 129 | 0 | 9457 | 504 | 0 | 0 | 11 | 0 | 205 | 201 | 11 | 0 | 12 | 11 | 0 | 0 | 0 | 317 | 52 | 245 |
| SRR4836504 | 24 | 26 | 1598 | 0 | 0 | 0 | 26 | 163 | 0 | 0 | 0 | 212 | 0 | 16 | 284 | 0 | 26 | 137 | 0 | 9938 | 517 | 0 | 0 | 0 | 0 | 39 | 31 | 0 | 0 | 13 | 0 | 0 | 0 | 18 | 61 | 52 | 36 |
| SRR4836505 | 21 | 23 | 1786 | 0 | 0 | 0 | 29 | 184 | 0 | 0 | 0 | 198 | 0 | 13 | 253 | 0 | 26 | 129 | 0 | 9416 | 409 | 0 | 0 | 15 | 0 | 29 | 30 | 0 | 0 | 0 | 0 | 0 | 0 | 11 | 68 | 54 | 34 |
| SRR4836507 | 21 | 129 | 1826 | 0 | 0 | 0 | 18 | 1249 | 0 | 0 | 0 | 217 | 0 | 10 | 248 | 0 | 26 | 156 | 0 | 8942 | 445 | 0 | 0 | 12 | 0 | 175 | 228 | 0 | 0 | 15 | 0 | 0 | 0 | 12 | 446 | 40 | 292 |
| SRR4836509 | 39 | 34 | 1537 | 0 | 0 | 0 | 88 | 52 | 0 | 0 | 0 | 103 | 0 | 0 | 74 | 0 | 42 | 576 | 32 | 5120 | 247 | 0 | 0 | 0 | 0 | 40 | 75 | 0 | 0 | 0 | 23 | 0 | 0 | 17 | 154 | 18 | 75 |
| SRR4836511 | 50 | 22 | 1525 | 0 | 0 | 0 | 71 | 140 | 0 | 0 | 0 | 110 | 0 | 0 | 84 | 0 | 43 | 616 | 26 | 5501 | 301 | 0 | 0 | 0 | 0 | 137 | 245 | 0 | 0 | 11 | 17 | 0 | 0 | 18 | 556 | 76 | 244 |
| SRR4836512 | 55 | 27 | 1650 | 0 | 0 | 11 | 89 | 51 | 0 | 0 | 0 | 125 | 0 | 13 | 70 | 0 | 45 | 545 | 21 | 5420 | 287 | 0 | 0 | 0 | 0 | 35 | 67 | 0 | 0 | 0 | 15 | 0 | 0 | 18 | 164 | 17 | 67 |
| SRR4836513 | 66 | 31 | 1614 | 0 | 0 | 14 | 98 | 59 | 0 | 0 | 0 | 106 | 0 | 0 | 78 | 0 | 54 | 600 | 20 | 4987 | 261 | 0 | 0 | 0 | 0 | 45 | 80 | 0 | 0 | 17 | 17 | 0 | 0 | 19 | 200 | 22 | 77 |
| SRR4836514 | 20 | 53 | 2131 | 0 | 0 | 19 | 25 | 1778 | 13 | 0 | 0 | 0 | 0 | 0 | 130 | 0 | 21 | 26 | 25 | 428 | 893 | 0 | 0 | 0 | 0 | 251 | 203 | 0 | 0 | 0 | 64 | 0 | 0 | 0 | 803 | 43 | 876 |
| SRR4836515 | 34 | 16 | 1961 | 0 | 0 | 0 | 42 | 477 | 12 | 0 | 0 | 0 | 0 | 0 | 208 | 0 | 12 | 33 | 0 | 408 | 1467 | 0 | 0 | 0 | 0 | 83 | 59 | 0 | 0 | 0 | 16 | 0 | 0 | 0 | 237 | 16 | 253 |
| SRR4836517 | 24 | 55 | 2348 | 0 | 0 | 19 | 0 | 1694 | 16 | 0 | 0 | 0 | 0 | 14 | 142 | 0 | 14 | 32 | 16 | 377 | 1235 | 0 | 0 | 0 | 0 | 263 | 205 | 0 | 0 | 0 | 62 | 0 | 0 | 0 | 758 | 48 | 872 |
| SRR4836519 | 22 | 52 | 2244 | 0 | 0 | 21 | 23 | 1961 | 0 | 0 | 0 | 0 | 0 | 0 | 175 | 0 | 14 | 19 | 18 | 480 | 1053 | 0 | 0 | 0 | 0 | 286 | 232 | 0 | 0 | 0 | 52 | 0 | 0 | 0 | 977 | 53 | 1089 |
| SRR4836521 | 38 | 27 | 3298 | 0 | 0 | 51 | 23 | 1200 | 11 | 0 | 0 | 242 | 0 | 11 | 12377 | 0 | 15 | 125 | 0 | 375 | 1500 | 0 | 0 | 16 | 0 | 495 | 366 | 0 | 0 | 0 | 74 | 0 | 0 | 0 | 482 | 22 | 194 |
| SRR4836522 | 36 | 51 | 3027 | 0 | 0 | 20 | 16 | 1140 | 34 | 0 | 0 | 231 | 0 | 0 | 14389 | 0 | 13 | 190 | 0 | 498 | 1664 | 0 | 0 | 14 | 0 | 541 | 346 | 0 | 0 | 0 | 65 | 0 | 0 | 0 | 493 | 26 | 289 |
| SRR4836523 | 34 | 46 | 3320 | 0 | 0 | 32 | 59 | 1184 | 27 | 0 | 0 | 262 | 0 | 11 | 12710 | 0 | 14 | 150 | 0 | 577 | 1687 | 0 | 0 | 0 | 0 | 517 | 363 | 0 | 0 | 0 | 62 | 0 | 0 | 0 | 398 | 16 | 261 |
| SRR4836524 | 37 | 30 | 3390 | 0 | 0 | 24 | 49 | 1328 | 15 | 0 | 0 | 239 | 0 | 0 | 12229 | 0 | 0 | 128 | 0 | 361 | 1350 | 0 | 0 | 17 | 0 | 566 | 388 | 0 | 0 | 0 | 59 | 0 | 0 | 0 | 511 | 59 | 293 |
| SRR4836525 | 15 | 57 | 1983 | 0 | 0 | 30 | 51 | 1098 | 15 | 0 | 0 | 0 | 0 | 0 | 182 | 0 | 19 | 41 | 0 | 530 | 821 | 0 | 0 | 0 | 0 | 301 | 218 | 0 | 0 | 0 | 65 | 0 | 0 | 0 | 947 | 47 | 883 |
| SRR4836526 | 18 | 44 | 1793 | 0 | 0 | 15 | 22 | 1016 | 17 | 0 | 0 | 0 | 0 | 14 | 133 | 0 | 16 | 66 | 0 | 494 | 1025 | 0 | 0 | 0 | 0 | 283 | 187 | 0 | 0 | 0 | 99 | 0 | 0 | 0 | 1034 | 38 | 973 |
| SRR4836528 | 31 | 15 | 2064 | 0 | 0 | 33 | 47 | 305 | 17 | 0 | 0 | 0 | 0 | 0 | 147 | 0 | 15 | 56 | 0 | 527 | 975 | 0 | 0 | 0 | 0 | 85 | 63 | 0 | 0 | 0 | 25 | 0 | 0 | 0 | 311 | 16 | 255 |
| SRR4836529 | 16 | 60 | 2184 | 0 | 0 | 20 | 68 | 1176 | 14 | 0 | 0 | 0 | 0 | 0 | 166 | 0 | 25 | 33 | 10 | 458 | 952 | 0 | 0 | 0 | 0 | 357 | 275 | 0 | 0 | 11 | 74 | 0 | 0 | 0 | 1156 | 49 | 994 |
| SRR4836531 | 13 | 37 | 4226 | 0 | 0 | 14 | 0 | 26 | 23 | 0 | 0 | 123 | 0 | 43 | 727 | 0 | 16 | 255 | 29 | 587 | 773 | 0 | 0 | 0 | 11 | 309 | 360 | 0 | 0 | 0 | 17 | 14 | 0 | 11 | 723 | 45 | 366 |
| SRR4836533 | 13 | 30 | 4244 | 0 | 11 | 14 | 23 | 19 | 0 | 0 | 0 | 135 | 0 | 39 | 712 | 0 | 10 | 206 | 79 | 582 | 802 | 0 | 0 | 0 | 14 | 322 | 346 | 11 | 0 | 0 | 17 | 0 | 0 | 22 | 723 | 58 | 457 |
| SRR4836534 | 16 | 47 | 3777 | 0 | 0 | 25 | 38 | 0 | 0 | 0 | 0 | 145 | 0 | 52 | 697 | 0 | 17 | 245 | 66 | 604 | 821 | 0 | 0 | 0 | 17 | 55 | 47 | 0 | 0 | 0 | 22 | 14 | 0 | 25 | 129 | 11 | 80 |
| SRR4836535 | 26 | 102 | 3719 | 0 | 0 | 0 | 37 | 0 | 0 | 0 | 0 | 290 | 0 | 0 | 1228 | 0 | 30 | 1593 | 11 | 4153 | 936 | 0 | 0 | 0 | 0 | 151 | 293 | 0 | 0 | 0 | 12 | 0 | 0 | 0 | 303 | 14 | 182 |
| SRR4836536 | 20 | 79 | 3905 | 0 | 0 | 20 | 32 | 10 | 0 | 0 | 0 | 281 | 0 | 0 | 1166 | 0 | 20 | 1626 | 0 | 3860 | 895 | 0 | 0 | 0 | 0 | 214 | 337 | 0 | 0 | 0 | 14 | 0 | 0 | 10 | 353 | 26 | 218 |
| SRR4836537 | 26 | 29 | 3461 | 0 | 0 | 12 | 18 | 0 | 0 | 0 | 0 | 282 | 0 | 14 | 1267 | 0 | 22 | 1525 | 0 | 3856 | 936 | 0 | 0 | 0 | 0 | 40 | 68 | 0 | 0 | 0 | 0 | 0 | 0 | 0 | 78 | 15 | 48 |
| SRR4836539 | 42 | 44 | 798 | 0 | 0 | 0 | 0 | 14 | 0 | 0 | 0 | 60 | 0 | 36 | 28 | 0 | 0 | 1433 | 0 | 4445 | 1658 | 0 | 0 | 0 | 0 | 98 | 97 | 0 | 0 | 0 | 0 | 0 | 0 | 25 | 68 | 11 | 39 |
| SRR4836541 | 11 | 61 | 755 | 0 | 0 | 12 | 0 | 13 | 0 | 0 | 0 | 77 | 0 | 34 | 46 | 0 | 0 | 1391 | 0 | 3850 | 1624 | 0 | 0 | 0 | 0 | 102 | 90 | 0 | 0 | 0 | 12 | 0 | 0 | 33 | 92 | 18 | 58 |
| SRR4836543 | 42 | 51 | 778 | 0 | 0 | 0 | 0 | 0 | 0 | 0 | 0 | 59 | 0 | 30 | 55 | 0 | 15 | 1533 | 0 | 4225 | 1611 | 0 | 0 | 0 | 0 | 107 | 92 | 0 | 0 | 0 | 0 | 0 | 0 | 25 | 72 | 0 | 40 |
| SRR4836544 | 55 | 38 | 2473 | 0 | 0 | 56 | 151 | 44 | 15 | 0 | 0 | 416 | 0 | 0 | 1873 | 0 | 0 | 350 | 0 | 604 | 4049 | 0 | 0 | 0 | 0 | 697 | 803 | 0 | 0 | 0 | 56 | 0 | 0 | 24 | 1492 | 168 | 883 |
| SRR4836545 | 62 | 55 | 2481 | 0 | 0 | 55 | 30 | 78 | 14 | 0 | 0 | 445 | 0 | 0 | 1625 | 0 | 0 | 298 | 0 | 680 | 3517 | 0 | 0 | 0 | 0 | 773 | 822 | 0 | 0 | 0 | 69 | 0 | 0 | 0 | 1661 | 208 | 1012 |
| SRR4836546 | 68 | 32 | 2221 | 0 | 0 | 44 | 36 | 13 | 23 | 0 | 0 | 429 | 0 | 12 | 1530 | 0 | 11 | 328 | 0 | 637 | 3824 | 0 | 0 | 0 | 0 | 164 | 170 | 0 | 0 | 0 | 13 | 0 | 0 | 0 | 338 | 36 | 207 |
| SRR4836548 | 35 | 15 | 3155 | 0 | 18 | 13 | 21 | 13 | 45 | 0 | 0 | 24 | 0 | 0 | 419 | 0 | 34 | 67 | 0 | 1180 | 664 | 0 | 0 | 0 | 0 | 189 | 81 | 0 | 0 | 0 | 17 | 13 | 13 | 13 | 199 | 17 | 77 |
| SRR4836550 | 26 | 10 | 3173 | 0 | 13 | 14 | 41 | 49 | 49 | 0 | 0 | 24 | 0 | 0 | 428 | 0 | 23 | 37 | 0 | 993 | 748 | 0 | 0 | 0 | 0 | 233 | 103 | 0 | 0 | 0 | 16 | 13 | 0 | 18 | 265 | 15 | 98 |
| SRR4836552 | 32 | 109 | 2952 | 0 | 11 | 85 | 17 | 57 | 41 | 0 | 0 | 24 | 0 | 14 | 391 | 0 | 30 | 46 | 0 | 1047 | 921 | 0 | 0 | 0 | 0 | 1720 | 855 | 0 | 0 | 0 | 137 | 22 | 0 | 0 | 1961 | 112 | 582 |
| SRR4836553 | 89 | 14 | 1274 | 0 | 0 | 12 | 29 | 52 | 17 | 0 | 0 | 108 | 0 | 30 | 2684 | 22 | 15 | 3116 | 0 | 230 | 4881 | 0 | 0 | 0 | 0 | 154 | 139 | 0 | 0 | 17 | 14 | 0 | 0 | 39 | 255 | 18 | 93 |
| SRR4836554 | 109 | 42 | 1400 | 0 | 13 | 68 | 0 | 25 | 47 | 0 | 0 | 110 | 0 | 13 | 2292 | 0 | 0 | 2863 | 0 | 171 | 4683 | 0 | 0 | 0 | 0 | 909 | 947 | 0 | 0 | 0 | 47 | 0 | 0 | 51 | 1662 | 110 | 520 |
| SRR4836555 | 82 | 11 | 1197 | 0 | 0 | 50 | 17 | 29 | 34 | 0 | 0 | 119 | 0 | 29 | 2294 | 13 | 0 | 3284 | 0 | 87 | 4780 | 0 | 0 | 0 | 0 | 157 | 163 | 0 | 0 | 0 | 50 | 0 | 0 | 34 | 288 | 11 | 104 |
| SRR4836556 | 37 | 97 | 1208 | 0 | 15 | 24 | 0 | 19 | 0 | 0 | 0 | 191 | 0 | 68 | 1833 | 15 | 26 | 1805 | 0 | 172 | 1337 | 0 | 0 | 0 | 0 | 592 | 165 | 0 | 0 | 0 | 102 | 53 | 0 | 24 | 2704 | 257 | 932 |
| SRR4836560 | 40 | 93 | 1168 | 0 | 0 | 19 | 17 | 19 | 15 | 0 | 0 | 187 | 0 | 112 | 1637 | 0 | 38 | 1725 | 0 | 121 | 1223 | 0 | 0 | 0 | 0 | 585 | 186 | 12 | 0 | 0 | 116 | 23 | 0 | 15 | 2522 | 225 | 860 |
| SRR4836561 | 26 | 102 | 1153 | 0 | 0 | 16 | 17 | 0 | 12 | 0 | 0 | 242 | 0 | 117 | 1511 | 0 | 29 | 1763 | 0 | 122 | 1303 | 0 | 0 | 0 | 12 | 568 | 196 | 12 | 0 | 0 | 110 | 47 | 0 | 12 | 2792 | 204 | 916 |
| SRR4836562 | 0 | 68 | 703 | 0 | 0 | 0 | 0 | 16 | 20 | 0 | 0 | 0 | 0 | 0 | 1173 | 0 | 22 | 138 | 0 | 70 | 309 | 0 | 0 | 0 | 16 | 43 | 11 | 0 | 0 | 0 | 16 | 0 | 0 | 0 | 197 | 56 | 95 |
| SRR4836563 | 0 | 18 | 693 | 12 | 12 | 12 | 0 | 0 | 0 | 0 | 0 | 0 | 0 | 0 | 1316 | 0 | 28 | 150 | 0 | 55 | 310 | 0 | 0 | 0 | 16 | 39 | 21 | 0 | 0 | 0 | 20 | 0 | 0 | 12 | 201 | 51 | 77 |
| SRR4836565 | 13 | 119 | 667 | 0 | 0 | 13 | 0 | 0 | 0 | 0 | 0 | 0 | 0 | 0 | 1145 | 0 | 0 | 182 | 0 | 53 | 340 | 0 | 0 | 0 | 0 | 277 | 90 | 0 | 0 | 0 | 32 | 0 | 0 | 0 | 1479 | 45 | 630 |
| SRR4836567 | 0 | 85 | 651 | 0 | 0 | 13 | 0 | 0 | 0 | 0 | 0 | 0 | 0 | 0 | 1136 | 0 | 24 | 128 | 12 | 57 | 353 | 0 | 0 | 0 | 0 | 291 | 137 | 0 | 0 | 0 | 0 | 0 | 0 | 0 | 1421 | 49 | 686 |
| SRR4836569 | 23 | 25 | 4295 | 0 | 0 | 76 | 11 | 11 | 47 | 0 | 0 | 377 | 0 | 44 | 8504 | 0 | 0 | 543 | 0 | 174 | 488 | 0 | 0 | 0 | 0 | 1216 | 302 | 0 | 0 | 0 | 120 | 0 | 0 | 11 | 3112 | 429 | 3825 |
| SRR4836570 | 27 | 47 | 4167 | 0 | 0 | 82 | 0 | 14 | 61 | 0 | 0 | 359 | 0 | 25 | 9014 | 0 | 0 | 366 | 0 | 224 | 352 | 0 | 0 | 0 | 32 | 60 | 14 | 0 | 0 | 0 | 143 | 0 | 0 | 18 | 182 | 23 | 200 |
| SRR4836571 | 27 | 29 | 4219 | 0 | 0 | 84 | 17 | 35 | 81 | 0 | 0 | 345 | 0 | 35 | 8604 | 0 | 0 | 478 | 0 | 189 | 361 | 0 | 0 | 0 | 38 | 68 | 12 | 0 | 0 | 0 | 113 | 17 | 0 | 20 | 172 | 19 | 213 |
| SRR4836572 | 28 | 41 | 4179 | 0 | 0 | 94 | 17 | 15 | 105 | 0 | 0 | 363 | 0 | 38 | 8424 | 0 | 0 | 452 | 0 | 137 | 435 | 0 | 0 | 0 | 26 | 66 | 12 | 0 | 0 | 0 | 114 | 12 | 0 | 23 | 168 | 26 | 209 |
| SRR4896694 | 16 | 12 | 138 | 0 | 0 | 17 | 51 | 0 | 17 | 0 | 0 | 0 | 17 | 65 | 67 | 0 | 0 | 0 | 1830 | 4020 | 579 | 0 | 0 | 10 | 146 | 385 | 148 | 31 | 12 | 0 | 36 | 0 | 0 | 244 | 301 | 46 | 237 |
| SRR4896695 | 15 | 0 | 59 | 0 | 0 | 92 | 28 | 0 | 92 | 0 | 0 | 0 | 0 | 16 | 0 | 0 | 0 | 0 | 591 | 141290 | 2855 | 0 | 0 | 20 | 141 | 22 | 149 | 24 | 0 | 0 | 104 | 0 | 0 | 80 | 13 | 36 | 313 |
| SRR4896698 | 14 | 16 | 184 | 0 | 0 | 70 | 22 | 0 | 54 | 0 | 0 | 0 | 27 | 11 | 46 | 0 | 0 | 0 | 960 | 8786 | 1028 | 0 | 11 | 54 | 220 | 1126 | 139 | 27 | 27 | 0 | 129 | 0 | 0 | 134 | 236 | 43 | 182 |
| SRR4896701 | 48 | 0 | 259 | 0 | 0 | 63 | 29 | 28 | 23 | 0 | 0 | 11 | 0 | 0 | 0 | 0 | 35 | 0 | 958 | 35491 | 243 | 0 | 0 | 250 | 93 | 724 | 278 | 20 | 48 | 0 | 178 | 0 | 0 | 130 | 706 | 103 | 353 |
| SRR4896708 | 12 | 0 | 472 | 0 | 0 | 37 | 21 | 10 | 12 | 0 | 0 | 44 | 0 | 15 | 13 | 0 | 22 | 0 | 99 | 1722 | 428 | 0 | 0 | 36 | 154 | 642 | 86 | 15 | 87 | 0 | 47 | 0 | 0 | 125 | 757 | 240 | 586 |
| SRR5022482 | 59 | 13 | 13617 | 0 | 0 | 61 | 342 | 27 | 49 | 0 | 0 | 26 | 22 | 0 | 23 | 0 | 139 | 0 | 30 | 1065 | 323 | 0 | 0 | 0 | 14 | 1209 | 638 | 0 | 20 | 0 | 123 | 0 | 0 | 0 | 496 | 146 | 437 |
| SRR5022483 | 115 | 0 | 16768 | 0 | 31 | 10 | 186 | 11 | 11 | 0 | 0 | 34 | 0 | 0 | 0 | 0 | 95 | 0 | 401 | 2490 | 144 | 0 | 0 | 0 | 0 | 16 | 338 | 0 | 0 | 0 | 0 | 0 | 0 | 40 | 0 | 0 | 13 |
| SRR5022484 | 25 | 24 | 18733 | 0 | 0 | 32 | 29 | 19 | 20 | 0 | 0 | 46 | 0 | 11 | 73 | 0 | 75 | 0 | 0 | 402 | 304 | 0 | 0 | 0 | 0 | 18 | 34 | 0 | 0 | 0 | 75 | 0 | 0 | 0 | 48 | 96 | 26 |
| SRR5022485 | 29 | 20 | 14480 | 25 | 35 | 0 | 134 | 0 | 0 | 0 | 0 | 53 | 0 | 0 | 0 | 0 | 59 | 0 | 924 | 2660 | 185 | 0 | 0 | 0 | 0 | 101 | 556 | 0 | 0 | 0 | 0 | 0 | 0 | 19 | 23 | 0 | 41 |
| SRR5029397 | 189 | 0 | 59 | 0 | 0 | 0 | 0 | 0 | 0 | 20 | 232 | 149 | 15 | 12 | 48 | 0 | 19 | 0 | 0 | 0 | 130 | 0 | 0 | 0 | 333 | 39 | 0 | 0 | 0 | 0 | 27 | 0 | 10 | 93 | 53 | 14 | 81 |
| SRR5029398 | 105 | 0 | 91 | 0 | 0 | 0 | 0 | 0 | 0 | 25 | 198 | 108 | 24 | 0 | 165 | 0 | 25 | 69 | 0 | 0 | 49 | 0 | 0 | 0 | 445 | 79 | 0 | 0 | 0 | 0 | 99 | 0 | 10 | 1000 | 32 | 14 | 37 |
| SRR5029399 | 21 | 0 | 41 | 0 | 0 | 15 | 0 | 0 | 24 | 15 | 26 | 43 | 111 | 0 | 1222 | 0 | 16 | 88 | 0 | 0 | 421 | 0 | 0 | 0 | 104 | 142 | 100 | 0 | 0 | 0 | 16 | 0 | 0 | 16 | 13 | 94 | 39 |
| SRR5029400 | 50 | 0 | 139 | 0 | 0 | 14 | 0 | 0 | 10 | 65 | 718 | 83 | 60 | 0 | 783 | 0 | 32 | 0 | 0 | 0 | 40 | 0 | 0 | 0 | 1398 | 157 | 0 | 0 | 0 | 0 | 338 | 0 | 13 | 182 | 270 | 52 | 324 |
| SRR5029401 | 182 | 0 | 158 | 0 | 0 | 0 | 0 | 0 | 0 | 27 | 482 | 72 | 35 | 11 | 64 | 0 | 13 | 0 | 0 | 13 | 115 | 0 | 0 | 0 | 1128 | 122 | 0 | 13 | 0 | 0 | 109 | 14 | 35 | 338 | 73 | 22 | 139 |
| SRR5029402 | 347 | 0 | 112 | 0 | 0 | 15 | 0 | 0 | 0 | 10 | 237 | 125 | 46 | 23 | 165 | 0 | 20 | 0 | 0 | 0 | 127 | 0 | 0 | 0 | 547 | 127 | 0 | 0 | 0 | 0 | 55 | 0 | 19 | 419 | 95 | 32 | 198 |
| SRR5029403 | 35 | 0 | 111 | 0 | 0 | 0 | 0 | 0 | 22 | 35 | 26 | 72 | 65 | 0 | 368 | 0 | 11 | 34 | 0 | 0 | 36 | 0 | 0 | 0 | 71 | 10 | 11 | 0 | 0 | 0 | 16 | 0 | 11 | 108 | 86 | 13 | 50 |
| SRR5029404 | 276 | 0 | 57 | 0 | 0 | 0 | 0 | 0 | 0 | 43 | 543 | 64 | 56 | 0 | 0 | 0 | 12 | 0 | 26 | 0 | 26 | 0 | 0 | 0 | 891 | 37 | 0 | 0 | 0 | 0 | 53 | 0 | 21 | 395 | 46 | 0 | 46 |
| SRR5029405 | 589 | 0 | 88 | 0 | 0 | 0 | 0 | 0 | 14 | 32 | 542 | 151 | 54 | 0 | 35 | 0 | 0 | 0 | 0 | 0 | 207 | 0 | 0 | 0 | 921 | 110 | 0 | 0 | 0 | 0 | 69 | 0 | 19 | 427 | 121 | 22 | 171 |
| SRR5029406 | 840 | 0 | 64 | 0 | 0 | 32 | 0 | 0 | 0 | 28 | 284 | 477 | 40 | 15 | 38 | 0 | 33 | 0 | 0 | 0 | 288 | 0 | 0 | 0 | 396 | 364 | 13 | 0 | 0 | 0 | 971 | 0 | 46 | 488 | 341 | 150 | 894 |
| SRR5029407 | 68 | 0 | 90 | 0 | 0 | 0 | 0 | 0 | 0 | 42 | 49 | 64 | 33 | 13 | 1343 | 0 | 0 | 57 | 0 | 22 | 0 | 0 | 0 | 0 | 241 | 39 | 0 | 0 | 0 | 0 | 25 | 0 | 0 | 13 | 36 | 11 | 33 |
| SRR5029408 | 83 | 0 | 86 | 0 | 0 | 0 | 0 | 0 | 0 | 12 | 101 | 72 | 14 | 0 | 193 | 0 | 22 | 0 | 0 | 11 | 0 | 0 | 0 | 0 | 294 | 16 | 0 | 0 | 0 | 0 | 35 | 0 | 0 | 131 | 11 | 0 | 14 |
| SRR5029409 | 50 | 0 | 86 | 0 | 0 | 28 | 0 | 0 | 0 | 20 | 12 | 86 | 17 | 11 | 115 | 0 | 0 | 35 | 0 | 0 | 66 | 0 | 0 | 0 | 91 | 21 | 0 | 0 | 0 | 0 | 20 | 0 | 20 | 13 | 98 | 67 | 12 |
| SRR5029410 | 69 | 0 | 34 | 0 | 0 | 19 | 0 | 0 | 56 | 0 | 79 | 80 | 13 | 0 | 41 | 0 | 10 | 0 | 0 | 0 | 538 | 0 | 0 | 0 | 36 | 153 | 90 | 0 | 0 | 0 | 24 | 26 | 0 | 33 | 19 | 95 | 48 |
| SRR5029411 | 311 | 0 | 43 | 0 | 0 | 12 | 0 | 0 | 30 | 0 | 650 | 298 | 111 | 0 | 189 | 0 | 25 | 0 | 0 | 0 | 165 | 0 | 0 | 0 | 781 | 123 | 0 | 0 | 0 | 0 | 42 | 0 | 0 | 871 | 105 | 15 | 144 |
| SRR5029412 | 40 | 0 | 54 | 0 | 0 | 41 | 0 | 0 | 50 | 32 | 379 | 79 | 111 | 0 | 1187 | 0 | 12 | 274 | 0 | 0 | 112 | 0 | 0 | 0 | 1142 | 562 | 36 | 0 | 0 | 0 | 160 | 0 | 0 | 262 | 181 | 52 | 178 |
| SRR5029414 | 589 | 0 | 280 | 0 | 0 | 0 | 0 | 0 | 0 | 18 | 134 | 192 | 16 | 14 | 0 | 0 | 0 | 0 | 0 | 0 | 16 | 0 | 0 | 0 | 260 | 10 | 0 | 0 | 0 | 0 | 20 | 0 | 14 | 785 | 20 | 0 | 25 |
| SRR5029415 | 197 | 19 | 85 | 0 | 0 | 0 | 0 | 0 | 0 | 19 | 180 | 103 | 13 | 0 | 133 | 0 | 0 | 0 | 0 | 0 | 0 | 0 | 0 | 19 | 296 | 19 | 0 | 19 | 0 | 0 | 26 | 0 | 0 | 168 | 16 | 0 | 16 |
| SRR5029416 | 62 | 0 | 103 | 0 | 0 | 50 | 0 | 0 | 57 | 0 | 59 | 107 | 33 | 0 | 104 | 0 | 33 | 19 | 0 | 15 | 671 | 0 | 0 | 0 | 122 | 128 | 57 | 0 | 0 | 0 | 22 | 0 | 0 | 55 | 23 | 13 | 58 |
| SRR5029417 | 52 | 0 | 87 | 0 | 0 | 0 | 0 | 0 | 0 | 42 | 554 | 70 | 22 | 10 | 186 | 0 | 18 | 0 | 0 | 0 | 0 | 0 | 0 | 0 | 1990 | 40 | 0 | 0 | 0 | 0 | 109 | 0 | 53 | 53 | 55 | 18 | 50 |
| SRR5029418 | 578 | 0 | 84 | 0 | 0 | 39 | 0 | 0 | 0 | 13 | 238 | 159 | 28 | 0 | 64 | 0 | 0 | 25 | 0 | 0 | 383 | 0 | 0 | 0 | 316 | 335 | 16 | 0 | 0 | 0 | 85 | 0 | 0 | 1448 | 99 | 54 | 306 |
| SRR5029419 | 134 | 0 | 81 | 0 | 0 | 0 | 0 | 0 | 10 | 16 | 309 | 99 | 30 | 0 | 159 | 0 | 43 | 38 | 0 | 0 | 126 | 0 | 0 | 0 | 410 | 46 | 0 | 0 | 0 | 0 | 44 | 14 | 0 | 1728 | 21 | 0 | 40 |
| SRR5029420 | 63 | 0 | 322 | 14 | 0 | 49 | 0 | 0 | 30 | 0 | 121 | 176 | 38 | 0 | 61 | 0 | 17 | 0 | 0 | 0 | 98 | 0 | 0 | 0 | 715 | 505 | 11 | 0 | 0 | 0 | 270 | 0 | 11 | 216 | 111 | 113 | 173 |
| SRR5029422 | 113 | 0 | 69 | 0 | 0 | 113 | 0 | 0 | 10 | 13 | 160 | 212 | 31 | 0 | 223 | 0 | 67 | 0 | 0 | 0 | 148 | 0 | 0 | 0 | 54 | 73 | 20 | 0 | 0 | 0 | 29 | 0 | 27 | 16 | 13 | 113 | 20 |
| SRR5029424 | 11 | 0 | 0 | 0 | 0 | 0 | 0 | 0 | 0 | 0 | 194 | 96 | 18 | 0 | 6961 | 0 | 53 | 0 | 0 | 0 | 0 | 0 | 0 | 0 | 50 | 23 | 0 | 0 | 0 | 0 | 14 | 18 | 0 | 243 | 23 | 0 | 41 |
| SRR5029425 | 87 | 0 | 118 | 0 | 0 | 0 | 0 | 0 | 0 | 71 | 443 | 95 | 21 | 0 | 364 | 0 | 19 | 13 | 0 | 0 | 0 | 0 | 0 | 0 | 1289 | 95 | 0 | 0 | 0 | 0 | 111 | 0 | 51 | 156 | 46 | 14 | 52 |
| SRR5029426 | 73 | 0 | 81 | 0 | 0 | 0 | 0 | 0 | 27 | 0 | 294 | 116 | 88 | 0 | 143 | 0 | 10 | 0 | 0 | 0 | 66 | 0 | 0 | 0 | 879 | 134 | 0 | 0 | 0 | 0 | 195 | 0 | 0 | 144 | 58 | 24 | 88 |
| SRR5029427 | 188 | 0 | 182 | 0 | 0 | 47 | 0 | 0 | 19 | 40 | 474 | 147 | 21 | 0 | 40 | 0 | 14 | 0 | 0 | 0 | 181 | 0 | 0 | 0 | 965 | 683 | 32 | 0 | 0 | 0 | 87 | 0 | 28 | 359 | 256 | 100 | 449 |
| SRR5029428 | 16 | 0 | 0 | 0 | 0 | 27 | 27 | 0 | 18 | 22 | 226 | 61 | 0 | 22 | 1460 | 0 | 49 | 0 | 0 | 0 | 27 | 0 | 0 | 0 | 53 | 80 | 0 | 0 | 0 | 0 | 22 | 22 | 0 | 67 | 107 | 133 | 26 |
| SRR5029429 | 15 | 0 | 26 | 0 | 0 | 26 | 0 | 0 | 31 | 0 | 222 | 29 | 98 | 0 | 3682 | 0 | 0 | 620 | 0 | 0 | 149 | 0 | 0 | 0 | 1406 | 450 | 21 | 0 | 0 | 0 | 155 | 0 | 0 | 176 | 57 | 72 | 160 |
| SRR5029430 | 28 | 0 | 92 | 0 | 0 | 63 | 0 | 0 | 21 | 0 | 16 | 185 | 143 | 0 | 144 | 0 | 18 | 0 | 0 | 18 | 144 | 0 | 0 | 0 | 65 | 31 | 21 | 0 | 0 | 0 | 14 | 0 | 37 | 16 | 12 | 69 | 17 |
| SRR5029431 | 346 | 0 | 94 | 0 | 0 | 0 | 0 | 0 | 0 | 12 | 190 | 216 | 20 | 0 | 122 | 0 | 10 | 0 | 0 | 0 | 0 | 0 | 0 | 0 | 242 | 21 | 0 | 0 | 0 | 0 | 32 | 0 | 12 | 593 | 13 | 0 | 14 |
| SRR5029432 | 28 | 0 | 104 | 0 | 0 | 19 | 0 | 0 | 47 | 0 | 35 | 108 | 12 | 0 | 1399 | 0 | 40 | 33 | 0 | 0 | 46 | 0 | 0 | 0 | 122 | 41 | 13 | 0 | 0 | 0 | 15 | 0 | 0 | 38 | 16 | 82 | 27 |
| SRR5029433 | 297 | 0 | 145 | 0 | 0 | 0 | 0 | 0 | 0 | 47 | 372 | 217 | 27 | 38 | 310 | 0 | 24 | 0 | 0 | 0 | 30 | 0 | 0 | 0 | 809 | 18 | 0 | 0 | 0 | 0 | 95 | 0 | 57 | 228 | 23 | 12 | 33 |
| SRR5029434 | 895 | 0 | 116 | 0 | 0 | 0 | 15 | 0 | 19 | 16 | 56 | 46 | 25 | 17 | 0 | 0 | 0 | 0 | 0 | 0 | 190 | 0 | 0 | 0 | 182 | 49 | 0 | 0 | 0 | 0 | 48 | 11 | 16 | 112 | 65 | 18 | 171 |
| SRR5029435 | 69 | 0 | 65 | 0 | 0 | 31 | 0 | 0 | 41 | 25 | 419 | 48 | 169 | 31 | 207 | 0 | 0 | 15 | 0 | 0 | 127 | 0 | 0 | 0 | 1165 | 431 | 16 | 0 | 0 | 0 | 206 | 0 | 0 | 216 | 87 | 53 | 247 |
| SRR5029436 | 41 | 0 | 193 | 0 | 11 | 13 | 0 | 0 | 49 | 23 | 47 | 117 | 20 | 0 | 87 | 0 | 17 | 67 | 0 | 0 | 190 | 0 | 0 | 0 | 68 | 61 | 23 | 0 | 29 | 0 | 100 | 0 | 26 | 32 | 44 | 92 | 66 |
| SRR5029437 | 20 | 0 | 28 | 0 | 0 | 0 | 23 | 0 | 0 | 12 | 60 | 193 | 0 | 0 | 1079 | 0 | 45 | 0 | 0 | 0 | 23 | 0 | 0 | 0 | 0 | 109 | 0 | 0 | 0 | 0 | 36 | 0 | 0 | 30 | 66 | 91 | 181 |
| SRR5029438 | 44 | 0 | 90 | 0 | 0 | 0 | 0 | 0 | 29 | 14 | 13 | 150 | 14 | 12 | 56 | 0 | 38 | 15 | 0 | 0 | 64 | 0 | 0 | 0 | 54 | 15 | 0 | 0 | 0 | 0 | 21 | 0 | 40 | 88 | 83 | 62 | 20 |
| SRR5029439 | 66 | 0 | 96 | 0 | 0 | 12 | 0 | 0 | 20 | 22 | 215 | 103 | 17 | 0 | 1771 | 0 | 52 | 1548 | 0 | 0 | 52 | 0 | 0 | 0 | 51 | 11 | 0 | 0 | 54 | 0 | 132 | 0 | 61 | 36 | 17 | 24 | 24 |
| SRR5029440 | 16 | 0 | 37 | 0 | 0 | 0 | 62 | 0 | 0 | 0 | 68 | 123 | 23 | 23 | 1430 | 0 | 32 | 0 | 0 | 0 | 0 | 0 | 0 | 0 | 10 | 63 | 0 | 0 | 0 | 0 | 0 | 0 | 0 | 29 | 78 | 57 | 13 |
| SRR5029441 | 171 | 0 | 193 | 12 | 0 | 16 | 0 | 0 | 0 | 44 | 217 | 242 | 16 | 23 | 209 | 0 | 19 | 0 | 0 | 0 | 12 | 0 | 0 | 0 | 562 | 53 | 0 | 0 | 0 | 0 | 98 | 0 | 21 | 60 | 53 | 34 | 92 |
| SRR5029442 | 104 | 0 | 152 | 0 | 0 | 19 | 0 | 0 | 0 | 31 | 229 | 303 | 130 | 0 | 151 | 0 | 46 | 0 | 0 | 0 | 75 | 0 | 0 | 0 | 1548 | 132 | 0 | 0 | 0 | 0 | 303 | 0 | 17 | 84 | 132 | 48 | 166 |
| SRR5029444 | 36 | 0 | 207 | 0 | 0 | 0 | 0 | 0 | 0 | 0 | 417 | 270 | 0 | 0 | 248 | 0 | 27 | 0 | 0 | 0 | 0 | 0 | 0 | 15 | 1515 | 31 | 0 | 0 | 0 | 0 | 278 | 0 | 15 | 216 | 108 | 31 | 62 |
| SRR5029446 | 14 | 0 | 0 | 0 | 0 | 0 | 0 | 0 | 11 | 18 | 17 | 139 | 15 | 0 | 16541 | 0 | 53 | 76 | 0 | 0 | 0 | 0 | 0 | 0 | 44 | 15 | 0 | 0 | 0 | 0 | 33 | 0 | 0 | 138 | 18 | 11 | 22 |
| SRR5029447 | 64 | 0 | 119 | 0 | 0 | 0 | 0 | 0 | 0 | 20 | 70 | 86 | 12 | 0 | 270 | 0 | 21 | 0 | 0 | 10 | 67 | 0 | 0 | 0 | 399 | 10 | 11 | 0 | 0 | 0 | 19 | 0 | 16 | 40 | 19 | 33 | 18 |
| SRR5029448 | 47 | 0 | 49 | 0 | 0 | 16 | 0 | 0 | 42 | 0 | 319 | 61 | 198 | 0 | 300 | 0 | 15 | 0 | 0 | 26 | 117 | 0 | 0 | 0 | 1264 | 211 | 13 | 0 | 0 | 0 | 150 | 0 | 0 | 245 | 174 | 26 | 145 |
| SRR5029452 | 19 | 0 | 0 | 0 | 0 | 15 | 15 | 0 | 26 | 0 | 49 | 130 | 23 | 0 | 256 | 0 | 45 | 0 | 0 | 0 | 15 | 0 | 0 | 0 | 0 | 196 | 0 | 0 | 0 | 0 | 30 | 0 | 0 | 30 | 34 | 23 | 147 |
| SRR5029455 | 38 | 0 | 131 | 0 | 0 | 13 | 0 | 0 | 19 | 21 | 178 | 151 | 25 | 0 | 1042 | 0 | 53 | 433 | 0 | 0 | 147 | 0 | 0 | 0 | 38 | 88 | 0 | 0 | 84 | 0 | 23 | 0 | 63 | 17 | 11 | 38 | 69 |
| SRR5029456 | 79 | 0 | 141 | 0 | 0 | 15 | 0 | 0 | 0 | 11 | 52 | 168 | 29 | 19 | 46 | 0 | 14 | 0 | 0 | 0 | 11 | 0 | 0 | 0 | 252 | 16 | 0 | 0 | 0 | 0 | 73 | 0 | 42 | 39 | 28 | 32 | 32 |

### Supp Table S8.2.Columns_AM_to_BW

| | | | | | | | | | | | | | | | | | | | | | | | | | | | | | | | | | | | | | |
|---|---|---|---|---|---|---|---|---|---|---|---|---|---|---|---|---|---|---|---|---|---|---|---|---|---|---|---|---|---|---|---|---|---|---|---|---|---|
| SRR5029457 | 54 | 0 | 90 | 0 | 13 | 10 | 0 | 0 | 0 | 0 | 435 | 76 | 50 | 0 | 445 | 0 | 20 | 177 | 0 | 0 | 44 | 0 | 0 | 0 | 1458 | 89 | 0 | 0 | 0 | 0 | 123 | 0 | 0 | 177 | 47 | 15 | 62 |
| SRR5029459 | 45 | 0 | 107 | 0 | 0 | 81 | 0 | 0 | 48 | 30 | 196 | 250 | 107 | 0 | 57 | 0 | 20 | 56 | 0 | 0 | 170 | 0 | 0 | 0 | 878 | 924 | 32 | 0 | 0 | 0 | 223 | 0 | 0 | 192 | 200 | 110 | 589 |
| SRR5029460 | 68 | 0 | 168 | 0 | 0 | 0 | 0 | 0 | 16 | 33 | 83 | 175 | 18 | 0 | 1488 | 0 | 16 | 138 | 0 | 0 | 0 | 0 | 0 | 0 | 260 | 13 | 0 | 0 | 0 | 0 | 37 | 0 | 0 | 37 | 12 | 22 | 11 |
| SRR5029462 | 92 | 0 | 72 | 0 | 0 | 16 | 0 | 0 | 10 | 0 | 19 | 124 | 23 | 11 | 108 | 0 | 0 | 23 | 0 | 18 | 398 | 0 | 0 | 0 | 78 | 134 | 12 | 0 | 0 | 0 | 21 | 0 | 0 | 33 | 16 | 78 | 34 |
| SRR5029463 | 111 | 0 | 161 | 0 | 0 | 18 | 0 | 0 | 0 | 12 | 241 | 125 | 18 | 0 | 339 | 0 | 0 | 0 | 0 | 0 | 145 | 0 | 0 | 0 | 1127 | 158 | 12 | 0 | 0 | 0 | 117 | 0 | 12 | 229 | 53 | 29 | 94 |
| SRR5029465 | 12 | 0 | 0 | 0 | 0 | 0 | 0 | 0 | 0 | 12 | 128 | 29 | 0 | 12 | 55 | 0 | 39 | 0 | 0 | 14 | 41 | 0 | 0 | 0 | 37 | 27 | 0 | 0 | 0 | 0 | 0 | 15 | 12 | 391 | 0 | 0 | 24 |
| SRR5029466 | 103 | 0 | 170 | 0 | 0 | 72 | 0 | 0 | 78 | 12 | 177 | 154 | 101 | 0 | 62 | 0 | 24 | 0 | 0 | 0 | 169 | 0 | 0 | 0 | 1074 | 790 | 19 | 0 | 0 | 0 | 163 | 0 | 14 | 625 | 267 | 29 | 362 |
| SRR5029467 | 205 | 0 | 103 | 0 | 0 | 0 | 0 | 0 | 0 | 21 | 58 | 165 | 14 | 0 | 109 | 0 | 16 | 0 | 0 | 0 | 11 | 0 | 0 | 0 | 309 | 24 | 0 | 0 | 0 | 0 | 48 | 0 | 25 | 77 | 10 | 11 | 13 |
| SRR5029469 | 112 | 0 | 103 | 0 | 0 | 0 | 0 | 0 | 0 | 0 | 127 | 138 | 87 | 0 | 58 | 0 | 19 | 27 | 0 | 0 | 12 | 0 | 0 | 0 | 644 | 50 | 0 | 0 | 0 | 0 | 459 | 0 | 15 | 207 | 66 | 0 | 59 |
| SRR5029471 | 34 | 0 | 137 | 0 | 0 | 0 | 0 | 0 | 36 | 14 | 25 | 419 | 13 | 0 | 1686 | 0 | 33 | 129 | 0 | 0 | 48 | 0 | 0 | 0 | 96 | 15 | 0 | 0 | 0 | 0 | 27 | 0 | 20 | 38 | 56 | 16 | 69 |
| SRR5029472 | 121 | 0 | 71 | 0 | 0 | 41 | 0 | 0 | 22 | 18 | 182 | 79 | 96 | 0 | 0 | 0 | 0 | 0 | 0 | 15 | 128 | 0 | 0 | 0 | 827 | 672 | 41 | 0 | 0 | 0 | 195 | 0 | 18 | 241 | 207 | 72 | 509 |
| SRR5029473 | 111 | 0 | 258 | 0 | 0 | 58 | 0 | 0 | 16 | 34 | 331 | 371 | 50 | 24 | 0 | 0 | 31 | 0 | 0 | 0 | 181 | 0 | 0 | 0 | 1561 | 766 | 25 | 0 | 0 | 0 | 168 | 0 | 25 | 127 | 204 | 142 | 663 |
| SRR5029474 | 139 | 0 | 115 | 0 | 0 | 58 | 0 | 0 | 36 | 22 | 112 | 245 | 67 | 0 | 108 | 0 | 27 | 0 | 0 | 0 | 163 | 0 | 0 | 0 | 916 | 579 | 18 | 0 | 0 | 0 | 1157 | 0 | 18 | 129 | 158 | 100 | 335 |
| SRR5029476 | 32 | 0 | 74 | 0 | 0 | 43 | 0 | 0 | 27 | 38 | 288 | 139 | 24 | 0 | 600 | 0 | 18 | 0 | 0 | 0 | 106 | 0 | 0 | 0 | 1322 | 697 | 31 | 0 | 0 | 0 | 446 | 0 | 16 | 199 | 214 | 101 | 458 |
| SRR5029477 | 231 | 0 | 184 | 0 | 0 | 11 | 0 | 0 | 18 | 21 | 453 | 124 | 71 | 0 | 240 | 0 | 25 | 32 | 0 | 0 | 50 | 0 | 0 | 0 | 1195 | 103 | 0 | 0 | 0 | 0 | 366 | 0 | 34 | 172 | 108 | 25 | 90 |
| SRR5029478 | 253 | 0 | 133 | 0 | 0 | 0 | 0 | 0 | 0 | 15 | 75 | 367 | 20 | 0 | 61 | 0 | 28 | 0 | 0 | 0 | 134 | 0 | 0 | 0 | 289 | 46 | 17 | 0 | 0 | 0 | 29 | 0 | 26 | 120 | 41 | 34 | 48 |
| SRR5029480 | 36 | 0 | 53 | 0 | 0 | 20 | 0 | 0 | 35 | 11 | 13 | 75 | 14 | 11 | 712 | 0 | 17 | 0 | 0 | 0 | 178 | 0 | 0 | 0 | 80 | 21 | 0 | 0 | 0 | 0 | 11 | 0 | 0 | 25 | 110 | 33 | 13 |
| SRR5029481 | 111 | 0 | 105 | 0 | 0 | 16 | 0 | 0 | 17 | 16 | 161 | 220 | 39 | 0 | 23 | 0 | 29 | 0 | 0 | 0 | 126 | 0 | 0 | 0 | 811 | 208 | 0 | 11 | 0 | 0 | 84 | 0 | 21 | 123 | 107 | 36 | 172 |
| SRR5029482 | 87 | 0 | 100 | 0 | 0 | 0 | 0 | 0 | 0 | 25 | 63 | 72 | 27 | 13 | 508 | 0 | 14 | 40 | 0 | 0 | 18 | 0 | 0 | 0 | 390 | 15 | 12 | 0 | 0 | 0 | 36 | 0 | 17 | 63 | 13 | 13 | 14 |
| SRR5029483 | 405 | 0 | 102 | 0 | 0 | 0 | 0 | 0 | 0 | 32 | 286 | 145 | 63 | 0 | 59 | 0 | 0 | 0 | 0 | 0 | 24 | 0 | 0 | 0 | 991 | 57 | 0 | 0 | 0 | 0 | 123 | 0 | 0 | 390 | 82 | 42 | 92 |
| SRR5029484 | 24 | 0 | 145 | 0 | 0 | 34 | 0 | 0 | 35 | 14 | 101 | 62 | 27 | 12 | 119 | 0 | 49 | 0 | 0 | 0 | 266 | 0 | 0 | 0 | 212 | 346 | 14 | 0 | 21 | 0 | 32 | 0 | 25 | 33 | 39 | 20 | 90 |
| SRR5029486 | 51 | 0 | 427 | 0 | 0 | 12 | 0 | 0 | 10 | 35 | 250 | 165 | 18 | 13 | 136 | 0 | 51 | 0 | 0 | 0 | 23 | 0 | 10 | 0 | 233 | 114 | 13 | 0 | 28 | 0 | 55 | 0 | 35 | 54 | 32 | 11 | 48 |
| SRR5029487 | 74 | 0 | 159 | 0 | 0 | 0 | 0 | 0 | 0 | 45 | 254 | 97 | 37 | 0 | 349 | 0 | 32 | 0 | 0 | 0 | 25 | 0 | 0 | 0 | 1559 | 42 | 0 | 0 | 0 | 0 | 96 | 0 | 25 | 89 | 61 | 19 | 61 |
| SRR5029488 | 11 | 0 | 107 | 0 | 0 | 15 | 0 | 0 | 15 | 29 | 191 | 241 | 44 | 15 | 77 | 0 | 28 | 0 | 0 | 0 | 0 | 0 | 0 | 0 | 927 | 221 | 44 | 0 | 0 | 15 | 280 | 0 | 0 | 88 | 191 | 0 | 177 |
| SRR5029489 | 63 | 0 | 99 | 0 | 0 | 13 | 0 | 0 | 21 | 10 | 24 | 134 | 19 | 11 | 182 | 0 | 44 | 0 | 0 | 0 | 211 | 0 | 0 | 0 | 226 | 111 | 44 | 0 | 0 | 0 | 29 | 10 | 10 | 32 | 34 | 19 | 72 |
| SRR5029490 | 37 | 0 | 75 | 0 | 0 | 35 | 0 | 0 | 35 | 33 | 372 | 56 | 75 | 0 | 152 | 0 | 31 | 0 | 0 | 0 | 96 | 0 | 0 | 0 | 679 | 435 | 11 | 0 | 18 | 0 | 125 | 0 | 38 | 556 | 82 | 36 | 167 |
| SRR5029491 | 55 | 0 | 110 | 0 | 0 | 30 | 0 | 0 | 37 | 0 | 133 | 173 | 242 | 14 | 234 | 0 | 22 | 138 | 0 | 0 | 99 | 0 | 0 | 0 | 995 | 340 | 21 | 0 | 0 | 0 | 312 | 0 | 19 | 282 | 205 | 65 | 214 |
| SRR5029492 | 48 | 0 | 280 | 0 | 0 | 14 | 16 | 0 | 27 | 0 | 294 | 173 | 53 | 0 | 177 | 0 | 19 | 0 | 0 | 0 | 48 | 0 | 0 | 0 | 1422 | 161 | 14 | 0 | 0 | 0 | 167 | 0 | 0 | 278 | 83 | 25 | 108 |
| SRR5029493 | 50 | 0 | 118 | 0 | 0 | 0 | 0 | 0 | 11 | 30 | 200 | 314 | 88 | 21 | 113 | 0 | 53 | 0 | 0 | 0 | 0 | 0 | 0 | 0 | 940 | 82 | 0 | 0 | 0 | 0 | 280 | 0 | 13 | 250 | 120 | 69 | 103 |
| SRR5029494 | 54 | 0 | 210 | 0 | 0 | 13 | 0 | 0 | 11 | 25 | 61 | 308 | 14 | 19 | 801 | 0 | 41 | 216 | 0 | 0 | 13 | 0 | 0 | 0 | 264 | 50 | 0 | 0 | 0 | 0 | 96 | 0 | 17 | 63 | 49 | 41 | 62 |
| SRR5029495 | 201 | 0 | 120 | 0 | 0 | 95 | 0 | 0 | 51 | 22 | 150 | 218 | 78 | 19 | 579 | 0 | 42 | 28 | 0 | 0 | 204 | 0 | 0 | 0 | 1177 | 849 | 90 | 0 | 0 | 0 | 220 | 0 | 0 | 464 | 352 | 96 | 446 |
| SRR5029496 | 65 | 0 | 114 | 0 | 0 | 16 | 0 | 0 | 0 | 13 | 28 | 191 | 20 | 0 | 20 | 0 | 31 | 0 | 0 | 0 | 34 | 0 | 0 | 0 | 226 | 32 | 0 | 0 | 0 | 0 | 15 | 0 | 16 | 83 | 23 | 16 | 24 |
| SRR5029497 | 74 | 0 | 93 | 0 | 0 | 13 | 0 | 0 | 22 | 27 | 174 | 98 | 46 | 0 | 189 | 0 | 0 | 21 | 0 | 0 | 27 | 0 | 0 | 0 | 1130 | 230 | 17 | 0 | 0 | 0 | 156 | 0 | 12 | 203 | 218 | 26 | 196 |
| SRR5029498 | 542 | 0 | 92 | 0 | 0 | 0 | 0 | 0 | 38 | 21 | 20 | 377 | 52 | 0 | 0 | 0 | 0 | 0 | 0 | 0 | 336 | 0 | 0 | 0 | 122 | 36 | 21 | 0 | 0 | 0 | 291 | 0 | 19 | 35 | 53 | 14 | 57 |
| SRR5029499 | 54 | 0 | 163 | 0 | 0 | 10 | 0 | 0 | 18 | 12 | 31 | 420 | 10 | 0 | 30 | 0 | 24 | 11 | 0 | 0 | 37 | 0 | 0 | 0 | 210 | 16 | 0 | 0 | 0 | 0 | 40 | 0 | 10 | 37 | 19 | 29 | 18 |
| SRR5029500 | 51 | 0 | 197 | 0 | 0 | 0 | 0 | 11 | 11 | 32 | 296 | 253 | 74 | 0 | 0 | 0 | 35 | 0 | 0 | 0 | 0 | 0 | 0 | 0 | 445 | 85 | 0 | 0 | 11 | 11 | 233 | 11 | 64 | 349 | 138 | 21 | 106 |
| SRR5029501 | 25 | 0 | 63 | 0 | 0 | 0 | 0 | 0 | 0 | 44 | 40 | 41 | 15 | 0 | 381 | 0 | 43 | 347 | 0 | 0 | 0 | 0 | 0 | 0 | 214 | 30 | 0 | 0 | 0 | 0 | 23 | 0 | 22 | 15 | 44 | 13 | 49 |
| SRR5029502 | 39 | 22 | 96 | 0 | 0 | 67 | 23 | 0 | 11 | 67 | 44 | 220 | 14 | 11 | 0 | 0 | 28 | 0 | 0 | 23 | 23 | 0 | 0 | 0 | 87 | 52 | 56 | 11 | 0 | 0 | 16 | 0 | 22 | 82 | 38 | 22 | 47 |
| SRR5029503 | 19 | 0 | 86 | 0 | 0 | 0 | 0 | 0 | 0 | 28 | 37 | 34 | 32 | 26 | 636 | 0 | 30 | 514 | 0 | 0 | 24 | 0 | 0 | 0 | 230 | 21 | 0 | 13 | 0 | 0 | 29 | 0 | 20 | 22 | 24 | 32 | 19 |
| SRR5029504 | 253 | 0 | 87 | 0 | 0 | 0 | 0 | 0 | 0 | 39 | 253 | 177 | 77 | 13 | 47 | 0 | 20 | 0 | 0 | 0 | 55 | 0 | 0 | 0 | 1019 | 84 | 0 | 0 | 0 | 0 | 158 | 0 | 24 | 266 | 131 | 35 | 137 |
| SRR5029505 | 71 | 0 | 212 | 0 | 0 | 0 | 0 | 0 | 0 | 21 | 246 | 152 | 77 | 27 | 129 | 0 | 21 | 0 | 0 | 0 | 35 | 0 | 0 | 0 | 1788 | 109 | 0 | 0 | 0 | 0 | 105 | 0 | 11 | 127 | 135 | 41 | 106 |
| SRR5029506 | 17 | 0 | 24 | 0 | 0 | 78 | 0 | 0 | 19 | 17 | 177 | 144 | 58 | 11 | 49 | 0 | 50 | 43 | 0 | 0 | 152 | 0 | 0 | 0 | 760 | 729 | 47 | 0 | 0 | 0 | 635 | 0 | 30 | 217 | 214 | 137 | 455 |
| SRR5029507 | 85 | 0 | 50 | 0 | 0 | 40 | 0 | 0 | 57 | 0 | 159 | 180 | 211 | 0 | 69 | 0 | 29 | 0 | 0 | 0 | 91 | 0 | 0 | 0 | 546 | 399 | 0 | 0 | 0 | 0 | 173 | 0 | 21 | 743 | 166 | 71 | 344 |
| SRR5029508 | 347 | 0 | 57 | 0 | 0 | 16 | 0 | 0 | 13 | 13 | 26 | 97 | 22 | 29 | 0 | 0 | 15 | 0 | 0 | 0 | 87 | 0 | 0 | 0 | 211 | 27 | 36 | 0 | 0 | 0 | 31 | 0 | 0 | 60 | 42 | 53 | 59 |
| SRR5029509 | 31 | 0 | 106 | 0 | 0 | 11 | 0 | 0 | 19 | 132 | 840 | 39 | 20 | 22 | 1077 | 0 | 54 | 86 | 0 | 0 | 29 | 0 | 0 | 0 | 519 | 179 | 0 | 14 | 24 | 0 | 110 | 0 | 120 | 224 | 90 | 17 | 153 |
| SRR5029510 | 34 | 0 | 112 | 0 | 0 | 18 | 0 | 0 | 24 | 22 | 157 | 111 | 58 | 14 | 29 | 0 | 28 | 0 | 0 | 0 | 52 | 0 | 0 | 0 | 986 | 240 | 0 | 0 | 0 | 0 | 154 | 0 | 12 | 155 | 116 | 73 | 232 |
| SRR5029511 | 81 | 0 | 59 | 0 | 0 | 54 | 0 | 0 | 56 | 14 | 126 | 212 | 94 | 0 | 90 | 0 | 42 | 0 | 0 | 0 | 264 | 0 | 0 | 0 | 1269 | 573 | 19 | 0 | 0 | 0 | 207 | 0 | 13 | 204 | 161 | 70 | 307 |
| SRR5029512 | 74 | 0 | 114 | 0 | 0 | 36 | 0 | 0 | 0 | 21 | 145 | 69 | 48 | 0 | 59 | 0 | 23 | 23 | 0 | 0 | 118 | 0 | 0 | 0 | 1484 | 579 | 60 | 0 | 0 | 0 | 111 | 0 | 11 | 178 | 149 | 84 | 253 |
| SRR5029513 | 24 | 0 | 95 | 0 | 0 | 10 | 0 | 0 | 10 | 33 | 167 | 461 | 86 | 0 | 1078 | 0 | 36 | 805 | 0 | 0 | 17 | 0 | 0 | 0 | 956 | 131 | 0 | 0 | 0 | 0 | 1272 | 0 | 14 | 166 | 109 | 28 | 138 |
| SRR5029514 | 23 | 0 | 104 | 0 | 0 | 0 | 0 | 0 | 0 | 43 | 67 | 73 | 62 | 20 | 74 | 0 | 14 | 0 | 0 | 0 | 11 | 0 | 0 | 0 | 154 | 12 | 0 | 0 | 0 | 0 | 23 | 0 | 36 | 16 | 10 | 12 | 55 |
| SRR5029515 | 49 | 0 | 64 | 0 | 0 | 0 | 0 | 0 | 0 | 25 | 21 | 210 | 16 | 0 | 165 | 0 | 28 | 12 | 0 | 0 | 0 | 0 | 0 | 0 | 151 | 39 | 0 | 0 | 0 | 0 | 20 | 0 | 47 | 22 | 50 | 17 | 35 |
| SRR5029516 | 91 | 0 | 83 | 0 | 0 | 31 | 12 | 0 | 18 | 15 | 13 | 139 | 13 | 0 | 75 | 0 | 76 | 0 | 0 | 0 | 624 | 0 | 0 | 0 | 152 | 222 | 20 | 0 | 0 | 0 | 23 | 0 | 0 | 34 | 18 | 68 | 32 |
| SRR5029870 | 133 | 0 | 26378 | 28 | 0 | 55 | 0 | 28 | 55 | 0 | 0 | 98 | 0 | 14 | 17 | 0 | 36 | 0 | 0 | 835 | 1113 | 0 | 0 | 0 | 14 | 60 | 67 | 0 | 0 | 0 | 42 | 0 | 0 | 14 | 320 | 22 | 153 |
| SRR5029871 | 98 | 0 | 23692 | 19 | 0 | 76 | 0 | 0 | 94 | 0 | 0 | 100 | 0 | 19 | 0 | 0 | 23 | 26 | 0 | 326 | 826 | 0 | 0 | 0 | 0 | 35 | 24 | 0 | 0 | 0 | 0 | 19 | 0 | 0 | 234 | 18 | 112 |
| SRR5029872 | 116 | 0 | 27173 | 0 | 0 | 46 | 0 | 61 | 77 | 0 | 0 | 91 | 0 | 15 | 0 | 15 | 34 | 0 | 0 | 448 | 895 | 0 | 0 | 0 | 0 | 55 | 78 | 15 | 0 | 0 | 0 | 0 | 0 | 15 | 313 | 22 | 148 |
| SRR5029873 | 116 | 20 | 24582 | 59 | 0 | 79 | 27 | 0 | 39 | 0 | 0 | 93 | 0 | 0 | 54 | 0 | 39 | 27 | 27 | 403 | 403 | 0 | 0 | 0 | 0 | 31 | 20 | 20 | 0 | 0 | 20 | 20 | 0 | 20 | 238 | 16 | 112 |
| SRR5029874 | 131 | 14 | 26557 | 0 | 14 | 100 | 18 | 43 | 57 | 0 | 0 | 96 | 0 | 0 | 18 | 0 | 30 | 0 | 0 | 586 | 1227 | 0 | 0 | 0 | 0 | 77 | 76 | 0 | 0 | 0 | 0 | 0 | 0 | 14 | 304 | 24 | 136 |
| SRR5029875 | 115 | 0 | 23748 | 39 | 0 | 77 | 0 | 0 | 19 | 0 | 0 | 86 | 0 | 19 | 0 | 19 | 33 | 13 | 0 | 260 | 832 | 19 | 0 | 0 | 0 | 34 | 24 | 0 | 0 | 0 | 19 | 0 | 0 | 0 | 231 | 13 | 111 |
| SRR5029876 | 126 | 0 | 26547 | 30 | 0 | 135 | 19 | 60 | 45 | 0 | 0 | 116 | 0 | 30 | 0 | 0 | 30 | 0 | 0 | 655 | 1011 | 15 | 0 | 0 | 0 | 1678 | 1843 | 15 | 0 | 0 | 15 | 15 | 0 | 0 | 7538 | 719 | 4106 |
| SRR5029877 | 97 | 0 | 24225 | 20 | 20 | 139 | 0 | 20 | 60 | 0 | 0 | 91 | 0 | 20 | 27 | 0 | 30 | 54 | 0 | 351 | 811 | 20 | 0 | 0 | 0 | 1131 | 913 | 0 | 0 | 0 | 40 | 20 | 0 | 0 | 8155 | 655 | 4246 |
| SRR5029878 | 110 | 0 | 11703 | 0 | 0 | 343 | 0 | 30 | 262 | 0 | 0 | 43 | 0 | 0 | 10 | 0 | 15 | 0 | 10 | 178 | 346 | 0 | 0 | 0 | 0 | 3966 | 1382 | 10 | 0 | 0 | 111 | 10 | 0 | 10 | 8678 | 616 | 4894 |
| SRR5029879 | 87 | 0 | 10014 | 0 | 0 | 454 | 0 | 0 | 151 | 0 | 0 | 26 | 0 | 0 | 55 | 0 | 12 | 0 | 0 | 55 | 277 | 0 | 0 | 0 | 0 | 3029 | 909 | 0 | 0 | 0 | 151 | 0 | 0 | 50 | 8532 | 555 | 4342 |
| SRR5029880 | 113 | 0 | 11950 | 0 | 0 | 313 | 0 | 11 | 173 | 0 | 0 | 34 | 0 | 0 | 33 | 0 | 20 | 0 | 11 | 211 | 422 | 0 | 0 | 11 | 0 | 3250 | 1425 | 0 | 0 | 0 | 130 | 0 | 0 | 22 | 7980 | 842 | 4449 |
| SRR5029882 | 132 | 0 | 11541 | 0 | 0 | 14 | 22 | 21 | 216 | 0 | 0 | 34 | 0 | 0 | 11 | 0 | 19 | 0 | 0 | 249 | 249 | 0 | 0 | 0 | 10 | 120 | 63 | 10 | 0 | 0 | 103 | 0 | 0 | 10 | 295 | 24 | 178 |
| SRR5029883 | 100 | 0 | 10180 | 0 | 0 | 305 | 0 | 0 | 102 | 0 | 0 | 23 | 0 | 0 | 0 | 0 | 15 | 0 | 0 | 167 | 500 | 0 | 0 | 0 | 0 | 88 | 24 | 0 | 0 | 0 | 204 | 0 | 0 | 0 | 226 | 21 | 158 |
| SRR5029884 | 113 | 0 | 11703 | 10 | 0 | 356 | 11 | 0 | 105 | 0 | 0 | 30 | 0 | 0 | 22 | 0 | 23 | 0 | 0 | 184 | 358 | 0 | 0 | 0 | 10 | 3503 | 1652 | 0 | 0 | 0 | 105 | 10 | 0 | 21 | 8272 | 847 | 4968 |
| SRR5029885 | 66 | 0 | 10249 | 0 | 0 | 262 | 0 | 0 | 314 | 0 | 0 | 18 | 0 | 0 | 114 | 0 | 21 | 0 | 0 | 342 | 513 | 0 | 0 | 0 | 0 | 2934 | 681 | 0 | 0 | 0 | 52 | 0 | 0 | 0 | 7859 | 838 | 4349 |
| SRR5029886 | 415 | 0 | 5121 | 10 | 0 | 12 | 0 | 0 | 146 | 0 | 0 | 231 | 0 | 0 | 16 | 0 | 49 | 0 | 0 | 435 | 516 | 10 | 0 | 0 | 0 | 248 | 194 | 0 | 0 | 0 | 83 | 0 | 0 | 0 | 1054 | 114 | 745 |
| SRR5029887 | 393 | 0 | 4660 | 0 | 0 | 129 | 0 | 0 | 83 | 0 | 0 | 208 | 0 | 0 | 18 | 0 | 57 | 0 | 0 | 222 | 406 | 0 | 0 | 0 | 0 | 127 | 61 | 18 | 0 | 0 | 64 | 0 | 0 | 0 | 852 | 86 | 507 |
| SRR5029888 | 418 | 0 | 5287 | 11 | 0 | 167 | 0 | 0 | 111 | 0 | 0 | 218 | 0 | 0 | 0 | 0 | 35 | 0 | 0 | 279 | 645 | 22 | 0 | 11 | 0 | 234 | 170 | 0 | 0 | 0 | 44 | 0 | 0 | 0 | 1059 | 105 | 718 |
| SRR5029889 | 397 | 0 | 4834 | 0 | 0 | 162 | 0 | 0 | 105 | 0 | 0 | 204 | 0 | 0 | 58 | 0 | 54 | 0 | 0 | 173 | 365 | 0 | 0 | 0 | 0 | 130 | 56 | 0 | 0 | 0 | 76 | 0 | 0 | 0 | 833 | 89 | 517 |
| SRR5029890 | 416 | 0 | 5245 | 21 | 0 | 244 | 0 | 0 | 223 | 0 | 0 | 192 | 0 | 0 | 33 | 0 | 41 | 0 | 0 | 297 | 659 | 0 | 0 | 0 | 0 | 4226 | 2845 | 11 | 0 | 0 | 96 | 0 | 0 | 0 | 17529 | 2187 | 13250 |
| SRR5029891 | 349 | 0 | 4732 | 19 | 0 | 142 | 0 | 0 | 113 | 0 | 0 | 215 | 0 | 0 | 0 | 0 | 42 | 0 | 0 | 187 | 467 | 0 | 0 | 0 | 0 | 2900 | 1200 | 0 | 0 | 0 | 132 | 19 | 0 | 0 | 17511 | 1625 | 10815 |
| SRR5029892 | 412 | 0 | 5214 | 11 | 0 | 174 | 0 | 0 | 98 | 0 | 0 | 226 | 0 | 0 | 0 | 0 | 64 | 0 | 0 | 473 | 560 | 0 | 0 | 0 | 0 | 4035 | 2926 | 11 | 0 | 0 | 33 | 11 | 0 | 0 | 17947 | 2110 | 12400 |
| SRR5029893 | 390 | 0 | 4889 | 19 | 0 | 10 | 0 | 0 | 86 | 0 | 0 | 194 | 0 | 0 | 58 | 0 | 55 | 0 | 0 | 233 | 467 | 19 | 0 | 0 | 0 | 137 | 59 | 19 | 0 | 0 | 77 | 0 | 0 | 0 | 845 | 88 | 504 |
| SRR5029894 | 108 | 12 | 19606 | 0 | 0 | 43 | 0 | 18 | 18 | 0 | 0 | 141 | 0 | 25 | 0 | 0 | 32 | 0 | 0 | 138 | 402 | 12 | 0 | 0 | 0 | 77 | 71 | 0 | 0 | 0 | 18 | 31 | 0 | 0 | 345 | 24 | 233 |
| SRR5029895 | 93 | 77 | 17915 | 15 | 15 | 77 | 0 | 0 | 0 | 0 | 0 | 120 | 0 | 77 | 0 | 0 | 24 | 0 | 0 | 138 | 184 | 0 | 0 | 0 | 0 | 712 | 428 | 23 | 0 | 0 | 23 | 61 | 0 | 0 | 5027 | 367 | 2701 |
| SRR5029896 | 101 | 26 | 20301 | 0 | 13 | 79 | 25 | 13 | 20 | 0 | 0 | 144 | 0 | 53 | 0 | 0 | 44 | 0 | 0 | 86 | 331 | 0 | 0 | 0 | 0 | 920 | 906 | 0 | 0 | 0 | 0 | 20 | 0 | 0 | 4089 | 357 | 2547 |
| SRR5029897 | 100 | 48 | 18821 | 0 | 0 | 56 | 16 | 0 | 24 | 0 | 0 | 129 | 0 | 80 | 0 | 0 | 46 | 0 | 0 | 96 | 272 | 0 | 0 | 0 | 0 | 887 | 464 | 0 | 0 | 0 | 0 | 56 | 0 | 0 | 4684 | 264 | 2606 |
| SRR5029898 | 85 | 0 | 19745 | 0 | 0 | 63 | 0 | 25 | 0 | 0 | 0 | 166 | 0 | 25 | 0 | 0 | 46 | 0 | 24 | 224 | 413 | 0 | 0 | 0 | 0 | 81 | 69 | 13 | 0 | 0 | 13 | 13 | 0 | 0 | 346 | 23 | 242 |
| SRR5029899 | 103 | 23 | 18240 | 16 | 0 | 23 | 0 | 0 | 16 | 0 | 0 | 135 | 0 | 86 | 16 | 0 | 32 | 0 | 0 | 79 | 204 | 0 | 0 | 0 | 0 | 679 | 406 | 0 | 0 | 0 | 31 | 39 | 0 | 0 | 4729 | 164 | 2630 |
| SRR5029900 | 93 | 0 | 19692 | 13 | 0 | 64 | 24 | 19 | 13 | 0 | 0 | 158 | 0 | 26 | 0 | 0 | 36 | 0 | 0 | 120 | 192 | 0 | 0 | 0 | 0 | 785 | 746 | 0 | 0 | 0 | 13 | 39 | 0 | 0 | 3880 | 290 | 2484 |
| SRR5029901 | 93 | 40 | 18536 | 16 | 0 | 0 | 0 | 0 | 32 | 0 | 0 | 123 | 0 | 72 | 16 | 0 | 32 | 0 | 0 | 81 | 243 | 0 | 0 | 0 | 0 | 941 | 455 | 0 | 0 | 0 | 24 | 40 | 0 | 0 | 4826 | 319 | 2505 |
| SRR5029902 | 301 | 0 | 9870 | 0 | 0 | 232 | 0 | 0 | 75 | 0 | 0 | 434 | 0 | 0 | 391 | 0 | 26 | 0 | 0 | 416 | 2250 | 0 | 0 | 0 | 0 | 3822 | 1260 | 0 | 0 | 0 | 75 | 0 | 0 | 0 | 4817 | 198 | 3263 |
| SRR5029904 | 294 | 0 | 10174 | 0 | 0 | 216 | 0 | 0 | 37 | 0 | 0 | 414 | 0 | 0 | 241 | 0 | 29 | 0 | 0 | 429 | 2038 | 0 | 0 | 0 | 0 | 3656 | 1229 | 0 | 0 | 0 | 45 | 0 | 0 | 0 | 4751 | 320 | 2949 |
| SRR5029905 | 234 | 0 | 9111 | 0 | 0 | 112 | 0 | 0 | 56 | 0 | 0 | 373 | 0 | 0 | 626 | 0 | 22 | 0 | 0 | 209 | 834 | 0 | 0 | 0 | 0 | 2677 | 502 | 56 | 0 | 0 | 56 | 56 | 0 | 0 | 4740 | 335 | 2788 |
| SRR5029906 | 298 | 0 | 9798 | 14 | 0 | 11 | 0 | 0 | 77 | 0 | 0 | 443 | 0 | 0 | 378 | 0 | 27 | 0 | 0 | 353 | 1614 | 21 | 0 | 0 | 0 | 226 | 73 | 14 | 0 | 0 | 92 | 0 | 0 | 0 | 279 | 21 | 202 |
| SRR5029908 | 298 | 0 | 10004 | 0 | 0 | 13 | 0 | 0 | 58 | 0 | 0 | 444 | 0 | 0 | 473 | 0 | 26 | 0 | 0 | 552 | 2602 | 0 | 0 | 0 | 0 | 213 | 80 | 0 | 0 | 0 | 109 | 0 | 0 | 22 | 272 | 25 | 180 |
| SRR5029910 | 156 | 0 | 13052 | 0 | 20 | 123 | 0 | 61 | 20 | 0 | 0 | 59 | 0 | 0 | 0 | 0 | 19 | 0 | 0 | 104 | 151 | 20 | 0 | 0 | 0 | 4097 | 1373 | 0 | 0 | 0 | 41 | 0 | 0 | 225 | 11861 | 574 | 2847 |
| SRR5029911 | 165 | 0 | 13391 | 0 | 21 | 212 | 0 | 21 | 127 | 0 | 0 | 40 | 0 | 0 | 13 | 0 | 22 | 0 | 0 | 53 | 145 | 21 | 0 | 0 | 0 | 115 | 360 | 0 | 0 | 0 | 106 | 0 | 0 | 85 | 397 | 13 | 75 |
| SRR5029912 | 152 | 0 | 13324 | 22 | 0 | 245 | 0 | 89 | 89 | 0 | 0 | 49 | 0 | 22 | 0 | 0 | 26 | 0 | 0 | 140 | 76 | 0 | 0 | 0 | 0 | 126 | 29 | 0 | 0 | 0 | 89 | 0 | 0 | 0 | 377 | 13 | 87 |
| SRR5029913 | 174 | 0 | 13749 | 22 | 22 | 133 | 0 | 67 | 111 | 0 | 0 | 63 | 0 | 22 | 0 | 0 | 34 | 0 | 0 | 55 | 109 | 0 | 0 | 0 | 0 | 117 | 13 | 0 | 0 | 0 | 67 | 0 | 0 | 111 | 376 | 16 | 80 |
| SRR5029914 | 169 | 21 | 12993 | 0 | 43 | 342 | 0 | 149 | 43 | 0 | 0 | 54 | 0 | 0 | 24 | 0 | 28 | 0 | 0 | 122 | 171 | 0 | 0 | 0 | 0 | 132 | 20 | 0 | 0 | 0 | 85 | 0 | 0 | 64 | 360 | 12 | 92 |
| SRR5029915 | 179 | 0 | 13466 | 0 | 65 | 131 | 0 | 44 | 22 | 0 | 0 | 46 | 0 | 44 | 27 | 0 | 24 | 0 | 0 | 81 | 95 | 22 | 0 | 0 | 0 | 113 | 15 | 0 | 0 | 0 | 22 | 0 | 0 | 65 | 388 | 18 | 89 |
| SRR5029916 | 169 | 0 | 13332 | 0 | 0 | 301 | 0 | 129 | 43 | 0 | 0 | 49 | 0 | 0 | 0 | 0 | 28 | 0 | 0 | 136 | 123 | 0 | 0 | 0 | 0 | 140 | 30 | 0 | 0 | 0 | 64 | 0 | 0 | 43 | 365 | 17 | 93 |
| SRR5029917 | 167 | 0 | 13673 | 44 | 66 | 240 | 0 | 44 | 87 | 0 | 0 | 37 | 0 | 22 | 0 | 0 | 35 | 0 | 0 | 41 | 55 | 0 | 0 | 0 | 0 | 3870 | 437 | 0 | 0 | 0 | 66 | 0 | 0 | 87 | 14102 | 612 | 3192 |
| SRR5029918 | 647 | 17 | 19029 | 0 | 0 | 415 | 0 | 0 | 33 | 0 | 0 | 117 | 0 | 0 | 117 | 0 | 28 | 0 | 0 | 281 | 2225 | 50 | 0 | 0 | 0 | 8475 | 8624 | 0 | 17 | 0 | 83 | 0 | 0 | 0 | 7295 | 349 | 4487 |
| SRR5029919 | 592 | 24 | 17634 | 0 | 0 | 186 | 0 | 0 | 0 | 0 | 0 | 104 | 0 | 0 | 228 | 0 | 25 | 33 | 15 | 76 | 1324 | 0 | 0 | 0 | 0 | 5867 | 4455 | 16 | 0 | 0 | 210 | 0 | 0 | 0 | 8288 | 444 | 4116 |
| SRR5029920 | 661 | 0 | 19313 | 0 | 0 | 283 | 0 | 0 | 53 | 0 | 0 | 89 | 0 | 0 | 155 | 0 | 20 | 0 | 26 | 259 | 1966 | 0 | 0 | 0 | 0 | 7492 | 8413 | 0 | 18 | 0 | 124 | 0 | 0 | 0 | 7173 | 319 | 4924 |
| SRR5029921 | 583 | 59 | 18238 | 0 | 0 | 227 | 0 | 17 | 25 | 0 | 0 | 86 | 0 | 0 | 143 | 25 | 22 | 0 | 0 | 174 | 1648 | 0 | 0 | 0 | 0 | 6061 | 3921 | 0 | 0 | 0 | 101 | 0 | 0 | 0 | 8026 | 453 | 3979 |
| SRR5029922 | 645 | 34 | 19063 | 0 | 0 | 390 | 0 | 0 | 34 | 0 | 0 | 84 | 0 | 17 | 24 | 0 | 17 | 0 | 0 | 245 | 1885 | 17 | 0 | 0 | 0 | 7493 | 8883 | 0 | 0 | 0 | 68 | 0 | 0 | 0 | 8222 | 542 | 4645 |
| SRR5029923 | 588 | 25 | 17665 | 33 | 0 | 180 | 0 | 0 | 25 | 0 | 0 | 93 | 0 | 0 | 123 | 0 | 23 | 0 | 15 | 153 | 1565 | 16 | 0 | 0 | 0 | 6302 | 4373 | 0 | 0 | 0 | 229 | 0 | 0 | 0 | 7855 | 392 | 4349 |
| SRR5029924 | 625 | 0 | 19667 | 0 | 35 | 13 | 0 | 0 | 35 | 0 | 0 | 117 | 0 | 0 | 76 | 0 | 18 | 0 | 0 | 202 | 1641 | 0 | 0 | 0 | 0 | 334 | 385 | 0 | 17 | 0 | 139 | 0 | 0 | 0 | 326 | 25 | 229 |
| SRR5029925 | 587 | 33 | 18108 | 0 | 0 | 276 | 0 | 0 | 17 | 0 | 0 | 80 | 0 | 0 | 112 | 0 | 22 | 0 | 0 | 48 | 1578 | 0 | 0 | 0 | 0 | 5845 | 4172 | 25 | 0 | 0 | 167 | 0 | 0 | 17 | 8395 | 435 | 4373 |
| SRR5029926 | 160 | 0 | 17696 | 79 | 16 | 40 | 0 | 24 | 16 | 0 | 0 | 146 | 0 | 0 | 132 | 0 | 34 | 0 | 0 | 369 | 1080 | 40 | 0 | 0 | 0 | 60 | 72 | 0 | 0 | 0 | 0 | 0 | 0 | 0 | 226 | 11 | 117 |
| SRR5029927 | 154 | 0 | 16540 | 106 | 44 | 22 | 0 | 0 | 28 | 0 | 0 | 126 | 0 | 22 | 460 | 0 | 23 | 0 | 0 | 181 | 525 | 17 | 0 | 0 | 0 | 505 | 355 | 0 | 0 | 0 | 22 | 0 | 0 | 0 | 3287 | 183 | 1471 |
| SRR5029928 | 146 | 0 | 18661 | 52 | 0 | 34 | 29 | 0 | 34 | 0 | 0 | 162 | 0 | 0 | 260 | 0 | 31 | 0 | 29 | 490 | 1414 | 43 | 0 | 0 | 0 | 671 | 852 | 0 | 0 | 0 | 17 | 0 | 0 | 0 | 2246 | 120 | 1300 |
| SRR5029929 | 159 | 0 | 16949 | 150 | 29 | 23 | 0 | 0 | 0 | 0 | 0 | 129 | 0 | 17 | 345 | 12 | 29 | 0 | 0 | 207 | 603 | 29 | 0 | 0 | 0 | 24 | 12 | 17 | 0 | 0 | 12 | 0 | 0 | 0 | 169 | 11 | 75 |
| SRR5029930 | 148 | 0 | 17685 | 73 | 49 | 16 | 0 | 0 | 41 | 0 | 0 | 154 | 0 | 0 | 112 | 0 | 23 | 0 | 56 | 670 | 977 | 49 | 0 | 0 | 0 | 576 | 779 | 0 | 0 | 0 | 16 | 0 | 0 | 0 | 2426 | 203 | 1201 |
| SRR5029931 | 169 | 11 | 16537 | 129 | 39 | 0 | 0 | 0 | 34 | 0 | 0 | 130 | 0 | 11 | 480 | 0 | 24 | 0 | 0 | 132 | 562 | 17 | 0 | 0 | 0 | 21 | 13 | 11 | 0 | 0 | 17 | 0 | 0 | 0 | 177 | 18 | 71 |
| SRR5029932 | 195 | 0 | 18293 | 75 | 58 | 0 | 28 | 0 | 25 | 0 | 0 | 136 | 0 | 0 | 168 | 0 | 31 | 0 | 0 | 477 | 1066 | 25 | 0 | 0 | 0 | 610 | 1035 | 17 | 0 | 0 | 25 | 0 | 0 | 0 | 2346 | 159 | 1136 |
| SRR5029933 | 156 | 0 | 16708 | 116 | 17 | 12 | 17 | 0 | 17 | 0 | 0 | 153 | 0 | 23 | 441 | 0 | 28 | 0 | 17 | 136 | 610 | 17 | 0 | 0 | 0 | 26 | 13 | 17 | 0 | 0 | 23 | 0 | 0 | 0 | 169 | 13 | 77 |
| SRR5029934 | 353 | 53 | 9725 | 0 | 0 | 119 | 184 | 318 | 93 | 0 | 0 | 41 | 0 | 0 | 257 | 0 | 47 | 0 | 0 | 31 | 1341 | 0 | 0 | 0 | 0 | 2119 | 1192 | 40 | 0 | 0 | 26 | 0 | 0 | 0 | 7696 | 543 | 4464 |
| SRR5029935 | 314 | 15 | 9071 | 0 | 0 | 60 | 112 | 75 | 90 | 0 | 0 | 41 | 0 | 30 | 642 | 0 | 48 | 0 | 0 | 61 | 1071 | 90 | 0 | 0 | 0 | 1392 | 404 | 0 | 0 | 0 | 90 | 0 | 0 | 0 | 7212 | 389 | 3082 |
| SRR5029936 | 400 | 0 | 10203 | 0 | 0 | 129 | 140 | 259 | 101 | 0 | 0 | 24 | 0 | 14 | 180 | 0 | 48 | 0 | 0 | 60 | 1544 | 43 | 0 | 0 | 14 | 1811 | 1222 | 0 | 0 | 0 | 29 | 0 | 0 | 0 | 7273 | 460 | 4212 |
| SRR5029937 | 363 | 31 | 9141 | 0 | 0 | 78 | 97 | 110 | 78 | 0 | 0 | 39 | 0 | 16 | 537 | 0 | 39 | 0 | 0 | 54 | 1053 | 16 | 0 | 0 | 0 | 1535 | 298 | 16 | 0 | 0 | 47 | 0 | 0 | 0 | 6733 | 485 | 3367 |
| SRR5029938 | 317 | 0 | 9623 | 0 | 0 | 123 | 127 | 232 | 96 | 0 | 0 | 48 | 0 | 0 | 197 | 0 | 36 | 0 | 0 | 83 | 1356 | 82 | 0 | 0 | 0 | 86 | 50 | 14 | 0 | 0 | 0 | 0 | 0 | 14 | 320 | 16 | 188 |
| SRR5029939 | 304 | 61 | 9192 | 0 | 0 | 92 | 147 | 92 | 61 | 0 | 0 | 23 | 0 | 31 | 599 | 0 | 25 | 0 | 0 | 63 | 978 | 46 | 0 | 0 | 0 | 52 | 26 | 15 | 0 | 0 | 61 | 0 | 0 | 0 | 325 | 23 | 159 |
| SRR5029940 | 318 | 0 | 9914 | 0 | 0 | 140 | 129 | 323 | 70 | 0 | 0 | 34 | 0 | 14 | 200 | 0 | 38 | 0 | 0 | 90 | 1496 | 70 | 0 | 0 | 0 | 1770 | 1222 | 0 | 0 | 0 | 0 | 0 | 0 | 28 | 7992 | 534 | 4537 |
| SRR5029941 | 291 | 31 | 9236 | 0 | 0 | 0 | 75 | 125 | 63 | 0 | 0 | 39 | 0 | 47 | 651 | 0 | 43 | 0 | 0 | 32 | 1110 | 16 | 0 | 0 | 0 | 1111 | 516 | 16 | 0 | 0 | 31 | 0 | 0 | 0 | 6585 | 438 | 3582 |
| SRR5029942 | 924 | 0 | 12946 | 0 | 0 | 182 | 37 | 28 | 28 | 0 | 0 | 491 | 0 | 0 | 0 | 0 | 25 | 0 | 0 | 478 | 1323 | 0 | 0 | 0 | 0 | 2284 | 995 | 14 | 0 | 0 | 42 | 28 | 0 | 14 | 1864 | 98 | 799 |
| SRR5029943 | 832 | 11 | 11543 | 0 | 22 | 99 | 0 | 0 | 44 | 0 | 0 | 430 | 0 | 28 | 76 | 0 | 27 | 0 | 19 | 114 | 705 | 0 | 0 | 0 | 0 | 1376 | 248 | 44 | 0 | 0 | 44 | 28 | 0 | 0 | 2004 | 116 | 694 |
| SRR5029944 | 915 | 0 | 13300 | 31 | 15 | 16 | 40 | 15 | 15 | 0 | 0 | 485 | 0 | 0 | 0 | 0 | 22 | 0 | 0 | 440 | 1000 | 0 | 0 | 0 | 0 | 150 | 62 | 15 | 0 | 0 | 0 | 31 | 0 | 0 | 156 | 93 | 70 |
| SRR5029945 | 818 | 12 | 11710 | 0 | 0 | 92 | 20 | 0 | 52 | 0 | 0 | 409 | 0 | 0 | 59 | 0 | 33 | 0 | 0 | 157 | 786 | 0 | 0 | 0 | 0 | 71 | 13 | 29 | 12 | 0 | 69 | 23 | 0 | 0 | 108 | 109 | 37 |

### Supp Table S8.2.Columns_AM_to_BW

| | | | | | | | | | | | | | | | | | | | | | | | | | | | | | | | | | | | | | |
|---|---|---|---|---|---|---|---|---|---|---|---|---|---|---|---|---|---|---|---|---|---|---|---|---|---|---|---|---|---|---|---|---|---|---|---|---|---|
| SRR5029946 | 914 | 0 | 12794 | 0 | 0 | 13 | 38 | 29 | 29 | 0 | 0 | 487 | 0 | 0 | 0 | 0 | 30 | 0 | 0 | 345 | 1417 | 15 | 0 | 0 | 0 | 158 | 67 | 0 | 0 | 0 | 15 | 15 | 0 | 29 | 162 | 73 | 68 |
| SRR5029947 | 825 | 17 | 11323 | 0 | 28 | 123 | 0 | 0 | 67 | 0 | 0 | 422 | 0 | 11 | 96 | 0 | 33 | 0 | 0 | 172 | 592 | 0 | 0 | 0 | 0 | 1381 | 223 | 22 | 0 | 0 | 61 | 22 | 0 | 0 | 1866 | 67 | 757 |
| SRR5029948 | 904 | 0 | 12641 | 0 | 0 | 212 | 39 | 0 | 15 | 0 | 0 | 503 | 0 | 0 | 0 | 0 | 19 | 0 | 0 | 507 | 1015 | 15 | 0 | 0 | 0 | 1880 | 879 | 15 | 0 | 0 | 91 | 15 | 0 | 0 | 1592 | 45 | 698 |
| SRR5029949 | 834 | 0 | 11694 | 17 | 40 | 121 | 0 | 0 | 75 | 0 | 0 | 411 | 0 | 0 | 60 | 0 | 29 | 0 | 0 | 100 | 702 | 0 | 0 | 0 | 0 | 73 | 14 | 23 | 0 | 0 | 75 | 40 | 0 | 0 | 123 | 63 | 37 |
| SRR5029950 | 83 | 0 | 9857 | 0 | 0 | 140 | 27 | 14 | 28 | 0 | 0 | 80 | 0 | 19 | 80 | 0 | 16 | 0 | 27 | 517 | 1126 | 0 | 0 | 0 | 0 | 2997 | 909 | 0 | 14 | 0 | 103 | 14 | 0 | 0 | 6531 | 312 | 2657 |
| SRR5029951 | 119 | 0 | 9654 | 0 | 0 | 171 | 0 | 0 | 0 | 0 | 0 | 84 | 0 | 0 | 0 | 0 | 0 | 0 | 0 | 126 | 1260 | 0 | 0 | 0 | 0 | 3034 | 855 | 43 | 43 | 0 | 43 | 0 | 0 | 0 | 6708 | 555 | 2393 |
| SRR5029952 | 106 | 15 | 10149 | 10 | 0 | 110 | 29 | 0 | 20 | 0 | 0 | 58 | 0 | 10 | 158 | 0 | 18 | 0 | 14 | 575 | 1193 | 0 | 0 | 0 | 0 | 274 | 77 | 15 | 15 | 0 | 85 | 0 | 0 | 35 | 508 | 29 | 245 |
| SRR5029953 | 88 | 0 | 10250 | 0 | 0 | 352 | 0 | 0 | 0 | 0 | 0 | 73 | 0 | 0 | 0 | 0 | 0 | 0 | 129 | 129 | 516 | 0 | 0 | 0 | 0 | 2203 | 881 | 0 | 0 | 0 | 44 | 44 | 0 | 0 | 6079 | 308 | 3216 |
| SRR5029954 | 92 | 29 | 9844 | 0 | 0 | 148 | 14 | 14 | 24 | 0 | 0 | 88 | 0 | 34 | 82 | 0 | 21 | 0 | 14 | 590 | 1126 | 0 | 0 | 0 | 0 | 3304 | 1053 | 0 | 0 | 0 | 105 | 0 | 0 | 19 | 5985 | 263 | 3074 |
| SRR5029955 | 78 | 0 | 10132 | 0 | 0 | 262 | 127 | 0 | 0 | 0 | 0 | 36 | 0 | 0 | 127 | 0 | 0 | 0 | 0 | 127 | 765 | 0 | 0 | 0 | 0 | 3270 | 436 | 0 | 0 | 0 | 87 | 0 | 0 | 0 | 6191 | 654 | 2354 |
| SRR5029956 | 88 | 24 | 9970 | 20 | 0 | 11 | 28 | 0 | 29 | 0 | 0 | 75 | 0 | 0 | 56 | 0 | 23 | 0 | 28 | 361 | 1001 | 0 | 0 | 0 | 0 | 232 | 84 | 0 | 29 | 0 | 112 | 0 | 0 | 15 | 502 | 27 | 229 |
| SRR5029957 | 94 | 44 | 10586 | 0 | 0 | 177 | 0 | 0 | 0 | 0 | 0 | 65 | 0 | 44 | 130 | 0 | 29 | 0 | 130 | 648 | 518 | 0 | 0 | 0 | 0 | 3180 | 707 | 0 | 44 | 0 | 88 | 0 | 0 | 0 | 5388 | 221 | 2826 |
| SRR5029958 | 193 | 0 | 10527 | 0 | 0 | 29 | 20 | 29 | 0 | 0 | 0 | 31 | 0 | 0 | 20 | 0 | 35 | 0 | 0 | 670 | 1104 | 0 | 0 | 0 | 0 | 41 | 67 | 0 | 0 | 15 | 29 | 15 | 0 | 59 | 609 | 48 | 424 |
| SRR5029959 | 161 | 0 | 9080 | 0 | 16 | 16 | 16 | 0 | 0 | 0 | 0 | 28 | 0 | 0 | 32 | 0 | 27 | 0 | 0 | 112 | 529 | 0 | 0 | 0 | 0 | 25 | 12 | 25 | 0 | 0 | 33 | 0 | 0 | 41 | 503 | 34 | 267 |
| SRR5029960 | 184 | 16 | 10756 | 0 | 0 | 63 | 0 | 63 | 16 | 0 | 0 | 31 | 0 | 0 | 0 | 0 | 27 | 0 | 22 | 700 | 1246 | 0 | 0 | 0 | 0 | 916 | 1152 | 16 | 0 | 0 | 63 | 16 | 0 | 32 | 11256 | 789 | 7388 |
| SRR5029961 | 159 | 34 | 9451 | 0 | 0 | 17 | 0 | 0 | 17 | 0 | 0 | 18 | 0 | 0 | 151 | 0 | 27 | 0 | 0 | 100 | 536 | 0 | 0 | 0 | 0 | 627 | 297 | 0 | 0 | 0 | 42 | 17 | 0 | 34 | 11363 | 763 | 6304 |
| SRR5029962 | 159 | 0 | 10495 | 0 | 0 | 15 | 0 | 60 | 15 | 0 | 0 | 28 | 0 | 0 | 0 | 0 | 28 | 0 | 0 | 546 | 944 | 0 | 0 | 0 | 0 | 58 | 55 | 15 | 30 | 0 | 15 | 15 | 0 | 15 | 608 | 46 | 436 |
| SRR5029963 | 165 | 0 | 9273 | 0 | 0 | 0 | 0 | 17 | 0 | 0 | 0 | 23 | 0 | 0 | 49 | 0 | 33 | 0 | 16 | 163 | 505 | 0 | 0 | 0 | 0 | 26 | 15 | 0 | 17 | 0 | 0 | 0 | 0 | 17 | 531 | 34 | 277 |
| SRR5029964 | 193 | 0 | 10470 | 0 | 15 | 46 | 0 | 31 | 0 | 0 | 0 | 32 | 0 | 0 | 0 | 0 | 30 | 0 | 21 | 566 | 838 | 0 | 0 | 15 | 0 | 35 | 74 | 0 | 46 | 0 | 15 | 15 | 0 | 15 | 662 | 40 | 413 |
| SRR5029965 | 171 | 0 | 9398 | 0 | 0 | 17 | 17 | 0 | 0 | 0 | 0 | 24 | 0 | 0 | 101 | 0 | 28 | 0 | 0 | 168 | 522 | 0 | 0 | 0 | 0 | 509 | 297 | 0 | 17 | 0 | 25 | 0 | 0 | 17 | 11526 | 797 | 6043 |
| SRR5029966 | 214 | 11 | 11280 | 0 | 0 | 95 | 32 | 32 | 64 | 0 | 0 | 202 | 0 | 21 | 16 | 0 | 58 | 0 | 0 | 347 | 853 | 11 | 0 | 0 | 0 | 1262 | 1453 | 0 | 0 | 0 | 21 | 0 | 0 | 11 | 13781 | 976 | 4986 |
| SRR5029967 | 171 | 17 | 10288 | 0 | 0 | 50 | 0 | 0 | 22 | 0 | 0 | 139 | 0 | 17 | 95 | 0 | 42 | 0 | 0 | 106 | 423 | 0 | 0 | 0 | 0 | 46 | 21 | 0 | 0 | 0 | 72 | 0 | 0 | 11 | 784 | 47 | 249 |
| SRR5029968 | 202 | 12 | 11758 | 0 | 0 | 58 | 35 | 47 | 58 | 0 | 0 | 158 | 0 | 0 | 35 | 0 | 37 | 0 | 0 | 435 | 731 | 12 | 0 | 0 | 0 | 87 | 102 | 0 | 0 | 0 | 0 | 0 | 0 | 23 | 1053 | 65 | 404 |
| SRR5029969 | 189 | 17 | 10583 | 0 | 0 | 29 | 0 | 17 | 23 | 0 | 0 | 148 | 0 | 23 | 124 | 0 | 35 | 0 | 0 | 79 | 416 | 0 | 0 | 0 | 0 | 841 | 354 | 0 | 0 | 0 | 35 | 12 | 0 | 0 | 15138 | 946 | 4688 |
| SRR5029970 | 218 | 0 | 11632 | 0 | 0 | 88 | 17 | 44 | 44 | 0 | 0 | 207 | 0 | 11 | 17 | 0 | 38 | 0 | 50 | 315 | 880 | 0 | 0 | 0 | 0 | 81 | 91 | 0 | 0 | 0 | 11 | 0 | 0 | 0 | 1085 | 77 | 390 |
| SRR5029971 | 165 | 0 | 10159 | 0 | 0 | 56 | 0 | 0 | 40 | 0 | 0 | 145 | 0 | 11 | 132 | 0 | 38 | 0 | 22 | 88 | 482 | 11 | 0 | 0 | 0 | 796 | 390 | 0 | 0 | 0 | 28 | 11 | 0 | 0 | 15200 | 904 | 4615 |
| SRR5029972 | 217 | 0 | 11637 | 0 | 0 | 45 | 0 | 23 | 0 | 0 | 0 | 169 | 0 | 11 | 69 | 0 | 51 | 0 | 0 | 397 | 1052 | 0 | 0 | 0 | 0 | 1157 | 1384 | 23 | 0 | 0 | 34 | 11 | 0 | 0 | 14346 | 1009 | 5875 |
| SRR5029973 | 170 | 12 | 10523 | 0 | 0 | 35 | 0 | 0 | 41 | 0 | 0 | 158 | 0 | 18 | 88 | 0 | 40 | 0 | 0 | 66 | 516 | 0 | 0 | 0 | 0 | 40 | 20 | 0 | 0 | 0 | 29 | 0 | 0 | 0 | 807 | 46 | 244 |
| SRR5029974 | 462 | 0 | 13785 | 0 | 0 | 25 | 0 | 0 | 58 | 0 | 0 | 206 | 0 | 0 | 215 | 0 | 34 | 0 | 0 | 979 | 1313 | 0 | 0 | 0 | 0 | 107 | 107 | 0 | 0 | 0 | 25 | 0 | 0 | 0 | 502 | 45 | 285 |
| SRR5029975 | 420 | 0 | 12754 | 0 | 0 | 24 | 0 | 0 | 36 | 0 | 0 | 183 | 0 | 12 | 541 | 0 | 29 | 0 | 0 | 130 | 447 | 0 | 0 | 0 | 0 | 850 | 285 | 0 | 0 | 0 | 24 | 0 | 0 | 0 | 5576 | 446 | 2586 |
| SRR5029976 | 419 | 0 | 13444 | 0 | 0 | 54 | 0 | 0 | 0 | 0 | 0 | 188 | 0 | 0 | 308 | 0 | 25 | 0 | 26 | 1334 | 1283 | 0 | 0 | 0 | 0 | 120 | 89 | 0 | 0 | 0 | 27 | 0 | 0 | 0 | 523 | 55 | 327 |
| SRR5029977 | 417 | 0 | 12844 | 0 | 0 | 31 | 0 | 0 | 25 | 0 | 0 | 168 | 0 | 0 | 551 | 0 | 33 | 0 | 0 | 216 | 394 | 0 | 0 | 0 | 0 | 53 | 21 | 0 | 0 | 0 | 31 | 0 | 0 | 0 | 405 | 29 | 198 |
| SRR5029978 | 466 | 0 | 13293 | 0 | 0 | 43 | 0 | 0 | 103 | 0 | 0 | 206 | 0 | 0 | 122 | 0 | 51 | 0 | 0 | 1196 | 1098 | 0 | 0 | 0 | 0 | 1188 | 1214 | 0 | 0 | 0 | 34 | 0 | 0 | 0 | 5899 | 611 | 3462 |
| SRR5029979 | 391 | 0 | 12425 | 0 | 0 | 43 | 0 | 0 | 12 | 0 | 0 | 185 | 0 | 0 | 596 | 0 | 31 | 0 | 0 | 168 | 317 | 0 | 0 | 0 | 0 | 54 | 21 | 12 | 0 | 0 | 43 | 0 | 0 | 0 | 389 | 30 | 195 |
| SRR5029980 | 453 | 0 | 13847 | 0 | 0 | 35 | 0 | 0 | 70 | 0 | 0 | 212 | 0 | 0 | 322 | 0 | 27 | 0 | 25 | 941 | 1387 | 0 | 0 | 0 | 0 | 113 | 99 | 0 | 0 | 0 | 0 | 0 | 0 | 0 | 569 | 42 | 344 |
| SRR5029981 | 417 | 0 | 12983 | 0 | 0 | 19 | 0 | 0 | 43 | 0 | 0 | 217 | 0 | 12 | 496 | 0 | 32 | 0 | 0 | 267 | 381 | 0 | 0 | 0 | 12 | 695 | 298 | 0 | 0 | 0 | 31 | 0 | 0 | 0 | 5656 | 584 | 3036 |
| SRR5029982 | 100 | 0 | 14974 | 0 | 0 | 55 | 35 | 41 | 0 | 0 | 0 | 29 | 0 | 0 | 127 | 0 | 16 | 0 | 0 | 78 | 1259 | 0 | 0 | 14 | 0 | 760 | 718 | 0 | 0 | 0 | 14 | 0 | 0 | 0 | 5636 | 511 | 3453 |
| SRR5029983 | 116 | 0 | 15373 | 0 | 0 | 94 | 0 | 0 | 0 | 0 | 0 | 15 | 0 | 0 | 204 | 0 | 0 | 0 | 0 | 0 | 1022 | 0 | 0 | 0 | 0 | 31 | 23 | 0 | 0 | 0 | 0 | 0 | 0 | 0 | 145 | 10 | 104 |
| SRR5029984 | 104 | 0 | 15379 | 0 | 0 | 30 | 38 | 60 | 15 | 0 | 0 | 38 | 0 | 15 | 153 | 0 | 14 | 0 | 0 | 38 | 1450 | 15 | 0 | 0 | 0 | 839 | 629 | 0 | 0 | 0 | 30 | 0 | 0 | 0 | 5406 | 434 | 2755 |
| SRR5029986 | 91 | 0 | 14764 | 0 | 14 | 71 | 37 | 42 | 0 | 0 | 0 | 41 | 0 | 0 | 146 | 0 | 18 | 0 | 0 | 51 | 1633 | 28 | 0 | 0 | 0 | 18 | 19 | 0 | 0 | 0 | 28 | 0 | 0 | 0 | 158 | 11 | 75 |
| SRR5029988 | 87 | 0 | 14997 | 15 | 0 | 44 | 22 | 44 | 15 | 0 | 0 | 38 | 0 | 0 | 141 | 0 | 18 | 0 | 0 | 112 | 1287 | 15 | 0 | 0 | 0 | 847 | 701 | 0 | 15 | 0 | 15 | 0 | 0 | 0 | 5681 | 424 | 3126 |
| SRR5029990 | 116 | 0 | 12356 | 11 | 0 | 80 | 0 | 110 | 15 | 0 | 0 | 104 | 0 | 0 | 0 | 0 | 63 | 0 | 0 | 106 | 515 | 0 | 0 | 0 | 0 | 2359 | 647 | 0 | 329 | 22 | 37 | 0 | 0 | 124 | 2092 | 121 | 1368 |
| SRR5029991 | 84 | 0 | 11313 | 0 | 0 | 70 | 0 | 40 | 25 | 0 | 0 | 91 | 0 | 0 | 0 | 0 | 48 | 0 | 0 | 60 | 388 | 0 | 0 | 0 | 0 | 201 | 31 | 0 | 52 | 0 | 65 | 0 | 0 | 15 | 296 | 55 | 141 |
| SRR5029992 | 112 | 0 | 12777 | 0 | 0 | 68 | 19 | 84 | 16 | 0 | 0 | 125 | 0 | 0 | 0 | 0 | 21 | 0 | 0 | 190 | 362 | 20 | 0 | 0 | 0 | 2107 | 581 | 12 | 321 | 36 | 44 | 0 | 0 | 120 | 2127 | 116 | 1438 |
| SRR5029993 | 92 | 0 | 11168 | 0 | 0 | 58 | 0 | 26 | 11 | 0 | 0 | 116 | 0 | 0 | 32 | 0 | 43 | 0 | 0 | 158 | 412 | 0 | 0 | 0 | 0 | 198 | 30 | 11 | 59 | 0 | 53 | 0 | 0 | 58 | 284 | 11 | 142 |
| SRR5029994 | 116 | 0 | 12670 | 0 | 0 | 13 | 0 | 16 | 23 | 0 | 0 | 124 | 0 | 0 | 0 | 0 | 40 | 0 | 0 | 167 | 632 | 11 | 0 | 0 | 0 | 313 | 77 | 0 | 59 | 19 | 45 | 0 | 0 | 17 | 320 | 14 | 193 |
| SRR5029995 | 115 | 0 | 11215 | 0 | 0 | 61 | 0 | 36 | 15 | 0 | 0 | 90 | 0 | 10 | 0 | 0 | 32 | 0 | 0 | 122 | 336 | 0 | 0 | 0 | 15 | 207 | 37 | 0 | 60 | 0 | 17 | 0 | 0 | 13 | 284 | 16 | 166 |
| SRR5029996 | 92 | 0 | 12598 | 0 | 0 | 11 | 37 | 10 | 19 | 0 | 0 | 117 | 0 | 0 | 0 | 0 | 51 | 0 | 0 | 315 | 371 | 16 | 0 | 0 | 0 | 291 | 74 | 0 | 55 | 27 | 62 | 0 | 0 | 17 | 326 | 15 | 183 |
| SRR5029997 | 55 | 10 | 11265 | 0 | 0 | 42 | 0 | 37 | 31 | 0 | 0 | 107 | 0 | 0 | 0 | 0 | 38 | 0 | 0 | 161 | 290 | 16 | 0 | 0 | 0 | 1449 | 230 | 0 | 440 | 10 | 42 | 0 | 0 | 110 | 1936 | 84 | 1125 |
| SRR5029998 | 148 | 0 | 12115 | 0 | 0 | 115 | 17 | 0 | 71 | 0 | 0 | 232 | 0 | 0 | 17 | 0 | 39 | 58 | 0 | 275 | 4243 | 0 | 0 | 0 | 0 | 1941 | 1338 | 0 | 18 | 0 | 27 | 0 | 0 | 0 | 2854 | 186 | 2021 |
| SRR5029999 | 92 | 0 | 10948 | 11 | 0 | 48 | 18 | 0 | 37 | 0 | 0 | 207 | 0 | 11 | 24 | 0 | 39 | 204 | 0 | 132 | 2639 | 16 | 0 | 0 | 0 | 1260 | 566 | 37 | 0 | 0 | 37 | 11 | 0 | 11 | 3065 | 235 | 2067 |
| SRR5030000 | 115 | 0 | 12397 | 0 | 0 | 123 | 18 | 0 | 47 | 0 | 0 | 255 | 0 | 0 | 18 | 0 | 51 | 35 | 0 | 237 | 3805 | 0 | 0 | 0 | 0 | 1745 | 1254 | 0 | 0 | 0 | 0 | 0 | 0 | 0 | 2810 | 198 | 2263 |
| SRR5030001 | 103 | 11 | 11110 | 0 | 0 | 100 | 19 | 0 | 22 | 0 | 0 | 200 | 0 | 11 | 49 | 0 | 50 | 222 | 0 | 142 | 2527 | 0 | 0 | 0 | 11 | 1371 | 585 | 22 | 0 | 0 | 33 | 11 | 0 | 0 | 3255 | 268 | 1683 |
| SRR5030002 | 98 | 0 | 12350 | 0 | 0 | 14 | 18 | 18 | 54 | 0 | 0 | 246 | 0 | 0 | 18 | 0 | 36 | 73 | 0 | 247 | 3831 | 0 | 0 | 0 | 0 | 142 | 97 | 27 | 0 | 0 | 27 | 0 | 0 | 0 | 242 | 19 | 174 |
| SRR5030003 | 105 | 0 | 11093 | 0 | 0 | 60 | 18 | 0 | 38 | 0 | 0 | 217 | 0 | 0 | 74 | 0 | 38 | 259 | 0 | 80 | 2568 | 0 | 0 | 0 | 11 | 1459 | 544 | 33 | 11 | 0 | 71 | 16 | 0 | 0 | 3048 | 250 | 1758 |
| SRR5030004 | 94 | 0 | 12247 | 0 | 0 | 12 | 18 | 0 | 37 | 0 | 0 | 232 | 0 | 0 | 18 | 0 | 38 | 54 | 27 | 222 | 4285 | 0 | 0 | 0 | 0 | 121 | 79 | 0 | 0 | 0 | 0 | 0 | 0 | 37 | 243 | 28 | 194 |
| SRR5030005 | 76 | 11 | 11359 | 0 | 0 | 89 | 37 | 0 | 33 | 0 | 0 | 205 | 0 | 11 | 56 | 0 | 39 | 250 | 0 | 99 | 2636 | 0 | 0 | 0 | 11 | 94 | 30 | 0 | 0 | 0 | 50 | 11 | 0 | 0 | 205 | 22 | 125 |
| SRR5030006 | 238 | 0 | 14691 | 0 | 0 | 24 | 0 | 0 | 117 | 0 | 0 | 196 | 0 | 0 | 107 | 0 | 28 | 0 | 13 | 615 | 1323 | 0 | 0 | 0 | 0 | 344 | 92 | 0 | 0 | 0 | 59 | 0 | 0 | 0 | 703 | 52 | 338 |
| SRR5030008 | 236 | 0 | 15143 | 0 | 18 | 22 | 0 | 0 | 108 | 0 | 0 | 191 | 0 | 0 | 93 | 0 | 29 | 0 | 0 | 496 | 1626 | 0 | 0 | 0 | 0 | 321 | 102 | 0 | 0 | 0 | 27 | 0 | 0 | 0 | 682 | 49 | 307 |
| SRR5030009 | 181 | 0 | 14008 | 0 | 0 | 23 | 0 | 0 | 157 | 0 | 0 | 236 | 0 | 0 | 280 | 0 | 17 | 0 | 0 | 421 | 1122 | 0 | 0 | 0 | 0 | 228 | 48 | 0 | 0 | 0 | 79 | 0 | 0 | 0 | 587 | 40 | 231 |
| SRR5030010 | 234 | 0 | 14759 | 0 | 0 | 567 | 29 | 0 | 112 | 0 | 0 | 181 | 0 | 0 | 73 | 0 | 26 | 0 | 0 | 513 | 1114 | 0 | 0 | 0 | 0 | 7751 | 2672 | 0 | 0 | 0 | 77 | 0 | 0 | 0 | 16524 | 1298 | 7562 |
| SRR5030011 | 219 | 0 | 13511 | 0 | 0 | 22 | 0 | 0 | 155 | 0 | 0 | 174 | 0 | 0 | 0 | 0 | 12 | 0 | 0 | 127 | 1393 | 0 | 0 | 0 | 0 | 263 | 50 | 0 | 0 | 0 | 77 | 0 | 0 | 0 | 567 | 30 | 246 |
| SRR5030012 | 233 | 0 | 14881 | 0 | 0 | 535 | 0 | 0 | 132 | 0 | 0 | 198 | 0 | 0 | 91 | 0 | 27 | 0 | 0 | 438 | 1541 | 0 | 0 | 0 | 0 | 7567 | 2359 | 0 | 0 | 0 | 96 | 0 | 0 | 0 | 16475 | 1403 | 7549 |
| SRR5030014 | 133 | 0 | 11417 | 0 | 0 | 372 | 11 | 0 | 152 | 0 | 0 | 124 | 0 | 0 | 163 | 0 | 55 | 0 | 0 | 207 | 2166 | 55 | 0 | 14 | 0 | 7053 | 5083 | 0 | 14 | 0 | 124 | 0 | 0 | 0 | 5125 | 730 | 4767 |
| SRR5030015 | 118 | 0 | 10134 | 14 | 0 | 123 | 0 | 0 | 41 | 0 | 0 | 101 | 0 | 0 | 681 | 0 | 30 | 0 | 0 | 83 | 1098 | 0 | 0 | 0 | 0 | 4604 | 1489 | 0 | 14 | 0 | 287 | 0 | 0 | 27 | 4673 | 533 | 4071 |
| SRR5030016 | 133 | 0 | 11690 | 0 | 0 | 243 | 24 | 0 | 61 | 0 | 0 | 122 | 0 | 0 | 167 | 0 | 32 | 0 | 0 | 227 | 1889 | 15 | 0 | 0 | 0 | 7401 | 5045 | 0 | 0 | 0 | 122 | 0 | 0 | 0 | 5045 | 547 | 4893 |
| SRR5030017 | 108 | 0 | 10378 | 0 | 0 | 215 | 0 | 0 | 100 | 0 | 0 | 107 | 0 | 0 | 580 | 0 | 37 | 15 | 0 | 43 | 1000 | 29 | 0 | 0 | 0 | 165 | 50 | 0 | 0 | 0 | 258 | 0 | 0 | 0 | 184 | 18 | 160 |
| SRR5030018 | 164 | 0 | 11503 | 0 | 0 | 14 | 0 | 29 | 57 | 0 | 0 | 145 | 0 | 0 | 146 | 0 | 27 | 0 | 0 | 236 | 2161 | 29 | 0 | 0 | 0 | 333 | 249 | 0 | 0 | 0 | 12 | 0 | 0 | 0 | 231 | 35 | 221 |
| SRR5030019 | 106 | 0 | 9962 | 0 | 42 | 195 | 14 | 0 | 111 | 0 | 0 | 112 | 0 | 0 | 683 | 0 | 33 | 0 | 0 | 28 | 1110 | 14 | 0 | 0 | 0 | 5068 | 1420 | 0 | 0 | 0 | 223 | 0 | 0 | 0 | 5276 | 473 | 3912 |
| SRR5030020 | 141 | 0 | 11519 | 0 | 0 | 18 | 12 | 0 | 103 | 0 | 0 | 126 | 0 | 15 | 220 | 0 | 48 | 0 | 0 | 232 | 2146 | 0 | 0 | 0 | 0 | 339 | 255 | 0 | 15 | 0 | 162 | 0 | 0 | 0 | 247 | 30 | 230 |
| SRR5030021 | 123 | 0 | 10172 | 0 | 0 | 187 | 0 | 0 | 115 | 0 | 0 | 97 | 0 | 0 | 849 | 0 | 43 | 0 | 0 | 15 | 1172 | 43 | 0 | 0 | 14 | 165 | 55 | 29 | 14 | 0 | 10 | 0 | 0 | 0 | 192 | 22 | 151 |
| SRR5030022 | 148 | 12 | 15968 | 0 | 0 | 23 | 0 | 12 | 73 | 0 | 0 | 210 | 0 | 12 | 2533 | 0 | 22 | 0 | 0 | 719 | 6264 | 0 | 0 | 0 | 0 | 483 | 169 | 0 | 0 | 0 | 11 | 0 | 0 | 12 | 66 | 109 | 35 |
| SRR5030023 | 174 | 0 | 16332 | 0 | 22 | 13 | 0 | 22 | 134 | 0 | 0 | 253 | 0 | 0 | 3576 | 0 | 25 | 0 | 0 | 337 | 4115 | 0 | 0 | 0 | 0 | 386 | 68 | 22 | 0 | 0 | 11 | 0 | 0 | 22 | 56 | 134 | 23 |
| SRR5030024 | 148 | 13 | 16354 | 13 | 0 | 17 | 37 | 26 | 65 | 0 | 0 | 234 | 0 | 0 | 2257 | 0 | 20 | 0 | 74 | 666 | 7179 | 0 | 0 | 0 | 0 | 456 | 158 | 39 | 0 | 0 | 12 | 0 | 0 | 0 | 58 | 104 | 32 |
| SRR5030025 | 166 | 0 | 16197 | 23 | 0 | 465 | 0 | 0 | 23 | 0 | 0 | 237 | 0 | 0 | 4273 | 0 | 18 | 0 | 0 | 678 | 4002 | 0 | 0 | 0 | 0 | 10186 | 1953 | 0 | 0 | 0 | 419 | 0 | 0 | 0 | 1628 | 47 | 930 |
| SRR5030026 | 152 | 0 | 16199 | 12 | 12 | 771 | 0 | 0 | 37 | 0 | 0 | 239 | 0 | 12 | 2218 | 0 | 22 | 0 | 35 | 669 | 5667 | 0 | 0 | 0 | 0 | 11663 | 3494 | 12 | 0 | 0 | 224 | 0 | 0 | 0 | 1343 | 87 | 758 |
| SRR5030027 | 160 | 0 | 16162 | 0 | 0 | 428 | 0 | 0 | 90 | 0 | 0 | 219 | 0 | 0 | 2972 | 0 | 12 | 0 | 68 | 270 | 4391 | 0 | 0 | 0 | 0 | 9902 | 2543 | 0 | 0 | 0 | 450 | 0 | 0 | 0 | 1620 | 158 | 810 |
| SRR5030028 | 137 | 13 | 16602 | 13 | 0 | 21 | 0 | 0 | 38 | 0 | 0 | 261 | 0 | 0 | 2902 | 0 | 14 | 0 | 37 | 478 | 6319 | 0 | 0 | 0 | 0 | 462 | 162 | 0 | 0 | 0 | 189 | 0 | 0 | 0 | 57 | 114 | 39 |
| SRR5030030 | 210 | 11 | 17992 | 0 | 0 | 67 | 0 | 0 | 67 | 0 | 0 | 54 | 0 | 0 | 0 | 0 | 28 | 0 | 0 | 316 | 1199 | 0 | 0 | 0 | 11 | 2070 | 2036 | 79 | 11 | 0 | 67 | 0 | 0 | 0 | 5883 | 416 | 4848 |
| SRR5030031 | 178 | 0 | 16313 | 12 | 0 | 12 | 0 | 0 | 23 | 0 | 0 | 60 | 0 | 0 | 84 | 12 | 25 | 0 | 0 | 42 | 674 | 0 | 0 | 0 | 0 | 1309 | 347 | 93 | 0 | 0 | 46 | 0 | 0 | 12 | 5836 | 475 | 3613 |
| SRR5030032 | 180 | 0 | 18520 | 0 | 12 | 73 | 0 | 12 | 24 | 0 | 0 | 63 | 0 | 12 | 28 | 0 | 18 | 0 | 0 | 324 | 1354 | 36 | 0 | 0 | 0 | 2107 | 1913 | 73 | 0 | 0 | 36 | 0 | 0 | 0 | 5704 | 521 | 4796 |
| SRR5030033 | 172 | 0 | 16783 | 12 | 0 | 24 | 0 | 12 | 36 | 0 | 0 | 55 | 0 | 0 | 90 | 0 | 21 | 0 | 0 | 67 | 560 | 0 | 0 | 0 | 0 | 1386 | 458 | 96 | 0 | 0 | 96 | 0 | 0 | 0 | 6121 | 301 | 3820 |
| SRR5030034 | 227 | 0 | 18313 | 12 | 0 | 92 | 14 | 12 | 69 | 0 | 0 | 47 | 0 | 0 | 0 | 0 | 31 | 0 | 0 | 246 | 1298 | 0 | 0 | 0 | 0 | 104 | 107 | 69 | 12 | 0 | 46 | 0 | 0 | 0 | 356 | 31 | 261 |
| SRR5030035 | 173 | 12 | 16375 | 35 | 12 | 47 | 0 | 0 | 24 | 0 | 0 | 50 | 0 | 0 | 21 | 0 | 24 | 0 | 0 | 64 | 678 | 12 | 0 | 0 | 0 | 1133 | 519 | 94 | 0 | 0 | 35 | 0 | 0 | 0 | 5583 | 295 | 3671 |
| SRR5030036 | 190 | 0 | 18488 | 24 | 12 | 24 | 14 | 0 | 24 | 0 | 0 | 58 | 0 | 0 | 28 | 0 | 21 | 0 | 0 | 279 | 1034 | 12 | 0 | 0 | 0 | 1929 | 1729 | 59 | 12 | 0 | 35 | 0 | 0 | 0 | 5280 | 447 | 5033 |
| SRR5030038 | 201 | 0 | 8945 | 0 | 0 | 76 | 0 | 0 | 13 | 0 | 0 | 301 | 0 | 25 | 82 | 0 | 30 | 0 | 0 | 177 | 1157 | 0 | 0 | 0 | 0 | 1206 | 889 | 13 | 0 | 0 | 89 | 0 | 0 | 0 | 14323 | 775 | 4825 |
| SRR5030039 | 199 | 0 | 9092 | 0 | 0 | 47 | 0 | 0 | 71 | 0 | 0 | 274 | 0 | 0 | 174 | 0 | 36 | 0 | 0 | 58 | 581 | 0 | 0 | 0 | 0 | 39 | 17 | 0 | 0 | 0 | 118 | 0 | 0 | 0 | 543 | 23 | 152 |
| SRR5030040 | 215 | 0 | 9313 | 0 | 0 | 94 | 0 | 0 | 13 | 0 | 0 | 284 | 0 | 13 | 102 | 0 | 31 | 0 | 0 | 319 | 1002 | 13 | 0 | 0 | 0 | 41 | 33 | 13 | 13 | 0 | 81 | 0 | 0 | 0 | 552 | 28 | 177 |
| SRR5030041 | 208 | 0 | 9628 | 0 | 0 | 73 | 0 | 0 | 24 | 0 | 0 | 288 | 0 | 0 | 59 | 0 | 33 | 0 | 0 | 235 | 499 | 0 | 0 | 0 | 0 | 1023 | 414 | 0 | 0 | 0 | 219 | 0 | 0 | 0 | 15727 | 803 | 4041 |
| SRR5030042 | 216 | 0 | 9033 | 13 | 0 | 116 | 0 | 0 | 0 | 0 | 0 | 260 | 0 | 0 | 42 | 13 | 29 | 0 | 0 | 250 | 1069 | 13 | 0 | 0 | 0 | 1448 | 698 | 13 | 0 | 0 | 65 | 0 | 0 | 13 | 13913 | 595 | 4797 |
| SRR5030043 | 228 | 0 | 9005 | 0 | 0 | 95 | 0 | 0 | 48 | 0 | 0 | 313 | 0 | 0 | 144 | 0 | 35 | 0 | 0 | 29 | 692 | 0 | 0 | 0 | 0 | 953 | 500 | 0 | 24 | 0 | 95 | 0 | 0 | 0 | 14749 | 739 | 4575 |
| SRR5030044 | 201 | 0 | 8901 | 13 | 0 | 79 | 0 | 0 | 105 | 0 | 0 | 297 | 0 | 53 | 87 | 0 | 46 | 17 | 0 | 219 | 932 | 0 | 0 | 0 | 0 | 1157 | 1026 | 0 | 13 | 0 | 66 | 0 | 0 | 13 | 14770 | 671 | 4722 |
| SRR5030045 | 213 | 0 | 9186 | 0 | 0 | 24 | 0 | 0 | 24 | 0 | 0 | 298 | 0 | 49 | 236 | 0 | 33 | 0 | 0 | 89 | 768 | 24 | 0 | 0 | 0 | 1073 | 561 | 0 | 24 | 0 | 122 | 0 | 0 | 0 | 14510 | 561 | 4585 |
| SRR5030046 | 248 | 0 | 7573 | 30 | 0 | 23 | 14 | 59 | 141 | 0 | 0 | 50 | 0 | 0 | 14 | 0 | 38 | 0 | 0 | 373 | 690 | 0 | 0 | 0 | 0 | 515 | 67 | 0 | 0 | 0 | 11 | 0 | 0 | 52 | 324 | 12 | 160 |
| SRR5030048 | 248 | 0 | 7753 | 16 | 0 | 434 | 0 | 80 | 177 | 0 | 0 | 44 | 0 | 0 | 0 | 0 | 36 | 0 | 0 | 370 | 785 | 16 | 0 | 0 | 0 | 9099 | 1569 | 0 | 0 | 0 | 298 | 24 | 0 | 32 | 5519 | 193 | 2623 |
| SRR5030050 | 222 | 0 | 7633 | 0 | 15 | 472 | 0 | 69 | 107 | 0 | 0 | 41 | 0 | 0 | 14 | 0 | 38 | 0 | 0 | 492 | 605 | 0 | 0 | 0 | 0 | 9031 | 1357 | 15 | 0 | 0 | 152 | 0 | 0 | 15 | 5548 | 251 | 2774 |
| SRR5030052 | 220 | 0 | 7507 | 0 | 0 | 402 | 14 | 79 | 220 | 0 | 0 | 39 | 0 | 0 | 29 | 0 | 35 | 0 | 0 | 343 | 515 | 0 | 0 | 0 | 0 | 9039 | 1283 | 0 | 0 | 0 | 283 | 0 | 0 | 31 | 5543 | 276 | 2929 |
| SRR5030053 | 225 | 0 | 6969 | 0 | 0 | 593 | 0 | 0 | 137 | 0 | 0 | 46 | 0 | 0 | 0 | 0 | 15 | 0 | 0 | 91 | 366 | 0 | 0 | 0 | 0 | 6528 | 548 | 46 | 0 | 0 | 137 | 0 | 0 | 91 | 5341 | 137 | 2922 |
| SRR5030054 | 146 | 0 | 17078 | 0 | 0 | 102 | 39 | 0 | 29 | 0 | 0 | 22 | 0 | 0 | 78 | 0 | 40 | 0 | 0 | 94 | 404 | 44 | 0 | 0 | 0 | 2432 | 3262 | 0 | 0 | 0 | 15 | 0 | 0 | 15 | 10514 | 714 | 6510 |
| SRR5030055 | 117 | 0 | 15484 | 0 | 0 | 0 | 21 | 0 | 0 | 0 | 0 | 26 | 0 | 48 | 192 | 0 | 49 | 0 | 0 | 85 | 449 | 0 | 0 | 0 | 0 | 48 | 38 | 48 | 0 | 0 | 48 | 0 | 0 | 0 | 247 | 19 | 132 |
| SRR5030056 | 143 | 16 | 17361 | 0 | 0 | 63 | 54 | 47 | 31 | 0 | 0 | 29 | 0 | 0 | 54 | 0 | 65 | 0 | 0 | 71 | 363 | 16 | 0 | 0 | 0 | 2368 | 3308 | 16 | 16 | 0 | 31 | 0 | 0 | 31 | 10568 | 674 | 6711 |
| SRR5030057 | 118 | 0 | 15868 | 0 | 0 | 0 | 22 | 0 | 0 | 0 | 0 | 25 | 0 | 0 | 133 | 0 | 69 | 0 | 0 | 89 | 378 | 51 | 0 | 0 | 0 | 1480 | 1377 | 0 | 0 | 0 | 153 | 0 | 0 | 0 | 10408 | 969 | 6275 |
| SRR5030058 | 154 | 0 | 16888 | 15 | 0 | 90 | 63 | 15 | 0 | 0 | 0 | 23 | 0 | 15 | 51 | 0 | 55 | 0 | 0 | 103 | 439 | 15 | 0 | 0 | 0 | 64 | 93 | 15 | 0 | 0 | 30 | 0 | 0 | 15 | 294 | 25 | 208 |
| SRR5030059 | 129 | 0 | 15608 | 0 | 0 | 49 | 22 | 49 | 0 | 0 | 0 | 19 | 0 | 0 | 174 | 0 | 60 | 0 | 0 | 22 | 326 | 0 | 0 | 0 | 0 | 52 | 31 | 0 | 0 | 0 | 0 | 0 | 0 | 0 | 300 | 22 | 162 |
| SRR5030060 | 147 | 0 | 17333 | 0 | 31 | 77 | 23 | 31 | 15 | 0 | 0 | 19 | 0 | 0 | 52 | 0 | 65 | 0 | 0 | 98 | 422 | 15 | 0 | 0 | 0 | 68 | 88 | 15 | 0 | 0 | 15 | 15 | 0 | 15 | 298 | 25 | 192 |
| SRR5030061 | 147 | 0 | 16379 | 0 | 0 | 0 | 45 | 0 | 0 | 0 | 0 | 12 | 0 | 0 | 90 | 0 | 49 | 0 | 0 | 22 | 314 | 0 | 0 | 0 | 0 | 40 | 43 | 0 | 0 | 0 | 0 | 0 | 0 | 0 | 283 | 27 | 167 |
| SRR5030062 | 253 | 26 | 16525 | 0 | 0 | 156 | 0 | 0 | 91 | 0 | 0 | 33 | 0 | 13 | 236 | 0 | 12 | 15 | 0 | 253 | 1096 | 0 | 0 | 0 | 0 | 84 | 59 | 39 | 0 | 0 | 26 | 0 | 0 | 0 | 104 | 104 | 24 |
| SRR5030064 | 231 | 0 | 17106 | 0 | 0 | 56 | 0 | 0 | 126 | 0 | 0 | 38 | 0 | 0 | 362 | 0 | 17 | 0 | 0 | 217 | 1394 | 14 | 0 | 0 | 0 | 2273 | 1757 | 0 | 0 | 0 | 28 | 0 | 0 | 0 | 2859 | 56 | 683 |
| SRR5030066 | 247 | 0 | 16648 | 0 | 0 | 147 | 0 | 0 | 120 | 0 | 0 | 33 | 0 | 0 | 224 | 0 | 20 | 0 | 0 | 259 | 1208 | 0 | 0 | 0 | 0 | 2085 | 1751 | 0 | 0 | 0 | 13 | 0 | 0 | 0 | 2994 | 67 | 936 |
| SRR5030068 | 223 | 0 | 16798 | 0 | 0 | 67 | 0 | 0 | 188 | 0 | 0 | 34 | 0 | 0 | 122 | 0 | 11 | 0 | 0 | 157 | 1343 | 0 | 0 | 0 | 0 | 1991 | 1655 | 13 | 0 | 0 | 27 | 0 | 0 | 13 | 3188 | 135 | 753 |
| SRR5030069 | 287 | 0 | 16700 | 0 | 0 | 125 | 0 | 0 | 0 | 0 | 0 | 25 | 0 | 0 | 639 | 0 | 0 | 0 | 0 | 0 | 1186 | 0 | 0 | 0 | 0 | 1938 | 875 | 0 | 0 | 0 | 125 | 0 | 0 | 0 | 2000 | 125 | 563 |
| SRR5030070 | 423 | 0 | 3553 | 0 | 0 | 209 | 0 | 0 | 129 | 0 | 0 | 79 | 0 | 0 | 600 | 0 | 37 | 0 | 0 | 378 | 2021 | 0 | 0 | 0 | 0 | 4883 | 2367 | 0 | 0 | 0 | 119 | 0 | 0 | 0 | 1999 | 259 | 2188 |
| SRR5030071 | 414 | 0 | 3316 | 0 | 0 | 123 | 30 | 21 | 82 | 0 | 0 | 65 | 0 | 21 | 1091 | 0 | 36 | 0 | 0 | 182 | 1758 | 0 | 0 | 0 | 0 | 3740 | 1356 | 0 | 0 | 0 | 185 | 0 | 0 | 21 | 2404 | 288 | 2178 |
| SRR5030072 | 413 | 0 | 3584 | 0 | 0 | 311 | 0 | 0 | 97 | 0 | 0 | 76 | 0 | 0 | 439 | 0 | 44 | 0 | 14 | 425 | 2479 | 0 | 0 | 0 | 0 | 4876 | 2534 | 11 | 11 | 0 | 86 | 0 | 0 | 0 | 2008 | 290 | 2116 |
| SRR5030073 | 406 | 0 | 3342 | 0 | 0 | 212 | 0 | 0 | 106 | 0 | 0 | 84 | 0 | 0 | 1058 | 0 | 34 | 0 | 0 | 96 | 1987 | 0 | 0 | 0 | 0 | 141 | 56 | 0 | 0 | 0 | 191 | 0 | 0 | 0 | 95 | 13 | 73 |
| SRR5030074 | 425 | 0 | 3483 | 20 | 0 | 15 | 14 | 0 | 174 | 0 | 0 | 91 | 0 | 10 | 449 | 0 | 43 | 0 | 0 | 476 | 2668 | 0 | 0 | 0 | 0 | 253 | 105 | 0 | 0 | 0 | 92 | 0 | 0 | 0 | 103 | 18 | 103 |
| SRR5030076 | 489 | 0 | 3579 | 10 | 10 | 271 | 0 | 21 | 167 | 0 | 0 | 89 | 0 | 10 | 576 | 0 | 36 | 0 | 0 | 316 | 2963 | 0 | 0 | 0 | 0 | 4376 | 2219 | 10 | 0 | 0 | 63 | 0 | 0 | 0 | 2282 | 417 | 2282 |
| SRR5030077 | 347 | 0 | 3333 | 21 | 21 | 233 | 0 | 0 | 85 | 0 | 0 | 78 | 0 | 42 | 1260 | 0 | 29 | 0 | 0 | 158 | 1732 | 0 | 0 | 0 | 0 | 148 | 48 | 0 | 0 | 0 | 127 | 0 | 0 | 0 | 85 | 170 | 93 |
| SRR5030078 | 104 | 0 | 15883 | 0 | 11 | 180 | 40 | 42 | 74 | 0 | 0 | 246 | 0 | 0 | 0 | 0 | 44 | 0 | 100 | 220 | 0 | 0 | 0 | 0 | 0 | 138 | 61 | 0 | 21 | 11 | 21 | 0 | 0 | 63 | 110 | 63 | 71 |
| SRR5030079 | 81 | 0 | 14848 | 0 | 17 | 240 | 0 | 0 | 52 | 0 | 0 | 241 | 0 | 17 | 40 | 0 | 44 | 0 | 119 | 119 | 0 | 0 | 0 | 0 | 0 | 2146 | 429 | 0 | 0 | 0 | 0 | 0 | 0 | 0 | 2146 | 172 | 1271 |
| SRR5030080 | 91 | 0 | 16350 | 0 | 0 | 160 | 43 | 0 | 114 | 0 | 0 | 262 | 0 | 0 | 0 | 0 | 44 | 0 | 86 | 215 | 0 | 0 | 0 | 0 | 0 | 3079 | 1305 | 11 | 11 | 11 | 34 | 11 | 0 | 34 | 2472 | 114 | 1431 |
| SRR5030081 | 84 | 0 | 15371 | 36 | 18 | 161 | 41 | 0 | 72 | 0 | 0 | 245 | 0 | 0 | 41 | 0 | 39 | 0 | 41 | 41 | 0 | 0 | 0 | 0 | 0 | 88 | 24 | 0 | 36 | 0 | 72 | 0 | 0 | 18 | 97 | 125 | 59 |
| SRR5030082 | 109 | 0 | 15974 | 0 | 0 | 164 | 20 | 44 | 44 | 0 | 0 | 248 | 0 | 0 | 0 | 0 | 43 | 0 | 41 | 245 | 0 | 0 | 0 | 0 | 11 | 120 | 68 | 11 | 22 | 0 | 33 | 33 | 0 | 33 | 107 | 131 | 74 |
| SRR5030083 | 70 | 0 | 14617 | 35 | 18 | 105 | 0 | 18 | 53 | 0 | 0 | 238 | 0 | 18 | 0 | 0 | 29 | 0 | 0 | 200 | 0 | 0 | 0 | 0 | 35 | 80 | 21 | 0 | 53 | 0 | 35 | 18 | 0 | 35 | 92 | 70 | 40 |

### Supp Table S8.2.Columns_AM_to_BW

| | | | | | | | | | | | | | | | | | | | | | | | | | | | | | | | | | | | | | |
|---|---|---|---|---|---|---|---|---|---|---|---|---|---|---|---|---|---|---|---|---|---|---|---|---|---|---|---|---|---|---|---|---|---|---|---|---|---|
| SRR5030084 | 112 | 0 | 16185 | 0 | 0 | 191 | 21 | 22 | 22 | 0 | 0 | 247 | 0 | 0 | 0 | 0 | 52 | 0 | 62 | 247 | 0 | 0 | 0 | 0 | 11 | 130 | 61 | 0 | 11 | 0 | 56 | 11 | 0 | 11 | 104 | 168 | 85 |
| SRR5030085 | 71 | 0 | 14970 | 0 | 18 | 178 | 0 | 36 | 36 | 0 | 0 | 238 | 0 | 0 | 0 | 0 | 41 | 0 | 0 | 160 | 0 | 0 | 0 | 0 | 0 | 95 | 29 | 0 | 36 | 0 | 53 | 0 | 0 | 36 | 94 | 125 | 55 |
| SRR5030086 | 115 | 0 | 13331 | 25 | 0 | 64 | 48 | 20 | 25 | 0 | 0 | 186 | 0 | 35 | 48 | 0 | 25 | 17 | 24 | 289 | 674 | 0 | 0 | 0 | 0 | 866 | 297 | 0 | 0 | 0 | 30 | 0 | 0 | 0 | 1837 | 124 | 931 |
| SRR5030087 | 139 | 0 | 13258 | 0 | 34 | 43 | 0 | 0 | 0 | 0 | 0 | 161 | 0 | 34 | 122 | 0 | 23 | 15 | 0 | 245 | 653 | 0 | 0 | 0 | 0 | 792 | 145 | 0 | 17 | 0 | 43 | 0 | 0 | 0 | 2232 | 153 | 1048 |
| SRR5030088 | 105 | 10 | 13786 | 16 | 10 | 42 | 0 | 0 | 26 | 0 | 0 | 152 | 0 | 16 | 51 | 0 | 25 | 0 | 0 | 257 | 1054 | 0 | 0 | 0 | 0 | 104 | 34 | 0 | 16 | 0 | 26 | 0 | 0 | 0 | 202 | 10 | 95 |
| SRR5030090 | 149 | 15 | 13504 | 15 | 25 | 35 | 0 | 20 | 20 | 0 | 0 | 168 | 0 | 0 | 0 | 0 | 17 | 18 | 0 | 469 | 1112 | 0 | 0 | 0 | 0 | 932 | 216 | 0 | 10 | 0 | 40 | 0 | 0 | 10 | 1810 | 85 | 917 |
| SRR5030091 | 154 | 0 | 13485 | 17 | 0 | 52 | 0 | 17 | 43 | 0 | 0 | 201 | 0 | 0 | 0 | 0 | 26 | 0 | 0 | 451 | 820 | 0 | 0 | 0 | 0 | 66 | 14 | 0 | 0 | 0 | 26 | 17 | 0 | 17 | 204 | 10 | 89 |
| SRR5030092 | 115 | 10 | 13199 | 25 | 10 | 31 | 25 | 25 | 20 | 0 | 0 | 162 | 0 | 10 | 50 | 0 | 28 | 18 | 0 | 477 | 879 | 0 | 0 | 0 | 0 | 99 | 24 | 0 | 10 | 0 | 10 | 0 | 0 | 20 | 205 | 16 | 94 |
| SRR5030093 | 144 | 27 | 13737 | 53 | 18 | 35 | 0 | 0 | 35 | 0 | 0 | 182 | 0 | 18 | 84 | 0 | 26 | 15 | 0 | 167 | 753 | 0 | 0 | 0 | 0 | 742 | 221 | 0 | 0 | 0 | 35 | 0 | 0 | 0 | 2554 | 124 | 866 |
| SRR5030094 | 144 | 0 | 15165 | 31 | 12 | 32 | 0 | 0 | 0 | 0 | 0 | 619 | 0 | 0 | 0 | 0 | 54 | 13 | 12 | 373 | 1154 | 0 | 0 | 0 | 0 | 463 | 295 | 0 | 0 | 0 | 20 | 0 | 0 | 0 | 1246 | 45 | 296 |
| SRR5030095 | 131 | 0 | 13436 | 0 | 0 | 0 | 0 | 0 | 41 | 0 | 0 | 587 | 0 | 0 | 0 | 0 | 44 | 0 | 0 | 149 | 891 | 0 | 0 | 0 | 21 | 63 | 50 | 0 | 0 | 0 | 13 | 21 | 0 | 0 | 255 | 41 | 67 |
| SRR5030096 | 152 | 0 | 15160 | 32 | 12 | 15 | 13 | 0 | 10 | 0 | 0 | 597 | 0 | 0 | 13 | 0 | 53 | 0 | 13 | 363 | 1101 | 0 | 0 | 0 | 0 | 102 | 89 | 0 | 0 | 0 | 20 | 0 | 0 | 0 | 323 | 13 | 95 |
| SRR5030097 | 129 | 0 | 13584 | 0 | 21 | 0 | 0 | 0 | 21 | 0 | 0 | 450 | 0 | 0 | 0 | 21 | 28 | 0 | 0 | 302 | 302 | 0 | 0 | 0 | 0 | 426 | 234 | 0 | 21 | 0 | 43 | 0 | 0 | 0 | 1597 | 64 | 447 |
| SRR5030098 | 125 | 0 | 15049 | 32 | 11 | 27 | 0 | 0 | 11 | 0 | 0 | 642 | 0 | 0 | 12 | 0 | 42 | 0 | 0 | 413 | 947 | 0 | 0 | 0 | 0 | 377 | 306 | 0 | 0 | 0 | 17 | 0 | 0 | 0 | 1254 | 63 | 331 |
| SRR5030100 | 151 | 0 | 15378 | 24 | 11 | 13 | 25 | 0 | 0 | 0 | 0 | 638 | 0 | 0 | 13 | 0 | 46 | 0 | 0 | 463 | 876 | 0 | 0 | 0 | 0 | 129 | 75 | 0 | 0 | 0 | 19 | 0 | 0 | 0 | 360 | 13 | 92 |
| SRR5030101 | 109 | 0 | 12792 | 21 | 21 | 21 | 0 | 0 | 0 | 0 | 0 | 599 | 0 | 21 | 0 | 0 | 36 | 0 | 0 | 0 | 1089 | 0 | 0 | 0 | 0 | 383 | 170 | 21 | 0 | 0 | 21 | 0 | 0 | 0 | 1577 | 64 | 426 |
| SRR5030102 | 483 | 0 | 5650 | 0 | 0 | 285 | 0 | 0 | 46 | 0 | 0 | 123 | 0 | 0 | 168 | 0 | 67 | 0 | 0 | 958 | 2323 | 15 | 0 | 0 | 0 | 3359 | 2088 | 15 | 15 | 0 | 39 | 0 | 0 | 0 | 9338 | 624 | 4477 |
| SRR5030103 | 408 | 0 | 5291 | 0 | 0 | 150 | 0 | 0 | 66 | 0 | 0 | 92 | 0 | 0 | 163 | 0 | 65 | 0 | 27 | 298 | 1463 | 0 | 0 | 0 | 0 | 2797 | 882 | 19 | 28 | 0 | 84 | 0 | 0 | 0 | 9844 | 488 | 3716 |
| SRR5030104 | 449 | 0 | 5915 | 0 | 0 | 252 | 25 | 0 | 34 | 0 | 0 | 116 | 0 | 0 | 124 | 0 | 58 | 0 | 25 | 939 | 2224 | 0 | 0 | 0 | 0 | 3640 | 2374 | 0 | 0 | 0 | 84 | 0 | 0 | 0 | 9436 | 646 | 4479 |
| SRR5030105 | 431 | 0 | 5510 | 0 | 0 | 10 | 0 | 0 | 59 | 0 | 0 | 94 | 0 | 0 | 171 | 0 | 62 | 0 | 29 | 257 | 1427 | 0 | 0 | 0 | 0 | 163 | 61 | 0 | 20 | 0 | 98 | 0 | 0 | 0 | 544 | 33 | 216 |
| SRR5030106 | 442 | 0 | 5713 | 0 | 0 | 13 | 0 | 0 | 56 | 0 | 0 | 107 | 0 | 0 | 117 | 0 | 61 | 0 | 0 | 865 | 2782 | 0 | 0 | 0 | 0 | 284 | 168 | 0 | 0 | 0 | 56 | 0 | 0 | 0 | 671 | 43 | 293 |
| SRR5030107 | 466 | 0 | 5349 | 0 | 0 | 143 | 0 | 0 | 0 | 0 | 0 | 100 | 0 | 0 | 186 | 0 | 52 | 0 | 0 | 477 | 1749 | 0 | 0 | 0 | 0 | 2576 | 1302 | 0 | 19 | 0 | 48 | 0 | 0 | 0 | 10084 | 523 | 3716 |
| SRR5030108 | 500 | 0 | 5727 | 0 | 0 | 156 | 24 | 0 | 41 | 0 | 0 | 126 | 0 | 0 | 72 | 0 | 66 | 0 | 48 | 1270 | 2109 | 0 | 0 | 0 | 0 | 3672 | 2090 | 0 | 0 | 0 | 33 | 16 | 0 | 0 | 9580 | 647 | 4483 |
| SRR5030109 | 401 | 0 | 5406 | 0 | 0 | 206 | 0 | 0 | 29 | 0 | 0 | 106 | 0 | 0 | 165 | 0 | 62 | 0 | 0 | 440 | 1514 | 0 | 0 | 0 | 0 | 3195 | 804 | 0 | 0 | 0 | 59 | 0 | 0 | 0 | 9300 | 441 | 4057 |
| SRR5030110 | 129 | 0 | 15706 | 20 | 0 | 60 | 24 | 30 | 0 | 0 | 0 | 30 | 0 | 40 | 24 | 0 | 37 | 0 | 0 | 364 | 995 | 0 | 0 | 0 | 0 | 92 | 54 | 0 | 20 | 20 | 20 | 0 | 0 | 80 | 508 | 25 | 191 |
| SRR5030111 | 161 | 0 | 13588 | 28 | 0 | 50 | 0 | 17 | 39 | 0 | 0 | 23 | 0 | 28 | 180 | 0 | 23 | 0 | 0 | 162 | 539 | 0 | 0 | 0 | 0 | 691 | 150 | 0 | 22 | 0 | 0 | 0 | 0 | 78 | 6228 | 329 | 2072 |
| SRR5030112 | 141 | 0 | 15753 | 21 | 0 | 74 | 0 | 43 | 53 | 0 | 0 | 28 | 0 | 0 | 52 | 0 | 24 | 0 | 0 | 498 | 995 | 53 | 0 | 0 | 0 | 103 | 54 | 0 | 0 | 0 | 0 | 0 | 0 | 43 | 516 | 22 | 167 |
| SRR5030113 | 125 | 12 | 13863 | 23 | 0 | 0 | 0 | 0 | 17 | 0 | 0 | 28 | 0 | 17 | 133 | 0 | 20 | 0 | 0 | 76 | 476 | 17 | 0 | 0 | 0 | 52 | 150 | 0 | 0 | 0 | 29 | 0 | 12 | 75 | 415 | 21 | 121 |
| SRR5030114 | 164 | 0 | 15631 | 51 | 10 | 61 | 25 | 10 | 10 | 0 | 10 | 35 | 0 | 0 | 25 | 0 | 15 | 0 | 0 | 718 | 1065 | 41 | 0 | 0 | 0 | 1069 | 652 | 0 | 0 | 0 | 0 | 0 | 0 | 20 | 6680 | 316 | 2739 |
| SRR5030115 | 126 | 0 | 13487 | 23 | 0 | 40 | 0 | 0 | 28 | 0 | 0 | 26 | 0 | 11 | 55 | 0 | 24 | 0 | 0 | 37 | 442 | 0 | 0 | 0 | 0 | 45 | 11 | 17 | 11 | 0 | 28 | 0 | 0 | 34 | 415 | 22 | 136 |
| SRR5030116 | 155 | 0 | 15572 | 10 | 0 | 21 | 0 | 41 | 31 | 0 | 0 | 33 | 0 | 10 | 0 | 0 | 18 | 0 | 0 | 581 | 1111 | 10 | 0 | 0 | 0 | 1048 | 695 | 0 | 0 | 21 | 0 | 0 | 0 | 41 | 6680 | 322 | 2375 |
| SRR5030117 | 140 | 0 | 13755 | 23 | 0 | 34 | 0 | 0 | 40 | 0 | 0 | 34 | 0 | 11 | 93 | 0 | 22 | 0 | 0 | 74 | 556 | 0 | 0 | 0 | 0 | 43 | 138 | 0 | 0 | 0 | 57 | 0 | 0 | 40 | 415 | 19 | 147 |
| SRR5030118 | 193 | 0 | 13591 | 24 | 21 | 38 | 0 | 0 | 21 | 0 | 0 | 52 | 0 | 17 | 0 | 0 | 21 | 11 | 23 | 266 | 1954 | 14 | 0 | 0 | 0 | 130 | 64 | 0 | 0 | 0 | 21 | 0 | 0 | 28 | 573 | 22 | 211 |
| SRR5030119 | 192 | 0 | 14029 | 38 | 19 | 25 | 0 | 0 | 0 | 0 | 0 | 48 | 0 | 22 | 23 | 0 | 23 | 21 | 12 | 104 | 1531 | 0 | 0 | 0 | 0 | 533 | 187 | 0 | 0 | 0 | 22 | 19 | 0 | 35 | 3479 | 193 | 1151 |
| SRR5030120 | 188 | 0 | 14044 | 22 | 0 | 34 | 0 | 0 | 15 | 0 | 0 | 55 | 0 | 22 | 62 | 0 | 29 | 12 | 0 | 322 | 2054 | 0 | 0 | 0 | 0 | 116 | 69 | 11 | 0 | 0 | 45 | 0 | 0 | 19 | 582 | 35 | 226 |
| SRR5030121 | 162 | 0 | 14198 | 36 | 23 | 30 | 0 | 13 | 0 | 0 | 0 | 46 | 0 | 20 | 48 | 13 | 29 | 22 | 0 | 107 | 1358 | 0 | 0 | 0 | 0 | 572 | 159 | 0 | 0 | 0 | 23 | 17 | 0 | 23 | 3524 | 185 | 1049 |
| SRR5030122 | 147 | 0 | 13740 | 36 | 11 | 29 | 0 | 0 | 0 | 0 | 0 | 39 | 0 | 0 | 24 | 0 | 21 | 0 | 0 | 201 | 1800 | 14 | 0 | 0 | 0 | 135 | 65 | 0 | 0 | 0 | 18 | 11 | 0 | 25 | 606 | 36 | 233 |
| SRR5030123 | 165 | 0 | 14005 | 45 | 16 | 35 | 0 | 0 | 0 | 0 | 0 | 60 | 0 | 19 | 24 | 13 | 17 | 21 | 0 | 118 | 1745 | 0 | 0 | 0 | 13 | 544 | 180 | 13 | 0 | 0 | 29 | 16 | 0 | 55 | 3632 | 199 | 1062 |
| SRR5030124 | 172 | 0 | 14064 | 15 | 22 | 18 | 0 | 0 | 0 | 0 | 0 | 48 | 0 | 18 | 24 | 0 | 15 | 0 | 12 | 181 | 1858 | 15 | 0 | 0 | 0 | 624 | 315 | 0 | 0 | 0 | 33 | 0 | 0 | 36 | 3121 | 185 | 1044 |
| SRR5030125 | 174 | 0 | 14081 | 43 | 20 | 26 | 0 | 0 | 0 | 0 | 0 | 51 | 0 | 16 | 24 | 0 | 19 | 22 | 24 | 84 | 1494 | 0 | 0 | 0 | 0 | 617 | 200 | 0 | 0 | 0 | 43 | 0 | 0 | 30 | 3503 | 197 | 1092 |
| SRR5030126 | 70 | 0 | 8353 | 0 | 0 | 29 | 106 | 0 | 0 | 0 | 0 | 146 | 0 | 22 | 53 | 0 | 13 | 0 | 318 | 689 | 636 | 0 | 0 | 0 | 0 | 31 | 19 | 15 | 0 | 0 | 0 | 0 | 0 | 59 | 419 | 20 | 173 |
| SRR5030127 | 66 | 14 | 8209 | 48 | 34 | 0 | 0 | 0 | 0 | 0 | 0 | 169 | 0 | 90 | 218 | 0 | 24 | 0 | 131 | 0 | 218 | 0 | 0 | 0 | 0 | 269 | 83 | 41 | 0 | 0 | 0 | 21 | 0 | 110 | 5431 | 310 | 1944 |
| SRR5030128 | 60 | 0 | 8413 | 39 | 16 | 0 | 0 | 16 | 0 | 0 | 0 | 148 | 0 | 39 | 231 | 0 | 14 | 0 | 58 | 347 | 289 | 16 | 0 | 0 | 0 | 274 | 196 | 23 | 0 | 0 | 0 | 0 | 0 | 94 | 3946 | 235 | 1675 |
| SRR5030130 | 63 | 0 | 8193 | 15 | 15 | 15 | 0 | 0 | 0 | 0 | 0 | 129 | 0 | 37 | 221 | 0 | 0 | 0 | 221 | 720 | 277 | 0 | 0 | 0 | 0 | 36 | 22 | 0 | 0 | 0 | 0 | 0 | 0 | 60 | 379 | 20 | 179 |
| SRR5030131 | 72 | 0 | 8371 | 28 | 0 | 0 | 0 | 0 | 0 | 0 | 0 | 133 | 0 | 70 | 266 | 0 | 16 | 0 | 44 | 44 | 133 | 0 | 0 | 0 | 0 | 21 | 91 | 28 | 0 | 0 | 14 | 0 | 0 | 140 | 366 | 19 | 137 |
| SRR5030132 | 50 | 0 | 8347 | 31 | 0 | 0 | 56 | 0 | 0 | 0 | 0 | 133 | 0 | 0 | 56 | 0 | 25 | 0 | 279 | 391 | 224 | 31 | 0 | 0 | 0 | 284 | 154 | 0 | 0 | 0 | 0 | 0 | 0 | 84 | 3593 | 123 | 1658 |
| SRR5030133 | 71 | 0 | 8340 | 29 | 0 | 14 | 45 | 0 | 0 | 0 | 0 | 134 | 0 | 72 | 45 | 0 | 15 | 0 | 134 | 45 | 134 | 0 | 0 | 0 | 0 | 18 | 50 | 0 | 0 | 0 | 0 | 22 | 0 | 72 | 353 | 23 | 151 |
| SRR5030134 | 219 | 0 | 7895 | 18 | 10 | 20 | 0 | 0 | 14 | 0 | 0 | 14 | 0 | 0 | 0 | 0 | 25 | 38 | 13 | 453 | 1785 | 0 | 0 | 0 | 0 | 105 | 69 | 0 | 0 | 0 | 14 | 0 | 0 | 21 | 598 | 23 | 267 |
| SRR5030135 | 205 | 0 | 7297 | 23 | 17 | 17 | 0 | 0 | 10 | 0 | 0 | 13 | 0 | 21 | 29 | 0 | 10 | 79 | 0 | 265 | 972 | 10 | 0 | 0 | 0 | 70 | 19 | 0 | 0 | 0 | 0 | 27 | 0 | 0 | 511 | 20 | 214 |
| SRR5030136 | 206 | 0 | 8180 | 17 | 24 | 31 | 0 | 0 | 15 | 0 | 0 | 16 | 0 | 0 | 14 | 0 | 29 | 40 | 0 | 430 | 2094 | 0 | 0 | 0 | 0 | 98 | 63 | 0 | 0 | 0 | 0 | 0 | 0 | 11 | 600 | 23 | 236 |
| SRR5030137 | 190 | 0 | 7364 | 64 | 42 | 14 | 0 | 0 | 28 | 0 | 0 | 19 | 0 | 0 | 123 | 0 | 26 | 124 | 0 | 216 | 1018 | 0 | 0 | 0 | 0 | 333 | 85 | 0 | 0 | 0 | 11 | 11 | 0 | 21 | 2126 | 71 | 817 |
| SRR5030138 | 207 | 0 | 7743 | 48 | 25 | 29 | 0 | 0 | 18 | 0 | 0 | 21 | 0 | 0 | 27 | 0 | 0 | 0 | 14 | 311 | 1554 | 11 | 0 | 0 | 0 | 321 | 221 | 0 | 0 | 0 | 0 | 0 | 0 | 14 | 1895 | 94 | 827 |
| SRR5030139 | 199 | 0 | 7197 | 104 | 42 | 38 | 0 | 0 | 0 | 0 | 0 | 0 | 0 | 17 | 30 | 0 | 19 | 0 | 0 | 90 | 1049 | 0 | 0 | 0 | 0 | 267 | 97 | 0 | 0 | 0 | 10 | 17 | 0 | 21 | 2037 | 55 | 748 |
| SRR5030140 | 184 | 0 | 7851 | 30 | 25 | 17 | 14 | 0 | 14 | 0 | 0 | 13 | 0 | 0 | 56 | 0 | 19 | 0 | 0 | 389 | 1446 | 0 | 0 | 0 | 0 | 116 | 64 | 0 | 0 | 0 | 0 | 0 | 0 | 0 | 593 | 25 | 279 |
| SRR5030141 | 136 | 0 | 7613 | 13 | 28 | 21 | 0 | 0 | 14 | 0 | 0 | 28 | 0 | 11 | 31 | 14 | 15 | 82 | 0 | 248 | 1086 | 0 | 0 | 0 | 0 | 77 | 27 | 0 | 0 | 0 | 11 | 0 | 0 | 11 | 551 | 23 | 199 |
| SRR5030142 | 126 | 10 | 8739 | 17 | 10 | 24 | 20 | 14 | 24 | 0 | 0 | 738 | 0 | 24 | 20 | 21 | 127 | 155 | 0 | 133 | 5241 | 17 | 0 | 0 | 0 | 39 | 61 | 0 | 0 | 0 | 14 | 31 | 0 | 17 | 535 | 41 | 220 |
| SRR5030143 | 182 | 30 | 8437 | 120 | 0 | 30 | 0 | 30 | 0 | 0 | 0 | 714 | 0 | 30 | 27 | 0 | 160 | 365 | 0 | 107 | 4193 | 0 | 0 | 0 | 0 | 330 | 330 | 0 | 0 | 0 | 30 | 0 | 0 | 0 | 5639 | 210 | 1830 |
| SRR5030144 | 144 | 15 | 8852 | 34 | 23 | 11 | 0 | 15 | 0 | 0 | 0 | 780 | 0 | 27 | 15 | 23 | 147 | 149 | 0 | 76 | 5290 | 15 | 0 | 0 | 0 | 34 | 56 | 0 | 0 | 0 | 0 | 27 | 0 | 15 | 520 | 35 | 203 |
| SRR5030145 | 191 | 64 | 7804 | 64 | 0 | 0 | 0 | 0 | 32 | 0 | 0 | 612 | 0 | 64 | 28 | 32 | 79 | 326 | 0 | 28 | 4639 | 0 | 0 | 0 | 0 | 415 | 574 | 0 | 0 | 0 | 96 | 0 | 0 | 32 | 4434 | 319 | 1882 |
| SRR5030146 | 150 | 22 | 8566 | 25 | 18 | 18 | 0 | 11 | 25 | 0 | 0 | 809 | 0 | 18 | 12 | 11 | 137 | 141 | 12 | 99 | 5239 | 0 | 0 | 0 | 0 | 41 | 58 | 0 | 0 | 0 | 14 | 32 | 0 | 18 | 524 | 37 | 207 |
| SRR5030147 | 131 | 62 | 7940 | 62 | 31 | 0 | 0 | 0 | 0 | 0 | 0 | 617 | 0 | 0 | 0 | 0 | 177 | 420 | 0 | 27 | 4475 | 0 | 0 | 0 | 31 | 30 | 18 | 31 | 0 | 0 | 0 | 0 | 0 | 62 | 432 | 30 | 175 |
| SRR5030148 | 145 | 15 | 8695 | 22 | 15 | 15 | 12 | 0 | 15 | 0 | 0 | 744 | 0 | 33 | 12 | 15 | 145 | 157 | 0 | 98 | 4946 | 19 | 0 | 0 | 0 | 419 | 590 | 0 | 0 | 0 | 11 | 45 | 0 | 22 | 5201 | 408 | 1936 |
| SRR5030149 | 151 | 157 | 7891 | 0 | 31 | 31 | 0 | 0 | 0 | 0 | 0 | 714 | 0 | 31 | 84 | 0 | 214 | 217 | 0 | 56 | 4052 | 31 | 0 | 0 | 0 | 409 | 157 | 0 | 0 | 0 | 0 | 31 | 0 | 0 | 5185 | 566 | 1446 |
| SRR5030150 | 93 | 0 | 21584 | 66 | 27 | 0 | 52 | 23 | 0 | 0 | 0 | 38 | 0 | 43 | 0 | 25 | 24 | 15 | 15 | 125 | 1068 | 46 | 0 | 0 | 0 | 176 | 334 | 16 | 0 | 0 | 0 | 41 | 0 | 62 | 2268 | 146 | 1026 |
| SRR5030151 | 169 | 0 | 22411 | 118 | 20 | 0 | 0 | 0 | 0 | 0 | 0 | 28 | 0 | 20 | 67 | 39 | 0 | 34 | 0 | 0 | 469 | 20 | 0 | 0 | 0 | 177 | 158 | 20 | 0 | 0 | 20 | 0 | 0 | 39 | 2128 | 197 | 1143 |
| SRR5030152 | 99 | 0 | 22315 | 20 | 27 | 0 | 72 | 0 | 0 | 0 | 0 | 41 | 0 | 37 | 24 | 15 | 15 | 0 | 16 | 135 | 977 | 12 | 0 | 0 | 0 | 32 | 53 | 0 | 0 | 0 | 0 | 25 | 0 | 13 | 412 | 26 | 189 |
| SRR5030153 | 107 | 0 | 22893 | 102 | 41 | 0 | 69 | 0 | 0 | 0 | 0 | 78 | 0 | 41 | 69 | 0 | 0 | 0 | 0 | 0 | 554 | 0 | 0 | 0 | 20 | 266 | 123 | 0 | 0 | 20 | 20 | 61 | 0 | 61 | 2847 | 164 | 963 |
| SRR5030154 | 95 | 0 | 21976 | 14 | 16 | 0 | 68 | 0 | 0 | 0 | 0 | 29 | 0 | 35 | 15 | 19 | 27 | 12 | 38 | 121 | 843 | 13 | 0 | 0 | 0 | 28 | 72 | 0 | 0 | 0 | 0 | 26 | 0 | 42 | 425 | 24 | 194 |
| SRR5030155 | 106 | 0 | 21088 | 26 | 11 | 0 | 0 | 0 | 0 | 0 | 0 | 19 | 0 | 15 | 0 | 20 | 0 | 0 | 0 | 136 | 476 | 15 | 0 | 0 | 0 | 26 | 44 | 0 | 0 | 0 | 0 | 11 | 0 | 18 | 445 | 18 | 215 |
| SRR5030156 | 95 | 0 | 21789 | 20 | 22 | 0 | 46 | 14 | 0 | 0 | 0 | 34 | 0 | 34 | 15 | 14 | 20 | 12 | 15 | 123 | 924 | 11 | 0 | 0 | 0 | 31 | 76 | 0 | 0 | 0 | 0 | 22 | 0 | 10 | 430 | 27 | 198 |
| SRR5030157 | 108 | 0 | 22265 | 166 | 21 | 21 | 69 | 0 | 0 | 0 | 0 | 20 | 0 | 41 | 0 | 0 | 39 | 0 | 0 | 0 | 556 | 0 | 0 | 0 | 0 | 166 | 228 | 0 | 0 | 0 | 0 | 41 | 0 | 104 | 2279 | 145 | 1512 |
| SRR5030158 | 164 | 0 | 15837 | 10 | 25 | 37 | 32 | 0 | 16 | 0 | 0 | 25 | 0 | 16 | 0 | 0 | 34 | 0 | 19 | 128 | 711 | 16 | 0 | 0 | 0 | 136 | 215 | 0 | 0 | 0 | 29 | 29 | 0 | 66 | 673 | 43 | 373 |
| SRR5030159 | 143 | 0 | 14348 | 41 | 47 | 29 | 12 | 0 | 12 | 0 | 0 | 31 | 0 | 64 | 0 | 29 | 31 | 0 | 0 | 60 | 602 | 18 | 0 | 0 | 0 | 95 | 88 | 0 | 0 | 0 | 18 | 53 | 0 | 53 | 653 | 24 | 265 |
| SRR5030160 | 165 | 0 | 16373 | 13 | 0 | 22 | 0 | 0 | 22 | 0 | 0 | 43 | 0 | 0 | 0 | 18 | 24 | 0 | 21 | 145 | 540 | 0 | 0 | 0 | 0 | 126 | 208 | 0 | 0 | 0 | 18 | 31 | 0 | 13 | 682 | 37 | 317 |
| SRR5030161 | 166 | 0 | 14786 | 98 | 37 | 31 | 12 | 12 | 0 | 0 | 0 | 31 | 0 | 37 | 25 | 12 | 32 | 18 | 12 | 87 | 659 | 18 | 0 | 0 | 0 | 827 | 796 | 0 | 0 | 0 | 18 | 49 | 0 | 37 | 5964 | 269 | 2247 |
| SRR5030162 | 143 | 0 | 15685 | 59 | 50 | 59 | 27 | 13 | 13 | 0 | 0 | 40 | 0 | 42 | 0 | 0 | 30 | 0 | 0 | 139 | 671 | 21 | 0 | 0 | 0 | 1156 | 1648 | 0 | 0 | 0 | 17 | 21 | 0 | 50 | 5843 | 324 | 2829 |
| SRR5030163 | 144 | 12 | 14819 | 59 | 18 | 30 | 24 | 0 | 18 | 0 | 0 | 26 | 0 | 77 | 0 | 18 | 33 | 0 | 24 | 36 | 413 | 12 | 0 | 0 | 12 | 823 | 746 | 0 | 0 | 0 | 41 | 41 | 0 | 65 | 5521 | 326 | 2553 |
| SRR5030164 | 184 | 0 | 16160 | 60 | 17 | 39 | 14 | 0 | 0 | 0 | 0 | 38 | 0 | 22 | 0 | 0 | 36 | 0 | 14 | 136 | 817 | 17 | 0 | 0 | 13 | 1153 | 1600 | 0 | 0 | 0 | 22 | 30 | 0 | 56 | 5837 | 301 | 2568 |
| SRR5030165 | 144 | 0 | 14850 | 79 | 18 | 43 | 0 | 0 | 0 | 0 | 0 | 19 | 0 | 55 | 0 | 18 | 26 | 18 | 13 | 76 | 567 | 31 | 0 | 0 | 12 | 100 | 85 | 0 | 0 | 0 | 37 | 24 | 0 | 55 | 626 | 36 | 243 |
| SRR5030166 | 190 | 13 | 6260 | 0 | 0 | 15 | 0 | 0 | 52 | 0 | 0 | 15 | 0 | 0 | 175 | 0 | 25 | 0 | 0 | 655 | 1878 | 0 | 0 | 13 | 0 | 201 | 78 | 0 | 0 | 0 | 91 | 0 | 0 | 13 | 602 | 45 | 442 |
| SRR5030167 | 168 | 0 | 5924 | 0 | 0 | 200 | 0 | 0 | 19 | 0 | 0 | 20 | 0 | 0 | 311 | 0 | 20 | 0 | 0 | 415 | 1245 | 0 | 0 | 0 | 0 | 121 | 33 | 19 | 0 | 0 | 86 | 0 | 0 | 0 | 461 | 36 | 346 |
| SRR5030168 | 163 | 0 | 6443 | 0 | 0 | 212 | 0 | 14 | 28 | 0 | 0 | 14 | 0 | 0 | 143 | 0 | 24 | 0 | 0 | 744 | 2122 | 0 | 0 | 0 | 0 | 3678 | 1683 | 0 | 0 | 0 | 57 | 0 | 0 | 0 | 11670 | 891 | 9704 |
| SRR5030169 | 172 | 0 | 6025 | 10 | 10 | 211 | 0 | 0 | 20 | 0 | 0 | 22 | 0 | 10 | 400 | 0 | 19 | 0 | 0 | 345 | 1283 | 0 | 0 | 0 | 0 | 2748 | 822 | 30 | 0 | 0 | 60 | 0 | 0 | 10 | 11823 | 782 | 8714 |
| SRR5030170 | 176 | 0 | 6188 | 0 | 0 | 11 | 0 | 0 | 54 | 0 | 0 | 21 | 0 | 13 | 138 | 0 | 25 | 0 | 0 | 719 | 2096 | 0 | 0 | 0 | 0 | 172 | 84 | 13 | 0 | 0 | 13 | 27 | 0 | 54 | 539 | 47 | 462 |
| SRR5030171 | 168 | 0 | 5904 | 29 | 0 | 185 | 0 | 0 | 88 | 0 | 0 | 16 | 0 | 0 | 295 | 0 | 27 | 0 | 0 | 295 | 1207 | 0 | 0 | 0 | 0 | 2825 | 799 | 29 | 0 | 0 | 39 | 0 | 0 | 0 | 11729 | 740 | 8992 |
| SRR5030172 | 172 | 0 | 6384 | 28 | 0 | 19 | 15 | 0 | 56 | 0 | 0 | 22 | 0 | 14 | 201 | 0 | 22 | 0 | 15 | 914 | 1937 | 0 | 0 | 0 | 0 | 194 | 56 | 14 | 0 | 0 | 42 | 14 | 0 | 0 | 528 | 44 | 444 |
| SRR5030173 | 164 | 0 | 6072 | 0 | 0 | 168 | 0 | 0 | 79 | 0 | 0 | 13 | 0 | 0 | 218 | 0 | 31 | 0 | 0 | 300 | 1255 | 0 | 0 | 0 | 0 | 3286 | 730 | 0 | 0 | 0 | 30 | 0 | 0 | 0 | 11515 | 918 | 8269 |
| SRR5030174 | 126 | 0 | 16722 | 56 | 17 | 11 | 127 | 17 | 0 | 0 | 0 | 33 | 0 | 0 | 45 | 0 | 20 | 37 | 0 | 149 | 2734 | 17 | 0 | 0 | 0 | 152 | 244 | 0 | 0 | 0 | 0 | 14 | 0 | 20 | 1207 | 48 | 499 |
| SRR5030175 | 124 | 0 | 15071 | 25 | 14 | 14 | 121 | 0 | 0 | 0 | 0 | 24 | 0 | 14 | 158 | 26 | 33 | 73 | 0 | 61 | 1578 | 24 | 0 | 0 | 0 | 37 | 34 | 14 | 0 | 0 | 0 | 21 | 0 | 10 | 355 | 19 | 140 |
| SRR5030176 | 109 | 0 | 17660 | 19 | 11 | 16 | 103 | 22 | 0 | 0 | 0 | 27 | 0 | 0 | 40 | 0 | 16 | 16 | 16 | 127 | 3086 | 14 | 0 | 0 | 0 | 50 | 82 | 0 | 0 | 0 | 0 | 0 | 0 | 24 | 368 | 20 | 169 |
| SRR5030177 | 87 | 0 | 15187 | 108 | 25 | 0 | 89 | 0 | 0 | 0 | 0 | 24 | 0 | 0 | 190 | 22 | 14 | 38 | 0 | 25 | 1888 | 27 | 0 | 0 | 0 | 108 | 145 | 0 | 0 | 0 | 0 | 15 | 0 | 17 | 1313 | 52 | 469 |
| SRR5030178 | 117 | 0 | 17040 | 18 | 20 | 0 | 123 | 15 | 0 | 0 | 0 | 35 | 0 | 0 | 54 | 12 | 27 | 53 | 0 | 139 | 3025 | 25 | 0 | 0 | 0 | 41 | 73 | 0 | 0 | 0 | 0 | 14 | 0 | 15 | 399 | 23 | 177 |
| SRR5030179 | 121 | 12 | 15241 | 27 | 11 | 0 | 50 | 19 | 0 | 0 | 0 | 32 | 0 | 31 | 223 | 12 | 16 | 87 | 0 | 50 | 1933 | 0 | 0 | 0 | 0 | 43 | 26 | 0 | 0 | 0 | 0 | 24 | 0 | 17 | 364 | 20 | 147 |
| SRR5030180 | 119 | 0 | 17325 | 20 | 22 | 11 | 62 | 20 | 0 | 0 | 0 | 36 | 0 | 14 | 62 | 0 | 18 | 23 | 16 | 132 | 2892 | 22 | 0 | 0 | 0 | 55 | 80 | 0 | 0 | 0 | 0 | 0 | 0 | 22 | 393 | 22 | 178 |
| SRR5030181 | 137 | 17 | 15310 | 33 | 24 | 17 | 50 | 0 | 0 | 0 | 0 | 34 | 0 | 12 | 226 | 17 | 18 | 25 | 13 | 13 | 1921 | 20 | 0 | 0 | 0 | 40 | 31 | 0 | 0 | 0 | 0 | 15 | 0 | 20 | 356 | 24 | 157 |
| SRR5030182 | 282 | 0 | 12145 | 0 | 27 | 82 | 0 | 0 | 96 | 0 | 0 | 41 | 0 | 0 | 103 | 0 | 48 | 0 | 0 | 494 | 1401 | 0 | 0 | 0 | 0 | 2677 | 2787 | 27 | 0 | 0 | 82 | 14 | 0 | 0 | 6946 | 645 | 4818 |
| SRR5030183 | 261 | 0 | 11086 | 0 | 23 | 150 | 0 | 0 | 46 | 0 | 0 | 43 | 0 | 12 | 39 | 0 | 38 | 0 | 0 | 137 | 784 | 0 | 0 | 0 | 0 | 1854 | 1359 | 12 | 0 | 0 | 138 | 0 | 12 | 0 | 7245 | 587 | 4573 |
| SRR5030184 | 248 | 0 | 12535 | 45 | 0 | 238 | 0 | 0 | 30 | 0 | 0 | 50 | 0 | 0 | 131 | 0 | 56 | 0 | 0 | 392 | 1589 | 45 | 0 | 15 | 0 | 2504 | 2548 | 0 | 0 | 0 | 75 | 0 | 0 | 0 | 6319 | 730 | 4873 |
| SRR5030185 | 266 | 0 | 11431 | 12 | 48 | 108 | 0 | 0 | 60 | 0 | 0 | 42 | 0 | 0 | 148 | 0 | 44 | 0 | 0 | 191 | 657 | 0 | 0 | 0 | 0 | 1960 | 1243 | 24 | 0 | 0 | 119 | 12 | 0 | 0 | 7134 | 526 | 5055 |
| SRR5030186 | 266 | 14 | 12151 | 14 | 0 | 198 | 0 | 0 | 43 | 0 | 0 | 39 | 0 | 0 | 82 | 0 | 47 | 0 | 21 | 515 | 1093 | 28 | 0 | 0 | 0 | 2295 | 2494 | 43 | 0 | 0 | 43 | 0 | 0 | 0 | 7184 | 652 | 5285 |
| SRR5030187 | 260 | 0 | 11116 | 12 | 12 | 128 | 20 | 0 | 70 | 0 | 0 | 39 | 0 | 0 | 200 | 0 | 38 | 0 | 20 | 240 | 819 | 0 | 0 | 0 | 12 | 1841 | 1235 | 35 | 0 | 0 | 93 | 0 | 0 | 0 | 7214 | 466 | 4732 |
| SRR5030188 | 284 | 0 | 12243 | 0 | 15 | 102 | 0 | 0 | 44 | 0 | 0 | 42 | 0 | 0 | 109 | 0 | 56 | 0 | 0 | 239 | 1238 | 15 | 0 | 0 | 0 | 111 | 136 | 15 | 0 | 0 | 87 | 0 | 0 | 0 | 307 | 27 | 212 |
| SRR5030189 | 267 | 12 | 11572 | 0 | 12 | 60 | 0 | 0 | 0 | 0 | 0 | 46 | 0 | 12 | 126 | 0 | 43 | 0 | 0 | 126 | 777 | 24 | 0 | 0 | 0 | 67 | 44 | 12 | 0 | 0 | 48 | 0 | 0 | 0 | 234 | 23 | 160 |
| SRR5030190 | 286 | 194 | 3384 | 0 | 16 | 33 | 19 | 16 | 0 | 0 | 0 | 19 | 0 | 204 | 169 | 0 | 39 | 0 | 0 | 282 | 1495 | 0 | 0 | 0 | 0 | 269 | 168 | 0 | 0 | 0 | 19 | 16 | 0 | 0 | 590 | 56 | 489 |
| SRR5030191 | 236 | 38 | 3397 | 0 | 18 | 35 | 0 | 0 | 0 | 0 | 0 | 22 | 0 | 38 | 404 | 0 | 49 | 0 | 0 | 184 | 1028 | 0 | 0 | 0 | 0 | 49 | 21 | 0 | 0 | 0 | 0 | 27 | 0 | 0 | 152 | 18 | 103 |
| SRR5030192 | 255 | 191 | 3468 | 0 | 0 | 25 | 40 | 0 | 0 | 0 | 0 | 15 | 0 | 213 | 243 | 0 | 41 | 0 | 0 | 232 | 1404 | 0 | 0 | 0 | 0 | 273 | 166 | 0 | 0 | 0 | 0 | 22 | 0 | 0 | 615 | 52 | 496 |
| SRR5030193 | 240 | 293 | 3282 | 0 | 0 | 0 | 0 | 0 | 0 | 0 | 0 | 0 | 0 | 229 | 150 | 18 | 42 | 0 | 0 | 113 | 789 | 0 | 0 | 0 | 0 | 193 | 101 | 0 | 0 | 0 | 0 | 46 | 0 | 0 | 734 | 64 | 605 |
| SRR5030194 | 265 | 167 | 3460 | 0 | 19 | 24 | 29 | 0 | 0 | 0 | 0 | 16 | 0 | 238 | 173 | 0 | 55 | 0 | 0 | 260 | 1481 | 0 | 0 | 0 | 0 | 305 | 136 | 0 | 0 | 0 | 0 | 19 | 0 | 0 | 682 | 36 | 484 |
| SRR5030196 | 257 | 37 | 3474 | 0 | 0 | 24 | 49 | 0 | 0 | 0 | 0 | 24 | 0 | 46 | 315 | 0 | 42 | 0 | 0 | 237 | 1242 | 0 | 0 | 0 | 0 | 52 | 26 | 0 | 0 | 0 | 12 | 15 | 0 | 0 | 133 | 41 | 112 |
| SRR5030197 | 284 | 257 | 3618 | 0 | 0 | 0 | 0 | 0 | 0 | 0 | 0 | 23 | 0 | 165 | 262 | 0 | 51 | 0 | 0 | 112 | 937 | 0 | 0 | 0 | 0 | 312 | 101 | 0 | 0 | 0 | 0 | 18 | 0 | 0 | 752 | 28 | 541 |
| SRR5030198 | 160 | 0 | 4635 | 32 | 23 | 41 | 67 | 0 | 14 | 0 | 0 | 87 | 0 | 0 | 94 | 0 | 26 | 0 | 0 | 629 | 2221 | 0 | 0 | 0 | 0 | 628 | 314 | 0 | 0 | 0 | 27 | 0 | 0 | 0 | 3078 | 296 | 2554 |
| SRR5030199 | 159 | 0 | 4622 | 58 | 22 | 65 | 0 | 0 | 0 | 0 | 0 | 114 | 0 | 0 | 209 | 0 | 35 | 0 | 0 | 394 | 1508 | 0 | 0 | 0 | 0 | 363 | 116 | 0 | 0 | 0 | 73 | 22 | 0 | 0 | 3365 | 240 | 2188 |
| SRR5030200 | 155 | 0 | 4806 | 0 | 15 | 39 | 29 | 0 | 15 | 0 | 0 | 117 | 0 | 0 | 72 | 0 | 43 | 29 | 0 | 558 | 1946 | 0 | 0 | 0 | 0 | 65 | 34 | 0 | 0 | 0 | 54 | 20 | 0 | 0 | 376 | 40 | 308 |
| SRR5030201 | 147 | 0 | 4995 | 23 | 15 | 30 | 24 | 0 | 15 | 0 | 0 | 80 | 0 | 15 | 218 | 0 | 33 | 0 | 0 | 460 | 1499 | 0 | 0 | 0 | 0 | 48 | 14 | 0 | 0 | 0 | 61 | 15 | 0 | 0 | 409 | 42 | 262 |
| SRR5030202 | 191 | 0 | 4771 | 14 | 14 | 0 | 14 | 0 | 0 | 0 | 0 | 79 | 0 | 0 | 193 | 0 | 28 | 19 | 14 | 592 | 1816 | 0 | 0 | 0 | 0 | 63 | 31 | 0 | 0 | 0 | 47 | 0 | 0 | 0 | 370 | 36 | 323 |
| SRR5030203 | 147 | 0 | 4586 | 22 | 45 | 37 | 0 | 0 | 15 | 0 | 0 | 103 | 0 | 0 | 97 | 0 | 22 | 16 | 0 | 145 | 1477 | 0 | 0 | 0 | 0 | 62 | 17 | 15 | 0 | 0 | 67 | 0 | 0 | 0 | 373 | 38 | 277 |
| SRR5030204 | 156 | 0 | 4751 | 24 | 24 | 43 | 28 | 0 | 47 | 0 | 0 | 100 | 0 | 0 | 98 | 0 | 21 | 0 | 28 | 477 | 2062 | 0 | 0 | 0 | 0 | 67 | 33 | 0 | 0 | 0 | 28 | 0 | 0 | 0 | 387 | 41 | 336 |
| SRR5030205 | 185 | 0 | 4678 | 15 | 23 | 23 | 0 | 0 | 15 | 0 | 0 | 98 | 0 | 15 | 196 | 23 | 23 | 0 | 0 | 245 | 1128 | 0 | 0 | 0 | 0 | 485 | 129 | 0 | 0 | 0 | 38 | 0 | 0 | 0 | 3478 | 349 | 2546 |
| SRR5030206 | 73 | 0 | 5761 | 19 | 12 | 19 | 0 | 0 | 25 | 0 | 0 | 43 | 0 | 0 | 28 | 0 | 38 | 19 | 0 | 235 | 3152 | 0 | 0 | 0 | 0 | 480 | 362 | 0 | 0 | 0 | 19 | 0 | 0 | 0 | 7119 | 299 | 1135 |
| SRR5030207 | 74 | 0 | 5965 | 0 | 40 | 0 | 0 | 0 | 0 | 0 | 0 | 21 | 0 | 0 | 0 | 0 | 37 | 0 | 0 | 191 | 1531 | 0 | 0 | 0 | 0 | 30 | 18 | 0 | 0 | 0 | 0 | 0 | 0 | 0 | 462 | 18 | 60 |
| SRR5030208 | 65 | 0 | 6142 | 27 | 13 | 27 | 15 | 0 | 0 | 0 | 0 | 46 | 0 | 20 | 90 | 0 | 32 | 40 | 0 | 347 | 3105 | 0 | 0 | 0 | 0 | 33 | 21 | 0 | 0 | 0 | 20 | 0 | 0 | 13 | 471 | 14 | 82 |
| SRR5030209 | 51 | 0 | 5890 | 0 | 122 | 41 | 0 | 0 | 0 | 0 | 0 | 26 | 0 | 0 | 305 | 0 | 21 | 124 | 0 | 0 | 1930 | 0 | 0 | 0 | 0 | 31 | 10 | 0 | 0 | 0 | 82 | 0 | 0 | 0 | 455 | 18 | 71 |
| SRR5030210 | 63 | 0 | 5751 | 19 | 19 | 26 | 0 | 0 | 19 | 0 | 0 | 26 | 0 | 19 | 42 | 0 | 29 | 19 | 0 | 354 | 2745 | 0 | 0 | 0 | 0 | 544 | 397 | 0 | 0 | 0 | 19 | 0 | 0 | 13 | 7192 | 243 | 1159 |
| SRR5030211 | 67 | 0 | 5839 | 0 | 41 | 0 | 0 | 0 | 0 | 0 | 0 | 42 | 0 | 0 | 0 | 0 | 17 | 0 | 0 | 193 | 2701 | 41 | 0 | 0 | 0 | 123 | 20 | 0 | 0 | 0 | 0 | 0 | 0 | 0 | 405 | 13 | 94 |

### Supp Table S8.2.Columns_AM_to_BW

| | | | | | | | | | | | | | | | | | | | | | | | | | | | | | | | | | | | | | |
|---|---|---|---|---|---|---|---|---|---|---|---|---|---|---|---|---|---|---|---|---|---|---|---|---|---|---|---|---|---|---|---|---|---|---|---|---|---|
| SRR5030212 | 64 | 0 | 5941 | 13 | 0 | 52 | 0 | 0 | 0 | 0 | 0 | 35 | 0 | 0 | 14 | 0 | 27 | 39 | 0 | 229 | 3222 | 0 | 0 | 0 | 13 | 31 | 22 | 0 | 0 | 0 | 19 | 0 | 0 | 0 | 439 | 21 | 80 |
| SRR5030213 | 56 | 0 | 6209 | 0 | 0 | 0 | 0 | 0 | 0 | 0 | 0 | 30 | 0 | 0 | 100 | 0 | 34 | 246 | 0 | 200 | 2704 | 0 | 0 | 0 | 0 | 416 | 83 | 0 | 0 | 0 | 0 | 0 | 0 | 0 | 7813 | 332 | 1081 |
| SRR5030214 | 241 | 0 | 5765 | 11 | 0 | 50 | 0 | 0 | 18 | 0 | 0 | 111 | 0 | 0 | 272 | 0 | 51 | 0 | 0 | 453 | 2231 | 11 | 0 | 0 | 0 | 89 | 28 | 25 | 0 | 0 | 43 | 0 | 0 | 29 | 189 | 10 | 76 |
| SRR5030216 | 261 | 0 | 6115 | 0 | 0 | 27 | 0 | 0 | 12 | 0 | 0 | 117 | 0 | 0 | 245 | 0 | 46 | 0 | 0 | 319 | 2169 | 0 | 0 | 0 | 0 | 95 | 27 | 12 | 12 | 0 | 31 | 23 | 0 | 0 | 194 | 77 | 72 |
| SRR5030217 | 255 | 0 | 6136 | 0 | 0 | 37 | 0 | 0 | 0 | 0 | 0 | 162 | 0 | 0 | 0 | 0 | 62 | 0 | 0 | 472 | 2005 | 37 | 0 | 0 | 0 | 96 | 24 | 37 | 0 | 0 | 37 | 0 | 0 | 0 | 232 | 74 | 88 |
| SRR5030218 | 206 | 0 | 5963 | 15 | 0 | 33 | 0 | 0 | 15 | 0 | 0 | 107 | 0 | 0 | 155 | 0 | 52 | 0 | 0 | 250 | 2181 | 18 | 0 | 0 | 0 | 107 | 27 | 0 | 0 | 0 | 22 | 0 | 0 | 11 | 187 | 15 | 71 |
| SRR5030220 | 250 | 0 | 5871 | 0 | 15 | 26 | 0 | 0 | 30 | 0 | 0 | 111 | 0 | 0 | 182 | 0 | 74 | 0 | 0 | 353 | 2274 | 26 | 0 | 0 | 0 | 972 | 284 | 11 | 0 | 0 | 45 | 11 | 0 | 11 | 1768 | 142 | 669 |
| SRR5030221 | 238 | 0 | 6024 | 0 | 37 | 0 | 0 | 0 | 0 | 0 | 0 | 123 | 0 | 0 | 470 | 0 | 84 | 0 | 0 | 235 | 1646 | 0 | 0 | 0 | 0 | 148 | 20 | 0 | 0 | 0 | 37 | 0 | 0 | 37 | 192 | 20 | 72 |
| SRR5030222 | 113 | 16 | 9025 | 33 | 0 | 33 | 0 | 0 | 0 | 0 | 0 | 37 | 0 | 16 | 43 | 0 | 27 | 0 | 0 | 320 | 1410 | 33 | 0 | 0 | 0 | 623 | 606 | 49 | 0 | 0 | 33 | 0 | 0 | 0 | 4636 | 508 | 2883 |
| SRR5030223 | 139 | 0 | 8829 | 21 | 0 | 17 | 0 | 0 | 25 | 0 | 0 | 25 | 0 | 0 | 113 | 17 | 32 | 0 | 0 | 106 | 807 | 25 | 0 | 0 | 0 | 34 | 16 | 47 | 0 | 0 | 17 | 0 | 0 | 0 | 479 | 41 | 203 |
| SRR5030224 | 134 | 35 | 9033 | 18 | 0 | 18 | 0 | 0 | 0 | 0 | 0 | 52 | 0 | 0 | 69 | 0 | 29 | 0 | 0 | 297 | 1509 | 35 | 0 | 0 | 0 | 37 | 58 | 0 | 0 | 0 | 18 | 0 | 18 | 0 | 459 | 41 | 254 |
| SRR5030225 | 150 | 0 | 8905 | 22 | 0 | 39 | 0 | 0 | 0 | 0 | 0 | 18 | 0 | 18 | 131 | 13 | 33 | 0 | 0 | 77 | 739 | 0 | 0 | 0 | 13 | 333 | 153 | 13 | 0 | 0 | 26 | 0 | 0 | 18 | 5914 | 465 | 2407 |
| SRR5030226 | 122 | 0 | 8639 | 17 | 0 | 0 | 0 | 0 | 17 | 0 | 0 | 16 | 0 | 0 | 45 | 0 | 25 | 0 | 0 | 334 | 892 | 17 | 0 | 0 | 17 | 583 | 417 | 0 | 0 | 0 | 33 | 17 | 0 | 0 | 5317 | 483 | 2534 |
| SRR5030227 | 129 | 0 | 8738 | 22 | 0 | 0 | 0 | 0 | 0 | 0 | 0 | 23 | 0 | 0 | 115 | 13 | 36 | 0 | 0 | 92 | 785 | 17 | 0 | 0 | 0 | 36 | 13 | 35 | 0 | 0 | 30 | 0 | 0 | 13 | 467 | 35 | 203 |
| SRR5030229 | 116 | 0 | 8798 | 18 | 0 | 0 | 0 | 0 | 0 | 0 | 0 | 19 | 0 | 0 | 110 | 0 | 29 | 0 | 0 | 102 | 767 | 35 | 0 | 0 | 0 | 347 | 206 | 22 | 0 | 0 | 0 | 0 | 0 | 0 | 5615 | 399 | 2558 |
| SRR5030230 | 124 | 0 | 14294 | 34 | 0 | 68 | 0 | 0 | 0 | 0 | 0 | 47 | 0 | 0 | 0 | 0 | 33 | 120 | 0 | 300 | 3746 | 0 | 0 | 0 | 0 | 745 | 610 | 0 | 0 | 0 | 68 | 0 | 0 | 0 | 12534 | 610 | 5826 |
| SRR5030231 | 66 | 0 | 12783 | 17 | 13 | 21 | 0 | 0 | 0 | 0 | 0 | 31 | 0 | 17 | 269 | 0 | 42 | 129 | 0 | 82 | 1965 | 0 | 0 | 0 | 0 | 20 | 156 | 0 | 0 | 0 | 21 | 0 | 0 | 34 | 526 | 30 | 205 |
| SRR5030232 | 74 | 0 | 14524 | 36 | 0 | 0 | 0 | 0 | 0 | 0 | 0 | 23 | 0 | 0 | 53 | 0 | 31 | 0 | 0 | 368 | 4422 | 0 | 0 | 0 | 0 | 51 | 33 | 0 | 0 | 0 | 0 | 0 | 0 | 0 | 800 | 42 | 362 |
| SRR5030233 | 64 | 0 | 12939 | 13 | 0 | 48 | 0 | 0 | 0 | 0 | 0 | 27 | 0 | 0 | 319 | 0 | 43 | 125 | 0 | 61 | 1815 | 0 | 0 | 0 | 0 | 471 | 148 | 0 | 0 | 0 | 48 | 0 | 0 | 17 | 11912 | 562 | 4629 |
| SRR5030234 | 134 | 0 | 14568 | 0 | 0 | 35 | 0 | 0 | 0 | 0 | 0 | 22 | 0 | 0 | 103 | 0 | 30 | 0 | 0 | 155 | 4227 | 35 | 0 | 0 | 0 | 828 | 414 | 0 | 0 | 0 | 0 | 0 | 0 | 0 | 11355 | 587 | 5937 |
| SRR5030235 | 69 | 0 | 12818 | 26 | 17 | 34 | 0 | 0 | 13 | 0 | 0 | 27 | 0 | 17 | 259 | 0 | 41 | 44 | 0 | 99 | 1912 | 0 | 0 | 0 | 0 | 19 | 133 | 0 | 0 | 0 | 17 | 0 | 0 | 0 | 506 | 28 | 212 |
| SRR5030236 | 79 | 0 | 14714 | 35 | 0 | 70 | 0 | 70 | 0 | 0 | 0 | 45 | 0 | 0 | 0 | 0 | 26 | 63 | 0 | 420 | 4415 | 0 | 0 | 0 | 0 | 34 | 41 | 0 | 0 | 0 | 0 | 0 | 0 | 35 | 761 | 44 | 277 |
| SRR5030237 | 71 | 13 | 12979 | 35 | 22 | 26 | 0 | 0 | 0 | 0 | 0 | 20 | 0 | 13 | 321 | 0 | 35 | 133 | 0 | 61 | 1934 | 0 | 0 | 0 | 0 | 441 | 131 | 0 | 0 | 0 | 13 | 0 | 0 | 0 | 11928 | 655 | 4890 |
| SRR5030238 | 349 | 0 | 1776 | 50 | 0 | 65 | 13 | 11 | 0 | 0 | 0 | 417 | 0 | 0 | 2600 | 0 | 44 | 74 | 13 | 325 | 12195 | 0 | 0 | 0 | 0 | 56 | 40 | 0 | 0 | 0 | 36 | 0 | 0 | 0 | 1112 | 61 | 282 |
| SRR5030239 | 349 | 0 | 1759 | 50 | 37 | 74 | 23 | 62 | 0 | 0 | 0 | 411 | 0 | 12 | 3243 | 0 | 45 | 121 | 0 | 140 | 9075 | 0 | 0 | 0 | 0 | 40 | 15 | 0 | 12 | 0 | 37 | 12 | 0 | 0 | 1100 | 59 | 238 |
| SRR5030240 | 333 | 0 | 1881 | 31 | 15 | 62 | 42 | 139 | 0 | 0 | 0 | 434 | 0 | 0 | 2911 | 0 | 37 | 27 | 14 | 224 | 13157 | 0 | 0 | 0 | 0 | 709 | 516 | 15 | 0 | 0 | 46 | 15 | 0 | 0 | 15420 | 647 | 3774 |
| SRR5030241 | 331 | 26 | 1919 | 77 | 13 | 39 | 25 | 77 | 0 | 0 | 0 | 489 | 0 | 0 | 3705 | 0 | 68 | 84 | 0 | 74 | 8834 | 0 | 0 | 0 | 0 | 644 | 219 | 26 | 13 | 0 | 26 | 26 | 0 | 26 | 17034 | 695 | 4159 |
| SRR5030242 | 318 | 22 | 1846 | 44 | 15 | 73 | 27 | 168 | 0 | 0 | 0 | 413 | 0 | 0 | 3130 | 0 | 45 | 50 | 14 | 325 | 12169 | 0 | 0 | 0 | 0 | 652 | 520 | 0 | 0 | 0 | 29 | 0 | 0 | 0 | 15784 | 769 | 3567 |
| SRR5030243 | 345 | 0 | 1845 | 25 | 13 | 38 | 0 | 76 | 0 | 0 | 0 | 435 | 0 | 13 | 3006 | 0 | 53 | 123 | 0 | 168 | 9043 | 0 | 0 | 0 | 0 | 46 | 20 | 51 | 0 | 0 | 63 | 51 | 0 | 38 | 1122 | 52 | 261 |
| SRR5030244 | 318 | 38 | 1900 | 23 | 0 | 53 | 0 | 173 | 0 | 0 | 0 | 383 | 0 | 0 | 2518 | 0 | 41 | 0 | 14 | 287 | 12250 | 0 | 0 | 0 | 0 | 713 | 428 | 0 | 0 | 0 | 53 | 15 | 0 | 15 | 15031 | 825 | 3638 |
| SRR5030245 | 341 | 39 | 1957 | 26 | 13 | 64 | 0 | 52 | 0 | 0 | 0 | 470 | 0 | 0 | 3325 | 0 | 52 | 0 | 0 | 146 | 8931 | 0 | 0 | 0 | 0 | 567 | 232 | 0 | 0 | 0 | 52 | 13 | 0 | 13 | 17505 | 889 | 3903 |
| SRR5030246 | 188 | 11 | 13777 | 43 | 11 | 0 | 0 | 0 | 0 | 0 | 0 | 137 | 0 | 11 | 17 | 0 | 128 | 11 | 33 | 818 | 1202 | 0 | 0 | 0 | 0 | 235 | 439 | 11 | 11 | 0 | 0 | 11 | 0 | 32 | 3949 | 246 | 1776 |
| SRR5030247 | 175 | 0 | 12928 | 14 | 21 | 14 | 0 | 0 | 14 | 0 | 0 | 124 | 0 | 35 | 145 | 0 | 136 | 48 | 15 | 290 | 595 | 14 | 0 | 0 | 0 | 138 | 145 | 28 | 0 | 0 | 0 | 28 | 0 | 90 | 247 | 166 | 89 |
| SRR5030248 | 228 | 0 | 14585 | 23 | 0 | 34 | 18 | 34 | 11 | 0 | 0 | 153 | 0 | 23 | 18 | 11 | 121 | 24 | 0 | 762 | 851 | 46 | 0 | 11 | 0 | 22 | 29 | 11 | 0 | 0 | 11 | 0 | 0 | 0 | 263 | 27 | 132 |
| SRR5030249 | 180 | 22 | 13271 | 14 | 0 | 14 | 0 | 0 | 0 | 0 | 0 | 129 | 0 | 14 | 76 | 0 | 110 | 20 | 15 | 183 | 701 | 0 | 0 | 0 | 0 | 194 | 79 | 36 | 0 | 0 | 0 | 14 | 0 | 14 | 4630 | 266 | 1632 |
| SRR5030250 | 193 | 0 | 14244 | 11 | 0 | 32 | 0 | 0 | 22 | 0 | 0 | 137 | 0 | 22 | 50 | 0 | 104 | 11 | 0 | 824 | 1296 | 11 | 0 | 0 | 0 | 281 | 335 | 11 | 0 | 0 | 0 | 0 | 0 | 11 | 4239 | 151 | 1795 |
| SRR5030251 | 171 | 0 | 12724 | 21 | 14 | 0 | 0 | 0 | 0 | 0 | 0 | 133 | 0 | 21 | 89 | 0 | 138 | 0 | 30 | 356 | 624 | 21 | 0 | 0 | 0 | 203 | 84 | 14 | 0 | 0 | 0 | 0 | 0 | 28 | 4627 | 273 | 1388 |
| SRR5030252 | 191 | 0 | 14292 | 33 | 0 | 0 | 0 | 22 | 11 | 0 | 0 | 119 | 0 | 11 | 0 | 11 | 100 | 23 | 17 | 715 | 1429 | 11 | 0 | 0 | 0 | 21 | 18 | 11 | 0 | 0 | 0 | 0 | 0 | 33 | 335 | 16 | 131 |
| SRR5030253 | 167 | 0 | 13092 | 22 | 14 | 0 | 0 | 0 | 0 | 0 | 0 | 120 | 0 | 14 | 31 | 14 | 132 | 0 | 0 | 305 | 611 | 0 | 0 | 0 | 0 | 251 | 86 | 14 | 14 | 0 | 0 | 14 | 0 | 65 | 4343 | 251 | 1741 |
| SRR5030254 | 75 | 0 | 23038 | 20 | 0 | 0 | 14 | 0 | 0 | 0 | 0 | 14 | 0 | 16 | 70 | 0 | 43 | 0 | 0 | 28 | 168 | 0 | 0 | 0 | 0 | 88 | 40 | 0 | 119 | 0 | 0 | 0 | 0 | 0 | 94 | 0 | 52 |
| SRR5030255 | 89 | 10 | 23148 | 17 | 0 | 0 | 0 | 0 | 0 | 0 | 0 | 13 | 0 | 13 | 86 | 0 | 41 | 0 | 0 | 57 | 158 | 0 | 0 | 0 | 0 | 96 | 62 | 0 | 139 | 0 | 0 | 0 | 0 | 0 | 108 | 0 | 38 |
| SRR5030256 | 75 | 0 | 23087 | 16 | 0 | 0 | 0 | 0 | 0 | 0 | 0 | 15 | 0 | 19 | 28 | 0 | 45 | 0 | 0 | 28 | 153 | 0 | 0 | 0 | 0 | 101 | 53 | 0 | 111 | 0 | 0 | 0 | 0 | 0 | 68 | 0 | 54 |
| SRR5030257 | 85 | 0 | 23191 | 10 | 0 | 0 | 14 | 0 | 0 | 0 | 0 | 15 | 0 | 18 | 43 | 0 | 47 | 0 | 0 | 14 | 170 | 0 | 0 | 0 | 0 | 108 | 46 | 0 | 123 | 0 | 0 | 0 | 0 | 10 | 62 | 0 | 54 |
| SRR5030258 | 35 | 0 | 10116 | 0 | 0 | 0 | 0 | 0 | 0 | 0 | 0 | 261 | 0 | 10 | 66 | 0 | 26 | 0 | 0 | 66 | 164 | 0 | 0 | 0 | 0 | 24 | 82 | 21 | 17 | 0 | 0 | 0 | 0 | 0 | 164 | 31 | 127 |
| SRR5030259 | 37 | 0 | 10077 | 0 | 0 | 0 | 0 | 0 | 0 | 0 | 0 | 265 | 0 | 11 | 69 | 0 | 28 | 0 | 0 | 17 | 224 | 0 | 0 | 0 | 0 | 43 | 65 | 11 | 14 | 0 | 18 | 0 | 0 | 0 | 105 | 0 | 119 |
| SRR5030260 | 34 | 10 | 10041 | 0 | 10 | 0 | 0 | 0 | 0 | 0 | 0 | 242 | 0 | 0 | 102 | 0 | 29 | 0 | 0 | 34 | 187 | 0 | 0 | 0 | 0 | 31 | 91 | 17 | 17 | 0 | 17 | 0 | 0 | 0 | 188 | 10 | 101 |
| SRR5030261 | 41 | 17 | 10106 | 17 | 0 | 0 | 0 | 0 | 0 | 0 | 0 | 259 | 0 | 0 | 85 | 0 | 27 | 0 | 0 | 17 | 239 | 0 | 0 | 0 | 10 | 45 | 70 | 0 | 14 | 0 | 10 | 0 | 0 | 0 | 101 | 0 | 94 |
| SRR5030262 | 260 | 36 | 15846 | 0 | 0 | 20 | 0 | 0 | 31 | 0 | 0 | 123 | 0 | 10 | 29 | 0 | 55 | 0 | 0 | 29 | 101 | 0 | 0 | 0 | 0 | 105 | 14 | 10 | 15 | 0 | 41 | 0 | 0 | 20 | 65 | 76 | 42 |
| SRR5030263 | 225 | 26 | 15933 | 11 | 0 | 42 | 0 | 11 | 26 | 0 | 0 | 150 | 0 | 11 | 60 | 0 | 53 | 0 | 0 | 15 | 104 | 0 | 0 | 0 | 0 | 1333 | 169 | 32 | 11 | 0 | 545 | 16 | 11 | 42 | 857 | 63 | 666 |
| SRR5030265 | 237 | 26 | 16004 | 15 | 0 | 52 | 0 | 0 | 31 | 0 | 0 | 129 | 0 | 0 | 57 | 0 | 52 | 0 | 0 | 0 | 14 | 0 | 0 | 0 | 0 | 101 | 18 | 10 | 10 | 0 | 39 | 31 | 0 | 31 | 73 | 36 | 49 |
| SRR5030266 | 72 | 0 | 15985 | 0 | 0 | 23 | 27 | 0 | 0 | 0 | 0 | 58 | 0 | 0 | 81 | 0 | 29 | 0 | 0 | 108 | 215 | 0 | 0 | 0 | 0 | 637 | 508 | 15 | 23 | 0 | 76 | 0 | 0 | 15 | 470 | 30 | 182 |
| SRR5030267 | 49 | 0 | 15828 | 0 | 0 | 24 | 0 | 0 | 0 | 0 | 0 | 61 | 0 | 0 | 112 | 0 | 39 | 0 | 0 | 28 | 140 | 16 | 0 | 0 | 0 | 701 | 441 | 31 | 24 | 0 | 110 | 0 | 0 | 24 | 386 | 31 | 102 |
| SRR5030268 | 54 | 0 | 15889 | 0 | 0 | 31 | 0 | 0 | 15 | 0 | 0 | 62 | 0 | 0 | 107 | 0 | 25 | 0 | 0 | 187 | 133 | 15 | 0 | 0 | 0 | 37 | 28 | 0 | 23 | 0 | 11 | 0 | 15 | 0 | 19 | 31 | 15 |
| SRR5030270 | 82 | 0 | 2729 | 20 | 0 | 72 | 0 | 0 | 0 | 0 | 0 | 38 | 13 | 26 | 22 | 0 | 69 | 0 | 0 | 11 | 66 | 0 | 0 | 0 | 0 | 46 | 243 | 26 | 0 | 0 | 138 | 0 | 0 | 0 | 36 | 53 | 14 |
| SRR5030271 | 80 | 0 | 2833 | 21 | 0 | 62 | 0 | 0 | 14 | 0 | 0 | 36 | 14 | 21 | 24 | 0 | 80 | 0 | 0 | 47 | 165 | 0 | 0 | 0 | 0 | 49 | 12 | 14 | 0 | 0 | 132 | 21 | 0 | 0 | 32 | 49 | 15 |
| SRR5030272 | 84 | 0 | 2837 | 33 | 0 | 94 | 0 | 0 | 47 | 0 | 0 | 46 | 0 | 20 | 44 | 0 | 64 | 14 | 0 | 11 | 78 | 0 | 0 | 0 | 0 | 48 | 10 | 0 | 0 | 0 | 181 | 13 | 0 | 0 | 38 | 40 | 15 |
| SRR5030273 | 79 | 0 | 2845 | 0 | 0 | 73 | 0 | 0 | 27 | 0 | 0 | 47 | 0 | 0 | 0 | 0 | 60 | 0 | 0 | 11 | 124 | 0 | 0 | 0 | 0 | 1309 | 407 | 0 | 13 | 0 | 200 | 20 | 0 | 0 | 948 | 67 | 341 |
| SRR5030274 | 82 | 0 | 11065 | 0 | 0 | 26 | 0 | 0 | 0 | 0 | 0 | 78 | 0 | 13 | 35 | 0 | 59 | 0 | 0 | 18 | 88 | 0 | 0 | 0 | 0 | 29 | 13 | 0 | 0 | 0 | 199 | 0 | 0 | 0 | 185 | 0 | 258 |
| SRR5030275 | 81 | 0 | 11152 | 14 | 0 | 42 | 0 | 0 | 14 | 0 | 0 | 70 | 0 | 0 | 37 | 0 | 49 | 0 | 0 | 19 | 93 | 0 | 0 | 0 | 0 | 30 | 17 | 28 | 0 | 0 | 146 | 0 | 0 | 14 | 194 | 35 | 12 |
| SRR5030276 | 82 | 0 | 11122 | 14 | 0 | 14 | 0 | 0 | 0 | 0 | 0 | 82 | 0 | 0 | 55 | 0 | 56 | 0 | 0 | 18 | 127 | 0 | 0 | 0 | 14 | 28 | 15 | 0 | 0 | 0 | 81 | 14 | 0 | 0 | 189 | 14 | 10 |
| SRR5030277 | 82 | 20 | 11056 | 0 | 0 | 34 | 0 | 0 | 20 | 0 | 0 | 73 | 0 | 0 | 107 | 0 | 63 | 0 | 0 | 18 | 178 | 0 | 0 | 0 | 0 | 663 | 379 | 0 | 0 | 0 | 135 | 0 | 0 | 20 | 176 | 14 | 257 |
| SRR5030278 | 96 | 0 | 12932 | 0 | 0 | 32 | 37 | 0 | 29 | 0 | 0 | 230 | 0 | 0 | 267 | 0 | 51 | 0 | 0 | 28 | 92 | 0 | 0 | 0 | 0 | 960 | 253 | 22 | 15 | 0 | 261 | 0 | 0 | 0 | 302 | 29 | 224 |
| SRR5030279 | 112 | 10 | 12741 | 0 | 0 | 41 | 30 | 0 | 15 | 0 | 0 | 237 | 0 | 13 | 257 | 0 | 51 | 0 | 0 | 49 | 158 | 0 | 0 | 0 | 0 | 62 | 18 | 13 | 15 | 0 | 20 | 0 | 0 | 18 | 22 | 18 | 13 |
| SRR5030280 | 111 | 0 | 12751 | 0 | 0 | 42 | 0 | 0 | 30 | 0 | 0 | 244 | 0 | 0 | 191 | 0 | 55 | 0 | 0 | 19 | 133 | 0 | 0 | 0 | 0 | 65 | 17 | 15 | 22 | 0 | 23 | 0 | 0 | 15 | 23 | 17 | 13 |
| SRR5030281 | 100 | 0 | 12880 | 12 | 0 | 45 | 57 | 0 | 30 | 0 | 0 | 231 | 0 | 0 | 228 | 0 | 56 | 0 | 0 | 19 | 104 | 0 | 0 | 0 | 0 | 66 | 18 | 25 | 17 | 0 | 21 | 0 | 0 | 17 | 23 | 27 | 17 |
| SRR5030282 | 140 | 47 | 15379 | 21 | 0 | 10 | 0 | 0 | 0 | 0 | 0 | 19 | 0 | 10 | 40 | 0 | 21 | 0 | 0 | 60 | 80 | 0 | 0 | 0 | 26 | 260 | 78 | 0 | 0 | 0 | 47 | 0 | 16 | 0 | 2602 | 99 | 623 |
| SRR5030283 | 131 | 16 | 15507 | 11 | 0 | 16 | 11 | 0 | 0 | 0 | 0 | 19 | 0 | 27 | 54 | 0 | 24 | 21 | 0 | 43 | 43 | 0 | 0 | 0 | 33 | 217 | 87 | 0 | 0 | 0 | 27 | 0 | 22 | 0 | 2535 | 60 | 619 |
| SRR5030284 | 144 | 16 | 15463 | 0 | 0 | 0 | 0 | 0 | 0 | 0 | 0 | 16 | 0 | 11 | 115 | 0 | 20 | 0 | 0 | 42 | 83 | 0 | 0 | 0 | 21 | 222 | 74 | 0 | 0 | 0 | 53 | 0 | 16 | 0 | 84 | 101 | 19 |
| SRR5030285 | 140 | 21 | 15542 | 0 | 0 | 11 | 0 | 0 | 16 | 0 | 0 | 18 | 0 | 0 | 105 | 0 | 20 | 0 | 0 | 63 | 21 | 0 | 0 | 0 | 26 | 264 | 53 | 0 | 0 | 0 | 16 | 0 | 37 | 0 | 76 | 116 | 19 |
| SRR5030287 | 149 | 0 | 18711 | 79 | 44 | 0 | 0 | 0 | 0 | 0 | 0 | 29 | 0 | 26 | 53 | 0 | 31 | 16 | 0 | 53 | 0 | 0 | 0 | 0 | 0 | 473 | 132 | 18 | 0 | 0 | 18 | 0 | 0 | 0 | 2893 | 123 | 491 |
| SRR5030289 | 151 | 0 | 18708 | 17 | 43 | 0 | 0 | 0 | 0 | 0 | 0 | 22 | 0 | 0 | 35 | 0 | 35 | 0 | 0 | 87 | 69 | 0 | 0 | 0 | 0 | 359 | 128 | 0 | 0 | 0 | 34 | 0 | 0 | 0 | 2735 | 85 | 752 |
| SRR5030290 | 140 | 0 | 12280 | 26 | 0 | 31 | 0 | 0 | 78 | 0 | 0 | 157 | 0 | 26 | 177 | 0 | 35 | 156 | 0 | 62 | 231 | 0 | 0 | 0 | 16 | 87 | 110 | 10 | 0 | 0 | 11 | 0 | 0 | 0 | 235 | 17 | 71 |
| SRR5030291 | 152 | 22 | 12405 | 11 | 0 | 65 | 0 | 0 | 38 | 0 | 0 | 165 | 0 | 22 | 71 | 0 | 39 | 195 | 0 | 47 | 245 | 0 | 0 | 0 | 0 | 80 | 87 | 0 | 0 | 0 | 11 | 0 | 0 | 0 | 247 | 20 | 73 |
| SRR5030292 | 149 | 11 | 12357 | 27 | 16 | 48 | 0 | 0 | 91 | 0 | 0 | 143 | 0 | 0 | 102 | 11 | 25 | 198 | 0 | 55 | 227 | 0 | 0 | 0 | 0 | 1368 | 69 | 0 | 0 | 0 | 176 | 0 | 0 | 0 | 4297 | 319 | 1342 |
| SRR5030293 | 154 | 21 | 12310 | 0 | 0 | 37 | 0 | 0 | 69 | 0 | 0 | 149 | 0 | 11 | 141 | 0 | 35 | 144 | 0 | 31 | 188 | 0 | 0 | 0 | 0 | 1479 | 69 | 16 | 0 | 0 | 180 | 0 | 0 | 0 | 4428 | 308 | 1331 |
| SRR5030294 | 137 | 13 | 14473 | 0 | 0 | 80 | 0 | 0 | 100 | 0 | 0 | 76 | 0 | 20 | 29 | 13 | 32 | 36 | 0 | 0 | 175 | 0 | 0 | 0 | 40 | 60 | 139 | 13 | 0 | 0 | 219 | 0 | 0 | 0 | 104 | 11 | 66 |
| SRR5030295 | 143 | 14 | 14614 | 0 | 0 | 56 | 0 | 0 | 35 | 0 | 0 | 89 | 0 | 0 | 40 | 0 | 31 | 50 | 0 | 60 | 100 | 0 | 0 | 0 | 28 | 68 | 202 | 21 | 0 | 0 | 139 | 0 | 0 | 0 | 112 | 14 | 59 |
| SRR5030296 | 144 | 27 | 14412 | 13 | 0 | 108 | 0 | 0 | 74 | 0 | 0 | 95 | 0 | 61 | 79 | 0 | 31 | 37 | 0 | 49 | 118 | 0 | 0 | 0 | 61 | 63 | 148 | 0 | 0 | 0 | 202 | 0 | 20 | 34 | 99 | 15 | 60 |
| SRR5030297 | 141 | 0 | 14441 | 0 | 0 | 47 | 0 | 0 | 47 | 0 | 0 | 89 | 0 | 20 | 0 | 0 | 26 | 42 | 0 | 30 | 99 | 0 | 0 | 0 | 20 | 1608 | 189 | 14 | 0 | 0 | 142 | 0 | 14 | 0 | 2837 | 270 | 1317 |
| SRR5034616 | 384 | 49 | 235 | 99 | 25 | 0 | 0 | 25 | 0 | 0 | 0 | 373 | 0 | 12 | 0 | 0 | 0 | 0 | 101 | 226 | 1433 | 12 | 0 | 0 | 0 | 0 | 420 | 0 | 0 | 0 | 0 | 0 | 0 | 25 | 0 | 0 | 0 |
| SRR5034617 | 266 | 0 | 164 | 19 | 13 | 0 | 143 | 228 | 0 | 0 | 0 | 416 | 0 | 0 | 11 | 0 | 0 | 0 | 50 | 250 | 624 | 16 | 0 | 0 | 0 | 79 | 455 | 0 | 0 | 15 | 0 | 0 | 0 | 19 | 69 | 0 | 48 |
| SRR5034618 | 711 | 23 | 496 | 10 | 11 | 0 | 206 | 3052 | 0 | 0 | 0 | 1449 | 0 | 0 | 15 | 0 | 0 | 12 | 32 | 272 | 1073 | 0 | 0 | 0 | 0 | 60 | 444 | 0 | 0 | 125 | 0 | 0 | 0 | 33 | 47 | 0 | 19 |
| SRR5034619 | 395 | 0 | 253 | 415 | 80 | 0 | 0 | 15 | 0 | 0 | 0 | 510 | 0 | 0 | 0 | 0 | 0 | 95 | 27 | 133 | 1357 | 0 | 0 | 0 | 0 | 0 | 226 | 15 | 0 | 0 | 0 | 29 | 0 | 0 | 0 | 0 | 15 |
| SRR5034620 | 106 | 18 | 94 | 31 | 0 | 0 | 36 | 1244 | 0 | 0 | 0 | 182 | 0 | 0 | 18 | 0 | 0 | 56 | 90 | 525 | 1538 | 0 | 0 | 0 | 0 | 25 | 727 | 18 | 0 | 25 | 0 | 0 | 0 | 12 | 43 | 0 | 99 |
| SRR5034621 | 125 | 54 | 105 | 0 | 16 | 0 | 36 | 557 | 0 | 0 | 0 | 212 | 0 | 0 | 14 | 0 | 0 | 21 | 115 | 324 | 748 | 0 | 0 | 0 | 0 | 44 | 351 | 0 | 0 | 32 | 0 | 0 | 0 | 13 | 73 | 0 | 82 |
| SRR5034622 | 789 | 0 | 748 | 16 | 16 | 0 | 159 | 2807 | 0 | 0 | 0 | 1394 | 0 | 0 | 16 | 0 | 10 | 18 | 21 | 274 | 1221 | 0 | 0 | 0 | 0 | 65 | 299 | 0 | 0 | 98 | 0 | 0 | 0 | 31 | 34 | 0 | 17 |
| SRR5034623 | 71 | 0 | 38 | 0 | 0 | 0 | 301 | 308 | 0 | 0 | 0 | 70 | 0 | 0 | 15 | 0 | 0 | 0 | 0 | 0 | 20 | 0 | 0 | 0 | 0 | 37 | 261 | 0 | 0 | 0 | 0 | 0 | 0 | 45 | 0 | 0 | 0 |
| SRR5034625 | 356 | 20 | 348 | 26 | 0 | 0 | 16 | 2837 | 13 | 0 | 0 | 309 | 0 | 0 | 0 | 0 | 0 | 0 | 0 | 0 | 138 | 20 | 0 | 0 | 0 | 20 | 626 | 0 | 0 | 98 | 0 | 0 | 0 | 52 | 0 | 0 | 0 |
| SRR5034626 | 105 | 11 | 175 | 0 | 0 | 15 | 708 | 1763 | 32 | 0 | 0 | 194 | 0 | 0 | 33 | 0 | 0 | 0 | 0 | 18 | 91 | 0 | 0 | 0 | 0 | 21 | 22 | 0 | 0 | 132 | 0 | 0 | 0 | 114 | 28 | 0 | 11 |
| SRR5034627 | 58 | 0 | 23 | 10 | 0 | 0 | 215 | 16 | 0 | 0 | 0 | 14 | 0 | 0 | 31 | 0 | 0 | 0 | 62 | 203 | 160 | 10 | 0 | 0 | 0 | 0 | 12 | 0 | 41 | 15 | 21 | 0 | 0 | 21 | 0 | 0 | 0 |
| SRR5034628 | 124 | 0 | 29 | 80 | 16 | 0 | 0 | 31 | 0 | 0 | 0 | 55 | 0 | 0 | 0 | 0 | 0 | 0 | 55 | 165 | 631 | 0 | 0 | 0 | 0 | 16 | 25 | 0 | 0 | 80 | 0 | 0 | 0 | 32 | 48 | 0 | 48 |
| SRR5034629 | 97 | 44 | 82 | 11 | 22 | 0 | 27 | 291 | 0 | 0 | 0 | 32 | 0 | 0 | 91 | 11 | 0 | 0 | 0 | 164 | 18 | 0 | 0 | 0 | 0 | 71 | 176 | 0 | 0 | 22 | 0 | 0 | 0 | 93 | 0 | 0 | 22 |
| SRR5034630 | 249 | 35 | 330 | 47 | 24 | 0 | 125 | 259 | 0 | 0 | 0 | 191 | 0 | 0 | 208 | 0 | 0 | 0 | 0 | 187 | 146 | 0 | 0 | 0 | 0 | 12 | 177 | 0 | 0 | 0 | 0 | 0 | 0 | 47 | 24 | 0 | 0 |
| SRR5034632 | 427 | 74 | 229 | 126 | 21 | 0 | 0 | 32 | 0 | 0 | 0 | 201 | 0 | 0 | 68 | 0 | 0 | 0 | 0 | 34 | 119 | 11 | 0 | 0 | 0 | 11 | 285 | 11 | 0 | 0 | 0 | 0 | 0 | 0 | 0 | 0 | 0 |
| SRR5034634 | 304 | 19 | 187 | 19 | 0 | 0 | 0 | 205 | 0 | 0 | 0 | 257 | 0 | 0 | 83 | 0 | 18 | 0 | 33 | 17 | 83 | 19 | 0 | 0 | 0 | 0 | 410 | 0 | 0 | 0 | 56 | 0 | 0 | 19 | 0 | 0 | 0 |
| SRR5034635 | 407 | 0 | 343 | 112 | 16 | 0 | 28 | 48 | 0 | 0 | 0 | 377 | 0 | 0 | 14 | 0 | 15 | 0 | 0 | 71 | 156 | 0 | 0 | 0 | 0 | 32 | 398 | 0 | 0 | 0 | 16 | 0 | 0 | 48 | 0 | 0 | 0 |
| SRR5034636 | 402 | 25 | 449 | 50 | 0 | 0 | 0 | 75 | 0 | 0 | 0 | 398 | 0 | 0 | 48 | 0 | 10 | 0 | 0 | 0 | 95 | 75 | 0 | 0 | 0 | 50 | 201 | 0 | 0 | 0 | 0 | 75 | 0 | 25 | 0 | 0 | 0 |
| SRR5034637 | 173 | 77 | 145 | 102 | 77 | 0 | 50 | 154 | 0 | 0 | 0 | 156 | 0 | 0 | 151 | 0 | 0 | 0 | 0 | 0 | 352 | 0 | 0 | 0 | 0 | 0 | 999 | 0 | 0 | 26 | 0 | 0 | 0 | 26 | 0 | 0 | 0 |
| SRR5034639 | 445 | 0 | 664 | 273 | 149 | 0 | 0 | 0 | 0 | 0 | 0 | 616 | 0 | 0 | 0 | 0 | 0 | 0 | 40 | 40 | 120 | 0 | 0 | 0 | 0 | 0 | 398 | 0 | 0 | 0 | 0 | 0 | 0 | 0 | 0 | 0 | 0 |
| SRR5034641 | 86 | 12 | 25 | 0 | 0 | 0 | 0 | 12 | 0 | 0 | 0 | 13 | 0 | 0 | 158 | 0 | 0 | 0 | 79 | 158 | 355 | 0 | 0 | 0 | 0 | 0 | 162 | 0 | 0 | 0 | 0 | 0 | 0 | 24 | 0 | 0 | 0 |
| SRR5034642 | 84 | 65 | 15 | 16 | 11 | 0 | 186 | 188 | 0 | 0 | 0 | 25 | 0 | 16 | 0 | 0 | 0 | 0 | 0 | 10 | 33 | 16 | 0 | 0 | 0 | 0 | 397 | 22 | 0 | 370 | 0 | 0 | 0 | 33 | 22 | 0 | 0 |
| SRR5034643 | 392 | 0 | 192 | 33 | 0 | 0 | 458 | 1405 | 0 | 0 | 0 | 222 | 0 | 0 | 0 | 0 | 0 | 0 | 0 | 12 | 84 | 0 | 0 | 0 | 0 | 15 | 198 | 0 | 0 | 0 | 0 | 0 | 0 | 29 | 26 | 0 | 29 |
| SRR5034644 | 789 | 17 | 525 | 40 | 29 | 17 | 231 | 63 | 12 | 12 | 0 | 330 | 0 | 0 | 50 | 0 | 0 | 0 | 14 | 68 | 77 | 23 | 0 | 0 | 0 | 115 | 19 | 17 | 12 | 87 | 0 | 0 | 0 | 29 | 23 | 0 | 0 |
| SRR5034645 | 59 | 29 | 22 | 19 | 19 | 38 | 0 | 333 | 0 | 0 | 0 | 16 | 0 | 0 | 0 | 0 | 0 | 0 | 0 | 146 | 92 | 0 | 0 | 0 | 0 | 48 | 228 | 0 | 0 | 48 | 0 | 0 | 0 | 48 | 0 | 0 | 0 |
| SRR5034646 | 287 | 49 | 224 | 97 | 49 | 0 | 126 | 12276 | 0 | 0 | 0 | 148 | 0 | 12 | 91 | 0 | 0 | 0 | 23 | 103 | 434 | 0 | 0 | 0 | 0 | 61 | 256 | 24 | 0 | 852 | 0 | 0 | 0 | 61 | 0 | 0 | 0 |
| SRR5034647 | 212 | 51 | 129 | 14 | 0 | 0 | 249 | 1805 | 0 | 0 | 0 | 70 | 0 | 0 | 0 | 0 | 0 | 0 | 0 | 13 | 14 | 0 | 0 | 0 | 0 | 27 | 297 | 0 | 0 | 95 | 0 | 0 | 0 | 24 | 0 | 0 | 0 |
| SRR5034648 | 274 | 0 | 194 | 61 | 15 | 0 | 122 | 1560 | 0 | 0 | 0 | 286 | 0 | 30 | 0 | 0 | 0 | 0 | 24 | 122 | 219 | 30 | 0 | 45 | 0 | 76 | 1273 | 15 | 0 | 364 | 0 | 0 | 0 | 45 | 0 | 0 | 0 |
| SRR5034649 | 435 | 56 | 360 | 56 | 0 | 0 | 14 | 235 | 0 | 0 | 0 | 246 | 0 | 0 | 0 | 0 | 0 | 0 | 11 | 14 | 36 | 0 | 0 | 0 | 0 | 28 | 713 | 28 | 0 | 19 | 0 | 0 | 0 | 19 | 0 | 0 | 0 |
| SRR5034650 | 115 | 30 | 103 | 75 | 60 | 65 | 105 | 558 | 20 | 0 | 0 | 69 | 0 | 0 | 75 | 0 | 0 | 0 | 30 | 301 | 286 | 0 | 0 | 0 | 0 | 21 | 56 | 0 | 15 | 97 | 0 | 0 | 0 | 30 | 13 | 0 | 20 |
| SRR5034651 | 256 | 35 | 140 | 14 | 42 | 0 | 393 | 673 | 0 | 0 | 0 | 104 | 0 | 0 | 0 | 0 | 0 | 0 | 27 | 27 | 352 | 0 | 0 | 0 | 0 | 0 | 24 | 0 | 0 | 81 | 0 | 0 | 0 | 21 | 0 | 0 | 0 |
| SRR5034652 | 103 | 145 | 65 | 16 | 0 | 0 | 478 | 14728 | 0 | 0 | 0 | 66 | 0 | 0 | 20 | 0 | 0 | 0 | 20 | 45 | 115 | 0 | 0 | 0 | 0 | 139 | 285 | 0 | 0 | 1015 | 0 | 0 | 0 | 51 | 0 | 0 | 13 |
| SRR5034653 | 97 | 132 | 56 | 25 | 0 | 12 | 731 | 29091 | 0 | 0 | 0 | 103 | 0 | 0 | 0 | 0 | 0 | 0 | 24 | 44 | 96 | 0 | 0 | 0 | 0 | 161 | 252 | 0 | 0 | 602 | 0 | 0 | 0 | 50 | 0 | 0 | 0 |
| SRR5034654 | 223 | 22 | 221 | 0 | 11 | 18 | 138 | 140 | 11 | 0 | 0 | 261 | 0 | 0 | 31 | 0 | 0 | 0 | 16 | 27 | 114 | 0 | 0 | 0 | 0 | 190 | 12 | 0 | 0 | 14 | 0 | 0 | 0 | 80 | 24 | 0 | 11 |
| SRR5034655 | 231 | 108 | 124 | 60 | 0 | 0 | 903 | 31563 | 0 | 0 | 0 | 111 | 0 | 0 | 0 | 0 | 0 | 0 | 0 | 26 | 38 | 13 | 0 | 0 | 0 | 22 | 138 | 13 | 0 | 1265 | 0 | 0 | 0 | 125 | 0 | 0 | 0 |
| SRR5034656 | 317 | 105 | 279 | 26 | 0 | 13 | 598 | 15648 | 0 | 0 | 0 | 181 | 0 | 0 | 0 | 0 | 0 | 0 | 0 | 10 | 92 | 0 | 0 | 0 | 0 | 0 | 183 | 0 | 0 | 681 | 0 | 0 | 0 | 69 | 0 | 0 | 0 |
| SRR5034657 | 195 | 32 | 105 | 13 | 32 | 0 | 50 | 818 | 0 | 0 | 0 | 78 | 0 | 0 | 0 | 0 | 11 | 0 | 0 | 19 | 31 | 0 | 0 | 0 | 0 | 13 | 331 | 0 | 0 | 52 | 19 | 0 | 0 | 175 | 26 | 0 | 32 |
| SRR5034658 | 230 | 17 | 125 | 0 | 11 | 28 | 213 | 492 | 0 | 0 | 0 | 124 | 0 | 0 | 47 | 0 | 0 | 0 | 16 | 27 | 144 | 0 | 0 | 0 | 0 | 15 | 18 | 0 | 0 | 61 | 0 | 0 | 0 | 88 | 0 | 0 | 0 |
| SRR5034659 | 137 | 37 | 147 | 11 | 0 | 18 | 873 | 13787 | 54 | 0 | 0 | 126 | 0 | 0 | 0 | 0 | 0 | 0 | 0 | 0 | 27 | 0 | 0 | 0 | 0 | 274 | 307 | 0 | 0 | 534 | 0 | 0 | 0 | 423 | 0 | 0 | 0 |
| SRR5034660 | 133 | 23 | 85 | 11 | 0 | 0 | 2148 | 89916 | 11 | 0 | 0 | 140 | 0 | 0 | 0 | 0 | 0 | 0 | 0 | 0 | 31 | 0 | 0 | 0 | 0 | 164 | 176 | 0 | 0 | 1205 | 0 | 0 | 0 | 191 | 0 | 0 | 0 |
| SRR5034661 | 246 | 0 | 79 | 14 | 0 | 0 | 0 | 426 | 0 | 0 | 0 | 0 | 0 | 0 | 0 | 0 | 0 | 0 | 0 | 70 | 45 | 0 | 0 | 0 | 0 | 25 | 40 | 0 | 25 | 0 | 0 | 0 | 0 | 72 | 11 | 0 | 11 |
| SRR5034662 | 341 | 12 | 279 | 10 | 10 | 0 | 1666 | 1593 | 0 | 0 | 0 | 414 | 0 | 0 | 15 | 0 | 0 | 0 | 0 | 12 | 59 | 0 | 0 | 0 | 0 | 93 | 17 | 0 | 0 | 26 | 0 | 0 | 0 | 72 | 0 | 0 | 0 |
| SRR5034663 | 195 | 42 | 169 | 90 | 0 | 0 | 145 | 1493 | 0 | 0 | 0 | 88 | 0 | 0 | 13 | 0 | 0 | 0 | 0 | 135 | 84 | 19 | 0 | 0 | 0 | 137 | 650 | 0 | 0 | 122 | 0 | 0 | 0 | 19 | 19 | 0 | 14 |
| SRR5054304 | 169 | 0 | 21 | 0 | 0 | 239 | 0 | 0 | 349 | 171 | 223 | 413 | 459 | 70 | 167 | 52 | 203 | 0 | 0 | 0 | 40 | 0 | 0 | 0 | 32 | 2049 | 20 | 29 | 25 | 0 | 2711 | 0 | 0 | 1894 | 9587 | 20460 | 39195 |
| SRR5054305 | 146 | 13 | 30 | 0 | 0 | 1694 | 0 | 0 | 734 | 0 | 55 | 461 | 23 | 65 | 123 | 13 | 163 | 0 | 0 | 18 | 335 | 159 | 0 | 21 | 18 | 10489 | 130 | 68 | 0 | 0 | 3325 | 36 | 31 | 234 | 18534 | 19627 | 102562 |

### Supp Table S8.2.Columns_AM_to_BW

| | | | | | | | | | | | | | | | | | | | | | | | | | | | | | | | | | | | | | |
|---|---|---|---|---|---|---|---|---|---|---|---|---|---|---|---|---|---|---|---|---|---|---|---|---|---|---|---|---|---|---|---|---|---|---|---|---|---|
| SRR5087508 | 47 | 22 | 487 | 0 | 0 | 25 | 30 | 0 | 70 | 0 | 0 | 89 | 0 | 64 | 66 | 0 | 0 | 0 | 0 | 47 | 127 | 0 | 0 | 0 | 15 | 66 | 23 | 46 | 18 | 0 | 72 | 0 | 0 | 32 | 15 | 11 | 69 |
| SRR5087510 | 48 | 15 | 432 | 0 | 0 | 31 | 40 | 10 | 63 | 0 | 0 | 97 | 0 | 67 | 63 | 0 | 11 | 0 | 0 | 73 | 98 | 0 | 0 | 0 | 10 | 75 | 37 | 36 | 21 | 0 | 75 | 0 | 0 | 36 | 16 | 16 | 80 |
| SRR5087512 | 47 | 14 | 447 | 0 | 0 | 32 | 58 | 0 | 68 | 0 | 0 | 102 | 0 | 61 | 74 | 0 | 0 | 0 | 0 | 51 | 82 | 0 | 0 | 0 | 12 | 77 | 33 | 37 | 19 | 0 | 87 | 0 | 0 | 28 | 16 | 11 | 72 |
| SRR5087516 | 64 | 18 | 599 | 0 | 0 | 41 | 42 | 18 | 77 | 0 | 0 | 133 | 0 | 41 | 21 | 0 | 11 | 0 | 0 | 32 | 94 | 0 | 0 | 0 | 15 | 76 | 41 | 34 | 19 | 0 | 59 | 0 | 0 | 63 | 13 | 0 | 52 |
| SRR5087517 | 61 | 16 | 562 | 0 | 0 | 50 | 34 | 18 | 87 | 0 | 0 | 116 | 0 | 42 | 24 | 0 | 0 | 0 | 0 | 37 | 97 | 0 | 0 | 0 | 17 | 87 | 45 | 45 | 15 | 0 | 74 | 0 | 0 | 60 | 11 | 0 | 50 |
| SRR5087519 | 54 | 29 | 562 | 0 | 0 | 40 | 40 | 22 | 10 | 0 | 0 | 125 | 0 | 35 | 25 | 0 | 10 | 0 | 0 | 31 | 82 | 0 | 0 | 0 | 0 | 85 | 45 | 40 | 29 | 0 | 65 | 0 | 0 | 57 | 13 | 13 | 41 |
| SRR5087522 | 42 | 11 | 452 | 0 | 0 | 66 | 28 | 0 | 36 | 0 | 0 | 63 | 0 | 0 | 51 | 0 | 0 | 0 | 0 | 192 | 462 | 0 | 0 | 0 | 18 | 1090 | 503 | 0 | 0 | 0 | 131 | 0 | 0 | 12 | 519 | 27 | 244 |
| SRR5087527 | 43 | 14 | 451 | 0 | 0 | 52 | 30 | 0 | 62 | 0 | 0 | 59 | 0 | 15 | 30 | 0 | 0 | 0 | 0 | 190 | 571 | 0 | 0 | 0 | 34 | 92 | 40 | 0 | 0 | 0 | 11 | 0 | 0 | 0 | 38 | 43 | 17 |
| SRR5087554 | 42 | 0 | 470 | 0 | 0 | 64 | 25 | 0 | 44 | 0 | 0 | 61 | 0 | 0 | 38 | 0 | 0 | 0 | 0 | 239 | 522 | 0 | 0 | 0 | 44 | 98 | 37 | 0 | 0 | 0 | 10 | 0 | 0 | 0 | 34 | 21 | 16 |
| SRR5090283 | 55 | 0 | 0 | 14 | 31 | 18 | 88 | 56 | 23 | 12 | 75 | 542 | 0 | 41 | 219 | 0 | 14 | 0 | 32 | 196 | 38 | 0 | 0 | 658 | 32 | 71 | 56 | 15 | 0 | 87 | 59 | 0 | 0 | 51 | 12 | 21 | 30 |
| SRR5092303 | 2620 | 0 | 0 | 0 | 39 | 0 | 0 | 20 | 20 | 20 | 39 | 17923 | 3368 | 20 | 2996 | 0 | 992 | 0 | 0 | 0 | 214 | 0 | 0 | 236 | 39 | 0 | 0 | 0 | 0 | 20 | 0 | 0 | 0 | 512 | 0 | 0 | 39 |
| SRR5092304 | 1842 | 0 | 0 | 0 | 0 | 0 | 0 | 47 | 0 | 0 | 0 | 12254 | 2215 | 0 | 1776 | 0 | 547 | 0 | 0 | 0 | 384 | 0 | 0 | 163 | 70 | 23 | 0 | 0 | 0 | 70 | 23 | 0 | 0 | 326 | 23 | 0 | 93 |
| SRR5092305 | 593 | 0 | 0 | 0 | 22 | 0 | 0 | 22 | 22 | 0 | 0 | 4216 | 925 | 88 | 1149 | 0 | 387 | 0 | 0 | 0 | 418 | 0 | 0 | 353 | 22 | 0 | 0 | 0 | 0 | 88 | 22 | 0 | 0 | 264 | 22 | 0 | 22 |
| SRR5092836 | 25 | 38 | 2496 | 0 | 0 | 38 | 601 | 114 | 38 | 0 | 0 | 310 | 0 | 38 | 0 | 0 | 15 | 0 | 651 | 2203 | 0 | 38 | 0 | 0 | 38 | 36 | 26 | 38 | 38 | 0 | 0 | 0 | 0 | 38 | 190 | 0 | 0 |
| SRR5100734 | 0 | 36 | 70 | 50 | 29 | 0 | 2219 | 78 | 13 | 0 | 12 | 5664 | 0 | 0 | 30 | 0 | 0 | 0 | 0 | 0 | 0 | 0 | 0 | 0 | 0 | 0 | 0 | 0 | 0 | 179 | 47 | 15 | 211 | 234 | 0 | 0 | 12 |
| SRR5100735 | 0 | 34 | 0 | 0 | 0 | 0 | 1535 | 128 | 92 | 0 | 15 | 1178 | 0 | 0 | 17 | 0 | 17 | 0 | 60 | 0 | 105 | 0 | 0 | 13 | 0 | 19 | 0 | 0 | 0 | 302 | 64 | 0 | 54 | 167 | 72 | 224 | 544 |
| SRR5104206 | 43 | 20 | 1250 | 0 | 0 | 17 | 0 | 0 | 17 | 0 | 0 | 183 | 0 | 16 | 378 | 0 | 0 | 14 | 0 | 402 | 0 | 0 | 0 | 0 | 0 | 71 | 24 | 19 | 0 | 0 | 24 | 0 | 0 | 0 | 30 | 12 | 19 |
| SRR5104207 | 36 | 21 | 2681 | 0 | 0 | 26 | 25 | 0 | 24 | 0 | 0 | 267 | 0 | 33 | 126 | 0 | 0 | 10 | 0 | 370 | 0 | 0 | 0 | 0 | 11 | 76 | 30 | 15 | 0 | 0 | 49 | 0 | 11 | 0 | 22 | 20 | 15 |
| SRR5104208 | 34 | 21 | 2216 | 0 | 0 | 16 | 0 | 0 | 31 | 0 | 0 | 219 | 0 | 14 | 405 | 0 | 0 | 21 | 0 | 336 | 0 | 0 | 0 | 0 | 24 | 78 | 33 | 11 | 0 | 0 | 35 | 0 | 11 | 0 | 17 | 21 | 15 |
| SRR5104209 | 38 | 18 | 2274 | 0 | 0 | 15 | 0 | 0 | 31 | 0 | 0 | 173 | 0 | 28 | 620 | 0 | 0 | 0 | 0 | 255 | 0 | 0 | 0 | 0 | 18 | 281 | 116 | 25 | 0 | 0 | 28 | 0 | 0 | 0 | 74 | 0 | 62 |
| SRR5104210 | 53 | 28 | 1780 | 0 | 0 | 21 | 0 | 0 | 17 | 0 | 0 | 186 | 0 | 22 | 433 | 0 | 0 | 0 | 0 | 208 | 0 | 0 | 0 | 0 | 11 | 351 | 63 | 0 | 0 | 0 | 29 | 0 | 0 | 0 | 118 | 21 | 64 |
| SRR5104211 | 64 | 15 | 1974 | 0 | 0 | 19 | 0 | 0 | 14 | 0 | 0 | 149 | 0 | 22 | 74 | 0 | 0 | 10 | 0 | 147 | 0 | 0 | 0 | 0 | 13 | 372 | 65 | 16 | 0 | 0 | 24 | 0 | 0 | 0 | 65 | 0 | 18 |
| SRR5104212 | 59 | 26 | 2596 | 0 | 0 | 16 | 0 | 0 | 0 | 0 | 0 | 121 | 0 | 20 | 755 | 0 | 0 | 0 | 0 | 185 | 0 | 0 | 0 | 0 | 12 | 350 | 97 | 0 | 0 | 0 | 48 | 0 | 16 | 0 | 55 | 12 | 39 |
| SRR5104213 | 66 | 52 | 1891 | 0 | 0 | 10 | 23 | 0 | 24 | 0 | 0 | 107 | 0 | 16 | 317 | 0 | 0 | 29 | 0 | 141 | 0 | 0 | 0 | 0 | 16 | 361 | 69 | 19 | 0 | 0 | 58 | 0 | 0 | 0 | 82 | 21 | 49 |
| SRR5104214 | 79 | 19 | 1553 | 0 | 0 | 17 | 23 | 0 | 33 | 0 | 0 | 53 | 0 | 14 | 245 | 0 | 0 | 0 | 0 | 152 | 0 | 0 | 0 | 0 | 16 | 345 | 61 | 0 | 0 | 0 | 45 | 0 | 0 | 0 | 31 | 0 | 0 |
| SRR5104215 | 51 | 19 | 1652 | 0 | 0 | 18 | 11 | 0 | 14 | 0 | 0 | 78 | 0 | 18 | 628 | 0 | 0 | 0 | 0 | 152 | 0 | 0 | 0 | 0 | 14 | 470 | 75 | 13 | 0 | 0 | 43 | 0 | 0 | 0 | 58 | 0 | 18 |
| SRR5104216 | 36 | 11 | 1445 | 0 | 0 | 17 | 0 | 0 | 30 | 0 | 0 | 91 | 0 | 16 | 83 | 0 | 0 | 0 | 0 | 249 | 0 | 0 | 0 | 0 | 15 | 47 | 57 | 15 | 0 | 0 | 74 | 0 | 12 | 0 | 90 | 11 | 39 |
| SRR5104217 | 53 | 20 | 1624 | 0 | 0 | 28 | 0 | 0 | 13 | 0 | 0 | 61 | 0 | 20 | 94 | 0 | 0 | 0 | 0 | 121 | 0 | 0 | 0 | 0 | 0 | 330 | 80 | 0 | 0 | 0 | 57 | 0 | 13 | 0 | 64 | 12 | 25 |
| SRR5104218 | 52 | 20 | 981 | 0 | 0 | 17 | 20 | 0 | 20 | 0 | 0 | 37 | 0 | 27 | 276 | 0 | 0 | 12 | 0 | 217 | 0 | 0 | 0 | 0 | 17 | 40 | 39 | 17 | 0 | 0 | 46 | 0 | 13 | 0 | 43 | 13 | 21 |
| SRR5104219 | 54 | 14 | 1109 | 0 | 0 | 32 | 0 | 0 | 20 | 0 | 0 | 49 | 0 | 19 | 111 | 0 | 0 | 0 | 0 | 282 | 0 | 0 | 0 | 0 | 14 | 50 | 66 | 17 | 0 | 0 | 69 | 0 | 0 | 0 | 60 | 16 | 23 |
| SRR5104220 | 43 | 22 | 827 | 0 | 0 | 28 | 42 | 0 | 20 | 0 | 0 | 50 | 0 | 43 | 192 | 0 | 0 | 0 | 0 | 186 | 0 | 0 | 0 | 0 | 14 | 497 | 49 | 26 | 0 | 0 | 54 | 0 | 0 | 0 | 94 | 18 | 36 |
| SRR5104221 | 29 | 0 | 572 | 0 | 0 | 17 | 18 | 0 | 16 | 0 | 0 | 29 | 0 | 0 | 150 | 0 | 0 | 0 | 0 | 176 | 0 | 0 | 0 | 0 | 0 | 311 | 43 | 11 | 0 | 0 | 16 | 0 | 0 | 0 | 60 | 11 | 16 |
| SRR5104222 | 45 | 15 | 819 | 0 | 0 | 18 | 0 | 0 | 13 | 0 | 0 | 29 | 0 | 24 | 128 | 0 | 0 | 0 | 0 | 163 | 0 | 0 | 0 | 0 | 10 | 303 | 54 | 10 | 0 | 0 | 35 | 0 | 0 | 0 | 46 | 0 | 28 |
| SRR5104223 | 46 | 14 | 280 | 0 | 0 | 13 | 0 | 0 | 16 | 0 | 0 | 32 | 0 | 10 | 53 | 0 | 0 | 14 | 0 | 190 | 0 | 0 | 0 | 0 | 0 | 39 | 25 | 12 | 0 | 0 | 48 | 0 | 0 | 0 | 35 | 0 | 13 |
| SRR5104224 | 60 | 11 | 818 | 0 | 0 | 13 | 0 | 0 | 0 | 0 | 0 | 18 | 0 | 23 | 159 | 0 | 0 | 0 | 0 | 51 | 0 | 0 | 0 | 0 | 0 | 31 | 25 | 0 | 0 | 0 | 34 | 0 | 11 | 0 | 20 | 0 | 0 |
| SRR5104225 | 44 | 25 | 837 | 0 | 0 | 28 | 0 | 0 | 17 | 0 | 0 | 32 | 0 | 13 | 213 | 0 | 0 | 0 | 0 | 205 | 0 | 0 | 0 | 0 | 13 | 43 | 55 | 18 | 0 | 0 | 45 | 0 | 0 | 0 | 68 | 0 | 35 |
| SRR5116489 | 73 | 0 | 5518 | 0 | 0 | 0 | 0 | 0 | 0 | 0 | 0 | 427 | 0 | 0 | 0 | 0 | 0 | 0 | 0 | 147 | 74 | 0 | 0 | 0 | 73 | 73 | 623 | 0 | 0 | 0 | 110 | 0 | 0 | 37 | 1247 | 73 | 660 |
| SRR5116490 | 183 | 0 | 3710 | 0 | 0 | 158 | 0 | 26 | 26 | 0 | 0 | 1032 | 0 | 0 | 0 | 13 | 0 | 0 | 0 | 4789 | 1906 | 13 | 0 | 0 | 119 | 843 | 9048 | 0 | 0 | 0 | 13 | 0 | 0 | 13 | 1093 | 53 | 777 |
| SRR5116491 | 85 | 0 | 4458 | 0 | 0 | 43 | 0 | 21 | 21 | 0 | 0 | 515 | 0 | 21 | 0 | 0 | 0 | 0 | 0 | 3524 | 611 | 0 | 0 | 21 | 43 | 490 | 7395 | 0 | 0 | 0 | 128 | 0 | 0 | 0 | 1151 | 43 | 831 |
| SRR5116492 | 116 | 0 | 3302 | 0 | 0 | 50 | 291 | 96 | 50 | 0 | 0 | 970 | 0 | 14 | 0 | 23 | 0 | 0 | 13 | 5162 | 6920 | 0 | 0 | 14 | 46 | 34 | 425 | 32 | 0 | 0 | 82 | 0 | 0 | 0 | 115 | 11 | 76 |
| SRR5116493 | 80 | 0 | 3964 | 0 | 0 | 16 | 0 | 16 | 47 | 0 | 0 | 395 | 0 | 0 | 0 | 0 | 13 | 0 | 0 | 1769 | 993 | 0 | 0 | 0 | 16 | 488 | 3935 | 16 | 0 | 0 | 110 | 0 | 0 | 0 | 1385 | 47 | 803 |
| SRR5116494 | 118 | 0 | 3209 | 0 | 0 | 34 | 0 | 0 | 67 | 0 | 0 | 477 | 0 | 11 | 0 | 0 | 0 | 46 | 0 | 755 | 1748 | 0 | 0 | 11 | 56 | 426 | 1369 | 0 | 0 | 0 | 146 | 0 | 0 | 0 | 2042 | 101 | 1010 |
| SRR5116495 | 117 | 0 | 4026 | 0 | 0 | 33 | 34 | 29 | 19 | 0 | 0 | 649 | 0 | 0 | 0 | 0 | 0 | 0 | 0 | 2254 | 2612 | 0 | 0 | 0 | 19 | 129 | 2448 | 14 | 0 | 0 | 48 | 0 | 0 | 0 | 1150 | 81 | 835 |
| SRR5116496 | 68 | 0 | 2651 | 0 | 0 | 20 | 44 | 0 | 69 | 0 | 0 | 801 | 0 | 20 | 0 | 0 | 0 | 0 | 0 | 1684 | 1507 | 0 | 0 | 0 | 0 | 177 | 1228 | 20 | 0 | 0 | 59 | 0 | 0 | 0 | 1620 | 59 | 697 |
| SRR5121477 | 32 | 0 | 2223 | 0 | 0 | 0 | 58 | 125 | 0 | 0 | 0 | 0 | 0 | 0 | 0 | 0 | 74 | 14 | 13 | 5243 | 923 | 0 | 0 | 19 | 0 | 0 | 0 | 0 | 62 | 13 | 0 | 0 | 0 | 26 | 76 | 10 | 42 |
| SRR5121478 | 39 | 0 | 294 | 0 | 0 | 0 | 48 | 926 | 0 | 0 | 0 | 0 | 0 | 0 | 0 | 0 | 19 | 0 | 29 | 9176 | 1631 | 0 | 0 | 123 | 0 | 0 | 0 | 0 | 0 | 61 | 0 | 0 | 0 | 0 | 370 | 65 | 770 |
| SRR5121479 | 51 | 0 | 2199 | 0 | 0 | 119 | 33 | 35 | 30 | 0 | 0 | 12 | 0 | 11 | 47 | 0 | 16 | 0 | 0 | 2597 | 957 | 0 | 0 | 0 | 0 | 1432 | 1938 | 0 | 11 | 0 | 184 | 0 | 0 | 37 | 4283 | 621 | 2397 |
| SRR5121480 | 0 | 0 | 3410 | 0 | 0 | 33 | 167 | 81 | 20 | 0 | 0 | 0 | 0 | 22 | 0 | 0 | 38 | 0 | 13 | 640 | 380 | 0 | 0 | 0 | 0 | 545 | 1214 | 75 | 41 | 11 | 222 | 0 | 0 | 27 | 483 | 154 | 1120 |
| SRR5121481 | 16 | 0 | 4248 | 0 | 11 | 17 | 0 | 163 | 19 | 0 | 0 | 0 | 0 | 17 | 15 | 0 | 0 | 0 | 0 | 1142 | 1093 | 0 | 0 | 0 | 0 | 402 | 656 | 0 | 55 | 22 | 17 | 0 | 0 | 41 | 1063 | 88 | 554 |
| SRR5121482 | 0 | 0 | 12988 | 0 | 0 | 58 | 21 | 13 | 24 | 0 | 0 | 0 | 0 | 13 | 0 | 0 | 17 | 0 | 0 | 707 | 1198 | 17 | 0 | 0 | 0 | 77 | 92 | 0 | 0 | 16 | 56 | 0 | 0 | 23 | 58 | 15 | 82 |
| SRR5121483 | 14 | 13 | 5712 | 0 | 0 | 0 | 230 | 4533 | 0 | 0 | 0 | 0 | 0 | 23 | 818 | 0 | 31 | 0 | 0 | 11520 | 8871 | 0 | 0 | 34 | 0 | 0 | 0 | 0 | 0 | 54 | 0 | 0 | 0 | 13 | 13 | 14 | 69 |
| SRR5121484 | 0 | 0 | 5329 | 0 | 0 | 78 | 22 | 0 | 20 | 0 | 0 | 0 | 0 | 0 | 40 | 0 | 16 | 0 | 22 | 8555 | 5690 | 0 | 0 | 12 | 0 | 63 | 222 | 72 | 0 | 0 | 58 | 0 | 0 | 29 | 110 | 14 | 137 |
| SRR5121485 | 0 | 0 | 2923 | 0 | 0 | 26 | 0 | 0 | 16 | 0 | 0 | 0 | 0 | 0 | 78 | 0 | 18 | 0 | 34 | 2486 | 3145 | 0 | 0 | 0 | 0 | 71 | 41 | 0 | 0 | 0 | 33 | 0 | 0 | 28 | 786 | 98 | 530 |
| SRR5121486 | 13 | 40 | 3452 | 0 | 0 | 0 | 76 | 696 | 12 | 0 | 0 | 0 | 0 | 35 | 111 | 0 | 19 | 0 | 0 | 377 | 9216 | 0 | 0 | 0 | 0 | 24 | 52 | 0 | 0 | 0 | 0 | 0 | 0 | 14 | 142 | 0 | 54 |
| SRR5121487 | 0 | 0 | 4354 | 0 | 11 | 71 | 0 | 55 | 13 | 0 | 0 | 0 | 0 | 0 | 92 | 0 | 25 | 0 | 98 | 6030 | 2605 | 0 | 0 | 0 | 11 | 1735 | 780 | 11 | 0 | 0 | 49 | 0 | 0 | 11 | 2428 | 326 | 2660 |
| SRR5127200 | 210 | 145 | 706 | 0 | 0 | 29 | 0 | 1046 | 0 | 0 | 0 | 0 | 0 | 10 | 51 | 0 | 12 | 0 | 0 | 664 | 441 | 0 | 0 | 0 | 10 | 439 | 376 | 0 | 0 | 0 | 0 | 0 | 0 | 0 | 923 | 51 | 390 |
| SRR5127201 | 24 | 0 | 4643 | 0 | 0 | 25 | 53 | 35 | 22 | 0 | 0 | 558 | 0 | 0 | 42 | 0 | 0 | 0 | 0 | 1018 | 242 | 0 | 0 | 0 | 0 | 569 | 1033 | 0 | 0 | 0 | 21 | 0 | 0 | 0 | 1326 | 236 | 917 |
| SRR5127202 | 48 | 0 | 4112 | 0 | 0 | 102 | 93 | 0 | 37 | 0 | 0 | 1422 | 0 | 0 | 36 | 0 | 0 | 0 | 14 | 1602 | 467 | 0 | 0 | 0 | 0 | 178 | 673 | 15 | 0 | 0 | 57 | 0 | 0 | 15 | 186 | 31 | 167 |
| SRR5127203 | 67 | 0 | 3038 | 0 | 0 | 166 | 15 | 19 | 134 | 0 | 0 | 643 | 14 | 0 | 31 | 0 | 0 | 0 | 12 | 2685 | 606 | 0 | 0 | 0 | 0 | 3150 | 5154 | 10 | 0 | 0 | 72 | 0 | 0 | 24 | 3031 | 610 | 3129 |
| SRR5127204 | 22 | 0 | 1108 | 0 | 0 | 79 | 0 | 0 | 34 | 0 | 0 | 228 | 0 | 0 | 85 | 0 | 0 | 0 | 0 | 2174 | 1140 | 0 | 0 | 0 | 0 | 2063 | 2671 | 0 | 0 | 0 | 54 | 0 | 0 | 0 | 674 | 129 | 787 |
| SRR5127205 | 41 | 0 | 1361 | 0 | 0 | 29 | 24 | 0 | 15 | 0 | 0 | 1626 | 0 | 0 | 128 | 0 | 0 | 21 | 18 | 3685 | 512 | 0 | 0 | 0 | 0 | 1357 | 2016 | 0 | 0 | 0 | 49 | 0 | 0 | 0 | 444 | 174 | 1247 |
| SRR5127206 | 26 | 0 | 4137 | 0 | 0 | 94 | 41 | 36 | 34 | 0 | 0 | 648 | 0 | 0 | 34 | 0 | 0 | 0 | 0 | 1650 | 424 | 0 | 0 | 0 | 0 | 189 | 290 | 0 | 0 | 0 | 44 | 0 | 0 | 19 | 89 | 20 | 93 |
| SRR5127207 | 14 | 0 | 635 | 0 | 0 | 10 | 60 | 0 | 0 | 0 | 0 | 901 | 0 | 0 | 0 | 0 | 0 | 0 | 50 | 411 | 882 | 0 | 0 | 0 | 0 | 75 | 75 | 0 | 0 | 0 | 0 | 0 | 0 | 0 | 22 | 0 | 11 |
| SRR5127208 | 78 | 0 | 3310 | 0 | 0 | 107 | 0 | 22 | 52 | 0 | 0 | 798 | 0 | 0 | 18 | 0 | 0 | 0 | 0 | 1066 | 834 | 0 | 0 | 0 | 0 | 135 | 309 | 0 | 0 | 0 | 22 | 0 | 0 | 0 | 114 | 39 | 276 |
| SRR5127209 | 162 | 11 | 15298 | 0 | 0 | 208 | 307 | 30 | 64 | 0 | 0 | 346 | 0 | 0 | 169 | 0 | 30 | 0 | 21 | 547 | 984 | 0 | 0 | 0 | 12 | 3183 | 2084 | 14 | 19 | 0 | 89 | 0 | 0 | 30 | 773 | 130 | 861 |
| SRR5127210 | 24 | 0 | 4218 | 0 | 0 | 96 | 0 | 0 | 44 | 0 | 0 | 883 | 0 | 0 | 35 | 0 | 0 | 0 | 52 | 3320 | 1103 | 0 | 0 | 0 | 0 | 194 | 576 | 0 | 0 | 0 | 44 | 0 | 0 | 0 | 204 | 38 | 277 |
| SRR5127211 | 33 | 0 | 2622 | 0 | 0 | 12 | 13 | 18 | 0 | 0 | 0 | 535 | 0 | 0 | 53 | 0 | 0 | 0 | 12 | 965 | 513 | 0 | 0 | 0 | 0 | 67 | 78 | 0 | 0 | 0 | 0 | 0 | 0 | 0 | 36 | 0 | 44 |
| SRR5127212 | 66 | 0 | 3346 | 0 | 0 | 63 | 0 | 0 | 25 | 0 | 0 | 346 | 0 | 0 | 15 | 0 | 0 | 0 | 0 | 895 | 472 | 0 | 0 | 0 | 0 | 1274 | 4356 | 0 | 0 | 0 | 34 | 0 | 0 | 0 | 1137 | 103 | 711 |
| SRR5127213 | 16 | 0 | 5677 | 0 | 0 | 41 | 22 | 0 | 39 | 0 | 0 | 356 | 0 | 0 | 11 | 0 | 0 | 0 | 17 | 3318 | 576 | 0 | 0 | 0 | 0 | 78 | 480 | 15 | 0 | 0 | 34 | 0 | 0 | 0 | 209 | 23 | 168 |
| SRR5127214 | 26 | 0 | 8742 | 0 | 0 | 125 | 43 | 0 | 63 | 0 | 0 | 553 | 0 | 0 | 26 | 0 | 0 | 0 | 22 | 2396 | 438 | 0 | 0 | 0 | 0 | 3354 | 3417 | 0 | 0 | 0 | 100 | 0 | 0 | 0 | 1125 | 278 | 1290 |
| SRR5127215 | 45 | 0 | 2141 | 0 | 0 | 106 | 0 | 0 | 54 | 0 | 0 | 568 | 0 | 0 | 0 | 0 | 0 | 0 | 0 | 1005 | 174 | 0 | 0 | 0 | 0 | 2450 | 2891 | 0 | 0 | 0 | 46 | 0 | 0 | 0 | 1455 | 291 | 1303 |
| SRR5127216 | 52 | 0 | 3508 | 0 | 0 | 106 | 0 | 0 | 56 | 0 | 0 | 641 | 0 | 0 | 0 | 0 | 0 | 0 | 0 | 881 | 482 | 0 | 0 | 0 | 0 | 1739 | 4620 | 0 | 0 | 0 | 51 | 0 | 0 | 17 | 810 | 153 | 1059 |
| SRR5127217 | 18 | 0 | 3650 | 0 | 0 | 60 | 10 | 0 | 36 | 0 | 0 | 541 | 0 | 0 | 12 | 0 | 0 | 0 | 0 | 499 | 164 | 0 | 0 | 0 | 0 | 96 | 172 | 0 | 0 | 0 | 47 | 0 | 0 | 0 | 37 | 108 | 50 |
| SRR5127218 | 21 | 0 | 2690 | 0 | 0 | 114 | 29 | 15 | 48 | 0 | 0 | 1332 | 0 | 0 | 115 | 0 | 0 | 13 | 0 | 3058 | 745 | 0 | 0 | 0 | 0 | 180 | 267 | 0 | 20 | 0 | 103 | 0 | 0 | 12 | 91 | 19 | 102 |
| SRR5127219 | 47 | 35 | 144 | 0 | 0 | 0 | 123 | 583 | 0 | 0 | 0 | 0 | 0 | 21 | 0 | 0 | 0 | 0 | 0 | 303 | 130 | 0 | 0 | 0 | 0 | 87 | 118 | 0 | 0 | 28 | 0 | 0 | 0 | 70 | 102 | 0 | 50 |
| SRR5127220 | 21 | 0 | 3592 | 0 | 0 | 42 | 11 | 49 | 23 | 0 | 0 | 767 | 0 | 0 | 101 | 0 | 0 | 0 | 0 | 695 | 615 | 0 | 0 | 0 | 0 | 964 | 3118 | 0 | 0 | 0 | 52 | 0 | 0 | 14 | 2265 | 486 | 2357 |
| SRR5127221 | 19 | 0 | 1804 | 0 | 0 | 78 | 77 | 19 | 32 | 0 | 0 | 1147 | 0 | 0 | 19 | 0 | 0 | 0 | 0 | 2043 | 192 | 0 | 0 | 14 | 0 | 93 | 290 | 11 | 0 | 0 | 39 | 0 | 0 | 24 | 134 | 31 | 161 |
| SRR5127222 | 31 | 0 | 2457 | 0 | 0 | 117 | 15 | 0 | 24 | 0 | 0 | 1096 | 0 | 0 | 55 | 0 | 0 | 0 | 0 | 1273 | 717 | 0 | 0 | 0 | 0 | 2442 | 6060 | 0 | 0 | 0 | 74 | 0 | 0 | 12 | 1300 | 294 | 1794 |
| SRR5127223 | 16 | 0 | 3648 | 0 | 0 | 66 | 14 | 42 | 19 | 0 | 0 | 943 | 0 | 0 | 30 | 0 | 0 | 0 | 0 | 638 | 440 | 0 | 0 | 0 | 0 | 1054 | 5705 | 0 | 0 | 0 | 55 | 0 | 0 | 27 | 1943 | 318 | 2018 |
| SRR5127224 | 102 | 0 | 2141 | 0 | 0 | 11 | 0 | 0 | 45 | 0 | 0 | 1044 | 0 | 0 | 42 | 0 | 0 | 21 | 11 | 2408 | 1146 | 0 | 0 | 0 | 0 | 178 | 350 | 0 | 0 | 0 | 52 | 0 | 0 | 0 | 207 | 24 | 147 |
| SRR5127225 | 43 | 0 | 1722 | 0 | 0 | 95 | 107 | 0 | 21 | 0 | 0 | 1622 | 0 | 0 | 21 | 0 | 0 | 0 | 11 | 3568 | 671 | 0 | 0 | 0 | 0 | 1815 | 12505 | 0 | 0 | 0 | 53 | 0 | 0 | 12 | 4732 | 543 | 3648 |
| SRR5127226 | 31 | 0 | 1596 | 0 | 0 | 89 | 15 | 15 | 31 | 0 | 0 | 1505 | 0 | 0 | 121 | 0 | 0 | 0 | 12 | 938 | 594 | 0 | 0 | 0 | 0 | 211 | 247 | 0 | 0 | 0 | 47 | 0 | 0 | 0 | 144 | 35 | 178 |
| SRR5127227 | 42 | 0 | 1789 | 0 | 0 | 59 | 54 | 25 | 29 | 0 | 0 | 2013 | 0 | 0 | 132 | 0 | 0 | 0 | 0 | 3996 | 667 | 0 | 0 | 0 | 0 | 1861 | 5892 | 0 | 0 | 0 | 50 | 0 | 0 | 0 | 1060 | 247 | 1343 |
| SRR5127228 | 36 | 0 | 2985 | 14 | 0 | 26 | 29 | 42 | 37 | 0 | 0 | 539 | 0 | 0 | 42 | 0 | 0 | 15 | 11 | 2062 | 1365 | 0 | 0 | 0 | 0 | 834 | 1841 | 13 | 0 | 24 | 52 | 0 | 0 | 0 | 1116 | 287 | 937 |
| SRR5127229 | 21 | 0 | 27 | 0 | 0 | 78 | 0 | 0 | 53 | 0 | 0 | 1429 | 0 | 0 | 40 | 0 | 0 | 0 | 0 | 1902 | 356 | 0 | 0 | 14 | 0 | 113 | 267 | 0 | 0 | 0 | 61 | 0 | 0 | 0 | 87 | 21 | 144 |
| SRR5127230 | 13 | 0 | 2018 | 0 | 0 | 97 | 0 | 0 | 61 | 0 | 0 | 277 | 0 | 0 | 97 | 0 | 0 | 0 | 0 | 1645 | 593 | 0 | 0 | 0 | 0 | 183 | 166 | 0 | 0 | 0 | 88 | 0 | 0 | 0 | 98 | 15 | 98 |
| SRR5127231 | 75 | 11 | 1115 | 0 | 0 | 0 | 13 | 0 | 0 | 0 | 0 | 987 | 0 | 0 | 102 | 0 | 0 | 0 | 40 | 5713 | 2063 | 0 | 0 | 0 | 0 | 134 | 470 | 0 | 0 | 0 | 0 | 0 | 0 | 0 | 96 | 12 | 64 |
| SRR5127232 | 15 | 0 | 25 | 0 | 0 | 0 | 0 | 0 | 0 | 0 | 0 | 0 | 0 | 0 | 18 | 0 | 0 | 0 | 0 | 567 | 71 | 0 | 0 | 14 | 0 | 76 | 56 | 0 | 0 | 0 | 0 | 0 | 0 | 118 | 25 | 0 | 14 |
| SRR5127233 | 30 | 0 | 1748 | 0 | 0 | 64 | 138 | 15 | 42 | 0 | 0 | 1038 | 0 | 0 | 23 | 0 | 0 | 0 | 0 | 811 | 251 | 0 | 0 | 0 | 0 | 132 | 176 | 0 | 0 | 11 | 37 | 0 | 0 | 63 | 45 | 12 | 55 |
| SRR5127234 | 160 | 117 | 8518 | 0 | 0 | 75 | 420 | 138 | 42 | 0 | 0 | 171 | 0 | 0 | 36 | 0 | 17 | 0 | 13 | 391 | 294 | 0 | 0 | 0 | 13 | 74 | 96 | 0 | 21 | 21 | 48 | 0 | 0 | 22 | 28 | 52 | 29 |
| SRR5127235 | 0 | 0 | 5315 | 0 | 0 | 82 | 77 | 19 | 51 | 0 | 0 | 207 | 0 | 0 | 19 | 0 | 0 | 0 | 0 | 1783 | 827 | 0 | 0 | 0 | 0 | 1922 | 7991 | 17 | 0 | 0 | 32 | 0 | 0 | 25 | 2392 | 504 | 2559 |
| SRR5127236 | 25 | 0 | 3624 | 0 | 0 | 0 | 40 | 0 | 0 | 0 | 0 | 462 | 0 | 0 | 30 | 0 | 0 | 0 | 0 | 1029 | 2683 | 0 | 0 | 0 | 0 | 112 | 234 | 0 | 0 | 0 | 0 | 0 | 0 | 0 | 59 | 16 | 127 |
| SRR5136285 | 0 | 0 | 1250 | 0 | 21 | 0 | 525 | 517 | 0 | 0 | 0 | 156 | 0 | 31 | 0 | 0 | 0 | 0 | 0 | 131 | 0 | 0 | 0 | 0 | 0 | 0 | 0 | 14 | 0 | 191 | 0 | 0 | 14 | 38 | 0 | 0 | 0 |
| SRR5136286 | 81 | 0 | 2036 | 0 | 19 | 0 | 128 | 198 | 0 | 0 | 0 | 0 | 0 | 17 | 128 | 0 | 0 | 0 | 0 | 514 | 0 | 0 | 0 | 0 | 0 | 0 | 0 | 0 | 0 | 56 | 0 | 0 | 0 | 13 | 0 | 0 | 0 |
| SRR5136287 | 0 | 0 | 1595 | 0 | 27 | 0 | 122 | 463 | 0 | 0 | 0 | 76 | 0 | 30 | 0 | 0 | 0 | 0 | 0 | 122 | 0 | 0 | 0 | 0 | 0 | 0 | 0 | 13 | 0 | 195 | 0 | 0 | 0 | 57 | 0 | 0 | 0 |
| SRR5136288 | 0 | 0 | 1409 | 14 | 14 | 0 | 249 | 522 | 0 | 0 | 0 | 201 | 0 | 30 | 0 | 0 | 0 | 0 | 0 | 0 | 0 | 0 | 0 | 0 | 0 | 0 | 11 | 0 | 0 | 201 | 0 | 0 | 0 | 52 | 0 | 0 | 0 |
| SRR5136289 | 0 | 0 | 1184 | 0 | 34 | 0 | 244 | 583 | 0 | 0 | 0 | 0 | 0 | 30 | 0 | 0 | 0 | 0 | 0 | 244 | 0 | 0 | 0 | 0 | 0 | 0 | 0 | 14 | 0 | 169 | 0 | 0 | 0 | 48 | 0 | 0 | 0 |
| SRR5136290 | 145 | 0 | 800 | 0 | 23 | 0 | 134 | 185 | 0 | 0 | 0 | 0 | 0 | 20 | 0 | 0 | 0 | 0 | 0 | 0 | 0 | 0 | 0 | 0 | 0 | 0 | 0 | 17 | 0 | 68 | 0 | 0 | 0 | 12 | 0 | 0 | 0 |
| SRR5136291 | 0 | 0 | 1633 | 0 | 25 | 0 | 253 | 560 | 0 | 0 | 0 | 74 | 0 | 19 | 0 | 0 | 0 | 0 | 0 | 127 | 0 | 0 | 0 | 0 | 0 | 0 | 14 | 0 | 0 | 218 | 0 | 0 | 0 | 55 | 0 | 0 | 0 |
| SRR5136292 | 93 | 0 | 1757 | 0 | 26 | 0 | 124 | 190 | 0 | 0 | 0 | 0 | 0 | 10 | 0 | 0 | 0 | 0 | 0 | 0 | 0 | 0 | 0 | 0 | 0 | 0 | 0 | 0 | 0 | 74 | 0 | 0 | 0 | 18 | 0 | 0 | 0 |
| SRR5136293 | 51 | 0 | 1022 | 10 | 23 | 0 | 145 | 2073 | 21 | 0 | 0 | 77 | 0 | 19 | 0 | 0 | 0 | 0 | 0 | 0 | 0 | 0 | 0 | 0 | 0 | 0 | 33 | 21 | 0 | 625 | 0 | 0 | 0 | 114 | 0 | 0 | 0 |
| SRR5136294 | 166 | 0 | 1853 | 0 | 15 | 0 | 156 | 429 | 30 | 0 | 0 | 138 | 0 | 17 | 0 | 0 | 0 | 0 | 0 | 0 | 0 | 0 | 0 | 0 | 0 | 0 | 15 | 0 | 0 | 131 | 0 | 0 | 0 | 19 | 0 | 0 | 0 |
| SRR5136295 | 77 | 0 | 1356 | 0 | 26 | 0 | 125 | 411 | 18 | 0 | 0 | 77 | 0 | 16 | 0 | 0 | 0 | 0 | 0 | 16 | 0 | 0 | 0 | 0 | 0 | 0 | 26 | 10 | 0 | 141 | 0 | 0 | 0 | 18 | 0 | 0 | 0 |
| SRR5136296 | 82 | 0 | 1292 | 0 | 15 | 0 | 98 | 430 | 32 | 0 | 0 | 82 | 0 | 21 | 0 | 0 | 0 | 0 | 0 | 16 | 0 | 0 | 0 | 0 | 0 | 0 | 19 | 11 | 0 | 149 | 0 | 0 | 0 | 22 | 0 | 0 | 0 |
| SRR5136297 | 105 | 0 | 1580 | 0 | 11 | 0 | 142 | 2009 | 17 | 0 | 0 | 132 | 0 | 0 | 0 | 0 | 0 | 0 | 0 | 0 | 0 | 0 | 0 | 0 | 0 | 0 | 24 | 19 | 0 | 693 | 0 | 0 | 0 | 115 | 0 | 0 | 0 |
| SRR5136298 | 54 | 0 | 1623 | 0 | 16 | 0 | 124 | 1923 | 27 | 0 | 0 | 54 | 0 | 16 | 0 | 0 | 0 | 0 | 0 | 0 | 0 | 0 | 0 | 0 | 0 | 0 | 22 | 23 | 0 | 711 | 0 | 0 | 0 | 119 | 0 | 0 | 0 |
| SRR5136299 | 103 | 0 | 1674 | 11 | 18 | 0 | 84 | 441 | 17 | 0 | 0 | 129 | 0 | 0 | 0 | 0 | 0 | 0 | 0 | 0 | 0 | 0 | 0 | 0 | 0 | 0 | 28 | 0 | 0 | 154 | 0 | 0 | 0 | 23 | 0 | 0 | 0 |
| SRR5136300 | 0 | 0 | 1217 | 0 | 14 | 0 | 116 | 2093 | 23 | 0 | 0 | 55 | 0 | 17 | 0 | 0 | 0 | 0 | 0 | 0 | 0 | 0 | 0 | 0 | 0 | 0 | 15 | 0 | 0 | 705 | 0 | 0 | 0 | 136 | 0 | 0 | 0 |
| SRR5139121 | 24 | 0 | 5569 | 0 | 0 | 100 | 17 | 0 | 48 | 0 | 0 | 364 | 0 | 0 | 51 | 0 | 0 | 0 | 28 | 2976 | 1335 | 0 | 0 | 0 | 0 | 189 | 287 | 0 | 0 | 0 | 43 | 0 | 0 | 36 | 181 | 21 | 114 |
| SRR5145238 | 274 | 0 | 866 | 0 | 10 | 61 | 117 | 64 | 28 | 83 | 104 | 2127 | 39 | 13 | 18 | 0 | 1143 | 0 | 34 | 390 | 186 | 0 | 0 | 55 | 57 | 338 | 863 | 0 | 68 | 35 | 15 | 0 | 0 | 190 | 116 | 70 | 142 |
| SRR5145239 | 297 | 0 | 350 | 0 | 36 | 286 | 0 | 118 | 87 | 0 | 15 | 4128 | 235 | 0 | 0 | 0 | 1745 | 0 | 0 | 84 | 246 | 0 | 0 | 551 | 186 | 1526 | 2139 | 0 | 0 | 82 | 103 | 0 | 0 | 236 | 922 | 1511 | 2126 |
| SRR5145240 | 139 | 0 | 716 | 12 | 17 | 47 | 305 | 279 | 18 | 1526 | 2314 | 1660 | 18 | 35 | 134 | 0 | 1305 | 111 | 0 | 0 | 1247 | 0 | 0 | 16 | 121 | 234 | 385 | 0 | 1098 | 16 | 40 | 0 | 19 | 14204 | 102 | 62 | 134 |
| SRR5145241 | 172 | 0 | 922 | 0 | 37 | 143 | 143 | 51 | 60 | 13 | 42 | 2891 | 27 | 38 | 49 | 0 | 1803 | 0 | 0 | 26 | 256 | 0 | 0 | 12 | 165 | 853 | 1432 | 19 | 68 | 10 | 94 | 0 | 0 | 2355 | 383 | 404 | 1077 |
| SRR5145242 | 139 | 0 | 753 | 0 | 14 | 131 | 1090 | 274 | 46 | 1768 | 2135 | 1601 | 0 | 31 | 56 | 0 | 1757 | 41 | 52 | 12 | 613 | 0 | 0 | 12 | 103 | 633 | 271 | 0 | 2085 | 54 | 41 | 0 | 0 | 11714 | 30 | 25 | 45 |
| SRR5145243 | 119 | 0 | 791 | 0 | 13 | 40 | 10 | 12 | 18 | 27 | 31 | 2238 | 19 | 22 | 83 | 0 | 1632 | 11 | 0 | 40 | 95 | 0 | 0 | 27 | 159 | 285 | 106 | 25 | 10 | 12 | 56 | 0 | 0 | 145 | 42 | 39 | 50 |
| SRR5145244 | 166 | 0 | 1236 | 0 | 14 | 171 | 1296 | 88 | 37 | 3808 | 4146 | 1304 | 20 | 52 | 44 | 0 | 1200 | 0 | 93 | 280 | 573 | 0 | 0 | 0 | 180 | 854 | 832 | 0 | 1138 | 16 | 44 | 0 | 0 | 1737 | 53 | 17 | 44 |
| SRR5145245 | 129 | 0 | 480 | 0 | 29 | 209 | 29 | 332 | 21 | 0 | 15 | 2641 | 90 | 0 | 18 | 0 | 1684 | 0 | 88 | 862 | 539 | 0 | 0 | 1602 | 125 | 1188 | 2268 | 0 | 0 | 176 | 32 | 0 | 0 | 262 | 339 | 365 | 553 |
| SRR5163971 | 49 | 0 | 32 | 0 | 0 | 34 | 1863 | 226 | 38 | 0 | 28 | 54 | 0 | 38 | 488 | 0 | 38 | 0 | 54 | 190 | 687 | 0 | 0 | 1236 | 71 | 204 | 238 | 0 | 0 | 30 | 44 | 0 | 0 | 232 | 115 | 71 | 331 |
| SRR5163972 | 54 | 12 | 24 | 0 | 0 | 35 | 2049 | 184 | 20 | 0 | 35 | 43 | 0 | 67 | 371 | 0 | 23 | 21 | 18 | 177 | 689 | 12 | 0 | 1235 | 63 | 231 | 259 | 20 | 0 | 24 | 35 | 12 | 0 | 165 | 122 | 106 | 314 |
| SRR5163973 | 42 | 0 | 41 | 0 | 0 | 110 | 353 | 22 | 49 | 0 | 45 | 89 | 0 | 25 | 82 | 0 | 136 | 0 | 0 | 86 | 287 | 30 | 0 | 86 | 64 | 43 | 25 | 13 | 21 | 12 | 51 | 13 | 0 | 102 | 10 | 13 | 58 |
| SRR5163974 | 37 | 0 | 43 | 0 | 0 | 165 | 399 | 271 | 58 | 0 | 34 | 103 | 0 | 48 | 171 | 15 | 138 | 0 | 0 | 57 | 257 | 15 | 0 | 1094 | 116 | 595 | 402 | 19 | 34 | 155 | 39 | 0 | 0 | 92 | 106 | 194 | 896 |
| SRR5163975 | 49 | 0 | 91 | 0 | 0 | 41 | 296 | 0 | 25 | 0 | 71 | 859 | 0 | 52 | 101 | 18 | 98 | 0 | 23 | 288 | 1651 | 27 | 0 | 470 | 83 | 224 | 187 | 38 | 0 | 0 | 80 | 29 | 12 | 146 | 241 | 456 | 1481 |
| SRR5163976 | 42 | 0 | 98 | 0 | 0 | 42 | 341 | 0 | 32 | 0 | 71 | 856 | 0 | 45 | 120 | 42 | 86 | 0 | 20 | 240 | 1944 | 42 | 0 | 415 | 81 | 279 | 178 | 29 | 16 | 0 | 97 | 29 | 0 | 139 | 194 | 398 | 1467 |
| SRR5163977 | 443 | 0 | 38 | 0 | 0 | 15 | 24 | 24 | 57 | 0 | 51 | 30 | 0 | 65 | 217 | 15 | 98 | 59 | 120 | 795 | 6358 | 18 | 0 | 204 | 11 | 50 | 26 | 50 | 0 | 51 | 42 | 22 | 0 | 56 | 34 | 38 | 166 |

### Supp Table S8.2.Columns_AM_to_BW

| | | | | | | | | | | | | | | | | | | |
|---|---|---|---|---|---|---|---|---|---|---|---|---|---|---|---|---|---|---|
| SRR5163978 | 438 | 0 | 30 | 0 | 0 | 98 | 122 | 40 | 79 | 0 | 28 | 35 | 0 | 46 | 30 | 0 | 101 | 0 |
| SRR5163979 | 31 | 0 | 29 | 0 | 0 | 40 | 186 | 17 | 13 | 0 | 23 | 52 | 0 | 36 | 321 | 0 | 58 | 0 |
| SRR5163980 | 28 | 0 | 32 | 0 | 0 | 46 | 269 | 0 | 35 | 0 | 16 | 54 | 0 | 32 | 319 | 0 | 51 | 0 |
| SRR5163981 | 32 | 0 | 16 | 0 | 0 | 83 | 1777 | 91 | 37 | 0 | 17 | 31 | 0 | 52 | 0 | 17 | 52 | 0 |
| SRR5163982 | 177 | 0 | 174 | 0 | 0 | 66 | 1222 | 29 | 24 | 0 | 37 | 22 | 0 | 40 | 42 | 0 | 52 | 0 |
| SRR5163983 | 54 | 0 | 13 | 0 | 0 | 49 | 2192 | 29 | 24 | 0 | 22 | 20 | 0 | 25 | 24 | 10 | 34 | 0 |
| SRR5163984 | 55 | 0 | 16 | 0 | 0 | 38 | 2174 | 29 | 34 | 0 | 21 | 20 | 0 | 34 | 20 | 0 | 38 | 0 |
| SRR5163985 | 15 | 0 | 0 | 0 | 0 | 53 | 435 | 18 | 12 | 0 | 12 | 0 | 0 | 29 | 762 | 0 | 0 | 0 |
| SRR5163986 | 12 | 0 | 0 | 0 | 0 | 33 | 420 | 14 | 0 | 0 | 28 | 0 | 0 | 28 | 168 | 0 | 0 | 0 |
| SRR5163987 | 54 | 0 | 14 | 0 | 0 | 60 | 1611 | 31 | 42 | 0 | 17 | 20 | 0 | 66 | 552 | 0 | 38 | 0 |
| SRR5168444 | 181 | 0 | 45 | 0 | 0 | 33 | 519 | 37 | 21 | 0 | 28 | 0 | 0 | 14 | 1533 | 0 | 18 | 148 |
| SRR5168445 | 270 | 0 | 640 | 0 | 0 | 74 | 589 | 13465 | 38 | 0 | 0 | 141 | 0 | 0 | 0 | 0 | 25 | 0 |
| SRR5168447 | 156 | 0 | 145 | 0 | 0 | 138 | 673 | 678 | 51 | 0 | 25 | 153 | 22 | 41 | 808 | 0 | 33 | 142 |
| SRR5168448 | 641 | 0 | 509 | 0 | 14 | 34 | 350 | 60 | 0 | 0 | 22 | 56 | 27 | 108 | 13 | 0 | 74 | 0 |
| SRR5168451 | 261 | 0 | 652 | 0 | 0 | 37 | 64 | 889 | 0 | 0 | 0 | 52 | 18 | 16 | 0 | 0 | 33 | 0 |
| SRR5168454 | 12 | 0 | 94 | 0 | 0 | 23 | 62 | 1154 | 14 | 12 | 16 | 66 | 0 | 0 | 18 | 0 | 22 | 0 |
| SRR5168458 | 118 | 0 | 185 | 0 | 0 | 53 | 506 | 614 | 25 | 0 | 25 | 123 | 0 | 0 | 186 | 0 | 60 | 15 |
| SRR5168460 | 52 | 0 | 351 | 0 | 0 | 17 | 754 | 556 | 12 | 65 | 11 | 31 | 0 | 14 | 0 | 0 | 47 | 0 |
| SRR5168465 | 14 | 0 | 875 | 0 | 0 | 20 | 629 | 77 | 94 | 0 | 15 | 161 | 28 | 0 | 93 | 0 | 67 | 0 |
| SRR5168467 | 136 | 0 | 339 | 0 | 0 | 20 | 272 | 88 | 23 | 12 | 43 | 224 | 0 | 12 | 0 | 0 | 75 | 0 |
| SRR5168471 | 90 | 0 | 131 | 0 | 0 | 191 | 181 | 1526 | 36 | 24 | 105 | 151 | 14 | 0 | 0 | 0 | 64 | 0 |
| SRR5168475 | 299 | 0 | 817 | 0 | 0 | 33 | 205 | 49 | 49 | 83 | 189 | 80 | 0 | 21 | 0 | 0 | 17 | 0 |
| SRR5171728 | 139 | 0 | 7119 | 0 | 0 | 19 | 0 | 13 | 0 | 0 | 0 | 980 | 0 | 0 | 0 | 0 | 99 | 0 |
| SRR5171729 | 226 | 0 | 6182 | 0 | 0 | 0 | 130 | 65 | 0 | 0 | 0 | 1034 | 0 | 0 | 65 | 0 | 143 | 0 |
| SRR5171730 | 102 | 0 | 12523 | 0 | 0 | 16 | 59 | 15 | 0 | 0 | 0 | 1078 | 0 | 0 | 0 | 0 | 131 | 0 |
| SRR5171731 | 81 | 0 | 9855 | 0 | 0 | 20 | 0 | 23 | 0 | 0 | 0 | 1175 | 0 | 0 | 0 | 0 | 128 | 0 |
| SRR5171732 | 200 | 0 | 14675 | 0 | 0 | 0 | 0 | 0 | 0 | 0 | 0 | 1644 | 0 | 0 | 0 | 0 | 40 | 0 |
| SRR5171733 | 174 | 0 | 8838 | 0 | 0 | 15 | 63 | 13 | 0 | 0 | 0 | 1807 | 0 | 0 | 0 | 0 | 65 | 0 |
| SRR5195030 | 32 | 0 | 1601 | 0 | 0 | 0 | 133 | 22 | 0 | 0 | 0 | 1147 | 0 | 0 | 29 | 0 | 0 | 0 |
| SRR5195032 | 85 | 0 | 615 | 0 | 0 | 0 | 77 | 0 | 0 | 0 | 0 | 1340 | 0 | 0 | 113 | 0 | 0 | 0 |
| SRR5195033 | 36 | 0 | 2305 | 0 | 0 | 0 | 114 | 0 | 0 | 0 | 0 | 1668 | 0 | 0 | 38 | 0 | 0 | 0 |
| SRR5195034 | 60 | 0 | 1365 | 0 | 0 | 16 | 144 | 160 | 0 | 0 | 0 | 1135 | 0 | 0 | 32 | 0 | 0 | 0 |
| SRR5224010 | 124 | 0 | 65 | 14 | 32 | 188 | 231 | 9655 | 180 | 30 | 175 | 94 | 18 | 23 | 48 | 0 | 12 | 13 |
| SRR5224011 | 128 | 0 | 64 | 0 | 14 | 38 | 457 | 4968 | 29 | 21 | 30 | 148 | 14 | 15 | 0 | 0 | 17 | 14 |
| SRR5224012 | 166 | 0 | 37 | 0 | 19 | 50 | 325 | 2080 | 56 | 17 | 17 | 147 | 18 | 24 | 37 | 0 | 19 | 0 |
| SRR5224013 | 52 | 0 | 140 | 17 | 18 | 30 | 16 | 38 | 25 | 0 | 12 | 807 | 24 | 15 | 0 | 0 | 45 | 0 |
| SRR5224014 | 146 | 0 | 60 | 26 | 14 | 25 | 15 | 222 | 22 | 0 | 24 | 130 | 0 | 23 | 77 | 0 | 0 | 0 |
| SRR5224015 | 61 | 0 | 151 | 18 | 17 | 42 | 12 | 29 | 30 | 0 | 26 | 946 | 16 | 11 | 17 | 0 | 22 | 0 |
| SRR5224016 | 95 | 0 | 103 | 21 | 15 | 47 | 25 | 33 | 33 | 0 | 10 | 667 | 17 | 12 | 40 | 0 | 22 | 0 |
| SRR5224017 | 29 | 0 | 79 | 40 | 48 | 43 | 18 | 145 | 74 | 0 | 25 | 380 | 11 | 37 | 0 | 0 | 10 | 0 |
| SRR5224018 | 77 | 0 | 162 | 15 | 39 | 137 | 14 | 79 | 115 | 0 | 36 | 920 | 0 | 33 | 70 | 0 | 21 | 14 |
| SRR5224019 | 49 | 0 | 188 | 19 | 19 | 27 | 0 | 31 | 15 | 0 | 12 | 1077 | 22 | 14 | 76 | 0 | 57 | 0 |
| SRR5224020 | 31 | 0 | 120 | 11 | 27 | 27 | 13 | 43 | 27 | 0 | 15 | 644 | 17 | 27 | 119 | 0 | 0 | 0 |
| SRR5224021 | 78 | 0 | 248 | 17 | 15 | 27 | 0 | 30 | 21 | 12 | 14 | 1125 | 24 | 17 | 0 | 0 | 58 | 0 |
| SRR5224022 | 46 | 0 | 139 | 24 | 24 | 38 | 0 | 39 | 26 | 0 | 11 | 1173 | 11 | 14 | 12 | 0 | 20 | 0 |
| SRR5224023 | 48 | 0 | 334 | 25 | 67 | 79 | 19 | 76 | 29 | 0 | 37 | 722 | 59 | 31 | 0 | 0 | 72 | 0 |
| SRR5224024 | 35 | 0 | 204 | 24 | 53 | 116 | 12 | 121 | 76 | 0 | 39 | 857 | 32 | 36 | 88 | 0 | 26 | 0 |
| SRR5230550 | 367 | 0 | 0 | 0 | 0 | 52 | 7821 | 2652 | 30 | 0 | 47 | 458 | 0 | 0 | 0 | 0 | 61 | 0 |
| SRR5230551 | 131 | 0 | 0 | 0 | 0 | 49 | 771 | 64 | 42 | 0 | 31 | 76 | 0 | 0 | 0 | 0 | 79 | 0 |
| SRR5230552 | 322 | 0 | 33 | 31 | 42 | 25 | 77 | 54 | 26 | 10 | 42 | 627 | 0 | 20 | 20 | 0 | 79 | 0 |
| SRR5230553 | 709 | 0 | 16 | 0 | 0 | 21 | 92 | 26 | 27 | 0 | 62 | 876 | 0 | 0 | 66 | 0 | 74 | 0 |
| SRR5230554 | 545 | 0 | 0 | 0 | 0 | 30 | 3236 | 860 | 48 | 19 | 105 | 1183 | 0 | 13 | 12 | 0 | 67 | 0 |
| SRR5230555 | 580 | 0 | 32 | 0 | 0 | 22 | 811 | 67 | 29 | 0 | 10 | 941 | 0 | 0 | 0 | 0 | 68 | 0 |
| SRR5230556 | 577 | 0 | 0 | 0 | 0 | 11 | 993 | 58 | 50 | 0 | 18 | 355 | 0 | 0 | 0 | 0 | 25 | 0 |
| SRR5230557 | 484 | 0 | 16 | 0 | 0 | 15 | 2109 | 315 | 30 | 0 | 61 | 614 | 0 | 0 | 33 | 0 | 20 | 0 |
| SRR5230558 | 328 | 0 | 11 | 0 | 0 | 17 | 3031 | 574 | 26 | 60 | 537 | 583 | 0 | 27 | 36 | 0 | 65 | 0 |
| SRR5230559 | 1242 | 0 | 41 | 0 | 0 | 35 | 484 | 114 | 44 | 14 | 49 | 1866 | 0 | 0 | 37 | 0 | 38 | 0 |
| SRR5230560 | 446 | 0 | 12 | 0 | 0 | 44 | 710 | 23 | 23 | 0 | 25 | 498 | 0 | 0 | 11 | 0 | 77 | 0 |
| SRR5230561 | 296 | 0 | 0 | 0 | 0 | 34 | 359 | 85 | 18 | 14 | 60 | 407 | 0 | 0 | 17 | 0 | 45 | 0 |
| SRR5230562 | 773 | 0 | 0 | 0 | 0 | 21 | 38 | 53 | 46 | 0 | 97 | 995 | 0 | 0 | 34 | 0 | 32 | 0 |
| SRR5230563 | 765 | 0 | 11 | 0 | 10 | 31 | 40 | 91 | 31 | 0 | 29 | 957 | 0 | 0 | 0 | 0 | 28 | 0 |
| SRR5230564 | 541 | 0 | 38 | 0 | 19 | 42 | 1401 | 486 | 34 | 12 | 43 | 655 | 0 | 0 | 55 | 0 | 44 | 0 |
| SRR5230565 | 79 | 0 | 0 | 15 | 14 | 19 | 570 | 45 | 31 | 73 | 496 | 35 | 0 | 0 | 0 | 0 | 42 | 0 |
| SRR5230566 | 670 | 0 | 53 | 0 | 0 | 32 | 6154 | 278 | 33 | 0 | 32 | 688 | 0 | 0 | 0 | 0 | 50 | 0 |
| SRR5230567 | 109 | 0 | 0 | 12 | 19 | 21 | 61 | 36 | 42 | 14 | 17 | 140 | 0 | 0 | 52 | 0 | 17 | 0 |
| SRR5230568 | 359 | 0 | 16 | 0 | 0 | 31 | 163 | 84 | 23 | 0 | 13 | 890 | 0 | 0 | 16 | 0 | 0 | 0 |
| SRR5230569 | 550 | 0 | 18 | 19 | 20 | 43 | 53 | 16 | 10 | 0 | 32 | 1150 | 0 | 0 | 15 | 0 | 43 | 0 |
| SRR5230570 | 347 | 0 | 10 | 0 | 0 | 38 | 1285 | 603 | 53 | 0 | 25 | 392 | 0 | 0 | 17 | 0 | 50 | 0 |
| SRR5230571 | 153 | 0 | 0 | 0 | 12 | 36 | 332 | 74 | 31 | 30 | 12 | 181 | 0 | 11 | 26 | 0 | 84 | 0 |
| SRR5230572 | 169 | 0 | 0 | 0 | 0 | 34 | 21762 | 1862 | 37 | 23 | 161 | 342 | 0 | 0 | 0 | 0 | 78 | 0 |
| SRR5230573 | 145 | 0 | 26 | 0 | 0 | 64 | 1864 | 337 | 21 | 0 | 26 | 172 | 0 | 0 | 75 | 0 | 100 | 0 |
| SRR5230574 | 406 | 0 | 15 | 18 | 16 | 50 | 182 | 19 | 38 | 27 | 16 | 499 | 0 | 24 | 65 | 0 | 30 | 0 |
| SRR5230575 | 478 | 0 | 15 | 16 | 28 | 37 | 173 | 0 | 33 | 19 | 109 | 621 | 0 | 14 | 14 | 0 | 33 | 0 |
| SRR5230576 | 521 | 0 | 14 | 10 | 0 | 34 | 123 | 15 | 52 | 33 | 117 | 631 | 0 | 11 | 80 | 0 | 135 | 0 |
| SRR5230577 | 617 | 0 | 0 | 0 | 22 | 49 | 120 | 33 | 55 | 31 | 108 | 724 | 0 | 15 | 55 | 0 | 116 | 0 |
| SRR5230578 | 138 | 0 | 0 | 0 | 0 | 37 | 532 | 116 | 53 | 106 | 52 | 203 | 0 | 17 | 12 | 0 | 63 | 0 |
| SRR5230579 | 163 | 0 | 0 | 0 | 0 | 32 | 638 | 101 | 64 | 75 | 45 | 227 | 0 | 27 | 13 | 0 | 65 | 0 |
| SRR5230580 | 217 | 0 | 0 | 0 | 10 | 34 | 255 | 32 | 30 | 0 | 46 | 146 | 0 | 24 | 69 | 0 | 64 | 0 |
| SRR5230581 | 230 | 0 | 0 | 0 | 0 | 44 | 216 | 297 | 22 | 0 | 36 | 119 | 0 | 15 | 130 | 0 | 35 | 0 |
| SRR5230582 | 338 | 0 | 0 | 0 | 0 | 50 | 556 | 1521 | 45 | 0 | 21 | 697 | 0 | 0 | 80 | 0 | 46 | 0 |
| SRR5230583 | 337 | 0 | 0 | 0 | 16 | 94 | 686 | 1415 | 31 | 0 | 0 | 841 | 0 | 0 | 39 | 0 | 53 | 0 |
| SRR5230584 | 499 | 0 | 0 | 0 | 0 | 31 | 118 | 82 | 62 | 12 | 63 | 791 | 0 | 24 | 301 | 0 | 18 | 0 |
| SRR5230585 | 554 | 0 | 22 | 0 | 0 | 22 | 104 | 77 | 73 | 0 | 77 | 854 | 17 | 26 | 207 | 0 | 0 | 0 |
| SRR5230586 | 302 | 0 | 14 | 0 | 0 | 42 | 194 | 167 | 33 | 19 | 42 | 232 | 0 | 0 | 0 | 0 | 59 | 0 |
| SRR5230587 | 269 | 0 | 13 | 0 | 0 | 56 | 277 | 150 | 30 | 12 | 41 | 286 | 0 | 0 | 0 | 0 | 56 | 0 |
| SRR5230588 | 477 | 0 | 13 | 0 | 0 | 50 | 198 | 80 | 44 | 15 | 45 | 517 | 0 | 0 | 0 | 0 | 42 | 0 |
| SRR5230589 | 474 | 0 | 0 | 0 | 0 | 38 | 103 | 46 | 38 | 0 | 15 | 627 | 0 | 0 | 69 | 0 | 30 | 0 |
| SRR5230590 | 187 | 0 | 13 | 0 | 0 | 0 | 0 | 0 | 55 | 96 | 110 | 354 | 0 | 0 | 0 | 0 | 40 | 0 |
| SRR5230591 | 230 | 0 | 14 | 0 | 0 | 12 | 17 | 0 | 15 | 37 | 114 | 412 | 0 | 0 | 106 | 0 | 51 | 0 |
| SRR5230592 | 68 | 0 | 0 | 0 | 0 | 0 | 0 | 0 | 0 | 0 | 97 | 55 | 0 | 0 | 0 | 0 | 147 | 0 |
| SRR5230593 | 67 | 0 | 0 | 0 | 0 | 0 | 67 | 0 | 0 | 29 | 55 | 58 | 0 | 0 | 29 | 0 | 100 | 0 |
| SRR5230594 | 272 | 0 | 0 | 0 | 0 | 63 | 39 | 35 | 36 | 0 | 23 | 314 | 0 | 0 | 26 | 0 | 86 | 0 |
| SRR5230595 | 280 | 0 | 11 | 0 | 17 | 11 | 81 | 34 | 40 | 17 | 23 | 369 | 0 | 0 | 54 | 0 | 97 | 0 |
| SRR5230596 | 613 | 0 | 36 | 0 | 0 | 29 | 646 | 26 | 45 | 0 | 31 | 795 | 0 | 0 | 35 | 0 | 42 | 0 |
| SRR5230597 | 616 | 0 | 39 | 0 | 0 | 33 | 665 | 162 | 41 | 0 | 28 | 882 | 0 | 13 | 44 | 0 | 64 | 0 |
| SRR5230600 | 362 | 0 | 15 | 0 | 0 | 22 | 20 | 0 | 29 | 14 | 55 | 412 | 0 | 0 | 0 | 0 | 52 | 0 |
| SRR5230601 | 403 | 10 | 18 | 10 | 24 | 13 | 0 | 0 | 24 | 20 | 67 | 486 | 0 | 17 | 32 | 0 | 44 | 0 |
| SRR5230602 | 372 | 0 | 16 | 0 | 0 | 32 | 169 | 33 | 16 | 0 | 34 | 449 | 0 | 0 | 86 | 0 | 67 | 0 |
| SRR5230603 | 372 | 0 | 20 | 0 | 0 | 44 | 237 | 368 | 15 | 0 | 22 | 543 | 0 | 0 | 130 | 0 | 72 | 0 |
| SRR5230604 | 1095 | 0 | 14 | 20 | 0 | 36 | 60 | 20 | 32 | 0 | 12 | 1479 | 0 | 0 | 0 | 0 | 54 | 0 |
| SRR5230605 | 1198 | 0 | 28 | 17 | 0 | 53 | 44 | 18 | 35 | 0 | 56 | 1559 | 0 | 0 | 13 | 0 | 62 | 0 |
| SRR5230606 | 690 | 0 | 19 | 13 | 0 | 10 | 254 | 650 | 55 | 17 | 100 | 1107 | 27 | 13 | 118 | 0 | 80 | 0 |
| SRR5230607 | 767 | 0 | 25 | 0 | 12 | 0 | 246 | 47 | 52 | 29 | 13 | 1123 | 17 | 0 | 130 | 0 | 86 | 0 |
| SRR5230608 | 196 | 0 | 0 | 0 | 0 | 22 | 125 | 550 | 28 | 0 | 14 | 181 | 0 | 0 | 0 | 0 | 56 | 0 |
| SRR5230609 | 218 | 0 | 0 | 12 | 0 | 12 | 30 | 539 | 24 | 0 | 18 | 208 | 0 | 0 | 0 | 0 | 50 | 0 |
| SRR5230610 | 259 | 0 | 0 | 0 | 0 | 0 | 0 | 20 | 0 | 29 | 49 | 601 | 0 | 0 | 120 | 0 | 91 | 16 |
| SRR5230611 | 354 | 0 | 0 | 0 | 11 | 11 | 50 | 20 | 17 | 0 | 53 | 610 | 0 | 14 | 107 | 0 | 70 | 0 |
| SRR5230613 | 240 | 0 | 0 | 0 | 0 | 13 | 0 | 0 | 25 | 10 | 56 | 504 | 0 | 13 | 25 | 0 | 42 | 0 |
| SRR5230614 | 168 | 20 | 0 | 53 | 27 | 0 | 47 | 10 | 10 | 27 | 97 | 128 | 0 | 57 | 0 | 0 | 89 | 0 |
| SRR5230615 | 198 | 24 | 0 | 35 | 19 | 0 | 0 | 0 | 16 | 19 | 92 | 155 | 0 | 59 | 23 | 0 | 94 | 0 |
| SRR5230617 | 596 | 0 | 0 | 0 | 14 | 0 | 0 | 0 | 11 | 0 | 76 | 1257 | 0 | 12 | 40 | 0 | 47 | 0 |
| SRR5230618 | 157 | 0 | 0 | 0 | 0 | 11 | 289 | 33 | 27 | 33 | 19 | 445 | 0 | 27 | 58 | 0 | 60 | 0 |
| SRR5230619 | 144 | 0 | 10 | 14 | 0 | 12 | 24 | 22 | 21 | 39 | 163 | 504 | 14 | 19 | 141 | 0 | 75 | 0 |
| SRR5230620 | 854 | 0 | 42 | 10 | 0 | 21 | 44 | 0 | 13 | 24 | 50 | 1172 | 0 | 11 | 64 | 0 | 77 | 0 |
| SRR5230621 | 847 | 0 | 71 | 0 | 0 | 31 | 46 | 0 | 0 | 15 | 38 | 1288 | 0 | 27 | 62 | 0 | 66 | 0 |
| SRR5230622 | 128 | 0 | 0 | 0 | 0 | 36 | 253 | 10 | 22 | 0 | 46 | 141 | 0 | 0 | 0 | 0 | 86 | 0 |
| SRR5230623 | 130 | 0 | 0 | 0 | 0 | 23 | 7138 | 9657 | 33 | 18 | 75 | 142 | 0 | 0 | 0 | 0 | 78 | 0 |
| SRR5230624 | 800 | 0 | 46 | 11 | 0 | 59 | 379 | 603 | 44 | 0 | 29 | 1787 | 0 | 0 | 19 | 0 | 28 | 0 |
| SRR5230625 | 606 | 0 | 25 | 0 | 0 | 15 | 408 | 115 | 13 | 0 | 25 | 934 | 0 | 0 | 0 | 0 | 41 | 0 |
| SRR5230626 | 149 | 0 | 0 | 0 | 13 | 74 | 563 | 829 | 49 | 0 | 26 | 85 | 0 | 0 | 0 | 0 | 35 | 0 |
| SRR5230627 | 407 | 0 | 21 | 0 | 0 | 55 | 1690 | 4014 | 40 | 12 | 63 | 480 | 0 | 0 | 0 | 0 | 41 | 0 |
| SRR5230628 | 256 | 0 | 14 | 0 | 0 | 47 | 161 | 75 | 48 | 12 | 24 | 345 | 0 | 0 | 0 | 0 | 33 | 0 |
| SRR5230629 | 89 | 0 | 14 | 0 | 0 | 22 | 7987 | 545 | 14 | 29 | 25 | 179 | 0 | 0 | 0 | 0 | 72 | 0 |
| SRR5230630 | 1039 | 0 | 25 | 0 | 13 | 34 | 1408 | 117 | 11 | 30 | 21 | 1473 | 0 | 0 | 18 | 0 | 60 | 0 |

| | | | | | | | | | | | | | | | | | | | |
|---|---|---|---|---|---|---|---|---|---|---|---|---|---|---|---|---|---|---|---|
| SRR5163978 | 61 | 699 | 6113 | 17 | 0 | 1183 | 81 | 318 | 207 | 38 | 0 | 22 | 43 | 18 | 0 | 397 | 223 | 220 | 1036 |
| SRR5163979 | 0 | 186 | 714 | 26 | 0 | 785 | 83 | 202 | 174 | 28 | 0 | 0 | 41 | 0 | 0 | 94 | 43 | 65 | 273 |
| SRR5163980 | 34 | 252 | 722 | 11 | 0 | 76 | 73 | 20 | 17 | 32 | 11 | 11 | 49 | 30 | 0 | 68 | 51 | 35 | 21 |
| SRR5163981 | 228 | 719 | 732 | 0 | 0 | 352 | 73 | 614 | 515 | 26 | 0 | 18 | 67 | 0 | 0 | 148 | 92 | 101 | 361 |
| SRR5163982 | 30 | 129 | 1192 | 24 | 0 | 51 | 47 | 15 | 15 | 31 | 0 | 35 | 68 | 16 | 0 | 11 | 87 | 143 | 37 |
| SRR5163983 | 71 | 276 | 694 | 0 | 0 | 831 | 78 | 214 | 231 | 37 | 0 | 75 | 49 | 12 | 0 | 158 | 66 | 76 | 478 |
| SRR5163984 | 20 | 353 | 646 | 17 | 0 | 68 | 88 | 23 | 19 | 29 | 0 | 46 | 34 | 0 | 0 | 16 | 92 | 67 | 38 |
| SRR5163985 | 0 | 435 | 2069 | 0 | 0 | 515 | 117 | 123 | 158 | 41 | 0 | 187 | 0 | 0 | 12 | 755 | 82 | 123 | 427 |
| SRR5163986 | 0 | 252 | 2602 | 19 | 0 | 587 | 93 | 163 | 182 | 23 | 0 | 154 | 0 | 19 | 0 | 563 | 140 | 88 | 438 |
| SRR5163987 | 81 | 620 | 1003 | 0 | 0 | 688 | 103 | 370 | 216 | 36 | 0 | 24 | 91 | 0 | 0 | 159 | 130 | 154 | 838 |
| SRR5168444 | 36 | 34 | 3427 | 0 | 0 | 188 | 0 | 146 | 20 | 0 | 0 | 17 | 23 | 0 | 19 | 121 | 89 | 44 | 134 |
| SRR5168445 | 28 | 18 | 88 | 0 | 0 | 626 | 0 | 297 | 328 | 0 | 11 | 2642 | 15 | 0 | 0 | 950 | 0 | 0 | 13 |
| SRR5168447 | 56 | 178 | 477 | 0 | 0 | 156 | 69 | 646 | 347 | 10 | 0 | 109 | 109 | 0 | 18 | 298 | 47 | 0 | 39 |
| SRR5168448 | 27 | 23 | 499 | 0 | 0 | 14 | 48 | 372 | 941 | 0 | 0 | 41 | 41 | 0 | 10 | 308 | 203 | 44 | 247 |
| SRR5168451 | 0 | 0 | 413 | 0 | 0 | 41 | 33 | 282 | 5076 | 14 | 0 | 198 | 15 | 0 | 0 | 392 | 15 | 0 | 19 |
| SRR5168454 | 0 | 0 | 108 | 0 | 0 | 125 | 41 | 127 | 231 | 0 | 0 | 382 | 0 | 0 | 0 | 190 | 22 | 20 | 26 |
| SRR5168458 | 0 | 0 | 138 | 0 | 0 | 15 | 24 | 12 | 105 | 0 | 0 | 210 | 19 | 0 | 0 | 72 | 16 | 0 | 17 |
| SRR5168460 | 0 | 0 | 15 | 0 | 0 | 18 | 18 | 35 | 33 | 0 | 0 | 210 | 53 | 0 | 11 | 402 | 24 | 14 | 30 |
| SRR5168465 | 14 | 19 | 128 | 0 | 0 | 27 | 22 | 100 | 135 | 0 | 0 | 18 | 61 | 0 | 0 | 105 | 74 | 22 | 71 |
| SRR5168467 | 0 | 0 | 0 | 0 | 0 | 16 | 12 | 12 | 35 | 0 | 0 | 45 | 35 | 0 | 0 | 509 | 0 | 0 | 14 |
| SRR5168471 | 0 | 0 | 50 | 0 | 0 | 68 | 52 | 1021 | 1372 | 0 | 0 | 347 | 30 | 0 | 0 | 600 | 30 | 28 | 34 |
| SRR5168475 | 0 | 0 | 24 | 0 | 0 | 0 | 150 | 338 | 466 | 13 | 0 | 26 | 81 | 0 | 10 | 292 | 0 | 0 | 0 |
| SRR5171728 | 38 | 8180 | 269 | 0 | 0 | 241 | 0 | 195 | 321 | 0 | 0 | 0 | 10 | 0 | 0 | 0 | 84 | 90 | 126 |
| SRR5171729 | 33 | 12099 | 65 | 0 | 0 | 212 | 25 | 273 | 470 | 0 | 0 | 0 | 11 | 0 | 0 | 0 | 92 | 119 | 170 |
| SRR5171730 | 0 | 8562 | 297 | 0 | 0 | 671 | 27 | 69 | 185 | 0 | 0 | 0 | 0 | 0 | 0 | 0 | 16 | 19 | 27 |
| SRR5171731 | 80 | 16429 | 438 | 0 | 0 | 237 | 18 | 80 | 257 | 0 | 0 | 0 | 13 | 0 | 0 | 0 | 28 | 34 | 60 |
| SRR5171732 | 0 | 8386 | 286 | 0 | 0 | 402 | 20 | 48 | 161 | 0 | 0 | 13 | 0 | 0 | 0 | 0 | 29 | 20 | 43 |
| SRR5171733 | 21 | 16677 | 375 | 0 | 0 | 176 | 0 | 114 | 300 | 0 | 0 | 0 | 0 | 0 | 0 | 0 | 50 | 59 | 87 |
| SRR5195030 | 270 | 8765 | 151 | 0 | 0 | 15 | 0 | 10 | 105 | 0 | 0 | 0 | 10 | 0 | 0 | 0 | 11 | 18 | 87 |
| SRR5195032 | 91 | 12129 | 549 | 0 | 0 | 0 | 0 | 91 | 506 | 0 | 0 | 0 | 0 | 0 | 0 | 0 | 45 | 0 | 29 |
| SRR5195033 | 360 | 22908 | 1691 | 0 | 0 | 0 | 0 | 87 | 481 | 0 | 0 | 0 | 0 | 0 | 0 | 0 | 23 | 0 | 26 |
| SRR5195034 | 320 | 25494 | 1285 | 0 | 0 | 130 | 0 | 248 | 1211 | 0 | 0 | 0 | 0 | 0 | 0 | 0 | 51 | 0 | 46 |
| SRR5224010 | 0 | 279 | 287 | 0 | 0 | 759 | 78 | 700 | 883 | 0 | 0 | 2660 | 147 | 0 | 0 | 1271 | 154 | 50 | 278 |
| SRR5224011 | 111 | 523 | 155 | 0 | 0 | 171 | 17 | 144 | 163 | 0 | 0 | 616 | 23 | 0 | 0 | 236 | 21 | 44 | 37 |
| SRR5224012 | 175 | 1512 | 248 | 0 | 0 | 133 | 22 | 161 | 90 | 11 | 10 | 138 | 25 | 0 | 0 | 176 | 31 | 21 | 88 |
| SRR5224013 | 22 | 143 | 77 | 0 | 0 | 309 | 20 | 102 | 349 | 0 | 0 | 48 | 27 | 0 | 0 | 88 | 67 | 15 | 87 |
| SRR5224014 | 15 | 62 | 114 | 0 | 0 | 177 | 25 | 82 | 129 | 0 | 0 | 456 | 51 | 0 | 0 | 139 | 45 | 17 | 97 |
| SRR5224015 | 38 | 315 | 102 | 0 | 0 | 296 | 39 | 139 | 341 | 0 | 0 | 31 | 38 | 0 | 0 | 72 | 48 | 13 | 61 |
| SRR5224016 | 25 | 301 | 89 | 0 | 0 | 210 | 49 | 129 | 311 | 12 | 0 | 42 | 67 | 0 | 0 | 71 | 76 | 18 | 87 |
| SRR5224017 | 0 | 86 | 50 | 0 | 0 | 254 | 10 | 130 | 336 | 0 | 0 | 229 | 154 | 0 | 0 | 340 | 274 | 106 | 773 |
| SRR5224018 | 21 | 238 | 119 | 0 | 0 | 704 | 169 | 453 | 1118 | 13 | 0 | 113 | 179 | 0 | 0 | 157 | 282 | 76 | 381 |
| SRR5224019 | 50 | 195 | 50 | 0 | 0 | 249 | 45 | 91 | 423 | 0 | 0 | 34 | 27 | 0 | 0 | 91 | 72 | 11 | 79 |
| SRR5224020 | 18 | 181 | 128 | 0 | 0 | 153 | 28 | 90 | 238 | 11 | 0 | 51 | 38 | 0 | 0 | 106 | 70 | 16 | 85 |
| SRR5224021 | 20 | 193 | 79 | 0 | 0 | 239 | 40 | 96 | 411 | 0 | 0 | 37 | 29 | 0 | 0 | 107 | 79 | 18 | 104 |
| SRR5224022 | 40 | 276 | 89 | 0 | 0 | 278 | 33 | 122 | 344 | 0 | 0 | 41 | 30 | 0 | 0 | 86 | 90 | 26 | 115 |
| SRR5224023 | 52 | 450 | 76 | 0 | 0 | 686 | 114 | 321 | 1723 | 15 | 0 | 65 | 69 | 0 | 0 | 197 | 320 | 36 | 367 |
| SRR5224024 | 36 | 247 | 48 | 0 | 0 | 680 | 163 | 353 | 1173 | 14 | 0 | 134 | 156 | 0 | 0 | 253 | 182 | 88 | 277 |
| SRR5230550 | 23 | 0 | 229 | 0 | 0 | 32 | 44 | 222 | 901 | 0 | 0 | 386 | 87 | 0 | 0 | 353 | 0 | 0 | 0 |
| SRR5230551 | 0 | 0 | 0 | 0 | 0 | 0 | 34 | 12 | 33 | 0 | 0 | 29 | 54 | 0 | 0 | 33 | 0 | 0 | 0 |
| SRR5230552 | 0 | 0 | 81 | 0 | 0 | 0 | 52 | 16 | 55 | 0 | 0 | 21 | 59 | 0 | 0 | 171 | 0 | 0 | 0 |
| SRR5230553 | 0 | 0 | 33 | 0 | 0 | 11 | 40 | 12 | 61 | 18 | 0 | 11 | 16 | 0 | 0 | 12 | 0 | 0 | 0 |
| SRR5230554 | 32 | 79 | 22 | 0 | 0 | 42 | 43 | 183 | 730 | 0 | 108 | 159 | 132 | 0 | 0 | 47 | 0 | 0 | 0 |
| SRR5230555 | 23 | 16 | 86 | 0 | 0 | 12 | 12 | 73 | 362 | 0 | 0 | 11 | 23 | 0 | 0 | 18 | 0 | 0 | 0 |
| SRR5230556 | 166 | 111 | 186 | 0 | 0 | 18 | 20 | 16 | 33 | 0 | 14 | 18 | 11 | 0 | 0 | 36 | 0 | 0 | 0 |
| SRR5230557 | 33 | 0 | 70 | 0 | 0 | 0 | 58 | 154 | 422 | 0 | 0 | 0 | 177 | 0 | 0 | 18 | 0 | 0 | 0 |
| SRR5230558 | 10 | 23 | 52 | 0 | 0 | 0 | 27 | 78 | 199 | 0 | 85 | 247 | 92 | 0 | 0 | 332 | 0 | 0 | 0 |
| SRR5230559 | 0 | 0 | 157 | 0 | 0 | 39 | 32 | 143 | 696 | 0 | 0 | 12 | 159 | 0 | 0 | 20 | 0 | 0 | 0 |
| SRR5230560 | 16 | 38 | 92 | 0 | 0 | 55 | 25 | 24 | 138 | 0 | 0 | 42 | 16 | 0 | 0 | 27 | 0 | 0 | 0 |
| SRR5230561 | 11 | 11 | 34 | 0 | 0 | 0 | 50 | 17 | 49 | 0 | 0 | 16 | 88 | 0 | 0 | 17 | 0 | 0 | 0 |
| SRR5230562 | 0 | 0 | 104 | 15 | 0 | 20 | 0 | 77 | 214 | 0 | 0 | 46 | 120 | 0 | 0 | 196 | 0 | 0 | 0 |
| SRR5230563 | 0 | 0 | 63 | 0 | 0 | 75 | 21 | 151 | 638 | 0 | 0 | 63 | 72 | 0 | 0 | 222 | 0 | 0 | 0 |
| SRR5230564 | 0 | 0 | 76 | 0 | 0 | 49 | 23 | 167 | 470 | 0 | 32 | 191 | 115 | 0 | 0 | 132 | 0 | 0 | 0 |
| SRR5230565 | 0 | 17 | 17 | 0 | 0 | 0 | 33 | 151 | 195 | 0 | 123 | 26 | 119 | 0 | 0 | 96 | 0 | 0 | 0 |
| SRR5230566 | 0 | 16 | 41 | 0 | 0 | 0 | 35 | 436 | 835 | 0 | 11 | 0 | 144 | 0 | 0 | 39 | 0 | 0 | 0 |
| SRR5230567 | 0 | 0 | 44 | 0 | 0 | 0 | 21 | 46 | 50 | 0 | 0 | 15 | 27 | 0 | 0 | 57 | 0 | 0 | 0 |
| SRR5230568 | 0 | 0 | 116 | 0 | 0 | 17 | 0 | 26 | 58 | 0 | 0 | 22 | 16 | 0 | 0 | 47 | 0 | 0 | 0 |
| SRR5230569 | 11 | 0 | 167 | 20 | 0 | 85 | 12 | 15 | 60 | 0 | 16 | 12 | 12 | 0 | 0 | 93 | 0 | 0 | 0 |
| SRR5230570 | 0 | 13 | 47 | 0 | 0 | 36 | 30 | 238 | 804 | 0 | 0 | 359 | 63 | 0 | 0 | 451 | 0 | 0 | 0 |
| SRR5230571 | 13 | 10 | 47 | 0 | 0 | 0 | 14 | 29 | 83 | 0 | 0 | 30 | 12 | 0 | 0 | 24 | 0 | 0 | 0 |
| SRR5230572 | 0 | 26 | 60 | 0 | 0 | 25 | 38 | 293 | 957 | 0 | 52 | 105 | 77 | 0 | 0 | 60 | 0 | 0 | 0 |
| SRR5230573 | 20 | 80 | 286 | 0 | 0 | 21 | 60 | 831 | 2122 | 0 | 17 | 46 | 72 | 0 | 0 | 49 | 0 | 0 | 0 |
| SRR5230574 | 30 | 12 | 60 | 0 | 0 | 0 | 36 | 35 | 54 | 0 | 0 | 0 | 24 | 0 | 0 | 49 | 0 | 0 | 0 |
| SRR5230575 | 0 | 0 | 116 | 0 | 0 | 12 | 49 | 156 | 328 | 14 | 14 | 14 | 156 | 0 | 0 | 47 | 0 | 0 | 0 |
| SRR5230576 | 14 | 24 | 121 | 0 | 0 | 0 | 101 | 223 | 520 | 10 | 0 | 0 | 234 | 0 | 0 | 195 | 0 | 0 | 0 |
| SRR5230577 | 11 | 22 | 76 | 0 | 0 | 0 | 110 | 214 | 526 | 0 | 0 | 0 | 285 | 0 | 0 | 197 | 0 | 0 | 0 |
| SRR5230578 | 33 | 23 | 17 | 0 | 0 | 11 | 19 | 12 | 35 | 0 | 82 | 94 | 35 | 0 | 0 | 14 | 0 | 0 | 0 |
| SRR5230579 | 13 | 26 | 13 | 0 | 0 | 0 | 0 | 17 | 30 | 11 | 107 | 89 | 37 | 0 | 0 | 15 | 0 | 0 | 0 |
| SRR5230580 | 14 | 14 | 103 | 0 | 0 | 46 | 24 | 28 | 95 | 0 | 0 | 21 | 12 | 0 | 0 | 25 | 0 | 0 | 0 |
| SRR5230581 | 0 | 0 | 303 | 0 | 0 | 51 | 15 | 152 | 631 | 0 | 0 | 189 | 44 | 0 | 0 | 218 | 0 | 0 | 0 |
| SRR5230582 | 0 | 0 | 111 | 0 | 0 | 61 | 21 | 328 | 747 | 0 | 0 | 144 | 131 | 0 | 0 | 223 | 0 | 0 | 32 |
| SRR5230583 | 0 | 0 | 98 | 0 | 0 | 79 | 47 | 291 | 660 | 0 | 0 | 189 | 63 | 0 | 0 | 220 | 0 | 0 | 47 |
| SRR5230584 | 0 | 0 | 107 | 0 | 0 | 0 | 15 | 79 | 110 | 0 | 0 | 32 | 569 | 0 | 0 | 174 | 0 | 0 | 0 |
| SRR5230585 | 17 | 0 | 104 | 0 | 0 | 0 | 0 | 60 | 103 | 0 | 0 | 26 | 525 | 0 | 13 | 129 | 0 | 0 | 0 |
| SRR5230586 | 0 | 18 | 15 | 0 | 0 | 27 | 24 | 172 | 931 | 0 | 0 | 47 | 39 | 0 | 0 | 159 | 0 | 0 | 0 |
| SRR5230587 | 0 | 16 | 0 | 0 | 0 | 0 | 44 | 215 | 959 | 0 | 0 | 41 | 47 | 0 | 0 | 168 | 0 | 0 | 0 |
| SRR5230588 | 0 | 11 | 72 | 0 | 0 | 69 | 48 | 288 | 1136 | 0 | 0 | 22 | 62 | 0 | 0 | 66 | 0 | 0 | 0 |
| SRR5230589 | 0 | 0 | 69 | 0 | 0 | 46 | 38 | 200 | 1284 | 0 | 0 | 31 | 69 | 0 | 0 | 85 | 0 | 0 | 15 |
| SRR5230590 | 0 | 0 | 163 | 0 | 0 | 28 | 41 | 83 | 83 | 0 | 0 | 0 | 14 | 0 | 0 | 96 | 0 | 0 | 0 |
| SRR5230591 | 0 | 0 | 39 | 0 | 0 | 0 | 95 | 73 | 51 | 0 | 0 | 0 | 103 | 0 | 0 | 137 | 0 | 0 | 0 |
| SRR5230592 | 0 | 0 | 0 | 0 | 0 | 0 | 145 | 48 | 194 | 0 | 0 | 0 | 97 | 0 | 0 | 0 | 0 | 0 | 0 |
| SRR5230593 | 0 | 0 | 0 | 0 | 0 | 0 | 123 | 57 | 60 | 0 | 0 | 0 | 72 | 0 | 0 | 52 | 0 | 0 | 0 |
| SRR5230594 | 0 | 13 | 34 | 0 | 0 | 11 | 35 | 42 | 189 | 0 | 0 | 23 | 55 | 0 | 0 | 66 | 0 | 0 | 0 |
| SRR5230595 | 0 | 54 | 134 | 0 | 0 | 13 | 45 | 37 | 192 | 0 | 0 | 0 | 12 | 0 | 0 | 16 | 0 | 0 | 17 |
| SRR5230596 | 0 | 0 | 81 | 0 | 0 | 0 | 53 | 26 | 74 | 0 | 0 | 20 | 39 | 0 | 0 | 122 | 0 | 0 | 0 |
| SRR5230597 | 0 | 0 | 79 | 0 | 0 | 0 | 61 | 128 | 381 | 0 | 0 | 26 | 229 | 0 | 0 | 673 | 0 | 0 | 0 |
| SRR5230600 | 0 | 0 | 15 | 0 | 0 | 0 | 30 | 11 | 39 | 0 | 0 | 0 | 16 | 0 | 0 | 18 | 0 | 0 | 0 |
| SRR5230601 | 0 | 0 | 0 | 0 | 0 | 0 | 30 | 87 | 310 | 0 | 0 | 0 | 108 | 0 | 0 | 188 | 0 | 0 | 0 |
| SRR5230602 | 0 | 0 | 76 | 0 | 0 | 10 | 22 | 16 | 84 | 0 | 0 | 14 | 53 | 0 | 0 | 22 | 0 | 0 | 14 |
| SRR5230603 | 0 | 22 | 216 | 0 | 0 | 96 | 15 | 81 | 780 | 0 | 0 | 96 | 66 | 0 | 0 | 15 | 0 | 0 | 0 |
| SRR5230604 | 0 | 0 | 35 | 0 | 0 | 0 | 39 | 29 | 70 | 0 | 0 | 0 | 24 | 0 | 0 | 18 | 0 | 0 | 0 |
| SRR5230605 | 0 | 0 | 13 | 0 | 0 | 0 | 42 | 145 | 349 | 0 | 0 | 0 | 138 | 0 | 0 | 97 | 0 | 0 | 0 |
| SRR5230606 | 0 | 0 | 74 | 0 | 0 | 0 | 20 | 100 | 193 | 0 | 0 | 37 | 107 | 0 | 0 | 335 | 0 | 0 | 0 |
| SRR5230607 | 0 | 0 | 87 | 0 | 0 | 0 | 41 | 11 | 99 | 0 | 0 | 17 | 11 | 0 | 0 | 33 | 0 | 0 | 0 |
| SRR5230608 | 0 | 0 | 14 | 0 | 0 | 55 | 12 | 162 | 623 | 0 | 0 | 159 | 75 | 0 | 0 | 131 | 0 | 0 | 0 |
| SRR5230609 | 0 | 0 | 0 | 0 | 0 | 77 | 18 | 148 | 617 | 0 | 0 | 190 | 77 | 0 | 0 | 160 | 0 | 0 | 0 |
| SRR5230610 | 0 | 0 | 0 | 0 | 0 | 0 | 20 | 59 | 68 | 20 | 0 | 0 | 117 | 0 | 0 | 166 | 0 | 0 | 0 |
| SRR5230611 | 0 | 0 | 0 | 0 | 0 | 0 | 61 | 62 | 55 | 11 | 0 | 0 | 18 | 0 | 0 | 19 | 0 | 0 | 0 |
| SRR5230613 | 0 | 0 | 12 | 0 | 0 | 0 | 63 | 91 | 58 | 13 | 0 | 0 | 177 | 0 | 0 | 88 | 0 | 0 | 0 |
| SRR5230614 | 0 | 0 | 0 | 0 | 13 | 0 | 30 | 50 | 30 | 0 | 0 | 23 | 107 | 0 | 0 | 193 | 0 | 0 | 0 |
| SRR5230615 | 0 | 0 | 0 | 0 | 0 | 0 | 25 | 33 | 22 | 16 | 0 | 29 | 133 | 0 | 0 | 189 | 0 | 0 | 0 |
| SRR5230617 | 0 | 0 | 0 | 0 | 0 | 0 | 31 | 70 | 30 | 0 | 0 | 0 | 10 | 0 | 0 | 39 | 0 | 0 | 0 |
| SRR5230618 | 0 | 0 | 0 | 0 | 0 | 0 | 15 | 82 | 66 | 11 | 11 | 0 | 26 | 0 | 0 | 28 | 0 | 0 | 0 |
| SRR5230619 | 0 | 0 | 18 | 0 | 0 | 0 | 88 | 92 | 51 | 0 | 0 | 0 | 180 | 0 | 0 | 250 | 0 | 0 | 0 |
| SRR5230620 | 0 | 0 | 127 | 0 | 0 | 0 | 18 | 186 | 183 | 0 | 0 | 0 | 50 | 0 | 0 | 103 | 0 | 0 | 0 |
| SRR5230621 | 0 | 31 | 62 | 0 | 0 | 0 | 31 | 28 | 15 | 15 | 0 | 0 | 61 | 0 | 0 | 11 | 0 | 0 | 0 |
| SRR5230622 | 19 | 54 | 78 | 0 | 0 | 0 | 25 | 46 | 134 | 0 | 0 | 48 | 31 | 0 | 0 | 21 | 0 | 0 | 0 |
| SRR5230623 | 0 | 17 | 32 | 0 | 0 | 58 | 14 | 144 | 573 | 0 | 18 | 2458 | 48 | 0 | 0 | 333 | 0 | 0 | 0 |
| SRR5230624 | 0 | 0 | 272 | 13 | 0 | 47 | 27 | 270 | 944 | 0 | 0 | 287 | 71 | 0 | 0 | 265 | 0 | 0 | 0 |
| SRR5230625 | 0 | 15 | 66 | 0 | 0 | 13 | 24 | 60 | 175 | 0 | 0 | 41 | 14 | 0 | 0 | 15 | 0 | 0 | 0 |
| SRR5230626 | 0 | 0 | 33 | 0 | 0 | 56 | 26 | 260 | 791 | 0 | 0 | 180 | 65 | 0 | 0 | 414 | 0 | 0 | 0 |
| SRR5230627 | 65 | 50 | 61 | 0 | 0 | 29 | 23 | 248 | 757 | 0 | 47 | 2843 | 67 | 0 | 0 | 141 | 0 | 0 | 0 |
| SRR5230628 | 0 | 15 | 153 | 0 | 0 | 52 | 16 | 31 | 67 | 0 | 0 | 28 | 24 | 0 | 0 | 28 | 0 | 0 | 0 |
| SRR5230629 | 0 | 20 | 37 | 0 | 0 | 33 | 12 | 80 | 291 | 0 | 0 | 11 | 14 | 0 | 0 | 11 | 0 | 0 | 0 |
| SRR5230630 | 0 | 39 | 38 | 15 | 0 | 26 | 23 | 55 | 175 | 0 | 20 | 12 | 22 | 0 | 0 | 13 | 0 | 0 | 0 |

### Supp Table S8.2.Columns_AM_to_BW

| | | | | | | | | | | | | | | | | | | | | | | | | | | | | | | | | | | | | | |
|---|---|---|---|---|---|---|---|---|---|---|---|---|---|---|---|---|---|---|---|---|---|---|---|---|---|---|---|---|---|---|---|---|---|---|---|---|---|
| SRR5230631 | 339 | 0 | 0 | 0 | 0 | 55 | 471 | 99 | 40 | 0 | 36 | 636 | 0 | 0 | 18 | 0 | 45 | 0 | 0 | 15 | 78 | 0 | 0 | 50 | 11 | 26 | 54 | 0 | 0 | 35 | 68 | 0 | 0 | 34 | 0 | 0 | 0 |
| SRR5230632 | 245 | 0 | 12 | 0 | 0 | 17 | 1235 | 378 | 36 | 0 | 23 | 485 | 0 | 0 | 0 | 0 | 62 | 0 | 19 | 16 | 122 | 0 | 0 | 30 | 29 | 64 | 267 | 0 | 15 | 171 | 20 | 0 | 0 | 70 | 0 | 0 | 0 |
| SRR5230633 | 358 | 0 | 17 | 0 | 11 | 31 | 3582 | 194 | 27 | 0 | 13 | 715 | 0 | 0 | 23 | 0 | 59 | 0 | 29 | 10 | 157 | 19 | 0 | 14 | 0 | 19 | 70 | 0 | 14 | 57 | 60 | 0 | 0 | 14 | 0 | 0 | 0 |
| SRR5230634 | 304 | 0 | 0 | 0 | 20 | 0 | 13 | 85 | 129 | 0 | 14 | 166 | 0 | 12 | 25 | 0 | 0 | 0 | 0 | 0 | 313 | 18 | 0 | 46 | 0 | 62 | 0 | 0 | 34 | 36 | 12 | 0 | 0 | 87 | 0 | 0 | 0 |
| SRR5230635 | 98 | 0 | 0 | 0 | 23 | 14 | 23 | 263 | 97 | 0 | 199 | 112 | 23 | 0 | 441 | 0 | 13 | 0 | 0 | 0 | 1709 | 0 | 0 | 79 | 0 | 0 | 14 | 18 | 0 | 323 | 0 | 0 | 0 | 148 | 0 | 0 | 0 |
| SRR5230636 | 152 | 0 | 0 | 15 | 16 | 0 | 126 | 119 | 66 | 41 | 27 | 137 | 0 | 0 | 512 | 0 | 0 | 0 | 0 | 0 | 325 | 0 | 0 | 35 | 0 | 0 | 49 | 0 | 19 | 150 | 0 | 15 | 0 | 27 | 0 | 0 | 0 |
| SRR5230637 | 168 | 0 | 0 | 0 | 32 | 12 | 172 | 150 | 47 | 0 | 24 | 107 | 0 | 44 | 2027 | 0 | 0 | 0 | 0 | 0 | 585 | 0 | 0 | 24 | 0 | 20 | 59 | 0 | 0 | 134 | 40 | 0 | 16 | 71 | 0 | 0 | 0 |
| SRR5230638 | 726 | 0 | 0 | 34 | 0 | 0 | 68 | 166 | 18 | 0 | 0 | 510 | 29 | 0 | 1007 | 0 | 13 | 0 | 0 | 0 | 3845 | 0 | 0 | 79 | 11 | 0 | 34 | 18 | 0 | 18 | 13 | 0 | 0 | 185 | 0 | 0 | 0 |
| SRR5230639 | 583 | 0 | 0 | 0 | 0 | 22 | 16 | 12 | 0 | 0 | 0 | 621 | 61 | 0 | 212 | 0 | 0 | 0 | 0 | 0 | 1560 | 15 | 0 | 177 | 0 | 22 | 0 | 0 | 16 | 29 | 0 | 0 | 0 | 17 | 0 | 0 | 0 |
| SRR5230640 | 256 | 0 | 0 | 11 | 17 | 0 | 35 | 69 | 17 | 0 | 0 | 189 | 0 | 0 | 579 | 0 | 0 | 0 | 0 | 0 | 1448 | 0 | 0 | 19 | 0 | 0 | 17 | 0 | 0 | 190 | 0 | 15 | 0 | 106 | 0 | 0 | 0 |
| SRR5230641 | 377 | 0 | 0 | 0 | 0 | 0 | 46 | 12 | 0 | 0 | 0 | 160 | 13 | 19 | 2675 | 0 | 0 | 0 | 21 | 0 | 1002 | 0 | 0 | 16 | 0 | 0 | 0 | 0 | 0 | 48 | 0 | 0 | 0 | 69 | 0 | 0 | 0 |
| SRR5230642 | 32 | 0 | 0 | 0 | 0 | 0 | 1150 | 187 | 13 | 0 | 21 | 63 | 0 | 0 | 53 | 0 | 34 | 0 | 27 | 59 | 225 | 0 | 0 | 37 | 0 | 27 | 69 | 0 | 29 | 24 | 19 | 0 | 0 | 56 | 0 | 0 | 0 |
| SRR5230643 | 437 | 0 | 0 | 0 | 0 | 0 | 107 | 14 | 0 | 0 | 0 | 449 | 14 | 0 | 1835 | 0 | 12 | 0 | 27 | 27 | 1701 | 0 | 0 | 72 | 0 | 0 | 34 | 16 | 68 | 22 | 14 | 0 | 0 | 109 | 0 | 0 | 0 |
| SRR5230644 | 268 | 0 | 0 | 0 | 0 | 0 | 77 | 58 | 12 | 0 | 0 | 507 | 48 | 0 | 2368 | 0 | 16 | 0 | 21 | 13 | 3233 | 0 | 0 | 120 | 0 | 30 | 0 | 0 | 14 | 377 | 18 | 0 | 0 | 172 | 0 | 0 | 0 |
| SRR5230645 | 258 | 0 | 0 | 0 | 0 | 0 | 100 | 409 | 0 | 0 | 0 | 602 | 31 | 0 | 717 | 0 | 13 | 0 | 65 | 105 | 1109 | 0 | 0 | 162 | 0 | 18 | 13 | 0 | 23 | 507 | 0 | 13 | 0 | 301 | 0 | 0 | 0 |
| SRR5230646 | 353 | 0 | 0 | 31 | 28 | 0 | 547 | 179 | 28 | 0 | 17 | 252 | 14 | 0 | 3625 | 0 | 15 | 0 | 16 | 25 | 1555 | 46 | 0 | 69 | 0 | 0 | 23 | 0 | 36 | 13 | 0 | 0 | 0 | 73 | 0 | 0 | 0 |
| SRR5230647 | 193 | 0 | 0 | 15 | 0 | 0 | 0 | 42 | 0 | 0 | 13 | 13 | 11 | 0 | 300 | 0 | 0 | 0 | 0 | 0 | 1327 | 0 | 0 | 0 | 0 | 0 | 0 | 0 | 0 | 37 | 0 | 0 | 0 | 23 | 0 | 0 | 0 |
| SRR5230648 | 113 | 0 | 0 | 0 | 28 | 0 | 352 | 38 | 37 | 0 | 132 | 299 | 10 | 0 | 267 | 0 | 25 | 0 | 52 | 38 | 874 | 0 | 0 | 69 | 0 | 27 | 23 | 37 | 75 | 39 | 15 | 0 | 0 | 13 | 0 | 0 | 15 |
| SRR5230649 | 276 | 0 | 0 | 0 | 0 | 0 | 44 | 112 | 24 | 0 | 14 | 316 | 19 | 37 | 2374 | 0 | 11 | 0 | 14 | 30 | 1122 | 10 | 0 | 53 | 0 | 17 | 72 | 0 | 0 | 281 | 20 | 0 | 0 | 70 | 0 | 0 | 0 |
| SRR5230650 | 98 | 0 | 0 | 0 | 0 | 0 | 26 | 21 | 0 | 0 | 0 | 0 | 0 | 0 | 149 | 0 | 12 | 0 | 0 | 0 | 1164 | 0 | 0 | 67 | 0 | 17 | 0 | 0 | 11 | 39 | 15 | 0 | 11 | 64 | 0 | 0 | 0 |
| SRR5230651 | 464 | 0 | 0 | 0 | 0 | 0 | 193 | 117 | 0 | 0 | 0 | 399 | 0 | 0 | 3470 | 0 | 49 | 0 | 37 | 77 | 487 | 0 | 0 | 197 | 12 | 0 | 14 | 12 | 56 | 267 | 0 | 0 | 0 | 91 | 0 | 0 | 0 |
| SRR5230652 | 139 | 0 | 0 | 21 | 33 | 0 | 301 | 391 | 41 | 0 | 19 | 115 | 0 | 193 | 1256 | 30 | 26 | 0 | 49 | 227 | 684 | 24 | 0 | 15 | 0 | 17 | 0 | 19 | 0 | 285 | 13 | 0 | 0 | 78 | 0 | 0 | 0 |
| SRR5230653 | 197 | 0 | 0 | 0 | 0 | 0 | 25 | 59 | 0 | 0 | 0 | 139 | 0 | 0 | 480 | 0 | 22 | 0 | 11 | 0 | 1391 | 0 | 0 | 64 | 0 | 12 | 40 | 0 | 0 | 187 | 0 | 0 | 0 | 26 | 0 | 0 | 0 |
| SRR5230654 | 167 | 14 | 0 | 13 | 36 | 0 | 39 | 75 | 53 | 0 | 0 | 58 | 14 | 47 | 9256 | 0 | 0 | 0 | 168 | 426 | 1203 | 13 | 0 | 35 | 0 | 0 | 17 | 68 | 0 | 95 | 0 | 0 | 0 | 59 | 0 | 0 | 0 |
| SRR5230655 | 419 | 0 | 0 | 0 | 0 | 0 | 61 | 154 | 0 | 0 | 0 | 288 | 0 | 0 | 1105 | 0 | 25 | 0 | 11 | 0 | 2236 | 11 | 0 | 113 | 0 | 0 | 20 | 0 | 22 | 23 | 0 | 0 | 0 | 176 | 0 | 0 | 0 |
| SRR5230656 | 413 | 0 | 0 | 0 | 14 | 0 | 46 | 136 | 19 | 0 | 0 | 288 | 0 | 33 | 55 | 0 | 14 | 0 | 21 | 13 | 571 | 0 | 0 | 70 | 0 | 23 | 19 | 23 | 19 | 323 | 0 | 0 | 0 | 243 | 0 | 0 | 19 |
| SRR5230657 | 397 | 0 | 0 | 0 | 0 | 0 | 468 | 256 | 48 | 0 | 18 | 227 | 0 | 0 | 4721 | 0 | 18 | 0 | 88 | 38 | 813 | 0 | 0 | 127 | 0 | 0 | 22 | 0 | 64 | 436 | 0 | 0 | 0 | 90 | 0 | 0 | 0 |
| SRR5230658 | 362 | 0 | 0 | 0 | 0 | 0 | 47 | 15 | 0 | 0 | 0 | 341 | 33 | 0 | 36 | 0 | 20 | 0 | 0 | 0 | 567 | 0 | 0 | 27 | 0 | 25 | 13 | 0 | 38 | 76 | 0 | 0 | 0 | 11 | 0 | 0 | 0 |
| SRR5230659 | 163 | 0 | 0 | 0 | 0 | 0 | 665 | 354 | 0 | 0 | 0 | 147 | 0 | 0 | 336 | 0 | 0 | 0 | 19 | 0 | 569 | 0 | 0 | 314 | 0 | 0 | 11 | 17 | 11 | 987 | 0 | 0 | 0 | 205 | 0 | 0 | 0 |
| SRR5230660 | 184 | 0 | 0 | 0 | 0 | 0 | 101 | 35 | 23 | 0 | 42 | 196 | 28 | 0 | 66 | 0 | 15 | 0 | 0 | 0 | 1013 | 23 | 0 | 290 | 0 | 0 | 23 | 19 | 98 | 61 | 0 | 0 | 0 | 15 | 0 | 0 | 14 |
| SRR5230661 | 102 | 0 | 0 | 44 | 0 | 0 | 585 | 2473 | 15 | 0 | 0 | 76 | 0 | 0 | 0 | 0 | 11 | 0 | 14 | 28 | 228 | 0 | 0 | 468 | 0 | 0 | 0 | 0 | 0 | 1657 | 0 | 0 | 0 | 436 | 0 | 0 | 0 |
| SRR5230662 | 92 | 0 | 0 | 0 | 0 | 0 | 55 | 12 | 0 | 0 | 0 | 48 | 0 | 0 | 118 | 0 | 19 | 0 | 0 | 24 | 693 | 0 | 0 | 76 | 0 | 0 | 0 | 0 | 16 | 31 | 0 | 0 | 0 | 12 | 0 | 0 | 0 |
| SRR5230663 | 477 | 0 | 0 | 0 | 0 | 0 | 473 | 2160 | 48 | 0 | 24 | 393 | 28 | 0 | 927 | 0 | 16 | 0 | 68 | 59 | 759 | 0 | 0 | 252 | 0 | 0 | 47 | 0 | 73 | 956 | 0 | 0 | 0 | 181 | 0 | 0 | 0 |
| SRR5230664 | 241 | 0 | 0 | 0 | 0 | 0 | 18 | 81 | 0 | 0 | 0 | 126 | 0 | 0 | 248 | 0 | 38 | 0 | 0 | 0 | 1609 | 0 | 0 | 58 | 0 | 0 | 10 | 0 | 0 | 166 | 0 | 0 | 0 | 102 | 0 | 0 | 0 |
| SRR5230665 | 295 | 0 | 0 | 0 | 0 | 0 | 285 | 481 | 32 | 0 | 14 | 493 | 12 | 86 | 242 | 0 | 23 | 0 | 79 | 206 | 1404 | 0 | 0 | 331 | 0 | 0 | 29 | 22 | 314 | 663 | 0 | 0 | 0 | 192 | 0 | 0 | 0 |
| SRR5230666 | 292 | 0 | 0 | 0 | 0 | 0 | 2263 | 34 | 36 | 0 | 43 | 127 | 17 | 33 | 6574 | 0 | 10 | 0 | 70 | 153 | 945 | 0 | 0 | 13 | 0 | 0 | 50 | 23 | 0 | 18 | 23 | 0 | 0 | 178 | 0 | 0 | 0 |
| SRR5230667 | 92 | 0 | 0 | 0 | 0 | 0 | 105 | 200 | 0 | 0 | 0 | 28 | 0 | 0 | 79 | 0 | 23 | 0 | 66 | 0 | 630 | 0 | 0 | 73 | 0 | 12 | 0 | 0 | 13 | 639 | 0 | 0 | 0 | 123 | 0 | 0 | 0 |
| SRR5230668 | 187 | 0 | 0 | 0 | 0 | 0 | 123 | 15 | 0 | 0 | 0 | 224 | 13 | 0 | 1086 | 0 | 19 | 0 | 54 | 126 | 1586 | 0 | 0 | 129 | 0 | 0 | 15 | 0 | 72 | 25 | 12 | 0 | 0 | 83 | 0 | 0 | 0 |
| SRR5230669 | 499 | 0 | 0 | 0 | 0 | 0 | 123 | 477 | 39 | 0 | 27 | 336 | 45 | 0 | 8164 | 0 | 0 | 0 | 12 | 14 | 4879 | 0 | 0 | 150 | 0 | 39 | 14 | 18 | 0 | 678 | 14 | 0 | 0 | 688 | 0 | 14 | 16 |
| SRR5230670 | 99 | 0 | 0 | 0 | 0 | 0 | 16 | 10 | 24 | 0 | 0 | 131 | 0 | 15 | 16 | 0 | 26 | 0 | 33 | 0 | 258 | 0 | 0 | 120 | 0 | 0 | 15 | 0 | 23 | 23 | 0 | 0 | 0 | 16 | 0 | 0 | 0 |
| SRR5230671 | 52 | 16 | 0 | 29 | 16 | 0 | 270 | 330 | 0 | 0 | 0 | 68 | 0 | 0 | 0 | 0 | 12 | 0 | 0 | 72 | 1487 | 0 | 0 | 404 | 0 | 0 | 25 | 0 | 29 | 573 | 0 | 0 | 0 | 675 | 0 | 0 | 0 |
| SRR5230673 | 202 | 0 | 0 | 0 | 0 | 0 | 163 | 444 | 0 | 0 | 16 | 139 | 21 | 0 | 746 | 0 | 0 | 0 | 48 | 17 | 5348 | 0 | 0 | 128 | 0 | 11 | 60 | 37 | 0 | 1096 | 0 | 0 | 0 | 302 | 0 | 0 | 0 |
| SRR5230674 | 257 | 0 | 0 | 13 | 0 | 0 | 28 | 731 | 0 | 0 | 0 | 208 | 0 | 0 | 61 | 0 | 22 | 0 | 41 | 17 | 760 | 13 | 0 | 509 | 0 | 13 | 0 | 11 | 37 | 1737 | 0 | 0 | 0 | 306 | 0 | 0 | 0 |
| SRR5230675 | 148 | 0 | 0 | 0 | 0 | 0 | 2381 | 2492 | 0 | 0 | 0 | 100 | 0 | 27 | 0 | 0 | 31 | 0 | 30 | 36 | 254 | 0 | 0 | 12 | 0 | 0 | 0 | 12 | 15 | 357 | 0 | 0 | 0 | 208 | 0 | 0 | 0 |
| SRR5230676 | 193 | 0 | 0 | 30 | 33 | 23 | 21 | 19 | 26 | 0 | 0 | 249 | 36 | 76 | 126 | 0 | 11 | 0 | 21 | 0 | 2683 | 46 | 0 | 212 | 0 | 23 | 0 | 36 | 0 | 40 | 0 | 0 | 0 | 16 | 0 | 0 | 0 |
| SRR5230677 | 256 | 0 | 0 | 0 | 0 | 0 | 58 | 97 | 30 | 0 | 11 | 134 | 26 | 0 | 40 | 0 | 14 | 0 | 0 | 0 | 789 | 0 | 0 | 202 | 0 | 0 | 13 | 0 | 24 | 398 | 0 | 0 | 0 | 271 | 0 | 0 | 0 |
| SRR5230678 | 225 | 14 | 0 | 0 | 0 | 0 | 18 | 15 | 28 | 0 | 0 | 306 | 75 | 19 | 69 | 0 | 0 | 0 | 0 | 0 | 1067 | 0 | 0 | 219 | 0 | 23 | 0 | 0 | 75 | 58 | 23 | 0 | 0 | 359 | 0 | 0 | 0 |
| SRR5230679 | 131 | 0 | 0 | 22 | 0 | 0 | 57 | 20 | 0 | 0 | 11 | 102 | 0 | 0 | 23 | 0 | 0 | 0 | 0 | 0 | 1422 | 29 | 0 | 42 | 0 | 0 | 74 | 0 | 40 | 57 | 45 | 0 | 0 | 92 | 0 | 0 | 0 |
| SRR5230680 | 72 | 0 | 0 | 0 | 0 | 0 | 459 | 170 | 54 | 0 | 0 | 27 | 0 | 0 | 57 | 0 | 27 | 0 | 34 | 29 | 843 | 0 | 0 | 116 | 0 | 0 | 14 | 25 | 57 | 13 | 0 | 0 | 0 | 37 | 0 | 0 | 0 |
| SRR5230681 | 152 | 0 | 0 | 0 | 0 | 0 | 30 | 34 | 18 | 0 | 0 | 82 | 0 | 0 | 417 | 0 | 0 | 0 | 0 | 0 | 771 | 0 | 0 | 21 | 0 | 18 | 25 | 0 | 0 | 147 | 18 | 0 | 0 | 39 | 0 | 0 | 0 |
| SRR5230682 | 734 | 0 | 0 | 0 | 0 | 0 | 105 | 609 | 0 | 0 | 17 | 516 | 29 | 0 | 94 | 0 | 31 | 0 | 20 | 0 | 708 | 0 | 0 | 221 | 0 | 0 | 32 | 0 | 585 | 1408 | 0 | 11 | 0 | 639 | 0 | 0 | 0 |
| SRR5230683 | 113 | 0 | 0 | 0 | 0 | 0 | 109 | 270 | 12 | 0 | 0 | 158 | 45 | 0 | 134 | 0 | 19 | 0 | 14 | 14 | 3850 | 0 | 0 | 126 | 0 | 0 | 16 | 0 | 0 | 728 | 0 | 0 | 12 | 490 | 0 | 0 | 0 |
| SRR5230684 | 346 | 0 | 0 | 18 | 0 | 0 | 175 | 387 | 36 | 0 | 0 | 197 | 0 | 0 | 31 | 0 | 17 | 0 | 0 | 0 | 323 | 22 | 0 | 324 | 0 | 17 | 102 | 0 | 57 | 664 | 0 | 20 | 0 | 105 | 0 | 0 | 0 |
| SRR5230685 | 253 | 0 | 0 | 19 | 0 | 0 | 35 | 444 | 35 | 0 | 0 | 219 | 31 | 0 | 46 | 0 | 19 | 0 | 26 | 29 | 1685 | 0 | 0 | 204 | 0 | 0 | 0 | 0 | 0 | 1184 | 15 | 0 | 0 | 845 | 0 | 0 | 0 |
| SRR5230686 | 390 | 0 | 0 | 51 | 0 | 0 | 65 | 20 | 23 | 0 | 14 | 357 | 94 | 0 | 98 | 0 | 11 | 0 | 10 | 0 | 2690 | 43 | 0 | 11 | 0 | 28 | 48 | 0 | 85 | 67 | 0 | 0 | 20 | 38 | 0 | 0 | 20 |
| SRR5230687 | 737 | 0 | 0 | 11 | 0 | 0 | 15 | 33 | 15 | 0 | 0 | 747 | 31 | 0 | 82 | 0 | 18 | 0 | 0 | 0 | 2272 | 0 | 0 | 13 | 0 | 0 | 81 | 0 | 95 | 85 | 0 | 0 | 0 | 68 | 0 | 0 | 0 |
| SRR5230688 | 186 | 0 | 0 | 0 | 0 | 0 | 138 | 255 | 0 | 0 | 0 | 76 | 22 | 0 | 85 | 0 | 20 | 0 | 18 | 13 | 1651 | 0 | 0 | 77 | 0 | 0 | 0 | 0 | 13 | 603 | 0 | 0 | 0 | 124 | 0 | 0 | 0 |
| SRR5230689 | 129 | 0 | 0 | 0 | 0 | 0 | 24 | 353 | 69 | 0 | 0 | 54 | 90 | 0 | 158 | 0 | 0 | 0 | 17 | 0 | 2841 | 0 | 0 | 180 | 0 | 36 | 48 | 0 | 36 | 1054 | 15 | 36 | 0 | 1033 | 0 | 0 | 0 |
| SRR5230690 | 540 | 0 | 0 | 0 | 0 | 0 | 59 | 512 | 24 | 0 | 0 | 604 | 29 | 0 | 102 | 0 | 31 | 0 | 22 | 0 | 3960 | 19 | 0 | 241 | 0 | 0 | 13 | 0 | 30 | 1366 | 0 | 0 | 0 | 746 | 0 | 0 | 0 |
| SRR5230691 | 336 | 0 | 0 | 0 | 0 | 0 | 42 | 136 | 11 | 0 | 0 | 60 | 0 | 0 | 127 | 0 | 0 | 0 | 26 | 0 | 2182 | 0 | 0 | 90 | 0 | 0 | 39 | 0 | 0 | 274 | 0 | 0 | 0 | 202 | 0 | 0 | 0 |
| SRR5230692 | 223 | 0 | 11 | 0 | 0 | 0 | 102 | 268 | 14 | 0 | 0 | 219 | 36 | 0 | 1235 | 0 | 15 | 0 | 26 | 71 | 2156 | 0 | 0 | 148 | 15 | 17 | 105 | 0 | 0 | 623 | 13 | 0 | 0 | 311 | 0 | 0 | 0 |
| SRR5230693 | 327 | 16 | 0 | 0 | 0 | 21 | 54 | 408 | 0 | 0 | 30 | 339 | 0 | 0 | 67 | 0 | 17 | 0 | 0 | 0 | 892 | 0 | 0 | 503 | 0 | 18 | 34 | 0 | 25 | 2006 | 14 | 0 | 0 | 561 | 0 | 0 | 0 |
| SRR5230694 | 116 | 0 | 0 | 0 | 0 | 0 | 993 | 5647 | 31 | 12 | 192 | 86 | 0 | 0 | 144 | 0 | 19 | 0 | 16 | 22 | 296 | 0 | 0 | 33 | 0 | 24 | 53 | 0 | 44 | 6252 | 0 | 0 | 0 | 1850 | 0 | 0 | 0 |
| SRR5230695 | 273 | 0 | 0 | 0 | 0 | 0 | 33 | 348 | 0 | 0 | 18 | 130 | 0 | 0 | 107 | 0 | 13 | 0 | 0 | 0 | 1772 | 0 | 0 | 313 | 0 | 20 | 26 | 0 | 0 | 1345 | 0 | 0 | 0 | 817 | 0 | 0 | 0 |
| SRR5230696 | 247 | 0 | 0 | 0 | 0 | 0 | 514 | 3648 | 21 | 19 | 119 | 151 | 32 | 0 | 169 | 0 | 17 | 0 | 43 | 40 | 1061 | 0 | 0 | 71 | 0 | 15 | 55 | 29 | 39 | 2455 | 0 | 0 | 0 | 767 | 0 | 0 | 0 |
| SRR5230697 | 353 | 0 | 0 | 0 | 0 | 0 | 45 | 359 | 13 | 0 | 0 | 180 | 23 | 0 | 32 | 0 | 0 | 0 | 11 | 0 | 2021 | 0 | 0 | 186 | 0 | 0 | 10 | 0 | 0 | 1366 | 0 | 0 | 0 | 635 | 0 | 0 | 0 |
| SRR5230698 | 816 | 12 | 0 | 0 | 0 | 0 | 19 | 508 | 20 | 0 | 0 | 635 | 27 | 0 | 50 | 0 | 10 | 0 | 0 | 0 | 348 | 0 | 0 | 114 | 0 | 0 | 64 | 0 | 1106 | 1312 | 0 | 0 | 0 | 384 | 0 | 0 | 0 |
| SRR5230699 | 340 | 0 | 0 | 0 | 0 | 0 | 1695 | 3615 | 11 | 70 | 1942 | 241 | 20 | 11 | 5346 | 0 | 28 | 0 | 65 | 90 | 2285 | 11 | 0 | 99 | 0 | 16 | 58 | 20 | 52 | 543 | 0 | 0 | 0 | 256 | 0 | 0 | 0 |
| SRR5230700 | 264 | 0 | 0 | 0 | 0 | 0 | 35 | 18 | 20 | 0 | 17 | 122 | 15 | 0 | 104 | 0 | 20 | 0 | 0 | 0 | 1565 | 0 | 0 | 19 | 0 | 0 | 19 | 12 | 22 | 83 | 0 | 0 | 0 | 25 | 0 | 0 | 0 |
| SRR5230701 | 218 | 0 | 0 | 0 | 0 | 0 | 53 | 30 | 0 | 0 | 0 | 88 | 0 | 0 | 579 | 0 | 0 | 0 | 22 | 0 | 790 | 0 | 0 | 136 | 0 | 17 | 30 | 0 | 0 | 65 | 0 | 0 | 0 | 15 | 0 | 0 | 0 |
| SRR5230702 | 293 | 0 | 0 | 13 | 0 | 0 | 27 | 130 | 15 | 0 | 0 | 297 | 0 | 47 | 941 | 0 | 0 | 0 | 0 | 0 | 0 | 0 | 0 | 82 | 0 | 27 | 124 | 0 | 0 | 27 | 0 | 0 | 0 | 79 | 0 | 0 | 0 |
| SRR5230703 | 139 | 0 | 0 | 0 | 0 | 0 | 301 | 190 | 16 | 0 | 118 | 172 | 12 | 0 | 68 | 0 | 19 | 0 | 31 | 43 | 839 | 0 | 0 | 105 | 0 | 10 | 46 | 0 | 130 | 225 | 15 | 0 | 0 | 55 | 0 | 0 | 0 |
| SRR5230704 | 364 | 0 | 0 | 0 | 0 | 15 | 118 | 249 | 0 | 0 | 0 | 75 | 0 | 0 | 148 | 0 | 0 | 0 | 0 | 22 | 553 | 0 | 0 | 51 | 0 | 0 | 0 | 0 | 0 | 332 | 15 | 0 | 0 | 161 | 0 | 0 | 0 |
| SRR5230705 | 54 | 0 | 0 | 0 | 0 | 0 | 445 | 57 | 11 | 0 | 0 | 17 | 0 | 0 | 35 | 0 | 0 | 0 | 0 | 0 | 231 | 0 | 0 | 14 | 0 | 0 | 0 | 0 | 10 | 11 | 0 | 0 | 0 | 37 | 0 | 0 | 0 |
| SRR5230706 | 210 | 0 | 0 | 0 | 15 | 0 | 1961 | 213 | 17 | 14 | 136 | 263 | 17 | 0 | 573 | 0 | 27 | 0 | 30 | 30 | 408 | 0 | 0 | 39 | 0 | 14 | 36 | 0 | 34 | 95 | 0 | 0 | 0 | 17 | 0 | 0 | 0 |
| SRR5230707 | 657 | 0 | 0 | 0 | 0 | 0 | 319 | 515 | 0 | 0 | 11 | 670 | 0 | 0 | 28 | 0 | 37 | 0 | 46 | 20 | 319 | 0 | 0 | 181 | 25 | 0 | 0 | 0 | 252 | 881 | 13 | 0 | 0 | 221 | 0 | 0 | 0 |
| SRR5230708 | 658 | 0 | 0 | 0 | 0 | 0 | 54 | 489 | 50 | 0 | 0 | 934 | 36 | 0 | 101 | 0 | 34 | 0 | 0 | 0 | 1453 | 0 | 0 | 36 | 0 | 22 | 22 | 14 | 94 | 15 | 50 | 0 | 0 | 295 | 0 | 0 | 0 |
| SRR5230709 | 433 | 0 | 0 | 0 | 0 | 0 | 211 | 16 | 0 | 0 | 21 | 311 | 21 | 17 | 0 | 17 | 32 | 0 | 30 | 20 | 627 | 17 | 0 | 81 | 0 | 34 | 21 | 0 | 0 | 20 | 30 | 0 | 0 | 247 | 0 | 0 | 0 |
| SRR5230710 | 383 | 0 | 0 | 0 | 0 | 0 | 27 | 59 | 30 | 0 | 0 | 232 | 0 | 0 | 102 | 0 | 0 | 0 | 0 | 0 | 485 | 0 | 0 | 20 | 0 | 17 | 0 | 0 | 0 | 127 | 0 | 0 | 0 | 136 | 0 | 0 | 0 |
| SRR5230711 | 239 | 0 | 0 | 21 | 0 | 0 | 15 | 100 | 21 | 0 | 0 | 71 | 0 | 0 | 0 | 0 | 0 | 0 | 0 | 0 | 204 | 42 | 0 | 107 | 0 | 21 | 0 | 0 | 0 | 405 | 0 | 0 | 0 | 357 | 0 | 0 | 0 |
| SRR5230712 | 222 | 0 | 0 | 0 | 0 | 0 | 75 | 123 | 21 | 0 | 0 | 163 | 0 | 0 | 135 | 0 | 31 | 0 | 10 | 45 | 1969 | 0 | 0 | 170 | 0 | 0 | 64 | 19 | 53 | 511 | 11 | 0 | 0 | 168 | 0 | 0 | 0 |
| SRR5230713 | 639 | 0 | 0 | 0 | 0 | 0 | 96 | 17 | 0 | 0 | 0 | 241 | 0 | 0 | 358 | 0 | 23 | 0 | 80 | 105 | 624 | 0 | 0 | 13 | 0 | 0 | 63 | 0 | 0 | 42 | 18 | 0 | 0 | 94 | 0 | 0 | 0 |
| SRR5230714 | 262 | 0 | 0 | 0 | 0 | 0 | 939 | 885 | 14 | 0 | 16 | 124 | 10 | 0 | 446 | 0 | 23 | 0 | 48 | 61 | 835 | 0 | 0 | 44 | 0 | 10 | 26 | 0 | 75 | 550 | 0 | 0 | 0 | 118 | 0 | 0 | 0 |
| SRR5230716 | 271 | 0 | 0 | 0 | 0 | 19 | 113 | 23 | 21 | 0 | 12 | 124 | 0 | 0 | 0 | 0 | 65 | 0 | 16 | 60 | 286 | 0 | 0 | 77 | 0 | 40 | 154 | 0 | 0 | 25 | 23 | 0 | 0 | 21 | 0 | 0 | 0 |
| SRR5230717 | 345 | 0 | 0 | 0 | 16 | 16 | 214 | 421 | 0 | 0 | 0 | 199 | 10 | 0 | 194 | 0 | 23 | 0 | 36 | 56 | 710 | 0 | 0 | 46 | 0 | 12 | 16 | 20 | 0 | 496 | 12 | 0 | 0 | 183 | 0 | 0 | 0 |
| SRR5230718 | 525 | 15 | 0 | 0 | 0 | 0 | 55 | 155 | 77 | 0 | 0 | 322 | 0 | 0 | 36 | 0 | 22 | 0 | 0 | 0 | 659 | 0 | 0 | 87 | 0 | 0 | 25 | 12 | 0 | 325 | 12 | 0 | 0 | 16 | 0 | 0 | 0 |
| SRR5230719 | 405 | 0 | 0 | 0 | 0 | 11 | 45 | 184 | 0 | 0 | 0 | 147 | 27 | 0 | 202 | 0 | 0 | 0 | 12 | 29 | 594 | 36 | 0 | 81 | 11 | 18 | 106 | 0 | 0 | 35 | 0 | 0 | 0 | 177 | 0 | 0 | 0 |
| SRR5230720 | 673 | 0 | 0 | 0 | 14 | 0 | 161 | 691 | 0 | 0 | 0 | 369 | 48 | 0 | 884 | 0 | 11 | 0 | 0 | 24 | 2467 | 0 | 0 | 121 | 0 | 0 | 26 | 0 | 48 | 1712 | 12 | 0 | 11 | 326 | 0 | 0 | 0 |
| SRR5230721 | 205 | 0 | 0 | 0 | 0 | 0 | 223 | 31 | 14 | 0 | 19 | 78 | 16 | 0 | 93 | 0 | 0 | 0 | 32 | 0 | 218 | 44 | 0 | 56 | 0 | 0 | 33 | 0 | 121 | 62 | 0 | 0 | 0 | 10 | 0 | 0 | 0 |
| SRR5230722 | 469 | 0 | 0 | 0 | 0 | 0 | 103 | 97 | 15 | 0 | 0 | 176 | 0 | 0 | 220 | 0 | 0 | 0 | 41 | 0 | 1176 | 0 | 0 | 85 | 0 | 12 | 63 | 0 | 0 | 21 | 12 | 0 | 0 | 24 | 0 | 0 | 0 |
| SRR5230723 | 287 | 0 | 0 | 0 | 0 | 0 | 79 | 220 | 65 | 0 | 12 | 492 | 37 | 0 | 60 | 0 | 47 | 0 | 0 | 0 | 483 | 0 | 0 | 77 | 0 | 24 | 41 | 29 | 73 | 16 | 12 | 0 | 0 | 11 | 0 | 0 | 0 |
| SRR5230724 | 155 | 10 | 0 | 0 | 0 | 0 | 389 | 109 | 0 | 0 | 0 | 63 | 0 | 15 | 11 | 0 | 22 | 0 | 0 | 0 | 138 | 0 | 0 | 57 | 0 | 10 | 12 | 10 | 0 | 162 | 0 | 0 | 0 | 113 | 0 | 0 | 0 |
| SRR5230725 | 310 | 0 | 0 | 12 | 0 | 0 | 68 | 169 | 0 | 0 | 0 | 262 | 51 | 0 | 22 | 0 | 21 | 0 | 0 | 0 | 994 | 0 | 0 | 55 | 0 | 16 | 0 | 0 | 0 | 10 | 0 | 0 | 0 | 83 | 0 | 0 | 0 |
| SRR5230726 | 210 | 0 | 0 | 0 | 17 | 0 | 421 | 292 | 41 | 0 | 23 | 128 | 0 | 0 | 16 | 0 | 21 | 0 | 23 | 26 | 1883 | 11 | 0 | 32 | 0 | 0 | 17 | 11 | 47 | 360 | 0 | 0 | 0 | 136 | 0 | 0 | 0 |
| SRR5230727 | 94 | 0 | 0 | 0 | 0 | 0 | 102 | 317 | 39 | 0 | 0 | 41 | 0 | 0 | 50 | 0 | 21 | 0 | 33 | 50 | 1212 | 0 | 0 | 77 | 0 | 0 | 15 | 0 | 62 | 731 | 0 | 0 | 0 | 284 | 0 | 0 | 0 |
| SRR5230728 | 360 | 0 | 0 | 13 | 0 | 0 | 566 | 1322 | 15 | 0 | 41 | 132 | 13 | 0 | 1427 | 0 | 15 | 0 | 88 | 80 | 634 | 0 | 0 | 32 | 0 | 0 | 32 | 15 | 0 | 178 | 11 | 0 | 0 | 62 | 0 | 0 | 0 |
| SRR5230729 | 441 | 0 | 0 | 0 | 0 | 0 | 284 | 640 | 0 | 0 | 26 | 476 | 0 | 0 | 345 | 0 | 28 | 0 | 10 | 0 | 1625 | 0 | 0 | 86 | 0 | 0 | 106 | 0 | 0 | 1036 | 29 | 0 | 0 | 336 | 0 | 0 | 0 |
| SRR5230732 | 412 | 0 | 0 | 32 | 0 | 0 | 121 | 139 | 26 | 0 | 12 | 320 | 0 | 20 | 76 | 0 | 23 | 0 | 38 | 21 | 323 | 0 | 0 | 44 | 0 | 0 | 49 | 23 | 162 | 113 | 0 | 12 | 0 | 200 | 0 | 0 | 0 |
| SRR5230733 | 195 | 0 | 0 | 0 | 0 | 0 | 72 | 85 | 33 | 0 | 0 | 50 | 0 | 0 | 39 | 0 | 15 | 0 | 11 | 0 | 160 | 0 | 0 | 12 | 0 | 0 | 12 | 0 | 0 | 74 | 11 | 0 | 0 | 15 | 0 | 0 | 0 |
| SRR5230734 | 247 | 27 | 0 | 0 | 0 | 0 | 245 | 177 | 15 | 0 | 27 | 95 | 0 | 88 | 150 | 15 | 0 | 0 | 42 | 0 | 262 | 0 | 0 | 12 | 0 | 50 | 15 | 35 | 35 | 138 | 0 | 0 | 0 | 38 | 0 | 0 | 0 |
| SRR5230735 | 396 | 0 | 0 | 0 | 0 | 28 | 58 | 133 | 13 | 0 | 11 | 239 | 11 | 0 | 20 | 0 | 0 | 0 | 20 | 15 | 379 | 35 | 0 | 65 | 0 | 11 | 65 | 37 | 35 | 13 | 0 | 0 | 0 | 18 | 0 | 0 | 0 |
| SRR5230736 | 254 | 0 | 0 | 0 | 0 | 0 | 1005 | 64 | 25 | 0 | 0 | 65 | 0 | 0 | 0 | 0 | 22 | 0 | 18 | 0 | 318 | 17 | 0 | 15 | 0 | 0 | 0 | 0 | 11 | 86 | 0 | 0 | 0 | 69 | 0 | 0 | 0 |
| SRR5230737 | 395 | 0 | 0 | 15 | 25 | 10 | 176 | 257 | 61 | 0 | 0 | 335 | 0 | 40 | 263 | 0 | 21 | 0 | 12 | 15 | 892 | 0 | 0 | 45 | 0 | 0 | 40 | 56 | 15 | 217 | 25 | 0 | 0 | 131 | 0 | 0 | 0 |
| SRR5230738 | 270 | 12 | 0 | 0 | 0 | 0 | 102 | 52 | 0 | 0 | 17 | 204 | 0 | 0 | 31 | 0 | 40 | 0 | 0 | 0 | 237 | 0 | 0 | 26 | 0 | 0 | 14 | 20 | 0 | 116 | 0 | 0 | 0 | 180 | 0 | 0 | 0 |
| SRR5230739 | 335 | 0 | 0 | 14 | 0 | 0 | 13 | 41 | 134 | 0 | 0 | 256 | 14 | 11 | 137 | 0 | 0 | 0 | 0 | 0 | 278 | 41 | 0 | 11 | 0 | 33 | 11 | 0 | 0 | 87 | 125 | 0 | 0 | 95 | 0 | 0 | 0 |
| SRR5230740 | 237 | 0 | 0 | 0 | 0 | 0 | 819 | 28 | 0 | 0 | 10 | 110 | 0 | 0 | 338 | 0 | 15 | 0 | 64 | 15 | 113 | 0 | 0 | 0 | 0 | 0 | 15 | 38 | 21 | 94 | 0 | 0 | 0 | 42 | 0 | 0 | 0 |
| SRR5230741 | 271 | 0 | 0 | 0 | 12 | 0 | 715 | 12 | 18 | 0 | 59 | 303 | 0 | 0 | 1248 | 0 | 0 | 0 | 51 | 30 | 441 | 53 | 0 | 0 | 0 | 0 | 21 | 0 | 12 | 88 | 14 | 0 | 0 | 90 | 0 | 0 | 0 |
| SRR5230742 | 704 | 0 | 0 | 0 | 14 | 0 | 46 | 126 | 21 | 0 | 0 | 684 | 49 | 0 | 72 | 0 | 37 | 0 | 0 | 0 | 1037 | 14 | 0 | 14 | 0 | 0 | 28 | 14 | 0 | 161 | 14 | 0 | 0 | 308 | 0 | 0 | 0 |
| SRR5230743 | 237 | 0 | 0 | 0 | 13 | 0 | 681 | 319 | 0 | 0 | 28 | 461 | 0 | 0 | 588 | 0 | 35 | 0 | 15 | 37 | 600 | 0 | 0 | 72 | 0 | 0 | 51 | 0 | 61 | 283 | 0 | 0 | 0 | 199 | 0 | 0 | 0 |
| SRR5230744 | 265 | 0 | 0 | 18 | 0 | 0 | 74 | 43 | 14 | 0 | 0 | 78 | 0 | 0 | 794 | 0 | 15 | 0 | 19 | 0 | 574 | 18 | 0 | 36 | 0 | 0 | 43 | 0 | 0 | 127 | 0 | 0 | 0 | 13 | 0 | 0 | 0 |
| SRR5230745 | 416 | 0 | 0 | 22 | 22 | 0 | 129 | 142 | 38 | 0 | 22 | 239 | 0 | 32 | 1128 | 0 | 40 | 0 | 67 | 36 | 948 | 0 | 0 | 0 | 0 | 16 | 19 | 13 | 0 | 290 | 32 | 0 | 0 | 129 | 0 | 0 | 0 |
| SRR5230746 | 426 | 0 | 0 | 0 | 32 | 0 | 95 | 324 | 18 | 0 | 0 | 250 | 0 | 0 | 49 | 0 | 25 | 0 | 16 | 46 | 1032 | 0 | 0 | 64 | 0 | 0 | 36 | 64 | 0 | 383 | 46 | 0 | 0 | 401 | 0 | 0 | 0 |
| SRR5230747 | 322 | 0 | 0 | 0 | 0 | 0 | 57 | 90 | 18 | 0 | 0 | 415 | 22 | 0 | 96 | 0 | 36 | 0 | 0 | 0 | 2125 | 0 | 0 | 27 | 0 | 0 | 0 | 31 | 31 | 242 | 0 | 0 | 0 | 287 | 0 | 0 | 0 |
| SRR5230748 | 558 | 0 | 0 | 0 | 0 | 0 | 102 | 227 | 32 | 0 | 0 | 680 | 27 | 16 | 169 | 0 | 31 | 0 | 14 | 11 | 2311 | 0 | 0 | 54 | 0 | 0 | 0 | 27 | 54 | 324 | 0 | 0 | 0 | 670 | 0 | 0 | 11 |
| SRR5230749 | 263 | 0 | 0 | 0 | 0 | 0 | 478 | 186 | 0 | 0 | 53 | 249 | 0 | 0 | 97 | 0 | 27 | 0 | 0 | 0 | 715 | 0 | 0 | 14 | 0 | 11 | 31 | 17 | 25 | 142 | 0 | 0 | 0 | 153 | 19 | 0 | 0 |
| SRR5230750 | 252 | 0 | 0 | 17 | 14 | 21 | 15 | 31 | 51 | 0 | 0 | 146 | 0 | 0 | 0 | 0 | 33 | 0 | 0 | 0 | 467 | 10 | 0 | 24 | 0 | 24 | 0 | 0 | 0 | 96 | 34 | 0 | 0 | 151 | 0 | 0 | 0 |
| SRR5230751 | 171 | 0 | 0 | 0 | 0 | 0 | 40 | 56 | 0 | 0 | 0 | 76 | 0 | 0 | 28 | 0 | 46 | 0 | 0 | 0 | 281 | 0 | 0 | 0 | 0 | 0 | 0 | 0 | 22 | 140 | 0 | 0 | 0 | 106 | 0 | 0 | 0 |
| SRR5230752 | 499 | 11 | 0 | 0 | 22 | 0 | 73 | 61 | 83 | 0 | 17 | 324 | 0 | 11 | 50 | 0 | 25 | 0 | 0 | 0 | 263 | 39 | 0 | 17 | 0 | 0 | 0 | 61 | 17 | 105 | 17 | 0 | 0 | 44 | 0 | 0 | 0 |
| SRR5230753 | 655 | 0 | 0 | 0 | 0 | 0 | 0 | 56 | 25 | 0 | 0 | 602 | 0 | 0 | 94 | 0 | 34 | 0 | 23 | 0 | 832 | 0 | 0 | 62 | 0 | 31 | 51 | 0 | 0 | 135 | 20 | 0 | 17 | 219 | 0 | 0 | 0 |
| SRR5231309 | 61 | 37 | 4152 | 27 | 14 | 0 | 0 | 0 | 0 | 0 | 0 | 287 | 0 | 51 | 0 | 32 | 0 | 0 | 0 | 22 | 27 | 157 | 29 | 0 | 15 | 139 | 143 | 55 | 98 | 0 | 0 | 0 | 31 | 33 | 31 | 0 | 17 |
| SRR5231310 | 72 | 44 | 3500 | 39 | 18 | 0 | 19 | 0 | 0 | 0 | 0 | 193 | 0 | 93 | 0 | 52 | 0 | 0 | 0 | 33 | 33 | 92 | 34 | 0 | 30 | 125 | 176 | 61 | 63 | 0 | 0 | 0 | 63 | 20 | 26 | 0 | 17 |
| SRR5231311 | 64 | 36 | 4022 | 55 | 17 | 0 | 22 | 0 | 0 | 0 | 0 | 153 | 0 | 63 | 0 | 30 | 24 | 0 | 0 | 35 | 31 | 31 | 27 | 0 | 38 | 343 | 187 | 82 | 28 | 0 | 0 | 0 | 57 | 29 | 10 | 0 | 0 |
| SRR5231312 | 30 | 87 | 3082 | 65 | 26 | 0 | 17 | 0 | 0 | 0 | 0 | 255 | 0 | 116 | 0 | 76 | 12 | 0 | 0 | 13 | 21 | 155 | 26 | 0 | 43 | 64 | 115 | 97 | 84 | 0 | 0 | 0 | 150 | 25 | 15 | 0 | 0 |
| SRR5231313 | 37 | 11 | 3773 | 14 | 18 | 0 | 23 | 0 | 0 | 0 | 0 | 260 | 0 | 21 | 0 | 21 | 22 | 0 | 0 | 32 | 39 | 46 | 14 | 0 | 10 | 47 | 101 | 19 | 25 | 0 | 0 | 0 | 12 | 15 | 15 | 0 | 16 |

### Supp Table S8.2.Columns_AM_to_BW

| | | | | | | | | | | | | | | | | | | | | | | | | | | | | | | | | | | | | | |
|---|---|---|---|---|---|---|---|---|---|---|---|---|---|---|---|---|---|---|---|---|---|---|---|---|---|---|---|---|---|---|---|---|---|---|---|---|---|
| SRR5231314 | 58 | 26 | 3940 | 55 | 16 | 0 | 53 | 0 | 12 | 0 | 0 | 194 | 0 | 45 | 0 | 37 | 0 | 0 | 0 | 51 | 45 | 38 | 24 | 0 | 28 | 316 | 254 | 84 | 41 | 0 | 0 | 0 | 28 | 55 | 14 | 0 | 0 |
| SRR5239182 | 0 | 0 | 43 | 0 | 0 | 0 | 66 | 121 | 0 | 0 | 0 | 129 | 0 | 0 | 109 | 0 | 18 | 33 | 35 | 3293 | 2177 | 0 | 0 | 0 | 0 | 10 | 0 | 0 | 0 | 0 | 0 | 0 | 0 | 0 | 128 | 11 | 57 |
| SRR5239183 | 0 | 0 | 14 | 0 | 0 | 0 | 37 | 113 | 0 | 0 | 0 | 14 | 0 | 0 | 0 | 0 | 0 | 0 | 73 | 1435 | 3126 | 0 | 0 | 0 | 0 | 16 | 0 | 0 | 0 | 18 | 0 | 0 | 0 | 0 | 22 | 0 | 18 |
| SRR5239184 | 0 | 0 | 41 | 15 | 0 | 0 | 37 | 133 | 0 | 0 | 0 | 63 | 0 | 0 | 79 | 0 | 0 | 0 | 58 | 694 | 4166 | 0 | 0 | 0 | 0 | 13 | 17 | 0 | 0 | 13 | 0 | 0 | 0 | 11 | 51 | 0 | 35 |
| SRR5239185 | 10 | 0 | 45 | 0 | 0 | 0 | 20 | 241 | 0 | 0 | 0 | 136 | 0 | 0 | 267 | 0 | 21 | 48 | 0 | 2307 | 1853 | 0 | 0 | 0 | 0 | 0 | 0 | 0 | 0 | 0 | 0 | 0 | 0 | 0 | 98 | 14 | 61 |
| SRR5239186 | 0 | 0 | 40 | 0 | 0 | 0 | 83 | 94 | 0 | 0 | 0 | 69 | 0 | 0 | 228 | 0 | 20 | 75 | 45 | 3905 | 2406 | 0 | 0 | 0 | 0 | 0 | 0 | 0 | 0 | 0 | 0 | 0 | 0 | 0 | 132 | 19 | 90 |
| SRR5239187 | 0 | 12 | 170 | 0 | 0 | 0 | 157 | 255 | 11 | 0 | 0 | 110 | 0 | 0 | 409 | 0 | 21 | 0 | 135 | 2761 | 3231 | 0 | 0 | 0 | 0 | 20 | 35 | 0 | 0 | 0 | 0 | 0 | 0 | 0 | 61 | 15 | 51 |
| SRR5239188 | 11 | 12 | 154 | 0 | 0 | 0 | 105 | 412 | 0 | 0 | 0 | 121 | 0 | 0 | 273 | 0 | 15 | 20 | 61 | 1937 | 3063 | 0 | 0 | 0 | 0 | 10 | 19 | 0 | 0 | 0 | 0 | 0 | 0 | 0 | 128 | 12 | 80 |
| SRR5275577 | 13 | 0 | 1292 | 35 | 58 | 0 | 27 | 0 | 0 | 0 | 0 | 641 | 0 | 554 | 135 | 49 | 10 | 12 | 0 | 93 | 999 | 55 | 0 | 0 | 54 | 75 | 269 | 294 | 0 | 0 | 0 | 0 | 33 | 10 | 262 | 59 | 468 |
| SRR5275578 | 15 | 0 | 1218 | 51 | 66 | 0 | 24 | 0 | 0 | 0 | 0 | 571 | 0 | 851 | 263 | 78 | 0 | 47 | 0 | 95 | 1050 | 51 | 0 | 0 | 82 | 62 | 238 | 350 | 0 | 0 | 0 | 0 | 52 | 0 | 224 | 52 | 393 |
| SRR5275579 | 20 | 0 | 1345 | 44 | 62 | 0 | 25 | 0 | 0 | 0 | 0 | 557 | 0 | 570 | 137 | 53 | 15 | 46 | 0 | 102 | 852 | 56 | 0 | 0 | 45 | 58 | 238 | 250 | 0 | 0 | 0 | 0 | 41 | 13 | 260 | 59 | 436 |
| SRR5275580 | 14 | 0 | 965 | 16 | 22 | 0 | 17 | 0 | 0 | 0 | 0 | 840 | 0 | 388 | 266 | 29 | 0 | 58 | 0 | 72 | 667 | 12 | 0 | 0 | 27 | 23 | 62 | 175 | 0 | 0 | 13 | 11 | 14 | 22 | 385 | 60 | 387 |
| SRR5275581 | 21 | 0 | 932 | 50 | 60 | 0 | 0 | 0 | 0 | 0 | 0 | 834 | 0 | 971 | 253 | 85 | 0 | 64 | 0 | 38 | 490 | 33 | 0 | 0 | 77 | 46 | 110 | 431 | 0 | 0 | 13 | 10 | 44 | 28 | 913 | 133 | 831 |
| SRR5275582 | 21 | 0 | 965 | 27 | 37 | 0 | 54 | 0 | 0 | 0 | 0 | 951 | 0 | 338 | 78 | 43 | 0 | 42 | 0 | 211 | 1092 | 73 | 0 | 0 | 31 | 105 | 438 | 186 | 0 | 0 | 0 | 0 | 22 | 32 | 936 | 172 | 1243 |
| SRR5284195 | 88 | 0 | 811 | 44 | 62 | 36 | 297 | 0 | 12 | 0 | 14 | 5537 | 10 | 18 | 301 | 0 | 14 | 18 | 35 | 297 | 346 | 0 | 0 | 0 | 23 | 259 | 159 | 0 | 0 | 0 | 13 | 0 | 0 | 20 | 17 | 62 | 89 |
| SRR5284196 | 87 | 0 | 605 | 14 | 25 | 64 | 174 | 15 | 21 | 0 | 12 | 5189 | 0 | 0 | 1613 | 0 | 0 | 32 | 0 | 254 | 973 | 0 | 0 | 13 | 16 | 329 | 433 | 0 | 0 | 11 | 15 | 0 | 0 | 14 | 22 | 35 | 44 |
| SRR5284197 | 84 | 0 | 931 | 34 | 52 | 36 | 163 | 12 | 21 | 0 | 15 | 12481 | 0 | 11 | 338 | 0 | 0 | 35 | 111 | 1176 | 376 | 0 | 0 | 20 | 23 | 205 | 392 | 11 | 0 | 14 | 35 | 0 | 0 | 57 | 23 | 73 | 172 |
| SRR5284198 | 42 | 0 | 602 | 36 | 232 | 143 | 257 | 0 | 21 | 12 | 66 | 9350 | 0 | 11 | 147 | 0 | 0 | 0 | 23 | 720 | 399 | 0 | 0 | 34 | 138 | 875 | 1010 | 11 | 0 | 27 | 26 | 0 | 0 | 63 | 53 | 119 | 158 |
| SRR5284199 | 69 | 0 | 803 | 28 | 145 | 162 | 86 | 0 | 37 | 0 | 24 | 2758 | 0 | 11 | 79 | 0 | 11 | 0 | 79 | 807 | 278 | 0 | 0 | 74 | 114 | 1051 | 1304 | 0 | 0 | 0 | 66 | 0 | 0 | 34 | 50 | 55 | 67 |
| SRR5284200 | 59 | 0 | 549 | 18 | 105 | 176 | 109 | 16 | 50 | 0 | 31 | 2052 | 0 | 0 | 412 | 0 | 0 | 0 | 49 | 978 | 422 | 0 | 0 | 129 | 69 | 1006 | 1148 | 0 | 0 | 27 | 43 | 0 | 0 | 44 | 40 | 68 | 118 |
| SRR5284201 | 110 | 0 | 1038 | 15 | 74 | 32 | 124 | 0 | 12 | 0 | 21 | 8894 | 0 | 13 | 89 | 0 | 16 | 0 | 17 | 869 | 196 | 0 | 0 | 20 | 45 | 217 | 416 | 11 | 0 | 15 | 28 | 0 | 0 | 13 | 19 | 17 | 44 |
| SRR5284202 | 79 | 0 | 399 | 40 | 215 | 96 | 88 | 0 | 19 | 0 | 63 | 10217 | 11 | 33 | 157 | 0 | 14 | 0 | 31 | 333 | 126 | 0 | 0 | 14 | 105 | 604 | 763 | 27 | 0 | 0 | 40 | 0 | 0 | 233 | 45 | 55 | 138 |
| SRR5284203 | 53 | 0 | 364 | 0 | 13 | 29 | 114 | 15 | 14 | 0 | 19 | 1955 | 0 | 13 | 67 | 0 | 14 | 18 | 43 | 1379 | 444 | 0 | 0 | 21 | 34 | 180 | 232 | 0 | 0 | 34 | 15 | 0 | 0 | 29 | 17 | 13 | 41 |
| SRR5284204 | 43 | 0 | 307 | 24 | 96 | 232 | 229 | 0 | 113 | 0 | 36 | 1372 | 0 | 16 | 427 | 0 | 0 | 20 | 48 | 837 | 922 | 0 | 0 | 0 | 90 | 1218 | 709 | 13 | 0 | 0 | 96 | 0 | 0 | 40 | 52 | 79 | 120 |
| SRR5284205 | 50 | 0 | 376 | 38 | 46 | 25 | 306 | 0 | 10 | 14 | 20 | 6356 | 0 | 21 | 522 | 0 | 11 | 69 | 81 | 1219 | 531 | 0 | 0 | 0 | 27 | 190 | 240 | 11 | 0 | 0 | 10 | 0 | 0 | 38 | 32 | 53 | 148 |
| SRR5284206 | 49 | 0 | 318 | 37 | 42 | 36 | 279 | 0 | 11 | 0 | 18 | 6738 | 0 | 13 | 207 | 0 | 13 | 38 | 62 | 765 | 528 | 0 | 0 | 12 | 24 | 228 | 294 | 14 | 0 | 0 | 34 | 0 | 0 | 28 | 20 | 49 | 137 |
| SRR5317818 | 12 | 0 | 816 | 0 | 0 | 0 | 68 | 0 | 0 | 0 | 0 | 161 | 0 | 14 | 794 | 0 | 0 | 0 | 0 | 90 | 8130 | 0 | 0 | 0 | 39 | 34 | 16 | 14 | 0 | 0 | 0 | 0 | 0 | 51 | 36 | 0 | 32 |
| SRR5317819 | 72 | 0 | 2044 | 0 | 0 | 17 | 84 | 39 | 0 | 0 | 0 | 23 | 0 | 0 | 103 | 0 | 11 | 0 | 11 | 755 | 5099 | 13 | 0 | 0 | 14 | 98 | 155 | 10 | 0 | 13 | 0 | 0 | 0 | 46 | 23 | 0 | 14 |
| SRR5317820 | 0 | 0 | 757 | 0 | 0 | 0 | 0 | 0 | 0 | 0 | 0 | 30 | 0 | 20 | 15190 | 0 | 0 | 0 | 33 | 69 | 22464 | 12 | 0 | 0 | 31 | 38 | 18 | 12 | 10 | 0 | 0 | 0 | 0 | 57 | 41 | 0 | 28 |
| SRR5317821 | 13 | 25 | 775 | 0 | 0 | 0 | 55 | 18 | 14 | 0 | 0 | 0 | 0 | 0 | 0 | 0 | 0 | 0 | 20 | 46 | 828 | 0 | 0 | 0 | 23 | 280 | 145 | 0 | 0 | 0 | 20 | 0 | 0 | 319 | 0 | 0 | 0 |
| SRR5317822 | 0 | 14 | 711 | 0 | 0 | 0 | 43 | 20 | 0 | 0 | 0 | 102 | 0 | 23 | 1061 | 0 | 0 | 85 | 0 | 14 | 3909 | 0 | 0 | 0 | 50 | 58 | 95 | 31 | 0 | 0 | 0 | 0 | 0 | 19 | 36 | 0 | 27 |
| SRR5317823 | 51 | 0 | 413 | 0 | 0 | 0 | 102 | 66 | 0 | 0 | 0 | 17 | 0 | 0 | 22 | 0 | 0 | 0 | 11 | 190 | 9292 | 13 | 0 | 0 | 18 | 159 | 235 | 22 | 13 | 32 | 0 | 0 | 0 | 130 | 20 | 0 | 18 |
| SRR5317824 | 15 | 0 | 1456 | 18 | 13 | 0 | 29 | 0 | 0 | 0 | 0 | 141 | 0 | 17 | 1777 | 0 | 0 | 0 | 0 | 45 | 7655 | 13 | 0 | 0 | 37 | 40 | 25 | 33 | 18 | 0 | 0 | 0 | 0 | 133 | 36 | 0 | 16 |
| SRR5317825 | 17 | 0 | 352 | 0 | 0 | 15 | 48 | 81 | 13 | 0 | 0 | 14 | 0 | 0 | 0 | 0 | 0 | 0 | 0 | 0 | 1894 | 0 | 0 | 0 | 0 | 355 | 575 | 0 | 27 | 23 | 0 | 0 | 0 | 523 | 25 | 0 | 15 |
| SRR5317826 | 34 | 0 | 1306 | 0 | 0 | 0 | 17 | 0 | 0 | 0 | 0 | 61 | 0 | 0 | 62 | 0 | 11 | 16 | 0 | 50 | 2693 | 11 | 0 | 0 | 55 | 85 | 128 | 33 | 19 | 0 | 0 | 0 | 0 | 156 | 40 | 0 | 35 |
| SRR5317827 | 90 | 0 | 747 | 0 | 0 | 0 | 76 | 14 | 0 | 0 | 0 | 16 | 0 | 0 | 15 | 0 | 0 | 0 | 48 | 199 | 9580 | 0 | 0 | 0 | 10 | 67 | 88 | 13 | 0 | 0 | 0 | 0 | 0 | 29 | 14 | 0 | 13 |
| SRR5387779 | 0 | 0 | 0 | 0 | 0 | 55 | 0 | 55 | 27 | 0 | 0 | 0 | 0 | 0 | 0 | 0 | 0 | 0 | 0 | 1186 | 237 | 0 | 0 | 55 | 0 | 522 | 412 | 0 | 0 | 0 | 0 | 0 | 0 | 14 | 0 | 0 | 0 |
| SRR5387780 | 0 | 0 | 0 | 0 | 0 | 0 | 0 | 53 | 0 | 0 | 0 | 0 | 0 | 0 | 0 | 0 | 0 | 0 | 0 | 1058 | 423 | 0 | 0 | 98 | 0 | 181 | 399 | 0 | 0 | 0 | 0 | 0 | 0 | 0 | 0 | 0 | 15 |
| SRR5387781 | 0 | 12 | 11 | 12 | 0 | 0 | 0 | 0 | 0 | 0 | 0 | 0 | 0 | 0 | 0 | 0 | 0 | 0 | 0 | 1348 | 0 | 0 | 0 | 49 | 0 | 203 | 246 | 0 | 0 | 0 | 0 | 0 | 0 | 0 | 0 | 0 | 0 |
| SRR5387782 | 0 | 0 | 16 | 0 | 0 | 29 | 0 | 14 | 0 | 0 | 0 | 0 | 0 | 0 | 65 | 0 | 0 | 0 | 0 | 1363 | 65 | 0 | 0 | 57 | 0 | 294 | 359 | 0 | 0 | 0 | 14 | 0 | 0 | 14 | 0 | 0 | 0 |
| SRR5387783 | 0 | 0 | 0 | 0 | 0 | 25 | 0 | 17 | 0 | 0 | 0 | 0 | 0 | 0 | 0 | 0 | 0 | 0 | 82 | 1642 | 164 | 0 | 0 | 100 | 0 | 276 | 335 | 0 | 0 | 0 | 0 | 0 | 0 | 17 | 0 | 0 | 0 |
| SRR5387784 | 0 | 0 | 10 | 0 | 0 | 0 | 0 | 0 | 0 | 0 | 0 | 0 | 0 | 0 | 0 | 0 | 0 | 0 | 0 | 1278 | 58 | 0 | 0 | 86 | 0 | 47 | 266 | 0 | 0 | 0 | 0 | 0 | 0 | 16 | 0 | 0 | 0 |
| SRR5387785 | 0 | 0 | 0 | 0 | 0 | 0 | 0 | 66 | 0 | 0 | 0 | 0 | 0 | 0 | 0 | 0 | 0 | 0 | 0 | 1314 | 227 | 0 | 0 | 46 | 0 | 79 | 99 | 0 | 0 | 0 | 0 | 0 | 0 | 40 | 13 | 0 | 0 |
| SRR5387786 | 0 | 0 | 0 | 0 | 0 | 0 | 0 | 32 | 0 | 0 | 0 | 0 | 0 | 16 | 0 | 0 | 0 | 0 | 0 | 1867 | 0 | 0 | 0 | 138 | 0 | 130 | 170 | 0 | 0 | 0 | 0 | 0 | 0 | 16 | 0 | 0 | 0 |
| SRR5387787 | 0 | 45 | 0 | 0 | 0 | 27 | 0 | 45 | 0 | 0 | 0 | 0 | 0 | 0 | 0 | 0 | 0 | 0 | 0 | 958 | 120 | 18 | 0 | 125 | 0 | 161 | 215 | 0 | 0 | 0 | 0 | 0 | 0 | 0 | 0 | 0 | 0 |
| SRR5387788 | 0 | 14 | 0 | 0 | 0 | 0 | 0 | 0 | 21 | 0 | 0 | 0 | 0 | 0 | 149 | 0 | 0 | 0 | 0 | 297 | 149 | 0 | 0 | 97 | 0 | 200 | 283 | 0 | 0 | 0 | 0 | 0 | 0 | 14 | 0 | 0 | 0 |
| SRR5387789 | 0 | 0 | 0 | 0 | 0 | 29 | 0 | 0 | 0 | 0 | 0 | 0 | 0 | 0 | 0 | 0 | 0 | 0 | 0 | 900 | 0 | 0 | 0 | 107 | 0 | 195 | 224 | 0 | 0 | 0 | 0 | 0 | 19 | 0 | 0 | 0 | 0 |
| SRR5387790 | 0 | 0 | 0 | 0 | 0 | 0 | 65 | 0 | 0 | 0 | 0 | 0 | 0 | 0 | 0 | 0 | 0 | 0 | 0 | 908 | 0 | 0 | 0 | 129 | 0 | 107 | 207 | 0 | 0 | 0 | 0 | 0 | 0 | 0 | 0 | 0 | 14 |
| SRR5387791 | 0 | 18 | 0 | 0 | 0 | 88 | 0 | 18 | 0 | 0 | 0 | 0 | 0 | 0 | 0 | 0 | 0 | 0 | 0 | 1077 | 196 | 0 | 0 | 70 | 0 | 405 | 599 | 0 | 0 | 0 | 0 | 0 | 0 | 0 | 18 | 0 | 0 |
| SRR5387792 | 0 | 0 | 0 | 10 | 0 | 10 | 0 | 0 | 0 | 0 | 0 | 0 | 0 | 0 | 0 | 0 | 0 | 0 | 0 | 805 | 0 | 0 | 0 | 51 | 0 | 307 | 379 | 0 | 0 | 0 | 0 | 0 | 0 | 20 | 0 | 0 | 10 |
| SRR5387793 | 0 | 29 | 0 | 0 | 0 | 0 | 0 | 43 | 14 | 0 | 14 | 0 | 0 | 0 | 0 | 0 | 0 | 0 | 0 | 998 | 125 | 0 | 0 | 72 | 0 | 331 | 489 | 0 | 0 | 0 | 0 | 0 | 0 | 0 | 14 | 0 | 0 |
| SRR5387794 | 0 | 0 | 20 | 0 | 0 | 0 | 0 | 50 | 0 | 0 | 0 | 0 | 0 | 0 | 252 | 0 | 0 | 0 | 0 | 945 | 315 | 0 | 0 | 75 | 0 | 181 | 200 | 0 | 0 | 0 | 0 | 25 | 0 | 0 | 0 | 0 | 0 |
| SRR5387795 | 0 | 11 | 0 | 0 | 0 | 22 | 0 | 11 | 11 | 0 | 0 | 0 | 0 | 0 | 0 | 0 | 0 | 0 | 114 | 1593 | 228 | 0 | 0 | 54 | 0 | 280 | 194 | 0 | 0 | 0 | 11 | 0 | 11 | 22 | 0 | 0 | 0 |
| SRR5387796 | 0 | 13 | 0 | 0 | 0 | 27 | 0 | 40 | 0 | 0 | 0 | 0 | 0 | 0 | 0 | 0 | 0 | 0 | 0 | 1242 | 414 | 0 | 0 | 80 | 0 | 241 | 322 | 0 | 0 | 13 | 0 | 13 | 0 | 13 | 0 | 0 | 0 |
| SRR5387797 | 0 | 0 | 0 | 0 | 0 | 0 | 0 | 41 | 16 | 0 | 0 | 0 | 0 | 0 | 56 | 0 | 0 | 0 | 0 | 2074 | 224 | 0 | 0 | 49 | 0 | 188 | 245 | 0 | 0 | 0 | 0 | 0 | 0 | 16 | 16 | 0 | 0 |
| SRR5387798 | 0 | 22 | 21 | 0 | 0 | 29 | 0 | 15 | 0 | 0 | 0 | 0 | 0 | 29 | 0 | 0 | 0 | 0 | 79 | 1101 | 314 | 0 | 0 | 80 | 0 | 204 | 460 | 0 | 0 | 0 | 0 | 0 | 0 | 0 | 0 | 0 | 0 |
| SRR5387799 | 0 | 16 | 0 | 0 | 0 | 23 | 0 | 23 | 0 | 0 | 0 | 0 | 0 | 0 | 164 | 0 | 0 | 0 | 0 | 1423 | 219 | 0 | 0 | 47 | 0 | 296 | 249 | 0 | 0 | 0 | 16 | 0 | 0 | 23 | 0 | 0 | 0 |
| SRR5387800 | 0 | 0 | 0 | 0 | 0 | 0 | 0 | 0 | 17 | 0 | 0 | 0 | 0 | 0 | 0 | 0 | 0 | 0 | 0 | 1368 | 526 | 0 | 0 | 157 | 0 | 323 | 481 | 0 | 0 | 0 | 25 | 0 | 0 | 0 | 0 | 0 | 0 |
| SRR5387801 | 0 | 28 | 0 | 0 | 0 | 0 | 0 | 0 | 0 | 0 | 0 | 0 | 0 | 0 | 103 | 0 | 0 | 0 | 0 | 928 | 103 | 0 | 0 | 160 | 0 | 123 | 189 | 0 | 0 | 0 | 0 | 0 | 0 | 0 | 0 | 0 | 0 |
| SRR5387802 | 0 | 26 | 17 | 0 | 0 | 0 | 0 | 0 | 0 | 0 | 0 | 0 | 0 | 0 | 0 | 0 | 0 | 0 | 0 | 1233 | 164 | 0 | 0 | 57 | 0 | 211 | 268 | 0 | 0 | 0 | 0 | 0 | 0 | 0 | 13 | 0 | 0 |
| SRR5387803 | 0 | 13 | 0 | 0 | 0 | 0 | 0 | 0 | 0 | 0 | 0 | 0 | 0 | 0 | 0 | 0 | 0 | 0 | 0 | 1215 | 243 | 0 | 0 | 124 | 0 | 72 | 274 | 0 | 0 | 0 | 0 | 0 | 0 | 0 | 0 | 0 | 0 |
| SRR5387804 | 0 | 0 | 10 | 0 | 0 | 0 | 0 | 0 | 0 | 0 | 0 | 0 | 0 | 0 | 0 | 0 | 0 | 0 | 0 | 572 | 245 | 0 | 0 | 165 | 0 | 228 | 338 | 0 | 0 | 0 | 16 | 0 | 0 | 0 | 0 | 0 | 0 |
| SRR5387805 | 0 | 0 | 10 | 0 | 0 | 28 | 0 | 0 | 19 | 0 | 0 | 0 | 0 | 0 | 0 | 0 | 0 | 0 | 0 | 1228 | 614 | 0 | 0 | 84 | 0 | 176 | 195 | 0 | 0 | 0 | 0 | 0 | 0 | 0 | 0 | 0 | 0 |
| SRR5387806 | 0 | 0 | 17 | 0 | 0 | 16 | 0 | 24 | 0 | 0 | 0 | 0 | 0 | 0 | 91 | 0 | 0 | 0 | 0 | 2545 | 273 | 0 | 0 | 72 | 0 | 128 | 215 | 0 | 0 | 0 | 0 | 0 | 0 | 0 | 0 | 0 | 0 |
| SRR5387807 | 0 | 12 | 0 | 0 | 0 | 24 | 0 | 12 | 12 | 0 | 0 | 0 | 0 | 0 | 0 | 0 | 0 | 0 | 64 | 955 | 127 | 0 | 0 | 71 | 12 | 367 | 391 | 0 | 0 | 0 | 0 | 0 | 0 | 0 | 0 | 0 | 0 |
| SRR5387808 | 0 | 0 | 0 | 0 | 0 | 20 | 0 | 0 | 0 | 0 | 0 | 0 | 0 | 0 | 0 | 0 | 0 | 0 | 0 | 608 | 182 | 0 | 0 | 199 | 0 | 457 | 427 | 0 | 0 | 0 | 0 | 0 | 0 | 0 | 0 | 0 | 0 |
| SRR5387809 | 0 | 18 | 0 | 18 | 0 | 18 | 0 | 18 | 0 | 0 | 0 | 0 | 0 | 0 | 152 | 0 | 0 | 0 | 0 | 1219 | 0 | 0 | 0 | 18 | 0 | 355 | 373 | 0 | 0 | 0 | 0 | 0 | 0 | 0 | 18 | 0 | 18 |
| SRR5387810 | 0 | 0 | 21 | 0 | 0 | 21 | 0 | 14 | 14 | 0 | 0 | 0 | 0 | 0 | 0 | 0 | 0 | 0 | 0 | 1357 | 522 | 0 | 0 | 63 | 0 | 153 | 181 | 0 | 0 | 0 | 0 | 0 | 0 | 14 | 0 | 0 | 0 |
| SRR5387811 | 0 | 23 | 0 | 0 | 0 | 0 | 0 | 16 | 0 | 0 | 0 | 0 | 0 | 16 | 0 | 0 | 0 | 0 | 0 | 554 | 370 | 0 | 0 | 86 | 0 | 125 | 179 | 0 | 0 | 0 | 0 | 0 | 0 | 0 | 0 | 0 | 0 |
| SRR5387812 | 0 | 0 | 0 | 0 | 0 | 14 | 0 | 14 | 0 | 0 | 0 | 0 | 0 | 21 | 67 | 0 | 0 | 0 | 0 | 674 | 67 | 0 | 0 | 167 | 0 | 167 | 313 | 0 | 0 | 0 | 0 | 0 | 0 | 0 | 0 | 0 | 0 |
| SRR5387813 | 0 | 0 | 0 | 0 | 0 | 15 | 0 | 15 | 15 | 0 | 0 | 0 | 0 | 15 | 0 | 0 | 0 | 0 | 0 | 1075 | 98 | 0 | 0 | 45 | 0 | 303 | 257 | 0 | 0 | 0 | 15 | 0 | 0 | 0 | 0 | 0 | 0 |
| SRR5387814 | 0 | 0 | 12 | 0 | 0 | 35 | 0 | 0 | 0 | 0 | 0 | 0 | 0 | 0 | 0 | 0 | 0 | 0 | 0 | 683 | 427 | 0 | 0 | 140 | 0 | 359 | 570 | 0 | 0 | 0 | 0 | 18 | 0 | 0 | 0 | 0 | 35 |
| SRR5387815 | 0 | 14 | 11 | 0 | 0 | 28 | 0 | 14 | 0 | 0 | 0 | 0 | 0 | 0 | 0 | 0 | 0 | 0 | 0 | 1018 | 424 | 0 | 0 | 174 | 0 | 313 | 341 | 0 | 0 | 0 | 0 | 0 | 0 | 0 | 0 | 0 | 0 |
| SRR5387816 | 0 | 0 | 0 | 0 | 0 | 37 | 0 | 0 | 0 | 0 | 0 | 0 | 0 | 0 | 104 | 0 | 0 | 0 | 0 | 2712 | 104 | 0 | 0 | 177 | 19 | 392 | 401 | 0 | 0 | 0 | 28 | 0 | 28 | 0 | 0 | 0 | 0 |
| SRR5387817 | 0 | 0 | 0 | 0 | 0 | 0 | 0 | 13 | 0 | 0 | 0 | 0 | 0 | 0 | 0 | 0 | 0 | 0 | 0 | 1020 | 78 | 0 | 0 | 183 | 13 | 248 | 301 | 0 | 13 | 0 | 0 | 0 | 0 | 0 | 13 | 0 | 0 |
| SRR5387818 | 0 | 17 | 0 | 0 | 0 | 17 | 0 | 17 | 0 | 0 | 0 | 0 | 0 | 0 | 0 | 0 | 0 | 0 | 0 | 505 | 126 | 0 | 0 | 174 | 0 | 165 | 383 | 0 | 0 | 0 | 0 | 0 | 0 | 0 | 0 | 0 | 0 |
| SRR5387819 | 0 | 0 | 0 | 0 | 0 | 17 | 0 | 0 | 0 | 0 | 0 | 0 | 0 | 0 | 0 | 0 | 0 | 0 | 45 | 893 | 223 | 0 | 0 | 42 | 0 | 395 | 479 | 0 | 0 | 0 | 0 | 0 | 0 | 0 | 34 | 0 | 0 |
| SRR5387820 | 0 | 0 | 0 | 0 | 0 | 29 | 0 | 0 | 0 | 0 | 0 | 0 | 0 | 0 | 0 | 0 | 0 | 0 | 0 | 565 | 188 | 0 | 0 | 58 | 0 | 623 | 662 | 0 | 0 | 0 | 0 | 0 | 0 | 0 | 0 | 0 | 0 |
| SRR5387821 | 0 | 0 | 15 | 0 | 0 | 35 | 0 | 0 | 0 | 0 | 0 | 0 | 0 | 0 | 0 | 0 | 0 | 0 | 102 | 1330 | 307 | 0 | 0 | 26 | 0 | 244 | 401 | 0 | 0 | 0 | 0 | 0 | 0 | 0 | 0 | 0 | 0 |
| SRR5387822 | 0 | 0 | 16 | 0 | 0 | 13 | 0 | 13 | 0 | 0 | 0 | 0 | 0 | 0 | 0 | 0 | 0 | 0 | 0 | 1213 | 324 | 0 | 0 | 94 | 0 | 202 | 255 | 0 | 0 | 0 | 13 | 0 | 0 | 0 | 0 | 0 | 13 |
| SRR5387823 | 0 | 0 | 12 | 0 | 0 | 0 | 0 | 16 | 0 | 0 | 0 | 0 | 0 | 0 | 0 | 0 | 0 | 0 | 71 | 1066 | 0 | 0 | 0 | 57 | 0 | 130 | 252 | 0 | 0 | 0 | 0 | 0 | 0 | 0 | 0 | 0 | 0 |
| SRR5387824 | 0 | 0 | 17 | 0 | 0 | 0 | 0 | 27 | 0 | 0 | 0 | 0 | 0 | 0 | 0 | 0 | 0 | 0 | 74 | 1626 | 148 | 0 | 0 | 73 | 0 | 136 | 227 | 0 | 0 | 0 | 0 | 0 | 0 | 0 | 0 | 0 | 0 |
| SRR5387825 | 0 | 18 | 10 | 0 | 0 | 28 | 0 | 37 | 0 | 0 | 0 | 0 | 0 | 0 | 0 | 0 | 0 | 0 | 0 | 1227 | 82 | 0 | 0 | 73 | 0 | 230 | 129 | 0 | 0 | 0 | 0 | 0 | 0 | 0 | 0 | 0 | 0 |
| SRR5387826 | 0 | 0 | 0 | 0 | 0 | 24 | 0 | 24 | 12 | 0 | 0 | 0 | 0 | 12 | 0 | 0 | 0 | 0 | 0 | 118 | 118 | 0 | 0 | 85 | 12 | 378 | 341 | 0 | 0 | 0 | 12 | 0 | 0 | 12 | 0 | 0 | 0 |
| SRR5387827 | 0 | 0 | 0 | 0 | 0 | 32 | 0 | 0 | 0 | 0 | 0 | 0 | 0 | 0 | 0 | 0 | 0 | 0 | 0 | 1144 | 654 | 0 | 0 | 16 | 0 | 426 | 205 | 0 | 0 | 0 | 16 | 0 | 0 | 0 | 16 | 0 | 16 |
| SRR5387828 | 0 | 0 | 0 | 0 | 0 | 0 | 0 | 0 | 0 | 0 | 0 | 0 | 0 | 0 | 0 | 0 | 0 | 0 | 84 | 503 | 168 | 0 | 0 | 64 | 0 | 366 | 403 | 0 | 0 | 0 | 0 | 0 | 0 | 0 | 0 | 0 | 0 |
| SRR5387829 | 0 | 10 | 0 | 0 | 0 | 20 | 0 | 10 | 0 | 0 | 0 | 0 | 0 | 0 | 0 | 0 | 0 | 0 | 0 | 665 | 39 | 0 | 0 | 105 | 0 | 347 | 357 | 0 | 0 | 0 | 0 | 0 | 0 | 0 | 0 | 0 | 0 |
| SRR5387830 | 0 | 13 | 15 | 0 | 0 | 13 | 0 | 0 | 0 | 0 | 0 | 0 | 0 | 0 | 0 | 0 | 0 | 0 | 0 | 1004 | 359 | 0 | 0 | 78 | 0 | 168 | 259 | 0 | 0 | 0 | 0 | 0 | 0 | 0 | 0 | 0 | 0 |
| SRR5387831 | 0 | 0 | 0 | 0 | 0 | 37 | 0 | 28 | 0 | 0 | 0 | 0 | 0 | 0 | 0 | 0 | 0 | 0 | 0 | 1147 | 115 | 0 | 0 | 139 | 0 | 120 | 176 | 0 | 0 | 0 | 0 | 0 | 0 | 0 | 0 | 0 | 0 |
| SRR5387832 | 0 | 21 | 0 | 0 | 0 | 21 | 119 | 0 | 0 | 0 | 0 | 0 | 0 | 10 | 0 | 0 | 0 | 0 | 0 | 595 | 119 | 0 | 0 | 104 | 0 | 290 | 248 | 0 | 0 | 0 | 0 | 0 | 0 | 0 | 10 | 0 | 0 |
| SRR5387833 | 0 | 12 | 0 | 0 | 0 | 35 | 0 | 12 | 12 | 0 | 0 | 0 | 0 | 0 | 0 | 0 | 0 | 0 | 0 | 1531 | 109 | 0 | 0 | 92 | 23 | 439 | 762 | 12 | 0 | 0 | 0 | 0 | 0 | 0 | 0 | 0 | 12 |
| SRR5387834 | 0 | 0 | 0 | 0 | 0 | 0 | 0 | 0 | 0 | 0 | 0 | 0 | 0 | 0 | 0 | 0 | 0 | 0 | 0 | 664 | 0 | 0 | 0 | 82 | 0 | 228 | 391 | 0 | 0 | 0 | 0 | 0 | 0 | 0 | 0 | 0 | 0 |
| SRR5387835 | 0 | 21 | 13 | 0 | 0 | 21 | 0 | 0 | 0 | 0 | 0 | 0 | 0 | 14 | 77 | 0 | 0 | 0 | 0 | 537 | 153 | 0 | 0 | 83 | 0 | 241 | 441 | 0 | 0 | 0 | 0 | 0 | 0 | 28 | 0 | 0 | 0 |
| SRR5387836 | 0 | 0 | 0 | 0 | 0 | 15 | 0 | 30 | 0 | 0 | 0 | 0 | 0 | 0 | 87 | 0 | 0 | 0 | 0 | 262 | 175 | 0 | 0 | 81 | 15 | 155 | 266 | 0 | 0 | 0 | 0 | 0 | 0 | 0 | 15 | 0 | 0 |
| SRR5387837 | 0 | 0 | 0 | 0 | 0 | 0 | 0 | 0 | 0 | 0 | 0 | 0 | 0 | 0 | 0 | 0 | 0 | 0 | 0 | 578 | 96 | 0 | 0 | 73 | 0 | 146 | 256 | 0 | 0 | 0 | 0 | 0 | 22 | 0 | 0 | 0 | 0 |
| SRR5387838 | 0 | 0 | 0 | 0 | 0 | 13 | 0 | 19 | 25 | 0 | 0 | 0 | 0 | 0 | 0 | 0 | 0 | 0 | 0 | 97 | 292 | 0 | 0 | 88 | 0 | 265 | 360 | 0 | 0 | 0 | 13 | 0 | 0 | 0 | 0 | 0 | 0 |
| SRR5387839 | 0 | 21 | 0 | 0 | 0 | 64 | 0 | 0 | 11 | 0 | 0 | 0 | 0 | 0 | 0 | 0 | 0 | 0 | 116 | 1041 | 0 | 0 | 0 | 139 | 0 | 610 | 236 | 0 | 0 | 11 | 11 | 0 | 0 | 0 | 0 | 0 | 0 |
| SRR5387840 | 0 | 14 | 0 | 0 | 0 | 72 | 0 | 0 | 43 | 0 | 0 | 0 | 0 | 0 | 0 | 0 | 0 | 0 | 0 | 707 | 0 | 0 | 0 | 143 | 0 | 11 | 443 | 43 | 0 | 0 | 0 | 0 | 0 | 0 | 14 | 0 | 0 |
| SRR5387841 | 0 | 16 | 0 | 0 | 0 | 25 | 0 | 0 | 0 | 0 | 0 | 0 | 0 | 0 | 0 | 0 | 0 | 0 | 84 | 2867 | 590 | 0 | 0 | 114 | 0 | 237 | 123 | 0 | 0 | 0 | 0 | 0 | 0 | 0 | 0 | 0 | 0 |
| SRR5387842 | 0 | 0 | 0 | 0 | 0 | 18 | 0 | 18 | 0 | 0 | 0 | 0 | 0 | 0 | 0 | 0 | 0 | 0 | 0 | 929 | 186 | 0 | 0 | 105 | 0 | 219 | 316 | 0 | 0 | 0 | 0 | 0 | 0 | 0 | 0 | 0 | 0 |
| SRR5387843 | 0 | 0 | 16 | 0 | 0 | 26 | 0 | 0 | 0 | 0 | 0 | 0 | 0 | 0 | 0 | 0 | 0 | 0 | 0 | 1447 | 145 | 0 | 0 | 164 | 0 | 249 | 315 | 0 | 0 | 0 | 0 | 0 | 0 | 0 | 0 | 0 | 13 |
| SRR5387844 | 11 | 18 | 0 | 0 | 0 | 0 | 0 | 0 | 0 | 0 | 0 | 0 | 0 | 18 | 0 | 0 | 0 | 0 | 78 | 704 | 235 | 0 | 0 | 119 | 0 | 192 | 449 | 0 | 0 | 0 | 0 | 0 | 0 | 0 | 0 | 0 | 0 |
| SRR5387845 | 0 | 10 | 0 | 0 | 0 | 31 | 0 | 31 | 0 | 0 | 0 | 0 | 0 | 20 | 62 | 0 | 0 | 0 | 0 | 1058 | 62 | 10 | 0 | 71 | 0 | 183 | 143 | 0 | 0 | 0 | 10 | 0 | 0 | 31 | 0 | 0 | 0 |
| SRR5387846 | 0 | 0 | 0 | 0 | 0 | 0 | 0 | 25 | 0 | 0 | 0 | 0 | 0 | 0 | 0 | 0 | 0 | 0 | 0 | 1069 | 164 | 0 | 0 | 127 | 0 | 296 | 330 | 0 | 0 | 0 | 25 | 0 | 0 | 0 | 0 | 0 | 0 |
| SRR5387847 | 0 | 33 | 0 | 0 | 0 | 22 | 0 | 33 | 0 | 0 | 0 | 0 | 0 | 0 | 0 | 0 | 0 | 0 | 64 | 1596 | 64 | 0 | 0 | 100 | 0 | 276 | 199 | 0 | 0 | 0 | 11 | 0 | 0 | 0 | 0 | 0 | 0 |
| SRR5387848 | 0 | 0 | 18 | 0 | 0 | 14 | 0 | 0 | 0 | 0 | 0 | 0 | 0 | 20 | 0 | 0 | 0 | 0 | 0 | 1779 | 123 | 0 | 0 | 136 | 0 | 143 | 224 | 0 | 0 | 0 | 0 | 0 | 0 | 0 | 0 | 0 | 0 |
| SRR5387849 | 0 | 0 | 0 | 0 | 0 | 0 | 0 | 15 | 0 | 0 | 0 | 0 | 0 | 0 | 55 | 0 | 0 | 0 | 0 | 872 | 55 | 0 | 0 | 170 | 0 | 170 | 378 | 0 | 0 | 0 | 0 | 0 | 0 | 0 | 0 | 0 | 0 |
| SRR5387850 | 0 | 18 | 0 | 0 | 0 | 27 | 0 | 18 | 0 | 0 | 0 | 0 | 0 | 0 | 0 | 0 | 0 | 0 | 80 | 957 | 80 | 0 | 0 | 156 | 0 | 256 | 275 | 0 | 0 | 0 | 0 | 0 | 0 | 0 | 0 | 0 | 0 |
| SRR5387851 | 0 | 0 | 0 | 0 | 0 | 0 | 0 | 0 | 14 | 0 | 0 | 0 | 0 | 0 | 0 | 0 | 0 | 0 | 0 | 1002 | 501 | 0 | 0 | 41 | 14 | 358 | 317 | 0 | 0 | 0 | 0 | 0 | 0 | 0 | 14 | 0 | 0 |
| SRR5387852 | 0 | 18 | 15 | 0 | 0 | 37 | 0 | 18 | 0 | 0 | 0 | 0 | 0 | 0 | 0 | 0 | 0 | 0 | 0 | 1819 | 79 | 0 | 0 | 119 | 0 | 247 | 247 | 0 | 0 | 0 | 0 | 0 | 0 | 0 | 0 | 0 | 18 |
| SRR5387853 | 0 | 0 | 10 | 0 | 0 | 62 | 0 | 0 | 0 | 0 | 0 | 0 | 0 | 0 | 126 | 0 | 0 | 0 | 0 | 758 | 126 | 0 | 0 | 177 | 10 | 260 | 468 | 0 | 0 | 0 | 0 | 0 | 0 | 10 | 0 | 0 | 0 |
| SRR5387854 | 0 | 0 | 17 | 0 | 0 | 21 | 0 | 0 | 14 | 0 | 0 | 0 | 0 | 0 | 0 | 0 | 0 | 0 | 0 | 993 | 132 | 0 | 0 | 71 | 14 | 226 | 424 | 0 | 0 | 0 | 0 | 0 | 0 | 0 | 0 | 0 | 0 |
| SRR5387855 | 0 | 18 | 0 | 0 | 0 | 0 | 0 | 18 | 0 | 0 | 0 | 0 | 0 | 0 | 0 | 0 | 0 | 0 | 73 | 2037 | 0 | 0 | 0 | 120 | 0 | 248 | 285 | 0 | 0 | 0 | 0 | 0 | 0 | 0 | 0 | 0 | 0 |
| SRR5387856 | 0 | 27 | 0 | 0 | 0 | 0 | 0 | 14 | 0 | 0 | 0 | 0 | 0 | 0 | 0 | 0 | 0 | 0 | 69 | 827 | 138 | 0 | 0 | 81 | 0 | 108 | 209 | 0 | 0 | 0 | 0 | 0 | 0 | 0 | 0 | 0 | 0 |
| SRR5387857 | 0 | 0 | 0 | 0 | 0 | 22 | 93 | 22 | 0 | 0 | 0 | 0 | 0 | 11 | 0 | 0 | 0 | 0 | 0 | 559 | 93 | 0 | 0 | 167 | 0 | 156 | 322 | 0 | 0 | 0 | 0 | 0 | 0 | 0 | 0 | 0 | 0 |
| SRR5387858 | 0 | 0 | 0 | 0 | 0 | 0 | 0 | 33 | 0 | 0 | 0 | 0 | 0 | 0 | 0 | 0 | 0 | 0 | 0 | 1025 | 205 | 0 | 0 | 90 | 0 | 65 | 278 | 0 | 0 | 0 | 0 | 0 | 0 | 0 | 0 | 0 | 0 |
| SRR5387859 | 0 | 13 | 0 | 0 | 0 | 13 | 0 | 13 | 0 | 0 | 0 | 0 | 0 | 0 | 0 | 0 | 0 | 0 | 0 | 1147 | 191 | 0 | 0 | 66 | 0 | 249 | 144 | 0 | 0 | 0 | 0 | 0 | 0 | 0 | 0 | 0 | 0 |
| SRR5387860 | 0 | 0 | 0 | 0 | 0 | 11 | 0 | 11 | 0 | 0 | 0 | 0 | 0 | 0 | 0 | 0 | 0 | 0 | 0 | 554 | 277 | 0 | 0 | 148 | 0 | 170 | 361 | 0 | 0 | 0 | 0 | 0 | 0 | 0 | 11 | 0 | 0 |
| SRR5387861 | 0 | 10 | 0 | 0 | 0 | 0 | 0 | 0 | 10 | 0 | 0 | 0 | 0 | 0 | 0 | 0 | 0 | 0 | 0 | 832 | 462 | 0 | 0 | 112 | 0 | 275 | 458 | 0 | 0 | 0 | 0 | 0 | 0 | 0 | 20 | 0 | 20 |
| SRR5387862 | 0 | 17 | 12 | 0 | 0 | 25 | 0 | 0 | 0 | 0 | 0 | 0 | 0 | 0 | 0 | 0 | 0 | 0 | 0 | 492 | 109 | 0 | 0 | 109 | 0 | 151 | 353 | 0 | 0 | 0 | 0 | 0 | 0 | 0 | 0 | 0 | 0 |
| SRR5387863 | 0 | 0 | 31 | 0 | 0 | 0 | 0 | 19 | 0 | 0 | 0 | 0 | 0 | 0 | 0 | 0 | 0 | 0 | 0 | 787 | 79 | 0 | 0 | 29 | 0 | 191 | 172 | 0 | 0 | 0 | 0 | 0 | 0 | 0 | 19 | 0 | 0 |
| SRR5387864 | 0 | 0 | 30 | 0 | 0 | 0 | 0 | 0 | 0 | 0 | 0 | 0 | 0 | 0 | 0 | 0 | 0 | 0 | 0 | 1032 | 147 | 0 | 0 | 91 | 0 | 114 | 289 | 0 | 0 | 0 | 0 | 0 | 0 | 0 | 0 | 0 | 0 |
| SRR5387865 | 11 | 0 | 16 | 0 | 0 | 20 | 0 | 13 | 0 | 0 | 0 | 0 | 0 | 0 | 0 | 0 | 0 | 0 | 102 | 922 | 410 | 0 | 0 | 79 | 0 | 53 | 151 | 0 | 0 | 0 | 0 | 0 | 0 | 0 | 0 | 0 | 0 |
| SRR5387866 | 0 | 22 | 0 | 22 | 0 | 0 | 0 | 0 | 0 | 0 | 0 | 0 | 0 | 0 | 0 | 24 | 0 | 0 | 0 | 644 | 176 | 0 | 0 | 20 | 11 | 34 | 177 | 23 | 0 | 0 | 0 | 13 | 0 | 0 | 0 | 0 | 0 |

### Supp Table S8.2.Columns_AM_to_BW

| | | | | | | | | | | | | | | | | | | | | | | | | | | | | | | | | | | | | | |
|---|---|---|---|---|---|---|---|---|---|---|---|---|---|---|---|---|---|---|---|---|---|---|---|---|---|---|---|---|---|---|---|---|---|---|---|---|---|
| SRR5387867 | 0 | 0 | 0 | 0 | 0 | 16 | 0 | 0 | 0 | 0 | 0 | 0 | 0 | 0 | 0 | 0 | 0 | 0 | 0 | 812 | 312 | 0 | 0 | 190 | 0 | 555 | 404 | 0 | 0 | 0 | 0 | 0 | 0 | 0 | 0 | 0 | 16 |
| SRR5387868 | 0 | 0 | 0 | 0 | 0 | 10 | 0 | 21 | 10 | 0 | 0 | 0 | 0 | 0 | 0 | 10 | 0 | 0 | 0 | 435 | 0 | 0 | 0 | 125 | 10 | 63 | 230 | 10 | 0 | 0 | 0 | 0 | 0 | 10 | 0 | 0 | 0 |
| SRR5387869 | 0 | 0 | 0 | 0 | 0 | 32 | 0 | 11 | 0 | 0 | 0 | 0 | 0 | 11 | 0 | 0 | 0 | 0 | 0 | 1240 | 338 | 0 | 0 | 151 | 0 | 258 | 355 | 0 | 0 | 0 | 11 | 0 | 0 | 11 | 0 | 0 | 0 |
| SRR5387870 | 0 | 0 | 14 | 0 | 0 | 0 | 0 | 0 | 0 | 0 | 0 | 0 | 0 | 0 | 0 | 0 | 0 | 0 | 0 | 390 | 65 | 0 | 0 | 108 | 0 | 161 | 215 | 0 | 0 | 0 | 0 | 0 | 0 | 0 | 0 | 0 | 0 |
| SRR5387871 | 0 | 0 | 0 | 0 | 0 | 0 | 0 | 12 | 0 | 0 | 0 | 0 | 0 | 0 | 0 | 0 | 0 | 0 | 0 | 582 | 233 | 0 | 0 | 107 | 0 | 166 | 249 | 0 | 0 | 0 | 0 | 0 | 0 | 12 | 0 | 0 | 0 |
| SRR5387872 | 0 | 0 | 13 | 0 | 0 | 20 | 0 | 0 | 0 | 0 | 0 | 0 | 0 | 20 | 0 | 0 | 0 | 0 | 0 | 625 | 521 | 0 | 0 | 179 | 0 | 239 | 249 | 0 | 0 | 0 | 0 | 0 | 0 | 0 | 20 | 0 | 0 |
| SRR5387873 | 0 | 0 | 0 | 0 | 0 | 54 | 0 | 18 | 0 | 0 | 0 | 0 | 0 | 54 | 134 | 0 | 0 | 0 | 0 | 1338 | 268 | 0 | 0 | 73 | 0 | 816 | 599 | 0 | 0 | 0 | 0 | 0 | 0 | 0 | 18 | 0 | 0 |
| SRR5387874 | 0 | 0 | 0 | 0 | 0 | 25 | 0 | 17 | 0 | 0 | 0 | 0 | 0 | 0 | 58 | 0 | 0 | 0 | 58 | 637 | 58 | 0 | 0 | 76 | 0 | 305 | 390 | 0 | 0 | 0 | 0 | 0 | 0 | 0 | 0 | 0 | 0 |
| SRR5387875 | 0 | 24 | 12 | 0 | 0 | 24 | 0 | 0 | 0 | 0 | 0 | 0 | 0 | 0 | 0 | 0 | 0 | 0 | 0 | 1324 | 722 | 0 | 0 | 49 | 0 | 243 | 260 | 0 | 0 | 0 | 0 | 0 | 0 | 0 | 0 | 0 | 0 |
| SRR5387876 | 0 | 31 | 0 | 0 | 0 | 0 | 0 | 0 | 0 | 0 | 0 | 0 | 0 | 0 | 0 | 0 | 0 | 0 | 0 | 952 | 0 | 0 | 0 | 46 | 0 | 162 | 208 | 0 | 0 | 0 | 0 | 0 | 0 | 0 | 0 | 0 | 0 |
| SRR5387877 | 0 | 23 | 0 | 0 | 0 | 23 | 0 | 68 | 11 | 0 | 0 | 0 | 0 | 23 | 140 | 0 | 0 | 0 | 0 | 1684 | 140 | 0 | 0 | 91 | 0 | 251 | 285 | 0 | 0 | 0 | 0 | 0 | 0 | 34 | 0 | 0 | 0 |
| SRR5387878 | 0 | 23 | 0 | 0 | 0 | 15 | 0 | 68 | 0 | 0 | 0 | 0 | 0 | 0 | 0 | 0 | 0 | 0 | 157 | 1414 | 157 | 0 | 0 | 167 | 0 | 274 | 10 | 0 | 0 | 0 | 0 | 0 | 0 | 0 | 0 | 0 | 0 |
| SRR5387879 | 0 | 0 | 0 | 0 | 0 | 19 | 0 | 12 | 0 | 0 | 0 | 0 | 0 | 0 | 0 | 0 | 0 | 0 | 86 | 1458 | 343 | 0 | 0 | 74 | 0 | 229 | 248 | 0 | 0 | 0 | 0 | 0 | 0 | 0 | 0 | 0 | 0 |
| SRR5387880 | 0 | 0 | 0 | 0 | 0 | 0 | 105 | 25 | 0 | 0 | 0 | 0 | 0 | 0 | 0 | 0 | 0 | 0 | 0 | 1991 | 0 | 0 | 0 | 74 | 0 | 288 | 271 | 0 | 0 | 0 | 0 | 0 | 0 | 0 | 0 | 0 | 0 |
| SRR5387881 | 0 | 15 | 0 | 0 | 0 | 15 | 0 | 46 | 0 | 0 | 0 | 0 | 0 | 0 | 0 | 0 | 0 | 0 | 0 | 1564 | 0 | 0 | 0 | 39 | 0 | 85 | 92 | 0 | 15 | 0 | 0 | 0 | 0 | 15 | 0 | 0 | 0 |
| SRR5387882 | 0 | 0 | 10 | 0 | 0 | 0 | 0 | 12 | 0 | 0 | 0 | 0 | 0 | 12 | 0 | 0 | 0 | 0 | 108 | 1569 | 108 | 0 | 0 | 61 | 12 | 103 | 164 | 0 | 0 | 0 | 0 | 0 | 0 | 0 | 0 | 0 | 0 |
| SRR5387883 | 0 | 0 | 18 | 0 | 0 | 0 | 0 | 15 | 0 | 0 | 0 | 13 | 0 | 0 | 159 | 0 | 0 | 0 | 0 | 713 | 396 | 0 | 0 | 90 | 0 | 53 | 158 | 0 | 0 | 0 | 0 | 0 | 0 | 0 | 15 | 0 | 0 |
| SRR5387884 | 0 | 0 | 11 | 0 | 0 | 0 | 0 | 0 | 0 | 0 | 0 | 0 | 0 | 0 | 0 | 0 | 0 | 0 | 0 | 1853 | 98 | 0 | 0 | 156 | 0 | 183 | 374 | 0 | 0 | 0 | 0 | 0 | 0 | 0 | 0 | 0 | 0 |
| SRR5387885 | 0 | 10 | 0 | 0 | 0 | 31 | 0 | 10 | 0 | 0 | 0 | 0 | 0 | 10 | 0 | 0 | 0 | 0 | 0 | 940 | 0 | 10 | 0 | 52 | 0 | 176 | 351 | 10 | 0 | 0 | 0 | 0 | 0 | 0 | 10 | 0 | 10 |
| SRR5387886 | 10 | 27 | 0 | 18 | 0 | 18 | 0 | 0 | 0 | 0 | 0 | 0 | 0 | 0 | 0 | 0 | 0 | 0 | 0 | 1052 | 191 | 0 | 0 | 55 | 0 | 274 | 338 | 0 | 0 | 0 | 0 | 0 | 0 | 0 | 0 | 0 | 0 |
| SRR5387887 | 0 | 11 | 0 | 0 | 0 | 11 | 0 | 22 | 11 | 0 | 0 | 0 | 0 | 0 | 0 | 0 | 0 | 0 | 0 | 1661 | 0 | 0 | 0 | 67 | 0 | 223 | 189 | 0 | 0 | 0 | 0 | 0 | 0 | 11 | 11 | 0 | 11 |
| SRR5387888 | 0 | 0 | 15 | 0 | 0 | 0 | 0 | 0 | 0 | 0 | 0 | 0 | 0 | 0 | 0 | 0 | 0 | 0 | 0 | 170 | 113 | 0 | 0 | 93 | 0 | 132 | 162 | 0 | 0 | 0 | 0 | 0 | 15 | 23 | 0 | 0 | 0 |
| SRR5387889 | 0 | 0 | 0 | 0 | 0 | 19 | 0 | 0 | 0 | 0 | 0 | 0 | 0 | 0 | 0 | 0 | 0 | 0 | 0 | 2298 | 306 | 0 | 0 | 0 | 0 | 87 | 154 | 0 | 0 | 0 | 0 | 0 | 0 | 0 | 0 | 0 | 0 |
| SRR5387890 | 0 | 0 | 15 | 0 | 0 | 0 | 0 | 0 | 0 | 0 | 0 | 0 | 0 | 0 | 0 | 0 | 0 | 0 | 0 | 1175 | 69 | 0 | 0 | 47 | 19 | 170 | 217 | 0 | 0 | 0 | 0 | 0 | 0 | 0 | 0 | 0 | 0 |
| SRR5387891 | 0 | 48 | 0 | 0 | 0 | 19 | 0 | 0 | 48 | 0 | 0 | 0 | 0 | 0 | 0 | 0 | 0 | 0 | 0 | 1052 | 55 | 0 | 0 | 67 | 0 | 468 | 373 | 0 | 0 | 0 | 0 | 0 | 0 | 0 | 19 | 0 | 0 |
| SRR5387892 | 0 | 0 | 15 | 0 | 0 | 14 | 0 | 0 | 0 | 0 | 0 | 0 | 0 | 0 | 0 | 0 | 0 | 0 | 54 | 1186 | 216 | 0 | 0 | 170 | 0 | 271 | 407 | 0 | 0 | 0 | 0 | 0 | 0 | 14 | 0 | 0 | 0 |
| SRR5387893 | 0 | 0 | 14 | 12 | 0 | 0 | 0 | 0 | 0 | 0 | 0 | 0 | 0 | 0 | 0 | 0 | 0 | 0 | 0 | 778 | 60 | 0 | 0 | 160 | 12 | 86 | 308 | 0 | 0 | 0 | 0 | 0 | 0 | 0 | 0 | 0 | 12 |
| SRR5387894 | 0 | 0 | 0 | 0 | 0 | 0 | 0 | 0 | 0 | 0 | 0 | 0 | 0 | 0 | 0 | 0 | 0 | 0 | 0 | 720 | 131 | 0 | 0 | 135 | 0 | 232 | 377 | 0 | 0 | 0 | 0 | 0 | 0 | 0 | 0 | 0 | 0 |
| SRR5387895 | 0 | 0 | 21 | 0 | 0 | 24 | 0 | 0 | 0 | 0 | 0 | 0 | 0 | 24 | 0 | 0 | 0 | 0 | 0 | 1018 | 712 | 0 | 0 | 100 | 0 | 330 | 11 | 0 | 0 | 0 | 0 | 0 | 0 | 12 | 0 | 0 | 0 |
| SRR5387896 | 0 | 0 | 18 | 0 | 0 | 13 | 0 | 13 | 0 | 0 | 0 | 0 | 0 | 0 | 0 | 0 | 0 | 0 | 0 | 1078 | 479 | 0 | 0 | 79 | 20 | 205 | 390 | 0 | 0 | 0 | 0 | 0 | 0 | 0 | 0 | 0 | 0 |
| SRR5387897 | 0 | 0 | 0 | 0 | 0 | 11 | 74 | 0 | 11 | 0 | 0 | 0 | 0 | 11 | 0 | 0 | 0 | 0 | 0 | 962 | 222 | 0 | 0 | 137 | 0 | 179 | 359 | 0 | 0 | 0 | 0 | 0 | 0 | 11 | 11 | 0 | 11 |
| SRR5387898 | 0 | 26 | 0 | 0 | 0 | 17 | 0 | 0 | 0 | 0 | 0 | 0 | 0 | 0 | 0 | 0 | 0 | 0 | 0 | 1177 | 69 | 0 | 0 | 138 | 0 | 258 | 327 | 0 | 0 | 0 | 34 | 0 | 0 | 0 | 0 | 0 | 0 |
| SRR5387899 | 0 | 18 | 0 | 0 | 0 | 0 | 0 | 0 | 0 | 0 | 0 | 0 | 0 | 0 | 0 | 0 | 0 | 0 | 0 | 372 | 93 | 0 | 0 | 83 | 0 | 119 | 413 | 0 | 0 | 0 | 0 | 0 | 0 | 18 | 0 | 0 | 0 |
| SRR5387900 | 0 | 0 | 0 | 0 | 0 | 0 | 0 | 0 | 0 | 0 | 0 | 0 | 0 | 0 | 0 | 0 | 0 | 0 | 0 | 1361 | 136 | 0 | 0 | 43 | 0 | 198 | 242 | 17 | 0 | 0 | 0 | 17 | 0 | 0 | 0 | 0 | 0 |
| SRR5387901 | 14 | 0 | 52 | 0 | 0 | 17 | 0 | 0 | 0 | 0 | 0 | 16 | 0 | 0 | 0 | 0 | 0 | 0 | 69 | 1180 | 69 | 0 | 0 | 46 | 0 | 133 | 11 | 0 | 0 | 0 | 0 | 0 | 0 | 25 | 0 | 0 | 0 |
| SRR5387902 | 15 | 0 | 84 | 0 | 0 | 0 | 0 | 0 | 0 | 0 | 0 | 27 | 0 | 0 | 433 | 0 | 0 | 0 | 72 | 1732 | 794 | 0 | 0 | 50 | 0 | 91 | 100 | 0 | 0 | 0 | 0 | 0 | 0 | 0 | 0 | 0 | 0 |
| SRR5387903 | 0 | 0 | 12 | 0 | 0 | 23 | 0 | 0 | 0 | 0 | 0 | 0 | 0 | 0 | 0 | 0 | 0 | 0 | 49 | 1970 | 99 | 0 | 0 | 62 | 0 | 480 | 356 | 0 | 0 | 0 | 0 | 0 | 0 | 0 | 0 | 0 | 0 |
| SRR5387904 | 0 | 18 | 0 | 0 | 0 | 27 | 0 | 0 | 0 | 0 | 0 | 0 | 0 | 0 | 0 | 0 | 0 | 0 | 0 | 597 | 239 | 0 | 0 | 64 | 0 | 191 | 155 | 0 | 0 | 0 | 0 | 0 | 18 | 0 | 0 | 0 | 0 |
| SRR5387905 | 0 | 0 | 0 | 0 | 0 | 0 | 0 | 12 | 0 | 0 | 0 | 0 | 0 | 0 | 0 | 0 | 0 | 0 | 0 | 541 | 120 | 0 | 0 | 94 | 0 | 112 | 224 | 0 | 0 | 0 | 0 | 0 | 0 | 0 | 0 | 0 | 12 |
| SRR5387906 | 0 | 0 | 0 | 0 | 0 | 11 | 0 | 11 | 0 | 0 | 0 | 0 | 0 | 0 | 0 | 0 | 0 | 0 | 0 | 488 | 70 | 0 | 0 | 101 | 11 | 169 | 220 | 0 | 0 | 0 | 0 | 0 | 0 | 0 | 0 | 0 | 0 |
| SRR5387907 | 0 | 154 | 0 | 0 | 0 | 0 | 0 | 0 | 0 | 0 | 0 | 0 | 0 | 0 | 60 | 23 | 0 | 0 | 0 | 598 | 0 | 0 | 0 | 40 | 0 | 54 | 233 | 117 | 0 | 0 | 0 | 145 | 0 | 16 | 0 | 0 | 0 |
| SRR5387908 | 0 | 0 | 0 | 0 | 0 | 0 | 0 | 0 | 0 | 0 | 0 | 0 | 0 | 0 | 0 | 0 | 0 | 0 | 0 | 780 | 142 | 0 | 0 | 81 | 0 | 146 | 293 | 0 | 0 | 0 | 0 | 0 | 0 | 0 | 0 | 0 | 0 |
| SRR5387909 | 0 | 0 | 0 | 0 | 0 | 27 | 0 | 36 | 18 | 0 | 0 | 0 | 0 | 0 | 0 | 0 | 0 | 0 | 0 | 765 | 765 | 0 | 0 | 27 | 0 | 224 | 206 | 0 | 0 | 0 | 0 | 0 | 0 | 0 | 0 | 0 | 18 |
| SRR5387910 | 0 | 0 | 12 | 0 | 0 | 0 | 0 | 0 | 17 | 0 | 0 | 0 | 0 | 0 | 0 | 0 | 0 | 0 | 0 | 847 | 242 | 0 | 0 | 87 | 0 | 251 | 268 | 0 | 0 | 0 | 0 | 0 | 0 | 0 | 0 | 0 | 0 |
| SRR5387911 | 0 | 25 | 0 | 0 | 0 | 25 | 0 | 25 | 0 | 0 | 0 | 0 | 0 | 17 | 0 | 0 | 0 | 0 | 0 | 674 | 300 | 0 | 0 | 17 | 0 | 252 | 328 | 0 | 0 | 0 | 0 | 0 | 0 | 0 | 0 | 0 | 0 |
| SRR5387912 | 0 | 0 | 0 | 0 | 0 | 23 | 0 | 0 | 0 | 0 | 0 | 0 | 0 | 0 | 100 | 0 | 0 | 0 | 0 | 1798 | 100 | 0 | 0 | 100 | 0 | 270 | 439 | 0 | 0 | 0 | 0 | 0 | 0 | 0 | 15 | 0 | 0 |
| SRR5387913 | 0 | 23 | 18 | 0 | 0 | 12 | 0 | 12 | 12 | 0 | 0 | 0 | 0 | 35 | 0 | 0 | 0 | 0 | 0 | 1669 | 196 | 0 | 0 | 104 | 0 | 485 | 566 | 0 | 0 | 0 | 0 | 0 | 0 | 0 | 12 | 0 | 12 |
| SRR5387914 | 0 | 0 | 12 | 0 | 0 | 29 | 0 | 0 | 0 | 0 | 0 | 0 | 0 | 0 | 0 | 0 | 0 | 0 | 0 | 407 | 244 | 0 | 0 | 107 | 0 | 381 | 596 | 0 | 0 | 0 | 0 | 0 | 0 | 20 | 0 | 0 | 0 |
| SRR5387915 | 0 | 15 | 0 | 0 | 0 | 15 | 0 | 0 | 0 | 0 | 0 | 0 | 0 | 0 | 0 | 0 | 0 | 0 | 0 | 1326 | 0 | 0 | 0 | 38 | 0 | 307 | 345 | 0 | 0 | 0 | 0 | 0 | 0 | 0 | 0 | 0 | 0 |
| SRR5387916 | 0 | 35 | 17 | 0 | 0 | 0 | 0 | 0 | 18 | 0 | 0 | 0 | 0 | 0 | 82 | 0 | 0 | 0 | 0 | 740 | 329 | 0 | 0 | 168 | 0 | 212 | 389 | 0 | 0 | 0 | 0 | 0 | 0 | 0 | 0 | 0 | 0 |
| SRR5387917 | 0 | 0 | 16 | 0 | 0 | 40 | 0 | 0 | 0 | 0 | 0 | 0 | 0 | 0 | 87 | 0 | 0 | 0 | 0 | 608 | 347 | 0 | 0 | 162 | 0 | 210 | 331 | 0 | 0 | 0 | 0 | 0 | 0 | 0 | 0 | 0 | 0 |
| SRR5387918 | 0 | 0 | 0 | 0 | 0 | 16 | 0 | 0 | 0 | 0 | 0 | 0 | 0 | 0 | 0 | 0 | 0 | 0 | 92 | 1010 | 92 | 0 | 0 | 98 | 16 | 270 | 343 | 0 | 0 | 16 | 0 | 0 | 0 | 16 | 0 | 0 | 0 |
| SRR5387919 | 0 | 0 | 10 | 0 | 0 | 0 | 0 | 20 | 0 | 0 | 0 | 0 | 0 | 0 | 0 | 0 | 0 | 0 | 0 | 863 | 308 | 0 | 0 | 80 | 0 | 80 | 60 | 0 | 0 | 0 | 0 | 0 | 0 | 0 | 20 | 0 | 0 |
| SRR5387920 | 0 | 0 | 11 | 0 | 0 | 12 | 0 | 0 | 12 | 0 | 0 | 0 | 0 | 0 | 0 | 0 | 0 | 0 | 0 | 1570 | 185 | 0 | 0 | 73 | 0 | 488 | 464 | 0 | 0 | 0 | 0 | 0 | 0 | 0 | 0 | 0 | 0 |
| SRR5387921 | 0 | 0 | 0 | 0 | 0 | 22 | 0 | 0 | 22 | 0 | 0 | 0 | 0 | 0 | 0 | 0 | 0 | 0 | 0 | 0 | 178 | 0 | 0 | 22 | 0 | 132 | 132 | 0 | 0 | 0 | 0 | 0 | 0 | 0 | 0 | 0 | 0 |
| SRR5387922 | 0 | 20 | 0 | 0 | 0 | 20 | 148 | 0 | 0 | 0 | 0 | 0 | 0 | 40 | 148 | 0 | 0 | 0 | 0 | 148 | 297 | 0 | 0 | 81 | 0 | 645 | 746 | 20 | 0 | 0 | 81 | 0 | 0 | 0 | 0 | 0 | 0 |
| SRR5387923 | 0 | 0 | 0 | 0 | 0 | 90 | 0 | 23 | 11 | 0 | 0 | 0 | 0 | 0 | 0 | 0 | 0 | 0 | 0 | 1460 | 487 | 0 | 0 | 79 | 0 | 543 | 305 | 0 | 0 | 11 | 0 | 0 | 0 | 0 | 11 | 0 | 0 |
| SRR5387924 | 0 | 0 | 0 | 0 | 0 | 16 | 0 | 0 | 0 | 0 | 0 | 0 | 0 | 0 | 0 | 0 | 0 | 0 | 0 | 499 | 250 | 0 | 0 | 39 | 0 | 273 | 444 | 0 | 0 | 0 | 0 | 0 | 0 | 0 | 0 | 0 | 0 |
| SRR5387925 | 0 | 10 | 13 | 0 | 0 | 0 | 0 | 0 | 0 | 0 | 0 | 0 | 0 | 0 | 35 | 0 | 0 | 0 | 70 | 800 | 104 | 0 | 0 | 129 | 0 | 61 | 149 | 0 | 0 | 0 | 0 | 0 | 0 | 10 | 0 | 0 | 0 |
| SRR5387926 | 0 | 0 | 11 | 0 | 0 | 10 | 0 | 16 | 10 | 0 | 0 | 0 | 0 | 10 | 0 | 0 | 0 | 0 | 0 | 1286 | 161 | 0 | 0 | 162 | 0 | 12 | 15 | 0 | 0 | 0 | 0 | 0 | 0 | 0 | 0 | 0 | 0 |
| SRR5387927 | 0 | 0 | 0 | 0 | 0 | 0 | 0 | 0 | 0 | 0 | 0 | 0 | 0 | 0 | 0 | 0 | 0 | 0 | 0 | 1787 | 357 | 0 | 0 | 85 | 0 | 247 | 436 | 0 | 0 | 0 | 0 | 0 | 0 | 0 | 38 | 0 | 0 |
| SRR5387928 | 0 | 0 | 18 | 0 | 0 | 24 | 0 | 16 | 0 | 0 | 0 | 0 | 0 | 0 | 84 | 0 | 0 | 0 | 0 | 1011 | 253 | 0 | 0 | 146 | 0 | 211 | 325 | 0 | 0 | 0 | 0 | 0 | 0 | 0 | 0 | 0 | 0 |
| SRR5387929 | 0 | 0 | 16 | 0 | 0 | 23 | 0 | 0 | 0 | 0 | 0 | 0 | 0 | 0 | 0 | 0 | 0 | 0 | 0 | 1611 | 201 | 0 | 0 | 115 | 0 | 368 | 391 | 0 | 0 | 0 | 0 | 0 | 0 | 0 | 0 | 0 | 0 |
| SRR5387930 | 0 | 0 | 0 | 0 | 0 | 0 | 0 | 0 | 0 | 0 | 0 | 0 | 0 | 0 | 0 | 0 | 0 | 0 | 0 | 972 | 108 | 0 | 0 | 147 | 0 | 184 | 491 | 0 | 0 | 0 | 0 | 0 | 0 | 0 | 0 | 0 | 0 |
| SRR5387931 | 0 | 0 | 0 | 0 | 0 | 26 | 0 | 0 | 0 | 0 | 0 | 0 | 0 | 0 | 104 | 0 | 0 | 0 | 0 | 829 | 207 | 0 | 0 | 128 | 0 | 451 | 358 | 0 | 0 | 0 | 0 | 26 | 0 | 0 | 0 | 0 | 0 |
| SRR5387932 | 0 | 16 | 0 | 0 | 0 | 39 | 0 | 0 | 24 | 0 | 0 | 0 | 0 | 0 | 0 | 0 | 0 | 0 | 0 | 535 | 0 | 0 | 0 | 181 | 0 | 197 | 441 | 0 | 0 | 0 | 0 | 0 | 0 | 16 | 0 | 0 | 0 |
| SRR5387933 | 0 | 20 | 13 | 0 | 0 | 20 | 0 | 0 | 0 | 0 | 0 | 0 | 0 | 14 | 76 | 14 | 0 | 0 | 0 | 1221 | 76 | 0 | 0 | 81 | 0 | 176 | 365 | 41 | 0 | 0 | 0 | 27 | 0 | 27 | 0 | 0 | 0 |
| SRR5387934 | 0 | 0 | 11 | 0 | 0 | 0 | 0 | 0 | 0 | 0 | 0 | 0 | 0 | 12 | 0 | 0 | 0 | 0 | 57 | 745 | 57 | 0 | 0 | 81 | 0 | 187 | 256 | 0 | 0 | 0 | 0 | 0 | 0 | 19 | 0 | 0 | 0 |
| SRR5387935 | 0 | 0 | 15 | 0 | 0 | 10 | 0 | 0 | 31 | 0 | 0 | 0 | 0 | 42 | 0 | 0 | 0 | 0 | 0 | 1579 | 226 | 0 | 0 | 21 | 10 | 251 | 251 | 0 | 0 | 0 | 10 | 0 | 0 | 10 | 10 | 0 | 21 |
| SRR5387936 | 0 | 0 | 14 | 0 | 0 | 0 | 0 | 25 | 0 | 0 | 0 | 0 | 0 | 0 | 0 | 0 | 0 | 0 | 0 | 1073 | 248 | 0 | 0 | 91 | 0 | 141 | 315 | 0 | 0 | 0 | 0 | 0 | 0 | 0 | 0 | 0 | 0 |
| SRR5387937 | 0 | 0 | 14 | 0 | 0 | 0 | 0 | 0 | 0 | 0 | 0 | 0 | 0 | 0 | 0 | 0 | 0 | 0 | 0 | 656 | 364 | 0 | 0 | 75 | 0 | 212 | 417 | 0 | 0 | 0 | 0 | 0 | 0 | 21 | 0 | 0 | 0 |
| SRR5387938 | 0 | 0 | 19 | 0 | 0 | 22 | 0 | 0 | 0 | 0 | 0 | 0 | 0 | 0 | 86 | 0 | 0 | 0 | 172 | 687 | 172 | 0 | 0 | 79 | 0 | 180 | 374 | 0 | 0 | 0 | 0 | 0 | 0 | 14 | 14 | 0 | 0 |
| SRR5387939 | 0 | 0 | 0 | 0 | 0 | 25 | 0 | 25 | 0 | 0 | 0 | 0 | 0 | 0 | 0 | 0 | 0 | 0 | 0 | 1743 | 152 | 0 | 0 | 118 | 0 | 395 | 303 | 0 | 0 | 17 | 0 | 0 | 0 | 0 | 0 | 0 | 0 |
| SRR5387940 | 0 | 15 | 0 | 12 | 19 | 23 | 0 | 0 | 0 | 0 | 0 | 0 | 0 | 0 | 72 | 0 | 0 | 0 | 0 | 719 | 288 | 0 | 0 | 77 | 15 | 54 | 11 | 12 | 0 | 0 | 0 | 0 | 0 | 0 | 0 | 0 | 0 |
| SRR5387941 | 0 | 0 | 12 | 15 | 0 | 29 | 0 | 0 | 15 | 0 | 0 | 0 | 0 | 0 | 0 | 0 | 0 | 0 | 0 | 998 | 399 | 0 | 0 | 103 | 0 | 278 | 432 | 0 | 0 | 0 | 0 | 0 | 15 | 0 | 0 | 0 | 0 |
| SRR5387942 | 0 | 143 | 0 | 0 | 0 | 0 | 0 | 0 | 0 | 0 | 0 | 0 | 0 | 0 | 247 | 16 | 0 | 0 | 0 | 742 | 371 | 12 | 0 | 15 | 0 | 38 | 166 | 163 | 0 | 20 | 0 | 124 | 0 | 13 | 0 | 0 | 0 |
| SRR5387943 | 0 | 16 | 12 | 0 | 0 | 0 | 0 | 16 | 0 | 0 | 0 | 0 | 0 | 0 | 0 | 0 | 0 | 0 | 108 | 970 | 108 | 0 | 0 | 16 | 0 | 413 | 334 | 0 | 16 | 0 | 0 | 0 | 0 | 0 | 0 | 0 | 0 |
| SRR5387944 | 0 | 0 | 0 | 0 | 0 | 16 | 0 | 0 | 0 | 0 | 0 | 0 | 0 | 0 | 0 | 0 | 0 | 0 | 87 | 782 | 174 | 0 | 0 | 137 | 0 | 241 | 338 | 0 | 0 | 0 | 0 | 0 | 0 | 0 | 0 | 0 | 0 |
| SRR5387945 | 0 | 19 | 11 | 0 | 0 | 0 | 0 | 19 | 0 | 0 | 0 | 0 | 0 | 0 | 0 | 0 | 0 | 0 | 0 | 902 | 69 | 0 | 0 | 115 | 0 | 221 | 192 | 0 | 0 | 0 | 0 | 0 | 0 | 19 | 0 | 0 | 0 |
| SRR5387946 | 0 | 16 | 14 | 0 | 0 | 11 | 0 | 11 | 0 | 0 | 0 | 0 | 0 | 0 | 0 | 0 | 0 | 0 | 0 | 1155 | 272 | 0 | 0 | 79 | 0 | 79 | 242 | 0 | 0 | 0 | 11 | 0 | 0 | 0 | 0 | 0 | 0 |
| SRR5387947 | 0 | 11 | 0 | 0 | 0 | 34 | 0 | 0 | 11 | 0 | 0 | 0 | 0 | 11 | 0 | 11 | 0 | 0 | 0 | 1099 | 220 | 0 | 0 | 90 | 0 | 314 | 527 | 0 | 0 | 0 | 0 | 0 | 0 | 11 | 0 | 0 | 11 |
| SRR5387948 | 0 | 32 | 14 | 0 | 0 | 0 | 0 | 0 | 0 | 0 | 0 | 0 | 0 | 0 | 0 | 0 | 0 | 0 | 0 | 1074 | 126 | 0 | 0 | 54 | 0 | 109 | 249 | 0 | 0 | 0 | 0 | 32 | 0 | 18 | 0 | 0 | 0 |
| SRR5387949 | 0 | 0 | 0 | 0 | 0 | 39 | 0 | 0 | 0 | 0 | 0 | 0 | 0 | 52 | 0 | 0 | 0 | 0 | 0 | 1613 | 302 | 0 | 0 | 142 | 0 | 464 | 555 | 0 | 0 | 0 | 0 | 0 | 13 | 13 | 0 | 0 | 0 |
| SRR5387950 | 0 | 0 | 11 | 0 | 0 | 0 | 0 | 0 | 15 | 0 | 0 | 0 | 0 | 0 | 0 | 0 | 0 | 0 | 99 | 1094 | 99 | 0 | 0 | 131 | 0 | 263 | 12 | 15 | 0 | 0 | 0 | 0 | 0 | 0 | 0 | 0 | 0 |
| SRR5387951 | 0 | 0 | 0 | 0 | 0 | 33 | 0 | 22 | 0 | 0 | 0 | 0 | 0 | 0 | 113 | 0 | 0 | 0 | 0 | 567 | 453 | 0 | 0 | 122 | 0 | 155 | 376 | 0 | 0 | 0 | 0 | 0 | 11 | 0 | 0 | 0 | 0 |
| SRR5387952 | 0 | 10 | 0 | 0 | 0 | 20 | 0 | 10 | 0 | 0 | 0 | 0 | 0 | 10 | 84 | 0 | 0 | 0 | 0 | 1597 | 84 | 0 | 0 | 122 | 0 | 122 | 254 | 0 | 0 | 0 | 10 | 0 | 0 | 0 | 0 | 0 | 0 |
| SRR5387953 | 0 | 0 | 0 | 0 | 0 | 0 | 0 | 29 | 0 | 0 | 0 | 0 | 0 | 0 | 0 | 0 | 0 | 0 | 0 | 780 | 390 | 0 | 0 | 48 | 0 | 115 | 384 | 0 | 0 | 0 | 29 | 0 | 0 | 0 | 19 | 0 | 0 |
| SRR5387954 | 0 | 0 | 19 | 0 | 0 | 12 | 0 | 0 | 0 | 0 | 0 | 0 | 0 | 0 | 0 | 0 | 0 | 0 | 0 | 1223 | 235 | 0 | 0 | 55 | 0 | 110 | 220 | 0 | 0 | 0 | 0 | 0 | 0 | 12 | 0 | 0 | 0 |
| SRR5387955 | 0 | 33 | 0 | 0 | 0 | 11 | 99 | 22 | 0 | 0 | 0 | 0 | 0 | 0 | 0 | 0 | 0 | 0 | 0 | 2463 | 99 | 0 | 0 | 77 | 0 | 154 | 319 | 0 | 0 | 0 | 0 | 0 | 0 | 11 | 22 | 0 | 22 |
| SRR5387956 | 0 | 0 | 11 | 0 | 0 | 23 | 0 | 15 | 0 | 0 | 0 | 0 | 0 | 0 | 0 | 0 | 0 | 0 | 0 | 1652 | 275 | 0 | 0 | 75 | 0 | 166 | 354 | 0 | 0 | 0 | 0 | 0 | 0 | 0 | 0 | 0 | 0 |
| SRR5387957 | 0 | 0 | 16 | 0 | 0 | 0 | 0 | 24 | 0 | 0 | 0 | 0 | 0 | 0 | 53 | 0 | 0 | 0 | 0 | 1440 | 160 | 0 | 0 | 73 | 0 | 113 | 194 | 0 | 0 | 0 | 0 | 0 | 0 | 24 | 0 | 0 | 0 |
| SRR5387958 | 0 | 0 | 14 | 0 | 0 | 10 | 0 | 21 | 0 | 0 | 0 | 0 | 0 | 0 | 0 | 0 | 0 | 0 | 0 | 575 | 48 | 0 | 0 | 88 | 0 | 93 | 124 | 0 | 0 | 0 | 10 | 0 | 0 | 0 | 0 | 0 | 0 |
| SRR5387959 | 0 | 0 | 0 | 0 | 0 | 0 | 0 | 0 | 0 | 0 | 0 | 0 | 0 | 0 | 0 | 0 | 0 | 0 | 0 | 1383 | 361 | 0 | 0 | 62 | 0 | 81 | 124 | 0 | 0 | 0 | 12 | 0 | 0 | 12 | 0 | 0 | 0 |
| SRR5387960 | 13 | 0 | 11 | 0 | 0 | 30 | 105 | 0 | 0 | 0 | 0 | 0 | 0 | 0 | 0 | 0 | 0 | 0 | 0 | 840 | 420 | 0 | 0 | 120 | 0 | 60 | 187 | 0 | 0 | 0 | 0 | 0 | 0 | 0 | 0 | 0 | 0 |
| SRR5387961 | 0 | 0 | 10 | 0 | 0 | 18 | 0 | 18 | 0 | 0 | 0 | 0 | 0 | 0 | 0 | 0 | 0 | 0 | 0 | 1675 | 447 | 0 | 0 | 45 | 0 | 299 | 345 | 0 | 0 | 0 | 0 | 0 | 0 | 18 | 0 | 0 | 0 |
| SRR5387962 | 0 | 0 | 23 | 0 | 0 | 0 | 0 | 19 | 0 | 0 | 0 | 0 | 0 | 0 | 0 | 0 | 0 | 0 | 0 | 739 | 164 | 0 | 0 | 124 | 0 | 168 | 255 | 0 | 0 | 0 | 0 | 0 | 0 | 25 | 0 | 0 | 12 |
| SRR5387963 | 0 | 0 | 14 | 0 | 0 | 0 | 0 | 0 | 0 | 0 | 0 | 0 | 0 | 17 | 0 | 0 | 0 | 0 | 63 | 1389 | 126 | 0 | 0 | 100 | 0 | 142 | 125 | 0 | 0 | 0 | 0 | 0 | 0 | 0 | 0 | 0 | 0 |
| SRR5387964 | 0 | 0 | 0 | 0 | 0 | 0 | 0 | 0 | 0 | 0 | 0 | 0 | 0 | 0 | 0 | 0 | 0 | 0 | 0 | 662 | 147 | 0 | 0 | 148 | 0 | 176 | 222 | 0 | 0 | 0 | 0 | 0 | 0 | 0 | 0 | 0 | 0 |
| SRR5387965 | 0 | 0 | 0 | 0 | 0 | 20 | 0 | 13 | 0 | 0 | 0 | 0 | 0 | 0 | 0 | 0 | 0 | 0 | 0 | 1577 | 0 | 0 | 0 | 179 | 0 | 113 | 285 | 0 | 0 | 0 | 0 | 0 | 0 | 0 | 0 | 0 | 0 |
| SRR5387966 | 0 | 0 | 11 | 0 | 0 | 0 | 0 | 24 | 0 | 0 | 0 | 0 | 0 | 0 | 0 | 0 | 0 | 0 | 0 | 595 | 85 | 0 | 0 | 181 | 16 | 173 | 260 | 0 | 0 | 0 | 0 | 0 | 0 | 0 | 31 | 0 | 0 |
| SRR5387967 | 0 | 0 | 0 | 0 | 0 | 0 | 0 | 14 | 0 | 0 | 0 | 0 | 0 | 0 | 57 | 0 | 0 | 0 | 0 | 802 | 0 | 0 | 0 | 137 | 0 | 108 | 259 | 0 | 0 | 0 | 0 | 0 | 14 | 0 | 0 | 0 | 22 |
| SRR5387968 | 0 | 0 | 0 | 0 | 0 | 23 | 0 | 0 | 0 | 0 | 0 | 0 | 0 | 0 | 0 | 0 | 0 | 0 | 0 | 288 | 96 | 0 | 0 | 101 | 0 | 257 | 250 | 0 | 0 | 0 | 0 | 0 | 0 | 0 | 0 | 0 | 0 |
| SRR5387969 | 0 | 24 | 15 | 0 | 0 | 24 | 0 | 0 | 0 | 0 | 0 | 0 | 0 | 0 | 0 | 0 | 0 | 0 | 0 | 1606 | 155 | 0 | 0 | 112 | 0 | 168 | 616 | 0 | 0 | 0 | 0 | 0 | 0 | 0 | 0 | 0 | 0 |
| SRR5387970 | 0 | 0 | 14 | 0 | 0 | 43 | 0 | 0 | 17 | 0 | 0 | 0 | 0 | 0 | 0 | 0 | 0 | 0 | 0 | 1453 | 0 | 0 | 0 | 77 | 0 | 240 | 402 | 0 | 0 | 0 | 0 | 0 | 0 | 0 | 0 | 0 | 0 |
| SRR5387971 | 0 | 10 | 16 | 0 | 0 | 21 | 0 | 0 | 0 | 0 | 0 | 0 | 0 | 0 | 0 | 0 | 0 | 0 | 59 | 1535 | 0 | 0 | 0 | 51 | 0 | 298 | 463 | 10 | 0 | 0 | 0 | 0 | 0 | 21 | 21 | 0 | 21 |
| SRR5387972 | 0 | 0 | 16 | 0 | 0 | 23 | 97 | 15 | 0 | 0 | 0 | 0 | 0 | 0 | 0 | 0 | 0 | 0 | 0 | 1453 | 0 | 0 | 0 | 123 | 0 | 169 | 231 | 0 | 0 | 0 | 0 | 0 | 0 | 0 | 0 | 0 | 15 |
| SRR5387973 | 0 | 0 | 11 | 10 | 0 | 0 | 0 | 10 | 10 | 0 | 0 | 0 | 0 | 51 | 0 | 0 | 0 | 0 | 0 | 1175 | 353 | 0 | 0 | 154 | 31 | 369 | 369 | 0 | 0 | 0 | 0 | 0 | 0 | 0 | 0 | 0 | 10 |
| SRR5387974 | 0 | 0 | 25 | 0 | 0 | 20 | 0 | 0 | 0 | 0 | 0 | 0 | 0 | 0 | 0 | 0 | 0 | 0 | 122 | 2683 | 244 | 0 | 0 | 199 | 0 | 398 | 508 | 0 | 20 | 0 | 0 | 0 | 0 | 0 | 0 | 0 | 0 |
| SRR5387975 | 0 | 0 | 0 | 0 | 0 | 18 | 0 | 55 | 18 | 0 | 0 | 0 | 0 | 0 | 0 | 0 | 0 | 0 | 0 | 953 | 0 | 18 | 0 | 146 | 0 | 310 | 182 | 0 | 0 | 0 | 0 | 0 | 0 | 0 | 0 | 0 | 0 |
| SRR5387976 | 10 | 12 | 35 | 0 | 0 | 12 | 0 | 29 | 0 | 0 | 0 | 0 | 0 | 0 | 162 | 0 | 0 | 0 | 0 | 487 | 893 | 0 | 0 | 58 | 12 | 173 | 253 | 0 | 0 | 0 | 0 | 0 | 0 | 0 | 0 | 0 | 0 |
| SRR5387977 | 0 | 0 | 16 | 0 | 0 | 0 | 0 | 19 | 0 | 0 | 0 | 0 | 0 | 0 | 0 | 0 | 0 | 0 | 75 | 2177 | 75 | 0 | 0 | 154 | 0 | 231 | 259 | 0 | 0 | 0 | 0 | 0 | 0 | 19 | 0 | 0 | 0 |
| SRR5387978 | 0 | 0 | 0 | 0 | 0 | 0 | 0 | 0 | 0 | 0 | 0 | 0 | 0 | 0 | 0 | 0 | 0 | 0 | 0 | 1994 | 153 | 0 | 0 | 123 | 0 | 104 | 406 | 0 | 0 | 0 | 0 | 0 | 0 | 0 | 0 | 0 | 28 |
| SRR5387979 | 0 | 0 | 0 | 0 | 0 | 24 | 0 | 12 | 12 | 0 | 0 | 0 | 0 | 12 | 0 | 0 | 0 | 0 | 0 | 1080 | 432 | 0 | 0 | 110 | 0 | 269 | 232 | 12 | 0 | 0 | 12 | 0 | 0 | 0 | 0 | 0 | 0 |
| SRR5387980 | 0 | 0 | 0 | 0 | 0 | 18 | 0 | 18 | 0 | 0 | 0 | 0 | 0 | 0 | 0 | 0 | 0 | 0 | 0 | 982 | 151 | 0 | 0 | 116 | 0 | 242 | 259 | 0 | 0 | 0 | 0 | 0 | 0 | 0 | 0 | 0 | 18 |
| SRR5387981 | 0 | 0 | 0 | 0 | 0 | 69 | 0 | 0 | 14 | 0 | 0 | 0 | 0 | 0 | 0 | 0 | 0 | 0 | 0 | 2290 | 135 | 0 | 0 | 97 | 0 | 375 | 333 | 0 | 0 | 0 | 0 | 0 | 0 | 0 | 14 | 0 | 0 |
| SRR5387982 | 0 | 11 | 0 | 0 | 0 | 11 | 0 | 0 | 0 | 0 | 0 | 0 | 0 | 0 | 0 | 0 | 0 | 0 | 0 | 684 | 0 | 0 | 0 | 79 | 0 | 293 | 349 | 0 | 0 | 0 | 0 | 0 | 0 | 0 | 0 | 0 | 11 |
| SRR5387983 | 0 | 0 | 0 | 0 | 0 | 16 | 0 | 0 | 0 | 0 | 0 | 0 | 0 | 24 | 0 | 0 | 0 | 0 | 0 | 887 | 118 | 0 | 0 | 118 | 0 | 395 | 434 | 0 | 0 | 0 | 0 | 0 | 0 | 0 | 0 | 0 | 0 |
| SRR5387984 | 0 | 11 | 13 | 11 | 0 | 0 | 0 | 0 | 11 | 0 | 0 | 0 | 0 | 16 | 0 | 0 | 0 | 0 | 0 | 379 | 0 | 0 | 0 | 65 | 0 | 142 | 212 | 11 | 0 | 0 | 0 | 11 | 0 | 16 | 0 | 0 | 0 |
| SRR5387985 | 0 | 0 | 11 | 0 | 0 | 13 | 0 | 0 | 0 | 0 | 0 | 0 | 0 | 0 | 0 | 0 | 0 | 0 | 25 | 1237 | 49 | 0 | 0 | 88 | 0 | 144 | 276 | 0 | 0 | 0 | 0 | 0 | 0 | 13 | 0 | 0 | 0 |
| SRR5387986 | 0 | 0 | 30 | 0 | 0 | 0 | 0 | 15 | 0 | 0 | 0 | 0 | 0 | 0 | 29 | 0 | 0 | 0 | 58 | 1442 | 87 | 0 | 0 | 74 | 0 | 188 | 271 | 0 | 0 | 0 | 0 | 0 | 0 | 15 | 0 | 0 | 0 |
| SRR5387987 | 0 | 10 | 11 | 0 | 0 | 10 | 0 | 20 | 0 | 0 | 0 | 0 | 0 | 0 | 0 | 0 | 0 | 295 | 103 | 1138 | 207 | 0 | 0 | 161 | 0 | 402 | 563 | 10 | 0 | 0 | 0 | 0 | 10 | 10 | 0 | 0 | 0 |
| SRR5387988 | 0 | 0 | 10 | 0 | 0 | 0 | 0 | 0 | 17 | 0 | 0 | 0 | 0 | 0 | 0 | 0 | 0 | 0 | 0 | 871 | 87 | 0 | 0 | 96 | 0 | 357 | 635 | 0 | 0 | 0 | 17 | 0 | 0 | 0 | 0 | 0 | 0 |
| SRR5387989 | 0 | 67 | 10 | 0 | 0 | 28 | 0 | 0 | 0 | 0 | 0 | 0 | 0 | 0 | 0 | 0 | 0 | 0 | 0 | 648 | 162 | 0 | 0 | 78 | 0 | 339 | 889 | 0 | 0 | 0 | 0 | 17 | 0 | 0 | 0 | 0 | 11 |
| SRR5387990 | 0 | 76 | 0 | 0 | 0 | 30 | 0 | 0 | 0 | 0 | 0 | 0 | 0 | 0 | 0 | 0 | 0 | 0 | 0 | 362 | 241 | 15 | 0 | 53 | 0 | 167 | 318 | 91 | 0 | 15 | 0 | 15 | 0 | 0 | 0 | 0 | 0 |

### Supp Table S8.2.Columns_AM_to_BW

| | | | | | | | | | | | | | | | | | | | | | | | | | | | | | | | | | | | | | |
|---|---|---|---|---|---|---|---|---|---|---|---|---|---|---|---|---|---|---|---|---|---|---|---|---|---|---|---|---|---|---|---|---|---|---|---|---|---|
| SRR5387991 | 0 | 0 | 12 | 0 | 0 | 23 | 0 | 31 | 0 | 0 | 0 | 0 | 0 | 0 | 57 | 0 | 0 | 0 | 0 | 1469 | 0 | 0 | 0 | 61 | 15 | 276 | 314 | 0 | 0 | 0 | 0 | 15 | 0 | 15 | 0 | 0 | 0 |
| SRR5387992 | 0 | 0 | 0 | 0 | 0 | 17 | 0 | 17 | 17 | 0 | 0 | 0 | 0 | 17 | 82 | 0 | 0 | 0 | 0 | 1062 | 0 | 0 | 0 | 122 | 0 | 480 | 585 | 0 | 0 | 0 | 0 | 0 | 0 | 26 | 0 | 0 | 0 |
| SRR5387993 | 0 | 26 | 11 | 0 | 0 | 34 | 0 | 0 | 0 | 0 | 0 | 0 | 0 | 0 | 0 | 0 | 0 | 0 | 0 | 177 | 266 | 0 | 0 | 103 | 0 | 15 | 14 | 17 | 0 | 0 | 0 | 0 | 0 | 17 | 0 | 0 | 0 |
| SRR5387994 | 0 | 0 | 0 | 15 | 0 | 0 | 0 | 0 | 0 | 0 | 0 | 0 | 0 | 0 | 0 | 0 | 0 | 0 | 0 | 1680 | 249 | 0 | 0 | 69 | 0 | 251 | 320 | 0 | 0 | 0 | 0 | 0 | 0 | 0 | 0 | 0 | 0 |
| SRR5387995 | 0 | 0 | 10 | 0 | 0 | 26 | 0 | 0 | 0 | 0 | 0 | 0 | 0 | 0 | 0 | 0 | 0 | 0 | 0 | 914 | 141 | 0 | 0 | 78 | 0 | 217 | 539 | 0 | 0 | 0 | 0 | 0 | 0 | 0 | 17 | 0 | 0 |
| SRR5387996 | 0 | 14 | 11 | 0 | 0 | 21 | 0 | 0 | 0 | 0 | 0 | 0 | 0 | 0 | 0 | 0 | 0 | 0 | 57 | 680 | 340 | 0 | 0 | 76 | 0 | 277 | 520 | 0 | 0 | 0 | 0 | 0 | 14 | 0 | 0 | 0 | 0 |
| SRR5387997 | 0 | 15 | 14 | 0 | 0 | 0 | 0 | 15 | 0 | 0 | 0 | 0 | 0 | 15 | 0 | 0 | 0 | 0 | 0 | 1312 | 131 | 0 | 0 | 156 | 0 | 126 | 245 | 0 | 0 | 0 | 0 | 0 | 0 | 0 | 0 | 0 | 0 |
| SRR5387998 | 0 | 0 | 17 | 0 | 0 | 11 | 0 | 23 | 0 | 0 | 0 | 0 | 0 | 0 | 0 | 0 | 0 | 0 | 0 | 841 | 70 | 0 | 0 | 73 | 0 | 146 | 225 | 0 | 0 | 0 | 0 | 0 | 0 | 28 | 0 | 0 | 0 |
| SRR5387999 | 0 | 0 | 12 | 0 | 0 | 23 | 0 | 18 | 0 | 0 | 0 | 0 | 0 | 0 | 83 | 0 | 0 | 0 | 0 | 1585 | 500 | 0 | 0 | 29 | 0 | 82 | 117 | 0 | 0 | 0 | 0 | 0 | 0 | 35 | 0 | 0 | 0 |
| SRR5388000 | 0 | 0 | 17 | 0 | 0 | 34 | 0 | 17 | 0 | 0 | 0 | 0 | 0 | 0 | 0 | 0 | 0 | 0 | 0 | 2046 | 241 | 0 | 0 | 117 | 0 | 117 | 319 | 0 | 0 | 0 | 0 | 0 | 0 | 0 | 0 | 0 | 0 |
| SRR5388001 | 0 | 15 | 12 | 0 | 0 | 22 | 0 | 15 | 0 | 0 | 0 | 0 | 0 | 0 | 0 | 0 | 0 | 0 | 0 | 712 | 0 | 0 | 0 | 90 | 0 | 142 | 291 | 0 | 0 | 0 | 0 | 0 | 0 | 0 | 0 | 0 | 0 |
| SRR5388002 | 0 | 0 | 14 | 0 | 0 | 14 | 0 | 77 | 0 | 0 | 0 | 0 | 0 | 14 | 0 | 0 | 0 | 0 | 94 | 1698 | 189 | 0 | 0 | 160 | 0 | 146 | 251 | 0 | 0 | 0 | 0 | 0 | 0 | 14 | 0 | 0 | 0 |
| SRR5388003 | 0 | 13 | 0 | 0 | 0 | 13 | 0 | 0 | 26 | 0 | 0 | 0 | 0 | 13 | 0 | 0 | 0 | 0 | 0 | 1650 | 733 | 0 | 0 | 66 | 0 | 434 | 420 | 0 | 13 | 0 | 0 | 13 | 0 | 13 | 0 | 0 | 0 |
| SRR5388004 | 0 | 0 | 0 | 0 | 0 | 0 | 0 | 0 | 0 | 0 | 0 | 0 | 0 | 13 | 0 | 0 | 0 | 0 | 0 | 881 | 196 | 0 | 0 | 90 | 0 | 116 | 149 | 0 | 0 | 0 | 13 | 0 | 0 | 0 | 0 | 0 | 0 |
| SRR5388005 | 0 | 0 | 18 | 0 | 0 | 0 | 0 | 21 | 0 | 0 | 0 | 0 | 0 | 0 | 49 | 0 | 0 | 0 | 49 | 2781 | 195 | 0 | 0 | 28 | 0 | 49 | 201 | 0 | 0 | 0 | 0 | 0 | 0 | 0 | 0 | 0 | 0 |
| SRR5388006 | 0 | 0 | 18 | 0 | 0 | 0 | 0 | 0 | 0 | 0 | 0 | 0 | 0 | 0 | 0 | 0 | 0 | 0 | 67 | 1415 | 202 | 0 | 0 | 24 | 0 | 130 | 252 | 0 | 0 | 0 | 0 | 0 | 0 | 24 | 0 | 0 | 0 |
| SRR5388007 | 0 | 13 | 21 | 0 | 0 | 0 | 0 | 18 | 0 | 0 | 0 | 0 | 0 | 18 | 50 | 0 | 0 | 0 | 50 | 942 | 248 | 0 | 0 | 36 | 0 | 53 | 58 | 0 | 0 | 0 | 0 | 0 | 0 | 0 | 0 | 0 | 0 |
| SRR5388008 | 0 | 10 | 41 | 0 | 0 | 0 | 0 | 0 | 0 | 0 | 0 | 0 | 0 | 0 | 0 | 0 | 0 | 0 | 0 | 612 | 245 | 0 | 0 | 97 | 0 | 55 | 128 | 0 | 0 | 0 | 0 | 0 | 0 | 0 | 0 | 0 | 0 |
| SRR5388009 | 0 | 0 | 0 | 0 | 0 | 10 | 0 | 10 | 0 | 0 | 0 | 0 | 0 | 0 | 0 | 0 | 0 | 0 | 0 | 518 | 207 | 0 | 0 | 103 | 0 | 123 | 298 | 0 | 0 | 0 | 0 | 0 | 0 | 10 | 10 | 0 | 0 |
| SRR5388010 | 0 | 0 | 11 | 0 | 0 | 15 | 0 | 0 | 0 | 0 | 0 | 0 | 0 | 15 | 332 | 0 | 0 | 0 | 0 | 886 | 222 | 0 | 0 | 128 | 0 | 279 | 422 | 0 | 0 | 0 | 0 | 0 | 0 | 15 | 0 | 0 | 0 |
| SRR5388011 | 0 | 0 | 0 | 0 | 0 | 11 | 0 | 0 | 11 | 0 | 0 | 0 | 0 | 23 | 0 | 0 | 0 | 0 | 0 | 1066 | 246 | 0 | 0 | 23 | 0 | 203 | 259 | 0 | 0 | 0 | 11 | 0 | 0 | 0 | 11 | 0 | 11 |
| SRR5388012 | 0 | 0 | 10 | 0 | 0 | 0 | 0 | 31 | 10 | 0 | 0 | 0 | 0 | 10 | 72 | 0 | 0 | 0 | 0 | 1512 | 360 | 0 | 0 | 63 | 10 | 293 | 168 | 10 | 0 | 0 | 0 | 0 | 0 | 10 | 10 | 0 | 10 |
| SRR5388013 | 0 | 0 | 16 | 0 | 0 | 0 | 0 | 18 | 0 | 0 | 0 | 0 | 0 | 0 | 38 | 0 | 0 | 0 | 38 | 1362 | 38 | 0 | 0 | 248 | 0 | 128 | 266 | 0 | 0 | 0 | 0 | 0 | 0 | 18 | 0 | 0 | 0 |
| SRR5388014 | 0 | 0 | 17 | 0 | 0 | 0 | 0 | 12 | 0 | 0 | 0 | 0 | 0 | 0 | 52 | 0 | 0 | 0 | 0 | 627 | 157 | 0 | 0 | 82 | 0 | 199 | 14 | 0 | 0 | 0 | 0 | 0 | 0 | 0 | 0 | 0 | 0 |
| SRR5388015 | 0 | 0 | 0 | 0 | 0 | 0 | 0 | 0 | 0 | 0 | 0 | 0 | 0 | 0 | 0 | 0 | 0 | 0 | 0 | 1013 | 0 | 0 | 0 | 104 | 0 | 235 | 270 | 0 | 0 | 0 | 0 | 0 | 0 | 17 | 0 | 0 | 0 |
| SRR5388016 | 0 | 29 | 33 | 0 | 0 | 23 | 0 | 53 | 0 | 0 | 0 | 16 | 0 | 0 | 47 | 0 | 0 | 0 | 0 | 699 | 233 | 0 | 0 | 129 | 0 | 158 | 479 | 12 | 0 | 0 | 12 | 12 | 0 | 18 | 0 | 0 | 12 |
| SRR5388017 | 0 | 0 | 0 | 0 | 0 | 29 | 0 | 38 | 0 | 0 | 0 | 0 | 0 | 0 | 0 | 0 | 0 | 0 | 0 | 708 | 59 | 0 | 0 | 96 | 0 | 201 | 297 | 0 | 0 | 0 | 0 | 0 | 0 | 19 | 0 | 0 | 0 |
| SRR5388018 | 0 | 44 | 0 | 0 | 0 | 0 | 0 | 0 | 0 | 0 | 0 | 0 | 0 | 0 | 0 | 0 | 0 | 0 | 0 | 596 | 85 | 0 | 0 | 66 | 0 | 132 | 257 | 51 | 0 | 0 | 0 | 59 | 0 | 0 | 0 | 0 | 0 |
| SRR5388019 | 0 | 11 | 12 | 0 | 11 | 11 | 0 | 0 | 11 | 0 | 0 | 0 | 0 | 0 | 180 | 0 | 0 | 0 | 0 | 1351 | 180 | 0 | 0 | 101 | 11 | 347 | 347 | 0 | 0 | 11 | 0 | 0 | 0 | 0 | 22 | 0 | 0 |
| SRR5388020 | 0 | 19 | 0 | 0 | 0 | 0 | 0 | 0 | 0 | 0 | 0 | 0 | 0 | 0 | 0 | 0 | 0 | 0 | 0 | 540 | 648 | 0 | 0 | 177 | 0 | 270 | 681 | 28 | 0 | 0 | 0 | 0 | 0 | 0 | 0 | 0 | 0 |
| SRR5388021 | 0 | 25 | 22 | 0 | 0 | 0 | 0 | 25 | 0 | 0 | 0 | 0 | 0 | 0 | 63 | 0 | 0 | 0 | 0 | 940 | 125 | 0 | 0 | 58 | 0 | 341 | 358 | 0 | 0 | 0 | 0 | 0 | 0 | 0 | 0 | 0 | 0 |
| SRR5388022 | 0 | 0 | 18 | 0 | 0 | 19 | 0 | 38 | 0 | 0 | 0 | 0 | 0 | 0 | 0 | 0 | 0 | 0 | 0 | 859 | 66 | 0 | 0 | 88 | 0 | 214 | 302 | 0 | 0 | 0 | 0 | 0 | 0 | 13 | 0 | 0 | 0 |
| SRR5388023 | 0 | 0 | 20 | 0 | 0 | 52 | 0 | 0 | 10 | 0 | 0 | 0 | 0 | 10 | 0 | 0 | 0 | 0 | 0 | 1639 | 0 | 0 | 0 | 73 | 0 | 514 | 608 | 10 | 0 | 0 | 31 | 0 | 0 | 0 | 0 | 0 | 0 |
| SRR5388024 | 0 | 0 | 11 | 0 | 0 | 16 | 0 | 0 | 0 | 0 | 0 | 0 | 0 | 0 | 234 | 0 | 0 | 0 | 0 | 1637 | 936 | 0 | 0 | 106 | 0 | 295 | 344 | 0 | 0 | 0 | 0 | 0 | 0 | 0 | 0 | 0 | 0 |
| SRR5388025 | 0 | 0 | 0 | 0 | 0 | 65 | 0 | 16 | 0 | 0 | 0 | 0 | 0 | 0 | 0 | 0 | 0 | 0 | 0 | 1515 | 95 | 0 | 0 | 163 | 0 | 506 | 441 | 0 | 0 | 0 | 0 | 0 | 0 | 0 | 16 | 0 | 0 |
| SRR5388026 | 0 | 15 | 0 | 0 | 0 | 46 | 0 | 31 | 46 | 0 | 0 | 0 | 0 | 15 | 0 | 0 | 0 | 0 | 0 | 1083 | 98 | 0 | 0 | 123 | 0 | 766 | 705 | 0 | 0 | 0 | 0 | 0 | 15 | 0 | 15 | 0 | 0 |
| SRR5388027 | 0 | 0 | 0 | 0 | 0 | 0 | 0 | 24 | 12 | 0 | 0 | 0 | 0 | 0 | 0 | 0 | 0 | 0 | 0 | 1430 | 191 | 0 | 0 | 72 | 12 | 346 | 299 | 0 | 12 | 0 | 0 | 0 | 0 | 0 | 0 | 0 | 12 |
| SRR5388028 | 0 | 0 | 14 | 0 | 0 | 0 | 0 | 19 | 0 | 0 | 0 | 0 | 0 | 0 | 111 | 0 | 0 | 0 | 0 | 1555 | 111 | 0 | 0 | 104 | 0 | 236 | 368 | 19 | 0 | 0 | 0 | 0 | 0 | 0 | 0 | 0 | 0 |
| SRR5388029 | 0 | 0 | 0 | 0 | 0 | 0 | 0 | 25 | 0 | 0 | 0 | 0 | 0 | 0 | 0 | 0 | 0 | 0 | 0 | 1301 | 65 | 0 | 0 | 141 | 17 | 158 | 299 | 17 | 0 | 0 | 0 | 0 | 0 | 0 | 0 | 0 | 0 |
| SRR5388030 | 0 | 0 | 11 | 0 | 0 | 34 | 0 | 17 | 0 | 0 | 0 | 0 | 0 | 0 | 0 | 0 | 0 | 0 | 0 | 242 | 161 | 0 | 0 | 127 | 0 | 161 | 424 | 0 | 0 | 0 | 0 | 0 | 0 | 0 | 17 | 0 | 0 |
| SRR5388031 | 0 | 16 | 0 | 0 | 0 | 0 | 0 | 0 | 0 | 0 | 0 | 0 | 0 | 0 | 59 | 0 | 0 | 0 | 0 | 614 | 117 | 0 | 0 | 36 | 0 | 80 | 164 | 12 | 0 | 0 | 0 | 0 | 0 | 0 | 0 | 0 | 0 |
| SRR5388032 | 0 | 105 | 0 | 0 | 0 | 0 | 0 | 11 | 0 | 0 | 0 | 0 | 0 | 0 | 0 | 0 | 0 | 0 | 0 | 750 | 0 | 0 | 0 | 39 | 0 | 127 | 226 | 72 | 0 | 0 | 0 | 55 | 11 | 0 | 0 | 0 | 0 |
| SRR5388033 | 0 | 20 | 0 | 20 | 0 | 29 | 0 | 0 | 0 | 0 | 0 | 0 | 0 | 0 | 0 | 0 | 0 | 0 | 64 | 1096 | 129 | 0 | 0 | 88 | 0 | 440 | 254 | 0 | 0 | 0 | 0 | 0 | 0 | 0 | 0 | 0 | 0 |
| SRR5388034 | 0 | 0 | 20 | 0 | 0 | 17 | 77 | 0 | 0 | 0 | 0 | 0 | 0 | 0 | 77 | 0 | 0 | 0 | 0 | 461 | 307 | 0 | 0 | 117 | 0 | 241 | 400 | 0 | 0 | 0 | 0 | 0 | 0 | 0 | 0 | 0 | 0 |
| SRR5388035 | 0 | 24 | 0 | 0 | 0 | 0 | 0 | 12 | 24 | 0 | 0 | 0 | 0 | 12 | 92 | 0 | 0 | 0 | 0 | 922 | 92 | 0 | 0 | 130 | 0 | 496 | 496 | 0 | 0 | 0 | 0 | 0 | 0 | 0 | 0 | 0 | 0 |
| SRR5388036 | 0 | 19 | 0 | 0 | 0 | 19 | 0 | 0 | 0 | 0 | 0 | 0 | 0 | 29 | 0 | 0 | 0 | 0 | 0 | 1274 | 85 | 0 | 0 | 145 | 0 | 280 | 473 | 0 | 0 | 0 | 19 | 0 | 0 | 0 | 0 | 0 | 0 |
| SRR5388037 | 0 | 0 | 0 | 0 | 0 | 16 | 41 | 0 | 0 | 0 | 0 | 0 | 0 | 0 | 82 | 0 | 0 | 0 | 82 | 1757 | 82 | 0 | 0 | 25 | 0 | 115 | 206 | 0 | 0 | 0 | 0 | 0 | 0 | 25 | 0 | 0 | 0 |
| SRR5388038 | 0 | 0 | 0 | 0 | 0 | 11 | 0 | 0 | 11 | 0 | 0 | 0 | 0 | 44 | 0 | 0 | 0 | 0 | 0 | 773 | 193 | 0 | 0 | 66 | 0 | 219 | 317 | 0 | 0 | 0 | 0 | 0 | 0 | 0 | 11 | 0 | 0 |
| SRR5388039 | 0 | 0 | 13 | 0 | 0 | 28 | 0 | 14 | 0 | 0 | 0 | 0 | 0 | 0 | 87 | 0 | 0 | 0 | 0 | 959 | 174 | 0 | 0 | 55 | 0 | 83 | 97 | 0 | 0 | 0 | 0 | 0 | 0 | 14 | 0 | 0 | 0 |
| SRR5388040 | 0 | 0 | 17 | 0 | 0 | 0 | 0 | 0 | 0 | 0 | 0 | 0 | 0 | 15 | 0 | 0 | 0 | 0 | 0 | 1467 | 140 | 0 | 0 | 67 | 0 | 164 | 335 | 0 | 0 | 0 | 0 | 0 | 0 | 0 | 0 | 0 | 15 |
| SRR5388041 | 0 | 0 | 0 | 0 | 0 | 18 | 0 | 0 | 0 | 0 | 0 | 0 | 0 | 0 | 0 | 0 | 0 | 0 | 0 | 1053 | 0 | 0 | 0 | 73 | 0 | 301 | 273 | 18 | 0 | 0 | 0 | 0 | 0 | 0 | 0 | 0 | 0 |
| SRR5388042 | 0 | 0 | 0 | 0 | 0 | 0 | 0 | 0 | 0 | 0 | 0 | 0 | 0 | 0 | 90 | 0 | 0 | 0 | 0 | 628 | 359 | 0 | 0 | 69 | 0 | 317 | 412 | 0 | 0 | 0 | 26 | 0 | 0 | 0 | 0 | 0 | 0 |
| SRR5388043 | 0 | 163 | 11 | 0 | 0 | 0 | 0 | 0 | 0 | 0 | 0 | 0 | 0 | 0 | 0 | 0 | 0 | 0 | 0 | 927 | 116 | 11 | 0 | 32 | 0 | 52 | 126 | 151 | 0 | 14 | 0 | 134 | 0 | 25 | 0 | 0 | 0 |
| SRR5388044 | 0 | 0 | 14 | 10 | 0 | 15 | 41 | 0 | 0 | 0 | 0 | 0 | 0 | 0 | 0 | 0 | 0 | 0 | 0 | 1100 | 245 | 0 | 0 | 166 | 0 | 166 | 257 | 0 | 0 | 10 | 0 | 0 | 0 | 15 | 0 | 0 | 0 |
| SRR5388045 | 0 | 16 | 30 | 0 | 0 | 16 | 0 | 0 | 0 | 0 | 0 | 0 | 0 | 0 | 0 | 0 | 0 | 0 | 0 | 1736 | 801 | 0 | 0 | 91 | 0 | 11 | 15 | 0 | 0 | 0 | 0 | 0 | 0 | 0 | 0 | 0 | 0 |
| SRR5388046 | 0 | 17 | 0 | 0 | 0 | 0 | 0 | 0 | 0 | 0 | 0 | 0 | 0 | 0 | 0 | 0 | 0 | 0 | 0 | 601 | 0 | 0 | 0 | 59 | 0 | 194 | 363 | 0 | 0 | 0 | 0 | 0 | 0 | 0 | 0 | 0 | 0 |
| SRR5388047 | 0 | 17 | 12 | 0 | 0 | 26 | 0 | 17 | 17 | 0 | 0 | 0 | 0 | 0 | 46 | 0 | 0 | 0 | 0 | 1386 | 92 | 0 | 0 | 149 | 0 | 271 | 341 | 0 | 0 | 0 | 0 | 0 | 0 | 0 | 0 | 0 | 0 |
| SRR5388048 | 0 | 16 | 10 | 0 | 0 | 25 | 0 | 0 | 0 | 0 | 0 | 0 | 0 | 0 | 0 | 0 | 0 | 0 | 0 | 750 | 115 | 16 | 0 | 106 | 0 | 262 | 335 | 0 | 0 | 0 | 0 | 0 | 0 | 16 | 0 | 0 | 16 |
| SRR5388049 | 0 | 0 | 13 | 0 | 0 | 0 | 0 | 0 | 0 | 0 | 0 | 0 | 0 | 0 | 139 | 0 | 0 | 0 | 0 | 487 | 348 | 0 | 0 | 67 | 0 | 335 | 447 | 0 | 0 | 0 | 0 | 0 | 0 | 0 | 0 | 0 | 0 |
| SRR5388050 | 0 | 12 | 14 | 0 | 0 | 19 | 0 | 0 | 0 | 0 | 0 | 0 | 0 | 0 | 0 | 0 | 0 | 0 | 0 | 651 | 195 | 0 | 0 | 93 | 12 | 236 | 11 | 0 | 0 | 0 | 0 | 0 | 0 | 0 | 12 | 0 | 12 |
| SRR5388051 | 0 | 0 | 18 | 0 | 0 | 20 | 0 | 14 | 0 | 0 | 0 | 0 | 0 | 0 | 99 | 0 | 0 | 0 | 0 | 991 | 297 | 0 | 0 | 75 | 0 | 54 | 95 | 0 | 0 | 0 | 0 | 0 | 0 | 0 | 0 | 0 | 0 |
| SRR5388052 | 0 | 0 | 15 | 0 | 0 | 0 | 0 | 17 | 0 | 0 | 0 | 0 | 0 | 0 | 0 | 0 | 0 | 0 | 0 | 803 | 0 | 0 | 0 | 91 | 0 | 103 | 177 | 0 | 0 | 0 | 0 | 0 | 0 | 17 | 29 | 0 | 0 |
| SRR5388053 | 0 | 0 | 0 | 0 | 0 | 19 | 0 | 12 | 0 | 0 | 0 | 0 | 0 | 12 | 49 | 0 | 0 | 0 | 0 | 394 | 197 | 0 | 0 | 99 | 0 | 253 | 284 | 0 | 0 | 0 | 0 | 0 | 0 | 31 | 0 | 0 | 12 |
| SRR5388054 | 0 | 0 | 0 | 0 | 0 | 0 | 0 | 0 | 0 | 0 | 0 | 0 | 0 | 0 | 0 | 0 | 0 | 0 | 0 | 624 | 104 | 15 | 0 | 113 | 0 | 302 | 257 | 0 | 0 | 0 | 0 | 0 | 0 | 0 | 0 | 0 | 0 |
| SRR5388055 | 0 | 0 | 0 | 0 | 0 | 28 | 0 | 0 | 0 | 0 | 0 | 0 | 0 | 0 | 82 | 0 | 0 | 0 | 0 | 657 | 246 | 0 | 0 | 171 | 0 | 209 | 446 | 0 | 0 | 0 | 0 | 0 | 0 | 0 | 0 | 0 | 0 |
| SRR5388056 | 13 | 15 | 0 | 0 | 0 | 39 | 0 | 0 | 15 | 0 | 0 | 0 | 0 | 0 | 0 | 0 | 0 | 0 | 0 | 1145 | 82 | 0 | 0 | 131 | 0 | 170 | 271 | 0 | 0 | 0 | 0 | 0 | 0 | 0 | 0 | 0 | 0 |
| SRR5388057 | 0 | 0 | 0 | 0 | 0 | 0 | 0 | 18 | 0 | 0 | 0 | 0 | 0 | 0 | 112 | 0 | 0 | 0 | 0 | 562 | 337 | 0 | 0 | 70 | 0 | 229 | 273 | 0 | 0 | 0 | 0 | 0 | 0 | 0 | 0 | 0 | 0 |
| SRR5388058 | 0 | 18 | 19 | 0 | 0 | 0 | 0 | 12 | 0 | 0 | 0 | 0 | 0 | 0 | 177 | 0 | 0 | 0 | 89 | 1152 | 177 | 0 | 0 | 85 | 12 | 146 | 267 | 0 | 0 | 0 | 0 | 0 | 0 | 0 | 0 | 0 | 12 |
| SRR5388059 | 0 | 14 | 0 | 0 | 0 | 22 | 0 | 0 | 0 | 0 | 0 | 0 | 0 | 29 | 90 | 0 | 0 | 0 | 0 | 1258 | 270 | 0 | 0 | 101 | 0 | 216 | 339 | 0 | 0 | 0 | 0 | 0 | 0 | 0 | 0 | 0 | 0 |
| SRR5388060 | 0 | 20 | 0 | 0 | 0 | 0 | 0 | 0 | 0 | 0 | 0 | 0 | 0 | 0 | 262 | 0 | 0 | 0 | 0 | 698 | 175 | 0 | 0 | 108 | 0 | 276 | 350 | 34 | 0 | 0 | 0 | 0 | 0 | 13 | 0 | 0 | 0 |
| SRR5388061 | 0 | 0 | 0 | 0 | 0 | 23 | 0 | 0 | 0 | 0 | 0 | 0 | 0 | 0 | 0 | 0 | 0 | 0 | 0 | 624 | 374 | 0 | 0 | 193 | 0 | 228 | 421 | 0 | 0 | 0 | 11 | 0 | 0 | 0 | 0 | 0 | 0 |
| SRR5388062 | 0 | 0 | 13 | 0 | 0 | 0 | 0 | 86 | 0 | 0 | 0 | 0 | 0 | 0 | 0 | 0 | 0 | 0 | 0 | 1202 | 100 | 0 | 0 | 133 | 0 | 180 | 274 | 0 | 0 | 0 | 0 | 0 | 0 | 16 | 0 | 0 | 0 |
| SRR5388063 | 0 | 11 | 0 | 0 | 0 | 53 | 0 | 43 | 11 | 0 | 0 | 0 | 0 | 0 | 0 | 0 | 0 | 0 | 0 | 702 | 70 | 0 | 0 | 75 | 0 | 256 | 384 | 0 | 0 | 0 | 0 | 0 | 0 | 0 | 0 | 0 | 0 |
| SRR5388064 | 0 | 0 | 0 | 0 | 0 | 0 | 0 | 27 | 0 | 0 | 0 | 0 | 0 | 0 | 239 | 0 | 0 | 0 | 0 | 718 | 160 | 0 | 0 | 124 | 0 | 133 | 274 | 0 | 0 | 0 | 0 | 0 | 0 | 18 | 18 | 0 | 0 |
| SRR5388065 | 0 | 0 | 0 | 0 | 0 | 28 | 0 | 0 | 0 | 0 | 0 | 0 | 0 | 0 | 0 | 0 | 0 | 0 | 0 | 2095 | 0 | 0 | 0 | 28 | 0 | 375 | 472 | 0 | 0 | 0 | 14 | 0 | 0 | 0 | 14 | 0 | 0 |
| SRR5388066 | 0 | 0 | 0 | 0 | 0 | 0 | 0 | 27 | 0 | 0 | 0 | 0 | 0 | 0 | 0 | 0 | 0 | 0 | 0 | 1283 | 112 | 0 | 0 | 37 | 0 | 302 | 192 | 27 | 0 | 0 | 0 | 0 | 0 | 18 | 0 | 0 | 0 |
| SRR5388067 | 11 | 0 | 27 | 0 | 0 | 0 | 0 | 0 | 0 | 0 | 0 | 0 | 0 | 0 | 0 | 0 | 0 | 0 | 0 | 1065 | 609 | 0 | 0 | 135 | 0 | 292 | 479 | 0 | 0 | 0 | 0 | 0 | 0 | 0 | 15 | 0 | 0 |
| SRR5388068 | 0 | 0 | 25 | 0 | 0 | 26 | 0 | 17 | 0 | 0 | 0 | 0 | 0 | 0 | 0 | 0 | 0 | 0 | 0 | 1003 | 0 | 0 | 0 | 86 | 0 | 120 | 351 | 0 | 0 | 17 | 0 | 0 | 0 | 0 | 0 | 0 | 0 |
| SRR5388069 | 0 | 24 | 11 | 0 | 0 | 0 | 0 | 16 | 0 | 0 | 0 | 0 | 0 | 0 | 0 | 0 | 0 | 0 | 0 | 1183 | 278 | 0 | 0 | 87 | 0 | 182 | 435 | 0 | 0 | 0 | 16 | 0 | 0 | 0 | 0 | 0 | 0 |
| SRR5388070 | 0 | 0 | 13 | 0 | 0 | 24 | 0 | 16 | 0 | 0 | 0 | 0 | 0 | 0 | 0 | 0 | 0 | 0 | 0 | 852 | 0 | 0 | 0 | 89 | 0 | 267 | 534 | 0 | 0 | 0 | 0 | 0 | 0 | 0 | 0 | 0 | 0 |
| SRR5388071 | 0 | 36 | 0 | 0 | 0 | 0 | 0 | 0 | 12 | 12 | 0 | 0 | 0 | 0 | 90 | 0 | 0 | 0 | 90 | 1537 | 271 | 0 | 0 | 60 | 0 | 72 | 120 | 0 | 12 | 0 | 0 | 0 | 0 | 12 | 12 | 0 | 0 |
| SRR5388072 | 0 | 0 | 0 | 0 | 0 | 0 | 0 | 0 | 0 | 0 | 0 | 0 | 0 | 0 | 138 | 0 | 0 | 0 | 0 | 963 | 0 | 0 | 0 | 99 | 0 | 208 | 326 | 0 | 20 | 0 | 0 | 0 | 0 | 0 | 0 | 0 | 0 |
| SRR5388073 | 0 | 0 | 0 | 0 | 0 | 27 | 0 | 36 | 0 | 0 | 0 | 0 | 0 | 0 | 65 | 0 | 0 | 0 | 0 | 3071 | 392 | 0 | 0 | 64 | 0 | 218 | 245 | 0 | 0 | 0 | 0 | 0 | 0 | 18 | 0 | 0 | 0 |
| SRR5388074 | 0 | 0 | 13 | 0 | 0 | 19 | 0 | 0 | 0 | 0 | 0 | 0 | 0 | 0 | 0 | 0 | 0 | 0 | 0 | 1138 | 152 | 0 | 0 | 173 | 0 | 221 | 337 | 0 | 0 | 0 | 0 | 0 | 0 | 0 | 0 | 0 | 0 |
| SRR5388075 | 0 | 15 | 27 | 0 | 0 | 0 | 0 | 15 | 0 | 0 | 0 | 17 | 0 | 0 | 156 | 0 | 0 | 0 | 0 | 730 | 209 | 0 | 0 | 59 | 0 | 198 | 337 | 0 | 0 | 0 | 0 | 0 | 0 | 0 | 0 | 0 | 0 |
| SRR5388076 | 0 | 25 | 12 | 17 | 0 | 0 | 0 | 0 | 0 | 0 | 0 | 0 | 0 | 0 | 0 | 0 | 0 | 0 | 0 | 925 | 77 | 0 | 0 | 51 | 0 | 263 | 441 | 0 | 0 | 0 | 0 | 0 | 0 | 0 | 0 | 0 | 25 |
| SRR5388077 | 0 | 0 | 0 | 0 | 0 | 31 | 0 | 10 | 0 | 0 | 0 | 0 | 0 | 0 | 0 | 0 | 0 | 0 | 0 | 1601 | 283 | 0 | 0 | 84 | 0 | 167 | 198 | 0 | 0 | 0 | 0 | 0 | 0 | 21 | 10 | 0 | 0 |
| SRR5388078 | 0 | 35 | 15 | 19 | 0 | 0 | 0 | 0 | 0 | 0 | 0 | 0 | 0 | 0 | 0 | 0 | 0 | 0 | 0 | 755 | 0 | 0 | 0 | 26 | 25 | 17 | 54 | 15 | 0 | 0 | 0 | 11 | 0 | 0 | 0 | 0 | 0 |
| SRR5388079 | 0 | 0 | 0 | 0 | 0 | 38 | 0 | 38 | 19 | 0 | 0 | 0 | 0 | 19 | 0 | 0 | 0 | 0 | 0 | 1146 | 164 | 0 | 0 | 125 | 0 | 375 | 298 | 0 | 0 | 0 | 19 | 0 | 0 | 0 | 0 | 0 | 0 |
| SRR5388080 | 0 | 0 | 12 | 0 | 0 | 43 | 0 | 17 | 0 | 0 | 0 | 0 | 0 | 0 | 98 | 0 | 0 | 0 | 0 | 1767 | 196 | 0 | 0 | 34 | 0 | 385 | 454 | 0 | 0 | 0 | 0 | 0 | 0 | 0 | 0 | 0 | 0 |
| SRR5388081 | 0 | 0 | 0 | 0 | 0 | 43 | 0 | 34 | 0 | 0 | 0 | 0 | 0 | 0 | 133 | 0 | 0 | 0 | 0 | 1259 | 331 | 0 | 0 | 86 | 0 | 163 | 120 | 0 | 0 | 0 | 0 | 0 | 0 | 0 | 0 | 0 | 0 |
| SRR5388082 | 0 | 0 | 13 | 13 | 0 | 0 | 0 | 0 | 0 | 0 | 0 | 0 | 0 | 0 | 0 | 0 | 0 | 0 | 0 | 1009 | 63 | 0 | 0 | 53 | 0 | 173 | 286 | 0 | 0 | 0 | 0 | 0 | 0 | 13 | 0 | 0 | 13 |
| SRR5388083 | 0 | 15 | 0 | 0 | 0 | 23 | 0 | 0 | 0 | 0 | 0 | 0 | 0 | 0 | 0 | 0 | 0 | 0 | 0 | 981 | 54 | 0 | 0 | 161 | 0 | 177 | 284 | 0 | 0 | 0 | 0 | 0 | 0 | 0 | 0 | 0 | 0 |
| SRR5388084 | 0 | 19 | 0 | 0 | 0 | 0 | 0 | 28 | 0 | 0 | 0 | 0 | 0 | 0 | 0 | 0 | 0 | 0 | 0 | 1892 | 146 | 0 | 0 | 74 | 0 | 195 | 372 | 0 | 0 | 0 | 0 | 0 | 0 | 0 | 0 | 0 | 0 |
| SRR5388085 | 0 | 10 | 0 | 0 | 0 | 0 | 0 | 10 | 0 | 0 | 0 | 0 | 0 | 0 | 0 | 0 | 0 | 0 | 0 | 1260 | 110 | 0 | 0 | 31 | 0 | 105 | 272 | 0 | 0 | 0 | 0 | 0 | 0 | 10 | 0 | 0 | 0 |
| SRR5388086 | 0 | 0 | 0 | 0 | 0 | 17 | 0 | 0 | 0 | 0 | 0 | 0 | 0 | 0 | 0 | 0 | 0 | 0 | 72 | 1302 | 0 | 0 | 0 | 78 | 0 | 157 | 383 | 0 | 0 | 0 | 0 | 0 | 0 | 0 | 17 | 0 | 0 |
| SRR5388087 | 0 | 23 | 0 | 0 | 0 | 15 | 30 | 30 | 0 | 0 | 0 | 0 | 0 | 0 | 0 | 0 | 0 | 0 | 0 | 1713 | 118 | 0 | 0 | 15 | 0 | 343 | 190 | 0 | 0 | 0 | 0 | 0 | 0 | 38 | 15 | 0 | 0 |
| SRR5388088 | 0 | 0 | 13 | 0 | 0 | 20 | 0 | 13 | 0 | 0 | 0 | 0 | 0 | 0 | 0 | 0 | 0 | 0 | 0 | 1212 | 0 | 0 | 0 | 98 | 0 | 16 | 18 | 13 | 0 | 0 | 20 | 0 | 0 | 0 | 13 | 0 | 0 |
| SRR5388089 | 0 | 0 | 11 | 0 | 0 | 0 | 63 | 38 | 0 | 0 | 0 | 0 | 0 | 23 | 0 | 0 | 0 | 0 | 0 | 1706 | 126 | 0 | 0 | 108 | 0 | 131 | 331 | 0 | 0 | 0 | 15 | 0 | 0 | 15 | 0 | 0 | 0 |
| SRR5388090 | 0 | 12 | 12 | 0 | 0 | 0 | 0 | 31 | 0 | 0 | 0 | 0 | 0 | 18 | 0 | 0 | 0 | 0 | 0 | 540 | 135 | 0 | 0 | 129 | 0 | 98 | 264 | 0 | 0 | 0 | 0 | 0 | 0 | 25 | 0 | 0 | 0 |
| SRR5388091 | 0 | 11 | 11 | 0 | 0 | 16 | 0 | 16 | 0 | 0 | 0 | 0 | 0 | 0 | 0 | 0 | 0 | 0 | 70 | 1047 | 209 | 0 | 0 | 91 | 0 | 107 | 209 | 0 | 0 | 0 | 0 | 0 | 0 | 0 | 0 | 0 | 0 |
| SRR5388092 | 0 | 0 | 0 | 0 | 0 | 0 | 0 | 0 | 0 | 0 | 0 | 0 | 0 | 0 | 0 | 0 | 0 | 0 | 0 | 1068 | 200 | 0 | 0 | 121 | 0 | 121 | 161 | 0 | 0 | 0 | 0 | 0 | 0 | 0 | 0 | 0 | 0 |
| SRR5388093 | 21 | 13 | 43 | 0 | 0 | 20 | 0 | 0 | 0 | 0 | 0 | 0 | 0 | 0 | 82 | 0 | 0 | 0 | 0 | 1230 | 410 | 0 | 0 | 10 | 24 | 31 | 54 | 0 | 0 | 0 | 0 | 0 | 0 | 0 | 0 | 0 | 0 |
| SRR5388094 | 0 | 17 | 19 | 15 | 0 | 0 | 0 | 0 | 0 | 0 | 0 | 0 | 0 | 0 | 0 | 12 | 0 | 0 | 0 | 702 | 255 | 0 | 0 | 31 | 0 | 52 | 102 | 19 | 0 | 0 | 0 | 15 | 0 | 0 | 0 | 0 | 0 |
| SRR5388095 | 0 | 0 | 0 | 0 | 0 | 12 | 105 | 0 | 0 | 0 | 0 | 0 | 0 | 12 | 0 | 0 | 0 | 0 | 0 | 1150 | 105 | 0 | 0 | 50 | 0 | 297 | 310 | 0 | 0 | 0 | 0 | 0 | 0 | 0 | 12 | 0 | 12 |
| SRR5388096 | 0 | 0 | 18 | 0 | 0 | 14 | 0 | 14 | 0 | 0 | 0 | 0 | 0 | 0 | 0 | 0 | 0 | 0 | 0 | 742 | 148 | 0 | 0 | 122 | 0 | 151 | 259 | 0 | 0 | 0 | 14 | 0 | 0 | 0 | 0 | 0 | 0 |
| SRR5388097 | 0 | 19 | 0 | 0 | 0 | 38 | 0 | 28 | 0 | 0 | 0 | 0 | 0 | 0 | 0 | 0 | 0 | 0 | 197 | 492 | 98 | 0 | 0 | 152 | 0 | 303 | 559 | 0 | 0 | 0 | 0 | 0 | 0 | 19 | 0 | 0 | 0 |
| SRR5388098 | 0 | 0 | 0 | 0 | 0 | 35 | 0 | 12 | 0 | 0 | 0 | 0 | 0 | 0 | 0 | 0 | 0 | 0 | 0 | 811 | 348 | 0 | 0 | 153 | 0 | 340 | 399 | 0 | 0 | 12 | 12 | 0 | 12 | 0 | 0 | 0 | 0 |
| SRR5388099 | 0 | 0 | 0 | 0 | 0 | 55 | 0 | 0 | 16 | 0 | 0 | 0 | 0 | 16 | 177 | 0 | 0 | 0 | 89 | 974 | 443 | 0 | 0 | 103 | 0 | 434 | 331 | 16 | 0 | 0 | 16 | 0 | 0 | 0 | 0 | 0 | 0 |
| SRR5388100 | 0 | 0 | 22 | 0 | 0 | 33 | 0 | 11 | 11 | 0 | 0 | 0 | 0 | 11 | 0 | 0 | 0 | 0 | 0 | 1118 | 112 | 0 | 0 | 76 | 0 | 327 | 360 | 0 | 0 | 0 | 22 | 0 | 0 | 11 | 0 | 0 | 0 |
| SRR5388101 | 0 | 0 | 0 | 0 | 0 | 0 | 0 | 0 | 0 | 0 | 0 | 0 | 0 | 0 | 75 | 0 | 0 | 0 | 150 | 1276 | 75 | 0 | 0 | 92 | 0 | 330 | 275 | 0 | 0 | 0 | 0 | 0 | 0 | 0 | 0 | 0 | 0 |
| SRR5388102 | 0 | 0 | 0 | 0 | 0 | 26 | 0 | 26 | 0 | 0 | 0 | 0 | 0 | 0 | 0 | 0 | 0 | 0 | 0 | 1273 | 0 | 0 | 0 | 121 | 0 | 121 | 311 | 0 | 0 | 0 | 0 | 0 | 0 | 0 | 0 | 0 | 0 |
| SRR5388103 | 0 | 11 | 0 | 0 | 0 | 21 | 0 | 21 | 0 | 0 | 0 | 0 | 0 | 11 | 0 | 0 | 0 | 0 | 0 | 629 | 236 | 0 | 0 | 85 | 0 | 309 | 234 | 21 | 0 | 0 | 0 | 0 | 21 | 0 | 0 | 0 | 0 |
| SRR5388104 | 0 | 0 | 0 | 0 | 0 | 0 | 0 | 0 | 0 | 0 | 0 | 0 | 0 | 0 | 384 | 0 | 0 | 0 | 0 | 1153 | 480 | 0 | 0 | 81 | 0 | 125 | 294 | 0 | 0 | 0 | 0 | 0 | 0 | 0 | 0 | 0 | 0 |
| SRR5388105 | 0 | 25 | 13 | 12 | 0 | 12 | 0 | 12 | 12 | 0 | 0 | 0 | 0 | 12 | 0 | 0 | 0 | 0 | 0 | 777 | 0 | 0 | 0 | 137 | 0 | 299 | 361 | 0 | 0 | 0 | 12 | 0 | 0 | 12 | 0 | 0 | 0 |
| SRR5388106 | 0 | 0 | 0 | 0 | 0 | 0 | 0 | 17 | 0 | 0 | 0 | 0 | 0 | 0 | 0 | 0 | 0 | 0 | 0 | 806 | 0 | 0 | 0 | 74 | 0 | 124 | 290 | 0 | 0 | 0 | 0 | 0 | 0 | 0 | 0 | 0 | 0 |
| SRR5388107 | 0 | 24 | 0 | 0 | 0 | 12 | 0 | 0 | 0 | 0 | 0 | 0 | 0 | 0 | 0 | 0 | 0 | 0 | 0 | 677 | 97 | 0 | 0 | 84 | 0 | 313 | 277 | 12 | 0 | 0 | 12 | 0 | 0 | 12 | 0 | 0 | 0 |
| SRR5388108 | 0 | 14 | 0 | 0 | 0 | 0 | 0 | 0 | 0 | 0 | 0 | 0 | 0 | 0 | 0 | 0 | 0 | 0 | 0 | 616 | 176 | 0 | 0 | 79 | 0 | 121 | 193 | 0 | 0 | 0 | 0 | 0 | 0 | 21 | 0 | 0 | 0 |
| SRR5388109 | 12 | 0 | 22 | 0 | 0 | 0 | 0 | 57 | 0 | 0 | 0 | 0 | 0 | 0 | 177 | 0 | 0 | 0 | 59 | 1358 | 708 | 0 | 0 | 74 | 0 | 154 | 11 | 0 | 0 | 0 | 0 | 0 | 17 | 23 | 0 | 0 | 23 |
| SRR5388110 | 0 | 0 | 31 | 0 | 0 | 0 | 0 | 24 | 0 | 0 | 0 | 11 | 0 | 14 | 158 | 0 | 0 | 0 | 0 | 1260 | 210 | 0 | 0 | 130 | 0 | 154 | 270 | 0 | 0 | 0 | 0 | 0 | 0 | 0 | 0 | 0 | 0 |
| SRR5388111 | 0 | 0 | 20 | 0 | 0 | 0 | 0 | 31 | 0 | 0 | 0 | 13 | 0 | 12 | 219 | 0 | 0 | 0 | 0 | 512 | 73 | 0 | 0 | 112 | 0 | 56 | 180 | 0 | 0 | 0 | 0 | 0 | 0 | 25 | 0 | 0 | 12 |
| SRR5388112 | 0 | 0 | 11 | 0 | 0 | 0 | 0 | 22 | 0 | 0 | 0 | 0 | 0 | 16 | 0 | 0 | 0 | 0 | 72 | 1293 | 790 | 11 | 0 | 115 | 0 | 93 | 274 | 0 | 0 | 0 | 0 | 0 | 0 | 0 | 0 | 0 | 16 |
| SRR5388113 | 0 | 0 | 0 | 0 | 0 | 0 | 0 | 15 | 0 | 0 | 0 | 0 | 0 | 0 | 0 | 0 | 0 | 0 | 0 | 732 | 651 | 0 | 0 | 77 | 0 | 116 | 77 | 0 | 0 | 0 | 0 | 0 | 0 | 0 | 0 | 0 | 0 |
| SRR5388114 | 0 | 0 | 33 | 0 | 0 | 12 | 0 | 0 | 0 | 0 | 0 | 0 | 0 | 17 | 0 | 0 | 0 | 0 | 0 | 731 | 0 | 0 | 0 | 104 | 12 | 196 | 242 | 0 | 0 | 0 | 12 | 0 | 0 | 0 | 0 | 0 | 0 |

### Supp Table S8.2.Columns_AM_to_BW

| | | | | | | | | | | | | | | | | | | | | | | | | | | | | | | | | | | | | | |
|---|---|---|---|---|---|---|---|---|---|---|---|---|---|---|---|---|---|---|---|---|---|---|---|---|---|---|---|---|---|---|---|---|---|---|---|---|---|
| SRR5388115 | 0 | 0 | 20 | 0 | 16 | 0 | 0 | 0 | 0 | 0 | 0 | 16 | 0 | 0 | 0 | 0 | 0 | 0 | 0 | 1300 | 200 | 0 | 0 | 180 | 0 | 401 | 12 | 0 | 0 | 0 | 0 | 0 | 0 | 25 | 16 | 0 | 0 |
| SRR5388116 | 0 | 0 | 11 | 0 | 0 | 29 | 0 | 0 | 0 | 0 | 0 | 0 | 0 | 0 | 0 | 0 | 0 | 0 | 0 | 809 | 81 | 0 | 0 | 135 | 0 | 231 | 298 | 0 | 0 | 0 | 19 | 0 | 0 | 0 | 0 | 0 | 0 |
| SRR5388117 | 0 | 0 | 0 | 0 | 0 | 0 | 0 | 0 | 0 | 0 | 0 | 0 | 0 | 0 | 0 | 0 | 0 | 0 | 35 | 1434 | 35 | 0 | 0 | 103 | 0 | 78 | 107 | 0 | 0 | 0 | 0 | 0 | 0 | 0 | 0 | 0 | 0 |
| SRR5388118 | 0 | 0 | 12 | 0 | 0 | 0 | 0 | 0 | 0 | 0 | 0 | 0 | 0 | 0 | 0 | 0 | 0 | 0 | 60 | 1266 | 60 | 0 | 0 | 117 | 0 | 170 | 117 | 0 | 0 | 0 | 0 | 0 | 0 | 0 | 0 | 0 | 0 |
| SRR5388119 | 0 | 0 | 23 | 0 | 0 | 18 | 0 | 0 | 0 | 0 | 0 | 0 | 0 | 18 | 0 | 0 | 0 | 0 | 174 | 2174 | 696 | 0 | 0 | 116 | 0 | 304 | 455 | 0 | 0 | 18 | 0 | 0 | 0 | 0 | 18 | 0 | 0 |
| SRR5388120 | 14 | 13 | 49 | 0 | 0 | 0 | 0 | 13 | 0 | 0 | 0 | 0 | 0 | 0 | 231 | 0 | 0 | 0 | 0 | 1620 | 555 | 0 | 0 | 59 | 0 | 139 | 264 | 0 | 0 | 0 | 0 | 0 | 0 | 0 | 0 | 0 | 0 |
| SRR5388121 | 0 | 0 | 11 | 0 | 0 | 0 | 0 | 0 | 0 | 0 | 0 | 0 | 0 | 0 | 0 | 0 | 0 | 0 | 0 | 1201 | 114 | 0 | 0 | 155 | 0 | 325 | 384 | 0 | 0 | 0 | 0 | 0 | 0 | 0 | 0 | 0 | 15 |
| SRR5388122 | 0 | 0 | 0 | 0 | 0 | 0 | 0 | 0 | 0 | 0 | 0 | 0 | 0 | 0 | 0 | 0 | 0 | 0 | 64 | 1020 | 128 | 0 | 0 | 160 | 0 | 388 | 480 | 0 | 0 | 0 | 34 | 0 | 0 | 0 | 0 | 0 | 0 |
| SRR5388123 | 0 | 12 | 14 | 0 | 0 | 12 | 0 | 12 | 12 | 0 | 0 | 0 | 0 | 0 | 0 | 0 | 0 | 0 | 0 | 890 | 0 | 0 | 0 | 61 | 0 | 98 | 172 | 0 | 0 | 0 | 0 | 0 | 0 | 0 | 12 | 0 | 0 |
| SRR5388124 | 0 | 0 | 14 | 0 | 0 | 35 | 0 | 18 | 0 | 0 | 0 | 0 | 0 | 0 | 61 | 0 | 0 | 0 | 0 | 549 | 0 | 0 | 0 | 150 | 0 | 106 | 273 | 0 | 0 | 0 | 0 | 0 | 0 | 0 | 0 | 0 | 0 |
| SRR5388125 | 0 | 11 | 10 | 33 | 0 | 0 | 0 | 0 | 0 | 0 | 0 | 0 | 0 | 0 | 0 | 0 | 0 | 0 | 0 | 1410 | 371 | 0 | 0 | 54 | 0 | 92 | 179 | 16 | 0 | 11 | 0 | 0 | 11 | 0 | 0 | 0 | 0 |
| SRR5388126 | 0 | 0 | 0 | 0 | 0 | 37 | 0 | 0 | 0 | 0 | 0 | 0 | 0 | 0 | 0 | 0 | 0 | 0 | 0 | 885 | 221 | 0 | 0 | 124 | 12 | 286 | 584 | 12 | 0 | 0 | 0 | 0 | 0 | 0 | 25 | 0 | 0 |
| SRR5388127 | 0 | 22 | 0 | 0 | 0 | 65 | 0 | 0 | 0 | 0 | 0 | 0 | 0 | 0 | 0 | 0 | 0 | 0 | 0 | 475 | 0 | 0 | 0 | 140 | 11 | 312 | 721 | 0 | 0 | 0 | 0 | 0 | 0 | 11 | 0 | 0 | 22 |
| SRR5388128 | 0 | 0 | 12 | 0 | 0 | 16 | 0 | 0 | 0 | 0 | 0 | 0 | 0 | 0 | 0 | 0 | 0 | 0 | 0 | 715 | 130 | 0 | 0 | 109 | 0 | 327 | 507 | 0 | 0 | 0 | 0 | 0 | 0 | 0 | 0 | 0 | 16 |
| SRR5388129 | 0 | 0 | 0 | 0 | 10 | 10 | 0 | 31 | 0 | 0 | 0 | 0 | 0 | 0 | 0 | 0 | 0 | 0 | 0 | 794 | 132 | 0 | 0 | 61 | 0 | 164 | 205 | 0 | 0 | 0 | 0 | 0 | 0 | 10 | 0 | 0 | 0 |
| SRR5388130 | 0 | 15 | 0 | 0 | 0 | 31 | 0 | 23 | 0 | 0 | 0 | 0 | 0 | 0 | 0 | 0 | 0 | 0 | 92 | 1283 | 92 | 0 | 0 | 100 | 0 | 483 | 398 | 0 | 0 | 0 | 0 | 0 | 0 | 23 | 0 | 0 | 0 |
| SRR5388131 | 0 | 0 | 0 | 0 | 0 | 29 | 0 | 0 | 0 | 0 | 0 | 0 | 0 | 0 | 66 | 0 | 0 | 0 | 66 | 1247 | 66 | 0 | 0 | 43 | 0 | 72 | 301 | 0 | 0 | 0 | 0 | 0 | 0 | 0 | 0 | 0 | 14 |
| SRR5388132 | 0 | 0 | 14 | 0 | 0 | 0 | 0 | 17 | 0 | 0 | 0 | 0 | 0 | 0 | 0 | 0 | 0 | 0 | 0 | 1632 | 131 | 0 | 0 | 71 | 0 | 125 | 379 | 0 | 0 | 0 | 0 | 0 | 0 | 0 | 0 | 0 | 0 |
| SRR5388133 | 0 | 20 | 0 | 0 | 0 | 20 | 0 | 69 | 0 | 0 | 0 | 0 | 0 | 0 | 0 | 0 | 0 | 0 | 42 | 754 | 293 | 0 | 0 | 109 | 0 | 79 | 79 | 0 | 0 | 0 | 0 | 0 | 0 | 0 | 0 | 0 | 0 |
| SRR5388134 | 12 | 0 | 30 | 0 | 0 | 0 | 65 | 17 | 0 | 0 | 0 | 10 | 0 | 0 | 0 | 0 | 0 | 0 | 0 | 916 | 262 | 0 | 0 | 112 | 0 | 133 | 129 | 0 | 0 | 0 | 0 | 0 | 0 | 0 | 0 | 0 | 0 |
| SRR5388135 | 0 | 16 | 17 | 0 | 0 | 16 | 0 | 16 | 0 | 0 | 0 | 0 | 0 | 16 | 0 | 0 | 0 | 0 | 0 | 1802 | 212 | 0 | 0 | 78 | 0 | 334 | 419 | 0 | 0 | 0 | 16 | 0 | 0 | 0 | 0 | 0 | 0 |
| SRR5388136 | 0 | 0 | 0 | 0 | 0 | 39 | 141 | 0 | 0 | 0 | 0 | 0 | 0 | 0 | 0 | 0 | 0 | 0 | 0 | 1267 | 141 | 0 | 0 | 116 | 0 | 406 | 367 | 0 | 0 | 0 | 0 | 0 | 0 | 0 | 0 | 0 | 0 |
| SRR5388137 | 0 | 37 | 0 | 0 | 0 | 0 | 0 | 19 | 0 | 0 | 0 | 0 | 0 | 0 | 0 | 0 | 0 | 0 | 0 | 1180 | 84 | 0 | 0 | 118 | 12 | 143 | 242 | 19 | 0 | 0 | 0 | 12 | 0 | 0 | 0 | 0 | 0 |
| SRR5388138 | 0 | 0 | 21 | 0 | 12 | 0 | 0 | 0 | 12 | 0 | 0 | 0 | 0 | 0 | 0 | 0 | 0 | 0 | 75 | 901 | 375 | 0 | 0 | 153 | 0 | 47 | 100 | 12 | 0 | 0 | 0 | 0 | 0 | 0 | 0 | 0 | 0 |
| SRR5388139 | 0 | 0 | 0 | 0 | 0 | 0 | 0 | 18 | 0 | 0 | 0 | 0 | 0 | 0 | 0 | 0 | 0 | 0 | 0 | 1089 | 91 | 0 | 0 | 65 | 0 | 379 | 517 | 0 | 0 | 0 | 0 | 0 | 0 | 0 | 0 | 0 | 0 |
| SRR5388140 | 0 | 0 | 25 | 0 | 0 | 0 | 39 | 0 | 0 | 0 | 0 | 0 | 0 | 0 | 39 | 0 | 0 | 0 | 0 | 1358 | 116 | 0 | 0 | 104 | 0 | 267 | 12 | 13 | 0 | 0 | 0 | 0 | 0 | 0 | 0 | 0 | 0 |
| SRR5388141 | 0 | 0 | 0 | 11 | 0 | 21 | 0 | 11 | 0 | 0 | 0 | 0 | 0 | 0 | 0 | 0 | 0 | 0 | 0 | 605 | 242 | 0 | 0 | 97 | 0 | 225 | 257 | 0 | 0 | 0 | 0 | 0 | 0 | 0 | 0 | 0 | 21 |
| SRR5388142 | 0 | 37 | 18 | 0 | 0 | 0 | 0 | 0 | 0 | 0 | 0 | 0 | 0 | 0 | 0 | 0 | 0 | 0 | 0 | 1223 | 102 | 0 | 0 | 22 | 0 | 200 | 318 | 37 | 0 | 0 | 0 | 52 | 0 | 22 | 0 | 0 | 0 |
| SRR5388143 | 0 | 10 | 0 | 0 | 0 | 31 | 0 | 0 | 20 | 0 | 0 | 0 | 0 | 0 | 87 | 0 | 0 | 0 | 87 | 524 | 0 | 0 | 0 | 184 | 0 | 429 | 511 | 0 | 0 | 0 | 20 | 0 | 0 | 0 | 0 | 0 | 10 |
| SRR5388144 | 0 | 0 | 0 | 0 | 0 | 36 | 0 | 0 | 0 | 0 | 0 | 0 | 0 | 0 | 0 | 0 | 0 | 0 | 0 | 495 | 0 | 0 | 0 | 168 | 0 | 324 | 575 | 0 | 0 | 0 | 0 | 0 | 0 | 0 | 0 | 0 | 36 |
| SRR5388145 | 0 | 11 | 0 | 0 | 0 | 11 | 80 | 33 | 0 | 0 | 0 | 0 | 0 | 11 | 0 | 0 | 0 | 0 | 0 | 1364 | 161 | 0 | 0 | 76 | 11 | 163 | 218 | 0 | 0 | 11 | 0 | 0 | 0 | 0 | 0 | 0 | 11 |
| SRR5388146 | 0 | 43 | 11 | 11 | 0 | 11 | 0 | 0 | 0 | 0 | 0 | 0 | 0 | 0 | 0 | 0 | 0 | 0 | 0 | 618 | 103 | 0 | 0 | 86 | 0 | 160 | 278 | 0 | 0 | 0 | 0 | 0 | 0 | 0 | 0 | 0 | 0 |
| SRR5388147 | 0 | 0 | 24 | 0 | 0 | 25 | 0 | 0 | 0 | 0 | 0 | 0 | 0 | 0 | 0 | 0 | 0 | 0 | 0 | 1113 | 257 | 0 | 0 | 58 | 0 | 359 | 359 | 0 | 0 | 0 | 0 | 0 | 0 | 0 | 0 | 0 | 17 |
| SRR5388148 | 0 | 22 | 18 | 0 | 0 | 22 | 0 | 0 | 0 | 0 | 0 | 0 | 0 | 0 | 0 | 0 | 0 | 0 | 0 | 670 | 201 | 22 | 0 | 103 | 0 | 140 | 436 | 0 | 0 | 0 | 15 | 0 | 0 | 0 | 0 | 0 | 0 |
| SRR5388149 | 0 | 0 | 12 | 0 | 0 | 26 | 0 | 18 | 0 | 0 | 0 | 0 | 0 | 0 | 72 | 0 | 0 | 0 | 143 | 1577 | 215 | 0 | 0 | 97 | 0 | 246 | 343 | 0 | 0 | 0 | 0 | 0 | 18 | 0 | 0 | 0 | 0 |
| SRR5388150 | 0 | 0 | 0 | 0 | 0 | 24 | 0 | 0 | 0 | 0 | 0 | 0 | 0 | 0 | 61 | 0 | 0 | 0 | 0 | 860 | 123 | 0 | 0 | 135 | 0 | 175 | 310 | 0 | 0 | 0 | 0 | 0 | 0 | 0 | 0 | 0 | 0 |
| SRR5388151 | 0 | 0 | 0 | 0 | 0 | 49 | 0 | 20 | 0 | 0 | 0 | 0 | 0 | 0 | 0 | 0 | 0 | 0 | 0 | 1649 | 82 | 0 | 0 | 79 | 0 | 296 | 266 | 0 | 0 | 0 | 0 | 0 | 0 | 0 | 0 | 0 | 0 |
| SRR5388152 | 0 | 0 | 10 | 0 | 0 | 62 | 0 | 35 | 35 | 0 | 0 | 0 | 0 | 26 | 0 | 0 | 0 | 0 | 0 | 1097 | 137 | 0 | 0 | 88 | 0 | 371 | 441 | 0 | 0 | 0 | 18 | 0 | 0 | 0 | 0 | 0 | 0 |
| SRR5388153 | 0 | 0 | 0 | 0 | 0 | 33 | 0 | 0 | 0 | 0 | 0 | 0 | 0 | 0 | 0 | 0 | 0 | 0 | 0 | 1419 | 0 | 0 | 0 | 50 | 0 | 367 | 383 | 0 | 0 | 0 | 0 | 0 | 0 | 0 | 0 | 0 | 0 |
| SRR5388154 | 0 | 0 | 13 | 0 | 0 | 15 | 0 | 0 | 0 | 0 | 0 | 0 | 0 | 0 | 193 | 0 | 0 | 0 | 0 | 1801 | 386 | 0 | 0 | 61 | 0 | 182 | 380 | 0 | 0 | 0 | 0 | 0 | 0 | 0 | 0 | 0 | 0 |
| SRR5388155 | 0 | 56 | 0 | 0 | 0 | 0 | 0 | 28 | 0 | 0 | 0 | 0 | 0 | 0 | 0 | 0 | 0 | 0 | 84 | 674 | 505 | 0 | 0 | 21 | 0 | 161 | 105 | 0 | 0 | 0 | 0 | 0 | 0 | 0 | 0 | 0 | 0 |
| SRR5388156 | 0 | 0 | 13 | 0 | 0 | 23 | 0 | 28 | 0 | 0 | 0 | 0 | 0 | 0 | 0 | 17 | 0 | 0 | 0 | 1438 | 240 | 0 | 0 | 125 | 0 | 436 | 300 | 0 | 0 | 0 | 0 | 0 | 11 | 11 | 11 | 0 | 0 |
| SRR5388157 | 0 | 12 | 0 | 0 | 0 | 12 | 0 | 35 | 12 | 0 | 0 | 0 | 0 | 0 | 0 | 0 | 0 | 0 | 0 | 1340 | 357 | 0 | 0 | 47 | 0 | 200 | 246 | 0 | 0 | 0 | 0 | 0 | 0 | 0 | 0 | 0 | 0 |
| SRR5388158 | 0 | 12 | 0 | 0 | 0 | 12 | 0 | 0 | 12 | 0 | 0 | 0 | 0 | 12 | 0 | 0 | 0 | 0 | 139 | 832 | 139 | 0 | 0 | 59 | 12 | 131 | 380 | 0 | 0 | 0 | 0 | 12 | 0 | 0 | 0 | 0 | 0 |
| SRR5388159 | 0 | 0 | 0 | 0 | 0 | 10 | 0 | 10 | 0 | 0 | 0 | 0 | 0 | 10 | 0 | 0 | 0 | 0 | 0 | 784 | 261 | 0 | 0 | 21 | 0 | 293 | 188 | 0 | 0 | 10 | 0 | 0 | 10 | 10 | 0 | 0 | 10 |
| SRR5388160 | 0 | 16 | 20 | 0 | 0 | 16 | 0 | 24 | 0 | 0 | 0 | 0 | 0 | 0 | 95 | 0 | 0 | 0 | 0 | 1902 | 475 | 0 | 0 | 118 | 0 | 182 | 355 | 0 | 0 | 0 | 0 | 0 | 0 | 0 | 0 | 0 | 0 |
| SRR5388161 | 0 | 0 | 0 | 0 | 0 | 0 | 0 | 19 | 0 | 0 | 0 | 0 | 0 | 12 | 55 | 0 | 0 | 0 | 0 | 1425 | 55 | 0 | 0 | 37 | 0 | 62 | 112 | 0 | 0 | 0 | 0 | 0 | 0 | 12 | 0 | 0 | 0 |
| SRR5388162 | 0 | 0 | 0 | 0 | 0 | 0 | 85 | 28 | 0 | 0 | 0 | 0 | 0 | 0 | 85 | 0 | 0 | 0 | 0 | 2473 | 171 | 0 | 0 | 65 | 0 | 158 | 195 | 0 | 0 | 0 | 0 | 0 | 0 | 0 | 0 | 0 | 0 |
| SRR5388163 | 0 | 13 | 0 | 0 | 0 | 0 | 0 | 19 | 13 | 0 | 0 | 0 | 0 | 0 | 0 | 0 | 0 | 0 | 0 | 995 | 51 | 0 | 0 | 96 | 0 | 199 | 302 | 0 | 0 | 0 | 13 | 0 | 0 | 26 | 0 | 0 | 13 |
| SRR5388164 | 0 | 37 | 10 | 0 | 0 | 75 | 0 | 0 | 0 | 0 | 0 | 0 | 0 | 0 | 0 | 0 | 0 | 0 | 0 | 543 | 241 | 0 | 0 | 50 | 0 | 150 | 237 | 0 | 0 | 12 | 12 | 12 | 0 | 12 | 0 | 12 | 0 |
| SRR5388165 | 0 | 0 | 0 | 0 | 0 | 16 | 0 | 11 | 11 | 0 | 0 | 0 | 0 | 11 | 0 | 0 | 0 | 0 | 0 | 1502 | 1093 | 0 | 0 | 37 | 0 | 277 | 112 | 0 | 0 | 0 | 16 | 0 | 11 | 0 | 0 | 0 | 0 |
| SRR5388166 | 0 | 0 | 0 | 11 | 0 | 44 | 0 | 0 | 11 | 0 | 0 | 0 | 0 | 33 | 0 | 11 | 0 | 0 | 0 | 415 | 83 | 11 | 0 | 88 | 0 | 318 | 307 | 11 | 0 | 0 | 0 | 0 | 0 | 0 | 0 | 0 | 0 |
| SRR5388167 | 0 | 0 | 0 | 0 | 0 | 0 | 0 | 0 | 0 | 0 | 0 | 0 | 0 | 13 | 0 | 0 | 0 | 0 | 0 | 1382 | 10782 | 0 | 0 | 97 | 0 | 15 | 20 | 0 | 0 | 0 | 0 | 0 | 0 | 0 | 0 | 0 | 0 |
| SRR5388168 | 0 | 0 | 12 | 0 | 0 | 0 | 0 | 16 | 0 | 0 | 0 | 0 | 0 | 20 | 51 | 0 | 0 | 0 | 0 | 967 | 153 | 0 | 0 | 67 | 0 | 83 | 177 | 0 | 0 | 0 | 0 | 0 | 0 | 0 | 0 | 0 | 0 |
| SRR5388169 | 14 | 12 | 22 | 0 | 0 | 0 | 0 | 12 | 0 | 0 | 0 | 0 | 0 | 0 | 0 | 0 | 0 | 0 | 0 | 435 | 62 | 0 | 0 | 131 | 0 | 41 | 126 | 0 | 0 | 0 | 0 | 0 | 0 | 0 | 0 | 0 | 0 |
| SRR5388170 | 0 | 0 | 0 | 0 | 0 | 0 | 0 | 0 | 0 | 0 | 0 | 0 | 0 | 0 | 0 | 0 | 0 | 0 | 0 | 245 | 82 | 0 | 0 | 88 | 0 | 29 | 58 | 0 | 0 | 0 | 0 | 0 | 0 | 0 | 0 | 0 | 0 |
| SRR5388171 | 0 | 20 | 0 | 0 | 0 | 0 | 0 | 20 | 0 | 0 | 0 | 0 | 0 | 0 | 40 | 0 | 0 | 0 | 0 | 160 | 0 | 0 | 0 | 45 | 0 | 20 | 73 | 0 | 0 | 0 | 0 | 0 | 0 | 0 | 0 | 0 | 0 |
| SRR5388172 | 0 | 0 | 0 | 0 | 0 | 0 | 0 | 62 | 0 | 0 | 0 | 0 | 0 | 0 | 0 | 0 | 0 | 0 | 0 | 210 | 0 | 0 | 0 | 97 | 0 | 0 | 97 | 0 | 0 | 0 | 0 | 0 | 0 | 0 | 0 | 0 | 0 |
| SRR5388173 | 0 | 0 | 13 | 0 | 0 | 20 | 0 | 0 | 0 | 0 | 0 | 0 | 0 | 0 | 59 | 0 | 0 | 0 | 0 | 1119 | 236 | 0 | 0 | 69 | 0 | 89 | 128 | 0 | 0 | 0 | 0 | 0 | 0 | 0 | 0 | 0 | 0 |
| SRR5388174 | 0 | 12 | 0 | 0 | 0 | 12 | 0 | 12 | 0 | 0 | 0 | 0 | 0 | 12 | 0 | 0 | 0 | 0 | 0 | 882 | 189 | 0 | 0 | 37 | 0 | 135 | 208 | 0 | 0 | 0 | 0 | 0 | 0 | 24 | 0 | 0 | 12 |
| SRR5388175 | 0 | 0 | 0 | 0 | 0 | 0 | 0 | 12 | 0 | 0 | 0 | 0 | 0 | 0 | 0 | 0 | 0 | 0 | 0 | 710 | 59 | 0 | 0 | 93 | 0 | 50 | 156 | 0 | 0 | 0 | 0 | 0 | 0 | 0 | 0 | 0 | 0 |
| SRR5388176 | 0 | 21 | 14 | 0 | 0 | 0 | 0 | 0 | 0 | 0 | 0 | 0 | 0 | 10 | 0 | 0 | 0 | 0 | 0 | 711 | 0 | 0 | 0 | 72 | 0 | 124 | 83 | 0 | 0 | 21 | 0 | 0 | 0 | 0 | 0 | 0 | 0 |
| SRR5388177 | 0 | 10 | 0 | 0 | 0 | 40 | 0 | 0 | 20 | 0 | 0 | 0 | 0 | 0 | 0 | 0 | 0 | 0 | 0 | 1678 | 148 | 0 | 0 | 20 | 0 | 445 | 202 | 0 | 0 | 0 | 20 | 10 | 0 | 0 | 0 | 0 | 0 |
| SRR5388178 | 0 | 22 | 0 | 0 | 0 | 22 | 0 | 0 | 0 | 0 | 0 | 0 | 0 | 22 | 0 | 0 | 0 | 0 | 45 | 903 | 181 | 0 | 0 | 76 | 0 | 164 | 218 | 0 | 0 | 0 | 33 | 0 | 0 | 0 | 0 | 0 | 0 |
| SRR5388179 | 0 | 0 | 12 | 0 | 0 | 0 | 0 | 15 | 0 | 0 | 0 | 0 | 0 | 0 | 0 | 0 | 0 | 0 | 49 | 1174 | 98 | 0 | 0 | 115 | 0 | 54 | 161 | 0 | 0 | 0 | 0 | 0 | 0 | 0 | 0 | 0 | 0 |
| SRR5388180 | 11 | 13 | 15 | 0 | 0 | 0 | 0 | 0 | 0 | 0 | 0 | 0 | 0 | 0 | 0 | 0 | 0 | 0 | 0 | 2073 | 194 | 0 | 0 | 91 | 0 | 72 | 85 | 0 | 0 | 0 | 0 | 0 | 0 | 0 | 0 | 0 | 0 |
| SRR5388181 | 0 | 10 | 13 | 0 | 0 | 0 | 0 | 10 | 0 | 0 | 0 | 0 | 0 | 0 | 0 | 0 | 0 | 0 | 0 | 515 | 74 | 0 | 0 | 92 | 0 | 185 | 287 | 0 | 0 | 0 | 0 | 0 | 10 | 31 | 0 | 0 | 0 |
| SRR5388182 | 0 | 0 | 13 | 0 | 0 | 14 | 0 | 0 | 0 | 0 | 0 | 0 | 0 | 0 | 0 | 0 | 0 | 0 | 0 | 653 | 137 | 0 | 0 | 96 | 0 | 192 | 323 | 0 | 0 | 0 | 0 | 0 | 0 | 0 | 0 | 0 | 0 |
| SRR5388183 | 0 | 0 | 0 | 0 | 0 | 14 | 0 | 0 | 0 | 0 | 0 | 0 | 0 | 0 | 0 | 0 | 0 | 0 | 0 | 1902 | 293 | 0 | 0 | 68 | 0 | 231 | 177 | 0 | 0 | 0 | 0 | 0 | 0 | 27 | 0 | 0 | 0 |
| SRR5388184 | 0 | 11 | 0 | 0 | 0 | 11 | 0 | 11 | 0 | 0 | 0 | 0 | 0 | 11 | 0 | 0 | 0 | 0 | 0 | 254 | 0 | 0 | 0 | 87 | 0 | 55 | 120 | 0 | 0 | 0 | 0 | 0 | 11 | 11 | 0 | 0 | 0 |
| SRR5388185 | 0 | 0 | 0 | 0 | 0 | 0 | 0 | 31 | 16 | 0 | 0 | 0 | 0 | 0 | 0 | 0 | 0 | 0 | 0 | 218 | 218 | 0 | 0 | 62 | 0 | 171 | 249 | 0 | 0 | 0 | 0 | 0 | 0 | 0 | 0 | 0 | 0 |
| SRR5388186 | 0 | 17 | 0 | 0 | 0 | 0 | 0 | 26 | 0 | 0 | 0 | 0 | 0 | 0 | 0 | 0 | 0 | 0 | 0 | 923 | 173 | 0 | 0 | 34 | 0 | 128 | 60 | 0 | 0 | 0 | 0 | 0 | 0 | 0 | 0 | 0 | 0 |
| SRR5388187 | 0 | 0 | 13 | 0 | 0 | 24 | 0 | 0 | 0 | 0 | 0 | 0 | 0 | 0 | 191 | 0 | 0 | 0 | 95 | 764 | 382 | 0 | 0 | 97 | 12 | 157 | 121 | 0 | 0 | 12 | 0 | 0 | 0 | 0 | 0 | 0 | 0 |
| SRR5388188 | 0 | 14 | 0 | 0 | 0 | 21 | 0 | 0 | 0 | 0 | 0 | 0 | 0 | 14 | 118 | 0 | 0 | 0 | 59 | 1655 | 473 | 0 | 0 | 85 | 0 | 170 | 332 | 0 | 0 | 0 | 0 | 0 | 0 | 14 | 0 | 0 | 0 |
| SRR5388189 | 0 | 12 | 0 | 0 | 0 | 0 | 0 | 0 | 0 | 0 | 0 | 0 | 0 | 0 | 27 | 0 | 0 | 0 | 53 | 559 | 293 | 0 | 0 | 68 | 0 | 126 | 151 | 0 | 0 | 0 | 0 | 0 | 0 | 0 | 0 | 0 | 0 |
| SRR5388190 | 0 | 0 | 12 | 0 | 0 | 14 | 0 | 14 | 0 | 0 | 0 | 0 | 0 | 23 | 52 | 0 | 0 | 0 | 0 | 982 | 103 | 0 | 0 | 72 | 0 | 95 | 168 | 0 | 0 | 0 | 0 | 0 | 0 | 0 | 0 | 0 | 0 |
| SRR5388191 | 0 | 35 | 0 | 0 | 0 | 88 | 0 | 0 | 0 | 0 | 0 | 0 | 0 | 0 | 0 | 0 | 0 | 0 | 0 | 789 | 99 | 0 | 0 | 70 | 18 | 455 | 508 | 0 | 0 | 18 | 35 | 0 | 0 | 18 | 0 | 0 | 0 |
| SRR5388192 | 0 | 0 | 17 | 0 | 0 | 17 | 0 | 0 | 0 | 0 | 0 | 0 | 0 | 0 | 151 | 0 | 0 | 0 | 0 | 954 | 301 | 0 | 0 | 46 | 0 | 137 | 200 | 0 | 0 | 0 | 0 | 0 | 0 | 0 | 0 | 0 | 0 |
| SRR5388193 | 0 | 17 | 13 | 0 | 0 | 0 | 0 | 0 | 0 | 0 | 0 | 0 | 0 | 0 | 0 | 0 | 0 | 0 | 51 | 967 | 178 | 0 | 0 | 78 | 0 | 111 | 219 | 0 | 0 | 0 | 0 | 0 | 0 | 0 | 0 | 0 | 0 |
| SRR5388194 | 11 | 0 | 15 | 0 | 0 | 0 | 0 | 19 | 0 | 0 | 0 | 0 | 0 | 0 | 0 | 0 | 0 | 0 | 52 | 1145 | 312 | 0 | 0 | 102 | 0 | 102 | 196 | 0 | 0 | 0 | 0 | 0 | 0 | 28 | 0 | 0 | 19 |
| SRR5388195 | 0 | 20 | 10 | 0 | 0 | 0 | 0 | 30 | 0 | 0 | 0 | 0 | 0 | 0 | 0 | 0 | 0 | 0 | 0 | 755 | 40 | 0 | 0 | 119 | 0 | 109 | 298 | 0 | 0 | 0 | 0 | 0 | 20 | 50 | 0 | 0 | 0 |
| SRR5388196 | 0 | 18 | 12 | 0 | 0 | 18 | 0 | 18 | 0 | 0 | 0 | 0 | 0 | 0 | 30 | 0 | 0 | 0 | 30 | 907 | 91 | 0 | 0 | 59 | 0 | 130 | 218 | 0 | 0 | 0 | 0 | 0 | 0 | 18 | 0 | 0 | 0 |
| SRR5388197 | 0 | 14 | 0 | 0 | 0 | 0 | 0 | 14 | 0 | 0 | 0 | 0 | 0 | 0 | 0 | 0 | 0 | 0 | 0 | 733 | 85 | 0 | 0 | 136 | 0 | 100 | 335 | 0 | 0 | 0 | 0 | 0 | 14 | 14 | 0 | 0 | 0 |
| SRR5388198 | 0 | 12 | 0 | 0 | 0 | 0 | 0 | 12 | 0 | 0 | 0 | 0 | 0 | 0 | 0 | 0 | 0 | 0 | 0 | 635 | 173 | 0 | 0 | 111 | 0 | 123 | 160 | 0 | 0 | 0 | 0 | 0 | 0 | 12 | 12 | 0 | 0 |
| SRR5388199 | 0 | 0 | 0 | 0 | 0 | 0 | 0 | 0 | 0 | 0 | 0 | 0 | 0 | 0 | 0 | 0 | 0 | 0 | 43 | 2560 | 4225 | 0 | 0 | 77 | 0 | 40 | 40 | 0 | 0 | 0 | 0 | 0 | 0 | 0 | 0 | 0 | 0 |
| SRR5388200 | 0 | 0 | 0 | 0 | 0 | 0 | 0 | 0 | 0 | 0 | 0 | 0 | 0 | 0 | 0 | 0 | 0 | 0 | 77 | 1003 | 154 | 0 | 0 | 60 | 0 | 180 | 114 | 0 | 0 | 0 | 0 | 0 | 0 | 0 | 0 | 0 | 0 |
| SRR5388201 | 0 | 16 | 0 | 0 | 0 | 0 | 47 | 16 | 16 | 0 | 0 | 0 | 0 | 0 | 47 | 0 | 0 | 0 | 0 | 378 | 236 | 0 | 0 | 73 | 0 | 113 | 178 | 0 | 0 | 0 | 0 | 0 | 0 | 0 | 0 | 0 | 0 |
| SRR5388202 | 0 | 0 | 13 | 0 | 0 | 22 | 0 | 22 | 11 | 0 | 0 | 0 | 0 | 0 | 0 | 0 | 0 | 0 | 0 | 788 | 72 | 0 | 0 | 76 | 0 | 65 | 185 | 0 | 0 | 0 | 11 | 0 | 0 | 11 | 0 | 0 | 0 |
| SRR5388203 | 0 | 0 | 0 | 0 | 0 | 0 | 0 | 0 | 0 | 0 | 0 | 0 | 0 | 0 | 105 | 0 | 0 | 0 | 0 | 575 | 157 | 0 | 0 | 76 | 0 | 161 | 246 | 0 | 0 | 0 | 0 | 0 | 0 | 25 | 0 | 0 | 0 |
| SRR5388204 | 0 | 24 | 0 | 0 | 0 | 24 | 0 | 12 | 12 | 0 | 0 | 0 | 0 | 12 | 0 | 0 | 0 | 0 | 0 | 664 | 83 | 0 | 0 | 94 | 0 | 153 | 212 | 0 | 0 | 0 | 0 | 0 | 0 | 0 | 12 | 0 | 12 |
| SRR5388205 | 0 | 0 | 0 | 0 | 0 | 0 | 0 | 0 | 0 | 0 | 0 | 0 | 0 | 0 | 0 | 0 | 0 | 0 | 0 | 681 | 120 | 0 | 0 | 97 | 0 | 62 | 204 | 0 | 0 | 0 | 0 | 0 | 0 | 0 | 0 | 0 | 0 |
| SRR5388206 | 0 | 0 | 0 | 0 | 0 | 0 | 0 | 0 | 0 | 0 | 0 | 0 | 0 | 0 | 0 | 0 | 0 | 0 | 0 | 410 | 27 | 0 | 0 | 134 | 16 | 63 | 182 | 0 | 0 | 0 | 0 | 0 | 0 | 0 | 0 | 0 | 0 |
| SRR5388207 | 0 | 0 | 0 | 0 | 0 | 32 | 0 | 16 | 0 | 0 | 0 | 0 | 0 | 0 | 0 | 0 | 0 | 0 | 0 | 909 | 165 | 0 | 0 | 89 | 16 | 259 | 299 | 0 | 0 | 0 | 0 | 0 | 0 | 0 | 0 | 0 | 0 |
| SRR5388208 | 0 | 0 | 0 | 0 | 0 | 0 | 0 | 0 | 0 | 0 | 0 | 0 | 0 | 0 | 0 | 0 | 0 | 0 | 0 | 435 | 0 | 0 | 0 | 111 | 0 | 139 | 223 | 0 | 0 | 0 | 0 | 0 | 0 | 0 | 0 | 0 | 0 |
| SRR5388209 | 0 | 0 | 0 | 0 | 0 | 28 | 0 | 14 | 0 | 0 | 0 | 0 | 0 | 0 | 0 | 0 | 0 | 0 | 0 | 395 | 36 | 0 | 0 | 85 | 0 | 71 | 85 | 0 | 0 | 0 | 28 | 0 | 0 | 0 | 0 | 0 | 0 |
| SRR5388210 | 0 | 22 | 0 | 0 | 0 | 22 | 0 | 43 | 0 | 0 | 0 | 0 | 0 | 0 | 0 | 0 | 0 | 0 | 0 | 223 | 89 | 0 | 0 | 65 | 0 | 22 | 86 | 22 | 0 | 0 | 0 | 0 | 0 | 0 | 0 | 0 | 0 |
| SRR5388211 | 0 | 101 | 0 | 0 | 0 | 0 | 0 | 12 | 0 | 0 | 0 | 0 | 0 | 0 | 0 | 0 | 0 | 0 | 51 | 714 | 0 | 0 | 0 | 37 | 0 | 71 | 144 | 104 | 0 | 0 | 0 | 101 | 0 | 0 | 0 | 0 | 0 |
| SRR5388212 | 0 | 0 | 0 | 0 | 0 | 0 | 0 | 0 | 0 | 0 | 0 | 0 | 0 | 0 | 85 | 0 | 0 | 0 | 0 | 677 | 0 | 0 | 0 | 100 | 0 | 127 | 273 | 0 | 0 | 0 | 0 | 0 | 0 | 0 | 0 | 0 | 0 |
| SRR5388213 | 0 | 24 | 0 | 0 | 0 | 0 | 0 | 39 | 0 | 0 | 0 | 0 | 0 | 0 | 0 | 0 | 0 | 0 | 0 | 503 | 77 | 0 | 0 | 87 | 0 | 55 | 95 | 0 | 0 | 0 | 0 | 0 | 0 | 0 | 0 | 0 | 0 |
| SRR5388214 | 0 | 11 | 0 | 0 | 0 | 11 | 0 | 63 | 0 | 0 | 0 | 0 | 0 | 0 | 0 | 0 | 0 | 0 | 0 | 110 | 74 | 0 | 0 | 63 | 0 | 95 | 158 | 0 | 0 | 0 | 0 | 11 | 0 | 0 | 11 | 0 | 0 |
| SRR5388215 | 0 | 10 | 0 | 0 | 0 | 21 | 0 | 73 | 0 | 0 | 0 | 0 | 0 | 0 | 0 | 0 | 0 | 0 | 0 | 1668 | 371 | 0 | 0 | 63 | 0 | 115 | 52 | 0 | 0 | 0 | 0 | 0 | 0 | 0 | 21 | 0 | 10 |
| SRR5388216 | 0 | 23 | 11 | 0 | 0 | 0 | 0 | 12 | 0 | 0 | 0 | 0 | 0 | 0 | 0 | 0 | 0 | 0 | 0 | 583 | 0 | 0 | 0 | 109 | 0 | 86 | 219 | 20 | 0 | 0 | 0 | 16 | 0 | 0 | 0 | 0 | 0 |
| SRR5388217 | 0 | 0 | 0 | 0 | 0 | 0 | 0 | 0 | 0 | 0 | 0 | 0 | 0 | 0 | 0 | 0 | 0 | 0 | 0 | 2636 | 264 | 0 | 0 | 18 | 0 | 0 | 144 | 0 | 0 | 0 | 18 | 0 | 0 | 0 | 18 | 0 | 0 |
| SRR5388218 | 0 | 0 | 0 | 0 | 0 | 19 | 0 | 58 | 0 | 0 | 0 | 0 | 0 | 0 | 0 | 0 | 0 | 0 | 0 | 1010 | 126 | 0 | 0 | 145 | 0 | 77 | 222 | 0 | 0 | 0 | 0 | 0 | 0 | 0 | 0 | 0 | 0 |
| SRR5388219 | 0 | 18 | 0 | 0 | 0 | 0 | 0 | 0 | 0 | 0 | 0 | 0 | 0 | 0 | 125 | 0 | 0 | 0 | 0 | 626 | 0 | 0 | 0 | 80 | 0 | 133 | 319 | 0 | 0 | 0 | 0 | 0 | 0 | 0 | 0 | 0 | 0 |
| SRR5388220 | 0 | 20 | 0 | 0 | 0 | 27 | 0 | 13 | 0 | 0 | 0 | 0 | 0 | 0 | 0 | 0 | 0 | 0 | 138 | 276 | 138 | 0 | 0 | 94 | 0 | 100 | 234 | 0 | 0 | 0 | 0 | 0 | 0 | 0 | 0 | 0 | 0 |
| SRR5388221 | 0 | 0 | 0 | 0 | 0 | 24 | 0 | 24 | 10 | 0 | 0 | 0 | 0 | 0 | 0 | 0 | 0 | 0 | 0 | 1318 | 107 | 0 | 0 | 169 | 0 | 236 | 320 | 0 | 0 | 0 | 0 | 0 | 0 | 0 | 0 | 0 | 0 |
| SRR5388222 | 0 | 0 | 0 | 0 | 0 | 31 | 0 | 0 | 0 | 0 | 0 | 0 | 0 | 0 | 0 | 0 | 0 | 0 | 0 | 459 | 0 | 10 | 0 | 51 | 0 | 153 | 254 | 0 | 0 | 0 | 0 | 0 | 0 | 0 | 0 | 0 | 0 |
| SRR5388223 | 0 | 11 | 0 | 0 | 0 | 22 | 0 | 11 | 0 | 0 | 0 | 0 | 0 | 0 | 0 | 0 | 0 | 0 | 70 | 1547 | 141 | 0 | 0 | 76 | 11 | 142 | 164 | 0 | 0 | 0 | 0 | 0 | 0 | 11 | 0 | 0 | 0 |
| SRR5388224 | 0 | 10 | 0 | 0 | 0 | 0 | 0 | 0 | 0 | 10 | 0 | 0 | 0 | 0 | 0 | 0 | 0 | 0 | 0 | 1373 | 0 | 0 | 0 | 164 | 0 | 72 | 245 | 0 | 0 | 0 | 0 | 0 | 0 | 0 | 10 | 0 | 0 |
| SRR5388225 | 0 | 0 | 0 | 0 | 0 | 0 | 0 | 0 | 0 | 0 | 0 | 0 | 0 | 0 | 0 | 0 | 0 | 0 | 60 | 300 | 180 | 0 | 0 | 81 | 0 | 162 | 189 | 0 | 0 | 0 | 0 | 0 | 18 | 0 | 0 | 0 | 0 |
| SRR5388226 | 0 | 0 | 0 | 0 | 0 | 0 | 0 | 0 | 0 | 0 | 0 | 0 | 0 | 0 | 111 | 0 | 0 | 0 | 0 | 612 | 278 | 0 | 0 | 114 | 19 | 66 | 265 | 0 | 0 | 0 | 0 | 0 | 0 | 0 | 0 | 0 | 0 |
| SRR5388227 | 0 | 0 | 0 | 0 | 0 | 27 | 0 | 27 | 0 | 0 | 0 | 0 | 0 | 0 | 0 | 0 | 0 | 0 | 0 | 849 | 319 | 0 | 0 | 53 | 0 | 146 | 27 | 0 | 0 | 0 | 0 | 0 | 0 | 13 | 0 | 0 | 0 |
| SRR5388228 | 0 | 0 | 12 | 0 | 0 | 11 | 0 | 0 | 0 | 0 | 0 | 0 | 0 | 0 | 0 | 0 | 0 | 0 | 53 | 425 | 212 | 0 | 0 | 38 | 0 | 77 | 131 | 0 | 0 | 0 | 0 | 0 | 0 | 0 | 11 | 0 | 0 |
| SRR5388229 | 0 | 11 | 0 | 0 | 11 | 0 | 0 | 0 | 0 | 0 | 0 | 0 | 0 | 0 | 0 | 0 | 0 | 0 | 0 | 1080 | 324 | 0 | 0 | 98 | 0 | 164 | 142 | 0 | 0 | 0 | 0 | 0 | 0 | 11 | 0 | 0 | 11 |
| SRR5388230 | 0 | 0 | 0 | 0 | 0 | 0 | 0 | 0 | 0 | 0 | 0 | 0 | 0 | 0 | 0 | 0 | 0 | 0 | 0 | 540 | 337 | 0 | 0 | 86 | 0 | 229 | 229 | 0 | 0 | 0 | 0 | 0 | 0 | 0 | 0 | 0 | 0 |
| SRR5388231 | 0 | 0 | 0 | 0 | 0 | 25 | 0 | 33 | 0 | 0 | 0 | 0 | 0 | 0 | 0 | 0 | 0 | 0 | 0 | 879 | 258 | 0 | 0 | 66 | 0 | 313 | 346 | 0 | 0 | 0 | 16 | 0 | 0 | 0 | 16 | 0 | 0 |
| SRR5388232 | 0 | 18 | 0 | 0 | 0 | 27 | 0 | 0 | 0 | 0 | 0 | 0 | 0 | 0 | 0 | 0 | 0 | 0 | 0 | 601 | 0 | 0 | 0 | 161 | 0 | 206 | 251 | 0 | 0 | 0 | 0 | 0 | 0 | 0 | 0 | 0 | 0 |
| SRR5388233 | 0 | 0 | 14 | 0 | 0 | 12 | 0 | 0 | 0 | 0 | 0 | 0 | 0 | 0 | 0 | 0 | 0 | 0 | 34 | 1019 | 68 | 0 | 0 | 45 | 0 | 138 | 65 | 0 | 0 | 0 | 0 | 0 | 0 | 0 | 0 | 0 | 0 |
| SRR5388234 | 0 | 0 | 0 | 0 | 0 | 32 | 0 | 0 | 0 | 0 | 0 | 0 | 0 | 0 | 0 | 0 | 0 | 0 | 0 | 726 | 81 | 0 | 0 | 96 | 0 | 304 | 288 | 0 | 0 | 0 | 0 | 0 | 0 | 0 | 0 | 0 | 0 |
| SRR5388235 | 10 | 14 | 13 | 0 | 0 | 0 | 0 | 0 | 0 | 0 | 0 | 0 | 0 | 0 | 0 | 0 | 0 | 0 | 0 | 507 | 362 | 0 | 0 | 50 | 0 | 110 | 151 | 0 | 0 | 0 | 0 | 0 | 0 | 14 | 14 | 0 | 0 |
| SRR5388236 | 0 | 0 | 0 | 0 | 0 | 0 | 0 | 23 | 0 | 0 | 0 | 0 | 0 | 0 | 0 | 0 | 0 | 0 | 0 | 601 | 86 | 0 | 0 | 82 | 0 | 129 | 152 | 0 | 0 | 0 | 0 | 0 | 0 | 12 | 0 | 0 | 0 |
| SRR5388237 | 0 | 13 | 21 | 0 | 0 | 0 | 0 | 0 | 0 | 0 | 0 | 0 | 0 | 0 | 66 | 0 | 0 | 0 | 0 | 1988 | 133 | 0 | 0 | 154 | 0 | 115 | 103 | 0 | 0 | 0 | 0 | 0 | 0 | 0 | 0 | 0 | 0 |
| SRR5388238 | 0 | 0 | 0 | 0 | 0 | 0 | 0 | 0 | 0 | 0 | 0 | 0 | 0 | 0 | 0 | 0 | 0 | 0 | 0 | 691 | 92 | 0 | 0 | 126 | 0 | 278 | 185 | 0 | 0 | 0 | 0 | 0 | 0 | 0 | 0 | 0 | 0 |

### Supp Table S8.2.Columns_AM_to_BW

| | | | | | | | | | | | | | | | | | | | | | | | | | | | | | | | | | | | | | |
|---|---|---|---|---|---|---|---|---|---|---|---|---|---|---|---|---|---|---|---|---|---|---|---|---|---|---|---|---|---|---|---|---|---|---|---|---|---|
| SRR5388239 | 0 | 0 | 12 | 0 | 0 | 0 | 0 | 0 | 0 | 0 | 0 | 0 | 0 | 0 | 0 | 0 | 0 | 0 | 0 | 965 | 161 | 0 | 0 | 108 | 0 | 185 | 223 | 0 | 0 | 0 | 0 | 0 | 0 | 13 | 0 | 0 | 0 |
| SRR5388240 | 0 | 0 | 0 | 0 | 0 | 0 | 0 | 0 | 0 | 0 | 0 | 0 | 0 | 0 | 0 | 0 | 0 | 0 | 0 | 791 | 352 | 0 | 0 | 97 | 0 | 176 | 220 | 0 | 0 | 0 | 0 | 0 | 0 | 0 | 0 | 0 | 0 |
| SRR5388241 | 11 | 22 | 37 | 0 | 0 | 0 | 0 | 0 | 0 | 0 | 0 | 0 | 0 | 11 | 0 | 0 | 0 | 0 | 0 | 1026 | 79 | 0 | 0 | 86 | 0 | 70 | 97 | 0 | 0 | 0 | 0 | 0 | 0 | 11 | 0 | 0 | 0 |
| SRR5388242 | 12 | 0 | 39 | 0 | 0 | 0 | 0 | 15 | 0 | 0 | 0 | 13 | 0 | 11 | 61 | 0 | 0 | 0 | 0 | 1706 | 61 | 0 | 0 | 64 | 0 | 60 | 56 | 0 | 0 | 0 | 0 | 0 | 0 | 11 | 0 | 0 | 0 |
| SRR5388243 | 0 | 0 | 0 | 0 | 0 | 14 | 0 | 20 | 0 | 0 | 0 | 0 | 0 | 0 | 0 | 0 | 0 | 0 | 0 | 1005 | 0 | 0 | 0 | 88 | 0 | 102 | 210 | 0 | 0 | 0 | 20 | 0 | 0 | 0 | 0 | 0 | 0 |
| SRR5388244 | 0 | 0 | 0 | 0 | 0 | 31 | 0 | 10 | 20 | 0 | 0 | 0 | 0 | 0 | 0 | 0 | 0 | 0 | 0 | 1039 | 283 | 0 | 0 | 122 | 0 | 357 | 531 | 0 | 0 | 0 | 10 | 10 | 10 | 0 | 0 | 0 | 0 |
| SRR5388245 | 0 | 0 | 0 | 0 | 0 | 0 | 0 | 15 | 0 | 0 | 0 | 0 | 0 | 0 | 0 | 0 | 0 | 0 | 0 | 960 | 160 | 0 | 0 | 69 | 0 | 23 | 84 | 0 | 0 | 0 | 0 | 0 | 0 | 0 | 0 | 0 | 0 |
| SRR5388246 | 0 | 0 | 0 | 0 | 0 | 0 | 0 | 17 | 0 | 0 | 0 | 0 | 0 | 0 | 0 | 0 | 0 | 0 | 0 | 581 | 508 | 0 | 0 | 99 | 0 | 23 | 70 | 0 | 0 | 0 | 0 | 0 | 0 | 0 | 0 | 0 | 0 |
| SRR5388247 | 0 | 0 | 0 | 0 | 0 | 0 | 0 | 15 | 0 | 0 | 0 | 0 | 0 | 0 | 0 | 0 | 0 | 0 | 0 | 286 | 95 | 0 | 0 | 67 | 0 | 165 | 262 | 0 | 0 | 0 | 0 | 0 | 15 | 0 | 0 | 0 | 0 |
| SRR5388248 | 0 | 0 | 0 | 0 | 0 | 0 | 0 | 0 | 0 | 0 | 0 | 0 | 0 | 0 | 0 | 0 | 0 | 0 | 0 | 390 | 49 | 0 | 0 | 51 | 13 | 129 | 328 | 0 | 0 | 0 | 0 | 0 | 0 | 0 | 0 | 0 | 0 |
| SRR5388249 | 0 | 0 | 0 | 0 | 0 | 0 | 0 | 0 | 0 | 0 | 0 | 0 | 0 | 0 | 0 | 0 | 0 | 0 | 0 | 1033 | 387 | 0 | 0 | 38 | 0 | 132 | 227 | 19 | 0 | 0 | 0 | 0 | 19 | 0 | 0 | 0 | 0 |
| SRR5388250 | 0 | 24 | 0 | 0 | 0 | 0 | 0 | 0 | 0 | 0 | 0 | 0 | 0 | 0 | 0 | 0 | 0 | 0 | 0 | 1464 | 105 | 0 | 0 | 43 | 0 | 109 | 189 | 61 | 0 | 12 | 0 | 43 | 0 | 0 | 0 | 0 | 0 |
| SRR5388251 | 0 | 35 | 0 | 0 | 0 | 35 | 0 | 0 | 0 | 0 | 0 | 0 | 0 | 0 | 0 | 0 | 0 | 0 | 95 | 473 | 378 | 0 | 0 | 88 | 0 | 131 | 166 | 26 | 0 | 0 | 0 | 0 | 0 | 0 | 0 | 0 | 0 |
| SRR5388252 | 0 | 0 | 0 | 0 | 0 | 17 | 0 | 0 | 0 | 0 | 0 | 0 | 0 | 0 | 0 | 0 | 0 | 0 | 0 | 746 | 166 | 0 | 0 | 185 | 0 | 244 | 303 | 0 | 0 | 0 | 0 | 0 | 0 | 0 | 0 | 0 | 0 |
| SRR5388253 | 0 | 0 | 12 | 0 | 0 | 0 | 0 | 0 | 0 | 0 | 0 | 0 | 0 | 0 | 0 | 0 | 0 | 0 | 0 | 2137 | 142 | 0 | 0 | 0 | 0 | 188 | 267 | 0 | 0 | 0 | 0 | 0 | 20 | 0 | 0 | 0 | 0 |
| SRR5388254 | 0 | 0 | 0 | 0 | 0 | 0 | 0 | 0 | 0 | 0 | 0 | 0 | 0 | 0 | 0 | 0 | 0 | 0 | 0 | 2100 | 191 | 0 | 0 | 115 | 0 | 186 | 186 | 18 | 0 | 0 | 0 | 0 | 0 | 0 | 0 | 0 | 0 |
| SRR5388255 | 0 | 0 | 14 | 0 | 0 | 0 | 0 | 0 | 0 | 0 | 0 | 0 | 0 | 0 | 0 | 0 | 0 | 0 | 88 | 1945 | 88 | 0 | 0 | 42 | 0 | 21 | 52 | 0 | 0 | 0 | 0 | 0 | 0 | 0 | 0 | 0 | 10 |
| SRR5388256 | 0 | 0 | 13 | 0 | 0 | 21 | 0 | 14 | 0 | 0 | 0 | 0 | 0 | 0 | 48 | 0 | 0 | 0 | 48 | 476 | 238 | 0 | 0 | 76 | 0 | 62 | 62 | 0 | 0 | 0 | 0 | 0 | 0 | 0 | 0 | 0 | 0 |
| SRR5388257 | 0 | 0 | 0 | 0 | 0 | 22 | 85 | 87 | 0 | 0 | 0 | 0 | 0 | 0 | 0 | 0 | 0 | 0 | 0 | 1438 | 0 | 0 | 0 | 76 | 0 | 162 | 97 | 0 | 0 | 0 | 0 | 0 | 0 | 11 | 11 | 0 | 0 |
| SRR5388258 | 0 | 23 | 0 | 0 | 0 | 35 | 0 | 23 | 0 | 0 | 0 | 0 | 0 | 0 | 0 | 0 | 0 | 0 | 0 | 194 | 65 | 0 | 0 | 58 | 12 | 534 | 302 | 0 | 12 | 0 | 0 | 0 | 23 | 12 | 12 | 0 | 0 |
| SRR5388259 | 0 | 0 | 0 | 0 | 0 | 14 | 0 | 14 | 20 | 0 | 0 | 0 | 0 | 0 | 0 | 0 | 0 | 0 | 0 | 642 | 198 | 0 | 0 | 75 | 0 | 96 | 287 | 0 | 0 | 0 | 0 | 0 | 0 | 0 | 0 | 0 | 0 |
| SRR5388260 | 0 | 0 | 15 | 0 | 0 | 0 | 0 | 0 | 19 | 0 | 0 | 0 | 0 | 0 | 0 | 0 | 0 | 0 | 83 | 998 | 83 | 0 | 0 | 28 | 0 | 312 | 340 | 0 | 0 | 0 | 0 | 0 | 0 | 0 | 0 | 0 | 0 |
| SRR5388261 | 0 | 14 | 0 | 0 | 0 | 14 | 0 | 0 | 0 | 0 | 0 | 0 | 0 | 0 | 0 | 0 | 0 | 0 | 0 | 676 | 608 | 0 | 0 | 74 | 0 | 149 | 290 | 0 | 0 | 0 | 0 | 0 | 0 | 0 | 0 | 0 | 0 |
| SRR5388262 | 0 | 0 | 14 | 0 | 0 | 13 | 0 | 0 | 0 | 0 | 0 | 0 | 0 | 0 | 0 | 0 | 0 | 0 | 0 | 795 | 217 | 0 | 0 | 120 | 13 | 171 | 190 | 0 | 0 | 0 | 0 | 0 | 0 | 0 | 0 | 0 | 0 |
| SRR5388263 | 0 | 0 | 0 | 0 | 0 | 29 | 0 | 0 | 20 | 0 | 0 | 0 | 0 | 0 | 0 | 0 | 0 | 0 | 0 | 423 | 265 | 0 | 0 | 78 | 0 | 528 | 782 | 0 | 0 | 0 | 29 | 0 | 0 | 0 | 0 | 0 | 0 |
| SRR5388264 | 0 | 0 | 0 | 0 | 0 | 11 | 0 | 45 | 0 | 0 | 0 | 0 | 0 | 0 | 0 | 0 | 0 | 0 | 0 | 1149 | 72 | 0 | 0 | 123 | 11 | 336 | 415 | 0 | 0 | 0 | 0 | 0 | 22 | 0 | 11 | 0 | 0 |
| SRR5388265 | 0 | 0 | 15 | 0 | 0 | 0 | 0 | 0 | 0 | 0 | 0 | 0 | 0 | 0 | 0 | 0 | 0 | 0 | 0 | 506 | 42 | 0 | 0 | 135 | 0 | 48 | 115 | 0 | 0 | 0 | 0 | 0 | 0 | 0 | 0 | 0 | 0 |
| SRR5388266 | 0 | 0 | 0 | 0 | 0 | 0 | 0 | 0 | 0 | 0 | 0 | 0 | 0 | 0 | 0 | 0 | 0 | 0 | 0 | 465 | 233 | 0 | 0 | 85 | 0 | 131 | 154 | 0 | 0 | 0 | 0 | 0 | 0 | 0 | 15 | 0 | 0 |
| SRR5388267 | 0 | 11 | 0 | 0 | 0 | 0 | 0 | 11 | 11 | 0 | 0 | 0 | 0 | 0 | 0 | 0 | 0 | 0 | 0 | 765 | 170 | 0 | 0 | 66 | 0 | 55 | 155 | 0 | 0 | 0 | 0 | 0 | 0 | 0 | 0 | 0 | 0 |
| SRR5388268 | 0 | 0 | 0 | 0 | 0 | 19 | 0 | 0 | 0 | 0 | 0 | 0 | 0 | 0 | 0 | 0 | 0 | 0 | 0 | 820 | 46 | 0 | 0 | 95 | 0 | 227 | 290 | 0 | 0 | 0 | 0 | 0 | 0 | 13 | 0 | 0 | 0 |
| SRR5388269 | 0 | 0 | 0 | 0 | 0 | 0 | 0 | 0 | 0 | 0 | 0 | 0 | 0 | 0 | 0 | 0 | 0 | 0 | 0 | 1392 | 371 | 0 | 0 | 83 | 0 | 101 | 258 | 28 | 0 | 0 | 0 | 0 | 0 | 0 | 0 | 0 | 0 |
| SRR5388270 | 0 | 19 | 0 | 0 | 0 | 19 | 0 | 0 | 0 | 0 | 0 | 0 | 0 | 0 | 0 | 0 | 0 | 0 | 0 | 426 | 160 | 0 | 0 | 139 | 0 | 121 | 278 | 0 | 0 | 0 | 0 | 0 | 0 | 0 | 0 | 0 | 28 |
| SRR5388271 | 0 | 0 | 0 | 0 | 0 | 119 | 0 | 30 | 0 | 0 | 0 | 0 | 0 | 0 | 0 | 0 | 0 | 0 | 0 | 0 | 218 | 0 | 0 | 59 | 0 | 386 | 267 | 0 | 0 | 0 | 0 | 0 | 0 | 0 | 0 | 0 | 0 |
| SRR5388272 | 0 | 0 | 10 | 0 | 0 | 0 | 0 | 0 | 0 | 0 | 0 | 0 | 0 | 0 | 100 | 0 | 0 | 0 | 0 | 701 | 100 | 0 | 0 | 213 | 0 | 301 | 456 | 0 | 0 | 0 | 0 | 0 | 0 | 0 | 0 | 0 | 19 |
| SRR5388273 | 0 | 0 | 0 | 0 | 0 | 0 | 0 | 16 | 0 | 0 | 0 | 0 | 0 | 0 | 68 | 0 | 0 | 0 | 0 | 272 | 204 | 0 | 0 | 72 | 0 | 145 | 249 | 0 | 0 | 0 | 0 | 0 | 0 | 0 | 0 | 0 | 24 |
| SRR5388274 | 0 | 24 | 16 | 0 | 0 | 0 | 0 | 30 | 0 | 0 | 0 | 0 | 0 | 0 | 0 | 0 | 0 | 0 | 0 | 683 | 152 | 0 | 0 | 35 | 0 | 106 | 213 | 0 | 0 | 0 | 0 | 0 | 12 | 12 | 0 | 0 | 0 |
| SRR5388275 | 0 | 0 | 0 | 0 | 0 | 0 | 0 | 56 | 0 | 0 | 0 | 0 | 0 | 0 | 0 | 0 | 0 | 0 | 0 | 1607 | 189 | 0 | 0 | 131 | 0 | 56 | 103 | 0 | 0 | 37 | 0 | 19 | 0 | 19 | 0 | 0 | 0 |
| SRR5388276 | 0 | 0 | 0 | 0 | 0 | 0 | 0 | 0 | 0 | 0 | 0 | 0 | 0 | 0 | 0 | 0 | 0 | 0 | 0 | 1083 | 125 | 0 | 0 | 130 | 0 | 79 | 119 | 0 | 0 | 0 | 0 | 0 | 0 | 23 | 0 | 0 | 0 |
| SRR5388277 | 0 | 0 | 0 | 0 | 0 | 0 | 0 | 0 | 0 | 0 | 0 | 0 | 0 | 0 | 0 | 0 | 0 | 0 | 0 | 557 | 93 | 0 | 0 | 41 | 0 | 184 | 306 | 0 | 0 | 10 | 0 | 0 | 0 | 10 | 0 | 0 | 0 |
| SRR5388278 | 0 | 0 | 0 | 0 | 0 | 17 | 0 | 0 | 0 | 0 | 0 | 0 | 0 | 0 | 0 | 0 | 0 | 0 | 121 | 668 | 121 | 0 | 0 | 50 | 0 | 167 | 192 | 0 | 0 | 0 | 0 | 0 | 0 | 0 | 0 | 0 | 0 |
| SRR5388279 | 0 | 0 | 0 | 0 | 0 | 0 | 0 | 18 | 0 | 0 | 0 | 0 | 0 | 18 | 0 | 0 | 0 | 0 | 0 | 657 | 0 | 0 | 0 | 53 | 0 | 0 | 98 | 0 | 0 | 0 | 0 | 0 | 0 | 0 | 18 | 0 | 0 |
| SRR5388280 | 0 | 0 | 0 | 0 | 0 | 0 | 0 | 0 | 0 | 0 | 0 | 0 | 0 | 0 | 0 | 0 | 0 | 0 | 0 | 480 | 60 | 0 | 0 | 87 | 0 | 94 | 121 | 0 | 0 | 0 | 0 | 0 | 0 | 0 | 0 | 0 | 0 |
| SRR5388281 | 0 | 0 | 0 | 0 | 0 | 24 | 0 | 169 | 0 | 0 | 0 | 0 | 0 | 24 | 0 | 0 | 0 | 0 | 0 | 1048 | 262 | 0 | 0 | 169 | 24 | 242 | 290 | 0 | 0 | 0 | 0 | 0 | 0 | 0 | 48 | 0 | 0 |
| SRR5388282 | 0 | 14 | 0 | 0 | 0 | 43 | 0 | 28 | 14 | 0 | 0 | 0 | 0 | 0 | 0 | 0 | 0 | 0 | 0 | 757 | 108 | 0 | 0 | 185 | 28 | 469 | 724 | 0 | 0 | 0 | 0 | 0 | 0 | 0 | 14 | 0 | 0 |
| SRR5388283 | 0 | 20 | 15 | 0 | 0 | 0 | 0 | 26 | 0 | 0 | 0 | 0 | 0 | 0 | 0 | 0 | 0 | 0 | 0 | 701 | 263 | 0 | 0 | 66 | 13 | 66 | 124 | 0 | 0 | 0 | 0 | 0 | 0 | 0 | 0 | 0 | 0 |
| SRR5388284 | 0 | 0 | 0 | 0 | 0 | 24 | 0 | 0 | 0 | 0 | 0 | 0 | 0 | 0 | 0 | 0 | 0 | 0 | 0 | 307 | 137 | 0 | 0 | 67 | 0 | 62 | 271 | 0 | 0 | 0 | 0 | 0 | 0 | 0 | 0 | 0 | 0 |
| SRR5388285 | 0 | 28 | 0 | 0 | 0 | 0 | 0 | 56 | 0 | 0 | 0 | 0 | 0 | 0 | 0 | 14 | 0 | 0 | 0 | 1221 | 333 | 0 | 0 | 56 | 0 | 56 | 112 | 0 | 0 | 0 | 0 | 0 | 0 | 14 | 14 | 0 | 0 |
| SRR5388286 | 0 | 0 | 0 | 0 | 0 | 0 | 0 | 44 | 0 | 0 | 0 | 0 | 0 | 0 | 0 | 0 | 0 | 0 | 80 | 879 | 80 | 15 | 0 | 30 | 0 | 192 | 163 | 0 | 0 | 15 | 0 | 15 | 15 | 0 | 0 | 0 | 15 |
| SRR5388287 | 0 | 0 | 0 | 0 | 0 | 0 | 0 | 11 | 0 | 0 | 0 | 0 | 0 | 11 | 0 | 0 | 0 | 0 | 0 | 1033 | 176 | 0 | 0 | 34 | 11 | 137 | 148 | 0 | 0 | 0 | 0 | 0 | 0 | 11 | 0 | 0 | 0 |
| SRR5388288 | 0 | 0 | 0 | 0 | 0 | 0 | 0 | 0 | 0 | 0 | 0 | 0 | 0 | 0 | 91 | 0 | 0 | 0 | 91 | 727 | 45 | 0 | 0 | 206 | 0 | 67 | 73 | 0 | 0 | 0 | 0 | 0 | 0 | 0 | 0 | 0 | 0 |
| SRR5388289 | 0 | 0 | 0 | 0 | 0 | 12 | 0 | 12 | 0 | 0 | 0 | 0 | 0 | 12 | 0 | 0 | 0 | 0 | 0 | 670 | 149 | 0 | 0 | 105 | 0 | 128 | 279 | 0 | 0 | 0 | 0 | 0 | 0 | 12 | 0 | 0 | 0 |
| SRR5388290 | 0 | 0 | 0 | 0 | 0 | 0 | 0 | 0 | 0 | 0 | 0 | 0 | 0 | 0 | 0 | 0 | 0 | 0 | 63 | 1007 | 126 | 0 | 0 | 123 | 0 | 199 | 304 | 0 | 0 | 0 | 0 | 0 | 0 | 0 | 0 | 0 | 0 |
| SRR5388291 | 0 | 0 | 10 | 0 | 0 | 0 | 0 | 30 | 0 | 0 | 0 | 0 | 0 | 0 | 26 | 0 | 0 | 0 | 26 | 761 | 79 | 0 | 0 | 85 | 0 | 73 | 177 | 0 | 0 | 0 | 0 | 0 | 0 | 0 | 0 | 0 | 0 |
| SRR5388292 | 0 | 0 | 0 | 0 | 0 | 0 | 0 | 0 | 15 | 0 | 0 | 0 | 0 | 0 | 0 | 0 | 0 | 0 | 0 | 947 | 59 | 0 | 0 | 135 | 15 | 128 | 369 | 0 | 0 | 0 | 15 | 0 | 0 | 0 | 0 | 0 | 0 |
| SRR5388293 | 0 | 0 | 10 | 0 | 0 | 0 | 0 | 21 | 0 | 0 | 0 | 0 | 0 | 21 | 0 | 0 | 0 | 0 | 0 | 645 | 176 | 0 | 0 | 134 | 0 | 78 | 98 | 0 | 0 | 0 | 0 | 10 | 0 | 16 | 16 | 0 | 0 |
| SRR5388294 | 0 | 0 | 0 | 0 | 0 | 0 | 0 | 13 | 0 | 0 | 0 | 0 | 0 | 13 | 0 | 0 | 0 | 0 | 0 | 900 | 69 | 0 | 0 | 138 | 0 | 118 | 184 | 0 | 0 | 0 | 0 | 0 | 0 | 0 | 0 | 0 | 0 |
| SRR5476966 | 0 | 0 | 192 | 0 | 0 | 0 | 0 | 0 | 0 | 0 | 0 | 64 | 0 | 0 | 0 | 0 | 0 | 0 | 0 | 0 | 0 | 0 | 0 | 0 | 0 | 21 | 0 | 0 | 0 | 0 | 0 | 0 | 0 | 0 | 21 | 0 | 21 |
| SRR5476967 | 0 | 0 | 327 | 0 | 0 | 0 | 0 | 0 | 0 | 0 | 0 | 187 | 0 | 0 | 0 | 0 | 0 | 0 | 0 | 0 | 0 | 0 | 0 | 0 | 0 | 301 | 0 | 0 | 0 | 0 | 0 | 0 | 0 | 301 | 0 | 0 | 0 |
| SRR5476969 | 0 | 0 | 0 | 0 | 0 | 0 | 0 | 0 | 0 | 0 | 0 | 0 | 0 | 0 | 0 | 0 | 0 | 0 | 0 | 0 | 0 | 0 | 0 | 0 | 0 | 1866 | 0 | 0 | 0 | 0 | 933 | 0 | 0 | 0 | 0 | 0 | 0 |
| SRR5517463 | 0 | 13 | 452 | 19 | 18 | 30 | 0 | 59 | 19 | 17 | 87 | 759 | 0 | 35 | 12779 | 0 | 37 | 6571 | 0 | 0 | 1568 | 0 | 0 | 18 | 20 | 17 | 15 | 0 | 0 | 0 | 14 | 0 | 30 | 39 | 24 | 22 | 19 |
| SRR5517464 | 0 | 53 | 601 | 0 | 21 | 86 | 88 | 21 | 26 | 0 | 64 | 1035 | 0 | 27 | 7338 | 0 | 115 | 10865 | 0 | 0 | 5521 | 0 | 0 | 53 | 36 | 46 | 16 | 0 | 0 | 0 | 83 | 0 | 21 | 47 | 114 | 81 | 95 |
| SRR5517465 | 0 | 54 | 421 | 12 | 12 | 81 | 82 | 342 | 392 | 11 | 47 | 848 | 0 | 20 | 6264 | 0 | 67 | 6940 | 0 | 0 | 6992 | 0 | 0 | 45 | 24 | 472 | 252 | 0 | 0 | 0 | 84 | 0 | 21 | 33 | 628 | 467 | 482 |
| SRR5517466 | 0 | 50 | 492 | 20 | 26 | 31 | 0 | 86 | 283 | 18 | 47 | 708 | 0 | 37 | 8670 | 17 | 66 | 8726 | 0 | 0 | 2597 | 0 | 0 | 19 | 26 | 235 | 59 | 0 | 24 | 0 | 88 | 0 | 27 | 39 | 256 | 132 | 141 |
| SRR5517467 | 21 | 67 | 900 | 0 | 0 | 485 | 91 | 141 | 372 | 16 | 40 | 1058 | 0 | 14 | 4051 | 0 | 80 | 564 | 0 | 0 | 2491 | 0 | 0 | 18 | 14 | 1794 | 390 | 0 | 17 | 0 | 165 | 0 | 21 | 24 | 577 | 359 | 413 |
| SRR5517468 | 13 | 50 | 818 | 0 | 0 | 587 | 72 | 228 | 389 | 17 | 72 | 1173 | 0 | 0 | 3609 | 0 | 90 | 652 | 0 | 0 | 3477 | 0 | 0 | 28 | 18 | 2640 | 490 | 0 | 25 | 0 | 147 | 0 | 26 | 29 | 1483 | 1025 | 996 |
| SRR5517469 | 11 | 94 | 505 | 0 | 0 | 20 | 33 | 96 | 34 | 11 | 43 | 799 | 0 | 16 | 6939 | 0 | 46 | 1258 | 0 | 0 | 2010 | 0 | 0 | 11 | 0 | 92 | 11 | 0 | 0 | 0 | 13 | 0 | 21 | 11 | 36 | 25 | 23 |
| SRR5517470 | 13 | 51 | 419 | 0 | 0 | 38 | 53 | 23 | 36 | 0 | 37 | 915 | 0 | 17 | 4449 | 0 | 59 | 712 | 0 | 11 | 3259 | 0 | 0 | 23 | 15 | 143 | 27 | 0 | 11 | 0 | 127 | 0 | 32 | 17 | 72 | 37 | 47 |
| SRR5575011 | 0 | 0 | 211 | 0 | 0 | 62 | 21 | 0 | 22 | 0 | 0 | 222 | 25 | 0 | 0 | 0 | 0 | 0 | 260 | 4019 | 145 | 0 | 0 | 20 | 22 | 71 | 15 | 0 | 0 | 0 | 17 | 0 | 35 | 0 | 17 | 30 | 20 |
| SRR5575012 | 0 | 0 | 274 | 0 | 0 | 56 | 0 | 0 | 11 | 0 | 0 | 118 | 22 | 0 | 0 | 0 | 0 | 0 | 233 | 2724 | 54 | 0 | 0 | 109 | 0 | 68 | 17 | 0 | 0 | 0 | 11 | 0 | 16 | 0 | 29 | 33 | 16 |
| SRR5575013 | 0 | 0 | 265 | 0 | 0 | 62 | 85 | 0 | 42 | 0 | 0 | 246 | 28 | 0 | 0 | 0 | 0 | 0 | 170 | 7139 | 68 | 0 | 0 | 149 | 24 | 952 | 273 | 0 | 0 | 0 | 21 | 0 | 10 | 0 | 31 | 24 | 17 |
| SRR5575014 | 0 | 0 | 291 | 0 | 0 | 70 | 66 | 0 | 22 | 0 | 0 | 338 | 28 | 0 | 0 | 0 | 0 | 0 | 262 | 5113 | 169 | 0 | 0 | 150 | 15 | 895 | 385 | 0 | 0 | 0 | 17 | 0 | 15 | 0 | 43 | 24 | 26 |
| SRR5575015 | 0 | 0 | 334 | 0 | 0 | 45 | 88 | 0 | 13 | 0 | 0 | 290 | 24 | 0 | 0 | 0 | 0 | 0 | 234 | 6383 | 154 | 0 | 0 | 11 | 13 | 56 | 23 | 0 | 0 | 0 | 0 | 0 | 0 | 0 | 86 | 80 | 56 |
| SRR5575016 | 0 | 0 | 251 | 0 | 0 | 60 | 44 | 0 | 39 | 0 | 0 | 233 | 48 | 0 | 0 | 0 | 0 | 0 | 353 | 6096 | 44 | 0 | 0 | 136 | 42 | 985 | 395 | 0 | 0 | 0 | 39 | 0 | 15 | 0 | 30 | 27 | 24 |
| SRR5575017 | 0 | 0 | 321 | 0 | 0 | 73 | 41 | 0 | 0 | 0 | 0 | 229 | 26 | 0 | 0 | 0 | 0 | 0 | 238 | 4232 | 107 | 0 | 0 | 108 | 24 | 48 | 18 | 0 | 0 | 0 | 22 | 0 | 20 | 0 | 29 | 59 | 18 |
| SRR5575018 | 0 | 0 | 367 | 0 | 0 | 67 | 77 | 0 | 0 | 0 | 0 | 433 | 41 | 0 | 0 | 0 | 0 | 0 | 319 | 7811 | 108 | 0 | 0 | 11 | 31 | 46 | 22 | 0 | 0 | 0 | 14 | 0 | 0 | 0 | 104 | 88 | 41 |
| SRR5575019 | 0 | 0 | 253 | 0 | 0 | 44 | 83 | 0 | 24 | 0 | 0 | 254 | 34 | 0 | 0 | 0 | 0 | 0 | 306 | 6077 | 56 | 0 | 24 | 146 | 39 | 846 | 269 | 0 | 0 | 0 | 13 | 0 | 21 | 0 | 24 | 47 | 16 |
| SRR5644067 | 27 | 0 | 126 | 0 | 0 | 175 | 689 | 454 | 15 | 0 | 0 | 27 | 0 | 0 | 522 | 0 | 0 | 221 | 25 | 654 | 1845 | 0 | 0 | 37 | 0 | 912 | 264 | 11 | 0 | 15 | 78 | 0 | 0 | 0 | 231 | 30 | 130 |
| SRR5644068 | 30 | 0 | 3314 | 0 | 0 | 46 | 282 | 213 | 0 | 0 | 0 | 213 | 0 | 0 | 0 | 0 | 43 | 0 | 32 | 92 | 54 | 0 | 0 | 76 | 0 | 609 | 426 | 15 | 0 | 0 | 0 | 0 | 0 | 15 | 396 | 0 | 243 |
| SRR5644069 | 39 | 0 | 96 | 0 | 0 | 16 | 2242 | 20 | 0 | 0 | 0 | 1312 | 0 | 0 | 44 | 0 | 11 | 0 | 135 | 899 | 171 | 0 | 0 | 55 | 47 | 314 | 362 | 0 | 0 | 0 | 0 | 0 | 0 | 31 | 94 | 0 | 47 |
| SRR5644070 | 12 | 13 | 289 | 0 | 0 | 30 | 389 | 11 | 15 | 0 | 0 | 0 | 0 | 0 | 197 | 0 | 0 | 306 | 17 | 237 | 1093 | 0 | 0 | 0 | 0 | 19 | 10 | 0 | 0 | 0 | 23 | 0 | 0 | 18 | 61 | 10 | 48 |
| SRR5644071 | 26 | 0 | 1227 | 0 | 0 | 62 | 489 | 379 | 31 | 0 | 0 | 229 | 0 | 49 | 11 | 0 | 70 | 0 | 0 | 232 | 180 | 18 | 0 | 45 | 18 | 21 | 11 | 49 | 27 | 0 | 22 | 0 | 0 | 18 | 138 | 0 | 76 |
| SRR5644072 | 22 | 0 | 2091 | 0 | 0 | 21 | 100 | 368 | 31 | 0 | 0 | 21 | 0 | 26 | 98 | 0 | 12 | 0 | 0 | 61 | 100 | 0 | 0 | 0 | 0 | 43 | 54 | 0 | 132 | 102 | 0 | 0 | 0 | 76 | 14 | 0 | 28 |
| SRR5644073 | 42 | 0 | 243 | 0 | 0 | 73 | 949 | 609 | 26 | 0 | 0 | 11 | 0 | 0 | 22 | 0 | 0 | 0 | 37 | 506 | 270 | 0 | 0 | 45 | 13 | 1024 | 641 | 0 | 0 | 0 | 41 | 0 | 0 | 54 | 54 | 0 | 32 |
| SRR5644074 | 62 | 14 | 343 | 0 | 0 | 19 | 714 | 173 | 12 | 0 | 0 | 12 | 0 | 58 | 22 | 0 | 0 | 0 | 28 | 539 | 118 | 39 | 0 | 0 | 0 | 294 | 420 | 21 | 0 | 0 | 0 | 0 | 0 | 79 | 112 | 0 | 51 |
| SRR5644075 | 28 | 24 | 340 | 0 | 0 | 48 | 1032 | 30 | 42 | 0 | 0 | 0 | 0 | 18 | 21 | 0 | 0 | 12 | 38 | 392 | 131 | 0 | 0 | 60 | 18 | 21 | 12 | 18 | 0 | 18 | 30 | 0 | 0 | 18 | 60 | 0 | 36 |
| SRR5644076 | 40 | 28 | 298 | 0 | 0 | 44 | 531 | 26 | 28 | 0 | 0 | 13 | 0 | 57 | 236 | 0 | 10 | 44 | 48 | 130 | 288 | 0 | 0 | 0 | 10 | 377 | 70 | 41 | 10 | 0 | 54 | 0 | 0 | 34 | 132 | 0 | 54 |
| SRR5644077 | 41 | 72 | 525 | 0 | 0 | 38 | 110 | 25 | 28 | 0 | 0 | 0 | 0 | 133 | 208 | 0 | 15 | 0 | 0 | 136 | 93 | 0 | 0 | 0 | 22 | 38 | 26 | 35 | 0 | 0 | 38 | 0 | 0 | 107 | 95 | 0 | 73 |
| SRR5644078 | 53 | 16 | 314 | 0 | 0 | 24 | 738 | 98 | 14 | 0 | 0 | 12 | 0 | 88 | 40 | 0 | 0 | 0 | 47 | 856 | 67 | 26 | 0 | 0 | 0 | 18 | 15 | 58 | 14 | 0 | 11 | 0 | 0 | 41 | 38 | 0 | 16 |
| SRR5644085 | 40 | 18 | 245 | 0 | 0 | 25 | 495 | 50 | 0 | 0 | 0 | 71 | 0 | 65 | 529 | 0 | 0 | 0 | 26 | 1007 | 145 | 14 | 0 | 47 | 22 | 133 | 12 | 25 | 18 | 0 | 0 | 0 | 0 | 32 | 12 | 11 | 108 |
| SRR5644086 | 30 | 11 | 288 | 0 | 0 | 0 | 162 | 113 | 0 | 0 | 0 | 45 | 0 | 42 | 0 | 0 | 17 | 18 | 14 | 115 | 851 | 18 | 0 | 49 | 11 | 70 | 123 | 43 | 0 | 17 | 0 | 0 | 0 | 31 | 33 | 0 | 38 |
| SRR5644087 | 33 | 11 | 403 | 0 | 0 | 55 | 192 | 49 | 33 | 0 | 0 | 22 | 0 | 71 | 41 | 0 | 0 | 0 | 17 | 33 | 211 | 11 | 0 | 22 | 0 | 17 | 15 | 0 | 0 | 44 | 33 | 0 | 0 | 214 | 143 | 16 | 121 |
| SRR5644088 | 32 | 0 | 229 | 0 | 0 | 48 | 444 | 626 | 10 | 0 | 0 | 16 | 10 | 20 | 17 | 0 | 13 | 0 | 73 | 116 | 341 | 0 | 0 | 14 | 0 | 810 | 477 | 0 | 0 | 44 | 24 | 0 | 0 | 531 | 82 | 10 | 71 |
| SRR5657537 | 82 | 0 | 1172 | 0 | 0 | 0 | 322 | 28 | 0 | 0 | 0 | 13 | 0 | 0 | 457 | 0 | 12 | 0 | 39 | 320 | 7365 | 0 | 0 | 0 | 39 | 87 | 163 | 19 | 0 | 0 | 0 | 0 | 0 | 135 | 120 | 10 | 88 |
| SRR5657538 | 40 | 10 | 2320 | 12 | 16 | 0 | 39 | 0 | 0 | 0 | 0 | 90 | 0 | 43 | 903 | 0 | 14 | 43 | 0 | 53 | 4422 | 0 | 0 | 0 | 85 | 38 | 63 | 57 | 10 | 0 | 0 | 0 | 0 | 193 | 34 | 0 | 22 |
| SRR5657539 | 36 | 0 | 677 | 12 | 11 | 0 | 18 | 0 | 0 | 0 | 0 | 17 | 0 | 13 | 631 | 0 | 0 | 0 | 0 | 153 | 7774 | 0 | 0 | 0 | 19 | 34 | 39 | 14 | 18 | 0 | 0 | 0 | 0 | 64 | 16 | 0 | 21 |
| SRR5657540 | 10 | 0 | 1325 | 0 | 0 | 0 | 30 | 0 | 0 | 0 | 0 | 35 | 0 | 10 | 3447 | 0 | 0 | 0 | 0 | 43 | 4827 | 0 | 0 | 0 | 82 | 25 | 25 | 24 | 0 | 0 | 0 | 0 | 0 | 265 | 68 | 0 | 38 |
| SRR5657541 | 12 | 0 | 196 | 0 | 13 | 0 | 73 | 0 | 0 | 0 | 0 | 0 | 0 | 32 | 508 | 0 | 0 | 0 | 22 | 137 | 4985 | 0 | 0 | 0 | 59 | 20 | 23 | 58 | 0 | 0 | 0 | 0 | 0 | 39 | 16 | 19 | 56 |
| SRR5657542 | 17 | 0 | 2313 | 0 | 0 | 0 | 81 | 0 | 0 | 0 | 0 | 0 | 0 | 27 | 698 | 10 | 0 | 11 | 0 | 45 | 1037 | 0 | 0 | 0 | 61 | 49 | 46 | 58 | 0 | 0 | 0 | 0 | 0 | 280 | 70 | 10 | 38 |
| SRR5657543 | 11 | 0 | 388 | 11 | 19 | 0 | 27 | 0 | 0 | 0 | 0 | 0 | 0 | 17 | 973 | 0 | 0 | 0 | 0 | 30 | 1953 | 0 | 0 | 0 | 37 | 24 | 18 | 13 | 0 | 0 | 0 | 0 | 0 | 68 | 11 | 0 | 20 |
| SRR5657544 | 18 | 0 | 1346 | 0 | 0 | 0 | 56 | 18 | 0 | 0 | 0 | 31 | 0 | 0 | 236 | 0 | 0 | 0 | 31 | 58 | 4639 | 0 | 0 | 0 | 23 | 14 | 21 | 46 | 0 | 0 | 0 | 0 | 0 | 22 | 24 | 0 | 24 |
| SRR5657545 | 0 | 11 | 509 | 0 | 0 | 0 | 0 | 0 | 0 | 0 | 0 | 11 | 20 | 11 | 12 | 0 | 11 | 0 | 0 | 0 | 369 | 0 | 0 | 0 | 580 | 252 | 118 | 0 | 0 | 0 | 28 | 0 | 0 | 33 | 24 | 0 | 13 |
| SRR5657546 | 0 | 0 | 1109 | 0 | 0 | 14 | 0 | 0 | 0 | 0 | 0 | 90 | 0 | 12 | 0 | 0 | 0 | 0 | 0 | 23 | 468 | 0 | 0 | 0 | 70 | 80 | 89 | 0 | 0 | 0 | 40 | 0 | 0 | 36 | 46 | 0 | 27 |
| SRR5658693 | 48 | 0 | 1117 | 0 | 0 | 13 | 0 | 0 | 17 | 0 | 0 | 23 | 0 | 0 | 54 | 0 | 12 | 0 | 15 | 1633 | 966 | 0 | 0 | 0 | 22 | 581 | 144 | 12 | 0 | 0 | 15 | 0 | 0 | 14 | 141 | 16 | 81 |
| SRR5658694 | 69 | 0 | 1798 | 0 | 0 | 46 | 22 | 25 | 28 | 0 | 0 | 26 | 0 | 0 | 39 | 0 | 14 | 0 | 17 | 1966 | 1344 | 0 | 0 | 0 | 29 | 967 | 224 | 0 | 15 | 0 | 14 | 0 | 0 | 0 | 310 | 21 | 223 |
| SRR5658695 | 43 | 0 | 924 | 0 | 0 | 39 | 30 | 0 | 28 | 0 | 0 | 26 | 0 | 0 | 60 | 0 | 12 | 0 | 0 | 2137 | 963 | 0 | 0 | 0 | 32 | 764 | 151 | 0 | 13 | 0 | 31 | 0 | 0 | 0 | 360 | 28 | 243 |
| SRR5658696 | 52 | 0 | 1520 | 0 | 0 | 41 | 39 | 0 | 28 | 0 | 0 | 32 | 0 | 0 | 94 | 0 | 12 | 0 | 11 | 2592 | 1268 | 0 | 0 | 0 | 23 | 900 | 230 | 0 | 13 | 0 | 29 | 0 | 0 | 0 | 375 | 41 | 263 |
| SRR5658697 | 74 | 0 | 1661 | 0 | 0 | 67 | 11 | 0 | 46 | 0 | 0 | 44 | 0 | 11 | 131 | 0 | 16 | 0 | 0 | 2042 | 1412 | 0 | 0 | 0 | 36 | 110 | 29 | 0 | 15 | 0 | 46 | 0 | 0 | 0 | 56 | 34 | 37 |
| SRR5687455 | 88 | 162 | 5340 | 0 | 14 | 0 | 31 | 121 | 0 | 0 | 0 | 269 | 0 | 90 | 4105 | 0 | 0 | 10 | 0 | 34 | 3231 | 0 | 0 | 0 | 0 | 58 | 140 | 0 | 0 | 0 | 0 | 0 | 0 | 25 | 802 | 83 | 1120 |
| SRR5687456 | 71 | 141 | 5412 | 0 | 21 | 0 | 28 | 132 | 0 | 0 | 0 | 252 | 0 | 88 | 4122 | 0 | 0 | 0 | 0 | 28 | 3092 | 0 | 0 | 0 | 0 | 88 | 187 | 0 | 0 | 0 | 0 | 0 | 0 | 49 | 1635 | 113 | 1413 |
| SRR5687457 | 96 | 29 | 6153 | 12 | 0 | 0 | 0 | 19 | 0 | 0 | 0 | 228 | 0 | 15 | 4823 | 0 | 0 | 0 | 0 | 12 | 3355 | 0 | 0 | 0 | 0 | 17 | 21 | 0 | 0 | 0 | 0 | 0 | 0 | 35 | 286 | 15 | 223 |
| SRR5687458 | 80 | 160 | 5582 | 0 | 14 | 0 | 16 | 66 | 0 | 0 | 0 | 253 | 0 | 96 | 4707 | 0 | 0 | 0 | 0 | 12 | 4041 | 0 | 0 | 0 | 0 | 75 | 168 | 0 | 0 | 0 | 0 | 0 | 0 | 20 | 1658 | 146 | 1427 |
| SRR5687459 | 86 | 23 | 5522 | 0 | 20 | 0 | 13 | 12 | 0 | 0 | 0 | 255 | 0 | 11 | 4319 | 0 | 0 | 0 | 0 | 22 | 2942 | 0 | 0 | 0 | 0 | 11 | 21 | 11 | 0 | 0 | 0 | 0 | 0 | 19 | 165 | 18 | 184 |
| SRR5687461 | 35 | 736 | 1799 | 0 | 0 | 0 | 19 | 72 | 0 | 0 | 0 | 303 | 0 | 459 | 2435 | 0 | 0 | 10 | 0 | 0 | 567 | 0 | 0 | 0 | 0 | 25 | 52 | 0 | 0 | 0 | 0 | 0 | 11 | 21 | 835 | 59 | 551 |
| SRR5687462 | 60 | 143 | 2130 | 0 | 15 | 0 | 16 | 27 | 0 | 0 | 0 | 391 | 0 | 79 | 2317 | 0 | 0 | 0 | 0 | 0 | 982 | 0 | 0 | 0 | 0 | 12 | 17 | 0 | 0 | 0 | 0 | 0 | 0 | 22 | 230 | 15 | 166 |
| SRR5687463 | 58 | 123 | 2006 | 0 | 0 | 0 | 20 | 20 | 0 | 0 | 0 | 361 | 0 | 68 | 3020 | 0 | 0 | 0 | 0 | 16 | 1066 | 0 | 0 | 0 | 0 | 13 | 15 | 13 | 0 | 0 | 0 | 0 | 0 | 21 | 292 | 21 | 207 |
| SRR5687464 | 46 | 518 | 1962 | 0 | 11 | 0 | 0 | 23 | 0 | 0 | 0 | 310 | 0 | 246 | 5510 | 0 | 0 | 28 | 0 | 0 | 477 | 0 | 0 | 0 | 0 | 17 | 15 | 0 | 0 | 0 | 0 | 0 | 0 | 23 | 516 | 35 | 351 |
| SRR5687465 | 82 | 85 | 2378 | 11 | 13 | 0 | 0 | 83 | 11 | 0 | 0 | 421 | 0 | 79 | 649 | 0 | 0 | 0 | 0 | 0 | 1492 | 0 | 0 | 0 | 0 | 253 | 248 | 0 | 0 | 0 | 0 | 0 | 0 | 27 | 1360 | 89 | 1011 |
| SRR5687466 | 56 | 12 | 2520 | 18 | 18 | 0 | 0 | 57 | 0 | 0 | 0 | 373 | 0 | 17 | 880 | 0 | 0 | 0 | 0 | 16 | 981 | 0 | 0 | 0 | 0 | 23 | 29 | 0 | 0 | 0 | 0 | 0 | 0 | 34 | 247 | 13 | 155 |
| SRR5687467 | 79 | 67 | 2626 | 15 | 21 | 0 | 0 | 56 | 10 | 0 | 0 | 425 | 0 | 89 | 838 | 0 | 0 | 0 | 0 | 0 | 1258 | 0 | 0 | 0 | 0 | 177 | 223 | 0 | 0 | 0 | 0 | 0 | 0 | 34 | 1510 | 85 | 995 |
| SRR5687468 | 26 | 75 | 2872 | 13 | 22 | 0 | 0 | 23 | 0 | 0 | 0 | 184 | 0 | 139 | 1279 | 0 | 0 | 0 | 0 | 0 | 422 | 0 | 0 | 0 | 0 | 56 | 75 | 0 | 0 | 0 | 0 | 0 | 0 | 28 | 1605 | 72 | 741 |
| SRR5687469 | 52 | 12 | 3131 | 11 | 18 | 0 | 0 | 37 | 0 | 0 | 0 | 381 | 0 | 17 | 1701 | 0 | 0 | 0 | 0 | 0 | 1404 | 0 | 0 | 0 | 0 | 22 | 25 | 0 | 0 | 0 | 0 | 0 | 0 | 27 | 294 | 18 | 167 |
| SRR5753916 | 72 | 0 | 43 | 0 | 0 | 35 | 87 | 70 | 13 | 0 | 15 | 398 | 26 | 0 | 69 | 0 | 339 | 0 | 54 | 360 | 359 | 0 | 0 | 33 | 17 | 158 | 182 | 0 | 0 | 23 | 40 | 20 | 0 | 61 | 1627 | 713 | 1032 |
| SRR5755813 | 48 | 0 | 861 | 0 | 0 | 126 | 13 | 126 | 20 | 0 | 0 | 0 | 0 | 10 | 567 | 0 | 16 | 23 | 13 | 607 | 675 | 0 | 0 | 0 | 0 | 1345 | 922 | 0 | 0 | 0 | 30 | 0 | 0 | 0 | 1173 | 35 | 267 |
| SRR5755814 | 83 | 0 | 1621 | 0 | 0 | 26 | 0 | 0 | 26 | 0 | 0 | 17 | 0 | 0 | 220 | 0 | 12 | 0 | 0 | 879 | 366 | 0 | 0 | 0 | 0 | 60 | 31 | 0 | 0 | 0 | 52 | 0 | 0 | 0 | 45 | 104 | 18 |

### Supp Table S8.2.Columns_AM_to_BW

| | | | | | | | | | | | | | | | | | | | | | | | | | | | | | | | | | | | | | |
|---|---|---|---|---|---|---|---|---|---|---|---|---|---|---|---|---|---|---|---|---|---|---|---|---|---|---|---|---|---|---|---|---|---|---|---|---|---|
| SRR5755815 | 48 | 0 | 3438 | 0 | 0 | 35 | 58 | 105 | 35 | 0 | 0 | 219 | 0 | 0 | 58 | 0 | 57 | 0 | 0 | 1336 | 116 | 0 | 0 | 0 | 0 | 13 | 209 | 0 | 35 | 0 | 0 | 0 | 0 | 35 | 113 | 175 | 26 |
| SRR5755818 | 32 | 0 | 8132 | 0 | 0 | 12 | 59 | 0 | 0 | 0 | 0 | 0 | 0 | 0 | 887 | 0 | 11 | 24 | 0 | 225 | 367 | 0 | 0 | 0 | 0 | 189 | 155 | 0 | 0 | 0 | 12 | 0 | 0 | 0 | 464 | 19 | 104 |
| SRR5755819 | 39 | 0 | 1087 | 0 | 0 | 78 | 0 | 11 | 45 | 0 | 0 | 0 | 0 | 0 | 522 | 0 | 17 | 0 | 0 | 384 | 412 | 0 | 0 | 0 | 11 | 62 | 17 | 0 | 0 | 0 | 56 | 0 | 0 | 0 | 40 | 11 | 202 |
| SRR5755820 | 52 | 0 | 1570 | 0 | 0 | 29 | 0 | 0 | 59 | 0 | 0 | 0 | 0 | 0 | 576 | 0 | 0 | 0 | 0 | 432 | 432 | 0 | 0 | 0 | 0 | 2532 | 707 | 0 | 0 | 0 | 206 | 0 | 0 | 0 | 1678 | 29 | 353 |
| SRR5755821 | 51 | 0 | 2448 | 0 | 0 | 30 | 180 | 20 | 20 | 0 | 0 | 297 | 0 | 60 | 80 | 0 | 84 | 23 | 20 | 100 | 40 | 0 | 0 | 0 | 20 | 37 | 25 | 0 | 0 | 0 | 30 | 0 | 0 | 0 | 88 | 100 | 17 |
| SRR5755824 | 36 | 0 | 8048 | 0 | 0 | 28 | 49 | 14 | 0 | 0 | 0 | 0 | 0 | 14 | 98 | 0 | 21 | 0 | 0 | 341 | 49 | 0 | 0 | 0 | 0 | 268 | 99 | 0 | 0 | 0 | 42 | 0 | 0 | 14 | 480 | 0 | 113 |
| SRR5755828 | 47 | 0 | 2270 | 0 | 0 | 25 | 662 | 10 | 10 | 0 | 0 | 341 | 0 | 25 | 31 | 0 | 79 | 16 | 61 | 785 | 41 | 0 | 0 | 0 | 0 | 354 | 348 | 0 | 0 | 0 | 17 | 0 | 0 | 0 | 1354 | 52 | 254 |
| SRR5755830 | 80 | 11 | 1438 | 0 | 0 | 119 | 0 | 43 | 0 | 0 | 0 | 0 | 0 | 0 | 49 | 0 | 16 | 0 | 97 | 534 | 194 | 0 | 0 | 11 | 33 | 2105 | 1139 | 11 | 11 | 0 | 54 | 0 | 0 | 0 | 4199 | 11 | 434 |
| SRR5755831 | 31 | 0 | 8230 | 0 | 0 | 17 | 468 | 38 | 0 | 0 | 0 | 0 | 0 | 0 | 134 | 0 | 31 | 0 | 22 | 22 | 45 | 0 | 0 | 0 | 0 | 49 | 20 | 0 | 0 | 0 | 24 | 0 | 0 | 0 | 53 | 0 | 12 |
| SRR5755833 | 66 | 0 | 791 | 0 | 0 | 129 | 0 | 17 | 53 | 0 | 0 | 0 | 0 | 12 | 531 | 0 | 0 | 0 | 0 | 292 | 265 | 0 | 0 | 18 | 0 | 114 | 68 | 0 | 0 | 0 | 18 | 0 | 0 | 12 | 64 | 29 | 25 |
| SRR5755835 | 53 | 19 | 2886 | 13 | 0 | 31 | 165 | 23 | 0 | 0 | 0 | 173 | 0 | 13 | 135 | 0 | 40 | 0 | 30 | 75 | 30 | 0 | 0 | 13 | 0 | 18 | 13 | 0 | 0 | 0 | 31 | 0 | 0 | 0 | 67 | 69 | 23 |
| SRR5755837 | 46 | 19 | 1393 | 23 | 0 | 80 | 18 | 30 | 0 | 0 | 0 | 0 | 0 | 0 | 1666 | 0 | 26 | 27 | 0 | 107 | 914 | 0 | 0 | 0 | 0 | 151 | 97 | 0 | 0 | 0 | 14 | 0 | 0 | 0 | 152 | 23 | 33 |
| SRR5755838 | 40 | 0 | 5518 | 0 | 0 | 33 | 84 | 59 | 46 | 0 | 0 | 0 | 0 | 0 | 292 | 0 | 0 | 0 | 0 | 71 | 80 | 0 | 0 | 0 | 0 | 21 | 12 | 0 | 0 | 0 | 50 | 0 | 0 | 0 | 35 | 13 | 197 |
| SRR5755842 | 50 | 30 | 2942 | 0 | 0 | 60 | 37 | 104 | 0 | 0 | 0 | 248 | 0 | 0 | 19 | 0 | 65 | 0 | 93 | 708 | 37 | 0 | 0 | 0 | 0 | 34 | 32 | 0 | 0 | 15 | 30 | 0 | 0 | 0 | 82 | 75 | 23 |
| SRR5755844 | 28 | 0 | 2229 | 0 | 0 | 17 | 82 | 333 | 17 | 0 | 0 | 0 | 0 | 0 | 247 | 0 | 0 | 0 | 412 | 577 | 824 | 0 | 0 | 17 | 0 | 1116 | 650 | 0 | 0 | 0 | 0 | 50 | 0 | 0 | 1466 | 0 | 300 |
| SRR5755847 | 74 | 0 | 1346 | 0 | 0 | 75 | 0 | 299 | 37 | 0 | 0 | 0 | 0 | 0 | 541 | 0 | 0 | 0 | 180 | 1983 | 90 | 0 | 0 | 75 | 0 | 2278 | 1008 | 0 | 0 | 0 | 112 | 0 | 0 | 0 | 1307 | 37 | 149 |
| SRR5755849 | 37 | 0 | 2599 | 16 | 0 | 0 | 87 | 31 | 0 | 0 | 0 | 257 | 16 | 16 | 43 | 0 | 81 | 0 | 11 | 54 | 54 | 0 | 0 | 31 | 16 | 24 | 27 | 31 | 31 | 0 | 0 | 0 | 0 | 0 | 61 | 47 | 21 |
| SRR5755851 | 75 | 0 | 1763 | 0 | 0 | 59 | 138 | 21 | 12 | 0 | 0 | 0 | 0 | 0 | 874 | 0 | 27 | 0 | 414 | 1012 | 690 | 0 | 0 | 0 | 0 | 71 | 29 | 0 | 0 | 0 | 0 | 0 | 0 | 0 | 73 | 24 | 15 |
| SRR5755854 | 62 | 0 | 699 | 0 | 0 | 205 | 46 | 205 | 85 | 0 | 0 | 0 | 0 | 0 | 291 | 0 | 21 | 0 | 0 | 368 | 276 | 0 | 0 | 34 | 0 | 1914 | 718 | 0 | 0 | 0 | 68 | 0 | 0 | 0 | 1316 | 17 | 342 |
| SRR5755855 | 101 | 0 | 1168 | 0 | 0 | 122 | 65 | 223 | 61 | 0 | 0 | 0 | 0 | 20 | 366 | 0 | 22 | 0 | 0 | 301 | 172 | 0 | 0 | 61 | 0 | 85 | 34 | 0 | 0 | 0 | 61 | 0 | 0 | 0 | 52 | 41 | 11 |
| SRR5755856 | 52 | 0 | 2660 | 0 | 0 | 68 | 78 | 190 | 14 | 0 | 0 | 294 | 0 | 27 | 52 | 0 | 83 | 0 | 44 | 497 | 35 | 0 | 0 | 0 | 0 | 1263 | 991 | 14 | 0 | 14 | 14 | 0 | 0 | 41 | 1969 | 109 | 489 |
| SRR5755858 | 36 | 0 | 1982 | 0 | 0 | 113 | 56 | 273 | 19 | 0 | 0 | 0 | 0 | 0 | 1181 | 0 | 39 | 51 | 14 | 436 | 1069 | 0 | 0 | 0 | 0 | 1381 | 865 | 0 | 0 | 0 | 28 | 0 | 0 | 0 | 1720 | 23 | 268 |
| SRR5755861 | 86 | 0 | 1281 | 0 | 0 | 132 | 88 | 143 | 44 | 0 | 0 | 0 | 0 | 0 | 326 | 0 | 19 | 25 | 0 | 514 | 796 | 0 | 0 | 11 | 11 | 2555 | 984 | 0 | 0 | 0 | 27 | 0 | 0 | 0 | 670 | 27 | 198 |
| SRR5755862 | 80 | 0 | 1460 | 0 | 0 | 11 | 73 | 13 | 34 | 0 | 0 | 0 | 0 | 0 | 260 | 0 | 23 | 0 | 0 | 390 | 511 | 0 | 0 | 25 | 0 | 96 | 43 | 0 | 0 | 17 | 25 | 0 | 0 | 0 | 38 | 17 | 143 |
| SRR5755863 | 41 | 0 | 3234 | 0 | 17 | 59 | 148 | 31 | 34 | 0 | 0 | 407 | 0 | 0 | 15 | 0 | 96 | 0 | 37 | 655 | 93 | 0 | 0 | 34 | 17 | 43 | 37 | 0 | 17 | 0 | 25 | 0 | 0 | 0 | 69 | 59 | 18 |
| SRR5755865 | 48 | 0 | 1652 | 0 | 0 | 45 | 39 | 27 | 26 | 0 | 0 | 0 | 0 | 13 | 157 | 0 | 15 | 20 | 20 | 372 | 1332 | 19 | 0 | 13 | 0 | 77 | 59 | 0 | 0 | 0 | 26 | 0 | 0 | 0 | 88 | 13 | 16 |
| SRR5755866 | 62 | 0 | 7744 | 0 | 0 | 65 | 81 | 106 | 24 | 0 | 0 | 11 | 0 | 0 | 71 | 0 | 19 | 0 | 0 | 323 | 337 | 0 | 0 | 0 | 0 | 54 | 38 | 0 | 0 | 0 | 47 | 0 | 0 | 0 | 37 | 24 | 15 |
| SRR5755868 | 102 | 0 | 732 | 0 | 0 | 52 | 21 | 118 | 39 | 0 | 0 | 0 | 0 | 0 | 342 | 0 | 0 | 0 | 0 | 427 | 427 | 0 | 0 | 0 | 13 | 1008 | 458 | 0 | 0 | 0 | 13 | 0 | 0 | 0 | 262 | 0 | 157 |
| SRR5755870 | 69 | 0 | 2301 | 0 | 0 | 33 | 124 | 432 | 13 | 0 | 0 | 418 | 0 | 26 | 32 | 0 | 66 | 0 | 16 | 434 | 80 | 0 | 0 | 39 | 13 | 674 | 608 | 0 | 0 | 0 | 26 | 0 | 0 | 0 | 1001 | 20 | 471 |
| SRR5755875 | 32 | 0 | 1529 | 0 | 0 | 202 | 35 | 104 | 97 | 0 | 0 | 0 | 0 | 0 | 124 | 0 | 31 | 0 | 0 | 230 | 124 | 0 | 0 | 0 | 0 | 146 | 49 | 0 | 0 | 0 | 52 | 0 | 0 | 0 | 38 | 0 | 12 |
| SRR5755876 | 36 | 0 | 2303 | 0 | 0 | 11 | 0 | 22 | 22 | 0 | 0 | 0 | 0 | 15 | 77 | 0 | 53 | 0 | 0 | 0 | 48 | 0 | 0 | 0 | 0 | 176 | 44 | 0 | 0 | 0 | 96 | 0 | 0 | 0 | 38 | 15 | 133 |
| SRR5755877 | 43 | 12 | 3047 | 0 | 0 | 51 | 46 | 636 | 27 | 0 | 0 | 354 | 0 | 30 | 17 | 0 | 57 | 0 | 0 | 723 | 61 | 0 | 0 | 27 | 21 | 857 | 1237 | 0 | 0 | 12 | 27 | 0 | 0 | 24 | 1485 | 84 | 630 |
| SRR5755879 | 43 | 17 | 2405 | 0 | 0 | 94 | 13 | 65 | 19 | 0 | 0 | 0 | 0 | 19 | 758 | 0 | 33 | 24 | 0 | 280 | 1026 | 0 | 0 | 0 | 0 | 117 | 121 | 0 | 0 | 0 | 24 | 0 | 0 | 0 | 71 | 22 | 14 |
| SRR5755880 | 23 | 0 | 9132 | 0 | 0 | 25 | 17 | 72 | 21 | 0 | 0 | 0 | 0 | 0 | 61 | 0 | 27 | 0 | 0 | 142 | 78 | 0 | 0 | 0 | 0 | 21 | 15 | 0 | 0 | 0 | 25 | 0 | 0 | 0 | 23 | 0 | 191 |
| SRR5755882 | 75 | 0 | 1499 | 0 | 0 | 102 | 27 | 166 | 13 | 0 | 0 | 0 | 0 | 13 | 121 | 0 | 16 | 0 | 0 | 135 | 67 | 0 | 0 | 26 | 0 | 66 | 24 | 0 | 13 | 0 | 26 | 0 | 0 | 0 | 26 | 26 | 229 |
| SRR5755884 | 46 | 13 | 2625 | 0 | 0 | 50 | 109 | 730 | 25 | 0 | 0 | 209 | 0 | 38 | 28 | 0 | 60 | 0 | 46 | 524 | 37 | 0 | 0 | 20 | 0 | 690 | 757 | 0 | 10 | 0 | 28 | 0 | 0 | 13 | 1843 | 75 | 642 |
| SRR5755887 | 20 | 0 | 9996 | 0 | 0 | 81 | 12 | 48 | 16 | 0 | 0 | 0 | 0 | 0 | 93 | 0 | 12 | 0 | 0 | 23 | 35 | 0 | 0 | 0 | 0 | 516 | 323 | 0 | 0 | 0 | 0 | 0 | 0 | 0 | 694 | 16 | 113 |
| SRR5755889 | 95 | 34 | 1016 | 0 | 0 | 109 | 91 | 1204 | 34 | 0 | 0 | 0 | 0 | 21 | 224 | 0 | 0 | 0 | 0 | 373 | 207 | 0 | 0 | 0 | 0 | 992 | 513 | 0 | 0 | 14 | 41 | 0 | 0 | 0 | 1135 | 68 | 486 |
| SRR5755890 | 81 | 15 | 970 | 0 | 0 | 85 | 87 | 495 | 0 | 0 | 0 | 0 | 0 | 0 | 87 | 0 | 0 | 0 | 0 | 87 | 87 | 0 | 0 | 0 | 0 | 1245 | 348 | 0 | 0 | 0 | 62 | 15 | 0 | 15 | 1570 | 46 | 364 |
| SRR5755891 | 82 | 0 | 2313 | 0 | 0 | 55 | 183 | 289 | 43 | 0 | 0 | 453 | 0 | 0 | 0 | 0 | 60 | 0 | 26 | 142 | 26 | 0 | 0 | 0 | 12 | 1187 | 744 | 0 | 0 | 0 | 55 | 0 | 0 | 0 | 1236 | 80 | 891 |
| SRR5755894 | 61 | 23 | 4497 | 0 | 0 | 37 | 145 | 138 | 23 | 0 | 0 | 12 | 0 | 0 | 102 | 0 | 0 | 11 | 0 | 327 | 348 | 0 | 0 | 0 | 0 | 29 | 10 | 0 | 0 | 0 | 51 | 0 | 0 | 0 | 23 | 37 | 20 |
| SRR5755896 | 23 | 0 | 963 | 0 | 0 | 199 | 33 | 47 | 88 | 0 | 0 | 0 | 0 | 12 | 151 | 0 | 19 | 0 | 0 | 118 | 63 | 0 | 0 | 0 | 0 | 77 | 25 | 0 | 0 | 0 | 47 | 0 | 0 | 0 | 54 | 41 | 11 |
| SRR5755897 | 26 | 0 | 1582 | 0 | 0 | 136 | 0 | 0 | 48 | 0 | 0 | 0 | 0 | 0 | 37 | 0 | 24 | 0 | 0 | 0 | 17 | 0 | 0 | 0 | 0 | 1868 | 527 | 0 | 0 | 0 | 40 | 0 | 0 | 0 | 1445 | 24 | 359 |
| SRR5755898 | 53 | 0 | 2236 | 14 | 0 | 72 | 143 | 19 | 0 | 0 | 0 | 269 | 0 | 14 | 158 | 0 | 53 | 0 | 43 | 416 | 57 | 0 | 0 | 14 | 0 | 43 | 65 | 0 | 14 | 0 | 57 | 0 | 0 | 0 | 182 | 129 | 48 |
| SRR5755901 | 15 | 0 | 8277 | 0 | 0 | 31 | 12 | 26 | 22 | 0 | 0 | 0 | 0 | 0 | 31 | 0 | 25 | 0 | 0 | 12 | 12 | 0 | 0 | 0 | 0 | 724 | 331 | 0 | 0 | 0 | 31 | 0 | 0 | 0 | 1298 | 57 | 406 |
| SRR5755903 | 32 | 12 | 935 | 0 | 0 | 160 | 105 | 141 | 42 | 0 | 0 | 0 | 0 | 0 | 37 | 0 | 18 | 0 | 0 | 125 | 68 | 0 | 0 | 0 | 0 | 61 | 24 | 0 | 0 | 0 | 54 | 0 | 0 | 0 | 29 | 26 | 286 |
| SRR5755905 | 37 | 0 | 2649 | 0 | 0 | 49 | 535 | 53 | 11 | 0 | 0 | 256 | 0 | 0 | 15 | 0 | 81 | 0 | 36 | 72 | 24 | 0 | 0 | 0 | 19 | 541 | 734 | 11 | 0 | 0 | 19 | 0 | 0 | 0 | 1526 | 83 | 519 |
| SRR5755908 | 27 | 0 | 7162 | 0 | 0 | 48 | 13 | 48 | 0 | 0 | 0 | 0 | 0 | 0 | 0 | 0 | 19 | 0 | 0 | 0 | 0 | 0 | 0 | 0 | 0 | 875 | 461 | 0 | 0 | 0 | 32 | 0 | 0 | 0 | 422 | 16 | 127 |
| SRR5755910 | 19 | 0 | 588 | 0 | 0 | 194 | 56 | 27 | 57 | 0 | 0 | 0 | 0 | 0 | 22 | 0 | 23 | 0 | 0 | 20 | 13 | 0 | 0 | 0 | 19 | 2930 | 782 | 0 | 0 | 0 | 46 | 0 | 0 | 0 | 660 | 16 | 194 |
| SRR5755911 | 13 | 0 | 774 | 0 | 0 | 235 | 0 | 63 | 23 | 0 | 0 | 0 | 0 | 0 | 0 | 0 | 15 | 0 | 0 | 0 | 14 | 0 | 0 | 0 | 0 | 88 | 18 | 0 | 0 | 0 | 47 | 0 | 0 | 23 | 18 | 47 | 243 |
| SRR5755912 | 28 | 0 | 2002 | 0 | 10 | 66 | 1001 | 107 | 35 | 0 | 0 | 216 | 0 | 0 | 17 | 0 | 69 | 0 | 35 | 199 | 30 | 0 | 0 | 0 | 0 | 21 | 21 | 0 | 14 | 0 | 24 | 0 | 0 | 17 | 49 | 86 | 16 |
| SRR5755914 | 57 | 13 | 666 | 0 | 0 | 67 | 20 | 63 | 13 | 0 | 0 | 0 | 0 | 0 | 572 | 0 | 29 | 56 | 0 | 257 | 869 | 0 | 0 | 0 | 0 | 1038 | 603 | 0 | 0 | 0 | 21 | 0 | 0 | 0 | 883 | 0 | 151 |
| SRR5755915 | 13 | 0 | 6306 | 0 | 0 | 46 | 12 | 63 | 0 | 0 | 0 | 12 | 0 | 0 | 0 | 0 | 28 | 0 | 0 | 0 | 0 | 0 | 0 | 0 | 0 | 1388 | 427 | 0 | 0 | 11 | 34 | 0 | 0 | 63 | 586 | 17 | 222 |
| SRR5755917 | 40 | 0 | 1692 | 0 | 0 | 165 | 122 | 12 | 50 | 0 | 0 | 110 | 0 | 14 | 366 | 0 | 40 | 0 | 10 | 732 | 203 | 0 | 0 | 22 | 0 | 89 | 81 | 0 | 0 | 0 | 36 | 0 | 0 | 0 | 101 | 136 | 38 |
| SRR5755918 | 69 | 0 | 1923 | 0 | 0 | 11 | 128 | 183 | 27 | 0 | 0 | 0 | 0 | 0 | 256 | 0 | 0 | 0 | 38 | 602 | 243 | 0 | 0 | 14 | 0 | 121 | 58 | 0 | 0 | 0 | 54 | 0 | 0 | 0 | 46 | 34 | 17 |
| SRR5755919 | 48 | 0 | 2309 | 19 | 0 | 97 | 88 | 204 | 0 | 0 | 0 | 367 | 0 | 0 | 69 | 0 | 53 | 0 | 31 | 663 | 69 | 0 | 0 | 19 | 0 | 25 | 20 | 0 | 39 | 0 | 0 | 0 | 0 | 19 | 57 | 78 | 21 |
| SRR5755922 | 32 | 0 | 6592 | 17 | 0 | 34 | 121 | 86 | 17 | 0 | 0 | 24 | 0 | 0 | 182 | 0 | 11 | 0 | 0 | 313 | 141 | 0 | 0 | 11 | 0 | 589 | 629 | 0 | 0 | 0 | 11 | 0 | 0 | 0 | 1063 | 29 | 337 |
| SRR5755924 | 62 | 0 | 1285 | 0 | 0 | 187 | 0 | 62 | 33 | 0 | 0 | 0 | 0 | 0 | 382 | 0 | 13 | 21 | 11 | 1293 | 495 | 0 | 0 | 24 | 0 | 2647 | 1126 | 0 | 0 | 0 | 42 | 0 | 0 | 0 | 1103 | 0 | 228 |
| SRR5755925 | 83 | 0 | 1776 | 11 | 0 | 205 | 0 | 233 | 11 | 0 | 0 | 0 | 0 | 0 | 362 | 0 | 11 | 0 | 0 | 1087 | 477 | 0 | 0 | 0 | 0 | 3634 | 1371 | 0 | 0 | 0 | 40 | 0 | 0 | 0 | 1263 | 0 | 324 |
| SRR5755926 | 34 | 0 | 2365 | 11 | 0 | 53 | 60 | 13 | 16 | 0 | 0 | 254 | 0 | 0 | 111 | 0 | 52 | 0 | 23 | 396 | 69 | 0 | 0 | 16 | 0 | 34 | 30 | 16 | 0 | 0 | 42 | 11 | 0 | 32 | 54 | 74 | 23 |
| SRR5755928 | 64 | 0 | 1367 | 0 | 0 | 129 | 23 | 81 | 38 | 0 | 0 | 25 | 0 | 0 | 822 | 0 | 23 | 57 | 0 | 502 | 799 | 0 | 0 | 16 | 0 | 105 | 40 | 0 | 0 | 0 | 48 | 0 | 0 | 0 | 43 | 11 | 11 |
| SRR5755929 | 49 | 0 | 7500 | 0 | 12 | 87 | 35 | 99 | 0 | 0 | 0 | 0 | 0 | 0 | 175 | 0 | 16 | 0 | 0 | 594 | 70 | 0 | 0 | 37 | 0 | 669 | 384 | 0 | 0 | 0 | 12 | 0 | 0 | 0 | 929 | 12 | 223 |
| SRR5755931 | 45 | 0 | 696 | 0 | 0 | 172 | 196 | 609 | 49 | 0 | 0 | 0 | 0 | 0 | 82 | 0 | 17 | 0 | 0 | 428 | 155 | 0 | 0 | 0 | 0 | 2497 | 1076 | 0 | 0 | 0 | 93 | 0 | 0 | 0 | 869 | 18 | 287 |
| SRR5755933 | 43 | 30 | 2260 | 0 | 40 | 151 | 236 | 312 | 20 | 0 | 0 | 247 | 0 | 0 | 15 | 10 | 61 | 0 | 15 | 186 | 95 | 0 | 0 | 0 | 0 | 23 | 18 | 0 | 40 | 0 | 71 | 10 | 0 | 10 | 37 | 20 | 11 |
| SRR5755936 | 47 | 0 | 7080 | 0 | 0 | 72 | 55 | 449 | 13 | 0 | 0 | 12 | 0 | 0 | 0 | 0 | 24 | 0 | 0 | 30 | 0 | 0 | 0 | 15 | 0 | 753 | 469 | 0 | 0 | 0 | 31 | 0 | 0 | 0 | 616 | 26 | 294 |
| SRR5755938 | 44 | 0 | 1618 | 0 | 0 | 191 | 41 | 211 | 26 | 0 | 0 | 0 | 0 | 0 | 290 | 0 | 15 | 0 | 0 | 373 | 282 | 0 | 0 | 0 | 0 | 2587 | 1023 | 0 | 0 | 0 | 46 | 0 | 0 | 0 | 1663 | 40 | 561 |
| SRR5755939 | 59 | 0 | 1628 | 0 | 0 | 254 | 66 | 314 | 29 | 0 | 0 | 0 | 0 | 0 | 53 | 0 | 17 | 0 | 0 | 192 | 94 | 0 | 0 | 0 | 0 | 3318 | 1090 | 0 | 0 | 0 | 36 | 0 | 0 | 0 | 1679 | 22 | 520 |
| SRR5755940 | 26 | 0 | 2405 | 0 | 24 | 63 | 253 | 53 | 31 | 0 | 0 | 269 | 0 | 0 | 139 | 0 | 83 | 0 | 12 | 361 | 78 | 0 | 0 | 0 | 0 | 49 | 55 | 0 | 16 | 0 | 39 | 0 | 0 | 16 | 136 | 180 | 58 |
| SRR5755943 | 36 | 17 | 7518 | 0 | 0 | 56 | 93 | 338 | 13 | 0 | 0 | 0 | 0 | 0 | 60 | 0 | 22 | 0 | 0 | 138 | 28 | 0 | 0 | 40 | 0 | 1024 | 703 | 0 | 0 | 0 | 40 | 0 | 0 | 0 | 1611 | 46 | 527 |
| SRR5755946 | 76 | 0 | 1206 | 0 | 0 | 213 | 63 | 12 | 61 | 0 | 0 | 0 | 0 | 0 | 738 | 0 | 0 | 0 | 0 | 675 | 0 | 0 | 0 | 15 | 0 | 170 | 43 | 0 | 0 | 0 | 99 | 0 | 0 | 0 | 73 | 38 | 13 |
| SRR5755947 | 76 | 0 | 2817 | 0 | 16 | 132 | 109 | 972 | 25 | 0 | 0 | 317 | 0 | 0 | 98 | 0 | 49 | 0 | 11 | 635 | 0 | 0 | 0 | 66 | 0 | 1376 | 1079 | 0 | 0 | 0 | 58 | 0 | 0 | 0 | 3254 | 124 | 1021 |
| SRR5755950 | 41 | 0 | 6166 | 0 | 10 | 72 | 54 | 10 | 21 | 0 | 0 | 0 | 0 | 0 | 305 | 0 | 0 | 36 | 18 | 323 | 0 | 0 | 0 | 10 | 0 | 35 | 15 | 0 | 0 | 0 | 41 | 0 | 0 | 0 | 51 | 21 | 11 |
| SRR5755952 | 32 | 0 | 1317 | 0 | 0 | 160 | 204 | 16 | 28 | 0 | 0 | 0 | 0 | 0 | 216 | 0 | 12 | 0 | 11 | 318 | 148 | 0 | 0 | 0 | 0 | 113 | 57 | 0 | 0 | 0 | 75 | 0 | 0 | 0 | 71 | 19 | 21 |
| SRR5755953 | 54 | 0 | 2035 | 0 | 10 | 114 | 163 | 989 | 31 | 0 | 0 | 0 | 0 | 0 | 66 | 0 | 16 | 0 | 0 | 166 | 46 | 0 | 0 | 26 | 0 | 1973 | 1025 | 0 | 0 | 0 | 21 | 0 | 0 | 0 | 1134 | 26 | 285 |
| SRR5755954 | 44 | 0 | 2444 | 0 | 0 | 66 | 898 | 731 | 0 | 0 | 0 | 325 | 0 | 17 | 86 | 0 | 65 | 11 | 86 | 295 | 37 | 0 | 0 | 50 | 0 | 382 | 615 | 0 | 0 | 0 | 17 | 0 | 0 | 17 | 1662 | 166 | 515 |
| SRR5755956 | 43 | 0 | 1412 | 0 | 0 | 78 | 43 | 375 | 19 | 0 | 0 | 0 | 0 | 0 | 486 | 0 | 27 | 14 | 14 | 229 | 614 | 0 | 0 | 26 | 0 | 1245 | 710 | 0 | 0 | 11 | 15 | 0 | 0 | 0 | 985 | 30 | 186 |
| SRR5755957 | 43 | 12 | 8615 | 0 | 12 | 36 | 90 | 18 | 24 | 0 | 0 | 28 | 0 | 0 | 43 | 0 | 19 | 0 | 0 | 43 | 14 | 0 | 0 | 24 | 0 | 17 | 13 | 0 | 12 | 0 | 0 | 0 | 0 | 0 | 21 | 49 | 122 |
| SRR5755959 | 51 | 0 | 1232 | 0 | 11 | 164 | 18 | 77 | 49 | 0 | 0 | 0 | 0 | 0 | 430 | 0 | 23 | 0 | 0 | 466 | 233 | 0 | 0 | 0 | 0 | 2763 | 1552 | 0 | 0 | 0 | 82 | 0 | 0 | 11 | 2133 | 82 | 724 |
| SRR5755960 | 87 | 0 | 1989 | 0 | 0 | 238 | 0 | 79 | 44 | 0 | 0 | 0 | 0 | 0 | 121 | 0 | 15 | 0 | 0 | 114 | 44 | 0 | 0 | 71 | 0 | 160 | 55 | 0 | 0 | 0 | 88 | 0 | 0 | 0 | 71 | 18 | 308 |
| SRR5755961 | 63 | 0 | 3153 | 0 | 0 | 117 | 231 | 176 | 13 | 0 | 0 | 284 | 0 | 21 | 97 | 0 | 68 | 0 | 66 | 982 | 75 | 0 | 0 | 88 | 0 | 53 | 52 | 0 | 13 | 0 | 17 | 0 | 0 | 0 | 132 | 142 | 36 |
| SRR5755963 | 72 | 0 | 1511 | 0 | 0 | 121 | 33 | 1106 | 20 | 0 | 0 | 0 | 0 | 0 | 366 | 0 | 25 | 57 | 0 | 732 | 599 | 0 | 0 | 0 | 0 | 1998 | 1274 | 0 | 0 | 0 | 40 | 0 | 0 | 0 | 2494 | 20 | 402 |
| SRR5755964 | 33 | 0 | 8219 | 0 | 0 | 125 | 32 | 152 | 0 | 0 | 0 | 53 | 0 | 18 | 77 | 0 | 21 | 0 | 0 | 81 | 18 | 0 | 0 | 27 | 0 | 40 | 20 | 0 | 0 | 0 | 107 | 0 | 0 | 0 | 52 | 80 | 10 |
| SRR5755966 | 68 | 0 | 1088 | 0 | 0 | 196 | 78 | 898 | 45 | 0 | 0 | 0 | 0 | 10 | 205 | 0 | 0 | 0 | 0 | 666 | 354 | 0 | 0 | 45 | 0 | 2695 | 1390 | 0 | 0 | 0 | 45 | 0 | 0 | 0 | 1215 | 25 | 351 |
| SRR5755967 | 93 | 0 | 1565 | 0 | 0 | 119 | 121 | 44 | 48 | 0 | 0 | 0 | 0 | 0 | 29 | 0 | 16 | 0 | 0 | 112 | 87 | 0 | 0 | 0 | 0 | 66 | 32 | 0 | 0 | 0 | 0 | 0 | 0 | 0 | 39 | 48 | 11 |
| SRR5755968 | 47 | 29 | 3243 | 0 | 0 | 0 | 144 | 1877 | 17 | 0 | 0 | 294 | 0 | 0 | 183 | 0 | 43 | 0 | 39 | 269 | 79 | 0 | 0 | 0 | 0 | 345 | 697 | 0 | 0 | 0 | 23 | 0 | 0 | 0 | 1612 | 75 | 403 |
| SRR5755971 | 30 | 0 | 8488 | 0 | 0 | 15 | 69 | 66 | 15 | 0 | 0 | 0 | 0 | 0 | 418 | 0 | 26 | 0 | 0 | 156 | 168 | 0 | 0 | 0 | 0 | 20 | 13 | 0 | 0 | 0 | 0 | 0 | 0 | 0 | 42 | 12 | 169 |
| SRR5755973 | 29 | 0 | 756 | 0 | 0 | 227 | 13 | 15 | 61 | 0 | 0 | 0 | 0 | 0 | 311 | 0 | 21 | 0 | 0 | 225 | 99 | 0 | 0 | 0 | 0 | 2600 | 1410 | 0 | 0 | 0 | 30 | 0 | 0 | 0 | 1220 | 38 | 258 |
| SRR5755974 | 45 | 0 | 2314 | 0 | 0 | 37 | 0 | 0 | 33 | 0 | 0 | 0 | 0 | 0 | 263 | 0 | 21 | 0 | 0 | 0 | 14 | 0 | 0 | 0 | 25 | 71 | 31 | 0 | 0 | 0 | 62 | 0 | 0 | 0 | 36 | 0 | 115 |
| SRR5755975 | 46 | 0 | 2666 | 0 | 0 | 45 | 95 | 198 | 16 | 0 | 0 | 318 | 0 | 12 | 280 | 0 | 72 | 0 | 51 | 590 | 47 | 0 | 0 | 0 | 0 | 28 | 31 | 12 | 0 | 0 | 29 | 0 | 0 | 0 | 73 | 95 | 30 |
| SRR5755978 | 34 | 0 | 7830 | 0 | 0 | 0 | 39 | 0 | 0 | 0 | 0 | 36 | 0 | 0 | 169 | 0 | 22 | 0 | 0 | 0 | 15 | 0 | 0 | 0 | 0 | 553 | 352 | 17 | 0 | 0 | 0 | 0 | 0 | 0 | 888 | 17 | 226 |
| SRR5755980 | 41 | 0 | 1044 | 0 | 0 | 134 | 32 | 15 | 39 | 0 | 0 | 0 | 0 | 0 | 2888 | 0 | 19 | 29 | 0 | 317 | 371 | 0 | 0 | 0 | 0 | 2084 | 934 | 0 | 0 | 0 | 49 | 0 | 0 | 0 | 1071 | 46 | 225 |
| SRR5755981 | 58 | 0 | 1334 | 0 | 0 | 164 | 0 | 0 | 40 | 0 | 0 | 0 | 0 | 0 | 1550 | 0 | 16 | 17 | 0 | 366 | 327 | 0 | 0 | 0 | 0 | 3270 | 1235 | 0 | 0 | 0 | 69 | 0 | 0 | 0 | 1335 | 0 | 203 |
| SRR5755982 | 47 | 0 | 2885 | 0 | 0 | 50 | 32 | 27 | 18 | 0 | 0 | 261 | 0 | 0 | 121 | 0 | 88 | 63 | 0 | 500 | 60 | 0 | 0 | 15 | 15 | 37 | 35 | 0 | 0 | 0 | 30 | 0 | 0 | 0 | 80 | 74 | 22 |
| SRR5755984 | 57 | 0 | 625 | 0 | 0 | 114 | 0 | 49 | 34 | 0 | 0 | 0 | 0 | 0 | 1556 | 0 | 11 | 95 | 0 | 217 | 485 | 0 | 0 | 0 | 0 | 118 | 51 | 0 | 0 | 0 | 49 | 0 | 0 | 0 | 77 | 28 | 11 |
| SRR5755985 | 33 | 0 | 6810 | 0 | 0 | 69 | 0 | 0 | 16 | 0 | 0 | 0 | 0 | 0 | 255 | 0 | 24 | 0 | 0 | 21 | 50 | 0 | 0 | 0 | 0 | 898 | 425 | 0 | 0 | 0 | 44 | 0 | 0 | 0 | 1023 | 20 | 226 |
| SRR5755987 | 32 | 0 | 1106 | 0 | 0 | 84 | 21 | 27 | 35 | 0 | 0 | 0 | 0 | 13 | 447 | 0 | 21 | 0 | 0 | 115 | 53 | 0 | 0 | 0 | 0 | 82 | 30 | 0 | 0 | 0 | 53 | 0 | 0 | 0 | 39 | 27 | 199 |
| SRR5755988 | 35 | 0 | 1626 | 0 | 0 | 73 | 24 | 20 | 73 | 0 | 0 | 0 | 0 | 0 | 228 | 0 | 18 | 0 | 0 | 0 | 12 | 0 | 0 | 0 | 0 | 1618 | 497 | 0 | 0 | 0 | 63 | 0 | 0 | 0 | 585 | 29 | 93 |
| SRR5755989 | 58 | 0 | 2000 | 0 | 0 | 65 | 182 | 15 | 28 | 0 | 0 | 304 | 0 | 28 | 88 | 0 | 75 | 0 | 35 | 481 | 47 | 0 | 0 | 18 | 0 | 41 | 50 | 28 | 0 | 0 | 55 | 0 | 0 | 18 | 110 | 148 | 39 |
| SRR5755991 | 46 | 0 | 2058 | 0 | 0 | 10 | 0 | 121 | 33 | 0 | 0 | 0 | 0 | 17 | 2031 | 0 | 27 | 0 | 0 | 262 | 575 | 0 | 0 | 0 | 0 | 102 | 52 | 0 | 0 | 0 | 40 | 0 | 0 | 0 | 73 | 0 | 11 |
| SRR5755992 | 23 | 0 | 7785 | 0 | 0 | 17 | 18 | 0 | 0 | 0 | 0 | 17 | 0 | 0 | 334 | 0 | 26 | 0 | 0 | 35 | 61 | 0 | 0 | 0 | 0 | 484 | 283 | 0 | 0 | 0 | 0 | 0 | 0 | 0 | 527 | 0 | 141 |
| SRR5755994 | 59 | 0 | 2677 | 0 | 11 | 106 | 0 | 34 | 67 | 0 | 0 | 77 | 0 | 0 | 299 | 0 | 45 | 18 | 0 | 466 | 111 | 0 | 0 | 17 | 17 | 1759 | 1167 | 11 | 22 | 0 | 34 | 0 | 0 | 0 | 2754 | 106 | 586 |
| SRR5755995 | 55 | 0 | 1944 | 0 | 0 | 200 | 0 | 28 | 56 | 0 | 0 | 0 | 0 | 0 | 657 | 0 | 0 | 0 | 0 | 241 | 154 | 0 | 0 | 0 | 0 | 115 | 42 | 0 | 0 | 0 | 65 | 0 | 0 | 0 | 68 | 23 | 12 |
| SRR5755996 | 65 | 0 | 3303 | 0 | 0 | 74 | 0 | 56 | 32 | 0 | 0 | 134 | 0 | 21 | 23 | 0 | 55 | 11 | 0 | 325 | 24 | 0 | 0 | 0 | 0 | 1049 | 1171 | 13 | 21 | 0 | 61 | 0 | 0 | 0 | 4187 | 146 | 977 |
| SRR5755997 | 88 | 0 | 308 | 0 | 0 | 0 | 28 | 0 | 0 | 0 | 0 | 181 | 0 | 47 | 138 | 0 | 78 | 0 | 0 | 551 | 55 | 0 | 0 | 0 | 19 | 56 | 13 | 0 | 0 | 0 | 0 | 0 | 0 | 0 | 28 | 0 | 0 |
| SRR5755998 | 60 | 19 | 1347 | 0 | 0 | 130 | 0 | 59 | 19 | 0 | 0 | 0 | 0 | 0 | 1721 | 0 | 15 | 0 | 0 | 201 | 306 | 0 | 0 | 0 | 0 | 1756 | 958 | 0 | 0 | 0 | 26 | 0 | 0 | 0 | 2269 | 19 | 304 |
| SRR5755999 | 29 | 0 | 9821 | 0 | 0 | 106 | 0 | 11 | 17 | 0 | 0 | 0 | 0 | 0 | 272 | 0 | 18 | 0 | 0 | 127 | 100 | 0 | 0 | 11 | 0 | 32 | 26 | 0 | 17 | 0 | 67 | 0 | 0 | 0 | 51 | 44 | 10 |
| SRR5756000 | 16 | 0 | 0 | 0 | 0 | 0 | 0 | 0 | 0 | 0 | 0 | 0 | 0 | 0 | 0 | 0 | 0 | 0 | 0 | 0 | 0 | 0 | 0 | 163 | 0 | 0 | 0 | 0 | 0 | 0 | 0 | 0 | 0 | 163 | 0 | 0 | 0 |
| SRR5756001 | 38 | 0 | 1181 | 0 | 0 | 12 | 50 | 12 | 85 | 0 | 0 | 13 | 0 | 0 | 1598 | 0 | 24 | 0 | 10 | 378 | 534 | 0 | 0 | 0 | 0 | 197 | 56 | 0 | 0 | 0 | 73 | 0 | 0 | 0 | 55 | 0 | 10 |
| SRR5756002 | 83 | 0 | 1158 | 0 | 0 | 14 | 0 | 0 | 89 | 0 | 0 | 0 | 0 | 0 | 638 | 0 | 19 | 0 | 0 | 360 | 196 | 0 | 0 | 0 | 16 | 247 | 57 | 0 | 0 | 0 | 57 | 0 | 0 | 24 | 81 | 40 | 12 |
| SRR5756003 | 55 | 0 | 2726 | 0 | 12 | 111 | 70 | 34 | 54 | 0 | 0 | 264 | 0 | 19 | 67 | 0 | 65 | 0 | 12 | 753 | 58 | 0 | 0 | 0 | 0 | 1839 | 1176 | 15 | 0 | 0 | 73 | 0 | 0 | 12 | 3127 | 115 | 870 |
| SRR5756005 | 71 | 0 | 780 | 0 | 0 | 17 | 16 | 45 | 45 | 0 | 0 | 0 | 0 | 0 | 2081 | 0 | 21 | 0 | 0 | 186 | 543 | 0 | 0 | 0 | 0 | 235 | 77 | 0 | 0 | 0 | 54 | 0 | 0 | 0 | 77 | 15 | 11 |
| SRR5756006 | 43 | 0 | 8112 | 0 | 0 | 133 | 0 | 13 | 33 | 0 | 0 | 15 | 0 | 0 | 308 | 0 | 21 | 0 | 0 | 248 | 173 | 0 | 0 | 0 | 0 | 69 | 29 | 0 | 0 | 0 | 66 | 0 | 0 | 0 | 66 | 73 | 16 |
| SRR5756007 | 0 | 0 | 0 | 0 | 0 | 0 | 0 | 0 | 0 | 0 | 0 | 0 | 0 | 0 | 0 | 0 | 0 | 0 | 0 | 909 | 0 | 0 | 0 | 0 | 0 | 0 | 0 | 0 | 0 | 0 | 0 | 0 | 0 | 0 | 0 | 0 | 0 |
| SRR5756008 | 35 | 0 | 1085 | 0 | 0 | 146 | 35 | 0 | 79 | 0 | 0 | 0 | 0 | 0 | 86 | 0 | 12 | 0 | 0 | 17 | 104 | 0 | 0 | 0 | 18 | 219 | 35 | 0 | 0 | 0 | 61 | 0 | 0 | 0 | 32 | 24 | 213 |
| SRR5756009 | 51 | 0 | 3319 | 0 | 0 | 135 | 32 | 11 | 49 | 0 | 0 | 348 | 0 | 32 | 108 | 0 | 42 | 0 | 23 | 796 | 85 | 0 | 0 | 0 | 0 | 65 | 39 | 0 | 32 | 0 | 70 | 0 | 0 | 0 | 47 | 70 | 20 |
| SRR5756011 | 64 | 12 | 2069 | 0 | 0 | 68 | 25 | 14 | 26 | 0 | 0 | 0 | 0 | 16 | 1121 | 0 | 30 | 36 | 0 | 177 | 590 | 0 | 0 | 0 | 0 | 122 | 82 | 0 | 0 | 0 | 26 | 0 | 0 | 0 | 78 | 21 | 14 |
| SRR5756012 | 22 | 0 | 7312 | 0 | 0 | 59 | 18 | 0 | 15 | 0 | 0 | 15 | 0 | 0 | 96 | 0 | 25 | 0 | 0 | 11 | 0 | 0 | 0 | 0 | 0 | 1511 | 496 | 0 | 0 | 0 | 28 | 0 | 0 | 0 | 496 | 12 | 155 |
| SRR5756014 | 35 | 0 | 1014 | 0 | 0 | 184 | 15 | 22 | 90 | 0 | 0 | 37 | 0 | 0 | 98 | 0 | 31 | 0 | 0 | 0 | 31 | 0 | 0 | 0 | 0 | 242 | 48 | 0 | 0 | 0 | 72 | 0 | 0 | 0 | 39 | 31 | 184 |
| SRR5756016 | 57 | 0 | 2486 | 0 | 0 | 53 | 96 | 192 | 27 | 0 | 0 | 379 | 0 | 16 | 67 | 0 | 84 | 0 | 43 | 659 | 86 | 0 | 0 | 0 | 0 | 653 | 669 | 0 | 11 | 0 | 18 | 0 | 0 | 0 | 1445 | 98 | 464 |
| SRR5756017 | 38 | 0 | 235 | 0 | 0 | 20 | 186 | 20 | 0 | 0 | 0 | 296 | 0 | 0 | 0 | 0 | 95 | 0 | 0 | 558 | 93 | 0 | 0 | 0 | 0 | 41 | 223 | 20 | 0 | 0 | 0 | 0 | 0 | 0 | 0 | 20 | 41 |
| SRR5756018 | 59 | 13 | 977 | 0 | 0 | 126 | 0 | 95 | 13 | 0 | 0 | 0 | 0 | 35 | 1163 | 0 | 17 | 53 | 0 | 314 | 676 | 0 | 0 | 0 | 0 | 1735 | 920 | 0 | 0 | 0 | 30 | 0 | 0 | 0 | 902 | 0 | 135 |
| SRR5756019 | 34 | 13 | 6433 | 0 | 0 | 52 | 24 | 0 | 26 | 0 | 0 | 100 | 0 | 0 | 49 | 0 | 41 | 0 | 0 | 24 | 24 | 0 | 0 | 0 | 0 | 1711 | 674 | 0 | 0 | 0 | 65 | 0 | 0 | 0 | 544 | 0 | 194 |
| SRR5756021 | 33 | 0 | 901 | 0 | 0 | 255 | 21 | 30 | 41 | 0 | 0 | 0 | 0 | 0 | 563 | 0 | 25 | 0 | 0 | 169 | 258 | 0 | 0 | 0 | 11 | 2804 | 1410 | 0 | 0 | 0 | 52 | 0 | 0 | 0 | 814 | 34 | 210 |
| SRR5756022 | 30 | 0 | 1244 | 0 | 0 | 190 | 40 | 31 | 68 | 0 | 0 | 0 | 0 | 0 | 357 | 0 | 17 | 0 | 0 | 0 | 26 | 0 | 0 | 0 | 0 | 2900 | 1075 | 0 | 0 | 0 | 61 | 0 | 0 | 0 | 522 | 12 | 178 |

### Supp Table S8.2.Columns_AM_to_BW

| | | | | | | | | | | | | | | | | | | | | | | | | | | | | | | | | | | | | | |
|---|---|---|---|---|---|---|---|---|---|---|---|---|---|---|---|---|---|---|---|---|---|---|---|---|---|---|---|---|---|---|---|---|---|---|---|---|---|
| SRR5756023 | 57 | 0 | 2692 | 0 | 0 | 68 | 46 | 250 | 15 | 0 | 0 | 334 | 0 | 53 | 35 | 0 | 68 | 29 | 0 | 970 | 92 | 0 | 0 | 0 | 0 | 992 | 1014 | 23 | 0 | 0 | 23 | 0 | 0 | 15 | 1438 | 45 | 454 |
| SRR5756025 | 55 | 0 | 1603 | 0 | 0 | 98 | 30 | 49 | 22 | 0 | 0 | 0 | 0 | 0 | 1656 | 0 | 25 | 82 | 0 | 343 | 597 | 0 | 0 | 0 | 0 | 92 | 49 | 0 | 0 | 0 | 0 | 0 | 0 | 0 | 45 | 27 | 161 |
| SRR5756026 | 22 | 0 | 6184 | 0 | 0 | 44 | 43 | 19 | 38 | 0 | 0 | 19 | 0 | 0 | 86 | 0 | 33 | 0 | 0 | 14 | 64 | 0 | 0 | 0 | 13 | 41 | 20 | 0 | 0 | 0 | 25 | 0 | 0 | 0 | 18 | 38 | 151 |
| SRR5756027 | 39 | 0 | 1118 | 0 | 0 | 108 | 20 | 34 | 24 | 0 | 0 | 0 | 0 | 10 | 361 | 0 | 24 | 0 | 0 | 128 | 199 | 0 | 0 | 0 | 17 | 1334 | 1104 | 0 | 0 | 0 | 20 | 0 | 0 | 0 | 415 | 24 | 101 |
| SRR5756028 | 36 | 0 | 1252 | 0 | 0 | 70 | 0 | 0 | 31 | 0 | 0 | 0 | 0 | 15 | 455 | 0 | 25 | 0 | 0 | 165 | 290 | 0 | 0 | 0 | 23 | 41 | 30 | 0 | 0 | 0 | 31 | 0 | 0 | 0 | 12 | 0 | 116 |
| SRR5756029 | 51 | 0 | 2575 | 0 | 0 | 36 | 57 | 106 | 0 | 0 | 0 | 264 | 0 | 28 | 100 | 0 | 73 | 58 | 24 | 775 | 91 | 0 | 0 | 0 | 0 | 481 | 873 | 0 | 0 | 0 | 20 | 0 | 0 | 17 | 601 | 53 | 285 |
| SRR5756031 | 34 | 0 | 1776 | 0 | 0 | 73 | 12 | 12 | 11 | 0 | 0 | 0 | 0 | 18 | 1577 | 0 | 22 | 111 | 0 | 312 | 954 | 0 | 0 | 0 | 0 | 56 | 84 | 0 | 0 | 0 | 11 | 0 | 0 | 0 | 38 | 11 | 121 |
| SRR5756032 | 28 | 0 | 6973 | 0 | 0 | 13 | 17 | 0 | 13 | 0 | 0 | 13 | 0 | 0 | 225 | 0 | 24 | 0 | 0 | 25 | 58 | 0 | 0 | 0 | 13 | 21 | 27 | 0 | 0 | 0 | 26 | 0 | 0 | 0 | 13 | 0 | 132 |
| SRR5756033 | 40 | 0 | 1145 | 0 | 0 | 174 | 0 | 120 | 74 | 0 | 0 | 0 | 0 | 0 | 1781 | 0 | 15 | 107 | 0 | 306 | 1032 | 0 | 0 | 20 | 0 | 147 | 72 | 0 | 0 | 0 | 74 | 0 | 0 | 0 | 69 | 27 | 17 |
| SRR5756034 | 63 | 0 | 1232 | 0 | 0 | 263 | 0 | 55 | 51 | 0 | 0 | 0 | 0 | 0 | 1754 | 0 | 11 | 17 | 0 | 443 | 899 | 0 | 0 | 0 | 0 | 3868 | 1409 | 0 | 0 | 0 | 115 | 0 | 0 | 0 | 1177 | 24 | 162 |
| SRR5756035 | 53 | 0 | 3121 | 0 | 0 | 76 | 22 | 84 | 20 | 0 | 0 | 279 | 0 | 12 | 116 | 0 | 58 | 23 | 0 | 733 | 152 | 0 | 0 | 12 | 0 | 818 | 862 | 0 | 20 | 0 | 48 | 0 | 0 | 0 | 1528 | 44 | 385 |
| SRR5756037 | 67 | 0 | 1270 | 0 | 0 | 13 | 0 | 57 | 45 | 0 | 0 | 0 | 0 | 0 | 1432 | 0 | 21 | 34 | 0 | 388 | 817 | 0 | 0 | 0 | 0 | 205 | 88 | 0 | 0 | 0 | 60 | 0 | 0 | 0 | 87 | 17 | 159 |
| SRR5756038 | 15 | 11 | 7130 | 0 | 0 | 85 | 69 | 11 | 11 | 0 | 0 | 0 | 0 | 0 | 764 | 0 | 17 | 335 | 0 | 317 | 377 | 0 | 0 | 0 | 0 | 648 | 393 | 0 | 0 | 0 | 16 | 0 | 0 | 0 | 626 | 11 | 42 |
| SRR5756040 | 45 | 0 | 638 | 0 | 0 | 141 | 21 | 68 | 21 | 0 | 0 | 0 | 0 | 0 | 1841 | 0 | 15 | 11 | 0 | 261 | 628 | 0 | 0 | 0 | 0 | 2158 | 1004 | 0 | 0 | 0 | 68 | 0 | 0 | 0 | 906 | 13 | 141 |
| SRR5756041 | 39 | 0 | 912 | 0 | 0 | 63 | 32 | 17 | 23 | 0 | 0 | 0 | 0 | 0 | 1301 | 0 | 24 | 37 | 0 | 183 | 258 | 0 | 0 | 0 | 0 | 57 | 26 | 0 | 0 | 0 | 34 | 0 | 0 | 0 | 22 | 11 | 97 |
| SRR5756042 | 35 | 0 | 3268 | 0 | 0 | 86 | 15 | 164 | 47 | 0 | 0 | 286 | 0 | 0 | 96 | 0 | 58 | 17 | 15 | 382 | 37 | 0 | 0 | 47 | 0 | 46 | 45 | 0 | 0 | 0 | 62 | 0 | 0 | 0 | 61 | 16 | 16 |
| SRR5756044 | 64 | 0 | 855 | 0 | 0 | 195 | 0 | 140 | 12 | 0 | 0 | 0 | 0 | 0 | 3013 | 0 | 23 | 50 | 0 | 260 | 499 | 0 | 0 | 43 | 0 | 2064 | 1325 | 0 | 0 | 0 | 37 | 0 | 0 | 0 | 1014 | 0 | 153 |
| SRR5756045 | 39 | 0 | 6446 | 0 | 0 | 70 | 107 | 0 | 35 | 0 | 0 | 17 | 0 | 0 | 514 | 0 | 38 | 31 | 0 | 150 | 107 | 0 | 0 | 0 | 21 | 469 | 280 | 0 | 0 | 0 | 14 | 0 | 0 | 0 | 455 | 21 | 161 |
| SRR5756047 | 47 | 0 | 1263 | 0 | 0 | 220 | 14 | 46 | 73 | 0 | 0 | 0 | 0 | 0 | 680 | 0 | 21 | 24 | 0 | 426 | 384 | 0 | 0 | 23 | 0 | 133 | 83 | 0 | 0 | 0 | 69 | 0 | 0 | 0 | 58 | 50 | 17 |
| SRR5756048 | 68 | 0 | 2334 | 0 | 0 | 15 | 28 | 46 | 74 | 0 | 0 | 0 | 0 | 0 | 303 | 0 | 19 | 0 | 0 | 168 | 65 | 0 | 0 | 14 | 0 | 208 | 68 | 0 | 0 | 0 | 125 | 0 | 0 | 0 | 38 | 19 | 12 |
| SRR5756049 | 51 | 0 | 3559 | 0 | 0 | 59 | 53 | 150 | 21 | 0 | 0 | 283 | 0 | 17 | 68 | 10 | 62 | 0 | 28 | 598 | 43 | 0 | 0 | 38 | 0 | 954 | 1073 | 10 | 0 | 0 | 28 | 0 | 0 | 14 | 1929 | 70 | 582 |
| SRR5756051 | 49 | 0 | 2069 | 0 | 0 | 221 | 0 | 151 | 39 | 0 | 0 | 0 | 0 | 0 | 1252 | 0 | 18 | 16 | 0 | 397 | 334 | 0 | 0 | 27 | 0 | 2480 | 1401 | 0 | 0 | 0 | 35 | 0 | 0 | 0 | 1797 | 19 | 268 |
| SRR5756052 | 36 | 0 | 11403 | 0 | 0 | 82 | 53 | 18 | 0 | 0 | 0 | 13 | 0 | 0 | 98 | 0 | 21 | 0 | 0 | 98 | 30 | 0 | 0 | 18 | 0 | 31 | 25 | 0 | 0 | 0 | 18 | 0 | 0 | 0 | 26 | 0 | 187 |
| SRR5756054 | 47 | 0 | 893 | 0 | 0 | 187 | 0 | 84 | 19 | 0 | 0 | 0 | 0 | 0 | 942 | 0 | 12 | 0 | 0 | 311 | 735 | 0 | 0 | 13 | 0 | 2498 | 1530 | 0 | 0 | 0 | 77 | 0 | 0 | 0 | 1659 | 39 | 336 |
| SRR5756055 | 80 | 0 | 1758 | 0 | 0 | 240 | 0 | 56 | 78 | 0 | 0 | 0 | 0 | 21 | 659 | 0 | 12 | 0 | 0 | 190 | 135 | 0 | 0 | 14 | 0 | 3035 | 1355 | 0 | 0 | 0 | 14 | 0 | 0 | 0 | 1384 | 28 | 212 |
| SRR5756056 | 84 | 0 | 3652 | 0 | 0 | 33 | 17 | 88 | 33 | 0 | 0 | 268 | 0 | 0 | 112 | 0 | 54 | 0 | 0 | 561 | 39 | 0 | 0 | 28 | 0 | 33 | 38 | 0 | 22 | 0 | 0 | 0 | 0 | 0 | 74 | 44 | 25 |
| SRR5756058 | 51 | 0 | 1323 | 0 | 0 | 219 | 12 | 119 | 24 | 0 | 0 | 0 | 0 | 0 | 1594 | 0 | 25 | 95 | 0 | 303 | 663 | 0 | 0 | 30 | 0 | 2873 | 1433 | 0 | 0 | 0 | 49 | 0 | 0 | 0 | 1138 | 18 | 189 |
| SRR5756059 | 58 | 0 | 9921 | 0 | 0 | 19 | 15 | 0 | 19 | 0 | 0 | 0 | 0 | 0 | 121 | 0 | 27 | 0 | 0 | 166 | 45 | 0 | 0 | 28 | 0 | 709 | 755 | 0 | 0 | 0 | 0 | 0 | 0 | 0 | 1138 | 19 | 261 |
| SRR5756061 | 37 | 13 | 1317 | 0 | 0 | 106 | 31 | 13 | 49 | 0 | 0 | 0 | 0 | 0 | 792 | 0 | 24 | 0 | 0 | 146 | 575 | 0 | 0 | 0 | 0 | 104 | 54 | 0 | 0 | 27 | 40 | 0 | 0 | 0 | 57 | 27 | 14 |
| SRR5756062 | 75 | 0 | 2554 | 0 | 0 | 72 | 34 | 17 | 31 | 0 | 0 | 0 | 0 | 0 | 303 | 0 | 20 | 0 | 0 | 67 | 135 | 0 | 0 | 24 | 0 | 100 | 38 | 0 | 0 | 0 | 0 | 0 | 0 | 0 | 40 | 14 | 96 |
| SRR5756063 | 63 | 0 | 3656 | 0 | 0 | 58 | 16 | 77 | 13 | 0 | 0 | 231 | 0 | 19 | 68 | 0 | 63 | 14 | 11 | 314 | 70 | 0 | 0 | 39 | 0 | 34 | 52 | 16 | 0 | 0 | 13 | 0 | 0 | 0 | 75 | 52 | 19 |
| SRR5756065 | 82 | 0 | 1651 | 0 | 0 | 83 | 0 | 55 | 23 | 0 | 0 | 0 | 0 | 0 | 1135 | 0 | 18 | 48 | 0 | 155 | 897 | 0 | 0 | 17 | 0 | 1236 | 773 | 0 | 0 | 0 | 14 | 0 | 0 | 0 | 1250 | 14 | 170 |
| SRR5756066 | 33 | 0 | 9072 | 0 | 0 | 49 | 49 | 18 | 21 | 0 | 0 | 0 | 0 | 0 | 275 | 0 | 25 | 11 | 0 | 108 | 138 | 0 | 0 | 18 | 0 | 33 | 31 | 0 | 0 | 0 | 31 | 0 | 0 | 0 | 55 | 20 | 13 |
| SRR5756068 | 59 | 0 | 1588 | 0 | 0 | 154 | 25 | 64 | 64 | 0 | 0 | 0 | 0 | 0 | 1415 | 0 | 13 | 0 | 0 | 197 | 516 | 0 | 0 | 26 | 0 | 2677 | 1432 | 0 | 0 | 0 | 23 | 0 | 0 | 0 | 718 | 13 | 193 |
| SRR5756069 | 80 | 0 | 1571 | 0 | 0 | 14 | 17 | 73 | 102 | 0 | 0 | 0 | 0 | 0 | 597 | 0 | 18 | 0 | 0 | 282 | 199 | 0 | 0 | 29 | 0 | 218 | 87 | 0 | 0 | 0 | 94 | 0 | 0 | 0 | 39 | 20 | 183 |
| SRR5756070 | 41 | 0 | 2789 | 0 | 0 | 73 | 84 | 106 | 41 | 0 | 0 | 310 | 0 | 0 | 105 | 0 | 59 | 0 | 0 | 414 | 77 | 0 | 0 | 16 | 0 | 1559 | 923 | 0 | 0 | 0 | 16 | 0 | 0 | 0 | 3225 | 171 | 865 |
| SRR5756072 | 53 | 12 | 1483 | 0 | 0 | 11 | 0 | 10 | 36 | 0 | 0 | 0 | 0 | 0 | 1888 | 0 | 24 | 0 | 0 | 199 | 385 | 0 | 0 | 18 | 0 | 188 | 92 | 0 | 0 | 0 | 18 | 0 | 0 | 0 | 47 | 0 | 93 |
| SRR5756073 | 76 | 22 | 7848 | 0 | 0 | 88 | 61 | 73 | 29 | 0 | 0 | 0 | 0 | 0 | 137 | 0 | 19 | 0 | 0 | 244 | 76 | 0 | 0 | 0 | 0 | 1649 | 708 | 0 | 0 | 0 | 36 | 0 | 0 | 0 | 781 | 0 | 285 |
| SRR5756075 | 32 | 0 | 906 | 0 | 0 | 169 | 21 | 144 | 31 | 0 | 0 | 0 | 0 | 13 | 1658 | 0 | 15 | 16 | 0 | 322 | 1108 | 0 | 0 | 25 | 0 | 2392 | 1359 | 0 | 0 | 0 | 25 | 0 | 0 | 0 | 745 | 13 | 207 |
| SRR5756076 | 67 | 0 | 1199 | 0 | 0 | 115 | 0 | 89 | 25 | 0 | 0 | 0 | 0 | 0 | 2191 | 0 | 12 | 0 | 0 | 194 | 498 | 0 | 0 | 26 | 0 | 2002 | 1113 | 0 | 0 | 0 | 28 | 0 | 0 | 0 | 403 | 0 | 91 |
| SRR5756077 | 42 | 0 | 2678 | 0 | 0 | 110 | 80 | 69 | 14 | 0 | 0 | 210 | 0 | 0 | 161 | 0 | 58 | 19 | 27 | 607 | 107 | 0 | 0 | 28 | 14 | 1821 | 1021 | 28 | 14 | 0 | 110 | 0 | 0 | 0 | 1917 | 97 | 717 |
| SRR5756079 | 105 | 0 | 1075 | 0 | 0 | 233 | 0 | 45 | 54 | 0 | 0 | 0 | 0 | 0 | 2072 | 0 | 18 | 0 | 0 | 134 | 585 | 0 | 0 | 13 | 0 | 3541 | 1230 | 0 | 0 | 0 | 85 | 0 | 0 | 0 | 880 | 13 | 202 |
| SRR5756080 | 42 | 0 | 6493 | 0 | 0 | 54 | 22 | 30 | 13 | 0 | 0 | 0 | 0 | 0 | 490 | 0 | 18 | 0 | 0 | 138 | 49 | 0 | 0 | 11 | 0 | 44 | 34 | 0 | 0 | 0 | 25 | 0 | 0 | 0 | 25 | 11 | 127 |
| SRR5756082 | 26 | 0 | 907 | 0 | 0 | 115 | 0 | 92 | 26 | 0 | 0 | 0 | 0 | 0 | 2248 | 0 | 18 | 19 | 0 | 432 | 609 | 0 | 0 | 20 | 0 | 97 | 69 | 0 | 0 | 0 | 15 | 0 | 0 | 0 | 26 | 15 | 97 |
| SRR5756083 | 40 | 0 | 1253 | 0 | 0 | 67 | 15 | 58 | 14 | 0 | 0 | 0 | 0 | 0 | 2551 | 0 | 15 | 77 | 0 | 355 | 464 | 0 | 0 | 19 | 0 | 1120 | 745 | 0 | 0 | 0 | 36 | 0 | 0 | 0 | 439 | 0 | 106 |
| SRR5756084 | 28 | 0 | 2290 | 0 | 0 | 62 | 89 | 22 | 31 | 0 | 0 | 258 | 0 | 23 | 193 | 0 | 84 | 0 | 0 | 491 | 164 | 0 | 0 | 15 | 0 | 43 | 47 | 0 | 0 | 0 | 39 | 0 | 0 | 23 | 73 | 39 | 25 |
| SRR5756086 | 36 | 0 | 870 | 0 | 0 | 11 | 0 | 49 | 41 | 0 | 0 | 0 | 0 | 0 | 1776 | 0 | 16 | 20 | 0 | 302 | 1044 | 0 | 0 | 15 | 0 | 197 | 113 | 0 | 0 | 0 | 37 | 0 | 0 | 0 | 49 | 0 | 11 |
| SRR5756087 | 19 | 0 | 8673 | 0 | 0 | 29 | 40 | 0 | 12 | 0 | 0 | 12 | 0 | 0 | 871 | 0 | 19 | 223 | 0 | 101 | 202 | 0 | 0 | 12 | 0 | 34 | 29 | 0 | 0 | 0 | 29 | 0 | 0 | 0 | 22 | 29 | 121 |
| SRR5756089 | 35 | 0 | 914 | 0 | 0 | 158 | 0 | 35 | 35 | 0 | 0 | 0 | 0 | 0 | 4179 | 0 | 18 | 10 | 0 | 442 | 1674 | 0 | 0 | 0 | 0 | 127 | 86 | 0 | 0 | 0 | 35 | 0 | 0 | 0 | 39 | 35 | 12 |
| SRR5756090 | 50 | 0 | 1448 | 0 | 0 | 208 | 23 | 28 | 36 | 0 | 0 | 0 | 0 | 0 | 2813 | 0 | 16 | 0 | 0 | 520 | 595 | 0 | 0 | 0 | 0 | 2818 | 1492 | 0 | 0 | 0 | 51 | 0 | 0 | 0 | 478 | 15 | 152 |
| SRR5756091 | 52 | 0 | 4306 | 0 | 0 | 47 | 26 | 51 | 25 | 0 | 0 | 224 | 0 | 0 | 159 | 0 | 68 | 19 | 0 | 555 | 87 | 0 | 0 | 34 | 0 | 1009 | 1004 | 0 | 0 | 0 | 38 | 0 | 0 | 0 | 1161 | 30 | 292 |
| SRR5756093 | 44 | 0 | 912 | 0 | 0 | 15 | 0 | 38 | 30 | 0 | 0 | 0 | 0 | 0 | 2090 | 0 | 34 | 32 | 0 | 260 | 443 | 0 | 0 | 22 | 0 | 182 | 97 | 0 | 0 | 0 | 24 | 0 | 0 | 0 | 47 | 11 | 122 |
| SRR5756094 | 23 | 0 | 9577 | 0 | 0 | 36 | 26 | 0 | 22 | 0 | 0 | 0 | 0 | 0 | 1255 | 0 | 19 | 71 | 0 | 244 | 176 | 0 | 0 | 0 | 0 | 604 | 712 | 0 | 0 | 0 | 36 | 0 | 0 | 0 | 492 | 0 | 137 |
| SRR5756096 | 25 | 0 | 1218 | 0 | 0 | 291 | 25 | 27 | 85 | 0 | 0 | 0 | 0 | 0 | 387 | 0 | 26 | 0 | 0 | 168 | 150 | 0 | 0 | 18 | 18 | 4273 | 1922 | 0 | 0 | 0 | 57 | 0 | 0 | 0 | 758 | 18 | 146 |
| SRR5756097 | 44 | 0 | 2321 | 0 | 0 | 13 | 0 | 0 | 90 | 0 | 0 | 10 | 0 | 0 | 43 | 0 | 28 | 0 | 0 | 216 | 216 | 0 | 0 | 0 | 0 | 279 | 102 | 0 | 0 | 0 | 81 | 0 | 0 | 0 | 38 | 18 | 127 |
| SRR5756098 | 37 | 0 | 3215 | 0 | 0 | 50 | 52 | 147 | 31 | 0 | 0 | 258 | 0 | 0 | 74 | 0 | 48 | 30 | 0 | 661 | 103 | 0 | 0 | 19 | 0 | 793 | 736 | 0 | 0 | 0 | 17 | 0 | 0 | 0 | 1588 | 71 | 371 |
| SRR5756100 | 50 | 12 | 2883 | 0 | 0 | 85 | 16 | 65 | 18 | 0 | 0 | 0 | 0 | 0 | 689 | 0 | 21 | 0 | 0 | 269 | 428 | 0 | 0 | 12 | 0 | 1787 | 1310 | 0 | 0 | 0 | 15 | 0 | 0 | 0 | 1088 | 17 | 174 |
| SRR5756101 | 31 | 0 | 11129 | 0 | 0 | 101 | 37 | 0 | 30 | 0 | 0 | 16 | 0 | 0 | 55 | 0 | 25 | 0 | 0 | 203 | 55 | 0 | 0 | 12 | 42 | 1009 | 1003 | 0 | 0 | 0 | 0 | 0 | 0 | 0 | 549 | 0 | 125 |
| SRR5756103 | 41 | 0 | 988 | 0 | 0 | 11 | 0 | 0 | 67 | 0 | 0 | 0 | 0 | 0 | 25 | 0 | 22 | 0 | 0 | 1311 | 101 | 0 | 0 | 22 | 0 | 180 | 95 | 0 | 0 | 0 | 82 | 0 | 0 | 0 | 76 | 89 | 21 |
| SRR5756104 | 49 | 0 | 1527 | 0 | 0 | 211 | 0 | 0 | 41 | 0 | 0 | 0 | 0 | 0 | 0 | 0 | 0 | 0 | 0 | 577 | 68 | 0 | 0 | 0 | 0 | 3170 | 1791 | 0 | 0 | 0 | 91 | 0 | 0 | 0 | 1784 | 46 | 444 |
| SRR5756105 | 38 | 0 | 3559 | 0 | 0 | 87 | 0 | 0 | 23 | 0 | 0 | 230 | 0 | 10 | 0 | 0 | 54 | 0 | 0 | 1101 | 45 | 0 | 0 | 27 | 10 | 34 | 31 | 0 | 20 | 0 | 57 | 0 | 0 | 0 | 91 | 120 | 24 |
| SRR5756106 | 33 | 0 | 411 | 0 | 18 | 0 | 0 | 0 | 0 | 0 | 0 | 230 | 0 | 0 | 0 | 0 | 77 | 0 | 0 | 1811 | 44 | 0 | 0 | 0 | 0 | 27 | 163 | 18 | 0 | 0 | 0 | 0 | 0 | 0 | 407 | 0 | 18 |
| SRR5756107 | 34 | 0 | 712 | 0 | 0 | 208 | 0 | 0 | 51 | 0 | 0 | 0 | 0 | 0 | 0 | 0 | 17 | 0 | 0 | 413 | 252 | 0 | 0 | 0 | 0 | 123 | 72 | 0 | 0 | 0 | 64 | 0 | 0 | 0 | 76 | 0 | 16 |
| SRR5756108 | 27 | 0 | 7081 | 0 | 0 | 67 | 0 | 0 | 0 | 0 | 0 | 16 | 0 | 10 | 19 | 0 | 14 | 0 | 0 | 412 | 28 | 0 | 0 | 0 | 0 | 30 | 16 | 0 | 0 | 0 | 83 | 0 | 0 | 0 | 48 | 36 | 12 |
| SRR5756110 | 0 | 35 | 0 | 0 | 0 | 106 | 0 | 0 | 35 | 0 | 0 | 0 | 0 | 0 | 0 | 0 | 0 | 0 | 0 | 138 | 138 | 0 | 0 | 0 | 0 | 637 | 425 | 0 | 35 | 0 | 35 | 0 | 0 | 0 | 0 | 0 | 0 |
| SRR5756114 | 0 | 0 | 0 | 0 | 0 | 95 | 0 | 0 | 24 | 0 | 0 | 0 | 0 | 0 | 0 | 0 | 0 | 0 | 0 | 340 | 0 | 0 | 0 | 0 | 0 | 474 | 261 | 0 | 0 | 0 | 0 | 0 | 0 | 71 | 24 | 0 | 0 |
| SRR5756115 | 0 | 0 | 0 | 0 | 0 | 0 | 0 | 0 | 0 | 0 | 0 | 0 | 0 | 0 | 0 | 0 | 0 | 0 | 0 | 331 | 0 | 0 | 0 | 0 | 0 | 421 | 281 | 0 | 0 | 0 | 0 | 0 | 0 | 70 | 0 | 0 | 0 |
| SRR5756116 | 0 | 68 | 0 | 0 | 0 | 68 | 0 | 0 | 0 | 0 | 0 | 0 | 0 | 0 | 252 | 0 | 0 | 0 | 0 | 0 | 0 | 0 | 0 | 0 | 0 | 1016 | 609 | 0 | 0 | 0 | 0 | 0 | 0 | 0 | 0 | 0 | 0 |
| SRR5756118 | 0 | 0 | 0 | 0 | 0 | 0 | 0 | 0 | 0 | 0 | 0 | 0 | 0 | 0 | 0 | 0 | 0 | 0 | 0 | 0 | 0 | 0 | 0 | 0 | 0 | 427 | 555 | 0 | 0 | 0 | 43 | 0 | 0 | 0 | 0 | 0 | 0 |
| SRR5756120 | 0 | 27 | 0 | 0 | 0 | 0 | 0 | 0 | 0 | 0 | 0 | 0 | 0 | 0 | 110 | 0 | 0 | 0 | 0 | 883 | 110 | 0 | 0 | 0 | 81 | 651 | 217 | 0 | 0 | 0 | 0 | 0 | 0 | 0 | 0 | 0 | 0 |
| SRR5756121 | 0 | 0 | 0 | 0 | 0 | 132 | 0 | 0 | 0 | 0 | 0 | 0 | 0 | 0 | 0 | 0 | 0 | 0 | 0 | 188 | 0 | 0 | 0 | 0 | 0 | 439 | 527 | 0 | 0 | 0 | 0 | 0 | 0 | 0 | 0 | 0 | 0 |
| SRR5756122 | 0 | 0 | 17 | 0 | 0 | 27 | 0 | 0 | 27 | 0 | 0 | 0 | 0 | 0 | 0 | 0 | 0 | 0 | 0 | 119 | 237 | 0 | 0 | 0 | 0 | 479 | 186 | 0 | 0 | 0 | 0 | 0 | 0 | 53 | 27 | 0 | 0 |
| SRR5756123 | 0 | 0 | 0 | 0 | 0 | 0 | 0 | 0 | 0 | 0 | 0 | 0 | 0 | 0 | 0 | 0 | 0 | 0 | 0 | 210 | 0 | 0 | 0 | 0 | 0 | 343 | 315 | 0 | 29 | 0 | 0 | 0 | 29 | 29 | 0 | 0 | 0 |
| SRR5756125 | 0 | 0 | 0 | 0 | 0 | 0 | 0 | 32 | 0 | 0 | 0 | 0 | 0 | 0 | 0 | 0 | 0 | 0 | 0 | 496 | 0 | 0 | 0 | 0 | 16 | 221 | 79 | 0 | 0 | 0 | 16 | 0 | 0 | 47 | 16 | 0 | 0 |
| SRR5756128 | 0 | 0 | 0 | 0 | 0 | 0 | 0 | 0 | 0 | 0 | 0 | 0 | 0 | 0 | 245 | 0 | 0 | 0 | 0 | 245 | 0 | 0 | 0 | 0 | 0 | 973 | 324 | 0 | 0 | 0 | 46 | 0 | 0 | 0 | 0 | 0 | 0 |
| SRR5756130 | 0 | 0 | 0 | 0 | 0 | 0 | 0 | 0 | 23 | 0 | 0 | 0 | 0 | 23 | 132 | 0 | 0 | 0 | 0 | 132 | 0 | 0 | 0 | 0 | 0 | 158 | 90 | 0 | 0 | 0 | 0 | 0 | 0 | 23 | 0 | 0 | 0 |
| SRR5756131 | 0 | 26 | 0 | 0 | 0 | 0 | 0 | 0 | 0 | 0 | 0 | 0 | 0 | 0 | 0 | 0 | 0 | 0 | 0 | 424 | 0 | 0 | 0 | 0 | 0 | 486 | 307 | 0 | 0 | 0 | 26 | 0 | 26 | 0 | 26 | 0 | 0 |
| SRR5756132 | 0 | 30 | 0 | 0 | 0 | 0 | 0 | 30 | 0 | 0 | 0 | 0 | 0 | 0 | 0 | 0 | 0 | 0 | 0 | 734 | 122 | 0 | 0 | 0 | 0 | 386 | 238 | 0 | 0 | 0 | 0 | 0 | 0 | 59 | 0 | 0 | 0 |
| SRR5756133 | 0 | 0 | 0 | 0 | 0 | 0 | 0 | 0 | 0 | 0 | 0 | 0 | 0 | 0 | 0 | 0 | 0 | 0 | 0 | 364 | 121 | 0 | 0 | 0 | 0 | 291 | 219 | 0 | 36 | 0 | 0 | 0 | 0 | 0 | 73 | 0 | 36 |
| SRR5756135 | 0 | 0 | 0 | 0 | 0 | 37 | 0 | 0 | 0 | 0 | 0 | 0 | 0 | 74 | 0 | 0 | 0 | 0 | 0 | 797 | 0 | 0 | 0 | 0 | 0 | 662 | 515 | 0 | 0 | 0 | 0 | 0 | 0 | 37 | 37 | 0 | 37 |
| SRR5756136 | 0 | 0 | 0 | 0 | 0 | 181 | 0 | 0 | 36 | 0 | 0 | 0 | 0 | 0 | 0 | 0 | 0 | 0 | 0 | 642 | 160 | 0 | 0 | 36 | 0 | 1301 | 1012 | 0 | 0 | 0 | 0 | 0 | 0 | 0 | 0 | 0 | 0 |
| SRR5756137 | 0 | 18 | 0 | 0 | 0 | 36 | 130 | 18 | 53 | 0 | 0 | 0 | 0 | 0 | 0 | 0 | 0 | 0 | 0 | 2610 | 783 | 0 | 0 | 0 | 0 | 16 | 535 | 0 | 18 | 0 | 18 | 0 | 18 | 53 | 18 | 0 | 36 |
| SRR5756138 | 0 | 0 | 0 | 0 | 0 | 0 | 0 | 0 | 29 | 0 | 0 | 0 | 0 | 0 | 0 | 0 | 0 | 0 | 0 | 1459 | 417 | 0 | 0 | 0 | 0 | 687 | 486 | 0 | 0 | 0 | 0 | 0 | 0 | 29 | 0 | 0 | 0 |
| SRR5756139 | 0 | 0 | 0 | 0 | 0 | 207 | 0 | 0 | 35 | 0 | 0 | 0 | 0 | 0 | 0 | 0 | 0 | 0 | 0 | 646 | 0 | 0 | 0 | 35 | 0 | 863 | 932 | 0 | 0 | 0 | 0 | 0 | 0 | 0 | 0 | 0 | 35 |
| SRR5756140 | 0 | 0 | 0 | 0 | 0 | 87 | 0 | 116 | 0 | 0 | 0 | 0 | 0 | 29 | 0 | 0 | 0 | 0 | 0 | 1920 | 213 | 0 | 0 | 0 | 0 | 839 | 810 | 0 | 0 | 0 | 0 | 0 | 0 | 0 | 0 | 0 | 29 |
| SRR5756141 | 0 | 16 | 0 | 0 | 0 | 16 | 0 | 0 | 16 | 0 | 0 | 0 | 0 | 0 | 0 | 0 | 0 | 0 | 122 | 1094 | 851 | 0 | 0 | 16 | 0 | 765 | 383 | 0 | 0 | 0 | 0 | 0 | 0 | 48 | 0 | 0 | 0 |
| SRR5756142 | 0 | 18 | 0 | 0 | 0 | 18 | 0 | 36 | 0 | 0 | 0 | 0 | 0 | 18 | 0 | 0 | 0 | 0 | 0 | 1465 | 0 | 0 | 0 | 0 | 0 | 745 | 600 | 0 | 0 | 0 | 0 | 0 | 0 | 73 | 36 | 0 | 0 |
| SRR5756143 | 0 | 0 | 0 | 0 | 0 | 0 | 0 | 18 | 18 | 0 | 0 | 0 | 0 | 0 | 0 | 0 | 0 | 0 | 0 | 959 | 137 | 0 | 0 | 0 | 0 | 810 | 576 | 0 | 0 | 0 | 0 | 18 | 0 | 18 | 0 | 0 | 0 |
| SRR5756144 | 0 | 0 | 0 | 0 | 0 | 20 | 0 | 0 | 20 | 0 | 0 | 0 | 0 | 20 | 0 | 0 | 0 | 0 | 0 | 1456 | 0 | 0 | 0 | 0 | 20 | 592 | 388 | 0 | 20 | 0 | 20 | 0 | 0 | 20 | 0 | 0 | 0 |
| SRR5756145 | 0 | 76 | 0 | 0 | 0 | 76 | 0 | 0 | 0 | 0 | 0 | 0 | 0 | 0 | 0 | 0 | 0 | 0 | 0 | 458 | 229 | 0 | 0 | 0 | 0 | 879 | 1070 | 0 | 38 | 0 | 0 | 0 | 0 | 0 | 0 | 0 | 0 |
| SRR5756146 | 0 | 0 | 0 | 20 | 0 | 79 | 0 | 0 | 20 | 0 | 0 | 0 | 0 | 20 | 0 | 0 | 0 | 0 | 0 | 1097 | 0 | 0 | 0 | 20 | 0 | 1028 | 870 | 0 | 0 | 0 | 0 | 0 | 0 | 119 | 0 | 0 | 0 |
| SRR5756147 | 0 | 0 | 0 | 0 | 0 | 195 | 0 | 0 | 0 | 0 | 0 | 0 | 0 | 0 | 184 | 0 | 0 | 0 | 0 | 736 | 368 | 0 | 0 | 0 | 0 | 702 | 1014 | 0 | 39 | 0 | 0 | 0 | 0 | 0 | 0 | 0 | 0 |
| SRR5756148 | 0 | 0 | 0 | 0 | 0 | 27 | 0 | 0 | 54 | 0 | 0 | 0 | 0 | 0 | 0 | 0 | 0 | 0 | 0 | 192 | 0 | 0 | 0 | 27 | 0 | 755 | 512 | 0 | 0 | 0 | 0 | 0 | 0 | 0 | 0 | 0 | 0 |
| SRR5756149 | 0 | 0 | 0 | 0 | 0 | 38 | 0 | 0 | 0 | 0 | 0 | 0 | 0 | 0 | 195 | 0 | 0 | 0 | 0 | 391 | 195 | 0 | 0 | 0 | 0 | 1133 | 831 | 0 | 0 | 0 | 0 | 0 | 0 | 0 | 0 | 0 | 0 |
| SRR5756150 | 0 | 0 | 0 | 0 | 0 | 39 | 0 | 58 | 0 | 0 | 0 | 0 | 0 | 0 | 0 | 0 | 0 | 0 | 0 | 1159 | 258 | 0 | 0 | 0 | 0 | 660 | 601 | 0 | 0 | 0 | 19 | 0 | 0 | 19 | 39 | 0 | 0 |
| SRR5756151 | 0 | 0 | 0 | 0 | 0 | 115 | 0 | 0 | 38 | 0 | 0 | 0 | 0 | 38 | 0 | 0 | 0 | 0 | 0 | 704 | 235 | 0 | 0 | 0 | 0 | 12 | 1186 | 0 | 0 | 0 | 38 | 0 | 38 | 0 | 0 | 0 | 0 |
| SRR5756152 | 0 | 17 | 0 | 0 | 0 | 35 | 0 | 885 | 0 | 0 | 0 | 0 | 0 | 0 | 0 | 0 | 0 | 0 | 0 | 805 | 345 | 0 | 0 | 0 | 0 | 364 | 469 | 0 | 0 | 0 | 17 | 0 | 0 | 69 | 17 | 0 | 0 |
| SRR5756153 | 0 | 47 | 0 | 0 | 0 | 0 | 0 | 0 | 24 | 0 | 0 | 0 | 0 | 24 | 0 | 0 | 0 | 0 | 0 | 158 | 158 | 0 | 0 | 0 | 0 | 757 | 757 | 0 | 0 | 0 | 0 | 0 | 0 | 47 | 0 | 0 | 0 |
| SRR5756154 | 0 | 0 | 0 | 0 | 0 | 45 | 0 | 0 | 0 | 0 | 0 | 0 | 0 | 0 | 0 | 0 | 0 | 0 | 0 | 604 | 0 | 0 | 0 | 0 | 0 | 1492 | 1311 | 0 | 45 | 0 | 0 | 0 | 0 | 0 | 0 | 0 | 45 |
| SRR5756155 | 0 | 0 | 0 | 0 | 0 | 14 | 0 | 28 | 28 | 0 | 0 | 0 | 0 | 0 | 0 | 0 | 0 | 0 | 0 | 953 | 136 | 0 | 0 | 0 | 0 | 363 | 335 | 0 | 14 | 0 | 28 | 0 | 14 | 0 | 14 | 0 | 0 |
| SRR5756156 | 0 | 0 | 0 | 0 | 0 | 41 | 0 | 62 | 0 | 0 | 0 | 0 | 0 | 21 | 0 | 0 | 0 | 0 | 0 | 1642 | 410 | 0 | 0 | 21 | 0 | 1029 | 761 | 0 | 0 | 0 | 0 | 0 | 0 | 0 | 21 | 0 | 21 |
| SRR5756157 | 0 | 0 | 0 | 0 | 0 | 0 | 0 | 38 | 19 | 0 | 0 | 0 | 0 | 19 | 0 | 0 | 0 | 0 | 0 | 144 | 288 | 0 | 0 | 0 | 0 | 874 | 646 | 0 | 0 | 0 | 0 | 0 | 0 | 76 | 0 | 0 | 0 |
| SRR5756158 | 0 | 0 | 0 | 0 | 0 | 23 | 0 | 0 | 0 | 0 | 0 | 0 | 0 | 0 | 0 | 0 | 0 | 0 | 0 | 774 | 0 | 0 | 0 | 0 | 0 | 927 | 927 | 0 | 0 | 0 | 0 | 0 | 0 | 45 | 23 | 0 | 0 |
| SRR5756159 | 0 | 0 | 0 | 21 | 0 | 21 | 0 | 42 | 0 | 0 | 0 | 0 | 0 | 0 | 180 | 0 | 0 | 0 | 0 | 1624 | 361 | 0 | 0 | 0 | 0 | 527 | 421 | 0 | 0 | 0 | 21 | 0 | 0 | 21 | 0 | 0 | 21 |
| SRR5756160 | 0 | 53 | 0 | 0 | 0 | 0 | 0 | 0 | 0 | 0 | 0 | 0 | 0 | 53 | 0 | 0 | 0 | 506 | 0 | 0 | 0 | 0 | 0 | 0 | 0 | 475 | 264 | 0 | 0 | 0 | 0 | 0 | 0 | 53 | 53 | 0 | 0 |
| SRR5756162 | 0 | 0 | 12 | 0 | 0 | 0 | 0 | 0 | 0 | 0 | 0 | 0 | 0 | 0 | 0 | 0 | 0 | 0 | 0 | 286 | 286 | 0 | 0 | 0 | 0 | 930 | 232 | 0 | 0 | 0 | 0 | 0 | 0 | 58 | 58 | 0 | 0 |
| SRR5756164 | 0 | 0 | 19 | 0 | 0 | 0 | 0 | 0 | 0 | 0 | 0 | 0 | 0 | 45 | 105 | 0 | 0 | 0 | 0 | 525 | 0 | 0 | 0 | 0 | 0 | 201 | 45 | 22 | 0 | 0 | 22 | 0 | 0 | 67 | 0 | 0 | 22 |
| SRR5756166 | 0 | 0 | 10 | 0 | 0 | 25 | 0 | 0 | 75 | 0 | 0 | 0 | 0 | 0 | 0 | 0 | 0 | 0 | 0 | 385 | 0 | 0 | 0 | 0 | 0 | 175 | 276 | 0 | 0 | 0 | 25 | 0 | 0 | 200 | 0 | 0 | 25 |
| SRR5756167 | 0 | 0 | 0 | 0 | 0 | 30 | 0 | 0 | 0 | 0 | 0 | 0 | 0 | 0 | 0 | 0 | 0 | 0 | 0 | 0 | 0 | 0 | 0 | 0 | 0 | 502 | 59 | 0 | 0 | 0 | 0 | 0 | 0 | 0 | 0 | 30 | 30 |
| SRR5756168 | 0 | 0 | 0 | 0 | 0 | 0 | 0 | 0 | 0 | 0 | 0 | 0 | 0 | 36 | 0 | 0 | 0 | 0 | 0 | 659 | 220 | 0 | 0 | 0 | 0 | 639 | 426 | 0 | 0 | 0 | 0 | 0 | 0 | 36 | 0 | 0 | 0 |
| SRR5756177 | 0 | 0 | 0 | 110 | 0 | 0 | 0 | 0 | 0 | 0 | 0 | 0 | 0 | 0 | 0 | 0 | 0 | 0 | 0 | 396 | 0 | 0 | 0 | 0 | 0 | 1206 | 768 | 0 | 0 | 0 | 0 | 0 | 0 | 0 | 0 | 0 | 0 |
| SRR5756178 | 0 | 0 | 0 | 0 | 0 | 0 | 0 | 0 | 0 | 0 | 0 | 0 | 0 | 0 | 0 | 0 | 0 | 0 | 0 | 784 | 0 | 0 | 0 | 0 | 0 | 625 | 268 | 89 | 0 | 0 | 0 | 0 | 0 | 0 | 0 | 0 | 0 |
| SRR5756179 | 0 | 0 | 0 | 0 | 0 | 0 | 0 | 0 | 83 | 0 | 0 | 0 | 0 | 0 | 0 | 0 | 0 | 0 | 349 | 0 | 0 | 0 | 0 | 0 | 0 | 1156 | 330 | 0 | 0 | 0 | 83 | 0 | 0 | 0 | 0 | 0 | 0 |
| SRR5756180 | 0 | 0 | 0 | 0 | 0 | 28 | 0 | 0 | 28 | 0 | 0 | 0 | 0 | 28 | 0 | 0 | 0 | 0 | 0 | 460 | 115 | 0 | 0 | 0 | 0 | 446 | 167 | 0 | 0 | 0 | 0 | 0 | 0 | 56 | 0 | 0 | 28 |
| SRR5756182 | 0 | 0 | 0 | 0 | 0 | 0 | 0 | 0 | 0 | 0 | 0 | 0 | 0 | 0 | 0 | 0 | 0 | 0 | 0 | 0 | 0 | 0 | 0 | 0 | 0 | 666 | 250 | 0 | 0 | 0 | 83 | 0 | 0 | 0 | 0 | 0 | 0 |
| SRR5756183 | 0 | 23 | 0 | 0 | 0 | 34 | 0 | 0 | 0 | 0 | 0 | 0 | 0 | 34 | 45 | 0 | 0 | 0 | 0 | 677 | 135 | 0 | 0 | 0 | 0 | 694 | 375 | 0 | 0 | 0 | 11 | 0 | 0 | 0 | 0 | 0 | 23 |
| SRR5756184 | 0 | 0 | 0 | 0 | 0 | 30 | 0 | 0 | 0 | 0 | 0 | 0 | 0 | 30 | 0 | 0 | 0 | 0 | 0 | 127 | 0 | 0 | 0 | 0 | 0 | 597 | 477 | 0 | 0 | 0 | 0 | 0 | 30 | 0 | 30 | 0 | 0 |
| SRR5756185 | 0 | 0 | 0 | 0 | 0 | 35 | 0 | 17 | 0 | 0 | 0 | 0 | 0 | 0 | 0 | 0 | 0 | 0 | 67 | 931 | 0 | 0 | 0 | 0 | 0 | 974 | 278 | 17 | 0 | 0 | 17 | 0 | 17 | 52 | 0 | 0 | 0 |
| SRR5756186 | 0 | 0 | 0 | 0 | 0 | 16 | 63 | 33 | 16 | 0 | 0 | 0 | 0 | 0 | 0 | 0 | 0 | 0 | 0 | 818 | 0 | 0 | 0 | 16 | 0 | 902 | 312 | 0 | 0 | 0 | 0 | 0 | 0 | 66 | 49 | 0 | 0 |
| SRR5804909 | 373 | 0 | 124 | 0 | 0 | 0 | 328 | 0 | 0 | 0 | 46 | 124 | 0 | 12 | 73 | 0 | 0 | 0 | 73 | 182 | 0 | 0 | 0 | 11 | 0 | 0 | 0 | 0 | 0 | 0 | 0 | 0 | 0 | 896 | 0 | 0 | 0 |
| SRR5804912 | 47 | 0 | 0 | 0 | 0 | 0 | 472 | 0 | 0 | 0 | 57 | 711 | 0 | 27 | 220 | 0 | 142 | 0 | 63 | 1573 | 0 | 0 | 0 | 11 | 0 | 0 | 0 | 0 | 0 | 0 | 0 | 0 | 0 | 773 | 0 | 0 | 0 |
| SRR5804914 | 0 | 0 | 0 | 0 | 13 | 0 | 0 | 0 | 0 | 0 | 71 | 271 | 0 | 0 | 566 | 0 | 0 | 0 | 113 | 226 | 0 | 0 | 0 | 17 | 0 | 0 | 0 | 0 | 0 | 0 | 0 | 0 | 0 | 9218 | 0 | 0 | 0 |
| SRR5817623 | 41 | 0 | 356 | 0 | 0 | 26 | 74 | 22 | 84 | 0 | 20 | 3186 | 0 | 11 | 0 | 0 | 26 | 11 | 394 | 20879 | 1598 | 0 | 0 | 0 | 15 | 471 | 1437 | 0 | 0 | 0 | 16 | 0 | 11 | 0 | 14 | 31 | 30 |
| SRR5817624 | 45 | 0 | 1531 | 0 | 0 | 28 | 152 | 46 | 94 | 0 | 20 | 2921 | 0 | 20 | 32 | 11 | 31 | 126 | 159 | 12976 | 1975 | 0 | 0 | 0 | 19 | 484 | 766 | 0 | 0 | 21 | 16 | 0 | 16 | 0 | 157 | 20 | 19 |

### Supp Table S8.2.Columns_AM_to_BW

| | | | | | | | | | | | | | | | | | | | | | | | | | | | | | | | | | | | | | |
|---|---|---|---|---|---|---|---|---|---|---|---|---|---|---|---|---|---|---|---|---|---|---|---|---|---|---|---|---|---|---|---|---|---|---|---|---|---|
| SRR5817625 | 47 | 0 | 281 | 0 | 0 | 511 | 76 | 10 | 100 | 0 | 15 | 3465 | 0 | 0 | 0 | 0 | 23 | 17 | 427 | 21571 | 1493 | 0 | 0 | 0 | 236 | 8115 | 24306 | 0 | 0 | 0 | 267 | 0 | 0 | 0 | 248 | 570 | 514 |
| SRR5817626 | 48 | 0 | 1271 | 0 | 0 | 547 | 176 | 85 | 101 | 0 | 18 | 3159 | 0 | 12 | 12 | 0 | 25 | 60 | 233 | 20377 | 2309 | 0 | 0 | 0 | 217 | 8162 | 13235 | 0 | 0 | 34 | 260 | 0 | 0 | 0 | 197 | 392 | 448 |
| SRR5821803 | 185 | 0 | 7109 | 0 | 0 | 12 | 13 | 0 | 14 | 0 | 0 | 446 | 0 | 0 | 40 | 0 | 0 | 0 | 0 | 1132 | 397 | 0 | 0 | 0 | 12 | 111 | 83 | 0 | 0 | 0 | 20 | 0 | 0 | 0 | 85 | 12 | 23 |
| SRR5821804 | 40 | 87 | 2018 | 0 | 49 | 0 | 16 | 0 | 0 | 0 | 0 | 580 | 0 | 34 | 233 | 0 | 0 | 0 | 0 | 133 | 49 | 0 | 20 | 0 | 0 | 77 | 58 | 36 | 0 | 0 | 42 | 0 | 0 | 0 | 120 | 19 | 65 |
| SRR5821805 | 40 | 14 | 1456 | 0 | 0 | 49 | 0 | 0 | 29 | 0 | 0 | 640 | 0 | 0 | 38 | 0 | 0 | 0 | 0 | 1813 | 626 | 0 | 0 | 0 | 0 | 906 | 984 | 45 | 0 | 0 | 275 | 0 | 0 | 28 | 1434 | 157 | 806 |
| SRR5832003 | 0 | 0 | 1233 | 0 | 0 | 37 | 41 | 37 | 0 | 0 | 0 | 332 | 0 | 0 | 0 | 0 | 0 | 0 | 326 | 407 | 9363 | 0 | 0 | 37 | 0 | 407 | 74 | 0 | 0 | 0 | 74 | 0 | 0 | 0 | 74 | 0 | 37 |
| SRR5832004 | 0 | 0 | 767 | 0 | 0 | 0 | 296 | 19 | 0 | 0 | 0 | 211 | 0 | 0 | 0 | 0 | 0 | 0 | 82 | 66 | 856 | 0 | 0 | 62 | 0 | 94 | 125 | 0 | 0 | 0 | 31 | 0 | 0 | 0 | 62 | 0 | 125 |
| SRR5832005 | 0 | 0 | 823 | 0 | 0 | 0 | 1072 | 224 | 0 | 0 | 0 | 155 | 0 | 0 | 0 | 0 | 0 | 22 | 193 | 64 | 343 | 0 | 0 | 28 | 0 | 0 | 14 | 0 | 0 | 42 | 14 | 0 | 0 | 28 | 70 | 0 | 0 |
| SRR5832006 | 0 | 0 | 767 | 0 | 0 | 0 | 190 | 86 | 0 | 0 | 0 | 318 | 0 | 0 | 0 | 0 | 14 | 0 | 48 | 95 | 8277 | 0 | 0 | 86 | 0 | 0 | 15 | 0 | 0 | 0 | 86 | 0 | 0 | 86 | 86 | 0 | 0 |
| SRR5832007 | 0 | 0 | 792 | 0 | 0 | 0 | 124 | 126 | 42 | 0 | 0 | 422 | 0 | 0 | 0 | 0 | 0 | 0 | 16 | 0 | 31 | 0 | 0 | 0 | 0 | 84 | 210 | 0 | 126 | 42 | 0 | 0 | 0 | 42 | 252 | 42 | 126 |
| SRR5832008 | 0 | 0 | 565 | 0 | 0 | 0 | 1590 | 53 | 0 | 0 | 0 | 273 | 0 | 0 | 0 | 0 | 0 | 0 | 89 | 0 | 89 | 0 | 0 | 12 | 0 | 12 | 12 | 0 | 37 | 11 | 0 | 0 | 0 | 49 | 18 | 0 | 12 |
| SRR5832009 | 0 | 0 | 684 | 0 | 0 | 0 | 382 | 143 | 0 | 0 | 0 | 171 | 0 | 0 | 0 | 0 | 0 | 0 | 215 | 334 | 1193 | 0 | 0 | 143 | 0 | 20 | 16 | 0 | 0 | 36 | 71 | 0 | 0 | 0 | 179 | 0 | 0 |
| SRR5832010 | 0 | 0 | 1041 | 0 | 0 | 0 | 450 | 195 | 0 | 0 | 0 | 165 | 0 | 0 | 0 | 0 | 0 | 0 | 937 | 337 | 1387 | 0 | 0 | 56 | 0 | 251 | 112 | 0 | 0 | 84 | 0 | 0 | 0 | 0 | 28 | 0 | 28 |
| SRR5832011 | 0 | 0 | 840 | 0 | 0 | 0 | 35 | 58 | 58 | 0 | 0 | 121 | 0 | 0 | 0 | 0 | 0 | 0 | 35 | 162 | 371 | 0 | 0 | 116 | 0 | 14 | 233 | 0 | 0 | 0 | 0 | 0 | 0 | 0 | 58 | 0 | 116 |
| SRR5832012 | 0 | 0 | 669 | 0 | 0 | 0 | 154 | 0 | 0 | 0 | 0 | 198 | 0 | 0 | 0 | 0 | 0 | 0 | 56 | 154 | 1952 | 0 | 0 | 260 | 0 | 325 | 325 | 0 | 0 | 0 | 0 | 0 | 0 | 0 | 65 | 0 | 130 |
| SRR5832013 | 0 | 0 | 535 | 0 | 0 | 42 | 481 | 84 | 0 | 0 | 0 | 180 | 0 | 0 | 0 | 0 | 0 | 0 | 336 | 144 | 1298 | 0 | 0 | 84 | 0 | 126 | 335 | 0 | 0 | 0 | 42 | 0 | 0 | 0 | 251 | 0 | 0 |
| SRR5832014 | 0 | 0 | 993 | 0 | 0 | 61 | 57 | 0 | 61 | 0 | 0 | 156 | 0 | 0 | 0 | 0 | 0 | 0 | 190 | 362 | 17149 | 0 | 0 | 61 | 0 | 15 | 17 | 0 | 0 | 0 | 0 | 0 | 0 | 0 | 122 | 0 | 61 |
| SRR5832015 | 14 | 0 | 2786 | 0 | 0 | 37 | 372 | 73 | 0 | 0 | 0 | 153 | 0 | 0 | 0 | 0 | 0 | 0 | 223 | 74 | 323 | 0 | 0 | 0 | 0 | 147 | 183 | 0 | 0 | 37 | 0 | 0 | 0 | 0 | 110 | 37 | 73 |
| SRR5832016 | 25 | 0 | 2733 | 0 | 0 | 0 | 825 | 200 | 0 | 0 | 0 | 159 | 0 | 0 | 0 | 29 | 0 | 0 | 127 | 0 | 222 | 0 | 0 | 29 | 0 | 172 | 57 | 0 | 0 | 29 | 0 | 0 | 0 | 0 | 143 | 0 | 29 |
| SRR5832017 | 0 | 0 | 5088 | 0 | 0 | 50 | 33 | 50 | 50 | 0 | 0 | 108 | 0 | 0 | 0 | 0 | 14 | 0 | 51 | 257 | 121 | 0 | 0 | 0 | 0 | 503 | 402 | 0 | 0 | 0 | 0 | 0 | 0 | 0 | 0 | 0 | 50 |
| SRR5832018 | 20 | 0 | 3691 | 0 | 0 | 25 | 96 | 25 | 0 | 0 | 0 | 149 | 0 | 0 | 0 | 0 | 0 | 0 | 144 | 344 | 1981 | 0 | 0 | 25 | 0 | 424 | 399 | 0 | 0 | 0 | 0 | 0 | 0 | 0 | 25 | 25 | 0 |
| SRR5832019 | 12 | 0 | 3747 | 0 | 0 | 73 | 13 | 0 | 73 | 0 | 0 | 118 | 0 | 0 | 0 | 0 | 12 | 0 | 25 | 376 | 413 | 0 | 0 | 36 | 0 | 36 | 23 | 0 | 0 | 0 | 36 | 0 | 0 | 0 | 36 | 0 | 36 |
| SRR5832020 | 31 | 0 | 3436 | 0 | 0 | 63 | 73 | 63 | 0 | 0 | 0 | 197 | 0 | 0 | 0 | 0 | 0 | 13 | 37 | 49 | 691 | 0 | 0 | 156 | 0 | 313 | 344 | 0 | 0 | 0 | 31 | 0 | 0 | 0 | 94 | 0 | 31 |
| SRR5832021 | 17 | 0 | 3786 | 0 | 0 | 19 | 537 | 37 | 19 | 0 | 0 | 231 | 0 | 0 | 0 | 0 | 17 | 0 | 313 | 201 | 1476 | 0 | 0 | 37 | 0 | 130 | 149 | 0 | 0 | 37 | 56 | 0 | 0 | 19 | 56 | 0 | 0 |
| SRR5832022 | 15 | 0 | 3686 | 0 | 0 | 0 | 132 | 21 | 0 | 0 | 0 | 137 | 0 | 0 | 0 | 0 | 0 | 0 | 25 | 38 | 227 | 0 | 0 | 0 | 0 | 208 | 130 | 26 | 0 | 52 | 26 | 0 | 0 | 26 | 52 | 0 | 0 |
| SRR5832023 | 45 | 0 | 3386 | 0 | 0 | 22 | 567 | 109 | 0 | 0 | 0 | 127 | 0 | 0 | 0 | 0 | 17 | 59 | 567 | 120 | 671 | 0 | 0 | 0 | 0 | 131 | 261 | 0 | 0 | 44 | 22 | 0 | 0 | 22 | 65 | 0 | 22 |
| SRR5832024 | 22 | 0 | 3680 | 0 | 0 | 56 | 37 | 0 | 0 | 0 | 0 | 149 | 0 | 0 | 0 | 0 | 11 | 0 | 18 | 256 | 493 | 0 | 0 | 56 | 0 | 391 | 335 | 0 | 0 | 0 | 0 | 0 | 0 | 0 | 223 | 0 | 0 |
| SRR5832025 | 26 | 0 | 3598 | 0 | 0 | 11 | 1199 | 249 | 0 | 0 | 0 | 106 | 0 | 0 | 0 | 0 | 0 | 16 | 107 | 43 | 193 | 0 | 0 | 0 | 0 | 136 | 159 | 0 | 0 | 57 | 0 | 0 | 0 | 57 | 57 | 0 | 45 |
| SRR5832026 | 0 | 0 | 3725 | 0 | 0 | 0 | 889 | 20 | 0 | 0 | 0 | 160 | 0 | 0 | 0 | 0 | 15 | 0 | 250 | 56 | 306 | 0 | 0 | 16 | 0 | 97 | 97 | 32 | 0 | 48 | 0 | 0 | 0 | 0 | 15 | 0 | 48 |
| SRR5832027 | 25 | 0 | 3902 | 0 | 0 | 0 | 204 | 33 | 33 | 0 | 0 | 168 | 0 | 0 | 0 | 0 | 16 | 0 | 450 | 347 | 2371 | 0 | 0 | 65 | 0 | 555 | 228 | 0 | 0 | 0 | 0 | 0 | 0 | 0 | 33 | 0 | 98 |
| SRR5832028 | 21 | 0 | 3796 | 0 | 0 | 14 | 828 | 150 | 0 | 0 | 0 | 224 | 0 | 0 | 0 | 0 | 0 | 39 | 349 | 239 | 920 | 14 | 0 | 0 | 0 | 82 | 177 | 0 | 14 | 0 | 0 | 0 | 0 | 0 | 82 | 0 | 55 |
| SRR5832029 | 0 | 0 | 4418 | 0 | 0 | 49 | 24 | 98 | 49 | 0 | 0 | 91 | 0 | 0 | 0 | 0 | 0 | 0 | 84 | 276 | 828 | 0 | 0 | 49 | 0 | 10 | 295 | 0 | 0 | 0 | 49 | 0 | 0 | 0 | 0 | 0 | 0 |
| SRR5868085 | 235 | 0 | 118 | 0 | 0 | 517 | 55 | 37 | 717 | 0 | 0 | 1074 | 774 | 0 | 293 | 0 | 89 | 17 | 24 | 142 | 12493 | 15 | 0 | 30 | 54 | 3327 | 333 | 0 | 0 | 72 | 1565 | 0 | 0 | 9702 | 1994 | 10503 | 22088 |
| SRR5901584 | 14 | 52 | 2021 | 0 | 41 | 16 | 215 | 25 | 14 | 0 | 0 | 2025 | 0 | 18 | 1557 | 0 | 10 | 0 | 17 | 1039 | 100 | 0 | 0 | 0 | 0 | 103 | 108 | 0 | 0 | 0 | 71 | 0 | 0 | 0 | 48 | 22 | 27 |
| SRR5901585 | 0 | 75 | 1967 | 0 | 51 | 12 | 185 | 37 | 26 | 0 | 0 | 1981 | 0 | 27 | 1800 | 0 | 0 | 14 | 25 | 1243 | 100 | 0 | 0 | 0 | 0 | 111 | 102 | 0 | 0 | 0 | 53 | 0 | 0 | 0 | 43 | 18 | 25 |
| SRR5901586 | 12 | 11 | 2147 | 0 | 68 | 15 | 209 | 43 | 27 | 0 | 0 | 2299 | 0 | 22 | 1808 | 0 | 0 | 0 | 29 | 1213 | 172 | 0 | 0 | 0 | 0 | 143 | 159 | 0 | 0 | 0 | 63 | 0 | 0 | 0 | 64 | 29 | 46 |
| SRR5901587 | 15 | 60 | 2969 | 0 | 17 | 55 | 69 | 108 | 11 | 0 | 0 | 2249 | 0 | 19 | 1208 | 0 | 0 | 0 | 0 | 247 | 72 | 0 | 0 | 0 | 0 | 361 | 389 | 0 | 0 | 0 | 47 | 0 | 0 | 0 | 580 | 28 | 345 |
| SRR5901588 | 14 | 52 | 2516 | 0 | 17 | 74 | 115 | 112 | 19 | 0 | 14 | 2504 | 0 | 15 | 731 | 0 | 12 | 0 | 17 | 585 | 42 | 0 | 0 | 0 | 0 | 631 | 486 | 0 | 0 | 0 | 59 | 0 | 0 | 0 | 569 | 35 | 358 |
| SRR5901589 | 14 | 75 | 3222 | 0 | 10 | 42 | 55 | 13 | 16 | 0 | 0 | 2037 | 0 | 21 | 1836 | 0 | 11 | 0 | 0 | 304 | 92 | 0 | 0 | 0 | 0 | 47 | 52 | 0 | 0 | 0 | 37 | 0 | 0 | 0 | 68 | 35 | 42 |
| SRR5901590 | 11 | 18 | 714 | 0 | 0 | 0 | 0 | 32 | 0 | 0 | 0 | 2109 | 0 | 14 | 1772 | 0 | 17 | 0 | 0 | 69 | 31 | 0 | 0 | 0 | 0 | 32 | 14 | 0 | 0 | 0 | 14 | 0 | 0 | 0 | 183 | 11 | 81 |
| SRR5901591 | 10 | 39 | 648 | 0 | 0 | 0 | 0 | 32 | 0 | 0 | 0 | 1369 | 0 | 17 | 894 | 0 | 11 | 0 | 0 | 78 | 29 | 0 | 0 | 0 | 0 | 42 | 303 | 13 | 0 | 0 | 35 | 0 | 0 | 0 | 1954 | 94 | 810 |
| SRR5901592 | 11 | 41 | 776 | 0 | 0 | 0 | 0 | 62 | 0 | 0 | 0 | 1376 | 0 | 24 | 738 | 0 | 11 | 0 | 0 | 43 | 16 | 0 | 0 | 0 | 0 | 17 | 18 | 0 | 0 | 0 | 0 | 0 | 0 | 0 | 151 | 70 | 72 |
| SRR5901593 | 0 | 0 | 1417 | 0 | 0 | 0 | 0 | 19 | 0 | 0 | 0 | 1745 | 0 | 27 | 361 | 0 | 19 | 0 | 0 | 53 | 29 | 0 | 0 | 0 | 0 | 19 | 40 | 0 | 0 | 0 | 13 | 0 | 0 | 0 | 103 | 96 | 67 |
| SRR5901594 | 15 | 18 | 1899 | 0 | 0 | 0 | 0 | 16 | 0 | 0 | 0 | 1470 | 0 | 26 | 605 | 0 | 15 | 0 | 0 | 52 | 29 | 0 | 0 | 0 | 0 | 43 | 57 | 13 | 0 | 0 | 17 | 0 | 0 | 0 | 97 | 57 | 64 |
| SRR5901595 | 0 | 12 | 1417 | 0 | 0 | 0 | 0 | 13 | 0 | 0 | 0 | 1880 | 0 | 19 | 707 | 0 | 14 | 0 | 0 | 39 | 40 | 0 | 0 | 0 | 0 | 40 | 38 | 23 | 0 | 0 | 17 | 0 | 0 | 0 | 105 | 52 | 68 |
| SRR5922753 | 62 | 0 | 2167 | 0 | 0 | 228 | 59 | 0 | 197 | 0 | 0 | 1298 | 485 | 22 | 13 | 0 | 0 | 0 | 17 | 247 | 80 | 0 | 0 | 0 | 516 | 6913 | 1540 | 0 | 0 | 0 | 331 | 0 | 0 | 11 | 398 | 308 | 320 |
| SRR5922754 | 35 | 0 | 1867 | 0 | 0 | 47 | 23 | 0 | 64 | 0 | 18 | 934 | 231 | 20 | 55 | 0 | 11 | 13 | 0 | 31 | 0 | 0 | 0 | 0 | 31 | 49 | 89 | 0 | 0 | 0 | 15 | 0 | 0 | 0 | 129 | 86 | 106 |
| SRR5922756 | 45 | 0 | 2743 | 0 | 0 | 0 | 0 | 0 | 0 | 34 | 44 | 934 | 11 | 17 | 94 | 0 | 17 | 19 | 0 | 0 | 0 | 0 | 20 | 0 | 36 | 19 | 0 | 14 | 0 | 0 | 18 | 0 | 0 | 29 | 14 | 0 | 0 |
| SRR5925650 | 46 | 16 | 2070 | 0 | 0 | 0 | 19 | 0 | 0 | 0 | 0 | 112 | 0 | 54 | 0 | 10 | 0 | 0 | 0 | 297 | 308 | 16 | 0 | 0 | 12 | 21 | 29 | 35 | 35 | 0 | 0 | 0 | 27 | 14 | 29 | 0 | 24 |
| SRR5925651 | 61 | 12 | 4998 | 0 | 0 | 0 | 63 | 0 | 0 | 12 | 0 | 110 | 0 | 86 | 35 | 37 | 0 | 0 | 0 | 645 | 223 | 23 | 0 | 0 | 46 | 17 | 101 | 118 | 14 | 0 | 0 | 0 | 71 | 39 | 50 | 0 | 23 |
| SRR5925652 | 38 | 0 | 1068 | 0 | 0 | 19 | 81 | 0 | 11 | 0 | 0 | 394 | 0 | 53 | 167 | 0 | 15 | 0 | 24 | 1690 | 3664 | 0 | 0 | 0 | 51 | 242 | 1473 | 37 | 0 | 0 | 26 | 0 | 30 | 20 | 218 | 46 | 175 |
| SRR5925665 | 55 | 0 | 869 | 0 | 0 | 50 | 36 | 317 | 0 | 0 | 0 | 62 | 0 | 0 | 84 | 0 | 12 | 0 | 48 | 2060 | 988 | 0 | 0 | 89 | 0 | 406 | 936 | 0 | 0 | 25 | 0 | 0 | 0 | 0 | 936 | 183 | 1075 |
| SRR5925666 | 59 | 0 | 935 | 0 | 0 | 16 | 0 | 14 | 0 | 0 | 0 | 66 | 0 | 0 | 17 | 0 | 0 | 0 | 0 | 1866 | 967 | 0 | 0 | 54 | 0 | 23 | 50 | 0 | 0 | 70 | 12 | 0 | 0 | 0 | 51 | 11 | 61 |
| SRR5925667 | 63 | 0 | 959 | 0 | 0 | 20 | 11 | 12 | 0 | 0 | 0 | 99 | 12 | 0 | 34 | 0 | 0 | 0 | 22 | 2526 | 1673 | 0 | 0 | 162 | 0 | 32 | 63 | 16 | 0 | 36 | 12 | 0 | 0 | 0 | 102 | 24 | 103 |
| SRR5925668 | 67 | 0 | 732 | 0 | 0 | 11 | 0 | 258 | 0 | 0 | 0 | 95 | 0 | 0 | 33 | 0 | 0 | 0 | 13 | 2249 | 1075 | 11 | 0 | 116 | 0 | 536 | 1109 | 13 | 0 | 24 | 11 | 0 | 0 | 0 | 1393 | 331 | 1430 |
| SRR5925669 | 54 | 0 | 863 | 0 | 0 | 20 | 0 | 12 | 0 | 0 | 0 | 101 | 0 | 12 | 170 | 0 | 0 | 0 | 24 | 2078 | 1081 | 0 | 0 | 131 | 0 | 31 | 54 | 24 | 0 | 20 | 0 | 0 | 0 | 0 | 101 | 20 | 100 |
| SRR5925670 | 66 | 0 | 802 | 0 | 0 | 39 | 23 | 793 | 0 | 0 | 0 | 91 | 0 | 0 | 46 | 0 | 0 | 0 | 34 | 1769 | 1149 | 0 | 0 | 105 | 0 | 522 | 1025 | 18 | 0 | 39 | 26 | 0 | 0 | 0 | 1617 | 364 | 1389 |
| SRR5925671 | 60 | 0 | 682 | 0 | 0 | 57 | 0 | 245 | 19 | 0 | 0 | 60 | 0 | 0 | 72 | 0 | 10 | 0 | 0 | 1428 | 1548 | 0 | 0 | 122 | 23 | 936 | 948 | 23 | 0 | 19 | 31 | 0 | 0 | 0 | 1609 | 271 | 1311 |
| SRR5925672 | 60 | 0 | 764 | 0 | 0 | 37 | 19 | 144 | 29 | 0 | 0 | 70 | 0 | 0 | 67 | 0 | 0 | 11 | 38 | 1750 | 1655 | 0 | 0 | 168 | 0 | 1329 | 1062 | 16 | 0 | 33 | 41 | 0 | 0 | 0 | 1833 | 361 | 1411 |
| SRR5925673 | 88 | 0 | 1143 | 0 | 0 | 24 | 0 | 109 | 0 | 0 | 0 | 55 | 0 | 0 | 39 | 0 | 0 | 0 | 39 | 3445 | 1905 | 0 | 0 | 44 | 0 | 28 | 52 | 0 | 0 | 62 | 15 | 0 | 0 | 0 | 40 | 10 | 44 |
| SRR5925674 | 92 | 0 | 845 | 0 | 0 | 15 | 44 | 96 | 0 | 0 | 0 | 62 | 0 | 0 | 22 | 0 | 0 | 0 | 66 | 3655 | 1598 | 0 | 0 | 74 | 0 | 40 | 61 | 0 | 0 | 59 | 30 | 0 | 0 | 0 | 31 | 177 | 32 |
| SRR5925675 | 92 | 0 | 1165 | 0 | 0 | 0 | 0 | 17 | 0 | 0 | 0 | 82 | 0 | 0 | 0 | 0 | 0 | 0 | 45 | 2567 | 1149 | 0 | 0 | 56 | 0 | 34 | 61 | 0 | 0 | 56 | 12 | 0 | 0 | 0 | 26 | 53 | 16 |
| SRR5925676 | 77 | 0 | 1087 | 0 | 0 | 24 | 24 | 92 | 21 | 0 | 0 | 86 | 0 | 0 | 0 | 0 | 0 | 0 | 32 | 2116 | 1062 | 0 | 0 | 86 | 0 | 34 | 58 | 12 | 0 | 24 | 18 | 0 | 0 | 0 | 34 | 89 | 25 |
| SRR5925677 | 54 | 0 | 1321 | 0 | 0 | 27 | 11 | 138 | 0 | 0 | 0 | 90 | 0 | 0 | 11 | 0 | 0 | 0 | 32 | 1894 | 1288 | 0 | 0 | 129 | 0 | 33 | 72 | 0 | 0 | 58 | 31 | 0 | 0 | 0 | 29 | 76 | 32 |
| SRR5925678 | 65 | 0 | 1326 | 0 | 0 | 0 | 0 | 141 | 0 | 0 | 0 | 117 | 0 | 0 | 0 | 0 | 0 | 0 | 44 | 2186 | 1182 | 0 | 0 | 108 | 0 | 32 | 77 | 0 | 0 | 37 | 24 | 0 | 0 | 0 | 51 | 155 | 44 |
| SRR5925679 | 66 | 0 | 726 | 0 | 0 | 19 | 20 | 146 | 0 | 0 | 0 | 92 | 0 | 0 | 49 | 0 | 0 | 0 | 39 | 2045 | 1868 | 0 | 0 | 92 | 0 | 34 | 49 | 0 | 0 | 23 | 38 | 0 | 0 | 0 | 32 | 73 | 26 |
| SRR5925680 | 63 | 0 | 561 | 0 | 0 | 24 | 31 | 79 | 12 | 0 | 0 | 96 | 0 | 12 | 31 | 0 | 0 | 0 | 31 | 1647 | 1696 | 0 | 0 | 105 | 14 | 502 | 921 | 12 | 0 | 24 | 29 | 0 | 0 | 0 | 743 | 100 | 498 |
| SRR5925681 | 95 | 0 | 301 | 0 | 17 | 25 | 0 | 126 | 0 | 0 | 0 | 54 | 0 | 0 | 22 | 0 | 0 | 0 | 11 | 1357 | 912 | 0 | 0 | 172 | 0 | 612 | 775 | 29 | 0 | 29 | 21 | 0 | 0 | 0 | 813 | 117 | 549 |
| SRR5925682 | 61 | 0 | 420 | 0 | 0 | 23 | 14 | 112 | 0 | 0 | 0 | 79 | 0 | 12 | 35 | 0 | 14 | 0 | 14 | 1225 | 616 | 0 | 0 | 162 | 0 | 22 | 35 | 12 | 0 | 42 | 0 | 0 | 0 | 0 | 51 | 138 | 39 |
| SRR5925683 | 65 | 13 | 590 | 0 | 0 | 63 | 0 | 11 | 0 | 0 | 0 | 85 | 0 | 0 | 28 | 0 | 0 | 0 | 0 | 1512 | 898 | 0 | 0 | 67 | 13 | 32 | 40 | 36 | 0 | 0 | 0 | 0 | 0 | 0 | 59 | 130 | 34 |
| SRR5925684 | 68 | 0 | 485 | 0 | 0 | 41 | 11 | 134 | 10 | 0 | 0 | 86 | 0 | 0 | 22 | 0 | 11 | 0 | 0 | 1445 | 1254 | 0 | 0 | 124 | 16 | 28 | 36 | 0 | 0 | 16 | 21 | 0 | 0 | 0 | 51 | 134 | 35 |
| SRR5925685 | 17 | 0 | 15 | 0 | 0 | 25 | 257 | 215 | 11 | 0 | 0 | 0 | 0 | 11 | 30 | 0 | 0 | 0 | 24 | 2215 | 0 | 0 | 0 | 32 | 0 | 34 | 41 | 14 | 0 | 185 | 60 | 0 | 0 | 11 | 33 | 57 | 27 |
| SRR5925686 | 17 | 0 | 23 | 0 | 0 | 19 | 143 | 3757 | 23 | 0 | 0 | 0 | 0 | 0 | 13 | 0 | 0 | 0 | 32 | 2164 | 0 | 0 | 0 | 42 | 0 | 654 | 609 | 11 | 0 | 148 | 34 | 0 | 0 | 11 | 730 | 83 | 416 |
| SRR5925687 | 12 | 0 | 17 | 0 | 0 | 44 | 212 | 4258 | 13 | 0 | 0 | 0 | 0 | 0 | 0 | 0 | 0 | 11 | 65 | 2290 | 0 | 0 | 0 | 26 | 0 | 725 | 629 | 0 | 0 | 201 | 66 | 0 | 0 | 31 | 935 | 105 | 734 |
| SRR5925688 | 12 | 16 | 14 | 0 | 0 | 24 | 228 | 215 | 0 | 0 | 0 | 0 | 0 | 0 | 0 | 0 | 0 | 10 | 41 | 2138 | 0 | 0 | 0 | 12 | 0 | 41 | 32 | 20 | 0 | 126 | 69 | 0 | 0 | 16 | 43 | 61 | 27 |
| SRR5925689 | 16 | 0 | 18 | 0 | 0 | 36 | 291 | 277 | 14 | 0 | 0 | 0 | 0 | 0 | 0 | 0 | 0 | 0 | 36 | 2930 | 0 | 0 | 0 | 40 | 0 | 45 | 40 | 0 | 0 | 148 | 61 | 0 | 0 | 18 | 39 | 65 | 27 |
| SRR5925690 | 13 | 0 | 19 | 0 | 0 | 46 | 331 | 255 | 15 | 0 | 0 | 0 | 0 | 0 | 20 | 0 | 0 | 0 | 13 | 2085 | 0 | 0 | 0 | 15 | 0 | 37 | 39 | 0 | 0 | 156 | 55 | 0 | 0 | 12 | 46 | 70 | 32 |
| SRR5925691 | 18 | 0 | 16 | 0 | 0 | 53 | 182 | 2744 | 11 | 0 | 0 | 0 | 0 | 11 | 27 | 0 | 0 | 0 | 40 | 2188 | 0 | 18 | 0 | 21 | 0 | 676 | 648 | 0 | 0 | 137 | 67 | 0 | 0 | 21 | 813 | 112 | 631 |
| SRR5925692 | 21 | 0 | 24 | 0 | 0 | 23 | 258 | 148 | 23 | 0 | 0 | 0 | 0 | 0 | 25 | 0 | 0 | 0 | 19 | 2191 | 0 | 0 | 0 | 26 | 0 | 47 | 38 | 0 | 0 | 108 | 69 | 0 | 0 | 26 | 43 | 75 | 30 |
| SRR5925693 | 17 | 0 | 23 | 0 | 0 | 42 | 258 | 211 | 23 | 0 | 0 | 0 | 0 | 26 | 31 | 0 | 0 | 0 | 19 | 3106 | 0 | 0 | 0 | 0 | 0 | 41 | 37 | 26 | 0 | 12 | 42 | 0 | 0 | 19 | 56 | 81 | 42 |
| SRR5925694 | 21 | 12 | 26 | 0 | 0 | 33 | 216 | 250 | 15 | 0 | 0 | 0 | 0 | 12 | 38 | 0 | 0 | 0 | 32 | 3234 | 0 | 0 | 0 | 45 | 0 | 31 | 45 | 0 | 0 | 16 | 36 | 0 | 0 | 15 | 57 | 93 | 41 |
| SRR5925695 | 12 | 0 | 15 | 0 | 0 | 24 | 143 | 2478 | 0 | 0 | 0 | 0 | 0 | 0 | 13 | 0 | 0 | 0 | 20 | 2596 | 13 | 0 | 0 | 28 | 0 | 953 | 1005 | 14 | 0 | 165 | 66 | 0 | 0 | 19 | 699 | 80 | 543 |
| SRR5925696 | 13 | 0 | 17 | 0 | 0 | 38 | 161 | 93 | 14 | 0 | 0 | 0 | 0 | 14 | 38 | 0 | 0 | 30 | 38 | 3401 | 0 | 0 | 0 | 0 | 0 | 44 | 38 | 0 | 0 | 183 | 66 | 0 | 0 | 14 | 33 | 178 | 41 |
| SRR5925697 | 13 | 0 | 19 | 0 | 0 | 55 | 171 | 3898 | 16 | 0 | 0 | 0 | 0 | 0 | 19 | 0 | 0 | 0 | 46 | 3322 | 0 | 0 | 0 | 29 | 0 | 808 | 1190 | 10 | 0 | 178 | 88 | 0 | 0 | 12 | 1057 | 101 | 749 |
| SRR5925698 | 11 | 0 | 13 | 0 | 0 | 35 | 237 | 3167 | 20 | 0 | 0 | 0 | 0 | 0 | 0 | 0 | 0 | 0 | 29 | 2710 | 0 | 0 | 0 | 30 | 0 | 854 | 1120 | 0 | 0 | 168 | 74 | 0 | 0 | 15 | 982 | 143 | 947 |
| SRR5925699 | 12 | 0 | 14 | 0 | 0 | 59 | 185 | 146 | 16 | 0 | 0 | 0 | 0 | 0 | 32 | 0 | 0 | 10 | 23 | 2265 | 0 | 0 | 0 | 39 | 0 | 47 | 40 | 0 | 0 | 139 | 70 | 0 | 0 | 13 | 38 | 68 | 29 |
| SRR5925700 | 14 | 0 | 15 | 0 | 0 | 42 | 215 | 2277 | 17 | 0 | 0 | 0 | 0 | 0 | 50 | 0 | 0 | 16 | 36 | 2377 | 0 | 0 | 0 | 21 | 0 | 808 | 724 | 0 | 0 | 126 | 114 | 0 | 0 | 21 | 838 | 84 | 711 |
| SRR5925701 | 21 | 0 | 20 | 0 | 0 | 41 | 194 | 2517 | 17 | 0 | 0 | 0 | 0 | 14 | 33 | 0 | 0 | 0 | 44 | 1911 | 0 | 0 | 0 | 39 | 0 | 596 | 598 | 0 | 0 | 275 | 57 | 0 | 0 | 38 | 698 | 72 | 569 |
| SRR5925702 | 24 | 0 | 19 | 0 | 0 | 31 | 178 | 2648 | 0 | 0 | 0 | 0 | 0 | 0 | 36 | 0 | 0 | 0 | 36 | 2174 | 0 | 0 | 0 | 39 | 0 | 530 | 814 | 27 | 0 | 187 | 62 | 0 | 0 | 0 | 752 | 86 | 619 |
| SRR5925703 | 16 | 11 | 25 | 0 | 0 | 42 | 223 | 204 | 11 | 0 | 0 | 0 | 0 | 19 | 39 | 0 | 0 | 0 | 20 | 1877 | 0 | 0 | 0 | 23 | 0 | 33 | 42 | 19 | 0 | 165 | 77 | 0 | 0 | 23 | 43 | 115 | 36 |
| SRR5925704 | 14 | 11 | 20 | 0 | 0 | 42 | 232 | 208 | 25 | 0 | 0 | 0 | 0 | 11 | 15 | 0 | 0 | 0 | 0 | 1573 | 12 | 0 | 0 | 42 | 0 | 39 | 32 | 0 | 0 | 189 | 53 | 0 | 0 | 24 | 37 | 137 | 40 |
| SRR5925705 | 102 | 0 | 969 | 0 | 0 | 0 | 0 | 52 | 0 | 0 | 0 | 97 | 0 | 0 | 4383 | 0 | 0 | 0 | 15 | 1052 | 49282 | 0 | 0 | 19 | 0 | 167 | 311 | 0 | 0 | 0 | 0 | 0 | 0 | 0 | 426 | 160 | 629 |
| SRR5925706 | 131 | 0 | 712 | 0 | 0 | 0 | 25 | 14 | 0 | 0 | 0 | 106 | 0 | 0 | 12389 | 0 | 0 | 11 | 0 | 922 | 39566 | 0 | 0 | 22 | 0 | 40 | 38 | 0 | 0 | 0 | 0 | 0 | 0 | 0 | 132 | 19 | 187 |
| SRR5925707 | 91 | 0 | 837 | 0 | 0 | 0 | 0 | 68 | 0 | 0 | 0 | 93 | 0 | 0 | 3941 | 0 | 0 | 0 | 0 | 776 | 39742 | 0 | 0 | 28 | 0 | 12 | 41 | 0 | 0 | 0 | 0 | 0 | 0 | 0 | 80 | 25 | 104 |
| SRR5925708 | 127 | 0 | 727 | 0 | 0 | 0 | 0 | 94 | 0 | 0 | 0 | 117 | 0 | 0 | 8784 | 0 | 0 | 16 | 13 | 838 | 38251 | 0 | 0 | 21 | 0 | 38 | 241 | 0 | 0 | 0 | 0 | 0 | 0 | 0 | 908 | 144 | 1048 |
| SRR5925709 | 105 | 0 | 64 | 0 | 0 | 101 | 0 | 303 | 74 | 0 | 0 | 38 | 0 | 0 | 605 | 0 | 0 | 0 | 0 | 464 | 2113 | 0 | 0 | 21 | 0 | 52 | 14 | 11 | 0 | 32 | 74 | 0 | 0 | 0 | 31 | 223 | 16 |
| SRR5925710 | 116 | 0 | 290 | 0 | 0 | 0 | 0 | 214 | 0 | 0 | 0 | 0 | 0 | 0 | 0 | 0 | 0 | 0 | 0 | 0 | 5906 | 0 | 0 | 0 | 0 | 0 | 0 | 0 | 0 | 0 | 0 | 0 | 0 | 0 | 53 | 27 | 0 |
| SRR5925711 | 199 | 0 | 376 | 0 | 0 | 0 | 0 | 95 | 0 | 0 | 0 | 56 | 0 | 0 | 2711 | 0 | 0 | 0 | 0 | 1135 | 12797 | 0 | 0 | 53 | 0 | 47 | 10 | 0 | 0 | 17 | 0 | 0 | 0 | 0 | 69 | 75 | 64 |
| SRR5925712 | 439 | 0 | 853 | 0 | 0 | 11 | 0 | 60 | 0 | 0 | 0 | 76 | 0 | 0 | 236 | 0 | 11 | 0 | 11 | 908 | 1950 | 0 | 0 | 19 | 0 | 262 | 91 | 0 | 0 | 0 | 0 | 0 | 0 | 0 | 239 | 97 | 232 |
| SRR5925713 | 162 | 0 | 617 | 0 | 0 | 0 | 0 | 70 | 0 | 0 | 0 | 80 | 0 | 0 | 4284 | 0 | 0 | 0 | 11 | 814 | 29135 | 0 | 0 | 17 | 0 | 20 | 38 | 0 | 0 | 0 | 0 | 0 | 0 | 0 | 56 | 20 | 78 |
| SRR5925714 | 177 | 0 | 661 | 0 | 0 | 0 | 0 | 63 | 0 | 0 | 0 | 86 | 0 | 0 | 3643 | 0 | 0 | 30 | 17 | 952 | 30091 | 0 | 0 | 24 | 0 | 26 | 44 | 0 | 0 | 0 | 15 | 0 | 0 | 0 | 61 | 20 | 85 |
| SRR5925715 | 133 | 0 | 529 | 0 | 0 | 0 | 0 | 18 | 10 | 0 | 0 | 36 | 0 | 0 | 10720 | 0 | 0 | 11 | 0 | 417 | 30747 | 0 | 0 | 15 | 0 | 99 | 15 | 0 | 0 | 10 | 10 | 0 | 0 | 0 | 76 | 11 | 102 |
| SRR5925716 | 130 | 0 | 521 | 0 | 0 | 0 | 0 | 84 | 0 | 0 | 0 | 43 | 0 | 0 | 8535 | 0 | 0 | 46 | 0 | 427 | 28575 | 0 | 0 | 17 | 0 | 70 | 93 | 0 | 0 | 0 | 0 | 0 | 0 | 0 | 529 | 101 | 673 |
| SRR5925717 | 201 | 0 | 386 | 0 | 0 | 0 | 0 | 71 | 0 | 0 | 0 | 45 | 0 | 0 | 14384 | 0 | 0 | 19 | 21 | 1192 | 32556 | 0 | 0 | 0 | 0 | 36 | 106 | 0 | 0 | 11 | 0 | 0 | 0 | 0 | 511 | 77 | 769 |
| SRR5925718 | 204 | 0 | 384 | 0 | 0 | 0 | 0 | 83 | 0 | 0 | 0 | 47 | 0 | 0 | 12801 | 0 | 0 | 14 | 18 | 1365 | 30433 | 0 | 0 | 0 | 0 | 36 | 121 | 0 | 0 | 14 | 0 | 0 | 0 | 0 | 554 | 92 | 821 |
| SRR5925719 | 31 | 93 | 16 | 0 | 0 | 0 | 573 | 2366 | 21 | 0 | 0 | 0 | 0 | 166 | 165 | 0 | 0 | 24 | 44 | 1368 | 116 | 0 | 0 | 0 | 0 | 16 | 18 | 18 | 0 | 107 | 42 | 0 | 0 | 0 | 45 | 55 | 40 |
| SRR5925720 | 32 | 88 | 15 | 0 | 0 | 0 | 765 | 2646 | 10 | 0 | 0 | 0 | 0 | 154 | 156 | 0 | 0 | 30 | 43 | 1719 | 71 | 0 | 0 | 0 | 0 | 21 | 18 | 16 | 0 | 108 | 34 | 0 | 0 | 20 | 43 | 56 | 42 |
| SRR5925721 | 41 | 29 | 20 | 11 | 0 | 0 | 456 | 16063 | 14 | 0 | 0 | 0 | 0 | 49 | 293 | 0 | 0 | 13 | 60 | 1379 | 49 | 0 | 0 | 0 | 0 | 267 | 174 | 0 | 19 | 182 | 26 | 0 | 0 | 19 | 389 | 27 | 328 |
| SRR5925722 | 40 | 29 | 22 | 0 | 10 | 0 | 506 | 16072 | 0 | 0 | 0 | 0 | 0 | 59 | 286 | 0 | 0 | 0 | 77 | 1518 | 46 | 0 | 0 | 0 | 0 | 288 | 206 | 0 | 28 | 183 | 34 | 0 | 0 | 17 | 389 | 28 | 290 |
| SRR5925723 | 39 | 19 | 32 | 10 | 0 | 0 | 138 | 7166 | 0 | 0 | 0 | 0 | 0 | 37 | 295 | 0 | 0 | 0 | 39 | 1302 | 74 | 0 | 0 | 0 | 0 | 93 | 121 | 0 | 33 | 224 | 0 | 0 | 0 | 10 | 209 | 36 | 392 |
| SRR5925724 | 46 | 18 | 33 | 0 | 0 | 0 | 173 | 1332 | 0 | 0 | 0 | 0 | 0 | 43 | 257 | 0 | 0 | 0 | 20 | 1642 | 79 | 0 | 0 | 0 | 0 | 15 | 26 | 0 | 32 | 39 | 11 | 0 | 0 | 17 | 33 | 38 | 60 |
| SRR5925725 | 46 | 19 | 25 | 0 | 0 | 0 | 316 | 10011 | 0 | 0 | 0 | 0 | 0 | 59 | 201 | 0 | 0 | 0 | 67 | 1571 | 72 | 0 | 0 | 0 | 0 | 151 | 169 | 0 | 21 | 381 | 18 | 0 | 0 | 23 | 160 | 28 | 315 |
| SRR5925726 | 38 | 13 | 29 | 0 | 0 | 0 | 226 | 1343 | 0 | 0 | 0 | 0 | 0 | 12 | 162 | 0 | 0 | 15 | 41 | 1350 | 35 | 0 | 0 | 0 | 0 | 20 | 23 | 0 | 34 | 56 | 17 | 0 | 0 | 24 | 25 | 27 | 41 |
| SRR5925727 | 0 | 0 | 0 | 0 | 0 | 0 | 0 | 1660 | 0 | 0 | 0 | 0 | 0 | 0 | 0 | 0 | 0 | 0 | 0 | 0 | 16260 | 0 | 0 | 0 | 0 | 0 | 0 | 0 | 0 | 208 | 0 | 0 | 0 | 0 | 0 | 0 | 0 |
| SRR5925728 | 47 | 46 | 23 | 0 | 0 | 0 | 648 | 31071 | 0 | 0 | 0 | 0 | 0 | 160 | 501 | 0 | 0 | 0 | 85 | 3425 | 39 | 0 | 0 | 0 | 0 | 80 | 210 | 0 | 0 | 207 | 0 | 0 | 0 | 10 | 272 | 39 | 480 |
| SRR5925729 | 41 | 56 | 18 | 0 | 0 | 14 | 272 | 26223 | 0 | 0 | 0 | 0 | 0 | 93 | 268 | 0 | 0 | 28 | 37 | 1343 | 54 | 0 | 0 | 0 | 0 | 184 | 192 | 11 | 0 | 163 | 14 | 0 | 0 | 0 | 585 | 22 | 469 |
| SRR5925730 | 41 | 61 | 21 | 0 | 0 | 0 | 222 | 26328 | 0 | 0 | 0 | 0 | 0 | 103 | 192 | 0 | 0 | 35 | 30 | 1426 | 47 | 0 | 0 | 0 | 0 | 155 | 204 | 0 | 0 | 165 | 27 | 0 | 0 | 22 | 465 | 30 | 410 |
| SRR5925731 | 36 | 112 | 20 | 0 | 0 | 0 | 351 | 42800 | 0 | 0 | 0 | 0 | 0 | 145 | 608 | 0 | 0 | 14 | 132 | 6174 | 113 | 0 | 0 | 0 | 0 | 87 | 517 | 0 | 0 | 211 | 21 | 0 | 0 | 17 | 815 | 54 | 932 |
| SRR5925732 | 39 | 86 | 23 | 10 | 0 | 0 | 202 | 2260 | 0 | 0 | 0 | 0 | 0 | 12 | 671 | 0 | 0 | 15 | 97 | 4468 | 145 | 0 | 0 | 13 | 0 | 81 | 25 | 10 | 0 | 14 | 0 | 0 | 0 | 10 | 47 | 84 | 66 |
| SRR5930976 | 19 | 0 | 2037 | 0 | 0 | 22 | 20 | 15 | 14 | 0 | 0 | 78 | 0 | 12 | 152 | 0 | 12 | 0 | 20 | 1558 | 4056 | 0 | 0 | 0 | 0 | 24 | 337 | 20 | 0 | 0 | 39 | 0 | 0 | 0 | 95 | 11 | 72 |
| SRR5930977 | 34 | 0 | 1650 | 0 | 0 | 0 | 0 | 0 | 0 | 0 | 0 | 82 | 0 | 12 | 15 | 13 | 0 | 0 | 0 | 1003 | 2846 | 0 | 0 | 0 | 0 | 53 | 62 | 49 | 0 | 0 | 0 | 0 | 12 | 12 | 16 | 0 | 11 |
| SRR5930978 | 50 | 48 | 3533 | 0 | 0 | 0 | 0 | 0 | 0 | 17 | 0 | 181 | 0 | 83 | 0 | 23 | 0 | 0 | 0 | 78 | 94 | 10 | 0 | 0 | 24 | 19 | 53 | 73 | 59 | 0 | 13 | 13 | 81 | 29 | 18 | 0 | 62 |
| SRR5936591 | 19 | 0 | 585 | 0 | 0 | 10 | 44 | 0 | 17 | 0 | 0 | 80 | 0 | 0 | 88 | 0 | 0 | 0 | 66 | 2996 | 4780 | 0 | 0 | 0 | 0 | 144 | 413 | 0 | 0 | 0 | 30 | 0 | 0 | 0 | 281 | 54 | 140 |
| SRR5936592 | 19 | 69 | 2278 | 0 | 0 | 0 | 205 | 0 | 0 | 15 | 0 | 69 | 0 | 185 | 0 | 39 | 0 | 0 | 0 | 162 | 226 | 23 | 11 | 0 | 111 | 153 | 48 | 208 | 18 | 0 | 19 | 0 | 59 | 12 | 31 | 0 | 23 |

### Supp Table S8.2.Columns_AM_to_BW

| | | | | | | | | | | | | | | | | | | | | | | | | | | | | | | | | | | | | | |
|---|---|---|---|---|---|---|---|---|---|---|---|---|---|---|---|---|---|---|---|---|---|---|---|---|---|---|---|---|---|---|---|---|---|---|---|---|---|
| SRR5951878 | 40 | 0 | 21 | 0 | 0 | 99 | 67 | 182 | 55 | 0 | 0 | 304 | 0 | 14 | 9060 | 0 | 242 | 91 | 16 | 361 | 574 | 0 | 0 | 13 | 21 | 18 | 18 | 0 | 0 | 12 | 64 | 0 | 55 | 21 | 20 | 94 | 84 |
| SRR5951879 | 25 | 0 | 231 | 0 | 0 | 78 | 90 | 10 | 37 | 0 | 0 | 416 | 0 | 0 | 8176 | 0 | 249 | 40 | 99 | 1109 | 688 | 0 | 0 | 62 | 22 | 30 | 48 | 0 | 0 | 34 | 35 | 0 | 0 | 19 | 16 | 12 | 14 |
| SRR5951880 | 0 | 0 | 0 | 0 | 0 | 0 | 0 | 0 | 0 | 0 | 0 | 0 | 0 | 0 | 50 | 0 | 0 | 0 | 0 | 326 | 100 | 0 | 0 | 0 | 452 | 0 | 0 | 0 | 0 | 0 | 0 | 0 | 0 | 0 | 0 | 0 | 0 |
| SRR5951881 | 21 | 0 | 13 | 0 | 0 | 219 | 63 | 10 | 203 | 0 | 16 | 265 | 0 | 0 | 10933 | 0 | 62 | 50 | 19 | 171 | 442 | 0 | 0 | 14 | 78 | 30 | 27 | 0 | 0 | 28 | 214 | 12 | 62 | 12 | 313 | 47 | 51 |
| SRR5951882 | 29 | 0 | 96 | 0 | 0 | 669 | 533 | 702 | 192 | 0 | 17 | 465 | 0 | 15 | 23423 | 0 | 140 | 38 | 100 | 783 | 1848 | 0 | 0 | 1059 | 40 | 4277 | 2253 | 0 | 0 | 30 | 767 | 0 | 20 | 25 | 194 | 231 | 515 |
| SRR5951883 | 0 | 0 | 0 | 0 | 0 | 16 | 0 | 0 | 0 | 0 | 48 | 0 | 0 | 0 | 0 | 0 | 0 | 12 | 0 | 212 | 124 | 0 | 0 | 0 | 302 | 0 | 0 | 0 | 0 | 0 | 0 | 0 | 16 | 0 | 0 | 0 | 32 |
| SRR5951884 | 22 | 0 | 37 | 0 | 0 | 255 | 30 | 648 | 93 | 0 | 13 | 210 | 13 | 0 | 1677 | 0 | 37 | 0 | 142 | 1063 | 659 | 0 | 0 | 2436 | 0 | 1205 | 265 | 0 | 0 | 81 | 197 | 0 | 64 | 66 | 1660 | 3452 | 4263 |
| SRR5951885 | 22 | 0 | 42 | 0 | 0 | 329 | 0 | 671 | 109 | 0 | 15 | 206 | 13 | 0 | 2001 | 0 | 33 | 0 | 131 | 911 | 662 | 0 | 0 | 2348 | 0 | 1352 | 253 | 0 | 0 | 126 | 169 | 0 | 86 | 57 | 1584 | 3481 | 3896 |
| SRR5951886 | 18 | 0 | 29 | 0 | 0 | 269 | 0 | 2016 | 89 | 0 | 14 | 254 | 13 | 0 | 1140 | 0 | 31 | 0 | 63 | 388 | 548 | 0 | 0 | 2185 | 0 | 1199 | 274 | 0 | 0 | 193 | 158 | 0 | 58 | 64 | 1482 | 2615 | 3667 |
| SRR5951887 | 13 | 0 | 15 | 0 | 0 | 312 | 15 | 3066 | 72 | 0 | 26 | 175 | 16 | 0 | 36535 | 0 | 21 | 0 | 45 | 400 | 407 | 0 | 0 | 1882 | 0 | 1422 | 312 | 0 | 0 | 198 | 150 | 10 | 48 | 83 | 4769 | 8413 | 8447 |
| SRR5951888 | 14 | 0 | 16 | 0 | 0 | 282 | 34 | 4346 | 88 | 0 | 32 | 162 | 0 | 0 | 26292 | 0 | 25 | 0 | 59 | 286 | 375 | 0 | 0 | 1592 | 0 | 1511 | 319 | 0 | 0 | 181 | 161 | 11 | 60 | 62 | 5667 | 9614 | 10533 |
| SRR5951889 | 12 | 0 | 14 | 0 | 0 | 14 | 73 | 309 | 81 | 0 | 35 | 200 | 12 | 0 | 23820 | 0 | 29 | 0 | 89 | 394 | 467 | 0 | 0 | 87 | 0 | 64 | 16 | 0 | 0 | 15 | 133 | 12 | 58 | 58 | 218 | 464 | 449 |
| SRR5951890 | 14 | 0 | 47 | 0 | 0 | 349 | 84 | 4253 | 132 | 0 | 25 | 216 | 0 | 0 | 20851 | 0 | 25 | 0 | 81 | 718 | 366 | 0 | 0 | 924 | 15 | 1454 | 326 | 0 | 0 | 441 | 114 | 18 | 24 | 45 | 3808 | 10248 | 9593 |
| SRR5951891 | 11 | 0 | 38 | 0 | 0 | 21 | 86 | 430 | 102 | 0 | 34 | 172 | 0 | 0 | 34235 | 0 | 29 | 0 | 37 | 574 | 431 | 0 | 0 | 48 | 0 | 87 | 15 | 0 | 0 | 27 | 128 | 27 | 33 | 63 | 212 | 493 | 491 |
| SRR5951892 | 0 | 0 | 30 | 0 | 0 | 20 | 53 | 205 | 116 | 0 | 39 | 179 | 0 | 0 | 44075 | 0 | 32 | 0 | 57 | 377 | 625 | 0 | 0 | 45 | 0 | 73 | 10 | 0 | 0 | 11 | 117 | 0 | 44 | 51 | 216 | 785 | 750 |
| SRR5951893 | 25 | 0 | 46 | 0 | 0 | 304 | 65 | 9025 | 99 | 0 | 20 | 253 | 16 | 0 | 70169 | 0 | 23 | 0 | 45 | 532 | 712 | 0 | 0 | 1543 | 0 | 1543 | 266 | 0 | 0 | 554 | 242 | 49 | 56 | 62 | 831 | 1579 | 2634 |
| SRR5951894 | 31 | 0 | 116 | 0 | 0 | 272 | 21 | 439 | 84 | 0 | 22 | 285 | 0 | 0 | 2902 | 0 | 28 | 0 | 78 | 681 | 504 | 0 | 0 | 1413 | 0 | 1016 | 833 | 0 | 0 | 102 | 250 | 0 | 150 | 81 | 1195 | 1618 | 2896 |
| SRR5951895 | 24 | 0 | 46 | 0 | 0 | 11 | 105 | 379 | 89 | 0 | 37 | 277 | 0 | 0 | 78527 | 0 | 25 | 0 | 61 | 588 | 842 | 0 | 0 | 52 | 0 | 51 | 219 | 0 | 0 | 27 | 160 | 87 | 84 | 67 | 28 | 69 | 110 |
| SRR5951896 | 0 | 0 | 195 | 0 | 0 | 83 | 52 | 192 | 43 | 0 | 36 | 663 | 0 | 12 | 7202 | 0 | 47 | 15 | 23 | 168 | 1079 | 0 | 0 | 271 | 106 | 354 | 332 | 0 | 0 | 14 | 39 | 0 | 43 | 25 | 313 | 463 | 424 |
| SRR5951897 | 13 | 0 | 838 | 0 | 0 | 71 | 91 | 117 | 36 | 0 | 22 | 642 | 0 | 0 | 3070 | 0 | 104 | 87 | 18 | 251 | 560 | 0 | 0 | 29 | 68 | 20 | 21 | 0 | 0 | 88 | 38 | 0 | 22 | 57 | 153 | 13 | 16 |
| SRR5951898 | 0 | 0 | 0 | 0 | 0 | 0 | 0 | 0 | 0 | 0 | 98 | 0 | 0 | 0 | 28 | 0 | 0 | 0 | 0 | 838 | 196 | 0 | 0 | 0 | 363 | 12 | 0 | 0 | 0 | 0 | 0 | 0 | 0 | 0 | 0 | 0 | 0 |
| SRR5951899 | 12 | 0 | 182 | 0 | 14 | 106 | 265 | 61 | 59 | 17 | 108 | 235 | 0 | 0 | 10026 | 0 | 30 | 24 | 41 | 299 | 597 | 0 | 0 | 294 | 79 | 484 | 457 | 0 | 0 | 23 | 31 | 0 | 93 | 21 | 245 | 665 | 615 |
| SRR5951900 | 15 | 0 | 916 | 0 | 0 | 197 | 232 | 105 | 117 | 12 | 44 | 402 | 0 | 0 | 1662 | 0 | 121 | 81 | 39 | 750 | 271 | 0 | 0 | 17 | 80 | 58 | 28 | 0 | 0 | 25 | 104 | 0 | 39 | 30 | 244 | 17 | 18 |
| SRR5951901 | 0 | 0 | 0 | 0 | 0 | 0 | 0 | 0 | 0 | 0 | 111 | 0 | 0 | 0 | 33 | 0 | 0 | 0 | 0 | 881 | 163 | 0 | 0 | 0 | 541 | 0 | 0 | 0 | 0 | 0 | 0 | 0 | 0 | 0 | 0 | 0 | 0 |
| SRR5951902 | 11 | 0 | 227 | 0 | 0 | 135 | 87 | 329 | 46 | 0 | 35 | 534 | 0 | 24 | 12562 | 0 | 30 | 79 | 25 | 194 | 1303 | 0 | 0 | 385 | 115 | 435 | 298 | 0 | 0 | 28 | 44 | 0 | 69 | 25 | 294 | 449 | 354 |
| SRR5951903 | 12 | 0 | 815 | 0 | 0 | 126 | 28 | 145 | 51 | 0 | 10 | 799 | 0 | 0 | 4163 | 0 | 62 | 65 | 40 | 451 | 703 | 0 | 0 | 34 | 93 | 17 | 14 | 0 | 0 | 10 | 40 | 0 | 32 | 25 | 11 | 214 | 11 |
| SRR5951904 | 0 | 0 | 0 | 0 | 0 | 0 | 0 | 0 | 0 | 0 | 120 | 0 | 0 | 0 | 85 | 0 | 0 | 0 | 0 | 2192 | 370 | 0 | 0 | 0 | 517 | 0 | 0 | 0 | 0 | 0 | 0 | 0 | 0 | 0 | 0 | 0 | 11 |
| SRR5951905 | 0 | 0 | 304 | 0 | 0 | 129 | 121 | 25 | 88 | 0 | 56 | 606 | 0 | 0 | 15144 | 0 | 20 | 48 | 0 | 130 | 1188 | 0 | 0 | 18 | 104 | 28 | 14 | 0 | 0 | 37 | 70 | 21 | 54 | 38 | 170 | 13 | 15 |
| SRR5951906 | 23 | 0 | 997 | 0 | 0 | 13 | 19 | 168 | 159 | 0 | 19 | 882 | 0 | 0 | 2307 | 0 | 20 | 274 | 49 | 419 | 736 | 0 | 0 | 58 | 162 | 52 | 27 | 0 | 0 | 73 | 93 | 0 | 16 | 47 | 14 | 11 | 16 |
| SRR5951907 | 0 | 0 | 0 | 0 | 0 | 0 | 0 | 0 | 0 | 0 | 61 | 0 | 0 | 0 | 108 | 0 | 0 | 0 | 0 | 235 | 199 | 0 | 0 | 0 | 127 | 0 | 0 | 0 | 0 | 0 | 0 | 0 | 0 | 0 | 0 | 0 | 0 |
| SRR5951908 | 21 | 0 | 185 | 0 | 0 | 127 | 60 | 8616 | 167 | 0 | 84 | 420 | 11 | 0 | 62186 | 0 | 0 | 0 | 64 | 466 | 684 | 0 | 0 | 414 | 33 | 809 | 90 | 0 | 0 | 1394 | 86 | 0 | 65 | 310 | 2436 | 3889 | 4602 |
| SRR5951909 | 21 | 0 | 252 | 0 | 0 | 141 | 37 | 10193 | 208 | 0 | 132 | 345 | 0 | 0 | 95125 | 0 | 0 | 12 | 51 | 352 | 441 | 0 | 0 | 453 | 18 | 937 | 152 | 0 | 0 | 989 | 88 | 0 | 54 | 86 | 1937 | 2473 | 3024 |
| SRR5951910 | 14 | 0 | 175 | 0 | 0 | 143 | 32 | 5637 | 147 | 0 | 118 | 288 | 0 | 0 | 50133 | 0 | 0 | 0 | 68 | 347 | 599 | 0 | 0 | 514 | 0 | 841 | 124 | 0 | 0 | 614 | 73 | 0 | 77 | 92 | 2297 | 3160 | 4453 |
| SRR5951911 | 24 | 0 | 245 | 0 | 0 | 104 | 30 | 8521 | 125 | 13 | 121 | 454 | 0 | 0 | 69412 | 0 | 0 | 0 | 69 | 664 | 841 | 0 | 0 | 308 | 0 | 557 | 116 | 0 | 0 | 585 | 62 | 17 | 34 | 72 | 521 | 702 | 1282 |
| SRR5951912 | 17 | 0 | 182 | 0 | 0 | 98 | 44 | 17008 | 97 | 0 | 120 | 379 | 0 | 0 | 69275 | 0 | 10 | 11 | 89 | 521 | 709 | 0 | 0 | 321 | 0 | 562 | 99 | 0 | 0 | 1049 | 55 | 14 | 61 | 50 | 437 | 440 | 900 |
| SRR5951913 | 19 | 0 | 163 | 0 | 0 | 97 | 36 | 443 | 108 | 0 | 137 | 347 | 0 | 0 | 78053 | 0 | 11 | 0 | 42 | 505 | 667 | 0 | 0 | 16 | 13 | 39 | 108 | 0 | 0 | 47 | 69 | 16 | 69 | 70 | 23 | 34 | 68 |
| SRR6020460 | 18 | 0 | 165 | 0 | 0 | 0 | 0 | 0 | 0 | 0 | 0 | 0 | 0 | 21 | 6779 | 16 | 0 | 0 | 0 | 170 | 1141 | 0 | 0 | 0 | 33 | 21 | 0 | 11 | 0 | 0 | 25 | 0 | 39 | 33 | 0 | 0 | 0 |
| SRR6020461 | 22 | 12 | 371 | 0 | 0 | 0 | 0 | 0 | 0 | 0 | 0 | 0 | 0 | 19 | 2348 | 0 | 0 | 23 | 0 | 1083 | 959 | 0 | 0 | 0 | 14 | 14 | 26 | 26 | 0 | 0 | 18 | 0 | 41 | 14 | 21 | 0 | 14 |
| SRR6020462 | 30 | 0 | 196 | 0 | 0 | 0 | 0 | 0 | 0 | 0 | 0 | 0 | 0 | 14 | 1469 | 12 | 0 | 0 | 0 | 362 | 2466 | 0 | 0 | 0 | 150 | 64 | 24 | 17 | 10 | 0 | 31 | 0 | 61 | 34 | 0 | 0 | 0 |
| SRR6020463 | 17 | 0 | 147 | 0 | 0 | 0 | 16 | 0 | 0 | 0 | 0 | 0 | 0 | 14 | 1551 | 18 | 0 | 0 | 0 | 410 | 3230 | 0 | 0 | 0 | 15 | 26 | 25 | 10 | 22 | 0 | 30 | 0 | 11 | 40 | 24 | 0 | 11 |
| SRR6020464 | 15 | 0 | 147 | 0 | 0 | 0 | 0 | 0 | 0 | 0 | 0 | 0 | 0 | 31 | 6670 | 27 | 0 | 0 | 0 | 67 | 1239 | 0 | 0 | 0 | 37 | 12 | 15 | 14 | 0 | 0 | 13 | 0 | 21 | 51 | 26 | 0 | 12 |
| SRR6020465 | 19 | 12 | 216 | 0 | 0 | 0 | 0 | 0 | 0 | 0 | 0 | 0 | 0 | 33 | 1834 | 20 | 0 | 33 | 0 | 123 | 778 | 0 | 0 | 0 | 184 | 18 | 11 | 16 | 0 | 0 | 26 | 0 | 176 | 28 | 12 | 0 | 0 |
| SRR6020466 | 23 | 10 | 207 | 0 | 0 | 0 | 0 | 0 | 0 | 0 | 0 | 0 | 0 | 19 | 1378 | 18 | 0 | 0 | 0 | 995 | 3835 | 0 | 0 | 0 | 113 | 49 | 26 | 16 | 11 | 0 | 26 | 0 | 80 | 59 | 11 | 0 | 0 |
| SRR6020467 | 13 | 0 | 264 | 0 | 0 | 0 | 0 | 0 | 0 | 0 | 0 | 0 | 0 | 19 | 4808 | 33 | 0 | 22 | 0 | 96 | 1557 | 0 | 0 | 0 | 413 | 50 | 15 | 14 | 0 | 0 | 23 | 0 | 218 | 39 | 0 | 0 | 0 |
| SRR6020468 | 18 | 11 | 214 | 0 | 0 | 0 | 0 | 0 | 0 | 0 | 0 | 0 | 0 | 19 | 1150 | 11 | 0 | 0 | 0 | 378 | 1759 | 0 | 0 | 0 | 287 | 51 | 15 | 0 | 0 | 0 | 34 | 0 | 182 | 30 | 0 | 0 | 0 |
| SRR6020469 | 21 | 0 | 203 | 0 | 0 | 0 | 0 | 0 | 0 | 0 | 0 | 0 | 0 | 19 | 1003 | 0 | 0 | 12 | 0 | 322 | 3162 | 0 | 0 | 0 | 36 | 12 | 35 | 0 | 10 | 0 | 23 | 0 | 27 | 12 | 11 | 0 | 0 |
| SRR6020470 | 14 | 0 | 191 | 0 | 0 | 0 | 0 | 0 | 0 | 0 | 0 | 0 | 0 | 25 | 6312 | 0 | 0 | 0 | 0 | 162 | 650 | 0 | 0 | 0 | 40 | 46 | 0 | 19 | 0 | 0 | 31 | 0 | 21 | 12 | 0 | 0 | 0 |
| SRR6020471 | 24 | 12 | 177 | 0 | 0 | 0 | 36 | 0 | 0 | 0 | 0 | 0 | 0 | 33 | 2489 | 0 | 0 | 31 | 0 | 711 | 1432 | 0 | 0 | 0 | 26 | 36 | 15 | 0 | 0 | 0 | 15 | 0 | 26 | 39 | 0 | 0 | 0 |
| SRR6020472 | 15 | 13 | 203 | 0 | 0 | 0 | 0 | 0 | 0 | 0 | 0 | 0 | 0 | 37 | 7154 | 18 | 0 | 0 | 0 | 41 | 453 | 0 | 0 | 0 | 26 | 32 | 0 | 17 | 0 | 0 | 10 | 0 | 36 | 24 | 0 | 0 | 0 |
| SRR6020473 | 20 | 10 | 237 | 0 | 0 | 0 | 0 | 0 | 0 | 0 | 0 | 0 | 0 | 27 | 1660 | 0 | 0 | 48 | 0 | 70 | 943 | 0 | 0 | 0 | 336 | 62 | 0 | 12 | 0 | 0 | 30 | 0 | 104 | 36 | 24 | 0 | 0 |
| SRR6020474 | 22 | 0 | 166 | 0 | 0 | 0 | 0 | 0 | 0 | 0 | 0 | 0 | 0 | 18 | 3182 | 0 | 0 | 0 | 0 | 184 | 2226 | 0 | 0 | 0 | 264 | 68 | 16 | 0 | 0 | 0 | 17 | 0 | 113 | 44 | 10 | 0 | 0 |
| SRR6020475 | 24 | 10 | 156 | 0 | 0 | 0 | 0 | 0 | 0 | 0 | 0 | 0 | 0 | 13 | 2578 | 14 | 0 | 18 | 0 | 211 | 1037 | 0 | 0 | 0 | 26 | 33 | 0 | 14 | 0 | 0 | 29 | 0 | 20 | 25 | 10 | 0 | 0 |
| SRR6020476 | 17 | 0 | 112 | 0 | 0 | 0 | 0 | 0 | 0 | 0 | 0 | 0 | 0 | 10 | 1343 | 12 | 0 | 0 | 0 | 1435 | 4919 | 0 | 0 | 0 | 19 | 13 | 42 | 0 | 0 | 0 | 17 | 0 | 45 | 45 | 10 | 0 | 0 |
| SRR6020477 | 28 | 0 | 168 | 0 | 0 | 0 | 0 | 0 | 0 | 0 | 0 | 0 | 0 | 32 | 7609 | 14 | 0 | 0 | 0 | 217 | 1767 | 0 | 0 | 0 | 39 | 13 | 14 | 10 | 0 | 0 | 35 | 0 | 24 | 29 | 12 | 0 | 0 |
| SRR6020478 | 18 | 12 | 240 | 0 | 0 | 0 | 0 | 0 | 0 | 0 | 0 | 0 | 0 | 18 | 5901 | 25 | 0 | 0 | 0 | 94 | 1403 | 0 | 0 | 0 | 186 | 23 | 12 | 15 | 0 | 0 | 32 | 0 | 176 | 23 | 0 | 0 | 0 |
| SRR6020479 | 17 | 0 | 211 | 0 | 0 | 0 | 0 | 0 | 0 | 0 | 0 | 0 | 0 | 15 | 2049 | 20 | 0 | 423 | 0 | 26 | 1642 | 0 | 0 | 0 | 63 | 16 | 12 | 10 | 0 | 0 | 42 | 0 | 47 | 41 | 18 | 0 | 0 |
| SRR6020480 | 35 | 0 | 239 | 0 | 0 | 0 | 0 | 0 | 0 | 0 | 0 | 0 | 0 | 14 | 458 | 28 | 0 | 215 | 0 | 354 | 2058 | 0 | 0 | 0 | 26 | 11 | 28 | 0 | 11 | 0 | 21 | 0 | 32 | 16 | 13 | 0 | 0 |
| SRR6020481 | 19 | 0 | 242 | 0 | 0 | 0 | 0 | 0 | 0 | 0 | 0 | 0 | 0 | 28 | 2763 | 12 | 0 | 20 | 0 | 233 | 830 | 0 | 0 | 0 | 33 | 25 | 0 | 13 | 0 | 0 | 21 | 0 | 29 | 28 | 16 | 0 | 0 |
| SRR6020482 | 23 | 0 | 214 | 0 | 0 | 0 | 24 | 0 | 0 | 0 | 0 | 0 | 0 | 18 | 1353 | 11 | 0 | 0 | 0 | 628 | 3257 | 0 | 0 | 0 | 173 | 144 | 51 | 0 | 10 | 0 | 31 | 0 | 57 | 51 | 11 | 0 | 0 |
| SRR6020483 | 22 | 0 | 222 | 0 | 0 | 0 | 0 | 0 | 0 | 0 | 0 | 0 | 0 | 30 | 7478 | 14 | 0 | 0 | 0 | 64 | 925 | 0 | 0 | 0 | 220 | 20 | 0 | 20 | 0 | 0 | 20 | 0 | 148 | 21 | 0 | 0 | 0 |
| SRR6020484 | 30 | 11 | 242 | 0 | 0 | 0 | 0 | 0 | 0 | 0 | 0 | 0 | 0 | 18 | 2558 | 27 | 0 | 18 | 0 | 238 | 1700 | 0 | 0 | 0 | 224 | 71 | 17 | 0 | 0 | 0 | 40 | 0 | 139 | 63 | 0 | 0 | 0 |
| SRR6020485 | 18 | 0 | 227 | 0 | 0 | 0 | 0 | 0 | 0 | 0 | 0 | 0 | 0 | 30 | 5078 | 16 | 0 | 0 | 0 | 332 | 2391 | 0 | 0 | 0 | 164 | 76 | 24 | 19 | 0 | 0 | 27 | 0 | 104 | 40 | 15 | 0 | 0 |
| SRR6020486 | 27 | 0 | 194 | 0 | 0 | 0 | 0 | 0 | 0 | 0 | 0 | 0 | 0 | 13 | 5891 | 12 | 0 | 65 | 0 | 122 | 793 | 0 | 0 | 0 | 152 | 22 | 0 | 0 | 0 | 0 | 25 | 0 | 232 | 28 | 0 | 0 | 0 |
| SRR6020487 | 23 | 20 | 226 | 0 | 0 | 0 | 0 | 0 | 0 | 0 | 0 | 0 | 0 | 17 | 7235 | 0 | 0 | 0 | 0 | 205 | 1083 | 0 | 0 | 0 | 26 | 38 | 0 | 19 | 0 | 0 | 17 | 0 | 24 | 20 | 11 | 0 | 0 |
| SRR6020488 | 16 | 12 | 206 | 0 | 0 | 0 | 13 | 0 | 0 | 0 | 0 | 0 | 0 | 21 | 2670 | 13 | 0 | 0 | 0 | 271 | 2058 | 0 | 0 | 0 | 227 | 45 | 20 | 16 | 0 | 0 | 27 | 0 | 104 | 44 | 0 | 0 | 0 |
| SRR6020489 | 0 | 16 | 165 | 0 | 0 | 0 | 0 | 0 | 0 | 0 | 0 | 0 | 0 | 16 | 4853 | 18 | 0 | 30 | 0 | 78 | 203 | 0 | 0 | 0 | 117 | 0 | 0 | 12 | 0 | 0 | 18 | 0 | 391 | 20 | 0 | 0 | 0 |
| SRR6020490 | 22 | 0 | 142 | 0 | 0 | 0 | 0 | 0 | 0 | 0 | 0 | 0 | 0 | 22 | 6060 | 20 | 0 | 15 | 0 | 123 | 1085 | 0 | 0 | 0 | 281 | 33 | 0 | 0 | 0 | 0 | 21 | 0 | 270 | 26 | 12 | 0 | 0 |
| SRR6020491 | 22 | 0 | 221 | 0 | 0 | 0 | 0 | 0 | 0 | 0 | 0 | 0 | 0 | 19 | 5418 | 10 | 0 | 0 | 0 | 134 | 933 | 0 | 0 | 0 | 333 | 15 | 0 | 12 | 0 | 0 | 19 | 0 | 230 | 19 | 0 | 0 | 0 |
| SRR6020492 | 30 | 0 | 206 | 0 | 0 | 15 | 20 | 0 | 0 | 0 | 0 | 0 | 0 | 19 | 1028 | 0 | 0 | 0 | 0 | 1377 | 8697 | 0 | 0 | 0 | 26 | 34 | 46 | 13 | 16 | 0 | 33 | 0 | 12 | 17 | 15 | 0 | 14 |
| SRR6020493 | 13 | 0 | 190 | 0 | 0 | 0 | 12 | 0 | 0 | 0 | 0 | 0 | 0 | 21 | 7202 | 24 | 0 | 0 | 0 | 72 | 1166 | 0 | 0 | 0 | 163 | 23 | 12 | 17 | 0 | 0 | 34 | 0 | 202 | 34 | 0 | 0 | 0 |
| SRR6020494 | 17 | 14 | 212 | 0 | 0 | 0 | 18 | 0 | 0 | 0 | 0 | 0 | 0 | 27 | 2010 | 13 | 0 | 11 | 0 | 188 | 305 | 0 | 0 | 0 | 40 | 25 | 0 | 12 | 0 | 0 | 27 | 0 | 57 | 22 | 0 | 0 | 0 |
| SRR6020495 | 29 | 0 | 37 | 0 | 0 | 0 | 0 | 0 | 0 | 0 | 0 | 0 | 0 | 15 | 6992 | 0 | 0 | 17 | 0 | 158 | 1838 | 0 | 0 | 0 | 27 | 10 | 0 | 0 | 0 | 0 | 11 | 0 | 32 | 18 | 0 | 0 | 0 |
| SRR6020496 | 24 | 0 | 179 | 0 | 0 | 0 | 0 | 0 | 0 | 0 | 0 | 0 | 0 | 15 | 8217 | 13 | 0 | 28 | 0 | 91 | 1126 | 0 | 0 | 0 | 33 | 16 | 0 | 0 | 0 | 0 | 21 | 0 | 48 | 20 | 0 | 0 | 0 |
| SRR6020497 | 18 | 0 | 332 | 0 | 0 | 0 | 0 | 0 | 0 | 0 | 0 | 0 | 0 | 29 | 1965 | 15 | 0 | 2371 | 0 | 46 | 213 | 0 | 0 | 0 | 33 | 0 | 0 | 0 | 0 | 0 | 37 | 0 | 51 | 23 | 0 | 0 | 0 |
| SRR6020498 | 39 | 0 | 162 | 0 | 0 | 0 | 29 | 0 | 0 | 0 | 0 | 0 | 0 | 0 | 337 | 0 | 0 | 0 | 0 | 2684 | 3656 | 0 | 0 | 0 | 59 | 200 | 70 | 12 | 0 | 0 | 12 | 0 | 74 | 63 | 40 | 0 | 34 |
| SRR6020499 | 30 | 10 | 288 | 0 | 0 | 0 | 0 | 0 | 0 | 0 | 0 | 0 | 0 | 23 | 630 | 15 | 0 | 0 | 0 | 246 | 1525 | 0 | 0 | 0 | 33 | 33 | 12 | 21 | 0 | 0 | 20 | 0 | 30 | 35 | 0 | 0 | 0 |
| SRR6020500 | 42 | 0 | 133 | 0 | 0 | 0 | 0 | 0 | 0 | 0 | 0 | 0 | 0 | 0 | 839 | 0 | 0 | 0 | 0 | 581 | 5105 | 0 | 0 | 0 | 15 | 37 | 30 | 13 | 20 | 0 | 0 | 0 | 12 | 21 | 17 | 0 | 0 |
| SRR6020501 | 23 | 0 | 157 | 0 | 0 | 0 | 0 | 0 | 0 | 0 | 0 | 0 | 0 | 22 | 3041 | 0 | 0 | 335 | 0 | 68 | 559 | 0 | 0 | 0 | 19 | 22 | 0 | 15 | 0 | 0 | 20 | 0 | 16 | 33 | 18 | 0 | 0 |
| SRR6020502 | 38 | 0 | 226 | 0 | 0 | 0 | 0 | 0 | 0 | 0 | 0 | 0 | 0 | 25 | 1214 | 0 | 0 | 44 | 0 | 1372 | 1521 | 0 | 0 | 0 | 12 | 39 | 19 | 15 | 0 | 0 | 11 | 0 | 14 | 11 | 0 | 0 | 0 |
| SRR6020503 | 14 | 0 | 181 | 0 | 0 | 0 | 0 | 0 | 0 | 0 | 0 | 0 | 0 | 25 | 5443 | 22 | 0 | 17 | 0 | 171 | 1914 | 0 | 0 | 0 | 248 | 37 | 20 | 15 | 0 | 0 | 20 | 0 | 171 | 34 | 10 | 0 | 0 |
| SRR6020504 | 23 | 14 | 263 | 0 | 0 | 0 | 0 | 0 | 0 | 0 | 0 | 0 | 0 | 28 | 2355 | 19 | 0 | 24 | 0 | 111 | 1214 | 0 | 0 | 0 | 45 | 34 | 13 | 16 | 0 | 0 | 24 | 0 | 32 | 32 | 23 | 0 | 0 |
| SRR6020505 | 24 | 10 | 189 | 0 | 0 | 0 | 0 | 0 | 0 | 0 | 0 | 0 | 0 | 22 | 4113 | 10 | 0 | 0 | 0 | 143 | 1580 | 0 | 0 | 0 | 217 | 66 | 20 | 16 | 0 | 0 | 26 | 0 | 151 | 38 | 15 | 0 | 0 |
| SRR6020506 | 24 | 0 | 190 | 0 | 0 | 0 | 0 | 0 | 0 | 0 | 0 | 0 | 0 | 23 | 5075 | 40 | 0 | 116 | 0 | 69 | 1517 | 0 | 0 | 0 | 38 | 31 | 11 | 14 | 0 | 0 | 30 | 0 | 27 | 30 | 0 | 0 | 0 |
| SRR6020507 | 20 | 16 | 201 | 0 | 0 | 0 | 0 | 0 | 0 | 0 | 0 | 0 | 0 | 16 | 2521 | 0 | 0 | 95 | 0 | 134 | 500 | 0 | 0 | 0 | 163 | 38 | 0 | 0 | 0 | 0 | 18 | 0 | 105 | 0 | 0 | 0 | 0 |
| SRR6020508 | 25 | 0 | 297 | 0 | 0 | 0 | 44 | 0 | 0 | 0 | 0 | 0 | 0 | 20 | 654 | 18 | 0 | 0 | 0 | 1050 | 2352 | 0 | 0 | 0 | 18 | 15 | 39 | 10 | 13 | 0 | 23 | 0 | 15 | 44 | 13 | 0 | 0 |
| SRR6020509 | 17 | 0 | 138 | 0 | 0 | 0 | 0 | 0 | 0 | 0 | 0 | 0 | 0 | 21 | 5263 | 14 | 0 | 29 | 0 | 154 | 987 | 0 | 0 | 0 | 125 | 26 | 0 | 0 | 0 | 0 | 15 | 0 | 143 | 16 | 11 | 0 | 0 |
| SRR6020510 | 18 | 14 | 178 | 0 | 0 | 0 | 0 | 0 | 0 | 0 | 0 | 0 | 0 | 19 | 3541 | 14 | 0 | 10 | 0 | 58 | 504 | 0 | 0 | 0 | 255 | 23 | 0 | 19 | 0 | 0 | 37 | 0 | 204 | 23 | 0 | 0 | 0 |
| SRR6020511 | 22 | 11 | 214 | 0 | 0 | 0 | 0 | 0 | 0 | 0 | 0 | 0 | 0 | 33 | 2614 | 24 | 0 | 0 | 0 | 168 | 2012 | 0 | 0 | 0 | 43 | 24 | 0 | 12 | 0 | 0 | 18 | 0 | 77 | 13 | 0 | 0 | 0 |
| SRR6020512 | 14 | 0 | 185 | 0 | 0 | 0 | 0 | 0 | 0 | 0 | 0 | 0 | 0 | 30 | 4901 | 17 | 0 | 0 | 0 | 119 | 843 | 0 | 0 | 0 | 260 | 22 | 0 | 11 | 0 | 0 | 22 | 0 | 215 | 18 | 0 | 0 | 0 |
| SRR6020513 | 21 | 11 | 208 | 0 | 0 | 0 | 0 | 0 | 0 | 0 | 0 | 0 | 0 | 22 | 5094 | 16 | 0 | 17 | 0 | 73 | 1468 | 0 | 0 | 0 | 30 | 30 | 0 | 13 | 0 | 0 | 27 | 0 | 26 | 14 | 0 | 0 | 0 |
| SRR6020514 | 36 | 0 | 155 | 0 | 0 | 0 | 0 | 0 | 0 | 0 | 0 | 0 | 0 | 19 | 4648 | 0 | 0 | 0 | 0 | 236 | 680 | 0 | 0 | 0 | 27 | 12 | 0 | 0 | 0 | 0 | 11 | 0 | 36 | 14 | 0 | 0 | 0 |
| SRR6020515 | 14 | 15 | 231 | 0 | 0 | 0 | 0 | 0 | 0 | 0 | 0 | 0 | 0 | 37 | 12230 | 22 | 0 | 41 | 0 | 13 | 692 | 0 | 0 | 0 | 253 | 37 | 0 | 15 | 0 | 0 | 30 | 0 | 129 | 22 | 13 | 0 | 0 |
| SRR6020516 | 32 | 0 | 155 | 0 | 0 | 0 | 31 | 0 | 0 | 0 | 0 | 0 | 0 | 24 | 1631 | 0 | 0 | 22 | 0 | 554 | 2431 | 0 | 0 | 0 | 173 | 82 | 31 | 0 | 0 | 0 | 20 | 0 | 69 | 44 | 13 | 0 | 0 |
| SRR6020517 | 28 | 0 | 214 | 0 | 0 | 0 | 0 | 0 | 0 | 0 | 0 | 0 | 0 | 18 | 1787 | 12 | 0 | 0 | 0 | 643 | 4436 | 0 | 0 | 0 | 100 | 86 | 40 | 18 | 0 | 0 | 14 | 0 | 73 | 72 | 11 | 0 | 0 |
| SRR6020518 | 28 | 10 | 142 | 0 | 0 | 0 | 0 | 0 | 0 | 0 | 0 | 0 | 0 | 12 | 3396 | 0 | 0 | 0 | 0 | 295 | 987 | 0 | 0 | 0 | 54 | 49 | 0 | 12 | 0 | 0 | 47 | 0 | 20 | 0 | 0 | 0 | 0 |
| SRR6020519 | 25 | 0 | 190 | 0 | 0 | 0 | 0 | 0 | 0 | 0 | 0 | 0 | 0 | 25 | 1971 | 0 | 0 | 39 | 0 | 170 | 1457 | 0 | 0 | 0 | 23 | 48 | 26 | 17 | 0 | 0 | 27 | 0 | 15 | 44 | 18 | 0 | 0 |
| SRR6020520 | 27 | 0 | 169 | 0 | 0 | 0 | 0 | 0 | 0 | 0 | 0 | 0 | 0 | 44 | 2099 | 22 | 0 | 91 | 0 | 222 | 1544 | 0 | 0 | 0 | 49 | 73 | 32 | 13 | 0 | 0 | 30 | 0 | 16 | 52 | 0 | 0 | 0 |
| SRR6020521 | 25 | 0 | 284 | 0 | 0 | 0 | 0 | 0 | 0 | 0 | 0 | 0 | 0 | 20 | 2812 | 24 | 0 | 0 | 0 | 393 | 2038 | 0 | 0 | 0 | 193 | 44 | 19 | 16 | 0 | 0 | 20 | 0 | 148 | 34 | 0 | 0 | 0 |
| SRR6020522 | 16 | 10 | 194 | 0 | 0 | 0 | 0 | 0 | 0 | 0 | 0 | 0 | 0 | 26 | 2133 | 0 | 0 | 0 | 0 | 135 | 2034 | 0 | 0 | 0 | 53 | 12 | 13 | 20 | 0 | 0 | 15 | 0 | 41 | 23 | 0 | 0 | 0 |
| SRR6020523 | 0 | 0 | 202 | 0 | 0 | 0 | 0 | 0 | 0 | 0 | 0 | 0 | 0 | 19 | 5676 | 28 | 0 | 15 | 0 | 204 | 2866 | 0 | 0 | 0 | 39 | 11 | 15 | 22 | 0 | 0 | 29 | 0 | 23 | 42 | 13 | 0 | 0 |
| SRR6020524 | 17 | 11 | 263 | 0 | 0 | 0 | 0 | 0 | 0 | 0 | 0 | 0 | 0 | 29 | 9870 | 0 | 0 | 0 | 0 | 172 | 664 | 0 | 0 | 0 | 34 | 18 | 0 | 0 | 0 | 0 | 18 | 0 | 38 | 19 | 0 | 0 | 0 |
| SRR6020525 | 23 | 0 | 231 | 0 | 0 | 0 | 0 | 0 | 0 | 0 | 0 | 0 | 0 | 20 | 4799 | 20 | 0 | 11 | 0 | 233 | 743 | 0 | 0 | 0 | 31 | 27 | 0 | 17 | 0 | 0 | 32 | 0 | 71 | 29 | 0 | 0 | 0 |
| SRR6020526 | 22 | 0 | 205 | 0 | 0 | 0 | 0 | 0 | 0 | 0 | 0 | 0 | 0 | 22 | 2380 | 26 | 0 | 11 | 0 | 130 | 1942 | 0 | 0 | 0 | 219 | 47 | 17 | 12 | 11 | 0 | 25 | 0 | 244 | 27 | 0 | 0 | 0 |
| SRR6020527 | 21 | 0 | 283 | 0 | 0 | 0 | 0 | 0 | 0 | 0 | 0 | 0 | 0 | 26 | 2749 | 14 | 0 | 40 | 0 | 617 | 971 | 0 | 0 | 0 | 84 | 65 | 24 | 17 | 0 | 0 | 28 | 0 | 169 | 45 | 14 | 0 | 0 |
| SRR6020528 | 45 | 0 | 99 | 0 | 0 | 0 | 25 | 0 | 0 | 0 | 0 | 0 | 0 | 0 | 345 | 0 | 0 | 20 | 0 | 2855 | 5100 | 0 | 0 | 0 | 12 | 30 | 16 | 0 | 0 | 0 | 11 | 0 | 12 | 10 | 12 | 0 | 0 |
| SRR6020529 | 20 | 13 | 182 | 0 | 0 | 0 | 0 | 0 | 0 | 0 | 0 | 0 | 0 | 26 | 5367 | 17 | 0 | 169 | 0 | 111 | 768 | 0 | 0 | 0 | 29 | 20 | 0 | 25 | 0 | 0 | 10 | 0 | 22 | 15 | 0 | 0 | 0 |
| SRR6020530 | 15 | 19 | 217 | 0 | 0 | 0 | 0 | 0 | 0 | 0 | 0 | 0 | 0 | 36 | 4965 | 11 | 0 | 13 | 0 | 83 | 426 | 0 | 0 | 0 | 39 | 0 | 0 | 22 | 0 | 0 | 10 | 0 | 38 | 0 | 0 | 0 | 0 |
| SRR6020531 | 15 | 0 | 244 | 0 | 0 | 0 | 0 | 0 | 0 | 0 | 0 | 0 | 0 | 25 | 6247 | 0 | 0 | 0 | 0 | 43 | 802 | 0 | 0 | 0 | 58 | 0 | 0 | 11 | 0 | 0 | 22 | 0 | 81 | 16 | 0 | 0 | 0 |
| SRR6020532 | 18 | 0 | 162 | 0 | 0 | 0 | 0 | 0 | 0 | 0 | 0 | 0 | 0 | 24 | 2077 | 16 | 0 | 48 | 0 | 297 | 2328 | 0 | 0 | 0 | 288 | 26 | 0 | 11 | 0 | 0 | 29 | 0 | 180 | 42 | 14 | 0 | 0 |
| SRR6025030 | 22 | 0 | 54 | 0 | 19 | 101 | 205 | 1977 | 18 | 12 | 167 | 15 | 89 | 0 | 180 | 0 | 23 | 0 | 63 | 18 | 91 | 0 | 0 | 355 | 287 | 538 | 654 | 0 | 0 | 781 | 19 | 0 | 0 | 1181 | 44 | 31 | 138 |
| SRR6025031 | 260 | 0 | 230 | 0 | 0 | 0 | 0 | 0 | 0 | 0 | 0 | 950 | 0 | 0 | 0 | 0 | 200 | 24 | 0 | 0 | 0 | 0 | 0 | 0 | 0 | 26 | 0 | 0 | 0 | 0 | 0 | 0 | 0 | 0 | 0 | 13 | 0 |
| SRR6025033 | 356 | 0 | 299 | 0 | 0 | 0 | 0 | 0 | 0 | 0 | 0 | 785 | 0 | 0 | 0 | 0 | 84 | 0 | 0 | 0 | 0 | 0 | 0 | 1002 | 463 | 1221 | 819 | 0 | 0 | 0 | 0 | 0 | 0 | 0 | 11 | 0 | 0 |
| SRR6025035 | 34 | 0 | 505 | 0 | 0 | 12 | 107 | 22 | 0 | 0 | 0 | 0 | 0 | 0 | 18 | 0 | 0 | 0 | 53 | 18 | 83 | 0 | 0 | 0 | 37 | 126 | 653 | 0 | 0 | 0 | 0 | 0 | 0 | 0 | 44 | 0 | 30 |
| SRR6025036 | 52 | 0 | 1808 | 0 | 0 | 0 | 10 | 0 | 0 | 0 | 0 | 289 | 0 | 0 | 437 | 0 | 0 | 0 | 0 | 182 | 380 | 0 | 0 | 0 | 0 | 15 | 147 | 0 | 0 | 0 | 0 | 0 | 0 | 0 | 40 | 0 | 18 |
| SRR6025037 | 47 | 0 | 556 | 0 | 0 | 11 | 116 | 52 | 0 | 0 | 0 | 0 | 0 | 0 | 0 | 0 | 0 | 0 | 55 | 22 | 110 | 0 | 0 | 35 | 13 | 115 | 603 | 0 | 0 | 0 | 0 | 0 | 0 | 0 | 40 | 0 | 28 |
| SRR6025038 | 220 | 0 | 252 | 0 | 0 | 661 | 0 | 0 | 0 | 0 | 0 | 1069 | 0 | 0 | 0 | 0 | 15 | 0 | 0 | 1458 | 0 | 0 | 0 | 1287 | 0 | 1720 | 0 | 0 | 0 | 0 | 0 | 0 | 0 | 0 | 0 | 0 | 0 |
| SRR6025041 | 100 | 0 | 101 | 18 | 15 | 15 | 65 | 490 | 19 | 0 | 40 | 66 | 14 | 15 | 5193 | 0 | 0 | 0 | 0 | 0 | 216 | 0 | 0 | 0 | 12 | 50 | 0 | 0 | 0 | 150 | 109 | 0 | 0 | 282 | 15 | 13 | 59 |
| SRR6025042 | 160 | 0 | 14 | 0 | 0 | 0 | 322 | 0 | 0 | 0 | 0 | 953 | 0 | 0 | 0 | 0 | 27 | 0 | 0 | 0 | 199 | 0 | 0 | 928 | 757 | 1810 | 38 | 0 | 0 | 0 | 0 | 0 | 0 | 0 | 0 | 0 | 10 |
| SRR6025043 | 37 | 0 | 486 | 0 | 11 | 18 | 96 | 0 | 0 | 0 | 0 | 13 | 0 | 0 | 0 | 0 | 0 | 0 | 42 | 10 | 84 | 0 | 0 | 0 | 25 | 22 | 94 | 0 | 0 | 0 | 0 | 0 | 0 | 0 | 71 | 0 | 35 |
| SRR6025044 | 70 | 0 | 5396 | 0 | 0 | 24 | 163 | 31 | 0 | 0 | 0 | 86 | 11 | 0 | 71 | 0 | 0 | 0 | 105 | 79 | 565 | 0 | 0 | 0 | 24 | 358 | 462 | 0 | 0 | 0 | 0 | 0 | 0 | 0 | 64 | 0 | 44 |
| SRR6025047 | 84 | 0 | 74 | 15 | 31 | 32 | 0 | 0 | 31 | 18 | 31 | 173 | 13 | 17 | 1086 | 0 | 15 | 0 | 0 | 0 | 155 | 0 | 0 | 0 | 44 | 18 | 31 | 0 | 0 | 0 | 12 | 0 | 0 | 158 | 41 | 18 | 15 |
| SRR6025048 | 74 | 0 | 33 | 0 | 0 | 15 | 0 | 91 | 23 | 0 | 14 | 327 | 11 | 0 | 754 | 0 | 0 | 0 | 0 | 0 | 348 | 0 | 0 | 199 | 18 | 17 | 17 | 0 | 0 | 169 | 27 | 0 | 0 | 134 | 11 | 15 | 44 |
| SRR6025050 | 15 | 0 | 18 | 0 | 0 | 10 | 91 | 263 | 13 | 0 | 19 | 0 | 50 | 0 | 518 | 0 | 0 | 0 | 38 | 0 | 114 | 0 | 0 | 11 | 52 | 34 | 12 | 0 | 0 | 93 | 26 | 0 | 11 | 123 | 24 | 36 | 21 |
| SRR6025051 | 43 | 0 | 1315 | 0 | 0 | 15 | 0 | 0 | 0 | 0 | 0 | 244 | 0 | 0 | 620 | 0 | 0 | 0 | 0 | 108 | 307 | 0 | 0 | 0 | 18 | 10 | 83 | 0 | 0 | 0 | 0 | 0 | 0 | 0 | 10 | 0 | 31 |

### Supp Table S8.2.Columns_AM_to_BW

| | | | | | | | | | | | | | | | | | | | | | | | | | | | | | | | | | | | | | |
|---|---|---|---|---|---|---|---|---|---|---|---|---|---|---|---|---|---|---|---|---|---|---|---|---|---|---|---|---|---|---|---|---|---|---|---|---|---|
| SRR6025053 | 56 | 0 | 4675 | 0 | 0 | 0 | 258 | 37 | 0 | 0 | 0 | 59 | 13 | 0 | 75 | 0 | 0 | 0 | 258 | 199 | 690 | 0 | 0 | 11 | 0 | 29 | 52 | 0 | 0 | 0 | 0 | 0 | 0 | 0 | 28 | 0 | 15 |
| SRR6025055 | 76 | 0 | 677 | 0 | 11 | 16 | 395 | 477 | 36 | 14 | 105 | 166 | 25 | 0 | 4044 | 0 | 20 | 58 | 294 | 181 | 727 | 0 | 0 | 77 | 31 | 81 | 46 | 0 | 0 | 162 | 44 | 0 | 0 | 67 | 61 | 28 | 12 |
| SRR6025056 | 40 | 0 | 172 | 0 | 0 | 38 | 31 | 21 | 0 | 0 | 32 | 460 | 156 | 0 | 12584 | 0 | 55 | 0 | 16 | 155 | 140 | 0 | 0 | 455 | 258 | 341 | 767 | 0 | 0 | 94 | 26 | 0 | 0 | 872 | 65 | 32 | 106 |
| SRR6025058 | 95 | 0 | 5333 | 0 | 0 | 30 | 257 | 180 | 11 | 0 | 0 | 81 | 0 | 0 | 26 | 0 | 0 | 0 | 286 | 219 | 820 | 0 | 0 | 19 | 0 | 298 | 497 | 0 | 0 | 0 | 0 | 0 | 0 | 0 | 31 | 0 | 24 |
| SRR6025061 | 39 | 0 | 4714 | 0 | 15 | 49 | 258 | 94 | 0 | 0 | 0 | 45 | 30 | 10 | 330 | 0 | 0 | 0 | 322 | 415 | 1684 | 0 | 0 | 43 | 197 | 518 | 488 | 0 | 0 | 13 | 13 | 0 | 0 | 0 | 40 | 0 | 23 |
| SRR6025062 | 46 | 0 | 4007 | 0 | 0 | 17 | 263 | 49 | 12 | 0 | 0 | 52 | 0 | 0 | 156 | 0 | 0 | 0 | 335 | 334 | 1636 | 0 | 0 | 0 | 32 | 293 | 188 | 0 | 0 | 13 | 16 | 0 | 0 | 13 | 61 | 0 | 38 |
| SRR6025065 | 16 | 0 | 367 | 0 | 0 | 31 | 104 | 25 | 0 | 0 | 0 | 0 | 0 | 0 | 23 | 28 | 0 | 0 | 113 | 47 | 236 | 0 | 0 | 30 | 165 | 20 | 34 | 0 | 0 | 0 | 18 | 0 | 0 | 0 | 72 | 0 | 34 |
| SRR6025066 | 242 | 0 | 3645 | 0 | 0 | 190 | 4221 | 15 | 116 | 0 | 0 | 108 | 28 | 0 | 633 | 0 | 62 | 23 | 1342 | 971 | 2068 | 0 | 0 | 1017 | 930 | 2822 | 392 | 0 | 0 | 0 | 52 | 0 | 0 | 0 | 46 | 16 | 35 |
| SRR6025067 | 32 | 0 | 349 | 0 | 0 | 14 | 91 | 17 | 0 | 0 | 0 | 0 | 0 | 0 | 0 | 0 | 0 | 0 | 86 | 31 | 174 | 0 | 0 | 0 | 18 | 13 | 31 | 0 | 0 | 0 | 0 | 0 | 0 | 0 | 78 | 0 | 51 |
| SRR6025068 | 23 | 0 | 1188 | 0 | 0 | 0 | 0 | 0 | 0 | 0 | 0 | 134 | 0 | 0 | 72 | 0 | 0 | 0 | 0 | 262 | 264 | 0 | 0 | 0 | 46 | 77 | 57 | 0 | 0 | 0 | 0 | 0 | 0 | 0 | 63 | 0 | 37 |
| SRR6025069 | 21 | 0 | 1051 | 0 | 0 | 31 | 0 | 0 | 0 | 0 | 0 | 99 | 11 | 0 | 166 | 31 | 13 | 0 | 0 | 360 | 395 | 0 | 0 | 53 | 12 | 20 | 42 | 0 | 0 | 0 | 0 | 0 | 0 | 0 | 73 | 0 | 29 |
| SRR6025070 | 40 | 0 | 11 | 13 | 43 | 57 | 37 | 85 | 37 | 25 | 23 | 0 | 57 | 19 | 82 | 0 | 0 | 0 | 0 | 0 | 48 | 0 | 0 | 19 | 32 | 21 | 30 | 0 | 0 | 22 | 12 | 0 | 0 | 216 | 61 | 12 | 19 |
| SRR6025071 | 12 | 0 | 0 | 0 | 0 | 27 | 45 | 3373 | 34 | 0 | 95 | 0 | 0 | 0 | 188 | 0 | 0 | 0 | 0 | 0 | 76 | 0 | 0 | 449 | 18 | 63 | 48 | 0 | 0 | 2072 | 109 | 0 | 0 | 767 | 16 | 13 | 133 |
| SRR6025072 | 219 | 0 | 3899 | 0 | 0 | 0 | 73 | 23 | 0 | 0 | 0 | 93 | 0 | 14 | 367 | 0 | 0 | 0 | 143 | 116 | 1126 | 0 | 0 | 24 | 13 | 83 | 19 | 0 | 0 | 0 | 0 | 0 | 0 | 0 | 33 | 0 | 23 |
| SRR6025073 | 175 | 0 | 1449 | 0 | 0 | 0 | 0 | 0 | 0 | 0 | 0 | 412 | 0 | 0 | 68 | 0 | 0 | 0 | 0 | 95 | 520 | 0 | 0 | 11 | 0 | 33 | 125 | 0 | 0 | 0 | 0 | 0 | 0 | 0 | 37 | 0 | 21 |
| SRR6025075 | 139 | 0 | 1384 | 0 | 0 | 0 | 0 | 0 | 0 | 0 | 0 | 174 | 0 | 0 | 184 | 0 | 0 | 0 | 0 | 104 | 247 | 0 | 0 | 0 | 15 | 25 | 45 | 0 | 0 | 0 | 0 | 0 | 0 | 0 | 48 | 0 | 40 |
| SRR6025079 | 65 | 0 | 0 | 0 | 0 | 49 | 42 | 7990 | 48 | 0 | 0 | 13 | 12 | 0 | 0 | 0 | 0 | 0 | 0 | 0 | 101 | 0 | 0 | 776 | 15 | 115 | 86 | 0 | 0 | 5189 | 17 | 0 | 0 | 1285 | 28 | 23 | 194 |
| SRR6025080 | 25 | 0 | 121 | 0 | 13 | 50 | 16 | 65 | 21 | 0 | 140 | 271 | 50 | 0 | 1294 | 0 | 35 | 0 | 0 | 57 | 283 | 0 | 0 | 305 | 94 | 240 | 955 | 0 | 0 | 136 | 37 | 0 | 0 | 710 | 41 | 46 | 129 |
| SRR6025082 | 57 | 0 | 367 | 0 | 17 | 76 | 455 | 1957 | 24 | 0 | 62 | 263 | 63 | 20 | 1311 | 0 | 0 | 36 | 280 | 87 | 804 | 0 | 0 | 2267 | 102 | 366 | 328 | 0 | 0 | 1960 | 45 | 0 | 0 | 284 | 24 | 24 | 70 |
| SRR6025083 | 16 | 0 | 34 | 0 | 13 | 115 | 312 | 1730 | 30 | 12 | 146 | 11 | 49 | 0 | 152 | 0 | 13 | 0 | 50 | 12 | 158 | 0 | 0 | 199 | 88 | 461 | 869 | 0 | 0 | 886 | 46 | 0 | 0 | 2422 | 72 | 43 | 187 |
| SRR6025084 | 50 | 0 | 419 | 0 | 13 | 16 | 383 | 240 | 52 | 11 | 91 | 128 | 14 | 16 | 2845 | 0 | 20 | 20 | 111 | 42 | 1056 | 0 | 0 | 85 | 11 | 71 | 60 | 0 | 0 | 91 | 89 | 0 | 0 | 158 | 51 | 42 | 16 |
| SRR6025085 | 19 | 0 | 33 | 0 | 19 | 14 | 250 | 71 | 13 | 19 | 41 | 21 | 19 | 20 | 81 | 0 | 0 | 0 | 78 | 11 | 108 | 0 | 0 | 208 | 20 | 60 | 79 | 0 | 0 | 96 | 34 | 0 | 0 | 116 | 25 | 22 | 14 |
| SRR6025086 | 219 | 0 | 1931 | 0 | 0 | 107 | 44 | 0 | 43 | 0 | 0 | 557 | 0 | 0 | 293 | 14 | 148 | 0 | 0 | 371 | 354 | 0 | 0 | 1074 | 1306 | 1269 | 704 | 0 | 0 | 0 | 45 | 0 | 0 | 0 | 95 | 31 | 64 |
| SRR6025087 | 134 | 0 | 453 | 0 | 0 | 13 | 917 | 0 | 102 | 0 | 0 | 18 | 0 | 0 | 27 | 12 | 49 | 0 | 220 | 50 | 170 | 0 | 0 | 39 | 82 | 136 | 32 | 0 | 0 | 0 | 72 | 0 | 0 | 0 | 48 | 18 | 39 |
| SRR6025088 | 31 | 0 | 116 | 0 | 15 | 44 | 24 | 13 | 11 | 0 | 20 | 451 | 24 | 12 | 700 | 0 | 17 | 0 | 0 | 80 | 287 | 0 | 0 | 299 | 20 | 54 | 96 | 0 | 0 | 44 | 25 | 0 | 0 | 23 | 16 | 14 | 11 |
| SRR6025091 | 157 | 0 | 153 | 0 | 15 | 63 | 52 | 442 | 33 | 17 | 13 | 40 | 14 | 22 | 1532 | 0 | 0 | 25 | 0 | 0 | 159 | 0 | 0 | 52 | 51 | 26 | 13 | 0 | 0 | 25 | 11 | 0 | 0 | 99 | 41 | 14 | 11 |
| SRR6025092 | 161 | 0 | 579 | 0 | 0 | 0 | 43 | 0 | 0 | 0 | 0 | 22 | 0 | 0 | 0 | 0 | 0 | 0 | 23 | 11 | 230 | 0 | 0 | 15 | 0 | 58 | 418 | 0 | 0 | 0 | 0 | 0 | 0 | 0 | 25 | 0 | 25 |
| SRR6025093 | 37 | 0 | 12 | 18 | 29 | 29 | 140 | 331 | 30 | 0 | 12 | 0 | 0 | 15 | 33 | 0 | 0 | 16 | 17 | 0 | 46 | 0 | 0 | 470 | 11 | 22 | 40 | 10 | 0 | 233 | 22 | 0 | 0 | 143 | 11 | 16 | 14 |
| SRR6025094 | 145 | 0 | 538 | 0 | 0 | 0 | 11 | 0 | 0 | 0 | 0 | 26 | 0 | 0 | 22 | 0 | 0 | 0 | 24 | 18 | 159 | 0 | 0 | 0 | 16 | 79 | 30 | 0 | 0 | 0 | 0 | 0 | 0 | 0 | 39 | 0 | 26 |
| SRR6025095 | 20 | 0 | 0 | 23 | 27 | 48 | 94 | 1560 | 65 | 11 | 191 | 0 | 29 | 0 | 317 | 0 | 0 | 0 | 0 | 0 | 86 | 0 | 0 | 26 | 19 | 175 | 48 | 0 | 0 | 2211 | 117 | 0 | 0 | 423 | 67 | 26 | 226 |
| SRR6025096 | 164 | 0 | 158 | 11 | 25 | 81 | 57 | 442 | 45 | 15 | 79 | 52 | 16 | 27 | 844 | 0 | 0 | 15 | 11 | 0 | 204 | 0 | 0 | 54 | 21 | 30 | 30 | 0 | 0 | 30 | 12 | 0 | 0 | 91 | 76 | 16 | 20 |
| SRR6025097 | 103 | 0 | 133 | 25 | 24 | 32 | 150 | 2922 | 28 | 0 | 55 | 103 | 17 | 23 | 602 | 0 | 0 | 61 | 62 | 16 | 275 | 0 | 0 | 407 | 44 | 95 | 104 | 11 | 0 | 1642 | 104 | 0 | 0 | 505 | 54 | 23 | 99 |
| SRR6025099 | 315 | 0 | 757 | 0 | 0 | 0 | 269 | 0 | 0 | 0 | 0 | 912 | 0 | 0 | 455 | 0 | 24 | 0 | 47 | 643 | 0 | 0 | 0 | 79 | 19 | 75 | 0 | 0 | 0 | 0 | 0 | 0 | 0 | 0 | 0 | 10 | 0 |
| SRR6025101 | 339 | 0 | 99 | 0 | 0 | 11 | 28 | 2225 | 13 | 0 | 0 | 174 | 0 | 0 | 116 | 0 | 0 | 0 | 0 | 0 | 359 | 0 | 0 | 304 | 15 | 38 | 16 | 0 | 0 | 850 | 24 | 0 | 0 | 58 | 23 | 25 | 43 |
| SRR6025103 | 53 | 0 | 24 | 15 | 0 | 16 | 0 | 16 | 24 | 0 | 25 | 299 | 0 | 11 | 1692 | 0 | 0 | 0 | 0 | 0 | 322 | 0 | 0 | 15 | 15 | 17 | 18 | 0 | 0 | 12 | 29 | 0 | 0 | 44 | 14 | 15 | 39 |
| SRR6025104 | 19 | 0 | 14 | 62 | 0 | 0 | 0 | 0 | 14 | 0 | 88 | 226 | 15 | 26 | 3714 | 0 | 0 | 0 | 0 | 0 | 139 | 0 | 0 | 0 | 20 | 17 | 0 | 0 | 0 | 0 | 173 | 0 | 0 | 312 | 0 | 0 | 38 |
| SRR6025105 | 179 | 0 | 3722 | 0 | 0 | 27 | 37 | 74 | 0 | 0 | 0 | 39 | 0 | 0 | 414 | 0 | 0 | 0 | 110 | 102 | 979 | 0 | 0 | 31 | 162 | 376 | 388 | 0 | 0 | 13 | 14 | 0 | 0 | 0 | 28 | 0 | 30 |
| SRR6025106 | 71 | 0 | 52 | 21 | 0 | 26 | 17 | 11953 | 25 | 0 | 32 | 78 | 24 | 0 | 2212 | 0 | 0 | 0 | 0 | 0 | 321 | 0 | 0 | 142 | 12 | 98 | 28 | 0 | 0 | 6052 | 126 | 0 | 0 | 593 | 20 | 16 | 122 |
| SRR6025108 | 257 | 0 | 50 | 0 | 0 | 11 | 0 | 13 | 0 | 0 | 0 | 1225 | 0 | 0 | 719 | 0 | 48 | 0 | 0 | 0 | 0 | 0 | 0 | 0 | 0 | 16 | 0 | 0 | 0 | 0 | 12 | 0 | 0 | 0 | 0 | 0 | 0 |
| SRR6025111 | 71 | 0 | 629 | 0 | 0 | 168 | 372 | 10348 | 34 | 25 | 101 | 168 | 508 | 0 | 3130 | 0 | 25 | 25 | 207 | 159 | 622 | 0 | 0 | 735 | 294 | 840 | 701 | 0 | 0 | 3624 | 38 | 13 | 0 | 1302 | 46 | 25 | 139 |
| SRR6025113 | 36 | 0 | 107 | 0 | 0 | 22 | 0 | 80 | 0 | 0 | 51 | 370 | 58 | 0 | 30770 | 0 | 25 | 0 | 0 | 229 | 381 | 0 | 0 | 75 | 29 | 22 | 23 | 0 | 0 | 80 | 0 | 0 | 0 | 59 | 29 | 18 | 11 |
| SRR6025114 | 34 | 0 | 94 | 0 | 0 | 47 | 0 | 54 | 0 | 14 | 47 | 333 | 72 | 0 | 36491 | 0 | 18 | 0 | 63 | 126 | 536 | 0 | 0 | 57 | 58 | 35 | 20 | 0 | 0 | 68 | 11 | 0 | 0 | 58 | 29 | 43 | 72 |
| SRR6025116 | 13 | 0 | 15 | 0 | 0 | 11 | 54 | 336 | 15 | 0 | 30 | 0 | 0 | 0 | 583 | 0 | 0 | 0 | 11 | 0 | 153 | 0 | 0 | 11 | 26 | 34 | 33 | 0 | 0 | 147 | 12 | 0 | 13 | 68 | 0 | 21 | 38 |
| SRR6025119 | 16 | 0 | 0 | 10 | 0 | 20 | 55 | 511 | 32 | 0 | 113 | 0 | 0 | 0 | 168 | 0 | 0 | 0 | 0 | 0 | 90 | 0 | 0 | 76 | 20 | 56 | 46 | 0 | 0 | 434 | 99 | 0 | 0 | 241 | 17 | 14 | 129 |
| SRR6025120 | 185 | 0 | 3784 | 0 | 0 | 0 | 48 | 41 | 0 | 0 | 0 | 70 | 0 | 0 | 355 | 0 | 0 | 0 | 106 | 90 | 873 | 0 | 0 | 0 | 0 | 108 | 95 | 0 | 0 | 0 | 0 | 0 | 0 | 0 | 36 | 0 | 41 |
| SRR6025121 | 105 | 0 | 1274 | 0 | 0 | 19 | 0 | 0 | 0 | 0 | 0 | 120 | 0 | 0 | 319 | 60 | 0 | 0 | 0 | 71 | 411 | 0 | 0 | 15 | 100 | 109 | 405 | 0 | 0 | 0 | 0 | 0 | 0 | 0 | 47 | 0 | 19 |
| SRR6025123 | 119 | 0 | 72 | 10 | 19 | 23 | 0 | 34 | 25 | 15 | 20 | 178 | 11 | 12 | 608 | 0 | 13 | 0 | 0 | 0 | 243 | 0 | 0 | 42 | 28 | 16 | 30 | 0 | 0 | 18 | 41 | 0 | 0 | 143 | 26 | 11 | 18 |
| SRR6025125 | 100 | 0 | 56 | 0 | 0 | 38 | 49 | 20364 | 26 | 0 | 27 | 89 | 19 | 12 | 2169 | 0 | 0 | 0 | 0 | 0 | 343 | 0 | 0 | 590 | 16 | 90 | 41 | 0 | 0 | 8245 | 103 | 0 | 0 | 607 | 14 | 17 | 99 |
| SRR6025126 | 60 | 0 | 32 | 12 | 0 | 16 | 0 | 158 | 17 | 0 | 73 | 277 | 0 | 0 | 1425 | 0 | 0 | 0 | 0 | 0 | 416 | 0 | 0 | 515 | 15 | 43 | 49 | 0 | 0 | 312 | 74 | 0 | 0 | 362 | 0 | 13 | 97 |
| SRR6025128 | 56 | 0 | 0 | 0 | 12 | 36 | 44 | 637 | 36 | 12 | 14 | 0 | 40 | 11 | 56 | 0 | 0 | 0 | 0 | 0 | 51 | 0 | 0 | 43 | 27 | 22 | 40 | 0 | 0 | 154 | 12 | 0 | 0 | 264 | 53 | 0 | 16 |
| SRR6040044 | 58 | 0 | 206 | 0 | 12 | 92 | 31 | 89 | 73 | 0 | 27 | 1531 | 24 | 0 | 67 | 0 | 1048 | 46 | 0 | 17 | 944 | 0 | 0 | 15 | 111 | 43 | 92 | 0 | 0 | 51 | 95 | 0 | 0 | 194 | 273 | 401 | 680 |
| SRR6040045 | 56 | 0 | 198 | 0 | 0 | 108 | 27 | 119 | 91 | 0 | 28 | 1622 | 304 | 0 | 87 | 0 | 893 | 58 | 0 | 0 | 727 | 0 | 0 | 16 | 162 | 581 | 72 | 0 | 0 | 37 | 118 | 0 | 0 | 3094 | 3958 | 6945 | 10543 |
| SRR6076852 | 109 | 55 | 0 | 0 | 39 | 0 | 0 | 0 | 47 | 31 | 219 | 54 | 0 | 47 | 0 | 0 | 32 | 203 | 0 | 0 | 39 | 16 | 0 | 0 | 0 | 23 | 0 | 16 | 39 | 0 | 23 | 16 | 16 | 141 | 0 | 31 | 39 |
| SRR6076853 | 72 | 76 | 0 | 21 | 35 | 0 | 23 | 28 | 28 | 0 | 159 | 53 | 0 | 21 | 161 | 0 | 17 | 0 | 0 | 0 | 138 | 0 | 0 | 0 | 21 | 21 | 0 | 0 | 21 | 90 | 28 | 0 | 21 | 672 | 21 | 0 | 55 |
| SRR6076854 | 69 | 245 | 0 | 0 | 21 | 11 | 0 | 43 | 11 | 0 | 160 | 20 | 11 | 32 | 54 | 0 | 0 | 0 | 72 | 36 | 127 | 0 | 0 | 11 | 0 | 64 | 11 | 0 | 128 | 53 | 11 | 11 | 11 | 479 | 43 | 64 | 96 |
| SRR6076855 | 100 | 30 | 0 | 22 | 30 | 0 | 0 | 0 | 52 | 0 | 518 | 103 | 15 | 0 | 565 | 0 | 17 | 0 | 0 | 0 | 85 | 0 | 0 | 0 | 0 | 30 | 0 | 15 | 37 | 0 | 44 | 0 | 0 | 214 | 0 | 22 | 22 |
| SRR6076856 | 120 | 134 | 0 | 17 | 22 | 11 | 22 | 0 | 50 | 0 | 73 | 110 | 0 | 22 | 223 | 0 | 16 | 0 | 22 | 22 | 111 | 0 | 0 | 0 | 0 | 67 | 0 | 34 | 39 | 28 | 285 | 22 | 11 | 285 | 17 | 0 | 22 |
| SRR6076858 | 82 | 61 | 0 | 35 | 22 | 0 | 0 | 0 | 22 | 13 | 236 | 47 | 0 | 17 | 1027 | 0 | 15 | 0 | 76 | 0 | 38 | 0 | 0 | 0 | 13 | 13 | 0 | 0 | 17 | 0 | 31 | 0 | 44 | 249 | 17 | 0 | 0 |
| SRR6076861 | 169 | 59 | 0 | 11 | 16 | 16 | 38 | 54 | 103 | 0 | 135 | 127 | 0 | 16 | 48 | 0 | 29 | 0 | 0 | 0 | 140 | 0 | 0 | 0 | 0 | 97 | 16 | 0 | 38 | 54 | 65 | 22 | 0 | 611 | 0 | 38 | 43 |
| SRR6153965 | 189 | 0 | 1058 | 0 | 0 | 0 | 444 | 0 | 0 | 0 | 0 | 255 | 0 | 0 | 30 | 0 | 20 | 18 | 0 | 537 | 294 | 0 | 0 | 0 | 0 | 135 | 68 | 0 | 0 | 0 | 0 | 0 | 0 | 0 | 0 | 0 | 0 |
| SRR6153968 | 819 | 14 | 118 | 0 | 0 | 19 | 794 | 17 | 0 | 0 | 0 | 82 | 0 | 0 | 0 | 0 | 0 | 0 | 21 | 4813 | 232 | 0 | 0 | 14 | 12 | 209 | 95 | 0 | 0 | 0 | 12 | 0 | 0 | 0 | 0 | 0 | 0 |
| SRR6153976 | 371 | 0 | 49 | 0 | 0 | 0 | 4061 | 43 | 0 | 0 | 0 | 323 | 0 | 0 | 0 | 0 | 0 | 0 | 0 | 80 | 530 | 0 | 0 | 0 | 0 | 54 | 35 | 0 | 0 | 0 | 0 | 0 | 0 | 0 | 0 | 0 | 0 |
| SRR6153984 | 981 | 0 | 0 | 0 | 0 | 12 | 1017 | 0 | 0 | 0 | 0 | 234 | 0 | 0 | 0 | 0 | 0 | 0 | 19 | 3738 | 958 | 0 | 0 | 0 | 18 | 198 | 76 | 0 | 0 | 0 | 0 | 0 | 0 | 16 | 0 | 0 | 0 |
| SRR6153998 | 593 | 15 | 178 | 0 | 0 | 0 | 512 | 21 | 0 | 0 | 0 | 390 | 0 | 11 | 0 | 0 | 0 | 0 | 0 | 984 | 377 | 0 | 0 | 0 | 0 | 131 | 111 | 0 | 0 | 0 | 10 | 0 | 0 | 17 | 0 | 0 | 0 |
| SRR6154002 | 220 | 0 | 419 | 0 | 0 | 44 | 580 | 0 | 0 | 0 | 0 | 90 | 0 | 0 | 0 | 0 | 41 | 24 | 17 | 324 | 1629 | 0 | 0 | 0 | 0 | 177 | 17 | 0 | 0 | 0 | 12 | 0 | 0 | 0 | 0 | 0 | 0 |
| SRR6154003 | 41 | 14 | 87 | 0 | 11 | 11 | 468 | 3452 | 19 | 12 | 0 | 95 | 0 | 0 | 70 | 0 | 20 | 0 | 27 | 99 | 73 | 0 | 0 | 0 | 12 | 86 | 368 | 0 | 24 | 526 | 0 | 0 | 0 | 121 | 0 | 0 | 0 |
| SRR6154004 | 1118 | 14 | 235 | 0 | 0 | 15 | 1904 | 0 | 0 | 0 | 0 | 477 | 0 | 17 | 0 | 0 | 26 | 0 | 0 | 1307 | 459 | 0 | 0 | 11 | 34 | 149 | 182 | 0 | 0 | 0 | 0 | 0 | 0 | 0 | 0 | 0 | 0 |
| SRR6154005 | 899 | 0 | 167 | 0 | 0 | 12 | 11766 | 20 | 0 | 0 | 0 | 580 | 0 | 0 | 12 | 0 | 23 | 13 | 17 | 602 | 710 | 0 | 0 | 0 | 0 | 156 | 170 | 0 | 0 | 0 | 0 | 0 | 0 | 14 | 0 | 0 | 0 |
| SRR6154006 | 238 | 18 | 64 | 0 | 0 | 0 | 685 | 1377 | 14 | 0 | 0 | 105 | 0 | 0 | 18 | 0 | 0 | 0 | 48 | 153 | 158 | 0 | 0 | 0 | 0 | 60 | 159 | 0 | 28 | 217 | 0 | 0 | 0 | 20 | 0 | 0 | 0 |
| SRR6203722 | 13 | 25 | 2636 | 87 | 212 | 0 | 15 | 50 | 0 | 0 | 0 | 18 | 0 | 0 | 786 | 0 | 0 | 0 | 44 | 30 | 59 | 0 | 0 | 12 | 200 | 62 | 162 | 0 | 0 | 0 | 0 | 0 | 0 | 0 | 12 | 0 | 12 |
| SRR6203757 | 49 | 0 | 2429 | 0 | 0 | 0 | 94 | 46 | 0 | 0 | 0 | 18 | 0 | 23 | 376 | 0 | 0 | 0 | 31 | 110 | 0 | 0 | 0 | 0 | 139 | 92 | 231 | 0 | 0 | 23 | 0 | 0 | 0 | 0 | 46 | 23 | 0 |
| SRR6203760 | 0 | 0 | 63 | 0 | 0 | 0 | 0 | 0 | 0 | 0 | 0 | 0 | 0 | 0 | 297 | 0 | 0 | 0 | 0 | 0 | 297 | 0 | 0 | 0 | 0 | 0 | 0 | 0 | 0 | 0 | 0 | 0 | 0 | 0 | 0 | 0 | 0 |
| SRR6203761 | 34 | 0 | 75 | 0 | 0 | 0 | 0 | 0 | 0 | 0 | 0 | 0 | 0 | 0 | 0 | 0 | 0 | 0 | 0 | 0 | 0 | 0 | 0 | 0 | 0 | 0 | 0 | 0 | 0 | 0 | 0 | 0 | 0 | 0 | 0 | 0 | 0 |
| SRR6203772 | 0 | 0 | 2168 | 66 | 0 | 0 | 106 | 47 | 0 | 0 | 0 | 12 | 0 | 0 | 395 | 0 | 12 | 0 | 219 | 148 | 0 | 0 | 0 | 0 | 284 | 104 | 171 | 0 | 0 | 0 | 0 | 0 | 0 | 0 | 28 | 0 | 0 |
| SRR6203807 | 39 | 0 | 1954 | 0 | 0 | 0 | 56 | 0 | 0 | 0 | 0 | 0 | 0 | 0 | 898 | 14 | 0 | 0 | 253 | 84 | 28 | 0 | 0 | 0 | 110 | 41 | 137 | 0 | 0 | 0 | 41 | 0 | 0 | 0 | 0 | 0 | 0 |
| SRR6224558 | 427 | 0 | 0 | 0 | 41 | 0 | 0 | 0 | 0 | 18 | 42 | 2815 | 0 | 18 | 116 | 0 | 0 | 0 | 0 | 0 | 19 | 0 | 0 | 0 | 0 | 0 | 0 | 13 | 0 | 0 | 0 | 0 | 0 | 17 | 0 | 0 | 0 |
| SRR6226650 | 41 | 16 | 13 | 0 | 0 | 0 | 12 | 0 | 13 | 0 | 39 | 54 | 10 | 47 | 1613 | 0 | 0 | 23 | 0 | 12 | 154 | 0 | 0 | 0 | 165 | 152 | 0 | 0 | 0 | 0 | 117 | 0 | 0 | 140 | 236 | 564 | 628 |
| SRR6226651 | 40 | 18 | 22 | 0 | 0 | 19 | 19 | 0 | 0 | 0 | 36 | 103 | 20 | 66 | 433 | 0 | 10 | 0 | 13 | 0 | 153 | 0 | 0 | 0 | 242 | 235 | 0 | 20 | 0 | 0 | 93 | 0 | 0 | 603 | 1003 | 3420 | 2864 |
| SRR6226652 | 82 | 0 | 10 | 0 | 18 | 23 | 20 | 0 | 18 | 16 | 88 | 31 | 16 | 21 | 1434 | 0 | 0 | 0 | 0 | 0 | 438 | 0 | 0 | 0 | 101 | 266 | 0 | 0 | 0 | 0 | 66 | 0 | 10 | 466 | 251 | 518 | 539 |
| SRR6226653 | 54 | 11 | 68 | 0 | 16 | 17 | 20 | 0 | 26 | 45 | 30 | 168 | 13 | 15 | 548 | 21 | 11 | 0 | 0 | 0 | 57 | 0 | 11 | 0 | 19 | 21 | 0 | 23 | 0 | 0 | 71 | 0 | 29 | 21 | 12 | 15 | 22 |
| SRR6226654 | 126 | 0 | 140 | 0 | 20 | 88 | 41 | 0 | 22 | 10 | 47 | 84 | 0 | 52 | 979 | 0 | 36 | 0 | 0 | 0 | 124 | 0 | 0 | 0 | 537 | 871 | 40 | 22 | 0 | 0 | 147 | 0 | 12 | 59 | 288 | 554 | 780 |
| SRR6226655 | 84 | 0 | 112 | 0 | 13 | 69 | 78 | 0 | 22 | 17 | 125 | 138 | 0 | 54 | 791 | 0 | 12 | 0 | 0 | 0 | 110 | 0 | 0 | 0 | 274 | 676 | 0 | 30 | 0 | 0 | 151 | 11 | 0 | 35 | 188 | 507 | 874 |
| SRR6226656 | 104 | 10 | 305 | 0 | 14 | 11 | 0 | 0 | 12 | 20 | 11 | 55 | 0 | 32 | 170 | 0 | 0 | 0 | 0 | 0 | 17 | 0 | 0 | 0 | 28 | 24 | 0 | 23 | 0 | 0 | 51 | 0 | 0 | 43 | 14 | 75 | 100 |
| SRR6226657 | 88 | 0 | 197 | 0 | 0 | 37 | 79 | 0 | 10 | 17 | 92 | 77 | 0 | 22 | 45 | 0 | 11 | 0 | 0 | 0 | 98 | 0 | 0 | 0 | 270 | 469 | 22 | 10 | 0 | 0 | 36 | 16 | 0 | 63 | 89 | 547 | 931 |
| SRR6230411 | 63 | 0 | 0 | 0 | 0 | 122 | 27 | 0 | 83 | 0 | 0 | 67 | 27 | 31 | 61 | 0 | 62 | 26 | 0 | 24 | 2048 | 0 | 0 | 0 | 0 | 1082 | 27 | 0 | 10 | 0 | 2982 | 0 | 0 | 0 | 6633 | 7876 | 21858 |
| SRR6230412 | 144 | 0 | 0 | 0 | 0 | 73 | 38 | 12 | 44 | 0 | 36 | 152 | 35 | 0 | 80 | 27 | 187 | 0 | 73 | 45 | 500 | 0 | 0 | 37 | 25 | 32 | 141 | 0 | 0 | 0 | 73 | 20 | 33 | 84 | 428 | 705 | 2743 |
| SRR6230413 | 126 | 0 | 11 | 0 | 0 | 92 | 45 | 0 | 94 | 0 | 49 | 188 | 43 | 12 | 133 | 12 | 177 | 0 | 17 | 117 | 451 | 0 | 0 | 10 | 65 | 780 | 190 | 10 | 0 | 0 | 2541 | 0 | 22 | 1812 | 6865 | 8671 | 21909 |
| SRR6246305 | 670 | 0 | 1971 | 0 | 0 | 12 | 27 | 0 | 23 | 0 | 0 | 904 | 11 | 0 | 22 | 0 | 25 | 34 | 0 | 18 | 106 | 0 | 0 | 0 | 12 | 315 | 90 | 0 | 0 | 0 | 0 | 0 | 0 | 24 | 43 | 32 | 47 |
| SRR6246306 | 774 | 0 | 1908 | 0 | 0 | 50 | 29 | 0 | 22 | 0 | 0 | 869 | 0 | 0 | 19 | 0 | 33 | 27 | 0 | 21 | 118 | 0 | 0 | 0 | 47 | 1335 | 347 | 0 | 0 | 0 | 12 | 0 | 0 | 18 | 182 | 41 | 180 |
| SRR6246307 | 727 | 0 | 1940 | 0 | 0 | 12 | 25 | 0 | 26 | 0 | 0 | 907 | 0 | 0 | 17 | 0 | 50 | 28 | 0 | 15 | 122 | 0 | 0 | 0 | 11 | 314 | 85 | 0 | 0 | 0 | 0 | 0 | 0 | 16 | 45 | 11 | 45 |
| SRR6246308 | 739 | 0 | 1883 | 0 | 0 | 54 | 21 | 0 | 24 | 0 | 0 | 926 | 10 | 0 | 16 | 0 | 37 | 21 | 0 | 20 | 123 | 0 | 0 | 0 | 47 | 1244 | 324 | 0 | 0 | 0 | 0 | 0 | 0 | 17 | 189 | 41 | 177 |
| SRR6246309 | 213 | 0 | 933 | 0 | 0 | 12 | 93 | 139 | 31 | 0 | 0 | 522 | 0 | 0 | 0 | 0 | 17 | 0 | 17 | 0 | 48 | 0 | 0 | 0 | 19 | 80 | 343 | 24 | 0 | 58 | 0 | 0 | 0 | 34 | 35 | 0 | 32 |
| SRR6246310 | 176 | 0 | 956 | 0 | 0 | 20 | 96 | 149 | 19 | 0 | 0 | 544 | 0 | 14 | 0 | 0 | 24 | 0 | 18 | 0 | 64 | 0 | 0 | 0 | 26 | 78 | 349 | 28 | 14 | 37 | 0 | 0 | 0 | 36 | 42 | 0 | 28 |
| SRR6246311 | 153 | 0 | 977 | 0 | 0 | 20 | 103 | 138 | 22 | 0 | 0 | 507 | 0 | 10 | 0 | 0 | 27 | 0 | 11 | 12 | 51 | 0 | 0 | 0 | 21 | 72 | 353 | 19 | 10 | 53 | 11 | 0 | 0 | 40 | 35 | 0 | 19 |
| SRR6246312 | 169 | 0 | 971 | 0 | 0 | 15 | 117 | 961 | 28 | 0 | 0 | 496 | 0 | 10 | 0 | 0 | 33 | 0 | 18 | 11 | 64 | 0 | 0 | 0 | 23 | 534 | 2338 | 17 | 0 | 49 | 0 | 0 | 0 | 261 | 36 | 0 | 21 |
| SRR6246313 | 257 | 10 | 1481 | 10 | 0 | 10 | 67 | 117 | 0 | 0 | 0 | 721 | 0 | 15 | 0 | 0 | 17 | 0 | 21 | 20 | 20 | 0 | 0 | 0 | 17 | 49 | 129 | 27 | 0 | 67 | 0 | 0 | 0 | 22 | 12 | 0 | 0 |
| SRR6246314 | 233 | 0 | 1598 | 0 | 0 | 0 | 60 | 1116 | 21 | 0 | 0 | 584 | 0 | 0 | 0 | 0 | 17 | 0 | 19 | 22 | 13 | 0 | 0 | 0 | 22 | 384 | 1076 | 31 | 0 | 65 | 0 | 0 | 0 | 181 | 0 | 0 | 0 |
| SRR6246315 | 210 | 0 | 1594 | 0 | 0 | 15 | 80 | 122 | 0 | 0 | 0 | 646 | 0 | 0 | 0 | 0 | 14 | 0 | 20 | 23 | 15 | 0 | 0 | 0 | 25 | 44 | 118 | 22 | 0 | 65 | 0 | 0 | 0 | 19 | 0 | 0 | 0 |
| SRR6246316 | 270 | 0 | 1485 | 0 | 10 | 20 | 77 | 1007 | 15 | 0 | 0 | 730 | 0 | 0 | 0 | 0 | 23 | 0 | 29 | 19 | 0 | 0 | 0 | 0 | 26 | 380 | 963 | 26 | 0 | 63 | 0 | 0 | 0 | 162 | 10 | 0 | 0 |
| SRR6246317 | 318 | 28 | 1454 | 0 | 0 | 0 | 156 | 13 | 0 | 0 | 0 | 828 | 0 | 0 | 39 | 0 | 67 | 0 | 273 | 2649 | 0 | 0 | 0 | 0 | 0 | 28 | 51 | 0 | 0 | 19 | 0 | 0 | 0 | 14 | 0 | 0 | 0 |
| SRR6246318 | 207 | 19 | 1531 | 0 | 0 | 0 | 79 | 15 | 0 | 0 | 0 | 652 | 0 | 0 | 0 | 0 | 0 | 0 | 432 | 2473 | 157 | 0 | 0 | 0 | 0 | 19 | 44 | 19 | 0 | 0 | 0 | 0 | 0 | 0 | 0 | 0 | 0 |
| SRR6246319 | 242 | 28 | 1598 | 0 | 0 | 0 | 77 | 47 | 0 | 0 | 0 | 746 | 0 | 0 | 39 | 0 | 19 | 0 | 270 | 2699 | 0 | 0 | 0 | 0 | 0 | 33 | 145 | 0 | 0 | 0 | 0 | 0 | 0 | 19 | 0 | 0 | 0 |
| SRR6246320 | 373 | 13 | 1523 | 0 | 0 | 0 | 117 | 12 | 0 | 0 | 0 | 776 | 0 | 0 | 39 | 0 | 29 | 0 | 195 | 2693 | 39 | 0 | 0 | 0 | 0 | 24 | 34 | 0 | 0 | 0 | 0 | 0 | 0 | 0 | 0 | 0 | 0 |
| SRR6246321 | 456 | 0 | 2636 | 0 | 0 | 0 | 11 | 98 | 0 | 0 | 0 | 515 | 0 | 0 | 106 | 0 | 0 | 438 | 96 | 659 | 999 | 0 | 0 | 0 | 11 | 88 | 117 | 0 | 0 | 14 | 0 | 0 | 0 | 23 | 59 | 0 | 20 |
| SRR6246322 | 442 | 0 | 2775 | 0 | 0 | 0 | 44 | 102 | 0 | 0 | 0 | 705 | 0 | 0 | 89 | 0 | 0 | 448 | 100 | 488 | 943 | 0 | 0 | 0 | 11 | 67 | 125 | 0 | 0 | 15 | 0 | 0 | 0 | 21 | 55 | 0 | 24 |
| SRR6246323 | 516 | 0 | 2513 | 0 | 0 | 0 | 74 | 177 | 0 | 0 | 0 | 531 | 0 | 0 | 95 | 0 | 0 | 455 | 95 | 686 | 1066 | 0 | 0 | 0 | 0 | 150 | 244 | 0 | 0 | 37 | 0 | 0 | 0 | 40 | 107 | 0 | 41 |
| SRR6246324 | 451 | 0 | 2624 | 0 | 0 | 0 | 22 | 217 | 0 | 0 | 0 | 520 | 0 | 0 | 142 | 0 | 23 | 474 | 55 | 612 | 994 | 0 | 0 | 0 | 0 | 155 | 228 | 0 | 0 | 43 | 0 | 0 | 0 | 35 | 97 | 0 | 50 |
| SRR6246325 | 385 | 0 | 899 | 0 | 0 | 16 | 62 | 1235 | 16 | 0 | 0 | 498 | 0 | 11 | 0 | 0 | 12 | 0 | 0 | 13 | 17 | 0 | 0 | 0 | 17 | 49 | 159 | 12 | 0 | 21 | 14 | 0 | 0 | 41 | 0 | 0 | 11 |
| SRR6246326 | 436 | 0 | 1014 | 0 | 0 | 24 | 58 | 9657 | 22 | 0 | 0 | 509 | 0 | 11 | 0 | 0 | 0 | 0 | 10 | 0 | 25 | 0 | 0 | 0 | 0 | 379 | 1153 | 16 | 0 | 187 | 0 | 0 | 0 | 308 | 16 | 0 | 0 |
| SRR6246327 | 419 | 0 | 830 | 0 | 0 | 12 | 59 | 1203 | 26 | 0 | 0 | 516 | 0 | 0 | 0 | 0 | 0 | 0 | 15 | 12 | 27 | 0 | 0 | 0 | 12 | 54 | 152 | 17 | 0 | 25 | 17 | 0 | 0 | 39 | 0 | 0 | 12 |
| SRR6246328 | 436 | 0 | 1048 | 0 | 0 | 19 | 56 | 1201 | 22 | 0 | 0 | 540 | 0 | 13 | 0 | 0 | 0 | 0 | 14 | 0 | 20 | 0 | 0 | 0 | 0 | 55 | 151 | 17 | 0 | 24 | 13 | 0 | 0 | 37 | 0 | 0 | 19 |
| SRR6246329 | 311 | 0 | 1099 | 0 | 0 | 0 | 58 | 1031 | 24 | 0 | 0 | 377 | 0 | 0 | 0 | 0 | 0 | 0 | 54 | 48 | 15 | 0 | 0 | 0 | 15 | 39 | 104 | 33 | 0 | 26 | 0 | 0 | 0 | 23 | 13 | 0 | 0 |
| SRR6246330 | 353 | 0 | 1175 | 0 | 0 | 11 | 56 | 8017 | 23 | 0 | 0 | 557 | 0 | 16 | 0 | 0 | 0 | 0 | 41 | 44 | 17 | 0 | 0 | 0 | 0 | 308 | 777 | 32 | 0 | 179 | 0 | 0 | 0 | 138 | 0 | 0 | 0 |
| SRR6246331 | 378 | 0 | 1282 | 0 | 0 | 0 | 65 | 7884 | 23 | 0 | 0 | 468 | 0 | 0 | 0 | 0 | 24 | 0 | 49 | 32 | 0 | 0 | 0 | 0 | 13 | 253 | 886 | 11 | 0 | 177 | 0 | 0 | 0 | 177 | 13 | 0 | 0 |
| SRR6246332 | 289 | 0 | 1186 | 11 | 0 | 11 | 40 | 7632 | 16 | 0 | 0 | 583 | 0 | 0 | 0 | 0 | 25 | 0 | 49 | 34 | 0 | 0 | 0 | 0 | 20 | 291 | 853 | 22 | 0 | 149 | 0 | 0 | 0 | 182 | 0 | 0 | 0 |
| SRR6246333 | 516 | 29 | 1296 | 0 | 0 | 0 | 31 | 175 | 0 | 0 | 0 | 604 | 0 | 48 | 0 | 0 | 0 | 0 | 153 | 2166 | 0 | 0 | 0 | 0 | 0 | 14 | 21 | 0 | 0 | 0 | 0 | 0 | 0 | 29 | 0 | 0 | 0 |
| SRR6246334 | 615 | 20 | 1430 | 0 | 0 | 0 | 128 | 205 | 0 | 0 | 0 | 584 | 0 | 20 | 0 | 0 | 15 | 0 | 288 | 2113 | 32 | 0 | 0 | 0 | 0 | 20 | 30 | 20 | 20 | 20 | 0 | 0 | 0 | 0 | 0 | 0 | 0 |
| SRR6246335 | 464 | 0 | 1521 | 0 | 0 | 0 | 0 | 930 | 0 | 0 | 0 | 666 | 0 | 28 | 0 | 0 | 0 | 0 | 150 | 1622 | 0 | 0 | 0 | 0 | 0 | 76 | 152 | 0 | 0 | 28 | 0 | 0 | 0 | 19 | 0 | 0 | 0 |
| SRR6246336 | 498 | 0 | 1328 | 0 | 0 | 0 | 31 | 1059 | 0 | 0 | 0 | 709 | 0 | 0 | 0 | 0 | 15 | 0 | 248 | 1797 | 0 | 0 | 0 | 0 | 0 | 59 | 177 | 20 | 0 | 39 | 0 | 0 | 0 | 0 | 0 | 0 | 0 |
| SRR6246337 | 183 | 0 | 2289 | 0 | 0 | 0 | 67 | 100 | 0 | 0 | 0 | 119 | 0 | 0 | 519 | 0 | 0 | 628 | 0 | 308 | 798 | 0 | 0 | 0 | 16 | 88 | 278 | 0 | 0 | 0 | 0 | 0 | 0 | 0 | 26 | 0 | 12 |
| SRR6246338 | 158 | 0 | 2414 | 0 | 0 | 0 | 147 | 72 | 0 | 0 | 0 | 105 | 0 | 0 | 460 | 0 | 19 | 769 | 29 | 274 | 949 | 0 | 0 | 0 | 11 | 52 | 171 | 0 | 0 | 11 | 0 | 0 | 0 | 0 | 18 | 0 | 0 |
| SRR6246339 | 171 | 0 | 2516 | 0 | 0 | 0 | 167 | 96 | 0 | 0 | 0 | 144 | 0 | 10 | 389 | 0 | 14 | 684 | 0 | 287 | 759 | 0 | 0 | 0 | 20 | 95 | 245 | 0 | 0 | 10 | 0 | 0 | 0 | 11 | 21 | 0 | 0 |
| SRR6246340 | 216 | 0 | 2241 | 0 | 0 | 0 | 96 | 111 | 0 | 0 | 0 | 108 | 0 | 0 | 489 | 0 | 0 | 676 | 38 | 336 | 758 | 0 | 0 | 0 | 18 | 86 | 243 | 0 | 0 | 12 | 0 | 0 | 0 | 11 | 26 | 0 | 12 |
| SRR6246341 | 100 | 0 | 1398 | 0 | 0 | 0 | 193 | 539 | 0 | 0 | 0 | 171 | 0 | 0 | 0 | 0 | 0 | 0 | 15 | 17 | 37 | 0 | 0 | 0 | 0 | 19 | 412 | 11 | 0 | 26 | 0 | 0 | 0 | 14 | 0 | 0 | 0 |

### Supp Table S8.2.Columns_AM_to_BW

| | | | | | | | | | | | | | | | | | | | | | | | | | | | | | | | | | | | | | |
|---|---|---|---|---|---|---|---|---|---|---|---|---|---|---|---|---|---|---|---|---|---|---|---|---|---|---|---|---|---|---|---|---|---|---|---|---|---|
| SRR6246342 | 177 | 0 | 2063 | 0 | 0 | 0 | 175 | 513 | 0 | 0 | 0 | 192 | 0 | 0 | 0 | 0 | 0 | 0 | 13 | 10 | 23 | 0 | 0 | 0 | 0 | 28 | 388 | 18 | 0 | 33 | 0 | 0 | 0 | 14 | 0 | 0 | 0 |
| SRR6246343 | 85 | 0 | 2073 | 0 | 0 | 0 | 199 | 528 | 0 | 0 | 0 | 156 | 0 | 0 | 0 | 0 | 0 | 0 | 15 | 12 | 27 | 0 | 0 | 0 | 0 | 33 | 420 | 0 | 0 | 29 | 0 | 0 | 0 | 21 | 0 | 0 | 0 |
| SRR6246344 | 145 | 0 | 1931 | 0 | 0 | 0 | 173 | 2002 | 0 | 0 | 0 | 174 | 0 | 0 | 0 | 0 | 0 | 0 | 0 | 20 | 18 | 0 | 0 | 0 | 0 | 39 | 1696 | 16 | 0 | 143 | 0 | 0 | 0 | 50 | 0 | 0 | 0 |
| SRR6246345 | 201 | 0 | 2161 | 0 | 0 | 0 | 86 | 1216 | 0 | 22 | 0 | 352 | 0 | 0 | 0 | 0 | 0 | 0 | 57 | 172 | 57 | 0 | 0 | 0 | 0 | 44 | 487 | 22 | 0 | 44 | 0 | 0 | 0 | 22 | 0 | 0 | 0 |
| SRR6246346 | 314 | 0 | 2459 | 23 | 23 | 0 | 153 | 216 | 0 | 0 | 0 | 314 | 0 | 0 | 0 | 0 | 0 | 0 | 61 | 153 | 0 | 0 | 0 | 0 | 0 | 23 | 127 | 0 | 0 | 46 | 0 | 0 | 0 | 23 | 0 | 0 | 0 |
| SRR6246347 | 399 | 0 | 2047 | 44 | 0 | 0 | 202 | 1140 | 0 | 0 | 0 | 150 | 0 | 0 | 29 | 0 | 0 | 0 | 29 | 173 | 29 | 0 | 0 | 0 | 0 | 22 | 285 | 22 | 0 | 66 | 0 | 0 | 0 | 44 | 0 | 0 | 0 |
| SRR6246348 | 306 | 23 | 2192 | 0 | 23 | 0 | 0 | 1558 | 0 | 0 | 0 | 255 | 0 | 23 | 0 | 0 | 0 | 18 | 89 | 30 | 30 | 0 | 0 | 0 | 0 | 69 | 504 | 69 | 0 | 115 | 0 | 0 | 0 | 46 | 0 | 0 | 0 |
| SRR6246349 | 204 | 0 | 1349 | 0 | 0 | 0 | 0 | 249 | 83 | 0 | 0 | 368 | 0 | 0 | 0 | 0 | 0 | 0 | 0 | 1334 | 0 | 41 | 0 | 0 | 0 | 41 | 332 | 0 | 0 | 124 | 0 | 0 | 0 | 41 | 0 | 0 | 0 |
| SRR6246350 | 255 | 43 | 893 | 0 | 0 | 0 | 0 | 107 | 20 | 0 | 0 | 298 | 0 | 0 | 0 | 0 | 0 | 0 | 452 | 679 | 0 | 43 | 0 | 0 | 0 | 20 | 98 | 43 | 0 | 0 | 0 | 0 | 0 | 0 | 0 | 0 | 0 |
| SRR6246351 | 202 | 0 | 1295 | 0 | 0 | 0 | 0 | 530 | 41 | 0 | 0 | 243 | 0 | 0 | 0 | 0 | 0 | 0 | 210 | 1262 | 0 | 0 | 0 | 0 | 0 | 41 | 245 | 0 | 0 | 163 | 0 | 0 | 0 | 0 | 0 | 0 | 0 |
| SRR6246352 | 250 | 0 | 1497 | 0 | 0 | 0 | 0 | 57 | 42 | 0 | 0 | 166 | 0 | 0 | 0 | 0 | 0 | 0 | 0 | 684 | 0 | 0 | 0 | 0 | 0 | 0 | 86 | 0 | 0 | 42 | 0 | 0 | 0 | 0 | 0 | 0 | 0 |
| SRR6246353 | 330 | 0 | 1052 | 0 | 0 | 0 | 177 | 21 | 0 | 0 | 0 | 583 | 18 | 20 | 57 | 0 | 0 | 0 | 0 | 0 | 68 | 0 | 22 | 0 | 41 | 111 | 257 | 0 | 0 | 0 | 0 | 0 | 0 | 21 | 10 | 15 | 12 |
| SRR6246354 | 336 | 0 | 1041 | 0 | 0 | 12 | 164 | 22 | 0 | 0 | 0 | 462 | 38 | 14 | 33 | 0 | 19 | 0 | 0 | 0 | 81 | 0 | 20 | 0 | 101 | 251 | 612 | 11 | 0 | 0 | 0 | 0 | 0 | 40 | 26 | 0 | 25 |
| SRR6246355 | 469 | 0 | 1236 | 0 | 0 | 0 | 169 | 10 | 0 | 0 | 0 | 469 | 18 | 23 | 43 | 0 | 14 | 0 | 0 | 0 | 75 | 0 | 21 | 0 | 42 | 105 | 272 | 11 | 0 | 0 | 0 | 0 | 0 | 15 | 22 | 13 | 10 |
| SRR6246356 | 410 | 0 | 1076 | 0 | 0 | 0 | 154 | 22 | 0 | 0 | 0 | 502 | 43 | 18 | 50 | 0 | 0 | 0 | 0 | 0 | 70 | 0 | 18 | 0 | 97 | 250 | 611 | 0 | 0 | 0 | 0 | 0 | 0 | 45 | 18 | 11 | 25 |
| SRR6246358 | 442 | 11 | 1106 | 0 | 0 | 11 | 61 | 444 | 11 | 0 | 0 | 221 | 0 | 11 | 0 | 0 | 0 | 0 | 0 | 0 | 15 | 0 | 0 | 0 | 0 | 11 | 29 | 11 | 0 | 18 | 0 | 0 | 0 | 44 | 0 | 0 | 0 |
| SRR6246359 | 0 | 0 | 1479 | 0 | 0 | 0 | 37 | 491 | 0 | 0 | 0 | 211 | 0 | 11 | 0 | 11 | 0 | 0 | 0 | 0 | 0 | 0 | 0 | 0 | 0 | 11 | 49 | 0 | 0 | 11 | 0 | 0 | 11 | 11 | 0 | 0 | 0 |
| SRR6246360 | 219 | 0 | 2186 | 0 | 0 | 0 | 46 | 2307 | 0 | 0 | 0 | 219 | 0 | 11 | 0 | 0 | 0 | 0 | 0 | 0 | 23 | 0 | 0 | 0 | 11 | 87 | 152 | 0 | 0 | 54 | 0 | 0 | 0 | 22 | 0 | 0 | 0 |
| SRR6246361 | 155 | 0 | 2123 | 0 | 12 | 0 | 104 | 362 | 0 | 0 | 0 | 570 | 0 | 0 | 0 | 0 | 104 | 0 | 166 | 250 | 83 | 0 | 0 | 0 | 0 | 48 | 145 | 36 | 0 | 24 | 0 | 0 | 12 | 12 | 0 | 12 | 0 |
| SRR6246362 | 266 | 0 | 1488 | 0 | 25 | 0 | 214 | 320 | 12 | 0 | 0 | 159 | 0 | 12 | 0 | 0 | 0 | 0 | 86 | 128 | 43 | 0 | 0 | 0 | 0 | 25 | 185 | 49 | 0 | 37 | 0 | 0 | 12 | 37 | 0 | 0 | 0 |
| SRR6246363 | 155 | 24 | 1652 | 24 | 12 | 12 | 272 | 140 | 0 | 12 | 0 | 465 | 0 | 12 | 0 | 0 | 0 | 0 | 105 | 105 | 0 | 0 | 0 | 0 | 12 | 14 | 32 | 11 | 12 | 11 | 0 | 0 | 0 | 12 | 0 | 0 | 12 |
| SRR6246364 | 264 | 0 | 1634 | 25 | 25 | 0 | 254 | 127 | 12 | 0 | 0 | 316 | 0 | 0 | 0 | 0 | 0 | 0 | 63 | 106 | 42 | 0 | 0 | 0 | 0 | 18 | 44 | 25 | 0 | 0 | 0 | 0 | 0 | 11 | 0 | 12 | 0 |
| SRR6246365 | 272 | 29 | 1177 | 0 | 0 | 0 | 77 | 29 | 0 | 0 | 0 | 498 | 0 | 0 | 0 | 0 | 0 | 0 | 0 | 618 | 0 | 0 | 0 | 0 | 0 | 29 | 88 | 0 | 0 | 0 | 0 | 0 | 0 | 29 | 0 | 0 | 29 |
| SRR6246366 | 413 | 150 | 1239 | 0 | 0 | 30 | 80 | 0 | 0 | 0 | 0 | 275 | 0 | 0 | 0 | 0 | 0 | 0 | 0 | 885 | 0 | 0 | 0 | 0 | 0 | 0 | 60 | 30 | 0 | 30 | 0 | 0 | 0 | 0 | 0 | 0 | 0 |
| SRR6246367 | 362 | 29 | 1086 | 0 | 0 | 0 | 239 | 0 | 0 | 0 | 0 | 226 | 0 | 0 | 0 | 29 | 0 | 0 | 0 | 319 | 0 | 0 | 0 | 0 | 0 | 0 | 116 | 29 | 0 | 0 | 0 | 0 | 0 | 0 | 0 | 0 | 0 |
| SRR6246368 | 183 | 60 | 1370 | 0 | 0 | 0 | 0 | 0 | 0 | 0 | 0 | 274 | 0 | 0 | 0 | 30 | 0 | 0 | 80 | 641 | 0 | 0 | 0 | 0 | 0 | 0 | 90 | 0 | 0 | 0 | 0 | 0 | 0 | 0 | 0 | 0 | 30 |
| SRR6246369 | 428 | 0 | 1141 | 0 | 0 | 0 | 50 | 24 | 0 | 0 | 0 | 29 | 13 | 18 | 0 | 0 | 0 | 0 | 0 | 0 | 36 | 0 | 0 | 0 | 0 | 75 | 429 | 0 | 0 | 0 | 16 | 0 | 0 | 46 | 27 | 0 | 28 |
| SRR6246370 | 235 | 0 | 1291 | 0 | 0 | 0 | 58 | 26 | 0 | 0 | 0 | 147 | 16 | 12 | 0 | 0 | 0 | 0 | 0 | 0 | 36 | 0 | 0 | 0 | 0 | 93 | 431 | 0 | 0 | 0 | 14 | 0 | 0 | 22 | 17 | 0 | 27 |
| SRR6246371 | 174 | 0 | 1247 | 0 | 0 | 0 | 50 | 31 | 0 | 0 | 0 | 145 | 0 | 13 | 0 | 0 | 0 | 0 | 0 | 0 | 23 | 0 | 0 | 0 | 0 | 24 | 113 | 11 | 0 | 0 | 17 | 0 | 0 | 30 | 24 | 0 | 32 |
| SRR6246372 | 292 | 0 | 1225 | 0 | 0 | 0 | 55 | 32 | 0 | 0 | 0 | 117 | 13 | 16 | 10 | 0 | 0 | 0 | 0 | 0 | 27 | 0 | 0 | 0 | 0 | 73 | 404 | 0 | 0 | 0 | 13 | 0 | 0 | 41 | 25 | 0 | 27 |
| SRR6246373 | 136 | 0 | 1773 | 0 | 0 | 0 | 33 | 401 | 0 | 0 | 0 | 546 | 0 | 0 | 0 | 0 | 0 | 0 | 0 | 0 | 0 | 0 | 0 | 0 | 15 | 23 | 53 | 0 | 0 | 38 | 0 | 0 | 0 | 15 | 0 | 0 | 0 |
| SRR6246374 | 421 | 0 | 2387 | 0 | 0 | 0 | 20 | 410 | 0 | 0 | 0 | 0 | 0 | 0 | 0 | 0 | 0 | 0 | 0 | 0 | 34 | 0 | 0 | 0 | 0 | 0 | 58 | 23 | 0 | 16 | 0 | 0 | 0 | 39 | 0 | 0 | 0 |
| SRR6246375 | 270 | 0 | 2162 | 0 | 0 | 0 | 80 | 400 | 0 | 0 | 0 | 135 | 0 | 0 | 0 | 0 | 0 | 0 | 0 | 0 | 20 | 0 | 0 | 0 | 0 | 30 | 67 | 23 | 0 | 21 | 0 | 0 | 0 | 30 | 0 | 0 | 0 |
| SRR6246376 | 427 | 0 | 2137 | 0 | 0 | 0 | 47 | 1930 | 0 | 0 | 0 | 0 | 0 | 0 | 0 | 0 | 0 | 0 | 0 | 0 | 34 | 0 | 0 | 0 | 0 | 31 | 285 | 23 | 0 | 85 | 0 | 0 | 0 | 46 | 0 | 0 | 0 |
| SRR6246379 | 0 | 0 | 0 | 0 | 0 | 0 | 0 | 1287 | 0 | 0 | 0 | 0 | 0 | 0 | 0 | 0 | 0 | 0 | 0 | 0 | 0 | 0 | 0 | 0 | 0 | 322 | 322 | 0 | 0 | 0 | 0 | 0 | 0 | 0 | 0 | 0 | 0 |
| SRR6246385 | 352 | 0 | 1339 | 0 | 0 | 12 | 509 | 64 | 0 | 0 | 0 | 207 | 0 | 44 | 75 | 0 | 12 | 0 | 20 | 27 | 79 | 0 | 0 | 0 | 62 | 269 | 322 | 0 | 0 | 0 | 0 | 0 | 0 | 38 | 37 | 22 | 34 |
| SRR6246386 | 411 | 0 | 1285 | 0 | 0 | 0 | 479 | 64 | 0 | 0 | 0 | 174 | 0 | 37 | 84 | 0 | 0 | 0 | 20 | 17 | 71 | 0 | 0 | 0 | 54 | 257 | 304 | 0 | 0 | 0 | 0 | 0 | 0 | 47 | 41 | 17 | 38 |
| SRR6246387 | 365 | 0 | 1263 | 0 | 0 | 0 | 505 | 59 | 0 | 0 | 0 | 202 | 0 | 38 | 81 | 0 | 14 | 0 | 18 | 22 | 80 | 0 | 0 | 0 | 63 | 281 | 324 | 0 | 0 | 0 | 0 | 0 | 0 | 43 | 38 | 16 | 32 |
| SRR6246388 | 363 | 0 | 1309 | 0 | 0 | 0 | 477 | 62 | 0 | 0 | 0 | 184 | 0 | 41 | 71 | 0 | 16 | 0 | 20 | 28 | 75 | 0 | 0 | 0 | 58 | 282 | 310 | 0 | 0 | 0 | 0 | 0 | 0 | 37 | 41 | 16 | 29 |
| SRR6246389 | 137 | 0 | 1375 | 0 | 0 | 0 | 45 | 338 | 0 | 0 | 0 | 309 | 0 | 0 | 0 | 0 | 0 | 0 | 0 | 17 | 68 | 0 | 0 | 0 | 0 | 52 | 69 | 60 | 0 | 60 | 0 | 0 | 0 | 52 | 0 | 0 | 0 |
| SRR6246390 | 107 | 0 | 1280 | 0 | 0 | 0 | 29 | 2100 | 0 | 0 | 0 | 284 | 0 | 0 | 0 | 0 | 0 | 0 | 0 | 0 | 75 | 0 | 0 | 0 | 0 | 62 | 335 | 44 | 0 | 62 | 35 | 0 | 0 | 62 | 0 | 0 | 0 |
| SRR6246391 | 173 | 34 | 1105 | 0 | 0 | 0 | 63 | 347 | 0 | 0 | 0 | 276 | 0 | 0 | 0 | 0 | 0 | 0 | 0 | 11 | 57 | 0 | 0 | 0 | 0 | 16 | 59 | 51 | 0 | 11 | 0 | 0 | 26 | 26 | 0 | 0 | 0 |
| SRR6246392 | 35 | 0 | 1500 | 0 | 17 | 0 | 68 | 394 | 0 | 0 | 0 | 244 | 0 | 0 | 0 | 0 | 0 | 0 | 0 | 0 | 62 | 0 | 0 | 0 | 0 | 11 | 77 | 17 | 0 | 10 | 0 | 0 | 0 | 10 | 0 | 0 | 0 |
| SRR6246393 | 61 | 0 | 790 | 0 | 0 | 0 | 348 | 58 | 0 | 14 | 0 | 365 | 0 | 0 | 0 | 0 | 0 | 0 | 44 | 65 | 87 | 0 | 0 | 0 | 0 | 18 | 0 | 13 | 0 | 13 | 0 | 0 | 14 | 27 | 0 | 0 | 0 |
| SRR6246394 | 251 | 30 | 627 | 0 | 0 | 0 | 312 | 254 | 0 | 0 | 0 | 188 | 0 | 0 | 0 | 0 | 0 | 0 | 156 | 45 | 89 | 0 | 0 | 0 | 0 | 15 | 15 | 75 | 0 | 15 | 0 | 0 | 0 | 75 | 0 | 0 | 0 |
| SRR6246395 | 239 | 27 | 717 | 0 | 0 | 0 | 326 | 104 | 0 | 0 | 0 | 358 | 0 | 0 | 0 | 0 | 0 | 0 | 130 | 174 | 87 | 0 | 0 | 15 | 0 | 29 | 14 | 0 | 0 | 15 | 0 | 0 | 0 | 14 | 0 | 0 | 0 |
| SRR6246396 | 63 | 0 | 500 | 0 | 0 | 0 | 240 | 115 | 0 | 0 | 0 | 188 | 0 | 0 | 0 | 0 | 0 | 0 | 196 | 44 | 87 | 0 | 0 | 0 | 0 | 14 | 14 | 0 | 0 | 30 | 0 | 0 | 30 | 23 | 0 | 0 | 0 |
| SRR6246397 | 340 | 321 | 1097 | 0 | 0 | 0 | 64 | 13 | 0 | 0 | 0 | 246 | 0 | 0 | 0 | 0 | 0 | 0 | 64 | 860 | 0 | 0 | 0 | 0 | 0 | 0 | 207 | 0 | 0 | 0 | 0 | 0 | 0 | 0 | 0 | 0 | 0 |
| SRR6246398 | 217 | 154 | 1084 | 0 | 0 | 0 | 0 | 16 | 0 | 0 | 0 | 197 | 0 | 0 | 0 | 0 | 0 | 0 | 162 | 844 | 0 | 0 | 0 | 0 | 0 | 13 | 113 | 13 | 0 | 0 | 0 | 0 | 0 | 0 | 0 | 0 | 0 |
| SRR6246399 | 231 | 165 | 904 | 0 | 0 | 0 | 224 | 10 | 0 | 0 | 0 | 115 | 0 | 0 | 0 | 0 | 19 | 0 | 128 | 1057 | 32 | 0 | 0 | 0 | 0 | 19 | 99 | 0 | 0 | 0 | 0 | 0 | 0 | 0 | 0 | 0 | 0 |
| SRR6246400 | 251 | 323 | 1180 | 0 | 0 | 0 | 65 | 0 | 0 | 0 | 0 | 193 | 0 | 0 | 0 | 0 | 0 | 0 | 163 | 781 | 0 | 0 | 0 | 0 | 0 | 16 | 206 | 13 | 0 | 0 | 0 | 0 | 0 | 0 | 0 | 0 | 0 |
| SRR6265480 | 0 | 10 | 6282 | 0 | 12 | 14 | 42 | 14 | 0 | 0 | 0 | 14 | 0 | 10 | 106 | 0 | 0 | 22 | 21 | 918 | 696 | 0 | 0 | 0 | 0 | 64 | 45 | 0 | 0 | 0 | 13 | 0 | 0 | 43 | 48 | 20 | 40 |
| SRR6265481 | 14 | 16 | 6571 | 0 | 0 | 0 | 132 | 172 | 0 | 0 | 0 | 20 | 0 | 0 | 83 | 0 | 14 | 19 | 14 | 1178 | 1012 | 0 | 0 | 0 | 0 | 222 | 290 | 0 | 0 | 32 | 0 | 0 | 0 | 35 | 76 | 0 | 53 |
| SRR6265482 | 15 | 12 | 3192 | 0 | 38 | 12 | 18 | 0 | 0 | 0 | 0 | 62 | 0 | 0 | 445 | 0 | 11 | 14 | 0 | 71 | 534 | 0 | 0 | 0 | 0 | 222 | 153 | 0 | 24 | 0 | 44 | 0 | 0 | 18 | 212 | 24 | 177 |
| SRR6265483 | 12 | 15 | 5052 | 0 | 35 | 17 | 64 | 0 | 0 | 0 | 0 | 81 | 0 | 17 | 222 | 0 | 0 | 41 | 0 | 636 | 254 | 0 | 0 | 0 | 0 | 66 | 34 | 0 | 25 | 0 | 25 | 0 | 0 | 37 | 36 | 30 | 26 |
| SRR6265484 | 15 | 0 | 3318 | 0 | 28 | 18 | 67 | 71 | 0 | 0 | 0 | 150 | 0 | 16 | 67 | 0 | 0 | 0 | 0 | 1036 | 713 | 0 | 0 | 0 | 0 | 274 | 360 | 0 | 17 | 14 | 12 | 0 | 0 | 13 | 83 | 13 | 87 |
| SRR6265485 | 11 | 0 | 3760 | 0 | 19 | 0 | 60 | 30 | 0 | 0 | 0 | 57 | 0 | 10 | 96 | 0 | 0 | 11 | 60 | 2932 | 769 | 0 | 0 | 10 | 0 | 46 | 84 | 0 | 0 | 10 | 13 | 0 | 0 | 10 | 43 | 13 | 20 |
| SRR6265486 | 0 | 0 | 4934 | 0 | 30 | 0 | 0 | 0 | 0 | 0 | 0 | 27 | 0 | 16 | 288 | 0 | 0 | 0 | 0 | 387 | 394 | 0 | 0 | 0 | 0 | 141 | 178 | 0 | 0 | 0 | 13 | 0 | 0 | 0 | 676 | 47 | 333 |
| SRR6265487 | 13 | 0 | 4687 | 0 | 24 | 14 | 68 | 0 | 0 | 0 | 0 | 0 | 0 | 0 | 194 | 0 | 14 | 0 | 39 | 426 | 1035 | 0 | 0 | 0 | 0 | 574 | 163 | 0 | 0 | 0 | 15 | 0 | 0 | 13 | 174 | 0 | 94 |
| SRR6265488 | 11 | 0 | 3873 | 0 | 22 | 29 | 42 | 0 | 20 | 0 | 0 | 25 | 0 | 0 | 110 | 0 | 10 | 18 | 11 | 703 | 1040 | 0 | 0 | 0 | 0 | 124 | 78 | 0 | 0 | 0 | 20 | 0 | 0 | 16 | 54 | 25 | 36 |
| SRR6265489 | 13 | 0 | 8069 | 0 | 0 | 0 | 20 | 0 | 0 | 0 | 0 | 0 | 0 | 0 | 39 | 0 | 0 | 0 | 20 | 2823 | 888 | 0 | 0 | 0 | 0 | 12 | 11 | 0 | 0 | 0 | 0 | 0 | 0 | 20 | 12 | 0 | 0 |
| SRR6265490 | 14 | 0 | 3049 | 0 | 16 | 16 | 122 | 111 | 0 | 0 | 0 | 109 | 0 | 0 | 35 | 0 | 0 | 15 | 20 | 2511 | 1332 | 0 | 0 | 0 | 0 | 267 | 319 | 0 | 14 | 16 | 0 | 0 | 0 | 0 | 239 | 35 | 210 |
| SRR6265491 | 15 | 15 | 3508 | 0 | 13 | 0 | 22 | 23 | 12 | 0 | 0 | 17 | 0 | 0 | 146 | 0 | 0 | 20 | 0 | 764 | 1169 | 0 | 0 | 0 | 0 | 40 | 22 | 0 | 17 | 0 | 18 | 0 | 0 | 22 | 25 | 0 | 22 |
| SRR6306014 | 167 | 25 | 1414 | 0 | 0 | 0 | 84 | 1222 | 0 | 0 | 0 | 450 | 0 | 228 | 3811 | 0 | 0 | 0 | 0 | 719 | 1856 | 0 | 0 | 0 | 0 | 77 | 2567 | 0 | 0 | 0 | 12 | 0 | 0 | 0 | 2500 | 257 | 1711 |
| SRR6306015 | 144 | 17 | 1703 | 0 | 0 | 0 | 71 | 513 | 0 | 0 | 0 | 484 | 0 | 54 | 3542 | 0 | 0 | 0 | 0 | 488 | 1407 | 0 | 0 | 0 | 0 | 84 | 2915 | 14 | 0 | 0 | 14 | 0 | 0 | 0 | 2701 | 198 | 1988 |
| SRR6313948 | 98 | 0 | 55 | 12 | 12 | 136 | 14 | 16 | 100 | 14 | 540 | 1088 | 223 | 171 | 126 | 30 | 97 | 0 | 0 | 23 | 1923 | 40 | 0 | 30 | 31 | 1265 | 151 | 65 | 20 | 0 | 502 | 59 | 131 | 20388 | 1086 | 2967 | 5637 |
| SRR6313949 | 110 | 0 | 38 | 37 | 0 | 63 | 0 | 0 | 93 | 0 | 249 | 1481 | 250 | 413 | 164 | 68 | 154 | 16 | 0 | 0 | 507 | 27 | 0 | 0 | 80 | 607 | 57 | 140 | 38 | 0 | 504 | 92 | 19 | 6004 | 752 | 1221 | 2456 |
| SRR6317622 | 67 | 14 | 1700 | 0 | 0 | 49 | 65 | 120 | 0 | 0 | 0 | 379 | 0 | 0 | 32 | 0 | 0 | 0 | 65 | 7295 | 1102 | 0 | 0 | 0 | 21 | 41 | 150 | 0 | 0 | 35 | 77 | 0 | 0 | 14 | 44 | 49 | 31 |
| SRR6317623 | 38 | 0 | 1653 | 0 | 0 | 0 | 0 | 0 | 0 | 0 | 0 | 350 | 0 | 24 | 163 | 0 | 0 | 0 | 47 | 3796 | 1258 | 0 | 0 | 0 | 85 | 33 | 73 | 0 | 0 | 0 | 76 | 0 | 0 | 0 | 41 | 66 | 27 |
| SRR6317624 | 54 | 0 | 2008 | 0 | 0 | 38 | 17 | 17 | 17 | 0 | 0 | 386 | 0 | 0 | 17 | 0 | 0 | 0 | 17 | 4554 | 851 | 0 | 0 | 0 | 25 | 45 | 157 | 0 | 0 | 13 | 80 | 0 | 0 | 25 | 35 | 88 | 33 |
| SRR6317625 | 66 | 14 | 1760 | 0 | 0 | 28 | 24 | 0 | 14 | 0 | 0 | 309 | 0 | 0 | 24 | 0 | 0 | 0 | 0 | 4387 | 498 | 0 | 0 | 0 | 0 | 41 | 106 | 0 | 0 | 0 | 78 | 0 | 0 | 0 | 37 | 135 | 41 |
| SRR6317626 | 29 | 14 | 1355 | 0 | 0 | 0 | 91 | 30 | 11 | 0 | 0 | 72 | 0 | 16 | 18 | 0 | 0 | 0 | 0 | 2138 | 380 | 0 | 0 | 0 | 0 | 34 | 55 | 0 | 0 | 11 | 46 | 0 | 0 | 0 | 14 | 16 | 97 |
| SRR6317627 | 29 | 15 | 1533 | 0 | 0 | 0 | 489 | 190 | 16 | 0 | 0 | 104 | 0 | 16 | 60 | 0 | 0 | 12 | 54 | 730 | 670 | 0 | 0 | 0 | 16 | 206 | 383 | 0 | 0 | 43 | 28 | 0 | 12 | 96 | 50 | 0 | 31 |
| SRR6317628 | 28 | 0 | 1151 | 0 | 0 | 27 | 26 | 0 | 13 | 0 | 0 | 201 | 0 | 0 | 78 | 0 | 0 | 0 | 26 | 5969 | 860 | 0 | 0 | 0 | 0 | 392 | 801 | 0 | 0 | 0 | 57 | 0 | 0 | 0 | 219 | 33 | 246 |
| SRR6317629 | 23 | 0 | 1325 | 0 | 0 | 0 | 89 | 0 | 0 | 0 | 0 | 79 | 0 | 0 | 30 | 0 | 0 | 0 | 30 | 3312 | 857 | 0 | 0 | 0 | 25 | 21 | 43 | 0 | 0 | 0 | 50 | 0 | 12 | 0 | 11 | 0 | 79 |
| SRR6317630 | 40 | 10 | 1830 | 0 | 0 | 48 | 0 | 20 | 34 | 0 | 0 | 393 | 0 | 0 | 15 | 0 | 0 | 0 | 31 | 5724 | 1270 | 0 | 0 | 0 | 0 | 51 | 200 | 0 | 0 | 0 | 82 | 0 | 0 | 14 | 53 | 31 | 32 |
| SRR6317631 | 53 | 0 | 1645 | 0 | 0 | 28 | 0 | 0 | 12 | 0 | 0 | 327 | 0 | 0 | 65 | 0 | 0 | 0 | 33 | 6187 | 944 | 0 | 0 | 0 | 53 | 433 | 1021 | 0 | 0 | 0 | 84 | 0 | 0 | 0 | 342 | 72 | 339 |
| SRR6317632 | 65 | 21 | 1939 | 0 | 0 | 53 | 0 | 0 | 25 | 0 | 0 | 368 | 0 | 0 | 16 | 0 | 0 | 0 | 48 | 4582 | 671 | 0 | 0 | 0 | 16 | 740 | 2043 | 0 | 0 | 0 | 99 | 0 | 0 | 0 | 617 | 78 | 440 |
| SRR6317633 | 78 | 0 | 2155 | 0 | 0 | 23 | 22 | 0 | 14 | 0 | 0 | 455 | 0 | 0 | 22 | 0 | 0 | 0 | 43 | 4028 | 517 | 0 | 0 | 0 | 0 | 37 | 107 | 0 | 0 | 0 | 84 | 0 | 0 | 0 | 32 | 56 | 28 |
| SRR6317634 | 29 | 15 | 1616 | 0 | 0 | 27 | 135 | 25 | 25 | 0 | 0 | 81 | 0 | 15 | 50 | 0 | 0 | 0 | 17 | 2288 | 303 | 0 | 0 | 0 | 15 | 413 | 591 | 17 | 0 | 0 | 42 | 0 | 12 | 0 | 178 | 15 | 94 |
| SRR6317635 | 37 | 21 | 1534 | 0 | 0 | 17 | 511 | 184 | 11 | 0 | 0 | 126 | 0 | 15 | 45 | 0 | 0 | 0 | 65 | 1656 | 1100 | 0 | 0 | 0 | 12 | 297 | 398 | 0 | 0 | 45 | 28 | 0 | 0 | 62 | 55 | 0 | 41 |
| SRR6317636 | 38 | 0 | 1231 | 0 | 0 | 40 | 0 | 0 | 14 | 0 | 0 | 209 | 0 | 0 | 52 | 0 | 0 | 0 | 52 | 5651 | 1029 | 0 | 0 | 0 | 20 | 28 | 64 | 0 | 0 | 0 | 71 | 0 | 0 | 0 | 16 | 45 | 18 |
| SRR6317637 | 23 | 15 | 1299 | 0 | 0 | 0 | 17 | 0 | 0 | 0 | 0 | 83 | 0 | 27 | 33 | 0 | 0 | 0 | 0 | 2368 | 834 | 0 | 0 | 0 | 30 | 196 | 466 | 38 | 0 | 0 | 35 | 0 | 0 | 25 | 83 | 0 | 65 |
| SRR6317638 | 38 | 0 | 1755 | 0 | 0 | 48 | 22 | 0 | 14 | 0 | 0 | 323 | 0 | 0 | 43 | 0 | 0 | 0 | 22 | 3891 | 1081 | 0 | 0 | 0 | 19 | 37 | 73 | 0 | 0 | 0 | 58 | 0 | 0 | 0 | 37 | 43 | 18 |
| SRR6317639 | 33 | 0 | 1672 | 0 | 0 | 11 | 97 | 34 | 30 | 0 | 0 | 411 | 0 | 0 | 155 | 0 | 0 | 0 | 19 | 5005 | 1086 | 0 | 0 | 0 | 45 | 512 | 1035 | 0 | 0 | 0 | 79 | 0 | 0 | 19 | 561 | 45 | 414 |
| SRR6317640 | 60 | 0 | 1847 | 0 | 0 | 35 | 38 | 32 | 29 | 0 | 0 | 416 | 0 | 0 | 0 | 0 | 0 | 0 | 25 | 4457 | 700 | 0 | 0 | 0 | 21 | 42 | 157 | 0 | 0 | 0 | 79 | 0 | 0 | 0 | 44 | 56 | 35 |
| SRR6317641 | 81 | 0 | 1942 | 0 | 0 | 23 | 20 | 0 | 14 | 0 | 0 | 381 | 0 | 0 | 61 | 0 | 0 | 0 | 41 | 3881 | 305 | 0 | 0 | 0 | 18 | 413 | 1082 | 0 | 0 | 0 | 50 | 0 | 0 | 0 | 316 | 83 | 275 |
| SRR6317642 | 21 | 12 | 1392 | 0 | 0 | 15 | 153 | 83 | 0 | 0 | 0 | 68 | 0 | 0 | 0 | 0 | 0 | 0 | 0 | 2703 | 479 | 0 | 0 | 0 | 15 | 33 | 70 | 0 | 0 | 15 | 52 | 0 | 15 | 25 | 13 | 12 | 89 |
| SRR6317643 | 44 | 18 | 1430 | 0 | 0 | 17 | 395 | 36 | 27 | 0 | 0 | 208 | 0 | 0 | 216 | 0 | 0 | 0 | 19 | 903 | 705 | 0 | 0 | 0 | 0 | 89 | 209 | 0 | 0 | 11 | 45 | 0 | 0 | 13 | 22 | 17 | 11 |
| SRR6317644 | 28 | 0 | 1206 | 0 | 0 | 33 | 0 | 0 | 14 | 0 | 0 | 189 | 0 | 0 | 52 | 0 | 0 | 0 | 52 | 8262 | 1016 | 0 | 0 | 0 | 19 | 568 | 1062 | 0 | 0 | 0 | 71 | 0 | 0 | 0 | 283 | 33 | 402 |
| SRR6317645 | 24 | 23 | 1369 | 0 | 0 | 0 | 33 | 0 | 14 | 0 | 0 | 89 | 0 | 11 | 0 | 0 | 0 | 0 | 49 | 4003 | 993 | 0 | 0 | 0 | 16 | 30 | 73 | 18 | 0 | 0 | 47 | 0 | 0 | 18 | 13 | 0 | 70 |
| SRR6324194 | 0 | 0 | 601 | 0 | 0 | 0 | 0 | 0 | 10 | 0 | 0 | 154 | 0 | 0 | 0 | 0 | 0 | 0 | 0 | 40 | 0 | 0 | 0 | 0 | 0 | 52 | 63 | 0 | 0 | 0 | 45 | 0 | 0 | 10 | 10 | 24 | 55 |
| SRR6324195 | 0 | 15 | 569 | 0 | 0 | 22 | 37 | 32 | 12 | 0 | 0 | 253 | 0 | 0 | 15 | 0 | 0 | 0 | 15 | 128 | 67 | 0 | 0 | 0 | 10 | 299 | 199 | 24 | 0 | 12 | 54 | 0 | 0 | 32 | 149 | 57 | 142 |
| SRR6363052 | 0 | 0 | 1692 | 0 | 0 | 67 | 0 | 0 | 47 | 0 | 0 | 123 | 0 | 70 | 514 | 0 | 0 | 67 | 27 | 460 | 4132 | 0 | 0 | 0 | 41 | 115 | 86 | 24 | 0 | 0 | 12 | 0 | 14 | 34 | 232 | 124 | 202 |
| SRR6363053 | 17 | 0 | 2127 | 0 | 0 | 53 | 0 | 0 | 30 | 0 | 0 | 112 | 0 | 24 | 406 | 0 | 0 | 11 | 12 | 271 | 2092 | 0 | 0 | 0 | 20 | 1492 | 976 | 22 | 0 | 0 | 180 | 0 | 0 | 25 | 3364 | 1368 | 2563 |
| SRR6363054 | 0 | 0 | 2311 | 0 | 0 | 26 | 25 | 0 | 26 | 0 | 0 | 96 | 0 | 37 | 234 | 0 | 0 | 29 | 31 | 197 | 2885 | 0 | 0 | 0 | 27 | 78 | 62 | 17 | 0 | 0 | 125 | 0 | 0 | 24 | 215 | 84 | 149 |
| SRR6370846 | 161 | 0 | 0 | 0 | 0 | 0 | 57 | 0 | 0 | 0 | 0 | 0 | 0 | 0 | 0 | 0 | 141 | 0 | 0 | 57 | 0 | 0 | 0 | 0 | 0 | 0 | 0 | 0 | 0 | 0 | 0 | 0 | 0 | 10 | 0 | 0 | 0 |
| SRR6370847 | 20 | 0 | 0 | 0 | 0 | 0 | 0 | 0 | 17 | 0 | 0 | 19 | 17 | 0 | 32 | 0 | 136 | 0 | 0 | 0 | 65 | 0 | 0 | 0 | 34 | 76 | 17 | 0 | 0 | 0 | 17 | 0 | 0 | 706 | 25 | 17 | 67 |
| SRR6370848 | 56 | 0 | 0 | 0 | 0 | 24 | 209 | 0 | 0 | 0 | 0 | 19 | 0 | 18 | 0 | 0 | 434 | 0 | 0 | 0 | 0 | 0 | 0 | 0 | 12 | 24 | 0 | 0 | 0 | 12 | 30 | 0 | 0 | 103 | 12 | 0 | 30 |
| SRR6370849 | 67 | 0 | 0 | 0 | 0 | 0 | 229 | 24 | 10 | 0 | 0 | 43 | 0 | 0 | 0 | 0 | 204 | 0 | 0 | 114 | 29 | 0 | 0 | 10 | 64 | 74 | 101 | 0 | 0 | 13 | 0 | 0 | 13 | 437 | 57 | 27 | 94 |
| SRR6370852 | 66 | 0 | 11 | 0 | 0 | 0 | 111 | 0 | 0 | 0 | 0 | 40 | 0 | 0 | 0 | 0 | 435 | 0 | 37 | 74 | 0 | 0 | 0 | 0 | 118 | 55 | 155 | 0 | 0 | 0 | 0 | 0 | 0 | 291 | 55 | 146 | 173 |
| SRR6370853 | 42 | 0 | 0 | 0 | 0 | 0 | 123 | 0 | 0 | 0 | 26 | 35 | 0 | 22 | 0 | 0 | 245 | 0 | 0 | 0 | 245 | 0 | 0 | 0 | 17 | 13 | 0 | 0 | 0 | 0 | 30 | 0 | 0 | 789 | 13 | 0 | 0 |
| SRR6370856 | 41 | 0 | 0 | 0 | 0 | 11 | 0 | 112 | 0 | 0 | 0 | 37 | 0 | 0 | 0 | 0 | 164 | 0 | 17 | 104 | 0 | 0 | 0 | 44 | 25 | 134 | 35 | 0 | 11 | 71 | 19 | 0 | 0 | 22 | 30 | 46 | 112 |
| SRR6372849 | 0 | 0 | 30 | 0 | 0 | 175 | 103 | 27 | 56 | 113 | 255 | 117 | 23 | 17 | 116 | 0 | 177 | 24 | 19 | 71 | 361 | 0 | 0 | 106 | 91 | 1053 | 1162 | 14 | 16 | 0 | 23 | 0 | 21 | 31 | 147 | 81 | 360 |
| SRR6372850 | 11 | 21 | 29 | 0 | 0 | 131 | 15 | 0 | 56 | 67 | 235 | 102 | 0 | 15 | 215 | 0 | 123 | 15 | 0 | 92 | 115 | 0 | 0 | 164 | 115 | 1068 | 391 | 0 | 0 | 0 | 115 | 0 | 0 | 0 | 128 | 108 | 149 |
| SRR6372851 | 0 | 13 | 22 | 0 | 0 | 55 | 19 | 0 | 17 | 86 | 30 | 67 | 0 | 14 | 394 | 0 | 24 | 0 | 11 | 34 | 1207 | 0 | 0 | 56 | 52 | 314 | 83 | 11 | 20 | 0 | 10 | 0 | 24 | 24 | 69 | 14 | 18 |
| SRR6372852 | 83 | 0 | 12 | 0 | 0 | 91 | 18 | 12 | 34 | 17 | 23 | 114 | 12 | 0 | 119 | 0 | 115 | 0 | 79 | 174 | 420 | 0 | 0 | 0 | 48 | 442 | 53 | 0 | 0 | 0 | 26 | 0 | 29 | 28 | 311 | 105 | 197 |
| SRR6372853 | 32 | 0 | 14 | 0 | 0 | 80 | 37 | 24 | 21 | 13 | 37 | 153 | 0 | 0 | 327 | 11 | 87 | 14 | 68 | 241 | 309 | 0 | 0 | 75 | 21 | 412 | 49 | 0 | 0 | 15 | 37 | 0 | 11 | 36 | 136 | 44 | 46 |
| SRR6372854 | 55 | 31 | 46 | 0 | 0 | 221 | 0 | 0 | 143 | 21 | 77 | 157 | 0 | 11 | 1016 | 0 | 68 | 12 | 0 | 0 | 153 | 0 | 0 | 30 | 65 | 1237 | 42 | 0 | 0 | 0 | 159 | 0 | 19 | 18 | 1526 | 336 | 733 |
| SRR6372855 | 36 | 0 | 54 | 0 | 0 | 0 | 21 | 0 | 0 | 24 | 80 | 39 | 14 | 15 | 228 | 0 | 0 | 34 | 0 | 0 | 42 | 0 | 0 | 25 | 22 | 18 | 0 | 0 | 0 | 0 | 18 | 0 | 0 | 10 | 11 | 0 | 0 |
| SRR6372856 | 70 | 0 | 218 | 0 | 0 | 62 | 87 | 0 | 47 | 22 | 79 | 62 | 26 | 0 | 220 | 0 | 0 | 37 | 20 | 107 | 511 | 0 | 0 | 16 | 53 | 398 | 65 | 0 | 0 | 0 | 82 | 0 | 0 | 41 | 39 | 16 | 34 |
| SRR6372857 | 246 | 0 | 695 | 0 | 0 | 35 | 48 | 0 | 15 | 16 | 25 | 111 | 20 | 0 | 216 | 0 | 0 | 82 | 0 | 0 | 153 | 0 | 0 | 0 | 62 | 217 | 68 | 10 | 0 | 0 | 30 | 0 | 19 | 31 | 16 | 10 | 23 |
| SRR6389795 | 429 | 0 | 556 | 0 | 0 | 33 | 41 | 75 | 0 | 0 | 0 | 694 | 0 | 0 | 16 | 0 | 0 | 24 | 30 | 1039 | 6200 | 12 | 0 | 0 | 0 | 48 | 631 | 0 | 0 | 0 | 0 | 20 | 0 | 0 | 734 | 58 | 637 |
| SRR6389796 | 481 | 0 | 1177 | 0 | 0 | 72 | 230 | 308 | 16 | 0 | 0 | 1455 | 0 | 0 | 39 | 0 | 0 | 0 | 28 | 547 | 3152 | 0 | 0 | 0 | 0 | 63 | 905 | 0 | 0 | 18 | 20 | 33 | 0 | 48 | 641 | 45 | 373 |
| SRR6389797 | 192 | 0 | 3674 | 0 | 0 | 11 | 187 | 50 | 15 | 0 | 0 | 1209 | 0 | 0 | 0 | 0 | 0 | 0 | 74 | 1601 | 6243 | 0 | 0 | 0 | 0 | 150 | 1955 | 0 | 0 | 15 | 30 | 0 | 0 | 24 | 1385 | 73 | 887 |
| SRR6389798 | 73 | 0 | 1188 | 0 | 0 | 31 | 14 | 78 | 0 | 0 | 0 | 0 | 0 | 0 | 44 | 0 | 28 | 0 | 0 | 257 | 663 | 0 | 0 | 0 | 0 | 372 | 1043 | 0 | 0 | 0 | 20 | 0 | 0 | 25 | 7722 | 434 | 3287 |
| SRR6389799 | 56 | 0 | 2993 | 0 | 0 | 21 | 42 | 289 | 0 | 0 | 0 | 980 | 0 | 33 | 40 | 0 | 11 | 0 | 16 | 656 | 2833 | 0 | 0 | 0 | 0 | 34 | 857 | 0 | 0 | 0 | 0 | 13 | 0 | 0 | 852 | 66 | 612 |
| SRR6389800 | 217 | 0 | 4077 | 0 | 0 | 41 | 166 | 682 | 25 | 0 | 0 | 710 | 0 | 0 | 32 | 0 | 16 | 0 | 0 | 928 | 1992 | 0 | 0 | 0 | 0 | 670 | 2558 | 0 | 0 | 0 | 15 | 17 | 0 | 0 | 14844 | 950 | 8573 |
| SRR6389801 | 104 | 12 | 5527 | 0 | 0 | 42 | 0 | 78 | 22 | 0 | 0 | 55 | 0 | 11 | 407 | 0 | 10 | 18 | 0 | 45 | 2123 | 0 | 0 | 0 | 0 | 104 | 41 | 0 | 0 | 0 | 42 | 0 | 0 | 0 | 1040 | 93 | 470 |
| SRR6389802 | 302 | 14 | 4197 | 18 | 11 | 229 | 0 | 129 | 105 | 0 | 0 | 298 | 0 | 15 | 1508 | 0 | 0 | 85 | 0 | 29 | 12954 | 0 | 0 | 0 | 0 | 4115 | 1432 | 0 | 0 | 0 | 158 | 0 | 0 | 0 | 33229 | 3858 | 20410 |
| SRR6389803 | 361 | 10 | 2357 | 0 | 0 | 226 | 36 | 14 | 122 | 0 | 0 | 680 | 0 | 19 | 57 | 0 | 20 | 31 | 0 | 129 | 1307 | 0 | 0 | 0 | 0 | 99 | 93 | 0 | 0 | 0 | 84 | 0 | 0 | 0 | 1404 | 119 | 894 |
| SRR6389804 | 142 | 0 | 5721 | 0 | 0 | 337 | 0 | 11 | 122 | 0 | 0 | 276 | 0 | 0 | 2779 | 0 | 18 | 0 | 0 | 50 | 1949 | 0 | 0 | 0 | 0 | 57 | 144 | 11 | 25 | 0 | 269 | 11 | 0 | 0 | 385 | 49 | 353 |
| SRR6389805 | 122 | 141 | 2486 | 0 | 0 | 68 | 13 | 142 | 38 | 0 | 0 | 472 | 0 | 25 | 172 | 0 | 16 | 121 | 0 | 45 | 3878 | 0 | 0 | 0 | 0 | 1187 | 958 | 0 | 17 | 0 | 28 | 0 | 0 | 17 | 9199 | 692 | 5863 |
| SRR6389806 | 124 | 75 | 1325 | 0 | 0 | 81 | 94 | 17736 | 26 | 0 | 0 | 960 | 0 | 28 | 1873 | 0 | 12 | 26 | 13 | 1478 | 8905 | 0 | 0 | 28 | 0 | 773 | 13275 | 0 | 0 | 17 | 35 | 0 | 0 | 0 | 14828 | 1634 | 11054 |

### Supp Table S8.2.Columns_AM_to_BW

| | | | | | | | | | | | | | | | | | | | | | | | | | | | | | | | | | | | | | |
|---|---|---|---|---|---|---|---|---|---|---|---|---|---|---|---|---|---|---|---|---|---|---|---|---|---|---|---|---|---|---|---|---|---|---|---|---|---|
| SRR6389807 | 65 | 22 | 1871 | 0 | 0 | 51 | 20 | 52 | 16 | 0 | 0 | 1083 | 0 | 11 | 119 | 0 | 32 | 109 | 0 | 625 | 1352 | 0 | 0 | 0 | 0 | 44 | 364 | 0 | 0 | 0 | 55 | 32 | 0 | 0 | 618 | 34 | 322 |
| SRR6389808 | 201 | 0 | 1733 | 0 | 0 | 26 | 171 | 33 | 59 | 0 | 0 | 341 | 0 | 0 | 84 | 0 | 0 | 0 | 19 | 490 | 6507 | 0 | 0 | 0 | 0 | 234 | 1405 | 10 | 0 | 0 | 68 | 19 | 0 | 0 | 2518 | 219 | 1416 |
| SRR6389809 | 141 | 0 | 4449 | 0 | 0 | 79 | 64 | 26 | 17 | 0 | 0 | 201 | 0 | 16 | 20 | 0 | 29 | 0 | 0 | 228 | 1652 | 0 | 0 | 0 | 0 | 82 | 181 | 0 | 14 | 0 | 38 | 80 | 0 | 15 | 705 | 37 | 365 |
| SRR6389810 | 300 | 38 | 2571 | 0 | 0 | 27 | 90 | 53 | 55 | 0 | 0 | 572 | 0 | 0 | 47 | 0 | 24 | 18 | 13 | 398 | 3093 | 0 | 0 | 0 | 0 | 352 | 181 | 0 | 0 | 0 | 21 | 15 | 0 | 12 | 1108 | 76 | 582 |
| SRR6389811 | 147 | 24 | 2763 | 0 | 0 | 79 | 156 | 2570 | 29 | 0 | 0 | 779 | 0 | 0 | 511 | 0 | 0 | 40 | 27 | 780 | 5924 | 0 | 0 | 12 | 0 | 1334 | 9505 | 0 | 0 | 0 | 21 | 0 | 0 | 0 | 3448 | 278 | 3084 |
| SRR6389812 | 250 | 12 | 4798 | 0 | 0 | 47 | 65 | 12 | 69 | 0 | 0 | 798 | 0 | 15 | 564 | 0 | 0 | 66 | 0 | 110 | 1377 | 0 | 0 | 0 | 0 | 21 | 76 | 0 | 0 | 0 | 52 | 20 | 0 | 15 | 181 | 14 | 130 |
| SRR6389813 | 224 | 0 | 5886 | 0 | 0 | 10 | 48 | 20 | 77 | 0 | 0 | 1842 | 0 | 0 | 73 | 0 | 0 | 12 | 19 | 197 | 2946 | 0 | 0 | 0 | 0 | 178 | 302 | 19 | 0 | 0 | 14 | 10 | 0 | 0 | 2195 | 166 | 1345 |
| SRR6389814 | 426 | 65 | 479 | 0 | 27 | 59 | 49 | 520 | 26 | 0 | 0 | 494 | 0 | 13 | 485 | 0 | 13 | 61 | 12 | 640 | 3155 | 0 | 0 | 0 | 0 | 699 | 1409 | 0 | 0 | 0 | 60 | 20 | 0 | 0 | 2393 | 228 | 2205 |
| SRR6389815 | 71 | 0 | 5869 | 0 | 0 | 0 | 0 | 0 | 0 | 0 | 0 | 0 | 0 | 0 | 104 | 0 | 32 | 312 | 0 | 32 | 2307 | 0 | 0 | 0 | 0 | 71 | 25 | 12 | 0 | 0 | 0 | 16 | 0 | 18 | 305 | 21 | 260 |
| SRR6389816 | 235 | 0 | 6358 | 0 | 0 | 12 | 123 | 426 | 16 | 0 | 0 | 2216 | 0 | 17 | 17 | 0 | 27 | 0 | 0 | 317 | 564 | 0 | 0 | 0 | 0 | 459 | 9575 | 0 | 0 | 0 | 15 | 0 | 0 | 42 | 23450 | 1431 | 13548 |
| SRR6389817 | 170 | 0 | 12330 | 0 | 0 | 307 | 11 | 196 | 88 | 0 | 0 | 0 | 0 | 0 | 860 | 0 | 0 | 10 | 11 | 312 | 10115 | 0 | 0 | 0 | 0 | 5215 | 1695 | 0 | 25 | 0 | 511 | 54 | 0 | 0 | 22013 | 1894 | 21995 |
| SRR6389818 | 340 | 0 | 5346 | 0 | 0 | 124 | 69 | 2776 | 48 | 0 | 0 | 2079 | 0 | 40 | 187 | 0 | 0 | 0 | 0 | 437 | 3446 | 0 | 0 | 0 | 0 | 1280 | 5746 | 0 | 0 | 0 | 37 | 14 | 0 | 15 | 11892 | 1851 | 10425 |
| SRR6389819 | 942 | 0 | 725 | 0 | 0 | 57 | 35 | 22 | 30 | 0 | 0 | 14 | 0 | 0 | 0 | 0 | 0 | 0 | 22 | 547 | 2465 | 0 | 0 | 0 | 0 | 1117 | 856 | 12 | 0 | 0 | 35 | 24 | 0 | 0 | 958 | 149 | 1034 |
| SRR6389820 | 336 | 0 | 5224 | 0 | 0 | 25 | 56 | 26 | 10 | 0 | 0 | 437 | 0 | 13 | 36 | 0 | 30 | 13 | 0 | 409 | 2712 | 0 | 0 | 0 | 0 | 372 | 185 | 11 | 0 | 0 | 70 | 0 | 0 | 21 | 979 | 57 | 463 |
| SRR6409751 | 0 | 0 | 55 | 15 | 0 | 0 | 0 | 0 | 0 | 0 | 0 | 0 | 0 | 0 | 0 | 0 | 0 | 0 | 0 | 0 | 422 | 0 | 0 | 0 | 0 | 0 | 0 | 11 | 0 | 13 | 0 | 0 | 0 | 0 | 0 | 0 | 0 |
| SRR6409760 | 0 | 0 | 0 | 0 | 0 | 0 | 0 | 0 | 0 | 0 | 0 | 0 | 0 | 0 | 0 | 0 | 0 | 0 | 0 | 0 | 2778 | 0 | 0 | 0 | 0 | 0 | 0 | 0 | 0 | 0 | 0 | 0 | 0 | 0 | 0 | 0 | 0 |
| SRR6409798 | 0 | 0 | 1855 | 0 | 0 | 0 | 0 | 0 | 0 | 0 | 0 | 0 | 0 | 0 | 0 | 0 | 0 | 0 | 85 | 0 | 2238 | 0 | 0 | 0 | 0 | 0 | 0 | 0 | 0 | 0 | 18 | 0 | 0 | 0 | 0 | 0 | 0 |
| SRR6409800 | 0 | 0 | 0 | 0 | 0 | 0 | 0 | 0 | 0 | 0 | 0 | 0 | 0 | 0 | 0 | 0 | 0 | 0 | 0 | 0 | 0 | 0 | 0 | 0 | 0 | 0 | 0 | 0 | 0 | 0 | 0 | 0 | 0 | 0 | 0 | 0 | 0 |
| SRR6409827 | 0 | 0 | 2042 | 0 | 0 | 0 | 487 | 0 | 0 | 0 | 0 | 0 | 0 | 0 | 0 | 0 | 0 | 0 | 0 | 224 | 3680 | 0 | 0 | 0 | 0 | 0 | 0 | 0 | 0 | 0 | 0 | 0 | 0 | 0 | 0 | 0 | 0 |
| SRR6409832 | 0 | 0 | 0 | 0 | 0 | 0 | 0 | 0 | 0 | 0 | 0 | 0 | 0 | 0 | 0 | 0 | 0 | 0 | 0 | 0 | 0 | 0 | 0 | 0 | 0 | 0 | 0 | 0 | 0 | 0 | 0 | 0 | 0 | 0 | 0 | 0 | 0 |
| SRR6409833 | 0 | 0 | 1473 | 0 | 0 | 0 | 0 | 0 | 0 | 0 | 0 | 0 | 0 | 0 | 283 | 0 | 50 | 0 | 0 | 0 | 1033 | 0 | 0 | 0 | 0 | 22 | 0 | 0 | 0 | 0 | 0 | 0 | 0 | 0 | 0 | 0 | 0 |
| SRR6409834 | 0 | 0 | 2453 | 0 | 0 | 0 | 0 | 0 | 0 | 0 | 0 | 0 | 0 | 0 | 482 | 0 | 0 | 0 | 283 | 0 | 154 | 0 | 0 | 0 | 0 | 2051 | 0 | 0 | 0 | 0 | 0 | 0 | 0 | 0 | 0 | 0 | 0 |
| SRR6409845 | 0 | 0 | 0 | 0 | 0 | 0 | 0 | 0 | 0 | 0 | 0 | 0 | 0 | 0 | 0 | 0 | 0 | 0 | 0 | 0 | 4488 | 0 | 0 | 0 | 0 | 0 | 0 | 0 | 0 | 0 | 0 | 0 | 0 | 0 | 0 | 0 | 0 |
| SRR6409851 | 37 | 0 | 2436 | 0 | 0 | 0 | 0 | 0 | 0 | 0 | 0 | 0 | 0 | 0 | 179 | 0 | 0 | 0 | 23 | 38 | 1624 | 0 | 0 | 0 | 0 | 268 | 0 | 0 | 0 | 0 | 368 | 0 | 0 | 0 | 0 | 0 | 526 |
| SRR6409853 | 80 | 0 | 2778 | 0 | 0 | 0 | 0 | 0 | 0 | 0 | 0 | 0 | 0 | 0 | 67 | 0 | 0 | 0 | 0 | 85 | 2174 | 0 | 0 | 0 | 0 | 0 | 0 | 0 | 0 | 0 | 0 | 0 | 0 | 0 | 0 | 0 | 0 |
| SRR6409859 | 87 | 0 | 2518 | 0 | 0 | 0 | 0 | 0 | 0 | 0 | 0 | 0 | 0 | 0 | 123 | 0 | 0 | 0 | 212 | 289 | 1697 | 0 | 0 | 0 | 0 | 35 | 0 | 0 | 0 | 0 | 0 | 0 | 0 | 0 | 0 | 0 | 0 |
| SRR6409860 | 0 | 0 | 2905 | 0 | 0 | 0 | 0 | 0 | 0 | 0 | 0 | 0 | 0 | 0 | 65 | 0 | 0 | 0 | 111 | 151 | 1586 | 0 | 0 | 0 | 0 | 0 | 0 | 0 | 0 | 0 | 0 | 0 | 0 | 0 | 0 | 0 | 0 |
| SRR6409863 | 0 | 0 | 857 | 0 | 0 | 1415 | 0 | 0 | 0 | 0 | 0 | 0 | 0 | 0 | 0 | 0 | 0 | 0 | 0 | 0 | 1921 | 0 | 0 | 0 | 0 | 0 | 0 | 0 | 0 | 0 | 0 | 0 | 0 | 0 | 0 | 0 | 0 |
| SRR6409866 | 0 | 0 | 685 | 0 | 0 | 0 | 0 | 0 | 0 | 0 | 0 | 0 | 0 | 0 | 532 | 0 | 0 | 0 | 0 | 0 | 369 | 0 | 0 | 0 | 0 | 1689 | 0 | 0 | 0 | 0 | 0 | 0 | 0 | 0 | 0 | 0 | 0 |
| SRR6409867 | 0 | 0 | 2387 | 0 | 0 | 0 | 0 | 0 | 0 | 0 | 0 | 0 | 0 | 0 | 252 | 0 | 0 | 0 | 0 | 100 | 2356 | 0 | 0 | 0 | 0 | 0 | 0 | 0 | 0 | 0 | 0 | 0 | 0 | 0 | 0 | 0 | 0 |
| SRR6409868 | 0 | 0 | 1447 | 0 | 0 | 0 | 0 | 0 | 0 | 0 | 0 | 0 | 0 | 0 | 0 | 0 | 0 | 0 | 0 | 0 | 1889 | 0 | 0 | 0 | 0 | 0 | 0 | 0 | 0 | 0 | 0 | 0 | 0 | 0 | 0 | 0 | 0 |
| SRR6450851 | 0 | 0 | 0 | 0 | 0 | 76 | 0 | 19 | 19 | 0 | 0 | 321 | 19 | 0 | 0 | 19 | 64 | 0 | 0 | 0 | 212 | 0 | 0 | 57 | 19 | 420 | 95 | 19 | 19 | 0 | 0 | 0 | 210 | 57 | 57 | 38 | 229 |
| SRR6450852 | 67 | 41 | 72 | 24 | 29 | 25 | 12 | 128 | 29 | 24 | 0 | 187 | 44 | 64 | 12 | 46 | 206 | 0 | 12 | 116 | 46 | 18 | 24 | 63 | 18 | 158 | 25 | 45 | 25 | 55 | 47 | 24 | 244 | 258 | 80 | 108 | 198 |
| SRR6450853 | 36 | 33 | 60 | 17 | 17 | 149 | 0 | 149 | 17 | 0 | 17 | 169 | 99 | 132 | 0 | 215 | 181 | 0 | 41 | 81 | 204 | 17 | 17 | 182 | 330 | 628 | 165 | 165 | 116 | 149 | 33 | 50 | 958 | 942 | 446 | 380 | 793 |
| SRR6450854 | 68 | 0 | 53 | 12 | 35 | 116 | 0 | 12 | 81 | 35 | 0 | 204 | 116 | 256 | 26 | 105 | 211 | 0 | 0 | 26 | 105 | 35 | 12 | 0 | 232 | 686 | 70 | 151 | 81 | 23 | 58 | 0 | 1453 | 837 | 244 | 546 | 814 |
| SRR6450855 | 128 | 25 | 43 | 0 | 0 | 25 | 171 | 51 | 0 | 0 | 13 | 43 | 51 | 38 | 0 | 38 | 213 | 0 | 0 | 0 | 171 | 0 | 13 | 13 | 13 | 76 | 64 | 25 | 38 | 38 | 0 | 13 | 369 | 127 | 76 | 38 | 114 |
| SRR6450856 | 46 | 0 | 123 | 0 | 36 | 28 | 0 | 87 | 24 | 12 | 24 | 162 | 48 | 35 | 0 | 15 | 223 | 0 | 44 | 65 | 131 | 0 | 24 | 113 | 22 | 203 | 92 | 41 | 20 | 46 | 11 | 48 | 111 | 196 | 107 | 85 | 155 |
| SRR6450857 | 60 | 0 | 60 | 0 | 0 | 35 | 0 | 52 | 0 | 0 | 0 | 240 | 35 | 35 | 0 | 52 | 180 | 0 | 0 | 0 | 376 | 17 | 0 | 87 | 17 | 174 | 70 | 35 | 0 | 17 | 0 | 0 | 191 | 209 | 70 | 17 | 139 |
| SRR6450858 | 0 | 23 | 77 | 0 | 12 | 128 | 0 | 383 | 23 | 35 | 23 | 123 | 232 | 116 | 0 | 174 | 84 | 0 | 52 | 52 | 260 | 0 | 12 | 337 | 116 | 685 | 290 | 174 | 81 | 209 | 46 | 23 | 929 | 1045 | 441 | 476 | 1301 |
| SRR6450859 | 34 | 0 | 34 | 0 | 55 | 221 | 0 | 332 | 55 | 0 | 111 | 171 | 221 | 0 | 0 | 111 | 376 | 0 | 0 | 332 | 221 | 0 | 0 | 609 | 277 | 996 | 609 | 332 | 55 | 111 | 55 | 111 | 387 | 553 | 332 | 609 | 664 |
| SRR6450860 | 27 | 0 | 55 | 0 | 46 | 35 | 108 | 53 | 18 | 46 | 0 | 110 | 27 | 18 | 0 | 0 | 247 | 0 | 0 | 0 | 325 | 0 | 0 | 71 | 44 | 177 | 44 | 46 | 27 | 62 | 18 | 46 | 53 | 186 | 106 | 89 | 133 |
| SRR6450861 | 79 | 31 | 39 | 0 | 31 | 31 | 0 | 31 | 0 | 31 | 0 | 177 | 28 | 50 | 63 | 11 | 157 | 0 | 0 | 0 | 252 | 31 | 31 | 11 | 61 | 121 | 33 | 55 | 22 | 31 | 33 | 31 | 237 | 171 | 72 | 66 | 121 |
| SRR6450862 | 64 | 12 | 60 | 27 | 30 | 96 | 0 | 25 | 30 | 39 | 20 | 166 | 103 | 224 | 178 | 143 | 182 | 79 | 0 | 24 | 196 | 17 | 20 | 12 | 234 | 716 | 204 | 187 | 94 | 12 | 81 | 49 | 1260 | 704 | 507 | 409 | 842 |
| SRR6450863 | 68 | 16 | 113 | 55 | 0 | 94 | 0 | 251 | 31 | 24 | 0 | 211 | 117 | 211 | 0 | 188 | 105 | 24 | 21 | 82 | 165 | 47 | 16 | 102 | 157 | 744 | 157 | 219 | 117 | 180 | 55 | 24 | 1379 | 1324 | 243 | 533 | 971 |
| SRR6450867 | 150 | 0 | 476 | 0 | 0 | 55 | 149 | 0 | 0 | 0 | 0 | 80 | 0 | 0 | 0 | 0 | 1945 | 0 | 0 | 1943 | 664 | 0 | 138 | 55 | 18 | 1253 | 4109 | 0 | 111 | 0 | 55 | 0 | 28 | 0 | 673 | 396 | 332 |
| SRR6450868 | 161 | 0 | 3681 | 0 | 0 | 0 | 97 | 0 | 0 | 0 | 0 | 0 | 0 | 26 | 0 | 0 | 1931 | 0 | 32 | 647 | 291 | 0 | 292 | 0 | 44 | 20 | 56 | 146 | 51 | 0 | 0 | 0 | 330 | 36 | 77 | 38 | 23 |
| SRR6450869 | 236 | 0 | 3907 | 0 | 0 | 0 | 187 | 0 | 0 | 0 | 0 | 0 | 0 | 30 | 93 | 0 | 1546 | 0 | 93 | 374 | 93 | 0 | 419 | 0 | 60 | 50 | 40 | 120 | 90 | 0 | 0 | 0 | 279 | 30 | 130 | 40 | 0 |
| SRR6453341 | 13 | 0 | 0 | 0 | 0 | 23 | 0 | 26 | 17 | 0 | 31 | 22 | 0 | 11 | 438 | 0 | 0 | 0 | 14 | 274 | 55 | 0 | 0 | 374 | 0 | 130 | 11 | 0 | 0 | 51 | 29 | 0 | 0 | 12 | 0 | 0 | 40 |
| SRR6453342 | 0 | 0 | 0 | 0 | 0 | 12 | 11 | 82 | 20 | 0 | 48 | 33 | 0 | 19 | 260 | 0 | 14 | 0 | 54 | 487 | 119 | 0 | 0 | 720 | 11 | 67 | 25 | 0 | 0 | 128 | 42 | 0 | 0 | 25 | 0 | 0 | 29 |
| SRR6453343 | 12 | 0 | 0 | 0 | 13 | 30 | 0 | 29 | 32 | 0 | 41 | 31 | 0 | 13 | 202 | 0 | 0 | 0 | 38 | 304 | 76 | 0 | 0 | 106 | 14 | 24 | 35 | 0 | 0 | 11 | 37 | 0 | 0 | 25 | 0 | 14 | 11 |
| SRR6453344 | 0 | 0 | 0 | 12 | 12 | 45 | 0 | 53 | 37 | 11 | 11 | 31 | 0 | 15 | 135 | 0 | 0 | 0 | 79 | 598 | 124 | 0 | 0 | 198 | 20 | 35 | 10 | 0 | 0 | 17 | 54 | 0 | 0 | 25 | 0 | 13 | 11 |
| SRR6453345 | 0 | 0 | 0 | 0 | 10 | 36 | 0 | 66 | 49 | 0 | 55 | 29 | 0 | 12 | 150 | 0 | 0 | 0 | 70 | 420 | 210 | 0 | 0 | 93 | 18 | 25 | 27 | 0 | 0 | 21 | 53 | 0 | 0 | 45 | 15 | 12 | 12 |
| SRR6453346 | 0 | 0 | 0 | 0 | 0 | 54 | 0 | 48 | 48 | 13 | 39 | 23 | 0 | 0 | 183 | 0 | 0 | 26 | 52 | 419 | 183 | 0 | 0 | 782 | 0 | 240 | 37 | 0 | 0 | 64 | 58 | 0 | 19 | 12 | 0 | 0 | 43 |
| SRR6453347 | 0 | 0 | 0 | 0 | 14 | 18 | 13 | 66 | 10 | 0 | 51 | 15 | 0 | 26 | 66 | 0 | 0 | 0 | 0 | 184 | 118 | 0 | 0 | 789 | 17 | 80 | 46 | 0 | 0 | 89 | 41 | 0 | 0 | 32 | 0 | 0 | 21 |
| SRR6453348 | 0 | 0 | 0 | 0 | 11 | 39 | 0 | 38 | 50 | 0 | 81 | 46 | 0 | 14 | 630 | 0 | 0 | 0 | 112 | 495 | 292 | 0 | 0 | 621 | 12 | 202 | 29 | 0 | 0 | 126 | 46 | 0 | 0 | 25 | 11 | 16 | 103 |
| SRR6453349 | 0 | 0 | 0 | 0 | 11 | 36 | 0 | 38 | 39 | 0 | 60 | 21 | 0 | 0 | 167 | 0 | 0 | 0 | 56 | 204 | 185 | 0 | 0 | 680 | 0 | 177 | 32 | 0 | 0 | 95 | 45 | 0 | 0 | 30 | 14 | 18 | 95 |
| SRR6453350 | 0 | 0 | 0 | 0 | 14 | 57 | 14 | 50 | 65 | 0 | 10 | 26 | 0 | 11 | 179 | 0 | 0 | 0 | 62 | 283 | 124 | 0 | 0 | 112 | 19 | 38 | 39 | 0 | 0 | 17 | 69 | 0 | 0 | 37 | 19 | 22 | 15 |
| SRR6453351 | 11 | 0 | 0 | 0 | 0 | 25 | 21 | 85 | 20 | 11 | 57 | 27 | 0 | 15 | 63 | 0 | 12 | 0 | 56 | 346 | 56 | 0 | 0 | 775 | 18 | 99 | 37 | 0 | 0 | 156 | 43 | 0 | 0 | 38 | 0 | 0 | 36 |
| SRR6453352 | 11 | 0 | 0 | 0 | 0 | 24 | 40 | 29 | 20 | 0 | 12 | 33 | 0 | 18 | 27 | 0 | 0 | 0 | 27 | 200 | 67 | 0 | 0 | 200 | 18 | 29 | 35 | 0 | 0 | 12 | 11 | 0 | 0 | 15 | 12 | 16 | 16 |
| SRR6453353 | 10 | 0 | 0 | 0 | 13 | 17 | 16 | 67 | 16 | 0 | 44 | 23 | 0 | 19 | 149 | 0 | 0 | 0 | 0 | 181 | 31 | 0 | 0 | 350 | 0 | 67 | 22 | 0 | 0 | 140 | 41 | 0 | 0 | 32 | 0 | 0 | 24 |
| SRR6453354 | 0 | 0 | 0 | 0 | 17 | 31 | 0 | 80 | 24 | 0 | 55 | 20 | 0 | 13 | 240 | 0 | 0 | 0 | 21 | 302 | 73 | 0 | 0 | 630 | 22 | 147 | 40 | 0 | 0 | 134 | 33 | 0 | 0 | 49 | 0 | 12 | 25 |
| SRR6453355 | 13 | 0 | 0 | 0 | 18 | 41 | 0 | 105 | 32 | 0 | 61 | 26 | 0 | 17 | 43 | 0 | 0 | 0 | 34 | 187 | 38 | 0 | 0 | 831 | 44 | 169 | 64 | 0 | 0 | 166 | 45 | 0 | 0 | 71 | 11 | 10 | 33 |
| SRR6453356 | 0 | 0 | 0 | 13 | 16 | 30 | 0 | 24 | 30 | 0 | 11 | 13 | 0 | 14 | 33 | 0 | 0 | 0 | 25 | 190 | 66 | 0 | 0 | 104 | 22 | 24 | 33 | 0 | 0 | 32 | 37 | 0 | 0 | 11 | 13 | 16 | 43 |
| SRR6453357 | 0 | 0 | 0 | 0 | 0 | 22 | 0 | 26 | 17 | 0 | 48 | 44 | 0 | 22 | 128 | 0 | 14 | 0 | 47 | 349 | 128 | 0 | 0 | 146 | 20 | 18 | 38 | 0 | 0 | 10 | 55 | 0 | 0 | 20 | 11 | 14 | 11 |
| SRR6453358 | 11 | 0 | 0 | 13 | 26 | 35 | 0 | 64 | 51 | 17 | 109 | 19 | 0 | 32 | 44 | 0 | 0 | 0 | 38 | 245 | 71 | 0 | 0 | 572 | 20 | 153 | 34 | 0 | 0 | 120 | 50 | 0 | 0 | 27 | 11 | 0 | 46 |
| SRR6453359 | 0 | 0 | 0 | 14 | 19 | 26 | 0 | 11 | 17 | 12 | 16 | 23 | 0 | 16 | 48 | 0 | 0 | 0 | 31 | 186 | 10 | 0 | 0 | 85 | 43 | 19 | 46 | 0 | 0 | 19 | 40 | 0 | 0 | 51 | 0 | 0 | 13 |
| SRR6453360 | 0 | 0 | 0 | 0 | 0 | 12 | 0 | 16 | 18 | 0 | 38 | 18 | 0 | 26 | 132 | 0 | 0 | 0 | 0 | 221 | 177 | 0 | 0 | 423 | 13 | 80 | 16 | 0 | 0 | 36 | 39 | 0 | 0 | 11 | 0 | 16 | 55 |
| SRR6453361 | 14 | 0 | 0 | 0 | 14 | 38 | 0 | 19 | 25 | 12 | 73 | 33 | 0 | 12 | 134 | 0 | 0 | 0 | 67 | 329 | 67 | 0 | 0 | 80 | 35 | 21 | 54 | 0 | 0 | 23 | 34 | 0 | 11 | 42 | 0 | 0 | 21 |
| SRR6453362 | 10 | 0 | 0 | 0 | 24 | 31 | 11 | 15 | 24 | 0 | 13 | 23 | 0 | 33 | 32 | 0 | 0 | 0 | 27 | 250 | 11 | 0 | 0 | 172 | 45 | 26 | 13 | 0 | 0 | 27 | 46 | 0 | 0 | 11 | 0 | 0 | 27 |
| SRR6453363 | 11 | 0 | 0 | 0 | 18 | 32 | 53 | 16 | 32 | 0 | 19 | 30 | 0 | 33 | 88 | 0 | 0 | 0 | 71 | 270 | 47 | 0 | 0 | 114 | 24 | 31 | 29 | 0 | 0 | 24 | 12 | 0 | 0 | 11 | 0 | 0 | 28 |
| SRR6453364 | 0 | 0 | 0 | 0 | 0 | 15 | 0 | 29 | 21 | 0 | 32 | 15 | 0 | 26 | 210 | 0 | 0 | 0 | 35 | 158 | 123 | 0 | 0 | 468 | 10 | 94 | 15 | 0 | 0 | 52 | 39 | 0 | 0 | 17 | 0 | 14 | 64 |
| SRR6453365 | 0 | 0 | 0 | 0 | 11 | 11 | 19 | 18 | 39 | 0 | 11 | 17 | 0 | 13 | 77 | 0 | 0 | 0 | 87 | 319 | 68 | 0 | 0 | 89 | 13 | 40 | 26 | 0 | 0 | 30 | 33 | 0 | 0 | 23 | 13 | 16 | 11 |
| SRR6453366 | 12 | 0 | 0 | 11 | 16 | 23 | 13 | 18 | 19 | 0 | 13 | 19 | 0 | 30 | 75 | 0 | 0 | 0 | 25 | 163 | 31 | 0 | 0 | 142 | 35 | 23 | 45 | 11 | 0 | 26 | 40 | 0 | 0 | 43 | 0 | 0 | 24 |
| SRR6453367 | 10 | 0 | 0 | 19 | 29 | 25 | 0 | 0 | 29 | 0 | 0 | 18 | 0 | 45 | 17 | 0 | 29 | 0 | 51 | 745 | 34 | 0 | 0 | 79 | 55 | 11 | 87 | 13 | 0 | 0 | 45 | 0 | 0 | 0 | 0 | 0 | 12 |
| SRR6453368 | 15 | 0 | 0 | 24 | 31 | 29 | 0 | 0 | 40 | 0 | 0 | 26 | 0 | 37 | 11 | 0 | 33 | 0 | 44 | 948 | 65 | 0 | 0 | 991 | 33 | 169 | 125 | 16 | 0 | 0 | 41 | 0 | 0 | 0 | 10 | 0 | 17 |
| SRR6453369 | 12 | 0 | 0 | 0 | 0 | 18 | 12 | 29 | 15 | 0 | 45 | 20 | 0 | 0 | 415 | 0 | 0 | 0 | 58 | 323 | 115 | 0 | 0 | 520 | 12 | 101 | 17 | 0 | 0 | 65 | 35 | 0 | 0 | 18 | 0 | 0 | 45 |
| SRR6453370 | 0 | 0 | 0 | 0 | 0 | 21 | 16 | 81 | 19 | 0 | 39 | 19 | 0 | 18 | 31 | 0 | 0 | 0 | 94 | 437 | 109 | 0 | 0 | 1116 | 17 | 83 | 40 | 0 | 0 | 186 | 44 | 0 | 0 | 24 | 0 | 0 | 48 |
| SRR6453371 | 11 | 0 | 0 | 0 | 0 | 16 | 26 | 500 | 13 | 0 | 43 | 27 | 0 | 20 | 91 | 0 | 0 | 0 | 26 | 234 | 39 | 0 | 0 | 305 | 0 | 55 | 18 | 0 | 0 | 435 | 43 | 0 | 0 | 130 | 0 | 0 | 37 |
| SRR6453372 | 14 | 0 | 0 | 0 | 12 | 0 | 14 | 155 | 0 | 0 | 13 | 12 | 0 | 10 | 82 | 0 | 0 | 0 | 0 | 218 | 0 | 0 | 0 | 121 | 10 | 12 | 17 | 13 | 0 | 219 | 30 | 0 | 0 | 60 | 0 | 0 | 24 |
| SRR6453373 | 12 | 0 | 0 | 0 | 13 | 22 | 0 | 367 | 19 | 0 | 44 | 25 | 13 | 16 | 251 | 0 | 0 | 0 | 0 | 151 | 117 | 0 | 0 | 401 | 14 | 108 | 20 | 0 | 0 | 520 | 54 | 0 | 0 | 231 | 11 | 17 | 69 |
| SRR6453374 | 16 | 0 | 0 | 0 | 0 | 16 | 0 | 384 | 11 | 0 | 41 | 25 | 0 | 12 | 83 | 0 | 0 | 0 | 31 | 114 | 93 | 0 | 0 | 367 | 12 | 72 | 15 | 0 | 0 | 627 | 25 | 0 | 0 | 224 | 0 | 0 | 33 |
| SRR6453375 | 0 | 0 | 0 | 0 | 0 | 23 | 12 | 435 | 27 | 0 | 41 | 27 | 0 | 28 | 193 | 0 | 0 | 0 | 12 | 108 | 145 | 0 | 0 | 406 | 0 | 106 | 13 | 0 | 0 | 544 | 44 | 0 | 0 | 203 | 0 | 0 | 30 |
| SRR6453376 | 0 | 0 | 0 | 0 | 34 | 48 | 26 | 61 | 32 | 0 | 48 | 12 | 18 | 15 | 77 | 0 | 0 | 0 | 26 | 359 | 154 | 0 | 0 | 78 | 14 | 17 | 104 | 0 | 0 | 83 | 41 | 0 | 0 | 26 | 0 | 0 | 25 |
| SRR6453377 | 0 | 0 | 0 | 0 | 12 | 16 | 0 | 194 | 11 | 0 | 38 | 0 | 0 | 16 | 32 | 0 | 0 | 0 | 21 | 264 | 53 | 0 | 0 | 316 | 13 | 71 | 23 | 0 | 0 | 388 | 31 | 0 | 0 | 116 | 11 | 10 | 49 |
| SRR6453378 | 0 | 0 | 0 | 0 | 0 | 16 | 0 | 72 | 18 | 0 | 41 | 15 | 0 | 26 | 102 | 0 | 0 | 0 | 85 | 271 | 68 | 0 | 0 | 89 | 12 | 16 | 20 | 0 | 0 | 138 | 38 | 0 | 0 | 36 | 0 | 10 | 11 |
| SRR6453379 | 13 | 0 | 0 | 0 | 0 | 19 | 13 | 195 | 13 | 0 | 33 | 19 | 10 | 31 | 396 | 0 | 0 | 25 | 26 | 488 | 158 | 0 | 0 | 137 | 20 | 21 | 31 | 0 | 0 | 181 | 12 | 0 | 0 | 51 | 0 | 13 | 18 |
| SRR6453380 | 10 | 0 | 0 | 0 | 0 | 0 | 0 | 206 | 0 | 0 | 40 | 20 | 0 | 16 | 151 | 0 | 0 | 0 | 13 | 189 | 63 | 0 | 0 | 337 | 12 | 39 | 22 | 0 | 0 | 445 | 37 | 0 | 0 | 123 | 0 | 0 | 20 |
| SRR6453381 | 13 | 0 | 0 | 15 | 17 | 17 | 0 | 40 | 10 | 0 | 12 | 15 | 0 | 20 | 90 | 0 | 0 | 0 | 18 | 145 | 54 | 0 | 0 | 98 | 12 | 15 | 23 | 0 | 0 | 11 | 23 | 0 | 0 | 19 | 0 | 11 | 38 |
| SRR6453382 | 10 | 0 | 0 | 13 | 21 | 16 | 11 | 54 | 15 | 11 | 110 | 43 | 0 | 29 | 74 | 0 | 13 | 0 | 42 | 191 | 32 | 0 | 0 | 734 | 27 | 75 | 48 | 11 | 0 | 73 | 35 | 0 | 0 | 31 | 0 | 0 | 37 |
| SRR6453383 | 0 | 0 | 0 | 0 | 0 | 24 | 0 | 10 | 19 | 0 | 10 | 15 | 0 | 19 | 128 | 0 | 0 | 0 | 0 | 255 | 69 | 0 | 0 | 71 | 17 | 13 | 22 | 0 | 0 | 19 | 33 | 0 | 0 | 25 | 0 | 14 | 54 |
| SRR6453384 | 0 | 0 | 0 | 10 | 0 | 16 | 0 | 12 | 19 | 0 | 13 | 16 | 0 | 33 | 129 | 0 | 0 | 0 | 53 | 144 | 45 | 0 | 0 | 88 | 11 | 15 | 17 | 0 | 0 | 16 | 10 | 0 | 0 | 20 | 0 | 0 | 40 |
| SRR6453385 | 15 | 0 | 0 | 11 | 15 | 11 | 33 | 11 | 12 | 11 | 14 | 49 | 0 | 22 | 67 | 0 | 0 | 0 | 45 | 871 | 156 | 0 | 0 | 86 | 11 | 12 | 14 | 0 | 0 | 20 | 31 | 0 | 0 | 10 | 0 | 0 | 24 |
| SRR6453386 | 1128 | 0 | 51 | 24 | 0 | 0 | 0 | 38 | 11 | 0 | 72 | 2560 | 11 | 17 | 228 | 0 | 0 | 0 | 91 | 91 | 592 | 0 | 0 | 324 | 0 | 56 | 0 | 11 | 0 | 36 | 42 | 0 | 0 | 29 | 0 | 0 | 25 |
| SRR6453387 | 1275 | 0 | 90 | 13 | 12 | 10 | 0 | 16 | 18 | 0 | 29 | 4070 | 20 | 12 | 88 | 0 | 25 | 31 | 53 | 528 | 194 | 0 | 0 | 344 | 0 | 22 | 15 | 0 | 0 | 22 | 18 | 0 | 0 | 25 | 0 | 0 | 13 |
| SRR6453388 | 2412 | 0 | 112 | 0 | 0 | 25 | 39 | 19 | 17 | 0 | 108 | 4971 | 35 | 49 | 13 | 0 | 40 | 70 | 39 | 656 | 2543 | 0 | 0 | 1092 | 36 | 107 | 74 | 14 | 0 | 38 | 97 | 0 | 29 | 44 | 0 | 17 | 41 |
| SRR6453389 | 1691 | 0 | 129 | 19 | 22 | 21 | 34 | 21 | 21 | 12 | 32 | 4015 | 16 | 13 | 124 | 0 | 28 | 79 | 11 | 191 | 2360 | 0 | 0 | 290 | 14 | 36 | 19 | 0 | 0 | 41 | 31 | 0 | 18 | 57 | 0 | 0 | 17 |
| SRR6453390 | 1779 | 0 | 433 | 0 | 17 | 16 | 30 | 14 | 10 | 13 | 33 | 4306 | 27 | 12 | 30 | 0 | 119 | 0 | 126 | 1933 | 2267 | 0 | 0 | 716 | 36 | 86 | 73 | 11 | 0 | 32 | 29 | 0 | 11 | 69 | 12 | 23 | 25 |
| SRR6453391 | 2553 | 0 | 225 | 0 | 11 | 16 | 179 | 24 | 16 | 0 | 61 | 3586 | 48 | 43 | 38 | 0 | 70 | 83 | 51 | 718 | 2973 | 0 | 0 | 840 | 30 | 106 | 56 | 14 | 0 | 42 | 41 | 0 | 13 | 44 | 0 | 13 | 56 |
| SRR6453392 | 1742 | 0 | 113 | 0 | 0 | 31 | 19 | 27 | 18 | 0 | 58 | 5403 | 49 | 41 | 45 | 0 | 108 | 57 | 64 | 526 | 1814 | 0 | 0 | 1278 | 54 | 153 | 120 | 10 | 0 | 46 | 70 | 0 | 14 | 74 | 0 | 11 | 33 |
| SRR6453393 | 2803 | 0 | 226 | 0 | 10 | 37 | 37 | 32 | 20 | 12 | 69 | 4804 | 97 | 25 | 28 | 0 | 104 | 27 | 93 | 972 | 2083 | 0 | 0 | 1149 | 102 | 187 | 169 | 10 | 0 | 79 | 102 | 0 | 25 | 148 | 0 | 12 | 37 |
| SRR6453394 | 1767 | 0 | 217 | 0 | 21 | 24 | 44 | 14 | 19 | 13 | 89 | 2843 | 22 | 35 | 87 | 0 | 36 | 368 | 22 | 763 | 807 | 0 | 0 | 1010 | 41 | 118 | 56 | 13 | 0 | 16 | 90 | 0 | 38 | 0 | 0 | 0 | 21 |
| SRR6453395 | 2335 | 0 | 142 | 0 | 11 | 25 | 10 | 23 | 22 | 0 | 88 | 6258 | 46 | 40 | 71 | 0 | 80 | 57 | 81 | 589 | 2620 | 0 | 0 | 1067 | 55 | 92 | 91 | 14 | 0 | 37 | 97 | 0 | 16 | 67 | 0 | 0 | 32 |
| SRR6453396 | 2628 | 0 | 256 | 12 | 15 | 17 | 12 | 14 | 14 | 11 | 56 | 3164 | 15 | 20 | 0 | 10 | 56 | 162 | 61 | 650 | 1349 | 0 | 0 | 518 | 35 | 59 | 44 | 14 | 0 | 27 | 42 | 0 | 19 | 43 | 0 | 0 | 12 |
| SRR6453397 | 2833 | 0 | 104 | 10 | 0 | 11 | 97 | 20 | 0 | 0 | 70 | 3520 | 31 | 36 | 121 | 0 | 36 | 0 | 24 | 315 | 1721 | 0 | 0 | 590 | 19 | 56 | 27 | 12 | 0 | 40 | 60 | 0 | 22 | 41 | 0 | 0 | 15 |
| SRR6453398 | 2358 | 0 | 258 | 10 | 10 | 15 | 0 | 13 | 14 | 0 | 37 | 3906 | 18 | 22 | 150 | 0 | 46 | 326 | 30 | 538 | 628 | 0 | 0 | 398 | 12 | 36 | 15 | 11 | 0 | 17 | 30 | 0 | 11 | 13 | 0 | 0 | 13 |
| SRR6453399 | 1372 | 0 | 57 | 17 | 12 | 25 | 0 | 15 | 17 | 0 | 23 | 2801 | 0 | 10 | 215 | 0 | 28 | 224 | 27 | 148 | 2622 | 0 | 0 | 231 | 23 | 34 | 30 | 0 | 0 | 21 | 46 | 0 | 38 | 14 | 0 | 13 | 29 |
| SRR6453400 | 2148 | 0 | 115 | 11 | 16 | 20 | 72 | 14 | 17 | 0 | 111 | 4992 | 17 | 54 | 129 | 0 | 60 | 160 | 0 | 229 | 2392 | 0 | 0 | 878 | 45 | 90 | 62 | 16 | 0 | 17 | 74 | 0 | 20 | 35 | 0 | 0 | 22 |
| SRR6453401 | 2499 | 0 | 232 | 0 | 0 | 11 | 21 | 16 | 22 | 0 | 24 | 6528 | 15 | 18 | 64 | 0 | 86 | 69 | 96 | 696 | 3340 | 0 | 0 | 372 | 19 | 48 | 34 | 12 | 0 | 10 | 25 | 0 | 12 | 19 | 0 | 15 | 17 |
| SRR6453402 | 1257 | 0 | 139 | 12 | 14 | 23 | 38 | 27 | 22 | 0 | 21 | 6745 | 21 | 18 | 76 | 0 | 42 | 19 | 76 | 1164 | 801 | 0 | 0 | 373 | 26 | 40 | 28 | 19 | 0 | 36 | 38 | 0 | 12 | 20 | 0 | 15 | 19 |
| SRR6453403 | 1269 | 0 | 89 | 0 | 10 | 18 | 43 | 59 | 26 | 0 | 42 | 4903 | 32 | 43 | 64 | 0 | 37 | 26 | 86 | 644 | 2693 | 0 | 0 | 865 | 24 | 97 | 42 | 10 | 0 | 128 | 91 | 0 | 17 | 68 | 0 | 0 | 35 |
| SRR6453404 | 1588 | 0 | 60 | 20 | 13 | 17 | 38 | 25 | 25 | 0 | 23 | 4111 | 19 | 31 | 164 | 0 | 11 | 32 | 13 | 429 | 3001 | 0 | 0 | 299 | 15 | 28 | 26 | 10 | 0 | 34 | 42 | 0 | 19 | 11 | 0 | 0 | 10 |
| SRR6453405 | 1739 | 0 | 86 | 0 | 13 | 14 | 0 | 11 | 16 | 0 | 21 | 4287 | 29 | 11 | 59 | 0 | 12 | 162 | 79 | 594 | 4121 | 0 | 0 | 264 | 18 | 32 | 30 | 0 | 0 | 15 | 21 | 0 | 13 | 28 | 0 | 24 | 21 |
| SRR6453406 | 1351 | 0 | 113 | 10 | 14 | 21 | 100 | 17 | 22 | 0 | 37 | 6371 | 20 | 22 | 131 | 0 | 37 | 78 | 46 | 761 | 1391 | 0 | 0 | 381 | 25 | 37 | 20 | 16 | 0 | 24 | 34 | 0 | 23 | 17 | 0 | 15 | 19 |
| SRR6453407 | 1603 | 0 | 91 | 12 | 11 | 17 | 19 | 54 | 15 | 0 | 49 | 3644 | 26 | 35 | 131 | 0 | 13 | 0 | 19 | 449 | 1608 | 0 | 0 | 770 | 18 | 85 | 28 | 0 | 0 | 80 | 92 | 0 | 48 | 32 | 0 | 0 | 17 |
| SRR6453408 | 1632 | 0 | 98 | 15 | 18 | 11 | 73 | 40 | 16 | 0 | 54 | 3549 | 11 | 39 | 316 | 0 | 14 | 0 | 24 | 607 | 3327 | 0 | 0 | 641 | 0 | 74 | 17 | 12 | 0 | 54 | 69 | 0 | 56 | 20 | 0 | 11 | 45 |
| SRR6453409 | 2159 | 0 | 68 | 0 | 0 | 19 | 29 | 14 | 30 | 0 | 18 | 4745 | 14 | 11 | 29 | 0 | 29 | 40 | 195 | 1562 | 2226 | 0 | 0 | 516 | 11 | 28 | 32 | 0 | 0 | 13 | 26 | 0 | 25 | 11 | 0 | 11 | 15 |
| SRR6453410 | 1187 | 0 | 107 | 12 | 13 | 14 | 32 | 29 | 25 | 0 | 19 | 4281 | 10 | 16 | 254 | 0 | 34 | 0 | 16 | 477 | 2321 | 0 | 0 | 363 | 16 | 28 | 25 | 13 | 0 | 38 | 40 | 0 | 18 | 15 | 0 | 0 | 12 |
| SRR6453411 | 1629 | 0 | 233 | 11 | 10 | 22 | 51 | 13 | 23 | 0 | 29 | 5231 | 28 | 12 | 51 | 0 | 18 | 18 | 51 | 669 | 2591 | 0 | 0 | 328 | 10 | 35 | 20 | 15 | 0 | 25 | 27 | 0 | 15 | 27 | 0 | 15 | 19 |
| SRR6453412 | 1844 | 0 | 37 | 0 | 0 | 0 | 36 | 18 | 15 | 0 | 23 | 1826 | 18 | 13 | 399 | 0 | 0 | 0 | 109 | 1271 | 617 | 0 | 0 | 195 | 0 | 30 | 10 | 0 | 0 | 19 | 29 | 0 | 0 | 21 | 0 | 0 | 14 |
| SRR6453413 | 1151 | 0 | 91 | 0 | 0 | 13 | 48 | 23 | 16 | 0 | 37 | 3062 | 11 | 17 | 48 | 0 | 28 | 51 | 119 | 882 | 357 | 0 | 0 | 409 | 0 | 43 | 11 | 0 | 0 | 32 | 23 | 0 | 0 | 37 | 0 | 0 | 18 |
| SRR6453414 | 1366 | 0 | 495 | 24 | 20 | 16 | 12 | 0 | 25 | 0 | 0 | 6095 | 15 | 61 | 0 | 0 | 181 | 72 | 184 | 1888 | 1728 | 0 | 0 | 788 | 68 | 105 | 126 | 22 | 0 | 0 | 63 | 0 | 15 | 0 | 0 | 0 | 0 |
| SRR6453415 | 1293 | 0 | 307 | 20 | 28 | 16 | 0 | 12 | 17 | 0 | 0 | 3347 | 12 | 50 | 28 | 0 | 109 | 62 | 112 | 1397 | 1216 | 0 | 0 | 1012 | 27 | 92 | 59 | 15 | 0 | 0 | 43 | 0 | 56 | 0 | 0 | 0 | 0 |
| SRR6453416 | 1091 | 0 | 1151 | 13 | 20 | 14 | 74 | 12 | 0 | 0 | 0 | 3534 | 23 | 33 | 0 | 0 | 237 | 0 | 262 | 1790 | 382 | 0 | 0 | 306 | 20 | 35 | 72 | 27 | 0 | 0 | 11 | 0 | 0 | 0 | 0 | 0 | 22 |
| SRR6453417 | 1432 | 0 | 280 | 32 | 34 | 23 | 0 | 16 | 33 | 0 | 0 | 5060 | 0 | 56 | 0 | 0 | 121 | 0 | 283 | 3113 | 940 | 0 | 0 | 209 | 52 | 20 | 26 | 23 | 0 | 0 | 12 | 0 | 22 | 0 | 10 | 0 | 12 |

### Supp Table S8.2.Columns_AM_to_BW

| | | | | | | | | | | | | | | | | | | | | | | | | | | | | | | | | | | | | | |
|---|---|---|---|---|---|---|---|---|---|---|---|---|---|---|---|---|---|---|---|---|---|---|---|---|---|---|---|---|---|---|---|---|---|---|---|---|---|
| SRR6453418 | 3025 | 0 | 226 | 13 | 27 | 30 | 75 | 88 | 33 | 0 | 83 | 5828 | 27 | 27 | 45 | 0 | 58 | 33 | 60 | 1239 | 3239 | 0 | 0 | 1174 | 45 | 169 | 64 | 0 | 0 | 155 | 70 | 0 | 20 | 123 | 0 | 0 | 30 |
| SRR6453419 | 954 | 0 | 223 | 17 | 25 | 20 | 0 | 12 | 27 | 0 | 39 | 3477 | 17 | 12 | 124 | 0 | 42 | 36 | 71 | 1262 | 1138 | 0 | 0 | 282 | 28 | 33 | 18 | 0 | 0 | 32 | 39 | 0 | 17 | 28 | 0 | 17 | 18 |
| SRR6453420 | 1681 | 0 | 118 | 20 | 11 | 12 | 25 | 20 | 16 | 0 | 43 | 4214 | 14 | 18 | 58 | 0 | 27 | 56 | 58 | 907 | 1113 | 0 | 0 | 262 | 16 | 24 | 12 | 11 | 0 | 41 | 36 | 0 | 17 | 31 | 0 | 0 | 11 |
| SRR6453421 | 683 | 0 | 111 | 15 | 18 | 20 | 324 | 271 | 28 | 11 | 105 | 4771 | 0 | 32 | 73 | 0 | 24 | 0 | 42 | 764 | 776 | 0 | 0 | 468 | 19 | 96 | 32 | 0 | 0 | 86 | 67 | 0 | 27 | 79 | 16 | 24 | 112 |
| SRR6453422 | 590 | 0 | 489 | 0 | 15 | 0 | 61 | 32 | 13 | 0 | 37 | 5940 | 13 | 35 | 68 | 0 | 36 | 0 | 61 | 943 | 376 | 0 | 0 | 346 | 0 | 33 | 13 | 18 | 0 | 37 | 33 | 0 | 0 | 48 | 0 | 10 | 37 |
| SRR6453423 | 910 | 0 | 299 | 13 | 25 | 0 | 55 | 51 | 11 | 13 | 113 | 8716 | 22 | 35 | 22 | 0 | 37 | 0 | 77 | 548 | 515 | 0 | 0 | 1038 | 12 | 48 | 15 | 10 | 0 | 82 | 28 | 0 | 0 | 204 | 0 | 0 | 18 |
| SRR6453424 | 1493 | 0 | 493 | 31 | 25 | 23 | 44 | 21 | 21 | 0 | 0 | 5388 | 19 | 43 | 0 | 0 | 157 | 22 | 220 | 2110 | 672 | 0 | 0 | 1022 | 52 | 140 | 154 | 18 | 0 | 0 | 67 | 0 | 16 | 0 | 11 | 0 | 0 |
| SRR6453425 | 1112 | 0 | 380 | 21 | 22 | 19 | 98 | 25 | 20 | 0 | 0 | 6422 | 13 | 46 | 0 | 0 | 78 | 28 | 204 | 1789 | 621 | 0 | 0 | 297 | 45 | 27 | 36 | 24 | 0 | 0 | 11 | 0 | 24 | 0 | 0 | 0 | 0 |
| SRR6453426 | 2060 | 0 | 1133 | 24 | 25 | 17 | 33 | 17 | 10 | 0 | 0 | 3683 | 12 | 69 | 0 | 0 | 143 | 29 | 172 | 1264 | 1921 | 0 | 0 | 727 | 26 | 109 | 54 | 16 | 0 | 0 | 39 | 0 | 12 | 0 | 0 | 0 | 20 |
| SRR6453427 | 1578 | 0 | 389 | 25 | 29 | 18 | 0 | 10 | 16 | 0 | 0 | 3838 | 11 | 66 | 0 | 0 | 150 | 79 | 159 | 1293 | 1886 | 0 | 0 | 1032 | 37 | 107 | 74 | 19 | 0 | 0 | 44 | 0 | 0 | 0 | 0 | 0 | 0 |
| SRR6453428 | 1684 | 0 | 762 | 23 | 25 | 18 | 11 | 10 | 17 | 0 | 12 | 4753 | 22 | 15 | 11 | 0 | 123 | 44 | 216 | 2672 | 1933 | 0 | 0 | 309 | 10 | 25 | 21 | 20 | 0 | 0 | 33 | 0 | 0 | 0 | 0 | 0 | 11 |
| SRR6454160 | 0 | 27 | 235 | 0 | 0 | 27 | 33 | 0 | 53 | 0 | 0 | 50 | 0 | 80 | 0 | 0 | 11 | 0 | 0 | 68 | 27 | 0 | 0 | 0 | 80 | 824 | 851 | 0 | 0 | 0 | 0 | 0 | 0 | 106 | 1223 | 160 | 558 |
| SRR6454161 | 15 | 23 | 338 | 0 | 0 | 0 | 12 | 14 | 0 | 0 | 0 | 126 | 0 | 33 | 12 | 0 | 0 | 0 | 0 | 47 | 0 | 0 | 0 | 0 | 14 | 194 | 199 | 84 | 262 | 0 | 0 | 0 | 0 | 73 | 314 | 15 | 107 |
| SRR6454162 | 0 | 0 | 0 | 0 | 0 | 0 | 0 | 0 | 0 | 0 | 0 | 0 | 0 | 0 | 0 | 0 | 0 | 0 | 0 | 155 | 464 | 0 | 0 | 0 | 0 | 827 | 337 | 0 | 0 | 0 | 0 | 0 | 0 | 0 | 0 | 0 | 31 |
| SRR6454163 | 19 | 0 | 596 | 0 | 0 | 10 | 12 | 268 | 10 | 0 | 0 | 208 | 0 | 72 | 0 | 0 | 0 | 0 | 0 | 77 | 11 | 0 | 0 | 0 | 0 | 423 | 826 | 10 | 10 | 10 | 62 | 0 | 0 | 10 | 4243 | 93 | 1662 |
| SRR6454164 | 0 | 0 | 0 | 0 | 0 | 123 | 0 | 0 | 31 | 0 | 0 | 0 | 0 | 0 | 0 | 0 | 0 | 0 | 0 | 755 | 1359 | 0 | 0 | 0 | 0 | 705 | 245 | 0 | 0 | 0 | 0 | 0 | 0 | 31 | 61 | 0 | 31 |
| SRR6454165 | 19 | 15 | 480 | 0 | 0 | 15 | 16 | 169 | 31 | 0 | 0 | 144 | 0 | 15 | 17 | 0 | 0 | 0 | 0 | 159 | 0 | 0 | 0 | 0 | 0 | 575 | 675 | 69 | 0 | 0 | 31 | 0 | 0 | 92 | 28 | 84 | 10 |
| SRR6454166 | 0 | 0 | 0 | 0 | 0 | 63 | 0 | 0 | 0 | 0 | 0 | 0 | 0 | 0 | 0 | 0 | 0 | 0 | 0 | 0 | 358 | 0 | 0 | 0 | 0 | 348 | 348 | 0 | 0 | 0 | 0 | 0 | 0 | 0 | 63 | 0 | 0 |
| SRR6454167 | 11 | 0 | 539 | 12 | 0 | 0 | 0 | 96 | 0 | 0 | 0 | 187 | 0 | 24 | 0 | 12 | 0 | 0 | 0 | 80 | 16 | 12 | 0 | 12 | 12 | 191 | 693 | 48 | 0 | 0 | 36 | 0 | 12 | 96 | 26 | 120 | 1399 |
| SRR6454169 | 11 | 0 | 295 | 0 | 10 | 10 | 20 | 328 | 41 | 10 | 0 | 132 | 0 | 62 | 0 | 10 | 0 | 0 | 0 | 40 | 0 | 0 | 0 | 0 | 21 | 503 | 287 | 0 | 0 | 0 | 21 | 10 | 10 | 51 | 29 | 123 | 1005 |
| SRR6454170 | 0 | 0 | 0 | 0 | 0 | 20 | 0 | 0 | 20 | 0 | 0 | 0 | 0 | 0 | 125 | 0 | 0 | 0 | 0 | 1374 | 1374 | 0 | 0 | 20 | 41 | 306 | 429 | 0 | 41 | 0 | 0 | 0 | 0 | 41 | 0 | 0 | 20 |
| SRR6454171 | 0 | 0 | 544 | 0 | 0 | 27 | 21 | 115 | 0 | 0 | 0 | 324 | 0 | 89 | 0 | 0 | 0 | 0 | 0 | 156 | 0 | 0 | 0 | 0 | 27 | 293 | 612 | 0 | 0 | 0 | 44 | 0 | 0 | 35 | 1047 | 44 | 550 |
| SRR6454172 | 0 | 0 | 0 | 0 | 0 | 33 | 88 | 0 | 0 | 0 | 0 | 0 | 0 | 16 | 0 | 0 | 0 | 0 | 88 | 971 | 706 | 0 | 0 | 0 | 0 | 590 | 459 | 0 | 0 | 0 | 0 | 0 | 0 | 49 | 0 | 0 | 0 |
| SRR6454173 | 16 | 0 | 799 | 0 | 0 | 0 | 22 | 203 | 0 | 0 | 0 | 102 | 0 | 16 | 19 | 0 | 0 | 0 | 0 | 230 | 0 | 0 | 0 | 0 | 16 | 187 | 242 | 0 | 0 | 0 | 16 | 0 | 0 | 0 | 1146 | 23 | 343 |
| SRR6454174 | 0 | 0 | 0 | 0 | 0 | 0 | 0 | 0 | 0 | 0 | 0 | 0 | 0 | 0 | 0 | 0 | 0 | 0 | 182 | 455 | 455 | 0 | 0 | 0 | 16 | 334 | 191 | 16 | 0 | 0 | 0 | 0 | 16 | 0 | 0 | 16 | 16 |
| SRR6454175 | 0 | 0 | 497 | 0 | 0 | 25 | 24 | 140 | 19 | 0 | 0 | 63 | 0 | 38 | 0 | 0 | 0 | 0 | 0 | 115 | 17 | 0 | 0 | 13 | 0 | 533 | 934 | 38 | 0 | 0 | 44 | 0 | 0 | 0 | 699 | 57 | 413 |
| SRR6454176 | 0 | 0 | 0 | 0 | 0 | 66 | 0 | 0 | 39 | 0 | 0 | 0 | 0 | 0 | 0 | 13 | 0 | 0 | 0 | 689 | 230 | 0 | 0 | 13 | 0 | 605 | 526 | 0 | 0 | 0 | 0 | 0 | 13 | 0 | 13 | 0 | 13 |
| SRR6454177 | 14 | 0 | 866 | 0 | 0 | 26 | 0 | 13 | 26 | 13 | 0 | 292 | 0 | 13 | 145 | 0 | 0 | 0 | 0 | 115 | 18 | 0 | 0 | 0 | 0 | 1003 | 2045 | 26 | 0 | 0 | 39 | 0 | 0 | 0 | 3244 | 78 | 1120 |
| SRR6454178 | 0 | 0 | 0 | 0 | 0 | 35 | 0 | 0 | 0 | 0 | 0 | 0 | 0 | 0 | 0 | 0 | 0 | 0 | 0 | 324 | 216 | 0 | 0 | 0 | 0 | 658 | 346 | 0 | 0 | 0 | 0 | 0 | 0 | 0 | 17 | 0 | 0 |
| SRR6454179 | 12 | 13 | 501 | 0 | 13 | 38 | 32 | 390 | 25 | 0 | 0 | 135 | 0 | 13 | 0 | 0 | 0 | 0 | 0 | 163 | 0 | 0 | 0 | 0 | 0 | 528 | 779 | 25 | 0 | 13 | 25 | 0 | 13 | 25 | 1508 | 0 | 440 |
| SRR6454180 | 0 | 0 | 0 | 0 | 0 | 24 | 0 | 0 | 0 | 0 | 0 | 0 | 0 | 0 | 0 | 0 | 0 | 0 | 53 | 1920 | 213 | 0 | 0 | 0 | 0 | 375 | 232 | 0 | 0 | 0 | 0 | 0 | 0 | 64 | 0 | 0 | 0 |
| SRR6454181 | 16 | 0 | 718 | 0 | 0 | 10 | 0 | 10 | 0 | 0 | 0 | 183 | 0 | 30 | 40 | 10 | 0 | 0 | 0 | 131 | 13 | 0 | 0 | 0 | 0 | 252 | 697 | 30 | 0 | 0 | 20 | 0 | 0 | 0 | 12 | 71 | 727 |
| SRR6454182 | 0 | 0 | 0 | 0 | 0 | 0 | 0 | 0 | 0 | 0 | 0 | 0 | 0 | 0 | 0 | 0 | 0 | 0 | 37 | 673 | 112 | 0 | 0 | 0 | 0 | 269 | 235 | 0 | 0 | 0 | 0 | 0 | 0 | 34 | 0 | 0 | 0 |
| SRR6454183 | 14 | 0 | 707 | 0 | 0 | 0 | 19 | 106 | 0 | 0 | 0 | 118 | 0 | 28 | 0 | 0 | 0 | 0 | 0 | 74 | 0 | 0 | 0 | 0 | 0 | 155 | 395 | 35 | 0 | 14 | 42 | 0 | 14 | 0 | 10 | 35 | 331 |
| SRR6454184 | 0 | 0 | 0 | 16 | 0 | 65 | 104 | 0 | 32 | 0 | 0 | 0 | 0 | 0 | 104 | 0 | 0 | 0 | 104 | 417 | 417 | 0 | 0 | 0 | 0 | 807 | 468 | 0 | 0 | 0 | 0 | 0 | 0 | 32 | 16 | 0 | 0 |
| SRR6454185 | 34 | 0 | 1106 | 0 | 0 | 16 | 100 | 190 | 0 | 0 | 0 | 119 | 0 | 0 | 13 | 0 | 0 | 0 | 13 | 206 | 22 | 0 | 0 | 0 | 0 | 252 | 16 | 0 | 0 | 0 | 20 | 0 | 0 | 0 | 18 | 23 | 10 |
| SRR6454186 | 0 | 0 | 0 | 0 | 0 | 0 | 0 | 0 | 16 | 0 | 0 | 0 | 0 | 0 | 0 | 0 | 0 | 0 | 120 | 1563 | 962 | 0 | 0 | 0 | 16 | 513 | 529 | 0 | 0 | 0 | 16 | 0 | 0 | 31 | 31 | 0 | 0 |
| SRR6454187 | 13 | 0 | 520 | 0 | 0 | 18 | 13 | 304 | 0 | 0 | 0 | 183 | 0 | 75 | 0 | 0 | 0 | 0 | 0 | 109 | 0 | 0 | 0 | 0 | 18 | 242 | 13 | 0 | 18 | 53 | 13 | 0 | 13 | 119 | 16 | 62 | 423 |
| SRR6454188 | 0 | 21 | 0 | 0 | 0 | 0 | 0 | 0 | 0 | 0 | 0 | 0 | 0 | 0 | 0 | 0 | 0 | 0 | 0 | 473 | 473 | 0 | 0 | 0 | 0 | 621 | 310 | 0 | 0 | 0 | 0 | 0 | 0 | 41 | 21 | 0 | 0 |
| SRR6454189 | 15 | 22 | 629 | 0 | 0 | 0 | 0 | 0 | 0 | 0 | 0 | 131 | 0 | 0 | 79 | 0 | 0 | 0 | 0 | 32 | 15 | 0 | 0 | 0 | 0 | 335 | 11 | 0 | 0 | 0 | 89 | 0 | 0 | 0 | 19 | 45 | 581 |
| SRR6454190 | 0 | 0 | 0 | 0 | 0 | 75 | 0 | 0 | 0 | 0 | 0 | 0 | 0 | 0 | 0 | 0 | 0 | 0 | 0 | 978 | 783 | 0 | 0 | 0 | 0 | 848 | 299 | 0 | 0 | 0 | 0 | 0 | 25 | 25 | 0 | 0 | 0 |
| SRR6454191 | 15 | 0 | 923 | 0 | 0 | 15 | 26 | 61 | 38 | 0 | 0 | 89 | 0 | 38 | 36 | 0 | 0 | 0 | 0 | 103 | 0 | 0 | 0 | 0 | 0 | 457 | 10 | 15 | 0 | 0 | 38 | 0 | 0 | 0 | 14 | 15 | 198 |
| SRR6454193 | 15 | 0 | 669 | 0 | 0 | 42 | 29 | 223 | 14 | 0 | 0 | 131 | 0 | 42 | 0 | 0 | 0 | 0 | 0 | 98 | 22 | 0 | 0 | 0 | 0 | 335 | 545 | 0 | 0 | 0 | 112 | 0 | 0 | 0 | 28 | 56 | 810 |
| SRR6454194 | 0 | 0 | 0 | 0 | 0 | 25 | 0 | 0 | 25 | 0 | 0 | 0 | 0 | 0 | 0 | 0 | 0 | 0 | 0 | 2059 | 588 | 25 | 0 | 0 | 25 | 838 | 345 | 0 | 25 | 0 | 0 | 0 | 25 | 25 | 0 | 0 | 0 |
| SRR6454195 | 0 | 0 | 522 | 0 | 0 | 71 | 13 | 89 | 36 | 0 | 0 | 96 | 0 | 0 | 0 | 0 | 0 | 0 | 0 | 67 | 11 | 0 | 0 | 0 | 0 | 1190 | 1297 | 0 | 0 | 0 | 160 | 0 | 0 | 18 | 2949 | 53 | 942 |
| SRR6454196 | 0 | 0 | 0 | 0 | 0 | 0 | 0 | 0 | 25 | 0 | 0 | 0 | 0 | 0 | 0 | 0 | 0 | 0 | 0 | 228 | 228 | 0 | 0 | 0 | 25 | 396 | 50 | 0 | 0 | 0 | 0 | 0 | 0 | 0 | 0 | 0 | 0 |
| SRR6454197 | 10 | 0 | 429 | 0 | 0 | 42 | 103 | 226 | 0 | 0 | 0 | 138 | 0 | 42 | 0 | 0 | 14 | 0 | 0 | 99 | 14 | 0 | 0 | 0 | 0 | 508 | 770 | 14 | 0 | 0 | 14 | 0 | 14 | 14 | 861 | 92 | 643 |
| SRR6454198 | 0 | 13 | 0 | 0 | 0 | 13 | 0 | 0 | 13 | 0 | 0 | 0 | 0 | 40 | 111 | 0 | 0 | 0 | 0 | 890 | 445 | 27 | 0 | 13 | 40 | 306 | 266 | 27 | 0 | 13 | 0 | 0 | 0 | 27 | 13 | 0 | 0 |
| SRR6454199 | 14 | 0 | 271 | 0 | 0 | 62 | 44 | 234 | 18 | 0 | 0 | 72 | 0 | 25 | 14 | 0 | 12 | 0 | 12 | 85 | 38 | 0 | 0 | 0 | 0 | 634 | 702 | 31 | 0 | 0 | 31 | 0 | 18 | 18 | 622 | 43 | 443 |
| SRR6454201 | 15 | 0 | 432 | 0 | 0 | 95 | 51 | 29 | 58 | 0 | 0 | 79 | 0 | 0 | 0 | 0 | 0 | 0 | 0 | 195 | 17 | 0 | 0 | 0 | 0 | 22 | 788 | 0 | 0 | 0 | 88 | 0 | 0 | 0 | 16 | 15 | 526 |
| SRR6454203 | 15 | 0 | 777 | 0 | 0 | 85 | 22 | 339 | 0 | 0 | 0 | 70 | 0 | 0 | 0 | 0 | 0 | 0 | 0 | 93 | 0 | 0 | 0 | 17 | 0 | 1101 | 1185 | 0 | 0 | 0 | 34 | 0 | 0 | 0 | 20 | 17 | 1236 |
| SRR6454205 | 16 | 12 | 585 | 0 | 0 | 49 | 25 | 123 | 12 | 0 | 0 | 157 | 0 | 61 | 0 | 0 | 0 | 0 | 0 | 104 | 30 | 12 | 0 | 12 | 0 | 258 | 799 | 0 | 0 | 0 | 25 | 0 | 0 | 37 | 2877 | 49 | 824 |
| SRR6454206 | 0 | 0 | 0 | 0 | 0 | 28 | 0 | 0 | 28 | 0 | 0 | 0 | 0 | 0 | 0 | 0 | 0 | 0 | 0 | 730 | 243 | 0 | 0 | 0 | 0 | 470 | 332 | 0 | 0 | 0 | 0 | 0 | 0 | 28 | 0 | 0 | 28 |
| SRR6454207 | 16 | 0 | 376 | 0 | 0 | 34 | 0 | 34 | 34 | 0 | 0 | 85 | 0 | 34 | 290 | 0 | 0 | 0 | 0 | 132 | 83 | 0 | 0 | 11 | 11 | 501 | 900 | 23 | 0 | 11 | 11 | 0 | 0 | 11 | 2983 | 91 | 1093 |
| SRR6454208 | 0 | 58 | 0 | 0 | 0 | 58 | 0 | 0 | 29 | 0 | 0 | 0 | 0 | 0 | 0 | 0 | 0 | 0 | 0 | 129 | 388 | 0 | 0 | 0 | 29 | 835 | 461 | 0 | 0 | 0 | 29 | 0 | 0 | 0 | 0 | 0 | 0 |
| SRR6454209 | 16 | 0 | 614 | 0 | 0 | 0 | 0 | 101 | 0 | 0 | 0 | 155 | 0 | 0 | 0 | 0 | 0 | 0 | 0 | 117 | 26 | 0 | 0 | 0 | 17 | 294 | 857 | 42 | 0 | 0 | 84 | 0 | 0 | 34 | 2513 | 67 | 681 |
| SRR6454211 | 20 | 0 | 443 | 0 | 0 | 33 | 21 | 141 | 16 | 0 | 0 | 121 | 0 | 33 | 0 | 11 | 0 | 0 | 0 | 70 | 0 | 0 | 0 | 0 | 0 | 515 | 732 | 11 | 0 | 16 | 49 | 0 | 11 | 43 | 1063 | 22 | 483 |
| SRR6454213 | 17 | 0 | 648 | 0 | 11 | 23 | 0 | 102 | 11 | 0 | 0 | 142 | 0 | 45 | 0 | 0 | 0 | 0 | 0 | 72 | 17 | 0 | 0 | 0 | 0 | 440 | 1287 | 23 | 11 | 0 | 23 | 0 | 11 | 11 | 2235 | 68 | 734 |
| SRR6454214 | 0 | 0 | 0 | 0 | 0 | 27 | 0 | 0 | 0 | 0 | 0 | 0 | 0 | 0 | 130 | 0 | 0 | 0 | 0 | 913 | 261 | 0 | 0 | 0 | 0 | 295 | 134 | 0 | 0 | 0 | 0 | 0 | 27 | 0 | 0 | 0 | 0 |
| SRR6454215 | 13 | 0 | 368 | 0 | 0 | 18 | 13 | 65 | 0 | 0 | 0 | 97 | 0 | 36 | 0 | 0 | 0 | 0 | 0 | 64 | 0 | 0 | 0 | 0 | 0 | 298 | 637 | 30 | 0 | 0 | 42 | 0 | 12 | 18 | 12 | 42 | 601 |
| SRR6454216 | 0 | 0 | 0 | 0 | 0 | 0 | 0 | 0 | 49 | 0 | 0 | 0 | 0 | 0 | 0 | 0 | 0 | 0 | 0 | 0 | 0 | 0 | 0 | 0 | 0 | 245 | 0 | 0 | 0 | 0 | 0 | 0 | 0 | 49 | 98 | 0 | 0 |
| SRR6454217 | 14 | 0 | 1162 | 0 | 0 | 17 | 27 | 25 | 0 | 0 | 0 | 75 | 0 | 42 | 17 | 0 | 0 | 0 | 0 | 72 | 58 | 0 | 0 | 0 | 0 | 415 | 798 | 0 | 0 | 0 | 42 | 0 | 17 | 0 | 20 | 108 | 1205 |
| SRR6454218 | 0 | 0 | 0 | 0 | 0 | 54 | 0 | 0 | 0 | 0 | 0 | 0 | 0 | 0 | 0 | 0 | 0 | 0 | 0 | 192 | 128 | 0 | 0 | 0 | 54 | 298 | 108 | 0 | 0 | 0 | 0 | 0 | 0 | 0 | 0 | 0 | 27 |
| SRR6454219 | 0 | 0 | 199 | 0 | 0 | 17 | 29 | 52 | 0 | 0 | 0 | 98 | 0 | 43 | 0 | 0 | 0 | 0 | 0 | 128 | 18 | 0 | 0 | 0 | 0 | 600 | 1147 | 35 | 0 | 0 | 17 | 0 | 52 | 0 | 991 | 61 | 504 |
| SRR6454220 | 0 | 22 | 0 | 0 | 0 | 22 | 0 | 0 | 0 | 0 | 0 | 0 | 0 | 0 | 0 | 0 | 0 | 0 | 0 | 103 | 103 | 0 | 0 | 0 | 45 | 357 | 67 | 0 | 0 | 0 | 89 | 0 | 0 | 45 | 22 | 0 | 0 |
| SRR6454221 | 11 | 0 | 343 | 0 | 0 | 11 | 26 | 68 | 0 | 0 | 0 | 114 | 0 | 45 | 23 | 0 | 15 | 0 | 0 | 186 | 27 | 0 | 0 | 11 | 0 | 632 | 1117 | 34 | 0 | 0 | 45 | 0 | 45 | 11 | 1906 | 135 | 1072 |
| SRR6454222 | 0 | 74 | 0 | 0 | 0 | 0 | 0 | 0 | 0 | 0 | 0 | 0 | 0 | 25 | 117 | 0 | 0 | 0 | 0 | 0 | 0 | 0 | 0 | 0 | 0 | 173 | 25 | 0 | 0 | 0 | 0 | 0 | 0 | 0 | 50 | 0 | 0 |
| SRR6454223 | 12 | 0 | 260 | 0 | 0 | 20 | 28 | 72 | 13 | 0 | 0 | 122 | 0 | 33 | 17 | 0 | 0 | 0 | 0 | 82 | 60 | 0 | 0 | 0 | 0 | 431 | 869 | 33 | 0 | 13 | 33 | 0 | 46 | 20 | 12 | 111 | 732 |
| SRR6454224 | 0 | 26 | 0 | 0 | 0 | 0 | 0 | 0 | 0 | 0 | 0 | 0 | 0 | 0 | 0 | 0 | 0 | 0 | 0 | 0 | 0 | 0 | 0 | 0 | 0 | 208 | 26 | 0 | 0 | 0 | 0 | 0 | 0 | 0 | 0 | 0 | 0 |
| SRR6454225 | 10 | 0 | 420 | 0 | 14 | 14 | 25 | 42 | 14 | 0 | 0 | 107 | 0 | 0 | 0 | 0 | 0 | 0 | 0 | 83 | 36 | 0 | 0 | 0 | 0 | 475 | 866 | 70 | 0 | 0 | 42 | 0 | 14 | 0 | 1061 | 126 | 698 |
| SRR6454226 | 0 | 24 | 0 | 0 | 0 | 0 | 0 | 0 | 0 | 0 | 0 | 0 | 0 | 0 | 104 | 0 | 0 | 0 | 0 | 104 | 208 | 0 | 0 | 0 | 0 | 166 | 0 | 24 | 0 | 0 | 0 | 0 | 0 | 0 | 0 | 0 | 0 |
| SRR6454227 | 13 | 0 | 235 | 0 | 0 | 49 | 54 | 320 | 12 | 0 | 0 | 77 | 0 | 53 | 11 | 0 | 0 | 0 | 0 | 248 | 34 | 0 | 0 | 0 | 12 | 14 | 14 | 20 | 0 | 16 | 49 | 0 | 20 | 0 | 904 | 57 | 426 |
| SRR6454228 | 0 | 35 | 0 | 0 | 17 | 35 | 0 | 0 | 0 | 0 | 0 | 0 | 0 | 0 | 0 | 0 | 0 | 0 | 0 | 0 | 233 | 0 | 0 | 0 | 0 | 260 | 52 | 0 | 0 | 0 | 0 | 0 | 0 | 35 | 0 | 0 | 0 |
| SRR6454229 | 0 | 0 | 466 | 0 | 0 | 17 | 0 | 0 | 33 | 0 | 0 | 119 | 0 | 17 | 10 | 17 | 0 | 0 | 0 | 77 | 20 | 0 | 0 | 0 | 0 | 433 | 1083 | 33 | 0 | 0 | 50 | 0 | 33 | 0 | 2016 | 100 | 1066 |
| SRR6454230 | 0 | 19 | 0 | 0 | 0 | 19 | 0 | 0 | 19 | 0 | 0 | 0 | 0 | 19 | 0 | 0 | 0 | 0 | 0 | 0 | 0 | 0 | 0 | 0 | 0 | 324 | 57 | 0 | 0 | 0 | 0 | 0 | 0 | 19 | 19 | 0 | 0 |
| SRR6454231 | 0 | 0 | 371 | 0 | 0 | 26 | 33 | 65 | 13 | 0 | 0 | 149 | 0 | 39 | 0 | 0 | 0 | 0 | 0 | 124 | 62 | 0 | 0 | 0 | 0 | 855 | 1256 | 0 | 0 | 0 | 52 | 0 | 26 | 0 | 1373 | 117 | 699 |
| SRR6454232 | 0 | 16 | 0 | 32 | 0 | 49 | 0 | 0 | 16 | 0 | 0 | 0 | 0 | 0 | 0 | 0 | 0 | 0 | 0 | 0 | 105 | 0 | 0 | 0 | 0 | 437 | 32 | 32 | 0 | 0 | 0 | 32 | 0 | 49 | 32 | 0 | 0 |
| SRR6454233 | 0 | 0 | 257 | 0 | 0 | 16 | 28 | 24 | 0 | 0 | 0 | 58 | 0 | 47 | 11 | 0 | 0 | 0 | 0 | 57 | 24 | 0 | 0 | 0 | 0 | 251 | 243 | 47 | 0 | 0 | 31 | 0 | 0 | 102 | 10 | 39 | 424 |
| SRR6454234 | 13 | 0 | 570 | 0 | 0 | 0 | 65 | 244 | 0 | 0 | 0 | 71 | 0 | 19 | 0 | 0 | 0 | 0 | 17 | 115 | 14 | 0 | 0 | 0 | 19 | 263 | 620 | 0 | 0 | 0 | 0 | 0 | 0 | 28 | 451 | 0 | 254 |
| SRR6454235 | 0 | 0 | 0 | 0 | 0 | 33 | 0 | 0 | 0 | 0 | 0 | 0 | 0 | 33 | 68 | 0 | 0 | 0 | 0 | 137 | 205 | 0 | 0 | 0 | 16 | 244 | 130 | 33 | 0 | 0 | 0 | 16 | 0 | 33 | 33 | 0 | 16 |
| SRR6454236 | 12 | 14 | 232 | 0 | 0 | 29 | 69 | 76 | 0 | 0 | 0 | 57 | 0 | 76 | 0 | 0 | 0 | 0 | 18 | 150 | 39 | 0 | 0 | 0 | 29 | 343 | 266 | 33 | 0 | 0 | 19 | 0 | 0 | 33 | 609 | 38 | 338 |
| SRR6454237 | 0 | 0 | 0 | 0 | 0 | 0 | 0 | 0 | 0 | 0 | 0 | 0 | 0 | 19 | 0 | 19 | 0 | 0 | 0 | 120 | 179 | 0 | 0 | 0 | 19 | 195 | 78 | 0 | 0 | 0 | 0 | 0 | 0 | 19 | 19 | 0 | 19 |
| SRR6454238 | 11 | 0 | 121 | 0 | 0 | 16 | 24 | 0 | 16 | 0 | 0 | 46 | 0 | 40 | 0 | 0 | 0 | 0 | 0 | 81 | 0 | 0 | 0 | 0 | 24 | 128 | 265 | 48 | 0 | 0 | 16 | 0 | 0 | 48 | 746 | 16 | 329 |
| SRR6454239 | 0 | 10 | 10 | 10 | 0 | 10 | 0 | 0 | 0 | 0 | 0 | 0 | 0 | 0 | 91 | 0 | 0 | 0 | 46 | 183 | 91 | 0 | 0 | 0 | 0 | 175 | 41 | 10 | 0 | 0 | 0 | 0 | 10 | 82 | 10 | 10 | 10 |
| SRR6454240 | 15 | 0 | 645 | 0 | 0 | 27 | 56 | 16 | 0 | 0 | 0 | 22 | 0 | 27 | 17 | 0 | 0 | 0 | 37 | 97 | 38 | 0 | 0 | 0 | 27 | 322 | 134 | 32 | 0 | 0 | 27 | 0 | 0 | 32 | 1177 | 59 | 484 |
| SRR6454241 | 0 | 12 | 0 | 0 | 0 | 0 | 0 | 0 | 12 | 0 | 0 | 0 | 0 | 23 | 0 | 0 | 0 | 0 | 0 | 54 | 109 | 0 | 0 | 0 | 12 | 139 | 35 | 0 | 0 | 0 | 0 | 0 | 0 | 46 | 0 | 0 | 0 |
| SRR6454242 | 12 | 0 | 172 | 0 | 0 | 10 | 60 | 10 | 0 | 0 | 0 | 50 | 0 | 70 | 13 | 0 | 0 | 0 | 27 | 144 | 10 | 0 | 0 | 0 | 15 | 155 | 255 | 25 | 0 | 0 | 10 | 0 | 0 | 55 | 11 | 50 | 346 |
| SRR6454244 | 0 | 0 | 223 | 0 | 0 | 28 | 14 | 0 | 0 | 0 | 0 | 67 | 0 | 41 | 105 | 0 | 0 | 0 | 0 | 101 | 31 | 0 | 0 | 0 | 0 | 633 | 661 | 14 | 0 | 0 | 34 | 0 | 21 | 21 | 12 | 117 | 723 |
| SRR6454245 | 0 | 27 | 0 | 0 | 0 | 0 | 0 | 0 | 0 | 0 | 0 | 0 | 0 | 0 | 0 | 0 | 0 | 0 | 0 | 0 | 0 | 0 | 0 | 0 | 0 | 220 | 55 | 0 | 0 | 0 | 0 | 0 | 0 | 0 | 27 | 27 | 0 |
| SRR6454246 | 0 | 0 | 596 | 0 | 0 | 50 | 43 | 75 | 0 | 0 | 0 | 68 | 0 | 33 | 14 | 0 | 0 | 0 | 0 | 126 | 0 | 0 | 0 | 0 | 0 | 606 | 656 | 25 | 0 | 0 | 66 | 0 | 25 | 0 | 1519 | 17 | 681 |
| SRR6454247 | 0 | 0 | 251 | 0 | 0 | 27 | 0 | 0 | 0 | 0 | 0 | 43 | 0 | 14 | 45 | 0 | 0 | 0 | 0 | 89 | 26 | 0 | 0 | 0 | 0 | 486 | 500 | 21 | 0 | 0 | 21 | 0 | 21 | 0 | 541 | 27 | 301 |
| SRR6454249 | 0 | 0 | 496 | 0 | 0 | 39 | 12 | 158 | 0 | 0 | 0 | 126 | 0 | 39 | 13 | 0 | 11 | 0 | 0 | 122 | 13 | 0 | 0 | 13 | 0 | 354 | 1838 | 39 | 13 | 0 | 0 | 0 | 53 | 13 | 971 | 39 | 564 |
| SRR6454251 | 0 | 0 | 1056 | 0 | 0 | 17 | 17 | 17 | 17 | 0 | 0 | 46 | 0 | 25 | 42 | 0 | 0 | 0 | 0 | 41 | 0 | 0 | 0 | 0 | 0 | 435 | 426 | 0 | 0 | 0 | 0 | 0 | 17 | 0 | 994 | 33 | 543 |
| SRR6454252 | 0 | 0 | 0 | 0 | 0 | 0 | 0 | 0 | 0 | 0 | 0 | 0 | 0 | 0 | 0 | 0 | 0 | 0 | 0 | 157 | 157 | 0 | 0 | 0 | 0 | 29 | 17 | 0 | 0 | 0 | 0 | 0 | 0 | 0 | 0 | 0 | 0 |
| SRR6454253 | 12 | 0 | 319 | 0 | 0 | 21 | 46 | 147 | 0 | 0 | 0 | 62 | 0 | 56 | 15 | 0 | 0 | 0 | 17 | 113 | 18 | 0 | 0 | 0 | 0 | 657 | 881 | 49 | 0 | 0 | 21 | 0 | 0 | 0 | 13 | 70 | 482 |
| SRR6454254 | 0 | 14 | 0 | 0 | 0 | 28 | 0 | 14 | 14 | 0 | 0 | 0 | 0 | 0 | 0 | 0 | 0 | 0 | 0 | 964 | 0 | 0 | 0 | 28 | 0 | 266 | 126 | 0 | 0 | 0 | 0 | 0 | 0 | 28 | 0 | 0 | 14 |
| SRR6454255 | 0 | 0 | 586 | 0 | 0 | 20 | 22 | 161 | 0 | 0 | 0 | 43 | 0 | 27 | 21 | 0 | 0 | 0 | 0 | 109 | 0 | 0 | 0 | 20 | 13 | 490 | 476 | 20 | 0 | 20 | 13 | 0 | 13 | 0 | 798 | 34 | 429 |
| SRR6454256 | 0 | 95 | 0 | 0 | 0 | 32 | 0 | 0 | 0 | 0 | 0 | 0 | 0 | 0 | 0 | 0 | 0 | 0 | 0 | 144 | 144 | 0 | 0 | 0 | 0 | 32 | 63 | 0 | 0 | 0 | 0 | 0 | 32 | 0 | 0 | 0 | 0 |
| SRR6454257 | 18 | 0 | 728 | 0 | 0 | 14 | 80 | 581 | 14 | 0 | 0 | 43 | 0 | 36 | 14 | 0 | 0 | 0 | 16 | 197 | 19 | 0 | 0 | 22 | 0 | 703 | 645 | 14 | 0 | 36 | 29 | 0 | 0 | 0 | 746 | 57 | 516 |
| SRR6454258 | 0 | 0 | 440 | 0 | 0 | 16 | 29 | 33 | 33 | 0 | 0 | 158 | 0 | 66 | 0 | 0 | 11 | 0 | 0 | 81 | 0 | 0 | 0 | 16 | 0 | 509 | 772 | 16 | 0 | 0 | 16 | 0 | 49 | 16 | 870 | 33 | 591 |
| SRR6454259 | 11 | 0 | 311 | 0 | 0 | 0 | 24 | 0 | 20 | 0 | 0 | 133 | 0 | 0 | 0 | 0 | 0 | 0 | 0 | 46 | 0 | 0 | 0 | 0 | 0 | 400 | 781 | 20 | 0 | 0 | 29 | 0 | 0 | 29 | 752 | 49 | 322 |
| SRR6454260 | 0 | 0 | 452 | 0 | 0 | 0 | 15 | 19 | 0 | 0 | 0 | 138 | 0 | 86 | 0 | 0 | 13 | 0 | 0 | 24 | 0 | 0 | 0 | 0 | 0 | 259 | 518 | 48 | 0 | 0 | 0 | 0 | 0 | 0 | 1045 | 48 | 546 |
| SRR6454261 | 0 | 0 | 322 | 0 | 0 | 13 | 52 | 51 | 19 | 0 | 0 | 122 | 0 | 102 | 0 | 0 | 0 | 0 | 0 | 74 | 0 | 0 | 0 | 0 | 0 | 441 | 504 | 13 | 0 | 0 | 13 | 0 | 13 | 13 | 536 | 26 | 300 |
| SRR6454262 | 0 | 13 | 195 | 0 | 0 | 13 | 31 | 13 | 0 | 0 | 0 | 76 | 0 | 76 | 0 | 0 | 0 | 0 | 0 | 22 | 0 | 0 | 0 | 0 | 13 | 139 | 304 | 0 | 0 | 0 | 38 | 0 | 13 | 0 | 443 | 25 | 177 |
| SRR6454263 | 16 | 13 | 618 | 0 | 0 | 19 | 47 | 0 | 0 | 0 | 0 | 48 | 0 | 25 | 18 | 0 | 0 | 0 | 0 | 62 | 34 | 0 | 0 | 0 | 0 | 348 | 202 | 19 | 0 | 0 | 0 | 0 | 13 | 0 | 594 | 70 | 291 |
| SRR6454264 | 0 | 18 | 217 | 0 | 0 | 0 | 35 | 18 | 0 | 18 | 0 | 64 | 0 | 18 | 0 | 0 | 0 | 0 | 0 | 47 | 10 | 0 | 0 | 0 | 0 | 178 | 250 | 54 | 0 | 0 | 18 | 0 | 0 | 18 | 571 | 89 | 303 |
| SRR6454265 | 0 | 0 | 116 | 0 | 0 | 15 | 20 | 0 | 0 | 0 | 0 | 35 | 0 | 30 | 0 | 0 | 0 | 0 | 0 | 92 | 0 | 0 | 0 | 0 | 15 | 180 | 135 | 15 | 0 | 0 | 15 | 0 | 15 | 0 | 706 | 45 | 345 |
| SRR6454266 | 0 | 0 | 349 | 0 | 0 | 0 | 29 | 13 | 0 | 0 | 0 | 64 | 0 | 63 | 12 | 0 | 0 | 0 | 0 | 13 | 10 | 0 | 0 | 0 | 0 | 331 | 291 | 18 | 0 | 0 | 22 | 0 | 0 | 13 | 654 | 40 | 233 |
| SRR6454267 | 0 | 0 | 781 | 0 | 0 | 21 | 49 | 24 | 0 | 0 | 0 | 59 | 0 | 18 | 0 | 0 | 0 | 0 | 0 | 71 | 27 | 0 | 0 | 0 | 0 | 300 | 237 | 16 | 0 | 0 | 18 | 0 | 0 | 0 | 770 | 18 | 407 |
| SRR6454268 | 0 | 0 | 257 | 0 | 0 | 0 | 14 | 0 | 0 | 0 | 0 | 168 | 0 | 0 | 0 | 0 | 0 | 0 | 0 | 70 | 19 | 0 | 0 | 0 | 0 | 375 | 589 | 0 | 0 | 0 | 27 | 0 | 0 | 0 | 696 | 54 | 401 |
| SRR6454269 | 0 | 0 | 223 | 0 | 0 | 106 | 28 | 26 | 0 | 0 | 0 | 100 | 0 | 53 | 0 | 0 | 0 | 0 | 0 | 41 | 0 | 26 | 0 | 0 | 0 | 714 | 608 | 79 | 26 | 0 | 0 | 0 | 0 | 0 | 688 | 26 | 132 |
| SRR6454270 | 0 | 0 | 178 | 0 | 0 | 0 | 11 | 28 | 0 | 0 | 0 | 51 | 14 | 56 | 12 | 0 | 0 | 0 | 0 | 70 | 12 | 0 | 0 | 0 | 14 | 139 | 209 | 0 | 0 | 0 | 14 | 0 | 0 | 28 | 376 | 14 | 293 |
| SRR6454271 | 11 | 0 | 401 | 0 | 0 | 32 | 120 | 150 | 22 | 0 | 0 | 109 | 0 | 0 | 55 | 0 | 0 | 0 | 0 | 337 | 273 | 0 | 0 | 41 | 0 | 541 | 805 | 13 | 0 | 0 | 29 | 0 | 16 | 29 | 11 | 162 | 891 |
| SRR6454272 | 16 | 0 | 407 | 0 | 0 | 103 | 61 | 460 | 24 | 0 | 0 | 86 | 0 | 32 | 0 | 0 | 15 | 0 | 16 | 206 | 106 | 0 | 0 | 40 | 16 | 1618 | 1801 | 0 | 16 | 111 | 63 | 0 | 0 | 190 | 1626 | 127 | 547 |
| SRR6454273 | 149 | 27 | 2433 | 0 | 0 | 44 | 383 | 333 | 53 | 0 | 0 | 966 | 0 | 0 | 52 | 0 | 28 | 0 | 46 | 190 | 335 | 0 | 0 | 44 | 0 | 12 | 10 | 0 | 0 | 0 | 18 | 0 | 0 | 35 | 26 | 124 | 14 |
| SRR6454274 | 208 | 0 | 1787 | 0 | 0 | 19 | 601 | 8084 | 57 | 0 | 0 | 1203 | 0 | 0 | 92 | 0 | 23 | 0 | 62 | 292 | 548 | 0 | 0 | 19 | 29 | 980 | 894 | 0 | 0 | 0 | 29 | 19 | 0 | 0 | 1550 | 57 | 361 |
| SRR6454275 | 20 | 0 | 744 | 0 | 0 | 32 | 102 | 202 | 48 | 0 | 0 | 143 | 0 | 43 | 0 | 0 | 0 | 0 | 0 | 166 | 39 | 11 | 0 | 27 | 0 | 874 | 815 | 48 | 32 | 0 | 48 | 0 | 11 | 16 | 12 | 37 | 469 |
| SRR6454276 | 27 | 0 | 1159 | 0 | 0 | 60 | 74 | 70 | 30 | 0 | 0 | 94 | 0 | 10 | 134 | 0 | 0 | 0 | 0 | 199 | 213 | 35 | 0 | 55 | 0 | 11 | 11 | 35 | 0 | 0 | 40 | 0 | 0 | 10 | 22 | 45 | 744 |
| SRR6454277 | 21 | 0 | 962 | 0 | 0 | 25 | 50 | 35 | 20 | 0 | 0 | 204 | 0 | 74 | 98 | 0 | 0 | 0 | 0 | 356 | 142 | 30 | 0 | 15 | 0 | 479 | 889 | 119 | 0 | 0 | 25 | 0 | 0 | 20 | 904 | 49 | 361 |
| SRR6454278 | 17 | 0 | 734 | 0 | 0 | 40 | 34 | 186 | 35 | 0 | 0 | 112 | 0 | 20 | 0 | 15 | 0 | 0 | 0 | 127 | 37 | 0 | 0 | 15 | 0 | 940 | 442 | 10 | 25 | 0 | 55 | 0 | 0 | 10 | 3185 | 55 | 1140 |
| SRR6454279 | 0 | 0 | 579 | 0 | 14 | 0 | 23 | 0 | 0 | 0 | 0 | 140 | 0 | 53 | 12 | 0 | 0 | 0 | 0 | 108 | 0 | 14 | 0 | 0 | 0 | 410 | 362 | 58 | 0 | 0 | 53 | 0 | 14 | 0 | 801 | 53 | 256 |
| SRR6454280 | 17 | 23 | 618 | 11 | 11 | 31 | 62 | 31 | 27 | 0 | 0 | 72 | 0 | 42 | 455 | 0 | 0 | 0 | 0 | 95 | 46 | 0 | 0 | 0 | 0 | 588 | 252 | 34 | 0 | 0 | 27 | 0 | 0 | 15 | 413 | 19 | 157 |
| SRR6454281 | 13 | 0 | 550 | 0 | 0 | 27 | 25 | 63 | 27 | 0 | 0 | 86 | 0 | 82 | 14 | 0 | 10 | 0 | 0 | 122 | 12 | 0 | 0 | 0 | 24 | 416 | 322 | 24 | 0 | 0 | 71 | 0 | 12 | 12 | 12 | 55 | 330 |
| SRR6454282 | 18 | 0 | 481 | 0 | 0 | 0 | 29 | 19 | 23 | 0 | 0 | 48 | 0 | 14 | 23 | 0 | 0 | 0 | 0 | 38 | 16 | 0 | 0 | 0 | 0 | 328 | 80 | 0 | 0 | 0 | 61 | 0 | 0 | 0 | 1310 | 14 | 370 |

### Supp Table S8.2.Columns_AM_to_BW

| | | | | | | | | | | | | | | | | | | | | | | | | | | | | | | | | | | | | | |
|---|---|---|---|---|---|---|---|---|---|---|---|---|---|---|---|---|---|---|---|---|---|---|---|---|---|---|---|---|---|---|---|---|---|---|---|---|---|
| SRR6454283 | 17 | 14 | 617 | 0 | 0 | 33 | 47 | 14 | 23 | 0 | 0 | 102 | 0 | 56 | 0 | 0 | 0 | 0 | 0 | 85 | 0 | 0 | 0 | 0 | 0 | 634 | 317 | 23 | 0 | 0 | 93 | 0 | 19 | 0 | 1889 | 61 | 513 |
| SRR6454284 | 11 | 0 | 980 | 0 | 0 | 32 | 35 | 22 | 0 | 0 | 0 | 39 | 0 | 22 | 25 | 0 | 0 | 0 | 0 | 89 | 24 | 0 | 0 | 0 | 16 | 594 | 297 | 22 | 11 | 0 | 86 | 0 | 11 | 0 | 2280 | 32 | 686 |
| SRR6454285 | 20 | 30 | 727 | 0 | 0 | 20 | 71 | 20 | 25 | 0 | 0 | 231 | 0 | 61 | 10 | 0 | 0 | 0 | 0 | 83 | 23 | 10 | 0 | 0 | 0 | 324 | 293 | 71 | 0 | 0 | 0 | 0 | 0 | 20 | 116 | 15 | 56 |
| SRR6454286 | 16 | 0 | 1086 | 0 | 0 | 39 | 88 | 84 | 39 | 0 | 0 | 29 | 0 | 18 | 12 | 0 | 0 | 0 | 0 | 171 | 79 | 0 | 0 | 0 | 0 | 840 | 354 | 15 | 0 | 0 | 63 | 0 | 21 | 15 | 11 | 93 | 699 |
| SRR6454287 | 234 | 0 | 2283 | 0 | 0 | 27 | 0 | 54 | 0 | 0 | 0 | 602 | 0 | 0 | 756 | 0 | 13 | 0 | 0 | 832 | 2269 | 0 | 0 | 27 | 0 | 21 | 16 | 0 | 0 | 0 | 0 | 0 | 0 | 0 | 30 | 54 | 12 |
| SRR6454288 | 251 | 41 | 1520 | 0 | 0 | 0 | 0 | 0 | 0 | 0 | 0 | 856 | 0 | 0 | 135 | 0 | 12 | 0 | 0 | 719 | 2517 | 0 | 0 | 41 | 0 | 702 | 1568 | 0 | 0 | 0 | 41 | 0 | 0 | 0 | 11 | 41 | 578 |
| SRR6454289 | 185 | 0 | 1776 | 0 | 0 | 17 | 32 | 118 | 0 | 0 | 0 | 1007 | 0 | 34 | 473 | 0 | 17 | 0 | 0 | 819 | 1008 | 0 | 0 | 34 | 34 | 824 | 706 | 0 | 0 | 0 | 0 | 0 | 0 | 0 | 1716 | 50 | 370 |
| SRR6454290 | 183 | 0 | 3536 | 0 | 0 | 123 | 21 | 145 | 33 | 0 | 0 | 584 | 0 | 22 | 646 | 11 | 35 | 0 | 0 | 1064 | 626 | 0 | 0 | 45 | 0 | 39 | 30 | 0 | 0 | 0 | 89 | 0 | 0 | 0 | 52 | 45 | 18 |
| SRR6454291 | 117 | 10 | 1588 | 0 | 20 | 50 | 22 | 10 | 0 | 0 | 0 | 705 | 0 | 0 | 1500 | 0 | 28 | 0 | 11 | 162 | 324 | 0 | 0 | 0 | 40 | 666 | 414 | 0 | 20 | 0 | 81 | 0 | 10 | 10 | 23 | 91 | 514 |
| SRR6454292 | 119 | 0 | 1492 | 0 | 0 | 59 | 43 | 0 | 0 | 0 | 0 | 721 | 0 | 20 | 414 | 0 | 21 | 0 | 0 | 157 | 357 | 0 | 0 | 0 | 20 | 16 | 255 | 0 | 0 | 0 | 29 | 0 | 49 | 0 | 32 | 88 | 412 |
| SRR6454293 | 89 | 0 | 1333 | 0 | 0 | 35 | 0 | 0 | 0 | 0 | 0 | 483 | 0 | 35 | 841 | 0 | 16 | 0 | 0 | 112 | 280 | 0 | 0 | 0 | 0 | 10 | 455 | 0 | 35 | 0 | 0 | 0 | 35 | 0 | 26 | 70 | 105 |
| SRR6454294 | 132 | 0 | 1664 | 0 | 0 | 24 | 23 | 24 | 0 | 0 | 0 | 676 | 0 | 49 | 871 | 0 | 17 | 0 | 0 | 252 | 458 | 0 | 0 | 24 | 0 | 854 | 513 | 24 | 0 | 0 | 98 | 0 | 24 | 24 | 3563 | 146 | 415 |
| SRR6454295 | 103 | 26 | 1886 | 0 | 0 | 39 | 0 | 0 | 13 | 0 | 0 | 488 | 0 | 13 | 1623 | 0 | 31 | 0 | 0 | 200 | 266 | 0 | 0 | 13 | 13 | 762 | 328 | 13 | 0 | 0 | 13 | 0 | 79 | 0 | 1865 | 105 | 722 |
| SRR6454296 | 106 | 0 | 1440 | 0 | 18 | 0 | 0 | 0 | 18 | 0 | 0 | 587 | 0 | 36 | 726 | 0 | 32 | 0 | 0 | 556 | 242 | 0 | 0 | 0 | 0 | 641 | 374 | 18 | 0 | 0 | 53 | 0 | 36 | 0 | 1692 | 178 | 374 |
| SRR6454298 | 11 | 0 | 1483 | 0 | 0 | 47 | 0 | 0 | 70 | 0 | 0 | 38 | 0 | 16 | 2173 | 0 | 0 | 0 | 0 | 0 | 259 | 0 | 0 | 0 | 16 | 613 | 47 | 62 | 0 | 0 | 85 | 0 | 0 | 85 | 2103 | 101 | 714 |
| SRR6454299 | 224 | 0 | 3142 | 0 | 0 | 0 | 0 | 0 | 14 | 0 | 0 | 696 | 0 | 21 | 508 | 0 | 16 | 0 | 0 | 10 | 100 | 0 | 0 | 0 | 0 | 136 | 0 | 0 | 0 | 0 | 0 | 0 | 0 | 0 | 19 | 14 | 221 |
| SRR6454300 | 127 | 0 | 2835 | 0 | 0 | 19 | 0 | 0 | 17 | 0 | 0 | 702 | 0 | 60 | 883 | 0 | 44 | 0 | 0 | 0 | 129 | 0 | 0 | 24 | 0 | 259 | 0 | 24 | 0 | 0 | 17 | 0 | 0 | 15 | 1729 | 63 | 227 |
| SRR6454301 | 20 | 0 | 724 | 0 | 0 | 17 | 58 | 17 | 13 | 0 | 0 | 151 | 0 | 47 | 64 | 0 | 0 | 0 | 0 | 196 | 77 | 26 | 0 | 0 | 0 | 601 | 759 | 81 | 17 | 0 | 64 | 0 | 17 | 0 | 2103 | 55 | 525 |
| SRR6454302 | 24 | 0 | 1419 | 0 | 0 | 27 | 104 | 0 | 0 | 0 | 0 | 64 | 0 | 23 | 46 | 0 | 0 | 0 | 49 | 221 | 152 | 12 | 0 | 0 | 0 | 595 | 462 | 31 | 12 | 0 | 47 | 0 | 27 | 0 | 1143 | 31 | 474 |
| SRR6454303 | 18 | 0 | 725 | 0 | 0 | 30 | 82 | 57 | 34 | 0 | 0 | 63 | 0 | 47 | 15 | 0 | 10 | 0 | 0 | 118 | 23 | 0 | 0 | 0 | 0 | 589 | 426 | 25 | 0 | 0 | 44 | 0 | 0 | 12 | 11 | 27 | 374 |
| SRR6454304 | 38 | 0 | 863 | 0 | 0 | 40 | 35 | 11 | 30 | 0 | 0 | 98 | 0 | 43 | 159 | 0 | 0 | 0 | 0 | 103 | 41 | 0 | 0 | 0 | 0 | 905 | 287 | 46 | 11 | 0 | 35 | 0 | 0 | 0 | 389 | 0 | 156 |
| SRR6454305 | 23 | 16 | 805 | 27 | 11 | 22 | 54 | 0 | 22 | 0 | 0 | 247 | 0 | 48 | 301 | 0 | 0 | 0 | 0 | 86 | 22 | 11 | 0 | 0 | 0 | 317 | 264 | 108 | 16 | 0 | 48 | 0 | 16 | 11 | 12 | 11 | 188 |
| SRR6454306 | 18 | 27 | 641 | 0 | 12 | 32 | 55 | 0 | 17 | 0 | 0 | 85 | 0 | 36 | 132 | 0 | 0 | 0 | 0 | 66 | 29 | 0 | 0 | 0 | 0 | 596 | 175 | 51 | 0 | 0 | 63 | 0 | 0 | 0 | 288 | 15 | 112 |
| SRR6454307 | 10 | 0 | 723 | 0 | 0 | 12 | 27 | 0 | 17 | 0 | 0 | 157 | 0 | 50 | 34 | 0 | 0 | 0 | 0 | 118 | 23 | 0 | 0 | 0 | 0 | 361 | 298 | 45 | 19 | 0 | 50 | 0 | 17 | 0 | 13 | 31 | 283 |
| SRR6454308 | 25 | 0 | 1146 | 0 | 0 | 30 | 58 | 19 | 28 | 0 | 0 | 45 | 0 | 19 | 45 | 0 | 0 | 0 | 0 | 92 | 34 | 0 | 0 | 0 | 11 | 566 | 155 | 25 | 30 | 0 | 83 | 0 | 0 | 0 | 740 | 44 | 248 |
| SRR6454309 | 68 | 13 | 829 | 0 | 0 | 89 | 143 | 16 | 79 | 0 | 0 | 160 | 0 | 48 | 518 | 0 | 15 | 0 | 0 | 358 | 568 | 0 | 0 | 13 | 0 | 1826 | 957 | 51 | 0 | 0 | 117 | 0 | 13 | 0 | 1138 | 41 | 469 |
| SRR6454310 | 29 | 0 | 1420 | 0 | 0 | 57 | 182 | 12 | 35 | 0 | 0 | 44 | 0 | 25 | 30 | 0 | 10 | 0 | 41 | 210 | 137 | 16 | 0 | 21 | 0 | 958 | 841 | 21 | 12 | 0 | 51 | 0 | 0 | 0 | 441 | 31 | 199 |
| SRR6454311 | 134 | 32 | 2008 | 0 | 0 | 65 | 0 | 87 | 22 | 0 | 0 | 486 | 0 | 54 | 4795 | 0 | 20 | 0 | 0 | 430 | 471 | 0 | 0 | 0 | 0 | 854 | 627 | 43 | 0 | 0 | 54 | 0 | 0 | 0 | 2574 | 54 | 627 |
| SRR6454312 | 124 | 29 | 1118 | 0 | 0 | 26 | 17 | 139 | 18 | 0 | 0 | 782 | 0 | 86 | 44 | 0 | 42 | 0 | 0 | 900 | 148 | 0 | 0 | 0 | 26 | 350 | 12 | 18 | 24 | 0 | 15 | 0 | 0 | 0 | 29 | 70 | 390 |
| SRR6454313 | 142 | 0 | 1726 | 0 | 0 | 128 | 0 | 0 | 46 | 0 | 0 | 587 | 0 | 35 | 3033 | 0 | 26 | 0 | 0 | 473 | 768 | 0 | 0 | 0 | 0 | 1161 | 557 | 0 | 12 | 0 | 12 | 0 | 12 | 0 | 1149 | 23 | 221 |
| SRR6454314 | 194 | 154 | 1406 | 0 | 15 | 46 | 18 | 11 | 0 | 0 | 0 | 661 | 0 | 69 | 42 | 23 | 21 | 0 | 0 | 378 | 84 | 0 | 0 | 0 | 0 | 378 | 11 | 0 | 0 | 0 | 0 | 0 | 0 | 0 | 17 | 69 | 216 |
| SRR6454315 | 90 | 0 | 1242 | 0 | 26 | 32 | 0 | 0 | 19 | 0 | 0 | 630 | 0 | 64 | 501 | 0 | 18 | 0 | 0 | 167 | 146 | 0 | 0 | 0 | 19 | 736 | 403 | 90 | 26 | 0 | 77 | 0 | 0 | 0 | 2881 | 102 | 493 |
| SRR6454316 | 101 | 0 | 1166 | 17 | 0 | 26 | 0 | 0 | 17 | 0 | 0 | 559 | 0 | 43 | 100 | 0 | 17 | 0 | 0 | 92 | 92 | 0 | 0 | 0 | 34 | 13 | 375 | 34 | 17 | 0 | 0 | 0 | 17 | 0 | 47 | 85 | 444 |
| SRR6454317 | 97 | 0 | 1349 | 0 | 0 | 73 | 0 | 0 | 33 | 0 | 0 | 444 | 0 | 13 | 532 | 0 | 19 | 0 | 0 | 225 | 148 | 0 | 0 | 0 | 0 | 1057 | 634 | 13 | 0 | 0 | 93 | 0 | 0 | 0 | 17 | 86 | 429 |
| SRR6454318 | 125 | 11 | 1595 | 11 | 21 | 54 | 0 | 0 | 11 | 0 | 0 | 482 | 0 | 11 | 183 | 0 | 21 | 0 | 0 | 287 | 69 | 0 | 0 | 0 | 11 | 868 | 439 | 21 | 11 | 0 | 32 | 0 | 32 | 0 | 15 | 32 | 450 |
| SRR6454319 | 101 | 0 | 1461 | 15 | 0 | 74 | 0 | 0 | 37 | 0 | 0 | 426 | 0 | 0 | 367 | 0 | 20 | 0 | 0 | 272 | 214 | 0 | 0 | 0 | 45 | 1236 | 492 | 15 | 0 | 0 | 37 | 0 | 22 | 0 | 4349 | 216 | 864 |
| SRR6454320 | 119 | 0 | 1834 | 23 | 34 | 46 | 0 | 0 | 23 | 0 | 0 | 508 | 0 | 11 | 654 | 0 | 17 | 0 | 0 | 164 | 227 | 0 | 0 | 0 | 0 | 14 | 11 | 0 | 11 | 0 | 57 | 0 | 46 | 0 | 65 | 320 | 12 |
| SRR6454321 | 167 | 15 | 1215 | 0 | 0 | 30 | 0 | 15 | 30 | 0 | 0 | 666 | 0 | 0 | 1217 | 0 | 18 | 0 | 0 | 330 | 811 | 0 | 0 | 0 | 15 | 831 | 634 | 0 | 15 | 0 | 15 | 0 | 15 | 0 | 1420 | 60 | 151 |
| SRR6454322 | 127 | 27 | 1479 | 0 | 0 | 66 | 0 | 27 | 53 | 0 | 0 | 629 | 0 | 40 | 1154 | 0 | 18 | 0 | 0 | 366 | 2027 | 0 | 0 | 13 | 0 | 771 | 572 | 0 | 0 | 0 | 13 | 0 | 0 | 0 | 705 | 53 | 253 |
| SRR6454323 | 174 | 107 | 1442 | 14 | 19 | 0 | 0 | 25 | 0 | 0 | 0 | 657 | 0 | 102 | 114 | 0 | 25 | 0 | 11 | 707 | 160 | 19 | 0 | 0 | 19 | 232 | 14 | 0 | 0 | 0 | 0 | 0 | 0 | 0 | 66 | 65 | 15 |
| SRR6454324 | 182 | 0 | 1908 | 0 | 0 | 85 | 0 | 12 | 36 | 0 | 0 | 588 | 0 | 0 | 16 | 0 | 22 | 0 | 0 | 323 | 189 | 0 | 0 | 12 | 0 | 1485 | 1207 | 18 | 0 | 0 | 18 | 0 | 18 | 0 | 3538 | 217 | 1189 |
| SRR6454325 | 12 | 0 | 806 | 0 | 0 | 78 | 107 | 140 | 55 | 0 | 0 | 119 | 0 | 20 | 28 | 0 | 12 | 0 | 0 | 255 | 39 | 0 | 0 | 16 | 0 | 16 | 15 | 23 | 0 | 0 | 117 | 0 | 0 | 0 | 1152 | 49 | 433 |
| SRR6454326 | 47 | 0 | 1005 | 0 | 0 | 41 | 70 | 0 | 28 | 0 | 0 | 52 | 0 | 37 | 687 | 0 | 18 | 0 | 0 | 500 | 320 | 13 | 0 | 16 | 0 | 749 | 500 | 31 | 16 | 0 | 51 | 0 | 0 | 0 | 704 | 39 | 276 |
| SRR6454331 | 102 | 0 | 1713 | 0 | 0 | 107 | 0 | 75 | 32 | 0 | 0 | 622 | 0 | 11 | 2052 | 0 | 23 | 0 | 0 | 477 | 927 | 0 | 0 | 11 | 0 | 1239 | 1068 | 0 | 0 | 0 | 53 | 0 | 0 | 21 | 1463 | 32 | 374 |
| SRR6454332 | 136 | 11 | 1378 | 0 | 0 | 34 | 0 | 102 | 11 | 0 | 0 | 862 | 0 | 34 | 660 | 0 | 25 | 0 | 0 | 227 | 873 | 0 | 0 | 0 | 56 | 564 | 745 | 0 | 11 | 0 | 0 | 0 | 0 | 0 | 17 | 113 | 373 |
| SRR6454335 | 182 | 0 | 1348 | 34 | 34 | 34 | 0 | 0 | 34 | 0 | 0 | 988 | 0 | 68 | 102 | 0 | 14 | 0 | 0 | 0 | 0 | 0 | 0 | 0 | 0 | 410 | 0 | 34 | 34 | 0 | 68 | 0 | 34 | 0 | 25 | 137 | 819 |
| SRR6454337 | 15 | 0 | 3383 | 0 | 0 | 0 | 13 | 0 | 0 | 0 | 0 | 67 | 0 | 59 | 3943 | 0 | 12 | 27 | 0 | 0 | 238 | 0 | 0 | 0 | 0 | 287 | 32 | 36 | 0 | 0 | 23 | 0 | 18 | 41 | 24 | 96 | 619 |
| SRR6454338 | 22 | 15 | 1162 | 0 | 0 | 10 | 14 | 0 | 0 | 0 | 0 | 43 | 0 | 21 | 1899 | 0 | 20 | 24 | 0 | 0 | 198 | 0 | 0 | 0 | 10 | 139 | 26 | 26 | 0 | 0 | 26 | 0 | 0 | 72 | 3874 | 170 | 783 |
| SRR6454339 | 27 | 0 | 1795 | 0 | 0 | 0 | 24 | 0 | 0 | 0 | 0 | 55 | 0 | 71 | 1632 | 0 | 17 | 116 | 0 | 0 | 146 | 0 | 0 | 0 | 0 | 86 | 12 | 34 | 0 | 0 | 31 | 0 | 18 | 46 | 1842 | 34 | 472 |
| SRR6454340 | 24 | 0 | 2012 | 0 | 0 | 17 | 40 | 0 | 17 | 0 | 0 | 40 | 0 | 22 | 40 | 0 | 13 | 0 | 0 | 17 | 69 | 0 | 0 | 0 | 11 | 269 | 28 | 34 | 0 | 0 | 174 | 0 | 28 | 0 | 331 | 34 | 213 |
| SRR6454341 | 15 | 0 | 1523 | 0 | 0 | 0 | 14 | 0 | 0 | 0 | 0 | 23 | 0 | 26 | 883 | 0 | 0 | 107 | 0 | 0 | 492 | 0 | 0 | 0 | 0 | 89 | 0 | 0 | 0 | 0 | 30 | 0 | 0 | 122 | 761 | 41 | 347 |
| SRR6454342 | 18 | 0 | 1229 | 0 | 0 | 0 | 36 | 0 | 14 | 0 | 0 | 17 | 0 | 17 | 530 | 0 | 0 | 169 | 0 | 0 | 375 | 0 | 0 | 0 | 0 | 40 | 0 | 0 | 0 | 0 | 12 | 0 | 12 | 132 | 283 | 42 | 141 |
| SRR6454343 | 16 | 0 | 1141 | 0 | 0 | 0 | 22 | 0 | 0 | 0 | 0 | 28 | 0 | 30 | 1942 | 0 | 10 | 78 | 0 | 0 | 890 | 0 | 0 | 0 | 0 | 212 | 37 | 0 | 0 | 0 | 44 | 0 | 17 | 27 | 741 | 53 | 419 |
| SRR6454345 | 287 | 0 | 2933 | 17 | 0 | 0 | 0 | 0 | 0 | 0 | 0 | 383 | 0 | 140 | 55 | 0 | 20 | 0 | 0 | 0 | 41 | 0 | 0 | 0 | 0 | 124 | 0 | 17 | 0 | 0 | 0 | 17 | 17 | 0 | 26 | 0 | 281 |
| SRR6454347 | 169 | 0 | 1804 | 26 | 19 | 49 | 0 | 0 | 24 | 0 | 0 | 608 | 0 | 161 | 28 | 0 | 16 | 0 | 0 | 0 | 0 | 0 | 0 | 0 | 0 | 1094 | 0 | 81 | 0 | 0 | 54 | 0 | 0 | 0 | 4572 | 139 | 968 |
| SRR6454348 | 108 | 0 | 1681 | 14 | 11 | 98 | 0 | 0 | 22 | 0 | 0 | 651 | 0 | 182 | 57 | 0 | 15 | 0 | 0 | 0 | 0 | 0 | 0 | 0 | 0 | 884 | 0 | 59 | 14 | 0 | 50 | 0 | 0 | 0 | 4355 | 154 | 714 |
| SRR6454349 | 212 | 11 | 1132 | 0 | 0 | 83 | 0 | 0 | 22 | 0 | 0 | 714 | 0 | 19 | 47 | 0 | 12 | 0 | 0 | 0 | 0 | 0 | 0 | 0 | 0 | 11 | 0 | 0 | 0 | 0 | 22 | 0 | 0 | 0 | 38 | 83 | 585 |
| SRR6454350 | 350 | 0 | 1254 | 66 | 19 | 19 | 0 | 0 | 19 | 0 | 0 | 786 | 0 | 47 | 28 | 0 | 0 | 0 | 0 | 0 | 0 | 0 | 0 | 0 | 0 | 266 | 0 | 38 | 0 | 0 | 19 | 0 | 0 | 0 | 19 | 47 | 199 |
| SRR6454351 | 23 | 0 | 762 | 0 | 0 | 0 | 0 | 0 | 20 | 0 | 0 | 29 | 0 | 15 | 221 | 0 | 0 | 22 | 0 | 0 | 0 | 0 | 0 | 0 | 24 | 390 | 0 | 0 | 20 | 0 | 15 | 0 | 0 | 0 | 439 | 20 | 171 |
| SRR6454352 | 11 | 0 | 1150 | 0 | 0 | 0 | 0 | 0 | 0 | 0 | 0 | 45 | 0 | 28 | 197 | 0 | 0 | 0 | 0 | 0 | 0 | 0 | 0 | 0 | 0 | 162 | 0 | 0 | 0 | 0 | 0 | 0 | 0 | 38 | 13 | 38 | 757 |
| SRR6454354 | 16 | 0 | 1497 | 0 | 0 | 0 | 0 | 0 | 0 | 0 | 0 | 54 | 0 | 27 | 55 | 0 | 0 | 30 | 0 | 0 | 0 | 0 | 0 | 0 | 0 | 143 | 0 | 0 | 0 | 0 | 0 | 0 | 0 | 66 | 10 | 19 | 418 |
| SRR6496456 | 35 | 0 | 577 | 0 | 0 | 0 | 23 | 0 | 0 | 23 | 28 | 3176 | 0 | 84 | 0 | 22 | 29 | 22 | 15 | 625 | 41 | 0 | 19 | 0 | 130 | 78 | 159 | 19 | 136 | 0 | 13 | 0 | 12 | 12 | 75 | 99 | 81 |
| SRR6496457 | 32 | 0 | 493 | 0 | 0 | 0 | 48 | 0 | 0 | 47 | 34 | 4664 | 0 | 84 | 0 | 16 | 27 | 14 | 34 | 1096 | 67 | 0 | 15 | 0 | 118 | 37 | 126 | 16 | 17 | 0 | 0 | 0 | 21 | 0 | 56 | 54 | 34 |
| SRR6496458 | 26 | 0 | 584 | 0 | 0 | 11 | 20 | 0 | 0 | 173 | 127 | 4924 | 0 | 116 | 0 | 20 | 29 | 15 | 24 | 703 | 82 | 0 | 10 | 0 | 182 | 131 | 507 | 13 | 196 | 0 | 0 | 0 | 29 | 12 | 126 | 151 | 117 |
| SRR6496459 | 29 | 0 | 667 | 0 | 0 | 0 | 0 | 0 | 0 | 113 | 79 | 2850 | 0 | 137 | 18 | 11 | 26 | 37 | 0 | 150 | 41 | 0 | 11 | 0 | 205 | 28 | 51 | 13 | 28 | 0 | 0 | 0 | 60 | 14 | 47 | 57 | 33 |
| SRR6496460 | 34 | 0 | 629 | 0 | 0 | 0 | 0 | 0 | 0 | 404 | 254 | 3241 | 0 | 149 | 0 | 21 | 22 | 15 | 0 | 389 | 42 | 0 | 10 | 0 | 182 | 113 | 204 | 0 | 234 | 0 | 12 | 0 | 45 | 32 | 109 | 182 | 124 |
| SRR6496461 | 111 | 0 | 1887 | 0 | 0 | 18 | 19 | 31 | 0 | 0 | 0 | 475 | 0 | 77 | 145 | 0 | 19 | 12 | 0 | 1522 | 1154 | 26 | 0 | 46 | 13 | 308 | 1612 | 0 | 19 | 0 | 13 | 0 | 0 | 13 | 61 | 0 | 38 |
| SRR6496462 | 85 | 0 | 1892 | 0 | 0 | 15 | 15 | 29 | 0 | 0 | 0 | 453 | 0 | 61 | 151 | 0 | 19 | 0 | 0 | 1533 | 1197 | 23 | 0 | 40 | 14 | 344 | 1526 | 11 | 15 | 0 | 0 | 0 | 0 | 14 | 55 | 0 | 40 |
| SRR6496463 | 69 | 0 | 1890 | 11 | 0 | 13 | 15 | 34 | 0 | 0 | 0 | 462 | 0 | 69 | 159 | 0 | 20 | 0 | 0 | 1514 | 1170 | 18 | 0 | 32 | 15 | 345 | 1537 | 0 | 13 | 0 | 0 | 0 | 0 | 13 | 57 | 0 | 30 |
| SRR6496464 | 71 | 0 | 1928 | 0 | 0 | 0 | 29 | 24 | 0 | 0 | 0 | 451 | 0 | 25 | 180 | 0 | 0 | 0 | 0 | 1719 | 1208 | 10 | 0 | 10 | 11 | 117 | 560 | 0 | 21 | 0 | 11 | 0 | 0 | 18 | 23 | 0 | 12 |
| SRR6496465 | 131 | 0 | 1765 | 0 | 0 | 24 | 0 | 21 | 0 | 0 | 0 | 539 | 0 | 79 | 201 | 0 | 20 | 34 | 14 | 1476 | 1060 | 17 | 0 | 31 | 0 | 366 | 1540 | 0 | 17 | 0 | 0 | 0 | 10 | 0 | 69 | 17 | 48 |
| SRR6498854 | 14 | 0 | 3509 | 0 | 0 | 10 | 22 | 37 | 20 | 0 | 0 | 165 | 0 | 10 | 142 | 0 | 0 | 23 | 0 | 3994 | 722 | 0 | 0 | 30 | 0 | 139 | 350 | 10 | 0 | 10 | 37 | 0 | 0 | 14 | 73 | 37 | 43 |
| SRR6498855 | 23 | 0 | 3931 | 0 | 0 | 17 | 35 | 29 | 12 | 0 | 0 | 438 | 0 | 0 | 104 | 0 | 0 | 0 | 0 | 3026 | 654 | 0 | 0 | 55 | 0 | 65 | 189 | 0 | 0 | 0 | 29 | 0 | 0 | 0 | 84 | 55 | 65 |
| SRR6498856 | 16 | 0 | 3863 | 0 | 0 | 23 | 48 | 36 | 16 | 0 | 0 | 228 | 0 | 0 | 126 | 0 | 0 | 0 | 23 | 3054 | 561 | 0 | 0 | 31 | 0 | 618 | 1816 | 0 | 0 | 14 | 35 | 0 | 0 | 20 | 439 | 22 | 235 |
| SRR6498857 | 21 | 0 | 3352 | 0 | 0 | 25 | 65 | 29 | 27 | 0 | 0 | 412 | 0 | 0 | 81 | 0 | 0 | 0 | 41 | 2548 | 885 | 0 | 0 | 10 | 0 | 73 | 203 | 0 | 12 | 12 | 37 | 0 | 0 | 39 | 73 | 39 | 68 |
| SRR6498858 | 17 | 0 | 3324 | 0 | 0 | 12 | 41 | 57 | 18 | 0 | 0 | 189 | 0 | 0 | 160 | 0 | 0 | 0 | 30 | 3385 | 1343 | 0 | 0 | 40 | 0 | 476 | 1248 | 0 | 12 | 20 | 24 | 0 | 0 | 44 | 650 | 40 | 325 |
| SRR6498860 | 16 | 0 | 3134 | 0 | 0 | 17 | 10 | 15 | 37 | 0 | 0 | 208 | 0 | 0 | 141 | 0 | 0 | 14 | 40 | 2855 | 293 | 0 | 0 | 19 | 0 | 650 | 1382 | 0 | 13 | 136 | 42 | 0 | 0 | 15 | 379 | 54 | 396 |
| SRR6498861 | 14 | 0 | 5066 | 0 | 0 | 20 | 36 | 0 | 16 | 0 | 0 | 190 | 0 | 13 | 85 | 0 | 0 | 0 | 0 | 690 | 436 | 0 | 0 | 16 | 0 | 373 | 406 | 0 | 36 | 0 | 23 | 0 | 0 | 33 | 916 | 26 | 684 |
| SRR6498862 | 12 | 0 | 2408 | 0 | 0 | 22 | 72 | 17 | 15 | 0 | 0 | 183 | 0 | 0 | 159 | 0 | 0 | 21 | 43 | 2520 | 753 | 0 | 0 | 12 | 0 | 115 | 234 | 0 | 12 | 0 | 11 | 0 | 0 | 10 | 85 | 30 | 64 |
| SRR6498863 | 13 | 0 | 3769 | 0 | 0 | 0 | 22 | 25 | 51 | 0 | 0 | 418 | 0 | 16 | 32 | 0 | 0 | 0 | 11 | 1366 | 430 | 0 | 0 | 0 | 0 | 388 | 931 | 14 | 25 | 0 | 11 | 0 | 0 | 110 | 556 | 46 | 559 |
| SRR6498864 | 12 | 0 | 3127 | 0 | 0 | 18 | 64 | 34 | 12 | 0 | 0 | 230 | 15 | 0 | 39 | 0 | 0 | 16 | 19 | 1579 | 1008 | 0 | 0 | 15 | 11 | 197 | 309 | 0 | 15 | 0 | 35 | 0 | 0 | 21 | 164 | 21 | 121 |
| SRR6498865 | 13 | 0 | 4274 | 18 | 26 | 0 | 112 | 13 | 18 | 0 | 0 | 366 | 0 | 26 | 0 | 0 | 0 | 0 | 56 | 1062 | 894 | 0 | 0 | 0 | 26 | 81 | 243 | 0 | 0 | 10 | 0 | 0 | 0 | 18 | 190 | 30 | 132 |
| SRR6498866 | 14 | 0 | 2857 | 0 | 0 | 22 | 52 | 17 | 21 | 0 | 0 | 213 | 0 | 0 | 155 | 0 | 0 | 0 | 26 | 4588 | 710 | 0 | 0 | 31 | 0 | 129 | 190 | 0 | 20 | 22 | 33 | 0 | 0 | 24 | 82 | 26 | 51 |
| SRR6498867 | 50 | 0 | 4032 | 0 | 0 | 0 | 40 | 49 | 0 | 0 | 0 | 206 | 0 | 0 | 167 | 0 | 0 | 0 | 72 | 3545 | 359 | 0 | 0 | 10 | 0 | 49 | 62 | 0 | 13 | 49 | 14 | 0 | 0 | 0 | 29 | 15 | 17 |
| SRR6498868 | 17 | 0 | 2630 | 0 | 0 | 15 | 175 | 90 | 15 | 0 | 0 | 167 | 0 | 0 | 128 | 0 | 0 | 11 | 105 | 5367 | 2130 | 0 | 0 | 32 | 0 | 458 | 569 | 0 | 21 | 87 | 19 | 0 | 0 | 15 | 258 | 24 | 139 |
| SRR6498869 | 32 | 0 | 3878 | 0 | 0 | 23 | 37 | 137 | 17 | 0 | 0 | 257 | 0 | 0 | 64 | 0 | 0 | 0 | 27 | 2980 | 549 | 0 | 0 | 23 | 0 | 428 | 702 | 0 | 13 | 157 | 25 | 0 | 0 | 43 | 398 | 23 | 240 |
| SRR6498870 | 26 | 25 | 3755 | 0 | 0 | 17 | 62 | 102 | 13 | 0 | 0 | 138 | 0 | 0 | 79 | 0 | 0 | 0 | 18 | 1910 | 519 | 0 | 0 | 14 | 0 | 336 | 303 | 0 | 69 | 38 | 30 | 0 | 0 | 17 | 486 | 19 | 175 |
| SRR6498871 | 55 | 20 | 2729 | 0 | 0 | 12 | 77 | 63 | 0 | 0 | 0 | 458 | 0 | 0 | 69 | 0 | 0 | 0 | 23 | 1810 | 990 | 0 | 0 | 11 | 0 | 265 | 414 | 0 | 11 | 145 | 21 | 0 | 0 | 55 | 298 | 40 | 203 |
| SRR6705804 | 59 | 0 | 243 | 0 | 0 | 0 | 31 | 0 | 0 | 0 | 0 | 163 | 0 | 0 | 23 | 0 | 0 | 43 | 162 | 85 | 39 | 0 | 0 | 0 | 16 | 11 | 0 | 0 | 0 | 0 | 0 | 0 | 0 | 0 | 12 | 0 | 0 |
| SRR6705805 | 15 | 0 | 107 | 0 | 0 | 0 | 0 | 0 | 0 | 0 | 0 | 92 | 0 | 0 | 39 | 0 | 0 | 191 | 52 | 117 | 39 | 0 | 0 | 0 | 27 | 0 | 0 | 0 | 0 | 0 | 0 | 0 | 0 | 0 | 0 | 0 | 0 |
| SRR6705806 | 0 | 0 | 0 | 0 | 0 | 0 | 0 | 0 | 0 | 0 | 0 | 0 | 0 | 0 | 0 | 0 | 0 | 0 | 0 | 0 | 0 | 0 | 0 | 0 | 0 | 0 | 0 | 0 | 0 | 0 | 0 | 0 | 0 | 0 | 0 | 0 | 0 |
| SRR6705808 | 268 | 0 | 205 | 0 | 0 | 43 | 0 | 0 | 75 | 0 | 0 | 164 | 0 | 0 | 406 | 0 | 0 | 0 | 0 | 0 | 45 | 0 | 0 | 0 | 54 | 41 | 0 | 0 | 0 | 0 | 0 | 0 | 0 | 96 | 123 | 43 | 86 |
| SRR6705809 | 76 | 0 | 409 | 0 | 0 | 0 | 0 | 0 | 0 | 0 | 0 | 398 | 0 | 0 | 1768 | 0 | 11 | 0 | 11 | 32 | 42 | 0 | 0 | 0 | 10 | 45 | 0 | 0 | 0 | 0 | 35 | 0 | 0 | 15 | 41 | 70 | 14 |
| SRR6705810 | 69 | 0 | 931 | 0 | 0 | 44 | 0 | 0 | 15 | 0 | 0 | 130 | 15 | 0 | 44 | 0 | 0 | 0 | 33 | 0 | 196 | 0 | 0 | 0 | 51 | 26 | 51 | 0 | 0 | 0 | 15 | 0 | 0 | 0 | 13 | 29 | 102 |
| SRR6705811 | 10 | 0 | 244 | 0 | 0 | 0 | 0 | 0 | 0 | 0 | 0 | 20 | 0 | 0 | 35 | 0 | 0 | 0 | 0 | 0 | 36 | 0 | 0 | 0 | 14 | 106 | 0 | 0 | 0 | 0 | 0 | 0 | 0 | 0 | 0 | 0 | 0 |
| SRR6705812 | 32 | 0 | 246 | 0 | 0 | 0 | 60 | 0 | 0 | 0 | 0 | 225 | 0 | 0 | 68 | 0 | 0 | 232 | 51 | 102 | 77 | 0 | 0 | 0 | 44 | 0 | 0 | 0 | 0 | 0 | 0 | 0 | 0 | 0 | 0 | 0 | 0 |
| SRR6705813 | 51 | 0 | 226 | 0 | 0 | 0 | 0 | 0 | 0 | 0 | 0 | 165 | 0 | 0 | 31 | 0 | 0 | 32 | 194 | 102 | 82 | 0 | 0 | 0 | 44 | 16 | 18 | 0 | 0 | 0 | 0 | 0 | 0 | 0 | 0 | 0 | 0 |
| SRR6705814 | 206 | 0 | 339 | 0 | 0 | 23 | 27 | 0 | 0 | 0 | 0 | 163 | 10 | 0 | 29 | 0 | 0 | 21 | 0 | 0 | 100 | 0 | 0 | 0 | 0 | 48 | 97 | 0 | 0 | 0 | 27 | 0 | 0 | 47 | 0 | 0 | 0 |
| SRR6705815 | 181 | 0 | 269 | 0 | 0 | 100 | 57 | 0 | 35 | 0 | 0 | 296 | 43 | 0 | 114 | 0 | 0 | 129 | 0 | 22 | 470 | 0 | 0 | 0 | 22 | 1237 | 174 | 0 | 0 | 0 | 17 | 0 | 0 | 65 | 117 | 48 | 87 |
| SRR6705816 | 91 | 0 | 551 | 0 | 0 | 0 | 41 | 0 | 0 | 0 | 0 | 99 | 12 | 0 | 78 | 0 | 0 | 67 | 0 | 20 | 322 | 0 | 0 | 0 | 0 | 198 | 51 | 0 | 0 | 0 | 0 | 0 | 0 | 0 | 44 | 14 | 39 |
| SRR6705817 | 52 | 0 | 1138 | 0 | 0 | 0 | 645 | 0 | 0 | 0 | 0 | 137 | 0 | 0 | 271 | 0 | 16 | 0 | 135 | 98 | 378 | 0 | 0 | 0 | 31 | 131 | 15 | 0 | 20 | 0 | 0 | 0 | 0 | 13 | 108 | 38 | 58 |
| SRR6705818 | 134 | 0 | 532 | 0 | 0 | 37 | 50 | 0 | 12 | 0 | 0 | 114 | 0 | 0 | 81 | 0 | 17 | 0 | 0 | 101 | 512 | 0 | 0 | 17 | 37 | 112 | 14 | 0 | 0 | 15 | 47 | 0 | 0 | 0 | 13 | 0 | 30 |
| SRR6705819 | 79 | 0 | 434 | 0 | 0 | 28 | 143 | 41 | 90 | 0 | 0 | 135 | 0 | 0 | 130 | 0 | 15 | 15 | 37 | 200 | 644 | 0 | 0 | 0 | 62 | 478 | 51 | 0 | 21 | 74 | 13 | 0 | 0 | 18 | 16 | 39 | 59 |
| SRR6705820 | 264 | 0 | 667 | 0 | 0 | 0 | 69 | 0 | 0 | 0 | 0 | 124 | 0 | 0 | 176 | 0 | 0 | 15 | 698 | 1333 | 1166 | 0 | 0 | 0 | 12 | 130 | 21 | 0 | 0 | 0 | 0 | 0 | 0 | 0 | 13 | 0 | 10 |
| SRR6705821 | 178 | 0 | 931 | 0 | 0 | 104 | 46 | 0 | 26 | 0 | 0 | 205 | 0 | 0 | 71 | 0 | 13 | 11 | 39 | 82 | 300 | 0 | 0 | 14 | 32 | 1507 | 92 | 0 | 0 | 0 | 43 | 0 | 0 | 0 | 63 | 35 | 66 |
| SRR6705822 | 313 | 0 | 87 | 0 | 0 | 10 | 270 | 0 | 0 | 0 | 0 | 111 | 14 | 0 | 984 | 0 | 0 | 20 | 111 | 611 | 1643 | 0 | 0 | 0 | 17 | 66 | 14 | 0 | 0 | 0 | 30 | 0 | 0 | 0 | 48 | 16 | 28 |
| SRR6705823 | 221 | 0 | 669 | 0 | 0 | 91 | 243 | 0 | 45 | 0 | 0 | 332 | 0 | 0 | 375 | 0 | 26 | 30 | 100 | 497 | 1966 | 0 | 0 | 0 | 13 | 1058 | 96 | 0 | 79 | 0 | 32 | 0 | 0 | 0 | 87 | 51 | 58 |
| SRR6705824 | 46 | 0 | 431 | 0 | 0 | 0 | 0 | 0 | 0 | 0 | 0 | 83 | 0 | 0 | 137 | 0 | 0 | 14 | 0 | 0 | 39 | 0 | 0 | 0 | 0 | 58 | 0 | 0 | 0 | 0 | 37 | 0 | 0 | 0 | 59 | 0 | 14 |
| SRR6705825 | 13 | 0 | 571 | 0 | 0 | 0 | 0 | 0 | 84 | 0 | 0 | 34 | 0 | 23 | 268 | 0 | 0 | 12 | 0 | 0 | 23 | 0 | 0 | 0 | 35 | 211 | 0 | 0 | 147 | 0 | 104 | 0 | 0 | 32 | 185 | 58 | 185 |
| SRR6705826 | 16 | 0 | 352 | 0 | 0 | 0 | 0 | 0 | 0 | 0 | 0 | 26 | 0 | 0 | 34 | 0 | 0 | 0 | 0 | 0 | 22 | 0 | 0 | 0 | 0 | 66 | 0 | 0 | 0 | 0 | 0 | 0 | 0 | 21 | 49 | 0 | 17 |
| SRR6705827 | 37 | 0 | 1945 | 0 | 0 | 0 | 0 | 0 | 201 | 0 | 0 | 102 | 0 | 0 | 16 | 0 | 0 | 31 | 0 | 0 | 13 | 0 | 0 | 0 | 25 | 19 | 0 | 0 | 17 | 0 | 76 | 0 | 0 | 34 | 185 | 101 | 59 |
| SRR6705828 | 27 | 0 | 184 | 0 | 0 | 0 | 0 | 0 | 22 | 0 | 0 | 72 | 0 | 0 | 1736 | 0 | 0 | 58 | 0 | 0 | 39 | 0 | 0 | 0 | 33 | 37 | 0 | 0 | 0 | 0 | 65 | 0 | 0 | 0 | 19 | 17 | 0 |
| SRR6705829 | 0 | 0 | 13420 | 0 | 0 | 0 | 0 | 0 | 28 | 0 | 0 | 125 | 0 | 14 | 0 | 0 | 14 | 45 | 0 | 0 | 0 | 0 | 0 | 0 | 0 | 18 | 0 | 0 | 0 | 0 | 14 | 0 | 0 | 0 | 101 | 99 | 73 |
| SRR6705830 | 133 | 0 | 273 | 0 | 0 | 12 | 32 | 0 | 0 | 0 | 0 | 328 | 0 | 0 | 0 | 0 | 0 | 0 | 28 | 52 | 42 | 0 | 0 | 0 | 39 | 42 | 17 | 0 | 0 | 0 | 0 | 0 | 0 | 14 | 0 | 0 | 0 |
| SRR6705831 | 109 | 0 | 349 | 0 | 0 | 11 | 253 | 0 | 0 | 0 | 0 | 310 | 0 | 0 | 0 | 0 | 0 | 41 | 253 | 981 | 238 | 0 | 0 | 0 | 28 | 162 | 37 | 0 | 0 | 0 | 11 | 0 | 0 | 0 | 12 | 14 | 15 |
| SRR6705832 | 121 | 0 | 340 | 0 | 0 | 0 | 30 | 0 | 0 | 0 | 0 | 137 | 0 | 0 | 0 | 0 | 0 | 24 | 315 | 305 | 108 | 0 | 0 | 0 | 11 | 30 | 0 | 0 | 0 | 0 | 0 | 0 | 0 | 0 | 0 | 0 | 0 |
| SRR6705833 | 49 | 0 | 334 | 0 | 0 | 0 | 36 | 0 | 0 | 0 | 0 | 203 | 0 | 0 | 0 | 0 | 0 | 11 | 127 | 155 | 79 | 0 | 0 | 0 | 34 | 106 | 10 | 0 | 0 | 0 | 0 | 0 | 0 | 0 | 19 | 0 | 13 |
| SRR6705834 | 55 | 0 | 717 | 0 | 0 | 34 | 53 | 0 | 55 | 0 | 0 | 430 | 0 | 0 | 86 | 0 | 45 | 342 | 47 | 76 | 792 | 0 | 0 | 0 | 29 | 378 | 19 | 0 | 48 | 0 | 11 | 0 | 0 | 0 | 22 | 10 | 44 |
| SRR6705835 | 160 | 0 | 563 | 0 | 0 | 66 | 41 | 0 | 23 | 0 | 0 | 70 | 26 | 0 | 124 | 0 | 0 | 12 | 18 | 20 | 252 | 0 | 0 | 0 | 37 | 89 | 13 | 0 | 0 | 0 | 23 | 0 | 0 | 0 | 13 | 34 | 46 |
| SRR6705837 | 21 | 0 | 155 | 0 | 0 | 0 | 25 | 0 | 0 | 0 | 0 | 49 | 0 | 0 | 259 | 0 | 0 | 13 | 15 | 13 | 361 | 0 | 0 | 0 | 0 | 62 | 16 | 0 | 0 | 0 | 0 | 0 | 0 | 0 | 15 | 0 | 0 |
| SRR6705838 | 33 | 0 | 929 | 0 | 0 | 18 | 53 | 0 | 0 | 0 | 0 | 100 | 0 | 0 | 35 | 0 | 15 | 0 | 80 | 38 | 443 | 0 | 0 | 0 | 16 | 42 | 63 | 0 | 0 | 0 | 13 | 0 | 0 | 0 | 30 | 21 | 21 |
| SRR6705839 | 173 | 0 | 470 | 0 | 0 | 0 | 53 | 0 | 0 | 0 | 0 | 64 | 0 | 0 | 133 | 0 | 0 | 10 | 121 | 125 | 501 | 0 | 0 | 0 | 0 | 96 | 15 | 0 | 0 | 0 | 0 | 0 | 0 | 0 | 29 | 16 | 19 |
| SRR6705840 | 96 | 0 | 862 | 0 | 0 | 62 | 34 | 0 | 14 | 0 | 0 | 311 | 0 | 16 | 69 | 0 | 25 | 253 | 13 | 13 | 565 | 0 | 0 | 0 | 22 | 613 | 22 | 0 | 0 | 0 | 24 | 0 | 0 | 0 | 59 | 0 | 14 |
| SRR6705841 | 124 | 0 | 925 | 0 | 0 | 11 | 0 | 0 | 0 | 0 | 0 | 139 | 0 | 0 | 13 | 0 | 0 | 0 | 0 | 0 | 36 | 0 | 0 | 11 | 0 | 205 | 18 | 0 | 0 | 0 | 0 | 0 | 0 | 0 | 43 | 29 | 36 |

Supp Table S8.2.Columns_AM_to_BW

| | | | | | | | | | | | | | | | | | | | | | | | | | | | | | | | | | | | | | |
|---|---|---|---|---|---|---|---|---|---|---|---|---|---|---|---|---|---|---|---|---|---|---|---|---|---|---|---|---|---|---|---|---|---|---|---|---|---|
| SRR6705842 | 32 | 0 | 461 | 0 | 0 | 61 | 48 | 0 | 0 | 0 | 0 | 156 | 0 | 0 | 32 | 0 | 0 | 0 | 47 | 0 | 83 | 0 | 0 | 0 | 20 | 90 | 39 | 0 | 0 | 0 | 29 | 0 | 0 | 0 | 63 | 22 | 37 |
| SRR6705843 | 63 | 0 | 776 | 0 | 0 | 0 | 45 | 0 | 0 | 0 | 0 | 298 | 0 | 0 | 0 | 0 | 0 | 34 | 0 | 0 | 73 | 0 | 0 | 0 | 0 | 28 | 29 | 0 | 0 | 0 | 0 | 0 | 0 | 0 | 60 | 0 | 29 |
| SRR6705844 | 308 | 0 | 421 | 0 | 0 | 11 | 291 | 35 | 0 | 0 | 0 | 180 | 0 | 0 | 0 | 0 | 15 | 0 | 0 | 41 | 26 | 0 | 0 | 47 | 0 | 160 | 48 | 0 | 0 | 0 | 0 | 0 | 0 | 13 | 0 | 0 | 0 |
| SRR6705845 | 192 | 0 | 297 | 0 | 0 | 15 | 404 | 54 | 16 | 0 | 0 | 212 | 0 | 0 | 199 | 0 | 35 | 47 | 284 | 1065 | 433 | 0 | 0 | 10 | 30 | 185 | 24 | 0 | 0 | 16 | 61 | 0 | 0 | 0 | 12 | 20 | 33 |
| SRR6705846 | 329 | 0 | 343 | 0 | 0 | 0 | 3437 | 18 | 0 | 0 | 0 | 164 | 0 | 0 | 134 | 0 | 0 | 18 | 5907 | 10267 | 798 | 0 | 0 | 0 | 11 | 208 | 68 | 0 | 0 | 0 | 12 | 0 | 0 | 0 | 33 | 18 | 18 |
| SRR6705847 | 198 | 0 | 239 | 0 | 0 | 59 | 918 | 73 | 29 | 0 | 0 | 166 | 0 | 0 | 244 | 0 | 30 | 24 | 748 | 2960 | 347 | 0 | 22 | 11 | 19 | 613 | 142 | 0 | 59 | 42 | 18 | 0 | 0 | 0 | 35 | 14 | 18 |
| SRR6705848 | 299 | 0 | 299 | 0 | 0 | 0 | 4441 | 0 | 0 | 0 | 0 | 159 | 0 | 0 | 64 | 0 | 0 | 0 | 9136 | 18971 | 1483 | 0 | 0 | 0 | 0 | 16 | 0 | 0 | 0 | 0 | 0 | 0 | 0 | 0 | 11 | 0 | 0 |
| SRR6705849 | 211 | 0 | 292 | 0 | 0 | 0 | 2826 | 0 | 0 | 0 | 0 | 374 | 0 | 0 | 0 | 0 | 33 | 0 | 1167 | 386 | 61 | 0 | 0 | 0 | 0 | 33 | 19 | 0 | 0 | 0 | 0 | 0 | 0 | 0 | 0 | 0 | 11 |
| SRR6705850 | 372 | 0 | 424 | 0 | 0 | 0 | 602 | 18 | 0 | 0 | 0 | 123 | 0 | 0 | 94 | 0 | 17 | 14 | 461 | 20423 | 1391 | 0 | 0 | 35 | 16 | 149 | 40 | 0 | 0 | 0 | 13 | 0 | 0 | 0 | 15 | 0 | 11 |
| SRR6705851 | 192 | 0 | 613 | 0 | 0 | 0 | 503 | 14 | 0 | 0 | 0 | 84 | 0 | 0 | 18 | 0 | 17 | 26 | 392 | 5682 | 767 | 0 | 0 | 0 | 0 | 99 | 42 | 0 | 0 | 0 | 0 | 0 | 0 | 0 | 21 | 0 | 0 |
| SRR6705852 | 386 | 0 | 263 | 0 | 0 | 0 | 686 | 28 | 0 | 0 | 0 | 107 | 0 | 0 | 165 | 0 | 18 | 0 | 642 | 1178 | 119 | 0 | 0 | 33 | 16 | 184 | 40 | 0 | 0 | 11 | 0 | 0 | 0 | 0 | 19 | 0 | 0 |
| SRR6705853 | 144 | 0 | 985 | 0 | 0 | 14 | 560 | 19 | 0 | 0 | 0 | 208 | 0 | 0 | 22 | 0 | 24 | 192 | 273 | 641 | 231 | 0 | 0 | 14 | 0 | 267 | 62 | 0 | 19 | 0 | 32 | 0 | 0 | 14 | 24 | 21 | 19 |
| SRR6705854 | 215 | 0 | 435 | 0 | 0 | 47 | 1499 | 45 | 40 | 0 | 13 | 137 | 0 | 11 | 19 | 0 | 30 | 68 | 7245 | 6449 | 265 | 0 | 40 | 0 | 24 | 65 | 15 | 0 | 48 | 21 | 21 | 0 | 0 | 11 | 40 | 13 | 0 |
| SRR6705855 | 441 | 0 | 341 | 0 | 0 | 24 | 1346 | 0 | 12 | 0 | 0 | 183 | 0 | 0 | 22 | 0 | 14 | 0 | 9749 | 1907 | 348 | 0 | 0 | 0 | 16 | 203 | 41 | 0 | 0 | 0 | 0 | 0 | 0 | 0 | 16 | 0 | 10 |
| SRR6705856 | 46 | 0 | 321 | 0 | 0 | 82 | 533 | 12 | 15 | 0 | 0 | 136 | 0 | 0 | 51 | 0 | 14 | 13 | 0 | 30 | 342 | 0 | 0 | 0 | 40 | 1009 | 137 | 0 | 15 | 0 | 43 | 0 | 0 | 24 | 243 | 152 | 155 |
| SRR6705857 | 104 | 0 | 408 | 0 | 0 | 39 | 180 | 13 | 0 | 0 | 0 | 100 | 0 | 0 | 52 | 0 | 16 | 0 | 0 | 14 | 264 | 0 | 0 | 0 | 23 | 39 | 81 | 0 | 0 | 13 | 0 | 0 | 0 | 0 | 71 | 48 | 64 |
| SRR6705859 | 62 | 0 | 488 | 0 | 0 | 25 | 49 | 0 | 0 | 0 | 0 | 97 | 0 | 0 | 21 | 0 | 15 | 29 | 0 | 0 | 240 | 0 | 0 | 0 | 20 | 395 | 69 | 0 | 0 | 0 | 0 | 0 | 0 | 30 | 54 | 15 | 0 |
| SRR6705860 | 316 | 0 | 664 | 0 | 0 | 34 | 25 | 0 | 45 | 0 | 0 | 477 | 0 | 0 | 56 | 0 | 0 | 184 | 0 | 0 | 378 | 0 | 0 | 0 | 17 | 39 | 10 | 0 | 28 | 0 | 38 | 0 | 0 | 14 | 55 | 10 | 21 |
| SRR6705861 | 233 | 0 | 488 | 0 | 0 | 0 | 17 | 0 | 0 | 0 | 0 | 135 | 0 | 0 | 0 | 0 | 0 | 0 | 0 | 14 | 124 | 0 | 0 | 0 | 0 | 21 | 0 | 0 | 0 | 0 | 0 | 0 | 0 | 17 | 11 | 0 | 19 |
| SRR6705862 | 45 | 0 | 520 | 0 | 0 | 17 | 51 | 0 | 0 | 0 | 0 | 150 | 0 | 0 | 19 | 0 | 0 | 0 | 17 | 11 | 178 | 0 | 0 | 0 | 0 | 35 | 40 | 0 | 16 | 0 | 19 | 0 | 0 | 0 | 20 | 0 | 14 |
| SRR6705863 | 187 | 0 | 449 | 0 | 0 | 0 | 141 | 0 | 0 | 0 | 0 | 124 | 27 | 0 | 19 | 0 | 0 | 27 | 11 | 0 | 128 | 0 | 0 | 0 | 0 | 162 | 47 | 0 | 0 | 0 | 0 | 0 | 0 | 36 | 41 | 27 | 30 |
| SRR6705866 | 187 | 0 | 703 | 0 | 0 | 0 | 160 | 25 | 0 | 0 | 0 | 187 | 0 | 0 | 107 | 0 | 47 | 101 | 107 | 8496 | 214 | 12 | 0 | 50 | 0 | 15 | 13 | 0 | 0 | 0 | 0 | 0 | 0 | 12 | 0 | 12 | 12 |
| SRR6705867 | 286 | 0 | 753 | 0 | 0 | 39 | 822 | 78 | 0 | 0 | 0 | 226 | 0 | 0 | 17 | 0 | 30 | 0 | 3289 | 3118 | 411 | 0 | 0 | 823 | 16 | 388 | 202 | 0 | 0 | 0 | 0 | 0 | 0 | 16 | 47 | 0 | 16 |
| SRR6705868 | 215 | 0 | 927 | 0 | 0 | 0 | 296 | 30 | 0 | 0 | 0 | 69 | 0 | 0 | 0 | 0 | 0 | 15 | 546 | 4787 | 425 | 0 | 0 | 78 | 19 | 58 | 58 | 0 | 0 | 0 | 0 | 0 | 0 | 0 | 0 | 0 | 0 |
| SRR6705869 | 193 | 0 | 502 | 0 | 0 | 11 | 317 | 36 | 31 | 0 | 0 | 283 | 0 | 0 | 22 | 0 | 26 | 0 | 1002 | 1334 | 184 | 0 | 0 | 17 | 19 | 108 | 37 | 0 | 19 | 25 | 31 | 0 | 0 | 13 | 38 | 19 | 31 |
| SRR6705870 | 204 | 0 | 551 | 0 | 0 | 0 | 653 | 11 | 0 | 0 | 0 | 127 | 0 | 0 | 74 | 0 | 23 | 25 | 193 | 1569 | 543 | 0 | 0 | 15 | 15 | 47 | 11 | 0 | 0 | 15 | 11 | 0 | 0 | 0 | 31 | 0 | 0 |
| SRR6705871 | 412 | 0 | 539 | 0 | 0 | 52 | 1075 | 75 | 24 | 0 | 0 | 400 | 0 | 0 | 46 | 0 | 54 | 0 | 2604 | 1913 | 451 | 0 | 0 | 16 | 21 | 100 | 35 | 0 | 0 | 31 | 16 | 0 | 0 | 12 | 35 | 12 | 28 |
| SRR6705874 | 38 | 0 | 76 | 0 | 0 | 0 | 75 | 0 | 0 | 0 | 0 | 0 | 0 | 0 | 402 | 0 | 0 | 0 | 537 | 909 | 149 | 0 | 0 | 0 | 0 | 0 | 0 | 0 | 0 | 0 | 0 | 0 | 0 | 0 | 0 | 0 | 0 |
| SRR6705875 | 79 | 0 | 776 | 0 | 0 | 0 | 69 | 0 | 0 | 0 | 0 | 125 | 0 | 0 | 41 | 0 | 0 | 0 | 1087 | 845 | 117 | 0 | 0 | 0 | 0 | 18 | 0 | 0 | 0 | 0 | 0 | 0 | 0 | 0 | 0 | 0 | 0 |
| SRR6705876 | 729 | 0 | 625 | 0 | 0 | 0 | 144 | 54 | 0 | 0 | 0 | 451 | 0 | 0 | 0 | 0 | 0 | 0 | 324 | 504 | 32 | 0 | 0 | 15 | 0 | 0 | 0 | 0 | 0 | 21 | 0 | 0 | 0 | 121 | 0 | 0 | 0 |
| SRR6705877 | 27 | 0 | 137 | 0 | 0 | 0 | 288 | 0 | 0 | 0 | 0 | 55 | 0 | 0 | 0 | 0 | 0 | 0 | 83 | 32 | 0 | 0 | 0 | 0 | 0 | 0 | 0 | 0 | 0 | 0 | 0 | 0 | 0 | 245 | 0 | 0 | 0 |
| SRR6705878 | 166 | 0 | 296 | 0 | 0 | 0 | 42 | 0 | 0 | 0 | 0 | 55 | 0 | 0 | 12 | 0 | 0 | 0 | 151 | 94 | 16 | 0 | 0 | 0 | 0 | 14 | 0 | 0 | 0 | 0 | 0 | 0 | 0 | 39 | 14 | 0 | 17 |
| SRR6705879 | 110 | 0 | 108 | 0 | 0 | 0 | 151 | 0 | 10 | 0 | 0 | 87 | 0 | 0 | 478 | 0 | 0 | 39 | 446 | 1107 | 239 | 0 | 0 | 0 | 0 | 15 | 0 | 0 | 15 | 0 | 0 | 0 | 0 | 0 | 11 | 0 | 0 |
| SRR6705880 | 11 | 0 | 284 | 0 | 0 | 0 | 0 | 0 | 0 | 0 | 0 | 67 | 0 | 0 | 26 | 0 | 13 | 97 | 0 | 26 | 77 | 0 | 0 | 0 | 16 | 0 | 0 | 0 | 0 | 0 | 0 | 0 | 0 | 0 | 0 | 0 | 0 |
| SRR6705881 | 11 | 0 | 454 | 0 | 0 | 0 | 0 | 0 | 0 | 0 | 0 | 207 | 0 | 0 | 88 | 0 | 0 | 143 | 0 | 0 | 33 | 0 | 0 | 0 | 13 | 16 | 0 | 0 | 11 | 0 | 12 | 0 | 0 | 0 | 0 | 0 | 0 |
| SRR6705882 | 0 | 0 | 602 | 0 | 0 | 0 | 0 | 0 | 0 | 0 | 0 | 58 | 0 | 0 | 115 | 0 | 0 | 62 | 0 | 0 | 58 | 0 | 0 | 0 | 20 | 0 | 0 | 0 | 0 | 0 | 0 | 0 | 0 | 0 | 0 | 0 | 0 |
| SRR6705883 | 11 | 0 | 530 | 0 | 0 | 0 | 0 | 0 | 0 | 0 | 0 | 253 | 0 | 0 | 13 | 0 | 0 | 21 | 0 | 0 | 54 | 0 | 0 | 0 | 13 | 36 | 0 | 0 | 0 | 0 | 21 | 0 | 0 | 0 | 0 | 0 | 0 |
| SRR6705884 | 160 | 0 | 410 | 0 | 0 | 16 | 790 | 20 | 0 | 0 | 0 | 132 | 0 | 0 | 58 | 0 | 11 | 28 | 627 | 732 | 155 | 0 | 0 | 37 | 0 | 209 | 69 | 0 | 0 | 0 | 0 | 0 | 0 | 0 | 19 | 0 | 0 |
| SRR6705885 | 242 | 0 | 359 | 0 | 0 | 33 | 2062 | 40 | 20 | 0 | 0 | 135 | 0 | 0 | 113 | 0 | 27 | 0 | 3408 | 5686 | 386 | 0 | 0 | 14 | 13 | 43 | 13 | 0 | 0 | 13 | 0 | 0 | 0 | 0 | 13 | 0 | 0 |
| SRR6705886 | 246 | 0 | 349 | 0 | 0 | 30 | 1476 | 15 | 0 | 0 | 0 | 142 | 0 | 0 | 27 | 0 | 0 | 52 | 3834 | 2367 | 189 | 0 | 0 | 70 | 0 | 200 | 94 | 0 | 21 | 0 | 12 | 0 | 0 | 0 | 12 | 12 | 0 |
| SRR6705887 | 163 | 0 | 539 | 0 | 0 | 18 | 683 | 33 | 0 | 0 | 0 | 84 | 0 | 0 | 0 | 0 | 25 | 15 | 1825 | 2048 | 228 | 0 | 17 | 17 | 28 | 44 | 18 | 0 | 15 | 0 | 17 | 0 | 0 | 0 | 30 | 10 | 17 |
| SRR6705888 | 90 | 0 | 429 | 0 | 0 | 0 | 2333 | 0 | 0 | 0 | 0 | 122 | 0 | 0 | 18 | 0 | 13 | 37 | 964 | 145 | 78 | 0 | 0 | 10 | 0 | 22 | 19 | 0 | 84 | 0 | 0 | 0 | 0 | 0 | 15 | 20 | 11 |
| SRR6705889 | 128 | 0 | 765 | 0 | 0 | 13 | 3750 | 18 | 0 | 0 | 0 | 54 | 0 | 0 | 13 | 0 | 0 | 20 | 4163 | 1628 | 120 | 0 | 14 | 86 | 0 | 243 | 83 | 0 | 21 | 0 | 0 | 0 | 0 | 0 | 17 | 0 | 0 |
| SRR6705890 | 342 | 0 | 171 | 0 | 0 | 231 | 1320 | 90 | 130 | 0 | 0 | 213 | 0 | 18 | 131 | 0 | 32 | 29 | 563 | 1283 | 669 | 0 | 25 | 0 | 14 | 2132 | 184 | 0 | 0 | 14 | 36 | 0 | 0 | 29 | 97 | 14 | 51 |
| SRR6705891 | 511 | 0 | 316 | 0 | 0 | 20 | 1121 | 0 | 10 | 0 | 0 | 113 | 0 | 0 | 253 | 0 | 14 | 37 | 788 | 5646 | 599 | 0 | 0 | 0 | 10 | 42 | 22 | 0 | 0 | 0 | 0 | 0 | 0 | 0 | 36 | 12 | 20 |
| SRR6705892 | 126 | 0 | 504 | 0 | 0 | 75 | 2079 | 0 | 25 | 0 | 0 | 157 | 0 | 0 | 10 | 0 | 77 | 0 | 617 | 389 | 73 | 0 | 0 | 0 | 16 | 983 | 60 | 0 | 0 | 0 | 217 | 0 | 0 | 0 | 22 | 0 | 47 |
| SRR6705893 | 400 | 0 | 491 | 0 | 0 | 12 | 1346 | 39 | 16 | 0 | 0 | 151 | 0 | 0 | 178 | 0 | 16 | 11 | 759 | 17831 | 1135 | 0 | 0 | 0 | 14 | 91 | 39 | 0 | 0 | 10 | 20 | 0 | 0 | 0 | 29 | 10 | 0 |
| SRR6705894 | 57 | 0 | 1098 | 0 | 0 | 0 | 0 | 0 | 0 | 0 | 0 | 165 | 0 | 47 | 203 | 0 | 0 | 0 | 22 | 0 | 29 | 0 | 23 | 0 | 31 | 55 | 0 | 0 | 0 | 0 | 0 | 0 | 0 | 0 | 45 | 31 | 49 |
| SRR6705895 | 55 | 0 | 946 | 0 | 33 | 0 | 0 | 0 | 0 | 0 | 0 | 61 | 0 | 0 | 305 | 0 | 0 | 0 | 12 | 0 | 0 | 13 | 0 | 0 | 65 | 104 | 20 | 0 | 0 | 0 | 26 | 0 | 0 | 0 | 248 | 202 | 195 |
| SRR6705897 | 28 | 0 | 310 | 0 | 0 | 0 | 0 | 0 | 13 | 0 | 0 | 31 | 0 | 0 | 440 | 0 | 0 | 0 | 15 | 29 | 24 | 0 | 19 | 0 | 13 | 153 | 13 | 0 | 0 | 0 | 25 | 0 | 0 | 0 | 312 | 293 | 395 |
| SRR6705898 | 36 | 0 | 76 | 0 | 0 | 0 | 29 | 0 | 0 | 0 | 0 | 0 | 0 | 0 | 29 | 0 | 0 | 0 | 175 | 67 | 41 | 0 | 0 | 0 | 0 | 0 | 0 | 0 | 0 | 0 | 0 | 0 | 0 | 0 | 0 | 0 | 0 |
| SRR6705899 | 22 | 0 | 0 | 0 | 0 | 0 | 39 | 0 | 0 | 0 | 0 | 0 | 0 | 0 | 888 | 0 | 0 | 0 | 425 | 174 | 135 | 0 | 0 | 0 | 0 | 45 | 0 | 0 | 0 | 0 | 0 | 0 | 0 | 0 | 11 | 0 | 0 |
| SRR6705900 | 218 | 0 | 62 | 0 | 0 | 0 | 57 | 0 | 0 | 0 | 0 | 0 | 0 | 0 | 28 | 0 | 0 | 0 | 469 | 341 | 14 | 0 | 0 | 0 | 0 | 0 | 0 | 0 | 0 | 0 | 0 | 0 | 0 | 0 | 0 | 0 | 0 |
| SRR6705901 | 0 | 0 | 98 | 0 | 0 | 0 | 13 | 0 | 0 | 0 | 0 | 0 | 0 | 0 | 10 | 0 | 0 | 0 | 55 | 0 | 10 | 0 | 0 | 0 | 0 | 13 | 0 | 0 | 0 | 0 | 0 | 0 | 0 | 0 | 12 | 17 | 51 |
| SRR6705902 | 229 | 0 | 406 | 0 | 0 | 53 | 60 | 0 | 14 | 0 | 0 | 165 | 14 | 0 | 0 | 0 | 0 | 0 | 0 | 10 | 265 | 0 | 0 | 18 | 0 | 643 | 109 | 0 | 0 | 0 | 0 | 0 | 0 | 18 | 21 | 14 | 14 |
| SRR6705903 | 98 | 0 | 379 | 0 | 0 | 51 | 58 | 0 | 34 | 0 | 0 | 137 | 0 | 0 | 31 | 0 | 29 | 163 | 46 | 42 | 859 | 0 | 0 | 0 | 20 | 63 | 37 | 0 | 23 | 0 | 19 | 0 | 0 | 19 | 26 | 11 | 22 |
| SRR6726498 | 0 | 0 | 12 | 0 | 0 | 0 | 0 | 0 | 0 | 0 | 0 | 113 | 0 | 0 | 0 | 0 | 0 | 0 | 0 | 1012 | 849 | 0 | 0 | 0 | 0 | 110 | 142 | 0 | 0 | 0 | 0 | 0 | 0 | 0 | 53 | 54 | 71 |
| SRR6726499 | 0 | 0 | 13 | 0 | 0 | 189 | 111 | 0 | 30 | 0 | 0 | 82 | 0 | 120 | 0 | 0 | 0 | 0 | 0 | 589 | 125 | 0 | 0 | 0 | 50 | 3239 | 578 | 0 | 0 | 0 | 718 | 0 | 0 | 0 | 837 | 0 | 209 |
| SRR6757230 | 16 | 46 | 3145 | 0 | 0 | 29 | 198 | 2270 | 29 | 0 | 0 | 15 | 0 | 17 | 145 | 0 | 20 | 0 | 112 | 2039 | 2491 | 0 | 0 | 53 | 11 | 565 | 1456 | 0 | 0 | 0 | 22 | 0 | 0 | 0 | 1021 | 151 | 750 |
| SRR6757231 | 17 | 43 | 2965 | 0 | 0 | 24 | 213 | 446 | 21 | 0 | 0 | 19 | 0 | 0 | 159 | 0 | 16 | 0 | 102 | 1945 | 2394 | 0 | 0 | 11 | 0 | 96 | 276 | 0 | 0 | 15 | 30 | 0 | 0 | 0 | 175 | 27 | 134 |
| SRR6757232 | 20 | 53 | 4140 | 0 | 0 | 29 | 201 | 151 | 16 | 0 | 0 | 43 | 0 | 36 | 227 | 0 | 29 | 25 | 20 | 1993 | 1270 | 0 | 0 | 0 | 20 | 92 | 202 | 0 | 0 | 0 | 18 | 0 | 0 | 0 | 131 | 13 | 93 |
| SRR6757233 | 23 | 46 | 3808 | 0 | 0 | 24 | 289 | 160 | 15 | 0 | 0 | 40 | 0 | 17 | 249 | 0 | 20 | 37 | 44 | 1942 | 1388 | 0 | 0 | 0 | 13 | 105 | 221 | 0 | 0 | 0 | 14 | 0 | 0 | 0 | 142 | 13 | 90 |
| SRR6757234 | 24 | 50 | 2833 | 0 | 0 | 17 | 408 | 1038 | 0 | 0 | 0 | 32 | 0 | 26 | 261 | 0 | 15 | 23 | 62 | 1410 | 1485 | 0 | 0 | 32 | 0 | 57 | 173 | 0 | 0 | 19 | 14 | 0 | 0 | 0 | 116 | 13 | 64 |
| SRR6757235 | 18 | 47 | 2674 | 0 | 0 | 18 | 445 | 1126 | 15 | 0 | 0 | 32 | 0 | 31 | 322 | 0 | 22 | 17 | 55 | 1368 | 1480 | 0 | 0 | 30 | 0 | 55 | 170 | 0 | 0 | 0 | 0 | 0 | 0 | 0 | 119 | 12 | 58 |
| SRR6757236 | 24 | 35 | 3175 | 0 | 0 | 20 | 328 | 23 | 23 | 0 | 0 | 34 | 0 | 20 | 345 | 0 | 11 | 0 | 34 | 1347 | 2404 | 0 | 0 | 0 | 18 | 91 | 216 | 0 | 0 | 0 | 22 | 0 | 0 | 0 | 253 | 32 | 191 |
| SRR6757237 | 23 | 165 | 2841 | 0 | 0 | 17 | 302 | 176 | 17 | 0 | 0 | 41 | 0 | 17 | 329 | 0 | 20 | 0 | 41 | 1231 | 2524 | 0 | 0 | 0 | 13 | 422 | 1074 | 0 | 0 | 0 | 19 | 0 | 0 | 0 | 1267 | 168 | 929 |
| SRR6757238 | 26 | 0 | 3679 | 0 | 0 | 11 | 29 | 76 | 16 | 0 | 0 | 54 | 0 | 0 | 37 | 0 | 14 | 0 | 45 | 2049 | 2837 | 0 | 0 | 19 | 0 | 292 | 459 | 0 | 0 | 0 | 0 | 0 | 0 | 0 | 342 | 54 | 313 |
| SRR6757239 | 25 | 0 | 3349 | 0 | 0 | 35 | 44 | 247 | 12 | 0 | 0 | 56 | 0 | 0 | 51 | 0 | 0 | 0 | 51 | 1888 | 2699 | 0 | 0 | 43 | 0 | 764 | 1237 | 0 | 0 | 0 | 0 | 0 | 0 | 0 | 884 | 146 | 745 |
| SRR6757240 | 39 | 0 | 6232 | 0 | 0 | 58 | 373 | 2137 | 49 | 0 | 0 | 39 | 0 | 0 | 12 | 0 | 29 | 0 | 71 | 3051 | 4195 | 0 | 0 | 117 | 0 | 1052 | 2862 | 0 | 0 | 19 | 16 | 0 | 0 | 0 | 1087 | 223 | 896 |
| SRR6757241 | 21 | 0 | 5860 | 0 | 0 | 13 | 462 | 546 | 15 | 0 | 0 | 26 | 0 | 0 | 21 | 0 | 12 | 0 | 87 | 3264 | 4308 | 0 | 0 | 29 | 0 | 266 | 703 | 0 | 0 | 16 | 13 | 0 | 0 | 0 | 283 | 50 | 234 |
| SRR6757242 | 28 | 0 | 4709 | 0 | 0 | 32 | 68 | 321 | 35 | 0 | 0 | 39 | 0 | 0 | 36 | 0 | 21 | 0 | 60 | 3527 | 2581 | 0 | 0 | 69 | 0 | 821 | 1571 | 0 | 0 | 10 | 22 | 0 | 0 | 0 | 517 | 92 | 561 |
| SRR6757243 | 17 | 0 | 4816 | 0 | 0 | 26 | 51 | 74 | 37 | 0 | 0 | 34 | 0 | 0 | 45 | 0 | 18 | 0 | 39 | 3631 | 2742 | 0 | 0 | 14 | 0 | 186 | 390 | 0 | 0 | 10 | 19 | 0 | 0 | 0 | 127 | 23 | 138 |
| SRR6757244 | 29 | 13 | 2644 | 0 | 0 | 30 | 127 | 861 | 20 | 0 | 0 | 59 | 0 | 0 | 65 | 0 | 24 | 0 | 93 | 2634 | 3675 | 0 | 0 | 109 | 0 | 673 | 2139 | 0 | 0 | 10 | 11 | 0 | 0 | 0 | 849 | 142 | 811 |
| SRR6757245 | 37 | 14 | 2580 | 0 | 0 | 31 | 84 | 923 | 24 | 0 | 0 | 58 | 0 | 0 | 51 | 0 | 25 | 0 | 72 | 2364 | 3072 | 0 | 0 | 92 | 0 | 629 | 2147 | 0 | 0 | 0 | 15 | 0 | 0 | 0 | 824 | 173 | 787 |
| SRR6757246 | 19 | 168 | 2913 | 0 | 0 | 20 | 384 | 1670 | 21 | 0 | 0 | 25 | 0 | 24 | 520 | 0 | 20 | 15 | 67 | 3689 | 0 | 0 | 0 | 45 | 15 | 353 | 1410 | 0 | 0 | 0 | 21 | 0 | 0 | 0 | 408 | 94 | 415 |
| SRR6757247 | 16 | 159 | 2926 | 0 | 0 | 11 | 304 | 1703 | 17 | 0 | 0 | 31 | 0 | 26 | 511 | 0 | 16 | 14 | 127 | 3448 | 0 | 0 | 0 | 34 | 12 | 352 | 1360 | 0 | 0 | 0 | 10 | 0 | 0 | 0 | 342 | 80 | 389 |
| SRR6757248 | 24 | 57 | 3940 | 0 | 0 | 26 | 326 | 933 | 23 | 0 | 0 | 27 | 0 | 17 | 175 | 0 | 11 | 42 | 75 | 3893 | 2764 | 0 | 0 | 48 | 10 | 538 | 1065 | 0 | 0 | 0 | 17 | 0 | 0 | 0 | 503 | 61 | 340 |
| SRR6757249 | 22 | 68 | 3953 | 0 | 0 | 15 | 314 | 948 | 18 | 0 | 0 | 31 | 0 | 12 | 187 | 0 | 13 | 32 | 57 | 3910 | 2778 | 0 | 0 | 48 | 15 | 516 | 1044 | 0 | 0 | 0 | 21 | 0 | 0 | 0 | 465 | 67 | 302 |
| SRR6757250 | 19 | 11 | 4093 | 0 | 0 | 22 | 206 | 144 | 26 | 0 | 0 | 21 | 0 | 20 | 379 | 0 | 22 | 0 | 59 | 1772 | 2834 | 0 | 0 | 57 | 22 | 80 | 235 | 0 | 0 | 0 | 43 | 0 | 0 | 0 | 88 | 12 | 69 |
| SRR6757251 | 22 | 56 | 4223 | 0 | 0 | 16 | 184 | 878 | 22 | 0 | 0 | 24 | 0 | 28 | 282 | 0 | 23 | 0 | 45 | 1811 | 2598 | 0 | 0 | 63 | 13 | 506 | 1309 | 0 | 0 | 0 | 40 | 0 | 0 | 0 | 529 | 72 | 453 |
| SRR6757252 | 18 | 11 | 4199 | 0 | 0 | 23 | 144 | 661 | 27 | 0 | 0 | 25 | 0 | 10 | 16 | 0 | 16 | 0 | 64 | 2715 | 2072 | 0 | 0 | 60 | 0 | 572 | 1286 | 0 | 0 | 0 | 18 | 0 | 0 | 0 | 665 | 111 | 573 |
| SRR6757253 | 20 | 15 | 4202 | 0 | 0 | 21 | 149 | 141 | 21 | 0 | 0 | 24 | 0 | 0 | 22 | 0 | 18 | 0 | 56 | 2764 | 1828 | 0 | 0 | 13 | 0 | 119 | 262 | 0 | 0 | 0 | 16 | 0 | 0 | 0 | 122 | 22 | 109 |
| SRR6757254 | 31 | 21 | 2312 | 0 | 0 | 23 | 42 | 71 | 23 | 0 | 0 | 24 | 0 | 0 | 415 | 0 | 20 | 0 | 34 | 1204 | 1638 | 0 | 0 | 44 | 16 | 103 | 220 | 0 | 0 | 0 | 38 | 0 | 0 | 0 | 267 | 23 | 179 |
| SRR6757255 | 30 | 16 | 2109 | 0 | 0 | 32 | 36 | 419 | 36 | 0 | 0 | 26 | 0 | 14 | 467 | 0 | 20 | 0 | 33 | 1260 | 1457 | 0 | 0 | 31 | 17 | 590 | 1179 | 0 | 0 | 0 | 26 | 0 | 0 | 0 | 1409 | 131 | 985 |
| SRR6757256 | 25 | 56 | 2214 | 0 | 0 | 19 | 152 | 1613 | 21 | 0 | 0 | 35 | 0 | 24 | 297 | 0 | 11 | 0 | 69 | 1639 | 2035 | 0 | 0 | 43 | 10 | 444 | 975 | 0 | 0 | 12 | 30 | 0 | 0 | 0 | 795 | 84 | 791 |
| SRR6757257 | 23 | 14 | 2125 | 0 | 0 | 18 | 116 | 384 | 19 | 0 | 0 | 32 | 0 | 16 | 222 | 0 | 0 | 0 | 88 | 1613 | 1927 | 0 | 0 | 11 | 0 | 112 | 225 | 0 | 0 | 14 | 14 | 0 | 0 | 0 | 199 | 22 | 192 |
| SRR6757258 | 34 | 68 | 3044 | 0 | 0 | 25 | 150 | 1704 | 35 | 0 | 0 | 11 | 0 | 18 | 80 | 0 | 13 | 0 | 35 | 278 | 614 | 0 | 0 | 51 | 0 | 488 | 943 | 12 | 0 | 20 | 18 | 0 | 0 | 0 | 695 | 70 | 430 |
| SRR6757259 | 30 | 61 | 2694 | 0 | 0 | 22 | 152 | 1608 | 22 | 0 | 0 | 12 | 0 | 17 | 95 | 0 | 11 | 0 | 49 | 344 | 641 | 0 | 0 | 51 | 12 | 481 | 947 | 0 | 0 | 25 | 18 | 0 | 0 | 0 | 673 | 61 | 434 |
| SRR6757260 | 28 | 17 | 3414 | 0 | 0 | 58 | 58 | 167 | 39 | 0 | 0 | 15 | 0 | 12 | 655 | 0 | 18 | 0 | 39 | 1176 | 2051 | 0 | 0 | 62 | 0 | 90 | 200 | 0 | 0 | 12 | 11 | 0 | 0 | 11 | 226 | 23 | 188 |
| SRR6757261 | 18 | 16 | 3244 | 0 | 0 | 40 | 23 | 176 | 26 | 0 | 0 | 21 | 0 | 14 | 654 | 0 | 13 | 0 | 38 | 1177 | 2039 | 0 | 0 | 12 | 16 | 89 | 193 | 0 | 0 | 0 | 61 | 0 | 0 | 0 | 215 | 24 | 173 |
| SRR6757262 | 24 | 0 | 1790 | 0 | 0 | 20 | 22 | 236 | 30 | 0 | 0 | 21 | 0 | 0 | 46 | 0 | 12 | 0 | 16 | 1265 | 2471 | 0 | 0 | 112 | 0 | 639 | 2075 | 0 | 0 | 0 | 29 | 0 | 0 | 0 | 2722 | 344 | 2543 |
| SRR6757263 | 21 | 0 | 1662 | 0 | 0 | 23 | 14 | 237 | 34 | 0 | 0 | 16 | 0 | 0 | 85 | 0 | 10 | 0 | 20 | 1284 | 2533 | 0 | 0 | 131 | 0 | 556 | 1890 | 0 | 0 | 0 | 22 | 0 | 0 | 0 | 2593 | 353 | 2389 |
| SRR6757264 | 32 | 0 | 1986 | 0 | 0 | 30 | 31 | 153 | 33 | 0 | 0 | 21 | 0 | 0 | 631 | 0 | 19 | 0 | 24 | 1372 | 1805 | 0 | 0 | 85 | 16 | 622 | 1417 | 0 | 0 | 0 | 30 | 0 | 0 | 0 | 1655 | 219 | 1721 |
| SRR6757265 | 32 | 0 | 1938 | 0 | 0 | 32 | 22 | 151 | 28 | 0 | 0 | 15 | 0 | 0 | 692 | 0 | 14 | 0 | 19 | 1429 | 1712 | 0 | 0 | 87 | 19 | 568 | 1395 | 0 | 0 | 0 | 35 | 0 | 0 | 0 | 1578 | 183 | 1672 |
| SRR6757266 | 26 | 0 | 2180 | 0 | 0 | 27 | 76 | 259 | 25 | 0 | 0 | 20 | 0 | 0 | 23 | 0 | 11 | 0 | 19 | 2110 | 2990 | 0 | 0 | 131 | 0 | 538 | 2584 | 0 | 0 | 0 | 18 | 0 | 0 | 0 | 1052 | 141 | 1148 |
| SRR6757267 | 25 | 0 | 2059 | 0 | 0 | 41 | 30 | 226 | 24 | 0 | 0 | 21 | 0 | 0 | 20 | 0 | 23 | 0 | 10 | 2034 | 2575 | 0 | 0 | 126 | 0 | 483 | 2386 | 0 | 0 | 0 | 18 | 0 | 0 | 0 | 858 | 117 | 1102 |
| SRR6757268 | 32 | 0 | 1610 | 0 | 0 | 28 | 14 | 312 | 15 | 0 | 0 | 24 | 0 | 0 | 154 | 0 | 11 | 0 | 35 | 1150 | 3487 | 0 | 0 | 99 | 0 | 530 | 1643 | 0 | 0 | 0 | 28 | 0 | 0 | 0 | 798 | 96 | 747 |
| SRR6757269 | 32 | 0 | 1582 | 0 | 0 | 22 | 21 | 51 | 23 | 0 | 0 | 21 | 0 | 0 | 150 | 0 | 13 | 0 | 49 | 940 | 3513 | 0 | 0 | 20 | 0 | 86 | 265 | 0 | 0 | 0 | 30 | 0 | 0 | 0 | 128 | 15 | 130 |
| SRR6757270 | 28 | 64 | 3405 | 0 | 0 | 15 | 179 | 127 | 21 | 0 | 0 | 27 | 0 | 12 | 320 | 0 | 16 | 0 | 84 | 1573 | 3529 | 0 | 0 | 15 | 0 | 61 | 197 | 0 | 0 | 0 | 18 | 0 | 0 | 0 | 78 | 11 | 88 |
| SRR6757271 | 24 | 56 | 3443 | 0 | 0 | 13 | 113 | 129 | 21 | 0 | 0 | 34 | 0 | 0 | 319 | 0 | 11 | 0 | 57 | 1692 | 3382 | 0 | 0 | 16 | 15 | 71 | 205 | 0 | 0 | 0 | 21 | 0 | 0 | 0 | 81 | 12 | 84 |
| SRR6757272 | 21 | 147 | 1746 | 0 | 0 | 23 | 149 | 1510 | 28 | 0 | 0 | 16 | 0 | 23 | 267 | 0 | 14 | 15 | 42 | 1710 | 2698 | 0 | 0 | 73 | 15 | 552 | 1300 | 0 | 0 | 0 | 43 | 0 | 0 | 0 | 527 | 64 | 519 |
| SRR6757273 | 21 | 163 | 1755 | 0 | 0 | 36 | 127 | 1377 | 24 | 0 | 0 | 10 | 0 | 22 | 264 | 0 | 0 | 16 | 48 | 1680 | 2616 | 0 | 0 | 60 | 16 | 489 | 1247 | 10 | 0 | 0 | 40 | 0 | 0 | 0 | 520 | 74 | 478 |
| SRR6757274 | 10 | 165 | 875 | 0 | 0 | 29 | 454 | 1815 | 40 | 0 | 0 | 11 | 0 | 30 | 382 | 0 | 0 | 0 | 72 | 1876 | 2282 | 0 | 0 | 38 | 0 | 479 | 1143 | 0 | 0 | 0 | 45 | 0 | 0 | 0 | 807 | 95 | 940 |
| SRR6757275 | 12 | 27 | 923 | 0 | 0 | 37 | 477 | 290 | 27 | 0 | 0 | 0 | 0 | 17 | 353 | 0 | 0 | 0 | 56 | 1972 | 2223 | 0 | 0 | 26 | 0 | 69 | 178 | 0 | 0 | 0 | 42 | 0 | 0 | 0 | 124 | 13 | 140 |
| SRR6757276 | 18 | 17 | 2014 | 0 | 0 | 39 | 73 | 96 | 27 | 0 | 0 | 13 | 0 | 0 | 84 | 0 | 15 | 0 | 65 | 1160 | 2353 | 0 | 0 | 12 | 0 | 61 | 172 | 10 | 0 | 0 | 46 | 0 | 0 | 0 | 73 | 75 | 60 |
| SRR6757277 | 18 | 28 | 2056 | 0 | 0 | 22 | 51 | 794 | 18 | 0 | 0 | 0 | 0 | 0 | 59 | 0 | 0 | 0 | 67 | 1195 | 2259 | 0 | 0 | 108 | 0 | 470 | 1438 | 0 | 0 | 0 | 37 | 0 | 0 | 0 | 577 | 60 | 537 |
| SRR6757373 | 19 | 0 | 752 | 0 | 0 | 0 | 0 | 0 | 0 | 0 | 0 | 22 | 0 | 0 | 2697 | 0 | 0 | 0 | 0 | 0 | 56 | 0 | 0 | 0 | 18 | 38 | 0 | 0 | 0 | 0 | 48 | 0 | 0 | 13 | 26 | 0 | 12 |
| SRR6757374 | 18 | 0 | 648 | 0 | 0 | 0 | 14 | 0 | 0 | 0 | 0 | 80 | 0 | 11 | 136 | 0 | 0 | 30 | 0 | 0 | 14 | 0 | 0 | 0 | 15 | 98 | 0 | 20 | 0 | 0 | 181 | 0 | 0 | 13 | 50 | 0 | 12 |
| SRR6757375 | 0 | 0 | 574 | 0 | 0 | 0 | 0 | 0 | 0 | 0 | 0 | 12 | 0 | 0 | 82 | 0 | 0 | 49 | 0 | 15 | 0 | 0 | 0 | 0 | 31 | 0 | 0 | 0 | 0 | 0 | 19 | 0 | 0 | 0 | 0 | 0 | 0 |
| SRR6757376 | 16 | 0 | 861 | 11 | 0 | 0 | 14 | 0 | 0 | 0 | 0 | 38 | 0 | 50 | 42 | 0 | 22 | 19 | 0 | 0 | 14 | 0 | 0 | 0 | 86 | 38 | 0 | 21 | 0 | 0 | 250 | 0 | 0 | 0 | 38 | 0 | 0 |
| SRR6757377 | 14 | 0 | 1132 | 0 | 0 | 0 | 0 | 0 | 0 | 0 | 0 | 16 | 0 | 0 | 1278 | 0 | 0 | 33 | 0 | 51 | 34 | 0 | 0 | 0 | 14 | 0 | 0 | 0 | 0 | 0 | 18 | 0 | 0 | 0 | 0 | 0 | 0 |
| SRR6757378 | 22 | 0 | 2274 | 0 | 0 | 13 | 39 | 0 | 19 | 0 | 0 | 17 | 0 | 0 | 133 | 10 | 16 | 0 | 0 | 12 | 27 | 0 | 0 | 0 | 36 | 290 | 37 | 36 | 0 | 0 | 97 | 0 | 0 | 37 | 166 | 0 | 36 |
| SRR6757379 | 24 | 0 | 2146 | 0 | 0 | 0 | 0 | 0 | 0 | 0 | 0 | 27 | 0 | 0 | 0 | 0 | 0 | 0 | 0 | 32 | 222 | 0 | 0 | 0 | 0 | 60 | 0 | 0 | 0 | 0 | 17 | 0 | 0 | 32 | 89 | 0 | 21 |
| SRR6757380 | 25 | 0 | 1864 | 0 | 0 | 0 | 0 | 0 | 13 | 0 | 0 | 36 | 0 | 0 | 0 | 0 | 25 | 0 | 0 | 29 | 393 | 0 | 0 | 0 | 0 | 120 | 14 | 0 | 0 | 0 | 36 | 0 | 0 | 0 | 126 | 0 | 31 |
| SRR6757381 | 36 | 0 | 1322 | 0 | 0 | 0 | 0 | 0 | 0 | 0 | 0 | 32 | 0 | 0 | 4184 | 0 | 0 | 0 | 0 | 35 | 2423 | 0 | 0 | 0 | 0 | 20 | 0 | 0 | 0 | 0 | 13 | 0 | 0 | 41 | 19 | 12 | 10 |
| SRR6757382 | 19 | 0 | 1811 | 0 | 0 | 0 | 0 | 0 | 0 | 0 | 0 | 34 | 0 | 0 | 7096 | 0 | 0 | 0 | 0 | 0 | 85 | 0 | 0 | 0 | 35 | 44 | 0 | 0 | 0 | 0 | 32 | 0 | 0 | 15 | 121 | 0 | 33 |
| SRR6757383 | 29 | 0 | 623 | 0 | 0 | 0 | 0 | 0 | 0 | 0 | 0 | 18 | 0 | 0 | 37 | 0 | 0 | 13 | 0 | 66 | 59 | 0 | 0 | 0 | 43 | 33 | 0 | 13 | 0 | 0 | 69 | 0 | 0 | 10 | 35 | 0 | 0 |
| SRR6757384 | 28 | 0 | 1395 | 0 | 0 | 0 | 10 | 0 | 0 | 0 | 0 | 30 | 0 | 15 | 4385 | 0 | 0 | 69 | 0 | 10 | 31 | 0 | 0 | 0 | 42 | 120 | 15 | 10 | 0 | 0 | 552 | 0 | 0 | 0 | 35 | 0 | 18 |
| SRR6757385 | 14 | 0 | 634 | 0 | 0 | 0 | 16 | 11 | 0 | 0 | 0 | 21 | 0 | 27 | 156 | 0 | 0 | 0 | 0 | 97 | 43 | 0 | 0 | 0 | 26 | 30 | 0 | 14 | 0 | 0 | 85 | 0 | 0 | 30 | 24 | 0 | 11 |
| SRR6757386 | 11 | 0 | 865 | 0 | 0 | 0 | 13 | 0 | 0 | 0 | 0 | 45 | 0 | 149 | 0 | 0 | 15 | 0 | 0 | 12 | 0 | 0 | 0 | 0 | 0 | 74 | 0 | 0 | 0 | 0 | 142 | 0 | 0 | 0 | 36 | 0 | 12 |
| SRR6757387 | 19 | 0 | 843 | 0 | 0 | 0 | 0 | 0 | 0 | 0 | 0 | 25 | 0 | 12 | 7624 | 0 | 10 | 27 | 0 | 0 | 45 | 0 | 0 | 0 | 51 | 23 | 0 | 0 | 0 | 0 | 70 | 0 | 0 | 0 | 27 | 0 | 0 |
| SRR6757388 | 0 | 0 | 2005 | 14 | 10 | 0 | 40 | 0 | 0 | 0 | 0 | 31 | 0 | 43 | 204 | 0 | 13 | 31 | 0 | 0 | 22 | 0 | 0 | 0 | 35 | 27 | 0 | 10 | 11 | 0 | 151 | 0 | 0 | 34 | 106 | 0 | 27 |
| SRR6757389 | 11 | 0 | 944 | 0 | 0 | 0 | 0 | 0 | 0 | 0 | 0 | 0 | 0 | 0 | 854 | 0 | 0 | 0 | 0 | 168 | 155 | 0 | 0 | 0 | 15 | 149 | 37 | 0 | 0 | 0 | 31 | 0 | 0 | 23 | 85 | 0 | 53 |
| SRR6757390 | 0 | 0 | 785 | 0 | 0 | 50 | 14 | 0 | 44 | 0 | 0 | 15 | 0 | 12 | 0 | 0 | 0 | 0 | 0 | 21 | 48 | 0 | 0 | 0 | 28 | 24 | 137 | 0 | 17 | 0 | 396 | 0 | 0 | 57 | 140 | 17 | 104 |

### Supp Table S8.2.Columns_AM_to_BW

| | | | | | | | | | | | | | | | | | | | | | | | | | | | | | | | | | | | | | |
|---|---|---|---|---|---|---|---|---|---|---|---|---|---|---|---|---|---|---|---|---|---|---|---|---|---|---|---|---|---|---|---|---|---|---|---|---|---|
| SRR6757391 | 18 | 0 | 985 | 0 | 0 | 0 | 0 | 0 | 0 | 0 | 0 | 33 | 0 | 0 | 94 | 0 | 11 | 32 | 0 | 23 | 133 | 0 | 0 | 0 | 0 | 52 | 0 | 0 | 0 | 0 | 47 | 0 | 0 | 10 | 48 | 0 | 0 |
| SRR6757392 | 13 | 0 | 852 | 0 | 0 | 0 | 28 | 0 | 0 | 0 | 0 | 42 | 0 | 35 | 19 | 0 | 13 | 0 | 0 | 0 | 18 | 0 | 0 | 0 | 0 | 98 | 21 | 0 | 0 | 0 | 196 | 0 | 0 | 0 | 93 | 0 | 18 |
| SRR6757393 | 11 | 0 | 633 | 0 | 0 | 0 | 0 | 0 | 0 | 0 | 0 | 14 | 0 | 24 | 57 | 0 | 0 | 46 | 0 | 50 | 164 | 0 | 0 | 0 | 16 | 30 | 0 | 0 | 0 | 0 | 81 | 0 | 0 | 0 | 0 | 0 | 0 |
| SRR6757394 | 0 | 0 | 499 | 0 | 0 | 0 | 33 | 0 | 0 | 0 | 0 | 0 | 0 | 33 | 99 | 0 | 0 | 17 | 0 | 23 | 52 | 0 | 0 | 0 | 11 | 42 | 0 | 0 | 0 | 0 | 101 | 0 | 0 | 0 | 15 | 0 | 0 |
| SRR6757395 | 30 | 0 | 978 | 0 | 0 | 20 | 0 | 0 | 21 | 0 | 0 | 32 | 0 | 0 | 0 | 0 | 12 | 0 | 0 | 179 | 1440 | 0 | 0 | 0 | 0 | 517 | 28 | 0 | 0 | 0 | 35 | 0 | 0 | 40 | 130 | 17 | 65 |
| SRR6757396 | 14 | 0 | 1103 | 0 | 0 | 23 | 0 | 0 | 32 | 0 | 0 | 35 | 0 | 16 | 0 | 0 | 0 | 0 | 0 | 51 | 258 | 0 | 0 | 0 | 0 | 426 | 32 | 14 | 0 | 0 | 35 | 0 | 0 | 24 | 95 | 0 | 36 |
| SRR6757397 | 23 | 0 | 1197 | 0 | 0 | 0 | 0 | 0 | 0 | 0 | 0 | 30 | 0 | 0 | 37 | 0 | 11 | 0 | 0 | 22 | 132 | 0 | 0 | 0 | 0 | 23 | 0 | 0 | 0 | 0 | 18 | 0 | 0 | 0 | 40 | 0 | 14 |
| SRR6757398 | 18 | 0 | 833 | 0 | 0 | 0 | 11 | 0 | 0 | 0 | 0 | 22 | 0 | 19 | 11 | 0 | 0 | 0 | 0 | 23 | 52 | 0 | 0 | 0 | 0 | 105 | 15 | 11 | 0 | 0 | 209 | 0 | 0 | 15 | 113 | 0 | 34 |
| SRR6757399 | 15 | 0 | 1270 | 0 | 0 | 0 | 0 | 0 | 0 | 0 | 0 | 17 | 0 | 0 | 380 | 0 | 0 | 0 | 0 | 49 | 442 | 0 | 0 | 0 | 0 | 25 | 0 | 0 | 0 | 0 | 17 | 0 | 0 | 25 | 37 | 0 | 0 |
| SRR6757400 | 13 | 0 | 1479 | 0 | 0 | 0 | 21 | 0 | 18 | 0 | 0 | 51 | 0 | 27 | 0 | 0 | 0 | 0 | 0 | 15 | 107 | 0 | 0 | 0 | 0 | 105 | 0 | 0 | 0 | 0 | 78 | 0 | 0 | 25 | 208 | 13 | 57 |
| SRR6757401 | 20 | 0 | 989 | 0 | 0 | 0 | 0 | 0 | 0 | 0 | 0 | 22 | 0 | 10 | 154 | 0 | 0 | 0 | 0 | 66 | 0 | 0 | 0 | 0 | 0 | 86 | 0 | 0 | 0 | 0 | 42 | 0 | 0 | 21 | 71 | 0 | 25 |
| SRR6757402 | 15 | 0 | 917 | 0 | 0 | 0 | 0 | 0 | 0 | 0 | 0 | 41 | 0 | 0 | 0 | 0 | 14 | 0 | 0 | 14 | 0 | 0 | 0 | 0 | 0 | 52 | 0 | 0 | 0 | 0 | 59 | 0 | 0 | 0 | 65 | 0 | 19 |
| SRR6757403 | 20 | 0 | 709 | 0 | 0 | 0 | 0 | 0 | 0 | 0 | 0 | 19 | 0 | 0 | 429 | 0 | 0 | 19 | 0 | 174 | 60 | 0 | 0 | 0 | 21 | 57 | 0 | 0 | 0 | 0 | 45 | 0 | 0 | 15 | 33 | 0 | 0 |
| SRR6757404 | 20 | 0 | 915 | 0 | 0 | 15 | 15 | 0 | 42 | 0 | 0 | 30 | 0 | 12 | 14 | 0 | 0 | 0 | 0 | 0 | 0 | 0 | 0 | 0 | 0 | 314 | 16 | 13 | 0 | 0 | 126 | 0 | 0 | 21 | 218 | 12 | 50 |
| SRR6757405 | 17 | 0 | 1324 | 0 | 0 | 0 | 0 | 0 | 0 | 0 | 0 | 21 | 0 | 0 | 45 | 0 | 0 | 0 | 0 | 82 | 609 | 0 | 0 | 0 | 0 | 60 | 0 | 0 | 0 | 0 | 25 | 0 | 0 | 0 | 63 | 0 | 19 |
| SRR6757406 | 14 | 0 | 1839 | 0 | 0 | 0 | 52 | 0 | 0 | 0 | 0 | 23 | 0 | 0 | 45 | 0 | 0 | 0 | 0 | 30 | 275 | 0 | 0 | 0 | 0 | 57 | 0 | 0 | 0 | 0 | 27 | 0 | 0 | 17 | 43 | 0 | 12 |
| SRR6757407 | 18 | 0 | 798 | 0 | 0 | 0 | 0 | 0 | 0 | 0 | 0 | 23 | 0 | 0 | 34 | 0 | 0 | 30 | 17 | 77 | 341 | 0 | 0 | 0 | 0 | 29 | 0 | 0 | 0 | 0 | 16 | 0 | 0 | 0 | 45 | 0 | 0 |
| SRR6757408 | 12 | 0 | 639 | 0 | 0 | 0 | 14 | 0 | 0 | 0 | 0 | 18 | 0 | 18 | 11 | 0 | 0 | 0 | 0 | 0 | 18 | 0 | 0 | 0 | 0 | 18 | 0 | 0 | 0 | 0 | 56 | 0 | 0 | 0 | 14 | 0 | 0 |
| SRR6757409 | 11 | 0 | 842 | 0 | 0 | 0 | 0 | 0 | 0 | 0 | 0 | 23 | 0 | 23 | 26 | 0 | 0 | 46 | 0 | 28 | 23 | 0 | 0 | 0 | 0 | 45 | 0 | 0 | 0 | 0 | 59 | 0 | 0 | 0 | 18 | 0 | 0 |
| SRR6757410 | 15 | 0 | 820 | 0 | 0 | 10 | 12 | 0 | 14 | 0 | 0 | 33 | 0 | 14 | 0 | 0 | 12 | 0 | 0 | 52 | 98 | 0 | 0 | 0 | 0 | 250 | 25 | 17 | 0 | 0 | 244 | 0 | 0 | 0 | 100 | 0 | 17 |
| SRR6757411 | 27 | 0 | 1201 | 0 | 0 | 0 | 0 | 0 | 0 | 0 | 0 | 20 | 0 | 0 | 100 | 0 | 0 | 0 | 0 | 23 | 77 | 0 | 0 | 0 | 15 | 42 | 0 | 0 | 0 | 0 | 18 | 0 | 0 | 39 | 70 | 0 | 24 |
| SRR6757412 | 16 | 0 | 1218 | 0 | 0 | 0 | 0 | 0 | 49 | 0 | 0 | 12 | 0 | 0 | 18 | 0 | 0 | 0 | 0 | 20 | 114 | 0 | 0 | 0 | 0 | 258 | 16 | 0 | 0 | 0 | 155 | 0 | 0 | 10 | 11 | 16 | 131 |
| SRR6757413 | 23 | 0 | 1290 | 0 | 0 | 15 | 0 | 0 | 33 | 0 | 0 | 26 | 0 | 11 | 16 | 0 | 0 | 0 | 0 | 21 | 582 | 0 | 0 | 0 | 0 | 19 | 16 | 0 | 0 | 0 | 134 | 0 | 0 | 60 | 229 | 17 | 78 |
| SRR6757414 | 17 | 0 | 976 | 0 | 0 | 0 | 0 | 0 | 0 | 0 | 0 | 21 | 0 | 26 | 33 | 0 | 0 | 0 | 0 | 24 | 54 | 0 | 0 | 0 | 0 | 66 | 0 | 0 | 0 | 0 | 460 | 0 | 0 | 0 | 40 | 0 | 0 |
| SRR6757415 | 22 | 0 | 758 | 0 | 0 | 0 | 0 | 0 | 0 | 0 | 0 | 21 | 0 | 0 | 119 | 0 | 0 | 11 | 0 | 0 | 60 | 0 | 0 | 0 | 28 | 25 | 0 | 0 | 0 | 0 | 25 | 0 | 0 | 11 | 24 | 0 | 0 |
| SRR6757416 | 15 | 0 | 780 | 0 | 0 | 0 | 13 | 0 | 0 | 0 | 0 | 36 | 0 | 18 | 214 | 0 | 13 | 0 | 0 | 0 | 22 | 0 | 0 | 0 | 47 | 18 | 0 | 0 | 0 | 0 | 41 | 0 | 0 | 0 | 16 | 0 | 0 |
| SRR6757417 | 18 | 0 | 1395 | 0 | 0 | 0 | 0 | 0 | 0 | 0 | 0 | 25 | 0 | 0 | 56 | 0 | 0 | 0 | 0 | 39 | 168 | 0 | 0 | 0 | 0 | 75 | 0 | 0 | 0 | 0 | 12 | 0 | 0 | 0 | 74 | 0 | 35 |
| SRR6757418 | 26 | 0 | 2323 | 0 | 0 | 19 | 0 | 0 | 36 | 0 | 0 | 41 | 0 | 0 | 0 | 0 | 19 | 0 | 0 | 0 | 28 | 0 | 0 | 0 | 0 | 14 | 28 | 12 | 0 | 0 | 34 | 0 | 0 | 16 | 14 | 18 | 124 |
| SRR6757419 | 18 | 0 | 1191 | 0 | 0 | 0 | 0 | 0 | 0 | 0 | 0 | 19 | 0 | 0 | 34 | 0 | 0 | 0 | 0 | 88 | 217 | 0 | 0 | 0 | 0 | 143 | 26 | 0 | 0 | 0 | 13 | 0 | 0 | 22 | 145 | 0 | 68 |
| SRR6757420 | 20 | 0 | 1500 | 0 | 0 | 18 | 0 | 0 | 20 | 0 | 0 | 41 | 0 | 0 | 0 | 0 | 12 | 0 | 0 | 0 | 42 | 0 | 0 | 0 | 0 | 17 | 35 | 12 | 0 | 0 | 38 | 0 | 0 | 11 | 140 | 12 | 82 |
| SRR6757421 | 16 | 0 | 1258 | 0 | 0 | 0 | 0 | 0 | 0 | 0 | 0 | 24 | 0 | 0 | 57 | 0 | 0 | 0 | 0 | 29 | 0 | 0 | 0 | 0 | 0 | 11 | 0 | 0 | 0 | 0 | 13 | 0 | 0 | 0 | 28 | 0 | 0 |
| SRR6757422 | 16 | 0 | 1285 | 0 | 0 | 0 | 0 | 0 | 0 | 0 | 0 | 15 | 0 | 0 | 0 | 0 | 11 | 0 | 0 | 18 | 0 | 0 | 0 | 0 | 0 | 41 | 0 | 0 | 0 | 0 | 34 | 0 | 0 | 0 | 25 | 0 | 0 |
| SRR6757423 | 14 | 0 | 807 | 0 | 0 | 0 | 50 | 0 | 0 | 0 | 0 | 19 | 0 | 0 | 12 | 0 | 0 | 0 | 0 | 263 | 192 | 0 | 0 | 0 | 0 | 12 | 36 | 0 | 0 | 0 | 17 | 0 | 0 | 0 | 23 | 0 | 11 |
| SRR6757424 | 18 | 0 | 1254 | 0 | 0 | 20 | 70 | 13 | 17 | 0 | 0 | 41 | 0 | 0 | 25 | 0 | 0 | 0 | 0 | 336 | 475 | 0 | 0 | 0 | 0 | 515 | 228 | 0 | 0 | 0 | 47 | 0 | 0 | 19 | 204 | 16 | 67 |
| SRR6757425 | 17 | 0 | 1239 | 0 | 0 | 0 | 22 | 11 | 0 | 0 | 0 | 26 | 0 | 0 | 55 | 0 | 0 | 0 | 11 | 340 | 132 | 0 | 0 | 0 | 0 | 47 | 11 | 0 | 0 | 0 | 15 | 0 | 0 | 14 | 42 | 0 | 18 |
| SRR6757426 | 24 | 0 | 1836 | 0 | 0 | 0 | 0 | 0 | 20 | 0 | 0 | 45 | 0 | 23 | 0 | 0 | 0 | 0 | 0 | 0 | 41 | 0 | 0 | 0 | 0 | 123 | 15 | 0 | 0 | 0 | 38 | 0 | 0 | 31 | 129 | 0 | 29 |
| SRR6757427 | 14 | 0 | 1124 | 0 | 0 | 13 | 35 | 0 | 10 | 0 | 0 | 22 | 0 | 0 | 27 | 0 | 0 | 0 | 12 | 398 | 101 | 0 | 0 | 0 | 0 | 248 | 64 | 0 | 0 | 10 | 25 | 0 | 0 | 13 | 87 | 11 | 31 |
| SRR6757428 | 20 | 0 | 1886 | 0 | 0 | 13 | 0 | 0 | 29 | 0 | 0 | 51 | 0 | 0 | 0 | 0 | 0 | 0 | 0 | 0 | 0 | 0 | 0 | 0 | 0 | 348 | 15 | 0 | 0 | 0 | 163 | 0 | 0 | 23 | 237 | 0 | 84 |
| SRR6757429 | 18 | 0 | 1136 | 0 | 0 | 0 | 0 | 0 | 0 | 0 | 0 | 12 | 0 | 0 | 56 | 0 | 0 | 16 | 0 | 93 | 297 | 0 | 0 | 0 | 0 | 13 | 0 | 0 | 0 | 0 | 13 | 0 | 0 | 0 | 26 | 0 | 0 |
| SRR6757430 | 23 | 0 | 1354 | 0 | 0 | 0 | 0 | 0 | 0 | 0 | 0 | 40 | 0 | 0 | 0 | 0 | 0 | 0 | 0 | 0 | 131 | 0 | 0 | 0 | 0 | 30 | 0 | 0 | 0 | 0 | 23 | 0 | 0 | 0 | 29 | 0 | 0 |
| SRR6757431 | 15 | 0 | 628 | 0 | 0 | 0 | 0 | 0 | 0 | 0 | 0 | 12 | 0 | 0 | 308 | 0 | 0 | 82 | 0 | 113 | 21 | 0 | 0 | 0 | 0 | 26 | 0 | 0 | 0 | 0 | 29 | 0 | 0 | 0 | 26 | 0 | 0 |
| SRR6757432 | 17 | 0 | 1751 | 0 | 0 | 14 | 16 | 0 | 34 | 0 | 0 | 46 | 0 | 20 | 0 | 0 | 0 | 0 | 0 | 0 | 0 | 0 | 0 | 0 | 0 | 380 | 11 | 0 | 0 | 0 | 294 | 0 | 0 | 0 | 350 | 16 | 83 |
| SRR6757433 | 15 | 0 | 788 | 0 | 0 | 0 | 0 | 0 | 0 | 0 | 0 | 32 | 0 | 0 | 23 | 0 | 0 | 0 | 0 | 46 | 99 | 0 | 0 | 0 | 0 | 19 | 0 | 0 | 0 | 0 | 20 | 0 | 0 | 0 | 17 | 0 | 0 |
| SRR6757434 | 21 | 0 | 2602 | 17 | 15 | 0 | 11 | 11 | 0 | 0 | 0 | 35 | 0 | 30 | 17 | 0 | 26 | 0 | 0 | 28 | 28 | 0 | 0 | 0 | 28 | 45 | 17 | 0 | 93 | 0 | 24 | 0 | 15 | 100 | 85 | 0 | 18 |
| SRR6757435 | 21 | 0 | 1678 | 0 | 0 | 0 | 0 | 0 | 0 | 0 | 0 | 26 | 0 | 0 | 24 | 0 | 0 | 0 | 0 | 47 | 214 | 0 | 0 | 0 | 0 | 113 | 0 | 0 | 0 | 0 | 32 | 0 | 0 | 23 | 121 | 0 | 38 |
| SRR6757436 | 19 | 0 | 2019 | 0 | 0 | 0 | 16 | 0 | 0 | 0 | 0 | 63 | 0 | 0 | 0 | 0 | 14 | 0 | 0 | 11 | 42 | 0 | 0 | 0 | 0 | 112 | 0 | 0 | 0 | 0 | 60 | 0 | 0 | 15 | 194 | 0 | 48 |
| SRR6757437 | 22 | 0 | 1530 | 0 | 0 | 0 | 0 | 0 | 16 | 0 | 0 | 26 | 0 | 16 | 0 | 0 | 0 | 0 | 0 | 15 | 183 | 0 | 0 | 0 | 0 | 19 | 0 | 0 | 0 | 0 | 31 | 0 | 0 | 16 | 11 | 0 | 39 |
| SRR6757438 | 20 | 0 | 3174 | 0 | 0 | 24 | 0 | 0 | 48 | 0 | 0 | 126 | 0 | 0 | 0 | 0 | 14 | 0 | 0 | 15 | 88 | 0 | 0 | 0 | 0 | 24 | 33 | 0 | 0 | 0 | 10 | 0 | 0 | 32 | 12 | 22 | 124 |
| SRR6757439 | 37 | 0 | 2109 | 0 | 0 | 18 | 0 | 0 | 22 | 0 | 0 | 33 | 0 | 0 | 0 | 0 | 11 | 0 | 0 | 87 | 597 | 0 | 0 | 0 | 0 | 376 | 66 | 0 | 0 | 0 | 17 | 0 | 0 | 21 | 245 | 18 | 113 |
| SRR6757440 | 32 | 0 | 2625 | 0 | 0 | 24 | 18 | 0 | 46 | 0 | 0 | 46 | 0 | 0 | 0 | 0 | 0 | 0 | 0 | 12 | 75 | 0 | 0 | 0 | 0 | 30 | 49 | 0 | 16 | 0 | 126 | 0 | 0 | 70 | 14 | 13 | 137 |
| SRR6757441 | 50 | 0 | 4372 | 0 | 0 | 14 | 0 | 0 | 20 | 0 | 0 | 34 | 0 | 0 | 16 | 0 | 12 | 0 | 0 | 30 | 0 | 0 | 0 | 0 | 0 | 341 | 25 | 0 | 0 | 0 | 15 | 0 | 0 | 37 | 257 | 16 | 127 |
| SRR6757442 | 13 | 0 | 2033 | 0 | 0 | 0 | 19 | 0 | 0 | 0 | 0 | 58 | 0 | 0 | 0 | 0 | 14 | 0 | 0 | 0 | 0 | 0 | 0 | 0 | 0 | 124 | 14 | 0 | 0 | 0 | 23 | 0 | 0 | 12 | 99 | 0 | 33 |
| SRR6757443 | 25 | 0 | 2047 | 0 | 0 | 0 | 0 | 0 | 0 | 0 | 0 | 20 | 0 | 0 | 17 | 0 | 0 | 0 | 0 | 43 | 137 | 0 | 0 | 0 | 0 | 66 | 0 | 17 | 0 | 0 | 14 | 0 | 0 | 31 | 11 | 0 | 19 |
| SRR6757444 | 23 | 0 | 2498 | 0 | 0 | 0 | 27 | 0 | 15 | 0 | 0 | 70 | 0 | 0 | 0 | 0 | 14 | 0 | 0 | 0 | 112 | 0 | 0 | 0 | 0 | 14 | 15 | 12 | 0 | 0 | 87 | 0 | 0 | 105 | 18 | 0 | 90 |
| SRR6757445 | 19 | 0 | 2160 | 0 | 0 | 0 | 0 | 0 | 0 | 0 | 0 | 37 | 0 | 0 | 12 | 0 | 11 | 0 | 0 | 50 | 256 | 0 | 0 | 0 | 0 | 99 | 0 | 0 | 0 | 0 | 18 | 0 | 0 | 16 | 130 | 0 | 38 |
| SRR6757446 | 20 | 0 | 1528 | 0 | 0 | 0 | 0 | 0 | 0 | 0 | 0 | 27 | 0 | 0 | 44 | 0 | 0 | 0 | 0 | 27 | 186 | 0 | 0 | 0 | 13 | 37 | 0 | 0 | 0 | 0 | 92 | 0 | 0 | 14 | 55 | 0 | 0 |
| SRR6757447 | 20 | 0 | 1878 | 0 | 0 | 0 | 0 | 0 | 0 | 0 | 0 | 22 | 0 | 0 | 33 | 0 | 10 | 0 | 0 | 49 | 311 | 0 | 0 | 0 | 0 | 35 | 0 | 0 | 0 | 0 | 12 | 0 | 0 | 17 | 10 | 0 | 25 |
| SRR6757448 | 22 | 0 | 1830 | 0 | 0 | 0 | 11 | 0 | 12 | 0 | 0 | 55 | 0 | 14 | 0 | 0 | 0 | 0 | 0 | 0 | 13 | 0 | 0 | 0 | 0 | 169 | 16 | 0 | 0 | 0 | 143 | 0 | 0 | 55 | 13 | 11 | 76 |
| SRR6757449 | 19 | 0 | 501 | 0 | 0 | 0 | 0 | 0 | 0 | 0 | 0 | 17 | 0 | 0 | 46 | 0 | 0 | 15 | 0 | 148 | 157 | 0 | 0 | 0 | 0 | 21 | 0 | 0 | 0 | 0 | 30 | 0 | 0 | 0 | 12 | 0 | 0 |
| SRR6757450 | 20 | 0 | 1537 | 0 | 0 | 0 | 34 | 0 | 0 | 0 | 0 | 31 | 0 | 20 | 25 | 0 | 21 | 15 | 0 | 159 | 311 | 0 | 0 | 0 | 21 | 38 | 31 | 27 | 0 | 0 | 57 | 0 | 0 | 11 | 69 | 0 | 11 |
| SRR6757451 | 14 | 0 | 1045 | 0 | 0 | 0 | 0 | 18 | 0 | 0 | 0 | 30 | 0 | 18 | 0 | 0 | 0 | 0 | 0 | 43 | 16 | 0 | 0 | 0 | 46 | 201 | 22 | 0 | 0 | 0 | 68 | 0 | 0 | 40 | 185 | 0 | 64 |
| SRR6757452 | 44 | 0 | 2678 | 0 | 0 | 39 | 0 | 11 | 20 | 0 | 0 | 137 | 0 | 15 | 0 | 0 | 26 | 0 | 0 | 0 | 0 | 0 | 0 | 0 | 38 | 31 | 25 | 21 | 11 | 0 | 23 | 0 | 0 | 121 | 36 | 47 | 11 |
| SRR6757453 | 11 | 0 | 1089 | 0 | 0 | 0 | 0 | 0 | 15 | 0 | 0 | 17 | 0 | 0 | 0 | 0 | 0 | 0 | 0 | 48 | 87 | 0 | 0 | 0 | 0 | 13 | 19 | 0 | 0 | 0 | 39 | 0 | 0 | 46 | 104 | 11 | 35 |
| SRR6757454 | 22 | 0 | 1687 | 0 | 0 | 36 | 0 | 0 | 25 | 0 | 0 | 36 | 0 | 10 | 0 | 0 | 0 | 0 | 0 | 73 | 427 | 0 | 0 | 0 | 0 | 51 | 42 | 0 | 0 | 0 | 85 | 0 | 0 | 16 | 132 | 13 | 51 |
| SRR6757455 | 16 | 0 | 794 | 0 | 0 | 0 | 0 | 0 | 0 | 0 | 0 | 14 | 0 | 11 | 0 | 0 | 0 | 0 | 0 | 47 | 75 | 0 | 0 | 0 | 27 | 35 | 0 | 11 | 0 | 0 | 58 | 0 | 0 | 45 | 29 | 0 | 0 |
| SRR6757456 | 11 | 0 | 2055 | 0 | 0 | 0 | 14 | 0 | 0 | 0 | 0 | 37 | 0 | 29 | 0 | 0 | 10 | 0 | 0 | 0 | 17 | 0 | 0 | 0 | 69 | 94 | 0 | 11 | 0 | 0 | 246 | 0 | 0 | 27 | 98 | 0 | 31 |
| SRR6757457 | 13 | 0 | 729 | 0 | 0 | 0 | 0 | 0 | 0 | 0 | 0 | 18 | 0 | 0 | 49 | 0 | 0 | 0 | 0 | 57 | 90 | 0 | 0 | 0 | 30 | 20 | 0 | 0 | 0 | 0 | 30 | 0 | 0 | 17 | 19 | 0 | 0 |
| SRR6757458 | 0 | 0 | 591 | 0 | 0 | 0 | 0 | 0 | 0 | 0 | 0 | 21 | 0 | 30 | 12 | 0 | 0 | 37 | 0 | 35 | 50 | 0 | 0 | 0 | 71 | 12 | 0 | 12 | 0 | 0 | 45 | 0 | 0 | 0 | 0 | 0 | 0 |
| SRR6757459 | 15 | 0 | 1322 | 0 | 0 | 0 | 0 | 0 | 0 | 0 | 0 | 35 | 0 | 0 | 0 | 0 | 10 | 0 | 0 | 44 | 59 | 0 | 0 | 0 | 0 | 72 | 0 | 0 | 0 | 0 | 22 | 0 | 0 | 21 | 42 | 0 | 18 |
| SRR6757460 | 12 | 0 | 611 | 0 | 0 | 0 | 0 | 0 | 0 | 0 | 0 | 48 | 0 | 21 | 28 | 0 | 12 | 0 | 0 | 0 | 42 | 0 | 0 | 0 | 62 | 112 | 0 | 0 | 0 | 0 | 306 | 0 | 0 | 32 | 34 | 0 | 0 |
| SRR6757461 | 0 | 0 | 665 | 0 | 0 | 0 | 0 | 0 | 0 | 0 | 0 | 15 | 0 | 35 | 16 | 0 | 0 | 43 | 0 | 51 | 28 | 0 | 0 | 0 | 55 | 20 | 0 | 10 | 0 | 0 | 17 | 0 | 0 | 25 | 15 | 0 | 0 |
| SRR6757462 | 11 | 14 | 714 | 0 | 0 | 0 | 12 | 0 | 0 | 0 | 0 | 12 | 0 | 20 | 0 | 0 | 19 | 0 | 0 | 12 | 0 | 0 | 0 | 0 | 73 | 18 | 0 | 15 | 0 | 0 | 30 | 0 | 0 | 23 | 24 | 0 | 0 |
| SRR6757463 | 17 | 0 | 1083 | 0 | 0 | 0 | 0 | 0 | 0 | 0 | 0 | 31 | 0 | 0 | 0 | 0 | 0 | 0 | 0 | 72 | 46 | 0 | 0 | 0 | 16 | 79 | 12 | 0 | 0 | 0 | 35 | 0 | 0 | 47 | 72 | 0 | 14 |
| SRR6757464 | 25 | 0 | 1819 | 12 | 10 | 18 | 41 | 31 | 10 | 0 | 0 | 77 | 0 | 27 | 24 | 0 | 20 | 0 | 0 | 12 | 14 | 0 | 0 | 0 | 36 | 32 | 29 | 12 | 0 | 0 | 59 | 0 | 0 | 139 | 14 | 0 | 56 |
| SRR6757465 | 27 | 0 | 1123 | 0 | 0 | 0 | 12 | 0 | 0 | 0 | 0 | 34 | 0 | 0 | 30 | 0 | 0 | 23 | 0 | 21 | 113 | 0 | 0 | 0 | 32 | 71 | 0 | 15 | 0 | 0 | 39 | 0 | 0 | 39 | 65 | 0 | 21 |
| SRR6757466 | 27 | 0 | 1573 | 0 | 11 | 0 | 0 | 0 | 0 | 0 | 0 | 39 | 0 | 13 | 17 | 0 | 14 | 0 | 0 | 0 | 27 | 0 | 0 | 0 | 22 | 135 | 23 | 0 | 12 | 0 | 135 | 0 | 0 | 49 | 322 | 20 | 86 |
| SRR6757467 | 12 | 10 | 639 | 0 | 0 | 0 | 11 | 0 | 0 | 0 | 0 | 12 | 0 | 77 | 65 | 0 | 0 | 71 | 0 | 38 | 27 | 0 | 0 | 0 | 70 | 29 | 0 | 0 | 0 | 0 | 53 | 0 | 0 | 0 | 21 | 0 | 0 |
| SRR6757468 | 11 | 0 | 714 | 0 | 0 | 0 | 16 | 0 | 0 | 0 | 0 | 26 | 0 | 32 | 95 | 0 | 0 | 38 | 0 | 36 | 16 | 0 | 0 | 0 | 71 | 23 | 0 | 16 | 0 | 0 | 88 | 0 | 0 | 0 | 26 | 0 | 0 |
| SRR6761159 | 54 | 0 | 44 | 0 | 170 | 0 | 861 | 256 | 0 | 0 | 0 | 44 | 0 | 0 | 0 | 0 | 0 | 0 | 0 | 0 | 1721 | 170 | 0 | 85 | 0 | 0 | 0 | 85 | 0 | 85 | 0 | 0 | 0 | 0 | 0 | 0 | 0 |
| SRR6761162 | 11 | 0 | 0 | 0 | 302 | 0 | 0 | 0 | 0 | 0 | 0 | 0 | 0 | 0 | 0 | 0 | 0 | 0 | 0 | 0 | 2183 | 0 | 0 | 0 | 0 | 0 | 0 | 0 | 0 | 302 | 0 | 0 | 0 | 0 | 0 | 0 | 0 |
| SRR6761163 | 20 | 0 | 0 | 0 | 216 | 0 | 0 | 324 | 0 | 0 | 0 | 30 | 0 | 0 | 0 | 0 | 0 | 0 | 0 | 0 | 0 | 0 | 0 | 0 | 108 | 0 | 0 | 0 | 0 | 0 | 0 | 0 | 0 | 0 | 0 | 0 | 0 |
| SRR6761165 | 0 | 0 | 29 | 90 | 179 | 0 | 0 | 90 | 0 | 0 | 0 | 38 | 0 | 0 | 0 | 0 | 0 | 0 | 0 | 0 | 0 | 0 | 0 | 0 | 0 | 0 | 0 | 0 | 0 | 0 | 0 | 0 | 0 | 0 | 0 | 0 | 0 |
| SRR6761167 | 51 | 0 | 0 | 0 | 0 | 0 | 0 | 0 | 0 | 0 | 0 | 23 | 0 | 0 | 0 | 0 | 0 | 0 | 0 | 0 | 0 | 384 | 0 | 384 | 0 | 0 | 0 | 0 | 0 | 0 | 0 | 0 | 0 | 0 | 0 | 0 | 0 |
| SRR6761175 | 28 | 0 | 11 | 0 | 0 | 0 | 0 | 258 | 0 | 0 | 0 | 0 | 0 | 0 | 0 | 0 | 0 | 0 | 1547 | 0 | 773 | 0 | 0 | 0 | 0 | 0 | 0 | 0 | 0 | 172 | 0 | 0 | 0 | 0 | 0 | 0 | 0 |
| SRR6761179 | 16 | 0 | 13 | 0 | 0 | 0 | 0 | 171 | 0 | 0 | 0 | 20 | 0 | 0 | 0 | 0 | 0 | 0 | 0 | 0 | 1969 | 0 | 0 | 0 | 0 | 0 | 0 | 0 | 0 | 0 | 0 | 0 | 0 | 0 | 0 | 0 | 0 |
| SRR6761181 | 24 | 0 | 15 | 56 | 0 | 0 | 0 | 167 | 0 | 0 | 0 | 23 | 0 | 0 | 0 | 0 | 0 | 0 | 0 | 585 | 9942 | 0 | 0 | 0 | 0 | 0 | 0 | 0 | 0 | 0 | 0 | 0 | 0 | 0 | 0 | 0 | 0 |
| SRR6761182 | 23 | 0 | 11 | 168 | 210 | 0 | 0 | 0 | 0 | 0 | 0 | 14 | 0 | 0 | 0 | 0 | 0 | 0 | 536 | 2144 | 1608 | 0 | 0 | 0 | 0 | 0 | 0 | 0 | 0 | 0 | 0 | 0 | 0 | 0 | 0 | 0 | 0 |
| SRR6761183 | 33 | 0 | 0 | 164 | 55 | 0 | 1206 | 55 | 0 | 0 | 0 | 24 | 0 | 0 | 0 | 0 | 0 | 0 | 603 | 1206 | 4219 | 0 | 0 | 0 | 0 | 0 | 0 | 0 | 0 | 55 | 0 | 0 | 0 | 0 | 0 | 0 | 0 |
| SRR6761184 | 19 | 0 | 11 | 0 | 0 | 0 | 0 | 32 | 0 | 0 | 0 | 23 | 0 | 0 | 0 | 0 | 0 | 0 | 768 | 768 | 2689 | 0 | 0 | 0 | 0 | 0 | 32 | 0 | 0 | 32 | 0 | 0 | 0 | 32 | 0 | 0 | 0 |
| SRR6761188 | 61 | 0 | 34 | 113 | 150 | 0 | 0 | 113 | 0 | 0 | 0 | 82 | 0 | 75 | 0 | 0 | 0 | 0 | 0 | 2493 | 5982 | 0 | 0 | 75 | 0 | 0 | 0 | 38 | 0 | 0 | 0 | 0 | 0 | 0 | 0 | 0 | 0 |
| SRR6761192 | 34 | 0 | 17 | 340 | 0 | 0 | 0 | 340 | 0 | 0 | 0 | 137 | 0 | 0 | 0 | 0 | 0 | 0 | 0 | 0 | 3175 | 0 | 0 | 0 | 0 | 0 | 0 | 0 | 0 | 0 | 0 | 0 | 0 | 0 | 0 | 0 | 0 |
| SRR6761198 | 69 | 0 | 80 | 18 | 18 | 0 | 2578 | 53 | 0 | 0 | 0 | 19 | 0 | 0 | 0 | 0 | 0 | 0 | 516 | 1805 | 1031 | 0 | 0 | 0 | 0 | 0 | 0 | 0 | 0 | 88 | 0 | 0 | 0 | 0 | 0 | 0 | 0 |
| SRR6767691 | 0 | 100 | 0 | 560 | 0 | 0 | 0 | 0 | 0 | 0 | 0 | 249 | 0 | 112 | 0 | 0 | 0 | 0 | 0 | 386 | 0 | 37 | 0 | 0 | 0 | 0 | 0 | 25 | 0 | 0 | 0 | 0 | 0 | 0 | 0 | 0 | 0 |
| SRR6767701 | 0 | 0 | 0 | 0 | 0 | 0 | 0 | 0 | 0 | 0 | 0 | 0 | 0 | 134 | 0 | 0 | 0 | 0 | 0 | 0 | 0 | 0 | 0 | 0 | 0 | 0 | 0 | 0 | 0 | 0 | 0 | 0 | 0 | 0 | 0 | 0 | 0 |
| SRR6767712 | 11 | 0 | 0 | 574 | 0 | 0 | 122 | 0 | 0 | 0 | 0 | 65 | 0 | 174 | 0 | 0 | 0 | 0 | 0 | 1219 | 0 | 27 | 0 | 0 | 0 | 0 | 0 | 0 | 0 | 0 | 0 | 0 | 0 | 0 | 0 | 0 | 0 |
| SRR6767713 | 15 | 0 | 0 | 801 | 0 | 0 | 0 | 0 | 0 | 0 | 0 | 89 | 0 | 95 | 0 | 0 | 0 | 0 | 0 | 0 | 0 | 0 | 0 | 0 | 0 | 0 | 0 | 0 | 0 | 0 | 0 | 0 | 0 | 0 | 0 | 0 | 0 |
| SRR6821085 | 19 | 17 | 1234 | 0 | 0 | 21 | 197 | 87 | 15 | 0 | 0 | 156 | 0 | 89 | 99 | 0 | 12 | 0 | 25 | 903 | 205 | 0 | 0 | 54 | 41 | 16 | 23 | 14 | 20 | 20 | 20 | 0 | 0 | 20 | 64 | 22 | 89 |
| SRR6821086 | 11 | 21 | 822 | 0 | 0 | 60 | 129 | 120 | 40 | 0 | 0 | 47 | 0 | 194 | 45 | 0 | 17 | 0 | 0 | 238 | 332 | 0 | 0 | 22 | 73 | 48 | 14 | 16 | 0 | 15 | 66 | 0 | 0 | 84 | 48 | 37 | 50 |
| SRR6821087 | 16 | 28 | 836 | 0 | 0 | 0 | 123 | 94 | 0 | 0 | 0 | 48 | 0 | 270 | 75 | 0 | 12 | 18 | 0 | 135 | 272 | 0 | 0 | 0 | 82 | 201 | 292 | 154 | 0 | 0 | 59 | 0 | 0 | 314 | 29 | 17 | 34 |
| SRR6821088 | 63 | 0 | 1339 | 0 | 0 | 26 | 213 | 19 | 0 | 0 | 0 | 0 | 0 | 0 | 951 | 0 | 0 | 0 | 71 | 83 | 344 | 0 | 0 | 16 | 0 | 308 | 88 | 0 | 0 | 0 | 13 | 0 | 0 | 0 | 182 | 16 | 123 |
| SRR6821089 | 42 | 0 | 433 | 0 | 0 | 10 | 113 | 0 | 0 | 0 | 0 | 248 | 0 | 0 | 3910 | 0 | 0 | 0 | 244 | 440 | 5609 | 0 | 0 | 19 | 0 | 107 | 23 | 0 | 0 | 0 | 0 | 0 | 0 | 0 | 19 | 0 | 26 |
| SRR6821090 | 73 | 0 | 1075 | 0 | 0 | 39 | 240 | 10 | 10 | 0 | 0 | 0 | 0 | 0 | 2847 | 0 | 0 | 23 | 99 | 44 | 1079 | 0 | 0 | 33 | 39 | 358 | 133 | 0 | 0 | 0 | 10 | 0 | 0 | 12 | 76 | 0 | 63 |
| SRR6823984 | 59 | 0 | 91 | 0 | 0 | 1488 | 31 | 86 | 539 | 0 | 10 | 470 | 14 | 0 | 719 | 0 | 339 | 961 | 78 | 641 | 1204 | 0 | 0 | 907 | 206 | 9126 | 1428 | 0 | 0 | 22 | 1570 | 0 | 0 | 22 | 1878 | 2565 | 9929 |
| SRR6823985 | 39 | 0 | 67 | 0 | 0 | 690 | 0 | 15 | 491 | 0 | 23 | 530 | 13 | 0 | 1975 | 0 | 163 | 2174 | 0 | 280 | 996 | 0 | 0 | 10 | 242 | 5048 | 374 | 0 | 0 | 0 | 2462 | 0 | 0 | 158 | 1530 | 2113 | 8398 |
| SRR6823986 | 114 | 0 | 282 | 0 | 0 | 63 | 24 | 0 | 44 | 0 | 13 | 149 | 42 | 0 | 533 | 0 | 130 | 44 | 65 | 291 | 1115 | 0 | 0 | 140 | 159 | 671 | 1100 | 0 | 0 | 0 | 1363 | 0 | 343 | 138 | 6699 | 3538 | 12793 |
| SRR6823987 | 104 | 0 | 390 | 0 | 0 | 54 | 12 | 0 | 56 | 0 | 11 | 189 | 38 | 16 | 668 | 40 | 161 | 20 | 35 | 311 | 1163 | 0 | 0 | 110 | 20 | 79 | 93 | 32 | 0 | 0 | 166 | 0 | 35 | 38 | 384 | 178 | 704 |
| SRR6823990 | 51 | 0 | 42 | 0 | 0 | 945 | 116 | 22 | 361 | 0 | 13 | 842 | 35 | 0 | 1804 | 0 | 255 | 0 | 182 | 1821 | 3029 | 0 | 0 | 289 | 91 | 5508 | 954 | 0 | 0 | 126 | 1221 | 0 | 0 | 163 | 2062 | 1190 | 5716 |
| SRR6823991 | 71 | 0 | 461 | 0 | 0 | 69 | 27 | 0 | 25 | 0 | 0 | 961 | 64 | 16 | 493 | 0 | 322 | 0 | 55 | 1451 | 3040 | 19 | 0 | 51 | 27 | 870 | 225 | 0 | 16 | 29 | 69 | 0 | 0 | 15 | 104 | 59 | 282 |
| SRR6829958 | 146 | 0 | 50 | 0 | 0 | 11 | 469 | 99 | 14 | 0 | 0 | 322 | 109 | 0 | 213 | 0 | 56 | 13 | 23 | 52 | 1170 | 0 | 0 | 36 | 24 | 79 | 117 | 0 | 0 | 43 | 18 | 50 | 0 | 121 | 202 | 1076 | 1410 |
| SRR6829959 | 273 | 0 | 20 | 0 | 0 | 13 | 152 | 14 | 23 | 0 | 0 | 552 | 35 | 12 | 17 | 28 | 56 | 0 | 43 | 195 | 605 | 34 | 0 | 15 | 28 | 91 | 142 | 39 | 17 | 18 | 10 | 46 | 0 | 117 | 56 | 153 | 369 |
| SRR6829960 | 153 | 0 | 68 | 0 | 0 | 359 | 349 | 41 | 344 | 0 | 14 | 282 | 226 | 12 | 55 | 0 | 71 | 0 | 22 | 83 | 1128 | 0 | 0 | 16 | 70 | 2512 | 196 | 17 | 0 | 22 | 473 | 42 | 0 | 3363 | 8753 | 48034 | 56513 |
| SRR6829961 | 198 | 0 | 15 | 0 | 17 | 258 | 127 | 0 | 13 | 0 | 0 | 372 | 54 | 47 | 71 | 0 | 28 | 12 | 37 | 85 | 267 | 0 | 0 | 0 | 109 | 44 | 215 | 17 | 0 | 0 | 10 | 35 | 0 | 24 | 42 | 113 | 224 |
| SRR6829962 | 124 | 0 | 39 | 0 | 0 | 235 | 196 | 26 | 13 | 0 | 0 | 310 | 11 | 18 | 1756 | 11 | 60 | 0 | 12 | 33 | 969 | 14 | 0 | 30 | 48 | 47 | 84 | 36 | 0 | 23 | 19 | 61 | 0 | 119 | 142 | 873 | 1013 |
| SRR6829963 | 94 | 0 | 0 | 0 | 19 | 169 | 149 | 13 | 466 | 0 | 44 | 188 | 61 | 19 | 3292 | 10 | 12 | 27 | 0 | 48 | 585 | 0 | 0 | 0 | 42 | 1140 | 10 | 19 | 0 | 13 | 779 | 36 | 0 | 750 | 773 | 4616 | 11709 |
| SRR6831609 | 449 | 0 | 39 | 0 | 22 | 0 | 0 | 0 | 22 | 0 | 43 | 2371 | 1340 | 22 | 935 | 0 | 131 | 27 | 0 | 0 | 2405 | 0 | 0 | 43 | 173 | 22 | 22 | 0 | 0 | 0 | 43 | 0 | 0 | 12 | 0 | 0 | 22 |
| SRR6831611 | 529 | 0 | 24 | 20 | 59 | 20 | 0 | 0 | 20 | 0 | 0 | 1697 | 5408 | 0 | 558 | 0 | 83 | 0 | 0 | 0 | 5843 | 0 | 20 | 20 | 59 | 98 | 20 | 0 | 0 | 0 | 20 | 0 | 20 | 3441 | 0 | 0 | 0 |
| SRR6831613 | 1076 | 0 | 41 | 0 | 57 | 29 | 0 | 0 | 29 | 57 | 29 | 1995 | 1261 | 0 | 434 | 0 | 104 | 0 | 0 | 0 | 5786 | 0 | 0 | 0 | 57 | 0 | 57 | 0 | 0 | 0 | 57 | 0 | 29 | 1634 | 0 | 0 | 0 |
| SRR6837863 | 862 | 0 | 14330 | 30 | 26 | 0 | 0 | 0 | 0 | 18 | 0 | 1151 | 0 | 171 | 77 | 11 | 19 | 0 | 0 | 50 | 0 | 0 | 0 | 0 | 22 | 20 | 12 | 47 | 127 | 0 | 32 | 0 | 0 | 39 | 438 | 33 | 90 |
| SRR6837867 | 679 | 0 | 13983 | 59 | 45 | 0 | 0 | 0 | 0 | 31 | 0 | 912 | 0 | 17 | 100 | 0 | 20 | 16 | 0 | 14 | 0 | 0 | 10 | 0 | 42 | 24 | 0 | 57 | 101 | 0 | 31 | 0 | 0 | 40 | 128 | 12 | 29 |
| SRR6837868 | 770 | 0 | 13677 | 44 | 46 | 0 | 0 | 0 | 0 | 19 | 0 | 1124 | 0 | 202 | 77 | 0 | 23 | 0 | 0 | 0 | 0 | 0 | 0 | 0 | 41 | 74 | 0 | 51 | 118 | 0 | 23 | 0 | 13 | 53 | 244 | 28 | 75 |
| SRR6880326 | 158 | 0 | 18 | 35 | 0 | 0 | 12 | 0 | 0 | 0 | 0 | 3148 | 0 | 76 | 0 | 0 | 0 | 66 | 0 | 12 | 211 | 0 | 0 | 0 | 0 | 0 | 0 | 35 | 0 | 0 | 14 | 0 | 0 | 0 | 0 | 0 | 0 |
| SRR6880328 | 10 | 0 | 0 | 11 | 0 | 0 | 44 | 0 | 0 | 30 | 0 | 22 | 0 | 55 | 0 | 0 | 0 | 0 | 11 | 2448 | 98 | 0 | 0 | 0 | 0 | 0 | 0 | 13 | 0 | 0 | 0 | 0 | 15 | 0 | 0 | 0 | 0 |
| SRR6880329 | 14 | 0 | 0 | 14 | 0 | 0 | 53 | 0 | 0 | 27 | 0 | 25 | 0 | 61 | 0 | 0 | 0 | 12 | 30 | 1915 | 83 | 11 | 0 | 22 | 0 | 0 | 0 | 14 | 0 | 0 | 0 | 0 | 15 | 0 | 0 | 0 | 0 |
| SRR6880330 | 0 | 0 | 0 | 13 | 0 | 0 | 73 | 0 | 0 | 33 | 12 | 25 | 0 | 47 | 0 | 0 | 0 | 0 | 0 | 1302 | 257 | 12 | 0 | 12 | 0 | 0 | 0 | 0 | 0 | 0 | 0 | 0 | 14 | 0 | 0 | 0 | 0 |

### Supp Table S8.2.Columns_AM_to_BW

| | | | | | | | | | | | | | | | | | | | | | | | | | | | | | | | | | | | | | |
|---|---|---|---|---|---|---|---|---|---|---|---|---|---|---|---|---|---|---|---|---|---|---|---|---|---|---|---|---|---|---|---|---|---|---|---|---|---|
| SRR6880331 | 14 | 0 | 0 | 0 | 0 | 0 | 108 | 0 | 0 | 55 | 0 | 15 | 0 | 43 | 0 | 0 | 0 | 0 | 86 | 1978 | 215 | 0 | 0 | 19 | 0 | 0 | 0 | 11 | 0 | 0 | 0 | 0 | 18 | 0 | 0 | 0 | 0 |
| SRR6880333 | 57 | 0 | 41 | 29 | 0 | 0 | 55 | 0 | 0 | 87 | 16 | 570 | 0 | 312 | 21 | 0 | 0 | 0 | 34 | 902 | 406 | 22 | 0 | 0 | 0 | 0 | 0 | 90 | 0 | 0 | 0 | 0 | 0 | 0 | 14 | 0 | 20 |
| SRR6880334 | 50 | 0 | 52 | 22 | 0 | 0 | 0 | 0 | 0 | 49 | 15 | 305 | 0 | 185 | 17 | 0 | 0 | 13 | 17 | 1421 | 503 | 10 | 0 | 0 | 0 | 0 | 0 | 43 | 0 | 0 | 0 | 0 | 0 | 0 | 10 | 0 | 11 |
| SRR6880335 | 62 | 0 | 29 | 33 | 0 | 0 | 34 | 0 | 0 | 74 | 16 | 348 | 0 | 270 | 0 | 0 | 0 | 0 | 101 | 1709 | 514 | 20 | 0 | 0 | 0 | 12 | 0 | 82 | 0 | 0 | 0 | 0 | 0 | 0 | 20 | 0 | 23 |
| SRR6880336 | 767 | 0 | 22 | 0 | 0 | 0 | 585 | 0 | 0 | 0 | 0 | 1929 | 0 | 16 | 0 | 0 | 0 | 100 | 0 | 81 | 1642 | 18 | 0 | 0 | 0 | 0 | 0 | 11 | 0 | 0 | 13 | 0 | 14 | 12 | 0 | 0 | 0 |
| SRR6880337 | 126 | 0 | 20 | 12 | 0 | 0 | 876 | 13 | 0 | 16 | 0 | 393 | 0 | 45 | 30 | 0 | 20 | 24 | 37 | 408 | 6003 | 11 | 0 | 40 | 0 | 0 | 20 | 13 | 0 | 25 | 11 | 0 | 0 | 0 | 0 | 0 | 0 |
| SRR6880338 | 138 | 0 | 0 | 46 | 0 | 0 | 12 | 0 | 0 | 21 | 0 | 2991 | 0 | 137 | 0 | 0 | 0 | 50 | 0 | 23 | 205 | 0 | 0 | 0 | 0 | 0 | 0 | 53 | 0 | 0 | 0 | 0 | 0 | 0 | 0 | 0 | 0 |
| SRR6880339 | 178 | 0 | 21 | 19 | 0 | 0 | 0 | 0 | 0 | 21 | 0 | 4762 | 0 | 109 | 0 | 0 | 0 | 163 | 0 | 0 | 186 | 19 | 0 | 0 | 0 | 0 | 0 | 66 | 0 | 0 | 0 | 0 | 0 | 0 | 0 | 0 | 0 |
| SRR6880341 | 32 | 0 | 0 | 82 | 0 | 0 | 25 | 0 | 0 | 20 | 0 | 1086 | 0 | 99 | 0 | 0 | 0 | 18 | 0 | 30 | 163 | 0 | 0 | 0 | 0 | 0 | 0 | 36 | 0 | 13 | 23 | 0 | 0 | 0 | 0 | 0 | 0 |
| SRR6880342 | 123 | 0 | 0 | 22 | 0 | 0 | 15 | 20 | 0 | 13 | 0 | 1072 | 0 | 16 | 0 | 0 | 0 | 10 | 0 | 23 | 94 | 0 | 0 | 0 | 0 | 0 | 0 | 18 | 14 | 0 | 20 | 0 | 0 | 0 | 0 | 0 | 0 |
| SRR6880343 | 58 | 0 | 0 | 39 | 0 | 0 | 33 | 20 | 0 | 15 | 15 | 1418 | 0 | 42 | 0 | 0 | 0 | 0 | 0 | 0 | 84 | 0 | 0 | 0 | 0 | 0 | 0 | 22 | 0 | 0 | 43 | 0 | 0 | 0 | 0 | 0 | 0 |
| SRR6880345 | 135 | 0 | 0 | 0 | 0 | 0 | 663 | 0 | 0 | 24 | 14 | 491 | 0 | 83 | 0 | 0 | 0 | 0 | 78 | 571 | 4198 | 13 | 0 | 11 | 15 | 22 | 17 | 106 | 0 | 0 | 0 | 0 | 12 | 0 | 0 | 0 | 0 |
| SRR6880351 | 0 | 0 | 0 | 40 | 0 | 0 | 3599 | 19 | 0 | 20 | 21 | 324 | 0 | 122 | 15 | 0 | 0 | 0 | 83 | 404 | 886 | 14 | 0 | 0 | 0 | 0 | 0 | 115 | 0 | 0 | 10 | 0 | 0 | 0 | 0 | 0 | 0 |
| SRR6880356 | 54 | 0 | 0 | 13 | 0 | 0 | 889 | 20 | 0 | 23 | 0 | 250 | 0 | 100 | 46 | 0 | 0 | 0 | 0 | 380 | 2408 | 18 | 0 | 13 | 0 | 0 | 0 | 73 | 0 | 0 | 19 | 0 | 0 | 0 | 0 | 0 | 0 |
| SRR6880357 | 76 | 0 | 0 | 29 | 0 | 0 | 376 | 0 | 0 | 26 | 0 | 295 | 0 | 34 | 0 | 0 | 0 | 0 | 29 | 96 | 1755 | 10 | 0 | 0 | 0 | 0 | 0 | 21 | 0 | 0 | 21 | 0 | 0 | 0 | 0 | 0 | 0 |
| SRR6898464 | 846 | 0 | 639 | 0 | 0 | 11 | 777 | 111 | 24 | 0 | 0 | 106 | 0 | 0 | 22 | 0 | 19 | 0 | 140 | 3760 | 334 | 0 | 0 | 18 | 18 | 153 | 202 | 0 | 11 | 43 | 11 | 0 | 0 | 19 | 35 | 12 | 18 |
| SRR6914752 | 13 | 0 | 64 | 0 | 0 | 147 | 25 | 21 | 84 | 0 | 63 | 326 | 0 | 21 | 0 | 0 | 88 | 0 | 62 | 9377 | 511 | 0 | 0 | 0 | 0 | 1470 | 420 | 42 | 0 | 0 | 336 | 0 | 0 | 0 | 3464 | 9784 | 25447 |
| SRR6914753 | 20 | 0 | 454 | 0 | 0 | 308 | 19 | 0 | 32 | 0 | 0 | 602 | 0 | 0 | 0 | 0 | 492 | 0 | 52 | 6741 | 989 | 0 | 0 | 51 | 461 | 32 | 20 | 0 | 0 | 0 | 240 | 0 | 0 | 0 | 66 | 171 | 143 |
| SRR6914754 | 22 | 13 | 156 | 0 | 0 | 93 | 165 | 213 | 60 | 0 | 126 | 214 | 0 | 0 | 18 | 0 | 36 | 31 | 142 | 6540 | 248 | 0 | 0 | 20 | 73 | 19 | 12 | 0 | 0 | 0 | 286 | 0 | 0 | 0 | 16 | 38 | 70 |
| SRR6914755 | 29 | 11 | 666 | 0 | 0 | 395 | 148 | 19 | 117 | 0 | 81 | 558 | 0 | 0 | 0 | 0 | 374 | 0 | 127 | 9692 | 762 | 0 | 0 | 39 | 10 | 69 | 35 | 0 | 14 | 0 | 331 | 0 | 0 | 0 | 25 | 86 | 76 |
| SRR6928219 | 114 | 0 | 204 | 0 | 0 | 137 | 55 | 47 | 50 | 0 | 19 | 351 | 40 | 0 | 114 | 0 | 63 | 0 | 67 | 480 | 1179 | 0 | 0 | 514 | 428 | 911 | 1932 | 0 | 0 | 43 | 170 | 0 | 0 | 69 | 1036 | 3364 | 3560 |
| SRR6928220 | 137 | 0 | 212 | 0 | 0 | 10 | 116 | 33 | 38 | 16 | 43 | 574 | 19 | 0 | 38 | 0 | 76 | 0 | 54 | 304 | 2028 | 0 | 0 | 28 | 32 | 90 | 169 | 0 | 0 | 34 | 38 | 0 | 0 | 48 | 135 | 376 | 404 |
| SRR6928221 | 63 | 0 | 165 | 0 | 0 | 152 | 50 | 156 | 71 | 0 | 29 | 266 | 47 | 12 | 31 | 0 | 70 | 0 | 48 | 389 | 2737 | 0 | 0 | 475 | 151 | 1303 | 1938 | 0 | 0 | 33 | 91 | 0 | 0 | 179 | 1035 | 2852 | 3928 |
| SRR6928222 | 68 | 0 | 122 | 0 | 0 | 562 | 84 | 76 | 124 | 19 | 42 | 442 | 26 | 0 | 18 | 0 | 34 | 0 | 50 | 433 | 4344 | 0 | 0 | 359 | 315 | 4332 | 1462 | 0 | 0 | 14 | 709 | 0 | 0 | 39 | 357 | 907 | 1224 |
| SRR6928223 | 99 | 0 | 122 | 0 | 0 | 97 | 37 | 36 | 55 | 0 | 15 | 511 | 34 | 0 | 98 | 0 | 74 | 0 | 31 | 295 | 965 | 0 | 0 | 575 | 271 | 765 | 1996 | 0 | 0 | 23 | 185 | 0 | 0 | 50 | 2445 | 11341 | 11512 |
| SRR6928224 | 220 | 0 | 429 | 0 | 0 | 17 | 152 | 22 | 41 | 16 | 44 | 663 | 19 | 0 | 18 | 0 | 93 | 0 | 45 | 323 | 1441 | 0 | 0 | 68 | 49 | 146 | 225 | 14 | 0 | 22 | 64 | 0 | 0 | 75 | 148 | 383 | 375 |
| SRR6928225 | 73 | 0 | 165 | 0 | 0 | 233 | 23 | 94 | 95 | 0 | 15 | 363 | 32 | 0 | 18 | 0 | 38 | 0 | 54 | 430 | 3394 | 0 | 0 | 467 | 184 | 1755 | 2201 | 0 | 0 | 18 | 138 | 0 | 0 | 169 | 1600 | 5071 | 5776 |
| SRR6928226 | 65 | 0 | 123 | 0 | 0 | 119 | 89 | 23 | 25 | 14 | 35 | 527 | 25 | 0 | 21 | 0 | 48 | 0 | 70 | 463 | 3673 | 0 | 0 | 92 | 68 | 825 | 340 | 0 | 0 | 19 | 140 | 0 | 0 | 42 | 81 | 186 | 251 |
| SRR6930936 | 102 | 0 | 128 | 0 | 14 | 40 | 157 | 0 | 17 | 0 | 36 | 290 | 14 | 32 | 2157 | 18 | 71 | 43 | 52 | 395 | 2535 | 0 | 0 | 271 | 142 | 229 | 223 | 21 | 0 | 0 | 25 | 0 | 0 | 12 | 75 | 377 | 423 |
| SRR6930937 | 117 | 0 | 197 | 0 | 13 | 48 | 112 | 0 | 22 | 0 | 31 | 555 | 46 | 50 | 1909 | 22 | 249 | 18 | 81 | 274 | 2132 | 0 | 0 | 349 | 133 | 270 | 221 | 22 | 0 | 0 | 33 | 0 | 0 | 16 | 191 | 268 | 545 |
| SRR6930939 | 90 | 0 | 569 | 0 | 0 | 36 | 260 | 50 | 0 | 0 | 0 | 582 | 14 | 30 | 676 | 0 | 113 | 0 | 0 | 104 | 728 | 0 | 0 | 1689 | 103 | 158 | 185 | 11 | 0 | 0 | 16 | 0 | 0 | 18 | 98 | 34 | 114 |
| SRR6930941 | 105 | 0 | 370 | 0 | 0 | 11 | 310 | 31 | 21 | 0 | 23 | 615 | 20 | 43 | 381 | 0 | 116 | 76 | 60 | 316 | 1388 | 0 | 0 | 110 | 25 | 51 | 80 | 20 | 0 | 0 | 27 | 0 | 0 | 36 | 54 | 97 | 195 |
| SRR6932941 | 81 | 0 | 662 | 0 | 0 | 0 | 0 | 20 | 0 | 0 | 0 | 1479 | 0 | 0 | 407 | 0 | 22 | 15 | 0 | 42 | 42 | 0 | 0 | 0 | 0 | 17 | 46 | 12 | 0 | 0 | 56 | 0 | 0 | 0 | 91 | 11 | 17 |
| SRR6932942 | 1828 | 0 | 753 | 0 | 0 | 0 | 0 | 0 | 0 | 0 | 0 | 332 | 0 | 30 | 415 | 0 | 22 | 40 | 0 | 126 | 18 | 0 | 0 | 0 | 30 | 30 | 50 | 0 | 0 | 0 | 50 | 0 | 0 | 0 | 187 | 50 | 29 |
| SRR6932943 | 180 | 67 | 2845 | 0 | 0 | 0 | 0 | 0 | 30 | 0 | 0 | 1288 | 0 | 15 | 243 | 0 | 22 | 0 | 0 | 17 | 17 | 0 | 0 | 0 | 112 | 0 | 0 | 0 | 15 | 0 | 37 | 0 | 0 | 0 | 83 | 17 | 49 |
| SRR6932946 | 888 | 60 | 1936 | 0 | 0 | 0 | 0 | 0 | 23 | 0 | 0 | 465 | 0 | 0 | 168 | 0 | 0 | 0 | 0 | 0 | 23 | 0 | 0 | 0 | 28 | 83 | 0 | 18 | 32 | 0 | 51 | 0 | 0 | 0 | 99 | 41 | 83 |
| SRR6932949 | 955 | 15 | 2633 | 0 | 0 | 0 | 0 | 0 | 0 | 0 | 0 | 951 | 0 | 0 | 441 | 0 | 58 | 0 | 0 | 23 | 0 | 0 | 0 | 0 | 24 | 0 | 0 | 0 | 0 | 0 | 39 | 0 | 0 | 0 | 0 | 0 | 0 |
| SRR6932950 | 609 | 12 | 2568 | 0 | 0 | 0 | 0 | 0 | 0 | 0 | 0 | 1317 | 0 | 0 | 1250 | 0 | 54 | 0 | 0 | 0 | 0 | 0 | 0 | 0 | 52 | 0 | 0 | 0 | 0 | 0 | 49 | 0 | 0 | 0 | 0 | 15 | 0 |
| SRR6932951 | 297 | 21 | 1895 | 0 | 0 | 0 | 0 | 0 | 0 | 0 | 0 | 872 | 0 | 0 | 706 | 0 | 60 | 0 | 0 | 13 | 13 | 0 | 0 | 0 | 73 | 0 | 14 | 0 | 0 | 0 | 38 | 0 | 0 | 0 | 0 | 10 | 0 |
| SRR6932952 | 1634 | 27 | 2661 | 0 | 0 | 965 | 74 | 0 | 76 | 0 | 0 | 1082 | 0 | 16 | 49 | 0 | 36 | 0 | 0 | 31 | 1130 | 0 | 0 | 0 | 11 | 5275 | 195 | 27 | 0 | 0 | 141 | 0 | 0 | 0 | 7178 | 2223 | 6230 |
| SRR6932953 | 1455 | 21 | 4188 | 0 | 0 | 64 | 76 | 0 | 67 | 0 | 0 | 852 | 0 | 21 | 83 | 0 | 34 | 0 | 0 | 38 | 545 | 0 | 0 | 15 | 15 | 367 | 29 | 21 | 10 | 0 | 108 | 0 | 0 | 0 | 487 | 69 | 297 |
| SRR6932954 | 1708 | 179 | 3659 | 0 | 0 | 14 | 22 | 0 | 125 | 0 | 0 | 819 | 0 | 27 | 594 | 0 | 27 | 0 | 0 | 0 | 327 | 0 | 0 | 0 | 11 | 144 | 70 | 32 | 0 | 0 | 87 | 0 | 0 | 0 | 405 | 18 | 125 |
| SRR6932955 | 1569 | 76 | 2010 | 0 | 0 | 0 | 0 | 0 | 20 | 0 | 0 | 1101 | 0 | 20 | 192 | 0 | 17 | 13 | 0 | 0 | 12 | 0 | 0 | 15 | 76 | 15 | 0 | 15 | 25 | 0 | 61 | 0 | 0 | 20 | 104 | 20 | 62 |
| SRR7062776 | 89 | 12 | 8417 | 0 | 0 | 106 | 1008 | 44 | 166 | 0 | 0 | 1009 | 12 | 24 | 0 | 0 | 168 | 0 | 50 | 482 | 26 | 0 | 0 | 122 | 113 | 1183 | 2768 | 19 | 0 | 24 | 25 | 0 | 0 | 27 | 94 | 31 | 74 |
| SRR7062777 | 86 | 14 | 8283 | 0 | 0 | 116 | 930 | 16 | 140 | 0 | 0 | 909 | 0 | 44 | 0 | 0 | 164 | 0 | 50 | 540 | 24 | 0 | 0 | 46 | 87 | 1067 | 2648 | 20 | 0 | 14 | 35 | 0 | 0 | 29 | 70 | 23 | 49 |
| SRR7062778 | 99 | 10 | 8839 | 0 | 0 | 124 | 822 | 17 | 158 | 0 | 0 | 955 | 0 | 34 | 0 | 0 | 168 | 0 | 58 | 555 | 31 | 0 | 0 | 58 | 126 | 1166 | 1369 | 26 | 52 | 0 | 49 | 0 | 0 | 15 | 66 | 15 | 53 |
| SRR7062779 | 99 | 0 | 8955 | 0 | 0 | 134 | 744 | 16 | 181 | 0 | 0 | 1008 | 0 | 43 | 0 | 0 | 155 | 0 | 43 | 592 | 35 | 0 | 0 | 43 | 122 | 1087 | 1284 | 16 | 45 | 13 | 36 | 0 | 0 | 23 | 75 | 23 | 45 |
| SRR7062780 | 92 | 0 | 8569 | 0 | 0 | 20 | 614 | 20 | 26 | 0 | 0 | 984 | 0 | 44 | 0 | 0 | 177 | 0 | 43 | 520 | 40 | 0 | 0 | 21 | 23 | 205 | 263 | 27 | 0 | 0 | 30 | 0 | 0 | 11 | 17 | 28 | 11 |
| SRR7062781 | 85 | 0 | 8992 | 0 | 0 | 83 | 499 | 15 | 128 | 0 | 0 | 956 | 0 | 47 | 0 | 0 | 145 | 0 | 44 | 476 | 30 | 0 | 0 | 61 | 118 | 921 | 1250 | 26 | 15 | 10 | 33 | 0 | 10 | 47 | 80 | 28 | 46 |
| SRR7081766 | 154 | 45 | 283 | 195 | 60 | 0 | 0 | 0 | 0 | 0 | 0 | 152 | 0 | 15 | 0 | 30 | 0 | 0 | 0 | 576 | 346 | 45 | 0 | 0 | 0 | 0 | 314 | 0 | 0 | 0 | 0 | 0 | 0 | 15 | 0 | 0 | 0 |
| SRR7081767 | 249 | 0 | 313 | 344 | 283 | 0 | 0 | 0 | 0 | 0 | 0 | 233 | 0 | 101 | 0 | 0 | 0 | 0 | 0 | 190 | 190 | 61 | 0 | 0 | 0 | 61 | 121 | 0 | 0 | 0 | 0 | 0 | 0 | 0 | 0 | 0 | 0 |
| SRR7081768 | 210 | 0 | 655 | 107 | 103 | 0 | 0 | 0 | 0 | 0 | 0 | 199 | 0 | 0 | 0 | 0 | 18 | 0 | 0 | 201 | 134 | 0 | 0 | 0 | 0 | 127 | 32 | 0 | 0 | 0 | 0 | 0 | 0 | 0 | 0 | 0 | 0 |
| SRR7081769 | 273 | 0 | 149 | 173 | 26 | 0 | 0 | 60 | 0 | 0 | 0 | 309 | 0 | 0 | 173 | 0 | 54 | 0 | 0 | 230 | 115 | 35 | 0 | 0 | 0 | 60 | 0 | 52 | 0 | 0 | 0 | 0 | 0 | 0 | 0 | 0 | 0 |
| SRR7081770 | 91 | 59 | 107 | 176 | 59 | 0 | 0 | 0 | 0 | 0 | 0 | 73 | 0 | 20 | 0 | 0 | 0 | 0 | 0 | 531 | 743 | 78 | 0 | 0 | 0 | 0 | 117 | 0 | 0 | 0 | 0 | 0 | 0 | 20 | 98 | 0 | 0 |
| SRR7081771 | 116 | 10 | 234 | 271 | 125 | 0 | 0 | 10 | 0 | 0 | 0 | 132 | 0 | 0 | 0 | 0 | 10 | 0 | 0 | 44 | 267 | 0 | 0 | 0 | 0 | 94 | 177 | 0 | 0 | 0 | 0 | 0 | 0 | 10 | 21 | 0 | 0 |
| SRR7081772 | 316 | 75 | 132 | 212 | 75 | 0 | 0 | 25 | 0 | 0 | 0 | 274 | 0 | 25 | 0 | 0 | 0 | 0 | 0 | 0 | 551 | 37 | 0 | 0 | 0 | 12 | 75 | 0 | 0 | 0 | 0 | 12 | 0 | 0 | 0 | 0 | 0 |
| SRR7081773 | 153 | 0 | 122 | 194 | 194 | 0 | 0 | 33 | 0 | 0 | 0 | 26 | 0 | 0 | 0 | 0 | 0 | 0 | 0 | 145 | 872 | 33 | 0 | 0 | 0 | 0 | 120 | 0 | 0 | 0 | 0 | 0 | 0 | 47 | 13 | 0 | 0 |
| SRR7081774 | 256 | 0 | 248 | 224 | 112 | 0 | 0 | 0 | 0 | 0 | 0 | 150 | 0 | 0 | 0 | 0 | 0 | 0 | 0 | 125 | 375 | 43 | 0 | 0 | 0 | 0 | 163 | 0 | 0 | 0 | 0 | 0 | 0 | 0 | 0 | 0 | 0 |
| SRR7081775 | 126 | 0 | 90 | 156 | 182 | 0 | 69 | 0 | 0 | 0 | 0 | 142 | 0 | 0 | 0 | 0 | 0 | 0 | 0 | 139 | 416 | 35 | 0 | 0 | 0 | 0 | 78 | 0 | 0 | 0 | 0 | 0 | 0 | 0 | 0 | 0 | 0 |
| SRR7081776 | 329 | 15 | 125 | 91 | 183 | 0 | 0 | 0 | 0 | 0 | 0 | 217 | 0 | 0 | 0 | 0 | 0 | 0 | 0 | 0 | 0 | 15 | 0 | 0 | 0 | 0 | 15 | 0 | 0 | 0 | 0 | 0 | 0 | 0 | 0 | 0 | 0 |
| SRR7081777 | 983 | 0 | 366 | 385 | 37 | 0 | 190 | 0 | 0 | 0 | 0 | 760 | 0 | 0 | 0 | 0 | 0 | 0 | 0 | 95 | 476 | 0 | 0 | 0 | 0 | 0 | 128 | 0 | 0 | 0 | 0 | 0 | 0 | 0 | 0 | 0 | 0 |
| SRR7107416 | 0 | 12 | 707 | 0 | 0 | 461 | 0 | 0 | 62 | 0 | 0 | 0 | 0 | 0 | 46 | 0 | 14 | 0 | 44 | 0 | 2784 | 0 | 0 | 0 | 13 | 4622 | 2524 | 0 | 0 | 0 | 214 | 0 | 0 | 0 | 0 | 2776 | 0 |
| SRR7107417 | 0 | 12 | 682 | 0 | 0 | 26 | 0 | 0 | 63 | 0 | 0 | 0 | 0 | 0 | 61 | 0 | 13 | 0 | 27 | 0 | 2563 | 0 | 0 | 0 | 0 | 287 | 91 | 0 | 0 | 0 | 14 | 0 | 0 | 0 | 0 | 204 | 0 |
| SRR7107418 | 167 | 0 | 1721 | 0 | 0 | 0 | 15 | 109 | 0 | 0 | 0 | 195 | 0 | 0 | 0 | 0 | 0 | 0 | 14 | 1545 | 703 | 20 | 0 | 13 | 0 | 0 | 669 | 19 | 0 | 13 | 0 | 0 | 0 | 0 | 88 | 105 | 69 |
| SRR7107419 | 204 | 0 | 1692 | 0 | 0 | 0 | 11 | 65 | 0 | 0 | 0 | 195 | 0 | 14 | 0 | 0 | 10 | 0 | 16 | 1479 | 493 | 27 | 0 | 0 | 0 | 0 | 9406 | 18 | 0 | 0 | 0 | 0 | 0 | 0 | 1122 | 109 | 832 |
| SRR7107420 | 0 | 15 | 0 | 0 | 0 | 450 | 0 | 0 | 60 | 0 | 0 | 0 | 0 | 0 | 29 | 0 | 12 | 0 | 36 | 5037 | 3923 | 0 | 0 | 0 | 0 | 4968 | 0 | 0 | 0 | 0 | 274 | 0 | 0 | 0 | 0 | 2693 | 0 |
| SRR7107421 | 0 | 15 | 0 | 0 | 0 | 26 | 0 | 0 | 53 | 0 | 0 | 0 | 0 | 0 | 45 | 0 | 13 | 0 | 29 | 4738 | 3605 | 0 | 0 | 0 | 0 | 247 | 0 | 0 | 0 | 0 | 12 | 0 | 0 | 0 | 0 | 121 | 0 |
| SRR7126083 | 511 | 0 | 0 | 0 | 15 | 0 | 133 | 1082 | 15 | 15 | 58 | 151 | 29 | 15 | 67 | 0 | 17 | 0 | 0 | 0 | 666 | 29 | 0 | 44 | 0 | 15 | 29 | 0 | 161 | 1009 | 117 | 0 | 29 | 395 | 0 | 29 | 44 |
| SRR7126084 | 162 | 19 | 200 | 0 | 0 | 70 | 2192 | 715 | 44 | 0 | 0 | 1070 | 55 | 14 | 1967 | 0 | 331 | 94 | 211 | 1261 | 2353 | 0 | 0 | 46 | 73 | 376 | 25 | 0 | 0 | 70 | 96 | 25 | 40 | 82 | 517 | 1097 | 1425 |
| SRR7126085 | 215 | 28 | 341 | 0 | 0 | 115 | 3871 | 594 | 45 | 0 | 14 | 955 | 38 | 0 | 3888 | 0 | 273 | 34 | 312 | 1533 | 3576 | 0 | 0 | 28 | 82 | 683 | 32 | 0 | 0 | 53 | 103 | 34 | 39 | 80 | 699 | 1810 | 2117 |
| SRR7126086 | 259 | 28 | 434 | 0 | 0 | 1309 | 4359 | 4035 | 530 | 0 | 28 | 831 | 160 | 11 | 3275 | 0 | 317 | 0 | 163 | 489 | 2522 | 0 | 0 | 173 | 43 | 8386 | 214 | 0 | 0 | 393 | 1324 | 11 | 56 | 981 | 5320 | 13090 | 15271 |
| SRR7126087 | 217 | 28 | 297 | 0 | 0 | 853 | 2774 | 7045 | 367 | 0 | 24 | 980 | 161 | 55 | 1220 | 0 | 421 | 82 | 182 | 1194 | 1703 | 0 | 0 | 127 | 42 | 5132 | 149 | 0 | 0 | 430 | 682 | 18 | 28 | 427 | 4839 | 5647 | 6147 |
| SRR7126088 | 204 | 27 | 224 | 0 | 0 | 1236 | 5679 | 3928 | 559 | 0 | 23 | 799 | 95 | 30 | 2855 | 0 | 238 | 28 | 182 | 455 | 2128 | 0 | 0 | 200 | 21 | 7073 | 131 | 0 | 0 | 342 | 1141 | 15 | 55 | 610 | 4373 | 10583 | 12885 |
| SRR7126089 | 154 | 30 | 151 | 0 | 0 | 140 | 3457 | 500 | 69 | 0 | 23 | 824 | 17 | 15 | 6737 | 0 | 185 | 13 | 76 | 360 | 4947 | 24 | 0 | 24 | 54 | 703 | 16 | 0 | 0 | 43 | 112 | 30 | 14 | 107 | 430 | 1243 | 1508 |
| SRR7126090 | 186 | 30 | 228 | 0 | 0 | 120 | 4692 | 1961 | 52 | 0 | 18 | 976 | 61 | 11 | 3807 | 0 | 234 | 39 | 253 | 1185 | 2712 | 13 | 0 | 28 | 68 | 667 | 25 | 0 | 0 | 138 | 123 | 62 | 25 | 145 | 626 | 1792 | 2495 |
| SRR7126091 | 143 | 39 | 258 | 0 | 0 | 75 | 2697 | 833 | 38 | 0 | 16 | 1116 | 28 | 13 | 1433 | 0 | 308 | 28 | 249 | 1323 | 1238 | 0 | 0 | 33 | 47 | 436 | 21 | 0 | 0 | 57 | 93 | 19 | 14 | 58 | 603 | 1377 | 1730 |
| SRR7126092 | 171 | 22 | 184 | 0 | 0 | 114 | 2051 | 972 | 58 | 0 | 29 | 783 | 15 | 41 | 1342 | 0 | 296 | 27 | 159 | 698 | 1385 | 0 | 0 | 26 | 27 | 647 | 20 | 0 | 0 | 39 | 94 | 35 | 10 | 82 | 534 | 1075 | 1271 |
| SRR7126093 | 190 | 39 | 396 | 0 | 0 | 119 | 3738 | 530 | 50 | 0 | 25 | 740 | 25 | 23 | 2447 | 0 | 277 | 0 | 141 | 615 | 2295 | 0 | 0 | 27 | 50 | 662 | 21 | 0 | 0 | 64 | 116 | 18 | 12 | 88 | 760 | 1754 | 1907 |
| SRR7153253 | 35 | 0 | 1063 | 0 | 0 | 14 | 0 | 16 | 14 | 0 | 0 | 169 | 0 | 12 | 0 | 0 | 0 | 0 | 0 | 89 | 32 | 0 | 0 | 0 | 0 | 10 | 68 | 0 | 0 | 0 | 0 | 0 | 0 | 29 | 83 | 0 | 34 |
| SRR7153254 | 39 | 0 | 945 | 0 | 0 | 19 | 0 | 11 | 0 | 0 | 0 | 167 | 0 | 0 | 18 | 0 | 11 | 0 | 0 | 136 | 75 | 0 | 0 | 0 | 0 | 155 | 41 | 11 | 0 | 0 | 0 | 0 | 0 | 0 | 138 | 0 | 68 |
| SRR7153255 | 46 | 0 | 1214 | 0 | 0 | 11 | 0 | 11 | 0 | 0 | 0 | 233 | 0 | 0 | 120 | 0 | 0 | 0 | 0 | 209 | 84 | 0 | 0 | 38 | 0 | 158 | 694 | 0 | 0 | 0 | 0 | 0 | 0 | 0 | 117 | 0 | 63 |
| SRR7153256 | 47 | 0 | 1125 | 0 | 0 | 25 | 0 | 50 | 10 | 0 | 0 | 264 | 0 | 0 | 19 | 0 | 11 | 0 | 0 | 302 | 64 | 0 | 0 | 61 | 0 | 294 | 905 | 10 | 0 | 12 | 11 | 0 | 0 | 0 | 238 | 0 | 118 |
| SRR7153257 | 51 | 0 | 955 | 0 | 0 | 0 | 0 | 75 | 0 | 0 | 0 | 86 | 0 | 0 | 12 | 0 | 11 | 0 | 0 | 106 | 50 | 0 | 0 | 52 | 0 | 129 | 733 | 0 | 0 | 0 | 0 | 0 | 0 | 0 | 54 | 0 | 22 |
| SRR7153258 | 30 | 0 | 925 | 0 | 0 | 17 | 0 | 26 | 0 | 0 | 0 | 115 | 0 | 0 | 0 | 0 | 12 | 0 | 11 | 205 | 34 | 0 | 0 | 78 | 0 | 10 | 61 | 0 | 0 | 0 | 0 | 0 | 0 | 0 | 101 | 0 | 53 |
| SRR7153259 | 42 | 0 | 1013 | 0 | 0 | 17 | 0 | 17 | 0 | 0 | 0 | 224 | 0 | 0 | 0 | 0 | 0 | 0 | 0 | 95 | 27 | 0 | 0 | 34 | 0 | 235 | 1034 | 0 | 0 | 0 | 0 | 0 | 0 | 0 | 74 | 0 | 44 |
| SRR7153260 | 32 | 0 | 1096 | 0 | 0 | 17 | 0 | 14 | 12 | 0 | 0 | 189 | 0 | 0 | 0 | 0 | 0 | 0 | 0 | 77 | 28 | 0 | 0 | 13 | 0 | 254 | 1206 | 0 | 0 | 0 | 0 | 0 | 0 | 0 | 100 | 0 | 52 |
| SRR7153261 | 55 | 0 | 1464 | 0 | 0 | 13 | 0 | 44 | 0 | 0 | 0 | 204 | 0 | 0 | 0 | 0 | 0 | 0 | 0 | 80 | 35 | 0 | 0 | 48 | 0 | 185 | 1490 | 0 | 0 | 21 | 0 | 0 | 0 | 13 | 81 | 0 | 38 |
| SRR7153262 | 41 | 0 | 1201 | 0 | 0 | 0 | 22 | 19 | 0 | 0 | 0 | 195 | 0 | 0 | 0 | 0 | 11 | 0 | 0 | 188 | 56 | 0 | 0 | 71 | 0 | 78 | 105 | 0 | 0 | 0 | 0 | 0 | 0 | 0 | 51 | 0 | 24 |
| SRR7153263 | 25 | 0 | 1449 | 0 | 0 | 17 | 13 | 0 | 10 | 0 | 0 | 241 | 0 | 0 | 30 | 0 | 0 | 0 | 0 | 228 | 73 | 0 | 0 | 25 | 0 | 21 | 115 | 0 | 0 | 0 | 0 | 0 | 0 | 0 | 11 | 0 | 49 |
| SRR7153264 | 38 | 0 | 1034 | 0 | 0 | 38 | 0 | 32 | 21 | 0 | 0 | 150 | 0 | 0 | 11 | 0 | 11 | 0 | 0 | 85 | 32 | 0 | 0 | 38 | 0 | 13 | 41 | 12 | 0 | 0 | 13 | 0 | 0 | 0 | 10 | 0 | 114 |
| SRR7153265 | 42 | 0 | 1175 | 0 | 0 | 20 | 0 | 15 | 0 | 0 | 0 | 216 | 0 | 0 | 20 | 0 | 0 | 0 | 0 | 247 | 58 | 0 | 0 | 50 | 0 | 227 | 1328 | 0 | 0 | 0 | 0 | 0 | 0 | 0 | 122 | 0 | 58 |
| SRR7153266 | 58 | 0 | 1070 | 0 | 0 | 22 | 11 | 30 | 0 | 0 | 0 | 228 | 0 | 0 | 0 | 0 | 0 | 0 | 0 | 255 | 33 | 0 | 0 | 54 | 0 | 17 | 151 | 0 | 0 | 0 | 0 | 0 | 0 | 0 | 69 | 0 | 44 |
| SRR7153267 | 49 | 0 | 1030 | 0 | 0 | 37 | 0 | 0 | 0 | 0 | 0 | 231 | 0 | 0 | 18 | 0 | 0 | 0 | 0 | 145 | 29 | 0 | 0 | 0 | 0 | 12 | 94 | 0 | 0 | 0 | 0 | 0 | 0 | 0 | 81 | 0 | 42 |
| SRR7153268 | 67 | 0 | 1586 | 0 | 0 | 34 | 0 | 11 | 0 | 0 | 0 | 278 | 0 | 0 | 0 | 0 | 13 | 0 | 0 | 115 | 14 | 0 | 0 | 55 | 0 | 223 | 1235 | 0 | 0 | 0 | 0 | 0 | 0 | 0 | 54 | 0 | 42 |
| SRR7153269 | 46 | 0 | 1073 | 0 | 0 | 11 | 0 | 0 | 15 | 0 | 0 | 146 | 0 | 0 | 12 | 0 | 0 | 0 | 0 | 173 | 36 | 0 | 0 | 0 | 0 | 235 | 801 | 0 | 0 | 0 | 0 | 0 | 0 | 0 | 83 | 0 | 51 |
| SRR7153270 | 42 | 0 | 1277 | 0 | 0 | 18 | 0 | 0 | 11 | 0 | 0 | 184 | 0 | 0 | 15 | 0 | 0 | 0 | 0 | 120 | 56 | 0 | 0 | 33 | 0 | 177 | 1089 | 0 | 0 | 0 | 0 | 0 | 0 | 0 | 54 | 0 | 47 |
| SRR7153271 | 44 | 0 | 972 | 0 | 0 | 0 | 0 | 14 | 0 | 0 | 0 | 172 | 0 | 0 | 33 | 0 | 0 | 0 | 0 | 394 | 165 | 0 | 0 | 14 | 0 | 13 | 159 | 0 | 0 | 0 | 0 | 0 | 0 | 0 | 40 | 0 | 32 |
| SRR7153272 | 40 | 0 | 864 | 0 | 0 | 0 | 12 | 0 | 0 | 0 | 0 | 133 | 0 | 0 | 11 | 0 | 0 | 0 | 0 | 137 | 50 | 0 | 0 | 55 | 0 | 79 | 85 | 0 | 0 | 0 | 0 | 0 | 0 | 0 | 60 | 0 | 23 |
| SRR7153273 | 37 | 0 | 1700 | 0 | 0 | 0 | 0 | 12 | 0 | 0 | 0 | 143 | 0 | 0 | 58 | 0 | 11 | 0 | 0 | 181 | 79 | 0 | 0 | 50 | 0 | 59 | 731 | 0 | 0 | 0 | 0 | 0 | 0 | 0 | 192 | 0 | 71 |
| SRR7153274 | 40 | 0 | 1048 | 0 | 0 | 0 | 0 | 21 | 0 | 0 | 0 | 129 | 0 | 0 | 0 | 0 | 0 | 0 | 0 | 377 | 68 | 0 | 0 | 58 | 0 | 133 | 1091 | 0 | 0 | 0 | 0 | 0 | 0 | 0 | 86 | 0 | 34 |
| SRR7153275 | 48 | 0 | 1263 | 0 | 0 | 10 | 0 | 18 | 0 | 0 | 0 | 121 | 0 | 0 | 0 | 0 | 11 | 0 | 0 | 197 | 76 | 0 | 0 | 38 | 0 | 221 | 1685 | 0 | 0 | 0 | 0 | 0 | 0 | 0 | 198 | 12 | 86 |
| SRR7153276 | 51 | 0 | 1140 | 0 | 0 | 14 | 21 | 15 | 0 | 0 | 0 | 133 | 0 | 0 | 68 | 0 | 0 | 0 | 0 | 1027 | 438 | 0 | 0 | 29 | 0 | 123 | 1352 | 0 | 0 | 0 | 0 | 0 | 0 | 0 | 62 | 0 | 29 |
| SRR7153277 | 35 | 0 | 975 | 0 | 0 | 11 | 0 | 25 | 0 | 0 | 0 | 70 | 0 | 0 | 22 | 0 | 0 | 0 | 0 | 164 | 0 | 0 | 0 | 11 | 0 | 81 | 967 | 0 | 0 | 0 | 0 | 0 | 0 | 0 | 44 | 0 | 18 |
| SRR7153278 | 58 | 0 | 1374 | 0 | 0 | 0 | 24 | 2695 | 0 | 0 | 0 | 147 | 0 | 0 | 74 | 0 | 12 | 0 | 0 | 248 | 110 | 0 | 0 | 12 | 0 | 69 | 486 | 0 | 20 | 661 | 0 | 0 | 0 | 20 | 38 | 0 | 23 |
| SRR7153279 | 37 | 0 | 2182 | 0 | 0 | 10 | 0 | 20 | 0 | 0 | 0 | 123 | 0 | 0 | 58 | 0 | 0 | 0 | 0 | 350 | 82 | 0 | 0 | 36 | 0 | 93 | 1412 | 0 | 0 | 0 | 0 | 0 | 0 | 15 | 115 | 0 | 52 |
| SRR7153280 | 39 | 0 | 1055 | 0 | 0 | 11 | 0 | 37 | 0 | 0 | 0 | 116 | 0 | 0 | 34 | 0 | 0 | 0 | 0 | 404 | 0 | 0 | 0 | 45 | 0 | 12 | 136 | 0 | 0 | 0 | 0 | 0 | 0 | 0 | 58 | 0 | 35 |
| SRR7153281 | 31 | 0 | 1075 | 0 | 0 | 13 | 0 | 64 | 0 | 0 | 0 | 93 | 0 | 0 | 28 | 0 | 11 | 0 | 0 | 192 | 47 | 0 | 0 | 48 | 0 | 12 | 61 | 0 | 0 | 16 | 0 | 0 | 0 | 0 | 106 | 0 | 42 |
| SRR7153282 | 53 | 0 | 979 | 0 | 0 | 0 | 14 | 0 | 0 | 0 | 0 | 115 | 0 | 0 | 10 | 0 | 0 | 0 | 0 | 219 | 109 | 0 | 0 | 45 | 0 | 57 | 663 | 0 | 0 | 0 | 0 | 0 | 0 | 0 | 44 | 0 | 27 |
| SRR7153283 | 54 | 0 | 868 | 0 | 0 | 17 | 0 | 12 | 13 | 0 | 0 | 116 | 0 | 0 | 117 | 0 | 0 | 0 | 0 | 234 | 67 | 0 | 0 | 27 | 0 | 13 | 83 | 0 | 0 | 0 | 0 | 0 | 0 | 0 | 118 | 0 | 50 |
| SRR7153284 | 52 | 0 | 1039 | 0 | 0 | 0 | 0 | 22 | 0 | 0 | 0 | 66 | 0 | 0 | 0 | 0 | 0 | 0 | 0 | 81 | 0 | 0 | 0 | 26 | 0 | 111 | 1411 | 0 | 0 | 0 | 0 | 0 | 0 | 0 | 23 | 0 | 16 |
| SRR7153285 | 41 | 0 | 925 | 0 | 0 | 12 | 0 | 0 | 0 | 0 | 0 | 117 | 0 | 0 | 37 | 0 | 0 | 0 | 0 | 93 | 25 | 0 | 0 | 0 | 0 | 135 | 1316 | 0 | 0 | 0 | 0 | 0 | 0 | 0 | 75 | 0 | 42 |
| SRR7153286 | 43 | 0 | 904 | 0 | 0 | 24 | 10 | 59 | 0 | 0 | 0 | 109 | 0 | 0 | 131 | 0 | 0 | 0 | 0 | 346 | 119 | 0 | 0 | 0 | 0 | 290 | 1124 | 0 | 0 | 19 | 0 | 0 | 0 | 0 | 75 | 0 | 30 |
| SRR7153287 | 44 | 0 | 1310 | 0 | 0 | 22 | 0 | 19 | 0 | 0 | 0 | 102 | 0 | 0 | 0 | 0 | 0 | 0 | 0 | 188 | 17 | 0 | 0 | 0 | 0 | 238 | 1526 | 0 | 0 | 0 | 0 | 0 | 0 | 0 | 25 | 0 | 15 |
| SRR7153288 | 52 | 0 | 961 | 0 | 0 | 20 | 12 | 15 | 0 | 0 | 0 | 91 | 0 | 0 | 23 | 0 | 0 | 0 | 0 | 167 | 28 | 0 | 0 | 0 | 0 | 161 | 2132 | 0 | 0 | 0 | 0 | 0 | 0 | 0 | 119 | 0 | 48 |
| SRR7153289 | 39 | 0 | 1150 | 0 | 0 | 0 | 0 | 30 | 0 | 0 | 0 | 85 | 0 | 0 | 13 | 0 | 11 | 0 | 0 | 141 | 42 | 0 | 0 | 39 | 0 | 95 | 91 | 0 | 0 | 0 | 0 | 0 | 0 | 0 | 51 | 0 | 30 |
| SRR7153290 | 55 | 0 | 1213 | 0 | 0 | 17 | 0 | 0 | 0 | 0 | 0 | 95 | 0 | 0 | 11 | 0 | 0 | 0 | 0 | 206 | 27 | 0 | 0 | 44 | 0 | 16 | 107 | 0 | 0 | 0 | 0 | 0 | 0 | 0 | 94 | 0 | 50 |
| SRR7153291 | 43 | 0 | 1920 | 0 | 0 | 10 | 15 | 0 | 0 | 0 | 0 | 120 | 0 | 0 | 101 | 0 | 0 | 0 | 0 | 261 | 84 | 0 | 0 | 0 | 0 | 102 | 140 | 0 | 0 | 0 | 0 | 0 | 0 | 0 | 102 | 0 | 67 |
| SRR7153292 | 43 | 0 | 1837 | 0 | 0 | 0 | 0 | 37 | 0 | 0 | 0 | 155 | 0 | 0 | 48 | 0 | 0 | 0 | 0 | 207 | 68 | 0 | 0 | 15 | 0 | 67 | 86 | 0 | 0 | 0 | 0 | 0 | 0 | 12 | 102 | 0 | 58 |
| SRR7153293 | 89 | 0 | 2434 | 0 | 0 | 0 | 28 | 43 | 0 | 0 | 0 | 109 | 0 | 0 | 40 | 0 | 13 | 0 | 0 | 1080 | 297 | 0 | 0 | 65 | 0 | 108 | 2500 | 0 | 0 | 0 | 0 | 0 | 0 | 0 | 82 | 0 | 55 |
| SRR7153294 | 33 | 0 | 877 | 0 | 0 | 0 | 16 | 28 | 0 | 0 | 0 | 105 | 0 | 0 | 37 | 0 | 0 | 0 | 0 | 241 | 42 | 0 | 0 | 63 | 0 | 84 | 1124 | 0 | 0 | 0 | 0 | 0 | 0 | 0 | 22 | 0 | 11 |
| SRR7153295 | 43 | 0 | 1024 | 0 | 0 | 0 | 0 | 90 | 0 | 0 | 0 | 169 | 0 | 0 | 0 | 0 | 12 | 0 | 0 | 145 | 68 | 0 | 0 | 44 | 0 | 128 | 83 | 0 | 0 | 15 | 0 | 0 | 0 | 0 | 35 | 0 | 29 |
| SRR7153296 | 36 | 0 | 814 | 0 | 0 | 12 | 0 | 47 | 0 | 0 | 0 | 119 | 0 | 0 | 22 | 0 | 12 | 0 | 0 | 488 | 110 | 0 | 0 | 70 | 0 | 68 | 931 | 0 | 0 | 14 | 0 | 0 | 0 | 0 | 27 | 0 | 19 |
| SRR7153297 | 51 | 0 | 904 | 0 | 0 | 0 | 71 | 590 | 0 | 0 | 0 | 104 | 0 | 27 | 52 | 0 | 0 | 0 | 0 | 295 | 186 | 0 | 0 | 0 | 0 | 43 | 496 | 0 | 12 | 215 | 0 | 0 | 0 | 29 | 12 | 0 | 0 |
| SRR7153298 | 28 | 0 | 1265 | 0 | 0 | 14 | 14 | 26 | 0 | 0 | 0 | 103 | 0 | 0 | 22 | 0 | 0 | 0 | 0 | 228 | 28 | 0 | 0 | 29 | 0 | 119 | 1619 | 0 | 0 | 0 | 0 | 0 | 0 | 0 | 59 | 0 | 21 |

### Supp Table S8.2.Columns_AM_to_BW

| | | | | | | | | | | | | | | | | | | | | | | | | | | | | | | | | | | | | | |
|---|---|---|---|---|---|---|---|---|---|---|---|---|---|---|---|---|---|---|---|---|---|---|---|---|---|---|---|---|---|---|---|---|---|---|---|---|---|
| SRR7153299 | 65 | 0 | 1389 | 0 | 0 | 18 | 0 | 0 | 0 | 0 | 0 | 134 | 0 | 0 | 0 | 0 | 12 | 0 | 0 | 105 | 45 | 0 | 0 | 0 | 0 | 170 | 1649 | 0 | 0 | 0 | 0 | 0 | 0 | 0 | 135 | 0 | 68 |
| SRR7153300 | 71 | 0 | 1084 | 0 | 0 | 13 | 0 | 41 | 0 | 0 | 0 | 103 | 0 | 0 | 14 | 0 | 0 | 0 | 0 | 186 | 39 | 0 | 0 | 0 | 0 | 15 | 84 | 0 | 0 | 17 | 0 | 0 | 0 | 0 | 82 | 0 | 54 |
| SRR7153301 | 38 | 0 | 1308 | 0 | 0 | 0 | 19 | 1982 | 13 | 0 | 0 | 109 | 0 | 0 | 246 | 0 | 13 | 0 | 0 | 201 | 73 | 0 | 0 | 19 | 0 | 124 | 578 | 11 | 15 | 483 | 0 | 0 | 0 | 12 | 30 | 0 | 14 |
| SRR7153302 | 38 | 0 | 1004 | 0 | 0 | 0 | 0 | 28 | 0 | 0 | 0 | 90 | 0 | 0 | 16 | 0 | 0 | 0 | 0 | 194 | 33 | 0 | 0 | 26 | 0 | 66 | 1230 | 0 | 0 | 0 | 0 | 0 | 0 | 0 | 40 | 0 | 25 |
| SRR7153303 | 64 | 0 | 1870 | 0 | 0 | 0 | 21 | 0 | 0 | 0 | 0 | 73 | 0 | 0 | 530 | 0 | 0 | 0 | 0 | 763 | 305 | 0 | 0 | 0 | 0 | 102 | 897 | 0 | 0 | 0 | 0 | 0 | 0 | 0 | 169 | 0 | 76 |
| SRR7153304 | 66 | 0 | 1347 | 0 | 0 | 24 | 12 | 20 | 0 | 0 | 0 | 165 | 0 | 0 | 0 | 0 | 10 | 0 | 0 | 285 | 38 | 0 | 0 | 0 | 0 | 183 | 1550 | 0 | 0 | 12 | 0 | 0 | 0 | 0 | 46 | 0 | 33 |
| SRR7153305 | 44 | 0 | 1290 | 0 | 0 | 0 | 11 | 25 | 0 | 0 | 0 | 103 | 0 | 0 | 12 | 0 | 0 | 0 | 0 | 427 | 45 | 0 | 0 | 42 | 0 | 13 | 141 | 0 | 0 | 0 | 0 | 0 | 0 | 0 | 60 | 0 | 29 |
| SRR7153306 | 55 | 0 | 1428 | 0 | 0 | 14 | 0 | 17 | 0 | 0 | 0 | 105 | 0 | 0 | 16 | 0 | 0 | 0 | 0 | 171 | 61 | 0 | 0 | 63 | 0 | 12 | 102 | 0 | 0 | 28 | 0 | 0 | 0 | 0 | 48 | 0 | 23 |
| SRR7153307 | 33 | 0 | 1132 | 0 | 0 | 0 | 0 | 19 | 0 | 0 | 0 | 133 | 0 | 0 | 10 | 0 | 0 | 0 | 0 | 177 | 25 | 0 | 0 | 45 | 0 | 132 | 995 | 0 | 0 | 0 | 0 | 0 | 0 | 0 | 93 | 0 | 40 |
| SRR7153308 | 57 | 0 | 1015 | 0 | 0 | 20 | 12 | 15 | 0 | 0 | 0 | 143 | 0 | 0 | 44 | 0 | 11 | 0 | 0 | 194 | 61 | 0 | 0 | 12 | 0 | 170 | 1096 | 0 | 0 | 0 | 0 | 0 | 0 | 0 | 71 | 0 | 29 |
| SRR7153309 | 37 | 0 | 1247 | 0 | 0 | 12 | 10 | 31 | 0 | 0 | 0 | 85 | 0 | 0 | 0 | 0 | 0 | 0 | 0 | 215 | 53 | 0 | 0 | 56 | 0 | 92 | 91 | 0 | 0 | 0 | 0 | 0 | 0 | 0 | 47 | 0 | 25 |
| SRR7153310 | 59 | 0 | 1080 | 0 | 0 | 27 | 0 | 17 | 0 | 0 | 0 | 100 | 0 | 11 | 14 | 0 | 11 | 0 | 0 | 16 | 28 | 0 | 0 | 0 | 0 | 19 | 60 | 0 | 23 | 0 | 0 | 0 | 0 | 13 | 121 | 0 | 55 |
| SRR7153311 | 57 | 0 | 1012 | 0 | 0 | 11 | 12 | 0 | 0 | 0 | 0 | 111 | 0 | 0 | 51 | 0 | 0 | 0 | 0 | 472 | 183 | 0 | 0 | 0 | 0 | 113 | 1246 | 0 | 0 | 0 | 0 | 0 | 0 | 0 | 66 | 0 | 30 |
| SRR7153312 | 63 | 0 | 2292 | 0 | 0 | 10 | 0 | 0 | 0 | 0 | 0 | 102 | 0 | 0 | 51 | 0 | 0 | 0 | 0 | 475 | 121 | 0 | 0 | 58 | 0 | 10 | 153 | 0 | 0 | 0 | 0 | 0 | 0 | 0 | 16 | 0 | 83 |
| SRR7153313 | 29 | 0 | 1845 | 0 | 0 | 16 | 21 | 21 | 0 | 0 | 0 | 70 | 0 | 0 | 38 | 0 | 0 | 0 | 0 | 308 | 35 | 0 | 0 | 25 | 0 | 150 | 68 | 0 | 0 | 34 | 0 | 0 | 0 | 20 | 126 | 11 | 55 |
| SRR7153314 | 46 | 0 | 988 | 0 | 0 | 11 | 10 | 26 | 0 | 0 | 0 | 81 | 0 | 0 | 11 | 0 | 0 | 0 | 0 | 175 | 26 | 0 | 0 | 0 | 0 | 90 | 92 | 0 | 0 | 0 | 0 | 0 | 0 | 0 | 56 | 0 | 35 |
| SRR7153315 | 50 | 0 | 960 | 0 | 0 | 15 | 0 | 30 | 0 | 0 | 0 | 214 | 0 | 0 | 29 | 0 | 0 | 0 | 0 | 229 | 193 | 0 | 0 | 45 | 0 | 15 | 110 | 11 | 0 | 15 | 0 | 0 | 0 | 19 | 50 | 0 | 31 |
| SRR7153316 | 57 | 0 | 1006 | 0 | 0 | 0 | 20 | 16 | 0 | 0 | 0 | 269 | 0 | 0 | 113 | 0 | 0 | 0 | 0 | 799 | 122 | 0 | 0 | 21 | 0 | 19 | 325 | 0 | 0 | 0 | 0 | 0 | 0 | 0 | 38 | 0 | 23 |
| SRR7153317 | 46 | 0 | 1921 | 0 | 0 | 11 | 0 | 11 | 0 | 0 | 0 | 97 | 0 | 0 | 26 | 0 | 0 | 0 | 0 | 52 | 115 | 0 | 0 | 0 | 0 | 148 | 787 | 0 | 0 | 0 | 0 | 0 | 0 | 110 | 190 | 0 | 95 |
| SRR7153318 | 25 | 11 | 1146 | 0 | 0 | 13 | 139 | 70 | 0 | 0 | 0 | 125 | 0 | 28 | 33 | 0 | 0 | 0 | 0 | 127 | 85 | 0 | 0 | 0 | 14 | 92 | 474 | 0 | 72 | 35 | 0 | 0 | 0 | 19 | 28 | 0 | 20 |
| SRR7153319 | 67 | 0 | 2691 | 0 | 0 | 0 | 77 | 0 | 0 | 0 | 0 | 110 | 0 | 26 | 148 | 0 | 0 | 0 | 0 | 626 | 190 | 0 | 0 | 0 | 0 | 34 | 1009 | 0 | 0 | 0 | 0 | 0 | 0 | 0 | 95 | 0 | 53 |
| SRR7153320 | 49 | 0 | 1913 | 0 | 0 | 13 | 0 | 0 | 0 | 0 | 0 | 138 | 0 | 0 | 138 | 0 | 0 | 0 | 0 | 556 | 141 | 0 | 0 | 36 | 0 | 101 | 127 | 0 | 0 | 0 | 0 | 0 | 0 | 0 | 15 | 0 | 82 |
| SRR7153321 | 100 | 0 | 2902 | 0 | 0 | 13 | 150 | 20 | 0 | 0 | 0 | 116 | 0 | 0 | 0 | 0 | 14 | 0 | 0 | 893 | 151 | 0 | 0 | 13 | 0 | 33 | 215 | 0 | 0 | 0 | 0 | 0 | 0 | 0 | 19 | 0 | 15 |
| SRR7153322 | 52 | 0 | 2280 | 0 | 0 | 10 | 0 | 0 | 0 | 0 | 0 | 82 | 0 | 0 | 39 | 0 | 0 | 0 | 0 | 29 | 61 | 0 | 0 | 0 | 0 | 198 | 34 | 0 | 0 | 0 | 13 | 0 | 0 | 23 | 14 | 11 | 180 |
| SRR7153323 | 62 | 0 | 2016 | 0 | 0 | 12 | 17 | 80 | 0 | 0 | 0 | 102 | 0 | 0 | 110 | 0 | 0 | 0 | 0 | 530 | 119 | 0 | 0 | 46 | 0 | 127 | 1475 | 0 | 0 | 0 | 0 | 0 | 0 | 15 | 94 | 0 | 46 |
| SRR7153324 | 35 | 0 | 1002 | 0 | 0 | 14 | 0 | 12 | 19 | 0 | 0 | 106 | 0 | 19 | 0 | 0 | 0 | 0 | 0 | 111 | 44 | 0 | 0 | 0 | 0 | 10 | 31 | 12 | 0 | 0 | 0 | 0 | 10 | 21 | 92 | 0 | 42 |
| SRR7153325 | 54 | 0 | 2266 | 0 | 0 | 0 | 0 | 0 | 0 | 0 | 0 | 195 | 0 | 0 | 25 | 0 | 12 | 0 | 0 | 241 | 135 | 0 | 0 | 0 | 0 | 61 | 641 | 0 | 0 | 0 | 0 | 0 | 0 | 16 | 49 | 0 | 20 |
| SRR7153326 | 69 | 0 | 2126 | 0 | 0 | 0 | 36 | 16 | 0 | 0 | 0 | 99 | 0 | 0 | 148 | 0 | 0 | 0 | 0 | 376 | 100 | 0 | 0 | 28 | 0 | 112 | 1115 | 0 | 0 | 0 | 0 | 0 | 0 | 0 | 135 | 0 | 58 |
| SRR7153327 | 93 | 0 | 1836 | 0 | 0 | 14 | 0 | 0 | 0 | 0 | 0 | 152 | 0 | 0 | 98 | 0 | 0 | 0 | 0 | 358 | 127 | 0 | 0 | 0 | 0 | 147 | 1743 | 0 | 0 | 0 | 0 | 0 | 0 | 0 | 206 | 0 | 100 |
| SRR7153328 | 52 | 0 | 1805 | 0 | 0 | 0 | 0 | 12 | 0 | 0 | 0 | 94 | 0 | 0 | 29 | 0 | 10 | 0 | 0 | 223 | 88 | 0 | 0 | 28 | 0 | 91 | 92 | 0 | 0 | 0 | 0 | 0 | 0 | 0 | 89 | 0 | 34 |
| SRR7153329 | 54 | 0 | 1655 | 0 | 0 | 0 | 12 | 11 | 0 | 0 | 0 | 155 | 0 | 0 | 204 | 0 | 0 | 0 | 0 | 378 | 135 | 0 | 0 | 0 | 0 | 84 | 83 | 0 | 0 | 0 | 0 | 0 | 0 | 27 | 10 | 0 | 71 |
| SRR7153330 | 58 | 0 | 2787 | 0 | 0 | 0 | 18 | 236 | 0 | 0 | 0 | 117 | 0 | 0 | 37 | 0 | 0 | 0 | 0 | 385 | 70 | 0 | 0 | 30 | 0 | 85 | 862 | 0 | 0 | 27 | 0 | 0 | 0 | 99 | 84 | 0 | 31 |
| SRR7153331 | 60 | 0 | 1879 | 0 | 0 | 0 | 24 | 58 | 0 | 0 | 0 | 88 | 0 | 17 | 139 | 0 | 11 | 0 | 0 | 674 | 282 | 0 | 0 | 38 | 0 | 11 | 181 | 0 | 0 | 12 | 0 | 0 | 0 | 33 | 13 | 0 | 49 |
| SRR7162919 | 126 | 29 | 454 | 0 | 0 | 0 | 15 | 0 | 29 | 0 | 0 | 1133 | 86 | 14 | 258 | 0 | 629 | 0 | 0 | 0 | 2294 | 0 | 0 | 0 | 302 | 165 | 0 | 0 | 0 | 0 | 14 | 0 | 0 | 0 | 237 | 43 | 72 |
| SRR7162920 | 38 | 106 | 316 | 0 | 14 | 0 | 0 | 0 | 23 | 0 | 0 | 513 | 81 | 0 | 44 | 0 | 677 | 41 | 0 | 0 | 1002 | 0 | 0 | 0 | 79 | 38 | 0 | 0 | 0 | 0 | 0 | 11 | 0 | 0 | 115 | 41 | 50 |
| SRR7162921 | 101 | 57 | 395 | 23 | 0 | 0 | 0 | 0 | 28 | 0 | 0 | 1234 | 28 | 11 | 108 | 0 | 625 | 24 | 0 | 0 | 998 | 0 | 0 | 0 | 23 | 120 | 0 | 23 | 0 | 0 | 0 | 11 | 0 | 0 | 205 | 114 | 51 |
| SRR7178585 | 32 | 0 | 401 | 0 | 0 | 14 | 24 | 18 | 0 | 0 | 0 | 23 | 0 | 35 | 13 | 0 | 0 | 0 | 0 | 151 | 65 | 0 | 0 | 14 | 46 | 14 | 12 | 0 | 0 | 0 | 11 | 0 | 0 | 0 | 74 | 0 | 67 |
| SRR7178586 | 34 | 0 | 423 | 0 | 0 | 20 | 24 | 0 | 0 | 0 | 0 | 27 | 0 | 30 | 0 | 0 | 0 | 0 | 0 | 134 | 70 | 0 | 0 | 13 | 20 | 18 | 15 | 0 | 0 | 0 | 13 | 0 | 0 | 0 | 82 | 0 | 106 |
| SRR7178587 | 22 | 0 | 279 | 0 | 0 | 16 | 21 | 32 | 0 | 0 | 0 | 15 | 0 | 64 | 0 | 0 | 0 | 0 | 0 | 175 | 37 | 0 | 0 | 16 | 16 | 350 | 238 | 0 | 16 | 16 | 32 | 0 | 0 | 16 | 32 | 16 | 16 |
| SRR7178588 | 16 | 0 | 215 | 0 | 0 | 18 | 0 | 18 | 0 | 0 | 0 | 16 | 0 | 0 | 0 | 0 | 0 | 0 | 0 | 98 | 64 | 0 | 0 | 0 | 35 | 442 | 389 | 0 | 0 | 0 | 0 | 0 | 0 | 0 | 18 | 0 | 0 |
| SRR7178589 | 30 | 0 | 758 | 0 | 0 | 0 | 18 | 0 | 0 | 0 | 0 | 16 | 0 | 0 | 0 | 0 | 0 | 0 | 0 | 626 | 152 | 0 | 0 | 0 | 18 | 151 | 314 | 11 | 0 | 0 | 0 | 0 | 0 | 0 | 93 | 0 | 45 |
| SRR7178590 | 37 | 0 | 816 | 0 | 0 | 0 | 13 | 0 | 13 | 0 | 0 | 23 | 0 | 11 | 0 | 0 | 0 | 0 | 0 | 528 | 186 | 0 | 0 | 0 | 19 | 162 | 232 | 13 | 0 | 0 | 15 | 0 | 0 | 0 | 114 | 0 | 59 |
| SRR7178591 | 26 | 0 | 472 | 0 | 0 | 0 | 16 | 0 | 47 | 0 | 0 | 14 | 0 | 0 | 16 | 0 | 0 | 0 | 0 | 447 | 64 | 0 | 0 | 0 | 28 | 253 | 10 | 0 | 0 | 0 | 0 | 0 | 0 | 0 | 56 | 0 | 37 |
| SRR7178592 | 17 | 0 | 340 | 0 | 0 | 11 | 0 | 0 | 11 | 0 | 0 | 15 | 0 | 22 | 16 | 0 | 0 | 0 | 0 | 536 | 142 | 0 | 0 | 22 | 33 | 196 | 229 | 0 | 11 | 0 | 0 | 0 | 0 | 11 | 44 | 0 | 44 |
| SRR7178593 | 40 | 0 | 529 | 0 | 0 | 49 | 0 | 49 | 27 | 0 | 0 | 19 | 0 | 0 | 0 | 0 | 0 | 0 | 0 | 290 | 162 | 0 | 0 | 62 | 0 | 455 | 910 | 0 | 0 | 18 | 0 | 0 | 0 | 0 | 177 | 13 | 80 |
| SRR7178594 | 37 | 0 | 510 | 0 | 0 | 13 | 13 | 43 | 17 | 0 | 0 | 16 | 0 | 0 | 0 | 0 | 0 | 0 | 0 | 270 | 175 | 0 | 0 | 111 | 13 | 17 | 38 | 0 | 0 | 13 | 21 | 0 | 0 | 0 | 137 | 0 | 116 |
| SRR7178595 | 28 | 0 | 462 | 0 | 0 | 69 | 0 | 35 | 0 | 0 | 0 | 11 | 0 | 17 | 0 | 0 | 0 | 0 | 0 | 224 | 62 | 0 | 0 | 121 | 35 | 311 | 865 | 0 | 0 | 0 | 0 | 0 | 0 | 0 | 104 | 0 | 35 |
| SRR7178596 | 17 | 0 | 346 | 0 | 0 | 24 | 0 | 72 | 0 | 0 | 0 | 0 | 0 | 0 | 0 | 0 | 0 | 0 | 0 | 260 | 65 | 0 | 0 | 48 | 72 | 193 | 1037 | 0 | 0 | 48 | 24 | 0 | 0 | 24 | 121 | 24 | 24 |
| SRR7178597 | 29 | 0 | 824 | 0 | 0 | 16 | 86 | 95 | 14 | 0 | 0 | 30 | 0 | 0 | 0 | 0 | 0 | 0 | 14 | 671 | 207 | 0 | 0 | 0 | 14 | 17 | 51 | 0 | 0 | 0 | 25 | 0 | 0 | 0 | 109 | 0 | 41 |
| SRR7178598 | 29 | 0 | 815 | 0 | 0 | 19 | 12 | 0 | 0 | 0 | 0 | 20 | 0 | 0 | 0 | 0 | 0 | 0 | 0 | 390 | 54 | 0 | 0 | 0 | 30 | 16 | 28 | 0 | 0 | 0 | 12 | 0 | 0 | 0 | 102 | 0 | 44 |
| SRR7178599 | 29 | 0 | 772 | 0 | 0 | 0 | 0 | 0 | 0 | 0 | 0 | 21 | 0 | 0 | 0 | 0 | 0 | 0 | 0 | 395 | 83 | 0 | 0 | 0 | 17 | 17 | 31 | 0 | 0 | 0 | 0 | 0 | 0 | 0 | 101 | 0 | 46 |
| SRR7178600 | 34 | 0 | 2276 | 0 | 0 | 0 | 27 | 14 | 0 | 0 | 0 | 18 | 0 | 0 | 0 | 0 | 0 | 0 | 0 | 103 | 107 | 0 | 0 | 0 | 12 | 19 | 10 | 0 | 0 | 0 | 0 | 0 | 0 | 0 | 33 | 0 | 20 |
| SRR7178601 | 27 | 0 | 2563 | 0 | 0 | 0 | 15 | 11 | 0 | 0 | 0 | 22 | 0 | 0 | 11 | 0 | 0 | 0 | 0 | 112 | 105 | 0 | 0 | 0 | 12 | 158 | 85 | 0 | 0 | 0 | 0 | 0 | 0 | 0 | 36 | 0 | 0 |
| SRR7178602 | 14 | 0 | 1525 | 0 | 0 | 0 | 15 | 16 | 0 | 0 | 0 | 20 | 0 | 0 | 30 | 0 | 0 | 0 | 0 | 136 | 90 | 0 | 0 | 0 | 0 | 101 | 75 | 0 | 0 | 0 | 0 | 0 | 0 | 0 | 53 | 0 | 16 |
| SRR7178603 | 10 | 0 | 1103 | 0 | 0 | 0 | 0 | 0 | 13 | 0 | 0 | 0 | 0 | 0 | 0 | 0 | 0 | 0 | 0 | 60 | 60 | 0 | 0 | 0 | 0 | 133 | 89 | 0 | 0 | 0 | 0 | 0 | 0 | 0 | 44 | 0 | 0 |
| SRR7178604 | 25 | 0 | 420 | 0 | 0 | 0 | 29 | 16 | 0 | 0 | 0 | 13 | 0 | 48 | 0 | 0 | 0 | 0 | 0 | 70 | 43 | 0 | 0 | 0 | 0 | 13 | 112 | 0 | 0 | 0 | 13 | 0 | 0 | 19 | 29 | 0 | 32 |
| SRR7178605 | 33 | 0 | 439 | 0 | 0 | 0 | 35 | 12 | 0 | 0 | 0 | 15 | 0 | 40 | 12 | 0 | 0 | 0 | 0 | 85 | 54 | 0 | 0 | 15 | 15 | 12 | 144 | 0 | 0 | 0 | 12 | 0 | 0 | 21 | 28 | 0 | 46 |
| SRR7178606 | 25 | 0 | 425 | 0 | 0 | 0 | 14 | 0 | 0 | 0 | 0 | 0 | 0 | 0 | 0 | 0 | 11 | 0 | 0 | 42 | 63 | 0 | 0 | 14 | 14 | 208 | 139 | 0 | 0 | 0 | 0 | 0 | 0 | 55 | 28 | 0 | 28 |
| SRR7178607 | 31 | 0 | 298 | 0 | 0 | 0 | 0 | 0 | 0 | 0 | 0 | 12 | 0 | 19 | 0 | 0 | 0 | 0 | 0 | 106 | 53 | 0 | 0 | 0 | 0 | 136 | 194 | 0 | 0 | 0 | 0 | 0 | 0 | 78 | 58 | 0 | 39 |
| SRR7178608 | 26 | 0 | 571 | 0 | 0 | 21 | 48 | 49 | 0 | 0 | 0 | 11 | 0 | 21 | 0 | 0 | 0 | 0 | 0 | 1284 | 268 | 0 | 0 | 16 | 23 | 156 | 452 | 0 | 0 | 0 | 0 | 0 | 0 | 23 | 75 | 0 | 47 |
| SRR7178609 | 28 | 0 | 631 | 0 | 0 | 20 | 45 | 57 | 0 | 0 | 0 | 11 | 0 | 13 | 0 | 0 | 0 | 0 | 0 | 1330 | 239 | 0 | 0 | 37 | 37 | 150 | 31 | 0 | 0 | 0 | 0 | 0 | 0 | 24 | 105 | 0 | 39 |
| SRR7178610 | 23 | 0 | 408 | 0 | 0 | 0 | 71 | 57 | 0 | 0 | 0 | 0 | 0 | 19 | 0 | 0 | 0 | 0 | 0 | 1232 | 212 | 0 | 0 | 28 | 47 | 142 | 332 | 0 | 0 | 19 | 19 | 0 | 0 | 19 | 0 | 0 | 0 |
| SRR7178611 | 0 | 0 | 286 | 0 | 0 | 0 | 28 | 43 | 11 | 0 | 0 | 0 | 0 | 0 | 0 | 0 | 0 | 0 | 0 | 897 | 117 | 0 | 0 | 22 | 22 | 119 | 10 | 0 | 0 | 11 | 0 | 0 | 0 | 11 | 43 | 0 | 33 |
| SRR7178612 | 26 | 0 | 531 | 0 | 0 | 25 | 50 | 123 | 14 | 0 | 0 | 12 | 0 | 0 | 19 | 0 | 0 | 0 | 0 | 1504 | 264 | 0 | 0 | 44 | 30 | 27 | 38 | 0 | 0 | 16 | 23 | 0 | 0 | 25 | 118 | 0 | 63 |
| SRR7178613 | 30 | 0 | 547 | 0 | 0 | 45 | 56 | 137 | 0 | 0 | 0 | 12 | 0 | 0 | 25 | 0 | 0 | 0 | 0 | 1669 | 339 | 0 | 0 | 49 | 22 | 409 | 510 | 0 | 0 | 25 | 34 | 0 | 0 | 29 | 119 | 0 | 54 |
| SRR7178614 | 19 | 0 | 449 | 0 | 0 | 29 | 21 | 105 | 0 | 0 | 0 | 0 | 0 | 19 | 0 | 0 | 0 | 0 | 0 | 1111 | 206 | 0 | 0 | 0 | 0 | 15 | 19 | 0 | 0 | 0 | 0 | 0 | 0 | 0 | 96 | 0 | 48 |
| SRR7178615 | 11 | 0 | 360 | 0 | 0 | 40 | 0 | 93 | 13 | 0 | 0 | 0 | 0 | 13 | 0 | 0 | 0 | 0 | 0 | 1233 | 322 | 0 | 0 | 53 | 27 | 332 | 531 | 0 | 0 | 13 | 27 | 0 | 13 | 0 | 27 | 0 | 40 |
| SRR7178616 | 25 | 0 | 781 | 0 | 0 | 0 | 34 | 0 | 14 | 0 | 0 | 20 | 0 | 18 | 0 | 0 | 0 | 0 | 0 | 425 | 169 | 12 | 0 | 18 | 12 | 23 | 32 | 36 | 0 | 0 | 0 | 0 | 0 | 18 | 59 | 0 | 27 |
| SRR7178617 | 27 | 0 | 868 | 0 | 0 | 10 | 20 | 0 | 11 | 0 | 0 | 21 | 0 | 19 | 0 | 0 | 0 | 0 | 0 | 423 | 134 | 0 | 0 | 20 | 17 | 29 | 33 | 29 | 0 | 0 | 11 | 0 | 0 | 14 | 58 | 0 | 32 |
| SRR7178618 | 16 | 0 | 634 | 0 | 0 | 0 | 0 | 0 | 12 | 0 | 0 | 20 | 0 | 165 | 0 | 0 | 12 | 0 | 0 | 302 | 95 | 0 | 0 | 24 | 18 | 306 | 283 | 0 | 0 | 0 | 12 | 0 | 12 | 12 | 53 | 0 | 29 |
| SRR7178619 | 0 | 0 | 395 | 0 | 0 | 14 | 0 | 0 | 0 | 0 | 0 | 17 | 0 | 88 | 0 | 0 | 0 | 0 | 0 | 305 | 74 | 0 | 0 | 20 | 0 | 10 | 204 | 0 | 14 | 0 | 14 | 0 | 0 | 20 | 48 | 0 | 14 |
| SRR7178620 | 23 | 0 | 1057 | 0 | 0 | 14 | 29 | 0 | 14 | 0 | 0 | 23 | 0 | 0 | 0 | 0 | 0 | 0 | 0 | 245 | 62 | 0 | 0 | 14 | 0 | 14 | 20 | 0 | 0 | 0 | 12 | 0 | 0 | 0 | 45 | 0 | 38 |
| SRR7178621 | 25 | 0 | 1093 | 0 | 0 | 18 | 30 | 16 | 11 | 0 | 0 | 21 | 0 | 0 | 0 | 0 | 0 | 0 | 0 | 260 | 40 | 0 | 0 | 18 | 0 | 13 | 25 | 0 | 0 | 0 | 0 | 0 | 0 | 0 | 50 | 0 | 0 |
| SRR7178622 | 19 | 11 | 966 | 0 | 0 | 22 | 45 | 0 | 0 | 0 | 0 | 14 | 0 | 11 | 0 | 0 | 0 | 0 | 11 | 224 | 34 | 0 | 0 | 33 | 0 | 176 | 15 | 0 | 0 | 0 | 0 | 0 | 0 | 0 | 11 | 0 | 33 |
| SRR7178623 | 13 | 0 | 746 | 0 | 0 | 30 | 22 | 0 | 0 | 0 | 0 | 10 | 0 | 0 | 0 | 0 | 0 | 0 | 0 | 158 | 14 | 0 | 0 | 0 | 0 | 182 | 10 | 0 | 0 | 0 | 0 | 0 | 0 | 0 | 76 | 0 | 15 |
| SRR7178624 | 29 | 0 | 1168 | 0 | 0 | 20 | 27 | 83 | 0 | 0 | 0 | 32 | 0 | 14 | 0 | 0 | 0 | 0 | 0 | 902 | 93 | 0 | 0 | 37 | 0 | 276 | 606 | 0 | 0 | 0 | 23 | 0 | 0 | 0 | 83 | 0 | 57 |
| SRR7178625 | 27 | 0 | 1295 | 0 | 0 | 16 | 23 | 61 | 29 | 0 | 0 | 23 | 0 | 11 | 11 | 0 | 0 | 0 | 0 | 894 | 111 | 0 | 0 | 58 | 21 | 26 | 53 | 0 | 0 | 11 | 0 | 0 | 0 | 0 | 103 | 0 | 58 |
| SRR7178626 | 20 | 13 | 918 | 0 | 0 | 0 | 30 | 63 | 0 | 0 | 0 | 28 | 0 | 0 | 10 | 0 | 0 | 0 | 0 | 855 | 40 | 0 | 0 | 51 | 63 | 14 | 29 | 13 | 0 | 0 | 0 | 0 | 0 | 0 | 114 | 0 | 63 |
| SRR7178627 | 11 | 0 | 585 | 0 | 0 | 14 | 19 | 69 | 28 | 0 | 0 | 26 | 0 | 0 | 0 | 0 | 0 | 0 | 19 | 646 | 84 | 0 | 0 | 14 | 28 | 264 | 23 | 0 | 0 | 14 | 14 | 0 | 0 | 0 | 125 | 0 | 83 |
| SRR7178628 | 41 | 0 | 992 | 0 | 0 | 13 | 19 | 21 | 0 | 0 | 0 | 29 | 0 | 0 | 0 | 0 | 0 | 0 | 0 | 319 | 58 | 0 | 0 | 23 | 16 | 19 | 24 | 0 | 0 | 15 | 16 | 0 | 0 | 0 | 29 | 0 | 21 |
| SRR7178629 | 31 | 0 | 990 | 0 | 0 | 0 | 10 | 30 | 0 | 0 | 0 | 35 | 0 | 0 | 10 | 0 | 0 | 0 | 0 | 358 | 29 | 0 | 0 | 19 | 0 | 14 | 20 | 0 | 0 | 17 | 0 | 0 | 0 | 0 | 15 | 0 | 22 |
| SRR7178630 | 28 | 0 | 907 | 0 | 0 | 0 | 17 | 20 | 0 | 0 | 0 | 34 | 0 | 0 | 0 | 0 | 0 | 0 | 0 | 270 | 27 | 0 | 0 | 14 | 16 | 175 | 211 | 0 | 0 | 20 | 0 | 0 | 0 | 0 | 27 | 0 | 20 |
| SRR7178631 | 43 | 0 | 876 | 0 | 0 | 12 | 20 | 0 | 0 | 0 | 0 | 23 | 0 | 0 | 41 | 0 | 0 | 0 | 0 | 279 | 41 | 0 | 0 | 16 | 14 | 228 | 237 | 0 | 0 | 0 | 0 | 0 | 0 | 0 | 39 | 0 | 16 |
| SRR7178632 | 36 | 0 | 987 | 0 | 0 | 15 | 25 | 11 | 0 | 0 | 0 | 20 | 0 | 0 | 42 | 0 | 0 | 0 | 0 | 305 | 59 | 0 | 0 | 26 | 15 | 247 | 275 | 0 | 0 | 0 | 17 | 0 | 0 | 0 | 41 | 0 | 13 |
| SRR7178633 | 26 | 0 | 630 | 0 | 0 | 0 | 20 | 30 | 0 | 0 | 0 | 11 | 0 | 0 | 20 | 0 | 0 | 0 | 0 | 180 | 30 | 0 | 0 | 50 | 30 | 15 | 11 | 0 | 0 | 10 | 0 | 0 | 0 | 0 | 30 | 0 | 10 |
| SRR7178634 | 17 | 0 | 473 | 0 | 0 | 0 | 0 | 0 | 11 | 0 | 0 | 18 | 0 | 0 | 48 | 0 | 0 | 0 | 0 | 228 | 38 | 0 | 0 | 11 | 0 | 194 | 171 | 0 | 0 | 11 | 11 | 0 | 0 | 0 | 23 | 0 | 34 |
| SRR7178635 | 35 | 0 | 652 | 0 | 0 | 24 | 16 | 35 | 14 | 0 | 0 | 12 | 0 | 0 | 0 | 0 | 0 | 0 | 0 | 786 | 96 | 0 | 0 | 19 | 16 | 303 | 558 | 0 | 0 | 0 | 17 | 0 | 0 | 0 | 88 | 0 | 41 |
| SRR7178636 | 28 | 0 | 642 | 0 | 0 | 15 | 27 | 29 | 14 | 0 | 0 | 11 | 0 | 0 | 0 | 0 | 0 | 0 | 16 | 826 | 131 | 0 | 0 | 15 | 11 | 21 | 37 | 0 | 11 | 0 | 20 | 0 | 0 | 0 | 105 | 0 | 43 |
| SRR7178637 | 32 | 0 | 574 | 0 | 0 | 20 | 15 | 33 | 0 | 0 | 0 | 0 | 0 | 0 | 0 | 0 | 0 | 0 | 0 | 748 | 114 | 0 | 0 | 27 | 27 | 12 | 17 | 0 | 0 | 0 | 0 | 0 | 0 | 0 | 27 | 0 | 40 |
| SRR7178638 | 23 | 0 | 429 | 0 | 0 | 28 | 0 | 74 | 18 | 0 | 0 | 12 | 0 | 0 | 0 | 0 | 0 | 0 | 0 | 719 | 97 | 0 | 0 | 0 | 0 | 324 | 453 | 0 | 0 | 0 | 0 | 0 | 0 | 0 | 83 | 0 | 18 |
| SRR7178639 | 30 | 0 | 938 | 0 | 0 | 34 | 32 | 21 | 21 | 0 | 0 | 14 | 0 | 0 | 0 | 0 | 0 | 0 | 28 | 2540 | 406 | 0 | 0 | 23 | 26 | 33 | 46 | 0 | 0 | 21 | 44 | 0 | 0 | 0 | 114 | 13 | 49 |
| SRR7178640 | 30 | 0 | 914 | 0 | 0 | 42 | 10 | 45 | 26 | 0 | 0 | 15 | 0 | 0 | 0 | 0 | 0 | 0 | 0 | 2278 | 295 | 0 | 0 | 13 | 23 | 24 | 40 | 0 | 0 | 16 | 42 | 0 | 0 | 0 | 123 | 0 | 68 |
| SRR7178641 | 31 | 0 | 842 | 0 | 0 | 30 | 57 | 54 | 17 | 0 | 0 | 14 | 0 | 0 | 0 | 0 | 0 | 0 | 16 | 1981 | 306 | 0 | 0 | 0 | 20 | 549 | 760 | 0 | 17 | 20 | 74 | 0 | 0 | 10 | 104 | 0 | 67 |
| SRR7178642 | 30 | 0 | 701 | 0 | 0 | 15 | 14 | 18 | 0 | 0 | 0 | 33 | 0 | 0 | 0 | 0 | 0 | 0 | 0 | 557 | 107 | 0 | 0 | 72 | 15 | 25 | 32 | 0 | 0 | 33 | 0 | 0 | 0 | 0 | 72 | 0 | 17 |
| SRR7178643 | 28 | 0 | 708 | 0 | 0 | 19 | 13 | 16 | 12 | 0 | 0 | 34 | 0 | 0 | 0 | 0 | 0 | 0 | 0 | 545 | 108 | 0 | 0 | 70 | 31 | 16 | 23 | 0 | 0 | 45 | 0 | 0 | 0 | 0 | 61 | 0 | 14 |
| SRR7178644 | 24 | 0 | 673 | 0 | 0 | 12 | 18 | 15 | 0 | 0 | 0 | 31 | 0 | 0 | 0 | 0 | 0 | 0 | 11 | 521 | 91 | 0 | 0 | 66 | 15 | 16 | 28 | 0 | 0 | 32 | 0 | 0 | 0 | 0 | 39 | 0 | 22 |
| SRR7178645 | 19 | 0 | 597 | 0 | 0 | 20 | 14 | 0 | 20 | 0 | 0 | 20 | 0 | 0 | 0 | 0 | 0 | 0 | 0 | 171 | 40 | 0 | 0 | 0 | 17 | 18 | 18 | 0 | 0 | 0 | 0 | 0 | 0 | 13 | 46 | 0 | 26 |
| SRR7178646 | 23 | 0 | 681 | 0 | 0 | 31 | 0 | 0 | 19 | 0 | 0 | 19 | 0 | 0 | 0 | 0 | 0 | 0 | 0 | 203 | 39 | 0 | 0 | 0 | 12 | 17 | 22 | 0 | 0 | 0 | 28 | 0 | 0 | 15 | 37 | 0 | 40 |
| SRR7178647 | 0 | 0 | 399 | 0 | 0 | 13 | 12 | 27 | 27 | 0 | 0 | 21 | 0 | 0 | 0 | 0 | 0 | 0 | 0 | 151 | 19 | 0 | 0 | 0 | 40 | 282 | 605 | 0 | 0 | 0 | 0 | 0 | 0 | 0 | 27 | 0 | 27 |
| SRR7178648 | 0 | 0 | 299 | 0 | 0 | 0 | 14 | 0 | 0 | 0 | 0 | 13 | 15 | 0 | 0 | 0 | 0 | 0 | 0 | 112 | 18 | 0 | 0 | 15 | 15 | 272 | 302 | 0 | 0 | 15 | 15 | 0 | 0 | 15 | 15 | 0 | 15 |
| SRR7178649 | 32 | 0 | 437 | 0 | 0 | 14 | 33 | 0 | 0 | 0 | 0 | 14 | 0 | 0 | 11 | 0 | 0 | 0 | 0 | 263 | 115 | 0 | 0 | 33 | 29 | 506 | 339 | 0 | 0 | 11 | 18 | 0 | 0 | 0 | 112 | 0 | 76 |
| SRR7178650 | 33 | 0 | 480 | 0 | 0 | 19 | 31 | 0 | 19 | 0 | 0 | 19 | 0 | 0 | 11 | 0 | 0 | 0 | 0 | 306 | 109 | 0 | 0 | 34 | 24 | 445 | 464 | 0 | 0 | 0 | 19 | 0 | 0 | 0 | 57 | 14 | 62 |
| SRR7178651 | 39 | 0 | 437 | 0 | 0 | 10 | 34 | 0 | 20 | 0 | 0 | 18 | 0 | 10 | 11 | 0 | 0 | 0 | 0 | 251 | 113 | 0 | 0 | 30 | 45 | 18 | 15 | 0 | 0 | 0 | 10 | 0 | 0 | 0 | 86 | 0 | 30 |
| SRR7178652 | 20 | 0 | 968 | 0 | 0 | 22 | 23 | 0 | 0 | 0 | 0 | 22 | 0 | 0 | 0 | 0 | 0 | 0 | 0 | 100 | 32 | 0 | 0 | 0 | 12 | 24 | 20 | 0 | 0 | 0 | 20 | 0 | 0 | 0 | 42 | 0 | 27 |
| SRR7178653 | 22 | 0 | 994 | 0 | 0 | 0 | 21 | 0 | 0 | 0 | 0 | 33 | 0 | 0 | 0 | 0 | 0 | 0 | 0 | 120 | 32 | 0 | 0 | 0 | 17 | 275 | 322 | 0 | 0 | 0 | 14 | 0 | 0 | 0 | 48 | 0 | 31 |
| SRR7178654 | 14 | 0 | 878 | 0 | 0 | 0 | 0 | 10 | 0 | 0 | 10 | 14 | 0 | 0 | 0 | 0 | 0 | 0 | 0 | 134 | 32 | 0 | 0 | 0 | 21 | 301 | 249 | 0 | 0 | 0 | 31 | 0 | 0 | 21 | 21 | 0 | 21 |
| SRR7178655 | 0 | 0 | 653 | 0 | 0 | 15 | 24 | 0 | 0 | 0 | 0 | 21 | 0 | 0 | 0 | 0 | 0 | 0 | 0 | 57 | 33 | 0 | 0 | 0 | 0 | 320 | 247 | 0 | 0 | 0 | 15 | 0 | 0 | 15 | 15 | 0 | 15 |
| SRR7178656 | 25 | 0 | 713 | 0 | 0 | 43 | 12 | 0 | 36 | 0 | 0 | 15 | 0 | 0 | 0 | 0 | 0 | 0 | 0 | 833 | 91 | 0 | 0 | 15 | 38 | 30 | 33 | 0 | 0 | 13 | 38 | 0 | 0 | 0 | 132 | 0 | 51 |
| SRR7178657 | 25 | 0 | 743 | 0 | 0 | 29 | 0 | 0 | 34 | 0 | 0 | 17 | 0 | 0 | 0 | 0 | 0 | 0 | 0 | 986 | 86 | 0 | 0 | 27 | 34 | 599 | 731 | 0 | 0 | 0 | 19 | 0 | 0 | 0 | 117 | 0 | 58 |
| SRR7178658 | 22 | 11 | 595 | 0 | 0 | 33 | 13 | 11 | 11 | 0 | 0 | 15 | 0 | 0 | 0 | 0 | 0 | 0 | 0 | 764 | 40 | 0 | 0 | 11 | 55 | 602 | 766 | 0 | 0 | 0 | 44 | 0 | 0 | 0 | 77 | 11 | 33 |
| SRR7178659 | 17 | 0 | 489 | 0 | 0 | 15 | 26 | 15 | 0 | 0 | 0 | 10 | 0 | 0 | 0 | 0 | 0 | 0 | 0 | 630 | 34 | 0 | 0 | 15 | 30 | 11 | 14 | 0 | 0 | 0 | 0 | 0 | 0 | 0 | 30 | 0 | 60 |
| SRR7178660 | 21 | 0 | 677 | 0 | 0 | 17 | 19 | 28 | 11 | 0 | 0 | 23 | 0 | 0 | 0 | 0 | 0 | 0 | 0 | 260 | 38 | 0 | 0 | 70 | 28 | 25 | 25 | 0 | 0 | 0 | 20 | 0 | 0 | 0 | 13 | 13 | 79 |
| SRR7178661 | 17 | 0 | 670 | 0 | 0 | 24 | 22 | 33 | 14 | 0 | 0 | 19 | 0 | 0 | 0 | 0 | 0 | 0 | 0 | 224 | 53 | 0 | 0 | 52 | 14 | 363 | 326 | 0 | 0 | 0 | 21 | 0 | 0 | 17 | 203 | 0 | 68 |
| SRR7178662 | 15 | 0 | 620 | 0 | 0 | 17 | 15 | 44 | 12 | 0 | 0 | 22 | 0 | 0 | 0 | 0 | 0 | 0 | 0 | 208 | 31 | 0 | 0 | 71 | 0 | 20 | 17 | 0 | 0 | 0 | 22 | 0 | 0 | 0 | 159 | 0 | 59 |
| SRR7178663 | 32 | 0 | 567 | 0 | 0 | 16 | 29 | 0 | 0 | 0 | 0 | 25 | 0 | 0 | 11 | 0 | 0 | 0 | 0 | 221 | 67 | 0 | 0 | 0 | 0 | 20 | 15 | 0 | 0 | 0 | 20 | 0 | 0 | 0 | 65 | 0 | 41 |
| SRR7178664 | 26 | 0 | 643 | 0 | 0 | 19 | 38 | 0 | 15 | 0 | 0 | 17 | 0 | 0 | 14 | 0 | 0 | 0 | 0 | 175 | 50 | 0 | 0 | 0 | 23 | 419 | 261 | 0 | 0 | 0 | 30 | 0 | 0 | 0 | 76 | 0 | 38 |
| SRR7178665 | 23 | 19 | 436 | 0 | 0 | 19 | 22 | 0 | 19 | 0 | 0 | 10 | 0 | 0 | 0 | 0 | 0 | 0 | 0 | 149 | 45 | 0 | 0 | 0 | 19 | 467 | 168 | 0 | 0 | 0 | 19 | 0 | 0 | 0 | 75 | 0 | 19 |
| SRR7178666 | 18 | 0 | 253 | 22 | 0 | 44 | 21 | 22 | 0 | 0 | 0 | 0 | 0 | 0 | 0 | 0 | 0 | 0 | 0 | 183 | 71 | 0 | 0 | 44 | 0 | 457 | 326 | 0 | 22 | 0 | 22 | 0 | 0 | 0 | 0 | 0 | 44 |
| SRR7178667 | 41 | 0 | 655 | 0 | 0 | 21 | 13 | 21 | 0 | 0 | 0 | 16 | 0 | 14 | 48 | 0 | 0 | 0 | 0 | 515 | 428 | 0 | 0 | 0 | 26 | 28 | 21 | 0 | 0 | 0 | 17 | 0 | 0 | 0 | 69 | 0 | 24 |
| SRR7178668 | 49 | 0 | 701 | 0 | 0 | 27 | 62 | 16 | 11 | 0 | 0 | 17 | 0 | 20 | 45 | 0 | 0 | 0 | 0 | 452 | 452 | 0 | 0 | 0 | 27 | 30 | 25 | 0 | 0 | 0 | 16 | 0 | 0 | 0 | 49 | 0 | 24 |
| SRR7178669 | 29 | 0 | 427 | 0 | 0 | 32 | 61 | 11 | 0 | 11 | 0 | 13 | 0 | 32 | 46 | 0 | 0 | 0 | 0 | 412 | 366 | 0 | 0 | 0 | 0 | 285 | 328 | 11 | 0 | 0 | 21 | 0 | 0 | 0 | 63 | 11 | 53 |
| SRR7178670 | 16 | 0 | 307 | 0 | 0 | 12 | 15 | 24 | 0 | 0 | 0 | 16 | 0 | 12 | 29 | 0 | 0 | 0 | 0 | 263 | 394 | 0 | 0 | 0 | 24 | 292 | 207 | 12 | 0 | 0 | 37 | 0 | 12 | 12 | 49 | 0 | 24 |
| SRR7178671 | 20 | 0 | 863 | 0 | 0 | 35 | 21 | 14 | 11 | 0 | 0 | 29 | 0 | 0 | 0 | 0 | 12 | 0 | 0 | 209 | 135 | 0 | 0 | 27 | 19 | 479 | 642 | 0 | 0 | 0 | 16 | 0 | 0 | 0 | 65 | 0 | 22 |
| SRR7178672 | 24 | 0 | 887 | 0 | 0 | 34 | 15 | 13 | 0 | 0 | 0 | 30 | 0 | 0 | 0 | 0 | 11 | 0 | 0 | 188 | 121 | 0 | 0 | 45 | 37 | 26 | 38 | 0 | 0 | 0 | 18 | 0 | 0 | 0 | 79 | 0 | 47 |

### Supp Table S8.2.Columns_AM_to_BW

| | | | | | | | | | | | | | | | | | | | | | | | | | | | | | | | | | | | | | |
|---|---|---|---|---|---|---|---|---|---|---|---|---|---|---|---|---|---|---|---|---|---|---|---|---|---|---|---|---|---|---|---|---|---|---|---|---|---|
| SRR7178673 | 14 | 0 | 800 | 0 | 0 | 0 | 16 | 11 | 33 | 0 | 0 | 15 | 0 | 0 | 0 | 0 | 0 | 0 | 0 | 173 | 115 | 0 | 0 | 44 | 11 | 522 | 729 | 0 | 11 | 22 | 33 | 0 | 0 | 0 | 76 | 0 | 11 |
| SRR7178674 | 19 | 0 | 600 | 0 | 0 | 0 | 13 | 0 | 15 | 0 | 0 | 13 | 0 | 0 | 0 | 0 | 0 | 0 | 0 | 121 | 34 | 0 | 0 | 73 | 15 | 11 | 17 | 0 | 0 | 0 | 15 | 0 | 15 | 0 | 44 | 0 | 15 |
| SRR7178675 | 10 | 0 | 753 | 0 | 0 | 18 | 0 | 32 | 0 | 0 | 0 | 15 | 0 | 14 | 0 | 0 | 0 | 0 | 0 | 368 | 192 | 0 | 0 | 23 | 18 | 349 | 477 | 0 | 0 | 0 | 0 | 0 | 0 | 55 | 106 | 0 | 46 |
| SRR7178676 | 11 | 0 | 813 | 0 | 0 | 21 | 13 | 38 | 0 | 0 | 0 | 15 | 0 | 0 | 0 | 0 | 0 | 0 | 0 | 348 | 221 | 0 | 0 | 0 | 0 | 14 | 23 | 0 | 0 | 13 | 17 | 0 | 17 | 94 | 90 | 0 | 30 |
| SRR7178677 | 0 | 0 | 516 | 0 | 0 | 19 | 17 | 39 | 0 | 0 | 0 | 0 | 0 | 0 | 0 | 0 | 0 | 0 | 0 | 215 | 232 | 0 | 19 | 39 | 0 | 350 | 10 | 0 | 0 | 19 | 19 | 0 | 19 | 58 | 78 | 0 | 39 |
| SRR7178678 | 11 | 0 | 347 | 0 | 0 | 0 | 47 | 0 | 22 | 0 | 0 | 13 | 0 | 22 | 0 | 0 | 0 | 0 | 0 | 221 | 126 | 0 | 0 | 22 | 44 | 288 | 488 | 0 | 0 | 0 | 0 | 0 | 0 | 44 | 44 | 0 | 22 |
| SRR7178679 | 30 | 0 | 622 | 0 | 0 | 54 | 24 | 50 | 19 | 0 | 0 | 28 | 0 | 0 | 0 | 0 | 0 | 0 | 0 | 529 | 65 | 0 | 0 | 32 | 16 | 35 | 39 | 0 | 19 | 0 | 44 | 0 | 0 | 16 | 13 | 13 | 69 |
| SRR7178680 | 29 | 0 | 633 | 0 | 0 | 27 | 23 | 50 | 15 | 0 | 0 | 22 | 0 | 0 | 0 | 0 | 0 | 0 | 0 | 490 | 62 | 0 | 0 | 46 | 15 | 24 | 28 | 0 | 0 | 0 | 31 | 0 | 0 | 19 | 10 | 15 | 108 |
| SRR7178681 | 26 | 0 | 573 | 0 | 0 | 44 | 30 | 76 | 28 | 0 | 0 | 25 | 0 | 0 | 0 | 0 | 0 | 0 | 0 | 444 | 60 | 0 | 0 | 20 | 12 | 613 | 829 | 0 | 0 | 0 | 20 | 0 | 0 | 12 | 244 | 20 | 100 |
| SRR7178682 | 29 | 0 | 403 | 0 | 0 | 35 | 22 | 15 | 20 | 0 | 0 | 19 | 0 | 0 | 0 | 0 | 0 | 0 | 0 | 694 | 132 | 0 | 0 | 29 | 35 | 39 | 40 | 0 | 0 | 11 | 39 | 0 | 0 | 0 | 103 | 11 | 44 |
| SRR7178683 | 30 | 0 | 441 | 0 | 0 | 41 | 22 | 0 | 21 | 0 | 0 | 17 | 0 | 0 | 0 | 0 | 0 | 0 | 0 | 706 | 130 | 0 | 0 | 21 | 28 | 38 | 45 | 0 | 0 | 17 | 27 | 0 | 0 | 0 | 133 | 0 | 39 |
| SRR7178684 | 21 | 0 | 316 | 0 | 0 | 29 | 10 | 0 | 22 | 0 | 0 | 14 | 0 | 0 | 0 | 0 | 0 | 0 | 0 | 683 | 132 | 0 | 0 | 22 | 44 | 14 | 24 | 0 | 0 | 15 | 22 | 0 | 0 | 0 | 80 | 0 | 15 |
| SRR7178685 | 15 | 0 | 235 | 0 | 0 | 25 | 0 | 0 | 17 | 0 | 0 | 13 | 0 | 0 | 0 | 0 | 0 | 0 | 0 | 489 | 122 | 0 | 0 | 51 | 17 | 14 | 19 | 0 | 0 | 25 | 0 | 0 | 0 | 0 | 76 | 0 | 25 |
| SRR7178686 | 38 | 0 | 422 | 0 | 0 | 18 | 0 | 0 | 0 | 0 | 0 | 17 | 0 | 0 | 112 | 0 | 0 | 0 | 0 | 158 | 408 | 0 | 0 | 0 | 14 | 304 | 153 | 0 | 0 | 0 | 0 | 0 | 0 | 0 | 112 | 16 | 53 |
| SRR7178687 | 40 | 0 | 415 | 0 | 0 | 0 | 0 | 0 | 17 | 0 | 0 | 18 | 0 | 0 | 61 | 0 | 0 | 0 | 0 | 228 | 462 | 0 | 0 | 0 | 27 | 19 | 167 | 0 | 0 | 0 | 0 | 0 | 0 | 0 | 119 | 0 | 53 |
| SRR7178688 | 33 | 0 | 347 | 0 | 0 | 0 | 0 | 0 | 17 | 0 | 0 | 0 | 0 | 0 | 107 | 0 | 0 | 0 | 0 | 161 | 287 | 0 | 0 | 0 | 0 | 322 | 110 | 0 | 0 | 0 | 17 | 0 | 0 | 0 | 76 | 0 | 51 |
| SRR7178689 | 30 | 0 | 265 | 0 | 0 | 0 | 0 | 0 | 0 | 0 | 0 | 0 | 0 | 0 | 23 | 0 | 0 | 0 | 0 | 69 | 229 | 0 | 0 | 0 | 0 | 246 | 129 | 0 | 0 | 0 | 0 | 0 | 0 | 0 | 59 | 0 | 12 |
| SRR7178690 | 16 | 0 | 876 | 0 | 0 | 0 | 14 | 13 | 15 | 0 | 0 | 19 | 0 | 0 | 0 | 0 | 0 | 0 | 0 | 224 | 108 | 0 | 0 | 41 | 21 | 23 | 29 | 0 | 0 | 0 | 0 | 0 | 0 | 0 | 72 | 0 | 26 |
| SRR7178691 | 24 | 0 | 777 | 0 | 0 | 12 | 98 | 96 | 0 | 0 | 0 | 30 | 0 | 12 | 0 | 0 | 0 | 0 | 0 | 670 | 166 | 0 | 0 | 15 | 12 | 209 | 668 | 0 | 0 | 23 | 17 | 0 | 0 | 0 | 99 | 0 | 49 |
| SRR7178692 | 20 | 0 | 717 | 0 | 0 | 15 | 90 | 51 | 0 | 0 | 0 | 22 | 0 | 15 | 0 | 0 | 0 | 0 | 0 | 632 | 174 | 0 | 0 | 18 | 15 | 210 | 674 | 0 | 0 | 12 | 0 | 0 | 0 | 0 | 87 | 0 | 27 |
| SRR7178693 | 33 | 0 | 630 | 0 | 0 | 30 | 16 | 23 | 20 | 0 | 0 | 19 | 0 | 16 | 0 | 0 | 0 | 0 | 0 | 792 | 0 | 0 | 0 | 13 | 11 | 24 | 31 | 0 | 0 | 0 | 11 | 0 | 0 | 11 | 79 | 0 | 56 |
| SRR7178694 | 27 | 0 | 629 | 0 | 0 | 16 | 18 | 29 | 0 | 0 | 0 | 19 | 0 | 14 | 0 | 0 | 0 | 0 | 0 | 824 | 0 | 0 | 0 | 12 | 21 | 384 | 560 | 0 | 0 | 0 | 14 | 0 | 0 | 0 | 67 | 12 | 41 |
| SRR7178695 | 24 | 0 | 584 | 0 | 0 | 0 | 17 | 0 | 0 | 0 | 0 | 10 | 0 | 17 | 0 | 0 | 0 | 0 | 0 | 844 | 0 | 0 | 0 | 0 | 0 | 11 | 15 | 0 | 0 | 0 | 0 | 0 | 0 | 17 | 68 | 0 | 26 |
| SRR7178696 | 15 | 0 | 510 | 0 | 0 | 12 | 11 | 24 | 0 | 0 | 0 | 0 | 0 | 0 | 0 | 0 | 0 | 0 | 0 | 739 | 0 | 12 | 0 | 12 | 0 | 295 | 16 | 0 | 0 | 24 | 12 | 0 | 0 | 24 | 35 | 0 | 35 |
| SRR7178697 | 30 | 0 | 854 | 0 | 0 | 18 | 0 | 0 | 0 | 0 | 0 | 23 | 0 | 0 | 0 | 0 | 0 | 0 | 0 | 489 | 92 | 0 | 0 | 18 | 14 | 249 | 497 | 0 | 0 | 0 | 0 | 0 | 0 | 0 | 98 | 0 | 30 |
| SRR7178698 | 19 | 0 | 917 | 0 | 0 | 14 | 13 | 0 | 0 | 0 | 0 | 23 | 0 | 0 | 0 | 0 | 0 | 0 | 0 | 225 | 105 | 0 | 0 | 33 | 21 | 16 | 25 | 0 | 0 | 0 | 21 | 0 | 0 | 0 | 60 | 0 | 21 |
| SRR7178699 | 17 | 0 | 821 | 0 | 0 | 25 | 14 | 0 | 15 | 0 | 0 | 27 | 0 | 0 | 0 | 0 | 0 | 0 | 0 | 252 | 121 | 0 | 0 | 23 | 23 | 17 | 22 | 0 | 0 | 0 | 13 | 0 | 0 | 0 | 68 | 0 | 25 |
| SRR7178700 | 19 | 0 | 573 | 0 | 0 | 17 | 18 | 0 | 0 | 0 | 0 | 29 | 0 | 0 | 0 | 0 | 0 | 0 | 0 | 153 | 137 | 0 | 0 | 0 | 38 | 384 | 333 | 0 | 0 | 0 | 19 | 0 | 0 | 0 | 132 | 0 | 36 |
| SRR7178701 | 22 | 0 | 581 | 0 | 0 | 21 | 18 | 0 | 12 | 0 | 0 | 24 | 0 | 0 | 0 | 0 | 0 | 0 | 0 | 138 | 128 | 0 | 0 | 0 | 28 | 25 | 26 | 0 | 0 | 0 | 14 | 0 | 0 | 0 | 124 | 0 | 25 |
| SRR7178702 | 18 | 0 | 478 | 0 | 0 | 32 | 18 | 11 | 0 | 0 | 0 | 23 | 0 | 0 | 18 | 0 | 0 | 0 | 0 | 177 | 71 | 0 | 0 | 32 | 32 | 296 | 253 | 0 | 0 | 0 | 0 | 0 | 11 | 0 | 74 | 0 | 42 |
| SRR7178703 | 10 | 0 | 357 | 0 | 0 | 0 | 0 | 0 | 15 | 0 | 0 | 12 | 0 | 15 | 0 | 0 | 0 | 0 | 0 | 148 | 46 | 0 | 0 | 0 | 88 | 280 | 11 | 0 | 0 | 0 | 44 | 0 | 0 | 0 | 44 | 0 | 29 |
| SRR7178704 | 41 | 0 | 479 | 0 | 0 | 18 | 0 | 12 | 12 | 0 | 0 | 20 | 0 | 0 | 124 | 0 | 0 | 0 | 0 | 435 | 398 | 0 | 0 | 28 | 18 | 27 | 13 | 0 | 0 | 0 | 0 | 0 | 0 | 0 | 50 | 0 | 42 |
| SRR7178705 | 52 | 0 | 472 | 0 | 0 | 0 | 32 | 25 | 12 | 0 | 0 | 20 | 0 | 0 | 101 | 0 | 0 | 0 | 0 | 336 | 349 | 0 | 0 | 17 | 22 | 21 | 10 | 0 | 0 | 0 | 22 | 0 | 0 | 0 | 32 | 0 | 27 |
| SRR7178706 | 43 | 0 | 419 | 0 | 0 | 18 | 24 | 26 | 10 | 0 | 0 | 16 | 0 | 0 | 115 | 0 | 0 | 0 | 0 | 439 | 363 | 0 | 0 | 16 | 31 | 21 | 12 | 0 | 0 | 16 | 10 | 0 | 0 | 0 | 49 | 10 | 26 |
| SRR7178707 | 15 | 0 | 538 | 0 | 0 | 19 | 20 | 0 | 19 | 0 | 0 | 13 | 0 | 0 | 0 | 0 | 0 | 0 | 0 | 74 | 127 | 0 | 0 | 0 | 13 | 17 | 12 | 0 | 19 | 0 | 25 | 0 | 0 | 0 | 89 | 0 | 13 |
| SRR7178708 | 10 | 0 | 570 | 0 | 0 | 31 | 23 | 0 | 31 | 0 | 0 | 0 | 0 | 0 | 0 | 0 | 10 | 0 | 0 | 102 | 112 | 0 | 0 | 0 | 43 | 12 | 11 | 0 | 0 | 0 | 18 | 0 | 0 | 0 | 86 | 0 | 68 |
| SRR7178709 | 14 | 0 | 505 | 0 | 0 | 25 | 0 | 0 | 0 | 0 | 0 | 0 | 0 | 0 | 0 | 0 | 11 | 0 | 0 | 25 | 74 | 0 | 0 | 25 | 0 | 277 | 176 | 0 | 0 | 0 | 0 | 0 | 0 | 0 | 25 | 0 | 0 |
| SRR7178710 | 0 | 0 | 405 | 0 | 0 | 34 | 0 | 0 | 0 | 0 | 0 | 0 | 0 | 0 | 0 | 0 | 0 | 0 | 0 | 63 | 94 | 0 | 0 | 0 | 34 | 238 | 375 | 0 | 0 | 0 | 0 | 0 | 0 | 68 | 34 | 0 | 68 |
| SRR7178711 | 23 | 0 | 428 | 0 | 0 | 43 | 38 | 25 | 40 | 0 | 0 | 0 | 0 | 0 | 19 | 0 | 0 | 0 | 19 | 685 | 599 | 0 | 0 | 0 | 58 | 899 | 637 | 0 | 11 | 0 | 50 | 0 | 0 | 0 | 288 | 0 | 122 |
| SRR7178712 | 26 | 0 | 457 | 0 | 0 | 62 | 21 | 13 | 62 | 0 | 0 | 0 | 0 | 0 | 11 | 0 | 0 | 0 | 0 | 495 | 505 | 0 | 0 | 22 | 35 | 873 | 567 | 0 | 0 | 0 | 18 | 0 | 0 | 0 | 257 | 0 | 128 |
| SRR7178713 | 25 | 0 | 416 | 0 | 0 | 46 | 0 | 32 | 55 | 0 | 0 | 11 | 0 | 0 | 22 | 0 | 0 | 0 | 11 | 627 | 436 | 0 | 0 | 0 | 37 | 767 | 553 | 0 | 0 | 0 | 23 | 0 | 0 | 0 | 251 | 23 | 105 |
| SRR7178714 | 20 | 0 | 616 | 0 | 0 | 15 | 0 | 11 | 0 | 0 | 0 | 0 | 0 | 0 | 0 | 0 | 0 | 0 | 11 | 504 | 131 | 0 | 0 | 11 | 15 | 15 | 31 | 0 | 0 | 0 | 11 | 0 | 0 | 0 | 86 | 15 | 71 |
| SRR7178715 | 17 | 0 | 677 | 0 | 0 | 11 | 0 | 0 | 11 | 0 | 0 | 0 | 0 | 0 | 0 | 0 | 0 | 0 | 0 | 502 | 92 | 0 | 0 | 11 | 18 | 275 | 567 | 0 | 0 | 0 | 0 | 0 | 0 | 0 | 116 | 0 | 35 |
| SRR7178716 | 10 | 0 | 483 | 0 | 0 | 31 | 0 | 0 | 16 | 0 | 0 | 0 | 0 | 0 | 0 | 0 | 0 | 0 | 0 | 468 | 87 | 0 | 0 | 16 | 16 | 249 | 16 | 0 | 0 | 0 | 0 | 0 | 0 | 0 | 16 | 0 | 31 |
| SRR7178717 | 10 | 0 | 288 | 0 | 0 | 0 | 0 | 18 | 0 | 0 | 0 | 0 | 0 | 0 | 0 | 0 | 0 | 0 | 16 | 273 | 40 | 0 | 0 | 0 | 18 | 245 | 350 | 0 | 0 | 0 | 0 | 0 | 0 | 18 | 88 | 0 | 35 |
| SRR7178718 | 16 | 0 | 500 | 0 | 0 | 32 | 28 | 19 | 20 | 0 | 0 | 19 | 0 | 0 | 0 | 0 | 0 | 0 | 0 | 304 | 124 | 0 | 0 | 30 | 24 | 522 | 488 | 0 | 0 | 19 | 20 | 0 | 0 | 0 | 105 | 0 | 37 |
| SRR7178719 | 17 | 0 | 523 | 0 | 0 | 19 | 35 | 21 | 0 | 0 | 0 | 11 | 0 | 0 | 0 | 0 | 0 | 0 | 0 | 323 | 127 | 0 | 0 | 19 | 35 | 24 | 21 | 0 | 0 | 16 | 0 | 0 | 0 | 14 | 106 | 0 | 42 |
| SRR7178720 | 15 | 0 | 493 | 0 | 0 | 22 | 24 | 14 | 19 | 0 | 0 | 12 | 0 | 0 | 0 | 0 | 0 | 0 | 0 | 293 | 119 | 0 | 0 | 17 | 0 | 479 | 467 | 0 | 0 | 0 | 24 | 0 | 0 | 0 | 123 | 0 | 43 |
| SRR7178721 | 20 | 0 | 353 | 0 | 0 | 36 | 37 | 12 | 24 | 0 | 0 | 11 | 0 | 20 | 0 | 0 | 0 | 0 | 0 | 216 | 253 | 0 | 0 | 20 | 0 | 24 | 23 | 0 | 16 | 0 | 12 | 0 | 0 | 40 | 11 | 0 | 84 |
| SRR7178722 | 15 | 0 | 346 | 0 | 0 | 31 | 35 | 31 | 16 | 0 | 0 | 13 | 0 | 0 | 12 | 0 | 0 | 0 | 0 | 301 | 262 | 0 | 0 | 0 | 31 | 571 | 633 | 0 | 0 | 0 | 27 | 0 | 0 | 31 | 241 | 12 | 93 |
| SRR7178723 | 11 | 0 | 317 | 0 | 0 | 47 | 15 | 16 | 31 | 0 | 0 | 0 | 0 | 0 | 0 | 0 | 0 | 0 | 0 | 225 | 173 | 0 | 0 | 16 | 0 | 420 | 435 | 16 | 0 | 0 | 31 | 0 | 0 | 31 | 249 | 0 | 78 |
| SRR7178724 | 19 | 0 | 245 | 0 | 0 | 43 | 19 | 0 | 0 | 0 | 0 | 0 | 0 | 0 | 0 | 0 | 0 | 0 | 0 | 95 | 76 | 22 | 0 | 22 | 22 | 517 | 495 | 22 | 0 | 0 | 22 | 0 | 0 | 22 | 194 | 22 | 43 |
| SRR7213733 | 2152 | 0 | 47 | 0 | 28 | 19 | 60 | 0 | 0 | 0 | 43 | 5026 | 33 | 24 | 0 | 0 | 35 | 0 | 0 | 0 | 1132 | 0 | 0 | 17 | 0 | 62 | 163 | 0 | 11 | 0 | 0 | 0 | 0 | 166 | 0 | 0 | 0 |
| SRR7213734 | 2112 | 0 | 28 | 32 | 36 | 15 | 0 | 0 | 0 | 0 | 43 | 4655 | 27 | 0 | 17 | 0 | 67 | 0 | 0 | 0 | 596 | 0 | 0 | 32 | 11 | 47 | 157 | 0 | 13 | 15 | 0 | 0 | 0 | 148 | 0 | 0 | 15 |
| SRR7213735 | 2057 | 0 | 20 | 22 | 20 | 13 | 11 | 0 | 11 | 0 | 35 | 4487 | 53 | 22 | 23 | 0 | 71 | 0 | 0 | 0 | 960 | 0 | 0 | 15 | 13 | 31 | 93 | 0 | 24 | 0 | 0 | 0 | 0 | 168 | 0 | 0 | 0 |
| SRR7226512 | 492 | 0 | 203 | 0 | 0 | 38 | 280 | 12 | 20 | 0 | 0 | 22 | 0 | 0 | 545 | 0 | 13 | 55 | 98 | 64 | 3110 | 0 | 0 | 37 | 278 | 610 | 1127 | 0 | 0 | 11 | 47 | 0 | 0 | 14 | 29 | 18 | 31 |
| SRR7226513 | 700 | 0 | 231 | 0 | 0 | 50 | 697 | 22 | 32 | 0 | 0 | 25 | 0 | 0 | 1083 | 0 | 22 | 41 | 279 | 155 | 2781 | 0 | 0 | 32 | 729 | 876 | 2774 | 0 | 0 | 0 | 52 | 0 | 0 | 0 | 20 | 15 | 17 |
| SRR7226514 | 458 | 0 | 7908 | 0 | 0 | 29 | 0 | 0 | 20 | 0 | 0 | 300 | 19 | 0 | 9336 | 0 | 29 | 215 | 37 | 82 | 9484 | 0 | 0 | 11 | 314 | 666 | 732 | 0 | 0 | 0 | 43 | 30 | 24 | 0 | 149 | 105 | 163 |
| SRR7226515 | 807 | 0 | 6145 | 0 | 0 | 50 | 13 | 0 | 42 | 0 | 0 | 417 | 0 | 24 | 2767 | 0 | 13 | 75 | 27 | 64 | 1757 | 0 | 0 | 0 | 1324 | 1189 | 1565 | 0 | 0 | 0 | 106 | 0 | 0 | 0 | 182 | 90 | 147 |
| SRR7226516 | 211 | 0 | 10064 | 0 | 11 | 114 | 25 | 0 | 79 | 0 | 0 | 95 | 13 | 10 | 13029 | 0 | 18 | 63 | 0 | 0 | 7890 | 0 | 0 | 11 | 1875 | 2519 | 7670 | 0 | 0 | 0 | 235 | 34 | 12 | 11 | 231 | 148 | 276 |
| SRR7226517 | 1426 | 44 | 1351 | 0 | 0 | 210 | 77 | 10 | 150 | 0 | 0 | 46 | 0 | 0 | 0 | 12 | 70 | 76 | 0 | 214 | 4372 | 0 | 0 | 0 | 1088 | 2945 | 4167 | 0 | 15 | 0 | 243 | 0 | 0 | 11 | 606 | 231 | 368 |
| SRR7226518 | 1421 | 12 | 1240 | 0 | 0 | 69 | 115 | 11 | 46 | 0 | 0 | 65 | 0 | 0 | 0 | 21 | 68 | 36 | 25 | 242 | 8279 | 0 | 0 | 20 | 120 | 917 | 314 | 0 | 12 | 12 | 70 | 0 | 0 | 10 | 142 | 40 | 76 |
| SRR7226519 | 1243 | 0 | 919 | 0 | 0 | 142 | 56 | 13 | 98 | 0 | 0 | 67 | 0 | 0 | 0 | 17 | 38 | 18 | 18 | 143 | 12321 | 0 | 0 | 18 | 165 | 2237 | 429 | 0 | 0 | 0 | 175 | 13 | 0 | 0 | 72 | 22 | 50 |
| SRR7226520 | 248 | 0 | 604 | 0 | 0 | 17 | 281 | 13 | 27 | 0 | 0 | 63 | 0 | 0 | 3601 | 0 | 0 | 206 | 44 | 710 | 6378 | 0 | 0 | 30 | 301 | 203 | 890 | 0 | 0 | 0 | 22 | 0 | 0 | 0 | 66 | 39 | 50 |
| SRR7226521 | 230 | 0 | 1429 | 0 | 0 | 80 | 97 | 20 | 59 | 0 | 0 | 111 | 0 | 0 | 2498 | 0 | 36 | 295 | 25 | 520 | 8417 | 0 | 0 | 13 | 1298 | 1145 | 5071 | 0 | 0 | 0 | 87 | 0 | 0 | 0 | 373 | 254 | 295 |
| SRR7226522 | 285 | 0 | 1428 | 0 | 0 | 81 | 160 | 11 | 61 | 0 | 0 | 61 | 0 | 0 | 8050 | 0 | 0 | 0 | 26 | 624 | 6478 | 0 | 0 | 17 | 1219 | 1258 | 1906 | 0 | 0 | 0 | 141 | 0 | 0 | 0 | 220 | 192 | 289 |
| SRR7226523 | 1230 | 17 | 5977 | 0 | 0 | 26 | 31 | 14 | 21 | 0 | 0 | 318 | 13 | 0 | 1978 | 0 | 31 | 465 | 47 | 315 | 5339 | 0 | 0 | 13 | 415 | 611 | 1662 | 0 | 0 | 0 | 37 | 0 | 0 | 0 | 78 | 95 | 136 |
| SRR7226524 | 462 | 0 | 7623 | 16 | 0 | 0 | 37 | 184 | 15 | 0 | 0 | 417 | 0 | 20 | 193 | 45 | 16 | 676 | 53 | 270 | 5952 | 0 | 0 | 33 | 156 | 193 | 284 | 0 | 0 | 48 | 0 | 18 | 0 | 79 | 48 | 29 | 53 |
| SRR7226525 | 479 | 11 | 8097 | 0 | 0 | 14 | 18 | 0 | 19 | 0 | 0 | 285 | 13 | 0 | 3822 | 0 | 17 | 33 | 18 | 112 | 3888 | 0 | 0 | 46 | 246 | 422 | 578 | 0 | 0 | 0 | 32 | 0 | 11 | 0 | 198 | 118 | 176 |
| SRR7226526 | 458 | 11 | 6533 | 0 | 0 | 33 | 82 | 25 | 28 | 0 | 0 | 847 | 0 | 11 | 7116 | 0 | 29 | 134 | 36 | 298 | 5650 | 0 | 0 | 12 | 431 | 795 | 1540 | 0 | 0 | 0 | 43 | 11 | 11 | 0 | 398 | 234 | 290 |
| SRR7233553 | 64 | 0 | 0 | 15 | 31 | 33 | 0 | 0 | 11 | 30 | 45 | 38 | 0 | 31 | 113 | 0 | 0 | 0 | 11 | 11 | 56 | 0 | 0 | 0 | 10 | 42 | 16 | 16 | 0 | 0 | 43 | 0 | 0 | 38 | 38 | 50 | 45 |
| SRR7233554 | 61 | 0 | 0 | 0 | 33 | 36 | 0 | 0 | 30 | 22 | 32 | 44 | 0 | 31 | 213 | 0 | 0 | 0 | 11 | 53 | 181 | 0 | 0 | 0 | 37 | 35 | 12 | 10 | 0 | 0 | 35 | 0 | 0 | 20 | 21 | 42 | 25 |
| SRR7233556 | 41 | 0 | 0 | 0 | 32 | 15 | 0 | 0 | 20 | 25 | 30 | 36 | 0 | 18 | 199 | 0 | 0 | 0 | 0 | 0 | 55 | 0 | 0 | 0 | 46 | 21 | 0 | 12 | 0 | 0 | 37 | 0 | 0 | 12 | 0 | 27 | 18 |
| SRR7233557 | 84 | 0 | 0 | 11 | 27 | 0 | 0 | 0 | 16 | 21 | 148 | 18 | 0 | 24 | 27 | 0 | 0 | 0 | 0 | 0 | 27 | 0 | 0 | 0 | 27 | 51 | 0 | 10 | 0 | 0 | 23 | 0 | 0 | 69 | 20 | 25 | 104 |
| SRR7233558 | 76 | 0 | 0 | 13 | 41 | 14 | 0 | 0 | 27 | 20 | 136 | 30 | 0 | 29 | 106 | 0 | 0 | 0 | 0 | 0 | 149 | 0 | 0 | 0 | 36 | 127 | 19 | 14 | 0 | 0 | 38 | 0 | 0 | 119 | 16 | 30 | 113 |
| SRR7233559 | 103 | 0 | 0 | 12 | 33 | 21 | 0 | 0 | 33 | 43 | 50 | 30 | 0 | 34 | 0 | 0 | 0 | 0 | 0 | 0 | 87 | 0 | 0 | 0 | 53 | 23 | 0 | 0 | 0 | 0 | 49 | 0 | 0 | 14 | 25 | 36 | 26 |
| SRR7233560 | 53 | 0 | 0 | 0 | 29 | 11 | 0 | 0 | 20 | 19 | 152 | 46 | 0 | 32 | 0 | 0 | 0 | 0 | 0 | 0 | 15 | 0 | 0 | 0 | 39 | 64 | 0 | 0 | 0 | 0 | 43 | 0 | 0 | 48 | 17 | 20 | 80 |
| SRR7233561 | 61 | 0 | 0 | 14 | 21 | 56 | 0 | 0 | 15 | 19 | 26 | 24 | 0 | 36 | 82 | 0 | 0 | 0 | 0 | 0 | 82 | 0 | 0 | 0 | 19 | 61 | 16 | 11 | 0 | 0 | 55 | 0 | 0 | 30 | 49 | 13 | 51 |
| SRR7233562 | 70 | 0 | 0 | 11 | 33 | 24 | 0 | 0 | 43 | 19 | 30 | 28 | 0 | 25 | 98 | 0 | 0 | 0 | 0 | 0 | 43 | 0 | 0 | 0 | 40 | 38 | 0 | 16 | 0 | 0 | 32 | 0 | 0 | 21 | 34 | 44 | 36 |
| SRR7233563 | 53 | 0 | 0 | 0 | 18 | 56 | 0 | 0 | 45 | 14 | 98 | 27 | 0 | 29 | 124 | 0 | 0 | 0 | 0 | 15 | 325 | 0 | 0 | 0 | 47 | 320 | 28 | 14 | 0 | 0 | 59 | 0 | 0 | 155 | 28 | 62 | 258 |
| SRR7233564 | 66 | 0 | 0 | 10 | 26 | 50 | 0 | 0 | 53 | 15 | 17 | 33 | 0 | 44 | 175 | 0 | 0 | 0 | 0 | 15 | 364 | 0 | 0 | 0 | 51 | 48 | 31 | 20 | 0 | 0 | 67 | 0 | 0 | 22 | 40 | 65 | 50 |
| SRR7233565 | 100 | 0 | 0 | 14 | 42 | 12 | 0 | 0 | 58 | 30 | 28 | 18 | 10 | 24 | 75 | 0 | 0 | 21 | 0 | 19 | 84 | 0 | 0 | 0 | 14 | 66 | 34 | 17 | 0 | 0 | 48 | 0 | 0 | 31 | 38 | 14 | 56 |
| SRR7233566 | 52 | 0 | 0 | 11 | 33 | 84 | 0 | 0 | 57 | 12 | 105 | 15 | 16 | 17 | 94 | 0 | 0 | 0 | 0 | 42 | 178 | 0 | 0 | 0 | 42 | 407 | 37 | 15 | 0 | 0 | 52 | 0 | 0 | 156 | 49 | 92 | 390 |
| SRR7233568 | 74 | 0 | 0 | 0 | 30 | 22 | 0 | 0 | 32 | 47 | 45 | 34 | 0 | 28 | 34 | 0 | 0 | 0 | 0 | 0 | 46 | 0 | 0 | 0 | 10 | 22 | 13 | 18 | 0 | 0 | 36 | 0 | 0 | 14 | 18 | 26 | 16 |
| SRR7233570 | 68 | 0 | 0 | 0 | 14 | 36 | 0 | 0 | 46 | 26 | 42 | 39 | 0 | 23 | 228 | 0 | 0 | 0 | 0 | 0 | 254 | 0 | 0 | 0 | 12 | 43 | 0 | 0 | 0 | 0 | 48 | 0 | 0 | 32 | 12 | 29 | 21 |
| SRR7233571 | 68 | 0 | 0 | 12 | 35 | 42 | 0 | 0 | 30 | 21 | 43 | 20 | 0 | 32 | 122 | 0 | 0 | 0 | 0 | 0 | 28 | 0 | 0 | 0 | 17 | 55 | 30 | 15 | 0 | 0 | 10 | 0 | 0 | 34 | 20 | 45 | 37 |
| SRR7233572 | 68 | 0 | 0 | 0 | 27 | 15 | 0 | 0 | 22 | 23 | 40 | 16 | 0 | 27 | 106 | 0 | 0 | 0 | 0 | 0 | 73 | 0 | 0 | 0 | 12 | 23 | 0 | 0 | 0 | 0 | 31 | 0 | 0 | 27 | 20 | 38 | 27 |
| SRR7262826 | 607 | 0 | 171 | 0 | 27 | 175 | 130 | 0 | 68 | 0 | 39 | 289 | 0 | 26 | 300 | 0 | 62 | 0 | 0 | 0 | 457 | 11 | 0 | 17 | 176 | 1167 | 79 | 49 | 12 | 0 | 81 | 50 | 12 | 142 | 77 | 74 | 218 |
| SRR7262827 | 371 | 0 | 250 | 0 | 24 | 139 | 319 | 0 | 52 | 0 | 39 | 283 | 13 | 29 | 275 | 0 | 151 | 0 | 0 | 0 | 597 | 17 | 0 | 13 | 203 | 1095 | 258 | 24 | 22 | 0 | 57 | 25 | 20 | 579 | 79 | 93 | 256 |
| SRR7262828 | 408 | 0 | 219 | 0 | 15 | 26 | 309 | 0 | 63 | 0 | 28 | 316 | 15 | 14 | 131 | 0 | 96 | 0 | 0 | 17 | 786 | 15 | 0 | 0 | 22 | 183 | 31 | 24 | 34 | 0 | 66 | 19 | 0 | 127 | 15 | 20 | 42 |
| SRR7262829 | 476 | 0 | 133 | 0 | 21 | 301 | 335 | 0 | 64 | 0 | 38 | 235 | 0 | 14 | 206 | 11 | 22 | 0 | 31 | 57 | 775 | 22 | 0 | 47 | 91 | 1713 | 121 | 33 | 12 | 12 | 68 | 69 | 0 | 321 | 46 | 96 | 206 |
| SRR7262830 | 444 | 0 | 264 | 0 | 18 | 20 | 133 | 0 | 61 | 0 | 32 | 323 | 16 | 11 | 231 | 0 | 178 | 0 | 0 | 0 | 494 | 12 | 0 | 0 | 24 | 148 | 16 | 27 | 21 | 0 | 79 | 41 | 13 | 84 | 14 | 14 | 35 |
| SRR7288164 | 97 | 0 | 11 | 552 | 245 | 0 | 0 | 0 | 0 | 0 | 0 | 55 | 0 | 0 | 0 | 0 | 0 | 0 | 0 | 0 | 0 | 0 | 0 | 0 | 0 | 0 | 0 | 0 | 0 | 0 | 0 | 0 | 0 | 0 | 0 | 0 | 0 |
| SRR7288166 | 97 | 0 | 0 | 19 | 199 | 0 | 0 | 0 | 0 | 0 | 0 | 55 | 0 | 0 | 0 | 0 | 0 | 0 | 0 | 0 | 0 | 46 | 0 | 0 | 0 | 0 | 31 | 0 | 0 | 0 | 0 | 0 | 0 | 0 | 0 | 0 | 0 |
| SRR7288167 | 97 | 0 | 0 | 13 | 126 | 0 | 0 | 0 | 0 | 0 | 0 | 43 | 0 | 0 | 0 | 0 | 0 | 0 | 0 | 0 | 0 | 16 | 0 | 0 | 0 | 0 | 0 | 0 | 0 | 0 | 0 | 0 | 0 | 0 | 0 | 0 | 0 |
| SRR7288236 | 31 | 0 | 0 | 17 | 94 | 0 | 0 | 31 | 0 | 0 | 0 | 0 | 0 | 0 | 0 | 0 | 0 | 0 | 0 | 0 | 0 | 0 | 0 | 0 | 0 | 0 | 0 | 0 | 0 | 0 | 0 | 0 | 0 | 0 | 0 | 0 | 0 |
| SRR7288237 | 24 | 0 | 0 | 433 | 177 | 0 | 0 | 0 | 0 | 0 | 0 | 13 | 0 | 0 | 0 | 0 | 0 | 0 | 0 | 0 | 0 | 0 | 0 | 0 | 0 | 0 | 0 | 0 | 0 | 0 | 0 | 0 | 0 | 0 | 0 | 0 | 0 |
| SRR7288238 | 17 | 15 | 0 | 18 | 62 | 0 | 0 | 0 | 0 | 0 | 0 | 0 | 0 | 0 | 0 | 0 | 0 | 0 | 0 | 0 | 0 | 0 | 0 | 0 | 0 | 0 | 0 | 0 | 0 | 0 | 0 | 0 | 0 | 0 | 0 | 0 | 0 |
| SRR7288239 | 13 | 0 | 0 | 711 | 108 | 0 | 0 | 31 | 0 | 0 | 0 | 0 | 0 | 0 | 0 | 0 | 0 | 0 | 0 | 0 | 0 | 0 | 0 | 0 | 0 | 0 | 0 | 0 | 0 | 0 | 0 | 0 | 0 | 0 | 0 | 0 | 0 |
| SRR7288244 | 55 | 0 | 0 | 853 | 0 | 0 | 0 | 0 | 0 | 0 | 0 | 0 | 0 | 0 | 0 | 0 | 0 | 0 | 0 | 0 | 0 | 0 | 0 | 0 | 0 | 0 | 0 | 0 | 0 | 0 | 0 | 0 | 0 | 0 | 0 | 0 | 0 |
| SRR7288264 | 86 | 0 | 33 | 359 | 157 | 0 | 0 | 0 | 0 | 0 | 0 | 89 | 0 | 0 | 0 | 0 | 24 | 0 | 0 | 0 | 0 | 0 | 0 | 0 | 0 | 0 | 0 | 45 | 0 | 0 | 0 | 0 | 0 | 0 | 0 | 0 | 0 |
| SRR7288265 | 124 | 0 | 21 | 14 | 160 | 0 | 0 | 0 | 0 | 0 | 0 | 85 | 0 | 0 | 0 | 0 | 0 | 0 | 0 | 0 | 0 | 0 | 0 | 0 | 0 | 0 | 0 | 0 | 0 | 0 | 0 | 0 | 0 | 0 | 0 | 0 | 0 |
| SRR7288266 | 112 | 0 | 21 | 16 | 133 | 0 | 0 | 0 | 0 | 0 | 0 | 77 | 0 | 0 | 0 | 0 | 0 | 0 | 0 | 0 | 0 | 22 | 0 | 0 | 0 | 0 | 0 | 22 | 0 | 0 | 0 | 0 | 0 | 0 | 0 | 0 | 0 |
| SRR7288267 | 105 | 0 | 0 | 364 | 46 | 0 | 0 | 0 | 0 | 0 | 0 | 87 | 0 | 0 | 0 | 0 | 0 | 0 | 0 | 0 | 0 | 23 | 0 | 0 | 0 | 0 | 0 | 0 | 0 | 0 | 0 | 0 | 0 | 0 | 0 | 0 | 0 |
| SRR7288277 | 161 | 0 | 11 | 490 | 140 | 0 | 0 | 140 | 0 | 0 | 0 | 23 | 0 | 0 | 0 | 0 | 0 | 0 | 0 | 0 | 0 | 0 | 0 | 0 | 0 | 0 | 210 | 0 | 0 | 0 | 0 | 0 | 0 | 0 | 0 | 0 | 0 |
| SRR7288300 | 165 | 0 | 25 | 378 | 327 | 0 | 0 | 0 | 0 | 0 | 0 | 121 | 0 | 0 | 0 | 0 | 0 | 0 | 0 | 0 | 174 | 25 | 0 | 0 | 0 | 0 | 25 | 0 | 0 | 0 | 0 | 0 | 0 | 0 | 0 | 0 | 0 |
| SRR7288301 | 110 | 0 | 32 | 309 | 283 | 0 | 0 | 0 | 0 | 0 | 0 | 146 | 0 | 0 | 0 | 0 | 0 | 0 | 0 | 0 | 0 | 77 | 0 | 0 | 0 | 0 | 0 | 0 | 0 | 0 | 0 | 0 | 0 | 0 | 0 | 0 | 0 |
| SRR7288302 | 136 | 0 | 35 | 12 | 276 | 0 | 0 | 0 | 0 | 0 | 0 | 146 | 0 | 0 | 0 | 0 | 0 | 0 | 0 | 0 | 0 | 25 | 0 | 0 | 0 | 0 | 0 | 0 | 0 | 0 | 0 | 0 | 0 | 0 | 0 | 0 | 0 |
| SRR7288303 | 104 | 0 | 49 | 389 | 181 | 0 | 0 | 0 | 0 | 0 | 0 | 163 | 0 | 0 | 0 | 0 | 0 | 0 | 0 | 0 | 178 | 26 | 0 | 0 | 0 | 0 | 26 | 0 | 0 | 0 | 0 | 0 | 0 | 0 | 0 | 0 | 0 |
| SRR7288308 | 133 | 0 | 63 | 13 | 251 | 0 | 0 | 0 | 0 | 0 | 0 | 108 | 0 | 0 | 0 | 0 | 0 | 0 | 0 | 594 | 594 | 0 | 0 | 0 | 0 | 0 | 36 | 72 | 0 | 0 | 0 | 0 | 0 | 0 | 0 | 0 | 0 |
| SRR7288309 | 128 | 75 | 33 | 12 | 149 | 0 | 0 | 0 | 0 | 0 | 0 | 121 | 0 | 0 | 0 | 0 | 0 | 0 | 0 | 620 | 0 | 0 | 0 | 0 | 0 | 0 | 37 | 0 | 0 | 0 | 0 | 0 | 0 | 0 | 0 | 0 | 0 |
| SRR7288310 | 106 | 0 | 59 | 674 | 213 | 0 | 0 | 0 | 0 | 0 | 0 | 140 | 0 | 0 | 0 | 0 | 0 | 0 | 0 | 0 | 0 | 0 | 0 | 0 | 0 | 0 | 35 | 0 | 0 | 0 | 0 | 0 | 0 | 0 | 0 | 0 | 0 |
| SRR7288311 | 103 | 0 | 55 | 297 | 223 | 0 | 0 | 0 | 0 | 0 | 0 | 123 | 0 | 0 | 0 | 0 | 0 | 0 | 0 | 0 | 0 | 0 | 0 | 0 | 0 | 0 | 74 | 37 | 0 | 0 | 0 | 0 | 0 | 0 | 0 | 0 | 0 |
| SRR7288320 | 171 | 49 | 105 | 320 | 98 | 0 | 0 | 0 | 0 | 0 | 0 | 258 | 0 | 0 | 534 | 0 | 0 | 0 | 0 | 0 | 0 | 25 | 0 | 0 | 0 | 0 | 0 | 0 | 0 | 0 | 0 | 0 | 0 | 0 | 0 | 0 | 0 |
| SRR7288321 | 210 | 26 | 111 | 261 | 52 | 0 | 0 | 0 | 0 | 0 | 0 | 282 | 0 | 0 | 0 | 0 | 11 | 0 | 0 | 0 | 0 | 26 | 0 | 0 | 0 | 0 | 0 | 26 | 0 | 0 | 0 | 0 | 0 | 0 | 0 | 0 | 0 |
| SRR7288322 | 206 | 0 | 122 | 341 | 97 | 0 | 0 | 24 | 0 | 0 | 0 | 260 | 0 | 49 | 0 | 0 | 0 | 0 | 0 | 0 | 0 | 49 | 0 | 0 | 0 | 0 | 0 | 0 | 0 | 0 | 0 | 0 | 0 | 0 | 0 | 0 | 0 |
| SRR7288323 | 149 | 0 | 119 | 179 | 26 | 0 | 0 | 26 | 0 | 0 | 0 | 231 | 0 | 0 | 549 | 0 | 11 | 0 | 0 | 0 | 0 | 26 | 0 | 0 | 0 | 0 | 26 | 0 | 0 | 0 | 0 | 0 | 0 | 0 | 0 | 0 | 0 |
| SRR7288376 | 166 | 0 | 0 | 11 | 118 | 0 | 0 | 0 | 0 | 0 | 0 | 48 | 0 | 0 | 0 | 0 | 0 | 0 | 0 | 0 | 0 | 59 | 0 | 0 | 0 | 0 | 0 | 0 | 0 | 0 | 0 | 0 | 0 | 0 | 0 | 0 | 0 |
| SRR7288377 | 143 | 0 | 15 | 611 | 183 | 0 | 0 | 0 | 0 | 0 | 0 | 85 | 0 | 0 | 0 | 0 | 0 | 0 | 0 | 0 | 0 | 61 | 0 | 0 | 0 | 0 | 61 | 0 | 0 | 0 | 0 | 0 | 0 | 0 | 0 | 0 | 0 |
| SRR7288378 | 172 | 0 | 18 | 411 | 118 | 0 | 0 | 0 | 0 | 0 | 0 | 73 | 0 | 0 | 0 | 0 | 0 | 0 | 0 | 0 | 0 | 59 | 0 | 0 | 0 | 0 | 0 | 0 | 0 | 59 | 0 | 0 | 0 | 0 | 0 | 0 | 0 |
| SRR7288379 | 140 | 0 | 42 | 490 | 245 | 0 | 0 | 0 | 0 | 0 | 0 | 102 | 0 | 0 | 0 | 0 | 0 | 0 | 0 | 0 | 0 | 61 | 0 | 0 | 0 | 0 | 61 | 0 | 0 | 61 | 0 | 0 | 0 | 0 | 0 | 0 | 0 |
| SRR7288380 | 78 | 0 | 0 | 486 | 88 | 0 | 0 | 0 | 0 | 0 | 0 | 46 | 0 | 0 | 0 | 0 | 0 | 0 | 0 | 0 | 0 | 66 | 0 | 0 | 0 | 0 | 0 | 0 | 0 | 0 | 0 | 0 | 0 | 0 | 0 | 0 | 0 |
| SRR7288381 | 75 | 23 | 0 | 25 | 113 | 0 | 0 | 0 | 0 | 0 | 0 | 55 | 0 | 0 | 0 | 0 | 0 | 0 | 0 | 0 | 0 | 0 | 0 | 0 | 0 | 0 | 45 | 0 | 0 | 0 | 0 | 0 | 0 | 0 | 0 | 0 | 0 |
| SRR7288382 | 70 | 0 | 20 | 21 | 173 | 0 | 0 | 0 | 0 | 0 | 0 | 56 | 0 | 0 | 0 | 0 | 0 | 0 | 0 | 0 | 0 | 22 | 0 | 0 | 0 | 0 | 0 | 0 | 0 | 0 | 0 | 0 | 0 | 0 | 0 | 0 | 0 |

### Supp Table S8.2.Columns_AM_to_BW

| | | | | | | | | | | | | | | | | | | | | | | | | | | | | | | | | | | | | | |
|---|---|---|---|---|---|---|---|---|---|---|---|---|---|---|---|---|---|---|---|---|---|---|---|---|---|---|---|---|---|---|---|---|---|---|---|---|---|
| SRR7288383 | 92 | 21 | 0 | 15 | 150 | 0 | 0 | 0 | 0 | 0 | 0 | 42 | 0 | 0 | 0 | 0 | 0 | 0 | 0 | 0 | 0 | 21 | 0 | 0 | 0 | 0 | 21 | 0 | 0 | 0 | 0 | 0 | 0 | 0 | 0 | 0 | 0 |
| SRR7288384 | 318 | 0 | 266 | 396 | 86 | 0 | 0 | 0 | 0 | 0 | 0 | 103 | 0 | 0 | 0 | 0 | 0 | 0 | 0 | 0 | 0 | 34 | 0 | 0 | 0 | 0 | 0 | 0 | 0 | 0 | 0 | 0 | 0 | 0 | 0 | 0 | 0 |
| SRR7288385 | 284 | 0 | 241 | 332 | 105 | 0 | 0 | 0 | 0 | 0 | 0 | 116 | 0 | 0 | 0 | 0 | 0 | 0 | 0 | 0 | 0 | 52 | 0 | 0 | 0 | 0 | 0 | 0 | 0 | 0 | 0 | 0 | 0 | 0 | 0 | 0 | 0 |
| SRR7288386 | 252 | 0 | 222 | 16 | 168 | 0 | 0 | 0 | 0 | 0 | 0 | 83 | 0 | 0 | 0 | 0 | 0 | 0 | 0 | 940 | 0 | 34 | 0 | 0 | 0 | 0 | 0 | 0 | 0 | 0 | 0 | 0 | 0 | 0 | 0 | 0 | 0 |
| SRR7288387 | 293 | 0 | 255 | 17 | 84 | 0 | 0 | 0 | 0 | 0 | 0 | 91 | 0 | 17 | 0 | 0 | 0 | 0 | 0 | 0 | 0 | 67 | 0 | 0 | 0 | 0 | 0 | 0 | 0 | 0 | 0 | 0 | 0 | 0 | 0 | 0 | 0 |
| SRR7288400 | 92 | 0 | 0 | 0 | 0 | 0 | 0 | 0 | 0 | 0 | 0 | 46 | 0 | 0 | 0 | 0 | 0 | 0 | 0 | 0 | 0 | 0 | 0 | 0 | 0 | 0 | 0 | 0 | 0 | 0 | 0 | 0 | 0 | 0 | 0 | 0 | 0 |
| SRR7288402 | 156 | 0 | 22 | 26 | 0 | 0 | 0 | 0 | 0 | 0 | 0 | 89 | 0 | 0 | 0 | 0 | 0 | 0 | 0 | 0 | 0 | 0 | 0 | 0 | 0 | 0 | 0 | 0 | 0 | 0 | 0 | 0 | 0 | 0 | 0 | 0 | 0 |
| SRR7288472 | 25 | 0 | 16 | 16 | 398 | 0 | 0 | 0 | 0 | 0 | 0 | 33 | 0 | 0 | 0 | 0 | 0 | 0 | 0 | 0 | 0 | 0 | 0 | 0 | 0 | 0 | 0 | 0 | 0 | 0 | 0 | 0 | 0 | 0 | 0 | 0 | 0 |
| SRR7288474 | 65 | 0 | 0 | 22 | 97 | 0 | 0 | 0 | 0 | 0 | 0 | 81 | 0 | 0 | 0 | 0 | 0 | 0 | 0 | 0 | 0 | 0 | 0 | 0 | 0 | 0 | 0 | 0 | 0 | 0 | 0 | 0 | 0 | 0 | 0 | 0 | 0 |
| SRR7288475 | 34 | 0 | 0 | 721 | 515 | 0 | 0 | 0 | 0 | 0 | 0 | 94 | 0 | 0 | 0 | 0 | 0 | 0 | 0 | 0 | 0 | 0 | 0 | 0 | 0 | 0 | 0 | 0 | 0 | 0 | 0 | 0 | 0 | 0 | 0 | 0 | 0 |
| SRR7288512 | 127 | 0 | 21 | 421 | 50 | 0 | 0 | 0 | 0 | 0 | 0 | 71 | 0 | 0 | 0 | 0 | 0 | 0 | 0 | 0 | 0 | 0 | 0 | 0 | 0 | 0 | 0 | 0 | 25 | 0 | 0 | 0 | 0 | 0 | 0 | 0 | 0 |
| SRR7288513 | 122 | 0 | 13 | 381 | 25 | 0 | 0 | 25 | 0 | 0 | 0 | 64 | 0 | 0 | 0 | 0 | 0 | 0 | 0 | 0 | 606 | 25 | 0 | 0 | 0 | 0 | 0 | 0 | 0 | 0 | 0 | 0 | 0 | 0 | 0 | 0 | 0 |
| SRR7288514 | 115 | 0 | 15 | 244 | 0 | 0 | 0 | 0 | 0 | 0 | 0 | 62 | 0 | 0 | 0 | 0 | 0 | 0 | 0 | 0 | 0 | 49 | 0 | 0 | 0 | 0 | 0 | 0 | 24 | 0 | 0 | 0 | 0 | 0 | 0 | 0 | 0 |
| SRR7288515 | 115 | 24 | 15 | 484 | 0 | 0 | 0 | 0 | 0 | 0 | 0 | 63 | 0 | 0 | 0 | 0 | 0 | 0 | 0 | 0 | 0 | 0 | 0 | 0 | 0 | 0 | 0 | 0 | 24 | 0 | 0 | 0 | 0 | 0 | 0 | 0 | 0 |
| SRR7293222 | 20 | 0 | 2581 | 0 | 0 | 0 | 0 | 82 | 0 | 0 | 0 | 443 | 0 | 0 | 3200 | 0 | 0 | 0 | 0 | 63 | 1310 | 0 | 0 | 0 | 0 | 34 | 570 | 16 | 0 | 16 | 0 | 0 | 0 | 14 | 1747 | 94 | 673 |
| SRR7293223 | 26 | 0 | 2631 | 0 | 0 | 0 | 0 | 52 | 0 | 0 | 0 | 418 | 0 | 0 | 2601 | 0 | 0 | 0 | 0 | 89 | 1076 | 0 | 0 | 0 | 0 | 27 | 357 | 20 | 0 | 14 | 0 | 0 | 0 | 0 | 1252 | 66 | 598 |
| SRR7293224 | 20 | 63 | 1733 | 0 | 0 | 0 | 26 | 163 | 0 | 0 | 0 | 247 | 0 | 43 | 9825 | 0 | 0 | 11 | 46 | 560 | 7462 | 0 | 0 | 0 | 0 | 51 | 31 | 0 | 0 | 0 | 0 | 0 | 0 | 0 | 94 | 109 | 66 |
| SRR7293225 | 14 | 0 | 2654 | 0 | 0 | 0 | 0 | 45 | 0 | 0 | 0 | 222 | 0 | 0 | 131 | 0 | 0 | 0 | 0 | 136 | 413 | 0 | 0 | 0 | 0 | 151 | 23 | 85 | 0 | 24 | 12 | 0 | 12 | 0 | 41 | 63 | 15 |
| SRR7293226 | 22 | 0 | 3616 | 0 | 0 | 0 | 0 | 156 | 0 | 0 | 0 | 305 | 0 | 0 | 484 | 0 | 0 | 0 | 0 | 105 | 662 | 0 | 0 | 0 | 0 | 91 | 50 | 40 | 0 | 58 | 0 | 0 | 0 | 43 | 58 | 74 | 22 |
| SRR7293227 | 14 | 13 | 2882 | 0 | 0 | 0 | 0 | 492 | 0 | 0 | 0 | 267 | 0 | 0 | 324 | 0 | 0 | 0 | 0 | 87 | 564 | 0 | 0 | 0 | 0 | 19 | 441 | 21 | 0 | 177 | 0 | 0 | 0 | 25 | 719 | 28 | 278 |
| SRR7293228 | 28 | 14 | 1520 | 0 | 0 | 0 | 0 | 17 | 0 | 0 | 0 | 470 | 0 | 14 | 32250 | 0 | 0 | 0 | 0 | 30 | 860 | 0 | 0 | 0 | 0 | 43 | 231 | 0 | 0 | 0 | 0 | 0 | 0 | 0 | 96 | 136 | 34 |
| SRR7293229 | 12 | 13 | 2328 | 0 | 0 | 0 | 0 | 0 | 0 | 0 | 0 | 323 | 0 | 0 | 3914 | 0 | 0 | 0 | 0 | 0 | 366 | 0 | 0 | 0 | 0 | 44 | 200 | 67 | 13 | 0 | 13 | 0 | 0 | 22 | 46 | 93 | 16 |
| SRR7293230 | 127 | 50 | 2934 | 20 | 38 | 12 | 0 | 495 | 0 | 0 | 0 | 67 | 0 | 0 | 209 | 0 | 17 | 0 | 20 | 866 | 606 | 0 | 0 | 0 | 0 | 292 | 1715 | 85 | 0 | 174 | 12 | 0 | 12 | 46 | 192 | 25 | 108 |
| SRR7293231 | 126 | 37 | 2966 | 20 | 19 | 0 | 0 | 162 | 0 | 0 | 0 | 59 | 0 | 0 | 165 | 0 | 19 | 0 | 17 | 344 | 462 | 0 | 0 | 0 | 0 | 160 | 665 | 50 | 0 | 59 | 16 | 0 | 0 | 88 | 170 | 16 | 106 |
| SRR7293232 | 167 | 69 | 2124 | 0 | 0 | 0 | 0 | 20 | 0 | 0 | 0 | 71 | 0 | 22 | 1597 | 0 | 20 | 292 | 0 | 412 | 730 | 0 | 0 | 0 | 0 | 18 | 35 | 0 | 0 | 0 | 11 | 0 | 0 | 0 | 48 | 66 | 75 |
| SRR7293233 | 96 | 22 | 2643 | 23 | 27 | 14 | 12 | 447 | 0 | 0 | 0 | 62 | 0 | 0 | 29 | 0 | 19 | 0 | 16 | 813 | 817 | 0 | 0 | 0 | 0 | 520 | 1604 | 69 | 0 | 143 | 21 | 0 | 0 | 61 | 161 | 0 | 74 |
| SRR7343626 | 0 | 0 | 0 | 0 | 0 | 0 | 0 | 0 | 0 | 0 | 0 | 0 | 0 | 41 | 0 | 0 | 0 | 0 | 0 | 0 | 0 | 0 | 0 | 0 | 41 | 62 | 0 | 0 | 0 | 0 | 0 | 0 | 0 | 21 | 0 | 0 | 0 |
| SRR7343627 | 0 | 0 | 0 | 0 | 0 | 0 | 0 | 0 | 0 | 0 | 0 | 0 | 0 | 0 | 0 | 0 | 0 | 0 | 0 | 0 | 0 | 0 | 0 | 0 | 0 | 149 | 0 | 0 | 0 | 0 | 0 | 0 | 0 | 0 | 50 | 0 | 0 |
| SRR7343629 | 0 | 0 | 0 | 0 | 0 | 0 | 0 | 0 | 0 | 0 | 0 | 0 | 0 | 0 | 0 | 0 | 0 | 0 | 0 | 0 | 0 | 0 | 0 | 0 | 50 | 846 | 50 | 0 | 0 | 0 | 0 | 0 | 0 | 0 | 0 | 0 | 0 |
| SRR7343630 | 0 | 0 | 0 | 0 | 0 | 0 | 0 | 0 | 0 | 0 | 0 | 0 | 0 | 0 | 0 | 0 | 0 | 0 | 0 | 0 | 0 | 0 | 0 | 0 | 0 | 0 | 0 | 0 | 0 | 0 | 0 | 0 | 0 | 36 | 0 | 0 | 0 |
| SRR7343631 | 0 | 0 | 0 | 0 | 0 | 0 | 0 | 21 | 0 | 0 | 0 | 0 | 0 | 0 | 0 | 0 | 0 | 0 | 0 | 0 | 0 | 0 | 0 | 0 | 0 | 21 | 0 | 0 | 0 | 0 | 0 | 0 | 0 | 0 | 0 | 0 | 0 |
| SRR7343632 | 0 | 0 | 0 | 0 | 0 | 41 | 0 | 0 | 0 | 0 | 0 | 0 | 0 | 0 | 0 | 0 | 0 | 0 | 0 | 0 | 0 | 0 | 0 | 0 | 41 | 290 | 41 | 41 | 0 | 0 | 0 | 0 | 0 | 41 | 0 | 0 | 0 |
| SRR7343633 | 0 | 0 | 0 | 0 | 0 | 0 | 0 | 0 | 21 | 10 | 0 | 0 | 0 | 10 | 0 | 0 | 0 | 0 | 0 | 0 | 0 | 0 | 0 | 0 | 0 | 134 | 41 | 0 | 0 | 0 | 31 | 0 | 0 | 10 | 0 | 0 | 0 |
| SRR7343635 | 0 | 0 | 0 | 0 | 0 | 0 | 0 | 0 | 0 | 0 | 0 | 0 | 0 | 39 | 0 | 0 | 0 | 0 | 0 | 0 | 249 | 0 | 0 | 0 | 0 | 39 | 0 | 0 | 0 | 0 | 39 | 0 | 0 | 0 | 0 | 0 | 39 |
| SRR7343636 | 0 | 0 | 0 | 0 | 0 | 12 | 0 | 0 | 12 | 0 | 0 | 0 | 0 | 12 | 0 | 0 | 0 | 0 | 0 | 78 | 0 | 24 | 0 | 0 | 0 | 329 | 0 | 0 | 0 | 0 | 0 | 0 | 0 | 12 | 0 | 0 | 24 |
| SRR7343637 | 0 | 0 | 0 | 0 | 0 | 0 | 0 | 0 | 25 | 0 | 0 | 0 | 0 | 0 | 0 | 0 | 0 | 0 | 0 | 0 | 116 | 0 | 0 | 0 | 0 | 51 | 0 | 0 | 0 | 0 | 25 | 0 | 0 | 0 | 0 | 0 | 0 |
| SRR7343638 | 0 | 0 | 0 | 0 | 0 | 0 | 0 | 0 | 0 | 0 | 0 | 0 | 0 | 0 | 0 | 0 | 0 | 0 | 0 | 0 | 0 | 0 | 0 | 0 | 78 | 78 | 0 | 0 | 0 | 0 | 0 | 0 | 0 | 0 | 0 | 0 | 0 |
| SRR7343639 | 0 | 0 | 0 | 0 | 0 | 0 | 0 | 0 | 0 | 0 | 0 | 0 | 0 | 0 | 0 | 0 | 0 | 0 | 0 | 0 | 265 | 0 | 0 | 0 | 0 | 133 | 0 | 44 | 0 | 0 | 0 | 0 | 0 | 0 | 44 | 0 | 44 |
| SRR7343640 | 0 | 0 | 0 | 0 | 0 | 0 | 0 | 39 | 0 | 0 | 0 | 0 | 0 | 0 | 0 | 0 | 0 | 0 | 0 | 0 | 0 | 0 | 0 | 0 | 0 | 96 | 0 | 0 | 0 | 0 | 0 | 0 | 0 | 0 | 0 | 0 | 0 |
| SRR7412243 | 100 | 0 | 668 | 0 | 0 | 14 | 0 | 43 | 36 | 0 | 0 | 107 | 0 | 0 | 30 | 0 | 22 | 0 | 0 | 2377 | 208 | 0 | 0 | 36 | 0 | 746 | 149 | 0 | 0 | 0 | 21 | 0 | 0 | 0 | 71 | 21 | 21 |
| SRR7412244 | 82 | 0 | 1462 | 0 | 0 | 18 | 0 | 0 | 18 | 0 | 0 | 67 | 0 | 18 | 0 | 0 | 40 | 0 | 43 | 2261 | 304 | 0 | 0 | 65 | 28 | 931 | 129 | 18 | 0 | 0 | 37 | 0 | 0 | 0 | 65 | 18 | 74 |
| SRR7412245 | 113 | 0 | 518 | 0 | 0 | 23 | 0 | 16 | 31 | 0 | 0 | 108 | 0 | 0 | 0 | 0 | 19 | 0 | 0 | 2801 | 423 | 0 | 0 | 39 | 16 | 81 | 20 | 0 | 0 | 0 | 23 | 0 | 0 | 0 | 31 | 31 | 11 |
| SRR7412246 | 90 | 0 | 513 | 0 | 0 | 19 | 0 | 0 | 32 | 0 | 0 | 85 | 0 | 0 | 0 | 0 | 37 | 0 | 0 | 3435 | 470 | 0 | 0 | 58 | 0 | 744 | 128 | 0 | 0 | 0 | 32 | 0 | 0 | 0 | 45 | 71 | 147 |
| SRR7412247 | 85 | 0 | 1216 | 0 | 0 | 16 | 0 | 16 | 0 | 0 | 0 | 78 | 0 | 12 | 0 | 0 | 24 | 0 | 17 | 1897 | 356 | 0 | 0 | 36 | 0 | 746 | 262 | 0 | 0 | 0 | 36 | 0 | 0 | 0 | 119 | 20 | 91 |
| SRR7412248 | 69 | 0 | 2058 | 0 | 0 | 0 | 0 | 28 | 0 | 0 | 0 | 56 | 0 | 23 | 0 | 0 | 32 | 0 | 0 | 1041 | 83 | 0 | 0 | 33 | 0 | 496 | 201 | 0 | 0 | 0 | 23 | 0 | 0 | 0 | 168 | 19 | 65 |
| SRR7412249 | 58 | 0 | 1718 | 0 | 0 | 23 | 0 | 14 | 0 | 0 | 0 | 51 | 0 | 0 | 0 | 0 | 17 | 0 | 0 | 2358 | 181 | 0 | 0 | 23 | 0 | 92 | 18 | 0 | 0 | 0 | 14 | 0 | 0 | 0 | 14 | 28 | 89 |
| SRR7412250 | 90 | 0 | 1550 | 0 | 0 | 14 | 0 | 0 | 41 | 0 | 0 | 112 | 0 | 0 | 0 | 0 | 19 | 0 | 0 | 1483 | 175 | 0 | 0 | 27 | 0 | 850 | 178 | 14 | 0 | 0 | 34 | 0 | 0 | 0 | 75 | 34 | 69 |
| SRR7412251 | 50 | 0 | 792 | 0 | 0 | 17 | 0 | 17 | 50 | 0 | 0 | 111 | 0 | 17 | 0 | 0 | 24 | 89 | 0 | 2162 | 254 | 0 | 0 | 0 | 0 | 859 | 215 | 17 | 0 | 0 | 17 | 0 | 0 | 0 | 248 | 0 | 116 |
| SRR7412252 | 92 | 0 | 1321 | 0 | 0 | 33 | 0 | 0 | 20 | 0 | 0 | 77 | 0 | 0 | 13 | 0 | 31 | 0 | 0 | 1402 | 135 | 0 | 0 | 29 | 0 | 835 | 209 | 0 | 0 | 0 | 20 | 0 | 0 | 0 | 29 | 0 | 29 |
| SRR7412253 | 102 | 0 | 1594 | 0 | 0 | 25 | 0 | 25 | 19 | 0 | 0 | 177 | 0 | 0 | 0 | 0 | 24 | 0 | 0 | 1555 | 329 | 0 | 0 | 37 | 37 | 1167 | 670 | 25 | 0 | 0 | 37 | 0 | 0 | 0 | 366 | 50 | 149 |
| SRR7412254 | 93 | 0 | 2336 | 0 | 0 | 53 | 0 | 0 | 53 | 0 | 0 | 154 | 0 | 0 | 0 | 0 | 14 | 0 | 0 | 1157 | 600 | 0 | 0 | 40 | 13 | 1214 | 567 | 0 | 0 | 0 | 53 | 0 | 0 | 0 | 158 | 79 | 79 |
| SRR7412255 | 107 | 0 | 1447 | 0 | 0 | 31 | 0 | 26 | 36 | 0 | 0 | 164 | 0 | 10 | 0 | 0 | 26 | 0 | 0 | 1088 | 297 | 0 | 0 | 0 | 10 | 863 | 196 | 0 | 0 | 0 | 16 | 0 | 0 | 0 | 83 | 36 | 72 |
| SRR7412256 | 148 | 0 | 376 | 0 | 0 | 21 | 0 | 0 | 62 | 0 | 0 | 167 | 0 | 0 | 0 | 0 | 36 | 0 | 0 | 2493 | 241 | 0 | 0 | 10 | 10 | 1174 | 291 | 0 | 0 | 0 | 42 | 0 | 0 | 0 | 135 | 31 | 62 |
| SRR7412257 | 71 | 0 | 1230 | 0 | 0 | 74 | 0 | 14 | 54 | 0 | 0 | 93 | 0 | 14 | 0 | 0 | 34 | 0 | 0 | 2109 | 412 | 0 | 0 | 20 | 14 | 196 | 28 | 0 | 0 | 0 | 68 | 0 | 0 | 0 | 15 | 27 | 95 |
| SRR7412258 | 88 | 0 | 778 | 0 | 0 | 0 | 0 | 0 | 0 | 36 | 0 | 165 | 0 | 0 | 0 | 0 | 15 | 0 | 0 | 2449 | 230 | 0 | 0 | 0 | 0 | 30 | 36 | 0 | 0 | 0 | 72 | 0 | 0 | 0 | 0 | 0 | 108 |
| SRR7412259 | 51 | 0 | 1736 | 0 | 0 | 38 | 0 | 13 | 38 | 0 | 0 | 143 | 0 | 0 | 26 | 0 | 22 | 0 | 0 | 1439 | 334 | 0 | 0 | 19 | 0 | 1264 | 314 | 0 | 13 | 0 | 77 | 0 | 0 | 0 | 289 | 19 | 51 |
| SRR7412260 | 105 | 0 | 490 | 0 | 0 | 0 | 48 | 0 | 0 | 0 | 0 | 185 | 0 | 0 | 0 | 0 | 0 | 0 | 0 | 774 | 97 | 0 | 0 | 36 | 0 | 1061 | 90 | 0 | 0 | 0 | 54 | 18 | 0 | 0 | 72 | 72 | 90 |
| SRR7412261 | 69 | 0 | 1499 | 0 | 0 | 42 | 25 | 21 | 49 | 0 | 0 | 32 | 0 | 0 | 0 | 0 | 20 | 0 | 0 | 1497 | 152 | 0 | 0 | 0 | 0 | 2227 | 271 | 0 | 0 | 0 | 76 | 0 | 0 | 0 | 97 | 49 | 62 |
| SRR7412262 | 103 | 0 | 1003 | 0 | 0 | 43 | 0 | 0 | 34 | 0 | 0 | 61 | 0 | 0 | 0 | 0 | 15 | 0 | 0 | 1044 | 261 | 0 | 0 | 17 | 0 | 112 | 13 | 17 | 0 | 0 | 102 | 0 | 0 | 0 | 11 | 17 | 60 |
| SRR7412263 | 96 | 0 | 1788 | 0 | 0 | 24 | 0 | 71 | 24 | 0 | 0 | 142 | 0 | 24 | 0 | 0 | 13 | 0 | 0 | 532 | 125 | 0 | 0 | 0 | 0 | 61 | 52 | 0 | 0 | 0 | 24 | 0 | 0 | 0 | 26 | 95 | 95 |
| SRR7412264 | 81 | 0 | 711 | 0 | 0 | 26 | 41 | 13 | 26 | 0 | 0 | 55 | 0 | 26 | 0 | 0 | 31 | 0 | 0 | 1666 | 0 | 0 | 0 | 39 | 13 | 1342 | 605 | 0 | 0 | 0 | 26 | 0 | 13 | 0 | 289 | 39 | 197 |
| SRR7412265 | 128 | 0 | 1815 | 0 | 0 | 25 | 0 | 0 | 34 | 0 | 0 | 84 | 0 | 0 | 0 | 0 | 25 | 0 | 0 | 418 | 154 | 0 | 0 | 25 | 0 | 120 | 46 | 0 | 0 | 0 | 42 | 0 | 0 | 0 | 19 | 25 | 12 |
| SRR7412266 | 64 | 0 | 2664 | 0 | 0 | 52 | 0 | 22 | 59 | 0 | 0 | 136 | 0 | 0 | 0 | 0 | 18 | 0 | 0 | 297 | 177 | 0 | 0 | 81 | 0 | 1617 | 532 | 0 | 0 | 0 | 44 | 0 | 0 | 0 | 502 | 30 | 140 |
| SRR7412267 | 99 | 0 | 1965 | 0 | 0 | 40 | 0 | 13 | 20 | 0 | 0 | 126 | 0 | 13 | 0 | 0 | 11 | 0 | 0 | 943 | 338 | 0 | 0 | 40 | 0 | 1388 | 407 | 0 | 0 | 0 | 20 | 0 | 13 | 0 | 133 | 0 | 80 |
| SRR7412268 | 102 | 0 | 2551 | 0 | 0 | 25 | 0 | 81 | 38 | 0 | 0 | 145 | 0 | 19 | 0 | 0 | 17 | 0 | 0 | 603 | 276 | 13 | 0 | 25 | 0 | 126 | 32 | 0 | 0 | 0 | 56 | 0 | 0 | 0 | 13 | 13 | 107 |
| SRR7412269 | 89 | 0 | 2066 | 0 | 12 | 18 | 0 | 18 | 18 | 0 | 0 | 98 | 0 | 0 | 0 | 0 | 22 | 0 | 0 | 846 | 185 | 0 | 0 | 24 | 12 | 137 | 86 | 0 | 0 | 0 | 60 | 0 | 0 | 0 | 29 | 24 | 16 |
| SRR7412270 | 82 | 0 | 1804 | 0 | 0 | 30 | 0 | 30 | 30 | 0 | 0 | 70 | 0 | 0 | 0 | 0 | 0 | 0 | 0 | 501 | 451 | 0 | 0 | 15 | 0 | 2201 | 907 | 15 | 0 | 0 | 89 | 0 | 0 | 15 | 268 | 15 | 134 |
| SRR7412271 | 131 | 0 | 1035 | 0 | 0 | 11 | 0 | 11 | 46 | 0 | 0 | 275 | 0 | 0 | 0 | 0 | 30 | 0 | 0 | 1738 | 541 | 0 | 0 | 23 | 11 | 959 | 343 | 11 | 0 | 0 | 57 | 0 | 11 | 23 | 137 | 46 | 80 |
| SRR7412272 | 86 | 0 | 1499 | 0 | 0 | 19 | 0 | 0 | 29 | 0 | 0 | 148 | 0 | 0 | 0 | 0 | 19 | 0 | 0 | 1067 | 103 | 0 | 0 | 29 | 0 | 791 | 328 | 0 | 0 | 0 | 0 | 0 | 0 | 0 | 116 | 19 | 39 |
| SRR7412273 | 63 | 0 | 1086 | 0 | 0 | 77 | 47 | 36 | 28 | 0 | 0 | 117 | 0 | 12 | 0 | 0 | 27 | 0 | 0 | 2483 | 609 | 0 | 0 | 0 | 20 | 1426 | 377 | 0 | 0 | 0 | 49 | 0 | 0 | 0 | 357 | 36 | 97 |
| SRR7412274 | 52 | 0 | 1391 | 0 | 0 | 68 | 0 | 0 | 59 | 0 | 0 | 129 | 0 | 0 | 37 | 0 | 38 | 0 | 0 | 1084 | 710 | 0 | 0 | 20 | 20 | 1845 | 596 | 0 | 0 | 0 | 49 | 0 | 0 | 0 | 244 | 49 | 146 |
| SRR7412275 | 96 | 0 | 1448 | 0 | 0 | 13 | 0 | 84 | 26 | 0 | 0 | 184 | 0 | 26 | 0 | 0 | 17 | 0 | 0 | 461 | 461 | 0 | 0 | 0 | 0 | 723 | 220 | 13 | 0 | 0 | 19 | 0 | 0 | 0 | 194 | 32 | 97 |
| SRR7412276 | 88 | 0 | 1673 | 0 | 0 | 18 | 0 | 0 | 0 | 0 | 0 | 67 | 0 | 0 | 0 | 0 | 21 | 0 | 0 | 2409 | 634 | 0 | 0 | 18 | 0 | 74 | 26 | 0 | 0 | 0 | 27 | 0 | 0 | 0 | 19 | 27 | 54 |
| SRR7412277 | 141 | 0 | 1297 | 0 | 0 | 40 | 0 | 16 | 40 | 0 | 0 | 225 | 0 | 0 | 33 | 0 | 22 | 0 | 0 | 2346 | 358 | 0 | 0 | 16 | 0 | 97 | 21 | 0 | 0 | 0 | 16 | 0 | 0 | 0 | 22 | 16 | 72 |
| SRR7412278 | 116 | 0 | 930 | 0 | 0 | 56 | 0 | 42 | 14 | 0 | 0 | 95 | 0 | 0 | 0 | 0 | 12 | 0 | 0 | 1287 | 0 | 0 | 0 | 14 | 0 | 159 | 42 | 0 | 0 | 0 | 71 | 0 | 0 | 0 | 42 | 71 | 0 |
| SRR7412279 | 102 | 0 | 794 | 0 | 0 | 16 | 0 | 25 | 0 | 0 | 0 | 190 | 0 | 0 | 0 | 0 | 20 | 0 | 0 | 1245 | 453 | 0 | 0 | 16 | 0 | 866 | 98 | 0 | 0 | 0 | 25 | 0 | 0 | 0 | 25 | 25 | 57 |
| SRR7412280 | 52 | 0 | 1654 | 0 | 0 | 18 | 0 | 18 | 36 | 0 | 0 | 60 | 0 | 0 | 0 | 0 | 12 | 0 | 0 | 565 | 246 | 0 | 0 | 0 | 0 | 134 | 36 | 0 | 0 | 0 | 18 | 0 | 0 | 0 | 11 | 55 | 73 |
| SRR7412281 | 107 | 0 | 1581 | 0 | 0 | 30 | 41 | 0 | 30 | 0 | 0 | 144 | 0 | 0 | 0 | 0 | 16 | 0 | 41 | 1433 | 205 | 0 | 0 | 20 | 0 | 76 | 11 | 0 | 0 | 0 | 39 | 0 | 0 | 0 | 12 | 49 | 79 |
| SRR7412282 | 97 | 0 | 1906 | 0 | 0 | 68 | 0 | 0 | 11 | 0 | 0 | 67 | 0 | 0 | 0 | 0 | 0 | 0 | 0 | 920 | 212 | 0 | 0 | 0 | 0 | 1939 | 342 | 0 | 0 | 0 | 34 | 0 | 0 | 0 | 103 | 11 | 80 |
| SRR7412283 | 141 | 0 | 1368 | 0 | 0 | 26 | 0 | 0 | 0 | 0 | 0 | 133 | 0 | 0 | 0 | 0 | 16 | 0 | 0 | 1671 | 134 | 0 | 0 | 18 | 0 | 84 | 16 | 0 | 0 | 0 | 18 | 0 | 0 | 0 | 114 | 53 | 114 |
| SRR7412284 | 92 | 0 | 1187 | 0 | 0 | 0 | 0 | 0 | 0 | 0 | 0 | 53 | 0 | 0 | 0 | 0 | 15 | 0 | 0 | 951 | 605 | 0 | 0 | 18 | 0 | 56 | 15 | 18 | 0 | 0 | 27 | 0 | 0 | 0 | 10 | 36 | 63 |
| SRR7412285 | 90 | 0 | 1102 | 0 | 0 | 47 | 37 | 41 | 12 | 0 | 29 | 108 | 0 | 0 | 0 | 0 | 32 | 0 | 0 | 1358 | 316 | 0 | 0 | 47 | 0 | 1244 | 298 | 0 | 0 | 0 | 64 | 0 | 0 | 0 | 99 | 47 | 82 |
| SRR7412286 | 108 | 0 | 1510 | 0 | 0 | 32 | 0 | 0 | 0 | 0 | 0 | 185 | 0 | 0 | 0 | 0 | 20 | 0 | 0 | 1258 | 314 | 0 | 0 | 0 | 0 | 110 | 14 | 0 | 0 | 0 | 0 | 0 | 0 | 0 | 32 | 24 | 40 |
| SRR7412287 | 81 | 0 | 935 | 0 | 0 | 13 | 0 | 0 | 0 | 0 | 0 | 123 | 0 | 0 | 0 | 0 | 14 | 0 | 0 | 1638 | 983 | 0 | 0 | 0 | 0 | 457 | 101 | 0 | 0 | 0 | 20 | 0 | 0 | 0 | 94 | 13 | 101 |
| SRR7412288 | 73 | 0 | 971 | 0 | 0 | 0 | 0 | 0 | 0 | 0 | 0 | 133 | 0 | 0 | 13 | 0 | 17 | 0 | 0 | 537 | 486 | 0 | 0 | 15 | 0 | 550 | 382 | 0 | 0 | 0 | 0 | 0 | 0 | 0 | 322 | 84 | 134 |
| SRR7412289 | 57 | 0 | 1094 | 0 | 0 | 61 | 0 | 0 | 10 | 0 | 0 | 79 | 0 | 0 | 0 | 0 | 12 | 0 | 0 | 1008 | 475 | 0 | 0 | 20 | 0 | 78 | 92 | 10 | 0 | 0 | 0 | 0 | 0 | 10 | 72 | 41 | 41 |
| SRR7412290 | 110 | 0 | 166 | 0 | 0 | 28 | 0 | 0 | 42 | 0 | 0 | 213 | 0 | 0 | 0 | 0 | 28 | 0 | 0 | 2269 | 454 | 0 | 0 | 14 | 0 | 958 | 28 | 0 | 0 | 0 | 69 | 0 | 0 | 14 | 56 | 42 | 42 |
| SRR7412291 | 98 | 0 | 1789 | 0 | 0 | 16 | 67 | 0 | 56 | 0 | 0 | 109 | 0 | 0 | 0 | 0 | 13 | 0 | 0 | 1281 | 405 | 0 | 0 | 24 | 0 | 749 | 207 | 0 | 0 | 0 | 24 | 0 | 0 | 0 | 48 | 16 | 32 |
| SRR7412292 | 199 | 0 | 689 | 0 | 0 | 0 | 0 | 0 | 27 | 0 | 0 | 168 | 0 | 0 | 0 | 0 | 18 | 0 | 0 | 2526 | 161 | 0 | 0 | 27 | 0 | 1061 | 101 | 0 | 0 | 0 | 0 | 0 | 0 | 0 | 55 | 27 | 46 |
| SRR7412293 | 72 | 0 | 1257 | 0 | 0 | 18 | 26 | 28 | 18 | 0 | 0 | 129 | 0 | 0 | 0 | 18 | 16 | 0 | 0 | 361 | 876 | 0 | 0 | 28 | 0 | 73 | 19 | 0 | 0 | 0 | 55 | 0 | 0 | 0 | 12 | 37 | 83 |
| SRR7412294 | 25 | 0 | 1075 | 0 | 0 | 29 | 0 | 21 | 14 | 0 | 0 | 113 | 0 | 0 | 0 | 0 | 30 | 0 | 0 | 2462 | 703 | 0 | 0 | 57 | 143 | 1040 | 235 | 0 | 0 | 0 | 21 | 0 | 0 | 0 | 78 | 29 | 36 |
| SRR7412295 | 220 | 0 | 1273 | 0 | 0 | 35 | 0 | 0 | 26 | 0 | 0 | 139 | 0 | 0 | 86 | 0 | 17 | 0 | 0 | 1946 | 601 | 0 | 0 | 26 | 18 | 123 | 39 | 0 | 0 | 0 | 0 | 0 | 0 | 0 | 22 | 122 | 11 |
| SRR7412296 | 18 | 0 | 535 | 0 | 0 | 24 | 0 | 12 | 49 | 0 | 0 | 27 | 0 | 0 | 0 | 0 | 19 | 0 | 0 | 575 | 679 | 0 | 0 | 12 | 24 | 523 | 170 | 0 | 12 | 0 | 37 | 0 | 0 | 0 | 402 | 49 | 97 |
| SRR7412297 | 59 | 0 | 1643 | 0 | 0 | 16 | 0 | 11 | 0 | 0 | 0 | 141 | 0 | 0 | 0 | 0 | 14 | 0 | 0 | 369 | 338 | 0 | 0 | 16 | 0 | 55 | 23 | 16 | 11 | 0 | 11 | 0 | 0 | 0 | 26 | 16 | 71 |
| SRR7412298 | 128 | 0 | 1128 | 0 | 0 | 17 | 0 | 17 | 0 | 0 | 0 | 78 | 0 | 0 | 24 | 0 | 20 | 0 | 24 | 968 | 0 | 0 | 0 | 17 | 0 | 66 | 19 | 0 | 0 | 0 | 0 | 0 | 0 | 0 | 75 | 42 | 50 |
| SRR7412299 | 104 | 0 | 974 | 0 | 0 | 53 | 0 | 53 | 53 | 0 | 0 | 204 | 0 | 0 | 46 | 0 | 12 | 0 | 0 | 1012 | 506 | 0 | 0 | 18 | 18 | 1212 | 160 | 0 | 0 | 0 | 0 | 0 | 0 | 0 | 160 | 18 | 36 |
| SRR7412300 | 112 | 0 | 992 | 0 | 0 | 45 | 0 | 33 | 22 | 0 | 0 | 250 | 0 | 11 | 0 | 0 | 13 | 0 | 0 | 377 | 237 | 0 | 0 | 45 | 11 | 1181 | 390 | 0 | 11 | 11 | 45 | 0 | 0 | 0 | 279 | 67 | 45 |
| SRR7412301 | 55 | 0 | 1239 | 0 | 0 | 11 | 0 | 11 | 0 | 0 | 0 | 57 | 0 | 0 | 0 | 0 | 0 | 0 | 0 | 1923 | 846 | 0 | 0 | 34 | 0 | 1112 | 227 | 11 | 0 | 0 | 45 | 0 | 0 | 11 | 136 | 45 | 79 |
| SRR7412302 | 53 | 0 | 1630 | 0 | 0 | 0 | 0 | 34 | 0 | 0 | 0 | 20 | 0 | 0 | 0 | 0 | 17 | 0 | 0 | 709 | 501 | 0 | 0 | 39 | 0 | 509 | 304 | 0 | 0 | 0 | 0 | 0 | 0 | 0 | 98 | 56 | 73 |
| SRR7412303 | 95 | 0 | 2044 | 0 | 0 | 12 | 0 | 0 | 0 | 0 | 0 | 34 | 0 | 12 | 0 | 0 | 12 | 0 | 0 | 541 | 418 | 0 | 0 | 18 | 0 | 99 | 19 | 0 | 0 | 0 | 31 | 0 | 0 | 0 | 30 | 67 | 13 |
| SRR7412304 | 79 | 0 | 2070 | 0 | 0 | 0 | 0 | 25 | 0 | 0 | 0 | 82 | 0 | 0 | 0 | 0 | 0 | 0 | 0 | 334 | 435 | 0 | 0 | 33 | 0 | 43 | 38 | 0 | 0 | 0 | 0 | 0 | 0 | 17 | 21 | 33 | 13 |
| SRR7412305 | 75 | 0 | 983 | 0 | 0 | 69 | 35 | 46 | 11 | 0 | 0 | 129 | 0 | 11 | 0 | 0 | 18 | 0 | 0 | 743 | 266 | 0 | 0 | 11 | 11 | 97 | 20 | 11 | 11 | 0 | 23 | 0 | 11 | 0 | 126 | 57 | 69 |
| SRR7412306 | 52 | 0 | 1695 | 0 | 0 | 61 | 0 | 0 | 10 | 0 | 0 | 52 | 0 | 0 | 0 | 0 | 23 | 0 | 0 | 843 | 222 | 0 | 0 | 41 | 0 | 1945 | 580 | 10 | 0 | 0 | 71 | 0 | 0 | 0 | 143 | 61 | 31 |
| SRR7412307 | 117 | 0 | 917 | 0 | 0 | 33 | 94 | 16 | 16 | 0 | 0 | 186 | 0 | 0 | 0 | 0 | 14 | 0 | 0 | 1603 | 189 | 0 | 0 | 16 | 0 | 1164 | 148 | 0 | 0 | 0 | 16 | 0 | 0 | 0 | 66 | 66 | 33 |
| SRR7412308 | 104 | 0 | 1542 | 0 | 0 | 19 | 0 | 0 | 29 | 0 | 0 | 138 | 0 | 0 | 0 | 0 | 15 | 0 | 0 | 1533 | 219 | 0 | 0 | 38 | 0 | 640 | 258 | 0 | 0 | 0 | 29 | 0 | 0 | 0 | 86 | 19 | 29 |
| SRR7412309 | 63 | 0 | 1045 | 0 | 0 | 25 | 33 | 0 | 17 | 0 | 0 | 40 | 0 | 0 | 0 | 0 | 32 | 0 | 0 | 1635 | 434 | 0 | 0 | 25 | 25 | 61 | 26 | 0 | 0 | 0 | 25 | 0 | 0 | 0 | 83 | 50 | 14 |
| SRR7412310 | 124 | 0 | 1526 | 0 | 0 | 19 | 0 | 0 | 11 | 0 | 0 | 77 | 0 | 0 | 0 | 0 | 30 | 0 | 0 | 1872 | 212 | 0 | 0 | 31 | 0 | 723 | 180 | 0 | 0 | 0 | 11 | 0 | 0 | 0 | 84 | 27 | 61 |
| SRR7412311 | 94 | 0 | 1411 | 0 | 0 | 25 | 0 | 0 | 16 | 0 | 0 | 101 | 0 | 0 | 0 | 0 | 26 | 0 | 0 | 1561 | 114 | 0 | 0 | 0 | 0 | 912 | 230 | 0 | 0 | 0 | 33 | 0 | 0 | 0 | 49 | 33 | 74 |
| SRR7412312 | 120 | 0 | 909 | 0 | 0 | 17 | 0 | 0 | 0 | 0 | 0 | 149 | 0 | 0 | 0 | 0 | 18 | 0 | 0 | 946 | 355 | 0 | 0 | 25 | 0 | 891 | 272 | 0 | 0 | 0 | 25 | 0 | 0 | 0 | 68 | 17 | 34 |
| SRR7412313 | 127 | 0 | 1403 | 0 | 0 | 19 | 0 | 105 | 19 | 0 | 0 | 158 | 0 | 0 | 0 | 0 | 11 | 0 | 0 | 427 | 197 | 0 | 0 | 0 | 0 | 860 | 267 | 0 | 0 | 0 | 76 | 0 | 0 | 0 | 67 | 0 | 86 |
| SRR7412314 | 60 | 0 | 354 | 0 | 0 | 0 | 0 | 0 | 15 | 0 | 0 | 204 | 0 | 0 | 0 | 45 | 24 | 0 | 0 | 1176 | 527 | 0 | 0 | 0 | 15 | 401 | 111 | 37 | 0 | 0 | 30 | 0 | 0 | 0 | 37 | 22 | 45 |
| SRR7412315 | 71 | 0 | 454 | 0 | 0 | 35 | 0 | 35 | 35 | 0 | 0 | 90 | 0 | 0 | 0 | 0 | 18 | 0 | 0 | 1875 | 110 | 0 | 0 | 35 | 0 | 905 | 223 | 0 | 0 | 0 | 35 | 0 | 0 | 12 | 188 | 59 | 59 |
| SRR7412316 | 120 | 0 | 2591 | 0 | 0 | 36 | 0 | 27 | 55 | 0 | 0 | 87 | 0 | 0 | 0 | 0 | 19 | 0 | 0 | 452 | 266 | 0 | 0 | 46 | 0 | 154 | 95 | 0 | 0 | 0 | 0 | 0 | 0 | 0 | 32 | 46 | 14 |
| SRR7412317 | 72 | 0 | 437 | 0 | 0 | 40 | 32 | 0 | 12 | 0 | 0 | 97 | 0 | 0 | 0 | 0 | 20 | 0 | 0 | 1675 | 741 | 0 | 0 | 17 | 23 | 57 | 15 | 23 | 0 | 0 | 23 | 0 | 0 | 0 | 23 | 12 | 17 |
| SRR7412318 | 59 | 0 | 703 | 0 | 0 | 25 | 0 | 0 | 0 | 0 | 0 | 86 | 0 | 0 | 0 | 0 | 11 | 0 | 0 | 1211 | 681 | 0 | 0 | 13 | 0 | 967 | 127 | 0 | 0 | 0 | 13 | 0 | 0 | 0 | 51 | 38 | 13 |
| SRR7412319 | 66 | 0 | 624 | 0 | 0 | 50 | 0 | 0 | 0 | 0 | 0 | 86 | 0 | 0 | 0 | 0 | 28 | 0 | 0 | 2209 | 363 | 0 | 0 | 28 | 0 | 63 | 12 | 21 | 0 | 0 | 64 | 0 | 0 | 0 | 12 | 28 | 10 |
| SRR7412320 | 98 | 0 | 888 | 0 | 0 | 60 | 0 | 65 | 40 | 0 | 0 | 86 | 0 | 0 | 0 | 0 | 31 | 0 | 0 | 588 | 235 | 0 | 0 | 20 | 10 | 1261 | 528 | 0 | 0 | 0 | 60 | 0 | 0 | 0 | 377 | 151 | 181 |
| SRR7412321 | 108 | 0 | 1836 | 0 | 0 | 0 | 0 | 0 | 0 | 0 | 0 | 76 | 0 | 0 | 0 | 0 | 0 | 0 | 0 | 319 | 228 | 0 | 0 | 0 | 0 | 853 | 275 | 0 | 0 | 0 | 19 | 0 | 0 | 0 | 209 | 28 | 161 |
| SRR7412322 | 130 | 0 | 1190 | 0 | 0 | 29 | 0 | 0 | 29 | 0 | 0 | 68 | 0 | 0 | 0 | 0 | 26 | 0 | 0 | 730 | 446 | 0 | 0 | 120 | 17 | 1128 | 269 | 11 | 0 | 0 | 17 | 0 | 0 | 0 | 149 | 29 | 109 |
| SRR7412323 | 240 | 0 | 2135 | 0 | 0 | 33 | 0 | 0 | 33 | 0 | 0 | 89 | 0 | 0 | 0 | 0 | 15 | 0 | 0 | 297 | 82 | 0 | 0 | 74 | 0 | 95 | 34 | 0 | 0 | 0 | 66 | 0 | 0 | 0 | 13 | 49 | 13 |
| SRR7412324 | 135 | 0 | 2825 | 0 | 0 | 27 | 0 | 0 | 82 | 0 | 0 | 71 | 0 | 0 | 0 | 0 | 16 | 0 | 0 | 502 | 281 | 0 | 0 | 82 | 0 | 882 | 291 | 0 | 0 | 0 | 18 | 0 | 0 | 0 | 109 | 73 | 100 |
| SRR7412325 | 100 | 0 | 1175 | 0 | 0 | 65 | 0 | 0 | 43 | 0 | 0 | 127 | 0 | 11 | 31 | 0 | 22 | 0 | 0 | 1462 | 405 | 0 | 0 | 54 | 0 | 1865 | 347 | 22 | 11 | 0 | 98 | 0 | 0 | 0 | 54 | 43 | 108 |
| SRR7412326 | 96 | 0 | 1305 | 0 | 0 | 0 | 22 | 16 | 20 | 0 | 0 | 90 | 0 | 12 | 0 | 0 | 35 | 0 | 0 | 1480 | 188 | 0 | 0 | 12 | 0 | 51 | 20 | 0 | 0 | 0 | 36 | 0 | 0 | 0 | 117 | 68 | 60 |
| SRR7412327 | 101 | 0 | 1895 | 0 | 0 | 26 | 25 | 26 | 26 | 0 | 0 | 69 | 0 | 0 | 0 | 0 | 29 | 0 | 25 | 1430 | 75 | 0 | 0 | 0 | 0 | 1323 | 229 | 0 | 0 | 0 | 59 | 0 | 0 | 0 | 111 | 33 | 144 |

### Supp Table S8.2.Columns_AM_to_BW

| | | | | | | | | | | | | | | | | | | | | | | | | | | | | | | | | | | | | | |
|---|---|---|---|---|---|---|---|---|---|---|---|---|---|---|---|---|---|---|---|---|---|---|---|---|---|---|---|---|---|---|---|---|---|---|---|---|---|
| SRR7412328 | 67 | 0 | 1765 | 0 | 0 | 35 | 0 | 46 | 35 | 0 | 0 | 63 | 0 | 0 | 0 | 0 | 10 | 0 | 0 | 1048 | 393 | 0 | 0 | 29 | 0 | 1039 | 219 | 12 | 12 | 0 | 35 | 0 | 0 | 0 | 473 | 81 | 237 |
| SRR7412329 | 43 | 0 | 3264 | 0 | 0 | 56 | 0 | 19 | 0 | 0 | 0 | 101 | 0 | 0 | 0 | 0 | 27 | 0 | 0 | 992 | 173 | 0 | 0 | 37 | 18 | 227 | 87 | 0 | 0 | 0 | 28 | 0 | 0 | 0 | 30 | 19 | 12 |
| SRR7412330 | 42 | 16 | 510 | 0 | 0 | 40 | 0 | 0 | 16 | 0 | 0 | 89 | 0 | 0 | 0 | 0 | 0 | 0 | 83 | 1835 | 1084 | 0 | 0 | 32 | 223 | 1340 | 662 | 16 | 0 | 0 | 40 | 0 | 24 | 0 | 359 | 24 | 112 |
| SRR7412331 | 88 | 0 | 1656 | 0 | 0 | 56 | 0 | 159 | 48 | 0 | 0 | 139 | 0 | 0 | 0 | 0 | 25 | 0 | 21 | 316 | 506 | 0 | 0 | 40 | 0 | 1442 | 757 | 0 | 0 | 0 | 24 | 0 | 0 | 24 | 430 | 24 | 135 |
| SRR7412332 | 72 | 0 | 1406 | 0 | 0 | 66 | 0 | 218 | 0 | 0 | 0 | 80 | 0 | 0 | 34 | 0 | 28 | 70 | 0 | 963 | 447 | 0 | 0 | 0 | 0 | 1202 | 322 | 0 | 0 | 0 | 76 | 0 | 0 | 0 | 199 | 76 | 66 |
| SRR7412333 | 86 | 0 | 1874 | 0 | 0 | 54 | 0 | 0 | 24 | 0 | 0 | 168 | 0 | 0 | 0 | 0 | 19 | 0 | 0 | 1001 | 476 | 0 | 0 | 0 | 0 | 1169 | 331 | 0 | 0 | 0 | 39 | 0 | 0 | 0 | 97 | 39 | 97 |
| SRR7412334 | 67 | 0 | 1307 | 0 | 0 | 21 | 29 | 14 | 21 | 0 | 0 | 83 | 0 | 0 | 0 | 0 | 19 | 0 | 0 | 902 | 407 | 0 | 0 | 56 | 0 | 607 | 293 | 0 | 0 | 0 | 35 | 0 | 0 | 0 | 356 | 0 | 126 |
| SRR7412335 | 84 | 0 | 1134 | 0 | 0 | 0 | 40 | 10 | 15 | 0 | 0 | 117 | 0 | 30 | 0 | 0 | 16 | 0 | 0 | 802 | 80 | 0 | 0 | 30 | 0 | 70 | 29 | 0 | 15 | 0 | 0 | 0 | 0 | 0 | 17 | 15 | 75 |
| SRR7412336 | 132 | 0 | 3172 | 0 | 0 | 19 | 0 | 28 | 28 | 0 | 0 | 161 | 0 | 19 | 0 | 0 | 13 | 0 | 0 | 509 | 254 | 0 | 0 | 38 | 0 | 872 | 216 | 0 | 0 | 0 | 66 | 0 | 0 | 0 | 300 | 38 | 159 |
| SRR7412337 | 134 | 0 | 565 | 0 | 0 | 40 | 0 | 0 | 0 | 0 | 0 | 231 | 0 | 0 | 0 | 0 | 26 | 0 | 0 | 2298 | 615 | 0 | 0 | 69 | 0 | 1324 | 237 | 0 | 0 | 0 | 30 | 0 | 0 | 0 | 395 | 79 | 198 |
| SRR7412338 | 67 | 0 | 2057 | 0 | 0 | 0 | 0 | 30 | 45 | 0 | 0 | 32 | 0 | 0 | 0 | 0 | 19 | 0 | 0 | 819 | 252 | 0 | 0 | 0 | 0 | 1444 | 542 | 0 | 0 | 0 | 45 | 0 | 0 | 0 | 406 | 60 | 226 |
| SRR7412339 | 142 | 0 | 1571 | 0 | 0 | 13 | 0 | 0 | 27 | 0 | 0 | 240 | 0 | 0 | 0 | 0 | 12 | 0 | 34 | 1014 | 203 | 0 | 0 | 54 | 13 | 49 | 13 | 0 | 13 | 0 | 40 | 0 | 0 | 0 | 148 | 0 | 94 |
| SRR7412340 | 223 | 0 | 786 | 0 | 0 | 12 | 0 | 0 | 48 | 0 | 0 | 123 | 0 | 0 | 0 | 24 | 11 | 0 | 0 | 735 | 678 | 12 | 0 | 36 | 0 | 938 | 36 | 0 | 12 | 0 | 72 | 0 | 0 | 0 | 96 | 72 | 96 |
| SRR7412341 | 272 | 0 | 464 | 0 | 0 | 19 | 0 | 0 | 28 | 0 | 0 | 98 | 0 | 0 | 0 | 0 | 0 | 0 | 0 | 828 | 773 | 0 | 0 | 0 | 0 | 740 | 103 | 0 | 0 | 0 | 75 | 0 | 0 | 0 | 112 | 75 | 253 |
| SRR7412342 | 46 | 0 | 497 | 0 | 0 | 30 | 0 | 0 | 0 | 0 | 0 | 96 | 0 | 0 | 0 | 0 | 15 | 0 | 0 | 2926 | 279 | 0 | 0 | 23 | 0 | 81 | 19 | 0 | 0 | 0 | 38 | 0 | 0 | 0 | 13 | 0 | 10 |
| SRR7412343 | 43 | 17 | 1031 | 0 | 0 | 0 | 0 | 33 | 25 | 0 | 0 | 70 | 0 | 0 | 0 | 0 | 39 | 0 | 50 | 1850 | 2101 | 0 | 0 | 42 | 0 | 37 | 28 | 25 | 0 | 0 | 0 | 0 | 0 | 0 | 17 | 33 | 92 |
| SRR7412344 | 111 | 0 | 2662 | 0 | 0 | 0 | 0 | 0 | 37 | 0 | 0 | 63 | 0 | 0 | 0 | 0 | 0 | 0 | 0 | 168 | 134 | 0 | 0 | 30 | 0 | 172 | 23 | 0 | 0 | 0 | 12 | 0 | 0 | 0 | 24 | 22 | 10 |
| SRR7412345 | 70 | 0 | 657 | 0 | 0 | 0 | 0 | 20 | 0 | 0 | 0 | 112 | 0 | 20 | 0 | 0 | 21 | 0 | 0 | 2340 | 312 | 0 | 0 | 20 | 0 | 41 | 21 | 20 | 0 | 0 | 39 | 0 | 0 | 0 | 17 | 20 | 117 |
| SRR7466734 | 20 | 0 | 4144 | 0 | 0 | 20 | 0 | 0 | 43 | 0 | 0 | 21 | 0 | 12 | 38 | 0 | 32 | 0 | 19 | 647 | 1693 | 0 | 0 | 0 | 0 | 319 | 862 | 12 | 0 | 0 | 22 | 0 | 0 | 0 | 1206 | 52 | 439 |
| SRR7466735 | 13 | 0 | 4238 | 0 | 0 | 15 | 20 | 0 | 36 | 0 | 0 | 26 | 0 | 0 | 0 | 0 | 22 | 0 | 0 | 684 | 1973 | 0 | 0 | 0 | 12 | 326 | 858 | 0 | 0 | 0 | 20 | 0 | 0 | 0 | 1220 | 43 | 448 |
| SRR7466736 | 32 | 0 | 4446 | 0 | 0 | 45 | 17 | 0 | 15 | 0 | 0 | 22 | 0 | 0 | 0 | 0 | 15 | 0 | 0 | 768 | 1553 | 0 | 0 | 0 | 27 | 873 | 2432 | 0 | 0 | 0 | 31 | 0 | 0 | 0 | 3681 | 119 | 1289 |
| SRR7466737 | 25 | 0 | 4416 | 0 | 0 | 40 | 0 | 0 | 11 | 0 | 0 | 31 | 0 | 0 | 34 | 0 | 18 | 0 | 17 | 1000 | 1724 | 0 | 0 | 0 | 27 | 830 | 2586 | 0 | 0 | 0 | 29 | 0 | 0 | 0 | 3826 | 97 | 1242 |
| SRR7466738 | 38 | 0 | 4318 | 0 | 0 | 12 | 0 | 0 | 0 | 0 | 0 | 15 | 0 | 0 | 16 | 0 | 15 | 0 | 0 | 952 | 1575 | 0 | 0 | 0 | 11 | 253 | 787 | 0 | 0 | 0 | 11 | 0 | 0 | 0 | 1166 | 34 | 379 |
| SRR7466739 | 33 | 0 | 4284 | 0 | 0 | 48 | 0 | 0 | 0 | 0 | 0 | 22 | 0 | 0 | 50 | 0 | 27 | 0 | 0 | 773 | 1546 | 0 | 0 | 0 | 39 | 874 | 2615 | 0 | 0 | 0 | 24 | 0 | 0 | 0 | 3904 | 125 | 1181 |
| SRR7466740 | 16 | 0 | 4576 | 0 | 0 | 41 | 40 | 0 | 23 | 0 | 0 | 16 | 0 | 0 | 20 | 0 | 25 | 0 | 61 | 727 | 1958 | 0 | 0 | 0 | 31 | 978 | 1499 | 0 | 0 | 0 | 34 | 0 | 0 | 0 | 5087 | 173 | 1950 |
| SRR7466741 | 30 | 0 | 4423 | 0 | 0 | 39 | 41 | 0 | 11 | 0 | 0 | 16 | 0 | 0 | 0 | 0 | 31 | 0 | 0 | 1170 | 2012 | 0 | 0 | 0 | 39 | 988 | 1662 | 11 | 0 | 0 | 34 | 0 | 0 | 0 | 5229 | 161 | 2129 |
| SRR7466742 | 21 | 0 | 4548 | 0 | 0 | 25 | 0 | 0 | 20 | 0 | 0 | 25 | 0 | 0 | 0 | 0 | 14 | 0 | 16 | 853 | 1986 | 0 | 0 | 0 | 36 | 258 | 423 | 0 | 0 | 0 | 37 | 0 | 0 | 0 | 1306 | 42 | 523 |
| SRR7466743 | 30 | 0 | 4673 | 0 | 0 | 40 | 20 | 0 | 11 | 0 | 0 | 12 | 0 | 0 | 0 | 0 | 27 | 0 | 40 | 786 | 2277 | 0 | 0 | 0 | 46 | 965 | 1643 | 0 | 0 | 0 | 29 | 0 | 0 | 0 | 5258 | 157 | 2034 |
| SRR7466744 | 32 | 0 | 4160 | 0 | 0 | 34 | 0 | 0 | 18 | 0 | 0 | 16 | 0 | 0 | 36 | 0 | 30 | 0 | 0 | 873 | 2055 | 0 | 0 | 0 | 32 | 298 | 453 | 0 | 0 | 0 | 17 | 0 | 0 | 0 | 1420 | 58 | 666 |
| SRR7466745 | 31 | 0 | 4287 | 0 | 0 | 52 | 0 | 0 | 16 | 0 | 0 | 31 | 0 | 0 | 66 | 0 | 24 | 0 | 22 | 1099 | 2814 | 0 | 0 | 0 | 24 | 1571 | 2285 | 0 | 0 | 0 | 73 | 0 | 0 | 0 | 7373 | 272 | 3694 |
| SRR7466746 | 21 | 0 | 4372 | 0 | 0 | 90 | 0 | 0 | 51 | 0 | 0 | 38 | 0 | 0 | 39 | 0 | 13 | 0 | 0 | 828 | 2542 | 0 | 0 | 0 | 36 | 2625 | 3559 | 18 | 0 | 0 | 159 | 0 | 0 | 0 | 10245 | 366 | 4280 |
| SRR7466747 | 24 | 0 | 4343 | 0 | 0 | 146 | 19 | 0 | 30 | 0 | 0 | 31 | 0 | 0 | 19 | 0 | 17 | 0 | 0 | 859 | 2977 | 0 | 0 | 0 | 33 | 2748 | 3656 | 0 | 0 | 0 | 155 | 0 | 0 | 0 | 9997 | 408 | 3938 |
| SRR7466748 | 28 | 0 | 4459 | 0 | 0 | 103 | 0 | 0 | 32 | 0 | 0 | 39 | 0 | 0 | 22 | 0 | 22 | 0 | 0 | 1155 | 2789 | 0 | 0 | 0 | 32 | 2739 | 3767 | 12 | 0 | 0 | 141 | 0 | 0 | 0 | 9829 | 370 | 4170 |
| SRR7466749 | 23 | 0 | 4485 | 0 | 0 | 18 | 21 | 0 | 43 | 0 | 0 | 26 | 0 | 0 | 64 | 0 | 19 | 0 | 21 | 1265 | 3622 | 0 | 0 | 0 | 46 | 388 | 520 | 0 | 0 | 0 | 15 | 0 | 0 | 0 | 1484 | 51 | 609 |
| SRR7466750 | 28 | 0 | 4354 | 0 | 0 | 22 | 0 | 0 | 0 | 0 | 0 | 22 | 0 | 0 | 0 | 0 | 34 | 0 | 0 | 704 | 2407 | 0 | 0 | 0 | 61 | 823 | 1508 | 0 | 0 | 0 | 32 | 0 | 0 | 0 | 4472 | 151 | 1722 |
| SRR7466751 | 26 | 0 | 4244 | 0 | 0 | 13 | 79 | 0 | 0 | 0 | 0 | 26 | 0 | 0 | 79 | 0 | 14 | 0 | 0 | 1079 | 2448 | 0 | 0 | 0 | 18 | 236 | 445 | 0 | 0 | 0 | 30 | 0 | 0 | 0 | 1305 | 42 | 504 |
| SRR7466752 | 28 | 0 | 4384 | 0 | 0 | 14 | 26 | 0 | 0 | 0 | 0 | 10 | 0 | 0 | 0 | 0 | 30 | 0 | 0 | 942 | 2564 | 0 | 0 | 0 | 15 | 249 | 447 | 0 | 0 | 0 | 20 | 0 | 0 | 0 | 1333 | 49 | 515 |
| SRR7466753 | 22 | 0 | 4219 | 0 | 0 | 14 | 0 | 0 | 0 | 0 | 0 | 16 | 0 | 0 | 0 | 0 | 22 | 0 | 0 | 966 | 2668 | 0 | 0 | 0 | 17 | 236 | 455 | 0 | 0 | 0 | 29 | 0 | 0 | 0 | 1334 | 46 | 506 |
| SRR7466754 | 30 | 0 | 4445 | 0 | 19 | 11 | 0 | 12 | 22 | 0 | 0 | 22 | 0 | 0 | 22 | 0 | 21 | 0 | 0 | 681 | 2548 | 0 | 0 | 0 | 15 | 297 | 822 | 0 | 0 | 0 | 14 | 0 | 0 | 0 | 1493 | 46 | 503 |
| SRR7466755 | 36 | 0 | 4265 | 0 | 0 | 16 | 0 | 0 | 22 | 0 | 0 | 32 | 0 | 0 | 22 | 0 | 0 | 0 | 22 | 936 | 2653 | 0 | 0 | 0 | 22 | 292 | 823 | 0 | 0 | 0 | 16 | 0 | 0 | 0 | 1538 | 46 | 519 |
| SRR7466756 | 41 | 0 | 4152 | 0 | 0 | 83 | 0 | 18 | 27 | 0 | 0 | 26 | 0 | 0 | 50 | 0 | 0 | 0 | 0 | 665 | 1528 | 0 | 0 | 0 | 24 | 1872 | 4702 | 0 | 0 | 0 | 95 | 0 | 0 | 0 | 8718 | 225 | 2806 |
| SRR7466757 | 25 | 0 | 4400 | 0 | 0 | 67 | 0 | 0 | 0 | 0 | 0 | 47 | 0 | 0 | 34 | 0 | 27 | 0 | 17 | 582 | 1558 | 0 | 0 | 0 | 15 | 1668 | 4841 | 12 | 0 | 0 | 82 | 0 | 0 | 0 | 8600 | 259 | 2902 |
| SRR7466758 | 27 | 0 | 3858 | 0 | 0 | 15 | 0 | 0 | 34 | 0 | 0 | 29 | 0 | 0 | 57 | 0 | 19 | 0 | 29 | 718 | 1923 | 0 | 0 | 0 | 0 | 259 | 770 | 0 | 0 | 0 | 67 | 0 | 0 | 0 | 1538 | 53 | 487 |
| SRR7466759 | 36 | 0 | 3883 | 14 | 0 | 12 | 37 | 14 | 0 | 0 | 0 | 25 | 0 | 0 | 37 | 0 | 15 | 0 | 0 | 853 | 2113 | 0 | 0 | 0 | 14 | 254 | 861 | 20 | 0 | 0 | 11 | 0 | 0 | 0 | 1496 | 41 | 516 |
| SRR7466760 | 30 | 0 | 3585 | 0 | 0 | 96 | 0 | 0 | 32 | 0 | 0 | 24 | 0 | 0 | 28 | 0 | 14 | 0 | 0 | 692 | 1439 | 13 | 0 | 0 | 0 | 1792 | 6701 | 13 | 0 | 0 | 84 | 0 | 0 | 0 | 11738 | 296 | 3990 |
| SRR7466761 | 25 | 0 | 3862 | 0 | 0 | 85 | 0 | 0 | 20 | 0 | 0 | 0 | 0 | 0 | 0 | 0 | 19 | 0 | 0 | 687 | 1633 | 0 | 0 | 0 | 0 | 2156 | 6107 | 0 | 0 | 0 | 99 | 0 | 0 | 0 | 11859 | 335 | 3589 |
| SRR7466762 | 24 | 0 | 4150 | 12 | 0 | 14 | 20 | 0 | 20 | 0 | 0 | 23 | 0 | 12 | 39 | 0 | 23 | 0 | 0 | 685 | 1469 | 12 | 0 | 0 | 0 | 319 | 841 | 0 | 0 | 0 | 19 | 0 | 0 | 0 | 1137 | 41 | 458 |
| SRR7466763 | 24 | 0 | 4158 | 0 | 0 | 15 | 0 | 17 | 74 | 0 | 0 | 24 | 0 | 0 | 0 | 0 | 22 | 0 | 0 | 723 | 2218 | 12 | 0 | 0 | 0 | 344 | 889 | 12 | 0 | 0 | 22 | 0 | 0 | 0 | 1195 | 41 | 459 |
| SRR7466764 | 47 | 10 | 3903 | 0 | 0 | 13 | 0 | 0 | 25 | 0 | 0 | 33 | 0 | 0 | 43 | 0 | 19 | 0 | 0 | 1195 | 2454 | 0 | 0 | 0 | 35 | 307 | 419 | 0 | 0 | 0 | 12 | 0 | 0 | 0 | 1401 | 66 | 669 |
| SRR7466765 | 30 | 0 | 4272 | 0 | 0 | 12 | 0 | 0 | 15 | 0 | 0 | 25 | 0 | 0 | 43 | 0 | 21 | 0 | 0 | 1018 | 2838 | 0 | 0 | 0 | 26 | 305 | 428 | 0 | 0 | 0 | 12 | 0 | 0 | 0 | 1407 | 62 | 651 |
| SRR7541164 | 46 | 0 | 1434 | 0 | 0 | 19 | 0 | 0 | 0 | 0 | 0 | 0 | 0 | 0 | 101 | 0 | 0 | 0 | 0 | 369 | 319 | 0 | 0 | 0 | 0 | 265 | 19 | 0 | 0 | 0 | 29 | 0 | 0 | 0 | 1376 | 54 | 391 |
| SRR7541165 | 371 | 0 | 468 | 0 | 0 | 0 | 12 | 0 | 0 | 0 | 0 | 0 | 0 | 0 | 24 | 0 | 10 | 0 | 12 | 545 | 711 | 0 | 0 | 0 | 11 | 118 | 190 | 0 | 15 | 0 | 0 | 0 | 0 | 0 | 61 | 0 | 20 |
| SRR7541166 | 42 | 0 | 1247 | 0 | 0 | 0 | 0 | 0 | 0 | 0 | 0 | 0 | 0 | 12 | 34 | 0 | 0 | 0 | 0 | 376 | 1093 | 0 | 0 | 0 | 0 | 78 | 17 | 0 | 0 | 0 | 0 | 0 | 0 | 0 | 2211 | 44 | 700 |
| SRR7541167 | 156 | 0 | 1095 | 0 | 0 | 0 | 34 | 0 | 0 | 0 | 0 | 0 | 0 | 41 | 13 | 0 | 0 | 0 | 0 | 294 | 602 | 0 | 0 | 0 | 0 | 28 | 20 | 0 | 0 | 0 | 0 | 0 | 0 | 0 | 43 | 0 | 10 |
| SRR7541168 | 16 | 0 | 1505 | 0 | 0 | 27 | 247 | 0 | 21 | 0 | 0 | 40 | 0 | 54 | 109 | 0 | 21 | 0 | 62 | 621 | 905 | 13 | 0 | 0 | 12 | 582 | 117 | 11 | 0 | 0 | 57 | 0 | 0 | 0 | 3973 | 232 | 1744 |
| SRR7541169 | 106 | 0 | 1379 | 0 | 0 | 23 | 21 | 0 | 22 | 0 | 0 | 0 | 0 | 23 | 243 | 0 | 16 | 0 | 22 | 152 | 331 | 0 | 0 | 0 | 13 | 52 | 28 | 0 | 0 | 0 | 0 | 0 | 0 | 0 | 427 | 47 | 110 |
| SRR7541170 | 188 | 13 | 819 | 0 | 0 | 0 | 0 | 0 | 0 | 0 | 0 | 0 | 0 | 22 | 42 | 0 | 0 | 0 | 0 | 206 | 617 | 0 | 0 | 0 | 0 | 21 | 0 | 0 | 14 | 0 | 0 | 0 | 0 | 10 | 202 | 18 | 44 |
| SRR7541171 | 127 | 17 | 2550 | 0 | 0 | 36 | 0 | 0 | 0 | 0 | 0 | 0 | 0 | 0 | 87 | 0 | 10 | 0 | 0 | 74 | 173 | 0 | 0 | 0 | 0 | 59 | 82 | 12 | 0 | 0 | 36 | 0 | 0 | 10 | 397 | 24 | 194 |
| SRR7541172 | 108 | 0 | 1495 | 0 | 0 | 27 | 0 | 0 | 22 | 0 | 0 | 17 | 0 | 0 | 51 | 0 | 0 | 0 | 0 | 133 | 255 | 0 | 0 | 0 | 0 | 69 | 30 | 0 | 0 | 0 | 0 | 0 | 0 | 0 | 390 | 71 | 127 |
| SRR7541173 | 118 | 0 | 333 | 0 | 0 | 61 | 0 | 0 | 67 | 0 | 0 | 0 | 0 | 16 | 37 | 0 | 0 | 11 | 0 | 120 | 695 | 0 | 0 | 0 | 0 | 1227 | 40 | 0 | 18 | 0 | 87 | 0 | 0 | 0 | 2890 | 125 | 603 |
| SRR7541174 | 188 | 0 | 2404 | 0 | 0 | 10 | 19 | 0 | 0 | 0 | 0 | 0 | 0 | 0 | 0 | 0 | 11 | 0 | 12 | 261 | 1232 | 0 | 0 | 0 | 11 | 21 | 23 | 0 | 0 | 0 | 0 | 0 | 0 | 0 | 333 | 12 | 102 |
| SRR7541175 | 89 | 24 | 556 | 0 | 0 | 18 | 25 | 0 | 0 | 0 | 0 | 0 | 0 | 0 | 0 | 0 | 0 | 0 | 0 | 248 | 176 | 0 | 0 | 0 | 0 | 65 | 38 | 0 | 0 | 0 | 23 | 0 | 0 | 0 | 441 | 17 | 103 |
| SRR7541176 | 83 | 0 | 792 | 0 | 0 | 30 | 15 | 0 | 14 | 0 | 0 | 0 | 0 | 0 | 19 | 0 | 0 | 0 | 0 | 170 | 1518 | 0 | 0 | 0 | 16 | 34 | 34 | 11 | 16 | 0 | 14 | 0 | 0 | 0 | 436 | 33 | 153 |
| SRR7541177 | 80 | 117 | 1601 | 0 | 0 | 49 | 0 | 0 | 36 | 0 | 0 | 11 | 0 | 12 | 82 | 0 | 10 | 0 | 0 | 92 | 203 | 0 | 0 | 0 | 13 | 758 | 196 | 0 | 12 | 0 | 128 | 0 | 0 | 0 | 10172 | 336 | 2612 |
| SRR7541178 | 52 | 0 | 778 | 0 | 0 | 11 | 0 | 0 | 51 | 0 | 0 | 0 | 0 | 11 | 0 | 0 | 17 | 0 | 0 | 169 | 0 | 0 | 0 | 0 | 17 | 210 | 64 | 0 | 11 | 0 | 32 | 0 | 0 | 0 | 317 | 85 | 62 |
| SRR7541179 | 59 | 0 | 448 | 0 | 0 | 15 | 22 | 0 | 15 | 0 | 0 | 0 | 0 | 21 | 92 | 0 | 0 | 0 | 0 | 103 | 491 | 0 | 0 | 0 | 0 | 288 | 13 | 0 | 18 | 0 | 0 | 0 | 0 | 0 | 1021 | 41 | 231 |
| SRR7541180 | 94 | 27 | 2848 | 0 | 0 | 86 | 0 | 0 | 56 | 0 | 0 | 0 | 0 | 12 | 0 | 0 | 34 | 0 | 0 | 209 | 327 | 0 | 0 | 0 | 19 | 136 | 98 | 0 | 32 | 0 | 104 | 0 | 0 | 12 | 697 | 26 | 143 |
| SRR7541181 | 73 | 16 | 2463 | 0 | 0 | 45 | 0 | 0 | 16 | 0 | 0 | 0 | 0 | 17 | 289 | 0 | 0 | 16 | 13 | 99 | 978 | 0 | 0 | 0 | 0 | 720 | 66 | 0 | 40 | 0 | 80 | 0 | 0 | 0 | 5073 | 225 | 804 |
| SRR7541182 | 60 | 16 | 1813 | 0 | 0 | 22 | 15 | 0 | 0 | 0 | 0 | 0 | 0 | 39 | 14 | 0 | 0 | 0 | 0 | 130 | 163 | 0 | 0 | 0 | 11 | 433 | 339 | 0 | 0 | 0 | 19 | 0 | 0 | 0 | 8363 | 359 | 2352 |
| SRR7541183 | 83 | 0 | 1213 | 0 | 0 | 31 | 0 | 0 | 29 | 0 | 0 | 0 | 0 | 14 | 12 | 0 | 0 | 0 | 0 | 204 | 117 | 0 | 0 | 0 | 0 | 654 | 39 | 0 | 33 | 0 | 39 | 0 | 0 | 0 | 3619 | 110 | 1062 |
| SRR7541184 | 57 | 11 | 1938 | 0 | 0 | 35 | 0 | 0 | 51 | 0 | 0 | 0 | 0 | 56 | 31 | 0 | 11 | 0 | 0 | 335 | 62 | 0 | 0 | 0 | 11 | 58 | 36 | 0 | 17 | 0 | 34 | 0 | 0 | 0 | 510 | 14 | 139 |
| SRR7541185 | 121 | 0 | 2827 | 0 | 0 | 11 | 0 | 0 | 77 | 0 | 0 | 0 | 0 | 22 | 13 | 0 | 15 | 0 | 0 | 57 | 73 | 0 | 0 | 0 | 16 | 58 | 189 | 0 | 18 | 0 | 76 | 0 | 0 | 0 | 378 | 11 | 96 |
| SRR7541186 | 54 | 0 | 1020 | 0 | 0 | 20 | 0 | 0 | 13 | 0 | 0 | 0 | 0 | 11 | 0 | 0 | 0 | 0 | 0 | 77 | 103 | 0 | 0 | 0 | 0 | 61 | 23 | 0 | 0 | 0 | 19 | 0 | 0 | 0 | 607 | 16 | 169 |
| SRR7541187 | 66 | 14 | 816 | 0 | 0 | 0 | 0 | 0 | 0 | 0 | 0 | 0 | 0 | 11 | 16 | 0 | 12 | 0 | 0 | 36 | 112 | 0 | 0 | 0 | 13 | 12 | 0 | 0 | 0 | 0 | 0 | 0 | 0 | 0 | 247 | 19 | 63 |
| SRR7541188 | 39 | 18 | 391 | 0 | 0 | 0 | 0 | 0 | 0 | 0 | 0 | 0 | 0 | 0 | 0 | 0 | 0 | 0 | 0 | 82 | 133 | 0 | 0 | 0 | 0 | 84 | 13 | 0 | 0 | 0 | 0 | 0 | 0 | 0 | 1997 | 61 | 395 |
| SRR7541189 | 46 | 0 | 917 | 0 | 0 | 30 | 0 | 0 | 0 | 0 | 0 | 0 | 0 | 0 | 60 | 0 | 0 | 0 | 0 | 58 | 960 | 0 | 0 | 0 | 18 | 435 | 408 | 0 | 44 | 0 | 37 | 0 | 0 | 0 | 5968 | 145 | 983 |
| SRR7541190 | 51 | 37 | 956 | 0 | 0 | 25 | 0 | 0 | 15 | 0 | 0 | 0 | 0 | 78 | 125 | 0 | 31 | 0 | 0 | 56 | 484 | 0 | 0 | 0 | 18 | 416 | 18 | 0 | 23 | 0 | 65 | 0 | 0 | 0 | 6323 | 214 | 1099 |
| SRR7541191 | 114 | 22 | 3872 | 0 | 0 | 34 | 0 | 0 | 30 | 0 | 0 | 0 | 0 | 37 | 13 | 0 | 0 | 0 | 0 | 113 | 307 | 0 | 0 | 0 | 0 | 28 | 20 | 0 | 10 | 0 | 17 | 0 | 0 | 0 | 159 | 59 | 25 |
| SRR7541192 | 75 | 55 | 158 | 0 | 0 | 0 | 43 | 0 | 0 | 0 | 0 | 0 | 0 | 29 | 20 | 0 | 0 | 0 | 0 | 125 | 118 | 0 | 0 | 0 | 0 | 268 | 35 | 0 | 29 | 0 | 24 | 0 | 0 | 0 | 2538 | 127 | 542 |
| SRR7541193 | 51 | 80 | 1218 | 0 | 0 | 14 | 28 | 0 | 0 | 0 | 0 | 0 | 0 | 28 | 20 | 0 | 0 | 51 | 0 | 83 | 336 | 0 | 0 | 0 | 0 | 198 | 18 | 0 | 0 | 0 | 13 | 0 | 0 | 0 | 2456 | 43 | 302 |
| SRR7547853 | 66 | 0 | 175 | 0 | 0 | 23 | 58 | 61 | 28 | 0 | 21 | 38 | 12 | 0 | 185 | 0 | 219 | 0 | 18 | 83 | 268 | 0 | 0 | 569 | 10 | 157 | 233 | 0 | 0 | 55 | 26 | 0 | 0 | 321 | 12 | 48 | 22 |
| SRR7547854 | 94 | 0 | 175 | 0 | 0 | 105 | 53 | 236 | 16 | 0 | 18 | 18 | 62 | 0 | 567 | 0 | 30 | 0 | 63 | 194 | 410 | 0 | 0 | 784 | 19 | 390 | 190 | 0 | 0 | 203 | 15 | 0 | 0 | 503 | 19 | 40 | 17 |
| SRR7547855 | 58 | 0 | 92 | 0 | 0 | 327 | 122 | 1540 | 109 | 0 | 14 | 85 | 80 | 0 | 863 | 0 | 15 | 0 | 77 | 306 | 519 | 0 | 0 | 3864 | 90 | 1622 | 1092 | 0 | 0 | 785 | 49 | 0 | 0 | 1860 | 19 | 18 | 24 |
| SRR7547856 | 184 | 0 | 259 | 0 | 0 | 138 | 159 | 1367 | 32 | 0 | 17 | 202 | 99 | 0 | 0 | 0 | 187 | 0 | 13 | 146 | 123 | 0 | 0 | 3239 | 82 | 558 | 977 | 0 | 0 | 638 | 25 | 0 | 0 | 2380 | 75 | 37 | 100 |
| SRR7547857 | 24 | 0 | 202 | 0 | 0 | 142 | 90 | 1200 | 44 | 0 | 10 | 12 | 141 | 0 | 0 | 0 | 49 | 0 | 0 | 32 | 15 | 0 | 0 | 3042 | 92 | 813 | 1149 | 0 | 0 | 792 | 36 | 0 | 0 | 2612 | 48 | 17 | 64 |
| SRR7547858 | 89 | 0 | 89 | 0 | 0 | 183 | 118 | 820 | 64 | 0 | 11 | 90 | 72 | 0 | 255 | 0 | 52 | 0 | 0 | 75 | 149 | 0 | 0 | 3892 | 57 | 1013 | 893 | 0 | 0 | 1110 | 49 | 0 | 0 | 2294 | 46 | 19 | 68 |
| SRR7547859 | 23 | 0 | 185 | 0 | 0 | 19 | 233 | 370 | 35 | 10 | 23 | 64 | 52 | 0 | 0 | 0 | 72 | 0 | 15 | 169 | 26 | 0 | 0 | 108 | 16 | 133 | 158 | 0 | 39 | 102 | 12 | 0 | 0 | 410 | 24 | 32 | 57 |
| SRR7547860 | 42 | 0 | 140 | 0 | 0 | 187 | 853 | 663 | 43 | 0 | 30 | 86 | 147 | 0 | 138 | 0 | 31 | 0 | 47 | 209 | 448 | 0 | 0 | 3895 | 145 | 959 | 1111 | 0 | 10 | 515 | 22 | 0 | 0 | 1614 | 35 | 15 | 35 |
| SRR7547861 | 42 | 0 | 143 | 0 | 0 | 81 | 116 | 638 | 39 | 23 | 10 | 117 | 37 | 0 | 119 | 0 | 58 | 0 | 23 | 71 | 326 | 0 | 0 | 728 | 23 | 398 | 142 | 0 | 17 | 312 | 19 | 0 | 0 | 543 | 36 | 31 | 58 |
| SRR7547862 | 67 | 0 | 243 | 0 | 0 | 19 | 274 | 93 | 22 | 0 | 0 | 63 | 0 | 0 | 1245 | 0 | 104 | 17 | 23 | 101 | 323 | 0 | 0 | 813 | 18 | 138 | 238 | 0 | 0 | 90 | 19 | 0 | 0 | 149 | 44 | 35 | 15 |
| SRR7547863 | 109 | 0 | 258 | 0 | 0 | 22 | 236 | 201 | 24 | 0 | 14 | 169 | 12 | 0 | 132 | 0 | 184 | 0 | 30 | 81 | 329 | 0 | 0 | 435 | 16 | 107 | 148 | 0 | 20 | 88 | 22 | 0 | 0 | 194 | 55 | 14 | 21 |
| SRR7547864 | 37 | 0 | 88 | 0 | 0 | 182 | 188 | 60 | 64 | 16 | 48 | 41 | 36 | 16 | 6572 | 0 | 28 | 0 | 18 | 162 | 363 | 0 | 0 | 758 | 333 | 966 | 629 | 0 | 0 | 73 | 134 | 0 | 0 | 681 | 47 | 27 | 55 |
| SRR7547865 | 106 | 0 | 215 | 0 | 0 | 31 | 135 | 359 | 36 | 0 | 0 | 94 | 13 | 0 | 97 | 0 | 88 | 0 | 16 | 130 | 125 | 0 | 0 | 420 | 62 | 194 | 173 | 0 | 0 | 81 | 34 | 0 | 0 | 188 | 10 | 36 | 13 |
| SRR7547866 | 81 | 0 | 305 | 0 | 0 | 22 | 195 | 16 | 28 | 0 | 0 | 136 | 10 | 0 | 931 | 0 | 188 | 0 | 24 | 215 | 403 | 0 | 0 | 599 | 28 | 145 | 280 | 0 | 0 | 36 | 54 | 0 | 0 | 133 | 15 | 17 | 59 |
| SRR7547867 | 149 | 0 | 242 | 0 | 0 | 83 | 69 | 988 | 22 | 0 | 11 | 1381 | 83 | 0 | 50 | 0 | 184 | 17 | 21 | 133 | 175 | 0 | 0 | 6894 | 59 | 416 | 871 | 0 | 0 | 650 | 32 | 0 | 0 | 2037 | 52 | 65 | 143 |
| SRR7547868 | 51 | 0 | 100 | 0 | 0 | 262 | 300 | 731 | 109 | 0 | 22 | 54 | 71 | 0 | 628 | 0 | 56 | 0 | 28 | 142 | 271 | 0 | 0 | 4463 | 78 | 1534 | 771 | 0 | 0 | 618 | 49 | 0 | 0 | 1525 | 44 | 30 | 89 |
| SRR7547869 | 58 | 0 | 216 | 0 | 0 | 122 | 492 | 2487 | 37 | 0 | 0 | 21 | 500 | 0 | 544 | 0 | 69 | 0 | 32 | 214 | 219 | 0 | 0 | 3422 | 50 | 798 | 1117 | 0 | 26 | 1227 | 19 | 0 | 0 | 2232 | 61 | 28 | 89 |
| SRR7547870 | 41 | 0 | 135 | 0 | 15 | 33 | 108 | 1358 | 16 | 0 | 19 | 28 | 36 | 0 | 86 | 0 | 100 | 0 | 25 | 65 | 209 | 0 | 0 | 761 | 10 | 151 | 168 | 0 | 17 | 744 | 22 | 0 | 0 | 811 | 11 | 16 | 18 |
| SRR7547871 | 29 | 0 | 310 | 0 | 0 | 238 | 74 | 713 | 73 | 0 | 0 | 52 | 172 | 0 | 973 | 0 | 57 | 0 | 11 | 76 | 621 | 0 | 0 | 3645 | 73 | 1402 | 862 | 0 | 0 | 788 | 188 | 0 | 0 | 1906 | 91 | 31 | 136 |
| SRR7547872 | 31 | 0 | 148 | 0 | 13 | 17 | 279 | 127 | 57 | 0 | 20 | 37 | 19 | 0 | 54 | 0 | 85 | 0 | 0 | 41 | 32 | 0 | 0 | 539 | 14 | 86 | 106 | 0 | 0 | 86 | 42 | 0 | 0 | 396 | 41 | 22 | 50 |
| SRR7547873 | 101 | 0 | 379 | 0 | 0 | 18 | 121 | 557 | 23 | 0 | 24 | 261 | 25 | 0 | 34 | 0 | 423 | 0 | 21 | 172 | 62 | 0 | 0 | 835 | 13 | 137 | 153 | 0 | 36 | 212 | 39 | 0 | 0 | 245 | 25 | 26 | 63 |
| SRR7547874 | 98 | 0 | 137 | 0 | 0 | 150 | 126 | 834 | 52 | 0 | 0 | 104 | 94 | 0 | 145 | 0 | 45 | 0 | 17 | 143 | 147 | 0 | 0 | 4093 | 53 | 897 | 889 | 0 | 0 | 1345 | 32 | 0 | 0 | 2414 | 137 | 48 | 164 |
| SRR7547875 | 41 | 0 | 119 | 0 | 0 | 163 | 422 | 1104 | 31 | 0 | 0 | 102 | 97 | 0 | 156 | 0 | 43 | 0 | 0 | 100 | 103 | 0 | 0 | 4532 | 76 | 831 | 932 | 0 | 0 | 957 | 20 | 0 | 0 | 1724 | 77 | 44 | 123 |
| SRR7547876 | 63 | 0 | 211 | 0 | 16 | 17 | 574 | 14 | 10 | 16 | 10 | 256 | 34 | 26 | 4711 | 0 | 165 | 0 | 33 | 250 | 136 | 0 | 0 | 364 | 83 | 92 | 187 | 13 | 0 | 21 | 11 | 0 | 0 | 97 | 22 | 17 | 32 |
| SRR7547877 | 110 | 0 | 124 | 0 | 0 | 108 | 134 | 1154 | 34 | 0 | 0 | 76 | 103 | 0 | 494 | 0 | 41 | 0 | 78 | 233 | 565 | 0 | 0 | 4506 | 81 | 532 | 696 | 0 | 0 | 1008 | 31 | 0 | 0 | 1242 | 117 | 70 | 209 |
| SRR7547878 | 11 | 0 | 126 | 0 | 13 | 25 | 108 | 73 | 36 | 29 | 19 | 43 | 17 | 0 | 420 | 0 | 78 | 0 | 0 | 56 | 89 | 0 | 0 | 172 | 80 | 128 | 97 | 0 | 0 | 39 | 19 | 0 | 0 | 349 | 36 | 22 | 11 |
| SRR7547879 | 43 | 0 | 158 | 0 | 19 | 13 | 116 | 100 | 25 | 43 | 21 | 17 | 22 | 18 | 660 | 0 | 51 | 0 | 16 | 103 | 198 | 0 | 0 | 116 | 47 | 77 | 69 | 0 | 21 | 58 | 52 | 0 | 0 | 271 | 44 | 35 | 10 |
| SRR7547880 | 35 | 0 | 97 | 0 | 0 | 95 | 139 | 147 | 30 | 52 | 127 | 58 | 31 | 18 | 1054 | 0 | 66 | 0 | 27 | 296 | 92 | 0 | 0 | 797 | 397 | 394 | 527 | 11 | 0 | 81 | 136 | 0 | 0 | 638 | 14 | 15 | 14 |
| SRR7547881 | 37 | 0 | 139 | 0 | 0 | 196 | 106 | 83 | 56 | 27 | 70 | 74 | 37 | 13 | 5975 | 0 | 23 | 0 | 25 | 119 | 96 | 0 | 0 | 874 | 382 | 938 | 463 | 0 | 0 | 201 | 230 | 0 | 0 | 744 | 39 | 20 | 38 |
| SRR7547882 | 477 | 0 | 77 | 0 | 0 | 94 | 328 | 457 | 19 | 0 | 10 | 120 | 36 | 0 | 0 | 0 | 12 | 0 | 33 | 320 | 1122 | 0 | 0 | 1938 | 38 | 408 | 826 | 0 | 10 | 639 | 19 | 0 | 0 | 844 | 13 | 17 | 33 |
| SRR7547883 | 1435 | 0 | 211 | 0 | 0 | 82 | 165 | 144 | 17 | 0 | 14 | 228 | 48 | 0 | 0 | 0 | 41 | 0 | 0 | 124 | 246 | 0 | 0 | 6765 | 90 | 461 | 2553 | 0 | 0 | 126 | 18 | 0 | 0 | 232 | 20 | 12 | 37 |
| SRR7547884 | 524 | 0 | 296 | 0 | 0 | 46 | 318 | 330 | 11 | 0 | 12 | 758 | 49 | 0 | 28 | 0 | 61 | 0 | 56 | 633 | 290 | 0 | 0 | 1297 | 44 | 284 | 1199 | 0 | 0 | 256 | 12 | 0 | 0 | 689 | 16 | 16 | 45 |
| SRR7547885 | 24 | 0 | 269 | 0 | 22 | 58 | 329 | 322 | 31 | 16 | 61 | 49 | 136 | 0 | 113 | 0 | 150 | 0 | 22 | 66 | 27 | 0 | 0 | 911 | 257 | 487 | 402 | 0 | 16 | 176 | 57 | 0 | 0 | 1849 | 86 | 44 | 87 |
| SRR7547886 | 74 | 0 | 163 | 0 | 22 | 33 | 53 | 124 | 11 | 22 | 13 | 29 | 18 | 12 | 773 | 0 | 67 | 724 | 0 | 20 | 348 | 0 | 0 | 98 | 47 | 155 | 84 | 0 | 0 | 65 | 13 | 0 | 0 | 239 | 18 | 36 | 14 |
| SRR7547887 | 33 | 0 | 444 | 0 | 23 | 14 | 92 | 25 | 30 | 39 | 17 | 59 | 17 | 16 | 901 | 0 | 142 | 0 | 0 | 51 | 119 | 0 | 0 | 219 | 73 | 78 | 105 | 0 | 0 | 18 | 13 | 0 | 0 | 209 | 40 | 27 | 40 |
| SRR7547888 | 52 | 0 | 171 | 0 | 22 | 196 | 61 | 388 | 69 | 46 | 105 | 251 | 27 | 24 | 184 | 0 | 162 | 0 | 0 | 32 | 77 | 0 | 0 | 620 | 327 | 853 | 424 | 0 | 26 | 219 | 70 | 0 | 0 | 1924 | 63 | 29 | 59 |
| SRR7547889 | 34 | 0 | 246 | 0 | 21 | 77 | 106 | 530 | 43 | 35 | 122 | 161 | 43 | 12 | 1747 | 0 | 209 | 63 | 53 | 183 | 425 | 0 | 0 | 718 | 226 | 343 | 399 | 0 | 140 | 485 | 72 | 0 | 0 | 2234 | 45 | 26 | 45 |
| SRR7547890 | 29 | 0 | 488 | 0 | 24 | 16 | 83 | 25 | 29 | 31 | 17 | 0 | 14 | 15 | 447 | 0 | 166 | 33 | 16 | 86 | 238 | 0 | 0 | 141 | 62 | 83 | 91 | 0 | 0 | 18 | 39 | 0 | 0 | 305 | 15 | 30 | 14 |
| SRR7547891 | 48 | 0 | 258 | 0 | 24 | 91 | 261 | 39 | 34 | 43 | 107 | 88 | 45 | 12 | 6024 | 0 | 78 | 0 | 34 | 433 | 146 | 0 | 0 | 898 | 290 | 439 | 594 | 0 | 0 | 80 | 88 | 0 | 0 | 778 | 25 | 15 | 27 |
| SRR7547892 | 34 | 0 | 245 | 0 | 14 | 25 | 41 | 30 | 30 | 23 | 12 | 39 | 39 | 28 | 236 | 0 | 137 | 0 | 52 | 60 | 143 | 0 | 0 | 225 | 51 | 103 | 163 | 0 | 0 | 39 | 27 | 0 | 0 | 168 | 32 | 25 | 10 |
| SRR7547893 | 21 | 0 | 227 | 0 | 26 | 16 | 164 | 52 | 45 | 27 | 61 | 0 | 59 | 12 | 942 | 0 | 49 | 43 | 56 | 104 | 407 | 0 | 0 | 109 | 44 | 89 | 83 | 0 | 0 | 37 | 51 | 0 | 0 | 236 | 50 | 25 | 47 |
| SRR7547894 | 34 | 0 | 256 | 0 | 16 | 114 | 63 | 125 | 31 | 19 | 41 | 24 | 43 | 15 | 0 | 0 | 169 | 0 | 0 | 25 | 12 | 0 | 0 | 844 | 304 | 492 | 989 | 0 | 0 | 230 | 59 | 0 | 0 | 1285 | 11 | 0 | 20 |
| SRR7547895 | 19 | 0 | 1235 | 0 | 33 | 11 | 88 | 16 | 26 | 32 | 16 | 87 | 18 | 11 | 2315 | 0 | 107 | 0 | 11 | 92 | 122 | 0 | 0 | 174 | 69 | 94 | 115 | 0 | 0 | 21 | 15 | 0 | 0 | 250 | 11 | 28 | 46 |
| SRR7547896 | 83 | 0 | 229 | 0 | 20 | 108 | 127 | 30 | 28 | 34 | 73 | 75 | 31 | 21 | 669 | 0 | 254 | 16 | 33 | 265 | 105 | 0 | 0 | 1300 | 505 | 729 | 1096 | 0 | 0 | 33 | 78 | 0 | 0 | 204 | 47 | 48 | 67 |

### Supp Table S8.2.Columns_AM_to_BW

| | | | | | | | | | | | | | | | | | | | | | | | | | | | | | | | | | | | | | |
|---|---|---|---|---|---|---|---|---|---|---|---|---|---|---|---|---|---|---|---|---|---|---|---|---|---|---|---|---|---|---|---|---|---|---|---|---|---|
| SRR7547897 | 47 | 0 | 173 | 0 | 11 | 34 | 94 | 49 | 13 | 22 | 52 | 29 | 60 | 0 | 9388 | 0 | 27 | 30 | 0 | 118 | 306 | 0 | 0 | 141 | 67 | 171 | 53 | 0 | 0 | 10 | 15 | 0 | 0 | 85 | 49 | 32 | 52 |
| SRR7547898 | 68 | 0 | 335 | 0 | 0 | 101 | 118 | 38 | 37 | 0 | 28 | 46 | 65 | 17 | 347 | 0 | 79 | 0 | 124 | 632 | 298 | 0 | 0 | 2058 | 286 | 560 | 866 | 0 | 0 | 77 | 126 | 0 | 0 | 425 | 70 | 68 | 122 |
| SRR7547899 | 54 | 0 | 154 | 0 | 0 | 102 | 123 | 81 | 44 | 0 | 30 | 364 | 60 | 24 | 1627 | 0 | 62 | 0 | 0 | 93 | 254 | 0 | 0 | 1808 | 210 | 469 | 688 | 0 | 0 | 167 | 121 | 0 | 0 | 378 | 60 | 26 | 50 |
| SRR7547900 | 23 | 0 | 99 | 0 | 15 | 72 | 73 | 146 | 35 | 40 | 84 | 78 | 52 | 16 | 1251 | 0 | 44 | 0 | 70 | 124 | 380 | 0 | 0 | 1080 | 292 | 378 | 403 | 0 | 10 | 141 | 85 | 0 | 0 | 1125 | 27 | 14 | 30 |
| SRR7547901 | 97 | 0 | 191 | 0 | 16 | 265 | 163 | 51 | 124 | 15 | 53 | 29 | 86 | 26 | 232 | 0 | 88 | 0 | 21 | 126 | 113 | 0 | 0 | 1399 | 399 | 1650 | 849 | 10 | 0 | 74 | 94 | 0 | 0 | 508 | 71 | 73 | 93 |
| SRR7547902 | 121 | 0 | 216 | 0 | 11 | 110 | 1033 | 3985 | 37 | 12 | 43 | 43 | 50 | 0 | 93 | 0 | 75 | 78 | 11 | 81 | 767 | 0 | 0 | 928 | 252 | 782 | 492 | 0 | 0 | 1577 | 72 | 0 | 0 | 498 | 34 | 28 | 44 |
| SRR7547903 | 46 | 0 | 354 | 0 | 12 | 16 | 147 | 60 | 29 | 0 | 49 | 23 | 56 | 18 | 233 | 0 | 94 | 0 | 35 | 104 | 225 | 0 | 0 | 164 | 55 | 104 | 123 | 0 | 0 | 14 | 16 | 0 | 0 | 84 | 35 | 28 | 49 |
| SRR7547904 | 49 | 0 | 177 | 0 | 12 | 91 | 805 | 53 | 28 | 22 | 69 | 74 | 46 | 22 | 380 | 0 | 82 | 11 | 25 | 184 | 50 | 0 | 0 | 1475 | 320 | 425 | 696 | 0 | 0 | 69 | 241 | 0 | 0 | 548 | 16 | 15 | 26 |
| SRR7547905 | 41 | 0 | 196 | 0 | 16 | 122 | 195 | 65 | 42 | 25 | 62 | 103 | 58 | 17 | 539 | 0 | 19 | 0 | 30 | 133 | 133 | 0 | 0 | 1122 | 301 | 567 | 387 | 0 | 0 | 86 | 103 | 0 | 0 | 569 | 35 | 29 | 25 |
| SRR7547906 | 142 | 0 | 142 | 0 | 12 | 84 | 66 | 124 | 33 | 14 | 38 | 153 | 80 | 16 | 47 | 0 | 43 | 81 | 38 | 141 | 780 | 0 | 0 | 2806 | 398 | 378 | 1330 | 0 | 0 | 55 | 51 | 0 | 0 | 3403 | 32 | 18 | 43 |
| SRR7547907 | 359 | 0 | 141 | 0 | 16 | 107 | 190 | 238 | 36 | 17 | 44 | 175 | 96 | 15 | 66 | 0 | 15 | 0 | 35 | 120 | 307 | 0 | 0 | 1235 | 329 | 469 | 1357 | 0 | 0 | 163 | 73 | 0 | 0 | 3635 | 41 | 31 | 56 |
| SRR7547908 | 317 | 0 | 236 | 0 | 0 | 285 | 314 | 39 | 73 | 34 | 67 | 105 | 69 | 0 | 51 | 0 | 62 | 0 | 59 | 424 | 636 | 0 | 0 | 1057 | 760 | 2089 | 1655 | 0 | 0 | 37 | 80 | 0 | 23 | 553 | 81 | 107 | 145 |
| SRR7547909 | 32 | 0 | 100 | 0 | 16 | 348 | 215 | 117 | 71 | 19 | 61 | 19 | 51 | 13 | 79 | 0 | 115 | 0 | 16 | 86 | 22 | 0 | 0 | 705 | 226 | 1725 | 510 | 0 | 0 | 84 | 472 | 0 | 0 | 1558 | 66 | 54 | 75 |
| SRR7547910 | 14 | 0 | 132 | 0 | 13 | 166 | 330 | 197 | 60 | 11 | 48 | 15 | 73 | 0 | 279 | 0 | 65 | 0 | 0 | 55 | 47 | 0 | 0 | 659 | 194 | 1068 | 329 | 0 | 0 | 177 | 207 | 0 | 0 | 2187 | 62 | 37 | 68 |
| SRR7547911 | 28 | 0 | 142 | 0 | 12 | 14 | 405 | 16 | 24 | 25 | 17 | 72 | 18 | 14 | 2663 | 0 | 50 | 0 | 0 | 69 | 314 | 0 | 0 | 221 | 48 | 84 | 55 | 0 | 0 | 13 | 17 | 0 | 0 | 313 | 31 | 21 | 37 |
| SRR7547912 | 32 | 0 | 133 | 0 | 12 | 189 | 235 | 119 | 61 | 13 | 35 | 80 | 74 | 0 | 679 | 0 | 35 | 0 | 17 | 150 | 148 | 0 | 0 | 921 | 226 | 980 | 384 | 0 | 0 | 181 | 88 | 0 | 0 | 1557 | 44 | 29 | 46 |
| SRR7547913 | 41 | 0 | 181 | 0 | 17 | 23 | 224 | 22 | 45 | 21 | 11 | 41 | 43 | 14 | 1767 | 0 | 42 | 0 | 18 | 143 | 170 | 0 | 0 | 211 | 57 | 134 | 65 | 0 | 0 | 30 | 18 | 0 | 0 | 251 | 46 | 28 | 44 |
| SRR7547914 | 27 | 0 | 101 | 0 | 21 | 132 | 69 | 32 | 58 | 27 | 82 | 39 | 45 | 13 | 1468 | 0 | 35 | 0 | 0 | 55 | 155 | 0 | 0 | 647 | 273 | 754 | 264 | 0 | 0 | 66 | 197 | 0 | 0 | 1258 | 28 | 16 | 36 |
| SRR7547915 | 45 | 0 | 550 | 0 | 14 | 17 | 72 | 15 | 35 | 22 | 63 | 40 | 18 | 11 | 436 | 0 | 135 | 0 | 0 | 29 | 90 | 0 | 0 | 203 | 48 | 109 | 108 | 0 | 0 | 12 | 13 | 0 | 0 | 177 | 48 | 35 | 54 |
| SRR7547916 | 67 | 0 | 451 | 0 | 15 | 78 | 316 | 15 | 15 | 30 | 93 | 107 | 44 | 13 | 316 | 0 | 207 | 12 | 23 | 125 | 64 | 0 | 0 | 1311 | 345 | 412 | 903 | 10 | 0 | 12 | 47 | 0 | 0 | 210 | 19 | 19 | 16 |
| SRR7547917 | 278 | 0 | 229 | 0 | 13 | 192 | 436 | 124 | 46 | 14 | 43 | 77 | 54 | 0 | 115 | 0 | 30 | 52 | 21 | 230 | 444 | 0 | 0 | 1297 | 501 | 1063 | 920 | 0 | 0 | 162 | 72 | 0 | 0 | 1475 | 50 | 57 | 122 |
| SRR7547918 | 281 | 0 | 308 | 0 | 12 | 110 | 129 | 59 | 29 | 39 | 108 | 110 | 52 | 25 | 134 | 0 | 50 | 37 | 14 | 48 | 585 | 0 | 0 | 813 | 451 | 638 | 1007 | 18 | 0 | 39 | 66 | 0 | 0 | 2096 | 18 | 18 | 28 |
| SRR7547919 | 376 | 0 | 224 | 0 | 12 | 76 | 172 | 236 | 27 | 27 | 73 | 103 | 96 | 22 | 125 | 0 | 49 | 0 | 21 | 312 | 405 | 0 | 0 | 1332 | 274 | 368 | 695 | 13 | 0 | 96 | 55 | 0 | 0 | 2907 | 21 | 13 | 32 |
| SRR7547920 | 129 | 0 | 234 | 0 | 0 | 60 | 135 | 21 | 12 | 14 | 15 | 427 | 22 | 11 | 14 | 0 | 39 | 0 | 155 | 797 | 3981 | 0 | 0 | 653 | 130 | 371 | 309 | 0 | 0 | 18 | 28 | 14 | 12 | 507 | 23 | 30 | 52 |
| SRR7585373 | 61 | 44 | 638 | 0 | 0 | 0 | 384 | 590 | 0 | 0 | 0 | 447 | 0 | 12 | 1763 | 0 | 14 | 28 | 36 | 10141 | 483 | 0 | 0 | 0 | 0 | 18 | 62 | 0 | 0 | 0 | 27 | 27 | 0 | 0 | 21 | 15 | 15 |
| SRR7586110 | 56 | 0 | 427 | 0 | 0 | 145 | 31 | 979 | 74 | 0 | 23 | 15 | 46 | 16 | 912 | 0 | 42 | 0 | 48 | 401 | 1234 | 0 | 0 | 5171 | 46 | 643 | 1075 | 0 | 0 | 530 | 117 | 0 | 0 | 191 | 299 | 131 | 298 |
| SRR7586111 | 51 | 0 | 427 | 0 | 14 | 54 | 31 | 334 | 32 | 0 | 27 | 19 | 20 | 12 | 710 | 0 | 39 | 0 | 26 | 342 | 1083 | 0 | 0 | 1800 | 22 | 248 | 420 | 0 | 0 | 189 | 44 | 0 | 0 | 61 | 106 | 48 | 108 |
| SRR7586113 | 68 | 0 | 234 | 0 | 0 | 137 | 21 | 818 | 71 | 0 | 35 | 0 | 59 | 19 | 136 | 0 | 54 | 0 | 39 | 443 | 476 | 0 | 0 | 3273 | 81 | 690 | 1504 | 0 | 0 | 975 | 128 | 0 | 0 | 596 | 370 | 181 | 435 |
| SRR7586114 | 61 | 0 | 253 | 0 | 19 | 143 | 24 | 799 | 86 | 0 | 36 | 0 | 55 | 18 | 88 | 0 | 40 | 0 | 72 | 357 | 405 | 0 | 0 | 3181 | 77 | 702 | 1637 | 0 | 0 | 965 | 120 | 0 | 0 | 576 | 356 | 185 | 445 |
| SRR7586116 | 20 | 0 | 458 | 0 | 0 | 42 | 21 | 94 | 21 | 0 | 28 | 12 | 16 | 11 | 1993 | 0 | 72 | 0 | 81 | 952 | 3395 | 0 | 0 | 1518 | 42 | 215 | 544 | 0 | 0 | 105 | 32 | 0 | 0 | 94 | 153 | 141 | 273 |
| SRR7586117 | 23 | 0 | 489 | 0 | 14 | 187 | 11 | 365 | 94 | 0 | 29 | 16 | 48 | 12 | 1841 | 0 | 70 | 0 | 74 | 971 | 3311 | 0 | 0 | 5554 | 164 | 907 | 2272 | 0 | 0 | 402 | 137 | 0 | 0 | 366 | 577 | 558 | 1057 |
| SRR7586118 | 21 | 0 | 464 | 0 | 17 | 51 | 18 | 93 | 25 | 0 | 28 | 12 | 14 | 13 | 2277 | 0 | 79 | 0 | 97 | 1186 | 4108 | 0 | 0 | 1431 | 40 | 237 | 597 | 0 | 0 | 100 | 34 | 0 | 0 | 92 | 153 | 149 | 280 |
| SRR7586119 | 79 | 0 | 421 | 0 | 0 | 72 | 16 | 95 | 33 | 0 | 13 | 12 | 25 | 0 | 404 | 0 | 131 | 0 | 72 | 1301 | 1746 | 0 | 0 | 1005 | 60 | 449 | 976 | 0 | 19 | 101 | 39 | 0 | 0 | 186 | 190 | 209 | 325 |
| SRR7586120 | 82 | 0 | 412 | 0 | 33 | 218 | 31 | 237 | 103 | 0 | 36 | 12 | 64 | 11 | 386 | 0 | 132 | 0 | 62 | 1152 | 1654 | 0 | 0 | 2659 | 154 | 1274 | 2790 | 0 | 16 | 269 | 122 | 0 | 0 | 458 | 497 | 550 | 885 |
| SRR7586122 | 25 | 0 | 666 | 0 | 0 | 196 | 12 | 1307 | 92 | 0 | 24 | 0 | 56 | 0 | 1262 | 0 | 21 | 0 | 105 | 808 | 2723 | 0 | 0 | 5276 | 58 | 1000 | 1466 | 0 | 0 | 1384 | 159 | 0 | 0 | 620 | 390 | 317 | 626 |
| SRR7586123 | 40 | 0 | 688 | 0 | 19 | 206 | 40 | 1265 | 97 | 0 | 23 | 0 | 48 | 0 | 1447 | 0 | 29 | 0 | 85 | 851 | 3220 | 0 | 0 | 5053 | 53 | 1008 | 1526 | 0 | 0 | 1330 | 155 | 0 | 0 | 611 | 352 | 306 | 617 |
| SRR7586125 | 49 | 0 | 461 | 0 | 0 | 104 | 23 | 239 | 53 | 0 | 19 | 0 | 35 | 11 | 993 | 0 | 39 | 0 | 70 | 450 | 2858 | 0 | 0 | 455 | 25 | 484 | 689 | 0 | 11 | 266 | 64 | 0 | 0 | 308 | 202 | 79 | 181 |
| SRR7586126 | 32 | 0 | 471 | 0 | 33 | 320 | 52 | 624 | 164 | 0 | 29 | 12 | 93 | 15 | 886 | 0 | 32 | 0 | 86 | 490 | 2761 | 0 | 0 | 1193 | 64 | 1423 | 2111 | 0 | 13 | 696 | 192 | 0 | 0 | 833 | 531 | 230 | 465 |
| SRR7586128 | 47 | 0 | 260 | 0 | 0 | 69 | 47 | 302 | 39 | 0 | 12 | 20 | 25 | 13 | 2283 | 0 | 30 | 0 | 86 | 565 | 5867 | 0 | 0 | 836 | 30 | 343 | 644 | 0 | 0 | 336 | 67 | 0 | 0 | 287 | 189 | 181 | 280 |
| SRR7586129 | 49 | 0 | 249 | 0 | 10 | 77 | 71 | 306 | 41 | 0 | 11 | 26 | 28 | 15 | 2559 | 0 | 35 | 0 | 78 | 626 | 6544 | 0 | 0 | 825 | 30 | 372 | 712 | 0 | 0 | 328 | 70 | 0 | 0 | 275 | 189 | 185 | 273 |
| SRR7586131 | 50 | 0 | 439 | 0 | 0 | 54 | 29 | 307 | 29 | 0 | 30 | 0 | 12 | 18 | 2665 | 0 | 15 | 0 | 72 | 375 | 2939 | 0 | 0 | 1358 | 12 | 267 | 270 | 0 | 0 | 490 | 64 | 0 | 0 | 197 | 119 | 125 | 233 |
| SRR7586132 | 42 | 0 | 433 | 0 | 16 | 174 | 18 | 976 | 101 | 0 | 32 | 0 | 39 | 14 | 2787 | 0 | 16 | 0 | 95 | 467 | 3394 | 0 | 0 | 4240 | 41 | 914 | 893 | 0 | 0 | 1486 | 197 | 0 | 0 | 577 | 368 | 406 | 772 |
| SRR7586134 | 63 | 0 | 480 | 0 | 0 | 94 | 39 | 263 | 51 | 0 | 10 | 18 | 25 | 13 | 956 | 0 | 31 | 0 | 74 | 532 | 3588 | 0 | 0 | 890 | 31 | 446 | 868 | 0 | 0 | 412 | 82 | 0 | 0 | 348 | 150 | 162 | 271 |
| SRR7586135 | 58 | 0 | 536 | 0 | 23 | 99 | 45 | 274 | 55 | 0 | 11 | 12 | 28 | 15 | 982 | 0 | 29 | 0 | 58 | 612 | 3397 | 0 | 0 | 868 | 31 | 465 | 953 | 0 | 0 | 398 | 77 | 0 | 0 | 338 | 149 | 168 | 276 |
| SRR7586137 | 58 | 0 | 334 | 0 | 0 | 117 | 47 | 1502 | 60 | 0 | 17 | 13 | 65 | 12 | 52 | 0 | 45 | 14 | 52 | 634 | 660 | 0 | 0 | 5573 | 33 | 563 | 1512 | 0 | 0 | 1680 | 74 | 0 | 0 | 573 | 264 | 131 | 442 |
| SRR7586138 | 51 | 0 | 403 | 0 | 22 | 32 | 43 | 378 | 19 | 0 | 17 | 17 | 14 | 14 | 43 | 0 | 38 | 0 | 72 | 459 | 596 | 0 | 0 | 1457 | 37 | 165 | 440 | 0 | 0 | 433 | 21 | 0 | 0 | 140 | 73 | 35 | 117 |
| SRR7586140 | 66 | 0 | 380 | 0 | 0 | 100 | 98 | 305 | 58 | 0 | 12 | 0 | 30 | 13 | 635 | 0 | 16 | 0 | 78 | 492 | 2665 | 0 | 0 | 515 | 20 | 448 | 605 | 0 | 0 | 359 | 62 | 0 | 0 | 301 | 110 | 110 | 219 |
| SRR7586141 | 66 | 0 | 392 | 0 | 12 | 114 | 73 | 292 | 65 | 0 | 13 | 0 | 29 | 14 | 570 | 0 | 16 | 0 | 82 | 541 | 2768 | 0 | 0 | 511 | 20 | 503 | 665 | 0 | 0 | 335 | 70 | 0 | 0 | 289 | 102 | 109 | 224 |
| SRR7586143 | 51 | 0 | 217 | 0 | 0 | 73 | 0 | 330 | 47 | 0 | 21 | 11 | 23 | 24 | 708 | 0 | 21 | 0 | 76 | 567 | 1660 | 0 | 0 | 864 | 14 | 336 | 397 | 0 | 0 | 477 | 81 | 0 | 0 | 337 | 187 | 145 | 264 |
| SRR7586144 | 41 | 0 | 215 | 0 | 27 | 204 | 0 | 815 | 120 | 0 | 23 | 0 | 57 | 26 | 715 | 0 | 25 | 0 | 95 | 544 | 1618 | 0 | 0 | 2119 | 40 | 935 | 1125 | 0 | 0 | 1152 | 198 | 0 | 0 | 800 | 470 | 380 | 674 |
| SRR7586146 | 87 | 0 | 489 | 0 | 0 | 100 | 16 | 31 | 74 | 0 | 15 | 41 | 28 | 17 | 3168 | 0 | 82 | 0 | 27 | 202 | 1645 | 0 | 0 | 30 | 46 | 531 | 378 | 11 | 0 | 13 | 94 | 0 | 0 | 137 | 130 | 94 | 208 |
| SRR7586147 | 73 | 0 | 503 | 0 | 27 | 355 | 14 | 38 | 262 | 0 | 46 | 33 | 89 | 17 | 3689 | 0 | 114 | 0 | 41 | 158 | 1982 | 0 | 0 | 110 | 142 | 1830 | 1387 | 0 | 0 | 42 | 307 | 0 | 0 | 428 | 422 | 281 | 665 |
| SRR7586149 | 21 | 0 | 18 | 0 | 0 | 119 | 18 | 354 | 85 | 0 | 47 | 0 | 99 | 38 | 488 | 0 | 41 | 0 | 41 | 337 | 888 | 0 | 0 | 567 | 60 | 610 | 1050 | 12 | 0 | 388 | 159 | 0 | 0 | 791 | 590 | 226 | 919 |
| SRR7586150 | 30 | 0 | 17 | 0 | 12 | 47 | 17 | 117 | 31 | 0 | 16 | 0 | 35 | 14 | 484 | 0 | 41 | 0 | 60 | 388 | 842 | 0 | 0 | 194 | 22 | 228 | 382 | 13 | 0 | 139 | 59 | 0 | 0 | 266 | 208 | 90 | 331 |
| SRR7586152 | 110 | 0 | 338 | 0 | 0 | 161 | 42 | 387 | 114 | 0 | 36 | 48 | 143 | 26 | 112 | 0 | 81 | 0 | 38 | 319 | 784 | 0 | 0 | 908 | 55 | 834 | 1467 | 0 | 0 | 398 | 155 | 0 | 0 | 855 | 557 | 426 | 1312 |
| SRR7586153 | 108 | 0 | 366 | 0 | 28 | 60 | 38 | 134 | 44 | 0 | 29 | 49 | 41 | 18 | 139 | 0 | 84 | 0 | 45 | 335 | 843 | 0 | 0 | 300 | 20 | 300 | 527 | 0 | 0 | 135 | 53 | 0 | 0 | 280 | 181 | 154 | 444 |
| SRR7586155 | 65 | 0 | 189 | 0 | 0 | 47 | 30 | 28 | 25 | 0 | 13 | 53 | 32 | 12 | 2891 | 0 | 77 | 0 | 20 | 76 | 1362 | 0 | 0 | 53 | 37 | 227 | 299 | 20 | 0 | 40 | 52 | 0 | 0 | 310 | 182 | 111 | 260 |
| SRR7586156 | 65 | 0 | 178 | 0 | 27 | 53 | 19 | 32 | 29 | 0 | 13 | 50 | 31 | 12 | 2566 | 0 | 97 | 0 | 29 | 90 | 1174 | 0 | 0 | 52 | 37 | 242 | 323 | 12 | 0 | 36 | 52 | 0 | 0 | 301 | 185 | 123 | 258 |
| SRR7586158 | 47 | 0 | 67 | 0 | 0 | 224 | 38 | 713 | 145 | 0 | 20 | 109 | 106 | 11 | 791 | 0 | 98 | 0 | 134 | 1044 | 2520 | 0 | 0 | 1219 | 43 | 1178 | 1750 | 0 | 0 | 903 | 124 | 0 | 0 | 1425 | 983 | 860 | 2429 |
| SRR7586159 | 47 | 0 | 79 | 0 | 16 | 253 | 33 | 680 | 151 | 0 | 19 | 101 | 91 | 14 | 752 | 0 | 99 | 0 | 135 | 927 | 2513 | 0 | 0 | 1194 | 46 | 1276 | 1905 | 0 | 0 | 905 | 120 | 0 | 0 | 1380 | 974 | 853 | 2454 |
| SRR7609258 | 37 | 0 | 31 | 0 | 0 | 15 | 15 | 11 | 11 | 0 | 17 | 130 | 123 | 17 | 17796 | 0 | 44 | 0 | 0 | 0 | 80 | 0 | 0 | 13 | 137 | 83 | 11 | 0 | 0 | 0 | 10 | 0 | 57 | 17 | 35 | 13 | 15 |
| SRR7609259 | 30 | 0 | 16 | 20 | 0 | 61 | 23 | 20 | 14 | 25 | 144 | 26 | 81 | 26 | 391 | 0 | 0 | 0 | 0 | 16 | 83 | 0 | 0 | 0 | 36 | 16 | 0 | 0 | 0 | 48 | 68 | 0 | 202 | 30 | 17 | 29 | 47 |
| SRR7609260 | 42 | 25 | 38 | 0 | 0 | 89 | 23 | 12 | 13 | 42 | 206 | 14 | 84 | 36 | 4689 | 0 | 16 | 0 | 0 | 0 | 55 | 0 | 0 | 18 | 50 | 21 | 0 | 0 | 0 | 37 | 88 | 0 | 104 | 22 | 151 | 160 | 13 |
| SRR7609261 | 37 | 55 | 0 | 0 | 0 | 0 | 130 | 1003 | 48 | 0 | 147 | 0 | 58 | 38 | 3951 | 0 | 13 | 85 | 0 | 0 | 87 | 0 | 0 | 17 | 97 | 37 | 0 | 0 | 0 | 514 | 199 | 0 | 67 | 270 | 42 | 66 | 111 |
| SRR7609262 | 12 | 34 | 0 | 0 | 0 | 0 | 56 | 28 | 36 | 40 | 111 | 80 | 20 | 61 | 6043 | 0 | 41 | 0 | 0 | 0 | 38 | 0 | 0 | 16 | 111 | 30 | 0 | 0 | 0 | 18 | 220 | 0 | 91 | 246 | 59 | 81 | 117 |
| SRR7609263 | 0 | 47 | 17 | 0 | 0 | 12 | 21 | 35 | 16 | 26 | 137 | 0 | 75 | 40 | 547 | 0 | 28 | 37 | 0 | 17 | 21 | 0 | 0 | 19 | 109 | 47 | 0 | 14 | 0 | 33 | 29 | 0 | 184 | 91 | 44 | 30 | 91 |
| SRR7609264 | 19 | 42 | 20 | 0 | 0 | 15 | 47 | 64 | 42 | 20 | 118 | 0 | 0 | 23 | 356 | 0 | 0 | 15 | 0 | 24 | 27 | 0 | 0 | 74 | 139 | 54 | 10 | 0 | 0 | 115 | 20 | 0 | 103 | 31 | 39 | 29 | 65 |
| SRR7609265 | 51 | 63 | 18 | 0 | 0 | 0 | 20 | 21 | 19 | 29 | 110 | 15 | 76 | 17 | 266 | 0 | 16 | 18 | 0 | 12 | 23 | 0 | 0 | 39 | 116 | 30 | 0 | 0 | 0 | 44 | 36 | 0 | 132 | 17 | 24 | 42 | 72 |
| SRR7609266 | 55 | 180 | 10 | 0 | 0 | 0 | 12 | 0 | 19 | 0 | 29 | 24 | 31 | 13 | 136 | 31 | 21 | 11 | 0 | 0 | 17 | 0 | 0 | 21 | 29 | 19 | 0 | 117 | 0 | 22 | 283 | 133 | 248 | 458 | 58 | 63 | 118 |
| SRR7609267 | 37 | 103 | 21 | 0 | 0 | 0 | 34 | 0 | 16 | 57 | 12 | 17 | 49 | 13 | 8695 | 11 | 16 | 0 | 0 | 0 | 33 | 0 | 0 | 12 | 87 | 39 | 0 | 48 | 0 | 15 | 25 | 21 | 18 | 90 | 61 | 53 | 71 |
| SRR7609268 | 43 | 107 | 0 | 0 | 0 | 0 | 228 | 101 | 51 | 15 | 95 | 21 | 0 | 72 | 1610 | 0 | 0 | 0 | 24 | 63 | 26 | 0 | 0 | 63 | 132 | 36 | 0 | 0 | 0 | 102 | 21 | 0 | 48 | 15 | 52 | 89 | 128 |
| SRR7609269 | 17 | 52 | 49 | 0 | 0 | 0 | 48 | 31 | 12 | 47 | 25 | 91 | 68 | 40 | 9452 | 0 | 52 | 0 | 0 | 0 | 0 | 0 | 0 | 0 | 10 | 29 | 0 | 0 | 0 | 0 | 15 | 0 | 69 | 93 | 45 | 51 | 81 |
| SRR7609270 | 0 | 101 | 0 | 0 | 0 | 0 | 18 | 109 | 20 | 93 | 844 | 11 | 53 | 65 | 430 | 0 | 0 | 0 | 0 | 0 | 17 | 0 | 0 | 11 | 0 | 30 | 0 | 0 | 0 | 49 | 364 | 0 | 232 | 355 | 127 | 186 | 220 |
| SRR7609271 | 56 | 41 | 56 | 0 | 0 | 0 | 57 | 23 | 11 | 19 | 54 | 38 | 142 | 24 | 174 | 0 | 12 | 0 | 0 | 0 | 0 | 0 | 0 | 22 | 121 | 22 | 0 | 0 | 0 | 24 | 14 | 0 | 59 | 113 | 42 | 68 | 92 |
| SRR7609272 | 64 | 29 | 0 | 0 | 0 | 0 | 41 | 12 | 65 | 0 | 43 | 15 | 50 | 29 | 13570 | 0 | 0 | 0 | 0 | 11 | 36 | 0 | 0 | 11 | 14 | 43 | 0 | 0 | 0 | 25 | 32 | 0 | 47 | 20 | 23 | 72 | 108 |
| SRR7609273 | 28 | 36 | 43 | 0 | 0 | 0 | 0 | 20 | 12 | 55 | 204 | 39 | 68 | 122 | 693 | 0 | 22 | 195 | 0 | 0 | 49 | 0 | 0 | 0 | 151 | 16 | 0 | 0 | 0 | 16 | 294 | 0 | 69 | 500 | 25 | 25 | 48 |
| SRR7609274 | 16 | 11 | 17 | 0 | 0 | 0 | 34 | 0 | 14 | 13 | 63 | 36 | 38 | 17 | 121 | 0 | 0 | 0 | 0 | 0 | 37 | 0 | 0 | 17 | 133 | 36 | 0 | 0 | 0 | 13 | 11 | 0 | 60 | 149 | 36 | 54 | 87 |
| SRR7609275 | 16 | 146 | 43 | 0 | 0 | 23 | 46 | 16 | 66 | 24 | 190 | 11 | 87 | 41 | 4464 | 0 | 15 | 0 | 0 | 0 | 27 | 0 | 0 | 26 | 61 | 244 | 0 | 0 | 0 | 26 | 5836 | 0 | 87 | 328 | 132 | 242 | 305 |
| SRR7609276 | 35 | 38 | 0 | 0 | 0 | 0 | 32 | 27 | 31 | 16 | 46 | 27 | 36 | 34 | 63073 | 0 | 0 | 82 | 0 | 0 | 40 | 0 | 0 | 0 | 189 | 21 | 0 | 0 | 0 | 52 | 510 | 0 | 85 | 238 | 0 | 0 | 0 |
| SRR7609277 | 10 | 41 | 32 | 0 | 0 | 0 | 98 | 33 | 16 | 44 | 12 | 19 | 62 | 101 | 2669 | 0 | 12 | 0 | 0 | 11 | 45 | 0 | 0 | 18 | 17 | 23 | 0 | 11 | 0 | 22 | 14 | 0 | 114 | 13 | 150 | 18 | 27 |
| SRR7609278 | 28 | 27 | 24 | 0 | 0 | 0 | 84 | 44 | 15 | 26 | 14 | 0 | 42 | 18 | 18463 | 0 | 15 | 63 | 0 | 15 | 128 | 0 | 0 | 57 | 69 | 20 | 0 | 0 | 0 | 71 | 12 | 0 | 119 | 58 | 25 | 20 | 72 |
| SRR7609279 | 16 | 22 | 16 | 0 | 0 | 0 | 18 | 0 | 18 | 17 | 98 | 65 | 43 | 21 | 33046 | 0 | 0 | 0 | 0 | 0 | 18 | 0 | 0 | 0 | 62 | 21 | 0 | 0 | 0 | 13 | 11 | 0 | 123 | 126 | 108 | 114 | 12 |
| SRR7609280 | 23 | 83 | 13 | 0 | 0 | 0 | 94 | 19 | 11 | 0 | 46 | 16 | 40 | 89 | 6565 | 0 | 0 | 0 | 0 | 0 | 15 | 0 | 0 | 0 | 99 | 0 | 0 | 0 | 0 | 22 | 18 | 0 | 67 | 77 | 35 | 78 | 72 |
| SRR7609281 | 39 | 83 | 26 | 12 | 0 | 24 | 228 | 63 | 42 | 0 | 66 | 22 | 37 | 39 | 11337 | 0 | 0 | 39 | 0 | 0 | 58 | 0 | 0 | 0 | 61 | 66 | 0 | 0 | 0 | 66 | 666 | 0 | 66 | 176 | 54 | 12 | 12 |
| SRR7609282 | 15 | 37 | 0 | 0 | 0 | 23 | 17 | 27 | 70 | 16 | 80 | 0 | 127 | 49 | 115 | 0 | 0 | 0 | 0 | 0 | 22 | 0 | 0 | 12 | 78 | 127 | 0 | 21 | 0 | 21 | 1609 | 0 | 133 | 583 | 291 | 287 | 468 |
| SRR7609283 | 43 | 12 | 0 | 0 | 0 | 20 | 0 | 0 | 114 | 30 | 149 | 65 | 87 | 21 | 188 | 0 | 0 | 0 | 0 | 0 | 61 | 0 | 0 | 0 | 51 | 146 | 0 | 13 | 0 | 0 | 765 | 0 | 107 | 475 | 137 | 163 | 324 |
| SRR7609284 | 34 | 26 | 0 | 0 | 0 | 15 | 25 | 24 | 46 | 0 | 53 | 0 | 95 | 30 | 308 | 0 | 14 | 0 | 0 | 0 | 48 | 0 | 0 | 29 | 185 | 71 | 0 | 0 | 0 | 101 | 33 | 0 | 69 | 28 | 98 | 113 | 10 |
| SRR7609285 | 55 | 13 | 0 | 0 | 0 | 10 | 13 | 23 | 75 | 21 | 91 | 92 | 92 | 22 | 258 | 0 | 17 | 0 | 0 | 0 | 40 | 0 | 0 | 0 | 59 | 84 | 0 | 0 | 0 | 28 | 569 | 0 | 76 | 487 | 94 | 117 | 230 |
| SRR7609286 | 28 | 26 | 11 | 0 | 0 | 26 | 0 | 15 | 82 | 15 | 103 | 111 | 108 | 29 | 1576 | 0 | 12 | 0 | 0 | 0 | 176 | 0 | 0 | 0 | 59 | 123 | 0 | 0 | 0 | 12 | 440 | 0 | 62 | 371 | 75 | 80 | 199 |
| SRR7609287 | 0 | 20 | 17 | 0 | 11 | 31 | 33 | 34 | 91 | 26 | 216 | 24 | 40 | 40 | 877 | 0 | 12 | 0 | 0 | 0 | 14 | 0 | 0 | 0 | 37 | 185 | 0 | 34 | 0 | 11 | 443 | 0 | 156 | 207 | 153 | 134 | 523 |
| SRR7609288 | 20 | 0 | 41 | 0 | 0 | 19 | 13 | 32 | 28 | 15 | 162 | 17 | 81 | 12 | 471 | 0 | 82 | 0 | 0 | 0 | 238 | 0 | 0 | 21 | 170 | 75 | 14 | 12 | 0 | 12 | 103 | 0 | 62 | 165 | 152 | 14 | 36 |
| SRR7609289 | 24 | 13 | 29 | 0 | 0 | 0 | 75 | 78 | 26 | 63 | 34 | 19 | 23 | 30 | 935 | 0 | 18 | 14 | 16 | 0 | 50 | 0 | 0 | 14 | 36 | 35 | 0 | 12 | 0 | 30 | 37 | 37 | 16 | 26 | 42 | 45 | 58 |
| SRR7609290 | 19 | 31 | 0 | 0 | 12 | 0 | 23 | 52 | 55 | 52 | 275 | 0 | 67 | 15 | 333 | 0 | 15 | 0 | 0 | 0 | 0 | 0 | 0 | 0 | 119 | 64 | 12 | 15 | 0 | 37 | 294 | 0 | 119 | 11 | 159 | 135 | 303 |
| SRR7609291 | 29 | 35 | 33 | 0 | 0 | 0 | 26 | 0 | 0 | 29 | 90 | 96 | 0 | 27 | 871 | 0 | 130 | 20 | 0 | 0 | 64 | 0 | 0 | 0 | 24 | 21 | 0 | 0 | 0 | 0 | 94 | 0 | 174 | 57 | 35 | 62 | 77 |
| SRR7609292 | 30 | 11 | 0 | 0 | 0 | 0 | 36 | 88 | 29 | 18 | 135 | 21 | 0 | 124 | 2672 | 0 | 0 | 0 | 0 | 26 | 19 | 0 | 0 | 23 | 85 | 36 | 0 | 0 | 0 | 25 | 17 | 0 | 53 | 12 | 69 | 125 | 10 |
| SRR7609293 | 17 | 10 | 20 | 0 | 0 | 24 | 23 | 32 | 56 | 24 | 84 | 19 | 69 | 28 | 7029 | 0 | 0 | 0 | 0 | 0 | 0 | 0 | 0 | 13 | 109 | 153 | 0 | 0 | 0 | 35 | 835 | 0 | 134 | 193 | 316 | 209 | 498 |
| SRR7609294 | 66 | 13 | 0 | 0 | 0 | 25 | 27 | 24 | 99 | 20 | 50 | 66 | 31 | 29 | 5303 | 0 | 0 | 110 | 0 | 0 | 107 | 0 | 0 | 0 | 19 | 11 | 0 | 0 | 0 | 15 | 44 | 13 | 60 | 12 | 125 | 10 | 24 |
| SRR7609295 | 0 | 104 | 15 | 0 | 0 | 0 | 72 | 215 | 0 | 144 | 870 | 25 | 12 | 28 | 1409 | 0 | 25 | 0 | 0 | 67 | 23 | 0 | 0 | 90 | 38 | 36 | 0 | 0 | 0 | 158 | 68 | 0 | 299 | 94 | 12 | 80 | 235 |
| SRR7609296 | 0 | 0 | 0 | 0 | 0 | 11 | 30 | 30 | 0 | 0 | 92 | 36 | 0 | 0 | 2644 | 0 | 0 | 72 | 0 | 10 | 119 | 0 | 0 | 0 | 63 | 25 | 0 | 0 | 0 | 15 | 131 | 0 | 68 | 15 | 0 | 16 | 23 |
| SRR7609297 | 11 | 0 | 0 | 0 | 0 | 0 | 135 | 295 | 15 | 20 | 141 | 0 | 20 | 17 | 23147 | 0 | 0 | 32 | 0 | 0 | 152 | 0 | 0 | 31 | 32 | 43 | 0 | 0 | 0 | 251 | 513 | 0 | 101 | 75 | 24 | 43 | 107 |
| SRR7609298 | 0 | 19 | 0 | 0 | 0 | 16 | 53 | 99 | 29 | 0 | 53 | 25 | 27 | 16 | 2345 | 0 | 0 | 0 | 0 | 0 | 221 | 0 | 0 | 99 | 66 | 58 | 0 | 0 | 0 | 175 | 194 | 0 | 35 | 33 | 19 | 19 | 70 |
| SRR7609299 | 13 | 165 | 0 | 0 | 0 | 12 | 151 | 177 | 14 | 32 | 249 | 0 | 20 | 50 | 302 | 0 | 0 | 0 | 16 | 37 | 21 | 0 | 0 | 52 | 66 | 24 | 0 | 0 | 0 | 167 | 193 | 0 | 155 | 28 | 40 | 103 | 129 |
| SRR7609300 | 0 | 0 | 0 | 0 | 0 | 50 | 43 | 79 | 23 | 26 | 111 | 0 | 15 | 20 | 7976 | 0 | 0 | 0 | 12 | 0 | 186 | 0 | 0 | 73 | 67 | 108 | 15 | 0 | 0 | 149 | 465 | 0 | 132 | 73 | 67 | 117 | 199 |
| SRR7609301 | 0 | 145 | 0 | 0 | 0 | 39 | 221 | 18 | 66 | 37 | 91 | 0 | 15 | 22 | 46 | 0 | 10 | 0 | 29 | 42 | 35 | 0 | 0 | 223 | 44 | 96 | 0 | 0 | 0 | 169 | 152 | 0 | 213 | 59 | 83 | 113 | 14 |
| SRR7609302 | 19 | 92 | 0 | 0 | 0 | 55 | 191 | 13 | 64 | 21 | 70 | 0 | 0 | 52 | 1023 | 0 | 0 | 92 | 18 | 64 | 55 | 0 | 0 | 175 | 0 | 80 | 0 | 0 | 0 | 18 | 169 | 0 | 101 | 92 | 34 | 43 | 110 |
| SRR7644105 | 200 | 0 | 18 | 240 | 120 | 0 | 0 | 0 | 0 | 0 | 0 | 0 | 0 | 0 | 0 | 0 | 0 | 0 | 0 | 0 | 1752 | 0 | 0 | 120 | 210 | 12 | 240 | 0 | 0 | 0 | 0 | 0 | 0 | 0 | 0 | 0 | 0 |
| SRR7644106 | 191 | 0 | 34 | 163 | 98 | 0 | 0 | 0 | 0 | 0 | 0 | 0 | 0 | 0 | 0 | 0 | 0 | 0 | 0 | 0 | 958 | 0 | 0 | 33 | 163 | 455 | 390 | 33 | 0 | 0 | 0 | 0 | 0 | 0 | 0 | 0 | 0 |
| SRR7644107 | 212 | 0 | 47 | 274 | 61 | 0 | 0 | 0 | 0 | 0 | 0 | 0 | 0 | 0 | 0 | 0 | 0 | 0 | 0 | 0 | 2207 | 0 | 0 | 0 | 152 | 487 | 274 | 91 | 0 | 0 | 0 | 0 | 0 | 0 | 0 | 0 | 0 |
| SRR7644108 | 167 | 0 | 34 | 225 | 64 | 0 | 0 | 0 | 0 | 0 | 0 | 0 | 0 | 0 | 0 | 0 | 0 | 0 | 0 | 0 | 2418 | 0 | 0 | 0 | 257 | 19 | 289 | 32 | 0 | 0 | 0 | 0 | 0 | 0 | 0 | 0 | 0 |
| SRR7644109 | 171 | 0 | 68 | 240 | 133 | 80 | 0 | 0 | 0 | 0 | 0 | 14 | 0 | 27 | 0 | 0 | 0 | 0 | 0 | 0 | 489 | 27 | 0 | 53 | 107 | 213 | 213 | 0 | 0 | 0 | 0 | 0 | 0 | 0 | 0 | 0 | 0 |
| SRR7644110 | 254 | 0 | 42 | 260 | 58 | 29 | 0 | 0 | 0 | 0 | 0 | 0 | 0 | 0 | 0 | 0 | 0 | 0 | 0 | 0 | 1047 | 29 | 0 | 0 | 29 | 260 | 260 | 0 | 0 | 0 | 0 | 0 | 0 | 0 | 0 | 0 | 0 |
| SRR7644111 | 211 | 0 | 26 | 299 | 54 | 82 | 0 | 0 | 0 | 0 | 0 | 15 | 0 | 27 | 0 | 0 | 0 | 0 | 0 | 0 | 0 | 27 | 0 | 0 | 82 | 299 | 272 | 0 | 0 | 0 | 54 | 0 | 0 | 0 | 0 | 0 | 0 |
| SRR7644112 | 223 | 0 | 51 | 256 | 57 | 57 | 0 | 0 | 0 | 0 | 0 | 23 | 0 | 29 | 0 | 0 | 0 | 0 | 0 | 0 | 2076 | 29 | 0 | 29 | 29 | 256 | 313 | 0 | 0 | 0 | 29 | 0 | 0 | 0 | 0 | 0 | 0 |
| SRR7644113 | 425 | 0 | 74 | 148 | 0 | 0 | 0 | 0 | 0 | 0 | 0 | 17 | 0 | 0 | 0 | 0 | 0 | 0 | 0 | 0 | 1167 | 30 | 0 | 0 | 30 | 10 | 12 | 0 | 0 | 0 | 0 | 0 | 0 | 0 | 0 | 30 | 0 |
| SRR7644114 | 393 | 0 | 89 | 252 | 0 | 0 | 320 | 0 | 0 | 0 | 0 | 26 | 0 | 0 | 0 | 0 | 0 | 0 | 0 | 0 | 1919 | 31 | 0 | 0 | 0 | 472 | 755 | 0 | 31 | 0 | 0 | 0 | 0 | 0 | 0 | 31 | 0 |
| SRR7644115 | 447 | 0 | 52 | 268 | 30 | 0 | 0 | 0 | 0 | 0 | 0 | 15 | 0 | 0 | 0 | 0 | 0 | 0 | 0 | 0 | 2489 | 0 | 0 | 0 | 119 | 10 | 13 | 0 | 0 | 0 | 0 | 0 | 0 | 0 | 0 | 0 | 0 |
| SRR7644116 | 399 | 0 | 81 | 156 | 94 | 31 | 0 | 0 | 0 | 0 | 0 | 16 | 0 | 0 | 0 | 0 | 0 | 0 | 0 | 0 | 1298 | 31 | 0 | 0 | 0 | 14 | 343 | 0 | 31 | 0 | 0 | 0 | 0 | 0 | 0 | 0 | 0 |
| SRR7644117 | 373 | 0 | 70 | 106 | 133 | 0 | 0 | 0 | 53 | 0 | 0 | 12 | 0 | 0 | 0 | 0 | 0 | 0 | 0 | 0 | 917 | 80 | 0 | 0 | 53 | 106 | 212 | 0 | 0 | 0 | 0 | 0 | 0 | 0 | 0 | 0 | 0 |
| SRR7644118 | 380 | 0 | 87 | 113 | 85 | 0 | 0 | 0 | 0 | 0 | 0 | 0 | 0 | 0 | 0 | 0 | 0 | 0 | 0 | 0 | 0 | 113 | 0 | 0 | 0 | 368 | 57 | 0 | 0 | 0 | 0 | 0 | 0 | 0 | 0 | 0 | 0 |
| SRR7644119 | 391 | 0 | 130 | 212 | 106 | 27 | 0 | 0 | 53 | 0 | 0 | 12 | 0 | 0 | 456 | 0 | 0 | 0 | 0 | 0 | 0 | 80 | 0 | 80 | 53 | 212 | 159 | 0 | 0 | 0 | 53 | 0 | 0 | 0 | 0 | 0 | 0 |
| SRR7644120 | 383 | 0 | 88 | 112 | 168 | 56 | 0 | 0 | 0 | 0 | 0 | 31 | 0 | 0 | 501 | 0 | 0 | 0 | 0 | 0 | 0 | 28 | 0 | 28 | 0 | 140 | 84 | 0 | 0 | 0 | 0 | 0 | 0 | 0 | 0 | 0 | 0 |
| SRR7644121 | 507 | 0 | 107 | 79 | 79 | 0 | 284 | 26 | 0 | 0 | 0 | 0 | 0 | 0 | 0 | 0 | 0 | 0 | 0 | 0 | 284 | 26 | 0 | 0 | 236 | 289 | 892 | 0 | 0 | 0 | 0 | 0 | 0 | 0 | 0 | 0 | 0 |
| SRR7644122 | 635 | 0 | 147 | 86 | 86 | 29 | 300 | 29 | 0 | 0 | 0 | 0 | 0 | 0 | 0 | 0 | 0 | 0 | 300 | 0 | 601 | 29 | 0 | 0 | 144 | 230 | 604 | 0 | 0 | 0 | 0 | 0 | 0 | 0 | 0 | 0 | 0 |
| SRR7644123 | 588 | 0 | 113 | 134 | 54 | 54 | 0 | 0 | 0 | 0 | 0 | 11 | 0 | 0 | 0 | 0 | 0 | 0 | 554 | 0 | 1108 | 0 | 0 | 27 | 161 | 214 | 1099 | 27 | 0 | 0 | 0 | 0 | 0 | 0 | 0 | 0 | 0 |

### Supp Table S8.2.Columns_AM_to_BW

| | | | | | | | | | | | | | | | | | | | | | | | | | | | | | | | | | | | | | |
|---|---|---|---|---|---|---|---|---|---|---|---|---|---|---|---|---|---|---|---|---|---|---|---|---|---|---|---|---|---|---|---|---|---|---|---|---|---|
| SRR7644124 | 582 | 0 | 128 | 141 | 141 | 28 | 296 | 57 | 0 | 0 | 0 | 16 | 0 | 0 | 0 | 0 | 0 | 0 | 296 | 0 | 1184 | 28 | 0 | 0 | 85 | 255 | 651 | 0 | 0 | 0 | 0 | 0 | 0 | 0 | 0 | 0 | 0 |
| SRR7644125 | 751 | 0 | 117 | 275 | 229 | 0 | 0 | 23 | 0 | 0 | 0 | 37 | 0 | 0 | 0 | 0 | 0 | 0 | 0 | 0 | 2593 | 0 | 0 | 23 | 46 | 389 | 343 | 0 | 0 | 0 | 23 | 0 | 0 | 0 | 0 | 0 | 0 |
| SRR7644126 | 677 | 0 | 103 | 99 | 74 | 0 | 0 | 25 | 50 | 0 | 0 | 33 | 0 | 0 | 0 | 0 | 0 | 0 | 0 | 0 | 3378 | 25 | 0 | 50 | 74 | 14 | 12 | 0 | 0 | 0 | 0 | 0 | 0 | 0 | 0 | 0 | 0 |
| SRR7644128 | 754 | 0 | 132 | 138 | 115 | 0 | 0 | 46 | 0 | 0 | 0 | 50 | 0 | 0 | 0 | 0 | 0 | 0 | 0 | 0 | 1570 | 0 | 0 | 23 | 23 | 322 | 322 | 0 | 0 | 0 | 46 | 0 | 0 | 0 | 0 | 0 | 0 |
| SRR7644129 | 713 | 0 | 121 | 25 | 198 | 0 | 0 | 0 | 25 | 0 | 0 | 47 | 0 | 25 | 0 | 0 | 0 | 0 | 0 | 0 | 2549 | 0 | 0 | 0 | 99 | 223 | 173 | 0 | 0 | 0 | 25 | 0 | 0 | 0 | 0 | 0 | 0 |
| SRR7663662 | 0 | 110 | 573 | 151 | 12 | 0 | 11 | 0 | 0 | 0 | 0 | 213 | 0 | 82 | 0 | 67 | 0 | 0 | 0 | 120 | 0 | 16 | 0 | 0 | 13 | 0 | 11 | 136 | 0 | 0 | 0 | 0 | 292 | 11 | 0 | 0 | 0 |
| SRR7663663 | 0 | 24 | 2193 | 37 | 0 | 0 | 171 | 14 | 0 | 0 | 0 | 392 | 0 | 24 | 0 | 37 | 0 | 0 | 31 | 389 | 0 | 24 | 0 | 0 | 12 | 18 | 261 | 51 | 0 | 0 | 0 | 0 | 83 | 12 | 0 | 0 | 0 |
| SRR7664346 | 29 | 0 | 4110 | 0 | 0 | 22 | 14 | 0 | 24 | 0 | 0 | 483 | 0 | 0 | 12 | 0 | 0 | 0 | 0 | 224 | 616 | 0 | 0 | 0 | 0 | 565 | 399 | 0 | 0 | 0 | 53 | 0 | 0 | 0 | 311 | 32 | 136 |
| SRR7664347 | 20 | 0 | 3929 | 0 | 0 | 12 | 0 | 0 | 19 | 0 | 0 | 317 | 0 | 0 | 58 | 0 | 0 | 0 | 0 | 78 | 250 | 0 | 0 | 0 | 11 | 372 | 218 | 0 | 0 | 0 | 85 | 0 | 0 | 12 | 284 | 22 | 102 |
| SRR7690697 | 174 | 0 | 1925 | 0 | 0 | 38 | 79 | 36 | 10 | 0 | 0 | 43 | 0 | 0 | 0 | 0 | 16 | 0 | 0 | 838 | 807 | 0 | 0 | 54 | 36 | 547 | 1105 | 0 | 0 | 0 | 18 | 0 | 0 | 15 | 55 | 0 | 24 |
| SRR7690698 | 411 | 0 | 1425 | 0 | 0 | 15 | 55 | 24 | 0 | 0 | 0 | 65 | 0 | 0 | 0 | 0 | 24 | 0 | 0 | 1210 | 592 | 0 | 0 | 27 | 12 | 156 | 415 | 0 | 0 | 16 | 10 | 0 | 0 | 0 | 21 | 0 | 11 |
| SRR7690699 | 139 | 0 | 801 | 0 | 0 | 48 | 71 | 51 | 26 | 0 | 0 | 43 | 0 | 0 | 0 | 0 | 15 | 0 | 14 | 1461 | 750 | 0 | 0 | 57 | 32 | 842 | 924 | 0 | 0 | 15 | 20 | 0 | 0 | 10 | 48 | 0 | 26 |
| SRR7690700 | 68 | 0 | 158 | 0 | 31 | 155 | 37 | 0 | 47 | 13 | 58 | 83 | 143 | 24 | 83 | 0 | 96 | 0 | 0 | 43 | 183 | 0 | 0 | 45 | 276 | 977 | 455 | 12 | 0 | 0 | 64 | 0 | 0 | 712 | 78 | 42 | 70 |
| SRR7690701 | 64 | 0 | 99 | 0 | 39 | 64 | 50 | 0 | 23 | 14 | 76 | 85 | 85 | 23 | 100 | 0 | 55 | 17 | 0 | 25 | 203 | 0 | 0 | 0 | 352 | 501 | 108 | 0 | 0 | 0 | 54 | 0 | 0 | 484 | 51 | 34 | 52 |
| SRR7690702 | 100 | 0 | 102 | 0 | 53 | 133 | 45 | 0 | 45 | 39 | 90 | 63 | 115 | 11 | 36 | 0 | 47 | 0 | 0 | 59 | 308 | 0 | 0 | 14 | 366 | 915 | 401 | 0 | 0 | 0 | 70 | 0 | 0 | 385 | 39 | 30 | 42 |
| SRR7690703 | 103 | 0 | 78 | 10 | 75 | 69 | 21 | 0 | 34 | 41 | 128 | 82 | 76 | 21 | 13 | 0 | 43 | 0 | 13 | 0 | 226 | 0 | 0 | 0 | 376 | 576 | 121 | 15 | 0 | 0 | 35 | 0 | 11 | 2083 | 17 | 14 | 19 |
| SRR7690704 | 88 | 0 | 101 | 0 | 41 | 20 | 25 | 0 | 13 | 16 | 15 | 70 | 26 | 15 | 61 | 0 | 75 | 0 | 0 | 0 | 418 | 0 | 0 | 0 | 43 | 141 | 24 | 0 | 0 | 0 | 12 | 0 | 15 | 149 | 21 | 39 | 21 |
| SRR7690705 | 85 | 0 | 79 | 0 | 60 | 45 | 0 | 0 | 30 | 18 | 100 | 63 | 81 | 20 | 43 | 0 | 59 | 20 | 0 | 0 | 98 | 0 | 0 | 0 | 289 | 307 | 60 | 0 | 0 | 0 | 67 | 0 | 0 | 695 | 22 | 17 | 26 |
| SRR7690706 | 142 | 0 | 250 | 0 | 31 | 96 | 18 | 0 | 22 | 62 | 209 | 89 | 352 | 14 | 90 | 0 | 79 | 28 | 0 | 12 | 294 | 0 | 0 | 0 | 507 | 658 | 377 | 12 | 0 | 0 | 34 | 0 | 49 | 2991 | 21 | 20 | 53 |
| SRR7690707 | 586 | 0 | 1369 | 0 | 0 | 29 | 33 | 14 | 14 | 0 | 0 | 50 | 0 | 0 | 0 | 0 | 10 | 0 | 14 | 2308 | 1362 | 0 | 0 | 152 | 35 | 631 | 1093 | 0 | 0 | 0 | 0 | 0 | 0 | 0 | 18 | 0 | 11 |
| SRR7690708 | 155 | 0 | 270 | 0 | 42 | 143 | 14 | 0 | 34 | 39 | 141 | 91 | 281 | 0 | 69 | 12 | 97 | 0 | 14 | 111 | 671 | 0 | 0 | 13 | 401 | 974 | 785 | 0 | 0 | 0 | 55 | 0 | 57 | 1526 | 46 | 30 | 70 |
| SRR7690713 | 119 | 0 | 192 | 0 | 30 | 214 | 30 | 0 | 67 | 26 | 113 | 94 | 290 | 0 | 135 | 0 | 100 | 0 | 15 | 115 | 648 | 0 | 0 | 0 | 406 | 1388 | 852 | 0 | 0 | 0 | 58 | 0 | 56 | 1447 | 75 | 38 | 106 |
| SRR7690714 | 241 | 0 | 273 | 0 | 36 | 229 | 11 | 0 | 64 | 59 | 171 | 115 | 252 | 16 | 44 | 0 | 113 | 0 | 0 | 60 | 753 | 0 | 0 | 0 | 295 | 1229 | 296 | 12 | 0 | 0 | 62 | 0 | 38 | 973 | 41 | 24 | 59 |
| SRR7690715 | 398 | 0 | 1094 | 0 | 0 | 12 | 45 | 12 | 12 | 0 | 0 | 30 | 0 | 0 | 15 | 0 | 0 | 0 | 0 | 1243 | 1580 | 0 | 0 | 44 | 12 | 167 | 312 | 0 | 0 | 0 | 12 | 0 | 0 | 0 | 12 | 0 | 0 |
| SRR7690716 | 253 | 11 | 666 | 0 | 0 | 10 | 69 | 12 | 11 | 0 | 0 | 33 | 0 | 0 | 0 | 0 | 0 | 0 | 0 | 1404 | 2330 | 0 | 0 | 32 | 25 | 133 | 243 | 0 | 0 | 27 | 10 | 0 | 0 | 14 | 26 | 0 | 11 |
| SRR7690717 | 166 | 0 | 89 | 0 | 66 | 129 | 13 | 0 | 75 | 15 | 73 | 156 | 251 | 14 | 87 | 0 | 76 | 0 | 0 | 0 | 404 | 0 | 0 | 0 | 266 | 1032 | 171 | 0 | 0 | 0 | 59 | 0 | 0 | 1795 | 44 | 34 | 67 |
| SRR7690718 | 132 | 0 | 46 | 12 | 91 | 78 | 41 | 0 | 18 | 23 | 129 | 120 | 0 | 19 | 82 | 0 | 71 | 63 | 0 | 19 | 450 | 0 | 0 | 12 | 222 | 427 | 244 | 15 | 0 | 0 | 28 | 0 | 0 | 366 | 0 | 0 | 16 |
| SRR7690719 | 142 | 0 | 152 | 0 | 33 | 39 | 21 | 0 | 13 | 18 | 18 | 118 | 37 | 14 | 33 | 0 | 139 | 0 | 0 | 21 | 258 | 0 | 0 | 0 | 118 | 291 | 85 | 0 | 0 | 0 | 17 | 0 | 26 | 138 | 14 | 11 | 21 |
| SRR7690720 | 115 | 0 | 271 | 0 | 31 | 33 | 30 | 0 | 37 | 22 | 16 | 76 | 47 | 10 | 37 | 0 | 147 | 0 | 0 | 37 | 573 | 0 | 0 | 0 | 91 | 255 | 132 | 0 | 0 | 0 | 21 | 0 | 13 | 235 | 16 | 47 | 18 |
| SRR7690721 | 164 | 0 | 369 | 0 | 16 | 57 | 56 | 0 | 26 | 15 | 40 | 97 | 22 | 13 | 19 | 0 | 158 | 0 | 25 | 119 | 393 | 0 | 0 | 0 | 118 | 503 | 324 | 14 | 10 | 0 | 38 | 0 | 23 | 290 | 14 | 11 | 13 |
| SRR7690722 | 60 | 13 | 23 | 0 | 0 | 21 | 170 | 0 | 21 | 0 | 75 | 76 | 13 | 0 | 256 | 52 | 15 | 0 | 10 | 40 | 1810 | 41 | 0 | 0 | 180 | 130 | 16 | 11 | 0 | 30 | 20 | 49 | 15 | 128 | 30 | 22 | 65 |
| SRR7690723 | 69 | 15 | 13 | 0 | 0 | 141 | 0 | 0 | 246 | 10 | 69 | 87 | 203 | 12 | 2597 | 59 | 21 | 0 | 14 | 14 | 2754 | 27 | 0 | 0 | 187 | 1006 | 85 | 18 | 0 | 0 | 266 | 77 | 241 | 1592 | 494 | 166 | 683 |
| SRR7690724 | 42 | 0 | 24 | 0 | 15 | 13 | 68 | 12 | 176 | 32 | 149 | 44 | 19 | 17 | 4464 | 37 | 28 | 0 | 0 | 0 | 769 | 20 | 0 | 0 | 136 | 76 | 107 | 15 | 30 | 15 | 206 | 42 | 11 | 259 | 28 | 23 | 71 |
| SRR7690726 | 60 | 16 | 34 | 0 | 10 | 302 | 101 | 0 | 295 | 14 | 67 | 132 | 708 | 10 | 90 | 30 | 34 | 0 | 0 | 0 | 890 | 10 | 0 | 10 | 253 | 1961 | 263 | 12 | 14 | 24 | 208 | 24 | 192 | 3594 | 569 | 342 | 1068 |
| SRR7690727 | 112 | 0 | 71 | 0 | 50 | 132 | 0 | 0 | 40 | 21 | 111 | 132 | 102 | 22 | 78 | 0 | 126 | 0 | 0 | 0 | 232 | 0 | 0 | 0 | 327 | 758 | 248 | 14 | 0 | 0 | 57 | 0 | 0 | 950 | 54 | 29 | 73 |
| SRR7690728 | 166 | 0 | 53 | 0 | 61 | 77 | 14 | 0 | 39 | 26 | 104 | 187 | 117 | 16 | 18 | 0 | 61 | 15 | 0 | 11 | 317 | 0 | 0 | 0 | 342 | 510 | 100 | 0 | 0 | 0 | 58 | 0 | 0 | 995 | 37 | 41 | 51 |
| SRR7690729 | 95 | 0 | 111 | 0 | 39 | 44 | 23 | 0 | 40 | 25 | 21 | 131 | 42 | 12 | 69 | 0 | 136 | 0 | 0 | 0 | 352 | 0 | 0 | 0 | 57 | 247 | 27 | 0 | 0 | 0 | 22 | 0 | 0 | 215 | 34 | 20 | 47 |
| SRR7690730 | 479 | 0 | 151 | 17 | 76 | 107 | 23 | 0 | 40 | 65 | 188 | 65 | 424 | 23 | 23 | 0 | 71 | 0 | 0 | 11 | 364 | 0 | 0 | 0 | 419 | 692 | 429 | 0 | 0 | 0 | 50 | 0 | 0 | 882 | 76 | 23 | 89 |
| SRR7690731 | 104 | 0 | 144 | 14 | 119 | 64 | 17 | 0 | 23 | 50 | 202 | 97 | 221 | 16 | 72 | 0 | 88 | 0 | 0 | 13 | 89 | 0 | 0 | 0 | 491 | 492 | 215 | 10 | 0 | 0 | 42 | 0 | 0 | 1777 | 35 | 32 | 69 |
| SRR7690732 | 146 | 0 | 193 | 0 | 11 | 30 | 22 | 0 | 16 | 37 | 38 | 82 | 25 | 11 | 44 | 0 | 92 | 0 | 27 | 16 | 223 | 0 | 0 | 0 | 91 | 217 | 195 | 0 | 0 | 0 | 17 | 0 | 34 | 105 | 39 | 19 | 42 |
| SRR7690733 | 102 | 0 | 211 | 0 | 11 | 27 | 18 | 0 | 11 | 27 | 23 | 59 | 31 | 21 | 72 | 0 | 134 | 74 | 36 | 36 | 306 | 0 | 0 | 11 | 91 | 164 | 78 | 11 | 0 | 0 | 19 | 0 | 11 | 112 | 20 | 36 | 17 |
| SRR7690734 | 64 | 0 | 25 | 0 | 0 | 135 | 26 | 0 | 229 | 14 | 36 | 121 | 88 | 25 | 634 | 29 | 40 | 0 | 0 | 13 | 3013 | 18 | 0 | 0 | 100 | 932 | 35 | 0 | 0 | 11 | 157 | 0 | 114 | 1588 | 483 | 142 | 541 |
| SRR7690736 | 108 | 0 | 42 | 0 | 11 | 656 | 162 | 0 | 888 | 0 | 20 | 176 | 182 | 0 | 44 | 34 | 19 | 0 | 0 | 96 | 6068 | 33 | 0 | 0 | 108 | 4071 | 79 | 0 | 0 | 13 | 380 | 21 | 66 | 1324 | 1042 | 564 | 1542 |
| SRR7690737 | 58 | 0 | 39 | 0 | 0 | 379 | 30 | 0 | 637 | 0 | 42 | 127 | 555 | 0 | 209 | 11 | 52 | 0 | 0 | 22 | 536 | 11 | 0 | 0 | 280 | 2778 | 54 | 25 | 20 | 20 | 919 | 0 | 175 | 3794 | 1273 | 586 | 1565 |
| SRR7690739 | 59 | 0 | 29 | 0 | 0 | 612 | 44 | 0 | 672 | 0 | 42 | 110 | 1263 | 0 | 310 | 27 | 41 | 0 | 0 | 13 | 1063 | 0 | 0 | 0 | 205 | 3268 | 253 | 12 | 12 | 48 | 573 | 21 | 106 | 3217 | 892 | 609 | 1827 |
| SRR7690740 | 74 | 0 | 22 | 0 | 0 | 767 | 84 | 10 | 671 | 0 | 30 | 126 | 1228 | 0 | 44 | 13 | 42 | 0 | 0 | 0 | 1165 | 10 | 0 | 0 | 338 | 3488 | 557 | 0 | 0 | 50 | 494 | 15 | 91 | 3278 | 973 | 580 | 1934 |
| SRR7690741 | 216 | 0 | 1037 | 0 | 0 | 37 | 10 | 0 | 18 | 0 | 0 | 35 | 0 | 0 | 17 | 0 | 20 | 0 | 0 | 895 | 563 | 0 | 0 | 17 | 25 | 132 | 189 | 0 | 0 | 0 | 13 | 0 | 0 | 0 | 14 | 0 | 31 |
| SRR7690742 | 226 | 0 | 1143 | 0 | 0 | 33 | 15 | 0 | 14 | 0 | 0 | 41 | 0 | 0 | 15 | 0 | 23 | 0 | 0 | 816 | 578 | 0 | 0 | 14 | 30 | 98 | 164 | 0 | 0 | 0 | 16 | 0 | 0 | 0 | 45 | 0 | 25 |
| SRR7690743 | 265 | 0 | 1122 | 0 | 0 | 35 | 17 | 0 | 18 | 0 | 0 | 41 | 0 | 0 | 34 | 0 | 21 | 0 | 0 | 1025 | 693 | 0 | 0 | 44 | 43 | 701 | 682 | 0 | 0 | 0 | 15 | 0 | 0 | 0 | 43 | 0 | 27 |
| SRR7690744 | 110 | 0 | 141 | 0 | 22 | 48 | 35 | 0 | 12 | 13 | 10 | 90 | 11 | 0 | 91 | 0 | 47 | 0 | 15 | 79 | 188 | 0 | 0 | 15 | 119 | 336 | 266 | 0 | 0 | 0 | 15 | 0 | 13 | 11 | 11 | 50 | 19 |
| SRR7690745 | 83 | 0 | 92 | 0 | 22 | 17 | 14 | 0 | 69 | 10 | 34 | 83 | 62 | 11 | 72 | 17 | 117 | 20 | 0 | 19 | 477 | 0 | 0 | 0 | 60 | 172 | 60 | 0 | 0 | 0 | 16 | 0 | 57 | 19 | 16 | 47 | 10 |
| SRR7690746 | 93 | 0 | 77 | 0 | 16 | 34 | 44 | 0 | 17 | 0 | 10 | 66 | 13 | 15 | 171 | 0 | 33 | 0 | 22 | 136 | 343 | 0 | 0 | 19 | 20 | 224 | 63 | 0 | 0 | 0 | 14 | 0 | 0 | 16 | 11 | 27 | 15 |
| SRR7690747 | 82 | 0 | 103 | 0 | 38 | 15 | 0 | 0 | 30 | 17 | 16 | 100 | 14 | 11 | 86 | 0 | 168 | 0 | 0 | 15 | 76 | 0 | 0 | 0 | 113 | 119 | 66 | 0 | 0 | 0 | 13 | 0 | 0 | 108 | 43 | 26 | 11 |
| SRR7690748 | 110 | 0 | 325 | 0 | 62 | 157 | 21 | 0 | 37 | 33 | 114 | 88 | 361 | 0 | 25 | 0 | 56 | 40 | 13 | 89 | 693 | 0 | 0 | 10 | 355 | 1116 | 930 | 0 | 0 | 0 | 39 | 0 | 26 | 1922 | 40 | 49 | 113 |
| SRR7690749 | 247 | 0 | 257 | 0 | 58 | 139 | 12 | 0 | 40 | 30 | 94 | 191 | 76 | 12 | 26 | 0 | 35 | 55 | 0 | 17 | 285 | 0 | 0 | 0 | 434 | 681 | 328 | 0 | 0 | 0 | 66 | 0 | 19 | 631 | 23 | 14 | 23 |
| SRR7690750 | 96 | 0 | 83 | 0 | 47 | 159 | 40 | 0 | 42 | 12 | 76 | 105 | 97 | 18 | 51 | 0 | 67 | 0 | 0 | 19 | 243 | 0 | 0 | 0 | 282 | 891 | 345 | 0 | 0 | 0 | 51 | 0 | 0 | 814 | 50 | 34 | 56 |
| SRR7690751 | 71 | 0 | 98 | 0 | 17 | 44 | 24 | 0 | 11 | 20 | 28 | 106 | 32 | 16 | 74 | 0 | 52 | 0 | 0 | 47 | 285 | 0 | 0 | 13 | 103 | 285 | 161 | 0 | 0 | 0 | 20 | 0 | 0 | 352 | 28 | 27 | 12 |
| SRR7690752 | 70 | 0 | 125 | 0 | 12 | 20 | 17 | 0 | 29 | 23 | 23 | 84 | 48 | 16 | 41 | 0 | 98 | 0 | 0 | 0 | 151 | 0 | 0 | 0 | 82 | 138 | 34 | 0 | 0 | 0 | 12 | 0 | 0 | 418 | 11 | 30 | 13 |
| SRR7690753 | 122 | 0 | 251 | 0 | 30 | 65 | 47 | 0 | 31 | 27 | 24 | 87 | 41 | 14 | 81 | 10 | 55 | 0 | 19 | 123 | 1043 | 0 | 0 | 17 | 147 | 537 | 498 | 0 | 0 | 0 | 18 | 0 | 20 | 368 | 18 | 12 | 21 |
| SRR7690754 | 185 | 0 | 241 | 0 | 17 | 58 | 32 | 0 | 13 | 24 | 12 | 87 | 98 | 15 | 77 | 15 | 100 | 0 | 0 | 83 | 849 | 0 | 0 | 0 | 96 | 372 | 197 | 13 | 11 | 0 | 20 | 0 | 13 | 393 | 21 | 14 | 32 |
| SRR7690755 | 270 | 0 | 532 | 0 | 44 | 208 | 71 | 0 | 22 | 35 | 165 | 70 | 82 | 17 | 14 | 0 | 144 | 0 | 43 | 206 | 768 | 0 | 0 | 0 | 391 | 2245 | 1458 | 0 | 12 | 0 | 188 | 0 | 167 | 924 | 68 | 55 | 97 |
| SRR7690756 | 120 | 0 | 363 | 0 | 23 | 20 | 10 | 0 | 37 | 18 | 14 | 78 | 64 | 12 | 51 | 0 | 128 | 12 | 10 | 31 | 225 | 0 | 0 | 11 | 85 | 243 | 67 | 0 | 0 | 0 | 23 | 0 | 44 | 212 | 12 | 47 | 14 |
| SRR7690757 | 65 | 0 | 222 | 0 | 33 | 126 | 34 | 0 | 21 | 20 | 85 | 39 | 245 | 12 | 28 | 0 | 118 | 0 | 45 | 214 | 304 | 0 | 0 | 17 | 480 | 1010 | 1930 | 0 | 0 | 0 | 59 | 0 | 36 | 1044 | 63 | 28 | 64 |
| SRR7690758 | 33 | 0 | 17 | 0 | 0 | 15 | 39 | 0 | 15 | 0 | 31 | 68 | 35 | 0 | 264 | 14 | 19 | 0 | 0 | 0 | 223 | 14 | 0 | 0 | 331 | 58 | 216 | 0 | 0 | 0 | 13 | 38 | 52 | 244 | 22 | 20 | 52 |
| SRR7690759 | 119 | 0 | 44 | 0 | 13 | 49 | 25 | 0 | 40 | 11 | 53 | 113 | 148 | 0 | 183 | 40 | 14 | 0 | 17 | 133 | 7967 | 32 | 0 | 0 | 72 | 297 | 88 | 24 | 14 | 13 | 33 | 21 | 138 | 105 | 71 | 41 | 141 |
| SRR7690762 | 60 | 20 | 21 | 0 | 0 | 16 | 30 | 0 | 16 | 0 | 60 | 103 | 47 | 0 | 129 | 37 | 36 | 0 | 0 | 0 | 1425 | 20 | 0 | 11 | 214 | 88 | 219 | 0 | 11 | 57 | 19 | 20 | 157 | 124 | 38 | 20 | 65 |
| SRR7690764 | 388 | 0 | 1216 | 10 | 0 | 13 | 12 | 21 | 15 | 0 | 0 | 34 | 0 | 0 | 0 | 0 | 16 | 0 | 0 | 1780 | 836 | 0 | 0 | 46 | 15 | 221 | 479 | 10 | 0 | 0 | 16 | 0 | 0 | 13 | 21 | 0 | 17 |
| SRR7690765 | 329 | 0 | 974 | 0 | 0 | 33 | 24 | 0 | 15 | 0 | 0 | 36 | 0 | 24 | 0 | 0 | 20 | 0 | 0 | 932 | 1010 | 0 | 0 | 114 | 44 | 600 | 924 | 0 | 0 | 0 | 13 | 0 | 0 | 0 | 15 | 0 | 0 |
| SRR7690766 | 96 | 0 | 92 | 0 | 42 | 22 | 20 | 0 | 30 | 27 | 18 | 118 | 34 | 13 | 333 | 0 | 78 | 0 | 0 | 57 | 430 | 0 | 0 | 17 | 81 | 148 | 108 | 0 | 0 | 0 | 41 | 0 | 0 | 360 | 39 | 24 | 11 |
| SRR7690767 | 92 | 0 | 69 | 0 | 46 | 101 | 48 | 0 | 39 | 18 | 75 | 94 | 0 | 24 | 157 | 0 | 83 | 12 | 0 | 22 | 377 | 0 | 0 | 0 | 264 | 628 | 202 | 10 | 0 | 0 | 55 | 0 | 0 | 514 | 29 | 32 | 42 |
| SRR7690768 | 132 | 0 | 244 | 0 | 31 | 24 | 13 | 0 | 11 | 28 | 12 | 71 | 36 | 0 | 45 | 0 | 86 | 0 | 0 | 58 | 391 | 0 | 0 | 0 | 80 | 188 | 73 | 0 | 0 | 0 | 12 | 0 | 12 | 186 | 12 | 43 | 12 |
| SRR7690769 | 57 | 0 | 17 | 0 | 0 | 16 | 29 | 0 | 15 | 0 | 50 | 83 | 87 | 11 | 3164 | 18 | 30 | 0 | 15 | 29 | 4549 | 57 | 0 | 0 | 100 | 83 | 130 | 0 | 0 | 0 | 14 | 32 | 11 | 34 | 30 | 13 | 61 |
| SRR7690770 | 85 | 11 | 32 | 0 | 0 | 654 | 32 | 0 | 559 | 11 | 60 | 87 | 333 | 16 | 216 | 38 | 25 | 0 | 0 | 74 | 1478 | 29 | 0 | 11 | 122 | 3408 | 211 | 16 | 0 | 24 | 293 | 44 | 180 | 1586 | 745 | 488 | 1706 |
| SRR7690773 | 47 | 0 | 28 | 0 | 11 | 247 | 0 | 0 | 285 | 15 | 49 | 60 | 17 | 0 | 12956 | 23 | 21 | 0 | 0 | 0 | 1058 | 11 | 0 | 0 | 13 | 65 | 57 | 19 | 0 | 0 | 307 | 15 | 205 | 70 | 24 | 23 | 47 |
| SRR7690775 | 155 | 0 | 247 | 0 | 21 | 196 | 30 | 0 | 56 | 27 | 76 | 94 | 75 | 0 | 71 | 0 | 302 | 0 | 0 | 61 | 420 | 0 | 0 | 0 | 932 | 2047 | 1623 | 0 | 0 | 0 | 144 | 0 | 29 | 47 | 68 | 31 | 59 |
| SRR7690777 | 46 | 0 | 112 | 0 | 23 | 158 | 27 | 0 | 67 | 0 | 30 | 79 | 66 | 13 | 19 | 0 | 190 | 0 | 0 | 49 | 353 | 0 | 0 | 0 | 521 | 1534 | 661 | 0 | 0 | 0 | 88 | 0 | 0 | 55 | 129 | 34 | 119 |
| SRR7690778 | 552 | 0 | 344 | 12 | 32 | 103 | 24 | 0 | 19 | 12 | 38 | 155 | 28 | 10 | 0 | 0 | 130 | 133 | 28 | 55 | 241 | 0 | 0 | 16 | 234 | 565 | 261 | 0 | 0 | 0 | 27 | 0 | 16 | 201 | 14 | 10 | 16 |
| SRR7690779 | 270 | 0 | 335 | 0 | 29 | 167 | 201 | 0 | 16 | 21 | 109 | 29 | 21 | 47 | 52 | 0 | 31 | 0 | 30 | 201 | 394 | 0 | 0 | 0 | 136 | 1912 | 783 | 0 | 10 | 0 | 266 | 0 | 142 | 925 | 44 | 49 | 45 |
| SRR7690780 | 220 | 0 | 309 | 0 | 17 | 39 | 26 | 0 | 35 | 16 | 14 | 125 | 102 | 0 | 48 | 0 | 210 | 10 | 13 | 52 | 361 | 0 | 0 | 0 | 139 | 389 | 181 | 0 | 0 | 0 | 20 | 0 | 31 | 188 | 14 | 38 | 16 |
| SRR7690781 | 105 | 0 | 192 | 0 | 27 | 64 | 25 | 0 | 13 | 21 | 53 | 109 | 58 | 0 | 53 | 0 | 100 | 0 | 0 | 56 | 359 | 0 | 0 | 0 | 63 | 443 | 254 | 0 | 0 | 0 | 14 | 0 | 34 | 440 | 18 | 11 | 26 |
| SRR7690783 | 57 | 0 | 23 | 0 | 0 | 398 | 81 | 0 | 307 | 0 | 32 | 80 | 155 | 0 | 392 | 0 | 21 | 0 | 14 | 41 | 4340 | 0 | 0 | 19 | 280 | 1821 | 443 | 0 | 0 | 23 | 383 | 0 | 30 | 2491 | 678 | 487 | 1385 |
| SRR7690788 | 116 | 0 | 53 | 0 | 0 | 39 | 55 | 16 | 24 | 0 | 36 | 139 | 80 | 0 | 45 | 0 | 62 | 0 | 0 | 32 | 1773 | 11 | 0 | 0 | 19 | 214 | 22 | 0 | 0 | 59 | 16 | 17 | 47 | 482 | 69 | 26 | 100 |
| SRR7690789 | 62 | 0 | 25 | 0 | 10 | 17 | 53 | 19 | 16 | 0 | 40 | 92 | 34 | 0 | 13 | 15 | 34 | 0 | 0 | 11 | 598 | 34 | 0 | 42 | 88 | 118 | 30 | 10 | 10 | 82 | 159 | 40 | 46 | 324 | 30 | 44 | 112 |
| SRR7690790 | 170 | 0 | 256 | 0 | 11 | 447 | 43 | 0 | 66 | 0 | 26 | 118 | 65 | 0 | 29 | 0 | 252 | 0 | 11 | 124 | 1086 | 0 | 0 | 10 | 559 | 3626 | 4172 | 0 | 0 | 0 | 69 | 0 | 14 | 52 | 53 | 46 | 102 |
| SRR7690791 | 151 | 0 | 90 | 0 | 22 | 225 | 0 | 0 | 52 | 0 | 41 | 141 | 0 | 0 | 79 | 0 | 159 | 0 | 0 | 56 | 530 | 0 | 0 | 26 | 611 | 1755 | 1179 | 0 | 0 | 0 | 99 | 0 | 0 | 94 | 56 | 57 | 80 |
| SRR7690792 | 53 | 0 | 92 | 0 | 29 | 161 | 13 | 0 | 32 | 16 | 41 | 82 | 40 | 10 | 241 | 0 | 116 | 0 | 0 | 29 | 349 | 0 | 0 | 0 | 502 | 1358 | 525 | 0 | 0 | 0 | 86 | 0 | 28 | 69 | 90 | 29 | 73 |
| SRR7690793 | 231 | 0 | 173 | 0 | 39 | 156 | 10 | 0 | 49 | 26 | 106 | 114 | 108 | 15 | 104 | 0 | 45 | 15 | 0 | 13 | 534 | 0 | 0 | 0 | 303 | 880 | 144 | 0 | 0 | 0 | 59 | 0 | 64 | 530 | 18 | 22 | 32 |
| SRR7690794 | 308 | 0 | 282 | 0 | 13 | 66 | 0 | 0 | 32 | 12 | 32 | 157 | 78 | 0 | 111 | 0 | 59 | 37 | 0 | 16 | 483 | 0 | 0 | 0 | 188 | 363 | 33 | 0 | 0 | 0 | 40 | 0 | 19 | 606 | 13 | 16 | 10 |
| SRR7690795 | 355 | 0 | 352 | 0 | 20 | 103 | 35 | 0 | 20 | 15 | 18 | 164 | 52 | 0 | 53 | 0 | 61 | 62 | 26 | 188 | 1409 | 0 | 0 | 32 | 206 | 554 | 362 | 0 | 0 | 14 | 20 | 0 | 12 | 368 | 14 | 0 | 0 |
| SRR7690796 | 143 | 0 | 184 | 0 | 29 | 217 | 32 | 0 | 40 | 21 | 64 | 108 | 361 | 0 | 187 | 17 | 100 | 0 | 11 | 49 | 649 | 0 | 0 | 0 | 427 | 1441 | 911 | 0 | 0 | 0 | 83 | 0 | 50 | 1658 | 83 | 33 | 103 |
| SRR7690797 | 42 | 0 | 12 | 0 | 0 | 31 | 79 | 0 | 39 | 0 | 11 | 80 | 44 | 0 | 2335 | 15 | 18 | 0 | 0 | 290 | 5541 | 11 | 0 | 22 | 251 | 126 | 137 | 11 | 26 | 0 | 36 | 0 | 77 | 196 | 23 | 27 | 65 |
| SRR7690798 | 60 | 0 | 29 | 0 | 0 | 14 | 53 | 0 | 13 | 0 | 47 | 104 | 11 | 0 | 7070 | 0 | 28 | 0 | 0 | 15 | 2498 | 28 | 0 | 11 | 201 | 81 | 170 | 11 | 0 | 25 | 14 | 39 | 31 | 70 | 28 | 17 | 69 |
| SRR7690800 | 61 | 14 | 19 | 0 | 0 | 17 | 43 | 10 | 14 | 0 | 106 | 70 | 22 | 19 | 496 | 34 | 20 | 0 | 0 | 97 | 4646 | 48 | 0 | 0 | 117 | 81 | 134 | 22 | 0 | 20 | 14 | 37 | 19 | 167 | 37 | 15 | 70 |
| SRR7690801 | 41 | 0 | 36 | 0 | 0 | 304 | 41 | 0 | 389 | 0 | 25 | 93 | 748 | 17 | 70 | 25 | 50 | 0 | 12 | 0 | 397 | 0 | 0 | 0 | 283 | 2237 | 169 | 17 | 0 | 0 | 499 | 0 | 140 | 2791 | 1091 | 643 | 1844 |
| SRR7690802 | 77 | 0 | 21 | 0 | 11 | 142 | 29 | 0 | 10 | 13 | 70 | 110 | 174 | 20 | 1129 | 70 | 16 | 14 | 15 | 15 | 1524 | 0 | 0 | 0 | 11 | 34 | 20 | 22 | 0 | 0 | 16 | 24 | 215 | 47 | 20 | 11 | 25 |
| SRR7690803 | 81 | 0 | 32 | 0 | 0 | 414 | 91 | 0 | 321 | 17 | 68 | 81 | 265 | 0 | 906 | 38 | 17 | 0 | 27 | 36 | 3506 | 0 | 0 | 0 | 316 | 2359 | 276 | 13 | 11 | 0 | 271 | 34 | 212 | 1239 | 415 | 269 | 667 |
| SRR7819267 | 75 | 77 | 0 | 26 | 21 | 0 | 0 | 0 | 0 | 36 | 421 | 172 | 0 | 21 | 2570 | 0 | 45 | 0 | 0 | 0 | 0 | 0 | 0 | 0 | 21 | 0 | 0 | 15 | 0 | 0 | 421 | 0 | 36 | 0 | 0 | 0 | 0 |
| SRR7819269 | 878 | 0 | 0 | 0 | 29 | 0 | 0 | 0 | 0 | 29 | 29 | 979 | 0 | 86 | 236 | 0 | 95 | 288 | 0 | 0 | 59 | 0 | 0 | 0 | 0 | 0 | 0 | 0 | 0 | 0 | 57 | 0 | 86 | 29 | 0 | 0 | 0 |
| SRR7819272 | 572 | 72 | 0 | 39 | 39 | 0 | 0 | 0 | 0 | 0 | 196 | 560 | 0 | 0 | 1405 | 20 | 92 | 0 | 0 | 0 | 83 | 0 | 0 | 13 | 33 | 0 | 0 | 13 | 0 | 0 | 131 | 0 | 13 | 33 | 13 | 20 | 0 |
| SRR7851651 | 56 | 15 | 724 | 33 | 16 | 0 | 0 | 0 | 0 | 188 | 91 | 1923 | 16 | 754 | 16 | 191 | 21 | 0 | 0 | 38 | 0 | 0 | 127 | 0 | 1775 | 11 | 0 | 145 | 12 | 0 | 28 | 0 | 0 | 0 | 0 | 0 | 0 |
| SRR7851652 | 46 | 11 | 711 | 11 | 0 | 0 | 0 | 0 | 0 | 85 | 36 | 1921 | 0 | 606 | 23 | 70 | 19 | 0 | 0 | 70 | 0 | 0 | 64 | 0 | 753 | 0 | 0 | 68 | 13 | 0 | 11 | 0 | 0 | 0 | 0 | 0 | 0 |
| SRR7851653 | 35 | 19 | 623 | 0 | 0 | 0 | 0 | 0 | 0 | 123 | 51 | 1060 | 0 | 905 | 120 | 72 | 20 | 30 | 0 | 41 | 0 | 0 | 53 | 0 | 999 | 0 | 0 | 77 | 0 | 0 | 0 | 0 | 0 | 0 | 0 | 0 | 0 |
| SRR7851655 | 52 | 24 | 728 | 21 | 0 | 0 | 0 | 0 | 0 | 173 | 85 | 2098 | 10 | 1364 | 11 | 188 | 22 | 0 | 0 | 79 | 0 | 0 | 189 | 0 | 1960 | 0 | 0 | 209 | 0 | 0 | 14 | 0 | 0 | 0 | 0 | 0 | 0 |
| SRR7851656 | 47 | 18 | 732 | 15 | 0 | 0 | 0 | 0 | 0 | 63 | 28 | 1626 | 0 | 328 | 46 | 44 | 21 | 19 | 0 | 144 | 0 | 0 | 44 | 0 | 758 | 13 | 0 | 49 | 0 | 0 | 23 | 0 | 0 | 0 | 0 | 0 | 0 |
| SRR7901057 | 685 | 0 | 0 | 0 | 0 | 22 | 0 | 357 | 23 | 0 | 27 | 221 | 62 | 15 | 6836 | 0 | 0 | 0 | 19 | 30 | 551 | 14 | 0 | 0 | 0 | 207 | 10 | 0 | 0 | 278 | 61 | 12 | 0 | 6147 | 16 | 14 | 44 |
| SRR7901058 | 677 | 0 | 0 | 0 | 17 | 10 | 0 | 253 | 52 | 13 | 91 | 254 | 237 | 28 | 5515 | 0 | 0 | 0 | 0 | 38 | 484 | 21 | 0 | 0 | 0 | 514 | 0 | 0 | 0 | 139 | 116 | 0 | 11 | 9166 | 45 | 48 | 107 |
| SRR7901059 | 842 | 0 | 0 | 0 | 0 | 21 | 0 | 129 | 31 | 0 | 19 | 228 | 49 | 11 | 3306 | 0 | 0 | 0 | 10 | 127 | 550 | 0 | 0 | 0 | 0 | 224 | 0 | 0 | 0 | 71 | 52 | 0 | 0 | 2125 | 21 | 16 | 41 |
| SRR7903701 | 54 | 0 | 64 | 0 | 0 | 1078 | 1709 | 21 | 267 | 0 | 30 | 152 | 106 | 0 | 2460 | 0 | 253 | 0 | 97 | 553 | 8652 | 0 | 0 | 24 | 133 | 5107 | 930 | 0 | 0 | 0 | 777 | 11 | 0 | 42 | 1609 | 1923 | 4325 |
| SRR7903702 | 49 | 0 | 51 | 0 | 0 | 1008 | 1875 | 28 | 365 | 0 | 21 | 158 | 82 | 0 | 2789 | 0 | 236 | 0 | 76 | 477 | 7824 | 0 | 0 | 38 | 123 | 4379 | 950 | 0 | 0 | 18 | 800 | 12 | 0 | 62 | 1841 | 1490 | 3924 |
| SRR7903703 | 53 | 0 | 62 | 0 | 0 | 941 | 1363 | 15 | 277 | 0 | 26 | 166 | 89 | 0 | 2441 | 0 | 281 | 0 | 80 | 492 | 7055 | 0 | 0 | 26 | 149 | 4451 | 1313 | 0 | 0 | 0 | 689 | 0 | 0 | 33 | 1780 | 1684 | 3760 |
| SRR7903704 | 611 | 0 | 0 | 0 | 0 | 166 | 59 | 243 | 992 | 0 | 0 | 59 | 18 | 12 | 110 | 0 | 93 | 0 | 0 | 41 | 3087 | 10 | 0 | 275 | 0 | 6016 | 81 | 11 | 0 | 169 | 585 | 32 | 0 | 970 | 258 | 125 | 413 |
| SRR7903705 | 678 | 0 | 0 | 0 | 0 | 166 | 192 | 167 | 858 | 0 | 0 | 71 | 25 | 11 | 93 | 0 | 93 | 0 | 11 | 101 | 3960 | 0 | 0 | 223 | 0 | 6049 | 49 | 0 | 0 | 107 | 599 | 18 | 0 | 404 | 288 | 139 | 392 |
| SRR7903706 | 927 | 0 | 0 | 0 | 0 | 144 | 85 | 165 | 631 | 0 | 0 | 57 | 18 | 0 | 53 | 0 | 115 | 0 | 0 | 61 | 2714 | 0 | 0 | 295 | 0 | 6852 | 21 | 0 | 0 | 113 | 422 | 0 | 0 | 909 | 281 | 120 | 428 |
| SRR8063607 | 134 | 0 | 64 | 0 | 0 | 288 | 0 | 0 | 171 | 0 | 0 | 11910 | 233 | 0 | 205 | 0 | 1092 | 32 | 60 | 72 | 1362 | 0 | 0 | 15 | 219 | 1451 | 908 | 18 | 0 | 0 | 550 | 0 | 0 | 918 | 10861 | 1764 | 9654 |
| SRR8063608 | 139 | 0 | 51 | 0 | 0 | 33 | 0 | 0 | 30 | 0 | 0 | 12473 | 16 | 0 | 147 | 0 | 1008 | 0 | 0 | 147 | 1588 | 0 | 0 | 0 | 25 | 159 | 86 | 39 | 0 | 19 | 57 | 0 | 0 | 71 | 1177 | 205 | 1025 |
| SRR8063609 | 163 | 0 | 70 | 0 | 0 | 41 | 0 | 0 | 18 | 0 | 0 | 12661 | 21 | 0 | 127 | 0 | 983 | 0 | 38 | 331 | 2139 | 0 | 0 | 0 | 22 | 189 | 253 | 22 | 0 | 0 | 70 | 0 | 0 | 88 | 1099 | 246 | 1009 |
| SRR8063610 | 131 | 29 | 51 | 0 | 0 | 18 | 100 | 15 | 332 | 0 | 46 | 8275 | 13 | 13 | 301 | 0 | 963 | 16 | 14 | 104 | 1723 | 0 | 0 | 0 | 12 | 139 | 20 | 0 | 0 | 21 | 11 | 17 | 0 | 154 | 101 | 47 | 200 |
| SRR8063611 | 141 | 16 | 53 | 0 | 0 | 21 | 103 | 32 | 323 | 0 | 37 | 8289 | 11 | 12 | 217 | 0 | 858 | 12 | 13 | 95 | 2098 | 0 | 0 | 0 | 48 | 133 | 19 | 0 | 0 | 0 | 11 | 16 | 0 | 118 | 97 | 44 | 179 |
| SRR8063612 | 132 | 13 | 61 | 0 | 0 | 20 | 113 | 20 | 299 | 0 | 50 | 9054 | 44 | 0 | 193 | 0 | 867 | 0 | 49 | 170 | 2494 | 0 | 0 | 13 | 44 | 136 | 32 | 0 | 0 | 13 | 10 | 19 | 0 | 163 | 85 | 44 | 157 |
| SRR8063613 | 206 | 50 | 0 | 0 | 34 | 0 | 192 | 34 | 754 | 0 | 0 | 15246 | 0 | 0 | 256 | 34 | 125 | 0 | 192 | 128 | 1215 | 34 | 0 | 0 | 50 | 117 | 117 | 0 | 0 | 0 | 67 | 34 | 0 | 1492 | 168 | 101 | 251 |
| SRR8063614 | 153 | 0 | 34 | 0 | 0 | 0 | 194 | 248 | 13 | 0 | 22 | 18028 | 0 | 0 | 141 | 0 | 264 | 703 | 317 | 581 | 968 | 0 | 0 | 60 | 14 | 18 | 61 | 0 | 0 | 106 | 14 | 0 | 0 | 32 | 18 | 22 | 11 |
| SRR8063615 | 162 | 16 | 44 | 0 | 25 | 16 | 114 | 0 | 16 | 0 | 16 | 18307 | 0 | 25 | 284 | 0 | 297 | 739 | 57 | 0 | 512 | 0 | 0 | 0 | 17 | 41 | 21 | 0 | 0 | 0 | 25 | 25 | 0 | 46 | 23 | 25 | 13 |
| SRR8074870 | 27 | 0 | 20 | 32 | 0 | 0 | 0 | 0 | 0 | 0 | 69 | 510 | 0 | 13 | 0 | 176 | 29 | 170 | 0 | 0 | 0 | 0 | 0 | 0 | 169 | 0 | 0 | 146 | 0 | 0 | 181 | 15 | 0 | 0 | 0 | 0 | 13 |
| SRR8074871 | 26 | 0 | 22 | 32 | 0 | 0 | 0 | 0 | 0 | 0 | 32 | 537 | 0 | 287 | 0 | 154 | 26 | 170 | 0 | 0 | 12 | 0 | 0 | 0 | 163 | 0 | 0 | 149 | 0 | 0 | 190 | 15 | 0 | 0 | 12 | 0 | 18 |

### Supp Table S8.2.Columns_AM_to_BW

| | | | | | | | | | | | | | | | | | | | | | | | | | | | | | | | | | | | | | |
|---|---|---|---|---|---|---|---|---|---|---|---|---|---|---|---|---|---|---|---|---|---|---|---|---|---|---|---|---|---|---|---|---|---|---|---|---|---|
| SRR8074873 | 31 | 0 | 16 | 17 | 0 | 0 | 0 | 0 | 0 | 0 | 37 | 677 | 0 | 174 | 0 | 127 | 27 | 80 | 0 | 0 | 0 | 0 | 0 | 0 | 88 | 0 | 0 | 96 | 0 | 0 | 103 | 21 | 0 | 0 | 0 | 0 | 48 |
| SRR8074874 | 25 | 0 | 0 | 11 | 0 | 0 | 12 | 0 | 23 | 0 | 29 | 763 | 0 | 115 | 0 | 71 | 28 | 46 | 0 | 39 | 42 | 0 | 0 | 0 | 52 | 0 | 0 | 55 | 0 | 0 | 70 | 25 | 0 | 0 | 14 | 0 | 30 |
| SRR8074875 | 26 | 0 | 0 | 14 | 0 | 0 | 0 | 0 | 11 | 0 | 29 | 759 | 0 | 103 | 0 | 67 | 31 | 67 | 0 | 26 | 35 | 0 | 0 | 0 | 77 | 0 | 0 | 58 | 0 | 0 | 91 | 31 | 0 | 0 | 0 | 0 | 27 |
| SRR8094756 | 161 | 34 | 560 | 14 | 21 | 18 | 133 | 0 | 11 | 43 | 144 | 6767 | 58 | 44 | 9587 | 103 | 76 | 0 | 42 | 233 | 2534 | 46 | 33 | 39 | 27 | 217 | 19 | 0 | 18 | 0 | 21 | 31 | 34 | 42 | 43 | 280 | 252 |
| SRR8094757 | 155 | 0 | 341 | 0 | 0 | 82 | 143 | 107 | 86 | 93 | 209 | 6799 | 15 | 61 | 1938 | 11 | 78 | 0 | 26 | 129 | 3623 | 0 | 0 | 144 | 74 | 674 | 170 | 27 | 604 | 62 | 28 | 0 | 0 | 7775 | 408 | 1807 | 3977 |
| SRR8094758 | 43 | 0 | 116 | 0 | 12 | 23 | 144 | 11 | 12 | 15 | 10 | 3907 | 26 | 70 | 28 | 18 | 0 | 0 | 66 | 169 | 511 | 39 | 42 | 17 | 36 | 203 | 50 | 51 | 15 | 44 | 26 | 63 | 33 | 43 | 44 | 306 | 585 |
| SRR8094759 | 182 | 0 | 99 | 0 | 15 | 44 | 5469 | 13 | 29 | 0 | 14 | 20 | 0 | 13 | 22 | 18 | 17 | 0 | 47 | 202 | 3804 | 12 | 0 | 47 | 29 | 408 | 28 | 13 | 14 | 0 | 15 | 26 | 0 | 97 | 14 | 54 | 98 |
| SRR8094760 | 60 | 0 | 42 | 0 | 65 | 34 | 1341 | 0 | 69 | 27 | 20 | 42 | 0 | 27 | 150 | 13 | 0 | 22 | 70 | 100 | 11096 | 11 | 0 | 0 | 23 | 222 | 30 | 29 | 0 | 0 | 20 | 65 | 38 | 17 | 31 | 82 | 159 |
| SRR8094761 | 125 | 0 | 0 | 0 | 0 | 44 | 294 | 39 | 29 | 0 | 24 | 38 | 255 | 12 | 570 | 12 | 18 | 1431 | 78 | 32 | 3943 | 0 | 0 | 12 | 24 | 316 | 95 | 0 | 0 | 59 | 46 | 0 | 0 | 31 | 39 | 103 | 198 |
| SRR8094762 | 118 | 0 | 87 | 0 | 15 | 1030 | 2913 | 74 | 408 | 0 | 71 | 90 | 20 | 18 | 94 | 13 | 20 | 199 | 1210 | 594 | 4651 | 0 | 0 | 64 | 35 | 6877 | 86 | 0 | 0 | 74 | 369 | 26 | 0 | 200 | 2506 | 7446 | 12077 |
| SRR8094763 | 210 | 0 | 317 | 0 | 0 | 551 | 106 | 22 | 180 | 31 | 81 | 19 | 270 | 11 | 15 | 10 | 22 | 0 | 30 | 211 | 373 | 0 | 0 | 384 | 35 | 3004 | 298 | 0 | 691 | 125 | 123 | 12 | 55 | 265 | 687 | 730 | 2453 |
| SRR8094764 | 401 | 14 | 128 | 0 | 0 | 73 | 18 | 0 | 45 | 0 | 0 | 53 | 34 | 35 | 0 | 32 | 67 | 0 | 26 | 48 | 69 | 13 | 24 | 32 | 69 | 375 | 18 | 0 | 0 | 13 | 16 | 115 | 0 | 30 | 103 | 122 | 326 |
| SRR8094765 | 276 | 31 | 75 | 0 | 0 | 18 | 27 | 0 | 148 | 0 | 36 | 20 | 100 | 22 | 12 | 38 | 65 | 0 | 48 | 203 | 417 | 29 | 15 | 107 | 62 | 116 | 14 | 0 | 46 | 51 | 97 | 49 | 10 | 33 | 35 | 33 | 91 |
| SRR8094766 | 426 | 0 | 292 | 0 | 0 | 62 | 39 | 0 | 47 | 19 | 101 | 30 | 36 | 59 | 0 | 29 | 57 | 0 | 25 | 44 | 51 | 25 | 21 | 17 | 23 | 386 | 118 | 17 | 44 | 61 | 21 | 74 | 17 | 51 | 89 | 78 | 270 |
| SRR8094767 | 338 | 0 | 125 | 17 | 29 | 21 | 50 | 0 | 12 | 76 | 81 | 126 | 19 | 35 | 29 | 64 | 37 | 0 | 14 | 128 | 178 | 0 | 17 | 47 | 134 | 99 | 128 | 12 | 81 | 35 | 11 | 0 | 477 | 15 | 26 | 22 | 58 |
| SRR8094768 | 693 | 0 | 201 | 0 | 0 | 157 | 46 | 0 | 84 | 0 | 0 | 21 | 38 | 18 | 0 | 16 | 64 | 0 | 65 | 116 | 105 | 0 | 13 | 31 | 52 | 731 | 12 | 0 | 0 | 13 | 32 | 70 | 0 | 19 | 110 | 132 | 340 |
| SRR8094769 | 404 | 0 | 335 | 0 | 0 | 1775 | 27 | 0 | 629 | 0 | 51 | 23 | 467 | 0 | 15 | 0 | 55 | 0 | 36 | 89 | 59 | 0 | 0 | 126 | 59 | 8102 | 199 | 14 | 0 | 24 | 467 | 0 | 43 | 402 | 3143 | 3110 | 9922 |
| SRR8094770 | 68 | 0 | 41 | 0 | 34 | 166 | 0 | 0 | 47 | 42 | 234 | 474 | 168 | 24 | 28 | 47 | 77 | 0 | 0 | 17 | 341 | 27 | 20 | 71 | 61 | 1327 | 208 | 14 | 0 | 0 | 166 | 52 | 0 | 315 | 2062 | 7521 | 11489 |
| SRR8137033 | 112 | 20 | 76 | 0 | 0 | 16 | 53 | 20 | 81 | 0 | 20 | 3360 | 20 | 0 | 160 | 20 | 439 | 0 | 0 | 213 | 372 | 20 | 0 | 40 | 27 | 78 | 36 | 20 | 0 | 0 | 89 | 0 | 0 | 41 | 51 | 78 | 306 |
| SRR8137034 | 85 | 0 | 19 | 0 | 0 | 70 | 288 | 160 | 36 | 0 | 0 | 2549 | 0 | 0 | 185 | 0 | 358 | 0 | 41 | 288 | 1029 | 0 | 0 | 654 | 24 | 365 | 272 | 0 | 0 | 31 | 252 | 0 | 0 | 25 | 167 | 266 | 627 |
| SRR8137035 | 28 | 0 | 0 | 0 | 0 | 11 | 0 | 45 | 33 | 0 | 17 | 1464 | 0 | 0 | 260 | 0 | 26 | 0 | 0 | 130 | 3288 | 12 | 0 | 65 | 0 | 37 | 42 | 0 | 0 | 36 | 75 | 0 | 0 | 25 | 18 | 100 | 493 |
| SRR8137036 | 92 | 0 | 24 | 0 | 0 | 131 | 0 | 101 | 50 | 0 | 20 | 5826 | 0 | 0 | 36 | 0 | 175 | 0 | 0 | 178 | 748 | 0 | 0 | 272 | 91 | 422 | 221 | 0 | 0 | 80 | 674 | 0 | 30 | 0 | 292 | 513 | 2766 |
| SRR8137037 | 158 | 0 | 226 | 0 | 0 | 39 | 413 | 18 | 16 | 0 | 0 | 967 | 0 | 0 | 330 | 0 | 151 | 0 | 248 | 1652 | 4543 | 0 | 0 | 546 | 0 | 227 | 71 | 13 | 0 | 20 | 22 | 0 | 0 | 0 | 152 | 158 | 629 |
| SRR8203922 | 19 | 11 | 173 | 0 | 0 | 0 | 48 | 11 | 0 | 0 | 0 | 821 | 0 | 0 | 0 | 0 | 0 | 0 | 305 | 4764 | 0 | 0 | 0 | 0 | 0 | 33 | 24 | 0 | 0 | 0 | 0 | 0 | 91 | 0 | 0 | 0 | 0 |
| SRR8203923 | 0 | 0 | 154 | 0 | 0 | 0 | 25 | 17 | 0 | 0 | 0 | 860 | 0 | 0 | 0 | 0 | 30 | 0 | 279 | 4781 | 0 | 0 | 0 | 0 | 0 | 37 | 29 | 0 | 0 | 0 | 0 | 0 | 82 | 0 | 11 | 11 | 0 |
| SRR8203924 | 0 | 0 | 171 | 0 | 0 | 0 | 50 | 25 | 0 | 0 | 0 | 935 | 0 | 0 | 0 | 0 | 13 | 0 | 232 | 4791 | 0 | 0 | 0 | 0 | 0 | 25 | 17 | 0 | 0 | 0 | 0 | 0 | 82 | 0 | 0 | 0 | 0 |
| SRR8203925 | 17 | 0 | 181 | 0 | 0 | 0 | 17 | 20 | 0 | 17 | 0 | 830 | 0 | 0 | 0 | 0 | 17 | 0 | 194 | 4621 | 0 | 0 | 0 | 0 | 0 | 20 | 17 | 0 | 0 | 0 | 0 | 0 | 590 | 11 | 0 | 0 | 0 |
| SRR8203926 | 11 | 0 | 203 | 0 | 0 | 0 | 58 | 15 | 13 | 0 | 0 | 806 | 0 | 0 | 40 | 0 | 35 | 0 | 240 | 3821 | 22 | 0 | 0 | 0 | 0 | 26 | 41 | 0 | 0 | 0 | 0 | 0 | 443 | 0 | 21 | 0 | 13 |
| SRR8203927 | 25 | 0 | 221 | 0 | 0 | 0 | 89 | 24 | 0 | 0 | 0 | 941 | 0 | 0 | 0 | 0 | 48 | 0 | 272 | 3784 | 28 | 0 | 0 | 0 | 0 | 38 | 35 | 0 | 0 | 0 | 0 | 0 | 507 | 0 | 11 | 0 | 16 |
| SRR8203928 | 16 | 0 | 227 | 0 | 0 | 0 | 50 | 13 | 16 | 0 | 0 | 769 | 0 | 0 | 18 | 0 | 41 | 0 | 292 | 3920 | 0 | 0 | 0 | 0 | 0 | 32 | 58 | 0 | 0 | 0 | 0 | 0 | 47 | 0 | 16 | 0 | 26 |
| SRR8203929 | 0 | 0 | 234 | 0 | 0 | 0 | 51 | 16 | 0 | 0 | 0 | 843 | 0 | 0 | 23 | 0 | 36 | 0 | 210 | 4159 | 0 | 0 | 0 | 0 | 0 | 40 | 38 | 0 | 0 | 0 | 13 | 0 | 470 | 0 | 19 | 0 | 0 |
| SRR8203930 | 20 | 0 | 225 | 0 | 0 | 0 | 34 | 31 | 0 | 0 | 0 | 830 | 0 | 0 | 0 | 0 | 28 | 0 | 326 | 8143 | 0 | 0 | 0 | 0 | 0 | 47 | 54 | 0 | 0 | 0 | 0 | 0 | 74 | 0 | 0 | 0 | 0 |
| SRR8203931 | 19 | 0 | 247 | 0 | 0 | 0 | 16 | 26 | 0 | 0 | 0 | 885 | 0 | 0 | 0 | 0 | 23 | 0 | 323 | 7542 | 0 | 0 | 0 | 0 | 0 | 50 | 33 | 0 | 0 | 0 | 0 | 0 | 72 | 0 | 10 | 0 | 0 |
| SRR8203932 | 24 | 0 | 234 | 0 | 0 | 0 | 27 | 25 | 0 | 0 | 0 | 809 | 0 | 0 | 0 | 0 | 31 | 0 | 307 | 7890 | 11 | 0 | 0 | 0 | 0 | 33 | 50 | 0 | 0 | 0 | 0 | 0 | 837 | 0 | 13 | 0 | 0 |
| SRR8203933 | 14 | 0 | 240 | 0 | 0 | 0 | 28 | 17 | 0 | 0 | 0 | 767 | 0 | 0 | 0 | 0 | 28 | 0 | 307 | 7788 | 0 | 0 | 0 | 0 | 0 | 26 | 55 | 0 | 0 | 0 | 0 | 0 | 82 | 0 | 10 | 0 | 0 |
| SRR8203934 | 23 | 0 | 358 | 0 | 0 | 0 | 29 | 14 | 0 | 0 | 0 | 1049 | 0 | 0 | 0 | 0 | 24 | 0 | 133 | 2286 | 0 | 0 | 0 | 0 | 0 | 37 | 56 | 0 | 0 | 0 | 0 | 0 | 531 | 0 | 0 | 0 | 0 |
| SRR8203935 | 17 | 0 | 349 | 0 | 0 | 0 | 41 | 0 | 15 | 0 | 0 | 1049 | 0 | 0 | 0 | 0 | 36 | 0 | 103 | 2240 | 0 | 0 | 0 | 0 | 0 | 24 | 68 | 0 | 0 | 0 | 0 | 0 | 49 | 0 | 11 | 0 | 0 |
| SRR8203936 | 27 | 0 | 353 | 0 | 0 | 0 | 37 | 0 | 15 | 0 | 0 | 1101 | 0 | 0 | 0 | 0 | 58 | 0 | 114 | 2271 | 0 | 0 | 0 | 0 | 0 | 33 | 71 | 0 | 0 | 0 | 0 | 0 | 575 | 0 | 12 | 0 | 0 |
| SRR8203937 | 26 | 0 | 311 | 0 | 0 | 0 | 33 | 12 | 11 | 0 | 0 | 1033 | 0 | 0 | 0 | 0 | 53 | 0 | 114 | 2277 | 0 | 0 | 0 | 0 | 0 | 35 | 62 | 0 | 0 | 0 | 0 | 0 | 48 | 0 | 0 | 0 | 0 |
| SRR8203938 | 38 | 0 | 476 | 0 | 0 | 0 | 31 | 19 | 0 | 0 | 0 | 889 | 0 | 0 | 0 | 0 | 30 | 0 | 173 | 3199 | 0 | 0 | 0 | 0 | 0 | 25 | 15 | 0 | 0 | 0 | 0 | 0 | 57 | 0 | 11 | 0 | 0 |
| SRR8203939 | 27 | 0 | 439 | 0 | 0 | 0 | 26 | 16 | 0 | 0 | 0 | 828 | 0 | 0 | 0 | 0 | 40 | 0 | 128 | 3201 | 0 | 0 | 0 | 0 | 0 | 25 | 115 | 0 | 0 | 0 | 0 | 0 | 460 | 0 | 13 | 0 | 0 |
| SRR8203940 | 20 | 0 | 435 | 0 | 0 | 0 | 32 | 11 | 0 | 0 | 0 | 808 | 0 | 0 | 0 | 0 | 39 | 0 | 167 | 3126 | 0 | 0 | 0 | 0 | 0 | 28 | 121 | 0 | 0 | 0 | 0 | 0 | 464 | 10 | 0 | 0 | 0 |
| SRR8203941 | 26 | 0 | 450 | 0 | 0 | 0 | 38 | 14 | 0 | 0 | 0 | 909 | 0 | 0 | 0 | 0 | 41 | 0 | 152 | 3108 | 0 | 0 | 0 | 0 | 0 | 28 | 160 | 0 | 0 | 0 | 0 | 0 | 465 | 12 | 0 | 0 | 0 |
| SRR8203942 | 26 | 0 | 444 | 0 | 0 | 0 | 29 | 15 | 0 | 0 | 0 | 1795 | 0 | 0 | 0 | 0 | 0 | 0 | 24 | 883 | 0 | 0 | 0 | 0 | 0 | 10 | 0 | 0 | 0 | 0 | 0 | 0 | 64 | 0 | 0 | 0 | 0 |
| SRR8203943 | 27 | 0 | 421 | 0 | 0 | 0 | 19 | 0 | 0 | 0 | 0 | 1771 | 0 | 0 | 0 | 0 | 0 | 0 | 25 | 977 | 0 | 0 | 0 | 0 | 0 | 27 | 0 | 0 | 0 | 0 | 0 | 0 | 805 | 0 | 0 | 0 | 0 |
| SRR8203944 | 10 | 0 | 398 | 0 | 0 | 0 | 12 | 21 | 10 | 0 | 0 | 1654 | 0 | 0 | 0 | 0 | 16 | 0 | 42 | 1079 | 0 | 0 | 0 | 0 | 0 | 10 | 21 | 0 | 0 | 0 | 0 | 0 | 728 | 0 | 0 | 0 | 0 |
| SRR8203945 | 27 | 16 | 472 | 0 | 0 | 0 | 12 | 21 | 0 | 0 | 0 | 1734 | 0 | 0 | 0 | 0 | 11 | 0 | 43 | 901 | 0 | 0 | 0 | 0 | 0 | 11 | 21 | 0 | 0 | 0 | 0 | 0 | 64 | 0 | 0 | 0 | 0 |
| SRR8203946 | 21 | 0 | 675 | 0 | 0 | 0 | 40 | 64 | 0 | 0 | 0 | 1163 | 0 | 0 | 0 | 0 | 23 | 0 | 30 | 1545 | 0 | 0 | 0 | 0 | 0 | 15 | 49 | 0 | 0 | 0 | 0 | 0 | 46 | 0 | 0 | 0 | 0 |
| SRR8203947 | 33 | 0 | 762 | 0 | 0 | 10 | 18 | 52 | 0 | 0 | 0 | 1141 | 0 | 0 | 0 | 0 | 11 | 0 | 50 | 1589 | 0 | 0 | 0 | 0 | 0 | 15 | 55 | 0 | 0 | 0 | 0 | 0 | 46 | 0 | 0 | 0 | 0 |
| SRR8203948 | 26 | 0 | 731 | 0 | 0 | 0 | 27 | 59 | 0 | 0 | 0 | 1101 | 0 | 0 | 0 | 0 | 13 | 0 | 51 | 1507 | 10 | 0 | 0 | 0 | 0 | 29 | 65 | 0 | 0 | 0 | 0 | 0 | 387 | 0 | 0 | 0 | 0 |
| SRR8203949 | 26 | 0 | 672 | 0 | 0 | 0 | 32 | 48 | 0 | 0 | 0 | 1139 | 0 | 0 | 0 | 0 | 11 | 0 | 63 | 1590 | 0 | 0 | 0 | 0 | 0 | 14 | 58 | 0 | 0 | 0 | 0 | 0 | 47 | 0 | 0 | 0 | 0 |
| SRR8248787 | 231 | 0 | 25 | 0 | 0 | 173 | 1067 | 949 | 317 | 0 | 27 | 4682 | 83 | 13 | 728 | 0 | 170 | 0 | 351 | 2282 | 1594 | 0 | 0 | 849 | 54 | 983 | 1175 | 0 | 0 | 4485 | 362 | 0 | 20 | 1016 | 1098 | 1537 | 3486 |
| SRR8248788 | 195 | 12 | 18 | 0 | 0 | 380 | 838 | 95 | 846 | 0 | 67 | 4356 | 247 | 24 | 2275 | 0 | 142 | 0 | 120 | 641 | 2324 | 14 | 0 | 67 | 112 | 2105 | 922 | 0 | 0 | 562 | 1005 | 0 | 35 | 2279 | 2177 | 3691 | 8342 |
| SRR8248789 | 200 | 0 | 17 | 0 | 0 | 264 | 717 | 318 | 634 | 0 | 54 | 3885 | 212 | 28 | 1812 | 0 | 120 | 0 | 175 | 778 | 2116 | 16 | 0 | 118 | 146 | 1560 | 860 | 0 | 0 | 1752 | 772 | 0 | 47 | 2135 | 1625 | 3000 | 7063 |
| SRR8248790 | 216 | 0 | 0 | 0 | 0 | 57 | 1106 | 2195 | 189 | 0 | 20 | 3838 | 37 | 0 | 1543 | 0 | 78 | 0 | 390 | 1768 | 2184 | 0 | 0 | 1970 | 24 | 328 | 229 | 0 | 0 | 8510 | 207 | 0 | 12 | 638 | 292 | 674 | 1526 |
| SRR8248791 | 172 | 11 | 16 | 0 | 0 | 16 | 364 | 29 | 53 | 0 | 84 | 3551 | 14 | 25 | 2231 | 0 | 107 | 0 | 61 | 167 | 1054 | 0 | 0 | 28 | 110 | 94 | 22 | 0 | 0 | 15 | 78 | 0 | 59 | 166 | 160 | 305 | 624 |
| SRR8248792 | 146 | 0 | 0 | 0 | 0 | 150 | 351 | 54 | 746 | 0 | 90 | 3438 | 107 | 65 | 2172 | 0 | 53 | 0 | 60 | 204 | 2429 | 18 | 0 | 15 | 74 | 949 | 254 | 12 | 0 | 187 | 1229 | 0 | 103 | 2594 | 1869 | 5483 | 13916 |
| SRR8272058 | 149 | 0 | 1042 | 60 | 0 | 0 | 0 | 0 | 0 | 0 | 0 | 97 | 0 | 74 | 43 | 14 | 0 | 0 | 0 | 0 | 0 | 0 | 0 | 0 | 10 | 0 | 0 | 33 | 14 | 0 | 11 | 0 | 11 | 0 | 28 | 0 | 0 |
| SRR8280405 | 27 | 87 | 14 | 44 | 0 | 0 | 730 | 36 | 0 | 0 | 0 | 0 | 0 | 0 | 0 | 0 | 0 | 0 | 0 | 0 | 0 | 0 | 0 | 0 | 0 | 0 | 0 | 0 | 44 | 0 | 0 | 0 | 0 | 219 | 0 | 0 | 0 |
| SRR8280408 | 48 | 0 | 48 | 0 | 0 | 0 | 551 | 1823 | 0 | 0 | 0 | 0 | 0 | 0 | 0 | 0 | 0 | 0 | 0 | 0 | 1102 | 0 | 0 | 0 | 0 | 0 | 110 | 166 | 0 | 55 | 0 | 0 | 0 | 0 | 0 | 0 | 0 |
| SRR8280414 | 11 | 0 | 72 | 246 | 112 | 0 | 0 | 112 | 0 | 0 | 0 | 16 | 0 | 45 | 0 | 0 | 0 | 0 | 0 | 219 | 0 | 134 | 0 | 0 | 0 | 0 | 0 | 0 | 0 | 0 | 0 | 0 | 0 | 0 | 0 | 0 | 0 |
| SRR8280415 | 59 | 0 | 51 | 467 | 611 | 0 | 460 | 899 | 0 | 0 | 0 | 0 | 0 | 0 | 0 | 0 | 0 | 0 | 0 | 0 | 92 | 288 | 0 | 0 | 0 | 0 | 0 | 0 | 0 | 0 | 0 | 0 | 0 | 0 | 0 | 0 | 0 |
| SRR8280416 | 25 | 0 | 44 | 486 | 11 | 0 | 0 | 583 | 0 | 0 | 0 | 25 | 0 | 0 | 0 | 0 | 0 | 0 | 0 | 0 | 0 | 825 | 0 | 0 | 0 | 0 | 0 | 0 | 0 | 0 | 0 | 0 | 0 | 0 | 0 | 0 | 0 |
| SRR8280417 | 15 | 0 | 85 | 154 | 325 | 0 | 0 | 0 | 0 | 0 | 0 | 10 | 0 | 0 | 0 | 0 | 0 | 0 | 0 | 49 | 97 | 51 | 0 | 0 | 0 | 0 | 34 | 0 | 0 | 51 | 0 | 0 | 0 | 0 | 0 | 0 | 0 |
| SRR8280423 | 18 | 0 | 0 | 290 | 32 | 0 | 1330 | 3420 | 0 | 0 | 0 | 0 | 0 | 0 | 0 | 0 | 0 | 0 | 0 | 0 | 363 | 0 | 0 | 0 | 0 | 0 | 0 | 0 | 0 | 129 | 0 | 0 | 0 | 97 | 0 | 0 | 0 |
| SRR8280425 | 0 | 0 | 28 | 203 | 68 | 0 | 898 | 1216 | 0 | 0 | 0 | 0 | 0 | 0 | 0 | 0 | 0 | 0 | 0 | 0 | 673 | 135 | 0 | 0 | 0 | 0 | 0 | 0 | 0 | 0 | 0 | 0 | 0 | 0 | 0 | 0 | 0 |
| SRR8280432 | 42 | 0 | 47 | 183 | 0 | 0 | 2048 | 1173 | 0 | 0 | 0 | 15 | 0 | 0 | 0 | 0 | 0 | 0 | 0 | 267 | 0 | 0 | 0 | 0 | 0 | 0 | 110 | 0 | 0 | 147 | 0 | 0 | 0 | 0 | 0 | 0 | 0 |
| SRR8280440 | 115 | 0 | 128 | 11 | 0 | 0 | 2001 | 19805 | 0 | 0 | 0 | 41 | 0 | 28 | 0 | 0 | 0 | 0 | 0 | 0 | 104 | 0 | 0 | 0 | 0 | 0 | 101 | 56 | 0 | 448 | 0 | 0 | 0 | 45 | 0 | 0 | 0 |
| SRR8280442 | 46 | 0 | 55 | 0 | 18 | 0 | 676 | 15961 | 0 | 0 | 0 | 15 | 0 | 0 | 0 | 0 | 0 | 0 | 0 | 0 | 108 | 0 | 0 | 0 | 0 | 0 | 18 | 0 | 0 | 931 | 0 | 0 | 0 | 73 | 0 | 0 | 0 |
| SRR8280452 | 12 | 0 | 46 | 0 | 0 | 0 | 0 | 2061 | 0 | 0 | 0 | 0 | 0 | 0 | 0 | 0 | 0 | 0 | 0 | 0 | 0 | 0 | 0 | 0 | 0 | 0 | 0 | 0 | 0 | 179 | 0 | 0 | 0 | 0 | 0 | 0 | 0 |
| SRR8280460 | 29 | 0 | 49 | 43 | 0 | 0 | 1679 | 59 | 0 | 0 | 0 | 0 | 0 | 0 | 0 | 0 | 0 | 0 | 133 | 88 | 177 | 57 | 0 | 0 | 0 | 43 | 0 | 0 | 0 | 14 | 0 | 0 | 0 | 0 | 0 | 0 | 0 |
| SRR8280461 | 0 | 0 | 0 | 0 | 0 | 0 | 1463 | 1921 | 0 | 0 | 0 | 0 | 0 | 0 | 0 | 0 | 0 | 0 | 0 | 0 | 0 | 60 | 0 | 0 | 0 | 0 | 0 | 0 | 0 | 0 | 0 | 0 | 0 | 0 | 0 | 0 | 0 |
| SRR8280463 | 0 | 0 | 0 | 0 | 0 | 0 | 875 | 4445 | 0 | 0 | 0 | 0 | 0 | 0 | 0 | 0 | 0 | 0 | 0 | 0 | 42 | 0 | 0 | 0 | 0 | 0 | 0 | 0 | 0 | 95 | 0 | 0 | 0 | 0 | 0 | 0 | 0 |
| SRR8280464 | 100 | 0 | 75 | 0 | 0 | 0 | 793 | 34 | 0 | 0 | 0 | 44 | 0 | 46 | 53 | 0 | 0 | 0 | 0 | 0 | 53 | 0 | 0 | 0 | 0 | 0 | 0 | 0 | 0 | 0 | 0 | 0 | 0 | 0 | 0 | 0 | 0 |
| SRR8280467 | 36 | 0 | 84 | 25 | 225 | 0 | 220 | 587 | 0 | 0 | 0 | 39 | 0 | 0 | 0 | 0 | 0 | 0 | 0 | 0 | 439 | 0 | 0 | 0 | 0 | 37 | 37 | 0 | 0 | 200 | 0 | 0 | 0 | 0 | 0 | 0 | 0 |
| SRR8280468 | 18 | 0 | 22 | 0 | 542 | 0 | 0 | 271 | 0 | 0 | 0 | 0 | 0 | 0 | 0 | 0 | 0 | 0 | 0 | 351 | 0 | 0 | 0 | 0 | 0 | 0 | 0 | 0 | 0 | 0 | 0 | 0 | 0 | 0 | 0 | 0 | 0 |
| SRR8280472 | 0 | 0 | 68 | 0 | 273 | 0 | 993 | 3691 | 0 | 0 | 0 | 0 | 0 | 0 | 0 | 0 | 0 | 0 | 0 | 0 | 0 | 0 | 0 | 0 | 0 | 0 | 0 | 0 | 0 | 547 | 0 | 0 | 0 | 137 | 0 | 0 | 0 |
| SRR8280473 | 29 | 0 | 46 | 48 | 0 | 0 | 597 | 862 | 0 | 0 | 0 | 23 | 0 | 0 | 199 | 0 | 0 | 0 | 0 | 0 | 1194 | 0 | 0 | 0 | 0 | 0 | 0 | 0 | 0 | 0 | 0 | 0 | 0 | 0 | 0 | 0 | 0 |
| SRR8280476 | 29 | 0 | 19 | 221 | 0 | 0 | 0 | 221 | 0 | 0 | 0 | 0 | 0 | 0 | 0 | 0 | 0 | 0 | 0 | 0 | 530 | 0 | 0 | 0 | 0 | 0 | 0 | 0 | 0 | 0 | 0 | 0 | 0 | 0 | 0 | 0 | 0 |
| SRR8280479 | 0 | 0 | 33 | 773 | 331 | 0 | 0 | 773 | 0 | 0 | 0 | 0 | 0 | 0 | 0 | 0 | 0 | 0 | 0 | 0 | 0 | 110 | 0 | 0 | 0 | 0 | 0 | 110 | 0 | 0 | 0 | 0 | 0 | 0 | 0 | 0 | 0 |
| SRR8280481 | 22 | 0 | 35 | 516 | 129 | 0 | 0 | 516 | 0 | 0 | 0 | 17 | 0 | 0 | 0 | 0 | 0 | 0 | 0 | 0 | 181 | 258 | 0 | 0 | 0 | 129 | 0 | 0 | 0 | 0 | 0 | 0 | 0 | 0 | 0 | 0 | 0 |
| SRR8280483 | 0 | 0 | 39 | 748 | 1293 | 0 | 547 | 0 | 0 | 0 | 0 | 0 | 0 | 0 | 0 | 0 | 0 | 0 | 0 | 0 | 0 | 204 | 0 | 0 | 0 | 0 | 136 | 0 | 0 | 0 | 0 | 0 | 0 | 0 | 0 | 0 | 0 |
| SRR8280485 | 0 | 0 | 17 | 475 | 855 | 0 | 0 | 0 | 0 | 0 | 0 | 0 | 0 | 27 | 0 | 0 | 0 | 0 | 0 | 0 | 0 | 122 | 0 | 0 | 0 | 0 | 0 | 0 | 0 | 0 | 0 | 0 | 0 | 0 | 0 | 0 | 0 |
| SRR8280487 | 81 | 0 | 93 | 0 | 119 | 0 | 299 | 714 | 0 | 0 | 0 | 41 | 0 | 0 | 0 | 0 | 0 | 0 | 0 | 0 | 0 | 0 | 0 | 0 | 0 | 0 | 0 | 0 | 0 | 0 | 0 | 0 | 0 | 0 | 0 | 0 | 0 |
| SRR8280490 | 67 | 0 | 146 | 142 | 0 | 0 | 1142 | 853 | 0 | 0 | 0 | 53 | 0 | 0 | 0 | 0 | 0 | 0 | 0 | 0 | 571 | 0 | 0 | 0 | 0 | 0 | 0 | 0 | 0 | 142 | 0 | 0 | 0 | 142 | 0 | 0 | 0 |
| SRR8280494 | 26 | 0 | 123 | 11 | 0 | 0 | 1008 | 362 | 0 | 0 | 0 | 0 | 0 | 0 | 0 | 0 | 0 | 0 | 0 | 0 | 0 | 121 | 0 | 0 | 0 | 0 | 0 | 0 | 0 | 121 | 0 | 0 | 0 | 0 | 0 | 0 | 0 |
| SRR8280497 | 22 | 0 | 62 | 43 | 32 | 0 | 90 | 259 | 0 | 0 | 0 | 14 | 0 | 0 | 0 | 0 | 0 | 0 | 0 | 90 | 270 | 0 | 0 | 0 | 0 | 22 | 11 | 0 | 0 | 0 | 0 | 0 | 0 | 22 | 0 | 0 | 0 |
| SRR8280498 | 39 | 0 | 110 | 311 | 509 | 0 | 0 | 57 | 0 | 0 | 0 | 0 | 0 | 0 | 0 | 0 | 0 | 0 | 0 | 0 | 0 | 28 | 0 | 0 | 0 | 0 | 0 | 0 | 0 | 0 | 0 | 0 | 0 | 0 | 0 | 0 | 0 |
| SRR8280500 | 36 | 0 | 107 | 229 | 229 | 0 | 33 | 31 | 0 | 0 | 0 | 16 | 0 | 0 | 0 | 0 | 0 | 0 | 0 | 0 | 0 | 0 | 0 | 0 | 0 | 0 | 15 | 0 | 0 | 46 | 0 | 0 | 0 | 0 | 0 | 0 | 0 |
| SRR8280501 | 85 | 0 | 163 | 0 | 0 | 0 | 66 | 79 | 0 | 0 | 0 | 333 | 13 | 0 | 0 | 0 | 0 | 0 | 0 | 201 | 571 | 0 | 0 | 0 | 0 | 18 | 16 | 0 | 0 | 252 | 0 | 0 | 0 | 0 | 0 | 0 | 0 |
| SRR8280502 | 23 | 0 | 70 | 622 | 1243 | 0 | 0 | 414 | 0 | 0 | 0 | 16 | 0 | 0 | 0 | 0 | 0 | 0 | 0 | 0 | 0 | 104 | 0 | 0 | 0 | 0 | 104 | 0 | 0 | 0 | 0 | 0 | 0 | 0 | 0 | 0 | 0 |
| SRR8280503 | 76 | 0 | 112 | 0 | 0 | 0 | 64 | 0 | 0 | 0 | 0 | 178 | 0 | 13 | 13 | 0 | 0 | 0 | 0 | 69 | 245 | 0 | 0 | 0 | 0 | 20 | 23 | 0 | 0 | 0 | 0 | 0 | 0 | 0 | 0 | 0 | 0 |
| SRR8280504 | 41 | 0 | 121 | 296 | 1103 | 0 | 176 | 995 | 0 | 0 | 0 | 36 | 0 | 0 | 0 | 0 | 0 | 0 | 0 | 0 | 527 | 54 | 0 | 0 | 0 | 0 | 0 | 0 | 0 | 0 | 0 | 0 | 0 | 0 | 0 | 0 | 0 |
| SRR8280505 | 114 | 0 | 139 | 0 | 0 | 0 | 194 | 248 | 0 | 0 | 0 | 479 | 0 | 90 | 0 | 0 | 0 | 0 | 0 | 26 | 259 | 0 | 0 | 0 | 0 | 20 | 81 | 0 | 0 | 13 | 0 | 0 | 0 | 0 | 0 | 0 | 0 |
| SRR8280506 | 20 | 0 | 36 | 214 | 214 | 0 | 103 | 171 | 0 | 0 | 0 | 0 | 0 | 43 | 0 | 0 | 0 | 0 | 0 | 0 | 0 | 43 | 0 | 0 | 0 | 0 | 0 | 0 | 0 | 0 | 0 | 0 | 0 | 0 | 0 | 0 | 0 |
| SRR8280507 | 117 | 0 | 151 | 0 | 0 | 0 | 91 | 0 | 0 | 0 | 0 | 309 | 11 | 0 | 17 | 0 | 0 | 0 | 0 | 180 | 370 | 0 | 0 | 0 | 0 | 22 | 17 | 0 | 0 | 0 | 0 | 0 | 0 | 0 | 0 | 0 | 0 |
| SRR8280509 | 164 | 0 | 159 | 0 | 0 | 0 | 300 | 15 | 0 | 0 | 0 | 518 | 19 | 24 | 17 | 0 | 0 | 0 | 0 | 229 | 288 | 0 | 0 | 0 | 0 | 13 | 42 | 0 | 0 | 0 | 0 | 0 | 0 | 0 | 0 | 0 | 0 |
| SRR8280510 | 52 | 0 | 101 | 59 | 11 | 0 | 0 | 59 | 0 | 0 | 0 | 0 | 0 | 0 | 0 | 0 | 0 | 0 | 0 | 0 | 0 | 237 | 0 | 0 | 0 | 0 | 0 | 0 | 0 | 0 | 0 | 0 | 0 | 0 | 0 | 0 | 0 |
| SRR8280511 | 104 | 0 | 165 | 0 | 0 | 0 | 126 | 45 | 0 | 0 | 0 | 284 | 10 | 19 | 0 | 0 | 0 | 0 | 0 | 81 | 227 | 0 | 0 | 0 | 0 | 0 | 29 | 0 | 0 | 0 | 0 | 0 | 0 | 13 | 0 | 0 | 0 |
| SRR8280512 | 17 | 46 | 79 | 139 | 371 | 0 | 416 | 648 | 0 | 0 | 0 | 12 | 0 | 0 | 0 | 0 | 0 | 0 | 208 | 0 | 624 | 648 | 0 | 0 | 0 | 0 | 0 | 0 | 0 | 0 | 0 | 0 | 0 | 0 | 0 | 0 | 0 |
| SRR8280513 | 148 | 0 | 86 | 0 | 0 | 0 | 96 | 26 | 0 | 0 | 0 | 330 | 0 | 0 | 26 | 0 | 0 | 0 | 0 | 1708 | 1353 | 0 | 0 | 0 | 0 | 18 | 36 | 0 | 0 | 0 | 0 | 0 | 0 | 0 | 0 | 0 | 0 |
| SRR8280515 | 21 | 0 | 67 | 0 | 613 | 0 | 188 | 123 | 0 | 0 | 0 | 12 | 0 | 0 | 0 | 0 | 0 | 0 | 0 | 0 | 188 | 306 | 0 | 123 | 0 | 0 | 0 | 0 | 0 | 0 | 0 | 0 | 0 | 0 | 0 | 0 | 0 |
| SRR8280519 | 17 | 0 | 35 | 146 | 146 | 0 | 1105 | 3984 | 0 | 0 | 0 | 16 | 0 | 0 | 0 | 0 | 0 | 0 | 0 | 0 | 0 | 0 | 0 | 0 | 0 | 0 | 0 | 0 | 0 | 292 | 0 | 0 | 0 | 146 | 0 | 0 | 0 |
| SRR8280521 | 210 | 0 | 697 | 25 | 59 | 0 | 657 | 418 | 0 | 0 | 0 | 121 | 0 | 0 | 56 | 0 | 0 | 0 | 19 | 0 | 282 | 0 | 0 | 0 | 0 | 0 | 0 | 25 | 0 | 229 | 0 | 0 | 0 | 17 | 0 | 0 | 0 |
| SRR8280525 | 42 | 0 | 120 | 470 | 564 | 0 | 1334 | 71 | 0 | 0 | 0 | 18 | 0 | 0 | 0 | 0 | 0 | 0 | 0 | 0 | 0 | 0 | 0 | 0 | 0 | 0 | 0 | 0 | 0 | 10 | 0 | 0 | 0 | 0 | 0 | 0 | 0 |
| SRR8280531 | 0 | 0 | 42 | 54 | 54 | 0 | 1876 | 80 | 0 | 0 | 0 | 26 | 0 | 0 | 0 | 0 | 0 | 0 | 0 | 0 | 395 | 54 | 0 | 0 | 0 | 0 | 0 | 0 | 0 | 0 | 0 | 0 | 0 | 0 | 0 | 0 | 0 |
| SRR8280533 | 18 | 0 | 39 | 60 | 0 | 0 | 165 | 1444 | 0 | 0 | 0 | 0 | 0 | 0 | 0 | 0 | 0 | 0 | 0 | 82 | 495 | 0 | 0 | 0 | 0 | 0 | 0 | 0 | 0 | 120 | 0 | 0 | 0 | 0 | 0 | 0 | 0 |
| SRR8280536 | 15 | 0 | 28 | 0 | 0 | 0 | 1280 | 3779 | 0 | 0 | 0 | 0 | 0 | 0 | 0 | 0 | 0 | 0 | 0 | 0 | 295 | 0 | 0 | 0 | 0 | 62 | 31 | 31 | 0 | 280 | 16 | 0 | 0 | 0 | 0 | 0 | 0 |
| SRR8280539 | 61 | 0 | 78 | 153 | 81 | 0 | 1719 | 95 | 0 | 0 | 0 | 38 | 0 | 0 | 0 | 0 | 0 | 0 | 20 | 0 | 263 | 0 | 0 | 0 | 0 | 0 | 0 | 0 | 0 | 54 | 0 | 0 | 0 | 36 | 0 | 0 | 0 |
| SRR8280541 | 79 | 0 | 102 | 73 | 0 | 0 | 620 | 1753 | 0 | 0 | 0 | 31 | 0 | 0 | 0 | 0 | 0 | 0 | 0 | 34 | 620 | 0 | 0 | 0 | 0 | 0 | 37 | 0 | 0 | 37 | 0 | 0 | 0 | 0 | 0 | 0 | 0 |
| SRR8280543 | 0 | 0 | 15 | 0 | 639 | 0 | 0 | 128 | 0 | 0 | 0 | 0 | 0 | 0 | 0 | 0 | 0 | 0 | 0 | 0 | 0 | 0 | 0 | 0 | 0 | 0 | 0 | 0 | 0 | 0 | 0 | 0 | 0 | 0 | 0 | 0 | 0 |
| SRR8280544 | 16 | 0 | 34 | 63 | 188 | 0 | 0 | 1129 | 0 | 0 | 0 | 0 | 0 | 0 | 0 | 0 | 0 | 0 | 0 | 0 | 270 | 0 | 0 | 0 | 0 | 0 | 63 | 0 | 0 | 125 | 0 | 0 | 0 | 0 | 0 | 0 | 0 |
| SRR8280546 | 11 | 0 | 17 | 118 | 0 | 0 | 0 | 235 | 0 | 0 | 0 | 0 | 0 | 0 | 0 | 0 | 0 | 0 | 0 | 0 | 0 | 0 | 0 | 0 | 0 | 0 | 0 | 0 | 0 | 118 | 0 | 0 | 0 | 0 | 0 | 0 | 0 |
| SRR8280548 | 40 | 0 | 52 | 387 | 0 | 0 | 0 | 0 | 0 | 0 | 0 | 17 | 0 | 0 | 0 | 0 | 0 | 0 | 0 | 0 | 1152 | 0 | 0 | 0 | 0 | 0 | 0 | 0 | 0 | 0 | 0 | 0 | 0 | 0 | 0 | 0 | 0 |
| SRR8280550 | 21 | 0 | 18 | 0 | 0 | 0 | 501 | 2313 | 0 | 0 | 0 | 11 | 0 | 0 | 0 | 0 | 0 | 0 | 0 | 0 | 215 | 0 | 0 | 0 | 0 | 0 | 0 | 0 | 0 | 31 | 0 | 0 | 0 | 0 | 0 | 0 | 0 |
| SRR8280551 | 15 | 0 | 14 | 632 | 456 | 0 | 578 | 19 | 0 | 0 | 0 | 0 | 0 | 0 | 0 | 0 | 0 | 0 | 0 | 0 | 0 | 0 | 0 | 0 | 0 | 0 | 0 | 0 | 0 | 70 | 0 | 0 | 0 | 0 | 0 | 0 | 0 |
| SRR8280552 | 56 | 0 | 63 | 99 | 0 | 0 | 911 | 13 | 0 | 0 | 0 | 17 | 0 | 0 | 0 | 0 | 0 | 0 | 0 | 0 | 0 | 0 | 0 | 0 | 0 | 0 | 0 | 0 | 0 | 199 | 0 | 0 | 0 | 298 | 0 | 0 | 0 |
| SRR8280554 | 63 | 0 | 141 | 0 | 0 | 0 | 1266 | 1808 | 0 | 0 | 0 | 52 | 0 | 0 | 0 | 0 | 0 | 0 | 0 | 0 | 1012 | 0 | 0 | 0 | 0 | 0 | 0 | 0 | 0 | 70 | 0 | 0 | 0 | 0 | 0 | 0 | 0 |
| SRR8280558 | 19 | 0 | 27 | 219 | 0 | 0 | 1303 | 1974 | 0 | 0 | 0 | 0 | 0 | 0 | 0 | 0 | 0 | 0 | 0 | 0 | 0 | 0 | 0 | 0 | 0 | 0 | 0 | 0 | 0 | 0 | 0 | 0 | 0 | 0 | 0 | 0 | 0 |
| SRR8280560 | 18 | 0 | 14 | 0 | 0 | 0 | 200 | 807 | 0 | 0 | 0 | 0 | 0 | 0 | 0 | 0 | 0 | 0 | 50 | 50 | 0 | 0 | 0 | 0 | 0 | 0 | 0 | 0 | 0 | 73 | 0 | 0 | 0 | 110 | 0 | 0 | 0 |
| SRR8280561 | 0 | 62 | 24 | 0 | 0 | 0 | 1610 | 2086 | 0 | 0 | 0 | 0 | 0 | 0 | 293 | 0 | 0 | 0 | 0 | 0 | 0 | 0 | 0 | 0 | 0 | 62 | 0 | 0 | 0 | 62 | 0 | 0 | 0 | 0 | 0 | 0 | 0 |
| SRR8280562 | 30 | 0 | 99 | 0 | 0 | 0 | 399 | 65 | 0 | 0 | 0 | 26 | 0 | 0 | 0 | 0 | 0 | 0 | 100 | 50 | 449 | 0 | 0 | 0 | 0 | 0 | 35 | 0 | 71 | 213 | 0 | 0 | 0 | 0 | 0 | 0 | 0 |
| SRR8280564 | 33 | 0 | 52 | 24 | 0 | 0 | 2562 | 395 | 0 | 0 | 0 | 0 | 0 | 0 | 51 | 0 | 0 | 0 | 0 | 154 | 307 | 0 | 0 | 0 | 0 | 0 | 73 | 0 | 0 | 194 | 24 | 0 | 0 | 0 | 0 | 0 | 0 |
| SRR8280566 | 28 | 0 | 86 | 26 | 78 | 0 | 1688 | 9010 | 0 | 0 | 0 | 12 | 0 | 0 | 0 | 0 | 0 | 0 | 38 | 0 | 113 | 0 | 0 | 0 | 0 | 0 | 0 | 0 | 0 | 182 | 52 | 0 | 0 | 26 | 0 | 0 | 0 |

### Supp Table S8.2.Columns_AM_to_BW

| | | | | | | | | | | | | | | | | | | | | | | | | | | | | | | | | | | | | | |
|---|---|---|---|---|---|---|---|---|---|---|---|---|---|---|---|---|---|---|---|---|---|---|---|---|---|---|---|---|---|---|---|---|---|---|---|---|---|
| SRR8280568 | 22 | 0 | 47 | 0 | 0 | 0 | 4646 | 101 | 0 | 0 | 0 | 11 | 0 | 0 | 0 | 0 | 0 | 0 | 101 | 101 | 303 | 0 | 0 | 0 | 0 | 0 | 0 | 0 | 0 | 170 | 0 | 0 | 0 | 0 | 0 | 0 | 0 |
| SRR8280572 | 14 | 0 | 39 | 128 | 171 | 0 | 0 | 157 | 0 | 0 | 0 | 0 | 0 | 0 | 0 | 0 | 0 | 0 | 0 | 61 | 183 | 29 | 0 | 0 | 0 | 0 | 0 | 29 | 0 | 0 | 0 | 0 | 0 | 0 | 0 | 0 | 43 |
| SRR8280573 | 64 | 0 | 147 | 346 | 499 | 0 | 0 | 2015 | 0 | 0 | 0 | 49 | 0 | 0 | 0 | 0 | 0 | 0 | 0 | 0 | 447 | 58 | 0 | 0 | 0 | 0 | 38 | 0 | 0 | 0 | 0 | 0 | 0 | 0 | 0 | 0 | 0 |
| SRR8280574 | 20 | 0 | 43 | 13 | 15 | 0 | 0 | 0 | 0 | 0 | 0 | 45 | 0 | 0 | 0 | 0 | 0 | 0 | 0 | 0 | 0 | 191 | 0 | 0 | 0 | 0 | 0 | 0 | 0 | 267 | 0 | 0 | 0 | 0 | 0 | 0 | 0 |
| SRR8280576 | 38 | 0 | 33 | 0 | 399 | 0 | 0 | 0 | 0 | 0 | 0 | 0 | 0 | 0 | 0 | 0 | 0 | 0 | 0 | 0 | 0 | 150 | 0 | 0 | 0 | 0 | 0 | 0 | 0 | 0 | 0 | 0 | 0 | 0 | 0 | 0 | 0 |
| SRR8280580 | 0 | 0 | 23 | 403 | 927 | 0 | 79 | 202 | 0 | 0 | 0 | 0 | 0 | 0 | 0 | 0 | 0 | 0 | 0 | 0 | 79 | 81 | 0 | 0 | 0 | 0 | 40 | 81 | 0 | 0 | 0 | 0 | 0 | 0 | 0 | 0 | 0 |
| SRR8280582 | 29 | 0 | 71 | 127 | 509 | 0 | 0 | 18 | 0 | 0 | 0 | 12 | 0 | 0 | 0 | 0 | 0 | 0 | 0 | 0 | 172 | 255 | 0 | 0 | 0 | 0 | 0 | 0 | 0 | 0 | 0 | 0 | 0 | 0 | 0 | 0 | 0 |
| SRR8280583 | 80 | 0 | 118 | 0 | 0 | 0 | 11 | 142 | 0 | 0 | 0 | 301 | 0 | 24 | 0 | 0 | 0 | 0 | 0 | 27 | 40 | 12 | 0 | 0 | 18 | 24 | 12 | 0 | 0 | 0 | 0 | 0 | 0 | 0 | 0 | 0 | 0 |
| SRR8280584 | 42 | 0 | 61 | 147 | 11 | 0 | 100 | 441 | 0 | 0 | 0 | 14 | 0 | 0 | 0 | 0 | 0 | 0 | 0 | 0 | 399 | 245 | 0 | 0 | 0 | 0 | 0 | 0 | 0 | 98 | 0 | 0 | 0 | 0 | 0 | 0 | 0 |
| SRR8280589 | 14 | 0 | 43 | 165 | 132 | 0 | 794 | 15 | 0 | 0 | 0 | 0 | 0 | 0 | 0 | 0 | 0 | 0 | 0 | 0 | 99 | 0 | 0 | 0 | 0 | 0 | 0 | 66 | 0 | 66 | 0 | 0 | 0 | 0 | 0 | 0 | 0 |
| SRR8280590 | 29 | 0 | 44 | 0 | 110 | 0 | 1217 | 69 | 0 | 0 | 0 | 27 | 0 | 0 | 0 | 0 | 0 | 0 | 0 | 0 | 0 | 0 | 0 | 0 | 0 | 0 | 0 | 0 | 0 | 478 | 0 | 0 | 0 | 74 | 0 | 0 | 0 |
| SRR8280592 | 52 | 0 | 105 | 388 | 659 | 0 | 115 | 427 | 0 | 0 | 0 | 0 | 0 | 0 | 0 | 0 | 0 | 0 | 229 | 0 | 0 | 155 | 0 | 0 | 0 | 0 | 0 | 0 | 0 | 116 | 0 | 0 | 0 | 0 | 0 | 0 | 0 |
| SRR8280593 | 75 | 0 | 40 | 0 | 133 | 0 | 4885 | 116 | 0 | 0 | 0 | 0 | 0 | 0 | 0 | 0 | 0 | 0 | 0 | 0 | 54 | 0 | 0 | 0 | 0 | 0 | 0 | 0 | 33 | 133 | 0 | 0 | 0 | 0 | 0 | 0 | 0 |
| SRR8311061 | 127 | 0 | 32 | 0 | 23 | 0 | 0 | 176 | 0 | 0 | 0 | 0 | 0 | 0 | 0 | 0 | 0 | 0 | 0 | 602 | 0 | 54 | 0 | 0 | 0 | 145 | 38 | 0 | 0 | 0 | 0 | 0 | 0 | 0 | 0 | 0 | 0 |
| SRR8311062 | 167 | 0 | 99 | 505 | 0 | 0 | 0 | 162 | 0 | 0 | 0 | 0 | 0 | 0 | 0 | 0 | 0 | 0 | 0 | 0 | 0 | 144 | 0 | 0 | 0 | 126 | 86 | 0 | 0 | 0 | 0 | 86 | 0 | 0 | 0 | 0 | 0 |
| SRR8311063 | 165 | 0 | 137 | 0 | 0 | 0 | 0 | 369 | 0 | 0 | 0 | 0 | 0 | 0 | 0 | 105 | 0 | 0 | 0 | 689 | 0 | 0 | 0 | 0 | 199 | 760 | 339 | 0 | 0 | 0 | 105 | 124 | 0 | 0 | 0 | 0 | 0 |
| SRR8311064 | 332 | 0 | 314 | 0 | 215 | 0 | 0 | 0 | 0 | 0 | 0 | 52 | 0 | 0 | 0 | 0 | 0 | 0 | 0 | 3062 | 0 | 168 | 0 | 33 | 332 | 346 | 190 | 0 | 0 | 0 | 0 | 303 | 0 | 0 | 0 | 0 | 0 |
| SRR8311070 | 274 | 0 | 532 | 0 | 0 | 0 | 0 | 165 | 0 | 0 | 0 | 0 | 0 | 0 | 0 | 0 | 0 | 0 | 0 | 0 | 0 | 0 | 0 | 0 | 55 | 1102 | 0 | 0 | 0 | 0 | 0 | 0 | 0 | 0 | 0 | 0 | 0 |
| SRR8311071 | 278 | 0 | 182 | 0 | 0 | 0 | 0 | 0 | 0 | 0 | 0 | 0 | 0 | 0 | 0 | 0 | 0 | 0 | 697 | 0 | 0 | 0 | 0 | 129 | 407 | 190 | 473 | 0 | 0 | 0 | 0 | 0 | 0 | 0 | 0 | 0 | 0 |
| SRR8311072 | 131 | 0 | 102 | 65 | 31 | 0 | 0 | 12 | 0 | 0 | 0 | 0 | 0 | 0 | 0 | 0 | 0 | 0 | 0 | 0 | 0 | 0 | 0 | 0 | 116 | 13 | 188 | 0 | 41 | 0 | 0 | 0 | 0 | 0 | 0 | 0 | 0 |
| SRR8311073 | 110 | 0 | 156 | 43 | 346 | 0 | 0 | 0 | 0 | 0 | 0 | 0 | 0 | 0 | 0 | 0 | 0 | 0 | 0 | 0 | 0 | 0 | 0 | 149 | 235 | 0 | 254 | 0 | 0 | 0 | 0 | 0 | 0 | 0 | 0 | 0 | 0 |
| SRR8311074 | 663 | 0 | 673 | 0 | 0 | 0 | 0 | 20 | 0 | 0 | 0 | 101 | 0 | 0 | 3336 | 0 | 0 | 0 | 0 | 0 | 2756 | 0 | 0 | 0 | 0 | 16 | 0 | 0 | 0 | 213 | 0 | 0 | 0 | 0 | 0 | 0 | 0 |
| SRR8311076 | 85 | 0 | 60 | 0 | 209 | 106 | 0 | 0 | 0 | 0 | 0 | 0 | 0 | 0 | 0 | 0 | 0 | 0 | 0 | 0 | 0 | 0 | 0 | 100 | 26 | 271 | 42 | 0 | 0 | 0 | 0 | 0 | 0 | 58 | 0 | 0 | 0 |
| SRR8311078 | 225 | 0 | 111 | 152 | 237 | 0 | 0 | 0 | 170 | 0 | 0 | 0 | 0 | 0 | 0 | 0 | 0 | 0 | 0 | 0 | 0 | 58 | 0 | 0 | 424 | 40 | 107 | 0 | 0 | 0 | 0 | 0 | 0 | 0 | 0 | 0 | 0 |
| SRR8311080 | 71 | 0 | 29 | 0 | 132 | 0 | 0 | 119 | 0 | 0 | 0 | 0 | 0 | 0 | 0 | 0 | 0 | 0 | 0 | 0 | 0 | 0 | 0 | 0 | 151 | 15 | 356 | 0 | 0 | 0 | 0 | 0 | 0 | 72 | 0 | 0 | 0 |
| SRR8311081 | 62 | 0 | 20 | 468 | 0 | 0 | 0 | 0 | 0 | 0 | 0 | 0 | 0 | 0 | 0 | 0 | 0 | 0 | 0 | 0 | 0 | 0 | 0 | 0 | 0 | 291 | 0 | 0 | 0 | 0 | 0 | 0 | 0 | 0 | 0 | 0 | 0 |
| SRR8311082 | 196 | 0 | 20 | 216 | 0 | 0 | 0 | 0 | 0 | 0 | 0 | 0 | 0 | 0 | 0 | 0 | 0 | 0 | 0 | 0 | 0 | 178 | 0 | 0 | 203 | 368 | 279 | 0 | 0 | 0 | 0 | 0 | 0 | 114 | 0 | 0 | 0 |
| SRR8311085 | 235 | 0 | 333 | 0 | 0 | 0 | 0 | 58 | 0 | 0 | 0 | 0 | 0 | 0 | 0 | 0 | 0 | 0 | 0 | 0 | 0 | 113 | 0 | 0 | 103 | 222 | 0 | 0 | 0 | 0 | 0 | 0 | 0 | 142 | 0 | 0 | 0 |
| SRR8311087 | 230 | 0 | 236 | 0 | 231 | 0 | 0 | 462 | 0 | 0 | 0 | 40 | 0 | 0 | 0 | 0 | 0 | 0 | 0 | 1658 | 0 | 0 | 0 | 0 | 0 | 190 | 0 | 0 | 0 | 0 | 0 | 0 | 0 | 0 | 0 | 0 | 0 |
| SRR8311089 | 248 | 0 | 136 | 0 | 0 | 0 | 0 | 0 | 0 | 0 | 0 | 11 | 0 | 0 | 0 | 0 | 0 | 0 | 0 | 0 | 0 | 240 | 0 | 0 | 240 | 0 | 0 | 0 | 0 | 0 | 0 | 0 | 0 | 0 | 0 | 0 | 0 |
| SRR8311092 | 668 | 0 | 427 | 0 | 0 | 0 | 0 | 229 | 0 | 0 | 0 | 40 | 0 | 0 | 0 | 0 | 0 | 0 | 0 | 0 | 0 | 0 | 0 | 0 | 0 | 0 | 0 | 0 | 0 | 0 | 0 | 0 | 0 | 0 | 0 | 0 | 0 |
| SRR8311093 | 113 | 0 | 62 | 99 | 85 | 0 | 0 | 112 | 59 | 0 | 0 | 0 | 0 | 0 | 0 | 0 | 0 | 0 | 0 | 0 | 0 | 43 | 0 | 0 | 46 | 0 | 99 | 0 | 0 | 0 | 0 | 0 | 0 | 0 | 0 | 0 | 0 |
| SRR8311095 | 221 | 0 | 117 | 93 | 294 | 0 | 0 | 131 | 0 | 0 | 0 | 0 | 0 | 0 | 0 | 0 | 0 | 0 | 0 | 0 | 0 | 125 | 0 | 0 | 0 | 245 | 38 | 0 | 120 | 0 | 0 | 0 | 0 | 0 | 0 | 0 | 0 |
| SRR8311098 | 225 | 0 | 290 | 0 | 0 | 0 | 0 | 319 | 0 | 0 | 0 | 0 | 0 | 0 | 0 | 0 | 37 | 0 | 0 | 0 | 0 | 0 | 0 | 0 | 0 | 0 | 148 | 0 | 0 | 0 | 0 | 0 | 0 | 0 | 0 | 0 | 0 |
| SRR8311105 | 285 | 0 | 90 | 417 | 93 | 0 | 0 | 338 | 0 | 0 | 0 | 15 | 0 | 0 | 0 | 0 | 0 | 0 | 0 | 0 | 0 | 172 | 0 | 0 | 0 | 152 | 0 | 0 | 0 | 0 | 0 | 0 | 0 | 0 | 0 | 0 | 0 |
| SRR8311106 | 447 | 0 | 249 | 0 | 227 | 16 | 0 | 0 | 0 | 0 | 0 | 0 | 0 | 0 | 0 | 0 | 0 | 0 | 0 | 0 | 0 | 0 | 0 | 0 | 0 | 0 | 0 | 0 | 174 | 0 | 0 | 0 | 0 | 231 | 0 | 0 | 0 |
| SRR835324 | 0 | 0 | 13 | 0 | 44 | 44 | 72 | 0 | 0 | 24 | 58 | 0 | 29 | 12 | 16 | 0 | 0 | 22 | 12 | 624 | 1999 | 0 | 0 | 0 | 130 | 104 | 222 | 0 | 0 | 18 | 45 | 0 | 0 | 120 | 11 | 20 | 15 |
| SRR835325 | 0 | 0 | 0 | 0 | 39 | 15 | 172 | 30 | 0 | 10 | 27 | 0 | 21 | 13 | 0 | 0 | 0 | 14 | 0 | 244 | 201 | 0 | 0 | 0 | 107 | 145 | 538 | 0 | 0 | 18 | 26 | 0 | 0 | 161 | 22 | 34 | 29 |
| SRR8354354 | 0 | 0 | 0 | 0 | 0 | 10 | 0 | 0 | 0 | 0 | 0 | 0 | 0 | 0 | 0 | 0 | 186 | 0 | 0 | 0 | 0 | 0 | 0 | 18 | 0 | 90 | 0 | 0 | 0 | 10 | 17 | 0 | 0 | 39 | 0 | 0 | 0 |
| SRR837775 | 0 | 0 | 4420 | 0 | 0 | 24 | 0 | 0 | 0 | 0 | 0 | 0 | 0 | 23 | 74 | 0 | 29 | 0 | 0 | 181 | 302 | 0 | 0 | 0 | 41 | 312 | 288 | 48 | 0 | 0 | 138 | 0 | 0 | 66 | 2036 | 310 | 1304 |
| SRR837776 | 23 | 0 | 1860 | 0 | 0 | 13 | 0 | 0 | 0 | 0 | 0 | 0 | 0 | 21 | 94 | 0 | 39 | 0 | 0 | 33 | 216 | 0 | 0 | 0 | 26 | 12 | 170 | 32 | 0 | 0 | 24 | 0 | 0 | 25 | 131 | 161 | 49 |
| SRR837777 | 31 | 0 | 3495 | 0 | 0 | 0 | 0 | 0 | 0 | 0 | 0 | 406 | 0 | 0 | 388 | 0 | 0 | 13 | 0 | 27 | 14 | 0 | 0 | 0 | 0 | 0 | 56 | 0 | 0 | 0 | 13 | 0 | 0 | 0 | 46 | 0 | 19 |
| SRR837778 | 0 | 0 | 1506 | 0 | 0 | 0 | 0 | 0 | 0 | 0 | 0 | 578 | 0 | 0 | 94 | 0 | 0 | 0 | 0 | 27 | 0 | 0 | 0 | 0 | 0 | 0 | 31 | 0 | 0 | 0 | 0 | 0 | 0 | 0 | 30 | 0 | 22 |
| SRR837779 | 23 | 0 | 1860 | 0 | 0 | 13 | 0 | 0 | 0 | 0 | 0 | 0 | 0 | 21 | 94 | 0 | 39 | 0 | 0 | 33 | 216 | 0 | 0 | 0 | 26 | 12 | 170 | 32 | 0 | 0 | 24 | 0 | 0 | 25 | 131 | 161 | 49 |
| SRR837780 | 0 | 0 | 4420 | 0 | 0 | 24 | 0 | 0 | 0 | 0 | 0 | 0 | 0 | 23 | 72 | 0 | 29 | 0 | 0 | 176 | 294 | 0 | 0 | 0 | 41 | 312 | 288 | 48 | 0 | 0 | 138 | 0 | 0 | 66 | 2036 | 310 | 1304 |
| SRR8380550 | 20 | 0 | 27 | 0 | 0 | 41 | 42 | 0 | 0 | 16 | 0 | 15 | 11 | 0 | 16 | 0 | 35 | 0 | 0 | 0 | 0 | 0 | 0 | 0 | 649 | 89 | 146 | 0 | 0 | 0 | 28 | 0 | 0 | 18 | 0 | 0 | 0 |
| SRR8380552 | 21 | 0 | 0 | 0 | 18 | 29 | 256 | 74 | 58 | 0 | 0 | 40 | 195 | 0 | 8274 | 0 | 0 | 567 | 0 | 0 | 1009 | 0 | 0 | 0 | 57 | 105 | 15 | 0 | 0 | 153 | 155 | 0 | 0 | 1375 | 143 | 80 | 293 |
| SRR8380553 | 10 | 0 | 20 | 0 | 0 | 0 | 18 | 303 | 16 | 0 | 15 | 301 | 244 | 0 | 370 | 0 | 0 | 174 | 0 | 0 | 561 | 0 | 0 | 0 | 60 | 44 | 17 | 0 | 0 | 459 | 39 | 0 | 0 | 1657 | 175 | 80 | 277 |
| SRR8380554 | 0 | 0 | 0 | 0 | 0 | 0 | 116 | 479 | 0 | 0 | 11 | 51 | 14 | 0 | 82 | 0 | 0 | 0 | 0 | 0 | 47 | 0 | 0 | 0 | 343 | 50 | 120 | 0 | 0 | 555 | 60 | 0 | 0 | 480 | 0 | 0 | 0 |
| SRR8380555 | 15 | 0 | 133 | 0 | 0 | 22 | 37 | 0 | 0 | 13 | 15 | 910 | 45 | 0 | 627 | 0 | 154 | 140 | 0 | 0 | 55 | 0 | 0 | 0 | 2821 | 216 | 433 | 0 | 0 | 0 | 38 | 0 | 0 | 88 | 20 | 0 | 15 |
| SRR8380722 | 64 | 0 | 416 | 0 | 12 | 230 | 462 | 30 | 50 | 0 | 14 | 352 | 112 | 34 | 81 | 0 | 59 | 0 | 32 | 68 | 883 | 12 | 0 | 23 | 183 | 1840 | 526 | 14 | 0 | 41 | 91 | 0 | 0 | 1225 | 124 | 312 | 1035 |
| SRR8380723 | 80 | 0 | 797 | 0 | 25 | 142 | 208 | 0 | 46 | 0 | 10 | 358 | 198 | 61 | 564 | 10 | 90 | 16 | 13 | 27 | 396 | 0 | 0 | 0 | 527 | 1409 | 380 | 35 | 0 | 0 | 193 | 0 | 0 | 644 | 132 | 167 | 502 |
| SRR8380724 | 132 | 0 | 449 | 0 | 0 | 432 | 947 | 16 | 80 | 0 | 0 | 259 | 67 | 0 | 557 | 0 | 64 | 0 | 98 | 203 | 858 | 0 | 0 | 301 | 83 | 2525 | 2416 | 0 | 0 | 48 | 26 | 22 | 0 | 208 | 115 | 173 | 458 |
| SRR8380725 | 62 | 0 | 1085 | 0 | 12 | 70 | 105 | 0 | 21 | 12 | 24 | 779 | 92 | 50 | 361 | 16 | 125 | 0 | 10 | 18 | 213 | 0 | 0 | 0 | 926 | 754 | 178 | 20 | 0 | 0 | 86 | 0 | 13 | 590 | 173 | 96 | 226 |
| SRR8380726 | 55 | 14 | 418 | 0 | 24 | 114 | 450 | 0 | 18 | 0 | 55 | 586 | 91 | 32 | 3813 | 28 | 40 | 12 | 16 | 14 | 791 | 0 | 0 | 0 | 13 | 43 | 84 | 22 | 0 | 0 | 117 | 0 | 16 | 59 | 114 | 11 | 24 |
| SRR8380727 | 104 | 0 | 645 | 0 | 20 | 112 | 103 | 0 | 21 | 0 | 22 | 346 | 78 | 40 | 122 | 20 | 72 | 16 | 0 | 0 | 234 | 0 | 0 | 0 | 329 | 870 | 86 | 26 | 0 | 0 | 121 | 0 | 0 | 812 | 92 | 99 | 267 |
| SRR8400006 | 38 | 22 | 668 | 19 | 30 | 78 | 546 | 0 | 19 | 18 | 0 | 32 | 55 | 0 | 20 | 0 | 140 | 27 | 0 | 0 | 49 | 0 | 0 | 0 | 15 | 141 | 44 | 0 | 0 | 0 | 12 | 0 | 11 | 0 | 44 | 13 | 14 |
| SRR8400007 | 21 | 23 | 278 | 21 | 14 | 0 | 20 | 0 | 0 | 22 | 14 | 13 | 20 | 13 | 3052 | 0 | 26 | 40 | 0 | 0 | 12 | 0 | 0 | 0 | 882 | 34 | 0 | 0 | 0 | 0 | 161 | 0 | 26 | 0 | 0 | 0 | 0 |
| SRR8400008 | 22 | 20 | 477 | 16 | 16 | 30 | 71 | 0 | 13 | 10 | 0 | 15 | 76 | 0 | 294 | 0 | 41 | 129 | 0 | 16 | 73 | 0 | 0 | 0 | 452 | 808 | 27 | 0 | 0 | 0 | 645 | 0 | 0 | 0 | 102 | 20 | 22 |
| SRR8400009 | 25 | 13 | 99 | 18 | 20 | 57 | 178 | 0 | 18 | 0 | 0 | 0 | 27 | 0 | 132 | 0 | 24 | 447 | 0 | 0 | 262 | 0 | 0 | 0 | 274 | 1365 | 15 | 0 | 0 | 0 | 100 | 0 | 0 | 0 | 18 | 0 | 0 |
| SRR8400010 | 37 | 11 | 95 | 0 | 0 | 17 | 75 | 0 | 22 | 0 | 0 | 22 | 44 | 0 | 2507 | 0 | 21 | 1036 | 0 | 0 | 410 | 0 | 0 | 0 | 294 | 21 | 0 | 0 | 0 | 0 | 260 | 0 | 0 | 0 | 55 | 26 | 23 |
| SRR8417876 | 15 | 0 | 1794 | 0 | 0 | 0 | 17 | 28 | 0 | 0 | 0 | 1625 | 0 | 0 | 56 | 0 | 0 | 0 | 85 | 4187 | 516 | 0 | 0 | 12 | 14 | 169 | 2998 | 0 | 0 | 0 | 18 | 0 | 0 | 0 | 882 | 81 | 563 |
| SRR8417877 | 11 | 0 | 1537 | 0 | 0 | 12 | 56 | 51 | 0 | 0 | 0 | 1319 | 0 | 0 | 70 | 0 | 0 | 0 | 95 | 3132 | 418 | 0 | 0 | 0 | 22 | 43 | 237 | 0 | 0 | 0 | 22 | 0 | 0 | 0 | 47 | 50 | 38 |
| SRR8417878 | 15 | 0 | 1663 | 0 | 0 | 43 | 36 | 13 | 11 | 0 | 0 | 1269 | 0 | 0 | 53 | 0 | 0 | 0 | 47 | 1653 | 412 | 0 | 0 | 0 | 220 | 1473 | 862 | 0 | 0 | 0 | 36 | 0 | 0 | 0 | 272 | 31 | 197 |
| SRR8435171 | 244 | 0 | 413 | 23 | 33 | 0 | 11 | 0 | 15 | 100 | 184 | 27 | 257 | 126 | 155 | 47 | 196 | 318 | 0 | 28 | 17 | 0 | 0 | 0 | 166 | 72 | 0 | 0 | 0 | 0 | 259 | 0 | 20 | 334 | 135 | 122 | 143 |
| SRR8435172 | 292 | 10 | 281 | 27 | 42 | 0 | 13 | 0 | 0 | 34 | 92 | 28 | 52 | 345 | 585 | 109 | 145 | 197 | 0 | 19 | 13 | 0 | 0 | 0 | 104 | 15 | 0 | 0 | 0 | 0 | 83 | 13 | 12 | 114 | 43 | 40 | 57 |
| SRR8481810 | 69 | 0 | 1363 | 0 | 0 | 19 | 0 | 0 | 21 | 0 | 0 | 2476 | 0 | 0 | 62 | 0 | 0 | 0 | 0 | 600 | 169 | 0 | 0 | 0 | 0 | 160 | 179 | 0 | 0 | 0 | 14 | 0 | 0 | 0 | 40 | 73 | 33 |
| SRR8481811 | 63 | 0 | 1506 | 0 | 0 | 48 | 0 | 0 | 11 | 0 | 0 | 3021 | 0 | 51 | 37 | 0 | 0 | 0 | 0 | 422 | 56 | 0 | 0 | 0 | 13 | 87 | 131 | 0 | 0 | 0 | 39 | 0 | 0 | 0 | 34 | 17 | 21 |
| SRR8481812 | 52 | 0 | 1314 | 0 | 0 | 113 | 0 | 0 | 30 | 0 | 0 | 2749 | 0 | 0 | 91 | 0 | 0 | 0 | 23 | 845 | 148 | 0 | 0 | 0 | 0 | 122 | 168 | 0 | 0 | 0 | 12 | 0 | 0 | 21 | 27 | 33 | 18 |
| SRR8481813 | 54 | 0 | 1731 | 0 | 0 | 0 | 27 | 0 | 0 | 0 | 0 | 2346 | 0 | 115 | 0 | 0 | 0 | 0 | 0 | 408 | 20 | 0 | 0 | 0 | 18 | 47 | 256 | 0 | 0 | 0 | 0 | 0 | 0 | 0 | 47 | 0 | 21 |
| SRR8481814 | 52 | 0 | 765 | 0 | 0 | 18 | 0 | 0 | 0 | 0 | 0 | 3393 | 0 | 0 | 48 | 0 | 0 | 0 | 0 | 317 | 125 | 0 | 0 | 0 | 0 | 22 | 124 | 0 | 0 | 0 | 19 | 0 | 0 | 0 | 31 | 35 | 23 |
| SRR8481815 | 55 | 0 | 660 | 0 | 0 | 29 | 11 | 0 | 0 | 0 | 0 | 3838 | 0 | 68 | 84 | 0 | 0 | 0 | 0 | 327 | 148 | 0 | 0 | 0 | 0 | 235 | 1214 | 10 | 0 | 0 | 29 | 0 | 0 | 0 | 268 | 27 | 158 |
| SRR8481816 | 58 | 0 | 1010 | 0 | 0 | 30 | 0 | 10 | 0 | 0 | 0 | 3314 | 0 | 0 | 61 | 0 | 0 | 0 | 17 | 1174 | 261 | 0 | 0 | 15 | 0 | 364 | 2278 | 0 | 0 | 0 | 22 | 0 | 0 | 0 | 568 | 61 | 497 |
| SRR8481817 | 62 | 0 | 984 | 0 | 0 | 0 | 49 | 0 | 0 | 0 | 0 | 2841 | 0 | 17 | 0 | 0 | 0 | 0 | 24 | 1030 | 42 | 0 | 0 | 0 | 16 | 26 | 140 | 0 | 0 | 0 | 0 | 0 | 0 | 0 | 21 | 0 | 13 |
| SRR8481818 | 45 | 0 | 766 | 0 | 0 | 30 | 0 | 0 | 14 | 0 | 0 | 2777 | 0 | 11 | 29 | 0 | 10 | 0 | 58 | 3758 | 1923 | 0 | 0 | 29 | 0 | 35 | 182 | 0 | 0 | 0 | 19 | 0 | 0 | 0 | 31 | 28 | 20 |
| SRR8481819 | 36 | 0 | 804 | 0 | 0 | 74 | 36 | 15 | 0 | 0 | 0 | 2424 | 0 | 18 | 109 | 0 | 0 | 0 | 36 | 3359 | 761 | 0 | 0 | 0 | 0 | 39 | 184 | 0 | 0 | 12 | 30 | 0 | 0 | 0 | 29 | 74 | 23 |
| SRR8481820 | 38 | 0 | 619 | 0 | 0 | 71 | 12 | 0 | 18 | 0 | 0 | 1944 | 0 | 0 | 12 | 0 | 0 | 0 | 61 | 5550 | 649 | 0 | 0 | 13 | 0 | 29 | 155 | 18 | 0 | 0 | 31 | 0 | 0 | 22 | 34 | 54 | 22 |
| SRR8481821 | 31 | 0 | 820 | 0 | 0 | 0 | 32 | 11 | 0 | 0 | 0 | 1623 | 0 | 29 | 0 | 0 | 0 | 0 | 47 | 898 | 90 | 0 | 0 | 0 | 26 | 34 | 76 | 0 | 0 | 0 | 0 | 0 | 0 | 0 | 13 | 0 | 26 |
| SRR8485055 | 297 | 0 | 1399 | 0 | 0 | 56 | 29 | 0 | 19 | 0 | 0 | 44 | 45 | 0 | 79 | 0 | 32 | 15 | 44 | 35 | 568 | 0 | 0 | 61 | 102 | 671 | 642 | 0 | 0 | 0 | 53 | 0 | 10 | 0 | 91 | 45 | 89 |
| SRR8485056 | 246 | 0 | 1923 | 0 | 0 | 366 | 147 | 0 | 194 | 0 | 0 | 151 | 0 | 0 | 2470 | 0 | 106 | 0 | 23 | 445 | 1038 | 0 | 0 | 555 | 1808 | 6170 | 13802 | 0 | 0 | 0 | 404 | 0 | 0 | 23 | 79 | 92 | 118 |
| SRR8485057 | 1014 | 12 | 2586 | 0 | 0 | 27 | 266 | 0 | 11 | 0 | 0 | 63 | 37 | 0 | 760 | 0 | 82 | 0 | 21 | 179 | 1466 | 0 | 0 | 141 | 369 | 332 | 1867 | 0 | 0 | 0 | 24 | 0 | 0 | 0 | 17 | 36 | 11 |
| SRR8485058 | 588 | 0 | 1801 | 0 | 0 | 343 | 611 | 12 | 212 | 0 | 0 | 46 | 12 | 0 | 1484 | 0 | 40 | 0 | 58 | 88 | 677 | 0 | 0 | 834 | 1357 | 4380 | 17124 | 0 | 0 | 10 | 311 | 0 | 0 | 0 | 47 | 22 | 28 |
| SRR8485059 | 282 | 19 | 1909 | 0 | 0 | 13 | 91 | 0 | 62 | 0 | 0 | 59 | 31 | 0 | 143 | 0 | 158 | 0 | 0 | 117 | 327 | 0 | 0 | 100 | 316 | 183 | 1297 | 0 | 0 | 0 | 17 | 0 | 0 | 0 | 20 | 10 | 13 |
| SRR8485060 | 1080 | 0 | 3297 | 0 | 0 | 72 | 50 | 0 | 54 | 0 | 0 | 177 | 22 | 0 | 11030 | 0 | 84 | 45 | 25 | 212 | 5920 | 0 | 0 | 290 | 636 | 899 | 3459 | 0 | 0 | 0 | 90 | 0 | 0 | 0 | 17 | 10 | 35 |
| SRR8485061 | 165 | 0 | 1892 | 0 | 0 | 384 | 149 | 16 | 205 | 0 | 0 | 183 | 14 | 10 | 930 | 0 | 109 | 0 | 29 | 478 | 1180 | 0 | 0 | 979 | 1238 | 6184 | 8775 | 0 | 0 | 0 | 413 | 0 | 15 | 0 | 108 | 42 | 97 |
| SRR8485062 | 445 | 0 | 919 | 0 | 0 | 28 | 193 | 0 | 16 | 0 | 0 | 76 | 0 | 0 | 2775 | 0 | 32 | 0 | 55 | 166 | 1377 | 0 | 0 | 105 | 161 | 329 | 927 | 0 | 0 | 0 | 30 | 0 | 0 | 0 | 11 | 37 | 75 |
| SRR8485063 | 380 | 0 | 989 | 0 | 0 | 185 | 94 | 0 | 74 | 0 | 0 | 118 | 49 | 0 | 4583 | 0 | 41 | 16 | 100 | 360 | 1234 | 0 | 0 | 579 | 1608 | 2321 | 8388 | 0 | 0 | 0 | 525 | 0 | 16 | 11 | 117 | 50 | 97 |
| SRR8485064 | 522 | 0 | 1321 | 0 | 0 | 327 | 77 | 0 | 155 | 0 | 0 | 46 | 39 | 0 | 1518 | 0 | 52 | 18 | 146 | 123 | 1782 | 0 | 0 | 1168 | 1699 | 4153 | 7442 | 0 | 0 | 0 | 626 | 0 | 13 | 20 | 168 | 66 | 111 |
| SRR8485065 | 235 | 0 | 1054 | 0 | 0 | 103 | 252 | 0 | 44 | 0 | 0 | 42 | 68 | 11 | 63 | 0 | 45 | 48 | 20 | 46 | 383 | 0 | 0 | 33 | 131 | 102 | 209 | 0 | 0 | 0 | 11 | 0 | 0 | 0 | 50 | 23 | 37 |
| SRR8485066 | 312 | 0 | 1086 | 0 | 0 | 130 | 256 | 0 | 57 | 0 | 0 | 56 | 15 | 0 | 621 | 0 | 29 | 0 | 17 | 107 | 1400 | 0 | 0 | 502 | 1366 | 1796 | 3804 | 0 | 0 | 0 | 212 | 0 | 0 | 0 | 60 | 29 | 48 |
| SRR8485067 | 163 | 0 | 798 | 0 | 0 | 16 | 65 | 0 | 109 | 0 | 0 | 36 | 43 | 0 | 63 | 0 | 30 | 0 | 132 | 28 | 393 | 0 | 0 | 18 | 20 | 145 | 28 | 0 | 0 | 0 | 24 | 0 | 22 | 0 | 203 | 46 | 143 |
| SRR8487217 | 178 | 0 | 3717 | 0 | 0 | 43 | 56 | 12 | 22 | 0 | 0 | 183 | 11 | 12 | 41 | 0 | 77 | 111 | 85 | 364 | 12114 | 0 | 0 | 52 | 224 | 498 | 745 | 0 | 0 | 0 | 50 | 11 | 0 | 0 | 205 | 54 | 125 |
| SRR8487218 | 221 | 0 | 1826 | 0 | 0 | 19 | 218 | 75 | 15 | 0 | 0 | 180 | 0 | 0 | 0 | 0 | 25 | 0 | 13 | 99 | 125 | 0 | 0 | 89 | 221 | 333 | 963 | 0 | 0 | 0 | 22 | 0 | 0 | 0 | 16 | 13 | 20 |
| SRR8487219 | 174 | 0 | 3712 | 0 | 0 | 66 | 1230 | 16 | 33 | 0 | 0 | 205 | 46 | 13 | 537 | 0 | 28 | 0 | 216 | 305 | 1320 | 0 | 0 | 47 | 155 | 77 | 508 | 0 | 0 | 0 | 110 | 0 | 0 | 0 | 36 | 21 | 28 |
| SRR8487220 | 275 | 0 | 2158 | 0 | 0 | 119 | 13 | 0 | 34 | 0 | 0 | 162 | 84 | 0 | 22 | 0 | 28 | 13 | 0 | 0 | 261 | 0 | 0 | 14 | 56 | 54 | 128 | 0 | 0 | 0 | 22 | 0 | 0 | 0 | 136 | 67 | 108 |
| SRR8487221 | 616 | 0 | 1874 | 0 | 13 | 96 | 336 | 0 | 16 | 0 | 0 | 87 | 36 | 12 | 449 | 0 | 30 | 0 | 33 | 56 | 903 | 0 | 0 | 27 | 263 | 945 | 1336 | 0 | 0 | 0 | 119 | 0 | 0 | 0 | 38 | 15 | 25 |
| SRR8487222 | 533 | 0 | 1034 | 0 | 18 | 328 | 2073 | 0 | 118 | 0 | 0 | 78 | 33 | 0 | 2389 | 0 | 16 | 0 | 216 | 900 | 1281 | 0 | 0 | 129 | 1044 | 4177 | 6055 | 0 | 0 | 0 | 565 | 0 | 0 | 0 | 160 | 80 | 111 |
| SRR8487223 | 428 | 23 | 2279 | 0 | 0 | 19 | 658 | 0 | 30 | 0 | 0 | 182 | 0 | 11 | 116 | 0 | 28 | 61 | 31 | 94 | 618 | 0 | 0 | 23 | 163 | 227 | 932 | 0 | 18 | 0 | 34 | 0 | 0 | 35 | 92 | 36 | 52 |
| SRR8487224 | 98 | 0 | 1459 | 0 | 0 | 88 | 841 | 0 | 52 | 0 | 0 | 113 | 34 | 18 | 66 | 0 | 39 | 38 | 63 | 82 | 1378 | 0 | 0 | 55 | 1015 | 1415 | 2082 | 0 | 0 | 0 | 167 | 0 | 0 | 41 | 27 | 16 | 35 |
| SRR8586304 | 61 | 0 | 39 | 0 | 0 | 0 | 653 | 46 | 0 | 0 | 0 | 72 | 0 | 0 | 0 | 0 | 0 | 0 | 3559 | 1136 | 0 | 0 | 0 | 0 | 14 | 0 | 0 | 0 | 14 | 16 | 0 | 0 | 0 | 54 | 0 | 0 | 0 |
| SRR871609 | 17 | 0 | 15 | 0 | 0 | 15 | 0 | 0 | 71 | 0 | 0 | 0 | 0 | 0 | 142 | 0 | 0 | 0 | 0 | 0 | 711 | 0 | 0 | 0 | 0 | 158 | 0 | 0 | 145 | 0 | 39 | 0 | 0 | 0 | 0 | 0 | 0 |
| SRR871671 | 22 | 15 | 0 | 30 | 0 | 15 | 0 | 0 | 23 | 0 | 0 | 0 | 0 | 0 | 1852 | 0 | 0 | 0 | 109 | 0 | 1253 | 0 | 0 | 0 | 0 | 23 | 0 | 0 | 45 | 0 | 76 | 0 | 0 | 0 | 61 | 38 | 0 |
| SRR873381 | 92 | 15 | 530 | 0 | 0 | 53 | 23 | 0 | 30 | 0 | 0 | 59 | 0 | 0 | 2032 | 0 | 0 | 0 | 0 | 228 | 1922 | 0 | 0 | 0 | 0 | 29 | 27 | 0 | 27 | 0 | 47 | 0 | 0 | 0 | 198 | 62 | 20 |
| SRR873401 | 106 | 0 | 86 | 0 | 0 | 36 | 0 | 0 | 0 | 0 | 0 | 34 | 0 | 0 | 897 | 0 | 0 | 0 | 18 | 364 | 4470 | 0 | 0 | 0 | 0 | 288 | 28 | 0 | 0 | 0 | 0 | 0 | 0 | 11 | 932 | 157 | 1029 |
| SRR873410 | 49 | 0 | 145 | 13 | 103 | 32 | 19 | 11 | 13 | 0 | 0 | 53 | 0 | 0 | 149 | 0 | 0 | 0 | 19 | 1971 | 2070 | 0 | 0 | 0 | 23 | 340 | 93 | 0 | 0 | 0 | 0 | 0 | 0 | 0 | 803 | 65 | 428 |
| SRR8834226 | 0 | 0 | 0 | 0 | 0 | 0 | 1179 | 0 | 0 | 0 | 0 | 418 | 0 | 0 | 0 | 0 | 0 | 0 | 144448 | 0 | 0 | 0 | 0 | 0 | 0 | 45 | 17 | 0 | 0 | 0 | 0 | 0 | 0 | 0 | 0 | 0 | 0 |
| SRR8834227 | 0 | 0 | 0 | 0 | 0 | 0 | 618 | 0 | 0 | 0 | 0 | 966 | 0 | 0 | 13 | 0 | 0 | 22 | 1582 | 0 | 0 | 0 | 0 | 17 | 0 | 12 | 11 | 0 | 0 | 0 | 0 | 0 | 0 | 0 | 0 | 0 | 0 |
| SRR8839811 | 77 | 0 | 12 | 0 | 0 | 0 | 156 | 37 | 13 | 48 | 1774 | 323 | 0 | 31 | 0 | 0 | 31 | 0 | 11 | 0 | 21 | 0 | 0 | 0 | 0 | 0 | 0 | 0 | 65 | 0 | 0 | 0 | 0 | 20 | 0 | 0 | 0 |
| SRR8839812 | 71 | 0 | 0 | 0 | 0 | 0 | 20 | 55 | 81 | 0 | 18 | 284 | 0 | 0 | 0 | 0 | 36 | 0 | 0 | 0 | 0 | 0 | 0 | 0 | 0 | 0 | 0 | 0 | 44 | 22 | 0 | 0 | 0 | 48 | 0 | 0 | 18 |
| SRR8850891 | 80 | 0 | 74 | 0 | 0 | 27 | 221 | 138 | 49 | 0 | 0 | 5299 | 306 | 0 | 999 | 0 | 108 | 0 | 987 | 2787 | 1870 | 0 | 0 | 866 | 150 | 425 | 548 | 0 | 0 | 961 | 80 | 0 | 0 | 458 | 250 | 412 | 602 |
| SRR8850892 | 86 | 0 | 63 | 0 | 0 | 88 | 135 | 11 | 10 | 0 | 0 | 4361 | 46 | 0 | 411 | 0 | 144 | 21 | 810 | 3169 | 1807 | 0 | 0 | 38 | 14 | 52 | 65 | 0 | 0 | 54 | 14 | 0 | 0 | 49 | 36 | 59 | 95 |
| SRR8850893 | 92 | 0 | 73 | 0 | 0 | 83 | 129 | 484 | 106 | 0 | 0 | 4562 | 599 | 0 | 3333 | 0 | 154 | 0 | 563 | 1714 | 6126 | 0 | 0 | 364 | 167 | 925 | 1519 | 0 | 0 | 549 | 138 | 0 | 0 | 546 | 899 | 2341 | 2703 |
| SRR8850894 | 59 | 0 | 61 | 0 | 0 | 61 | 124 | 44 | 48 | 0 | 0 | 3467 | 50 | 0 | 6594 | 0 | 138 | 0 | 458 | 1525 | 5984 | 0 | 0 | 49 | 28 | 60 | 128 | 0 | 0 | 61 | 94 | 0 | 0 | 48 | 72 | 170 | 193 |
| SRR8850895 | 109 | 0 | 119 | 0 | 0 | 60 | 91 | 18 | 34 | 0 | 0 | 5464 | 32 | 0 | 129 | 0 | 147 | 13 | 819 | 2821 | 872 | 0 | 0 | 110 | 16 | 41 | 82 | 0 | 0 | 68 | 78 | 0 | 0 | 28 | 30 | 40 | 69 |
| SRR8850896 | 116 | 0 | 109 | 0 | 0 | 36 | 271 | 349 | 20 | 11 | 0 | 4732 | 475 | 0 | 157 | 0 | 107 | 0 | 841 | 3661 | 983 | 0 | 0 | 1253 | 46 | 288 | 456 | 0 | 0 | 1233 | 42 | 0 | 0 | 304 | 176 | 251 | 483 |
| SRR8850897 | 102 | 0 | 148 | 0 | 0 | 48 | 58 | 3274 | 27 | 0 | 0 | 4581 | 433 | 0 | 1796 | 0 | 126 | 17 | 321 | 1482 | 2547 | 0 | 0 | 1457 | 96 | 353 | 455 | 0 | 0 | 4392 | 59 | 0 | 0 | 674 | 286 | 1184 | 1308 |
| SRR8850898 | 45 | 0 | 144 | 0 | 0 | 65 | 67 | 43 | 28 | 0 | 11 | 4811 | 39 | 0 | 1151 | 0 | 201 | 0 | 338 | 1207 | 1494 | 0 | 0 | 83 | 52 | 97 | 243 | 0 | 0 | 58 | 18 | 0 | 0 | 54 | 64 | 154 | 166 |
| SRR8850899 | 86 | 0 | 115 | 0 | 11 | 44 | 61 | 90 | 66 | 0 | 16 | 6475 | 95 | 32 | 158 | 15 | 89 | 0 | 232 | 912 | 657 | 0 | 0 | 57 | 17 | 35 | 84 | 0 | 0 | 47 | 89 | 0 | 14 | 154 | 49 | 50 | 110 |
| SRR8850900 | 97 | 0 | 135 | 0 | 17 | 21 | 90 | 40 | 23 | 0 | 0 | 6680 | 578 | 0 | 60 | 0 | 162 | 0 | 386 | 1060 | 590 | 0 | 0 | 49 | 224 | 292 | 1387 | 0 | 0 | 224 | 57 | 0 | 0 | 746 | 656 | 328 | 1214 |
| SRR8850901 | 69 | 0 | 134 | 0 | 23 | 25 | 29 | 435 | 45 | 0 | 0 | 5830 | 998 | 14 | 2789 | 0 | 97 | 0 | 251 | 677 | 2024 | 0 | 0 | 42 | 239 | 408 | 1046 | 0 | 0 | 523 | 57 | 0 | 0 | 1096 | 687 | 1029 | 1683 |
| SRR8850902 | 74 | 0 | 153 | 0 | 19 | 45 | 55 | 78 | 37 | 0 | 12 | 6062 | 73 | 0 | 2056 | 0 | 106 | 0 | 86 | 258 | 1473 | 0 | 0 | 64 | 24 | 38 | 87 | 0 | 0 | 87 | 57 | 15 | 0 | 89 | 71 | 90 | 157 |
| SRR8858246 | 208 | 0 | 219 | 29 | 0 | 0 | 0 | 0 | 0 | 0 | 0 | 126 | 0 | 34 | 422 | 0 | 0 | 0 | 0 | 0 | 0 | 0 | 0 | 0 | 0 | 29 | 0 | 39 | 0 | 0 | 151 | 0 | 0 | 0 | 3879 | 0 | 16 |
| SRR8858247 | 301 | 0 | 369 | 33 | 0 | 0 | 0 | 0 | 0 | 0 | 0 | 257 | 0 | 30 | 225 | 0 | 0 | 0 | 0 | 0 | 0 | 0 | 0 | 0 | 0 | 0 | 0 | 0 | 0 | 0 | 93 | 21 | 0 | 17 | 275 | 92 | 33 |
| SRR8858248 | 440 | 0 | 378 | 53 | 0 | 0 | 0 | 0 | 0 | 0 | 0 | 196 | 0 | 89 | 0 | 33 | 18 | 0 | 0 | 0 | 0 | 0 | 0 | 0 | 0 | 0 | 0 | 0 | 0 | 0 | 57 | 0 | 0 | 0 | 44 | 0 | 34 |

### Supp Table S8.2.Columns_AM_to_BW

| | | | | | | | | | | | | | | | | | | | | | | | | | | | | | | | | | | | | | |
|---|---|---|---|---|---|---|---|---|---|---|---|---|---|---|---|---|---|---|---|---|---|---|---|---|---|---|---|---|---|---|---|---|---|---|---|---|---|
| SRR8858254 | 1456 | 0 | 1943 | 16 | 0 | 0 | 0 | 0 | 0 | 0 | 0 | 69 | 0 | 0 | 1171 | 0 | 0 | 0 | 0 | 0 | 0 | 0 | 0 | 0 | 0 | 0 | 0 | 0 | 0 | 0 | 82 | 0 | 0 | 0 | 0 | 0 | 0 |
| SRR8858258 | 520 | 0 | 537 | 15 | 0 | 0 | 0 | 0 | 0 | 0 | 0 | 50 | 0 | 40 | 1180 | 0 | 13 | 0 | 0 | 0 | 0 | 0 | 0 | 0 | 0 | 0 | 0 | 0 | 0 | 0 | 0 | 0 | 0 | 0 | 82 | 0 | 0 |
| SRR8907824 | 18 | 21 | 0 | 20 | 0 | 0 | 54 | 0 | 39 | 0 | 12 | 1297 | 0 | 0 | 261 | 12 | 0 | 0 | 0 | 0 | 20 | 0 | 0 | 0 | 0 | 0 | 0 | 17 | 0 | 23 | 17 | 66 | 34 | 27 | 0 | 15 | 11 |
| SRR8932122 | 11 | 0 | 1063 | 0 | 0 | 0 | 28 | 46 | 0 | 0 | 0 | 19 | 0 | 27 | 0 | 0 | 0 | 0 | 0 | 35 | 101 | 0 | 0 | 0 | 11 | 15 | 19 | 15 | 0 | 0 | 0 | 0 | 0 | 65 | 42 | 0 | 20 |
| SRR8932123 | 0 | 0 | 61 | 0 | 0 | 0 | 74 | 0 | 0 | 0 | 0 | 0 | 0 | 36 | 1132 | 0 | 0 | 0 | 26 | 249 | 745 | 0 | 0 | 0 | 15 | 45 | 63 | 23 | 0 | 0 | 0 | 0 | 0 | 24 | 10 | 0 | 0 |
| SRR8932124 | 11 | 0 | 369 | 0 | 11 | 0 | 29 | 0 | 0 | 0 | 0 | 11 | 0 | 36 | 0 | 0 | 0 | 0 | 0 | 78 | 879 | 0 | 0 | 0 | 59 | 14 | 12 | 28 | 0 | 0 | 0 | 0 | 10 | 67 | 20 | 0 | 12 |
| SRR8932125 | 24 | 12 | 487 | 11 | 11 | 0 | 18 | 0 | 0 | 0 | 0 | 0 | 0 | 30 | 89 | 0 | 0 | 0 | 0 | 199 | 2300 | 0 | 0 | 0 | 159 | 52 | 63 | 23 | 0 | 0 | 0 | 0 | 0 | 124 | 20 | 0 | 10 |
| SRR8944248 | 0 | 0 | 0 | 0 | 0 | 0 | 0 | 0 | 0 | 0 | 0 | 20 | 0 | 0 | 0 | 0 | 0 | 0 | 0 | 0 | 1451 | 0 | 0 | 0 | 0 | 140 | 0 | 0 | 0 | 0 | 0 | 0 | 0 | 0 | 0 | 0 | 0 |
| SRR8944250 | 34 | 0 | 34 | 0 | 0 | 0 | 0 | 0 | 0 | 0 | 0 | 35 | 0 | 0 | 1751 | 0 | 0 | 0 | 0 | 0 | 16813 | 0 | 0 | 0 | 0 | 0 | 0 | 0 | 0 | 0 | 0 | 0 | 0 | 0 | 0 | 0 | 0 |
| SRR8944251 | 22 | 0 | 60 | 0 | 0 | 0 | 0 | 0 | 0 | 0 | 0 | 17 | 0 | 0 | 0 | 0 | 0 | 0 | 0 | 0 | 7501 | 0 | 0 | 0 | 0 | 0 | 0 | 0 | 0 | 0 | 0 | 0 | 0 | 0 | 0 | 0 | 0 |
| SRR8944252 | 58 | 0 | 86 | 0 | 0 | 0 | 0 | 0 | 0 | 0 | 0 | 56 | 0 | 0 | 0 | 0 | 0 | 0 | 0 | 0 | 14744 | 0 | 0 | 0 | 0 | 0 | 0 | 0 | 0 | 0 | 0 | 0 | 0 | 0 | 0 | 0 | 0 |
| SRR8944257 | 24 | 0 | 43 | 0 | 0 | 0 | 0 | 0 | 0 | 0 | 0 | 37 | 0 | 0 | 0 | 0 | 0 | 0 | 0 | 0 | 7095 | 0 | 0 | 0 | 0 | 0 | 0 | 0 | 0 | 0 | 0 | 0 | 0 | 0 | 0 | 0 | 0 |
| SRR8944259 | 20 | 0 | 109 | 0 | 0 | 0 | 0 | 0 | 0 | 0 | 0 | 52 | 0 | 0 | 0 | 0 | 0 | 0 | 0 | 0 | 6693 | 0 | 0 | 0 | 0 | 0 | 0 | 0 | 0 | 0 | 0 | 0 | 0 | 0 | 0 | 0 | 0 |
| SRR8956086 | 52 | 0 | 66 | 0 | 0 | 13 | 189 | 14 | 22 | 0 | 0 | 1289 | 10 | 0 | 0 | 0 | 80 | 0 | 0 | 0 | 151 | 0 | 0 | 0 | 0 | 29 | 35 | 0 | 0 | 12 | 16 | 0 | 0 | 133 | 16 | 0 | 22 |
| SRR8956087 | 70 | 0 | 68 | 0 | 0 | 0 | 124 | 15 | 46 | 0 | 0 | 1695 | 20 | 0 | 0 | 0 | 101 | 0 | 13 | 0 | 180 | 0 | 0 | 0 | 0 | 68 | 112 | 0 | 0 | 0 | 33 | 0 | 0 | 321 | 18 | 0 | 22 |
| SRR8956089 | 92 | 0 | 98 | 0 | 0 | 13 | 275 | 0 | 19 | 0 | 13 | 1462 | 27 | 0 | 0 | 0 | 119 | 0 | 23 | 0 | 134 | 0 | 0 | 0 | 0 | 69 | 18 | 0 | 0 | 0 | 14 | 0 | 0 | 137 | 11 | 20 | 14 |
| SRR8956090 | 50 | 0 | 82 | 0 | 0 | 55 | 424 | 0 | 79 | 0 | 13 | 1040 | 14 | 0 | 0 | 0 | 86 | 0 | 27 | 0 | 260 | 0 | 0 | 0 | 0 | 276 | 60 | 0 | 0 | 0 | 47 | 0 | 0 | 451 | 34 | 47 | 121 |
| SRR8956091 | 68 | 0 | 101 | 0 | 0 | 45 | 269 | 13 | 86 | 0 | 13 | 1261 | 31 | 0 | 0 | 0 | 111 | 0 | 21 | 0 | 185 | 0 | 0 | 0 | 0 | 266 | 78 | 0 | 0 | 0 | 61 | 0 | 0 | 562 | 30 | 27 | 93 |
| SRR8992368 | 789 | 0 | 0 | 20 | 31 | 12 | 70 | 26 | 59 | 51 | 31 | 210 | 0 | 12 | 0 | 0 | 10 | 0 | 38 | 0 | 0 | 0 | 0 | 0 | 0 | 35 | 16 | 0 | 0 | 44 | 16 | 0 | 31 | 159 | 0 | 0 | 0 |
| SRR8992369 | 517 | 0 | 15 | 0 | 16 | 36 | 867 | 64 | 32 | 42 | 76 | 322 | 0 | 17 | 0 | 0 | 12 | 0 | 19 | 0 | 0 | 0 | 0 | 0 | 18 | 52 | 11 | 0 | 0 | 63 | 16 | 0 | 10 | 63 | 0 | 0 | 0 |
| SRR8992370 | 306 | 0 | 11 | 0 | 18 | 18 | 1027 | 676 | 39 | 42 | 91 | 285 | 0 | 14 | 0 | 0 | 0 | 0 | 28 | 0 | 0 | 0 | 0 | 0 | 0 | 44 | 0 | 0 | 0 | 901 | 11 | 0 | 101 | 793 | 0 | 0 | 0 |
| SRR8992371 | 415 | 0 | 22 | 0 | 16 | 41 | 764 | 1284 | 59 | 49 | 68 | 366 | 0 | 17 | 0 | 0 | 0 | 0 | 37 | 0 | 0 | 0 | 0 | 0 | 20 | 67 | 15 | 0 | 0 | 1791 | 0 | 0 | 99 | 1899 | 0 | 0 | 0 |
| SRR8992378 | 511 | 0 | 0 | 0 | 22 | 21 | 756 | 131 | 63 | 45 | 11 | 354 | 0 | 0 | 0 | 0 | 15 | 0 | 31 | 0 | 0 | 0 | 0 | 11 | 0 | 52 | 10 | 0 | 0 | 203 | 10 | 0 | 54 | 170 | 0 | 0 | 0 |
| SRR8992379 | 515 | 0 | 0 | 11 | 33 | 28 | 116 | 24 | 105 | 39 | 77 | 170 | 0 | 17 | 0 | 0 | 0 | 0 | 0 | 0 | 0 | 0 | 0 | 0 | 0 | 39 | 17 | 0 | 0 | 50 | 28 | 0 | 39 | 77 | 0 | 0 | 0 |
| SRR8992380 | 410 | 0 | 14 | 0 | 29 | 48 | 864 | 1415 | 60 | 35 | 75 | 463 | 0 | 17 | 0 | 0 | 0 | 0 | 32 | 0 | 18 | 0 | 0 | 0 | 12 | 69 | 14 | 0 | 0 | 1952 | 17 | 0 | 70 | 1597 | 0 | 0 | 0 |
| SRR8992381 | 653 | 0 | 0 | 10 | 18 | 20 | 87 | 40 | 71 | 16 | 41 | 249 | 0 | 0 | 0 | 0 | 12 | 0 | 0 | 0 | 0 | 0 | 0 | 0 | 0 | 24 | 14 | 0 | 0 | 51 | 12 | 0 | 45 | 54 | 0 | 0 | 0 |
| SRR8992382 | 788 | 0 | 0 | 0 | 32 | 31 | 147 | 54 | 10 | 45 | 52 | 155 | 0 | 0 | 0 | 0 | 0 | 0 | 24 | 0 | 0 | 0 | 0 | 0 | 0 | 50 | 16 | 0 | 0 | 85 | 22 | 0 | 23 | 99 | 0 | 0 | 0 |
| SRR8992383 | 689 | 0 | 0 | 0 | 29 | 27 | 160 | 68 | 77 | 28 | 38 | 172 | 0 | 0 | 0 | 0 | 0 | 0 | 17 | 0 | 0 | 10 | 0 | 0 | 0 | 50 | 32 | 0 | 0 | 95 | 17 | 0 | 28 | 82 | 0 | 0 | 0 |
| SRR8992384 | 741 | 0 | 10 | 0 | 28 | 31 | 883 | 55 | 61 | 14 | 56 | 349 | 0 | 12 | 0 | 0 | 10 | 0 | 27 | 0 | 0 | 0 | 0 | 0 | 0 | 43 | 0 | 0 | 0 | 74 | 0 | 0 | 46 | 90 | 0 | 0 | 0 |
| SRR8992385 | 433 | 0 | 0 | 0 | 22 | 20 | 821 | 320 | 53 | 31 | 93 | 274 | 0 | 21 | 0 | 0 | 0 | 0 | 27 | 0 | 0 | 0 | 0 | 0 | 0 | 27 | 0 | 0 | 0 | 447 | 14 | 0 | 87 | 648 | 0 | 0 | 0 |
| SRR8992387 | 477 | 0 | 0 | 13 | 36 | 18 | 876 | 121 | 54 | 47 | 79 | 306 | 0 | 0 | 0 | 0 | 0 | 0 | 29 | 0 | 0 | 0 | 0 | 0 | 0 | 54 | 10 | 0 | 0 | 224 | 21 | 0 | 88 | 657 | 0 | 0 | 0 |
| SRR8992388 | 455 | 0 | 19 | 0 | 19 | 27 | 919 | 906 | 41 | 48 | 90 | 339 | 0 | 16 | 0 | 0 | 10 | 0 | 31 | 0 | 0 | 0 | 0 | 0 | 11 | 34 | 11 | 0 | 0 | 1100 | 13 | 0 | 119 | 751 | 0 | 0 | 0 |
| SRR8992389 | 381 | 0 | 11 | 0 | 22 | 39 | 899 | 86 | 55 | 45 | 69 | 356 | 0 | 18 | 0 | 0 | 16 | 0 | 38 | 0 | 0 | 0 | 0 | 14 | 19 | 47 | 12 | 0 | 0 | 95 | 16 | 0 | 11 | 59 | 0 | 0 | 0 |
| SRR8992390 | 537 | 0 | 0 | 0 | 35 | 28 | 863 | 540 | 76 | 21 | 54 | 305 | 0 | 20 | 0 | 0 | 0 | 0 | 36 | 0 | 0 | 0 | 0 | 0 | 0 | 58 | 17 | 0 | 0 | 733 | 0 | 0 | 53 | 621 | 0 | 0 | 0 |
| SRR8992391 | 517 | 0 | 0 | 0 | 0 | 23 | 1730 | 146 | 31 | 52 | 87 | 99 | 0 | 0 | 0 | 0 | 0 | 0 | 31 | 0 | 0 | 0 | 0 | 0 | 0 | 38 | 0 | 0 | 0 | 191 | 0 | 0 | 102 | 301 | 0 | 0 | 0 |
| SRR9003330 | 0 | 0 | 25 | 0 | 0 | 31 | 95 | 0 | 74 | 0 | 15 | 0 | 0 | 16 | 211 | 0 | 41 | 0 | 72 | 390 | 9464 | 0 | 0 | 0 | 59 | 71 | 30 | 0 | 15 | 0 | 183 | 0 | 0 | 11 | 115 | 89 | 157 |
| SRR9003332 | 0 | 15 | 86 | 0 | 0 | 0 | 0 | 0 | 0 | 21 | 106 | 0 | 0 | 26 | 341 | 0 | 41 | 12 | 0 | 0 | 0 | 0 | 0 | 0 | 34 | 0 | 0 | 0 | 43 | 0 | 154 | 0 | 11 | 0 | 35 | 0 | 0 |
| SRR9003333 | 0 | 0 | 27 | 0 | 0 | 583 | 102 | 0 | 1053 | 0 | 22 | 0 | 0 | 12 | 229 | 0 | 57 | 0 | 55 | 359 | 11614 | 0 | 0 | 0 | 67 | 1392 | 370 | 0 | 59 | 0 | 3041 | 0 | 12 | 15 | 1889 | 1931 | 3039 |
| SRR9003334 | 0 | 26 | 101 | 0 | 13 | 0 | 0 | 0 | 0 | 26 | 63 | 0 | 0 | 133 | 407 | 0 | 48 | 0 | 0 | 0 | 0 | 0 | 0 | 0 | 61 | 0 | 0 | 19 | 0 | 0 | 2848 | 10 | 24 | 0 | 0 | 0 | 0 |
| SRR9003336 | 0 | 0 | 31 | 0 | 0 | 495 | 99 | 0 | 1190 | 0 | 19 | 0 | 0 | 18 | 272 | 0 | 43 | 0 | 40 | 330 | 12371 | 0 | 0 | 0 | 60 | 1225 | 511 | 0 | 63 | 0 | 3242 | 0 | 0 | 16 | 2685 | 2504 | 3684 |
| SRR9003338 | 0 | 13 | 96 | 0 | 17 | 0 | 0 | 0 | 0 | 15 | 20 | 0 | 0 | 53 | 304 | 0 | 43 | 20 | 0 | 0 | 0 | 0 | 0 | 0 | 44 | 0 | 0 | 0 | 0 | 0 | 5274 | 0 | 18 | 0 | 13 | 0 | 0 |
| SRR9003339 | 0 | 0 | 24 | 0 | 0 | 26 | 119 | 0 | 56 | 0 | 19 | 0 | 11 | 13 | 168 | 0 | 66 | 0 | 74 | 508 | 13508 | 0 | 0 | 52 | 63 | 68 | 33 | 0 | 55 | 14 | 166 | 0 | 0 | 10 | 140 | 137 | 207 |
| SRR9003341 | 0 | 23 | 132 | 0 | 21 | 0 | 0 | 0 | 15 | 0 | 27 | 0 | 10 | 56 | 319 | 0 | 98 | 0 | 0 | 0 | 0 | 0 | 0 | 0 | 160 | 31 | 0 | 12 | 0 | 0 | 3397 | 0 | 0 | 0 | 46 | 46 | 46 |
| SRR9003342 | 0 | 0 | 26 | 0 | 0 | 34 | 69 | 0 | 69 | 0 | 13 | 0 | 0 | 17 | 281 | 0 | 49 | 0 | 50 | 300 | 13679 | 0 | 0 | 0 | 88 | 82 | 29 | 0 | 77 | 0 | 237 | 0 | 0 | 17 | 115 | 109 | 180 |
| SRR9018258 | 145 | 0 | 64 | 0 | 13 | 69 | 16 | 0 | 62 | 0 | 0 | 10951 | 149 | 29 | 18 | 0 | 615 | 0 | 0 | 77 | 171 | 0 | 0 | 44 | 260 | 379 | 455 | 43 | 0 | 0 | 333 | 0 | 219 | 709 | 6635 | 1425 | 5002 |
| SRR9018259 | 82 | 0 | 88 | 0 | 15 | 80 | 71 | 27 | 91 | 0 | 13 | 2676 | 121 | 0 | 180 | 0 | 496 | 0 | 95 | 551 | 256 | 0 | 0 | 36 | 289 | 423 | 628 | 0 | 0 | 0 | 229 | 0 | 14 | 1071 | 8075 | 730 | 6846 |
| SRR9018260 | 78 | 27 | 0 | 80 | 10 | 13 | 405 | 5716 | 55 | 0 | 0 | 4279 | 0 | 58 | 10965 | 36 | 214 | 0 | 850 | 389 | 850 | 0 | 0 | 1386 | 100 | 45 | 215 | 38 | 0 | 891 | 114 | 23 | 0 | 763 | 1147 | 2345 | 4103 |
| SRR9018261 | 71 | 0 | 25 | 0 | 0 | 24 | 137 | 475 | 20 | 0 | 0 | 3255 | 13 | 41 | 186 | 58 | 194 | 10 | 66 | 368 | 128 | 0 | 0 | 1169 | 22 | 80 | 26 | 57 | 0 | 213 | 40 | 0 | 327 | 445 | 132 | 186 | 432 |
| SRR952248 | 38 | 26 | 1531 | 0 | 0 | 49 | 171 | 268 | 19 | 0 | 0 | 33 | 0 | 14 | 343 | 0 | 11 | 22 | 26 | 3333 | 4112 | 0 | 0 | 63 | 0 | 74 | 258 | 0 | 0 | 21 | 76 | 0 | 0 | 19 | 69 | 13 | 67 |
| SRR952249 | 34 | 70 | 1451 | 0 | 0 | 39 | 363 | 5227 | 0 | 0 | 0 | 39 | 0 | 10 | 1094 | 0 | 12 | 55 | 31 | 2813 | 11830 | 0 | 0 | 19 | 0 | 599 | 1905 | 0 | 0 | 21 | 47 | 0 | 0 | 13 | 619 | 125 | 554 |
| SRR952288 | 35 | 66 | 1465 | 0 | 0 | 38 | 402 | 683 | 13 | 0 | 0 | 36 | 0 | 16 | 1010 | 0 | 12 | 64 | 37 | 2830 | 11668 | 0 | 0 | 17 | 0 | 80 | 249 | 0 | 0 | 20 | 47 | 0 | 0 | 13 | 80 | 18 | 72 |
| SRR952289 | 38 | 63 | 1643 | 0 | 0 | 56 | 152 | 50 | 15 | 0 | 0 | 46 | 0 | 16 | 1103 | 0 | 16 | 23 | 0 | 741 | 9135 | 0 | 0 | 0 | 0 | 76 | 156 | 0 | 0 | 0 | 53 | 0 | 0 | 28 | 102 | 21 | 109 |
| SRR952290 | 36 | 57 | 1645 | 0 | 0 | 54 | 150 | 51 | 0 | 0 | 0 | 47 | 0 | 18 | 1018 | 0 | 15 | 19 | 12 | 759 | 8932 | 0 | 0 | 0 | 0 | 77 | 157 | 0 | 0 | 0 | 57 | 0 | 0 | 29 | 101 | 20 | 112 |
| SRR952309 | 40 | 0 | 2179 | 0 | 0 | 62 | 13 | 18 | 17 | 0 | 0 | 22 | 0 | 0 | 354 | 0 | 15 | 0 | 0 | 663 | 646 | 0 | 0 | 0 | 0 | 55 | 138 | 0 | 0 | 0 | 10 | 0 | 0 | 0 | 126 | 11 | 75 |
| SRR952310 | 42 | 0 | 2224 | 0 | 0 | 68 | 19 | 43 | 15 | 0 | 0 | 25 | 0 | 0 | 228 | 0 | 20 | 0 | 0 | 1365 | 1227 | 0 | 0 | 0 | 0 | 752 | 3474 | 0 | 0 | 0 | 81 | 0 | 0 | 0 | 1129 | 109 | 774 |
| SRR952311 | 57 | 28 | 2199 | 0 | 0 | 66 | 122 | 198 | 16 | 0 | 0 | 31 | 0 | 13 | 4156 | 0 | 19 | 0 | 11 | 741 | 3213 | 0 | 0 | 0 | 0 | 760 | 1978 | 0 | 0 | 0 | 71 | 0 | 0 | 0 | 1465 | 93 | 825 |
| SRR952312 | 13 | 14 | 542 | 0 | 0 | 53 | 81 | 266 | 32 | 0 | 0 | 0 | 0 | 0 | 778 | 0 | 0 | 0 | 27 | 1041 | 12426 | 0 | 0 | 0 | 0 | 1061 | 1369 | 11 | 0 | 0 | 133 | 0 | 0 | 14 | 686 | 186 | 434 |
| SRR952313 | 11 | 0 | 761 | 0 | 0 | 28 | 54 | 442 | 12 | 0 | 0 | 0 | 0 | 0 | 970 | 0 | 0 | 0 | 20 | 1379 | 18012 | 0 | 0 | 11 | 0 | 1011 | 1384 | 0 | 10 | 11 | 132 | 0 | 0 | 0 | 444 | 151 | 445 |
| SRR952314 | 28 | 0 | 1234 | 0 | 0 | 134 | 0 | 76 | 47 | 0 | 0 | 0 | 0 | 17 | 10 | 0 | 0 | 11 | 23 | 1459 | 4645 | 0 | 0 | 0 | 0 | 1659 | 753 | 0 | 0 | 0 | 217 | 0 | 0 | 0 | 918 | 343 | 918 |
| SRR952315 | 30 | 0 | 1026 | 0 | 0 | 103 | 0 | 166 | 38 | 0 | 0 | 0 | 0 | 12 | 0 | 0 | 0 | 0 | 71 | 2024 | 4160 | 0 | 0 | 20 | 0 | 75 | 38 | 13 | 0 | 0 | 12 | 0 | 0 | 0 | 32 | 11 | 37 |
| SRR952316 | 1006 | 0 | 233 | 0 | 0 | 96 | 174 | 263 | 84 | 0 | 0 | 1323 | 0 | 0 | 0 | 0 | 0 | 0 | 201 | 3876 | 106 | 0 | 0 | 0 | 0 | 1719 | 3662 | 0 | 0 | 0 | 205 | 0 | 0 | 0 | 0 | 19 | 32 |
| SRR952317 | 865 | 0 | 132 | 0 | 0 | 75 | 379 | 963 | 119 | 0 | 0 | 882 | 0 | 0 | 27 | 0 | 0 | 0 | 44 | 6682 | 142 | 0 | 0 | 0 | 0 | 3870 | 9108 | 0 | 0 | 0 | 469 | 0 | 0 | 0 | 12 | 0 | 0 |
| SRR952318 | 2158 | 0 | 275 | 0 | 0 | 337 | 0 | 149 | 194 | 0 | 0 | 1025 | 0 | 0 | 0 | 0 | 0 | 0 | 124 | 3928 | 0 | 0 | 0 | 0 | 0 | 6673 | 5247 | 0 | 0 | 0 | 940 | 0 | 13 | 0 | 0 | 0 | 0 |
| SRR952319 | 2435 | 0 | 221 | 0 | 0 | 728 | 0 | 173 | 389 | 0 | 0 | 0 | 0 | 0 | 0 | 0 | 0 | 0 | 82 | 4520 | 0 | 0 | 0 | 0 | 0 | 6189 | 4761 | 0 | 0 | 0 | 1085 | 0 | 0 | 0 | 0 | 0 | 0 |
| SRR9588764 | 67 | 0 | 852 | 0 | 48 | 38 | 55 | 13 | 0 | 0 | 22 | 85 | 23 | 49 | 1195 | 0 | 29 | 0 | 18 | 100 | 52 | 0 | 0 | 1012 | 61 | 364 | 776 | 32 | 0 | 0 | 42 | 0 | 0 | 35 | 0 | 0 | 0 |
| SRR9588765 | 80 | 0 | 967 | 0 | 49 | 37 | 27 | 15 | 0 | 0 | 38 | 306 | 92 | 26 | 8164 | 0 | 31 | 30 | 42 | 96 | 342 | 0 | 0 | 2245 | 93 | 233 | 810 | 0 | 0 | 0 | 15 | 0 | 0 | 15 | 15 | 0 | 0 |
| SRR9588766 | 85 | 0 | 817 | 14 | 23 | 28 | 49 | 0 | 0 | 0 | 17 | 494 | 20 | 15 | 0 | 0 | 63 | 0 | 0 | 23 | 189 | 0 | 0 | 29 | 51 | 91 | 350 | 23 | 20 | 0 | 11 | 0 | 13 | 45 | 0 | 0 | 0 |
| SRR9588767 | 94 | 0 | 795 | 0 | 34 | 112 | 28 | 0 | 15 | 0 | 11 | 73 | 87 | 39 | 1153 | 0 | 56 | 0 | 0 | 31 | 155 | 0 | 0 | 607 | 161 | 666 | 1225 | 13 | 0 | 0 | 35 | 0 | 0 | 30 | 43 | 0 | 20 |
| SRR9618218 | 69 | 0 | 3351 | 0 | 0 | 35 | 0 | 0 | 14 | 0 | 0 | 242 | 0 | 0 | 13 | 0 | 79 | 0 | 13 | 17461 | 297 | 0 | 0 | 35 | 0 | 65 | 44 | 14 | 0 | 0 | 14 | 0 | 0 | 0 | 72 | 24 | 28 |
| SRR9618219 | 144 | 0 | 5496 | 0 | 0 | 46 | 0 | 0 | 25 | 0 | 0 | 586 | 0 | 18 | 11 | 0 | 119 | 0 | 0 | 257 | 53 | 0 | 0 | 0 | 18 | 940 | 236 | 11 | 25 | 0 | 34 | 0 | 0 | 0 | 2141 | 396 | 958 |
| SRR9618220 | 51 | 0 | 3181 | 0 | 0 | 43 | 11 | 20 | 0 | 0 | 0 | 244 | 0 | 0 | 34 | 0 | 60 | 0 | 11 | 28762 | 405 | 0 | 0 | 124 | 17 | 605 | 581 | 10 | 0 | 0 | 33 | 0 | 0 | 0 | 1176 | 114 | 408 |
| SRR9618221 | 53 | 0 | 3621 | 0 | 0 | 24 | 0 | 0 | 0 | 0 | 0 | 222 | 0 | 0 | 0 | 0 | 60 | 0 | 11 | 12060 | 857 | 0 | 0 | 0 | 0 | 43 | 27 | 0 | 0 | 0 | 0 | 0 | 0 | 0 | 76 | 45 | 25 |
| SRR9618222 | 127 | 0 | 5049 | 0 | 0 | 13 | 13 | 0 | 30 | 0 | 0 | 337 | 0 | 39 | 40 | 0 | 60 | 0 | 0 | 480 | 293 | 0 | 0 | 0 | 0 | 544 | 167 | 26 | 0 | 0 | 47 | 0 | 0 | 21 | 1267 | 235 | 813 |
| SRR9627609 | 190 | 0 | 970 | 0 | 0 | 20 | 54 | 0 | 27 | 0 | 0 | 52 | 0 | 0 | 380 | 0 | 0 | 0 | 0 | 145 | 1267 | 0 | 0 | 0 | 10 | 919 | 10 | 0 | 14 | 0 | 41 | 0 | 0 | 0 | 16 | 14 | 28 |
| SRR9627610 | 181 | 0 | 877 | 0 | 0 | 13 | 18 | 0 | 20 | 0 | 0 | 64 | 0 | 0 | 527 | 0 | 0 | 0 | 0 | 45 | 679 | 0 | 0 | 0 | 11 | 473 | 16 | 0 | 10 | 0 | 40 | 0 | 0 | 0 | 50 | 11 | 23 |
| SRR9627611 | 209 | 0 | 1005 | 0 | 0 | 89 | 0 | 0 | 103 | 0 | 0 | 72 | 0 | 0 | 399 | 0 | 0 | 0 | 17 | 83 | 831 | 0 | 0 | 0 | 49 | 3204 | 22 | 0 | 12 | 0 | 181 | 0 | 0 | 0 | 39 | 0 | 16 |
| SRR9627612 | 207 | 0 | 967 | 0 | 0 | 23 | 0 | 0 | 24 | 0 | 0 | 43 | 0 | 0 | 967 | 0 | 0 | 0 | 0 | 101 | 1006 | 0 | 0 | 0 | 30 | 775 | 27 | 0 | 0 | 0 | 35 | 0 | 0 | 0 | 11 | 0 | 17 |
| SRR9627613 | 173 | 0 | 859 | 0 | 0 | 21 | 14 | 0 | 26 | 0 | 0 | 39 | 0 | 0 | 1173 | 0 | 0 | 0 | 0 | 90 | 888 | 0 | 0 | 0 | 39 | 685 | 27 | 0 | 0 | 0 | 32 | 0 | 0 | 0 | 32 | 0 | 11 |
| SRR9627614 | 185 | 0 | 929 | 0 | 0 | 12 | 0 | 0 | 19 | 0 | 0 | 44 | 0 | 0 | 1376 | 0 | 0 | 74 | 0 | 13 | 538 | 0 | 0 | 0 | 11 | 549 | 0 | 0 | 0 | 0 | 46 | 0 | 0 | 0 | 11 | 0 | 19 |
| SRR9627651 | 878 | 0 | 1765 | 0 | 0 | 61 | 0 | 0 | 24 | 0 | 0 | 177 | 0 | 0 | 1326 | 0 | 88 | 0 | 0 | 204 | 212 | 0 | 0 | 0 | 102 | 1121 | 104 | 0 | 0 | 0 | 41 | 0 | 0 | 0 | 12 | 60 | 68 |
| SRR9627653 | 660 | 0 | 1660 | 0 | 0 | 12 | 0 | 0 | 10 | 0 | 0 | 139 | 0 | 0 | 1599 | 0 | 88 | 0 | 0 | 23 | 15 | 0 | 0 | 0 | 119 | 309 | 19 | 0 | 0 | 0 | 47 | 0 | 0 | 0 | 136 | 44 | 49 |
| SRR9627654 | 645 | 0 | 1700 | 0 | 0 | 403 | 0 | 0 | 201 | 0 | 0 | 97 | 0 | 0 | 2811 | 0 | 108 | 31 | 0 | 65 | 49 | 0 | 0 | 0 | 1591 | 8500 | 56 | 0 | 0 | 0 | 406 | 0 | 0 | 0 | 95 | 43 | 41 |
| SRR9627655 | 548 | 0 | 1484 | 0 | 0 | 177 | 0 | 0 | 125 | 0 | 0 | 81 | 0 | 0 | 3407 | 0 | 98 | 0 | 0 | 45 | 54 | 0 | 0 | 0 | 1796 | 4406 | 23 | 0 | 0 | 0 | 499 | 0 | 17 | 0 | 76 | 28 | 43 |
| SRR9627656 | 754 | 0 | 1586 | 0 | 0 | 64 | 13 | 0 | 26 | 0 | 0 | 101 | 0 | 0 | 1939 | 0 | 101 | 13 | 0 | 180 | 253 | 0 | 0 | 0 | 114 | 1331 | 11 | 0 | 0 | 0 | 37 | 0 | 0 | 0 | 15 | 49 | 59 |
| SRR9627693 | 856 | 0 | 1289 | 0 | 0 | 21 | 208 | 11 | 23 | 0 | 0 | 128 | 0 | 18 | 776 | 0 | 12 | 35 | 18 | 668 | 1120 | 0 | 0 | 11 | 57 | 512 | 357 | 0 | 0 | 0 | 18 | 0 | 0 | 0 | 57 | 22 | 27 |
| SRR9627694 | 815 | 0 | 1312 | 0 | 0 | 21 | 172 | 10 | 24 | 0 | 0 | 126 | 0 | 13 | 516 | 0 | 0 | 13 | 0 | 717 | 1500 | 0 | 0 | 13 | 60 | 524 | 426 | 0 | 0 | 0 | 19 | 0 | 0 | 0 | 62 | 19 | 41 |
| SRR9627695 | 776 | 0 | 1334 | 0 | 0 | 21 | 112 | 11 | 21 | 0 | 0 | 146 | 0 | 0 | 733 | 0 | 11 | 0 | 48 | 629 | 1243 | 0 | 0 | 15 | 63 | 512 | 509 | 0 | 0 | 0 | 19 | 0 | 0 | 0 | 48 | 26 | 29 |
| SRR9627696 | 781 | 0 | 1098 | 0 | 0 | 13 | 442 | 40 | 13 | 0 | 0 | 70 | 0 | 14 | 895 | 0 | 0 | 0 | 84 | 1189 | 1452 | 0 | 0 | 50 | 27 | 343 | 327 | 0 | 0 | 0 | 59 | 0 | 0 | 0 | 34 | 0 | 12 |
| SRR9627697 | 873 | 0 | 1242 | 0 | 0 | 85 | 245 | 386 | 95 | 0 | 0 | 80 | 0 | 0 | 1153 | 0 | 0 | 0 | 43 | 730 | 1300 | 0 | 0 | 47 | 134 | 2450 | 1565 | 0 | 0 | 0 | 49 | 0 | 0 | 0 | 29 | 11 | 15 |
| SRR9627698 | 910 | 0 | 1152 | 0 | 0 | 14 | 364 | 88 | 16 | 0 | 0 | 65 | 0 | 12 | 1220 | 0 | 0 | 33 | 0 | 705 | 1750 | 0 | 0 | 45 | 21 | 415 | 213 | 0 | 0 | 11 | 69 | 0 | 0 | 0 | 41 | 0 | 0 |
| SRR9648387 | 0 | 0 | 0 | 0 | 0 | 0 | 0 | 0 | 35 | 0 | 0 | 110 | 0 | 0 | 0 | 0 | 0 | 0 | 0 | 0 | 0 | 0 | 0 | 0 | 0 | 0 | 0 | 0 | 0 | 0 | 11 | 0 | 0 | 31 | 0 | 0 | 11 |
| SRR9648388 | 0 | 0 | 0 | 0 | 0 | 0 | 0 | 0 | 21 | 0 | 10 | 0 | 0 | 0 | 0 | 0 | 0 | 0 | 0 | 0 | 22 | 0 | 29 | 0 | 0 | 21 | 0 | 0 | 134 | 0 | 38 | 0 | 0 | 86 | 0 | 0 | 0 |
| SRR9648389 | 0 | 0 | 0 | 0 | 0 | 0 | 0 | 0 | 13 | 0 | 13 | 187 | 0 | 0 | 0 | 0 | 0 | 0 | 0 | 0 | 0 | 0 | 60 | 0 | 11 | 0 | 0 | 0 | 29 | 0 | 23 | 0 | 24 | 17 | 0 | 0 | 0 |
| SRR965653 | 291 | 119 | 12567 | 0 | 0 | 0 | 19 | 18 | 0 | 0 | 0 | 0 | 0 | 0 | 2315 | 0 | 25 | 0 | 19 | 1309 | 13926 | 0 | 0 | 0 | 0 | 18 | 10 | 0 | 0 | 18 | 0 | 0 | 0 | 39 | 27 | 0 | 18 |
| SRR965655 | 19 | 0 | 8447 | 0 | 0 | 40 | 36 | 20 | 22 | 0 | 0 | 0 | 0 | 0 | 47 | 0 | 34 | 0 | 14 | 3630 | 1369 | 0 | 0 | 16 | 0 | 47 | 304 | 0 | 37 | 30 | 182 | 0 | 0 | 42 | 70 | 149 | 44 |
| SRR9833150 | 59 | 32 | 342 | 0 | 0 | 0 | 77 | 0 | 40 | 0 | 0 | 125 | 0 | 12 | 292 | 0 | 47 | 159 | 0 | 194 | 476 | 0 | 0 | 0 | 0 | 115 | 33 | 0 | 0 | 0 | 0 | 0 | 0 | 0 | 604 | 34 | 150 |
| SRR9833151 | 65 | 52 | 314 | 0 | 0 | 0 | 0 | 0 | 25 | 0 | 0 | 167 | 0 | 11 | 507 | 0 | 42 | 85 | 0 | 106 | 134 | 0 | 0 | 0 | 0 | 78 | 0 | 0 | 0 | 0 | 12 | 0 | 0 | 0 | 228 | 18 | 102 |
| SRR9833152 | 62 | 11 | 332 | 0 | 0 | 0 | 29 | 0 | 18 | 0 | 0 | 133 | 0 | 12 | 319 | 0 | 43 | 110 | 0 | 209 | 354 | 0 | 0 | 0 | 10 | 47 | 20 | 0 | 0 | 0 | 0 | 0 | 0 | 0 | 171 | 12 | 50 |
| SRR9833153 | 57 | 83 | 358 | 0 | 0 | 0 | 37 | 0 | 19 | 0 | 0 | 181 | 0 | 16 | 381 | 0 | 31 | 39 | 0 | 68 | 123 | 0 | 0 | 0 | 0 | 35 | 0 | 0 | 0 | 0 | 10 | 0 | 0 | 0 | 842 | 44 | 365 |
| SRR9833154 | 73 | 21 | 335 | 0 | 0 | 0 | 16 | 0 | 18 | 0 | 0 | 186 | 0 | 0 | 152 | 0 | 46 | 43 | 0 | 71 | 82 | 0 | 0 | 0 | 0 | 27 | 0 | 0 | 0 | 0 | 24 | 0 | 0 | 0 | 170 | 12 | 68 |
| SRR9833155 | 53 | 54 | 430 | 0 | 0 | 0 | 222 | 0 | 52 | 0 | 0 | 183 | 0 | 18 | 222 | 0 | 45 | 45 | 29 | 240 | 291 | 0 | 0 | 0 | 0 | 137 | 20 | 0 | 0 | 0 | 16 | 0 | 0 | 0 | 785 | 71 | 729 |
| SRR9833156 | 68 | 40 | 516 | 0 | 0 | 11 | 191 | 0 | 36 | 0 | 0 | 272 | 0 | 0 | 277 | 0 | 31 | 40 | 0 | 490 | 740 | 0 | 0 | 0 | 0 | 236 | 56 | 0 | 0 | 0 | 0 | 0 | 0 | 0 | 289 | 28 | 254 |
| SRR9833157 | 62 | 12 | 427 | 0 | 0 | 15 | 97 | 23 | 20 | 0 | 0 | 170 | 0 | 0 | 344 | 0 | 35 | 71 | 0 | 2058 | 1390 | 0 | 0 | 0 | 0 | 139 | 36 | 0 | 0 | 0 | 14 | 0 | 0 | 0 | 110 | 10 | 67 |
| SRR9833158 | 58 | 21 | 418 | 0 | 0 | 17 | 199 | 528 | 35 | 0 | 0 | 167 | 0 | 13 | 361 | 0 | 50 | 126 | 19 | 3091 | 2702 | 0 | 0 | 0 | 0 | 225 | 204 | 0 | 0 | 0 | 0 | 0 | 0 | 0 | 263 | 16 | 144 |
| SRR9833159 | 41 | 23 | 653 | 18 | 23 | 11 | 191 | 516 | 22 | 0 | 0 | 85 | 0 | 33 | 1066 | 14 | 26 | 999 | 30 | 1579 | 3515 | 0 | 0 | 0 | 21 | 67 | 173 | 0 | 0 | 23 | 0 | 0 | 0 | 12 | 19 | 0 | 16 |
| SRR9833160 | 64 | 29 | 942 | 12 | 12 | 21 | 106 | 2273 | 28 | 0 | 0 | 97 | 0 | 57 | 1799 | 17 | 30 | 362 | 15 | 1268 | 2999 | 0 | 0 | 0 | 13 | 210 | 174 | 0 | 0 | 46 | 0 | 11 | 0 | 13 | 127 | 0 | 55 |
| SRR9833161 | 93 | 17 | 719 | 10 | 11 | 0 | 57 | 437 | 16 | 0 | 0 | 40 | 0 | 21 | 1562 | 20 | 19 | 520 | 23 | 722 | 2570 | 0 | 0 | 0 | 20 | 45 | 26 | 0 | 0 | 15 | 0 | 13 | 0 | 17 | 37 | 0 | 23 |
| SRR9833162 | 54 | 63 | 873 | 18 | 11 | 21 | 81 | 1527 | 26 | 0 | 0 | 81 | 0 | 83 | 2204 | 30 | 20 | 495 | 19 | 1163 | 2310 | 0 | 0 | 0 | 17 | 310 | 208 | 0 | 0 | 19 | 11 | 0 | 0 | 0 | 172 | 0 | 90 |
| SRR9833163 | 57 | 24 | 785 | 29 | 15 | 13 | 276 | 1059 | 33 | 0 | 0 | 66 | 0 | 80 | 822 | 28 | 35 | 985 | 33 | 1427 | 2791 | 0 | 0 | 0 | 21 | 217 | 240 | 0 | 0 | 25 | 0 | 0 | 0 | 16 | 80 | 0 | 61 |
| SRR9833164 | 73 | 55 | 661 | 0 | 0 | 26 | 154 | 2335 | 33 | 0 | 0 | 73 | 0 | 31 | 2021 | 22 | 50 | 921 | 0 | 2809 | 2478 | 0 | 0 | 0 | 15 | 432 | 183 | 0 | 0 | 20 | 0 | 0 | 0 | 0 | 135 | 0 | 65 |
| SRR9856157 | 41 | 0 | 67 | 0 | 0 | 118 | 164 | 0 | 23 | 12 | 60 | 162 | 198 | 27 | 109049 | 18 | 0 | 17 | 28 | 388 | 27722 | 0 | 0 | 11 | 11 | 16 | 153 | 0 | 0 | 0 | 44 | 11 | 12 | 11 | 48 | 107 | 167 |
| SRR9856158 | 46 | 0 | 121 | 0 | 0 | 95 | 320 | 0 | 18 | 0 | 37 | 216 | 190 | 28 | 42289 | 0 | 28 | 82 | 0 | 108 | 10096 | 0 | 0 | 18 | 542 | 518 | 178 | 0 | 15 | 0 | 40 | 0 | 0 | 294 | 211 | 162 | 551 |
| SRR9856159 | 50 | 0 | 103 | 0 | 0 | 118 | 274 | 0 | 71 | 0 | 42 | 125 | 169 | 22 | 120518 | 23 | 20 | 0 | 12 | 224 | 15435 | 0 | 0 | 53 | 16 | 38 | 12 | 0 | 11 | 0 | 98 | 0 | 16 | 170 | 95 | 147 | 14 |
| SRR9856160 | 12 | 0 | 11 | 0 | 0 | 100 | 354 | 0 | 26 | 0 | 61 | 38 | 112 | 72 | 194750 | 0 | 0 | 0 | 16 | 128 | 20405 | 19 | 0 | 0 | 19 | 366 | 37 | 26 | 0 | 0 | 37 | 23 | 0 | 144 | 19 | 86 | 13 |
| SRR9856161 | 64 | 0 | 128 | 0 | 0 | 121 | 526 | 0 | 30 | 0 | 24 | 211 | 456 | 0 | 19438 | 0 | 25 | 139 | 13 | 114 | 7506 | 0 | 0 | 0 | 290 | 546 | 232 | 0 | 0 | 0 | 91 | 0 | 0 | 317 | 121 | 97 | 305 |
| SRR9856162 | 52 | 0 | 160 | 0 | 0 | 131 | 258 | 0 | 14 | 0 | 23 | 528 | 580 | 0 | 3781 | 14 | 32 | 89 | 0 | 163 | 4338 | 0 | 0 | 0 | 220 | 651 | 229 | 0 | 0 | 0 | 117 | 0 | 0 | 267 | 159 | 98 | 398 |
| SRR9856163 | 62 | 0 | 78 | 0 | 0 | 153 | 544 | 0 | 61 | 0 | 12 | 128 | 336 | 15 | 10459 | 0 | 37 | 1452 | 0 | 300 | 11478 | 0 | 0 | 18 | 345 | 584 | 385 | 0 | 0 | 0 | 67 | 0 | 0 | 150 | 131 | 70 | 287 |
| SRR9856164 | 41 | 0 | 104 | 0 | 0 | 81 | 252 | 0 | 38 | 0 | 42 | 147 | 275 | 29 | 89488 | 0 | 33 | 0 | 0 | 79 | 16058 | 0 | 0 | 20 | 12 | 13 | 11 | 0 | 0 | 11 | 52 | 13 | 0 | 347 | 69 | 38 | 210 |
| SRR9856165 | 38 | 0 | 63 | 0 | 0 | 136 | 310 | 0 | 78 | 10 | 54 | 272 | 614 | 34 | 95108 | 0 | 15 | 0 | 44 | 524 | 14325 | 0 | 0 | 27 | 295 | 533 | 292 | 20 | 0 | 10 | 81 | 0 | 0 | 478 | 78 | 112 | 360 |
| SRR9856166 | 34 | 0 | 33 | 0 | 0 | 84 | 466 | 0 | 25 | 0 | 28 | 120 | 252 | 22 | 52222 | 0 | 20 | 126 | 27 | 351 | 12019 | 0 | 0 | 16 | 539 | 343 | 268 | 16 | 0 | 0 | 69 | 0 | 0 | 121 | 78 | 41 | 100 |
| SRR9856167 | 32 | 0 | 63 | 0 | 0 | 56 | 199 | 17 | 18 | 0 | 26 | 178 | 27 | 47 | 124512 | 0 | 10 | 143 | 77 | 1021 | 11324 | 0 | 0 | 72 | 99 | 12 | 243 | 15 | 0 | 0 | 35 | 0 | 0 | 16 | 56 | 68 | 11 |
| SRR9856168 | 67 | 0 | 71 | 0 | 0 | 73 | 871 | 114 | 31 | 0 | 28 | 197 | 331 | 62 | 95886 | 12 | 29 | 40 | 26 | 574 | 9612 | 0 | 0 | 137 | 272 | 431 | 393 | 19 | 0 | 0 | 59 | 12 | 0 | 163 | 106 | 99 | 478 |
| SRR9856169 | 87 | 0 | 115 | 0 | 0 | 68 | 257 | 74 | 26 | 0 | 53 | 505 | 18 | 24 | 62564 | 0 | 27 | 17 | 14 | 291 | 14720 | 0 | 0 | 108 | 14 | 13 | 17 | 0 | 0 | 20 | 47 | 0 | 0 | 19 | 125 | 56 | 13 |
| SRR9856170 | 83 | 0 | 186 | 0 | 0 | 37 | 123 | 0 | 40 | 47 | 127 | 222 | 442 | 35 | 179266 | 10 | 31 | 54 | 0 | 34 | 6960 | 0 | 0 | 0 | 677 | 472 | 103 | 19 | 0 | 0 | 80 | 0 | 0 | 310 | 261 | 195 | 841 |

### Supp Table S8.2.Columns_AM_to_BW

| | | | | | | | | | | | | | | | | | | | | | | | | | | | | | | | | | | | | | |
|---|---|---|---|---|---|---|---|---|---|---|---|---|---|---|---|---|---|---|---|---|---|---|---|---|---|---|---|---|---|---|---|---|---|---|---|---|---|
| SRR9856171 | 55 | 0 | 137 | 0 | 0 | 13 | 18 | 0 | 0 | 30 | 105 | 1149 | 23 | 42 | 154593 | 21 | 39 | 604 | 0 | 13 | 4966 | 0 | 0 | 0 | 52 | 10 | 18 | 29 | 0 | 0 | 30 | 0 | 20 | 34 | 49 | 49 | 11 |
| SRR9856172 | 96 | 0 | 162 | 0 | 0 | 81 | 500 | 0 | 41 | 12 | 24 | 269 | 13 | 26 | 95043 | 0 | 50 | 155 | 54 | 500 | 4549 | 0 | 0 | 19 | 12 | 11 | 297 | 12 | 0 | 0 | 108 | 0 | 0 | 263 | 194 | 96 | 441 |
| SRR9856173 | 93 | 0 | 145 | 0 | 0 | 38 | 169 | 0 | 19 | 13 | 65 | 637 | 148 | 51 | 21712 | 29 | 39 | 23 | 22 | 226 | 14871 | 0 | 0 | 0 | 238 | 243 | 164 | 36 | 16 | 0 | 43 | 28 | 0 | 234 | 44 | 41 | 160 |
| SRR9856174 | 60 | 0 | 100 | 0 | 0 | 51 | 788 | 0 | 22 | 0 | 61 | 157 | 342 | 72 | 120254 | 0 | 14 | 11 | 26 | 263 | 14097 | 0 | 0 | 67 | 153 | 13 | 150 | 25 | 0 | 0 | 64 | 0 | 0 | 261 | 194 | 12 | 53 |
| SRR9856175 | 63 | 0 | 97 | 0 | 0 | 58 | 306 | 134 | 22 | 0 | 50 | 246 | 492 | 27 | 37330 | 0 | 23 | 458 | 22 | 357 | 14613 | 0 | 0 | 115 | 244 | 367 | 439 | 0 | 0 | 53 | 52 | 0 | 0 | 590 | 84 | 122 | 500 |
| SRR9856176 | 49 | 0 | 73 | 0 | 0 | 75 | 473 | 0 | 33 | 12 | 63 | 209 | 118 | 27 | 42221 | 27 | 20 | 33 | 50 | 663 | 10231 | 0 | 0 | 40 | 32 | 30 | 12 | 0 | 14 | 0 | 70 | 11 | 0 | 23 | 41 | 94 | 13 |
| SRR9856177 | 82 | 0 | 126 | 0 | 0 | 36 | 614 | 10 | 0 | 0 | 48 | 143 | 0 | 30 | 21727 | 25 | 28 | 60 | 22 | 247 | 5121 | 0 | 0 | 32 | 370 | 314 | 137 | 22 | 11 | 0 | 33 | 0 | 13 | 319 | 28 | 63 | 103 |
| SRR9856178 | 10 | 0 | 17 | 0 | 0 | 59 | 124 | 16 | 0 | 0 | 49 | 107 | 319 | 53 | 44983 | 0 | 0 | 0 | 96 | 651 | 6065 | 12 | 0 | 18 | 39 | 237 | 71 | 23 | 0 | 36 | 35 | 0 | 0 | 327 | 59 | 215 | 21 |
| SRR9856179 | 58 | 0 | 112 | 0 | 0 | 52 | 453 | 0 | 29 | 17 | 19 | 473 | 349 | 30 | 7085 | 0 | 25 | 56 | 61 | 749 | 10323 | 0 | 0 | 48 | 447 | 327 | 219 | 17 | 0 | 0 | 40 | 0 | 0 | 657 | 51 | 46 | 167 |
| SRR9856180 | 47 | 0 | 86 | 0 | 0 | 71 | 206 | 0 | 36 | 0 | 16 | 480 | 10 | 0 | 66898 | 0 | 30 | 39 | 14 | 588 | 17446 | 0 | 0 | 48 | 375 | 320 | 320 | 19 | 0 | 0 | 61 | 0 | 0 | 453 | 181 | 207 | 517 |
| SRR9856181 | 40 | 0 | 89 | 0 | 0 | 160 | 303 | 0 | 42 | 0 | 46 | 440 | 413 | 0 | 94023 | 13 | 19 | 0 | 17 | 286 | 11362 | 0 | 0 | 0 | 308 | 522 | 291 | 0 | 0 | 0 | 139 | 0 | 0 | 333 | 160 | 265 | 788 |
| SRR9856182 | 44 | 0 | 130 | 0 | 0 | 81 | 249 | 0 | 55 | 0 | 23 | 459 | 11 | 16 | 77113 | 0 | 17 | 39 | 28 | 446 | 20912 | 0 | 0 | 19 | 203 | 300 | 387 | 0 | 0 | 0 | 110 | 0 | 0 | 300 | 52 | 87 | 277 |
| SRR9856183 | 37 | 0 | 71 | 0 | 0 | 19 | 844 | 0 | 17 | 0 | 26 | 735 | 23 | 29 | 36010 | 10 | 21 | 130 | 0 | 155 | 12719 | 0 | 0 | 0 | 13 | 121 | 171 | 18 | 0 | 0 | 35 | 45 | 0 | 158 | 27 | 34 | 118 |
| SRR9856184 | 50 | 0 | 63 | 0 | 0 | 62 | 978 | 89 | 25 | 0 | 37 | 820 | 17 | 21 | 91614 | 0 | 16 | 0 | 65 | 1170 | 44187 | 0 | 0 | 268 | 205 | 10 | 14 | 15 | 0 | 55 | 42 | 0 | 0 | 267 | 79 | 93 | 12 |
| SRR9856185 | 18 | 0 | 20 | 0 | 0 | 65 | 540 | 0 | 44 | 0 | 36 | 101 | 319 | 51 | 15370 | 15 | 0 | 0 | 203 | 1520 | 9087 | 15 | 0 | 73 | 189 | 522 | 102 | 44 | 0 | 0 | 36 | 0 | 15 | 131 | 44 | 254 | 769 |
| SRR9856186 | 32 | 0 | 34 | 0 | 0 | 53 | 2941 | 29 | 17 | 17 | 138 | 331 | 959 | 34 | 213434 | 0 | 0 | 0 | 44 | 431 | 22650 | 0 | 0 | 82 | 220 | 375 | 148 | 22 | 0 | 39 | 68 | 17 | 17 | 896 | 196 | 889 | 1758 |
| SRR990360 | 0 | 0 | 93 | 0 | 0 | 0 | 0 | 0 | 0 | 0 | 0 | 0 | 0 | 32 | 523 | 0 | 0 | 277 | 0 | 0 | 0 | 0 | 0 | 0 | 80 | 0 | 0 | 0 | 0 | 0 | 0 | 0 | 0 | 23 | 0 | 0 | 0 |
| SRR9924500 | 67 | 0 | 4291 | 0 | 0 | 56 | 0 | 0 | 23 | 0 | 0 | 97 | 0 | 0 | 18 | 0 | 23 | 0 | 68 | 6103 | 562 | 0 | 0 | 0 | 15 | 902 | 5350 | 0 | 0 | 0 | 32 | 0 | 0 | 25 | 350 | 24 | 212 |
| SRR9924502 | 50 | 21 | 4783 | 0 | 0 | 31 | 12 | 0 | 15 | 0 | 0 | 102 | 0 | 0 | 72 | 0 | 24 | 0 | 12 | 3541 | 674 | 13 | 0 | 0 | 17 | 82 | 348 | 10 | 0 | 0 | 38 | 11 | 13 | 22 | 27 | 0 | 19 |
| SRR9924511 | 69 | 17 | 5553 | 0 | 0 | 21 | 24 | 77 | 13 | 0 | 0 | 89 | 0 | 0 | 47 | 0 | 20 | 0 | 74 | 6712 | 925 | 12 | 0 | 11 | 0 | 521 | 4920 | 19 | 0 | 16 | 21 | 0 | 0 | 11 | 257 | 26 | 159 |
| SRR9924512 | 73 | 0 | 6769 | 0 | 0 | 38 | 24 | 44 | 15 | 0 | 0 | 70 | 0 | 0 | 29 | 0 | 18 | 0 | 29 | 5091 | 1014 | 0 | 0 | 0 | 14 | 589 | 3250 | 12 | 14 | 0 | 16 | 0 | 16 | 14 | 267 | 18 | 125 |
| SRR9924514 | 83 | 13 | 5567 | 0 | 0 | 25 | 0 | 25 | 0 | 0 | 0 | 91 | 0 | 17 | 55 | 0 | 23 | 13 | 55 | 8577 | 1116 | 25 | 0 | 11 | 0 | 60 | 602 | 38 | 0 | 0 | 10 | 0 | 0 | 34 | 31 | 22 | 23 |
| SRR9924515 | 60 | 11 | 5329 | 0 | 0 | 18 | 13 | 29 | 0 | 0 | 0 | 78 | 0 | 0 | 35 | 22 | 26 | 0 | 57 | 4900 | 716 | 0 | 0 | 0 | 11 | 47 | 417 | 0 | 0 | 0 | 13 | 11 | 0 | 11 | 27 | 10 | 16 |
| SRR9924516 | 73 | 17 | 5407 | 0 | 0 | 22 | 0 | 0 | 0 | 0 | 0 | 83 | 0 | 23 | 56 | 0 | 14 | 12 | 19 | 1312 | 479 | 0 | 0 | 0 | 15 | 39 | 194 | 69 | 0 | 0 | 17 | 0 | 0 | 26 | 35 | 16 | 17 |
| SRR9924517 | 73 | 10 | 5656 | 0 | 0 | 21 | 0 | 0 | 0 | 0 | 0 | 90 | 0 | 16 | 43 | 0 | 25 | 32 | 22 | 5627 | 1207 | 27 | 0 | 0 | 18 | 55 | 543 | 46 | 0 | 0 | 19 | 0 | 0 | 40 | 41 | 22 | 23 |
| SRR9924518 | 77 | 0 | 5318 | 0 | 0 | 40 | 0 | 11 | 0 | 0 | 0 | 97 | 0 | 0 | 23 | 0 | 25 | 0 | 13 | 2397 | 281 | 0 | 0 | 0 | 16 | 81 | 368 | 11 | 0 | 0 | 30 | 0 | 10 | 15 | 22 | 0 | 13 |
| SRR9924519 | 84 | 0 | 6184 | 0 | 0 | 21 | 17 | 32 | 0 | 0 | 0 | 120 | 0 | 12 | 23 | 0 | 19 | 0 | 25 | 4433 | 505 | 0 | 0 | 11 | 17 | 90 | 439 | 19 | 0 | 0 | 15 | 0 | 38 | 30 | 23 | 0 | 14 |
| SRR9924520 | 68 | 0 | 5882 | 0 | 0 | 25 | 13 | 19 | 14 | 0 | 0 | 115 | 0 | 11 | 0 | 0 | 17 | 0 | 35 | 9434 | 1287 | 16 | 0 | 61 | 0 | 523 | 3977 | 0 | 0 | 0 | 25 | 0 | 0 | 27 | 184 | 14 | 143 |
| SRR9924521 | 83 | 0 | 5690 | 0 | 0 | 26 | 13 | 13 | 0 | 0 | 0 | 121 | 0 | 0 | 22 | 0 | 19 | 0 | 17 | 3017 | 357 | 11 | 0 | 0 | 10 | 81 | 370 | 0 | 0 | 0 | 25 | 0 | 11 | 25 | 20 | 12 | 13 |
| SRR9924522 | 70 | 13 | 5284 | 0 | 0 | 19 | 0 | 0 | 0 | 0 | 0 | 106 | 0 | 26 | 18 | 0 | 15 | 0 | 12 | 2931 | 743 | 11 | 0 | 0 | 18 | 293 | 1434 | 24 | 0 | 0 | 22 | 0 | 0 | 32 | 150 | 0 | 104 |
| SRR9924523 | 77 | 15 | 5928 | 0 | 0 | 24 | 0 | 0 | 0 | 0 | 0 | 119 | 0 | 0 | 48 | 0 | 19 | 0 | 27 | 5201 | 631 | 0 | 0 | 10 | 16 | 58 | 404 | 0 | 0 | 0 | 21 | 0 | 0 | 0 | 24 | 0 | 14 |
| SRR9924524 | 90 | 0 | 5386 | 0 | 0 | 11 | 26 | 42 | 18 | 0 | 0 | 110 | 0 | 0 | 20 | 12 | 24 | 0 | 66 | 6334 | 734 | 0 | 0 | 0 | 34 | 235 | 430 | 18 | 11 | 0 | 38 | 0 | 26 | 11 | 23 | 0 | 11 |
| SRR9924525 | 45 | 0 | 3848 | 0 | 0 | 50 | 20 | 27 | 17 | 0 | 0 | 71 | 0 | 0 | 41 | 0 | 20 | 0 | 29 | 8702 | 760 | 0 | 0 | 66 | 0 | 888 | 5805 | 0 | 0 | 0 | 31 | 0 | 16 | 10 | 242 | 25 | 141 |
| SRR9924526 | 90 | 0 | 5014 | 0 | 0 | 11 | 39 | 26 | 18 | 0 | 0 | 99 | 0 | 0 | 0 | 0 | 20 | 0 | 48 | 7221 | 525 | 0 | 0 | 0 | 17 | 230 | 476 | 0 | 11 | 0 | 32 | 0 | 19 | 47 | 20 | 14 | 11 |
| SRR9924527 | 82 | 13 | 5631 | 0 | 0 | 63 | 17 | 19 | 18 | 0 | 0 | 118 | 0 | 0 | 0 | 0 | 19 | 0 | 48 | 5068 | 947 | 0 | 0 | 0 | 14 | 1344 | 2561 | 0 | 0 | 0 | 32 | 0 | 20 | 26 | 118 | 0 | 74 |
