## Supplementary material for "Single-nucleotide Differences and Cell Type Decide the Subcellular Localization of miRNA Isoforms (isomiRs), tRNA-derived Fragments (tRFs) and rRNA-derived Fragments (rRFs)": Supp. Table S8 (3 of 5)

### Supp Table S8.3.Columns_BX_to_DH

**Supplemental Table S8:** The abundances

**111 Short RNA (Figure 5)**

| | >hg38_21_+_8433172_8446622_RNA45SN1_45S_50ntFlanks@475.506.32\|CTTCGTGATCGATGTGGTGACGTCGTGCTCTC | >hg38_21_+_8433172_8446622_RNA45SN1_45S_50ntFlanks@475.507.33\|CTTCGTGATCGATGTGGTGACGTCGTGCTCTCC | >hg38_21_+_8433172_8446622_RNA45SN1_45S_50ntFlanks@475.514.40\|CTTCGTGATCGATGTGGTGACGTCGTGCTCTCCCGGGCCG | >hg38_21_+_8433172_8446622_RNA45SN1_45S_50ntFlanks@475.515.41\|CTTCGTGATCGATGTGGTGACGTCGTGCTCTCCCGGGCCGG | >hg38_21_+_8433172_8446622_RNA45SN1_45S_50ntFlanks@6652.6683.32\|GACTCTTAGCGGTGGATCACTCGGCTCGTGCG | >trnaMT_SerGCT_MT_+_12207_12265@1.19.19\|GAGAAAGCTCACAAGAACT | >trnaMT_SerGCT_MT_+_12207_12265@1.21.21\|GAGAAAGCTCACAAGAACTGC | >trnaMT_SerGCT_MT_+_12207_12265@1.22.22\|GAGAAAGCTCACAAGAACTGCT | >trnaMT_SerGCT_MT_+_12207_12265@1.25.25\|GAGAAAGCTCACAAGAACTGCTAAC | >hg38_21_+_8433172_8446622_RNA45SN1_45S_50ntFlanks@5491.5516.26\|GAGAAGACGGTCGAACTTGACTATCT | >hg38_1_-_228634819_228635039_RNA5S12_5S_50ntFlanks@132.169.38\|GATGGGAGACCGCCTGGGAATACCGGGTGCTGTAGGCT | >hg38_21_+_8433172_8446622_RNA45SN1_45S_50ntFlanks@12419.12448.30\|GATGTCGGCTCTTCCTATCATTGTGAAGCA | >hg38_21_+_8433172_8446622_RNA45SN1_45S_50ntFlanks@485.505.21\|GATGTGGTGACGTCGTGCTCT | >hg38_21_+_8433172_8446622_RNA45SN1_45S_50ntFlanks@485.506.22\|GATGTGGTGACGTCGTGCTCTC | >trna2_GlyGCC_21_-_18827107_18827177@1.32.32,trna35_GlyGCC_1_+_161413094_161413164@1.32.32,trna37_GlyGCC_1_+_161420467_161420537@1.32.32,trna39_GlyGCC_1_+_161427898_161427968@1.32.32,trna41_GlyGCC_1_+_161435258_161435328@1.32.32\|GCATGGGTGGTTCAGTGGTAGAATTCTC | >am_trna128_GlyGCC_6_-_27870686_27870756@1.31.31,trna133_GlyCCC_1_-_16872434_16872504@1.31.31,trna18_GlyGCC_16_+_70822597_70822667@1.31.31,trna19_GlyGCC_16_+_70823410_70823480@1.31.31,trna19_GlyGCC_2_-_157257659_157257729@1.31.31,trna24_GlyGCC_16_- | >am_trna128_GlyGCC_6_-_27870686_27870756@1.32.32,trna133_GlyCCC_1_-_16872434_16872504@1.32.32,trna18_GlyGCC_16_+_70822597_70822667@1.32.32,trna19_GlyGCC_16_+_70823410_70823480@1.32.32,trna19_GlyGCC_2_-_157257659_157257729@1.32.32,trna24_GlyGCC_16_- | >trna119_LysCTT_1_-_145395522_145395594@1.29.29,trna11_LysCTT_5_-_180648979_180649051@1.29.29,trna13_LysCTT_6_+_26556774_26556846@1.29.29,trna32_LysCTT_16_-_3207406_3207478@1.29.29,trna7_LysCTT_16_+_3225692_3225764@1.29.29,trna9_LysCTT_5_+_180634755_180 | >trna119_LysCTT_1_-_145395522_145395594@1.35.35,trna11_LysCTT_5_-_180648979_180649051@1.35.35,trna13_LysCTT_6_+_26556774_26556846@1.35.35,trna7_LysCTT_16_+_3225692_3225764@1.35.35,trna9_LysCTT_5_+_180634755_180634827@1.35.35\|GCCCGGCTAGCTCAGTCGGTAGAGCAT | >trna10_LysCTT_16_+_3241501_3241573@1.34.34,trna13_LysCTT_14_-_58706613_58706685@1.34.34,trna2_LysCTT_15_+_79152904_79152976@1.34.34\|GCCCGGCTAGCTCAGTCGGTAGAGCATGGGACTC | >trna111_HisGTG_1_-_147774845_147774916@1.33.33,trna118_HisGTG_1_-_145396881_145396952@1.33.33,trna16_HisGTG_1_+_146544773_146544844@1.33.33,trna1_HisGTG_15_+_45493349_45493420@1.33.33,trna21_HisGTG_1_+_147753471_147753542@1.33.33,trna33_HisGTG_6_+_27125906_2712597 | >hg38_21_+_8433172_8446622_RNA45SN1_45S_50ntFlanks@8114.8140.27\|GCGCGGGACATGTGGCGTACGGAAGAC | >hg38_21_+_8433172_8446622_RNA45SN1_45S_50ntFlanks@8114.8141.28\|GCGCGGGACATGTGGCGTACGGAAGACC | >multi-hg38_MT_+_1621_3279_RNR2_16S_50ntFlanks@51.74.24\|GCTAAACCTAGCCCCAAACCCACT | >multi-hg38_MT_+_1621_3279_RNR2_16S_50ntFlanks@51.84.34\|GCTAAACCTAGCCCCAAACCCACTCCACCTTACT | >multi-hg38_MT_+_1621_3279_RNR2_16S_50ntFlanks@51.87.37\|GCTAAACCTAGCCCCAAACCCACTCCACCTTACTACC | >multi-hg38_MT_+_1621_3279_RNR2_16S_50ntFlanks@51.88.38\|GCTAAACCTAGCCCCAAACCCACTCCACCTTACTACCA | >hg38_MT_+_1621_3279_RNR2_16S_50ntFlanks@51.95.45\|GCTAAACCTAGCCCCAAACCCACTCCACCTTACTACCAGACAACC | >hg38_21_+_8433172_8446622_RNA45SN1_45S_50ntFlanks@12990.13013.24\|GCTCCCTCGCTGCGATCTATTGAA | >hg38_1_-_228634819_228635039_RNA5S12_5S_50ntFlanks@136.170.35\|GGAGACCGCCTGGGAATACCGGGTGCTGTAGGCTT | >hg38_21_+_8433172_8446622_RNA45SN1_45S_50ntFlanks@8113.8138.26\|GGCGCGGGACATGTGGCGTACGGAAG | >trna16_GlnTTG_17_+_47269890_47269961@1.29.29,trna64_GlnTTG_6_+_28557156_28557227@1.29.29,trna84_GlnTTG_6_+_145503859_145503930@1.29.29\|GGTCCCATGGTGTAATGGTTAGCACTCTG | >hg38_21_+_8433172_8446622_RNA45SN1_45S_50ntFlanks@9545.9566.22\|GTGGAGGTCCGTAGCGGTCCTG | >hg38_21_+_8433172_8446622_RNA45SN1_45S_50ntFlanks@9545.9570.26\|GTGGAGGTCCGTAGCGGTCCTGACGT | >am_trna10_ValCAC_5_-_180649395_180649467@1.23.23,trna12_ValAAC_5_-_180645270_180645342@1.23.23,trna132_ValAAC_6_-_27721179_27721251@1.23.23,trna136_ValAAC_6_-_27648885_27648957@1.23.23,trna139_ValAAC_6_-_27618707_27618779@1.23.23,trna18_ValCAC | >MI0000737\|hsa-mir-200a&WithFlank&1\|+\|1167857\|1167958@60.82.23&[MIMAT0000682&hsa-miR-200a-3p&offsets\|0\|+1;m-142&1\|+\|1167916\|1167938&offsets\|0\|0]\|TAACACTGTCTGGTAACGATGT | >hg38_21_+_8433172_8446622_RNA45SN1_45S_50ntFlanks@11729.11753.25\|TAACTATGACTCTCTTAAGGTAGCC |
|---|---|---|---|---|---|---|---|---|---|---|---|---|---|---|---|---|---|---|---|---|---|---|---|---|---|---|---|---|---|---|---|---|---|---|---|---|---|
| SRR10024150 | 634 | 485 | 66 | 102 | 501 | 383 | 35 | 221 | 581 | 250 | 859 | 0 | 0 | 0 | 906 | 142230 | 22091 | 12 | 93 | 558 | 163 | 0 | 0 | 332 | 87 | 107 | 220 | 0 | 0 | 342 | 0 | 23 | 0 | 0 | 7706 | 0 | 0 |
| SRR10024151 | 24 | 27 | 124 | 133 | 45 | 29 | 0 | 126 | 766 | 342 | 15 | 0 | 0 | 0 | 4846 | 292677 | 66064 | 29 | 136 | 417 | 998 | 0 | 0 | 364 | 83 | 113 | 163 | 0 | 0 | 243 | 0 | 107 | 0 | 0 | 14373 | 0 | 0 |
| SRR10024152 | 406 | 576 | 31 | 46 | 488 | 244 | 23 | 47 | 70 | 247 | 874 | 21 | 0 | 0 | 605 | 109784 | 12426 | 12 | 93 | 419 | 442 | 0 | 0 | 123 | 36 | 41 | 123 | 0 | 10 | 211 | 0 | 0 | 0 | 0 | 12688 | 0 | 0 |
| SRR10024153 | 597 | 707 | 0 | 25 | 1095 | 339 | 31 | 339 | 941 | 244 | 214 | 30 | 0 | 0 | 509 | 175650 | 25480 | 31 | 93 | 386 | 524 | 0 | 0 | 408 | 50 | 25 | 70 | 0 | 0 | 179 | 0 | 31 | 0 | 0 | 9753 | 0 | 0 |
| SRR10024154 | 168 | 142 | 13 | 41 | 304 | 96 | 16 | 193 | 818 | 164 | 310 | 0 | 0 | 0 | 706 | 126332 | 28429 | 0 | 48 | 401 | 273 | 0 | 0 | 76 | 25 | 28 | 66 | 0 | 0 | 60 | 0 | 16 | 0 | 0 | 12785 | 0 | 0 |
| SRR10024155 | 222 | 259 | 0 | 0 | 457 | 230 | 13 | 204 | 676 | 88 | 347 | 230 | 0 | 0 | 574 | 116137 | 29124 | 38 | 344 | 1326 | 2410 | 0 | 0 | 112 | 16 | 18 | 22 | 0 | 94 | 62 | 0 | 38 | 0 | 0 | 7382 | 0 | 0 |
| SRR10024156 | 12 | 15 | 30 | 23 | 21 | 280 | 16 | 156 | 607 | 356 | 25 | 25 | 0 | 0 | 1090 | 174688 | 36047 | 0 | 78 | 529 | 1448 | 0 | 0 | 262 | 155 | 104 | 117 | 0 | 0 | 391 | 0 | 0 | 0 | 0 | 26144 | 0 | 0 |
| SRR10024158 | 133 | 150 | 0 | 0 | 345 | 201 | 29 | 373 | 1651 | 213 | 82 | 0 | 0 | 0 | 631 | 81902 | 20878 | 0 | 43 | 258 | 330 | 0 | 0 | 116 | 58 | 17 | 14 | 0 | 0 | 51 | 0 | 29 | 0 | 0 | 13959 | 0 | 0 |
| SRR10024159 | 380 | 489 | 0 | 0 | 640 | 127 | 0 | 85 | 579 | 151 | 53 | 101 | 0 | 0 | 282 | 141323 | 17367 | 28 | 56 | 268 | 790 | 0 | 0 | 154 | 11 | 0 | 0 | 0 | 53 | 78 | 0 | 56 | 0 | 0 | 25483 | 0 | 0 |
| SRR10024161 | 244 | 256 | 19 | 15 | 542 | 94 | 0 | 214 | 617 | 95 | 321 | 115 | 0 | 0 | 1063 | 98972 | 26999 | 17 | 94 | 334 | 677 | 0 | 0 | 221 | 34 | 46 | 61 | 0 | 42 | 156 | 0 | 26 | 0 | 0 | 3637 | 0 | 0 |
| SRR10024162 | 18 | 28 | 34 | 53 | 30 | 179 | 11 | 137 | 580 | 280 | 424 | 12 | 0 | 0 | 949 | 210314 | 26711 | 21 | 116 | 285 | 295 | 0 | 0 | 299 | 84 | 84 | 146 | 0 | 0 | 215 | 0 | 21 | 0 | 0 | 14446 | 0 | 0 |
| SRR10024163 | 15 | 12 | 98 | 98 | 24 | 98 | 14 | 84 | 112 | 191 | 11 | 28 | 0 | 0 | 1772 | 177521 | 39413 | 14 | 49 | 293 | 251 | 0 | 0 | 252 | 47 | 61 | 56 | 0 | 0 | 266 | 0 | 14 | 0 | 0 | 9267 | 0 | 0 |
| SRR10024164 | 702 | 260 | 0 | 35 | 347 | 171 | 29 | 143 | 913 | 217 | 121 | 26 | 0 | 0 | 1569 | 178962 | 32056 | 29 | 143 | 913 | 285 | 0 | 0 | 78 | 0 | 0 | 26 | 0 | 0 | 130 | 0 | 114 | 0 | 0 | 6965 | 0 | 0 |
| SRR10024166 | 21 | 19 | 0 | 0 | 12 | 84 | 17 | 17 | 218 | 384 | 165 | 12 | 0 | 0 | 445 | 114638 | 18110 | 17 | 0 | 218 | 311 | 0 | 0 | 508 | 25 | 12 | 33 | 0 | 0 | 78 | 0 | 25 | 0 | 0 | 21158 | 0 | 0 |
| SRR10024167 | 92 | 171 | 0 | 0 | 232 | 240 | 32 | 144 | 624 | 212 | 27 | 0 | 0 | 0 | 256 | 97549 | 14196 | 16 | 48 | 256 | 112 | 0 | 0 | 154 | 14 | 0 | 0 | 0 | 0 | 24 | 0 | 16 | 0 | 0 | 8666 | 0 | 0 |
| SRR10024168 | 237 | 97 | 0 | 15 | 407 | 177 | 32 | 145 | 563 | 58 | 10 | 0 | 0 | 0 | 1319 | 247605 | 23909 | 113 | 241 | 1834 | 434 | 0 | 0 | 194 | 15 | 29 | 39 | 0 | 0 | 276 | 0 | 97 | 0 | 0 | 17950 | 0 | 0 |
| SRR10024169 | 15 | 18 | 11 | 13 | 30 | 34 | 11 | 149 | 480 | 312 | 69 | 0 | 0 | 0 | 434 | 112977 | 25413 | 23 | 80 | 263 | 1737 | 0 | 0 | 261 | 27 | 11 | 18 | 0 | 0 | 55 | 0 | 46 | 0 | 0 | 26371 | 0 | 0 |
| SRR10024170 | 525 | 382 | 22 | 30 | 633 | 91 | 23 | 68 | 285 | 221 | 299 | 13 | 0 | 0 | 854 | 118400 | 17757 | 11 | 23 | 251 | 399 | 0 | 0 | 174 | 48 | 35 | 26 | 0 | 0 | 108 | 0 | 11 | 0 | 0 | 5849 | 0 | 0 |
| SRR10024171 | 294 | 286 | 49 | 47 | 11 | 49 | 25 | 111 | 590 | 200 | 12 | 0 | 0 | 0 | 775 | 131195 | 24683 | 49 | 123 | 209 | 357 | 0 | 0 | 67 | 44 | 57 | 69 | 0 | 0 | 136 | 0 | 37 | 0 | 0 | 14721 | 0 | 0 |
| SRR10024173 | 545 | 252 | 23 | 23 | 329 | 240 | 34 | 445 | 2157 | 99 | 698 | 14 | 0 | 0 | 1900 | 176474 | 46370 | 0 | 154 | 993 | 719 | 0 | 0 | 90 | 68 | 32 | 50 | 0 | 0 | 167 | 0 | 68 | 0 | 0 | 4381 | 0 | 0 |
| SRR10024176 | 67 | 81 | 0 | 0 | 219 | 48 | 0 | 101 | 564 | 336 | 101 | 0 | 0 | 0 | 335 | 60462 | 15752 | 0 | 0 | 57 | 75 | 0 | 0 | 82 | 34 | 0 | 0 | 0 | 0 | 58 | 0 | 0 | 0 | 0 | 6070 | 0 | 0 |
| SRR10024178 | 214 | 323 | 0 | 17 | 17 | 198 | 0 | 64 | 92 | 231 | 10 | 0 | 0 | 0 | 454 | 78931 | 14735 | 43 | 35 | 248 | 333 | 0 | 0 | 270 | 22 | 22 | 45 | 0 | 11 | 161 | 0 | 35 | 0 | 0 | 32596 | 0 | 0 |
| SRR10024179 | 89 | 95 | 0 | 0 | 469 | 83 | 33 | 183 | 766 | 49 | 55 | 161 | 0 | 0 | 683 | 96899 | 18651 | 17 | 133 | 183 | 2599 | 0 | 0 | 40 | 0 | 0 | 13 | 0 | 201 | 17 | 0 | 67 | 0 | 15 | 14347 | 0 | 0 |
| SRR10024180 | 272 | 320 | 0 | 0 | 456 | 223 | 14 | 223 | 877 | 217 | 92 | 0 | 0 | 0 | 292 | 103135 | 14029 | 0 | 28 | 195 | 153 | 0 | 0 | 66 | 11 | 0 | 15 | 0 | 0 | 29 | 0 | 0 | 0 | 0 | 10741 | 0 | 0 |
| SRR10024181 | 367 | 590 | 19 | 75 | 1180 | 144 | 0 | 65 | 405 | 279 | 421 | 0 | 0 | 0 | 1501 | 130664 | 18823 | 13 | 13 | 183 | 183 | 0 | 0 | 66 | 28 | 94 | 176 | 0 | 0 | 129 | 0 | 13 | 0 | 0 | 16632 | 0 | 0 |
| SRR10024182 | 13 | 16 | 43 | 48 | 22 | 129 | 0 | 97 | 612 | 439 | 584 | 43 | 0 | 0 | 1837 | 175151 | 29361 | 16 | 16 | 226 | 97 | 0 | 0 | 154 | 48 | 43 | 82 | 0 | 63 | 179 | 0 | 64 | 0 | 0 | 21036 | 0 | 0 |
| SRR10024183 | 348 | 382 | 40 | 46 | 730 | 38 | 0 | 64 | 480 | 134 | 641 | 67 | 0 | 0 | 493 | 100752 | 21686 | 0 | 58 | 346 | 724 | 0 | 0 | 61 | 12 | 37 | 79 | 0 | 12 | 137 | 0 | 0 | 0 | 12 | 10863 | 0 | 0 |
| SRR10024185 | 602 | 563 | 27 | 48 | 506 | 63 | 0 | 125 | 601 | 2654 | 1431 | 15 | 0 | 0 | 576 | 180478 | 22432 | 38 | 13 | 451 | 338 | 0 | 0 | 328 | 105 | 24 | 66 | 0 | 0 | 768 | 0 | 38 | 0 | 0 | 6861 | 0 | 0 |
| SRR10024186 | 149 | 182 | 17 | 11 | 179 | 69 | 0 | 77 | 405 | 127 | 113 | 0 | 0 | 0 | 456 | 84558 | 15339 | 0 | 0 | 164 | 69 | 0 | 0 | 118 | 14 | 11 | 19 | 0 | 0 | 61 | 0 | 0 | 0 | 0 | 13132 | 0 | 0 |
| SRR10024187 | 13 | 359 | 44 | 31 | 290 | 44 | 0 | 79 | 421 | 31 | 913 | 31 | 0 | 0 | 1166 | 66730 | 24131 | 18 | 70 | 824 | 473 | 0 | 0 | 14 | 44 | 113 | 176 | 0 | 0 | 384 | 0 | 0 | 0 | 0 | 719 | 0 | 0 |
| SRR10024188 | 782 | 782 | 0 | 16 | 335 | 55 | 0 | 37 | 65 | 351 | 185 | 16 | 0 | 0 | 461 | 92026 | 12775 | 0 | 46 | 138 | 877 | 0 | 0 | 141 | 0 | 0 | 12 | 0 | 0 | 85 | 0 | 55 | 0 | 0 | 9685 | 0 | 0 |
| SRR10024189 | 813 | 575 | 0 | 0 | 967 | 59 | 59 | 188 | 377 | 163 | 121 | 21 | 0 | 0 | 1531 | 223674 | 37363 | 47 | 130 | 636 | 1213 | 0 | 0 | 167 | 63 | 0 | 25 | 0 | 0 | 104 | 0 | 24 | 0 | 0 | 10681 | 0 | 0 |
| SRR10024190 | 40 | 39 | 0 | 12 | 494 | 59 | 0 | 59 | 139 | 555 | 121 | 133 | 0 | 0 | 753 | 176452 | 22096 | 0 | 0 | 178 | 2438 | 0 | 0 | 76 | 12 | 0 | 12 | 0 | 0 | 48 | 0 | 79 | 0 | 0 | 51213 | 0 | 0 |
| SRR10024191 | 434 | 130 | 43 | 33 | 586 | 79 | 0 | 209 | 890 | 33 | 293 | 0 | 0 | 0 | 2198 | 134049 | 25887 | 79 | 157 | 1099 | 183 | 0 | 0 | 293 | 54 | 87 | 108 | 11 | 0 | 98 | 0 | 0 | 0 | 11 | 8943 | 0 | 0 |
| SRR10024192 | 1399 | 972 | 32 | 64 | 1335 | 141 | 35 | 141 | 528 | 32 | 288 | 21 | 0 | 0 | 1548 | 219959 | 20464 | 35 | 106 | 493 | 457 | 0 | 0 | 448 | 53 | 53 | 107 | 11 | 0 | 139 | 0 | 70 | 0 | 11 | 7849 | 0 | 0 |
| SRR10024193 | 547 | 11 | 18 | 69 | 27 | 327 | 55 | 454 | 2617 | 199 | 391 | 0 | 0 | 0 | 2817 | 205961 | 40397 | 18 | 164 | 582 | 273 | 0 | 0 | 323 | 91 | 47 | 72 | 0 | 0 | 185 | 0 | 18 | 0 | 0 | 5854 | 0 | 0 |
| SRR10024194 | 173 | 149 | 0 | 0 | 284 | 150 | 30 | 150 | 800 | 229 | 199 | 0 | 0 | 0 | 770 | 98727 | 21327 | 30 | 70 | 450 | 50 | 0 | 0 | 50 | 32 | 15 | 21 | 0 | 0 | 70 | 0 | 30 | 0 | 0 | 11228 | 0 | 0 |
| SRR10024195 | 397 | 837 | 24 | 31 | 1167 | 150 | 12 | 35 | 69 | 377 | 892 | 0 | 0 | 0 | 647 | 150545 | 15637 | 12 | 46 | 150 | 358 | 0 | 0 | 216 | 31 | 47 | 67 | 0 | 0 | 149 | 0 | 23 | 0 | 0 | 15098 | 0 | 0 |
| SRR10024196 | 884 | 438 | 0 | 31 | 665 | 71 | 0 | 18 | 248 | 524 | 203 | 0 | 0 | 0 | 656 | 194844 | 25778 | 0 | 106 | 319 | 603 | 0 | 0 | 376 | 16 | 31 | 23 | 0 | 0 | 78 | 0 | 0 | 0 | 16 | 11377 | 0 | 0 |
| SRR10024197 | 195 | 199 | 22 | 30 | 173 | 139 | 46 | 201 | 788 | 177 | 579 | 0 | 0 | 0 | 432 | 118195 | 13781 | 0 | 62 | 247 | 154 | 0 | 0 | 125 | 30 | 39 | 78 | 0 | 0 | 86 | 0 | 31 | 0 | 0 | 7013 | 0 | 0 |
| SRR10024198 | 165 | 159 | 11 | 13 | 446 | 157 | 13 | 92 | 616 | 101 | 239 | 0 | 0 | 0 | 1232 | 137462 | 33049 | 26 | 39 | 393 | 655 | 0 | 0 | 98 | 24 | 27 | 35 | 0 | 35 | 69 | 0 | 39 | 0 | 0 | 12659 | 0 | 0 |
| SRR10024199 | 626 | 420 | 19 | 26 | 659 | 196 | 18 | 71 | 178 | 497 | 697 | 19 | 0 | 0 | 1371 | 266552 | 32905 | 89 | 125 | 855 | 499 | 0 | 0 | 161 | 26 | 65 | 52 | 0 | 0 | 168 | 0 | 0 | 0 | 0 | 10991 | 0 | 0 |
| SRR10024201 | 429 | 541 | 0 | 23 | 1159 | 140 | 53 | 316 | 860 | 230 | 144 | 14 | 0 | 0 | 263 | 117881 | 15360 | 35 | 53 | 211 | 158 | 0 | 0 | 185 | 14 | 23 | 27 | 0 | 0 | 45 | 0 | 53 | 0 | 0 | 14853 | 0 | 0 |
| SRR10024202 | 1204 | 622 | 32 | 16 | 486 | 68 | 17 | 68 | 102 | 407 | 997 | 16 | 0 | 0 | 1654 | 198938 | 38289 | 68 | 136 | 921 | 699 | 0 | 0 | 207 | 40 | 32 | 72 | 0 | 0 | 391 | 0 | 34 | 0 | 0 | 10369 | 0 | 0 |
| SRR10024203 | 506 | 12 | 14 | 28 | 24 | 218 | 0 | 139 | 1291 | 221 | 156 | 0 | 0 | 0 | 2185 | 143213 | 25705 | 79 | 60 | 258 | 556 | 0 | 0 | 221 | 83 | 60 | 120 | 0 | 0 | 87 | 0 | 79 | 0 | 0 | 8943 | 0 | 0 |
| SRR10024204 | 306 | 12 | 0 | 0 | 13 | 293 | 0 | 70 | 153 | 163 | 94 | 0 | 0 | 0 | 474 | 75376 | 8478 | 14 | 28 | 167 | 209 | 0 | 0 | 85 | 0 | 0 | 0 | 0 | 0 | 59 | 0 | 28 | 0 | 0 | 14673 | 0 | 0 |
| SRR10024205 | 1649 | 1103 | 0 | 23 | 1533 | 153 | 76 | 191 | 458 | 197 | 163 | 35 | 0 | 0 | 420 | 187861 | 18990 | 0 | 76 | 458 | 1031 | 0 | 0 | 407 | 12 | 58 | 58 | 0 | 23 | 128 | 0 | 76 | 0 | 0 | 14303 | 0 | 12 |
| SRR10024206 | 316 | 12 | 31 | 19 | 32 | 32 | 0 | 169 | 1014 | 238 | 274 | 0 | 0 | 0 | 1848 | 218699 | 61406 | 53 | 95 | 475 | 2482 | 0 | 0 | 190 | 67 | 38 | 42 | 0 | 0 | 71 | 0 | 0 | 0 | 0 | 8295 | 0 | 0 |
| SRR10024207 | 265 | 250 | 0 | 0 | 265 | 47 | 16 | 63 | 598 | 178 | 92 | 15 | 0 | 0 | 929 | 122394 | 27378 | 63 | 16 | 567 | 252 | 0 | 0 | 163 | 0 | 0 | 0 | 0 | 15 | 31 | 0 | 47 | 0 | 0 | 8410 | 0 | 0 |
| SRR10024208 | 375 | 227 | 12 | 0 | 463 | 409 | 47 | 251 | 550 | 144 | 263 | 80 | 0 | 0 | 346 | 182725 | 13933 | 16 | 141 | 440 | 660 | 0 | 0 | 64 | 12 | 44 | 44 | 0 | 36 | 152 | 0 | 47 | 0 | 0 | 11793 | 0 | 0 |
| SRR1003025 | 85 | 37 | 106 | 13 | 13 | 168 | 208 | 327 | 50 | 13 | 25 | 0 | 0 | 0 | 6573 | 54024 | 67760 | 0 | 364 | 32959 | 2499 | 0 | 0 | 16 | 28 | 47 | 21 | 0 | 0 | 68 | 0 | 0 | 0 | 0 | 749 | 0 | 0 |
| SRR1005875 | 0 | 0 | 0 | 0 | 19 | 0 | 0 | 91 | 0 | 19 | 29 | 0 | 0 | 0 | 4009 | 189877 | 117466 | 91 | 0 | 6742 | 547 | 0 | 0 | 126 | 0 | 0 | 48 | 0 | 0 | 39 | 0 | 0 | 0 | 0 | 2487 | 0 | 58 |
| SRR10072377 | 16 | 0 | 0 | 0 | 146 | 15065 | 8712 | 48139 | 50572 | 147 | 363 | 0 | 0 | 0 | 22966 | 172542 | 269176 | 146 | 2317 | 6091 | 1488 | 0 | 0 | 224 | 490 | 257 | 286 | 81 | 0 | 2259 | 0 | 0 | 0 | 11 | 2625 | 0 | 0 |
| SRR10072378 | 0 | 0 | 0 | 0 | 21 | 15143 | 8723 | 46817 | 50468 | 15 | 54 | 14 | 0 | 0 | 24607 | 175791 | 274443 | 128 | 2321 | 6292 | 1720 | 0 | 0 | 30 | 67 | 40 | 38 | 12 | 0 | 314 | 0 | 0 | 0 | 15 | 2564 | 0 | 0 |
| SRR10072379 | 22 | 10 | 0 | 13 | 134 | 45335 | 22080 | 114411 | 96720 | 16 | 373 | 19 | 0 | 0 | 1661 | 67580 | 64174 | 267 | 2608 | 974 | 153 | 0 | 0 | 705 | 943 | 334 | 334 | 41 | 0 | 1960 | 0 | 0 | 0 | 0 | 1930 | 15 | 0 |
| SRR10072380 | 28 | 0 | 0 | 12 | 122 | 45250 | 22276 | 116152 | 98440 | 18 | 379 | 25 | 0 | 0 | 1815 | 69447 | 67004 | 274 | 2429 | 973 | 150 | 0 | 0 | 707 | 1001 | 385 | 303 | 43 | 0 | 2167 | 0 | 0 | 0 | 0 | 1816 | 14 | 0 |
| SRR10072381 | 0 | 0 | 0 | 17 | 24 | 24218 | 15878 | 75367 | 74233 | 11 | 42 | 13 | 0 | 0 | 10274 | 76364 | 320721 | 319 | 3699 | 5895 | 2189 | 0 | 0 | 84 | 141 | 62 | 69 | 15 | 0 | 181 | 0 | 0 | 0 | 16 | 2909 | 0 | 0 |
| SRR10072382 | 13 | 0 | 0 | 24 | 23 | 24995 | 16085 | 78155 | 75122 | 12 | 38 | 26 | 0 | 0 | 10249 | 77440 | 330357 | 396 | 3629 | 5815 | 2158 | 0 | 0 | 77 | 142 | 60 | 67 | 17 | 0 | 198 | 0 | 0 | 0 | 10 | 2897 | 0 | 0 |
| SRR10072383 | 45 | 30 | 12 | 21 | 47 | 34388 | 19410 | 81630 | 61385 | 0 | 34 | 0 | 0 | 0 | 20863 | 144807 | 355971 | 173 | 1907 | 3002 | 813 | 0 | 0 | 101 | 55 | 24 | 34 | 12 | 0 | 72 | 0 | 0 | 0 | 0 | 537 | 0 | 0 |
| SRR10072384 | 72 | 28 | 16 | 21 | 69 | 34603 | 20606 | 80237 | 59821 | 0 | 33 | 12 | 0 | 0 | 20838 | 149521 | 361494 | 211 | 1932 | 2924 | 1056 | 0 | 0 | 94 | 64 | 28 | 35 | 11 | 0 | 65 | 0 | 0 | 0 | 0 | 657 | 0 | 0 |
| SRR10072385 | 30 | 0 | 0 | 22 | 17 | 11147 | 5981 | 30608 | 24077 | 48 | 36 | 19 | 0 | 0 | 38387 | 273956 | 330460 | 269 | 2841 | 5828 | 1622 | 0 | 0 | 49 | 56 | 30 | 36 | 16 | 0 | 117 | 0 | 0 | 0 | 12 | 1024 | 0 | 0 |
| SRR10072386 | 26 | 0 | 0 | 20 | 156 | 11081 | 5985 | 29787 | 24257 | 48 | 328 | 26 | 0 | 0 | 38731 | 277721 | 333676 | 325 | 2743 | 5757 | 1489 | 0 | 0 | 444 | 561 | 325 | 412 | 149 | 0 | 1078 | 0 | 0 | 0 | 12 | 1016 | 0 | 0 |
| SRR10072387 | 12 | 0 | 0 | 0 | 136 | 11884 | 7374 | 37254 | 30301 | 79 | 186 | 14 | 0 | 0 | 13909 | 61562 | 370910 | 456 | 5225 | 27219 | 4828 | 0 | 0 | 645 | 935 | 369 | 345 | 104 | 0 | 1233 | 0 | 0 | 0 | 0 | 2147 | 0 | 0 |
| SRR10072388 | 0 | 0 | 0 | 15 | 19 | 12237 | 7155 | 37235 | 30531 | 69 | 25 | 12 | 0 | 0 | 14161 | 62689 | 375803 | 535 | 5288 | 27882 | 4910 | 0 | 0 | 88 | 136 | 52 | 51 | 17 | 0 | 194 | 0 | 0 | 0 | 13 | 2042 | 0 | 0 |
| SRR10072389 | 24 | 59 | 44 | 61 | 37 | 32949 | 10138 | 30221 | 49902 | 22 | 35 | 0 | 0 | 0 | 8819 | 44029 | 75782 | 26 | 1616 | 2677 | 569 | 0 | 0 | 69 | 79 | 22 | 42 | 11 | 0 | 60 | 0 | 0 | 0 | 15 | 479 | 0 | 0 |
| SRR10072390 | 255 | 95 | 29 | 63 | 34 | 32762 | 9862 | 30504 | 50799 | 27 | 471 | 0 | 0 | 0 | 8237 | 45406 | 78582 | 0 | 1588 | 2221 | 670 | 0 | 0 | 889 | 960 | 243 | 578 | 136 | 0 | 853 | 0 | 0 | 0 | 0 | 442 | 0 | 0 |
| SRR10072391 | 35 | 0 | 13 | 17 | 52 | 31590 | 16103 | 62760 | 69862 | 13 | 44 | 0 | 0 | 0 | 12026 | 92559 | 306064 | 444 | 2466 | 5187 | 1800 | 0 | 0 | 142 | 99 | 48 | 77 | 35 | 0 | 117 | 0 | 0 | 0 | 0 | 766 | 15 | 0 |
| SRR10072392 | 34 | 13 | 18 | 11 | 73 | 32133 | 15590 | 64056 | 70878 | 0 | 384 | 13 | 0 | 0 | 13321 | 95740 | 312801 | 452 | 2543 | 5643 | 1469 | 0 | 0 | 1101 | 846 | 387 | 576 | 323 | 0 | 1063 | 0 | 0 | 0 | 0 | 710 | 15 | 0 |
| SRR10072393 | 30 | 0 | 11 | 21 | 169 | 25313 | 8299 | 48646 | 39057 | 51 | 366 | 16 | 0 | 0 | 40155 | 169589 | 457259 | 152 | 2733 | 2075 | 698 | 0 | 0 | 341 | 511 | 320 | 343 | 119 | 0 | 1067 | 0 | 0 | 0 | 18 | 590 | 0 | 0 |
| SRR10072394 | 44 | 0 | 16 | 21 | 130 | 25068 | 8428 | 48768 | 38864 | 65 | 426 | 11 | 0 | 0 | 41222 | 173602 | 463585 | 180 | 3230 | 2012 | 799 | 0 | 0 | 323 | 551 | 314 | 395 | 140 | 0 | 1164 | 0 | 0 | 0 | 18 | 641 | 0 | 0 |
| SRR10072395 | 429 | 95 | 41 | 48 | 48 | 10121 | 16341 | 31541 | 19064 | 0 | 197 | 14 | 0 | 0 | 11413 | 83893 | 199435 | 25 | 823 | 1786 | 1685 | 0 | 0 | 14 | 0 | 20 | 0 | 0 | 0 | 225 | 0 | 0 | 0 | 0 | 87 | 0 | 0 |
| SRR10072396 | 342 | 114 | 40 | 27 | 54 | 11246 | 17093 | 30463 | 18743 | 0 | 188 | 0 | 0 | 0 | 12620 | 87067 | 207246 | 0 | 850 | 1762 | 1487 | 0 | 0 | 13 | 27 | 0 | 0 | 0 | 0 | 262 | 0 | 0 | 0 | 0 | 132 | 0 | 0 |
| SRR10072397 | 79 | 15 | 17 | 12 | 145 | 18288 | 6548 | 35768 | 39508 | 73 | 636 | 0 | 0 | 0 | 35675 | 241216 | 424698 | 190 | 2622 | 3374 | 2849 | 0 | 0 | 751 | 762 | 376 | 459 | 110 | 0 | 1842 | 0 | 0 | 0 | 0 | 574 | 21 | 0 |
| SRR10072398 | 60 | 12 | 17 | 15 | 15 | 18285 | 6609 | 35085 | 39578 | 10 | 75 | 0 | 0 | 0 | 37496 | 244675 | 429926 | 174 | 2799 | 3418 | 3053 | 0 | 0 | 84 | 89 | 36 | 52 | 15 | 0 | 222 | 0 | 0 | 0 | 0 | 514 | 24 | 0 |
| SRR10072399 | 13 | 0 | 0 | 14 | 205 | 17067 | 11722 | 50214 | 58270 | 60 | 268 | 17 | 0 | 0 | 28217 | 95524 | 545299 | 332 | 4528 | 3283 | 1955 | 0 | 0 | 531 | 708 | 412 | 377 | 145 | 0 | 1146 | 0 | 0 | 0 | 13 | 2399 | 0 | 0 |
| SRR10072400 | 0 | 0 | 0 | 14 | 246 | 16725 | 12200 | 49906 | 59769 | 81 | 246 | 15 | 0 | 0 | 29959 | 96254 | 554404 | 355 | 4404 | 3375 | 1865 | 0 | 0 | 551 | 703 | 349 | 328 | 123 | 0 | 1327 | 0 | 0 | 0 | 10 | 2460 | 0 | 0 |
| SRR10077412 | 29 | 0 | 0 | 19 | 77 | 191 | 287 | 574 | 134 | 0 | 0 | 0 | 0 | 0 | 4704 | 10953 | 12546 | 0 | 19 | 822 | 229 | 0 | 0 | 753 | 29 | 106 | 193 | 106 | 0 | 29 | 0 | 19 | 0 | 0 | 8421 | 0 | 0 |
| SRR10077413 | 20 | 30 | 0 | 15 | 54 | 195 | 507 | 429 | 97 | 0 | 0 | 0 | 0 | 0 | 4969 | 10446 | 12663 | 58 | 0 | 643 | 117 | 0 | 0 | 32 | 35 | 59 | 10 | 99 | 0 | 39 | 0 | 19 | 0 | 0 | 10067 | 0 | 15 |
| SRR10077414 | 26 | 11 | 0 | 21 | 79 | 162 | 404 | 505 | 121 | 0 | 0 | 11 | 0 | 0 | 4485 | 11340 | 11741 | 20 | 20 | 849 | 182 | 0 | 0 | 854 | 26 | 37 | 316 | 84 | 0 | 26 | 0 | 0 | 0 | 0 | 9177 | 0 | 0 |
| SRR10077415 | 0 | 0 | 0 | 15 | 80 | 59 | 491 | 530 | 157 | 0 | 0 | 0 | 0 | 0 | 5043 | 11418 | 11940 | 20 | 20 | 922 | 157 | 0 | 0 | 716 | 45 | 70 | 224 | 90 | 0 | 20 | 0 | 39 | 0 | 0 | 9273 | 0 | 0 |
| SRR10077416 | 0 | 0 | 0 | 14 | 14 | 547 | 634 | 897 | 175 | 0 | 0 | 0 | 0 | 0 | 1247 | 2290 | 2987 | 44 | 0 | 241 | 0 | 0 | 0 | 30 | 0 | 0 | 77 | 24 | 0 | 0 | 0 | 0 | 0 | 0 | 10655 | 0 | 0 |
| SRR10077417 | 0 | 0 | 0 | 0 | 0 | 418 | 462 | 857 | 66 | 0 | 0 | 0 | 0 | 0 | 1011 | 2010 | 2680 | 44 | 22 | 242 | 88 | 0 | 0 | 27 | 0 | 29 | 64 | 29 | 0 | 0 | 0 | 0 | 0 | 0 | 10753 | 0 | 0 |
| SRR10077418 | 0 | 0 | 0 | 0 | 10 | 335 | 551 | 790 | 144 | 0 | 0 | 0 | 0 | 0 | 1174 | 2882 | 2593 | 0 | 0 | 335 | 0 | 0 | 0 | 25 | 10 | 26 | 52 | 26 | 0 | 10 | 0 | 0 | 0 | 0 | 9726 | 0 | 0 |
| SRR10077419 | 0 | 0 | 0 | 0 | 30 | 444 | 422 | 688 | 155 | 0 | 0 | 0 | 0 | 0 | 1242 | 2705 | 3550 | 44 | 0 | 222 | 0 | 0 | 0 | 731 | 0 | 15 | 69 | 0 | 0 | 0 | 0 | 0 | 0 | 0 | 10245 | 0 | 0 |
| SRR10077420 | 0 | 0 | 0 | 0 | 0 | 335 | 553 | 687 | 84 | 0 | 0 | 0 | 0 | 0 | 1357 | 3503 | 3804 | 67 | 0 | 335 | 50 | 0 | 0 | 393 | 0 | 17 | 87 | 41 | 0 | 0 | 0 | 0 | 0 | 0 | 9659 | 0 | 0 |
| SRR10077421 | 0 | 0 | 0 | 0 | 0 | 293 | 552 | 673 | 224 | 0 | 0 | 0 | 0 | 0 | 1552 | 3406 | 4174 | 0 | 0 | 207 | 69 | 0 | 0 | 27 | 12 | 12 | 72 | 39 | 0 | 0 | 0 | 0 | 0 | 0 | 9756 | 0 | 0 |
| SRR10077422 | 0 | 0 | 0 | 0 | 0 | 220 | 661 | 844 | 220 | 0 | 0 | 0 | 0 | 0 | 1394 | 3919 | 3429 | 18 | 0 | 239 | 92 | 0 | 0 | 22 | 13 | 13 | 70 | 19 | 0 | 16 | 0 | 0 | 0 | 0 | 9498 | 0 | 0 |
| SRR10077423 | 0 | 0 | 0 | 0 | 27 | 244 | 470 | 679 | 209 | 0 | 0 | 0 | 0 | 0 | 1445 | 3183 | 3747 | 0 | 0 | 279 | 104 | 0 | 0 | 22 | 0 | 24 | 66 | 33 | 0 | 0 | 0 | 17 | 0 | 0 | 9317 | 0 | 12 |
| SRR10077424 | 18 | 0 | 0 | 0 | 29 | 240 | 576 | 352 | 144 | 0 | 0 | 0 | 0 | 0 | 18795 | 8158 | 35877 | 104 | 24 | 648 | 176 | 0 | 0 | 823 | 14 | 29 | 123 | 51 | 0 | 14 | 0 | 0 | 0 | 0 | 105241 | 0 | 43 |
| SRR10077425 | 0 | 0 | 0 | 0 | 26 | 172 | 441 | 433 | 49 | 0 | 0 | 0 | 0 | 0 | 19632 | 8793 | 35633 | 82 | 41 | 548 | 196 | 0 | 0 | 855 | 26 | 29 | 122 | 33 | 0 | 11 | 0 | 16 | 0 | 0 | 106576 | 0 | 37 |
| SRR10077426 | 0 | 0 | 0 | 0 | 0 | 184 | 596 | 447 | 105 | 0 | 0 | 0 | 0 | 0 | 19273 | 8843 | 34301 | 114 | 44 | 526 | 184 | 0 | 0 | 837 | 0 | 39 | 118 | 35 | 0 | 31 | 0 | 0 | 0 | 0 | 106544 | 0 | 39 |
| SRR10077427 | 0 | 0 | 0 | 0 | 33 | 125 | 535 | 343 | 159 | 0 | 0 | 0 | 0 | 0 | 19810 | 8782 | 35993 | 167 | 17 | 493 | 134 | 0 | 0 | 33 | 11 | 48 | 103 | 33 | 0 | 0 | 0 | 0 | 0 | 0 | 106578 | 0 | 30 |

### Supp Table S8.3.Columns_BX_to_DH

| | | | | | | | | | | | | | | | | | | | | | | | | | | | | | | | | | | | | | |
|---|---|---|---|---|---|---|---|---|---|---|---|---|---|---|---|---|---|---|---|---|---|---|---|---|---|---|---|---|---|---|---|---|---|---|---|---|---|
| SRR10077428 | 71 | 50 | 0 | 0 | 113 | 128 | 360 | 348 | 116 | 0 | 0 | 0 | 0 | 0 | 95942 | 16611 | 110846 | 93 | 174 | 3932 | 1241 | 0 | 0 | 34 | 170 | 227 | 20 | 92 | 0 | 120 | 0 | 0 | 0 | 0 | 45728 | 0 | 28 |
| SRR10077429 | 22 | 58 | 15 | 0 | 161 | 130 | 354 | 248 | 130 | 0 | 15 | 0 | 0 | 0 | 92306 | 17746 | 110399 | 118 | 154 | 4064 | 1158 | 0 | 0 | 868 | 109 | 255 | 642 | 117 | 0 | 88 | 0 | 0 | 0 | 0 | 45719 | 0 | 51 |
| SRR10077430 | 47 | 55 | 0 | 16 | 219 | 75 | 350 | 225 | 62 | 0 | 0 | 0 | 0 | 0 | 93063 | 16525 | 110103 | 137 | 225 | 3886 | 950 | 0 | 0 | 899 | 70 | 172 | 539 | 125 | 0 | 109 | 0 | 25 | 0 | 0 | 45519 | 0 | 63 |
| SRR10077431 | 87 | 29 | 15 | 0 | 109 | 47 | 318 | 318 | 71 | 0 | 0 | 0 | 0 | 0 | 94549 | 17494 | 110074 | 129 | 176 | 4070 | 1235 | 0 | 0 | 27 | 116 | 124 | 21 | 116 | 0 | 116 | 0 | 24 | 0 | 0 | 45551 | 0 | 51 |
| SRR10077432 | 21 | 0 | 0 | 0 | 32 | 148 | 531 | 488 | 136 | 0 | 0 | 0 | 0 | 0 | 34215 | 14184 | 61453 | 49 | 62 | 895 | 327 | 0 | 0 | 902 | 25 | 81 | 211 | 60 | 0 | 18 | 0 | 12 | 0 | 0 | 110989 | 0 | 42 |
| SRR10077433 | 33 | 0 | 0 | 0 | 40 | 115 | 633 | 460 | 109 | 0 | 0 | 0 | 0 | 0 | 34049 | 14062 | 62336 | 51 | 51 | 933 | 358 | 0 | 0 | 31 | 25 | 87 | 243 | 69 | 0 | 33 | 0 | 19 | 0 | 0 | 109948 | 0 | 58 |
| SRR10077434 | 12 | 0 | 0 | 0 | 62 | 181 | 586 | 446 | 126 | 0 | 0 | 0 | 0 | 0 | 35335 | 13180 | 61201 | 91 | 63 | 781 | 349 | 0 | 0 | 27 | 19 | 81 | 216 | 65 | 0 | 19 | 0 | 0 | 0 | 0 | 108739 | 0 | 23 |
| SRR10077435 | 29 | 0 | 0 | 0 | 84 | 197 | 558 | 414 | 105 | 11 | 0 | 0 | 0 | 0 | 35094 | 13877 | 60931 | 39 | 39 | 1018 | 328 | 0 | 0 | 1088 | 26 | 77 | 190 | 106 | 0 | 33 | 0 | 0 | 0 | 0 | 111340 | 0 | 37 |
| SRR10116271 | 0 | 0 | 0 | 0 | 145 | 5582 | 640 | 2244 | 821 | 63 | 0 | 0 | 0 | 0 | 74362 | 26860 | 709132 | 50 | 118 | 448 | 68 | 102 | 252 | 231 | 0 | 0 | 16 | 0 | 0 | 42 | 131 | 0 | 0 | 0 | 2246 | 3666 | 0 |
| SRR10116272 | 0 | 0 | 0 | 0 | 49 | 34764 | 2588 | 19112 | 2341 | 513 | 0 | 0 | 0 | 0 | 102168 | 69028 | 376543 | 139 | 117 | 512 | 436 | 14 | 28 | 812 | 0 | 0 | 0 | 0 | 0 | 71 | 25 | 0 | 0 | 0 | 4539 | 4393 | 0 |
| SRR10116273 | 0 | 12 | 0 | 0 | 14 | 9611 | 679 | 2422 | 508 | 31 | 0 | 0 | 0 | 0 | 26892 | 11971 | 72911 | 83 | 18 | 39 | 48 | 78 | 42 | 67 | 0 | 0 | 0 | 0 | 0 | 11 | 23 | 0 | 0 | 0 | 2092 | 13824 | 0 |
| SRR10116274 | 14 | 0 | 0 | 0 | 77 | 47801 | 2579 | 20863 | 2669 | 116 | 0 | 0 | 0 | 0 | 167274 | 52755 | 441718 | 120 | 269 | 428 | 269 | 0 | 11 | 60 | 0 | 0 | 0 | 0 | 0 | 34 | 0 | 0 | 0 | 0 | 4014 | 9250 | 0 |
| SRR10116275 | 0 | 18 | 0 | 0 | 146 | 3243 | 633 | 2332 | 968 | 35 | 14 | 0 | 0 | 0 | 83446 | 53459 | 423296 | 151 | 118 | 1532 | 1099 | 19 | 79 | 331 | 13 | 14 | 19 | 0 | 0 | 38 | 23 | 0 | 0 | 0 | 3698 | 3396 | 12 |
| SRR10119011 | 0 | 0 | 0 | 0 | 0 | 26 | 0 | 0 | 0 | 0 | 88 | 0 | 0 | 0 | 5037 | 28092 | 73844 | 0 | 45 | 9306 | 628 | 0 | 0 | 0 | 13 | 224 | 2759 | 724 | 0 | 490 | 0 | 0 | 0 | 0 | 706 | 0 | 0 |
| SRR10119012 | 0 | 0 | 0 | 0 | 0 | 0 | 0 | 0 | 0 | 0 | 0 | 104 | 0 | 0 | 0 | 15763 | 69648 | 0 | 98 | 0 | 11253 | 0 | 0 | 0 | 0 | 0 | 49 | 0 | 0 | 79 | 0 | 0 | 0 | 0 | 9164 | 0 | 293 |
| SRR10119014 | 0 | 0 | 0 | 0 | 0 | 0 | 0 | 0 | 0 | 0 | 16 | 0 | 0 | 0 | 4204 | 5197 | 19967 | 0 | 0 | 0 | 4729 | 0 | 0 | 0 | 0 | 0 | 0 | 0 | 0 | 0 | 0 | 0 | 0 | 0 | 3556 | 0 | 0 |
| SRR10119022 | 0 | 0 | 0 | 16 | 11 | 18 | 0 | 77 | 22 | 0 | 30 | 0 | 0 | 0 | 6534 | 15447 | 103918 | 0 | 36 | 14490 | 823 | 0 | 0 | 19 | 27 | 357 | 4162 | 1290 | 0 | 192 | 0 | 0 | 0 | 0 | 583 | 0 | 0 |
| SRR10119033 | 0 | 0 | 0 | 0 | 0 | 0 | 0 | 0 | 0 | 0 | 42 | 0 | 0 | 0 | 397 | 22474 | 49575 | 838 | 0 | 1808 | 3924 | 0 | 0 | 42 | 39 | 58 | 506 | 133 | 0 | 156 | 0 | 0 | 0 | 0 | 8687 | 0 | 64 |
| SRR10119034 | 0 | 0 | 0 | 0 | 0 | 0 | 0 | 0 | 0 | 0 | 0 | 390 | 0 | 0 | 0 | 13841 | 39474 | 0 | 0 | 2529 | 0 | 0 | 0 | 0 | 0 | 0 | 81 | 27 | 0 | 188 | 0 | 0 | 0 | 0 | 6152 | 0 | 108 |
| SRR10119035 | 0 | 0 | 0 | 23 | 20 | 0 | 0 | 18 | 0 | 0 | 22 | 0 | 0 | 0 | 9561 | 25777 | 66811 | 0 | 73 | 9849 | 189 | 0 | 0 | 54 | 51 | 482 | 5038 | 1130 | 0 | 121 | 0 | 12 | 0 | 0 | 220 | 0 | 0 |
| SRR10119037 | 0 | 0 | 0 | 0 | 0 | 0 | 3868 | 0 | 0 | 0 | 0 | 88 | 0 | 0 | 0 | 228 | 35332 | 0 | 0 | 0 | 6682 | 0 | 0 | 0 | 0 | 0 | 56 | 28 | 0 | 0 | 0 | 0 | 0 | 0 | 8662 | 0 | 163 |
| SRR10119040 | 0 | 0 | 0 | 0 | 0 | 0 | 0 | 0 | 0 | 0 | 0 | 215 | 0 | 0 | 804 | 11197 | 53370 | 0 | 0 | 1293 | 7865 | 0 | 0 | 0 | 0 | 46 | 10 | 77 | 0 | 0 | 0 | 0 | 0 | 0 | 3872 | 0 | 0 |
| SRR10119044 | 0 | 0 | 0 | 0 | 0 | 0 | 0 | 0 | 0 | 0 | 95 | 0 | 0 | 0 | 1662 | 21027 | 90605 | 0 | 0 | 5686 | 3761 | 0 | 0 | 0 | 0 | 132 | 0 | 34 | 0 | 0 | 0 | 2712 | 0 | 0 | 11373 | 0 | 122 |
| SRR10119046 | 0 | 0 | 0 | 11 | 14 | 23 | 0 | 15 | 0 | 0 | 10 | 0 | 0 | 0 | 9445 | 27522 | 67806 | 0 | 56 | 9568 | 253 | 0 | 0 | 12 | 10 | 128 | 1228 | 546 | 0 | 42 | 0 | 0 | 0 | 0 | 266 | 0 | 0 |
| SRR10119048 | 0 | 0 | 0 | 0 | 0 | 0 | 0 | 792 | 0 | 0 | 75 | 146 | 0 | 0 | 0 | 20590 | 106692 | 0 | 0 | 604 | 1471 | 0 | 0 | 93 | 22 | 296 | 446 | 252 | 0 | 84 | 0 | 0 | 0 | 0 | 3120 | 0 | 128 |
| SRR10119051 | 0 | 0 | 0 | 0 | 0 | 0 | 0 | 0 | 0 | 0 | 0 | 326 | 0 | 0 | 0 | 6020 | 40875 | 0 | 0 | 7555 | 5288 | 0 | 0 | 0 | 0 | 18 | 31 | 0 | 0 | 0 | 0 | 0 | 0 | 0 | 317 | 0 | 197 |
| SRR10119052 | 0 | 0 | 0 | 0 | 0 | 0 | 0 | 0 | 0 | 0 | 23 | 80 | 0 | 0 | 626 | 22532 | 44897 | 0 | 0 | 3380 | 3630 | 0 | 0 | 0 | 0 | 132 | 201 | 34 | 0 | 144 | 0 | 0 | 0 | 0 | 11431 | 0 | 155 |
| SRR10119056 | 0 | 0 | 0 | 0 | 0 | 0 | 0 | 0 | 0 | 0 | 42 | 99 | 0 | 0 | 0 | 16068 | 49947 | 0 | 0 | 4926 | 8303 | 0 | 0 | 0 | 0 | 87 | 311 | 24 | 0 | 146 | 0 | 0 | 0 | 0 | 3301 | 0 | 285 |
| SRR10119057 | 0 | 0 | 0 | 17 | 12 | 12 | 0 | 0 | 0 | 0 | 28 | 0 | 0 | 0 | 399 | 15980 | 23575 | 0 | 0 | 1591 | 95 | 0 | 0 | 12 | 17 | 85 | 812 | 411 | 0 | 186 | 0 | 0 | 0 | 0 | 1502 | 0 | 0 |
| SRR10119058 | 0 | 0 | 0 | 25 | 15 | 0 | 0 | 42 | 0 | 0 | 54 | 0 | 0 | 0 | 422 | 16600 | 20457 | 0 | 0 | 1310 | 21 | 0 | 0 | 19 | 16 | 182 | 1713 | 1066 | 0 | 521 | 0 | 0 | 0 | 0 | 1351 | 0 | 0 |
| SRR10119059 | 0 | 0 | 13 | 11 | 16 | 0 | 0 | 0 | 0 | 0 | 44 | 0 | 0 | 0 | 8090 | 16141 | 137618 | 0 | 0 | 17501 | 1515 | 0 | 0 | 0 | 12 | 21 | 213 | 149 | 0 | 65 | 0 | 0 | 0 | 0 | 495 | 0 | 0 |
| SRR10119063 | 0 | 0 | 0 | 0 | 0 | 0 | 0 | 1020 | 0 | 0 | 0 | 0 | 0 | 0 | 32162 | 7237 | 199857 | 0 | 0 | 2589 | 0 | 0 | 0 | 0 | 0 | 0 | 0 | 0 | 0 | 15 | 0 | 0 | 0 | 0 | 510 | 0 | 0 |
| SRR10119066 | 0 | 0 | 0 | 33 | 24 | 0 | 13 | 12 | 0 | 0 | 49 | 0 | 0 | 0 | 1849 | 24448 | 38820 | 0 | 14 | 12070 | 214 | 0 | 0 | 40 | 39 | 257 | 2503 | 1204 | 0 | 437 | 0 | 0 | 0 | 0 | 187 | 0 | 0 |
| SRR10119067 | 0 | 0 | 0 | 17 | 20 | 0 | 0 | 15 | 0 | 0 | 18 | 0 | 0 | 0 | 2078 | 18289 | 43498 | 0 | 22 | 12190 | 234 | 0 | 0 | 16 | 14 | 138 | 976 | 689 | 0 | 104 | 0 | 0 | 0 | 0 | 276 | 0 | 0 |
| SRR10119068 | 0 | 0 | 0 | 22 | 18 | 0 | 0 | 0 | 0 | 0 | 19 | 0 | 0 | 0 | 6274 | 14229 | 73160 | 0 | 16 | 7974 | 1673 | 0 | 0 | 12 | 25 | 178 | 1568 | 437 | 0 | 110 | 0 | 0 | 0 | 0 | 402 | 0 | 0 |
| SRR10119069 | 0 | 0 | 0 | 20 | 0 | 0 | 13 | 61 | 0 | 0 | 17 | 0 | 0 | 0 | 5618 | 15158 | 70808 | 0 | 27 | 7418 | 1114 | 0 | 0 | 11 | 14 | 130 | 1016 | 424 | 0 | 145 | 0 | 0 | 0 | 0 | 424 | 0 | 0 |
| SRR10119070 | 0 | 0 | 0 | 0 | 0 | 0 | 0 | 0 | 0 | 0 | 45 | 0 | 0 | 0 | 3928 | 20412 | 63490 | 0 | 0 | 11562 | 544 | 0 | 0 | 0 | 11 | 129 | 868 | 513 | 0 | 331 | 0 | 0 | 0 | 0 | 1080 | 0 | 0 |
| SRR10119071 | 0 | 0 | 0 | 10 | 0 | 0 | 0 | 11 | 0 | 0 | 57 | 0 | 0 | 0 | 799 | 8203 | 23028 | 0 | 0 | 1289 | 22 | 0 | 0 | 0 | 31 | 268 | 1506 | 970 | 0 | 371 | 0 | 0 | 0 | 0 | 738 | 0 | 0 |
| SRR10119072 | 0 | 0 | 0 | 0 | 0 | 0 | 0 | 0 | 0 | 0 | 151 | 0 | 0 | 0 | 1687 | 13810 | 45876 | 0 | 0 | 1953 | 46 | 0 | 0 | 0 | 0 | 107 | 578 | 439 | 0 | 679 | 0 | 0 | 0 | 0 | 496 | 0 | 0 |
| SRR10119073 | 0 | 0 | 0 | 0 | 0 | 0 | 0 | 0 | 14 | 0 | 115 | 0 | 0 | 0 | 410 | 4992 | 10553 | 0 | 0 | 650 | 64 | 0 | 0 | 0 | 0 | 70 | 707 | 306 | 0 | 941 | 0 | 0 | 0 | 0 | 823 | 0 | 0 |
| SRR10119074 | 0 | 0 | 0 | 0 | 0 | 0 | 0 | 0 | 0 | 0 | 49 | 0 | 0 | 0 | 378 | 6555 | 13770 | 0 | 0 | 526 | 83 | 0 | 0 | 0 | 12 | 46 | 435 | 259 | 0 | 391 | 0 | 0 | 0 | 0 | 1079 | 0 | 0 |
| SRR10119075 | 0 | 0 | 0 | 15 | 14 | 0 | 0 | 0 | 0 | 0 | 22 | 0 | 0 | 0 | 4283 | 24278 | 48758 | 0 | 0 | 6361 | 396 | 0 | 0 | 10 | 22 | 103 | 1217 | 656 | 0 | 171 | 0 | 0 | 0 | 0 | 842 | 0 | 0 |
| SRR10119076 | 0 | 0 | 0 | 19 | 0 | 0 | 24 | 0 | 36 | 0 | 32 | 0 | 0 | 0 | 5238 | 26907 | 60052 | 0 | 16 | 7078 | 548 | 0 | 0 | 24 | 15 | 60 | 820 | 420 | 0 | 191 | 0 | 0 | 0 | 0 | 1230 | 0 | 0 |
| SRR10119077 | 0 | 0 | 0 | 24 | 27 | 0 | 23 | 28 | 0 | 0 | 42 | 0 | 0 | 0 | 4023 | 22801 | 29266 | 0 | 44 | 4207 | 194 | 0 | 0 | 53 | 39 | 264 | 1676 | 972 | 0 | 202 | 0 | 0 | 0 | 0 | 261 | 0 | 0 |
| SRR10119078 | 0 | 0 | 0 | 11 | 24 | 0 | 10 | 22 | 0 | 0 | 14 | 0 | 0 | 0 | 3723 | 21116 | 28014 | 0 | 27 | 6022 | 235 | 0 | 0 | 22 | 12 | 72 | 325 | 205 | 0 | 83 | 0 | 0 | 0 | 0 | 289 | 0 | 0 |
| SRR10119079 | 0 | 0 | 0 | 25 | 12 | 16 | 0 | 0 | 0 | 0 | 90 | 0 | 0 | 0 | 11301 | 33772 | 55772 | 0 | 14 | 10233 | 987 | 0 | 0 | 17 | 43 | 340 | 3194 | 1199 | 0 | 417 | 0 | 13 | 0 | 0 | 355 | 0 | 0 |
| SRR10119080 | 0 | 0 | 0 | 14 | 0 | 17 | 0 | 37 | 0 | 0 | 14 | 0 | 0 | 0 | 12924 | 34255 | 66600 | 0 | 34 | 11112 | 927 | 0 | 0 | 14 | 12 | 122 | 724 | 401 | 0 | 138 | 0 | 0 | 0 | 0 | 451 | 0 | 0 |
| SRR10119081 | 0 | 0 | 0 | 0 | 0 | 0 | 0 | 23 | 0 | 0 | 76 | 0 | 0 | 0 | 261 | 5041 | 10095 | 0 | 16 | 943 | 0 | 0 | 0 | 13 | 65 | 508 | 2336 | 1214 | 0 | 437 | 0 | 0 | 0 | 0 | 510 | 0 | 0 |
| SRR10119082 | 0 | 0 | 0 | 0 | 0 | 0 | 0 | 0 | 0 | 0 | 129 | 0 | 0 | 0 | 542 | 7350 | 15133 | 0 | 15 | 1308 | 11 | 0 | 0 | 0 | 34 | 289 | 2298 | 464 | 0 | 530 | 0 | 0 | 0 | 0 | 847 | 0 | 0 |
| SRR1014666 | 0 | 0 | 0 | 0 | 0 | 2005 | 2310 | 11896 | 218 | 50 | 0 | 0 | 0 | 0 | 22 | 159 | 1128 | 22 | 0 | 6438 | 33 | 0 | 0 | 22 | 0 | 0 | 0 | 0 | 0 | 0 | 0 | 0 | 0 | 0 | 6998 | 279 | 0 |
| SRR1014667 | 0 | 0 | 0 | 0 | 0 | 416 | 3184 | 11395 | 405 | 20 | 0 | 0 | 0 | 0 | 18837 | 116053 | 120770 | 29 | 12 | 38911 | 1267 | 0 | 0 | 55 | 0 | 0 | 0 | 0 | 0 | 0 | 0 | 18 | 0 | 0 | 10196 | 379 | 0 |
| SRR1014669 | 0 | 0 | 0 | 0 | 0 | 192 | 128 | 460 | 0 | 29 | 0 | 0 | 0 | 0 | 19224 | 7355 | 159455 | 11 | 0 | 9489 | 1763 | 0 | 12 | 0 | 0 | 0 | 0 | 0 | 0 | 0 | 0 | 21 | 0 | 0 | 28372 | 480 | 0 |
| SRR10197184 | 37 | 0 | 0 | 0 | 28 | 9406 | 2183 | 10333 | 5705 | 6517 | 19 | 0 | 0 | 0 | 5055 | 29062 | 21712 | 134 | 154 | 202 | 1435 | 0 | 0 | 304 | 15 | 0 | 0 | 0 | 0 | 48 | 0 | 0 | 0 | 0 | 7152 | 0 | 0 |
| SRR10197185 | 41 | 83 | 0 | 0 | 0 | 222 | 0 | 0 | 296 | 4916 | 0 | 32 | 24 | 0 | 259 | 8425 | 1256 | 0 | 37 | 0 | 16465 | 0 | 0 | 0 | 0 | 0 | 0 | 0 | 12 | 0 | 0 | 37 | 0 | 0 | 35060 | 0 | 0 |
| SRR10197186 | 13 | 0 | 0 | 0 | 13 | 2451 | 558 | 2451 | 7641 | 2434 | 0 | 0 | 0 | 0 | 2181 | 36354 | 11094 | 0 | 44 | 32 | 5701 | 0 | 0 | 79 | 0 | 0 | 0 | 0 | 0 | 0 | 0 | 16 | 0 | 0 | 38954 | 0 | 0 |
| SRR10197187 | 0 | 0 | 0 | 0 | 0 | 2039 | 623 | 2820 | 5913 | 2358 | 10 | 0 | 0 | 0 | 1876 | 11996 | 3458 | 55 | 38 | 17 | 1276 | 209 | 167 | 126 | 0 | 0 | 0 | 0 | 49 | 0 | 36 | 139 | 0 | 0 | 55385 | 0 | 0 |
| SRR10197188 | 0 | 0 | 0 | 0 | 0 | 2894 | 1031 | 3728 | 4169 | 2532 | 11 | 0 | 0 | 0 | 2242 | 8701 | 3846 | 153 | 64 | 56 | 641 | 227 | 184 | 135 | 0 | 0 | 0 | 0 | 58 | 19 | 36 | 83 | 0 | 0 | 62511 | 0 | 0 |
| SRR10197189 | 13 | 0 | 0 | 0 | 0 | 1740 | 292 | 1208 | 2877 | 4745 | 0 | 0 | 0 | 0 | 923 | 13270 | 3247 | 0 | 27 | 14 | 4264 | 266 | 313 | 121 | 0 | 0 | 0 | 0 | 31 | 0 | 27 | 216 | 0 | 0 | 55792 | 0 | 0 |
| SRR10197190 | 0 | 0 | 0 | 0 | 0 | 3552 | 1318 | 4575 | 6117 | 303 | 0 | 0 | 0 | 0 | 3354 | 17499 | 10506 | 88 | 72 | 110 | 648 | 205 | 172 | 19 | 0 | 0 | 0 | 0 | 11 | 11 | 40 | 66 | 0 | 0 | 36619 | 0 | 0 |
| SRR10197191 | 0 | 0 | 0 | 0 | 0 | 3007 | 1457 | 3729 | 7537 | 369 | 11 | 0 | 0 | 0 | 3387 | 19349 | 10356 | 56 | 84 | 89 | 778 | 171 | 150 | 31 | 0 | 0 | 0 | 0 | 22 | 16 | 32 | 42 | 0 | 0 | 44023 | 0 | 0 |
| SRR10197192 | 0 | 0 | 0 | 0 | 0 | 2169 | 967 | 3140 | 4185 | 929 | 11 | 0 | 0 | 0 | 4866 | 25451 | 17756 | 66 | 87 | 145 | 908 | 247 | 232 | 22 | 0 | 0 | 0 | 0 | 15 | 16 | 32 | 41 | 0 | 0 | 38471 | 0 | 0 |
| SRR10225092 | 88 | 0 | 2190 | 2278 | 350 | 0 | 0 | 166 | 166 | 0 | 0 | 0 | 0 | 0 | 52246 | 10338 | 77068 | 0 | 0 | 1997 | 333 | 0 | 0 | 175 | 263 | 876 | 4030 | 1840 | 0 | 175 | 0 | 0 | 0 | 0 | 0 | 0 | 0 |
| SRR10225093 | 0 | 0 | 0 | 0 | 28 | 0 | 0 | 0 | 0 | 110 | 87 | 28 | 0 | 0 | 257402 | 25208 | 276425 | 0 | 321 | 2843 | 28016 | 0 | 0 | 0 | 0 | 28 | 0 | 14 | 0 | 266 | 0 | 18 | 0 | 0 | 615 | 0 | 0 |
| SRR10225094 | 11 | 11 | 101 | 101 | 18 | 0 | 38 | 38 | 0 | 0 | 20 | 44 | 0 | 0 | 15252 | 32129 | 89357 | 38 | 38 | 1287 | 530 | 0 | 0 | 13 | 20 | 78 | 211 | 202 | 0 | 31 | 0 | 0 | 0 | 0 | 251 | 0 | 0 |
| SRR10225096 | 0 | 29 | 15 | 58 | 52 | 0 | 0 | 46 | 0 | 0 | 17 | 0 | 0 | 0 | 11123 | 74080 | 16634 | 46 | 46 | 415 | 508 | 39 | 42 | 10 | 54 | 123 | 350 | 60 | 0 | 27 | 39 | 138 | 0 | 0 | 441 | 0 | 0 |
| SRR10282881 | 0 | 0 | 0 | 0 | 29 | 238 | 46 | 258 | 0 | 13 | 0 | 0 | 0 | 0 | 159129 | 4694 | 234281 | 0 | 56 | 2938 | 45514 | 0 | 0 | 0 | 21 | 11 | 0 | 0 | 0 | 0 | 0 | 0 | 0 | 0 | 59 | 0 | 23 |
| SRR10282882 | 0 | 0 | 0 | 0 | 19 | 268 | 61 | 328 | 0 | 16 | 0 | 0 | 0 | 0 | 168526 | 5263 | 265374 | 0 | 61 | 2989 | 46411 | 0 | 0 | 13 | 27 | 14 | 0 | 0 | 0 | 14 | 0 | 0 | 0 | 0 | 41 | 0 | 23 |
| SRR10282883 | 0 | 0 | 0 | 0 | 30 | 283 | 37 | 299 | 0 | 12 | 0 | 0 | 0 | 0 | 77765 | 7745 | 333459 | 0 | 173 | 2704 | 69797 | 0 | 0 | 14 | 33 | 66 | 85 | 0 | 0 | 20 | 0 | 0 | 0 | 0 | 58 | 0 | 0 |
| SRR10282884 | 0 | 0 | 0 | 0 | 19 | 345 | 45 | 362 | 0 | 0 | 0 | 0 | 0 | 0 | 80943 | 8446 | 371522 | 0 | 144 | 2646 | 69946 | 0 | 0 | 22 | 36 | 62 | 84 | 0 | 0 | 187 | 0 | 0 | 0 | 0 | 50 | 0 | 14 |
| SRR10282885 | 0 | 0 | 0 | 0 | 39 | 130 | 17 | 152 | 0 | 11 | 0 | 0 | 0 | 0 | 84997 | 6221 | 454804 | 0 | 177 | 2385 | 98352 | 0 | 0 | 49 | 54 | 98 | 110 | 0 | 0 | 71 | 0 | 0 | 0 | 0 | 35 | 0 | 18 |
| SRR10282886 | 0 | 0 | 0 | 0 | 39 | 158 | 25 | 135 | 0 | 17 | 0 | 0 | 0 | 0 | 89179 | 6621 | 498188 | 0 | 191 | 2244 | 99398 | 0 | 0 | 50 | 76 | 98 | 126 | 0 | 0 | 54 | 0 | 0 | 0 | 0 | 37 | 0 | 17 |
| SRR10282890 | 0 | 0 | 0 | 0 | 66 | 127 | 64 | 515 | 26 | 18 | 0 | 0 | 0 | 0 | 114552 | 8082 | 329162 | 0 | 160 | 3514 | 52068 | 0 | 0 | 22 | 29 | 28 | 12 | 0 | 0 | 74 | 0 | 0 | 0 | 0 | 188 | 0 | 0 |
| SRR10282891 | 0 | 0 | 0 | 0 | 53 | 175 | 95 | 581 | 22 | 15 | 0 | 0 | 0 | 0 | 118981 | 8779 | 362608 | 0 | 130 | 3467 | 52627 | 0 | 0 | 20 | 37 | 31 | 13 | 0 | 0 | 64 | 0 | 0 | 0 | 0 | 170 | 0 | 0 |
| SRR10282892 | 0 | 0 | 0 | 0 | 23 | 26 | 17 | 131 | 0 | 36 | 0 | 0 | 0 | 0 | 87904 | 6538 | 442160 | 0 | 14 | 1664 | 221276 | 0 | 0 | 25 | 23 | 12 | 0 | 0 | 0 | 0 | 0 | 0 | 0 | 0 | 31 | 0 | 0 |
| SRR10282893 | 0 | 0 | 0 | 0 | 20 | 42 | 17 | 129 | 0 | 34 | 0 | 0 | 0 | 0 | 94189 | 7030 | 482434 | 0 | 13 | 1722 | 225230 | 0 | 0 | 22 | 34 | 16 | 0 | 0 | 0 | 0 | 0 | 0 | 0 | 0 | 24 | 0 | 0 |
| SRR10282894 | 0 | 0 | 0 | 0 | 0 | 11 | 0 | 55 | 0 | 0 | 0 | 0 | 0 | 0 | 59901 | 7713 | 376252 | 0 | 18 | 853 | 139610 | 0 | 0 | 0 | 22 | 32 | 13 | 0 | 0 | 0 | 0 | 0 | 0 | 0 | 23 | 0 | 0 |
| SRR10282895 | 0 | 0 | 0 | 0 | 0 | 17 | 17 | 75 | 0 | 0 | 0 | 0 | 0 | 0 | 61374 | 8262 | 413370 | 0 | 13 | 842 | 139934 | 0 | 0 | 0 | 28 | 39 | 16 | 0 | 0 | 0 | 0 | 0 | 0 | 0 | 20 | 0 | 0 |
| SRR10323931 | 85 | 12 | 31 | 0 | 13 | 3350 | 596 | 2599 | 5946 | 34 | 0 | 16 | 13 | 0 | 3032 | 212501 | 10304 | 102 | 82 | 490 | 1020 | 13 | 21 | 45 | 21 | 0 | 21 | 0 | 0 | 0 | 0 | 16 | 0 | 11 | 16296 | 191 | 0 |
| SRR10323932 | 2192 | 353 | 0 | 0 | 193 | 3043 | 825 | 2757 | 6165 | 1793 | 0 | 42 | 0 | 0 | 3923 | 435241 | 12385 | 47 | 318 | 1465 | 5450 | 36 | 105 | 357 | 0 | 0 | 0 | 0 | 0 | 17 | 30 | 35 | 0 | 39 | 7373 | 43 | 0 |
| SRR10323933 | 185 | 44 | 131 | 48 | 71 | 517 | 306 | 787 | 2323 | 60 | 30 | 11 | 0 | 0 | 6553 | 324164 | 21327 | 39 | 431 | 1711 | 3435 | 12 | 47 | 38 | 125 | 82 | 12 | 0 | 0 | 71 | 12 | 0 | 0 | 14 | 2439 | 87 | 0 |
| SRR10323939 | 599 | 79 | 0 | 0 | 202 | 5762 | 1019 | 4512 | 10013 | 621 | 0 | 11 | 0 | 0 | 1109 | 105318 | 6623 | 65 | 45 | 316 | 683 | 0 | 27 | 1003 | 12 | 0 | 11 | 0 | 0 | 0 | 16 | 0 | 0 | 14 | 23726 | 106 | 0 |
| SRR10323940 | 1309 | 213 | 0 | 0 | 695 | 3181 | 606 | 2687 | 5494 | 391 | 11 | 11 | 0 | 0 | 13975 | 276823 | 61765 | 200 | 1037 | 3225 | 13687 | 30 | 63 | 941 | 47 | 16 | 31 | 0 | 0 | 38 | 17 | 56 | 0 | 28 | 7290 | 10 | 0 |
| SRR10323941 | 714 | 116 | 0 | 0 | 806 | 2299 | 377 | 2405 | 3843 | 209 | 0 | 0 | 0 | 0 | 16404 | 360686 | 66179 | 135 | 861 | 2882 | 9650 | 39 | 101 | 472 | 19 | 0 | 21 | 0 | 0 | 22 | 108 | 64 | 15 | 28 | 5275 | 0 | 0 |
| SRR10385125 | 43 | 13 | 52 | 0 | 303 | 6222 | 359 | 2644 | 1358 | 176 | 0 | 0 | 36 | 14 | 3872 | 7985 | 14784 | 36 | 1387 | 295 | 1315 | 0 | 0 | 39 | 18 | 14 | 0 | 0 | 0 | 25 | 0 | 0 | 0 | 0 | 5267 | 16 | 0 |
| SRR10385126 | 1032 | 468 | 123 | 32 | 6899 | 1524 | 193 | 1797 | 1989 | 672 | 0 | 35 | 17 | 0 | 1315 | 7736 | 9919 | 199 | 568 | 136 | 774 | 0 | 0 | 481 | 23 | 0 | 0 | 0 | 78 | 53 | 0 | 0 | 0 | 14 | 5833 | 0 | 0 |
| SRR10416439 | 227 | 686 | 0 | 0 | 3255 | 345 | 80 | 737 | 1342 | 433 | 89 | 13 | 125 | 15 | 27 | 2331 | 2628 | 166 | 20 | 372 | 578 | 0 | 0 | 62 | 105 | 43 | 83 | 0 | 0 | 1130 | 0 | 20 | 0 | 0 | 4324 | 16 | 0 |
| SRR10416440 | 33 | 83 | 0 | 0 | 434 | 282 | 86 | 688 | 782 | 240 | 11 | 0 | 12 | 0 | 931 | 2000 | 7254 | 141 | 16 | 328 | 673 | 0 | 0 | 0 | 29 | 16 | 31 | 0 | 0 | 168 | 0 | 78 | 0 | 0 | 1637 | 40 | 0 |
| SRR10416441 | 36 | 102 | 0 | 0 | 651 | 274 | 84 | 916 | 1725 | 158 | 12 | 0 | 17 | 14 | 179 | 1909 | 6365 | 125 | 11 | 585 | 539 | 0 | 0 | 29 | 17 | 40 | 22 | 0 | 0 | 199 | 0 | 19 | 0 | 0 | 4548 | 32 | 0 |
| SRR10416442 | 285 | 403 | 0 | 0 | 3825 | 282 | 551 | 3126 | 5756 | 372 | 115 | 31 | 53 | 12 | 14 | 3503 | 6727 | 213 | 28 | 709 | 627 | 0 | 0 | 137 | 175 | 63 | 254 | 0 | 0 | 1418 | 0 | 34 | 0 | 22 | 6508 | 17 | 0 |
| SRR10416496 | 0 | 0 | 0 | 0 | 0 | 0 | 0 | 0 | 0 | 0 | 585 | 0 | 0 | 0 | 529 | 26209 | 41351 | 265 | 265 | 12434 | 0 | 0 | 0 | 0 | 0 | 0 | 21 | 10 | 0 | 1252 | 0 | 0 | 0 | 0 | 4659 | 0 | 0 |
| SRR10416497 | 0 | 0 | 0 | 0 | 18 | 717 | 0 | 717 | 0 | 0 | 23 | 0 | 0 | 0 | 0 | 18683 | 24021 | 0 | 359 | 6812 | 0 | 0 | 0 | 0 | 0 | 0 | 72 | 18 | 0 | 51 | 0 | 0 | 0 | 0 | 4448 | 0 | 36 |
| SRR10416498 | 0 | 0 | 0 | 0 | 0 | 0 | 251 | 754 | 0 | 0 | 35 | 0 | 0 | 0 | 754 | 6547 | 19640 | 0 | 1256 | 4521 | 0 | 0 | 0 | 18 | 0 | 0 | 82 | 63 | 0 | 98 | 0 | 0 | 0 | 0 | 9820 | 0 | 0 |
| SRR10416499 | 0 | 0 | 0 | 0 | 0 | 792 | 396 | 0 | 0 | 0 | 36 | 0 | 0 | 0 | 1189 | 27746 | 17341 | 396 | 0 | 9905 | 0 | 0 | 0 | 23 | 0 | 23 | 23 | 23 | 0 | 87 | 0 | 0 | 0 | 0 | 5780 | 0 | 0 |
| SRR10416500 | 0 | 0 | 0 | 15 | 0 | 287 | 0 | 287 | 575 | 0 | 15 | 0 | 0 | 0 | 287 | 22209 | 37815 | 287 | 287 | 4884 | 0 | 0 | 0 | 0 | 0 | 0 | 149 | 30 | 0 | 31 | 0 | 0 | 0 | 0 | 18607 | 0 | 0 |
| SRR10416501 | 0 | 0 | 0 | 0 | 30 | 0 | 0 | 0 | 0 | 0 | 40 | 0 | 0 | 0 | 2188 | 27354 | 47710 | 0 | 625 | 10316 | 0 | 0 | 0 | 0 | 15 | 0 | 13 | 15 | 0 | 87 | 0 | 0 | 0 | 0 | 8906 | 0 | 15 |
| SRR10416502 | 0 | 0 | 0 | 20 | 0 | 0 | 460 | 0 | 0 | 0 | 258 | 0 | 0 | 0 | 2301 | 19792 | 26042 | 0 | 0 | 5983 | 0 | 0 | 0 | 0 | 0 | 20 | 20 | 0 | 0 | 953 | 0 | 0 | 0 | 0 | 13542 | 0 | 0 |
| SRR10416503 | 0 | 0 | 0 | 0 | 0 | 0 | 0 | 416 | 0 | 0 | 15 | 0 | 0 | 0 | 832 | 23256 | 29070 | 0 | 416 | 4158 | 0 | 0 | 0 | 0 | 0 | 22 | 44 | 0 | 0 | 60 | 0 | 0 | 0 | 0 | 7752 | 0 | 0 |
| SRR10416504 | 0 | 0 | 0 | 0 | 0 | 574 | 0 | 574 | 0 | 0 | 22 | 0 | 0 | 0 | 574 | 14389 | 26722 | 574 | 0 | 4016 | 0 | 0 | 0 | 21 | 21 | 21 | 148 | 42 | 0 | 28 | 0 | 0 | 0 | 0 | 17472 | 0 | 0 |
| SRR10416505 | 0 | 0 | 0 | 0 | 0 | 0 | 0 | 787 | 0 | 0 | 1102 | 0 | 0 | 0 | 1890 | 17485 | 32473 | 0 | 630 | 4882 | 0 | 0 | 0 | 0 | 0 | 0 | 26 | 15 | 0 | 1678 | 0 | 0 | 0 | 0 | 8326 | 0 | 0 |
| SRR10416506 | 35 | 0 | 0 | 0 | 18 | 0 | 0 | 386 | 0 | 0 | 16 | 0 | 0 | 0 | 1544 | 29683 | 35824 | 386 | 772 | 7333 | 0 | 0 | 0 | 0 | 0 | 0 | 18 | 0 | 0 | 50 | 0 | 0 | 0 | 0 | 19447 | 0 | 0 |
| SRR10416507 | 0 | 0 | 0 | 0 | 0 | 0 | 0 | 461 | 0 | 0 | 20 | 0 | 0 | 0 | 1382 | 13514 | 24570 | 0 | 0 | 6912 | 0 | 0 | 0 | 15 | 0 | 0 | 15 | 15 | 0 | 27 | 0 | 0 | 0 | 0 | 8600 | 0 | 0 |
| SRR10416508 | 0 | 0 | 0 | 23 | 0 | 758 | 758 | 0 | 0 | 0 | 10 | 0 | 0 | 0 | 379 | 16411 | 13129 | 0 | 379 | 2273 | 0 | 0 | 0 | 0 | 0 | 0 | 68 | 45 | 0 | 28 | 0 | 0 | 0 | 0 | 15317 | 0 | 0 |
| SRR10416509 | 0 | 0 | 0 | 0 | 0 | 431 | 0 | 0 | 0 | 0 | 57 | 0 | 0 | 0 | 216 | 14862 | 20170 | 216 | 647 | 3666 | 216 | 0 | 0 | 0 | 0 | 0 | 82 | 15 | 0 | 109 | 0 | 0 | 0 | 0 | 12208 | 0 | 0 |
| SRR10416510 | 0 | 0 | 31 | 0 | 0 | 0 | 0 | 0 | 0 | 0 | 41 | 0 | 0 | 0 | 1178 | 11321 | 3774 | 0 | 0 | 0 | 0 | 0 | 0 | 0 | 0 | 0 | 31 | 94 | 0 | 26 | 0 | 0 | 0 | 0 | 7547 | 0 | 0 |
| SRR10416513 | 0 | 0 | 0 | 0 | 0 | 133 | 0 | 665 | 133 | 0 | 711 | 0 | 0 | 0 | 0 | 7894 | 11633 | 133 | 2262 | 9050 | 133 | 0 | 0 | 0 | 0 | 0 | 14 | 0 | 0 | 1847 | 0 | 0 | 0 | 0 | 8309 | 0 | 0 |
| SRR10416514 | 0 | 0 | 0 | 0 | 23 | 0 | 0 | 513 | 513 | 0 | 468 | 0 | 0 | 0 | 1539 | 23897 | 36765 | 0 | 0 | 9236 | 0 | 0 | 0 | 0 | 0 | 23 | 70 | 47 | 0 | 1077 | 0 | 0 | 0 | 0 | 9191 | 0 | 23 |
| SRR10416515 | 0 | 0 | 0 | 0 | 23 | 0 | 0 | 0 | 0 | 0 | 1171 | 0 | 0 | 0 | 3017 | 28011 | 51821 | 0 | 0 | 9483 | 0 | 0 | 0 | 0 | 0 | 0 | 68 | 45 | 0 | 1734 | 0 | 0 | 0 | 0 | 9804 | 0 | 0 |
| SRR10416516 | 0 | 0 | 0 | 0 | 0 | 400 | 400 | 400 | 0 | 0 | 14 | 0 | 0 | 0 | 1200 | 29376 | 25704 | 0 | 0 | 10004 | 0 | 0 | 0 | 0 | 0 | 0 | 0 | 0 | 0 | 41 | 0 | 0 | 0 | 0 | 4896 | 0 | 0 |
| SRR10416517 | 0 | 0 | 0 | 46 | 0 | 0 | 0 | 0 | 358 | 0 | 43 | 0 | 0 | 0 | 4656 | 40758 | 47393 | 0 | 716 | 7521 | 358 | 0 | 0 | 0 | 0 | 15 | 0 | 62 | 0 | 61 | 0 | 0 | 0 | 0 | 5687 | 0 | 0 |
| SRR10416518 | 0 | 0 | 0 | 0 | 0 | 0 | 323 | 0 | 0 | 28 | 38 | 0 | 0 | 0 | 2581 | 10669 | 32978 | 0 | 0 | 8390 | 323 | 0 | 0 | 0 | 0 | 0 | 111 | 28 | 0 | 41 | 0 | 0 | 0 | 0 | 31038 | 0 | 0 |
| SRR10416519 | 0 | 0 | 0 | 32 | 16 | 0 | 0 | 0 | 0 | 32 | 2208 | 0 | 0 | 0 | 0 | 19332 | 14060 | 0 | 0 | 4169 | 0 | 0 | 0 | 32 | 0 | 16 | 177 | 32 | 0 | 2111 | 0 | 0 | 0 | 0 | 14060 | 0 | 0 |
| SRR10416520 | 0 | 0 | 0 | 39 | 0 | 0 | 0 | 0 | 0 | 0 | 503 | 0 | 0 | 0 | 649 | 23758 | 45356 | 0 | 0 | 2596 | 0 | 0 | 0 | 0 | 77 | 77 | 15 | 155 | 0 | 348 | 0 | 0 | 0 | 0 | 8639 | 0 | 0 |
| SRR10416521 | 0 | 0 | 0 | 0 | 0 | 0 | 485 | 485 | 0 | 0 | 1553 | 0 | 0 | 0 | 1455 | 27496 | 37627 | 0 | 0 | 5820 | 0 | 0 | 0 | 0 | 74 | 148 | 518 | 222 | 0 | 592 | 0 | 0 | 0 | 0 | 18813 | 0 | 0 |
| SRR10416522 | 0 | 0 | 0 | 0 | 0 | 0 | 0 | 0 | 0 | 0 | 20 | 0 | 0 | 0 | 527 | 10130 | 11577 | 0 | 0 | 527 | 0 | 0 | 0 | 0 | 0 | 45 | 361 | 45 | 0 | 11 | 0 | 0 | 0 | 0 | 15919 | 0 | 0 |
| SRR10416523 | 0 | 0 | 0 | 0 | 0 | 0 | 0 | 0 | 0 | 0 | 1027 | 0 | 0 | 0 | 409 | 18576 | 19608 | 0 | 0 | 1226 | 0 | 0 | 0 | 0 | 63 | 210 | 1006 | 293 | 0 | 650 | 0 | 0 | 0 | 0 | 13416 | 0 | 0 |
| SRR10416524 | 0 | 0 | 0 | 30 | 30 | 0 | 0 | 663 | 0 | 0 | 13 | 0 | 0 | 0 | 0 | 26201 | 8734 | 0 | 0 | 663 | 663 | 0 | 0 | 30 | 89 | 148 | 33 | 266 | 0 | 12 | 0 | 0 | 0 | 0 | 21834 | 0 | 30 |
| SRR10416525 | 0 | 0 | 0 | 0 | 0 | 0 | 0 | 0 | 0 | 0 | 2369 | 0 | 0 | 0 | 1453 | 26816 | 34637 | 0 | 0 | 1453 | 0 | 0 | 0 | 0 | 0 | 86 | 258 | 129 | 0 | 2929 | 0 | 0 | 0 | 0 | 12291 | 0 | 0 |
| SRR10416526 | 0 | 0 | 0 | 39 | 0 | 0 | 0 | 0 | 0 | 0 | 1853 | 39 | 0 | 0 | 2028 | 13289 | 19934 | 0 | 0 | 2705 | 0 | 0 | 0 | 0 | 0 | 39 | 849 | 154 | 0 | 618 | 0 | 0 | 0 | 0 | 6645 | 0 | 0 |
| SRR10416527 | 0 | 47 | 0 | 0 | 0 | 0 | 0 | 0 | 0 | 0 | 22 | 0 | 0 | 0 | 736 | 25281 | 33708 | 0 | 0 | 2209 | 0 | 0 | 0 | 47 | 0 | 0 | 0 | 0 | 0 | 21 | 0 | 0 | 0 | 0 | 33708 | 0 | 0 |
| SRR10416528 | 0 | 0 | 0 | 81 | 41 | 0 | 0 | 0 | 0 | 0 | 1262 | 0 | 0 | 0 | 0 | 21127 | 16432 | 0 | 0 | 2530 | 0 | 0 | 0 | 81 | 122 | 326 | 1099 | 41 | 0 | 529 | 0 | 0 | 0 | 0 | 24648 | 0 | 0 |
| SRR10416529 | 0 | 0 | 0 | 0 | 0 | 0 | 0 | 0 | 0 | 0 | 273 | 0 | 0 | 0 | 0 | 7335 | 22005 | 0 | 0 | 2708 | 0 | 0 | 0 | 55 | 0 | 109 | 382 | 218 | 55 | 327 | 0 | 0 | 0 | 0 | 19560 | 0 | 0 |
| SRR10416530 | 0 | 0 | 0 | 60 | 0 | 0 | 749 | 1497 | 0 | 0 | 13 | 0 | 0 | 0 | 749 | 10256 | 19231 | 0 | 0 | 2246 | 0 | 0 | 0 | 0 | 0 | 119 | 476 | 60 | 0 | 655 | 0 | 0 | 0 | 0 | 10256 | 0 | 0 |
| SRR10416531 | 0 | 0 | 0 | 0 | 0 | 0 | 0 | 0 | 0 | 0 | 1489 | 0 | 0 | 0 | 2210 | 20790 | 16632 | 0 | 0 | 1105 | 0 | 0 | 0 | 0 | 0 | 199 | 695 | 99 | 0 | 1390 | 0 | 0 | 0 | 0 | 16632 | 0 | 0 |
| SRR10416532 | 0 | 0 | 0 | 0 | 0 | 0 | 0 | 0 | 0 | 0 | 2211 | 0 | 0 | 0 | 1540 | 20767 | 43131 | 0 | 0 | 1540 | 0 | 0 | 0 | 0 | 69 | 345 | 138 | 207 | 0 | 1520 | 0 | 0 | 0 | 0 | 14377 | 0 | 0 |

### Supp Table S8.3.Columns_BX_to_DH

| | | | | | | | | | | | | | | | | | | | | | | | | | | | | | | | | | | | | | |
|---|---|---|---|---|---|---|---|---|---|---|---|---|---|---|---|---|---|---|---|---|---|---|---|---|---|---|---|---|---|---|---|---|---|---|---|---|---|
| SRR10416533 | 0 | 0 | 0 | 44 | 0 | 0 | 0 | 0 | 0 | 0 | 14 | 0 | 0 | 0 | 0 | 19868 | 33113 | 0 | 0 | 4020 | 2010 | 0 | 0 | 0 | 0 | 44 | 87 | 44 | 0 | 350 | 0 | 0 | 0 | 0 | 33113 | 0 | 0 |
| SRR10416534 | 0 | 0 | 0 | 0 | 85 | 0 | 0 | 0 | 0 | 0 | 2469 | 0 | 0 | 0 | 2312 | 17036 | 30664 | 0 | 0 | 0 | 0 | 0 | 0 | 0 | 0 | 0 | 0 | 85 | 0 | 1873 | 0 | 0 | 0 | 0 | 6814 | 0 | 0 |
| SRR10416535 | 0 | 0 | 0 | 0 | 0 | 0 | 0 | 0 | 0 | 0 | 42 | 0 | 0 | 0 | 840 | 27219 | 40237 | 0 | 0 | 3359 | 0 | 0 | 0 | 0 | 0 | 137 | 502 | 91 | 0 | 39 | 0 | 0 | 0 | 0 | 10651 | 0 | 0 |
| SRR10416536 | 0 | 0 | 0 | 0 | 0 | 0 | 0 | 0 | 0 | 44 | 62 | 0 | 0 | 0 | 1442 | 20588 | 23529 | 0 | 0 | 6489 | 0 | 0 | 0 | 0 | 0 | 0 | 310 | 177 | 0 | 30 | 0 | 0 | 0 | 0 | 5882 | 0 | 0 |
| SRR10416537 | 0 | 0 | 0 | 0 | 0 | 0 | 548 | 0 | 0 | 0 | 36 | 0 | 0 | 0 | 4384 | 27322 | 49180 | 0 | 0 | 3836 | 0 | 0 | 0 | 0 | 0 | 179 | 224 | 179 | 0 | 24 | 0 | 0 | 0 | 0 | 3643 | 0 | 0 |
| SRR10416538 | 0 | 0 | 0 | 0 | 0 | 0 | 0 | 0 | 0 | 0 | 16 | 0 | 0 | 0 | 661 | 18610 | 23573 | 0 | 0 | 1321 | 661 | 0 | 0 | 0 | 137 | 102 | 580 | 171 | 0 | 444 | 0 | 0 | 0 | 0 | 13648 | 0 | 0 |
| SRR10416539 | 0 | 0 | 0 | 0 | 0 | 0 | 0 | 704 | 0 | 0 | 2342 | 0 | 0 | 0 | 1407 | 9950 | 31095 | 0 | 0 | 2111 | 0 | 0 | 0 | 0 | 0 | 0 | 134 | 0 | 0 | 1004 | 0 | 0 | 0 | 0 | 23632 | 0 | 0 |
| SRR10416540 | 0 | 0 | 0 | 0 | 0 | 0 | 0 | 0 | 0 | 0 | 34 | 0 | 0 | 0 | 0 | 19028 | 35941 | 0 | 0 | 3226 | 0 | 0 | 0 | 0 | 0 | 151 | 16 | 303 | 0 | 22 | 0 | 0 | 0 | 0 | 16913 | 0 | 0 |
| SRR10416541 | 0 | 0 | 0 | 0 | 0 | 0 | 0 | 0 | 0 | 0 | 11 | 0 | 0 | 0 | 4292 | 17167 | 17167 | 0 | 0 | 2146 | 0 | 0 | 0 | 0 | 114 | 0 | 228 | 456 | 0 | 684 | 0 | 0 | 0 | 0 | 17167 | 0 | 0 |
| SRR10416542 | 0 | 0 | 0 | 0 | 0 | 0 | 3017 | 1508 | 0 | 0 | 13 | 0 | 0 | 0 | 3017 | 10274 | 20548 | 0 | 0 | 3017 | 0 | 0 | 0 | 0 | 0 | 271 | 542 | 452 | 0 | 722 | 0 | 0 | 0 | 0 | 6849 | 0 | 0 |
| SRR10416543 | 0 | 0 | 0 | 0 | 0 | 0 | 263 | 263 | 0 | 0 | 863 | 0 | 0 | 0 | 1052 | 47170 | 58176 | 0 | 2105 | 14470 | 0 | 0 | 0 | 0 | 0 | 0 | 0 | 0 | 0 | 2299 | 0 | 0 | 0 | 0 | 5503 | 0 | 0 |
| SRR10416544 | 0 | 0 | 0 | 0 | 0 | 0 | 0 | 0 | 0 | 0 | 546 | 0 | 0 | 0 | 693 | 33537 | 9146 | 0 | 0 | 4161 | 0 | 0 | 0 | 0 | 0 | 78 | 11 | 624 | 0 | 312 | 0 | 0 | 0 | 0 | 51829 | 0 | 0 |
| SRR10416545 | 0 | 0 | 0 | 38 | 38 | 0 | 0 | 1251 | 0 | 0 | 639 | 19 | 0 | 0 | 834 | 15737 | 20744 | 0 | 417 | 1667 | 0 | 0 | 0 | 0 | 0 | 0 | 150 | 75 | 0 | 676 | 0 | 0 | 0 | 0 | 18598 | 0 | 0 |
| SRR10416546 | 0 | 0 | 0 | 0 | 0 | 0 | 0 | 0 | 612 | 0 | 11 | 0 | 0 | 0 | 1224 | 10329 | 21596 | 0 | 0 | 3060 | 0 | 0 | 0 | 0 | 0 | 0 | 0 | 0 | 0 | 396 | 0 | 0 | 0 | 0 | 16901 | 0 | 33 |
| SRR10416547 | 0 | 0 | 0 | 0 | 0 | 0 | 0 | 0 | 0 | 0 | 387 | 0 | 0 | 0 | 0 | 3125 | 9375 | 0 | 0 | 3155 | 0 | 0 | 0 | 0 | 0 | 48 | 242 | 48 | 0 | 435 | 0 | 0 | 0 | 0 | 12500 | 0 | 0 |
| SRR10416549 | 0 | 0 | 0 | 45 | 23 | 280 | 0 | 280 | 0 | 0 | 11 | 0 | 0 | 0 | 841 | 22121 | 20169 | 0 | 0 | 2524 | 0 | 0 | 0 | 23 | 23 | 114 | 30 | 159 | 0 | 15 | 0 | 0 | 0 | 0 | 16266 | 0 | 0 |
| SRR10416550 | 0 | 0 | 0 | 21 | 0 | 0 | 0 | 0 | 0 | 0 | 17 | 0 | 0 | 0 | 986 | 27429 | 26286 | 0 | 0 | 2464 | 0 | 0 | 0 | 42 | 83 | 42 | 14 | 167 | 0 | 11 | 0 | 0 | 0 | 0 | 5714 | 0 | 0 |
| SRR10416551 | 0 | 0 | 0 | 27 | 27 | 0 | 0 | 0 | 471 | 27 | 901 | 0 | 0 | 0 | 941 | 20656 | 20656 | 0 | 0 | 1882 | 0 | 0 | 0 | 0 | 0 | 106 | 398 | 27 | 0 | 1034 | 0 | 471 | 0 | 0 | 8505 | 0 | 0 |
| SRR10416553 | 0 | 0 | 0 | 130 | 0 | 0 | 0 | 727 | 0 | 0 | 843 | 0 | 0 | 0 | 727 | 8982 | 22455 | 0 | 0 | 0 | 0 | 0 | 0 | 0 | 0 | 130 | 519 | 0 | 0 | 713 | 0 | 0 | 0 | 0 | 13473 | 0 | 0 |
| SRR10416554 | 0 | 0 | 0 | 0 | 0 | 0 | 0 | 0 | 0 | 0 | 18 | 0 | 0 | 0 | 0 | 45704 | 51188 | 0 | 0 | 3492 | 0 | 0 | 0 | 0 | 0 | 71 | 142 | 71 | 0 | 708 | 0 | 0 | 0 | 0 | 14625 | 0 | 0 |
| SRR10416555 | 0 | 0 | 0 | 0 | 114 | 0 | 0 | 1168 | 0 | 0 | 1828 | 0 | 0 | 0 | 2336 | 9506 | 19011 | 0 | 0 | 1168 | 1168 | 0 | 0 | 0 | 0 | 114 | 0 | 228 | 0 | 457 | 0 | 0 | 0 | 0 | 19011 | 0 | 0 |
| SRR10416557 | 0 | 0 | 0 | 115 | 0 | 0 | 0 | 0 | 0 | 0 | 461 | 0 | 0 | 0 | 0 | 15625 | 12500 | 0 | 0 | 0 | 0 | 0 | 0 | 0 | 0 | 0 | 231 | 231 | 0 | 346 | 0 | 0 | 0 | 0 | 0 | 0 | 0 |
| SRR10416558 | 0 | 0 | 0 | 0 | 0 | 0 | 673 | 0 | 0 | 0 | 30 | 0 | 0 | 0 | 2020 | 29178 | 34483 | 0 | 0 | 2020 | 0 | 0 | 0 | 0 | 51 | 152 | 380 | 228 | 0 | 405 | 0 | 0 | 0 | 0 | 23873 | 0 | 0 |
| SRR10416559 | 0 | 0 | 0 | 0 | 0 | 0 | 0 | 545 | 0 | 0 | 1713 | 0 | 0 | 0 | 2181 | 13889 | 18056 | 0 | 0 | 1091 | 0 | 0 | 0 | 0 | 0 | 0 | 0 | 0 | 0 | 467 | 0 | 0 | 0 | 0 | 11111 | 0 | 0 |
| SRR10416560 | 0 | 0 | 0 | 0 | 0 | 0 | 0 | 0 | 0 | 0 | 1035 | 0 | 0 | 0 | 1302 | 13720 | 25915 | 0 | 0 | 2604 | 0 | 0 | 0 | 23 | 46 | 184 | 897 | 230 | 0 | 575 | 0 | 0 | 0 | 0 | 15244 | 0 | 0 |
| SRR10416562 | 0 | 0 | 0 | 0 | 0 | 0 | 0 | 0 | 0 | 0 | 11 | 0 | 0 | 0 | 0 | 15411 | 22260 | 0 | 0 | 1443 | 0 | 0 | 0 | 0 | 0 | 0 | 277 | 35 | 0 | 243 | 0 | 0 | 0 | 0 | 23973 | 0 | 0 |
| SRR10416563 | 0 | 0 | 15 | 15 | 0 | 0 | 0 | 0 | 0 | 0 | 2355 | 0 | 0 | 0 | 0 | 24775 | 15766 | 0 | 0 | 1095 | 0 | 0 | 0 | 0 | 0 | 75 | 285 | 75 | 0 | 1935 | 0 | 0 | 0 | 0 | 6757 | 0 | 0 |
| SRR10416564 | 0 | 0 | 0 | 0 | 43 | 0 | 0 | 1398 | 0 | 0 | 560 | 0 | 0 | 0 | 699 | 8242 | 8242 | 0 | 0 | 0 | 0 | 0 | 0 | 43 | 0 | 0 | 689 | 86 | 0 | 646 | 0 | 0 | 0 | 0 | 9615 | 0 | 0 |
| SRR10416565 | 0 | 0 | 31 | 0 | 0 | 0 | 0 | 632 | 632 | 0 | 467 | 0 | 0 | 0 | 0 | 17258 | 9588 | 0 | 0 | 3793 | 0 | 0 | 0 | 31 | 31 | 93 | 249 | 62 | 0 | 529 | 0 | 0 | 0 | 0 | 21093 | 0 | 0 |
| SRR10416566 | 0 | 0 | 0 | 72 | 0 | 0 | 0 | 0 | 0 | 0 | 360 | 0 | 0 | 0 | 0 | 16529 | 22039 | 0 | 0 | 0 | 0 | 0 | 0 | 0 | 0 | 72 | 72 | 216 | 0 | 216 | 0 | 0 | 0 | 0 | 2755 | 0 | 0 |
| SRR10416567 | 0 | 0 | 0 | 0 | 0 | 0 | 0 | 1055 | 0 | 0 | 642 | 0 | 0 | 0 | 528 | 4335 | 8671 | 0 | 0 | 4749 | 0 | 0 | 0 | 0 | 0 | 0 | 214 | 107 | 0 | 749 | 0 | 0 | 0 | 0 | 21676 | 0 | 0 |
| SRR10416568 | 0 | 0 | 0 | 0 | 79 | 0 | 0 | 0 | 0 | 0 | 867 | 0 | 0 | 0 | 2527 | 26846 | 20134 | 0 | 0 | 2527 | 0 | 0 | 0 | 0 | 79 | 0 | 236 | 315 | 0 | 473 | 0 | 0 | 0 | 0 | 17897 | 0 | 0 |
| SRR10416569 | 0 | 0 | 0 | 0 | 0 | 0 | 0 | 0 | 637 | 0 | 858 | 0 | 0 | 0 | 0 | 5148 | 7722 | 0 | 0 | 0 | 0 | 0 | 0 | 0 | 0 | 0 | 334 | 0 | 0 | 572 | 0 | 0 | 0 | 0 | 6435 | 0 | 0 |
| SRR10416570 | 0 | 0 | 0 | 0 | 0 | 0 | 0 | 0 | 0 | 0 | 15 | 0 | 0 | 0 | 0 | 28409 | 26515 | 0 | 0 | 644 | 0 | 0 | 0 | 0 | 0 | 0 | 158 | 106 | 0 | 21 | 0 | 0 | 0 | 0 | 17046 | 0 | 0 |
| SRR10416571 | 0 | 0 | 0 | 61 | 0 | 0 | 0 | 0 | 0 | 0 | 734 | 0 | 0 | 0 | 0 | 11719 | 21484 | 0 | 0 | 1541 | 0 | 0 | 0 | 61 | 0 | 245 | 734 | 795 | 0 | 367 | 0 | 770 | 0 | 0 | 5859 | 0 | 0 |
| SRR10416572 | 0 | 0 | 0 | 131 | 0 | 0 | 0 | 546 | 0 | 0 | 1874 | 0 | 0 | 0 | 1638 | 37594 | 40602 | 0 | 0 | 2729 | 0 | 0 | 0 | 0 | 44 | 44 | 392 | 87 | 0 | 785 | 0 | 0 | 0 | 0 | 13534 | 0 | 0 |
| SRR10416573 | 0 | 0 | 0 | 204 | 102 | 0 | 0 | 0 | 0 | 0 | 511 | 0 | 0 | 0 | 1701 | 15038 | 18797 | 0 | 0 | 1701 | 0 | 0 | 0 | 0 | 0 | 102 | 716 | 511 | 0 | 204 | 0 | 0 | 0 | 0 | 15038 | 0 | 0 |
| SRR10416574 | 0 | 0 | 0 | 0 | 0 | 752 | 0 | 0 | 0 | 0 | 766 | 0 | 0 | 0 | 1505 | 6623 | 11038 | 0 | 0 | 0 | 0 | 0 | 0 | 96 | 96 | 96 | 287 | 287 | 0 | 670 | 0 | 0 | 0 | 0 | 4415 | 0 | 0 |
| SRR10416575 | 0 | 0 | 0 | 0 | 0 | 0 | 0 | 0 | 0 | 0 | 19 | 0 | 0 | 0 | 0 | 21053 | 10526 | 0 | 0 | 2594 | 0 | 0 | 0 | 0 | 0 | 0 | 0 | 0 | 0 | 29 | 0 | 0 | 0 | 0 | 21053 | 0 | 0 |
| SRR10416576 | 0 | 0 | 0 | 0 | 0 | 0 | 0 | 0 | 0 | 0 | 2768 | 0 | 0 | 0 | 2210 | 37975 | 21097 | 0 | 0 | 1105 | 0 | 0 | 0 | 0 | 0 | 56 | 198 | 113 | 0 | 2344 | 0 | 0 | 0 | 0 | 12658 | 0 | 0 |
| SRR10416577 | 0 | 0 | 0 | 0 | 71 | 0 | 682 | 0 | 0 | 0 | 712 | 0 | 0 | 0 | 1363 | 16839 | 7772 | 0 | 0 | 2727 | 0 | 0 | 0 | 0 | 0 | 71 | 783 | 427 | 0 | 427 | 0 | 0 | 0 | 0 | 19430 | 0 | 0 |
| SRR10416578 | 0 | 0 | 0 | 0 | 0 | 746 | 0 | 0 | 0 | 0 | 877 | 0 | 0 | 0 | 746 | 8929 | 10714 | 0 | 0 | 1491 | 0 | 0 | 0 | 146 | 0 | 0 | 292 | 0 | 0 | 730 | 0 | 0 | 0 | 0 | 3571 | 0 | 0 |
| SRR10416579 | 0 | 0 | 0 | 0 | 68 | 0 | 0 | 0 | 0 | 0 | 2595 | 0 | 0 | 0 | 3371 | 17391 | 60870 | 0 | 0 | 0 | 0 | 0 | 0 | 0 | 0 | 137 | 137 | 0 | 0 | 1161 | 0 | 0 | 0 | 0 | 11594 | 0 | 0 |
| SRR10416580 | 0 | 0 | 0 | 0 | 0 | 0 | 0 | 0 | 0 | 0 | 519 | 0 | 0 | 0 | 772 | 27273 | 14546 | 0 | 0 | 3086 | 0 | 0 | 0 | 0 | 52 | 208 | 363 | 52 | 0 | 415 | 0 | 0 | 0 | 0 | 3636 | 0 | 0 |
| SRR10416581 | 0 | 0 | 0 | 20 | 20 | 0 | 0 | 746 | 0 | 40 | 788 | 0 | 0 | 0 | 0 | 55000 | 42500 | 0 | 0 | 0 | 746 | 0 | 0 | 0 | 0 | 0 | 283 | 40 | 0 | 1151 | 0 | 0 | 0 | 0 | 12500 | 0 | 0 |
| SRR10416582 | 0 | 0 | 0 | 0 | 92 | 0 | 935 | 0 | 0 | 0 | 12 | 0 | 0 | 0 | 935 | 7874 | 15748 | 0 | 0 | 1869 | 0 | 0 | 0 | 92 | 0 | 184 | 920 | 92 | 0 | 644 | 0 | 0 | 0 | 0 | 5249 | 0 | 0 |
| SRR10416583 | 0 | 0 | 0 | 0 | 0 | 0 | 0 | 0 | 0 | 0 | 1925 | 0 | 0 | 0 | 0 | 22727 | 7576 | 0 | 0 | 3810 | 0 | 0 | 0 | 0 | 0 | 84 | 0 | 84 | 0 | 1088 | 0 | 0 | 0 | 0 | 3788 | 0 | 0 |
| SRR10416584 | 0 | 0 | 0 | 0 | 44 | 0 | 0 | 0 | 0 | 0 | 1245 | 0 | 0 | 0 | 0 | 30340 | 25485 | 0 | 0 | 2787 | 0 | 0 | 0 | 0 | 44 | 133 | 1245 | 133 | 0 | 489 | 0 | 0 | 0 | 0 | 13350 | 0 | 0 |
| SRR10416585 | 0 | 0 | 0 | 0 | 0 | 0 | 0 | 0 | 0 | 0 | 988 | 0 | 0 | 0 | 2028 | 26316 | 11278 | 0 | 0 | 2028 | 0 | 0 | 0 | 0 | 165 | 0 | 165 | 0 | 0 | 329 | 0 | 0 | 0 | 0 | 3759 | 0 | 0 |
| SRR10416586 | 0 | 0 | 0 | 0 | 0 | 0 | 0 | 0 | 0 | 0 | 1975 | 0 | 0 | 0 | 0 | 10152 | 17767 | 0 | 0 | 0 | 0 | 0 | 0 | 0 | 0 | 0 | 188 | 94 | 0 | 188 | 0 | 0 | 0 | 0 | 7614 | 0 | 0 |
| SRR10416587 | 0 | 0 | 0 | 0 | 0 | 0 | 0 | 0 | 0 | 0 | 2001 | 0 | 0 | 0 | 1002 | 24324 | 40541 | 0 | 0 | 0 | 0 | 0 | 0 | 0 | 65 | 65 | 323 | 129 | 0 | 516 | 0 | 0 | 0 | 0 | 32432 | 0 | 0 |
| SRR10416588 | 0 | 0 | 0 | 0 | 0 | 0 | 0 | 991 | 0 | 0 | 1088 | 0 | 0 | 0 | 991 | 5291 | 19400 | 0 | 0 | 0 | 0 | 0 | 0 | 0 | 78 | 78 | 622 | 155 | 0 | 1010 | 0 | 0 | 0 | 0 | 8818 | 0 | 0 |
| SRR10416589 | 0 | 0 | 0 | 0 | 0 | 0 | 0 | 0 | 0 | 0 | 3492 | 0 | 0 | 0 | 1587 | 16166 | 32333 | 0 | 0 | 6349 | 0 | 0 | 0 | 0 | 0 | 0 | 0 | 49 | 0 | 3319 | 0 | 0 | 0 | 0 | 5774 | 0 | 0 |
| SRR10416590 | 0 | 0 | 0 | 0 | 0 | 0 | 0 | 0 | 0 | 0 | 1683 | 0 | 0 | 0 | 0 | 14377 | 19169 | 0 | 0 | 3968 | 0 | 0 | 0 | 0 | 0 | 0 | 269 | 471 | 0 | 1414 | 0 | 0 | 0 | 0 | 20767 | 0 | 0 |
| SRR10416591 | 0 | 0 | 0 | 0 | 0 | 0 | 1511 | 1511 | 0 | 0 | 18 | 0 | 0 | 0 | 0 | 39514 | 24316 | 0 | 0 | 4532 | 1511 | 0 | 0 | 0 | 90 | 90 | 179 | 359 | 0 | 12 | 0 | 0 | 0 | 0 | 6079 | 0 | 0 |
| SRR10416592 | 0 | 0 | 0 | 0 | 0 | 0 | 371 | 371 | 0 | 45 | 30 | 0 | 0 | 0 | 1114 | 8065 | 21889 | 0 | 0 | 2972 | 0 | 0 | 0 | 45 | 0 | 0 | 227 | 91 | 0 | 21 | 0 | 0 | 0 | 0 | 5760 | 0 | 0 |
| SRR10416593 | 0 | 0 | 0 | 140 | 0 | 0 | 0 | 0 | 0 | 0 | 3226 | 0 | 0 | 0 | 1871 | 24336 | 33186 | 0 | 0 | 5613 | 0 | 0 | 0 | 0 | 47 | 140 | 421 | 327 | 0 | 2057 | 0 | 0 | 0 | 0 | 11062 | 0 | 0 |
| SRR10416594 | 0 | 0 | 0 | 0 | 0 | 0 | 0 | 0 | 0 | 0 | 1051 | 0 | 0 | 0 | 931 | 31031 | 20020 | 0 | 0 | 1862 | 0 | 0 | 0 | 75 | 75 | 75 | 300 | 225 | 0 | 825 | 0 | 0 | 0 | 0 | 9009 | 0 | 0 |
| SRR10416595 | 0 | 0 | 0 | 42 | 0 | 0 | 0 | 0 | 0 | 0 | 586 | 84 | 0 | 0 | 492 | 26506 | 38554 | 0 | 0 | 985 | 985 | 0 | 0 | 0 | 84 | 84 | 879 | 209 | 0 | 628 | 0 | 0 | 0 | 0 | 19277 | 0 | 168 |
| SRR10416596 | 0 | 0 | 0 | 39 | 39 | 0 | 0 | 0 | 0 | 0 | 887 | 0 | 0 | 0 | 0 | 20690 | 20690 | 0 | 0 | 1048 | 0 | 0 | 0 | 0 | 0 | 19 | 116 | 173 | 0 | 1157 | 0 | 0 | 0 | 0 | 0 | 0 | 0 |
| SRR10416597 | 0 | 0 | 0 | 27 | 0 | 0 | 0 | 0 | 0 | 0 | 2825 | 27 | 0 | 0 | 548 | 52555 | 43796 | 0 | 0 | 3836 | 0 | 0 | 0 | 0 | 0 | 0 | 247 | 110 | 0 | 1289 | 0 | 0 | 0 | 27 | 24818 | 0 | 0 |
| SRR10416598 | 0 | 0 | 0 | 0 | 0 | 0 | 0 | 1182 | 0 | 0 | 385 | 28 | 0 | 0 | 0 | 9677 | 19355 | 0 | 0 | 0 | 0 | 0 | 0 | 0 | 0 | 0 | 83 | 83 | 0 | 991 | 0 | 0 | 0 | 0 | 0 | 0 | 28 |
| SRR10416599 | 0 | 0 | 0 | 0 | 0 | 0 | 0 | 0 | 0 | 0 | 819 | 0 | 0 | 0 | 888 | 23876 | 9831 | 0 | 0 | 2221 | 0 | 0 | 0 | 63 | 0 | 0 | 63 | 0 | 0 | 504 | 0 | 0 | 0 | 0 | 2809 | 0 | 0 |
| SRR10416600 | 0 | 0 | 0 | 20 | 0 | 0 | 0 | 0 | 0 | 0 | 168 | 20 | 0 | 0 | 2094 | 22796 | 22796 | 0 | 0 | 2931 | 419 | 0 | 0 | 20 | 0 | 0 | 80 | 20 | 0 | 99 | 0 | 0 | 0 | 0 | 15198 | 0 | 0 |
| SRR10428553 | 0 | 0 | 0 | 0 | 52 | 106 | 66 | 268 | 13 | 40 | 128 | 0 | 0 | 0 | 43578 | 254765 | 104078 | 17 | 0 | 806 | 1133 | 0 | 0 | 117 | 0 | 13 | 47 | 13 | 0 | 434 | 0 | 0 | 0 | 0 | 58 | 0 | 0 |
| SRR10428554 | 0 | 0 | 0 | 14 | 43 | 536 | 301 | 2272 | 137 | 19 | 62 | 0 | 0 | 0 | 124570 | 160603 | 392487 | 0 | 51 | 4868 | 1747 | 0 | 0 | 69 | 0 | 54 | 45 | 21 | 0 | 190 | 0 | 0 | 0 | 0 | 586 | 0 | 12 |
| SRR10428555 | 0 | 0 | 0 | 0 | 15 | 88 | 77 | 364 | 18 | 18 | 56 | 15 | 0 | 0 | 43424 | 315105 | 176307 | 14 | 0 | 1491 | 503 | 0 | 0 | 38 | 0 | 0 | 23 | 0 | 0 | 48 | 0 | 0 | 0 | 0 | 288 | 0 | 16 |
| SRR10428556 | 0 | 0 | 0 | 12 | 36 | 170 | 258 | 1314 | 53 | 11 | 46 | 0 | 0 | 0 | 32175 | 78623 | 87202 | 16 | 60 | 14697 | 1304 | 0 | 0 | 40 | 0 | 21 | 75 | 15 | 0 | 134 | 0 | 11 | 0 | 0 | 363 | 0 | 0 |
| SRR10428557 | 0 | 0 | 0 | 0 | 0 | 528 | 257 | 2429 | 92 | 17 | 57 | 14 | 0 | 0 | 43656 | 64993 | 272451 | 0 | 42 | 6079 | 2301 | 0 | 0 | 27 | 0 | 13 | 52 | 0 | 0 | 300 | 0 | 0 | 0 | 0 | 571 | 0 | 11 |
| SRR10428558 | 0 | 0 | 0 | 0 | 22 | 142 | 197 | 889 | 45 | 17 | 28 | 0 | 0 | 0 | 35403 | 95754 | 171168 | 0 | 69 | 29786 | 905 | 0 | 0 | 28 | 0 | 23 | 50 | 12 | 0 | 93 | 0 | 0 | 0 | 0 | 242 | 0 | 0 |
| SRR10428559 | 0 | 0 | 0 | 0 | 27 | 371 | 272 | 1414 | 47 | 27 | 40 | 18 | 0 | 0 | 83087 | 119842 | 240231 | 0 | 58 | 12492 | 681 | 0 | 0 | 87 | 0 | 25 | 62 | 13 | 0 | 132 | 0 | 0 | 0 | 0 | 203 | 0 | 11 |
| SRR10428560 | 0 | 0 | 0 | 0 | 19 | 42 | 42 | 588 | 0 | 25 | 21 | 0 | 0 | 0 | 82549 | 88041 | 298127 | 0 | 32 | 12423 | 4214 | 0 | 0 | 58 | 0 | 26 | 53 | 0 | 0 | 42 | 0 | 0 | 0 | 0 | 93 | 0 | 0 |
| SRR10446207 | 97 | 34 | 26 | 11 | 608 | 644 | 804 | 6141 | 2351 | 0 | 11 | 0 | 0 | 0 | 196340 | 56575 | 274840 | 0 | 63 | 1689 | 992 | 0 | 0 | 125 | 26 | 31 | 105 | 105 | 0 | 23 | 0 | 0 | 0 | 0 | 1270 | 129 | 0 |
| SRR10446208 | 48 | 32 | 19 | 0 | 39 | 2006 | 2276 | 11821 | 3989 | 0 | 74 | 0 | 0 | 0 | 103000 | 91772 | 331304 | 0 | 180 | 9184 | 3685 | 0 | 0 | 16 | 80 | 99 | 21 | 16 | 0 | 58 | 0 | 0 | 0 | 0 | 1758 | 213 | 0 |
| SRR10446209 | 128 | 54 | 25 | 39 | 1322 | 4037 | 4646 | 20285 | 10264 | 0 | 108 | 0 | 0 | 0 | 39475 | 160128 | 195783 | 0 | 146 | 5594 | 3551 | 0 | 0 | 324 | 69 | 98 | 236 | 290 | 0 | 69 | 0 | 0 | 0 | 0 | 1757 | 69 | 0 |
| SRR10446210 | 131 | 92 | 0 | 26 | 85 | 2936 | 2443 | 13780 | 6301 | 0 | 72 | 0 | 0 | 0 | 11530 | 224560 | 318668 | 0 | 21 | 1329 | 2400 | 0 | 0 | 48 | 118 | 243 | 28 | 14 | 0 | 171 | 0 | 0 | 0 | 0 | 3969 | 33 | 0 |
| SRR10446211 | 23 | 18 | 0 | 0 | 429 | 674 | 595 | 2077 | 743 | 13 | 101 | 16 | 0 | 0 | 96530 | 109078 | 510567 | 0 | 378 | 3106 | 3092 | 16 | 12 | 17 | 17 | 28 | 32 | 30 | 0 | 158 | 0 | 0 | 12 | 0 | 5584 | 113 | 0 |
| SRR10446212 | 95 | 73 | 13 | 13 | 2024 | 820 | 2328 | 5766 | 2605 | 0 | 274 | 0 | 0 | 0 | 18938 | 125377 | 492043 | 0 | 331 | 6639 | 1256 | 0 | 0 | 137 | 127 | 293 | 283 | 426 | 0 | 465 | 0 | 0 | 0 | 0 | 3310 | 11 | 0 |
| SRR10531363 | 0 | 0 | 0 | 0 | 0 | 0 | 0 | 0 | 0 | 0 | 0 | 221 | 0 | 0 | 0 | 345967 | 8586 | 0 | 0 | 0 | 19668 | 0 | 0 | 0 | 0 | 0 | 0 | 0 | 0 | 0 | 377 | 0 | 0 | 0 | 2921 | 0 | 0 |
| SRR10531364 | 0 | 0 | 0 | 0 | 0 | 0 | 0 | 0 | 0 | 151 | 52 | 186 | 0 | 0 | 578 | 436517 | 19426 | 0 | 0 | 0 | 18726 | 46 | 0 | 0 | 0 | 0 | 0 | 0 | 0 | 0 | 0 | 0 | 0 | 0 | 3275 | 0 | 0 |
| SRR10531365 | 0 | 0 | 0 | 0 | 0 | 0 | 0 | 0 | 0 | 0 | 0 | 526 | 0 | 0 | 64 | 545915 | 8940 | 0 | 0 | 0 | 11818 | 139 | 0 | 0 | 0 | 0 | 0 | 0 | 0 | 0 | 0 | 0 | 0 | 0 | 2646 | 0 | 0 |
| SRR10531367 | 0 | 0 | 0 | 0 | 0 | 0 | 0 | 0 | 0 | 0 | 0 | 108 | 0 | 0 | 0 | 721624 | 16643 | 0 | 0 | 0 | 1994 | 0 | 0 | 0 | 0 | 0 | 131 | 0 | 0 | 0 | 0 | 0 | 0 | 0 | 1705 | 0 | 0 |
| SRR10531368 | 0 | 0 | 0 | 0 | 0 | 0 | 0 | 0 | 0 | 0 | 0 | 31 | 0 | 0 | 14 | 556090 | 11473 | 0 | 0 | 0 | 8200 | 0 | 0 | 0 | 0 | 0 | 0 | 0 | 0 | 0 | 0 | 0 | 0 | 0 | 6705 | 0 | 0 |
| SRR10531369 | 0 | 0 | 0 | 0 | 0 | 0 | 0 | 0 | 0 | 361 | 0 | 268 | 0 | 0 | 1748 | 630442 | 10936 | 0 | 11 | 0 | 6315 | 0 | 0 | 0 | 0 | 0 | 0 | 0 | 0 | 0 | 0 | 0 | 0 | 0 | 2426 | 0 | 0 |
| SRR10531370 | 0 | 104 | 0 | 0 | 101 | 0 | 0 | 0 | 0 | 231 | 0 | 88 | 0 | 0 | 0 | 360165 | 11255 | 0 | 477 | 0 | 6496 | 0 | 0 | 0 | 0 | 0 | 0 | 88 | 0 | 0 | 89 | 0 | 0 | 0 | 3639 | 0 | 0 |
| SRR10544939 | 0 | 0 | 0 | 0 | 14 | 22 | 19 | 22 | 0 | 0 | 139 | 0 | 0 | 0 | 4846 | 2975 | 20675 | 19 | 27 | 1417 | 0 | 0 | 0 | 0 | 0 | 0 | 27 | 15 | 0 | 1383 | 0 | 0 | 0 | 0 | 312 | 0 | 0 |
| SRR10544940 | 0 | 0 | 0 | 0 | 16 | 0 | 0 | 0 | 0 | 0 | 37 | 0 | 0 | 0 | 1501 | 4170 | 29386 | 12 | 0 | 515 | 0 | 0 | 0 | 13 | 0 | 0 | 10 | 14 | 0 | 145 | 0 | 0 | 0 | 0 | 602 | 0 | 0 |
| SRR10544941 | 24 | 0 | 14 | 0 | 39 | 0 | 0 | 0 | 0 | 0 | 175 | 0 | 0 | 0 | 2463 | 4190 | 25164 | 21 | 20 | 1075 | 0 | 0 | 0 | 27 | 0 | 14 | 82 | 36 | 0 | 333 | 0 | 0 | 0 | 0 | 370 | 0 | 0 |
| SRR10544942 | 0 | 0 | 0 | 0 | 0 | 0 | 10 | 0 | 0 | 0 | 102 | 0 | 0 | 0 | 957 | 2909 | 14589 | 15 | 25 | 2577 | 0 | 0 | 0 | 17 | 0 | 0 | 51 | 18 | 0 | 823 | 0 | 0 | 0 | 0 | 263 | 0 | 0 |
| SRR10544943 | 0 | 0 | 0 | 0 | 11 | 0 | 0 | 11 | 0 | 0 | 19 | 0 | 0 | 0 | 1661 | 4428 | 25044 | 14 | 17 | 632 | 0 | 0 | 0 | 0 | 0 | 0 | 12 | 17 | 0 | 73 | 0 | 0 | 0 | 0 | 369 | 0 | 0 |
| SRR10544944 | 0 | 0 | 0 | 0 | 0 | 30 | 23 | 27 | 14 | 0 | 50 | 0 | 0 | 0 | 1858 | 2420 | 21652 | 36 | 43 | 1723 | 10 | 0 | 0 | 15 | 0 | 0 | 28 | 17 | 0 | 157 | 0 | 0 | 0 | 0 | 462 | 0 | 0 |
| SRR10544945 | 0 | 0 | 0 | 0 | 0 | 18 | 0 | 14 | 0 | 0 | 0 | 0 | 0 | 0 | 2707 | 674 | 6185 | 12 | 23 | 443 | 0 | 0 | 0 | 0 | 0 | 0 | 0 | 0 | 0 | 26 | 0 | 0 | 0 | 0 | 51 | 0 | 0 |
| SRR10544946 | 0 | 0 | 0 | 0 | 0 | 14 | 18 | 16 | 0 | 0 | 20 | 0 | 0 | 0 | 2690 | 555 | 4805 | 0 | 29 | 245 | 20 | 0 | 0 | 0 | 0 | 0 | 0 | 0 | 0 | 62 | 0 | 0 | 0 | 0 | 26 | 0 | 0 |
| SRR10544947 | 0 | 0 | 0 | 0 | 0 | 37 | 20 | 0 | 0 | 0 | 13 | 0 | 0 | 0 | 4383 | 1262 | 11929 | 0 | 64 | 416 | 34 | 0 | 0 | 0 | 0 | 0 | 0 | 0 | 0 | 20 | 0 | 0 | 0 | 0 | 231 | 0 | 0 |
| SRR10544948 | 0 | 0 | 0 | 0 | 0 | 0 | 13 | 15 | 11 | 0 | 16 | 0 | 0 | 0 | 3257 | 953 | 5950 | 11 | 39 | 713 | 31 | 0 | 0 | 0 | 0 | 0 | 0 | 0 | 0 | 82 | 0 | 0 | 0 | 0 | 97 | 0 | 0 |
| SRR10544949 | 0 | 0 | 0 | 0 | 0 | 17 | 11 | 17 | 22 | 0 | 16 | 0 | 0 | 0 | 4419 | 2344 | 9215 | 17 | 150 | 835 | 11 | 0 | 0 | 0 | 0 | 0 | 0 | 0 | 0 | 23 | 0 | 0 | 0 | 0 | 53 | 0 | 0 |
| SRR10544950 | 0 | 0 | 0 | 0 | 0 | 26 | 0 | 36 | 0 | 0 | 15 | 0 | 0 | 0 | 4488 | 854 | 9817 | 21 | 57 | 285 | 36 | 0 | 0 | 0 | 0 | 0 | 0 | 0 | 0 | 23 | 0 | 0 | 0 | 0 | 52 | 0 | 0 |
| SRR10756575 | 56 | 18 | 176 | 133 | 664 | 631 | 195 | 917 | 885 | 199 | 122 | 149 | 0 | 0 | 992 | 209878 | 215977 | 55 | 185 | 742 | 1909 | 0 | 0 | 25 | 27 | 66 | 106 | 0 | 17 | 127 | 0 | 42 | 0 | 12 | 8661 | 0 | 0 |
| SRR10756576 | 77 | 25 | 148 | 66 | 542 | 8880 | 1452 | 5947 | 5048 | 78 | 40 | 102 | 0 | 0 | 701 | 163725 | 74608 | 74 | 1006 | 1419 | 9288 | 0 | 0 | 17 | 15 | 20 | 31 | 0 | 0 | 82 | 0 | 41 | 0 | 24 | 18563 | 0 | 0 |
| SRR10756577 | 1009 | 262 | 28 | 15 | 7294 | 1131 | 457 | 1592 | 1307 | 2499 | 33 | 106 | 0 | 10 | 293 | 73467 | 68261 | 60 | 72 | 754 | 1127 | 0 | 0 | 335 | 0 | 0 | 0 | 0 | 58 | 33 | 0 | 88 | 0 | 15 | 11297 | 0 | 0 |
| SRR10756578 | 889 | 255 | 126 | 126 | 4035 | 1684 | 687 | 2738 | 2771 | 1930 | 79 | 63 | 0 | 0 | 1731 | 265086 | 503173 | 13 | 159 | 1156 | 4526 | 0 | 0 | 626 | 50 | 51 | 95 | 0 | 0 | 71 | 0 | 12 | 0 | 17 | 2071 | 0 | 0 |
| SRR10756579 | 58 | 17 | 104 | 81 | 278 | 1462 | 469 | 1726 | 2368 | 299 | 67 | 104 | 0 | 0 | 1152 | 196954 | 233340 | 77 | 223 | 2217 | 3105 | 0 | 12 | 18 | 55 | 23 | 90 | 0 | 0 | 107 | 0 | 191 | 0 | 0 | 7533 | 0 | 0 |
| SRR10756581 | 160 | 33 | 11 | 171 | 212 | 955 | 397 | 2607 | 1071 | 65 | 66 | 43 | 0 | 0 | 1628 | 260643 | 335299 | 34 | 283 | 4655 | 5277 | 0 | 0 | 55 | 73 | 100 | 201 | 0 | 0 | 109 | 0 | 26 | 0 | 0 | 9188 | 0 | 0 |
| SRR10756582 | 4044 | 1153 | 444 | 384 | 19733 | 177 | 54 | 408 | 254 | 2041 | 81 | 74 | 15 | 0 | 1023 | 318101 | 382977 | 0 | 135 | 1589 | 11811 | 0 | 15 | 1405 | 22 | 140 | 237 | 0 | 15 | 155 | 0 | 0 | 0 | 0 | 6691 | 0 | 0 |
| SRR10756583 | 1215 | 486 | 117 | 151 | 10303 | 669 | 96 | 995 | 406 | 695 | 25 | 17 | 0 | 0 | 422 | 182326 | 256528 | 16 | 80 | 931 | 3057 | 34 | 0 | 2161 | 67 | 84 | 159 | 0 | 0 | 67 | 0 | 32 | 0 | 0 | 13673 | 0 | 0 |
| SRR10875785 | 3582 | 1161 | 95 | 76 | 209 | 2347 | 346 | 665 | 1558 | 192 | 70 | 67 | 37 | 19 | 2562 | 105642 | 32017 | 3162 | 2106 | 6083 | 38659 | 122 | 46 | 310 | 0 | 12 | 19 | 0 | 44 | 102 | 253 | 3104 | 18 | 112 | 10520 | 2359 | 30 |
| SRR10875786 | 129 | 45 | 0 | 0 | 18 | 1315 | 84 | 382 | 457 | 0 | 40 | 0 | 0 | 0 | 690 | 21779 | 4695 | 2435 | 373 | 1017 | 2948 | 123 | 41 | 15 | 0 | 0 | 0 | 0 | 13 | 41 | 146 | 4599 | 30 | 52 | 57584 | 2112 | 64 |
| SRR10953234 | 225 | 43 | 148 | 210 | 532 | 540 | 342 | 1488 | 9025 | 74 | 698 | 23 | 0 | 0 | 9761 | 89265 | 125104 | 266 | 698 | 7659 | 5877 | 0 | 0 | 22 | 215 | 221 | 165 | 145 | 0 | 474 | 0 | 12 | 0 | 0 | 1717 | 0 | 0 |
| SRR10953235 | 226 | 44 | 120 | 172 | 513 | 421 | 268 | 1273 | 6382 | 73 | 639 | 18 | 0 | 0 | 7257 | 80936 | 108747 | 191 | 543 | 6137 | 3804 | 0 | 0 | 19 | 148 | 155 | 138 | 157 | 0 | 401 | 0 | 0 | 0 | 0 | 2118 | 0 | 0 |
| SRR10953236 | 189 | 35 | 99 | 148 | 556 | 610 | 409 | 1863 | 9357 | 176 | 619 | 17 | 0 | 0 | 8745 | 104345 | 120062 | 203 | 516 | 4672 | 5735 | 0 | 0 | 28 | 162 | 166 | 138 | 151 | 0 | 428 | 0 | 0 | 0 | 0 | 1986 | 0 | 0 |
| SRR10953237 | 94 | 14 | 39 | 49 | 113 | 497 | 273 | 1199 | 5675 | 32 | 304 | 14 | 0 | 0 | 10532 | 114717 | 123401 | 225 | 777 | 5065 | 8496 | 0 | 0 | 15 | 85 | 82 | 80 | 100 | 0 | 173 | 0 | 0 | 0 | 0 | 1935 | 12 | 0 |
| SRR10953238 | 70 | 13 | 52 | 55 | 132 | 520 | 394 | 1828 | 9722 | 47 | 252 | 24 | 0 | 0 | 10728 | 59362 | 105988 | 239 | 565 | 6622 | 3465 | 0 | 0 | 26 | 72 | 69 | 57 | 51 | 0 | 174 | 0 | 0 | 0 | 0 | 1601 | 0 | 0 |
| SRR10953239 | 246 | 42 | 108 | 118 | 470 | 733 | 450 | 1977 | 11309 | 269 | 780 | 32 | 0 | 0 | 14535 | 67313 | 102264 | 262 | 614 | 7349 | 3800 | 0 | 16 | 32 | 172 | 157 | 130 | 115 | 0 | 512 | 0 | 0 | 0 | 0 | 1730 | 0 | 0 |
| SRR1107492 | 73 | 67 | 62 | 12 | 0 | 613 | 520 | 1653 | 149 | 15 | 19 | 10 | 0 | 0 | 9024 | 385 | 1770 | 0 | 0 | 0 | 27091 | 0 | 0 | 217 | 248 | 429 | 847 | 50 | 0 | 229 | 0 | 19 | 0 | 0 | 1263 | 0 | 0 |
| SRR1107493 | 14 | 0 | 0 | 0 | 0 | 540 | 308 | 1311 | 154 | 24 | 32 | 14 | 0 | 0 | 22131 | 31286 | 222613 | 0 | 19 | 1388 | 19336 | 0 | 14 | 18 | 17 | 45 | 71 | 26 | 0 | 43 | 0 | 19 | 0 | 0 | 524 | 0 | 14 |
| SRR1107494 | 28 | 20 | 47 | 13 | 24 | 590 | 389 | 1205 | 50 | 12 | 18 | 0 | 0 | 0 | 16640 | 21671 | 181529 | 13 | 75 | 4681 | 24646 | 0 | 0 | 14 | 20 | 34 | 61 | 26 | 0 | 74 | 0 | 0 | 0 | 0 | 786 | 0 | 0 |
| SRR1107495 | 32 | 18 | 64 | 14 | 20 | 559 | 191 | 927 | 82 | 13 | 16 | 0 | 0 | 0 | 18208 | 21912 | 199387 | 14 | 14 | 4774 | 24795 | 0 | 0 | 14 | 19 | 30 | 52 | 15 | 0 | 60 | 0 | 0 | 0 | 0 | 827 | 0 | 0 |
| SRR1107496 | 25 | 26 | 83 | 148 | 190 | 650 | 214 | 992 | 151 | 15 | 114 | 0 | 0 | 0 | 24211 | 30907 | 252740 | 0 | 87 | 8210 | 21085 | 0 | 0 | 137 | 171 | 325 | 502 | 26 | 0 | 403 | 0 | 16 | 0 | 0 | 358 | 0 | 0 |
| SRR1107497 | 47 | 21 | 72 | 17 | 13 | 404 | 224 | 1347 | 45 | 17 | 13 | 0 | 0 | 0 | 23212 | 32096 | 247146 | 0 | 0 | 6196 | 21775 | 0 | 0 | 16 | 13 | 33 | 51 | 0 | 0 | 41 | 0 | 45 | 0 | 0 | 396 | 0 | 0 |
| SRR1159512 | 14 | 13 | 82 | 17 | 21 | 155 | 162 | 182 | 37 | 0 | 40 | 0 | 0 | 0 | 67717 | 11907 | 160832 | 0 | 71 | 2537 | 760 | 0 | 0 | 27 | 36 | 55 | 145 | 42 | 0 | 38 | 0 | 571 | 0 | 0 | 2845 | 13 | 0 |
| SRR1207050 | 0 | 0 | 0 | 0 | 176 | 0 | 0 | 0 | 0 | 0 | 0 | 0 | 0 | 0 | 0 | 114 | 85 | 108 | 0 | 108 | 0 | 517 | 93 | 15 | 0 | 41 | 39 | 32 | 0 | 0 | 94 | 62 | 0 | 0 | 199 | 0 | 0 |
| SRR1207051 | 25 | 31 | 15 | 15 | 33 | 0 | 12 | 0 | 202 | 0 | 0 | 0 | 0 | 0 | 40 | 12 | 30 | 127 | 16 | 127 | 0 | 188 | 15 | 15 | 55 | 329 | 59 | 0 | 49 | 0 | 49 | 75 | 0 | 0 | 392 | 0 | 0 |
| SRR1207053 | 14 | 0 | 0 | 0 | 1179 | 0 | 0 | 0 | 28 | 0 | 0 | 0 | 0 | 0 | 15 | 17 | 29 | 267 | 0 | 11 | 0 | 2862 | 77 | 304 | 0 | 26 | 108 | 0 | 20 | 0 | 654 | 138 | 0 | 0 | 294 | 0 | 0 |
| SRR1207054 | 0 | 0 | 0 | 0 | 855 | 0 | 0 | 0 | 0 | 0 | 0 | 0 | 0 | 0 | 0 | 65 | 102 | 181 | 26 | 90 | 0 | 27208 | 3177 | 11 | 0 | 0 | 14 | 0 | 0 | 0 | 5655 | 58 | 0 | 0 | 223 | 0 | 0 |

### Supp Table S8.3.Columns_BX_to_DH

| | | | | | | | | | | | | | | | | | | | | | | | | | | | | | | | | | | | | | |
|---|---|---|---|---|---|---|---|---|---|---|---|---|---|---|---|---|---|---|---|---|---|---|---|---|---|---|---|---|---|---|---|---|---|---|---|---|---|
| SRR1207055 | 0 | 0 | 0 | 0 | 31 | 0 | 0 | 0 | 11 | 0 | 0 | 0 | 0 | 0 | 0 | 17 | 0 | 315 | 0 | 23 | 0 | 272 | 86 | 79 | 16 | 0 | 11 | 0 | 23 | 0 | 14 | 84 | 0 | 0 | 440 | 0 | 45 |
| SRR1232072 | 0 | 0 | 0 | 0 | 18 | 308 | 77 | 605 | 35 | 11 | 130 | 0 | 0 | 0 | 223215 | 41096 | 425258 | 0 | 17 | 1886 | 11904 | 0 | 0 | 13 | 13 | 30 | 23 | 15 | 0 | 1520 | 0 | 29 | 0 | 0 | 777 | 123 | 0 |
| SRR1232073 | 0 | 0 | 0 | 11 | 24 | 207 | 51 | 527 | 23 | 0 | 243 | 0 | 0 | 0 | 201976 | 27463 | 303662 | 0 | 0 | 3293 | 5881 | 0 | 0 | 12 | 24 | 38 | 18 | 0 | 0 | 2610 | 0 | 49 | 0 | 0 | 790 | 42 | 0 |
| SRR1240812 | 27 | 27 | 11 | 78 | 33 | 260 | 512 | 450 | 114 | 0 | 91 | 0 | 0 | 0 | 10258 | 4284 | 28242 | 18 | 20 | 6444 | 682 | 0 | 0 | 61 | 26 | 93 | 341 | 104 | 0 | 34 | 0 | 0 | 0 | 0 | 4840 | 482 | 0 |
| SRR1240813 | 21 | 20 | 15 | 99 | 24 | 245 | 279 | 440 | 97 | 0 | 53 | 0 | 0 | 0 | 9323 | 3007 | 21661 | 0 | 25 | 6638 | 964 | 0 | 0 | 61 | 24 | 102 | 270 | 100 | 0 | 31 | 0 | 0 | 0 | 0 | 4610 | 176 | 0 |
| SRR1240814 | 117 | 61 | 100 | 357 | 143 | 278 | 232 | 284 | 146 | 0 | 40 | 0 | 0 | 0 | 14163 | 3571 | 18541 | 0 | 60 | 4379 | 807 | 0 | 0 | 327 | 94 | 409 | 1481 | 582 | 0 | 259 | 0 | 0 | 0 | 0 | 4885 | 29 | 0 |
| SRR1240815 | 106 | 71 | 111 | 328 | 142 | 302 | 356 | 348 | 139 | 0 | 37 | 0 | 0 | 0 | 15118 | 3871 | 23691 | 0 | 31 | 4766 | 720 | 0 | 0 | 360 | 103 | 385 | 1557 | 504 | 0 | 142 | 0 | 0 | 0 | 0 | 4873 | 37 | 0 |
| SRR1240816 | 26 | 20 | 21 | 97 | 12 | 169 | 129 | 181 | 58 | 0 | 74 | 0 | 0 | 0 | 11669 | 2634 | 12406 | 16 | 22 | 6777 | 356 | 0 | 0 | 247 | 37 | 113 | 351 | 142 | 0 | 47 | 0 | 0 | 0 | 0 | 2732 | 0 | 0 |
| SRR1240817 | 240 | 156 | 233 | 943 | 98 | 120 | 113 | 204 | 62 | 0 | 82 | 0 | 0 | 0 | 8751 | 2612 | 10720 | 12 | 19 | 7900 | 348 | 0 | 0 | 1106 | 317 | 1040 | 2975 | 1407 | 0 | 435 | 0 | 0 | 0 | 0 | 3170 | 0 | 0 |
| SRR1240818 | 22 | 0 | 0 | 32 | 0 | 288 | 541 | 1071 | 222 | 23 | 153 | 147 | 0 | 0 | 43161 | 44442 | 68573 | 10 | 586 | 9071 | 4334 | 0 | 0 | 55 | 49 | 141 | 285 | 88 | 0 | 355 | 0 | 15 | 0 | 0 | 1100 | 80 | 0 |
| SRR1241600 | 0 | 0 | 25 | 0 | 36 | 93 | 187 | 83 | 164 | 0 | 18 | 0 | 0 | 21 | 10025 | 4379 | 35767 | 68 | 0 | 1670 | 31 | 0 | 24 | 10 | 0 | 0 | 0 | 0 | 0 | 17 | 0 | 0 | 0 | 0 | 367 | 0 | 0 |
| SRR1241601 | 42 | 45 | 110 | 36 | 426 | 44 | 86 | 52 | 65 | 0 | 22 | 0 | 0 | 0 | 42535 | 26569 | 82192 | 23 | 0 | 1228 | 57 | 14 | 21 | 22 | 0 | 14 | 24 | 0 | 0 | 159 | 0 | 0 | 0 | 0 | 106 | 0 | 0 |
| SRR1241602 | 13 | 11 | 18 | 0 | 152 | 52 | 193 | 65 | 100 | 0 | 15 | 0 | 0 | 15 | 36122 | 3193 | 53079 | 13 | 0 | 765 | 89 | 38 | 22 | 28 | 0 | 0 | 0 | 0 | 0 | 50 | 0 | 0 | 0 | 0 | 157 | 0 | 0 |
| SRR1241603 | 290 | 300 | 45 | 94 | 85 | 155 | 273 | 380 | 79 | 14 | 938 | 0 | 0 | 0 | 35597 | 9062 | 40951 | 12 | 52 | 2907 | 7744 | 0 | 0 | 39 | 0 | 16 | 48 | 0 | 0 | 3275 | 0 | 0 | 0 | 0 | 18410 | 0 | 0 |
| SRR1241604 | 199 | 245 | 68 | 160 | 104 | 122 | 459 | 288 | 51 | 15 | 526 | 0 | 0 | 0 | 42235 | 15978 | 49798 | 0 | 61 | 2702 | 9227 | 0 | 0 | 39 | 13 | 19 | 55 | 0 | 0 | 1602 | 0 | 0 | 0 | 0 | 14471 | 0 | 0 |
| SRR1241605 | 304 | 304 | 81 | 154 | 94 | 120 | 252 | 366 | 74 | 14 | 601 | 0 | 0 | 0 | 38626 | 21482 | 47760 | 0 | 73 | 2546 | 13730 | 0 | 0 | 73 | 16 | 29 | 55 | 0 | 0 | 1488 | 0 | 0 | 0 | 0 | 10370 | 0 | 0 |
| SRR1264353 | 13 | 16 | 0 | 0 | 12 | 1795 | 1250 | 1923 | 737 | 0 | 15 | 0 | 0 | 0 | 43293 | 119116 | 131864 | 993 | 192 | 1250 | 6890 | 0 | 0 | 18 | 17 | 0 | 0 | 0 | 0 | 152 | 0 | 96 | 0 | 0 | 1567 | 0 | 0 |
| SRR1264354 | 22 | 23 | 0 | 0 | 0 | 1022 | 317 | 670 | 282 | 0 | 12 | 0 | 0 | 0 | 11280 | 71060 | 18453 | 1163 | 71 | 1234 | 1234 | 0 | 0 | 15 | 11 | 0 | 0 | 0 | 0 | 23 | 0 | 71 | 0 | 0 | 2519 | 19 | 0 |
| SRR1270153 | 0 | 0 | 0 | 0 | 237 | 0 | 17 | 12 | 0 | 0 | 0 | 0 | 0 | 0 | 11664 | 216821 | 138803 | 243 | 31 | 41 | 202 | 0 | 0 | 17 | 0 | 22 | 86 | 0 | 0 | 22 | 0 | 1323 | 0 | 0 | 1537 | 0 | 0 |
| SRR1290784 | 16 | 57 | 39 | 40 | 26 | 229 | 145 | 245 | 68 | 0 | 30 | 0 | 32 | 0 | 60 | 7274 | 19246 | 64 | 28 | 5806 | 1038 | 0 | 0 | 11 | 13 | 28 | 49 | 82 | 0 | 71 | 0 | 0 | 0 | 0 | 2908 | 0 | 0 |
| SRR1290785 | 12 | 20 | 49 | 32 | 18 | 495 | 230 | 362 | 151 | 0 | 69 | 0 | 44 | 0 | 79 | 5770 | 24182 | 30 | 30 | 6681 | 961 | 0 | 0 | 15 | 13 | 31 | 42 | 28 | 0 | 123 | 0 | 0 | 0 | 0 | 3674 | 0 | 0 |
| SRR1290786 | 113 | 21 | 38 | 24 | 274 | 373 | 189 | 368 | 96 | 0 | 41 | 0 | 29 | 0 | 61 | 5828 | 16976 | 61 | 22 | 5258 | 820 | 0 | 0 | 121 | 115 | 254 | 371 | 417 | 0 | 721 | 0 | 0 | 0 | 0 | 3274 | 0 | 0 |
| SRR1290787 | 131 | 23 | 18 | 16 | 305 | 412 | 279 | 434 | 69 | 0 | 47 | 0 | 28 | 0 | 133 | 9543 | 41067 | 28 | 77 | 9425 | 3119 | 0 | 0 | 112 | 242 | 387 | 525 | 0 | 0 | 2855 | 0 | 0 | 0 | 0 | 1940 | 0 | 0 |
| SRR1290788 | 106 | 20 | 16 | 24 | 222 | 423 | 462 | 617 | 136 | 0 | 38 | 0 | 39 | 0 | 136 | 8861 | 32726 | 27 | 62 | 7764 | 2628 | 0 | 0 | 119 | 204 | 423 | 584 | 22 | 0 | 2180 | 0 | 0 | 0 | 0 | 3259 | 0 | 0 |
| SRR1290789 | 164 | 38 | 0 | 0 | 255 | 495 | 266 | 378 | 75 | 0 | 14 | 0 | 32 | 0 | 85 | 5473 | 18959 | 64 | 43 | 6102 | 1693 | 0 | 0 | 149 | 63 | 67 | 113 | 16 | 0 | 833 | 0 | 0 | 0 | 0 | 2602 | 0 | 0 |
| SRR1290790 | 95 | 29 | 20 | 26 | 207 | 403 | 119 | 314 | 47 | 0 | 51 | 0 | 44 | 0 | 17 | 4580 | 12744 | 30 | 30 | 9226 | 891 | 0 | 0 | 128 | 157 | 320 | 455 | 36 | 0 | 811 | 0 | 0 | 0 | 0 | 2656 | 0 | 0 |
| SRR1290791 | 25 | 81 | 50 | 42 | 19 | 212 | 66 | 208 | 26 | 0 | 46 | 0 | 28 | 0 | 106 | 7771 | 26019 | 40 | 40 | 11228 | 1607 | 0 | 0 | 11 | 10 | 19 | 31 | 28 | 0 | 62 | 0 | 0 | 0 | 0 | 2902 | 0 | 0 |
| SRR1290792 | 18 | 20 | 45 | 54 | 25 | 351 | 177 | 354 | 87 | 0 | 16 | 0 | 23 | 0 | 49 | 6436 | 20003 | 45 | 49 | 6439 | 1143 | 0 | 0 | 13 | 10 | 20 | 28 | 109 | 0 | 56 | 0 | 0 | 0 | 0 | 2742 | 0 | 0 |
| SRR1290793 | 192 | 28 | 36 | 32 | 273 | 609 | 290 | 505 | 106 | 0 | 21 | 0 | 38 | 0 | 41 | 4002 | 16269 | 26 | 10 | 5342 | 839 | 0 | 0 | 189 | 110 | 277 | 231 | 26 | 0 | 659 | 0 | 0 | 0 | 0 | 3608 | 0 | 0 |
| SRR1290794 | 131 | 15 | 17 | 11 | 165 | 647 | 201 | 517 | 125 | 0 | 25 | 0 | 29 | 0 | 44 | 1496 | 10484 | 60 | 22 | 7352 | 506 | 0 | 0 | 131 | 73 | 157 | 127 | 10 | 0 | 1622 | 0 | 0 | 0 | 0 | 3343 | 0 | 0 |
| SRR1290795 | 12 | 22 | 17 | 0 | 26 | 386 | 170 | 367 | 163 | 0 | 44 | 0 | 14 | 0 | 30 | 1649 | 8882 | 27 | 0 | 8505 | 609 | 0 | 0 | 12 | 12 | 23 | 16 | 18 | 0 | 231 | 0 | 0 | 0 | 0 | 3334 | 0 | 0 |
| SRR1290796 | 49 | 12 | 0 | 11 | 265 | 378 | 250 | 449 | 119 | 0 | 25 | 0 | 26 | 0 | 67 | 3328 | 21090 | 26 | 10 | 5363 | 2274 | 0 | 0 | 59 | 65 | 117 | 157 | 285 | 0 | 1445 | 0 | 0 | 0 | 0 | 3847 | 0 | 0 |
| SRR1290797 | 84 | 16 | 35 | 41 | 37 | 284 | 267 | 415 | 96 | 0 | 57 | 0 | 30 | 0 | 111 | 8849 | 45133 | 12 | 17 | 9835 | 4297 | 0 | 0 | 76 | 23 | 56 | 90 | 67 | 0 | 164 | 0 | 0 | 0 | 0 | 3953 | 0 | 0 |
| SRR1290798 | 201 | 44 | 101 | 89 | 440 | 131 | 121 | 216 | 51 | 0 | 56 | 0 | 17 | 0 | 118 | 17022 | 72593 | 0 | 0 | 8330 | 3324 | 0 | 0 | 144 | 233 | 570 | 1073 | 651 | 0 | 571 | 0 | 0 | 0 | 0 | 5207 | 0 | 0 |
| SRR1290799 | 11 | 29 | 21 | 65 | 68 | 385 | 79 | 425 | 36 | 0 | 55 | 53 | 0 | 0 | 547 | 29454 | 210156 | 0 | 36 | 14747 | 8607 | 0 | 0 | 22 | 116 | 137 | 131 | 22 | 0 | 65 | 0 | 0 | 0 | 0 | 1108 | 0 | 0 |
| SRR1290800 | 43 | 15 | 29 | 42 | 219 | 301 | 92 | 356 | 42 | 0 | 74 | 22 | 0 | 0 | 415 | 27883 | 179288 | 0 | 33 | 13522 | 6283 | 0 | 0 | 109 | 665 | 887 | 1053 | 863 | 0 | 240 | 0 | 0 | 0 | 0 | 1246 | 0 | 0 |
| SRR1290801 | 50 | 0 | 27 | 45 | 36 | 482 | 194 | 736 | 102 | 18 | 15 | 26 | 0 | 0 | 353 | 8626 | 86122 | 16 | 89 | 24262 | 6754 | 0 | 0 | 27 | 97 | 117 | 84 | 78 | 0 | 126 | 0 | 0 | 0 | 0 | 1210 | 0 | 0 |
| SRR1290802 | 133 | 16 | 55 | 55 | 16 | 380 | 188 | 392 | 81 | 0 | 58 | 0 | 10 | 0 | 54 | 8144 | 19340 | 27 | 31 | 3562 | 265 | 0 | 0 | 14 | 19 | 51 | 58 | 37 | 0 | 74 | 0 | 0 | 0 | 0 | 3659 | 0 | 0 |
| SRR1290803 | 91 | 15 | 63 | 67 | 153 | 342 | 183 | 280 | 124 | 0 | 73 | 0 | 19 | 0 | 19 | 7626 | 17441 | 31 | 39 | 5892 | 353 | 0 | 0 | 128 | 159 | 535 | 606 | 1035 | 0 | 1138 | 0 | 0 | 0 | 0 | 4804 | 0 | 0 |
| SRR1290804 | 18 | 25 | 90 | 85 | 11 | 515 | 463 | 678 | 127 | 0 | 67 | 0 | 40 | 0 | 46 | 11643 | 29680 | 20 | 91 | 5843 | 545 | 0 | 0 | 23 | 16 | 56 | 82 | 34 | 0 | 198 | 0 | 0 | 0 | 0 | 4915 | 0 | 0 |
| SRR1290805 | 19 | 11 | 72 | 69 | 21 | 381 | 363 | 514 | 79 | 0 | 15 | 0 | 33 | 0 | 54 | 9913 | 26425 | 30 | 175 | 10436 | 1033 | 0 | 0 | 16 | 16 | 57 | 89 | 12 | 0 | 427 | 0 | 0 | 0 | 0 | 3840 | 0 | 0 |
| SRR1290806 | 17 | 28 | 34 | 31 | 16 | 309 | 269 | 505 | 98 | 0 | 40 | 0 | 40 | 0 | 73 | 10775 | 25744 | 16 | 41 | 10268 | 1059 | 0 | 0 | 16 | 13 | 19 | 22 | 40 | 0 | 165 | 0 | 0 | 0 | 0 | 4518 | 0 | 0 |
| SRR1290807 | 241 | 47 | 52 | 44 | 220 | 249 | 145 | 276 | 110 | 0 | 76 | 0 | 42 | 0 | 55 | 10006 | 26816 | 21 | 28 | 10738 | 939 | 0 | 0 | 209 | 149 | 290 | 280 | 42 | 0 | 1649 | 0 | 0 | 0 | 0 | 4613 | 0 | 0 |
| SRR1290808 | 321 | 45 | 100 | 53 | 240 | 241 | 190 | 235 | 101 | 0 | 53 | 0 | 14 | 0 | 89 | 12797 | 48188 | 38 | 63 | 14607 | 1598 | 0 | 0 | 237 | 167 | 309 | 360 | 47 | 0 | 1789 | 0 | 0 | 0 | 0 | 5233 | 0 | 0 |
| SRR1290809 | 14 | 65 | 140 | 114 | 14 | 405 | 160 | 399 | 79 | 0 | 98 | 0 | 20 | 0 | 106 | 12371 | 35932 | 38 | 77 | 11868 | 857 | 0 | 0 | 15 | 23 | 63 | 86 | 167 | 0 | 154 | 0 | 0 | 0 | 0 | 4553 | 0 | 0 |
| SRR1292461 | 15 | 18 | 0 | 0 | 47 | 0 | 33 | 26 | 37 | 0 | 0 | 0 | 29 | 0 | 0 | 0 | 19 | 40 | 0 | 1415 | 0 | 307 | 146 | 24 | 15 | 0 | 0 | 0 | 0 | 0 | 77 | 0 | 0 | 0 | 4035 | 0 | 0 |
| SRR1304304 | 61 | 20 | 0 | 0 | 0 | 165 | 187 | 497 | 0 | 0 | 0 | 0 | 27 | 0 | 90 | 110 | 482 | 0 | 0 | 487 | 11 | 0 | 0 | 221 | 0 | 0 | 0 | 0 | 0 | 25 | 0 | 0 | 0 | 0 | 2074 | 0 | 0 |
| SRR1304307 | 0 | 0 | 0 | 0 | 0 | 202 | 306 | 571 | 69 | 76 | 0 | 18 | 0 | 0 | 225 | 1760 | 4724 | 0 | 0 | 8692 | 190 | 0 | 0 | 203 | 0 | 0 | 0 | 0 | 0 | 0 | 0 | 0 | 0 | 0 | 5022 | 0 | 0 |
| SRR1304309 | 13 | 0 | 0 | 0 | 0 | 15 | 205 | 163 | 27 | 0 | 0 | 55 | 15 | 0 | 242 | 1600 | 2619 | 0 | 0 | 546 | 174 | 0 | 0 | 983 | 0 | 0 | 0 | 0 | 0 | 25 | 0 | 0 | 0 | 0 | 6235 | 0 | 0 |
| SRR1304313 | 102 | 73 | 188 | 349 | 115 | 245 | 382 | 460 | 66 | 40 | 730 | 15 | 0 | 0 | 7010 | 123815 | 85754 | 0 | 74 | 5715 | 1182 | 0 | 0 | 136 | 0 | 33 | 35 | 0 | 0 | 1381 | 0 | 0 | 0 | 0 | 2536 | 0 | 0 |
| SRR1393733 | 512 | 100 | 0 | 0 | 283 | 377 | 1479 | 2542 | 135 | 0 | 0 | 36 | 0 | 0 | 4889 | 10145 | 14919 | 10 | 22 | 412 | 28 | 0 | 0 | 113 | 0 | 0 | 0 | 0 | 0 | 0 | 0 | 0 | 0 | 0 | 8740 | 12 | 113 |
| SRR1510077 | 0 | 0 | 0 | 21 | 23 | 43 | 444 | 507 | 287 | 27 | 246 | 0 | 0 | 0 | 42932 | 21457 | 183051 | 0 | 153 | 6884 | 3433 | 0 | 0 | 252 | 220 | 998 | 4212 | 1354 | 0 | 869 | 0 | 35 | 0 | 0 | 371 | 0 | 0 |
| SRR1510078 | 0 | 0 | 0 | 0 | 0 | 98 | 611 | 841 | 297 | 42 | 20 | 11 | 0 | 0 | 47813 | 45777 | 142222 | 0 | 190 | 16812 | 7153 | 0 | 0 | 12 | 0 | 68 | 38 | 74 | 0 | 90 | 0 | 162 | 0 | 0 | 1160 | 0 | 13 |
| SRR1510079 | 0 | 0 | 0 | 64 | 105 | 0 | 118 | 101 | 44 | 14 | 132 | 0 | 0 | 0 | 174721 | 12636 | 236461 | 0 | 50 | 5916 | 10800 | 0 | 0 | 429 | 420 | 1615 | 4789 | 2048 | 0 | 333 | 0 | 38 | 0 | 0 | 21 | 0 | 0 |
| SRR1510080 | 0 | 0 | 0 | 14 | 23 | 90 | 437 | 615 | 225 | 14 | 109 | 18 | 0 | 0 | 46687 | 58509 | 221002 | 0 | 106 | 12530 | 11478 | 0 | 0 | 94 | 109 | 570 | 2392 | 596 | 0 | 247 | 0 | 417 | 0 | 0 | 133 | 0 | 58 |
| SRR1510081 | 0 | 0 | 0 | 21 | 23 | 177 | 393 | 1146 | 137 | 21 | 114 | 0 | 0 | 0 | 74088 | 41007 | 251721 | 12 | 170 | 8751 | 11420 | 0 | 0 | 83 | 36 | 20 | 94 | 78 | 0 | 51 | 0 | 398 | 0 | 0 | 119 | 0 | 13 |
| SRR1510082 | 0 | 0 | 0 | 17 | 26 | 45 | 221 | 322 | 182 | 0 | 207 | 22 | 0 | 0 | 19558 | 55326 | 157510 | 0 | 350 | 10345 | 7703 | 0 | 0 | 496 | 245 | 1169 | 5701 | 4511 | 0 | 695 | 0 | 37 | 0 | 0 | 132 | 0 | 0 |
| SRR1510083 | 0 | 0 | 0 | 23 | 19 | 34 | 496 | 468 | 306 | 0 | 104 | 0 | 0 | 0 | 16634 | 17641 | 143610 | 0 | 364 | 15216 | 28016 | 0 | 0 | 306 | 270 | 1518 | 10390 | 6199 | 0 | 344 | 0 | 35 | 0 | 0 | 124 | 0 | 0 |
| SRR1510084 | 0 | 0 | 0 | 25 | 0 | 23 | 119 | 225 | 68 | 14 | 263 | 0 | 0 | 0 | 63676 | 37131 | 269103 | 0 | 183 | 8790 | 16163 | 0 | 0 | 209 | 206 | 627 | 3395 | 1110 | 0 | 833 | 0 | 144 | 0 | 0 | 107 | 0 | 11 |
| SRR1510085 | 0 | 0 | 0 | 0 | 0 | 52 | 591 | 692 | 293 | 10 | 87 | 10 | 0 | 0 | 61803 | 21029 | 238743 | 0 | 334 | 13016 | 9479 | 0 | 0 | 462 | 325 | 923 | 4287 | 1245 | 0 | 376 | 0 | 0 | 0 | 0 | 161 | 0 | 0 |
| SRR1510086 | 0 | 0 | 0 | 10 | 48 | 39 | 110 | 220 | 41 | 86 | 19 | 0 | 0 | 0 | 108708 | 43219 | 203218 | 0 | 308 | 11112 | 11500 | 0 | 0 | 43 | 129 | 29 | 86 | 24 | 0 | 18 | 0 | 63 | 0 | 0 | 73 | 0 | 0 |
| SRR1510087 | 0 | 18 | 15 | 53 | 29 | 21 | 167 | 222 | 184 | 0 | 16 | 0 | 0 | 0 | 4126 | 33539 | 42900 | 0 | 333 | 17701 | 10765 | 0 | 0 | 35 | 90 | 350 | 1759 | 469 | 0 | 21 | 0 | 38 | 0 | 0 | 226 | 0 | 0 |
| SRR1510088 | 0 | 0 | 0 | 10 | 0 | 112 | 923 | 1347 | 415 | 32 | 78 | 0 | 0 | 0 | 7557 | 56018 | 140216 | 0 | 385 | 20309 | 16133 | 0 | 12 | 102 | 89 | 319 | 1743 | 402 | 0 | 459 | 0 | 751 | 0 | 0 | 460 | 0 | 0 |
| SRR1510089 | 0 | 0 | 0 | 23 | 34 | 12 | 145 | 220 | 167 | 0 | 15 | 19 | 0 | 0 | 27972 | 12172 | 105774 | 0 | 175 | 13676 | 12520 | 0 | 0 | 16 | 18 | 56 | 291 | 81 | 0 | 43 | 0 | 22 | 0 | 0 | 162 | 0 | 0 |
| SRR1510090 | 0 | 0 | 0 | 52 | 31 | 25 | 395 | 386 | 111 | 18 | 83 | 0 | 0 | 0 | 22342 | 103941 | 365807 | 0 | 926 | 76556 | 15953 | 0 | 0 | 525 | 260 | 1204 | 6226 | 1750 | 0 | 569 | 0 | 0 | 0 | 0 | 234 | 0 | 0 |
| SRR1510091 | 0 | 12 | 0 | 12 | 19 | 32 | 182 | 228 | 146 | 0 | 158 | 34 | 0 | 0 | 73045 | 45644 | 217022 | 12 | 122 | 12365 | 16810 | 0 | 0 | 130 | 183 | 902 | 4722 | 2400 | 0 | 456 | 0 | 322 | 0 | 0 | 74 | 0 | 0 |
| SRR1510092 | 0 | 0 | 0 | 10 | 26 | 0 | 62 | 75 | 33 | 0 | 128 | 0 | 0 | 0 | 2953 | 14994 | 57284 | 0 | 403 | 17501 | 4247 | 0 | 0 | 283 | 364 | 1448 | 6855 | 6051 | 0 | 212 | 0 | 0 | 0 | 0 | 32 | 0 | 0 |
| SRR1510093 | 0 | 0 | 0 | 14 | 0 | 25 | 578 | 544 | 282 | 0 | 63 | 0 | 0 | 0 | 10979 | 57326 | 181599 | 0 | 285 | 27147 | 17336 | 0 | 0 | 341 | 350 | 1733 | 8456 | 4082 | 0 | 341 | 0 | 14 | 0 | 0 | 93 | 0 | 0 |
| SRR1510094 | 0 | 0 | 0 | 17 | 31 | 43 | 324 | 362 | 160 | 10 | 238 | 20 | 0 | 0 | 24787 | 37010 | 241806 | 0 | 148 | 12525 | 35926 | 0 | 0 | 191 | 41 | 341 | 1083 | 637 | 0 | 484 | 0 | 226 | 0 | 0 | 102 | 0 | 0 |
| SRR1510095 | 0 | 0 | 11 | 71 | 37 | 42 | 235 | 319 | 105 | 11 | 100 | 0 | 0 | 0 | 123028 | 16755 | 280402 | 0 | 194 | 15417 | 19984 | 0 | 0 | 178 | 156 | 591 | 2278 | 1264 | 0 | 439 | 0 | 81 | 0 | 0 | 66 | 0 | 0 |
| SRR1510096 | 0 | 0 | 0 | 30 | 0 | 107 | 1061 | 916 | 475 | 0 | 100 | 0 | 0 | 0 | 85178 | 17481 | 390068 | 0 | 453 | 27195 | 3057 | 0 | 0 | 272 | 223 | 806 | 3733 | 3661 | 0 | 589 | 0 | 22 | 0 | 0 | 360 | 0 | 0 |
| SRR1510097 | 0 | 0 | 14 | 45 | 36 | 16 | 133 | 127 | 23 | 23 | 172 | 32 | 0 | 0 | 15132 | 24123 | 163992 | 0 | 305 | 28440 | 3222 | 0 | 0 | 113 | 14 | 127 | 263 | 100 | 0 | 458 | 0 | 36 | 0 | 0 | 566 | 0 | 0 |
| SRR1510098 | 0 | 0 | 0 | 78 | 20 | 64 | 286 | 482 | 95 | 15 | 14 | 0 | 0 | 0 | 16024 | 11179 | 77139 | 0 | 101 | 12896 | 4299 | 0 | 0 | 14 | 12 | 71 | 347 | 137 | 0 | 45 | 0 | 64 | 0 | 0 | 626 | 0 | 0 |
| SRR1510099 | 0 | 11 | 0 | 16 | 14 | 32 | 36 | 149 | 0 | 18 | 34 | 63 | 0 | 0 | 3686 | 12529 | 24345 | 0 | 113 | 6947 | 9192 | 0 | 0 | 61 | 29 | 22 | 133 | 131 | 0 | 56 | 0 | 0 | 0 | 0 | 164 | 0 | 0 |
| SRR1510100 | 0 | 0 | 0 | 14 | 0 | 80 | 153 | 511 | 29 | 0 | 11 | 11 | 0 | 0 | 2562 | 11560 | 26762 | 0 | 88 | 8407 | 5087 | 0 | 0 | 23 | 25 | 77 | 404 | 181 | 0 | 22 | 0 | 29 | 0 | 0 | 297 | 0 | 16 |
| SRR1510101 | 16 | 24 | 0 | 33 | 19 | 29 | 336 | 550 | 110 | 0 | 54 | 11 | 0 | 0 | 6546 | 39287 | 158517 | 0 | 1337 | 48071 | 40476 | 0 | 0 | 492 | 696 | 4453 | 13059 | 1949 | 0 | 139 | 0 | 0 | 0 | 0 | 300 | 0 | 0 |
| SRR1510102 | 0 | 0 | 0 | 15 | 14 | 65 | 851 | 993 | 263 | 11 | 121 | 0 | 0 | 0 | 10422 | 59958 | 214720 | 0 | 669 | 29585 | 24017 | 0 | 0 | 212 | 256 | 919 | 4533 | 1139 | 0 | 771 | 0 | 30 | 0 | 0 | 1075 | 0 | 0 |
| SRR1510103 | 0 | 11 | 0 | 45 | 19 | 54 | 317 | 452 | 114 | 13 | 94 | 11 | 0 | 0 | 66484 | 23847 | 211543 | 0 | 205 | 14916 | 8389 | 0 | 0 | 358 | 276 | 956 | 4769 | 1176 | 0 | 340 | 0 | 380 | 0 | 0 | 114 | 0 | 0 |
| SRR1510104 | 0 | 0 | 0 | 23 | 22 | 12 | 56 | 108 | 38 | 0 | 204 | 0 | 0 | 0 | 22806 | 16324 | 48106 | 0 | 103 | 4505 | 5205 | 0 | 0 | 265 | 284 | 962 | 4256 | 1533 | 0 | 326 | 0 | 75 | 0 | 0 | 102 | 0 | 0 |
| SRR1510105 | 0 | 0 | 0 | 24 | 44 | 139 | 1427 | 1750 | 574 | 0 | 52 | 0 | 0 | 0 | 8536 | 32255 | 87297 | 0 | 116 | 13982 | 8004 | 0 | 0 | 31 | 209 | 44 | 263 | 132 | 0 | 12 | 0 | 23 | 0 | 0 | 1323 | 0 | 0 |
| SRR1510106 | 0 | 0 | 0 | 19 | 19 | 14 | 178 | 244 | 87 | 16 | 19 | 14 | 0 | 0 | 17452 | 46717 | 196476 | 12 | 190 | 17270 | 23828 | 0 | 0 | 15 | 99 | 35 | 174 | 56 | 0 | 42 | 0 | 439 | 0 | 0 | 135 | 0 | 0 |
| SRR1510107 | 0 | 0 | 19 | 14 | 24 | 26 | 413 | 404 | 122 | 19 | 58 | 24 | 0 | 0 | 135126 | 30406 | 363128 | 0 | 310 | 31719 | 15239 | 0 | 0 | 17 | 247 | 39 | 172 | 70 | 0 | 10 | 0 | 0 | 0 | 0 | 103 | 0 | 34 |
| SRR1510108 | 0 | 0 | 0 | 17 | 17 | 72 | 206 | 215 | 62 | 17 | 181 | 11 | 0 | 0 | 23296 | 9790 | 41007 | 0 | 134 | 7262 | 10300 | 0 | 0 | 167 | 224 | 977 | 4168 | 2588 | 0 | 526 | 0 | 98 | 0 | 0 | 154 | 0 | 0 |
| SRR1510109 | 0 | 0 | 0 | 34 | 38 | 77 | 440 | 640 | 256 | 15 | 132 | 11 | 0 | 0 | 30056 | 33280 | 175259 | 0 | 280 | 18227 | 14357 | 0 | 0 | 14 | 92 | 25 | 147 | 51 | 0 | 24 | 0 | 108 | 0 | 0 | 318 | 0 | 0 |
| SRR1510110 | 0 | 0 | 0 | 24 | 13 | 32 | 106 | 199 | 54 | 0 | 14 | 15 | 0 | 0 | 1966 | 31054 | 37106 | 0 | 235 | 4366 | 2097 | 0 | 0 | 37 | 23 | 104 | 1023 | 403 | 0 | 38 | 0 | 167 | 0 | 0 | 244 | 0 | 0 |
| SRR1510111 | 0 | 0 | 0 | 43 | 17 | 111 | 360 | 705 | 97 | 0 | 124 | 0 | 0 | 0 | 108357 | 26997 | 263523 | 0 | 338 | 18472 | 12510 | 0 | 0 | 367 | 128 | 409 | 1470 | 857 | 0 | 281 | 0 | 115 | 0 | 0 | 273 | 0 | 0 |
| SRR1510112 | 11 | 16 | 19 | 40 | 32 | 41 | 337 | 348 | 714 | 16 | 210 | 27 | 0 | 0 | 44889 | 26461 | 256488 | 0 | 401 | 16034 | 10759 | 0 | 0 | 11 | 232 | 33 | 231 | 43 | 0 | 21 | 0 | 301 | 0 | 0 | 88 | 0 | 0 |
| SRR1510113 | 0 | 0 | 11 | 46 | 26 | 117 | 863 | 1135 | 266 | 0 | 19 | 0 | 0 | 0 | 13059 | 36734 | 245646 | 0 | 462 | 49780 | 29587 | 0 | 0 | 211 | 15 | 52 | 223 | 96 | 0 | 144 | 0 | 64 | 0 | 0 | 193 | 0 | 0 |
| SRR1510114 | 0 | 0 | 0 | 12 | 18 | 98 | 529 | 913 | 378 | 26 | 13 | 18 | 0 | 0 | 44326 | 52109 | 200031 | 0 | 143 | 8939 | 12408 | 0 | 0 | 24 | 95 | 34 | 159 | 124 | 0 | 63 | 0 | 432 | 0 | 0 | 328 | 0 | 0 |
| SRR1510115 | 0 | 0 | 0 | 11 | 0 | 75 | 540 | 754 | 307 | 0 | 95 | 0 | 0 | 0 | 29138 | 82210 | 200883 | 18 | 384 | 25724 | 19376 | 0 | 0 | 12 | 93 | 47 | 274 | 124 | 0 | 29 | 0 | 1362 | 0 | 0 | 183 | 0 | 0 |
| SRR1510116 | 0 | 0 | 0 | 37 | 35 | 89 | 195 | 345 | 97 | 0 | 123 | 21 | 0 | 0 | 7402 | 13985 | 42389 | 0 | 131 | 5583 | 9777 | 0 | 0 | 288 | 0 | 0 | 0 | 0 | 0 | 218 | 0 | 44 | 0 | 0 | 92 | 0 | 0 |
| SRR1510117 | 0 | 0 | 16 | 57 | 26 | 20 | 95 | 136 | 39 | 11 | 29 | 0 | 0 | 0 | 2113 | 13344 | 34800 | 0 | 301 | 8702 | 8313 | 0 | 0 | 23 | 42 | 204 | 1112 | 583 | 0 | 74 | 0 | 34 | 0 | 0 | 76 | 0 | 0 |
| SRR1510118 | 0 | 0 | 0 | 43 | 27 | 50 | 309 | 754 | 288 | 0 | 206 | 19 | 0 | 0 | 9052 | 27221 | 93075 | 0 | 217 | 8075 | 11085 | 0 | 0 | 382 | 234 | 1209 | 5159 | 2087 | 0 | 314 | 0 | 101 | 0 | 0 | 226 | 0 | 0 |
| SRR1510119 | 0 | 0 | 12 | 24 | 35 | 85 | 619 | 829 | 405 | 10 | 115 | 0 | 0 | 0 | 47987 | 50390 | 241990 | 0 | 417 | 33986 | 22710 | 0 | 0 | 26 | 11 | 61 | 423 | 107 | 0 | 26 | 0 | 137 | 0 | 0 | 176 | 0 | 0 |
| SRR1510120 | 0 | 0 | 0 | 23 | 38 | 150 | 783 | 1343 | 590 | 11 | 98 | 16 | 0 | 0 | 17998 | 42078 | 212466 | 0 | 175 | 13766 | 14295 | 0 | 0 | 29 | 11 | 44 | 295 | 107 | 0 | 26 | 0 | 188 | 0 | 0 | 156 | 0 | 0 |
| SRR1510121 | 0 | 16 | 19 | 60 | 37 | 195 | 540 | 1231 | 212 | 16 | 77 | 0 | 0 | 0 | 16244 | 42147 | 250486 | 0 | 148 | 23063 | 30829 | 0 | 0 | 174 | 95 | 26 | 100 | 59 | 0 | 25 | 0 | 139 | 0 | 0 | 415 | 0 | 0 |
| SRR1519906 | 0 | 0 | 0 | 0 | 0 | 2615 | 785 | 2746 | 523 | 14 | 0 | 22 | 0 | 0 | 7061 | 51736 | 36853 | 0 | 0 | 2615 | 1700 | 0 | 0 | 0 | 0 | 0 | 0 | 0 | 0 | 14 | 0 | 0 | 0 | 0 | 1367 | 0 | 231 |
| SRR1519919 | 0 | 0 | 0 | 0 | 0 | 844 | 352 | 1230 | 0 | 32 | 0 | 11 | 0 | 0 | 31324 | 30701 | 134419 | 0 | 35 | 2953 | 45140 | 0 | 0 | 32 | 0 | 21 | 21 | 0 | 0 | 32 | 0 | 0 | 0 | 0 | 125 | 0 | 117 |
| SRR1525198 | 353 | 180 | 95 | 106 | 60 | 129 | 154 | 606 | 22 | 11 | 155 | 18 | 0 | 0 | 30170 | 5956 | 52047 | 639 | 115 | 1442 | 1198 | 0 | 0 | 0 | 0 | 0 | 0 | 0 | 0 | 470 | 0 | 11 | 0 | 0 | 5895 | 0 | 11 |
| SRR1525199 | 226 | 181 | 148 | 123 | 47 | 117 | 216 | 593 | 36 | 0 | 92 | 0 | 0 | 0 | 34049 | 5899 | 46801 | 114 | 140 | 1964 | 1648 | 0 | 0 | 11 | 0 | 0 | 14 | 0 | 0 | 600 | 0 | 15 | 0 | 0 | 4901 | 0 | 11 |
| SRR1525200 | 307 | 225 | 157 | 82 | 41 | 175 | 168 | 504 | 29 | 0 | 130 | 27 | 0 | 0 | 63487 | 7470 | 91165 | 281 | 175 | 1903 | 1297 | 0 | 0 | 14 | 0 | 0 | 14 | 0 | 0 | 738 | 0 | 0 | 0 | 0 | 5185 | 0 | 0 |
| SRR1525201 | 291 | 158 | 158 | 145 | 60 | 211 | 291 | 771 | 54 | 0 | 148 | 20 | 0 | 0 | 52472 | 9968 | 72956 | 197 | 151 | 2750 | 1682 | 0 | 0 | 10 | 0 | 18 | 25 | 10 | 0 | 674 | 0 | 0 | 0 | 0 | 4951 | 0 | 43 |
| SRR1525202 | 29 | 33 | 15 | 15 | 0 | 179 | 222 | 816 | 39 | 0 | 12 | 0 | 0 | 0 | 24660 | 4676 | 37275 | 1291 | 131 | 1666 | 973 | 0 | 0 | 0 | 0 | 0 | 32 | 20 | 0 | 25 | 0 | 0 | 0 | 0 | 5929 | 0 | 0 |
| SRR1525203 | 11 | 271 | 118 | 121 | 43 | 208 | 357 | 857 | 71 | 0 | 180 | 0 | 0 | 0 | 25529 | 7383 | 36849 | 185 | 198 | 2236 | 1905 | 0 | 0 | 13 | 0 | 0 | 11 | 0 | 0 | 25 | 0 | 0 | 0 | 0 | 5479 | 0 | 13 |
| SRR1563015 | 123 | 42 | 0 | 0 | 61 | 482 | 464 | 1157 | 184 | 165 | 710 | 22 | 0 | 0 | 14863 | 7045 | 51073 | 34 | 129 | 6019 | 16362 | 0 | 33 | 192 | 0 | 0 | 0 | 0 | 0 | 2631 | 0 | 220 | 0 | 0 | 3052 | 0 | 0 |
| SRR1563018 | 26 | 11 | 66 | 105 | 33 | 123 | 210 | 376 | 99 | 92 | 14 | 0 | 0 | 0 | 1117 | 3138 | 29911 | 0 | 28 | 1874 | 9418 | 0 | 13 | 15 | 0 | 0 | 13 | 0 | 0 | 22 | 0 | 0 | 0 | 0 | 1367 | 0 | 0 |
| SRR1563057 | 26 | 16 | 332 | 41 | 156 | 274 | 75 | 316 | 0 | 52 | 124 | 0 | 31 | 0 | 1188 | 2268 | 21091 | 42 | 17 | 3474 | 6299 | 0 | 0 | 166 | 0 | 0 | 31 | 0 | 0 | 218 | 0 | 0 | 0 | 0 | 2396 | 0 | 10 |
| SRR1563058 | 0 | 0 | 0 | 0 | 15 | 164 | 13 | 221 | 0 | 27 | 177 | 13 | 0 | 0 | 8910 | 17353 | 128682 | 38 | 189 | 3093 | 18754 | 0 | 0 | 0 | 0 | 0 | 0 | 0 | 0 | 647 | 0 | 28 | 0 | 0 | 8241 | 0 | 17 |
| SRR1563059 | 371 | 209 | 280 | 233 | 1368 | 192 | 277 | 441 | 66 | 27 | 489 | 13 | 20 | 0 | 4130 | 4581 | 44388 | 28 | 66 | 7383 | 1214 | 0 | 0 | 694 | 17 | 0 | 57 | 0 | 0 | 1197 | 0 | 0 | 0 | 0 | 4255 | 0 | 0 |
| SRR1563060 | 0 | 0 | 0 | 0 | 21 | 216 | 100 | 255 | 23 | 58 | 15 | 11 | 0 | 0 | 67605 | 33271 | 205663 | 23 | 247 | 14098 | 11887 | 0 | 11 | 0 | 0 | 0 | 0 | 0 | 0 | 126 | 0 | 124 | 0 | 0 | 8094 | 0 | 20 |
| SRR1563061 | 0 | 16 | 0 | 0 | 30 | 130 | 137 | 267 | 69 | 17 | 480 | 21 | 0 | 0 | 15097 | 2990 | 43954 | 0 | 0 | 2097 | 5902 | 0 | 0 | 14 | 0 | 0 | 0 | 0 | 0 | 719 | 0 | 76 | 0 | 0 | 2356 | 0 | 0 |
| SRR1563062 | 0 | 0 | 18 | 11 | 0 | 15 | 45 | 74 | 45 | 21 | 15 | 0 | 0 | 0 | 237348 | 31672 | 365452 | 0 | 193 | 5012 | 37580 | 0 | 0 | 0 | 0 | 0 | 0 | 0 | 0 | 29 | 0 | 0 | 0 | 0 | 7885 | 0 | 14 |
| SRR1563063 | 801 | 428 | 223 | 192 | 1037 | 97 | 189 | 456 | 58 | 11 | 108 | 0 | 18 | 0 | 7396 | 10218 | 88274 | 43 | 43 | 8119 | 13602 | 0 | 0 | 691 | 11 | 11 | 39 | 0 | 0 | 730 | 0 | 0 | 0 | 0 | 2063 | 0 | 0 |
| SRR1563064 | 0 | 0 | 0 | 0 | 18 | 129 | 37 | 104 | 0 | 125 | 119 | 46 | 0 | 0 | 22443 | 31636 | 193026 | 0 | 356 | 14034 | 24857 | 0 | 0 | 0 | 0 | 0 | 0 | 0 | 0 | 542 | 0 | 49 | 0 | 0 | 8905 | 0 | 22 |
| SRR1575598 | 0 | 0 | 0 | 0 | 0 | 43 | 85 | 171 | 0 | 0 | 0 | 0 | 0 | 0 | 43 | 82 | 82 | 64 | 0 | 0 | 0 | 0 | 0 | 0 | 0 | 0 | 0 | 0 | 0 | 0 | 0 | 790 | 16 | 0 | 15505 | 0 | 16 |
| SRR1575599 | 0 | 0 | 0 | 0 | 0 | 30 | 15 | 101 | 0 | 0 | 0 | 0 | 0 | 0 | 109 | 3246 | 153 | 828 | 0 | 0 | 0 | 18 | 0 | 0 | 0 | 0 | 0 | 0 | 0 | 0 | 0 | 32261 | 0 | 0 | 15412 | 0 | 55 |
| SRR1575601 | 0 | 0 | 0 | 0 | 0 | 19 | 35 | 135 | 0 | 0 | 0 | 0 | 0 | 0 | 32 | 678 | 25 | 29 | 0 | 0 | 0 | 0 | 0 | 0 | 0 | 0 | 0 | 0 | 0 | 0 | 0 | 516 | 0 | 0 | 100226 | 0 | 0 |
| SRR1575602 | 18 | 25 | 0 | 0 | 0 | 88 | 91 | 274 | 15 | 0 | 0 | 0 | 0 | 0 | 1753 | 4746 | 1223 | 725 | 0 | 494 | 24 | 0 | 0 | 18 | 0 | 0 | 0 | 0 | 0 | 0 | 0 | 28632 | 0 | 0 | 43338 | 0 | 57 |
| SRR1592244 | 0 | 0 | 48 | 96 | 22 | 40 | 92 | 169 | 33 | 0 | 24 | 0 | 0 | 0 | 1764 | 23682 | 108058 | 0 | 0 | 99 | 414 | 0 | 0 | 25 | 44 | 128 | 305 | 89 | 0 | 46 | 0 | 0 | 0 | 0 | 252 | 0 | 0 |
| SRR1592245 | 0 | 0 | 0 | 0 | 0 | 30 | 66 | 113 | 15 | 0 | 30 | 0 | 0 | 0 | 2552 | 188047 | 131013 | 0 | 15 | 2495 | 381 | 0 | 0 | 12 | 33 | 115 | 327 | 81 | 0 | 60 | 0 | 0 | 0 | 0 | 218 | 0 | 0 |
| SRR1592246 | 0 | 0 | 15 | 0 | 0 | 0 | 21 | 40 | 0 | 12 | 229 | 0 | 0 | 0 | 2333 | 556568 | 132912 | 0 | 20 | 2655 | 435 | 0 | 0 | 93 | 311 | 1072 | 2510 | 615 | 0 | 543 | 0 | 0 | 0 | 0 | 35 | 0 | 0 |
| SRR1592247 | 0 | 0 | 0 | 12 | 0 | 0 | 13 | 19 | 0 | 0 | 21 | 0 | 0 | 0 | 2186 | 356317 | 232660 | 0 | 0 | 868 | 705 | 0 | 0 | 16 | 25 | 100 | 383 | 331 | 0 | 37 | 0 | 0 | 0 | 0 | 174 | 0 | 0 |
| SRR1592248 | 0 | 0 | 0 | 25 | 0 | 11 | 33 | 49 | 0 | 18 | 182 | 0 | 0 | 0 | 2196 | 419616 | 177880 | 0 | 0 | 1570 | 986 | 0 | 0 | 28 | 126 | 453 | 1502 | 975 | 0 | 519 | 0 | 0 | 0 | 0 | 98 | 0 | 0 |
| SRR1596226 | 0 | 0 | 0 | 0 | 13 | 87 | 130 | 165 | 26 | 0 | 59 | 0 | 0 | 0 | 234 | 19562 | 14980 | 78 | 35 | 2175 | 121 | 0 | 0 | 23 | 0 | 16 | 16 | 0 | 0 | 1498 | 0 | 0 | 0 | 0 | 21170 | 134 | 0 |

### Supp Table S8.3.Columns_BX_to_DH

| | | | | | | | | | | | | | | | | | | | | | | | | | | | | | | | | | | | | | |
|---|---|---|---|---|---|---|---|---|---|---|---|---|---|---|---|---|---|---|---|---|---|---|---|---|---|---|---|---|---|---|---|---|---|---|---|---|---|
| SRR1596227 | 0 | 0 | 11 | 22 | 12 | 113 | 88 | 50 | 25 | 0 | 62 | 0 | 0 | 0 | 126 | 15304 | 9702 | 25 | 13 | 1858 | 163 | 0 | 0 | 22 | 0 | 14 | 27 | 0 | 0 | 1192 | 0 | 13 | 0 | 0 | 22754 | 222 | 0 |
| SRR1596228 | 0 | 0 | 0 | 0 | 80 | 168 | 273 | 337 | 126 | 13 | 374 | 0 | 0 | 0 | 1704 | 139492 | 36373 | 0 | 137 | 7426 | 736 | 0 | 0 | 94 | 20 | 63 | 94 | 0 | 0 | 4899 | 0 | 53 | 0 | 0 | 12627 | 273 | 0 |
| SRR1596229 | 0 | 0 | 0 | 0 | 14 | 174 | 501 | 305 | 22 | 0 | 74 | 0 | 0 | 0 | 1873 | 147934 | 36664 | 22 | 131 | 6924 | 414 | 0 | 0 | 14 | 13 | 11 | 25 | 0 | 0 | 1282 | 0 | 22 | 0 | 0 | 12044 | 306 | 0 |
| SRR1596230 | 24 | 10 | 0 | 20 | 81 | 350 | 467 | 593 | 153 | 27 | 91 | 0 | 0 | 0 | 18856 | 121718 | 275791 | 0 | 270 | 5058 | 970 | 0 | 0 | 95 | 30 | 44 | 54 | 0 | 0 | 2292 | 0 | 0 | 0 | 0 | 10595 | 174 | 0 |
| SRR1596231 | 0 | 16 | 12 | 33 | 122 | 264 | 518 | 687 | 74 | 0 | 98 | 0 | 0 | 0 | 19473 | 124207 | 277284 | 0 | 148 | 5022 | 877 | 0 | 12 | 90 | 16 | 57 | 77 | 0 | 0 | 1780 | 0 | 0 | 0 | 0 | 10083 | 160 | 16 |
| SRR1608128 | 0 | 0 | 0 | 0 | 0 | 139 | 666 | 124 | 418 | 0 | 0 | 0 | 0 | 0 | 170 | 6677 | 2245 | 93 | 0 | 0 | 0 | 0 | 0 | 44 | 0 | 0 | 0 | 0 | 0 | 0 | 0 | 139 | 0 | 0 | 1511 | 39 | 0 |
| SRR1608138 | 0 | 0 | 0 | 0 | 18 | 842 | 5420 | 755 | 2574 | 0 | 0 | 0 | 0 | 0 | 408 | 2149 | 726 | 25 | 0 | 25 | 0 | 0 | 0 | 80 | 0 | 0 | 0 | 0 | 0 | 0 | 0 | 161 | 0 | 0 | 1189 | 82 | 0 |
| SRR1608143 | 0 | 0 | 0 | 0 | 0 | 188 | 1119 | 131 | 425 | 0 | 0 | 0 | 0 | 0 | 684 | 15786 | 4198 | 67 | 0 | 0 | 28 | 0 | 0 | 0 | 0 | 0 | 0 | 0 | 0 | 0 | 11 | 227 | 0 | 0 | 506 | 49 | 0 |
| SRR1608150 | 0 | 0 | 0 | 0 | 0 | 326 | 1224 | 109 | 721 | 0 | 0 | 0 | 0 | 0 | 558 | 5420 | 1959 | 54 | 0 | 27 | 14 | 0 | 0 | 29 | 0 | 0 | 0 | 0 | 0 | 0 | 0 | 190 | 0 | 0 | 960 | 26 | 0 |
| SRR1608158 | 0 | 0 | 0 | 0 | 0 | 460 | 3395 | 386 | 1720 | 0 | 0 | 0 | 0 | 0 | 15 | 5427 | 3263 | 0 | 0 | 0 | 15 | 0 | 0 | 44 | 0 | 0 | 0 | 0 | 0 | 0 | 0 | 104 | 0 | 0 | 1238 | 107 | 0 |
| SRR1608160 | 0 | 0 | 0 | 0 | 0 | 454 | 2034 | 420 | 1412 | 0 | 0 | 0 | 0 | 0 | 34 | 3810 | 1526 | 34 | 0 | 0 | 0 | 0 | 0 | 69 | 19 | 39 | 11 | 0 | 0 | 0 | 0 | 185 | 0 | 0 | 893 | 49 | 0 |
| SRR1608163 | 0 | 0 | 0 | 0 | 0 | 184 | 800 | 64 | 377 | 0 | 0 | 0 | 0 | 0 | 395 | 12530 | 3227 | 46 | 0 | 0 | 0 | 0 | 0 | 24 | 12 | 0 | 0 | 0 | 0 | 0 | 0 | 129 | 0 | 0 | 981 | 20 | 0 |
| SRR1608165 | 0 | 0 | 0 | 0 | 0 | 32 | 318 | 65 | 175 | 0 | 0 | 0 | 0 | 0 | 2113 | 12714 | 4710 | 19 | 0 | 0 | 0 | 0 | 0 | 12 | 0 | 0 | 0 | 0 | 0 | 0 | 0 | 123 | 0 | 0 | 274 | 32 | 0 |
| SRR1608166 | 0 | 0 | 0 | 0 | 0 | 350 | 2051 | 250 | 826 | 0 | 0 | 0 | 0 | 0 | 550 | 3731 | 1336 | 0 | 0 | 0 | 0 | 0 | 0 | 76 | 0 | 26 | 0 | 0 | 0 | 0 | 0 | 100 | 0 | 0 | 1206 | 19 | 0 |
| SRR1608167 | 0 | 0 | 0 | 0 | 0 | 15 | 531 | 106 | 167 | 0 | 0 | 0 | 0 | 0 | 91 | 1582 | 690 | 0 | 0 | 0 | 0 | 0 | 0 | 22 | 0 | 0 | 0 | 0 | 0 | 0 | 0 | 76 | 0 | 0 | 943 | 12 | 0 |
| SRR1608168 | 0 | 0 | 0 | 0 | 0 | 97 | 469 | 70 | 431 | 0 | 0 | 0 | 0 | 0 | 2794 | 15635 | 9318 | 32 | 0 | 27 | 421 | 0 | 0 | 28 | 0 | 0 | 0 | 0 | 0 | 0 | 0 | 54 | 0 | 0 | 317 | 91 | 0 |
| SRR1608169 | 0 | 0 | 0 | 0 | 0 | 727 | 2919 | 338 | 707 | 0 | 0 | 0 | 0 | 0 | 307 | 2291 | 1211 | 10 | 0 | 10 | 20 | 0 | 0 | 35 | 35 | 59 | 11 | 0 | 0 | 11 | 0 | 72 | 0 | 0 | 624 | 28 | 0 |
| SRR1608170 | 0 | 0 | 0 | 0 | 0 | 187 | 818 | 108 | 488 | 0 | 0 | 0 | 0 | 0 | 115 | 1912 | 818 | 14 | 0 | 0 | 0 | 0 | 0 | 0 | 0 | 0 | 0 | 0 | 0 | 0 | 0 | 57 | 0 | 0 | 711 | 53 | 0 |
| SRR1608171 | 0 | 0 | 0 | 0 | 0 | 153 | 955 | 34 | 332 | 0 | 0 | 0 | 0 | 0 | 588 | 5788 | 1952 | 26 | 0 | 17 | 26 | 0 | 0 | 29 | 0 | 0 | 0 | 0 | 0 | 0 | 0 | 94 | 0 | 0 | 386 | 67 | 0 |
| SRR1608172 | 0 | 0 | 0 | 0 | 0 | 196 | 1121 | 116 | 338 | 0 | 0 | 0 | 0 | 0 | 614 | 2783 | 1625 | 0 | 0 | 0 | 0 | 0 | 0 | 18 | 0 | 0 | 0 | 0 | 0 | 0 | 0 | 125 | 0 | 0 | 579 | 35 | 0 |
| SRR1608174 | 0 | 0 | 0 | 0 | 0 | 117 | 898 | 143 | 573 | 0 | 0 | 0 | 0 | 0 | 39 | 6095 | 1879 | 0 | 0 | 0 | 0 | 0 | 0 | 26 | 0 | 0 | 0 | 0 | 0 | 0 | 0 | 65 | 0 | 0 | 1090 | 21 | 0 |
| SRR1608175 | 0 | 0 | 0 | 0 | 0 | 22 | 456 | 45 | 120 | 0 | 0 | 0 | 0 | 0 | 2063 | 37254 | 6642 | 0 | 0 | 15 | 60 | 0 | 0 | 23 | 39 | 58 | 0 | 0 | 0 | 0 | 0 | 67 | 0 | 0 | 883 | 36 | 0 |
| SRR1608177 | 0 | 0 | 0 | 0 | 0 | 201 | 1296 | 104 | 700 | 0 | 0 | 0 | 0 | 0 | 665 | 4458 | 1542 | 42 | 0 | 0 | 0 | 0 | 0 | 18 | 0 | 0 | 0 | 0 | 0 | 0 | 0 | 69 | 0 | 0 | 1495 | 11 | 0 |
| SRR1608178 | 0 | 0 | 0 | 0 | 19 | 254 | 3139 | 308 | 1052 | 0 | 0 | 0 | 0 | 0 | 1071 | 2896 | 1573 | 18 | 0 | 0 | 18 | 0 | 0 | 81 | 0 | 0 | 0 | 0 | 0 | 0 | 0 | 272 | 0 | 0 | 952 | 53 | 0 |
| SRR1608179 | 0 | 0 | 0 | 0 | 0 | 217 | 1847 | 163 | 598 | 0 | 11 | 0 | 0 | 0 | 964 | 6767 | 2540 | 14 | 41 | 14 | 27 | 0 | 0 | 65 | 54 | 69 | 11 | 0 | 0 | 0 | 0 | 95 | 0 | 0 | 1187 | 35 | 0 |
| SRR1608180 | 0 | 0 | 0 | 0 | 0 | 291 | 1909 | 393 | 1311 | 0 | 0 | 0 | 0 | 0 | 510 | 6148 | 3721 | 44 | 0 | 0 | 0 | 0 | 0 | 114 | 11 | 0 | 11 | 0 | 0 | 0 | 0 | 291 | 0 | 0 | 1187 | 14 | 0 |
| SRR1608181 | 0 | 0 | 0 | 0 | 0 | 360 | 2060 | 360 | 1749 | 0 | 0 | 0 | 0 | 0 | 408 | 5418 | 2006 | 29 | 0 | 0 | 0 | 0 | 0 | 30 | 0 | 13 | 0 | 0 | 0 | 0 | 0 | 165 | 0 | 0 | 1177 | 37 | 0 |
| SRR1608182 | 0 | 0 | 0 | 0 | 0 | 74 | 429 | 67 | 207 | 0 | 0 | 0 | 0 | 0 | 2558 | 20485 | 6460 | 52 | 0 | 15 | 30 | 0 | 0 | 27 | 11 | 0 | 0 | 0 | 0 | 0 | 0 | 104 | 0 | 0 | 604 | 51 | 0 |
| SRR1608184 | 0 | 0 | 0 | 0 | 0 | 378 | 3482 | 318 | 1134 | 0 | 0 | 0 | 0 | 0 | 199 | 2590 | 1190 | 0 | 0 | 0 | 0 | 0 | 0 | 94 | 0 | 14 | 0 | 0 | 0 | 0 | 0 | 239 | 0 | 0 | 693 | 30 | 0 |
| SRR1608185 | 0 | 0 | 0 | 0 | 0 | 304 | 2829 | 291 | 820 | 0 | 0 | 0 | 0 | 0 | 225 | 7195 | 3306 | 13 | 0 | 0 | 0 | 0 | 0 | 65 | 0 | 0 | 0 | 0 | 0 | 0 | 0 | 93 | 0 | 0 | 925 | 31 | 0 |
| SRR1608186 | 0 | 0 | 0 | 0 | 0 | 238 | 3135 | 253 | 966 | 0 | 0 | 0 | 0 | 0 | 758 | 2663 | 879 | 0 | 0 | 0 | 0 | 0 | 0 | 48 | 24 | 10 | 0 | 0 | 0 | 10 | 0 | 45 | 0 | 0 | 2075 | 50 | 0 |
| SRR1608189 | 0 | 0 | 0 | 0 | 0 | 548 | 2294 | 270 | 869 | 0 | 0 | 0 | 0 | 0 | 1138 | 11400 | 6235 | 51 | 0 | 0 | 0 | 0 | 0 | 57 | 0 | 0 | 0 | 0 | 0 | 0 | 0 | 143 | 0 | 0 | 945 | 42 | 0 |
| SRR1608190 | 0 | 0 | 0 | 0 | 0 | 292 | 2519 | 432 | 1081 | 0 | 0 | 0 | 0 | 0 | 54 | 2474 | 539 | 43 | 0 | 0 | 11 | 0 | 0 | 87 | 0 | 0 | 0 | 0 | 0 | 0 | 0 | 184 | 0 | 0 | 1052 | 43 | 0 |
| SRR1608191 | 0 | 0 | 0 | 0 | 0 | 1018 | 5855 | 775 | 2203 | 0 | 0 | 0 | 0 | 0 | 44 | 742 | 177 | 22 | 0 | 0 | 0 | 0 | 0 | 162 | 12 | 0 | 0 | 0 | 0 | 0 | 0 | 144 | 0 | 0 | 1263 | 49 | 0 |
| SRR1608192 | 0 | 0 | 0 | 0 | 0 | 561 | 2389 | 305 | 1540 | 0 | 0 | 0 | 0 | 0 | 449 | 11494 | 3892 | 0 | 0 | 16 | 0 | 0 | 0 | 38 | 0 | 0 | 0 | 0 | 0 | 0 | 0 | 257 | 0 | 0 | 1337 | 31 | 0 |
| SRR1608193 | 0 | 0 | 0 | 0 | 0 | 354 | 2509 | 354 | 1045 | 0 | 0 | 0 | 0 | 0 | 261 | 4784 | 1701 | 0 | 0 | 0 | 0 | 0 | 0 | 52 | 13 | 0 | 0 | 0 | 0 | 0 | 0 | 56 | 0 | 0 | 1236 | 28 | 0 |
| SRR1608194 | 0 | 0 | 0 | 0 | 15 | 253 | 2038 | 208 | 1547 | 0 | 0 | 0 | 0 | 0 | 535 | 2237 | 1050 | 30 | 0 | 0 | 15 | 0 | 0 | 56 | 11 | 26 | 0 | 0 | 0 | 0 | 0 | 431 | 0 | 0 | 1315 | 55 | 0 |
| SRR1608196 | 0 | 0 | 0 | 0 | 0 | 497 | 2254 | 307 | 529 | 0 | 0 | 0 | 0 | 0 | 275 | 5599 | 2487 | 53 | 0 | 21 | 0 | 0 | 0 | 27 | 0 | 0 | 0 | 0 | 0 | 0 | 0 | 85 | 0 | 0 | 705 | 20 | 0 |
| SRR1608201 | 0 | 0 | 0 | 0 | 0 | 388 | 1077 | 141 | 618 | 0 | 0 | 0 | 0 | 0 | 971 | 5492 | 2619 | 71 | 0 | 0 | 0 | 0 | 0 | 23 | 0 | 0 | 0 | 0 | 0 | 0 | 0 | 159 | 0 | 0 | 1101 | 45 | 0 |
| SRR1608208 | 0 | 0 | 0 | 0 | 0 | 294 | 1784 | 200 | 775 | 0 | 0 | 0 | 0 | 0 | 1221 | 6234 | 2749 | 47 | 0 | 23 | 23 | 0 | 0 | 89 | 11 | 0 | 0 | 0 | 0 | 0 | 0 | 129 | 0 | 0 | 1024 | 184 | 0 |
| SRR1608213 | 0 | 0 | 0 | 0 | 0 | 343 | 4709 | 438 | 1449 | 0 | 0 | 0 | 0 | 0 | 515 | 1697 | 860 | 0 | 0 | 0 | 0 | 0 | 0 | 38 | 0 | 0 | 0 | 0 | 0 | 0 | 0 | 210 | 0 | 0 | 715 | 38 | 0 |
| SRR1608214 | 0 | 0 | 0 | 0 | 0 | 236 | 1221 | 69 | 361 | 0 | 0 | 0 | 0 | 0 | 180 | 4765 | 1884 | 28 | 0 | 14 | 0 | 0 | 0 | 12 | 0 | 0 | 0 | 0 | 0 | 0 | 0 | 194 | 0 | 0 | 435 | 44 | 0 |
| SRR1608222 | 0 | 0 | 0 | 0 | 0 | 0 | 192 | 64 | 96 | 0 | 0 | 0 | 0 | 0 | 32 | 3898 | 1157 | 0 | 0 | 32 | 0 | 0 | 0 | 0 | 0 | 0 | 0 | 0 | 0 | 0 | 0 | 96 | 0 | 0 | 386 | 0 | 0 |
| SRR1608232 | 0 | 0 | 0 | 0 | 0 | 98 | 846 | 33 | 228 | 0 | 0 | 0 | 0 | 0 | 716 | 15060 | 13924 | 0 | 0 | 33 | 65 | 0 | 0 | 42 | 0 | 10 | 0 | 0 | 0 | 0 | 0 | 163 | 0 | 0 | 624 | 0 | 0 |
| SRR1608237 | 0 | 0 | 0 | 0 | 0 | 77 | 351 | 33 | 88 | 0 | 0 | 0 | 0 | 0 | 3833 | 49720 | 31098 | 44 | 0 | 0 | 44 | 0 | 0 | 13 | 0 | 0 | 0 | 0 | 0 | 0 | 0 | 99 | 0 | 0 | 119 | 0 | 0 |
| SRR1608254 | 0 | 0 | 0 | 0 | 0 | 242 | 1556 | 89 | 319 | 0 | 0 | 0 | 0 | 0 | 89 | 8037 | 3554 | 26 | 0 | 0 | 0 | 0 | 0 | 10 | 0 | 0 | 0 | 0 | 0 | 0 | 0 | 38 | 0 | 0 | 516 | 0 | 0 |
| SRR1608255 | 0 | 0 | 0 | 0 | 0 | 223 | 2605 | 213 | 679 | 0 | 0 | 0 | 0 | 0 | 122 | 6716 | 3454 | 10 | 0 | 0 | 0 | 0 | 0 | 18 | 0 | 0 | 0 | 0 | 0 | 0 | 0 | 91 | 0 | 0 | 645 | 0 | 0 |
| SRR1608256 | 0 | 0 | 0 | 0 | 0 | 144 | 1602 | 106 | 230 | 0 | 0 | 0 | 0 | 0 | 163 | 10429 | 4005 | 0 | 0 | 0 | 0 | 0 | 0 | 0 | 0 | 0 | 0 | 0 | 0 | 0 | 0 | 0 | 0 | 0 | 312 | 0 | 0 |
| SRR1608257 | 0 | 0 | 0 | 0 | 0 | 269 | 2520 | 343 | 636 | 0 | 0 | 0 | 0 | 0 | 563 | 9899 | 5088 | 0 | 0 | 37 | 0 | 0 | 0 | 0 | 0 | 0 | 0 | 0 | 0 | 0 | 0 | 110 | 0 | 0 | 698 | 0 | 0 |
| SRR1608260 | 0 | 0 | 0 | 0 | 0 | 209 | 2322 | 190 | 362 | 0 | 0 | 0 | 0 | 0 | 114 | 7174 | 2042 | 0 | 0 | 0 | 0 | 0 | 0 | 10 | 0 | 0 | 0 | 0 | 0 | 0 | 0 | 57 | 0 | 0 | 541 | 13 | 0 |
| SRR1608263 | 0 | 0 | 0 | 0 | 0 | 164 | 753 | 98 | 458 | 0 | 0 | 0 | 0 | 0 | 65 | 9181 | 4629 | 0 | 0 | 0 | 33 | 0 | 0 | 20 | 0 | 0 | 0 | 0 | 0 | 0 | 0 | 164 | 0 | 0 | 311 | 0 | 0 |
| SRR1608264 | 0 | 0 | 0 | 0 | 0 | 52 | 207 | 52 | 155 | 0 | 0 | 0 | 0 | 0 | 594 | 19832 | 13312 | 0 | 0 | 0 | 0 | 0 | 0 | 0 | 0 | 0 | 0 | 0 | 0 | 0 | 0 | 181 | 0 | 0 | 245 | 0 | 0 |
| SRR1608266 | 0 | 0 | 0 | 0 | 0 | 46 | 458 | 46 | 137 | 0 | 0 | 0 | 0 | 0 | 92 | 3008 | 1143 | 0 | 0 | 0 | 0 | 0 | 0 | 18 | 0 | 0 | 0 | 0 | 0 | 0 | 0 | 92 | 0 | 0 | 0 | 0 | 0 |
| SRR1608267 | 0 | 0 | 0 | 0 | 0 | 54 | 298 | 0 | 108 | 0 | 0 | 0 | 0 | 0 | 189 | 11043 | 7420 | 0 | 0 | 0 | 0 | 0 | 0 | 22 | 0 | 0 | 0 | 0 | 0 | 0 | 0 | 108 | 0 | 0 | 29 | 0 | 0 |
| SRR1608269 | 0 | 0 | 0 | 0 | 0 | 0 | 505 | 34 | 202 | 0 | 0 | 0 | 0 | 0 | 1044 | 50736 | 23421 | 0 | 0 | 0 | 34 | 0 | 0 | 40 | 0 | 0 | 0 | 0 | 0 | 0 | 0 | 135 | 0 | 0 | 329 | 0 | 0 |
| SRR1608270 | 0 | 0 | 0 | 0 | 0 | 78 | 466 | 0 | 214 | 0 | 0 | 0 | 0 | 0 | 2155 | 45033 | 30284 | 58 | 0 | 19 | 58 | 0 | 0 | 83 | 0 | 0 | 0 | 0 | 0 | 0 | 0 | 214 | 0 | 0 | 231 | 0 | 0 |
| SRR1608271 | 0 | 0 | 0 | 0 | 0 | 80 | 598 | 0 | 239 | 0 | 0 | 0 | 0 | 0 | 518 | 17571 | 3735 | 40 | 0 | 0 | 40 | 0 | 0 | 33 | 0 | 0 | 0 | 0 | 0 | 0 | 0 | 398 | 0 | 0 | 552 | 0 | 0 |
| SRR1608274 | 0 | 0 | 0 | 0 | 0 | 189 | 1303 | 126 | 420 | 0 | 0 | 0 | 0 | 0 | 714 | 20879 | 8835 | 21 | 0 | 0 | 42 | 0 | 0 | 40 | 0 | 0 | 0 | 0 | 0 | 0 | 0 | 210 | 0 | 0 | 426 | 0 | 0 |
| SRR1608276 | 0 | 0 | 0 | 0 | 0 | 88 | 852 | 59 | 323 | 0 | 14 | 0 | 0 | 0 | 1851 | 40269 | 20282 | 59 | 0 | 29 | 59 | 0 | 0 | 28 | 0 | 80 | 0 | 0 | 0 | 0 | 0 | 59 | 0 | 0 | 689 | 0 | 0 |
| SRR1608277 | 0 | 0 | 0 | 0 | 0 | 115 | 1177 | 172 | 775 | 0 | 0 | 0 | 0 | 0 | 431 | 15653 | 7125 | 0 | 0 | 0 | 29 | 0 | 0 | 30 | 0 | 0 | 0 | 0 | 0 | 0 | 0 | 345 | 0 | 0 | 617 | 0 | 0 |
| SRR1608278 | 0 | 0 | 0 | 0 | 0 | 33 | 598 | 66 | 266 | 0 | 0 | 0 | 0 | 0 | 365 | 15198 | 7817 | 0 | 0 | 0 | 0 | 0 | 0 | 0 | 0 | 0 | 0 | 0 | 0 | 0 | 0 | 33 | 0 | 0 | 145 | 0 | 0 |
| SRR1608281 | 0 | 0 | 0 | 0 | 0 | 304 | 1381 | 138 | 470 | 0 | 0 | 0 | 0 | 0 | 304 | 13188 | 7051 | 0 | 0 | 0 | 0 | 0 | 0 | 13 | 0 | 0 | 0 | 0 | 0 | 0 | 0 | 249 | 0 | 0 | 398 | 0 | 0 |
| SRR1608282 | 0 | 0 | 0 | 0 | 0 | 38 | 649 | 76 | 293 | 0 | 0 | 0 | 0 | 0 | 420 | 17238 | 6918 | 13 | 0 | 0 | 0 | 0 | 0 | 15 | 0 | 0 | 0 | 0 | 0 | 0 | 0 | 25 | 0 | 0 | 754 | 0 | 0 |
| SRR1608283 | 0 | 0 | 0 | 0 | 0 | 65 | 240 | 87 | 87 | 0 | 0 | 0 | 0 | 0 | 1919 | 56652 | 33835 | 0 | 0 | 44 | 44 | 0 | 0 | 18 | 0 | 0 | 0 | 0 | 0 | 0 | 0 | 174 | 0 | 0 | 226 | 0 | 0 |
| SRR1608286 | 0 | 0 | 0 | 0 | 0 | 0 | 79 | 39 | 158 | 0 | 0 | 0 | 0 | 0 | 394 | 13278 | 4313 | 0 | 0 | 0 | 0 | 0 | 0 | 11 | 0 | 0 | 0 | 0 | 0 | 0 | 0 | 118 | 0 | 0 | 48 | 0 | 0 |
| SRR1608287 | 0 | 0 | 0 | 0 | 0 | 195 | 779 | 167 | 139 | 0 | 0 | 0 | 0 | 0 | 696 | 31751 | 13315 | 0 | 0 | 0 | 0 | 0 | 0 | 42 | 0 | 0 | 0 | 0 | 0 | 0 | 0 | 83 | 0 | 0 | 151 | 0 | 0 |
| SRR1608289 | 0 | 0 | 0 | 0 | 0 | 0 | 427 | 78 | 194 | 0 | 0 | 0 | 0 | 0 | 155 | 14291 | 4921 | 0 | 0 | 0 | 0 | 0 | 0 | 39 | 0 | 0 | 0 | 0 | 0 | 0 | 0 | 39 | 0 | 0 | 314 | 0 | 0 |
| SRR1608290 | 0 | 0 | 0 | 0 | 0 | 34 | 67 | 34 | 67 | 0 | 0 | 0 | 0 | 0 | 168 | 13833 | 7834 | 101 | 0 | 0 | 34 | 0 | 0 | 22 | 0 | 0 | 0 | 0 | 0 | 0 | 0 | 101 | 0 | 0 | 313 | 0 | 0 |
| SRR1608302 | 0 | 0 | 0 | 0 | 0 | 59 | 589 | 0 | 118 | 0 | 0 | 0 | 0 | 0 | 295 | 7587 | 3794 | 0 | 0 | 0 | 0 | 0 | 0 | 31 | 0 | 0 | 0 | 0 | 0 | 0 | 0 | 59 | 0 | 0 | 131 | 0 | 0 |
| SRR1608307 | 0 | 0 | 0 | 0 | 0 | 99 | 644 | 50 | 248 | 0 | 0 | 0 | 0 | 0 | 297 | 7563 | 4291 | 0 | 0 | 0 | 0 | 0 | 0 | 19 | 0 | 0 | 0 | 0 | 0 | 0 | 0 | 173 | 0 | 0 | 255 | 0 | 0 |
| SRR1608308 | 0 | 0 | 0 | 0 | 0 | 0 | 1046 | 37 | 710 | 0 | 0 | 0 | 0 | 0 | 523 | 16810 | 10408 | 0 | 0 | 37 | 0 | 0 | 0 | 28 | 0 | 11 | 0 | 0 | 0 | 0 | 0 | 299 | 0 | 0 | 612 | 0 | 0 |
| SRR1634379 | 0 | 15 | 0 | 0 | 38 | 165 | 486 | 651 | 138 | 0 | 44 | 0 | 0 | 0 | 5189 | 106012 | 100437 | 37 | 536 | 14942 | 880 | 0 | 0 | 65 | 25 | 39 | 24 | 0 | 0 | 105 | 0 | 23 | 0 | 0 | 12671 | 26 | 12 |
| SRR1634381 | 15 | 14 | 0 | 0 | 82 | 210 | 493 | 576 | 165 | 13 | 75 | 0 | 0 | 0 | 9451 | 47948 | 67668 | 34 | 440 | 6559 | 1211 | 0 | 0 | 77 | 31 | 47 | 39 | 22 | 0 | 203 | 0 | 13 | 0 | 0 | 25050 | 293 | 0 |
| SRR1634384 | 0 | 0 | 0 | 0 | 75 | 159 | 514 | 499 | 256 | 0 | 85 | 0 | 0 | 0 | 6881 | 37195 | 52516 | 24 | 448 | 5516 | 559 | 0 | 0 | 134 | 66 | 90 | 115 | 51 | 0 | 240 | 0 | 0 | 0 | 0 | 8283 | 62 | 0 |
| SRR1634388 | 0 | 0 | 0 | 0 | 46 | 114 | 286 | 314 | 78 | 0 | 56 | 0 | 0 | 0 | 6493 | 36545 | 69232 | 17 | 498 | 19044 | 550 | 0 | 0 | 77 | 28 | 42 | 61 | 23 | 0 | 122 | 0 | 0 | 0 | 0 | 8065 | 88 | 0 |
| SRR1634390 | 0 | 0 | 0 | 0 | 39 | 378 | 665 | 828 | 198 | 26 | 81 | 0 | 0 | 0 | 74227 | 38532 | 285118 | 24 | 300 | 4194 | 986 | 0 | 0 | 98 | 26 | 43 | 50 | 17 | 0 | 353 | 0 | 0 | 0 | 0 | 34013 | 573 | 0 |
| SRR1634392 | 10 | 17 | 0 | 0 | 64 | 230 | 403 | 388 | 165 | 0 | 42 | 0 | 0 | 0 | 7795 | 32690 | 50774 | 24 | 459 | 6131 | 451 | 0 | 0 | 114 | 46 | 57 | 52 | 23 | 0 | 172 | 0 | 0 | 0 | 0 | 14581 | 258 | 0 |
| SRR1634394 | 0 | 0 | 0 | 0 | 62 | 190 | 462 | 510 | 128 | 15 | 48 | 0 | 0 | 0 | 19061 | 42655 | 75055 | 46 | 632 | 12886 | 809 | 0 | 0 | 82 | 35 | 50 | 40 | 14 | 0 | 120 | 0 | 0 | 0 | 0 | 19503 | 96 | 0 |
| SRR1634395 | 21 | 15 | 0 | 0 | 21 | 590 | 1216 | 1543 | 256 | 12 | 42 | 0 | 0 | 0 | 29657 | 58056 | 166639 | 52 | 232 | 7356 | 1017 | 0 | 0 | 19 | 20 | 13 | 12 | 11 | 0 | 190 | 0 | 0 | 0 | 0 | 65610 | 249 | 0 |
| SRR1634398 | 15 | 12 | 0 | 0 | 44 | 339 | 555 | 750 | 142 | 89 | 100 | 0 | 0 | 0 | 57714 | 44104 | 185468 | 42 | 411 | 11763 | 3038 | 0 | 0 | 41 | 27 | 35 | 27 | 13 | 0 | 470 | 0 | 0 | 0 | 0 | 70943 | 540 | 0 |
| SRR1634399 | 11 | 13 | 0 | 0 | 19 | 203 | 493 | 617 | 161 | 13 | 38 | 0 | 0 | 0 | 22222 | 51963 | 140920 | 35 | 328 | 10782 | 745 | 0 | 0 | 30 | 11 | 13 | 16 | 16 | 0 | 130 | 0 | 0 | 0 | 0 | 30744 | 163 | 0 |
| SRR1634400 | 11 | 14 | 0 | 0 | 44 | 218 | 458 | 622 | 167 | 0 | 101 | 0 | 0 | 0 | 22704 | 52685 | 141137 | 28 | 355 | 10864 | 714 | 0 | 0 | 91 | 18 | 41 | 46 | 11 | 0 | 387 | 0 | 0 | 0 | 0 | 31140 | 172 | 0 |
| SRR1634406 | 13 | 19 | 0 | 0 | 69 | 170 | 382 | 451 | 133 | 11 | 83 | 0 | 0 | 0 | 11110 | 31681 | 49865 | 22 | 387 | 5832 | 683 | 0 | 0 | 106 | 30 | 42 | 62 | 17 | 0 | 228 | 0 | 13 | 0 | 0 | 13786 | 98 | 0 |
| SRR1634409 | 21 | 15 | 0 | 0 | 19 | 644 | 1039 | 1241 | 258 | 15 | 32 | 0 | 0 | 0 | 11885 | 42260 | 85252 | 48 | 248 | 4647 | 690 | 0 | 0 | 19 | 17 | 10 | 16 | 0 | 0 | 142 | 0 | 16 | 0 | 0 | 91538 | 348 | 0 |
| SRR1634411 | 15 | 12 | 0 | 0 | 14 | 396 | 707 | 806 | 208 | 12 | 78 | 0 | 0 | 0 | 19753 | 35185 | 147872 | 25 | 276 | 17325 | 755 | 0 | 0 | 43 | 18 | 28 | 21 | 18 | 0 | 240 | 0 | 0 | 0 | 0 | 15257 | 114 | 0 |
| SRR1634412 | 21 | 12 | 0 | 12 | 14 | 315 | 707 | 819 | 211 | 12 | 78 | 0 | 0 | 0 | 19511 | 36502 | 147092 | 0 | 240 | 16927 | 826 | 0 | 0 | 38 | 14 | 10 | 18 | 12 | 0 | 241 | 0 | 0 | 0 | 0 | 16052 | 128 | 0 |
| SRR1634414 | 11 | 0 | 0 | 0 | 28 | 502 | 832 | 931 | 188 | 15 | 40 | 0 | 0 | 0 | 20048 | 39365 | 93297 | 30 | 292 | 3575 | 689 | 0 | 0 | 20 | 20 | 11 | 20 | 0 | 0 | 179 | 0 | 0 | 0 | 0 | 65703 | 347 | 0 |
| SRR1634415 | 16 | 17 | 0 | 0 | 26 | 407 | 913 | 774 | 210 | 19 | 49 | 0 | 0 | 0 | 9769 | 45487 | 70839 | 22 | 284 | 8096 | 893 | 0 | 0 | 30 | 11 | 15 | 17 | 0 | 0 | 123 | 0 | 13 | 0 | 0 | 27519 | 241 | 15 |
| SRR1634417 | 15 | 13 | 0 | 0 | 44 | 664 | 1066 | 1338 | 266 | 37 | 95 | 0 | 0 | 0 | 2328 | 41338 | 86252 | 16 | 127 | 4572 | 552 | 0 | 0 | 75 | 23 | 32 | 34 | 0 | 0 | 376 | 0 | 0 | 0 | 0 | 48416 | 988 | 0 |
| SRR1636959 | 0 | 0 | 14 | 43 | 36 | 1097 | 981 | 4270 | 538 | 22 | 38 | 150 | 0 | 0 | 1271 | 6147 | 35801 | 116 | 44 | 12770 | 1925 | 0 | 0 | 213 | 85 | 729 | 7022 | 2756 | 0 | 87 | 0 | 0 | 0 | 0 | 2909 | 0 | 0 |
| SRR1636960 | 0 | 0 | 28 | 49 | 24 | 2948 | 1287 | 12212 | 349 | 12 | 21 | 59 | 0 | 0 | 647 | 2089 | 15664 | 0 | 33 | 6707 | 853 | 0 | 0 | 240 | 130 | 594 | 8208 | 4644 | 0 | 47 | 0 | 0 | 0 | 0 | 688 | 0 | 0 |
| SRR1636961 | 11 | 0 | 14 | 28 | 0 | 3516 | 1449 | 14526 | 343 | 11 | 47 | 36 | 0 | 0 | 611 | 2625 | 20196 | 0 | 24 | 4975 | 992 | 0 | 0 | 327 | 152 | 614 | 7153 | 4226 | 0 | 28 | 0 | 14 | 0 | 0 | 618 | 0 | 11 |
| SRR1636962 | 47 | 28 | 28 | 115 | 109 | 134 | 208 | 383 | 74 | 0 | 140 | 0 | 0 | 0 | 609 | 6105 | 35491 | 0 | 55 | 2286 | 2198 | 0 | 0 | 342 | 90 | 696 | 5618 | 2665 | 0 | 65 | 0 | 0 | 0 | 0 | 2345 | 0 | 0 |
| SRR1636963 | 59 | 40 | 36 | 244 | 82 | 148 | 115 | 304 | 21 | 0 | 147 | 0 | 0 | 0 | 548 | 9604 | 34971 | 21 | 33 | 2571 | 2362 | 0 | 0 | 362 | 101 | 725 | 5684 | 2210 | 0 | 171 | 0 | 16 | 0 | 0 | 3548 | 0 | 0 |
| SRR1636964 | 48 | 57 | 36 | 11 | 102 | 88 | 88 | 195 | 51 | 0 | 144 | 0 | 0 | 0 | 561 | 6144 | 23142 | 0 | 32 | 2570 | 2264 | 0 | 0 | 22 | 72 | 38 | 302 | 119 | 0 | 63 | 0 | 23 | 0 | 0 | 6659 | 0 | 0 |
| SRR1636965 | 0 | 0 | 16 | 18 | 0 | 115 | 79 | 491 | 20 | 60 | 197 | 112 | 0 | 0 | 67163 | 8990 | 345092 | 50 | 124 | 12709 | 16693 | 0 | 0 | 25 | 0 | 33 | 168 | 173 | 0 | 139 | 0 | 27 | 0 | 0 | 551 | 0 | 14 |
| SRR1636966 | 0 | 0 | 0 | 11 | 0 | 50 | 17 | 233 | 17 | 26 | 38 | 42 | 0 | 0 | 30834 | 8887 | 162793 | 50 | 100 | 8945 | 5864 | 0 | 0 | 0 | 0 | 0 | 10 | 20 | 0 | 82 | 0 | 100 | 0 | 0 | 281 | 0 | 20 |
| SRR1636967 | 0 | 14 | 0 | 83 | 18 | 2284 | 776 | 7950 | 194 | 46 | 65 | 106 | 0 | 0 | 11138 | 10919 | 110404 | 22 | 108 | 7583 | 6657 | 0 | 0 | 18 | 18 | 157 | 212 | 475 | 0 | 129 | 0 | 129 | 0 | 0 | 117 | 0 | 14 |
| SRR1636968 | 131 | 40 | 57 | 116 | 52 | 1060 | 579 | 3956 | 73 | 471 | 66 | 0 | 0 | 0 | 22529 | 8408 | 179735 | 15 | 27 | 68456 | 30615 | 0 | 0 | 1420 | 112 | 574 | 1315 | 2617 | 0 | 60 | 0 | 15 | 0 | 0 | 353 | 40 | 0 |
| SRR1636969 | 164 | 52 | 84 | 121 | 22 | 950 | 346 | 2903 | 57 | 1761 | 60 | 23 | 0 | 0 | 16381 | 10722 | 186396 | 0 | 42 | 81796 | 20106 | 0 | 0 | 887 | 84 | 515 | 1648 | 1678 | 0 | 41 | 0 | 0 | 0 | 0 | 635 | 12 | 0 |
| SRR1636970 | 183 | 65 | 151 | 179 | 31 | 503 | 274 | 1894 | 50 | 1685 | 53 | 22 | 0 | 0 | 19268 | 12445 | 191916 | 0 | 56 | 89109 | 30077 | 0 | 0 | 637 | 125 | 1191 | 5238 | 1473 | 0 | 47 | 0 | 0 | 0 | 0 | 499 | 15 | 0 |
| SRR1642941 | 0 | 0 | 0 | 0 | 79 | 9348 | 5896 | 29790 | 66309 | 55 | 321 | 97 | 0 | 0 | 14946 | 140165 | 143889 | 670 | 521 | 2044 | 196 | 0 | 0 | 27 | 90 | 73 | 22 | 19 | 0 | 1664 | 0 | 0 | 0 | 27 | 7131 | 578 | 0 |
| SRR1642942 | 13 | 0 | 0 | 0 | 93 | 9620 | 5309 | 25793 | 60591 | 105 | 410 | 73 | 0 | 0 | 8187 | 29746 | 24292 | 1087 | 465 | 685 | 1370 | 0 | 0 | 81 | 246 | 143 | 88 | 52 | 0 | 1397 | 0 | 0 | 0 | 19 | 11793 | 0 | 0 |
| SRR1642943 | 0 | 0 | 0 | 0 | 13 | 9911 | 3819 | 17345 | 45820 | 46 | 118 | 28 | 0 | 0 | 12797 | 173786 | 309977 | 457 | 577 | 14268 | 1775 | 0 | 0 | 12 | 30 | 20 | 11 | 16 | 0 | 712 | 0 | 0 | 0 | 11 | 10566 | 123 | 0 |
| SRR1642944 | 10 | 0 | 0 | 0 | 87 | 16314 | 12691 | 59424 | 136529 | 44 | 353 | 115 | 10 | 0 | 433 | 31597 | 23100 | 23 | 114 | 980 | 319 | 0 | 0 | 53 | 129 | 93 | 36 | 40 | 0 | 1269 | 0 | 0 | 0 | 61 | 3160 | 47 | 0 |
| SRR1642945 | 0 | 0 | 0 | 0 | 43 | 4614 | 2140 | 9871 | 19986 | 113 | 377 | 96 | 0 | 0 | 15163 | 82616 | 474328 | 227 | 523 | 7586 | 957 | 0 | 0 | 23 | 59 | 53 | 17 | 15 | 0 | 1906 | 0 | 0 | 0 | 21 | 7812 | 126 | 0 |
| SRR1642946 | 0 | 0 | 0 | 0 | 23 | 8166 | 3881 | 18424 | 38372 | 17 | 220 | 29 | 0 | 0 | 5707 | 128775 | 151342 | 1242 | 151 | 6274 | 2139 | 0 | 0 | 19 | 51 | 43 | 18 | 14 | 0 | 811 | 0 | 0 | 0 | 13 | 5089 | 12 | 0 |
| SRR1642947 | 0 | 0 | 0 | 0 | 23 | 16226 | 8572 | 38440 | 87969 | 14 | 175 | 59 | 0 | 0 | 2173 | 98311 | 87333 | 181 | 237 | 2968 | 237 | 0 | 0 | 15 | 40 | 35 | 11 | 21 | 0 | 800 | 0 | 0 | 0 | 12 | 5555 | 127 | 0 |
| SRR1642948 | 0 | 0 | 0 | 0 | 36 | 11654 | 6864 | 32756 | 69309 | 16 | 149 | 44 | 0 | 0 | 1015 | 78293 | 113571 | 584 | 285 | 1322 | 500 | 0 | 0 | 17 | 40 | 35 | 18 | 25 | 0 | 630 | 0 | 0 | 0 | 16 | 8081 | 68 | 0 |
| SRR1642949 | 0 | 0 | 0 | 0 | 57 | 13137 | 8837 | 41332 | 97815 | 17 | 375 | 164 | 0 | 0 | 913 | 38389 | 45120 | 135 | 156 | 2407 | 230 | 0 | 0 | 24 | 68 | 67 | 17 | 14 | 0 | 1629 | 0 | 0 | 0 | 33 | 2283 | 160 | 0 |
| SRR1642950 | 0 | 0 | 0 | 0 | 42 | 18392 | 11005 | 51203 | 111121 | 11 | 401 | 157 | 0 | 0 | 384 | 57992 | 49162 | 196 | 119 | 2269 | 293 | 0 | 0 | 49 | 113 | 83 | 18 | 21 | 0 | 1868 | 0 | 0 | 0 | 35 | 2728 | 25 | 0 |
| SRR1642951 | 0 | 0 | 0 | 0 | 20 | 13347 | 8298 | 38916 | 82224 | 12 | 102 | 41 | 0 | 0 | 835 | 40113 | 49977 | 722 | 266 | 2208 | 355 | 0 | 0 | 11 | 37 | 34 | 19 | 15 | 0 | 509 | 0 | 0 | 0 | 23 | 11155 | 413 | 0 |
| SRR1642952 | 0 | 0 | 0 | 0 | 18 | 9450 | 6483 | 32561 | 62775 | 15 | 221 | 62 | 0 | 0 | 589 | 94340 | 191091 | 1011 | 319 | 2747 | 1119 | 0 | 0 | 13 | 32 | 22 | 12 | 18 | 0 | 824 | 0 | 0 | 0 | 18 | 7270 | 79 | 0 |
| SRR1642953 | 0 | 0 | 0 | 0 | 33 | 9084 | 6096 | 30742 | 64311 | 124 | 68 | 22 | 0 | 0 | 5563 | 208598 | 176091 | 273 | 207 | 13880 | 1241 | 0 | 0 | 15 | 0 | 0 | 0 | 0 | 0 | 566 | 0 | 0 | 0 | 17 | 5009 | 176 | 0 |
| SRR1642954 | 13 | 0 | 0 | 0 | 93 | 8487 | 6114 | 34001 | 66095 | 87 | 71 | 19 | 0 | 0 | 7609 | 261349 | 179118 | 1439 | 442 | 5148 | 904 | 0 | 0 | 34 | 0 | 0 | 0 | 0 | 0 | 440 | 0 | 0 | 0 | 14 | 2392 | 149 | 0 |
| SRR1642955 | 0 | 0 | 0 | 0 | 58 | 1557 | 792 | 4822 | 13685 | 2638 | 144 | 53 | 0 | 0 | 23178 | 248659 | 477524 | 130 | 449 | 10149 | 4192 | 0 | 0 | 30 | 129 | 48 | 24 | 0 | 0 | 1500 | 0 | 0 | 0 | 24 | 1552 | 85 | 0 |
| SRR1642956 | 21 | 10 | 0 | 0 | 94 | 2452 | 1805 | 9483 | 22060 | 807 | 373 | 71 | 0 | 0 | 118821 | 168005 | 196686 | 2461 | 296 | 2896 | 2564 | 0 | 0 | 30 | 102 | 43 | 0 | 0 | 0 | 2046 | 0 | 0 | 0 | 35 | 4245 | 616 | 0 |
| SRR1642957 | 0 | 0 | 0 | 0 | 354 | 6030 | 4282 | 19963 | 47993 | 156 | 288 | 59 | 0 | 0 | 9529 | 84472 | 183988 | 709 | 783 | 4568 | 1198 | 0 | 0 | 94 | 239 | 136 | 60 | 0 | 0 | 1987 | 0 | 0 | 0 | 36 | 4036 | 223 | 0 |
| SRR1642958 | 0 | 0 | 0 | 0 | 36 | 9654 | 7044 | 35855 | 84374 | 120 | 88 | 21 | 0 | 0 | 3997 | 54873 | 66639 | 722 | 393 | 3154 | 694 | 0 | 0 | 18 | 78 | 30 | 22 | 0 | 0 | 556 | 0 | 0 | 0 | 14 | 7665 | 317 | 0 |
| SRR1642959 | 0 | 0 | 0 | 0 | 89 | 13376 | 11229 | 49688 | 118994 | 138 | 246 | 112 | 0 | 0 | 1389 | 46839 | 65557 | 649 | 325 | 3139 | 289 | 0 | 0 | 39 | 130 | 69 | 13 | 0 | 0 | 1612 | 0 | 0 | 0 | 64 | 4017 | 238 | 0 |
| SRR1642960 | 0 | 0 | 0 | 0 | 48 | 13271 | 11120 | 56426 | 132307 | 56 | 131 | 57 | 0 | 0 | 1064 | 68544 | 77117 | 1554 | 980 | 3444 | 490 | 0 | 0 | 17 | 55 | 27 | 11 | 0 | 0 | 709 | 0 | 0 | 0 | 30 | 8061 | 10 | 0 |
| SRR1642961 | 0 | 0 | 0 | 0 | 25 | 2094 | 1449 | 6561 | 15274 | 200 | 80 | 30 | 0 | 0 | 6180 | 296101 | 336016 | 145 | 383 | 5570 | 1122 | 0 | 0 | 31 | 35 | 17 | 11 | 0 | 0 | 642 | 0 | 0 | 0 | 14 | 3295 | 83 | 0 |
| SRR1642962 | 0 | 0 | 0 | 0 | 26 | 11544 | 8767 | 39203 | 64921 | 24 | 55 | 31 | 0 | 0 | 1733 | 78696 | 114349 | 585 | 251 | 4676 | 699 | 0 | 0 | 23 | 83 | 40 | 10 | 0 | 0 | 543 | 0 | 0 | 0 | 14 | 2890 | 20 | 0 |
| SRR1642963 | 0 | 0 | 0 | 0 | 83 | 12444 | 9394 | 49976 | 119913 | 272 | 179 | 88 | 0 | 0 | 2650 | 145406 | 143894 | 348 | 109 | 4501 | 239 | 0 | 0 | 146 | 519 | 211 | 51 | 0 | 0 | 1874 | 0 | 0 | 0 | 63 | 6738 | 756 | 0 |

### Supp Table S8.3.Columns_BX_to_DH

| | | | | | | | | | | | | | | | | | | | | | | | | | | | | | | | | | | | | | |
|---|---|---|---|---|---|---|---|---|---|---|---|---|---|---|---|---|---|---|---|---|---|---|---|---|---|---|---|---|---|---|---|---|---|---|---|---|---|
| SRR1642964 | 16 | 0 | 0 | 0 | 64 | 17575 | 13434 | 62154 | 128536 | 73 | 115 | 17 | 0 | 0 | 1341 | 105431 | 121407 | 369 | 292 | 2032 | 690 | 0 | 0 | 34 | 97 | 50 | 68 | 0 | 0 | 844 | 0 | 0 | 0 | 13 | 6219 | 35 | 0 |
| SRR1642965 | 0 | 0 | 0 | 0 | 13 | 15578 | 13363 | 70429 | 145349 | 93 | 15 | 31 | 0 | 0 | 1192 | 48553 | 27872 | 944 | 149 | 1182 | 288 | 0 | 0 | 15 | 17 | 0 | 0 | 0 | 0 | 74 | 0 | 0 | 0 | 20 | 11198 | 380 | 0 |
| SRR1642966 | 0 | 0 | 0 | 0 | 30 | 10882 | 6979 | 35451 | 66252 | 138 | 30 | 98 | 0 | 0 | 22460 | 54448 | 75507 | 1097 | 163 | 697 | 678 | 0 | 0 | 47 | 24 | 0 | 0 | 0 | 0 | 339 | 0 | 0 | 0 | 57 | 3907 | 98 | 0 |
| SRR1642967 | 0 | 0 | 0 | 0 | 12 | 11024 | 9002 | 44259 | 84456 | 109 | 16 | 20 | 0 | 0 | 13306 | 61837 | 185420 | 751 | 306 | 1582 | 315 | 0 | 0 | 20 | 15 | 0 | 0 | 0 | 0 | 78 | 0 | 0 | 0 | 23 | 9389 | 927 | 0 |
| SRR1642968 | 0 | 0 | 0 | 0 | 42 | 15651 | 12036 | 63797 | 132708 | 218 | 16 | 75 | 0 | 0 | 3761 | 144475 | 61253 | 430 | 49 | 932 | 276 | 0 | 0 | 66 | 41 | 0 | 0 | 0 | 0 | 247 | 0 | 0 | 0 | 66 | 6193 | 532 | 0 |
| SRR1642969 | 0 | 0 | 0 | 0 | 30 | 8369 | 3803 | 17082 | 33750 | 162 | 13 | 12 | 0 | 0 | 7687 | 410241 | 176407 | 254 | 45 | 2330 | 339 | 0 | 0 | 22 | 11 | 0 | 0 | 0 | 0 | 58 | 0 | 0 | 0 | 31 | 3451 | 357 | 0 |
| SRR1642970 | 0 | 0 | 0 | 0 | 23 | 8857 | 5157 | 27650 | 54448 | 86 | 11 | 12 | 0 | 0 | 21377 | 190325 | 360662 | 734 | 167 | 1575 | 841 | 0 | 0 | 25 | 15 | 13 | 0 | 0 | 0 | 60 | 0 | 0 | 0 | 11 | 4535 | 211 | 0 |
| SRR1642971 | 0 | 0 | 0 | 0 | 20 | 9531 | 6124 | 29630 | 70994 | 104 | 12 | 22 | 0 | 0 | 15876 | 315589 | 106849 | 547 | 47 | 1243 | 187 | 0 | 0 | 20 | 12 | 0 | 0 | 0 | 0 | 79 | 0 | 0 | 0 | 18 | 2936 | 284 | 0 |
| SRR1642972 | 0 | 0 | 0 | 0 | 16 | 9458 | 7604 | 36132 | 85040 | 70 | 22 | 20 | 0 | 0 | 2349 | 101668 | 85315 | 979 | 93 | 1463 | 1339 | 0 | 0 | 22 | 11 | 0 | 0 | 0 | 0 | 76 | 0 | 0 | 0 | 18 | 10058 | 1046 | 0 |
| SRR1642973 | 0 | 0 | 0 | 0 | 19 | 16583 | 12934 | 64211 | 135164 | 94 | 15 | 22 | 0 | 0 | 1685 | 49153 | 43766 | 399 | 105 | 1603 | 158 | 0 | 0 | 26 | 13 | 0 | 0 | 0 | 0 | 76 | 0 | 0 | 0 | 18 | 7254 | 1028 | 0 |
| SRR1642974 | 0 | 0 | 0 | 0 | 68 | 11019 | 9927 | 51323 | 110460 | 427 | 12 | 41 | 0 | 0 | 794 | 27745 | 28098 | 539 | 85 | 638 | 128 | 0 | 0 | 48 | 35 | 0 | 0 | 0 | 0 | 220 | 0 | 0 | 0 | 51 | 19794 | 139 | 0 |
| SRR1642975 | 0 | 0 | 0 | 0 | 23 | 20071 | 14226 | 60549 | 124598 | 57 | 11 | 41 | 0 | 0 | 3309 | 85890 | 45425 | 482 | 59 | 1024 | 125 | 0 | 0 | 30 | 12 | 14 | 0 | 0 | 0 | 60 | 0 | 0 | 0 | 24 | 4785 | 254 | 0 |
| SRR1642976 | 0 | 0 | 0 | 0 | 42 | 11836 | 10050 | 52112 | 109353 | 98 | 12 | 15 | 0 | 0 | 1710 | 250817 | 212210 | 1523 | 81 | 1025 | 340 | 0 | 0 | 17 | 34 | 10 | 0 | 0 | 0 | 69 | 0 | 0 | 0 | 34 | 6049 | 44 | 0 |
| SRR1642977 | 16 | 0 | 0 | 0 | 30 | 4384 | 3011 | 15594 | 38054 | 77 | 27 | 17 | 0 | 0 | 16797 | 129805 | 311909 | 403 | 1236 | 1811 | 511 | 0 | 0 | 26 | 88 | 22 | 13 | 0 | 0 | 542 | 0 | 0 | 0 | 16 | 6127 | 365 | 0 |
| SRR1642978 | 0 | 0 | 0 | 0 | 98 | 6545 | 3830 | 18151 | 36251 | 225 | 93 | 81 | 0 | 0 | 2742 | 147065 | 316657 | 278 | 1080 | 7710 | 1373 | 0 | 0 | 86 | 210 | 76 | 15 | 0 | 0 | 1082 | 0 | 0 | 0 | 56 | 3475 | 36 | 0 |
| SRR1642979 | 11 | 0 | 0 | 0 | 141 | 12377 | 8914 | 42787 | 88097 | 183 | 126 | 66 | 0 | 0 | 2040 | 77104 | 60133 | 707 | 206 | 3096 | 358 | 0 | 0 | 49 | 133 | 51 | 0 | 0 | 0 | 1482 | 0 | 0 | 0 | 44 | 6879 | 833 | 0 |
| SRR1642980 | 18 | 15 | 0 | 0 | 99 | 17653 | 11714 | 59781 | 163274 | 618 | 152 | 107 | 20 | 0 | 565 | 83553 | 72298 | 469 | 220 | 1612 | 1736 | 0 | 0 | 77 | 123 | 55 | 14 | 0 | 0 | 1171 | 0 | 0 | 0 | 108 | 11654 | 11 | 0 |
| SRR1642981 | 0 | 0 | 0 | 0 | 88 | 8042 | 5927 | 26897 | 54606 | 273 | 130 | 103 | 0 | 0 | 11886 | 105461 | 139726 | 504 | 1434 | 2065 | 381 | 0 | 0 | 39 | 70 | 35 | 0 | 0 | 0 | 1564 | 0 | 0 | 0 | 36 | 9650 | 145 | 0 |
| SRR1642982 | 14 | 0 | 0 | 0 | 156 | 2901 | 2414 | 14766 | 32904 | 650 | 126 | 72 | 0 | 0 | 3570 | 136227 | 527375 | 171 | 451 | 2466 | 591 | 0 | 0 | 107 | 220 | 96 | 32 | 0 | 0 | 1065 | 0 | 0 | 0 | 54 | 3851 | 36 | 0 |
| SRR1642983 | 0 | 0 | 0 | 0 | 99 | 5190 | 3936 | 18199 | 43374 | 406 | 52 | 44 | 0 | 0 | 6769 | 78161 | 274293 | 214 | 615 | 15006 | 3340 | 0 | 0 | 68 | 204 | 35 | 11 | 0 | 0 | 764 | 0 | 0 | 0 | 35 | 14794 | 145 | 0 |
| SRR1642984 | 12 | 0 | 0 | 0 | 48 | 9612 | 7855 | 38854 | 88931 | 122 | 40 | 11 | 0 | 0 | 3119 | 113319 | 332846 | 412 | 428 | 3881 | 1003 | 0 | 0 | 63 | 102 | 21 | 11 | 0 | 0 | 388 | 0 | 0 | 0 | 23 | 6771 | 506 | 0 |
| SRR1642985 | 0 | 0 | 0 | 0 | 39 | 7014 | 6246 | 29790 | 63641 | 189 | 48 | 39 | 0 | 0 | 3509 | 95925 | 152519 | 322 | 521 | 5215 | 906 | 0 | 0 | 27 | 56 | 26 | 17 | 0 | 0 | 468 | 0 | 0 | 0 | 13 | 6973 | 177 | 0 |
| SRR1642986 | 0 | 0 | 0 | 0 | 35 | 13971 | 10406 | 52155 | 120094 | 175 | 45 | 40 | 0 | 0 | 1884 | 34324 | 57099 | 346 | 258 | 1181 | 412 | 0 | 0 | 37 | 105 | 37 | 11 | 0 | 0 | 504 | 0 | 0 | 0 | 18 | 12219 | 165 | 0 |
| SRR1642987 | 0 | 0 | 0 | 0 | 33 | 19027 | 14521 | 70059 | 132607 | 90 | 46 | 53 | 0 | 0 | 1152 | 60891 | 65758 | 142 | 92 | 7310 | 1093 | 0 | 0 | 17 | 34 | 14 | 0 | 0 | 0 | 527 | 0 | 0 | 0 | 24 | 5189 | 41 | 0 |
| SRR1642988 | 17 | 0 | 0 | 0 | 88 | 6477 | 6602 | 36930 | 87683 | 576 | 127 | 60 | 0 | 0 | 1832 | 196198 | 81896 | 594 | 119 | 2501 | 2351 | 0 | 0 | 96 | 236 | 112 | 16 | 0 | 0 | 1024 | 0 | 0 | 0 | 36 | 10136 | 141 | 0 |
| SRR1657194 | 69 | 106 | 22 | 65 | 22 | 1275 | 478 | 2719 | 133 | 0 | 45 | 0 | 0 | 0 | 26650 | 10682 | 52307 | 0 | 35 | 11636 | 659 | 0 | 0 | 48 | 139 | 19 | 54 | 82 | 0 | 31 | 0 | 17 | 0 | 0 | 2645 | 0 | 0 |
| SRR1657195 | 29 | 36 | 17 | 12 | 27 | 1490 | 632 | 3802 | 249 | 20 | 0 | 0 | 13 | 0 | 32430 | 10162 | 56884 | 19 | 30 | 11205 | 1737 | 0 | 0 | 85 | 16 | 20 | 44 | 73 | 0 | 24 | 0 | 11 | 0 | 0 | 2543 | 0 | 0 |
| SRR1657196 | 139 | 166 | 21 | 56 | 143 | 1126 | 536 | 2744 | 132 | 0 | 17 | 0 | 0 | 0 | 16745 | 8715 | 31678 | 18 | 47 | 15143 | 1398 | 0 | 0 | 530 | 105 | 212 | 425 | 77 | 0 | 295 | 0 | 18 | 0 | 0 | 2741 | 0 | 0 |
| SRR1657197 | 180 | 41 | 111 | 235 | 111 | 589 | 177 | 1665 | 162 | 0 | 111 | 0 | 0 | 0 | 22352 | 9964 | 42321 | 0 | 0 | 8060 | 1532 | 0 | 0 | 553 | 332 | 788 | 2432 | 1119 | 0 | 732 | 0 | 0 | 0 | 0 | 2491 | 0 | 0 |
| SRR1657198 | 21 | 25 | 11 | 0 | 38 | 1448 | 628 | 2456 | 316 | 27 | 0 | 0 | 0 | 0 | 44907 | 4696 | 44657 | 0 | 0 | 1076 | 2038 | 0 | 0 | 47 | 34 | 45 | 11 | 0 | 0 | 18 | 0 | 0 | 0 | 0 | 2542 | 0 | 0 |
| SRR1686806 | 0 | 0 | 0 | 0 | 0 | 29795 | 831 | 6470 | 125 | 0 | 79 | 24 | 0 | 0 | 0 | 0 | 0 | 46 | 145 | 3702 | 5604 | 0 | 0 | 0 | 12 | 48 | 32 | 0 | 0 | 107 | 0 | 0 | 0 | 0 | 1932 | 11 | 0 |
| SRR1707574 | 0 | 0 | 16 | 86 | 0 | 397 | 31 | 74 | 13 | 0 | 0 | 0 | 0 | 0 | 21698 | 30712 | 36955 | 19259 | 0 | 57 | 39 | 0 | 0 | 24 | 0 | 48 | 101 | 77 | 0 | 0 | 0 | 11 | 0 | 0 | 0 | 0 | 92 |
| SRR1707575 | 13 | 54 | 0 | 0 | 133 | 0 | 127 | 0 | 245 | 0 | 208 | 0 | 0 | 0 | 978 | 206331 | 1624 | 37 | 14 | 76 | 11 | 0 | 0 | 0 | 0 | 26 | 13 | 37 | 0 | 60 | 0 | 10 | 0 | 0 | 2069 | 0 | 67 |
| SRR1708906 | 220 | 58 | 81 | 18 | 65 | 306 | 225 | 702 | 81 | 16 | 102 | 10 | 0 | 0 | 1584 | 23528 | 19683 | 36 | 0 | 6550 | 1323 | 0 | 0 | 348 | 204 | 633 | 2045 | 144 | 0 | 659 | 0 | 171 | 0 | 0 | 5252 | 0 | 31 |
| SRR1708907 | 106 | 44 | 0 | 0 | 56 | 204 | 182 | 623 | 97 | 0 | 0 | 15 | 12 | 0 | 1106 | 15784 | 10019 | 129 | 32 | 3011 | 1685 | 0 | 0 | 31 | 131 | 128 | 28 | 0 | 0 | 12 | 0 | 242 | 0 | 0 | 70808 | 0 | 0 |
| SRR1708908 | 50 | 29 | 0 | 0 | 38 | 267 | 216 | 642 | 123 | 0 | 0 | 0 | 0 | 0 | 657 | 35138 | 5018 | 0 | 0 | 1162 | 1075 | 0 | 0 | 106 | 58 | 84 | 182 | 0 | 0 | 43 | 0 | 72 | 0 | 0 | 65639 | 0 | 0 |
| SRR1708909 | 37 | 10 | 14 | 65 | 56 | 192 | 216 | 533 | 88 | 0 | 80 | 0 | 0 | 0 | 924 | 5415 | 12990 | 17 | 0 | 4987 | 2495 | 0 | 0 | 28 | 78 | 24 | 85 | 58 | 0 | 63 | 0 | 121 | 0 | 0 | 68930 | 0 | 0 |
| SRR1708910 | 57 | 11 | 0 | 0 | 56 | 169 | 188 | 335 | 96 | 11 | 0 | 13 | 0 | 0 | 328 | 31131 | 4304 | 122 | 0 | 954 | 1264 | 0 | 0 | 21 | 56 | 72 | 119 | 17 | 0 | 54 | 0 | 317 | 0 | 0 | 79536 | 0 | 0 |
| SRR1708911 | 45 | 0 | 0 | 0 | 20 | 421 | 239 | 569 | 68 | 24 | 0 | 12 | 0 | 0 | 159 | 20671 | 3795 | 46 | 0 | 569 | 546 | 0 | 0 | 16 | 16 | 53 | 110 | 12 | 0 | 45 | 0 | 91 | 0 | 0 | 54700 | 0 | 0 |
| SRR1708912 | 64 | 20 | 0 | 0 | 40 | 494 | 202 | 494 | 61 | 0 | 0 | 0 | 12 | 0 | 918 | 5792 | 10788 | 40 | 10 | 3086 | 1170 | 0 | 0 | 375 | 35 | 58 | 572 | 23 | 0 | 245 | 0 | 61 | 0 | 0 | 7726 | 0 | 0 |
| SRR1708913 | 42 | 12 | 0 | 0 | 12 | 408 | 160 | 461 | 68 | 0 | 0 | 0 | 0 | 0 | 287 | 2420 | 3196 | 97 | 0 | 898 | 481 | 0 | 0 | 15 | 16 | 47 | 150 | 0 | 0 | 94 | 0 | 146 | 0 | 0 | 67772 | 0 | 0 |
| SRR1708914 | 104 | 15 | 0 | 0 | 75 | 405 | 217 | 405 | 29 | 0 | 0 | 35 | 0 | 0 | 723 | 1725 | 10583 | 14 | 0 | 2995 | 1360 | 0 | 0 | 134 | 79 | 308 | 15 | 25 | 0 | 179 | 0 | 0 | 0 | 0 | 10312 | 0 | 0 |
| SRR1708915 | 85 | 30 | 22 | 19 | 48 | 342 | 279 | 475 | 56 | 0 | 24 | 0 | 0 | 0 | 1068 | 3289 | 18826 | 77 | 35 | 6152 | 5768 | 0 | 0 | 320 | 63 | 139 | 836 | 19 | 0 | 383 | 0 | 28 | 0 | 0 | 13046 | 0 | 0 |
| SRR1708916 | 46 | 14 | 0 | 0 | 62 | 314 | 188 | 458 | 63 | 0 | 0 | 0 | 0 | 0 | 206 | 2231 | 4269 | 0 | 0 | 1382 | 1095 | 0 | 0 | 482 | 14 | 43 | 95 | 11 | 0 | 106 | 0 | 18 | 0 | 0 | 11751 | 0 | 0 |
| SRR1708917 | 71 | 0 | 0 | 0 | 80 | 251 | 173 | 494 | 102 | 0 | 17 | 0 | 0 | 0 | 267 | 2446 | 6418 | 16 | 0 | 4636 | 588 | 0 | 0 | 262 | 47 | 311 | 629 | 26 | 0 | 445 | 0 | 16 | 0 | 0 | 11363 | 0 | 0 |
| SRR1708918 | 26 | 0 | 0 | 0 | 39 | 354 | 243 | 619 | 55 | 0 | 0 | 0 | 0 | 0 | 2221 | 6400 | 16221 | 44 | 0 | 1404 | 1105 | 0 | 0 | 451 | 26 | 16 | 65 | 0 | 0 | 78 | 0 | 0 | 0 | 0 | 4159 | 0 | 0 |
| SRR1708919 | 69 | 14 | 0 | 0 | 94 | 485 | 457 | 840 | 93 | 0 | 0 | 0 | 0 | 0 | 4058 | 9411 | 50849 | 0 | 0 | 6185 | 4039 | 0 | 0 | 13 | 69 | 105 | 11 | 14 | 0 | 207 | 0 | 47 | 0 | 0 | 5151 | 0 | 0 |
| SRR1708920 | 32 | 12 | 0 | 0 | 87 | 540 | 230 | 685 | 127 | 0 | 0 | 0 | 0 | 0 | 3068 | 6566 | 31665 | 61 | 18 | 4971 | 3892 | 0 | 0 | 148 | 105 | 215 | 239 | 19 | 0 | 128 | 0 | 12 | 0 | 0 | 10516 | 0 | 0 |
| SRR1708921 | 47 | 0 | 0 | 16 | 111 | 426 | 234 | 625 | 122 | 12 | 0 | 45 | 0 | 0 | 4629 | 12434 | 53338 | 87 | 0 | 7043 | 7981 | 0 | 0 | 530 | 117 | 162 | 516 | 24 | 0 | 196 | 0 | 0 | 0 | 0 | 5706 | 0 | 0 |
| SRR1708922 | 49 | 0 | 0 | 0 | 100 | 361 | 280 | 582 | 103 | 0 | 0 | 0 | 0 | 0 | 1260 | 5276 | 21707 | 0 | 0 | 4760 | 3419 | 0 | 0 | 20 | 72 | 267 | 22 | 56 | 0 | 12 | 0 | 22 | 0 | 0 | 5477 | 0 | 0 |
| SRR1708923 | 76 | 33 | 0 | 12 | 152 | 1216 | 501 | 775 | 215 | 0 | 0 | 0 | 0 | 0 | 3361 | 19973 | 50949 | 72 | 48 | 13290 | 6663 | 0 | 0 | 16 | 149 | 11 | 34 | 15 | 0 | 17 | 0 | 12 | 0 | 0 | 7009 | 0 | 0 |
| SRR1708924 | 10 | 70 | 14 | 0 | 61 | 234 | 220 | 846 | 126 | 0 | 15 | 19 | 0 | 0 | 2716 | 26305 | 21563 | 30 | 37 | 14200 | 1537 | 0 | 0 | 28 | 31 | 28 | 57 | 34 | 0 | 20 | 0 | 74 | 0 | 0 | 42231 | 0 | 0 |
| SRR1708925 | 75 | 23 | 0 | 11 | 75 | 124 | 261 | 655 | 110 | 0 | 37 | 0 | 0 | 0 | 5746 | 49524 | 57263 | 37 | 55 | 19566 | 6232 | 0 | 0 | 29 | 13 | 31 | 47 | 29 | 0 | 24 | 0 | 243 | 0 | 0 | 29115 | 0 | 11 |
| SRR1708926 | 133 | 34 | 36 | 73 | 52 | 83 | 138 | 317 | 48 | 0 | 107 | 0 | 0 | 0 | 1817 | 35665 | 13269 | 89 | 62 | 6334 | 3826 | 0 | 0 | 144 | 27 | 132 | 248 | 27 | 0 | 31 | 0 | 200 | 0 | 0 | 51550 | 0 | 0 |
| SRR1708927 | 116 | 50 | 18 | 16 | 93 | 258 | 409 | 766 | 205 | 0 | 70 | 11 | 0 | 0 | 3798 | 33517 | 42692 | 61 | 30 | 8992 | 8097 | 0 | 0 | 471 | 188 | 390 | 1910 | 45 | 0 | 646 | 0 | 136 | 0 | 0 | 11703 | 0 | 0 |
| SRR1708928 | 112 | 93 | 0 | 0 | 62 | 452 | 331 | 1220 | 158 | 12 | 14 | 14 | 0 | 0 | 2298 | 93437 | 23511 | 0 | 0 | 6132 | 2960 | 0 | 0 | 301 | 234 | 163 | 483 | 0 | 0 | 277 | 0 | 0 | 0 | 0 | 6756 | 0 | 19 |
| SRR1708929 | 82 | 34 | 0 | 0 | 65 | 292 | 264 | 639 | 182 | 0 | 14 | 0 | 0 | 0 | 793 | 39146 | 12082 | 33 | 0 | 4048 | 1432 | 0 | 0 | 11 | 139 | 160 | 24 | 13 | 0 | 14 | 0 | 61 | 0 | 0 | 24331 | 0 | 0 |
| SRR1708930 | 56 | 13 | 35 | 0 | 49 | 730 | 158 | 561 | 37 | 10 | 22 | 10 | 0 | 0 | 2548 | 9747 | 23478 | 44 | 0 | 4784 | 2346 | 0 | 0 | 19 | 116 | 15 | 48 | 40 | 0 | 20 | 0 | 81 | 0 | 0 | 10388 | 0 | 15 |
| SRR1708931 | 67 | 15 | 21 | 0 | 33 | 392 | 110 | 491 | 44 | 0 | 31 | 13 | 0 | 0 | 1447 | 6371 | 16290 | 33 | 11 | 5285 | 3114 | 0 | 0 | 356 | 167 | 792 | 2527 | 90 | 0 | 500 | 0 | 110 | 0 | 0 | 21019 | 0 | 21 |
| SRR1708932 | 48 | 0 | 0 | 0 | 23 | 595 | 191 | 825 | 112 | 0 | 0 | 0 | 0 | 0 | 601 | 4416 | 6293 | 152 | 0 | 1269 | 904 | 0 | 0 | 184 | 45 | 107 | 13 | 11 | 0 | 102 | 0 | 11 | 0 | 0 | 9167 | 0 | 0 |
| SRR1708933 | 57 | 70 | 15 | 0 | 42 | 297 | 279 | 496 | 108 | 0 | 0 | 0 | 0 | 0 | 975 | 3112 | 12463 | 11 | 0 | 2081 | 1620 | 0 | 0 | 707 | 76 | 115 | 453 | 0 | 0 | 212 | 0 | 23 | 0 | 0 | 11245 | 0 | 0 |
| SRR1708934 | 102 | 57 | 19 | 24 | 59 | 152 | 117 | 270 | 86 | 0 | 26 | 0 | 0 | 0 | 514 | 3038 | 7990 | 41 | 16 | 5590 | 1006 | 0 | 0 | 423 | 146 | 610 | 4314 | 169 | 0 | 582 | 0 | 29 | 0 | 0 | 16389 | 0 | 0 |
| SRR1708935 | 31 | 0 | 0 | 0 | 42 | 412 | 133 | 383 | 64 | 0 | 0 | 0 | 0 | 0 | 273 | 2870 | 4395 | 23 | 0 | 2048 | 290 | 0 | 0 | 253 | 12 | 29 | 96 | 12 | 0 | 62 | 0 | 29 | 0 | 0 | 8928 | 0 | 0 |
| SRR1708936 | 43 | 0 | 32 | 43 | 156 | 490 | 198 | 714 | 76 | 0 | 24 | 0 | 0 | 0 | 8376 | 11194 | 53124 | 45 | 22 | 4365 | 8223 | 0 | 0 | 381 | 261 | 561 | 1447 | 129 | 0 | 341 | 0 | 54 | 0 | 0 | 3267 | 0 | 0 |
| SRR1708937 | 0 | 0 | 0 | 0 | 24 | 720 | 265 | 749 | 83 | 10 | 0 | 0 | 0 | 0 | 560 | 1444 | 5020 | 41 | 0 | 442 | 236 | 0 | 0 | 22 | 18 | 18 | 80 | 14 | 0 | 10 | 0 | 29 | 0 | 0 | 11976 | 0 | 0 |
| SRR1708938 | 19 | 19 | 0 | 0 | 59 | 1093 | 511 | 1352 | 150 | 0 | 0 | 0 | 0 | 0 | 2859 | 6875 | 18520 | 30 | 0 | 799 | 2457 | 0 | 0 | 235 | 81 | 87 | 142 | 0 | 0 | 34 | 0 | 24 | 0 | 0 | 7398 | 0 | 0 |
| SRR1708939 | 39 | 0 | 0 | 0 | 101 | 810 | 486 | 1134 | 155 | 0 | 0 | 0 | 0 | 0 | 5521 | 9337 | 40509 | 15 | 15 | 2120 | 5992 | 0 | 0 | 20 | 111 | 59 | 114 | 0 | 0 | 107 | 0 | 15 | 0 | 0 | 15456 | 0 | 0 |
| SRR1708940 | 33 | 0 | 0 | 0 | 110 | 684 | 419 | 717 | 135 | 0 | 0 | 0 | 0 | 0 | 5608 | 10247 | 41634 | 39 | 0 | 3423 | 5170 | 0 | 0 | 448 | 112 | 39 | 141 | 12 | 0 | 141 | 0 | 0 | 0 | 0 | 7985 | 0 | 0 |
| SRR1708941 | 41 | 11 | 0 | 0 | 114 | 785 | 317 | 838 | 171 | 0 | 0 | 0 | 0 | 0 | 3611 | 17903 | 43515 | 39 | 0 | 3407 | 4513 | 0 | 0 | 15 | 71 | 54 | 148 | 0 | 0 | 114 | 0 | 49 | 0 | 0 | 14704 | 0 | 0 |
| SRR1708942 | 158 | 60 | 0 | 0 | 60 | 559 | 427 | 1290 | 214 | 0 | 11 | 23 | 11 | 0 | 3336 | 60946 | 29414 | 41 | 16 | 8637 | 748 | 0 | 0 | 48 | 24 | 14 | 26 | 17 | 0 | 10 | 0 | 148 | 0 | 0 | 22507 | 0 | 17 |
| SRR1708943 | 31 | 0 | 0 | 0 | 23 | 172 | 188 | 394 | 61 | 0 | 0 | 0 | 0 | 0 | 626 | 12302 | 6252 | 81 | 0 | 1167 | 1329 | 0 | 0 | 40 | 27 | 29 | 71 | 0 | 0 | 30 | 0 | 114 | 0 | 0 | 54058 | 0 | 0 |
| SRR1708944 | 99 | 19 | 0 | 0 | 51 | 141 | 194 | 433 | 80 | 0 | 0 | 32 | 0 | 0 | 1778 | 37604 | 14251 | 15 | 11 | 3997 | 1657 | 0 | 0 | 228 | 199 | 234 | 331 | 16 | 0 | 75 | 0 | 205 | 0 | 0 | 53383 | 0 | 13 |
| SRR1708945 | 101 | 29 | 21 | 32 | 98 | 279 | 457 | 807 | 127 | 0 | 69 | 19 | 0 | 0 | 2356 | 35501 | 21246 | 10 | 41 | 11119 | 4778 | 0 | 0 | 26 | 23 | 46 | 104 | 50 | 0 | 28 | 0 | 310 | 0 | 0 | 53637 | 0 | 0 |
| SRR1708946 | 66 | 23 | 0 | 0 | 63 | 624 | 492 | 1834 | 252 | 23 | 0 | 0 | 0 | 0 | 4258 | 116434 | 31366 | 40 | 13 | 9613 | 1973 | 0 | 0 | 25 | 19 | 12 | 212 | 0 | 0 | 181 | 0 | 80 | 0 | 0 | 13157 | 0 | 0 |
| SRR1708947 | 83 | 27 | 35 | 20 | 86 | 133 | 188 | 415 | 94 | 0 | 69 | 15 | 0 | 0 | 1322 | 31701 | 14257 | 107 | 50 | 7505 | 2163 | 0 | 0 | 469 | 824 | 1879 | 2473 | 36 | 0 | 547 | 0 | 183 | 0 | 0 | 47679 | 0 | 0 |
| SRR1708948 | 30 | 0 | 0 | 0 | 36 | 1034 | 331 | 984 | 89 | 0 | 0 | 11 | 0 | 0 | 3433 | 15358 | 27784 | 54 | 0 | 3339 | 1835 | 0 | 0 | 334 | 90 | 54 | 137 | 0 | 0 | 146 | 0 | 84 | 0 | 0 | 13773 | 0 | 17 |
| SRR1708949 | 65 | 20 | 0 | 0 | 51 | 328 | 209 | 532 | 67 | 0 | 14 | 0 | 0 | 0 | 1903 | 6884 | 15536 | 35 | 40 | 3097 | 2946 | 0 | 0 | 20 | 148 | 13 | 38 | 20 | 0 | 14 | 0 | 40 | 0 | 0 | 56665 | 0 | 0 |
| SRR1708950 | 43 | 0 | 0 | 0 | 21 | 767 | 203 | 661 | 123 | 0 | 0 | 0 | 0 | 0 | 2646 | 11677 | 23021 | 123 | 0 | 4696 | 5578 | 0 | 0 | 101 | 106 | 115 | 114 | 0 | 0 | 98 | 0 | 62 | 0 | 0 | 23098 | 0 | 0 |
| SRR1708951 | 54 | 0 | 41 | 14 | 41 | 636 | 280 | 1133 | 130 | 0 | 23 | 16 | 0 | 0 | 3259 | 10758 | 39283 | 19 | 0 | 4358 | 6185 | 0 | 0 | 484 | 310 | 637 | 1379 | 84 | 0 | 316 | 0 | 24 | 0 | 0 | 9494 | 0 | 0 |
| SRR1708952 | 37 | 0 | 0 | 0 | 59 | 565 | 200 | 1104 | 108 | 15 | 0 | 0 | 0 | 0 | 1331 | 8974 | 17636 | 19 | 12 | 3870 | 2716 | 0 | 0 | 16 | 255 | 10 | 11 | 29 | 0 | 191 | 0 | 35 | 0 | 0 | 5374 | 0 | 15 |
| SRR1708953 | 39 | 18 | 0 | 0 | 49 | 918 | 309 | 918 | 150 | 0 | 14 | 0 | 0 | 0 | 1363 | 13545 | 22033 | 62 | 18 | 4006 | 1915 | 0 | 0 | 288 | 208 | 231 | 612 | 25 | 0 | 292 | 0 | 44 | 0 | 0 | 3641 | 0 | 0 |
| SRR1708954 | 29 | 11 | 0 | 0 | 66 | 814 | 262 | 956 | 106 | 0 | 0 | 0 | 0 | 0 | 5058 | 6761 | 34901 | 18 | 0 | 1729 | 2561 | 0 | 0 | 333 | 101 | 69 | 128 | 11 | 0 | 73 | 0 | 32 | 0 | 0 | 3107 | 0 | 0 |
| SRR1708955 | 24 | 0 | 0 | 0 | 114 | 582 | 263 | 566 | 116 | 0 | 0 | 0 | 0 | 0 | 2635 | 6393 | 28957 | 45 | 0 | 2538 | 1353 | 0 | 0 | 141 | 53 | 52 | 107 | 0 | 0 | 192 | 0 | 35 | 0 | 0 | 4945 | 0 | 0 |
| SRR1708956 | 17 | 0 | 15 | 0 | 101 | 930 | 190 | 1155 | 99 | 0 | 13 | 0 | 0 | 0 | 3293 | 7628 | 30313 | 28 | 0 | 2616 | 3138 | 0 | 0 | 96 | 266 | 603 | 686 | 51 | 0 | 160 | 0 | 39 | 0 | 0 | 5563 | 0 | 0 |
| SRR1708957 | 30 | 14 | 25 | 51 | 163 | 783 | 255 | 707 | 63 | 0 | 46 | 0 | 0 | 0 | 4888 | 6469 | 44251 | 13 | 0 | 3133 | 5743 | 0 | 0 | 20 | 13 | 22 | 57 | 62 | 0 | 15 | 0 | 22 | 0 | 0 | 3595 | 0 | 0 |
| SRR1708958 | 0 | 0 | 0 | 0 | 74 | 926 | 293 | 1261 | 178 | 0 | 0 | 0 | 0 | 0 | 2613 | 5674 | 25536 | 52 | 17 | 1970 | 2627 | 0 | 0 | 260 | 140 | 96 | 209 | 25 | 0 | 113 | 0 | 17 | 0 | 0 | 5914 | 0 | 0 |
| SRR1708959 | 32 | 14 | 0 | 0 | 130 | 573 | 156 | 898 | 117 | 0 | 0 | 0 | 0 | 0 | 2793 | 7097 | 33987 | 21 | 21 | 3604 | 2075 | 0 | 0 | 214 | 181 | 154 | 343 | 23 | 0 | 360 | 0 | 14 | 0 | 0 | 4538 | 0 | 0 |
| SRR1708960 | 159 | 35 | 39 | 27 | 49 | 345 | 282 | 1130 | 62 | 18 | 45 | 33 | 0 | 0 | 4947 | 41123 | 49232 | 28 | 83 | 14262 | 1695 | 0 | 0 | 404 | 497 | 879 | 1364 | 65 | 0 | 292 | 0 | 48 | 0 | 0 | 4004 | 0 | 10 |
| SRR1708961 | 171 | 75 | 14 | 25 | 54 | 200 | 225 | 900 | 136 | 23 | 51 | 18 | 0 | 0 | 4545 | 35159 | 57995 | 25 | 57 | 19030 | 8336 | 0 | 0 | 575 | 429 | 880 | 2996 | 59 | 0 | 826 | 0 | 57 | 0 | 0 | 14237 | 0 | 20 |
| SRR1708962 | 80 | 20 | 0 | 0 | 47 | 549 | 319 | 1115 | 324 | 14 | 0 | 10 | 10 | 0 | 5923 | 89841 | 34161 | 24 | 18 | 7751 | 3711 | 0 | 0 | 12 | 15 | 18 | 27 | 22 | 0 | 164 | 0 | 59 | 0 | 0 | 12217 | 0 | 0 |
| SRR1708963 | 99 | 54 | 0 | 0 | 29 | 216 | 211 | 639 | 86 | 0 | 21 | 12 | 0 | 0 | 2513 | 26722 | 19286 | 34 | 0 | 4209 | 3541 | 0 | 0 | 31 | 11 | 182 | 22 | 10 | 0 | 12 | 0 | 178 | 0 | 0 | 54206 | 0 | 21 |
| SRR1708964 | 192 | 41 | 15 | 0 | 87 | 371 | 274 | 1090 | 96 | 23 | 11 | 49 | 0 | 0 | 5635 | 94307 | 44998 | 15 | 22 | 12108 | 3470 | 0 | 0 | 14 | 11 | 12 | 16 | 23 | 0 | 233 | 0 | 96 | 0 | 0 | 17718 | 0 | 0 |
| SRR1708965 | 45 | 0 | 0 | 0 | 71 | 890 | 496 | 1460 | 219 | 0 | 0 | 0 | 0 | 0 | 2832 | 105680 | 32363 | 15 | 0 | 6408 | 1730 | 0 | 0 | 256 | 77 | 65 | 96 | 14 | 0 | 63 | 0 | 0 | 0 | 0 | 8833 | 0 | 22 |
| SRR1708966 | 65 | 11 | 44 | 15 | 33 | 863 | 181 | 878 | 91 | 11 | 33 | 17 | 0 | 0 | 4202 | 16784 | 35874 | 48 | 14 | 5771 | 2251 | 0 | 0 | 293 | 312 | 561 | 1651 | 59 | 0 | 487 | 0 | 48 | 0 | 0 | 5654 | 0 | 11 |
| SRR1708967 | 23 | 10 | 0 | 0 | 15 | 699 | 282 | 975 | 141 | 27 | 0 | 10 | 0 | 0 | 938 | 8122 | 9654 | 166 | 0 | 1104 | 975 | 0 | 0 | 17 | 46 | 31 | 136 | 0 | 0 | 79 | 0 | 67 | 0 | 0 | 18024 | 0 | 15 |
| SRR1708968 | 16 | 19 | 0 | 0 | 25 | 1223 | 376 | 1348 | 288 | 0 | 0 | 0 | 0 | 0 | 1342 | 9171 | 12758 | 19 | 0 | 1141 | 1411 | 0 | 0 | 291 | 87 | 103 | 197 | 0 | 0 | 62 | 0 | 19 | 0 | 0 | 6824 | 0 | 0 |
| SRR1708969 | 49 | 25 | 0 | 0 | 19 | 601 | 338 | 1001 | 124 | 0 | 0 | 0 | 0 | 0 | 2962 | 6902 | 31667 | 14 | 21 | 3183 | 3038 | 0 | 0 | 451 | 142 | 80 | 328 | 0 | 0 | 182 | 0 | 0 | 0 | 0 | 10220 | 0 | 0 |
| SRR1708970 | 35 | 0 | 0 | 0 | 30 | 962 | 305 | 1126 | 137 | 0 | 0 | 0 | 0 | 0 | 1172 | 9104 | 15607 | 34 | 0 | 2416 | 1214 | 0 | 0 | 29 | 61 | 30 | 40 | 0 | 0 | 58 | 0 | 31 | 0 | 0 | 7459 | 0 | 0 |
| SRR1708971 | 22 | 0 | 0 | 0 | 25 | 874 | 667 | 1334 | 214 | 0 | 0 | 0 | 0 | 0 | 824 | 11332 | 13274 | 0 | 0 | 1151 | 340 | 0 | 0 | 235 | 30 | 19 | 64 | 25 | 0 | 58 | 0 | 50 | 0 | 0 | 9217 | 0 | 0 |
| SRR1708972 | 18 | 0 | 0 | 0 | 52 | 1658 | 392 | 1540 | 164 | 15 | 0 | 0 | 0 | 0 | 6009 | 9855 | 41269 | 109 | 0 | 1335 | 1640 | 0 | 0 | 23 | 87 | 20 | 44 | 0 | 0 | 62 | 0 | 36 | 0 | 0 | 6665 | 0 | 18 |
| SRR1708973 | 0 | 0 | 0 | 0 | 12 | 1356 | 267 | 1561 | 156 | 11 | 0 | 0 | 0 | 0 | 863 | 2485 | 8036 | 12 | 0 | 218 | 111 | 0 | 0 | 325 | 12 | 0 | 12 | 0 | 0 | 14 | 0 | 0 | 0 | 0 | 4588 | 0 | 0 |
| SRR1708974 | 17 | 0 | 0 | 0 | 74 | 1195 | 248 | 927 | 136 | 0 | 0 | 0 | 0 | 0 | 2616 | 4482 | 19166 | 20 | 0 | 1498 | 2226 | 0 | 0 | 80 | 106 | 67 | 89 | 19 | 0 | 79 | 0 | 12 | 0 | 0 | 3426 | 0 | 15 |
| SRR1708975 | 32 | 0 | 0 | 0 | 106 | 1146 | 238 | 981 | 170 | 0 | 0 | 0 | 0 | 0 | 3975 | 5489 | 31427 | 60 | 0 | 2095 | 2659 | 0 | 0 | 14 | 73 | 53 | 69 | 0 | 0 | 106 | 0 | 0 | 0 | 0 | 5980 | 0 | 11 |
| SRR1708976 | 17 | 0 | 0 | 0 | 87 | 707 | 257 | 1161 | 140 | 15 | 0 | 14 | 14 | 0 | 4581 | 8754 | 36584 | 54 | 0 | 2271 | 2636 | 0 | 0 | 356 | 163 | 92 | 97 | 0 | 0 | 172 | 0 | 13 | 0 | 0 | 4922 | 0 | 0 |
| SRR1708977 | 31 | 0 | 18 | 0 | 108 | 656 | 181 | 766 | 102 | 0 | 55 | 0 | 0 | 0 | 5297 | 14244 | 60565 | 16 | 0 | 3902 | 2587 | 0 | 0 | 10 | 12 | 26 | 54 | 20 | 0 | 32 | 0 | 22 | 0 | 0 | 4737 | 0 | 0 |
| SRR1708978 | 103 | 34 | 0 | 0 | 107 | 377 | 286 | 594 | 65 | 0 | 0 | 21 | 0 | 0 | 1764 | 79585 | 11598 | 39 | 17 | 2102 | 520 | 0 | 0 | 31 | 23 | 13 | 14 | 0 | 0 | 10 | 0 | 91 | 0 | 0 | 61927 | 0 | 0 |
| SRR1708979 | 123 | 75 | 0 | 0 | 43 | 80 | 111 | 296 | 38 | 0 | 13 | 17 | 0 | 0 | 828 | 18009 | 7125 | 17 | 31 | 2368 | 1158 | 0 | 0 | 491 | 248 | 469 | 1137 | 30 | 0 | 254 | 0 | 247 | 0 | 0 | 51812 | 0 | 0 |
| SRR1708980 | 65 | 0 | 0 | 0 | 25 | 620 | 417 | 898 | 241 | 0 | 0 | 0 | 0 | 0 | 1973 | 48238 | 15123 | 19 | 0 | 2834 | 2500 | 0 | 0 | 71 | 67 | 123 | 129 | 0 | 0 | 163 | 0 | 83 | 0 | 0 | 16419 | 0 | 0 |
| SRR1708981 | 63 | 32 | 0 | 0 | 52 | 218 | 205 | 430 | 91 | 0 | 0 | 0 | 0 | 0 | 418 | 7140 | 4269 | 108 | 19 | 1880 | 1434 | 0 | 0 | 27 | 52 | 65 | 18 | 25 | 0 | 10 | 0 | 68 | 0 | 0 | 70223 | 0 | 0 |
| SRR1708982 | 95 | 15 | 0 | 0 | 76 | 1009 | 513 | 1690 | 151 | 24 | 15 | 27 | 0 | 0 | 2161 | 107984 | 14708 | 17 | 0 | 3558 | 866 | 0 | 0 | 381 | 230 | 242 | 239 | 37 | 0 | 308 | 0 | 50 | 0 | 0 | 9790 | 0 | 0 |
| SRR1708983 | 79 | 31 | 14 | 11 | 92 | 200 | 139 | 361 | 67 | 11 | 39 | 0 | 0 | 0 | 433 | 18896 | 4743 | 178 | 44 | 2958 | 910 | 0 | 0 | 269 | 175 | 423 | 1204 | 43 | 0 | 413 | 0 | 264 | 0 | 0 | 69713 | 0 | 0 |
| SRR1708984 | 85 | 38 | 0 | 0 | 27 | 387 | 114 | 364 | 61 | 0 | 0 | 0 | 0 | 0 | 1100 | 3747 | 9789 | 15 | 23 | 2853 | 895 | 0 | 0 | 222 | 33 | 99 | 18 | 25 | 0 | 189 | 0 | 175 | 0 | 0 | 15760 | 0 | 14 |
| SRR1708985 | 60 | 13 | 32 | 36 | 52 | 340 | 172 | 325 | 47 | 0 | 48 | 13 | 0 | 0 | 1560 | 7492 | 17299 | 113 | 31 | 6421 | 3203 | 0 | 0 | 213 | 99 | 536 | 2175 | 114 | 0 | 631 | 0 | 117 | 0 | 0 | 22783 | 0 | 20 |
| SRR1708986 | 84 | 20 | 0 | 10 | 71 | 270 | 270 | 525 | 93 | 0 | 27 | 0 | 0 | 0 | 718 | 1890 | 5588 | 15 | 15 | 4839 | 1304 | 0 | 0 | 81 | 128 | 39 | 77 | 47 | 0 | 15 | 0 | 62 | 0 | 0 | 15414 | 0 | 0 |
| SRR1708987 | 79 | 34 | 0 | 0 | 71 | 191 | 108 | 172 | 32 | 0 | 0 | 0 | 0 | 0 | 992 | 2490 | 9124 | 19 | 13 | 3668 | 2028 | 0 | 0 | 14 | 116 | 266 | 36 | 60 | 0 | 12 | 0 | 25 | 0 | 0 | 8655 | 0 | 11 |
| SRR1708988 | 26 | 14 | 0 | 0 | 39 | 898 | 318 | 831 | 124 | 14 | 11 | 26 | 0 | 0 | 1919 | 5179 | 15742 | 85 | 0 | 2576 | 1191 | 0 | 0 | 419 | 90 | 72 | 269 | 20 | 0 | 238 | 0 | 89 | 0 | 0 | 12910 | 0 | 13 |
| SRR1708989 | 51 | 13 | 0 | 0 | 28 | 956 | 326 | 892 | 240 | 0 | 13 | 0 | 0 | 0 | 2559 | 6094 | 19530 | 26 | 0 | 2199 | 2255 | 0 | 0 | 16 | 53 | 42 | 215 | 0 | 0 | 148 | 0 | 26 | 0 | 0 | 18941 | 0 | 25 |
| SRR1708990 | 60 | 0 | 45 | 18 | 136 | 487 | 163 | 506 | 93 | 0 | 18 | 0 | 0 | 0 | 16654 | 21873 | 90098 | 70 | 56 | 7907 | 15289 | 0 | 0 | 28 | 177 | 15 | 39 | 27 | 0 | 181 | 0 | 107 | 0 | 0 | 2478 | 0 | 0 |
| SRR1708991 | 14 | 10 | 0 | 0 | 108 | 673 | 417 | 1043 | 94 | 0 | 0 | 14 | 0 | 0 | 9239 | 15549 | 61789 | 21 | 0 | 4528 | 11279 | 0 | 0 | 15 | 189 | 10 | 13 | 14 | 0 | 173 | 0 | 57 | 0 | 0 | 2112 | 0 | 16 |
| SRR1708992 | 52 | 0 | 39 | 58 | 175 | 1089 | 395 | 1302 | 175 | 0 | 13 | 0 | 0 | 0 | 9343 | 12936 | 57978 | 32 | 39 | 4710 | 12019 | 0 | 0 | 15 | 24 | 51 | 76 | 68 | 0 | 16 | 0 | 19 | 0 | 0 | 5163 | 0 | 0 |
| SRR1708993 | 12 | 0 | 0 | 0 | 147 | 1524 | 637 | 1755 | 217 | 16 | 0 | 0 | 0 | 0 | 7344 | 14932 | 42028 | 88 | 0 | 2242 | 12134 | 0 | 0 | 26 | 206 | 97 | 12 | 16 | 0 | 162 | 0 | 20 | 0 | 0 | 14965 | 0 | 0 |
| SRR1708994 | 17 | 0 | 0 | 0 | 48 | 725 | 294 | 1098 | 149 | 0 | 29 | 15 | 0 | 0 | 3492 | 4535 | 23945 | 61 | 0 | 1998 | 1761 | 0 | 0 | 13 | 26 | 23 | 59 | 0 | 0 | 11 | 0 | 650 | 0 | 0 | 5777 | 0 | 25 |
| SRR1708995 | 38 | 15 | 0 | 0 | 98 | 375 | 117 | 416 | 70 | 0 | 40 | 0 | 0 | 0 | 2727 | 2789 | 20724 | 53 | 0 | 3243 | 1120 | 0 | 0 | 166 | 130 | 136 | 584 | 0 | 0 | 286 | 0 | 73 | 0 | 0 | 7958 | 0 | 14 |
| SRR1708996 | 98 | 19 | 10 | 0 | 83 | 589 | 496 | 985 | 159 | 16 | 0 | 49 | 0 | 0 | 3081 | 72109 | 19214 | 0 | 13 | 9429 | 1488 | 0 | 0 | 258 | 295 | 271 | 369 | 24 | 0 | 273 | 0 | 185 | 0 | 0 | 8973 | 0 | 18 |

### Supp Table S8.3.Columns_BX_to_DH

| | | | | | | | | | | | | | | | | | | | | | | | | | | | | | | | | | | | | | |
|---|---|---|---|---|---|---|---|---|---|---|---|---|---|---|---|---|---|---|---|---|---|---|---|---|---|---|---|---|---|---|---|---|---|---|---|---|---|
| SRR1708997 | 92 | 23 | 31 | 50 | 21 | 131 | 247 | 701 | 131 | 0 | 141 | 11 | 0 | 0 | 3667 | 42008 | 30595 | 31 | 54 | 12404 | 9854 | 0 | 0 | 254 | 153 | 808 | 1628 | 153 | 0 | 556 | 0 | 231 | 0 | 0 | 20049 | 0 | 19 |
| SRR1708998 | 103 | 15 | 0 | 18 | 59 | 311 | 285 | 691 | 147 | 0 | 21 | 10 | 0 | 0 | 2229 | 75703 | 19544 | 17 | 26 | 4510 | 2868 | 0 | 0 | 168 | 186 | 553 | 615 | 44 | 0 | 369 | 0 | 69 | 0 | 0 | 12162 | 0 | 13 |
| SRR1708999 | 33 | 19 | 0 | 0 | 39 | 231 | 231 | 534 | 68 | 0 | 0 | 0 | 0 | 0 | 293 | 6068 | 3584 | 42 | 0 | 660 | 906 | 0 | 0 | 470 | 21 | 29 | 153 | 0 | 0 | 66 | 0 | 100 | 0 | 0 | 85139 | 0 | 0 |
| SRR1709000 | 142 | 14 | 15 | 0 | 57 | 228 | 115 | 372 | 56 | 11 | 16 | 0 | 0 | 0 | 1525 | 4286 | 13174 | 43 | 27 | 5342 | 1392 | 0 | 0 | 397 | 85 | 266 | 850 | 64 | 0 | 443 | 0 | 99 | 0 | 0 | 39757 | 0 | 18 |
| SRR1709001 | 73 | 0 | 0 | 0 | 32 | 354 | 139 | 534 | 69 | 0 | 0 | 0 | 0 | 0 | 617 | 2799 | 6081 | 111 | 14 | 2053 | 1172 | 0 | 0 | 334 | 41 | 64 | 235 | 15 | 0 | 165 | 0 | 111 | 0 | 0 | 14985 | 0 | 0 |
| SRR1709002 | 75 | 31 | 63 | 38 | 41 | 364 | 154 | 767 | 108 | 0 | 51 | 0 | 0 | 0 | 544 | 1540 | 4554 | 23 | 16 | 4733 | 1046 | 0 | 0 | 88 | 142 | 71 | 306 | 20 | 0 | 24 | 0 | 0 | 0 | 0 | 11209 | 0 | 0 |
| SRR1709003 | 59 | 0 | 0 | 0 | 53 | 274 | 187 | 454 | 114 | 0 | 0 | 0 | 0 | 0 | 821 | 2191 | 6581 | 47 | 0 | 3031 | 1015 | 0 | 0 | 484 | 50 | 104 | 395 | 0 | 0 | 246 | 0 | 0 | 0 | 0 | 8207 | 0 | 0 |
| SRR1709004 | 192 | 72 | 0 | 0 | 62 | 130 | 211 | 559 | 93 | 13 | 0 | 0 | 0 | 0 | 671 | 15015 | 6233 | 43 | 19 | 1696 | 304 | 0 | 0 | 28 | 137 | 75 | 15 | 16 | 0 | 153 | 0 | 205 | 0 | 0 | 73727 | 0 | 0 |
| SRR1709005 | 137 | 17 | 26 | 24 | 85 | 345 | 257 | 515 | 99 | 0 | 24 | 24 | 0 | 0 | 1352 | 32221 | 13052 | 44 | 33 | 6258 | 1949 | 0 | 0 | 24 | 74 | 22 | 57 | 18 | 0 | 28 | 0 | 214 | 0 | 0 | 42346 | 0 | 0 |
| SRR1709006 | 99 | 0 | 33 | 65 | 86 | 231 | 357 | 682 | 100 | 0 | 50 | 0 | 0 | 0 | 3353 | 90137 | 24685 | 13 | 19 | 4954 | 3847 | 0 | 0 | 117 | 11 | 55 | 90 | 99 | 0 | 25 | 0 | 88 | 0 | 0 | 30655 | 0 | 0 |
| SRR1709007 | 110 | 27 | 24 | 27 | 51 | 298 | 260 | 654 | 144 | 0 | 53 | 0 | 11 | 0 | 1865 | 17457 | 19265 | 0 | 19 | 4950 | 8055 | 0 | 0 | 574 | 147 | 489 | 2110 | 120 | 0 | 537 | 0 | 77 | 0 | 0 | 39306 | 0 | 0 |
| SRR1709008 | 79 | 17 | 0 | 0 | 63 | 334 | 367 | 799 | 131 | 0 | 0 | 0 | 0 | 0 | 947 | 83537 | 7624 | 115 | 0 | 1724 | 892 | 0 | 0 | 19 | 151 | 103 | 13 | 11 | 0 | 160 | 0 | 175 | 0 | 0 | 53845 | 0 | 0 |
| SRR1709009 | 99 | 77 | 0 | 0 | 53 | 356 | 310 | 817 | 197 | 0 | 0 | 0 | 0 | 0 | 1407 | 64287 | 13984 | 30 | 15 | 3147 | 1422 | 0 | 0 | 248 | 246 | 320 | 741 | 24 | 0 | 360 | 0 | 61 | 0 | 0 | 14057 | 0 | 0 |
| SRR1709010 | 61 | 13 | 0 | 0 | 71 | 460 | 391 | 718 | 52 | 0 | 0 | 0 | 0 | 0 | 1000 | 4999 | 10414 | 17 | 0 | 3040 | 557 | 0 | 0 | 11 | 15 | 46 | 152 | 13 | 0 | 206 | 0 | 40 | 0 | 0 | 6656 | 0 | 13 |
| SRR1709011 | 96 | 22 | 17 | 15 | 58 | 196 | 134 | 238 | 62 | 0 | 14 | 0 | 0 | 0 | 720 | 3085 | 8300 | 62 | 28 | 4128 | 850 | 0 | 0 | 552 | 81 | 407 | 2722 | 60 | 0 | 745 | 0 | 28 | 0 | 0 | 27179 | 0 | 12 |
| SRR1709012 | 76 | 0 | 29 | 15 | 39 | 244 | 144 | 559 | 52 | 0 | 37 | 0 | 0 | 0 | 620 | 1786 | 6907 | 26 | 13 | 4075 | 1780 | 0 | 0 | 125 | 196 | 47 | 104 | 81 | 0 | 26 | 0 | 13 | 0 | 0 | 10877 | 0 | 0 |
| SRR1709013 | 36 | 27 | 0 | 0 | 48 | 349 | 250 | 402 | 38 | 0 | 0 | 0 | 0 | 0 | 674 | 1641 | 6237 | 53 | 0 | 2121 | 1076 | 0 | 0 | 17 | 42 | 60 | 19 | 0 | 0 | 14 | 0 | 23 | 0 | 0 | 18720 | 0 | 0 |
| SRR1709014 | 83 | 53 | 0 | 0 | 53 | 209 | 197 | 310 | 45 | 0 | 18 | 0 | 0 | 0 | 694 | 1733 | 6024 | 11 | 11 | 5444 | 643 | 0 | 0 | 473 | 62 | 192 | 830 | 41 | 0 | 369 | 0 | 17 | 0 | 0 | 21801 | 0 | 0 |
| SRR1709015 | 73 | 37 | 0 | 0 | 37 | 310 | 90 | 264 | 60 | 0 | 0 | 0 | 0 | 0 | 350 | 1497 | 3058 | 30 | 0 | 2565 | 190 | 0 | 0 | 165 | 47 | 120 | 21 | 23 | 0 | 12 | 0 | 0 | 0 | 0 | 4918 | 0 | 0 |
| SRR1709016 | 65 | 0 | 31 | 0 | 140 | 540 | 234 | 782 | 110 | 0 | 24 | 0 | 0 | 0 | 6166 | 11391 | 40670 | 53 | 23 | 3571 | 5392 | 0 | 0 | 434 | 159 | 285 | 672 | 18 | 0 | 230 | 0 | 38 | 0 | 0 | 2648 | 0 | 12 |
| SRR1709017 | 34 | 0 | 13 | 0 | 143 | 231 | 397 | 600 | 50 | 0 | 0 | 0 | 0 | 0 | 3796 | 3544 | 25622 | 11 | 0 | 3019 | 309 | 0 | 0 | 151 | 47 | 132 | 19 | 0 | 0 | 16 | 0 | 33 | 0 | 0 | 4884 | 0 | 0 |
| SRR1709018 | 16 | 12 | 0 | 0 | 110 | 877 | 297 | 866 | 95 | 16 | 0 | 0 | 0 | 0 | 2477 | 5300 | 14249 | 154 | 0 | 1435 | 2216 | 0 | 0 | 153 | 118 | 118 | 233 | 0 | 0 | 147 | 0 | 33 | 0 | 0 | 11590 | 0 | 0 |
| SRR1709019 | 42 | 13 | 0 | 0 | 94 | 300 | 332 | 502 | 111 | 0 | 0 | 0 | 0 | 0 | 2444 | 1637 | 12801 | 46 | 0 | 1662 | 241 | 0 | 0 | 12 | 42 | 84 | 11 | 0 | 0 | 118 | 0 | 0 | 0 | 0 | 8704 | 0 | 21 |
| SRR1709020 | 36 | 13 | 0 | 16 | 92 | 470 | 146 | 563 | 116 | 0 | 13 | 0 | 0 | 0 | 1873 | 2498 | 12905 | 30 | 11 | 4470 | 1765 | 0 | 0 | 14 | 111 | 16 | 94 | 39 | 0 | 25 | 0 | 0 | 0 | 0 | 6472 | 0 | 11 |
| SRR1709021 | 29 | 0 | 0 | 0 | 11 | 514 | 464 | 742 | 76 | 0 | 0 | 0 | 0 | 0 | 2209 | 5207 | 24626 | 17 | 0 | 4127 | 405 | 0 | 0 | 136 | 54 | 88 | 14 | 0 | 0 | 12 | 0 | 17 | 0 | 0 | 6992 | 0 | 0 |
| SRR1732015 | 17 | 11 | 0 | 12 | 0 | 103 | 130 | 377 | 101 | 0 | 23 | 0 | 0 | 0 | 4216 | 8681 | 7178 | 170 | 29 | 3508 | 533 | 0 | 11 | 0 | 0 | 0 | 12 | 0 | 0 | 83 | 0 | 907 | 21 | 0 | 21591 | 99 | 56 |
| SRR1732016 | 0 | 0 | 0 | 0 | 0 | 38 | 84 | 252 | 28 | 0 | 105 | 0 | 0 | 0 | 14387 | 38479 | 25974 | 1288 | 30 | 1137 | 1356 | 15 | 0 | 0 | 0 | 0 | 0 | 0 | 0 | 772 | 11 | 4201 | 79 | 0 | 39335 | 138 | 463 |
| SRR1748132 | 0 | 0 | 0 | 0 | 55 | 51 | 70 | 253 | 14 | 100 | 17 | 12 | 0 | 0 | 132826 | 99278 | 266919 | 11 | 225 | 12241 | 15601 | 0 | 0 | 118 | 11 | 13 | 15 | 0 | 0 | 201 | 0 | 0 | 0 | 0 | 243 | 41 | 0 |
| SRR1748133 | 0 | 0 | 0 | 0 | 16 | 65 | 106 | 710 | 45 | 55 | 14 | 21 | 0 | 0 | 100134 | 102433 | 262940 | 0 | 260 | 23113 | 2236 | 0 | 0 | 23 | 14 | 21 | 23 | 0 | 0 | 164 | 0 | 0 | 0 | 0 | 2154 | 65 | 16 |
| SRR1759212 | 0 | 0 | 0 | 16 | 64 | 280 | 943 | 1616 | 519 | 0 | 80 | 0 | 0 | 0 | 28533 | 21260 | 400873 | 0 | 678 | 20064 | 2133 | 0 | 0 | 226 | 282 | 513 | 1641 | 1200 | 0 | 877 | 0 | 25 | 0 | 0 | 604 | 0 | 0 |
| SRR1759213 | 0 | 0 | 0 | 0 | 24 | 593 | 2199 | 4847 | 1053 | 0 | 85 | 0 | 0 | 0 | 2389 | 19579 | 108504 | 20 | 757 | 16660 | 1347 | 0 | 0 | 30 | 22 | 37 | 126 | 83 | 0 | 124 | 0 | 83 | 0 | 0 | 2464 | 0 | 0 |
| SRR1759214 | 0 | 0 | 0 | 0 | 32 | 345 | 825 | 1704 | 625 | 0 | 84 | 0 | 0 | 0 | 3431 | 20818 | 117500 | 0 | 374 | 15172 | 1497 | 0 | 0 | 15 | 76 | 19 | 74 | 40 | 0 | 128 | 0 | 68 | 0 | 0 | 1694 | 0 | 0 |
| SRR1759215 | 0 | 0 | 0 | 0 | 19 | 668 | 790 | 1882 | 631 | 0 | 112 | 0 | 0 | 0 | 9518 | 21256 | 189826 | 0 | 326 | 9030 | 2175 | 0 | 0 | 250 | 205 | 379 | 710 | 432 | 0 | 1384 | 0 | 241 | 0 | 0 | 1279 | 0 | 0 |
| SRR1759216 | 0 | 0 | 0 | 18 | 37 | 374 | 888 | 2130 | 610 | 0 | 41 | 0 | 0 | 0 | 6945 | 19225 | 206289 | 0 | 415 | 11358 | 3723 | 0 | 0 | 234 | 132 | 377 | 1487 | 643 | 0 | 592 | 0 | 61 | 0 | 0 | 1258 | 0 | 0 |
| SRR1759217 | 0 | 0 | 0 | 13 | 25 | 401 | 632 | 1170 | 442 | 0 | 402 | 0 | 0 | 0 | 9166 | 27391 | 216794 | 0 | 344 | 8165 | 804 | 0 | 0 | 339 | 59 | 130 | 406 | 369 | 0 | 2744 | 0 | 24 | 0 | 0 | 664 | 0 | 0 |
| SRR1759218 | 0 | 0 | 0 | 11 | 48 | 666 | 663 | 2171 | 714 | 0 | 120 | 0 | 0 | 0 | 6327 | 22762 | 140708 | 0 | 418 | 11017 | 1572 | 0 | 0 | 396 | 84 | 158 | 601 | 182 | 0 | 1514 | 0 | 215 | 0 | 0 | 1303 | 0 | 0 |
| SRR1759219 | 0 | 0 | 0 | 19 | 50 | 502 | 1019 | 2361 | 1471 | 14 | 70 | 18 | 0 | 0 | 3659 | 23503 | 110848 | 13 | 936 | 28978 | 1127 | 0 | 0 | 15 | 13 | 35 | 108 | 44 | 0 | 58 | 0 | 111 | 0 | 0 | 1489 | 0 | 0 |
| SRR1759220 | 0 | 0 | 0 | 19 | 36 | 571 | 985 | 2412 | 1159 | 0 | 82 | 0 | 0 | 0 | 11473 | 18620 | 173146 | 0 | 271 | 3354 | 504 | 0 | 0 | 383 | 68 | 214 | 767 | 160 | 0 | 1072 | 0 | 40 | 0 | 0 | 936 | 0 | 0 |
| SRR1759221 | 0 | 0 | 0 | 62 | 38 | 356 | 358 | 1022 | 127 | 11 | 86 | 0 | 0 | 0 | 72532 | 24517 | 340492 | 0 | 148 | 13622 | 10202 | 0 | 0 | 12 | 112 | 22 | 130 | 32 | 0 | 15 | 0 | 50 | 0 | 0 | 151 | 0 | 0 |
| SRR1759222 | 0 | 0 | 0 | 0 | 54 | 531 | 1753 | 4024 | 998 | 0 | 0 | 0 | 0 | 0 | 7928 | 25259 | 119291 | 0 | 162 | 20271 | 16883 | 0 | 0 | 159 | 26 | 57 | 188 | 11 | 0 | 85 | 0 | 77 | 0 | 0 | 1137 | 0 | 0 |
| SRR1759223 | 0 | 0 | 14 | 56 | 66 | 157 | 645 | 959 | 532 | 0 | 89 | 0 | 0 | 0 | 6605 | 40427 | 200683 | 0 | 470 | 21378 | 6039 | 0 | 0 | 18 | 184 | 44 | 199 | 123 | 0 | 19 | 0 | 121 | 0 | 0 | 341 | 0 | 0 |
| SRR1759224 | 0 | 0 | 0 | 33 | 38 | 167 | 544 | 1229 | 187 | 0 | 50 | 11 | 0 | 0 | 33293 | 22233 | 243855 | 0 | 99 | 17499 | 10151 | 0 | 0 | 650 | 153 | 1029 | 3176 | 320 | 0 | 302 | 0 | 35 | 0 | 0 | 125 | 0 | 0 |
| SRR1759225 | 0 | 0 | 0 | 0 | 13 | 67 | 442 | 650 | 230 | 18 | 26 | 10 | 0 | 0 | 5658 | 89889 | 129814 | 0 | 383 | 32723 | 44841 | 0 | 26 | 127 | 43 | 173 | 790 | 45 | 0 | 194 | 0 | 153 | 0 | 0 | 370 | 0 | 0 |
| SRR1759226 | 0 | 0 | 0 | 0 | 36 | 70 | 257 | 521 | 142 | 0 | 29 | 0 | 0 | 0 | 8021 | 26946 | 193194 | 0 | 75 | 51035 | 12558 | 0 | 0 | 15 | 70 | 19 | 77 | 73 | 0 | 16 | 0 | 48 | 0 | 0 | 315 | 0 | 0 |
| SRR1759227 | 0 | 0 | 14 | 31 | 42 | 289 | 422 | 824 | 266 | 0 | 53 | 0 | 0 | 0 | 59214 | 32777 | 326002 | 0 | 189 | 10976 | 8152 | 0 | 0 | 259 | 66 | 350 | 1775 | 218 | 0 | 305 | 0 | 67 | 0 | 0 | 117 | 0 | 0 |
| SRR1759228 | 0 | 0 | 12 | 21 | 33 | 171 | 303 | 603 | 151 | 21 | 124 | 14 | 0 | 0 | 73161 | 39690 | 245017 | 0 | 234 | 15688 | 12670 | 0 | 0 | 222 | 62 | 189 | 24 | 56 | 0 | 15 | 0 | 257 | 0 | 0 | 117 | 0 | 0 |
| SRR1759229 | 0 | 0 | 12 | 42 | 52 | 230 | 352 | 820 | 242 | 19 | 57 | 0 | 0 | 0 | 16748 | 61202 | 243775 | 0 | 427 | 38385 | 8807 | 0 | 0 | 234 | 103 | 418 | 2494 | 1436 | 0 | 318 | 0 | 122 | 0 | 0 | 183 | 0 | 0 |
| SRR1759230 | 0 | 0 | 0 | 0 | 0 | 192 | 371 | 950 | 153 | 12 | 59 | 12 | 0 | 0 | 29340 | 39491 | 197441 | 0 | 163 | 22621 | 29834 | 0 | 13 | 73 | 46 | 172 | 592 | 42 | 0 | 231 | 0 | 461 | 0 | 0 | 181 | 0 | 0 |
| SRR1759231 | 0 | 0 | 0 | 24 | 112 | 110 | 637 | 1028 | 123 | 0 | 33 | 0 | 0 | 0 | 10510 | 47014 | 214025 | 0 | 505 | 56678 | 19395 | 0 | 0 | 13 | 75 | 12 | 49 | 151 | 0 | 229 | 0 | 14 | 0 | 0 | 361 | 0 | 0 |
| SRR1759232 | 0 | 11 | 0 | 41 | 67 | 230 | 764 | 1420 | 480 | 0 | 37 | 13 | 0 | 0 | 13137 | 51578 | 162974 | 0 | 223 | 19103 | 15655 | 0 | 0 | 18 | 82 | 15 | 61 | 52 | 0 | 165 | 0 | 71 | 0 | 0 | 365 | 0 | 0 |
| SRR1759233 | 0 | 0 | 0 | 0 | 43 | 231 | 560 | 1165 | 137 | 11 | 21 | 11 | 0 | 0 | 68529 | 39094 | 262672 | 0 | 92 | 13561 | 30479 | 0 | 0 | 19 | 48 | 186 | 23 | 45 | 0 | 120 | 0 | 283 | 0 | 0 | 168 | 0 | 0 |
| SRR1759234 | 0 | 0 | 15 | 25 | 51 | 85 | 405 | 519 | 191 | 18 | 36 | 0 | 0 | 0 | 16598 | 52734 | 278684 | 0 | 755 | 43691 | 15964 | 0 | 0 | 13 | 10 | 45 | 212 | 44 | 0 | 28 | 0 | 57 | 0 | 0 | 148 | 0 | 0 |
| SRR1759235 | 0 | 0 | 0 | 14 | 62 | 597 | 604 | 2262 | 565 | 13 | 113 | 0 | 0 | 0 | 5220 | 42590 | 105322 | 17 | 345 | 19369 | 7564 | 0 | 0 | 32 | 0 | 0 | 0 | 0 | 0 | 73 | 0 | 791 | 0 | 0 | 942 | 0 | 0 |
| SRR1759236 | 0 | 0 | 11 | 73 | 44 | 138 | 632 | 942 | 352 | 17 | 28 | 0 | 0 | 0 | 9536 | 38489 | 198821 | 0 | 219 | 19008 | 13354 | 0 | 0 | 283 | 137 | 489 | 2916 | 1834 | 0 | 232 | 0 | 70 | 0 | 0 | 382 | 0 | 0 |
| SRR1759237 | 0 | 0 | 0 | 107 | 44 | 206 | 794 | 1321 | 417 | 14 | 45 | 0 | 0 | 0 | 12864 | 32042 | 241659 | 0 | 251 | 11918 | 11119 | 0 | 0 | 386 | 143 | 588 | 3000 | 2047 | 0 | 306 | 0 | 272 | 0 | 0 | 515 | 0 | 0 |
| SRR1759238 | 0 | 0 | 13 | 113 | 76 | 92 | 600 | 745 | 272 | 17 | 15 | 0 | 0 | 0 | 25216 | 27419 | 304388 | 0 | 308 | 26618 | 6325 | 0 | 0 | 20 | 176 | 45 | 284 | 131 | 0 | 12 | 0 | 40 | 0 | 0 | 288 | 0 | 0 |
| SRR1759239 | 11 | 0 | 0 | 0 | 14 | 1301 | 4356 | 5804 | 1957 | 27 | 71 | 0 | 0 | 0 | 32382 | 112478 | 228392 | 23 | 1287 | 21904 | 3478 | 11 | 10 | 51 | 16 | 46 | 10 | 42 | 0 | 44 | 0 | 0 | 0 | 0 | 154 | 0 | 0 |
| SRR1759240 | 0 | 11 | 0 | 48 | 47 | 191 | 837 | 1726 | 686 | 13 | 61 | 0 | 0 | 0 | 7408 | 37618 | 159900 | 0 | 291 | 15440 | 3457 | 0 | 0 | 32 | 12 | 55 | 325 | 177 | 0 | 17 | 0 | 195 | 0 | 0 | 342 | 0 | 0 |
| SRR1759241 | 0 | 0 | 0 | 50 | 49 | 340 | 1041 | 2671 | 1000 | 12 | 63 | 13 | 0 | 0 | 6313 | 49044 | 107988 | 0 | 205 | 12810 | 11767 | 0 | 0 | 41 | 12 | 39 | 196 | 63 | 0 | 21 | 0 | 300 | 0 | 0 | 387 | 0 | 0 |
| SRR1759242 | 15 | 16 | 33 | 99 | 144 | 35 | 140 | 267 | 87 | 39 | 157 | 0 | 0 | 0 | 81044 | 29463 | 335796 | 0 | 162 | 10935 | 14734 | 0 | 0 | 451 | 196 | 647 | 2383 | 778 | 0 | 447 | 0 | 206 | 0 | 0 | 101 | 0 | 0 |
| SRR1759243 | 0 | 0 | 0 | 30 | 44 | 95 | 400 | 603 | 292 | 32 | 23 | 0 | 0 | 0 | 7955 | 87655 | 247763 | 0 | 552 | 62998 | 30780 | 0 | 0 | 466 | 184 | 400 | 1620 | 342 | 0 | 745 | 0 | 19 | 0 | 0 | 340 | 0 | 0 |
| SRR1759244 | 0 | 0 | 0 | 0 | 33 | 1058 | 2240 | 4589 | 715 | 0 | 246 | 0 | 0 | 0 | 14436 | 29380 | 219234 | 0 | 1173 | 20536 | 1253 | 0 | 0 | 873 | 0 | 0 | 14 | 0 | 0 | 1991 | 0 | 32 | 0 | 0 | 1106 | 0 | 0 |
| SRR1759245 | 0 | 0 | 0 | 18 | 33 | 292 | 641 | 1536 | 371 | 15 | 88 | 0 | 0 | 0 | 21606 | 28993 | 337764 | 0 | 291 | 13442 | 3973 | 0 | 0 | 425 | 175 | 396 | 1399 | 1175 | 0 | 768 | 0 | 100 | 0 | 0 | 475 | 0 | 0 |
| SRR1759246 | 0 | 0 | 15 | 39 | 75 | 1012 | 1198 | 1850 | 467 | 0 | 24 | 0 | 0 | 0 | 36530 | 68589 | 423191 | 0 | 882 | 32097 | 5102 | 0 | 0 | 27 | 293 | 41 | 224 | 44 | 0 | 209 | 0 | 0 | 0 | 0 | 228 | 0 | 0 |
| SRR1759247 | 0 | 0 | 0 | 33 | 37 | 149 | 293 | 701 | 102 | 11 | 67 | 15 | 0 | 0 | 63555 | 52029 | 242934 | 0 | 165 | 14614 | 15061 | 0 | 0 | 249 | 80 | 356 | 1312 | 154 | 0 | 212 | 0 | 156 | 0 | 0 | 99 | 0 | 0 |
| SRR1759248 | 0 | 10 | 0 | 89 | 59 | 714 | 887 | 2430 | 452 | 10 | 102 | 0 | 0 | 0 | 9245 | 36345 | 200956 | 0 | 511 | 18801 | 8429 | 0 | 0 | 14 | 11 | 49 | 230 | 118 | 0 | 16 | 0 | 139 | 0 | 0 | 248 | 0 | 0 |
| SRR1759249 | 0 | 0 | 0 | 42 | 47 | 233 | 1881 | 2074 | 386 | 0 | 13 | 0 | 0 | 0 | 11439 | 46356 | 241156 | 0 | 1093 | 39787 | 1375 | 0 | 0 | 20 | 210 | 44 | 162 | 58 | 0 | 151 | 0 | 13 | 0 | 0 | 480 | 0 | 0 |
| SRR1759250 | 0 | 0 | 0 | 0 | 75 | 258 | 1404 | 1767 | 702 | 0 | 49 | 0 | 0 | 0 | 14014 | 40099 | 243255 | 0 | 1157 | 45556 | 5307 | 0 | 0 | 17 | 242 | 29 | 101 | 12 | 0 | 13 | 0 | 0 | 0 | 0 | 786 | 0 | 0 |
| SRR1759251 | 0 | 0 | 0 | 24 | 27 | 129 | 508 | 584 | 228 | 0 | 38 | 0 | 0 | 0 | 34561 | 13628 | 479226 | 0 | 468 | 18050 | 3763 | 0 | 0 | 240 | 147 | 24 | 128 | 121 | 0 | 176 | 0 | 0 | 0 | 0 | 62 | 0 | 0 |
| SRR1759252 | 0 | 0 | 0 | 69 | 74 | 268 | 445 | 832 | 213 | 16 | 48 | 12 | 0 | 0 | 38376 | 39849 | 414458 | 0 | 437 | 10739 | 4590 | 0 | 0 | 502 | 216 | 981 | 4988 | 3452 | 0 | 332 | 0 | 11 | 0 | 0 | 119 | 26 | 0 |
| SRR1759253 | 0 | 0 | 0 | 30 | 32 | 67 | 493 | 677 | 314 | 0 | 36 | 0 | 0 | 0 | 6599 | 20718 | 167162 | 0 | 298 | 9842 | 7799 | 0 | 0 | 149 | 188 | 505 | 2886 | 2285 | 0 | 233 | 0 | 36 | 0 | 0 | 552 | 0 | 0 |
| SRR1759254 | 0 | 0 | 0 | 38 | 33 | 134 | 954 | 907 | 228 | 0 | 35 | 0 | 0 | 0 | 21625 | 55849 | 366784 | 0 | 1149 | 31751 | 6144 | 0 | 0 | 345 | 372 | 1500 | 6904 | 4587 | 0 | 305 | 0 | 11 | 0 | 0 | 298 | 0 | 0 |
| SRR1759255 | 0 | 0 | 0 | 0 | 18 | 187 | 2034 | 2598 | 468 | 0 | 35 | 0 | 0 | 0 | 9361 | 15601 | 174941 | 0 | 660 | 16452 | 2356 | 0 | 0 | 27 | 17 | 84 | 494 | 274 | 0 | 12 | 0 | 0 | 0 | 0 | 397 | 0 | 0 |
| SRR1759256 | 0 | 0 | 0 | 38 | 47 | 273 | 1814 | 2259 | 983 | 12 | 28 | 0 | 0 | 0 | 7963 | 66173 | 143939 | 0 | 179 | 22056 | 13166 | 0 | 0 | 754 | 148 | 546 | 3008 | 697 | 0 | 290 | 0 | 162 | 0 | 0 | 1227 | 0 | 0 |
| SRR1759257 | 0 | 0 | 14 | 0 | 28 | 26 | 306 | 263 | 61 | 39 | 17 | 0 | 0 | 0 | 43074 | 41668 | 561103 | 0 | 813 | 71259 | 13377 | 0 | 0 | 498 | 197 | 495 | 2081 | 383 | 0 | 262 | 0 | 0 | 0 | 0 | 81 | 0 | 0 |
| SRR1759258 | 28 | 29 | 52 | 475 | 403 | 3755 | 1996 | 6672 | 216 | 0 | 213 | 88 | 0 | 0 | 121324 | 27523 | 344472 | 21 | 457 | 6800 | 5282 | 0 | 0 | 861 | 550 | 1390 | 7947 | 2141 | 0 | 549 | 0 | 48 | 0 | 0 | 52 | 0 | 14 |
| SRR1759259 | 25 | 31 | 17 | 13 | 11 | 1007 | 1305 | 3954 | 656 | 0 | 68 | 0 | 0 | 0 | 30261 | 25773 | 222875 | 0 | 228 | 4765 | 2179 | 0 | 0 | 34 | 184 | 26 | 127 | 30 | 0 | 14 | 0 | 70 | 0 | 0 | 272 | 0 | 0 |
| SRR1759260 | 0 | 0 | 0 | 0 | 15 | 181 | 329 | 773 | 291 | 39 | 26 | 0 | 0 | 0 | 3164 | 72126 | 106887 | 13 | 651 | 77236 | 21906 | 0 | 29 | 53 | 53 | 100 | 313 | 206 | 0 | 642 | 0 | 245 | 0 | 0 | 633 | 0 | 0 |
| SRR1759261 | 0 | 10 | 18 | 36 | 24 | 427 | 671 | 1448 | 273 | 12 | 139 | 17 | 0 | 0 | 55098 | 35362 | 306835 | 0 | 340 | 11190 | 7128 | 0 | 0 | 297 | 206 | 790 | 3752 | 1808 | 0 | 278 | 0 | 100 | 0 | 0 | 205 | 0 | 0 |
| SRR1759262 | 0 | 0 | 0 | 15 | 16 | 346 | 201 | 1330 | 244 | 13 | 69 | 0 | 0 | 0 | 31263 | 37505 | 337523 | 18 | 588 | 32755 | 8833 | 0 | 0 | 14 | 70 | 17 | 52 | 27 | 0 | 48 | 0 | 245 | 0 | 0 | 324 | 0 | 0 |
| SRR1759263 | 0 | 0 | 0 | 0 | 55 | 513 | 2373 | 2788 | 877 | 11 | 0 | 0 | 0 | 0 | 5818 | 70531 | 161054 | 0 | 603 | 37618 | 6836 | 0 | 0 | 164 | 53 | 232 | 888 | 65 | 0 | 184 | 0 | 23 | 0 | 0 | 1335 | 0 | 0 |
| SRR1759264 | 0 | 0 | 0 | 25 | 23 | 285 | 1378 | 2000 | 339 | 0 | 0 | 0 | 0 | 0 | 27855 | 17636 | 239671 | 0 | 193 | 30245 | 5265 | 0 | 0 | 289 | 183 | 995 | 5240 | 404 | 0 | 119 | 0 | 11 | 0 | 0 | 281 | 0 | 0 |
| SRR1759265 | 0 | 0 | 0 | 0 | 59 | 83 | 311 | 380 | 52 | 0 | 0 | 0 | 0 | 0 | 31985 | 52471 | 590592 | 0 | 667 | 53919 | 10464 | 0 | 0 | 251 | 59 | 98 | 302 | 0 | 0 | 70 | 0 | 0 | 0 | 0 | 95 | 0 | 0 |
| SRR1759266 | 0 | 0 | 0 | 18 | 20 | 120 | 255 | 446 | 175 | 0 | 17 | 0 | 0 | 0 | 29919 | 90943 | 443665 | 0 | 1223 | 103604 | 3368 | 0 | 0 | 14 | 147 | 16 | 62 | 36 | 0 | 13 | 0 | 15 | 0 | 0 | 116 | 0 | 0 |
| SRR1759267 | 0 | 11 | 0 | 0 | 47 | 1741 | 931 | 2633 | 659 | 50 | 11 | 0 | 0 | 0 | 40289 | 79281 | 483301 | 0 | 372 | 35289 | 5930 | 0 | 0 | 26 | 63 | 117 | 210 | 14 | 0 | 126 | 0 | 77 | 0 | 0 | 529 | 0 | 0 |
| SRR1759268 | 0 | 0 | 0 | 22 | 127 | 292 | 973 | 1460 | 361 | 0 | 19 | 0 | 0 | 0 | 57455 | 70906 | 496515 | 0 | 412 | 22761 | 6874 | 0 | 0 | 1519 | 231 | 794 | 3660 | 212 | 0 | 127 | 0 | 0 | 0 | 0 | 446 | 0 | 0 |
| SRR1759269 | 0 | 0 | 0 | 18 | 42 | 441 | 2140 | 3923 | 412 | 0 | 31 | 0 | 0 | 0 | 18280 | 43437 | 382790 | 0 | 842 | 25909 | 7217 | 0 | 0 | 18 | 199 | 33 | 113 | 31 | 0 | 143 | 0 | 0 | 0 | 0 | 941 | 0 | 0 |
| SRR1759270 | 0 | 0 | 0 | 23 | 45 | 207 | 517 | 1007 | 188 | 0 | 23 | 0 | 0 | 0 | 15248 | 30430 | 358671 | 0 | 953 | 32037 | 4546 | 0 | 0 | 472 | 161 | 341 | 1137 | 1461 | 0 | 404 | 0 | 22 | 0 | 0 | 757 | 0 | 0 |
| SRR1759271 | 0 | 0 | 0 | 10 | 12 | 243 | 1294 | 2147 | 365 | 0 | 16 | 10 | 0 | 0 | 9791 | 33839 | 187908 | 0 | 337 | 35377 | 22406 | 0 | 0 | 178 | 44 | 228 | 30 | 40 | 0 | 161 | 0 | 209 | 0 | 0 | 602 | 0 | 0 |
| SRR1759272 | 0 | 0 | 0 | 23 | 23 | 154 | 743 | 1113 | 188 | 0 | 0 | 0 | 0 | 0 | 32166 | 22531 | 294392 | 0 | 306 | 35886 | 3041 | 0 | 0 | 365 | 131 | 420 | 1245 | 100 | 0 | 201 | 0 | 0 | 0 | 0 | 267 | 0 | 0 |
| SRR1759273 | 11 | 0 | 11 | 22 | 55 | 299 | 771 | 1142 | 165 | 0 | 111 | 0 | 0 | 0 | 46465 | 157883 | 389818 | 0 | 563 | 45136 | 22631 | 0 | 0 | 71 | 20 | 39 | 122 | 51 | 0 | 13 | 0 | 31 | 0 | 0 | 479 | 0 | 0 |
| SRR1759274 | 0 | 0 | 0 | 0 | 0 | 366 | 350 | 1200 | 273 | 63 | 95 | 0 | 0 | 0 | 11933 | 38319 | 174306 | 0 | 405 | 27295 | 18093 | 0 | 18 | 153 | 182 | 442 | 681 | 335 | 0 | 609 | 0 | 506 | 0 | 0 | 501 | 0 | 0 |
| SRR1759275 | 0 | 0 | 0 | 33 | 31 | 382 | 330 | 1123 | 213 | 0 | 420 | 0 | 0 | 0 | 8667 | 52503 | 136899 | 14 | 683 | 31170 | 9276 | 0 | 0 | 246 | 75 | 180 | 517 | 371 | 0 | 1531 | 0 | 553 | 0 | 0 | 641 | 0 | 0 |
| SRR1777167 | 17 | 10 | 0 | 0 | 290 | 84 | 84 | 301 | 736 | 758 | 37 | 34 | 0 | 0 | 10290 | 22810 | 53049 | 17 | 134 | 1799 | 15042 | 26 | 40 | 44 | 50 | 48 | 13 | 29 | 15 | 69 | 14 | 1623 | 0 | 44 | 7240 | 0 | 0 |
| SRR1820679 | 20 | 11 | 0 | 0 | 209 | 8628 | 12011 | 55972 | 96949 | 34 | 124 | 10 | 0 | 0 | 3403 | 84340 | 75547 | 200 | 480 | 3303 | 140 | 0 | 0 | 50 | 140 | 151 | 225 | 157 | 0 | 1446 | 0 | 0 | 0 | 0 | 21043 | 0 | 0 |
| SRR1820680 | 0 | 0 | 0 | 0 | 105 | 55971 | 39424 | 147197 | 175525 | 45 | 0 | 0 | 0 | 0 | 128 | 47417 | 30216 | 128 | 64 | 299 | 0 | 0 | 0 | 227 | 57 | 0 | 0 | 13 | 0 | 136 | 0 | 0 | 0 | 0 | 12762 | 0 | 0 |
| SRR1820681 | 0 | 0 | 0 | 0 | 201 | 8507 | 9121 | 41824 | 72753 | 54 | 105 | 12 | 0 | 0 | 1774 | 73517 | 93259 | 166 | 1161 | 11542 | 348 | 0 | 0 | 106 | 346 | 355 | 539 | 526 | 0 | 1099 | 0 | 0 | 0 | 0 | 10240 | 0 | 0 |
| SRR1820682 | 0 | 0 | 0 | 0 | 126 | 49632 | 29890 | 124550 | 115773 | 60 | 0 | 0 | 0 | 0 | 1473 | 93154 | 82739 | 430 | 123 | 2373 | 20 | 0 | 0 | 252 | 103 | 11 | 14 | 31 | 0 | 133 | 0 | 0 | 0 | 0 | 15708 | 0 | 0 |
| SRR1820683 | 12 | 0 | 0 | 0 | 210 | 5898 | 8508 | 38577 | 58726 | 32 | 98 | 0 | 0 | 0 | 2533 | 82631 | 107110 | 155 | 851 | 9436 | 174 | 0 | 0 | 72 | 195 | 212 | 469 | 220 | 0 | 1081 | 0 | 0 | 0 | 0 | 14208 | 0 | 0 |
| SRR1820684 | 20 | 0 | 0 | 0 | 28 | 24123 | 22438 | 82553 | 101442 | 98 | 0 | 0 | 0 | 0 | 3063 | 68628 | 113626 | 204 | 0 | 1072 | 77 | 0 | 0 | 21 | 49 | 0 | 0 | 0 | 0 | 20 | 0 | 0 | 0 | 0 | 22978 | 0 | 0 |
| SRR1820685 | 0 | 0 | 0 | 0 | 66 | 9423 | 12115 | 54005 | 97337 | 24 | 47 | 13 | 0 | 0 | 2051 | 69465 | 81901 | 32 | 192 | 1346 | 256 | 0 | 0 | 27 | 91 | 101 | 171 | 75 | 0 | 433 | 0 | 0 | 0 | 0 | 16475 | 0 | 0 |
| SRR1820686 | 0 | 0 | 0 | 0 | 23 | 42010 | 44201 | 169350 | 176176 | 44 | 0 | 0 | 0 | 0 | 427 | 46941 | 35206 | 199 | 28 | 0 | 85 | 0 | 0 | 41 | 12 | 31 | 22 | 13 | 0 | 20 | 0 | 0 | 0 | 0 | 18610 | 0 | 0 |
| SRR1820687 | 10 | 0 | 0 | 0 | 98 | 25174 | 18466 | 74618 | 162782 | 75 | 0 | 13 | 0 | 0 | 1116 | 49198 | 34891 | 372 | 73 | 1110 | 282 | 0 | 0 | 76 | 103 | 18 | 11 | 20 | 0 | 91 | 0 | 0 | 0 | 0 | 52525 | 0 | 0 |
| SRR1820688 | 0 | 0 | 0 | 0 | 72 | 20347 | 13736 | 58641 | 112220 | 74 | 0 | 35 | 0 | 0 | 660 | 67500 | 39107 | 438 | 202 | 6577 | 210 | 0 | 0 | 47 | 61 | 11 | 28 | 12 | 0 | 64 | 0 | 0 | 0 | 0 | 53216 | 0 | 0 |
| SRR1820689 | 0 | 0 | 0 | 0 | 113 | 17573 | 11922 | 54736 | 101997 | 916 | 0 | 20 | 0 | 0 | 1899 | 49046 | 65841 | 418 | 143 | 3645 | 559 | 0 | 0 | 94 | 21 | 0 | 0 | 0 | 0 | 38 | 0 | 0 | 0 | 0 | 70338 | 0 | 0 |
| SRR1820690 | 0 | 0 | 0 | 0 | 76 | 29501 | 19570 | 78319 | 176620 | 315 | 0 | 36 | 0 | 0 | 1074 | 66246 | 53754 | 324 | 80 | 3418 | 142 | 0 | 0 | 150 | 54 | 15 | 0 | 21 | 0 | 69 | 0 | 0 | 0 | 12 | 39666 | 0 | 0 |
| SRR1853804 | 0 | 15 | 15 | 19 | 51 | 0 | 0 | 103 | 0 | 0 | 0 | 0 | 0 | 0 | 103 | 1211 | 1561 | 0 | 0 | 1402 | 1060 | 0 | 0 | 41 | 12 | 37 | 152 | 39 | 0 | 51 | 0 | 34 | 0 | 0 | 646 | 0 | 0 |
| SRR1853805 | 0 | 22 | 0 | 53 | 61 | 0 | 0 | 155 | 0 | 0 | 18 | 13 | 0 | 0 | 104 | 757 | 1570 | 104 | 0 | 1035 | 1449 | 0 | 0 | 57 | 31 | 180 | 759 | 250 | 0 | 79 | 0 | 0 | 0 | 0 | 364 | 0 | 0 |
| SRR1853806 | 17 | 32 | 11 | 40 | 32 | 0 | 0 | 17 | 0 | 0 | 29 | 0 | 0 | 0 | 216 | 6899 | 18694 | 17 | 0 | 2098 | 24009 | 0 | 0 | 57 | 11 | 43 | 213 | 41 | 0 | 13 | 0 | 133 | 0 | 0 | 834 | 0 | 0 |
| SRR1853807 | 17 | 23 | 0 | 24 | 46 | 917 | 164 | 2947 | 33 | 0 | 21 | 11 | 0 | 0 | 66 | 4982 | 20621 | 196 | 98 | 491 | 10119 | 0 | 0 | 16 | 21 | 63 | 42 | 0 | 0 | 53 | 0 | 229 | 0 | 0 | 1677 | 0 | 0 |
| SRR1853808 | 48 | 74 | 67 | 888 | 122 | 301 | 0 | 752 | 150 | 0 | 55 | 29 | 0 | 0 | 451 | 24493 | 40532 | 150 | 301 | 2104 | 13378 | 0 | 0 | 103 | 45 | 125 | 417 | 32 | 0 | 77 | 0 | 301 | 0 | 0 | 3730 | 0 | 0 |
| SRR1853809 | 15 | 42 | 40 | 580 | 55 | 1159 | 901 | 2189 | 0 | 0 | 45 | 20 | 0 | 0 | 343 | 30537 | 78514 | 472 | 515 | 3220 | 38034 | 0 | 0 | 32 | 22 | 92 | 396 | 0 | 0 | 135 | 0 | 644 | 0 | 0 | 1815 | 0 | 0 |
| SRR1917323 | 263 | 77 | 98 | 52 | 275 | 2903 | 735 | 3031 | 4315 | 323 | 58 | 28 | 36 | 31 | 28881 | 77527 | 93126 | 183 | 505 | 517 | 13676 | 0 | 0 | 97 | 20 | 58 | 99 | 0 | 0 | 177 | 0 | 35 | 0 | 16 | 5968 | 12087 | 0 |
| SRR1917324 | 4488 | 849 | 56 | 24 | 7107 | 1853 | 485 | 2351 | 4224 | 2986 | 713 | 250 | 82 | 55 | 20797 | 47177 | 63926 | 105 | 302 | 339 | 8790 | 0 | 0 | 58 | 25 | 53 | 90 | 0 | 0 | 2125 | 0 | 101 | 0 | 0 | 4752 | 8763 | 0 |
| SRR1917325 | 319 | 86 | 126 | 32 | 327 | 2923 | 1219 | 6850 | 11814 | 251 | 50 | 18 | 39 | 29 | 48978 | 79416 | 160973 | 191 | 809 | 669 | 16387 | 0 | 0 | 154 | 48 | 66 | 110 | 0 | 0 | 248 | 0 | 31 | 0 | 14 | 3794 | 11485 | 0 |
| SRR1917326 | 0 | 0 | 0 | 0 | 1327 | 0 | 0 | 0 | 0 | 349 | 32 | 0 | 0 | 0 | 277916 | 192617 | 254307 | 43 | 1118 | 1648 | 11853 | 235 | 546 | 0 | 0 | 0 | 0 | 0 | 0 | 353 | 120 | 18 | 0 | 0 | 13198 | 2831 | 0 |
| SRR1917327 | 0 | 0 | 0 | 0 | 1290 | 0 | 0 | 12 | 16 | 147 | 18 | 0 | 0 | 0 | 237554 | 266606 | 263762 | 41 | 1874 | 1366 | 12819 | 54 | 101 | 0 | 0 | 0 | 0 | 0 | 0 | 76 | 16 | 10 | 0 | 0 | 8641 | 2516 | 0 |
| SRR1917328 | 0 | 0 | 0 | 0 | 214 | 18 | 0 | 0 | 18 | 20 | 0 | 0 | 0 | 0 | 41529 | 276630 | 59565 | 46 | 416 | 453 | 25488 | 0 | 0 | 0 | 0 | 0 | 0 | 0 | 0 | 0 | 0 | 0 | 0 | 0 | 5000 | 327 | 0 |
| SRR1917329 | 162 | 47 | 84 | 57 | 352 | 5893 | 2302 | 12286 | 17068 | 112 | 60 | 69 | 24 | 20 | 44634 | 76158 | 83272 | 285 | 1329 | 1112 | 13967 | 0 | 0 | 173 | 99 | 108 | 11 | 0 | 0 | 227 | 0 | 36 | 0 | 12 | 5574 | 15618 | 0 |
| SRR1917330 | 243 | 70 | 97 | 68 | 403 | 6708 | 1703 | 11574 | 12277 | 164 | 79 | 17 | 34 | 20 | 45636 | 81549 | 81690 | 231 | 1419 | 1174 | 18241 | 0 | 0 | 150 | 62 | 60 | 117 | 0 | 0 | 197 | 0 | 107 | 0 | 12 | 4093 | 14818 | 0 |
| SRR1917331 | 1721 | 801 | 201 | 167 | 2747 | 6793 | 1990 | 12572 | 14476 | 3664 | 1124 | 175 | 27 | 17 | 38658 | 85230 | 72636 | 345 | 1841 | 1428 | 28776 | 0 | 0 | 129 | 40 | 65 | 138 | 31 | 0 | 3323 | 0 | 94 | 0 | 14 | 4770 | 16774 | 0 |
| SRR1917332 | 0 | 0 | 0 | 0 | 429 | 24 | 24 | 0 | 118 | 55 | 20 | 0 | 17 | 0 | 142776 | 189161 | 359034 | 24 | 1395 | 14089 | 19691 | 162 | 340 | 0 | 0 | 0 | 0 | 0 | 0 | 123 | 45 | 189 | 0 | 0 | 8449 | 490 | 0 |
| SRR1917333 | 0 | 0 | 0 | 0 | 870 | 60 | 0 | 20 | 40 | 58 | 13 | 0 | 15 | 0 | 159738 | 209224 | 351399 | 181 | 1526 | 5299 | 12164 | 180 | 352 | 0 | 0 | 0 | 0 | 0 | 0 | 71 | 39 | 80 | 0 | 0 | 11219 | 428 | 0 |
| SRR1917334 | 0 | 0 | 0 | 0 | 947 | 0 | 0 | 19 | 0 | 581 | 82 | 0 | 35 | 0 | 141631 | 141230 | 330286 | 127 | 1752 | 6108 | 18248 | 468 | 1630 | 0 | 0 | 0 | 0 | 0 | 0 | 652 | 148 | 83 | 0 | 0 | 8351 | 565 | 0 |

### Supp Table S8.3.Columns_BX_to_DH

| | | | | | | | | | | | | | | | | | | | | | | | | | | | | | | | | | | | | | |
|---|---|---|---|---|---|---|---|---|---|---|---|---|---|---|---|---|---|---|---|---|---|---|---|---|---|---|---|---|---|---|---|---|---|---|---|---|---|
| SRR1917335 | 244 | 78 | 110 | 49 | 337 | 4113 | 1056 | 5518 | 8960 | 414 | 66 | 14 | 34 | 35 | 31310 | 57212 | 89987 | 259 | 838 | 937 | 13985 | 0 | 0 | 115 | 36 | 59 | 109 | 0 | 0 | 301 | 0 | 53 | 0 | 14 | 4457 | 11967 | 0 |
| SRR1917336 | 5540 | 1774 | 48 | 30 | 5130 | 6990 | 884 | 6326 | 5774 | 4238 | 642 | 388 | 66 | 42 | 30273 | 37810 | 74247 | 238 | 1044 | 341 | 13669 | 0 | 0 | 136 | 36 | 40 | 59 | 0 | 0 | 2365 | 0 | 76 | 0 | 16 | 5398 | 13900 | 0 |
| SRR1917337 | 199 | 79 | 74 | 27 | 236 | 10585 | 1673 | 15052 | 12210 | 308 | 67 | 50 | 47 | 32 | 41283 | 56924 | 86125 | 447 | 1687 | 742 | 11730 | 0 | 0 | 116 | 45 | 48 | 92 | 0 | 13 | 288 | 0 | 82 | 0 | 21 | 3457 | 19964 | 0 |
| SRR1917338 | 0 | 0 | 0 | 0 | 1189 | 0 | 0 | 0 | 0 | 79 | 16 | 0 | 0 | 0 | 20418 | 276388 | 79376 | 43 | 663 | 289 | 9141 | 117 | 244 | 0 | 0 | 0 | 0 | 0 | 0 | 30 | 23 | 0 | 0 | 0 | 33837 | 361 | 0 |
| SRR1917339 | 15 | 11 | 0 | 0 | 230 | 0 | 0 | 0 | 0 | 27 | 21 | 0 | 0 | 0 | 34952 | 320726 | 157829 | 21 | 319 | 226 | 18999 | 18 | 40 | 0 | 0 | 0 | 0 | 0 | 0 | 13 | 17 | 21 | 0 | 0 | 14217 | 337 | 0 |
| SRR1917340 | 14 | 0 | 0 | 0 | 462 | 0 | 0 | 0 | 0 | 191 | 31 | 0 | 10 | 0 | 42057 | 237432 | 178054 | 0 | 604 | 336 | 30484 | 97 | 155 | 0 | 0 | 0 | 0 | 0 | 0 | 29 | 21 | 0 | 0 | 0 | 9024 | 295 | 0 |
| SRR1922516 | 0 | 0 | 0 | 0 | 0 | 133 | 241 | 512 | 34 | 0 | 0 | 0 | 0 | 0 | 10724 | 6195 | 17556 | 34 | 26 | 564 | 1218 | 0 | 0 | 0 | 0 | 0 | 0 | 0 | 0 | 52 | 0 | 34 | 0 | 0 | 848 | 34 | 0 |
| SRR1922517 | 0 | 0 | 0 | 0 | 0 | 133 | 162 | 389 | 32 | 0 | 0 | 0 | 0 | 0 | 6964 | 4000 | 11271 | 14 | 36 | 381 | 1540 | 0 | 0 | 0 | 0 | 0 | 0 | 0 | 0 | 63 | 0 | 14 | 0 | 0 | 241 | 22 | 0 |
| SRR1922518 | 0 | 0 | 0 | 0 | 0 | 167 | 225 | 494 | 52 | 0 | 0 | 0 | 0 | 0 | 7987 | 5851 | 15005 | 15 | 20 | 362 | 928 | 0 | 0 | 0 | 0 | 0 | 0 | 0 | 0 | 101 | 0 | 0 | 0 | 0 | 466 | 18 | 0 |
| SRR1922519 | 0 | 0 | 0 | 0 | 0 | 145 | 148 | 298 | 25 | 0 | 0 | 0 | 0 | 0 | 4750 | 3333 | 8379 | 16 | 0 | 151 | 1018 | 0 | 0 | 15 | 0 | 0 | 11 | 0 | 0 | 33 | 0 | 0 | 0 | 0 | 258 | 16 | 0 |
| SRR1922520 | 0 | 0 | 0 | 0 | 0 | 191 | 237 | 516 | 37 | 0 | 0 | 0 | 0 | 0 | 10076 | 7631 | 19807 | 0 | 0 | 172 | 1208 | 0 | 0 | 0 | 0 | 0 | 0 | 0 | 0 | 45 | 0 | 0 | 0 | 0 | 188 | 22 | 0 |
| SRR1922521 | 0 | 0 | 0 | 0 | 0 | 220 | 256 | 537 | 25 | 0 | 0 | 0 | 0 | 0 | 8321 | 4519 | 12494 | 18 | 40 | 573 | 1376 | 0 | 0 | 0 | 0 | 0 | 0 | 0 | 0 | 43 | 0 | 54 | 0 | 0 | 244 | 15 | 0 |
| SRR1922522 | 0 | 0 | 0 | 0 | 0 | 220 | 297 | 569 | 24 | 0 | 0 | 0 | 0 | 0 | 9964 | 6751 | 19242 | 0 | 14 | 297 | 713 | 0 | 0 | 0 | 0 | 0 | 0 | 0 | 0 | 64 | 0 | 0 | 0 | 0 | 342 | 38 | 0 |
| SRR1922523 | 0 | 0 | 0 | 0 | 0 | 169 | 277 | 487 | 51 | 0 | 0 | 0 | 0 | 0 | 7158 | 3605 | 9689 | 21 | 31 | 441 | 1009 | 0 | 0 | 0 | 0 | 0 | 0 | 0 | 0 | 24 | 0 | 10 | 0 | 0 | 329 | 22 | 0 |
| SRR1922524 | 0 | 0 | 0 | 0 | 0 | 254 | 257 | 879 | 54 | 0 | 0 | 0 | 0 | 0 | 14469 | 8651 | 22034 | 18 | 44 | 524 | 1204 | 0 | 0 | 0 | 0 | 0 | 0 | 0 | 0 | 77 | 0 | 145 | 0 | 0 | 312 | 21 | 0 |
| SRR1922525 | 0 | 0 | 0 | 0 | 0 | 199 | 338 | 521 | 11 | 0 | 0 | 0 | 0 | 0 | 8827 | 6214 | 15161 | 11 | 22 | 183 | 925 | 0 | 0 | 0 | 0 | 0 | 0 | 0 | 0 | 18 | 0 | 44 | 0 | 0 | 422 | 41 | 0 |
| SRR1922526 | 0 | 0 | 0 | 0 | 0 | 232 | 304 | 819 | 43 | 0 | 0 | 0 | 0 | 0 | 18482 | 14600 | 29351 | 0 | 0 | 268 | 913 | 0 | 0 | 11 | 0 | 0 | 0 | 0 | 0 | 95 | 0 | 14 | 0 | 0 | 324 | 76 | 0 |
| SRR1922527 | 0 | 0 | 0 | 0 | 0 | 227 | 383 | 721 | 52 | 0 | 0 | 0 | 0 | 0 | 5918 | 3300 | 9702 | 11 | 19 | 227 | 613 | 0 | 0 | 11 | 0 | 0 | 0 | 0 | 0 | 28 | 0 | 22 | 0 | 0 | 469 | 39 | 0 |
| SRR1973407 | 21 | 32 | 53 | 106 | 127 | 0 | 136 | 746 | 136 | 11 | 32 | 11 | 0 | 0 | 32569 | 41957 | 256560 | 0 | 68 | 6107 | 8617 | 0 | 11 | 223 | 318 | 403 | 1197 | 201 | 0 | 583 | 0 | 68 | 0 | 0 | 3039 | 0 | 0 |
| SRR1973408 | 47 | 24 | 35 | 35 | 189 | 0 | 77 | 542 | 0 | 0 | 106 | 12 | 0 | 0 | 52615 | 37414 | 325678 | 0 | 77 | 8834 | 9609 | 0 | 12 | 248 | 260 | 225 | 1017 | 106 | 0 | 461 | 0 | 77 | 0 | 0 | 3396 | 0 | 0 |
| SRR1973409 | 59 | 0 | 39 | 118 | 177 | 0 | 189 | 520 | 47 | 59 | 79 | 0 | 0 | 0 | 57906 | 25416 | 311920 | 47 | 189 | 8556 | 9076 | 0 | 0 | 158 | 138 | 207 | 896 | 108 | 0 | 335 | 0 | 95 | 0 | 0 | 2855 | 0 | 0 |
| SRR1973410 | 40 | 0 | 60 | 139 | 268 | 0 | 460 | 1437 | 57 | 20 | 79 | 0 | 0 | 0 | 115639 | 50538 | 559032 | 0 | 57 | 6782 | 5862 | 0 | 0 | 377 | 189 | 328 | 1093 | 139 | 0 | 298 | 0 | 0 | 0 | 0 | 2619 | 14 | 0 |
| SRR1973411 | 0 | 0 | 0 | 11 | 18 | 0 | 483 | 1448 | 0 | 31 | 62 | 31 | 0 | 0 | 83273 | 53774 | 513774 | 0 | 483 | 4586 | 6276 | 0 | 31 | 93 | 22 | 18 | 70 | 93 | 0 | 20 | 31 | 0 | 0 | 0 | 3962 | 0 | 0 |
| SRR1973412 | 30 | 60 | 15 | 12 | 15 | 0 | 737 | 1014 | 0 | 0 | 0 | 0 | 0 | 0 | 138881 | 50661 | 604279 | 0 | 92 | 5161 | 5437 | 0 | 0 | 14 | 12 | 19 | 44 | 150 | 0 | 150 | 15 | 0 | 0 | 0 | 2758 | 0 | 0 |
| SRR1973413 | 22 | 22 | 56 | 112 | 349 | 0 | 465 | 542 | 0 | 11 | 0 | 11 | 0 | 0 | 125978 | 51583 | 617292 | 0 | 77 | 2867 | 4804 | 0 | 0 | 135 | 180 | 259 | 843 | 180 | 0 | 169 | 0 | 0 | 0 | 0 | 4917 | 0 | 0 |
| SRR1973414 | 15 | 29 | 15 | 293 | 293 | 0 | 188 | 754 | 188 | 15 | 29 | 0 | 0 | 0 | 151766 | 74637 | 660762 | 0 | 0 | 1507 | 4522 | 0 | 0 | 264 | 205 | 425 | 982 | 205 | 0 | 29 | 0 | 0 | 0 | 0 | 4379 | 0 | 29 |
| SRR1973415 | 32 | 0 | 129 | 129 | 274 | 0 | 605 | 847 | 0 | 32 | 0 | 16 | 0 | 0 | 219102 | 50666 | 765959 | 0 | 0 | 484 | 2300 | 0 | 32 | 226 | 161 | 387 | 1031 | 258 | 0 | 32 | 0 | 0 | 0 | 0 | 2873 | 0 | 0 |
| SRR1973952 | 8757 | 4965 | 2386 | 1496 | 52 | 1362 | 730 | 1216 | 4476 | 64 | 684 | 116 | 168 | 39 | 2968 | 5655 | 11576 | 0 | 341 | 1897 | 5595 | 0 | 0 | 555 | 155 | 168 | 258 | 103 | 0 | 890 | 0 | 0 | 0 | 13 | 533 | 1943 | 0 |
| SRR1973953 | 108 | 25 | 70 | 67 | 22 | 4504 | 875 | 3052 | 5468 | 146 | 882 | 10 | 0 | 0 | 20987 | 21140 | 74703 | 805 | 2694 | 3022 | 1541 | 0 | 0 | 214 | 392 | 78 | 73 | 155 | 0 | 3678 | 0 | 0 | 0 | 0 | 18776 | 5468 | 0 |
| SRR1973956 | 0 | 0 | 0 | 0 | 15 | 0 | 0 | 0 | 0 | 231 | 51 | 12 | 0 | 0 | 9382 | 415133 | 46935 | 0 | 110 | 0 | 165 | 15 | 25 | 0 | 0 | 0 | 0 | 0 | 38 | 34 | 25 | 0 | 0 | 13 | 13555 | 0 | 0 |
| SRR1973958 | 0 | 0 | 0 | 0 | 35 | 0 | 0 | 0 | 0 | 19 | 31 | 0 | 0 | 0 | 3649 | 422312 | 58434 | 0 | 58 | 170 | 389 | 0 | 0 | 0 | 0 | 0 | 0 | 0 | 0 | 26 | 0 | 13 | 0 | 0 | 1157 | 0 | 0 |
| SRR1973964 | 0 | 0 | 0 | 0 | 0 | 0 | 0 | 0 | 0 | 27 | 635 | 0 | 0 | 0 | 940 | 200337 | 2207 | 0 | 0 | 18 | 121 | 0 | 0 | 0 | 0 | 0 | 0 | 0 | 0 | 65 | 11 | 18 | 0 | 0 | 207 | 365 | 0 |
| SRR1973966 | 33 | 10 | 13 | 20 | 170 | 6468 | 2424 | 6208 | 8187 | 818 | 276 | 12 | 0 | 0 | 11736 | 65311 | 45335 | 952 | 2016 | 1657 | 482 | 0 | 0 | 56 | 75 | 42 | 26 | 65 | 0 | 2737 | 0 | 0 | 0 | 0 | 78843 | 0 | 0 |
| SRR1973967 | 31 | 12 | 18 | 35 | 38 | 10635 | 1402 | 2916 | 5527 | 281 | 299 | 0 | 0 | 0 | 6526 | 42655 | 15489 | 2256 | 1337 | 983 | 661 | 0 | 0 | 48 | 26 | 12 | 0 | 0 | 0 | 922 | 0 | 0 | 0 | 16 | 53772 | 12 | 0 |
| SRR1979367 | 33 | 0 | 0 | 0 | 25 | 1463 | 694 | 1119 | 665 | 17 | 0 | 0 | 0 | 0 | 105 | 1447 | 1301 | 0 | 12 | 490 | 58 | 0 | 0 | 39 | 41 | 33 | 149 | 17 | 0 | 33 | 0 | 12 | 0 | 0 | 838 | 7105 | 17 |
| SRR1988280 | 16 | 37 | 24 | 21 | 26 | 153 | 213 | 1221 | 89 | 0 | 13 | 0 | 0 | 0 | 6656 | 124803 | 22465 | 0 | 59 | 2512 | 475 | 0 | 0 | 30 | 14 | 37 | 29 | 0 | 0 | 41 | 0 | 119 | 0 | 0 | 1293 | 0 | 0 |
| SRR1988281 | 51 | 44 | 39 | 25 | 29 | 110 | 219 | 1165 | 57 | 0 | 0 | 0 | 0 | 0 | 6960 | 172406 | 29376 | 0 | 84 | 2299 | 758 | 0 | 0 | 51 | 16 | 45 | 170 | 0 | 0 | 47 | 0 | 115 | 0 | 0 | 1166 | 0 | 0 |
| SRR1988282 | 13 | 10 | 16 | 18 | 27 | 321 | 239 | 2046 | 144 | 0 | 11 | 0 | 0 | 0 | 6898 | 125969 | 22526 | 0 | 82 | 2930 | 498 | 0 | 0 | 11 | 13 | 34 | 39 | 0 | 0 | 19 | 0 | 173 | 0 | 0 | 1131 | 0 | 0 |
| SRR1988283 | 43 | 12 | 25 | 25 | 13 | 697 | 453 | 1879 | 201 | 14 | 0 | 0 | 0 | 0 | 8342 | 100222 | 29648 | 23 | 849 | 5386 | 252 | 0 | 0 | 21 | 22 | 13 | 38 | 0 | 0 | 18 | 0 | 116 | 0 | 0 | 1893 | 0 | 0 |
| SRR1988284 | 35 | 71 | 12 | 25 | 12 | 623 | 429 | 1554 | 162 | 0 | 12 | 0 | 0 | 0 | 6286 | 87519 | 21828 | 0 | 597 | 5135 | 304 | 0 | 0 | 88 | 15 | 61 | 244 | 0 | 0 | 80 | 0 | 84 | 0 | 0 | 2182 | 0 | 0 |
| SRR1988285 | 38 | 78 | 20 | 21 | 0 | 646 | 427 | 2332 | 229 | 13 | 10 | 12 | 0 | 0 | 6285 | 80415 | 22512 | 0 | 512 | 4129 | 293 | 0 | 0 | 79 | 14 | 72 | 214 | 10 | 0 | 79 | 0 | 48 | 0 | 0 | 1626 | 0 | 0 |
| SRR1988286 | 42 | 11 | 18 | 27 | 0 | 799 | 473 | 1865 | 178 | 14 | 12 | 0 | 0 | 0 | 6574 | 142621 | 23048 | 0 | 1404 | 8795 | 310 | 0 | 0 | 19 | 16 | 16 | 32 | 0 | 0 | 23 | 0 | 86 | 0 | 0 | 1813 | 0 | 0 |
| SRR1988287 | 20 | 23 | 30 | 24 | 11 | 239 | 192 | 476 | 77 | 25 | 0 | 0 | 0 | 0 | 165342 | 23139 | 146571 | 0 | 102 | 25381 | 4918 | 0 | 0 | 29 | 31 | 32 | 40 | 0 | 0 | 27 | 0 | 82 | 0 | 0 | 416 | 13 | 0 |
| SRR1988288 | 77 | 65 | 26 | 26 | 41 | 328 | 283 | 911 | 85 | 21 | 0 | 0 | 0 | 0 | 173161 | 26140 | 153420 | 0 | 152 | 23857 | 4873 | 0 | 0 | 67 | 111 | 96 | 156 | 0 | 0 | 36 | 0 | 88 | 0 | 0 | 561 | 15 | 0 |
| SRR1988289 | 60 | 55 | 20 | 20 | 21 | 1212 | 404 | 2937 | 176 | 14 | 0 | 0 | 0 | 0 | 138375 | 26882 | 126490 | 17 | 153 | 21808 | 4120 | 0 | 0 | 53 | 70 | 64 | 86 | 0 | 0 | 36 | 0 | 162 | 0 | 0 | 482 | 18 | 0 |
| SRR1988290 | 23 | 17 | 10 | 17 | 24 | 855 | 246 | 2074 | 128 | 22 | 0 | 0 | 0 | 0 | 151137 | 23103 | 145628 | 11 | 173 | 22712 | 4742 | 0 | 0 | 17 | 20 | 20 | 30 | 0 | 0 | 16 | 0 | 179 | 0 | 0 | 441 | 21 | 0 |
| SRR1988291 | 45 | 49 | 15 | 19 | 76 | 377 | 383 | 5200 | 167 | 13 | 0 | 10 | 0 | 0 | 4041 | 16335 | 14887 | 0 | 30 | 1638 | 532 | 0 | 0 | 33 | 21 | 56 | 203 | 0 | 0 | 69 | 0 | 72 | 0 | 0 | 1168 | 0 | 0 |
| SRR1988292 | 46 | 59 | 17 | 28 | 71 | 409 | 500 | 7667 | 165 | 27 | 11 | 0 | 0 | 0 | 4446 | 16852 | 17061 | 12 | 24 | 1665 | 531 | 0 | 0 | 51 | 25 | 42 | 151 | 10 | 0 | 90 | 0 | 79 | 0 | 0 | 1505 | 0 | 0 |
| SRR1988293 | 44 | 50 | 21 | 35 | 77 | 365 | 397 | 6063 | 199 | 16 | 0 | 0 | 0 | 0 | 3888 | 17050 | 16557 | 11 | 43 | 1976 | 483 | 0 | 0 | 41 | 16 | 69 | 189 | 0 | 0 | 84 | 0 | 118 | 0 | 0 | 1134 | 0 | 0 |
| SRR1988294 | 47 | 10 | 20 | 20 | 14 | 333 | 398 | 4904 | 149 | 13 | 18 | 11 | 0 | 0 | 3668 | 16008 | 15397 | 12 | 36 | 1956 | 535 | 0 | 0 | 46 | 15 | 11 | 37 | 11 | 0 | 16 | 0 | 77 | 0 | 0 | 1484 | 0 | 0 |
| SRR1988295 | 65 | 91 | 0 | 15 | 93 | 94 | 129 | 805 | 79 | 33 | 23 | 0 | 0 | 0 | 44609 | 33462 | 85392 | 119 | 233 | 26561 | 954 | 0 | 0 | 21 | 27 | 42 | 187 | 0 | 0 | 49 | 0 | 1271 | 0 | 0 | 806 | 0 | 0 |
| SRR1988296 | 74 | 67 | 16 | 22 | 83 | 115 | 106 | 653 | 55 | 28 | 10 | 0 | 0 | 0 | 54752 | 31625 | 114639 | 124 | 193 | 37154 | 1342 | 0 | 0 | 22 | 20 | 45 | 133 | 0 | 0 | 61 | 0 | 1714 | 0 | 0 | 772 | 0 | 0 |
| SRR1988297 | 66 | 70 | 16 | 24 | 70 | 121 | 121 | 513 | 27 | 34 | 11 | 0 | 0 | 0 | 50321 | 27443 | 102641 | 98 | 149 | 35679 | 1128 | 0 | 0 | 29 | 19 | 39 | 150 | 0 | 0 | 41 | 0 | 1371 | 0 | 0 | 660 | 0 | 0 |
| SRR1988298 | 10 | 12 | 12 | 12 | 13 | 63 | 66 | 497 | 36 | 26 | 0 | 0 | 0 | 0 | 134283 | 24562 | 212336 | 53 | 342 | 19479 | 1730 | 0 | 0 | 22 | 22 | 33 | 22 | 0 | 0 | 37 | 0 | 875 | 0 | 0 | 540 | 0 | 0 |
| SRR1988299 | 52 | 42 | 34 | 83 | 75 | 359 | 529 | 3634 | 90 | 31 | 13 | 0 | 0 | 0 | 4902 | 22642 | 30304 | 0 | 50 | 2256 | 1008 | 0 | 0 | 60 | 21 | 59 | 392 | 17 | 0 | 52 | 0 | 40 | 0 | 0 | 1360 | 0 | 0 |
| SRR1988300 | 77 | 74 | 24 | 40 | 11 | 611 | 683 | 5168 | 230 | 75 | 19 | 0 | 0 | 0 | 3506 | 15610 | 22463 | 0 | 33 | 2338 | 637 | 0 | 0 | 51 | 15 | 72 | 35 | 12 | 0 | 21 | 0 | 289 | 0 | 0 | 3189 | 0 | 0 |
| SRR1988301 | 10 | 86 | 35 | 11 | 13 | 265 | 510 | 4392 | 119 | 48 | 26 | 0 | 0 | 0 | 3518 | 21810 | 28092 | 0 | 40 | 2199 | 490 | 0 | 0 | 37 | 11 | 31 | 74 | 0 | 0 | 15 | 0 | 172 | 0 | 0 | 1958 | 0 | 0 |
| SRR1988302 | 63 | 50 | 19 | 64 | 77 | 370 | 450 | 4986 | 120 | 36 | 18 | 0 | 0 | 0 | 3784 | 15665 | 24203 | 0 | 46 | 2251 | 701 | 0 | 0 | 38 | 14 | 103 | 323 | 19 | 0 | 117 | 0 | 234 | 0 | 0 | 1596 | 0 | 0 |
| SRR1988303 | 58 | 53 | 20 | 51 | 75 | 258 | 484 | 3519 | 186 | 43 | 12 | 0 | 0 | 0 | 4837 | 15386 | 23241 | 0 | 86 | 2015 | 696 | 0 | 0 | 43 | 0 | 61 | 277 | 11 | 0 | 118 | 0 | 252 | 0 | 0 | 2415 | 0 | 0 |
| SRR1988304 | 67 | 68 | 17 | 66 | 72 | 203 | 519 | 3341 | 102 | 34 | 24 | 11 | 0 | 0 | 4921 | 12576 | 19847 | 0 | 56 | 2110 | 925 | 0 | 0 | 21 | 13 | 42 | 216 | 16 | 0 | 86 | 0 | 90 | 0 | 0 | 1828 | 0 | 0 |
| SRR1988305 | 41 | 61 | 28 | 54 | 73 | 242 | 457 | 2812 | 161 | 52 | 16 | 0 | 0 | 0 | 5362 | 15765 | 28173 | 0 | 27 | 2025 | 780 | 0 | 0 | 43 | 15 | 58 | 310 | 16 | 0 | 78 | 0 | 161 | 0 | 0 | 1309 | 0 | 0 |
| SRR1988306 | 53 | 75 | 16 | 38 | 49 | 231 | 429 | 2926 | 110 | 13 | 11 | 0 | 0 | 0 | 2981 | 8122 | 14643 | 0 | 55 | 1540 | 506 | 0 | 0 | 38 | 0 | 51 | 39 | 15 | 0 | 87 | 0 | 11 | 0 | 0 | 1778 | 0 | 15 |
| SRR1988307 | 10 | 11 | 41 | 51 | 86 | 399 | 570 | 3608 | 120 | 51 | 17 | 20 | 0 | 0 | 2964 | 10935 | 13604 | 0 | 46 | 3813 | 416 | 0 | 0 | 76 | 26 | 11 | 58 | 15 | 0 | 80 | 0 | 245 | 0 | 0 | 1665 | 0 | 0 |
| SRR1988308 | 67 | 13 | 28 | 11 | 11 | 451 | 451 | 4512 | 137 | 73 | 24 | 24 | 0 | 0 | 2562 | 21250 | 16187 | 0 | 46 | 5175 | 435 | 0 | 0 | 75 | 13 | 62 | 37 | 19 | 0 | 12 | 0 | 192 | 0 | 0 | 2061 | 0 | 0 |
| SRR1988309 | 10 | 89 | 37 | 72 | 11 | 330 | 303 | 2120 | 128 | 44 | 35 | 11 | 0 | 0 | 3588 | 8522 | 10439 | 0 | 64 | 2817 | 349 | 0 | 0 | 74 | 0 | 10 | 60 | 28 | 0 | 12 | 0 | 128 | 0 | 0 | 979 | 0 | 0 |
| SRR1988310 | 99 | 59 | 27 | 50 | 91 | 201 | 358 | 1862 | 131 | 34 | 19 | 0 | 0 | 0 | 4706 | 9588 | 16932 | 0 | 48 | 2473 | 545 | 0 | 0 | 54 | 21 | 96 | 523 | 22 | 0 | 106 | 0 | 92 | 0 | 0 | 1633 | 0 | 0 |
| SRR1988311 | 72 | 68 | 33 | 55 | 155 | 240 | 304 | 2159 | 64 | 129 | 16 | 14 | 0 | 0 | 5334 | 55994 | 27004 | 0 | 56 | 6629 | 712 | 0 | 0 | 143 | 18 | 98 | 440 | 45 | 0 | 84 | 0 | 264 | 0 | 0 | 1601 | 0 | 0 |
| SRR1988312 | 77 | 107 | 19 | 46 | 104 | 64 | 241 | 891 | 48 | 71 | 27 | 13 | 0 | 0 | 4642 | 56191 | 24319 | 0 | 32 | 7493 | 634 | 0 | 0 | 125 | 17 | 69 | 22 | 44 | 0 | 11 | 0 | 217 | 0 | 0 | 1649 | 0 | 10 |
| SRR2052873 | 242 | 289 | 30 | 110 | 565 | 3483 | 2099 | 17853 | 755 | 35 | 56 | 272 | 0 | 0 | 11953 | 17452 | 59088 | 54 | 101 | 1716 | 269 | 0 | 0 | 201 | 42 | 177 | 501 | 35 | 0 | 576 | 0 | 22 | 0 | 0 | 7355 | 0 | 17 |
| SRR2052874 | 179 | 147 | 23 | 102 | 773 | 3338 | 2236 | 18054 | 675 | 34 | 74 | 252 | 0 | 0 | 19840 | 16208 | 82305 | 23 | 65 | 1586 | 341 | 0 | 0 | 147 | 38 | 178 | 328 | 33 | 0 | 649 | 0 | 14 | 0 | 0 | 6791 | 0 | 18 |
| SRR2052875 | 178 | 200 | 31 | 109 | 983 | 2525 | 1906 | 13702 | 679 | 49 | 55 | 340 | 0 | 0 | 14948 | 30653 | 67474 | 0 | 113 | 1753 | 143 | 0 | 0 | 83 | 26 | 127 | 486 | 30 | 0 | 408 | 0 | 14 | 0 | 0 | 8437 | 0 | 20 |
| SRR2052877 | 140 | 70 | 10 | 77 | 310 | 2460 | 1374 | 16699 | 580 | 19 | 76 | 235 | 0 | 0 | 6866 | 16656 | 46882 | 32 | 107 | 1162 | 111 | 0 | 0 | 55 | 17 | 83 | 175 | 20 | 0 | 319 | 0 | 13 | 0 | 0 | 6671 | 0 | 0 |
| SRR2052878 | 32 | 11 | 14 | 13 | 54 | 2322 | 1203 | 13269 | 638 | 30 | 42 | 40 | 0 | 0 | 8569 | 11200 | 48228 | 34 | 51 | 1887 | 164 | 0 | 0 | 18 | 44 | 17 | 47 | 28 | 0 | 45 | 0 | 56 | 0 | 0 | 7820 | 0 | 11 |
| SRR2053027 | 137 | 112 | 0 | 68 | 586 | 1113 | 1012 | 10750 | 385 | 12 | 125 | 17 | 0 | 0 | 8525 | 40284 | 48069 | 11 | 88 | 1502 | 135 | 0 | 0 | 154 | 75 | 365 | 835 | 62 | 0 | 832 | 0 | 11 | 0 | 0 | 3513 | 0 | 0 |
| SRR2053028 | 167 | 110 | 18 | 78 | 476 | 1455 | 1024 | 11848 | 474 | 47 | 109 | 26 | 0 | 0 | 6388 | 28313 | 45070 | 75 | 121 | 2173 | 77 | 0 | 0 | 139 | 73 | 260 | 586 | 42 | 0 | 679 | 0 | 21 | 0 | 0 | 5500 | 0 | 0 |
| SRR2053029 | 185 | 195 | 20 | 106 | 968 | 927 | 1099 | 9620 | 414 | 15 | 141 | 49 | 0 | 0 | 4998 | 26578 | 37566 | 70 | 107 | 2118 | 84 | 0 | 0 | 243 | 117 | 511 | 1409 | 135 | 0 | 862 | 0 | 37 | 0 | 0 | 4284 | 0 | 0 |
| SRR2061931 | 0 | 0 | 0 | 67 | 50 | 277 | 169 | 952 | 60 | 17 | 248 | 42 | 0 | 0 | 15590 | 11440 | 51537 | 0 | 229 | 44865 | 1096 | 0 | 0 | 50 | 105 | 441 | 2917 | 122 | 0 | 621 | 0 | 24 | 0 | 0 | 635 | 599 | 0 |
| SRR2061932 | 0 | 0 | 0 | 14 | 68 | 401 | 445 | 757 | 134 | 14 | 162 | 14 | 0 | 0 | 14603 | 14219 | 61749 | 0 | 267 | 41405 | 712 | 0 | 0 | 0 | 41 | 311 | 1269 | 365 | 0 | 365 | 0 | 0 | 0 | 0 | 482 | 275 | 0 |
| SRR2061933 | 0 | 0 | 0 | 30 | 17 | 480 | 680 | 1661 | 260 | 0 | 59 | 0 | 0 | 0 | 7923 | 10049 | 40791 | 0 | 280 | 39416 | 540 | 0 | 0 | 22 | 20 | 76 | 716 | 48 | 0 | 127 | 0 | 0 | 0 | 0 | 731 | 674 | 0 |
| SRR2061934 | 0 | 0 | 0 | 46 | 46 | 1028 | 93 | 1495 | 467 | 0 | 27 | 0 | 0 | 0 | 17101 | 28775 | 69610 | 0 | 280 | 28596 | 1028 | 0 | 0 | 0 | 46 | 54 | 260 | 81 | 0 | 67 | 0 | 0 | 0 | 0 | 763 | 351 | 0 |
| SRR2061935 | 0 | 0 | 0 | 51 | 25 | 743 | 531 | 1698 | 106 | 0 | 14 | 25 | 0 | 0 | 26112 | 32196 | 139829 | 0 | 106 | 25793 | 1911 | 0 | 0 | 51 | 51 | 49 | 216 | 36 | 0 | 38 | 0 | 0 | 0 | 0 | 445 | 444 | 0 |
| SRR2061936 | 0 | 0 | 0 | 48 | 102 | 238 | 442 | 1088 | 204 | 16 | 417 | 11 | 0 | 0 | 5575 | 9339 | 42388 | 0 | 68 | 32771 | 544 | 0 | 0 | 11 | 11 | 107 | 1159 | 53 | 0 | 1015 | 0 | 34 | 0 | 0 | 2671 | 1295 | 0 |
| SRR2061937 | 0 | 0 | 0 | 0 | 159 | 482 | 764 | 1648 | 523 | 0 | 289 | 0 | 0 | 0 | 2010 | 6331 | 25251 | 0 | 322 | 45506 | 925 | 0 | 0 | 14 | 0 | 43 | 902 | 130 | 0 | 786 | 0 | 0 | 0 | 0 | 1210 | 1107 | 14 |
| SRR2061938 | 0 | 0 | 0 | 0 | 83 | 685 | 657 | 1589 | 356 | 0 | 426 | 15 | 0 | 0 | 4629 | 5014 | 25910 | 55 | 164 | 39881 | 383 | 0 | 0 | 29 | 39 | 132 | 2594 | 117 | 0 | 1126 | 0 | 27 | 0 | 0 | 1533 | 1155 | 0 |
| SRR2061940 | 0 | 0 | 0 | 0 | 24 | 292 | 292 | 1020 | 292 | 24 | 212 | 0 | 0 | 0 | 3352 | 14630 | 58136 | 0 | 292 | 50722 | 1020 | 0 | 0 | 71 | 71 | 71 | 1841 | 472 | 0 | 354 | 0 | 0 | 0 | 0 | 958 | 577 | 0 |
| SRR2061941 | 24 | 73 | 36 | 219 | 1117 | 51 | 342 | 164 | 0 | 24 | 255 | 0 | 0 | 0 | 45679 | 267298 | 41415 | 0 | 63 | 16825 | 354 | 0 | 0 | 73 | 36 | 121 | 874 | 121 | 0 | 2028 | 0 | 0 | 0 | 0 | 1541 | 1218 | 12 |
| SRR2061942 | 22 | 30 | 37 | 21 | 51 | 82 | 321 | 250 | 65 | 30 | 36 | 0 | 0 | 0 | 44756 | 234247 | 26142 | 0 | 60 | 10619 | 806 | 0 | 0 | 41 | 48 | 29 | 116 | 28 | 0 | 148 | 0 | 0 | 0 | 0 | 1069 | 529 | 18 |
| SRR2061943 | 31 | 28 | 46 | 14 | 21 | 145 | 222 | 565 | 54 | 15 | 23 | 0 | 0 | 0 | 51847 | 167475 | 37294 | 0 | 113 | 11438 | 529 | 0 | 0 | 46 | 31 | 35 | 114 | 11 | 0 | 123 | 0 | 0 | 0 | 0 | 1781 | 410 | 0 |
| SRR2061944 | 15 | 0 | 49 | 89 | 213 | 73 | 97 | 121 | 24 | 0 | 277 | 0 | 0 | 0 | 36518 | 120945 | 16673 | 0 | 85 | 12732 | 863 | 0 | 0 | 35 | 35 | 119 | 737 | 84 | 0 | 1800 | 0 | 12 | 0 | 0 | 952 | 346 | 0 |
| SRR2061945 | 10 | 0 | 52 | 111 | 301 | 71 | 355 | 254 | 35 | 21 | 252 | 0 | 0 | 0 | 54597 | 208089 | 29482 | 12 | 136 | 10724 | 1076 | 0 | 0 | 73 | 42 | 173 | 806 | 176 | 0 | 799 | 0 | 0 | 0 | 0 | 1007 | 456 | 0 |
| SRR2061946 | 0 | 0 | 16 | 55 | 210 | 90 | 138 | 244 | 40 | 88 | 177 | 0 | 0 | 0 | 153198 | 133341 | 98302 | 0 | 185 | 8346 | 667 | 0 | 0 | 45 | 21 | 149 | 517 | 56 | 0 | 473 | 0 | 13 | 0 | 0 | 862 | 514 | 0 |
| SRR2061947 | 51 | 42 | 56 | 93 | 321 | 113 | 291 | 184 | 28 | 37 | 270 | 0 | 0 | 0 | 48036 | 202949 | 25985 | 0 | 128 | 20157 | 844 | 0 | 0 | 84 | 33 | 191 | 815 | 98 | 0 | 1435 | 0 | 0 | 0 | 0 | 1546 | 590 | 0 |
| SRR2061948 | 12 | 16 | 39 | 75 | 247 | 86 | 214 | 293 | 57 | 16 | 212 | 0 | 0 | 0 | 49269 | 185202 | 24460 | 0 | 136 | 9110 | 1063 | 0 | 0 | 55 | 59 | 263 | 867 | 220 | 0 | 812 | 0 | 0 | 0 | 0 | 1062 | 364 | 0 |
| SRR2061949 | 17 | 0 | 17 | 53 | 27 | 230 | 124 | 473 | 12 | 39 | 18 | 0 | 0 | 0 | 85977 | 170079 | 61255 | 0 | 95 | 11049 | 609 | 0 | 0 | 59 | 36 | 25 | 83 | 45 | 0 | 64 | 0 | 12 | 0 | 0 | 1145 | 432 | 0 |
| SRR2061950 | 54 | 54 | 59 | 31 | 116 | 25 | 255 | 74 | 25 | 11 | 47 | 0 | 0 | 0 | 8624 | 236674 | 15138 | 0 | 82 | 3871 | 2273 | 0 | 0 | 12 | 19 | 42 | 160 | 65 | 0 | 212 | 0 | 0 | 0 | 0 | 1554 | 0 | 0 |
| SRR2061951 | 45 | 38 | 66 | 287 | 1038 | 30 | 347 | 67 | 30 | 86 | 225 | 0 | 0 | 0 | 8371 | 310870 | 16045 | 0 | 85 | 3395 | 1709 | 0 | 0 | 69 | 69 | 332 | 855 | 630 | 0 | 920 | 0 | 0 | 0 | 0 | 1536 | 0 | 0 |
| SRR2061952 | 40 | 58 | 38 | 30 | 101 | 101 | 355 | 153 | 77 | 13 | 27 | 0 | 0 | 0 | 13142 | 354516 | 24478 | 10 | 91 | 3568 | 1107 | 0 | 0 | 12 | 11 | 46 | 120 | 53 | 0 | 44 | 0 | 31 | 0 | 0 | 1836 | 0 | 0 |
| SRR2061953 | 11 | 15 | 56 | 23 | 208 | 39 | 206 | 116 | 19 | 15 | 54 | 0 | 0 | 0 | 10743 | 415818 | 23026 | 0 | 97 | 5085 | 1076 | 0 | 0 | 12 | 97 | 27 | 130 | 52 | 0 | 164 | 0 | 0 | 0 | 0 | 2404 | 0 | 0 |
| SRR2061954 | 44 | 40 | 24 | 28 | 140 | 39 | 162 | 154 | 23 | 21 | 42 | 0 | 0 | 0 | 10205 | 305748 | 18057 | 0 | 31 | 2780 | 2357 | 0 | 0 | 69 | 19 | 51 | 111 | 104 | 0 | 108 | 0 | 31 | 0 | 0 | 2075 | 0 | 0 |
| SRR2061955 | 53 | 53 | 39 | 26 | 94 | 96 | 289 | 165 | 23 | 17 | 78 | 0 | 0 | 0 | 11864 | 285526 | 22488 | 11 | 45 | 3489 | 1532 | 0 | 0 | 15 | 11 | 55 | 114 | 58 | 0 | 18 | 0 | 17 | 0 | 0 | 1975 | 0 | 0 |
| SRR2061956 | 91 | 111 | 49 | 281 | 1438 | 37 | 110 | 78 | 21 | 199 | 480 | 0 | 0 | 0 | 10285 | 319488 | 18558 | 0 | 52 | 4625 | 1910 | 0 | 0 | 105 | 88 | 343 | 1147 | 451 | 0 | 2006 | 0 | 16 | 0 | 0 | 1964 | 0 | 0 |
| SRR2061957 | 75 | 71 | 60 | 33 | 119 | 33 | 190 | 85 | 33 | 29 | 49 | 0 | 0 | 0 | 10031 | 297023 | 18760 | 0 | 59 | 3163 | 2620 | 0 | 0 | 12 | 12 | 40 | 110 | 89 | 0 | 109 | 0 | 0 | 0 | 0 | 2290 | 0 | 0 |
| SRR2061958 | 52 | 39 | 33 | 212 | 856 | 65 | 375 | 245 | 55 | 212 | 192 | 0 | 0 | 0 | 15739 | 343993 | 23477 | 10 | 15 | 2944 | 1684 | 0 | 0 | 107 | 127 | 472 | 960 | 381 | 0 | 511 | 0 | 15 | 0 | 0 | 1542 | 0 | 0 |
| SRR2069629 | 0 | 0 | 0 | 0 | 117 | 15893 | 6734 | 15950 | 7255 | 109 | 305 | 0 | 0 | 0 | 14722 | 5702 | 8829 | 732 | 192 | 494 | 273 | 0 | 0 | 0 | 0 | 0 | 0 | 0 | 0 | 2259 | 0 | 404 | 0 | 0 | 4224 | 0 | 0 |
| SRR2069630 | 0 | 0 | 0 | 0 | 21 | 20365 | 5681 | 11443 | 2074 | 109 | 219 | 0 | 0 | 0 | 3209 | 657 | 2246 | 180 | 143 | 265 | 164 | 0 | 0 | 44 | 0 | 0 | 0 | 0 | 0 | 681 | 0 | 308 | 0 | 17 | 999 | 0 | 0 |
| SRR2084358 | 392 | 98 | 0 | 392 | 11 | 957 | 34 | 308 | 410 | 0 | 0 | 0 | 0 | 0 | 10048 | 14143 | 30714 | 1435 | 376 | 3828 | 11484 | 0 | 0 | 49 | 49 | 196 | 15 | 98 | 0 | 294 | 0 | 3008 | 49 | 0 | 7867 | 804 | 98 |
| SRR2084359 | 317 | 13 | 35 | 211 | 11 | 3716 | 169 | 2051 | 893 | 0 | 20 | 0 | 0 | 0 | 19813 | 43659 | 51540 | 1593 | 434 | 4658 | 10449 | 0 | 0 | 247 | 211 | 21 | 55 | 35 | 0 | 20 | 0 | 3330 | 35 | 0 | 26721 | 1223 | 0 |
| SRR2085173 | 0 | 0 | 0 | 0 | 0 | 0 | 28 | 137 | 14 | 0 | 0 | 0 | 0 | 0 | 30 | 1245 | 81 | 488 | 0 | 0 | 0 | 64 | 79 | 18 | 0 | 0 | 0 | 0 | 0 | 0 | 0 | 2449 | 0 | 0 | 166 | 466 | 20 |
| SRR2103379 | 74 | 24 | 23 | 37 | 80 | 1104 | 196 | 3111 | 146 | 177 | 115 | 16 | 0 | 0 | 17928 | 1890 | 21428 | 252 | 123 | 2302 | 6895 | 0 | 0 | 74 | 100 | 134 | 99 | 23 | 0 | 1366 | 0 | 13 | 0 | 0 | 658 | 0 | 0 |
| SRR2103380 | 63 | 21 | 32 | 54 | 97 | 980 | 225 | 3534 | 137 | 123 | 118 | 0 | 0 | 0 | 19083 | 2404 | 27382 | 337 | 129 | 2211 | 6818 | 0 | 0 | 57 | 96 | 151 | 113 | 30 | 0 | 1296 | 0 | 17 | 0 | 0 | 550 | 0 | 0 |
| SRR2103381 | 17 | 18 | 37 | 11 | 20 | 1006 | 211 | 3462 | 121 | 38 | 24 | 0 | 0 | 0 | 20232 | 2025 | 26592 | 268 | 92 | 2558 | 7405 | 0 | 0 | 15 | 20 | 30 | 23 | 27 | 0 | 283 | 0 | 30 | 0 | 0 | 569 | 0 | 0 |
| SRR2103382 | 96 | 23 | 52 | 123 | 116 | 4015 | 949 | 14544 | 215 | 105 | 37 | 0 | 0 | 0 | 7267 | 6186 | 39831 | 48 | 251 | 3389 | 3914 | 0 | 0 | 367 | 50 | 205 | 448 | 105 | 0 | 268 | 0 | 61 | 0 | 0 | 1641 | 0 | 0 |
| SRR2103383 | 100 | 19 | 52 | 95 | 103 | 5022 | 1019 | 15944 | 276 | 510 | 30 | 20 | 0 | 0 | 7126 | 6438 | 37869 | 27 | 254 | 3854 | 3315 | 0 | 0 | 409 | 50 | 179 | 370 | 70 | 0 | 322 | 0 | 55 | 0 | 0 | 1800 | 0 | 0 |
| SRR2103384 | 92 | 18 | 48 | 126 | 132 | 1091 | 364 | 5253 | 181 | 40 | 51 | 29 | 0 | 0 | 7561 | 6539 | 32212 | 19 | 203 | 4003 | 3823 | 0 | 0 | 150 | 69 | 229 | 261 | 46 | 0 | 410 | 0 | 56 | 0 | 0 | 1879 | 0 | 0 |
| SRR2105940 | 0 | 0 | 0 | 0 | 0 | 152 | 195 | 1235 | 0 | 0 | 0 | 0 | 0 | 0 | 0 | 0 | 4569 | 117 | 0 | 17340 | 152 | 0 | 0 | 238 | 0 | 0 | 0 | 0 | 0 | 0 | 0 | 0 | 0 | 0 | 5953 | 24 | 0 |
| SRR2105943 | 0 | 0 | 0 | 0 | 0 | 67 | 188 | 1256 | 272 | 0 | 0 | 22 | 0 | 0 | 0 | 0 | 1816 | 0 | 0 | 2074 | 0 | 0 | 0 | 51 | 0 | 0 | 0 | 0 | 0 | 0 | 0 | 0 | 0 | 0 | 25442 | 14 | 0 |
| SRR2105944 | 0 | 0 | 0 | 0 | 0 | 81 | 88 | 930 | 139 | 0 | 0 | 15 | 0 | 0 | 0 | 69 | 2001 | 55 | 0 | 3628 | 0 | 0 | 0 | 16 | 0 | 0 | 0 | 0 | 0 | 0 | 0 | 0 | 0 | 0 | 6599 | 34 | 0 |
| SRR2105949 | 0 | 0 | 0 | 0 | 0 | 957 | 1217 | 5130 | 817 | 0 | 0 | 0 | 0 | 0 | 0 | 452 | 1202 | 0 | 0 | 397 | 0 | 0 | 0 | 55 | 0 | 0 | 0 | 0 | 0 | 0 | 0 | 0 | 0 | 0 | 5712 | 54 | 0 |
| SRR2105951 | 0 | 0 | 0 | 0 | 0 | 0 | 0 | 1408 | 0 | 0 | 0 | 13 | 0 | 0 | 270 | 787 | 15640 | 0 | 0 | 2051 | 0 | 0 | 0 | 12 | 0 | 0 | 0 | 0 | 0 | 0 | 0 | 0 | 0 | 0 | 5601 | 0 | 0 |
| SRR2106873 | 88 | 90 | 0 | 18 | 81 | 268 | 169 | 361 | 128 | 0 | 14 | 0 | 21 | 0 | 5530 | 2638 | 7982 | 47 | 17 | 3298 | 1026 | 0 | 0 | 196 | 23 | 44 | 196 | 32 | 0 | 194 | 0 | 0 | 0 | 0 | 2702 | 492 | 0 |
| SRR2106874 | 33 | 15 | 0 | 0 | 11 | 248 | 248 | 399 | 54 | 15 | 33 | 0 | 15 | 0 | 8634 | 8261 | 28255 | 43 | 32 | 1889 | 1176 | 0 | 0 | 21 | 24 | 24 | 94 | 0 | 0 | 57 | 0 | 11 | 0 | 0 | 3097 | 419 | 0 |
| SRR2106875 | 70 | 46 | 31 | 48 | 17 | 186 | 165 | 244 | 93 | 0 | 46 | 0 | 12 | 0 | 7068 | 12699 | 30138 | 29 | 65 | 3283 | 1283 | 0 | 0 | 11 | 75 | 16 | 85 | 27 | 0 | 46 | 0 | 22 | 0 | 0 | 2176 | 532 | 0 |
| SRR2106876 | 12 | 16 | 31 | 93 | 260 | 401 | 465 | 739 | 127 | 0 | 77 | 0 | 0 | 0 | 8952 | 19293 | 11152 | 0 | 21 | 5426 | 887 | 0 | 0 | 128 | 124 | 523 | 2565 | 938 | 0 | 957 | 0 | 0 | 0 | 0 | 1893 | 588 | 0 |
| SRR2106877 | 98 | 73 | 116 | 14 | 25 | 179 | 179 | 482 | 89 | 55 | 37 | 0 | 0 | 0 | 20169 | 167604 | 51475 | 54 | 36 | 4020 | 4055 | 0 | 0 | 19 | 146 | 45 | 169 | 129 | 0 | 25 | 0 | 0 | 0 | 0 | 1947 | 404 | 0 |

### Supp Table S8.3.Columns_BX_to_DH

| | | | | | | | | | | | | | | | | | | | | | | | | | | | | | | | | | | | | | |
|---|---|---|---|---|---|---|---|---|---|---|---|---|---|---|---|---|---|---|---|---|---|---|---|---|---|---|---|---|---|---|---|---|---|---|---|---|---|
| SRR2106878 | 49 | 31 | 154 | 362 | 289 | 62 | 227 | 372 | 62 | 0 | 203 | 0 | 0 | 0 | 10939 | 40408 | 169428 | 62 | 103 | 2745 | 2188 | 0 | 0 | 209 | 129 | 350 | 1395 | 1745 | 0 | 1075 | 0 | 0 | 0 | 0 | 6106 | 455 | 0 |
| SRR2106879 | 15 | 15 | 18 | 18 | 32 | 269 | 313 | 496 | 119 | 0 | 24 | 0 | 0 | 0 | 8923 | 16428 | 74853 | 11 | 65 | 1875 | 3944 | 0 | 0 | 12 | 104 | 13 | 53 | 83 | 0 | 32 | 0 | 22 | 0 | 0 | 2437 | 521 | 0 |
| SRR2106880 | 29 | 22 | 0 | 17 | 17 | 478 | 478 | 805 | 218 | 0 | 37 | 0 | 0 | 0 | 6508 | 2303 | 8257 | 50 | 50 | 4646 | 813 | 0 | 0 | 32 | 51 | 17 | 62 | 15 | 0 | 163 | 0 | 25 | 0 | 0 | 5075 | 562 | 0 |
| SRR2106881 | 24 | 36 | 0 | 24 | 17 | 1159 | 1043 | 1405 | 449 | 12 | 0 | 0 | 0 | 0 | 4475 | 5205 | 46253 | 87 | 14 | 3867 | 1970 | 0 | 0 | 34 | 57 | 11 | 47 | 57 | 0 | 35 | 0 | 0 | 0 | 0 | 2752 | 695 | 0 |
| SRR2106882 | 43 | 95 | 0 | 0 | 45 | 595 | 564 | 1270 | 231 | 0 | 56 | 0 | 13 | 0 | 9528 | 3593 | 15224 | 44 | 53 | 4102 | 608 | 0 | 0 | 180 | 31 | 60 | 400 | 17 | 0 | 1689 | 0 | 111 | 0 | 0 | 2876 | 728 | 0 |
| SRR2106883 | 46 | 65 | 42 | 106 | 69 | 361 | 404 | 559 | 248 | 0 | 187 | 0 | 0 | 0 | 12148 | 7123 | 73154 | 14 | 64 | 3058 | 3625 | 0 | 0 | 238 | 152 | 397 | 3488 | 332 | 0 | 1489 | 0 | 446 | 0 | 0 | 3735 | 644 | 0 |
| SRR2106884 | 43 | 77 | 20 | 49 | 49 | 512 | 492 | 988 | 198 | 0 | 98 | 0 | 0 | 0 | 9901 | 5034 | 35121 | 25 | 41 | 3587 | 1703 | 0 | 0 | 197 | 80 | 223 | 1536 | 146 | 0 | 1545 | 0 | 233 | 0 | 0 | 3446 | 669 | 0 |
| SRR2124760 | 0 | 0 | 0 | 22 | 0 | 284 | 380 | 947 | 168 | 0 | 150 | 0 | 0 | 0 | 726 | 11102 | 11580 | 0 | 101 | 20507 | 62 | 0 | 0 | 0 | 0 | 26 | 117 | 90 | 0 | 422 | 0 | 43 | 0 | 0 | 5568 | 54 | 0 |
| SRR2124761 | 0 | 0 | 0 | 24 | 20 | 66 | 143 | 260 | 69 | 0 | 48 | 0 | 0 | 0 | 2336 | 10062 | 29192 | 0 | 82 | 41841 | 395 | 0 | 0 | 22 | 0 | 34 | 84 | 99 | 0 | 161 | 0 | 15 | 0 | 0 | 853 | 27 | 0 |
| SRR2124762 | 0 | 0 | 0 | 15 | 28 | 41 | 66 | 161 | 18 | 0 | 202 | 0 | 0 | 0 | 1848 | 8684 | 31613 | 0 | 108 | 59166 | 700 | 0 | 0 | 19 | 0 | 50 | 618 | 371 | 0 | 718 | 0 | 0 | 0 | 0 | 498 | 79 | 0 |
| SRR2124763 | 0 | 0 | 0 | 46 | 59 | 28 | 63 | 153 | 31 | 0 | 26 | 11 | 0 | 0 | 3702 | 20353 | 65824 | 0 | 76 | 59633 | 1161 | 0 | 0 | 35 | 0 | 49 | 66 | 61 | 0 | 91 | 0 | 0 | 0 | 0 | 453 | 74 | 0 |
| SRR2124764 | 0 | 0 | 0 | 0 | 12 | 176 | 289 | 743 | 126 | 0 | 90 | 0 | 0 | 0 | 1817 | 12726 | 19765 | 0 | 114 | 41281 | 199 | 0 | 0 | 18 | 0 | 15 | 144 | 70 | 0 | 261 | 0 | 15 | 0 | 0 | 1528 | 10 | 0 |
| SRR2124765 | 0 | 0 | 0 | 11 | 20 | 190 | 267 | 602 | 82 | 0 | 263 | 0 | 0 | 0 | 867 | 12822 | 15430 | 0 | 99 | 60840 | 97 | 0 | 0 | 17 | 0 | 15 | 188 | 89 | 0 | 891 | 0 | 14 | 0 | 0 | 1425 | 14 | 0 |
| SRR2124766 | 0 | 0 | 0 | 16 | 0 | 155 | 292 | 626 | 107 | 0 | 436 | 0 | 0 | 0 | 391 | 5588 | 8886 | 0 | 103 | 75779 | 65 | 0 | 0 | 10 | 0 | 25 | 253 | 193 | 0 | 1315 | 0 | 44 | 0 | 0 | 2900 | 12 | 0 |
| SRR2124767 | 0 | 0 | 0 | 23 | 13 | 58 | 118 | 258 | 43 | 0 | 80 | 11 | 0 | 0 | 1414 | 9978 | 23831 | 0 | 129 | 116627 | 105 | 0 | 0 | 20 | 0 | 0 | 34 | 26 | 0 | 272 | 0 | 0 | 0 | 0 | 377 | 0 | 0 |
| SRR2124768 | 0 | 0 | 0 | 26 | 17 | 44 | 76 | 185 | 38 | 0 | 158 | 0 | 0 | 0 | 1981 | 8249 | 25670 | 0 | 79 | 77290 | 127 | 0 | 0 | 13 | 0 | 0 | 106 | 79 | 0 | 597 | 0 | 0 | 0 | 0 | 280 | 94 | 0 |
| SRR2124769 | 0 | 0 | 0 | 21 | 18 | 256 | 402 | 967 | 192 | 0 | 61 | 0 | 0 | 0 | 1072 | 13001 | 15234 | 0 | 58 | 52055 | 47 | 0 | 0 | 10 | 0 | 15 | 40 | 39 | 0 | 236 | 0 | 17 | 0 | 0 | 1715 | 171 | 0 |
| SRR2124770 | 0 | 0 | 0 | 0 | 0 | 85 | 130 | 269 | 48 | 0 | 214 | 0 | 0 | 0 | 2147 | 5187 | 35357 | 0 | 74 | 51491 | 395 | 0 | 0 | 20 | 0 | 25 | 219 | 126 | 0 | 889 | 0 | 0 | 0 | 0 | 967 | 90 | 0 |
| SRR2124771 | 0 | 0 | 0 | 0 | 13 | 180 | 197 | 348 | 114 | 0 | 224 | 0 | 0 | 0 | 1724 | 13875 | 25574 | 0 | 67 | 62774 | 190 | 0 | 0 | 17 | 0 | 11 | 244 | 130 | 0 | 852 | 0 | 0 | 0 | 0 | 1052 | 73 | 0 |
| SRR2124772 | 0 | 0 | 0 | 0 | 19 | 96 | 178 | 363 | 43 | 0 | 210 | 14 | 0 | 0 | 643 | 4738 | 7014 | 0 | 116 | 88280 | 45 | 0 | 0 | 0 | 0 | 0 | 87 | 79 | 0 | 673 | 0 | 0 | 0 | 0 | 822 | 0 | 0 |
| SRR2124773 | 0 | 0 | 0 | 13 | 13 | 391 | 558 | 1252 | 199 | 0 | 138 | 0 | 0 | 0 | 367 | 10021 | 9681 | 0 | 72 | 28886 | 40 | 0 | 0 | 16 | 0 | 10 | 117 | 115 | 0 | 468 | 0 | 72 | 0 | 0 | 5570 | 30 | 0 |
| SRR2124774 | 0 | 0 | 0 | 34 | 29 | 65 | 100 | 212 | 46 | 0 | 55 | 12 | 0 | 0 | 968 | 7187 | 19551 | 0 | 139 | 104038 | 121 | 0 | 0 | 11 | 0 | 11 | 22 | 19 | 0 | 206 | 0 | 11 | 0 | 0 | 279 | 53 | 0 |
| SRR2124775 | 0 | 0 | 0 | 47 | 30 | 98 | 183 | 330 | 69 | 0 | 156 | 11 | 0 | 0 | 1078 | 8414 | 22291 | 0 | 104 | 102293 | 196 | 0 | 0 | 19 | 0 | 0 | 104 | 84 | 0 | 677 | 0 | 11 | 0 | 0 | 750 | 38 | 0 |
| SRR2124776 | 0 | 0 | 0 | 41 | 22 | 65 | 129 | 283 | 52 | 0 | 262 | 0 | 0 | 0 | 391 | 6528 | 8241 | 0 | 93 | 63248 | 49 | 0 | 0 | 12 | 0 | 0 | 175 | 67 | 0 | 767 | 0 | 0 | 0 | 0 | 922 | 15 | 0 |
| SRR2124777 | 0 | 0 | 0 | 50 | 47 | 88 | 145 | 305 | 64 | 0 | 168 | 12 | 0 | 0 | 1708 | 14795 | 29458 | 0 | 65 | 44100 | 230 | 0 | 0 | 27 | 0 | 13 | 299 | 149 | 0 | 578 | 0 | 0 | 0 | 0 | 662 | 15 | 0 |
| SRR2124778 | 0 | 0 | 0 | 14 | 12 | 42 | 68 | 157 | 24 | 0 | 58 | 0 | 0 | 0 | 356 | 6262 | 6543 | 0 | 155 | 72827 | 86 | 0 | 0 | 12 | 0 | 11 | 74 | 38 | 0 | 185 | 0 | 0 | 0 | 0 | 1544 | 50 | 0 |
| SRR2124779 | 0 | 0 | 0 | 20 | 20 | 55 | 116 | 220 | 43 | 0 | 243 | 0 | 0 | 0 | 2107 | 9634 | 37097 | 0 | 86 | 52538 | 605 | 0 | 0 | 20 | 0 | 28 | 400 | 526 | 0 | 763 | 0 | 0 | 0 | 0 | 903 | 46 | 0 |
| SRR2126527 | 44 | 100 | 52 | 232 | 18 | 36 | 53 | 130 | 35 | 0 | 24 | 0 | 0 | 0 | 42737 | 18014 | 55804 | 30 | 68 | 6173 | 6942 | 0 | 0 | 33 | 66 | 325 | 1466 | 788 | 0 | 94 | 0 | 68 | 0 | 0 | 5665 | 680 | 0 |
| SRR2126528 | 352 | 513 | 333 | 1492 | 106 | 48 | 47 | 133 | 17 | 0 | 111 | 0 | 0 | 0 | 59136 | 17868 | 66415 | 30 | 78 | 10458 | 9058 | 0 | 0 | 171 | 431 | 1900 | 8137 | 4402 | 0 | 421 | 0 | 24 | 0 | 0 | 4229 | 743 | 0 |
| SRR2126529 | 321 | 654 | 361 | 1325 | 98 | 37 | 61 | 124 | 25 | 0 | 123 | 0 | 0 | 0 | 44357 | 22624 | 63808 | 23 | 67 | 5548 | 7876 | 0 | 0 | 203 | 498 | 1995 | 8872 | 4241 | 0 | 503 | 0 | 31 | 0 | 0 | 5788 | 660 | 0 |
| SRR2126530 | 37 | 84 | 46 | 226 | 17 | 43 | 38 | 101 | 21 | 0 | 35 | 0 | 0 | 0 | 46379 | 30143 | 64248 | 16 | 83 | 6147 | 12063 | 0 | 0 | 25 | 101 | 414 | 2144 | 1130 | 0 | 125 | 0 | 49 | 0 | 0 | 5181 | 711 | 0 |
| SRR2126531 | 181 | 378 | 232 | 1216 | 84 | 25 | 33 | 127 | 24 | 0 | 96 | 0 | 0 | 0 | 43065 | 25162 | 62407 | 14 | 96 | 10182 | 11576 | 0 | 0 | 105 | 410 | 1784 | 9390 | 4765 | 0 | 360 | 0 | 68 | 0 | 0 | 5692 | 727 | 0 |
| SRR2126532 | 67 | 130 | 69 | 315 | 29 | 46 | 51 | 78 | 25 | 0 | 24 | 0 | 0 | 0 | 40241 | 20146 | 57046 | 19 | 82 | 5336 | 4580 | 0 | 0 | 25 | 88 | 397 | 1820 | 682 | 0 | 98 | 0 | 41 | 0 | 0 | 5329 | 710 | 0 |
| SRR2135530 | 65 | 23 | 0 | 0 | 87 | 2989 | 480 | 13717 | 6641 | 585 | 19 | 20 | 35 | 22 | 9298 | 59033 | 69434 | 243 | 2749 | 5455 | 26288 | 0 | 0 | 26 | 19 | 46 | 27 | 0 | 0 | 179 | 0 | 61 | 0 | 0 | 13350 | 23 | 0 |
| SRR2135531 | 438 | 128 | 116 | 46 | 494 | 3156 | 1657 | 23622 | 9997 | 1178 | 151 | 0 | 0 | 0 | 8310 | 84272 | 75947 | 340 | 4862 | 9286 | 10481 | 0 | 0 | 178 | 139 | 229 | 316 | 123 | 0 | 1205 | 0 | 37 | 0 | 0 | 3680 | 31 | 0 |
| SRR2135532 | 67 | 17 | 34 | 0 | 88 | 3326 | 2212 | 25371 | 11712 | 261 | 47 | 0 | 0 | 0 | 7778 | 64511 | 61015 | 481 | 5996 | 15654 | 8303 | 0 | 0 | 30 | 12 | 14 | 16 | 0 | 0 | 153 | 0 | 51 | 0 | 0 | 1818 | 22 | 0 |
| SRR2229584 | 0 | 11 | 0 | 26 | 11 | 18 | 35 | 53 | 35 | 0 | 727 | 0 | 0 | 0 | 39333 | 12241 | 69451 | 0 | 0 | 1228 | 9930 | 0 | 0 | 0 | 0 | 32 | 13 | 22 | 0 | 1317 | 0 | 0 | 0 | 0 | 1667 | 0 | 0 |
| SRR2229585 | 10 | 0 | 31 | 14 | 17 | 43 | 85 | 85 | 43 | 0 | 126 | 0 | 0 | 0 | 18175 | 13161 | 43596 | 0 | 43 | 1451 | 6528 | 0 | 0 | 0 | 0 | 23 | 14 | 26 | 0 | 252 | 0 | 0 | 0 | 0 | 1880 | 0 | 0 |
| SRR2229586 | 0 | 0 | 0 | 18 | 359 | 0 | 0 | 0 | 0 | 0 | 154 | 0 | 0 | 0 | 0 | 0 | 0 | 0 | 0 | 0 | 8686 | 0 | 0 | 0 | 0 | 0 | 0 | 0 | 0 | 671 | 0 | 0 | 0 | 0 | 0 | 0 | 0 |
| SRR2229587 | 0 | 0 | 0 | 0 | 289 | 0 | 0 | 0 | 0 | 0 | 206 | 0 | 0 | 0 | 0 | 0 | 0 | 0 | 0 | 0 | 13861 | 0 | 0 | 0 | 0 | 0 | 0 | 0 | 0 | 739 | 0 | 0 | 0 | 0 | 0 | 0 | 0 |
| SRR2229588 | 29 | 54 | 39 | 24 | 15 | 104 | 104 | 701 | 26 | 0 | 224 | 0 | 0 | 0 | 158562 | 111269 | 238722 | 26 | 0 | 363 | 8568 | 0 | 0 | 0 | 15 | 11 | 83 | 37 | 0 | 332 | 0 | 26 | 0 | 0 | 24144 | 0 | 0 |
| SRR2229589 | 37 | 57 | 37 | 24 | 0 | 48 | 191 | 550 | 96 | 0 | 129 | 0 | 0 | 0 | 196351 | 115962 | 260921 | 24 | 0 | 335 | 6217 | 0 | 0 | 0 | 20 | 28 | 85 | 24 | 0 | 329 | 0 | 48 | 0 | 0 | 16177 | 0 | 0 |
| SRR2287868 | 0 | 15 | 19 | 119 | 40 | 152 | 277 | 1131 | 473 | 11 | 58 | 0 | 0 | 0 | 37842 | 27062 | 396936 | 0 | 2362 | 77864 | 7133 | 0 | 0 | 299 | 157 | 586 | 1360 | 799 | 0 | 77 | 0 | 0 | 0 | 0 | 0 | 0 | 0 |
| SRR2289624 | 0 | 0 | 0 | 0 | 14 | 72 | 646 | 574 | 293 | 25 | 162 | 14 | 0 | 0 | 104888 | 69364 | 449695 | 15 | 762 | 48022 | 9210 | 0 | 0 | 52 | 128 | 190 | 277 | 31 | 0 | 863 | 0 | 14 | 0 | 0 | 42 | 0 | 0 |
| SRR2289625 | 0 | 0 | 0 | 18 | 88 | 153 | 886 | 1391 | 963 | 0 | 20 | 12 | 0 | 0 | 56033 | 30923 | 295299 | 11 | 751 | 11317 | 4172 | 0 | 0 | 23 | 193 | 18 | 62 | 166 | 0 | 144 | 0 | 23 | 0 | 0 | 47 | 0 | 0 |
| SRR2289626 | 0 | 0 | 0 | 0 | 11 | 33 | 403 | 321 | 60 | 10 | 19 | 0 | 0 | 0 | 126791 | 25807 | 727030 | 0 | 380 | 17624 | 5176 | 0 | 0 | 457 | 228 | 597 | 1262 | 316 | 0 | 187 | 0 | 0 | 0 | 0 | 75 | 0 | 0 |
| SRR2294377 | 0 | 13 | 12 | 44 | 34 | 114 | 1487 | 1813 | 222 | 0 | 19 | 0 | 0 | 0 | 69937 | 58833 | 446235 | 0 | 771 | 47079 | 10317 | 0 | 0 | 913 | 127 | 680 | 2294 | 143 | 0 | 99 | 0 | 0 | 0 | 0 | 130 | 0 | 0 |
| SRR2294380 | 30 | 23 | 17 | 40 | 94 | 50 | 117 | 194 | 23 | 393 | 127 | 0 | 0 | 0 | 278643 | 103545 | 676981 | 0 | 776 | 13130 | 8803 | 0 | 0 | 232 | 251 | 562 | 948 | 50 | 0 | 299 | 0 | 0 | 0 | 0 | 16 | 0 | 0 |
| SRR2294739 | 11 | 14 | 18 | 80 | 129 | 192 | 329 | 1022 | 103 | 12 | 111 | 0 | 0 | 0 | 151308 | 24151 | 610766 | 0 | 1308 | 45399 | 9582 | 0 | 0 | 186 | 94 | 366 | 493 | 417 | 0 | 83 | 0 | 0 | 0 | 0 | 12 | 0 | 0 |
| SRR2294742 | 14 | 35 | 31 | 109 | 128 | 31 | 272 | 285 | 77 | 36 | 88 | 0 | 0 | 0 | 61581 | 36327 | 696160 | 0 | 895 | 22686 | 16204 | 0 | 0 | 490 | 121 | 492 | 20 | 194 | 0 | 355 | 0 | 0 | 0 | 0 | 45 | 0 | 0 |
| SRR2294746 | 11 | 11 | 11 | 48 | 122 | 25 | 71 | 224 | 32 | 48 | 111 | 0 | 0 | 0 | 102877 | 32515 | 516078 | 0 | 309 | 9465 | 14277 | 0 | 0 | 385 | 63 | 270 | 929 | 144 | 0 | 392 | 0 | 33 | 0 | 0 | 41 | 0 | 0 |
| SRR2294747 | 0 | 0 | 0 | 0 | 15 | 0 | 80 | 64 | 0 | 12 | 60 | 0 | 0 | 0 | 111384 | 8108 | 794936 | 0 | 252 | 9125 | 3451 | 0 | 0 | 142 | 160 | 544 | 10 | 605 | 0 | 232 | 0 | 0 | 0 | 0 | 40 | 0 | 0 |
| SRR2294748 | 0 | 11 | 29 | 70 | 66 | 32 | 209 | 350 | 88 | 0 | 16 | 0 | 0 | 0 | 54235 | 56108 | 498263 | 0 | 324 | 8029 | 36194 | 0 | 0 | 236 | 177 | 537 | 1411 | 119 | 0 | 73 | 0 | 0 | 0 | 0 | 80 | 0 | 0 |
| SRR2294820 | 15 | 54 | 25 | 81 | 88 | 77 | 153 | 405 | 44 | 14 | 260 | 0 | 0 | 0 | 206972 | 47608 | 791249 | 0 | 446 | 20612 | 6296 | 0 | 0 | 504 | 263 | 529 | 24 | 365 | 0 | 112 | 0 | 0 | 0 | 0 | 33 | 0 | 0 |
| SRR2296788 | 24 | 16 | 12 | 64 | 65 | 203 | 152 | 355 | 76 | 0 | 23 | 0 | 0 | 0 | 47577 | 26446 | 98075 | 76 | 76 | 6362 | 1724 | 0 | 0 | 16 | 34 | 108 | 459 | 376 | 0 | 131 | 0 | 0 | 0 | 0 | 18663 | 0 | 0 |
| SRR2296789 | 76 | 22 | 31 | 104 | 61 | 38 | 76 | 113 | 0 | 0 | 50 | 0 | 0 | 0 | 22895 | 7827 | 46639 | 38 | 38 | 4836 | 1662 | 0 | 0 | 107 | 30 | 110 | 586 | 517 | 0 | 172 | 0 | 0 | 0 | 0 | 19447 | 0 | 0 |
| SRR2296790 | 17 | 50 | 50 | 349 | 50 | 240 | 400 | 400 | 80 | 0 | 199 | 17 | 0 | 0 | 48747 | 12340 | 92843 | 0 | 80 | 3442 | 1121 | 0 | 0 | 83 | 149 | 697 | 2971 | 2988 | 0 | 830 | 0 | 0 | 0 | 0 | 14573 | 0 | 0 |
| SRR2296791 | 40 | 66 | 11 | 67 | 53 | 83 | 42 | 125 | 83 | 0 | 59 | 0 | 0 | 0 | 52344 | 15212 | 112477 | 0 | 42 | 4884 | 1211 | 0 | 0 | 92 | 11 | 93 | 450 | 611 | 0 | 112 | 0 | 0 | 0 | 0 | 13263 | 0 | 0 |
| SRR2296792 | 87 | 24 | 19 | 76 | 13 | 129 | 129 | 215 | 0 | 0 | 90 | 0 | 0 | 0 | 19586 | 11383 | 46312 | 0 | 215 | 8695 | 947 | 0 | 0 | 17 | 38 | 129 | 640 | 553 | 0 | 372 | 0 | 0 | 0 | 0 | 20199 | 0 | 0 |
| SRR2296793 | 27 | 21 | 48 | 48 | 41 | 163 | 489 | 285 | 163 | 0 | 26 | 0 | 0 | 0 | 33993 | 16366 | 76417 | 81 | 41 | 5537 | 651 | 0 | 0 | 12 | 21 | 73 | 390 | 272 | 0 | 122 | 0 | 0 | 0 | 0 | 20213 | 0 | 0 |
| SRR2296794 | 46 | 33 | 93 | 457 | 106 | 152 | 274 | 305 | 30 | 0 | 490 | 0 | 0 | 0 | 48434 | 61943 | 138159 | 0 | 122 | 4843 | 1097 | 0 | 0 | 119 | 219 | 1100 | 4266 | 3398 | 0 | 1537 | 0 | 0 | 0 | 0 | 14501 | 0 | 0 |
| SRR2296795 | 36 | 27 | 99 | 40 | 63 | 179 | 393 | 321 | 107 | 0 | 47 | 0 | 0 | 0 | 6036 | 10432 | 24184 | 36 | 36 | 4858 | 679 | 0 | 0 | 54 | 19 | 69 | 322 | 250 | 0 | 125 | 0 | 36 | 0 | 0 | 50265 | 0 | 0 |
| SRR2296797 | 24 | 24 | 75 | 50 | 13 | 62 | 172 | 94 | 47 | 19 | 24 | 0 | 0 | 0 | 23886 | 10940 | 70379 | 16 | 78 | 3604 | 780 | 0 | 0 | 15 | 33 | 93 | 378 | 318 | 0 | 120 | 0 | 0 | 0 | 0 | 15043 | 0 | 0 |
| SRR2296798 | 25 | 0 | 51 | 481 | 38 | 132 | 795 | 331 | 0 | 0 | 76 | 0 | 0 | 0 | 48666 | 10131 | 102864 | 0 | 66 | 4635 | 1390 | 0 | 0 | 63 | 114 | 455 | 1808 | 1783 | 0 | 733 | 0 | 0 | 0 | 0 | 18118 | 0 | 13 |
| SRR2296799 | 31 | 57 | 52 | 416 | 177 | 137 | 86 | 291 | 69 | 0 | 245 | 0 | 0 | 0 | 35167 | 22874 | 52757 | 34 | 120 | 3189 | 1800 | 0 | 0 | 260 | 422 | 1359 | 6075 | 1052 | 0 | 1390 | 0 | 0 | 0 | 0 | 20935 | 0 | 0 |
| SRR2296800 | 70 | 18 | 21 | 128 | 12 | 45 | 316 | 361 | 0 | 14 | 66 | 0 | 0 | 0 | 34223 | 22913 | 80047 | 90 | 0 | 6953 | 2754 | 0 | 0 | 31 | 60 | 151 | 680 | 629 | 0 | 252 | 0 | 0 | 0 | 0 | 20848 | 0 | 0 |
| SRR2296801 | 0 | 14 | 104 | 360 | 66 | 117 | 117 | 195 | 117 | 0 | 778 | 0 | 0 | 0 | 15654 | 4816 | 27889 | 39 | 195 | 4094 | 1384 | 0 | 0 | 138 | 270 | 996 | 4685 | 1062 | 0 | 3727 | 0 | 0 | 0 | 0 | 27032 | 0 | 0 |
| SRR2296802 | 24 | 48 | 48 | 435 | 169 | 195 | 325 | 325 | 130 | 0 | 555 | 0 | 0 | 0 | 9418 | 5103 | 26621 | 0 | 65 | 3637 | 585 | 0 | 0 | 169 | 459 | 918 | 3912 | 676 | 0 | 2270 | 0 | 0 | 0 | 0 | 38897 | 0 | 24 |
| SRR2296805 | 45 | 42 | 92 | 58 | 104 | 187 | 152 | 223 | 45 | 0 | 36 | 0 | 0 | 0 | 35029 | 7222 | 51005 | 18 | 71 | 5260 | 1970 | 0 | 0 | 17 | 27 | 79 | 354 | 50 | 0 | 184 | 0 | 0 | 0 | 0 | 18061 | 0 | 0 |
| SRR2296806 | 13 | 105 | 171 | 698 | 66 | 86 | 43 | 257 | 86 | 0 | 804 | 0 | 13 | 0 | 30573 | 6503 | 53596 | 0 | 43 | 9262 | 1415 | 0 | 0 | 250 | 211 | 843 | 3281 | 3519 | 0 | 2201 | 0 | 0 | 0 | 0 | 29825 | 0 | 0 |
| SRR2296807 | 36 | 24 | 14 | 67 | 16 | 202 | 101 | 202 | 87 | 0 | 51 | 0 | 0 | 0 | 46983 | 24947 | 76857 | 14 | 130 | 4853 | 1762 | 0 | 0 | 27 | 50 | 133 | 457 | 434 | 0 | 232 | 0 | 14 | 0 | 0 | 24748 | 0 | 12 |
| SRR2296808 | 69 | 52 | 14 | 64 | 16 | 55 | 55 | 330 | 55 | 0 | 63 | 0 | 0 | 0 | 34420 | 41528 | 74489 | 0 | 0 | 5883 | 1430 | 0 | 0 | 29 | 54 | 111 | 350 | 334 | 0 | 236 | 0 | 0 | 0 | 0 | 20260 | 0 | 0 |
| SRR2296809 | 0 | 15 | 11 | 32 | 13 | 237 | 162 | 409 | 151 | 0 | 93 | 0 | 0 | 0 | 17216 | 3286 | 30832 | 43 | 65 | 3327 | 1217 | 0 | 0 | 24 | 33 | 136 | 385 | 359 | 0 | 425 | 0 | 11 | 0 | 0 | 26889 | 0 | 0 |
| SRR2296810 | 44 | 17 | 19 | 89 | 22 | 136 | 136 | 227 | 0 | 0 | 86 | 0 | 0 | 0 | 17438 | 7626 | 48867 | 0 | 0 | 3968 | 1474 | 0 | 0 | 20 | 45 | 114 | 428 | 411 | 0 | 298 | 0 | 23 | 0 | 0 | 28428 | 0 | 0 |
| SRR2353417 | 0 | 0 | 0 | 0 | 43 | 436 | 1366 | 2462 | 2010 | 0 | 55 | 0 | 0 | 0 | 12716 | 17233 | 214059 | 0 | 1336 | 14547 | 1204 | 0 | 0 | 28 | 16 | 38 | 144 | 64 | 0 | 56 | 0 | 23 | 0 | 0 | 637 | 0 | 0 |
| SRR2353418 | 0 | 0 | 0 | 0 | 0 | 364 | 1169 | 2140 | 586 | 0 | 69 | 0 | 0 | 0 | 19582 | 19423 | 260172 | 0 | 1180 | 20434 | 8909 | 0 | 0 | 174 | 202 | 430 | 1333 | 725 | 0 | 693 | 0 | 22 | 0 | 0 | 1296 | 0 | 0 |
| SRR2353419 | 0 | 0 | 0 | 35 | 42 | 402 | 958 | 1969 | 992 | 0 | 57 | 0 | 0 | 0 | 8854 | 9180 | 175993 | 0 | 331 | 8726 | 537 | 0 | 0 | 14 | 103 | 29 | 127 | 52 | 0 | 58 | 0 | 90 | 0 | 0 | 515 | 0 | 0 |
| SRR2353420 | 0 | 0 | 0 | 0 | 37 | 541 | 943 | 2350 | 1254 | 0 | 54 | 0 | 0 | 0 | 1979 | 17960 | 64606 | 0 | 435 | 9005 | 723 | 0 | 0 | 119 | 40 | 155 | 760 | 325 | 0 | 881 | 0 | 245 | 0 | 0 | 1203 | 0 | 0 |
| SRR2353421 | 0 | 0 | 0 | 11 | 41 | 393 | 779 | 1951 | 811 | 0 | 59 | 0 | 0 | 0 | 4716 | 18426 | 145189 | 0 | 496 | 26181 | 1093 | 0 | 0 | 11 | 41 | 17 | 62 | 29 | 0 | 109 | 0 | 150 | 0 | 0 | 805 | 0 | 0 |
| SRR2353422 | 0 | 0 | 0 | 0 | 19 | 399 | 1258 | 2388 | 814 | 0 | 49 | 0 | 0 | 0 | 4529 | 26802 | 148975 | 0 | 977 | 34265 | 3063 | 0 | 0 | 10 | 12 | 27 | 91 | 40 | 0 | 54 | 0 | 52 | 0 | 0 | 1234 | 0 | 0 |
| SRR2353423 | 0 | 0 | 0 | 11 | 40 | 539 | 908 | 2129 | 1157 | 0 | 81 | 0 | 0 | 0 | 4726 | 16773 | 185397 | 0 | 432 | 11479 | 627 | 0 | 0 | 269 | 79 | 197 | 773 | 315 | 0 | 1365 | 0 | 53 | 0 | 0 | 849 | 0 | 0 |
| SRR2353424 | 0 | 0 | 0 | 0 | 43 | 218 | 1089 | 2038 | 429 | 0 | 73 | 0 | 0 | 0 | 2833 | 18941 | 227038 | 0 | 1129 | 25761 | 2708 | 0 | 0 | 10 | 139 | 19 | 63 | 30 | 0 | 92 | 0 | 24 | 0 | 0 | 1035 | 0 | 0 |
| SRR2353425 | 0 | 0 | 0 | 19 | 23 | 245 | 487 | 1050 | 595 | 12 | 79 | 0 | 0 | 0 | 7197 | 46045 | 159905 | 19 | 383 | 46550 | 6307 | 0 | 0 | 17 | 10 | 23 | 78 | 86 | 0 | 97 | 0 | 637 | 0 | 0 | 725 | 0 | 0 |
| SRR2353426 | 0 | 0 | 0 | 12 | 34 | 419 | 870 | 2079 | 815 | 0 | 49 | 0 | 0 | 0 | 2660 | 22750 | 91745 | 0 | 473 | 27623 | 1893 | 0 | 0 | 13 | 122 | 32 | 146 | 37 | 0 | 52 | 0 | 67 | 0 | 0 | 1452 | 0 | 0 |
| SRR2353427 | 0 | 0 | 0 | 0 | 24 | 470 | 871 | 2117 | 800 | 18 | 29 | 0 | 0 | 0 | 2772 | 24994 | 127916 | 0 | 485 | 32947 | 6495 | 0 | 0 | 126 | 70 | 17 | 51 | 83 | 0 | 44 | 0 | 182 | 0 | 0 | 1011 | 0 | 0 |
| SRR2353428 | 0 | 0 | 0 | 0 | 18 | 233 | 445 | 968 | 519 | 46 | 100 | 0 | 0 | 0 | 3830 | 45060 | 237465 | 0 | 751 | 51705 | 7280 | 0 | 0 | 16 | 94 | 14 | 31 | 68 | 0 | 85 | 0 | 124 | 0 | 0 | 1131 | 0 | 0 |
| SRR2353429 | 0 | 0 | 0 | 0 | 20 | 438 | 421 | 1481 | 467 | 28 | 67 | 0 | 0 | 0 | 6936 | 28649 | 138873 | 12 | 574 | 37202 | 9144 | 0 | 11 | 16 | 10 | 27 | 81 | 28 | 0 | 66 | 0 | 408 | 0 | 0 | 552 | 0 | 0 |
| SRR2353430 | 0 | 0 | 0 | 0 | 24 | 233 | 401 | 733 | 308 | 0 | 68 | 0 | 0 | 0 | 9334 | 43099 | 231104 | 0 | 1007 | 63839 | 7749 | 0 | 0 | 16 | 12 | 25 | 73 | 51 | 0 | 45 | 0 | 53 | 0 | 0 | 1041 | 0 | 0 |
| SRR2353431 | 0 | 0 | 0 | 12 | 43 | 639 | 656 | 2199 | 672 | 11 | 54 | 0 | 0 | 0 | 5100 | 27177 | 164325 | 0 | 257 | 12022 | 13664 | 0 | 0 | 22 | 14 | 42 | 103 | 43 | 0 | 59 | 0 | 253 | 0 | 0 | 814 | 0 | 0 |
| SRR2353432 | 0 | 0 | 0 | 0 | 29 | 289 | 697 | 1139 | 496 | 14 | 35 | 0 | 0 | 0 | 4357 | 62071 | 195452 | 0 | 880 | 67720 | 4140 | 0 | 0 | 290 | 179 | 356 | 1292 | 365 | 0 | 517 | 0 | 53 | 0 | 0 | 973 | 0 | 0 |
| SRR2353433 | 0 | 0 | 0 | 0 | 22 | 146 | 247 | 464 | 261 | 0 | 51 | 0 | 0 | 0 | 6490 | 71120 | 255300 | 0 | 676 | 70871 | 12348 | 0 | 0 | 262 | 307 | 452 | 1238 | 641 | 0 | 541 | 0 | 23 | 0 | 0 | 770 | 0 | 0 |
| SRR2353434 | 0 | 0 | 0 | 10 | 54 | 430 | 558 | 1508 | 588 | 14 | 62 | 0 | 0 | 0 | 3792 | 26723 | 180319 | 11 | 723 | 27470 | 1973 | 0 | 0 | 11 | 105 | 17 | 63 | 40 | 0 | 52 | 0 | 145 | 0 | 0 | 1067 | 0 | 0 |
| SRR2353435 | 0 | 0 | 0 | 19 | 52 | 492 | 792 | 1951 | 692 | 0 | 61 | 0 | 0 | 0 | 7909 | 23821 | 192222 | 0 | 320 | 15146 | 3517 | 0 | 0 | 14 | 104 | 19 | 59 | 51 | 0 | 63 | 0 | 44 | 0 | 0 | 1988 | 0 | 0 |
| SRR2353436 | 0 | 0 | 0 | 0 | 0 | 116 | 370 | 636 | 125 | 0 | 69 | 0 | 0 | 0 | 27143 | 38928 | 498029 | 0 | 1856 | 59235 | 19579 | 0 | 0 | 185 | 174 | 230 | 801 | 692 | 0 | 956 | 0 | 16 | 0 | 0 | 324 | 0 | 0 |
| SRR2353437 | 0 | 0 | 0 | 0 | 41 | 520 | 1993 | 4163 | 853 | 0 | 160 | 0 | 0 | 0 | 3568 | 23023 | 124488 | 12 | 913 | 24571 | 1345 | 0 | 0 | 263 | 111 | 353 | 1110 | 653 | 0 | 1233 | 0 | 51 | 0 | 0 | 1193 | 0 | 0 |
| SRR2353438 | 0 | 0 | 0 | 0 | 50 | 349 | 1072 | 2600 | 554 | 0 | 104 | 0 | 0 | 0 | 11790 | 43808 | 264566 | 0 | 931 | 25944 | 3985 | 0 | 0 | 14 | 19 | 51 | 123 | 22 | 0 | 93 | 0 | 28 | 0 | 0 | 1263 | 0 | 0 |
| SRR2353439 | 0 | 0 | 0 | 0 | 17 | 129 | 410 | 688 | 182 | 0 | 55 | 0 | 0 | 0 | 13903 | 44354 | 315201 | 0 | 1729 | 65698 | 16227 | 0 | 0 | 131 | 127 | 324 | 676 | 403 | 0 | 674 | 0 | 12 | 0 | 0 | 585 | 0 | 0 |
| SRR2353440 | 0 | 0 | 0 | 14 | 47 | 258 | 673 | 1259 | 382 | 0 | 38 | 0 | 0 | 0 | 6173 | 43749 | 200122 | 0 | 1447 | 60312 | 2730 | 0 | 0 | 13 | 120 | 19 | 57 | 27 | 0 | 44 | 0 | 36 | 0 | 0 | 569 | 0 | 0 |
| SRR2353441 | 0 | 0 | 0 | 22 | 27 | 140 | 282 | 636 | 171 | 0 | 82 | 0 | 0 | 0 | 5534 | 28962 | 339497 | 0 | 771 | 56724 | 8994 | 0 | 0 | 175 | 77 | 12 | 41 | 26 | 0 | 30 | 0 | 16 | 0 | 0 | 437 | 0 | 0 |
| SRR2353442 | 0 | 0 | 0 | 0 | 33 | 294 | 743 | 1395 | 338 | 0 | 62 | 0 | 0 | 0 | 10054 | 40205 | 312845 | 0 | 1493 | 53152 | 5535 | 0 | 0 | 207 | 183 | 287 | 733 | 298 | 0 | 694 | 0 | 23 | 0 | 0 | 656 | 0 | 0 |
| SRR2353443 | 0 | 0 | 0 | 26 | 28 | 423 | 240 | 1000 | 178 | 0 | 86 | 0 | 0 | 0 | 38016 | 28721 | 371914 | 0 | 178 | 7872 | 4334 | 0 | 0 | 348 | 152 | 370 | 1413 | 502 | 0 | 1118 | 0 | 45 | 0 | 0 | 237 | 0 | 0 |
| SRR2353444 | 0 | 0 | 0 | 0 | 14 | 207 | 441 | 992 | 310 | 0 | 88 | 0 | 0 | 0 | 26903 | 34505 | 318927 | 0 | 1041 | 35610 | 2113 | 0 | 0 | 155 | 213 | 270 | 911 | 776 | 0 | 956 | 0 | 36 | 0 | 0 | 391 | 0 | 0 |
| SRR2353445 | 0 | 0 | 0 | 13 | 35 | 200 | 735 | 1486 | 324 | 0 | 66 | 0 | 0 | 0 | 7411 | 55485 | 213795 | 0 | 1625 | 56929 | 5570 | 0 | 0 | 19 | 127 | 18 | 77 | 37 | 0 | 73 | 0 | 24 | 0 | 0 | 816 | 0 | 0 |
| SRR2422969 | 191 | 96 | 96 | 32 | 223 | 876 | 803 | 1095 | 146 | 32 | 255 | 0 | 0 | 0 | 57393 | 90015 | 98320 | 0 | 1533 | 2848 | 1825 | 0 | 0 | 64 | 32 | 287 | 318 | 605 | 32 | 891 | 0 | 0 | 0 | 0 | 286 | 0 | 0 |
| SRR2422970 | 133 | 33 | 166 | 166 | 200 | 1736 | 710 | 1578 | 316 | 0 | 366 | 0 | 0 | 0 | 58007 | 87957 | 94893 | 0 | 2289 | 3551 | 2131 | 0 | 0 | 100 | 67 | 333 | 233 | 566 | 0 | 999 | 0 | 0 | 0 | 0 | 302 | 14 | 0 |
| SRR2422971 | 91 | 91 | 30 | 60 | 242 | 487 | 975 | 1462 | 696 | 30 | 726 | 0 | 0 | 0 | 60224 | 89183 | 103514 | 0 | 1044 | 3899 | 2158 | 0 | 0 | 60 | 151 | 30 | 181 | 635 | 0 | 605 | 0 | 0 | 0 | 0 | 365 | 0 | 0 |
| SRR2422972 | 62 | 31 | 31 | 62 | 186 | 871 | 1016 | 2178 | 363 | 62 | 22 | 0 | 0 | 0 | 59097 | 97150 | 97802 | 0 | 1379 | 2904 | 2541 | 0 | 0 | 155 | 62 | 13 | 248 | 11 | 0 | 26 | 0 | 73 | 0 | 0 | 652 | 0 | 0 |
| SRR2454328 | 44 | 24 | 0 | 0 | 195 | 566 | 265 | 669 | 104 | 264 | 0 | 0 | 0 | 0 | 87187 | 48537 | 98224 | 19 | 26 | 2423 | 28645 | 0 | 0 | 345 | 24 | 14 | 37 | 0 | 0 | 67 | 0 | 0 | 0 | 0 | 2200 | 0 | 0 |
| SRR2454329 | 35 | 29 | 0 | 0 | 19 | 542 | 292 | 737 | 112 | 28 | 0 | 0 | 0 | 0 | 102601 | 49741 | 111180 | 24 | 36 | 2282 | 31944 | 0 | 0 | 40 | 30 | 13 | 21 | 0 | 0 | 67 | 0 | 0 | 0 | 0 | 2199 | 0 | 0 |
| SRR2454330 | 38 | 30 | 0 | 0 | 212 | 603 | 241 | 881 | 144 | 183 | 0 | 0 | 0 | 0 | 111766 | 62228 | 160026 | 11 | 25 | 1554 | 22120 | 0 | 0 | 435 | 24 | 17 | 31 | 0 | 0 | 65 | 0 | 0 | 0 | 0 | 2964 | 0 | 0 |
| SRR2454331 | 50 | 20 | 0 | 0 | 226 | 697 | 270 | 869 | 119 | 212 | 0 | 0 | 0 | 0 | 113806 | 53996 | 153102 | 24 | 38 | 1820 | 25983 | 0 | 0 | 494 | 17 | 17 | 39 | 11 | 0 | 86 | 0 | 0 | 0 | 0 | 2954 | 0 | 0 |
| SRR2454332 | 165 | 97 | 0 | 17 | 31 | 305 | 203 | 750 | 114 | 142 | 13 | 0 | 0 | 0 | 50738 | 53064 | 46816 | 21 | 79 | 2902 | 19262 | 0 | 0 | 36 | 34 | 36 | 127 | 0 | 0 | 14 | 0 | 0 | 0 | 0 | 1680 | 0 | 0 |
| SRR2454333 | 150 | 86 | 0 | 12 | 36 | 331 | 198 | 742 | 122 | 102 | 0 | 0 | 0 | 0 | 62809 | 74598 | 62436 | 13 | 36 | 2261 | 17003 | 0 | 0 | 42 | 60 | 70 | 160 | 0 | 0 | 15 | 0 | 11 | 0 | 0 | 1513 | 0 | 0 |
| SRR2454334 | 115 | 93 | 0 | 14 | 699 | 310 | 231 | 670 | 115 | 55 | 11 | 0 | 0 | 0 | 59143 | 73378 | 66977 | 0 | 36 | 1777 | 12644 | 0 | 0 | 1210 | 55 | 98 | 227 | 0 | 0 | 369 | 0 | 0 | 0 | 0 | 1475 | 0 | 25 |
| SRR2454335 | 64 | 30 | 0 | 11 | 235 | 806 | 502 | 1915 | 193 | 105 | 0 | 0 | 0 | 0 | 24871 | 57752 | 49487 | 18 | 0 | 607 | 4413 | 0 | 0 | 1147 | 11 | 37 | 67 | 0 | 0 | 64 | 0 | 0 | 0 | 0 | 6116 | 0 | 19 |
| SRR2454336 | 59 | 21 | 0 | 0 | 206 | 664 | 535 | 1841 | 198 | 122 | 10 | 0 | 0 | 0 | 32964 | 67206 | 72351 | 27 | 11 | 632 | 5074 | 0 | 0 | 1042 | 0 | 21 | 77 | 10 | 0 | 94 | 0 | 37 | 0 | 0 | 7407 | 0 | 14 |
| SRR2454337 | 21 | 14 | 0 | 0 | 184 | 1011 | 574 | 1940 | 274 | 104 | 0 | 0 | 0 | 0 | 32964 | 74058 | 74842 | 10 | 15 | 635 | 4281 | 0 | 0 | 1122 | 14 | 10 | 35 | 0 | 0 | 49 | 0 | 20 | 0 | 0 | 7753 | 0 | 14 |
| SRR2487412 | 0 | 0 | 0 | 11 | 79 | 737 | 1666 | 4724 | 1311 | 11 | 0 | 16 | 0 | 0 | 29817 | 61476 | 202917 | 0 | 82 | 8929 | 437 | 0 | 0 | 26 | 0 | 0 | 16 | 0 | 0 | 26 | 0 | 0 | 0 | 0 | 3448 | 45 | 0 |
| SRR2487413 | 0 | 0 | 0 | 0 | 91 | 797 | 1564 | 4660 | 1288 | 0 | 0 | 0 | 0 | 0 | 30047 | 61021 | 201638 | 0 | 123 | 8156 | 429 | 0 | 0 | 12 | 0 | 0 | 0 | 0 | 0 | 49 | 0 | 0 | 0 | 0 | 3846 | 40 | 0 |
| SRR2487414 | 0 | 10 | 0 | 0 | 67 | 491 | 1747 | 4695 | 1447 | 16 | 10 | 0 | 0 | 0 | 30708 | 59841 | 201170 | 0 | 136 | 7179 | 300 | 0 | 0 | 62 | 0 | 0 | 16 | 0 | 0 | 72 | 0 | 0 | 0 | 0 | 2952 | 40 | 0 |
| SRR2487415 | 14 | 69 | 0 | 42 | 139 | 1365 | 2345 | 10990 | 2555 | 14 | 0 | 28 | 0 | 0 | 18550 | 59211 | 155180 | 0 | 140 | 6370 | 1505 | 0 | 0 | 14 | 42 | 83 | 18 | 69 | 0 | 180 | 0 | 0 | 0 | 0 | 4761 | 294 | 14 |
| SRR2487416 | 0 | 64 | 0 | 0 | 177 | 957 | 2472 | 11445 | 2113 | 16 | 0 | 64 | 0 | 0 | 16589 | 59123 | 159316 | 0 | 0 | 6460 | 1715 | 0 | 0 | 15 | 32 | 145 | 19 | 96 | 0 | 161 | 0 | 0 | 0 | 16 | 4590 | 284 | 0 |
| SRR2487417 | 0 | 41 | 0 | 27 | 233 | 984 | 2355 | 10437 | 2460 | 0 | 0 | 96 | 0 | 0 | 18485 | 59839 | 155744 | 0 | 0 | 6396 | 2179 | 0 | 0 | 19 | 41 | 123 | 15 | 27 | 0 | 219 | 0 | 0 | 0 | 0 | 4404 | 296 | 0 |
| SRR2487418 | 0 | 20 | 0 | 20 | 141 | 207 | 538 | 1159 | 621 | 20 | 40 | 40 | 0 | 0 | 64678 | 44309 | 397471 | 0 | 0 | 6538 | 290 | 0 | 0 | 141 | 0 | 20 | 141 | 40 | 0 | 141 | 0 | 0 | 0 | 0 | 4532 | 57 | 0 |

### Supp Table S8.3.Columns_BX_to_DH

| | | | | | | | | | | | | | | | | | | | | | | | | | | | | | | | | | | | | | |
|---|---|---|---|---|---|---|---|---|---|---|---|---|---|---|---|---|---|---|---|---|---|---|---|---|---|---|---|---|---|---|---|---|---|---|---|---|---|
| SRR2487419 | 0 | 0 | 0 | 23 | 207 | 139 | 372 | 976 | 651 | 23 | 23 | 46 | 0 | 0 | 63871 | 44673 | 396696 | 0 | 139 | 5578 | 372 | 0 | 0 | 138 | 23 | 23 | 69 | 0 | 0 | 161 | 0 | 0 | 0 | 0 | 3155 | 47 | 0 |
| SRR2487420 | 20 | 20 | 0 | 0 | 159 | 82 | 371 | 906 | 453 | 40 | 20 | 20 | 0 | 0 | 65009 | 45883 | 392067 | 0 | 82 | 5105 | 288 | 0 | 0 | 219 | 0 | 0 | 60 | 0 | 0 | 199 | 0 | 0 | 0 | 0 | 3726 | 55 | 20 |
| SRR2487422 | 49 | 0 | 0 | 0 | 98 | 740 | 1283 | 4194 | 1135 | 25 | 25 | 49 | 0 | 0 | 19194 | 50880 | 291760 | 0 | 0 | 4835 | 395 | 0 | 0 | 14 | 0 | 196 | 490 | 74 | 0 | 147 | 0 | 0 | 0 | 0 | 4962 | 256 | 0 |
| SRR2487423 | 28 | 0 | 28 | 0 | 221 | 113 | 1974 | 4400 | 1297 | 28 | 0 | 83 | 0 | 0 | 20139 | 47850 | 296843 | 0 | 0 | 5133 | 395 | 0 | 0 | 12 | 28 | 83 | 552 | 110 | 0 | 221 | 0 | 0 | 0 | 0 | 4627 | 253 | 28 |
| SRR2487424 | 0 | 0 | 0 | 24 | 191 | 351 | 703 | 4267 | 1607 | 0 | 0 | 48 | 0 | 0 | 18324 | 52140 | 293608 | 0 | 151 | 4619 | 753 | 0 | 0 | 17 | 24 | 143 | 430 | 48 | 0 | 215 | 0 | 0 | 0 | 0 | 3797 | 272 | 24 |
| SRR2487426 | 0 | 35 | 0 | 0 | 10 | 167 | 895 | 1644 | 541 | 35 | 47 | 24 | 0 | 0 | 52219 | 29789 | 215660 | 21 | 271 | 7222 | 1769 | 0 | 0 | 130 | 24 | 0 | 130 | 12 | 0 | 201 | 0 | 0 | 0 | 0 | 2089 | 157 | 0 |
| SRR2487427 | 0 | 0 | 0 | 0 | 323 | 310 | 668 | 1671 | 788 | 27 | 27 | 40 | 0 | 0 | 52775 | 31444 | 220056 | 0 | 215 | 7256 | 2268 | 0 | 0 | 121 | 0 | 54 | 108 | 13 | 0 | 202 | 0 | 0 | 0 | 0 | 1980 | 147 | 0 |
| SRR2487428 | 0 | 12 | 0 | 0 | 421 | 210 | 884 | 1810 | 1094 | 47 | 0 | 70 | 0 | 0 | 52481 | 30033 | 217855 | 0 | 231 | 6818 | 2062 | 0 | 0 | 152 | 23 | 23 | 152 | 12 | 0 | 199 | 0 | 0 | 0 | 0 | 2039 | 131 | 23 |
| SRR2487430 | 0 | 16 | 0 | 0 | 214 | 98 | 337 | 870 | 163 | 0 | 66 | 16 | 0 | 0 | 42769 | 53055 | 235382 | 33 | 65 | 8473 | 2154 | 0 | 0 | 724 | 16 | 33 | 379 | 16 | 0 | 181 | 0 | 0 | 0 | 0 | 664 | 276 | 0 |
| SRR2487431 | 38 | 19 | 0 | 0 | 225 | 123 | 382 | 1023 | 222 | 19 | 56 | 0 | 0 | 0 | 43552 | 52319 | 235182 | 49 | 37 | 8471 | 2688 | 0 | 0 | 600 | 19 | 19 | 169 | 19 | 19 | 244 | 0 | 0 | 19 | 0 | 758 | 279 | 0 |
| SRR2487432 | 0 | 0 | 0 | 16 | 211 | 76 | 489 | 771 | 185 | 0 | 16 | 16 | 0 | 0 | 43949 | 52607 | 236000 | 22 | 43 | 8523 | 2389 | 0 | 0 | 504 | 0 | 16 | 146 | 146 | 0 | 211 | 0 | 0 | 0 | 0 | 657 | 270 | 16 |
| SRR2487434 | 0 | 0 | 0 | 0 | 108 | 33 | 48 | 244 | 76 | 27 | 0 | 0 | 0 | 0 | 107849 | 24852 | 345709 | 0 | 177 | 3086 | 468 | 0 | 0 | 27 | 0 | 20 | 14 | 0 | 0 | 88 | 0 | 0 | 0 | 0 | 496 | 77 | 0 |
| SRR2487435 | 0 | 0 | 0 | 0 | 109 | 22 | 38 | 229 | 76 | 0 | 16 | 0 | 0 | 0 | 106208 | 24803 | 347029 | 0 | 202 | 3239 | 464 | 0 | 0 | 47 | 0 | 0 | 0 | 0 | 0 | 132 | 0 | 0 | 0 | 0 | 447 | 97 | 0 |
| SRR2487436 | 0 | 0 | 0 | 0 | 113 | 33 | 57 | 219 | 86 | 27 | 0 | 0 | 0 | 0 | 107042 | 24926 | 348074 | 0 | 210 | 3101 | 538 | 0 | 0 | 27 | 0 | 0 | 33 | 0 | 0 | 93 | 0 | 0 | 0 | 0 | 476 | 84 | 0 |
| SRR2487438 | 11 | 0 | 0 | 11 | 312 | 178 | 356 | 2120 | 498 | 45 | 22 | 11 | 11 | 0 | 72122 | 54954 | 353131 | 25 | 51 | 6019 | 6304 | 0 | 0 | 36 | 134 | 134 | 14 | 67 | 0 | 134 | 0 | 15 | 0 | 0 | 328 | 311 | 0 |
| SRR2487439 | 0 | 0 | 0 | 0 | 152 | 87 | 272 | 2075 | 428 | 76 | 13 | 25 | 0 | 0 | 72747 | 54781 | 352975 | 17 | 58 | 6495 | 6443 | 0 | 0 | 1880 | 38 | 76 | 927 | 127 | 0 | 152 | 0 | 0 | 0 | 0 | 320 | 278 | 25 |
| SRR2487441 | 0 | 0 | 13 | 26 | 78 | 41 | 326 | 1591 | 775 | 13 | 118 | 0 | 0 | 0 | 13914 | 23802 | 125695 | 0 | 163 | 6569 | 367 | 0 | 0 | 52 | 0 | 39 | 209 | 13 | 0 | 1150 | 0 | 41 | 0 | 0 | 2497 | 96 | 0 |
| SRR2487443 | 0 | 40 | 0 | 67 | 40 | 167 | 500 | 1249 | 666 | 0 | 228 | 0 | 0 | 0 | 14451 | 23783 | 121434 | 0 | 250 | 6996 | 541 | 0 | 0 | 67 | 13 | 0 | 255 | 13 | 0 | 1128 | 0 | 42 | 0 | 0 | 2354 | 91 | 0 |
| SRR2487445 | 0 | 17 | 0 | 34 | 67 | 317 | 529 | 1640 | 635 | 17 | 270 | 17 | 0 | 0 | 14446 | 22958 | 116507 | 0 | 212 | 6456 | 423 | 0 | 0 | 34 | 0 | 34 | 202 | 0 | 0 | 1078 | 0 | 0 | 0 | 0 | 1766 | 85 | 0 |
| SRR2487447 | 0 | 0 | 17 | 33 | 67 | 51 | 618 | 1287 | 824 | 0 | 166 | 0 | 0 | 0 | 12719 | 22782 | 112568 | 51 | 51 | 6591 | 360 | 0 | 0 | 67 | 17 | 33 | 183 | 17 | 0 | 1180 | 0 | 0 | 0 | 0 | 1723 | 88 | 0 |
| SRR2487449 | 11 | 56 | 11 | 112 | 426 | 67 | 176 | 1548 | 297 | 0 | 90 | 11 | 0 | 0 | 12215 | 36213 | 152546 | 12 | 158 | 11128 | 1111 | 0 | 0 | 1087 | 157 | 381 | 3271 | 190 | 0 | 717 | 0 | 0 | 0 | 0 | 511 | 220 | 0 |
| SRR2487451 | 11 | 11 | 0 | 57 | 13 | 74 | 223 | 1551 | 205 | 0 | 92 | 0 | 0 | 0 | 11603 | 37087 | 151902 | 0 | 192 | 10654 | 1142 | 0 | 0 | 32 | 103 | 298 | 79 | 287 | 11 | 21 | 0 | 25 | 0 | 0 | 464 | 207 | 0 |
| SRR2487453 | 0 | 57 | 14 | 72 | 12 | 86 | 195 | 1498 | 250 | 0 | 101 | 0 | 0 | 0 | 11504 | 35730 | 148862 | 0 | 164 | 11379 | 1155 | 0 | 0 | 31 | 14 | 373 | 63 | 244 | 0 | 21 | 0 | 0 | 0 | 0 | 774 | 204 | 14 |
| SRR2487455 | 14 | 14 | 14 | 113 | 227 | 46 | 154 | 1430 | 231 | 0 | 99 | 14 | 0 | 0 | 11619 | 34981 | 148319 | 0 | 115 | 10773 | 1330 | 0 | 0 | 24 | 85 | 369 | 56 | 255 | 0 | 23 | 0 | 0 | 0 | 0 | 586 | 213 | 0 |
| SRR2487457 | 0 | 14 | 0 | 27 | 190 | 145 | 1241 | 3414 | 1966 | 0 | 95 | 0 | 0 | 0 | 17133 | 21099 | 85941 | 0 | 848 | 11918 | 310 | 0 | 0 | 27 | 14 | 81 | 366 | 27 | 0 | 352 | 0 | 41 | 0 | 0 | 1420 | 169 | 0 |
| SRR2487459 | 0 | 14 | 0 | 14 | 128 | 153 | 1271 | 2762 | 1731 | 57 | 28 | 0 | 14 | 0 | 16153 | 23256 | 87685 | 0 | 811 | 13260 | 482 | 0 | 0 | 43 | 0 | 28 | 271 | 0 | 0 | 356 | 0 | 22 | 0 | 0 | 1000 | 159 | 0 |
| SRR2487461 | 0 | 18 | 0 | 36 | 180 | 29 | 1434 | 2868 | 2107 | 18 | 18 | 0 | 0 | 0 | 16098 | 20622 | 79733 | 0 | 585 | 12644 | 702 | 0 | 0 | 18 | 18 | 90 | 234 | 0 | 0 | 433 | 0 | 0 | 0 | 0 | 797 | 152 | 0 |
| SRR2487463 | 0 | 35 | 0 | 0 | 124 | 303 | 1265 | 2393 | 1898 | 0 | 71 | 18 | 0 | 0 | 17300 | 20864 | 81777 | 28 | 798 | 12157 | 468 | 0 | 0 | 18 | 18 | 35 | 159 | 0 | 0 | 530 | 0 | 28 | 0 | 0 | 1425 | 172 | 0 |
| SRR2487465 | 33 | 33 | 0 | 83 | 199 | 153 | 212 | 1542 | 541 | 17 | 33 | 0 | 0 | 0 | 5283 | 61328 | 60316 | 44 | 22 | 8440 | 2024 | 0 | 0 | 598 | 50 | 249 | 1229 | 83 | 0 | 565 | 0 | 15 | 0 | 0 | 461 | 313 | 0 |
| SRR2487467 | 34 | 34 | 0 | 69 | 240 | 141 | 208 | 1522 | 408 | 0 | 69 | 0 | 0 | 0 | 5034 | 61570 | 60883 | 37 | 59 | 9281 | 2086 | 0 | 0 | 463 | 51 | 137 | 20 | 51 | 0 | 11 | 0 | 0 | 0 | 0 | 392 | 313 | 0 |
| SRR2487469 | 43 | 21 | 0 | 43 | 215 | 115 | 269 | 1228 | 527 | 0 | 21 | 21 | 0 | 0 | 5083 | 59361 | 59355 | 38 | 96 | 9466 | 2321 | 0 | 0 | 451 | 43 | 107 | 1419 | 64 | 0 | 580 | 0 | 0 | 0 | 0 | 401 | 311 | 0 |
| SRR2487471 | 21 | 0 | 43 | 21 | 64 | 142 | 246 | 1543 | 672 | 0 | 85 | 0 | 0 | 0 | 5046 | 60298 | 59908 | 28 | 66 | 9525 | 2036 | 0 | 0 | 492 | 21 | 107 | 15 | 43 | 0 | 13 | 0 | 0 | 0 | 0 | 488 | 343 | 0 |
| SRR2487473 | 17 | 0 | 0 | 51 | 475 | 787 | 1242 | 4555 | 1201 | 0 | 17 | 0 | 0 | 0 | 2484 | 30740 | 65462 | 41 | 704 | 4803 | 1366 | 0 | 0 | 102 | 0 | 34 | 119 | 0 | 0 | 14 | 0 | 41 | 0 | 0 | 17520 | 101 | 0 |
| SRR2487475 | 17 | 17 | 0 | 104 | 470 | 503 | 1173 | 4482 | 1089 | 0 | 104 | 0 | 0 | 0 | 2304 | 30002 | 65812 | 0 | 335 | 4942 | 1801 | 0 | 0 | 87 | 0 | 0 | 122 | 35 | 0 | 1045 | 0 | 0 | 17 | 0 | 15808 | 98 | 0 |
| SRR2487477 | 22 | 0 | 0 | 88 | 525 | 536 | 1930 | 4825 | 1019 | 22 | 175 | 0 | 0 | 0 | 1823 | 29370 | 61959 | 0 | 429 | 5254 | 1876 | 0 | 0 | 66 | 0 | 0 | 66 | 22 | 0 | 854 | 0 | 54 | 0 | 0 | 16697 | 83 | 0 |
| SRR2487479 | 0 | 0 | 0 | 43 | 304 | 730 | 1251 | 5107 | 1355 | 65 | 130 | 0 | 0 | 0 | 2345 | 29052 | 65765 | 0 | 365 | 4743 | 2189 | 0 | 0 | 0 | 0 | 0 | 130 | 22 | 0 | 978 | 0 | 0 | 0 | 0 | 19302 | 105 | 0 |
| SRR2487481 | 28 | 28 | 14 | 112 | 12 | 77 | 308 | 1606 | 450 | 0 | 56 | 0 | 0 | 0 | 3481 | 49330 | 105479 | 0 | 128 | 8555 | 861 | 0 | 0 | 23 | 112 | 266 | 113 | 210 | 0 | 20 | 0 | 0 | 0 | 0 | 1066 | 243 | 14 |
| SRR2487483 | 0 | 43 | 14 | 57 | 10 | 26 | 219 | 1777 | 554 | 0 | 86 | 14 | 0 | 0 | 3270 | 49532 | 99621 | 0 | 77 | 8998 | 978 | 0 | 0 | 23 | 101 | 302 | 93 | 187 | 0 | 20 | 0 | 0 | 0 | 0 | 914 | 281 | 0 |
| SRR2487485 | 0 | 54 | 0 | 90 | 361 | 120 | 256 | 1571 | 393 | 0 | 126 | 18 | 0 | 0 | 3124 | 47320 | 100576 | 0 | 137 | 8997 | 905 | 0 | 0 | 595 | 36 | 180 | 3101 | 234 | 0 | 703 | 0 | 0 | 0 | 0 | 1090 | 255 | 0 |
| SRR2487487 | 0 | 0 | 18 | 36 | 305 | 66 | 397 | 2084 | 480 | 0 | 72 | 36 | 0 | 0 | 3605 | 49366 | 100092 | 0 | 165 | 9542 | 926 | 0 | 0 | 520 | 36 | 287 | 2617 | 215 | 0 | 627 | 0 | 0 | 0 | 0 | 1083 | 262 | 0 |
| SRR2487489 | 0 | 0 | 0 | 0 | 103 | 17 | 70 | 250 | 58 | 47 | 34 | 28 | 0 | 0 | 13823 | 22498 | 123316 | 0 | 686 | 9419 | 3062 | 0 | 0 | 13 | 0 | 62 | 62 | 0 | 0 | 369 | 0 | 0 | 0 | 0 | 111 | 247 | 0 |
| SRR2487491 | 0 | 0 | 0 | 0 | 11 | 0 | 90 | 174 | 84 | 39 | 48 | 19 | 0 | 0 | 13933 | 21539 | 123659 | 0 | 618 | 9492 | 3132 | 0 | 0 | 13 | 19 | 52 | 64 | 0 | 0 | 52 | 0 | 18 | 0 | 0 | 127 | 232 | 0 |
| SRR2487493 | 0 | 0 | 0 | 0 | 59 | 29 | 74 | 169 | 66 | 35 | 55 | 31 | 0 | 0 | 13100 | 20016 | 115789 | 0 | 640 | 9540 | 3119 | 0 | 12 | 0 | 0 | 43 | 39 | 0 | 0 | 46 | 0 | 0 | 0 | 0 | 108 | 213 | 0 |
| SRR2487495 | 0 | 0 | 0 | 0 | 64 | 0 | 93 | 186 | 79 | 56 | 48 | 24 | 0 | 0 | 13701 | 20553 | 116726 | 0 | 666 | 9327 | 3400 | 0 | 0 | 20 | 0 | 40 | 32 | 0 | 0 | 348 | 0 | 14 | 0 | 0 | 183 | 190 | 0 |
| SRR2487497 | 0 | 17 | 0 | 34 | 19 | 95 | 296 | 3539 | 243 | 26 | 60 | 52 | 0 | 0 | 16783 | 17297 | 180479 | 0 | 225 | 21170 | 2976 | 0 | 0 | 22 | 121 | 11 | 72 | 155 | 0 | 17 | 0 | 0 | 0 | 0 | 628 | 252 | 0 |
| SRR2487499 | 0 | 18 | 0 | 18 | 18 | 117 | 227 | 3367 | 301 | 0 | 27 | 53 | 0 | 0 | 16957 | 16481 | 179904 | 12 | 252 | 21184 | 3133 | 0 | 0 | 22 | 71 | 14 | 65 | 186 | 0 | 17 | 0 | 12 | 0 | 0 | 563 | 268 | 0 |
| SRR2487501 | 0 | 22 | 0 | 44 | 498 | 61 | 222 | 3270 | 229 | 22 | 100 | 55 | 0 | 0 | 16233 | 16661 | 174249 | 0 | 229 | 21359 | 3231 | 0 | 11 | 653 | 44 | 332 | 1416 | 144 | 0 | 520 | 0 | 0 | 0 | 0 | 610 | 263 | 0 |
| SRR2487503 | 0 | 0 | 22 | 11 | 528 | 75 | 253 | 3034 | 253 | 11 | 88 | 33 | 0 | 0 | 16631 | 16108 | 175192 | 15 | 142 | 21088 | 3466 | 0 | 11 | 462 | 66 | 286 | 1563 | 132 | 0 | 363 | 0 | 0 | 0 | 0 | 610 | 277 | 0 |
| SRR2487504 | 60 | 0 | 0 | 0 | 240 | 68 | 136 | 476 | 340 | 120 | 300 | 0 | 0 | 0 | 24010 | 49949 | 34236 | 204 | 680 | 6530 | 476 | 0 | 0 | 120 | 0 | 120 | 120 | 60 | 0 | 1319 | 0 | 0 | 0 | 0 | 1405 | 161 | 0 |
| SRR2487505 | 0 | 0 | 0 | 41 | 372 | 53 | 318 | 1006 | 318 | 83 | 124 | 41 | 0 | 0 | 18630 | 46466 | 35027 | 0 | 635 | 8204 | 423 | 0 | 0 | 41 | 0 | 41 | 331 | 0 | 0 | 33 | 0 | 0 | 0 | 0 | 684 | 206 | 0 |
| SRR2487507 | 0 | 83 | 0 | 0 | 207 | 153 | 153 | 510 | 153 | 0 | 165 | 41 | 0 | 0 | 19084 | 48708 | 35519 | 153 | 1021 | 7195 | 459 | 0 | 0 | 83 | 0 | 41 | 165 | 0 | 0 | 1529 | 0 | 0 | 0 | 0 | 993 | 151 | 0 |
| SRR2487508 | 0 | 0 | 0 | 63 | 506 | 214 | 214 | 784 | 499 | 0 | 190 | 0 | 0 | 0 | 26447 | 51896 | 34474 | 0 | 713 | 6202 | 713 | 0 | 0 | 190 | 0 | 63 | 190 | 0 | 0 | 1454 | 0 | 0 | 0 | 0 | 1295 | 178 | 0 |
| SRR2487509 | 0 | 0 | 0 | 0 | 11 | 75 | 225 | 1126 | 526 | 0 | 132 | 0 | 0 | 0 | 24476 | 50422 | 33237 | 75 | 450 | 6232 | 375 | 0 | 0 | 66 | 0 | 66 | 198 | 0 | 0 | 36 | 0 | 0 | 0 | 0 | 1135 | 169 | 0 |
| SRR2487510 | 0 | 68 | 0 | 0 | 405 | 149 | 224 | 895 | 373 | 0 | 203 | 0 | 0 | 0 | 25511 | 49635 | 33491 | 0 | 522 | 5967 | 224 | 0 | 0 | 68 | 68 | 68 | 11 | 0 | 0 | 28 | 0 | 0 | 0 | 0 | 1175 | 206 | 0 |
| SRR2487512 | 0 | 0 | 0 | 64 | 11 | 39 | 117 | 782 | 508 | 96 | 288 | 0 | 0 | 0 | 19158 | 50140 | 37655 | 39 | 704 | 7155 | 665 | 0 | 0 | 96 | 0 | 96 | 320 | 0 | 0 | 30 | 0 | 0 | 0 | 0 | 799 | 197 | 0 |
| SRR2487514 | 33 | 33 | 0 | 33 | 233 | 41 | 286 | 980 | 327 | 67 | 67 | 0 | 0 | 0 | 19321 | 51486 | 36089 | 41 | 1185 | 7884 | 572 | 0 | 0 | 133 | 0 | 33 | 233 | 33 | 0 | 47 | 0 | 0 | 0 | 0 | 586 | 174 | 0 |
| SRR2487516 | 0 | 50 | 0 | 151 | 50 | 44 | 348 | 522 | 305 | 0 | 0 | 0 | 0 | 0 | 5656 | 24798 | 148855 | 0 | 131 | 4089 | 653 | 0 | 0 | 603 | 50 | 201 | 955 | 101 | 0 | 402 | 0 | 0 | 0 | 0 | 1312 | 339 | 50 |
| SRR2487517 | 65 | 33 | 0 | 33 | 131 | 29 | 349 | 553 | 116 | 0 | 33 | 0 | 0 | 0 | 4950 | 24609 | 145891 | 0 | 29 | 5561 | 670 | 0 | 0 | 10 | 131 | 131 | 29 | 98 | 0 | 13 | 0 | 0 | 0 | 0 | 1090 | 315 | 0 |
| SRR2487519 | 0 | 0 | 0 | 32 | 259 | 86 | 316 | 776 | 230 | 0 | 162 | 0 | 0 | 0 | 5804 | 25808 | 143416 | 0 | 144 | 6321 | 603 | 0 | 0 | 550 | 32 | 194 | 1198 | 194 | 0 | 615 | 0 | 0 | 0 | 0 | 1219 | 314 | 0 |
| SRR2487520 | 51 | 51 | 0 | 103 | 359 | 136 | 317 | 905 | 362 | 0 | 0 | 0 | 0 | 0 | 5473 | 26414 | 150578 | 0 | 45 | 4975 | 407 | 0 | 0 | 308 | 154 | 154 | 924 | 205 | 0 | 565 | 0 | 0 | 0 | 0 | 1419 | 329 | 0 |
| SRR2487521 | 56 | 56 | 0 | 0 | 223 | 0 | 251 | 954 | 201 | 0 | 0 | 0 | 0 | 0 | 4820 | 21815 | 144608 | 0 | 0 | 4820 | 653 | 0 | 0 | 12 | 0 | 56 | 18 | 0 | 0 | 334 | 0 | 0 | 0 | 0 | 1751 | 375 | 0 |
| SRR2487522 | 0 | 0 | 0 | 0 | 281 | 145 | 387 | 823 | 484 | 0 | 0 | 0 | 0 | 0 | 5570 | 24717 | 141962 | 0 | 0 | 4747 | 436 | 0 | 0 | 787 | 0 | 169 | 787 | 0 | 0 | 393 | 0 | 0 | 0 | 0 | 2738 | 318 | 0 |
| SRR2487524 | 0 | 76 | 0 | 76 | 152 | 65 | 260 | 629 | 217 | 0 | 76 | 0 | 0 | 0 | 5032 | 23653 | 149033 | 22 | 174 | 4902 | 781 | 0 | 0 | 355 | 102 | 102 | 1066 | 51 | 0 | 584 | 0 | 0 | 0 | 0 | 1152 | 312 | 0 |
| SRR2487526 | 26 | 0 | 0 | 52 | 182 | 156 | 179 | 647 | 156 | 26 | 0 | 0 | 0 | 0 | 4844 | 23549 | 142174 | 0 | 89 | 5648 | 580 | 0 | 0 | 339 | 78 | 156 | 1146 | 104 | 0 | 443 | 0 | 0 | 0 | 0 | 1175 | 337 | 0 |
| SRR2487529 | 0 | 18 | 0 | 44 | 98 | 312 | 637 | 2603 | 1003 | 18 | 62 | 18 | 0 | 0 | 15146 | 32304 | 123200 | 0 | 176 | 5654 | 312 | 0 | 0 | 133 | 27 | 27 | 204 | 0 | 0 | 328 | 0 | 0 | 0 | 0 | 1663 | 101 | 0 |
| SRR2487531 | 27 | 27 | 0 | 37 | 137 | 265 | 671 | 2320 | 866 | 18 | 55 | 0 | 0 | 0 | 15063 | 32502 | 121050 | 14 | 265 | 5394 | 363 | 0 | 0 | 137 | 0 | 64 | 302 | 0 | 0 | 11 | 0 | 28 | 0 | 0 | 1592 | 112 | 0 |
| SRR2487533 | 23 | 23 | 0 | 34 | 125 | 459 | 548 | 2315 | 795 | 0 | 68 | 11 | 0 | 0 | 15974 | 30226 | 115860 | 0 | 247 | 6414 | 495 | 0 | 0 | 113 | 11 | 34 | 340 | 11 | 0 | 726 | 0 | 0 | 0 | 0 | 1834 | 94 | 0 |
| SRR2487535 | 11 | 0 | 0 | 22 | 112 | 399 | 676 | 2254 | 1214 | 11 | 34 | 11 | 0 | 0 | 15119 | 30523 | 116353 | 0 | 225 | 6051 | 347 | 0 | 0 | 101 | 0 | 22 | 258 | 11 | 0 | 459 | 0 | 0 | 0 | 0 | 1827 | 122 | 0 |
| SRR2487537 | 15 | 30 | 0 | 30 | 258 | 163 | 263 | 2409 | 302 | 0 | 46 | 0 | 0 | 0 | 10501 | 29976 | 153787 | 0 | 93 | 19260 | 2470 | 0 | 0 | 1685 | 15 | 379 | 2550 | 91 | 0 | 607 | 0 | 0 | 0 | 0 | 393 | 417 | 15 |
| SRR2487539 | 0 | 0 | 0 | 0 | 252 | 229 | 260 | 2357 | 363 | 0 | 79 | 0 | 0 | 0 | 10483 | 30161 | 154429 | 0 | 158 | 20335 | 2428 | 0 | 0 | 1544 | 16 | 425 | 2301 | 63 | 0 | 425 | 0 | 0 | 0 | 0 | 412 | 373 | 0 |
| SRR2487541 | 20 | 0 | 0 | 0 | 293 | 159 | 358 | 2355 | 288 | 0 | 39 | 0 | 0 | 0 | 9925 | 28392 | 146683 | 0 | 179 | 19563 | 2434 | 0 | 0 | 1486 | 39 | 274 | 1623 | 78 | 0 | 371 | 0 | 0 | 0 | 0 | 408 | 393 | 0 |
| SRR2487543 | 39 | 58 | 0 | 0 | 213 | 204 | 272 | 2333 | 214 | 0 | 39 | 0 | 0 | 0 | 10088 | 29498 | 148820 | 0 | 136 | 20245 | 2634 | 0 | 0 | 18 | 19 | 155 | 38 | 116 | 0 | 13 | 0 | 0 | 0 | 0 | 368 | 396 | 0 |
| SRR2487545 | 122 | 0 | 0 | 122 | 245 | 0 | 0 | 27 | 10 | 0 | 183 | 0 | 0 | 0 | 43455 | 63761 | 122025 | 17 | 399 | 10196 | 686 | 0 | 0 | 183 | 61 | 61 | 306 | 0 | 0 | 887 | 0 | 0 | 0 | 0 | 85 | 142 | 0 |
| SRR2487547 | 31 | 31 | 0 | 93 | 62 | 0 | 0 | 52 | 10 | 31 | 93 | 0 | 0 | 0 | 43184 | 63488 | 121849 | 0 | 457 | 10335 | 879 | 0 | 0 | 0 | 0 | 31 | 125 | 0 | 0 | 841 | 0 | 0 | 0 | 0 | 74 | 174 | 0 |
| SRR2487549 | 78 | 39 | 0 | 0 | 196 | 0 | 0 | 13 | 0 | 0 | 78 | 0 | 0 | 0 | 41431 | 60731 | 115280 | 13 | 451 | 10448 | 779 | 0 | 0 | 118 | 0 | 0 | 39 | 0 | 0 | 785 | 0 | 0 | 0 | 0 | 63 | 168 | 0 |
| SRR2487551 | 0 | 0 | 0 | 78 | 117 | 0 | 0 | 50 | 0 | 117 | 78 | 0 | 0 | 0 | 42343 | 60721 | 115239 | 25 | 359 | 9788 | 685 | 0 | 0 | 0 | 0 | 0 | 156 | 0 | 0 | 896 | 0 | 0 | 0 | 0 | 72 | 165 | 39 |
| SRR2487553 | 20 | 61 | 20 | 283 | 344 | 0 | 0 | 45 | 16 | 0 | 162 | 20 | 0 | 0 | 24736 | 54371 | 203563 | 0 | 133 | 25827 | 4172 | 0 | 0 | 344 | 101 | 12 | 99 | 48 | 0 | 15 | 0 | 0 | 0 | 0 | 10 | 390 | 0 |
| SRR2487555 | 41 | 41 | 21 | 145 | 186 | 0 | 0 | 48 | 24 | 0 | 103 | 21 | 0 | 0 | 25348 | 54048 | 201818 | 0 | 126 | 26703 | 4015 | 0 | 0 | 290 | 41 | 497 | 90 | 41 | 0 | 17 | 0 | 0 | 0 | 0 | 12 | 423 | 0 |
| SRR2487560 | 0 | 0 | 0 | 0 | 623 | 0 | 0 | 142 | 0 | 0 | 138 | 0 | 0 | 0 | 24853 | 23980 | 93730 | 0 | 213 | 29404 | 907 | 0 | 0 | 138 | 0 | 69 | 692 | 0 | 0 | 623 | 0 | 0 | 0 | 0 | 254 | 269 | 0 |
| SRR2487561 | 0 | 0 | 0 | 38 | 246 | 0 | 0 | 0 | 0 | 0 | 246 | 0 | 0 | 0 | 18347 | 21418 | 89810 | 16 | 198 | 35483 | 1034 | 0 | 0 | 0 | 38 | 151 | 33 | 227 | 0 | 33 | 0 | 0 | 0 | 0 | 24 | 221 | 0 |
| SRR2487563 | 0 | 39 | 39 | 233 | 14 | 0 | 0 | 0 | 31 | 19 | 194 | 19 | 0 | 0 | 18304 | 20853 | 91249 | 10 | 253 | 35370 | 893 | 0 | 0 | 19 | 19 | 97 | 31 | 78 | 0 | 29 | 0 | 0 | 0 | 0 | 32 | 248 | 0 |
| SRR2487564 | 0 | 0 | 0 | 0 | 1042 | 0 | 37 | 56 | 19 | 0 | 223 | 0 | 0 | 0 | 24744 | 25019 | 95138 | 0 | 225 | 30551 | 749 | 0 | 0 | 298 | 0 | 74 | 670 | 0 | 0 | 447 | 0 | 0 | 0 | 0 | 203 | 266 | 0 |
| SRR2487565 | 77 | 0 | 0 | 230 | 921 | 0 | 0 | 59 | 20 | 0 | 77 | 0 | 0 | 0 | 24724 | 23281 | 90167 | 20 | 235 | 29213 | 1000 | 0 | 0 | 230 | 0 | 153 | 614 | 0 | 0 | 307 | 0 | 0 | 0 | 0 | 309 | 194 | 0 |
| SRR2487566 | 79 | 0 | 0 | 0 | 550 | 20 | 20 | 158 | 20 | 0 | 0 | 0 | 0 | 0 | 25602 | 23759 | 93745 | 39 | 178 | 28087 | 730 | 0 | 0 | 236 | 79 | 79 | 314 | 0 | 0 | 14 | 0 | 0 | 0 | 0 | 299 | 213 | 0 |
| SRR2487568 | 0 | 15 | 15 | 147 | 514 | 0 | 0 | 0 | 0 | 0 | 191 | 0 | 0 | 0 | 18905 | 21931 | 95973 | 0 | 240 | 34768 | 918 | 0 | 0 | 0 | 59 | 132 | 1191 | 132 | 0 | 1102 | 0 | 0 | 0 | 0 | 22 | 254 | 0 |
| SRR2487570 | 0 | 15 | 30 | 76 | 16 | 0 | 0 | 0 | 0 | 0 | 152 | 0 | 0 | 0 | 18949 | 22628 | 94854 | 13 | 239 | 35927 | 968 | 0 | 0 | 30 | 46 | 76 | 46 | 11 | 0 | 32 | 0 | 0 | 0 | 0 | 32 | 303 | 0 |
| SRR2487572 | 0 | 32 | 32 | 0 | 714 | 49 | 229 | 1759 | 106 | 0 | 32 | 0 | 0 | 0 | 23479 | 58611 | 145143 | 57 | 65 | 33558 | 3420 | 0 | 0 | 1428 | 97 | 454 | 1720 | 97 | 0 | 357 | 0 | 0 | 0 | 0 | 197 | 269 | 0 |
| SRR2487573 | 30 | 30 | 30 | 268 | 537 | 0 | 31 | 191 | 0 | 30 | 298 | 0 | 0 | 0 | 18343 | 50937 | 140242 | 15 | 153 | 43695 | 4599 | 0 | 0 | 89 | 60 | 746 | 3042 | 1253 | 0 | 895 | 0 | 0 | 0 | 0 | 38 | 254 | 0 |
| SRR2487575 | 0 | 60 | 60 | 210 | 240 | 0 | 0 | 189 | 38 | 0 | 330 | 0 | 0 | 0 | 18724 | 52109 | 141955 | 38 | 83 | 44140 | 4511 | 0 | 0 | 300 | 240 | 570 | 3389 | 1560 | 0 | 960 | 0 | 0 | 0 | 0 | 26 | 338 | 0 |
| SRR2487576 | 34 | 0 | 0 | 34 | 369 | 93 | 178 | 2075 | 110 | 0 | 67 | 0 | 0 | 0 | 24856 | 59477 | 145921 | 68 | 76 | 35307 | 3523 | 0 | 0 | 1240 | 101 | 402 | 1742 | 134 | 0 | 503 | 0 | 0 | 0 | 0 | 177 | 242 | 0 |
| SRR2487577 | 36 | 0 | 0 | 36 | 507 | 119 | 146 | 2067 | 229 | 0 | 217 | 0 | 0 | 0 | 24065 | 56138 | 140443 | 46 | 101 | 33724 | 4043 | 0 | 0 | 1087 | 36 | 507 | 2030 | 72 | 0 | 471 | 0 | 0 | 0 | 0 | 210 | 257 | 0 |
| SRR2487578 | 0 | 36 | 0 | 36 | 511 | 101 | 119 | 2190 | 110 | 0 | 73 | 0 | 0 | 0 | 24692 | 57523 | 141394 | 27 | 119 | 32268 | 3573 | 0 | 0 | 1350 | 146 | 219 | 1678 | 146 | 0 | 511 | 0 | 0 | 0 | 0 | 150 | 282 | 0 |
| SRR2487580 | 23 | 0 | 45 | 136 | 204 | 0 | 24 | 212 | 53 | 0 | 272 | 0 | 0 | 0 | 19635 | 53318 | 150344 | 24 | 100 | 42204 | 4379 | 0 | 0 | 159 | 249 | 24 | 147 | 62 | 0 | 25 | 0 | 0 | 0 | 0 | 0 | 266 | 0 |
| SRR2487582 | 70 | 23 | 0 | 116 | 603 | 0 | 0 | 265 | 36 | 0 | 185 | 0 | 0 | 0 | 18520 | 54295 | 149306 | 18 | 103 | 44181 | 4209 | 0 | 0 | 301 | 162 | 603 | 4172 | 1715 | 0 | 950 | 0 | 0 | 0 | 0 | 18 | 289 | 0 |
| SRR2487584 | 0 | 0 | 0 | 0 | 29 | 216 | 540 | 1026 | 108 | 0 | 40 | 0 | 0 | 0 | 12308 | 18552 | 86154 | 108 | 54 | 10041 | 324 | 0 | 0 | 0 | 11 | 0 | 16 | 0 | 0 | 94 | 0 | 0 | 0 | 0 | 1027 | 110 | 0 |
| SRR2487585 | 0 | 0 | 0 | 10 | 12 | 0 | 154 | 154 | 0 | 0 | 35 | 0 | 0 | 0 | 10526 | 15145 | 79234 | 0 | 461 | 13368 | 384 | 0 | 0 | 0 | 0 | 14 | 50 | 18 | 0 | 157 | 0 | 0 | 0 | 0 | 41 | 67 | 0 |
| SRR2487587 | 0 | 0 | 0 | 37 | 23 | 76 | 0 | 0 | 0 | 0 | 65 | 0 | 0 | 0 | 10522 | 14623 | 78314 | 0 | 303 | 12945 | 530 | 0 | 0 | 0 | 14 | 14 | 171 | 19 | 0 | 615 | 0 | 0 | 0 | 0 | 41 | 21 | 0 |
| SRR2487588 | 0 | 0 | 0 | 0 | 31 | 393 | 281 | 1234 | 112 | 0 | 35 | 0 | 0 | 0 | 12175 | 18870 | 83023 | 56 | 56 | 11277 | 337 | 0 | 0 | 23 | 0 | 12 | 62 | 0 | 0 | 384 | 0 | 0 | 0 | 0 | 1170 | 90 | 0 |
| SRR2487589 | 0 | 0 | 0 | 0 | 20 | 180 | 480 | 360 | 180 | 0 | 57 | 0 | 0 | 0 | 11525 | 17653 | 81640 | 60 | 60 | 10265 | 360 | 0 | 0 | 24 | 16 | 0 | 73 | 0 | 0 | 434 | 0 | 0 | 0 | 0 | 761 | 78 | 0 |
| SRR2487590 | 0 | 0 | 0 | 0 | 58 | 412 | 766 | 648 | 295 | 0 | 41 | 0 | 0 | 0 | 12021 | 17657 | 79154 | 59 | 59 | 10430 | 648 | 0 | 0 | 21 | 0 | 17 | 62 | 0 | 0 | 381 | 0 | 0 | 0 | 0 | 1025 | 61 | 0 |
| SRR2487592 | 0 | 0 | 0 | 32 | 28 | 0 | 60 | 60 | 0 | 0 | 74 | 0 | 0 | 0 | 11042 | 16638 | 85858 | 60 | 241 | 13456 | 181 | 0 | 0 | 0 | 0 | 11 | 148 | 28 | 0 | 620 | 0 | 0 | 0 | 0 | 36 | 125 | 0 |
| SRR2487594 | 0 | 0 | 0 | 26 | 15 | 0 | 122 | 61 | 61 | 0 | 87 | 0 | 0 | 0 | 11509 | 16251 | 83598 | 61 | 184 | 15549 | 490 | 0 | 0 | 0 | 0 | 40 | 142 | 22 | 0 | 645 | 0 | 0 | 0 | 0 | 144 | 166 | 0 |
| SRR2487597 | 32 | 21 | 0 | 42 | 264 | 64 | 399 | 1789 | 463 | 11 | 95 | 0 | 0 | 0 | 6421 | 56174 | 132453 | 0 | 103 | 7077 | 232 | 0 | 0 | 22 | 85 | 14 | 75 | 180 | 0 | 19 | 0 | 0 | 0 | 0 | 1710 | 305 | 0 |
| SRR2487598 | 0 | 23 | 0 | 56 | 13 | 141 | 353 | 2177 | 311 | 0 | 34 | 0 | 0 | 0 | 5782 | 53556 | 129191 | 0 | 99 | 6517 | 396 | 0 | 0 | 22 | 23 | 237 | 58 | 147 | 0 | 11 | 0 | 0 | 0 | 0 | 2154 | 298 | 11 |
| SRR2487599 | 11 | 0 | 0 | 23 | 330 | 125 | 390 | 2019 | 446 | 0 | 23 | 11 | 0 | 0 | 5975 | 53528 | 128933 | 0 | 84 | 6657 | 334 | 0 | 0 | 660 | 80 | 433 | 1890 | 114 | 0 | 421 | 0 | 0 | 0 | 0 | 1951 | 297 | 23 |
| SRR2487600 | 0 | 16 | 0 | 13 | 28 | 0 | 0 | 130 | 216 | 0 | 11 | 0 | 0 | 0 | 5009 | 51280 | 131951 | 0 | 108 | 8808 | 561 | 0 | 0 | 95 | 47 | 55 | 456 | 220 | 0 | 69 | 0 | 0 | 0 | 0 | 102 | 253 | 0 |
| SRR2487602 | 32 | 0 | 0 | 19 | 21 | 0 | 0 | 331 | 154 | 0 | 11 | 0 | 0 | 0 | 5333 | 48372 | 132671 | 22 | 198 | 8573 | 397 | 0 | 0 | 16 | 16 | 63 | 425 | 204 | 0 | 66 | 0 | 0 | 0 | 0 | 137 | 359 | 0 |
| SRR2487604 | 20 | 0 | 0 | 18 | 22 | 58 | 29 | 234 | 58 | 0 | 0 | 0 | 0 | 0 | 5810 | 46652 | 123244 | 0 | 88 | 10364 | 496 | 0 | 0 | 11 | 41 | 45 | 406 | 149 | 0 | 45 | 0 | 0 | 0 | 0 | 258 | 237 | 0 |
| SRR2487606 | 0 | 41 | 0 | 22 | 27 | 0 | 29 | 172 | 143 | 0 | 104 | 0 | 0 | 0 | 5633 | 48333 | 123734 | 0 | 200 | 9778 | 400 | 0 | 0 | 62 | 104 | 36 | 364 | 152 | 0 | 49 | 0 | 0 | 0 | 0 | 339 | 257 | 0 |
| SRR2487607 | 10 | 20 | 10 | 40 | 392 | 50 | 450 | 1814 | 563 | 10 | 60 | 0 | 0 | 0 | 6366 | 54713 | 130739 | 0 | 100 | 6454 | 488 | 0 | 0 | 583 | 50 | 251 | 1961 | 151 | 0 | 493 | 0 | 13 | 0 | 0 | 1707 | 301 | 10 |
| SRR2487609 | 23 | 0 | 0 | 35 | 10 | 24 | 56 | 207 | 64 | 0 | 16 | 0 | 0 | 0 | 61743 | 19356 | 141088 | 16 | 207 | 11258 | 741 | 0 | 0 | 311 | 12 | 35 | 15 | 23 | 0 | 28 | 0 | 0 | 0 | 0 | 394 | 100 | 12 |
| SRR2487610 | 0 | 0 | 12 | 12 | 208 | 25 | 33 | 141 | 50 | 0 | 10 | 0 | 0 | 0 | 59111 | 18355 | 133319 | 0 | 315 | 11308 | 871 | 0 | 0 | 301 | 12 | 23 | 278 | 12 | 0 | 32 | 0 | 0 | 0 | 0 | 443 | 108 | 12 |
| SRR2487611 | 12 | 12 | 12 | 0 | 200 | 25 | 67 | 200 | 108 | 0 | 516 | 0 | 0 | 0 | 60516 | 17953 | 134684 | 0 | 217 | 10557 | 909 | 0 | 0 | 399 | 12 | 35 | 317 | 35 | 0 | 951 | 0 | 0 | 0 | 0 | 547 | 101 | 12 |
| SRR2487612 | 0 | 0 | 34 | 201 | 302 | 0 | 0 | 66 | 0 | 0 | 873 | 0 | 0 | 0 | 48346 | 16174 | 140407 | 0 | 420 | 15040 | 1038 | 0 | 0 | 67 | 0 | 34 | 1175 | 235 | 0 | 1846 | 0 | 0 | 0 | 0 | 17 | 39 | 0 |
| SRR2487614 | 0 | 0 | 0 | 70 | 12 | 0 | 0 | 0 | 0 | 0 | 51 | 0 | 0 | 0 | 46551 | 18354 | 140789 | 0 | 433 | 15084 | 1481 | 0 | 0 | 70 | 0 | 0 | 63 | 19 | 0 | 143 | 0 | 0 | 0 | 0 | 17 | 82 | 0 |
| SRR2487616 | 0 | 0 | 0 | 136 | 14 | 0 | 28 | 28 | 0 | 45 | 32 | 0 | 0 | 0 | 45675 | 16656 | 131802 | 28 | 365 | 15000 | 1067 | 0 | 0 | 136 | 0 | 45 | 38 | 91 | 0 | 121 | 0 | 0 | 0 | 0 | 61 | 50 | 0 |
| SRR2487618 | 0 | 46 | 0 | 137 | 13 | 0 | 28 | 0 | 28 | 0 | 67 | 0 | 0 | 0 | 46817 | 16434 | 129156 | 0 | 563 | 13561 | 1013 | 0 | 0 | 46 | 0 | 46 | 49 | 16 | 0 | 119 | 0 | 0 | 0 | 0 | 0 | 121 | 0 |
| SRR2487619 | 0 | 0 | 11 | 11 | 286 | 0 | 45 | 174 | 60 | 11 | 15 | 32 | 0 | 0 | 59061 | 18846 | 137410 | 0 | 324 | 11325 | 822 | 0 | 0 | 11 | 0 | 21 | 14 | 32 | 0 | 30 | 0 | 15 | 0 | 0 | 366 | 104 | 11 |
| SRR2487621 | 0 | 0 | 0 | 0 | 51 | 130 | 183 | 600 | 417 | 0 | 102 | 0 | 0 | 0 | 22825 | 46977 | 127123 | 78 | 78 | 16199 | 2009 | 0 | 0 | 743 | 26 | 102 | 743 | 77 | 0 | 794 | 0 | 0 | 0 | 0 | 640 | 552 | 0 |
| SRR2487622 | 0 | 27 | 0 | 0 | 82 | 84 | 139 | 1032 | 112 | 0 | 55 | 0 | 0 | 0 | 22637 | 44462 | 122565 | 28 | 84 | 16643 | 1979 | 0 | 0 | 654 | 0 | 55 | 627 | 55 | 0 | 872 | 0 | 0 | 0 | 0 | 802 | 562 | 0 |
| SRR2487623 | 0 | 55 | 0 | 28 | 111 | 196 | 196 | 897 | 168 | 0 | 55 | 0 | 0 | 0 | 23857 | 44568 | 123720 | 56 | 84 | 16400 | 2046 | 0 | 0 | 12 | 55 | 28 | 581 | 55 | 0 | 15 | 0 | 0 | 0 | 0 | 716 | 561 | 0 |
| SRR2487624 | 0 | 46 | 15 | 107 | 46 | 0 | 17 | 120 | 51 | 0 | 138 | 0 | 0 | 0 | 17980 | 43305 | 131907 | 17 | 137 | 20275 | 2757 | 0 | 0 | 77 | 15 | 107 | 70 | 33 | 0 | 75 | 0 | 0 | 0 | 0 | 55 | 414 | 0 |
| SRR2487626 | 0 | 0 | 0 | 79 | 142 | 35 | 35 | 123 | 35 | 0 | 11 | 0 | 0 | 0 | 16960 | 42098 | 130653 | 18 | 123 | 21521 | 2501 | 0 | 0 | 13 | 16 | 12 | 71 | 45 | 0 | 75 | 0 | 0 | 0 | 0 | 95 | 457 | 0 |
| SRR2487628 | 0 | 20 | 0 | 60 | 40 | 0 | 22 | 130 | 22 | 0 | 14 | 0 | 0 | 0 | 17961 | 38783 | 124456 | 65 | 152 | 21214 | 2603 | 0 | 0 | 40 | 40 | 119 | 43 | 31 | 0 | 66 | 0 | 0 | 0 | 0 | 48 | 532 | 20 |

### Supp Table S8.3.Columns_BX_to_DH

| | | | | | | | | | | | | | | | | | | | | | | | | | | | | | | | | | | | | | |
|---|---|---|---|---|---|---|---|---|---|---|---|---|---|---|---|---|---|---|---|---|---|---|---|---|---|---|---|---|---|---|---|---|---|---|---|---|---|
| SRR2487630 | 0 | 0 | 0 | 60 | 99 | 44 | 0 | 65 | 87 | 0 | 159 | 0 | 0 | 0 | 18375 | 39785 | 125134 | 109 | 175 | 21845 | 2684 | 0 | 0 | 119 | 20 | 79 | 1152 | 437 | 0 | 1350 | 0 | 0 | 0 | 0 | 160 | 566 | 0 |
| SRR2487631 | 0 | 0 | 0 | 48 | 121 | 124 | 124 | 695 | 124 | 0 | 72 | 0 | 0 | 0 | 21789 | 45867 | 128078 | 25 | 74 | 17297 | 2283 | 0 | 0 | 14 | 24 | 24 | 531 | 24 | 0 | 12 | 0 | 0 | 0 | 0 | 672 | 501 | 0 |
| SRR2487633 | 37 | 18 | 18 | 221 | 55 | 0 | 0 | 25 | 0 | 0 | 221 | 0 | 0 | 0 | 30220 | 12708 | 148846 | 25 | 338 | 15529 | 4175 | 0 | 0 | 0 | 0 | 55 | 553 | 37 | 0 | 646 | 0 | 0 | 0 | 0 | 16 | 97 | 0 |
| SRR2487635 | 77 | 19 | 39 | 193 | 77 | 0 | 0 | 20 | 0 | 19 | 97 | 19 | 0 | 0 | 30019 | 12435 | 147824 | 36 | 392 | 16055 | 4214 | 0 | 0 | 135 | 0 | 77 | 13 | 193 | 0 | 13 | 0 | 0 | 0 | 0 | 23 | 167 | 0 |
| SRR2487637 | 24 | 0 | 24 | 95 | 24 | 0 | 0 | 32 | 16 | 0 | 71 | 0 | 0 | 0 | 28246 | 11997 | 140616 | 16 | 351 | 15962 | 4288 | 0 | 0 | 0 | 0 | 48 | 381 | 190 | 0 | 619 | 0 | 0 | 0 | 0 | 24 | 195 | 0 |
| SRR2487639 | 48 | 0 | 0 | 48 | 24 | 0 | 0 | 0 | 12 | 0 | 121 | 24 | 0 | 0 | 28681 | 11769 | 140500 | 24 | 402 | 15980 | 4456 | 0 | 0 | 97 | 0 | 24 | 12 | 97 | 0 | 13 | 0 | 0 | 0 | 0 | 20 | 188 | 0 |
| SRR2487641 | 0 | 23 | 23 | 229 | 274 | 0 | 0 | 143 | 22 | 0 | 183 | 0 | 0 | 0 | 29608 | 39081 | 173914 | 30 | 100 | 33425 | 3204 | 0 | 0 | 366 | 69 | 274 | 100 | 31 | 0 | 12 | 0 | 0 | 0 | 0 | 21 | 242 | 0 |
| SRR2487643 | 71 | 24 | 95 | 142 | 190 | 0 | 13 | 130 | 45 | 0 | 190 | 0 | 0 | 0 | 29849 | 38958 | 172284 | 36 | 112 | 34417 | 3124 | 0 | 0 | 404 | 71 | 427 | 3800 | 1876 | 0 | 594 | 0 | 0 | 0 | 0 | 14 | 337 | 0 |
| SRR2487645 | 0 | 30 | 0 | 242 | 182 | 0 | 22 | 94 | 55 | 0 | 151 | 0 | 0 | 0 | 28754 | 37141 | 165735 | 28 | 116 | 34144 | 3436 | 0 | 0 | 333 | 30 | 212 | 65 | 31 | 0 | 18 | 0 | 0 | 0 | 0 | 12 | 285 | 0 |
| SRR2487647 | 31 | 0 | 31 | 307 | 154 | 0 | 22 | 127 | 39 | 31 | 61 | 0 | 0 | 0 | 28721 | 37781 | 166421 | 33 | 88 | 34036 | 3131 | 0 | 0 | 215 | 123 | 276 | 73 | 25 | 0 | 10 | 0 | 0 | 0 | 0 | 0 | 213 | 0 |
| SRR2487648 | 21 | 0 | 0 | 0 | 272 | 25 | 38 | 228 | 25 | 0 | 147 | 42 | 0 | 0 | 47087 | 18416 | 106769 | 25 | 228 | 17797 | 633 | 0 | 0 | 209 | 0 | 126 | 335 | 21 | 0 | 544 | 0 | 0 | 0 | 0 | 914 | 234 | 0 |
| SRR2487649 | 0 | 0 | 24 | 72 | 13 | 0 | 15 | 15 | 15 | 0 | 19 | 0 | 0 | 0 | 37109 | 16558 | 102616 | 29 | 290 | 22082 | 581 | 0 | 0 | 191 | 24 | 119 | 41 | 143 | 0 | 58 | 0 | 0 | 0 | 0 | 81 | 358 | 0 |
| SRR2487651 | 48 | 24 | 24 | 214 | 238 | 14 | 0 | 43 | 0 | 0 | 309 | 0 | 0 | 0 | 38580 | 16798 | 102382 | 28 | 284 | 22793 | 568 | 0 | 0 | 48 | 0 | 95 | 523 | 24 | 0 | 1283 | 0 | 0 | 0 | 0 | 79 | 385 | 0 |
| SRR2487652 | 42 | 21 | 0 | 42 | 316 | 39 | 26 | 235 | 26 | 21 | 358 | 0 | 0 | 0 | 48269 | 18689 | 106740 | 13 | 274 | 18989 | 626 | 0 | 0 | 105 | 0 | 84 | 443 | 0 | 0 | 569 | 0 | 0 | 0 | 0 | 883 | 252 | 0 |
| SRR2487653 | 23 | 23 | 0 | 23 | 229 | 28 | 41 | 235 | 41 | 0 | 92 | 46 | 0 | 0 | 46508 | 17689 | 101433 | 0 | 235 | 16369 | 415 | 0 | 0 | 160 | 0 | 69 | 206 | 0 | 0 | 17 | 0 | 0 | 0 | 0 | 876 | 237 | 0 |
| SRR2487654 | 0 | 0 | 0 | 0 | 459 | 56 | 28 | 307 | 28 | 0 | 229 | 0 | 0 | 0 | 47451 | 17631 | 102625 | 28 | 140 | 17359 | 643 | 0 | 0 | 207 | 46 | 46 | 321 | 0 | 0 | 13 | 0 | 0 | 0 | 0 | 932 | 271 | 0 |
| SRR2487656 | 19 | 0 | 0 | 93 | 14 | 0 | 0 | 35 | 0 | 0 | 17 | 0 | 0 | 0 | 38230 | 16706 | 108918 | 0 | 301 | 21682 | 682 | 0 | 0 | 37 | 0 | 148 | 50 | 111 | 0 | 74 | 0 | 0 | 0 | 0 | 72 | 274 | 0 |
| SRR2487658 | 0 | 0 | 19 | 76 | 17 | 0 | 0 | 24 | 0 | 19 | 33 | 19 | 0 | 0 | 39126 | 16966 | 107990 | 12 | 306 | 22675 | 847 | 0 | 0 | 19 | 19 | 11 | 49 | 115 | 0 | 76 | 0 | 0 | 0 | 0 | 101 | 318 | 0 |
| SRR2487661 | 0 | 0 | 28 | 57 | 141 | 26 | 22 | 166 | 81 | 0 | 57 | 0 | 0 | 0 | 33789 | 19906 | 159007 | 15 | 63 | 14511 | 6527 | 0 | 0 | 57 | 57 | 254 | 1554 | 396 | 0 | 593 | 0 | 11 | 0 | 0 | 0 | 765 | 0 |
| SRR2487663 | 0 | 0 | 0 | 0 | 58 | 38 | 15 | 175 | 72 | 29 | 175 | 0 | 0 | 0 | 33675 | 19705 | 156621 | 0 | 61 | 15370 | 6617 | 0 | 0 | 87 | 0 | 175 | 1866 | 467 | 0 | 583 | 0 | 0 | 0 | 0 | 0 | 790 | 0 |
| SRR2487665 | 37 | 37 | 0 | 148 | 111 | 0 | 28 | 117 | 52 | 0 | 74 | 0 | 0 | 0 | 32296 | 18615 | 148133 | 0 | 61 | 14769 | 6964 | 0 | 0 | 74 | 0 | 297 | 18 | 371 | 0 | 667 | 0 | 0 | 0 | 0 | 0 | 793 | 0 |
| SRR2487667 | 37 | 0 | 0 | 75 | 37 | 23 | 19 | 102 | 42 | 37 | 112 | 0 | 0 | 0 | 33031 | 18668 | 149999 | 19 | 46 | 14766 | 6612 | 0 | 0 | 75 | 37 | 150 | 1122 | 860 | 0 | 636 | 0 | 0 | 0 | 0 | 10 | 1001 | 0 |
| SRR2487668 | 36 | 0 | 0 | 0 | 401 | 167 | 153 | 1124 | 97 | 0 | 109 | 0 | 0 | 0 | 52457 | 41968 | 184291 | 14 | 139 | 35966 | 597 | 0 | 0 | 73 | 0 | 0 | 36 | 0 | 0 | 14 | 0 | 0 | 0 | 0 | 502 | 122 | 36 |
| SRR2487669 | 66 | 0 | 0 | 165 | 463 | 0 | 10 | 135 | 10 | 0 | 199 | 0 | 0 | 0 | 36874 | 35842 | 177586 | 10 | 228 | 44048 | 559 | 0 | 0 | 99 | 0 | 0 | 232 | 33 | 0 | 1092 | 0 | 0 | 0 | 0 | 76 | 79 | 0 |
| SRR2487671 | 133 | 33 | 0 | 133 | 534 | 21 | 10 | 113 | 41 | 0 | 133 | 0 | 0 | 0 | 37251 | 35596 | 179833 | 0 | 175 | 43397 | 576 | 0 | 0 | 33 | 0 | 67 | 133 | 67 | 0 | 1434 | 0 | 0 | 0 | 0 | 50 | 92 | 0 |
| SRR2487672 | 38 | 0 | 0 | 0 | 13 | 158 | 286 | 1060 | 43 | 38 | 113 | 0 | 0 | 0 | 52705 | 42372 | 183698 | 14 | 72 | 38340 | 559 | 0 | 0 | 150 | 0 | 0 | 150 | 0 | 0 | 12 | 0 | 0 | 0 | 0 | 418 | 104 | 0 |
| SRR2487673 | 40 | 0 | 0 | 0 | 474 | 122 | 122 | 1240 | 77 | 40 | 40 | 0 | 0 | 0 | 51272 | 39158 | 180609 | 0 | 77 | 34620 | 735 | 0 | 0 | 158 | 0 | 0 | 158 | 0 | 0 | 790 | 0 | 0 | 0 | 0 | 410 | 118 | 0 |
| SRR2487674 | 0 | 0 | 0 | 40 | 13 | 139 | 185 | 1279 | 108 | 0 | 40 | 0 | 0 | 0 | 51462 | 40509 | 182711 | 0 | 92 | 34945 | 647 | 0 | 0 | 279 | 0 | 0 | 0 | 0 | 0 | 10 | 0 | 0 | 0 | 0 | 490 | 113 | 0 |
| SRR2487676 | 26 | 26 | 0 | 78 | 28 | 75 | 0 | 166 | 17 | 26 | 104 | 0 | 0 | 0 | 39288 | 36276 | 184174 | 0 | 191 | 41672 | 639 | 0 | 0 | 104 | 52 | 78 | 183 | 52 | 0 | 51 | 0 | 0 | 0 | 0 | 68 | 115 | 0 |
| SRR2487678 | 27 | 81 | 54 | 188 | 942 | 17 | 42 | 144 | 17 | 27 | 108 | 0 | 0 | 0 | 39128 | 36824 | 182408 | 0 | 220 | 42668 | 652 | 0 | 0 | 81 | 0 | 54 | 135 | 108 | 0 | 1022 | 0 | 0 | 0 | 0 | 48 | 118 | 0 |
| SRR2487688 | 0 | 0 | 0 | 68 | 906 | 634 | 852 | 4087 | 656 | 0 | 68 | 0 | 0 | 0 | 18163 | 23203 | 213823 | 0 | 87 | 16458 | 2076 | 0 | 0 | 793 | 45 | 227 | 1314 | 68 | 0 | 408 | 0 | 0 | 0 | 0 | 1915 | 421 | 0 |
| SRR2487689 | 0 | 59 | 0 | 118 | 1237 | 112 | 168 | 558 | 56 | 0 | 59 | 0 | 0 | 0 | 14855 | 19704 | 199739 | 0 | 56 | 24126 | 2848 | 0 | 0 | 294 | 177 | 412 | 2885 | 1531 | 0 | 59 | 0 | 56 | 0 | 0 | 130 | 466 | 0 |
| SRR2487691 | 0 | 0 | 60 | 299 | 1075 | 111 | 278 | 444 | 167 | 0 | 60 | 0 | 0 | 0 | 16499 | 18666 | 202714 | 0 | 56 | 25665 | 3055 | 0 | 0 | 179 | 179 | 179 | 3046 | 896 | 0 | 418 | 0 | 0 | 0 | 0 | 257 | 439 | 0 |
| SRR2487692 | 0 | 46 | 0 | 23 | 18 | 341 | 1183 | 4778 | 956 | 0 | 46 | 23 | 0 | 0 | 17565 | 23765 | 214660 | 0 | 0 | 16632 | 2002 | 0 | 0 | 16 | 115 | 69 | 29 | 0 | 0 | 299 | 0 | 0 | 0 | 0 | 2035 | 443 | 0 |
| SRR2487693 | 26 | 51 | 0 | 26 | 1002 | 446 | 1290 | 4167 | 769 | 26 | 26 | 0 | 0 | 0 | 17584 | 22834 | 212509 | 0 | 99 | 15179 | 2455 | 0 | 0 | 1027 | 26 | 180 | 1130 | 77 | 0 | 282 | 0 | 0 | 0 | 0 | 2041 | 396 | 0 |
| SRR2487694 | 0 | 76 | 0 | 76 | 943 | 399 | 1121 | 3936 | 922 | 0 | 25 | 0 | 0 | 0 | 18311 | 22508 | 214676 | 0 | 75 | 16468 | 2167 | 0 | 0 | 1070 | 0 | 178 | 1070 | 153 | 0 | 331 | 0 | 0 | 0 | 0 | 2072 | 438 | 0 |
| SRR2487696 | 0 | 0 | 0 | 228 | 1139 | 0 | 170 | 766 | 426 | 0 | 0 | 0 | 0 | 0 | 14430 | 18162 | 208643 | 0 | 255 | 23242 | 2767 | 0 | 0 | 46 | 137 | 319 | 3326 | 1549 | 0 | 410 | 0 | 0 | 0 | 0 | 102 | 400 | 0 |
| SRR2487698 | 0 | 46 | 0 | 14 | 62 | 0 | 218 | 522 | 131 | 0 | 184 | 0 | 0 | 0 | 15229 | 20101 | 204844 | 0 | 348 | 24453 | 2872 | 0 | 0 | 92 | 92 | 29 | 143 | 69 | 0 | 17 | 0 | 0 | 0 | 0 | 35 | 419 | 0 |
| SRR2487700 | 26 | 13 | 0 | 13 | 313 | 146 | 292 | 748 | 292 | 104 | 26 | 0 | 0 | 0 | 35632 | 18904 | 128879 | 0 | 109 | 6528 | 474 | 0 | 0 | 52 | 0 | 26 | 104 | 26 | 0 | 248 | 0 | 0 | 0 | 0 | 1223 | 108 | 0 |
| SRR2487701 | 24 | 0 | 0 | 13 | 22 | 0 | 33 | 66 | 33 | 24 | 49 | 0 | 0 | 0 | 26892 | 15842 | 124344 | 0 | 99 | 9271 | 953 | 0 | 0 | 0 | 24 | 98 | 32 | 19 | 0 | 32 | 0 | 0 | 0 | 0 | 46 | 126 | 0 |
| SRR2487703 | 0 | 0 | 24 | 122 | 170 | 32 | 32 | 160 | 0 | 73 | 49 | 24 | 0 | 0 | 28045 | 15293 | 126119 | 0 | 191 | 9572 | 670 | 0 | 0 | 24 | 0 | 49 | 389 | 195 | 0 | 681 | 0 | 0 | 0 | 0 | 136 | 141 | 0 |
| SRR2487704 | 39 | 26 | 0 | 26 | 406 | 187 | 225 | 600 | 337 | 52 | 52 | 0 | 0 | 0 | 33599 | 19219 | 130081 | 0 | 169 | 7008 | 506 | 0 | 0 | 52 | 0 | 66 | 170 | 0 | 0 | 288 | 0 | 19 | 0 | 0 | 1242 | 129 | 13 |
| SRR2487705 | 29 | 86 | 0 | 43 | 229 | 100 | 220 | 740 | 300 | 157 | 43 | 14 | 0 | 0 | 35868 | 18336 | 127567 | 0 | 140 | 6818 | 640 | 0 | 0 | 57 | 14 | 43 | 72 | 14 | 0 | 300 | 0 | 20 | 0 | 0 | 1298 | 139 | 0 |
| SRR2487706 | 14 | 100 | 0 | 43 | 315 | 121 | 141 | 707 | 485 | 57 | 43 | 0 | 0 | 0 | 36050 | 18234 | 127329 | 0 | 182 | 6830 | 667 | 0 | 0 | 43 | 14 | 72 | 272 | 0 | 0 | 358 | 0 | 0 | 0 | 0 | 1266 | 148 | 0 |
| SRR2487708 | 19 | 19 | 19 | 75 | 15 | 0 | 0 | 156 | 78 | 38 | 75 | 0 | 0 | 0 | 28790 | 16617 | 133260 | 0 | 287 | 8936 | 469 | 0 | 0 | 19 | 0 | 19 | 41 | 13 | 0 | 25 | 0 | 0 | 0 | 0 | 134 | 176 | 0 |
| SRR2487710 | 39 | 19 | 0 | 58 | 58 | 0 | 26 | 79 | 132 | 0 | 78 | 0 | 0 | 0 | 26976 | 16967 | 134702 | 0 | 158 | 8474 | 816 | 0 | 0 | 0 | 19 | 39 | 581 | 155 | 0 | 465 | 0 | 0 | 0 | 0 | 115 | 187 | 0 |
| SRR2529523 | 0 | 0 | 0 | 0 | 49 | 135 | 255 | 270 | 60 | 0 | 11 | 0 | 0 | 0 | 2704 | 78648 | 60881 | 0 | 120 | 12320 | 3561 | 0 | 0 | 23 | 0 | 55 | 138 | 52 | 0 | 139 | 0 | 30 | 0 | 0 | 6815 | 0 | 0 |
| SRR2529524 | 0 | 0 | 0 | 0 | 48 | 124 | 185 | 324 | 170 | 0 | 60 | 0 | 0 | 0 | 2517 | 77766 | 60077 | 0 | 139 | 11287 | 3134 | 0 | 0 | 19 | 0 | 60 | 131 | 53 | 0 | 136 | 0 | 31 | 0 | 0 | 5989 | 0 | 0 |
| SRR2529525 | 14 | 34 | 14 | 20 | 19 | 238 | 146 | 215 | 31 | 0 | 24 | 0 | 0 | 0 | 2522 | 34483 | 33309 | 0 | 69 | 2599 | 415 | 0 | 0 | 37 | 0 | 25 | 92 | 46 | 0 | 396 | 0 | 0 | 0 | 0 | 4505 | 0 | 0 |
| SRR2529526 | 21 | 40 | 10 | 21 | 12 | 237 | 90 | 228 | 33 | 0 | 144 | 0 | 0 | 0 | 2430 | 34149 | 33684 | 16 | 41 | 2749 | 375 | 0 | 0 | 52 | 0 | 42 | 546 | 28 | 0 | 2081 | 0 | 33 | 0 | 0 | 4387 | 0 | 0 |
| SRR2529527 | 25 | 35 | 16 | 13 | 0 | 169 | 95 | 204 | 18 | 0 | 10 | 0 | 0 | 0 | 35947 | 148299 | 157005 | 14 | 271 | 15340 | 5782 | 0 | 0 | 11 | 17 | 56 | 82 | 41 | 0 | 202 | 0 | 60 | 0 | 0 | 1890 | 0 | 0 |
| SRR2529528 | 19 | 43 | 16 | 14 | 13 | 173 | 123 | 177 | 47 | 0 | 78 | 0 | 0 | 0 | 36049 | 148087 | 155881 | 22 | 274 | 15255 | 5453 | 0 | 0 | 74 | 18 | 61 | 529 | 38 | 0 | 1306 | 0 | 54 | 0 | 0 | 2041 | 0 | 0 |
| SRR2529529 | 0 | 11 | 0 | 0 | 78 | 242 | 323 | 929 | 202 | 0 | 53 | 0 | 0 | 0 | 1635 | 62925 | 30626 | 0 | 81 | 20653 | 5108 | 0 | 0 | 57 | 23 | 66 | 1162 | 57 | 0 | 365 | 0 | 61 | 0 | 0 | 6721 | 0 | 11 |
| SRR2529530 | 0 | 0 | 0 | 0 | 75 | 311 | 249 | 787 | 145 | 0 | 47 | 0 | 0 | 0 | 1554 | 62085 | 28912 | 0 | 124 | 19636 | 5758 | 0 | 0 | 87 | 0 | 11 | 117 | 52 | 0 | 38 | 0 | 41 | 0 | 0 | 8039 | 0 | 0 |
| SRR2529531 | 12 | 0 | 23 | 17 | 49 | 178 | 178 | 261 | 119 | 0 | 43 | 0 | 0 | 0 | 1624 | 48423 | 20763 | 0 | 154 | 17134 | 5490 | 0 | 0 | 201 | 0 | 70 | 1058 | 70 | 0 | 409 | 0 | 12 | 0 | 0 | 6879 | 0 | 0 |
| SRR2529532 | 13 | 14 | 16 | 24 | 57 | 171 | 232 | 342 | 98 | 0 | 39 | 0 | 0 | 0 | 1842 | 47269 | 22065 | 0 | 98 | 17850 | 5503 | 0 | 0 | 203 | 11 | 64 | 1038 | 66 | 0 | 454 | 0 | 61 | 0 | 0 | 7071 | 0 | 0 |
| SRR2529533 | 12 | 13 | 12 | 0 | 69 | 143 | 215 | 573 | 172 | 0 | 42 | 0 | 0 | 0 | 4540 | 83335 | 77388 | 0 | 143 | 13276 | 2721 | 0 | 0 | 23 | 0 | 50 | 128 | 66 | 0 | 48 | 0 | 43 | 0 | 0 | 9026 | 0 | 0 |
| SRR2529534 | 11 | 0 | 11 | 17 | 91 | 278 | 322 | 439 | 176 | 0 | 44 | 0 | 0 | 0 | 4741 | 77882 | 76837 | 29 | 102 | 13273 | 2737 | 0 | 0 | 189 | 0 | 55 | 954 | 54 | 0 | 319 | 0 | 44 | 0 | 0 | 8976 | 0 | 0 |
| SRR2529535 | 0 | 0 | 0 | 0 | 28 | 558 | 180 | 968 | 83 | 0 | 149 | 0 | 0 | 0 | 2937 | 25510 | 56199 | 0 | 77 | 16003 | 26505 | 0 | 0 | 47 | 54 | 64 | 199 | 11 | 0 | 670 | 0 | 204 | 0 | 0 | 3478 | 0 | 0 |
| SRR2529536 | 0 | 0 | 0 | 0 | 38 | 527 | 202 | 978 | 110 | 0 | 105 | 0 | 0 | 0 | 2996 | 25155 | 56846 | 0 | 89 | 15592 | 26106 | 0 | 0 | 52 | 46 | 54 | 18 | 0 | 0 | 64 | 0 | 236 | 0 | 0 | 3491 | 0 | 0 |
| SRR2529537 | 11 | 0 | 12 | 36 | 19 | 555 | 98 | 436 | 60 | 0 | 19 | 0 | 0 | 0 | 5629 | 34355 | 35036 | 21 | 56 | 5150 | 12812 | 0 | 0 | 57 | 19 | 32 | 52 | 34 | 0 | 111 | 0 | 17 | 0 | 0 | 4199 | 0 | 0 |
| SRR2529538 | 0 | 0 | 12 | 29 | 17 | 610 | 134 | 410 | 67 | 0 | 115 | 0 | 0 | 0 | 5689 | 34336 | 35780 | 13 | 93 | 5346 | 12659 | 0 | 0 | 79 | 26 | 36 | 368 | 29 | 0 | 807 | 0 | 40 | 0 | 0 | 3826 | 0 | 0 |
| SRR2529539 | 0 | 0 | 13 | 33 | 0 | 346 | 118 | 289 | 122 | 0 | 75 | 0 | 0 | 0 | 2296 | 27468 | 13839 | 0 | 127 | 5323 | 3866 | 0 | 0 | 55 | 12 | 39 | 296 | 22 | 0 | 898 | 0 | 109 | 0 | 0 | 4298 | 0 | 0 |
| SRR2529540 | 0 | 0 | 13 | 34 | 14 | 433 | 207 | 332 | 83 | 0 | 98 | 0 | 0 | 0 | 2197 | 27386 | 13871 | 0 | 101 | 5545 | 4361 | 0 | 0 | 63 | 25 | 39 | 296 | 16 | 0 | 941 | 0 | 111 | 0 | 0 | 4202 | 0 | 0 |
| SRR2529541 | 0 | 0 | 0 | 19 | 38 | 164 | 107 | 549 | 95 | 0 | 70 | 0 | 0 | 0 | 3153 | 21783 | 43873 | 0 | 63 | 17438 | 11251 | 0 | 0 | 43 | 32 | 60 | 464 | 25 | 0 | 373 | 0 | 19 | 0 | 0 | 4094 | 0 | 0 |
| SRR2529542 | 0 | 0 | 0 | 21 | 45 | 223 | 217 | 613 | 140 | 0 | 68 | 0 | 0 | 0 | 2865 | 22855 | 44825 | 0 | 77 | 16918 | 11564 | 0 | 0 | 31 | 37 | 62 | 417 | 21 | 0 | 434 | 0 | 57 | 0 | 0 | 3787 | 0 | 0 |
| SRR2529543 | 0 | 0 | 0 | 23 | 53 | 271 | 170 | 774 | 217 | 0 | 67 | 0 | 0 | 0 | 3597 | 24131 | 44491 | 0 | 54 | 6034 | 17280 | 0 | 0 | 91 | 70 | 101 | 293 | 27 | 0 | 364 | 0 | 81 | 0 | 0 | 2059 | 0 | 0 |
| SRR2529544 | 0 | 0 | 0 | 25 | 35 | 314 | 251 | 969 | 167 | 0 | 64 | 0 | 0 | 0 | 3549 | 24366 | 44710 | 0 | 42 | 6401 | 16803 | 0 | 0 | 10 | 68 | 15 | 42 | 14 | 0 | 48 | 0 | 91 | 0 | 0 | 2083 | 0 | 0 |
| SRR2529545 | 0 | 0 | 0 | 20 | 54 | 412 | 212 | 725 | 62 | 11 | 65 | 0 | 0 | 0 | 3161 | 87220 | 51394 | 0 | 100 | 18941 | 5735 | 0 | 0 | 85 | 18 | 63 | 491 | 27 | 0 | 338 | 0 | 25 | 0 | 0 | 5821 | 0 | 0 |
| SRR2529546 | 0 | 0 | 0 | 23 | 53 | 369 | 305 | 840 | 89 | 0 | 69 | 0 | 0 | 0 | 3462 | 87370 | 49701 | 0 | 178 | 18593 | 5510 | 0 | 0 | 67 | 18 | 44 | 436 | 25 | 0 | 308 | 0 | 38 | 0 | 0 | 5249 | 0 | 0 |
| SRR2529547 | 0 | 0 | 0 | 0 | 84 | 818 | 491 | 1172 | 354 | 0 | 64 | 0 | 0 | 0 | 327 | 19496 | 13279 | 0 | 82 | 8423 | 491 | 0 | 0 | 93 | 0 | 38 | 604 | 52 | 0 | 602 | 0 | 136 | 0 | 0 | 10446 | 0 | 0 |
| SRR2529548 | 0 | 0 | 0 | 12 | 74 | 585 | 251 | 1143 | 251 | 0 | 83 | 0 | 0 | 0 | 446 | 19768 | 13278 | 0 | 28 | 7275 | 474 | 0 | 0 | 60 | 0 | 21 | 628 | 48 | 0 | 420 | 0 | 56 | 0 | 0 | 8795 | 0 | 0 |
| SRR2529549 | 18 | 13 | 13 | 31 | 20 | 175 | 187 | 325 | 75 | 0 | 140 | 0 | 0 | 0 | 250 | 10049 | 4457 | 50 | 75 | 2597 | 100 | 0 | 0 | 90 | 0 | 37 | 466 | 24 | 0 | 1450 | 0 | 37 | 0 | 0 | 9611 | 0 | 0 |
| SRR2529550 | 0 | 34 | 0 | 18 | 16 | 192 | 154 | 384 | 77 | 0 | 25 | 0 | 0 | 0 | 320 | 10002 | 3953 | 26 | 90 | 2699 | 64 | 0 | 0 | 41 | 0 | 18 | 82 | 38 | 0 | 234 | 0 | 38 | 0 | 0 | 9265 | 0 | 0 |
| SRR2529551 | 19 | 28 | 0 | 17 | 17 | 183 | 237 | 429 | 91 | 0 | 12 | 0 | 0 | 0 | 657 | 17889 | 8810 | 18 | 100 | 4053 | 785 | 0 | 0 | 21 | 0 | 41 | 102 | 54 | 0 | 184 | 0 | 100 | 0 | 0 | 6000 | 0 | 0 |
| SRR2529552 | 27 | 23 | 0 | 17 | 0 | 115 | 183 | 452 | 106 | 0 | 61 | 0 | 0 | 0 | 827 | 17781 | 8926 | 19 | 96 | 3594 | 875 | 0 | 0 | 118 | 0 | 42 | 695 | 54 | 0 | 999 | 0 | 96 | 0 | 0 | 6305 | 0 | 0 |
| SRR2529553 | 10 | 10 | 0 | 16 | 73 | 222 | 254 | 731 | 476 | 13 | 70 | 0 | 0 | 0 | 476 | 22128 | 18780 | 0 | 64 | 5114 | 1302 | 0 | 0 | 10 | 23 | 57 | 110 | 70 | 0 | 41 | 0 | 0 | 0 | 0 | 8620 | 0 | 10 |
| SRR2529554 | 0 | 0 | 0 | 0 | 51 | 355 | 548 | 1000 | 323 | 0 | 51 | 0 | 0 | 0 | 548 | 21213 | 17098 | 0 | 0 | 5323 | 1000 | 0 | 0 | 69 | 29 | 77 | 1037 | 74 | 0 | 396 | 0 | 65 | 0 | 0 | 8150 | 0 | 0 |
| SRR2529555 | 16 | 26 | 15 | 21 | 68 | 255 | 196 | 804 | 78 | 0 | 44 | 0 | 0 | 0 | 470 | 14518 | 12496 | 20 | 20 | 5312 | 490 | 0 | 0 | 215 | 0 | 83 | 1179 | 90 | 0 | 413 | 0 | 59 | 0 | 0 | 10307 | 0 | 0 |
| SRR2529556 | 0 | 17 | 15 | 38 | 60 | 130 | 168 | 596 | 75 | 0 | 54 | 0 | 0 | 0 | 354 | 13255 | 12975 | 56 | 0 | 4712 | 447 | 0 | 0 | 211 | 13 | 94 | 1151 | 79 | 0 | 383 | 0 | 37 | 0 | 0 | 10256 | 0 | 0 |
| SRR2529557 | 0 | 19 | 0 | 39 | 95 | 264 | 343 | 659 | 132 | 0 | 33 | 0 | 0 | 0 | 211 | 12934 | 9690 | 0 | 26 | 3244 | 211 | 0 | 0 | 149 | 21 | 93 | 1277 | 116 | 0 | 334 | 0 | 26 | 0 | 0 | 12613 | 0 | 0 |
| SRR2529558 | 12 | 17 | 14 | 41 | 12 | 270 | 189 | 595 | 81 | 0 | 48 | 0 | 0 | 0 | 325 | 13147 | 8293 | 0 | 0 | 4246 | 27 | 0 | 0 | 28 | 22 | 67 | 163 | 11 | 0 | 41 | 0 | 27 | 0 | 0 | 12823 | 0 | 0 |
| SRR2529559 | 0 | 0 | 0 | 0 | 34 | 1350 | 798 | 1391 | 307 | 0 | 106 | 0 | 0 | 0 | 511 | 62445 | 10847 | 61 | 102 | 10204 | 593 | 0 | 0 | 155 | 13 | 32 | 418 | 21 | 0 | 549 | 0 | 82 | 0 | 0 | 20264 | 0 | 0 |
| SRR2529560 | 0 | 0 | 0 | 0 | 41 | 1403 | 784 | 1320 | 268 | 0 | 107 | 0 | 0 | 0 | 330 | 62518 | 11403 | 21 | 103 | 10230 | 639 | 0 | 0 | 165 | 15 | 26 | 455 | 30 | 0 | 626 | 0 | 144 | 0 | 0 | 18705 | 0 | 0 |
| SRR2529561 | 0 | 17 | 0 | 23 | 14 | 214 | 158 | 563 | 101 | 0 | 79 | 0 | 0 | 0 | 259 | 18747 | 5005 | 23 | 68 | 2567 | 68 | 0 | 0 | 105 | 0 | 51 | 784 | 48 | 0 | 1165 | 0 | 23 | 0 | 0 | 8416 | 0 | 0 |
| SRR2529562 | 0 | 22 | 0 | 20 | 17 | 58 | 266 | 427 | 104 | 0 | 86 | 0 | 0 | 0 | 381 | 17529 | 4619 | 23 | 81 | 2436 | 23 | 0 | 0 | 86 | 0 | 33 | 703 | 43 | 0 | 1149 | 0 | 58 | 0 | 0 | 8047 | 0 | 0 |
| SRR2529563 | 0 | 11 | 0 | 11 | 17 | 235 | 310 | 556 | 182 | 0 | 62 | 0 | 0 | 0 | 225 | 30462 | 4097 | 118 | 139 | 4397 | 267 | 0 | 0 | 71 | 0 | 28 | 514 | 43 | 0 | 1561 | 0 | 96 | 0 | 0 | 7559 | 0 | 0 |
| SRR2529564 | 0 | 12 | 0 | 14 | 0 | 264 | 297 | 451 | 88 | 0 | 40 | 0 | 0 | 0 | 429 | 30847 | 4171 | 121 | 99 | 3407 | 319 | 0 | 0 | 68 | 0 | 40 | 478 | 35 | 0 | 1453 | 0 | 44 | 0 | 0 | 6955 | 0 | 0 |
| SRR2529565 | 0 | 0 | 0 | 0 | 65 | 279 | 677 | 1036 | 398 | 0 | 77 | 0 | 0 | 0 | 359 | 37250 | 8571 | 0 | 40 | 5299 | 717 | 0 | 0 | 87 | 0 | 59 | 764 | 53 | 0 | 495 | 0 | 120 | 0 | 0 | 12787 | 0 | 0 |
| SRR2529566 | 0 | 0 | 0 | 0 | 51 | 539 | 622 | 1077 | 622 | 0 | 70 | 0 | 0 | 0 | 580 | 38029 | 9459 | 0 | 83 | 5677 | 912 | 0 | 0 | 57 | 13 | 77 | 881 | 42 | 0 | 543 | 0 | 0 | 0 | 0 | 12757 | 0 | 0 |
| SRR2529567 | 0 | 0 | 0 | 21 | 96 | 308 | 560 | 448 | 280 | 0 | 68 | 0 | 0 | 0 | 784 | 85171 | 13154 | 0 | 56 | 6832 | 252 | 0 | 0 | 344 | 17 | 72 | 814 | 62 | 0 | 365 | 0 | 28 | 0 | 0 | 3537 | 0 | 0 |
| SRR2529568 | 0 | 0 | 0 | 15 | 15 | 198 | 255 | 708 | 255 | 0 | 14 | 0 | 0 | 0 | 708 | 83848 | 11475 | 0 | 28 | 6937 | 227 | 0 | 0 | 71 | 0 | 15 | 159 | 14 | 0 | 73 | 0 | 0 | 0 | 0 | 3891 | 0 | 0 |
| SRR2529569 | 0 | 0 | 0 | 0 | 12 | 520 | 378 | 1938 | 520 | 0 | 10 | 0 | 0 | 0 | 473 | 40824 | 9051 | 0 | 0 | 4586 | 189 | 0 | 0 | 17 | 0 | 28 | 135 | 11 | 0 | 62 | 0 | 95 | 0 | 0 | 13046 | 0 | 0 |
| SRR2529570 | 0 | 0 | 0 | 0 | 68 | 550 | 650 | 2250 | 400 | 0 | 49 | 0 | 0 | 0 | 600 | 34298 | 9800 | 0 | 150 | 4851 | 350 | 0 | 0 | 120 | 0 | 36 | 902 | 74 | 0 | 414 | 0 | 50 | 0 | 0 | 11900 | 0 | 0 |
| SRR2529571 | 0 | 0 | 0 | 12 | 57 | 275 | 353 | 393 | 98 | 0 | 47 | 0 | 0 | 0 | 510 | 39828 | 21320 | 0 | 39 | 6459 | 1198 | 0 | 0 | 20 | 28 | 12 | 146 | 13 | 0 | 54 | 0 | 39 | 0 | 0 | 6595 | 0 | 0 |
| SRR2529572 | 0 | 0 | 0 | 20 | 50 | 80 | 321 | 421 | 200 | 0 | 42 | 0 | 0 | 0 | 741 | 38564 | 18680 | 0 | 20 | 5829 | 1042 | 0 | 0 | 23 | 18 | 11 | 136 | 75 | 0 | 43 | 0 | 40 | 0 | 0 | 6366 | 0 | 0 |
| SRR2529573 | 13 | 27 | 0 | 27 | 16 | 122 | 287 | 209 | 235 | 0 | 114 | 0 | 0 | 0 | 443 | 22136 | 7144 | 0 | 156 | 5675 | 200 | 0 | 0 | 58 | 11 | 29 | 489 | 45 | 0 | 1495 | 0 | 130 | 0 | 0 | 6460 | 0 | 0 |
| SRR2529574 | 0 | 28 | 0 | 22 | 13 | 133 | 204 | 355 | 89 | 0 | 17 | 0 | 0 | 0 | 523 | 21895 | 6714 | 0 | 62 | 5273 | 186 | 0 | 0 | 59 | 0 | 30 | 69 | 33 | 0 | 232 | 0 | 53 | 0 | 0 | 6304 | 0 | 0 |
| SRR2529575 | 28 | 36 | 16 | 12 | 0 | 708 | 281 | 962 | 118 | 0 | 71 | 0 | 0 | 0 | 1652 | 26538 | 19296 | 0 | 154 | 3694 | 1488 | 0 | 0 | 79 | 55 | 87 | 590 | 55 | 0 | 959 | 0 | 64 | 0 | 0 | 5254 | 0 | 0 |
| SRR2529576 | 35 | 34 | 18 | 15 | 18 | 605 | 298 | 740 | 106 | 0 | 55 | 0 | 0 | 0 | 1652 | 25832 | 18874 | 58 | 115 | 3891 | 1739 | 0 | 0 | 49 | 40 | 90 | 566 | 41 | 0 | 924 | 0 | 67 | 0 | 0 | 5745 | 0 | 0 |
| SRR2529577 | 0 | 0 | 0 | 0 | 63 | 365 | 521 | 937 | 417 | 0 | 35 | 0 | 0 | 0 | 1458 | 35412 | 22733 | 0 | 104 | 10338 | 2396 | 0 | 0 | 57 | 0 | 85 | 100 | 55 | 0 | 37 | 0 | 0 | 0 | 0 | 7052 | 0 | 0 |
| SRR2529578 | 0 | 0 | 0 | 0 | 64 | 517 | 419 | 1010 | 197 | 0 | 30 | 0 | 0 | 0 | 1330 | 35831 | 23040 | 0 | 49 | 9950 | 2217 | 0 | 0 | 83 | 0 | 72 | 984 | 36 | 0 | 356 | 0 | 0 | 0 | 0 | 7349 | 0 | 0 |
| SRR2529579 | 15 | 12 | 16 | 31 | 37 | 160 | 288 | 640 | 144 | 0 | 56 | 0 | 0 | 0 | 368 | 13358 | 8144 | 0 | 16 | 5089 | 224 | 0 | 0 | 29 | 10 | 10 | 106 | 83 | 0 | 45 | 0 | 32 | 0 | 0 | 9534 | 0 | 0 |
| SRR2529580 | 12 | 0 | 0 | 20 | 50 | 147 | 228 | 814 | 130 | 0 | 52 | 0 | 0 | 0 | 309 | 11494 | 8103 | 0 | 16 | 4804 | 277 | 0 | 0 | 222 | 20 | 82 | 895 | 76 | 0 | 351 | 0 | 65 | 0 | 0 | 8726 | 0 | 0 |
| SRR2529581 | 0 | 0 | 0 | 0 | 63 | 223 | 610 | 792 | 305 | 0 | 31 | 0 | 0 | 0 | 528 | 19626 | 10356 | 0 | 41 | 5364 | 345 | 0 | 0 | 25 | 13 | 63 | 154 | 13 | 0 | 48 | 0 | 20 | 0 | 0 | 8742 | 0 | 0 |
| SRR2529582 | 0 | 0 | 0 | 19 | 61 | 478 | 395 | 706 | 104 | 0 | 35 | 0 | 0 | 0 | 519 | 19101 | 9097 | 0 | 21 | 5398 | 249 | 0 | 0 | 181 | 22 | 83 | 1186 | 85 | 0 | 331 | 0 | 42 | 0 | 0 | 8221 | 0 | 0 |
| SRR2529583 | 0 | 0 | 0 | 0 | 35 | 577 | 319 | 1166 | 270 | 0 | 78 | 0 | 0 | 0 | 650 | 50075 | 10593 | 0 | 37 | 9941 | 1191 | 0 | 0 | 65 | 12 | 23 | 39 | 16 | 0 | 50 | 0 | 196 | 0 | 0 | 10203 | 0 | 0 |
| SRR2529584 | 0 | 0 | 0 | 0 | 29 | 530 | 313 | 1204 | 277 | 0 | 53 | 0 | 0 | 0 | 554 | 50623 | 11074 | 0 | 0 | 9160 | 1396 | 0 | 0 | 64 | 0 | 25 | 384 | 22 | 0 | 482 | 0 | 120 | 0 | 0 | 10183 | 0 | 0 |
| SRR2529585 | 0 | 0 | 0 | 17 | 20 | 116 | 178 | 256 | 31 | 0 | 14 | 0 | 0 | 0 | 1124 | 18814 | 7380 | 16 | 47 | 4108 | 132 | 0 | 0 | 44 | 0 | 34 | 62 | 34 | 0 | 138 | 0 | 140 | 0 | 0 | 7758 | 0 | 0 |
| SRR2529586 | 0 | 0 | 0 | 22 | 13 | 115 | 138 | 262 | 23 | 0 | 125 | 0 | 0 | 0 | 1062 | 18114 | 7469 | 23 | 31 | 3769 | 169 | 0 | 0 | 66 | 0 | 36 | 418 | 31 | 0 | 932 | 0 | 38 | 0 | 0 | 7689 | 0 | 0 |
| SRR2529587 | 0 | 0 | 13 | 13 | 14 | 69 | 96 | 144 | 41 | 0 | 86 | 0 | 0 | 0 | 733 | 19392 | 4623 | 34 | 69 | 3488 | 377 | 0 | 0 | 48 | 0 | 30 | 336 | 0 | 0 | 850 | 0 | 48 | 0 | 0 | 3915 | 0 | 0 |
| SRR2529588 | 0 | 0 | 0 | 21 | 0 | 112 | 91 | 196 | 49 | 0 | 98 | 0 | 0 | 0 | 714 | 18669 | 4560 | 35 | 49 | 3624 | 273 | 0 | 0 | 49 | 0 | 0 | 319 | 0 | 0 | 955 | 0 | 63 | 0 | 0 | 4377 | 0 | 0 |
| SRR2529589 | 0 | 0 | 0 | 11 | 47 | 193 | 309 | 849 | 154 | 0 | 60 | 0 | 0 | 0 | 598 | 33168 | 12256 | 0 | 19 | 6559 | 887 | 0 | 0 | 51 | 0 | 35 | 87 | 44 | 0 | 42 | 0 | 58 | 0 | 0 | 6358 | 0 | 0 |
| SRR2529590 | 0 | 0 | 0 | 0 | 30 | 236 | 276 | 1024 | 177 | 0 | 52 | 0 | 0 | 0 | 689 | 32379 | 11627 | 0 | 0 | 6104 | 1024 | 0 | 0 | 48 | 0 | 41 | 78 | 30 | 0 | 42 | 0 | 39 | 0 | 0 | 5491 | 0 | 0 |
| SRR2529591 | 0 | 0 | 13 | 21 | 32 | 204 | 87 | 581 | 116 | 0 | 67 | 0 | 0 | 0 | 581 | 26518 | 7714 | 0 | 29 | 4186 | 494 | 0 | 0 | 56 | 11 | 35 | 586 | 43 | 0 | 312 | 0 | 29 | 0 | 0 | 4757 | 0 | 0 |
| SRR2529592 | 0 | 0 | 0 | 30 | 58 | 142 | 227 | 739 | 57 | 0 | 60 | 0 | 0 | 0 | 767 | 23993 | 6153 | 0 | 0 | 4008 | 256 | 0 | 0 | 44 | 0 | 41 | 603 | 60 | 0 | 379 | 0 | 28 | 0 | 0 | 4981 | 0 | 0 |
| SRR2529593 | 0 | 0 | 0 | 15 | 53 | 58 | 260 | 549 | 144 | 0 | 27 | 0 | 0 | 0 | 433 | 32389 | 7368 | 0 | 0 | 3726 | 173 | 0 | 0 | 104 | 0 | 56 | 893 | 50 | 0 | 361 | 0 | 29 | 0 | 0 | 7474 | 0 | 0 |
| SRR2529594 | 0 | 0 | 0 | 18 | 33 | 337 | 245 | 704 | 214 | 0 | 67 | 0 | 0 | 0 | 520 | 32571 | 6966 | 0 | 0 | 4623 | 337 | 0 | 0 | 104 | 0 | 43 | 944 | 58 | 0 | 265 | 0 | 0 | 0 | 0 | 6966 | 0 | 0 |
| SRR2648184 | 0 | 0 | 0 | 0 | 52 | 1562 | 2318 | 4814 | 305 | 22 | 0 | 0 | 0 | 0 | 9269 | 46691 | 126569 | 0 | 0 | 1239 | 373 | 0 | 14 | 160 | 0 | 0 | 28 | 0 | 0 | 88 | 0 | 21 | 0 | 0 | 4403 | 32 | 0 |
| SRR2648185 | 0 | 0 | 0 | 0 | 33 | 1675 | 2420 | 4936 | 310 | 18 | 0 | 0 | 0 | 0 | 9442 | 50431 | 119098 | 14 | 0 | 1502 | 500 | 0 | 0 | 18 | 0 | 0 | 18 | 0 | 0 | 12 | 0 | 18 | 0 | 0 | 5057 | 32 | 11 |
| SRR2648186 | 0 | 0 | 0 | 0 | 37 | 1586 | 2432 | 4969 | 285 | 19 | 0 | 0 | 0 | 0 | 8522 | 57377 | 113192 | 0 | 0 | 1856 | 465 | 0 | 11 | 116 | 0 | 0 | 21 | 0 | 0 | 78 | 0 | 21 | 0 | 0 | 3613 | 28 | 0 |
| SRR2648187 | 25 | 37 | 0 | 19 | 15 | 598 | 3717 | 1458 | 225 | 0 | 0 | 0 | 0 | 0 | 2727 | 203179 | 38473 | 0 | 0 | 5708 | 305 | 0 | 0 | 24 | 0 | 11 | 68 | 25 | 0 | 29 | 0 | 33 | 0 | 0 | 5048 | 36 | 0 |
| SRR2648188 | 31 | 63 | 0 | 20 | 386 | 572 | 3514 | 1431 | 277 | 0 | 0 | 0 | 0 | 0 | 2729 | 182895 | 37929 | 0 | 0 | 5800 | 200 | 0 | 0 | 631 | 0 | 13 | 61 | 28 | 0 | 55 | 0 | 14 | 0 | 0 | 6793 | 34 | 0 |
| SRR2648189 | 35 | 43 | 0 | 14 | 341 | 618 | 3110 | 1558 | 219 | 0 | 0 | 0 | 0 | 0 | 2973 | 174032 | 39004 | 0 | 0 | 4703 | 265 | 0 | 0 | 435 | 0 | 26 | 77 | 27 | 0 | 42 | 0 | 19 | 0 | 0 | 4315 | 30 | 0 |

### Supp Table S8.3.Columns_BX_to_DH

| | | | | | | | | | | | | | | | | | | | | | | | | | | | | | | | | | | | | | |
|---|---|---|---|---|---|---|---|---|---|---|---|---|---|---|---|---|---|---|---|---|---|---|---|---|---|---|---|---|---|---|---|---|---|---|---|---|---|
| SRR2728221 | 0 | 0 | 0 | 0 | 0 | 1606 | 1338 | 3747 | 0 | 0 | 0 | 0 | 0 | 0 | 0 | 2498 | 1665 | 0 | 0 | 0 | 535 | 0 | 0 | 0 | 0 | 0 | 0 | 0 | 0 | 0 | 0 | 268 | 0 | 0 | 833 | 0 | 0 |
| SRR2731278 | 0 | 0 | 0 | 0 | 207 | 102 | 55 | 515 | 2590 | 130 | 1147 | 0 | 0 | 0 | 33920 | 406122 | 211473 | 139 | 5450 | 23791 | 3167 | 0 | 0 | 22 | 74 | 33 | 23 | 0 | 0 | 10216 | 0 | 0 | 0 | 0 | 1300 | 672 | 0 |
| SRR2731279 | 24 | 17 | 0 | 0 | 94 | 176 | 60 | 321 | 1284 | 14 | 188 | 28 | 0 | 0 | 60621 | 131035 | 228825 | 480 | 2579 | 17312 | 1505 | 0 | 0 | 14 | 32 | 23 | 11 | 0 | 0 | 2050 | 0 | 0 | 0 | 0 | 9058 | 278 | 0 |
| SRR2731280 | 19 | 20 | 0 | 0 | 202 | 250 | 107 | 428 | 1250 | 126 | 109 | 20 | 0 | 0 | 2404 | 22577 | 32899 | 452 | 649 | 4315 | 1030 | 0 | 0 | 25 | 14 | 21 | 19 | 0 | 0 | 628 | 0 | 18 | 0 | 11 | 70383 | 386 | 0 |
| SRR2731281 | 26 | 16 | 0 | 12 | 789 | 730 | 566 | 2837 | 7262 | 134 | 541 | 24 | 0 | 0 | 6004 | 137622 | 88595 | 777 | 4711 | 8718 | 5717 | 0 | 0 | 82 | 200 | 140 | 113 | 0 | 0 | 9285 | 0 | 69 | 0 | 0 | 26321 | 1180 | 0 |
| SRR2731282 | 29 | 23 | 12 | 12 | 392 | 291 | 62 | 522 | 1235 | 46 | 1020 | 13 | 0 | 0 | 20861 | 135451 | 269092 | 597 | 11313 | 10323 | 2794 | 0 | 0 | 75 | 175 | 142 | 83 | 19 | 0 | 11159 | 0 | 19 | 0 | 0 | 8154 | 1247 | 0 |
| SRR2731283 | 15 | 0 | 0 | 0 | 108 | 210 | 117 | 644 | 2463 | 61 | 230 | 12 | 0 | 0 | 21832 | 80633 | 64410 | 1045 | 4891 | 4525 | 1280 | 0 | 0 | 44 | 116 | 110 | 67 | 17 | 0 | 2659 | 0 | 48 | 0 | 0 | 27993 | 2720 | 0 |
| SRR2731284 | 13 | 0 | 0 | 0 | 106 | 291 | 243 | 1290 | 4263 | 37 | 127 | 11 | 0 | 0 | 7674 | 60783 | 78030 | 825 | 3980 | 5521 | 1266 | 0 | 0 | 19 | 66 | 51 | 34 | 20 | 0 | 2565 | 0 | 42 | 0 | 0 | 17040 | 1065 | 0 |
| SRR2731285 | 13 | 0 | 0 | 0 | 82 | 1297 | 598 | 3743 | 6566 | 50 | 140 | 10 | 0 | 0 | 17149 | 40592 | 98087 | 860 | 4622 | 4714 | 1221 | 0 | 0 | 35 | 75 | 31 | 22 | 0 | 0 | 2474 | 0 | 69 | 0 | 12 | 40204 | 2342 | 0 |
| SRR2731286 | 13 | 0 | 0 | 0 | 108 | 376 | 183 | 843 | 2228 | 110 | 351 | 17 | 0 | 0 | 66873 | 216233 | 224858 | 299 | 8535 | 21643 | 2536 | 0 | 0 | 20 | 0 | 0 | 0 | 0 | 0 | 3454 | 0 | 27 | 0 | 18 | 13202 | 869 | 0 |
| SRR2731287 | 14 | 10 | 0 | 0 | 77 | 123 | 24 | 242 | 625 | 170 | 228 | 28 | 0 | 0 | 77627 | 198028 | 215192 | 188 | 3939 | 3597 | 3607 | 0 | 0 | 27 | 90 | 48 | 25 | 13 | 0 | 2588 | 0 | 53 | 0 | 0 | 17548 | 2488 | 0 |
| SRR2731288 | 26 | 18 | 0 | 0 | 327 | 1461 | 348 | 1978 | 3047 | 171 | 711 | 22 | 0 | 0 | 77480 | 61126 | 274678 | 531 | 5130 | 7827 | 3964 | 0 | 0 | 249 | 476 | 162 | 95 | 14 | 0 | 9634 | 0 | 22 | 0 | 0 | 16564 | 4271 | 0 |
| SRR2731289 | 24 | 20 | 11 | 0 | 218 | 101 | 18 | 185 | 840 | 191 | 319 | 86 | 0 | 0 | 14123 | 141291 | 122423 | 143 | 2811 | 4131 | 25055 | 0 | 0 | 22 | 105 | 114 | 23 | 23 | 0 | 2204 | 0 | 134 | 0 | 0 | 32478 | 1085 | 0 |
| SRR2731290 | 26 | 11 | 0 | 0 | 187 | 82 | 17 | 159 | 464 | 124 | 262 | 84 | 0 | 0 | 11099 | 84178 | 115240 | 96 | 2049 | 5835 | 19420 | 0 | 0 | 14 | 55 | 50 | 19 | 10 | 0 | 1596 | 0 | 166 | 0 | 11 | 36524 | 617 | 0 |
| SRR2731291 | 43 | 24 | 0 | 0 | 385 | 104 | 11 | 240 | 488 | 361 | 528 | 105 | 0 | 0 | 23212 | 49841 | 168430 | 173 | 3076 | 2167 | 34529 | 0 | 0 | 54 | 195 | 147 | 15 | 0 | 0 | 6896 | 0 | 110 | 0 | 0 | 35882 | 308 | 0 |
| SRR2731292 | 103 | 63 | 0 | 11 | 617 | 99 | 0 | 216 | 783 | 302 | 736 | 381 | 0 | 0 | 9287 | 72731 | 128428 | 70 | 1157 | 594 | 23453 | 0 | 0 | 36 | 147 | 158 | 21 | 13 | 0 | 3803 | 0 | 136 | 0 | 0 | 37710 | 479 | 0 |
| SRR2731293 | 12 | 0 | 0 | 0 | 87 | 423 | 213 | 1502 | 3580 | 64 | 189 | 11 | 0 | 0 | 37927 | 100392 | 255761 | 544 | 10860 | 9821 | 1152 | 0 | 0 | 22 | 69 | 28 | 18 | 0 | 0 | 2916 | 0 | 20 | 0 | 0 | 14797 | 2183 | 0 |
| SRR2731294 | 27 | 17 | 0 | 0 | 412 | 445 | 289 | 1141 | 3481 | 80 | 482 | 37 | 0 | 0 | 8586 | 88151 | 53895 | 808 | 3127 | 3754 | 644 | 0 | 0 | 121 | 221 | 146 | 88 | 0 | 0 | 5450 | 0 | 23 | 0 | 0 | 47691 | 1601 | 0 |
| SRR2731295 | 17 | 45 | 15 | 16 | 68 | 1760 | 373 | 1555 | 3190 | 38 | 70 | 16 | 0 | 0 | 13132 | 30442 | 80586 | 332 | 1964 | 1760 | 1079 | 0 | 0 | 34 | 50 | 40 | 36 | 17 | 0 | 1601 | 0 | 36 | 0 | 0 | 29633 | 541 | 0 |
| SRR2731296 | 11 | 0 | 0 | 13 | 385 | 491 | 119 | 852 | 2008 | 22 | 729 | 15 | 0 | 0 | 11066 | 69696 | 144508 | 902 | 5629 | 6520 | 1371 | 0 | 0 | 65 | 180 | 133 | 65 | 18 | 0 | 11622 | 0 | 0 | 0 | 0 | 16892 | 2098 | 0 |
| SRR2731297 | 0 | 0 | 0 | 0 | 193 | 363 | 148 | 985 | 2431 | 84 | 562 | 34 | 0 | 0 | 19546 | 98714 | 155256 | 595 | 9121 | 10089 | 2158 | 0 | 0 | 56 | 123 | 58 | 37 | 0 | 0 | 9061 | 0 | 43 | 0 | 0 | 15472 | 2469 | 0 |
| SRR2731298 | 13 | 55 | 16 | 19 | 30 | 142 | 28 | 138 | 287 | 16 | 25 | 0 | 0 | 0 | 7399 | 42517 | 80719 | 99 | 1439 | 9020 | 17227 | 104 | 22 | 19 | 0 | 0 | 32 | 0 | 0 | 332 | 0 | 11 | 0 | 23 | 36323 | 772 | 0 |
| SRR2731299 | 15 | 0 | 0 | 0 | 98 | 347 | 82 | 865 | 2823 | 16 | 185 | 0 | 0 | 0 | 62229 | 107440 | 277039 | 377 | 16919 | 9801 | 3420 | 0 | 0 | 14 | 73 | 48 | 32 | 15 | 0 | 2467 | 0 | 0 | 0 | 0 | 4889 | 693 | 0 |
| SRR2731300 | 14 | 0 | 0 | 0 | 93 | 1323 | 698 | 3957 | 7431 | 71 | 120 | 12 | 0 | 0 | 4727 | 57169 | 62412 | 758 | 3818 | 4252 | 1390 | 0 | 0 | 32 | 61 | 23 | 16 | 0 | 0 | 1745 | 0 | 64 | 0 | 14 | 24662 | 1579 | 0 |
| SRR2731301 | 21 | 11 | 0 | 0 | 426 | 258 | 199 | 814 | 2449 | 194 | 362 | 33 | 0 | 0 | 21575 | 148538 | 242019 | 564 | 12432 | 16781 | 1639 | 0 | 0 | 98 | 171 | 93 | 71 | 0 | 0 | 7181 | 0 | 27 | 0 | 0 | 13267 | 1470 | 0 |
| SRR2731302 | 28 | 14 | 0 | 15 | 454 | 161 | 70 | 745 | 1990 | 411 | 632 | 17 | 0 | 0 | 72813 | 75365 | 383199 | 410 | 22521 | 16136 | 3907 | 0 | 0 | 165 | 523 | 287 | 144 | 49 | 0 | 8076 | 0 | 0 | 0 | 0 | 4516 | 2015 | 0 |
| SRR2731303 | 37 | 15 | 0 | 0 | 428 | 177 | 97 | 290 | 755 | 319 | 780 | 16 | 0 | 0 | 9421 | 70431 | 181341 | 243 | 2531 | 12126 | 11531 | 0 | 0 | 72 | 42 | 47 | 103 | 0 | 0 | 3452 | 0 | 0 | 0 | 0 | 16507 | 525 | 0 |
| SRR2731304 | 24 | 13 | 0 | 11 | 600 | 446 | 290 | 1136 | 3636 | 39 | 568 | 20 | 0 | 0 | 12725 | 85336 | 80559 | 933 | 10738 | 9197 | 1127 | 0 | 0 | 57 | 113 | 88 | 45 | 0 | 0 | 7069 | 0 | 0 | 0 | 0 | 38225 | 1495 | 0 |
| SRR2731305 | 10 | 0 | 0 | 0 | 319 | 629 | 289 | 2240 | 6628 | 124 | 531 | 32 | 0 | 0 | 75128 | 76710 | 221174 | 443 | 8558 | 3707 | 634 | 0 | 0 | 103 | 361 | 223 | 154 | 25 | 0 | 7653 | 0 | 16 | 0 | 0 | 11315 | 1588 | 0 |
| SRR2731306 | 14 | 0 | 0 | 0 | 328 | 136 | 60 | 460 | 1615 | 186 | 845 | 33 | 0 | 0 | 24242 | 110443 | 208839 | 345 | 10461 | 8429 | 2885 | 0 | 0 | 76 | 340 | 185 | 78 | 27 | 0 | 7333 | 0 | 34 | 0 | 11 | 11387 | 641 | 0 |
| SRR2731307 | 21 | 11 | 0 | 0 | 87 | 1240 | 231 | 1790 | 4384 | 20 | 151 | 16 | 0 | 0 | 5574 | 99934 | 110911 | 154 | 1569 | 3057 | 1393 | 0 | 0 | 31 | 115 | 71 | 39 | 31 | 0 | 2038 | 0 | 14 | 0 | 0 | 13140 | 2711 | 0 |
| SRR2731308 | 20 | 0 | 0 | 0 | 206 | 268 | 146 | 1149 | 2675 | 34 | 171 | 18 | 0 | 0 | 32640 | 119207 | 191178 | 628 | 8722 | 8416 | 2190 | 0 | 0 | 18 | 75 | 49 | 29 | 26 | 0 | 2222 | 0 | 59 | 0 | 0 | 11777 | 532 | 0 |
| SRR2731309 | 11 | 0 | 0 | 0 | 297 | 424 | 173 | 1634 | 3937 | 138 | 441 | 25 | 0 | 0 | 7070 | 61253 | 108457 | 462 | 5000 | 6301 | 2402 | 0 | 0 | 92 | 354 | 145 | 112 | 28 | 0 | 7195 | 0 | 27 | 0 | 0 | 20383 | 2258 | 0 |
| SRR2731310 | 30 | 12 | 11 | 17 | 926 | 463 | 112 | 721 | 1804 | 25 | 688 | 14 | 0 | 0 | 15782 | 85771 | 196230 | 318 | 10210 | 5930 | 1189 | 0 | 0 | 113 | 239 | 198 | 121 | 25 | 0 | 8619 | 0 | 30 | 0 | 0 | 18781 | 951 | 0 |
| SRR2731311 | 11 | 0 | 0 | 0 | 89 | 177 | 63 | 371 | 807 | 78 | 189 | 27 | 0 | 0 | 41625 | 222510 | 249101 | 260 | 10102 | 15304 | 3458 | 0 | 0 | 23 | 77 | 13 | 15 | 0 | 0 | 2973 | 0 | 19 | 0 | 13 | 5814 | 2401 | 0 |
| SRR2731312 | 21 | 0 | 0 | 0 | 122 | 287 | 75 | 535 | 1013 | 31 | 195 | 11 | 0 | 0 | 24130 | 112311 | 162017 | 344 | 10341 | 7968 | 5067 | 0 | 0 | 12 | 24 | 26 | 17 | 0 | 0 | 3274 | 0 | 48 | 0 | 0 | 14710 | 1298 | 0 |
| SRR2731313 | 16 | 0 | 0 | 0 | 383 | 186 | 109 | 697 | 1181 | 310 | 463 | 26 | 0 | 0 | 73758 | 88881 | 266762 | 248 | 7749 | 7314 | 3083 | 0 | 0 | 67 | 148 | 56 | 42 | 0 | 0 | 5946 | 0 | 20 | 0 | 14 | 14346 | 897 | 0 |
| SRR2731314 | 0 | 0 | 0 | 0 | 64 | 1593 | 624 | 4383 | 6307 | 30 | 102 | 13 | 0 | 0 | 2947 | 47043 | 52108 | 1036 | 4771 | 5492 | 1357 | 0 | 0 | 24 | 40 | 17 | 31 | 0 | 0 | 2306 | 0 | 28 | 0 | 11 | 43158 | 1218 | 0 |
| SRR2731315 | 32 | 15 | 0 | 18 | 200 | 390 | 142 | 638 | 2643 | 28 | 106 | 18 | 0 | 0 | 36858 | 129972 | 261764 | 550 | 16746 | 10698 | 1836 | 0 | 0 | 20 | 59 | 47 | 26 | 0 | 0 | 2282 | 0 | 0 | 0 | 0 | 6781 | 1065 | 0 |
| SRR2731316 | 15 | 0 | 0 | 0 | 364 | 766 | 242 | 1284 | 4134 | 61 | 409 | 29 | 0 | 0 | 7298 | 71932 | 62279 | 693 | 5975 | 4881 | 1714 | 0 | 0 | 108 | 271 | 97 | 48 | 0 | 0 | 6849 | 0 | 18 | 0 | 0 | 26020 | 539 | 0 |
| SRR2731317 | 16 | 0 | 0 | 0 | 485 | 443 | 95 | 761 | 1889 | 103 | 473 | 29 | 0 | 0 | 39642 | 58170 | 283068 | 750 | 9524 | 3792 | 2343 | 0 | 0 | 79 | 221 | 75 | 44 | 0 | 0 | 7647 | 0 | 32 | 0 | 0 | 13802 | 625 | 0 |
| SRR2731318 | 40 | 32 | 13 | 18 | 50 | 243 | 55 | 239 | 885 | 20 | 172 | 12 | 0 | 0 | 11075 | 108730 | 76358 | 586 | 6836 | 8607 | 1262 | 0 | 0 | 14 | 18 | 25 | 0 | 0 | 0 | 1902 | 0 | 18 | 0 | 10 | 37102 | 724 | 0 |
| SRR2731319 | 17 | 10 | 0 | 0 | 300 | 409 | 171 | 749 | 2897 | 32 | 181 | 12 | 0 | 0 | 8622 | 79263 | 112708 | 575 | 5563 | 9006 | 1565 | 0 | 0 | 28 | 79 | 50 | 31 | 0 | 0 | 2535 | 0 | 51 | 0 | 0 | 17983 | 594 | 0 |
| SRR2731320 | 58 | 41 | 0 | 0 | 576 | 2561 | 136 | 848 | 1464 | 22 | 681 | 14 | 0 | 0 | 7319 | 76033 | 80587 | 223 | 3466 | 3562 | 1613 | 0 | 0 | 107 | 90 | 37 | 24 | 0 | 0 | 9811 | 0 | 17 | 0 | 13 | 26231 | 2439 | 0 |
| SRR2731321 | 24 | 16 | 0 | 0 | 31 | 800 | 198 | 717 | 2329 | 27 | 63 | 29 | 0 | 0 | 7867 | 84045 | 59525 | 631 | 5240 | 3545 | 3266 | 0 | 0 | 23 | 25 | 19 | 29 | 0 | 0 | 783 | 0 | 95 | 0 | 31 | 28098 | 1371 | 15 |
| SRR2731322 | 30 | 19 | 0 | 0 | 659 | 1017 | 274 | 1063 | 3358 | 44 | 324 | 33 | 0 | 0 | 4526 | 71599 | 47356 | 1101 | 4849 | 7812 | 779 | 0 | 0 | 158 | 205 | 119 | 169 | 0 | 0 | 2815 | 0 | 41 | 0 | 20 | 36811 | 1290 | 13 |
| SRR2731323 | 29 | 18 | 0 | 0 | 275 | 811 | 182 | 784 | 2525 | 118 | 535 | 20 | 0 | 0 | 5308 | 38870 | 39673 | 724 | 11555 | 12705 | 1357 | 0 | 0 | 65 | 85 | 0 | 0 | 0 | 0 | 7867 | 0 | 45 | 0 | 33 | 16284 | 661 | 0 |
| SRR2731324 | 0 | 0 | 0 | 0 | 182 | 1475 | 664 | 4398 | 6964 | 79 | 382 | 33 | 0 | 0 | 21306 | 54413 | 241625 | 1175 | 10235 | 10900 | 1078 | 0 | 0 | 51 | 76 | 35 | 26 | 0 | 0 | 7810 | 0 | 22 | 0 | 14 | 17848 | 827 | 0 |
| SRR2731325 | 12 | 0 | 0 | 0 | 67 | 1311 | 568 | 3082 | 5384 | 22 | 129 | 22 | 0 | 0 | 10609 | 75897 | 110020 | 1132 | 6277 | 7790 | 1438 | 0 | 0 | 34 | 0 | 0 | 0 | 0 | 0 | 2574 | 0 | 25 | 0 | 11 | 29836 | 2244 | 0 |
| SRR2731326 | 15 | 0 | 0 | 0 | 430 | 460 | 192 | 961 | 2325 | 105 | 574 | 33 | 0 | 0 | 13322 | 69546 | 99352 | 690 | 5218 | 7603 | 1552 | 0 | 0 | 58 | 176 | 77 | 32 | 0 | 0 | 7064 | 0 | 47 | 0 | 0 | 12944 | 298 | 0 |
| SRR2731327 | 13 | 0 | 0 | 0 | 123 | 580 | 292 | 1595 | 2426 | 62 | 103 | 13 | 0 | 0 | 13516 | 42776 | 84955 | 706 | 5229 | 4806 | 1796 | 0 | 0 | 33 | 62 | 24 | 18 | 0 | 0 | 1731 | 0 | 36 | 0 | 15 | 21637 | 1786 | 0 |
| SRR2731328 | 11 | 0 | 0 | 0 | 115 | 385 | 94 | 463 | 1163 | 17 | 236 | 11 | 0 | 0 | 124274 | 161539 | 347031 | 339 | 6700 | 8678 | 2138 | 0 | 0 | 12 | 34 | 23 | 18 | 0 | 0 | 3429 | 0 | 0 | 0 | 0 | 6062 | 740 | 0 |
| SRR2731329 | 56 | 53 | 22 | 23 | 382 | 196 | 58 | 419 | 1295 | 436 | 506 | 21 | 0 | 0 | 89413 | 39504 | 226217 | 716 | 5612 | 3266 | 6046 | 0 | 0 | 169 | 774 | 254 | 68 | 15 | 0 | 7145 | 0 | 76 | 0 | 0 | 6856 | 308 | 0 |
| SRR2731330 | 199 | 117 | 40 | 96 | 212 | 113 | 29 | 115 | 574 | 53 | 787 | 0 | 0 | 0 | 95706 | 249533 | 384348 | 112 | 10534 | 31181 | 5039 | 0 | 0 | 12 | 18 | 0 | 0 | 0 | 0 | 6754 | 0 | 0 | 0 | 25 | 3109 | 165 | 0 |
| SRR2731331 | 61 | 52 | 14 | 19 | 117 | 768 | 259 | 883 | 2330 | 14 | 193 | 18 | 0 | 0 | 8787 | 186891 | 129876 | 72 | 1159 | 154 | 400 | 0 | 0 | 43 | 37 | 27 | 30 | 0 | 0 | 731 | 0 | 0 | 0 | 12 | 34084 | 1002 | 11 |
| SRR2731332 | 39 | 25 | 0 | 0 | 559 | 261 | 82 | 411 | 1695 | 28 | 951 | 17 | 0 | 0 | 15694 | 119677 | 179435 | 280 | 6461 | 2765 | 1854 | 0 | 0 | 116 | 380 | 160 | 114 | 0 | 0 | 5850 | 0 | 0 | 0 | 0 | 8901 | 1141 | 0 |
| SRR2731333 | 126 | 82 | 13 | 20 | 274 | 107 | 30 | 136 | 473 | 61 | 1777 | 16 | 0 | 0 | 137566 | 189749 | 380017 | 385 | 16396 | 10485 | 5132 | 0 | 0 | 202 | 376 | 122 | 70 | 0 | 0 | 9860 | 0 | 0 | 0 | 11 | 6985 | 281 | 0 |
| SRR2731335 | 14 | 0 | 0 | 0 | 89 | 204 | 94 | 414 | 1267 | 12 | 159 | 23 | 0 | 0 | 9796 | 160717 | 155445 | 550 | 9391 | 11085 | 2380 | 0 | 0 | 10 | 24 | 22 | 14 | 0 | 0 | 2958 | 0 | 17 | 0 | 0 | 12486 | 1240 | 0 |
| SRR2731336 | 59 | 30 | 0 | 18 | 160 | 537 | 110 | 465 | 1876 | 20 | 584 | 12 | 0 | 0 | 55184 | 268381 | 196935 | 69 | 4389 | 1232 | 1040 | 0 | 0 | 39 | 83 | 18 | 13 | 0 | 0 | 1955 | 0 | 0 | 0 | 15 | 6038 | 759 | 0 |
| SRR2731337 | 11 | 23 | 0 | 12 | 159 | 1282 | 569 | 2507 | 7800 | 13 | 104 | 0 | 0 | 0 | 11255 | 122498 | 228661 | 151 | 3469 | 1000 | 689 | 0 | 0 | 30 | 78 | 43 | 28 | 18 | 0 | 1179 | 0 | 0 | 0 | 0 | 15767 | 1173 | 0 |
| SRR2731338 | 91 | 70 | 11 | 22 | 41 | 573 | 66 | 238 | 742 | 0 | 157 | 15 | 0 | 0 | 10840 | 223119 | 115943 | 40 | 2429 | 640 | 567 | 0 | 0 | 33 | 50 | 31 | 34 | 0 | 0 | 824 | 0 | 0 | 0 | 17 | 15715 | 1003 | 0 |
| SRR2731339 | 23 | 12 | 0 | 0 | 88 | 1148 | 478 | 2428 | 3480 | 24 | 100 | 0 | 0 | 0 | 8049 | 26742 | 30768 | 383 | 2978 | 2476 | 1029 | 40 | 56 | 45 | 15 | 0 | 0 | 0 | 11 | 492 | 26 | 24 | 27 | 204 | 46799 | 540 | 11 |
| SRR2731340 | 105 | 80 | 18 | 26 | 603 | 193 | 42 | 152 | 614 | 14 | 999 | 19 | 0 | 0 | 203968 | 116312 | 366787 | 334 | 1885 | 2504 | 1883 | 0 | 0 | 74 | 147 | 111 | 53 | 0 | 0 | 6702 | 0 | 0 | 0 | 0 | 4403 | 318 | 0 |
| SRR2731341 | 11 | 0 | 0 | 0 | 94 | 960 | 268 | 1209 | 2803 | 14 | 133 | 12 | 0 | 0 | 5934 | 62158 | 73563 | 473 | 9181 | 5898 | 1278 | 0 | 0 | 20 | 47 | 22 | 12 | 0 | 0 | 2573 | 0 | 14 | 0 | 0 | 19531 | 1121 | 0 |
| SRR2731342 | 0 | 0 | 0 | 0 | 363 | 822 | 463 | 2520 | 4022 | 130 | 347 | 48 | 0 | 0 | 4491 | 63494 | 66635 | 1094 | 8373 | 5927 | 2847 | 0 | 0 | 39 | 83 | 40 | 19 | 0 | 0 | 4933 | 0 | 45 | 0 | 11 | 28201 | 1154 | 0 |
| SRR2731343 | 0 | 0 | 0 | 0 | 89 | 623 | 268 | 1263 | 2582 | 31 | 171 | 15 | 0 | 0 | 26248 | 59959 | 50947 | 472 | 5024 | 7101 | 807 | 0 | 0 | 28 | 46 | 18 | 23 | 0 | 0 | 1884 | 0 | 10 | 0 | 23 | 35002 | 1271 | 0 |
| SRR2731344 | 23 | 15 | 0 | 14 | 172 | 1013 | 400 | 2424 | 2771 | 24 | 126 | 22 | 0 | 0 | 18944 | 55382 | 145900 | 939 | 5733 | 3754 | 554 | 0 | 0 | 49 | 63 | 28 | 22 | 0 | 0 | 2076 | 0 | 23 | 0 | 0 | 41513 | 1589 | 0 |
| SRR2731345 | 19 | 19 | 0 | 0 | 651 | 1443 | 357 | 1291 | 1761 | 104 | 618 | 0 | 0 | 0 | 3157 | 18844 | 65293 | 106 | 1575 | 1754 | 702 | 0 | 0 | 46 | 54 | 0 | 35 | 0 | 0 | 4549 | 0 | 0 | 0 | 12 | 20960 | 627 | 0 |
| SRR2731346 | 16 | 0 | 0 | 0 | 76 | 621 | 196 | 1020 | 1391 | 39 | 145 | 22 | 0 | 0 | 26291 | 92637 | 224246 | 434 | 14535 | 14256 | 2232 | 0 | 0 | 27 | 41 | 28 | 12 | 0 | 0 | 2222 | 0 | 0 | 0 | 0 | 13372 | 680 | 0 |
| SRR2731347 | 17 | 11 | 0 | 0 | 309 | 1159 | 457 | 2440 | 7277 | 48 | 659 | 31 | 0 | 0 | 17000 | 77257 | 143327 | 370 | 6667 | 4281 | 1655 | 0 | 0 | 145 | 297 | 146 | 89 | 0 | 0 | 6351 | 0 | 0 | 0 | 15 | 24682 | 764 | 0 |
| SRR2731348 | 12 | 0 | 0 | 0 | 244 | 579 | 414 | 1274 | 3581 | 14 | 122 | 30 | 0 | 0 | 48849 | 114602 | 192837 | 783 | 10407 | 9393 | 1257 | 0 | 0 | 29 | 59 | 67 | 22 | 0 | 0 | 2074 | 0 | 0 | 0 | 0 | 18226 | 1142 | 0 |
| SRR2731349 | 35 | 11 | 0 | 0 | 36 | 579 | 220 | 674 | 1444 | 27 | 102 | 15 | 0 | 0 | 55441 | 53573 | 372851 | 175 | 7347 | 30772 | 3844 | 0 | 0 | 17 | 14 | 47 | 44 | 0 | 0 | 734 | 0 | 0 | 0 | 0 | 14793 | 1091 | 0 |
| SRR2731350 | 15 | 0 | 0 | 0 | 854 | 1237 | 583 | 3648 | 7429 | 122 | 522 | 0 | 0 | 0 | 10099 | 50252 | 131404 | 969 | 13306 | 10297 | 1371 | 0 | 0 | 135 | 325 | 226 | 147 | 0 | 0 | 8061 | 0 | 23 | 0 | 0 | 16516 | 812 | 0 |
| SRR2910706 | 26 | 17 | 89 | 123 | 182 | 419 | 187 | 915 | 102 | 12 | 149 | 0 | 0 | 0 | 19869 | 20543 | 239555 | 0 | 33 | 5510 | 22386 | 0 | 0 | 134 | 179 | 271 | 436 | 26 | 0 | 619 | 0 | 12 | 0 | 0 | 838 | 0 | 0 |
| SRR2910737 | 34 | 14 | 36 | 73 | 199 | 607 | 330 | 1331 | 138 | 14 | 77 | 0 | 0 | 0 | 18631 | 20470 | 200306 | 0 | 11 | 6092 | 30009 | 0 | 0 | 119 | 74 | 108 | 177 | 0 | 0 | 652 | 0 | 39 | 0 | 0 | 1173 | 0 | 0 |
| SRR2910738 | 28 | 0 | 48 | 19 | 33 | 596 | 242 | 1202 | 81 | 14 | 16 | 0 | 0 | 0 | 21251 | 21179 | 202763 | 10 | 30 | 6937 | 31015 | 0 | 0 | 18 | 20 | 29 | 49 | 38 | 0 | 103 | 0 | 10 | 0 | 0 | 486 | 0 | 0 |
| SRR2912507 | 72 | 15 | 0 | 0 | 11 | 0 | 13 | 16 | 22 | 0 | 121 | 0 | 0 | 0 | 10396 | 18817 | 17741 | 434 | 2484 | 1137 | 532 | 0 | 0 | 0 | 0 | 16 | 0 | 0 | 0 | 32 | 0 | 0 | 0 | 0 | 876 | 0 | 0 |
| SRR2912508 | 259 | 54 | 0 | 0 | 61 | 0 | 13 | 21 | 61 | 0 | 1298 | 0 | 0 | 0 | 14416 | 14736 | 27180 | 925 | 3232 | 1051 | 161 | 0 | 0 | 0 | 0 | 13 | 0 | 0 | 0 | 284 | 0 | 0 | 0 | 0 | 1781 | 0 | 0 |
| SRR2912510 | 0 | 0 | 0 | 0 | 87 | 0 | 50 | 21 | 88 | 0 | 271 | 0 | 0 | 0 | 15641 | 49713 | 24644 | 3408 | 4407 | 485 | 183 | 0 | 0 | 0 | 0 | 13 | 0 | 0 | 0 | 310 | 0 | 0 | 0 | 0 | 807 | 0 | 0 |
| SRR2912511 | 0 | 0 | 0 | 0 | 137 | 0 | 42 | 20 | 72 | 0 | 257 | 0 | 0 | 0 | 13110 | 52122 | 23242 | 2124 | 3321 | 417 | 127 | 0 | 0 | 0 | 0 | 19 | 0 | 0 | 0 | 351 | 0 | 0 | 0 | 0 | 455 | 0 | 0 |
| SRR2912513 | 0 | 0 | 0 | 0 | 38 | 0 | 38 | 14 | 82 | 0 | 51 | 0 | 0 | 0 | 10742 | 70587 | 23195 | 656 | 3067 | 556 | 330 | 0 | 0 | 0 | 0 | 10 | 0 | 0 | 0 | 72 | 0 | 0 | 0 | 0 | 490 | 0 | 0 |
| SRR2912514 | 66 | 13 | 0 | 0 | 16 | 0 | 12 | 0 | 52 | 0 | 247 | 0 | 0 | 0 | 18014 | 22429 | 32144 | 853 | 3765 | 1556 | 345 | 0 | 0 | 0 | 0 | 28 | 0 | 0 | 0 | 59 | 0 | 0 | 0 | 0 | 2698 | 0 | 0 |
| SRR2912515 | 0 | 0 | 0 | 0 | 122 | 0 | 45 | 20 | 112 | 0 | 186 | 0 | 0 | 0 | 17678 | 42213 | 29923 | 993 | 4593 | 626 | 240 | 0 | 0 | 0 | 0 | 18 | 0 | 0 | 0 | 263 | 0 | 0 | 0 | 0 | 517 | 0 | 0 |
| SRR2913202 | 0 | 0 | 0 | 18 | 21 | 344 | 580 | 1198 | 226 | 0 | 392 | 0 | 0 | 0 | 250 | 5226 | 6941 | 19 | 141 | 28004 | 61 | 0 | 0 | 15 | 0 | 49 | 604 | 262 | 0 | 1389 | 0 | 33 | 0 | 0 | 3858 | 52 | 0 |
| SRR2913203 | 0 | 0 | 0 | 27 | 11 | 108 | 171 | 477 | 77 | 0 | 67 | 0 | 0 | 0 | 891 | 7079 | 11905 | 0 | 29 | 45100 | 128 | 0 | 0 | 27 | 0 | 46 | 89 | 55 | 0 | 234 | 0 | 11 | 0 | 0 | 561 | 13 | 0 |
| SRR2913204 | 0 | 0 | 0 | 21 | 11 | 62 | 126 | 231 | 64 | 0 | 45 | 0 | 0 | 0 | 1013 | 9800 | 17340 | 0 | 114 | 27906 | 169 | 0 | 0 | 26 | 0 | 27 | 139 | 86 | 0 | 143 | 0 | 0 | 0 | 0 | 812 | 38 | 0 |
| SRR2913205 | 0 | 0 | 0 | 15 | 29 | 139 | 280 | 462 | 52 | 0 | 63 | 0 | 0 | 0 | 685 | 5357 | 14239 | 0 | 82 | 26519 | 84 | 0 | 0 | 15 | 0 | 12 | 186 | 107 | 0 | 225 | 0 | 0 | 0 | 0 | 1669 | 39 | 0 |
| SRR2913206 | 0 | 0 | 0 | 0 | 16 | 345 | 634 | 1082 | 210 | 0 | 106 | 0 | 0 | 0 | 186 | 4459 | 4985 | 0 | 51 | 26770 | 37 | 0 | 0 | 0 | 0 | 14 | 186 | 89 | 0 | 330 | 0 | 19 | 0 | 0 | 3928 | 31 | 0 |
| SRR2913207 | 0 | 0 | 0 | 29 | 36 | 11 | 13 | 62 | 0 | 0 | 133 | 0 | 0 | 0 | 1588 | 8534 | 38875 | 0 | 196 | 77040 | 153 | 0 | 0 | 28 | 15 | 81 | 704 | 306 | 0 | 272 | 0 | 0 | 0 | 0 | 93 | 56 | 0 |
| SRR2913208 | 0 | 0 | 0 | 13 | 13 | 472 | 858 | 1463 | 374 | 0 | 60 | 0 | 0 | 0 | 806 | 10749 | 15498 | 12 | 20 | 10986 | 94 | 0 | 0 | 12 | 0 | 19 | 358 | 145 | 0 | 187 | 0 | 24 | 0 | 0 | 3970 | 48 | 0 |
| SRR2913209 | 0 | 0 | 0 | 19 | 12 | 303 | 524 | 1146 | 192 | 0 | 114 | 0 | 0 | 0 | 1224 | 17163 | 37832 | 0 | 61 | 5974 | 315 | 0 | 0 | 21 | 11 | 22 | 269 | 240 | 0 | 294 | 0 | 53 | 0 | 0 | 6116 | 52 | 0 |
| SRR2913210 | 0 | 0 | 0 | 14 | 20 | 306 | 482 | 1039 | 155 | 0 | 227 | 0 | 0 | 0 | 1325 | 5266 | 23177 | 0 | 34 | 11525 | 812 | 0 | 0 | 0 | 0 | 56 | 455 | 216 | 0 | 721 | 0 | 21 | 0 | 0 | 2575 | 141 | 0 |
| SRR2913211 | 0 | 0 | 0 | 13 | 20 | 583 | 682 | 935 | 264 | 0 | 69 | 0 | 0 | 0 | 418 | 6866 | 10007 | 0 | 44 | 11678 | 132 | 0 | 0 | 0 | 0 | 17 | 173 | 63 | 0 | 220 | 0 | 44 | 0 | 0 | 7889 | 112 | 0 |
| SRR2913212 | 0 | 0 | 0 | 11 | 16 | 264 | 496 | 897 | 168 | 0 | 103 | 0 | 0 | 0 | 2820 | 11836 | 24056 | 0 | 39 | 17358 | 339 | 0 | 0 | 20 | 12 | 26 | 208 | 261 | 0 | 255 | 0 | 22 | 0 | 0 | 1503 | 30 | 0 |
| SRR2913213 | 0 | 0 | 0 | 13 | 12 | 364 | 1173 | 2310 | 318 | 0 | 352 | 0 | 0 | 0 | 509 | 5884 | 18121 | 0 | 55 | 14135 | 155 | 0 | 0 | 0 | 0 | 58 | 320 | 405 | 0 | 1309 | 0 | 55 | 0 | 0 | 10551 | 45 | 0 |
| SRR2913214 | 0 | 0 | 0 | 16 | 21 | 623 | 895 | 1681 | 366 | 0 | 208 | 0 | 0 | 0 | 125 | 8125 | 10869 | 0 | 54 | 20886 | 47 | 0 | 0 | 0 | 0 | 30 | 344 | 410 | 0 | 824 | 0 | 23 | 0 | 0 | 6192 | 17 | 0 |
| SRR2913215 | 0 | 0 | 0 | 13 | 15 | 192 | 187 | 566 | 114 | 0 | 121 | 0 | 0 | 0 | 447 | 9902 | 13322 | 0 | 91 | 48491 | 196 | 0 | 0 | 18 | 0 | 17 | 224 | 170 | 0 | 390 | 0 | 0 | 0 | 0 | 1665 | 19 | 0 |
| SRR2913216 | 0 | 0 | 0 | 39 | 21 | 79 | 118 | 270 | 41 | 0 | 231 | 0 | 0 | 0 | 308 | 3698 | 9329 | 0 | 76 | 38375 | 176 | 0 | 0 | 21 | 0 | 75 | 628 | 686 | 0 | 684 | 0 | 0 | 0 | 0 | 1333 | 21 | 0 |
| SRR2913217 | 0 | 0 | 0 | 17 | 16 | 281 | 394 | 914 | 232 | 0 | 65 | 0 | 0 | 0 | 752 | 7704 | 15135 | 0 | 119 | 28496 | 105 | 0 | 0 | 0 | 0 | 17 | 114 | 128 | 0 | 169 | 0 | 14 | 0 | 0 | 3086 | 31 | 0 |
| SRR2913218 | 0 | 0 | 0 | 0 | 0 | 52 | 91 | 195 | 46 | 0 | 32 | 0 | 0 | 0 | 416 | 4931 | 15103 | 0 | 99 | 48177 | 155 | 0 | 0 | 0 | 0 | 28 | 64 | 50 | 0 | 120 | 0 | 0 | 0 | 0 | 981 | 0 | 0 |
| SRR2913219 | 0 | 0 | 0 | 13 | 0 | 282 | 440 | 811 | 213 | 0 | 100 | 11 | 0 | 0 | 275 | 5298 | 7735 | 0 | 83 | 32897 | 34 | 0 | 0 | 14 | 0 | 29 | 175 | 142 | 0 | 296 | 0 | 14 | 0 | 0 | 3901 | 12 | 0 |
| SRR2913220 | 0 | 0 | 0 | 15 | 17 | 55 | 84 | 220 | 41 | 0 | 47 | 0 | 0 | 0 | 998 | 8470 | 12427 | 0 | 120 | 35686 | 104 | 0 | 0 | 11 | 0 | 26 | 69 | 36 | 0 | 111 | 0 | 0 | 0 | 0 | 1117 | 49 | 0 |
| SRR2913221 | 0 | 0 | 0 | 11 | 13 | 18 | 21 | 61 | 0 | 0 | 68 | 15 | 0 | 0 | 1322 | 6994 | 14618 | 0 | 205 | 59871 | 167 | 0 | 0 | 10 | 0 | 13 | 86 | 53 | 0 | 242 | 0 | 0 | 0 | 0 | 380 | 96 | 0 |
| SRR2913222 | 0 | 0 | 0 | 17 | 22 | 49 | 114 | 209 | 30 | 0 | 58 | 0 | 0 | 0 | 1266 | 7575 | 17547 | 0 | 74 | 40555 | 540 | 0 | 0 | 0 | 0 | 11 | 87 | 32 | 0 | 153 | 0 | 0 | 0 | 0 | 941 | 11 | 0 |
| SRR2913223 | 0 | 0 | 0 | 17 | 12 | 48 | 81 | 202 | 35 | 0 | 39 | 0 | 0 | 0 | 649 | 3104 | 8002 | 0 | 98 | 49189 | 234 | 0 | 0 | 11 | 0 | 32 | 29 | 38 | 0 | 105 | 0 | 11 | 0 | 0 | 715 | 0 | 0 |
| SRR2913224 | 0 | 0 | 0 | 23 | 20 | 60 | 92 | 192 | 35 | 0 | 111 | 0 | 0 | 0 | 602 | 5618 | 18079 | 0 | 100 | 75810 | 145 | 0 | 0 | 26 | 0 | 31 | 254 | 258 | 0 | 350 | 0 | 0 | 0 | 0 | 974 | 28 | 0 |
| SRR2913225 | 0 | 0 | 0 | 25 | 22 | 69 | 141 | 267 | 38 | 0 | 54 | 10 | 0 | 0 | 1906 | 18387 | 34945 | 0 | 89 | 57064 | 344 | 0 | 0 | 29 | 0 | 20 | 67 | 79 | 0 | 158 | 0 | 0 | 0 | 0 | 666 | 33 | 0 |
| SRR2915342 | 25 | 34 | 0 | 0 | 130 | 2470 | 1599 | 6844 | 18779 | 197 | 65 | 25 | 26 | 0 | 56093 | 33274 | 75452 | 926 | 410 | 1685 | 1722 | 0 | 0 | 23 | 40 | 27 | 19 | 11 | 0 | 360 | 0 | 26 | 0 | 0 | 34699 | 0 | 0 |
| SRR2915343 | 32 | 42 | 23 | 13 | 138 | 3401 | 2039 | 8850 | 18834 | 258 | 50 | 20 | 27 | 11 | 74041 | 58931 | 84422 | 1134 | 506 | 1885 | 3443 | 0 | 0 | 18 | 34 | 23 | 17 | 10 | 0 | 288 | 0 | 26 | 0 | 0 | 40172 | 0 | 0 |
| SRR2915344 | 95 | 172 | 18 | 10 | 326 | 2940 | 2459 | 10054 | 25831 | 430 | 117 | 47 | 18 | 0 | 77232 | 50878 | 108146 | 1464 | 616 | 1786 | 2467 | 0 | 0 | 34 | 79 | 54 | 42 | 0 | 0 | 649 | 0 | 34 | 0 | 0 | 40337 | 0 | 0 |
| SRR2915345 | 316 | 67 | 56 | 23 | 907 | 4394 | 2958 | 11899 | 30658 | 423 | 618 | 76 | 28 | 16 | 6211 | 29716 | 26916 | 1744 | 792 | 3582 | 1210 | 0 | 0 | 72 | 88 | 51 | 35 | 0 | 0 | 2646 | 0 | 32 | 0 | 0 | 14821 | 0 | 0 |
| SRR2915346 | 56 | 14 | 20 | 21 | 332 | 4803 | 3671 | 13410 | 30577 | 227 | 280 | 29 | 20 | 10 | 6135 | 27462 | 27360 | 1628 | 720 | 4057 | 731 | 0 | 0 | 28 | 53 | 38 | 22 | 16 | 0 | 1061 | 0 | 37 | 0 | 0 | 11476 | 0 | 0 |
| SRR2915347 | 71 | 15 | 28 | 11 | 290 | 5008 | 3511 | 12633 | 24138 | 159 | 251 | 16 | 15 | 0 | 13412 | 32051 | 51590 | 1916 | 952 | 4185 | 1678 | 0 | 0 | 33 | 57 | 46 | 33 | 14 | 0 | 1000 | 0 | 15 | 0 | 11 | 16780 | 10 | 0 |
| SRR2915348 | 31 | 18 | 11 | 27 | 222 | 7001 | 5798 | 18548 | 41680 | 80 | 90 | 10 | 17 | 0 | 1176 | 30695 | 33641 | 1781 | 756 | 4811 | 217 | 0 | 0 | 30 | 47 | 40 | 40 | 11 | 0 | 526 | 0 | 56 | 0 | 0 | 8460 | 0 | 0 |
| SRR2915349 | 19 | 46 | 14 | 0 | 267 | 11480 | 10431 | 33146 | 57031 | 74 | 43 | 11 | 11 | 0 | 968 | 26091 | 24208 | 2526 | 987 | 5863 | 289 | 0 | 0 | 25 | 24 | 13 | 15 | 0 | 0 | 302 | 0 | 50 | 0 | 12 | 7888 | 0 | 0 |
| SRR2915350 | 37 | 19 | 11 | 18 | 291 | 9737 | 8375 | 28834 | 52538 | 46 | 56 | 11 | 16 | 0 | 1211 | 34668 | 31818 | 1446 | 996 | 4276 | 210 | 0 | 0 | 17 | 22 | 17 | 15 | 0 | 0 | 384 | 0 | 31 | 0 | 11 | 7960 | 0 | 0 |
| SRR2915905 | 0 | 0 | 0 | 34 | 117 | 34 | 60 | 217 | 49 | 130 | 33 | 0 | 0 | 0 | 136704 | 59690 | 409497 | 0 | 1069 | 22855 | 27872 | 0 | 18 | 85 | 26 | 72 | 254 | 38 | 0 | 10 | 0 | 96 | 0 | 0 | 191 | 75 | 0 |
| SRR2915915 | 0 | 31 | 0 | 20 | 44 | 112 | 49 | 340 | 29 | 21 | 16 | 0 | 0 | 0 | 69471 | 71692 | 434789 | 32 | 240 | 6606 | 29280 | 0 | 0 | 222 | 25 | 71 | 187 | 24 | 0 | 179 | 0 | 38 | 0 | 0 | 152 | 0 | 0 |
| SRR2915918 | 10 | 21 | 12 | 72 | 89 | 11 | 41 | 76 | 15 | 89 | 137 | 0 | 0 | 0 | 31100 | 27465 | 254394 | 29 | 215 | 61216 | 3612 | 0 | 0 | 131 | 92 | 210 | 25 | 17 | 0 | 12 | 0 | 0 | 0 | 0 | 110 | 0 | 0 |
| SRR2915921 | 0 | 0 | 0 | 0 | 24 | 204 | 103 | 763 | 39 | 166 | 327 | 39 | 0 | 0 | 71093 | 38557 | 242732 | 0 | 201 | 25036 | 11408 | 0 | 15 | 0 | 44 | 41 | 41 | 0 | 0 | 1606 | 0 | 85 | 0 | 0 | 644 | 0 | 22 |
| SRR2915923 | 0 | 0 | 0 | 0 | 49 | 0 | 19 | 47 | 0 | 57 | 173 | 14 | 0 | 0 | 96608 | 38869 | 531189 | 0 | 821 | 39864 | 721 | 0 | 0 | 111 | 18 | 39 | 135 | 21 | 0 | 22 | 0 | 0 | 0 | 0 | 61 | 0 | 0 |
| SRR2915926 | 0 | 0 | 0 | 18 | 39 | 12 | 15 | 55 | 0 | 203 | 109 | 0 | 0 | 0 | 51902 | 62224 | 378779 | 0 | 343 | 46105 | 3404 | 0 | 0 | 139 | 114 | 259 | 14 | 168 | 0 | 18 | 0 | 0 | 0 | 0 | 34 | 0 | 0 |
| SRR2915928 | 0 | 0 | 0 | 0 | 12 | 585 | 136 | 689 | 51 | 15 | 74 | 0 | 0 | 0 | 24518 | 46903 | 415115 | 0 | 317 | 24640 | 3928 | 0 | 0 | 47 | 41 | 85 | 19 | 12 | 0 | 32 | 0 | 19 | 0 | 0 | 918 | 0 | 13 |
| SRR2960039 | 0 | 0 | 0 | 0 | 0 | 3687 | 15940 | 29110 | 16041 | 0 | 0 | 0 | 0 | 0 | 0 | 2189 | 606 | 18 | 0 | 32 | 0 | 50 | 56 | 436 | 0 | 0 | 0 | 0 | 0 | 0 | 0 | 58 | 11 | 0 | 435 | 0 | 17 |

### Supp Table S8.3.Columns_BX_to_DH

| | | | | | | | | | | | | | | | | | | | | | | | | | | | | | | | | | | | | | |
|---|---|---|---|---|---|---|---|---|---|---|---|---|---|---|---|---|---|---|---|---|---|---|---|---|---|---|---|---|---|---|---|---|---|---|---|---|---|
| SRR2976280 | 0 | 0 | 0 | 0 | 0 | 240 | 0 | 959 | 229 | 16 | 0 | 0 | 0 | 0 | 0 | 10094 | 2782 | 125 | 0 | 17277 | 459 | 0 | 0 | 140 | 0 | 0 | 0 | 0 | 0 | 0 | 0 | 3689 | 0 | 0 | 2137 | 177 | 0 |
| SRR2976305 | 0 | 0 | 0 | 0 | 0 | 312 | 251 | 2676 | 242 | 0 | 0 | 0 | 0 | 0 | 0 | 1311 | 20690 | 0 | 0 | 0 | 1230 | 0 | 0 | 34 | 0 | 0 | 0 | 0 | 0 | 0 | 0 | 0 | 0 | 0 | 5802 | 181 | 0 |
| SRR3092114 | 190 | 39 | 0 | 0 | 23 | 440 | 714 | 540 | 540 | 0 | 0 | 0 | 0 | 0 | 44192 | 11876 | 62719 | 0 | 0 | 1456 | 165 | 0 | 0 | 14 | 0 | 0 | 0 | 0 | 0 | 17 | 0 | 18 | 0 | 0 | 896 | 0 | 0 |
| SRR3092115 | 68 | 20 | 0 | 0 | 0 | 505 | 1169 | 638 | 810 | 0 | 0 | 0 | 0 | 0 | 26711 | 11249 | 27315 | 27 | 0 | 1780 | 93 | 0 | 0 | 12 | 0 | 0 | 0 | 0 | 0 | 13 | 0 | 0 | 0 | 0 | 819 | 0 | 0 |
| SRR3103222 | 10 | 0 | 0 | 10 | 73 | 379 | 727 | 2665 | 281 | 16 | 0 | 0 | 0 | 0 | 42570 | 80681 | 97121 | 131 | 30 | 9494 | 2110 | 0 | 0 | 138 | 0 | 0 | 13 | 0 | 0 | 104 | 0 | 259 | 0 | 0 | 1411 | 93 | 0 |
| SRR3103223 | 0 | 0 | 0 | 28 | 28 | 15 | 54 | 133 | 44 | 14 | 114 | 28 | 0 | 0 | 88656 | 24483 | 158564 | 0 | 0 | 320 | 1436 | 0 | 0 | 43 | 0 | 85 | 142 | 0 | 0 | 611 | 0 | 0 | 0 | 0 | 97 | 128 | 0 |
| SRR3103224 | 0 | 0 | 0 | 13 | 26 | 262 | 360 | 655 | 229 | 0 | 13 | 0 | 0 | 0 | 4778 | 13115 | 60500 | 0 | 0 | 360 | 425 | 0 | 0 | 39 | 0 | 0 | 45 | 0 | 0 | 84 | 0 | 65 | 0 | 0 | 980 | 18 | 0 |
| SRR3103225 | 0 | 0 | 0 | 61 | 66 | 0 | 14 | 32 | 0 | 24 | 28 | 0 | 0 | 0 | 37363 | 17286 | 263414 | 16 | 97 | 32362 | 11826 | 0 | 0 | 71 | 24 | 213 | 889 | 217 | 0 | 246 | 0 | 16 | 0 | 0 | 38 | 18 | 0 |
| SRR3103226 | 0 | 0 | 21 | 0 | 42 | 37 | 37 | 662 | 74 | 0 | 106 | 21 | 0 | 0 | 195846 | 29248 | 73962 | 0 | 0 | 110 | 4044 | 0 | 0 | 106 | 21 | 106 | 338 | 21 | 0 | 254 | 0 | 0 | 0 | 0 | 297 | 89 | 0 |
| SRR3103227 | 0 | 10 | 0 | 71 | 121 | 26 | 34 | 112 | 0 | 0 | 20 | 0 | 0 | 0 | 19493 | 51972 | 262514 | 14 | 72 | 73356 | 3946 | 0 | 0 | 30 | 0 | 10 | 111 | 0 | 0 | 384 | 0 | 12 | 0 | 0 | 82 | 65 | 0 |
| SRR3103228 | 0 | 10 | 0 | 17 | 77 | 110 | 84 | 273 | 44 | 0 | 27 | 0 | 0 | 0 | 55727 | 64950 | 118743 | 26 | 79 | 36860 | 4541 | 0 | 0 | 0 | 0 | 20 | 34 | 0 | 0 | 215 | 0 | 22 | 0 | 0 | 184 | 19 | 0 |
| SRR3103229 | 11 | 14 | 0 | 56 | 49 | 33 | 53 | 219 | 0 | 11 | 28 | 0 | 0 | 0 | 26805 | 25332 | 347464 | 0 | 61 | 8358 | 1708 | 0 | 0 | 39 | 11 | 25 | 91 | 49 | 0 | 144 | 0 | 13 | 0 | 0 | 90 | 23 | 0 |
| SRR3103230 | 17 | 0 | 17 | 42 | 118 | 0 | 12 | 30 | 18 | 0 | 17 | 0 | 0 | 0 | 29327 | 26455 | 383447 | 0 | 56 | 29194 | 4145 | 0 | 0 | 67 | 0 | 42 | 135 | 0 | 0 | 295 | 0 | 0 | 0 | 0 | 63 | 66 | 0 |
| SRR3103231 | 0 | 0 | 18 | 46 | 46 | 22 | 28 | 125 | 0 | 14 | 27 | 0 | 0 | 0 | 18729 | 43524 | 306313 | 0 | 86 | 31027 | 8469 | 0 | 0 | 14 | 0 | 23 | 105 | 36 | 0 | 168 | 0 | 44 | 0 | 0 | 57 | 12 | 0 |
| SRR3103232 | 0 | 0 | 18 | 18 | 71 | 96 | 144 | 648 | 192 | 0 | 54 | 0 | 0 | 0 | 38083 | 61886 | 153878 | 0 | 96 | 39714 | 3429 | 0 | 0 | 36 | 0 | 0 | 71 | 0 | 0 | 590 | 0 | 72 | 0 | 0 | 536 | 80 | 0 |
| SRR3103233 | 0 | 0 | 0 | 12 | 30 | 66 | 137 | 472 | 48 | 0 | 0 | 0 | 0 | 0 | 24182 | 54993 | 253705 | 24 | 66 | 37026 | 6712 | 0 | 0 | 59 | 0 | 18 | 24 | 0 | 0 | 107 | 0 | 18 | 0 | 0 | 442 | 60 | 0 |
| SRR3103234 | 0 | 15 | 0 | 56 | 59 | 0 | 33 | 49 | 0 | 0 | 30 | 0 | 0 | 0 | 23150 | 29067 | 263629 | 0 | 49 | 29358 | 3855 | 0 | 0 | 48 | 0 | 26 | 152 | 41 | 0 | 163 | 0 | 0 | 0 | 0 | 43 | 0 | 0 |
| SRR3103235 | 0 | 0 | 0 | 48 | 87 | 12 | 35 | 138 | 10 | 13 | 58 | 0 | 0 | 0 | 17422 | 55802 | 261106 | 15 | 44 | 60018 | 9554 | 0 | 0 | 52 | 0 | 45 | 193 | 16 | 0 | 193 | 0 | 10 | 0 | 0 | 72 | 21 | 0 |
| SRR3103236 | 0 | 10 | 0 | 40 | 44 | 24 | 85 | 214 | 26 | 0 | 24 | 0 | 0 | 0 | 26254 | 119973 | 194644 | 0 | 0 | 7918 | 2897 | 0 | 0 | 67 | 13 | 40 | 299 | 30 | 0 | 57 | 0 | 50 | 0 | 0 | 47 | 25 | 0 |
| SRR3103237 | 0 | 0 | 0 | 67 | 124 | 0 | 13 | 70 | 0 | 0 | 0 | 0 | 0 | 0 | 23963 | 27404 | 366638 | 0 | 34 | 24289 | 9239 | 0 | 0 | 62 | 14 | 67 | 247 | 24 | 0 | 162 | 0 | 0 | 0 | 0 | 30 | 0 | 0 |
| SRR3103238 | 29 | 18 | 29 | 50 | 76 | 28 | 33 | 149 | 17 | 79 | 32 | 0 | 0 | 0 | 120639 | 41820 | 479603 | 0 | 42 | 35371 | 8140 | 0 | 0 | 111 | 0 | 26 | 140 | 15 | 0 | 143 | 0 | 22 | 0 | 0 | 29 | 11 | 0 |
| SRR3103239 | 0 | 0 | 0 | 79 | 14 | 206 | 265 | 884 | 118 | 0 | 0 | 0 | 0 | 0 | 11317 | 73486 | 129171 | 0 | 29 | 14382 | 1680 | 0 | 0 | 28 | 0 | 26 | 114 | 109 | 0 | 51 | 0 | 74 | 0 | 0 | 832 | 11 | 0 |
| SRR3103240 | 0 | 0 | 0 | 72 | 25 | 13 | 42 | 96 | 14 | 0 | 21 | 0 | 0 | 0 | 83883 | 19954 | 175201 | 0 | 164 | 10240 | 3126 | 0 | 0 | 65 | 30 | 233 | 914 | 145 | 0 | 97 | 0 | 32 | 0 | 0 | 99 | 32 | 0 |
| SRR3103241 | 0 | 0 | 19 | 101 | 0 | 56 | 167 | 234 | 45 | 0 | 57 | 0 | 0 | 0 | 53022 | 21977 | 120659 | 0 | 111 | 7116 | 1804 | 0 | 0 | 44 | 25 | 183 | 31 | 107 | 0 | 107 | 0 | 22 | 0 | 0 | 315 | 36 | 0 |
| SRR3103242 | 19 | 32 | 23 | 97 | 37 | 158 | 252 | 733 | 236 | 32 | 13 | 23 | 0 | 0 | 97412 | 14960 | 91399 | 0 | 79 | 11320 | 622 | 0 | 0 | 121 | 42 | 176 | 27 | 125 | 0 | 45 | 0 | 55 | 0 | 0 | 285 | 22 | 28 |
| SRR3103243 | 0 | 0 | 0 | 0 | 0 | 107 | 107 | 392 | 89 | 31 | 60 | 73 | 0 | 0 | 18595 | 8647 | 66277 | 0 | 0 | 0 | 374 | 0 | 0 | 98 | 17 | 19 | 31 | 0 | 0 | 333 | 0 | 178 | 0 | 0 | 1569 | 10 | 108 |
| SRR3103244 | 0 | 18 | 24 | 53 | 18 | 55 | 65 | 271 | 12 | 12 | 208 | 0 | 0 | 0 | 135176 | 28470 | 180083 | 0 | 0 | 246 | 6750 | 0 | 0 | 65 | 30 | 119 | 202 | 12 | 0 | 249 | 0 | 0 | 0 | 0 | 72 | 36 | 12 |
| SRR3103245 | 12 | 0 | 0 | 36 | 185 | 111 | 383 | 675 | 195 | 0 | 36 | 0 | 0 | 0 | 32161 | 16676 | 92181 | 0 | 0 | 195 | 7200 | 0 | 0 | 68 | 18 | 116 | 307 | 0 | 0 | 369 | 0 | 77 | 0 | 0 | 595 | 15 | 18 |
| SRR3103246 | 12 | 23 | 12 | 86 | 0 | 89 | 84 | 404 | 71 | 38 | 1210 | 66 | 0 | 0 | 247233 | 4240 | 48348 | 0 | 0 | 53 | 2686 | 0 | 0 | 12 | 17 | 64 | 76 | 10 | 0 | 2051 | 0 | 113 | 0 | 0 | 156 | 64 | 22 |
| SRR3103247 | 15 | 15 | 0 | 78 | 69 | 117 | 433 | 686 | 307 | 0 | 34 | 0 | 0 | 0 | 36613 | 33660 | 173211 | 18 | 162 | 12502 | 1795 | 0 | 0 | 59 | 0 | 64 | 132 | 24 | 0 | 269 | 0 | 27 | 0 | 0 | 962 | 17 | 0 |
| SRR3103248 | 0 | 0 | 0 | 13 | 118 | 31 | 35 | 92 | 0 | 11 | 122 | 19 | 0 | 0 | 126078 | 20988 | 157518 | 0 | 88 | 8439 | 9657 | 0 | 0 | 17 | 21 | 67 | 84 | 0 | 0 | 53 | 0 | 24 | 0 | 0 | 123 | 19 | 0 |
| SRR3103249 | 0 | 14 | 0 | 27 | 21 | 24 | 18 | 129 | 0 | 110 | 377 | 19 | 0 | 0 | 306003 | 19247 | 399042 | 0 | 80 | 2039 | 3970 | 0 | 0 | 13 | 0 | 31 | 66 | 21 | 0 | 1145 | 0 | 0 | 0 | 0 | 79 | 18 | 0 |
| SRR3103250 | 11 | 0 | 0 | 33 | 79 | 29 | 133 | 285 | 59 | 0 | 36 | 14 | 0 | 0 | 25391 | 30600 | 190851 | 20 | 49 | 15100 | 7344 | 0 | 0 | 22 | 14 | 36 | 77 | 0 | 0 | 20 | 0 | 103 | 0 | 0 | 456 | 206 | 0 |
| SRR3103251 | 0 | 0 | 0 | 72 | 168 | 48 | 60 | 290 | 36 | 0 | 14 | 0 | 0 | 0 | 25498 | 18509 | 152820 | 36 | 72 | 7791 | 2718 | 0 | 0 | 90 | 43 | 168 | 10 | 25 | 0 | 136 | 0 | 12 | 0 | 0 | 984 | 29 | 0 |
| SRR3103252 | 0 | 0 | 0 | 46 | 46 | 32 | 146 | 211 | 16 | 0 | 31 | 31 | 0 | 0 | 46370 | 20962 | 116724 | 49 | 49 | 10521 | 6998 | 0 | 0 | 46 | 0 | 77 | 108 | 0 | 0 | 123 | 0 | 114 | 0 | 0 | 883 | 364 | 0 |
| SRR3103253 | 17 | 26 | 0 | 0 | 78 | 55 | 167 | 344 | 30 | 0 | 17 | 0 | 0 | 0 | 53244 | 62376 | 342289 | 0 | 55 | 18134 | 3529 | 0 | 0 | 160 | 13 | 22 | 113 | 0 | 0 | 151 | 0 | 0 | 0 | 0 | 453 | 42 | 0 |
| SRR3103254 | 23 | 29 | 32 | 87 | 19 | 0 | 46 | 81 | 12 | 0 | 39 | 0 | 0 | 0 | 9940 | 31006 | 219932 | 0 | 48 | 23241 | 6196 | 0 | 0 | 100 | 0 | 110 | 248 | 174 | 0 | 103 | 0 | 0 | 0 | 0 | 73 | 0 | 0 |
| SRR3103255 | 21 | 23 | 14 | 30 | 21 | 178 | 382 | 494 | 93 | 0 | 19 | 10 | 0 | 0 | 56350 | 36228 | 98500 | 0 | 15 | 8455 | 111805 | 0 | 0 | 63 | 36 | 21 | 31 | 19 | 0 | 75 | 0 | 41 | 0 | 0 | 332 | 0 | 0 |
| SRR3103256 | 14 | 22 | 0 | 30 | 66 | 37 | 98 | 210 | 37 | 25 | 101 | 22 | 0 | 0 | 172826 | 20045 | 198320 | 0 | 69 | 5887 | 1305 | 0 | 0 | 164 | 25 | 156 | 22 | 41 | 0 | 188 | 0 | 97 | 0 | 0 | 33 | 0 | 0 |
| SRR3103257 | 0 | 0 | 10 | 19 | 101 | 37 | 122 | 220 | 24 | 12 | 48 | 17 | 0 | 0 | 20310 | 27260 | 198768 | 0 | 52 | 7479 | 27225 | 0 | 0 | 70 | 10 | 147 | 358 | 26 | 0 | 155 | 0 | 27 | 0 | 0 | 588 | 41 | 0 |
| SRR3103258 | 86 | 136 | 101 | 19 | 131 | 10 | 69 | 142 | 14 | 0 | 60 | 0 | 0 | 0 | 186741 | 27635 | 158846 | 14 | 35 | 4913 | 9228 | 0 | 0 | 136 | 20 | 232 | 43 | 186 | 0 | 151 | 0 | 0 | 0 | 0 | 268 | 15 | 10 |
| SRR3103259 | 0 | 0 | 0 | 31 | 18 | 38 | 60 | 251 | 23 | 46 | 54 | 0 | 0 | 0 | 17654 | 52941 | 223949 | 0 | 40 | 15198 | 2184 | 0 | 0 | 31 | 0 | 80 | 208 | 82 | 0 | 80 | 0 | 21 | 0 | 0 | 66 | 0 | 0 |
| SRR3103260 | 11 | 20 | 17 | 51 | 15 | 76 | 51 | 228 | 0 | 27 | 278 | 15 | 0 | 0 | 83212 | 14676 | 131666 | 0 | 58 | 4734 | 12049 | 0 | 0 | 19 | 31 | 88 | 130 | 33 | 0 | 666 | 0 | 23 | 0 | 0 | 233 | 0 | 0 |
| SRR3103261 | 31 | 35 | 32 | 107 | 18 | 163 | 155 | 330 | 13 | 111 | 226 | 23 | 0 | 0 | 97287 | 35234 | 121029 | 17 | 54 | 7867 | 5617 | 0 | 0 | 32 | 62 | 174 | 158 | 59 | 0 | 571 | 0 | 17 | 0 | 0 | 227 | 0 | 0 |
| SRR3103262 | 19 | 27 | 12 | 39 | 35 | 46 | 76 | 131 | 61 | 46 | 10 | 62 | 0 | 0 | 155548 | 10786 | 128389 | 0 | 90 | 14824 | 793 | 0 | 0 | 62 | 69 | 151 | 17 | 104 | 0 | 49 | 0 | 21 | 0 | 0 | 82 | 14 | 15 |
| SRR3103263 | 23 | 19 | 0 | 0 | 14 | 26 | 157 | 157 | 26 | 120 | 60 | 60 | 0 | 0 | 50209 | 7624 | 115550 | 0 | 0 | 105 | 1897 | 0 | 0 | 69 | 32 | 23 | 32 | 0 | 0 | 222 | 0 | 44 | 0 | 0 | 456 | 138 | 79 |
| SRR3103264 | 0 | 0 | 0 | 41 | 30 | 0 | 23 | 59 | 0 | 62 | 1098 | 77 | 0 | 0 | 159262 | 9458 | 190449 | 0 | 0 | 225 | 3550 | 0 | 0 | 11 | 26 | 49 | 63 | 28 | 0 | 772 | 0 | 11 | 0 | 0 | 24 | 0 | 42 |
| SRR3103265 | 0 | 17 | 17 | 46 | 252 | 20 | 53 | 66 | 19 | 23 | 34 | 26 | 0 | 0 | 67987 | 19137 | 298000 | 11 | 126 | 13691 | 8413 | 0 | 0 | 54 | 40 | 143 | 556 | 83 | 0 | 152 | 0 | 19 | 0 | 0 | 82 | 0 | 0 |
| SRR3103266 | 57 | 67 | 57 | 153 | 48 | 0 | 20 | 47 | 0 | 57 | 41 | 48 | 0 | 0 | 457435 | 8599 | 164772 | 0 | 0 | 32 | 5534 | 0 | 0 | 38 | 29 | 172 | 163 | 29 | 0 | 62 | 0 | 0 | 0 | 0 | 40 | 22 | 57 |
| SRR3103267 | 24 | 21 | 0 | 33 | 58 | 27 | 39 | 72 | 0 | 61 | 21 | 12 | 0 | 0 | 138087 | 12187 | 301664 | 12 | 60 | 7232 | 4208 | 0 | 0 | 0 | 15 | 21 | 36 | 0 | 0 | 236 | 0 | 0 | 0 | 0 | 195 | 114 | 0 |
| SRR3103268 | 20 | 0 | 26 | 26 | 215 | 24 | 12 | 57 | 0 | 78 | 179 | 26 | 0 | 0 | 187750 | 34547 | 219866 | 17 | 48 | 10597 | 7499 | 0 | 0 | 49 | 39 | 144 | 117 | 0 | 0 | 42 | 0 | 0 | 0 | 0 | 160 | 20 | 16 |
| SRR3103269 | 28 | 28 | 46 | 186 | 102 | 0 | 0 | 25 | 0 | 14 | 121 | 23 | 0 | 0 | 21129 | 14599 | 306511 | 0 | 25 | 2717 | 1310 | 0 | 0 | 65 | 121 | 418 | 502 | 116 | 0 | 11 | 0 | 0 | 0 | 0 | 55 | 54 | 0 |
| SRR3103270 | 24 | 36 | 24 | 108 | 227 | 0 | 0 | 54 | 0 | 12 | 48 | 0 | 0 | 0 | 61245 | 33871 | 382514 | 0 | 53 | 31554 | 7080 | 0 | 0 | 120 | 12 | 12 | 191 | 24 | 0 | 478 | 0 | 0 | 0 | 0 | 69 | 52 | 0 |
| SRR3103271 | 13 | 0 | 19 | 45 | 149 | 0 | 0 | 24 | 0 | 13 | 65 | 19 | 0 | 0 | 124848 | 32858 | 269870 | 42 | 166 | 18443 | 7429 | 0 | 0 | 71 | 19 | 317 | 944 | 272 | 0 | 181 | 0 | 0 | 0 | 0 | 117 | 29 | 0 |
| SRR3103272 | 0 | 0 | 0 | 19 | 19 | 0 | 0 | 39 | 0 | 19 | 19 | 0 | 19 | 0 | 105690 | 14243 | 163054 | 16 | 70 | 11309 | 6590 | 0 | 0 | 19 | 0 | 19 | 114 | 19 | 0 | 267 | 0 | 0 | 0 | 0 | 82 | 368 | 0 |
| SRR3103273 | 0 | 0 | 0 | 69 | 95 | 18 | 20 | 74 | 15 | 0 | 95 | 0 | 0 | 0 | 76074 | 41975 | 176846 | 0 | 203 | 18970 | 3102 | 0 | 0 | 104 | 43 | 424 | 1090 | 26 | 0 | 346 | 0 | 0 | 0 | 0 | 60 | 0 | 0 |
| SRR3103274 | 19 | 52 | 73 | 60 | 11 | 0 | 17 | 40 | 0 | 16 | 35 | 0 | 0 | 0 | 3719 | 27193 | 243609 | 13 | 31 | 26929 | 8286 | 0 | 0 | 114 | 24 | 136 | 312 | 33 | 0 | 128 | 0 | 0 | 0 | 0 | 100 | 0 | 0 |
| SRR3103275 | 20 | 20 | 26 | 42 | 13 | 0 | 10 | 39 | 0 | 117 | 104 | 50 | 0 | 0 | 174831 | 13961 | 235377 | 0 | 30 | 16696 | 121051 | 0 | 0 | 17 | 17 | 50 | 46 | 0 | 0 | 343 | 0 | 0 | 0 | 0 | 53 | 243 | 22 |
| SRR3103276 | 12 | 0 | 0 | 21 | 51 | 10 | 41 | 95 | 16 | 26 | 33 | 21 | 0 | 0 | 106579 | 19195 | 177943 | 0 | 79 | 16840 | 1964 | 0 | 0 | 138 | 35 | 98 | 14 | 21 | 0 | 173 | 0 | 41 | 0 | 0 | 56 | 0 | 23 |
| SRR3103277 | 0 | 12 | 12 | 24 | 77 | 0 | 11 | 32 | 0 | 12 | 18 | 0 | 0 | 0 | 30286 | 26341 | 252082 | 0 | 25 | 30715 | 26901 | 0 | 0 | 167 | 18 | 89 | 203 | 0 | 0 | 66 | 0 | 0 | 0 | 0 | 54 | 29 | 0 |
| SRR3103278 | 198 | 328 | 190 | 11 | 95 | 0 | 0 | 26 | 0 | 43 | 60 | 0 | 0 | 0 | 283142 | 34848 | 260794 | 37 | 19 | 8633 | 12071 | 0 | 0 | 147 | 69 | 319 | 13 | 86 | 0 | 147 | 0 | 0 | 0 | 0 | 49 | 0 | 0 |
| SRR3103279 | 10 | 0 | 10 | 44 | 25 | 0 | 10 | 56 | 0 | 0 | 0 | 0 | 0 | 0 | 12366 | 30968 | 283333 | 17 | 65 | 67397 | 5309 | 0 | 0 | 10 | 0 | 27 | 111 | 29 | 0 | 69 | 0 | 0 | 0 | 0 | 41 | 14 | 0 |
| SRR3103280 | 28 | 33 | 45 | 28 | 16 | 0 | 34 | 78 | 0 | 0 | 18 | 0 | 0 | 0 | 4014 | 19952 | 159413 | 0 | 97 | 9428 | 6552 | 0 | 0 | 16 | 0 | 59 | 129 | 19 | 0 | 63 | 0 | 0 | 0 | 0 | 126 | 0 | 11 |
| SRR3103281 | 56 | 105 | 62 | 738 | 553 | 12 | 30 | 72 | 0 | 0 | 24 | 0 | 0 | 0 | 4436 | 20952 | 198443 | 33 | 77 | 8640 | 5238 | 0 | 0 | 17 | 13 | 84 | 151 | 49 | 0 | 84 | 0 | 0 | 0 | 0 | 340 | 0 | 0 |
| SRR3103386 | 0 | 65 | 32 | 65 | 194 | 0 | 49 | 9267 | 49 | 97 | 0 | 0 | 0 | 0 | 31478 | 5906 | 624583 | 0 | 0 | 3481 | 1814 | 0 | 0 | 227 | 65 | 453 | 1198 | 3724 | 0 | 0 | 0 | 0 | 0 | 0 | 272 | 0 | 0 |
| SRR3103387 | 32 | 32 | 0 | 32 | 126 | 0 | 456 | 8117 | 182 | 63 | 0 | 0 | 0 | 0 | 9668 | 7513 | 454150 | 0 | 0 | 3740 | 1094 | 0 | 0 | 252 | 0 | 442 | 1388 | 3565 | 0 | 0 | 0 | 0 | 0 | 0 | 453 | 0 | 0 |
| SRR3103388 | 44 | 0 | 0 | 0 | 44 | 0 | 272 | 2138 | 170 | 0 | 0 | 0 | 0 | 0 | 25896 | 13320 | 589643 | 0 | 136 | 7093 | 2478 | 0 | 0 | 613 | 219 | 1707 | 3852 | 9147 | 0 | 0 | 0 | 0 | 0 | 0 | 118 | 0 | 0 |
| SRR3103389 | 0 | 0 | 32 | 63 | 32 | 0 | 205 | 9754 | 82 | 159 | 63 | 0 | 0 | 0 | 10246 | 5348 | 490532 | 0 | 82 | 10655 | 4057 | 0 | 0 | 13 | 127 | 25 | 59 | 137 | 0 | 63 | 0 | 0 | 0 | 0 | 217 | 0 | 0 |
| SRR3103390 | 0 | 0 | 0 | 28 | 56 | 0 | 74 | 12265 | 222 | 0 | 0 | 0 | 0 | 0 | 3621 | 4484 | 282685 | 0 | 74 | 3399 | 1330 | 0 | 0 | 251 | 56 | 1087 | 3400 | 7329 | 0 | 56 | 0 | 0 | 0 | 0 | 854 | 0 | 0 |
| SRR3103391 | 0 | 21 | 21 | 83 | 145 | 0 | 515 | 14797 | 47 | 21 | 42 | 0 | 0 | 0 | 20837 | 5848 | 364520 | 0 | 47 | 6228 | 2060 | 0 | 0 | 11 | 125 | 25 | 79 | 186 | 0 | 21 | 0 | 0 | 0 | 0 | 238 | 0 | 0 |
| SRR3103392 | 77 | 0 | 0 | 12 | 252 | 0 | 184 | 6023 | 52 | 0 | 0 | 0 | 0 | 0 | 68126 | 11279 | 649315 | 0 | 262 | 31845 | 12938 | 0 | 0 | 33 | 39 | 19 | 51 | 146 | 0 | 58 | 0 | 0 | 0 | 0 | 68 | 0 | 0 |
| SRR3103393 | 86 | 0 | 0 | 172 | 0 | 0 | 315 | 8194 | 0 | 0 | 0 | 0 | 0 | 0 | 15443 | 14386 | 677488 | 0 | 0 | 5358 | 2836 | 0 | 0 | 429 | 257 | 26 | 39 | 89 | 0 | 0 | 0 | 0 | 0 | 0 | 778 | 0 | 0 |
| SRR3103394 | 0 | 0 | 0 | 75 | 276 | 0 | 461 | 18702 | 276 | 17 | 0 | 0 | 0 | 0 | 32479 | 7894 | 583845 | 0 | 0 | 5385 | 3526 | 0 | 0 | 276 | 142 | 528 | 1206 | 3040 | 0 | 25 | 0 | 0 | 0 | 0 | 99 | 0 | 0 |
| SRR3103395 | 42 | 0 | 21 | 169 | 0 | 0 | 181 | 7966 | 121 | 42 | 0 | 0 | 0 | 0 | 43935 | 5245 | 633893 | 0 | 60 | 12191 | 6639 | 0 | 0 | 14 | 317 | 27 | 79 | 158 | 0 | 42 | 0 | 0 | 0 | 0 | 91 | 0 | 0 |
| SRR3103396 | 48 | 0 | 48 | 71 | 0 | 0 | 405 | 9238 | 442 | 24 | 24 | 0 | 0 | 0 | 18145 | 9074 | 483827 | 0 | 258 | 18844 | 6993 | 0 | 0 | 310 | 310 | 17 | 60 | 116 | 0 | 0 | 0 | 0 | 0 | 0 | 44 | 0 | 0 |
| SRR3103397 | 23 | 0 | 23 | 116 | 70 | 0 | 105 | 4222 | 174 | 0 | 0 | 0 | 0 | 0 | 28753 | 6358 | 629206 | 0 | 0 | 4990 | 6037 | 0 | 0 | 418 | 116 | 907 | 2952 | 7811 | 0 | 23 | 0 | 0 | 0 | 0 | 139 | 0 | 0 |
| SRR3103400 | 41 | 0 | 41 | 41 | 207 | 0 | 360 | 9061 | 216 | 83 | 83 | 0 | 0 | 0 | 19798 | 7762 | 493136 | 10 | 288 | 12794 | 9328 | 0 | 0 | 21 | 166 | 20 | 38 | 71 | 0 | 166 | 0 | 21 | 0 | 0 | 71 | 0 | 0 |
| SRR3103401 | 0 | 0 | 0 | 437 | 328 | 0 | 337 | 23584 | 270 | 109 | 0 | 0 | 0 | 0 | 24932 | 6454 | 496264 | 101 | 101 | 8996 | 8760 | 0 | 0 | 1422 | 328 | 1312 | 3171 | 7654 | 0 | 219 | 0 | 0 | 0 | 0 | 159 | 0 | 0 |
| SRR3103402 | 0 | 0 | 81 | 81 | 325 | 0 | 278 | 17842 | 139 | 0 | 0 | 0 | 0 | 0 | 28029 | 5716 | 520536 | 0 | 111 | 20597 | 4342 | 0 | 0 | 26 | 325 | 14 | 56 | 67 | 0 | 244 | 0 | 0 | 0 | 0 | 253 | 0 | 0 |
| SRR3103403 | 54 | 54 | 54 | 270 | 216 | 0 | 320 | 18573 | 200 | 323 | 0 | 0 | 0 | 0 | 16694 | 6388 | 362904 | 20 | 80 | 18553 | 5538 | 0 | 0 | 18 | 323 | 18 | 52 | 54 | 0 | 485 | 0 | 0 | 0 | 0 | 443 | 0 | 0 |
| SRR3103404 | 0 | 0 | 0 | 171 | 114 | 0 | 219 | 9242 | 122 | 0 | 0 | 0 | 0 | 0 | 41394 | 6525 | 589714 | 12 | 438 | 16003 | 16806 | 0 | 0 | 19 | 342 | 18 | 53 | 94 | 0 | 0 | 0 | 0 | 0 | 0 | 91 | 0 | 0 |
| SRR3103405 | 214 | 0 | 0 | 161 | 214 | 0 | 432 | 11343 | 84 | 0 | 54 | 0 | 0 | 0 | 81156 | 8190 | 652023 | 0 | 530 | 18910 | 19383 | 0 | 0 | 22 | 268 | 19 | 40 | 74 | 0 | 107 | 0 | 0 | 0 | 0 | 47 | 0 | 0 |
| SRR3103406 | 0 | 0 | 0 | 0 | 0 | 0 | 1081 | 29926 | 701 | 17 | 67 | 0 | 0 | 0 | 7452 | 3108 | 131726 | 0 | 175 | 7189 | 2367 | 0 | 0 | 17 | 200 | 27 | 96 | 109 | 0 | 33 | 0 | 0 | 33 | 0 | 252 | 0 | 0 |
| SRR3103407 | 117 | 0 | 0 | 0 | 292 | 0 | 80 | 5192 | 171 | 350 | 0 | 0 | 0 | 0 | 24672 | 14306 | 539161 | 10 | 321 | 10895 | 8345 | 0 | 0 | 21 | 233 | 14 | 36 | 51 | 0 | 350 | 0 | 0 | 0 | 0 | 58 | 0 | 0 |
| SRR3103408 | 114 | 0 | 0 | 0 | 0 | 0 | 360 | 19210 | 300 | 0 | 0 | 0 | 0 | 0 | 10385 | 5097 | 272399 | 0 | 240 | 6964 | 8344 | 0 | 0 | 799 | 685 | 1485 | 5254 | 4797 | 0 | 114 | 0 | 0 | 0 | 0 | 209 | 0 | 0 |
| SRR3103409 | 0 | 0 | 0 | 62 | 186 | 0 | 119 | 11613 | 357 | 10 | 62 | 0 | 0 | 0 | 6849 | 4312 | 223032 | 119 | 298 | 3633 | 2442 | 0 | 0 | 311 | 249 | 25 | 63 | 94 | 0 | 0 | 0 | 0 | 0 | 0 | 479 | 0 | 0 |
| SRR3138807 | 0 | 0 | 0 | 0 | 181 | 21 | 64 | 0 | 67 | 24 | 43447 | 0 | 0 | 0 | 11718 | 303616 | 11815 | 2665 | 571 | 2173 | 772 | 72 | 35 | 0 | 0 | 0 | 0 | 0 | 0 | 1206 | 10 | 574 | 0 | 14 | 87 | 676 | 0 |
| SRR3138808 | 0 | 0 | 0 | 0 | 37 | 13 | 35 | 0 | 76 | 16 | 23076 | 0 | 0 | 0 | 5180 | 243379 | 7774 | 4680 | 409 | 898 | 947 | 31 | 16 | 0 | 0 | 0 | 0 | 0 | 0 | 730 | 13 | 652 | 0 | 18 | 292 | 449 | 0 |
| SRR3138809 | 0 | 0 | 0 | 0 | 45 | 23 | 51 | 0 | 90 | 0 | 25732 | 0 | 0 | 0 | 4796 | 203232 | 7053 | 6360 | 440 | 784 | 847 | 38 | 20 | 0 | 0 | 0 | 0 | 0 | 0 | 862 | 17 | 380 | 0 | 19 | 353 | 467 | 0 |
| SRR3138810 | 0 | 0 | 0 | 0 | 82 | 43 | 122 | 21 | 204 | 36 | 15280 | 0 | 0 | 0 | 30636 | 195261 | 20210 | 2618 | 619 | 2095 | 617 | 14 | 0 | 0 | 0 | 0 | 0 | 0 | 0 | 735 | 0 | 498 | 0 | 22 | 89 | 713 | 0 |
| SRR3199254 | 286 | 75 | 0 | 0 | 233 | 1292 | 583 | 485 | 1589 | 0 | 911 | 0 | 18 | 18 | 124543 | 141183 | 156829 | 2294 | 82 | 41492 | 120 | 0 | 11 | 188 | 66 | 0 | 0 | 0 | 0 | 2083 | 0 | 0 | 0 | 0 | 3541 | 72 | 0 |
| SRR3199255 | 16 | 47 | 11 | 18 | 31 | 470 | 132 | 239 | 673 | 29 | 27 | 0 | 13 | 0 | 209914 | 217566 | 307096 | 1092 | 25 | 14099 | 94 | 0 | 29 | 73 | 20 | 0 | 0 | 0 | 0 | 56 | 0 | 0 | 0 | 0 | 907 | 50 | 0 |
| SRR3199256 | 232 | 70 | 0 | 0 | 58 | 467 | 101 | 115 | 647 | 17 | 141 | 0 | 25 | 21 | 447010 | 168769 | 534708 | 700 | 31 | 12641 | 391 | 0 | 12 | 203 | 79 | 174 | 141 | 29 | 0 | 1418 | 0 | 0 | 0 | 0 | 1611 | 0 | 0 |
| SRR3199257 | 471 | 80 | 0 | 10 | 171 | 741 | 434 | 930 | 535 | 30 | 80 | 0 | 10 | 0 | 79088 | 58034 | 257736 | 2395 | 28 | 8861 | 123 | 0 | 100 | 70 | 0 | 0 | 0 | 0 | 0 | 160 | 10 | 39 | 0 | 0 | 4143 | 0 | 20 |
| SRR3199258 | 215 | 45 | 0 | 0 | 59 | 393 | 202 | 173 | 601 | 59 | 48 | 0 | 0 | 0 | 225609 | 161528 | 605966 | 317 | 85 | 83591 | 270 | 36 | 277 | 53 | 11 | 0 | 0 | 0 | 0 | 12 | 0 | 0 | 0 | 0 | 2385 | 0 | 50 |
| SRR3199259 | 89 | 37 | 0 | 0 | 42 | 271 | 53 | 201 | 100 | 105 | 89 | 21 | 0 | 0 | 316703 | 109593 | 603737 | 683 | 176 | 21551 | 466 | 56 | 397 | 35 | 12 | 0 | 0 | 0 | 0 | 337 | 18 | 25 | 0 | 0 | 731 | 0 | 32 |
| SRR3199260 | 157 | 36 | 0 | 0 | 27 | 364 | 52 | 167 | 125 | 59 | 108 | 0 | 0 | 0 | 331981 | 55468 | 598506 | 1066 | 139 | 20571 | 303 | 13 | 100 | 43 | 0 | 0 | 0 | 0 | 0 | 717 | 0 | 0 | 0 | 0 | 745 | 0 | 48 |
| SRR3199261 | 64 | 27 | 0 | 0 | 27 | 551 | 154 | 652 | 202 | 43 | 107 | 0 | 0 | 0 | 207473 | 216314 | 592373 | 1039 | 192 | 29999 | 932 | 21 | 171 | 49 | 12 | 0 | 0 | 0 | 0 | 314 | 0 | 57 | 0 | 0 | 1359 | 0 | 24 |
| SRR3199262 | 230 | 43 | 0 | 0 | 39 | 519 | 190 | 355 | 315 | 39 | 172 | 16 | 12 | 0 | 142405 | 156217 | 554771 | 929 | 160 | 16166 | 904 | 12 | 97 | 55 | 0 | 0 | 0 | 0 | 0 | 300 | 20 | 25 | 0 | 0 | 2413 | 0 | 23 |
| SRR3199263 | 178 | 0 | 0 | 0 | 400 | 1078 | 283 | 763 | 566 | 0 | 311 | 0 | 178 | 44 | 7270 | 10881 | 12720 | 2018 | 27 | 9362 | 0 | 0 | 0 | 89 | 44 | 0 | 0 | 0 | 0 | 1246 | 0 | 0 | 0 | 0 | 7590 | 0 | 0 |
| SRR3235777 | 35 | 0 | 0 | 0 | 1687 | 20395 | 4555 | 11447 | 17013 | 1259 | 62 | 0 | 162 | 89 | 18950 | 66737 | 32135 | 204 | 1411 | 2949 | 484 | 0 | 0 | 54 | 81 | 0 | 23 | 0 | 0 | 328 | 0 | 25 | 0 | 0 | 5490 | 634 | 0 |
| SRR3235778 | 73 | 31 | 0 | 0 | 6155 | 2956 | 426 | 2571 | 1952 | 1843 | 54 | 0 | 103 | 38 | 31088 | 28907 | 28755 | 69 | 1031 | 2887 | 935 | 0 | 0 | 107 | 12 | 0 | 0 | 0 | 0 | 134 | 0 | 55 | 0 | 0 | 12573 | 2789 | 0 |
| SRR3235779 | 290 | 68 | 0 | 0 | 943 | 5371 | 1068 | 3458 | 3909 | 276 | 230 | 12 | 44 | 53 | 5771 | 22265 | 10418 | 248 | 2256 | 1036 | 858 | 0 | 0 | 152 | 0 | 0 | 0 | 0 | 0 | 194 | 0 | 13 | 0 | 0 | 4837 | 3514 | 0 |
| SRR3235780 | 80 | 11 | 0 | 0 | 1682 | 2568 | 931 | 4229 | 6467 | 773 | 917 | 19 | 32 | 19 | 6255 | 41824 | 14435 | 1413 | 1790 | 1166 | 825 | 19 | 0 | 53 | 11 | 0 | 0 | 0 | 16 | 619 | 159 | 141 | 0 | 48 | 4609 | 0 | 0 |
| SRR3235781 | 13 | 87 | 35 | 31 | 56 | 4723 | 1536 | 3243 | 3035 | 27 | 160 | 42 | 21 | 49 | 3016 | 17990 | 14947 | 275 | 835 | 1195 | 891 | 0 | 0 | 146 | 0 | 0 | 0 | 0 | 0 | 16 | 0 | 19 | 0 | 0 | 4651 | 317 | 0 |
| SRR3279553 | 0 | 0 | 0 | 0 | 0 | 0 | 0 | 300 | 0 | 0 | 36 | 0 | 0 | 0 | 17697 | 15050 | 82502 | 0 | 0 | 8098 | 2100 | 0 | 0 | 0 | 0 | 0 | 0 | 0 | 0 | 18 | 0 | 0 | 0 | 0 | 9066 | 710 | 0 |
| SRR3279554 | 0 | 0 | 0 | 0 | 0 | 49 | 0 | 98 | 0 | 0 | 14 | 0 | 0 | 0 | 63512 | 42861 | 210664 | 0 | 49 | 4878 | 18195 | 0 | 0 | 0 | 0 | 0 | 0 | 0 | 0 | 11 | 0 | 0 | 0 | 0 | 3502 | 176 | 0 |
| SRR3279555 | 0 | 0 | 0 | 0 | 0 | 58 | 58 | 232 | 0 | 0 | 28 | 0 | 0 | 0 | 33486 | 38386 | 168722 | 58 | 232 | 3824 | 2086 | 0 | 0 | 0 | 0 | 0 | 0 | 0 | 0 | 93 | 0 | 58 | 0 | 0 | 37145 | 168 | 0 |
| SRR3279556 | 10 | 10 | 0 | 0 | 0 | 327 | 303 | 350 | 70 | 24 | 47 | 0 | 0 | 0 | 31666 | 47480 | 225720 | 0 | 70 | 11831 | 443 | 0 | 0 | 14 | 0 | 0 | 0 | 0 | 0 | 128 | 0 | 70 | 0 | 0 | 46350 | 442 | 14 |
| SRR3279557 | 31 | 0 | 0 | 0 | 0 | 114 | 179 | 209 | 0 | 37 | 210 | 0 | 0 | 0 | 260955 | 76705 | 503216 | 14 | 103 | 4941 | 10327 | 0 | 0 | 0 | 0 | 0 | 0 | 0 | 0 | 443 | 0 | 14 | 0 | 0 | 7514 | 313 | 0 |
| SRR3279558 | 0 | 19 | 0 | 0 | 0 | 181 | 453 | 363 | 0 | 29 | 87 | 0 | 0 | 0 | 168676 | 83937 | 406385 | 0 | 272 | 16587 | 2538 | 0 | 0 | 0 | 0 | 0 | 0 | 0 | 0 | 250 | 0 | 0 | 0 | 0 | 29309 | 113 | 0 |
| SRR3279559 | 18 | 12 | 0 | 0 | 0 | 43 | 72 | 187 | 29 | 0 | 33 | 0 | 0 | 0 | 120880 | 56260 | 298354 | 201 | 72 | 15123 | 11820 | 0 | 0 | 0 | 0 | 0 | 0 | 0 | 0 | 75 | 0 | 14 | 0 | 0 | 4278 | 33 | 0 |
| SRR3279560 | 0 | 0 | 0 | 0 | 0 | 0 | 38 | 95 | 19 | 0 | 37 | 0 | 0 | 0 | 78130 | 21750 | 118253 | 57 | 19 | 8808 | 14680 | 0 | 0 | 0 | 0 | 0 | 0 | 0 | 0 | 16 | 0 | 38 | 0 | 0 | 1146 | 158 | 0 |
| SRR3279561 | 16 | 0 | 0 | 0 | 0 | 34 | 25 | 49 | 15 | 0 | 64 | 0 | 0 | 0 | 97510 | 27975 | 286197 | 69 | 59 | 21620 | 26098 | 0 | 0 | 0 | 0 | 0 | 0 | 0 | 0 | 550 | 0 | 29 | 0 | 0 | 1717 | 366 | 0 |
| SRR3279562 | 0 | 0 | 0 | 0 | 0 | 50 | 101 | 184 | 17 | 0 | 13 | 0 | 0 | 0 | 50349 | 66016 | 98566 | 34 | 50 | 5564 | 14364 | 0 | 0 | 0 | 0 | 0 | 0 | 0 | 0 | 61 | 0 | 84 | 0 | 0 | 7682 | 398 | 0 |
| SRR3279563 | 0 | 23 | 0 | 0 | 0 | 77 | 77 | 230 | 0 | 0 | 12 | 0 | 0 | 0 | 61638 | 25524 | 61503 | 153 | 230 | 9966 | 19703 | 0 | 0 | 0 | 0 | 0 | 0 | 0 | 0 | 22 | 0 | 0 | 0 | 0 | 4833 | 216 | 0 |
| SRR3279564 | 14 | 0 | 0 | 0 | 0 | 20 | 26 | 90 | 12 | 14 | 85 | 0 | 0 | 0 | 187868 | 18302 | 268466 | 0 | 0 | 4378 | 12610 | 0 | 0 | 0 | 0 | 0 | 0 | 0 | 0 | 177 | 0 | 32 | 0 | 0 | 635 | 189 | 0 |
| SRR3279565 | 0 | 17 | 0 | 0 | 0 | 115 | 104 | 253 | 0 | 0 | 125 | 0 | 0 | 0 | 44172 | 30584 | 250806 | 0 | 23 | 16083 | 8099 | 0 | 0 | 0 | 0 | 0 | 0 | 0 | 0 | 392 | 0 | 104 | 0 | 0 | 14702 | 433 | 0 |
| SRR3279566 | 0 | 0 | 0 | 0 | 0 | 0 | 167 | 209 | 0 | 0 | 25 | 0 | 0 | 0 | 46176 | 35854 | 226071 | 42 | 126 | 3224 | 4354 | 0 | 0 | 0 | 0 | 0 | 0 | 0 | 0 | 23 | 0 | 42 | 0 | 0 | 15309 | 246 | 0 |
| SRR3279567 | 11 | 24 | 0 | 0 | 0 | 81 | 127 | 226 | 18 | 27 | 15 | 0 | 0 | 0 | 219315 | 33969 | 572259 | 18 | 54 | 26397 | 9037 | 0 | 0 | 0 | 0 | 0 | 0 | 0 | 0 | 194 | 0 | 27 | 0 | 0 | 24858 | 280 | 0 |
| SRR3279568 | 0 | 0 | 0 | 0 | 0 | 49 | 25 | 123 | 0 | 0 | 195 | 0 | 0 | 0 | 48151 | 11957 | 112209 | 74 | 25 | 16961 | 10192 | 0 | 0 | 0 | 0 | 0 | 0 | 0 | 0 | 181 | 0 | 0 | 0 | 0 | 1683 | 119 | 0 |
| SRR3279569 | 0 | 32 | 0 | 0 | 0 | 18 | 37 | 110 | 0 | 0 | 10 | 0 | 0 | 0 | 75626 | 12162 | 110472 | 37 | 0 | 8934 | 28101 | 0 | 0 | 0 | 0 | 0 | 0 | 0 | 0 | 13 | 0 | 0 | 0 | 0 | 435 | 115 | 0 |
| SRR3279570 | 0 | 0 | 0 | 0 | 0 | 201 | 201 | 251 | 0 | 0 | 84 | 0 | 0 | 0 | 4575 | 19462 | 66038 | 0 | 50 | 35246 | 2263 | 0 | 0 | 0 | 0 | 0 | 0 | 0 | 0 | 217 | 0 | 0 | 0 | 0 | 10757 | 84 | 0 |
| SRR3279571 | 0 | 27 | 0 | 0 | 0 | 50 | 50 | 100 | 0 | 14 | 68 | 0 | 0 | 14 | 144925 | 42100 | 311391 | 17 | 17 | 6876 | 17267 | 0 | 0 | 0 | 0 | 0 | 0 | 0 | 0 | 68 | 0 | 0 | 0 | 0 | 1127 | 66 | 0 |

### Supp Table S8.3.Columns_BX_to_DH

| | | | | | | | | | | | | | | | | | | | | | | | | | | | | | | | | | | | | | |
|---|---|---|---|---|---|---|---|---|---|---|---|---|---|---|---|---|---|---|---|---|---|---|---|---|---|---|---|---|---|---|---|---|---|---|---|---|---|
| SRR3279572 | 17 | 0 | 0 | 0 | 0 | 45 | 81 | 179 | 36 | 11 | 13 | 0 | 0 | 0 | 107498 | 33380 | 292709 | 206 | 18 | 10774 | 43296 | 0 | 0 | 0 | 0 | 0 | 0 | 0 | 0 | 73 | 0 | 27 | 0 | 0 | 2033 | 235 | 0 |
| SRR3279573 | 0 | 14 | 0 | 0 | 0 | 18 | 128 | 128 | 0 | 0 | 12 | 0 | 0 | 0 | 128878 | 35468 | 270604 | 0 | 18 | 10500 | 3488 | 0 | 0 | 0 | 0 | 0 | 0 | 0 | 0 | 13 | 0 | 0 | 0 | 0 | 9492 | 307 | 0 |
| SRR3279574 | 0 | 20 | 0 | 0 | 0 | 70 | 211 | 164 | 0 | 0 | 16 | 0 | 0 | 0 | 56307 | 38472 | 227443 | 0 | 47 | 12591 | 13202 | 0 | 0 | 0 | 0 | 0 | 0 | 0 | 0 | 37 | 0 | 23 | 0 | 0 | 6642 | 351 | 0 |
| SRR3279575 | 11 | 13 | 0 | 0 | 0 | 71 | 102 | 173 | 10 | 13 | 15 | 0 | 0 | 0 | 158745 | 55750 | 463055 | 36 | 71 | 11504 | 10251 | 0 | 0 | 0 | 0 | 0 | 0 | 0 | 0 | 41 | 0 | 41 | 0 | 0 | 4918 | 381 | 11 |
| SRR3279576 | 0 | 0 | 0 | 0 | 0 | 45 | 83 | 70 | 0 | 27 | 76 | 0 | 0 | 0 | 218070 | 32947 | 394801 | 0 | 243 | 10538 | 7634 | 0 | 0 | 0 | 0 | 0 | 0 | 0 | 0 | 145 | 0 | 32 | 0 | 0 | 11509 | 324 | 0 |
| SRR3279577 | 0 | 20 | 0 | 0 | 0 | 24 | 49 | 130 | 24 | 12 | 66 | 12 | 0 | 0 | 82725 | 16156 | 180509 | 0 | 178 | 5216 | 11340 | 0 | 0 | 0 | 0 | 0 | 0 | 0 | 0 | 186 | 0 | 24 | 0 | 0 | 1677 | 316 | 0 |
| SRR3279578 | 0 | 0 | 0 | 0 | 0 | 68 | 68 | 287 | 14 | 0 | 123 | 0 | 0 | 0 | 32304 | 42346 | 243861 | 96 | 151 | 20784 | 21933 | 0 | 0 | 0 | 0 | 0 | 0 | 0 | 0 | 373 | 0 | 27 | 0 | 0 | 5702 | 141 | 0 |
| SRR3279579 | 0 | 15 | 0 | 0 | 0 | 0 | 41 | 205 | 41 | 0 | 24 | 0 | 0 | 0 | 67264 | 26341 | 79220 | 41 | 82 | 3573 | 11744 | 0 | 0 | 0 | 0 | 0 | 0 | 0 | 0 | 12 | 0 | 82 | 0 | 0 | 1854 | 209 | 0 |
| SRR3279580 | 0 | 0 | 0 | 0 | 0 | 134 | 106 | 236 | 31 | 42 | 15 | 0 | 0 | 0 | 144825 | 33169 | 450100 | 12 | 114 | 11015 | 5140 | 0 | 0 | 0 | 0 | 0 | 0 | 0 | 0 | 125 | 0 | 20 | 0 | 0 | 7867 | 147 | 18 |
| SRR3279581 | 0 | 0 | 0 | 0 | 0 | 0 | 16 | 28 | 0 | 34 | 249 | 0 | 0 | 0 | 107136 | 25710 | 473289 | 32 | 21 | 7322 | 90081 | 0 | 0 | 0 | 0 | 0 | 0 | 0 | 0 | 841 | 0 | 0 | 0 | 0 | 1213 | 839 | 0 |
| SRR3279582 | 0 | 0 | 0 | 0 | 0 | 12 | 12 | 12 | 0 | 0 | 65 | 0 | 0 | 0 | 211267 | 24177 | 357147 | 45 | 25 | 1944 | 30855 | 0 | 0 | 0 | 0 | 0 | 0 | 0 | 0 | 90 | 0 | 0 | 0 | 0 | 1008 | 315 | 0 |
| SRR3279583 | 0 | 14 | 0 | 0 | 0 | 48 | 161 | 321 | 16 | 0 | 138 | 0 | 0 | 0 | 71041 | 30925 | 249924 | 32 | 80 | 19749 | 16631 | 0 | 0 | 0 | 0 | 0 | 0 | 0 | 0 | 520 | 0 | 48 | 0 | 0 | 7296 | 277 | 0 |
| SRR3279584 | 0 | 0 | 0 | 12 | 0 | 44 | 97 | 173 | 17 | 13 | 73 | 0 | 0 | 0 | 73255 | 117811 | 419605 | 48 | 99 | 8411 | 9266 | 0 | 0 | 37 | 0 | 0 | 0 | 0 | 0 | 393 | 0 | 26 | 0 | 0 | 3437 | 859 | 0 |
| SRR3279585 | 0 | 0 | 0 | 0 | 0 | 32 | 97 | 259 | 65 | 0 | 26 | 0 | 0 | 0 | 40213 | 46420 | 146091 | 32 | 65 | 8211 | 15419 | 0 | 0 | 0 | 0 | 0 | 0 | 0 | 0 | 34 | 0 | 0 | 0 | 0 | 4229 | 128 | 0 |
| SRR3279586 | 0 | 0 | 0 | 0 | 0 | 0 | 41 | 41 | 0 | 0 | 10 | 0 | 0 | 0 | 49533 | 33613 | 235059 | 0 | 122 | 10402 | 15360 | 0 | 0 | 0 | 0 | 0 | 0 | 0 | 0 | 15 | 0 | 0 | 0 | 0 | 4073 | 256 | 0 |
| SRR3317221 | 179 | 0 | 0 | 0 | 251 | 8965 | 4890 | 26080 | 18745 | 18 | 0 | 36 | 36 | 0 | 4890 | 28591 | 8169 | 815 | 408 | 2037 | 6520 | 0 | 0 | 36 | 0 | 0 | 0 | 0 | 0 | 0 | 0 | 0 | 0 | 0 | 19741 | 2514 | 0 |
| SRR3477269 | 0 | 0 | 0 | 0 | 0 | 0 | 0 | 0 | 0 | 340 | 0 | 0 | 0 | 0 | 1007 | 2959 | 27792 | 0 | 0 | 45 | 437 | 0 | 0 | 0 | 0 | 0 | 0 | 0 | 0 | 0 | 0 | 0 | 0 | 0 | 2114 | 0 | 0 |
| SRR3477271 | 0 | 0 | 0 | 0 | 0 | 0 | 0 | 0 | 0 | 349 | 0 | 0 | 0 | 0 | 698 | 2514 | 21651 | 0 | 0 | 27 | 400 | 0 | 0 | 0 | 0 | 0 | 0 | 0 | 0 | 0 | 0 | 0 | 0 | 0 | 1050 | 0 | 0 |
| SRR3490374 | 63 | 0 | 0 | 0 | 189 | 0 | 148 | 740 | 0 | 0 | 63 | 0 | 0 | 0 | 48535 | 47819 | 431120 | 0 | 296 | 11986 | 13170 | 0 | 0 | 0 | 0 | 0 | 0 | 0 | 0 | 63 | 0 | 0 | 0 | 0 | 537 | 0 | 0 |
| SRR3490375 | 0 | 0 | 290 | 0 | 580 | 0 | 296 | 889 | 0 | 0 | 290 | 0 | 0 | 0 | 65185 | 51852 | 590123 | 0 | 593 | 29037 | 12148 | 0 | 0 | 0 | 0 | 1160 | 11 | 580 | 0 | 0 | 0 | 0 | 0 | 0 | 1152 | 0 | 0 |
| SRR3490376 | 15 | 0 | 0 | 30 | 74 | 0 | 260 | 1353 | 156 | 0 | 59 | 0 | 0 | 0 | 37261 | 22829 | 207907 | 0 | 312 | 28934 | 2238 | 0 | 0 | 55 | 15 | 18 | 55 | 11 | 0 | 35 | 0 | 0 | 0 | 0 | 129 | 0 | 0 |
| SRR3490377 | 0 | 0 | 0 | 0 | 0 | 0 | 248 | 2730 | 99 | 0 | 18 | 0 | 0 | 0 | 40897 | 11863 | 182152 | 0 | 298 | 13500 | 4219 | 0 | 0 | 16 | 27 | 0 | 0 | 0 | 0 | 20 | 0 | 0 | 0 | 0 | 268 | 0 | 0 |
| SRR3490378 | 0 | 0 | 24 | 0 | 24 | 0 | 430 | 2752 | 430 | 0 | 71 | 24 | 0 | 0 | 3440 | 36085 | 129984 | 0 | 602 | 33196 | 344 | 0 | 0 | 214 | 24 | 262 | 548 | 143 | 0 | 119 | 0 | 0 | 0 | 0 | 1794 | 0 | 0 |
| SRR3490379 | 0 | 0 | 0 | 275 | 92 | 0 | 1176 | 2548 | 980 | 0 | 0 | 0 | 0 | 0 | 2352 | 30220 | 156899 | 0 | 0 | 33908 | 2156 | 0 | 0 | 36 | 92 | 10 | 19 | 458 | 0 | 0 | 0 | 0 | 0 | 0 | 3358 | 0 | 0 |
| SRR3490380 | 0 | 0 | 0 | 0 | 171 | 0 | 934 | 4515 | 934 | 0 | 43 | 128 | 0 | 0 | 1557 | 39751 | 93870 | 0 | 156 | 29426 | 467 | 0 | 0 | 38 | 0 | 19 | 21 | 23 | 0 | 128 | 0 | 0 | 0 | 0 | 4310 | 0 | 0 |
| SRR3490382 | 0 | 0 | 0 | 202 | 202 | 0 | 569 | 569 | 0 | 0 | 0 | 0 | 0 | 0 | 80296 | 71994 | 439698 | 0 | 0 | 9681 | 2278 | 0 | 0 | 11 | 0 | 0 | 605 | 11 | 0 | 0 | 0 | 0 | 0 | 0 | 0 | 234 | 0 |
| SRR3490383 | 0 | 0 | 0 | 0 | 25 | 0 | 0 | 1289 | 0 | 13 | 25 | 20 | 0 | 0 | 8812 | 42290 | 160044 | 0 | 0 | 4943 | 645 | 0 | 0 | 11 | 0 | 13 | 50 | 38 | 25 | 75 | 0 | 0 | 0 | 0 | 18228 | 198 | 50 |
| SRR3490384 | 0 | 0 | 93 | 0 | 326 | 0 | 91 | 998 | 0 | 0 | 47 | 0 | 0 | 0 | 40907 | 40031 | 317506 | 0 | 272 | 12245 | 3991 | 0 | 0 | 512 | 0 | 140 | 838 | 279 | 0 | 326 | 0 | 0 | 0 | 0 | 468 | 316 | 0 |
| SRR3490385 | 51 | 0 | 152 | 101 | 101 | 0 | 329 | 658 | 329 | 51 | 0 | 0 | 0 | 0 | 17111 | 136276 | 208733 | 0 | 329 | 3949 | 987 | 0 | 0 | 16 | 101 | 0 | 203 | 101 | 0 | 304 | 0 | 0 | 0 | 0 | 12956 | 171 | 0 |
| SRR3490386 | 0 | 50 | 0 | 0 | 0 | 0 | 132 | 2245 | 0 | 0 | 0 | 0 | 0 | 0 | 117771 | 51686 | 447993 | 0 | 132 | 11751 | 3697 | 0 | 0 | 399 | 50 | 100 | 250 | 349 | 0 | 200 | 0 | 0 | 0 | 0 | 784 | 186 | 0 |
| SRR3490387 | 0 | 0 | 36 | 145 | 327 | 0 | 227 | 453 | 76 | 0 | 145 | 36 | 0 | 0 | 80420 | 32716 | 445269 | 0 | 76 | 5361 | 4531 | 0 | 0 | 399 | 0 | 145 | 109 | 399 | 0 | 254 | 0 | 0 | 0 | 0 | 307 | 156 | 36 |
| SRR3490388 | 0 | 200 | 14 | 200 | 0 | 0 | 0 | 0 | 0 | 0 | 0 | 0 | 0 | 0 | 228429 | 70201 | 653490 | 0 | 454 | 6812 | 2271 | 0 | 0 | 27 | 0 | 200 | 36 | 200 | 0 | 0 | 0 | 0 | 0 | 0 | 529 | 15 | 0 |
| SRR3490389 | 77 | 0 | 0 | 306 | 383 | 0 | 165 | 329 | 0 | 0 | 0 | 0 | 0 | 0 | 142292 | 142454 | 670575 | 0 | 0 | 3788 | 2470 | 0 | 0 | 536 | 0 | 460 | 996 | 613 | 0 | 536 | 0 | 0 | 0 | 0 | 1362 | 89 | 0 |
| SRR3490390 | 0 | 65 | 0 | 65 | 18 | 0 | 0 | 0 | 0 | 0 | 65 | 0 | 0 | 0 | 28763 | 36253 | 311061 | 0 | 0 | 11368 | 959 | 0 | 0 | 35 | 0 | 18 | 165 | 35 | 0 | 12 | 0 | 0 | 0 | 0 | 284 | 0 | 0 |
| SRR3490391 | 0 | 0 | 0 | 0 | 242 | 0 | 0 | 907 | 0 | 0 | 242 | 0 | 0 | 0 | 141561 | 45799 | 466154 | 0 | 0 | 6806 | 6352 | 0 | 0 | 0 | 0 | 242 | 59 | 13 | 0 | 0 | 0 | 0 | 0 | 0 | 0 | 12 | 0 |
| SRR3490392 | 17 | 17 | 17 | 34 | 429 | 0 | 105 | 891 | 0 | 0 | 34 | 17 | 0 | 0 | 267634 | 22410 | 803909 | 0 | 105 | 1887 | 9747 | 0 | 0 | 69 | 0 | 17 | 223 | 51 | 0 | 51 | 0 | 0 | 0 | 0 | 980 | 36 | 0 |
| SRR3490393 | 0 | 23 | 141 | 70 | 399 | 0 | 174 | 927 | 116 | 23 | 47 | 0 | 23 | 0 | 55964 | 36816 | 450344 | 0 | 521 | 14020 | 5330 | 0 | 0 | 493 | 141 | 141 | 1057 | 188 | 0 | 235 | 0 | 0 | 0 | 0 | 1168 | 145 | 47 |
| SRR3490394 | 0 | 0 | 0 | 266 | 531 | 0 | 498 | 456 | 249 | 0 | 0 | 0 | 0 | 0 | 39778 | 21717 | 238045 | 0 | 83 | 12900 | 4687 | 0 | 0 | 332 | 33 | 133 | 498 | 100 | 0 | 166 | 0 | 0 | 0 | 0 | 1166 | 91 | 0 |
| SRR3493372 | 0 | 0 | 0 | 0 | 29 | 36 | 33 | 73 | 119 | 2190 | 30 | 0 | 0 | 0 | 563022 | 32438 | 360037 | 0 | 53 | 363 | 946 | 0 | 0 | 0 | 0 | 0 | 0 | 0 | 0 | 12 | 0 | 0 | 0 | 53 | 703 | 5217 | 0 |
| SRR3493373 | 0 | 0 | 0 | 0 | 14 | 229 | 96 | 265 | 456 | 1671 | 18 | 0 | 0 | 0 | 247955 | 37198 | 139263 | 67 | 80 | 670 | 2242 | 0 | 0 | 0 | 0 | 0 | 0 | 0 | 0 | 43 | 0 | 57 | 0 | 14 | 2280 | 6940 | 0 |
| SRR3534933 | 89 | 126 | 35 | 54 | 20 | 121 | 128 | 460 | 102 | 109 | 25 | 0 | 0 | 0 | 6957 | 2216 | 10878 | 1608 | 101 | 2326 | 761 | 0 | 0 | 100 | 10 | 21 | 59 | 134 | 0 | 13 | 0 | 3801 | 42 | 0 | 1760 | 996 | 71 |
| SRR3534934 | 95 | 89 | 21 | 37 | 37 | 198 | 197 | 538 | 78 | 42 | 22 | 0 | 0 | 0 | 31237 | 3731 | 42438 | 1307 | 295 | 4251 | 2398 | 0 | 0 | 151 | 11 | 20 | 48 | 148 | 0 | 12 | 0 | 1626 | 11 | 0 | 2496 | 981 | 11 |
| SRR3534935 | 62 | 85 | 31 | 43 | 75 | 212 | 291 | 583 | 183 | 367 | 18 | 0 | 0 | 0 | 11201 | 5216 | 21934 | 246 | 176 | 4987 | 326 | 0 | 0 | 143 | 159 | 430 | 1573 | 178 | 0 | 220 | 0 | 356 | 0 | 0 | 3225 | 714 | 0 |
| SRR3534936 | 110 | 62 | 24 | 42 | 110 | 169 | 341 | 551 | 126 | 0 | 20 | 0 | 0 | 0 | 4954 | 4628 | 10310 | 288 | 221 | 4019 | 249 | 0 | 0 | 166 | 170 | 22 | 68 | 12 | 0 | 22 | 0 | 247 | 0 | 0 | 4560 | 750 | 0 |
| SRR3534937 | 46 | 35 | 53 | 72 | 11 | 692 | 1083 | 2862 | 377 | 0 | 81 | 0 | 0 | 0 | 19457 | 11179 | 37704 | 523 | 139 | 5465 | 1295 | 0 | 0 | 13 | 32 | 88 | 330 | 28 | 0 | 28 | 0 | 961 | 19 | 0 | 2272 | 774 | 30 |
| SRR3534938 | 89 | 71 | 48 | 39 | 150 | 483 | 863 | 1966 | 437 | 0 | 32 | 0 | 0 | 0 | 13122 | 7506 | 22134 | 316 | 155 | 5570 | 1129 | 0 | 0 | 186 | 383 | 817 | 2877 | 221 | 0 | 376 | 0 | 436 | 69 | 0 | 3109 | 748 | 32 |
| SRR3566024 | 11 | 21 | 0 | 0 | 0 | 239 | 96 | 430 | 0 | 0 | 46 | 0 | 0 | 0 | 10029 | 33672 | 15049 | 96 | 0 | 0 | 239 | 0 | 0 | 18 | 11 | 21 | 28 | 17 | 0 | 284 | 0 | 0 | 0 | 0 | 1505 | 0 | 0 |
| SRR3566026 | 26 | 26 | 28 | 0 | 47 | 62 | 187 | 218 | 0 | 0 | 63 | 0 | 0 | 0 | 9517 | 36786 | 17969 | 0 | 0 | 31 | 250 | 0 | 0 | 23 | 24 | 45 | 42 | 15 | 0 | 153 | 0 | 0 | 0 | 0 | 2314 | 0 | 0 |
| SRR3566150 | 21 | 47 | 0 | 10 | 0 | 243 | 223 | 465 | 0 | 0 | 784 | 0 | 0 | 0 | 5989 | 46253 | 14478 | 61 | 0 | 0 | 61 | 0 | 0 | 115 | 188 | 319 | 235 | 37 | 0 | 2316 | 0 | 0 | 0 | 0 | 1344 | 0 | 26 |
| SRR3566151 | 22 | 44 | 0 | 0 | 0 | 389 | 412 | 800 | 114 | 0 | 504 | 0 | 0 | 0 | 7364 | 33885 | 14867 | 46 | 0 | 0 | 91 | 0 | 0 | 88 | 137 | 361 | 186 | 22 | 0 | 2047 | 0 | 0 | 0 | 0 | 1103 | 0 | 38 |
| SRR3566152 | 16 | 39 | 0 | 0 | 0 | 191 | 447 | 351 | 191 | 0 | 412 | 0 | 0 | 0 | 7049 | 19095 | 11707 | 0 | 0 | 0 | 0 | 0 | 0 | 101 | 0 | 272 | 280 | 16 | 0 | 2652 | 0 | 0 | 0 | 0 | 1816 | 0 | 47 |
| SRR3566153 | 0 | 0 | 0 | 0 | 27 | 250 | 556 | 1168 | 139 | 0 | 144 | 0 | 0 | 0 | 4782 | 21764 | 8729 | 28 | 0 | 0 | 56 | 20 | 0 | 15 | 44 | 40 | 31 | 20 | 0 | 598 | 0 | 0 | 0 | 0 | 3834 | 0 | 16 |
| SRR3566154 | 0 | 0 | 0 | 0 | 0 | 0 | 903 | 903 | 0 | 0 | 149 | 0 | 0 | 0 | 2409 | 10801 | 4447 | 0 | 0 | 0 | 903 | 0 | 0 | 0 | 0 | 30 | 74 | 0 | 0 | 313 | 0 | 0 | 0 | 0 | 2541 | 0 | 0 |
| SRR3566155 | 0 | 0 | 0 | 0 | 53 | 0 | 701 | 701 | 0 | 0 | 212 | 0 | 0 | 0 | 1870 | 15681 | 13889 | 0 | 0 | 0 | 234 | 0 | 0 | 0 | 318 | 0 | 53 | 0 | 0 | 2068 | 0 | 0 | 0 | 0 | 4480 | 134 | 0 |
| SRR3582392 | 248 | 106 | 0 | 0 | 0 | 141 | 211 | 70 | 141 | 0 | 212 | 0 | 0 | 0 | 18215 | 38546 | 26589 | 0 | 0 | 0 | 352 | 0 | 0 | 194 | 53 | 159 | 194 | 71 | 0 | 583 | 0 | 0 | 0 | 0 | 2517 | 0 | 0 |
| SRR3582393 | 29 | 34 | 0 | 0 | 39 | 396 | 316 | 791 | 79 | 0 | 32 | 0 | 0 | 0 | 18989 | 39179 | 23740 | 0 | 0 | 158 | 396 | 0 | 0 | 24 | 58 | 18 | 24 | 39 | 0 | 100 | 0 | 0 | 0 | 0 | 2185 | 0 | 0 |
| SRR3623374 | 0 | 0 | 0 | 0 | 0 | 2586 | 476 | 813 | 138 | 0 | 0 | 0 | 0 | 0 | 0 | 336 | 702 | 0 | 0 | 26 | 0 | 0 | 0 | 446 | 73 | 52 | 218 | 0 | 0 | 41 | 0 | 0 | 0 | 0 | 2564 | 0 | 0 |
| SRR3624069 | 150 | 16 | 25 | 0 | 66 | 6860 | 2649 | 14275 | 21836 | 397 | 444 | 19 | 11 | 0 | 40259 | 31288 | 38165 | 10017 | 799 | 2129 | 2794 | 0 | 0 | 45 | 64 | 35 | 17 | 20 | 0 | 2011 | 0 | 1570 | 0 | 58 | 22641 | 12227 | 0 |
| SRR3624070 | 226 | 19 | 23 | 11 | 56 | 5148 | 1934 | 10601 | 15091 | 314 | 235 | 26 | 11 | 21 | 49292 | 119567 | 77537 | 3090 | 466 | 2048 | 5484 | 0 | 0 | 23 | 24 | 18 | 14 | 42 | 0 | 822 | 0 | 466 | 0 | 19 | 9207 | 11327 | 0 |
| SRR3624071 | 175 | 20 | 14 | 0 | 38 | 11110 | 4864 | 23508 | 27499 | 171 | 147 | 23 | 15 | 20 | 53915 | 30979 | 116832 | 2136 | 648 | 2272 | 4140 | 0 | 0 | 24 | 29 | 14 | 31 | 0 | 0 | 1001 | 0 | 107 | 0 | 15 | 7804 | 12969 | 0 |
| SRR3624072 | 299 | 25 | 40 | 15 | 39 | 4383 | 1097 | 6385 | 8370 | 159 | 145 | 20 | 15 | 23 | 92796 | 42952 | 185477 | 1825 | 581 | 1306 | 5906 | 0 | 0 | 16 | 20 | 16 | 38 | 11 | 0 | 426 | 0 | 191 | 0 | 12 | 7499 | 9361 | 0 |
| SRR3624073 | 104 | 11 | 26 | 0 | 31 | 10683 | 4067 | 22140 | 33805 | 201 | 66 | 17 | 11 | 19 | 93575 | 51110 | 225538 | 1616 | 659 | 2158 | 4741 | 0 | 0 | 26 | 30 | 13 | 25 | 0 | 0 | 458 | 0 | 48 | 0 | 31 | 4927 | 11255 | 0 |
| SRR3624074 | 1072 | 103 | 70 | 0 | 149 | 14460 | 5401 | 26312 | 33460 | 872 | 275 | 20 | 61 | 23 | 97914 | 33737 | 198837 | 1551 | 761 | 2326 | 3597 | 0 | 0 | 111 | 106 | 50 | 25 | 0 | 0 | 2307 | 0 | 20 | 0 | 16 | 4464 | 11567 | 0 |
| SRR3675003 | 0 | 0 | 0 | 0 | 25 | 193 | 609 | 967 | 197 | 19 | 117 | 0 | 0 | 0 | 80850 | 34629 | 214776 | 31 | 2261 | 23930 | 31701 | 0 | 0 | 0 | 0 | 0 | 0 | 0 | 0 | 219 | 0 | 0 | 0 | 0 | 132 | 0 | 0 |
| SRR3675004 | 0 | 0 | 0 | 12 | 28 | 191 | 361 | 880 | 152 | 22 | 57 | 0 | 0 | 0 | 229557 | 27593 | 326616 | 26 | 1736 | 10716 | 25266 | 0 | 0 | 0 | 0 | 0 | 0 | 0 | 0 | 41 | 0 | 0 | 0 | 0 | 67 | 0 | 0 |
| SRR3675005 | 231 | 110 | 51 | 86 | 342 | 149 | 609 | 420 | 139 | 28 | 137 | 0 | 12 | 0 | 23781 | 209702 | 25150 | 0 | 27 | 3330 | 1622 | 0 | 0 | 197 | 0 | 21 | 15 | 24 | 0 | 413 | 0 | 0 | 0 | 0 | 2696 | 0 | 0 |
| SRR3675006 | 0 | 0 | 0 | 14 | 22 | 155 | 448 | 734 | 157 | 28 | 250 | 0 | 0 | 0 | 180844 | 35502 | 282436 | 23 | 1456 | 15829 | 33629 | 0 | 0 | 0 | 0 | 0 | 0 | 0 | 0 | 171 | 0 | 0 | 0 | 0 | 77 | 0 | 0 |
| SRR3675007 | 12 | 0 | 0 | 15 | 28 | 158 | 523 | 743 | 134 | 15 | 26 | 0 | 0 | 0 | 299176 | 35779 | 372956 | 0 | 1996 | 13165 | 21806 | 0 | 0 | 0 | 0 | 0 | 0 | 0 | 0 | 37 | 0 | 0 | 0 | 0 | 58 | 0 | 0 |
| SRR3675008 | 0 | 0 | 0 | 18 | 27 | 181 | 326 | 806 | 120 | 13 | 61 | 0 | 0 | 0 | 206779 | 34908 | 298120 | 15 | 1417 | 10957 | 30610 | 0 | 0 | 0 | 0 | 11 | 0 | 0 | 0 | 42 | 0 | 0 | 0 | 0 | 67 | 0 | 0 |
| SRR3713941 | 15 | 11 | 0 | 0 | 41 | 372 | 227 | 1092 | 2028 | 47 | 24 | 19 | 0 | 0 | 912 | 724 | 439 | 1389 | 0 | 76 | 46 | 39 | 10 | 28 | 30 | 0 | 0 | 0 | 28 | 95 | 55 | 14171 | 45 | 18 | 6768 | 46 | 46 |
| SRR3713942 | 0 | 0 | 0 | 0 | 73 | 797 | 299 | 698 | 1744 | 11 | 144 | 0 | 0 | 0 | 49278 | 62674 | 201573 | 100 | 1196 | 0 | 18535 | 2331 | 670 | 18 | 11 | 18 | 0 | 0 | 0 | 25 | 1114 | 50 | 15 | 12 | 5082 | 0 | 0 |
| SRR3713943 | 25 | 0 | 0 | 0 | 34 | 566 | 400 | 1266 | 2764 | 34 | 40 | 0 | 0 | 0 | 649 | 1703 | 345 | 2231 | 50 | 416 | 100 | 123 | 49 | 0 | 0 | 0 | 0 | 0 | 159 | 102 | 162 | 29592 | 1617 | 312 | 16329 | 2770 | 247 |
| SRR3713945 | 30 | 0 | 0 | 0 | 291 | 2465 | 2707 | 8031 | 22972 | 141 | 608 | 40 | 0 | 0 | 4912 | 29028 | 39338 | 851 | 681 | 1560 | 968 | 244 | 76 | 21 | 20 | 28 | 50 | 0 | 40 | 2076 | 144 | 4006 | 151 | 71 | 5059 | 0 | 32 |
| SRR3713946 | 29 | 0 | 0 | 14 | 44 | 3609 | 2269 | 8352 | 17619 | 23 | 200 | 13 | 0 | 0 | 12726 | 55656 | 77111 | 902 | 1066 | 1804 | 2419 | 138 | 48 | 24 | 10 | 32 | 20 | 0 | 0 | 427 | 83 | 957 | 44 | 37 | 7345 | 0 | 18 |
| SRR3879751 | 0 | 0 | 0 | 0 | 70 | 406 | 0 | 186 | 41 | 177 | 0 | 34 | 0 | 0 | 311517 | 61170 | 440338 | 72 | 576 | 1669 | 28262 | 12 | 20 | 34 | 20 | 0 | 0 | 0 | 0 | 19 | 78 | 47 | 0 | 15 | 6646 | 4516 | 0 |
| SRR3879752 | 0 | 0 | 0 | 0 | 287 | 165 | 22 | 152 | 52 | 2910 | 0 | 19 | 0 | 0 | 292626 | 55790 | 402761 | 70 | 896 | 1400 | 31406 | 14 | 57 | 32 | 16 | 0 | 0 | 0 | 12 | 85 | 223 | 22 | 0 | 0 | 8744 | 5031 | 0 |
| SRR3879754 | 0 | 0 | 0 | 0 | 182 | 326 | 49 | 160 | 49 | 4077 | 0 | 0 | 0 | 0 | 294552 | 48550 | 318834 | 55 | 985 | 2247 | 5888 | 0 | 11 | 29 | 15 | 0 | 0 | 0 | 0 | 179 | 0 | 14 | 0 | 0 | 11139 | 6945 | 0 |
| SRR3879755 | 0 | 0 | 0 | 0 | 67 | 593 | 36 | 126 | 90 | 1953 | 0 | 0 | 0 | 0 | 300383 | 44900 | 303937 | 144 | 764 | 2525 | 5940 | 0 | 0 | 18 | 12 | 0 | 0 | 0 | 0 | 56 | 19 | 36 | 0 | 0 | 11456 | 6911 | 0 |
| SRR3879756 | 0 | 0 | 0 | 0 | 216 | 425 | 0 | 187 | 57 | 1405 | 0 | 0 | 0 | 0 | 243831 | 47729 | 260159 | 125 | 935 | 2442 | 7785 | 0 | 14 | 16 | 21 | 0 | 0 | 0 | 0 | 160 | 0 | 28 | 0 | 0 | 10980 | 5426 | 0 |
| SRR3879757 | 0 | 0 | 0 | 0 | 104 | 2341 | 177 | 2783 | 707 | 6570 | 0 | 0 | 0 | 0 | 187006 | 38697 | 202686 | 177 | 1369 | 2385 | 3622 | 0 | 14 | 27 | 16 | 0 | 0 | 0 | 0 | 183 | 0 | 0 | 0 | 0 | 12899 | 3374 | 0 |
| SRR3879758 | 0 | 0 | 0 | 0 | 26 | 2204 | 261 | 1980 | 560 | 2074 | 0 | 0 | 0 | 0 | 186800 | 33144 | 204053 | 37 | 896 | 1830 | 4071 | 0 | 0 | 35 | 0 | 0 | 0 | 0 | 0 | 31 | 16 | 0 | 0 | 0 | 11580 | 2763 | 0 |
| SRR3879759 | 0 | 0 | 0 | 0 | 180 | 2532 | 219 | 2701 | 338 | 4121 | 10 | 0 | 0 | 0 | 174400 | 44389 | 234239 | 101 | 1114 | 1739 | 4693 | 0 | 0 | 75 | 22 | 0 | 0 | 0 | 0 | 160 | 30 | 68 | 0 | 0 | 7079 | 2488 | 0 |
| SRR3879760 | 0 | 0 | 0 | 0 | 217 | 387 | 0 | 710 | 129 | 4939 | 0 | 0 | 0 | 0 | 195470 | 26739 | 118380 | 452 | 1033 | 839 | 4195 | 0 | 0 | 0 | 0 | 0 | 0 | 0 | 0 | 151 | 19 | 323 | 0 | 0 | 17913 | 4428 | 0 |
| SRR3879761 | 0 | 0 | 0 | 0 | 199 | 420 | 105 | 263 | 53 | 4111 | 0 | 0 | 0 | 0 | 134128 | 22215 | 86550 | 525 | 473 | 893 | 4098 | 25 | 16 | 0 | 0 | 0 | 0 | 0 | 0 | 124 | 21 | 53 | 0 | 0 | 18146 | 3670 | 0 |
| SRR3879766 | 153 | 35 | 0 | 0 | 1646 | 353 | 265 | 946 | 1324 | 267 | 0 | 0 | 10 | 0 | 3806 | 806 | 3361 | 174 | 30 | 141 | 298 | 0 | 0 | 393 | 35 | 0 | 18 | 0 | 0 | 156 | 0 | 0 | 0 | 27 | 12254 | 8701 | 0 |
| SRR3879767 | 34 | 0 | 0 | 0 | 35 | 0 | 0 | 629 | 5718 | 5221 | 0 | 14 | 105 | 20 | 858 | 1327 | 1695 | 0 | 0 | 172 | 343 | 0 | 0 | 0 | 0 | 0 | 0 | 0 | 0 | 10 | 0 | 0 | 0 | 0 | 12603 | 851 | 0 |
| SRR3879768 | 16 | 24 | 0 | 0 | 46 | 272 | 75 | 437 | 834 | 226 | 0 | 0 | 19 | 18 | 2197 | 725 | 1075 | 326 | 14 | 208 | 190 | 0 | 0 | 16 | 12 | 0 | 0 | 0 | 0 | 10 | 0 | 14 | 0 | 34 | 9264 | 2516 | 0 |
| SRR3879769 | 19 | 15 | 0 | 0 | 201 | 185 | 40 | 198 | 1412 | 290 | 0 | 0 | 17 | 31 | 1690 | 574 | 651 | 132 | 13 | 13 | 145 | 0 | 0 | 16 | 0 | 0 | 0 | 0 | 0 | 34 | 0 | 26 | 0 | 18 | 9859 | 1882 | 0 |
| SRR3879770 | 195 | 30 | 0 | 0 | 135 | 0 | 0 | 262 | 1392 | 5151 | 0 | 18 | 251 | 43 | 7625 | 1541 | 4297 | 20 | 0 | 61 | 3187 | 0 | 0 | 0 | 0 | 0 | 0 | 0 | 0 | 32 | 0 | 40 | 0 | 0 | 33492 | 16 | 0 |
| SRR3923813 | 0 | 0 | 0 | 0 | 0 | 0 | 0 | 0 | 0 | 0 | 0 | 0 | 0 | 0 | 1228 | 702 | 20645 | 24 | 0 | 5387 | 644 | 0 | 0 | 0 | 0 | 0 | 0 | 0 | 0 | 0 | 0 | 0 | 0 | 0 | 17038 | 21 | 0 |
| SRR3931357 | 113 | 120 | 30 | 113 | 251 | 509 | 501 | 1038 | 214 | 0 | 28 | 0 | 0 | 0 | 1387 | 687 | 5829 | 14 | 23 | 1317 | 723 | 0 | 0 | 673 | 347 | 875 | 2340 | 130 | 0 | 294 | 0 | 14 | 0 | 0 | 2757 | 2184 | 0 |
| SRR3931358 | 146 | 125 | 45 | 176 | 160 | 335 | 242 | 591 | 59 | 0 | 0 | 0 | 0 | 0 | 1486 | 873 | 6031 | 0 | 0 | 785 | 781 | 0 | 0 | 982 | 379 | 937 | 2416 | 118 | 0 | 126 | 0 | 0 | 0 | 0 | 2270 | 1872 | 0 |
| SRR3931359 | 62 | 49 | 47 | 197 | 20 | 2579 | 2639 | 7177 | 621 | 10 | 13 | 75 | 0 | 0 | 3931 | 19435 | 30946 | 0 | 906 | 14216 | 290 | 0 | 0 | 29 | 21 | 81 | 57 | 41 | 0 | 198 | 0 | 18 | 0 | 0 | 661 | 1174 | 11 |
| SRR3931360 | 184 | 177 | 171 | 843 | 62 | 2132 | 2377 | 6012 | 530 | 26 | 59 | 273 | 0 | 0 | 3595 | 18432 | 29826 | 13 | 602 | 14082 | 311 | 0 | 0 | 29 | 86 | 309 | 233 | 86 | 0 | 713 | 0 | 0 | 0 | 0 | 760 | 1138 | 35 |
| SRR3931984 | 0 | 0 | 0 | 0 | 0 | 1261 | 607 | 3377 | 133 | 73 | 0 | 0 | 0 | 0 | 137 | 2512 | 984 | 37 | 0 | 866 | 0 | 0 | 17 | 1079 | 0 | 0 | 0 | 0 | 0 | 0 | 0 | 340 | 0 | 0 | 18492 | 17 | 15 |
| SRR3931987 | 0 | 0 | 0 | 0 | 0 | 689 | 390 | 1296 | 127 | 64 | 0 | 0 | 0 | 0 | 37 | 3433 | 520 | 45 | 22 | 1596 | 0 | 0 | 0 | 198 | 0 | 0 | 0 | 0 | 0 | 0 | 0 | 637 | 0 | 0 | 13090 | 38 | 17 |
| SRR3931988 | 17 | 0 | 0 | 0 | 38 | 749 | 628 | 1986 | 248 | 84 | 0 | 0 | 0 | 0 | 1288 | 5725 | 4665 | 47 | 0 | 1920 | 16 | 0 | 39 | 60 | 0 | 0 | 0 | 0 | 0 | 0 | 0 | 469 | 0 | 0 | 22444 | 41 | 17 |
| SRR3992213 | 233 | 115 | 0 | 0 | 790 | 300 | 100 | 1448 | 2048 | 274 | 0 | 21 | 0 | 0 | 12437 | 34696 | 64926 | 0 | 50 | 250 | 1399 | 0 | 0 | 68 | 100 | 32 | 59 | 0 | 0 | 27 | 0 | 0 | 0 | 27 | 4294 | 0 | 0 |
| SRR3992215 | 29 | 0 | 0 | 0 | 239 | 622 | 848 | 6109 | 14085 | 43 | 0 | 15 | 12 | 0 | 16234 | 11356 | 45186 | 0 | 113 | 226 | 0 | 0 | 0 | 20 | 15 | 0 | 0 | 0 | 0 | 30 | 0 | 0 | 0 | 0 | 1419 | 0 | 0 |
| SRR3992474 | 0 | 0 | 0 | 0 | 762 | 0 | 0 | 0 | 0 | 0 | 304 | 0 | 0 | 0 | 95 | 4293 | 56244 | 63 | 20 | 2376 | 1132 | 0 | 0 | 0 | 0 | 0 | 0 | 0 | 0 | 540 | 0 | 1137 | 0 | 0 | 9537 | 0 | 17 |
| SRR3992475 | 0 | 0 | 0 | 0 | 522 | 0 | 0 | 0 | 0 | 0 | 642 | 0 | 0 | 0 | 48 | 3852 | 25707 | 96 | 19 | 1949 | 624 | 0 | 0 | 0 | 0 | 0 | 0 | 0 | 0 | 527 | 0 | 1089 | 0 | 0 | 7082 | 0 | 20 |
| SRR3992476 | 29 | 21 | 40 | 15 | 49 | 193 | 335 | 525 | 35 | 34 | 24 | 0 | 0 | 0 | 7938 | 32860 | 29872 | 511 | 32 | 1820 | 2189 | 0 | 0 | 16 | 11 | 32 | 72 | 130 | 0 | 371 | 0 | 2947 | 0 | 0 | 28268 | 0 | 0 |
| SRR3992477 | 47 | 56 | 73 | 232 | 62 | 218 | 168 | 292 | 75 | 0 | 93 | 0 | 0 | 0 | 17700 | 11030 | 70915 | 75 | 44 | 3083 | 4500 | 0 | 0 | 167 | 126 | 449 | 1304 | 1793 | 0 | 435 | 0 | 957 | 0 | 0 | 6471 | 0 | 0 |
| SRR3996364 | 34 | 54 | 0 | 13 | 0 | 0 | 1617 | 1775 | 158 | 0 | 382 | 0 | 0 | 0 | 32744 | 115469 | 42584 | 0 | 20 | 1400 | 11026 | 0 | 0 | 20 | 0 | 0 | 0 | 0 | 0 | 1575 | 0 | 0 | 0 | 0 | 7371 | 0 | 0 |
| SRR3996365 | 0 | 70 | 0 | 15 | 0 | 0 | 1270 | 1557 | 51 | 0 | 317 | 0 | 0 | 0 | 27442 | 126404 | 45851 | 0 | 34 | 1591 | 6044 | 0 | 0 | 23 | 0 | 0 | 0 | 0 | 0 | 1176 | 0 | 0 | 0 | 0 | 8957 | 0 | 0 |
| SRR3996366 | 34 | 67 | 0 | 13 | 21 | 0 | 1971 | 2439 | 176 | 0 | 26 | 0 | 0 | 0 | 53907 | 146030 | 110775 | 10 | 127 | 1980 | 6087 | 0 | 0 | 30 | 0 | 0 | 0 | 0 | 0 | 73 | 0 | 0 | 0 | 0 | 17606 | 0 | 0 |
| SRR3996367 | 35 | 76 | 14 | 28 | 14 | 0 | 2237 | 3191 | 100 | 0 | 16 | 0 | 0 | 0 | 77067 | 130255 | 119668 | 14 | 100 | 2650 | 8362 | 0 | 0 | 48 | 0 | 0 | 0 | 0 | 0 | 64 | 0 | 14 | 0 | 0 | 10725 | 0 | 0 |
| SRR3996368 | 21 | 21 | 0 | 0 | 0 | 0 | 1561 | 1340 | 39 | 0 | 154 | 0 | 0 | 0 | 60140 | 203157 | 110364 | 0 | 78 | 4554 | 3084 | 0 | 0 | 29 | 0 | 0 | 0 | 0 | 0 | 408 | 0 | 13 | 0 | 0 | 9917 | 0 | 0 |
| SRR3996369 | 0 | 0 | 0 | 0 | 0 | 0 | 1490 | 1063 | 85 | 0 | 153 | 0 | 0 | 0 | 67654 | 225612 | 145982 | 0 | 114 | 4413 | 2695 | 0 | 0 | 36 | 0 | 0 | 0 | 0 | 0 | 614 | 0 | 0 | 0 | 0 | 9934 | 0 | 0 |
| SRR4000625 | 42 | 0 | 0 | 0 | 219 | 6411 | 5155 | 24583 | 58457 | 221 | 0 | 78 | 0 | 0 | 2367 | 151277 | 5031 | 144 | 257 | 772 | 854 | 0 | 0 | 129 | 29 | 0 | 0 | 0 | 0 | 127 | 0 | 0 | 0 | 0 | 12842 | 761 | 0 |
| SRR4000626 | 34 | 0 | 0 | 0 | 208 | 6685 | 5296 | 25373 | 58280 | 209 | 0 | 87 | 0 | 0 | 2052 | 150055 | 4644 | 145 | 249 | 829 | 1078 | 0 | 0 | 124 | 29 | 0 | 0 | 0 | 0 | 122 | 0 | 41 | 0 | 0 | 12994 | 850 | 0 |
| SRR4000627 | 0 | 0 | 0 | 0 | 13 | 9137 | 9214 | 43983 | 74561 | 141 | 0 | 0 | 0 | 0 | 193 | 2658 | 454 | 0 | 97 | 0 | 116 | 0 | 0 | 89 | 0 | 0 | 0 | 0 | 0 | 16 | 0 | 0 | 0 | 0 | 28782 | 708 | 0 |
| SRR4000628 | 0 | 0 | 0 | 0 | 14 | 9102 | 8676 | 44426 | 73242 | 124 | 0 | 0 | 0 | 0 | 213 | 2300 | 263 | 58 | 174 | 0 | 77 | 0 | 0 | 89 | 0 | 0 | 0 | 0 | 0 | 12 | 0 | 0 | 0 | 0 | 29441 | 729 | 0 |
| SRR4000629 | 68 | 0 | 0 | 0 | 150 | 5580 | 3686 | 16984 | 38347 | 162 | 0 | 76 | 0 | 0 | 1317 | 27223 | 5989 | 69 | 335 | 300 | 335 | 0 | 0 | 115 | 20 | 0 | 0 | 0 | 0 | 69 | 0 | 0 | 0 | 0 | 19056 | 538 | 0 |
| SRR4000630 | 47 | 0 | 0 | 0 | 107 | 4958 | 3887 | 17342 | 38466 | 103 | 0 | 44 | 0 | 0 | 1525 | 27920 | 6033 | 47 | 396 | 268 | 466 | 0 | 0 | 74 | 11 | 0 | 0 | 0 | 0 | 42 | 0 | 0 | 0 | 0 | 18707 | 557 | 0 |
| SRR4000631 | 53 | 0 | 0 | 0 | 231 | 5128 | 4124 | 19569 | 37106 | 152 | 0 | 79 | 0 | 0 | 1028 | 35317 | 4877 | 191 | 873 | 347 | 526 | 0 | 0 | 151 | 37 | 0 | 0 | 0 | 0 | 91 | 0 | 0 | 0 | 0 | 9885 | 601 | 0 |
| SRR4000632 | 38 | 0 | 0 | 0 | 155 | 5464 | 3274 | 20276 | 39806 | 103 | 0 | 50 | 0 | 0 | 1070 | 34748 | 4914 | 174 | 934 | 349 | 548 | 0 | 0 | 88 | 27 | 0 | 0 | 0 | 0 | 58 | 0 | 0 | 0 | 0 | 11347 | 695 | 0 |
| SRR4000633 | 26 | 0 | 0 | 0 | 60 | 4613 | 4532 | 20479 | 49229 | 113 | 0 | 32 | 0 | 0 | 1852 | 35870 | 2394 | 140 | 233 | 210 | 897 | 0 | 0 | 96 | 17 | 0 | 0 | 0 | 0 | 32 | 0 | 0 | 0 | 0 | 20623 | 273 | 0 |
| SRR4000634 | 39 | 0 | 0 | 0 | 104 | 5149 | 4486 | 20705 | 49139 | 190 | 0 | 52 | 0 | 0 | 1990 | 35606 | 2460 | 205 | 374 | 60 | 977 | 0 | 0 | 137 | 21 | 0 | 0 | 0 | 0 | 54 | 0 | 24 | 0 | 0 | 22029 | 348 | 0 |
| SRR4000635 | 10 | 0 | 0 | 0 | 43 | 7754 | 9762 | 48324 | 118746 | 120 | 0 | 29 | 0 | 0 | 762 | 46552 | 1099 | 165 | 103 | 113 | 175 | 0 | 0 | 106 | 12 | 0 | 0 | 0 | 0 | 27 | 0 | 31 | 0 | 10 | 21108 | 420 | 0 |
| SRR4000636 | 12 | 0 | 0 | 0 | 46 | 7532 | 9966 | 47607 | 114592 | 120 | 0 | 35 | 0 | 0 | 704 | 46568 | 1191 | 101 | 91 | 171 | 101 | 0 | 0 | 103 | 16 | 0 | 0 | 0 | 0 | 24 | 0 | 10 | 0 | 0 | 19890 | 525 | 0 |
| SRR4000638 | 0 | 0 | 0 | 0 | 14 | 15175 | 8916 | 63672 | 77091 | 19 | 0 | 0 | 0 | 0 | 293 | 10587 | 2041 | 90 | 203 | 90 | 180 | 0 | 0 | 161 | 13 | 0 | 0 | 0 | 0 | 38 | 0 | 23 | 0 | 0 | 12914 | 443 | 0 |
| SRR4000639 | 0 | 0 | 0 | 0 | 0 | 22376 | 12060 | 75042 | 38251 | 21 | 0 | 0 | 0 | 0 | 188 | 3523 | 488 | 0 | 118 | 94 | 212 | 0 | 0 | 150 | 16 | 0 | 0 | 0 | 0 | 50 | 0 | 0 | 0 | 0 | 16966 | 557 | 0 |
| SRR4000640 | 0 | 0 | 0 | 0 | 0 | 22392 | 11124 | 75006 | 38458 | 23 | 0 | 0 | 0 | 0 | 95 | 3057 | 322 | 24 | 119 | 48 | 48 | 0 | 0 | 155 | 10 | 0 | 0 | 0 | 0 | 49 | 0 | 0 | 0 | 0 | 16360 | 502 | 0 |

### Supp Table S8.3.Columns_BX_to_DH

| | | | | | | | | | | | | | | | | | | | | | | | | | | | | | | | | | | | | | |
|---|---|---|---|---|---|---|---|---|---|---|---|---|---|---|---|---|---|---|---|---|---|---|---|---|---|---|---|---|---|---|---|---|---|---|---|---|---|
| SRR4000641 | 0 | 0 | 0 | 0 | 0 | 13036 | 7661 | 44022 | 60528 | 55 | 0 | 0 | 0 | 0 | 361 | 6181 | 628 | 20 | 281 | 120 | 140 | 0 | 0 | 172 | 20 | 0 | 0 | 0 | 0 | 61 | 0 | 0 | 0 | 0 | 18252 | 400 | 0 |
| SRR4000642 | 0 | 0 | 0 | 0 | 17 | 15008 | 7626 | 49924 | 65442 | 56 | 0 | 0 | 0 | 0 | 266 | 6137 | 641 | 67 | 222 | 44 | 200 | 0 | 0 | 167 | 15 | 0 | 0 | 0 | 0 | 71 | 0 | 0 | 0 | 0 | 15387 | 455 | 0 |
| SRR4000643 | 0 | 0 | 0 | 0 | 11 | 17496 | 11460 | 70041 | 128069 | 72 | 14 | 13 | 0 | 0 | 846 | 19697 | 5918 | 87 | 321 | 292 | 117 | 0 | 0 | 158 | 30 | 17 | 0 | 14 | 0 | 78 | 0 | 29 | 0 | 12 | 21895 | 591 | 0 |
| SRR4000644 | 0 | 0 | 0 | 0 | 31 | 19881 | 12891 | 82655 | 146550 | 122 | 14 | 23 | 0 | 0 | 890 | 15994 | 5486 | 132 | 297 | 264 | 231 | 0 | 0 | 275 | 54 | 18 | 0 | 12 | 0 | 143 | 0 | 0 | 0 | 0 | 21635 | 691 | 0 |
| SRR4000645 | 0 | 0 | 0 | 0 | 12 | 11865 | 6591 | 48225 | 91361 | 20 | 0 | 0 | 0 | 0 | 92 | 5016 | 330 | 31 | 92 | 31 | 0 | 0 | 0 | 166 | 0 | 0 | 0 | 0 | 0 | 11 | 0 | 0 | 0 | 19 | 11549 | 460 | 0 |
| SRR4000649 | 0 | 0 | 0 | 0 | 103 | 10439 | 5107 | 35135 | 80846 | 73 | 0 | 46 | 0 | 0 | 2673 | 73080 | 5883 | 623 | 229 | 417 | 266 | 0 | 0 | 112 | 12 | 0 | 0 | 0 | 0 | 71 | 0 | 0 | 0 | 0 | 23040 | 1857 | 0 |
| SRR4000650 | 0 | 0 | 0 | 0 | 99 | 10392 | 5528 | 35455 | 78210 | 72 | 0 | 50 | 0 | 0 | 2954 | 72784 | 5633 | 614 | 211 | 426 | 307 | 0 | 0 | 105 | 14 | 0 | 0 | 0 | 0 | 65 | 0 | 18 | 0 | 0 | 22977 | 1849 | 0 |
| SRR4011144 | 822 | 0 | 0 | 0 | 68 | 0 | 0 | 0 | 0 | 0 | 0 | 322 | 0 | 0 | 6183 | 15511 | 96510 | 0 | 0 | 0 | 0 | 0 | 0 | 19 | 107 | 71 | 858 | 0 | 0 | 0 | 0 | 0 | 0 | 0 | 0 | 0 | 0 |
| SRR4044175 | 0 | 0 | 0 | 0 | 0 | 0 | 0 | 21 | 0 | 14 | 45 | 0 | 0 | 0 | 510 | 6484 | 1388 | 219 | 0 | 125 | 110 | 0 | 0 | 0 | 0 | 0 | 0 | 0 | 0 | 98 | 0 | 12706 | 13 | 0 | 17747 | 0 | 208 |
| SRR4044176 | 0 | 0 | 0 | 0 | 0 | 0 | 0 | 30 | 0 | 0 | 148 | 0 | 0 | 0 | 566 | 5102 | 1128 | 185 | 0 | 83 | 367 | 19 | 0 | 0 | 0 | 0 | 0 | 0 | 0 | 153 | 0 | 24179 | 29 | 0 | 16352 | 0 | 1195 |
| SRR4044177 | 12 | 0 | 0 | 12 | 0 | 30 | 59 | 188 | 18 | 0 | 291 | 0 | 0 | 0 | 7396 | 8230 | 10756 | 68 | 42 | 713 | 73342 | 19 | 0 | 0 | 24 | 23 | 15 | 0 | 0 | 184 | 0 | 15242 | 21 | 0 | 17068 | 0 | 351 |
| SRR4044178 | 0 | 0 | 0 | 0 | 0 | 0 | 13 | 80 | 0 | 12 | 31 | 0 | 0 | 0 | 586 | 6772 | 1847 | 309 | 0 | 129 | 118 | 0 | 0 | 0 | 0 | 0 | 0 | 0 | 0 | 73 | 0 | 15994 | 14 | 0 | 13525 | 0 | 208 |
| SRR4044179 | 0 | 0 | 0 | 13 | 0 | 0 | 0 | 24 | 0 | 0 | 351 | 0 | 0 | 0 | 563 | 5191 | 1344 | 259 | 0 | 117 | 271 | 13 | 0 | 0 | 0 | 10 | 30 | 0 | 0 | 265 | 0 | 27380 | 27 | 0 | 11614 | 0 | 950 |
| SRR4044180 | 0 | 0 | 0 | 0 | 25 | 50 | 75 | 295 | 27 | 0 | 492 | 0 | 0 | 0 | 3054 | 4046 | 3386 | 120 | 37 | 425 | 31556 | 24 | 11 | 0 | 0 | 0 | 0 | 0 | 0 | 833 | 0 | 23291 | 25 | 0 | 11616 | 0 | 607 |
| SRR4044206 | 0 | 0 | 0 | 0 | 23 | 702 | 0 | 238 | 119 | 77 | 290 | 0 | 0 | 0 | 13977 | 247163 | 94881 | 220 | 1867 | 14304 | 14488 | 0 | 18 | 0 | 0 | 0 | 0 | 0 | 0 | 414 | 0 | 0 | 0 | 31 | 3117 | 4913 | 0 |
| SRR4044210 | 0 | 0 | 0 | 0 | 88 | 249 | 83 | 332 | 290 | 11737 | 38 | 22 | 0 | 0 | 14024 | 70643 | 25213 | 0 | 0 | 83 | 3195 | 0 | 16 | 0 | 0 | 0 | 0 | 0 | 0 | 353 | 0 | 290 | 0 | 0 | 59641 | 5068 | 0 |
| SRR4044211 | 0 | 0 | 0 | 0 | 187 | 1032 | 95 | 1207 | 619 | 1072 | 44 | 17 | 0 | 0 | 3350 | 83452 | 74093 | 79 | 191 | 127 | 889 | 39 | 44 | 22 | 17 | 20 | 11 | 11 | 0 | 220 | 72 | 286 | 0 | 0 | 47570 | 3853 | 0 |
| SRR4270461 | 208 | 79 | 52 | 71 | 496 | 3253 | 2413 | 10931 | 23977 | 890 | 625 | 33 | 0 | 0 | 21200 | 54773 | 48165 | 668 | 1341 | 4239 | 1862 | 274 | 82 | 178 | 207 | 156 | 130 | 157 | 0 | 997 | 144 | 212 | 0 | 28 | 8847 | 0 | 0 |
| SRR4270462 | 299 | 107 | 65 | 65 | 636 | 3357 | 2829 | 14451 | 29782 | 561 | 423 | 47 | 19 | 0 | 16315 | 41204 | 41953 | 550 | 681 | 1728 | 1322 | 345 | 115 | 195 | 239 | 151 | 108 | 93 | 0 | 727 | 199 | 140 | 0 | 19 | 7549 | 0 | 0 |
| SRR4292754 | 0 | 22 | 0 | 0 | 0 | 1517 | 637 | 1276 | 171 | 0 | 0 | 0 | 0 | 0 | 417 | 205 | 131 | 866 | 31 | 311 | 80 | 0 | 0 | 84 | 0 | 18 | 73 | 43 | 0 | 29 | 0 | 21676 | 0 | 0 | 4188 | 1673 | 0 |
| SRR4292755 | 0 | 17 | 0 | 10 | 0 | 1072 | 641 | 992 | 151 | 0 | 0 | 0 | 0 | 0 | 1103 | 227 | 215 | 325 | 14 | 399 | 77 | 0 | 0 | 1100 | 0 | 0 | 49 | 38 | 0 | 34 | 0 | 21208 | 0 | 0 | 6435 | 2806 | 0 |
| SRR4292756 | 11 | 14 | 0 | 11 | 0 | 1750 | 1208 | 1538 | 281 | 0 | 0 | 0 | 0 | 0 | 1148 | 5782 | 634 | 89 | 20 | 565 | 209 | 0 | 0 | 248 | 12 | 30 | 12 | 44 | 0 | 17 | 0 | 37 | 0 | 0 | 7839 | 1795 | 0 |
| SRR4292757 | 0 | 16 | 0 | 11 | 0 | 1415 | 717 | 840 | 167 | 0 | 0 | 0 | 0 | 0 | 1439 | 5029 | 565 | 114 | 35 | 633 | 320 | 0 | 0 | 390 | 0 | 29 | 11 | 30 | 0 | 17 | 0 | 53 | 0 | 0 | 5427 | 1490 | 0 |
| SRR4296311 | 2176 | 1007 | 547 | 197 | 1655 | 2589 | 1187 | 5340 | 6364 | 44 | 118 | 0 | 26 | 13 | 47786 | 16946 | 58022 | 701 | 1996 | 3560 | 11596 | 3013 | 709 | 333 | 127 | 123 | 219 | 118 | 0 | 114 | 1427 | 1079 | 13 | 18 | 12525 | 20 | 18 |
| SRR4296312 | 2095 | 721 | 462 | 192 | 1757 | 3221 | 1540 | 4061 | 5461 | 34 | 90 | 0 | 11 | 0 | 49993 | 18720 | 63033 | 280 | 2661 | 3501 | 15964 | 2861 | 687 | 417 | 68 | 225 | 101 | 158 | 0 | 90 | 1735 | 840 | 0 | 23 | 12796 | 31 | 0 |
| SRR4296313 | 125 | 63 | 31 | 25 | 157 | 2890 | 1870 | 8104 | 5894 | 57 | 12 | 0 | 12 | 12 | 63074 | 18356 | 69987 | 793 | 3740 | 3570 | 8671 | 240 | 53 | 31 | 14 | 69 | 17 | 97 | 0 | 105 | 117 | 737 | 0 | 0 | 6655 | 35 | 12 |
| SRR4296314 | 964 | 562 | 273 | 187 | 1370 | 2610 | 1789 | 6487 | 6487 | 80 | 59 | 0 | 11 | 0 | 66657 | 16377 | 70066 | 149 | 3952 | 4250 | 11930 | 2323 | 578 | 337 | 91 | 161 | 134 | 102 | 0 | 32 | 1290 | 596 | 32 | 0 | 7548 | 45 | 0 |
| SRR4296315 | 2550 | 1097 | 690 | 278 | 1169 | 4546 | 2013 | 7274 | 3897 | 78 | 56 | 11 | 0 | 0 | 96772 | 22344 | 94993 | 390 | 3182 | 4222 | 20394 | 3646 | 902 | 301 | 134 | 122 | 150 | 33 | 0 | 72 | 2399 | 65 | 0 | 0 | 6902 | 35 | 0 |
| SRR4296316 | 2438 | 1130 | 643 | 222 | 1341 | 3943 | 2390 | 6930 | 6094 | 89 | 66 | 0 | 0 | 0 | 88183 | 20693 | 88584 | 119 | 2987 | 5616 | 22464 | 3790 | 787 | 321 | 122 | 144 | 233 | 44 | 0 | 55 | 2106 | 239 | 11 | 0 | 5580 | 38 | 11 |
| SRR4296317 | 3969 | 1358 | 850 | 315 | 1104 | 2925 | 773 | 4305 | 3698 | 177 | 72 | 0 | 28 | 17 | 77046 | 27870 | 120442 | 110 | 1932 | 4415 | 28313 | 3660 | 949 | 348 | 121 | 110 | 221 | 149 | 0 | 77 | 2263 | 55 | 0 | 0 | 9576 | 32 | 0 |
| SRR4296318 | 195 | 76 | 41 | 15 | 55 | 4365 | 794 | 4286 | 3889 | 87 | 79 | 0 | 0 | 0 | 79371 | 25974 | 114662 | 0 | 1508 | 4286 | 29050 | 200 | 50 | 23 | 103 | 119 | 158 | 40 | 0 | 71 | 108 | 159 | 24 | 16 | 9569 | 46 | 0 |
| SRR4296319 | 81 | 17 | 18 | 57 | 26 | 4902 | 210 | 2031 | 2241 | 53 | 72 | 0 | 29 | 0 | 106730 | 44538 | 299500 | 210 | 2171 | 3151 | 71714 | 429 | 108 | 15 | 10 | 38 | 11 | 33 | 0 | 38 | 260 | 0 | 0 | 53 | 10489 | 23 | 0 |
| SRR4296320 | 882 | 290 | 246 | 81 | 388 | 3714 | 455 | 2729 | 2678 | 45 | 115 | 0 | 14 | 0 | 104753 | 37925 | 294582 | 76 | 2148 | 3411 | 71098 | 4866 | 1251 | 251 | 112 | 53 | 72 | 22 | 0 | 101 | 3252 | 354 | 0 | 62 | 8588 | 24 | 0 |
| SRR4296321 | 823 | 307 | 186 | 94 | 441 | 2682 | 530 | 2775 | 2994 | 38 | 142 | 0 | 0 | 10 | 112916 | 42441 | 270209 | 125 | 3118 | 3649 | 49956 | 4472 | 976 | 224 | 104 | 74 | 80 | 44 | 0 | 82 | 2911 | 405 | 12 | 32 | 6993 | 32 | 0 |
| SRR4296322 | 792 | 228 | 219 | 73 | 447 | 3492 | 380 | 2657 | 2277 | 24 | 155 | 0 | 0 | 0 | 115758 | 42032 | 279455 | 152 | 2505 | 3340 | 50175 | 4431 | 1117 | 194 | 151 | 78 | 102 | 58 | 0 | 58 | 2750 | 0 | 0 | 39 | 8763 | 34 | 0 |
| SRR4296979 | 26 | 40 | 20 | 29 | 0 | 280 | 173 | 547 | 200 | 26 | 323 | 0 | 0 | 0 | 7119 | 16196 | 28460 | 0 | 120 | 1853 | 1853 | 0 | 0 | 20 | 17 | 23 | 315 | 159 | 0 | 2032 | 0 | 13 | 0 | 0 | 6041 | 0 | 0 |
| SRR4296985 | 0 | 0 | 0 | 0 | 69 | 0 | 0 | 0 | 0 | 0 | 108 | 0 | 0 | 0 | 111912 | 103651 | 134681 | 0 | 541 | 4445 | 2703 | 30 | 28 | 15 | 0 | 0 | 16 | 0 | 0 | 3022 | 0 | 0 | 0 | 0 | 3042 | 0 | 0 |
| SRR4296986 | 0 | 0 | 0 | 0 | 40 | 30 | 0 | 0 | 0 | 0 | 61 | 0 | 0 | 0 | 82887 | 54650 | 100749 | 0 | 1149 | 5231 | 2994 | 0 | 12 | 0 | 0 | 0 | 0 | 0 | 0 | 2072 | 0 | 30 | 0 | 0 | 2651 | 0 | 21 |
| SRR4296987 | 0 | 0 | 0 | 0 | 0 | 27 | 0 | 31 | 0 | 17 | 11 | 0 | 0 | 0 | 32875 | 9879 | 87402 | 0 | 344 | 5353 | 1978 | 0 | 0 | 10 | 37 | 41 | 18 | 14 | 0 | 127 | 0 | 0 | 0 | 0 | 541 | 0 | 0 |
| SRR4296988 | 0 | 0 | 0 | 0 | 0 | 43 | 16 | 37 | 0 | 18 | 10 | 0 | 0 | 0 | 28868 | 12068 | 90542 | 0 | 423 | 4498 | 1701 | 0 | 0 | 13 | 32 | 12 | 27 | 16 | 0 | 113 | 0 | 0 | 0 | 0 | 468 | 0 | 0 |
| SRR4328127 | 0 | 0 | 0 | 0 | 0 | 67 | 0 | 301 | 0 | 122 | 0 | 0 | 0 | 0 | 2644 | 325735 | 78048 | 0 | 0 | 67 | 870 | 0 | 0 | 0 | 0 | 0 | 0 | 0 | 0 | 0 | 0 | 0 | 0 | 0 | 1602 | 0 | 0 |
| SRR4328130 | 0 | 0 | 0 | 0 | 0 | 0 | 0 | 247 | 495 | 24 | 25 | 0 | 0 | 0 | 19792 | 93273 | 184482 | 0 | 0 | 0 | 3216 | 0 | 0 | 0 | 0 | 0 | 15 | 0 | 0 | 0 | 0 | 0 | 0 | 0 | 4953 | 0 | 0 |
| SRR4328135 | 0 | 0 | 0 | 0 | 0 | 154 | 0 | 231 | 0 | 44 | 0 | 0 | 0 | 0 | 1696 | 420051 | 69066 | 0 | 77 | 0 | 3006 | 0 | 0 | 0 | 0 | 0 | 0 | 0 | 0 | 0 | 0 | 0 | 0 | 0 | 1028 | 0 | 0 |
| SRR4328136 | 0 | 0 | 0 | 0 | 0 | 380 | 20 | 880 | 320 | 43 | 0 | 0 | 0 | 0 | 1341 | 83190 | 66164 | 0 | 0 | 80 | 600 | 0 | 0 | 0 | 0 | 0 | 0 | 0 | 0 | 0 | 0 | 0 | 0 | 0 | 1796 | 57 | 0 |
| SRR4328137 | 0 | 0 | 0 | 0 | 0 | 23 | 0 | 554 | 231 | 149 | 0 | 0 | 0 | 0 | 3046 | 282457 | 96175 | 0 | 0 | 254 | 623 | 0 | 0 | 0 | 0 | 0 | 0 | 0 | 0 | 0 | 0 | 0 | 0 | 0 | 306 | 0 | 0 |
| SRR4409201 | 12 | 0 | 13 | 60 | 66 | 100 | 87 | 1336 | 24 | 0 | 75 | 0 | 0 | 0 | 39915 | 17640 | 318086 | 0 | 44 | 4690 | 5941 | 0 | 0 | 34 | 28 | 55 | 24 | 44 | 0 | 21 | 0 | 0 | 0 | 0 | 36 | 195 | 0 |
| SRR4409202 | 0 | 0 | 70 | 106 | 75 | 0 | 10 | 76 | 0 | 0 | 39 | 0 | 0 | 0 | 29150 | 27753 | 114200 | 10 | 0 | 611 | 1779 | 0 | 0 | 0 | 17 | 192 | 501 | 825 | 0 | 69 | 0 | 0 | 0 | 0 | 374 | 205 | 0 |
| SRR4409203 | 0 | 0 | 13 | 59 | 53 | 12 | 19 | 244 | 12 | 0 | 47 | 0 | 0 | 0 | 73931 | 19180 | 407892 | 0 | 44 | 5826 | 7499 | 0 | 0 | 25 | 13 | 55 | 14 | 13 | 0 | 13 | 0 | 0 | 0 | 0 | 18 | 41 | 0 |
| SRR4409204 | 0 | 0 | 0 | 0 | 0 | 0 | 0 | 148 | 70 | 0 | 43 | 27 | 0 | 0 | 83221 | 26539 | 217260 | 0 | 87 | 2287 | 15803 | 0 | 0 | 0 | 0 | 0 | 0 | 12 | 0 | 99 | 0 | 0 | 0 | 0 | 97 | 186 | 0 |
| SRR4417245 | 0 | 0 | 0 | 0 | 26 | 11471 | 6379 | 21969 | 14139 | 21 | 0 | 0 | 0 | 0 | 33 | 5636 | 2718 | 160 | 0 | 0 | 0 | 0 | 0 | 691 | 44 | 10 | 0 | 0 | 0 | 0 | 10 | 0 | 0 | 0 | 9172 | 17 | 0 |
| SRR4417246 | 0 | 0 | 0 | 0 | 0 | 8405 | 4740 | 17150 | 7239 | 0 | 0 | 0 | 0 | 0 | 0 | 1754 | 362 | 78 | 0 | 0 | 0 | 0 | 0 | 13 | 0 | 13 | 0 | 0 | 0 | 0 | 0 | 0 | 0 | 0 | 2450 | 19 | 0 |
| SRR4417247 | 0 | 0 | 0 | 0 | 21 | 858 | 1769 | 10171 | 13098 | 55 | 0 | 0 | 0 | 0 | 26 | 31664 | 4537 | 343 | 0 | 0 | 0 | 0 | 0 | 28 | 52 | 15 | 16 | 0 | 0 | 0 | 18 | 0 | 0 | 0 | 6070 | 13 | 0 |
| SRR4417248 | 0 | 0 | 0 | 0 | 0 | 2917 | 3801 | 19526 | 21288 | 53 | 0 | 0 | 0 | 0 | 33 | 65556 | 1213 | 474 | 0 | 0 | 0 | 0 | 0 | 18 | 17 | 0 | 0 | 0 | 0 | 0 | 0 | 0 | 0 | 0 | 4749 | 18 | 0 |
| SRR4417249 | 0 | 0 | 0 | 0 | 0 | 3978 | 5433 | 28435 | 13289 | 15 | 0 | 0 | 0 | 0 | 0 | 645 | 103 | 50 | 0 | 0 | 0 | 0 | 0 | 433 | 0 | 0 | 0 | 0 | 0 | 0 | 0 | 0 | 0 | 0 | 4488 | 17 | 0 |
| SRR4417250 | 0 | 0 | 0 | 0 | 11 | 2074 | 2155 | 15037 | 19300 | 81 | 0 | 0 | 0 | 0 | 39 | 43219 | 8584 | 549 | 12 | 15 | 26 | 0 | 0 | 557 | 61 | 51 | 44 | 0 | 0 | 0 | 0 | 0 | 0 | 0 | 6435 | 19 | 0 |
| SRR4423247 | 114 | 75 | 166 | 559 | 117 | 128 | 184 | 446 | 42 | 96 | 57 | 0 | 0 | 0 | 16156 | 12008 | 57168 | 39 | 176 | 58459 | 1187 | 0 | 0 | 102 | 68 | 340 | 1156 | 106 | 0 | 185 | 0 | 0 | 0 | 0 | 3516 | 1289 | 0 |
| SRR4423248 | 82 | 54 | 88 | 401 | 33 | 282 | 239 | 712 | 37 | 35 | 77 | 28 | 0 | 0 | 44092 | 36535 | 99378 | 58 | 133 | 66077 | 414 | 0 | 0 | 95 | 19 | 95 | 395 | 28 | 0 | 227 | 0 | 11 | 0 | 0 | 2891 | 607 | 11 |
| SRR4423250 | 364 | 85 | 213 | 327 | 152 | 564 | 379 | 1714 | 62 | 0 | 122 | 17 | 19 | 0 | 10535 | 25312 | 43484 | 131 | 249 | 8018 | 1258 | 0 | 0 | 164 | 109 | 494 | 1167 | 1660 | 0 | 768 | 0 | 38 | 0 | 0 | 2287 | 0 | 0 |
| SRR4423251 | 619 | 163 | 355 | 386 | 78 | 470 | 723 | 1496 | 74 | 13 | 80 | 0 | 15 | 0 | 32614 | 14363 | 53221 | 37 | 290 | 7243 | 564 | 0 | 0 | 224 | 201 | 1001 | 1467 | 1550 | 0 | 353 | 0 | 0 | 0 | 0 | 3638 | 0 | 0 |
| SRR4473670 | 48 | 47 | 43 | 10 | 20 | 378 | 325 | 2973 | 140 | 13 | 72 | 0 | 0 | 0 | 8281 | 25110 | 23787 | 53 | 521 | 5579 | 6794 | 0 | 0 | 32 | 37 | 17 | 57 | 16 | 0 | 10 | 0 | 28 | 0 | 0 | 2315 | 0 | 12 |
| SRR4734676 | 13 | 10 | 0 | 0 | 72 | 340 | 316 | 1282 | 87 | 16 | 0 | 20 | 0 | 0 | 1059 | 9228 | 104264 | 0 | 0 | 500 | 369 | 0 | 0 | 1146 | 24 | 62 | 196 | 0 | 11 | 149 | 0 | 0 | 92 | 0 | 3013 | 0 | 0 |
| SRR4734677 | 17 | 50 | 0 | 17 | 201 | 111 | 138 | 525 | 28 | 0 | 17 | 0 | 0 | 0 | 1989 | 20684 | 165261 | 0 | 0 | 1602 | 912 | 0 | 0 | 2079 | 17 | 151 | 419 | 34 | 0 | 34 | 0 | 0 | 0 | 0 | 1767 | 0 | 0 |
| SRR4734678 | 13 | 0 | 0 | 40 | 106 | 0 | 43 | 260 | 87 | 0 | 0 | 0 | 0 | 0 | 87 | 3530 | 11390 | 0 | 0 | 260 | 87 | 0 | 0 | 78 | 53 | 159 | 23 | 13 | 0 | 27 | 0 | 0 | 0 | 0 | 630 | 0 | 0 |
| SRR4734679 | 27 | 41 | 0 | 14 | 123 | 142 | 142 | 522 | 0 | 27 | 14 | 14 | 0 | 0 | 237 | 3782 | 11280 | 0 | 0 | 285 | 664 | 0 | 0 | 2750 | 137 | 397 | 1573 | 41 | 0 | 14 | 0 | 0 | 0 | 0 | 697 | 0 | 14 |
| SRR4734680 | 34 | 17 | 17 | 17 | 118 | 68 | 68 | 473 | 0 | 17 | 0 | 34 | 0 | 0 | 68 | 4212 | 11344 | 0 | 0 | 473 | 203 | 0 | 0 | 124 | 101 | 202 | 16 | 17 | 0 | 34 | 0 | 0 | 34 | 0 | 611 | 0 | 0 |
| SRR4734681 | 38 | 31 | 0 | 23 | 199 | 71 | 189 | 661 | 24 | 23 | 0 | 0 | 0 | 0 | 189 | 4646 | 15711 | 0 | 0 | 472 | 331 | 0 | 0 | 1764 | 77 | 130 | 537 | 0 | 0 | 69 | 0 | 0 | 0 | 0 | 3599 | 0 | 0 |
| SRR4734682 | 0 | 0 | 0 | 0 | 186 | 170 | 170 | 424 | 0 | 0 | 0 | 0 | 0 | 0 | 212 | 5613 | 15708 | 0 | 0 | 212 | 255 | 0 | 0 | 1802 | 40 | 81 | 307 | 16 | 0 | 113 | 0 | 0 | 0 | 0 | 1340 | 0 | 0 |
| SRR4734683 | 13 | 0 | 0 | 11 | 83 | 81 | 101 | 195 | 20 | 16 | 0 | 0 | 0 | 0 | 34 | 863 | 2392 | 0 | 0 | 94 | 101 | 0 | 0 | 90 | 85 | 12 | 56 | 36 | 0 | 104 | 0 | 0 | 0 | 0 | 450 | 0 | 0 |
| SRR4734684 | 49 | 39 | 0 | 0 | 192 | 67 | 146 | 517 | 16 | 0 | 12 | 0 | 0 | 0 | 203 | 3936 | 17796 | 0 | 0 | 584 | 412 | 0 | 0 | 77 | 85 | 127 | 13 | 15 | 0 | 112 | 0 | 0 | 13 | 0 | 1360 | 0 | 0 |
| SRR4734685 | 26 | 24 | 0 | 0 | 64 | 17 | 33 | 226 | 17 | 19 | 33 | 0 | 0 | 0 | 75 | 2213 | 5902 | 0 | 0 | 142 | 301 | 0 | 0 | 41 | 95 | 10 | 39 | 0 | 0 | 20 | 0 | 0 | 0 | 0 | 284 | 0 | 0 |
| SRR4734686 | 63 | 38 | 0 | 13 | 152 | 123 | 123 | 548 | 0 | 21 | 0 | 13 | 0 | 0 | 771 | 7309 | 47556 | 0 | 0 | 483 | 101 | 0 | 0 | 35 | 67 | 126 | 506 | 21 | 0 | 76 | 0 | 0 | 0 | 0 | 2415 | 0 | 0 |
| SRR4787084 | 0 | 0 | 0 | 0 | 69 | 8520 | 13952 | 18891 | 9260 | 0 | 0 | 0 | 0 | 0 | 10619 | 47526 | 107848 | 741 | 7408 | 8767 | 2469 | 0 | 0 | 1036 | 345 | 829 | 3385 | 2418 | 0 | 69 | 0 | 247 | 0 | 0 | 1828 | 10 | 0 |
| SRR4787085 | 0 | 0 | 0 | 36 | 0 | 446 | 3301 | 3390 | 625 | 0 | 36 | 0 | 0 | 0 | 13471 | 22191 | 165615 | 0 | 625 | 16594 | 10795 | 0 | 0 | 36 | 71 | 214 | 40 | 47 | 0 | 143 | 0 | 714 | 0 | 0 | 814 | 13 | 0 |
| SRR4787086 | 129 | 0 | 0 | 0 | 0 | 11586 | 14516 | 24371 | 12252 | 0 | 64 | 0 | 0 | 0 | 12385 | 50200 | 110053 | 266 | 7724 | 5061 | 1598 | 0 | 0 | 37 | 258 | 46 | 84 | 68 | 0 | 0 | 0 | 133 | 0 | 0 | 1931 | 25 | 0 |
| SRR4787087 | 0 | 0 | 0 | 0 | 90 | 9473 | 14874 | 18327 | 11155 | 0 | 0 | 45 | 0 | 0 | 10182 | 44882 | 102975 | 443 | 6994 | 9031 | 2922 | 0 | 0 | 806 | 672 | 1433 | 2284 | 1836 | 0 | 45 | 0 | 266 | 0 | 0 | 2450 | 10 | 0 |
| SRR4787088 | 0 | 0 | 0 | 0 | 132 | 8917 | 7410 | 12183 | 5401 | 0 | 0 | 0 | 0 | 0 | 6280 | 31581 | 81304 | 377 | 6154 | 8541 | 1507 | 0 | 0 | 927 | 662 | 993 | 3045 | 1721 | 0 | 66 | 0 | 251 | 0 | 0 | 3024 | 25 | 0 |
| SRR4787089 | 0 | 0 | 0 | 0 | 0 | 11146 | 9684 | 14983 | 5116 | 0 | 89 | 0 | 0 | 0 | 5847 | 32439 | 88681 | 0 | 4385 | 6395 | 1827 | 0 | 0 | 23 | 23 | 40 | 123 | 73 | 0 | 0 | 0 | 183 | 0 | 0 | 3034 | 16 | 0 |
| SRR4787090 | 13 | 31 | 0 | 0 | 184 | 5856 | 3004 | 8471 | 30231 | 627 | 148 | 54 | 0 | 0 | 5316 | 73072 | 109910 | 691 | 6072 | 7909 | 8622 | 0 | 0 | 85 | 228 | 188 | 278 | 313 | 0 | 340 | 0 | 43 | 0 | 18 | 11145 | 0 | 0 |
| SRR4787091 | 157 | 0 | 0 | 0 | 0 | 9455 | 12514 | 12792 | 5562 | 0 | 0 | 0 | 0 | 0 | 6118 | 35398 | 99558 | 278 | 5284 | 6118 | 1390 | 0 | 0 | 91 | 18 | 55 | 68 | 119 | 0 | 0 | 0 | 278 | 0 | 0 | 1475 | 21 | 0 |
| SRR4787092 | 19 | 0 | 14 | 0 | 135 | 6473 | 3088 | 12512 | 29828 | 611 | 111 | 29 | 0 | 0 | 4712 | 72885 | 107196 | 778 | 6679 | 7915 | 8532 | 0 | 0 | 111 | 154 | 197 | 236 | 313 | 0 | 476 | 0 | 183 | 0 | 0 | 12480 | 13 | 0 |
| SRR4787093 | 23 | 23 | 0 | 0 | 147 | 5985 | 2762 | 10436 | 29597 | 676 | 124 | 46 | 0 | 0 | 5569 | 72890 | 111647 | 1250 | 6818 | 9383 | 8375 | 0 | 0 | 74 | 184 | 244 | 290 | 368 | 0 | 377 | 0 | 88 | 0 | 14 | 13247 | 0 | 0 |
| SRR4787095 | 26 | 64 | 0 | 17 | 357 | 1700 | 4970 | 5942 | 3333 | 21 | 455 | 0 | 0 | 0 | 111519 | 77363 | 544140 | 0 | 4511 | 41183 | 8946 | 0 | 17 | 142 | 120 | 271 | 550 | 670 | 0 | 271 | 0 | 0 | 0 | 0 | 816 | 22 | 0 |
| SRR4787096 | 82 | 92 | 0 | 10 | 348 | 1133 | 5036 | 5429 | 3458 | 0 | 461 | 0 | 0 | 0 | 109966 | 73377 | 509707 | 0 | 5453 | 48781 | 10003 | 0 | 10 | 236 | 97 | 143 | 610 | 702 | 0 | 287 | 0 | 0 | 0 | 0 | 479 | 22 | 0 |
| SRR4787097 | 26 | 40 | 0 | 0 | 143 | 116 | 439 | 607 | 224 | 0 | 460 | 0 | 0 | 0 | 20812 | 44150 | 396499 | 0 | 323 | 1260 | 28650 | 15 | 79 | 24 | 66 | 119 | 156 | 310 | 0 | 269 | 31 | 0 | 0 | 0 | 262 | 712 | 0 |
| SRR4787098 | 0 | 39 | 0 | 0 | 134 | 21 | 512 | 787 | 127 | 0 | 477 | 0 | 0 | 0 | 18627 | 39046 | 377303 | 0 | 235 | 1354 | 28238 | 17 | 112 | 12 | 41 | 57 | 132 | 431 | 0 | 186 | 28 | 0 | 0 | 0 | 338 | 620 | 0 |
| SRR4787099 | 13 | 25 | 0 | 0 | 97 | 34 | 367 | 768 | 187 | 0 | 460 | 0 | 0 | 0 | 20708 | 37780 | 379394 | 0 | 333 | 1661 | 29570 | 14 | 70 | 0 | 0 | 95 | 190 | 353 | 0 | 146 | 30 | 0 | 0 | 0 | 569 | 696 | 0 |
| SRR4787100 | 46 | 49 | 12 | 0 | 277 | 330 | 2499 | 3557 | 1091 | 0 | 243 | 0 | 0 | 0 | 46222 | 68557 | 413214 | 27 | 3213 | 6399 | 22188 | 55 | 92 | 135 | 104 | 258 | 651 | 618 | 0 | 246 | 28 | 13 | 0 | 0 | 646 | 2184 | 0 |
| SRR4787101 | 40 | 43 | 0 | 0 | 19 | 239 | 2043 | 3161 | 779 | 13 | 17 | 0 | 0 | 0 | 48082 | 67277 | 425461 | 0 | 2388 | 6595 | 28719 | 24 | 128 | 88 | 88 | 14 | 35 | 39 | 0 | 12 | 29 | 0 | 0 | 0 | 595 | 1794 | 0 |
| SRR4787102 | 39 | 51 | 0 | 0 | 19 | 636 | 2835 | 4416 | 1455 | 0 | 21 | 0 | 0 | 0 | 46351 | 69799 | 416994 | 0 | 2841 | 5848 | 20996 | 39 | 58 | 60 | 56 | 15 | 33 | 29 | 0 | 17 | 44 | 23 | 0 | 0 | 870 | 2272 | 0 |
| SRR4787103 | 91 | 157 | 0 | 0 | 142 | 914 | 4546 | 5727 | 2192 | 10 | 66 | 0 | 0 | 0 | 39595 | 71020 | 369368 | 16 | 5961 | 9464 | 17456 | 20 | 35 | 20 | 172 | 14 | 42 | 37 | 0 | 177 | 10 | 32 | 0 | 0 | 713 | 7107 | 0 |
| SRR4787104 | 85 | 85 | 0 | 0 | 11 | 1614 | 5018 | 6385 | 2452 | 11 | 113 | 0 | 0 | 0 | 41749 | 69138 | 364527 | 0 | 6403 | 10252 | 17812 | 28 | 49 | 13 | 116 | 16 | 44 | 28 | 0 | 10 | 32 | 30 | 0 | 0 | 1049 | 6974 | 11 |
| SRR4787105 | 0 | 0 | 0 | 0 | 37 | 35 | 352 | 599 | 94 | 46 | 108 | 54 | 0 | 0 | 8854 | 17809 | 186378 | 0 | 124 | 9720 | 30923 | 0 | 0 | 37 | 29 | 75 | 18 | 0 | 0 | 22 | 0 | 84 | 0 | 0 | 994 | 407 | 0 |
| SRR4787106 | 13 | 0 | 0 | 0 | 19 | 69 | 413 | 672 | 122 | 0 | 10 | 38 | 0 | 0 | 8643 | 11970 | 173872 | 0 | 107 | 10500 | 29833 | 0 | 0 | 70 | 0 | 102 | 16 | 64 | 0 | 23 | 0 | 176 | 0 | 0 | 981 | 535 | 0 |
| SRR4787107 | 0 | 0 | 0 | 0 | 67 | 99 | 457 | 1227 | 191 | 0 | 94 | 39 | 0 | 0 | 7665 | 8760 | 131281 | 0 | 242 | 15429 | 29361 | 0 | 0 | 103 | 39 | 179 | 391 | 61 | 0 | 882 | 0 | 318 | 0 | 0 | 1138 | 969 | 12 |
| SRR4787108 | 0 | 0 | 0 | 0 | 34 | 40 | 390 | 894 | 181 | 0 | 14 | 94 | 0 | 0 | 4502 | 8739 | 113396 | 0 | 170 | 24229 | 19416 | 0 | 0 | 11 | 63 | 13 | 31 | 50 | 0 | 166 | 0 | 198 | 0 | 0 | 2134 | 568 | 0 |
| SRR4787109 | 0 | 0 | 0 | 18 | 18 | 48 | 386 | 625 | 121 | 18 | 93 | 25 | 0 | 0 | 8372 | 10477 | 145907 | 0 | 126 | 10467 | 30555 | 0 | 0 | 50 | 32 | 79 | 11 | 22 | 0 | 32 | 0 | 121 | 0 | 0 | 867 | 563 | 0 |
| SRR4787110 | 0 | 0 | 0 | 0 | 27 | 71 | 552 | 838 | 159 | 18 | 137 | 21 | 0 | 0 | 9959 | 11250 | 148702 | 0 | 170 | 12321 | 31129 | 0 | 0 | 84 | 69 | 119 | 349 | 36 | 0 | 415 | 0 | 216 | 0 | 0 | 1106 | 819 | 12 |
| SRR4787118 | 0 | 0 | 0 | 0 | 0 | 0 | 0 | 0 | 0 | 0 | 0 | 0 | 0 | 0 | 25504 | 27224 | 716575 | 0 | 0 | 3196 | 0 | 0 | 0 | 0 | 0 | 0 | 0 | 0 | 0 | 0 | 0 | 0 | 0 | 0 | 1931 | 0 | 0 |
| SRR4787119 | 0 | 0 | 0 | 653 | 0 | 0 | 0 | 0 | 0 | 0 | 0 | 0 | 0 | 0 | 14581 | 35631 | 713200 | 0 | 0 | 11010 | 1765 | 0 | 0 | 0 | 0 | 0 | 0 | 0 | 0 | 0 | 0 | 0 | 0 | 0 | 2976 | 0 | 0 |
| SRR4787120 | 101 | 28 | 0 | 0 | 267 | 3628 | 8799 | 11497 | 7311 | 18 | 239 | 0 | 0 | 0 | 82312 | 88638 | 496044 | 0 | 6853 | 44670 | 4901 | 0 | 0 | 13 | 138 | 13 | 26 | 20 | 0 | 179 | 0 | 0 | 0 | 0 | 782 | 36 | 0 |
| SRR4787121 | 11 | 10 | 0 | 0 | 283 | 369 | 1139 | 2132 | 1420 | 0 | 950 | 0 | 0 | 0 | 25320 | 69449 | 583931 | 0 | 633 | 16113 | 1105 | 0 | 11 | 0 | 12 | 27 | 70 | 83 | 0 | 662 | 0 | 0 | 0 | 0 | 4850 | 0 | 11 |
| SRR4787122 | 18 | 0 | 0 | 0 | 337 | 206 | 1013 | 1527 | 1255 | 0 | 1089 | 15 | 0 | 0 | 24629 | 64641 | 628942 | 0 | 578 | 18741 | 1094 | 0 | 75 | 0 | 26 | 48 | 75 | 101 | 0 | 666 | 0 | 0 | 0 | 0 | 1316 | 0 | 0 |
| SRR4787123 | 0 | 27 | 0 | 0 | 28 | 116 | 540 | 1043 | 740 | 0 | 84 | 0 | 0 | 0 | 22219 | 60235 | 633737 | 0 | 308 | 17188 | 1170 | 0 | 20 | 0 | 14 | 34 | 10 | 105 | 0 | 48 | 0 | 0 | 0 | 0 | 1188 | 0 | 0 |
| SRR4787124 | 0 | 0 | 0 | 0 | 494 | 0 | 0 | 184 | 201 | 0 | 389 | 0 | 0 | 0 | 15614 | 39522 | 691655 | 0 | 81 | 7289 | 1070 | 0 | 0 | 0 | 0 | 0 | 0 | 0 | 0 | 177 | 0 | 0 | 0 | 0 | 1158 | 0 | 0 |
| SRR4787125 | 0 | 0 | 0 | 0 | 520 | 0 | 144 | 273 | 271 | 0 | 1048 | 0 | 0 | 0 | 16574 | 48420 | 679354 | 0 | 81 | 9328 | 870 | 103 | 0 | 0 | 0 | 0 | 60 | 110 | 0 | 372 | 0 | 0 | 0 | 0 | 800 | 0 | 0 |
| SRR4787126 | 0 | 0 | 0 | 0 | 104 | 0 | 0 | 0 | 97 | 0 | 111 | 0 | 0 | 0 | 18643 | 41983 | 712773 | 0 | 93 | 6401 | 1430 | 0 | 0 | 0 | 0 | 0 | 32 | 18 | 0 | 107 | 0 | 0 | 0 | 0 | 404 | 0 | 0 |
| SRR4787127 | 0 | 0 | 0 | 0 | 0 | 0 | 0 | 0 | 0 | 0 | 0 | 0 | 0 | 0 | 13576 | 34544 | 743648 | 0 | 2262 | 6668 | 4583 | 0 | 0 | 0 | 0 | 0 | 0 | 0 | 0 | 0 | 0 | 0 | 0 | 0 | 0 | 0 | 0 |
| SRR4787132 | 0 | 0 | 0 | 47 | 47 | 282 | 2760 | 2985 | 676 | 0 | 0 | 0 | 0 | 0 | 12785 | 24783 | 163970 | 56 | 563 | 17066 | 7941 | 0 | 0 | 165 | 47 | 212 | 45 | 35 | 0 | 189 | 0 | 676 | 0 | 0 | 518 | 0 | 0 |
| SRR4787133 | 0 | 0 | 0 | 0 | 33 | 397 | 3570 | 3649 | 397 | 0 | 67 | 33 | 0 | 0 | 13645 | 25021 | 171682 | 0 | 397 | 17215 | 8965 | 0 | 0 | 33 | 67 | 233 | 48 | 35 | 0 | 100 | 0 | 476 | 0 | 0 | 592 | 0 | 0 |
| SRR4787134 | 0 | 0 | 0 | 0 | 44 | 22 | 28 | 285 | 12 | 28 | 18 | 0 | 0 | 0 | 20837 | 24308 | 375986 | 0 | 111 | 935 | 38652 | 0 | 41 | 0 | 41 | 19 | 0 | 35 | 0 | 27 | 0 | 0 | 0 | 0 | 1126 | 626 | 0 |
| SRR4787135 | 79 | 65 | 0 | 0 | 143 | 1049 | 4280 | 4787 | 1691 | 0 | 97 | 14 | 0 | 0 | 33878 | 66675 | 339173 | 21 | 6499 | 9538 | 16315 | 28 | 14 | 342 | 157 | 388 | 998 | 592 | 0 | 185 | 18 | 86 | 0 | 0 | 848 | 6340 | 18 |
| SRR4787136 | 11 | 69 | 11 | 0 | 35 | 4112 | 1809 | 6916 | 3232 | 58 | 15 | 0 | 0 | 0 | 26752 | 66631 | 347111 | 16 | 4786 | 19166 | 13964 | 45 | 77 | 18 | 93 | 14 | 12 | 13 | 0 | 207 | 29 | 58 | 0 | 0 | 6373 | 3139 | 0 |
| SRR4787137 | 0 | 56 | 13 | 0 | 215 | 20 | 72 | 498 | 153 | 0 | 258 | 0 | 0 | 0 | 27035 | 34527 | 413020 | 0 | 84 | 904 | 34452 | 0 | 56 | 0 | 99 | 163 | 39 | 17 | 0 | 142 | 0 | 0 | 0 | 0 | 1948 | 653 | 0 |
| SRR4787138 | 42 | 32 | 0 | 0 | 21 | 348 | 509 | 1930 | 288 | 0 | 13 | 0 | 0 | 0 | 37166 | 47649 | 449571 | 0 | 1053 | 2094 | 30277 | 29 | 89 | 25 | 63 | 10 | 38 | 35 | 0 | 38 | 12 | 0 | 0 | 0 | 5697 | 2215 | 0 |
| SRR4787139 | 26 | 25 | 0 | 0 | 317 | 723 | 1141 | 3811 | 838 | 20 | 200 | 0 | 0 | 0 | 42054 | 58676 | 438180 | 0 | 1534 | 3496 | 25319 | 24 | 90 | 31 | 105 | 165 | 63 | 27 | 0 | 471 | 29 | 0 | 0 | 0 | 6254 | 2496 | 0 |
| SRR4787140 | 45 | 68 | 0 | 0 | 348 | 1828 | 2804 | 7478 | 2150 | 34 | 111 | 0 | 0 | 0 | 54753 | 68044 | 396398 | 0 | 4215 | 5125 | 37912 | 28 | 83 | 122 | 267 | 355 | 171 | 68 | 0 | 340 | 28 | 0 | 0 | 0 | 5149 | 5267 | 13 |
| SRR4787141 | 28 | 38 | 0 | 0 | 33 | 1096 | 1513 | 4564 | 1498 | 10 | 39 | 0 | 0 | 0 | 38478 | 62094 | 430257 | 0 | 1220 | 3386 | 24000 | 19 | 66 | 48 | 15 | 21 | 38 | 47 | 0 | 63 | 30 | 0 | 0 | 0 | 6162 | 2138 | 0 |
| SRR4787142 | 105 | 84 | 12 | 0 | 358 | 1847 | 3414 | 8002 | 2536 | 23 | 183 | 0 | 0 | 0 | 61446 | 73101 | 405043 | 0 | 4788 | 6287 | 36419 | 28 | 67 | 107 | 167 | 353 | 195 | 63 | 0 | 255 | 28 | 24 | 0 | 0 | 3564 | 5701 | 0 |
| SRR4787143 | 88 | 48 | 0 | 0 | 36 | 1378 | 2408 | 6247 | 1656 | 13 | 112 | 0 | 0 | 0 | 63965 | 73695 | 460643 | 0 | 3165 | 3517 | 33822 | 28 | 112 | 11 | 16 | 28 | 14 | 37 | 0 | 29 | 42 | 22 | 0 | 0 | 4458 | 4388 | 11 |
| SRR4836291 | 10 | 15 | 10 | 15 | 10 | 389 | 698 | 885 | 295 | 0 | 0 | 0 | 0 | 0 | 1583 | 8382 | 4519 | 40 | 389 | 27189 | 496 | 0 | 0 | 12 | 14 | 68 | 243 | 165 | 0 | 11 | 0 | 13 | 0 | 0 | 859 | 0 | 0 |
| SRR4836293 | 25 | 19 | 13 | 16 | 25 | 296 | 774 | 889 | 148 | 0 | 0 | 0 | 0 | 0 | 1564 | 8455 | 4323 | 0 | 329 | 28746 | 329 | 0 | 0 | 22 | 35 | 145 | 662 | 546 | 0 | 47 | 0 | 16 | 0 | 0 | 684 | 0 | 0 |
| SRR4836294 | 26 | 15 | 23 | 30 | 13 | 355 | 769 | 976 | 276 | 0 | 0 | 0 | 0 | 0 | 1578 | 8926 | 5084 | 20 | 375 | 25886 | 286 | 0 | 0 | 30 | 30 | 67 | 234 | 157 | 0 | 12 | 0 | 0 | 0 | 0 | 638 | 0 | 0 |
| SRR4836295 | 129 | 26 | 57 | 36 | 31 | 688 | 934 | 2383 | 393 | 57 | 93 | 0 | 0 | 0 | 11203 | 16983 | 22394 | 49 | 762 | 27246 | 2236 | 0 | 0 | 118 | 52 | 129 | 191 | 201 | 0 | 227 | 0 | 25 | 0 | 0 | 1661 | 0 | 0 |
| SRR4836297 | 157 | 38 | 63 | 94 | 19 | 824 | 1098 | 1983 | 183 | 69 | 82 | 0 | 0 | 0 | 11165 | 21186 | 23690 | 244 | 580 | 29164 | 2196 | 0 | 0 | 126 | 38 | 94 | 126 | 132 | 0 | 176 | 0 | 61 | 0 | 0 | 1541 | 0 | 0 |

Supp Table S8.3.Columns_BX_to_DH

| | | | | | | | | | | | | | | | | | | | | | | | | | | | | | | | | | | | | | |
|---|---|---|---|---|---|---|---|---|---|---|---|---|---|---|---|---|---|---|---|---|---|---|---|---|---|---|---|---|---|---|---|---|---|---|---|---|---|
| SRR4836299 | 28 | 22 | 13 | 13 | 0 | 980 | 641 | 2192 | 374 | 10 | 19 | 0 | 0 | 0 | 11529 | 19387 | 24674 | 71 | 784 | 27636 | 2049 | 0 | 0 | 18 | 13 | 14 | 37 | 42 | 0 | 49 | 0 | 36 | 0 | 0 | 1997 | 0 | 0 |
| SRR4836301 | 23 | 15 | 16 | 12 | 10 | 2209 | 2061 | 7409 | 678 | 27 | 12 | 14 | 0 | 0 | 8079 | 27286 | 25592 | 70 | 1165 | 49857 | 339 | 0 | 0 | 16 | 12 | 62 | 160 | 146 | 0 | 36 | 0 | 78 | 0 | 0 | 988 | 0 | 0 |
| SRR4836302 | 33 | 50 | 15 | 24 | 11 | 2290 | 1922 | 7123 | 546 | 11 | 26 | 15 | 0 | 0 | 8509 | 29315 | 26464 | 74 | 1366 | 49785 | 378 | 0 | 0 | 39 | 28 | 139 | 367 | 402 | 0 | 83 | 0 | 84 | 0 | 0 | 1059 | 0 | 0 |
| SRR4836303 | 44 | 45 | 22 | 26 | 21 | 2450 | 2214 | 7275 | 921 | 12 | 31 | 13 | 0 | 0 | 9092 | 28950 | 27681 | 90 | 1190 | 47934 | 294 | 0 | 0 | 38 | 32 | 185 | 424 | 365 | 0 | 83 | 0 | 32 | 0 | 0 | 898 | 0 | 0 |
| SRR4836305 | 25 | 20 | 11 | 44 | 49 | 1162 | 1233 | 3237 | 925 | 25 | 12 | 11 | 0 | 0 | 2294 | 38181 | 13191 | 124 | 1233 | 61906 | 4286 | 0 | 0 | 19 | 40 | 31 | 166 | 158 | 0 | 38 | 0 | 24 | 0 | 0 | 2406 | 0 | 0 |
| SRR4836307 | 18 | 18 | 42 | 72 | 42 | 1174 | 1233 | 3196 | 635 | 18 | 139 | 36 | 0 | 0 | 2569 | 38892 | 12921 | 66 | 1144 | 64508 | 3632 | 0 | 0 | 48 | 24 | 151 | 1049 | 862 | 0 | 199 | 0 | 52 | 0 | 0 | 2410 | 0 | 0 |
| SRR4836308 | 47 | 51 | 36 | 54 | 11 | 1220 | 1251 | 3124 | 875 | 18 | 18 | 58 | 0 | 0 | 2673 | 40142 | 13401 | 71 | 1061 | 60203 | 3964 | 0 | 0 | 18 | 29 | 30 | 174 | 156 | 0 | 34 | 0 | 27 | 0 | 0 | 1893 | 0 | 0 |
| SRR4836310 | 51 | 55 | 58 | 86 | 43 | 528 | 749 | 1104 | 268 | 16 | 0 | 47 | 0 | 0 | 2382 | 18238 | 6026 | 39 | 1782 | 30459 | 1215 | 0 | 0 | 113 | 62 | 304 | 674 | 904 | 0 | 113 | 0 | 47 | 0 | 0 | 1205 | 0 | 0 |
| SRR4836311 | 33 | 57 | 14 | 12 | 38 | 373 | 837 | 928 | 282 | 0 | 19 | 24 | 0 | 0 | 3056 | 19531 | 6594 | 30 | 1634 | 31086 | 1281 | 0 | 0 | 18 | 11 | 50 | 119 | 148 | 0 | 16 | 0 | 20 | 0 | 0 | 1669 | 0 | 0 |
| SRR4836312 | 43 | 14 | 34 | 17 | 43 | 452 | 821 | 1097 | 411 | 0 | 26 | 14 | 0 | 0 | 2693 | 20960 | 6409 | 35 | 1490 | 29438 | 1150 | 0 | 0 | 19 | 15 | 56 | 127 | 176 | 0 | 14 | 0 | 29 | 0 | 0 | 1305 | 0 | 0 |
| SRR4836313 | 133 | 136 | 80 | 59 | 26 | 1255 | 632 | 2633 | 301 | 13 | 50 | 29 | 0 | 0 | 8927 | 6564 | 6185 | 249 | 176 | 3328 | 705 | 0 | 0 | 19 | 0 | 39 | 189 | 19 | 0 | 123 | 0 | 31 | 0 | 0 | 514 | 0 | 0 |
| SRR4836314 | 38 | 40 | 11 | 14 | 31 | 1425 | 792 | 2475 | 287 | 12 | 13 | 26 | 0 | 0 | 8374 | 7436 | 6185 | 287 | 148 | 3553 | 574 | 0 | 0 | 33 | 0 | 31 | 50 | 21 | 0 | 35 | 0 | 119 | 0 | 0 | 647 | 0 | 0 |
| SRR4836315 | 169 | 131 | 52 | 61 | 25 | 1518 | 792 | 2706 | 297 | 20 | 30 | 30 | 0 | 0 | 7655 | 7884 | 6035 | 190 | 99 | 3259 | 742 | 0 | 0 | 20 | 0 | 44 | 180 | 27 | 0 | 127 | 0 | 58 | 0 | 0 | 547 | 0 | 0 |
| SRR4836316 | 52 | 15 | 14 | 12 | 0 | 217 | 412 | 613 | 182 | 0 | 16 | 0 | 0 | 0 | 4179 | 9660 | 7264 | 23 | 386 | 27179 | 1317 | 0 | 0 | 11 | 23 | 17 | 47 | 28 | 0 | 17 | 0 | 0 | 0 | 0 | 680 | 0 | 0 |
| SRR4836317 | 47 | 65 | 16 | 25 | 0 | 245 | 444 | 652 | 193 | 0 | 29 | 0 | 0 | 0 | 4279 | 8994 | 6909 | 43 | 447 | 26966 | 1199 | 0 | 0 | 45 | 16 | 115 | 278 | 34 | 0 | 101 | 0 | 0 | 0 | 0 | 712 | 0 | 0 |
| SRR4836318 | 46 | 12 | 15 | 29 | 0 | 230 | 506 | 707 | 177 | 13 | 25 | 0 | 0 | 0 | 4444 | 9429 | 7558 | 13 | 421 | 25900 | 1091 | 0 | 0 | 12 | 32 | 20 | 56 | 50 | 0 | 19 | 0 | 0 | 0 | 0 | 611 | 0 | 0 |
| SRR4836320 | 83 | 37 | 70 | 10 | 83 | 95 | 472 | 271 | 162 | 0 | 0 | 0 | 0 | 0 | 1835 | 111221 | 17869 | 40 | 499 | 38450 | 13870 | 0 | 0 | 41 | 11 | 28 | 53 | 12 | 0 | 25 | 0 | 0 | 0 | 0 | 990 | 0 | 0 |
| SRR4836321 | 12 | 84 | 44 | 84 | 56 | 138 | 553 | 301 | 156 | 12 | 16 | 0 | 0 | 0 | 1904 | 110654 | 16754 | 30 | 515 | 37783 | 13664 | 0 | 0 | 60 | 72 | 29 | 68 | 28 | 0 | 22 | 0 | 0 | 0 | 0 | 878 | 0 | 0 |
| SRR4836323 | 68 | 47 | 78 | 74 | 58 | 143 | 594 | 334 | 135 | 0 | 20 | 0 | 0 | 0 | 1862 | 109430 | 18286 | 32 | 504 | 37115 | 13081 | 0 | 0 | 91 | 91 | 32 | 59 | 0 | 0 | 21 | 0 | 0 | 0 | 0 | 962 | 0 | 0 |
| SRR4836325 | 40 | 26 | 54 | 43 | 0 | 1464 | 900 | 4344 | 228 | 11 | 22 | 0 | 0 | 0 | 4056 | 8665 | 5637 | 48 | 204 | 3444 | 540 | 0 | 0 | 19 | 38 | 23 | 28 | 16 | 0 | 15 | 0 | 36 | 0 | 0 | 1064 | 0 | 0 |
| SRR4836326 | 46 | 27 | 49 | 47 | 0 | 1471 | 942 | 4655 | 138 | 0 | 31 | 0 | 0 | 0 | 4402 | 8313 | 5767 | 103 | 218 | 3391 | 529 | 0 | 0 | 23 | 39 | 29 | 25 | 18 | 0 | 16 | 0 | 92 | 0 | 0 | 905 | 0 | 0 |
| SRR4836327 | 251 | 179 | 31 | 55 | 0 | 1728 | 1053 | 4390 | 189 | 11 | 24 | 0 | 0 | 0 | 4301 | 8551 | 5374 | 30 | 258 | 3476 | 368 | 0 | 0 | 24 | 42 | 170 | 216 | 24 | 0 | 95 | 0 | 79 | 0 | 0 | 848 | 0 | 0 |
| SRR4836328 | 12 | 23 | 16 | 15 | 25 | 1229 | 838 | 3084 | 279 | 15 | 23 | 0 | 0 | 0 | 7018 | 17470 | 18681 | 22 | 268 | 19680 | 1486 | 0 | 0 | 19 | 25 | 24 | 33 | 15 | 0 | 35 | 0 | 0 | 0 | 0 | 946 | 0 | 0 |
| SRR4836330 | 13 | 18 | 23 | 11 | 14 | 1132 | 748 | 3748 | 352 | 21 | 37 | 0 | 0 | 0 | 6962 | 17071 | 18003 | 53 | 267 | 18517 | 1484 | 0 | 0 | 18 | 0 | 68 | 145 | 18 | 0 | 153 | 0 | 11 | 0 | 0 | 805 | 0 | 0 |
| SRR4836332 | 33 | 25 | 19 | 16 | 13 | 1307 | 957 | 3497 | 267 | 12 | 34 | 13 | 0 | 0 | 7399 | 17350 | 18961 | 0 | 267 | 16989 | 1334 | 0 | 0 | 15 | 16 | 68 | 178 | 13 | 0 | 149 | 0 | 28 | 0 | 0 | 1061 | 0 | 0 |
| SRR4836334 | 21 | 57 | 17 | 34 | 32 | 2014 | 1417 | 5138 | 273 | 11 | 23 | 27 | 0 | 0 | 13928 | 14692 | 10966 | 51 | 188 | 5513 | 1161 | 0 | 0 | 19 | 0 | 17 | 42 | 11 | 0 | 28 | 0 | 154 | 0 | 0 | 1060 | 0 | 0 |
| SRR4836335 | 18 | 49 | 16 | 11 | 26 | 2250 | 1208 | 5310 | 397 | 22 | 11 | 28 | 0 | 0 | 14110 | 15048 | 11138 | 99 | 215 | 5012 | 1191 | 0 | 0 | 14 | 0 | 16 | 45 | 26 | 0 | 33 | 0 | 165 | 0 | 0 | 965 | 0 | 0 |
| SRR4836336 | 71 | 162 | 17 | 50 | 19 | 2025 | 1419 | 5344 | 413 | 26 | 35 | 10 | 0 | 0 | 13621 | 15850 | 11461 | 55 | 124 | 4586 | 909 | 0 | 0 | 12 | 17 | 53 | 148 | 14 | 0 | 91 | 0 | 138 | 0 | 0 | 881 | 0 | 0 |
| SRR4836337 | 12 | 17 | 20 | 17 | 17 | 1398 | 1364 | 4432 | 548 | 22 | 42 | 13 | 0 | 0 | 3514 | 10794 | 4167 | 106 | 756 | 24359 | 280 | 0 | 0 | 12 | 21 | 132 | 254 | 295 | 0 | 35 | 0 | 170 | 0 | 0 | 905 | 0 | 0 |
| SRR4836339 | 12 | 26 | 19 | 32 | 30 | 1537 | 1287 | 4903 | 574 | 18 | 44 | 28 | 0 | 0 | 3857 | 10408 | 3675 | 97 | 681 | 22896 | 259 | 0 | 0 | 60 | 89 | 539 | 1111 | 1299 | 0 | 119 | 0 | 185 | 0 | 0 | 711 | 0 | 0 |
| SRR4836341 | 21 | 18 | 30 | 25 | 28 | 1482 | 1330 | 4674 | 594 | 18 | 53 | 45 | 0 | 0 | 3672 | 11268 | 4035 | 107 | 616 | 23065 | 284 | 0 | 0 | 66 | 89 | 595 | 1344 | 1343 | 0 | 139 | 0 | 149 | 0 | 0 | 824 | 0 | 0 |
| SRR4836343 | 24 | 40 | 17 | 28 | 30 | 1618 | 1421 | 5321 | 733 | 15 | 14 | 44 | 0 | 0 | 3907 | 11285 | 4188 | 69 | 674 | 22534 | 221 | 0 | 0 | 13 | 19 | 143 | 262 | 308 | 0 | 31 | 0 | 114 | 0 | 0 | 660 | 0 | 0 |
| SRR4836344 | 11 | 16 | 11 | 49 | 0 | 374 | 584 | 1434 | 262 | 0 | 10 | 15 | 0 | 0 | 3060 | 5608 | 5550 | 14 | 173 | 7175 | 276 | 0 | 0 | 11 | 63 | 43 | 172 | 171 | 0 | 24 | 0 | 10 | 0 | 0 | 938 | 0 | 0 |
| SRR4836345 | 92 | 183 | 51 | 81 | 20 | 389 | 656 | 1525 | 240 | 0 | 46 | 0 | 0 | 0 | 3125 | 5092 | 4889 | 21 | 144 | 6896 | 197 | 0 | 0 | 56 | 76 | 326 | 1338 | 1302 | 0 | 193 | 0 | 11 | 0 | 0 | 1018 | 0 | 0 |
| SRR4836346 | 84 | 134 | 73 | 65 | 27 | 362 | 652 | 1607 | 282 | 15 | 57 | 0 | 0 | 0 | 3089 | 5466 | 5189 | 26 | 172 | 7060 | 253 | 0 | 0 | 115 | 76 | 359 | 1670 | 1506 | 0 | 206 | 0 | 11 | 0 | 0 | 860 | 0 | 0 |
| SRR4836347 | 66 | 155 | 43 | 43 | 0 | 427 | 749 | 1768 | 270 | 12 | 58 | 0 | 0 | 0 | 3222 | 5471 | 5246 | 16 | 153 | 6662 | 197 | 0 | 0 | 128 | 58 | 383 | 1524 | 1574 | 0 | 190 | 0 | 0 | 0 | 0 | 821 | 0 | 0 |
| SRR4836348 | 17 | 29 | 13 | 49 | 41 | 1312 | 822 | 3705 | 159 | 16 | 15 | 0 | 0 | 0 | 1745 | 2805 | 2564 | 43 | 14 | 1932 | 519 | 0 | 0 | 37 | 33 | 41 | 105 | 145 | 0 | 23 | 0 | 29 | 0 | 0 | 787 | 0 | 0 |
| SRR4836349 | 10 | 18 | 43 | 30 | 30 | 1521 | 729 | 3430 | 233 | 0 | 56 | 13 | 0 | 0 | 1769 | 2740 | 2923 | 31 | 78 | 1924 | 559 | 0 | 0 | 13 | 34 | 46 | 107 | 150 | 0 | 31 | 0 | 16 | 0 | 0 | 809 | 0 | 0 |
| SRR4836350 | 16 | 18 | 10 | 45 | 48 | 1373 | 627 | 3363 | 303 | 0 | 13 | 13 | 0 | 0 | 2033 | 3067 | 2839 | 65 | 43 | 1860 | 562 | 0 | 0 | 42 | 35 | 47 | 120 | 156 | 0 | 19 | 0 | 43 | 0 | 0 | 762 | 0 | 0 |
| SRR4836351 | 21 | 22 | 53 | 13 | 53 | 1300 | 716 | 3978 | 242 | 0 | 12 | 0 | 0 | 0 | 2083 | 3126 | 2906 | 33 | 77 | 1917 | 507 | 0 | 0 | 17 | 40 | 43 | 124 | 193 | 0 | 20 | 0 | 11 | 0 | 0 | 641 | 0 | 0 |
| SRR4836352 | 15 | 19 | 12 | 11 | 28 | 728 | 1566 | 2618 | 894 | 15 | 32 | 0 | 0 | 0 | 2127 | 10681 | 4642 | 55 | 514 | 14141 | 419 | 0 | 0 | 13 | 0 | 30 | 105 | 93 | 0 | 57 | 0 | 47 | 0 | 0 | 1029 | 0 | 0 |
| SRR4836353 | 13 | 17 | 13 | 22 | 12 | 764 | 1799 | 2555 | 764 | 0 | 19 | 10 | 0 | 0 | 2105 | 9260 | 4865 | 85 | 356 | 13428 | 416 | 0 | 0 | 12 | 14 | 32 | 128 | 70 | 0 | 52 | 0 | 68 | 0 | 0 | 1070 | 0 | 0 |
| SRR4836354 | 11 | 23 | 11 | 15 | 27 | 731 | 1529 | 2561 | 933 | 0 | 34 | 16 | 0 | 0 | 2094 | 10324 | 4727 | 74 | 442 | 14694 | 491 | 0 | 0 | 22 | 14 | 31 | 108 | 94 | 0 | 58 | 0 | 86 | 0 | 0 | 1222 | 0 | 0 |
| SRR4836355 | 32 | 68 | 32 | 49 | 24 | 784 | 1569 | 2400 | 855 | 11 | 78 | 0 | 0 | 0 | 1952 | 11038 | 4891 | 53 | 472 | 12094 | 289 | 0 | 0 | 31 | 16 | 119 | 423 | 343 | 0 | 162 | 0 | 106 | 0 | 0 | 984 | 0 | 0 |
| SRR4836356 | 23 | 32 | 32 | 18 | 23 | 3553 | 2463 | 11683 | 1083 | 14 | 50 | 0 | 0 | 0 | 9606 | 13260 | 15960 | 89 | 401 | 14865 | 215 | 0 | 0 | 28 | 11 | 50 | 173 | 271 | 0 | 114 | 0 | 45 | 0 | 0 | 1923 | 0 | 0 |
| SRR4836357 | 24 | 21 | 28 | 21 | 25 | 3571 | 2414 | 13111 | 914 | 11 | 17 | 0 | 0 | 0 | 8876 | 12929 | 15421 | 94 | 359 | 13541 | 172 | 0 | 0 | 13 | 14 | 21 | 54 | 81 | 0 | 35 | 0 | 39 | 0 | 0 | 1459 | 0 | 0 |
| SRR4836359 | 29 | 23 | 35 | 28 | 14 | 3225 | 2463 | 12428 | 951 | 0 | 51 | 0 | 0 | 0 | 9331 | 13907 | 15100 | 61 | 417 | 14909 | 228 | 0 | 0 | 34 | 0 | 68 | 195 | 295 | 0 | 94 | 0 | 33 | 0 | 0 | 1760 | 0 | 0 |
| SRR4836361 | 27 | 29 | 21 | 18 | 21 | 3610 | 2917 | 14032 | 1031 | 0 | 50 | 0 | 0 | 0 | 8981 | 13719 | 14943 | 54 | 416 | 14213 | 157 | 0 | 0 | 30 | 12 | 66 | 181 | 311 | 0 | 105 | 0 | 78 | 0 | 0 | 1784 | 0 | 0 |
| SRR4836362 | 28 | 62 | 48 | 27 | 60 | 1106 | 731 | 3635 | 250 | 22 | 175 | 79 | 0 | 0 | 9654 | 12036 | 26542 | 99 | 191 | 2180 | 3747 | 0 | 0 | 22 | 39 | 168 | 793 | 433 | 0 | 277 | 0 | 151 | 0 | 0 | 1152 | 0 | 0 |
| SRR4836364 | 29 | 12 | 33 | 22 | 15 | 1197 | 746 | 3773 | 275 | 29 | 29 | 11 | 0 | 0 | 9298 | 12049 | 24664 | 92 | 169 | 1774 | 3365 | 0 | 0 | 26 | 26 | 38 | 163 | 99 | 0 | 53 | 0 | 127 | 0 | 0 | 772 | 0 | 0 |
| SRR4836366 | 32 | 55 | 43 | 33 | 54 | 1360 | 783 | 4155 | 238 | 28 | 183 | 68 | 0 | 0 | 9946 | 12033 | 25788 | 69 | 196 | 2276 | 4007 | 0 | 0 | 23 | 25 | 183 | 812 | 452 | 0 | 242 | 0 | 159 | 0 | 0 | 1021 | 0 | 0 |
| SRR4836368 | 32 | 71 | 39 | 36 | 59 | 1390 | 797 | 3941 | 198 | 27 | 166 | 67 | 0 | 0 | 9860 | 12281 | 27327 | 64 | 198 | 1759 | 3561 | 0 | 0 | 21 | 35 | 188 | 843 | 507 | 0 | 251 | 0 | 160 | 0 | 0 | 820 | 0 | 0 |
| SRR4836369 | 103 | 93 | 32 | 22 | 19 | 1783 | 868 | 5896 | 387 | 0 | 106 | 27 | 0 | 0 | 7037 | 4967 | 6974 | 104 | 170 | 4198 | 557 | 0 | 0 | 32 | 17 | 139 | 427 | 543 | 0 | 184 | 0 | 47 | 0 | 0 | 607 | 0 | 0 |
| SRR4836370 | 102 | 64 | 29 | 26 | 29 | 1366 | 861 | 5682 | 253 | 0 | 119 | 15 | 0 | 0 | 7059 | 5200 | 7622 | 69 | 69 | 3604 | 413 | 0 | 0 | 14 | 14 | 108 | 364 | 423 | 0 | 178 | 0 | 80 | 0 | 0 | 559 | 0 | 0 |
| SRR4836371 | 79 | 109 | 21 | 33 | 42 | 1513 | 855 | 5482 | 219 | 0 | 112 | 25 | 0 | 0 | 6930 | 4850 | 7690 | 66 | 121 | 3838 | 373 | 0 | 0 | 22 | 16 | 97 | 368 | 455 | 0 | 148 | 0 | 66 | 0 | 0 | 581 | 0 | 0 |
| SRR4836373 | 103 | 90 | 29 | 30 | 24 | 1557 | 726 | 5345 | 315 | 16 | 111 | 34 | 0 | 0 | 7628 | 4858 | 8218 | 114 | 175 | 3228 | 420 | 0 | 0 | 27 | 16 | 138 | 435 | 492 | 0 | 196 | 0 | 70 | 0 | 0 | 645 | 0 | 0 |
| SRR4836375 | 12 | 19 | 11 | 13 | 25 | 1803 | 1174 | 5700 | 293 | 0 | 11 | 12 | 0 | 0 | 2606 | 8085 | 11129 | 20 | 160 | 4834 | 3636 | 0 | 0 | 17 | 47 | 57 | 121 | 123 | 0 | 35 | 0 | 16 | 0 | 0 | 1064 | 0 | 0 |
| SRR4836376 | 48 | 75 | 56 | 54 | 25 | 1701 | 915 | 5470 | 293 | 10 | 65 | 21 | 0 | 0 | 2661 | 8187 | 11191 | 35 | 149 | 4158 | 3531 | 0 | 0 | 31 | 33 | 226 | 577 | 550 | 0 | 159 | 0 | 25 | 0 | 0 | 997 | 0 | 0 |
| SRR4836378 | 52 | 70 | 26 | 50 | 16 | 1709 | 1016 | 5545 | 347 | 0 | 56 | 16 | 0 | 0 | 2411 | 8238 | 11187 | 0 | 101 | 4890 | 3509 | 0 | 0 | 30 | 36 | 260 | 612 | 608 | 0 | 199 | 0 | 14 | 0 | 0 | 1030 | 0 | 0 |
| SRR4836380 | 18 | 18 | 14 | 11 | 13 | 1703 | 1076 | 5393 | 465 | 11 | 11 | 28 | 0 | 0 | 2753 | 8316 | 12089 | 30 | 155 | 4230 | 3335 | 0 | 0 | 23 | 10 | 61 | 141 | 130 | 0 | 40 | 0 | 34 | 0 | 0 | 1076 | 0 | 0 |
| SRR4836382 | 19 | 25 | 11 | 18 | 15 | 718 | 869 | 2171 | 386 | 0 | 11 | 0 | 0 | 0 | 2852 | 29180 | 8544 | 32 | 429 | 14079 | 3817 | 0 | 0 | 17 | 17 | 116 | 282 | 247 | 0 | 26 | 0 | 97 | 0 | 0 | 717 | 0 | 0 |
| SRR4836383 | 12 | 22 | 0 | 21 | 21 | 581 | 728 | 1943 | 534 | 0 | 0 | 0 | 0 | 0 | 2958 | 29296 | 9193 | 60 | 347 | 13230 | 3486 | 0 | 0 | 10 | 17 | 109 | 249 | 226 | 0 | 31 | 0 | 53 | 0 | 0 | 1056 | 0 | 0 |
| SRR4836384 | 16 | 18 | 10 | 19 | 21 | 511 | 803 | 2117 | 440 | 0 | 14 | 0 | 0 | 0 | 3101 | 29944 | 9068 | 71 | 382 | 14322 | 3574 | 0 | 0 | 11 | 21 | 92 | 226 | 199 | 0 | 27 | 0 | 58 | 0 | 0 | 889 | 0 | 0 |
| SRR4836386 | 13 | 25 | 0 | 13 | 16 | 602 | 867 | 2123 | 457 | 0 | 0 | 0 | 0 | 0 | 2969 | 30832 | 10008 | 93 | 384 | 13446 | 3655 | 0 | 0 | 13 | 20 | 120 | 301 | 224 | 0 | 32 | 0 | 47 | 0 | 0 | 832 | 0 | 0 |
| SRR4836388 | 14 | 14 | 11 | 14 | 32 | 3208 | 1290 | 9597 | 574 | 15 | 11 | 11 | 0 | 0 | 5349 | 11079 | 11055 | 68 | 162 | 2553 | 412 | 0 | 0 | 19 | 41 | 57 | 189 | 79 | 0 | 25 | 0 | 41 | 0 | 0 | 878 | 0 | 0 |
| SRR4836390 | 69 | 77 | 67 | 89 | 24 | 2701 | 1048 | 8973 | 469 | 24 | 26 | 18 | 0 | 0 | 5650 | 11337 | 11731 | 60 | 102 | 2582 | 366 | 0 | 0 | 20 | 38 | 212 | 620 | 222 | 0 | 129 | 0 | 94 | 0 | 0 | 857 | 0 | 0 |
| SRR4836391 | 58 | 71 | 52 | 81 | 21 | 2873 | 1298 | 8814 | 449 | 23 | 14 | 15 | 0 | 0 | 5835 | 11383 | 11354 | 73 | 147 | 2718 | 310 | 0 | 0 | 17 | 27 | 202 | 646 | 276 | 0 | 129 | 0 | 90 | 0 | 0 | 986 | 0 | 0 |
| SRR4836392 | 17 | 14 | 16 | 15 | 27 | 2967 | 1110 | 9618 | 523 | 13 | 25 | 17 | 0 | 0 | 6180 | 11306 | 12857 | 97 | 116 | 2606 | 316 | 0 | 0 | 16 | 41 | 58 | 162 | 75 | 0 | 28 | 0 | 84 | 0 | 0 | 710 | 0 | 0 |
| SRR4836394 | 19 | 15 | 17 | 18 | 48 | 2118 | 1238 | 4594 | 506 | 61 | 23 | 12 | 0 | 0 | 1452 | 12545 | 10171 | 42 | 196 | 8301 | 345 | 0 | 0 | 30 | 16 | 90 | 399 | 316 | 0 | 71 | 0 | 48 | 0 | 0 | 2862 | 0 | 0 |
| SRR4836396 | 16 | 11 | 18 | 17 | 10 | 1843 | 1068 | 4198 | 421 | 15 | 12 | 11 | 0 | 0 | 1542 | 12735 | 10251 | 38 | 203 | 8524 | 263 | 0 | 0 | 24 | 16 | 28 | 110 | 93 | 0 | 22 | 0 | 45 | 0 | 0 | 2807 | 0 | 0 |
| SRR4836398 | 10 | 12 | 19 | 15 | 41 | 1906 | 1201 | 4409 | 396 | 52 | 19 | 13 | 0 | 0 | 1474 | 12933 | 10286 | 108 | 209 | 9048 | 324 | 0 | 0 | 24 | 12 | 106 | 368 | 316 | 0 | 77 | 0 | 72 | 0 | 0 | 3049 | 0 | 0 |
| SRR4836399 | 0 | 19 | 22 | 20 | 12 | 1941 | 1264 | 4503 | 368 | 18 | 17 | 11 | 0 | 0 | 1490 | 13556 | 11372 | 44 | 176 | 7768 | 247 | 0 | 0 | 34 | 17 | 28 | 121 | 103 | 0 | 16 | 0 | 82 | 0 | 0 | 2990 | 0 | 0 |
| SRR4836400 | 138 | 133 | 40 | 196 | 150 | 1183 | 572 | 3337 | 259 | 14 | 17 | 12 | 0 | 0 | 3862 | 7281 | 6778 | 24 | 188 | 3807 | 376 | 0 | 0 | 52 | 29 | 216 | 335 | 943 | 0 | 75 | 0 | 0 | 0 | 0 | 1650 | 0 | 0 |
| SRR4836402 | 26 | 24 | 13 | 41 | 24 | 993 | 482 | 2829 | 351 | 14 | 29 | 14 | 0 | 0 | 4244 | 6645 | 6966 | 30 | 231 | 3180 | 522 | 0 | 0 | 11 | 11 | 26 | 56 | 154 | 0 | 17 | 0 | 0 | 0 | 0 | 1621 | 0 | 0 |
| SRR4836404 | 155 | 152 | 52 | 149 | 121 | 1250 | 481 | 2895 | 192 | 0 | 38 | 10 | 0 | 0 | 4174 | 7305 | 6645 | 48 | 183 | 3991 | 490 | 0 | 0 | 41 | 45 | 131 | 356 | 756 | 0 | 111 | 0 | 0 | 0 | 0 | 1627 | 0 | 0 |
| SRR4836406 | 25 | 25 | 23 | 37 | 24 | 1171 | 727 | 3091 | 306 | 0 | 14 | 0 | 0 | 0 | 4269 | 7725 | 6920 | 23 | 214 | 3535 | 497 | 0 | 0 | 11 | 42 | 37 | 68 | 163 | 0 | 23 | 0 | 0 | 0 | 0 | 1745 | 0 | 0 |
| SRR4836407 | 23 | 18 | 13 | 11 | 21 | 851 | 420 | 2288 | 215 | 10 | 11 | 0 | 0 | 0 | 11637 | 5657 | 16480 | 39 | 117 | 2494 | 1457 | 0 | 0 | 27 | 0 | 0 | 0 | 0 | 0 | 40 | 0 | 29 | 0 | 0 | 1604 | 0 | 0 |
| SRR4836408 | 27 | 23 | 15 | 13 | 22 | 713 | 504 | 2338 | 162 | 0 | 25 | 0 | 0 | 0 | 12458 | 6292 | 17949 | 95 | 162 | 2167 | 1587 | 0 | 0 | 22 | 0 | 0 | 0 | 0 | 0 | 32 | 0 | 95 | 0 | 0 | 1365 | 0 | 0 |
| SRR4836410 | 92 | 56 | 43 | 39 | 33 | 836 | 521 | 2453 | 158 | 14 | 38 | 0 | 0 | 0 | 12106 | 5970 | 18286 | 71 | 71 | 2342 | 1538 | 0 | 0 | 22 | 0 | 0 | 0 | 0 | 0 | 101 | 0 | 47 | 0 | 0 | 1351 | 0 | 0 |
| SRR4836412 | 17 | 17 | 23 | 23 | 25 | 95 | 765 | 264 | 314 | 10 | 10 | 0 | 0 | 0 | 4291 | 63249 | 17958 | 24 | 1107 | 25839 | 2283 | 0 | 0 | 36 | 13 | 92 | 343 | 149 | 0 | 48 | 0 | 19 | 0 | 0 | 1489 | 0 | 0 |
| SRR4836414 | 23 | 21 | 29 | 33 | 21 | 96 | 728 | 236 | 314 | 27 | 14 | 0 | 0 | 0 | 4866 | 69466 | 18950 | 30 | 1261 | 25035 | 2531 | 0 | 0 | 37 | 35 | 97 | 346 | 132 | 0 | 45 | 0 | 21 | 0 | 0 | 1309 | 0 | 0 |
| SRR4836415 | 12 | 10 | 27 | 27 | 15 | 120 | 805 | 336 | 378 | 17 | 26 | 0 | 0 | 0 | 4753 | 68425 | 19400 | 27 | 1121 | 23559 | 2374 | 0 | 0 | 29 | 31 | 29 | 96 | 55 | 0 | 15 | 0 | 0 | 0 | 0 | 1176 | 0 | 0 |
| SRR4836416 | 22 | 32 | 19 | 20 | 11 | 462 | 384 | 689 | 61 | 0 | 22 | 0 | 0 | 0 | 7572 | 3025 | 6061 | 35 | 70 | 3001 | 890 | 0 | 0 | 13 | 0 | 18 | 98 | 43 | 0 | 36 | 0 | 0 | 0 | 0 | 704 | 0 | 0 |
| SRR4836417 | 20 | 23 | 20 | 25 | 17 | 508 | 379 | 714 | 95 | 0 | 10 | 0 | 0 | 0 | 7615 | 2955 | 7117 | 26 | 95 | 3098 | 792 | 0 | 0 | 13 | 0 | 19 | 97 | 48 | 0 | 27 | 0 | 0 | 0 | 0 | 791 | 0 | 0 |
| SRR4836419 | 41 | 68 | 40 | 47 | 13 | 411 | 482 | 667 | 50 | 0 | 19 | 0 | 0 | 0 | 7696 | 2489 | 6313 | 92 | 21 | 2901 | 787 | 0 | 0 | 13 | 0 | 44 | 250 | 119 | 0 | 71 | 0 | 0 | 0 | 0 | 624 | 0 | 0 |
| SRR4836421 | 31 | 53 | 29 | 12 | 21 | 5148 | 1784 | 13939 | 510 | 39 | 37 | 14 | 0 | 0 | 2154 | 30274 | 10949 | 75 | 161 | 5407 | 2934 | 0 | 0 | 39 | 31 | 51 | 137 | 60 | 0 | 70 | 0 | 105 | 0 | 0 | 1096 | 0 | 21 |
| SRR4836422 | 32 | 61 | 42 | 65 | 30 | 5232 | 1986 | 13237 | 539 | 44 | 202 | 91 | 0 | 0 | 2029 | 30345 | 12379 | 93 | 187 | 5380 | 3577 | 0 | 0 | 23 | 48 | 285 | 887 | 316 | 0 | 379 | 0 | 126 | 0 | 0 | 1188 | 0 | 15 |
| SRR4836423 | 43 | 12 | 38 | 11 | 28 | 4988 | 1965 | 13882 | 564 | 11 | 35 | 15 | 0 | 0 | 2082 | 30772 | 12478 | 58 | 140 | 4909 | 3061 | 0 | 0 | 41 | 44 | 54 | 160 | 71 | 0 | 68 | 0 | 50 | 0 | 0 | 990 | 0 | 11 |
| SRR4836425 | 51 | 82 | 51 | 43 | 47 | 994 | 721 | 3456 | 167 | 31 | 145 | 59 | 0 | 0 | 4845 | 16062 | 10759 | 114 | 167 | 6182 | 387 | 0 | 0 | 35 | 31 | 348 | 923 | 759 | 0 | 297 | 0 | 26 | 0 | 0 | 1768 | 0 | 0 |
| SRR4836427 | 34 | 68 | 57 | 42 | 26 | 1255 | 658 | 3255 | 175 | 45 | 159 | 83 | 0 | 0 | 5281 | 17190 | 11478 | 44 | 228 | 6220 | 483 | 0 | 0 | 72 | 45 | 321 | 933 | 751 | 0 | 283 | 0 | 70 | 0 | 0 | 1722 | 0 | 0 |
| SRR4836428 | 38 | 75 | 41 | 38 | 25 | 973 | 588 | 3695 | 140 | 16 | 194 | 94 | 0 | 0 | 5004 | 17310 | 10466 | 70 | 182 | 5984 | 413 | 0 | 0 | 22 | 38 | 347 | 1097 | 885 | 0 | 275 | 0 | 77 | 0 | 0 | 1536 | 0 | 19 |
| SRR4836429 | 37 | 43 | 81 | 55 | 12 | 531 | 945 | 1078 | 563 | 12 | 14 | 0 | 0 | 0 | 3890 | 6635 | 5491 | 101 | 241 | 12244 | 314 | 0 | 0 | 89 | 86 | 317 | 680 | 585 | 0 | 127 | 0 | 28 | 0 | 0 | 1260 | 0 | 0 |
| SRR4836431 | 42 | 59 | 11 | 10 | 40 | 557 | 878 | 1006 | 518 | 14 | 25 | 11 | 0 | 0 | 3946 | 7694 | 6007 | 101 | 186 | 11966 | 340 | 0 | 0 | 17 | 15 | 48 | 94 | 81 | 0 | 20 | 0 | 19 | 0 | 0 | 1248 | 0 | 0 |
| SRR4836433 | 40 | 50 | 11 | 50 | 28 | 556 | 808 | 1115 | 540 | 19 | 43 | 0 | 0 | 0 | 3924 | 7580 | 6677 | 65 | 226 | 11203 | 271 | 0 | 0 | 17 | 11 | 49 | 94 | 80 | 0 | 22 | 0 | 0 | 0 | 0 | 1130 | 0 | 0 |
| SRR4836434 | 24 | 11 | 19 | 11 | 24 | 1327 | 948 | 3775 | 414 | 125 | 35 | 16 | 0 | 0 | 7705 | 15519 | 9864 | 34 | 138 | 3982 | 776 | 0 | 0 | 21 | 16 | 32 | 122 | 11 | 0 | 85 | 0 | 17 | 0 | 0 | 1603 | 0 | 0 |
| SRR4836435 | 16 | 10 | 0 | 0 | 18 | 1411 | 963 | 4168 | 432 | 33 | 15 | 11 | 0 | 0 | 7754 | 15014 | 9843 | 149 | 133 | 4549 | 648 | 0 | 0 | 26 | 18 | 17 | 45 | 0 | 0 | 23 | 0 | 66 | 0 | 0 | 1683 | 0 | 0 |
| SRR4836437 | 19 | 15 | 17 | 17 | 15 | 1490 | 1088 | 3679 | 470 | 38 | 11 | 21 | 0 | 0 | 7734 | 15701 | 9679 | 94 | 40 | 3921 | 859 | 0 | 0 | 10 | 13 | 22 | 44 | 0 | 0 | 26 | 0 | 107 | 0 | 0 | 1446 | 0 | 0 |
| SRR4836439 | 21 | 12 | 17 | 17 | 30 | 1758 | 999 | 3822 | 573 | 34 | 17 | 23 | 0 | 0 | 7350 | 16230 | 9925 | 40 | 226 | 3755 | 879 | 0 | 0 | 28 | 15 | 17 | 58 | 13 | 0 | 17 | 0 | 67 | 0 | 0 | 1398 | 0 | 0 |
| SRR4836440 | 22 | 15 | 11 | 58 | 48 | 637 | 347 | 1795 | 232 | 0 | 77 | 19 | 0 | 0 | 1698 | 2898 | 3167 | 97 | 174 | 2413 | 502 | 0 | 0 | 96 | 15 | 19 | 41 | 58 | 0 | 22 | 0 | 39 | 0 | 0 | 584 | 0 | 0 |
| SRR4836441 | 272 | 216 | 56 | 28 | 38 | 846 | 508 | 1975 | 75 | 19 | 141 | 47 | 0 | 0 | 1768 | 3164 | 3054 | 132 | 94 | 2313 | 639 | 0 | 0 | 94 | 103 | 281 | 562 | 84 | 0 | 347 | 0 | 19 | 0 | 0 | 761 | 0 | 0 |
| SRR4836443 | 17 | 15 | 92 | 61 | 54 | 639 | 312 | 1502 | 178 | 0 | 84 | 46 | 0 | 0 | 1621 | 3468 | 3030 | 119 | 178 | 1948 | 535 | 0 | 0 | 61 | 18 | 26 | 45 | 61 | 0 | 36 | 0 | 30 | 0 | 0 | 715 | 0 | 0 |
| SRR4836445 | 306 | 168 | 92 | 53 | 46 | 679 | 498 | 1583 | 90 | 23 | 130 | 31 | 0 | 0 | 1312 | 3272 | 3263 | 106 | 211 | 2518 | 483 | 0 | 0 | 76 | 153 | 290 | 520 | 46 | 0 | 290 | 0 | 60 | 0 | 0 | 739 | 0 | 0 |
| SRR4836446 | 43 | 27 | 27 | 24 | 27 | 473 | 661 | 1839 | 429 | 19 | 70 | 0 | 0 | 0 | 15998 | 10317 | 24240 | 55 | 297 | 10327 | 1817 | 0 | 0 | 0 | 48 | 108 | 145 | 0 | 0 | 170 | 0 | 77 | 0 | 0 | 1312 | 0 | 0 |
| SRR4836447 | 66 | 48 | 32 | 48 | 45 | 569 | 903 | 1837 | 591 | 21 | 26 | 0 | 0 | 0 | 16193 | 10913 | 24505 | 150 | 290 | 10401 | 1977 | 0 | 0 | 0 | 45 | 95 | 151 | 0 | 0 | 164 | 0 | 54 | 0 | 0 | 1222 | 0 | 0 |
| SRR4836448 | 60 | 47 | 62 | 30 | 56 | 550 | 763 | 1641 | 568 | 28 | 47 | 0 | 0 | 0 | 16477 | 10333 | 23847 | 124 | 328 | 11224 | 2023 | 0 | 0 | 13 | 75 | 105 | 169 | 0 | 0 | 201 | 0 | 124 | 0 | 0 | 1584 | 0 | 0 |
| SRR4836450 | 31 | 12 | 13 | 13 | 14 | 580 | 870 | 1915 | 597 | 20 | 14 | 0 | 0 | 0 | 16926 | 11591 | 25409 | 88 | 272 | 10637 | 1959 | 0 | 0 | 0 | 24 | 40 | 67 | 0 | 0 | 79 | 0 | 97 | 0 | 0 | 1317 | 0 | 0 |
| SRR4836452 | 13 | 32 | 16 | 0 | 0 | 1686 | 948 | 4636 | 316 | 0 | 32 | 12 | 0 | 0 | 14013 | 10124 | 17559 | 26 | 316 | 3898 | 527 | 0 | 0 | 0 | 20 | 25 | 42 | 0 | 0 | 14 | 0 | 0 | 0 | 0 | 796 | 164 | 0 |
| SRR4836453 | 15 | 12 | 16 | 12 | 0 | 1132 | 824 | 5045 | 541 | 0 | 16 | 16 | 0 | 0 | 14619 | 9811 | 16200 | 26 | 154 | 3809 | 489 | 0 | 0 | 12 | 16 | 17 | 42 | 0 | 0 | 23 | 0 | 51 | 0 | 0 | 1043 | 182 | 0 |
| SRR4836454 | 51 | 13 | 22 | 13 | 0 | 1611 | 909 | 4401 | 537 | 0 | 25 | 13 | 0 | 0 | 12520 | 11030 | 17440 | 41 | 124 | 3554 | 599 | 0 | 0 | 16 | 0 | 57 | 210 | 0 | 0 | 86 | 0 | 62 | 0 | 0 | 928 | 142 | 0 |
| SRR4836456 | 13 | 17 | 13 | 0 | 0 | 1438 | 833 | 5438 | 625 | 0 | 19 | 0 | 0 | 0 | 14626 | 10934 | 16600 | 21 | 250 | 4146 | 667 | 0 | 0 | 13 | 19 | 20 | 42 | 13 | 0 | 20 | 0 | 21 | 0 | 0 | 864 | 105 | 0 |
| SRR4836458 | 16 | 16 | 0 | 0 | 17 | 1224 | 762 | 5305 | 218 | 0 | 20 | 0 | 0 | 0 | 6149 | 4352 | 10149 | 109 | 82 | 2449 | 299 | 0 | 0 | 0 | 0 | 23 | 21 | 0 | 0 | 12 | 0 | 0 | 0 | 0 | 675 | 0 | 0 |
| SRR4836459 | 13 | 10 | 0 | 11 | 0 | 1057 | 898 | 5046 | 291 | 0 | 23 | 0 | 0 | 0 | 5865 | 4609 | 8923 | 132 | 132 | 2219 | 528 | 0 | 0 | 0 | 0 | 28 | 19 | 0 | 0 | 24 | 0 | 53 | 0 | 0 | 940 | 0 | 0 |
| SRR4836461 | 64 | 59 | 14 | 0 | 11 | 1139 | 746 | 4971 | 249 | 0 | 21 | 0 | 0 | 0 | 5882 | 4705 | 9777 | 83 | 62 | 2320 | 394 | 0 | 0 | 0 | 0 | 23 | 55 | 0 | 0 | 80 | 0 | 83 | 0 | 0 | 706 | 0 | 0 |
| SRR4836463 | 72 | 51 | 12 | 14 | 14 | 1023 | 901 | 5240 | 327 | 0 | 25 | 12 | 0 | 0 | 5465 | 4985 | 9881 | 102 | 164 | 2538 | 307 | 0 | 0 | 0 | 0 | 28 | 90 | 0 | 0 | 69 | 0 | 102 | 0 | 0 | 697 | 0 | 0 |
| SRR4836464 | 23 | 27 | 23 | 68 | 305 | 1445 | 2489 | 3395 | 420 | 0 | 15 | 12 | 0 | 0 | 15031 | 82157 | 43304 | 112 | 696 | 11360 | 144 | 0 | 0 | 82 | 0 | 83 | 283 | 11 | 0 | 80 | 0 | 46 | 0 | 0 | 1825 | 0 | 0 |
| SRR4836466 | 13 | 0 | 30 | 55 | 268 | 1385 | 2532 | 3699 | 513 | 0 | 22 | 10 | 0 | 0 | 14770 | 84466 | 43575 | 180 | 776 | 11398 | 103 | 0 | 0 | 115 | 24 | 95 | 294 | 18 | 0 | 80 | 0 | 19 | 0 | 0 | 1808 | 0 | 0 |
| SRR4836468 | 23 | 12 | 28 | 15 | 67 | 1517 | 2810 | 3462 | 563 | 0 | 12 | 0 | 0 | 0 | 15616 | 83401 | 44154 | 177 | 709 | 10658 | 78 | 0 | 0 | 24 | 30 | 24 | 74 | 12 | 0 | 20 | 0 | 42 | 0 | 0 | 1812 | 0 | 0 |
| SRR4836469 | 34 | 19 | 33 | 71 | 253 | 1388 | 2849 | 3707 | 550 | 11 | 16 | 0 | 0 | 0 | 15133 | 83359 | 42791 | 175 | 756 | 10413 | 57 | 0 | 0 | 133 | 18 | 97 | 290 | 13 | 0 | 78 | 0 | 51 | 0 | 0 | 1497 | 0 | 0 |
| SRR4836471 | 12 | 14 | 30 | 46 | 21 | 515 | 188 | 1252 | 172 | 24 | 62 | 16 | 0 | 0 | 9614 | 8192 | 5760 | 123 | 155 | 6881 | 1882 | 0 | 0 | 46 | 0 | 46 | 19 | 14 | 0 | 25 | 0 | 180 | 0 | 0 | 937 | 0 | 0 |
| SRR4836473 | 101 | 127 | 46 | 28 | 165 | 578 | 214 | 1573 | 173 | 22 | 89 | 14 | 0 | 0 | 10010 | 8582 | 6424 | 156 | 104 | 6697 | 2145 | 0 | 0 | 66 | 0 | 28 | 173 | 0 | 0 | 153 | 0 | 121 | 0 | 0 | 1042 | 0 | 0 |
| SRR4836474 | 11 | 16 | 49 | 56 | 21 | 518 | 274 | 1374 | 151 | 0 | 11 | 21 | 0 | 0 | 10546 | 9092 | 6499 | 163 | 157 | 7432 | 1990 | 0 | 0 | 43 | 0 | 52 | 31 | 0 | 0 | 23 | 0 | 163 | 0 | 0 | 986 | 0 | 0 |
| SRR4836475 | 63 | 58 | 21 | 21 | 75 | 1847 | 796 | 4305 | 369 | 17 | 0 | 29 | 0 | 0 | 5690 | 2316 | 4092 | 104 | 104 | 1951 | 369 | 0 | 0 | 54 | 0 | 121 | 330 | 25 | 0 | 75 | 0 | 12 | 0 | 0 | 602 | 0 | 0 |
| SRR4836477 | 52 | 13 | 0 | 0 | 13 | 1792 | 876 | 4018 | 524 | 0 | 15 | 37 | 0 | 0 | 6211 | 2409 | 4268 | 65 | 98 | 2005 | 556 | 0 | 0 | 37 | 37 | 15 | 61 | 0 | 0 | 12 | 0 | 0 | 0 | 0 | 681 | 0 | 0 |
| SRR4836479 | 66 | 119 | 34 | 13 | 78 | 1979 | 964 | 4431 | 615 | 0 | 0 | 22 | 0 | 0 | 6493 | 2441 | 4492 | 42 | 83 | 2220 | 449 | 0 | 0 | 47 | 25 | 153 | 385 | 28 | 0 | 91 | 0 | 17 | 0 | 0 | 819 | 0 | 0 |

### Supp Table S8.3.Columns_BX_to_DH

| | | | | | | | | | | | | | | | | | | | | | | | | | | | | | | | | | | | | | |
|---|---|---|---|---|---|---|---|---|---|---|---|---|---|---|---|---|---|---|---|---|---|---|---|---|---|---|---|---|---|---|---|---|---|---|---|---|---|
| SRR4836481 | 15 | 26 | 25 | 21 | 17 | 292 | 273 | 565 | 156 | 0 | 0 | 0 | 0 | 0 | 3916 | 18050 | 5560 | 19 | 175 | 11164 | 4520 | 0 | 0 | 16 | 0 | 21 | 32 | 0 | 0 | 25 | 0 | 0 | 0 | 0 | 1092 | 0 | 0 |
| SRR4836482 | 61 | 147 | 0 | 24 | 52 | 251 | 306 | 641 | 195 | 0 | 0 | 0 | 0 | 0 | 4498 | 17980 | 6622 | 28 | 278 | 11362 | 5055 | 0 | 0 | 12 | 12 | 28 | 92 | 0 | 0 | 21 | 0 | 56 | 0 | 0 | 1461 | 0 | 0 |
| SRR4836483 | 20 | 40 | 0 | 13 | 12 | 307 | 321 | 697 | 293 | 0 | 0 | 0 | 0 | 0 | 3834 | 19192 | 6853 | 0 | 279 | 10513 | 5020 | 0 | 0 | 0 | 0 | 22 | 24 | 0 | 0 | 22 | 0 | 56 | 0 | 0 | 1592 | 0 | 0 |
| SRR4836484 | 251 | 343 | 80 | 69 | 46 | 931 | 2460 | 3750 | 525 | 0 | 80 | 0 | 0 | 0 | 6401 | 2977 | 7522 | 48 | 96 | 1266 | 478 | 0 | 0 | 103 | 91 | 217 | 846 | 11 | 0 | 606 | 0 | 0 | 0 | 0 | 1170 | 0 | 11 |
| SRR4836485 | 15 | 29 | 103 | 52 | 103 | 529 | 2235 | 3378 | 273 | 0 | 43 | 0 | 0 | 0 | 6586 | 2825 | 8415 | 68 | 119 | 1280 | 512 | 0 | 0 | 103 | 94 | 18 | 80 | 0 | 0 | 35 | 0 | 17 | 17 | 0 | 687 | 0 | 0 |
| SRR4836487 | 16 | 27 | 52 | 35 | 78 | 762 | 2199 | 3307 | 416 | 0 | 87 | 0 | 0 | 0 | 6978 | 2958 | 8855 | 52 | 208 | 1403 | 710 | 0 | 0 | 104 | 35 | 19 | 73 | 17 | 0 | 39 | 0 | 0 | 0 | 0 | 1469 | 0 | 0 |
| SRR4836489 | 28 | 35 | 17 | 0 | 10 | 927 | 757 | 3094 | 563 | 0 | 0 | 22 | 0 | 0 | 2466 | 9837 | 6275 | 80 | 917 | 34931 | 399 | 0 | 0 | 26 | 14 | 20 | 27 | 12 | 0 | 14 | 0 | 65 | 0 | 0 | 657 | 21 | 0 |
| SRR4836491 | 10 | 13 | 20 | 13 | 10 | 934 | 803 | 3055 | 500 | 14 | 0 | 32 | 0 | 0 | 2500 | 10834 | 6818 | 55 | 1062 | 36758 | 352 | 0 | 0 | 24 | 26 | 20 | 28 | 0 | 0 | 10 | 0 | 59 | 0 | 0 | 782 | 22 | 0 |
| SRR4836492 | 32 | 11 | 0 | 0 | 0 | 925 | 847 | 3223 | 560 | 0 | 0 | 11 | 0 | 0 | 2535 | 11463 | 7166 | 43 | 1149 | 36117 | 397 | 0 | 0 | 30 | 19 | 23 | 28 | 0 | 0 | 14 | 0 | 96 | 0 | 0 | 635 | 30 | 0 |
| SRR4836493 | 65 | 51 | 11 | 12 | 62 | 849 | 584 | 2155 | 265 | 0 | 0 | 0 | 0 | 0 | 7295 | 7250 | 16660 | 149 | 117 | 2793 | 2198 | 0 | 0 | 17 | 0 | 23 | 65 | 0 | 0 | 57 | 0 | 85 | 0 | 0 | 1294 | 15 | 0 |
| SRR4836494 | 76 | 60 | 17 | 16 | 66 | 805 | 447 | 2148 | 291 | 0 | 11 | 0 | 0 | 0 | 7375 | 8006 | 16481 | 127 | 104 | 2774 | 2558 | 0 | 0 | 19 | 0 | 15 | 84 | 0 | 0 | 60 | 0 | 97 | 0 | 0 | 1257 | 18 | 0 |
| SRR4836495 | 67 | 63 | 25 | 19 | 63 | 961 | 465 | 2522 | 263 | 0 | 0 | 0 | 0 | 0 | 7618 | 7761 | 17182 | 105 | 90 | 2807 | 2454 | 0 | 0 | 24 | 0 | 22 | 89 | 0 | 0 | 62 | 0 | 173 | 0 | 0 | 1364 | 21 | 0 |
| SRR4836496 | 17 | 16 | 10 | 29 | 12 | 418 | 332 | 1746 | 183 | 23 | 18 | 0 | 0 | 0 | 7355 | 6849 | 14089 | 126 | 309 | 8528 | 1190 | 0 | 0 | 11 | 18 | 19 | 70 | 91 | 0 | 30 | 0 | 103 | 0 | 0 | 767 | 0 | 0 |
| SRR4836498 | 66 | 59 | 29 | 29 | 0 | 512 | 351 | 1787 | 176 | 32 | 52 | 18 | 0 | 0 | 7944 | 7688 | 15144 | 92 | 229 | 8128 | 978 | 0 | 0 | 12 | 21 | 59 | 233 | 335 | 0 | 88 | 0 | 69 | 0 | 0 | 677 | 0 | 0 |
| SRR4836500 | 14 | 13 | 28 | 29 | 22 | 510 | 417 | 1582 | 180 | 13 | 13 | 11 | 0 | 0 | 8053 | 7453 | 13799 | 50 | 266 | 8628 | 1150 | 0 | 0 | 0 | 12 | 18 | 60 | 89 | 0 | 26 | 0 | 86 | 0 | 0 | 853 | 0 | 0 |
| SRR4836502 | 20 | 18 | 13 | 25 | 11 | 481 | 323 | 1661 | 159 | 16 | 17 | 14 | 0 | 0 | 8225 | 8121 | 15066 | 63 | 233 | 8648 | 1391 | 0 | 0 | 15 | 16 | 21 | 68 | 111 | 0 | 18 | 0 | 116 | 0 | 0 | 743 | 0 | 0 |
| SRR4836503 | 21 | 18 | 28 | 20 | 30 | 235 | 691 | 882 | 350 | 0 | 16 | 0 | 0 | 0 | 1939 | 87854 | 25795 | 70 | 1274 | 40498 | 1827 | 0 | 0 | 87 | 96 | 436 | 1039 | 1236 | 0 | 96 | 0 | 33 | 0 | 0 | 440 | 0 | 0 |
| SRR4836504 | 11 | 16 | 18 | 11 | 27 | 273 | 707 | 842 | 230 | 13 | 20 | 13 | 0 | 0 | 2054 | 90460 | 26294 | 54 | 1312 | 39006 | 1765 | 0 | 0 | 11 | 13 | 70 | 142 | 182 | 0 | 20 | 0 | 34 | 0 | 0 | 323 | 0 | 0 |
| SRR4836505 | 19 | 15 | 17 | 0 | 30 | 256 | 671 | 809 | 294 | 0 | 17 | 11 | 0 | 0 | 2053 | 89671 | 27354 | 43 | 1296 | 41397 | 1783 | 0 | 0 | 11 | 15 | 64 | 160 | 198 | 0 | 18 | 0 | 16 | 0 | 0 | 415 | 0 | 0 |
| SRR4836507 | 22 | 13 | 25 | 10 | 15 | 288 | 667 | 820 | 361 | 0 | 18 | 18 | 0 | 0 | 2242 | 96735 | 28738 | 41 | 1337 | 39455 | 1884 | 0 | 0 | 72 | 76 | 498 | 1136 | 1306 | 0 | 102 | 0 | 36 | 0 | 0 | 434 | 0 | 0 |
| SRR4836509 | 37 | 15 | 30 | 37 | 16 | 251 | 688 | 790 | 236 | 0 | 23 | 10 | 0 | 0 | 5317 | 53347 | 19499 | 35 | 1411 | 57363 | 5966 | 0 | 0 | 20 | 11 | 39 | 84 | 126 | 0 | 24 | 0 | 0 | 0 | 0 | 747 | 0 | 0 |
| SRR4836511 | 37 | 71 | 27 | 29 | 11 | 296 | 660 | 791 | 207 | 0 | 19 | 12 | 0 | 0 | 5573 | 55176 | 20452 | 32 | 1373 | 55036 | 5759 | 0 | 0 | 60 | 32 | 140 | 248 | 436 | 0 | 99 | 0 | 0 | 0 | 0 | 838 | 0 | 0 |
| SRR4836512 | 12 | 16 | 31 | 36 | 0 | 265 | 622 | 709 | 217 | 10 | 19 | 14 | 0 | 0 | 5650 | 54966 | 20352 | 42 | 1421 | 58481 | 5628 | 0 | 0 | 16 | 30 | 40 | 76 | 110 | 0 | 24 | 0 | 0 | 0 | 0 | 997 | 0 | 0 |
| SRR4836513 | 15 | 19 | 35 | 11 | 10 | 276 | 672 | 809 | 246 | 0 | 22 | 19 | 0 | 0 | 5844 | 58826 | 21036 | 37 | 1388 | 56771 | 6333 | 0 | 0 | 20 | 11 | 40 | 83 | 131 | 0 | 24 | 0 | 10 | 0 | 0 | 835 | 0 | 0 |
| SRR4836514 | 125 | 64 | 79 | 60 | 25 | 347 | 378 | 1426 | 167 | 0 | 62 | 0 | 0 | 0 | 10907 | 5254 | 21908 | 50 | 74 | 2071 | 1922 | 0 | 0 | 0 | 11 | 91 | 418 | 107 | 0 | 120 | 0 | 31 | 0 | 0 | 847 | 0 | 0 |
| SRR4836515 | 42 | 21 | 25 | 16 | 17 | 350 | 267 | 1317 | 175 | 0 | 14 | 0 | 0 | 0 | 11492 | 6046 | 21593 | 67 | 67 | 2375 | 1875 | 0 | 0 | 0 | 0 | 25 | 95 | 36 | 0 | 31 | 0 | 25 | 0 | 0 | 783 | 0 | 0 |
| SRR4836517 | 110 | 68 | 51 | 62 | 22 | 220 | 228 | 1022 | 87 | 0 | 61 | 0 | 0 | 0 | 11803 | 5751 | 21963 | 94 | 16 | 2595 | 1801 | 0 | 0 | 13 | 25 | 98 | 346 | 116 | 0 | 104 | 0 | 0 | 0 | 0 | 867 | 0 | 0 |
| SRR4836519 | 144 | 80 | 77 | 76 | 20 | 339 | 287 | 1363 | 105 | 0 | 60 | 0 | 0 | 0 | 11780 | 6458 | 24862 | 58 | 35 | 2351 | 2082 | 0 | 0 | 0 | 15 | 90 | 433 | 126 | 0 | 107 | 0 | 0 | 0 | 0 | 832 | 0 | 0 |
| SRR4836521 | 25 | 40 | 0 | 40 | 38 | 434 | 873 | 1166 | 709 | 0 | 29 | 0 | 0 | 0 | 2731 | 14096 | 11363 | 0 | 1155 | 50324 | 1049 | 0 | 0 | 31 | 178 | 649 | 1974 | 578 | 0 | 76 | 0 | 0 | 0 | 0 | 1259 | 0 | 0 |
| SRR4836522 | 11 | 34 | 0 | 43 | 43 | 490 | 836 | 1141 | 667 | 0 | 31 | 0 | 0 | 0 | 2902 | 13986 | 11113 | 0 | 1214 | 48425 | 1125 | 0 | 0 | 51 | 207 | 513 | 1766 | 612 | 0 | 71 | 0 | 0 | 0 | 0 | 1444 | 0 | 0 |
| SRR4836523 | 19 | 46 | 16 | 30 | 27 | 592 | 658 | 1125 | 629 | 0 | 22 | 0 | 0 | 0 | 2900 | 14140 | 9941 | 37 | 1036 | 52304 | 1051 | 0 | 0 | 48 | 213 | 546 | 1913 | 476 | 0 | 89 | 0 | 0 | 0 | 0 | 1664 | 0 | 0 |
| SRR4836524 | 26 | 37 | 20 | 46 | 41 | 361 | 852 | 1180 | 754 | 11 | 37 | 0 | 0 | 0 | 2836 | 15235 | 11589 | 0 | 994 | 49406 | 1197 | 0 | 0 | 39 | 204 | 709 | 2178 | 570 | 0 | 89 | 0 | 0 | 0 | 0 | 1582 | 0 | 0 |
| SRR4836525 | 128 | 70 | 86 | 77 | 30 | 325 | 188 | 1037 | 85 | 0 | 65 | 0 | 0 | 0 | 7773 | 5225 | 16079 | 28 | 51 | 2234 | 1356 | 0 | 0 | 0 | 16 | 106 | 337 | 105 | 0 | 133 | 0 | 11 | 0 | 0 | 999 | 0 | 0 |
| SRR4836526 | 142 | 58 | 119 | 72 | 34 | 310 | 170 | 1047 | 44 | 0 | 67 | 0 | 0 | 0 | 7868 | 5055 | 15908 | 59 | 52 | 2389 | 1136 | 0 | 0 | 11 | 11 | 70 | 291 | 96 | 0 | 147 | 0 | 15 | 0 | 0 | 876 | 0 | 0 |
| SRR4836528 | 41 | 23 | 23 | 25 | 19 | 407 | 120 | 874 | 67 | 0 | 23 | 0 | 0 | 0 | 7423 | 5058 | 15327 | 53 | 73 | 2430 | 1188 | 0 | 0 | 0 | 17 | 28 | 85 | 27 | 0 | 43 | 0 | 27 | 0 | 0 | 1024 | 0 | 0 |
| SRR4836529 | 189 | 76 | 104 | 112 | 21 | 354 | 172 | 1176 | 68 | 0 | 88 | 0 | 0 | 0 | 8000 | 5494 | 18375 | 57 | 36 | 2387 | 1436 | 0 | 0 | 0 | 18 | 90 | 328 | 92 | 0 | 116 | 0 | 10 | 0 | 0 | 1033 | 0 | 0 |
| SRR4836531 | 51 | 113 | 11 | 20 | 48 | 512 | 215 | 1273 | 157 | 20 | 57 | 68 | 0 | 0 | 5837 | 9805 | 7909 | 52 | 291 | 10673 | 7400 | 0 | 0 | 68 | 26 | 82 | 238 | 11 | 0 | 173 | 0 | 23 | 0 | 0 | 875 | 0 | 11 |
| SRR4836533 | 61 | 114 | 19 | 25 | 58 | 610 | 254 | 1502 | 181 | 17 | 33 | 67 | 0 | 0 | 6037 | 10454 | 8611 | 45 | 215 | 10821 | 7861 | 0 | 0 | 50 | 17 | 75 | 266 | 14 | 0 | 166 | 0 | 40 | 0 | 0 | 1028 | 0 | 0 |
| SRR4836534 | 11 | 23 | 15 | 41 | 30 | 583 | 231 | 1373 | 176 | 17 | 49 | 10 | 0 | 0 | 6059 | 10404 | 8262 | 45 | 235 | 9848 | 7574 | 0 | 0 | 46 | 24 | 18 | 49 | 19 | 0 | 27 | 0 | 41 | 0 | 0 | 837 | 0 | 0 |
| SRR4836535 | 39 | 41 | 0 | 0 | 10 | 366 | 1335 | 1217 | 895 | 0 | 14 | 0 | 0 | 0 | 3051 | 16640 | 7145 | 48 | 991 | 41121 | 999 | 0 | 0 | 119 | 0 | 0 | 0 | 0 | 0 | 96 | 0 | 15 | 0 | 0 | 1044 | 0 | 0 |
| SRR4836536 | 26 | 42 | 26 | 0 | 12 | 485 | 1244 | 1215 | 1076 | 12 | 16 | 0 | 0 | 0 | 3400 | 16899 | 6626 | 68 | 905 | 40761 | 1041 | 0 | 0 | 95 | 0 | 0 | 0 | 0 | 0 | 69 | 0 | 14 | 0 | 0 | 958 | 0 | 0 |
| SRR4836537 | 33 | 12 | 27 | 12 | 14 | 404 | 1356 | 1300 | 1038 | 0 | 30 | 0 | 0 | 0 | 3457 | 17698 | 7321 | 31 | 865 | 38288 | 943 | 0 | 0 | 28 | 0 | 0 | 0 | 0 | 0 | 19 | 0 | 22 | 0 | 0 | 822 | 0 | 0 |
| SRR4836539 | 0 | 0 | 0 | 0 | 13 | 1256 | 1328 | 5673 | 399 | 25 | 37 | 23 | 0 | 0 | 5529 | 88032 | 19733 | 155 | 1933 | 33736 | 991 | 0 | 0 | 65 | 56 | 209 | 122 | 0 | 0 | 216 | 0 | 93 | 0 | 0 | 1433 | 0 | 0 |
| SRR4836541 | 11 | 0 | 0 | 0 | 13 | 1321 | 1482 | 5754 | 415 | 19 | 36 | 15 | 0 | 0 | 5879 | 92354 | 20819 | 148 | 1960 | 33240 | 1044 | 0 | 0 | 47 | 45 | 251 | 137 | 14 | 0 | 182 | 0 | 105 | 0 | 0 | 1360 | 0 | 0 |
| SRR4836543 | 0 | 0 | 0 | 0 | 13 | 1485 | 1566 | 6419 | 461 | 16 | 28 | 22 | 0 | 0 | 5777 | 91908 | 20438 | 130 | 1959 | 31329 | 1011 | 0 | 0 | 46 | 53 | 228 | 133 | 12 | 0 | 232 | 0 | 90 | 0 | 0 | 1436 | 0 | 0 |
| SRR4836544 | 118 | 147 | 12 | 12 | 106 | 1904 | 1360 | 4381 | 453 | 12 | 32 | 21 | 0 | 0 | 1330 | 9887 | 11239 | 60 | 121 | 635 | 302 | 0 | 0 | 18 | 0 | 24 | 74 | 0 | 0 | 82 | 0 | 30 | 0 | 0 | 1602 | 0 | 0 |
| SRR4836545 | 84 | 164 | 12 | 20 | 58 | 1862 | 1064 | 3871 | 266 | 0 | 23 | 20 | 0 | 0 | 1152 | 11481 | 12275 | 30 | 207 | 296 | 236 | 0 | 0 | 23 | 0 | 14 | 89 | 0 | 0 | 78 | 0 | 30 | 0 | 0 | 1339 | 0 | 0 |
| SRR4836546 | 13 | 36 | 27 | 25 | 22 | 1566 | 1311 | 5590 | 382 | 0 | 0 | 11 | 0 | 0 | 1111 | 11166 | 11374 | 36 | 237 | 273 | 237 | 0 | 0 | 0 | 0 | 0 | 19 | 0 | 0 | 11 | 0 | 36 | 0 | 0 | 1535 | 0 | 0 |
| SRR4836548 | 85 | 67 | 27 | 49 | 19 | 217 | 133 | 559 | 42 | 0 | 0 | 31 | 0 | 0 | 2570 | 4594 | 4488 | 56 | 105 | 7173 | 363 | 0 | 0 | 58 | 36 | 17 | 52 | 18 | 0 | 17 | 0 | 14 | 0 | 0 | 1573 | 0 | 0 |
| SRR4836550 | 49 | 79 | 40 | 49 | 17 | 156 | 150 | 619 | 136 | 22 | 27 | 31 | 0 | 0 | 2843 | 5804 | 4467 | 14 | 197 | 6896 | 347 | 0 | 0 | 79 | 35 | 19 | 56 | 0 | 0 | 18 | 0 | 0 | 0 | 0 | 1690 | 0 | 0 |
| SRR4836552 | 85 | 66 | 16 | 33 | 194 | 206 | 185 | 647 | 76 | 0 | 19 | 25 | 0 | 0 | 2917 | 5162 | 4516 | 38 | 198 | 6393 | 425 | 0 | 0 | 76 | 16 | 104 | 451 | 11 | 0 | 142 | 0 | 0 | 0 | 0 | 1635 | 0 | 0 |
| SRR4836553 | 10 | 21 | 17 | 35 | 19 | 2211 | 2182 | 7062 | 1034 | 26 | 39 | 15 | 0 | 0 | 4708 | 28615 | 19643 | 172 | 459 | 18216 | 258 | 0 | 0 | 26 | 35 | 25 | 80 | 0 | 0 | 30 | 0 | 57 | 0 | 0 | 1077 | 0 | 0 |
| SRR4836554 | 68 | 76 | 13 | 47 | 68 | 2391 | 1737 | 6832 | 1039 | 30 | 42 | 144 | 0 | 0 | 5366 | 31672 | 19893 | 114 | 427 | 17934 | 441 | 0 | 0 | 42 | 34 | 110 | 482 | 0 | 0 | 148 | 0 | 28 | 0 | 0 | 1021 | 0 | 21 |
| SRR4836555 | 12 | 20 | 0 | 34 | 19 | 2512 | 1710 | 7947 | 1090 | 47 | 47 | 24 | 0 | 0 | 4327 | 30559 | 20661 | 122 | 497 | 16208 | 366 | 0 | 0 | 47 | 34 | 29 | 95 | 0 | 0 | 27 | 0 | 17 | 0 | 0 | 796 | 0 | 18 |
| SRR4836556 | 92 | 150 | 19 | 0 | 49 | 1423 | 1121 | 5476 | 474 | 0 | 44 | 19 | 0 | 0 | 6382 | 9292 | 9328 | 86 | 0 | 2005 | 86 | 0 | 0 | 68 | 0 | 15 | 29 | 0 | 0 | 87 | 0 | 22 | 0 | 0 | 1787 | 0 | 0 |
| SRR4836560 | 89 | 159 | 15 | 0 | 27 | 1499 | 1189 | 5600 | 345 | 0 | 23 | 31 | 0 | 0 | 6634 | 9340 | 9626 | 103 | 52 | 2120 | 138 | 0 | 0 | 77 | 0 | 0 | 31 | 12 | 0 | 89 | 0 | 17 | 0 | 0 | 1640 | 0 | 0 |
| SRR4836561 | 102 | 149 | 20 | 0 | 59 | 1285 | 1181 | 6287 | 417 | 12 | 35 | 20 | 0 | 0 | 6704 | 9995 | 8445 | 69 | 35 | 2032 | 69 | 0 | 0 | 67 | 0 | 0 | 12 | 0 | 0 | 117 | 0 | 0 | 0 | 0 | 1777 | 0 | 0 |
| SRR4836562 | 56 | 11 | 44 | 0 | 44 | 604 | 464 | 1148 | 434 | 0 | 0 | 0 | 0 | 0 | 2645 | 5504 | 4584 | 45 | 175 | 14501 | 7550 | 0 | 0 | 14 | 20 | 16 | 33 | 0 | 0 | 48 | 0 | 10 | 0 | 0 | 1839 | 0 | 0 |
| SRR4836563 | 36 | 55 | 16 | 0 | 20 | 585 | 565 | 1266 | 505 | 12 | 0 | 12 | 0 | 0 | 2607 | 5212 | 4658 | 55 | 165 | 14490 | 7185 | 0 | 0 | 18 | 32 | 12 | 53 | 0 | 0 | 59 | 0 | 15 | 0 | 0 | 1354 | 0 | 0 |
| SRR4836565 | 61 | 42 | 23 | 0 | 42 | 526 | 538 | 1302 | 461 | 19 | 0 | 13 | 0 | 0 | 2763 | 5274 | 3924 | 69 | 214 | 14513 | 8179 | 0 | 0 | 109 | 45 | 74 | 338 | 0 | 0 | 58 | 0 | 28 | 0 | 0 | 1681 | 0 | 0 |
| SRR4836567 | 59 | 72 | 29 | 0 | 29 | 582 | 533 | 1267 | 529 | 26 | 0 | 0 | 0 | 0 | 2813 | 5809 | 4545 | 53 | 238 | 13860 | 7299 | 0 | 0 | 114 | 36 | 124 | 447 | 0 | 0 | 49 | 0 | 12 | 0 | 0 | 1799 | 0 | 0 |
| SRR4836569 | 211 | 87 | 15 | 0 | 142 | 6454 | 1768 | 13765 | 597 | 66 | 0 | 51 | 0 | 0 | 14850 | 11057 | 15750 | 54 | 76 | 2083 | 2679 | 0 | 0 | 400 | 120 | 175 | 309 | 11 | 0 | 80 | 0 | 54 | 0 | 0 | 1354 | 0 | 15 |
| SRR4836570 | 16 | 79 | 0 | 0 | 143 | 6987 | 1653 | 14134 | 501 | 104 | 0 | 100 | 0 | 0 | 14870 | 9900 | 15865 | 21 | 64 | 2272 | 2720 | 0 | 0 | 24 | 165 | 10 | 24 | 21 | 0 | 32 | 0 | 53 | 0 | 0 | 1219 | 0 | 14 |
| SRR4836571 | 14 | 87 | 12 | 0 | 139 | 6043 | 2063 | 13624 | 713 | 78 | 0 | 75 | 0 | 0 | 14939 | 10612 | 16183 | 34 | 103 | 1865 | 2991 | 0 | 0 | 24 | 98 | 179 | 19 | 35 | 0 | 52 | 0 | 60 | 0 | 0 | 1474 | 0 | 23 |
| SRR4836572 | 15 | 88 | 20 | 0 | 120 | 6606 | 2424 | 14611 | 597 | 117 | 12 | 102 | 0 | 0 | 14884 | 10547 | 16195 | 0 | 128 | 1997 | 2705 | 0 | 0 | 23 | 135 | 14 | 17 | 38 | 0 | 44 | 0 | 85 | 0 | 0 | 1428 | 0 | 0 |
| SRR4896694 | 101 | 31 | 19 | 34 | 70 | 244 | 499 | 1400 | 179 | 170 | 165 | 0 | 0 | 0 | 125774 | 66509 | 452350 | 0 | 215 | 49197 | 1325 | 0 | 0 | 103 | 34 | 38 | 17 | 0 | 0 | 242 | 0 | 0 | 0 | 0 | 129 | 0 | 0 |
| SRR4896695 | 24 | 24 | 40 | 36 | 193 | 185 | 351 | 610 | 139 | 116 | 36 | 0 | 0 | 0 | 74000 | 150091 | 339139 | 0 | 37 | 29219 | 3336 | 0 | 0 | 241 | 44 | 44 | 209 | 0 | 0 | 321 | 0 | 65 | 0 | 0 | 1604 | 0 | 0 |
| SRR4896698 | 64 | 48 | 0 | 21 | 209 | 68 | 169 | 188 | 74 | 107 | 70 | 0 | 0 | 0 | 209336 | 109328 | 421080 | 0 | 68 | 7145 | 5277 | 0 | 0 | 284 | 43 | 27 | 107 | 0 | 0 | 354 | 0 | 25 | 0 | 0 | 576 | 0 | 11 |
| SRR4896701 | 80 | 58 | 48 | 158 | 63 | 0 | 30 | 37 | 11 | 178 | 346 | 0 | 0 | 0 | 141952 | 134891 | 294478 | 0 | 219 | 40292 | 7330 | 0 | 0 | 113 | 33 | 58 | 223 | 10 | 0 | 967 | 0 | 0 | 0 | 0 | 469 | 0 | 0 |
| SRR4896708 | 180 | 199 | 23 | 18 | 68 | 24 | 57 | 73 | 29 | 199 | 59 | 0 | 0 | 0 | 269931 | 45858 | 625881 | 0 | 155 | 10483 | 18867 | 0 | 0 | 150 | 33 | 34 | 61 | 0 | 0 | 203 | 0 | 13 | 0 | 0 | 143 | 147 | 0 |
| SRR5022482 | 262 | 164 | 68 | 208 | 229 | 275 | 124 | 204 | 68 | 22 | 0 | 0 | 0 | 0 | 25418 | 18934 | 232861 | 0 | 78 | 12569 | 10077 | 0 | 0 | 1723 | 515 | 594 | 334 | 166 | 0 | 147 | 0 | 0 | 0 | 0 | 400 | 0 | 0 |
| SRR5022483 | 0 | 0 | 0 | 0 | 55 | 0 | 0 | 16 | 0 | 12 | 38 | 0 | 0 | 0 | 11958 | 20852 | 201664 | 87 | 465 | 10325 | 3047 | 0 | 0 | 14 | 53 | 39 | 26 | 0 | 0 | 34 | 0 | 0 | 0 | 0 | 4215 | 0 | 0 |
| SRR5022484 | 70 | 44 | 0 | 26 | 72 | 661 | 355 | 891 | 182 | 21 | 0 | 0 | 0 | 0 | 8816 | 11995 | 93957 | 0 | 77 | 4844 | 15415 | 0 | 0 | 13 | 141 | 184 | 83 | 117 | 0 | 13 | 0 | 0 | 0 | 0 | 1943 | 0 | 0 |
| SRR5022485 | 0 | 0 | 0 | 0 | 31 | 0 | 0 | 0 | 0 | 26 | 41 | 0 | 0 | 0 | 8337 | 25135 | 112501 | 127 | 332 | 11305 | 4010 | 0 | 0 | 0 | 0 | 0 | 0 | 0 | 0 | 420 | 0 | 67 | 0 | 0 | 5963 | 0 | 0 |
| SRR5029397 | 11 | 23 | 0 | 0 | 22 | 1578 | 335 | 1353 | 1496 | 282 | 0 | 0 | 16 | 16 | 4707 | 17582 | 117810 | 540 | 0 | 1851 | 792 | 0 | 0 | 45 | 0 | 0 | 0 | 0 | 0 | 14 | 0 | 0 | 0 | 27 | 83561 | 512 | 0 |
| SRR5029398 | 0 | 0 | 0 | 0 | 39 | 1457 | 387 | 1145 | 1910 | 707 | 12 | 11 | 0 | 0 | 99 | 1896 | 5780 | 156 | 0 | 519 | 66 | 0 | 0 | 41 | 0 | 0 | 0 | 0 | 0 | 117 | 0 | 0 | 0 | 0 | 49008 | 3093 | 0 |
| SRR5029399 | 14 | 44 | 0 | 0 | 37 | 2063 | 614 | 5379 | 5651 | 37 | 59 | 65 | 13 | 0 | 12613 | 71923 | 360277 | 386 | 89 | 5117 | 3315 | 0 | 0 | 36 | 105 | 33 | 41 | 0 | 0 | 68 | 0 | 99 | 0 | 15 | 24859 | 545 | 0 |
| SRR5029400 | 42 | 12 | 0 | 0 | 43 | 6578 | 944 | 8907 | 5119 | 952 | 0 | 15 | 12 | 0 | 16918 | 24188 | 187158 | 442 | 94 | 8526 | 7903 | 15 | 0 | 122 | 0 | 0 | 0 | 0 | 0 | 26 | 0 | 67 | 0 | 21 | 37923 | 3159 | 0 |
| SRR5029401 | 35 | 0 | 0 | 0 | 46 | 2670 | 524 | 2044 | 4855 | 616 | 0 | 0 | 0 | 0 | 243 | 3246 | 4379 | 460 | 26 | 1661 | 102 | 0 | 0 | 68 | 0 | 0 | 0 | 0 | 0 | 58 | 0 | 51 | 0 | 37 | 122258 | 1971 | 0 |
| SRR5029402 | 39 | 0 | 0 | 0 | 33 | 4130 | 608 | 1672 | 4294 | 479 | 12 | 0 | 13 | 0 | 1659 | 6505 | 34830 | 76 | 13 | 4370 | 253 | 0 | 0 | 171 | 0 | 0 | 0 | 0 | 0 | 118 | 0 | 38 | 0 | 0 | 72250 | 3975 | 0 |
| SRR5029403 | 0 | 0 | 0 | 0 | 69 | 2519 | 487 | 1771 | 3589 | 34 | 0 | 0 | 0 | 13 | 9269 | 24198 | 157741 | 559 | 36 | 1390 | 2472 | 0 | 0 | 21 | 0 | 0 | 0 | 0 | 0 | 61 | 0 | 95 | 0 | 0 | 59901 | 881 | 0 |
| SRR5029404 | 0 | 0 | 0 | 0 | 0 | 8991 | 1207 | 2132 | 4059 | 519 | 0 | 0 | 18 | 0 | 77 | 1969 | 2777 | 128 | 0 | 899 | 0 | 0 | 0 | 78 | 0 | 0 | 0 | 0 | 0 | 40 | 0 | 0 | 0 | 11 | 54787 | 2510 | 0 |
| SRR5029405 | 27 | 17 | 0 | 0 | 50 | 10281 | 933 | 2851 | 6704 | 875 | 0 | 11 | 0 | 0 | 553 | 6305 | 26841 | 346 | 17 | 3698 | 138 | 0 | 0 | 208 | 19 | 0 | 0 | 0 | 0 | 135 | 0 | 0 | 0 | 0 | 34807 | 2474 | 0 |
| SRR5029406 | 190 | 77 | 0 | 0 | 155 | 4419 | 801 | 1702 | 5746 | 693 | 14 | 20 | 10 | 0 | 2216 | 9695 | 44235 | 313 | 100 | 19390 | 213 | 0 | 0 | 106 | 0 | 0 | 0 | 0 | 0 | 417 | 0 | 138 | 0 | 31 | 114592 | 4434 | 0 |
| SRR5029407 | 10 | 0 | 0 | 0 | 18 | 5652 | 878 | 3092 | 3822 | 58 | 0 | 0 | 0 | 0 | 826 | 9705 | 15897 | 66 | 15 | 125 | 103 | 0 | 0 | 34 | 0 | 0 | 0 | 0 | 0 | 17 | 0 | 15 | 0 | 0 | 94470 | 772 | 0 |
| SRR5029408 | 0 | 0 | 0 | 0 | 0 | 2229 | 625 | 1421 | 3025 | 238 | 0 | 11 | 0 | 0 | 68 | 858 | 1994 | 114 | 0 | 273 | 34 | 0 | 0 | 52 | 0 | 0 | 0 | 0 | 0 | 24 | 0 | 0 | 0 | 0 | 62560 | 2497 | 0 |
| SRR5029409 | 36 | 0 | 0 | 0 | 75 | 3866 | 494 | 1563 | 3389 | 25 | 0 | 11 | 0 | 0 | 11582 | 17697 | 99747 | 66 | 33 | 1925 | 1135 | 0 | 0 | 25 | 0 | 0 | 0 | 0 | 0 | 84 | 0 | 247 | 0 | 0 | 59961 | 482 | 0 |
| SRR5029410 | 14 | 42 | 0 | 0 | 44 | 1607 | 228 | 1256 | 2740 | 72 | 69 | 42 | 0 | 0 | 25038 | 72109 | 506937 | 139 | 49 | 15586 | 2642 | 0 | 0 | 23 | 53 | 24 | 61 | 0 | 0 | 74 | 0 | 0 | 0 | 11 | 17355 | 959 | 0 |
| SRR5029411 | 12 | 0 | 0 | 0 | 129 | 3000 | 827 | 1654 | 3425 | 997 | 21 | 36 | 0 | 0 | 94 | 4245 | 10738 | 118 | 24 | 1039 | 118 | 0 | 0 | 371 | 0 | 0 | 12 | 0 | 0 | 186 | 0 | 0 | 0 | 12 | 43370 | 1710 | 0 |
| SRR5029412 | 59 | 14 | 0 | 0 | 226 | 4198 | 797 | 8070 | 7787 | 1693 | 0 | 20 | 14 | 0 | 35180 | 51284 | 576160 | 407 | 201 | 9346 | 8531 | 0 | 0 | 438 | 27 | 16 | 11 | 0 | 0 | 217 | 0 | 136 | 0 | 0 | 9855 | 698 | 0 |
| SRR5029414 | 0 | 0 | 0 | 0 | 0 | 6848 | 638 | 1556 | 3533 | 302 | 11 | 0 | 0 | 0 | 171 | 982 | 1394 | 202 | 16 | 1914 | 109 | 0 | 0 | 60 | 0 | 0 | 0 | 0 | 0 | 63 | 0 | 0 | 0 | 10 | 27018 | 1922 | 0 |
| SRR5029415 | 0 | 0 | 0 | 0 | 19 | 4380 | 531 | 3849 | 2787 | 280 | 23 | 0 | 0 | 0 | 6902 | 4335 | 121387 | 133 | 0 | 2389 | 1195 | 0 | 0 | 71 | 0 | 0 | 0 | 0 | 0 | 26 | 0 | 0 | 0 | 0 | 23121 | 2678 | 0 |
| SRR5029416 | 18 | 57 | 0 | 0 | 45 | 821 | 336 | 1141 | 2544 | 138 | 63 | 52 | 0 | 0 | 45038 | 36752 | 446718 | 1000 | 90 | 2440 | 2014 | 0 | 0 | 38 | 17 | 10 | 39 | 0 | 0 | 167 | 0 | 187 | 0 | 0 | 15579 | 3067 | 0 |
| SRR5029417 | 10 | 0 | 0 | 0 | 11 | 4494 | 894 | 2707 | 4371 | 285 | 0 | 0 | 0 | 0 | 991 | 3406 | 12743 | 425 | 18 | 416 | 230 | 0 | 0 | 125 | 0 | 0 | 0 | 0 | 0 | 11 | 0 | 53 | 0 | 13 | 75173 | 2070 | 0 |
| SRR5029418 | 70 | 24 | 0 | 0 | 95 | 3365 | 295 | 1555 | 3309 | 836 | 49 | 24 | 0 | 0 | 933 | 9091 | 30321 | 207 | 16 | 19558 | 303 | 0 | 0 | 95 | 15 | 0 | 0 | 0 | 0 | 645 | 0 | 0 | 0 | 16 | 52809 | 1713 | 0 |
| SRR5029419 | 0 | 0 | 0 | 0 | 34 | 683 | 239 | 696 | 1930 | 800 | 27 | 28 | 0 | 0 | 1167 | 5716 | 37569 | 212 | 0 | 1201 | 199 | 0 | 0 | 80 | 0 | 0 | 0 | 0 | 0 | 218 | 0 | 0 | 0 | 25 | 24125 | 2526 | 0 |
| SRR5029420 | 62 | 0 | 0 | 0 | 181 | 1585 | 147 | 848 | 1930 | 572 | 11 | 27 | 0 | 0 | 53867 | 37710 | 389187 | 160 | 61 | 4855 | 1536 | 0 | 0 | 65 | 0 | 0 | 0 | 0 | 0 | 373 | 0 | 49 | 0 | 0 | 62607 | 0 | 0 |
| SRR5029422 | 113 | 33 | 0 | 0 | 48 | 5886 | 841 | 2597 | 8779 | 36 | 0 | 20 | 20 | 0 | 1039 | 19575 | 47163 | 173 | 0 | 890 | 321 | 0 | 0 | 33 | 60 | 0 | 0 | 0 | 0 | 19 | 0 | 25 | 0 | 0 | 58014 | 4592 | 0 |
| SRR5029424 | 0 | 0 | 0 | 0 | 18 | 3112 | 170 | 3226 | 3735 | 20 | 0 | 0 | 0 | 0 | 13129 | 24471 | 269802 | 113 | 0 | 792 | 509 | 0 | 0 | 99 | 0 | 0 | 0 | 0 | 0 | 0 | 0 | 0 | 0 | 0 | 9571 | 0 | 0 |
| SRR5029425 | 12 | 0 | 0 | 0 | 22 | 4152 | 861 | 3884 | 8266 | 339 | 0 | 11 | 14 | 0 | 555 | 2141 | 10571 | 287 | 0 | 804 | 105 | 0 | 0 | 58 | 0 | 0 | 0 | 0 | 0 | 27 | 0 | 124 | 0 | 40 | 99843 | 3137 | 0 |
| SRR5029426 | 26 | 0 | 0 | 0 | 117 | 2823 | 736 | 1812 | 4613 | 513 | 0 | 0 | 0 | 13 | 1197 | 22832 | 40272 | 297 | 0 | 340 | 154 | 0 | 0 | 317 | 0 | 0 | 0 | 0 | 0 | 48 | 0 | 55 | 0 | 0 | 51730 | 881 | 0 |
| SRR5029427 | 121 | 45 | 0 | 0 | 163 | 2760 | 489 | 1789 | 3738 | 1040 | 15 | 26 | 0 | 0 | 2452 | 13357 | 63645 | 610 | 47 | 12279 | 824 | 0 | 0 | 110 | 10 | 0 | 0 | 0 | 0 | 248 | 0 | 54 | 0 | 30 | 80189 | 1594 | 0 |
| SRR5029428 | 11 | 22 | 0 | 0 | 129 | 811 | 352 | 3434 | 4408 | 218 | 0 | 0 | 13 | 22 | 36833 | 30992 | 638548 | 54 | 54 | 13332 | 9600 | 0 | 0 | 36 | 0 | 0 | 0 | 0 | 0 | 18 | 0 | 0 | 0 | 0 | 1494 | 0 | 0 |
| SRR5029429 | 41 | 10 | 0 | 0 | 284 | 5225 | 962 | 9222 | 8559 | 661 | 10 | 26 | 0 | 10 | 16122 | 59727 | 368128 | 630 | 100 | 4230 | 2189 | 0 | 0 | 434 | 10 | 10 | 0 | 0 | 0 | 134 | 0 | 116 | 0 | 0 | 28641 | 455 | 0 |
| SRR5029430 | 69 | 21 | 0 | 0 | 23 | 1672 | 252 | 1079 | 2607 | 35 | 21 | 37 | 11 | 0 | 47413 | 24264 | 197224 | 1115 | 36 | 2014 | 1906 | 0 | 0 | 18 | 11 | 0 | 0 | 0 | 0 | 15 | 0 | 54 | 0 | 0 | 40843 | 935 | 0 |
| SRR5029431 | 0 | 0 | 0 | 0 | 14 | 3471 | 642 | 1391 | 2202 | 341 | 0 | 0 | 0 | 0 | 229 | 1666 | 6359 | 122 | 0 | 688 | 31 | 0 | 0 | 29 | 0 | 0 | 0 | 0 | 0 | 29 | 0 | 0 | 0 | 12 | 40603 | 1279 | 0 |
| SRR5029432 | 82 | 16 | 0 | 0 | 26 | 2885 | 640 | 6229 | 3009 | 130 | 13 | 63 | 0 | 25 | 71759 | 72191 | 441811 | 784 | 67 | 1388 | 11948 | 0 | 0 | 30 | 22 | 0 | 0 | 0 | 0 | 12 | 0 | 52 | 0 | 0 | 8211 | 1258 | 0 |
| SRR5029433 | 0 | 0 | 0 | 0 | 0 | 8226 | 797 | 5065 | 5257 | 558 | 0 | 0 | 17 | 19 | 44 | 1942 | 2936 | 74 | 0 | 236 | 30 | 0 | 0 | 154 | 0 | 0 | 0 | 0 | 0 | 14 | 0 | 15 | 0 | 31 | 76220 | 2972 | 0 |
| SRR5029434 | 24 | 19 | 0 | 0 | 24 | 6989 | 438 | 2656 | 3298 | 231 | 0 | 0 | 0 | 0 | 3429 | 9333 | 93915 | 204 | 15 | 5982 | 350 | 0 | 0 | 53 | 0 | 0 | 0 | 0 | 0 | 40 | 0 | 44 | 0 | 0 | 80000 | 3743 | 0 |
| SRR5029435 | 50 | 16 | 0 | 0 | 153 | 1815 | 414 | 1114 | 3343 | 675 | 0 | 44 | 0 | 0 | 17000 | 56728 | 245609 | 605 | 32 | 4361 | 1544 | 0 | 0 | 184 | 0 | 0 | 0 | 0 | 0 | 212 | 0 | 111 | 0 | 16 | 28163 | 607 | 0 |
| SRR5029436 | 16 | 40 | 0 | 0 | 20 | 4391 | 734 | 2377 | 5188 | 221 | 29 | 34 | 29 | 0 | 42010 | 12447 | 210845 | 742 | 71 | 7518 | 6223 | 40 | 17 | 59 | 86 | 17 | 20 | 0 | 0 | 45 | 37 | 24 | 0 | 0 | 18795 | 3368 | 0 |
| SRR5029437 | 24 | 0 | 0 | 0 | 30 | 844 | 47 | 797 | 680 | 139 | 0 | 0 | 0 | 0 | 503636 | 17950 | 921723 | 23 | 23 | 2181 | 1759 | 0 | 0 | 115 | 0 | 0 | 0 | 0 | 0 | 0 | 0 | 0 | 0 | 0 | 221 | 0 | 0 |
| SRR5029438 | 55 | 0 | 0 | 0 | 11 | 1581 | 248 | 903 | 1853 | 76 | 12 | 17 | 0 | 0 | 24896 | 25229 | 284200 | 399 | 16 | 1941 | 2795 | 40 | 29 | 20 | 14 | 0 | 17 | 0 | 0 | 12 | 12 | 112 | 0 | 0 | 13285 | 3132 | 0 |
| SRR5029439 | 27 | 0 | 0 | 0 | 61 | 5441 | 1301 | 10695 | 15569 | 598 | 0 | 15 | 0 | 0 | 13335 | 19041 | 290149 | 161 | 64 | 2840 | 4346 | 61 | 12 | 45 | 10 | 0 | 0 | 0 | 0 | 11 | 14 | 71 | 0 | 13 | 16186 | 2070 | 0 |
| SRR5029440 | 42 | 0 | 0 | 0 | 31 | 1866 | 124 | 1928 | 1461 | 107 | 0 | 0 | 10 | 0 | 128798 | 19496 | 706501 | 62 | 0 | 4291 | 2736 | 0 | 0 | 102 | 0 | 0 | 0 | 0 | 0 | 0 | 0 | 0 | 0 | 0 | 1941 | 0 | 0 |
| SRR5029441 | 17 | 0 | 0 | 0 | 17 | 2967 | 849 | 2885 | 7050 | 141 | 0 | 0 | 13 | 0 | 209 | 2557 | 4241 | 70 | 12 | 1035 | 105 | 0 | 0 | 40 | 0 | 0 | 0 | 0 | 0 | 20 | 0 | 58 | 0 | 23 | 135803 | 2496 | 0 |
| SRR5029442 | 53 | 0 | 0 | 0 | 27 | 4282 | 989 | 2212 | 5924 | 246 | 0 | 15 | 0 | 0 | 3209 | 7993 | 43458 | 260 | 0 | 545 | 193 | 0 | 0 | 422 | 11 | 0 | 0 | 0 | 0 | 53 | 0 | 50 | 0 | 0 | 92136 | 5333 | 0 |
| SRR5029444 | 0 | 0 | 0 | 0 | 31 | 1651 | 0 | 4623 | 1651 | 912 | 15 | 0 | 0 | 0 | 45897 | 31077 | 366822 | 330 | 165 | 9906 | 6356 | 0 | 0 | 62 | 0 | 0 | 0 | 15 | 0 | 15 | 0 | 0 | 0 | 0 | 18936 | 0 | 0 |
| SRR5029446 | 11 | 0 | 0 | 0 | 29 | 1676 | 108 | 3568 | 5730 | 13 | 0 | 0 | 0 | 0 | 1676 | 10709 | 100858 | 108 | 0 | 108 | 486 | 0 | 0 | 116 | 15 | 0 | 0 | 0 | 0 | 0 | 0 | 0 | 0 | 0 | 16101 | 0 | 0 |
| SRR5029447 | 0 | 0 | 0 | 0 | 16 | 4288 | 973 | 2996 | 6137 | 112 | 0 | 0 | 11 | 0 | 877 | 4192 | 24759 | 385 | 39 | 626 | 405 | 0 | 0 | 66 | 0 | 0 | 0 | 0 | 10 | 37 | 0 | 48 | 0 | 14 | 71603 | 3601 | 0 |
| SRR5029448 | 24 | 11 | 0 | 0 | 172 | 2048 | 652 | 1317 | 3234 | 657 | 0 | 21 | 0 | 0 | 6143 | 23542 | 101727 | 887 | 13 | 430 | 1239 | 0 | 0 | 467 | 18 | 0 | 0 | 0 | 0 | 79 | 0 | 26 | 0 | 0 | 43160 | 595 | 0 |
| SRR5029452 | 23 | 0 | 0 | 0 | 68 | 646 | 45 | 706 | 526 | 128 | 0 | 0 | 34 | 0 | 361073 | 26632 | 880589 | 15 | 0 | 797 | 1353 | 0 | 0 | 98 | 19 | 0 | 0 | 0 | 0 | 19 | 0 | 0 | 0 | 0 | 1050 | 0 | 0 |
| SRR5029455 | 22 | 0 | 0 | 0 | 75 | 5384 | 525 | 6259 | 7578 | 102 | 0 | 47 | 0 | 0 | 46774 | 12278 | 117337 | 415 | 0 | 295 | 765 | 22 | 28 | 30 | 19 | 0 | 0 | 0 | 0 | 14 | 19 | 28 | 0 | 0 | 25221 | 2135 | 0 |
| SRR5029456 | 22 | 0 | 0 | 0 | 21 | 3970 | 562 | 2238 | 2662 | 110 | 0 | 16 | 22 | 0 | 4774 | 9979 | 46814 | 287 | 23 | 1664 | 2157 | 0 | 0 | 41 | 0 | 0 | 0 | 0 | 0 | 35 | 0 | 34 | 0 | 25 | 79782 | 907 | 0 |

### Supp Table S8.3.Columns_BX_to_DH

| | | | | | | | | | | | | | | | | | | | | | | | | | | | | | | | | | | | | | |
|---|---|---|---|---|---|---|---|---|---|---|---|---|---|---|---|---|---|---|---|---|---|---|---|---|---|---|---|---|---|---|---|---|---|---|---|---|---|
| SRR5029457 | 16 | 0 | 0 | 0 | 74 | 1729 | 699 | 1554 | 3301 | 553 | 0 | 19 | 0 | 0 | 1703 | 8774 | 32956 | 856 | 0 | 210 | 384 | 0 | 0 | 112 | 0 | 0 | 0 | 0 | 0 | 41 | 0 | 44 | 0 | 0 | 60756 | 0 | 0 |
| SRR5029459 | 160 | 69 | 0 | 0 | 286 | 1531 | 249 | 1111 | 2132 | 614 | 19 | 21 | 13 | 0 | 23957 | 25823 | 233068 | 567 | 62 | 2347 | 1548 | 0 | 0 | 160 | 37 | 0 | 0 | 0 | 0 | 417 | 0 | 79 | 0 | 0 | 56991 | 1425 | 0 |
| SRR5029460 | 10 | 0 | 0 | 0 | 49 | 6400 | 1251 | 9424 | 8447 | 168 | 0 | 0 | 0 | 0 | 11832 | 19227 | 170966 | 664 | 19 | 4228 | 1166 | 0 | 0 | 57 | 0 | 0 | 0 | 0 | 0 | 31 | 0 | 0 | 0 | 0 | 41564 | 929 | 0 |
| SRR5029462 | 81 | 42 | 0 | 0 | 72 | 2006 | 641 | 1816 | 6857 | 80 | 60 | 40 | 16 | 0 | 1771 | 27615 | 94285 | 1075 | 99 | 2213 | 723 | 0 | 0 | 53 | 17 | 36 | 89 | 0 | 0 | 104 | 0 | 27 | 0 | 0 | 64326 | 765 | 0 |
| SRR5029463 | 0 | 0 | 0 | 0 | 70 | 2177 | 581 | 968 | 2371 | 810 | 12 | 0 | 0 | 0 | 9096 | 16770 | 109385 | 629 | 0 | 1210 | 1306 | 0 | 0 | 123 | 0 | 0 | 0 | 0 | 12 | 88 | 0 | 48 | 0 | 0 | 38717 | 0 | 0 |
| SRR5029465 | 15 | 0 | 0 | 0 | 27 | 123 | 0 | 288 | 370 | 974 | 113 | 0 | 0 | 0 | 97667 | 24040 | 702182 | 27 | 123 | 8954 | 58376 | 0 | 0 | 0 | 0 | 0 | 0 | 0 | 0 | 1962 | 0 | 0 | 0 | 0 | 2234 | 0 | 0 |
| SRR5029466 | 93 | 12 | 0 | 0 | 296 | 4158 | 832 | 1663 | 6268 | 2197 | 31 | 31 | 0 | 0 | 15447 | 23737 | 299461 | 400 | 123 | 6422 | 1247 | 0 | 0 | 335 | 41 | 0 | 10 | 0 | 0 | 413 | 0 | 108 | 0 | 14 | 51709 | 2412 | 0 |
| SRR5029467 | 0 | 0 | 0 | 0 | 20 | 6506 | 1028 | 3237 | 5216 | 272 | 0 | 0 | 11 | 0 | 219 | 1435 | 3681 | 98 | 0 | 623 | 11 | 0 | 0 | 44 | 0 | 0 | 0 | 0 | 0 | 18 | 0 | 22 | 0 | 0 | 93840 | 3368 | 0 |
| SRR5029469 | 0 | 0 | 0 | 0 | 35 | 3878 | 380 | 1761 | 2716 | 400 | 0 | 0 | 17 | 0 | 20138 | 6487 | 65960 | 196 | 0 | 771 | 541 | 26 | 11 | 432 | 0 | 0 | 0 | 0 | 0 | 35 | 31 | 138 | 0 | 0 | 65974 | 4484 | 0 |
| SRR5029471 | 16 | 0 | 0 | 0 | 15 | 2572 | 623 | 3963 | 8500 | 318 | 0 | 13 | 15 | 0 | 32204 | 17555 | 221147 | 711 | 96 | 4050 | 6631 | 0 | 11 | 30 | 15 | 0 | 0 | 0 | 0 | 12 | 0 | 72 | 0 | 0 | 27628 | 2870 | 0 |
| SRR5029472 | 124 | 27 | 0 | 0 | 247 | 5115 | 377 | 1245 | 3342 | 659 | 21 | 24 | 35 | 0 | 21214 | 20769 | 196410 | 385 | 60 | 5673 | 5334 | 13 | 0 | 199 | 40 | 0 | 0 | 0 | 0 | 453 | 19 | 15 | 0 | 0 | 30815 | 2282 | 0 |
| SRR5029473 | 194 | 77 | 0 | 0 | 198 | 2956 | 791 | 2318 | 4822 | 391 | 11 | 37 | 31 | 18 | 3460 | 5317 | 29529 | 390 | 45 | 4104 | 904 | 0 | 0 | 246 | 21 | 0 | 0 | 0 | 0 | 203 | 0 | 170 | 0 | 30 | 138112 | 6116 | 0 |
| SRR5029474 | 123 | 22 | 0 | 0 | 212 | 1994 | 440 | 1356 | 3211 | 350 | 18 | 26 | 22 | 0 | 14424 | 23500 | 183310 | 223 | 24 | 2561 | 1795 | 0 | 0 | 129 | 21 | 0 | 0 | 0 | 0 | 352 | 0 | 66 | 0 | 0 | 94214 | 1920 | 0 |
| SRR5029476 | 118 | 36 | 0 | 0 | 235 | 1482 | 453 | 2065 | 2841 | 1020 | 11 | 42 | 0 | 0 | 47528 | 31907 | 462050 | 604 | 253 | 16824 | 11628 | 0 | 0 | 232 | 27 | 0 | 0 | 0 | 0 | 244 | 0 | 29 | 0 | 0 | 16434 | 1763 | 0 |
| SRR5029477 | 0 | 0 | 0 | 0 | 72 | 6147 | 669 | 3067 | 3206 | 663 | 0 | 0 | 34 | 16 | 6879 | 9966 | 158653 | 265 | 0 | 2158 | 4178 | 0 | 0 | 545 | 0 | 0 | 0 | 0 | 0 | 21 | 0 | 13 | 0 | 13 | 38146 | 3346 | 0 |
| SRR5029478 | 11 | 10 | 0 | 0 | 17 | 6799 | 631 | 2040 | 5330 | 438 | 0 | 13 | 23 | 0 | 5488 | 7814 | 59807 | 449 | 0 | 3133 | 401 | 0 | 0 | 54 | 20 | 0 | 0 | 0 | 0 | 26 | 0 | 97 | 0 | 16 | 51243 | 3006 | 0 |
| SRR5029480 | 29 | 13 | 0 | 0 | 17 | 2167 | 722 | 2355 | 7154 | 77 | 0 | 48 | 0 | 0 | 6303 | 26589 | 199000 | 772 | 30 | 801 | 851 | 0 | 0 | 36 | 44 | 0 | 0 | 0 | 0 | 13 | 0 | 89 | 0 | 11 | 43955 | 1476 | 0 |
| SRR5029481 | 54 | 0 | 0 | 0 | 58 | 6937 | 1045 | 2309 | 5823 | 262 | 0 | 0 | 24 | 0 | 115 | 2944 | 5517 | 57 | 34 | 494 | 230 | 0 | 0 | 222 | 0 | 0 | 0 | 0 | 0 | 125 | 0 | 92 | 0 | 0 | 94241 | 7008 | 0 |
| SRR5029482 | 0 | 0 | 0 | 0 | 0 | 7698 | 1206 | 12316 | 6291 | 229 | 0 | 0 | 0 | 0 | 19316 | 16177 | 170286 | 225 | 35 | 4600 | 5333 | 0 | 0 | 66 | 0 | 0 | 0 | 0 | 0 | 35 | 0 | 106 | 0 | 0 | 29812 | 4572 | 0 |
| SRR5029483 | 22 | 0 | 0 | 0 | 23 | 6576 | 1080 | 3335 | 4795 | 1458 | 0 | 0 | 0 | 0 | 3134 | 7008 | 56088 | 309 | 12 | 8522 | 831 | 0 | 0 | 172 | 0 | 0 | 0 | 0 | 0 | 35 | 0 | 12 | 0 | 0 | 23472 | 1687 | 0 |
| SRR5029484 | 50 | 11 | 0 | 0 | 94 | 1117 | 421 | 1419 | 3785 | 235 | 36 | 24 | 0 | 0 | 7724 | 14479 | 79296 | 437 | 131 | 2715 | 1332 | 10 | 11 | 20 | 40 | 0 | 18 | 0 | 0 | 149 | 28 | 187 | 0 | 16 | 146150 | 1859 | 0 |
| SRR5029486 | 10 | 11 | 0 | 0 | 47 | 1564 | 327 | 909 | 1596 | 625 | 23 | 36 | 10 | 0 | 59464 | 6457 | 57179 | 596 | 68 | 2000 | 1691 | 16 | 10 | 13 | 0 | 0 | 0 | 0 | 0 | 107 | 25 | 277 | 0 | 11 | 48281 | 4366 | 0 |
| SRR5029487 | 14 | 0 | 0 | 0 | 13 | 8689 | 1763 | 3900 | 6972 | 283 | 0 | 0 | 25 | 0 | 101 | 1183 | 2501 | 182 | 0 | 667 | 51 | 0 | 0 | 195 | 0 | 0 | 0 | 0 | 0 | 22 | 0 | 0 | 0 | 16 | 81230 | 3540 | 0 |
| SRR5029488 | 15 | 0 | 0 | 0 | 44 | 2223 | 153 | 2146 | 2376 | 706 | 0 | 15 | 0 | 15 | 3372 | 5984 | 35048 | 153 | 0 | 2146 | 537 | 0 | 15 | 162 | 0 | 0 | 15 | 0 | 0 | 147 | 0 | 77 | 0 | 15 | 67246 | 4187 | 0 |
| SRR5029489 | 20 | 35 | 0 | 0 | 34 | 5077 | 1235 | 3232 | 8221 | 80 | 19 | 16 | 11 | 0 | 1411 | 12285 | 66626 | 111 | 53 | 1786 | 281 | 0 | 0 | 75 | 49 | 10 | 13 | 0 | 0 | 37 | 0 | 64 | 0 | 0 | 93262 | 6351 | 0 |
| SRR5029490 | 36 | 15 | 0 | 0 | 199 | 1398 | 327 | 1376 | 2650 | 2967 | 12 | 62 | 0 | 0 | 33344 | 16174 | 340002 | 457 | 107 | 6337 | 13672 | 21 | 51 | 73 | 21 | 0 | 0 | 0 | 0 | 343 | 17 | 203 | 0 | 0 | 23726 | 6545 | 0 |
| SRR5029491 | 61 | 0 | 0 | 0 | 203 | 5449 | 575 | 3588 | 4235 | 799 | 0 | 28 | 0 | 21 | 18243 | 12208 | 140010 | 593 | 27 | 1636 | 1520 | 0 | 0 | 217 | 0 | 0 | 0 | 0 | 0 | 193 | 0 | 63 | 0 | 0 | 26852 | 3937 | 0 |
| SRR5029492 | 18 | 12 | 0 | 0 | 145 | 1602 | 354 | 2021 | 2423 | 766 | 0 | 12 | 0 | 0 | 55420 | 26139 | 401948 | 499 | 40 | 6915 | 14910 | 0 | 0 | 140 | 0 | 0 | 0 | 0 | 0 | 145 | 0 | 16 | 0 | 0 | 21320 | 774 | 0 |
| SRR5029493 | 28 | 13 | 0 | 0 | 45 | 2027 | 432 | 1468 | 2578 | 1268 | 0 | 13 | 0 | 13 | 46998 | 11135 | 120164 | 771 | 27 | 1322 | 6326 | 19 | 0 | 78 | 0 | 0 | 0 | 0 | 0 | 54 | 0 | 80 | 0 | 15 | 44222 | 5530 | 0 |
| SRR5029494 | 11 | 20 | 0 | 0 | 18 | 4115 | 935 | 5959 | 6334 | 636 | 0 | 0 | 0 | 0 | 76616 | 21983 | 297161 | 254 | 146 | 10157 | 4948 | 0 | 0 | 43 | 17 | 0 | 0 | 0 | 0 | 19 | 12 | 51 | 0 | 0 | 35085 | 2385 | 0 |
| SRR5029495 | 88 | 29 | 0 | 0 | 296 | 4761 | 720 | 4933 | 6037 | 1070 | 21 | 40 | 0 | 0 | 7102 | 17187 | 186813 | 298 | 86 | 8723 | 1738 | 0 | 0 | 374 | 105 | 0 | 0 | 0 | 0 | 473 | 0 | 70 | 0 | 0 | 40348 | 6013 | 0 |
| SRR5029496 | 26 | 0 | 0 | 0 | 12 | 5224 | 1042 | 2507 | 4652 | 104 | 0 | 22 | 0 | 0 | 1383 | 4635 | 36159 | 143 | 20 | 354 | 354 | 0 | 0 | 74 | 27 | 0 | 0 | 0 | 0 | 25 | 0 | 0 | 0 | 13 | 64775 | 6519 | 0 |
| SRR5029497 | 39 | 0 | 0 | 0 | 61 | 4456 | 440 | 2524 | 4438 | 572 | 0 | 0 | 11 | 0 | 11651 | 12064 | 154218 | 404 | 18 | 3899 | 5399 | 0 | 0 | 153 | 0 | 0 | 0 | 0 | 0 | 119 | 0 | 135 | 0 | 13 | 49488 | 2131 | 0 |
| SRR5029498 | 52 | 19 | 0 | 0 | 17 | 9396 | 257 | 2329 | 4501 | 249 | 0 | 0 | 36 | 0 | 4876 | 5346 | 53175 | 217 | 0 | 651 | 454 | 0 | 0 | 56 | 12 | 0 | 0 | 0 | 0 | 25 | 0 | 0 | 21 | 12 | 49392 | 2465 | 0 |
| SRR5029499 | 25 | 0 | 0 | 0 | 49 | 2061 | 523 | 1113 | 2397 | 127 | 0 | 0 | 0 | 12 | 76462 | 20174 | 255645 | 463 | 22 | 5488 | 7803 | 0 | 0 | 36 | 0 | 0 | 0 | 0 | 0 | 13 | 0 | 30 | 0 | 0 | 19792 | 4400 | 0 |
| SRR5029500 | 11 | 0 | 0 | 0 | 42 | 1169 | 53 | 1116 | 797 | 2128 | 21 | 0 | 0 | 0 | 34857 | 1596 | 16650 | 2125 | 0 | 1116 | 2869 | 42 | 85 | 106 | 0 | 0 | 0 | 0 | 0 | 116 | 53 | 3241 | 0 | 0 | 6641 | 1561 | 0 |
| SRR5029501 | 17 | 0 | 0 | 0 | 17 | 3999 | 568 | 3937 | 4442 | 181 | 0 | 13 | 0 | 0 | 12774 | 10043 | 106777 | 202 | 31 | 2015 | 3361 | 0 | 0 | 19 | 0 | 0 | 0 | 0 | 0 | 20 | 0 | 31 | 0 | 0 | 98177 | 3668 | 0 |
| SRR5029502 | 13 | 11 | 0 | 0 | 30 | 909 | 204 | 841 | 931 | 310 | 22 | 22 | 11 | 11 | 150405 | 15975 | 360769 | 1499 | 45 | 3521 | 7587 | 11 | 0 | 17 | 11 | 22 | 11 | 0 | 0 | 60 | 11 | 68 | 0 | 22 | 12879 | 1375 | 0 |
| SRR5029503 | 21 | 0 | 0 | 0 | 34 | 3604 | 777 | 5129 | 6077 | 197 | 0 | 13 | 0 | 0 | 13095 | 10691 | 122672 | 901 | 47 | 1819 | 4334 | 0 | 0 | 16 | 0 | 0 | 0 | 0 | 0 | 11 | 0 | 112 | 0 | 15 | 51362 | 3309 | 0 |
| SRR5029504 | 24 | 0 | 0 | 0 | 48 | 8001 | 736 | 2200 | 4812 | 754 | 0 | 0 | 38 | 21 | 2556 | 4539 | 33295 | 261 | 0 | 633 | 404 | 0 | 0 | 288 | 0 | 0 | 0 | 0 | 0 | 37 | 0 | 119 | 0 | 0 | 99015 | 5006 | 0 |
| SRR5029505 | 20 | 0 | 0 | 0 | 40 | 5384 | 1172 | 2806 | 6638 | 492 | 0 | 11 | 0 | 13 | 451 | 5568 | 18327 | 270 | 0 | 451 | 182 | 0 | 0 | 337 | 13 | 0 | 0 | 0 | 0 | 33 | 0 | 88 | 0 | 0 | 76608 | 6471 | 0 |
| SRR5029506 | 184 | 41 | 0 | 0 | 216 | 1453 | 411 | 1310 | 4095 | 1010 | 30 | 80 | 12 | 0 | 16585 | 13142 | 97010 | 550 | 139 | 2325 | 3554 | 0 | 10 | 153 | 47 | 12 | 19 | 0 | 0 | 644 | 10 | 134 | 0 | 20 | 110475 | 7528 | 0 |
| SRR5029507 | 43 | 12 | 0 | 0 | 299 | 2697 | 309 | 1257 | 4000 | 1104 | 0 | 17 | 0 | 0 | 14241 | 6948 | 116356 | 1154 | 80 | 2332 | 1966 | 90 | 28 | 209 | 26 | 0 | 0 | 0 | 0 | 252 | 472 | 960 | 0 | 0 | 16875 | 4854 | 0 |
| SRR5029508 | 29 | 11 | 0 | 0 | 15 | 10830 | 902 | 3726 | 7831 | 275 | 0 | 11 | 20 | 0 | 7162 | 9645 | 127251 | 378 | 0 | 5269 | 1135 | 0 | 0 | 61 | 0 | 0 | 0 | 0 | 0 | 16 | 0 | 0 | 0 | 13 | 40486 | 5884 | 0 |
| SRR5029509 | 32 | 12 | 0 | 0 | 78 | 2630 | 677 | 4746 | 5070 | 3513 | 0 | 38 | 0 | 0 | 79877 | 12410 | 167277 | 1134 | 48 | 2773 | 3440 | 35 | 71 | 85 | 11 | 0 | 0 | 0 | 0 | 89 | 69 | 467 | 13 | 22 | 34317 | 3217 | 0 |
| SRR5029510 | 51 | 0 | 0 | 0 | 116 | 2766 | 553 | 1268 | 3688 | 567 | 0 | 19 | 14 | 0 | 3071 | 8501 | 58015 | 346 | 17 | 870 | 830 | 0 | 0 | 130 | 11 | 0 | 0 | 0 | 0 | 82 | 0 | 127 | 0 | 0 | 76250 | 4195 | 0 |
| SRR5029511 | 60 | 13 | 0 | 0 | 167 | 4739 | 820 | 2946 | 7269 | 839 | 24 | 59 | 0 | 0 | 1369 | 13148 | 40970 | 160 | 42 | 1835 | 945 | 0 | 0 | 430 | 57 | 0 | 0 | 0 | 0 | 209 | 0 | 327 | 0 | 0 | 61351 | 6277 | 0 |
| SRR5029512 | 88 | 30 | 0 | 0 | 174 | 3205 | 570 | 2409 | 3849 | 463 | 21 | 31 | 16 | 0 | 2153 | 15574 | 55890 | 383 | 49 | 2964 | 821 | 0 | 0 | 239 | 46 | 11 | 0 | 0 | 0 | 338 | 0 | 54 | 0 | 0 | 79838 | 4911 | 0 |
| SRR5029513 | 12 | 0 | 0 | 0 | 35 | 8782 | 1143 | 12801 | 7280 | 845 | 0 | 0 | 0 | 28 | 25524 | 6373 | 74503 | 903 | 39 | 2491 | 4588 | 0 | 0 | 361 | 0 | 0 | 0 | 0 | 0 | 64 | 0 | 0 | 0 | 0 | 25091 | 4168 | 0 |
| SRR5029514 | 14 | 0 | 0 | 0 | 12 | 3493 | 836 | 2074 | 4181 | 104 | 0 | 18 | 12 | 0 | 3525 | 5329 | 36468 | 413 | 0 | 233 | 233 | 0 | 0 | 20 | 0 | 0 | 0 | 0 | 0 | 23 | 0 | 42 | 0 | 0 | 54796 | 1038 | 0 |
| SRR5029515 | 12 | 0 | 0 | 0 | 19 | 4229 | 573 | 2431 | 2813 | 219 | 0 | 0 | 12 | 0 | 13094 | 2978 | 34855 | 469 | 43 | 1198 | 747 | 0 | 0 | 29 | 0 | 0 | 0 | 0 | 0 | 23 | 0 | 87 | 0 | 16 | 33132 | 2314 | 0 |
| SRR5029516 | 11 | 28 | 0 | 0 | 106 | 2570 | 911 | 2794 | 6848 | 72 | 25 | 53 | 0 | 0 | 3480 | 29492 | 207199 | 200 | 137 | 1672 | 574 | 0 | 0 | 85 | 35 | 68 | 65 | 0 | 0 | 242 | 0 | 12 | 0 | 0 | 50564 | 9040 | 0 |
| SRR5029870 | 208 | 10 | 69 | 0 | 166 | 122 | 174 | 1304 | 17 | 14 | 42 | 28 | 0 | 0 | 8661 | 13878 | 287376 | 0 | 70 | 53951 | 13896 | 0 | 0 | 20 | 69 | 180 | 66 | 42 | 0 | 30 | 0 | 17 | 0 | 0 | 724 | 0 | 28 |
| SRR5029871 | 151 | 359 | 0 | 57 | 208 | 275 | 300 | 1828 | 175 | 19 | 57 | 19 | 0 | 0 | 7312 | 14999 | 277771 | 0 | 25 | 42792 | 11368 | 0 | 0 | 19 | 76 | 227 | 25 | 19 | 0 | 15 | 0 | 0 | 0 | 0 | 1326 | 0 | 0 |
| SRR5029872 | 123 | 214 | 31 | 107 | 138 | 195 | 58 | 1128 | 78 | 31 | 92 | 0 | 0 | 0 | 9242 | 13809 | 283709 | 0 | 19 | 55295 | 13172 | 0 | 0 | 14 | 123 | 230 | 65 | 46 | 0 | 28 | 0 | 19 | 0 | 0 | 892 | 0 | 31 |
| SRR5029873 | 157 | 118 | 20 | 39 | 216 | 457 | 295 | 1692 | 54 | 20 | 20 | 59 | 0 | 0 | 7977 | 14351 | 277161 | 0 | 0 | 45470 | 10797 | 0 | 0 | 16 | 20 | 79 | 26 | 39 | 0 | 15 | 0 | 27 | 0 | 0 | 1068 | 0 | 0 |
| SRR5029874 | 100 | 11 | 71 | 57 | 100 | 165 | 37 | 1025 | 37 | 28 | 100 | 57 | 0 | 0 | 8844 | 14229 | 285900 | 0 | 37 | 54803 | 12616 | 0 | 0 | 18 | 114 | 171 | 61 | 57 | 0 | 33 | 0 | 0 | 0 | 0 | 568 | 0 | 0 |
| SRR5029875 | 135 | 270 | 19 | 19 | 231 | 260 | 286 | 1976 | 104 | 39 | 19 | 0 | 0 | 0 | 7564 | 14630 | 277320 | 0 | 26 | 41591 | 11671 | 0 | 0 | 18 | 77 | 135 | 22 | 58 | 0 | 16 | 0 | 26 | 0 | 0 | 1355 | 0 | 0 |
| SRR5029876 | 240 | 225 | 45 | 75 | 120 | 131 | 94 | 1460 | 37 | 15 | 150 | 30 | 0 | 0 | 8837 | 14302 | 286670 | 0 | 19 | 55137 | 13461 | 0 | 0 | 510 | 45 | 240 | 1454 | 90 | 0 | 629 | 0 | 0 | 0 | 0 | 946 | 0 | 0 |
| SRR5029877 | 159 | 238 | 20 | 20 | 119 | 351 | 216 | 2405 | 108 | 40 | 60 | 79 | 0 | 0 | 7971 | 14455 | 276751 | 0 | 0 | 45043 | 10997 | 0 | 0 | 675 | 79 | 139 | 873 | 60 | 0 | 496 | 0 | 135 | 0 | 0 | 1193 | 0 | 20 |
| SRR5029878 | 444 | 605 | 10 | 81 | 474 | 21 | 0 | 168 | 21 | 0 | 40 | 10 | 0 | 0 | 9544 | 11554 | 622673 | 10 | 31 | 12082 | 24678 | 0 | 0 | 272 | 10 | 50 | 1080 | 50 | 0 | 313 | 0 | 10 | 0 | 0 | 178 | 0 | 0 |
| SRR5029879 | 202 | 505 | 0 | 151 | 505 | 0 | 0 | 111 | 0 | 0 | 0 | 0 | 0 | 0 | 9578 | 11869 | 617144 | 0 | 0 | 10962 | 23752 | 0 | 0 | 151 | 0 | 0 | 606 | 50 | 0 | 303 | 0 | 0 | 0 | 0 | 231 | 0 | 0 |
| SRR5029880 | 292 | 497 | 22 | 86 | 410 | 11 | 67 | 244 | 22 | 0 | 65 | 0 | 0 | 0 | 9231 | 11441 | 620902 | 0 | 67 | 12449 | 24033 | 0 | 0 | 205 | 22 | 65 | 983 | 11 | 0 | 302 | 0 | 0 | 0 | 0 | 156 | 0 | 0 |
| SRR5029882 | 14 | 20 | 21 | 103 | 13 | 11 | 65 | 97 | 0 | 0 | 82 | 10 | 0 | 0 | 9701 | 11386 | 622887 | 0 | 65 | 12276 | 24162 | 0 | 0 | 226 | 21 | 41 | 40 | 31 | 0 | 257 | 0 | 0 | 0 | 0 | 191 | 0 | 0 |
| SRR5029883 | 102 | 11 | 0 | 153 | 13 | 0 | 222 | 167 | 56 | 0 | 0 | 0 | 0 | 0 | 9270 | 10923 | 616602 | 0 | 0 | 11434 | 26477 | 0 | 0 | 153 | 51 | 0 | 16 | 0 | 0 | 16 | 0 | 0 | 0 | 0 | 296 | 0 | 0 |
| SRR5029884 | 366 | 596 | 21 | 126 | 335 | 22 | 54 | 206 | 22 | 0 | 94 | 10 | 0 | 0 | 9223 | 11274 | 622728 | 0 | 65 | 13005 | 24406 | 0 | 0 | 230 | 21 | 52 | 952 | 31 | 0 | 324 | 0 | 0 | 0 | 0 | 205 | 0 | 0 |
| SRR5029885 | 262 | 262 | 0 | 52 | 157 | 0 | 114 | 399 | 0 | 0 | 52 | 0 | 0 | 0 | 9240 | 11149 | 615708 | 0 | 57 | 11864 | 23785 | 0 | 0 | 52 | 0 | 0 | 786 | 52 | 0 | 367 | 0 | 0 | 0 | 0 | 302 | 0 | 0 |
| SRR5029886 | 32 | 48 | 42 | 125 | 28 | 0 | 48 | 65 | 32 | 0 | 42 | 0 | 0 | 0 | 452 | 18153 | 69885 | 0 | 0 | 3145 | 597 | 0 | 0 | 73 | 10 | 73 | 47 | 52 | 0 | 30 | 0 | 16 | 0 | 0 | 7523 | 0 | 0 |
| SRR5029887 | 25 | 25 | 0 | 0 | 17 | 74 | 74 | 185 | 18 | 0 | 0 | 0 | 0 | 0 | 554 | 10695 | 41358 | 18 | 18 | 2974 | 646 | 0 | 0 | 83 | 0 | 18 | 13 | 0 | 0 | 18 | 0 | 18 | 0 | 0 | 8512 | 0 | 0 |
| SRR5029888 | 29 | 51 | 67 | 56 | 36 | 52 | 52 | 122 | 17 | 0 | 89 | 0 | 0 | 0 | 383 | 17612 | 73549 | 17 | 35 | 3329 | 662 | 0 | 0 | 78 | 0 | 89 | 34 | 22 | 0 | 28 | 0 | 0 | 0 | 0 | 8307 | 0 | 0 |
| SRR5029889 | 20 | 30 | 0 | 0 | 17 | 58 | 19 | 212 | 38 | 0 | 0 | 0 | 0 | 0 | 519 | 10721 | 41870 | 38 | 0 | 2693 | 712 | 0 | 0 | 86 | 0 | 29 | 13 | 0 | 0 | 16 | 0 | 58 | 0 | 0 | 8904 | 0 | 0 |
| SRR5029890 | 584 | 828 | 32 | 53 | 478 | 66 | 16 | 165 | 49 | 0 | 64 | 0 | 11 | 0 | 313 | 17333 | 68766 | 16 | 49 | 3098 | 659 | 0 | 0 | 117 | 21 | 42 | 775 | 0 | 0 | 669 | 0 | 33 | 0 | 0 | 7715 | 0 | 0 |
| SRR5029891 | 378 | 623 | 0 | 28 | 453 | 37 | 19 | 187 | 19 | 0 | 19 | 0 | 0 | 0 | 449 | 11767 | 42260 | 0 | 0 | 2766 | 579 | 0 | 0 | 85 | 0 | 28 | 236 | 0 | 0 | 349 | 0 | 37 | 0 | 0 | 9113 | 0 | 0 |
| SRR5029892 | 555 | 718 | 54 | 76 | 555 | 18 | 35 | 53 | 18 | 0 | 65 | 0 | 0 | 0 | 543 | 16880 | 69606 | 18 | 18 | 2941 | 630 | 0 | 0 | 54 | 33 | 44 | 620 | 54 | 0 | 424 | 0 | 0 | 0 | 0 | 7522 | 0 | 0 |
| SRR5029893 | 26 | 33 | 29 | 38 | 19 | 78 | 19 | 195 | 19 | 0 | 38 | 0 | 0 | 0 | 331 | 11933 | 46619 | 19 | 19 | 2529 | 661 | 0 | 0 | 77 | 38 | 29 | 17 | 0 | 0 | 18 | 0 | 58 | 0 | 0 | 8956 | 0 | 0 |
| SRR5029894 | 20 | 26 | 12 | 12 | 98 | 103 | 138 | 436 | 57 | 0 | 111 | 12 | 0 | 0 | 3579 | 2955 | 90745 | 0 | 11 | 65502 | 4497 | 0 | 0 | 74 | 0 | 0 | 0 | 0 | 0 | 62 | 0 | 57 | 0 | 0 | 474 | 0 | 0 |
| SRR5029895 | 260 | 306 | 0 | 15 | 107 | 307 | 107 | 920 | 61 | 0 | 69 | 0 | 0 | 0 | 4019 | 3336 | 91870 | 0 | 31 | 57134 | 4141 | 0 | 0 | 184 | 0 | 0 | 0 | 0 | 0 | 474 | 0 | 92 | 0 | 0 | 909 | 0 | 0 |
| SRR5029896 | 251 | 371 | 26 | 20 | 93 | 196 | 49 | 625 | 61 | 0 | 119 | 33 | 0 | 0 | 3653 | 2928 | 90532 | 0 | 37 | 65606 | 4315 | 0 | 0 | 73 | 0 | 0 | 0 | 0 | 0 | 688 | 0 | 37 | 0 | 0 | 466 | 0 | 0 |
| SRR5029897 | 176 | 376 | 0 | 16 | 88 | 256 | 96 | 879 | 32 | 0 | 48 | 32 | 0 | 0 | 3420 | 3230 | 93781 | 0 | 0 | 55353 | 4330 | 0 | 0 | 152 | 0 | 0 | 0 | 0 | 0 | 560 | 0 | 96 | 0 | 0 | 843 | 0 | 0 |
| SRR5029898 | 18 | 23 | 19 | 19 | 63 | 189 | 106 | 696 | 59 | 0 | 113 | 19 | 0 | 0 | 3622 | 2903 | 90976 | 0 | 47 | 64860 | 4637 | 0 | 0 | 113 | 0 | 0 | 0 | 0 | 0 | 59 | 0 | 24 | 0 | 0 | 472 | 0 | 0 |
| SRR5029899 | 218 | 218 | 0 | 31 | 109 | 346 | 188 | 817 | 31 | 0 | 39 | 39 | 0 | 0 | 3722 | 3363 | 92684 | 0 | 31 | 56262 | 3942 | 0 | 0 | 117 | 0 | 0 | 0 | 0 | 0 | 476 | 0 | 16 | 0 | 0 | 857 | 0 | 0 |
| SRR5029900 | 238 | 232 | 13 | 19 | 77 | 120 | 60 | 517 | 48 | 0 | 58 | 51 | 0 | 0 | 3881 | 2977 | 90878 | 0 | 12 | 67043 | 4710 | 0 | 0 | 97 | 0 | 0 | 0 | 0 | 0 | 643 | 0 | 36 | 0 | 0 | 557 | 0 | 0 |
| SRR5029901 | 160 | 255 | 0 | 16 | 96 | 340 | 226 | 1100 | 32 | 0 | 56 | 0 | 0 | 0 | 3558 | 3306 | 94577 | 0 | 32 | 57373 | 4545 | 0 | 0 | 199 | 0 | 0 | 0 | 0 | 0 | 526 | 0 | 49 | 0 | 0 | 851 | 0 | 0 |
| SRR5029902 | 164 | 129 | 14 | 48 | 402 | 24 | 171 | 538 | 73 | 14 | 143 | 0 | 0 | 0 | 1712 | 13357 | 55750 | 24 | 0 | 2617 | 1027 | 0 | 0 | 95 | 0 | 20 | 491 | 20 | 0 | 1274 | 0 | 0 | 0 | 0 | 8242 | 0 | 20 |
| SRR5029904 | 201 | 119 | 15 | 37 | 410 | 0 | 295 | 322 | 0 | 37 | 119 | 15 | 0 | 0 | 1689 | 14339 | 56488 | 27 | 0 | 2467 | 1234 | 0 | 0 | 89 | 0 | 60 | 387 | 22 | 0 | 1333 | 0 | 0 | 0 | 0 | 7348 | 0 | 0 |
| SRR5029905 | 335 | 56 | 0 | 0 | 335 | 0 | 0 | 417 | 0 | 56 | 112 | 0 | 0 | 0 | 834 | 10354 | 44422 | 0 | 0 | 2294 | 1043 | 0 | 0 | 167 | 0 | 0 | 279 | 56 | 0 | 836 | 56 | 0 | 0 | 0 | 8016 | 0 | 0 |
| SRR5029906 | 17 | 169 | 21 | 42 | 20 | 0 | 202 | 454 | 50 | 28 | 155 | 0 | 0 | 0 | 2118 | 12897 | 56226 | 25 | 0 | 2320 | 1109 | 0 | 0 | 70 | 14 | 35 | 25 | 14 | 0 | 86 | 0 | 0 | 0 | 0 | 7439 | 0 | 35 |
| SRR5029908 | 12 | 101 | 22 | 58 | 21 | 53 | 263 | 552 | 184 | 0 | 12 | 0 | 0 | 0 | 1445 | 13240 | 53955 | 0 | 53 | 2155 | 1288 | 0 | 0 | 116 | 0 | 29 | 23 | 36 | 0 | 82 | 0 | 0 | 0 | 0 | 6869 | 0 | 29 |
| SRR5029910 | 61 | 102 | 41 | 102 | 287 | 0 | 0 | 0 | 0 | 41 | 61 | 0 | 0 | 0 | 672 | 22648 | 130930 | 0 | 0 | 3059 | 19909 | 0 | 0 | 328 | 0 | 41 | 471 | 0 | 0 | 369 | 0 | 12 | 0 | 0 | 452 | 0 | 0 |
| SRR5029911 | 148 | 85 | 0 | 42 | 11 | 13 | 26 | 26 | 39 | 64 | 42 | 0 | 0 | 0 | 802 | 24336 | 131407 | 0 | 0 | 2788 | 18343 | 0 | 0 | 360 | 0 | 42 | 11 | 0 | 0 | 233 | 0 | 0 | 0 | 0 | 650 | 0 | 0 |
| SRR5029912 | 156 | 111 | 22 | 134 | 10 | 25 | 13 | 64 | 25 | 67 | 111 | 0 | 0 | 0 | 737 | 22357 | 129597 | 0 | 0 | 3534 | 20100 | 0 | 0 | 178 | 0 | 67 | 18 | 0 | 0 | 290 | 0 | 13 | 0 | 0 | 403 | 0 | 0 |
| SRR5029913 | 111 | 89 | 0 | 0 | 13 | 0 | 0 | 68 | 14 | 177 | 44 | 0 | 22 | 0 | 780 | 23918 | 132428 | 0 | 0 | 2655 | 18532 | 0 | 0 | 355 | 0 | 67 | 155 | 0 | 0 | 10 | 0 | 0 | 0 | 0 | 653 | 0 | 0 |
| SRR5029914 | 107 | 149 | 43 | 85 | 11 | 0 | 12 | 49 | 0 | 107 | 43 | 0 | 0 | 0 | 683 | 22104 | 130024 | 0 | 0 | 3271 | 20088 | 0 | 0 | 214 | 0 | 85 | 13 | 21 | 0 | 235 | 0 | 12 | 0 | 0 | 481 | 0 | 0 |
| SRR5029915 | 109 | 152 | 22 | 0 | 11 | 0 | 27 | 95 | 27 | 65 | 44 | 0 | 22 | 0 | 719 | 23954 | 131517 | 14 | 0 | 2726 | 18854 | 0 | 0 | 12 | 0 | 44 | 283 | 22 | 0 | 305 | 0 | 0 | 0 | 0 | 717 | 0 | 22 |
| SRR5029916 | 64 | 86 | 86 | 64 | 14 | 0 | 25 | 111 | 12 | 172 | 43 | 0 | 0 | 0 | 654 | 22569 | 129235 | 0 | 0 | 3034 | 19401 | 0 | 0 | 13 | 0 | 64 | 13 | 21 | 0 | 11 | 0 | 25 | 0 | 0 | 446 | 0 | 0 |
| SRR5029917 | 131 | 175 | 0 | 22 | 306 | 28 | 28 | 55 | 0 | 66 | 22 | 0 | 0 | 0 | 619 | 23848 | 131688 | 0 | 0 | 3136 | 18749 | 0 | 0 | 437 | 0 | 0 | 415 | 0 | 0 | 284 | 0 | 14 | 0 | 0 | 710 | 0 | 22 |
| SRR5029918 | 216 | 399 | 50 | 17 | 764 | 23 | 0 | 0 | 47 | 17 | 33 | 0 | 0 | 0 | 2670 | 37420 | 85531 | 0 | 23 | 2225 | 16465 | 0 | 0 | 100 | 0 | 50 | 465 | 17 | 0 | 532 | 0 | 0 | 0 | 0 | 1812 | 0 | 17 |
| SRR5029919 | 315 | 299 | 16 | 0 | 767 | 0 | 15 | 15 | 0 | 0 | 24 | 0 | 0 | 0 | 2678 | 28390 | 54580 | 15 | 15 | 1841 | 17211 | 0 | 0 | 73 | 0 | 0 | 153 | 0 | 0 | 339 | 0 | 15 | 0 | 0 | 3465 | 0 | 0 |
| SRR5029920 | 478 | 283 | 106 | 53 | 762 | 52 | 0 | 0 | 0 | 0 | 18 | 18 | 0 | 0 | 2535 | 39953 | 89433 | 0 | 0 | 2095 | 16140 | 0 | 0 | 124 | 0 | 53 | 425 | 18 | 0 | 620 | 0 | 0 | 0 | 0 | 2333 | 0 | 0 |
| SRR5029921 | 285 | 361 | 0 | 0 | 730 | 32 | 0 | 63 | 32 | 17 | 25 | 0 | 0 | 0 | 2853 | 26950 | 57061 | 16 | 16 | 1743 | 15372 | 0 | 0 | 126 | 0 | 0 | 210 | 0 | 0 | 269 | 0 | 16 | 0 | 0 | 2744 | 0 | 0 |
| SRR5029922 | 237 | 475 | 34 | 17 | 797 | 0 | 0 | 0 | 0 | 0 | 102 | 0 | 0 | 0 | 2350 | 36699 | 91840 | 0 | 0 | 2056 | 15861 | 0 | 0 | 85 | 0 | 34 | 390 | 17 | 0 | 542 | 0 | 0 | 0 | 0 | 1752 | 0 | 17 |
| SRR5029923 | 458 | 384 | 16 | 0 | 613 | 31 | 61 | 77 | 0 | 16 | 25 | 0 | 0 | 0 | 2456 | 25523 | 56313 | 31 | 15 | 2026 | 15808 | 0 | 0 | 82 | 0 | 0 | 229 | 0 | 0 | 327 | 0 | 46 | 0 | 0 | 2991 | 0 | 0 |
| SRR5029924 | 18 | 22 | 87 | 17 | 40 | 0 | 0 | 25 | 50 | 0 | 35 | 0 | 0 | 0 | 2121 | 44838 | 85720 | 0 | 0 | 1919 | 16358 | 0 | 0 | 87 | 0 | 0 | 26 | 17 | 0 | 26 | 0 | 25 | 0 | 0 | 2449 | 0 | 0 |
| SRR5029925 | 276 | 293 | 25 | 0 | 652 | 16 | 16 | 80 | 32 | 0 | 25 | 0 | 0 | 0 | 2454 | 27282 | 53937 | 0 | 16 | 2215 | 16957 | 0 | 0 | 100 | 0 | 0 | 226 | 0 | 0 | 418 | 0 | 16 | 0 | 0 | 2927 | 0 | 0 |
| SRR5029926 | 63 | 19 | 0 | 24 | 32 | 26 | 158 | 580 | 105 | 0 | 16 | 32 | 0 | 0 | 3714 | 24170 | 46657 | 0 | 26 | 8377 | 5005 | 0 | 0 | 32 | 0 | 0 | 39 | 16 | 0 | 41 | 0 | 0 | 0 | 0 | 3995 | 0 | 0 |
| SRR5029927 | 56 | 172 | 0 | 0 | 39 | 148 | 164 | 591 | 49 | 0 | 17 | 33 | 0 | 0 | 2839 | 19828 | 34764 | 0 | 0 | 4956 | 3479 | 0 | 0 | 39 | 0 | 0 | 106 | 0 | 0 | 222 | 0 | 16 | 0 | 0 | 6465 | 0 | 0 |
| SRR5029928 | 60 | 232 | 0 | 17 | 43 | 87 | 87 | 548 | 144 | 0 | 95 | 52 | 0 | 0 | 4010 | 22764 | 45467 | 29 | 58 | 8424 | 4443 | 0 | 0 | 17 | 17 | 60 | 353 | 0 | 0 | 284 | 0 | 0 | 0 | 0 | 3666 | 0 | 0 |
| SRR5029929 | 75 | 150 | 0 | 0 | 23 | 207 | 207 | 706 | 121 | 0 | 12 | 29 | 0 | 0 | 2981 | 18371 | 34493 | 17 | 17 | 5583 | 3412 | 0 | 0 | 69 | 0 | 12 | 98 | 0 | 0 | 11 | 0 | 17 | 0 | 0 | 6101 | 0 | 0 |
| SRR5029930 | 65 | 146 | 0 | 0 | 41 | 195 | 112 | 530 | 28 | 0 | 65 | 49 | 0 | 0 | 4048 | 24689 | 48191 | 0 | 84 | 7844 | 4857 | 0 | 0 | 0 | 16 | 41 | 292 | 24 | 0 | 381 | 0 | 0 | 0 | 0 | 3763 | 0 | 0 |
| SRR5029931 | 101 | 13 | 0 | 0 | 45 | 165 | 215 | 612 | 99 | 11 | 11 | 34 | 0 | 0 | 2712 | 18045 | 33979 | 33 | 50 | 5407 | 3373 | 0 | 0 | 51 | 11 | 0 | 95 | 0 | 0 | 12 | 0 | 0 | 0 | 0 | 6140 | 0 | 0 |
| SRR5029932 | 109 | 217 | 17 | 33 | 58 | 84 | 28 | 645 | 112 | 0 | 50 | 58 | 0 | 0 | 3788 | 23839 | 46130 | 0 | 84 | 8615 | 4686 | 0 | 0 | 33 | 0 | 42 | 367 | 0 | 0 | 392 | 0 | 0 | 0 | 0 | 4792 | 0 | 0 |
| SRR5029933 | 98 | 145 | 0 | 0 | 29 | 153 | 136 | 509 | 102 | 0 | 0 | 75 | 0 | 0 | 2763 | 19468 | 34091 | 34 | 0 | 5493 | 3323 | 0 | 0 | 75 | 0 | 29 | 69 | 12 | 0 | 10 | 0 | 0 | 0 | 0 | 6188 | 0 | 0 |
| SRR5029934 | 212 | 437 | 40 | 40 | 212 | 37 | 43 | 2309 | 37 | 424 | 0 | 0 | 13 | 0 | 4201 | 5161 | 74154 | 12 | 12 | 19488 | 2964 | 0 | 0 | 40 | 265 | 291 | 821 | 26 | 0 | 331 | 0 | 0 | 13 | 0 | 134 | 0 | 0 |
| SRR5029935 | 180 | 299 | 15 | 15 | 224 | 112 | 143 | 4608 | 41 | 494 | 45 | 30 | 0 | 0 | 4425 | 5752 | 79462 | 0 | 10 | 19066 | 3089 | 0 | 0 | 120 | 45 | 120 | 254 | 30 | 0 | 239 | 0 | 0 | 0 | 0 | 274 | 0 | 15 |
| SRR5029936 | 244 | 402 | 14 | 58 | 259 | 40 | 87 | 2270 | 27 | 446 | 14 | 0 | 0 | 0 | 4427 | 5040 | 73825 | 0 | 27 | 20455 | 3049 | 0 | 0 | 29 | 144 | 316 | 762 | 29 | 0 | 316 | 0 | 0 | 0 | 0 | 139 | 0 | 0 |
| SRR5029937 | 125 | 235 | 0 | 31 | 125 | 150 | 129 | 4632 | 32 | 485 | 47 | 16 | 16 | 0 | 4729 | 5808 | 78280 | 0 | 32 | 19346 | 2687 | 0 | 0 | 125 | 31 | 125 | 266 | 16 | 0 | 251 | 0 | 0 | 0 | 0 | 287 | 0 | 0 |
| SRR5029938 | 13 | 13 | 68 | 0 | 12 | 51 | 64 | 2407 | 51 | 17 | 41 | 0 | 0 | 0 | 4279 | 4959 | 74374 | 0 | 38 | 20773 | 3076 | 0 | 0 | 55 | 123 | 12 | 30 | 14 | 0 | 17 | 0 | 0 | 0 | 0 | 150 | 0 | 0 |
| SRR5029939 | 107 | 92 | 0 | 15 | 214 | 189 | 305 | 4657 | 53 | 19 | 61 | 0 | 0 | 0 | 4352 | 5758 | 79608 | 0 | 42 | 19364 | 2954 | 0 | 0 | 92 | 76 | 122 | 214 | 15 | 0 | 183 | 0 | 21 | 0 | 0 | 274 | 0 | 15 |
| SRR5029940 | 211 | 351 | 28 | 0 | 183 | 52 | 58 | 2528 | 26 | 337 | 112 | 0 | 0 | 0 | 4257 | 5023 | 73848 | 13 | 45 | 20046 | 3206 | 0 | 0 | 84 | 140 | 281 | 885 | 0 | 0 | 421 | 0 | 0 | 0 | 0 | 154 | 0 | 0 |
| SRR5029941 | 156 | 172 | 16 | 0 | 235 | 117 | 171 | 4313 | 32 | 516 | 63 | 0 | 0 | 0 | 4591 | 5894 | 79626 | 0 | 21 | 18834 | 3267 | 0 | 0 | 125 | 47 | 203 | 250 | 0 | 0 | 297 | 0 | 11 | 0 | 0 | 288 | 0 | 16 |
| SRR5029942 | 14 | 112 | 0 | 70 | 378 | 184 | 257 | 1469 | 551 | 42 | 182 | 14 | 0 | 0 | 7421 | 23713 | 100481 | 37 | 514 | 14217 | 5584 | 0 | 0 | 70 | 0 | 28 | 84 | 14 | 0 | 659 | 0 | 37 | 0 | 0 | 1163 | 0 | 0 |
| SRR5029943 | 44 | 72 | 0 | 0 | 182 | 419 | 839 | 2592 | 438 | 0 | 39 | 0 | 0 | 0 | 7281 | 23307 | 96805 | 0 | 438 | 10178 | 5241 | 0 | 0 | 44 | 0 | 11 | 28 | 17 | 0 | 358 | 0 | 95 | 0 | 0 | 1980 | 0 | 0 |
| SRR5029944 | 46 | 62 | 0 | 31 | 14 | 160 | 360 | 1000 | 240 | 0 | 108 | 0 | 0 | 0 | 6919 | 24366 | 101458 | 40 | 400 | 14279 | 5880 | 0 | 0 | 62 | 0 | 93 | 124 | 0 | 0 | 57 | 0 | 0 | 0 | 0 | 1227 | 0 | 0 |
| SRR5029945 | 46 | 40 | 0 | 34 | 172 | 432 | 766 | 2476 | 786 | 0 | 17 | 0 | 0 | 0 | 6524 | 24211 | 96786 | 59 | 216 | 9609 | 4971 | 0 | 0 | 46 | 12 | 0 | 34 | 0 | 0 | 17 | 0 | 138 | 0 | 0 | 2545 | 0 | 0 |

### Supp Table S8.3.Columns_BX_to_DH

| | | | | | | | | | | | | | | | | | | | | | | | | | | | | | | | | | | | | |
|---|---|---|---|---|---|---|---|---|---|---|---|---|---|---|---|---|---|---|---|---|---|---|---|---|---|---|---|---|---|---|---|---|---|---|---|---|---|
| SRR5029946 | 58 | 102 | 0 | 102 | 18 | 115 | 498 | 1034 | 345 | 0 | 15 | 0 | 0 | 0 | 7430 | 24601 | 104343 | 0 | 613 | 13787 | 5859 | 0 | 0 | 87 | 15 | 29 | 16 | 0 | 0 | 47 | 0 | 38 | 0 | 0 | 1117 | 0 | 0 |
| SRR5029947 | 50 | 56 | 0 | 11 | 178 | 554 | 783 | 2273 | 592 | 28 | 45 | 0 | 0 | 0 | 6514 | 24365 | 95083 | 57 | 325 | 8768 | 5119 | 0 | 0 | 28 | 0 | 0 | 50 | 0 | 0 | 334 | 0 | 96 | 0 | 0 | 2514 | 0 | 0 |
| SRR5029948 | 45 | 76 | 30 | 45 | 212 | 234 | 390 | 1015 | 312 | 0 | 121 | 0 | 0 | 0 | 7649 | 23224 | 102827 | 0 | 429 | 12489 | 5425 | 0 | 0 | 30 | 15 | 15 | 212 | 0 | 0 | 667 | 0 | 0 | 0 | 0 | 888 | 0 | 0 |
| SRR5029949 | 58 | 98 | 0 | 17 | 161 | 381 | 642 | 2488 | 722 | 17 | 46 | 0 | 0 | 0 | 6401 | 25666 | 96611 | 0 | 161 | 10735 | 5337 | 0 | 0 | 63 | 0 | 0 | 63 | 12 | 0 | 19 | 0 | 100 | 0 | 0 | 2257 | 0 | 0 |
| SRR5029950 | 89 | 140 | 28 | 121 | 354 | 239 | 172 | 981 | 146 | 19 | 228 | 47 | 0 | 0 | 1550 | 30584 | 119703 | 27 | 40 | 3684 | 5115 | 0 | 0 | 93 | 0 | 47 | 364 | 33 | 0 | 778 | 0 | 40 | 0 | 0 | 2596 | 0 | 0 |
| SRR5029951 | 0 | 0 | 0 | 0 | 256 | 252 | 252 | 1260 | 0 | 43 | 43 | 43 | 0 | 0 | 2015 | 27036 | 124523 | 0 | 0 | 2015 | 4787 | 0 | 0 | 171 | 0 | 85 | 299 | 0 | 0 | 641 | 0 | 0 | 0 | 0 | 3022 | 0 | 0 |
| SRR5029952 | 70 | 105 | 35 | 90 | 26 | 230 | 201 | 949 | 101 | 25 | 20 | 25 | 0 | 0 | 1509 | 30114 | 120200 | 0 | 29 | 3622 | 5087 | 0 | 0 | 65 | 10 | 55 | 31 | 20 | 0 | 72 | 0 | 72 | 0 | 0 | 2933 | 0 | 0 |
| SRR5029953 | 44 | 132 | 0 | 0 | 396 | 258 | 0 | 516 | 0 | 0 | 44 | 44 | 0 | 0 | 2064 | 31621 | 120389 | 0 | 0 | 2322 | 4127 | 0 | 44 | 44 | 0 | 0 | 264 | 0 | 0 | 573 | 0 | 0 | 0 | 0 | 2964 | 0 | 44 |
| SRR5029954 | 81 | 96 | 14 | 91 | 340 | 137 | 288 | 1139 | 110 | 0 | 273 | 29 | 0 | 0 | 1428 | 29662 | 120532 | 0 | 0 | 2897 | 5628 | 0 | 14 | 77 | 0 | 38 | 441 | 14 | 0 | 809 | 0 | 27 | 0 | 0 | 2927 | 11 | 0 |
| SRR5029955 | 87 | 44 | 0 | 87 | 349 | 127 | 892 | 765 | 255 | 87 | 44 | 0 | 0 | 0 | 765 | 29922 | 123447 | 0 | 0 | 4205 | 5479 | 0 | 44 | 87 | 0 | 0 | 218 | 44 | 0 | 698 | 0 | 127 | 0 | 0 | 3107 | 0 | 44 |
| SRR5029956 | 68 | 68 | 20 | 13 | 24 | 209 | 236 | 751 | 97 | 0 | 17 | 49 | 0 | 0 | 1765 | 31735 | 123476 | 14 | 42 | 3378 | 5143 | 0 | 0 | 93 | 0 | 59 | 38 | 0 | 0 | 61 | 0 | 42 | 0 | 0 | 2892 | 0 | 0 |
| SRR5029957 | 44 | 221 | 0 | 88 | 397 | 0 | 1037 | 907 | 0 | 0 | 309 | 88 | 0 | 0 | 648 | 33018 | 129584 | 0 | 0 | 3499 | 5183 | 0 | 0 | 44 | 0 | 44 | 221 | 0 | 0 | 883 | 0 | 130 | 0 | 0 | 2655 | 14 | 0 |
| SRR5029958 | 25 | 26 | 29 | 44 | 133 | 39 | 39 | 197 | 0 | 44 | 177 | 0 | 0 | 0 | 2583 | 57129 | 118653 | 20 | 0 | 5226 | 7435 | 0 | 0 | 12 | 15 | 44 | 23 | 15 | 0 | 44 | 0 | 59 | 0 | 0 | 996 | 0 | 0 |
| SRR5029959 | 15 | 17 | 0 | 16 | 82 | 112 | 48 | 384 | 64 | 57 | 49 | 0 | 0 | 0 | 2114 | 57701 | 103164 | 0 | 0 | 3476 | 6487 | 0 | 0 | 14 | 0 | 25 | 114 | 0 | 0 | 19 | 0 | 0 | 0 | 0 | 2143 | 0 | 0 |
| SRR5029960 | 521 | 647 | 0 | 111 | 79 | 22 | 0 | 131 | 0 | 47 | 63 | 16 | 0 | 0 | 3017 | 58032 | 115075 | 0 | 0 | 4809 | 8110 | 0 | 0 | 205 | 0 | 111 | 489 | 47 | 0 | 647 | 0 | 22 | 0 | 0 | 1257 | 0 | 0 |
| SRR5029961 | 314 | 398 | 0 | 0 | 102 | 100 | 50 | 285 | 17 | 59 | 34 | 0 | 0 | 0 | 2393 | 56711 | 103330 | 0 | 0 | 3682 | 6962 | 0 | 0 | 271 | 17 | 42 | 102 | 17 | 0 | 314 | 0 | 17 | 0 | 0 | 2237 | 0 | 0 |
| SRR5029962 | 29 | 27 | 0 | 60 | 180 | 63 | 21 | 105 | 0 | 90 | 180 | 30 | 0 | 0 | 2707 | 59777 | 117721 | 21 | 0 | 4932 | 7744 | 0 | 0 | 13 | 30 | 10 | 17 | 75 | 0 | 42 | 0 | 21 | 0 | 0 | 1011 | 0 | 0 |
| SRR5029963 | 10 | 14 | 0 | 17 | 91 | 114 | 98 | 228 | 33 | 83 | 50 | 25 | 0 | 0 | 2542 | 57026 | 104262 | 0 | 0 | 3145 | 6616 | 0 | 0 | 216 | 0 | 17 | 75 | 0 | 0 | 19 | 0 | 16 | 0 | 0 | 2166 | 0 | 0 |
| SRR5029964 | 23 | 32 | 31 | 46 | 123 | 21 | 21 | 84 | 63 | 77 | 139 | 15 | 0 | 0 | 2704 | 59144 | 117625 | 0 | 0 | 4820 | 7754 | 0 | 0 | 14 | 15 | 77 | 27 | 46 | 0 | 48 | 0 | 21 | 0 | 0 | 1114 | 0 | 0 |
| SRR5029965 | 305 | 415 | 17 | 25 | 59 | 118 | 34 | 404 | 17 | 68 | 25 | 0 | 0 | 0 | 2643 | 56537 | 101021 | 0 | 0 | 3367 | 6886 | 0 | 0 | 212 | 0 | 34 | 76 | 0 | 0 | 356 | 0 | 67 | 0 | 0 | 2607 | 0 | 34 |
| SRR5029966 | 223 | 170 | 42 | 202 | 32 | 47 | 32 | 584 | 63 | 11 | 106 | 11 | 0 | 0 | 2163 | 26709 | 63752 | 0 | 0 | 7578 | 3994 | 0 | 0 | 0 | 0 | 0 | 0 | 0 | 0 | 721 | 0 | 16 | 0 | 0 | 1981 | 0 | 11 |
| SRR5029967 | 160 | 188 | 17 | 44 | 61 | 74 | 74 | 1384 | 85 | 17 | 28 | 0 | 17 | 0 | 2292 | 26492 | 55148 | 32 | 21 | 5208 | 3138 | 0 | 0 | 0 | 0 | 0 | 0 | 0 | 0 | 20 | 0 | 32 | 0 | 0 | 4037 | 0 | 0 |
| SRR5029968 | 16 | 23 | 35 | 12 | 47 | 122 | 17 | 662 | 17 | 12 | 117 | 12 | 0 | 0 | 2664 | 27389 | 65930 | 35 | 0 | 8080 | 3361 | 0 | 12 | 0 | 0 | 0 | 0 | 0 | 0 | 55 | 0 | 17 | 0 | 0 | 2083 | 0 | 0 |
| SRR5029969 | 186 | 174 | 0 | 17 | 52 | 203 | 56 | 1328 | 23 | 29 | 12 | 41 | 0 | 0 | 2138 | 25648 | 55111 | 23 | 0 | 5412 | 2824 | 0 | 0 | 0 | 0 | 0 | 0 | 0 | 0 | 319 | 0 | 56 | 0 | 0 | 4015 | 0 | 0 |
| SRR5029970 | 21 | 20 | 33 | 13 | 55 | 33 | 17 | 647 | 33 | 0 | 99 | 33 | 0 | 0 | 2274 | 26893 | 64206 | 33 | 0 | 7670 | 3370 | 0 | 0 | 0 | 0 | 0 | 0 | 0 | 0 | 52 | 0 | 50 | 0 | 0 | 2119 | 0 | 0 |
| SRR5029971 | 141 | 237 | 0 | 23 | 28 | 153 | 88 | 1644 | 66 | 28 | 34 | 28 | 0 | 17 | 2060 | 25121 | 55934 | 33 | 0 | 5173 | 3145 | 0 | 0 | 0 | 0 | 0 | 0 | 0 | 0 | 305 | 0 | 11 | 0 | 0 | 3798 | 0 | 0 |
| SRR5029972 | 250 | 272 | 45 | 181 | 57 | 52 | 0 | 552 | 17 | 34 | 57 | 11 | 0 | 0 | 2363 | 26478 | 64778 | 52 | 0 | 8297 | 3502 | 0 | 0 | 0 | 0 | 0 | 0 | 0 | 0 | 839 | 0 | 17 | 0 | 0 | 2015 | 0 | 0 |
| SRR5029973 | 11 | 169 | 0 | 41 | 64 | 132 | 77 | 1372 | 44 | 18 | 29 | 12 | 0 | 0 | 2217 | 26186 | 55325 | 11 | 11 | 5609 | 2876 | 0 | 18 | 12 | 0 | 0 | 0 | 0 | 0 | 19 | 0 | 0 | 0 | 0 | 3922 | 0 | 0 |
| SRR5029974 | 23 | 28 | 16 | 41 | 21 | 48 | 239 | 358 | 24 | 0 | 23 | 0 | 0 | 0 | 334 | 16206 | 48927 | 0 | 24 | 4940 | 1790 | 0 | 0 | 66 | 0 | 0 | 0 | 0 | 0 | 394 | 0 | 0 | 0 | 0 | 3930 | 0 | 0 |
| SRR5029975 | 125 | 196 | 0 | 0 | 89 | 168 | 242 | 373 | 37 | 0 | 95 | 0 | 0 | 0 | 317 | 9867 | 23885 | 0 | 0 | 2610 | 969 | 0 | 0 | 95 | 0 | 0 | 0 | 0 | 0 | 1855 | 0 | 37 | 12 | 0 | 5254 | 0 | 0 |
| SRR5029976 | 15 | 34 | 0 | 36 | 11 | 77 | 205 | 334 | 0 | 0 | 23 | 0 | 0 | 0 | 308 | 20219 | 52102 | 0 | 26 | 5619 | 1360 | 0 | 0 | 36 | 0 | 0 | 0 | 0 | 0 | 394 | 0 | 103 | 0 | 0 | 4714 | 0 | 0 |
| SRR5029977 | 136 | 13 | 0 | 0 | 81 | 138 | 335 | 394 | 39 | 0 | 68 | 0 | 0 | 0 | 256 | 11119 | 24725 | 0 | 20 | 2932 | 1102 | 0 | 0 | 99 | 0 | 0 | 0 | 0 | 0 | 143 | 0 | 20 | 0 | 0 | 5082 | 0 | 12 |
| SRR5029978 | 233 | 362 | 0 | 52 | 129 | 98 | 122 | 317 | 49 | 0 | 388 | 0 | 0 | 0 | 561 | 17399 | 47893 | 0 | 24 | 5685 | 1196 | 0 | 0 | 112 | 0 | 0 | 0 | 0 | 0 | 4685 | 0 | 122 | 0 | 0 | 4443 | 0 | 0 |
| SRR5029979 | 11 | 14 | 0 | 0 | 116 | 168 | 428 | 428 | 19 | 12 | 92 | 0 | 0 | 0 | 373 | 9851 | 26008 | 0 | 19 | 2757 | 857 | 0 | 0 | 80 | 0 | 0 | 0 | 0 | 0 | 124 | 0 | 0 | 0 | 0 | 5096 | 0 | 0 |
| SRR5029980 | 19 | 35 | 44 | 26 | 12 | 124 | 322 | 198 | 50 | 0 | 31 | 0 | 17 | 0 | 396 | 18637 | 48294 | 0 | 25 | 4632 | 1561 | 0 | 0 | 26 | 0 | 0 | 0 | 0 | 0 | 410 | 0 | 25 | 0 | 0 | 6133 | 0 | 0 |
| SRR5029981 | 149 | 248 | 0 | 0 | 124 | 191 | 248 | 133 | 0 | 0 | 124 | 0 | 0 | 0 | 248 | 10329 | 24918 | 0 | 0 | 3049 | 858 | 0 | 0 | 81 | 0 | 0 | 0 | 0 | 0 | 1813 | 0 | 57 | 0 | 0 | 5852 | 0 | 0 |
| SRR5029982 | 124 | 166 | 14 | 55 | 83 | 134 | 148 | 6632 | 71 | 166 | 124 | 28 | 0 | 0 | 6088 | 2785 | 115818 | 0 | 14 | 15103 | 24174 | 0 | 0 | 83 | 83 | 373 | 580 | 0 | 0 | 456 | 0 | 28 | 0 | 0 | 240 | 0 | 14 |
| SRR5029983 | 94 | 13 | 94 | 0 | 0 | 358 | 204 | 6900 | 102 | 187 | 187 | 0 | 0 | 0 | 6900 | 3036 | 120111 | 0 | 204 | 16355 | 24686 | 0 | 0 | 281 | 0 | 187 | 187 | 0 | 0 | 94 | 0 | 0 | 0 | 0 | 271 | 0 | 0 |
| SRR5029984 | 120 | 150 | 0 | 45 | 45 | 99 | 168 | 6624 | 84 | 105 | 90 | 0 | 0 | 0 | 5746 | 2540 | 116474 | 0 | 15 | 15942 | 24511 | 0 | 0 | 105 | 30 | 270 | 494 | 15 | 0 | 464 | 0 | 0 | 0 | 0 | 206 | 0 | 15 |
| SRR5029986 | 198 | 170 | 14 | 71 | 85 | 103 | 183 | 6350 | 73 | 85 | 142 | 0 | 0 | 0 | 5896 | 2668 | 114496 | 0 | 15 | 15089 | 24391 | 0 | 0 | 71 | 99 | 326 | 11 | 0 | 0 | 13 | 0 | 0 | 0 | 0 | 229 | 0 | 0 |
| SRR5029988 | 131 | 234 | 15 | 29 | 15 | 171 | 193 | 6495 | 89 | 204 | 161 | 29 | 0 | 0 | 5796 | 2673 | 116060 | 0 | 22 | 15007 | 23786 | 0 | 0 | 73 | 88 | 307 | 570 | 29 | 0 | 380 | 0 | 15 | 0 | 0 | 203 | 0 | 15 |
| SRR5029990 | 91 | 95 | 22 | 102 | 245 | 319 | 142 | 710 | 18 | 1309 | 161 | 18 | 0 | 0 | 6940 | 18234 | 104110 | 53 | 0 | 4721 | 40786 | 0 | 62 | 165 | 29 | 69 | 110 | 0 | 0 | 1262 | 0 | 36 | 0 | 0 | 281 | 0 | 0 |
| SRR5029991 | 70 | 40 | 0 | 50 | 31 | 836 | 90 | 1522 | 30 | 202 | 45 | 0 | 0 | 0 | 5582 | 19490 | 103978 | 60 | 30 | 4000 | 35612 | 0 | 13 | 21 | 25 | 20 | 14 | 20 | 0 | 89 | 0 | 90 | 0 | 0 | 547 | 0 | 0 |
| SRR5029992 | 60 | 88 | 28 | 124 | 264 | 362 | 38 | 875 | 0 | 1535 | 124 | 0 | 0 | 0 | 8011 | 18708 | 104120 | 38 | 76 | 4643 | 41351 | 12 | 76 | 124 | 40 | 48 | 180 | 16 | 0 | 1194 | 0 | 95 | 0 | 0 | 294 | 0 | 0 |
| SRR5029993 | 63 | 13 | 0 | 21 | 22 | 919 | 317 | 1362 | 63 | 193 | 53 | 11 | 0 | 0 | 6811 | 20471 | 104488 | 32 | 32 | 3485 | 40013 | 0 | 11 | 23 | 26 | 32 | 68 | 0 | 0 | 117 | 0 | 95 | 0 | 0 | 469 | 0 | 0 |
| SRR5029994 | 12 | 68 | 11 | 22 | 31 | 409 | 74 | 818 | 0 | 213 | 24 | 15 | 0 | 0 | 7756 | 17419 | 103343 | 56 | 37 | 4632 | 42372 | 11 | 14 | 19 | 45 | 38 | 19 | 11 | 0 | 180 | 0 | 93 | 0 | 0 | 297 | 0 | 0 |
| SRR5029995 | 11 | 11 | 15 | 41 | 19 | 1220 | 275 | 1465 | 0 | 213 | 66 | 15 | 0 | 0 | 6712 | 19032 | 103121 | 31 | 0 | 3478 | 35912 | 0 | 61 | 18 | 20 | 15 | 71 | 0 | 0 | 105 | 0 | 92 | 0 | 0 | 599 | 0 | 0 |
| SRR5029996 | 12 | 66 | 27 | 25 | 45 | 352 | 167 | 742 | 0 | 200 | 20 | 0 | 0 | 0 | 7420 | 17456 | 103902 | 37 | 37 | 4415 | 41236 | 0 | 66 | 17 | 62 | 47 | 18 | 23 | 0 | 189 | 0 | 130 | 0 | 0 | 266 | 0 | 12 |
| SRR5029997 | 37 | 58 | 16 | 31 | 173 | 1675 | 161 | 1546 | 0 | 1502 | 78 | 0 | 0 | 0 | 6539 | 19771 | 103575 | 97 | 0 | 4059 | 36657 | 0 | 68 | 162 | 31 | 47 | 84 | 16 | 0 | 764 | 0 | 32 | 0 | 0 | 624 | 0 | 0 |
| SRR5029998 | 319 | 505 | 35 | 44 | 248 | 438 | 404 | 3908 | 94 | 0 | 27 | 35 | 0 | 0 | 16896 | 6689 | 280172 | 0 | 0 | 27504 | 2697 | 0 | 0 | 585 | 213 | 727 | 1507 | 35 | 0 | 293 | 0 | 0 | 0 | 0 | 271 | 0 | 0 |
| SRR5029999 | 235 | 326 | 21 | 16 | 171 | 1136 | 962 | 6680 | 198 | 0 | 21 | 64 | 0 | 0 | 15067 | 6797 | 272403 | 0 | 24 | 23436 | 2110 | 0 | 0 | 780 | 150 | 320 | 753 | 0 | 0 | 166 | 0 | 18 | 0 | 0 | 641 | 0 | 11 |
| SRR5030000 | 330 | 481 | 57 | 38 | 302 | 456 | 356 | 3413 | 146 | 0 | 38 | 57 | 0 | 0 | 16653 | 6428 | 280652 | 0 | 0 | 27474 | 2062 | 0 | 0 | 500 | 255 | 905 | 1707 | 66 | 0 | 198 | 0 | 0 | 0 | 0 | 276 | 0 | 0 |
| SRR5030001 | 223 | 340 | 11 | 0 | 145 | 1094 | 785 | 6692 | 222 | 0 | 17 | 72 | 0 | 0 | 14805 | 7179 | 270108 | 0 | 19 | 22677 | 2169 | 0 | 0 | 758 | 178 | 334 | 702 | 33 | 0 | 150 | 0 | 25 | 0 | 0 | 510 | 0 | 0 |
| SRR5030002 | 21 | 24 | 36 | 72 | 13 | 273 | 449 | 3690 | 53 | 18 | 0 | 27 | 0 | 0 | 16769 | 6674 | 279820 | 0 | 44 | 26784 | 2528 | 0 | 0 | 37 | 16 | 62 | 129 | 45 | 0 | 21 | 0 | 0 | 0 | 0 | 286 | 0 | 27 |
| SRR5030003 | 234 | 310 | 11 | 33 | 142 | 1053 | 868 | 6687 | 179 | 0 | 11 | 76 | 0 | 0 | 15332 | 6585 | 271203 | 0 | 18 | 22542 | 2260 | 0 | 0 | 615 | 114 | 387 | 626 | 27 | 0 | 131 | 0 | 12 | 0 | 0 | 571 | 0 | 11 |
| SRR5030004 | 15 | 32 | 46 | 82 | 16 | 346 | 506 | 3371 | 80 | 0 | 37 | 18 | 0 | 0 | 16793 | 6560 | 279398 | 0 | 0 | 28299 | 2369 | 0 | 0 | 44 | 17 | 69 | 123 | 119 | 0 | 18 | 0 | 18 | 0 | 0 | 290 | 0 | 0 |
| SRR5030005 | 13 | 19 | 17 | 17 | 15 | 1163 | 866 | 6695 | 229 | 11 | 11 | 44 | 11 | 0 | 14577 | 6725 | 269008 | 0 | 0 | 22398 | 2339 | 0 | 0 | 45 | 11 | 18 | 48 | 17 | 0 | 14 | 0 | 12 | 0 | 0 | 577 | 0 | 11 |
| SRR5030006 | 234 | 17 | 17 | 42 | 23 | 13 | 0 | 267 | 27 | 0 | 84 | 33 | 0 | 0 | 4210 | 27645 | 98137 | 27 | 67 | 6175 | 3208 | 0 | 0 | 50 | 0 | 33 | 29 | 33 | 0 | 27 | 0 | 53 | 0 | 0 | 2617 | 0 | 0 |
| SRR5030008 | 198 | 19 | 18 | 45 | 23 | 15 | 31 | 201 | 46 | 0 | 90 | 36 | 0 | 0 | 4289 | 28168 | 96592 | 0 | 62 | 6751 | 4150 | 0 | 0 | 36 | 0 | 45 | 21 | 36 | 0 | 31 | 0 | 31 | 0 | 0 | 2500 | 0 | 0 |
| SRR5030009 | 236 | 26 | 0 | 0 | 17 | 0 | 140 | 280 | 0 | 0 | 157 | 0 | 0 | 0 | 4487 | 22374 | 82663 | 0 | 140 | 6170 | 2384 | 0 | 0 | 157 | 0 | 0 | 23 | 0 | 0 | 20 | 0 | 0 | 0 | 0 | 3858 | 0 | 0 |
| SRR5030010 | 249 | 344 | 26 | 69 | 490 | 29 | 44 | 352 | 44 | 0 | 60 | 0 | 0 | 0 | 4148 | 28038 | 94721 | 0 | 44 | 7211 | 3341 | 0 | 0 | 17 | 0 | 43 | 696 | 26 | 0 | 636 | 0 | 73 | 0 | 0 | 2573 | 0 | 0 |
| SRR5030011 | 155 | 14 | 0 | 232 | 14 | 0 | 0 | 380 | 0 | 0 | 77 | 0 | 77 | 0 | 4304 | 23328 | 83960 | 0 | 127 | 4811 | 2279 | 0 | 0 | 77 | 0 | 0 | 11 | 77 | 0 | 19 | 0 | 0 | 0 | 0 | 4193 | 0 | 0 |
| SRR5030012 | 263 | 342 | 18 | 26 | 447 | 0 | 30 | 257 | 45 | 0 | 79 | 0 | 0 | 0 | 4745 | 27366 | 97286 | 0 | 45 | 7102 | 4125 | 0 | 0 | 18 | 0 | 70 | 491 | 35 | 0 | 649 | 0 | 45 | 0 | 0 | 2422 | 0 | 26 |
| SRR5030014 | 303 | 372 | 28 | 152 | 964 | 33 | 120 | 294 | 44 | 0 | 138 | 28 | 0 | 0 | 642 | 11350 | 301489 | 44 | 44 | 4724 | 18276 | 0 | 0 | 179 | 41 | 69 | 758 | 41 | 0 | 441 | 0 | 33 | 0 | 0 | 556 | 0 | 0 |
| SRR5030015 | 150 | 205 | 0 | 27 | 615 | 56 | 195 | 848 | 14 | 14 | 14 | 14 | 0 | 0 | 640 | 12334 | 286331 | 56 | 83 | 3448 | 14363 | 0 | 0 | 478 | 27 | 14 | 205 | 27 | 0 | 424 | 0 | 0 | 0 | 0 | 1036 | 0 | 0 |
| SRR5030016 | 456 | 304 | 61 | 106 | 1018 | 12 | 96 | 383 | 24 | 0 | 46 | 0 | 0 | 0 | 586 | 11620 | 301937 | 24 | 12 | 5417 | 18462 | 0 | 0 | 289 | 122 | 61 | 547 | 46 | 0 | 729 | 0 | 48 | 0 | 0 | 470 | 0 | 0 |
| SRR5030017 | 10 | 215 | 29 | 0 | 22 | 87 | 159 | 869 | 87 | 29 | 29 | 0 | 0 | 0 | 594 | 12528 | 285257 | 101 | 14 | 3550 | 13562 | 0 | 0 | 15 | 0 | 0 | 129 | 0 | 0 | 13 | 0 | 0 | 0 | 0 | 1228 | 0 | 0 |
| SRR5030018 | 13 | 23 | 72 | 86 | 45 | 23 | 101 | 462 | 34 | 0 | 115 | 0 | 0 | 0 | 642 | 11374 | 299957 | 68 | 23 | 4886 | 17595 | 0 | 0 | 16 | 86 | 57 | 36 | 72 | 0 | 32 | 0 | 79 | 0 | 0 | 509 | 0 | 0 |
| SRR5030019 | 209 | 237 | 14 | 14 | 724 | 43 | 171 | 783 | 57 | 28 | 28 | 0 | 0 | 0 | 541 | 12199 | 285690 | 43 | 14 | 3260 | 14023 | 0 | 0 | 418 | 0 | 28 | 209 | 0 | 0 | 306 | 0 | 43 | 0 | 0 | 1083 | 0 | 0 |
| SRR5030020 | 14 | 12 | 74 | 103 | 36 | 23 | 128 | 452 | 46 | 0 | 74 | 15 | 0 | 0 | 603 | 11704 | 299512 | 12 | 81 | 4918 | 17771 | 0 | 0 | 16 | 44 | 88 | 38 | 44 | 0 | 30 | 0 | 46 | 0 | 0 | 457 | 0 | 0 |
| SRR5030021 | 216 | 172 | 0 | 57 | 22 | 59 | 161 | 835 | 29 | 29 | 43 | 0 | 0 | 0 | 425 | 12324 | 287054 | 29 | 44 | 3720 | 14397 | 0 | 0 | 15 | 14 | 14 | 158 | 0 | 0 | 12 | 0 | 73 | 0 | 0 | 900 | 0 | 0 |
| SRR5030022 | 97 | 109 | 0 | 36 | 37 | 34 | 205 | 342 | 34 | 24 | 36 | 12 | 0 | 0 | 787 | 29661 | 74362 | 68 | 0 | 11707 | 1472 | 0 | 0 | 73 | 0 | 48 | 14 | 12 | 0 | 206 | 0 | 0 | 0 | 0 | 7152 | 0 | 0 |
| SRR5030023 | 179 | 22 | 0 | 22 | 28 | 67 | 270 | 472 | 0 | 22 | 0 | 45 | 0 | 0 | 540 | 30221 | 63903 | 67 | 0 | 8568 | 1889 | 0 | 0 | 90 | 0 | 22 | 157 | 22 | 0 | 11 | 0 | 67 | 0 | 0 | 8327 | 0 | 22 |
| SRR5030024 | 39 | 130 | 13 | 26 | 34 | 37 | 148 | 333 | 37 | 65 | 0 | 0 | 0 | 0 | 444 | 29368 | 71030 | 37 | 37 | 10731 | 1850 | 0 | 0 | 104 | 0 | 13 | 14 | 13 | 0 | 16 | 0 | 0 | 0 | 0 | 6318 | 0 | 0 |
| SRR5030025 | 23 | 140 | 23 | 0 | 884 | 0 | 68 | 271 | 0 | 70 | 23 | 0 | 0 | 0 | 339 | 29635 | 64145 | 0 | 0 | 11192 | 1357 | 0 | 0 | 163 | 0 | 0 | 256 | 0 | 0 | 93 | 0 | 68 | 0 | 0 | 9164 | 0 | 0 |
| SRR5030026 | 99 | 162 | 12 | 50 | 933 | 0 | 70 | 141 | 35 | 12 | 62 | 12 | 0 | 0 | 880 | 30960 | 72947 | 0 | 0 | 10982 | 1338 | 0 | 0 | 99 | 0 | 37 | 435 | 25 | 0 | 286 | 0 | 35 | 0 | 0 | 6627 | 0 | 0 |
| SRR5030027 | 90 | 180 | 0 | 0 | 833 | 0 | 203 | 405 | 0 | 23 | 23 | 0 | 0 | 0 | 608 | 28841 | 70594 | 0 | 0 | 9728 | 1756 | 0 | 0 | 113 | 23 | 0 | 293 | 0 | 0 | 293 | 0 | 0 | 0 | 0 | 8483 | 0 | 0 |
| SRR5030028 | 101 | 139 | 0 | 25 | 31 | 37 | 147 | 110 | 37 | 38 | 76 | 0 | 0 | 0 | 772 | 28358 | 71609 | 0 | 37 | 10067 | 2021 | 0 | 0 | 50 | 0 | 38 | 18 | 38 | 0 | 11 | 0 | 0 | 0 | 0 | 7144 | 0 | 25 |
| SRR5030030 | 112 | 360 | 23 | 90 | 225 | 0 | 13 | 53 | 0 | 0 | 79 | 23 | 0 | 0 | 382 | 16792 | 50375 | 0 | 13 | 3360 | 6786 | 0 | 0 | 45 | 11 | 23 | 439 | 0 | 0 | 641 | 0 | 13 | 0 | 0 | 8965 | 0 | 23 |
| SRR5030031 | 127 | 197 | 0 | 23 | 174 | 63 | 21 | 189 | 21 | 0 | 12 | 23 | 0 | 0 | 611 | 9413 | 24566 | 0 | 21 | 2653 | 6042 | 0 | 0 | 58 | 0 | 12 | 69 | 0 | 0 | 278 | 0 | 21 | 0 | 0 | 9513 | 0 | 0 |
| SRR5030032 | 230 | 339 | 0 | 12 | 145 | 28 | 28 | 113 | 14 | 0 | 133 | 12 | 0 | 0 | 480 | 17502 | 51049 | 0 | 0 | 3640 | 6630 | 0 | 0 | 85 | 0 | 24 | 412 | 0 | 0 | 545 | 0 | 14 | 0 | 0 | 9003 | 0 | 0 |
| SRR5030033 | 60 | 205 | 0 | 0 | 169 | 22 | 45 | 224 | 45 | 36 | 12 | 0 | 0 | 0 | 605 | 9095 | 23440 | 22 | 0 | 3047 | 6274 | 0 | 0 | 108 | 0 | 12 | 72 | 0 | 0 | 217 | 0 | 45 | 0 | 0 | 10467 | 0 | 12 |
| SRR5030034 | 10 | 23 | 12 | 35 | 161 | 0 | 55 | 41 | 27 | 0 | 92 | 23 | 0 | 0 | 505 | 18552 | 52699 | 0 | 14 | 3292 | 6625 | 0 | 0 | 69 | 35 | 46 | 23 | 12 | 0 | 28 | 0 | 55 | 0 | 0 | 8441 | 0 | 0 |
| SRR5030035 | 94 | 142 | 0 | 0 | 94 | 21 | 106 | 212 | 42 | 12 | 24 | 24 | 0 | 0 | 445 | 8189 | 22391 | 42 | 21 | 3136 | 5848 | 0 | 0 | 94 | 0 | 0 | 71 | 12 | 0 | 271 | 0 | 21 | 0 | 0 | 9684 | 0 | 35 |
| SRR5030036 | 176 | 365 | 24 | 59 | 223 | 14 | 0 | 42 | 0 | 12 | 94 | 12 | 0 | 0 | 363 | 18240 | 52655 | 0 | 28 | 3660 | 7390 | 0 | 0 | 59 | 0 | 35 | 188 | 24 | 0 | 623 | 0 | 28 | 0 | 0 | 7719 | 0 | 0 |
| SRR5030038 | 203 | 419 | 13 | 51 | 114 | 54 | 68 | 368 | 0 | 25 | 102 | 13 | 0 | 0 | 422 | 48371 | 40926 | 27 | 0 | 4724 | 9828 | 0 | 0 | 216 | 25 | 51 | 863 | 38 | 0 | 432 | 0 | 27 | 0 | 0 | 3735 | 0 | 0 |
| SRR5030039 | 141 | 212 | 0 | 0 | 212 | 29 | 58 | 378 | 58 | 0 | 71 | 0 | 0 | 0 | 349 | 48866 | 39733 | 29 | 0 | 3343 | 9041 | 0 | 0 | 11 | 0 | 0 | 14 | 0 | 0 | 165 | 0 | 0 | 0 | 0 | 4723 | 0 | 24 |
| SRR5030040 | 188 | 11 | 13 | 13 | 175 | 44 | 87 | 261 | 29 | 0 | 67 | 0 | 13 | 0 | 407 | 47790 | 40618 | 0 | 0 | 3905 | 10395 | 0 | 0 | 11 | 0 | 27 | 30 | 13 | 0 | 14 | 0 | 44 | 0 | 0 | 3464 | 0 | 0 |
| SRR5030041 | 170 | 316 | 0 | 24 | 122 | 0 | 88 | 381 | 59 | 49 | 73 | 24 | 24 | 0 | 323 | 46785 | 39748 | 59 | 0 | 3608 | 9827 | 0 | 0 | 146 | 24 | 0 | 390 | 24 | 0 | 292 | 0 | 0 | 0 | 0 | 4927 | 0 | 0 |
| SRR5030042 | 259 | 168 | 26 | 52 | 103 | 28 | 56 | 291 | 42 | 26 | 103 | 0 | 0 | 0 | 458 | 47543 | 40819 | 14 | 0 | 4164 | 10410 | 0 | 0 | 194 | 13 | 26 | 698 | 0 | 0 | 388 | 0 | 0 | 0 | 0 | 3401 | 0 | 0 |
| SRR5030043 | 238 | 334 | 0 | 71 | 143 | 0 | 0 | 288 | 29 | 0 | 24 | 0 | 0 | 0 | 692 | 47648 | 38091 | 86 | 0 | 3602 | 8357 | 0 | 0 | 238 | 0 | 95 | 357 | 24 | 0 | 191 | 0 | 86 | 0 | 0 | 4367 | 0 | 24 |
| SRR5030044 | 237 | 224 | 26 | 105 | 184 | 44 | 29 | 437 | 44 | 26 | 53 | 0 | 0 | 0 | 539 | 47866 | 40843 | 15 | 29 | 4414 | 9687 | 0 | 0 | 158 | 0 | 26 | 513 | 13 | 0 | 381 | 0 | 73 | 0 | 0 | 3485 | 0 | 0 |
| SRR5030045 | 195 | 195 | 0 | 0 | 98 | 59 | 59 | 295 | 0 | 0 | 49 | 0 | 0 | 0 | 325 | 47681 | 38798 | 30 | 0 | 3189 | 9006 | 0 | 0 | 439 | 0 | 0 | 390 | 0 | 0 | 317 | 0 | 30 | 0 | 0 | 4553 | 0 | 0 |
| SRR5030046 | 111 | 14 | 15 | 119 | 73 | 83 | 55 | 648 | 110 | 163 | 163 | 22 | 0 | 0 | 924 | 133239 | 114287 | 0 | 124 | 7064 | 8734 | 0 | 0 | 10 | 22 | 45 | 24 | 15 | 0 | 57 | 0 | 207 | 0 | 0 | 665 | 0 | 15 |
| SRR5030048 | 137 | 201 | 48 | 80 | 1191 | 178 | 74 | 489 | 118 | 64 | 153 | 16 | 0 | 0 | 978 | 130908 | 115622 | 0 | 133 | 7036 | 8532 | 0 | 0 | 161 | 24 | 40 | 459 | 16 | 0 | 1054 | 0 | 104 | 0 | 0 | 819 | 0 | 0 |
| SRR5030050 | 107 | 267 | 15 | 122 | 1189 | 155 | 84 | 408 | 42 | 130 | 152 | 30 | 0 | 0 | 717 | 131514 | 113633 | 28 | 98 | 6696 | 9186 | 0 | 0 | 221 | 23 | 30 | 335 | 0 | 0 | 937 | 0 | 141 | 0 | 0 | 913 | 0 | 0 |
| SRR5030052 | 181 | 346 | 39 | 87 | 1102 | 129 | 86 | 429 | 43 | 157 | 134 | 31 | 0 | 0 | 1001 | 131489 | 116091 | 14 | 43 | 7368 | 9171 | 0 | 0 | 110 | 24 | 31 | 457 | 47 | 0 | 1220 | 0 | 114 | 0 | 0 | 532 | 0 | 0 |
| SRR5030053 | 46 | 274 | 0 | 46 | 776 | 183 | 183 | 731 | 91 | 91 | 0 | 46 | 0 | 0 | 640 | 132226 | 109553 | 0 | 0 | 7493 | 8132 | 0 | 0 | 137 | 0 | 0 | 320 | 0 | 0 | 822 | 0 | 548 | 0 | 0 | 1089 | 0 | 46 |
| SRR5030054 | 539 | 699 | 58 | 87 | 233 | 11 | 50 | 89 | 22 | 0 | 218 | 0 | 0 | 0 | 6604 | 12263 | 294014 | 22 | 17 | 3247 | 53349 | 0 | 0 | 29 | 29 | 204 | 1078 | 73 | 0 | 1034 | 0 | 17 | 0 | 0 | 318 | 0 | 29 |
| SRR5030055 | 11 | 14 | 48 | 48 | 240 | 43 | 43 | 214 | 0 | 0 | 48 | 0 | 0 | 0 | 7647 | 13890 | 291967 | 21 | 43 | 2948 | 51268 | 0 | 0 | 48 | 0 | 96 | 336 | 0 | 0 | 20 | 0 | 0 | 0 | 0 | 464 | 0 | 48 |
| SRR5030056 | 314 | 815 | 0 | 31 | 235 | 12 | 24 | 107 | 24 | 0 | 172 | 16 | 0 | 0 | 6631 | 12534 | 292409 | 18 | 24 | 3125 | 54293 | 0 | 0 | 47 | 78 | 204 | 1348 | 78 | 0 | 988 | 0 | 0 | 0 | 0 | 326 | 0 | 0 |
| SRR5030057 | 204 | 765 | 0 | 0 | 408 | 67 | 67 | 156 | 67 | 0 | 0 | 0 | 0 | 0 | 5739 | 13103 | 289946 | 44 | 0 | 3292 | 50984 | 0 | 0 | 51 | 0 | 102 | 1326 | 0 | 0 | 663 | 0 | 0 | 0 | 0 | 564 | 0 | 51 |
| SRR5030058 | 13 | 24 | 60 | 60 | 165 | 29 | 46 | 51 | 0 | 0 | 30 | 45 | 0 | 0 | 6650 | 12942 | 292564 | 40 | 11 | 3228 | 53467 | 0 | 0 | 15 | 30 | 135 | 35 | 75 | 0 | 31 | 0 | 0 | 0 | 0 | 302 | 0 | 30 |
| SRR5030059 | 295 | 17 | 0 | 148 | 246 | 44 | 22 | 131 | 44 | 0 | 0 | 0 | 0 | 0 | 6592 | 13536 | 290531 | 65 | 0 | 2850 | 52450 | 0 | 0 | 49 | 0 | 0 | 18 | 0 | 0 | 17 | 0 | 0 | 0 | 0 | 564 | 0 | 49 |
| SRR5030060 | 13 | 19 | 46 | 77 | 215 | 0 | 29 | 98 | 29 | 15 | 61 | 0 | 0 | 0 | 6283 | 12806 | 290614 | 29 | 12 | 3246 | 54434 | 0 | 0 | 15 | 0 | 230 | 40 | 0 | 0 | 30 | 0 | 17 | 0 | 0 | 273 | 0 | 15 |
| SRR5030061 | 302 | 16 | 0 | 0 | 101 | 45 | 112 | 202 | 22 | 0 | 0 | 50 | 0 | 0 | 7060 | 13379 | 290330 | 22 | 0 | 2958 | 50941 | 0 | 0 | 151 | 0 | 101 | 13 | 0 | 0 | 13 | 0 | 0 | 0 | 0 | 516 | 0 | 101 |
| SRR5030062 | 65 | 91 | 13 | 26 | 286 | 51 | 135 | 388 | 34 | 0 | 91 | 0 | 0 | 0 | 540 | 38274 | 48415 | 34 | 17 | 3239 | 7490 | 0 | 0 | 169 | 13 | 39 | 19 | 39 | 0 | 26 | 0 | 51 | 0 | 0 | 721 | 0 | 0 |
| SRR5030064 | 70 | 126 | 0 | 42 | 237 | 36 | 91 | 398 | 0 | 0 | 84 | 14 | 0 | 0 | 471 | 37754 | 48558 | 0 | 54 | 3259 | 7605 | 0 | 0 | 195 | 0 | 42 | 516 | 14 | 0 | 586 | 0 | 0 | 0 | 0 | 687 | 0 | 14 |
| SRR5030066 | 94 | 107 | 13 | 13 | 227 | 35 | 86 | 224 | 86 | 13 | 94 | 27 | 0 | 0 | 604 | 37824 | 48409 | 17 | 17 | 3089 | 8473 | 0 | 0 | 227 | 13 | 27 | 508 | 40 | 0 | 561 | 0 | 35 | 0 | 0 | 504 | 0 | 13 |
| SRR5030068 | 108 | 121 | 27 | 27 | 229 | 105 | 122 | 262 | 17 | 0 | 67 | 0 | 0 | 0 | 523 | 37342 | 49549 | 0 | 17 | 3384 | 8076 | 0 | 0 | 215 | 13 | 27 | 552 | 13 | 0 | 619 | 0 | 17 | 0 | 0 | 690 | 0 | 13 |
| SRR5030069 | 250 | 63 | 0 | 0 | 250 | 0 | 0 | 91 | 0 | 0 | 125 | 0 | 0 | 0 | 639 | 37943 | 47575 | 0 | 0 | 2737 | 6752 | 0 | 0 | 375 | 0 | 0 | 313 | 0 | 0 | 250 | 0 | 182 | 0 | 0 | 879 | 0 | 0 |
| SRR5030070 | 129 | 199 | 0 | 149 | 676 | 0 | 78 | 196 | 39 | 0 | 119 | 0 | 0 | 0 | 17303 | 19491 | 86077 | 13 | 26 | 4538 | 2230 | 0 | 0 | 169 | 20 | 0 | 497 | 0 | 0 | 567 | 0 | 13 | 0 | 0 | 1028 | 0 | 0 |
| SRR5030071 | 185 | 144 | 0 | 21 | 575 | 0 | 182 | 546 | 91 | 41 | 82 | 0 | 0 | 0 | 17244 | 19327 | 85238 | 0 | 0 | 3728 | 2091 | 0 | 0 | 226 | 0 | 0 | 144 | 0 | 0 | 432 | 0 | 0 | 0 | 0 | 1589 | 0 | 0 |
| SRR5030072 | 193 | 118 | 21 | 43 | 580 | 14 | 127 | 184 | 85 | 0 | 86 | 0 | 0 | 0 | 17692 | 18322 | 84857 | 85 | 71 | 4051 | 2309 | 0 | 0 | 75 | 11 | 86 | 397 | 32 | 0 | 537 | 0 | 28 | 0 | 0 | 880 | 0 | 11 |
| SRR5030073 | 170 | 85 | 0 | 21 | 23 | 64 | 288 | 609 | 0 | 21 | 106 | 0 | 0 | 0 | 17212 | 19631 | 85453 | 32 | 32 | 4103 | 2596 | 0 | 0 | 212 | 0 | 0 | 127 | 43 | 0 | 15 | 0 | 0 | 0 | 0 | 1553 | 0 | 0 |
| SRR5030074 | 10 | 194 | 51 | 31 | 30 | 27 | 54 | 327 | 41 | 0 | 41 | 0 | 0 | 0 | 18484 | 18571 | 86356 | 14 | 41 | 4519 | 2545 | 0 | 0 | 194 | 0 | 31 | 16 | 61 | 0 | 33 | 0 | 27 | 0 | 0 | 884 | 0 | 10 |
| SRR5030076 | 198 | 156 | 42 | 21 | 625 | 14 | 123 | 316 | 55 | 0 | 63 | 0 | 0 | 0 | 18273 | 18425 | 85363 | 27 | 27 | 4760 | 2414 | 0 | 0 | 198 | 10 | 31 | 563 | 42 | 0 | 521 | 0 | 14 | 0 | 0 | 796 | 0 | 0 |
| SRR5030077 | 149 | 85 | 21 | 0 | 12 | 63 | 189 | 315 | 63 | 0 | 21 | 0 | 0 | 0 | 16348 | 19965 | 84429 | 0 | 0 | 3717 | 2142 | 0 | 0 | 212 | 0 | 21 | 12 | 21 | 0 | 11 | 0 | 0 | 0 | 0 | 1642 | 0 | 0 |
| SRR5030078 | 180 | 12 | 0 | 32 | 19 | 40 | 0 | 320 | 40 | 106 | 11 | 21 | 0 | 0 | 260 | 16459 | 133223 | 0 | 100 | 5299 | 11718 | 0 | 32 | 53 | 0 | 11 | 21 | 21 | 0 | 167 | 0 | 40 | 0 | 0 | 1838 | 0 | 0 |
| SRR5030079 | 17 | 223 | 0 | 17 | 223 | 119 | 79 | 514 | 198 | 52 | 69 | 0 | 0 | 0 | 316 | 12754 | 104729 | 0 | 40 | 4668 | 11394 | 0 | 17 | 86 | 0 | 0 | 206 | 17 | 0 | 2163 | 0 | 40 | 0 | 0 | 3498 | 0 | 17 |
| SRR5030080 | 126 | 160 | 11 | 114 | 412 | 64 | 0 | 451 | 21 | 23 | 252 | 23 | 11 | 0 | 451 | 16893 | 135813 | 0 | 21 | 5738 | 12357 | 0 | 23 | 34 | 11 | 23 | 366 | 23 | 0 | 3445 | 0 | 43 | 0 | 0 | 1790 | 0 | 23 |
| SRR5030081 | 89 | 10 | 0 | 18 | 12 | 123 | 41 | 327 | 82 | 72 | 36 | 36 | 0 | 0 | 491 | 14496 | 107066 | 0 | 0 | 4254 | 10593 | 0 | 18 | 89 | 0 | 54 | 89 | 0 | 0 | 95 | 0 | 41 | 0 | 0 | 3076 | 0 | 0 |
| SRR5030082 | 55 | 10 | 0 | 98 | 24 | 61 | 20 | 225 | 143 | 55 | 11 | 11 | 0 | 0 | 225 | 16945 | 134592 | 20 | 41 | 4822 | 11748 | 0 | 0 | 98 | 0 | 11 | 15 | 11 | 0 | 165 | 0 | 41 | 0 | 0 | 1555 | 0 | 0 |
| SRR5030083 | 70 | 211 | 0 | 35 | 18 | 80 | 120 | 681 | 200 | 70 | 18 | 35 | 0 | 0 | 120 | 13493 | 104143 | 0 | 0 | 4124 | 11571 | 0 | 35 | 35 | 0 | 18 | 158 | 35 | 0 | 81 | 0 | 160 | 0 | 0 | 2400 | 0 | 0 |

Supp Table S8.3.Columns_BX_to_DH

| | | | | | | | | | | | | | | | | | | | | | | | | | | | | | | | | | | | | | |
|---|---|---|---|---|---|---|---|---|---|---|---|---|---|---|---|---|---|---|---|---|---|---|---|---|---|---|---|---|---|---|---|---|---|---|---|---|---|
| SRR5030084 | 168 | 17 | 11 | 34 | 18 | 21 | 41 | 268 | 41 | 79 | 213 | 22 | 0 | 0 | 309 | 15752 | 133852 | 0 | 82 | 5460 | 11869 | 0 | 11 | 79 | 11 | 56 | 18 | 11 | 0 | 156 | 0 | 21 | 0 | 0 | 1925 | 0 | 11 |
| SRR5030085 | 53 | 214 | 0 | 0 | 12 | 120 | 40 | 441 | 40 | 36 | 89 | 18 | 0 | 0 | 241 | 14321 | 108169 | 0 | 80 | 4130 | 12110 | 0 | 53 | 36 | 0 | 18 | 125 | 0 | 0 | 84 | 0 | 40 | 0 | 0 | 2441 | 0 | 18 |
| SRR5030086 | 59 | 104 | 15 | 15 | 25 | 601 | 168 | 2574 | 144 | 0 | 40 | 25 | 0 | 0 | 35335 | 6419 | 202454 | 0 | 72 | 14288 | 4426 | 0 | 0 | 40 | 0 | 20 | 238 | 0 | 0 | 347 | 0 | 0 | 0 | 0 | 523 | 0 | 0 |
| SRR5030087 | 119 | 68 | 0 | 0 | 145 | 286 | 449 | 2695 | 0 | 0 | 51 | 34 | 0 | 0 | 35277 | 6290 | 207534 | 0 | 0 | 12698 | 3716 | 0 | 0 | 43 | 0 | 0 | 77 | 0 | 0 | 366 | 0 | 41 | 0 | 0 | 930 | 0 | 0 |
| SRR5030088 | 63 | 78 | 0 | 16 | 42 | 180 | 488 | 2210 | 77 | 0 | 42 | 52 | 0 | 0 | 35209 | 5774 | 204555 | 0 | 51 | 15163 | 4240 | 0 | 0 | 58 | 16 | 42 | 23 | 0 | 0 | 40 | 0 | 0 | 0 | 0 | 430 | 0 | 0 |
| SRR5030090 | 55 | 75 | 10 | 20 | 75 | 395 | 296 | 2372 | 74 | 0 | 60 | 30 | 0 | 0 | 36664 | 6150 | 205985 | 0 | 25 | 13959 | 3928 | 0 | 0 | 80 | 25 | 45 | 145 | 0 | 0 | 351 | 0 | 49 | 0 | 0 | 399 | 0 | 0 |
| SRR5030091 | 103 | 103 | 0 | 26 | 69 | 328 | 287 | 2870 | 164 | 0 | 17 | 34 | 0 | 0 | 33255 | 6748 | 205163 | 0 | 82 | 13040 | 3978 | 0 | 0 | 86 | 0 | 0 | 103 | 0 | 0 | 18 | 0 | 82 | 0 | 0 | 859 | 0 | 0 |
| SRR5030092 | 81 | 10 | 20 | 41 | 76 | 352 | 251 | 2410 | 75 | 0 | 36 | 41 | 0 | 0 | 35954 | 6016 | 204360 | 0 | 25 | 14361 | 4670 | 0 | 0 | 25 | 0 | 25 | 29 | 20 | 10 | 32 | 0 | 25 | 0 | 0 | 591 | 0 | 0 |
| SRR5030093 | 88 | 106 | 0 | 0 | 106 | 460 | 544 | 3390 | 293 | 18 | 53 | 62 | 0 | 0 | 34574 | 6390 | 210105 | 0 | 42 | 12641 | 3265 | 0 | 0 | 88 | 0 | 18 | 71 | 0 | 18 | 247 | 0 | 42 | 0 | 0 | 719 | 0 | 0 |
| SRR5030094 | 12 | 11 | 0 | 0 | 48 | 96 | 84 | 697 | 24 | 11 | 79 | 79 | 0 | 0 | 19635 | 9076 | 220737 | 0 | 156 | 69610 | 22987 | 0 | 0 | 73 | 0 | 19 | 131 | 0 | 0 | 360 | 0 | 132 | 0 | 0 | 544 | 0 | 0 |
| SRR5030095 | 21 | 0 | 0 | 0 | 17 | 446 | 594 | 1486 | 0 | 0 | 41 | 29 | 0 | 0 | 16937 | 10358 | 217016 | 0 | 0 | 60912 | 20057 | 0 | 0 | 41 | 21 | 21 | 41 | 0 | 0 | 54 | 0 | 0 | 0 | 0 | 986 | 0 | 0 |
| SRR5030096 | 18 | 15 | 0 | 0 | 11 | 65 | 117 | 622 | 13 | 0 | 22 | 23 | 0 | 0 | 19529 | 9281 | 219777 | 0 | 117 | 72536 | 23026 | 0 | 0 | 14 | 0 | 25 | 33 | 0 | 0 | 110 | 0 | 26 | 0 | 0 | 734 | 0 | 0 |
| SRR5030097 | 43 | 64 | 0 | 0 | 43 | 0 | 151 | 1510 | 0 | 21 | 21 | 43 | 0 | 0 | 18119 | 9201 | 221418 | 0 | 0 | 60396 | 18421 | 0 | 0 | 85 | 0 | 21 | 64 | 0 | 0 | 213 | 0 | 151 | 0 | 0 | 1193 | 0 | 21 |
| SRR5030098 | 27 | 19 | 0 | 0 | 49 | 85 | 61 | 874 | 49 | 0 | 76 | 71 | 0 | 0 | 19899 | 9287 | 221790 | 12 | 170 | 71399 | 21574 | 0 | 0 | 52 | 0 | 16 | 154 | 0 | 0 | 370 | 0 | 36 | 0 | 0 | 695 | 0 | 0 |
| SRR5030100 | 19 | 24 | 0 | 0 | 10 | 75 | 50 | 726 | 0 | 0 | 20 | 28 | 0 | 0 | 19199 | 9306 | 218409 | 13 | 88 | 72254 | 21790 | 0 | 0 | 13 | 0 | 11 | 44 | 0 | 0 | 97 | 0 | 63 | 0 | 0 | 684 | 0 | 0 |
| SRR5030101 | 85 | 0 | 0 | 0 | 85 | 0 | 0 | 1245 | 0 | 0 | 21 | 85 | 0 | 0 | 19608 | 9857 | 219208 | 0 | 0 | 61158 | 18052 | 0 | 0 | 170 | 21 | 21 | 107 | 0 | 0 | 277 | 0 | 0 | 0 | 0 | 960 | 0 | 0 |
| SRR5030102 | 193 | 331 | 23 | 62 | 439 | 96 | 72 | 551 | 144 | 0 | 100 | 15 | 0 | 0 | 958 | 14083 | 40536 | 0 | 48 | 6584 | 2323 | 0 | 0 | 23 | 23 | 54 | 462 | 15 | 0 | 678 | 0 | 0 | 0 | 0 | 10059 | 0 | 0 |
| SRR5030103 | 235 | 235 | 28 | 0 | 235 | 217 | 271 | 894 | 0 | 0 | 103 | 0 | 0 | 0 | 379 | 11160 | 25060 | 0 | 27 | 3847 | 1598 | 0 | 0 | 0 | 0 | 38 | 244 | 0 | 0 | 610 | 0 | 0 | 0 | 0 | 10207 | 0 | 0 |
| SRR5030104 | 159 | 369 | 42 | 34 | 453 | 124 | 124 | 593 | 124 | 0 | 92 | 0 | 0 | 0 | 643 | 13523 | 40993 | 25 | 74 | 6500 | 2348 | 0 | 0 | 34 | 17 | 67 | 428 | 34 | 0 | 612 | 0 | 0 | 0 | 0 | 10225 | 0 | 0 |
| SRR5030105 | 156 | 14 | 0 | 29 | 19 | 200 | 257 | 599 | 86 | 0 | 49 | 0 | 0 | 0 | 685 | 8927 | 26740 | 0 | 57 | 3510 | 1541 | 0 | 0 | 29 | 0 | 29 | 14 | 0 | 0 | 25 | 0 | 0 | 0 | 0 | 10730 | 0 | 0 |
| SRR5030106 | 15 | 20 | 56 | 40 | 21 | 47 | 187 | 725 | 23 | 0 | 79 | 0 | 0 | 0 | 631 | 14878 | 43010 | 0 | 23 | 5984 | 2127 | 0 | 0 | 71 | 40 | 63 | 37 | 16 | 0 | 49 | 0 | 23 | 0 | 0 | 9558 | 13 | 0 |
| SRR5030107 | 133 | 200 | 0 | 29 | 475 | 133 | 133 | 583 | 27 | 0 | 48 | 0 | 0 | 0 | 636 | 9946 | 24664 | 0 | 0 | 3710 | 1325 | 0 | 0 | 67 | 0 | 19 | 209 | 0 | 0 | 513 | 0 | 0 | 0 | 0 | 11550 | 0 | 0 |
| SRR5030108 | 180 | 418 | 49 | 66 | 352 | 144 | 72 | 336 | 24 | 0 | 82 | 0 | 0 | 0 | 935 | 13664 | 41173 | 0 | 96 | 6832 | 1774 | 0 | 0 | 33 | 25 | 41 | 557 | 33 | 0 | 680 | 0 | 0 | 0 | 0 | 9838 | 0 | 0 |
| SRR5030109 | 137 | 225 | 29 | 20 | 343 | 193 | 165 | 578 | 28 | 0 | 78 | 0 | 0 | 0 | 633 | 10455 | 27867 | 28 | 28 | 4652 | 1211 | 0 | 0 | 78 | 39 | 29 | 206 | 0 | 0 | 461 | 0 | 0 | 0 | 0 | 10835 | 0 | 0 |
| SRR5030110 | 13 | 28 | 0 | 20 | 99 | 0 | 73 | 170 | 24 | 13 | 20 | 30 | 0 | 0 | 4321 | 41947 | 109227 | 24 | 24 | 5607 | 17404 | 0 | 0 | 0 | 50 | 30 | 14 | 0 | 0 | 63 | 0 | 146 | 0 | 0 | 2214 | 0 | 0 |
| SRR5030111 | 106 | 184 | 0 | 0 | 78 | 108 | 54 | 485 | 0 | 145 | 72 | 45 | 0 | 0 | 3701 | 38924 | 94104 | 72 | 0 | 3773 | 13133 | 0 | 0 | 39 | 17 | 22 | 50 | 0 | 0 | 407 | 0 | 287 | 0 | 0 | 4714 | 0 | 0 |
| SRR5030112 | 17 | 22 | 21 | 64 | 74 | 52 | 52 | 105 | 26 | 117 | 20 | 53 | 0 | 0 | 4244 | 41130 | 111550 | 0 | 52 | 6260 | 18127 | 0 | 0 | 64 | 11 | 11 | 16 | 11 | 0 | 77 | 0 | 288 | 0 | 0 | 1985 | 0 | 0 |
| SRR5030113 | 121 | 138 | 0 | 0 | 98 | 76 | 76 | 571 | 38 | 11 | 52 | 46 | 0 | 0 | 4000 | 38261 | 93016 | 19 | 19 | 4038 | 13868 | 17 | 17 | 46 | 29 | 0 | 58 | 12 | 0 | 19 | 0 | 210 | 0 | 0 | 4052 | 0 | 0 |
| SRR5030114 | 143 | 265 | 20 | 61 | 41 | 99 | 74 | 272 | 50 | 153 | 265 | 102 | 0 | 0 | 4655 | 40811 | 113163 | 0 | 25 | 6191 | 16963 | 0 | 0 | 51 | 20 | 31 | 244 | 10 | 0 | 866 | 0 | 198 | 0 | 0 | 1713 | 0 | 10 |
| SRR5030115 | 85 | 12 | 0 | 0 | 91 | 74 | 147 | 405 | 0 | 113 | 40 | 51 | 0 | 0 | 4103 | 39538 | 93663 | 0 | 18 | 3422 | 13726 | 0 | 0 | 79 | 17 | 17 | 74 | 0 | 0 | 26 | 0 | 331 | 0 | 0 | 4077 | 0 | 11 |
| SRR5030116 | 114 | 280 | 10 | 83 | 62 | 0 | 50 | 252 | 50 | 124 | 197 | 73 | 0 | 0 | 4645 | 42266 | 113018 | 25 | 25 | 5882 | 17443 | 0 | 10 | 73 | 41 | 31 | 270 | 10 | 0 | 1017 | 0 | 227 | 0 | 0 | 1737 | 0 | 10 |
| SRR5030117 | 132 | 115 | 11 | 0 | 80 | 111 | 37 | 463 | 56 | 15 | 29 | 57 | 0 | 0 | 3852 | 40969 | 92582 | 0 | 19 | 4186 | 13298 | 0 | 11 | 46 | 17 | 0 | 34 | 0 | 0 | 21 | 0 | 315 | 0 | 0 | 4466 | 0 | 0 |
| SRR5030118 | 16 | 15 | 0 | 14 | 13 | 717 | 682 | 3805 | 69 | 0 | 49 | 62 | 0 | 0 | 5412 | 14886 | 174314 | 23 | 12 | 117676 | 12038 | 0 | 0 | 10 | 0 | 14 | 13 | 0 | 0 | 311 | 0 | 185 | 0 | 0 | 744 | 0 | 17 |
| SRR5030119 | 73 | 41 | 0 | 16 | 57 | 898 | 932 | 5259 | 92 | 0 | 149 | 270 | 0 | 0 | 5846 | 16614 | 180471 | 12 | 46 | 107877 | 11669 | 0 | 0 | 63 | 16 | 16 | 54 | 0 | 0 | 1173 | 0 | 230 | 0 | 0 | 1094 | 0 | 32 |
| SRR5030120 | 45 | 12 | 0 | 15 | 11 | 693 | 557 | 3983 | 111 | 0 | 53 | 57 | 0 | 0 | 5418 | 13701 | 173502 | 12 | 49 | 119030 | 11468 | 0 | 0 | 48 | 0 | 41 | 22 | 0 | 0 | 291 | 0 | 99 | 0 | 0 | 633 | 0 | 41 |
| SRR5030121 | 73 | 89 | 13 | 0 | 63 | 1001 | 691 | 5194 | 107 | 26 | 165 | 285 | 0 | 0 | 5289 | 16562 | 178467 | 0 | 36 | 107945 | 11531 | 0 | 0 | 79 | 0 | 17 | 50 | 0 | 0 | 1175 | 0 | 203 | 0 | 0 | 1046 | 0 | 40 |
| SRR5030122 | 16 | 16 | 25 | 14 | 50 | 675 | 450 | 3706 | 118 | 0 | 58 | 49 | 0 | 0 | 5304 | 15025 | 170086 | 12 | 12 | 119604 | 11473 | 0 | 0 | 50 | 14 | 14 | 19 | 0 | 0 | 343 | 0 | 201 | 0 | 0 | 642 | 0 | 32 |
| SRR5030123 | 80 | 61 | 16 | 0 | 71 | 837 | 849 | 5283 | 94 | 23 | 174 | 306 | 0 | 0 | 5625 | 16223 | 174946 | 24 | 59 | 108686 | 10731 | 0 | 0 | 61 | 0 | 13 | 48 | 0 | 0 | 1123 | 0 | 318 | 0 | 0 | 1092 | 0 | 42 |
| SRR5030124 | 73 | 80 | 18 | 36 | 91 | 688 | 615 | 4222 | 121 | 11 | 272 | 297 | 0 | 0 | 5344 | 14882 | 174470 | 24 | 48 | 118445 | 11737 | 0 | 0 | 54 | 0 | 25 | 83 | 0 | 0 | 1686 | 0 | 217 | 0 | 0 | 851 | 0 | 29 |
| SRR5030125 | 66 | 72 | 16 | 16 | 89 | 860 | 872 | 5425 | 131 | 0 | 115 | 298 | 0 | 0 | 5520 | 15505 | 176015 | 0 | 24 | 107570 | 11530 | 0 | 0 | 43 | 13 | 0 | 52 | 0 | 0 | 1177 | 0 | 215 | 0 | 0 | 1112 | 0 | 13 |
| SRR5030126 | 88 | 10 | 0 | 15 | 37 | 212 | 53 | 477 | 53 | 0 | 11 | 37 | 0 | 0 | 3019 | 22672 | 133845 | 0 | 0 | 2648 | 5032 | 0 | 0 | 0 | 0 | 0 | 51 | 0 | 0 | 42 | 0 | 0 | 0 | 0 | 4310 | 0 | 0 |
| SRR5030127 | 117 | 41 | 0 | 0 | 0 | 305 | 174 | 1089 | 44 | 41 | 48 | 21 | 0 | 0 | 2789 | 23043 | 110519 | 0 | 0 | 1917 | 3791 | 0 | 0 | 69 | 0 | 0 | 34 | 0 | 0 | 255 | 0 | 44 | 0 | 0 | 4357 | 0 | 0 |
| SRR5030128 | 70 | 78 | 0 | 0 | 16 | 173 | 173 | 1041 | 0 | 23 | 125 | 0 | 0 | 0 | 3757 | 24371 | 129491 | 0 | 0 | 4278 | 4971 | 0 | 0 | 16 | 0 | 0 | 47 | 0 | 0 | 431 | 0 | 0 | 0 | 0 | 4051 | 0 | 0 |
| SRR5030130 | 90 | 52 | 0 | 15 | 0 | 388 | 55 | 720 | 55 | 22 | 90 | 0 | 0 | 0 | 2381 | 23177 | 132147 | 0 | 0 | 3599 | 5038 | 0 | 0 | 60 | 0 | 0 | 75 | 0 | 0 | 48 | 0 | 0 | 0 | 0 | 3009 | 0 | 0 |
| SRR5030131 | 91 | 112 | 0 | 0 | 35 | 310 | 133 | 1018 | 89 | 63 | 35 | 14 | 0 | 0 | 2656 | 21376 | 111769 | 0 | 0 | 2258 | 3276 | 0 | 0 | 77 | 0 | 0 | 21 | 0 | 0 | 20 | 0 | 0 | 0 | 0 | 5283 | 0 | 14 |
| SRR5030132 | 46 | 69 | 0 | 15 | 0 | 279 | 112 | 279 | 0 | 31 | 123 | 15 | 0 | 0 | 3521 | 22653 | 136110 | 0 | 56 | 3968 | 4639 | 0 | 0 | 54 | 0 | 0 | 69 | 0 | 0 | 345 | 0 | 0 | 0 | 0 | 4235 | 0 | 0 |
| SRR5030133 | 122 | 86 | 0 | 0 | 50 | 267 | 356 | 1069 | 45 | 79 | 36 | 36 | 0 | 0 | 2850 | 20840 | 112000 | 0 | 0 | 2806 | 3830 | 0 | 0 | 65 | 0 | 0 | 29 | 0 | 0 | 23 | 0 | 0 | 0 | 0 | 5487 | 0 | 0 |
| SRR5030134 | 12 | 27 | 0 | 0 | 17 | 266 | 813 | 3904 | 386 | 0 | 39 | 66 | 0 | 0 | 2305 | 23284 | 94546 | 0 | 120 | 5503 | 26248 | 0 | 0 | 0 | 0 | 12 | 39 | 0 | 0 | 222 | 0 | 240 | 0 | 0 | 1241 | 0 | 12 |
| SRR5030135 | 10 | 23 | 0 | 0 | 17 | 589 | 1472 | 6537 | 589 | 0 | 14 | 56 | 0 | 0 | 1944 | 20469 | 79064 | 0 | 29 | 4800 | 21879 | 0 | 0 | 24 | 0 | 0 | 13 | 0 | 0 | 122 | 0 | 177 | 0 | 0 | 2138 | 0 | 14 |
| SRR5030136 | 18 | 28 | 11 | 0 | 24 | 287 | 688 | 3169 | 301 | 0 | 36 | 65 | 0 | 0 | 2209 | 23069 | 97096 | 14 | 43 | 5894 | 24982 | 0 | 0 | 0 | 0 | 23 | 30 | 0 | 0 | 221 | 0 | 272 | 0 | 0 | 1371 | 0 | 11 |
| SRR5030137 | 57 | 81 | 0 | 0 | 11 | 833 | 895 | 6326 | 525 | 0 | 78 | 145 | 0 | 0 | 1790 | 19563 | 83893 | 0 | 62 | 5154 | 19380 | 0 | 0 | 46 | 0 | 0 | 28 | 0 | 0 | 552 | 0 | 185 | 0 | 0 | 1236 | 0 | 0 |
| SRR5030138 | 49 | 72 | 0 | 0 | 25 | 216 | 648 | 3837 | 405 | 0 | 123 | 209 | 0 | 0 | 2337 | 22154 | 94972 | 27 | 176 | 5809 | 24534 | 0 | 0 | 23 | 0 | 17 | 108 | 0 | 0 | 744 | 0 | 216 | 0 | 0 | 1333 | 0 | 0 |
| SRR5030139 | 87 | 107 | 0 | 0 | 31 | 630 | 989 | 6836 | 390 | 0 | 52 | 263 | 0 | 0 | 1649 | 19635 | 85017 | 0 | 60 | 4468 | 20868 | 0 | 0 | 10 | 0 | 14 | 69 | 0 | 0 | 544 | 0 | 180 | 0 | 0 | 1903 | 0 | 14 |
| SRR5030140 | 15 | 27 | 0 | 0 | 19 | 389 | 654 | 3602 | 362 | 0 | 42 | 72 | 0 | 0 | 2364 | 23417 | 94770 | 14 | 70 | 5563 | 25299 | 0 | 0 | 16 | 0 | 0 | 42 | 0 | 0 | 243 | 0 | 153 | 0 | 0 | 1141 | 0 | 0 |
| SRR5030141 | 13 | 19 | 0 | 0 | 21 | 589 | 869 | 4592 | 589 | 0 | 19 | 47 | 0 | 0 | 2389 | 20052 | 81880 | 0 | 31 | 4685 | 20477 | 0 | 0 | 35 | 0 | 11 | 35 | 0 | 0 | 134 | 0 | 186 | 0 | 0 | 2160 | 0 | 11 |
| SRR5030142 | 90 | 14 | 28 | 10 | 42 | 178 | 311 | 2524 | 96 | 0 | 23 | 26 | 0 | 0 | 1115 | 6506 | 115789 | 0 | 20 | 30077 | 914 | 0 | 0 | 17 | 63 | 31 | 79 | 0 | 0 | 170 | 0 | 57 | 0 | 0 | 535 | 0 | 21 |
| SRR5030143 | 150 | 90 | 0 | 60 | 0 | 481 | 481 | 4247 | 27 | 30 | 30 | 210 | 0 | 0 | 1095 | 7611 | 105411 | 0 | 27 | 28178 | 962 | 0 | 0 | 0 | 30 | 180 | 510 | 30 | 0 | 1380 | 0 | 134 | 0 | 0 | 886 | 0 | 120 |
| SRR5030144 | 69 | 13 | 11 | 23 | 50 | 197 | 252 | 2463 | 137 | 15 | 84 | 26 | 0 | 0 | 1102 | 6559 | 114917 | 0 | 24 | 29717 | 838 | 0 | 0 | 23 | 69 | 28 | 70 | 46 | 0 | 160 | 0 | 73 | 0 | 0 | 517 | 0 | 27 |
| SRR5030145 | 64 | 128 | 32 | 0 | 128 | 531 | 531 | 3745 | 140 | 0 | 64 | 255 | 0 | 0 | 1202 | 6576 | 109136 | 0 | 0 | 27807 | 699 | 0 | 0 | 32 | 0 | 255 | 542 | 0 | 0 | 1435 | 0 | 112 | 0 | 0 | 1033 | 0 | 32 |
| SRR5030146 | 57 | 13 | 22 | 43 | 54 | 204 | 248 | 2624 | 126 | 25 | 10 | 24 | 0 | 0 | 1059 | 6818 | 116408 | 0 | 29 | 29870 | 835 | 0 | 0 | 32 | 50 | 25 | 74 | 43 | 0 | 165 | 0 | 79 | 0 | 0 | 559 | 0 | 25 |
| SRR5030147 | 12 | 31 | 0 | 0 | 31 | 604 | 604 | 4090 | 110 | 0 | 93 | 24 | 0 | 0 | 1290 | 5670 | 107418 | 0 | 0 | 28550 | 631 | 0 | 0 | 0 | 93 | 15 | 45 | 62 | 0 | 91 | 0 | 55 | 0 | 0 | 1155 | 0 | 31 |
| SRR5030148 | 108 | 111 | 11 | 30 | 41 | 166 | 252 | 2623 | 101 | 26 | 100 | 286 | 0 | 0 | 1079 | 6672 | 116753 | 0 | 27 | 29932 | 877 | 0 | 0 | 37 | 59 | 245 | 805 | 26 | 0 | 1473 | 0 | 74 | 0 | 0 | 452 | 0 | 26 |
| SRR5030149 | 126 | 31 | 31 | 31 | 31 | 475 | 559 | 4052 | 168 | 31 | 63 | 189 | 0 | 0 | 1481 | 7312 | 110925 | 0 | 28 | 26157 | 1034 | 0 | 0 | 0 | 0 | 251 | 377 | 0 | 0 | 1634 | 0 | 140 | 0 | 0 | 1029 | 0 | 31 |
| SRR5030150 | 48 | 37 | 11 | 0 | 0 | 427 | 346 | 2459 | 37 | 23 | 139 | 450 | 0 | 0 | 18399 | 5479 | 246224 | 0 | 15 | 64510 | 13481 | 0 | 0 | 80 | 21 | 46 | 178 | 0 | 0 | 752 | 0 | 81 | 0 | 0 | 316 | 0 | 91 |
| SRR5030151 | 20 | 99 | 20 | 0 | 39 | 536 | 335 | 3151 | 0 | 20 | 59 | 709 | 0 | 0 | 17567 | 6299 | 249966 | 0 | 0 | 61620 | 14081 | 0 | 0 | 59 | 0 | 20 | 59 | 0 | 0 | 611 | 0 | 268 | 0 | 0 | 409 | 0 | 59 |
| SRR5030152 | 12 | 11 | 0 | 0 | 27 | 500 | 302 | 2582 | 24 | 15 | 27 | 88 | 0 | 0 | 17811 | 6101 | 243222 | 0 | 0 | 66031 | 14188 | 0 | 0 | 13 | 0 | 25 | 18 | 0 | 0 | 149 | 0 | 40 | 0 | 0 | 339 | 0 | 16 |
| SRR5030153 | 41 | 61 | 0 | 20 | 20 | 554 | 485 | 3739 | 69 | 82 | 41 | 492 | 0 | 0 | 15371 | 5905 | 250123 | 0 | 0 | 60098 | 13640 | 0 | 0 | 61 | 20 | 61 | 82 | 0 | 0 | 778 | 0 | 208 | 0 | 0 | 492 | 0 | 82 |
| SRR5030154 | 40 | 11 | 0 | 14 | 19 | 357 | 425 | 2202 | 30 | 21 | 24 | 89 | 0 | 0 | 18694 | 5621 | 244303 | 30 | 15 | 66674 | 13758 | 0 | 0 | 52 | 12 | 38 | 32 | 0 | 0 | 159 | 0 | 121 | 0 | 0 | 337 | 0 | 13 |
| SRR5030155 | 15 | 0 | 0 | 0 | 0 | 340 | 340 | 3329 | 0 | 15 | 15 | 88 | 0 | 0 | 19636 | 5732 | 250955 | 0 | 0 | 61286 | 12026 | 0 | 0 | 18 | 0 | 15 | 41 | 0 | 0 | 110 | 0 | 136 | 0 | 0 | 313 | 0 | 15 |
| SRR5030156 | 14 | 31 | 0 | 0 | 12 | 455 | 370 | 2450 | 54 | 14 | 24 | 87 | 0 | 0 | 17665 | 5808 | 244043 | 15 | 39 | 65697 | 13851 | 0 | 0 | 11 | 12 | 46 | 24 | 12 | 0 | 154 | 0 | 123 | 0 | 0 | 310 | 0 | 12 |
| SRR5030157 | 41 | 62 | 0 | 21 | 41 | 347 | 208 | 2986 | 69 | 0 | 166 | 456 | 0 | 0 | 17293 | 6021 | 247459 | 0 | 0 | 62991 | 13820 | 0 | 0 | 83 | 0 | 0 | 21 | 0 | 0 | 559 | 0 | 278 | 0 | 0 | 638 | 0 | 62 |
| SRR5030158 | 20 | 20 | 37 | 21 | 13 | 160 | 122 | 1013 | 51 | 37 | 15 | 56 | 0 | 0 | 10807 | 9104 | 256835 | 13 | 32 | 113309 | 29869 | 0 | 0 | 19 | 54 | 10 | 57 | 0 | 0 | 147 | 0 | 205 | 0 | 0 | 301 | 0 | 15 |
| SRR5030159 | 16 | 17 | 0 | 23 | 82 | 421 | 181 | 1950 | 36 | 12 | 12 | 50 | 0 | 0 | 11831 | 9648 | 265836 | 24 | 24 | 101809 | 28753 | 0 | 0 | 16 | 47 | 82 | 20 | 18 | 0 | 67 | 0 | 229 | 0 | 0 | 545 | 0 | 11 |
| SRR5030160 | 25 | 29 | 13 | 18 | 13 | 277 | 90 | 886 | 55 | 31 | 20 | 59 | 0 | 0 | 10878 | 8451 | 256125 | 0 | 21 | 114629 | 30265 | 0 | 0 | 12 | 75 | 71 | 54 | 31 | 13 | 137 | 0 | 138 | 0 | 0 | 320 | 0 | 11 |
| SRR5030161 | 104 | 141 | 24 | 24 | 92 | 510 | 249 | 1703 | 12 | 67 | 67 | 435 | 0 | 0 | 11799 | 9595 | 265472 | 25 | 12 | 102633 | 28335 | 0 | 0 | 135 | 12 | 67 | 196 | 12 | 0 | 576 | 0 | 137 | 0 | 0 | 607 | 0 | 178 |
| SRR5030162 | 185 | 193 | 0 | 29 | 88 | 246 | 120 | 936 | 27 | 25 | 172 | 479 | 0 | 0 | 10637 | 8308 | 255767 | 0 | 33 | 114168 | 30106 | 0 | 0 | 160 | 42 | 118 | 538 | 21 | 21 | 1240 | 0 | 173 | 0 | 0 | 336 | 0 | 105 |
| SRR5030163 | 154 | 124 | 0 | 18 | 71 | 425 | 267 | 1834 | 49 | 47 | 53 | 438 | 0 | 0 | 11527 | 9335 | 265936 | 0 | 12 | 101164 | 28434 | 0 | 0 | 184 | 36 | 59 | 225 | 0 | 12 | 640 | 0 | 243 | 0 | 0 | 539 | 0 | 124 |
| SRR5030164 | 176 | 219 | 0 | 30 | 129 | 191 | 157 | 844 | 20 | 0 | 159 | 503 | 0 | 0 | 10624 | 8355 | 256239 | 20 | 34 | 114497 | 29919 | 0 | 0 | 116 | 65 | 103 | 477 | 13 | 0 | 1148 | 0 | 143 | 0 | 0 | 335 | 0 | 108 |
| SRR5030165 | 16 | 12 | 18 | 12 | 73 | 479 | 252 | 1839 | 63 | 31 | 73 | 44 | 0 | 0 | 11339 | 9721 | 264601 | 25 | 38 | 102389 | 28284 | 0 | 0 | 12 | 67 | 43 | 27 | 18 | 18 | 65 | 0 | 252 | 0 | 0 | 494 | 0 | 13 |
| SRR5030166 | 21 | 29 | 39 | 65 | 17 | 116 | 102 | 480 | 146 | 13 | 156 | 13 | 0 | 0 | 175 | 31545 | 141191 | 0 | 29 | 3552 | 8633 | 0 | 0 | 52 | 13 | 0 | 16 | 13 | 0 | 69 | 0 | 291 | 0 | 0 | 2063 | 0 | 0 |
| SRR5030167 | 15 | 16 | 0 | 0 | 248 | 233 | 220 | 882 | 104 | 0 | 76 | 0 | 0 | 0 | 311 | 27127 | 115864 | 0 | 26 | 3228 | 8673 | 0 | 0 | 67 | 0 | 38 | 124 | 0 | 0 | 37 | 0 | 233 | 0 | 0 | 3435 | 0 | 0 |
| SRR5030168 | 566 | 637 | 14 | 14 | 325 | 63 | 111 | 412 | 95 | 14 | 311 | 28 | 0 | 0 | 222 | 29782 | 138040 | 16 | 0 | 4132 | 8312 | 0 | 0 | 28 | 14 | 71 | 269 | 28 | 0 | 1457 | 0 | 238 | 0 | 0 | 2266 | 0 | 0 |
| SRR5030169 | 381 | 311 | 20 | 30 | 201 | 207 | 248 | 828 | 193 | 20 | 110 | 20 | 10 | 0 | 166 | 28246 | 113896 | 0 | 14 | 3229 | 8513 | 0 | 0 | 40 | 0 | 30 | 150 | 0 | 0 | 963 | 0 | 442 | 0 | 0 | 3183 | 0 | 0 |
| SRR5030170 | 26 | 33 | 27 | 27 | 18 | 153 | 107 | 520 | 61 | 40 | 10 | 13 | 0 | 0 | 321 | 33420 | 135700 | 31 | 31 | 4116 | 9624 | 0 | 0 | 67 | 13 | 13 | 19 | 67 | 0 | 81 | 0 | 306 | 0 | 0 | 2423 | 0 | 0 |
| SRR5030171 | 429 | 399 | 0 | 0 | 302 | 308 | 228 | 872 | 134 | 0 | 117 | 29 | 19 | 0 | 349 | 27977 | 117417 | 0 | 27 | 3205 | 8275 | 0 | 0 | 58 | 19 | 29 | 224 | 0 | 0 | 974 | 0 | 402 | 0 | 0 | 3096 | 0 | 0 |
| SRR5030172 | 24 | 29 | 0 | 83 | 16 | 93 | 155 | 511 | 77 | 0 | 16 | 28 | 0 | 0 | 279 | 32656 | 138868 | 0 | 46 | 3935 | 9946 | 0 | 0 | 56 | 14 | 28 | 16 | 28 | 0 | 65 | 0 | 279 | 0 | 0 | 2844 | 0 | 0 |
| SRR5030173 | 296 | 414 | 20 | 30 | 257 | 218 | 327 | 532 | 123 | 20 | 109 | 0 | 0 | 0 | 218 | 27394 | 114875 | 14 | 68 | 3561 | 7503 | 0 | 0 | 109 | 0 | 30 | 187 | 20 | 0 | 888 | 0 | 423 | 0 | 0 | 3517 | 0 | 0 |
| SRR5030174 | 47 | 45 | 0 | 0 | 23 | 261 | 209 | 2890 | 67 | 11 | 355 | 260 | 0 | 0 | 10987 | 9627 | 237069 | 15 | 15 | 31062 | 13460 | 0 | 0 | 17 | 0 | 18 | 56 | 0 | 0 | 1925 | 0 | 156 | 0 | 0 | 279 | 0 | 23 |
| SRR5030175 | 18 | 13 | 0 | 0 | 17 | 825 | 668 | 4927 | 36 | 0 | 51 | 82 | 0 | 0 | 10620 | 9752 | 230527 | 0 | 12 | 25827 | 10984 | 0 | 0 | 12 | 0 | 14 | 11 | 0 | 0 | 344 | 0 | 255 | 0 | 0 | 648 | 0 | 31 |
| SRR5030176 | 13 | 20 | 0 | 0 | 27 | 255 | 262 | 2657 | 80 | 13 | 117 | 78 | 0 | 0 | 10802 | 9726 | 235268 | 16 | 16 | 31355 | 13196 | 0 | 0 | 16 | 0 | 29 | 26 | 0 | 0 | 651 | 0 | 88 | 0 | 0 | 250 | 0 | 21 |
| SRR5030177 | 37 | 32 | 0 | 0 | 15 | 773 | 595 | 4853 | 63 | 20 | 155 | 255 | 0 | 0 | 9401 | 10032 | 230117 | 38 | 25 | 24427 | 11732 | 0 | 0 | 27 | 0 | 29 | 22 | 0 | 0 | 1376 | 0 | 152 | 0 | 0 | 536 | 0 | 37 |
| SRR5030178 | 16 | 17 | 0 | 0 | 23 | 339 | 231 | 2664 | 77 | 0 | 118 | 86 | 0 | 0 | 10608 | 9796 | 237339 | 15 | 15 | 32087 | 12841 | 0 | 0 | 15 | 18 | 23 | 29 | 0 | 0 | 644 | 0 | 131 | 0 | 0 | 385 | 0 | 12 |
| SRR5030179 | 29 | 14 | 0 | 0 | 14 | 719 | 582 | 4561 | 50 | 0 | 43 | 70 | 0 | 0 | 10175 | 10608 | 224432 | 0 | 25 | 25369 | 11167 | 0 | 0 | 34 | 0 | 0 | 22 | 0 | 0 | 389 | 0 | 161 | 0 | 0 | 672 | 0 | 10 |
| SRR5030180 | 13 | 19 | 0 | 0 | 16 | 302 | 264 | 2916 | 78 | 17 | 120 | 94 | 0 | 0 | 10554 | 9339 | 238511 | 23 | 31 | 31142 | 13408 | 0 | 0 | 19 | 11 | 25 | 16 | 0 | 0 | 636 | 0 | 116 | 0 | 0 | 361 | 0 | 11 |
| SRR5030181 | 34 | 16 | 0 | 0 | 15 | 716 | 552 | 4884 | 50 | 0 | 52 | 72 | 0 | 0 | 10333 | 9774 | 229224 | 0 | 0 | 26041 | 11438 | 0 | 0 | 34 | 0 | 15 | 15 | 0 | 0 | 349 | 0 | 213 | 0 | 0 | 671 | 0 | 10 |
| SRR5030182 | 384 | 192 | 27 | 82 | 275 | 0 | 0 | 124 | 0 | 0 | 96 | 41 | 0 | 0 | 185 | 29884 | 39763 | 0 | 41 | 3152 | 2823 | 0 | 0 | 69 | 0 | 110 | 645 | 41 | 0 | 865 | 0 | 0 | 0 | 0 | 6190 | 0 | 14 |
| SRR5030183 | 161 | 438 | 12 | 46 | 219 | 0 | 0 | 137 | 20 | 12 | 23 | 12 | 12 | 0 | 98 | 24214 | 28567 | 0 | 20 | 2726 | 2608 | 0 | 0 | 150 | 23 | 12 | 230 | 0 | 0 | 438 | 0 | 0 | 0 | 0 | 8049 | 0 | 12 |
| SRR5030184 | 298 | 313 | 15 | 134 | 134 | 0 | 22 | 174 | 22 | 0 | 89 | 45 | 0 | 0 | 109 | 32210 | 40263 | 0 | 22 | 3439 | 2982 | 0 | 0 | 30 | 15 | 89 | 671 | 30 | 0 | 775 | 0 | 0 | 0 | 0 | 5292 | 0 | 0 |
| SRR5030185 | 275 | 275 | 0 | 12 | 251 | 21 | 0 | 21 | 42 | 12 | 60 | 36 | 0 | 0 | 127 | 24704 | 26808 | 0 | 0 | 3349 | 2904 | 0 | 0 | 48 | 0 | 72 | 323 | 0 | 0 | 562 | 0 | 21 | 0 | 0 | 8156 | 0 | 48 |
| SRR5030186 | 255 | 425 | 28 | 99 | 227 | 21 | 21 | 82 | 21 | 0 | 113 | 14 | 0 | 0 | 186 | 31312 | 38123 | 0 | 0 | 3773 | 2907 | 0 | 0 | 28 | 14 | 57 | 765 | 57 | 0 | 850 | 0 | 0 | 0 | 0 | 5487 | 0 | 0 |
| SRR5030187 | 186 | 338 | 0 | 23 | 163 | 60 | 20 | 200 | 60 | 0 | 70 | 23 | 0 | 0 | 260 | 25251 | 28891 | 0 | 0 | 2956 | 2596 | 0 | 0 | 82 | 0 | 12 | 245 | 12 | 0 | 478 | 0 | 20 | 0 | 0 | 8144 | 0 | 12 |
| SRR5030188 | 13 | 18 | 44 | 58 | 189 | 0 | 43 | 43 | 0 | 0 | 102 | 15 | 15 | 0 | 196 | 30251 | 38422 | 0 | 22 | 3997 | 3389 | 0 | 0 | 44 | 0 | 58 | 31 | 15 | 0 | 26 | 0 | 0 | 0 | 0 | 6024 | 0 | 0 |
| SRR5030189 | 179 | 10 | 12 | 12 | 227 | 42 | 21 | 84 | 21 | 0 | 60 | 24 | 0 | 0 | 42 | 25854 | 28159 | 0 | 0 | 3253 | 2456 | 0 | 0 | 84 | 0 | 24 | 263 | 12 | 0 | 11 | 0 | 0 | 0 | 0 | 7723 | 0 | 0 |
| SRR5030190 | 56 | 61 | 0 | 0 | 35 | 348 | 451 | 2162 | 103 | 0 | 23 | 26 | 0 | 0 | 3272 | 2346 | 120652 | 0 | 113 | 38378 | 7860 | 0 | 0 | 80 | 26 | 66 | 197 | 0 | 0 | 133 | 0 | 0 | 0 | 0 | 181 | 0 | 0 |
| SRR5030191 | 16 | 18 | 0 | 0 | 27 | 587 | 294 | 2791 | 294 | 0 | 18 | 27 | 0 | 0 | 3158 | 2771 | 125046 | 37 | 0 | 34257 | 7674 | 0 | 0 | 44 | 18 | 0 | 29 | 0 | 0 | 24 | 0 | 37 | 0 | 0 | 302 | 0 | 0 |
| SRR5030192 | 40 | 35 | 0 | 0 | 42 | 364 | 333 | 1970 | 141 | 0 | 32 | 12 | 0 | 0 | 3223 | 2525 | 119204 | 0 | 61 | 39336 | 7942 | 0 | 0 | 50 | 12 | 42 | 233 | 0 | 0 | 166 | 0 | 20 | 0 | 0 | 230 | 0 | 0 |
| SRR5030193 | 55 | 0 | 0 | 0 | 18 | 451 | 300 | 3230 | 75 | 0 | 28 | 37 | 0 | 0 | 4169 | 2923 | 126646 | 0 | 0 | 36169 | 7286 | 0 | 0 | 55 | 18 | 37 | 211 | 0 | 0 | 174 | 0 | 0 | 0 | 0 | 430 | 0 | 0 |
| SRR5030194 | 60 | 50 | 0 | 0 | 41 | 394 | 385 | 2308 | 135 | 0 | 31 | 38 | 0 | 0 | 3626 | 2682 | 118778 | 0 | 87 | 39765 | 8078 | 0 | 0 | 79 | 12 | 41 | 217 | 0 | 0 | 148 | 0 | 0 | 0 | 0 | 196 | 0 | 12 |
| SRR5030196 | 12 | 39 | 0 | 0 | 32 | 355 | 306 | 2070 | 138 | 0 | 32 | 24 | 0 | 0 | 3499 | 2308 | 118715 | 0 | 99 | 38074 | 7402 | 0 | 0 | 15 | 34 | 13 | 47 | 0 | 0 | 35 | 0 | 20 | 0 | 0 | 294 | 0 | 0 |
| SRR5030197 | 101 | 28 | 0 | 0 | 28 | 675 | 337 | 2249 | 375 | 0 | 18 | 37 | 0 | 0 | 3299 | 2876 | 125461 | 0 | 150 | 34491 | 7385 | 0 | 0 | 73 | 28 | 28 | 119 | 0 | 0 | 165 | 0 | 37 | 0 | 0 | 441 | 0 | 0 |
| SRR5030198 | 123 | 182 | 23 | 0 | 105 | 415 | 749 | 2971 | 268 | 0 | 96 | 87 | 0 | 0 | 9540 | 15690 | 173516 | 0 | 161 | 15789 | 26761 | 0 | 0 | 123 | 0 | 0 | 0 | 0 | 0 | 455 | 0 | 54 | 0 | 0 | 793 | 0 | 14 |
| SRR5030199 | 153 | 167 | 15 | 0 | 109 | 418 | 325 | 3597 | 325 | 0 | 58 | 51 | 0 | 0 | 10674 | 16787 | 178448 | 23 | 116 | 14595 | 25200 | 0 | 0 | 116 | 0 | 0 | 0 | 0 | 0 | 320 | 0 | 46 | 0 | 0 | 1272 | 0 | 15 |
| SRR5030200 | 25 | 29 | 39 | 0 | 78 | 343 | 644 | 2818 | 300 | 0 | 15 | 49 | 0 | 0 | 10172 | 16050 | 170705 | 0 | 100 | 15294 | 26953 | 0 | 0 | 13 | 0 | 0 | 0 | 0 | 0 | 42 | 0 | 129 | 0 | 0 | 806 | 0 | 0 |
| SRR5030201 | 17 | 13 | 15 | 0 | 61 | 411 | 798 | 3604 | 363 | 0 | 61 | 68 | 0 | 0 | 8658 | 16694 | 174105 | 48 | 73 | 13858 | 23846 | 0 | 0 | 83 | 0 | 0 | 0 | 0 | 0 | 43 | 0 | 242 | 0 | 0 | 1197 | 0 | 0 |
| SRR5030202 | 21 | 24 | 42 | 0 | 75 | 578 | 757 | 2751 | 289 | 0 | 17 | 0 | 0 | 0 | 9629 | 14935 | 172408 | 0 | 179 | 16425 | 26756 | 0 | 0 | 10 | 0 | 0 | 0 | 0 | 0 | 54 | 0 | 138 | 0 | 0 | 831 | 0 | 0 |
| SRR5030203 | 18 | 17 | 0 | 0 | 16 | 678 | 968 | 3438 | 266 | 15 | 52 | 52 | 0 | 0 | 9006 | 17404 | 173469 | 0 | 97 | 13291 | 26460 | 0 | 0 | 22 | 0 | 0 | 0 | 0 | 0 | 33 | 0 | 121 | 0 | 0 | 771 | 0 | 0 |
| SRR5030204 | 25 | 21 | 24 | 19 | 62 | 463 | 631 | 2735 | 182 | 0 | 15 | 57 | 0 | 0 | 9342 | 15547 | 172803 | 0 | 126 | 16791 | 27648 | 0 | 0 | 76 | 0 | 0 | 0 | 0 | 0 | 58 | 0 | 112 | 0 | 0 | 670 | 0 | 0 |
| SRR5030205 | 152 | 167 | 15 | 0 | 83 | 343 | 1153 | 3654 | 270 | 23 | 0 | 23 | 0 | 0 | 9171 | 15743 | 179529 | 25 | 74 | 14689 | 25283 | 0 | 0 | 144 | 0 | 0 | 0 | 0 | 0 | 242 | 0 | 123 | 0 | 0 | 924 | 0 | 0 |
| SRR5030206 | 81 | 100 | 0 | 0 | 31 | 124 | 180 | 1327 | 138 | 19 | 62 | 12 | 0 | 0 | 2834 | 11753 | 165614 | 28 | 55 | 19133 | 7092 | 0 | 0 | 156 | 25 | 150 | 505 | 12 | 0 | 181 | 0 | 41 | 0 | 0 | 1593 | 0 | 0 |
| SRR5030207 | 79 | 40 | 0 | 0 | 0 | 191 | 383 | 1531 | 287 | 0 | 119 | 40 | 0 | 0 | 2583 | 12753 | 166867 | 96 | 96 | 17603 | 6218 | 0 | 0 | 79 | 0 | 119 | 18 | 0 | 0 | 40 | 0 | 96 | 0 | 0 | 2286 | 0 | 0 |
| SRR5030208 | 86 | 12 | 13 | 13 | 40 | 136 | 136 | 1402 | 75 | 0 | 60 | 0 | 0 | 0 | 3060 | 12393 | 170881 | 15 | 30 | 20545 | 6753 | 0 | 0 | 146 | 13 | 80 | 38 | 13 | 0 | 11 | 0 | 30 | 0 | 0 | 1489 | 0 | 0 |
| SRR5030209 | 122 | 122 | 0 | 41 | 0 | 203 | 305 | 2234 | 406 | 0 | 41 | 0 | 0 | 0 | 3047 | 11664 | 162179 | 0 | 0 | 19600 | 8734 | 0 | 0 | 41 | 0 | 0 | 26 | 0 | 0 | 13 | 0 | 0 | 0 | 0 | 1613 | 0 | 41 |
| SRR5030210 | 51 | 115 | 13 | 0 | 38 | 255 | 255 | 1627 | 142 | 0 | 26 | 19 | 0 | 0 | 2915 | 11935 | 166878 | 14 | 28 | 21340 | 6779 | 0 | 0 | 141 | 13 | 128 | 423 | 0 | 0 | 231 | 0 | 28 | 0 | 0 | 1887 | 0 | 0 |
| SRR5030211 | 41 | 123 | 0 | 0 | 82 | 289 | 289 | 1833 | 289 | 0 | 41 | 0 | 0 | 0 | 2026 | 11899 | 164036 | 0 | 0 | 18329 | 6271 | 0 | 0 | 15 | 41 | 123 | 123 | 0 | 0 | 123 | 0 | 0 | 0 | 0 | 2186 | 0 | 0 |

### Supp Table S8.3.Columns_BX_to_DH

| | | | | | | | | | | | | | | | | | | | | | | | | | | | | | | | | | | | | | |
|---|---|---|---|---|---|---|---|---|---|---|---|---|---|---|---|---|---|---|---|---|---|---|---|---|---|---|---|---|---|---|---|---|---|---|---|---|---|
| SRR5030212 | 52 | 97 | 19 | 19 | 71 | 143 | 329 | 1446 | 158 | 13 | 65 | 0 | 0 | 0 | 2492 | 11125 | 167339 | 0 | 72 | 21423 | 6702 | 0 | 0 | 148 | 19 | 84 | 32 | 0 | 0 | 16 | 0 | 29 | 0 | 0 | 1665 | 0 | 0 |
| SRR5030213 | 42 | 291 | 0 | 42 | 83 | 401 | 501 | 2003 | 100 | 0 | 42 | 0 | 0 | 0 | 3104 | 12174 | 150394 | 0 | 0 | 20827 | 7109 | 0 | 0 | 83 | 0 | 83 | 291 | 83 | 0 | 83 | 0 | 0 | 0 | 0 | 2705 | 0 | 0 |
| SRR5030214 | 25 | 25 | 0 | 14 | 16 | 408 | 634 | 1823 | 419 | 11 | 49 | 0 | 0 | 0 | 3148 | 17438 | 159689 | 0 | 91 | 56655 | 7927 | 0 | 0 | 43 | 0 | 32 | 27 | 11 | 0 | 330 | 0 | 23 | 0 | 0 | 4335 | 0 | 0 |
| SRR5030216 | 31 | 54 | 0 | 27 | 12 | 429 | 846 | 1765 | 221 | 12 | 47 | 0 | 0 | 0 | 3174 | 17133 | 160160 | 0 | 74 | 57225 | 7879 | 0 | 0 | 50 | 0 | 31 | 26 | 0 | 0 | 313 | 0 | 0 | 0 | 0 | 4664 | 0 | 0 |
| SRR5030217 | 0 | 37 | 0 | 0 | 20 | 472 | 472 | 2005 | 236 | 0 | 28 | 37 | 0 | 0 | 2948 | 14482 | 157054 | 0 | 0 | 54009 | 7665 | 0 | 0 | 0 | 0 | 74 | 16 | 0 | 0 | 320 | 0 | 118 | 0 | 0 | 4120 | 0 | 37 |
| SRR5030218 | 30 | 52 | 0 | 26 | 13 | 393 | 763 | 2217 | 322 | 0 | 45 | 0 | 0 | 0 | 3278 | 17189 | 159258 | 12 | 131 | 59214 | 7473 | 0 | 0 | 41 | 18 | 55 | 28 | 15 | 0 | 318 | 0 | 0 | 0 | 0 | 4339 | 0 | 0 |
| SRR5030220 | 34 | 56 | 0 | 19 | 101 | 413 | 584 | 1726 | 316 | 11 | 374 | 11 | 0 | 0 | 3246 | 17229 | 159794 | 0 | 97 | 59758 | 7575 | 0 | 0 | 49 | 0 | 41 | 217 | 0 | 0 | 2933 | 0 | 12 | 0 | 0 | 4435 | 0 | 0 |
| SRR5030221 | 74 | 74 | 0 | 0 | 74 | 353 | 118 | 1646 | 353 | 0 | 24 | 37 | 0 | 0 | 2940 | 18875 | 160005 | 0 | 118 | 57738 | 6820 | 0 | 0 | 37 | 37 | 37 | 24 | 0 | 0 | 296 | 0 | 0 | 0 | 0 | 5058 | 0 | 0 |
| SRR5030222 | 147 | 377 | 49 | 0 | 147 | 0 | 21 | 107 | 0 | 16 | 180 | 49 | 0 | 0 | 64 | 20751 | 37473 | 0 | 43 | 42955 | 6878 | 0 | 0 | 49 | 16 | 16 | 180 | 0 | 0 | 1278 | 0 | 21 | 0 | 0 | 6689 | 0 | 0 |
| SRR5030223 | 119 | 17 | 0 | 0 | 93 | 15 | 136 | 173 | 30 | 0 | 59 | 47 | 0 | 0 | 106 | 18070 | 31153 | 23 | 60 | 35821 | 6348 | 0 | 0 | 68 | 0 | 13 | 47 | 0 | 0 | 59 | 0 | 0 | 0 | 0 | 8656 | 0 | 0 |
| SRR5030224 | 105 | 27 | 18 | 88 | 35 | 23 | 114 | 69 | 69 | 0 | 20 | 35 | 0 | 0 | 160 | 17907 | 36640 | 23 | 91 | 43291 | 6011 | 0 | 0 | 105 | 0 | 35 | 17 | 18 | 0 | 119 | 0 | 0 | 0 | 0 | 6024 | 0 | 0 |
| SRR5030225 | 180 | 245 | 0 | 0 | 70 | 0 | 108 | 115 | 0 | 0 | 70 | 39 | 0 | 0 | 77 | 16919 | 29443 | 0 | 38 | 35791 | 6363 | 0 | 0 | 57 | 0 | 0 | 61 | 0 | 0 | 710 | 0 | 0 | 0 | 0 | 8191 | 0 | 0 |
| SRR5030226 | 217 | 233 | 33 | 33 | 100 | 0 | 67 | 67 | 22 | 0 | 183 | 50 | 0 | 0 | 67 | 19379 | 37025 | 0 | 67 | 43506 | 6846 | 0 | 0 | 17 | 17 | 33 | 150 | 0 | 0 | 1600 | 0 | 22 | 0 | 0 | 5751 | 0 | 0 |
| SRR5030227 | 13 | 17 | 0 | 0 | 65 | 15 | 77 | 108 | 0 | 0 | 56 | 26 | 0 | 0 | 123 | 16268 | 29415 | 23 | 23 | 35440 | 6380 | 0 | 0 | 65 | 0 | 0 | 30 | 0 | 0 | 57 | 0 | 0 | 0 | 0 | 7536 | 0 | 0 |
| SRR5030229 | 167 | 215 | 18 | 0 | 48 | 0 | 94 | 164 | 31 | 0 | 61 | 31 | 0 | 0 | 149 | 17537 | 30083 | 23 | 31 | 36315 | 6201 | 0 | 0 | 53 | 0 | 0 | 66 | 0 | 0 | 601 | 0 | 0 | 0 | 0 | 8707 | 0 | 0 |
| SRR5030230 | 305 | 508 | 68 | 68 | 102 | 0 | 250 | 1598 | 100 | 0 | 136 | 68 | 0 | 0 | 1898 | 12454 | 179592 | 0 | 0 | 21580 | 3097 | 0 | 0 | 34 | 0 | 0 | 508 | 68 | 0 | 542 | 0 | 0 | 0 | 0 | 1023 | 0 | 34 |
| SRR5030231 | 11 | 16 | 13 | 34 | 55 | 314 | 381 | 3369 | 105 | 0 | 0 | 34 | 0 | 0 | 1987 | 11348 | 150881 | 15 | 22 | 14507 | 2062 | 0 | 0 | 110 | 0 | 51 | 93 | 17 | 0 | 12 | 0 | 22 | 0 | 0 | 2228 | 0 | 13 |
| SRR5030232 | 23 | 26 | 36 | 145 | 16 | 53 | 263 | 1526 | 158 | 0 | 145 | 36 | 0 | 0 | 1526 | 12969 | 177203 | 0 | 53 | 22371 | 2053 | 0 | 0 | 36 | 145 | 73 | 19 | 0 | 0 | 21 | 0 | 0 | 0 | 0 | 1303 | 0 | 0 |
| SRR5030233 | 296 | 375 | 0 | 22 | 48 | 243 | 380 | 3327 | 175 | 0 | 44 | 31 | 0 | 0 | 1405 | 11527 | 149818 | 0 | 23 | 14605 | 1921 | 0 | 0 | 118 | 0 | 22 | 126 | 0 | 0 | 244 | 0 | 0 | 0 | 0 | 2378 | 0 | 0 |
| SRR5030234 | 380 | 449 | 0 | 138 | 35 | 155 | 309 | 1186 | 155 | 0 | 104 | 0 | 0 | 0 | 1856 | 12345 | 177005 | 0 | 0 | 21394 | 2938 | 0 | 0 | 69 | 0 | 69 | 414 | 0 | 0 | 587 | 0 | 0 | 0 | 0 | 983 | 0 | 0 |
| SRR5030235 | 13 | 17 | 13 | 30 | 90 | 366 | 381 | 3192 | 152 | 0 | 22 | 0 | 17 | 0 | 1668 | 11753 | 149208 | 15 | 0 | 14925 | 1981 | 0 | 0 | 120 | 13 | 39 | 142 | 0 | 0 | 13 | 0 | 15 | 0 | 0 | 2678 | 0 | 22 |
| SRR5030236 | 11 | 32 | 70 | 141 | 70 | 53 | 105 | 1787 | 0 | 35 | 70 | 35 | 0 | 0 | 1892 | 12007 | 176349 | 0 | 0 | 20603 | 2681 | 0 | 0 | 35 | 0 | 11 | 41 | 35 | 0 | 37 | 0 | 0 | 0 | 0 | 813 | 0 | 35 |
| SRR5030237 | 249 | 393 | 0 | 22 | 65 | 237 | 405 | 3249 | 168 | 0 | 35 | 17 | 17 | 0 | 1521 | 11876 | 149961 | 31 | 0 | 14469 | 2010 | 0 | 0 | 83 | 0 | 35 | 131 | 0 | 0 | 288 | 0 | 0 | 0 | 0 | 2587 | 0 | 13 |
| SRR5030238 | 18 | 35 | 29 | 43 | 22 | 52 | 364 | 1352 | 208 | 86 | 108 | 43 | 0 | 0 | 559 | 8617 | 89269 | 39 | 26 | 7917 | 1755 | 0 | 0 | 0 | 36 | 65 | 13 | 0 | 0 | 58 | 0 | 13 | 0 | 0 | 4653 | 0 | 0 |
| SRR5030239 | 14 | 31 | 50 | 25 | 25 | 70 | 467 | 2216 | 140 | 62 | 74 | 25 | 0 | 0 | 723 | 7809 | 75434 | 0 | 23 | 6462 | 1703 | 0 | 0 | 25 | 0 | 87 | 149 | 0 | 0 | 49 | 0 | 23 | 0 | 0 | 5998 | 0 | 0 |
| SRR5030240 | 223 | 339 | 46 | 54 | 39 | 70 | 322 | 1414 | 98 | 62 | 131 | 15 | 0 | 0 | 574 | 7958 | 88384 | 42 | 42 | 7796 | 1554 | 0 | 0 | 0 | 0 | 77 | 123 | 0 | 0 | 662 | 0 | 14 | 0 | 0 | 4562 | 0 | 0 |
| SRR5030241 | 219 | 348 | 39 | 64 | 52 | 123 | 515 | 1448 | 98 | 77 | 64 | 52 | 0 | 0 | 736 | 8040 | 78427 | 0 | 49 | 7263 | 1816 | 0 | 0 | 13 | 39 | 77 | 90 | 13 | 0 | 515 | 0 | 49 | 0 | 0 | 4522 | 0 | 0 |
| SRR5030242 | 198 | 461 | 0 | 22 | 51 | 68 | 312 | 1315 | 95 | 44 | 212 | 73 | 0 | 0 | 772 | 7946 | 89629 | 0 | 54 | 8754 | 1531 | 0 | 0 | 0 | 15 | 88 | 205 | 22 | 0 | 718 | 0 | 0 | 0 | 0 | 4288 | 0 | 0 |
| SRR5030243 | 16 | 26 | 0 | 25 | 63 | 120 | 313 | 1539 | 72 | 51 | 88 | 13 | 0 | 0 | 553 | 6906 | 78008 | 24 | 24 | 7552 | 1467 | 0 | 0 | 13 | 13 | 0 | 126 | 0 | 0 | 34 | 0 | 0 | 0 | 0 | 5190 | 0 | 0 |
| SRR5030244 | 308 | 390 | 53 | 45 | 45 | 41 | 328 | 1506 | 192 | 83 | 135 | 30 | 0 | 0 | 534 | 8463 | 86967 | 27 | 27 | 7569 | 1779 | 0 | 0 | 23 | 30 | 135 | 165 | 0 | 0 | 878 | 0 | 14 | 0 | 0 | 3847 | 0 | 0 |
| SRR5030245 | 335 | 464 | 0 | 26 | 26 | 49 | 461 | 1893 | 170 | 116 | 90 | 64 | 0 | 0 | 510 | 7739 | 77473 | 24 | 0 | 6528 | 1602 | 0 | 0 | 0 | 0 | 64 | 77 | 0 | 0 | 567 | 0 | 0 | 0 | 0 | 6024 | 0 | 0 |
| SRR5030246 | 64 | 107 | 32 | 43 | 0 | 234 | 100 | 1352 | 17 | 0 | 289 | 21 | 0 | 0 | 16292 | 8925 | 194776 | 0 | 33 | 44719 | 7295 | 0 | 0 | 182 | 32 | 128 | 567 | 11 | 0 | 2483 | 0 | 0 | 0 | 0 | 871 | 0 | 0 |
| SRR5030247 | 97 | 110 | 0 | 0 | 21 | 624 | 450 | 3061 | 102 | 14 | 62 | 21 | 0 | 0 | 14973 | 9638 | 194551 | 0 | 0 | 35531 | 6079 | 0 | 0 | 13 | 21 | 48 | 152 | 14 | 0 | 55 | 0 | 29 | 0 | 0 | 1621 | 0 | 35 |
| SRR5030248 | 114 | 114 | 34 | 57 | 57 | 106 | 213 | 1595 | 18 | 0 | 14 | 68 | 0 | 0 | 15276 | 9017 | 191646 | 0 | 106 | 48168 | 6628 | 0 | 0 | 68 | 46 | 91 | 54 | 23 | 0 | 177 | 0 | 0 | 0 | 0 | 685 | 0 | 23 |
| SRR5030249 | 72 | 129 | 22 | 0 | 29 | 442 | 457 | 2849 | 152 | 22 | 43 | 29 | 14 | 0 | 15161 | 9969 | 193453 | 0 | 76 | 36616 | 5760 | 0 | 0 | 180 | 22 | 43 | 151 | 14 | 0 | 978 | 0 | 0 | 0 | 0 | 1603 | 0 | 22 |
| SRR5030250 | 151 | 76 | 11 | 32 | 32 | 135 | 118 | 1699 | 17 | 32 | 195 | 11 | 0 | 0 | 15312 | 9323 | 195287 | 0 | 17 | 47247 | 6882 | 0 | 0 | 205 | 43 | 108 | 606 | 32 | 0 | 2347 | 0 | 0 | 0 | 0 | 812 | 0 | 22 |
| SRR5030251 | 56 | 140 | 0 | 14 | 21 | 431 | 327 | 3964 | 104 | 0 | 70 | 21 | 0 | 0 | 15469 | 9737 | 192849 | 0 | 59 | 37159 | 6636 | 0 | 0 | 168 | 21 | 70 | 210 | 0 | 0 | 1080 | 0 | 0 | 0 | 0 | 1636 | 0 | 0 |
| SRR5030252 | 111 | 13 | 22 | 22 | 22 | 87 | 174 | 1882 | 87 | 11 | 19 | 33 | 0 | 0 | 16313 | 8647 | 192079 | 0 | 35 | 46274 | 6902 | 0 | 0 | 111 | 45 | 89 | 42 | 33 | 0 | 154 | 0 | 0 | 0 | 0 | 817 | 0 | 0 |
| SRR5030253 | 86 | 86 | 0 | 14 | 50 | 519 | 382 | 3434 | 153 | 57 | 79 | 36 | 0 | 0 | 16194 | 9676 | 194486 | 0 | 76 | 35714 | 5296 | 0 | 0 | 179 | 22 | 57 | 172 | 14 | 0 | 1046 | 0 | 15 | 0 | 0 | 1638 | 0 | 36 |
| SRR5030254 | 0 | 15 | 0 | 0 | 11 | 56 | 70 | 253 | 14 | 32 | 0 | 12 | 0 | 0 | 84 | 1590 | 5727 | 14 | 0 | 561 | 281 | 0 | 43 | 28 | 0 | 0 | 0 | 0 | 0 | 129 | 0 | 14 | 0 | 0 | 4747 | 0 | 0 |
| SRR5030255 | 0 | 0 | 0 | 0 | 10 | 100 | 115 | 258 | 14 | 32 | 0 | 10 | 0 | 0 | 72 | 1867 | 5660 | 14 | 0 | 645 | 287 | 0 | 31 | 16 | 0 | 0 | 0 | 0 | 0 | 132 | 0 | 29 | 0 | 0 | 5187 | 0 | 0 |
| SRR5030256 | 0 | 0 | 0 | 0 | 10 | 139 | 139 | 139 | 42 | 26 | 11 | 10 | 0 | 0 | 28 | 1919 | 5464 | 0 | 28 | 669 | 167 | 0 | 29 | 20 | 0 | 0 | 0 | 0 | 0 | 152 | 0 | 14 | 0 | 0 | 4618 | 0 | 0 |
| SRR5030257 | 0 | 10 | 0 | 0 | 0 | 213 | 85 | 57 | 0 | 54 | 0 | 13 | 0 | 0 | 85 | 1950 | 5663 | 28 | 0 | 483 | 156 | 0 | 34 | 31 | 0 | 0 | 0 | 0 | 0 | 152 | 0 | 0 | 0 | 0 | 4954 | 0 | 0 |
| SRR5030258 | 14 | 10 | 0 | 0 | 10 | 345 | 230 | 870 | 82 | 24 | 0 | 0 | 0 | 0 | 33 | 19574 | 14192 | 0 | 16 | 3775 | 722 | 0 | 0 | 48 | 45 | 10 | 34 | 0 | 0 | 110 | 0 | 0 | 0 | 0 | 5009 | 21 | 0 |
| SRR5030259 | 0 | 22 | 0 | 0 | 29 | 362 | 224 | 982 | 155 | 11 | 0 | 0 | 0 | 0 | 34 | 19069 | 14169 | 0 | 0 | 3565 | 654 | 0 | 0 | 25 | 14 | 11 | 11 | 0 | 0 | 101 | 0 | 0 | 0 | 0 | 4717 | 12 | 11 |
| SRR5030260 | 0 | 21 | 0 | 0 | 10 | 494 | 255 | 1056 | 204 | 0 | 0 | 0 | 0 | 0 | 17 | 19617 | 14420 | 0 | 0 | 3287 | 664 | 0 | 0 | 59 | 49 | 0 | 38 | 0 | 0 | 101 | 0 | 0 | 0 | 0 | 5268 | 15 | 0 |
| SRR5030261 | 0 | 24 | 0 | 0 | 0 | 632 | 290 | 957 | 120 | 0 | 0 | 0 | 0 | 0 | 17 | 18529 | 15450 | 0 | 0 | 3314 | 683 | 0 | 0 | 63 | 52 | 0 | 24 | 0 | 0 | 80 | 0 | 0 | 0 | 0 | 4959 | 18 | 10 |
| SRR5030262 | 87 | 97 | 0 | 0 | 16 | 145 | 72 | 347 | 29 | 71 | 0 | 0 | 0 | 0 | 4353 | 6205 | 50607 | 0 | 43 | 2025 | 4252 | 0 | 0 | 81 | 10 | 56 | 41 | 0 | 0 | 12 | 0 | 29 | 0 | 0 | 3123 | 0 | 0 |
| SRR5030263 | 53 | 122 | 0 | 0 | 201 | 164 | 45 | 268 | 30 | 48 | 16 | 0 | 0 | 0 | 4128 | 6005 | 49781 | 15 | 45 | 1982 | 3979 | 0 | 0 | 122 | 63 | 21 | 48 | 0 | 0 | 132 | 0 | 15 | 0 | 0 | 3265 | 0 | 0 |
| SRR5030265 | 62 | 67 | 0 | 0 | 19 | 185 | 142 | 370 | 14 | 21 | 15 | 0 | 0 | 0 | 4017 | 6255 | 49959 | 0 | 57 | 2137 | 4060 | 0 | 0 | 83 | 41 | 15 | 52 | 0 | 0 | 108 | 0 | 57 | 0 | 0 | 2856 | 0 | 0 |
| SRR5030266 | 0 | 38 | 0 | 0 | 83 | 134 | 27 | 511 | 0 | 0 | 15 | 0 | 0 | 0 | 12553 | 2084 | 18199 | 0 | 27 | 1828 | 2796 | 0 | 0 | 114 | 0 | 0 | 30 | 0 | 0 | 167 | 0 | 0 | 0 | 0 | 6784 | 0 | 0 |
| SRR5030267 | 24 | 16 | 0 | 0 | 134 | 112 | 28 | 392 | 112 | 63 | 0 | 16 | 0 | 0 | 13346 | 2230 | 18158 | 28 | 0 | 2126 | 2602 | 0 | 0 | 79 | 0 | 0 | 47 | 0 | 0 | 94 | 0 | 0 | 0 | 0 | 6133 | 0 | 39 |
| SRR5030268 | 31 | 23 | 0 | 0 | 147 | 160 | 107 | 373 | 27 | 23 | 0 | 0 | 0 | 0 | 12265 | 1605 | 17416 | 53 | 80 | 1920 | 2560 | 0 | 0 | 62 | 23 | 31 | 31 | 0 | 0 | 15 | 0 | 0 | 0 | 0 | 5546 | 0 | 15 |
| SRR5030270 | 33 | 0 | 0 | 0 | 85 | 143 | 77 | 230 | 55 | 0 | 0 | 0 | 0 | 0 | 22 | 20229 | 18051 | 22 | 11 | 2544 | 800 | 0 | 0 | 79 | 46 | 0 | 46 | 0 | 0 | 210 | 0 | 22 | 0 | 0 | 4716 | 0 | 13 |
| SRR5030271 | 21 | 0 | 0 | 0 | 111 | 47 | 130 | 177 | 71 | 14 | 0 | 28 | 0 | 0 | 24 | 21840 | 17924 | 0 | 0 | 3043 | 755 | 0 | 0 | 97 | 42 | 0 | 28 | 0 | 0 | 222 | 0 | 24 | 0 | 0 | 4806 | 0 | 14 |
| SRR5030272 | 27 | 54 | 0 | 0 | 114 | 122 | 67 | 234 | 56 | 27 | 13 | 0 | 13 | 0 | 22 | 19812 | 17730 | 22 | 0 | 2536 | 868 | 0 | 0 | 47 | 67 | 20 | 33 | 0 | 0 | 160 | 0 | 0 | 0 | 0 | 5143 | 0 | 0 |
| SRR5030273 | 13 | 13 | 0 | 0 | 140 | 101 | 146 | 258 | 45 | 27 | 0 | 27 | 0 | 0 | 22 | 20956 | 17700 | 0 | 56 | 2404 | 809 | 0 | 0 | 60 | 53 | 0 | 33 | 0 | 0 | 187 | 0 | 11 | 0 | 0 | 5328 | 0 | 13 |
| SRR5030274 | 20 | 66 | 0 | 0 | 93 | 298 | 70 | 280 | 35 | 26 | 0 | 0 | 20 | 0 | 0 | 6991 | 14082 | 18 | 0 | 2854 | 718 | 0 | 0 | 79 | 40 | 0 | 79 | 0 | 0 | 86 | 0 | 0 | 0 | 0 | 5751 | 0 | 0 |
| SRR5030275 | 28 | 42 | 0 | 0 | 111 | 224 | 75 | 299 | 37 | 0 | 0 | 0 | 0 | 0 | 0 | 8013 | 12672 | 56 | 0 | 2484 | 766 | 0 | 0 | 118 | 28 | 42 | 111 | 0 | 0 | 111 | 0 | 0 | 0 | 0 | 6025 | 0 | 0 |
| SRR5030276 | 34 | 61 | 0 | 0 | 81 | 146 | 73 | 455 | 73 | 20 | 0 | 14 | 0 | 0 | 0 | 7463 | 12451 | 18 | 0 | 2786 | 510 | 0 | 0 | 129 | 27 | 20 | 34 | 0 | 0 | 74 | 0 | 0 | 0 | 0 | 5672 | 0 | 0 |
| SRR5030277 | 27 | 34 | 0 | 0 | 68 | 284 | 36 | 498 | 53 | 27 | 0 | 14 | 0 | 0 | 0 | 7047 | 13234 | 0 | 0 | 2524 | 604 | 0 | 0 | 122 | 27 | 20 | 81 | 0 | 0 | 88 | 0 | 18 | 0 | 0 | 5225 | 0 | 0 |
| SRR5030278 | 32 | 98 | 0 | 0 | 132 | 1327 | 295 | 2286 | 55 | 56 | 17 | 0 | 0 | 0 | 0 | 1266 | 6703 | 0 | 0 | 2664 | 1899 | 0 | 0 | 105 | 122 | 37 | 24 | 0 | 0 | 154 | 0 | 0 | 0 | 0 | 712 | 0 | 0 |
| SRR5030279 | 21 | 82 | 0 | 0 | 110 | 1205 | 336 | 2321 | 99 | 41 | 15 | 0 | 0 | 0 | 20 | 1206 | 6737 | 0 | 0 | 2864 | 2024 | 0 | 0 | 82 | 108 | 18 | 46 | 0 | 0 | 12 | 0 | 30 | 0 | 0 | 670 | 0 | 0 |
| SRR5030280 | 37 | 104 | 0 | 0 | 132 | 1325 | 238 | 2554 | 76 | 52 | 20 | 0 | 0 | 0 | 0 | 1174 | 6680 | 0 | 0 | 2792 | 1830 | 0 | 0 | 82 | 122 | 32 | 52 | 0 | 0 | 132 | 0 | 38 | 0 | 0 | 697 | 0 | 0 |
| SRR5030281 | 17 | 89 | 0 | 0 | 97 | 1434 | 313 | 2440 | 161 | 57 | 20 | 0 | 0 | 0 | 0 | 1261 | 6646 | 0 | 0 | 2640 | 1880 | 0 | 0 | 67 | 114 | 27 | 32 | 0 | 0 | 124 | 0 | 28 | 0 | 0 | 681 | 0 | 0 |
| SRR5030282 | 36 | 26 | 0 | 0 | 26 | 60 | 50 | 289 | 20 | 21 | 16 | 10 | 0 | 0 | 358 | 1783 | 4675 | 0 | 0 | 358 | 378 | 0 | 0 | 88 | 0 | 0 | 42 | 0 | 0 | 244 | 0 | 0 | 0 | 0 | 2292 | 0 | 0 |
| SRR5030283 | 27 | 16 | 0 | 0 | 16 | 11 | 65 | 237 | 43 | 16 | 27 | 0 | 0 | 0 | 367 | 2090 | 4933 | 0 | 0 | 421 | 378 | 0 | 0 | 98 | 11 | 16 | 54 | 0 | 0 | 125 | 0 | 0 | 0 | 0 | 2325 | 0 | 0 |
| SRR5030284 | 21 | 0 | 0 | 0 | 32 | 10 | 73 | 271 | 10 | 11 | 0 | 0 | 0 | 0 | 344 | 1658 | 4522 | 0 | 0 | 616 | 417 | 0 | 0 | 106 | 0 | 0 | 48 | 0 | 0 | 127 | 0 | 0 | 0 | 0 | 2530 | 0 | 0 |
| SRR5030285 | 16 | 21 | 0 | 0 | 26 | 52 | 42 | 241 | 31 | 0 | 11 | 0 | 0 | 0 | 262 | 1870 | 4636 | 0 | 0 | 460 | 419 | 0 | 0 | 111 | 11 | 0 | 42 | 0 | 0 | 169 | 0 | 0 | 0 | 0 | 2423 | 0 | 0 |
| SRR5030287 | 0 | 35 | 0 | 0 | 44 | 53 | 89 | 249 | 36 | 0 | 18 | 0 | 0 | 0 | 125 | 1705 | 2510 | 0 | 18 | 1388 | 160 | 0 | 0 | 79 | 0 | 0 | 35 | 0 | 0 | 193 | 0 | 0 | 0 | 0 | 6961 | 0 | 35 |
| SRR5030289 | 0 | 43 | 0 | 0 | 43 | 35 | 121 | 225 | 87 | 26 | 17 | 0 | 17 | 0 | 87 | 1464 | 2256 | 0 | 0 | 1334 | 260 | 0 | 0 | 103 | 0 | 0 | 17 | 0 | 0 | 171 | 0 | 17 | 0 | 0 | 7211 | 0 | 26 |
| SRR5030290 | 89 | 52 | 0 | 0 | 17 | 1687 | 909 | 8927 | 123 | 16 | 0 | 0 | 16 | 0 | 1286 | 6639 | 14038 | 15 | 0 | 716 | 770 | 0 | 0 | 20 | 0 | 0 | 84 | 10 | 0 | 73 | 0 | 0 | 0 | 0 | 2744 | 12 | 0 |
| SRR5030291 | 87 | 60 | 0 | 0 | 13 | 1733 | 910 | 9069 | 127 | 16 | 16 | 16 | 11 | 0 | 1377 | 7088 | 14102 | 16 | 0 | 704 | 586 | 0 | 0 | 17 | 22 | 0 | 60 | 0 | 0 | 76 | 0 | 0 | 0 | 0 | 2762 | 11 | 0 |
| SRR5030292 | 59 | 69 | 0 | 0 | 250 | 1490 | 879 | 9138 | 126 | 0 | 0 | 0 | 21 | 0 | 1443 | 7051 | 14261 | 0 | 0 | 596 | 518 | 0 | 0 | 341 | 21 | 21 | 48 | 0 | 0 | 117 | 0 | 0 | 0 | 0 | 2196 | 0 | 0 |
| SRR5030293 | 90 | 64 | 0 | 0 | 244 | 1724 | 932 | 9043 | 188 | 16 | 0 | 0 | 21 | 0 | 1175 | 6831 | 14114 | 0 | 16 | 588 | 682 | 0 | 0 | 339 | 11 | 11 | 95 | 0 | 0 | 74 | 0 | 0 | 0 | 0 | 2526 | 0 | 11 |
| SRR5030294 | 113 | 100 | 0 | 0 | 11 | 603 | 389 | 3035 | 97 | 0 | 0 | 0 | 0 | 0 | 3181 | 8722 | 29018 | 0 | 0 | 506 | 798 | 0 | 0 | 13 | 0 | 13 | 106 | 0 | 0 | 106 | 0 | 0 | 0 | 0 | 1145 | 0 | 0 |
| SRR5030295 | 70 | 90 | 0 | 0 | 10 | 429 | 350 | 3006 | 140 | 0 | 35 | 0 | 14 | 0 | 3036 | 8431 | 29683 | 0 | 0 | 569 | 879 | 0 | 0 | 12 | 14 | 28 | 63 | 0 | 0 | 97 | 0 | 0 | 0 | 0 | 1388 | 0 | 0 |
| SRR5030296 | 88 | 67 | 13 | 13 | 236 | 658 | 403 | 3292 | 69 | 0 | 0 | 0 | 13 | 0 | 2880 | 8567 | 29209 | 0 | 0 | 678 | 688 | 0 | 20 | 10 | 27 | 0 | 88 | 0 | 0 | 101 | 0 | 0 | 0 | 0 | 1206 | 0 | 0 |
| SRR5030297 | 101 | 88 | 0 | 14 | 358 | 586 | 258 | 3416 | 129 | 0 | 0 | 0 | 14 | 0 | 3009 | 9064 | 28699 | 20 | 0 | 705 | 745 | 0 | 0 | 277 | 14 | 20 | 74 | 0 | 0 | 128 | 0 | 0 | 0 | 0 | 1157 | 0 | 0 |
| SRR5034616 | 0 | 0 | 0 | 0 | 0 | 302 | 0 | 905 | 25 | 12 | 49 | 0 | 0 | 0 | 528 | 6156 | 27860 | 0 | 0 | 2715 | 2841 | 12 | 0 | 432 | 49 | 321 | 864 | 25 | 0 | 160 | 0 | 75 | 0 | 0 | 1605 | 0 | 25 |
| SRR5034617 | 0 | 0 | 0 | 17 | 0 | 474 | 157 | 927 | 53 | 26 | 74 | 0 | 0 | 0 | 831 | 12904 | 41112 | 14 | 61 | 1943 | 4317 | 0 | 0 | 491 | 93 | 255 | 600 | 30 | 0 | 143 | 0 | 210 | 0 | 0 | 794 | 0 | 0 |
| SRR5034618 | 0 | 0 | 11 | 15 | 0 | 211 | 69 | 305 | 54 | 0 | 150 | 0 | 0 | 0 | 775 | 11464 | 41580 | 0 | 32 | 2239 | 4223 | 0 | 0 | 486 | 95 | 424 | 2591 | 482 | 0 | 192 | 0 | 107 | 0 | 0 | 763 | 0 | 13 |
| SRR5034619 | 0 | 0 | 0 | 0 | 0 | 160 | 106 | 878 | 80 | 0 | 0 | 0 | 0 | 0 | 1144 | 8163 | 36663 | 0 | 27 | 2022 | 2422 | 0 | 0 | 255 | 29 | 160 | 350 | 0 | 0 | 29 | 0 | 106 | 0 | 0 | 2276 | 0 | 299 |
| SRR5034620 | 0 | 0 | 18 | 12 | 0 | 90 | 18 | 326 | 0 | 43 | 117 | 0 | 0 | 0 | 1774 | 6674 | 46040 | 0 | 18 | 2660 | 6443 | 0 | 0 | 92 | 18 | 179 | 382 | 154 | 0 | 68 | 0 | 72 | 0 | 0 | 1092 | 0 | 37 |
| SRR5034621 | 0 | 0 | 0 | 32 | 0 | 58 | 36 | 230 | 29 | 16 | 142 | 0 | 0 | 0 | 705 | 8341 | 24401 | 0 | 22 | 5069 | 3603 | 0 | 0 | 206 | 13 | 155 | 586 | 152 | 0 | 161 | 0 | 223 | 0 | 0 | 694 | 0 | 16 |
| SRR5034622 | 0 | 0 | 0 | 15 | 0 | 146 | 50 | 391 | 23 | 36 | 172 | 18 | 0 | 0 | 1065 | 12556 | 54589 | 13 | 21 | 2986 | 4625 | 0 | 0 | 579 | 45 | 372 | 1054 | 307 | 0 | 201 | 0 | 164 | 0 | 0 | 1363 | 0 | 27 |
| SRR5034623 | 0 | 0 | 0 | 0 | 0 | 100 | 18 | 156 | 13 | 62 | 44 | 0 | 0 | 0 | 23397 | 44683 | 124118 | 0 | 0 | 912 | 2071 | 0 | 0 | 79 | 0 | 24 | 160 | 0 | 0 | 78 | 0 | 10 | 0 | 0 | 109 | 0 | 0 |
| SRR5034625 | 0 | 0 | 0 | 0 | 13 | 181 | 39 | 279 | 12 | 124 | 280 | 13 | 0 | 0 | 17874 | 34114 | 135009 | 0 | 0 | 433 | 987 | 0 | 0 | 228 | 39 | 137 | 1552 | 176 | 0 | 196 | 0 | 0 | 0 | 0 | 190 | 0 | 20 |
| SRR5034626 | 0 | 0 | 0 | 0 | 185 | 35 | 23 | 139 | 0 | 90 | 139 | 0 | 0 | 0 | 6881 | 120103 | 54983 | 0 | 0 | 256 | 8493 | 0 | 0 | 115 | 11 | 47 | 122 | 73 | 0 | 143 | 0 | 0 | 0 | 0 | 82 | 0 | 0 |
| SRR5034627 | 0 | 0 | 0 | 0 | 0 | 523 | 197 | 474 | 92 | 41 | 46 | 0 | 0 | 0 | 2025 | 30402 | 31534 | 12 | 0 | 345 | 4026 | 0 | 0 | 41 | 15 | 26 | 252 | 0 | 0 | 67 | 0 | 0 | 0 | 0 | 1386 | 0 | 0 |
| SRR5034628 | 0 | 0 | 0 | 0 | 0 | 686 | 110 | 631 | 0 | 64 | 177 | 0 | 0 | 0 | 7050 | 21215 | 58107 | 0 | 192 | 576 | 2277 | 0 | 0 | 80 | 48 | 112 | 24 | 112 | 0 | 209 | 0 | 0 | 0 | 0 | 546 | 0 | 129 |
| SRR5034629 | 0 | 0 | 16 | 22 | 22 | 566 | 109 | 1697 | 36 | 71 | 11 | 16 | 0 | 0 | 9697 | 21970 | 67340 | 0 | 0 | 620 | 1843 | 0 | 0 | 82 | 60 | 148 | 21 | 0 | 0 | 286 | 0 | 27 | 0 | 0 | 857 | 0 | 16 |
| SRR5034630 | 0 | 0 | 0 | 0 | 0 | 1062 | 229 | 3102 | 167 | 0 | 47 | 12 | 0 | 0 | 7431 | 31818 | 61985 | 0 | 0 | 1582 | 2498 | 0 | 0 | 247 | 71 | 82 | 801 | 12 | 0 | 235 | 0 | 83 | 0 | 0 | 1125 | 0 | 12 |
| SRR5034632 | 0 | 0 | 0 | 0 | 0 | 238 | 51 | 255 | 34 | 42 | 179 | 0 | 0 | 0 | 3742 | 44734 | 29979 | 0 | 0 | 850 | 952 | 0 | 0 | 0 | 11 | 63 | 295 | 0 | 0 | 242 | 0 | 0 | 0 | 0 | 574 | 0 | 105 |
| SRR5034634 | 0 | 0 | 0 | 0 | 75 | 199 | 0 | 315 | 17 | 0 | 93 | 0 | 0 | 0 | 6226 | 39953 | 39872 | 0 | 0 | 299 | 747 | 0 | 0 | 37 | 0 | 37 | 298 | 0 | 0 | 186 | 0 | 33 | 0 | 0 | 166 | 0 | 19 |
| SRR5034635 | 0 | 0 | 0 | 0 | 32 | 297 | 28 | 198 | 14 | 16 | 32 | 32 | 0 | 0 | 8470 | 46556 | 77854 | 0 | 14 | 269 | 1161 | 0 | 0 | 96 | 0 | 175 | 18 | 32 | 0 | 287 | 0 | 28 | 0 | 0 | 1689 | 0 | 32 |
| SRR5034636 | 0 | 0 | 0 | 0 | 0 | 222 | 48 | 175 | 32 | 0 | 25 | 25 | 0 | 0 | 11816 | 33848 | 79170 | 0 | 0 | 572 | 1938 | 0 | 0 | 126 | 0 | 201 | 979 | 50 | 0 | 25 | 0 | 32 | 0 | 0 | 268 | 0 | 226 |
| SRR5034637 | 0 | 0 | 0 | 0 | 0 | 201 | 101 | 352 | 0 | 26 | 0 | 0 | 0 | 0 | 6593 | 64033 | 33312 | 0 | 0 | 705 | 3271 | 0 | 0 | 77 | 51 | 51 | 282 | 128 | 0 | 102 | 0 | 0 | 0 | 0 | 471 | 0 | 77 |
| SRR5034639 | 0 | 0 | 0 | 0 | 0 | 321 | 0 | 241 | 0 | 0 | 25 | 0 | 0 | 0 | 12556 | 32455 | 62518 | 0 | 40 | 1524 | 3289 | 0 | 0 | 25 | 0 | 0 | 273 | 0 | 0 | 99 | 0 | 0 | 0 | 0 | 191 | 0 | 75 |
| SRR5034641 | 0 | 0 | 0 | 0 | 0 | 1458 | 177 | 1379 | 118 | 0 | 78 | 0 | 0 | 0 | 4414 | 29544 | 47677 | 0 | 0 | 2049 | 1537 | 0 | 0 | 24 | 0 | 30 | 150 | 0 | 0 | 54 | 0 | 20 | 0 | 0 | 368 | 0 | 12 |
| SRR5034642 | 0 | 0 | 0 | 0 | 0 | 23 | 0 | 53 | 0 | 98 | 76 | 0 | 0 | 0 | 678 | 15524 | 7726 | 0 | 0 | 79 | 842 | 0 | 0 | 65 | 27 | 38 | 375 | 0 | 0 | 98 | 0 | 0 | 0 | 0 | 125 | 0 | 16 |
| SRR5034643 | 0 | 0 | 0 | 0 | 0 | 61 | 0 | 119 | 14 | 15 | 81 | 29 | 0 | 0 | 3698 | 48483 | 48117 | 0 | 12 | 287 | 9135 | 0 | 0 | 318 | 29 | 110 | 673 | 0 | 0 | 88 | 0 | 19 | 0 | 0 | 360 | 0 | 15 |
| SRR5034644 | 0 | 0 | 0 | 0 | 35 | 190 | 50 | 281 | 14 | 98 | 202 | 0 | 0 | 0 | 8030 | 37040 | 105807 | 0 | 18 | 906 | 4783 | 0 | 0 | 266 | 92 | 98 | 16 | 0 | 0 | 254 | 0 | 18 | 0 | 0 | 784 | 0 | 46 |
| SRR5034645 | 0 | 0 | 0 | 0 | 0 | 84 | 15 | 138 | 46 | 0 | 266 | 0 | 0 | 0 | 15938 | 26275 | 131716 | 0 | 0 | 720 | 1931 | 0 | 0 | 86 | 48 | 133 | 542 | 114 | 0 | 95 | 0 | 0 | 0 | 0 | 92 | 0 | 38 |
| SRR5034646 | 0 | 0 | 0 | 0 | 12 | 171 | 34 | 263 | 34 | 24 | 97 | 134 | 0 | 0 | 13973 | 37816 | 101013 | 0 | 0 | 731 | 2434 | 0 | 0 | 85 | 85 | 219 | 1302 | 231 | 0 | 122 | 0 | 0 | 0 | 0 | 318 | 0 | 146 |
| SRR5034647 | 0 | 0 | 0 | 0 | 67 | 56 | 13 | 83 | 12 | 54 | 26 | 0 | 0 | 0 | 1940 | 75457 | 66701 | 0 | 0 | 78 | 1317 | 0 | 0 | 139 | 27 | 102 | 433 | 46 | 0 | 37 | 0 | 0 | 0 | 0 | 159 | 0 | 0 |
| SRR5034648 | 0 | 0 | 0 | 0 | 0 | 901 | 170 | 633 | 49 | 121 | 379 | 0 | 0 | 0 | 8425 | 40991 | 74209 | 0 | 73 | 1193 | 2118 | 0 | 0 | 0 | 0 | 76 | 682 | 182 | 0 | 924 | 0 | 0 | 0 | 0 | 330 | 0 | 30 |
| SRR5034649 | 0 | 0 | 0 | 0 | 0 | 52 | 14 | 109 | 14 | 84 | 113 | 38 | 0 | 0 | 3211 | 38275 | 67823 | 0 | 0 | 172 | 734 | 0 | 0 | 56 | 19 | 0 | 385 | 0 | 0 | 338 | 0 | 0 | 0 | 0 | 189 | 0 | 19 |
| SRR5034650 | 0 | 0 | 0 | 70 | 45 | 188 | 105 | 444 | 38 | 45 | 15 | 0 | 0 | 0 | 5145 | 38319 | 84096 | 23 | 0 | 851 | 8565 | 0 | 0 | 11 | 0 | 0 | 0 | 0 | 0 | 22 | 0 | 15 | 0 | 0 | 278 | 0 | 45 |
| SRR5034651 | 0 | 0 | 0 | 0 | 28 | 68 | 27 | 108 | 41 | 21 | 118 | 35 | 0 | 0 | 2874 | 30993 | 51643 | 0 | 27 | 285 | 5342 | 0 | 0 | 49 | 70 | 132 | 53 | 146 | 0 | 12 | 0 | 0 | 0 | 0 | 659 | 0 | 49 |
| SRR5034652 | 0 | 0 | 0 | 22 | 41 | 20 | 55 | 120 | 20 | 63 | 291 | 0 | 0 | 0 | 7272 | 42382 | 42499 | 0 | 15 | 379 | 5637 | 0 | 0 | 111 | 25 | 82 | 449 | 98 | 0 | 149 | 0 | 0 | 0 | 0 | 249 | 0 | 0 |
| SRR5034653 | 0 | 0 | 0 | 0 | 58 | 84 | 0 | 221 | 24 | 37 | 157 | 12 | 0 | 0 | 4821 | 56187 | 52202 | 0 | 20 | 293 | 3852 | 0 | 0 | 223 | 62 | 144 | 586 | 91 | 0 | 120 | 0 | 0 | 0 | 0 | 266 | 0 | 17 |
| SRR5034654 | 0 | 0 | 0 | 0 | 66 | 80 | 40 | 190 | 18 | 117 | 166 | 0 | 0 | 0 | 5623 | 47714 | 54698 | 0 | 0 | 352 | 2592 | 0 | 0 | 12 | 86 | 111 | 28 | 22 | 0 | 12 | 0 | 0 | 0 | 0 | 117 | 0 | 11 |
| SRR5034655 | 0 | 0 | 0 | 0 | 47 | 21 | 0 | 35 | 0 | 78 | 212 | 0 | 0 | 0 | 6051 | 51387 | 67480 | 0 | 0 | 120 | 4228 | 0 | 0 | 30 | 30 | 73 | 475 | 112 | 0 | 212 | 0 | 0 | 0 | 0 | 186 | 0 | 0 |
| SRR5034656 | 0 | 0 | 0 | 0 | 282 | 21 | 18 | 49 | 0 | 69 | 141 | 0 | 0 | 0 | 1723 | 108098 | 44398 | 0 | 15 | 141 | 3875 | 0 | 0 | 88 | 0 | 39 | 138 | 23 | 0 | 124 | 0 | 0 | 0 | 0 | 1943 | 0 | 16 |
| SRR5034657 | 0 | 0 | 0 | 0 | 110 | 135 | 15 | 244 | 0 | 143 | 97 | 0 | 0 | 0 | 3460 | 59219 | 49806 | 0 | 0 | 240 | 634 | 0 | 0 | 58 | 19 | 91 | 253 | 0 | 0 | 104 | 0 | 0 | 0 | 0 | 264 | 0 | 58 |
| SRR5034658 | 0 | 0 | 0 | 0 | 138 | 109 | 0 | 163 | 0 | 176 | 77 | 28 | 0 | 0 | 3200 | 52478 | 45720 | 0 | 0 | 272 | 1179 | 0 | 0 | 220 | 165 | 275 | 22 | 165 | 0 | 154 | 0 | 0 | 0 | 0 | 433 | 0 | 33 |
| SRR5034659 | 0 | 0 | 0 | 0 | 316 | 14 | 0 | 28 | 0 | 488 | 26 | 11 | 0 | 0 | 3183 | 60776 | 42379 | 0 | 0 | 79 | 3918 | 0 | 0 | 144 | 32 | 61 | 318 | 0 | 0 | 97 | 0 | 0 | 0 | 0 | 135 | 0 | 0 |
| SRR5034660 | 0 | 0 | 0 | 0 | 136 | 20 | 0 | 26 | 0 | 238 | 35 | 0 | 0 | 0 | 5120 | 81873 | 57119 | 0 | 0 | 32 | 2817 | 0 | 0 | 210 | 34 | 56 | 596 | 101 | 0 | 51 | 0 | 0 | 0 | 0 | 73 | 0 | 0 |
| SRR5034661 | 0 | 0 | 0 | 0 | 11 | 0 | 0 | 25 | 0 | 134 | 238 | 0 | 0 | 0 | 4879 | 99679 | 148256 | 0 | 0 | 105 | 451 | 0 | 0 | 76 | 0 | 40 | 227 | 58 | 0 | 321 | 0 | 0 | 0 | 0 | 3018 | 0 | 0 |
| SRR5034662 | 0 | 0 | 0 | 0 | 35 | 53 | 0 | 71 | 12 | 70 | 117 | 0 | 0 | 0 | 3974 | 48459 | 69966 | 0 | 0 | 146 | 3044 | 0 | 0 | 10 | 76 | 208 | 84 | 177 | 0 | 72 | 0 | 0 | 0 | 0 | 161 | 0 | 0 |
| SRR5034663 | 0 | 0 | 0 | 0 | 47 | 184 | 35 | 512 | 0 | 52 | 127 | 0 | 0 | 0 | 3664 | 39397 | 49934 | 0 | 0 | 309 | 1228 | 0 | 0 | 71 | 42 | 94 | 448 | 0 | 0 | 184 | 0 | 0 | 0 | 0 | 159 | 0 | 80 |
| SRR5054304 | 12280 | 2365 | 52 | 29 | 1000 | 560 | 58 | 710 | 3377 | 4015 | 0 | 16 | 32 | 23 | 271 | 3322 | 3454 | 69 | 40 | 58 | 306 | 3531 | 763 | 745 | 0 | 0 | 0 | 0 | 11 | 0 | 937 | 46 | 45 | 63 | 19226 | 0 | 0 |
| SRR5054305 | 20111 | 5113 | 156 | 125 | 3510 | 488 | 65 | 335 | 1199 | 494 | 36 | 0 | 34 | 0 | 3915 | 17280 | 38507 | 59 | 570 | 811 | 3433 | 16 | 0 | 861 | 39 | 10 | 13 | 0 | 0 | 68 | 18 | 1099 | 10 | 57 | 17602 | 0 | 0 |

### Supp Table S8.3.Columns_BX_to_DH

| | | | | | | | | | | | | | | | | | | | | | | | | | | | | | | | | | | | | | |
|---|---|---|---|---|---|---|---|---|---|---|---|---|---|---|---|---|---|---|---|---|---|---|---|---|---|---|---|---|---|---|---|---|---|---|---|---|---|
| SRR5087508 | 14 | 0 | 0 | 22 | 14 | 1358 | 313 | 1022 | 76 | 25 | 0 | 0 | 0 | 0 | 14414 | 50322 | 200090 | 0 | 65 | 6001 | 14238 | 0 | 0 | 19 | 99 | 12 | 15 | 0 | 0 | 75 | 0 | 29 | 0 | 0 | 304 | 0 | 0 |
| SRR5087510 | 17 | 13 | 0 | 25 | 16 | 1822 | 318 | 1157 | 85 | 28 | 0 | 0 | 0 | 0 | 13597 | 55030 | 199867 | 0 | 113 | 7013 | 14441 | 0 | 0 | 26 | 11 | 16 | 25 | 0 | 0 | 93 | 0 | 23 | 0 | 0 | 277 | 0 | 0 |
| SRR5087512 | 13 | 15 | 0 | 22 | 17 | 1849 | 323 | 1187 | 100 | 34 | 12 | 0 | 0 | 0 | 13171 | 44415 | 193792 | 0 | 102 | 6570 | 15063 | 0 | 0 | 24 | 11 | 15 | 26 | 0 | 0 | 99 | 0 | 41 | 0 | 0 | 267 | 0 | 0 |
| SRR5087516 | 11 | 15 | 0 | 11 | 18 | 1816 | 184 | 958 | 43 | 71 | 11 | 0 | 0 | 0 | 20515 | 48199 | 269295 | 0 | 58 | 6123 | 18285 | 0 | 0 | 18 | 12 | 14 | 14 | 0 | 0 | 75 | 0 | 12 | 0 | 0 | 196 | 0 | 0 |
| SRR5087517 | 0 | 0 | 0 | 0 | 22 | 1602 | 138 | 814 | 39 | 72 | 15 | 0 | 0 | 0 | 21106 | 37350 | 286443 | 0 | 50 | 5700 | 23730 | 0 | 0 | 16 | 12 | 13 | 10 | 0 | 0 | 78 | 0 | 15 | 0 | 0 | 187 | 0 | 0 |
| SRR5087519 | 11 | 0 | 0 | 12 | 21 | 1843 | 182 | 892 | 46 | 63 | 10 | 0 | 0 | 0 | 23046 | 37393 | 307809 | 0 | 67 | 5933 | 23705 | 0 | 0 | 18 | 12 | 16 | 13 | 0 | 0 | 56 | 0 | 11 | 0 | 0 | 175 | 0 | 0 |
| SRR5087522 | 13 | 21 | 19 | 117 | 195 | 1539 | 378 | 1550 | 124 | 0 | 12 | 0 | 0 | 0 | 5598 | 35906 | 115429 | 0 | 85 | 6483 | 4279 | 0 | 0 | 325 | 50 | 201 | 678 | 18 | 0 | 191 | 0 | 28 | 0 | 0 | 1604 | 0 | 0 |
| SRR5087527 | 30 | 15 | 18 | 74 | 19 | 1196 | 506 | 1684 | 226 | 14 | 0 | 0 | 0 | 0 | 7196 | 37056 | 135790 | 0 | 36 | 7166 | 4625 | 0 | 0 | 25 | 62 | 14 | 56 | 10 | 0 | 12 | 0 | 65 | 0 | 0 | 1512 | 0 | 0 |
| SRR5087554 | 24 | 21 | 11 | 114 | 15 | 1157 | 553 | 1553 | 245 | 0 | 16 | 0 | 0 | 0 | 6603 | 34214 | 128145 | 13 | 63 | 7251 | 4710 | 0 | 0 | 29 | 73 | 14 | 42 | 16 | 0 | 18 | 0 | 44 | 0 | 0 | 1861 | 0 | 0 |
| SRR5090283 | 17 | 0 | 0 | 0 | 124 | 5159 | 3507 | 12984 | 52476 | 183 | 93 | 51 | 0 | 0 | 3465 | 46996 | 58964 | 190 | 213 | 2607 | 5139 | 0 | 0 | 58 | 78 | 46 | 19 | 22 | 0 | 230 | 0 | 21 | 0 | 17 | 7051 | 0 | 0 |
| SRR5092303 | 0 | 0 | 0 | 0 | 59 | 13609 | 984 | 12197 | 7104 | 256 | 0 | 39 | 0 | 0 | 942 | 24822 | 5788 | 0 | 43 | 0 | 942 | 0 | 0 | 6185 | 39 | 0 | 0 | 20 | 0 | 39 | 0 | 0 | 0 | 0 | 6300 | 0 | 0 |
| SRR5092304 | 0 | 0 | 0 | 0 | 0 | 20066 | 1776 | 12145 | 9361 | 210 | 23 | 0 | 0 | 0 | 96 | 15362 | 3561 | 0 | 0 | 0 | 240 | 23 | 0 | 5293 | 0 | 0 | 0 | 0 | 0 | 0 | 23 | 0 | 0 | 0 | 9648 | 0 | 0 |
| SRR5092305 | 0 | 0 | 0 | 0 | 22 | 22972 | 1775 | 13679 | 7622 | 727 | 22 | 0 | 0 | 0 | 627 | 13268 | 3371 | 0 | 0 | 0 | 940 | 0 | 0 | 11 | 0 | 0 | 0 | 0 | 0 | 22 | 0 | 0 | 0 | 44 | 15443 | 0 | 0 |
| SRR5092836 | 0 | 0 | 38 | 38 | 114 | 0 | 0 | 0 | 0 | 76 | 38 | 0 | 0 | 0 | 149071 | 50878 | 431615 | 0 | 601 | 3454 | 25529 | 0 | 38 | 0 | 0 | 0 | 0 | 0 | 0 | 16 | 0 | 100 | 0 | 0 | 2326 | 30 | 0 |
| SRR5100734 | 0 | 0 | 0 | 0 | 10 | 0 | 0 | 19 | 32 | 15 | 6005 | 51 | 0 | 0 | 13 | 232 | 16 | 2645 | 0 | 0 | 0 | 13 | 0 | 0 | 0 | 0 | 0 | 0 | 25 | 97 | 0 | 35090 | 0 | 0 | 2192 | 909 | 0 |
| SRR5100735 | 304 | 285 | 17 | 10 | 68 | 0 | 0 | 0 | 66 | 104 | 7529 | 372 | 0 | 0 | 162 | 86801 | 2865 | 3207 | 215 | 193 | 1129 | 0 | 0 | 23 | 0 | 0 | 0 | 0 | 63 | 274 | 0 | 2439 | 0 | 0 | 334 | 0 | 0 |
| SRR5104206 | 0 | 0 | 0 | 32 | 13 | 475 | 329 | 1316 | 110 | 0 | 28 | 0 | 0 | 0 | 6544 | 5244 | 8989 | 12 | 49 | 2315 | 1816 | 0 | 0 | 48 | 16 | 19 | 36 | 21 | 0 | 22 | 0 | 37 | 0 | 0 | 1498 | 0 | 0 |
| SRR5104207 | 0 | 0 | 0 | 18 | 16 | 134 | 151 | 353 | 25 | 0 | 18 | 0 | 0 | 0 | 5543 | 4068 | 6351 | 0 | 17 | 1377 | 2091 | 0 | 0 | 45 | 18 | 13 | 36 | 13 | 0 | 22 | 0 | 25 | 0 | 0 | 1855 | 0 | 0 |
| SRR5104208 | 13 | 0 | 0 | 19 | 14 | 313 | 168 | 527 | 31 | 0 | 18 | 0 | 0 | 0 | 19120 | 4063 | 22772 | 0 | 53 | 3236 | 2168 | 0 | 0 | 36 | 0 | 15 | 44 | 17 | 0 | 19 | 0 | 0 | 0 | 0 | 1028 | 0 | 0 |
| SRR5104209 | 0 | 0 | 0 | 18 | 49 | 246 | 82 | 702 | 36 | 0 | 21 | 0 | 0 | 0 | 10478 | 4532 | 9398 | 0 | 0 | 2770 | 2661 | 0 | 0 | 156 | 0 | 90 | 158 | 11 | 0 | 106 | 0 | 18 | 0 | 0 | 949 | 0 | 0 |
| SRR5104210 | 0 | 0 | 0 | 0 | 68 | 133 | 200 | 416 | 92 | 0 | 54 | 0 | 0 | 0 | 3128 | 4205 | 12848 | 0 | 17 | 2878 | 1747 | 0 | 0 | 232 | 0 | 33 | 166 | 26 | 0 | 419 | 0 | 0 | 0 | 0 | 1528 | 0 | 0 |
| SRR5104211 | 0 | 0 | 0 | 12 | 61 | 37 | 206 | 191 | 81 | 0 | 31 | 0 | 0 | 0 | 2009 | 2330 | 7070 | 15 | 22 | 1052 | 1457 | 0 | 0 | 260 | 15 | 35 | 156 | 30 | 0 | 211 | 0 | 15 | 0 | 0 | 2037 | 0 | 0 |
| SRR5104212 | 0 | 0 | 0 | 0 | 40 | 247 | 208 | 377 | 69 | 0 | 26 | 11 | 0 | 0 | 3166 | 3301 | 9698 | 0 | 0 | 1695 | 847 | 0 | 0 | 247 | 12 | 47 | 174 | 28 | 0 | 226 | 0 | 0 | 0 | 0 | 999 | 0 | 0 |
| SRR5104213 | 0 | 0 | 0 | 0 | 72 | 117 | 223 | 317 | 94 | 0 | 18 | 0 | 0 | 0 | 3512 | 3204 | 12083 | 0 | 70 | 3888 | 987 | 0 | 0 | 243 | 12 | 43 | 190 | 22 | 0 | 202 | 0 | 0 | 0 | 0 | 991 | 0 | 0 |
| SRR5104214 | 0 | 0 | 0 | 0 | 78 | 70 | 58 | 117 | 12 | 0 | 37 | 0 | 0 | 0 | 1017 | 1631 | 5995 | 0 | 0 | 2594 | 374 | 0 | 0 | 331 | 11 | 47 | 188 | 28 | 0 | 447 | 0 | 0 | 0 | 0 | 560 | 0 | 0 |
| SRR5104215 | 0 | 0 | 0 | 15 | 71 | 11 | 108 | 173 | 76 | 0 | 21 | 0 | 0 | 0 | 2805 | 3865 | 13442 | 0 | 22 | 1018 | 758 | 0 | 0 | 259 | 0 | 50 | 178 | 25 | 0 | 206 | 0 | 0 | 0 | 0 | 1051 | 0 | 0 |
| SRR5104216 | 0 | 0 | 0 | 15 | 71 | 23 | 128 | 158 | 45 | 0 | 33 | 0 | 0 | 0 | 2613 | 6229 | 13290 | 0 | 23 | 9067 | 1250 | 0 | 0 | 22 | 16 | 51 | 15 | 12 | 0 | 24 | 0 | 0 | 0 | 0 | 2912 | 0 | 0 |
| SRR5104217 | 0 | 0 | 0 | 0 | 60 | 13 | 60 | 67 | 27 | 0 | 35 | 0 | 0 | 0 | 2574 | 3804 | 22161 | 0 | 74 | 32900 | 1240 | 0 | 0 | 273 | 10 | 32 | 172 | 16 | 0 | 334 | 0 | 0 | 0 | 0 | 867 | 0 | 0 |
| SRR5104218 | 0 | 0 | 0 | 0 | 65 | 79 | 118 | 118 | 0 | 0 | 52 | 0 | 0 | 0 | 3305 | 14269 | 17552 | 0 | 59 | 1648 | 996 | 0 | 0 | 38 | 0 | 31 | 12 | 20 | 0 | 37 | 0 | 20 | 0 | 0 | 1546 | 0 | 0 |
| SRR5104219 | 0 | 0 | 0 | 10 | 107 | 20 | 81 | 111 | 20 | 0 | 33 | 0 | 0 | 0 | 1641 | 3804 | 7672 | 20 | 30 | 1440 | 826 | 0 | 0 | 24 | 0 | 43 | 18 | 24 | 0 | 29 | 0 | 10 | 0 | 0 | 1813 | 0 | 0 |
| SRR5104220 | 0 | 0 | 0 | 0 | 89 | 54 | 84 | 204 | 24 | 0 | 20 | 0 | 0 | 0 | 4035 | 7003 | 19670 | 0 | 30 | 5489 | 1431 | 0 | 0 | 282 | 0 | 29 | 102 | 10 | 0 | 140 | 0 | 0 | 0 | 0 | 1101 | 0 | 0 |
| SRR5104221 | 0 | 0 | 0 | 0 | 49 | 53 | 35 | 88 | 0 | 0 | 43 | 0 | 0 | 0 | 2223 | 4634 | 14019 | 0 | 44 | 5399 | 979 | 0 | 0 | 261 | 14 | 34 | 179 | 46 | 0 | 314 | 0 | 0 | 0 | 0 | 1840 | 0 | 0 |
| SRR5104222 | 0 | 0 | 0 | 10 | 56 | 25 | 25 | 35 | 0 | 0 | 50 | 0 | 0 | 0 | 3740 | 4367 | 22870 | 0 | 69 | 20913 | 2087 | 0 | 0 | 344 | 13 | 51 | 148 | 22 | 0 | 351 | 0 | 0 | 0 | 0 | 355 | 0 | 0 |
| SRR5104223 | 0 | 0 | 0 | 0 | 38 | 53 | 21 | 21 | 0 | 0 | 46 | 0 | 0 | 0 | 2320 | 3295 | 8583 | 0 | 21 | 3521 | 1972 | 0 | 0 | 34 | 26 | 52 | 25 | 30 | 0 | 45 | 0 | 11 | 0 | 0 | 707 | 0 | 0 |
| SRR5104224 | 0 | 0 | 0 | 0 | 52 | 0 | 0 | 82 | 0 | 0 | 43 | 0 | 0 | 0 | 2182 | 3451 | 17217 | 0 | 13 | 9709 | 2645 | 0 | 0 | 49 | 11 | 13 | 15 | 16 | 0 | 51 | 0 | 0 | 0 | 0 | 402 | 0 | 0 |
| SRR5104225 | 0 | 0 | 0 | 0 | 83 | 24 | 39 | 79 | 0 | 0 | 63 | 0 | 0 | 0 | 4320 | 4271 | 18389 | 0 | 16 | 3025 | 1556 | 0 | 0 | 52 | 18 | 43 | 20 | 27 | 0 | 44 | 0 | 0 | 0 | 0 | 1016 | 0 | 0 |
| SRR5116489 | 73 | 37 | 0 | 0 | 0 | 0 | 956 | 6323 | 882 | 37 | 37 | 0 | 37 | 0 | 21028 | 25422 | 64096 | 147 | 882 | 10808 | 1397 | 0 | 0 | 367 | 0 | 0 | 0 | 0 | 0 | 880 | 0 | 74 | 0 | 0 | 59759 | 459 | 0 |
| SRR5116490 | 26 | 158 | 13 | 26 | 158 | 0 | 424 | 1222 | 244 | 0 | 777 | 0 | 0 | 0 | 56769 | 76209 | 87664 | 33 | 3339 | 18489 | 3502 | 0 | 0 | 290 | 26 | 13 | 184 | 0 | 0 | 3648 | 0 | 33 | 13 | 0 | 5403 | 334 | 0 |
| SRR5116491 | 149 | 85 | 43 | 85 | 64 | 0 | 390 | 2067 | 195 | 21 | 171 | 0 | 0 | 0 | 53815 | 42180 | 115171 | 0 | 4109 | 26173 | 1690 | 0 | 0 | 426 | 0 | 0 | 107 | 0 | 0 | 4753 | 0 | 13 | 0 | 0 | 6907 | 499 | 0 |
| SRR5116492 | 12 | 16 | 11 | 55 | 27 | 0 | 569 | 1607 | 493 | 18 | 26 | 0 | 0 | 0 | 40268 | 64153 | 67104 | 0 | 455 | 8109 | 2973 | 0 | 0 | 17 | 0 | 0 | 46 | 0 | 0 | 120 | 0 | 13 | 0 | 0 | 8273 | 316 | 0 |
| SRR5116493 | 63 | 16 | 0 | 0 | 47 | 0 | 621 | 1925 | 497 | 31 | 205 | 0 | 0 | 0 | 34612 | 66259 | 73281 | 62 | 1397 | 19029 | 2825 | 0 | 0 | 252 | 0 | 0 | 31 | 0 | 0 | 2660 | 0 | 62 | 0 | 0 | 12967 | 490 | 0 |
| SRR5116494 | 34 | 101 | 34 | 0 | 45 | 0 | 477 | 993 | 159 | 0 | 56 | 0 | 0 | 0 | 16128 | 100958 | 26142 | 0 | 199 | 6356 | 2900 | 0 | 0 | 213 | 0 | 0 | 0 | 0 | 0 | 752 | 0 | 0 | 0 | 0 | 9948 | 379 | 11 |
| SRR5116495 | 62 | 162 | 29 | 24 | 14 | 0 | 461 | 1400 | 290 | 29 | 157 | 0 | 0 | 0 | 25164 | 104088 | 39010 | 17 | 256 | 9577 | 2953 | 0 | 0 | 153 | 0 | 0 | 0 | 0 | 0 | 1737 | 0 | 17 | 0 | 0 | 6295 | 484 | 0 |
| SRR5116496 | 49 | 69 | 0 | 0 | 20 | 0 | 709 | 2127 | 266 | 59 | 59 | 0 | 0 | 0 | 21451 | 64591 | 38755 | 44 | 89 | 5540 | 1906 | 0 | 0 | 167 | 0 | 0 | 0 | 0 | 0 | 648 | 0 | 0 | 0 | 0 | 13849 | 432 | 0 |
| SRR5121477 | 0 | 13 | 20 | 21 | 0 | 564 | 320 | 872 | 237 | 29 | 0 | 0 | 0 | 0 | 107003 | 5660 | 125631 | 0 | 0 | 32 | 385 | 13 | 48 | 11 | 32 | 48 | 63 | 22 | 0 | 426 | 0 | 122 | 0 | 0 | 1619 | 0 | 0 |
| SRR5121478 | 86 | 59 | 51 | 31 | 0 | 372 | 286 | 601 | 191 | 12 | 0 | 0 | 0 | 0 | 12057 | 11496 | 46478 | 0 | 0 | 10 | 477 | 0 | 0 | 53 | 139 | 374 | 558 | 1007 | 0 | 922 | 0 | 29 | 0 | 0 | 2338 | 0 | 0 |
| SRR5121479 | 439 | 381 | 87 | 225 | 162 | 215 | 145 | 420 | 61 | 13 | 17 | 24 | 0 | 0 | 14767 | 3107 | 36031 | 33 | 42 | 3942 | 13030 | 0 | 0 | 143 | 78 | 221 | 430 | 0 | 0 | 123 | 0 | 0 | 0 | 0 | 930 | 0 | 0 |
| SRR5121480 | 197 | 91 | 194 | 83 | 88 | 127 | 90 | 260 | 40 | 22 | 41 | 0 | 0 | 0 | 62935 | 5113 | 99824 | 17 | 50 | 1179 | 8296 | 0 | 0 | 58 | 72 | 368 | 414 | 69 | 0 | 42 | 0 | 30 | 0 | 0 | 2232 | 0 | 0 |
| SRR5121481 | 63 | 55 | 89 | 175 | 34 | 204 | 84 | 273 | 47 | 30 | 324 | 50 | 0 | 0 | 1629 | 1130 | 10861 | 27 | 29 | 3764 | 8389 | 0 | 0 | 48 | 150 | 230 | 209 | 24 | 0 | 1129 | 0 | 15 | 0 | 0 | 807 | 0 | 10 |
| SRR5121482 | 17 | 19 | 168 | 14 | 12 | 302 | 59 | 225 | 19 | 35 | 12 | 0 | 0 | 0 | 2442 | 2457 | 43123 | 23 | 86 | 8154 | 17782 | 0 | 0 | 14 | 22 | 42 | 54 | 121 | 0 | 37 | 0 | 0 | 0 | 0 | 798 | 0 | 0 |
| SRR5121483 | 48 | 22 | 0 | 0 | 0 | 390 | 677 | 1096 | 812 | 316 | 0 | 0 | 0 | 0 | 17572 | 22634 | 95982 | 0 | 0 | 195 | 173 | 0 | 0 | 26 | 54 | 0 | 0 | 0 | 0 | 83 | 0 | 16 | 0 | 0 | 614 | 0 | 0 |
| SRR5121484 | 18 | 19 | 26 | 30 | 133 | 119 | 370 | 392 | 145 | 14 | 19 | 37 | 0 | 0 | 12195 | 8583 | 73580 | 0 | 277 | 1826 | 5946 | 0 | 0 | 89 | 60 | 27 | 39 | 18 | 0 | 19 | 0 | 0 | 0 | 0 | 2367 | 0 | 0 |
| SRR5121485 | 25 | 22 | 37 | 97 | 53 | 39 | 55 | 60 | 0 | 0 | 63 | 0 | 0 | 0 | 17516 | 15603 | 95281 | 0 | 10 | 1716 | 6724 | 0 | 0 | 68 | 13 | 19 | 95 | 92 | 0 | 43 | 0 | 0 | 0 | 0 | 1223 | 0 | 0 |
| SRR5121486 | 0 | 0 | 0 | 0 | 0 | 273 | 708 | 2978 | 524 | 14 | 93 | 35 | 0 | 0 | 7948 | 3484 | 29300 | 0 | 65 | 3507 | 3909 | 0 | 0 | 11 | 0 | 37 | 58 | 0 | 0 | 51 | 0 | 0 | 0 | 0 | 208 | 0 | 0 |
| SRR5121487 | 166 | 128 | 374 | 1327 | 91 | 0 | 68 | 40 | 12 | 0 | 219 | 22 | 0 | 0 | 37421 | 43822 | 163134 | 0 | 0 | 1422 | 11869 | 0 | 0 | 168 | 58 | 472 | 3057 | 233 | 0 | 222 | 0 | 49 | 0 | 0 | 902 | 0 | 11 |
| SRR5127200 | 16 | 21 | 44 | 61 | 58 | 26 | 99 | 176 | 56 | 0 | 560 | 0 | 0 | 0 | 1640 | 2553 | 33070 | 0 | 0 | 2484 | 5163 | 0 | 0 | 112 | 12 | 38 | 240 | 60 | 0 | 1014 | 0 | 53 | 0 | 0 | 1476 | 0 | 0 |
| SRR5127201 | 115 | 85 | 24 | 16 | 77 | 179 | 240 | 575 | 145 | 0 | 62 | 0 | 0 | 0 | 20606 | 85248 | 46195 | 20 | 234 | 19769 | 2312 | 0 | 0 | 218 | 93 | 177 | 845 | 44 | 0 | 326 | 0 | 0 | 0 | 0 | 1488 | 0 | 0 |
| SRR5127202 | 11 | 15 | 43 | 86 | 13 | 47 | 72 | 111 | 50 | 0 | 22 | 0 | 0 | 0 | 57791 | 76797 | 115686 | 36 | 144 | 17996 | 11751 | 0 | 0 | 97 | 102 | 21 | 70 | 68 | 0 | 15 | 0 | 0 | 0 | 0 | 1154 | 0 | 0 |
| SRR5127203 | 262 | 113 | 157 | 129 | 284 | 77 | 84 | 199 | 35 | 17 | 45 | 10 | 0 | 0 | 12308 | 159616 | 17010 | 0 | 92 | 26545 | 951 | 0 | 0 | 450 | 159 | 492 | 1767 | 126 | 0 | 365 | 0 | 0 | 0 | 0 | 5215 | 577 | 21 |
| SRR5127204 | 68 | 78 | 28 | 54 | 248 | 725 | 870 | 1470 | 443 | 0 | 23 | 0 | 0 | 0 | 5660 | 221530 | 13566 | 52 | 55 | 4778 | 397 | 0 | 0 | 534 | 147 | 429 | 2941 | 174 | 0 | 383 | 0 | 0 | 0 | 0 | 1115 | 447 | 0 |
| SRR5127205 | 77 | 29 | 22 | 72 | 73 | 189 | 140 | 1060 | 158 | 0 | 15 | 0 | 0 | 0 | 7967 | 223136 | 27574 | 30 | 152 | 30753 | 408 | 0 | 0 | 182 | 49 | 177 | 1249 | 69 | 0 | 201 | 0 | 12 | 0 | 0 | 1410 | 0 | 0 |
| SRR5127206 | 77 | 58 | 40 | 57 | 17 | 115 | 78 | 328 | 62 | 14 | 21 | 0 | 0 | 0 | 31405 | 184071 | 56187 | 31 | 100 | 18807 | 755 | 0 | 0 | 94 | 47 | 10 | 47 | 26 | 0 | 17 | 0 | 0 | 0 | 0 | 628 | 0 | 0 |
| SRR5127207 | 0 | 0 | 0 | 0 | 15 | 90 | 842 | 150 | 140 | 0 | 0 | 0 | 0 | 0 | 51625 | 36736 | 99150 | 70 | 140 | 10447 | 9465 | 0 | 0 | 22 | 16 | 17 | 74 | 0 | 0 | 15 | 0 | 10 | 0 | 0 | 275 | 0 | 0 |
| SRR5127208 | 20 | 13 | 52 | 28 | 118 | 46 | 41 | 113 | 50 | 24 | 23 | 0 | 0 | 0 | 19142 | 185874 | 24571 | 0 | 131 | 18776 | 769 | 0 | 0 | 21 | 21 | 35 | 131 | 93 | 0 | 19 | 0 | 0 | 0 | 0 | 2863 | 621 | 0 |
| SRR5127209 | 85 | 76 | 29 | 83 | 382 | 305 | 120 | 918 | 54 | 20 | 69 | 0 | 0 | 0 | 9189 | 27192 | 106243 | 34 | 274 | 7514 | 4195 | 0 | 0 | 238 | 129 | 253 | 1283 | 92 | 0 | 316 | 0 | 50 | 0 | 0 | 5222 | 0 | 0 |
| SRR5127210 | 22 | 26 | 80 | 18 | 16 | 133 | 186 | 284 | 99 | 0 | 27 | 0 | 0 | 0 | 10737 | 109483 | 33961 | 17 | 46 | 7684 | 754 | 0 | 0 | 18 | 10 | 36 | 180 | 91 | 0 | 22 | 0 | 0 | 0 | 0 | 1664 | 0 | 0 |
| SRR5127211 | 0 | 0 | 15 | 11 | 13 | 137 | 673 | 436 | 122 | 0 | 33 | 0 | 0 | 0 | 28747 | 43441 | 107253 | 0 | 120 | 16717 | 3251 | 0 | 0 | 10 | 0 | 48 | 17 | 46 | 0 | 12 | 0 | 15 | 0 | 0 | 6641 | 0 | 0 |
| SRR5127212 | 57 | 98 | 25 | 30 | 123 | 287 | 934 | 559 | 295 | 11 | 28 | 0 | 0 | 0 | 16067 | 189069 | 19401 | 37 | 106 | 28100 | 3668 | 0 | 0 | 187 | 86 | 223 | 1045 | 55 | 0 | 431 | 0 | 0 | 0 | 0 | 3403 | 538 | 0 |
| SRR5127213 | 15 | 11 | 79 | 131 | 86 | 50 | 116 | 150 | 83 | 0 | 58 | 0 | 0 | 0 | 10706 | 51552 | 31433 | 17 | 210 | 13924 | 2941 | 0 | 0 | 118 | 102 | 18 | 59 | 63 | 0 | 19 | 0 | 0 | 0 | 0 | 2401 | 0 | 0 |
| SRR5127214 | 68 | 76 | 32 | 52 | 252 | 87 | 654 | 420 | 113 | 0 | 43 | 0 | 0 | 0 | 24874 | 100664 | 55094 | 17 | 156 | 18087 | 5404 | 0 | 0 | 109 | 74 | 238 | 1272 | 69 | 0 | 297 | 0 | 0 | 0 | 0 | 1437 | 0 | 0 |
| SRR5127215 | 71 | 51 | 44 | 56 | 225 | 68 | 194 | 171 | 85 | 0 | 20 | 0 | 0 | 0 | 10333 | 234637 | 13766 | 0 | 60 | 22363 | 236 | 0 | 0 | 61 | 113 | 156 | 515 | 30 | 0 | 170 | 0 | 0 | 0 | 0 | 212 | 0 | 0 |
| SRR5127216 | 102 | 85 | 43 | 18 | 194 | 174 | 78 | 399 | 72 | 12 | 39 | 0 | 0 | 0 | 8876 | 142908 | 12214 | 33 | 182 | 21744 | 588 | 0 | 0 | 287 | 133 | 215 | 1028 | 74 | 0 | 315 | 0 | 0 | 0 | 0 | 4426 | 498 | 0 |
| SRR5127217 | 52 | 38 | 33 | 58 | 10 | 45 | 277 | 99 | 62 | 0 | 22 | 0 | 0 | 0 | 7345 | 213134 | 11492 | 19 | 63 | 26375 | 115 | 0 | 0 | 113 | 46 | 104 | 29 | 23 | 0 | 15 | 0 | 0 | 0 | 0 | 172 | 0 | 0 |
| SRR5127218 | 98 | 13 | 24 | 36 | 15 | 224 | 182 | 556 | 112 | 22 | 28 | 0 | 0 | 0 | 10954 | 71434 | 52727 | 32 | 174 | 22821 | 1077 | 0 | 0 | 13 | 66 | 14 | 113 | 73 | 0 | 28 | 0 | 0 | 0 | 0 | 1224 | 0 | 0 |
| SRR5127219 | 0 | 0 | 0 | 25 | 18 | 950 | 1057 | 2497 | 253 | 56 | 213 | 0 | 0 | 0 | 20561 | 24081 | 212384 | 10 | 55 | 16704 | 15741 | 0 | 0 | 141 | 31 | 113 | 188 | 58 | 0 | 601 | 0 | 2749 | 0 | 0 | 822 | 754 | 0 |
| SRR5127220 | 156 | 255 | 29 | 47 | 110 | 76 | 62 | 135 | 50 | 19 | 31 | 0 | 0 | 0 | 45985 | 35204 | 60098 | 41 | 64 | 13873 | 1105 | 0 | 0 | 217 | 155 | 325 | 1361 | 67 | 0 | 342 | 0 | 0 | 0 | 0 | 861 | 0 | 0 |
| SRR5127221 | 12 | 18 | 70 | 71 | 135 | 38 | 135 | 112 | 54 | 28 | 24 | 0 | 0 | 0 | 62968 | 66965 | 92292 | 54 | 208 | 8801 | 7261 | 0 | 0 | 142 | 120 | 16 | 73 | 84 | 0 | 13 | 0 | 0 | 0 | 0 | 546 | 0 | 0 |
| SRR5127222 | 93 | 123 | 24 | 48 | 287 | 154 | 482 | 588 | 151 | 0 | 22 | 0 | 0 | 0 | 16357 | 124091 | 21047 | 37 | 77 | 9965 | 526 | 0 | 0 | 214 | 101 | 254 | 1427 | 74 | 0 | 254 | 0 | 0 | 0 | 0 | 484 | 0 | 0 |
| SRR5127223 | 159 | 357 | 85 | 72 | 145 | 90 | 353 | 203 | 87 | 47 | 23 | 0 | 0 | 0 | 65888 | 50785 | 74343 | 33 | 109 | 9898 | 2046 | 0 | 0 | 203 | 100 | 196 | 1025 | 100 | 0 | 396 | 0 | 0 | 0 | 0 | 527 | 0 | 0 |
| SRR5127224 | 13 | 27 | 21 | 37 | 23 | 344 | 197 | 922 | 106 | 0 | 84 | 0 | 0 | 0 | 7550 | 98433 | 20823 | 26 | 49 | 6185 | 1217 | 0 | 0 | 18 | 10 | 35 | 222 | 11 | 0 | 93 | 0 | 0 | 0 | 0 | 3712 | 33 | 0 |
| SRR5127225 | 297 | 302 | 111 | 166 | 132 | 107 | 325 | 266 | 138 | 0 | 52 | 15 | 0 | 0 | 22717 | 118583 | 73810 | 11 | 117 | 15986 | 1267 | 0 | 0 | 246 | 123 | 367 | 2062 | 114 | 0 | 324 | 0 | 0 | 0 | 0 | 2002 | 0 | 0 |
| SRR5127226 | 15 | 16 | 46 | 52 | 24 | 214 | 251 | 529 | 149 | 0 | 23 | 0 | 0 | 0 | 19103 | 164893 | 38502 | 25 | 93 | 9479 | 737 | 0 | 0 | 18 | 70 | 26 | 132 | 81 | 0 | 26 | 0 | 0 | 0 | 0 | 835 | 0 | 0 |
| SRR5127227 | 98 | 91 | 45 | 77 | 146 | 140 | 85 | 419 | 132 | 0 | 31 | 0 | 0 | 0 | 12407 | 141265 | 35480 | 31 | 140 | 14894 | 345 | 0 | 0 | 206 | 102 | 253 | 1251 | 69 | 0 | 209 | 0 | 12 | 0 | 0 | 1289 | 0 | 0 |
| SRR5127228 | 104 | 60 | 286 | 201 | 159 | 308 | 474 | 1102 | 94 | 0 | 192 | 0 | 0 | 0 | 34624 | 18713 | 108046 | 0 | 103 | 2332 | 5552 | 0 | 0 | 434 | 43 | 118 | 287 | 94 | 0 | 390 | 0 | 34 | 0 | 0 | 5808 | 0 | 11 |
| SRR5127229 | 138 | 23 | 43 | 68 | 15 | 46 | 74 | 188 | 50 | 0 | 53 | 0 | 0 | 0 | 6766 | 159197 | 15273 | 50 | 64 | 5513 | 2487 | 0 | 0 | 135 | 86 | 20 | 121 | 83 | 0 | 56 | 0 | 0 | 0 | 0 | 1128 | 0 | 0 |
| SRR5127230 | 110 | 116 | 57 | 86 | 20 | 315 | 458 | 723 | 240 | 0 | 23 | 0 | 0 | 0 | 10196 | 301369 | 14060 | 22 | 97 | 9268 | 505 | 0 | 0 | 26 | 129 | 26 | 141 | 126 | 0 | 19 | 0 | 0 | 0 | 0 | 1382 | 462 | 0 |
| SRR5127231 | 0 | 0 | 12 | 17 | 21 | 93 | 353 | 318 | 66 | 0 | 117 | 0 | 0 | 0 | 9177 | 13195 | 66540 | 22 | 84 | 6296 | 548 | 0 | 0 | 19 | 22 | 217 | 738 | 189 | 0 | 415 | 0 | 1003 | 0 | 0 | 5888 | 0 | 0 |
| SRR5127232 | 0 | 0 | 0 | 0 | 14 | 0 | 0 | 18 | 0 | 0 | 609 | 0 | 0 | 0 | 11589 | 19065 | 58819 | 0 | 0 | 2357 | 868 | 0 | 0 | 0 | 0 | 0 | 0 | 0 | 0 | 461 | 0 | 35 | 0 | 0 | 8972 | 0 | 0 |
| SRR5127233 | 53 | 45 | 27 | 37 | 10 | 75 | 115 | 159 | 70 | 98 | 24 | 0 | 0 | 0 | 81954 | 125421 | 124787 | 17 | 215 | 14681 | 7058 | 0 | 0 | 103 | 85 | 12 | 55 | 38 | 0 | 15 | 0 | 0 | 0 | 0 | 201 | 0 | 0 |
| SRR5127234 | 36 | 40 | 11 | 56 | 129 | 94 | 57 | 222 | 25 | 21 | 49 | 11 | 0 | 0 | 35570 | 52115 | 224669 | 0 | 274 | 6978 | 13227 | 0 | 0 | 120 | 11 | 17 | 76 | 39 | 0 | 22 | 0 | 21 | 0 | 0 | 1457 | 104 | 0 |
| SRR5127235 | 190 | 220 | 106 | 136 | 123 | 122 | 83 | 269 | 32 | 0 | 33 | 0 | 0 | 0 | 16493 | 67988 | 30559 | 26 | 83 | 7233 | 1481 | 0 | 0 | 220 | 194 | 603 | 2788 | 73 | 0 | 269 | 0 | 0 | 0 | 0 | 1167 | 976 | 0 |
| SRR5127236 | 11 | 11 | 0 | 0 | 11 | 99 | 164 | 328 | 113 | 0 | 98 | 0 | 0 | 0 | 7007 | 68957 | 21677 | 26 | 115 | 7611 | 158 | 0 | 0 | 11 | 0 | 18 | 95 | 0 | 0 | 1339 | 0 | 0 | 0 | 0 | 679 | 0 | 0 |
| SRR5136285 | 0 | 0 | 0 | 0 | 0 | 0 | 131 | 131 | 0 | 850 | 17 | 0 | 0 | 0 | 480179 | 152457 | 617480 | 0 | 0 | 525 | 9976 | 0 | 0 | 0 | 0 | 0 | 0 | 0 | 0 | 29 | 0 | 0 | 0 | 0 | 612 | 0 | 14 |
| SRR5136286 | 0 | 0 | 0 | 0 | 0 | 0 | 257 | 257 | 0 | 293 | 11 | 0 | 0 | 0 | 450225 | 159206 | 611576 | 0 | 0 | 514 | 10533 | 0 | 0 | 0 | 0 | 0 | 0 | 0 | 0 | 11 | 0 | 0 | 0 | 0 | 1148 | 0 | 0 |
| SRR5136287 | 0 | 0 | 0 | 0 | 0 | 243 | 122 | 243 | 122 | 855 | 10 | 0 | 0 | 0 | 458141 | 156125 | 612465 | 0 | 0 | 973 | 6936 | 0 | 0 | 0 | 0 | 0 | 0 | 0 | 0 | 16 | 0 | 0 | 0 | 0 | 926 | 0 | 13 |
| SRR5136288 | 0 | 0 | 0 | 0 | 0 | 124 | 249 | 124 | 249 | 880 | 16 | 0 | 0 | 0 | 449304 | 156869 | 620494 | 0 | 124 | 249 | 9319 | 0 | 0 | 0 | 0 | 0 | 0 | 0 | 0 | 24 | 0 | 0 | 0 | 0 | 650 | 0 | 0 |
| SRR5136289 | 0 | 0 | 0 | 0 | 0 | 122 | 0 | 365 | 122 | 780 | 13 | 0 | 0 | 0 | 430381 | 67871 | 254777 | 0 | 122 | 609 | 9380 | 0 | 0 | 0 | 0 | 0 | 0 | 0 | 0 | 19 | 0 | 0 | 0 | 0 | 247 | 0 | 11 |
| SRR5136290 | 0 | 0 | 0 | 0 | 0 | 134 | 0 | 0 | 0 | 272 | 13 | 0 | 0 | 0 | 471557 | 69505 | 255188 | 0 | 0 | 669 | 8700 | 0 | 0 | 0 | 0 | 0 | 0 | 0 | 0 | 23 | 0 | 0 | 0 | 0 | 470 | 0 | 0 |
| SRR5136291 | 0 | 0 | 0 | 0 | 0 | 0 | 380 | 127 | 127 | 823 | 0 | 0 | 0 | 0 | 482204 | 157451 | 607683 | 0 | 0 | 760 | 10133 | 0 | 0 | 0 | 0 | 0 | 0 | 0 | 0 | 25 | 0 | 0 | 0 | 0 | 1892 | 0 | 0 |
| SRR5136292 | 0 | 0 | 0 | 0 | 0 | 247 | 0 | 247 | 124 | 294 | 20 | 0 | 0 | 0 | 452625 | 163841 | 598954 | 0 | 124 | 618 | 10006 | 0 | 0 | 0 | 0 | 0 | 0 | 0 | 0 | 15 | 0 | 0 | 0 | 0 | 1268 | 0 | 15 |
| SRR5136293 | 0 | 0 | 0 | 0 | 27 | 16 | 0 | 24 | 0 | 165 | 23 | 0 | 0 | 0 | 604055 | 235782 | 607728 | 0 | 0 | 105 | 2474 | 0 | 0 | 0 | 0 | 0 | 0 | 0 | 0 | 60 | 0 | 0 | 0 | 0 | 243 | 0 | 0 |
| SRR5136294 | 0 | 0 | 0 | 0 | 13 | 0 | 0 | 0 | 16 | 38 | 32 | 0 | 0 | 0 | 604421 | 227998 | 599957 | 0 | 0 | 107 | 2114 | 0 | 0 | 0 | 0 | 0 | 0 | 0 | 0 | 24 | 0 | 0 | 0 | 0 | 90 | 0 | 0 |
| SRR5136295 | 0 | 0 | 0 | 0 | 12 | 0 | 0 | 31 | 0 | 37 | 12 | 0 | 0 | 0 | 600778 | 233759 | 609259 | 0 | 16 | 109 | 2359 | 0 | 0 | 0 | 0 | 0 | 0 | 0 | 0 | 36 | 0 | 0 | 0 | 0 | 195 | 0 | 0 |
| SRR5136296 | 0 | 0 | 0 | 0 | 23 | 0 | 0 | 16 | 0 | 42 | 13 | 0 | 0 | 0 | 604737 | 233412 | 611050 | 0 | 0 | 82 | 2416 | 0 | 0 | 0 | 0 | 0 | 0 | 0 | 0 | 47 | 0 | 0 | 0 | 0 | 138 | 0 | 0 |
| SRR5136297 | 0 | 0 | 0 | 0 | 26 | 14 | 0 | 14 | 0 | 180 | 17 | 0 | 0 | 0 | 601295 | 229343 | 617001 | 0 | 0 | 71 | 2724 | 0 | 0 | 0 | 0 | 0 | 0 | 0 | 0 | 28 | 0 | 0 | 0 | 0 | 145 | 0 | 0 |
| SRR5136298 | 0 | 0 | 0 | 0 | 29 | 22 | 0 | 0 | 0 | 172 | 22 | 0 | 0 | 0 | 605823 | 227696 | 604979 | 0 | 0 | 65 | 2383 | 0 | 0 | 0 | 0 | 0 | 0 | 0 | 0 | 39 | 0 | 0 | 0 | 0 | 154 | 0 | 0 |
| SRR5136299 | 0 | 0 | 0 | 0 | 28 | 21 | 0 | 0 | 0 | 44 | 31 | 0 | 0 | 0 | 605078 | 225845 | 604508 | 0 | 0 | 49 | 2382 | 0 | 0 | 0 | 0 | 0 | 0 | 0 | 0 | 33 | 0 | 0 | 0 | 0 | 178 | 26 | 0 |
| SRR5136300 | 0 | 0 | 0 | 0 | 27 | 14 | 14 | 14 | 0 | 186 | 17 | 0 | 0 | 0 | 602913 | 232046 | 614856 | 0 | 0 | 29 | 2686 | 0 | 0 | 0 | 0 | 0 | 0 | 0 | 0 | 41 | 0 | 0 | 0 | 0 | 205 | 0 | 0 |
| SRR5139121 | 11 | 64 | 64 | 104 | 13 | 317 | 226 | 1601 | 266 | 14 | 15 | 0 | 0 | 0 | 3814 | 12719 | 15446 | 68 | 85 | 3723 | 419 | 0 | 0 | 11 | 90 | 24 | 82 | 83 | 0 | 41 | 0 | 34 | 0 | 0 | 2997 | 0 | 0 |
| SRR5145238 | 51 | 18 | 21 | 12 | 41 | 4748 | 1778 | 12302 | 18846 | 2707 | 194 | 21 | 0 | 0 | 23493 | 16939 | 23473 | 1578 | 1493 | 3363 | 1360 | 38 | 16 | 103 | 71 | 34 | 27 | 39 | 0 | 580 | 117 | 514 | 0 | 17 | 7792 | 33 | 0 |
| SRR5145239 | 1050 | 255 | 270 | 95 | 420 | 1423 | 811 | 5128 | 10068 | 816 | 1197 | 34 | 15 | 0 | 3365 | 8297 | 5912 | 339 | 200 | 1053 | 1028 | 0 | 0 | 528 | 352 | 279 | 332 | 84 | 0 | 2265 | 0 | 97 | 0 | 0 | 2675 | 0 | 0 |
| SRR5145240 | 82 | 23 | 0 | 0 | 74 | 2361 | 621 | 3437 | 6633 | 27028 | 269 | 0 | 0 | 0 | 195730 | 29427 | 135652 | 927 | 225 | 836 | 81050 | 1941 | 493 | 47 | 64 | 23 | 19 | 0 | 0 | 2245 | 5194 | 1027 | 11 | 63 | 4360 | 15 | 0 |
| SRR5145241 | 773 | 193 | 43 | 36 | 195 | 763 | 410 | 2405 | 5692 | 48011 | 689 | 13 | 23 | 26 | 386549 | 29485 | 344115 | 469 | 794 | 1423 | 26844 | 0 | 0 | 116 | 204 | 110 | 70 | 0 | 0 | 1729 | 0 | 0 | 0 | 74 | 2651 | 0 | 0 |
| SRR5145242 | 26 | 0 | 0 | 0 | 157 | 3848 | 1110 | 5984 | 12665 | 46009 | 329 | 0 | 0 | 0 | 204440 | 33262 | 215364 | 131 | 645 | 1198 | 47571 | 2658 | 663 | 32 | 58 | 21 | 20 | 0 | 0 | 2247 | 7596 | 633 | 14 | 43 | 6521 | 41 | 0 |
| SRR5145243 | 23 | 15 | 0 | 0 | 79 | 10636 | 6067 | 30819 | 48324 | 1234 | 39 | 14 | 15 | 0 | 8829 | 6316 | 5557 | 1758 | 450 | 1213 | 643 | 0 | 0 | 49 | 18 | 0 | 12 | 0 | 0 | 142 | 0 | 15 | 0 | 12 | 7324 | 97 | 0 |
| SRR5145244 | 18 | 10 | 0 | 0 | 146 | 9154 | 2106 | 12600 | 14369 | 6495 | 548 | 0 | 0 | 0 | 175228 | 18319 | 127653 | 798 | 1651 | 1595 | 41818 | 1486 | 364 | 92 | 110 | 41 | 43 | 0 | 0 | 2810 | 4998 | 393 | 10 | 36 | 9236 | 87 | 0 |
| SRR5145245 | 225 | 85 | 92 | 66 | 290 | 2929 | 1963 | 13032 | 28727 | 1679 | 1349 | 22 | 0 | 0 | 35812 | 86151 | 58291 | 1041 | 1167 | 4454 | 4003 | 0 | 0 | 202 | 328 | 209 | 305 | 140 | 0 | 2162 | 0 | 20 | 0 | 0 | 5464 | 40 | 0 |
| SRR5163971 | 85 | 36 | 16 | 71 | 75 | 298 | 72 | 534 | 1601 | 61 | 503 | 26 | 0 | 0 | 18154 | 206559 | 514267 | 109 | 1366 | 16209 | 6558 | 0 | 0 | 127 | 36 | 26 | 73 | 48 | 0 | 156 | 0 | 18 | 0 | 24 | 409 | 82 | 0 |
| SRR5163972 | 118 | 51 | 67 | 82 | 82 | 283 | 53 | 689 | 1343 | 78 | 431 | 16 | 0 | 0 | 16058 | 206194 | 511007 | 106 | 1590 | 16905 | 6147 | 0 | 0 | 118 | 59 | 59 | 98 | 63 | 0 | 157 | 0 | 18 | 0 | 24 | 433 | 81 | 0 |
| SRR5163973 | 18 | 42 | 64 | 38 | 96 | 197 | 16 | 119 | 390 | 74 | 161 | 87 | 0 | 0 | 26400 | 193881 | 623510 | 1005 | 1005 | 4249 | 14404 | 0 | 11 | 72 | 30 | 40 | 68 | 11 | 0 | 27 | 0 | 0 | 0 | 47 | 416 | 393 | 0 |
| SRR5163974 | 281 | 87 | 48 | 39 | 111 | 143 | 19 | 162 | 314 | 58 | 2130 | 53 | 0 | 0 | 25720 | 194064 | 625148 | 998 | 894 | 4554 | 15175 | 0 | 0 | 48 | 39 | 29 | 68 | 15 | 0 | 411 | 0 | 0 | 19 | 24 | 433 | 377 | 0 |
| SRR5163975 | 547 | 175 | 52 | 44 | 83 | 343 | 31 | 327 | 802 | 61 | 868 | 110 | 0 | 0 | 82182 | 187292 | 419612 | 1246 | 888 | 7751 | 8553 | 0 | 13 | 62 | 14 | 26 | 52 | 19 | 0 | 374 | 0 | 0 | 0 | 58 | 3776 | 33 | 0 |
| SRR5163976 | 667 | 181 | 52 | 65 | 91 | 321 | 80 | 441 | 1062 | 62 | 836 | 84 | 0 | 0 | 84395 | 184787 | 419221 | 1282 | 942 | 7354 | 8716 | 0 | 0 | 55 | 29 | 0 | 39 | 0 | 0 | 327 | 0 | 0 | 0 | 97 | 3606 | 32 | 0 |
| SRR5163977 | 44 | 15 | 50 | 10 | 18 | 963 | 24 | 1373 | 3468 | 28 | 276 | 15 | 0 | 0 | 50959 | 154329 | 375559 | 482 | 1108 | 5057 | 8092 | 0 | 0 | 16 | 63 | 10 | 24 | 36 | 0 | 60 | 0 | 0 | 0 | 63 | 4682 | 282 | 0 |

### Supp Table S8.3.Columns_BX_to_DH

| | | | | | | | | | | | | | | | | | | | | | | | | | | | | | | | | | | | | | |
|---|---|---|---|---|---|---|---|---|---|---|---|---|---|---|---|---|---|---|---|---|---|---|---|---|---|---|---|---|---|---|---|---|---|---|---|---|---|
| SRR5163978 | 251 | 98 | 41 | 71 | 99 | 1095 | 182 | 1156 | 3163 | 159 | 1785 | 79 | 0 | 0 | 54712 | 155901 | 369401 | 274 | 760 | 5201 | 8759 | 0 | 0 | 91 | 45 | 60 | 162 | 18 | 0 | 385 | 0 | 0 | 0 | 36 | 4483 | 292 | 0 |
| SRR5163979 | 53 | 41 | 12 | 33 | 76 | 435 | 41 | 248 | 642 | 38 | 634 | 78 | 0 | 0 | 56453 | 139491 | 635147 | 352 | 497 | 5321 | 5466 | 0 | 0 | 36 | 33 | 25 | 35 | 0 | 0 | 197 | 0 | 21 | 0 | 38 | 1616 | 62 | 0 |
| SRR5163980 | 70 | 49 | 0 | 22 | 92 | 336 | 0 | 218 | 739 | 22 | 58 | 57 | 0 | 0 | 55756 | 139418 | 629773 | 235 | 655 | 4718 | 5608 | 0 | 0 | 65 | 22 | 22 | 46 | 19 | 0 | 24 | 0 | 0 | 0 | 65 | 1510 | 70 | 0 |
| SRR5163981 | 138 | 41 | 38 | 83 | 93 | 1202 | 117 | 830 | 1445 | 55 | 739 | 26 | 0 | 0 | 19880 | 265442 | 376841 | 243 | 719 | 60465 | 10177 | 0 | 0 | 110 | 47 | 64 | 221 | 54 | 0 | 299 | 0 | 34 | 0 | 36 | 204 | 75 | 0 |
| SRR5163982 | 180 | 50 | 108 | 74 | 130 | 257 | 23 | 174 | 507 | 74 | 71 | 66 | 0 | 0 | 39893 | 285052 | 423507 | 378 | 1143 | 14951 | 6388 | 0 | 0 | 27 | 150 | 103 | 20 | 45 | 0 | 17 | 0 | 11 | 0 | 52 | 822 | 1402 | 0 |
| SRR5163983 | 137 | 36 | 20 | 80 | 115 | 292 | 118 | 615 | 1743 | 56 | 271 | 32 | 0 | 0 | 18150 | 193693 | 604904 | 39 | 804 | 26500 | 3580 | 0 | 0 | 108 | 37 | 46 | 103 | 69 | 0 | 105 | 0 | 0 | 0 | 29 | 518 | 64 | 0 |
| SRR5163984 | 11 | 25 | 34 | 75 | 13 | 294 | 39 | 646 | 1782 | 80 | 28 | 25 | 0 | 0 | 16783 | 196120 | 602388 | 78 | 744 | 27162 | 3388 | 0 | 0 | 113 | 34 | 25 | 63 | 92 | 0 | 80 | 0 | 0 | 0 | 17 | 505 | 62 | 0 |
| SRR5163985 | 117 | 76 | 18 | 18 | 35 | 218 | 0 | 109 | 980 | 281 | 702 | 99 | 0 | 0 | 13827 | 494486 | 221928 | 109 | 2177 | 5335 | 8383 | 0 | 0 | 41 | 12 | 12 | 12 | 0 | 0 | 181 | 0 | 0 | 0 | 35 | 2386 | 62 | 0 |
| SRR5163986 | 140 | 56 | 0 | 19 | 28 | 420 | 84 | 504 | 1427 | 261 | 591 | 37 | 0 | 0 | 15778 | 494711 | 220414 | 84 | 839 | 5875 | 12421 | 0 | 0 | 56 | 0 | 14 | 23 | 0 | 0 | 182 | 0 | 168 | 0 | 33 | 1511 | 70 | 0 |
| SRR5163987 | 196 | 96 | 42 | 123 | 130 | 632 | 77 | 564 | 1289 | 55 | 797 | 56 | 0 | 0 | 23308 | 239924 | 397766 | 242 | 1595 | 45456 | 7740 | 0 | 0 | 130 | 49 | 34 | 56 | 32 | 0 | 224 | 0 | 0 | 0 | 43 | 400 | 65 | 0 |
| SRR5168444 | 44 | 10 | 0 | 0 | 65 | 4904 | 594 | 3991 | 2883 | 34 | 29 | 17 | 0 | 0 | 4528 | 567671 | 107984 | 223 | 17783 | 54780 | 8865 | 0 | 0 | 16 | 0 | 0 | 15 | 0 | 0 | 18 | 0 | 53 | 0 | 0 | 487 | 0 | 0 |
| SRR5168445 | 11 | 0 | 0 | 0 | 310 | 121 | 30 | 341 | 423 | 2271 | 562 | 50 | 0 | 0 | 19677 | 176280 | 150316 | 269 | 13273 | 36654 | 2176 | 0 | 0 | 23 | 49 | 43 | 22 | 29 | 12 | 2018 | 0 | 16 | 0 | 16 | 1162 | 12 | 0 |
| SRR5168447 | 11 | 0 | 0 | 0 | 381 | 8629 | 985 | 9080 | 11043 | 808 | 380 | 0 | 0 | 0 | 15120 | 67813 | 307782 | 576 | 11981 | 41309 | 3765 | 0 | 0 | 93 | 55 | 28 | 19 | 0 | 0 | 3792 | 0 | 70 | 0 | 17 | 16447 | 15 | 0 |
| SRR5168448 | 58 | 29 | 0 | 0 | 66 | 489 | 89 | 764 | 902 | 657 | 637 | 15 | 0 | 0 | 10449 | 395579 | 242670 | 28 | 5368 | 92013 | 10828 | 0 | 0 | 46 | 52 | 18 | 45 | 0 | 0 | 373 | 0 | 0 | 0 | 47 | 660 | 0 | 0 |
| SRR5168451 | 0 | 0 | 0 | 0 | 75 | 540 | 44 | 718 | 625 | 1794 | 516 | 18 | 0 | 0 | 46416 | 28016 | 641352 | 273 | 4489 | 83189 | 8212 | 0 | 0 | 30 | 77 | 45 | 24 | 22 | 0 | 1293 | 0 | 25 | 0 | 166 | 2620 | 0 | 0 |
| SRR5168454 | 16 | 14 | 0 | 0 | 69 | 1072 | 276 | 1263 | 694 | 400 | 51 | 0 | 0 | 0 | 9704 | 48689 | 83321 | 793 | 427 | 7401 | 8253 | 36 | 107 | 0 | 0 | 0 | 0 | 13 | 0 | 1841 | 0 | 11 | 0 | 0 | 1291 | 10 | 0 |
| SRR5168458 | 10 | 0 | 0 | 0 | 76 | 425 | 183 | 780 | 1898 | 672 | 46 | 13 | 0 | 0 | 31784 | 89091 | 189096 | 589 | 9012 | 131619 | 2214 | 19 | 68 | 13 | 12 | 15 | 0 | 10 | 0 | 143 | 0 | 12 | 0 | 0 | 833 | 0 | 0 |
| SRR5168460 | 0 | 0 | 0 | 0 | 36 | 82 | 14 | 103 | 70 | 500 | 13 | 0 | 0 | 0 | 9027 | 225209 | 152235 | 178 | 1166 | 14252 | 3936 | 34 | 21 | 0 | 0 | 0 | 0 | 0 | 0 | 79 | 0 | 17 | 17 | 40 | 1076 | 0 | 0 |
| SRR5168465 | 22 | 11 | 0 | 0 | 45 | 1824 | 763 | 2492 | 6974 | 299 | 11 | 19 | 0 | 0 | 10058 | 164414 | 204976 | 601 | 1469 | 28862 | 1831 | 0 | 0 | 69 | 65 | 50 | 55 | 0 | 0 | 204 | 0 | 0 | 0 | 0 | 7030 | 10 | 0 |
| SRR5168467 | 0 | 0 | 0 | 0 | 82 | 217 | 14 | 318 | 123 | 1011 | 90 | 0 | 0 | 0 | 195955 | 120698 | 742113 | 122 | 1868 | 7651 | 1127 | 0 | 18 | 0 | 11 | 0 | 0 | 0 | 0 | 85 | 0 | 0 | 0 | 45 | 823 | 14 | 0 |
| SRR5168471 | 12 | 0 | 0 | 0 | 340 | 268 | 67 | 441 | 687 | 1492 | 34 | 0 | 0 | 0 | 31457 | 332248 | 389830 | 32 | 232 | 4516 | 5501 | 0 | 20 | 18 | 57 | 52 | 0 | 33 | 0 | 761 | 0 | 0 | 0 | 0 | 1208 | 0 | 0 |
| SRR5168475 | 0 | 0 | 0 | 0 | 181 | 1272 | 126 | 674 | 595 | 1014 | 56 | 0 | 0 | 0 | 6245 | 258939 | 88711 | 51 | 349 | 11384 | 1206 | 0 | 0 | 19 | 17 | 0 | 0 | 0 | 0 | 684 | 0 | 16 | 0 | 27 | 5007 | 0 | 0 |
| SRR5171728 | 73 | 25 | 103 | 97 | 0 | 77 | 192 | 384 | 115 | 0 | 860 | 0 | 0 | 0 | 5377 | 13376 | 34065 | 38 | 0 | 3879 | 1920 | 0 | 0 | 0 | 0 | 0 | 10 | 42 | 0 | 1160 | 0 | 0 | 0 | 0 | 3121 | 0 | 0 |
| SRR5171729 | 96 | 40 | 172 | 168 | 29 | 130 | 228 | 260 | 65 | 0 | 1040 | 0 | 0 | 0 | 25238 | 15528 | 123665 | 0 | 423 | 8651 | 3252 | 0 | 0 | 0 | 0 | 0 | 0 | 49 | 0 | 1593 | 0 | 65 | 0 | 0 | 1405 | 0 | 0 |
| SRR5171730 | 13 | 32 | 25 | 27 | 0 | 59 | 297 | 119 | 0 | 0 | 136 | 0 | 0 | 0 | 9810 | 18963 | 85536 | 0 | 59 | 3567 | 2913 | 0 | 0 | 22 | 11 | 18 | 22 | 100 | 0 | 222 | 0 | 0 | 0 | 0 | 3026 | 0 | 0 |
| SRR5171731 | 25 | 18 | 55 | 60 | 20 | 159 | 119 | 398 | 80 | 0 | 573 | 0 | 0 | 0 | 5649 | 22733 | 49911 | 0 | 40 | 6404 | 2864 | 0 | 0 | 0 | 0 | 25 | 20 | 56 | 0 | 904 | 0 | 40 | 0 | 0 | 2032 | 0 | 0 |
| SRR5171732 | 36 | 27 | 41 | 46 | 13 | 95 | 0 | 191 | 0 | 0 | 433 | 0 | 0 | 0 | 3907 | 16854 | 36517 | 0 | 95 | 2668 | 1525 | 0 | 0 | 0 | 0 | 0 | 13 | 41 | 0 | 609 | 0 | 0 | 0 | 0 | 2809 | 0 | 0 |
| SRR5171733 | 66 | 30 | 86 | 108 | 13 | 42 | 104 | 63 | 0 | 0 | 1227 | 0 | 0 | 0 | 9610 | 19768 | 80209 | 21 | 208 | 3544 | 2939 | 0 | 0 | 0 | 0 | 11 | 11 | 20 | 0 | 1290 | 0 | 42 | 0 | 0 | 1969 | 0 | 0 |
| SRR5195030 | 0 | 10 | 0 | 0 | 15 | 122 | 68 | 525 | 29 | 0 | 31 | 11 | 0 | 0 | 103780 | 12639 | 146916 | 0 | 159 | 3400 | 1139 | 0 | 0 | 62 | 0 | 0 | 0 | 0 | 0 | 66 | 0 | 16 | 0 | 0 | 547 | 0 | 0 |
| SRR5195032 | 0 | 0 | 0 | 0 | 17 | 118 | 113 | 798 | 122 | 0 | 15 | 0 | 0 | 0 | 23165 | 20669 | 85511 | 0 | 64 | 3207 | 708 | 0 | 0 | 94 | 11 | 38 | 197 | 0 | 0 | 44 | 0 | 18 | 0 | 0 | 975 | 0 | 0 |
| SRR5195033 | 0 | 0 | 0 | 0 | 15 | 103 | 188 | 570 | 76 | 0 | 28 | 0 | 0 | 0 | 36841 | 20572 | 78771 | 0 | 262 | 7429 | 5054 | 0 | 0 | 50 | 14 | 52 | 118 | 12 | 0 | 122 | 0 | 36 | 0 | 0 | 2317 | 0 | 0 |
| SRR5195034 | 0 | 0 | 0 | 16 | 38 | 47 | 72 | 475 | 61 | 0 | 64 | 0 | 0 | 0 | 53267 | 5846 | 112754 | 0 | 93 | 4354 | 1821 | 0 | 0 | 91 | 82 | 260 | 872 | 204 | 0 | 111 | 0 | 0 | 0 | 0 | 347 | 0 | 0 |
| SRR5224010 | 80 | 76 | 11 | 10 | 1253 | 3756 | 1762 | 9299 | 22114 | 6753 | 382 | 27 | 14 | 0 | 1483 | 31411 | 48915 | 1356 | 199 | 861 | 19642 | 0 | 0 | 60 | 118 | 87 | 87 | 19 | 0 | 1150 | 12 | 215 | 0 | 0 | 12128 | 0 | 0 |
| SRR5224011 | 13 | 10 | 0 | 0 | 257 | 2676 | 1187 | 5684 | 12099 | 858 | 87 | 25 | 0 | 0 | 4984 | 34510 | 73524 | 981 | 88 | 582 | 29978 | 0 | 0 | 38 | 18 | 19 | 21 | 22 | 0 | 276 | 10 | 125 | 0 | 0 | 15992 | 0 | 0 |
| SRR5224012 | 32 | 17 | 21 | 29 | 347 | 1395 | 756 | 3221 | 5518 | 736 | 52 | 29 | 0 | 0 | 9371 | 76794 | 241877 | 460 | 117 | 708 | 13717 | 0 | 0 | 11 | 28 | 20 | 23 | 18 | 0 | 209 | 16 | 58 | 0 | 0 | 10217 | 0 | 0 |
| SRR5224013 | 22 | 33 | 0 | 0 | 116 | 1967 | 1062 | 4725 | 9326 | 147 | 102 | 17 | 15 | 0 | 10771 | 61452 | 23496 | 1671 | 427 | 1467 | 584 | 0 | 0 | 12 | 25 | 13 | 12 | 16 | 0 | 827 | 0 | 175 | 0 | 0 | 9605 | 0 | 0 |
| SRR5224014 | 25 | 28 | 0 | 0 | 149 | 1387 | 492 | 2337 | 4730 | 620 | 164 | 17 | 23 | 0 | 10403 | 21416 | 19869 | 2844 | 81 | 374 | 4579 | 0 | 0 | 15 | 11 | 28 | 18 | 0 | 0 | 553 | 0 | 228 | 0 | 0 | 7211 | 0 | 0 |
| SRR5224015 | 24 | 34 | 0 | 0 | 176 | 1699 | 1395 | 3797 | 11672 | 124 | 139 | 20 | 0 | 0 | 10084 | 52174 | 15448 | 2396 | 352 | 1261 | 481 | 0 | 0 | 17 | 24 | 15 | 17 | 28 | 0 | 1111 | 0 | 264 | 0 | 0 | 11421 | 0 | 0 |
| SRR5224016 | 31 | 44 | 10 | 12 | 166 | 3281 | 1917 | 5362 | 13704 | 127 | 124 | 20 | 17 | 0 | 8821 | 34805 | 15093 | 2229 | 104 | 766 | 395 | 0 | 0 | 26 | 17 | 16 | 17 | 19 | 0 | 469 | 0 | 133 | 0 | 0 | 12906 | 0 | 0 |
| SRR5224017 | 163 | 183 | 0 | 0 | 261 | 1675 | 171 | 1468 | 2850 | 793 | 532 | 74 | 73 | 16 | 15356 | 19900 | 16610 | 1220 | 153 | 180 | 1062 | 0 | 0 | 21 | 26 | 23 | 0 | 0 | 0 | 2368 | 0 | 324 | 0 | 0 | 8297 | 0 | 0 |
| SRR5224018 | 152 | 159 | 13 | 13 | 550 | 4349 | 3167 | 9760 | 23883 | 337 | 426 | 51 | 20 | 0 | 6635 | 54027 | 15533 | 1384 | 91 | 1028 | 357 | 0 | 0 | 16 | 39 | 31 | 15 | 0 | 0 | 1520 | 0 | 126 | 0 | 0 | 25028 | 0 | 0 |
| SRR5224019 | 22 | 34 | 0 | 0 | 96 | 3166 | 1529 | 5815 | 9856 | 164 | 109 | 15 | 23 | 0 | 16345 | 27040 | 17333 | 2222 | 371 | 913 | 856 | 0 | 0 | 13 | 29 | 14 | 18 | 17 | 0 | 799 | 0 | 145 | 0 | 0 | 11770 | 0 | 0 |
| SRR5224020 | 34 | 54 | 0 | 0 | 124 | 2163 | 991 | 3158 | 8881 | 197 | 92 | 25 | 10 | 12 | 10323 | 29179 | 14382 | 2158 | 155 | 756 | 792 | 0 | 0 | 11 | 19 | 11 | 12 | 14 | 0 | 621 | 0 | 305 | 0 | 0 | 11612 | 0 | 0 |
| SRR5224021 | 32 | 32 | 12 | 0 | 92 | 3607 | 1727 | 6872 | 11698 | 196 | 125 | 13 | 14 | 0 | 11016 | 55695 | 22976 | 1691 | 292 | 871 | 1064 | 0 | 0 | 28 | 22 | 12 | 12 | 14 | 0 | 872 | 0 | 162 | 0 | 0 | 15942 | 0 | 0 |
| SRR5224022 | 47 | 66 | 12 | 10 | 137 | 1797 | 1342 | 4271 | 13667 | 139 | 110 | 25 | 17 | 0 | 10752 | 36690 | 15079 | 2654 | 326 | 1239 | 792 | 0 | 0 | 19 | 16 | 12 | 13 | 16 | 0 | 729 | 0 | 309 | 0 | 0 | 12869 | 0 | 0 |
| SRR5224023 | 74 | 117 | 0 | 0 | 176 | 3961 | 1744 | 7704 | 11239 | 363 | 552 | 39 | 25 | 0 | 23061 | 28356 | 21708 | 2435 | 384 | 1227 | 1815 | 0 | 0 | 86 | 155 | 61 | 19 | 34 | 0 | 2643 | 0 | 213 | 0 | 0 | 9440 | 0 | 0 |
| SRR5224024 | 115 | 100 | 17 | 12 | 396 | 2199 | 1564 | 4709 | 11856 | 426 | 408 | 56 | 16 | 0 | 12631 | 31662 | 16043 | 2558 | 227 | 1117 | 654 | 0 | 0 | 35 | 63 | 59 | 15 | 20 | 0 | 1263 | 0 | 180 | 0 | 0 | 12516 | 0 | 0 |
| SRR5230550 | 0 | 0 | 0 | 0 | 729 | 107 | 0 | 107 | 121 | 1091 | 777 | 0 | 0 | 0 | 12410 | 150906 | 98894 | 33 | 0 | 332 | 2806 | 17 | 107 | 0 | 25 | 16 | 0 | 0 | 0 | 1643 | 30 | 23 | 0 | 0 | 97446 | 0 | 0 |
| SRR5230551 | 0 | 0 | 0 | 0 | 17 | 161 | 18 | 63 | 54 | 114 | 23 | 10 | 0 | 0 | 29328 | 133771 | 50651 | 72 | 18 | 278 | 673 | 0 | 14 | 0 | 0 | 0 | 0 | 0 | 0 | 62 | 11 | 0 | 0 | 0 | 21497 | 0 | 0 |
| SRR5230552 | 0 | 0 | 0 | 0 | 31 | 360 | 28 | 230 | 259 | 536 | 46 | 45 | 0 | 0 | 11751 | 44162 | 40965 | 287 | 0 | 194 | 1621 | 0 | 18 | 25 | 23 | 23 | 0 | 0 | 0 | 136 | 0 | 101 | 0 | 0 | 191630 | 0 | 0 |
| SRR5230553 | 0 | 0 | 0 | 0 | 63 | 231 | 0 | 171 | 217 | 62 | 77 | 0 | 0 | 0 | 33134 | 75599 | 51741 | 79 | 99 | 323 | 909 | 15 | 14 | 12 | 36 | 12 | 0 | 0 | 0 | 241 | 27 | 20 | 0 | 0 | 48957 | 0 | 0 |
| SRR5230554 | 0 | 0 | 0 | 0 | 622 | 249 | 27 | 99 | 84 | 134 | 1102 | 13 | 0 | 0 | 95757 | 179030 | 459691 | 30 | 67 | 303 | 4559 | 162 | 434 | 0 | 0 | 0 | 0 | 0 | 0 | 3058 | 265 | 20 | 0 | 0 | 18659 | 0 | 0 |
| SRR5230555 | 0 | 0 | 0 | 0 | 153 | 140 | 0 | 164 | 86 | 10 | 210 | 12 | 0 | 0 | 22302 | 136575 | 119031 | 16 | 47 | 359 | 2027 | 0 | 14 | 0 | 19 | 16 | 0 | 0 | 0 | 765 | 0 | 31 | 0 | 0 | 43470 | 0 | 0 |
| SRR5230556 | 0 | 0 | 0 | 0 | 73 | 60 | 10 | 0 | 0 | 53 | 99 | 36 | 0 | 0 | 5064 | 156835 | 41105 | 111 | 373 | 141 | 1391 | 25 | 39 | 0 | 0 | 0 | 0 | 0 | 0 | 162 | 11 | 30 | 0 | 0 | 40559 | 0 | 0 |
| SRR5230557 | 0 | 0 | 0 | 0 | 534 | 393 | 45 | 215 | 120 | 59 | 413 | 0 | 0 | 0 | 29344 | 86837 | 70676 | 54 | 124 | 157 | 2912 | 14 | 53 | 0 | 0 | 0 | 0 | 0 | 0 | 1763 | 30 | 12 | 0 | 0 | 29955 | 0 | 0 |
| SRR5230558 | 0 | 0 | 0 | 0 | 351 | 398 | 16 | 237 | 276 | 1765 | 232 | 11 | 0 | 0 | 27608 | 148715 | 229607 | 31 | 36 | 200 | 3831 | 190 | 481 | 0 | 0 | 0 | 0 | 0 | 0 | 836 | 316 | 83 | 0 | 0 | 83329 | 0 | 0 |
| SRR5230559 | 0 | 0 | 0 | 0 | 497 | 289 | 21 | 268 | 115 | 103 | 785 | 0 | 0 | 0 | 53909 | 101790 | 261431 | 41 | 33 | 99 | 1455 | 14 | 71 | 22 | 26 | 32 | 0 | 0 | 0 | 1903 | 28 | 24 | 0 | 0 | 46746 | 0 | 0 |
| SRR5230560 | 0 | 0 | 0 | 0 | 86 | 49 | 11 | 65 | 0 | 60 | 156 | 0 | 0 | 0 | 28999 | 107167 | 65902 | 103 | 43 | 108 | 1734 | 14 | 12 | 0 | 12 | 0 | 0 | 0 | 0 | 390 | 49 | 0 | 0 | 0 | 43765 | 0 | 0 |
| SRR5230561 | 0 | 0 | 0 | 0 | 38 | 74 | 28 | 23 | 11 | 74 | 39 | 0 | 0 | 0 | 8428 | 67590 | 18668 | 51 | 0 | 63 | 775 | 0 | 23 | 0 | 0 | 11 | 0 | 0 | 0 | 128 | 13 | 0 | 0 | 0 | 62928 | 0 | 0 |
| SRR5230562 | 0 | 0 | 0 | 0 | 295 | 268 | 0 | 206 | 166 | 1004 | 1681 | 21 | 14 | 0 | 26309 | 52840 | 110083 | 53 | 23 | 51 | 2208 | 14 | 117 | 12 | 14 | 0 | 0 | 0 | 0 | 1440 | 18 | 0 | 0 | 0 | 145307 | 0 | 0 |
| SRR5230563 | 0 | 0 | 0 | 0 | 312 | 57 | 0 | 63 | 55 | 1496 | 584 | 20 | 0 | 0 | 3838 | 41950 | 10963 | 406 | 0 | 109 | 1225 | 0 | 34 | 0 | 0 | 0 | 0 | 0 | 0 | 1400 | 0 | 0 | 0 | 0 | 113795 | 0 | 0 |
| SRR5230564 | 0 | 0 | 0 | 0 | 433 | 307 | 47 | 207 | 113 | 501 | 502 | 14 | 0 | 0 | 29634 | 92906 | 180729 | 55 | 45 | 125 | 792 | 44 | 277 | 13 | 19 | 0 | 0 | 0 | 0 | 1113 | 68 | 0 | 0 | 0 | 122237 | 0 | 0 |
| SRR5230565 | 0 | 0 | 0 | 0 | 378 | 26 | 0 | 85 | 85 | 474 | 757 | 58 | 15 | 0 | 31978 | 136396 | 172710 | 17 | 43 | 162 | 1862 | 214 | 590 | 0 | 0 | 0 | 0 | 0 | 0 | 1791 | 268 | 26 | 0 | 0 | 52417 | 0 | 0 |
| SRR5230566 | 0 | 0 | 0 | 0 | 384 | 201 | 23 | 222 | 84 | 186 | 703 | 13 | 0 | 0 | 13656 | 110312 | 75037 | 51 | 39 | 74 | 1239 | 27 | 95 | 0 | 10 | 0 | 0 | 0 | 0 | 2101 | 36 | 0 | 0 | 0 | 126210 | 0 | 0 |
| SRR5230567 | 0 | 0 | 0 | 0 | 71 | 184 | 12 | 134 | 93 | 256 | 54 | 17 | 0 | 0 | 11217 | 29639 | 31206 | 99 | 70 | 114 | 1516 | 12 | 19 | 0 | 0 | 0 | 0 | 0 | 0 | 139 | 35 | 1667 | 0 | 0 | 226724 | 0 | 0 |
| SRR5230568 | 0 | 0 | 0 | 0 | 54 | 32 | 0 | 58 | 11 | 260 | 206 | 34 | 0 | 0 | 3834 | 47442 | 22544 | 11 | 42 | 42 | 2093 | 0 | 11 | 0 | 12 | 0 | 0 | 0 | 0 | 286 | 0 | 63 | 0 | 0 | 127838 | 168 | 0 |
| SRR5230569 | 0 | 0 | 0 | 0 | 76 | 107 | 0 | 122 | 72 | 21 | 57 | 33 | 0 | 0 | 12506 | 80572 | 35717 | 30 | 61 | 160 | 1906 | 0 | 79 | 0 | 33 | 21 | 19 | 11 | 0 | 117 | 20 | 38 | 0 | 0 | 49958 | 0 | 0 |
| SRR5230570 | 0 | 0 | 0 | 0 | 469 | 142 | 0 | 112 | 64 | 2410 | 896 | 34 | 0 | 0 | 33773 | 82775 | 74399 | 107 | 56 | 206 | 898 | 0 | 28 | 0 | 14 | 18 | 0 | 0 | 0 | 2327 | 0 | 17 | 0 | 0 | 63359 | 0 | 0 |
| SRR5230571 | 0 | 0 | 0 | 0 | 42 | 371 | 37 | 267 | 204 | 64 | 30 | 28 | 0 | 0 | 64865 | 43843 | 94772 | 460 | 21 | 238 | 1012 | 0 | 60 | 0 | 16 | 0 | 0 | 0 | 0 | 109 | 0 | 29 | 0 | 0 | 196017 | 20 | 0 |
| SRR5230572 | 0 | 0 | 0 | 0 | 641 | 210 | 17 | 103 | 73 | 99 | 653 | 0 | 0 | 0 | 44798 | 117933 | 229452 | 47 | 34 | 210 | 4142 | 50 | 161 | 0 | 16 | 10 | 0 | 0 | 0 | 1480 | 88 | 17 | 0 | 0 | 42567 | 14 | 0 |
| SRR5230573 | 0 | 0 | 0 | 0 | 593 | 371 | 40 | 391 | 165 | 96 | 807 | 0 | 0 | 0 | 39594 | 113920 | 97644 | 205 | 0 | 476 | 5187 | 0 | 57 | 0 | 24 | 19 | 0 | 0 | 0 | 1348 | 19 | 55 | 0 | 0 | 76929 | 0 | 0 |
| SRR5230574 | 0 | 0 | 0 | 0 | 80 | 152 | 94 | 88 | 120 | 39 | 30 | 0 | 0 | 0 | 10876 | 117001 | 47060 | 74 | 0 | 94 | 597 | 0 | 10 | 0 | 0 | 0 | 0 | 0 | 0 | 189 | 20 | 0 | 0 | 0 | 92805 | 0 | 0 |
| SRR5230575 | 0 | 0 | 0 | 0 | 494 | 231 | 43 | 101 | 217 | 228 | 163 | 0 | 0 | 0 | 11475 | 118829 | 46767 | 101 | 0 | 116 | 462 | 0 | 47 | 0 | 0 | 0 | 0 | 0 | 0 | 1032 | 0 | 0 | 0 | 16 | 88816 | 0 | 0 |
| SRR5230576 | 0 | 0 | 0 | 0 | 565 | 1565 | 137 | 676 | 428 | 647 | 294 | 15 | 0 | 0 | 17262 | 106083 | 48311 | 99 | 12 | 539 | 799 | 14 | 43 | 14 | 11 | 13 | 0 | 0 | 0 | 1700 | 48 | 0 | 0 | 0 | 100104 | 0 | 0 |
| SRR5230577 | 0 | 0 | 0 | 0 | 583 | 1397 | 109 | 709 | 469 | 630 | 265 | 20 | 0 | 0 | 17592 | 108021 | 49171 | 120 | 33 | 437 | 709 | 22 | 69 | 13 | 0 | 0 | 0 | 0 | 0 | 1648 | 44 | 0 | 0 | 0 | 95097 | 0 | 0 |
| SRR5230578 | 0 | 0 | 0 | 0 | 39 | 107 | 0 | 87 | 83 | 75 | 69 | 0 | 0 | 0 | 85997 | 313230 | 392848 | 14 | 89 | 512 | 3980 | 118 | 20 | 0 | 0 | 0 | 0 | 0 | 0 | 208 | 12 | 0 | 0 | 0 | 9169 | 0 | 0 |
| SRR5230579 | 11 | 0 | 0 | 0 | 41 | 52 | 13 | 65 | 65 | 79 | 74 | 11 | 11 | 0 | 90478 | 315686 | 399092 | 0 | 52 | 378 | 4402 | 91 | 18 | 0 | 16 | 0 | 0 | 0 | 0 | 178 | 11 | 39 | 0 | 0 | 8758 | 0 | 0 |
| SRR5230580 | 0 | 0 | 0 | 0 | 61 | 882 | 76 | 255 | 193 | 91 | 60 | 26 | 0 | 0 | 14670 | 85074 | 33110 | 96 | 110 | 358 | 2109 | 0 | 26 | 0 | 0 | 18 | 10 | 0 | 0 | 236 | 0 | 90 | 0 | 0 | 64063 | 71 | 0 |
| SRR5230581 | 0 | 0 | 0 | 0 | 457 | 562 | 130 | 303 | 433 | 624 | 500 | 36 | 0 | 0 | 13885 | 84801 | 32479 | 0 | 0 | 303 | 2206 | 0 | 22 | 0 | 0 | 0 | 15 | 0 | 0 | 1821 | 0 | 0 | 0 | 0 | 61317 | 66 | 0 |
| SRR5230582 | 0 | 0 | 0 | 0 | 616 | 117 | 12 | 93 | 90 | 516 | 609 | 15 | 0 | 0 | 13683 | 176464 | 115447 | 0 | 123 | 494 | 1726 | 0 | 21 | 0 | 15 | 35 | 20 | 0 | 0 | 1566 | 16 | 12 | 0 | 0 | 27954 | 41 | 0 |
| SRR5230583 | 0 | 0 | 0 | 0 | 558 | 157 | 20 | 59 | 78 | 582 | 707 | 16 | 0 | 0 | 14218 | 177435 | 116816 | 0 | 78 | 490 | 1549 | 16 | 16 | 0 | 0 | 16 | 31 | 0 | 0 | 1722 | 16 | 20 | 0 | 0 | 26452 | 43 | 0 |
| SRR5230584 | 0 | 0 | 0 | 0 | 268 | 1092 | 55 | 304 | 172 | 359 | 440 | 11 | 0 | 0 | 8806 | 118132 | 125275 | 57 | 11 | 523 | 1814 | 18 | 88 | 0 | 0 | 0 | 0 | 0 | 0 | 1603 | 17 | 0 | 0 | 0 | 51718 | 15 | 0 |
| SRR5230585 | 0 | 0 | 0 | 0 | 250 | 1227 | 86 | 346 | 156 | 340 | 387 | 0 | 0 | 0 | 9055 | 121893 | 127601 | 17 | 17 | 588 | 1711 | 30 | 108 | 0 | 0 | 0 | 0 | 0 | 0 | 1592 | 17 | 0 | 0 | 0 | 49824 | 29 | 0 |
| SRR5230586 | 0 | 0 | 0 | 0 | 403 | 33 | 0 | 41 | 54 | 609 | 899 | 0 | 0 | 0 | 14523 | 46838 | 30124 | 238 | 10 | 312 | 2610 | 0 | 50 | 0 | 0 | 16 | 0 | 0 | 0 | 2516 | 0 | 61 | 0 | 0 | 203563 | 0 | 0 |
| SRR5230587 | 0 | 0 | 0 | 0 | 448 | 33 | 0 | 114 | 33 | 593 | 909 | 0 | 0 | 0 | 14753 | 49073 | 31886 | 212 | 0 | 326 | 2657 | 0 | 47 | 0 | 12 | 18 | 0 | 0 | 0 | 2487 | 0 | 98 | 0 | 0 | 198688 | 0 | 0 |
| SRR5230588 | 0 | 0 | 0 | 0 | 700 | 143 | 22 | 204 | 287 | 192 | 1077 | 0 | 0 | 0 | 46149 | 65961 | 108688 | 77 | 28 | 413 | 866 | 16 | 61 | 0 | 39 | 28 | 0 | 0 | 0 | 2241 | 16 | 28 | 0 | 0 | 36115 | 0 | 0 |
| SRR5230589 | 0 | 0 | 0 | 0 | 484 | 206 | 0 | 34 | 206 | 177 | 1031 | 0 | 0 | 0 | 47429 | 66420 | 109393 | 69 | 0 | 412 | 343 | 0 | 38 | 15 | 69 | 38 | 0 | 0 | 0 | 2061 | 0 | 0 | 0 | 0 | 35136 | 0 | 0 |
| SRR5230590 | 0 | 0 | 0 | 0 | 220 | 327 | 490 | 327 | 490 | 661 | 41 | 0 | 0 | 0 | 10449 | 37581 | 22565 | 163 | 0 | 0 | 0 | 0 | 28 | 0 | 14 | 0 | 0 | 0 | 0 | 207 | 0 | 0 | 0 | 0 | 176358 | 0 | 0 |
| SRR5230591 | 0 | 0 | 0 | 0 | 188 | 531 | 123 | 592 | 357 | 555 | 15 | 0 | 0 | 0 | 9613 | 35927 | 20054 | 123 | 11 | 61 | 218 | 0 | 16 | 13 | 0 | 0 | 0 | 0 | 0 | 135 | 0 | 0 | 0 | 0 | 175850 | 0 | 0 |
| SRR5230592 | 0 | 0 | 0 | 0 | 0 | 273 | 0 | 0 | 273 | 484 | 290 | 0 | 0 | 0 | 11454 | 76962 | 32152 | 0 | 0 | 0 | 273 | 0 | 0 | 0 | 0 | 0 | 0 | 0 | 0 | 242 | 0 | 0 | 0 | 0 | 141941 | 0 | 0 |
| SRR5230593 | 0 | 0 | 0 | 0 | 136 | 230 | 19 | 211 | 67 | 11 | 148 | 13 | 0 | 0 | 13159 | 73127 | 28053 | 134 | 0 | 48 | 239 | 0 | 0 | 10 | 12 | 0 | 0 | 0 | 0 | 20 | 0 | 0 | 0 | 0 | 144331 | 0 | 0 |
| SRR5230594 | 0 | 0 | 0 | 0 | 91 | 215 | 0 | 133 | 60 | 37 | 66 | 0 | 0 | 0 | 12314 | 82092 | 30329 | 95 | 26 | 211 | 486 | 12 | 25 | 11 | 58 | 31 | 16 | 0 | 0 | 236 | 0 | 34 | 0 | 0 | 105437 | 0 | 0 |
| SRR5230595 | 0 | 0 | 0 | 0 | 100 | 188 | 0 | 54 | 134 | 36 | 58 | 11 | 0 | 0 | 12511 | 85000 | 30004 | 268 | 27 | 161 | 483 | 0 | 0 | 11 | 57 | 34 | 11 | 0 | 0 | 192 | 0 | 0 | 0 | 0 | 99301 | 0 | 0 |
| SRR5230596 | 0 | 0 | 0 | 0 | 65 | 410 | 38 | 264 | 141 | 630 | 38 | 20 | 0 | 0 | 19380 | 67517 | 65215 | 43 | 32 | 152 | 929 | 0 | 27 | 33 | 29 | 21 | 0 | 0 | 0 | 187 | 0 | 24 | 0 | 0 | 133324 | 0 | 0 |
| SRR5230597 | 0 | 0 | 0 | 0 | 379 | 402 | 61 | 394 | 87 | 3478 | 214 | 13 | 0 | 0 | 19066 | 68433 | 66640 | 17 | 61 | 184 | 822 | 0 | 33 | 32 | 32 | 20 | 0 | 0 | 0 | 1018 | 0 | 35 | 0 | 0 | 129325 | 0 | 0 |
| SRR5230600 | 0 | 0 | 0 | 0 | 41 | 86 | 0 | 76 | 43 | 163 | 16 | 0 | 0 | 0 | 8271 | 33187 | 8191 | 177 | 18 | 48 | 235 | 0 | 50 | 10 | 0 | 0 | 0 | 0 | 0 | 45 | 0 | 58 | 0 | 0 | 194291 | 0 | 0 |
| SRR5230601 | 0 | 0 | 0 | 0 | 310 | 96 | 0 | 48 | 48 | 1228 | 161 | 13 | 13 | 0 | 8932 | 34219 | 8206 | 144 | 0 | 48 | 208 | 10 | 47 | 20 | 10 | 0 | 0 | 0 | 0 | 370 | 0 | 16 | 0 | 0 | 186829 | 0 | 0 |
| SRR5230602 | 0 | 0 | 0 | 0 | 47 | 256 | 10 | 266 | 121 | 12 | 69 | 22 | 0 | 0 | 56407 | 35731 | 112065 | 66 | 48 | 304 | 2629 | 11 | 39 | 22 | 46 | 35 | 12 | 0 | 0 | 168 | 11 | 17 | 0 | 0 | 63808 | 0 | 0 |
| SRR5230603 | 0 | 0 | 0 | 0 | 346 | 324 | 0 | 389 | 237 | 118 | 743 | 0 | 0 | 0 | 57587 | 34941 | 113065 | 86 | 43 | 151 | 2612 | 15 | 51 | 15 | 37 | 29 | 22 | 0 | 0 | 1655 | 0 | 43 | 0 | 0 | 61659 | 14 | 0 |
| SRR5230604 | 0 | 0 | 0 | 0 | 60 | 173 | 22 | 107 | 68 | 86 | 32 | 28 | 0 | 0 | 32430 | 52066 | 47984 | 114 | 18 | 113 | 570 | 0 | 40 | 11 | 16 | 0 | 0 | 0 | 0 | 170 | 11 | 0 | 0 | 0 | 137559 | 0 | 0 |
| SRR5230605 | 0 | 0 | 0 | 0 | 313 | 176 | 25 | 176 | 94 | 446 | 189 | 23 | 0 | 0 | 32278 | 53588 | 48943 | 145 | 50 | 94 | 510 | 0 | 49 | 0 | 14 | 11 | 0 | 0 | 0 | 920 | 11 | 0 | 0 | 0 | 133018 | 0 | 0 |
| SRR5230606 | 0 | 0 | 0 | 0 | 273 | 233 | 21 | 238 | 150 | 1483 | 192 | 0 | 0 | 0 | 29710 | 103892 | 122597 | 32 | 16 | 76 | 959 | 0 | 49 | 23 | 11 | 12 | 0 | 0 | 0 | 1009 | 20 | 21 | 0 | 0 | 58499 | 11 | 0 |
| SRR5230607 | 0 | 0 | 0 | 0 | 27 | 159 | 14 | 333 | 101 | 114 | 17 | 35 | 0 | 0 | 32088 | 106126 | 123735 | 43 | 0 | 72 | 825 | 12 | 46 | 23 | 0 | 0 | 0 | 0 | 0 | 75 | 12 | 29 | 0 | 0 | 55344 | 0 | 0 |
| SRR5230608 | 0 | 0 | 0 | 0 | 285 | 97 | 0 | 82 | 53 | 620 | 2455 | 18 | 0 | 0 | 28664 | 56037 | 50790 | 106 | 68 | 290 | 2316 | 0 | 18 | 0 | 0 | 17 | 0 | 0 | 0 | 4268 | 0 | 72 | 0 | 0 | 109459 | 0 | 0 |
| SRR5230609 | 0 | 0 | 0 | 0 | 320 | 120 | 0 | 90 | 30 | 605 | 2371 | 0 | 0 | 0 | 30834 | 57716 | 53004 | 90 | 150 | 240 | 2277 | 0 | 30 | 12 | 0 | 18 | 24 | 0 | 0 | 4310 | 0 | 90 | 0 | 0 | 102818 | 10 | 0 |
| SRR5230610 | 0 | 0 | 0 | 0 | 137 | 422 | 60 | 422 | 241 | 1054 | 39 | 0 | 0 | 0 | 10056 | 56209 | 33957 | 60 | 0 | 0 | 1144 | 0 | 20 | 20 | 0 | 0 | 0 | 0 | 0 | 156 | 0 | 120 | 0 | 0 | 144570 | 11 | 0 |
| SRR5230611 | 0 | 0 | 0 | 0 | 24 | 705 | 28 | 456 | 147 | 147 | 31 | 0 | 0 | 0 | 9412 | 56395 | 34240 | 65 | 0 | 111 | 949 | 0 | 23 | 18 | 0 | 0 | 0 | 0 | 0 | 22 | 0 | 22 | 0 | 0 | 138761 | 0 | 0 |
| SRR5230613 | 0 | 0 | 0 | 0 | 200 | 245 | 18 | 100 | 85 | 453 | 22 | 0 | 0 | 0 | 2433 | 50313 | 7168 | 87 | 0 | 36 | 102 | 0 | 12 | 0 | 0 | 0 | 0 | 0 | 0 | 239 | 0 | 159 | 0 | 0 | 243298 | 0 | 0 |
| SRR5230614 | 0 | 0 | 0 | 0 | 143 | 70 | 0 | 70 | 70 | 847 | 13 | 40 | 0 | 0 | 1052 | 65423 | 13092 | 257 | 351 | 234 | 257 | 13 | 140 | 13 | 0 | 0 | 0 | 0 | 10 | 117 | 0 | 257 | 0 | 23 | 187565 | 0 | 0 |
| SRR5230615 | 0 | 0 | 0 | 0 | 128 | 129 | 0 | 87 | 47 | 865 | 11 | 34 | 0 | 0 | 871 | 67017 | 12241 | 191 | 296 | 311 | 222 | 23 | 128 | 0 | 0 | 0 | 0 | 0 | 22 | 111 | 14 | 306 | 0 | 15 | 186631 | 0 | 0 |
| SRR5230617 | 0 | 0 | 0 | 0 | 16 | 386 | 19 | 181 | 129 | 21 | 21 | 10 | 0 | 0 | 3327 | 65958 | 7427 | 56 | 0 | 32 | 210 | 0 | 24 | 25 | 0 | 0 | 0 | 0 | 0 | 11 | 0 | 85 | 0 | 0 | 138802 | 12 | 0 |
| SRR5230618 | 0 | 0 | 0 | 0 | 24 | 1592 | 58 | 723 | 550 | 125 | 60 | 0 | 11 | 0 | 20806 | 47519 | 80094 | 174 | 29 | 174 | 1476 | 0 | 22 | 71 | 0 | 0 | 0 | 0 | 0 | 33 | 11 | 87 | 0 | 0 | 200395 | 16 | 0 |
| SRR5230619 | 0 | 0 | 0 | 0 | 195 | 1272 | 158 | 766 | 653 | 1185 | 67 | 12 | 0 | 0 | 21072 | 45891 | 84193 | 130 | 0 | 187 | 1437 | 0 | 33 | 42 | 0 | 0 | 0 | 0 | 0 | 275 | 17 | 42 | 0 | 0 | 201107 | 21 | 0 |
| SRR5230620 | 0 | 0 | 0 | 0 | 258 | 301 | 24 | 159 | 208 | 411 | 231 | 14 | 0 | 0 | 14532 | 71321 | 51051 | 54 | 0 | 71 | 1315 | 0 | 38 | 23 | 29 | 0 | 11 | 0 | 0 | 931 | 0 | 37 | 0 | 0 | 125688 | 24 | 0 |
| SRR5230621 | 0 | 0 | 0 | 0 | 26 | 323 | 46 | 200 | 154 | 43 | 20 | 0 | 0 | 0 | 15012 | 72250 | 51639 | 62 | 0 | 77 | 1369 | 0 | 42 | 27 | 23 | 0 | 15 | 0 | 0 | 82 | 0 | 62 | 0 | 0 | 122059 | 21 | 0 |
| SRR5230622 | 0 | 0 | 0 | 0 | 63 | 107 | 0 | 29 | 88 | 41 | 162 | 0 | 0 | 0 | 39943 | 93046 | 70487 | 44 | 73 | 1159 | 1368 | 0 | 18 | 0 | 0 | 0 | 0 | 0 | 0 | 501 | 0 | 0 | 0 | 0 | 24647 | 0 | 0 |
| SRR5230623 | 0 | 0 | 0 | 0 | 401 | 68 | 0 | 58 | 42 | 1214 | 772 | 12 | 0 | 0 | 53989 | 99429 | 214793 | 71 | 37 | 264 | 4869 | 25 | 94 | 0 | 0 | 12 | 0 | 0 | 0 | 1773 | 44 | 29 | 0 | 0 | 41883 | 0 | 0 |
| SRR5230624 | 0 | 0 | 0 | 0 | 531 | 114 | 37 | 170 | 129 | 982 | 419 | 32 | 0 | 0 | 14810 | 172633 | 54928 | 62 | 35 | 214 | 1059 | 0 | 29 | 11 | 0 | 0 | 0 | 0 | 0 | 1099 | 0 | 0 | 0 | 0 | 92867 | 0 | 0 |
| SRR5230625 | 0 | 0 | 0 | 0 | 184 | 106 | 11 | 57 | 109 | 79 | 130 | 17 | 0 | 0 | 34696 | 115554 | 77128 | 102 | 160 | 248 | 1063 | 0 | 10 | 0 | 41 | 40 | 0 | 0 | 0 | 307 | 17 | 45 | 0 | 0 | 59384 | 857 | 0 |
| SRR5230626 | 0 | 0 | 0 | 0 | 410 | 86 | 12 | 57 | 68 | 1603 | 863 | 25 | 0 | 0 | 87656 | 111428 | 104246 | 103 | 22 | 313 | 1222 | 0 | 14 | 0 | 16 | 25 | 0 | 0 | 0 | 1117 | 0 | 48 | 0 | 0 | 102637 | 0 | 0 |
| SRR5230627 | 0 | 0 | 0 | 0 | 726 | 86 | 13 | 67 | 103 | 709 | 708 | 0 | 0 | 0 | 17736 | 125748 | 118071 | 17 | 71 | 318 | 6414 | 79 | 325 | 0 | 0 | 12 | 0 | 0 | 0 | 1480 | 98 | 71 | 0 | 0 | 77285 | 96 | 0 |
| SRR5230628 | 0 | 0 | 0 | 0 | 36 | 27 | 0 | 29 | 63 | 112 | 53 | 0 | 0 | 0 | 45229 | 152697 | 55493 | 97 | 0 | 170 | 261 | 0 | 0 | 20 | 34 | 49 | 10 | 0 | 0 | 90 | 0 | 0 | 0 | 0 | 48885 | 0 | 0 |
| SRR5230629 | 0 | 0 | 0 | 0 | 81 | 98 | 20 | 73 | 177 | 32 | 242 | 27 | 0 | 0 | 12195 | 186825 | 127323 | 132 | 0 | 492 | 1677 | 0 | 10 | 22 | 11 | 18 | 0 | 0 | 0 | 569 | 0 | 22 | 0 | 0 | 77531 | 0 | 0 |
| SRR5230630 | 0 | 0 | 0 | 0 | 131 | 41 | 25 | 69 | 65 | 51 | 244 | 18 | 0 | 0 | 8994 | 149530 | 62183 | 18 | 0 | 460 | 2392 | 19 | 20 | 0 | 0 | 0 | 0 | 0 | 0 | 850 | 11 | 0 | 0 | 0 | 31464 | 0 | 0 |

### Supp Table S8.3.Columns_BX_to_DH

| | | | | | | | | | | | | | | | | | | | | | | | | | | | | | | | | | | | | | |
|---|---|---|---|---|---|---|---|---|---|---|---|---|---|---|---|---|---|---|---|---|---|---|---|---|---|---|---|---|---|---|---|---|---|---|---|---|---|
| SRR5230631 | 0 | 0 | 0 | 0 | 70 | 143 | 21 | 64 | 55 | 79 | 76 | 0 | 0 | 0 | 63556 | 107292 | 242545 | 32 | 53 | 172 | 1345 | 0 | 32 | 20 | 51 | 58 | 17 | 11 | 0 | 197 | 0 | 24 | 0 | 0 | 42488 | 0 | 0 |
| SRR5230632 | 0 | 0 | 0 | 0 | 141 | 108 | 0 | 87 | 90 | 260 | 133 | 16 | 0 | 0 | 15610 | 75222 | 66284 | 188 | 64 | 482 | 3068 | 17 | 26 | 0 | 14 | 16 | 0 | 0 | 0 | 464 | 11 | 60 | 0 | 0 | 188534 | 18 | 0 |
| SRR5230633 | 0 | 0 | 0 | 0 | 78 | 116 | 0 | 85 | 188 | 39 | 74 | 49 | 0 | 0 | 47379 | 104210 | 143868 | 99 | 93 | 528 | 1666 | 25 | 24 | 0 | 0 | 12 | 0 | 0 | 0 | 152 | 13 | 39 | 0 | 0 | 94967 | 0 | 0 |
| SRR5230634 | 0 | 0 | 0 | 0 | 391 | 150 | 31 | 44 | 31 | 179 | 336 | 22 | 0 | 0 | 5388 | 220467 | 62490 | 63 | 56 | 125 | 1666 | 175 | 149 | 0 | 0 | 0 | 0 | 0 | 0 | 457 | 411 | 382 | 0 | 0 | 5470 | 0 | 0 |
| SRR5230635 | 0 | 0 | 0 | 0 | 69 | 964 | 86 | 77 | 59 | 199 | 60 | 60 | 0 | 0 | 2800 | 481419 | 14172 | 0 | 41 | 132 | 1614 | 55 | 83 | 0 | 0 | 14 | 69 | 42 | 0 | 60 | 134 | 282 | 0 | 0 | 1465 | 0 | 0 |
| SRR5230636 | 0 | 0 | 0 | 0 | 105 | 461 | 16 | 40 | 72 | 57 | 29 | 12 | 0 | 0 | 3587 | 445242 | 63529 | 0 | 32 | 99 | 4403 | 105 | 111 | 0 | 0 | 0 | 14 | 0 | 14 | 51 | 26 | 773 | 0 | 0 | 7221 | 0 | 0 |
| SRR5230637 | 0 | 0 | 0 | 0 | 221 | 546 | 13 | 76 | 122 | 154 | 67 | 111 | 0 | 0 | 1227 | 614210 | 3689 | 17 | 93 | 30 | 2450 | 0 | 16 | 0 | 0 | 51 | 83 | 16 | 0 | 12 | 20 | 66 | 0 | 0 | 1618 | 0 | 0 |
| SRR5230638 | 0 | 0 | 0 | 0 | 74 | 322 | 12 | 24 | 42 | 74 | 77 | 106 | 0 | 0 | 2049 | 565869 | 18926 | 0 | 160 | 24 | 4758 | 34 | 40 | 0 | 26 | 140 | 153 | 0 | 0 | 0 | 100 | 21 | 0 | 0 | 4326 | 0 | 0 |
| SRR5230639 | 0 | 0 | 0 | 0 | 70 | 125 | 0 | 0 | 39 | 17 | 179 | 27 | 0 | 0 | 2920 | 650864 | 38478 | 16 | 65 | 0 | 1739 | 162 | 14 | 12 | 65 | 56 | 150 | 17 | 17 | 116 | 24 | 0 | 0 | 0 | 3441 | 0 | 0 |
| SRR5230640 | 0 | 0 | 0 | 0 | 63 | 180 | 0 | 19 | 40 | 110 | 41 | 50 | 0 | 0 | 1556 | 563108 | 10991 | 0 | 19 | 0 | 1180 | 13 | 0 | 0 | 0 | 28 | 82 | 0 | 0 | 0 | 26 | 16 | 0 | 0 | 6539 | 0 | 0 |
| SRR5230641 | 0 | 0 | 0 | 0 | 17 | 248 | 0 | 0 | 18 | 21 | 48 | 13 | 0 | 0 | 1870 | 307308 | 81140 | 0 | 133 | 18 | 1391 | 26 | 0 | 0 | 0 | 12 | 28 | 0 | 0 | 14 | 22 | 14 | 0 | 0 | 12088 | 0 | 0 |
| SRR5230642 | 0 | 0 | 0 | 0 | 120 | 305 | 0 | 70 | 48 | 0 | 115 | 29 | 0 | 0 | 8556 | 329958 | 180143 | 0 | 59 | 262 | 3535 | 107 | 93 | 0 | 0 | 0 | 19 | 0 | 0 | 19 | 91 | 176 | 0 | 0 | 4685 | 0 | 0 |
| SRR5230643 | 0 | 0 | 0 | 0 | 68 | 468 | 0 | 80 | 154 | 135 | 58 | 0 | 0 | 0 | 2717 | 454163 | 31320 | 0 | 22 | 12 | 1750 | 18 | 42 | 0 | 44 | 0 | 107 | 0 | 0 | 20 | 103 | 12 | 0 | 0 | 2976 | 0 | 0 |
| SRR5230644 | 0 | 0 | 0 | 0 | 50 | 861 | 24 | 60 | 129 | 68 | 32 | 14 | 0 | 0 | 1623 | 547068 | 35657 | 0 | 54 | 43 | 1690 | 28 | 34 | 0 | 42 | 68 | 134 | 10 | 0 | 40 | 68 | 28 | 0 | 10 | 4914 | 0 | 0 |
| SRR5230645 | 0 | 0 | 0 | 0 | 46 | 772 | 32 | 152 | 185 | 136 | 93 | 49 | 0 | 0 | 4636 | 414211 | 50458 | 20 | 162 | 75 | 2615 | 93 | 152 | 21 | 10 | 95 | 185 | 0 | 0 | 15 | 190 | 20 | 0 | 0 | 4819 | 20 | 0 |
| SRR5230646 | 0 | 0 | 0 | 0 | 68 | 843 | 48 | 126 | 124 | 110 | 60 | 25 | 0 | 0 | 1990 | 454829 | 19648 | 0 | 78 | 64 | 2521 | 85 | 130 | 11 | 19 | 19 | 123 | 0 | 0 | 0 | 18 | 60 | 0 | 15 | 4865 | 0 | 0 |
| SRR5230647 | 0 | 0 | 0 | 0 | 0 | 142 | 0 | 0 | 0 | 22 | 22 | 0 | 0 | 0 | 5016 | 249317 | 14418 | 0 | 127 | 0 | 2036 | 0 | 0 | 0 | 0 | 0 | 0 | 0 | 0 | 0 | 0 | 64 | 0 | 0 | 1478 | 0 | 0 |
| SRR5230648 | 0 | 0 | 0 | 0 | 84 | 778 | 31 | 101 | 143 | 107 | 95 | 152 | 0 | 0 | 8027 | 482161 | 109287 | 0 | 110 | 122 | 4483 | 16 | 14 | 0 | 0 | 0 | 0 | 0 | 0 | 33 | 49 | 166 | 0 | 0 | 6392 | 19 | 0 |
| SRR5230649 | 0 | 0 | 0 | 0 | 17 | 567 | 0 | 106 | 25 | 208 | 40 | 107 | 0 | 0 | 2373 | 387557 | 32084 | 0 | 85 | 0 | 1157 | 0 | 0 | 0 | 0 | 0 | 67 | 0 | 0 | 29 | 56 | 35 | 0 | 31 | 5503 | 46 | 0 |
| SRR5230650 | 0 | 0 | 0 | 0 | 52 | 45 | 0 | 84 | 0 | 46 | 70 | 0 | 0 | 0 | 4444 | 266854 | 24513 | 104 | 71 | 78 | 1126 | 18 | 13 | 0 | 0 | 0 | 0 | 0 | 0 | 14 | 73 | 0 | 0 | 0 | 3231 | 0 | 0 |
| SRR5230651 | 0 | 0 | 0 | 0 | 87 | 560 | 13 | 117 | 200 | 108 | 54 | 19 | 0 | 0 | 4203 | 391602 | 37082 | 0 | 80 | 193 | 1480 | 169 | 148 | 12 | 0 | 30 | 94 | 0 | 0 | 21 | 380 | 0 | 0 | 0 | 2276 | 0 | 0 |
| SRR5230652 | 0 | 0 | 0 | 0 | 25 | 1678 | 44 | 178 | 176 | 683 | 100 | 566 | 0 | 0 | 5312 | 218222 | 68205 | 0 | 258 | 104 | 1881 | 24 | 32 | 0 | 0 | 0 | 30 | 0 | 0 | 20 | 44 | 466 | 0 | 30 | 3566 | 110 | 0 |
| SRR5230653 | 0 | 0 | 0 | 0 | 15 | 1229 | 0 | 91 | 105 | 13 | 61 | 18 | 0 | 0 | 4098 | 214252 | 29938 | 0 | 50 | 21 | 1880 | 0 | 11 | 0 | 0 | 17 | 25 | 0 | 0 | 28 | 12 | 73 | 0 | 0 | 3007 | 0 | 0 |
| SRR5230654 | 0 | 0 | 0 | 0 | 36 | 2393 | 131 | 184 | 236 | 52 | 0 | 20 | 0 | 0 | 5139 | 253549 | 80306 | 0 | 100 | 79 | 2651 | 0 | 0 | 0 | 0 | 19 | 23 | 0 | 0 | 0 | 92 | 131 | 0 | 0 | 20968 | 0 | 0 |
| SRR5230655 | 0 | 0 | 0 | 0 | 52 | 169 | 0 | 38 | 30 | 85 | 65 | 41 | 0 | 0 | 2602 | 570741 | 19409 | 0 | 66 | 17 | 881 | 0 | 15 | 0 | 0 | 50 | 183 | 17 | 0 | 28 | 83 | 0 | 0 | 0 | 7099 | 0 | 0 |
| SRR5230656 | 0 | 0 | 0 | 0 | 37 | 88 | 0 | 25 | 88 | 136 | 61 | 0 | 0 | 0 | 1545 | 293943 | 4419 | 21 | 76 | 101 | 2188 | 14 | 37 | 0 | 19 | 23 | 70 | 23 | 0 | 23 | 37 | 0 | 0 | 0 | 4337 | 0 | 0 |
| SRR5230657 | 0 | 0 | 0 | 0 | 66 | 456 | 0 | 80 | 68 | 160 | 53 | 15 | 0 | 0 | 2801 | 382930 | 10946 | 0 | 30 | 35 | 2095 | 50 | 110 | 0 | 0 | 0 | 28 | 0 | 0 | 64 | 230 | 0 | 0 | 0 | 2772 | 12 | 0 |
| SRR5230658 | 0 | 0 | 0 | 0 | 151 | 166 | 0 | 25 | 20 | 139 | 134 | 28 | 0 | 0 | 2582 | 401717 | 23071 | 0 | 81 | 31 | 1241 | 66 | 129 | 0 | 0 | 45 | 33 | 0 | 10 | 13 | 15 | 11 | 0 | 0 | 4665 | 0 | 0 |
| SRR5230659 | 0 | 0 | 0 | 0 | 43 | 112 | 0 | 0 | 0 | 152 | 39 | 28 | 0 | 0 | 2602 | 361553 | 24504 | 0 | 53 | 32 | 1511 | 22 | 22 | 0 | 0 | 0 | 11 | 0 | 0 | 28 | 50 | 182 | 0 | 0 | 2546 | 0 | 0 |
| SRR5230660 | 0 | 0 | 0 | 0 | 145 | 146 | 0 | 41 | 32 | 347 | 122 | 14 | 0 | 0 | 4080 | 447189 | 13731 | 0 | 89 | 22 | 1573 | 56 | 108 | 0 | 14 | 75 | 70 | 14 | 14 | 33 | 206 | 0 | 0 | 0 | 1641 | 0 | 0 |
| SRR5230661 | 0 | 0 | 0 | 0 | 124 | 23 | 0 | 0 | 0 | 130 | 241 | 18 | 0 | 0 | 740 | 393838 | 5889 | 14 | 45 | 53 | 1376 | 12 | 15 | 0 | 0 | 0 | 12 | 0 | 27 | 91 | 41 | 23 | 0 | 0 | 2570 | 0 | 0 |
| SRR5230662 | 0 | 0 | 0 | 0 | 32 | 118 | 0 | 0 | 47 | 78 | 103 | 37 | 0 | 0 | 1157 | 242005 | 7960 | 16 | 157 | 110 | 480 | 10 | 21 | 0 | 0 | 0 | 10 | 0 | 0 | 0 | 32 | 0 | 0 | 0 | 3415 | 0 | 0 |
| SRR5230663 | 0 | 0 | 0 | 0 | 321 | 665 | 0 | 135 | 130 | 214 | 95 | 0 | 0 | 0 | 2704 | 190429 | 26783 | 0 | 151 | 273 | 1478 | 76 | 82 | 0 | 0 | 24 | 82 | 0 | 0 | 60 | 268 | 0 | 0 | 0 | 1568 | 0 | 0 |
| SRR5230664 | 0 | 0 | 0 | 0 | 28 | 210 | 0 | 18 | 29 | 61 | 49 | 0 | 0 | 0 | 2398 | 558059 | 5351 | 11 | 16 | 20 | 2643 | 18 | 0 | 0 | 0 | 27 | 66 | 0 | 0 | 16 | 19 | 0 | 0 | 0 | 4368 | 0 | 0 |
| SRR5230665 | 0 | 0 | 0 | 0 | 44 | 909 | 0 | 140 | 75 | 414 | 194 | 101 | 0 | 0 | 6678 | 535359 | 97523 | 0 | 154 | 136 | 2534 | 317 | 303 | 0 | 0 | 19 | 57 | 12 | 16 | 27 | 1122 | 66 | 0 | 21 | 4115 | 165 | 0 |
| SRR5230666 | 0 | 0 | 0 | 17 | 106 | 1203 | 83 | 115 | 144 | 116 | 142 | 20 | 0 | 0 | 3207 | 537276 | 12246 | 22 | 83 | 233 | 3641 | 251 | 198 | 0 | 0 | 96 | 248 | 17 | 0 | 26 | 172 | 19 | 0 | 0 | 6584 | 0 | 0 |
| SRR5230667 | 0 | 0 | 0 | 0 | 96 | 92 | 0 | 26 | 0 | 92 | 88 | 0 | 0 | 0 | 1482 | 275998 | 5856 | 0 | 39 | 105 | 1312 | 0 | 0 | 0 | 0 | 0 | 0 | 0 | 0 | 19 | 31 | 0 | 0 | 0 | 4442 | 0 | 0 |
| SRR5230668 | 0 | 0 | 0 | 0 | 24 | 1385 | 17 | 136 | 143 | 60 | 86 | 0 | 0 | 0 | 6102 | 458804 | 59921 | 17 | 95 | 95 | 2399 | 78 | 80 | 0 | 0 | 13 | 57 | 0 | 0 | 35 | 20 | 0 | 0 | 0 | 2623 | 0 | 0 |
| SRR5230669 | 0 | 0 | 0 | 0 | 426 | 1247 | 17 | 174 | 124 | 397 | 324 | 37 | 0 | 0 | 1665 | 448337 | 11233 | 12 | 49 | 92 | 1351 | 16 | 0 | 10 | 0 | 57 | 174 | 12 | 0 | 78 | 47 | 0 | 0 | 0 | 5351 | 0 | 0 |
| SRR5230670 | 0 | 0 | 0 | 0 | 41 | 76 | 0 | 0 | 0 | 116 | 169 | 11 | 0 | 0 | 1205 | 377531 | 8879 | 0 | 16 | 17 | 901 | 0 | 15 | 0 | 0 | 0 | 17 | 0 | 0 | 26 | 11 | 0 | 0 | 0 | 5422 | 10 | 0 |
| SRR5230671 | 0 | 0 | 0 | 0 | 111 | 24 | 0 | 0 | 0 | 334 | 2035 | 0 | 0 | 0 | 2052 | 357671 | 12133 | 32 | 119 | 16 | 1805 | 16 | 0 | 0 | 0 | 0 | 0 | 0 | 0 | 391 | 25 | 0 | 0 | 0 | 2925 | 0 | 0 |
| SRR5230673 | 0 | 0 | 0 | 0 | 11 | 330 | 0 | 27 | 102 | 194 | 200 | 19 | 0 | 0 | 1127 | 548944 | 14424 | 0 | 51 | 24 | 1593 | 0 | 18 | 0 | 0 | 19 | 79 | 27 | 0 | 81 | 36 | 0 | 0 | 0 | 2344 | 0 | 0 |
| SRR5230674 | 0 | 0 | 0 | 0 | 64 | 83 | 0 | 0 | 17 | 352 | 134 | 48 | 0 | 0 | 2980 | 445436 | 25983 | 0 | 345 | 85 | 1414 | 70 | 55 | 0 | 0 | 15 | 66 | 0 | 15 | 40 | 178 | 17 | 0 | 0 | 3610 | 160 | 0 |
| SRR5230675 | 0 | 0 | 0 | 0 | 55 | 85 | 0 | 0 | 24 | 104 | 144 | 12 | 0 | 0 | 2702 | 303002 | 9518 | 0 | 54 | 502 | 32196 | 140 | 153 | 0 | 0 | 0 | 0 | 0 | 0 | 86 | 220 | 12 | 0 | 0 | 473 | 0 | 0 |
| SRR5230676 | 0 | 0 | 0 | 0 | 89 | 274 | 0 | 0 | 81 | 12 | 185 | 46 | 0 | 0 | 2491 | 421706 | 26940 | 0 | 39 | 150 | 972 | 30 | 30 | 0 | 0 | 76 | 126 | 23 | 17 | 129 | 96 | 42 | 0 | 0 | 2116 | 0 | 0 |
| SRR5230677 | 0 | 0 | 0 | 0 | 174 | 37 | 0 | 0 | 13 | 161 | 166 | 15 | 0 | 0 | 854 | 454434 | 11201 | 0 | 12 | 17 | 531 | 15 | 11 | 0 | 0 | 15 | 52 | 0 | 0 | 47 | 47 | 0 | 0 | 0 | 6871 | 0 | 0 |
| SRR5230678 | 0 | 0 | 0 | 0 | 214 | 92 | 0 | 0 | 13 | 238 | 228 | 93 | 0 | 0 | 1897 | 651163 | 23731 | 0 | 21 | 15 | 621 | 33 | 70 | 61 | 33 | 65 | 228 | 75 | 0 | 47 | 116 | 0 | 0 | 0 | 5092 | 0 | 0 |
| SRR5230679 | 0 | 0 | 0 | 0 | 83 | 59 | 0 | 0 | 0 | 43 | 77 | 65 | 0 | 0 | 960 | 373535 | 9481 | 33 | 15 | 0 | 625 | 0 | 16 | 0 | 0 | 0 | 0 | 0 | 11 | 26 | 18 | 0 | 0 | 0 | 5469 | 0 | 0 |
| SRR5230680 | 0 | 0 | 0 | 0 | 82 | 470 | 0 | 0 | 29 | 34 | 105 | 0 | 0 | 0 | 3372 | 402841 | 8783 | 0 | 23 | 92 | 2409 | 45 | 68 | 0 | 0 | 0 | 0 | 0 | 0 | 54 | 142 | 0 | 0 | 0 | 2610 | 0 | 0 |
| SRR5230681 | 0 | 0 | 0 | 0 | 15 | 163 | 0 | 0 | 17 | 23 | 24 | 12 | 0 | 0 | 1775 | 325573 | 27298 | 0 | 38 | 44 | 711 | 0 | 0 | 0 | 0 | 0 | 26 | 0 | 0 | 17 | 0 | 0 | 0 | 0 | 4226 | 125 | 0 |
| SRR5230682 | 0 | 0 | 0 | 0 | 33 | 81 | 0 | 12 | 28 | 351 | 225 | 0 | 0 | 0 | 3623 | 719478 | 52995 | 0 | 37 | 28 | 1124 | 352 | 635 | 0 | 0 | 19 | 94 | 15 | 0 | 79 | 1782 | 0 | 0 | 0 | 2351 | 0 | 0 |
| SRR5230683 | 0 | 0 | 0 | 0 | 30 | 182 | 0 | 34 | 30 | 283 | 271 | 37 | 0 | 0 | 2846 | 398441 | 21032 | 0 | 43 | 55 | 1145 | 0 | 16 | 0 | 0 | 26 | 124 | 18 | 0 | 112 | 12 | 0 | 0 | 0 | 6262 | 0 | 0 |
| SRR5230684 | 0 | 0 | 0 | 0 | 35 | 31 | 0 | 0 | 0 | 310 | 228 | 18 | 0 | 0 | 2022 | 363528 | 40311 | 0 | 340 | 45 | 784 | 46 | 45 | 0 | 0 | 13 | 87 | 0 | 21 | 157 | 38 | 16 | 0 | 0 | 6584 | 328 | 0 |
| SRR5230685 | 0 | 0 | 0 | 0 | 162 | 90 | 0 | 29 | 11 | 212 | 343 | 58 | 0 | 0 | 3468 | 445080 | 22867 | 0 | 64 | 15 | 751 | 12 | 12 | 0 | 0 | 89 | 39 | 15 | 12 | 89 | 66 | 0 | 0 | 0 | 3884 | 0 | 15 |
| SRR5230686 | 14 | 11 | 0 | 0 | 148 | 112 | 0 | 18 | 16 | 23 | 18 | 23 | 0 | 0 | 2247 | 480182 | 44494 | 0 | 57 | 22 | 1543 | 20 | 11 | 0 | 20 | 0 | 94 | 17 | 0 | 165 | 97 | 0 | 0 | 0 | 6443 | 0 | 0 |
| SRR5230687 | 0 | 0 | 0 | 0 | 53 | 85 | 0 | 0 | 24 | 18 | 14 | 97 | 0 | 0 | 5210 | 615536 | 60852 | 0 | 83 | 16 | 952 | 88 | 88 | 39 | 44 | 81 | 218 | 28 | 0 | 92 | 167 | 0 | 0 | 0 | 3140 | 0 | 0 |
| SRR5230688 | 0 | 0 | 0 | 0 | 36 | 384 | 0 | 18 | 49 | 75 | 75 | 0 | 0 | 0 | 6951 | 452351 | 48018 | 0 | 116 | 223 | 1847 | 25 | 45 | 0 | 0 | 13 | 21 | 0 | 0 | 42 | 45 | 0 | 0 | 0 | 3991 | 0 | 0 |
| SRR5230689 | 0 | 0 | 0 | 0 | 168 | 235 | 0 | 64 | 54 | 455 | 545 | 24 | 0 | 0 | 2253 | 135507 | 5539 | 44 | 60 | 151 | 1424 | 36 | 63 | 0 | 0 | 21 | 135 | 15 | 0 | 183 | 81 | 0 | 0 | 0 | 678 | 0 | 0 |
| SRR5230690 | 0 | 0 | 0 | 0 | 61 | 187 | 0 | 37 | 58 | 377 | 244 | 80 | 0 | 0 | 3670 | 452693 | 38505 | 24 | 67 | 32 | 759 | 130 | 146 | 19 | 29 | 106 | 278 | 45 | 0 | 95 | 262 | 19 | 0 | 0 | 11043 | 0 | 0 |
| SRR5230691 | 0 | 0 | 0 | 0 | 32 | 69 | 0 | 0 | 53 | 123 | 128 | 21 | 0 | 0 | 1226 | 460606 | 17675 | 39 | 183 | 29 | 953 | 0 | 0 | 18 | 0 | 30 | 114 | 13 | 0 | 63 | 0 | 0 | 0 | 0 | 2516 | 0 | 0 |
| SRR5230692 | 0 | 0 | 0 | 0 | 47 | 1295 | 15 | 130 | 143 | 173 | 126 | 36 | 0 | 0 | 5094 | 352199 | 50234 | 0 | 88 | 75 | 2032 | 0 | 0 | 11 | 0 | 42 | 147 | 14 | 0 | 45 | 27 | 17 | 0 | 0 | 5333 | 0 | 0 |
| SRR5230693 | 0 | 0 | 0 | 0 | 29 | 51 | 0 | 0 | 13 | 532 | 272 | 129 | 0 | 0 | 1445 | 455929 | 19336 | 0 | 57 | 0 | 660 | 68 | 118 | 16 | 0 | 0 | 0 | 0 | 25 | 143 | 134 | 13 | 0 | 11 | 11288 | 0 | 0 |
| SRR5230694 | 0 | 0 | 0 | 0 | 78 | 374 | 0 | 39 | 30 | 1482 | 1249 | 175 | 0 | 0 | 938 | 721193 | 12514 | 16 | 23 | 134 | 5195 | 87 | 96 | 0 | 0 | 0 | 21 | 0 | 0 | 232 | 100 | 62 | 0 | 0 | 3232 | 0 | 0 |
| SRR5230695 | 0 | 0 | 0 | 0 | 111 | 63 | 0 | 0 | 17 | 344 | 325 | 20 | 17 | 0 | 1492 | 360962 | 21559 | 15 | 32 | 54 | 445 | 45 | 26 | 0 | 19 | 22 | 118 | 17 | 32 | 75 | 64 | 11 | 0 | 0 | 10097 | 0 | 0 |
| SRR5230696 | 0 | 0 | 0 | 0 | 127 | 198 | 13 | 45 | 29 | 388 | 423 | 76 | 0 | 0 | 2914 | 404478 | 8879 | 0 | 95 | 119 | 7540 | 71 | 68 | 0 | 0 | 35 | 57 | 0 | 0 | 55 | 173 | 20 | 0 | 0 | 1642 | 0 | 0 |
| SRR5230697 | 0 | 0 | 0 | 0 | 37 | 18 | 0 | 11 | 16 | 216 | 237 | 22 | 0 | 0 | 2011 | 463076 | 27701 | 0 | 35 | 22 | 1055 | 31 | 46 | 0 | 18 | 31 | 109 | 20 | 36 | 42 | 36 | 14 | 0 | 0 | 8686 | 0 | 0 |
| SRR5230698 | 0 | 0 | 0 | 0 | 79 | 43 | 0 | 0 | 19 | 231 | 130 | 20 | 0 | 0 | 2632 | 667916 | 52511 | 0 | 48 | 17 | 840 | 73 | 141 | 0 | 20 | 55 | 74 | 12 | 17 | 22 | 293 | 0 | 0 | 0 | 1813 | 0 | 0 |
| SRR5230699 | 0 | 0 | 0 | 0 | 108 | 1258 | 17 | 151 | 200 | 307 | 142 | 25 | 0 | 0 | 3907 | 316259 | 42087 | 0 | 179 | 263 | 3091 | 166 | 60 | 0 | 0 | 106 | 252 | 18 | 0 | 58 | 298 | 0 | 0 | 0 | 2645 | 0 | 0 |
| SRR5230700 | 0 | 0 | 0 | 0 | 29 | 54 | 0 | 0 | 28 | 21 | 15 | 29 | 0 | 0 | 3048 | 312822 | 23838 | 17 | 45 | 52 | 655 | 14 | 29 | 0 | 12 | 29 | 32 | 0 | 22 | 164 | 105 | 0 | 0 | 0 | 4065 | 0 | 0 |
| SRR5230701 | 0 | 0 | 0 | 0 | 17 | 149 | 0 | 30 | 55 | 12 | 12 | 16 | 0 | 0 | 2219 | 339121 | 24708 | 16 | 33 | 23 | 1029 | 40 | 41 | 0 | 0 | 14 | 56 | 14 | 0 | 37 | 54 | 0 | 0 | 0 | 15573 | 0 | 0 |
| SRR5230702 | 0 | 0 | 0 | 0 | 55 | 662 | 0 | 59 | 36 | 24 | 157 | 104 | 0 | 0 | 4411 | 373071 | 31589 | 0 | 71 | 54 | 1607 | 24 | 0 | 0 | 0 | 29 | 51 | 0 | 0 | 92 | 0 | 15 | 0 | 0 | 4240 | 18 | 0 |
| SRR5230703 | 0 | 0 | 0 | 0 | 130 | 204 | 0 | 15 | 43 | 51 | 17 | 93 | 0 | 0 | 4525 | 618103 | 79320 | 0 | 100 | 69 | 12978 | 13 | 18 | 0 | 0 | 0 | 32 | 0 | 25 | 121 | 48 | 68 | 0 | 0 | 3619 | 0 | 0 |
| SRR5230704 | 0 | 0 | 0 | 0 | 18 | 30 | 0 | 0 | 0 | 179 | 118 | 42 | 0 | 0 | 862 | 521519 | 30846 | 0 | 76 | 33 | 919 | 0 | 90 | 0 | 0 | 0 | 0 | 0 | 16 | 0 | 105 | 0 | 0 | 0 | 4244 | 0 | 0 |
| SRR5230705 | 0 | 0 | 0 | 0 | 11 | 26 | 0 | 0 | 0 | 21 | 77 | 16 | 0 | 0 | 3010 | 533619 | 6505 | 0 | 44 | 0 | 1492 | 16 | 53 | 0 | 0 | 0 | 0 | 0 | 0 | 0 | 17 | 122 | 0 | 0 | 2471 | 0 | 0 |
| SRR5230706 | 0 | 0 | 0 | 0 | 68 | 741 | 21 | 135 | 165 | 20 | 18 | 25 | 0 | 0 | 5601 | 338331 | 174917 | 27 | 84 | 234 | 3604 | 11 | 102 | 0 | 0 | 10 | 24 | 0 | 0 | 129 | 32 | 42 | 0 | 0 | 7285 | 0 | 0 |
| SRR5230707 | 0 | 0 | 0 | 0 | 96 | 255 | 0 | 40 | 28 | 168 | 59 | 23 | 0 | 0 | 5700 | 451000 | 70350 | 13 | 51 | 116 | 1613 | 425 | 549 | 0 | 0 | 13 | 39 | 0 | 0 | 80 | 1301 | 0 | 0 | 0 | 3583 | 0 | 0 |
| SRR5230708 | 0 | 0 | 0 | 0 | 302 | 505 | 10 | 41 | 65 | 122 | 165 | 43 | 0 | 0 | 6222 | 399769 | 42648 | 0 | 60 | 80 | 1547 | 22 | 65 | 0 | 0 | 137 | 115 | 22 | 0 | 58 | 101 | 0 | 0 | 0 | 2662 | 0 | 0 |
| SRR5230709 | 0 | 0 | 0 | 0 | 34 | 188 | 0 | 23 | 80 | 115 | 187 | 77 | 0 | 0 | 4068 | 299582 | 13848 | 27 | 188 | 241 | 3197 | 77 | 30 | 0 | 43 | 38 | 17 | 13 | 0 | 128 | 77 | 0 | 0 | 0 | 4696 | 0 | 0 |
| SRR5230710 | 0 | 0 | 0 | 0 | 307 | 153 | 0 | 33 | 53 | 86 | 20 | 0 | 0 | 0 | 2826 | 280792 | 30412 | 0 | 84 | 0 | 1076 | 12 | 0 | 0 | 12 | 19 | 32 | 0 | 0 | 40 | 15 | 0 | 0 | 0 | 2091 | 0 | 0 |
| SRR5230711 | 0 | 0 | 0 | 0 | 24 | 133 | 0 | 0 | 0 | 242 | 353 | 97 | 0 | 0 | 1409 | 240956 | 10807 | 0 | 41 | 0 | 554 | 10 | 0 | 0 | 0 | 0 | 0 | 0 | 0 | 66 | 21 | 33 | 0 | 0 | 1094 | 0 | 0 |
| SRR5230712 | 0 | 0 | 0 | 0 | 77 | 950 | 0 | 105 | 105 | 64 | 89 | 19 | 0 | 0 | 6187 | 227553 | 26974 | 10 | 135 | 70 | 1064 | 134 | 147 | 0 | 0 | 43 | 136 | 19 | 0 | 57 | 355 | 0 | 0 | 0 | 3003 | 0 | 0 |
| SRR5230713 | 0 | 0 | 0 | 0 | 21 | 224 | 0 | 50 | 73 | 108 | 82 | 13 | 0 | 0 | 6161 | 250861 | 57586 | 25 | 346 | 78 | 850 | 32 | 46 | 0 | 30 | 0 | 62 | 0 | 0 | 32 | 73 | 11 | 0 | 0 | 4568 | 71 | 0 |
| SRR5230714 | 0 | 0 | 0 | 0 | 15 | 1510 | 26 | 247 | 203 | 100 | 241 | 46 | 0 | 0 | 4555 | 467727 | 68332 | 39 | 147 | 350 | 5127 | 166 | 155 | 0 | 0 | 35 | 90 | 0 | 0 | 121 | 534 | 74 | 0 | 0 | 2287 | 0 | 0 |
| SRR5230716 | 0 | 0 | 0 | 0 | 142 | 85 | 0 | 28 | 0 | 20 | 40 | 0 | 0 | 0 | 5449 | 248400 | 13081 | 16 | 202 | 129 | 1307 | 0 | 0 | 0 | 0 | 0 | 0 | 0 | 0 | 121 | 0 | 0 | 0 | 0 | 2745 | 0 | 0 |
| SRR5230717 | 0 | 0 | 0 | 0 | 58 | 345 | 0 | 0 | 62 | 125 | 81 | 0 | 0 | 0 | 979 | 215994 | 12567 | 0 | 92 | 53 | 2077 | 69 | 52 | 0 | 0 | 0 | 16 | 14 | 0 | 50 | 165 | 39 | 0 | 0 | 4131 | 0 | 0 |
| SRR5230718 | 0 | 0 | 0 | 0 | 176 | 126 | 0 | 0 | 37 | 241 | 59 | 0 | 0 | 0 | 3607 | 386089 | 24019 | 34 | 189 | 45 | 743 | 0 | 0 | 0 | 0 | 0 | 84 | 0 | 0 | 96 | 40 | 0 | 0 | 0 | 12558 | 25 | 0 |
| SRR5230719 | 0 | 0 | 0 | 0 | 45 | 88 | 0 | 0 | 19 | 99 | 43 | 13 | 0 | 0 | 3077 | 223531 | 14083 | 0 | 97 | 0 | 924 | 0 | 0 | 11 | 0 | 0 | 49 | 0 | 0 | 16 | 11 | 0 | 0 | 0 | 3417 | 0 | 0 |
| SRR5230720 | 0 | 0 | 0 | 0 | 0 | 225 | 0 | 0 | 37 | 193 | 99 | 11 | 0 | 0 | 3214 | 390782 | 31458 | 0 | 75 | 36 | 723 | 93 | 87 | 0 | 0 | 30 | 113 | 0 | 0 | 12 | 228 | 24 | 0 | 0 | 3612 | 0 | 0 |
| SRR5230721 | 0 | 0 | 0 | 0 | 112 | 407 | 0 | 25 | 66 | 184 | 37 | 16 | 0 | 0 | 1815 | 245860 | 9083 | 0 | 108 | 83 | 1641 | 212 | 156 | 0 | 0 | 0 | 21 | 0 | 12 | 0 | 21 | 0 | 0 | 0 | 5026 | 0 | 0 |
| SRR5230722 | 0 | 0 | 0 | 0 | 48 | 209 | 0 | 35 | 19 | 63 | 50 | 0 | 0 | 0 | 2778 | 257877 | 28869 | 11 | 117 | 0 | 1089 | 0 | 0 | 0 | 0 | 14 | 62 | 0 | 0 | 31 | 32 | 0 | 0 | 0 | 2993 | 14 | 0 |
| SRR5230723 | 0 | 0 | 0 | 0 | 21 | 153 | 0 | 0 | 32 | 179 | 183 | 0 | 0 | 0 | 1242 | 449313 | 10033 | 0 | 18 | 45 | 476 | 204 | 200 | 0 | 0 | 41 | 200 | 12 | 12 | 45 | 293 | 0 | 0 | 0 | 4801 | 0 | 0 |
| SRR5230724 | 0 | 0 | 0 | 0 | 0 | 56 | 0 | 0 | 32 | 153 | 86 | 24 | 0 | 0 | 1187 | 364077 | 6580 | 16 | 19 | 35 | 1290 | 0 | 13 | 0 | 0 | 0 | 13 | 0 | 0 | 24 | 0 | 0 | 0 | 0 | 3722 | 21 | 0 |
| SRR5230725 | 0 | 0 | 0 | 0 | 165 | 310 | 11 | 16 | 55 | 272 | 189 | 28 | 0 | 0 | 9297 | 368666 | 29296 | 0 | 110 | 41 | 1156 | 55 | 130 | 0 | 39 | 63 | 201 | 43 | 0 | 75 | 189 | 16 | 0 | 0 | 2581 | 18 | 0 |
| SRR5230726 | 0 | 0 | 0 | 0 | 62 | 36 | 0 | 23 | 101 | 89 | 47 | 0 | 0 | 0 | 1250 | 417331 | 6473 | 0 | 20 | 193 | 2722 | 83 | 74 | 0 | 15 | 0 | 28 | 0 | 0 | 28 | 281 | 0 | 0 | 0 | 1912 | 0 | 0 |
| SRR5230727 | 0 | 0 | 0 | 0 | 62 | 198 | 0 | 20 | 17 | 153 | 112 | 13 | 0 | 0 | 1688 | 315989 | 10813 | 46 | 50 | 86 | 1116 | 130 | 202 | 0 | 0 | 0 | 30 | 0 | 13 | 58 | 416 | 13 | 0 | 0 | 3980 | 0 | 0 |
| SRR5230728 | 0 | 0 | 0 | 0 | 17 | 1061 | 0 | 147 | 65 | 37 | 65 | 11 | 0 | 0 | 3490 | 209674 | 32918 | 0 | 133 | 357 | 1327 | 0 | 0 | 0 | 0 | 22 | 84 | 11 | 0 | 19 | 0 | 68 | 0 | 0 | 2013 | 20 | 0 |
| SRR5230729 | 0 | 0 | 0 | 0 | 29 | 1186 | 0 | 76 | 108 | 161 | 169 | 26 | 0 | 0 | 6074 | 394961 | 31031 | 0 | 104 | 98 | 2799 | 32 | 34 | 0 | 23 | 66 | 175 | 20 | 0 | 89 | 92 | 0 | 0 | 0 | 1373 | 0 | 0 |
| SRR5230732 | 0 | 0 | 0 | 0 | 78 | 742 | 0 | 17 | 73 | 189 | 125 | 136 | 0 | 0 | 7623 | 556797 | 49268 | 10 | 160 | 108 | 1141 | 22 | 27 | 17 | 23 | 32 | 55 | 0 | 0 | 52 | 71 | 62 | 0 | 15 | 4326 | 12 | 0 |
| SRR5230733 | 0 | 0 | 0 | 0 | 49 | 135 | 0 | 0 | 19 | 30 | 87 | 72 | 0 | 0 | 3769 | 229683 | 19528 | 0 | 64 | 17 | 1016 | 13 | 12 | 0 | 0 | 0 | 0 | 0 | 0 | 25 | 38 | 30 | 0 | 0 | 3001 | 31 | 0 |
| SRR5230734 | 0 | 0 | 0 | 0 | 0 | 179 | 0 | 0 | 0 | 223 | 42 | 250 | 0 | 0 | 4573 | 474833 | 30951 | 0 | 399 | 87 | 1823 | 0 | 19 | 0 | 0 | 0 | 0 | 0 | 0 | 15 | 0 | 415 | 0 | 0 | 3396 | 69 | 12 |
| SRR5230735 | 0 | 0 | 0 | 0 | 22 | 255 | 0 | 28 | 0 | 189 | 89 | 39 | 0 | 0 | 3827 | 245401 | 122284 | 0 | 98 | 50 | 924 | 122 | 83 | 0 | 0 | 0 | 50 | 0 | 0 | 52 | 198 | 38 | 0 | 0 | 7260 | 0 | 0 |
| SRR5230736 | 0 | 0 | 0 | 0 | 115 | 73 | 0 | 0 | 32 | 146 | 94 | 0 | 0 | 0 | 3118 | 277930 | 16082 | 123 | 59 | 73 | 1223 | 0 | 28 | 0 | 0 | 0 | 0 | 0 | 0 | 52 | 17 | 0 | 0 | 0 | 3708 | 0 | 0 |
| SRR5230737 | 0 | 0 | 0 | 0 | 414 | 430 | 0 | 42 | 57 | 86 | 40 | 15 | 0 | 0 | 4232 | 370843 | 17641 | 0 | 96 | 93 | 2680 | 56 | 61 | 10 | 0 | 15 | 40 | 0 | 15 | 15 | 141 | 12 | 0 | 0 | 1549 | 16 | 0 |
| SRR5230738 | 0 | 0 | 0 | 0 | 52 | 79 | 0 | 0 | 43 | 154 | 122 | 20 | 0 | 0 | 5001 | 302760 | 25944 | 43 | 54 | 61 | 498 | 0 | 0 | 0 | 32 | 0 | 35 | 0 | 0 | 72 | 0 | 13 | 0 | 0 | 7449 | 0 | 0 |
| SRR5230739 | 0 | 0 | 0 | 0 | 2677 | 111 | 0 | 10 | 29 | 194 | 41 | 49 | 0 | 0 | 386 | 318834 | 7736 | 0 | 0 | 0 | 519 | 0 | 0 | 0 | 0 | 14 | 63 | 0 | 0 | 0 | 0 | 11 | 0 | 11 | 2950 | 0 | 0 |
| SRR5230740 | 0 | 0 | 0 | 0 | 0 | 275 | 0 | 0 | 34 | 60 | 27 | 0 | 0 | 0 | 2870 | 421071 | 23586 | 0 | 113 | 191 | 7417 | 40 | 60 | 0 | 0 | 0 | 0 | 0 | 0 | 23 | 14 | 59 | 0 | 0 | 1849 | 0 | 0 |
| SRR5230741 | 0 | 0 | 0 | 0 | 27 | 813 | 0 | 89 | 57 | 70 | 64 | 76 | 0 | 0 | 4115 | 310787 | 81573 | 0 | 179 | 45 | 1549 | 96 | 53 | 0 | 0 | 0 | 23 | 0 | 0 | 12 | 18 | 30 | 0 | 0 | 11840 | 96 | 0 |
| SRR5230742 | 0 | 0 | 0 | 0 | 70 | 488 | 0 | 38 | 11 | 168 | 98 | 70 | 0 | 0 | 4981 | 372066 | 28169 | 23 | 38 | 61 | 694 | 21 | 0 | 0 | 28 | 21 | 49 | 0 | 0 | 84 | 28 | 61 | 0 | 0 | 4290 | 0 | 0 |
| SRR5230743 | 0 | 0 | 0 | 0 | 59 | 163 | 0 | 0 | 37 | 182 | 135 | 36 | 0 | 0 | 3866 | 301052 | 31229 | 0 | 86 | 77 | 3984 | 190 | 150 | 0 | 0 | 19 | 25 | 0 | 0 | 74 | 580 | 32 | 0 | 0 | 3522 | 0 | 0 |
| SRR5230744 | 0 | 0 | 0 | 0 | 29 | 100 | 0 | 23 | 42 | 53 | 65 | 14 | 0 | 0 | 3042 | 182556 | 23712 | 26 | 129 | 0 | 545 | 0 | 0 | 0 | 0 | 0 | 25 | 0 | 0 | 49 | 20 | 0 | 0 | 0 | 4845 | 0 | 0 |
| SRR5230745 | 13 | 0 | 0 | 0 | 929 | 3348 | 21 | 309 | 201 | 277 | 120 | 101 | 0 | 0 | 4075 | 147317 | 17777 | 46 | 149 | 144 | 1427 | 0 | 25 | 0 | 28 | 41 | 142 | 0 | 0 | 16 | 60 | 196 | 0 | 16 | 2103 | 0 | 0 |
| SRR5230746 | 0 | 0 | 0 | 0 | 296 | 312 | 0 | 0 | 23 | 374 | 292 | 18 | 0 | 0 | 4767 | 373982 | 32547 | 0 | 36 | 23 | 1907 | 73 | 36 | 0 | 27 | 0 | 87 | 0 | 18 | 141 | 141 | 0 | 0 | 0 | 3941 | 0 | 0 |
| SRR5230747 | 0 | 0 | 0 | 0 | 85 | 303 | 0 | 36 | 87 | 121 | 45 | 22 | 0 | 0 | 3616 | 475746 | 29399 | 0 | 60 | 41 | 907 | 81 | 36 | 0 | 0 | 0 | 36 | 22 | 0 | 31 | 58 | 0 | 0 | 0 | 5309 | 0 | 0 |
| SRR5230748 | 0 | 0 | 0 | 0 | 76 | 524 | 0 | 70 | 176 | 205 | 135 | 22 | 0 | 0 | 7911 | 450234 | 45224 | 28 | 88 | 32 | 1354 | 162 | 108 | 27 | 16 | 81 | 265 | 22 | 0 | 43 | 303 | 18 | 0 | 0 | 1835 | 0 | 0 |
| SRR5230749 | 0 | 0 | 0 | 0 | 36 | 400 | 0 | 40 | 57 | 11 | 70 | 0 | 0 | 0 | 4574 | 364091 | 33197 | 0 | 94 | 89 | 3046 | 47 | 14 | 0 | 0 | 0 | 0 | 0 | 0 | 33 | 125 | 14 | 0 | 0 | 4334 | 0 | 0 |
| SRR5230750 | 0 | 0 | 0 | 0 | 11 | 38 | 0 | 0 | 0 | 99 | 21 | 0 | 0 | 0 | 2669 | 329955 | 15400 | 34 | 91 | 38 | 771 | 0 | 31 | 0 | 0 | 0 | 0 | 0 | 0 | 65 | 31 | 19 | 0 | 0 | 7051 | 0 | 0 |
| SRR5230751 | 0 | 0 | 0 | 0 | 69 | 124 | 12 | 32 | 24 | 72 | 31 | 25 | 0 | 0 | 3282 | 298827 | 5475 | 36 | 36 | 108 | 377 | 56 | 65 | 0 | 0 | 0 | 0 | 0 | 0 | 31 | 156 | 0 | 0 | 0 | 5195 | 0 | 0 |
| SRR5230752 | 0 | 0 | 0 | 0 | 226 | 300 | 0 | 34 | 29 | 166 | 138 | 105 | 0 | 0 | 4350 | 401098 | 20702 | 10 | 216 | 99 | 693 | 28 | 105 | 39 | 0 | 22 | 83 | 11 | 22 | 94 | 66 | 60 | 0 | 0 | 3201 | 45 | 0 |
| SRR5230753 | 0 | 0 | 0 | 0 | 129 | 259 | 0 | 69 | 50 | 157 | 84 | 42 | 0 | 0 | 3417 | 243762 | 22162 | 0 | 115 | 22 | 1504 | 14 | 17 | 31 | 28 | 0 | 0 | 0 | 20 | 84 | 28 | 32 | 0 | 11 | 3361 | 0 | 0 |
| SRR5231309 | 0 | 0 | 0 | 11 | 12 | 0 | 14 | 19 | 0 | 0 | 446 | 0 | 0 | 0 | 3154 | 30237 | 4238 | 528 | 0 | 907 | 861 | 0 | 0 | 0 | 0 | 0 | 0 | 0 | 0 | 655 | 0 | 5333 | 21 | 0 | 738 | 741 | 371 |
| SRR5231310 | 0 | 0 | 0 | 0 | 0 | 11 | 37 | 32 | 0 | 0 | 391 | 0 | 0 | 0 | 4104 | 19357 | 3861 | 474 | 0 | 769 | 3532 | 0 | 0 | 0 | 0 | 0 | 0 | 0 | 0 | 557 | 0 | 5082 | 28 | 0 | 1592 | 594 | 477 |
| SRR5231311 | 0 | 0 | 0 | 0 | 30 | 18 | 32 | 49 | 0 | 0 | 463 | 0 | 0 | 0 | 11029 | 31121 | 7565 | 355 | 0 | 806 | 5709 | 0 | 0 | 0 | 0 | 0 | 0 | 0 | 0 | 638 | 0 | 6239 | 12 | 0 | 551 | 566 | 223 |
| SRR5231312 | 0 | 0 | 0 | 0 | 0 | 0 | 16 | 13 | 0 | 0 | 184 | 0 | 0 | 0 | 2310 | 17273 | 1886 | 635 | 0 | 617 | 768 | 0 | 0 | 0 | 0 | 0 | 0 | 0 | 0 | 379 | 0 | 7401 | 56 | 0 | 3503 | 644 | 1320 |
| SRR5231313 | 0 | 0 | 0 | 0 | 0 | 0 | 13 | 23 | 0 | 0 | 329 | 0 | 0 | 0 | 3124 | 16780 | 2307 | 346 | 0 | 516 | 2292 | 0 | 0 | 0 | 0 | 0 | 0 | 0 | 0 | 363 | 0 | 6182 | 10 | 0 | 1374 | 586 | 176 |

### Supp Table S8.3.Columns_BX_to_DH

| | | | | | | | | | | | | | | | | | | | | | | | | | | | | | | | | | | | | | |
|---|---|---|---|---|---|---|---|---|---|---|---|---|---|---|---|---|---|---|---|---|---|---|---|---|---|---|---|---|---|---|---|---|---|---|---|---|---|
| SRR5231314 | 0 | 0 | 0 | 0 | 36 | 25 | 57 | 85 | 13 | 0 | 480 | 0 | 0 | 0 | 25373 | 23421 | 13310 | 369 | 16 | 1106 | 13464 | 0 | 0 | 0 | 0 | 0 | 0 | 0 | 0 | 917 | 0 | 5472 | 0 | 0 | 986 | 533 | 212 |
| SRR5239182 | 0 | 0 | 0 | 0 | 0 | 210 | 187 | 1516 | 105 | 0 | 38 | 17 | 0 | 0 | 14927 | 12031 | 19647 | 70 | 117 | 5261 | 4682 | 0 | 0 | 0 | 0 | 0 | 0 | 0 | 0 | 123 | 0 | 124 | 0 | 0 | 27833 | 22 | 0 |
| SRR5239183 | 0 | 23 | 10 | 16 | 0 | 29 | 66 | 95 | 15 | 0 | 66 | 0 | 0 | 0 | 6933 | 4936 | 6767 | 22 | 44 | 549 | 3419 | 0 | 0 | 0 | 0 | 15 | 20 | 26 | 0 | 32 | 0 | 146 | 0 | 0 | 26643 | 0 | 0 |
| SRR5239184 | 10 | 17 | 11 | 15 | 0 | 121 | 105 | 721 | 37 | 0 | 141 | 17 | 0 | 0 | 6233 | 5734 | 6900 | 26 | 26 | 910 | 4224 | 0 | 0 | 0 | 0 | 0 | 0 | 0 | 0 | 107 | 0 | 110 | 0 | 0 | 23665 | 0 | 0 |
| SRR5239185 | 0 | 0 | 0 | 0 | 0 | 393 | 327 | 2547 | 167 | 0 | 78 | 13 | 0 | 0 | 8901 | 5498 | 10891 | 153 | 47 | 3280 | 4914 | 0 | 0 | 0 | 0 | 0 | 0 | 0 | 0 | 142 | 0 | 700 | 0 | 0 | 26490 | 0 | 0 |
| SRR5239186 | 0 | 0 | 0 | 0 | 0 | 793 | 699 | 4604 | 339 | 0 | 61 | 22 | 0 | 0 | 15762 | 19368 | 28858 | 142 | 201 | 9319 | 7010 | 0 | 0 | 0 | 0 | 0 | 0 | 0 | 0 | 141 | 0 | 582 | 0 | 0 | 16812 | 18 | 0 |
| SRR5239187 | 11 | 10 | 18 | 18 | 0 | 125 | 89 | 676 | 78 | 0 | 316 | 13 | 0 | 0 | 9176 | 7748 | 11373 | 157 | 135 | 1498 | 2818 | 0 | 0 | 0 | 0 | 0 | 0 | 0 | 0 | 113 | 0 | 363 | 0 | 0 | 23377 | 12 | 0 |
| SRR5239188 | 0 | 13 | 11 | 11 | 0 | 308 | 288 | 2272 | 148 | 0 | 279 | 36 | 0 | 0 | 10131 | 8705 | 19874 | 398 | 87 | 3080 | 2842 | 0 | 0 | 0 | 0 | 0 | 0 | 0 | 0 | 207 | 0 | 1088 | 0 | 0 | 23436 | 22 | 0 |
| SRR5275577 | 31 | 0 | 0 | 0 | 0 | 34 | 68 | 469 | 38 | 0 | 43 | 30 | 0 | 0 | 4040 | 5345 | 12150 | 1020 | 0 | 205 | 2000 | 0 | 0 | 12 | 21 | 61 | 34 | 0 | 14 | 62 | 0 | 22673 | 78 | 0 | 1281 | 0 | 109 |
| SRR5275578 | 23 | 0 | 0 | 0 | 0 | 61 | 122 | 811 | 47 | 0 | 33 | 39 | 0 | 0 | 4054 | 4566 | 11638 | 798 | 0 | 188 | 2116 | 0 | 0 | 15 | 16 | 45 | 28 | 0 | 16 | 52 | 0 | 22232 | 116 | 0 | 1577 | 0 | 119 |
| SRR5275579 | 26 | 0 | 0 | 0 | 0 | 49 | 94 | 663 | 44 | 0 | 49 | 54 | 0 | 0 | 2757 | 4559 | 8382 | 934 | 0 | 190 | 1502 | 0 | 0 | 12 | 20 | 56 | 31 | 0 | 15 | 65 | 0 | 24511 | 77 | 0 | 1020 | 0 | 117 |
| SRR5275580 | 31 | 11 | 11 | 16 | 0 | 185 | 234 | 1419 | 80 | 0 | 10 | 41 | 0 | 0 | 5786 | 5364 | 9006 | 1375 | 11 | 253 | 2827 | 0 | 0 | 15 | 16 | 13 | 18 | 0 | 20 | 24 | 0 | 23023 | 43 | 0 | 1525 | 0 | 44 |
| SRR5275581 | 63 | 16 | 0 | 0 | 0 | 204 | 277 | 1751 | 84 | 0 | 24 | 130 | 0 | 0 | 4580 | 4373 | 7036 | 1200 | 0 | 175 | 2188 | 0 | 0 | 15 | 13 | 25 | 0 | 0 | 20 | 55 | 0 | 26644 | 172 | 0 | 1608 | 0 | 121 |
| SRR5275582 | 107 | 36 | 27 | 45 | 0 | 78 | 102 | 699 | 56 | 0 | 115 | 229 | 0 | 0 | 5779 | 6577 | 12211 | 1071 | 10 | 326 | 2724 | 0 | 0 | 11 | 31 | 95 | 51 | 0 | 15 | 229 | 0 | 20007 | 77 | 0 | 846 | 0 | 82 |
| SRR5284195 | 61 | 27 | 17 | 0 | 127 | 1369 | 1538 | 4803 | 11281 | 24 | 32 | 25 | 0 | 0 | 9252 | 119065 | 70230 | 372 | 3394 | 3288 | 1790 | 0 | 0 | 14 | 15 | 0 | 0 | 0 | 0 | 571 | 0 | 44 | 0 | 19 | 7548 | 0 | 0 |
| SRR5284196 | 38 | 18 | 35 | 21 | 194 | 875 | 945 | 2892 | 8121 | 18 | 127 | 24 | 0 | 0 | 36412 | 94051 | 223797 | 174 | 3988 | 3155 | 1571 | 0 | 0 | 28 | 17 | 13 | 21 | 0 | 0 | 1110 | 0 | 52 | 0 | 11 | 4517 | 0 | 0 |
| SRR5284197 | 73 | 45 | 44 | 24 | 95 | 1394 | 847 | 4384 | 6771 | 49 | 100 | 18 | 0 | 0 | 24899 | 150129 | 129828 | 415 | 3495 | 1694 | 1711 | 0 | 0 | 27 | 11 | 11 | 14 | 0 | 0 | 964 | 0 | 30 | 0 | 20 | 5987 | 0 | 0 |
| SRR5284198 | 153 | 72 | 81 | 13 | 418 | 1871 | 2352 | 7299 | 17069 | 66 | 230 | 61 | 0 | 0 | 29975 | 123237 | 134417 | 440 | 3122 | 2421 | 1591 | 0 | 0 | 34 | 61 | 31 | 52 | 0 | 0 | 2345 | 0 | 69 | 0 | 13 | 9706 | 0 | 0 |
| SRR5284199 | 51 | 24 | 50 | 10 | 475 | 1039 | 1158 | 3261 | 7144 | 34 | 462 | 76 | 0 | 0 | 15738 | 175579 | 104296 | 311 | 2243 | 2878 | 1389 | 0 | 0 | 37 | 60 | 43 | 55 | 0 | 0 | 5099 | 0 | 46 | 0 | 10 | 2534 | 0 | 0 |
| SRR5284200 | 119 | 39 | 63 | 20 | 519 | 679 | 792 | 2519 | 6883 | 38 | 516 | 66 | 0 | 0 | 15254 | 194710 | 105694 | 246 | 2277 | 2516 | 943 | 0 | 0 | 23 | 70 | 51 | 68 | 0 | 0 | 4587 | 0 | 25 | 0 | 0 | 2405 | 0 | 0 |
| SRR5284201 | 30 | 18 | 16 | 12 | 77 | 1325 | 1223 | 6508 | 8037 | 14 | 108 | 22 | 0 | 0 | 27684 | 239885 | 132777 | 482 | 3755 | 1351 | 1683 | 0 | 0 | 10 | 21 | 16 | 10 | 0 | 0 | 1066 | 0 | 13 | 0 | 19 | 4316 | 0 | 0 |
| SRR5284202 | 53 | 13 | 0 | 0 | 465 | 2341 | 1519 | 8757 | 12812 | 243 | 117 | 65 | 0 | 0 | 16861 | 176229 | 113861 | 345 | 2341 | 1142 | 1977 | 0 | 0 | 46 | 64 | 0 | 0 | 0 | 0 | 2282 | 0 | 13 | 0 | 24 | 6905 | 0 | 0 |
| SRR5284203 | 14 | 30 | 23 | 25 | 145 | 817 | 365 | 2758 | 3795 | 30 | 233 | 40 | 0 | 0 | 32235 | 109434 | 236804 | 212 | 2220 | 1654 | 2070 | 0 | 0 | 31 | 15 | 16 | 22 | 0 | 0 | 1785 | 0 | 20 | 0 | 10 | 8260 | 0 | 0 |
| SRR5284204 | 135 | 56 | 39 | 17 | 946 | 965 | 1237 | 4079 | 11203 | 34 | 284 | 104 | 0 | 0 | 8771 | 122164 | 87218 | 261 | 1749 | 3242 | 805 | 0 | 0 | 37 | 40 | 11 | 12 | 0 | 0 | 3572 | 0 | 43 | 0 | 0 | 4835 | 0 | 0 |
| SRR5284205 | 84 | 38 | 12 | 12 | 91 | 2981 | 2170 | 10328 | 12892 | 49 | 72 | 18 | 15 | 0 | 35118 | 76315 | 208701 | 238 | 4365 | 3784 | 1236 | 0 | 0 | 17 | 17 | 13 | 12 | 0 | 0 | 776 | 0 | 17 | 0 | 13 | 6207 | 0 | 0 |
| SRR5284206 | 70 | 25 | 11 | 0 | 126 | 3879 | 2762 | 12578 | 18981 | 33 | 65 | 17 | 16 | 0 | 31508 | 120571 | 162901 | 414 | 3900 | 2803 | 2545 | 0 | 0 | 24 | 22 | 14 | 13 | 0 | 0 | 687 | 0 | 31 | 0 | 19 | 5734 | 0 | 0 |
| SRR5317818 | 0 | 0 | 0 | 0 | 0 | 79 | 180 | 366 | 107 | 27 | 10 | 10 | 0 | 0 | 94178 | 184998 | 482256 | 23 | 141 | 14773 | 4958 | 0 | 0 | 39 | 0 | 0 | 11 | 0 | 0 | 156 | 0 | 34 | 0 | 0 | 1863 | 121 | 20 |
| SRR5317819 | 12 | 14 | 14 | 17 | 15 | 34 | 57 | 87 | 0 | 16 | 58 | 13 | 0 | 0 | 112391 | 250383 | 364000 | 10 | 39 | 8401 | 7337 | 0 | 0 | 19 | 16 | 39 | 75 | 12 | 0 | 153 | 0 | 23 | 0 | 0 | 241 | 66 | 0 |
| SRR5317820 | 0 | 0 | 0 | 0 | 0 | 253 | 1399 | 1785 | 234 | 21 | 21 | 12 | 0 | 0 | 17769 | 140975 | 241118 | 0 | 43 | 1768 | 931 | 0 | 0 | 49 | 14 | 37 | 24 | 0 | 0 | 73 | 0 | 14 | 0 | 0 | 1981 | 254 | 11 |
| SRR5317821 | 0 | 0 | 0 | 0 | 61 | 52 | 11 | 41 | 0 | 72 | 0 | 0 | 0 | 0 | 135457 | 631722 | 275406 | 0 | 11 | 1462 | 11267 | 0 | 0 | 66 | 0 | 12 | 26 | 0 | 0 | 65 | 0 | 26 | 0 | 0 | 51 | 134 | 0 |
| SRR5317822 | 0 | 0 | 0 | 0 | 0 | 1240 | 753 | 5172 | 377 | 91 | 57 | 30 | 0 | 0 | 71501 | 200903 | 349065 | 24 | 84 | 9355 | 5173 | 0 | 0 | 35 | 46 | 93 | 13 | 0 | 0 | 57 | 0 | 46 | 0 | 0 | 144 | 261 | 0 |
| SRR5317823 | 0 | 0 | 0 | 0 | 28 | 45 | 55 | 300 | 11 | 37 | 209 | 13 | 0 | 0 | 81916 | 422866 | 250974 | 11 | 41 | 3076 | 10394 | 0 | 0 | 48 | 20 | 41 | 52 | 0 | 0 | 598 | 0 | 37 | 0 | 0 | 41 | 110 | 0 |
| SRR5317824 | 0 | 0 | 0 | 0 | 0 | 107 | 219 | 480 | 81 | 112 | 156 | 69 | 0 | 0 | 96986 | 56217 | 514264 | 0 | 108 | 25355 | 5858 | 0 | 0 | 19 | 0 | 0 | 27 | 0 | 0 | 1589 | 0 | 26 | 0 | 0 | 1888 | 136 | 19 |
| SRR5317825 | 0 | 0 | 0 | 11 | 159 | 30 | 52 | 160 | 0 | 134 | 348 | 0 | 0 | 0 | 226337 | 251597 | 640318 | 13 | 22 | 6740 | 24733 | 0 | 0 | 186 | 64 | 126 | 84 | 0 | 0 | 1900 | 0 | 0 | 0 | 0 | 0 | 210 | 0 |
| SRR5317826 | 0 | 0 | 0 | 0 | 14 | 115 | 187 | 597 | 40 | 48 | 177 | 31 | 0 | 0 | 76041 | 95821 | 495672 | 0 | 201 | 24958 | 17506 | 0 | 0 | 48 | 0 | 29 | 35 | 0 | 0 | 823 | 0 | 45 | 0 | 0 | 224 | 128 | 0 |
| SRR5317827 | 0 | 0 | 0 | 17 | 25 | 80 | 118 | 540 | 14 | 23 | 52 | 16 | 0 | 0 | 55210 | 263639 | 421200 | 0 | 150 | 10406 | 13404 | 0 | 0 | 20 | 13 | 44 | 39 | 0 | 0 | 68 | 0 | 0 | 0 | 0 | 292 | 65 | 0 |
| SRR5387779 | 0 | 0 | 0 | 14 | 82 | 0 | 0 | 0 | 0 | 0 | 233 | 0 | 0 | 0 | 3913 | 38201 | 52680 | 0 | 0 | 1423 | 0 | 0 | 0 | 14 | 14 | 27 | 124 | 69 | 0 | 124 | 0 | 0 | 0 | 0 | 1232 | 0 | 0 |
| SRR5387780 | 0 | 0 | 0 | 0 | 38 | 0 | 0 | 71 | 0 | 0 | 361 | 0 | 0 | 0 | 1481 | 11311 | 23065 | 0 | 0 | 1340 | 141 | 0 | 0 | 15 | 15 | 30 | 211 | 68 | 0 | 248 | 0 | 0 | 0 | 0 | 4214 | 0 | 0 |
| SRR5387781 | 0 | 0 | 0 | 0 | 31 | 0 | 0 | 67 | 0 | 0 | 290 | 74 | 0 | 0 | 2090 | 8416 | 17954 | 0 | 0 | 1348 | 202 | 0 | 0 | 0 | 0 | 25 | 105 | 74 | 0 | 333 | 0 | 0 | 0 | 0 | 5050 | 0 | 0 |
| SRR5387782 | 0 | 0 | 0 | 0 | 14 | 0 | 0 | 65 | 65 | 0 | 136 | 0 | 0 | 0 | 1947 | 12733 | 27662 | 0 | 0 | 973 | 130 | 0 | 0 | 0 | 14 | 29 | 179 | 108 | 0 | 143 | 0 | 0 | 0 | 0 | 1976 | 0 | 0 |
| SRR5387783 | 0 | 0 | 0 | 0 | 25 | 0 | 0 | 0 | 0 | 0 | 226 | 0 | 0 | 0 | 985 | 21155 | 16086 | 0 | 0 | 1560 | 82 | 0 | 0 | 0 | 25 | 42 | 234 | 125 | 0 | 151 | 0 | 0 | 0 | 0 | 2204 | 0 | 17 |
| SRR5387784 | 0 | 0 | 0 | 0 | 0 | 0 | 0 | 58 | 0 | 0 | 149 | 0 | 0 | 0 | 639 | 11852 | 14329 | 0 | 58 | 2092 | 58 | 0 | 0 | 0 | 0 | 31 | 188 | 47 | 0 | 266 | 0 | 0 | 0 | 0 | 2300 | 0 | 0 |
| SRR5387785 | 0 | 0 | 0 | 0 | 13 | 45 | 0 | 91 | 0 | 0 | 534 | 0 | 0 | 0 | 5346 | 21245 | 28594 | 0 | 45 | 1133 | 136 | 0 | 0 | 0 | 20 | 20 | 125 | 53 | 0 | 237 | 0 | 0 | 0 | 0 | 3474 | 0 | 0 |
| SRR5387786 | 0 | 0 | 0 | 0 | 32 | 0 | 0 | 78 | 0 | 0 | 365 | 0 | 0 | 0 | 1945 | 13042 | 14672 | 0 | 0 | 1245 | 0 | 0 | 0 | 24 | 0 | 32 | 113 | 57 | 0 | 308 | 0 | 0 | 0 | 0 | 3959 | 0 | 0 |
| SRR5387787 | 0 | 0 | 0 | 0 | 0 | 0 | 0 | 0 | 0 | 18 | 215 | 0 | 0 | 0 | 3115 | 28554 | 31968 | 0 | 0 | 1078 | 240 | 0 | 0 | 0 | 36 | 36 | 90 | 54 | 0 | 161 | 0 | 0 | 0 | 0 | 10553 | 0 | 0 |
| SRR5387788 | 0 | 0 | 0 | 14 | 0 | 74 | 0 | 0 | 0 | 0 | 214 | 0 | 0 | 0 | 1485 | 13089 | 18979 | 0 | 0 | 1114 | 149 | 0 | 0 | 0 | 14 | 0 | 166 | 28 | 0 | 138 | 0 | 0 | 0 | 0 | 1091 | 0 | 0 |
| SRR5387789 | 0 | 0 | 0 | 0 | 0 | 0 | 0 | 0 | 0 | 0 | 78 | 0 | 0 | 0 | 2025 | 14520 | 17256 | 0 | 0 | 975 | 0 | 0 | 0 | 19 | 0 | 0 | 0 | 29 | 0 | 117 | 0 | 0 | 0 | 0 | 2946 | 0 | 0 |
| SRR5387790 | 0 | 0 | 0 | 0 | 0 | 0 | 0 | 0 | 0 | 0 | 164 | 0 | 0 | 0 | 973 | 11823 | 10235 | 0 | 0 | 1687 | 0 | 0 | 0 | 0 | 0 | 21 | 178 | 79 | 0 | 207 | 0 | 0 | 0 | 0 | 1765 | 0 | 0 |
| SRR5387791 | 0 | 18 | 0 | 0 | 18 | 98 | 0 | 0 | 0 | 0 | 493 | 0 | 0 | 0 | 1566 | 19327 | 17570 | 0 | 0 | 1566 | 98 | 0 | 0 | 18 | 18 | 88 | 546 | 194 | 0 | 282 | 0 | 0 | 0 | 0 | 2928 | 0 | 0 |
| SRR5387792 | 0 | 0 | 0 | 31 | 51 | 89 | 0 | 0 | 0 | 0 | 287 | 0 | 0 | 0 | 715 | 11437 | 13783 | 0 | 0 | 1431 | 0 | 0 | 0 | 10 | 0 | 82 | 184 | 113 | 0 | 348 | 0 | 0 | 0 | 0 | 7038 | 0 | 0 |
| SRR5387793 | 0 | 0 | 0 | 29 | 101 | 0 | 0 | 0 | 0 | 0 | 259 | 0 | 0 | 0 | 1746 | 11920 | 14175 | 0 | 0 | 1247 | 125 | 0 | 0 | 14 | 14 | 29 | 129 | 43 | 0 | 144 | 0 | 0 | 0 | 0 | 6121 | 0 | 0 |
| SRR5387794 | 0 | 0 | 0 | 0 | 13 | 0 | 0 | 0 | 0 | 0 | 475 | 0 | 0 | 0 | 1260 | 12114 | 10443 | 0 | 0 | 1764 | 126 | 0 | 0 | 0 | 13 | 25 | 244 | 119 | 0 | 475 | 0 | 0 | 0 | 0 | 5848 | 0 | 0 |
| SRR5387795 | 0 | 0 | 0 | 22 | 11 | 0 | 0 | 114 | 0 | 0 | 183 | 0 | 0 | 0 | 1706 | 28197 | 22836 | 0 | 0 | 1593 | 228 | 0 | 0 | 22 | 11 | 75 | 205 | 97 | 0 | 118 | 0 | 0 | 0 | 0 | 1191 | 0 | 0 |
| SRR5387796 | 0 | 0 | 0 | 0 | 0 | 0 | 0 | 0 | 0 | 0 | 80 | 0 | 0 | 0 | 1656 | 22798 | 21904 | 0 | 0 | 2070 | 207 | 0 | 0 | 0 | 27 | 80 | 134 | 40 | 0 | 147 | 0 | 0 | 0 | 0 | 2682 | 0 | 0 |
| SRR5387797 | 0 | 0 | 16 | 0 | 0 | 0 | 0 | 112 | 0 | 0 | 318 | 0 | 0 | 0 | 8015 | 33976 | 33257 | 0 | 0 | 785 | 336 | 0 | 0 | 33 | 0 | 24 | 65 | 41 | 0 | 245 | 0 | 56 | 0 | 0 | 4175 | 0 | 0 |
| SRR5387798 | 0 | 0 | 0 | 44 | 15 | 0 | 0 | 79 | 0 | 0 | 161 | 15 | 0 | 0 | 2987 | 15506 | 17138 | 0 | 0 | 1572 | 1101 | 0 | 0 | 0 | 0 | 0 | 80 | 109 | 0 | 409 | 0 | 0 | 0 | 0 | 5713 | 0 | 0 |
| SRR5387799 | 0 | 0 | 0 | 0 | 23 | 0 | 0 | 0 | 0 | 0 | 109 | 0 | 0 | 0 | 821 | 26929 | 15498 | 0 | 0 | 1040 | 0 | 0 | 0 | 39 | 16 | 47 | 226 | 62 | 0 | 140 | 0 | 0 | 0 | 0 | 3078 | 0 | 0 |
| SRR5387800 | 0 | 0 | 0 | 0 | 33 | 0 | 0 | 105 | 0 | 0 | 174 | 0 | 0 | 0 | 947 | 14584 | 7292 | 0 | 0 | 210 | 105 | 0 | 0 | 33 | 17 | 66 | 323 | 149 | 0 | 207 | 0 | 0 | 0 | 0 | 3772 | 0 | 0 |
| SRR5387801 | 0 | 0 | 0 | 19 | 0 | 0 | 0 | 0 | 0 | 0 | 274 | 0 | 0 | 0 | 1134 | 11130 | 12243 | 0 | 0 | 1134 | 309 | 0 | 0 | 0 | 0 | 47 | 245 | 94 | 0 | 236 | 0 | 0 | 0 | 0 | 3339 | 0 | 0 |
| SRR5387802 | 0 | 0 | 0 | 13 | 19 | 0 | 0 | 0 | 0 | 0 | 223 | 0 | 0 | 0 | 740 | 10774 | 16491 | 0 | 0 | 986 | 247 | 0 | 0 | 0 | 13 | 19 | 121 | 45 | 0 | 153 | 0 | 0 | 0 | 0 | 1759 | 0 | 0 |
| SRR5387803 | 0 | 0 | 0 | 20 | 0 | 0 | 0 | 0 | 0 | 0 | 235 | 0 | 0 | 0 | 810 | 10540 | 16231 | 0 | 0 | 1701 | 81 | 0 | 0 | 0 | 0 | 13 | 209 | 59 | 0 | 10 | 0 | 0 | 0 | 0 | 4427 | 0 | 0 |
| SRR5387804 | 0 | 0 | 0 | 16 | 16 | 0 | 0 | 0 | 0 | 0 | 165 | 16 | 0 | 0 | 490 | 9021 | 18494 | 0 | 0 | 1796 | 82 | 0 | 0 | 16 | 0 | 39 | 196 | 94 | 0 | 220 | 0 | 0 | 0 | 0 | 3157 | 0 | 0 |
| SRR5387805 | 0 | 0 | 0 | 0 | 28 | 102 | 0 | 102 | 0 | 0 | 1392 | 0 | 0 | 0 | 5934 | 16208 | 28479 | 0 | 0 | 716 | 205 | 0 | 0 | 0 | 0 | 56 | 260 | 139 | 0 | 659 | 0 | 0 | 0 | 0 | 2778 | 0 | 0 |
| SRR5387806 | 0 | 0 | 0 | 32 | 32 | 0 | 0 | 0 | 0 | 0 | 375 | 0 | 0 | 0 | 2363 | 12665 | 33774 | 0 | 0 | 1636 | 545 | 0 | 0 | 0 | 16 | 56 | 239 | 120 | 0 | 207 | 0 | 0 | 0 | 0 | 5348 | 0 | 0 |
| SRR5387807 | 0 | 0 | 0 | 59 | 47 | 0 | 64 | 127 | 0 | 0 | 178 | 0 | 0 | 0 | 3693 | 21989 | 35521 | 0 | 0 | 1592 | 64 | 0 | 0 | 36 | 0 | 47 | 36 | 59 | 0 | 260 | 0 | 0 | 0 | 0 | 2537 | 0 | 0 |
| SRR5387808 | 0 | 0 | 0 | 0 | 30 | 0 | 0 | 61 | 0 | 0 | 209 | 0 | 0 | 0 | 1946 | 29570 | 22545 | 0 | 0 | 2007 | 0 | 0 | 0 | 20 | 0 | 0 | 129 | 60 | 0 | 258 | 0 | 0 | 0 | 0 | 2287 | 0 | 20 |
| SRR5387809 | 0 | 0 | 0 | 18 | 18 | 0 | 0 | 0 | 0 | 0 | 160 | 0 | 0 | 0 | 1828 | 10499 | 22497 | 0 | 0 | 762 | 0 | 0 | 0 | 0 | 0 | 89 | 142 | 177 | 0 | 71 | 0 | 0 | 0 | 0 | 1500 | 0 | 18 |
| SRR5387810 | 0 | 0 | 0 | 0 | 21 | 0 | 0 | 0 | 0 | 0 | 279 | 0 | 0 | 0 | 2818 | 21739 | 47315 | 0 | 0 | 1461 | 209 | 0 | 0 | 14 | 0 | 14 | 84 | 35 | 0 | 119 | 0 | 0 | 0 | 0 | 2877 | 0 | 0 |
| SRR5387811 | 0 | 0 | 0 | 31 | 0 | 0 | 0 | 0 | 0 | 0 | 218 | 0 | 0 | 0 | 554 | 7185 | 10159 | 0 | 0 | 1109 | 277 | 0 | 0 | 0 | 16 | 39 | 109 | 39 | 0 | 226 | 0 | 0 | 0 | 0 | 4460 | 0 | 0 |
| SRR5387812 | 0 | 0 | 14 | 0 | 14 | 0 | 0 | 0 | 0 | 0 | 174 | 0 | 0 | 0 | 742 | 8537 | 12110 | 0 | 0 | 1483 | 135 | 0 | 0 | 0 | 14 | 42 | 153 | 63 | 0 | 251 | 0 | 67 | 0 | 0 | 2184 | 0 | 0 |
| SRR5387813 | 0 | 0 | 0 | 30 | 15 | 0 | 98 | 0 | 0 | 0 | 303 | 0 | 0 | 0 | 1662 | 24021 | 21352 | 0 | 0 | 587 | 98 | 0 | 0 | 0 | 15 | 45 | 61 | 91 | 0 | 167 | 0 | 0 | 0 | 0 | 1779 | 0 | 15 |
| SRR5387814 | 0 | 0 | 0 | 0 | 35 | 0 | 0 | 85 | 0 | 0 | 324 | 0 | 0 | 0 | 1281 | 12356 | 13440 | 0 | 0 | 1196 | 171 | 0 | 0 | 0 | 0 | 35 | 289 | 105 | 0 | 430 | 0 | 0 | 0 | 0 | 3902 | 0 | 0 |
| SRR5387815 | 0 | 0 | 0 | 0 | 35 | 0 | 0 | 85 | 0 | 0 | 153 | 0 | 0 | 0 | 763 | 10115 | 18113 | 0 | 0 | 1442 | 170 | 0 | 0 | 28 | 0 | 49 | 279 | 174 | 0 | 174 | 0 | 0 | 0 | 0 | 1176 | 0 | 0 |
| SRR5387816 | 0 | 0 | 0 | 19 | 37 | 104 | 0 | 0 | 0 | 0 | 103 | 0 | 0 | 0 | 2504 | 17014 | 38090 | 0 | 0 | 2191 | 313 | 0 | 0 | 28 | 0 | 0 | 84 | 28 | 0 | 261 | 0 | 0 | 0 | 0 | 8380 | 0 | 0 |
| SRR5387817 | 0 | 0 | 0 | 0 | 0 | 0 | 78 | 0 | 0 | 0 | 183 | 0 | 0 | 0 | 1177 | 15243 | 14813 | 0 | 0 | 2354 | 78 | 0 | 0 | 26 | 0 | 26 | 222 | 131 | 0 | 314 | 0 | 0 | 0 | 0 | 1503 | 0 | 0 |
| SRR5387818 | 0 | 0 | 0 | 0 | 26 | 0 | 0 | 0 | 0 | 0 | 1193 | 0 | 0 | 0 | 821 | 8147 | 10421 | 0 | 0 | 2399 | 189 | 0 | 0 | 17 | 0 | 35 | 122 | 78 | 0 | 1028 | 0 | 0 | 0 | 0 | 4358 | 0 | 0 |
| SRR5387819 | 0 | 0 | 0 | 0 | 25 | 45 | 45 | 134 | 0 | 0 | 193 | 0 | 0 | 0 | 3976 | 21554 | 28140 | 0 | 0 | 938 | 134 | 0 | 0 | 17 | 17 | 50 | 227 | 160 | 0 | 176 | 0 | 0 | 0 | 0 | 1676 | 0 | 0 |
| SRR5387820 | 0 | 0 | 0 | 19 | 127 | 188 | 0 | 94 | 0 | 0 | 165 | 0 | 0 | 0 | 1130 | 12460 | 13193 | 0 | 0 | 1413 | 0 | 0 | 0 | 39 | 0 | 19 | 165 | 175 | 0 | 302 | 0 | 0 | 0 | 0 | 2688 | 0 | 0 |
| SRR5387821 | 0 | 0 | 0 | 0 | 26 | 0 | 102 | 102 | 0 | 0 | 165 | 17 | 0 | 0 | 4194 | 24308 | 23408 | 0 | 0 | 1841 | 409 | 0 | 0 | 26 | 35 | 35 | 226 | 61 | 0 | 270 | 0 | 102 | 0 | 0 | 4727 | 0 | 0 |
| SRR5387822 | 0 | 0 | 0 | 0 | 13 | 0 | 0 | 0 | 0 | 0 | 188 | 0 | 0 | 0 | 2993 | 11274 | 9998 | 0 | 0 | 1537 | 324 | 0 | 0 | 0 | 0 | 47 | 235 | 74 | 0 | 275 | 0 | 0 | 0 | 0 | 2978 | 0 | 0 |
| SRR5387823 | 0 | 0 | 0 | 0 | 16 | 0 | 0 | 71 | 0 | 0 | 228 | 0 | 0 | 0 | 4550 | 11092 | 18985 | 0 | 0 | 1280 | 284 | 0 | 0 | 16 | 16 | 41 | 49 | 49 | 0 | 212 | 0 | 0 | 0 | 0 | 1280 | 0 | 0 |
| SRR5387824 | 0 | 0 | 0 | 0 | 0 | 0 | 74 | 148 | 0 | 0 | 227 | 0 | 0 | 0 | 2438 | 14143 | 22411 | 0 | 0 | 443 | 0 | 0 | 0 | 0 | 0 | 27 | 91 | 27 | 0 | 318 | 0 | 74 | 0 | 0 | 3699 | 0 | 0 |
| SRR5387825 | 0 | 0 | 0 | 0 | 46 | 82 | 0 | 82 | 0 | 0 | 165 | 0 | 0 | 0 | 1800 | 13445 | 13648 | 0 | 0 | 2291 | 164 | 0 | 0 | 0 | 0 | 37 | 64 | 28 | 0 | 275 | 0 | 0 | 0 | 0 | 6722 | 0 | 0 |
| SRR5387826 | 0 | 0 | 0 | 0 | 24 | 0 | 0 | 0 | 0 | 0 | 183 | 0 | 0 | 0 | 707 | 13228 | 10251 | 0 | 0 | 1884 | 0 | 0 | 0 | 12 | 0 | 37 | 317 | 244 | 0 | 256 | 0 | 0 | 0 | 0 | 992 | 0 | 0 |
| SRR5387827 | 0 | 0 | 0 | 16 | 47 | 0 | 0 | 0 | 0 | 0 | 63 | 0 | 0 | 0 | 3596 | 36575 | 47332 | 0 | 0 | 817 | 0 | 0 | 0 | 16 | 16 | 0 | 174 | 142 | 0 | 158 | 0 | 0 | 0 | 0 | 1721 | 0 | 0 |
| SRR5387828 | 0 | 18 | 0 | 0 | 18 | 0 | 0 | 0 | 0 | 0 | 101 | 0 | 0 | 0 | 1340 | 10043 | 13239 | 0 | 0 | 2094 | 84 | 0 | 0 | 0 | 0 | 0 | 110 | 46 | 0 | 192 | 0 | 0 | 0 | 0 | 5706 | 0 | 0 |
| SRR5387829 | 0 | 0 | 0 | 30 | 15 | 0 | 0 | 0 | 0 | 0 | 236 | 0 | 0 | 0 | 274 | 6158 | 7969 | 0 | 0 | 939 | 157 | 0 | 0 | 0 | 10 | 0 | 90 | 45 | 0 | 321 | 0 | 0 | 0 | 0 | 2656 | 0 | 0 |
| SRR5387830 | 0 | 0 | 0 | 32 | 13 | 0 | 0 | 72 | 0 | 0 | 142 | 0 | 0 | 0 | 1076 | 16069 | 17947 | 0 | 0 | 1434 | 143 | 0 | 0 | 0 | 0 | 19 | 78 | 45 | 0 | 149 | 0 | 0 | 0 | 0 | 3339 | 0 | 0 |
| SRR5387831 | 0 | 0 | 0 | 0 | 0 | 0 | 0 | 0 | 0 | 0 | 398 | 0 | 0 | 0 | 1147 | 12978 | 14381 | 0 | 0 | 1261 | 115 | 0 | 0 | 0 | 19 | 19 | 157 | 111 | 0 | 259 | 0 | 0 | 0 | 0 | 3858 | 0 | 0 |
| SRR5387832 | 0 | 0 | 0 | 31 | 10 | 0 | 0 | 0 | 0 | 0 | 207 | 0 | 0 | 0 | 1905 | 9795 | 11374 | 0 | 0 | 1667 | 0 | 0 | 0 | 0 | 0 | 0 | 83 | 41 | 0 | 207 | 0 | 0 | 0 | 0 | 5055 | 0 | 0 |
| SRR5387833 | 0 | 0 | 0 | 0 | 81 | 109 | 0 | 219 | 0 | 12 | 185 | 0 | 0 | 0 | 2734 | 20584 | 31711 | 0 | 0 | 1203 | 219 | 0 | 0 | 35 | 0 | 35 | 23 | 81 | 0 | 196 | 0 | 0 | 0 | 0 | 9179 | 0 | 0 |
| SRR5387834 | 0 | 0 | 0 | 0 | 27 | 83 | 0 | 0 | 0 | 0 | 137 | 0 | 0 | 0 | 1826 | 9321 | 17686 | 0 | 0 | 498 | 0 | 0 | 0 | 18 | 0 | 27 | 91 | 46 | 0 | 218 | 0 | 0 | 0 | 0 | 5736 | 0 | 0 |
| SRR5387835 | 0 | 0 | 0 | 0 | 0 | 0 | 0 | 153 | 0 | 0 | 221 | 0 | 0 | 0 | 1074 | 15343 | 18276 | 0 | 0 | 1612 | 153 | 0 | 0 | 21 | 21 | 28 | 48 | 21 | 0 | 152 | 0 | 0 | 0 | 0 | 2256 | 0 | 0 |
| SRR5387836 | 0 | 0 | 0 | 0 | 0 | 0 | 0 | 0 | 175 | 0 | 192 | 0 | 0 | 0 | 874 | 8137 | 11934 | 0 | 0 | 699 | 0 | 0 | 0 | 22 | 0 | 30 | 192 | 59 | 0 | 148 | 0 | 0 | 0 | 0 | 4340 | 0 | 0 |
| SRR5387837 | 0 | 0 | 0 | 0 | 0 | 0 | 0 | 0 | 0 | 0 | 270 | 0 | 0 | 0 | 1253 | 11696 | 16244 | 0 | 0 | 1832 | 96 | 0 | 0 | 22 | 0 | 0 | 88 | 58 | 0 | 183 | 0 | 0 | 0 | 0 | 4982 | 0 | 0 |
| SRR5387838 | 0 | 0 | 0 | 19 | 0 | 0 | 0 | 0 | 97 | 0 | 189 | 0 | 0 | 0 | 972 | 6867 | 10987 | 0 | 97 | 1264 | 486 | 0 | 0 | 32 | 25 | 51 | 177 | 76 | 0 | 177 | 0 | 0 | 0 | 0 | 2289 | 0 | 0 |
| SRR5387839 | 0 | 11 | 0 | 32 | 11 | 0 | 0 | 0 | 0 | 0 | 182 | 11 | 0 | 0 | 1851 | 15296 | 14340 | 0 | 0 | 2430 | 0 | 0 | 0 | 32 | 11 | 54 | 21 | 21 | 0 | 193 | 0 | 0 | 0 | 0 | 2231 | 0 | 0 |
| SRR5387840 | 14 | 0 | 0 | 29 | 129 | 0 | 0 | 177 | 0 | 0 | 114 | 14 | 0 | 0 | 2297 | 24380 | 18392 | 0 | 0 | 883 | 353 | 0 | 0 | 0 | 0 | 0 | 29 | 0 | 0 | 57 | 0 | 0 | 0 | 0 | 4277 | 0 | 0 |
| SRR5387841 | 0 | 0 | 0 | 16 | 65 | 0 | 0 | 0 | 0 | 0 | 384 | 16 | 0 | 0 | 2698 | 29299 | 29512 | 0 | 0 | 253 | 169 | 0 | 0 | 33 | 25 | 82 | 139 | 90 | 0 | 196 | 0 | 0 | 0 | 0 | 2566 | 0 | 0 |
| SRR5387842 | 0 | 0 | 0 | 0 | 18 | 0 | 0 | 93 | 0 | 0 | 307 | 0 | 0 | 0 | 836 | 9611 | 13634 | 0 | 0 | 1301 | 186 | 0 | 0 | 18 | 0 | 18 | 114 | 88 | 0 | 359 | 0 | 93 | 0 | 0 | 4023 | 0 | 0 |
| SRR5387843 | 0 | 0 | 0 | 13 | 20 | 0 | 72 | 145 | 72 | 0 | 210 | 0 | 0 | 0 | 869 | 11117 | 15600 | 0 | 0 | 1882 | 72 | 0 | 0 | 20 | 13 | 33 | 243 | 79 | 0 | 361 | 0 | 0 | 0 | 0 | 3945 | 0 | 0 |
| SRR5387844 | 0 | 0 | 0 | 0 | 0 | 0 | 0 | 0 | 0 | 0 | 174 | 0 | 0 | 0 | 704 | 12893 | 12685 | 0 | 0 | 1487 | 156 | 0 | 0 | 0 | 0 | 55 | 293 | 101 | 0 | 293 | 0 | 0 | 0 | 0 | 2911 | 0 | 0 |
| SRR5387845 | 0 | 0 | 0 | 0 | 0 | 0 | 0 | 0 | 0 | 0 | 244 | 0 | 0 | 0 | 1556 | 30780 | 47981 | 0 | 0 | 934 | 249 | 0 | 0 | 0 | 10 | 0 | 51 | 61 | 0 | 193 | 0 | 0 | 0 | 0 | 3802 | 0 | 0 |
| SRR5387846 | 0 | 0 | 0 | 25 | 0 | 0 | 0 | 0 | 0 | 0 | 211 | 0 | 0 | 0 | 905 | 16755 | 15432 | 0 | 0 | 1316 | 247 | 0 | 0 | 0 | 17 | 25 | 144 | 84 | 0 | 279 | 0 | 0 | 0 | 0 | 1543 | 0 | 0 |
| SRR5387847 | 0 | 0 | 0 | 0 | 55 | 0 | 0 | 0 | 0 | 0 | 365 | 0 | 0 | 0 | 2617 | 33109 | 21555 | 0 | 0 | 1468 | 64 | 0 | 0 | 0 | 22 | 66 | 321 | 88 | 0 | 144 | 0 | 0 | 0 | 0 | 2587 | 0 | 0 |
| SRR5387848 | 0 | 0 | 0 | 34 | 27 | 0 | 0 | 123 | 0 | 0 | 306 | 0 | 0 | 0 | 2576 | 17107 | 13649 | 0 | 0 | 2638 | 0 | 0 | 0 | 116 | 27 | 82 | 102 | 136 | 0 | 10 | 0 | 0 | 0 | 0 | 7461 | 0 | 0 |
| SRR5387849 | 0 | 0 | 0 | 0 | 23 | 0 | 0 | 55 | 0 | 0 | 401 | 0 | 0 | 0 | 1254 | 12711 | 11383 | 0 | 0 | 927 | 109 | 0 | 0 | 15 | 0 | 39 | 131 | 100 | 0 | 278 | 0 | 0 | 0 | 0 | 3794 | 0 | 0 |
| SRR5387850 | 0 | 0 | 0 | 0 | 18 | 80 | 0 | 0 | 0 | 0 | 220 | 0 | 0 | 0 | 1276 | 9613 | 10912 | 0 | 80 | 2712 | 160 | 0 | 0 | 0 | 0 | 37 | 37 | 55 | 0 | 156 | 0 | 0 | 0 | 0 | 2078 | 0 | 0 |
| SRR5387851 | 0 | 0 | 0 | 0 | 69 | 0 | 0 | 0 | 0 | 0 | 152 | 0 | 0 | 0 | 5679 | 27948 | 21132 | 0 | 0 | 668 | 334 | 0 | 0 | 14 | 14 | 69 | 110 | 55 | 0 | 289 | 0 | 0 | 0 | 0 | 10566 | 0 | 0 |
| SRR5387852 | 0 | 0 | 0 | 18 | 27 | 79 | 0 | 0 | 0 | 0 | 91 | 0 | 0 | 0 | 5062 | 30635 | 27952 | 79 | 0 | 1424 | 158 | 0 | 0 | 0 | 0 | 27 | 64 | 55 | 0 | 146 | 0 | 0 | 0 | 0 | 5590 | 0 | 0 |
| SRR5387853 | 0 | 0 | 0 | 10 | 31 | 126 | 0 | 253 | 0 | 0 | 229 | 0 | 0 | 0 | 758 | 14326 | 16422 | 0 | 0 | 1516 | 126 | 0 | 0 | 10 | 31 | 21 | 135 | 73 | 0 | 166 | 0 | 0 | 0 | 0 | 1398 | 0 | 21 |
| SRR5387854 | 0 | 0 | 0 | 0 | 49 | 0 | 0 | 0 | 0 | 0 | 191 | 0 | 0 | 0 | 397 | 8769 | 6343 | 0 | 0 | 1258 | 331 | 0 | 0 | 14 | 42 | 35 | 198 | 56 | 0 | 452 | 0 | 0 | 0 | 0 | 5597 | 0 | 0 |
| SRR5387855 | 0 | 0 | 0 | 28 | 28 | 73 | 0 | 0 | 0 | 0 | 276 | 0 | 0 | 0 | 2037 | 14414 | 22352 | 0 | 0 | 1819 | 0 | 0 | 0 | 0 | 0 | 18 | 83 | 83 | 0 | 230 | 0 | 0 | 0 | 0 | 3551 | 0 | 0 |
| SRR5387856 | 0 | 0 | 0 | 14 | 0 | 0 | 0 | 69 | 0 | 0 | 371 | 0 | 0 | 0 | 1724 | 9424 | 17070 | 0 | 0 | 1448 | 138 | 0 | 0 | 0 | 0 | 27 | 88 | 61 | 0 | 344 | 0 | 0 | 0 | 0 | 5334 | 0 | 0 |
| SRR5387857 | 0 | 0 | 0 | 11 | 22 | 0 | 0 | 0 | 0 | 0 | 178 | 0 | 0 | 0 | 745 | 9002 | 9002 | 0 | 0 | 1863 | 93 | 0 | 0 | 11 | 0 | 11 | 56 | 22 | 0 | 300 | 0 | 0 | 0 | 0 | 5251 | 0 | 0 |
| SRR5387858 | 0 | 0 | 0 | 25 | 0 | 0 | 0 | 0 | 0 | 0 | 376 | 0 | 0 | 0 | 1094 | 13670 | 13670 | 0 | 0 | 1641 | 68 | 0 | 0 | 16 | 0 | 41 | 302 | 82 | 0 | 245 | 0 | 0 | 0 | 0 | 2071 | 0 | 0 |
| SRR5387859 | 0 | 0 | 0 | 0 | 26 | 0 | 0 | 191 | 0 | 0 | 223 | 0 | 0 | 0 | 3250 | 29468 | 35171 | 0 | 0 | 574 | 574 | 0 | 0 | 0 | 26 | 79 | 171 | 184 | 0 | 118 | 0 | 0 | 0 | 0 | 4278 | 0 | 0 |
| SRR5387860 | 0 | 0 | 11 | 11 | 0 | 0 | 0 | 0 | 0 | 0 | 286 | 0 | 0 | 0 | 1201 | 6698 | 12365 | 0 | 0 | 1293 | 92 | 0 | 0 | 0 | 0 | 21 | 95 | 32 | 0 | 180 | 0 | 0 | 0 | 0 | 1288 | 0 | 0 |
| SRR5387861 | 0 | 0 | 0 | 20 | 20 | 0 | 0 | 92 | 0 | 10 | 153 | 0 | 0 | 0 | 370 | 5259 | 9016 | 0 | 0 | 1017 | 185 | 0 | 0 | 0 | 0 | 41 | 183 | 183 | 0 | 224 | 0 | 0 | 0 | 0 | 4007 | 0 | 0 |
| SRR5387862 | 0 | 0 | 0 | 17 | 0 | 0 | 0 | 0 | 0 | 0 | 319 | 0 | 0 | 0 | 382 | 10123 | 14417 | 0 | 0 | 1366 | 164 | 0 | 0 | 0 | 0 | 50 | 67 | 50 | 0 | 227 | 0 | 55 | 0 | 0 | 920 | 0 | 0 |
| SRR5387863 | 0 | 0 | 0 | 0 | 0 | 0 | 0 | 79 | 0 | 0 | 95 | 0 | 0 | 0 | 2124 | 31516 | 30067 | 0 | 79 | 2361 | 157 | 0 | 0 | 0 | 19 | 48 | 105 | 48 | 0 | 162 | 0 | 0 | 0 | 0 | 3441 | 0 | 0 |
| SRR5387864 | 0 | 0 | 0 | 15 | 15 | 0 | 0 | 147 | 0 | 0 | 213 | 0 | 0 | 0 | 737 | 11186 | 11765 | 0 | 0 | 2875 | 0 | 0 | 0 | 15 | 0 | 76 | 183 | 91 | 0 | 190 | 0 | 0 | 0 | 0 | 3279 | 0 | 0 |
| SRR5387865 | 0 | 0 | 0 | 0 | 0 | 0 | 0 | 0 | 0 | 0 | 296 | 0 | 0 | 0 | 1537 | 22065 | 25460 | 0 | 0 | 2869 | 410 | 0 | 0 | 0 | 0 | 20 | 99 | 79 | 0 | 118 | 0 | 0 | 0 | 0 | 3112 | 0 | 0 |
| SRR5387866 | 0 | 0 | 0 | 0 | 0 | 0 | 0 | 59 | 0 | 0 | 175 | 116 | 0 | 0 | 1112 | 8323 | 16059 | 0 | 0 | 2574 | 351 | 0 | 0 | 0 | 0 | 0 | 13 | 0 | 11 | 621 | 0 | 59 | 0 | 0 | 8674 | 0 | 39 |

### Supp Table S8.3.Columns_BX_to_DH

| | | | | | | | | | | | | | | | | | | | | | | | | | | | | | | | | | | | | | |
|---|---|---|---|---|---|---|---|---|---|---|---|---|---|---|---|---|---|---|---|---|---|---|---|---|---|---|---|---|---|---|---|---|---|---|---|---|---|
| SRR5387867 | 0 | 0 | 0 | 0 | 32 | 0 | 0 | 0 | 0 | 0 | 325 | 0 | 0 | 0 | 1624 | 16164 | 16810 | 0 | 0 | 999 | 0 | 0 | 0 | 24 | 16 | 32 | 151 | 103 | 0 | 341 | 0 | 0 | 0 | 0 | 2802 | 0 | 0 |
| SRR5387868 | 0 | 0 | 0 | 10 | 0 | 0 | 0 | 0 | 0 | 0 | 209 | 0 | 0 | 0 | 684 | 6725 | 6553 | 0 | 0 | 1306 | 0 | 0 | 0 | 10 | 10 | 31 | 250 | 31 | 0 | 261 | 0 | 0 | 0 | 0 | 2069 | 0 | 0 |
| SRR5387869 | 0 | 11 | 0 | 0 | 54 | 0 | 0 | 0 | 0 | 0 | 161 | 11 | 0 | 0 | 1014 | 16369 | 11905 | 0 | 0 | 1353 | 0 | 0 | 0 | 0 | 11 | 43 | 194 | 97 | 0 | 183 | 0 | 0 | 0 | 0 | 5952 | 0 | 0 |
| SRR5387870 | 0 | 0 | 0 | 15 | 23 | 0 | 0 | 0 | 0 | 0 | 353 | 0 | 0 | 0 | 1365 | 15273 | 21617 | 0 | 0 | 1105 | 0 | 0 | 0 | 38 | 15 | 38 | 299 | 146 | 0 | 299 | 0 | 0 | 0 | 0 | 4464 | 0 | 15 |
| SRR5387871 | 0 | 0 | 12 | 12 | 12 | 0 | 0 | 0 | 0 | 0 | 142 | 0 | 0 | 0 | 349 | 9964 | 11172 | 0 | 0 | 1513 | 116 | 0 | 0 | 0 | 24 | 47 | 166 | 59 | 0 | 178 | 0 | 0 | 0 | 0 | 4529 | 0 | 0 |
| SRR5387872 | 0 | 0 | 0 | 0 | 20 | 0 | 0 | 0 | 104 | 0 | 119 | 0 | 0 | 0 | 625 | 11335 | 11618 | 0 | 0 | 1147 | 104 | 0 | 0 | 0 | 0 | 30 | 119 | 40 | 0 | 279 | 0 | 0 | 0 | 0 | 3400 | 0 | 0 |
| SRR5387873 | 0 | 0 | 0 | 36 | 145 | 0 | 0 | 0 | 0 | 0 | 163 | 0 | 0 | 0 | 2677 | 18157 | 27572 | 0 | 0 | 402 | 134 | 0 | 0 | 0 | 0 | 36 | 181 | 218 | 0 | 200 | 0 | 0 | 0 | 0 | 3026 | 0 | 0 |
| SRR5387874 | 0 | 0 | 0 | 0 | 17 | 0 | 0 | 0 | 0 | 0 | 221 | 0 | 0 | 0 | 1448 | 13547 | 13195 | 0 | 0 | 1738 | 0 | 0 | 0 | 17 | 0 | 34 | 68 | 76 | 0 | 365 | 0 | 58 | 0 | 0 | 3519 | 0 | 0 |
| SRR5387875 | 0 | 0 | 0 | 0 | 32 | 0 | 0 | 0 | 0 | 0 | 170 | 0 | 0 | 0 | 1083 | 20906 | 21539 | 0 | 0 | 1204 | 481 | 0 | 0 | 0 | 0 | 32 | 57 | 16 | 0 | 162 | 0 | 0 | 0 | 0 | 2217 | 0 | 0 |
| SRR5387876 | 0 | 0 | 0 | 0 | 0 | 0 | 0 | 238 | 0 | 0 | 170 | 0 | 0 | 0 | 952 | 11677 | 14508 | 0 | 0 | 833 | 0 | 0 | 0 | 0 | 0 | 46 | 270 | 69 | 0 | 278 | 0 | 0 | 0 | 0 | 7431 | 0 | 0 |
| SRR5387877 | 0 | 0 | 0 | 23 | 46 | 0 | 0 | 140 | 0 | 0 | 114 | 0 | 0 | 0 | 561 | 15496 | 14775 | 0 | 0 | 1123 | 140 | 0 | 0 | 11 | 0 | 46 | 228 | 68 | 0 | 171 | 0 | 0 | 0 | 0 | 3604 | 0 | 0 |
| SRR5387878 | 0 | 0 | 0 | 0 | 30 | 0 | 79 | 0 | 0 | 0 | 350 | 0 | 0 | 0 | 393 | 10191 | 11077 | 0 | 0 | 1414 | 0 | 0 | 0 | 0 | 46 | 129 | 144 | 114 | 0 | 12 | 0 | 0 | 0 | 0 | 1772 | 0 | 0 |
| SRR5387879 | 0 | 0 | 0 | 0 | 12 | 86 | 0 | 86 | 86 | 0 | 242 | 0 | 0 | 0 | 1115 | 14154 | 24615 | 0 | 0 | 1544 | 172 | 0 | 0 | 31 | 37 | 112 | 453 | 186 | 0 | 267 | 0 | 0 | 0 | 0 | 6564 | 0 | 0 |
| SRR5387880 | 0 | 0 | 0 | 0 | 66 | 0 | 0 | 105 | 105 | 0 | 164 | 0 | 0 | 0 | 2200 | 23354 | 26449 | 0 | 105 | 943 | 210 | 0 | 0 | 0 | 25 | 66 | 99 | 49 | 0 | 238 | 0 | 0 | 0 | 0 | 5627 | 0 | 0 |
| SRR5387881 | 0 | 0 | 0 | 0 | 23 | 0 | 52 | 0 | 0 | 0 | 11 | 0 | 0 | 0 | 3961 | 26605 | 46246 | 0 | 0 | 365 | 156 | 0 | 0 | 0 | 0 | 54 | 177 | 116 | 0 | 239 | 0 | 0 | 0 | 0 | 3204 | 0 | 0 |
| SRR5387882 | 0 | 0 | 0 | 0 | 18 | 0 | 0 | 108 | 0 | 0 | 309 | 0 | 0 | 0 | 3137 | 16158 | 34578 | 0 | 0 | 1893 | 325 | 0 | 0 | 36 | 24 | 55 | 188 | 91 | 0 | 218 | 0 | 0 | 0 | 0 | 6948 | 0 | 0 |
| SRR5387883 | 0 | 0 | 0 | 0 | 0 | 0 | 0 | 0 | 79 | 0 | 105 | 0 | 0 | 0 | 2140 | 25308 | 23107 | 0 | 0 | 1189 | 238 | 0 | 0 | 0 | 0 | 23 | 135 | 30 | 0 | 255 | 0 | 0 | 0 | 0 | 4401 | 0 | 0 |
| SRR5387884 | 0 | 0 | 0 | 0 | 0 | 98 | 0 | 0 | 0 | 0 | 148 | 0 | 0 | 0 | 1365 | 21435 | 16672 | 0 | 0 | 2243 | 0 | 0 | 0 | 52 | 0 | 17 | 165 | 113 | 0 | 165 | 0 | 0 | 0 | 0 | 2084 | 0 | 0 |
| SRR5387885 | 0 | 0 | 0 | 0 | 0 | 188 | 94 | 0 | 0 | 0 | 269 | 0 | 0 | 0 | 2915 | 20228 | 21006 | 0 | 94 | 1787 | 188 | 0 | 0 | 10 | 0 | 10 | 62 | 21 | 0 | 310 | 0 | 0 | 0 | 0 | 7261 | 0 | 0 |
| SRR5387886 | 0 | 0 | 0 | 18 | 0 | 0 | 0 | 0 | 0 | 0 | 174 | 0 | 0 | 0 | 574 | 19287 | 9059 | 0 | 0 | 1339 | 0 | 0 | 0 | 0 | 0 | 82 | 256 | 110 | 0 | 201 | 0 | 0 | 0 | 0 | 3507 | 0 | 0 |
| SRR5387887 | 0 | 0 | 0 | 0 | 0 | 79 | 0 | 0 | 0 | 11 | 401 | 0 | 0 | 0 | 1819 | 25000 | 31500 | 0 | 0 | 1740 | 79 | 0 | 0 | 33 | 0 | 11 | 145 | 78 | 0 | 223 | 0 | 0 | 0 | 0 | 5000 | 0 | 0 |
| SRR5387888 | 0 | 0 | 0 | 0 | 15 | 0 | 0 | 57 | 0 | 15 | 317 | 23 | 0 | 0 | 396 | 8349 | 15679 | 0 | 0 | 679 | 113 | 0 | 0 | 0 | 0 | 23 | 209 | 70 | 0 | 232 | 0 | 0 | 0 | 0 | 2443 | 0 | 0 |
| SRR5387889 | 0 | 0 | 0 | 0 | 0 | 77 | 0 | 0 | 0 | 0 | 366 | 0 | 0 | 0 | 4060 | 34355 | 30645 | 0 | 0 | 919 | 77 | 0 | 0 | 19 | 0 | 48 | 48 | 58 | 0 | 193 | 0 | 0 | 0 | 0 | 4355 | 0 | 0 |
| SRR5387890 | 0 | 0 | 0 | 0 | 19 | 69 | 0 | 0 | 0 | 0 | 179 | 0 | 0 | 0 | 2350 | 19730 | 33240 | 0 | 0 | 1382 | 276 | 0 | 0 | 0 | 19 | 28 | 57 | 76 | 0 | 246 | 0 | 69 | 0 | 0 | 3431 | 0 | 0 |
| SRR5387891 | 0 | 0 | 0 | 0 | 48 | 0 | 111 | 111 | 0 | 0 | 344 | 0 | 0 | 0 | 1717 | 14411 | 21788 | 0 | 0 | 1108 | 222 | 0 | 0 | 29 | 0 | 38 | 57 | 38 | 0 | 220 | 0 | 0 | 0 | 0 | 2230 | 0 | 0 |
| SRR5387892 | 0 | 0 | 0 | 0 | 14 | 0 | 0 | 108 | 0 | 0 | 360 | 0 | 0 | 0 | 1456 | 9763 | 14296 | 0 | 0 | 970 | 108 | 0 | 0 | 14 | 0 | 14 | 34 | 68 | 0 | 394 | 0 | 0 | 0 | 0 | 2789 | 0 | 0 |
| SRR5387893 | 0 | 0 | 0 | 0 | 0 | 0 | 0 | 60 | 0 | 0 | 185 | 0 | 0 | 0 | 1197 | 19734 | 12995 | 0 | 0 | 1437 | 0 | 0 | 0 | 12 | 0 | 12 | 62 | 12 | 0 | 357 | 0 | 0 | 0 | 0 | 6899 | 0 | 0 |
| SRR5387894 | 0 | 0 | 0 | 0 | 0 | 0 | 0 | 0 | 0 | 0 | 203 | 0 | 0 | 0 | 654 | 17955 | 15909 | 0 | 0 | 1505 | 65 | 0 | 0 | 0 | 0 | 29 | 174 | 106 | 0 | 155 | 0 | 65 | 0 | 0 | 7273 | 0 | 0 |
| SRR5387895 | 0 | 0 | 0 | 18 | 35 | 0 | 0 | 102 | 0 | 0 | 265 | 0 | 0 | 0 | 1425 | 15625 | 12292 | 0 | 0 | 2239 | 305 | 0 | 0 | 24 | 41 | 88 | 330 | 94 | 0 | 253 | 0 | 0 | 0 | 0 | 4583 | 0 | 0 |
| SRR5387896 | 0 | 0 | 0 | 33 | 20 | 0 | 120 | 0 | 0 | 0 | 225 | 33 | 0 | 0 | 839 | 8252 | 10905 | 0 | 0 | 479 | 0 | 0 | 0 | 0 | 0 | 26 | 132 | 66 | 0 | 363 | 0 | 0 | 0 | 0 | 5600 | 0 | 0 |
| SRR5387897 | 0 | 0 | 0 | 32 | 21 | 0 | 0 | 74 | 0 | 0 | 116 | 11 | 0 | 0 | 3330 | 11848 | 11119 | 0 | 0 | 2146 | 74 | 0 | 0 | 53 | 11 | 63 | 264 | 42 | 0 | 348 | 0 | 0 | 0 | 0 | 4557 | 0 | 11 |
| SRR5387898 | 0 | 0 | 0 | 0 | 17 | 0 | 0 | 69 | 0 | 0 | 198 | 0 | 0 | 0 | 1662 | 10927 | 10146 | 0 | 0 | 1523 | 0 | 0 | 0 | 17 | 0 | 52 | 215 | 26 | 0 | 172 | 0 | 0 | 0 | 0 | 2732 | 0 | 0 |
| SRR5387899 | 0 | 0 | 0 | 28 | 0 | 0 | 0 | 186 | 0 | 0 | 73 | 0 | 0 | 0 | 557 | 10612 | 9483 | 0 | 0 | 1300 | 186 | 0 | 0 | 18 | 0 | 0 | 73 | 28 | 0 | 174 | 0 | 93 | 0 | 0 | 1580 | 0 | 0 |
| SRR5387900 | 0 | 0 | 0 | 17 | 17 | 136 | 0 | 136 | 0 | 0 | 319 | 0 | 0 | 0 | 2859 | 17341 | 26333 | 0 | 0 | 1089 | 272 | 0 | 0 | 0 | 0 | 78 | 233 | 52 | 0 | 311 | 0 | 0 | 0 | 0 | 6744 | 0 | 0 |
| SRR5387901 | 0 | 0 | 0 | 0 | 0 | 0 | 0 | 208 | 0 | 0 | 141 | 0 | 0 | 0 | 2083 | 15058 | 34241 | 0 | 0 | 1736 | 208 | 0 | 0 | 0 | 0 | 33 | 112 | 54 | 0 | 129 | 0 | 139 | 0 | 0 | 3919 | 0 | 0 |
| SRR5387902 | 0 | 0 | 0 | 0 | 0 | 0 | 72 | 577 | 0 | 0 | 283 | 0 | 0 | 0 | 2093 | 19751 | 42610 | 0 | 0 | 577 | 72 | 0 | 0 | 0 | 0 | 18 | 50 | 88 | 0 | 209 | 0 | 0 | 0 | 0 | 9987 | 0 | 0 |
| SRR5387903 | 0 | 0 | 0 | 0 | 39 | 0 | 99 | 0 | 0 | 0 | 286 | 15 | 0 | 0 | 2561 | 22543 | 27217 | 0 | 0 | 1428 | 99 | 0 | 0 | 0 | 23 | 46 | 162 | 93 | 0 | 255 | 0 | 0 | 0 | 0 | 2612 | 0 | 0 |
| SRR5387904 | 0 | 0 | 0 | 18 | 18 | 0 | 0 | 0 | 0 | 0 | 491 | 0 | 0 | 0 | 1791 | 9797 | 22297 | 0 | 0 | 1313 | 239 | 0 | 0 | 0 | 0 | 0 | 36 | 45 | 0 | 418 | 0 | 0 | 0 | 0 | 2703 | 0 | 0 |
| SRR5387905 | 0 | 0 | 0 | 0 | 0 | 0 | 0 | 0 | 0 | 0 | 325 | 0 | 0 | 0 | 1082 | 17036 | 13347 | 0 | 0 | 1443 | 0 | 0 | 0 | 18 | 12 | 59 | 189 | 53 | 0 | 325 | 0 | 0 | 0 | 0 | 4215 | 0 | 0 |
| SRR5387906 | 0 | 0 | 0 | 0 | 17 | 0 | 0 | 70 | 0 | 0 | 226 | 0 | 0 | 0 | 1324 | 12556 | 12556 | 0 | 0 | 1045 | 279 | 0 | 0 | 0 | 0 | 23 | 197 | 56 | 0 | 231 | 0 | 0 | 0 | 0 | 3038 | 0 | 0 |
| SRR5387907 | 0 | 0 | 0 | 0 | 0 | 0 | 0 | 299 | 0 | 54 | 117 | 0 | 0 | 0 | 957 | 11162 | 11510 | 0 | 0 | 598 | 359 | 0 | 0 | 0 | 0 | 0 | 51 | 16 | 0 | 77 | 0 | 0 | 0 | 0 | 3139 | 0 | 0 |
| SRR5387908 | 0 | 0 | 0 | 0 | 15 | 0 | 0 | 71 | 0 | 0 | 168 | 0 | 0 | 0 | 1134 | 13404 | 11170 | 0 | 0 | 2126 | 0 | 0 | 0 | 0 | 0 | 0 | 73 | 44 | 0 | 227 | 0 | 0 | 0 | 0 | 3249 | 0 | 0 |
| SRR5387909 | 0 | 0 | 0 | 18 | 45 | 0 | 0 | 255 | 0 | 0 | 90 | 18 | 0 | 0 | 1656 | 52524 | 43478 | 0 | 0 | 1656 | 637 | 0 | 0 | 0 | 54 | 27 | 108 | 54 | 0 | 188 | 0 | 127 | 0 | 0 | 6128 | 0 | 0 |
| SRR5387910 | 0 | 0 | 0 | 0 | 35 | 0 | 0 | 0 | 0 | 0 | 216 | 0 | 0 | 0 | 363 | 8197 | 7514 | 0 | 0 | 847 | 121 | 0 | 0 | 0 | 52 | 52 | 199 | 87 | 0 | 234 | 0 | 0 | 0 | 0 | 3415 | 0 | 0 |
| SRR5387911 | 0 | 0 | 0 | 17 | 17 | 0 | 0 | 0 | 0 | 0 | 118 | 0 | 0 | 0 | 2172 | 26728 | 27532 | 0 | 0 | 1123 | 300 | 0 | 0 | 25 | 0 | 34 | 67 | 25 | 0 | 236 | 0 | 0 | 0 | 0 | 5024 | 0 | 0 |
| SRR5387912 | 0 | 0 | 0 | 15 | 23 | 0 | 0 | 0 | 0 | 0 | 185 | 0 | 0 | 0 | 799 | 14387 | 12817 | 0 | 0 | 1299 | 200 | 0 | 0 | 15 | 31 | 15 | 154 | 15 | 0 | 200 | 0 | 0 | 0 | 0 | 1831 | 0 | 0 |
| SRR5387913 | 0 | 0 | 0 | 23 | 81 | 0 | 0 | 0 | 0 | 0 | 197 | 0 | 0 | 0 | 1375 | 31073 | 23237 | 0 | 0 | 1571 | 98 | 0 | 0 | 12 | 12 | 104 | 243 | 81 | 0 | 173 | 0 | 0 | 0 | 0 | 2162 | 0 | 0 |
| SRR5387914 | 0 | 0 | 0 | 59 | 39 | 0 | 0 | 81 | 0 | 20 | 332 | 0 | 0 | 0 | 326 | 13185 | 12170 | 0 | 0 | 1304 | 163 | 0 | 0 | 0 | 0 | 59 | 146 | 127 | 0 | 322 | 0 | 0 | 0 | 0 | 1521 | 0 | 0 |
| SRR5387915 | 0 | 0 | 0 | 23 | 15 | 55 | 0 | 111 | 0 | 0 | 130 | 0 | 0 | 0 | 1658 | 36707 | 23778 | 0 | 0 | 1658 | 55 | 0 | 0 | 0 | 0 | 0 | 92 | 61 | 0 | 215 | 0 | 0 | 0 | 0 | 2972 | 0 | 0 |
| SRR5387916 | 0 | 0 | 0 | 27 | 27 | 0 | 0 | 0 | 0 | 0 | 239 | 0 | 0 | 0 | 1070 | 12494 | 11337 | 0 | 0 | 823 | 165 | 0 | 0 | 18 | 0 | 27 | 177 | 159 | 0 | 194 | 0 | 0 | 0 | 0 | 2314 | 0 | 0 |
| SRR5387917 | 0 | 0 | 0 | 0 | 24 | 0 | 0 | 87 | 0 | 0 | 97 | 0 | 0 | 0 | 1042 | 14468 | 14178 | 0 | 0 | 1650 | 87 | 0 | 0 | 0 | 0 | 40 | 65 | 16 | 0 | 137 | 0 | 0 | 0 | 0 | 6366 | 0 | 0 |
| SRR5387918 | 0 | 0 | 0 | 0 | 25 | 0 | 0 | 92 | 0 | 0 | 204 | 0 | 0 | 0 | 918 | 10054 | 17788 | 0 | 0 | 1101 | 642 | 0 | 0 | 16 | 25 | 33 | 155 | 74 | 0 | 531 | 0 | 0 | 0 | 0 | 6187 | 0 | 0 |
| SRR5387919 | 0 | 0 | 0 | 20 | 20 | 0 | 0 | 0 | 0 | 0 | 290 | 0 | 0 | 0 | 3330 | 25181 | 38401 | 0 | 0 | 1110 | 0 | 0 | 0 | 30 | 0 | 30 | 80 | 20 | 0 | 299 | 0 | 0 | 0 | 0 | 2046 | 0 | 0 |
| SRR5387920 | 0 | 0 | 0 | 12 | 37 | 0 | 92 | 92 | 0 | 0 | 159 | 0 | 0 | 0 | 2678 | 19568 | 38012 | 0 | 0 | 1570 | 185 | 0 | 0 | 12 | 12 | 0 | 122 | 24 | 0 | 195 | 0 | 0 | 0 | 0 | 2924 | 0 | 0 |
| SRR5387921 | 0 | 0 | 0 | 0 | 22 | 0 | 0 | 178 | 0 | 0 | 110 | 0 | 0 | 0 | 1421 | 41103 | 19342 | 0 | 0 | 178 | 0 | 0 | 0 | 22 | 0 | 22 | 88 | 44 | 0 | 198 | 0 | 0 | 0 | 0 | 2901 | 0 | 0 |
| SRR5387922 | 0 | 0 | 0 | 20 | 81 | 0 | 0 | 297 | 0 | 0 | 60 | 0 | 0 | 0 | 148 | 19178 | 9247 | 0 | 0 | 1188 | 0 | 0 | 0 | 40 | 60 | 121 | 81 | 141 | 0 | 202 | 0 | 0 | 0 | 0 | 4795 | 0 | 0 |
| SRR5387923 | 11 | 0 | 11 | 0 | 147 | 0 | 0 | 0 | 0 | 0 | 113 | 0 | 0 | 0 | 973 | 23146 | 13724 | 0 | 97 | 1071 | 97 | 0 | 0 | 23 | 34 | 34 | 294 | 226 | 0 | 237 | 0 | 0 | 0 | 0 | 1434 | 0 | 0 |
| SRR5387924 | 0 | 0 | 0 | 0 | 0 | 0 | 0 | 83 | 0 | 0 | 117 | 0 | 0 | 0 | 250 | 12160 | 14545 | 0 | 0 | 1663 | 166 | 0 | 0 | 0 | 0 | 31 | 148 | 55 | 0 | 335 | 0 | 166 | 0 | 0 | 3100 | 0 | 0 |
| SRR5387925 | 0 | 0 | 0 | 0 | 0 | 0 | 0 | 0 | 0 | 0 | 169 | 0 | 0 | 0 | 1217 | 11289 | 25635 | 0 | 0 | 1530 | 174 | 0 | 0 | 0 | 0 | 10 | 91 | 37 | 0 | 129 | 0 | 0 | 0 | 0 | 1176 | 0 | 0 |
| SRR5387926 | 0 | 0 | 0 | 0 | 10 | 54 | 0 | 0 | 0 | 0 | 10 | 0 | 0 | 0 | 857 | 8088 | 17613 | 0 | 0 | 1393 | 214 | 0 | 0 | 0 | 10 | 10 | 104 | 104 | 0 | 11 | 0 | 0 | 0 | 0 | 1977 | 0 | 0 |
| SRR5387927 | 0 | 0 | 0 | 28 | 28 | 0 | 0 | 0 | 0 | 0 | 237 | 0 | 0 | 0 | 2145 | 14011 | 22560 | 0 | 0 | 804 | 0 | 0 | 0 | 0 | 0 | 38 | 152 | 161 | 0 | 294 | 0 | 0 | 0 | 0 | 5462 | 0 | 0 |
| SRR5387928 | 0 | 0 | 0 | 0 | 16 | 84 | 84 | 0 | 0 | 0 | 18 | 0 | 0 | 0 | 3117 | 16011 | 18756 | 0 | 0 | 2106 | 337 | 0 | 0 | 0 | 16 | 33 | 73 | 106 | 0 | 11 | 0 | 0 | 0 | 0 | 5489 | 0 | 0 |
| SRR5387929 | 0 | 0 | 0 | 38 | 38 | 0 | 50 | 101 | 0 | 0 | 161 | 0 | 0 | 0 | 1309 | 13103 | 12292 | 0 | 0 | 1611 | 101 | 0 | 0 | 23 | 23 | 38 | 123 | 146 | 0 | 245 | 0 | 0 | 0 | 0 | 5268 | 0 | 0 |
| SRR5387930 | 0 | 0 | 12 | 37 | 0 | 0 | 0 | 108 | 0 | 0 | 197 | 0 | 0 | 0 | 972 | 27267 | 18859 | 0 | 0 | 1729 | 108 | 0 | 0 | 0 | 0 | 12 | 98 | 25 | 0 | 270 | 0 | 0 | 0 | 0 | 1591 | 0 | 0 |
| SRR5387931 | 0 | 0 | 0 | 0 | 34 | 0 | 207 | 0 | 0 | 0 | 170 | 0 | 0 | 0 | 1761 | 12400 | 15317 | 0 | 0 | 1243 | 104 | 0 | 0 | 17 | 0 | 85 | 247 | 145 | 0 | 187 | 0 | 0 | 0 | 0 | 4863 | 0 | 0 |
| SRR5387932 | 0 | 0 | 0 | 0 | 16 | 0 | 0 | 76 | 0 | 0 | 173 | 0 | 0 | 0 | 1147 | 15243 | 10358 | 0 | 0 | 1147 | 76 | 0 | 0 | 0 | 0 | 47 | 150 | 71 | 0 | 276 | 0 | 0 | 0 | 0 | 4104 | 0 | 0 |
| SRR5387933 | 0 | 0 | 0 | 20 | 0 | 0 | 76 | 76 | 0 | 0 | 230 | 0 | 0 | 0 | 992 | 21832 | 20171 | 0 | 0 | 1831 | 305 | 0 | 0 | 14 | 14 | 34 | 81 | 81 | 0 | 88 | 0 | 0 | 0 | 0 | 2610 | 0 | 0 |
| SRR5387934 | 0 | 0 | 0 | 0 | 19 | 0 | 0 | 0 | 0 | 0 | 131 | 0 | 0 | 0 | 803 | 11432 | 16647 | 0 | 0 | 2695 | 115 | 0 | 0 | 0 | 0 | 19 | 50 | 37 | 0 | 193 | 0 | 0 | 0 | 0 | 1203 | 0 | 0 |
| SRR5387935 | 0 | 0 | 0 | 0 | 0 | 0 | 0 | 0 | 0 | 0 | 84 | 0 | 0 | 0 | 789 | 25778 | 17882 | 0 | 0 | 2368 | 226 | 0 | 0 | 21 | 10 | 52 | 240 | 94 | 0 | 136 | 0 | 0 | 0 | 0 | 9057 | 0 | 0 |
| SRR5387936 | 0 | 0 | 0 | 17 | 25 | 0 | 0 | 0 | 0 | 0 | 240 | 0 | 0 | 0 | 1073 | 15250 | 13578 | 0 | 0 | 2229 | 165 | 0 | 0 | 0 | 25 | 50 | 149 | 66 | 0 | 348 | 0 | 0 | 0 | 0 | 3760 | 0 | 0 |
| SRR5387937 | 0 | 0 | 0 | 14 | 14 | 0 | 0 | 0 | 0 | 0 | 123 | 0 | 0 | 0 | 1821 | 14187 | 14954 | 0 | 0 | 1457 | 0 | 0 | 0 | 21 | 14 | 68 | 185 | 41 | 0 | 308 | 0 | 0 | 0 | 0 | 5368 | 0 | 0 |
| SRR5387938 | 0 | 0 | 0 | 29 | 0 | 0 | 0 | 0 | 0 | 0 | 137 | 0 | 0 | 0 | 1031 | 14107 | 13294 | 0 | 0 | 1203 | 86 | 0 | 0 | 0 | 14 | 72 | 172 | 122 | 0 | 244 | 0 | 0 | 0 | 0 | 1356 | 0 | 0 |
| SRR5387939 | 0 | 0 | 0 | 0 | 17 | 0 | 0 | 76 | 0 | 0 | 135 | 0 | 0 | 0 | 2349 | 24425 | 30378 | 0 | 0 | 1288 | 227 | 0 | 0 | 0 | 0 | 17 | 126 | 76 | 0 | 261 | 0 | 0 | 0 | 0 | 2874 | 0 | 0 |
| SRR5387940 | 0 | 0 | 0 | 0 | 0 | 0 | 0 | 0 | 0 | 0 | 11 | 46 | 0 | 0 | 863 | 14193 | 13667 | 0 | 0 | 2301 | 144 | 0 | 0 | 0 | 0 | 0 | 31 | 12 | 0 | 12 | 0 | 72 | 0 | 0 | 7885 | 0 | 31 |
| SRR5387941 | 0 | 0 | 0 | 37 | 44 | 0 | 0 | 67 | 0 | 0 | 557 | 0 | 0 | 0 | 1730 | 14824 | 15409 | 0 | 0 | 1464 | 0 | 0 | 0 | 51 | 22 | 81 | 227 | 103 | 0 | 506 | 0 | 0 | 0 | 0 | 1560 | 0 | 0 |
| SRR5387942 | 0 | 0 | 0 | 0 | 0 | 0 | 0 | 619 | 0 | 79 | 35 | 0 | 0 | 0 | 866 | 11720 | 16471 | 0 | 0 | 1608 | 990 | 0 | 0 | 0 | 0 | 0 | 22 | 17 | 0 | 39 | 0 | 0 | 0 | 0 | 3168 | 0 | 0 |
| SRR5387943 | 0 | 0 | 0 | 0 | 64 | 108 | 0 | 108 | 0 | 0 | 365 | 0 | 0 | 0 | 7867 | 31707 | 39478 | 0 | 0 | 647 | 108 | 0 | 0 | 64 | 0 | 64 | 175 | 32 | 0 | 254 | 0 | 0 | 0 | 0 | 4041 | 0 | 0 |
| SRR5387944 | 0 | 0 | 0 | 24 | 24 | 0 | 0 | 0 | 0 | 0 | 289 | 0 | 0 | 0 | 5038 | 24673 | 18630 | 0 | 0 | 1042 | 0 | 0 | 0 | 24 | 0 | 16 | 113 | 40 | 0 | 129 | 0 | 0 | 0 | 0 | 4028 | 0 | 0 |
| SRR5387945 | 0 | 0 | 0 | 0 | 38 | 0 | 139 | 0 | 0 | 19 | 346 | 0 | 0 | 0 | 2568 | 15813 | 31826 | 0 | 0 | 1805 | 139 | 0 | 0 | 0 | 19 | 38 | 106 | 48 | 0 | 202 | 0 | 0 | 0 | 0 | 2002 | 0 | 0 |
| SRR5387946 | 0 | 0 | 0 | 0 | 0 | 0 | 0 | 0 | 0 | 0 | 13 | 11 | 0 | 0 | 1019 | 11947 | 20331 | 0 | 0 | 2039 | 0 | 0 | 0 | 0 | 0 | 11 | 63 | 37 | 0 | 12 | 0 | 0 | 0 | 0 | 3982 | 0 | 0 |
| SRR5387947 | 0 | 0 | 0 | 11 | 22 | 0 | 0 | 0 | 0 | 0 | 247 | 0 | 0 | 0 | 3188 | 25872 | 13954 | 0 | 0 | 1649 | 110 | 0 | 0 | 11 | 0 | 45 | 224 | 67 | 0 | 269 | 0 | 0 | 0 | 0 | 6105 | 0 | 0 |
| SRR5387948 | 0 | 0 | 0 | 0 | 0 | 63 | 0 | 0 | 0 | 0 | 109 | 14 | 0 | 0 | 1705 | 16848 | 17245 | 0 | 0 | 1263 | 379 | 0 | 0 | 14 | 0 | 18 | 104 | 45 | 0 | 145 | 0 | 0 | 0 | 0 | 2577 | 0 | 18 |
| SRR5387949 | 0 | 0 | 0 | 0 | 39 | 101 | 0 | 0 | 0 | 0 | 129 | 0 | 0 | 0 | 2722 | 23232 | 26625 | 0 | 0 | 1915 | 0 | 0 | 0 | 13 | 13 | 64 | 52 | 39 | 0 | 206 | 0 | 0 | 0 | 0 | 3393 | 0 | 0 |
| SRR5387950 | 0 | 0 | 0 | 0 | 23 | 0 | 0 | 0 | 0 | 0 | 217 | 0 | 0 | 0 | 1691 | 14766 | 12346 | 0 | 0 | 597 | 99 | 0 | 0 | 0 | 31 | 31 | 278 | 108 | 0 | 155 | 0 | 0 | 0 | 0 | 6052 | 0 | 0 |
| SRR5387951 | 0 | 0 | 0 | 22 | 11 | 0 | 0 | 0 | 0 | 0 | 354 | 11 | 0 | 0 | 793 | 8956 | 7578 | 0 | 0 | 1360 | 0 | 0 | 0 | 11 | 11 | 55 | 221 | 177 | 0 | 376 | 0 | 0 | 0 | 0 | 6889 | 0 | 0 |
| SRR5387952 | 0 | 0 | 0 | 0 | 20 | 0 | 0 | 168 | 0 | 0 | 203 | 0 | 0 | 0 | 756 | 9046 | 9046 | 0 | 0 | 1260 | 84 | 0 | 0 | 41 | 0 | 20 | 193 | 61 | 0 | 447 | 0 | 0 | 0 | 0 | 2934 | 0 | 0 |
| SRR5387953 | 0 | 0 | 0 | 0 | 0 | 0 | 0 | 0 | 0 | 0 | 182 | 0 | 0 | 0 | 1716 | 10002 | 13753 | 0 | 0 | 702 | 0 | 0 | 0 | 0 | 0 | 19 | 58 | 19 | 0 | 240 | 0 | 78 | 0 | 0 | 2292 | 0 | 0 |
| SRR5387954 | 0 | 0 | 0 | 0 | 0 | 0 | 0 | 0 | 0 | 0 | 342 | 0 | 0 | 0 | 2775 | 14636 | 18055 | 0 | 0 | 753 | 94 | 0 | 0 | 0 | 30 | 79 | 159 | 73 | 0 | 287 | 0 | 0 | 0 | 0 | 1505 | 0 | 0 |
| SRR5387955 | 0 | 0 | 0 | 22 | 55 | 0 | 0 | 296 | 0 | 0 | 363 | 0 | 0 | 0 | 985 | 11878 | 12563 | 0 | 0 | 1576 | 0 | 0 | 0 | 11 | 33 | 66 | 231 | 33 | 0 | 198 | 0 | 0 | 0 | 0 | 5939 | 0 | 0 |
| SRR5387956 | 0 | 0 | 0 | 15 | 0 | 0 | 0 | 55 | 0 | 0 | 437 | 0 | 0 | 0 | 2038 | 8573 | 14821 | 0 | 0 | 1322 | 110 | 0 | 0 | 0 | 0 | 30 | 143 | 38 | 0 | 211 | 0 | 55 | 0 | 0 | 3342 | 0 | 0 |
| SRR5387957 | 0 | 0 | 0 | 0 | 16 | 0 | 0 | 53 | 0 | 0 | 444 | 16 | 0 | 0 | 2027 | 15878 | 17482 | 0 | 0 | 1547 | 53 | 0 | 0 | 0 | 0 | 24 | 89 | 48 | 0 | 452 | 0 | 0 | 0 | 0 | 3849 | 0 | 0 |
| SRR5387958 | 0 | 0 | 0 | 10 | 0 | 48 | 0 | 0 | 0 | 0 | 57 | 10 | 0 | 0 | 959 | 2630 | 3558 | 48 | 0 | 480 | 48 | 0 | 0 | 0 | 10 | 31 | 213 | 78 | 0 | 34 | 0 | 0 | 0 | 0 | 4486 | 0 | 0 |
| SRR5387959 | 0 | 0 | 0 | 0 | 0 | 0 | 0 | 60 | 0 | 0 | 13 | 0 | 0 | 0 | 5113 | 19842 | 34241 | 0 | 0 | 722 | 120 | 0 | 0 | 0 | 12 | 74 | 118 | 74 | 0 | 416 | 0 | 0 | 0 | 0 | 4390 | 0 | 12 |
| SRR5387960 | 0 | 0 | 0 | 15 | 0 | 0 | 0 | 0 | 0 | 0 | 10 | 0 | 0 | 0 | 1155 | 4886 | 14046 | 0 | 0 | 1365 | 315 | 0 | 0 | 0 | 30 | 45 | 112 | 60 | 0 | 321 | 0 | 0 | 0 | 0 | 2443 | 0 | 0 |
| SRR5387961 | 0 | 0 | 0 | 27 | 18 | 0 | 0 | 112 | 0 | 0 | 172 | 0 | 0 | 0 | 2010 | 21171 | 23262 | 0 | 0 | 1452 | 0 | 0 | 0 | 0 | 73 | 100 | 100 | 109 | 0 | 163 | 0 | 0 | 0 | 0 | 2352 | 0 | 0 |
| SRR5387962 | 12 | 0 | 0 | 0 | 0 | 0 | 0 | 82 | 0 | 0 | 124 | 0 | 0 | 0 | 1888 | 21463 | 28683 | 0 | 0 | 2135 | 82 | 0 | 0 | 0 | 0 | 19 | 62 | 43 | 0 | 106 | 0 | 0 | 0 | 0 | 3902 | 0 | 0 |
| SRR5387963 | 0 | 0 | 0 | 25 | 17 | 0 | 0 | 126 | 0 | 0 | 384 | 0 | 0 | 0 | 2714 | 26558 | 25516 | 0 | 0 | 758 | 126 | 0 | 0 | 0 | 0 | 25 | 142 | 100 | 0 | 284 | 0 | 0 | 0 | 0 | 2083 | 0 | 0 |
| SRR5387964 | 0 | 0 | 0 | 0 | 0 | 74 | 0 | 74 | 0 | 0 | 176 | 0 | 0 | 0 | 883 | 7389 | 12244 | 0 | 0 | 1545 | 74 | 0 | 0 | 0 | 0 | 0 | 120 | 37 | 0 | 259 | 0 | 0 | 0 | 0 | 2955 | 0 | 0 |
| SRR5387965 | 0 | 0 | 0 | 0 | 0 | 0 | 0 | 0 | 0 | 13 | 206 | 0 | 0 | 0 | 1351 | 15459 | 15266 | 0 | 0 | 2252 | 225 | 0 | 0 | 0 | 27 | 60 | 153 | 80 | 0 | 179 | 0 | 0 | 0 | 0 | 5797 | 0 | 0 |
| SRR5387966 | 0 | 0 | 0 | 0 | 31 | 0 | 0 | 0 | 0 | 0 | 173 | 0 | 0 | 0 | 1276 | 12650 | 15267 | 0 | 85 | 2211 | 85 | 0 | 0 | 0 | 0 | 47 | 87 | 55 | 0 | 338 | 0 | 0 | 0 | 0 | 6543 | 0 | 0 |
| SRR5387967 | 0 | 0 | 0 | 14 | 0 | 0 | 57 | 0 | 0 | 0 | 446 | 0 | 0 | 0 | 1318 | 10021 | 10197 | 0 | 0 | 1490 | 115 | 0 | 0 | 22 | 22 | 14 | 245 | 108 | 0 | 367 | 0 | 0 | 0 | 0 | 6329 | 0 | 0 |
| SRR5387968 | 0 | 0 | 0 | 0 | 31 | 96 | 192 | 0 | 0 | 0 | 164 | 0 | 0 | 0 | 1918 | 7057 | 9409 | 0 | 0 | 1726 | 96 | 0 | 0 | 0 | 16 | 23 | 211 | 62 | 0 | 195 | 0 | 0 | 0 | 0 | 3659 | 0 | 0 |
| SRR5387969 | 0 | 0 | 0 | 24 | 16 | 0 | 0 | 0 | 0 | 0 | 240 | 32 | 0 | 0 | 5232 | 16373 | 25189 | 0 | 0 | 1865 | 52 | 0 | 0 | 0 | 0 | 16 | 224 | 72 | 0 | 384 | 0 | 0 | 0 | 0 | 5038 | 0 | 0 |
| SRR5387970 | 0 | 0 | 0 | 34 | 17 | 0 | 0 | 0 | 0 | 0 | 163 | 0 | 0 | 0 | 4360 | 11581 | 24233 | 0 | 0 | 1615 | 0 | 0 | 0 | 51 | 43 | 60 | 257 | 197 | 0 | 265 | 0 | 0 | 0 | 0 | 1501 | 0 | 0 |
| SRR5387971 | 0 | 0 | 0 | 21 | 10 | 0 | 0 | 0 | 59 | 10 | 309 | 0 | 0 | 0 | 7969 | 14667 | 22578 | 0 | 0 | 1417 | 59 | 0 | 0 | 21 | 21 | 21 | 237 | 93 | 0 | 257 | 0 | 0 | 0 | 0 | 1648 | 0 | 10 |
| SRR5387972 | 0 | 0 | 0 | 0 | 0 | 0 | 0 | 0 | 0 | 0 | 254 | 0 | 0 | 0 | 5907 | 17209 | 23034 | 0 | 0 | 1743 | 291 | 0 | 0 | 0 | 15 | 0 | 54 | 54 | 0 | 161 | 0 | 0 | 0 | 0 | 3442 | 0 | 0 |
| SRR5387973 | 0 | 0 | 0 | 21 | 21 | 0 | 0 | 0 | 0 | 0 | 113 | 10 | 0 | 0 | 705 | 20843 | 20370 | 0 | 0 | 1763 | 118 | 0 | 0 | 10 | 62 | 82 | 185 | 164 | 0 | 185 | 0 | 0 | 0 | 0 | 3790 | 0 | 0 |
| SRR5387974 | 0 | 0 | 0 | 0 | 40 | 122 | 0 | 122 | 0 | 0 | 129 | 0 | 0 | 0 | 732 | 26905 | 20957 | 0 | 0 | 2805 | 610 | 0 | 0 | 0 | 40 | 70 | 139 | 80 | 0 | 269 | 0 | 122 | 0 | 0 | 4531 | 0 | 0 |
| SRR5387975 | 0 | 0 | 0 | 18 | 0 | 0 | 0 | 0 | 0 | 0 | 291 | 0 | 0 | 0 | 636 | 31847 | 23779 | 0 | 0 | 477 | 159 | 0 | 0 | 36 | 73 | 73 | 219 | 109 | 0 | 255 | 0 | 0 | 0 | 0 | 2972 | 0 | 0 |
| SRR5387976 | 0 | 0 | 0 | 17 | 0 | 0 | 0 | 0 | 0 | 0 | 345 | 0 | 0 | 0 | 974 | 12097 | 15373 | 0 | 0 | 2517 | 0 | 0 | 0 | 29 | 0 | 23 | 288 | 127 | 0 | 357 | 0 | 0 | 0 | 0 | 3276 | 0 | 0 |
| SRR5387977 | 0 | 0 | 0 | 0 | 19 | 0 | 0 | 0 | 0 | 0 | 298 | 0 | 0 | 0 | 2702 | 21077 | 33000 | 0 | 0 | 2177 | 75 | 0 | 0 | 0 | 0 | 19 | 48 | 19 | 0 | 221 | 0 | 0 | 0 | 0 | 1703 | 0 | 0 |
| SRR5387978 | 0 | 0 | 0 | 76 | 47 | 0 | 0 | 0 | 0 | 0 | 416 | 0 | 0 | 0 | 767 | 16938 | 21933 | 0 | 0 | 1534 | 77 | 0 | 0 | 19 | 19 | 47 | 208 | 104 | 0 | 236 | 0 | 0 | 0 | 0 | 1737 | 0 | 0 |
| SRR5387979 | 0 | 0 | 0 | 12 | 37 | 108 | 0 | 108 | 0 | 0 | 318 | 0 | 0 | 0 | 1728 | 39354 | 23209 | 0 | 108 | 2268 | 324 | 0 | 0 | 0 | 24 | 61 | 208 | 110 | 0 | 269 | 0 | 0 | 0 | 0 | 2523 | 0 | 0 |
| SRR5387980 | 0 | 0 | 0 | 45 | 18 | 0 | 0 | 151 | 0 | 0 | 376 | 0 | 0 | 0 | 1360 | 14414 | 15250 | 0 | 0 | 831 | 0 | 0 | 0 | 18 | 0 | 27 | 367 | 125 | 0 | 358 | 0 | 0 | 0 | 0 | 2925 | 0 | 0 |
| SRR5387981 | 0 | 0 | 0 | 0 | 42 | 269 | 0 | 0 | 0 | 0 | 208 | 0 | 0 | 0 | 1482 | 27020 | 18663 | 0 | 0 | 1347 | 135 | 0 | 0 | 14 | 83 | 139 | 444 | 278 | 0 | 167 | 0 | 0 | 0 | 0 | 6964 | 0 | 0 |
| SRR5387982 | 0 | 0 | 0 | 11 | 23 | 0 | 0 | 114 | 0 | 0 | 158 | 11 | 0 | 0 | 1482 | 11299 | 11299 | 0 | 0 | 3535 | 114 | 0 | 0 | 0 | 23 | 11 | 113 | 34 | 0 | 371 | 0 | 0 | 0 | 0 | 2959 | 0 | 0 |
| SRR5387983 | 16 | 0 | 0 | 16 | 24 | 0 | 0 | 0 | 0 | 0 | 134 | 0 | 0 | 0 | 1832 | 25536 | 25536 | 0 | 0 | 1832 | 118 | 0 | 0 | 16 | 0 | 32 | 158 | 39 | 0 | 182 | 0 | 0 | 0 | 0 | 2455 | 0 | 0 |
| SRR5387984 | 0 | 0 | 0 | 16 | 16 | 0 | 54 | 0 | 0 | 11 | 218 | 54 | 0 | 0 | 649 | 9861 | 15672 | 0 | 0 | 1082 | 0 | 0 | 0 | 16 | 11 | 0 | 60 | 33 | 11 | 294 | 0 | 0 | 0 | 0 | 704 | 0 | 0 |
| SRR5387985 | 0 | 0 | 0 | 19 | 0 | 0 | 0 | 0 | 0 | 0 | 320 | 0 | 0 | 0 | 2424 | 14629 | 30543 | 0 | 0 | 1212 | 223 | 0 | 0 | 0 | 0 | 38 | 119 | 13 | 0 | 264 | 0 | 0 | 0 | 0 | 2570 | 0 | 0 |
| SRR5387986 | 0 | 0 | 0 | 15 | 15 | 0 | 0 | 29 | 0 | 0 | 237 | 0 | 0 | 0 | 3547 | 14615 | 44744 | 0 | 0 | 1038 | 144 | 0 | 0 | 0 | 0 | 20 | 188 | 44 | 0 | 183 | 0 | 0 | 0 | 0 | 2586 | 0 | 0 |
| SRR5387987 | 0 | 0 | 0 | 20 | 20 | 0 | 0 | 0 | 0 | 0 | 241 | 0 | 0 | 0 | 1241 | 9453 | 15657 | 0 | 0 | 1552 | 310 | 0 | 0 | 0 | 10 | 40 | 181 | 191 | 0 | 261 | 0 | 0 | 0 | 0 | 2363 | 0 | 0 |
| SRR5387988 | 0 | 0 | 0 | 26 | 87 | 0 | 0 | 0 | 0 | 0 | 243 | 0 | 0 | 0 | 1045 | 10029 | 14433 | 0 | 0 | 436 | 0 | 0 | 0 | 17 | 17 | 35 | 374 | 252 | 0 | 339 | 0 | 0 | 0 | 0 | 4403 | 0 | 0 |
| SRR5387989 | 0 | 0 | 0 | 17 | 56 | 81 | 0 | 243 | 0 | 11 | 578 | 0 | 0 | 0 | 324 | 8366 | 12085 | 0 | 81 | 810 | 324 | 0 | 0 | 11 | 28 | 44 | 233 | 72 | 0 | 1506 | 0 | 0 | 0 | 0 | 2789 | 0 | 0 |
| SRR5387990 | 0 | 0 | 0 | 0 | 23 | 0 | 0 | 0 | 0 | 15 | 121 | 0 | 0 | 0 | 483 | 10882 | 7732 | 0 | 0 | 1328 | 362 | 0 | 0 | 0 | 0 | 0 | 38 | 23 | 0 | 288 | 0 | 0 | 0 | 0 | 6300 | 0 | 0 |

### Supp Table S8.3.Columns_BX_to_DH

| | | | | | | | | | | | | | | | | | | | | | | | | | | | | | | | | | | | | | |
|---|---|---|---|---|---|---|---|---|---|---|---|---|---|---|---|---|---|---|---|---|---|---|---|---|---|---|---|---|---|---|---|---|---|---|---|---|---|
| SRR5387991 | 0 | 0 | 0 | 15 | 31 | 0 | 170 | 57 | 0 | 0 | 360 | 15 | 0 | 0 | 1978 | 23288 | 21475 | 0 | 0 | 1413 | 113 | 0 | 0 | 15 | 15 | 54 | 107 | 69 | 0 | 460 | 0 | 0 | 0 | 0 | 6694 | 0 | 0 |
| SRR5387992 | 0 | 0 | 0 | 17 | 17 | 82 | 0 | 0 | 0 | 0 | 201 | 0 | 0 | 0 | 1308 | 13158 | 16917 | 0 | 0 | 1308 | 163 | 0 | 0 | 17 | 44 | 61 | 367 | 157 | 0 | 262 | 0 | 0 | 0 | 0 | 1253 | 0 | 0 |
| SRR5387993 | 0 | 0 | 0 | 43 | 17 | 0 | 0 | 177 | 0 | 0 | 86 | 0 | 0 | 0 | 1775 | 15969 | 14349 | 0 | 0 | 1331 | 0 | 0 | 0 | 0 | 17 | 34 | 292 | 172 | 0 | 224 | 0 | 0 | 0 | 0 | 463 | 0 | 0 |
| SRR5387994 | 0 | 0 | 0 | 20 | 25 | 0 | 62 | 62 | 0 | 0 | 197 | 49 | 0 | 0 | 1867 | 12084 | 15018 | 0 | 0 | 1742 | 62 | 0 | 0 | 0 | 0 | 15 | 167 | 89 | 0 | 177 | 0 | 0 | 0 | 0 | 1726 | 0 | 0 |
| SRR5387995 | 0 | 0 | 0 | 17 | 0 | 0 | 0 | 0 | 0 | 0 | 261 | 0 | 0 | 0 | 633 | 18347 | 11871 | 0 | 0 | 1054 | 70 | 0 | 0 | 35 | 0 | 52 | 304 | 139 | 0 | 209 | 0 | 0 | 0 | 0 | 3669 | 0 | 0 |
| SRR5387996 | 0 | 0 | 0 | 0 | 28 | 57 | 0 | 0 | 0 | 0 | 222 | 0 | 0 | 0 | 624 | 10906 | 9418 | 0 | 57 | 1644 | 227 | 0 | 0 | 21 | 0 | 0 | 90 | 111 | 0 | 589 | 0 | 0 | 0 | 0 | 6114 | 0 | 0 |
| SRR5387997 | 0 | 0 | 0 | 0 | 22 | 131 | 0 | 0 | 0 | 0 | 238 | 0 | 0 | 0 | 2361 | 30291 | 38395 | 0 | 0 | 1181 | 0 | 0 | 0 | 0 | 0 | 37 | 74 | 59 | 0 | 119 | 0 | 0 | 0 | 0 | 3666 | 0 | 0 |
| SRR5387998 | 0 | 0 | 0 | 0 | 17 | 0 | 0 | 70 | 0 | 0 | 372 | 0 | 0 | 0 | 1332 | 16068 | 28922 | 0 | 0 | 841 | 140 | 0 | 0 | 45 | 51 | 34 | 208 | 85 | 0 | 180 | 0 | 0 | 0 | 0 | 3816 | 0 | 0 |
| SRR5387999 | 0 | 0 | 0 | 0 | 23 | 0 | 250 | 167 | 0 | 0 | 269 | 23 | 0 | 0 | 7840 | 30539 | 42793 | 0 | 0 | 500 | 83 | 0 | 0 | 18 | 12 | 47 | 216 | 123 | 0 | 275 | 0 | 0 | 0 | 0 | 9337 | 0 | 0 |
| SRR5388000 | 0 | 0 | 0 | 0 | 0 | 120 | 0 | 0 | 0 | 0 | 151 | 0 | 0 | 0 | 3370 | 16837 | 33950 | 0 | 0 | 3250 | 120 | 0 | 0 | 34 | 0 | 42 | 84 | 59 | 0 | 168 | 0 | 0 | 0 | 0 | 8280 | 0 | 0 |
| SRR5388001 | 0 | 0 | 0 | 0 | 22 | 0 | 119 | 119 | 0 | 0 | 284 | 0 | 0 | 0 | 594 | 12612 | 8128 | 0 | 0 | 2968 | 0 | 0 | 0 | 0 | 0 | 0 | 0 | 0 | 0 | 291 | 0 | 0 | 0 | 0 | 3924 | 0 | 0 |
| SRR5388002 | 0 | 0 | 0 | 0 | 0 | 94 | 0 | 0 | 0 | 0 | 223 | 0 | 0 | 0 | 283 | 19793 | 9443 | 0 | 0 | 3585 | 94 | 0 | 0 | 35 | 0 | 0 | 0 | 0 | 0 | 251 | 0 | 0 | 0 | 0 | 6537 | 0 | 0 |
| SRR5388003 | 0 | 0 | 0 | 53 | 26 | 0 | 0 | 367 | 0 | 0 | 184 | 0 | 0 | 0 | 1466 | 38622 | 18441 | 0 | 0 | 1283 | 0 | 0 | 0 | 13 | 13 | 53 | 118 | 92 | 13 | 145 | 0 | 0 | 0 | 0 | 8351 | 0 | 0 |
| SRR5388004 | 0 | 0 | 0 | 0 | 0 | 0 | 98 | 98 | 0 | 0 | 175 | 0 | 0 | 0 | 783 | 5810 | 13640 | 0 | 0 | 685 | 0 | 0 | 0 | 0 | 19 | 32 | 97 | 39 | 0 | 175 | 0 | 0 | 0 | 0 | 3536 | 0 | 0 |
| SRR5388005 | 0 | 14 | 0 | 0 | 0 | 49 | 49 | 49 | 0 | 0 | 208 | 0 | 0 | 0 | 6879 | 38304 | 39545 | 0 | 0 | 342 | 293 | 0 | 0 | 28 | 0 | 0 | 49 | 14 | 0 | 250 | 0 | 0 | 0 | 0 | 4680 | 0 | 0 |
| SRR5388006 | 0 | 0 | 0 | 24 | 24 | 0 | 202 | 0 | 0 | 0 | 122 | 0 | 0 | 0 | 4111 | 23730 | 34723 | 0 | 0 | 1078 | 135 | 0 | 0 | 24 | 0 | 24 | 98 | 41 | 0 | 187 | 0 | 0 | 0 | 0 | 8044 | 0 | 0 |
| SRR5388007 | 0 | 0 | 0 | 0 | 13 | 0 | 0 | 50 | 0 | 0 | 12 | 0 | 0 | 0 | 6195 | 35978 | 129685 | 0 | 0 | 694 | 644 | 0 | 0 | 0 | 0 | 53 | 133 | 62 | 0 | 12 | 0 | 50 | 0 | 0 | 1642 | 0 | 13 |
| SRR5388008 | 0 | 0 | 0 | 0 | 14 | 41 | 0 | 41 | 0 | 0 | 417 | 0 | 0 | 0 | 1796 | 9333 | 37705 | 0 | 0 | 1265 | 571 | 0 | 0 | 0 | 0 | 41 | 217 | 38 | 0 | 358 | 0 | 0 | 0 | 0 | 1618 | 0 | 0 |
| SRR5388009 | 0 | 0 | 0 | 0 | 10 | 0 | 0 | 0 | 0 | 0 | 144 | 0 | 0 | 0 | 725 | 14539 | 6688 | 104 | 0 | 1656 | 0 | 0 | 0 | 0 | 0 | 0 | 92 | 0 | 0 | 175 | 0 | 0 | 0 | 0 | 4362 | 0 | 0 |
| SRR5388010 | 0 | 0 | 0 | 0 | 15 | 0 | 0 | 0 | 0 | 0 | 105 | 0 | 0 | 0 | 997 | 14269 | 19620 | 0 | 0 | 2216 | 111 | 0 | 0 | 0 | 0 | 15 | 105 | 75 | 0 | 136 | 0 | 0 | 0 | 0 | 5648 | 0 | 0 |
| SRR5388011 | 0 | 0 | 0 | 0 | 23 | 0 | 0 | 0 | 0 | 0 | 135 | 0 | 0 | 0 | 2215 | 36173 | 33460 | 0 | 0 | 1148 | 492 | 0 | 0 | 11 | 101 | 158 | 282 | 124 | 0 | 135 | 0 | 0 | 0 | 0 | 3075 | 0 | 11 |
| SRR5388012 | 0 | 0 | 0 | 10 | 42 | 72 | 72 | 72 | 0 | 0 | 241 | 0 | 0 | 0 | 2735 | 47303 | 42926 | 0 | 0 | 2663 | 432 | 0 | 0 | 10 | 63 | 126 | 220 | 105 | 0 | 325 | 0 | 0 | 0 | 0 | 3954 | 0 | 0 |
| SRR5388013 | 0 | 0 | 0 | 14 | 14 | 38 | 0 | 0 | 0 | 0 | 225 | 0 | 0 | 0 | 794 | 6538 | 11636 | 0 | 0 | 4539 | 151 | 0 | 0 | 0 | 0 | 23 | 83 | 28 | 0 | 266 | 0 | 0 | 0 | 0 | 1662 | 0 | 0 |
| SRR5388014 | 0 | 0 | 0 | 47 | 23 | 0 | 0 | 104 | 0 | 0 | 193 | 0 | 0 | 0 | 1515 | 9097 | 25285 | 0 | 0 | 2298 | 209 | 0 | 0 | 18 | 18 | 76 | 10 | 99 | 0 | 333 | 0 | 0 | 0 | 0 | 2621 | 0 | 0 |
| SRR5388015 | 0 | 0 | 0 | 0 | 26 | 0 | 0 | 0 | 0 | 0 | 183 | 0 | 0 | 0 | 1554 | 5063 | 11697 | 0 | 0 | 1081 | 68 | 0 | 0 | 0 | 17 | 35 | 235 | 157 | 0 | 200 | 0 | 0 | 0 | 0 | 1746 | 0 | 17 |
| SRR5388016 | 0 | 0 | 0 | 0 | 23 | 0 | 0 | 0 | 0 | 0 | 204 | 0 | 0 | 0 | 2050 | 9125 | 30113 | 0 | 0 | 1864 | 47 | 0 | 0 | 23 | 29 | 35 | 415 | 111 | 0 | 269 | 0 | 0 | 0 | 0 | 3650 | 0 | 12 |
| SRR5388017 | 0 | 0 | 0 | 0 | 29 | 0 | 0 | 0 | 0 | 0 | 191 | 0 | 0 | 0 | 1534 | 19930 | 18255 | 0 | 0 | 1416 | 0 | 0 | 0 | 48 | 0 | 48 | 201 | 124 | 0 | 201 | 0 | 59 | 0 | 0 | 2010 | 0 | 0 |
| SRR5388018 | 0 | 0 | 0 | 0 | 15 | 85 | 0 | 85 | 0 | 22 | 176 | 0 | 0 | 0 | 767 | 17018 | 16478 | 0 | 0 | 1874 | 256 | 0 | 0 | 0 | 0 | 15 | 73 | 59 | 0 | 169 | 0 | 0 | 0 | 0 | 3512 | 0 | 0 |
| SRR5388019 | 0 | 0 | 0 | 11 | 34 | 0 | 0 | 90 | 0 | 0 | 224 | 0 | 0 | 0 | 1982 | 29599 | 14909 | 0 | 0 | 2252 | 180 | 0 | 0 | 34 | 11 | 45 | 235 | 179 | 0 | 146 | 0 | 0 | 0 | 0 | 1754 | 0 | 0 |
| SRR5388020 | 0 | 0 | 0 | 28 | 37 | 0 | 0 | 0 | 0 | 0 | 140 | 0 | 0 | 0 | 108 | 15139 | 11836 | 0 | 108 | 1297 | 108 | 0 | 0 | 28 | 0 | 37 | 214 | 159 | 0 | 177 | 0 | 0 | 0 | 0 | 1927 | 0 | 0 |
| SRR5388021 | 0 | 0 | 0 | 0 | 67 | 0 | 0 | 0 | 0 | 0 | 275 | 0 | 0 | 0 | 2320 | 23505 | 20666 | 0 | 0 | 1254 | 125 | 0 | 0 | 0 | 33 | 50 | 192 | 75 | 0 | 308 | 0 | 0 | 0 | 0 | 6783 | 0 | 0 |
| SRR5388022 | 0 | 0 | 0 | 19 | 32 | 0 | 0 | 66 | 0 | 0 | 132 | 0 | 0 | 0 | 727 | 12597 | 12412 | 0 | 0 | 1058 | 66 | 0 | 0 | 0 | 0 | 19 | 139 | 50 | 0 | 164 | 0 | 0 | 0 | 0 | 2223 | 0 | 0 |
| SRR5388023 | 0 | 0 | 0 | 0 | 21 | 193 | 0 | 193 | 0 | 0 | 63 | 0 | 0 | 0 | 1735 | 27074 | 19444 | 0 | 0 | 1253 | 96 | 0 | 0 | 10 | 10 | 10 | 42 | 52 | 0 | 189 | 0 | 0 | 0 | 0 | 7384 | 0 | 0 |
| SRR5388024 | 0 | 0 | 0 | 16 | 25 | 0 | 0 | 117 | 0 | 0 | 74 | 0 | 0 | 0 | 468 | 8921 | 8633 | 0 | 0 | 1520 | 117 | 0 | 0 | 33 | 16 | 25 | 147 | 90 | 0 | 147 | 0 | 0 | 0 | 0 | 3165 | 0 | 0 |
| SRR5388025 | 0 | 0 | 0 | 16 | 33 | 0 | 0 | 0 | 95 | 0 | 147 | 0 | 0 | 0 | 2461 | 18315 | 17752 | 0 | 95 | 1609 | 189 | 0 | 0 | 0 | 0 | 16 | 131 | 131 | 0 | 229 | 0 | 0 | 0 | 0 | 3381 | 0 | 0 |
| SRR5388026 | 0 | 0 | 0 | 46 | 31 | 98 | 0 | 0 | 0 | 0 | 230 | 0 | 0 | 0 | 1280 | 18544 | 17453 | 0 | 0 | 2657 | 0 | 0 | 0 | 31 | 31 | 0 | 77 | 61 | 0 | 291 | 0 | 0 | 0 | 0 | 3818 | 0 | 0 |
| SRR5388027 | 0 | 0 | 0 | 0 | 12 | 0 | 95 | 95 | 0 | 0 | 191 | 0 | 0 | 0 | 1716 | 18791 | 13713 | 0 | 0 | 1144 | 191 | 0 | 0 | 12 | 0 | 36 | 143 | 96 | 0 | 167 | 0 | 0 | 0 | 0 | 2793 | 0 | 0 |
| SRR5388028 | 0 | 0 | 0 | 0 | 0 | 111 | 0 | 111 | 0 | 0 | 378 | 0 | 0 | 0 | 2777 | 15720 | 23579 | 0 | 0 | 1999 | 0 | 0 | 0 | 28 | 0 | 19 | 123 | 76 | 0 | 255 | 0 | 0 | 0 | 0 | 5441 | 0 | 0 |
| SRR5388029 | 0 | 0 | 0 | 0 | 0 | 0 | 0 | 0 | 0 | 0 | 232 | 0 | 0 | 0 | 1431 | 7266 | 11090 | 0 | 0 | 1756 | 260 | 0 | 0 | 0 | 0 | 17 | 83 | 33 | 0 | 390 | 0 | 65 | 0 | 0 | 2677 | 0 | 0 |
| SRR5388030 | 0 | 0 | 0 | 0 | 34 | 0 | 0 | 81 | 0 | 0 | 221 | 0 | 0 | 0 | 966 | 5584 | 12867 | 0 | 0 | 1852 | 403 | 0 | 0 | 0 | 0 | 17 | 288 | 127 | 0 | 212 | 0 | 0 | 0 | 0 | 3884 | 0 | 0 |
| SRR5388031 | 0 | 0 | 0 | 0 | 16 | 0 | 0 | 29 | 0 | 0 | 192 | 0 | 0 | 0 | 3305 | 15893 | 15268 | 0 | 0 | 1404 | 234 | 0 | 0 | 12 | 12 | 32 | 56 | 16 | 0 | 216 | 0 | 29 | 0 | 0 | 7148 | 0 | 0 |
| SRR5388032 | 0 | 0 | 0 | 0 | 11 | 0 | 0 | 94 | 0 | 33 | 105 | 0 | 0 | 0 | 938 | 9192 | 8708 | 0 | 0 | 1782 | 281 | 0 | 0 | 0 | 11 | 11 | 116 | 55 | 11 | 215 | 0 | 0 | 0 | 0 | 2177 | 0 | 17 |
| SRR5388033 | 0 | 0 | 0 | 39 | 39 | 0 | 0 | 193 | 0 | 0 | 244 | 0 | 0 | 0 | 1289 | 30308 | 13604 | 0 | 0 | 709 | 129 | 0 | 0 | 29 | 0 | 49 | 372 | 137 | 0 | 205 | 0 | 0 | 0 | 0 | 3789 | 0 | 0 |
| SRR5388034 | 0 | 0 | 0 | 0 | 50 | 0 | 0 | 0 | 0 | 0 | 200 | 17 | 0 | 0 | 1075 | 7646 | 10333 | 77 | 0 | 1767 | 0 | 0 | 0 | 0 | 17 | 17 | 133 | 33 | 0 | 216 | 0 | 0 | 0 | 0 | 3720 | 0 | 0 |
| SRR5388035 | 0 | 0 | 0 | 35 | 47 | 0 | 0 | 92 | 0 | 0 | 165 | 0 | 0 | 0 | 2857 | 33981 | 17106 | 0 | 0 | 1659 | 92 | 0 | 0 | 12 | 24 | 24 | 213 | 260 | 0 | 189 | 0 | 0 | 0 | 0 | 2080 | 0 | 0 |
| SRR5388036 | 0 | 0 | 0 | 29 | 0 | 0 | 0 | 0 | 0 | 0 | 212 | 0 | 0 | 0 | 510 | 17269 | 7044 | 0 | 0 | 1019 | 0 | 0 | 0 | 0 | 29 | 68 | 193 | 77 | 0 | 270 | 0 | 0 | 0 | 0 | 1363 | 0 | 0 |
| SRR5388037 | 0 | 0 | 0 | 0 | 33 | 0 | 41 | 123 | 0 | 0 | 289 | 0 | 0 | 0 | 3882 | 43059 | 27239 | 0 | 0 | 817 | 82 | 0 | 0 | 58 | 58 | 74 | 346 | 124 | 0 | 181 | 0 | 0 | 0 | 0 | 3667 | 0 | 0 |
| SRR5388038 | 0 | 0 | 11 | 33 | 11 | 0 | 0 | 0 | 0 | 0 | 273 | 0 | 0 | 0 | 1740 | 21532 | 12170 | 0 | 0 | 2578 | 322 | 0 | 0 | 22 | 22 | 66 | 98 | 98 | 0 | 437 | 0 | 0 | 0 | 0 | 6397 | 0 | 0 |
| SRR5388039 | 0 | 0 | 0 | 0 | 14 | 87 | 0 | 87 | 0 | 0 | 359 | 0 | 0 | 0 | 1656 | 33001 | 31894 | 0 | 0 | 872 | 0 | 0 | 0 | 14 | 14 | 55 | 138 | 41 | 0 | 470 | 0 | 0 | 0 | 0 | 5316 | 0 | 0 |
| SRR5388040 | 0 | 0 | 0 | 22 | 30 | 0 | 0 | 0 | 0 | 0 | 134 | 0 | 0 | 0 | 838 | 9247 | 15663 | 0 | 0 | 978 | 140 | 0 | 0 | 0 | 15 | 0 | 60 | 67 | 0 | 335 | 0 | 0 | 0 | 0 | 3019 | 0 | 0 |
| SRR5388041 | 0 | 0 | 0 | 0 | 55 | 96 | 96 | 191 | 0 | 0 | 182 | 0 | 0 | 0 | 1722 | 13707 | 18846 | 0 | 0 | 2009 | 96 | 0 | 0 | 0 | 27 | 27 | 164 | 173 | 0 | 200 | 0 | 0 | 0 | 0 | 5997 | 0 | 0 |
| SRR5388042 | 17 | 0 | 0 | 17 | 43 | 0 | 90 | 90 | 0 | 0 | 274 | 0 | 0 | 0 | 808 | 15381 | 21729 | 0 | 0 | 1885 | 0 | 0 | 0 | 34 | 17 | 60 | 111 | 103 | 0 | 197 | 0 | 0 | 0 | 0 | 2686 | 0 | 0 |
| SRR5388043 | 0 | 0 | 0 | 0 | 0 | 0 | 0 | 58 | 0 | 35 | 71 | 0 | 0 | 0 | 1622 | 9173 | 20718 | 0 | 0 | 1506 | 1158 | 0 | 0 | 0 | 0 | 18 | 35 | 29 | 0 | 52 | 0 | 0 | 0 | 0 | 1107 | 0 | 0 |
| SRR5388044 | 0 | 0 | 0 | 20 | 40 | 41 | 0 | 0 | 0 | 0 | 308 | 0 | 0 | 0 | 937 | 8337 | 15980 | 41 | 0 | 2730 | 245 | 0 | 0 | 20 | 10 | 30 | 192 | 66 | 0 | 11 | 0 | 0 | 0 | 0 | 5790 | 0 | 0 |
| SRR5388045 | 0 | 0 | 16 | 16 | 41 | 0 | 134 | 134 | 0 | 0 | 247 | 0 | 0 | 0 | 4941 | 20026 | 22933 | 0 | 0 | 267 | 0 | 0 | 0 | 16 | 0 | 25 | 58 | 49 | 0 | 181 | 0 | 0 | 0 | 0 | 20349 | 0 | 0 |
| SRR5388046 | 0 | 0 | 0 | 17 | 25 | 0 | 0 | 0 | 0 | 0 | 169 | 0 | 0 | 0 | 1602 | 10120 | 11297 | 0 | 0 | 3004 | 100 | 0 | 0 | 0 | 25 | 0 | 118 | 25 | 0 | 380 | 0 | 0 | 0 | 0 | 5884 | 0 | 0 |
| SRR5388047 | 0 | 0 | 0 | 0 | 0 | 0 | 0 | 92 | 0 | 0 | 323 | 0 | 0 | 0 | 2864 | 10163 | 24330 | 0 | 0 | 1617 | 46 | 0 | 0 | 0 | 17 | 17 | 35 | 44 | 0 | 297 | 0 | 0 | 0 | 0 | 2156 | 0 | 0 |
| SRR5388048 | 0 | 0 | 0 | 25 | 33 | 0 | 0 | 0 | 0 | 0 | 123 | 0 | 0 | 0 | 2828 | 13527 | 24579 | 0 | 0 | 2366 | 0 | 0 | 0 | 33 | 0 | 49 | 123 | 41 | 0 | 180 | 0 | 173 | 0 | 0 | 2145 | 0 | 0 |
| SRR5388049 | 0 | 0 | 0 | 30 | 75 | 0 | 0 | 70 | 0 | 0 | 306 | 0 | 0 | 0 | 765 | 10004 | 10540 | 0 | 0 | 1183 | 70 | 0 | 0 | 60 | 60 | 60 | 298 | 52 | 0 | 462 | 0 | 0 | 0 | 0 | 6967 | 0 | 0 |
| SRR5388050 | 0 | 0 | 0 | 19 | 12 | 0 | 65 | 0 | 0 | 0 | 255 | 37 | 0 | 0 | 521 | 7020 | 8816 | 0 | 0 | 1693 | 65 | 0 | 0 | 19 | 19 | 19 | 292 | 68 | 0 | 13 | 0 | 0 | 0 | 0 | 2449 | 0 | 0 |
| SRR5388051 | 0 | 0 | 0 | 0 | 0 | 0 | 0 | 0 | 0 | 0 | 170 | 0 | 0 | 0 | 2280 | 14230 | 16818 | 0 | 0 | 1190 | 99 | 0 | 0 | 0 | 0 | 14 | 197 | 48 | 0 | 204 | 0 | 0 | 0 | 0 | 5175 | 0 | 0 |
| SRR5388052 | 0 | 0 | 0 | 17 | 23 | 0 | 0 | 0 | 62 | 0 | 234 | 0 | 0 | 0 | 432 | 10851 | 10518 | 0 | 0 | 1421 | 62 | 0 | 0 | 0 | 0 | 23 | 74 | 17 | 0 | 229 | 0 | 0 | 0 | 0 | 2504 | 0 | 0 |
| SRR5388053 | 0 | 0 | 0 | 12 | 43 | 0 | 148 | 99 | 99 | 0 | 204 | 0 | 0 | 0 | 2758 | 20978 | 29601 | 0 | 0 | 936 | 197 | 0 | 0 | 0 | 0 | 37 | 167 | 167 | 0 | 179 | 0 | 0 | 0 | 0 | 1931 | 0 | 0 |
| SRR5388054 | 0 | 0 | 15 | 0 | 23 | 0 | 0 | 208 | 0 | 0 | 264 | 0 | 0 | 0 | 1247 | 8421 | 8158 | 0 | 104 | 1559 | 104 | 0 | 0 | 0 | 15 | 45 | 121 | 143 | 0 | 189 | 0 | 0 | 0 | 0 | 4211 | 0 | 0 |
| SRR5388055 | 0 | 0 | 0 | 38 | 0 | 0 | 0 | 0 | 0 | 0 | 294 | 0 | 0 | 0 | 1231 | 22419 | 17030 | 82 | 0 | 2134 | 0 | 0 | 0 | 0 | 0 | 66 | 104 | 66 | 0 | 266 | 0 | 0 | 0 | 0 | 2587 | 0 | 0 |
| SRR5388056 | 0 | 0 | 0 | 31 | 15 | 0 | 0 | 0 | 0 | 0 | 201 | 0 | 0 | 0 | 409 | 11108 | 8935 | 0 | 0 | 1226 | 0 | 0 | 0 | 23 | 23 | 39 | 356 | 162 | 0 | 201 | 0 | 0 | 0 | 0 | 1207 | 0 | 0 |
| SRR5388057 | 0 | 0 | 0 | 0 | 0 | 0 | 0 | 112 | 0 | 0 | 167 | 0 | 0 | 0 | 1237 | 11434 | 11761 | 0 | 0 | 1237 | 112 | 0 | 0 | 18 | 18 | 0 | 185 | 62 | 0 | 220 | 0 | 0 | 0 | 0 | 4247 | 0 | 0 |
| SRR5388058 | 0 | 0 | 0 | 0 | 18 | 0 | 0 | 0 | 0 | 0 | 212 | 0 | 0 | 0 | 1152 | 10858 | 14755 | 0 | 0 | 975 | 177 | 0 | 0 | 12 | 18 | 42 | 103 | 55 | 0 | 194 | 0 | 0 | 0 | 0 | 5568 | 0 | 0 |
| SRR5388059 | 0 | 0 | 0 | 0 | 36 | 0 | 90 | 90 | 0 | 0 | 187 | 0 | 0 | 0 | 1258 | 12987 | 13967 | 0 | 0 | 1977 | 90 | 0 | 0 | 14 | 14 | 29 | 65 | 43 | 0 | 238 | 0 | 0 | 0 | 0 | 9556 | 0 | 14 |
| SRR5388060 | 0 | 0 | 0 | 13 | 0 | 0 | 0 | 87 | 0 | 20 | 188 | 0 | 0 | 0 | 436 | 6239 | 6738 | 0 | 0 | 1658 | 349 | 0 | 0 | 0 | 20 | 27 | 128 | 47 | 0 | 229 | 0 | 0 | 0 | 0 | 3494 | 0 | 0 |
| SRR5388061 | 0 | 0 | 0 | 0 | 0 | 0 | 0 | 250 | 0 | 0 | 137 | 0 | 0 | 0 | 499 | 15424 | 13175 | 0 | 125 | 1872 | 0 | 0 | 0 | 0 | 0 | 46 | 80 | 46 | 0 | 137 | 0 | 0 | 0 | 0 | 1285 | 0 | 0 |
| SRR5388062 | 0 | 0 | 0 | 0 | 0 | 0 | 0 | 0 | 0 | 0 | 258 | 0 | 0 | 0 | 1302 | 7955 | 12464 | 0 | 0 | 1703 | 0 | 0 | 0 | 23 | 0 | 55 | 117 | 117 | 0 | 125 | 0 | 0 | 0 | 0 | 2121 | 0 | 0 |
| SRR5388063 | 0 | 0 | 11 | 32 | 21 | 0 | 0 | 0 | 0 | 11 | 245 | 0 | 0 | 0 | 983 | 10616 | 10616 | 0 | 0 | 2316 | 0 | 0 | 0 | 32 | 21 | 75 | 85 | 117 | 0 | 298 | 0 | 0 | 0 | 0 | 2548 | 0 | 0 |
| SRR5388064 | 0 | 0 | 0 | 0 | 0 | 0 | 0 | 160 | 0 | 0 | 247 | 0 | 0 | 0 | 638 | 12026 | 11790 | 0 | 0 | 2473 | 0 | 0 | 0 | 0 | 0 | 0 | 159 | 27 | 0 | 327 | 0 | 0 | 0 | 0 | 2358 | 0 | 0 |
| SRR5388065 | 0 | 0 | 0 | 0 | 69 | 0 | 72 | 72 | 72 | 0 | 167 | 0 | 0 | 0 | 5851 | 37084 | 37511 | 0 | 0 | 1373 | 0 | 0 | 0 | 0 | 14 | 14 | 69 | 83 | 0 | 111 | 0 | 0 | 0 | 0 | 3623 | 0 | 0 |
| SRR5388066 | 0 | 0 | 0 | 0 | 27 | 0 | 0 | 56 | 0 | 0 | 256 | 27 | 0 | 0 | 2902 | 13175 | 20952 | 0 | 0 | 1507 | 223 | 0 | 0 | 0 | 0 | 27 | 201 | 101 | 0 | 330 | 0 | 0 | 0 | 0 | 2540 | 0 | 0 |
| SRR5388067 | 0 | 0 | 0 | 15 | 22 | 0 | 76 | 152 | 76 | 0 | 195 | 0 | 0 | 0 | 1065 | 10602 | 12407 | 0 | 0 | 837 | 76 | 0 | 0 | 15 | 15 | 0 | 187 | 82 | 0 | 172 | 0 | 0 | 0 | 0 | 6091 | 0 | 0 |
| SRR5388068 | 0 | 0 | 0 | 17 | 17 | 0 | 0 | 143 | 0 | 0 | 359 | 0 | 0 | 0 | 1075 | 12749 | 15774 | 0 | 0 | 1792 | 0 | 0 | 0 | 43 | 17 | 34 | 239 | 188 | 0 | 10 | 0 | 0 | 0 | 0 | 1513 | 0 | 0 |
| SRR5388069 | 0 | 0 | 0 | 24 | 32 | 0 | 0 | 0 | 0 | 0 | 277 | 0 | 0 | 0 | 418 | 8502 | 7085 | 0 | 0 | 1322 | 209 | 0 | 0 | 0 | 0 | 55 | 214 | 55 | 0 | 293 | 0 | 0 | 0 | 0 | 4251 | 0 | 0 |
| SRR5388070 | 0 | 0 | 0 | 40 | 32 | 0 | 0 | 0 | 0 | 0 | 259 | 0 | 0 | 0 | 1022 | 13345 | 9255 | 0 | 0 | 767 | 170 | 0 | 0 | 0 | 0 | 57 | 251 | 105 | 0 | 275 | 0 | 0 | 0 | 0 | 2583 | 0 | 0 |
| SRR5388071 | 0 | 0 | 0 | 0 | 60 | 90 | 90 | 90 | 0 | 0 | 300 | 0 | 0 | 0 | 2532 | 22957 | 25727 | 0 | 0 | 452 | 271 | 0 | 0 | 24 | 0 | 12 | 132 | 72 | 0 | 541 | 0 | 0 | 0 | 0 | 7520 | 0 | 0 |
| SRR5388072 | 0 | 0 | 0 | 20 | 40 | 0 | 0 | 138 | 69 | 0 | 158 | 0 | 0 | 0 | 1170 | 19221 | 18720 | 0 | 69 | 2684 | 0 | 0 | 0 | 20 | 20 | 69 | 287 | 69 | 0 | 277 | 0 | 0 | 0 | 0 | 4346 | 0 | 0 |
| SRR5388073 | 0 | 0 | 0 | 18 | 45 | 65 | 0 | 65 | 0 | 0 | 454 | 0 | 0 | 0 | 2287 | 22829 | 23426 | 0 | 0 | 719 | 131 | 0 | 0 | 36 | 0 | 100 | 173 | 118 | 0 | 391 | 0 | 65 | 0 | 0 | 3730 | 0 | 0 |
| SRR5388074 | 0 | 0 | 0 | 0 | 0 | 76 | 0 | 0 | 0 | 0 | 144 | 0 | 0 | 0 | 911 | 4584 | 10836 | 0 | 0 | 1821 | 228 | 0 | 0 | 0 | 48 | 48 | 106 | 48 | 0 | 270 | 0 | 76 | 0 | 0 | 2501 | 0 | 0 |
| SRR5388075 | 0 | 15 | 0 | 0 | 0 | 0 | 0 | 52 | 52 | 0 | 227 | 0 | 0 | 0 | 1408 | 9169 | 10878 | 0 | 0 | 1200 | 52 | 0 | 0 | 0 | 0 | 37 | 220 | 51 | 0 | 176 | 0 | 0 | 0 | 0 | 2642 | 0 | 0 |
| SRR5388076 | 0 | 0 | 0 | 34 | 0 | 77 | 0 | 77 | 0 | 0 | 203 | 0 | 0 | 0 | 3085 | 10514 | 22780 | 0 | 0 | 771 | 77 | 0 | 0 | 17 | 0 | 0 | 161 | 93 | 0 | 263 | 0 | 0 | 0 | 0 | 3310 | 0 | 0 |
| SRR5388077 | 0 | 0 | 0 | 10 | 42 | 0 | 0 | 0 | 94 | 0 | 84 | 0 | 0 | 0 | 2826 | 39135 | 41967 | 0 | 0 | 2637 | 283 | 0 | 0 | 0 | 10 | 52 | 104 | 42 | 0 | 84 | 0 | 0 | 0 | 0 | 3605 | 0 | 0 |
| SRR5388078 | 0 | 0 | 0 | 0 | 0 | 0 | 0 | 432 | 0 | 0 | 109 | 11 | 0 | 0 | 701 | 8495 | 24946 | 0 | 0 | 2050 | 378 | 0 | 0 | 0 | 0 | 0 | 19 | 0 | 0 | 377 | 0 | 0 | 0 | 0 | 8280 | 0 | 0 |
| SRR5388079 | 0 | 0 | 0 | 19 | 58 | 0 | 0 | 0 | 0 | 0 | 279 | 0 | 0 | 0 | 1882 | 23885 | 18927 | 0 | 0 | 737 | 82 | 0 | 0 | 19 | 0 | 87 | 423 | 106 | 0 | 289 | 0 | 0 | 0 | 0 | 4507 | 0 | 0 |
| SRR5388080 | 0 | 0 | 0 | 17 | 26 | 0 | 0 | 0 | 0 | 0 | 146 | 0 | 0 | 0 | 883 | 16515 | 23121 | 0 | 0 | 1276 | 98 | 0 | 0 | 34 | 51 | 77 | 248 | 111 | 0 | 180 | 0 | 0 | 0 | 0 | 5780 | 0 | 0 |
| SRR5388081 | 0 | 0 | 0 | 0 | 0 | 0 | 0 | 0 | 0 | 0 | 223 | 0 | 0 | 0 | 928 | 21991 | 17671 | 0 | 0 | 2120 | 66 | 0 | 0 | 17 | 17 | 0 | 240 | 60 | 0 | 282 | 0 | 0 | 0 | 0 | 5301 | 0 | 0 |
| SRR5388082 | 0 | 0 | 0 | 20 | 0 | 0 | 0 | 63 | 0 | 0 | 273 | 0 | 0 | 0 | 1577 | 11765 | 15043 | 0 | 0 | 315 | 126 | 0 | 0 | 0 | 0 | 0 | 146 | 47 | 0 | 339 | 0 | 0 | 0 | 0 | 5400 | 0 | 0 |
| SRR5388083 | 0 | 0 | 0 | 0 | 15 | 54 | 0 | 54 | 0 | 0 | 238 | 84 | 0 | 0 | 2397 | 14408 | 22282 | 0 | 0 | 2506 | 54 | 0 | 0 | 15 | 23 | 46 | 177 | 54 | 0 | 284 | 0 | 54 | 0 | 0 | 2848 | 0 | 15 |
| SRR5388084 | 0 | 0 | 0 | 0 | 19 | 0 | 73 | 73 | 0 | 0 | 298 | 0 | 0 | 0 | 2619 | 10961 | 22351 | 0 | 0 | 1964 | 0 | 0 | 0 | 0 | 0 | 28 | 195 | 47 | 0 | 344 | 0 | 0 | 0 | 0 | 3224 | 0 | 0 |
| SRR5388085 | 0 | 0 | 0 | 0 | 0 | 0 | 0 | 0 | 0 | 10 | 335 | 10 | 0 | 0 | 7121 | 31292 | 30083 | 0 | 0 | 1369 | 55 | 0 | 0 | 10 | 10 | 52 | 178 | 105 | 0 | 325 | 0 | 0 | 0 | 0 | 4470 | 0 | 0 |
| SRR5388086 | 0 | 0 | 0 | 17 | 0 | 0 | 0 | 72 | 0 | 0 | 174 | 0 | 0 | 0 | 2169 | 14939 | 12024 | 0 | 0 | 2531 | 0 | 0 | 0 | 0 | 0 | 0 | 35 | 44 | 0 | 322 | 0 | 0 | 0 | 0 | 2186 | 0 | 0 |
| SRR5388087 | 0 | 0 | 0 | 0 | 15 | 0 | 0 | 0 | 0 | 0 | 350 | 15 | 0 | 0 | 7177 | 32261 | 41708 | 0 | 0 | 857 | 30 | 0 | 0 | 0 | 0 | 53 | 107 | 76 | 0 | 327 | 0 | 0 | 0 | 0 | 2674 | 0 | 0 |
| SRR5388088 | 0 | 0 | 0 | 13 | 13 | 0 | 0 | 0 | 0 | 20 | 176 | 0 | 0 | 0 | 1684 | 14895 | 14487 | 0 | 0 | 808 | 67 | 0 | 0 | 13 | 0 | 0 | 143 | 33 | 0 | 293 | 0 | 0 | 0 | 0 | 1020 | 0 | 20 |
| SRR5388089 | 0 | 0 | 0 | 0 | 23 | 0 | 0 | 63 | 0 | 0 | 185 | 0 | 0 | 0 | 2022 | 18968 | 23577 | 0 | 0 | 1580 | 379 | 0 | 0 | 23 | 15 | 123 | 192 | 131 | 0 | 185 | 0 | 0 | 0 | 0 | 3900 | 0 | 15 |
| SRR5388090 | 0 | 0 | 0 | 0 | 0 | 0 | 0 | 68 | 0 | 0 | 166 | 0 | 0 | 0 | 1283 | 10609 | 11609 | 0 | 0 | 2701 | 135 | 0 | 0 | 0 | 12 | 55 | 61 | 61 | 0 | 221 | 0 | 0 | 0 | 0 | 4404 | 0 | 0 |
| SRR5388091 | 0 | 0 | 0 | 0 | 16 | 0 | 0 | 0 | 0 | 0 | 327 | 0 | 0 | 0 | 1187 | 10862 | 19644 | 0 | 0 | 1117 | 0 | 0 | 0 | 16 | 21 | 21 | 182 | 75 | 0 | 214 | 0 | 0 | 0 | 0 | 1618 | 0 | 11 |
| SRR5388092 | 0 | 0 | 0 | 0 | 0 | 0 | 0 | 0 | 0 | 0 | 107 | 0 | 0 | 0 | 1402 | 14852 | 20233 | 0 | 0 | 734 | 67 | 0 | 0 | 0 | 0 | 20 | 94 | 40 | 0 | 175 | 0 | 0 | 0 | 0 | 4090 | 0 | 0 |
| SRR5388093 | 0 | 0 | 0 | 11 | 49 | 82 | 0 | 164 | 0 | 0 | 68 | 0 | 0 | 0 | 656 | 5576 | 14073 | 0 | 0 | 246 | 164 | 0 | 0 | 0 | 33 | 49 | 27 | 15 | 0 | 168 | 0 | 0 | 0 | 0 | 3452 | 0 | 0 |
| SRR5388094 | 0 | 0 | 0 | 0 | 0 | 64 | 0 | 0 | 0 | 0 | 79 | 320 | 0 | 0 | 447 | 11100 | 21505 | 0 | 0 | 3192 | 1788 | 0 | 0 | 0 | 0 | 12 | 31 | 19 | 0 | 155 | 0 | 0 | 0 | 0 | 5897 | 0 | 63 |
| SRR5388095 | 0 | 0 | 0 | 37 | 0 | 105 | 105 | 105 | 0 | 0 | 248 | 0 | 0 | 0 | 1569 | 37135 | 36176 | 0 | 0 | 1360 | 105 | 0 | 0 | 12 | 0 | 25 | 322 | 260 | 0 | 198 | 0 | 0 | 0 | 0 | 3833 | 0 | 0 |
| SRR5388096 | 0 | 0 | 0 | 0 | 14 | 148 | 0 | 0 | 0 | 0 | 295 | 14 | 0 | 0 | 1112 | 10722 | 10309 | 0 | 0 | 964 | 74 | 0 | 0 | 0 | 0 | 43 | 151 | 79 | 0 | 345 | 0 | 0 | 0 | 0 | 2062 | 0 | 0 |
| SRR5388097 | 0 | 0 | 0 | 0 | 28 | 0 | 0 | 0 | 0 | 0 | 256 | 0 | 0 | 0 | 2163 | 12471 | 12211 | 0 | 0 | 1868 | 0 | 0 | 0 | 0 | 19 | 0 | 47 | 19 | 0 | 227 | 0 | 0 | 0 | 0 | 3897 | 0 | 0 |
| SRR5388098 | 0 | 0 | 12 | 0 | 35 | 116 | 0 | 116 | 0 | 0 | 258 | 12 | 0 | 0 | 1159 | 7855 | 9221 | 0 | 0 | 1507 | 0 | 0 | 0 | 0 | 12 | 23 | 223 | 82 | 0 | 82 | 0 | 0 | 0 | 0 | 3757 | 0 | 0 |
| SRR5388099 | 0 | 0 | 0 | 0 | 55 | 0 | 0 | 89 | 0 | 0 | 174 | 0 | 0 | 0 | 1152 | 17477 | 14319 | 0 | 0 | 3012 | 89 | 0 | 0 | 0 | 0 | 63 | 276 | 126 | 0 | 229 | 0 | 0 | 16 | 0 | 3790 | 0 | 0 |
| SRR5388100 | 0 | 0 | 0 | 0 | 22 | 112 | 0 | 0 | 0 | 0 | 175 | 0 | 0 | 0 | 1677 | 26720 | 19841 | 0 | 0 | 2013 | 335 | 0 | 0 | 44 | 22 | 22 | 98 | 44 | 0 | 251 | 0 | 0 | 0 | 0 | 1852 | 0 | 22 |
| SRR5388101 | 0 | 0 | 0 | 0 | 18 | 0 | 0 | 0 | 0 | 0 | 147 | 0 | 0 | 0 | 1276 | 8015 | 13288 | 75 | 0 | 2627 | 0 | 0 | 0 | 28 | 28 | 73 | 165 | 73 | 0 | 248 | 0 | 0 | 0 | 0 | 2531 | 0 | 0 |
| SRR5388102 | 0 | 0 | 0 | 0 | 26 | 0 | 0 | 0 | 0 | 0 | 181 | 0 | 0 | 0 | 973 | 12724 | 11681 | 0 | 0 | 1572 | 0 | 0 | 0 | 17 | 0 | 17 | 86 | 52 | 0 | 164 | 0 | 0 | 0 | 0 | 2712 | 0 | 0 |
| SRR5388103 | 0 | 0 | 0 | 0 | 43 | 79 | 0 | 79 | 0 | 0 | 544 | 0 | 0 | 0 | 1731 | 36172 | 29669 | 0 | 0 | 2596 | 79 | 0 | 0 | 192 | 53 | 213 | 298 | 234 | 0 | 352 | 0 | 0 | 0 | 0 | 3048 | 0 | 0 |
| SRR5388104 | 0 | 0 | 0 | 0 | 0 | 96 | 0 | 0 | 0 | 0 | 235 | 22 | 0 | 0 | 288 | 13847 | 9139 | 0 | 0 | 1249 | 96 | 0 | 0 | 0 | 51 | 88 | 125 | 66 | 0 | 272 | 0 | 0 | 0 | 0 | 3600 | 0 | 0 |
| SRR5388105 | 0 | 0 | 0 | 25 | 62 | 0 | 0 | 52 | 0 | 0 | 311 | 0 | 0 | 0 | 2902 | 26136 | 36364 | 0 | 0 | 1399 | 0 | 0 | 0 | 37 | 12 | 62 | 162 | 125 | 0 | 336 | 0 | 0 | 0 | 0 | 2597 | 0 | 0 |
| SRR5388106 | 0 | 0 | 0 | 0 | 0 | 0 | 0 | 0 | 0 | 0 | 372 | 0 | 0 | 0 | 1567 | 12569 | 18770 | 0 | 0 | 1030 | 45 | 0 | 0 | 0 | 0 | 25 | 132 | 50 | 0 | 281 | 0 | 0 | 0 | 0 | 1676 | 0 | 0 |
| SRR5388107 | 0 | 0 | 0 | 12 | 48 | 0 | 0 | 0 | 0 | 0 | 217 | 12 | 0 | 0 | 677 | 16878 | 17815 | 0 | 0 | 774 | 0 | 0 | 0 | 24 | 12 | 24 | 108 | 48 | 0 | 241 | 0 | 0 | 0 | 0 | 1172 | 0 | 12 |
| SRR5388108 | 0 | 0 | 0 | 0 | 14 | 0 | 0 | 0 | 0 | 0 | 250 | 0 | 0 | 0 | 704 | 6944 | 10058 | 0 | 0 | 1496 | 88 | 0 | 0 | 0 | 21 | 0 | 64 | 50 | 0 | 257 | 0 | 0 | 0 | 0 | 2634 | 0 | 0 |
| SRR5388109 | 0 | 0 | 0 | 11 | 11 | 0 | 59 | 236 | 59 | 0 | 20 | 0 | 0 | 0 | 2598 | 21342 | 32013 | 0 | 0 | 2007 | 295 | 0 | 0 | 23 | 0 | 29 | 183 | 74 | 0 | 17 | 0 | 0 | 0 | 0 | 8246 | 0 | 0 |
| SRR5388110 | 0 | 0 | 0 | 14 | 48 | 0 | 0 | 105 | 0 | 0 | 478 | 0 | 0 | 0 | 1733 | 11248 | 21944 | 0 | 0 | 1260 | 53 | 0 | 0 | 24 | 19 | 39 | 125 | 121 | 0 | 415 | 0 | 0 | 0 | 0 | 8667 | 0 | 0 |
| SRR5388111 | 0 | 0 | 0 | 12 | 0 | 0 | 0 | 219 | 0 | 0 | 174 | 0 | 0 | 0 | 1317 | 15890 | 18509 | 0 | 0 | 2561 | 146 | 0 | 0 | 0 | 25 | 50 | 211 | 136 | 0 | 229 | 0 | 0 | 0 | 0 | 6635 | 0 | 0 |
| SRR5388112 | 0 | 0 | 0 | 11 | 11 | 0 | 72 | 72 | 0 | 0 | 269 | 0 | 0 | 0 | 934 | 17992 | 19300 | 0 | 0 | 3232 | 72 | 0 | 0 | 33 | 16 | 44 | 115 | 115 | 0 | 351 | 0 | 0 | 0 | 0 | 5070 | 0 | 0 |
| SRR5388113 | 0 | 0 | 0 | 23 | 0 | 0 | 0 | 81 | 0 | 0 | 255 | 0 | 0 | 0 | 1626 | 23563 | 22266 | 0 | 0 | 1545 | 407 | 0 | 0 | 0 | 31 | 62 | 263 | 178 | 0 | 131 | 0 | 0 | 0 | 0 | 2594 | 0 | 15 |
| SRR5388114 | 0 | 0 | 0 | 12 | 17 | 0 | 52 | 0 | 0 | 0 | 305 | 0 | 0 | 0 | 1933 | 15014 | 20397 | 0 | 0 | 1829 | 313 | 0 | 0 | 0 | 0 | 46 | 52 | 46 | 0 | 259 | 0 | 0 | 0 | 0 | 3399 | 0 | 0 |

### Supp Table S8.3.Columns_BX_to_DH

| | | | | | | | | | | | | | | | | | | | | | | | | | | | | | | | | | | | | | |
|---|---|---|---|---|---|---|---|---|---|---|---|---|---|---|---|---|---|---|---|---|---|---|---|---|---|---|---|---|---|---|---|---|---|---|---|---|---|
| SRR5388115 | 0 | 0 | 0 | 33 | 57 | 0 | 0 | 0 | 0 | 0 | 221 | 16 | 0 | 0 | 3550 | 8782 | 14852 | 0 | 0 | 2350 | 100 | 0 | 0 | 16 | 0 | 41 | 123 | 66 | 0 | 254 | 0 | 0 | 0 | 0 | 2841 | 0 | 0 |
| SRR5388116 | 0 | 0 | 0 | 38 | 0 | 0 | 0 | 0 | 81 | 0 | 164 | 0 | 0 | 0 | 1942 | 12032 | 12726 | 0 | 0 | 809 | 243 | 0 | 0 | 29 | 0 | 19 | 144 | 77 | 0 | 212 | 0 | 0 | 0 | 0 | 2082 | 0 | 0 |
| SRR5388117 | 0 | 0 | 0 | 0 | 0 | 0 | 0 | 0 | 0 | 0 | 12 | 0 | 0 | 0 | 1189 | 10020 | 23241 | 0 | 0 | 2064 | 35 | 0 | 0 | 0 | 0 | 25 | 214 | 66 | 0 | 296 | 0 | 0 | 0 | 0 | 1549 | 0 | 0 |
| SRR5388118 | 0 | 0 | 0 | 12 | 0 | 0 | 0 | 0 | 0 | 0 | 211 | 0 | 0 | 0 | 784 | 10434 | 22359 | 0 | 0 | 1085 | 181 | 0 | 0 | 0 | 0 | 29 | 88 | 53 | 0 | 223 | 0 | 0 | 0 | 0 | 2609 | 0 | 0 |
| SRR5388119 | 0 | 0 | 0 | 0 | 27 | 0 | 0 | 0 | 0 | 0 | 330 | 0 | 0 | 0 | 5825 | 10834 | 23316 | 0 | 0 | 435 | 174 | 0 | 0 | 0 | 0 | 27 | 134 | 63 | 0 | 357 | 0 | 0 | 0 | 0 | 942 | 0 | 0 |
| SRR5388120 | 0 | 0 | 0 | 0 | 20 | 46 | 93 | 93 | 0 | 0 | 614 | 0 | 0 | 0 | 6062 | 9664 | 69770 | 0 | 0 | 1296 | 278 | 0 | 0 | 0 | 0 | 23 | 284 | 92 | 0 | 1162 | 0 | 0 | 0 | 0 | 3182 | 0 | 0 |
| SRR5388121 | 0 | 0 | 0 | 0 | 37 | 0 | 57 | 57 | 0 | 0 | 584 | 0 | 0 | 0 | 2231 | 9002 | 10602 | 0 | 0 | 1373 | 0 | 0 | 0 | 0 | 15 | 22 | 118 | 111 | 0 | 754 | 0 | 0 | 0 | 0 | 2200 | 0 | 0 |
| SRR5388122 | 0 | 0 | 0 | 0 | 34 | 0 | 0 | 0 | 0 | 0 | 185 | 0 | 0 | 0 | 2040 | 11797 | 11023 | 0 | 0 | 1148 | 0 | 0 | 0 | 0 | 0 | 34 | 278 | 152 | 0 | 244 | 0 | 0 | 0 | 0 | 3094 | 0 | 0 |
| SRR5388123 | 0 | 0 | 0 | 12 | 25 | 0 | 0 | 52 | 52 | 0 | 209 | 0 | 0 | 0 | 4501 | 32277 | 26702 | 0 | 0 | 837 | 0 | 0 | 0 | 49 | 0 | 49 | 393 | 135 | 0 | 405 | 0 | 0 | 0 | 0 | 3961 | 0 | 0 |
| SRR5388124 | 0 | 0 | 0 | 0 | 0 | 61 | 0 | 0 | 0 | 0 | 238 | 0 | 0 | 0 | 1342 | 11390 | 11980 | 0 | 0 | 2501 | 0 | 0 | 0 | 18 | 0 | 26 | 141 | 35 | 0 | 299 | 0 | 0 | 0 | 0 | 2749 | 0 | 0 |
| SRR5388125 | 0 | 0 | 0 | 0 | 0 | 0 | 0 | 0 | 0 | 0 | 694 | 157 | 0 | 0 | 2894 | 7114 | 12769 | 0 | 0 | 1039 | 74 | 0 | 0 | 0 | 0 | 27 | 87 | 33 | 0 | 515 | 0 | 0 | 0 | 0 | 2554 | 0 | 49 |
| SRR5388126 | 0 | 0 | 0 | 37 | 25 | 0 | 0 | 0 | 111 | 0 | 187 | 0 | 0 | 0 | 3763 | 16772 | 22363 | 0 | 0 | 2103 | 111 | 0 | 0 | 0 | 0 | 12 | 373 | 124 | 0 | 87 | 0 | 0 | 0 | 0 | 1864 | 0 | 0 |
| SRR5388127 | 0 | 0 | 0 | 32 | 22 | 0 | 0 | 0 | 0 | 0 | 151 | 0 | 0 | 0 | 665 | 12121 | 8970 | 0 | 0 | 1140 | 190 | 0 | 0 | 11 | 11 | 11 | 118 | 22 | 0 | 215 | 0 | 0 | 0 | 0 | 1939 | 0 | 11 |
| SRR5388128 | 0 | 0 | 0 | 0 | 39 | 0 | 0 | 195 | 0 | 0 | 226 | 0 | 0 | 0 | 845 | 10293 | 11713 | 0 | 0 | 1885 | 130 | 0 | 0 | 31 | 0 | 47 | 374 | 140 | 0 | 343 | 0 | 0 | 0 | 0 | 2130 | 0 | 0 |
| SRR5388129 | 0 | 0 | 0 | 10 | 31 | 0 | 0 | 0 | 0 | 0 | 123 | 0 | 0 | 0 | 662 | 21625 | 16742 | 0 | 0 | 1324 | 265 | 0 | 0 | 10 | 10 | 0 | 51 | 51 | 0 | 153 | 0 | 0 | 0 | 0 | 2093 | 0 | 0 |
| SRR5388130 | 0 | 0 | 0 | 54 | 61 | 0 | 0 | 0 | 183 | 0 | 146 | 0 | 0 | 0 | 1283 | 26962 | 16440 | 0 | 0 | 1466 | 92 | 0 | 0 | 31 | 31 | 38 | 329 | 92 | 0 | 191 | 0 | 92 | 0 | 0 | 4165 | 0 | 0 |
| SRR5388131 | 0 | 0 | 0 | 29 | 14 | 0 | 0 | 131 | 0 | 14 | 244 | 0 | 0 | 0 | 3806 | 27716 | 42969 | 0 | 0 | 787 | 131 | 0 | 0 | 0 | 57 | 57 | 186 | 100 | 0 | 201 | 0 | 0 | 0 | 0 | 3534 | 0 | 14 |
| SRR5388132 | 0 | 0 | 0 | 0 | 13 | 65 | 0 | 33 | 33 | 0 | 429 | 0 | 0 | 0 | 2121 | 12655 | 30815 | 0 | 33 | 1599 | 131 | 0 | 0 | 0 | 13 | 33 | 187 | 25 | 0 | 646 | 0 | 0 | 0 | 0 | 6955 | 0 | 0 |
| SRR5388133 | 0 | 0 | 0 | 0 | 0 | 0 | 0 | 84 | 0 | 0 | 821 | 0 | 0 | 0 | 3479 | 24622 | 45117 | 0 | 0 | 880 | 0 | 0 | 0 | 30 | 0 | 0 | 0 | 0 | 0 | 475 | 0 | 0 | 0 | 0 | 3164 | 0 | 0 |
| SRR5388134 | 0 | 0 | 0 | 13 | 17 | 0 | 0 | 196 | 0 | 0 | 36 | 0 | 0 | 0 | 1832 | 10919 | 20190 | 0 | 65 | 1243 | 65 | 0 | 0 | 0 | 0 | 0 | 0 | 0 | 0 | 18 | 0 | 0 | 0 | 0 | 2678 | 0 | 0 |
| SRR5388135 | 0 | 0 | 0 | 23 | 54 | 0 | 0 | 318 | 0 | 0 | 147 | 0 | 0 | 0 | 530 | 15408 | 13897 | 0 | 0 | 1484 | 106 | 0 | 0 | 0 | 31 | 39 | 202 | 178 | 0 | 240 | 0 | 0 | 0 | 0 | 3927 | 0 | 0 |
| SRR5388136 | 0 | 0 | 0 | 0 | 48 | 0 | 0 | 282 | 0 | 0 | 241 | 0 | 0 | 0 | 422 | 11881 | 13861 | 0 | 0 | 845 | 282 | 0 | 0 | 0 | 0 | 0 | 39 | 19 | 0 | 232 | 0 | 0 | 0 | 0 | 2772 | 0 | 0 |
| SRR5388137 | 0 | 0 | 0 | 0 | 0 | 0 | 0 | 84 | 0 | 0 | 168 | 25 | 0 | 0 | 1180 | 13714 | 13489 | 0 | 0 | 1770 | 169 | 0 | 0 | 0 | 0 | 43 | 298 | 99 | 12 | 217 | 0 | 0 | 0 | 0 | 2923 | 0 | 0 |
| SRR5388138 | 0 | 0 | 0 | 0 | 12 | 0 | 0 | 0 | 0 | 0 | 282 | 12 | 0 | 0 | 2102 | 12826 | 17743 | 0 | 0 | 1501 | 300 | 0 | 0 | 18 | 0 | 35 | 153 | 76 | 0 | 199 | 0 | 0 | 0 | 0 | 1924 | 0 | 0 |
| SRR5388139 | 0 | 0 | 0 | 55 | 18 | 0 | 0 | 0 | 0 | 0 | 166 | 0 | 0 | 0 | 3266 | 25041 | 34727 | 0 | 0 | 907 | 0 | 0 | 0 | 0 | 18 | 46 | 194 | 65 | 0 | 148 | 0 | 0 | 0 | 0 | 3780 | 0 | 0 |
| SRR5388140 | 0 | 0 | 0 | 0 | 26 | 0 | 0 | 0 | 0 | 13 | 10 | 0 | 0 | 0 | 6051 | 17396 | 36233 | 0 | 0 | 1590 | 0 | 0 | 0 | 52 | 0 | 39 | 202 | 124 | 0 | 11 | 0 | 0 | 0 | 0 | 3435 | 0 | 0 |
| SRR5388141 | 0 | 0 | 0 | 32 | 43 | 0 | 0 | 0 | 0 | 0 | 139 | 11 | 0 | 0 | 968 | 14907 | 10952 | 0 | 0 | 1694 | 121 | 0 | 0 | 0 | 11 | 32 | 300 | 64 | 0 | 290 | 0 | 0 | 0 | 0 | 5172 | 0 | 11 |
| SRR5388142 | 0 | 0 | 0 | 0 | 15 | 0 | 0 | 102 | 0 | 30 | 81 | 0 | 0 | 0 | 815 | 10986 | 11535 | 0 | 0 | 2241 | 102 | 0 | 0 | 0 | 0 | 22 | 52 | 0 | 0 | 178 | 0 | 0 | 0 | 0 | 7965 | 0 | 0 |
| SRR5388143 | 0 | 0 | 0 | 20 | 31 | 0 | 0 | 0 | 0 | 0 | 266 | 0 | 0 | 0 | 1486 | 12110 | 11260 | 0 | 0 | 1486 | 0 | 0 | 0 | 20 | 20 | 41 | 153 | 174 | 0 | 398 | 0 | 0 | 0 | 0 | 2762 | 0 | 0 |
| SRR5388144 | 0 | 0 | 0 | 12 | 12 | 99 | 0 | 0 | 0 | 0 | 264 | 0 | 0 | 0 | 1978 | 16507 | 16241 | 99 | 0 | 692 | 0 | 0 | 0 | 48 | 0 | 36 | 60 | 48 | 0 | 216 | 0 | 0 | 0 | 0 | 2662 | 0 | 0 |
| SRR5388145 | 0 | 0 | 0 | 33 | 22 | 0 | 0 | 80 | 0 | 0 | 185 | 0 | 0 | 0 | 1284 | 27422 | 22852 | 0 | 0 | 1284 | 241 | 0 | 0 | 11 | 76 | 87 | 185 | 109 | 0 | 65 | 0 | 0 | 0 | 0 | 2925 | 0 | 11 |
| SRR5388146 | 0 | 0 | 0 | 11 | 11 | 0 | 0 | 309 | 0 | 0 | 246 | 0 | 0 | 0 | 206 | 24457 | 12352 | 0 | 0 | 2783 | 103 | 0 | 0 | 21 | 21 | 21 | 107 | 43 | 0 | 321 | 0 | 103 | 0 | 0 | 3459 | 0 | 0 |
| SRR5388147 | 0 | 0 | 0 | 17 | 58 | 0 | 86 | 0 | 0 | 0 | 175 | 0 | 0 | 0 | 2567 | 20608 | 17846 | 0 | 0 | 1797 | 86 | 0 | 0 | 17 | 33 | 33 | 83 | 67 | 0 | 250 | 0 | 0 | 0 | 0 | 8710 | 0 | 0 |
| SRR5388148 | 0 | 0 | 0 | 15 | 22 | 0 | 67 | 0 | 0 | 0 | 192 | 0 | 0 | 0 | 939 | 12505 | 14679 | 0 | 0 | 2279 | 268 | 0 | 0 | 0 | 0 | 30 | 103 | 74 | 0 | 347 | 0 | 0 | 0 | 0 | 1087 | 0 | 0 |
| SRR5388149 | 0 | 0 | 0 | 0 | 35 | 0 | 72 | 0 | 0 | 0 | 211 | 0 | 0 | 0 | 4015 | 23152 | 28894 | 0 | 0 | 2366 | 215 | 0 | 0 | 18 | 0 | 18 | 18 | 44 | 0 | 211 | 0 | 72 | 0 | 0 | 2778 | 0 | 0 |
| SRR5388150 | 0 | 0 | 0 | 0 | 24 | 0 | 0 | 0 | 0 | 0 | 342 | 0 | 0 | 0 | 1844 | 14320 | 15513 | 0 | 0 | 2335 | 246 | 0 | 0 | 16 | 40 | 56 | 135 | 64 | 0 | 239 | 0 | 0 | 0 | 0 | 1875 | 0 | 0 |
| SRR5388151 | 0 | 0 | 0 | 0 | 0 | 0 | 0 | 0 | 0 | 0 | 266 | 0 | 0 | 0 | 3629 | 17626 | 19699 | 0 | 0 | 1072 | 165 | 0 | 0 | 0 | 0 | 69 | 59 | 30 | 0 | 128 | 0 | 0 | 0 | 0 | 4147 | 0 | 0 |
| SRR5388152 | 0 | 0 | 0 | 18 | 71 | 69 | 0 | 69 | 0 | 0 | 335 | 0 | 0 | 0 | 3977 | 23471 | 23669 | 0 | 0 | 1371 | 274 | 0 | 0 | 0 | 18 | 62 | 194 | 141 | 0 | 238 | 0 | 0 | 0 | 0 | 4734 | 0 | 0 |
| SRR5388153 | 0 | 0 | 0 | 33 | 100 | 0 | 0 | 0 | 0 | 0 | 317 | 0 | 0 | 0 | 4139 | 42149 | 37645 | 0 | 0 | 709 | 0 | 0 | 0 | 0 | 0 | 0 | 183 | 83 | 0 | 267 | 0 | 0 | 0 | 0 | 6113 | 0 | 0 |
| SRR5388154 | 0 | 0 | 0 | 38 | 23 | 0 | 64 | 64 | 0 | 0 | 463 | 30 | 0 | 0 | 2894 | 21584 | 19375 | 0 | 64 | 1029 | 257 | 0 | 0 | 0 | 15 | 30 | 129 | 122 | 0 | 403 | 0 | 0 | 0 | 0 | 5438 | 0 | 15 |
| SRR5388155 | 0 | 0 | 0 | 0 | 49 | 84 | 84 | 0 | 0 | 0 | 245 | 49 | 0 | 0 | 2274 | 15005 | 27081 | 0 | 0 | 505 | 168 | 0 | 0 | 0 | 0 | 28 | 35 | 35 | 0 | 280 | 0 | 0 | 0 | 0 | 9332 | 0 | 21 |
| SRR5388156 | 0 | 0 | 0 | 0 | 23 | 0 | 0 | 60 | 0 | 0 | 430 | 17 | 0 | 0 | 1558 | 7896 | 17403 | 0 | 0 | 1318 | 539 | 0 | 0 | 28 | 57 | 57 | 210 | 91 | 11 | 379 | 0 | 0 | 0 | 0 | 4834 | 0 | 0 |
| SRR5388157 | 0 | 0 | 0 | 0 | 23 | 0 | 0 | 0 | 0 | 12 | 200 | 0 | 0 | 0 | 6966 | 31911 | 25529 | 0 | 0 | 1072 | 89 | 0 | 0 | 12 | 23 | 12 | 47 | 12 | 0 | 153 | 0 | 0 | 0 | 0 | 6183 | 0 | 0 |
| SRR5388158 | 0 | 0 | 0 | 0 | 24 | 0 | 0 | 139 | 0 | 0 | 214 | 0 | 0 | 0 | 1803 | 9204 | 15221 | 0 | 0 | 555 | 0 | 0 | 0 | 0 | 0 | 24 | 226 | 83 | 0 | 249 | 0 | 0 | 0 | 0 | 4248 | 0 | 0 |
| SRR5388159 | 0 | 0 | 0 | 31 | 42 | 87 | 87 | 87 | 0 | 0 | 230 | 10 | 0 | 0 | 2874 | 28885 | 23825 | 0 | 0 | 784 | 0 | 0 | 0 | 0 | 21 | 0 | 157 | 52 | 0 | 209 | 0 | 0 | 0 | 0 | 2530 | 0 | 0 |
| SRR5388160 | 0 | 0 | 0 | 0 | 24 | 0 | 0 | 0 | 0 | 0 | 269 | 0 | 0 | 0 | 2472 | 12386 | 9353 | 0 | 0 | 1426 | 190 | 0 | 0 | 0 | 0 | 32 | 150 | 103 | 0 | 276 | 0 | 0 | 0 | 0 | 3539 | 0 | 0 |
| SRR5388161 | 0 | 0 | 0 | 0 | 0 | 55 | 0 | 0 | 0 | 0 | 137 | 0 | 0 | 0 | 2138 | 18187 | 23566 | 0 | 0 | 822 | 55 | 0 | 0 | 19 | 0 | 37 | 50 | 44 | 0 | 87 | 0 | 0 | 0 | 0 | 2433 | 0 | 0 |
| SRR5388162 | 0 | 0 | 0 | 28 | 0 | 0 | 0 | 85 | 0 | 0 | 353 | 0 | 0 | 0 | 2046 | 9648 | 18581 | 0 | 0 | 767 | 85 | 0 | 0 | 0 | 56 | 19 | 167 | 65 | 0 | 260 | 0 | 0 | 0 | 0 | 9827 | 0 | 0 |
| SRR5388163 | 0 | 0 | 0 | 19 | 77 | 0 | 0 | 51 | 26 | 0 | 327 | 0 | 0 | 0 | 5845 | 33921 | 48669 | 0 | 0 | 766 | 179 | 0 | 0 | 19 | 19 | 32 | 289 | 269 | 0 | 231 | 0 | 26 | 0 | 0 | 898 | 0 | 0 |
| SRR5388164 | 0 | 0 | 0 | 25 | 100 | 0 | 0 | 0 | 0 | 0 | 687 | 0 | 0 | 0 | 1630 | 13402 | 20929 | 0 | 0 | 785 | 241 | 0 | 0 | 0 | 50 | 100 | 400 | 175 | 0 | 437 | 0 | 0 | 0 | 0 | 5691 | 0 | 0 |
| SRR5388165 | 0 | 0 | 0 | 27 | 16 | 0 | 0 | 68 | 0 | 0 | 437 | 0 | 0 | 0 | 1229 | 26898 | 25515 | 0 | 68 | 1298 | 137 | 0 | 0 | 53 | 11 | 53 | 155 | 85 | 0 | 389 | 0 | 0 | 0 | 0 | 1076 | 0 | 0 |
| SRR5388166 | 0 | 0 | 0 | 22 | 44 | 0 | 0 | 0 | 0 | 0 | 208 | 0 | 0 | 0 | 249 | 7291 | 7077 | 0 | 0 | 581 | 83 | 0 | 0 | 0 | 33 | 77 | 197 | 143 | 0 | 208 | 0 | 0 | 0 | 0 | 1072 | 0 | 11 |
| SRR5388167 | 0 | 0 | 0 | 0 | 0 | 0 | 0 | 0 | 0 | 0 | 1009 | 0 | 0 | 0 | 2765 | 28982 | 31119 | 0 | 0 | 553 | 0 | 0 | 0 | 0 | 0 | 15 | 57 | 37 | 0 | 1040 | 0 | 0 | 0 | 0 | 1624 | 0 | 0 |
| SRR5388168 | 0 | 0 | 0 | 0 | 0 | 0 | 0 | 102 | 0 | 16 | 150 | 0 | 0 | 0 | 1934 | 18644 | 27825 | 0 | 51 | 1782 | 153 | 0 | 0 | 0 | 16 | 67 | 205 | 67 | 0 | 165 | 0 | 51 | 0 | 0 | 4661 | 0 | 0 |
| SRR5388169 | 0 | 0 | 0 | 0 | 0 | 0 | 31 | 31 | 0 | 0 | 1187 | 0 | 0 | 0 | 1119 | 9603 | 11883 | 0 | 0 | 1337 | 31 | 0 | 0 | 33 | 0 | 41 | 453 | 86 | 0 | 526 | 0 | 0 | 0 | 0 | 2881 | 0 | 0 |
| SRR5388170 | 0 | 0 | 0 | 0 | 0 | 82 | 0 | 41 | 0 | 0 | 795 | 0 | 0 | 0 | 816 | 9322 | 11376 | 0 | 0 | 285 | 0 | 0 | 0 | 0 | 0 | 51 | 409 | 168 | 0 | 328 | 0 | 0 | 0 | 0 | 2528 | 0 | 0 |
| SRR5388171 | 0 | 0 | 0 | 0 | 0 | 20 | 0 | 0 | 0 | 0 | 26 | 0 | 0 | 0 | 360 | 6337 | 7279 | 0 | 0 | 380 | 0 | 0 | 0 | 12 | 0 | 53 | 11 | 73 | 0 | 16 | 0 | 0 | 0 | 0 | 4282 | 0 | 0 |
| SRR5388172 | 0 | 0 | 0 | 0 | 0 | 0 | 42 | 0 | 0 | 0 | 23 | 0 | 0 | 0 | 378 | 16491 | 14451 | 0 | 0 | 252 | 0 | 0 | 0 | 18 | 18 | 53 | 264 | 53 | 0 | 17 | 0 | 0 | 0 | 0 | 2210 | 0 | 0 |
| SRR5388173 | 0 | 0 | 0 | 0 | 0 | 0 | 0 | 59 | 0 | 0 | 612 | 0 | 0 | 0 | 1825 | 16021 | 22395 | 0 | 0 | 236 | 118 | 0 | 0 | 0 | 0 | 49 | 128 | 30 | 0 | 474 | 0 | 0 | 0 | 0 | 1723 | 0 | 0 |
| SRR5388174 | 0 | 0 | 0 | 24 | 12 | 63 | 63 | 126 | 0 | 0 | 648 | 0 | 0 | 0 | 2142 | 16419 | 23952 | 0 | 0 | 441 | 126 | 0 | 0 | 37 | 24 | 98 | 257 | 110 | 0 | 355 | 0 | 0 | 0 | 0 | 4056 | 0 | 0 |
| SRR5388175 | 0 | 0 | 0 | 0 | 0 | 0 | 0 | 59 | 0 | 0 | 255 | 0 | 0 | 0 | 473 | 9717 | 13999 | 0 | 0 | 887 | 118 | 0 | 0 | 0 | 0 | 0 | 106 | 37 | 0 | 286 | 0 | 0 | 0 | 0 | 3129 | 0 | 0 |
| SRR5388176 | 0 | 0 | 0 | 10 | 0 | 0 | 0 | 79 | 0 | 0 | 352 | 0 | 0 | 0 | 1026 | 16719 | 17610 | 0 | 0 | 474 | 158 | 0 | 0 | 10 | 10 | 104 | 456 | 83 | 0 | 466 | 0 | 0 | 0 | 0 | 1783 | 0 | 0 |
| SRR5388177 | 0 | 0 | 0 | 10 | 61 | 0 | 0 | 49 | 0 | 0 | 344 | 0 | 0 | 0 | 4245 | 43820 | 38093 | 0 | 0 | 888 | 49 | 0 | 0 | 0 | 0 | 30 | 132 | 71 | 0 | 223 | 0 | 0 | 0 | 0 | 1465 | 0 | 0 |
| SRR5388178 | 0 | 0 | 0 | 11 | 11 | 0 | 0 | 0 | 0 | 0 | 895 | 0 | 0 | 0 | 1942 | 14911 | 18208 | 0 | 0 | 542 | 135 | 0 | 0 | 0 | 0 | 0 | 33 | 11 | 0 | 480 | 0 | 0 | 0 | 0 | 2668 | 0 | 0 |
| SRR5388179 | 0 | 0 | 0 | 0 | 0 | 0 | 0 | 49 | 0 | 0 | 20 | 0 | 0 | 0 | 294 | 12650 | 17155 | 0 | 0 | 783 | 98 | 0 | 0 | 0 | 0 | 23 | 176 | 38 | 0 | 11 | 0 | 0 | 0 | 0 | 1213 | 0 | 0 |
| SRR5388180 | 0 | 0 | 0 | 0 | 20 | 0 | 65 | 65 | 0 | 0 | 905 | 0 | 0 | 0 | 907 | 18270 | 25208 | 0 | 0 | 1555 | 453 | 0 | 0 | 13 | 0 | 20 | 143 | 39 | 0 | 365 | 0 | 0 | 0 | 0 | 1156 | 0 | 0 |
| SRR5388181 | 0 | 0 | 0 | 10 | 21 | 0 | 0 | 0 | 0 | 0 | 195 | 0 | 0 | 0 | 883 | 10294 | 15546 | 0 | 0 | 1471 | 221 | 0 | 0 | 10 | 0 | 10 | 10 | 62 | 0 | 154 | 0 | 0 | 0 | 0 | 1471 | 0 | 0 |
| SRR5388182 | 0 | 0 | 0 | 0 | 27 | 0 | 0 | 0 | 0 | 0 | 426 | 0 | 0 | 0 | 893 | 12354 | 11479 | 0 | 0 | 790 | 69 | 0 | 0 | 41 | 14 | 0 | 378 | 103 | 0 | 323 | 0 | 0 | 0 | 0 | 984 | 0 | 0 |
| SRR5388183 | 0 | 0 | 0 | 14 | 27 | 0 | 0 | 293 | 73 | 0 | 381 | 0 | 0 | 0 | 2195 | 25141 | 20467 | 73 | 0 | 1756 | 146 | 0 | 0 | 27 | 14 | 68 | 245 | 184 | 0 | 170 | 0 | 0 | 0 | 0 | 2417 | 0 | 0 |
| SRR5388184 | 11 | 0 | 0 | 11 | 0 | 0 | 0 | 85 | 0 | 11 | 459 | 0 | 0 | 0 | 846 | 12961 | 12302 | 0 | 0 | 169 | 0 | 0 | 0 | 0 | 22 | 55 | 98 | 87 | 0 | 502 | 0 | 0 | 0 | 0 | 2856 | 0 | 0 |
| SRR5388185 | 0 | 0 | 0 | 0 | 16 | 218 | 0 | 0 | 0 | 0 | 257 | 0 | 0 | 0 | 819 | 11053 | 10063 | 0 | 0 | 1092 | 0 | 0 | 0 | 23 | 0 | 55 | 312 | 39 | 0 | 226 | 0 | 0 | 0 | 0 | 1815 | 0 | 0 |
| SRR5388186 | 0 | 0 | 0 | 17 | 0 | 0 | 58 | 0 | 0 | 0 | 486 | 0 | 0 | 0 | 1847 | 20844 | 20681 | 0 | 0 | 750 | 231 | 0 | 0 | 17 | 17 | 51 | 162 | 85 | 0 | 290 | 0 | 0 | 0 | 0 | 1954 | 0 | 0 |
| SRR5388187 | 12 | 12 | 12 | 36 | 12 | 286 | 0 | 0 | 0 | 12 | 678 | 0 | 0 | 0 | 1145 | 13579 | 6547 | 0 | 0 | 668 | 0 | 0 | 0 | 61 | 12 | 97 | 557 | 230 | 0 | 351 | 0 | 95 | 0 | 0 | 8002 | 0 | 0 |
| SRR5388188 | 0 | 0 | 0 | 28 | 0 | 59 | 0 | 0 | 0 | 0 | 559 | 0 | 0 | 0 | 1359 | 14645 | 9403 | 0 | 0 | 1359 | 0 | 0 | 0 | 64 | 14 | 85 | 523 | 191 | 0 | 424 | 0 | 0 | 0 | 0 | 4625 | 0 | 0 |
| SRR5388189 | 0 | 0 | 0 | 18 | 22 | 0 | 27 | 27 | 0 | 0 | 317 | 0 | 0 | 0 | 879 | 11811 | 11895 | 0 | 0 | 959 | 53 | 0 | 0 | 25 | 25 | 52 | 388 | 105 | 0 | 280 | 0 | 0 | 0 | 0 | 2345 | 0 | 0 |
| SRR5388190 | 0 | 0 | 0 | 14 | 18 | 0 | 0 | 0 | 0 | 0 | 390 | 0 | 0 | 0 | 1757 | 18036 | 16462 | 0 | 52 | 1395 | 52 | 0 | 0 | 27 | 18 | 45 | 163 | 59 | 0 | 362 | 0 | 0 | 0 | 0 | 4867 | 0 | 0 |
| SRR5388191 | 0 | 0 | 0 | 35 | 35 | 0 | 99 | 0 | 0 | 0 | 175 | 0 | 0 | 0 | 3353 | 31722 | 35499 | 0 | 0 | 888 | 99 | 0 | 0 | 0 | 18 | 0 | 70 | 88 | 0 | 88 | 0 | 0 | 0 | 0 | 755 | 0 | 0 |
| SRR5388192 | 0 | 0 | 0 | 0 | 11 | 100 | 50 | 201 | 0 | 0 | 361 | 0 | 0 | 0 | 753 | 11500 | 16341 | 0 | 0 | 201 | 0 | 0 | 0 | 23 | 57 | 92 | 452 | 126 | 0 | 298 | 0 | 0 | 0 | 0 | 1967 | 0 | 0 |
| SRR5388193 | 0 | 0 | 0 | 25 | 21 | 0 | 0 | 51 | 0 | 0 | 16 | 0 | 0 | 0 | 1043 | 11257 | 16886 | 25 | 0 | 407 | 102 | 0 | 0 | 0 | 33 | 87 | 388 | 107 | 0 | 437 | 0 | 0 | 0 | 0 | 4243 | 0 | 0 |
| SRR5388194 | 0 | 0 | 0 | 37 | 0 | 0 | 0 | 0 | 0 | 0 | 18 | 0 | 0 | 0 | 1197 | 12204 | 20036 | 0 | 0 | 625 | 52 | 0 | 0 | 0 | 19 | 28 | 12 | 214 | 0 | 12 | 0 | 0 | 0 | 0 | 3279 | 0 | 0 |
| SRR5388195 | 0 | 0 | 0 | 20 | 20 | 0 | 0 | 0 | 0 | 0 | 466 | 0 | 0 | 0 | 1907 | 17062 | 19283 | 0 | 0 | 556 | 119 | 0 | 0 | 0 | 20 | 20 | 248 | 99 | 0 | 437 | 0 | 0 | 0 | 0 | 3506 | 0 | 0 |
| SRR5388196 | 0 | 0 | 0 | 0 | 12 | 60 | 0 | 30 | 0 | 0 | 289 | 0 | 0 | 0 | 2116 | 15718 | 23485 | 0 | 30 | 1451 | 60 | 0 | 0 | 0 | 12 | 35 | 183 | 77 | 0 | 365 | 0 | 0 | 0 | 0 | 3381 | 0 | 0 |
| SRR5388197 | 0 | 0 | 0 | 21 | 0 | 28 | 28 | 0 | 28 | 0 | 492 | 0 | 0 | 0 | 930 | 11050 | 12355 | 0 | 0 | 1156 | 0 | 0 | 0 | 14 | 0 | 36 | 392 | 128 | 0 | 428 | 0 | 0 | 0 | 0 | 1305 | 0 | 0 |
| SRR5388198 | 0 | 0 | 0 | 37 | 37 | 173 | 0 | 115 | 0 | 0 | 578 | 0 | 0 | 0 | 1039 | 15573 | 16098 | 0 | 0 | 693 | 58 | 0 | 0 | 37 | 37 | 25 | 713 | 172 | 0 | 578 | 0 | 0 | 0 | 0 | 1225 | 0 | 12 |
| SRR5388199 | 0 | 0 | 0 | 0 | 0 | 43 | 0 | 128 | 0 | 0 | 2472 | 0 | 0 | 0 | 3499 | 37344 | 40891 | 0 | 0 | 811 | 85 | 0 | 0 | 0 | 0 | 0 | 30 | 16 | 0 | 1773 | 0 | 0 | 0 | 0 | 887 | 0 | 0 |
| SRR5388200 | 0 | 0 | 0 | 0 | 20 | 0 | 0 | 77 | 0 | 0 | 568 | 93 | 0 | 0 | 2545 | 16644 | 18893 | 0 | 0 | 926 | 77 | 0 | 0 | 13 | 0 | 33 | 140 | 47 | 0 | 240 | 0 | 0 | 0 | 0 | 3374 | 0 | 13 |
| SRR5388201 | 0 | 0 | 0 | 0 | 0 | 0 | 47 | 0 | 47 | 0 | 355 | 0 | 0 | 0 | 661 | 11660 | 10018 | 0 | 0 | 566 | 47 | 0 | 0 | 24 | 16 | 24 | 275 | 24 | 0 | 371 | 0 | 0 | 0 | 0 | 1642 | 0 | 0 |
| SRR5388202 | 0 | 0 | 11 | 0 | 11 | 72 | 143 | 0 | 0 | 0 | 240 | 0 | 0 | 0 | 788 | 16143 | 13417 | 0 | 0 | 358 | 0 | 0 | 0 | 22 | 11 | 87 | 360 | 131 | 0 | 261 | 0 | 0 | 0 | 0 | 1468 | 0 | 0 |
| SRR5388203 | 0 | 0 | 0 | 17 | 34 | 0 | 0 | 0 | 0 | 0 | 152 | 0 | 0 | 0 | 680 | 21658 | 21102 | 0 | 0 | 1098 | 209 | 0 | 0 | 42 | 0 | 68 | 178 | 76 | 0 | 229 | 0 | 52 | 0 | 0 | 1111 | 0 | 0 |
| SRR5388204 | 0 | 0 | 0 | 24 | 12 | 0 | 0 | 0 | 0 | 0 | 189 | 0 | 0 | 0 | 664 | 10016 | 8386 | 0 | 0 | 498 | 332 | 0 | 0 | 12 | 35 | 35 | 106 | 83 | 0 | 377 | 0 | 0 | 0 | 0 | 1398 | 0 | 12 |
| SRR5388205 | 0 | 0 | 0 | 0 | 0 | 0 | 0 | 0 | 0 | 0 | 708 | 0 | 0 | 0 | 1203 | 10028 | 18426 | 0 | 0 | 481 | 80 | 0 | 0 | 27 | 0 | 18 | 381 | 80 | 0 | 567 | 0 | 0 | 0 | 0 | 3134 | 0 | 0 |
| SRR5388206 | 0 | 0 | 0 | 24 | 0 | 0 | 0 | 109 | 0 | 0 | 601 | 0 | 0 | 0 | 1149 | 15427 | 18474 | 0 | 0 | 602 | 55 | 0 | 0 | 24 | 16 | 16 | 245 | 95 | 0 | 530 | 0 | 0 | 0 | 0 | 1619 | 0 | 0 |
| SRR5388207 | 0 | 0 | 0 | 16 | 32 | 83 | 0 | 83 | 0 | 0 | 461 | 0 | 0 | 0 | 2726 | 34784 | 28228 | 0 | 0 | 1239 | 83 | 0 | 0 | 24 | 0 | 40 | 194 | 105 | 0 | 202 | 0 | 0 | 0 | 0 | 4917 | 0 | 24 |
| SRR5388208 | 0 | 0 | 0 | 0 | 37 | 0 | 0 | 145 | 0 | 0 | 520 | 0 | 0 | 0 | 942 | 11535 | 7970 | 0 | 0 | 362 | 0 | 0 | 0 | 0 | 0 | 37 | 204 | 84 | 0 | 483 | 0 | 0 | 0 | 0 | 2936 | 0 | 0 |
| SRR5388209 | 0 | 0 | 0 | 28 | 0 | 0 | 0 | 0 | 0 | 0 | 13 | 0 | 0 | 0 | 575 | 5644 | 10372 | 0 | 0 | 180 | 0 | 0 | 0 | 71 | 0 | 43 | 470 | 71 | 0 | 655 | 0 | 0 | 0 | 0 | 915 | 0 | 0 |
| SRR5388210 | 0 | 0 | 0 | 22 | 22 | 0 | 0 | 45 | 0 | 0 | 1078 | 0 | 0 | 0 | 980 | 9619 | 13627 | 0 | 0 | 45 | 0 | 0 | 0 | 0 | 0 | 0 | 129 | 43 | 0 | 755 | 0 | 0 | 0 | 0 | 1603 | 0 | 0 |
| SRR5388211 | 0 | 0 | 0 | 0 | 0 | 0 | 0 | 357 | 0 | 46 | 175 | 0 | 0 | 0 | 1072 | 10510 | 12828 | 0 | 0 | 816 | 51 | 0 | 0 | 0 | 0 | 15 | 98 | 46 | 0 | 147 | 0 | 0 | 0 | 0 | 1082 | 0 | 0 |
| SRR5388212 | 0 | 0 | 0 | 0 | 18 | 85 | 0 | 85 | 0 | 0 | 346 | 0 | 0 | 0 | 677 | 9394 | 10099 | 0 | 0 | 1863 | 0 | 0 | 0 | 0 | 27 | 64 | 173 | 127 | 0 | 373 | 0 | 85 | 0 | 0 | 2114 | 0 | 0 |
| SRR5388213 | 0 | 0 | 0 | 0 | 0 | 77 | 0 | 39 | 0 | 0 | 12 | 0 | 0 | 0 | 2053 | 10378 | 25204 | 0 | 0 | 387 | 0 | 0 | 0 | 0 | 0 | 32 | 103 | 63 | 0 | 450 | 0 | 0 | 0 | 0 | 1668 | 0 | 0 |
| SRR5388214 | 0 | 11 | 0 | 21 | 0 | 0 | 0 | 37 | 0 | 0 | 507 | 0 | 0 | 0 | 368 | 6939 | 6395 | 0 | 0 | 368 | 37 | 0 | 0 | 0 | 32 | 32 | 53 | 32 | 0 | 528 | 0 | 0 | 0 | 0 | 952 | 0 | 0 |
| SRR5388215 | 0 | 0 | 0 | 0 | 0 | 93 | 0 | 0 | 93 | 0 | 365 | 0 | 0 | 0 | 2224 | 33881 | 25411 | 0 | 0 | 927 | 93 | 0 | 0 | 10 | 0 | 31 | 125 | 31 | 0 | 229 | 0 | 0 | 0 | 0 | 3337 | 0 | 0 |
| SRR5388216 | 0 | 0 | 0 | 0 | 16 | 0 | 0 | 34 | 0 | 12 | 12 | 0 | 0 | 0 | 721 | 7477 | 11704 | 0 | 0 | 1201 | 137 | 0 | 0 | 12 | 12 | 47 | 234 | 86 | 0 | 11 | 0 | 0 | 0 | 0 | 3901 | 0 | 0 |
| SRR5388217 | 0 | 0 | 0 | 18 | 36 | 0 | 0 | 0 | 0 | 0 | 1458 | 0 | 0 | 0 | 4086 | 38861 | 58292 | 0 | 0 | 527 | 132 | 0 | 0 | 72 | 0 | 18 | 270 | 198 | 0 | 576 | 0 | 0 | 0 | 0 | 670 | 0 | 18 |
| SRR5388218 | 0 | 0 | 0 | 0 | 0 | 0 | 0 | 63 | 0 | 0 | 647 | 0 | 0 | 0 | 821 | 12057 | 13941 | 63 | 0 | 1768 | 0 | 0 | 0 | 0 | 0 | 0 | 68 | 29 | 0 | 11 | 0 | 0 | 0 | 0 | 942 | 0 | 0 |
| SRR5388219 | 0 | 0 | 0 | 18 | 0 | 0 | 0 | 0 | 0 | 0 | 169 | 0 | 0 | 0 | 1001 | 16104 | 8210 | 0 | 0 | 1752 | 0 | 0 | 0 | 0 | 0 | 27 | 80 | 98 | 0 | 266 | 0 | 0 | 0 | 0 | 3158 | 0 | 0 |
| SRR5388220 | 0 | 0 | 0 | 0 | 13 | 138 | 0 | 0 | 0 | 0 | 160 | 0 | 0 | 0 | 552 | 19870 | 10116 | 0 | 0 | 828 | 0 | 0 | 0 | 0 | 0 | 27 | 154 | 114 | 0 | 174 | 0 | 0 | 0 | 0 | 5780 | 0 | 0 |
| SRR5388221 | 0 | 0 | 0 | 0 | 10 | 0 | 0 | 107 | 0 | 0 | 135 | 0 | 0 | 0 | 1211 | 17653 | 14905 | 0 | 0 | 1425 | 36 | 0 | 0 | 10 | 13 | 57 | 219 | 132 | 0 | 199 | 0 | 0 | 0 | 0 | 846 | 0 | 0 |
| SRR5388222 | 0 | 0 | 0 | 0 | 10 | 0 | 0 | 0 | 0 | 0 | 153 | 0 | 0 | 0 | 1493 | 13184 | 10681 | 0 | 0 | 1034 | 115 | 0 | 0 | 10 | 20 | 41 | 61 | 41 | 0 | 315 | 0 | 0 | 0 | 0 | 1502 | 0 | 0 |
| SRR5388223 | 0 | 0 | 0 | 22 | 0 | 70 | 0 | 141 | 0 | 0 | 240 | 0 | 0 | 0 | 3095 | 27702 | 18954 | 0 | 0 | 563 | 211 | 0 | 0 | 11 | 0 | 33 | 55 | 66 | 0 | 186 | 0 | 0 | 0 | 0 | 2369 | 0 | 0 |
| SRR5388224 | 0 | 0 | 0 | 0 | 10 | 0 | 0 | 0 | 0 | 0 | 419 | 0 | 0 | 0 | 1857 | 21035 | 19651 | 0 | 0 | 1938 | 0 | 0 | 0 | 20 | 10 | 31 | 256 | 82 | 0 | 296 | 0 | 0 | 0 | 0 | 554 | 0 | 0 |
| SRR5388225 | 0 | 0 | 0 | 18 | 18 | 120 | 60 | 60 | 0 | 0 | 459 | 0 | 0 | 0 | 600 | 9751 | 7078 | 0 | 0 | 1080 | 0 | 0 | 0 | 0 | 18 | 45 | 189 | 99 | 0 | 666 | 0 | 0 | 0 | 0 | 4718 | 0 | 0 |
| SRR5388226 | 0 | 0 | 0 | 28 | 0 | 0 | 0 | 167 | 56 | 0 | 474 | 0 | 0 | 0 | 390 | 15157 | 12432 | 0 | 0 | 835 | 56 | 0 | 0 | 0 | 28 | 19 | 189 | 85 | 0 | 815 | 0 | 0 | 0 | 0 | 4087 | 0 | 0 |
| SRR5388227 | 0 | 0 | 0 | 0 | 40 | 0 | 0 | 0 | 0 | 0 | 451 | 0 | 0 | 0 | 1593 | 28546 | 28841 | 0 | 0 | 319 | 106 | 0 | 0 | 13 | 0 | 53 | 186 | 133 | 0 | 226 | 0 | 0 | 0 | 0 | 1766 | 0 | 0 |
| SRR5388228 | 0 | 0 | 0 | 0 | 0 | 106 | 0 | 53 | 0 | 0 | 262 | 0 | 0 | 0 | 637 | 11416 | 12157 | 0 | 0 | 903 | 106 | 0 | 0 | 11 | 0 | 22 | 49 | 33 | 0 | 279 | 0 | 0 | 0 | 0 | 2372 | 0 | 0 |
| SRR5388229 | 0 | 0 | 0 | 22 | 11 | 0 | 108 | 324 | 0 | 0 | 558 | 0 | 0 | 0 | 1728 | 34257 | 29812 | 0 | 0 | 864 | 0 | 0 | 0 | 120 | 11 | 55 | 306 | 153 | 0 | 361 | 0 | 0 | 0 | 0 | 3138 | 0 | 0 |
| SRR5388230 | 0 | 0 | 0 | 0 | 0 | 0 | 0 | 0 | 0 | 0 | 386 | 0 | 0 | 0 | 472 | 12086 | 7407 | 0 | 0 | 1754 | 0 | 0 | 0 | 0 | 0 | 57 | 93 | 43 | 0 | 501 | 0 | 0 | 0 | 0 | 4289 | 0 | 0 |
| SRR5388231 | 0 | 0 | 0 | 0 | 16 | 52 | 0 | 0 | 0 | 0 | 461 | 0 | 0 | 0 | 930 | 17619 | 14866 | 0 | 52 | 1344 | 0 | 0 | 0 | 0 | 0 | 33 | 91 | 66 | 0 | 280 | 0 | 0 | 0 | 0 | 3964 | 0 | 0 |
| SRR5388232 | 0 | 0 | 0 | 0 | 0 | 0 | 0 | 0 | 0 | 0 | 304 | 0 | 0 | 0 | 1002 | 14207 | 8187 | 0 | 0 | 1803 | 100 | 0 | 0 | 0 | 0 | 27 | 81 | 27 | 0 | 313 | 0 | 0 | 0 | 0 | 3853 | 0 | 0 |
| SRR5388233 | 0 | 0 | 0 | 0 | 16 | 0 | 34 | 34 | 102 | 0 | 240 | 0 | 0 | 0 | 2004 | 29996 | 37756 | 0 | 0 | 1324 | 204 | 0 | 0 | 0 | 0 | 12 | 85 | 77 | 0 | 154 | 0 | 0 | 0 | 0 | 1514 | 0 | 0 |
| SRR5388234 | 0 | 0 | 0 | 0 | 0 | 0 | 0 | 81 | 0 | 0 | 176 | 0 | 0 | 0 | 807 | 16181 | 12945 | 0 | 0 | 1936 | 0 | 0 | 0 | 16 | 16 | 40 | 248 | 256 | 0 | 280 | 0 | 0 | 0 | 0 | 693 | 0 | 0 |
| SRR5388235 | 0 | 0 | 0 | 0 | 14 | 0 | 0 | 72 | 72 | 0 | 252 | 0 | 0 | 0 | 724 | 14416 | 25381 | 0 | 0 | 1230 | 72 | 0 | 0 | 18 | 0 | 50 | 302 | 133 | 0 | 233 | 0 | 72 | 0 | 0 | 3249 | 0 | 0 |
| SRR5388236 | 0 | 0 | 0 | 12 | 0 | 0 | 0 | 0 | 0 | 0 | 258 | 0 | 0 | 0 | 858 | 14604 | 16337 | 0 | 0 | 1373 | 172 | 0 | 0 | 18 | 23 | 53 | 182 | 100 | 0 | 293 | 0 | 0 | 0 | 0 | 3465 | 0 | 0 |
| SRR5388237 | 0 | 0 | 0 | 0 | 0 | 0 | 0 | 0 | 0 | 0 | 17 | 0 | 0 | 0 | 331 | 11140 | 14853 | 0 | 0 | 1789 | 265 | 0 | 0 | 13 | 13 | 32 | 122 | 71 | 0 | 21 | 0 | 0 | 0 | 0 | 3528 | 0 | 0 |
| SRR5388238 | 0 | 0 | 0 | 0 | 0 | 0 | 0 | 46 | 0 | 0 | 412 | 17 | 0 | 0 | 737 | 13372 | 13081 | 0 | 0 | 1335 | 0 | 0 | 0 | 0 | 0 | 59 | 236 | 101 | 0 | 345 | 0 | 46 | 0 | 0 | 1890 | 0 | 0 |

### Supp Table S8.3.Columns_BX_to_DH

| | | | | | | | | | | | | | | | | | | | | | | | | | | | | | | | | | | | | | |
|---|---|---|---|---|---|---|---|---|---|---|---|---|---|---|---|---|---|---|---|---|---|---|---|---|---|---|---|---|---|---|---|---|---|---|---|---|---|
| SRR5388239 | 0 | 0 | 0 | 19 | 19 | 80 | 0 | 161 | 0 | 0 | 274 | 0 | 0 | 0 | 1046 | 17613 | 15192 | 0 | 0 | 1931 | 80 | 0 | 0 | 13 | 13 | 19 | 57 | 96 | 0 | 447 | 0 | 0 | 0 | 0 | 2862 | 0 | 0 |
| SRR5388240 | 0 | 0 | 0 | 26 | 0 | 0 | 0 | 88 | 0 | 0 | 290 | 0 | 0 | 0 | 791 | 20357 | 17191 | 0 | 0 | 2726 | 88 | 0 | 0 | 0 | 0 | 53 | 211 | 149 | 0 | 193 | 0 | 0 | 0 | 0 | 2036 | 0 | 0 |
| SRR5388241 | 0 | 0 | 0 | 0 | 0 | 0 | 79 | 79 | 0 | 0 | 432 | 0 | 0 | 0 | 947 | 16690 | 19703 | 0 | 79 | 1579 | 316 | 0 | 0 | 0 | 22 | 49 | 81 | 86 | 0 | 232 | 0 | 0 | 0 | 0 | 2782 | 0 | 0 |
| SRR5388242 | 0 | 0 | 0 | 0 | 0 | 0 | 0 | 61 | 0 | 0 | 305 | 0 | 0 | 0 | 3351 | 18285 | 29824 | 0 | 0 | 2620 | 244 | 0 | 0 | 0 | 11 | 15 | 45 | 26 | 0 | 275 | 0 | 0 | 0 | 0 | 2663 | 0 | 0 |
| SRR5388243 | 0 | 0 | 0 | 0 | 14 | 0 | 0 | 0 | 0 | 0 | 379 | 0 | 0 | 0 | 1194 | 12692 | 16812 | 0 | 0 | 754 | 188 | 0 | 0 | 14 | 27 | 54 | 217 | 81 | 0 | 298 | 0 | 0 | 0 | 0 | 4121 | 0 | 0 |
| SRR5388244 | 0 | 0 | 0 | 41 | 41 | 94 | 0 | 0 | 0 | 10 | 347 | 0 | 0 | 0 | 1228 | 18468 | 14592 | 0 | 0 | 756 | 0 | 0 | 0 | 31 | 10 | 41 | 204 | 51 | 0 | 327 | 0 | 0 | 0 | 0 | 10716 | 0 | 0 |
| SRR5388245 | 0 | 0 | 0 | 0 | 15 | 0 | 107 | 0 | 53 | 0 | 548 | 0 | 0 | 0 | 3361 | 22868 | 26521 | 0 | 0 | 800 | 53 | 0 | 0 | 0 | 0 | 61 | 107 | 84 | 0 | 259 | 0 | 0 | 0 | 0 | 1906 | 0 | 0 |
| SRR5388246 | 0 | 0 | 0 | 0 | 0 | 0 | 0 | 0 | 73 | 0 | 17 | 0 | 0 | 0 | 1379 | 12961 | 32294 | 0 | 0 | 2177 | 218 | 0 | 0 | 0 | 0 | 23 | 168 | 46 | 0 | 15 | 0 | 73 | 0 | 0 | 7689 | 0 | 0 |
| SRR5388247 | 0 | 0 | 0 | 0 | 15 | 0 | 95 | 48 | 48 | 0 | 15 | 0 | 0 | 0 | 715 | 13022 | 8286 | 0 | 0 | 906 | 48 | 0 | 0 | 0 | 30 | 67 | 10 | 127 | 0 | 10 | 0 | 0 | 0 | 0 | 2664 | 0 | 0 |
| SRR5388248 | 0 | 0 | 0 | 0 | 13 | 0 | 0 | 49 | 0 | 0 | 10 | 0 | 0 | 0 | 243 | 9324 | 14486 | 0 | 0 | 390 | 49 | 0 | 0 | 0 | 0 | 19 | 238 | 122 | 0 | 315 | 0 | 0 | 0 | 0 | 2997 | 0 | 0 |
| SRR5388249 | 0 | 19 | 0 | 0 | 0 | 0 | 129 | 129 | 0 | 0 | 359 | 0 | 0 | 0 | 13685 | 54911 | 49454 | 0 | 0 | 516 | 129 | 0 | 0 | 0 | 0 | 57 | 170 | 246 | 0 | 113 | 0 | 0 | 0 | 0 | 1705 | 0 | 0 |
| SRR5388250 | 0 | 0 | 0 | 0 | 0 | 0 | 0 | 105 | 0 | 24 | 195 | 18 | 0 | 0 | 1987 | 22836 | 22495 | 0 | 0 | 2824 | 523 | 0 | 0 | 18 | 12 | 43 | 152 | 61 | 0 | 268 | 0 | 0 | 0 | 0 | 3067 | 0 | 0 |
| SRR5388251 | 0 | 0 | 0 | 18 | 0 | 0 | 0 | 189 | 0 | 0 | 219 | 0 | 0 | 0 | 1135 | 14698 | 12074 | 0 | 0 | 1514 | 95 | 0 | 0 | 0 | 0 | 44 | 201 | 53 | 0 | 350 | 0 | 0 | 0 | 0 | 2362 | 0 | 18 |
| SRR5388252 | 0 | 0 | 0 | 0 | 0 | 166 | 0 | 249 | 0 | 0 | 244 | 0 | 0 | 0 | 1078 | 10560 | 12626 | 0 | 0 | 1824 | 166 | 0 | 0 | 25 | 0 | 0 | 168 | 109 | 0 | 353 | 0 | 0 | 0 | 0 | 1607 | 0 | 0 |
| SRR5388253 | 0 | 20 | 0 | 30 | 20 | 0 | 0 | 142 | 0 | 0 | 218 | 0 | 0 | 0 | 712 | 13294 | 14955 | 0 | 0 | 997 | 285 | 0 | 0 | 0 | 0 | 59 | 178 | 20 | 0 | 139 | 0 | 0 | 0 | 0 | 7644 | 0 | 0 |
| SRR5388254 | 0 | 0 | 0 | 0 | 0 | 95 | 0 | 191 | 0 | 0 | 257 | 0 | 0 | 0 | 1146 | 13504 | 11048 | 0 | 0 | 1719 | 95 | 0 | 0 | 0 | 0 | 0 | 213 | 44 | 0 | 213 | 0 | 0 | 0 | 0 | 1473 | 0 | 0 |
| SRR5388255 | 0 | 0 | 0 | 0 | 10 | 0 | 0 | 88 | 0 | 0 | 23 | 0 | 0 | 0 | 6101 | 34643 | 40757 | 0 | 0 | 1061 | 0 | 0 | 0 | 10 | 31 | 31 | 21 | 94 | 0 | 366 | 0 | 0 | 0 | 0 | 2620 | 0 | 0 |
| SRR5388256 | 0 | 0 | 0 | 21 | 0 | 0 | 0 | 0 | 0 | 0 | 684 | 0 | 0 | 0 | 1667 | 9236 | 16874 | 0 | 0 | 1048 | 48 | 0 | 0 | 14 | 0 | 28 | 131 | 48 | 0 | 504 | 0 | 0 | 0 | 0 | 1421 | 0 | 0 |
| SRR5388257 | 0 | 0 | 0 | 22 | 22 | 0 | 0 | 0 | 0 | 0 | 823 | 0 | 0 | 0 | 6514 | 44829 | 45369 | 85 | 0 | 677 | 85 | 0 | 0 | 22 | 11 | 43 | 162 | 130 | 0 | 216 | 0 | 0 | 0 | 0 | 270 | 0 | 0 |
| SRR5388258 | 0 | 0 | 0 | 12 | 81 | 0 | 0 | 65 | 0 | 0 | 500 | 0 | 0 | 0 | 4265 | 39896 | 33930 | 0 | 0 | 905 | 129 | 0 | 0 | 0 | 23 | 46 | 198 | 221 | 0 | 232 | 0 | 0 | 0 | 0 | 3169 | 0 | 0 |
| SRR5388259 | 0 | 0 | 0 | 41 | 14 | 0 | 0 | 49 | 0 | 0 | 471 | 0 | 0 | 0 | 1730 | 14672 | 10189 | 0 | 0 | 1186 | 99 | 0 | 0 | 14 | 0 | 34 | 239 | 130 | 0 | 437 | 0 | 0 | 0 | 0 | 3804 | 0 | 0 |
| SRR5388260 | 0 | 0 | 0 | 28 | 28 | 0 | 0 | 0 | 0 | 0 | 208 | 0 | 0 | 0 | 3245 | 15733 | 18460 | 0 | 0 | 749 | 83 | 0 | 0 | 28 | 47 | 28 | 123 | 94 | 0 | 227 | 0 | 83 | 0 | 0 | 1678 | 0 | 0 |
| SRR5388261 | 0 | 0 | 0 | 0 | 20 | 0 | 68 | 203 | 0 | 0 | 385 | 0 | 0 | 0 | 540 | 9241 | 6685 | 0 | 0 | 946 | 68 | 0 | 0 | 0 | 20 | 95 | 358 | 95 | 0 | 270 | 0 | 0 | 0 | 0 | 1573 | 0 | 0 |
| SRR5388262 | 0 | 0 | 0 | 0 | 32 | 72 | 72 | 145 | 0 | 0 | 266 | 0 | 0 | 0 | 361 | 11241 | 12322 | 0 | 72 | 1879 | 145 | 0 | 0 | 0 | 51 | 51 | 298 | 171 | 0 | 12 | 0 | 0 | 0 | 0 | 3243 | 0 | 13 |
| SRR5388263 | 0 | 0 | 0 | 0 | 59 | 53 | 0 | 53 | 0 | 0 | 303 | 0 | 0 | 0 | 2063 | 18765 | 31275 | 0 | 0 | 794 | 0 | 0 | 0 | 0 | 0 | 39 | 244 | 176 | 0 | 323 | 0 | 0 | 0 | 0 | 2887 | 0 | 0 |
| SRR5388264 | 0 | 0 | 0 | 0 | 34 | 0 | 0 | 0 | 0 | 0 | 426 | 0 | 0 | 0 | 2299 | 20987 | 24116 | 0 | 0 | 647 | 144 | 0 | 0 | 0 | 11 | 67 | 325 | 168 | 0 | 370 | 0 | 0 | 0 | 0 | 3314 | 0 | 0 |
| SRR5388265 | 0 | 0 | 0 | 0 | 0 | 0 | 0 | 0 | 0 | 0 | 510 | 0 | 0 | 0 | 758 | 8464 | 15921 | 0 | 0 | 1222 | 0 | 0 | 0 | 29 | 0 | 67 | 67 | 77 | 0 | 702 | 0 | 0 | 0 | 0 | 2217 | 0 | 0 |
| SRR5388266 | 0 | 0 | 0 | 0 | 0 | 78 | 78 | 0 | 0 | 0 | 363 | 0 | 0 | 0 | 465 | 10154 | 9864 | 0 | 0 | 310 | 0 | 0 | 0 | 0 | 0 | 15 | 54 | 46 | 0 | 432 | 0 | 0 | 0 | 0 | 1741 | 0 | 0 |
| SRR5388267 | 0 | 0 | 0 | 0 | 0 | 0 | 0 | 0 | 0 | 0 | 398 | 0 | 0 | 0 | 1786 | 16388 | 20314 | 0 | 0 | 2381 | 85 | 0 | 0 | 11 | 22 | 121 | 309 | 121 | 0 | 353 | 0 | 0 | 0 | 0 | 2561 | 0 | 0 |
| SRR5388268 | 0 | 0 | 0 | 0 | 25 | 0 | 0 | 0 | 0 | 0 | 297 | 0 | 0 | 0 | 1093 | 7398 | 14674 | 0 | 0 | 1595 | 0 | 0 | 0 | 19 | 0 | 50 | 139 | 82 | 0 | 335 | 0 | 0 | 0 | 0 | 1334 | 0 | 0 |
| SRR5388269 | 0 | 0 | 0 | 18 | 0 | 0 | 0 | 93 | 0 | 0 | 488 | 0 | 0 | 0 | 1299 | 19584 | 22063 | 0 | 0 | 1670 | 93 | 0 | 0 | 37 | 0 | 28 | 194 | 46 | 0 | 350 | 0 | 0 | 0 | 0 | 2231 | 0 | 0 |
| SRR5388270 | 0 | 0 | 0 | 0 | 0 | 213 | 0 | 53 | 0 | 0 | 380 | 0 | 0 | 0 | 798 | 11370 | 10820 | 0 | 0 | 1011 | 53 | 0 | 0 | 19 | 0 | 0 | 46 | 65 | 0 | 575 | 0 | 0 | 0 | 0 | 2751 | 0 | 0 |
| SRR5388271 | 0 | 0 | 0 | 0 | 119 | 0 | 0 | 218 | 0 | 0 | 297 | 0 | 0 | 0 | 1310 | 75177 | 27245 | 0 | 0 | 218 | 0 | 0 | 0 | 0 | 0 | 89 | 148 | 148 | 0 | 237 | 0 | 0 | 0 | 0 | 1514 | 0 | 0 |
| SRR5388272 | 0 | 0 | 0 | 0 | 0 | 100 | 0 | 100 | 0 | 0 | 330 | 0 | 0 | 0 | 400 | 19268 | 10332 | 0 | 0 | 1602 | 0 | 0 | 0 | 0 | 0 | 19 | 58 | 39 | 0 | 543 | 0 | 0 | 0 | 0 | 4747 | 0 | 0 |
| SRR5388273 | 0 | 0 | 0 | 0 | 16 | 68 | 68 | 136 | 0 | 0 | 394 | 0 | 0 | 0 | 340 | 11961 | 13405 | 0 | 0 | 1224 | 136 | 0 | 0 | 0 | 0 | 32 | 129 | 48 | 0 | 458 | 0 | 0 | 0 | 0 | 1856 | 0 | 0 |
| SRR5388274 | 0 | 0 | 0 | 12 | 12 | 0 | 76 | 0 | 0 | 0 | 154 | 0 | 0 | 0 | 758 | 13812 | 12431 | 0 | 0 | 1669 | 76 | 0 | 0 | 0 | 12 | 12 | 41 | 41 | 0 | 354 | 0 | 76 | 0 | 0 | 2762 | 0 | 0 |
| SRR5388275 | 0 | 0 | 0 | 0 | 0 | 0 | 0 | 0 | 0 | 0 | 768 | 0 | 0 | 0 | 1512 | 23190 | 24444 | 0 | 0 | 378 | 284 | 0 | 0 | 19 | 28 | 56 | 450 | 169 | 0 | 169 | 0 | 0 | 0 | 0 | 627 | 0 | 0 |
| SRR5388276 | 0 | 0 | 0 | 0 | 0 | 0 | 0 | 0 | 0 | 0 | 17 | 0 | 0 | 0 | 791 | 8112 | 18977 | 42 | 0 | 1208 | 208 | 0 | 0 | 28 | 17 | 74 | 96 | 57 | 0 | 11 | 0 | 42 | 0 | 0 | 2028 | 0 | 0 |
| SRR5388277 | 0 | 0 | 0 | 10 | 20 | 0 | 0 | 0 | 0 | 0 | 235 | 0 | 0 | 0 | 1579 | 15381 | 16919 | 0 | 0 | 1301 | 0 | 0 | 0 | 10 | 0 | 20 | 41 | 10 | 0 | 214 | 0 | 93 | 0 | 0 | 2307 | 0 | 0 |
| SRR5388278 | 0 | 0 | 0 | 25 | 0 | 61 | 0 | 0 | 61 | 0 | 441 | 0 | 0 | 0 | 1822 | 19078 | 18879 | 0 | 0 | 1762 | 182 | 0 | 0 | 0 | 0 | 0 | 100 | 75 | 0 | 275 | 0 | 0 | 0 | 0 | 1987 | 0 | 0 |
| SRR5388279 | 0 | 0 | 0 | 0 | 0 | 55 | 0 | 55 | 0 | 0 | 667 | 0 | 0 | 0 | 1369 | 24940 | 27135 | 0 | 55 | 931 | 0 | 0 | 0 | 18 | 18 | 27 | 222 | 107 | 0 | 498 | 0 | 0 | 0 | 0 | 998 | 0 | 0 |
| SRR5388280 | 0 | 0 | 0 | 0 | 0 | 0 | 0 | 180 | 0 | 0 | 10 | 0 | 0 | 0 | 540 | 10801 | 9497 | 0 | 0 | 1080 | 60 | 0 | 0 | 13 | 0 | 20 | 114 | 47 | 0 | 12 | 0 | 0 | 0 | 0 | 3166 | 0 | 0 |
| SRR5388281 | 0 | 0 | 0 | 24 | 24 | 0 | 262 | 0 | 0 | 0 | 557 | 0 | 0 | 0 | 1573 | 42245 | 17502 | 0 | 0 | 262 | 0 | 0 | 0 | 48 | 97 | 339 | 508 | 411 | 0 | 363 | 0 | 0 | 0 | 0 | 604 | 0 | 0 |
| SRR5388282 | 0 | 0 | 14 | 28 | 43 | 0 | 0 | 0 | 0 | 0 | 213 | 0 | 0 | 0 | 866 | 15748 | 11982 | 0 | 0 | 649 | 108 | 0 | 0 | 14 | 0 | 0 | 355 | 241 | 0 | 341 | 0 | 0 | 0 | 0 | 5135 | 0 | 0 |
| SRR5388283 | 0 | 0 | 0 | 0 | 0 | 0 | 88 | 88 | 0 | 0 | 596 | 0 | 0 | 0 | 2324 | 33869 | 49622 | 0 | 0 | 614 | 0 | 0 | 0 | 13 | 13 | 33 | 79 | 85 | 0 | 354 | 0 | 0 | 0 | 0 | 630 | 0 | 0 |
| SRR5388284 | 0 | 0 | 0 | 0 | 0 | 0 | 0 | 34 | 0 | 0 | 186 | 0 | 0 | 0 | 854 | 16887 | 22883 | 0 | 0 | 615 | 34 | 0 | 0 | 14 | 0 | 33 | 162 | 57 | 0 | 300 | 0 | 0 | 0 | 0 | 1591 | 0 | 0 |
| SRR5388285 | 0 | 0 | 14 | 0 | 14 | 0 | 0 | 0 | 0 | 0 | 673 | 14 | 0 | 0 | 5882 | 37742 | 38422 | 0 | 0 | 1554 | 111 | 0 | 0 | 14 | 14 | 70 | 322 | 182 | 0 | 378 | 0 | 0 | 0 | 0 | 1360 | 0 | 0 |
| SRR5388286 | 0 | 0 | 0 | 0 | 15 | 0 | 0 | 80 | 0 | 0 | 651 | 0 | 0 | 0 | 4636 | 42062 | 29028 | 0 | 0 | 1279 | 80 | 0 | 0 | 15 | 15 | 15 | 177 | 148 | 0 | 222 | 0 | 80 | 0 | 0 | 5529 | 0 | 0 |
| SRR5388287 | 0 | 0 | 0 | 17 | 23 | 0 | 25 | 50 | 0 | 17 | 643 | 0 | 0 | 0 | 2519 | 22416 | 21980 | 0 | 0 | 428 | 0 | 0 | 0 | 23 | 28 | 85 | 296 | 193 | 0 | 336 | 0 | 0 | 0 | 0 | 2256 | 0 | 0 |
| SRR5388288 | 0 | 0 | 0 | 0 | 0 | 0 | 0 | 0 | 0 | 0 | 552 | 0 | 0 | 0 | 1227 | 12452 | 12613 | 0 | 45 | 1181 | 91 | 0 | 0 | 0 | 0 | 13 | 93 | 53 | 0 | 512 | 0 | 0 | 0 | 0 | 2587 | 0 | 0 |
| SRR5388289 | 0 | 0 | 0 | 47 | 0 | 0 | 0 | 37 | 0 | 0 | 849 | 0 | 0 | 0 | 1415 | 11827 | 18227 | 0 | 0 | 968 | 0 | 0 | 0 | 35 | 47 | 35 | 291 | 105 | 0 | 733 | 0 | 0 | 0 | 0 | 1252 | 0 | 0 |
| SRR5388290 | 0 | 0 | 0 | 38 | 0 | 0 | 0 | 0 | 0 | 0 | 294 | 0 | 0 | 0 | 1826 | 15965 | 14191 | 0 | 0 | 944 | 63 | 0 | 0 | 28 | 0 | 57 | 256 | 57 | 0 | 266 | 0 | 0 | 0 | 0 | 4032 | 0 | 0 |
| SRR5388291 | 0 | 0 | 0 | 0 | 0 | 0 | 0 | 26 | 0 | 0 | 621 | 0 | 0 | 0 | 1180 | 13037 | 18909 | 0 | 0 | 1364 | 52 | 0 | 0 | 0 | 24 | 104 | 554 | 104 | 0 | 420 | 0 | 0 | 0 | 0 | 2189 | 0 | 0 |
| SRR5388292 | 0 | 0 | 0 | 0 | 23 | 0 | 0 | 0 | 0 | 0 | 323 | 0 | 0 | 0 | 1361 | 14605 | 15276 | 0 | 0 | 2249 | 178 | 0 | 0 | 38 | 38 | 135 | 135 | 98 | 0 | 331 | 0 | 0 | 0 | 0 | 3693 | 0 | 0 |
| SRR5388293 | 0 | 0 | 0 | 0 | 0 | 59 | 0 | 293 | 0 | 0 | 491 | 0 | 0 | 0 | 1232 | 15668 | 14939 | 0 | 0 | 1173 | 117 | 0 | 0 | 0 | 21 | 57 | 414 | 119 | 0 | 222 | 0 | 0 | 0 | 0 | 2186 | 0 | 0 |
| SRR5388294 | 0 | 0 | 0 | 13 | 0 | 0 | 0 | 69 | 138 | 0 | 217 | 0 | 0 | 0 | 1038 | 13256 | 12401 | 0 | 0 | 1454 | 0 | 0 | 0 | 0 | 0 | 0 | 33 | 46 | 0 | 210 | 0 | 0 | 0 | 0 | 855 | 0 | 0 |
| SRR5476966 | 0 | 0 | 0 | 43 | 0 | 0 | 0 | 0 | 0 | 0 | 0 | 0 | 0 | 0 | 2967 | 0 | 0 | 0 | 0 | 0 | 0 | 0 | 0 | 21 | 21 | 0 | 86 | 0 | 0 | 0 | 0 | 5935 | 0 | 0 | 0 | 0 | 0 |
| SRR5476967 | 0 | 0 | 0 | 0 | 0 | 0 | 0 | 0 | 0 | 0 | 0 | 0 | 0 | 0 | 27982 | 2519 | 12595 | 0 | 0 | 2946 | 32401 | 0 | 0 | 301 | 0 | 301 | 0 | 301 | 0 | 301 | 0 | 4418 | 0 | 0 | 0 | 0 | 0 |
| SRR5476969 | 0 | 933 | 0 | 0 | 933 | 0 | 0 | 2915 | 0 | 0 | 0 | 0 | 0 | 0 | 58309 | 3175 | 25397 | 0 | 0 | 5831 | 8746 | 0 | 0 | 933 | 1866 | 1866 | 5597 | 3731 | 0 | 0 | 0 | 0 | 0 | 0 | 0 | 0 | 0 |
| SRR5517463 | 51 | 11 | 0 | 0 | 20 | 19494 | 1706 | 16867 | 9628 | 13 | 36 | 87 | 29 | 14 | 4760 | 19144 | 7875 | 33 | 33 | 108 | 338 | 0 | 0 | 49 | 0 | 0 | 0 | 0 | 31 | 28 | 0 | 19 | 0 | 53 | 14171 | 2137 | 0 |
| SRR5517464 | 23 | 39 | 0 | 0 | 25 | 17672 | 1950 | 14833 | 6819 | 90 | 18 | 15 | 21 | 70 | 14178 | 55664 | 36158 | 60 | 334 | 398 | 1078 | 0 | 0 | 80 | 21 | 0 | 0 | 0 | 24 | 102 | 0 | 0 | 0 | 56 | 4100 | 2299 | 0 |
| SRR5517465 | 149 | 45 | 0 | 0 | 346 | 10906 | 953 | 9864 | 5700 | 72 | 476 | 191 | 153 | 57 | 15814 | 59247 | 36612 | 44 | 383 | 424 | 1140 | 0 | 0 | 36 | 0 | 0 | 0 | 0 | 19 | 1474 | 0 | 16 | 0 | 43 | 5441 | 2129 | 0 |
| SRR5517466 | 31 | 13 | 0 | 0 | 190 | 19079 | 1354 | 18124 | 7893 | 94 | 83 | 155 | 163 | 78 | 10572 | 30266 | 17064 | 67 | 125 | 187 | 557 | 0 | 0 | 32 | 0 | 0 | 0 | 0 | 30 | 471 | 0 | 0 | 0 | 43 | 5290 | 2363 | 0 |
| SRR5517467 | 109 | 30 | 0 | 0 | 1078 | 5510 | 618 | 4488 | 2914 | 597 | 231 | 48 | 243 | 101 | 58866 | 92748 | 57762 | 94 | 474 | 493 | 2017 | 0 | 0 | 46 | 16 | 0 | 0 | 0 | 0 | 1754 | 0 | 31 | 0 | 52 | 4406 | 2733 | 0 |
| SRR5517468 | 248 | 57 | 0 | 0 | 1114 | 5062 | 804 | 3910 | 3437 | 667 | 244 | 65 | 224 | 107 | 51419 | 124923 | 66092 | 55 | 548 | 801 | 2348 | 0 | 14 | 71 | 17 | 0 | 0 | 0 | 0 | 1430 | 0 | 11 | 0 | 50 | 3557 | 2321 | 0 |
| SRR5517469 | 60 | 11 | 0 | 0 | 79 | 5028 | 401 | 4544 | 3142 | 51 | 64 | 38 | 25 | 105 | 16323 | 104426 | 25771 | 69 | 214 | 415 | 1280 | 0 | 12 | 27 | 0 | 0 | 0 | 0 | 0 | 53 | 0 | 23 | 0 | 41 | 6031 | 2048 | 0 |
| SRR5517470 | 11 | 30 | 0 | 0 | 89 | 2534 | 238 | 2601 | 1757 | 31 | 25 | 41 | 119 | 53 | 49202 | 115930 | 64545 | 63 | 438 | 700 | 2349 | 0 | 0 | 33 | 0 | 0 | 0 | 0 | 0 | 89 | 0 | 25 | 0 | 38 | 2992 | 1901 | 0 |
| SRR5575011 | 0 | 17 | 57 | 60 | 26 | 145 | 228 | 1039 | 644 | 0 | 55 | 0 | 0 | 0 | 1350 | 6785 | 4749 | 156 | 0 | 291 | 405 | 0 | 0 | 67 | 0 | 57 | 22 | 20 | 0 | 14 | 0 | 145 | 0 | 0 | 5108 | 81 | 0 |
| SRR5575012 | 0 | 0 | 51 | 38 | 18 | 233 | 81 | 1550 | 851 | 0 | 58 | 0 | 0 | 0 | 1640 | 5289 | 4066 | 90 | 0 | 385 | 582 | 0 | 0 | 24 | 0 | 33 | 58 | 10 | 0 | 20 | 0 | 99 | 0 | 0 | 5289 | 86 | 0 |
| SRR5575013 | 10 | 0 | 69 | 73 | 225 | 256 | 102 | 1670 | 1261 | 0 | 69 | 0 | 0 | 0 | 1568 | 4702 | 5010 | 119 | 51 | 358 | 494 | 0 | 0 | 76 | 10 | 48 | 204 | 228 | 0 | 194 | 0 | 204 | 0 | 0 | 6321 | 63 | 0 |
| SRR5575014 | 11 | 20 | 104 | 70 | 239 | 243 | 159 | 1302 | 768 | 0 | 59 | 0 | 0 | 0 | 2004 | 6630 | 6466 | 56 | 19 | 609 | 337 | 0 | 0 | 109 | 24 | 102 | 265 | 396 | 0 | 165 | 0 | 169 | 0 | 0 | 5198 | 113 | 0 |
| SRR5575015 | 32 | 17 | 80 | 60 | 19 | 439 | 175 | 1630 | 833 | 0 | 30 | 0 | 0 | 0 | 1572 | 8288 | 6033 | 80 | 44 | 1031 | 534 | 0 | 0 | 10 | 15 | 67 | 20 | 27 | 0 | 15 | 0 | 205 | 0 | 0 | 4510 | 117 | 0 |
| SRR5575016 | 15 | 15 | 102 | 48 | 229 | 426 | 250 | 1939 | 999 | 0 | 24 | 0 | 0 | 0 | 2453 | 3830 | 6114 | 44 | 44 | 323 | 837 | 0 | 0 | 69 | 24 | 102 | 238 | 75 | 0 | 217 | 0 | 118 | 0 | 0 | 6786 | 92 | 0 |
| SRR5575017 | 29 | 11 | 62 | 59 | 15 | 493 | 164 | 2473 | 1027 | 0 | 44 | 0 | 0 | 0 | 2506 | 6485 | 6578 | 66 | 16 | 526 | 551 | 0 | 0 | 59 | 11 | 99 | 14 | 23 | 0 | 17 | 0 | 279 | 0 | 0 | 6763 | 147 | 0 |
| SRR5575018 | 24 | 37 | 86 | 11 | 16 | 485 | 364 | 1787 | 874 | 0 | 53 | 0 | 0 | 0 | 1742 | 9374 | 8049 | 121 | 77 | 1244 | 453 | 0 | 0 | 16 | 25 | 11 | 38 | 43 | 0 | 23 | 0 | 121 | 0 | 0 | 3825 | 110 | 0 |
| SRR5575019 | 10 | 10 | 39 | 73 | 206 | 292 | 209 | 1724 | 960 | 0 | 57 | 0 | 0 | 0 | 1071 | 2672 | 6089 | 14 | 56 | 403 | 793 | 0 | 0 | 50 | 0 | 44 | 180 | 183 | 0 | 170 | 0 | 250 | 0 | 0 | 4846 | 134 | 0 |
| SRR5644067 | 37 | 22 | 15 | 74 | 71 | 3284 | 1279 | 15600 | 1165 | 19 | 283 | 0 | 0 | 0 | 15504 | 31743 | 104258 | 0 | 238 | 30183 | 10780 | 0 | 0 | 346 | 205 | 1403 | 3135 | 3458 | 0 | 625 | 0 | 614 | 0 | 0 | 326 | 0 | 0 |
| SRR5644068 | 61 | 0 | 61 | 61 | 61 | 45 | 17 | 184 | 15 | 46 | 563 | 0 | 0 | 0 | 103881 | 11024 | 283461 | 12 | 23 | 25230 | 88730 | 0 | 0 | 152 | 167 | 517 | 761 | 1232 | 0 | 1567 | 0 | 19 | 0 | 0 | 37 | 459 | 15 |
| SRR5644069 | 0 | 0 | 0 | 55 | 24 | 103 | 117 | 603 | 191 | 0 | 16 | 0 | 0 | 0 | 40986 | 30371 | 305081 | 0 | 39 | 45944 | 22376 | 0 | 0 | 503 | 424 | 27 | 68 | 51 | 0 | 307 | 0 | 13 | 0 | 0 | 28 | 0 | 0 |
| SRR5644070 | 13 | 0 | 0 | 18 | 43 | 6219 | 1994 | 26143 | 1122 | 18 | 46 | 10 | 0 | 0 | 4324 | 22094 | 99121 | 0 | 13 | 8694 | 3061 | 0 | 0 | 10 | 66 | 32 | 83 | 27 | 0 | 188 | 0 | 1646 | 0 | 0 | 359 | 11 | 0 |
| SRR5644071 | 0 | 0 | 0 | 67 | 98 | 81 | 34 | 383 | 41 | 45 | 31 | 31 | 0 | 0 | 73986 | 30053 | 331103 | 0 | 28 | 3413 | 8343 | 0 | 0 | 12 | 236 | 55 | 174 | 20 | 0 | 272 | 0 | 0 | 0 | 0 | 106 | 462 | 31 |
| SRR5644072 | 0 | 0 | 0 | 13 | 36 | 216 | 80 | 1681 | 70 | 202 | 147 | 0 | 0 | 0 | 94252 | 46540 | 376154 | 25 | 41 | 8667 | 5612 | 0 | 11 | 22 | 34 | 12 | 15 | 24 | 0 | 761 | 0 | 18 | 0 | 0 | 233 | 0 | 17 |
| SRR5644073 | 0 | 0 | 16 | 57 | 80 | 231 | 229 | 2063 | 281 | 32 | 344 | 0 | 0 | 0 | 21299 | 53850 | 265697 | 0 | 164 | 43409 | 7838 | 0 | 0 | 83 | 83 | 373 | 504 | 580 | 0 | 1225 | 0 | 847 | 0 | 0 | 40 | 35 | 0 |
| SRR5644074 | 0 | 0 | 0 | 23 | 56 | 146 | 177 | 919 | 132 | 82 | 37 | 23 | 0 | 0 | 20055 | 58307 | 290185 | 0 | 48 | 16535 | 4782 | 0 | 0 | 67 | 23 | 12 | 116 | 11 | 0 | 485 | 0 | 140 | 0 | 0 | 296 | 32 | 58 |
| SRR5644075 | 0 | 0 | 12 | 48 | 66 | 198 | 158 | 1217 | 125 | 0 | 120 | 0 | 0 | 0 | 44283 | 31868 | 490350 | 0 | 275 | 30152 | 8004 | 0 | 0 | 150 | 84 | 276 | 13 | 23 | 0 | 22 | 0 | 265 | 0 | 0 | 73 | 164 | 12 |
| SRR5644076 | 0 | 0 | 0 | 0 | 41 | 2784 | 1611 | 17371 | 1320 | 70 | 28 | 0 | 0 | 0 | 16759 | 37323 | 304434 | 0 | 59 | 26805 | 5488 | 0 | 0 | 90 | 13 | 28 | 34 | 0 | 0 | 475 | 0 | 671 | 0 | 0 | 387 | 50 | 23 |
| SRR5644077 | 13 | 0 | 0 | 0 | 102 | 294 | 222 | 1318 | 122 | 110 | 77 | 0 | 0 | 0 | 96621 | 49373 | 510479 | 0 | 103 | 12782 | 6369 | 0 | 0 | 13 | 33 | 32 | 50 | 0 | 0 | 25 | 0 | 67 | 0 | 0 | 916 | 195 | 27 |
| SRR5644078 | 0 | 0 | 0 | 0 | 34 | 83 | 100 | 407 | 63 | 51 | 26 | 36 | 0 | 0 | 23569 | 69426 | 318272 | 0 | 60 | 32839 | 4002 | 0 | 0 | 16 | 0 | 0 | 33 | 0 | 0 | 28 | 0 | 56 | 0 | 0 | 273 | 31 | 46 |
| SRR5644085 | 18 | 0 | 0 | 112 | 50 | 137 | 188 | 632 | 85 | 22 | 65 | 11 | 0 | 0 | 65868 | 33389 | 394047 | 0 | 43 | 7317 | 948 | 0 | 0 | 79 | 40 | 97 | 13 | 10 | 0 | 27 | 0 | 26 | 0 | 0 | 539 | 266 | 47 |
| SRR5644086 | 0 | 0 | 0 | 28 | 0 | 385 | 184 | 2048 | 120 | 36 | 17 | 20 | 0 | 0 | 142441 | 33811 | 250798 | 28 | 90 | 15330 | 18791 | 0 | 0 | 105 | 84 | 23 | 31 | 29 | 0 | 55 | 0 | 202 | 0 | 0 | 92 | 83 | 39 |
| SRR5644087 | 49 | 0 | 27 | 82 | 77 | 342 | 182 | 3915 | 218 | 66 | 330 | 0 | 0 | 0 | 23050 | 32590 | 505517 | 0 | 63 | 21482 | 16021 | 0 | 0 | 55 | 49 | 313 | 11 | 10 | 0 | 18 | 0 | 191 | 0 | 11 | 59 | 95 | 38 |
| SRR5644088 | 27 | 10 | 10 | 51 | 78 | 157 | 97 | 1121 | 102 | 174 | 262 | 0 | 0 | 0 | 22601 | 62680 | 409317 | 0 | 49 | 25795 | 7490 | 0 | 0 | 78 | 17 | 191 | 252 | 187 | 0 | 735 | 0 | 105 | 0 | 0 | 47 | 105 | 0 |
| SRR5657537 | 0 | 13 | 0 | 24 | 11 | 333 | 229 | 2011 | 94 | 63 | 73 | 22 | 0 | 0 | 29976 | 178213 | 255456 | 26 | 110 | 9244 | 4638 | 0 | 0 | 69 | 0 | 41 | 126 | 0 | 0 | 427 | 0 | 40 | 0 | 0 | 425 | 151 | 0 |
| SRR5657538 | 0 | 0 | 0 | 0 | 0 | 3024 | 568 | 7305 | 481 | 123 | 34 | 44 | 0 | 0 | 54157 | 51466 | 486248 | 15 | 109 | 9403 | 3136 | 0 | 0 | 101 | 0 | 12 | 22 | 0 | 0 | 821 | 0 | 12 | 0 | 0 | 2086 | 0 | 14 |
| SRR5657539 | 0 | 0 | 0 | 0 | 13 | 110 | 128 | 396 | 39 | 40 | 43 | 13 | 0 | 0 | 18818 | 210776 | 120377 | 11 | 23 | 3739 | 2506 | 0 | 0 | 32 | 0 | 0 | 0 | 0 | 0 | 467 | 0 | 446 | 0 | 0 | 518 | 230 | 0 |
| SRR5657540 | 0 | 0 | 0 | 0 | 0 | 900 | 450 | 3172 | 326 | 136 | 0 | 47 | 0 | 0 | 71049 | 82639 | 497934 | 43 | 34 | 47410 | 3022 | 0 | 0 | 121 | 0 | 0 | 0 | 0 | 0 | 205 | 0 | 0 | 0 | 0 | 1280 | 134 | 13 |
| SRR5657541 | 14 | 20 | 0 | 17 | 15 | 740 | 279 | 1467 | 117 | 27 | 29 | 19 | 0 | 0 | 49542 | 331126 | 373009 | 25 | 114 | 7027 | 15257 | 0 | 0 | 25 | 0 | 17 | 29 | 0 | 0 | 73 | 0 | 0 | 0 | 0 | 317 | 260 | 17 |
| SRR5657542 | 0 | 0 | 0 | 0 | 0 | 636 | 490 | 2935 | 518 | 165 | 18 | 38 | 0 | 0 | 103393 | 47266 | 774117 | 0 | 215 | 20705 | 5593 | 0 | 0 | 103 | 0 | 0 | 0 | 0 | 0 | 378 | 0 | 0 | 0 | 0 | 762 | 0 | 22 |
| SRR5657543 | 0 | 0 | 0 | 0 | 0 | 541 | 416 | 1120 | 117 | 75 | 11 | 19 | 0 | 0 | 47809 | 194197 | 532964 | 40 | 47 | 6531 | 7352 | 0 | 0 | 16 | 0 | 0 | 0 | 0 | 0 | 108 | 0 | 90 | 0 | 0 | 1280 | 314 | 25 |
| SRR5657544 | 11 | 16 | 0 | 11 | 0 | 0 | 22 | 29 | 0 | 14 | 19 | 12 | 0 | 0 | 251066 | 154152 | 565773 | 0 | 12 | 1074 | 33615 | 0 | 0 | 10 | 24 | 44 | 21 | 0 | 0 | 47 | 0 | 12 | 0 | 0 | 46 | 53 | 0 |
| SRR5657545 | 0 | 0 | 0 | 0 | 40 | 355 | 623 | 670 | 139 | 32 | 0 | 0 | 0 | 0 | 3542 | 178250 | 79603 | 14 | 0 | 1018 | 478 | 0 | 0 | 164 | 0 | 0 | 13 | 0 | 0 | 232 | 0 | 227 | 0 | 0 | 24216 | 470 | 0 |
| SRR5657546 | 12 | 10 | 0 | 0 | 63 | 290 | 411 | 511 | 73 | 17 | 0 | 0 | 0 | 0 | 22571 | 56411 | 197488 | 11 | 57 | 4509 | 901 | 0 | 0 | 11 | 0 | 0 | 0 | 0 | 0 | 43 | 0 | 135 | 0 | 0 | 67112 | 446 | 0 |
| SRR5658693 | 0 | 18 | 15 | 91 | 55 | 20 | 93 | 392 | 0 | 0 | 19 | 0 | 0 | 0 | 22971 | 49541 | 559487 | 0 | 0 | 1334 | 4302 | 0 | 0 | 35 | 24 | 131 | 141 | 0 | 0 | 56 | 0 | 0 | 0 | 0 | 1012 | 0 | 0 |
| SRR5658694 | 14 | 0 | 43 | 196 | 106 | 89 | 144 | 217 | 33 | 0 | 64 | 0 | 0 | 0 | 3327 | 55671 | 165213 | 0 | 0 | 2300 | 9687 | 0 | 0 | 31 | 36 | 214 | 332 | 32 | 0 | 100 | 0 | 11 | 0 | 0 | 4023 | 0 | 0 |
| SRR5658695 | 17 | 21 | 27 | 82 | 111 | 30 | 229 | 409 | 30 | 0 | 13 | 0 | 0 | 0 | 1607 | 25518 | 100274 | 0 | 12 | 1276 | 9533 | 0 | 0 | 60 | 41 | 124 | 144 | 0 | 0 | 65 | 0 | 0 | 0 | 0 | 2567 | 0 | 0 |
| SRR5658696 | 19 | 11 | 20 | 113 | 80 | 100 | 216 | 277 | 44 | 0 | 25 | 0 | 0 | 0 | 2958 | 37854 | 150493 | 17 | 22 | 1772 | 11404 | 0 | 0 | 59 | 33 | 145 | 190 | 0 | 0 | 88 | 0 | 17 | 0 | 0 | 3248 | 0 | 0 |
| SRR5658697 | 30 | 14 | 24 | 100 | 15 | 95 | 231 | 515 | 47 | 0 | 20 | 0 | 0 | 0 | 2100 | 26248 | 106576 | 0 | 0 | 2142 | 9660 | 0 | 0 | 110 | 47 | 10 | 15 | 0 | 0 | 97 | 0 | 11 | 0 | 0 | 3461 | 0 | 0 |
| SRR5687455 | 86 | 12 | 0 | 0 | 0 | 1389 | 1502 | 7747 | 785 | 175 | 41 | 39 | 0 | 0 | 47415 | 18363 | 112615 | 0 | 92 | 3244 | 1892 | 0 | 0 | 16 | 0 | 0 | 0 | 0 | 0 | 240 | 0 | 58 | 0 | 0 | 7610 | 61 | 0 |
| SRR5687456 | 86 | 16 | 0 | 0 | 0 | 1957 | 1655 | 10660 | 821 | 181 | 70 | 86 | 0 | 0 | 59548 | 19965 | 123209 | 31 | 100 | 3092 | 1400 | 0 | 0 | 17 | 0 | 0 | 0 | 0 | 0 | 381 | 0 | 50 | 0 | 0 | 6512 | 64 | 0 |
| SRR5687457 | 13 | 23 | 0 | 0 | 0 | 1233 | 1789 | 10891 | 642 | 23 | 58 | 12 | 0 | 0 | 54039 | 15674 | 129593 | 12 | 54 | 2280 | 1983 | 0 | 0 | 34 | 0 | 0 | 0 | 0 | 0 | 35 | 0 | 31 | 0 | 0 | 6955 | 52 | 0 |
| SRR5687458 | 87 | 15 | 0 | 0 | 0 | 1662 | 1782 | 9567 | 786 | 169 | 31 | 33 | 0 | 0 | 41991 | 15806 | 101861 | 12 | 49 | 3020 | 1691 | 0 | 0 | 18 | 0 | 0 | 0 | 0 | 0 | 218 | 0 | 29 | 0 | 0 | 8166 | 58 | 0 |
| SRR5687459 | 47 | 0 | 0 | 0 | 0 | 1502 | 1677 | 7811 | 738 | 28 | 32 | 30 | 0 | 0 | 46457 | 18383 | 113472 | 22 | 49 | 2973 | 1659 | 0 | 0 | 13 | 0 | 0 | 0 | 0 | 0 | 28 | 0 | 76 | 0 | 0 | 8848 | 61 | 0 |
| SRR5687461 | 26 | 0 | 0 | 0 | 0 | 1297 | 1044 | 6232 | 295 | 434 | 0 | 0 | 0 | 0 | 216966 | 19074 | 255353 | 23 | 53 | 3169 | 5532 | 0 | 0 | 0 | 0 | 0 | 0 | 0 | 0 | 105 | 0 | 23 | 0 | 0 | 6445 | 20 | 0 |
| SRR5687462 | 10 | 0 | 0 | 0 | 0 | 1295 | 959 | 7420 | 423 | 59 | 0 | 19 | 0 | 0 | 160217 | 23094 | 203451 | 16 | 27 | 3080 | 6813 | 0 | 0 | 0 | 0 | 0 | 0 | 0 | 0 | 26 | 0 | 27 | 0 | 0 | 5813 | 17 | 0 |
| SRR5687463 | 37 | 0 | 0 | 0 | 0 | 1112 | 1020 | 6569 | 387 | 50 | 0 | 15 | 0 | 0 | 151713 | 21941 | 197678 | 16 | 39 | 2729 | 6658 | 0 | 0 | 0 | 0 | 0 | 0 | 0 | 0 | 17 | 0 | 46 | 0 | 0 | 6335 | 15 | 0 |
| SRR5687464 | 20 | 0 | 0 | 0 | 0 | 2095 | 1592 | 10773 | 487 | 121 | 0 | 11 | 0 | 0 | 64512 | 12195 | 100566 | 0 | 0 | 1294 | 2158 | 0 | 0 | 0 | 0 | 0 | 0 | 0 | 0 | 24 | 0 | 52 | 0 | 0 | 10775 | 14 | 0 |
| SRR5687465 | 61 | 13 | 0 | 0 | 39 | 528 | 466 | 3604 | 223 | 205 | 70 | 28 | 0 | 0 | 73834 | 74987 | 79522 | 0 | 43 | 1964 | 1535 | 0 | 0 | 35 | 0 | 0 | 0 | 0 | 0 | 207 | 0 | 19 | 0 | 0 | 1821 | 15 | 0 |
| SRR5687466 | 40 | 0 | 0 | 0 | 20 | 908 | 847 | 6408 | 344 | 34 | 10 | 32 | 0 | 0 | 65844 | 65785 | 78414 | 0 | 29 | 1714 | 1212 | 0 | 0 | 15 | 0 | 0 | 0 | 0 | 0 | 31 | 0 | 14 | 0 | 0 | 2737 | 16 | 0 |
| SRR5687467 | 58 | 14 | 0 | 0 | 18 | 731 | 582 | 4877 | 259 | 154 | 50 | 39 | 0 | 0 | 53992 | 70195 | 69651 | 0 | 38 | 1803 | 1468 | 0 | 0 | 26 | 0 | 0 | 0 | 0 | 0 | 193 | 0 | 20 | 0 | 0 | 2266 | 17 | 0 |
| SRR5687468 | 21 | 0 | 0 | 0 | 0 | 1430 | 1216 | 10132 | 360 | 283 | 38 | 14 | 0 | 0 | 94505 | 49531 | 123790 | 11 | 0 | 1730 | 843 | 0 | 0 | 0 | 0 | 0 | 0 | 0 | 0 | 218 | 0 | 23 | 0 | 0 | 5247 | 18 | 0 |
| SRR5687469 | 41 | 0 | 0 | 0 | 15 | 1487 | 1083 | 9532 | 374 | 31 | 32 | 21 | 0 | 0 | 67341 | 66095 | 82527 | 0 | 36 | 1822 | 1706 | 0 | 0 | 21 | 0 | 0 | 0 | 0 | 0 | 26 | 0 | 23 | 0 | 0 | 3026 | 20 | 0 |
| SRR5753916 | 258 | 62 | 81 | 40 | 71 | 1871 | 379 | 2183 | 2982 | 149 | 32 | 56 | 32 | 11 | 33660 | 338968 | 99726 | 297 | 2009 | 1783 | 3472 | 0 | 0 | 34 | 14 | 12 | 12 | 49 | 0 | 48 | 0 | 73 | 0 | 0 | 11496 | 0 | 0 |
| SRR5755813 | 0 | 10 | 20 | 71 | 171 | 121 | 364 | 1444 | 648 | 0 | 50 | 0 | 0 | 0 | 580 | 8396 | 34598 | 13 | 81 | 12671 | 1309 | 0 | 0 | 55 | 71 | 171 | 1304 | 171 | 0 | 116 | 0 | 0 | 0 | 0 | 1978 | 0 | 0 |
| SRR5755814 | 26 | 0 | 0 | 0 | 12 | 220 | 439 | 2050 | 293 | 0 | 52 | 0 | 0 | 0 | 439 | 7876 | 40403 | 0 | 0 | 1611 | 293 | 0 | 0 | 130 | 26 | 130 | 34 | 26 | 0 | 52 | 0 | 0 | 0 | 0 | 4522 | 0 | 0 |

### Supp Table S8.3.Columns_BX_to_DH

| | | | | | | | | | | | | | | | | | | | | | | | | | | | | | | | | | | | | | |
|---|---|---|---|---|---|---|---|---|---|---|---|---|---|---|---|---|---|---|---|---|---|---|---|---|---|---|---|---|---|---|---|---|---|---|---|---|---|
| SRR5755815 | 0 | 0 | 0 | 105 | 35 | 174 | 349 | 2847 | 116 | 35 | 0 | 35 | 0 | 0 | 697 | 10027 | 51307 | 0 | 174 | 17894 | 465 | 0 | 0 | 140 | 35 | 175 | 32 | 209 | 0 | 105 | 0 | 0 | 0 | 0 | 4373 | 0 | 0 |
| SRR5755818 | 0 | 19 | 0 | 23 | 23 | 509 | 1171 | 5571 | 639 | 0 | 43 | 27 | 0 | 0 | 118 | 25986 | 31494 | 0 | 83 | 19920 | 745 | 0 | 0 | 0 | 0 | 19 | 201 | 15 | 0 | 101 | 0 | 47 | 0 | 0 | 1078 | 0 | 0 |
| SRR5755819 | 0 | 0 | 11 | 67 | 190 | 192 | 631 | 769 | 27 | 0 | 0 | 0 | 0 | 0 | 247 | 2418 | 24964 | 0 | 0 | 1949 | 933 | 0 | 0 | 56 | 22 | 90 | 18 | 45 | 0 | 157 | 0 | 0 | 0 | 0 | 1707 | 0 | 0 |
| SRR5755820 | 29 | 59 | 29 | 0 | 294 | 144 | 216 | 144 | 72 | 0 | 0 | 0 | 0 | 0 | 865 | 5340 | 20023 | 0 | 0 | 360 | 865 | 0 | 0 | 471 | 29 | 265 | 854 | 88 | 0 | 88 | 0 | 0 | 0 | 0 | 1335 | 0 | 0 |
| SRR5755821 | 50 | 20 | 0 | 129 | 70 | 661 | 341 | 6011 | 80 | 0 | 50 | 0 | 0 | 0 | 8195 | 6750 | 210401 | 0 | 321 | 60208 | 8034 | 0 | 0 | 30 | 0 | 129 | 24 | 139 | 0 | 90 | 0 | 0 | 0 | 0 | 4198 | 0 | 0 |
| SRR5755824 | 0 | 0 | 0 | 42 | 28 | 146 | 293 | 2341 | 146 | 14 | 28 | 0 | 0 | 0 | 195 | 10810 | 31912 | 0 | 0 | 2292 | 1219 | 0 | 0 | 71 | 14 | 14 | 197 | 0 | 0 | 42 | 0 | 0 | 0 | 0 | 1384 | 0 | 0 |
| SRR5755828 | 40 | 23 | 19 | 86 | 52 | 397 | 295 | 5899 | 61 | 0 | 13 | 0 | 0 | 0 | 12746 | 40597 | 380047 | 0 | 102 | 16383 | 4799 | 0 | 0 | 323 | 10 | 67 | 180 | 65 | 0 | 159 | 0 | 326 | 0 | 0 | 2384 | 16 | 0 |
| SRR5755830 | 11 | 11 | 0 | 33 | 282 | 243 | 632 | 3449 | 97 | 11 | 22 | 0 | 0 | 0 | 5636 | 5270 | 69503 | 0 | 0 | 2138 | 777 | 0 | 0 | 250 | 33 | 217 | 423 | 22 | 0 | 130 | 0 | 0 | 0 | 0 | 1069 | 0 | 0 |
| SRR5755831 | 0 | 0 | 0 | 62 | 10 | 334 | 490 | 4345 | 223 | 24 | 28 | 0 | 0 | 0 | 4590 | 29134 | 397219 | 0 | 89 | 17066 | 19005 | 0 | 10 | 19 | 10 | 28 | 14 | 24 | 0 | 23 | 0 | 111 | 0 | 0 | 1278 | 0 | 0 |
| SRR5755833 | 18 | 18 | 35 | 88 | 14 | 451 | 327 | 6862 | 141 | 0 | 65 | 0 | 0 | 0 | 964 | 3104 | 59906 | 0 | 88 | 4820 | 1716 | 0 | 0 | 53 | 0 | 100 | 43 | 106 | 0 | 147 | 0 | 18 | 0 | 0 | 883 | 0 | 0 |
| SRR5755835 | 38 | 0 | 0 | 150 | 44 | 293 | 319 | 10718 | 94 | 19 | 88 | 31 | 0 | 0 | 8211 | 4350 | 153409 | 0 | 56 | 22236 | 7794 | 0 | 0 | 250 | 25 | 156 | 21 | 119 | 0 | 175 | 0 | 15 | 0 | 0 | 310 | 0 | 0 |
| SRR5755837 | 23 | 14 | 19 | 33 | 80 | 1021 | 1272 | 36201 | 412 | 0 | 52 | 23 | 0 | 0 | 1451 | 5350 | 58713 | 0 | 0 | 7487 | 90 | 0 | 0 | 75 | 37 | 24 | 75 | 47 | 0 | 61 | 0 | 18 | 0 | 0 | 548 | 0 | 0 |
| SRR5755838 | 17 | 0 | 0 | 33 | 100 | 344 | 386 | 9455 | 180 | 0 | 46 | 0 | 0 | 0 | 3836 | 7881 | 216092 | 0 | 67 | 12574 | 8539 | 0 | 0 | 63 | 13 | 92 | 17 | 42 | 0 | 109 | 0 | 13 | 0 | 0 | 192 | 0 | 0 |
| SRR5755842 | 15 | 15 | 45 | 135 | 135 | 224 | 261 | 7173 | 130 | 0 | 45 | 15 | 0 | 0 | 7564 | 4564 | 205285 | 0 | 37 | 13936 | 7713 | 0 | 0 | 14 | 0 | 12 | 34 | 18 | 0 | 75 | 0 | 0 | 0 | 0 | 422 | 0 | 0 |
| SRR5755844 | 50 | 33 | 0 | 50 | 50 | 247 | 742 | 12693 | 165 | 0 | 17 | 33 | 0 | 0 | 9396 | 10333 | 265790 | 0 | 0 | 7253 | 907 | 0 | 0 | 67 | 33 | 167 | 233 | 83 | 0 | 83 | 0 | 0 | 0 | 0 | 550 | 0 | 0 |
| SRR5755847 | 0 | 0 | 37 | 75 | 187 | 0 | 90 | 2073 | 0 | 0 | 75 | 37 | 0 | 0 | 2254 | 3006 | 107921 | 0 | 90 | 5409 | 1082 | 0 | 0 | 0 | 0 | 75 | 710 | 112 | 0 | 149 | 0 | 0 | 0 | 0 | 859 | 0 | 0 |
| SRR5755849 | 62 | 62 | 0 | 171 | 93 | 163 | 196 | 6165 | 43 | 0 | 16 | 0 | 0 | 0 | 18885 | 6317 | 319552 | 0 | 54 | 12612 | 10187 | 0 | 0 | 217 | 31 | 171 | 31 | 16 | 0 | 171 | 0 | 33 | 0 | 0 | 388 | 0 | 0 |
| SRR5755851 | 0 | 12 | 0 | 24 | 130 | 368 | 690 | 20647 | 184 | 12 | 47 | 0 | 0 | 0 | 9932 | 9392 | 257685 | 0 | 0 | 7955 | 138 | 0 | 0 | 59 | 0 | 107 | 17 | 12 | 0 | 11 | 0 | 0 | 0 | 0 | 340 | 0 | 12 |
| SRR5755854 | 0 | 0 | 0 | 34 | 273 | 199 | 1057 | 6326 | 934 | 0 | 34 | 0 | 0 | 0 | 2344 | 3637 | 108912 | 0 | 77 | 20923 | 1716 | 0 | 0 | 273 | 171 | 359 | 1795 | 581 | 0 | 137 | 0 | 15 | 0 | 0 | 304 | 0 | 0 |
| SRR5755855 | 41 | 0 | 41 | 0 | 11 | 151 | 624 | 7149 | 861 | 0 | 41 | 0 | 0 | 0 | 3316 | 4491 | 186433 | 0 | 65 | 16301 | 2240 | 0 | 0 | 12 | 203 | 11 | 39 | 12 | 0 | 162 | 0 | 22 | 0 | 0 | 607 | 0 | 0 |
| SRR5755856 | 54 | 14 | 14 | 272 | 244 | 305 | 218 | 5363 | 131 | 27 | 81 | 0 | 0 | 0 | 7133 | 19735 | 208834 | 0 | 235 | 59679 | 9949 | 0 | 0 | 1277 | 41 | 312 | 1480 | 557 | 0 | 177 | 0 | 0 | 0 | 0 | 718 | 0 | 0 |
| SRR5755858 | 0 | 0 | 33 | 52 | 141 | 577 | 1420 | 28462 | 1167 | 0 | 14 | 0 | 0 | 0 | 703 | 7815 | 56901 | 0 | 84 | 25846 | 155 | 0 | 0 | 150 | 61 | 244 | 587 | 127 | 0 | 113 | 0 | 14 | 0 | 0 | 840 | 0 | 0 |
| SRR5755861 | 16 | 0 | 22 | 77 | 396 | 94 | 1266 | 3328 | 1454 | 0 | 60 | 0 | 0 | 0 | 545 | 13554 | 63612 | 0 | 232 | 37408 | 595 | 0 | 0 | 440 | 66 | 236 | 1637 | 654 | 0 | 247 | 0 | 13 | 0 | 0 | 728 | 0 | 0 |
| SRR5755862 | 0 | 0 | 0 | 76 | 14 | 154 | 958 | 5082 | 560 | 17 | 84 | 0 | 0 | 0 | 3125 | 11128 | 249742 | 0 | 260 | 23656 | 3012 | 0 | 0 | 21 | 59 | 10 | 37 | 14 | 0 | 14 | 0 | 0 | 0 | 0 | 240 | 0 | 0 |
| SRR5755863 | 67 | 0 | 25 | 218 | 168 | 126 | 178 | 4664 | 159 | 0 | 168 | 17 | 0 | 0 | 1188 | 20654 | 126013 | 0 | 237 | 113229 | 5397 | 0 | 0 | 45 | 34 | 12 | 49 | 28 | 0 | 14 | 0 | 26 | 0 | 0 | 1033 | 0 | 0 |
| SRR5755865 | 0 | 0 | 26 | 39 | 84 | 196 | 744 | 7756 | 470 | 0 | 14 | 0 | 0 | 0 | 5601 | 11649 | 137817 | 0 | 235 | 36056 | 1097 | 0 | 0 | 58 | 0 | 39 | 12 | 45 | 0 | 35 | 0 | 39 | 0 | 0 | 967 | 0 | 0 |
| SRR5755866 | 0 | 18 | 18 | 141 | 135 | 85 | 474 | 3092 | 417 | 0 | 59 | 24 | 0 | 0 | 379 | 26520 | 116283 | 0 | 199 | 34272 | 1935 | 0 | 0 | 10 | 18 | 177 | 32 | 18 | 0 | 11 | 0 | 24 | 0 | 0 | 435 | 0 | 0 |
| SRR5755868 | 0 | 0 | 0 | 26 | 327 | 64 | 449 | 727 | 470 | 0 | 92 | 0 | 0 | 0 | 556 | 7215 | 21881 | 0 | 64 | 10238 | 299 | 0 | 0 | 366 | 52 | 222 | 1426 | 406 | 0 | 236 | 0 | 0 | 0 | 0 | 3372 | 0 | 0 |
| SRR5755870 | 20 | 0 | 13 | 268 | 105 | 67 | 147 | 2576 | 185 | 13 | 236 | 20 | 0 | 0 | 1262 | 19041 | 88705 | 0 | 325 | 115880 | 3663 | 0 | 0 | 628 | 33 | 196 | 752 | 530 | 0 | 347 | 0 | 83 | 0 | 0 | 1759 | 0 | 0 |
| SRR5755875 | 15 | 0 | 30 | 45 | 24 | 130 | 507 | 6789 | 537 | 0 | 75 | 0 | 0 | 0 | 21174 | 13177 | 620813 | 0 | 324 | 56880 | 7874 | 0 | 0 | 34 | 172 | 16 | 44 | 18 | 0 | 13 | 0 | 24 | 0 | 0 | 278 | 0 | 0 |
| SRR5755876 | 22 | 0 | 29 | 110 | 25 | 183 | 337 | 7765 | 164 | 0 | 88 | 0 | 0 | 0 | 13789 | 12738 | 618090 | 0 | 289 | 58542 | 4676 | 0 | 0 | 29 | 59 | 19 | 45 | 13 | 0 | 15 | 0 | 19 | 0 | 0 | 470 | 0 | 0 |
| SRR5755877 | 51 | 24 | 18 | 224 | 206 | 151 | 217 | 3295 | 161 | 24 | 164 | 0 | 0 | 0 | 1238 | 37984 | 107679 | 0 | 385 | 107431 | 4037 | 0 | 0 | 974 | 90 | 254 | 1043 | 801 | 0 | 365 | 0 | 43 | 0 | 0 | 1244 | 13 | 0 |
| SRR5755879 | 0 | 0 | 0 | 41 | 18 | 599 | 1402 | 24013 | 1070 | 0 | 53 | 0 | 0 | 0 | 5384 | 17108 | 101367 | 13 | 166 | 27823 | 229 | 0 | 0 | 19 | 36 | 11 | 63 | 14 | 0 | 18 | 0 | 45 | 0 | 0 | 760 | 0 | 0 |
| SRR5755880 | 0 | 0 | 51 | 59 | 110 | 123 | 347 | 3360 | 269 | 0 | 81 | 0 | 0 | 0 | 10495 | 12549 | 564136 | 0 | 246 | 39028 | 9156 | 0 | 0 | 293 | 13 | 64 | 11 | 195 | 0 | 280 | 0 | 21 | 0 | 0 | 117 | 0 | 0 |
| SRR5755882 | 13 | 13 | 26 | 64 | 204 | 108 | 121 | 4824 | 0 | 0 | 38 | 0 | 0 | 0 | 10834 | 2846 | 254329 | 0 | 81 | 17559 | 3248 | 0 | 0 | 217 | 102 | 204 | 28 | 229 | 0 | 229 | 0 | 13 | 0 | 0 | 265 | 0 | 0 |
| SRR5755884 | 33 | 20 | 13 | 243 | 130 | 220 | 205 | 10482 | 69 | 23 | 63 | 0 | 0 | 0 | 1395 | 3825 | 115648 | 0 | 42 | 8278 | 3563 | 0 | 0 | 439 | 30 | 233 | 750 | 727 | 0 | 276 | 0 | 18 | 0 | 0 | 673 | 0 | 0 |
| SRR5755887 | 0 | 0 | 32 | 0 | 81 | 209 | 46 | 6640 | 58 | 0 | 65 | 0 | 0 | 0 | 9543 | 5940 | 435728 | 0 | 70 | 19782 | 6014 | 0 | 0 | 177 | 16 | 194 | 419 | 290 | 0 | 145 | 0 | 0 | 0 | 0 | 94 | 0 | 0 |
| SRR5755889 | 27 | 14 | 21 | 27 | 144 | 66 | 83 | 2289 | 83 | 0 | 103 | 21 | 0 | 0 | 1534 | 1882 | 39873 | 0 | 100 | 8136 | 1650 | 0 | 0 | 34 | 34 | 55 | 185 | 68 | 0 | 335 | 0 | 25 | 0 | 0 | 178 | 0 | 0 |
| SRR5755890 | 31 | 0 | 0 | 31 | 317 | 56 | 137 | 1218 | 44 | 0 | 46 | 0 | 0 | 0 | 1641 | 2873 | 73227 | 0 | 25 | 3500 | 535 | 0 | 0 | 101 | 54 | 70 | 348 | 23 | 0 | 70 | 0 | 0 | 0 | 0 | 255 | 0 | 0 |
| SRR5755891 | 68 | 18 | 37 | 160 | 264 | 32 | 43 | 1024 | 38 | 0 | 92 | 0 | 0 | 0 | 5012 | 2516 | 87017 | 0 | 73 | 8452 | 20227 | 0 | 0 | 934 | 25 | 191 | 652 | 117 | 0 | 209 | 0 | 19 | 0 | 0 | 315 | 0 | 0 |
| SRR5755894 | 46 | 0 | 23 | 46 | 152 | 109 | 189 | 2822 | 123 | 0 | 78 | 14 | 0 | 0 | 181 | 10092 | 33613 | 0 | 73 | 4136 | 3011 | 0 | 0 | 42 | 14 | 14 | 115 | 14 | 0 | 78 | 0 | 36 | 0 | 0 | 535 | 0 | 0 |
| SRR5755896 | 23 | 18 | 0 | 106 | 13 | 146 | 148 | 4670 | 38 | 12 | 41 | 0 | 0 | 0 | 22222 | 7710 | 444232 | 0 | 136 | 14671 | 7289 | 0 | 0 | 270 | 82 | 375 | 21 | 53 | 0 | 106 | 0 | 0 | 0 | 0 | 150 | 0 | 0 |
| SRR5755897 | 16 | 56 | 48 | 72 | 240 | 128 | 103 | 2550 | 0 | 0 | 32 | 0 | 0 | 0 | 25325 | 5204 | 404411 | 0 | 165 | 23951 | 2757 | 0 | 0 | 216 | 64 | 120 | 303 | 32 | 0 | 88 | 0 | 12 | 0 | 0 | 87 | 0 | 0 |
| SRR5755898 | 72 | 57 | 14 | 16 | 86 | 244 | 459 | 10603 | 129 | 14 | 43 | 0 | 0 | 0 | 2522 | 2896 | 77639 | 0 | 86 | 10059 | 960 | 0 | 0 | 12 | 0 | 17 | 72 | 29 | 0 | 15 | 0 | 0 | 0 | 0 | 796 | 0 | 0 |
| SRR5755901 | 0 | 0 | 0 | 62 | 119 | 79 | 54 | 1972 | 19 | 13 | 22 | 0 | 0 | 0 | 16756 | 6938 | 474317 | 0 | 129 | 18776 | 4716 | 0 | 0 | 102 | 13 | 119 | 340 | 49 | 0 | 62 | 0 | 0 | 0 | 0 | 74 | 0 | 0 |
| SRR5755903 | 16 | 0 | 16 | 35 | 272 | 64 | 99 | 1840 | 28 | 12 | 12 | 0 | 0 | 0 | 12902 | 5710 | 331962 | 0 | 80 | 9968 | 7985 | 0 | 0 | 183 | 23 | 134 | 17 | 38 | 0 | 110 | 0 | 0 | 0 | 0 | 99 | 0 | 0 |
| SRR5755905 | 79 | 27 | 30 | 170 | 155 | 181 | 162 | 4609 | 79 | 23 | 49 | 0 | 0 | 0 | 23951 | 6065 | 330531 | 0 | 130 | 17526 | 7909 | 0 | 0 | 602 | 53 | 231 | 632 | 95 | 0 | 148 | 0 | 10 | 0 | 0 | 208 | 0 | 0 |
| SRR5755908 | 0 | 16 | 0 | 95 | 159 | 56 | 78 | 1862 | 24 | 24 | 119 | 0 | 0 | 0 | 19989 | 7097 | 549402 | 0 | 182 | 28469 | 5352 | 0 | 0 | 95 | 40 | 56 | 199 | 0 | 0 | 119 | 0 | 0 | 0 | 0 | 79 | 11 | 0 |
| SRR5755910 | 33 | 0 | 35 | 41 | 514 | 81 | 67 | 2341 | 18 | 0 | 21 | 0 | 0 | 0 | 22427 | 12757 | 630682 | 0 | 152 | 34651 | 18864 | 0 | 0 | 428 | 44 | 143 | 282 | 11 | 0 | 108 | 0 | 0 | 0 | 0 | 51 | 0 | 0 |
| SRR5755911 | 0 | 16 | 16 | 78 | 11 | 66 | 96 | 2243 | 0 | 23 | 23 | 0 | 0 | 0 | 24641 | 13543 | 703008 | 0 | 164 | 47887 | 5616 | 0 | 0 | 399 | 47 | 235 | 352 | 47 | 0 | 157 | 0 | 0 | 0 | 0 | 39 | 0 | 0 |
| SRR5755912 | 41 | 0 | 21 | 138 | 166 | 165 | 119 | 4128 | 57 | 21 | 80 | 0 | 0 | 0 | 12431 | 4099 | 250466 | 0 | 120 | 23152 | 5778 | 0 | 0 | 21 | 28 | 111 | 17 | 62 | 0 | 145 | 0 | 0 | 0 | 0 | 276 | 0 | 0 |
| SRR5755914 | 0 | 17 | 0 | 33 | 134 | 1145 | 1086 | 20983 | 355 | 0 | 46 | 59 | 0 | 0 | 2408 | 6400 | 78852 | 0 | 20 | 4876 | 217 | 0 | 0 | 188 | 29 | 180 | 749 | 33 | 0 | 142 | 0 | 39 | 0 | 0 | 640 | 0 | 0 |
| SRR5755915 | 28 | 0 | 28 | 74 | 284 | 54 | 33 | 829 | 0 | 74 | 51 | 0 | 0 | 0 | 16697 | 16437 | 764711 | 0 | 169 | 60677 | 8907 | 0 | 0 | 364 | 40 | 154 | 239 | 23 | 0 | 148 | 0 | 0 | 0 | 0 | 24 | 0 | 0 |
| SRR5755917 | 79 | 65 | 43 | 13 | 14 | 193 | 254 | 7603 | 213 | 14 | 14 | 0 | 0 | 0 | 1362 | 3889 | 80071 | 0 | 51 | 7553 | 2704 | 0 | 0 | 12 | 86 | 16 | 59 | 19 | 0 | 179 | 0 | 0 | 0 | 0 | 979 | 0 | 0 |
| SRR5755918 | 20 | 27 | 20 | 88 | 14 | 128 | 269 | 4382 | 90 | 0 | 81 | 0 | 0 | 0 | 1640 | 2915 | 87931 | 0 | 0 | 5022 | 4971 | 0 | 0 | 54 | 54 | 210 | 30 | 142 | 0 | 237 | 0 | 26 | 0 | 0 | 369 | 0 | 0 |
| SRR5755919 | 49 | 78 | 39 | 126 | 107 | 138 | 238 | 8122 | 38 | 29 | 126 | 0 | 0 | 0 | 1389 | 2205 | 68457 | 0 | 31 | 7490 | 2134 | 0 | 0 | 243 | 87 | 223 | 27 | 389 | 0 | 14 | 0 | 0 | 0 | 0 | 803 | 0 | 0 |
| SRR5755922 | 17 | 0 | 0 | 109 | 86 | 313 | 293 | 4581 | 111 | 0 | 63 | 0 | 0 | 0 | 444 | 13224 | 77473 | 20 | 0 | 3865 | 5439 | 0 | 0 | 23 | 23 | 86 | 978 | 131 | 0 | 223 | 0 | 30 | 0 | 0 | 580 | 0 | 0 |
| SRR5755924 | 0 | 12 | 0 | 83 | 406 | 304 | 337 | 1810 | 169 | 0 | 56 | 0 | 0 | 0 | 427 | 3325 | 41992 | 0 | 67 | 2574 | 461 | 0 | 0 | 62 | 0 | 104 | 783 | 133 | 0 | 451 | 0 | 0 | 0 | 0 | 2892 | 0 | 0 |
| SRR5755925 | 34 | 0 | 23 | 91 | 353 | 172 | 439 | 3415 | 286 | 0 | 102 | 0 | 0 | 0 | 324 | 5217 | 54174 | 0 | 38 | 2404 | 172 | 0 | 0 | 51 | 23 | 57 | 745 | 102 | 0 | 353 | 0 | 0 | 0 | 0 | 1621 | 0 | 0 |
| SRR5755926 | 53 | 32 | 26 | 257 | 210 | 175 | 290 | 12036 | 106 | 26 | 126 | 16 | 0 | 0 | 1659 | 1892 | 76735 | 0 | 37 | 11593 | 678 | 0 | 0 | 131 | 0 | 137 | 36 | 221 | 0 | 29 | 0 | 0 | 0 | 0 | 821 | 0 | 11 |
| SRR5755928 | 0 | 0 | 11 | 38 | 15 | 594 | 548 | 16536 | 411 | 0 | 32 | 11 | 0 | 0 | 1462 | 4886 | 55445 | 23 | 23 | 3472 | 320 | 0 | 0 | 64 | 0 | 129 | 37 | 59 | 0 | 11 | 0 | 23 | 0 | 0 | 1207 | 13 | 0 |
| SRR5755929 | 25 | 25 | 25 | 12 | 74 | 140 | 105 | 2968 | 105 | 0 | 62 | 12 | 0 | 0 | 384 | 7921 | 92888 | 0 | 105 | 2654 | 454 | 0 | 0 | 0 | 0 | 12 | 446 | 173 | 0 | 260 | 0 | 35 | 0 | 0 | 708 | 0 | 0 |
| SRR5755931 | 22 | 0 | 0 | 44 | 357 | 132 | 182 | 2525 | 77 | 0 | 79 | 18 | 0 | 0 | 2940 | 2406 | 93559 | 0 | 32 | 5707 | 775 | 0 | 0 | 62 | 13 | 176 | 622 | 110 | 0 | 176 | 0 | 18 | 0 | 0 | 101 | 0 | 0 |
| SRR5755933 | 0 | 0 | 30 | 191 | 111 | 206 | 266 | 7274 | 90 | 0 | 111 | 10 | 0 | 0 | 6216 | 2354 | 105453 | 0 | 40 | 7866 | 371 | 0 | 0 | 312 | 50 | 222 | 24 | 191 | 0 | 181 | 0 | 30 | 0 | 0 | 257 | 14 | 0 |
| SRR5755936 | 23 | 18 | 23 | 70 | 139 | 30 | 43 | 891 | 10 | 13 | 70 | 15 | 0 | 0 | 5640 | 5223 | 311507 | 0 | 89 | 14937 | 3720 | 0 | 0 | 255 | 44 | 150 | 753 | 173 | 0 | 247 | 0 | 0 | 0 | 0 | 49 | 0 | 0 |
| SRR5755938 | 0 | 0 | 20 | 152 | 455 | 141 | 215 | 3812 | 133 | 0 | 73 | 0 | 0 | 0 | 6987 | 2661 | 175858 | 0 | 41 | 8205 | 2544 | 0 | 0 | 152 | 13 | 165 | 891 | 165 | 0 | 211 | 0 | 17 | 0 | 0 | 202 | 0 | 0 |
| SRR5755939 | 27 | 0 | 45 | 111 | 482 | 102 | 66 | 2635 | 26 | 0 | 62 | 27 | 0 | 0 | 6200 | 3441 | 231020 | 0 | 114 | 14751 | 3239 | 0 | 0 | 261 | 40 | 140 | 734 | 109 | 0 | 243 | 0 | 0 | 0 | 0 | 117 | 0 | 0 |
| SRR5755940 | 118 | 71 | 55 | 14 | 172 | 349 | 325 | 8673 | 120 | 16 | 31 | 0 | 0 | 0 | 2506 | 4912 | 103242 | 0 | 66 | 11907 | 2271 | 0 | 0 | 24 | 39 | 165 | 53 | 21 | 0 | 133 | 0 | 30 | 0 | 0 | 926 | 0 | 0 |
| SRR5755943 | 13 | 20 | 40 | 119 | 123 | 104 | 72 | 1645 | 42 | 0 | 63 | 23 | 0 | 0 | 1884 | 6921 | 198074 | 0 | 74 | 7647 | 2917 | 0 | 0 | 123 | 0 | 83 | 901 | 703 | 0 | 146 | 0 | 0 | 0 | 0 | 130 | 0 | 0 |
| SRR5755946 | 0 | 0 | 15 | 69 | 18 | 253 | 274 | 4051 | 84 | 0 | 61 | 0 | 0 | 0 | 591 | 4722 | 61752 | 0 | 63 | 2363 | 464 | 0 | 0 | 38 | 23 | 53 | 25 | 15 | 0 | 190 | 0 | 0 | 0 | 0 | 549 | 0 | 0 |
| SRR5755947 | 25 | 49 | 33 | 453 | 280 | 361 | 241 | 14638 | 98 | 25 | 82 | 0 | 0 | 0 | 2396 | 3177 | 116156 | 0 | 33 | 6263 | 1362 | 0 | 0 | 379 | 58 | 362 | 1590 | 1952 | 0 | 231 | 0 | 0 | 0 | 0 | 734 | 0 | 16 |
| SRR5755950 | 52 | 0 | 0 | 72 | 83 | 359 | 179 | 4932 | 54 | 0 | 41 | 0 | 0 | 0 | 1417 | 7968 | 139181 | 0 | 0 | 2314 | 735 | 0 | 0 | 31 | 21 | 52 | 18 | 20 | 0 | 114 | 0 | 0 | 0 | 0 | 304 | 0 | 0 |
| SRR5755952 | 19 | 19 | 38 | 75 | 12 | 182 | 148 | 2997 | 68 | 0 | 19 | 0 | 0 | 0 | 3746 | 2767 | 79196 | 0 | 23 | 5130 | 2316 | 0 | 0 | 66 | 56 | 188 | 30 | 85 | 0 | 151 | 0 | 0 | 0 | 0 | 177 | 0 | 0 |
| SRR5755953 | 26 | 10 | 0 | 124 | 285 | 66 | 57 | 2039 | 26 | 10 | 36 | 16 | 0 | 0 | 8014 | 4462 | 241228 | 0 | 46 | 9239 | 2139 | 0 | 0 | 155 | 109 | 321 | 1119 | 539 | 0 | 161 | 0 | 0 | 0 | 0 | 80 | 0 | 0 |
| SRR5755954 | 83 | 17 | 50 | 183 | 166 | 455 | 172 | 4464 | 49 | 17 | 50 | 0 | 0 | 0 | 8042 | 4766 | 141653 | 0 | 61 | 14904 | 7009 | 0 | 0 | 366 | 83 | 166 | 864 | 316 | 17 | 66 | 0 | 12 | 0 | 0 | 463 | 0 | 0 |
| SRR5755956 | 0 | 0 | 15 | 85 | 134 | 728 | 771 | 17410 | 286 | 0 | 30 | 59 | 0 | 0 | 5570 | 10145 | 140581 | 0 | 57 | 4613 | 243 | 0 | 0 | 52 | 37 | 193 | 925 | 379 | 0 | 82 | 0 | 0 | 0 | 0 | 443 | 0 | 0 |
| SRR5755957 | 0 | 0 | 36 | 109 | 170 | 52 | 43 | 1404 | 0 | 12 | 0 | 12 | 0 | 0 | 10375 | 5008 | 394484 | 0 | 66 | 13768 | 2808 | 0 | 0 | 134 | 24 | 195 | 12 | 11 | 0 | 122 | 0 | 0 | 0 | 0 | 82 | 0 | 0 |
| SRR5755959 | 22 | 0 | 27 | 82 | 313 | 278 | 269 | 5030 | 143 | 0 | 22 | 0 | 0 | 0 | 4582 | 3622 | 157090 | 0 | 63 | 6958 | 9541 | 0 | 0 | 143 | 38 | 192 | 1058 | 208 | 0 | 126 | 0 | 0 | 0 | 0 | 594 | 0 | 0 |
| SRR5755960 | 0 | 0 | 0 | 62 | 21 | 121 | 152 | 5018 | 13 | 0 | 44 | 0 | 0 | 0 | 12133 | 4712 | 349192 | 0 | 64 | 9465 | 3017 | 0 | 0 | 291 | 150 | 11 | 64 | 26 | 0 | 159 | 0 | 0 | 0 | 0 | 195 | 0 | 0 |
| SRR5755961 | 42 | 0 | 17 | 11 | 10 | 297 | 216 | 9163 | 94 | 17 | 88 | 0 | 0 | 0 | 9714 | 4676 | 195318 | 0 | 66 | 7596 | 3281 | 0 | 0 | 29 | 75 | 23 | 95 | 109 | 0 | 188 | 0 | 19 | 0 | 0 | 525 | 0 | 0 |
| SRR5755963 | 0 | 0 | 13 | 74 | 268 | 399 | 499 | 9749 | 133 | 0 | 40 | 13 | 0 | 0 | 4824 | 14486 | 52775 | 0 | 0 | 1730 | 266 | 0 | 0 | 74 | 54 | 268 | 892 | 557 | 0 | 114 | 0 | 67 | 0 | 0 | 2216 | 0 | 0 |
| SRR5755964 | 0 | 36 | 62 | 187 | 303 | 108 | 77 | 4070 | 32 | 0 | 36 | 0 | 0 | 0 | 11980 | 6669 | 444946 | 0 | 131 | 15445 | 5940 | 0 | 0 | 330 | 0 | 321 | 36 | 31 | 0 | 134 | 0 | 0 | 0 | 0 | 155 | 0 | 0 |
| SRR5755966 | 15 | 25 | 10 | 80 | 331 | 85 | 177 | 4150 | 64 | 0 | 15 | 0 | 0 | 0 | 5836 | 2668 | 135982 | 0 | 42 | 8718 | 5779 | 0 | 0 | 75 | 15 | 251 | 1104 | 627 | 0 | 95 | 0 | 0 | 0 | 0 | 196 | 0 | 0 |
| SRR5755967 | 24 | 0 | 40 | 87 | 11 | 29 | 87 | 2696 | 19 | 0 | 56 | 0 | 0 | 0 | 5314 | 2785 | 183149 | 0 | 68 | 6351 | 829 | 0 | 0 | 182 | 40 | 246 | 41 | 29 | 0 | 119 | 0 | 0 | 0 | 0 | 123 | 0 | 0 |
| SRR5755968 | 35 | 0 | 0 | 86 | 75 | 459 | 367 | 20941 | 105 | 17 | 0 | 0 | 0 | 0 | 2215 | 2736 | 87439 | 0 | 0 | 6985 | 1553 | 0 | 0 | 403 | 17 | 144 | 478 | 305 | 0 | 75 | 0 | 0 | 0 | 0 | 675 | 0 | 0 |
| SRR5755971 | 18 | 0 | 12 | 45 | 27 | 312 | 387 | 14386 | 156 | 0 | 15 | 0 | 0 | 0 | 1890 | 6656 | 168189 | 0 | 44 | 9102 | 5715 | 0 | 0 | 60 | 12 | 42 | 13 | 57 | 0 | 66 | 0 | 0 | 0 | 0 | 220 | 0 | 0 |
| SRR5755973 | 23 | 0 | 38 | 114 | 379 | 258 | 318 | 6397 | 73 | 0 | 23 | 0 | 0 | 0 | 14752 | 6827 | 377378 | 0 | 86 | 18641 | 18099 | 0 | 0 | 227 | 61 | 288 | 963 | 144 | 0 | 159 | 0 | 13 | 0 | 0 | 321 | 0 | 0 |
| SRR5755974 | 0 | 0 | 0 | 66 | 13 | 268 | 188 | 7196 | 24 | 0 | 16 | 0 | 0 | 0 | 24383 | 7001 | 521868 | 0 | 198 | 36481 | 5724 | 0 | 0 | 18 | 12 | 82 | 16 | 53 | 0 | 53 | 0 | 0 | 0 | 0 | 278 | 0 | 0 |
| SRR5755975 | 78 | 0 | 33 | 14 | 161 | 659 | 495 | 18547 | 76 | 29 | 41 | 0 | 0 | 0 | 9527 | 4276 | 210160 | 0 | 47 | 10761 | 7517 | 0 | 0 | 37 | 41 | 169 | 29 | 10 | 0 | 107 | 0 | 15 | 0 | 0 | 880 | 0 | 0 |
| SRR5755978 | 0 | 17 | 0 | 50 | 59 | 246 | 169 | 6230 | 46 | 0 | 0 | 0 | 0 | 0 | 11074 | 12049 | 526814 | 0 | 231 | 34730 | 12067 | 0 | 0 | 184 | 17 | 84 | 318 | 75 | 0 | 59 | 0 | 0 | 0 | 0 | 338 | 0 | 0 |
| SRR5755980 | 12 | 0 | 21 | 52 | 337 | 512 | 1947 | 6338 | 783 | 0 | 27 | 0 | 0 | 0 | 616 | 9348 | 59079 | 0 | 77 | 6858 | 385 | 0 | 0 | 361 | 85 | 282 | 1083 | 176 | 0 | 182 | 0 | 14 | 0 | 0 | 2381 | 0 | 0 |
| SRR5755981 | 15 | 0 | 15 | 65 | 531 | 327 | 1232 | 3312 | 270 | 15 | 0 | 0 | 0 | 0 | 1252 | 7544 | 75394 | 0 | 106 | 9580 | 356 | 0 | 0 | 357 | 84 | 288 | 1126 | 189 | 0 | 129 | 0 | 19 | 0 | 0 | 2610 | 0 | 0 |
| SRR5755982 | 39 | 15 | 0 | 10 | 113 | 269 | 429 | 7765 | 156 | 0 | 39 | 12 | 0 | 0 | 188 | 7610 | 24407 | 0 | 181 | 39865 | 730 | 0 | 0 | 31 | 39 | 190 | 47 | 21 | 0 | 160 | 0 | 0 | 0 | 0 | 4118 | 10 | 0 |
| SRR5755984 | 0 | 0 | 15 | 77 | 13 | 619 | 1204 | 10924 | 1489 | 0 | 12 | 12 | 0 | 0 | 4834 | 5723 | 35384 | 0 | 117 | 9644 | 217 | 0 | 0 | 10 | 34 | 99 | 39 | 117 | 0 | 117 | 0 | 42 | 0 | 0 | 1261 | 0 | 0 |
| SRR5755985 | 20 | 12 | 16 | 109 | 158 | 220 | 376 | 3949 | 121 | 0 | 20 | 0 | 0 | 0 | 851 | 6808 | 172549 | 0 | 177 | 29355 | 957 | 0 | 0 | 445 | 24 | 146 | 570 | 137 | 0 | 89 | 0 | 28 | 0 | 0 | 1964 | 0 | 0 |
| SRR5755987 | 27 | 0 | 0 | 57 | 12 | 267 | 185 | 6886 | 98 | 0 | 27 | 0 | 0 | 0 | 17772 | 4788 | 361851 | 0 | 90 | 18756 | 9589 | 0 | 0 | 234 | 35 | 137 | 16 | 49 | 0 | 71 | 0 | 0 | 0 | 0 | 253 | 0 | 0 |
| SRR5755988 | 15 | 0 | 0 | 58 | 292 | 324 | 270 | 11889 | 72 | 0 | 0 | 0 | 0 | 0 | 29211 | 6251 | 489247 | 0 | 114 | 45105 | 2434 | 0 | 0 | 195 | 15 | 97 | 346 | 68 | 0 | 68 | 0 | 30 | 0 | 0 | 213 | 0 | 0 |
| SRR5755989 | 74 | 46 | 55 | 21 | 176 | 264 | 170 | 7279 | 182 | 0 | 0 | 0 | 0 | 0 | 3789 | 5347 | 147266 | 0 | 135 | 35321 | 5085 | 0 | 0 | 21 | 46 | 10 | 45 | 15 | 0 | 92 | 0 | 0 | 0 | 0 | 735 | 0 | 0 |
| SRR5755991 | 0 | 0 | 0 | 44 | 15 | 511 | 1175 | 30497 | 933 | 0 | 17 | 0 | 0 | 0 | 1271 | 6779 | 36446 | 0 | 32 | 8164 | 109 | 0 | 0 | 85 | 0 | 75 | 25 | 116 | 0 | 71 | 0 | 13 | 0 | 0 | 902 | 0 | 0 |
| SRR5755992 | 0 | 0 | 0 | 73 | 86 | 404 | 259 | 10380 | 140 | 0 | 21 | 0 | 0 | 0 | 6869 | 5571 | 426149 | 0 | 158 | 29336 | 5517 | 0 | 0 | 129 | 0 | 60 | 146 | 86 | 0 | 73 | 0 | 0 | 0 | 0 | 230 | 0 | 0 |
| SRR5755994 | 45 | 0 | 50 | 285 | 251 | 170 | 239 | 1381 | 134 | 0 | 39 | 0 | 0 | 0 | 876 | 5993 | 80001 | 0 | 30 | 4195 | 424 | 0 | 0 | 614 | 78 | 201 | 1072 | 385 | 0 | 251 | 0 | 0 | 0 | 0 | 2360 | 0 | 0 |
| SRR5755995 | 0 | 14 | 19 | 88 | 16 | 116 | 204 | 2263 | 62 | 0 | 56 | 0 | 0 | 0 | 1231 | 9042 | 106465 | 0 | 0 | 1826 | 312 | 0 | 0 | 237 | 84 | 149 | 26 | 149 | 0 | 11 | 0 | 0 | 0 | 0 | 2355 | 0 | 0 |
| SRR5755996 | 64 | 13 | 37 | 287 | 146 | 87 | 86 | 1474 | 32 | 0 | 88 | 0 | 0 | 0 | 102 | 4953 | 43332 | 0 | 0 | 8305 | 51 | 0 | 0 | 635 | 66 | 122 | 528 | 374 | 0 | 348 | 0 | 0 | 0 | 0 | 2550 | 0 | 0 |
| SRR5755997 | 0 | 0 | 0 | 0 | 0 | 1707 | 881 | 3524 | 55 | 0 | 19 | 0 | 0 | 0 | 9664 | 11606 | 628692 | 0 | 28 | 12225 | 2808 | 0 | 0 | 18 | 37 | 56 | 140 | 47 | 0 | 158 | 0 | 0 | 0 | 0 | 3793 | 0 | 0 |
| SRR5755998 | 0 | 0 | 0 | 45 | 175 | 288 | 349 | 3171 | 341 | 0 | 41 | 0 | 0 | 0 | 2787 | 6830 | 30946 | 0 | 0 | 1599 | 341 | 0 | 0 | 130 | 45 | 115 | 342 | 111 | 0 | 230 | 0 | 52 | 0 | 0 | 2271 | 0 | 0 |
| SRR5755999 | 11 | 0 | 11 | 128 | 111 | 37 | 148 | 754 | 32 | 0 | 56 | 0 | 0 | 0 | 111 | 8348 | 55682 | 0 | 0 | 957 | 150 | 0 | 0 | 245 | 17 | 72 | 15 | 200 | 0 | 295 | 0 | 0 | 0 | 0 | 1762 | 0 | 0 |
| SRR5756000 | 0 | 0 | 0 | 0 | 0 | 730 | 0 | 0 | 0 | 0 | 163 | 0 | 0 | 0 | 730 | 0 | 40788 | 0 | 0 | 2922 | 0 | 0 | 0 | 0 | 0 | 0 | 0 | 0 | 0 | 0 | 0 | 0 | 0 | 0 | 4219 | 0 | 0 |
| SRR5756001 | 16 | 16 | 32 | 121 | 28 | 257 | 781 | 8000 | 459 | 0 | 44 | 0 | 0 | 0 | 4406 | 7439 | 382585 | 0 | 40 | 12109 | 1447 | 0 | 0 | 28 | 101 | 202 | 57 | 121 | 0 | 10 | 0 | 0 | 0 | 0 | 1307 | 0 | 0 |
| SRR5756002 | 0 | 0 | 24 | 81 | 36 | 147 | 295 | 3240 | 16 | 16 | 24 | 0 | 0 | 0 | 1571 | 10016 | 263564 | 0 | 49 | 5073 | 393 | 0 | 0 | 33 | 146 | 186 | 38 | 113 | 0 | 10 | 0 | 16 | 0 | 0 | 3537 | 0 | 0 |
| SRR5756003 | 96 | 31 | 42 | 402 | 287 | 324 | 242 | 7250 | 96 | 0 | 65 | 0 | 0 | 0 | 1673 | 6480 | 161365 | 0 | 55 | 28056 | 905 | 0 | 0 | 874 | 42 | 146 | 839 | 215 | 0 | 222 | 0 | 0 | 0 | 0 | 2177 | 0 | 0 |
| SRR5756005 | 0 | 0 | 0 | 45 | 30 | 357 | 978 | 10155 | 373 | 0 | 30 | 0 | 0 | 0 | 1040 | 6127 | 45763 | 16 | 16 | 2515 | 47 | 0 | 0 | 158 | 49 | 143 | 36 | 114 | 0 | 168 | 0 | 16 | 0 | 0 | 1865 | 0 | 0 |
| SRR5756006 | 33 | 0 | 33 | 192 | 10 | 195 | 225 | 2237 | 38 | 20 | 13 | 0 | 0 | 0 | 781 | 14800 | 187333 | 0 | 15 | 3708 | 428 | 0 | 0 | 20 | 20 | 153 | 32 | 139 | 0 | 146 | 0 | 0 | 0 | 0 | 2346 | 0 | 0 |
| SRR5756007 | 0 | 0 | 0 | 0 | 0 | 0 | 0 | 0 | 0 | 0 | 0 | 0 | 0 | 0 | 3636 | 1988 | 33797 | 0 | 0 | 909 | 0 | 0 | 0 | 0 | 0 | 0 | 0 | 0 | 0 | 260 | 0 | 0 | 0 | 0 | 1988 | 0 | 0 |
| SRR5756008 | 0 | 0 | 30 | 110 | 30 | 294 | 104 | 7777 | 52 | 18 | 12 | 0 | 0 | 0 | 18232 | 11550 | 563955 | 0 | 156 | 69335 | 4632 | 0 | 0 | 16 | 55 | 134 | 16 | 67 | 0 | 10 | 0 | 17 | 0 | 0 | 709 | 0 | 0 |
| SRR5756009 | 32 | 0 | 16 | 227 | 11 | 297 | 355 | 11932 | 139 | 0 | 22 | 0 | 0 | 0 | 2011 | 7579 | 128998 | 0 | 126 | 32373 | 1233 | 0 | 0 | 22 | 38 | 10 | 32 | 13 | 0 | 217 | 0 | 0 | 0 | 0 | 2092 | 0 | 0 |
| SRR5756011 | 0 | 0 | 26 | 52 | 14 | 497 | 1348 | 29414 | 674 | 0 | 23 | 0 | 0 | 0 | 1677 | 8906 | 123327 | 0 | 51 | 13936 | 101 | 0 | 0 | 17 | 49 | 16 | 39 | 103 | 0 | 13 | 0 | 0 | 0 | 0 | 1310 | 0 | 0 |
| SRR5756012 | 0 | 0 | 12 | 90 | 238 | 191 | 180 | 6017 | 66 | 0 | 25 | 0 | 0 | 0 | 12482 | 14987 | 751242 | 0 | 184 | 89541 | 6918 | 0 | 0 | 291 | 25 | 50 | 146 | 22 | 0 | 155 | 0 | 11 | 0 | 0 | 232 | 0 | 0 |
| SRR5756014 | 0 | 13 | 18 | 58 | 40 | 314 | 237 | 12372 | 93 | 0 | 40 | 0 | 0 | 0 | 44764 | 19138 | 678954 | 0 | 175 | 77589 | 12413 | 0 | 0 | 25 | 81 | 11 | 19 | 58 | 0 | 193 | 0 | 0 | 0 | 0 | 279 | 0 | 0 |
| SRR5756016 | 75 | 11 | 21 | 267 | 114 | 266 | 273 | 7696 | 94 | 0 | 57 | 16 | 0 | 0 | 5145 | 10919 | 230631 | 0 | 103 | 68454 | 6638 | 0 | 0 | 822 | 48 | 169 | 575 | 217 | 0 | 224 | 0 | 0 | 0 | 0 | 1472 | 0 | 0 |
| SRR5756017 | 0 | 20 | 20 | 0 | 20 | 1489 | 558 | 5771 | 93 | 0 | 41 | 0 | 0 | 0 | 8470 | 14241 | 582848 | 0 | 0 | 44676 | 6888 | 0 | 0 | 406 | 20 | 142 | 548 | 20 | 0 | 81 | 0 | 93 | 0 | 0 | 4084 | 0 | 0 |
| SRR5756018 | 0 | 0 | 0 | 26 | 295 | 723 | 1477 | 29273 | 629 | 0 | 22 | 17 | 0 | 0 | 1320 | 13587 | 49640 | 0 | 31 | 6756 | 126 | 0 | 0 | 174 | 13 | 148 | 659 | 113 | 0 | 148 | 0 | 16 | 0 | 0 | 1364 | 0 | 0 |
| SRR5756019 | 13 | 13 | 26 | 181 | 298 | 195 | 122 | 3395 | 0 | 0 | 65 | 0 | 0 | 0 | 15632 | 13301 | 721131 | 0 | 122 | 80237 | 18930 | 0 | 0 | 596 | 39 | 39 | 194 | 104 | 13 | 259 | 0 | 24 | 0 | 0 | 514 | 0 | 0 |
| SRR5756021 | 11 | 0 | 41 | 45 | 382 | 430 | 791 | 11735 | 655 | 0 | 26 | 0 | 0 | 0 | 18519 | 9873 | 469135 | 0 | 131 | 17158 | 5939 | 0 | 0 | 379 | 150 | 502 | 1188 | 135 | 0 | 157 | 0 | 0 | 0 | 0 | 317 | 0 | 0 |
| SRR5756022 | 0 | 0 | 18 | 55 | 356 | 543 | 675 | 18583 | 146 | 0 | 43 | 0 | 0 | 0 | 16929 | 13203 | 585992 | 0 | 212 | 28655 | 3693 | 0 | 0 | 424 | 104 | 258 | 479 | 117 | 0 | 98 | 0 | 13 | 0 | 0 | 735 | 0 | 0 |

### Supp Table S8.3.Columns_BX_to_DH

| | | | | | | | | | | | | | | | | | | | | | | | | | | | | | | | | | | | | | |
|---|---|---|---|---|---|---|---|---|---|---|---|---|---|---|---|---|---|---|---|---|---|---|---|---|---|---|---|---|---|---|---|---|---|---|---|---|---|
| SRR5756023 | 15 | 15 | 30 | 151 | 189 | 387 | 346 | 8468 | 127 | 0 | 23 | 0 | 0 | 0 | 2234 | 9536 | 129927 | 0 | 40 | 22605 | 375 | 0 | 0 | 901 | 61 | 212 | 1363 | 431 | 0 | 174 | 0 | 0 | 0 | 0 | 2056 | 11 | 0 |
| SRR5756025 | 0 | 0 | 13 | 0 | 11 | 552 | 1163 | 16751 | 731 | 0 | 13 | 0 | 0 | 0 | 6250 | 12388 | 68997 | 0 | 30 | 5877 | 298 | 0 | 0 | 85 | 45 | 170 | 27 | 85 | 0 | 165 | 0 | 30 | 0 | 0 | 1805 | 0 | 0 |
| SRR5756026 | 19 | 13 | 0 | 69 | 213 | 350 | 235 | 9730 | 86 | 0 | 0 | 0 | 0 | 0 | 10636 | 14473 | 658820 | 0 | 157 | 31357 | 6049 | 0 | 0 | 14 | 13 | 94 | 12 | 107 | 0 | 132 | 0 | 0 | 0 | 0 | 319 | 0 | 0 |
| SRR5756027 | 0 | 0 | 20 | 24 | 206 | 449 | 628 | 10372 | 358 | 0 | 24 | 0 | 0 | 0 | 32173 | 12412 | 602544 | 0 | 264 | 38290 | 9349 | 0 | 0 | 446 | 34 | 186 | 486 | 125 | 0 | 145 | 0 | 0 | 0 | 0 | 263 | 0 | 0 |
| SRR5756028 | 0 | 0 | 23 | 0 | 186 | 372 | 290 | 9720 | 41 | 0 | 31 | 0 | 0 | 0 | 5088 | 13079 | 308956 | 0 | 331 | 12988 | 4136 | 0 | 0 | 18 | 46 | 101 | 17 | 186 | 0 | 93 | 0 | 83 | 0 | 0 | 3270 | 0 | 0 |
| SRR5756029 | 28 | 0 | 17 | 53 | 109 | 450 | 329 | 8859 | 166 | 28 | 28 | 11 | 0 | 0 | 3567 | 12337 | 173352 | 0 | 163 | 57355 | 2399 | 0 | 0 | 674 | 67 | 182 | 674 | 277 | 0 | 187 | 0 | 57 | 0 | 0 | 1733 | 0 | 0 |
| SRR5756031 | 0 | 0 | 0 | 0 | 87 | 754 | 1490 | 21576 | 1134 | 0 | 12 | 12 | 0 | 0 | 1035 | 9148 | 54456 | 0 | 50 | 11455 | 150 | 0 | 0 | 78 | 32 | 121 | 48 | 91 | 0 | 103 | 0 | 12 | 0 | 0 | 1237 | 0 | 0 |
| SRR5756032 | 0 | 13 | 0 | 46 | 86 | 433 | 391 | 9592 | 67 | 0 | 33 | 0 | 0 | 0 | 11389 | 14411 | 667747 | 0 | 341 | 44967 | 8012 | 0 | 0 | 18 | 26 | 126 | 11 | 66 | 0 | 93 | 0 | 25 | 0 | 0 | 468 | 0 | 0 |
| SRR5756033 | 13 | 20 | 27 | 53 | 14 | 500 | 895 | 3691 | 1652 | 0 | 33 | 0 | 0 | 0 | 338 | 5758 | 38866 | 0 | 48 | 6278 | 532 | 0 | 0 | 53 | 47 | 13 | 50 | 160 | 0 | 167 | 0 | 32 | 0 | 0 | 1259 | 0 | 0 |
| SRR5756034 | 0 | 0 | 16 | 61 | 512 | 554 | 1072 | 4214 | 858 | 0 | 36 | 0 | 0 | 0 | 789 | 7682 | 83380 | 0 | 38 | 4947 | 439 | 0 | 0 | 57 | 45 | 154 | 721 | 156 | 0 | 209 | 0 | 10 | 0 | 0 | 835 | 0 | 0 |
| SRR5756035 | 20 | 0 | 0 | 160 | 200 | 253 | 289 | 6771 | 253 | 0 | 40 | 12 | 0 | 0 | 704 | 12091 | 55836 | 0 | 83 | 42675 | 718 | 0 | 0 | 184 | 80 | 289 | 1159 | 429 | 0 | 176 | 0 | 25 | 0 | 0 | 2294 | 0 | 0 |
| SRR5756037 | 12 | 0 | 30 | 32 | 21 | 354 | 1020 | 4778 | 632 | 0 | 27 | 0 | 0 | 0 | 1955 | 22522 | 23773 | 0 | 34 | 3185 | 295 | 0 | 0 | 70 | 35 | 10 | 31 | 90 | 0 | 137 | 0 | 25 | 0 | 0 | 1708 | 14 | 0 |
| SRR5756038 | 0 | 11 | 0 | 48 | 74 | 516 | 1190 | 9074 | 694 | 0 | 16 | 11 | 0 | 0 | 198 | 13786 | 36117 | 0 | 0 | 4165 | 238 | 0 | 0 | 16 | 27 | 53 | 297 | 101 | 0 | 154 | 0 | 20 | 0 | 0 | 957 | 0 | 0 |
| SRR5756040 | 0 | 13 | 0 | 26 | 321 | 442 | 1074 | 8722 | 919 | 0 | 30 | 17 | 0 | 0 | 813 | 3901 | 100711 | 0 | 46 | 8229 | 527 | 0 | 0 | 120 | 51 | 175 | 671 | 120 | 0 | 111 | 0 | 13 | 0 | 0 | 564 | 0 | 0 |
| SRR5756041 | 0 | 0 | 0 | 23 | 205 | 387 | 419 | 10085 | 280 | 0 | 45 | 0 | 0 | 0 | 1516 | 6980 | 139014 | 0 | 43 | 6225 | 323 | 0 | 0 | 11 | 51 | 205 | 25 | 205 | 0 | 148 | 0 | 0 | 0 | 0 | 2152 | 0 | 0 |
| SRR5756042 | 23 | 39 | 0 | 55 | 266 | 257 | 301 | 5167 | 132 | 16 | 39 | 0 | 16 | 0 | 573 | 6705 | 52944 | 0 | 59 | 28523 | 404 | 0 | 0 | 11 | 39 | 148 | 53 | 24 | 0 | 11 | 0 | 0 | 0 | 0 | 3642 | 0 | 0 |
| SRR5756044 | 0 | 0 | 0 | 18 | 293 | 303 | 780 | 10556 | 1322 | 0 | 24 | 0 | 0 | 0 | 672 | 6637 | 15470 | 0 | 43 | 7196 | 65 | 0 | 0 | 92 | 24 | 122 | 776 | 128 | 0 | 189 | 0 | 0 | 0 | 0 | 1198 | 0 | 0 |
| SRR5756045 | 0 | 0 | 0 | 63 | 56 | 385 | 171 | 9095 | 43 | 0 | 28 | 0 | 0 | 0 | 1241 | 9343 | 186803 | 0 | 0 | 4901 | 1070 | 0 | 0 | 273 | 14 | 105 | 357 | 112 | 0 | 84 | 0 | 0 | 0 | 0 | 2797 | 0 | 0 |
| SRR5756047 | 23 | 23 | 28 | 147 | 15 | 481 | 501 | 5121 | 446 | 0 | 18 | 0 | 0 | 0 | 5725 | 4134 | 145099 | 0 | 89 | 6638 | 8210 | 0 | 0 | 151 | 32 | 225 | 65 | 211 | 0 | 138 | 0 | 34 | 0 | 0 | 384 | 0 | 0 |
| SRR5756048 | 0 | 19 | 19 | 60 | 40 | 285 | 187 | 5047 | 70 | 0 | 19 | 0 | 0 | 0 | 17534 | 6080 | 352408 | 0 | 56 | 8014 | 2598 | 0 | 0 | 227 | 37 | 245 | 32 | 259 | 0 | 180 | 0 | 0 | 0 | 0 | 617 | 0 | 0 |
| SRR5756049 | 42 | 24 | 21 | 146 | 251 | 242 | 152 | 3774 | 87 | 0 | 52 | 0 | 0 | 0 | 2807 | 4662 | 103470 | 0 | 40 | 13019 | 1218 | 0 | 0 | 495 | 45 | 219 | 1181 | 613 | 0 | 320 | 0 | 28 | 0 | 0 | 1985 | 0 | 0 |
| SRR5756051 | 0 | 0 | 16 | 66 | 501 | 522 | 918 | 11229 | 741 | 0 | 35 | 0 | 0 | 0 | 18847 | 13432 | 75593 | 21 | 21 | 3632 | 877 | 0 | 0 | 101 | 31 | 109 | 601 | 213 | 0 | 272 | 0 | 52 | 0 | 0 | 986 | 10 | 0 |
| SRR5756052 | 0 | 12 | 0 | 64 | 134 | 151 | 151 | 3052 | 91 | 0 | 29 | 0 | 0 | 0 | 3453 | 4926 | 282681 | 0 | 68 | 13215 | 4165 | 0 | 0 | 181 | 12 | 76 | 16 | 193 | 0 | 198 | 0 | 15 | 0 | 0 | 689 | 0 | 0 |
| SRR5756054 | 26 | 13 | 19 | 58 | 329 | 490 | 546 | 5671 | 622 | 0 | 26 | 0 | 0 | 0 | 2770 | 3614 | 85223 | 0 | 66 | 8507 | 3862 | 0 | 0 | 148 | 39 | 219 | 1614 | 142 | 0 | 116 | 0 | 0 | 0 | 0 | 469 | 0 | 0 |
| SRR5756055 | 0 | 0 | 0 | 0 | 586 | 334 | 424 | 5279 | 126 | 14 | 71 | 0 | 0 | 0 | 4485 | 3714 | 169068 | 0 | 27 | 8429 | 1309 | 0 | 0 | 127 | 21 | 162 | 798 | 78 | 0 | 275 | 0 | 0 | 0 | 0 | 608 | 0 | 0 |
| SRR5756056 | 33 | 11 | 11 | 105 | 149 | 252 | 123 | 3286 | 112 | 11 | 100 | 0 | 0 | 0 | 2461 | 4103 | 89846 | 0 | 45 | 19815 | 1912 | 0 | 0 | 26 | 44 | 149 | 45 | 22 | 0 | 14 | 0 | 0 | 0 | 0 | 2474 | 0 | 0 |
| SRR5756058 | 0 | 0 | 0 | 43 | 487 | 640 | 838 | 13849 | 756 | 0 | 40 | 0 | 0 | 0 | 1234 | 7605 | 25319 | 0 | 35 | 4306 | 151 | 0 | 0 | 88 | 18 | 97 | 578 | 122 | 0 | 173 | 0 | 0 | 0 | 0 | 1347 | 10 | 0 |
| SRR5756059 | 19 | 0 | 0 | 28 | 65 | 75 | 90 | 1763 | 75 | 0 | 19 | 0 | 0 | 0 | 1311 | 4658 | 116565 | 0 | 0 | 4911 | 1100 | 0 | 0 | 224 | 37 | 140 | 877 | 308 | 0 | 205 | 0 | 15 | 0 | 0 | 2268 | 0 | 0 |
| SRR5756061 | 0 | 13 | 18 | 102 | 14 | 377 | 683 | 6028 | 925 | 0 | 49 | 0 | 0 | 0 | 6514 | 6861 | 211558 | 0 | 85 | 15575 | 5953 | 0 | 0 | 13 | 35 | 11 | 67 | 11 | 0 | 164 | 0 | 14 | 0 | 0 | 277 | 0 | 0 |
| SRR5756062 | 0 | 0 | 0 | 58 | 25 | 202 | 193 | 5517 | 193 | 0 | 79 | 0 | 0 | 0 | 8309 | 9225 | 308257 | 0 | 135 | 27457 | 1867 | 0 | 0 | 10 | 61 | 48 | 34 | 120 | 0 | 22 | 0 | 50 | 0 | 0 | 1213 | 0 | 0 |
| SRR5756063 | 13 | 0 | 13 | 167 | 126 | 285 | 278 | 5332 | 269 | 0 | 119 | 16 | 0 | 0 | 842 | 12267 | 73493 | 0 | 79 | 58410 | 705 | 0 | 0 | 29 | 68 | 13 | 63 | 33 | 0 | 21 | 0 | 16 | 0 | 0 | 2447 | 0 | 0 |
| SRR5756065 | 0 | 0 | 12 | 34 | 227 | 338 | 665 | 4724 | 986 | 0 | 63 | 0 | 0 | 0 | 5428 | 13030 | 52947 | 0 | 28 | 10373 | 216 | 0 | 0 | 69 | 23 | 101 | 647 | 118 | 0 | 250 | 0 | 66 | 0 | 0 | 1258 | 0 | 0 |
| SRR5756066 | 0 | 11 | 0 | 57 | 131 | 321 | 470 | 4183 | 371 | 0 | 75 | 0 | 0 | 0 | 3697 | 18023 | 264212 | 0 | 68 | 17592 | 2660 | 0 | 0 | 84 | 11 | 79 | 23 | 82 | 0 | 22 | 0 | 25 | 0 | 0 | 514 | 11 | 0 |
| SRR5756068 | 0 | 0 | 0 | 48 | 451 | 225 | 744 | 4930 | 1176 | 0 | 51 | 0 | 0 | 0 | 1647 | 3012 | 120054 | 0 | 74 | 14338 | 1563 | 0 | 0 | 151 | 42 | 148 | 788 | 267 | 0 | 235 | 0 | 14 | 0 | 0 | 677 | 0 | 0 |
| SRR5756069 | 12 | 0 | 0 | 98 | 38 | 133 | 249 | 3718 | 224 | 0 | 37 | 0 | 0 | 0 | 2962 | 3625 | 181479 | 0 | 66 | 12273 | 1834 | 0 | 0 | 155 | 37 | 183 | 40 | 16 | 0 | 11 | 0 | 0 | 0 | 0 | 1016 | 0 | 0 |
| SRR5756070 | 65 | 73 | 24 | 351 | 196 | 203 | 182 | 5655 | 154 | 0 | 33 | 0 | 0 | 0 | 2175 | 7731 | 84453 | 0 | 91 | 34921 | 1375 | 0 | 0 | 825 | 65 | 269 | 1674 | 767 | 0 | 188 | 0 | 14 | 0 | 0 | 2088 | 0 | 0 |
| SRR5756072 | 0 | 0 | 0 | 21 | 29 | 559 | 1006 | 12883 | 1404 | 15 | 21 | 0 | 0 | 0 | 2187 | 9206 | 31859 | 0 | 0 | 12473 | 87 | 0 | 0 | 93 | 75 | 11 | 42 | 126 | 0 | 16 | 0 | 0 | 0 | 0 | 1371 | 0 | 0 |
| SRR5756073 | 15 | 0 | 0 | 73 | 307 | 61 | 198 | 1801 | 137 | 0 | 29 | 0 | 0 | 0 | 534 | 12241 | 100696 | 0 | 0 | 3723 | 351 | 0 | 0 | 350 | 15 | 88 | 905 | 277 | 0 | 270 | 0 | 0 | 0 | 0 | 4512 | 0 | 0 |
| SRR5756075 | 0 | 0 | 31 | 75 | 263 | 393 | 1230 | 3717 | 1666 | 0 | 56 | 0 | 0 | 0 | 1015 | 4009 | 83768 | 0 | 64 | 6999 | 1673 | 0 | 0 | 81 | 100 | 282 | 920 | 257 | 0 | 244 | 0 | 0 | 0 | 0 | 629 | 0 | 0 |
| SRR5756076 | 0 | 0 | 0 | 42 | 261 | 502 | 676 | 5561 | 964 | 0 | 87 | 0 | 0 | 0 | 802 | 4214 | 75131 | 0 | 36 | 7201 | 721 | 0 | 0 | 55 | 83 | 161 | 643 | 187 | 0 | 340 | 0 | 0 | 0 | 0 | 1006 | 0 | 0 |
| SRR5756077 | 69 | 14 | 41 | 276 | 152 | 330 | 152 | 7187 | 161 | 14 | 41 | 14 | 14 | 0 | 1357 | 6636 | 64662 | 0 | 62 | 28935 | 1669 | 0 | 0 | 483 | 110 | 676 | 2276 | 938 | 0 | 221 | 0 | 0 | 0 | 0 | 1436 | 0 | 0 |
| SRR5756079 | 0 | 0 | 0 | 13 | 601 | 468 | 1019 | 11478 | 1103 | 0 | 36 | 0 | 0 | 0 | 1571 | 9272 | 27786 | 0 | 33 | 10008 | 117 | 0 | 0 | 112 | 45 | 135 | 462 | 121 | 0 | 202 | 0 | 17 | 0 | 0 | 1298 | 0 | 0 |
| SRR5756080 | 0 | 0 | 0 | 41 | 109 | 183 | 294 | 2869 | 267 | 0 | 59 | 0 | 0 | 0 | 989 | 8177 | 118832 | 0 | 0 | 7373 | 1822 | 0 | 0 | 45 | 20 | 82 | 29 | 136 | 0 | 19 | 0 | 18 | 0 | 0 | 642 | 0 | 0 |
| SRR5756082 | 0 | 0 | 0 | 41 | 11 | 556 | 911 | 4923 | 1538 | 0 | 38 | 0 | 0 | 0 | 444 | 4127 | 58482 | 18 | 59 | 8496 | 373 | 0 | 0 | 79 | 38 | 166 | 45 | 105 | 0 | 15 | 0 | 18 | 0 | 0 | 1117 | 0 | 0 |
| SRR5756083 | 0 | 0 | 0 | 25 | 225 | 834 | 892 | 7255 | 1326 | 0 | 42 | 0 | 0 | 0 | 210 | 7603 | 33892 | 0 | 22 | 5523 | 196 | 0 | 0 | 39 | 22 | 78 | 356 | 75 | 0 | 214 | 0 | 22 | 0 | 0 | 1579 | 0 | 0 |
| SRR5756084 | 39 | 23 | 23 | 13 | 139 | 409 | 379 | 14010 | 216 | 46 | 62 | 0 | 0 | 0 | 2105 | 6173 | 86189 | 0 | 104 | 45548 | 1919 | 0 | 0 | 26 | 23 | 11 | 52 | 21 | 0 | 93 | 0 | 15 | 0 | 0 | 1290 | 0 | 0 |
| SRR5756086 | 15 | 0 | 0 | 24 | 30 | 419 | 595 | 10497 | 946 | 0 | 32 | 0 | 0 | 0 | 546 | 7815 | 25285 | 0 | 0 | 9482 | 49 | 0 | 0 | 94 | 28 | 18 | 61 | 11 | 0 | 15 | 0 | 20 | 0 | 0 | 1555 | 0 | 0 |
| SRR5756087 | 0 | 12 | 0 | 17 | 69 | 790 | 425 | 5386 | 607 | 0 | 35 | 0 | 0 | 0 | 304 | 8925 | 33155 | 0 | 0 | 4130 | 466 | 0 | 0 | 29 | 0 | 29 | 13 | 75 | 0 | 13 | 0 | 20 | 0 | 0 | 1919 | 11 | 0 |
| SRR5756089 | 18 | 0 | 12 | 70 | 16 | 817 | 1005 | 5826 | 1406 | 0 | 18 | 0 | 0 | 0 | 1982 | 4635 | 81932 | 0 | 40 | 3697 | 1179 | 0 | 0 | 47 | 0 | 10 | 41 | 135 | 0 | 111 | 0 | 0 | 0 | 0 | 1086 | 0 | 0 |
| SRR5756090 | 0 | 0 | 0 | 62 | 458 | 687 | 999 | 5321 | 763 | 0 | 56 | 0 | 0 | 0 | 2409 | 6598 | 126191 | 0 | 35 | 4165 | 485 | 0 | 0 | 60 | 30 | 84 | 405 | 128 | 0 | 238 | 0 | 12 | 0 | 0 | 1395 | 0 | 0 |
| SRR5756091 | 17 | 0 | 0 | 123 | 203 | 288 | 283 | 3300 | 118 | 0 | 59 | 25 | 0 | 0 | 822 | 4404 | 43391 | 0 | 15 | 9093 | 262 | 0 | 0 | 284 | 25 | 106 | 644 | 263 | 0 | 220 | 0 | 0 | 0 | 0 | 2693 | 0 | 0 |
| SRR5756093 | 0 | 0 | 0 | 59 | 26 | 549 | 761 | 8833 | 877 | 0 | 27 | 0 | 0 | 0 | 1907 | 7533 | 22727 | 0 | 29 | 3179 | 29 | 0 | 0 | 81 | 32 | 132 | 51 | 130 | 0 | 12 | 0 | 19 | 0 | 0 | 1272 | 0 | 0 |
| SRR5756094 | 0 | 0 | 0 | 70 | 110 | 430 | 368 | 3402 | 301 | 0 | 31 | 0 | 0 | 0 | 503 | 11733 | 78501 | 0 | 0 | 2992 | 498 | 0 | 0 | 34 | 0 | 24 | 314 | 82 | 0 | 201 | 0 | 0 | 0 | 0 | 1160 | 0 | 0 |
| SRR5756096 | 0 | 0 | 23 | 53 | 739 | 181 | 447 | 4626 | 505 | 0 | 30 | 0 | 0 | 0 | 25727 | 9559 | 434977 | 0 | 125 | 26941 | 4104 | 0 | 0 | 350 | 48 | 206 | 1007 | 238 | 0 | 215 | 0 | 0 | 0 | 0 | 501 | 0 | 0 |
| SRR5756097 | 0 | 0 | 45 | 72 | 46 | 346 | 173 | 7949 | 0 | 0 | 18 | 0 | 0 | 0 | 12571 | 10841 | 413931 | 0 | 130 | 20951 | 778 | 0 | 0 | 33 | 72 | 172 | 27 | 13 | 0 | 13 | 0 | 0 | 0 | 0 | 3905 | 0 | 0 |
| SRR5756098 | 36 | 0 | 0 | 227 | 117 | 280 | 237 | 6750 | 144 | 13 | 44 | 17 | 0 | 0 | 1018 | 6883 | 81213 | 0 | 82 | 40304 | 688 | 0 | 0 | 484 | 42 | 231 | 782 | 434 | 0 | 124 | 0 | 17 | 0 | 0 | 2461 | 0 | 0 |
| SRR5756100 | 0 | 0 | 0 | 28 | 183 | 174 | 459 | 6255 | 261 | 0 | 13 | 20 | 0 | 0 | 491 | 4257 | 11194 | 0 | 0 | 4189 | 55 | 0 | 0 | 81 | 18 | 121 | 316 | 93 | 0 | 126 | 0 | 40 | 0 | 0 | 2284 | 0 | 0 |
| SRR5756101 | 12 | 0 | 12 | 48 | 221 | 277 | 184 | 2010 | 55 | 0 | 36 | 0 | 0 | 0 | 1402 | 13635 | 201892 | 0 | 0 | 5809 | 479 | 0 | 0 | 501 | 18 | 90 | 454 | 179 | 0 | 179 | 0 | 0 | 0 | 0 | 5020 | 12 | 0 |
| SRR5756103 | 59 | 15 | 15 | 104 | 24 | 76 | 50 | 277 | 25 | 0 | 0 | 0 | 0 | 0 | 655 | 2323 | 21921 | 25 | 0 | 1664 | 403 | 0 | 0 | 89 | 37 | 74 | 30 | 164 | 0 | 186 | 0 | 50 | 0 | 0 | 2626 | 0 | 0 |
| SRR5756104 | 0 | 0 | 46 | 91 | 375 | 12 | 135 | 338 | 0 | 0 | 29 | 0 | 0 | 0 | 491 | 3624 | 15952 | 0 | 0 | 854 | 258 | 0 | 0 | 77 | 34 | 158 | 507 | 214 | 0 | 211 | 0 | 0 | 0 | 0 | 2100 | 0 | 0 |
| SRR5756105 | 53 | 40 | 37 | 13 | 287 | 162 | 98 | 307 | 14 | 0 | 30 | 0 | 0 | 0 | 117 | 1469 | 11318 | 0 | 0 | 610 | 393 | 0 | 0 | 12 | 17 | 270 | 18 | 13 | 0 | 210 | 0 | 0 | 0 | 0 | 2266 | 0 | 0 |
| SRR5756106 | 0 | 0 | 0 | 18 | 0 | 287 | 221 | 1524 | 22 | 0 | 27 | 0 | 0 | 0 | 2076 | 2860 | 262929 | 0 | 0 | 4042 | 1679 | 0 | 0 | 136 | 0 | 181 | 533 | 72 | 0 | 99 | 0 | 22 | 0 | 0 | 4167 | 0 | 0 |
| SRR5756107 | 0 | 0 | 27 | 76 | 15 | 90 | 126 | 404 | 0 | 0 | 20 | 0 | 0 | 0 | 2867 | 2816 | 17305 | 0 | 0 | 683 | 153 | 0 | 0 | 162 | 29 | 181 | 25 | 140 | 0 | 191 | 0 | 0 | 0 | 0 | 1760 | 0 | 0 |
| SRR5756108 | 26 | 0 | 41 | 212 | 207 | 28 | 0 | 141 | 0 | 0 | 26 | 0 | 0 | 0 | 159 | 4498 | 15437 | 0 | 0 | 1190 | 937 | 0 | 0 | 41 | 0 | 52 | 274 | 98 | 0 | 139 | 0 | 0 | 0 | 0 | 1981 | 0 | 0 |
| SRR5756110 | 0 | 0 | 0 | 35 | 142 | 414 | 0 | 276 | 0 | 0 | 106 | 0 | 0 | 0 | 1931 | 26242 | 18744 | 0 | 0 | 1517 | 0 | 0 | 0 | 71 | 35 | 35 | 778 | 0 | 0 | 71 | 0 | 0 | 0 | 0 | 625 | 0 | 0 |
| SRR5756114 | 0 | 0 | 0 | 47 | 71 | 85 | 170 | 255 | 0 | 0 | 166 | 0 | 0 | 0 | 5779 | 23343 | 46150 | 0 | 0 | 2380 | 85 | 0 | 0 | 71 | 47 | 71 | 356 | 0 | 0 | 95 | 0 | 0 | 0 | 0 | 537 | 0 | 0 |
| SRR5756115 | 0 | 0 | 0 | 0 | 105 | 110 | 0 | 221 | 110 | 0 | 246 | 0 | 0 | 0 | 3422 | 20180 | 28829 | 0 | 0 | 1766 | 0 | 0 | 0 | 70 | 0 | 0 | 211 | 0 | 0 | 316 | 0 | 0 | 0 | 0 | 360 | 0 | 0 |
| SRR5756116 | 0 | 0 | 0 | 0 | 406 | 0 | 252 | 252 | 0 | 0 | 339 | 0 | 0 | 0 | 2019 | 21210 | 14348 | 0 | 0 | 3280 | 0 | 0 | 0 | 68 | 135 | 0 | 609 | 0 | 0 | 203 | 0 | 0 | 0 | 0 | 0 | 0 | 0 |
| SRR5756118 | 0 | 0 | 0 | 0 | 43 | 0 | 0 | 0 | 0 | 0 | 85 | 0 | 0 | 0 | 670 | 15436 | 12679 | 0 | 0 | 893 | 223 | 0 | 0 | 171 | 0 | 85 | 342 | 0 | 0 | 385 | 0 | 0 | 0 | 0 | 551 | 0 | 0 |
| SRR5756120 | 0 | 0 | 0 | 0 | 163 | 110 | 110 | 0 | 0 | 0 | 244 | 0 | 0 | 0 | 3200 | 23722 | 35120 | 0 | 0 | 2979 | 221 | 0 | 0 | 136 | 108 | 81 | 353 | 0 | 0 | 434 | 0 | 0 | 0 | 0 | 1232 | 0 | 27 |
| SRR5756121 | 0 | 0 | 0 | 0 | 176 | 188 | 0 | 188 | 0 | 0 | 0 | 0 | 0 | 0 | 2442 | 11742 | 28180 | 0 | 0 | 2442 | 188 | 0 | 0 | 176 | 44 | 88 | 220 | 44 | 0 | 308 | 0 | 0 | 0 | 0 | 0 | 0 | 0 |
| SRR5756122 | 0 | 0 | 0 | 0 | 160 | 237 | 475 | 119 | 0 | 0 | 80 | 0 | 0 | 0 | 7717 | 18636 | 30190 | 0 | 0 | 2256 | 475 | 0 | 0 | 0 | 27 | 27 | 53 | 0 | 0 | 266 | 0 | 0 | 0 | 0 | 745 | 0 | 0 |
| SRR5756123 | 0 | 0 | 0 | 57 | 86 | 315 | 0 | 315 | 0 | 0 | 86 | 0 | 0 | 0 | 3679 | 21821 | 18621 | 0 | 0 | 2207 | 105 | 0 | 0 | 143 | 0 | 57 | 315 | 0 | 0 | 229 | 0 | 0 | 0 | 0 | 873 | 0 | 0 |
| SRR5756125 | 0 | 0 | 0 | 0 | 63 | 99 | 198 | 198 | 0 | 0 | 206 | 0 | 0 | 0 | 4265 | 34678 | 41613 | 0 | 0 | 3769 | 198 | 0 | 0 | 47 | 47 | 16 | 95 | 0 | 0 | 206 | 0 | 0 | 0 | 0 | 771 | 0 | 0 |
| SRR5756128 | 0 | 0 | 0 | 46 | 93 | 245 | 0 | 245 | 0 | 0 | 185 | 0 | 0 | 0 | 1467 | 29499 | 26057 | 0 | 0 | 1711 | 0 | 0 | 0 | 46 | 0 | 139 | 232 | 0 | 0 | 232 | 0 | 0 | 0 | 0 | 1967 | 0 | 0 |
| SRR5756130 | 0 | 0 | 0 | 0 | 23 | 0 | 132 | 132 | 0 | 0 | 90 | 0 | 0 | 0 | 4610 | 24131 | 49681 | 0 | 0 | 2634 | 263 | 0 | 0 | 23 | 23 | 23 | 158 | 0 | 0 | 226 | 0 | 0 | 0 | 0 | 710 | 0 | 0 |
| SRR5756131 | 26 | 0 | 0 | 0 | 77 | 0 | 0 | 170 | 0 | 0 | 102 | 0 | 0 | 0 | 4496 | 16539 | 36896 | 0 | 0 | 1782 | 170 | 0 | 0 | 0 | 51 | 26 | 204 | 0 | 0 | 179 | 0 | 0 | 0 | 0 | 763 | 0 | 0 |
| SRR5756132 | 0 | 0 | 0 | 0 | 119 | 0 | 122 | 122 | 122 | 0 | 59 | 0 | 0 | 0 | 2325 | 13693 | 28560 | 0 | 0 | 2080 | 122 | 0 | 0 | 30 | 0 | 0 | 89 | 0 | 0 | 178 | 0 | 0 | 0 | 0 | 782 | 0 | 0 |
| SRR5756133 | 0 | 0 | 0 | 0 | 109 | 0 | 121 | 0 | 0 | 0 | 146 | 0 | 0 | 0 | 4130 | 30941 | 26122 | 0 | 0 | 2187 | 121 | 0 | 0 | 109 | 182 | 109 | 473 | 0 | 0 | 291 | 0 | 0 | 0 | 0 | 761 | 0 | 0 |
| SRR5756135 | 0 | 0 | 0 | 37 | 110 | 0 | 0 | 597 | 398 | 0 | 184 | 0 | 0 | 0 | 4581 | 21039 | 36258 | 0 | 0 | 1992 | 199 | 0 | 0 | 110 | 0 | 147 | 294 | 0 | 0 | 331 | 0 | 0 | 0 | 0 | 448 | 0 | 0 |
| SRR5756136 | 0 | 36 | 0 | 0 | 217 | 0 | 160 | 321 | 0 | 0 | 398 | 0 | 0 | 0 | 5453 | 41787 | 46930 | 0 | 0 | 7378 | 160 | 0 | 0 | 36 | 0 | 36 | 578 | 36 | 0 | 325 | 0 | 0 | 0 | 0 | 1929 | 0 | 36 |
| SRR5756137 | 0 | 0 | 0 | 36 | 160 | 0 | 0 | 522 | 0 | 0 | 250 | 0 | 0 | 0 | 17223 | 41295 | 37981 | 0 | 0 | 5611 | 1044 | 0 | 0 | 107 | 71 | 232 | 891 | 53 | 0 | 517 | 0 | 0 | 0 | 0 | 510 | 0 | 0 |
| SRR5756138 | 0 | 0 | 0 | 0 | 172 | 0 | 0 | 417 | 0 | 0 | 258 | 0 | 0 | 0 | 7918 | 36614 | 72753 | 0 | 208 | 3959 | 208 | 0 | 0 | 229 | 86 | 229 | 601 | 29 | 0 | 401 | 0 | 0 | 0 | 0 | 1902 | 0 | 0 |
| SRR5756139 | 0 | 0 | 0 | 35 | 242 | 323 | 0 | 0 | 0 | 0 | 414 | 0 | 0 | 0 | 3394 | 26925 | 28062 | 0 | 0 | 3394 | 0 | 0 | 0 | 276 | 69 | 311 | 1450 | 69 | 0 | 345 | 0 | 0 | 0 | 0 | 0 | 0 | 0 |
| SRR5756140 | 0 | 0 | 29 | 29 | 116 | 0 | 0 | 213 | 0 | 29 | 260 | 0 | 0 | 0 | 2347 | 40469 | 45261 | 0 | 0 | 6187 | 427 | 0 | 0 | 174 | 203 | 405 | 695 | 58 | 0 | 232 | 0 | 0 | 0 | 0 | 1597 | 0 | 0 |
| SRR5756141 | 0 | 16 | 0 | 16 | 144 | 0 | 0 | 122 | 122 | 0 | 239 | 0 | 0 | 0 | 7175 | 46843 | 47732 | 0 | 0 | 4621 | 608 | 0 | 0 | 112 | 128 | 303 | 590 | 0 | 0 | 351 | 0 | 0 | 0 | 0 | 889 | 0 | 16 |
| SRR5756142 | 0 | 18 | 0 | 55 | 73 | 0 | 0 | 133 | 133 | 36 | 564 | 0 | 0 | 0 | 5991 | 35554 | 56054 | 0 | 0 | 4394 | 666 | 0 | 0 | 73 | 73 | 164 | 491 | 0 | 0 | 345 | 0 | 133 | 0 | 0 | 961 | 0 | 0 |
| SRR5756143 | 0 | 0 | 0 | 0 | 270 | 0 | 137 | 274 | 0 | 0 | 288 | 0 | 0 | 0 | 8903 | 36906 | 56483 | 0 | 0 | 4794 | 822 | 0 | 0 | 126 | 54 | 144 | 540 | 18 | 0 | 234 | 0 | 0 | 0 | 0 | 642 | 0 | 0 |
| SRR5756144 | 0 | 0 | 0 | 61 | 122 | 291 | 0 | 291 | 0 | 0 | 245 | 0 | 0 | 0 | 7863 | 35361 | 33460 | 0 | 0 | 5387 | 146 | 0 | 0 | 0 | 61 | 61 | 449 | 41 | 0 | 326 | 0 | 0 | 0 | 0 | 1141 | 0 | 0 |
| SRR5756145 | 0 | 0 | 0 | 76 | 153 | 229 | 0 | 229 | 0 | 0 | 153 | 0 | 0 | 0 | 2978 | 32761 | 26853 | 0 | 0 | 2978 | 0 | 0 | 0 | 76 | 76 | 153 | 841 | 38 | 0 | 459 | 0 | 0 | 0 | 0 | 1611 | 0 | 0 |
| SRR5756146 | 0 | 0 | 0 | 40 | 158 | 219 | 219 | 658 | 0 | 0 | 178 | 0 | 0 | 0 | 5487 | 41391 | 57616 | 0 | 0 | 5926 | 219 | 0 | 0 | 59 | 99 | 237 | 573 | 40 | 0 | 217 | 0 | 0 | 0 | 0 | 0 | 0 | 0 |
| SRR5756147 | 0 | 0 | 0 | 78 | 234 | 0 | 184 | 184 | 0 | 0 | 312 | 39 | 0 | 0 | 4415 | 47510 | 48271 | 0 | 0 | 4967 | 0 | 0 | 0 | 117 | 39 | 117 | 663 | 39 | 0 | 429 | 0 | 0 | 0 | 0 | 1140 | 0 | 0 |
| SRR5756148 | 0 | 0 | 0 | 0 | 162 | 192 | 192 | 0 | 0 | 0 | 270 | 0 | 0 | 0 | 6136 | 41015 | 28330 | 0 | 0 | 3835 | 192 | 0 | 0 | 162 | 162 | 162 | 1051 | 0 | 0 | 404 | 0 | 0 | 0 | 0 | 1691 | 0 | 0 |
| SRR5756149 | 0 | 0 | 0 | 38 | 302 | 0 | 391 | 391 | 195 | 0 | 264 | 0 | 0 | 0 | 9578 | 43195 | 59902 | 0 | 0 | 4887 | 0 | 0 | 0 | 113 | 76 | 189 | 453 | 113 | 0 | 189 | 0 | 0 | 0 | 0 | 2037 | 0 | 0 |
| SRR5756150 | 0 | 0 | 0 | 0 | 78 | 0 | 258 | 258 | 0 | 0 | 621 | 0 | 0 | 0 | 5796 | 27805 | 50000 | 0 | 0 | 6440 | 258 | 0 | 0 | 116 | 39 | 136 | 718 | 0 | 0 | 504 | 0 | 0 | 0 | 0 | 1707 | 0 | 0 |
| SRR5756151 | 0 | 0 | 0 | 0 | 306 | 0 | 0 | 0 | 0 | 0 | 153 | 0 | 0 | 0 | 4926 | 61399 | 45916 | 0 | 0 | 3519 | 235 | 0 | 0 | 38 | 0 | 306 | 536 | 77 | 0 | 268 | 0 | 0 | 0 | 0 | 1068 | 0 | 0 |
| SRR5756152 | 0 | 17 | 0 | 17 | 87 | 0 | 0 | 230 | 0 | 0 | 417 | 0 | 0 | 0 | 12647 | 60112 | 61976 | 0 | 0 | 4599 | 345 | 0 | 0 | 104 | 156 | 226 | 902 | 52 | 0 | 451 | 0 | 0 | 0 | 0 | 1165 | 0 | 0 |
| SRR5756153 | 0 | 0 | 0 | 0 | 118 | 0 | 0 | 316 | 158 | 0 | 260 | 0 | 0 | 0 | 3003 | 34388 | 33216 | 0 | 0 | 3635 | 0 | 0 | 0 | 213 | 71 | 213 | 828 | 95 | 0 | 307 | 0 | 158 | 0 | 0 | 782 | 0 | 0 |
| SRR5756154 | 45 | 0 | 0 | 90 | 181 | 0 | 201 | 403 | 0 | 0 | 226 | 0 | 0 | 0 | 3221 | 43111 | 28741 | 0 | 0 | 4227 | 201 | 0 | 0 | 181 | 136 | 181 | 588 | 0 | 0 | 678 | 0 | 201 | 0 | 0 | 423 | 0 | 0 |
| SRR5756155 | 0 | 0 | 0 | 0 | 56 | 0 | 0 | 136 | 0 | 0 | 181 | 0 | 0 | 0 | 11436 | 60237 | 122255 | 0 | 0 | 4084 | 136 | 0 | 0 | 70 | 28 | 112 | 377 | 0 | 0 | 223 | 0 | 0 | 0 | 0 | 0 | 0 | 0 |
| SRR5756156 | 0 | 0 | 0 | 41 | 185 | 137 | 137 | 137 | 137 | 0 | 350 | 0 | 0 | 0 | 6293 | 42199 | 49629 | 0 | 0 | 8208 | 274 | 0 | 0 | 62 | 144 | 267 | 412 | 62 | 0 | 514 | 0 | 137 | 0 | 0 | 892 | 0 | 21 |
| SRR5756157 | 0 | 0 | 0 | 19 | 95 | 0 | 144 | 288 | 0 | 0 | 228 | 0 | 0 | 0 | 3894 | 49017 | 38402 | 0 | 0 | 7356 | 577 | 0 | 0 | 95 | 19 | 114 | 494 | 190 | 0 | 475 | 0 | 0 | 0 | 0 | 937 | 0 | 0 |
| SRR5756158 | 0 | 0 | 23 | 45 | 90 | 0 | 0 | 194 | 194 | 0 | 294 | 0 | 0 | 0 | 3291 | 28293 | 36642 | 0 | 0 | 6196 | 194 | 0 | 0 | 158 | 181 | 294 | 1108 | 158 | 0 | 294 | 0 | 0 | 0 | 0 | 1855 | 0 | 0 |
| SRR5756159 | 0 | 0 | 0 | 0 | 84 | 0 | 1082 | 541 | 180 | 0 | 190 | 0 | 0 | 0 | 2706 | 45241 | 49936 | 0 | 0 | 5232 | 361 | 0 | 0 | 42 | 21 | 42 | 274 | 21 | 0 | 400 | 0 | 0 | 0 | 0 | 2561 | 0 | 0 |
| SRR5756160 | 0 | 0 | 0 | 0 | 53 | 231 | 0 | 926 | 0 | 0 | 53 | 0 | 0 | 0 | 3471 | 18725 | 26316 | 0 | 0 | 1157 | 0 | 0 | 0 | 53 | 53 | 53 | 0 | 53 | 0 | 53 | 0 | 0 | 0 | 0 | 2530 | 0 | 0 |
| SRR5756162 | 0 | 0 | 0 | 0 | 116 | 0 | 0 | 286 | 0 | 0 | 116 | 0 | 0 | 0 | 2862 | 21372 | 20248 | 0 | 0 | 2576 | 286 | 0 | 0 | 58 | 232 | 116 | 407 | 0 | 0 | 174 | 0 | 0 | 0 | 0 | 0 | 0 | 0 |
| SRR5756164 | 0 | 0 | 0 | 0 | 111 | 210 | 105 | 210 | 0 | 0 | 22 | 0 | 0 | 0 | 5563 | 19790 | 49165 | 0 | 0 | 1784 | 0 | 0 | 0 | 45 | 0 | 0 | 45 | 0 | 0 | 156 | 0 | 0 | 0 | 0 | 1237 | 0 | 0 |
| SRR5756166 | 0 | 0 | 0 | 0 | 75 | 96 | 0 | 0 | 0 | 0 | 100 | 0 | 0 | 0 | 3174 | 13402 | 49954 | 0 | 0 | 866 | 0 | 0 | 0 | 75 | 0 | 25 | 125 | 0 | 0 | 175 | 0 | 0 | 0 | 0 | 305 | 0 | 0 |
| SRR5756167 | 0 | 0 | 0 | 0 | 89 | 0 | 0 | 0 | 0 | 0 | 30 | 0 | 0 | 0 | 1296 | 22112 | 20453 | 0 | 0 | 648 | 0 | 0 | 0 | 30 | 30 | 89 | 148 | 0 | 0 | 30 | 0 | 0 | 0 | 0 | 1658 | 0 | 30 |
| SRR5756168 | 0 | 36 | 0 | 0 | 178 | 220 | 0 | 439 | 0 | 0 | 71 | 0 | 0 | 0 | 4613 | 11799 | 34907 | 0 | 0 | 2856 | 439 | 0 | 0 | 142 | 71 | 0 | 142 | 0 | 0 | 284 | 0 | 0 | 0 | 0 | 492 | 0 | 0 |
| SRR5756177 | 110 | 0 | 0 | 0 | 110 | 0 | 0 | 0 | 0 | 0 | 219 | 0 | 0 | 0 | 1189 | 11228 | 28443 | 0 | 0 | 1585 | 0 | 0 | 0 | 0 | 0 | 0 | 219 | 0 | 0 | 439 | 0 | 0 | 0 | 0 | 1497 | 0 | 0 |
| SRR5756178 | 0 | 0 | 89 | 89 | 0 | 392 | 0 | 1176 | 0 | 0 | 0 | 0 | 0 | 0 | 784 | 16461 | 13169 | 0 | 0 | 784 | 0 | 0 | 0 | 268 | 0 | 0 | 268 | 0 | 0 | 178 | 0 | 0 | 0 | 0 | 1646 | 0 | 0 |
| SRR5756179 | 0 | 0 | 0 | 0 | 248 | 0 | 0 | 0 | 0 | 0 | 83 | 0 | 0 | 0 | 1744 | 12262 | 15668 | 0 | 0 | 1744 | 349 | 0 | 0 | 165 | 0 | 248 | 330 | 0 | 0 | 165 | 0 | 0 | 0 | 0 | 681 | 0 | 0 |
| SRR5756180 | 0 | 0 | 0 | 0 | 139 | 115 | 115 | 0 | 0 | 0 | 139 | 0 | 0 | 0 | 6446 | 19499 | 52678 | 0 | 0 | 1727 | 0 | 0 | 0 | 56 | 0 | 0 | 167 | 0 | 0 | 223 | 0 | 0 | 0 | 0 | 0 | 0 | 0 |
| SRR5756182 | 0 | 0 | 0 | 0 | 83 | 0 | 0 | 0 | 0 | 0 | 0 | 0 | 0 | 0 | 1667 | 21295 | 10222 | 0 | 0 | 1250 | 833 | 0 | 0 | 0 | 0 | 83 | 0 | 0 | 0 | 83 | 0 | 0 | 0 | 0 | 852 | 0 | 0 |
| SRR5756183 | 11 | 0 | 11 | 34 | 91 | 181 | 271 | 316 | 0 | 0 | 125 | 0 | 0 | 0 | 6593 | 44268 | 60677 | 0 | 0 | 2529 | 181 | 0 | 0 | 148 | 34 | 136 | 739 | 0 | 0 | 171 | 0 | 0 | 0 | 0 | 583 | 0 | 11 |
| SRR5756184 | 0 | 0 | 0 | 30 | 60 | 127 | 0 | 0 | 0 | 0 | 60 | 0 | 0 | 0 | 2658 | 21457 | 28386 | 0 | 0 | 2405 | 253 | 0 | 0 | 0 | 60 | 90 | 448 | 30 | 0 | 209 | 0 | 0 | 0 | 0 | 447 | 0 | 30 |
| SRR5756185 | 0 | 0 | 0 | 70 | 226 | 0 | 0 | 67 | 0 | 0 | 296 | 0 | 0 | 0 | 7650 | 25566 | 52913 | 0 | 0 | 2794 | 133 | 0 | 0 | 52 | 35 | 52 | 174 | 35 | 0 | 209 | 0 | 0 | 0 | 0 | 485 | 0 | 0 |
| SRR5756186 | 0 | 0 | 0 | 82 | 197 | 63 | 0 | 63 | 0 | 16 | 377 | 0 | 0 | 0 | 8117 | 26300 | 57588 | 0 | 0 | 2769 | 315 | 0 | 0 | 33 | 16 | 33 | 180 | 16 | 0 | 295 | 0 | 0 | 0 | 0 | 151 | 0 | 0 |
| SRR5804909 | 0 | 0 | 0 | 0 | 0 | 0 | 0 | 0 | 73 | 3973 | 15 | 51 | 0 | 0 | 36 | 1395 | 688 | 36 | 109 | 0 | 292 | 0 | 0 | 0 | 0 | 0 | 0 | 0 | 0 | 0 | 0 | 437 | 0 | 0 | 320154 | 0 | 0 |
| SRR5804912 | 0 | 0 | 0 | 0 | 0 | 315 | 0 | 189 | 566 | 5198 | 23 | 0 | 0 | 0 | 220 | 5412 | 5224 | 0 | 63 | 252 | 283 | 0 | 0 | 0 | 0 | 0 | 0 | 0 | 0 | 11 | 0 | 692 | 0 | 0 | 110432 | 0 | 0 |
| SRR5804914 | 0 | 0 | 0 | 0 | 0 | 0 | 0 | 0 | 339 | 39951 | 30 | 22 | 0 | 0 | 0 | 7808 | 4505 | 0 | 0 | 0 | 1697 | 0 | 0 | 0 | 0 | 0 | 0 | 0 | 0 | 0 | 0 | 3167 | 0 | 0 | 27327 | 0 | 0 |
| SRR5817623 | 24 | 87 | 155 | 52 | 66 | 2291 | 1128 | 1088 | 930 | 0 | 22 | 0 | 0 | 0 | 28237 | 10912 | 10931 | 4635 | 150 | 2353 | 158 | 0 | 0 | 16 | 84 | 12 | 46 | 17 | 0 | 206 | 0 | 56 | 0 | 0 | 8336 | 2647 | 0 |
| SRR5817624 | 15 | 71 | 121 | 48 | 85 | 4214 | 1352 | 3713 | 1185 | 0 | 28 | 0 | 0 | 0 | 55222 | 11927 | 16576 | 3303 | 147 | 1906 | 382 | 0 | 0 | 20 | 155 | 13 | 42 | 174 | 0 | 216 | 0 | 60 | 0 | 0 | 7521 | 2757 | 0 |

### Supp Table S8.3.Columns_BX_to_DH

| | | | | | | | | | | | | | | | | | | | | | | | | | | | | | | | | | | | | | |
|---|---|---|---|---|---|---|---|---|---|---|---|---|---|---|---|---|---|---|---|---|---|---|---|---|---|---|---|---|---|---|---|---|---|---|---|---|---|
| SRR5817625 | 398 | 82 | 154 | 38 | 1273 | 2015 | 1051 | 983 | 805 | 0 | 309 | 0 | 0 | 0 | 23477 | 10707 | 10693 | 4318 | 116 | 1344 | 165 | 0 | 0 | 291 | 87 | 243 | 944 | 641 | 0 | 3039 | 0 | 35 | 0 | 0 | 7694 | 2244 | 0 |
| SRR5817626 | 324 | 61 | 133 | 58 | 1486 | 2645 | 1014 | 1763 | 882 | 0 | 551 | 0 | 0 | 0 | 39781 | 12185 | 14678 | 3316 | 132 | 1929 | 479 | 0 | 0 | 277 | 87 | 248 | 1006 | 382 | 0 | 4051 | 0 | 61 | 0 | 0 | 8269 | 2317 | 0 |
| SRR5821803 | 11 | 0 | 0 | 0 | 52 | 120 | 36 | 201 | 0 | 0 | 101 | 0 | 0 | 0 | 4597 | 12504 | 92840 | 0 | 0 | 1106 | 419 | 0 | 0 | 40 | 28 | 14 | 34 | 0 | 0 | 398 | 0 | 18 | 0 | 0 | 7359 | 18 | 0 |
| SRR5821804 | 0 | 0 | 0 | 0 | 14 | 214 | 515 | 411 | 418 | 15 | 0 | 0 | 0 | 0 | 6168 | 36040 | 194918 | 52 | 42 | 2441 | 1470 | 0 | 0 | 64 | 0 | 0 | 21 | 0 | 0 | 0 | 0 | 23 | 0 | 0 | 549 | 21 | 11 |
| SRR5821805 | 116 | 183 | 0 | 22 | 102 | 42 | 56 | 108 | 10 | 0 | 0 | 0 | 0 | 0 | 1106 | 3176 | 10811 | 0 | 0 | 574 | 379 | 0 | 0 | 132 | 59 | 189 | 570 | 18 | 0 | 47 | 0 | 0 | 0 | 0 | 707 | 11 | 0 |
| SRR5832003 | 0 | 37 | 74 | 37 | 37 | 0 | 855 | 896 | 285 | 0 | 37 | 0 | 0 | 0 | 4641 | 67292 | 89244 | 0 | 407 | 18359 | 13352 | 0 | 0 | 185 | 259 | 370 | 3293 | 1813 | 0 | 148 | 0 | 163 | 0 | 0 | 507 | 0 | 0 |
| SRR5832004 | 0 | 0 | 0 | 94 | 31 | 0 | 230 | 642 | 214 | 0 | 0 | 0 | 0 | 0 | 5350 | 58265 | 215386 | 0 | 1185 | 46060 | 42356 | 0 | 31 | 94 | 15 | 52 | 272 | 171 | 0 | 19 | 0 | 66 | 0 | 0 | 294 | 0 | 0 |
| SRR5832005 | 0 | 0 | 14 | 0 | 14 | 0 | 172 | 794 | 193 | 0 | 14 | 0 | 0 | 0 | 8773 | 96940 | 148392 | 21 | 686 | 25010 | 31616 | 0 | 56 | 42 | 126 | 140 | 993 | 462 | 0 | 280 | 0 | 579 | 0 | 0 | 262 | 0 | 0 |
| SRR5832006 | 0 | 0 | 0 | 172 | 0 | 0 | 428 | 761 | 333 | 0 | 0 | 0 | 0 | 0 | 7278 | 38409 | 164282 | 0 | 190 | 26878 | 6279 | 0 | 0 | 172 | 0 | 0 | 86 | 172 | 0 | 86 | 0 | 95 | 0 | 0 | 238 | 0 | 0 |
| SRR5832007 | 42 | 0 | 0 | 0 | 0 | 0 | 0 | 31 | 23 | 0 | 0 | 0 | 0 | 0 | 220196 | 52832 | 348336 | 0 | 39 | 3266 | 40247 | 0 | 0 | 210 | 42 | 252 | 35 | 14 | 0 | 84 | 0 | 23 | 0 | 0 | 21 | 0 | 0 |
| SRR5832008 | 12 | 25 | 0 | 25 | 0 | 0 | 161 | 536 | 54 | 12 | 25 | 0 | 0 | 0 | 18990 | 52004 | 207189 | 36 | 786 | 41553 | 26029 | 12 | 61 | 37 | 25 | 11 | 51 | 40 | 0 | 29 | 0 | 286 | 0 | 0 | 63 | 0 | 0 |
| SRR5832009 | 0 | 36 | 0 | 0 | 0 | 0 | 215 | 549 | 48 | 0 | 107 | 0 | 0 | 0 | 26804 | 36117 | 233451 | 0 | 740 | 21410 | 20861 | 0 | 0 | 143 | 179 | 32 | 146 | 133 | 0 | 143 | 0 | 95 | 0 | 0 | 195 | 0 | 0 |
| SRR5832010 | 0 | 0 | 0 | 28 | 0 | 0 | 412 | 675 | 75 | 0 | 84 | 0 | 0 | 0 | 14321 | 72544 | 123765 | 0 | 787 | 19682 | 21669 | 0 | 0 | 84 | 28 | 307 | 2709 | 1731 | 0 | 391 | 0 | 375 | 0 | 0 | 319 | 0 | 0 |
| SRR5832011 | 0 | 0 | 0 | 58 | 0 | 0 | 197 | 278 | 58 | 0 | 0 | 0 | 0 | 0 | 7461 | 101888 | 234480 | 0 | 1008 | 72360 | 13508 | 0 | 0 | 174 | 58 | 37 | 208 | 161 | 0 | 174 | 0 | 0 | 0 | 0 | 433 | 0 | 0 |
| SRR5832012 | 0 | 0 | 65 | 0 | 0 | 0 | 393 | 463 | 84 | 0 | 0 | 0 | 0 | 0 | 5463 | 98093 | 177878 | 14 | 786 | 62324 | 9367 | 0 | 0 | 195 | 130 | 1430 | 7216 | 4811 | 0 | 195 | 0 | 28 | 0 | 0 | 207 | 0 | 0 |
| SRR5832013 | 0 | 42 | 0 | 167 | 42 | 0 | 336 | 865 | 288 | 0 | 0 | 0 | 0 | 0 | 8363 | 51420 | 145371 | 0 | 529 | 21773 | 20090 | 0 | 0 | 293 | 418 | 1130 | 4352 | 3557 | 0 | 209 | 0 | 48 | 0 | 0 | 1193 | 0 | 0 |
| SRR5832014 | 0 | 122 | 0 | 61 | 61 | 0 | 362 | 742 | 305 | 0 | 0 | 0 | 0 | 0 | 7499 | 61160 | 163193 | 0 | 1389 | 36010 | 13247 | 0 | 0 | 243 | 61 | 27 | 133 | 121 | 0 | 243 | 0 | 57 | 0 | 0 | 632 | 0 | 0 |
| SRR5832015 | 0 | 37 | 0 | 0 | 0 | 0 | 471 | 993 | 248 | 0 | 37 | 0 | 0 | 0 | 6352 | 101329 | 163942 | 25 | 596 | 37616 | 54687 | 0 | 110 | 147 | 183 | 404 | 2642 | 1541 | 0 | 330 | 0 | 99 | 0 | 0 | 62 | 0 | 0 |
| SRR5832016 | 29 | 0 | 0 | 29 | 29 | 0 | 492 | 777 | 111 | 0 | 29 | 0 | 0 | 0 | 27749 | 39865 | 254905 | 16 | 1349 | 50944 | 35221 | 0 | 86 | 143 | 143 | 1144 | 2802 | 1916 | 0 | 229 | 0 | 48 | 0 | 0 | 71 | 0 | 0 |
| SRR5832017 | 0 | 0 | 50 | 0 | 201 | 0 | 140 | 149 | 23 | 0 | 50 | 0 | 0 | 0 | 21390 | 141932 | 354476 | 0 | 467 | 70168 | 16555 | 0 | 0 | 453 | 0 | 0 | 101 | 0 | 0 | 201 | 0 | 0 | 0 | 0 | 72 | 19 | 0 |
| SRR5832018 | 0 | 25 | 0 | 25 | 25 | 0 | 765 | 1129 | 517 | 0 | 0 | 0 | 0 | 0 | 4832 | 87509 | 160882 | 0 | 1033 | 66946 | 7769 | 0 | 0 | 499 | 275 | 1472 | 10532 | 4817 | 0 | 200 | 0 | 0 | 0 | 0 | 346 | 0 | 0 |
| SRR5832019 | 36 | 36 | 0 | 146 | 109 | 0 | 1465 | 1966 | 100 | 0 | 36 | 0 | 0 | 0 | 29222 | 97034 | 278596 | 25 | 1527 | 98822 | 1377 | 0 | 0 | 29 | 255 | 38 | 155 | 150 | 0 | 109 | 0 | 0 | 0 | 0 | 370 | 16 | 0 |
| SRR5832020 | 0 | 31 | 0 | 63 | 0 | 0 | 453 | 600 | 232 | 31 | 94 | 0 | 0 | 0 | 5438 | 82567 | 179793 | 0 | 801 | 71273 | 6998 | 0 | 0 | 281 | 125 | 876 | 5661 | 2596 | 0 | 188 | 0 | 18 | 0 | 0 | 256 | 0 | 0 |
| SRR5832021 | 0 | 37 | 0 | 56 | 56 | 0 | 492 | 1118 | 693 | 0 | 130 | 37 | 0 | 0 | 9908 | 82531 | 129406 | 0 | 537 | 28002 | 15768 | 0 | 0 | 223 | 56 | 577 | 2772 | 949 | 0 | 205 | 0 | 760 | 0 | 0 | 620 | 0 | 0 |
| SRR5832022 | 0 | 0 | 0 | 0 | 0 | 0 | 63 | 145 | 19 | 104 | 26 | 0 | 0 | 0 | 128227 | 92123 | 304587 | 0 | 511 | 30688 | 57241 | 0 | 0 | 156 | 32 | 67 | 73 | 35 | 0 | 52 | 0 | 32 | 0 | 0 | 55 | 0 | 0 |
| SRR5832023 | 0 | 0 | 22 | 22 | 65 | 0 | 602 | 1685 | 378 | 44 | 44 | 0 | 0 | 0 | 9733 | 73958 | 149762 | 0 | 619 | 42286 | 29956 | 65 | 44 | 131 | 196 | 674 | 3371 | 1414 | 0 | 283 | 0 | 206 | 0 | 0 | 432 | 0 | 0 |
| SRR5832024 | 0 | 0 | 0 | 0 | 56 | 0 | 1040 | 1004 | 73 | 0 | 0 | 0 | 0 | 0 | 64795 | 22661 | 292343 | 18 | 529 | 47364 | 1405 | 0 | 0 | 447 | 279 | 1061 | 4298 | 2512 | 0 | 56 | 0 | 0 | 0 | 0 | 231 | 0 | 0 |
| SRR5832025 | 0 | 0 | 0 | 11 | 11 | 0 | 514 | 749 | 214 | 11 | 102 | 0 | 0 | 0 | 7044 | 85412 | 136868 | 21 | 364 | 31536 | 20767 | 23 | 0 | 23 | 57 | 193 | 861 | 680 | 0 | 419 | 0 | 728 | 0 | 0 | 294 | 0 | 0 |
| SRR5832026 | 0 | 16 | 0 | 16 | 0 | 0 | 236 | 570 | 194 | 16 | 64 | 0 | 0 | 0 | 14045 | 50961 | 214495 | 0 | 903 | 64863 | 44010 | 64 | 81 | 32 | 11 | 38 | 162 | 74 | 0 | 36 | 0 | 69 | 0 | 0 | 148 | 0 | 0 |
| SRR5832027 | 0 | 0 | 33 | 33 | 65 | 0 | 1063 | 2105 | 797 | 0 | 33 | 0 | 0 | 0 | 8238 | 39196 | 144622 | 20 | 368 | 32869 | 12040 | 0 | 0 | 294 | 98 | 1697 | 8811 | 1468 | 0 | 294 | 0 | 82 | 0 | 0 | 305 | 0 | 0 |
| SRR5832028 | 14 | 14 | 0 | 0 | 0 | 0 | 846 | 1582 | 478 | 27 | 14 | 0 | 0 | 0 | 9731 | 39103 | 154107 | 18 | 699 | 28805 | 39308 | 41 | 68 | 136 | 68 | 218 | 2698 | 286 | 0 | 422 | 0 | 221 | 0 | 0 | 233 | 0 | 0 |
| SRR5832029 | 0 | 0 | 0 | 0 | 98 | 0 | 984 | 1200 | 252 | 0 | 49 | 0 | 0 | 0 | 30090 | 29201 | 293707 | 0 | 528 | 57060 | 5279 | 0 | 0 | 14 | 295 | 22 | 110 | 15 | 0 | 197 | 0 | 0 | 0 | 0 | 202 | 0 | 0 |
| SRR5868085 | 6516 | 1077 | 148 | 107 | 1942 | 4185 | 285 | 2025 | 7326 | 7952 | 366 | 13 | 0 | 0 | 10269 | 102597 | 41866 | 119 | 910 | 1171 | 11765 | 0 | 0 | 1172 | 76 | 78 | 54 | 0 | 0 | 469 | 0 | 214 | 0 | 13 | 9580 | 609 | 0 |
| SRR5901584 | 27 | 24 | 0 | 20 | 71 | 1017 | 244 | 1012 | 137 | 27 | 39 | 0 | 0 | 0 | 8856 | 3212 | 28810 | 147 | 513 | 15583 | 589 | 0 | 0 | 13 | 20 | 19 | 49 | 0 | 0 | 340 | 0 | 15 | 0 | 0 | 1509 | 0 | 0 |
| SRR5901585 | 22 | 0 | 0 | 14 | 56 | 760 | 148 | 848 | 103 | 42 | 19 | 0 | 0 | 0 | 8534 | 3389 | 27474 | 90 | 415 | 15923 | 705 | 0 | 0 | 15 | 0 | 14 | 33 | 0 | 0 | 104 | 0 | 25 | 0 | 0 | 1840 | 0 | 0 |
| SRR5901586 | 30 | 26 | 18 | 20 | 57 | 844 | 260 | 927 | 137 | 47 | 27 | 0 | 0 | 0 | 8680 | 7424 | 31906 | 92 | 715 | 17434 | 727 | 0 | 0 | 13 | 0 | 13 | 27 | 0 | 0 | 205 | 0 | 14 | 0 | 0 | 2161 | 0 | 0 |
| SRR5901587 | 32 | 15 | 0 | 0 | 346 | 1069 | 284 | 1155 | 187 | 557 | 187 | 0 | 0 | 0 | 7876 | 2991 | 24777 | 99 | 187 | 8398 | 614 | 0 | 0 | 134 | 0 | 0 | 21 | 0 | 0 | 2993 | 0 | 18 | 0 | 0 | 2224 | 11 | 0 |
| SRR5901588 | 36 | 15 | 0 | 0 | 376 | 945 | 245 | 962 | 177 | 747 | 186 | 0 | 0 | 0 | 9326 | 2280 | 23378 | 125 | 332 | 9939 | 688 | 0 | 0 | 176 | 14 | 0 | 29 | 0 | 0 | 2556 | 0 | 26 | 0 | 0 | 1713 | 14 | 0 |
| SRR5901589 | 28 | 15 | 0 | 0 | 34 | 1099 | 357 | 1417 | 210 | 72 | 21 | 0 | 15 | 0 | 8348 | 2890 | 25422 | 72 | 171 | 10085 | 622 | 0 | 0 | 18 | 13 | 0 | 16 | 0 | 0 | 314 | 0 | 29 | 0 | 0 | 2188 | 12 | 0 |
| SRR5901590 | 59 | 34 | 0 | 0 | 0 | 1000 | 162 | 756 | 129 | 34 | 12 | 0 | 13 | 0 | 4187 | 532 | 4745 | 201 | 232 | 62380 | 59 | 0 | 0 | 81 | 0 | 0 | 0 | 0 | 0 | 140 | 0 | 10 | 0 | 0 | 1300 | 0 | 0 |
| SRR5901591 | 64 | 11 | 0 | 0 | 0 | 711 | 95 | 564 | 73 | 405 | 123 | 0 | 0 | 0 | 3650 | 692 | 5838 | 275 | 192 | 75855 | 62 | 0 | 0 | 99 | 0 | 23 | 25 | 0 | 0 | 929 | 0 | 17 | 0 | 0 | 1153 | 0 | 0 |
| SRR5901592 | 27 | 17 | 0 | 0 | 0 | 499 | 65 | 387 | 43 | 74 | 50 | 0 | 0 | 0 | 1660 | 431 | 3410 | 287 | 251 | 89584 | 45 | 0 | 0 | 59 | 0 | 0 | 0 | 0 | 0 | 112 | 0 | 0 | 0 | 0 | 1137 | 0 | 0 |
| SRR5901593 | 50 | 17 | 0 | 0 | 0 | 467 | 157 | 560 | 115 | 13 | 16 | 0 | 24 | 0 | 3152 | 657 | 4492 | 136 | 184 | 100020 | 50 | 0 | 0 | 124 | 0 | 0 | 0 | 0 | 0 | 256 | 0 | 17 | 0 | 0 | 2092 | 11 | 0 |
| SRR5901594 | 34 | 21 | 0 | 0 | 0 | 357 | 94 | 333 | 108 | 18 | 14 | 0 | 0 | 0 | 2387 | 578 | 3767 | 183 | 162 | 104182 | 44 | 0 | 0 | 82 | 0 | 0 | 0 | 0 | 0 | 191 | 0 | 13 | 0 | 0 | 841 | 0 | 0 |
| SRR5901595 | 29 | 0 | 0 | 0 | 0 | 401 | 140 | 498 | 136 | 18 | 18 | 0 | 35 | 0 | 1616 | 541 | 2307 | 163 | 226 | 96868 | 63 | 0 | 0 | 83 | 0 | 0 | 0 | 0 | 0 | 239 | 0 | 0 | 0 | 0 | 942 | 0 | 0 |
| SRR5922753 | 159 | 103 | 18 | 16 | 1412 | 3088 | 3830 | 6214 | 10735 | 0 | 31 | 0 | 0 | 0 | 11594 | 132163 | 96714 | 432 | 955 | 7978 | 193 | 0 | 0 | 3548 | 748 | 691 | 657 | 14 | 0 | 286 | 0 | 0 | 0 | 0 | 4728 | 12 | 11 |
| SRR5922754 | 21 | 16 | 0 | 0 | 11 | 12564 | 6535 | 13133 | 15173 | 0 | 0 | 0 | 0 | 0 | 4907 | 32388 | 30246 | 374 | 265 | 2212 | 70 | 0 | 0 | 79 | 77 | 47 | 45 | 13 | 0 | 16 | 0 | 0 | 0 | 0 | 6893 | 12 | 0 |
| SRR5922756 | 0 | 0 | 0 | 0 | 30 | 29969 | 5206 | 12368 | 10033 | 0 | 0 | 0 | 0 | 0 | 734 | 6870 | 10871 | 6165 | 89 | 641 | 98 | 263 | 45 | 95 | 0 | 0 | 0 | 0 | 0 | 11 | 131 | 61 | 0 | 0 | 17302 | 14 | 18 |
| SRR5925650 | 0 | 0 | 0 | 12 | 11 | 0 | 0 | 108 | 0 | 0 | 19 | 0 | 0 | 0 | 1146 | 7312 | 4637 | 254 | 0 | 79 | 223 | 0 | 0 | 45 | 0 | 21 | 17 | 30 | 0 | 82 | 0 | 7281 | 19 | 0 | 3341 | 48 | 53 |
| SRR5925651 | 0 | 0 | 0 | 0 | 15 | 24 | 37 | 73 | 13 | 0 | 12 | 0 | 0 | 0 | 3807 | 1766 | 2998 | 227 | 0 | 253 | 546 | 23 | 0 | 33 | 0 | 30 | 34 | 21 | 0 | 16 | 19 | 6656 | 24 | 0 | 11117 | 92 | 221 |
| SRR5925652 | 27 | 34 | 18 | 28 | 33 | 543 | 430 | 1168 | 209 | 17 | 114 | 18 | 0 | 0 | 23641 | 6806 | 21823 | 83 | 226 | 3699 | 7584 | 0 | 0 | 282 | 125 | 251 | 761 | 68 | 0 | 685 | 0 | 588 | 0 | 0 | 3927 | 0 | 20 |
| SRR5925665 | 134 | 114 | 99 | 74 | 40 | 385 | 554 | 2662 | 72 | 0 | 15 | 0 | 0 | 0 | 38341 | 36017 | 59883 | 0 | 60 | 2180 | 2493 | 0 | 0 | 253 | 327 | 1233 | 2338 | 2427 | 0 | 79 | 0 | 0 | 0 | 0 | 6027 | 307 | 0 |
| SRR5925666 | 116 | 74 | 82 | 70 | 35 | 390 | 466 | 2383 | 42 | 35 | 27 | 0 | 0 | 0 | 26127 | 35287 | 50041 | 0 | 42 | 1577 | 1815 | 0 | 0 | 16 | 18 | 65 | 113 | 131 | 0 | 109 | 0 | 0 | 0 | 0 | 5905 | 296 | 0 |
| SRR5925667 | 16 | 12 | 134 | 130 | 49 | 483 | 831 | 2739 | 79 | 0 | 28 | 0 | 0 | 0 | 35208 | 31690 | 56219 | 0 | 67 | 1448 | 2346 | 0 | 0 | 16 | 28 | 70 | 120 | 171 | 0 | 130 | 0 | 0 | 0 | 0 | 6497 | 268 | 0 |
| SRR5925668 | 223 | 129 | 126 | 105 | 21 | 317 | 765 | 2354 | 59 | 0 | 18 | 0 | 0 | 0 | 35455 | 35079 | 52442 | 0 | 53 | 1556 | 2255 | 0 | 0 | 242 | 271 | 1012 | 1453 | 2617 | 0 | 110 | 0 | 0 | 0 | 0 | 5703 | 351 | 0 |
| SRR5925669 | 12 | 135 | 69 | 118 | 82 | 255 | 705 | 2831 | 61 | 24 | 0 | 0 | 0 | 0 | 36550 | 28312 | 50143 | 0 | 24 | 1203 | 2673 | 0 | 0 | 13 | 16 | 65 | 112 | 192 | 0 | 69 | 0 | 0 | 0 | 0 | 6661 | 333 | 0 |
| SRR5925670 | 202 | 118 | 105 | 114 | 35 | 333 | 689 | 2653 | 80 | 44 | 22 | 0 | 0 | 0 | 35170 | 32651 | 55345 | 0 | 57 | 1252 | 2102 | 0 | 0 | 219 | 307 | 929 | 1586 | 2721 | 0 | 96 | 0 | 0 | 0 | 0 | 5808 | 316 | 0 |
| SRR5925671 | 126 | 180 | 134 | 172 | 76 | 287 | 766 | 2115 | 80 | 23 | 23 | 0 | 0 | 0 | 27259 | 32498 | 53483 | 0 | 32 | 1772 | 1556 | 0 | 0 | 386 | 275 | 910 | 1647 | 3416 | 0 | 107 | 0 | 0 | 0 | 0 | 5219 | 344 | 0 |
| SRR5925672 | 217 | 160 | 103 | 189 | 107 | 342 | 713 | 2396 | 57 | 0 | 0 | 0 | 0 | 0 | 24237 | 46911 | 47581 | 0 | 29 | 1274 | 1426 | 0 | 0 | 344 | 271 | 1029 | 1985 | 4096 | 0 | 127 | 0 | 0 | 0 | 0 | 6064 | 316 | 0 |
| SRR5925673 | 118 | 59 | 127 | 121 | 47 | 760 | 1066 | 5320 | 30 | 0 | 29 | 0 | 0 | 0 | 29304 | 24795 | 51258 | 0 | 20 | 1244 | 2823 | 0 | 0 | 20 | 23 | 80 | 149 | 165 | 0 | 121 | 0 | 0 | 0 | 0 | 7287 | 223 | 0 |
| SRR5925674 | 118 | 89 | 111 | 52 | 59 | 985 | 1313 | 4837 | 109 | 0 | 37 | 0 | 0 | 0 | 33376 | 37198 | 57227 | 0 | 0 | 1007 | 2429 | 0 | 0 | 20 | 12 | 61 | 124 | 161 | 0 | 148 | 0 | 0 | 0 | 0 | 8165 | 327 | 0 |
| SRR5925675 | 23 | 35 | 73 | 67 | 20 | 692 | 982 | 5524 | 112 | 0 | 26 | 0 | 0 | 0 | 54090 | 47642 | 89445 | 0 | 45 | 2500 | 3069 | 0 | 0 | 21 | 19 | 95 | 181 | 225 | 0 | 14 | 0 | 0 | 0 | 0 | 5214 | 224 | 0 |
| SRR5925676 | 77 | 44 | 142 | 101 | 59 | 620 | 772 | 3186 | 129 | 0 | 12 | 0 | 0 | 0 | 50894 | 48020 | 75011 | 0 | 24 | 1408 | 2309 | 0 | 0 | 16 | 16 | 71 | 150 | 166 | 0 | 104 | 0 | 0 | 0 | 0 | 7877 | 240 | 0 |
| SRR5925677 | 94 | 103 | 80 | 120 | 62 | 260 | 736 | 2122 | 22 | 0 | 0 | 0 | 0 | 0 | 23447 | 40985 | 37416 | 0 | 22 | 823 | 1635 | 0 | 0 | 13 | 25 | 84 | 154 | 164 | 0 | 112 | 0 | 0 | 0 | 0 | 6531 | 311 | 0 |
| SRR5925678 | 118 | 111 | 108 | 128 | 64 | 275 | 542 | 1981 | 80 | 0 | 30 | 0 | 0 | 0 | 24142 | 47058 | 39687 | 18 | 27 | 835 | 1875 | 0 | 0 | 13 | 35 | 105 | 193 | 174 | 0 | 98 | 0 | 0 | 0 | 0 | 6620 | 193 | 0 |
| SRR5925679 | 100 | 100 | 77 | 169 | 65 | 718 | 1269 | 3904 | 187 | 0 | 31 | 0 | 0 | 0 | 21427 | 44472 | 42848 | 0 | 59 | 1111 | 1986 | 0 | 0 | 20 | 20 | 82 | 141 | 181 | 0 | 61 | 0 | 0 | 0 | 0 | 6990 | 315 | 0 |
| SRR5925680 | 126 | 93 | 102 | 126 | 50 | 551 | 1028 | 2953 | 142 | 0 | 36 | 0 | 0 | 0 | 22341 | 46677 | 43301 | 0 | 43 | 960 | 1653 | 0 | 0 | 288 | 329 | 1159 | 1962 | 3012 | 0 | 105 | 0 | 0 | 0 | 0 | 6000 | 310 | 0 |
| SRR5925681 | 117 | 176 | 130 | 96 | 42 | 313 | 555 | 2264 | 71 | 25 | 17 | 0 | 0 | 0 | 21214 | 27121 | 42497 | 11 | 27 | 1270 | 1303 | 0 | 0 | 654 | 323 | 1307 | 1709 | 3858 | 0 | 163 | 0 | 0 | 0 | 0 | 5667 | 346 | 0 |
| SRR5925682 | 138 | 150 | 65 | 92 | 46 | 315 | 574 | 1756 | 28 | 0 | 12 | 0 | 0 | 0 | 30250 | 36316 | 50961 | 0 | 42 | 1232 | 1211 | 0 | 0 | 22 | 18 | 66 | 85 | 200 | 0 | 100 | 0 | 0 | 0 | 0 | 6111 | 357 | 0 |
| SRR5925683 | 143 | 170 | 72 | 130 | 80 | 293 | 775 | 2173 | 19 | 49 | 18 | 0 | 0 | 0 | 29035 | 30565 | 50415 | 0 | 0 | 1389 | 2277 | 0 | 0 | 16 | 16 | 75 | 127 | 211 | 0 | 103 | 0 | 0 | 0 | 0 | 4568 | 327 | 0 |
| SRR5925684 | 13 | 109 | 109 | 103 | 52 | 302 | 560 | 2094 | 45 | 67 | 10 | 0 | 0 | 0 | 36883 | 36279 | 64992 | 0 | 11 | 2744 | 1949 | 0 | 0 | 21 | 13 | 66 | 82 | 148 | 0 | 67 | 0 | 0 | 0 | 0 | 5217 | 398 | 0 |
| SRR5925685 | 82 | 153 | 99 | 167 | 71 | 215 | 322 | 776 | 12 | 21 | 89 | 0 | 0 | 0 | 96447 | 15165 | 131860 | 0 | 84 | 1093 | 49543 | 0 | 0 | 15 | 13 | 20 | 20 | 29 | 0 | 39 | 0 | 0 | 0 | 0 | 2438 | 0 | 0 |
| SRR5925686 | 79 | 110 | 102 | 151 | 53 | 227 | 246 | 887 | 13 | 19 | 95 | 0 | 0 | 0 | 84011 | 13875 | 118956 | 0 | 58 | 1049 | 44107 | 0 | 0 | 250 | 257 | 371 | 435 | 685 | 0 | 791 | 0 | 0 | 0 | 0 | 2467 | 0 | 0 |
| SRR5925687 | 79 | 118 | 74 | 144 | 39 | 196 | 98 | 562 | 81 | 26 | 96 | 0 | 0 | 0 | 77486 | 10313 | 105530 | 0 | 57 | 2493 | 50492 | 0 | 0 | 166 | 284 | 520 | 485 | 948 | 0 | 874 | 0 | 0 | 0 | 0 | 2522 | 0 | 0 |
| SRR5925688 | 98 | 163 | 85 | 126 | 33 | 81 | 122 | 415 | 24 | 0 | 65 | 0 | 0 | 0 | 82172 | 9119 | 106680 | 0 | 73 | 2504 | 54161 | 0 | 0 | 10 | 14 | 21 | 21 | 42 | 0 | 30 | 0 | 0 | 0 | 0 | 2095 | 0 | 0 |
| SRR5925689 | 97 | 119 | 86 | 148 | 65 | 160 | 276 | 662 | 0 | 11 | 50 | 0 | 0 | 0 | 94088 | 9475 | 108668 | 0 | 87 | 2632 | 51794 | 0 | 0 | 13 | 11 | 23 | 22 | 44 | 0 | 51 | 0 | 0 | 0 | 0 | 2220 | 0 | 0 |
| SRR5925690 | 55 | 113 | 46 | 113 | 40 | 185 | 199 | 516 | 33 | 0 | 67 | 0 | 0 | 0 | 92964 | 9538 | 108456 | 0 | 46 | 2455 | 56370 | 0 | 0 | 153 | 14 | 26 | 19 | 47 | 0 | 45 | 0 | 0 | 0 | 0 | 2162 | 0 | 0 |
| SRR5925691 | 112 | 186 | 119 | 123 | 56 | 283 | 256 | 868 | 20 | 14 | 102 | 0 | 0 | 0 | 93139 | 10902 | 107549 | 0 | 101 | 1696 | 58848 | 0 | 0 | 301 | 389 | 424 | 441 | 876 | 0 | 774 | 0 | 0 | 0 | 0 | 1693 | 0 | 0 |
| SRR5925692 | 144 | 167 | 11 | 128 | 59 | 214 | 214 | 862 | 76 | 16 | 92 | 0 | 0 | 0 | 102375 | 12570 | 126339 | 0 | 82 | 1794 | 56107 | 0 | 0 | 18 | 15 | 23 | 21 | 47 | 0 | 39 | 0 | 0 | 0 | 0 | 1861 | 0 | 0 |
| SRR5925693 | 97 | 13 | 97 | 12 | 65 | 252 | 226 | 874 | 13 | 0 | 110 | 0 | 0 | 0 | 88902 | 11020 | 114294 | 0 | 69 | 1352 | 51179 | 0 | 0 | 15 | 19 | 26 | 25 | 40 | 0 | 52 | 0 | 0 | 0 | 0 | 2070 | 0 | 0 |
| SRR5925694 | 99 | 13 | 105 | 132 | 57 | 374 | 292 | 1040 | 57 | 15 | 93 | 0 | 0 | 0 | 93479 | 13055 | 105999 | 0 | 82 | 1440 | 57967 | 0 | 0 | 17 | 15 | 24 | 25 | 43 | 0 | 56 | 0 | 0 | 0 | 0 | 1069 | 0 | 0 |
| SRR5925695 | 90 | 160 | 179 | 132 | 38 | 325 | 325 | 865 | 39 | 0 | 90 | 0 | 0 | 0 | 62462 | 15787 | 99194 | 0 | 39 | 4736 | 25353 | 0 | 0 | 227 | 227 | 349 | 430 | 1057 | 0 | 609 | 0 | 0 | 0 | 0 | 2893 | 0 | 0 |
| SRR5925696 | 103 | 160 | 225 | 146 | 33 | 437 | 322 | 1233 | 0 | 19 | 108 | 0 | 0 | 0 | 60879 | 19139 | 92107 | 0 | 46 | 3570 | 24774 | 0 | 0 | 221 | 174 | 22 | 19 | 48 | 0 | 37 | 0 | 0 | 0 | 0 | 3450 | 0 | 0 |
| SRR5925697 | 88 | 193 | 133 | 230 | 45 | 201 | 224 | 611 | 11 | 16 | 53 | 0 | 0 | 0 | 79352 | 11907 | 109293 | 0 | 83 | 3072 | 43059 | 0 | 0 | 142 | 261 | 476 | 451 | 886 | 0 | 763 | 0 | 0 | 0 | 0 | 1410 | 0 | 0 |
| SRR5925698 | 84 | 252 | 237 | 183 | 54 | 244 | 259 | 661 | 36 | 0 | 74 | 0 | 0 | 0 | 70025 | 11661 | 104650 | 0 | 65 | 1574 | 32586 | 0 | 0 | 133 | 321 | 553 | 395 | 1036 | 0 | 755 | 0 | 0 | 0 | 0 | 3191 | 0 | 0 |
| SRR5925699 | 124 | 173 | 99 | 174 | 45 | 218 | 302 | 718 | 47 | 0 | 68 | 0 | 0 | 0 | 67078 | 10282 | 92659 | 0 | 70 | 2182 | 40163 | 0 | 0 | 14 | 14 | 26 | 26 | 51 | 0 | 41 | 0 | 0 | 0 | 0 | 2989 | 0 | 0 |
| SRR5925700 | 109 | 253 | 265 | 173 | 59 | 336 | 279 | 959 | 0 | 0 | 105 | 0 | 0 | 0 | 77109 | 13169 | 103476 | 0 | 115 | 1976 | 40241 | 0 | 0 | 320 | 345 | 577 | 539 | 1023 | 0 | 964 | 0 | 0 | 0 | 0 | 2992 | 0 | 0 |
| SRR5925701 | 54 | 157 | 110 | 116 | 59 | 353 | 320 | 1140 | 38 | 17 | 151 | 0 | 0 | 0 | 89497 | 18467 | 115982 | 0 | 104 | 1490 | 45343 | 0 | 0 | 207 | 249 | 417 | 364 | 604 | 0 | 939 | 0 | 0 | 0 | 0 | 2737 | 0 | 0 |
| SRR5925702 | 117 | 121 | 202 | 97 | 39 | 234 | 206 | 625 | 28 | 12 | 58 | 0 | 0 | 0 | 68069 | 9689 | 93394 | 0 | 78 | 1030 | 42574 | 0 | 0 | 202 | 343 | 604 | 533 | 911 | 0 | 876 | 0 | 0 | 0 | 0 | 2921 | 0 | 0 |
| SRR5925703 | 92 | 11 | 119 | 146 | 61 | 276 | 394 | 1103 | 53 | 23 | 111 | 0 | 0 | 0 | 76461 | 15021 | 102596 | 0 | 79 | 2054 | 45636 | 0 | 0 | 13 | 14 | 23 | 19 | 37 | 0 | 52 | 0 | 0 | 0 | 0 | 2571 | 0 | 0 |
| SRR5925704 | 122 | 197 | 11 | 10 | 66 | 140 | 172 | 473 | 15 | 12 | 74 | 0 | 0 | 0 | 86743 | 9493 | 122740 | 0 | 59 | 1925 | 46359 | 0 | 0 | 15 | 15 | 25 | 22 | 39 | 0 | 45 | 0 | 0 | 0 | 0 | 2522 | 0 | 0 |
| SRR5925705 | 93 | 105 | 86 | 20 | 18 | 354 | 1865 | 7186 | 197 | 0 | 256 | 0 | 0 | 0 | 20142 | 81429 | 61840 | 0 | 100 | 1045 | 1791 | 0 | 0 | 100 | 0 | 0 | 0 | 0 | 0 | 808 | 0 | 0 | 0 | 0 | 20174 | 46 | 0 |
| SRR5925706 | 57 | 18 | 0 | 0 | 0 | 682 | 3169 | 8101 | 257 | 0 | 65 | 0 | 0 | 0 | 12381 | 54565 | 28088 | 0 | 80 | 1149 | 1023 | 0 | 0 | 45 | 0 | 0 | 0 | 0 | 0 | 161 | 0 | 0 | 0 | 0 | 28017 | 34 | 0 |
| SRR5925707 | 14 | 17 | 76 | 13 | 12 | 670 | 2760 | 11987 | 330 | 0 | 28 | 0 | 0 | 0 | 20967 | 77570 | 57280 | 0 | 105 | 1177 | 1816 | 0 | 0 | 83 | 0 | 0 | 0 | 0 | 0 | 85 | 0 | 0 | 0 | 0 | 34251 | 53 | 0 |
| SRR5925708 | 41 | 100 | 0 | 0 | 0 | 784 | 3633 | 11532 | 317 | 0 | 526 | 0 | 0 | 0 | 14280 | 59280 | 29874 | 0 | 100 | 1073 | 1070 | 0 | 0 | 39 | 0 | 0 | 0 | 0 | 0 | 1284 | 0 | 0 | 0 | 0 | 33595 | 44 | 0 |
| SRR5925709 | 149 | 431 | 250 | 80 | 229 | 189 | 868 | 2885 | 29 | 27 | 101 | 0 | 0 | 0 | 23682 | 92692 | 127246 | 0 | 0 | 1229 | 3883 | 0 | 0 | 25 | 16 | 69 | 43 | 0 | 0 | 191 | 0 | 0 | 0 | 0 | 10428 | 99 | 0 |
| SRR5925710 | 0 | 0 | 0 | 27 | 0 | 0 | 3937 | 2953 | 0 | 0 | 0 | 0 | 0 | 0 | 22638 | 64193 | 86259 | 0 | 0 | 0 | 7874 | 0 | 0 | 0 | 0 | 0 | 0 | 0 | 0 | 107 | 0 | 0 | 0 | 0 | 13039 | 58 | 0 |
| SRR5925711 | 35 | 103 | 14 | 0 | 0 | 399 | 2505 | 3777 | 173 | 21 | 56 | 0 | 0 | 0 | 32004 | 34289 | 46974 | 0 | 89 | 1127 | 1392 | 0 | 0 | 100 | 0 | 11 | 29 | 18 | 0 | 130 | 0 | 0 | 0 | 0 | 24126 | 38 | 0 |
| SRR5925712 | 60 | 140 | 165 | 25 | 40 | 63 | 544 | 737 | 20 | 10 | 437 | 0 | 0 | 0 | 22184 | 6111 | 17527 | 17 | 40 | 367 | 1597 | 0 | 0 | 86 | 0 | 0 | 0 | 0 | 0 | 1841 | 0 | 0 | 0 | 0 | 5208 | 50 | 0 |
| SRR5925713 | 12 | 14 | 64 | 15 | 18 | 829 | 2981 | 12112 | 223 | 0 | 26 | 0 | 0 | 0 | 32423 | 65816 | 77563 | 0 | 92 | 1148 | 2390 | 0 | 0 | 20 | 0 | 0 | 0 | 0 | 0 | 83 | 0 | 0 | 0 | 0 | 27867 | 37 | 0 |
| SRR5925714 | 14 | 15 | 55 | 21 | 18 | 794 | 3260 | 11451 | 289 | 0 | 30 | 0 | 0 | 0 | 28704 | 78232 | 82500 | 0 | 64 | 846 | 1898 | 0 | 0 | 21 | 0 | 19 | 0 | 0 | 0 | 99 | 0 | 0 | 0 | 0 | 21662 | 40 | 0 |
| SRR5925715 | 60 | 11 | 0 | 0 | 0 | 731 | 3492 | 9811 | 394 | 0 | 42 | 0 | 0 | 0 | 13604 | 64035 | 41850 | 0 | 66 | 1017 | 883 | 0 | 0 | 113 | 0 | 0 | 11 | 0 | 0 | 112 | 0 | 0 | 0 | 0 | 37711 | 26 | 0 |
| SRR5925716 | 27 | 66 | 0 | 0 | 10 | 1021 | 3202 | 10128 | 292 | 0 | 608 | 0 | 0 | 0 | 12640 | 63819 | 37812 | 0 | 60 | 875 | 640 | 0 | 0 | 60 | 0 | 0 | 0 | 0 | 0 | 1400 | 0 | 0 | 0 | 0 | 36876 | 25 | 0 |
| SRR5925717 | 37 | 56 | 0 | 0 | 0 | 913 | 5800 | 11550 | 450 | 0 | 581 | 0 | 0 | 0 | 20723 | 38859 | 52837 | 0 | 129 | 1950 | 1289 | 0 | 0 | 81 | 0 | 0 | 11 | 0 | 0 | 1370 | 0 | 0 | 0 | 0 | 30183 | 32 | 0 |
| SRR5925718 | 31 | 63 | 0 | 0 | 0 | 1333 | 6451 | 15153 | 502 | 0 | 621 | 0 | 0 | 0 | 21833 | 38999 | 53842 | 0 | 215 | 2199 | 1824 | 0 | 0 | 77 | 0 | 0 | 13 | 0 | 0 | 1389 | 0 | 0 | 0 | 0 | 26848 | 24 | 0 |
| SRR5925719 | 63 | 10 | 10 | 0 | 12 | 253 | 483 | 3340 | 41 | 37 | 15 | 0 | 0 | 0 | 23659 | 16902 | 69316 | 0 | 44 | 883 | 23158 | 0 | 0 | 81 | 0 | 0 | 0 | 0 | 0 | 36 | 0 | 0 | 0 | 0 | 14724 | 0 | 0 |
| SRR5925720 | 56 | 11 | 10 | 0 | 36 | 170 | 345 | 3442 | 80 | 46 | 15 | 0 | 0 | 0 | 26089 | 17883 | 72188 | 0 | 71 | 935 | 23823 | 0 | 0 | 66 | 0 | 0 | 0 | 0 | 0 | 32 | 0 | 0 | 0 | 0 | 9991 | 0 | 0 |
| SRR5925721 | 35 | 102 | 0 | 0 | 15 | 277 | 570 | 3796 | 76 | 27 | 554 | 0 | 0 | 0 | 26857 | 7184 | 33486 | 0 | 60 | 722 | 32641 | 0 | 0 | 91 | 0 | 0 | 0 | 0 | 0 | 1097 | 0 | 0 | 0 | 0 | 4568 | 0 | 0 |
| SRR5925722 | 38 | 113 | 0 | 0 | 23 | 276 | 608 | 3676 | 82 | 30 | 492 | 0 | 0 | 0 | 27571 | 7746 | 34198 | 0 | 41 | 690 | 32316 | 0 | 0 | 96 | 0 | 0 | 0 | 0 | 0 | 1114 | 0 | 0 | 0 | 0 | 5121 | 0 | 0 |
| SRR5925723 | 45 | 68 | 0 | 0 | 0 | 192 | 472 | 3784 | 59 | 0 | 435 | 0 | 0 | 0 | 11247 | 8926 | 23598 | 0 | 20 | 1145 | 5263 | 0 | 0 | 29 | 0 | 0 | 0 | 0 | 0 | 1083 | 0 | 0 | 0 | 0 | 9980 | 0 | 0 |
| SRR5925724 | 26 | 10 | 0 | 0 | 0 | 193 | 435 | 4111 | 84 | 17 | 83 | 0 | 0 | 0 | 13574 | 10221 | 27265 | 0 | 45 | 1257 | 5773 | 0 | 0 | 30 | 0 | 0 | 0 | 0 | 0 | 197 | 0 | 0 | 0 | 0 | 9629 | 0 | 0 |
| SRR5925725 | 29 | 70 | 0 | 0 | 11 | 129 | 421 | 2687 | 43 | 22 | 835 | 0 | 0 | 0 | 18383 | 13845 | 44875 | 0 | 53 | 1480 | 7889 | 0 | 0 | 32 | 0 | 0 | 0 | 0 | 0 | 1592 | 0 | 0 | 0 | 0 | 10954 | 0 | 0 |
| SRR5925726 | 33 | 50 | 0 | 0 | 14 | 184 | 416 | 3208 | 60 | 22 | 151 | 0 | 0 | 0 | 23253 | 17488 | 53208 | 10 | 32 | 1353 | 8853 | 0 | 0 | 26 | 0 | 0 | 0 | 0 | 0 | 286 | 0 | 0 | 0 | 0 | 14009 | 0 | 0 |
| SRR5925727 | 0 | 0 | 0 | 0 | 0 | 0 | 0 | 16260 | 0 | 0 | 0 | 0 | 0 | 0 | 24390 | 33149 | 49724 | 0 | 0 | 0 | 8130 | 0 | 0 | 0 | 0 | 0 | 0 | 0 | 0 | 208 | 0 | 0 | 0 | 0 | 5525 | 0 | 0 |
| SRR5925728 | 69 | 152 | 0 | 0 | 0 | 365 | 555 | 3646 | 78 | 12 | 414 | 0 | 0 | 0 | 54875 | 9530 | 56005 | 0 | 66 | 1188 | 54164 | 0 | 0 | 94 | 0 | 0 | 0 | 0 | 0 | 927 | 0 | 0 | 0 | 0 | 8873 | 0 | 0 |
| SRR5925729 | 35 | 124 | 0 | 0 | 0 | 185 | 396 | 2242 | 45 | 37 | 407 | 0 | 0 | 0 | 12634 | 18430 | 41149 | 0 | 37 | 1043 | 7137 | 0 | 0 | 45 | 0 | 0 | 0 | 0 | 0 | 931 | 0 | 0 | 0 | 0 | 20002 | 0 | 0 |
| SRR5925730 | 22 | 89 | 0 | 0 | 16 | 141 | 367 | 1922 | 68 | 28 | 405 | 0 | 0 | 0 | 14216 | 19712 | 47819 | 0 | 51 | 1004 | 7101 | 0 | 0 | 57 | 0 | 0 | 0 | 0 | 0 | 966 | 0 | 0 | 0 | 0 | 19973 | 0 | 0 |
| SRR5925731 | 84 | 246 | 0 | 13 | 17 | 702 | 1222 | 11307 | 182 | 17 | 307 | 0 | 0 | 0 | 35332 | 8677 | 39776 | 0 | 94 | 621 | 41099 | 0 | 0 | 93 | 0 | 0 | 0 | 0 | 0 | 815 | 0 | 13 | 0 | 0 | 4815 | 0 | 0 |
| SRR5925732 | 66 | 15 | 18 | 0 | 0 | 856 | 1131 | 11464 | 202 | 28 | 19 | 0 | 0 | 0 | 39498 | 9252 | 46417 | 0 | 81 | 566 | 39619 | 0 | 0 | 111 | 0 | 0 | 10 | 0 | 0 | 46 | 0 | 0 | 0 | 0 | 6405 | 0 | 0 |
| SRR5930976 | 102 | 137 | 57 | 80 | 50 | 268 | 200 | 510 | 113 | 0 | 26 | 0 | 0 | 0 | 28296 | 3808 | 14193 | 21 | 37 | 2733 | 2963 | 0 | 0 | 32 | 20 | 53 | 157 | 122 | 0 | 16 | 0 | 21 | 0 | 0 | 1340 | 0 | 15 |
| SRR5930977 | 0 | 0 | 0 | 25 | 0 | 51 | 81 | 367 | 12 | 0 | 26 | 12 | 0 | 0 | 547 | 43481 | 5573 | 185 | 21 | 639 | 424 | 0 | 0 | 0 | 0 | 21 | 41 | 0 | 0 | 63 | 0 | 738 | 0 | 0 | 18632 | 0 | 26 |
| SRR5930978 | 0 | 0 | 0 | 0 | 22 | 56 | 72 | 262 | 14 | 0 | 44 | 0 | 0 | 0 | 587 | 1753 | 704 | 358 | 0 | 67 | 67 | 15 | 0 | 22 | 0 | 53 | 42 | 41 | 0 | 43 | 0 | 13142 | 52 | 0 | 4320 | 121 | 100 |
| SRR5936591 | 30 | 80 | 18 | 36 | 18 | 62 | 97 | 132 | 13 | 0 | 0 | 0 | 0 | 0 | 28432 | 5140 | 22260 | 22 | 18 | 2167 | 4427 | 0 | 0 | 93 | 16 | 86 | 213 | 35 | 0 | 70 | 0 | 48 | 0 | 0 | 1045 | 0 | 0 |
| SRR5936592 | 0 | 0 | 0 | 0 | 30 | 0 | 0 | 29 | 0 | 0 | 15 | 0 | 0 | 0 | 482 | 1865 | 475 | 180 | 0 | 125 | 86 | 0 | 0 | 0 | 0 | 0 | 14 | 0 | 0 | 50 | 0 | 7737 | 62 | 0 | 6540 | 231 | 585 |

Supp Table S8.3.Columns_BX_to_DH

| | | | | | | | | | | | | | | | | | | | | | | | | | | | | | | | | | | | | | |
|---|---|---|---|---|---|---|---|---|---|---|---|---|---|---|---|---|---|---|---|---|---|---|---|---|---|---|---|---|---|---|---|---|---|---|---|---|---|
| SRR5951878 | 66 | 161 | 17 | 0 | 162 | 16218 | 9456 | 25015 | 25050 | 122 | 19 | 0 | 156 | 166 | 3937 | 15518 | 6974 | 935 | 850 | 1040 | 288 | 0 | 0 | 176 | 0 | 0 | 13 | 0 | 0 | 69 | 0 | 16 | 0 | 0 | 2942 | 3597 | 0 |
| SRR5951879 | 143 | 31 | 0 | 0 | 154 | 18949 | 11315 | 28356 | 14042 | 54 | 34 | 0 | 14 | 11 | 30540 | 36264 | 89448 | 812 | 7680 | 4250 | 513 | 0 | 0 | 130 | 14 | 20 | 35 | 0 | 0 | 284 | 0 | 0 | 0 | 0 | 3168 | 9192 | 0 |
| SRR5951880 | 0 | 0 | 0 | 0 | 32 | 0 | 0 | 0 | 0 | 0 | 32 | 65 | 0 | 0 | 0 | 0 | 0 | 4984 | 0 | 0 | 25 | 0 | 0 | 0 | 0 | 0 | 65 | 0 | 0 | 0 | 0 | 0 | 0 | 0 | 0 | 0 | 0 |
| SRR5951881 | 46 | 147 | 14 | 0 | 13 | 18025 | 7756 | 20968 | 18455 | 65 | 10 | 0 | 76 | 78 | 2949 | 37269 | 6114 | 1389 | 2242 | 669 | 259 | 0 | 0 | 127 | 0 | 0 | 0 | 0 | 0 | 87 | 0 | 25 | 0 | 0 | 8126 | 1345 | 0 |
| SRR5951882 | 311 | 66 | 0 | 0 | 1582 | 13052 | 7972 | 21746 | 11536 | 101 | 233 | 0 | 14 | 0 | 104885 | 38400 | 191480 | 701 | 6180 | 2522 | 369 | 0 | 0 | 269 | 12 | 27 | 39 | 11 | 0 | 3153 | 0 | 0 | 0 | 0 | 4992 | 3381 | 0 |
| SRR5951883 | 0 | 0 | 0 | 0 | 32 | 0 | 0 | 0 | 0 | 0 | 0 | 811 | 0 | 0 | 0 | 0 | 0 | 5686 | 35 | 0 | 0 | 0 | 0 | 0 | 0 | 0 | 48 | 0 | 127 | 0 | 0 | 0 | 0 | 0 | 12 | 0 | 0 |
| SRR5951884 | 1767 | 131 | 23 | 18 | 1680 | 2141 | 457 | 1744 | 2380 | 1423 | 95 | 0 | 57 | 28 | 13586 | 11298 | 17994 | 52 | 105 | 419 | 3077 | 0 | 0 | 91 | 0 | 0 | 0 | 0 | 0 | 380 | 0 | 0 | 0 | 0 | 8299 | 1965 | 0 |
| SRR5951885 | 1570 | 145 | 24 | 13 | 2046 | 1822 | 442 | 1767 | 2636 | 1522 | 90 | 0 | 36 | 25 | 10531 | 16624 | 13573 | 69 | 104 | 352 | 1981 | 0 | 0 | 94 | 0 | 0 | 19 | 0 | 0 | 384 | 0 | 0 | 0 | 0 | 9370 | 1999 | 0 |
| SRR5951886 | 1714 | 136 | 21 | 0 | 1665 | 941 | 310 | 795 | 1280 | 3050 | 111 | 0 | 49 | 32 | 12319 | 9830 | 15730 | 82 | 92 | 441 | 3497 | 0 | 0 | 81 | 0 | 0 | 11 | 0 | 0 | 376 | 0 | 19 | 0 | 0 | 7452 | 1738 | 0 |
| SRR5951887 | 3731 | 262 | 39 | 23 | 1980 | 879 | 374 | 1064 | 1615 | 2327 | 103 | 19 | 136 | 79 | 10371 | 2864 | 10263 | 87 | 196 | 404 | 2245 | 0 | 0 | 97 | 0 | 0 | 14 | 0 | 0 | 374 | 0 | 0 | 0 | 0 | 4648 | 1937 | 0 |
| SRR5951888 | 4053 | 297 | 44 | 27 | 2035 | 616 | 207 | 792 | 1002 | 2238 | 122 | 10 | 157 | 76 | 8625 | 3010 | 9912 | 62 | 172 | 344 | 2338 | 0 | 0 | 83 | 0 | 0 | 11 | 0 | 0 | 348 | 0 | 10 | 0 | 0 | 5278 | 1650 | 0 |
| SRR5951889 | 199 | 13 | 43 | 33 | 74 | 749 | 147 | 776 | 1196 | 80 | 110 | 31 | 108 | 66 | 13504 | 4213 | 16621 | 73 | 394 | 679 | 2856 | 0 | 0 | 63 | 0 | 0 | 21 | 0 | 0 | 14 | 0 | 12 | 0 | 0 | 5214 | 1705 | 0 |
| SRR5951890 | 4829 | 331 | 61 | 24 | 1414 | 1118 | 524 | 1334 | 2162 | 1062 | 157 | 14 | 123 | 82 | 6497 | 4563 | 10646 | 92 | 399 | 1044 | 718 | 0 | 0 | 64 | 0 | 0 | 0 | 0 | 0 | 688 | 0 | 0 | 0 | 0 | 3995 | 2216 | 0 |
| SRR5951891 | 223 | 15 | 48 | 20 | 90 | 800 | 246 | 907 | 1399 | 92 | 99 | 24 | 98 | 81 | 8410 | 3570 | 11864 | 94 | 287 | 903 | 948 | 0 | 0 | 66 | 0 | 0 | 15 | 0 | 0 | 18 | 0 | 16 | 0 | 0 | 3430 | 2103 | 0 |
| SRR5951892 | 348 | 22 | 55 | 24 | 67 | 789 | 324 | 877 | 1727 | 73 | 85 | 21 | 116 | 85 | 12542 | 3411 | 16576 | 65 | 286 | 736 | 1395 | 0 | 0 | 74 | 0 | 0 | 0 | 0 | 0 | 19 | 0 | 15 | 0 | 0 | 3330 | 1935 | 0 |
| SRR5951893 | 1222 | 135 | 24 | 0 | 2110 | 921 | 303 | 1093 | 2341 | 1769 | 81 | 0 | 24 | 18 | 20161 | 5167 | 16910 | 70 | 479 | 945 | 2140 | 0 | 0 | 108 | 0 | 12 | 37 | 0 | 0 | 348 | 0 | 12 | 0 | 0 | 3777 | 1572 | 0 |
| SRR5951894 | 1522 | 232 | 29 | 0 | 1744 | 1653 | 518 | 1852 | 2498 | 1138 | 114 | 0 | 40 | 55 | 15285 | 7787 | 18826 | 248 | 64 | 355 | 1121 | 0 | 0 | 146 | 0 | 17 | 38 | 0 | 0 | 726 | 0 | 14 | 0 | 0 | 9654 | 1589 | 0 |
| SRR5951895 | 52 | 111 | 19 | 20 | 55 | 1158 | 417 | 1421 | 2838 | 89 | 129 | 11 | 23 | 18 | 14818 | 4066 | 14644 | 66 | 662 | 1658 | 1921 | 0 | 0 | 128 | 0 | 0 | 13 | 0 | 0 | 18 | 0 | 13 | 0 | 0 | 3529 | 1667 | 0 |
| SRR5951896 | 304 | 20 | 28 | 11 | 86 | 13923 | 8323 | 20029 | 18143 | 54 | 338 | 0 | 98 | 48 | 2107 | 53228 | 5924 | 679 | 784 | 696 | 192 | 0 | 0 | 36 | 0 | 0 | 0 | 0 | 0 | 2619 | 0 | 23 | 0 | 0 | 8898 | 0 | 0 |
| SRR5951897 | 14 | 41 | 36 | 40 | 122 | 36526 | 21481 | 50874 | 33099 | 10 | 29 | 0 | 48 | 62 | 13303 | 73844 | 51835 | 309 | 1077 | 899 | 257 | 0 | 0 | 89 | 22 | 21 | 26 | 0 | 0 | 184 | 0 | 0 | 0 | 0 | 6327 | 0 | 0 |
| SRR5951898 | 0 | 0 | 0 | 0 | 0 | 28 | 0 | 0 | 0 | 0 | 0 | 328 | 0 | 0 | 28 | 0 | 19 | 5755 | 0 | 0 | 0 | 0 | 0 | 0 | 0 | 0 | 12 | 0 | 81 | 0 | 0 | 0 | 0 | 0 | 0 | 0 | 0 |
| SRR5951899 | 436 | 48 | 29 | 59 | 102 | 19918 | 12142 | 26299 | 25799 | 65 | 537 | 14 | 62 | 53 | 1578 | 54464 | 5384 | 619 | 929 | 1149 | 892 | 0 | 0 | 58 | 0 | 0 | 0 | 0 | 0 | 1933 | 0 | 0 | 0 | 0 | 6665 | 0 | 0 |
| SRR5951900 | 13 | 63 | 24 | 25 | 14 | 27493 | 14157 | 37399 | 21880 | 133 | 36 | 0 | 57 | 52 | 12755 | 43130 | 36359 | 562 | 1666 | 1053 | 1021 | 0 | 0 | 154 | 0 | 11 | 31 | 0 | 0 | 129 | 0 | 0 | 0 | 0 | 6206 | 0 | 0 |
| SRR5951901 | 0 | 0 | 0 | 0 | 0 | 0 | 0 | 0 | 0 | 0 | 0 | 287 | 0 | 0 | 65 | 22 | 0 | 6005 | 0 | 0 | 0 | 0 | 0 | 0 | 0 | 0 | 13 | 0 | 39 | 0 | 0 | 0 | 0 | 0 | 0 | 0 | 0 |
| SRR5951902 | 221 | 25 | 28 | 28 | 109 | 24098 | 13881 | 30170 | 35349 | 46 | 361 | 0 | 92 | 61 | 1781 | 64998 | 6501 | 367 | 899 | 849 | 544 | 0 | 0 | 41 | 0 | 0 | 0 | 0 | 0 | 2844 | 0 | 12 | 0 | 0 | 5893 | 0 | 0 |
| SRR5951903 | 169 | 17 | 15 | 13 | 148 | 51099 | 24632 | 59154 | 39891 | 31 | 20 | 0 | 39 | 50 | 6860 | 80469 | 34231 | 116 | 751 | 575 | 188 | 0 | 0 | 121 | 40 | 39 | 36 | 0 | 0 | 167 | 0 | 0 | 0 | 0 | 4064 | 0 | 0 |
| SRR5951904 | 0 | 0 | 0 | 0 | 0 | 0 | 0 | 0 | 0 | 0 | 0 | 582 | 0 | 0 | 28 | 39 | 0 | 3046 | 0 | 0 | 0 | 0 | 0 | 0 | 0 | 0 | 16 | 0 | 76 | 0 | 0 | 0 | 0 | 0 | 0 | 0 | 0 |
| SRR5951905 | 124 | 23 | 32 | 23 | 13 | 20790 | 9445 | 23063 | 19971 | 134 | 17 | 0 | 64 | 31 | 3188 | 43434 | 5242 | 317 | 1253 | 624 | 334 | 0 | 0 | 72 | 12 | 0 | 0 | 0 | 0 | 113 | 0 | 0 | 0 | 0 | 11701 | 0 | 0 |
| SRR5951906 | 165 | 24 | 17 | 10 | 25 | 23694 | 11526 | 29914 | 17285 | 61 | 13 | 0 | 83 | 49 | 5495 | 85825 | 13942 | 181 | 2054 | 695 | 147 | 0 | 0 | 90 | 37 | 33 | 34 | 0 | 0 | 142 | 0 | 0 | 0 | 0 | 4241 | 0 | 0 |
| SRR5951907 | 0 | 0 | 0 | 0 | 0 | 0 | 0 | 0 | 36 | 0 | 0 | 519 | 0 | 0 | 0 | 0 | 26 | 1859 | 18 | 0 | 0 | 0 | 0 | 0 | 0 | 0 | 17 | 0 | 201 | 0 | 0 | 0 | 0 | 0 | 0 | 0 | 0 |
| SRR5951908 | 1757 | 170 | 117 | 84 | 1266 | 1668 | 898 | 2083 | 3096 | 1403 | 324 | 64 | 119 | 69 | 3177 | 4884 | 7824 | 51 | 231 | 603 | 2634 | 0 | 0 | 48 | 0 | 0 | 28 | 11 | 18 | 956 | 0 | 17 | 0 | 0 | 4051 | 0 | 0 |
| SRR5951909 | 1128 | 125 | 47 | 36 | 1560 | 828 | 139 | 1073 | 1298 | 882 | 215 | 26 | 61 | 30 | 2897 | 2647 | 4268 | 80 | 133 | 465 | 1151 | 0 | 0 | 77 | 0 | 0 | 21 | 0 | 0 | 545 | 0 | 0 | 0 | 0 | 2588 | 0 | 0 |
| SRR5951910 | 1510 | 149 | 32 | 46 | 1329 | 730 | 176 | 830 | 1082 | 984 | 243 | 37 | 60 | 26 | 3129 | 3879 | 5774 | 72 | 180 | 802 | 1740 | 0 | 0 | 62 | 0 | 0 | 15 | 0 | 0 | 399 | 0 | 12 | 0 | 0 | 4361 | 0 | 0 |
| SRR5951911 | 519 | 45 | 12 | 10 | 1099 | 896 | 197 | 1107 | 1688 | 452 | 167 | 134 | 27 | 22 | 6702 | 6208 | 9244 | 59 | 408 | 654 | 1757 | 0 | 0 | 30 | 0 | 0 | 0 | 0 | 16 | 573 | 0 | 0 | 0 | 0 | 3524 | 0 | 0 |
| SRR5951912 | 300 | 35 | 13 | 14 | 879 | 842 | 216 | 1048 | 1330 | 456 | 167 | 47 | 23 | 13 | 3963 | 4901 | 5889 | 72 | 560 | 809 | 1580 | 0 | 0 | 27 | 0 | 0 | 0 | 0 | 14 | 378 | 0 | 11 | 0 | 0 | 4218 | 0 | 0 |
| SRR5951913 | 24 | 62 | 23 | 33 | 54 | 980 | 282 | 1208 | 1701 | 32 | 211 | 42 | 47 | 10 | 1935 | 4297 | 3572 | 24 | 252 | 709 | 1244 | 0 | 0 | 45 | 0 | 0 | 0 | 0 | 0 | 17 | 0 | 30 | 0 | 0 | 3949 | 0 | 0 |
| SRR6020460 | 0 | 0 | 0 | 0 | 0 | 5450 | 601 | 11531 | 401 | 30 | 0 | 0 | 0 | 0 | 2834 | 13164 | 104015 | 12 | 0 | 15657 | 194 | 0 | 0 | 43 | 0 | 0 | 16 | 0 | 0 | 30 | 0 | 0 | 0 | 0 | 7763 | 0 | 0 |
| SRR6020461 | 0 | 0 | 0 | 0 | 22 | 8068 | 1121 | 13996 | 908 | 18 | 11 | 0 | 0 | 0 | 10953 | 27386 | 162111 | 0 | 47 | 57149 | 196 | 0 | 0 | 177 | 0 | 12 | 18 | 0 | 0 | 31 | 0 | 0 | 0 | 0 | 1572 | 0 | 0 |
| SRR6020462 | 0 | 0 | 0 | 0 | 26 | 5608 | 555 | 15180 | 448 | 39 | 0 | 0 | 0 | 0 | 3884 | 24654 | 174668 | 0 | 14 | 18060 | 248 | 0 | 0 | 191 | 0 | 17 | 93 | 0 | 0 | 437 | 0 | 0 | 0 | 0 | 4612 | 0 | 0 |
| SRR6020463 | 0 | 0 | 0 | 0 | 40 | 2796 | 502 | 6421 | 418 | 52 | 0 | 17 | 0 | 0 | 6360 | 26302 | 204931 | 0 | 12 | 57067 | 189 | 0 | 0 | 72 | 16 | 38 | 36 | 0 | 0 | 122 | 0 | 0 | 0 | 0 | 3038 | 0 | 10 |
| SRR6020464 | 0 | 0 | 0 | 0 | 28 | 7149 | 1832 | 19976 | 773 | 51 | 0 | 0 | 0 | 0 | 1026 | 12826 | 75684 | 0 | 0 | 3272 | 47 | 0 | 0 | 59 | 0 | 0 | 0 | 0 | 0 | 36 | 0 | 0 | 0 | 0 | 20002 | 0 | 0 |
| SRR6020465 | 0 | 0 | 0 | 0 | 11 | 10635 | 880 | 25210 | 864 | 58 | 0 | 0 | 0 | 0 | 954 | 9818 | 42061 | 11 | 0 | 10316 | 37 | 0 | 0 | 207 | 0 | 0 | 0 | 0 | 0 | 158 | 0 | 0 | 0 | 0 | 11341 | 0 | 0 |
| SRR6020466 | 0 | 0 | 0 | 0 | 13 | 1330 | 491 | 4135 | 246 | 45 | 12 | 13 | 0 | 0 | 2607 | 20761 | 128513 | 0 | 0 | 44793 | 162 | 0 | 0 | 138 | 0 | 13 | 120 | 0 | 0 | 700 | 0 | 12 | 0 | 0 | 8538 | 0 | 10 |
| SRR6020467 | 0 | 0 | 0 | 0 | 14 | 4420 | 738 | 9812 | 334 | 53 | 0 | 0 | 0 | 0 | 538 | 8535 | 45959 | 0 | 0 | 7763 | 58 | 0 | 0 | 176 | 0 | 0 | 27 | 0 | 0 | 270 | 0 | 0 | 0 | 0 | 13821 | 0 | 0 |
| SRR6020468 | 0 | 0 | 0 | 0 | 25 | 2575 | 438 | 3740 | 223 | 46 | 0 | 0 | 0 | 0 | 2790 | 16891 | 142034 | 0 | 22 | 43554 | 215 | 0 | 0 | 276 | 0 | 10 | 82 | 0 | 0 | 271 | 0 | 0 | 0 | 0 | 5221 | 0 | 11 |
| SRR6020469 | 0 | 0 | 0 | 0 | 22 | 7825 | 727 | 19887 | 464 | 13 | 15 | 0 | 0 | 0 | 2339 | 31374 | 177091 | 0 | 20 | 25530 | 170 | 0 | 0 | 59 | 0 | 46 | 38 | 0 | 0 | 149 | 0 | 0 | 0 | 0 | 3412 | 0 | 0 |
| SRR6020470 | 0 | 0 | 0 | 0 | 12 | 7890 | 719 | 16128 | 557 | 28 | 0 | 0 | 0 | 0 | 719 | 7134 | 43033 | 0 | 0 | 5569 | 70 | 0 | 0 | 35 | 0 | 0 | 19 | 0 | 0 | 18 | 0 | 0 | 0 | 0 | 7721 | 0 | 0 |
| SRR6020471 | 0 | 0 | 0 | 0 | 12 | 4823 | 693 | 14169 | 538 | 41 | 0 | 0 | 0 | 0 | 4960 | 24133 | 165802 | 0 | 18 | 23424 | 264 | 0 | 0 | 115 | 0 | 0 | 46 | 0 | 0 | 34 | 0 | 0 | 0 | 0 | 3314 | 0 | 0 |
| SRR6020472 | 0 | 0 | 0 | 0 | 18 | 6619 | 755 | 11988 | 522 | 29 | 0 | 0 | 0 | 0 | 7113 | 26542 | 196017 | 0 | 0 | 6976 | 124 | 0 | 0 | 42 | 0 | 0 | 0 | 0 | 0 | 44 | 0 | 0 | 0 | 0 | 7108 | 0 | 0 |
| SRR6020473 | 0 | 0 | 0 | 0 | 21 | 11313 | 1380 | 33292 | 642 | 24 | 0 | 0 | 0 | 0 | 1083 | 14476 | 104188 | 0 | 0 | 7672 | 62 | 0 | 0 | 349 | 0 | 10 | 73 | 0 | 0 | 218 | 0 | 0 | 0 | 0 | 6543 | 0 | 11 |
| SRR6020474 | 0 | 0 | 0 | 0 | 0 | 2454 | 369 | 5260 | 195 | 45 | 0 | 0 | 0 | 0 | 3735 | 20370 | 178074 | 0 | 31 | 44705 | 297 | 0 | 0 | 424 | 0 | 23 | 101 | 0 | 0 | 528 | 0 | 0 | 0 | 0 | 2795 | 0 | 0 |
| SRR6020475 | 0 | 0 | 0 | 0 | 0 | 4119 | 367 | 10550 | 257 | 38 | 0 | 0 | 0 | 0 | 2798 | 13770 | 142208 | 0 | 0 | 10193 | 165 | 0 | 0 | 29 | 0 | 0 | 22 | 0 | 0 | 34 | 0 | 0 | 0 | 0 | 11960 | 0 | 11 |
| SRR6020476 | 0 | 0 | 0 | 0 | 15 | 1335 | 338 | 4712 | 253 | 37 | 20 | 13 | 0 | 0 | 2164 | 30216 | 145936 | 0 | 15 | 27618 | 177 | 0 | 0 | 88 | 13 | 28 | 89 | 0 | 0 | 137 | 0 | 0 | 0 | 0 | 5485 | 0 | 0 |
| SRR6020477 | 0 | 0 | 0 | 0 | 35 | 7301 | 1363 | 16574 | 772 | 32 | 0 | 10 | 0 | 0 | 3292 | 13765 | 104768 | 12 | 0 | 6427 | 247 | 0 | 0 | 106 | 0 | 10 | 11 | 0 | 0 | 47 | 0 | 0 | 0 | 0 | 9077 | 0 | 0 |
| SRR6020478 | 0 | 0 | 0 | 0 | 0 | 4962 | 998 | 11885 | 399 | 36 | 0 | 0 | 0 | 0 | 1656 | 15303 | 105324 | 0 | 0 | 19536 | 288 | 0 | 0 | 111 | 0 | 0 | 14 | 0 | 0 | 247 | 0 | 0 | 0 | 0 | 13608 | 0 | 0 |
| SRR6020479 | 0 | 0 | 0 | 0 | 23 | 7803 | 1815 | 25684 | 642 | 37 | 0 | 0 | 0 | 0 | 1068 | 15735 | 71768 | 0 | 0 | 2755 | 75 | 0 | 0 | 43 | 0 | 15 | 30 | 0 | 0 | 63 | 0 | 0 | 0 | 0 | 12824 | 0 | 0 |
| SRR6020480 | 0 | 0 | 0 | 0 | 21 | 6170 | 1054 | 21869 | 1156 | 15 | 13 | 0 | 0 | 0 | 1232 | 32413 | 106627 | 20 | 11 | 26721 | 260 | 0 | 0 | 148 | 0 | 26 | 26 | 0 | 0 | 72 | 0 | 0 | 0 | 0 | 5920 | 0 | 12 |
| SRR6020481 | 0 | 0 | 0 | 0 | 11 | 6168 | 655 | 11908 | 305 | 44 | 0 | 0 | 0 | 0 | 2049 | 12569 | 123479 | 13 | 13 | 30256 | 195 | 0 | 0 | 36 | 0 | 0 | 27 | 0 | 0 | 34 | 0 | 0 | 0 | 0 | 8376 | 0 | 14 |
| SRR6020482 | 0 | 0 | 0 | 0 | 54 | 3354 | 624 | 8942 | 342 | 45 | 0 | 20 | 0 | 0 | 4014 | 23151 | 180516 | 0 | 0 | 61440 | 334 | 0 | 0 | 403 | 0 | 51 | 348 | 0 | 0 | 609 | 0 | 0 | 0 | 0 | 3484 | 0 | 0 |
| SRR6020483 | 0 | 0 | 0 | 0 | 13 | 3369 | 796 | 7864 | 289 | 28 | 0 | 0 | 0 | 0 | 740 | 6793 | 61360 | 16 | 0 | 9730 | 48 | 0 | 0 | 195 | 0 | 0 | 12 | 0 | 0 | 108 | 0 | 0 | 0 | 0 | 11033 | 0 | 11 |
| SRR6020484 | 0 | 0 | 0 | 0 | 31 | 4020 | 907 | 12045 | 605 | 66 | 0 | 0 | 0 | 0 | 2949 | 25884 | 186634 | 0 | 0 | 27192 | 213 | 0 | 0 | 124 | 0 | 12 | 58 | 0 | 0 | 213 | 0 | 0 | 0 | 0 | 12302 | 0 | 11 |
| SRR6020485 | 0 | 0 | 0 | 0 | 23 | 4881 | 832 | 13437 | 437 | 48 | 0 | 0 | 0 | 0 | 3449 | 18585 | 143457 | 14 | 0 | 45177 | 106 | 0 | 0 | 71 | 0 | 17 | 64 | 0 | 0 | 328 | 0 | 0 | 0 | 0 | 6087 | 0 | 13 |
| SRR6020486 | 0 | 0 | 0 | 0 | 0 | 11192 | 1123 | 28507 | 793 | 33 | 0 | 0 | 0 | 0 | 1931 | 16841 | 90829 | 0 | 0 | 15847 | 61 | 0 | 0 | 172 | 0 | 0 | 16 | 0 | 0 | 194 | 0 | 0 | 0 | 0 | 4570 | 0 | 0 |
| SRR6020487 | 0 | 0 | 0 | 0 | 12 | 8120 | 812 | 17939 | 435 | 31 | 0 | 0 | 0 | 0 | 3650 | 16550 | 169742 | 0 | 0 | 21769 | 148 | 0 | 0 | 28 | 0 | 0 | 26 | 0 | 0 | 31 | 0 | 0 | 0 | 0 | 5679 | 0 | 0 |
| SRR6020488 | 0 | 0 | 0 | 0 | 18 | 2919 | 489 | 8007 | 278 | 53 | 0 | 0 | 0 | 0 | 2967 | 18273 | 143295 | 0 | 16 | 61699 | 448 | 0 | 0 | 177 | 0 | 14 | 88 | 0 | 0 | 392 | 0 | 25 | 0 | 0 | 3199 | 0 | 0 |
| SRR6020489 | 0 | 0 | 0 | 0 | 0 | 5478 | 703 | 10088 | 453 | 21 | 0 | 0 | 0 | 0 | 524 | 3364 | 14371 | 0 | 16 | 7869 | 211 | 0 | 0 | 214 | 0 | 0 | 0 | 0 | 0 | 13 | 0 | 0 | 0 | 0 | 5230 | 0 | 0 |
| SRR6020490 | 0 | 0 | 0 | 0 | 14 | 5952 | 819 | 10768 | 430 | 47 | 0 | 0 | 0 | 0 | 1453 | 12738 | 77685 | 0 | 15 | 43185 | 97 | 0 | 0 | 405 | 0 | 0 | 29 | 0 | 0 | 132 | 0 | 0 | 0 | 0 | 6181 | 0 | 0 |
| SRR6020491 | 0 | 0 | 0 | 0 | 0 | 3654 | 372 | 6027 | 324 | 22 | 0 | 0 | 0 | 0 | 791 | 6855 | 46713 | 0 | 16 | 8669 | 166 | 0 | 0 | 208 | 0 | 0 | 23 | 0 | 0 | 112 | 0 | 0 | 0 | 0 | 5627 | 0 | 0 |
| SRR6020492 | 0 | 0 | 0 | 10 | 13 | 1428 | 673 | 4722 | 267 | 16 | 23 | 15 | 0 | 0 | 2277 | 28520 | 148752 | 0 | 37 | 55022 | 244 | 0 | 0 | 120 | 38 | 17 | 195 | 13 | 0 | 146 | 0 | 10 | 0 | 0 | 2813 | 0 | 0 |
| SRR6020493 | 0 | 0 | 0 | 0 | 0 | 5104 | 619 | 13623 | 445 | 37 | 0 | 0 | 0 | 0 | 733 | 8922 | 69385 | 0 | 12 | 5735 | 84 | 0 | 0 | 120 | 0 | 0 | 12 | 0 | 0 | 198 | 0 | 12 | 0 | 0 | 14800 | 0 | 0 |
| SRR6020494 | 0 | 0 | 0 | 0 | 0 | 5501 | 359 | 8614 | 413 | 41 | 0 | 0 | 0 | 0 | 1409 | 6224 | 40523 | 0 | 0 | 38594 | 72 | 0 | 0 | 96 | 0 | 0 | 0 | 0 | 0 | 15 | 0 | 0 | 0 | 0 | 3423 | 0 | 0 |
| SRR6020495 | 0 | 0 | 0 | 0 | 0 | 3527 | 563 | 8813 | 176 | 13 | 0 | 0 | 0 | 0 | 1267 | 9549 | 77370 | 0 | 18 | 6034 | 79 | 0 | 0 | 17 | 0 | 0 | 0 | 0 | 0 | 104 | 0 | 0 | 0 | 0 | 5617 | 0 | 0 |
| SRR6020496 | 0 | 0 | 0 | 0 | 0 | 4329 | 921 | 14403 | 289 | 19 | 0 | 0 | 0 | 0 | 1255 | 10624 | 77291 | 0 | 0 | 3294 | 99 | 0 | 0 | 32 | 0 | 0 | 17 | 0 | 0 | 60 | 0 | 0 | 0 | 0 | 7520 | 0 | 0 |
| SRR6020497 | 0 | 0 | 0 | 0 | 0 | 10326 | 1225 | 32923 | 842 | 40 | 0 | 0 | 0 | 0 | 398 | 5787 | 20088 | 0 | 0 | 8456 | 72 | 0 | 0 | 23 | 0 | 0 | 13 | 0 | 0 | 46 | 0 | 11 | 0 | 0 | 2084 | 0 | 0 |
| SRR6020498 | 0 | 0 | 0 | 25 | 68 | 1247 | 284 | 3432 | 323 | 74 | 22 | 15 | 0 | 0 | 12778 | 45781 | 309679 | 0 | 49 | 51053 | 601 | 0 | 0 | 853 | 29 | 31 | 296 | 0 | 0 | 450 | 0 | 0 | 0 | 0 | 2160 | 0 | 17 |
| SRR6020499 | 0 | 0 | 0 | 0 | 0 | 3405 | 364 | 5766 | 177 | 45 | 0 | 0 | 0 | 0 | 2696 | 20292 | 122604 | 0 | 0 | 27453 | 295 | 0 | 0 | 30 | 0 | 0 | 18 | 0 | 0 | 38 | 0 | 0 | 0 | 0 | 10024 | 0 | 0 |
| SRR6020500 | 0 | 0 | 0 | 0 | 13 | 3071 | 508 | 12969 | 416 | 18 | 16 | 13 | 0 | 0 | 1738 | 20475 | 138483 | 0 | 18 | 17972 | 139 | 0 | 0 | 32 | 10 | 15 | 63 | 0 | 0 | 596 | 0 | 12 | 0 | 0 | 7821 | 0 | 0 |
| SRR6020501 | 0 | 0 | 0 | 0 | 0 | 13974 | 1215 | 40649 | 716 | 35 | 0 | 0 | 0 | 0 | 521 | 8546 | 55189 | 0 | 0 | 3480 | 45 | 0 | 0 | 32 | 0 | 0 | 25 | 0 | 0 | 34 | 0 | 0 | 0 | 0 | 5345 | 0 | 0 |
| SRR6020502 | 0 | 0 | 0 | 0 | 18 | 7825 | 1099 | 22251 | 1075 | 12 | 0 | 0 | 0 | 0 | 3747 | 16604 | 82501 | 0 | 38 | 36308 | 43 | 0 | 0 | 124 | 0 | 0 | 20 | 0 | 0 | 38 | 0 | 0 | 0 | 0 | 914 | 0 | 0 |
| SRR6020503 | 0 | 0 | 0 | 0 | 0 | 4105 | 704 | 9500 | 336 | 47 | 0 | 0 | 0 | 0 | 1717 | 10161 | 90186 | 0 | 0 | 11249 | 139 | 0 | 0 | 135 | 0 | 0 | 38 | 0 | 0 | 355 | 0 | 11 | 0 | 0 | 13733 | 0 | 0 |
| SRR6020504 | 0 | 0 | 0 | 0 | 19 | 13011 | 1257 | 31805 | 943 | 47 | 0 | 0 | 0 | 0 | 561 | 11294 | 59651 | 0 | 0 | 3496 | 97 | 0 | 0 | 91 | 0 | 24 | 12 | 0 | 0 | 41 | 0 | 0 | 0 | 0 | 8018 | 0 | 0 |
| SRR6020505 | 0 | 0 | 0 | 0 | 20 | 8401 | 814 | 21113 | 713 | 58 | 0 | 0 | 0 | 0 | 1411 | 13790 | 88974 | 0 | 0 | 22778 | 111 | 0 | 0 | 396 | 0 | 16 | 72 | 0 | 0 | 375 | 0 | 0 | 0 | 0 | 4211 | 0 | 10 |
| SRR6020506 | 0 | 0 | 0 | 0 | 0 | 10947 | 1430 | 33952 | 737 | 43 | 0 | 0 | 0 | 0 | 581 | 12758 | 79067 | 0 | 0 | 3458 | 82 | 0 | 0 | 18 | 0 | 15 | 30 | 0 | 0 | 54 | 0 | 0 | 0 | 0 | 9045 | 0 | 0 |
| SRR6020507 | 0 | 0 | 0 | 0 | 19 | 5197 | 768 | 11964 | 697 | 11 | 0 | 0 | 0 | 0 | 3549 | 9005 | 77620 | 14 | 0 | 14499 | 92 | 0 | 0 | 409 | 0 | 0 | 41 | 0 | 0 | 22 | 0 | 28 | 0 | 0 | 3960 | 0 | 0 |
| SRR6020508 | 0 | 0 | 0 | 0 | 24 | 1509 | 340 | 4321 | 270 | 13 | 0 | 0 | 0 | 0 | 5881 | 29060 | 189145 | 13 | 19 | 53549 | 465 | 0 | 0 | 39 | 0 | 17 | 21 | 0 | 0 | 83 | 0 | 0 | 0 | 0 | 6308 | 0 | 0 |
| SRR6020509 | 0 | 0 | 0 | 0 | 11 | 3786 | 806 | 9511 | 236 | 30 | 0 | 0 | 0 | 0 | 3198 | 11131 | 110573 | 18 | 0 | 16731 | 145 | 0 | 0 | 117 | 0 | 0 | 21 | 0 | 0 | 128 | 0 | 18 | 0 | 0 | 9017 | 0 | 0 |
| SRR6020510 | 0 | 0 | 0 | 0 | 15 | 7832 | 547 | 16822 | 468 | 31 | 0 | 0 | 0 | 0 | 2404 | 16391 | 138138 | 14 | 0 | 18362 | 331 | 0 | 0 | 234 | 0 | 0 | 0 | 0 | 0 | 114 | 0 | 0 | 0 | 0 | 9265 | 0 | 0 |
| SRR6020511 | 0 | 0 | 0 | 0 | 0 | 1529 | 597 | 3181 | 168 | 18 | 0 | 0 | 0 | 0 | 893 | 7820 | 52028 | 0 | 0 | 13250 | 89 | 0 | 0 | 46 | 0 | 0 | 20 | 0 | 0 | 40 | 0 | 0 | 0 | 0 | 11282 | 0 | 0 |
| SRR6020512 | 0 | 0 | 0 | 0 | 0 | 3696 | 551 | 7228 | 226 | 28 | 0 | 0 | 0 | 0 | 2993 | 13714 | 120287 | 0 | 0 | 32161 | 189 | 0 | 0 | 134 | 0 | 0 | 12 | 0 | 0 | 160 | 0 | 0 | 0 | 0 | 10111 | 0 | 0 |
| SRR6020513 | 0 | 0 | 0 | 0 | 0 | 3979 | 653 | 9716 | 281 | 18 | 0 | 0 | 0 | 0 | 1060 | 8545 | 70406 | 0 | 0 | 5012 | 91 | 0 | 0 | 18 | 0 | 0 | 29 | 0 | 0 | 21 | 0 | 0 | 0 | 0 | 11929 | 0 | 0 |
| SRR6020514 | 0 | 0 | 0 | 0 | 0 | 3373 | 642 | 7492 | 189 | 30 | 0 | 0 | 0 | 0 | 850 | 6397 | 43315 | 0 | 0 | 11440 | 47 | 0 | 0 | 80 | 0 | 0 | 26 | 0 | 0 | 26 | 0 | 0 | 0 | 0 | 4319 | 0 | 0 |
| SRR6020515 | 0 | 0 | 0 | 0 | 13 | 13728 | 1384 | 31281 | 759 | 41 | 0 | 0 | 0 | 0 | 519 | 5825 | 34922 | 0 | 0 | 5556 | 20 | 0 | 0 | 121 | 0 | 0 | 17 | 0 | 0 | 139 | 0 | 0 | 0 | 0 | 8941 | 0 | 12 |
| SRR6020516 | 0 | 0 | 0 | 0 | 30 | 3108 | 468 | 7847 | 339 | 47 | 13 | 0 | 0 | 0 | 4271 | 33147 | 243942 | 0 | 0 | 22804 | 172 | 0 | 0 | 475 | 0 | 0 | 99 | 0 | 0 | 682 | 0 | 0 | 0 | 0 | 6496 | 0 | 11 |
| SRR6020517 | 0 | 0 | 0 | 0 | 30 | 4601 | 970 | 17710 | 687 | 71 | 23 | 13 | 0 | 0 | 2641 | 33932 | 155355 | 0 | 22 | 38951 | 205 | 0 | 0 | 212 | 0 | 28 | 88 | 0 | 0 | 868 | 0 | 0 | 0 | 0 | 4271 | 0 | 12 |
| SRR6020518 | 0 | 0 | 0 | 0 | 36 | 2187 | 166 | 4633 | 157 | 0 | 0 | 0 | 0 | 0 | 2344 | 15412 | 116647 | 0 | 0 | 13335 | 138 | 0 | 0 | 166 | 0 | 10 | 52 | 0 | 0 | 245 | 0 | 0 | 0 | 0 | 3937 | 0 | 0 |
| SRR6020519 | 0 | 0 | 0 | 0 | 19 | 6898 | 535 | 16864 | 578 | 61 | 0 | 0 | 0 | 0 | 1298 | 13318 | 100415 | 0 | 0 | 5860 | 127 | 0 | 0 | 42 | 0 | 0 | 54 | 0 | 0 | 49 | 0 | 11 | 0 | 0 | 11508 | 0 | 0 |
| SRR6020520 | 0 | 0 | 0 | 0 | 12 | 8609 | 1253 | 21903 | 862 | 69 | 0 | 0 | 0 | 0 | 1771 | 16371 | 86970 | 0 | 21 | 37677 | 254 | 0 | 0 | 32 | 0 | 25 | 60 | 0 | 0 | 47 | 0 | 16 | 0 | 0 | 5913 | 0 | 13 |
| SRR6020521 | 0 | 0 | 0 | 0 | 14 | 7301 | 801 | 17557 | 485 | 41 | 0 | 0 | 0 | 0 | 2604 | 14283 | 138294 | 0 | 15 | 13246 | 170 | 0 | 0 | 219 | 0 | 14 | 64 | 0 | 0 | 392 | 0 | 0 | 0 | 0 | 6149 | 0 | 0 |
| SRR6020522 | 0 | 0 | 0 | 0 | 0 | 1728 | 351 | 3484 | 261 | 14 | 0 | 0 | 0 | 0 | 1746 | 10726 | 88302 | 0 | 0 | 20839 | 99 | 0 | 0 | 26 | 0 | 0 | 21 | 0 | 0 | 61 | 0 | 18 | 0 | 0 | 9935 | 0 | 13 |
| SRR6020523 | 0 | 0 | 0 | 0 | 25 | 3320 | 1187 | 11742 | 464 | 11 | 15 | 0 | 0 | 0 | 3914 | 33983 | 187377 | 0 | 0 | 10963 | 139 | 0 | 0 | 40 | 0 | 0 | 20 | 0 | 0 | 85 | 0 | 19 | 0 | 0 | 12654 | 0 | 12 |
| SRR6020524 | 0 | 0 | 0 | 0 | 0 | 6740 | 575 | 12491 | 362 | 21 | 0 | 0 | 0 | 0 | 759 | 10002 | 51735 | 0 | 18 | 18715 | 77 | 0 | 0 | 21 | 0 | 0 | 11 | 0 | 0 | 15 | 0 | 0 | 0 | 0 | 10104 | 0 | 0 |
| SRR6020525 | 0 | 0 | 0 | 0 | 11 | 3130 | 875 | 6865 | 737 | 42 | 0 | 0 | 0 | 0 | 1467 | 8551 | 46222 | 0 | 25 | 14347 | 176 | 0 | 0 | 80 | 0 | 0 | 0 | 0 | 0 | 11 | 0 | 0 | 0 | 0 | 7927 | 0 | 0 |
| SRR6020526 | 0 | 0 | 0 | 0 | 13 | 2777 | 438 | 6470 | 164 | 43 | 0 | 0 | 0 | 0 | 1772 | 13952 | 85456 | 21 | 14 | 16634 | 109 | 0 | 0 | 106 | 0 | 13 | 48 | 0 | 0 | 361 | 0 | 0 | 0 | 0 | 16728 | 0 | 14 |
| SRR6020527 | 0 | 0 | 0 | 0 | 22 | 7636 | 1317 | 20471 | 1105 | 60 | 11 | 0 | 0 | 0 | 1250 | 8385 | 41962 | 0 | 20 | 24458 | 114 | 0 | 0 | 430 | 0 | 0 | 48 | 0 | 0 | 125 | 0 | 0 | 0 | 0 | 5157 | 0 | 10 |
| SRR6020528 | 0 | 0 | 0 | 0 | 11 | 1354 | 359 | 5700 | 463 | 38 | 27 | 0 | 0 | 0 | 4613 | 21254 | 155967 | 0 | 25 | 32318 | 231 | 0 | 0 | 101 | 11 | 35 | 97 | 0 | 0 | 218 | 0 | 0 | 0 | 0 | 3933 | 0 | 0 |
| SRR6020529 | 0 | 0 | 0 | 0 | 0 | 15561 | 1314 | 31306 | 782 | 23 | 0 | 0 | 0 | 0 | 1033 | 8727 | 68264 | 0 | 0 | 13738 | 81 | 0 | 0 | 29 | 0 | 0 | 31 | 0 | 0 | 30 | 0 | 0 | 0 | 0 | 4900 | 0 | 0 |
| SRR6020530 | 0 | 0 | 0 | 0 | 0 | 14505 | 537 | 20007 | 445 | 25 | 0 | 0 | 0 | 0 | 1380 | 6391 | 46368 | 0 | 0 | 4826 | 130 | 0 | 0 | 21 | 0 | 0 | 0 | 0 | 0 | 13 | 0 | 0 | 0 | 0 | 6246 | 0 | 0 |
| SRR6020531 | 0 | 0 | 0 | 0 | 0 | 6963 | 1052 | 13014 | 520 | 24 | 0 | 0 | 0 | 0 | 202 | 3354 | 15347 | 0 | 0 | 1725 | 24 | 0 | 0 | 32 | 0 | 0 | 0 | 0 | 0 | 30 | 0 | 0 | 0 | 0 | 13302 | 0 | 0 |
| SRR6020532 | 0 | 0 | 0 | 0 | 10 | 3062 | 684 | 7768 | 302 | 44 | 0 | 0 | 0 | 0 | 1896 | 14624 | 119808 | 0 | 15 | 18282 | 246 | 0 | 0 | 205 | 0 | 0 | 73 | 0 | 0 | 165 | 0 | 15 | 0 | 0 | 8354 | 0 | 0 |
| SRR6025030 | 48 | 26 | 0 | 0 | 86 | 428 | 165 | 894 | 1004 | 21438 | 365 | 29 | 0 | 0 | 152467 | 91702 | 521151 | 253 | 983 | 7213 | 29920 | 0 | 0 | 104 | 51 | 63 | 173 | 72 | 0 | 1916 | 0 | 0 | 0 | 0 | 629 | 151 | 0 |
| SRR6025031 | 0 | 0 | 0 | 0 | 145 | 0 | 0 | 0 | 0 | 0 | 71 | 0 | 0 | 0 | 8893 | 12548 | 97710 | 0 | 0 | 928 | 1596 | 0 | 0 | 0 | 0 | 15 | 23 | 89 | 0 | 23 | 0 | 0 | 0 | 0 | 0 | 0 | 0 |
| SRR6025033 | 0 | 0 | 14 | 0 | 0 | 43 | 0 | 85 | 21 | 0 | 1957 | 0 | 0 | 0 | 49947 | 13948 | 173647 | 0 | 0 | 0 | 0 | 0 | 0 | 11 | 0 | 427 | 5153 | 1763 | 0 | 0 | 0 | 0 | 0 | 0 | 110 | 0 | 0 |
| SRR6025035 | 0 | 0 | 0 | 12 | 15 | 111 | 94 | 171 | 30 | 19 | 146 | 0 | 0 | 0 | 88965 | 33261 | 319299 | 18 | 82 | 11365 | 8777 | 0 | 0 | 95 | 70 | 96 | 307 | 72 | 0 | 1052 | 0 | 34 | 0 | 0 | 1000 | 15 | 0 |
| SRR6025036 | 0 | 0 | 0 | 15 | 18 | 797 | 548 | 2249 | 454 | 11 | 27 | 0 | 0 | 0 | 26992 | 71809 | 187192 | 131 | 1066 | 25311 | 1718 | 0 | 0 | 16 | 26 | 49 | 66 | 15 | 0 | 267 | 0 | 64 | 0 | 0 | 17821 | 0 | 0 |
| SRR6025037 | 0 | 0 | 0 | 13 | 21 | 46 | 63 | 135 | 23 | 18 | 143 | 0 | 0 | 0 | 89311 | 35682 | 331656 | 20 | 81 | 11812 | 9031 | 0 | 0 | 85 | 89 | 121 | 386 | 299 | 0 | 1119 | 0 | 39 | 0 | 0 | 750 | 18 | 0 |
| SRR6025038 | 0 | 0 | 0 | 0 | 439 | 0 | 0 | 0 | 0 | 0 | 0 | 0 | 0 | 0 | 23391 | 4814 | 218161 | 0 | 0 | 0 | 0 | 0 | 0 | 0 | 1159 | 0 | 465 | 341 | 0 | 570 | 0 | 0 | 0 | 0 | 21 | 43 | 0 |
| SRR6025041 | 13 | 0 | 0 | 0 | 194 | 845 | 140 | 682 | 1281 | 10639 | 0 | 31 | 11 | 0 | 964 | 25201 | 33965 | 11 | 230 | 310 | 5471 | 0 | 0 | 19 | 0 | 0 | 0 | 0 | 0 | 100 | 0 | 213 | 0 | 0 | 120256 | 13 | 0 |
| SRR6025042 | 0 | 0 | 0 | 10 | 17 | 0 | 385 | 0 | 0 | 0 | 505 | 0 | 0 | 0 | 90521 | 8807 | 211533 | 0 | 0 | 0 | 0 | 0 | 0 | 772 | 0 | 699 | 991 | 1416 | 0 | 288 | 0 | 0 | 0 | 0 | 29 | 0 | 121 |
| SRR6025043 | 0 | 0 | 0 | 0 | 35 | 91 | 111 | 191 | 37 | 20 | 17 | 0 | 0 | 0 | 97663 | 38044 | 332432 | 29 | 99 | 12726 | 10687 | 0 | 0 | 18 | 11 | 11 | 31 | 25 | 0 | 142 | 0 | 47 | 0 | 0 | 767 | 21 | 0 |
| SRR6025044 | 0 | 0 | 0 | 0 | 73 | 103 | 283 | 396 | 142 | 41 | 27 | 0 | 0 | 0 | 12291 | 31321 | 201935 | 22 | 375 | 19028 | 8885 | 0 | 29 | 231 | 214 | 331 | 837 | 40 | 0 | 1026 | 0 | 123 | 0 | 0 | 1760 | 0 | 0 |
| SRR6025047 | 34 | 18 | 0 | 0 | 36 | 1542 | 132 | 1057 | 962 | 1248 | 40 | 12 | 0 | 0 | 19788 | 18858 | 68899 | 474 | 2050 | 1207 | 5758 | 0 | 0 | 11 | 0 | 0 | 0 | 0 | 0 | 92 | 0 | 53 | 0 | 50 | 153276 | 0 | 0 |
| SRR6025048 | 11 | 13 | 0 | 0 | 63 | 505 | 48 | 342 | 819 | 1684 | 171 | 15 | 0 | 0 | 13560 | 51818 | 157599 | 118 | 1084 | 765 | 4953 | 0 | 0 | 11 | 14 | 11 | 13 | 16 | 0 | 318 | 0 | 89 | 0 | 0 | 149999 | 0 | 0 |
| SRR6025050 | 54 | 21 | 0 | 15 | 10 | 440 | 58 | 275 | 581 | 2402 | 29 | 50 | 0 | 0 | 121979 | 54281 | 482785 | 104 | 210 | 4266 | 48777 | 0 | 0 | 60 | 50 | 69 | 92 | 21 | 0 | 64 | 0 | 0 | 0 | 0 | 3973 | 79 | 0 |
| SRR6025051 | 0 | 0 | 0 | 12 | 19 | 911 | 755 | 2334 | 394 | 0 | 11 | 0 | 0 | 0 | 27050 | 84218 | 200118 | 135 | 1143 | 29211 | 2178 | 0 | 0 | 18 | 16 | 26 | 24 | 30 | 0 | 154 | 0 | 75 | 0 | 0 | 17051 | 0 | 0 |

### Supp Table S8.3.Columns_BX_to_DH

| | | | | | | | | | | | | | | | | | | | | | | | | | | | | | | | | | | | | | |
|---|---|---|---|---|---|---|---|---|---|---|---|---|---|---|---|---|---|---|---|---|---|---|---|---|---|---|---|---|---|---|---|---|---|---|---|---|---|
| SRR6025053 | 0 | 0 | 0 | 10 | 38 | 108 | 220 | 282 | 129 | 23 | 15 | 0 | 0 | 0 | 9764 | 21235 | 157497 | 12 | 490 | 19756 | 9548 | 0 | 0 | 15 | 25 | 51 | 188 | 130 | 0 | 97 | 0 | 129 | 0 | 0 | 1498 | 0 | 0 |
| SRR6025055 | 83 | 44 | 11 | 11 | 27 | 731 | 681 | 2177 | 4594 | 2684 | 12 | 61 | 0 | 0 | 10803 | 103024 | 265557 | 172 | 4351 | 11749 | 24090 | 17 | 88 | 10 | 61 | 14 | 58 | 19 | 0 | 72 | 0 | 25 | 0 | 0 | 2348 | 15 | 0 |
| SRR6025056 | 62 | 21 | 0 | 18 | 106 | 4630 | 1973 | 11994 | 12382 | 3936 | 332 | 12 | 0 | 0 | 40441 | 177423 | 265700 | 1383 | 13050 | 12243 | 3698 | 0 | 0 | 120 | 85 | 85 | 253 | 68 | 0 | 2464 | 0 | 0 | 0 | 0 | 11553 | 0 | 0 |
| SRR6025058 | 0 | 0 | 0 | 19 | 56 | 57 | 125 | 210 | 96 | 17 | 44 | 0 | 0 | 0 | 10655 | 19826 | 151821 | 19 | 504 | 20640 | 10281 | 0 | 11 | 123 | 231 | 525 | 1716 | 523 | 0 | 955 | 0 | 112 | 0 | 0 | 1025 | 0 | 0 |
| SRR6025061 | 0 | 0 | 0 | 42 | 62 | 41 | 326 | 305 | 109 | 12 | 22 | 13 | 0 | 0 | 22915 | 45097 | 268342 | 21 | 450 | 23460 | 15644 | 0 | 0 | 125 | 97 | 416 | 2012 | 1534 | 0 | 370 | 0 | 45 | 0 | 0 | 843 | 0 | 0 |
| SRR6025062 | 0 | 0 | 0 | 23 | 34 | 33 | 280 | 310 | 85 | 28 | 64 | 16 | 0 | 0 | 26202 | 29264 | 194972 | 23 | 457 | 24091 | 18722 | 0 | 10 | 157 | 128 | 588 | 2130 | 295 | 0 | 520 | 0 | 265 | 0 | 0 | 374 | 0 | 0 |
| SRR6025065 | 0 | 0 | 14 | 19 | 31 | 15 | 103 | 95 | 40 | 19 | 135 | 0 | 0 | 0 | 192659 | 68549 | 459464 | 29 | 104 | 15242 | 15390 | 0 | 0 | 116 | 33 | 122 | 33 | 36 | 0 | 13 | 0 | 14 | 0 | 0 | 287 | 15 | 0 |
| SRR6025066 | 26 | 21 | 76 | 113 | 807 | 44 | 453 | 243 | 272 | 0 | 66 | 0 | 0 | 0 | 9393 | 81836 | 172982 | 0 | 238 | 122 | 189 | 10 | 31 | 107 | 38 | 200 | 1522 | 1386 | 0 | 103 | 0 | 0 | 0 | 0 | 75 | 0 | 0 |
| SRR6025067 | 0 | 0 | 13 | 36 | 22 | 10 | 48 | 67 | 14 | 0 | 19 | 0 | 0 | 0 | 180175 | 50264 | 391669 | 25 | 77 | 13871 | 13421 | 0 | 0 | 97 | 40 | 11 | 44 | 25 | 0 | 37 | 0 | 51 | 0 | 0 | 256 | 11 | 0 |
| SRR6025068 | 0 | 0 | 0 | 30 | 17 | 90 | 262 | 459 | 54 | 0 | 27 | 12 | 0 | 0 | 112303 | 48498 | 411047 | 111 | 1433 | 33246 | 10878 | 0 | 0 | 15 | 41 | 12 | 36 | 22 | 0 | 45 | 0 | 21 | 0 | 0 | 847 | 0 | 0 |
| SRR6025069 | 11 | 0 | 11 | 41 | 20 | 108 | 267 | 383 | 74 | 0 | 154 | 0 | 0 | 0 | 139627 | 66279 | 463189 | 101 | 1785 | 33236 | 18875 | 0 | 0 | 128 | 23 | 105 | 19 | 29 | 0 | 138 | 0 | 0 | 0 | 0 | 677 | 0 | 0 |
| SRR6025070 | 25 | 36 | 0 | 0 | 36 | 274 | 48 | 231 | 218 | 2070 | 32 | 53 | 0 | 0 | 49041 | 45462 | 165494 | 62 | 133 | 434 | 5888 | 0 | 0 | 13 | 18 | 14 | 0 | 0 | 0 | 84 | 0 | 33 | 0 | 38 | 29516 | 53 | 0 |
| SRR6025071 | 44 | 21 | 0 | 0 | 189 | 145 | 12 | 72 | 241 | 7691 | 218 | 28 | 0 | 0 | 44404 | 72940 | 193147 | 35 | 53 | 308 | 3719 | 0 | 0 | 15 | 23 | 46 | 27 | 26 | 0 | 539 | 0 | 26 | 0 | 0 | 33582 | 15 | 0 |
| SRR6025072 | 0 | 11 | 0 | 10 | 0 | 27 | 185 | 296 | 98 | 0 | 19 | 14 | 0 | 0 | 5110 | 22561 | 95461 | 22 | 405 | 10005 | 6548 | 0 | 0 | 51 | 16 | 10 | 68 | 18 | 0 | 108 | 0 | 146 | 0 | 0 | 490 | 0 | 0 |
| SRR6025073 | 0 | 0 | 0 | 0 | 0 | 40 | 146 | 405 | 55 | 0 | 171 | 0 | 0 | 0 | 24054 | 18794 | 128463 | 153 | 1551 | 19766 | 6095 | 0 | 0 | 40 | 12 | 18 | 40 | 14 | 0 | 400 | 0 | 26 | 0 | 0 | 351 | 0 | 0 |
| SRR6025075 | 0 | 0 | 0 | 11 | 0 | 61 | 228 | 515 | 58 | 0 | 53 | 0 | 0 | 0 | 26561 | 23758 | 162403 | 90 | 973 | 15575 | 7673 | 0 | 0 | 42 | 20 | 34 | 30 | 10 | 0 | 45 | 0 | 20 | 0 | 0 | 537 | 0 | 0 |
| SRR6025079 | 61 | 40 | 12 | 11 | 276 | 11 | 0 | 24 | 114 | 10552 | 552 | 35 | 0 | 0 | 59063 | 75882 | 234946 | 27 | 55 | 342 | 4312 | 0 | 0 | 31 | 89 | 113 | 74 | 85 | 0 | 1202 | 0 | 25 | 0 | 0 | 6402 | 27 | 0 |
| SRR6025080 | 60 | 18 | 16 | 25 | 84 | 2522 | 402 | 1769 | 3472 | 3749 | 804 | 55 | 0 | 0 | 81271 | 104298 | 415054 | 1100 | 14996 | 12518 | 20208 | 0 | 0 | 58 | 47 | 58 | 85 | 35 | 0 | 1859 | 0 | 41 | 0 | 20 | 33332 | 0 | 0 |
| SRR6025082 | 36 | 0 | 0 | 0 | 282 | 1808 | 1575 | 4796 | 15104 | 2710 | 139 | 31 | 0 | 0 | 8432 | 90130 | 311011 | 73 | 2042 | 7358 | 10934 | 0 | 15 | 71 | 153 | 266 | 181 | 231 | 0 | 455 | 0 | 148 | 0 | 0 | 5017 | 11 | 0 |
| SRR6025083 | 65 | 32 | 14 | 17 | 96 | 712 | 153 | 600 | 1063 | 17642 | 581 | 36 | 0 | 0 | 162052 | 125630 | 407608 | 263 | 782 | 5902 | 22871 | 0 | 0 | 61 | 59 | 71 | 130 | 34 | 0 | 1912 | 0 | 51 | 0 | 12 | 4199 | 116 | 0 |
| SRR6025084 | 62 | 29 | 0 | 0 | 41 | 1679 | 555 | 2118 | 4130 | 2788 | 17 | 53 | 0 | 0 | 12234 | 111856 | 232934 | 88 | 4398 | 5921 | 23452 | 13 | 59 | 12 | 88 | 87 | 14 | 17 | 0 | 86 | 0 | 265 | 0 | 13 | 16493 | 17 | 0 |
| SRR6025085 | 33 | 15 | 0 | 0 | 19 | 798 | 530 | 1689 | 5013 | 402 | 55 | 27 | 0 | 0 | 139406 | 91533 | 420781 | 382 | 429 | 5201 | 9895 | 0 | 0 | 44 | 27 | 31 | 14 | 20 | 0 | 151 | 0 | 12 | 0 | 0 | 2499 | 80 | 0 |
| SRR6025086 | 32 | 31 | 109 | 146 | 325 | 135 | 404 | 342 | 162 | 0 | 159 | 0 | 0 | 0 | 46919 | 67599 | 250777 | 94 | 1577 | 667 | 147 | 0 | 0 | 57 | 12 | 47 | 269 | 509 | 0 | 78 | 0 | 0 | 0 | 0 | 219 | 0 | 0 |
| SRR6025087 | 30 | 16 | 67 | 58 | 30 | 24 | 108 | 62 | 39 | 0 | 85 | 0 | 0 | 0 | 79013 | 87171 | 267841 | 13 | 68 | 186 | 108 | 0 | 0 | 74 | 24 | 94 | 37 | 63 | 0 | 71 | 0 | 0 | 0 | 0 | 46 | 35 | 0 |
| SRR6025088 | 21 | 0 | 0 | 0 | 21 | 2339 | 1101 | 3967 | 13782 | 124 | 91 | 42 | 0 | 0 | 52898 | 116037 | 443978 | 968 | 6779 | 14479 | 12280 | 0 | 0 | 37 | 20 | 26 | 11 | 20 | 0 | 155 | 0 | 17 | 0 | 0 | 11745 | 0 | 0 |
| SRR6025091 | 19 | 29 | 0 | 0 | 70 | 519 | 141 | 514 | 685 | 3720 | 10 | 46 | 0 | 0 | 3689 | 22314 | 61992 | 22 | 377 | 527 | 4565 | 0 | 18 | 19 | 16 | 19 | 37 | 0 | 0 | 31 | 0 | 146 | 0 | 39 | 38578 | 0 | 0 |
| SRR6025092 | 0 | 0 | 0 | 0 | 0 | 14 | 57 | 77 | 14 | 0 | 387 | 0 | 0 | 0 | 41438 | 22655 | 153209 | 29 | 78 | 8688 | 7016 | 0 | 0 | 46 | 0 | 33 | 261 | 110 | 0 | 1036 | 0 | 39 | 0 | 0 | 218 | 37 | 0 |
| SRR6025093 | 23 | 15 | 0 | 0 | 42 | 353 | 69 | 295 | 467 | 889 | 99 | 24 | 0 | 0 | 55929 | 55845 | 214764 | 57 | 176 | 504 | 2361 | 0 | 0 | 18 | 13 | 16 | 19 | 37 | 0 | 222 | 0 | 32 | 0 | 0 | 19287 | 42 | 0 |
| SRR6025094 | 0 | 0 | 0 | 10 | 14 | 0 | 62 | 62 | 0 | 11 | 25 | 12 | 0 | 0 | 47847 | 24621 | 178322 | 30 | 72 | 8297 | 8131 | 0 | 0 | 50 | 11 | 50 | 36 | 53 | 0 | 31 | 0 | 54 | 0 | 0 | 265 | 11 | 0 |
| SRR6025095 | 65 | 60 | 0 | 0 | 339 | 169 | 21 | 116 | 199 | 14873 | 111 | 24 | 0 | 0 | 34120 | 60417 | 183299 | 46 | 48 | 390 | 8594 | 0 | 0 | 18 | 15 | 24 | 20 | 0 | 0 | 347 | 0 | 57 | 0 | 0 | 47077 | 22 | 0 |
| SRR6025096 | 35 | 51 | 0 | 0 | 76 | 285 | 63 | 352 | 434 | 3379 | 18 | 73 | 0 | 0 | 3613 | 28211 | 82049 | 16 | 476 | 518 | 4802 | 0 | 19 | 23 | 37 | 43 | 48 | 0 | 0 | 50 | 0 | 102 | 0 | 64 | 31899 | 0 | 0 |
| SRR6025097 | 37 | 30 | 0 | 0 | 343 | 493 | 140 | 419 | 1157 | 4921 | 223 | 40 | 0 | 0 | 2145 | 39320 | 111091 | 15 | 472 | 657 | 2138 | 0 | 0 | 29 | 80 | 124 | 46 | 19 | 0 | 354 | 0 | 86 | 0 | 13 | 24443 | 0 | 0 |
| SRR6025099 | 0 | 0 | 0 | 0 | 20 | 713 | 0 | 0 | 140 | 0 | 24 | 0 | 0 | 0 | 5597 | 11532 | 109803 | 0 | 0 | 0 | 457 | 0 | 0 | 0 | 0 | 0 | 165 | 61 | 0 | 0 | 0 | 0 | 0 | 0 | 14 | 0 | 0 |
| SRR6025101 | 14 | 34 | 0 | 0 | 140 | 14 | 0 | 43 | 314 | 880 | 38 | 14 | 0 | 0 | 1622 | 67404 | 142076 | 0 | 160 | 300 | 3512 | 0 | 10 | 22 | 21 | 45 | 24 | 28 | 0 | 105 | 0 | 119 | 0 | 0 | 17445 | 0 | 0 |
| SRR6025103 | 12 | 15 | 0 | 0 | 55 | 853 | 23 | 331 | 731 | 336 | 122 | 13 | 0 | 0 | 11580 | 50449 | 138819 | 163 | 983 | 802 | 4396 | 0 | 0 | 14 | 20 | 28 | 11 | 0 | 0 | 245 | 0 | 66 | 0 | 0 | 164040 | 0 | 0 |
| SRR6025104 | 14 | 0 | 0 | 0 | 70 | 907 | 69 | 622 | 1483 | 4385 | 47 | 58 | 12 | 0 | 1914 | 20928 | 34925 | 124 | 275 | 342 | 925 | 0 | 0 | 0 | 0 | 0 | 0 | 0 | 0 | 222 | 0 | 77 | 0 | 0 | 195507 | 0 | 0 |
| SRR6025105 | 0 | 0 | 0 | 0 | 33 | 0 | 169 | 149 | 50 | 13 | 137 | 20 | 0 | 0 | 8480 | 33048 | 162646 | 0 | 203 | 7591 | 6964 | 0 | 0 | 23 | 16 | 74 | 751 | 489 | 0 | 336 | 0 | 191 | 0 | 0 | 874 | 0 | 0 |
| SRR6025106 | 35 | 25 | 0 | 0 | 409 | 236 | 42 | 155 | 578 | 15563 | 82 | 24 | 0 | 0 | 1380 | 49430 | 86371 | 0 | 206 | 248 | 3069 | 0 | 11 | 22 | 34 | 76 | 37 | 14 | 0 | 253 | 0 | 233 | 0 | 0 | 48553 | 0 | 0 |
| SRR6025108 | 0 | 0 | 0 | 0 | 0 | 0 | 0 | 1581 | 0 | 0 | 0 | 0 | 0 | 0 | 106194 | 7905 | 205909 | 0 | 0 | 0 | 0 | 0 | 0 | 0 | 45 | 0 | 132 | 43 | 0 | 0 | 0 | 0 | 0 | 0 | 17 | 0 | 0 |
| SRR6025111 | 76 | 50 | 0 | 17 | 315 | 958 | 458 | 2764 | 3069 | 54565 | 160 | 46 | 0 | 0 | 11661 | 90182 | 330963 | 110 | 5052 | 7835 | 26098 | 29 | 76 | 139 | 84 | 92 | 554 | 151 | 0 | 878 | 0 | 31 | 0 | 0 | 1928 | 19 | 0 |
| SRR6025113 | 26 | 26 | 0 | 0 | 69 | 3508 | 839 | 4652 | 10524 | 404 | 16 | 0 | 0 | 0 | 38624 | 118407 | 254651 | 686 | 7702 | 9990 | 8884 | 0 | 0 | 55 | 51 | 13 | 26 | 14 | 0 | 67 | 0 | 0 | 0 | 0 | 34751 | 0 | 0 |
| SRR6025114 | 43 | 11 | 0 | 11 | 12 | 3277 | 599 | 5199 | 11912 | 393 | 13 | 14 | 0 | 0 | 30913 | 129522 | 197495 | 630 | 5483 | 13645 | 7342 | 0 | 0 | 11 | 83 | 14 | 30 | 12 | 0 | 59 | 0 | 0 | 0 | 11 | 39108 | 0 | 0 |
| SRR6025116 | 62 | 41 | 0 | 0 | 23 | 158 | 0 | 91 | 585 | 2220 | 29 | 14 | 0 | 0 | 61362 | 41213 | 366611 | 19 | 16 | 1174 | 49365 | 0 | 0 | 18 | 56 | 10 | 52 | 28 | 0 | 62 | 0 | 0 | 0 | 0 | 15526 | 29 | 0 |
| SRR6025119 | 33 | 19 | 0 | 0 | 187 | 196 | 0 | 90 | 229 | 2503 | 222 | 28 | 0 | 0 | 46299 | 70477 | 193412 | 41 | 42 | 290 | 3450 | 0 | 0 | 12 | 25 | 37 | 18 | 0 | 0 | 641 | 0 | 27 | 0 | 0 | 36781 | 10 | 0 |
| SRR6025120 | 0 | 0 | 0 | 11 | 14 | 0 | 130 | 201 | 58 | 0 | 173 | 11 | 0 | 0 | 8420 | 19724 | 115193 | 30 | 177 | 8305 | 8192 | 0 | 0 | 49 | 24 | 129 | 1241 | 175 | 0 | 469 | 0 | 147 | 0 | 0 | 714 | 0 | 11 |
| SRR6025121 | 0 | 0 | 0 | 12 | 18 | 87 | 198 | 365 | 39 | 0 | 199 | 18 | 0 | 0 | 50993 | 37244 | 248360 | 67 | 933 | 12111 | 9233 | 0 | 0 | 38 | 0 | 36 | 164 | 250 | 0 | 121 | 0 | 21 | 0 | 0 | 701 | 0 | 0 |
| SRR6025123 | 24 | 16 | 0 | 0 | 35 | 1022 | 126 | 601 | 552 | 944 | 92 | 36 | 0 | 0 | 32383 | 27681 | 125560 | 303 | 2253 | 1239 | 5152 | 0 | 0 | 11 | 14 | 21 | 17 | 0 | 0 | 153 | 0 | 48 | 0 | 14 | 119125 | 0 | 0 |
| SRR6025125 | 29 | 22 | 0 | 0 | 400 | 197 | 26 | 121 | 541 | 14150 | 122 | 22 | 0 | 0 | 1426 | 54156 | 96062 | 0 | 203 | 293 | 2879 | 0 | 10 | 12 | 45 | 73 | 59 | 72 | 0 | 232 | 0 | 184 | 0 | 0 | 43720 | 0 | 0 |
| SRR6025126 | 32 | 15 | 0 | 0 | 154 | 624 | 77 | 297 | 799 | 2994 | 463 | 39 | 0 | 0 | 12230 | 46518 | 146364 | 166 | 813 | 693 | 4260 | 0 | 0 | 12 | 14 | 43 | 12 | 19 | 0 | 715 | 0 | 104 | 0 | 0 | 152199 | 0 | 0 |
| SRR6025128 | 22 | 39 | 0 | 0 | 36 | 217 | 24 | 166 | 158 | 2172 | 51 | 32 | 0 | 0 | 56597 | 46215 | 169612 | 60 | 132 | 440 | 3955 | 0 | 0 | 0 | 24 | 29 | 27 | 0 | 0 | 117 | 0 | 21 | 0 | 21 | 29577 | 63 | 0 |
| SRR6040044 | 310 | 74 | 49 | 28 | 15 | 370 | 131 | 459 | 1378 | 1844 | 34 | 0 | 14 | 0 | 20220 | 319202 | 52531 | 83 | 1433 | 1702 | 49228 | 0 | 0 | 27 | 14 | 34 | 28 | 0 | 0 | 47 | 0 | 67 | 0 | 0 | 2285 | 0 | 0 |
| SRR6040045 | 5501 | 1253 | 29 | 20 | 258 | 298 | 114 | 528 | 1398 | 26459 | 17 | 0 | 14 | 13 | 15855 | 323109 | 42898 | 81 | 1008 | 1383 | 47715 | 0 | 0 | 375 | 107 | 20 | 17 | 0 | 0 | 56 | 0 | 61 | 0 | 0 | 2627 | 0 | 0 |
| SRR6076852 | 0 | 0 | 0 | 0 | 39 | 39 | 0 | 39 | 0 | 164 | 0 | 0 | 0 | 0 | 549 | 336366 | 12707 | 353 | 118 | 0 | 2472 | 16 | 55 | 0 | 0 | 0 | 0 | 0 | 0 | 0 | 102 | 196 | 0 | 0 | 19670 | 11 | 0 |
| SRR6076853 | 0 | 0 | 0 | 0 | 76 | 207 | 0 | 92 | 92 | 575 | 0 | 49 | 0 | 0 | 645 | 440263 | 19386 | 184 | 69 | 115 | 3387 | 49 | 14 | 35 | 0 | 0 | 0 | 0 | 0 | 0 | 118 | 230 | 0 | 0 | 13674 | 0 | 0 |
| SRR6076854 | 64 | 21 | 11 | 0 | 149 | 36 | 0 | 36 | 109 | 650 | 85 | 53 | 0 | 0 | 1121 | 499825 | 57645 | 109 | 127 | 54 | 3002 | 43 | 11 | 21 | 11 | 0 | 0 | 0 | 0 | 11 | 75 | 235 | 11 | 0 | 10565 | 14 | 0 |
| SRR6076855 | 0 | 0 | 0 | 0 | 148 | 452 | 85 | 311 | 254 | 392 | 0 | 15 | 0 | 22 | 734 | 268110 | 24888 | 311 | 56 | 113 | 2457 | 30 | 52 | 15 | 0 | 0 | 0 | 0 | 0 | 0 | 96 | 0 | 0 | 0 | 28754 | 0 | 0 |
| SRR6076856 | 0 | 0 | 0 | 0 | 129 | 290 | 0 | 89 | 178 | 397 | 0 | 50 | 0 | 0 | 535 | 245629 | 30265 | 0 | 67 | 45 | 2809 | 11 | 56 | 28 | 0 | 0 | 0 | 0 | 73 | 0 | 84 | 111 | 0 | 0 | 48720 | 0 | 0 |
| SRR6076858 | 0 | 0 | 0 | 0 | 0 | 571 | 76 | 381 | 38 | 502 | 0 | 57 | 0 | 0 | 266 | 105042 | 7331 | 266 | 76 | 38 | 2207 | 0 | 31 | 0 | 0 | 0 | 0 | 0 | 0 | 0 | 61 | 609 | 0 | 0 | 46057 | 16 | 0 |
| SRR6076861 | 16 | 0 | 0 | 0 | 130 | 97 | 0 | 65 | 134 | 616 | 38 | 49 | 0 | 0 | 898 | 433950 | 56432 | 81 | 75 | 32 | 2963 | 27 | 11 | 32 | 0 | 0 | 0 | 0 | 59 | 27 | 32 | 65 | 0 | 11 | 12482 | 37 | 0 |
| SRR6153965 | 0 | 0 | 0 | 0 | 23 | 72 | 528 | 1311 | 120 | 21 | 120 | 0 | 0 | 0 | 131467 | 16276 | 219732 | 0 | 426 | 9897 | 1401 | 0 | 0 | 58 | 25 | 112 | 148 | 33 | 0 | 531 | 0 | 0 | 0 | 0 | 65014 | 691 | 0 |
| SRR6153968 | 0 | 0 | 0 | 0 | 31 | 96 | 238 | 467 | 71 | 16 | 394 | 0 | 0 | 0 | 149066 | 15245 | 181899 | 35 | 1343 | 9088 | 3102 | 0 | 0 | 190 | 58 | 253 | 446 | 439 | 0 | 1047 | 0 | 13 | 0 | 0 | 11272 | 135 | 0 |
| SRR6153976 | 0 | 0 | 0 | 0 | 24 | 21 | 371 | 120 | 28 | 19 | 27 | 0 | 0 | 0 | 234144 | 23889 | 356022 | 0 | 231 | 4451 | 13221 | 0 | 0 | 32 | 35 | 149 | 125 | 70 | 0 | 187 | 0 | 0 | 0 | 0 | 3229 | 37 | 0 |
| SRR6153984 | 0 | 0 | 0 | 0 | 31 | 151 | 367 | 393 | 85 | 33 | 69 | 0 | 0 | 0 | 19392 | 6233 | 11631 | 21 | 212 | 2506 | 1119 | 0 | 0 | 102 | 44 | 256 | 368 | 33 | 0 | 158 | 0 | 0 | 0 | 0 | 1899 | 0 | 0 |
| SRR6153998 | 0 | 0 | 0 | 0 | 29 | 17 | 77 | 170 | 26 | 24 | 267 | 0 | 0 | 0 | 378110 | 21538 | 591806 | 0 | 341 | 4432 | 6731 | 0 | 0 | 107 | 43 | 145 | 250 | 63 | 0 | 623 | 0 | 0 | 0 | 0 | 32671 | 254 | 0 |
| SRR6154002 | 0 | 0 | 0 | 0 | 24 | 159 | 497 | 829 | 163 | 17 | 12 | 0 | 0 | 0 | 29848 | 29715 | 156138 | 0 | 1172 | 84181 | 9547 | 0 | 0 | 22 | 30 | 17 | 21 | 0 | 0 | 271 | 0 | 17 | 0 | 0 | 62555 | 299 | 0 |
| SRR6154003 | 0 | 0 | 0 | 0 | 53 | 219 | 391 | 1635 | 76 | 470 | 63 | 0 | 0 | 0 | 142367 | 28823 | 266792 | 0 | 210 | 11662 | 14868 | 0 | 0 | 13 | 23 | 37 | 20 | 15 | 0 | 242 | 0 | 11 | 0 | 0 | 139 | 908 | 0 |
| SRR6154004 | 0 | 0 | 0 | 0 | 18 | 96 | 298 | 587 | 88 | 48 | 389 | 0 | 0 | 0 | 116506 | 21327 | 126479 | 0 | 1037 | 10838 | 2450 | 0 | 0 | 120 | 46 | 175 | 248 | 64 | 0 | 2066 | 0 | 0 | 0 | 0 | 35720 | 138 | 0 |
| SRR6154005 | 0 | 0 | 0 | 0 | 34 | 82 | 217 | 720 | 82 | 85 | 141 | 0 | 0 | 0 | 148387 | 31065 | 267703 | 0 | 854 | 14483 | 7054 | 0 | 0 | 121 | 115 | 418 | 390 | 40 | 0 | 703 | 0 | 0 | 0 | 0 | 50679 | 293 | 0 |
| SRR6154006 | 0 | 0 | 0 | 0 | 14 | 35 | 61 | 296 | 44 | 159 | 17 | 0 | 0 | 0 | 170531 | 42344 | 260500 | 0 | 184 | 9462 | 60452 | 11 | 16 | 23 | 13 | 32 | 31 | 19 | 0 | 79 | 0 | 0 | 0 | 0 | 30 | 144 | 0 |
| SRR6203722 | 0 | 0 | 0 | 12 | 0 | 0 | 0 | 0 | 0 | 0 | 50 | 0 | 0 | 0 | 59207 | 52426 | 518014 | 15 | 356 | 29530 | 20319 | 0 | 0 | 37 | 37 | 100 | 500 | 362 | 0 | 162 | 0 | 30 | 0 | 0 | 122 | 0 | 75 |
| SRR6203757 | 0 | 0 | 0 | 23 | 23 | 0 | 0 | 0 | 0 | 23 | 46 | 115 | 0 | 0 | 72714 | 107467 | 512570 | 16 | 313 | 32284 | 20787 | 0 | 0 | 185 | 46 | 185 | 993 | 208 | 0 | 254 | 0 | 31 | 0 | 0 | 113 | 0 | 0 |
| SRR6203760 | 0 | 0 | 0 | 0 | 0 | 593 | 0 | 593 | 0 | 120 | 120 | 0 | 0 | 0 | 11862 | 336415 | 55019 | 0 | 0 | 1779 | 0 | 0 | 0 | 0 | 0 | 0 | 0 | 0 | 0 | 0 | 0 | 0 | 0 | 0 | 1150 | 0 | 0 |
| SRR6203761 | 0 | 0 | 0 | 0 | 0 | 1172 | 0 | 782 | 0 | 0 | 0 | 0 | 0 | 0 | 8597 | 319331 | 60381 | 0 | 0 | 2345 | 391 | 0 | 0 | 0 | 0 | 0 | 0 | 0 | 0 | 0 | 0 | 0 | 0 | 0 | 1595 | 0 | 0 |
| SRR6203772 | 0 | 0 | 0 | 0 | 0 | 0 | 0 | 177 | 0 | 28 | 208 | 0 | 0 | 0 | 85708 | 70147 | 553256 | 0 | 459 | 39408 | 25114 | 0 | 0 | 95 | 47 | 171 | 720 | 237 | 0 | 161 | 0 | 0 | 0 | 0 | 192 | 0 | 0 |
| SRR6203807 | 0 | 0 | 0 | 0 | 41 | 814 | 590 | 5082 | 28 | 0 | 14 | 0 | 0 | 0 | 56490 | 55864 | 500669 | 0 | 197 | 28807 | 16734 | 0 | 0 | 55 | 14 | 27 | 178 | 370 | 0 | 41 | 0 | 0 | 14 | 0 | 3335 | 0 | 0 |
| SRR6224558 | 0 | 0 | 0 | 0 | 0 | 39931 | 8052 | 10994 | 31744 | 46 | 0 | 0 | 0 | 0 | 387 | 65357 | 13813 | 0 | 19 | 39 | 0 | 0 | 20 | 26 | 0 | 0 | 0 | 0 | 0 | 0 | 17 | 0 | 0 | 0 | 6616 | 0 | 0 |
| SRR6226650 | 163 | 15 | 0 | 0 | 38 | 10959 | 949 | 3689 | 1317 | 367 | 0 | 16 | 0 | 0 | 3001 | 169043 | 13644 | 47 | 83 | 5681 | 7591 | 0 | 0 | 80 | 0 | 0 | 0 | 0 | 0 | 0 | 0 | 12 | 0 | 11 | 5751 | 0 | 0 |
| SRR6226651 | 884 | 49 | 0 | 0 | 40 | 4348 | 452 | 1712 | 751 | 1283 | 0 | 14 | 0 | 0 | 7995 | 201610 | 23660 | 121 | 159 | 12757 | 9822 | 0 | 0 | 72 | 0 | 0 | 0 | 0 | 0 | 0 | 0 | 32 | 0 | 40 | 4951 | 0 | 0 |
| SRR6226652 | 204 | 31 | 0 | 0 | 64 | 3116 | 357 | 1784 | 626 | 734 | 0 | 55 | 0 | 0 | 1501 | 414129 | 9978 | 34 | 249 | 9753 | 2605 | 0 | 0 | 62 | 0 | 0 | 0 | 0 | 0 | 0 | 0 | 20 | 0 | 43 | 3555 | 0 | 0 |
| SRR6226653 | 55 | 0 | 0 | 0 | 64 | 1265 | 172 | 702 | 1038 | 43 | 0 | 20 | 0 | 0 | 2638 | 540327 | 12987 | 217 | 104 | 2824 | 702 | 0 | 0 | 57 | 0 | 0 | 0 | 0 | 0 | 0 | 0 | 23 | 31 | 18 | 1142 | 0 | 0 |
| SRR6226654 | 317 | 84 | 0 | 0 | 121 | 1306 | 343 | 893 | 1434 | 42 | 0 | 32 | 0 | 0 | 6092 | 410335 | 54440 | 273 | 227 | 5418 | 2393 | 0 | 10 | 175 | 0 | 0 | 0 | 0 | 0 | 0 | 0 | 0 | 0 | 25 | 648 | 0 | 0 |
| SRR6226655 | 667 | 218 | 0 | 0 | 99 | 470 | 51 | 364 | 634 | 43 | 0 | 65 | 0 | 0 | 6521 | 472489 | 42405 | 247 | 274 | 4376 | 2576 | 0 | 0 | 266 | 0 | 0 | 0 | 0 | 0 | 0 | 0 | 16 | 0 | 19 | 754 | 0 | 0 |
| SRR6226656 | 47 | 24 | 0 | 0 | 27 | 598 | 66 | 215 | 252 | 36 | 0 | 45 | 0 | 0 | 1324 | 126926 | 4433 | 124 | 66 | 2346 | 1479 | 0 | 0 | 12 | 0 | 0 | 0 | 0 | 0 | 0 | 0 | 0 | 0 | 18 | 178 | 0 | 0 |
| SRR6226657 | 501 | 346 | 0 | 0 | 55 | 88 | 18 | 85 | 104 | 54 | 0 | 246 | 0 | 0 | 6578 | 322022 | 20441 | 78 | 93 | 4972 | 2068 | 0 | 0 | 49 | 0 | 0 | 0 | 0 | 0 | 15 | 0 | 0 | 0 | 60 | 75 | 0 | 0 |
| SRR6230411 | 5509 | 871 | 151 | 56 | 186 | 1557 | 44 | 361 | 1155 | 0 | 201 | 21 | 0 | 0 | 4958 | 97114 | 32202 | 126 | 1370 | 487 | 3408 | 0 | 0 | 820 | 33 | 12 | 25 | 0 | 0 | 213 | 0 | 242 | 0 | 10 | 5411 | 393 | 0 |
| SRR6230412 | 883 | 220 | 35 | 23 | 76 | 475 | 52 | 374 | 1535 | 21 | 90 | 55 | 0 | 0 | 6802 | 73507 | 50349 | 73 | 1821 | 951 | 2894 | 0 | 0 | 35 | 44 | 33 | 52 | 0 | 49 | 15 | 0 | 294 | 0 | 36 | 1781 | 595 | 0 |
| SRR6230413 | 8413 | 2324 | 204 | 76 | 147 | 701 | 100 | 640 | 2030 | 286 | 1648 | 20 | 12 | 0 | 9661 | 34436 | 49487 | 178 | 868 | 2280 | 3749 | 0 | 0 | 676 | 57 | 18 | 45 | 0 | 0 | 280 | 0 | 172 | 0 | 22 | 5087 | 803 | 0 |
| SRR6246305 | 11 | 28 | 12 | 67 | 58 | 913 | 158 | 1538 | 39 | 11 | 203 | 0 | 0 | 0 | 314161 | 11178 | 411473 | 0 | 830 | 2518 | 804 | 0 | 0 | 71 | 60 | 113 | 140 | 40 | 0 | 189 | 0 | 0 | 0 | 0 | 235 | 502 | 0 |
| SRR6246306 | 53 | 16 | 39 | 226 | 241 | 931 | 181 | 1499 | 54 | 36 | 799 | 0 | 0 | 0 | 316951 | 11179 | 410752 | 0 | 822 | 2588 | 755 | 0 | 0 | 263 | 221 | 463 | 593 | 53 | 0 | 724 | 0 | 0 | 0 | 0 | 248 | 483 | 0 |
| SRR6246307 | 11 | 23 | 11 | 59 | 61 | 908 | 166 | 1593 | 54 | 40 | 203 | 0 | 0 | 0 | 315614 | 11016 | 410729 | 0 | 841 | 2600 | 772 | 0 | 0 | 73 | 57 | 106 | 149 | 33 | 0 | 190 | 0 | 0 | 0 | 0 | 282 | 533 | 0 |
| SRR6246308 | 53 | 17 | 53 | 246 | 235 | 946 | 168 | 1530 | 47 | 32 | 805 | 0 | 0 | 0 | 316347 | 10816 | 411860 | 0 | 784 | 2548 | 774 | 0 | 0 | 280 | 241 | 450 | 555 | 43 | 0 | 768 | 0 | 0 | 0 | 0 | 273 | 591 | 0 |
| SRR6246309 | 0 | 0 | 0 | 11 | 33 | 79 | 12 | 56 | 0 | 45 | 68 | 0 | 0 | 0 | 548362 | 14362 | 483797 | 0 | 262 | 6691 | 34964 | 0 | 0 | 10 | 30 | 29 | 17 | 23 | 0 | 66 | 0 | 0 | 0 | 0 | 142 | 850 | 0 |
| SRR6246310 | 10 | 0 | 0 | 12 | 34 | 55 | 0 | 59 | 0 | 41 | 69 | 0 | 0 | 0 | 549094 | 14397 | 484222 | 0 | 242 | 6603 | 34619 | 0 | 0 | 12 | 34 | 27 | 13 | 22 | 0 | 62 | 0 | 0 | 0 | 0 | 150 | 841 | 0 |
| SRR6246311 | 0 | 0 | 0 | 12 | 32 | 84 | 0 | 52 | 0 | 47 | 67 | 0 | 0 | 0 | 548640 | 14420 | 485018 | 0 | 255 | 6695 | 34636 | 0 | 0 | 52 | 29 | 28 | 15 | 19 | 0 | 68 | 0 | 0 | 0 | 0 | 140 | 846 | 0 |
| SRR6246312 | 0 | 0 | 0 | 10 | 209 | 80 | 13 | 52 | 0 | 244 | 472 | 0 | 0 | 0 | 549213 | 14362 | 483697 | 0 | 257 | 6773 | 34392 | 0 | 0 | 77 | 210 | 170 | 97 | 22 | 0 | 436 | 0 | 0 | 0 | 0 | 147 | 883 | 0 |
| SRR6246313 | 0 | 0 | 0 | 0 | 18 | 0 | 0 | 0 | 0 | 17 | 19 | 0 | 0 | 0 | 638398 | 15177 | 705158 | 0 | 267 | 4702 | 38605 | 0 | 0 | 12 | 23 | 32 | 12 | 0 | 0 | 28 | 0 | 0 | 0 | 0 | 366 | 651 | 0 |
| SRR6246314 | 0 | 0 | 0 | 0 | 169 | 0 | 0 | 0 | 0 | 181 | 172 | 0 | 0 | 0 | 639767 | 15341 | 705627 | 0 | 241 | 4671 | 38597 | 0 | 0 | 14 | 19 | 31 | 24 | 0 | 0 | 255 | 0 | 0 | 0 | 0 | 392 | 656 | 12 |
| SRR6246315 | 0 | 0 | 0 | 0 | 17 | 0 | 0 | 0 | 0 | 22 | 22 | 0 | 0 | 0 | 638780 | 15253 | 705691 | 0 | 297 | 4820 | 38629 | 0 | 0 | 17 | 35 | 30 | 12 | 0 | 0 | 27 | 0 | 0 | 0 | 0 | 352 | 621 | 0 |
| SRR6246316 | 0 | 0 | 0 | 0 | 165 | 12 | 0 | 0 | 0 | 175 | 152 | 0 | 0 | 0 | 639862 | 15300 | 705805 | 0 | 264 | 4709 | 38640 | 0 | 0 | 15 | 46 | 36 | 15 | 0 | 0 | 220 | 0 | 0 | 0 | 0 | 380 | 698 | 19 |
| SRR6246317 | 0 | 0 | 0 | 0 | 0 | 0 | 39 | 0 | 0 | 0 | 18 | 0 | 0 | 0 | 116029 | 28190 | 185791 | 0 | 273 | 3273 | 13598 | 0 | 0 | 0 | 0 | 0 | 0 | 0 | 0 | 61 | 0 | 0 | 0 | 0 | 1402 | 510 | 0 |
| SRR6246318 | 0 | 0 | 0 | 0 | 0 | 39 | 0 | 39 | 0 | 0 | 34 | 0 | 0 | 0 | 121173 | 27385 | 188577 | 0 | 471 | 3022 | 12247 | 0 | 0 | 0 | 0 | 0 | 0 | 0 | 0 | 84 | 0 | 0 | 0 | 0 | 1515 | 366 | 0 |
| SRR6246319 | 0 | 0 | 0 | 0 | 0 | 39 | 0 | 0 | 0 | 0 | 56 | 0 | 0 | 0 | 115365 | 28065 | 184860 | 0 | 270 | 2776 | 12531 | 0 | 0 | 0 | 0 | 0 | 0 | 0 | 0 | 187 | 0 | 0 | 0 | 0 | 1421 | 416 | 0 |
| SRR6246320 | 0 | 0 | 0 | 0 | 14 | 0 | 0 | 78 | 0 | 0 | 22 | 0 | 0 | 0 | 118678 | 27744 | 191408 | 0 | 312 | 3786 | 12254 | 0 | 0 | 0 | 0 | 0 | 0 | 0 | 0 | 74 | 0 | 0 | 0 | 0 | 1415 | 570 | 0 |
| SRR6246321 | 0 | 0 | 12 | 56 | 12 | 6926 | 1636 | 20673 | 382 | 19 | 232 | 17 | 0 | 0 | 77018 | 88621 | 255097 | 21 | 457 | 2656 | 1987 | 0 | 0 | 25 | 33 | 134 | 449 | 101 | 0 | 158 | 0 | 21 | 0 | 0 | 577 | 344 | 0 |
| SRR6246322 | 0 | 0 | 12 | 33 | 13 | 7100 | 1509 | 21900 | 488 | 28 | 233 | 13 | 0 | 0 | 79846 | 88586 | 254664 | 0 | 288 | 2740 | 1709 | 0 | 0 | 24 | 34 | 129 | 470 | 89 | 0 | 138 | 0 | 11 | 0 | 0 | 514 | 364 | 0 |
| SRR6246323 | 0 | 10 | 20 | 97 | 31 | 7357 | 1393 | 21975 | 496 | 53 | 412 | 29 | 0 | 0 | 78695 | 86951 | 252981 | 0 | 369 | 2354 | 1689 | 0 | 0 | 61 | 56 | 252 | 783 | 155 | 0 | 230 | 0 | 11 | 0 | 0 | 497 | 273 | 0 |
| SRR6246324 | 0 | 0 | 12 | 74 | 25 | 6380 | 1376 | 21335 | 492 | 43 | 417 | 19 | 0 | 0 | 79366 | 87495 | 253290 | 11 | 350 | 2097 | 1923 | 0 | 0 | 37 | 72 | 269 | 832 | 147 | 0 | 233 | 0 | 11 | 0 | 0 | 571 | 321 | 0 |
| SRR6246325 | 0 | 0 | 0 | 11 | 17 | 144 | 14 | 45 | 0 | 70 | 42 | 0 | 0 | 0 | 498762 | 15828 | 431940 | 0 | 202 | 2124 | 24162 | 0 | 0 | 25 | 74 | 10 | 42 | 16 | 0 | 21 | 0 | 0 | 0 | 0 | 28 | 655 | 0 |
| SRR6246326 | 0 | 0 | 0 | 0 | 168 | 137 | 10 | 44 | 0 | 679 | 387 | 0 | 0 | 0 | 499293 | 15616 | 429839 | 0 | 250 | 2234 | 24223 | 0 | 0 | 22 | 85 | 94 | 54 | 112 | 0 | 185 | 0 | 0 | 0 | 0 | 28 | 569 | 0 |
| SRR6246327 | 0 | 0 | 0 | 0 | 20 | 130 | 0 | 32 | 0 | 79 | 44 | 0 | 0 | 0 | 498298 | 15704 | 431586 | 0 | 234 | 2311 | 23730 | 0 | 0 | 23 | 65 | 70 | 37 | 16 | 0 | 21 | 0 | 0 | 0 | 0 | 33 | 547 | 0 |
| SRR6246328 | 0 | 0 | 0 | 0 | 22 | 148 | 13 | 45 | 0 | 80 | 43 | 0 | 0 | 0 | 499036 | 15564 | 429010 | 0 | 203 | 2169 | 23892 | 0 | 0 | 28 | 11 | 11 | 38 | 13 | 0 | 22 | 0 | 0 | 0 | 0 | 27 | 604 | 0 |
| SRR6246329 | 0 | 0 | 0 | 0 | 15 | 18 | 0 | 16 | 0 | 43 | 17 | 0 | 0 | 0 | 375173 | 10857 | 314912 | 0 | 511 | 1981 | 27513 | 0 | 0 | 0 | 0 | 13 | 0 | 13 | 0 | 20 | 0 | 0 | 0 | 0 | 110 | 449 | 11 |
| SRR6246330 | 0 | 0 | 0 | 0 | 122 | 19 | 0 | 12 | 0 | 326 | 140 | 0 | 0 | 0 | 376975 | 10906 | 314063 | 0 | 511 | 2071 | 27264 | 0 | 0 | 0 | 0 | 0 | 0 | 0 | 0 | 140 | 0 | 0 | 0 | 0 | 109 | 483 | 14 |
| SRR6246331 | 0 | 0 | 0 | 0 | 96 | 28 | 0 | 19 | 0 | 353 | 102 | 0 | 0 | 0 | 374956 | 11043 | 314563 | 0 | 485 | 2209 | 27752 | 0 | 0 | 0 | 0 | 11 | 0 | 15 | 0 | 164 | 0 | 0 | 0 | 0 | 106 | 492 | 15 |
| SRR6246332 | 0 | 0 | 0 | 0 | 109 | 18 | 0 | 13 | 0 | 333 | 151 | 0 | 0 | 0 | 377067 | 10997 | 313131 | 0 | 505 | 2139 | 27686 | 0 | 0 | 0 | 0 | 0 | 0 | 11 | 0 | 122 | 0 | 0 | 0 | 0 | 114 | 495 | 0 |
| SRR6246333 | 0 | 0 | 0 | 0 | 29 | 732 | 397 | 2501 | 183 | 29 | 39 | 0 | 0 | 0 | 181924 | 13888 | 167027 | 0 | 183 | 1159 | 18119 | 0 | 0 | 0 | 0 | 0 | 0 | 0 | 0 | 29 | 0 | 0 | 0 | 0 | 240 | 412 | 0 |
| SRR6246334 | 0 | 0 | 0 | 0 | 0 | 768 | 416 | 2241 | 128 | 30 | 18 | 0 | 0 | 0 | 184432 | 14123 | 166293 | 0 | 256 | 1345 | 19052 | 0 | 0 | 0 | 0 | 0 | 0 | 0 | 0 | 48 | 0 | 0 | 0 | 0 | 222 | 461 | 20 |
| SRR6246335 | 0 | 0 | 0 | 0 | 28 | 991 | 300 | 2493 | 210 | 47 | 57 | 0 | 0 | 0 | 184614 | 13852 | 166761 | 0 | 360 | 1171 | 16431 | 0 | 0 | 0 | 0 | 0 | 0 | 0 | 0 | 142 | 0 | 0 | 0 | 0 | 247 | 522 | 0 |
| SRR6246336 | 0 | 0 | 0 | 0 | 39 | 991 | 341 | 2323 | 186 | 49 | 49 | 0 | 0 | 0 | 186780 | 13565 | 165367 | 0 | 155 | 1146 | 17315 | 0 | 0 | 0 | 0 | 0 | 0 | 0 | 0 | 157 | 0 | 0 | 0 | 0 | 153 | 483 | 20 |
| SRR6246337 | 0 | 0 | 0 | 12 | 13 | 7214 | 1472 | 20257 | 500 | 13 | 74 | 13 | 0 | 0 | 181412 | 23810 | 246462 | 0 | 183 | 1020 | 7782 | 0 | 0 | 41 | 56 | 301 | 990 | 148 | 0 | 96 | 0 | 0 | 0 | 0 | 747 | 353 | 0 |
| SRR6246338 | 0 | 0 | 0 | 13 | 15 | 6625 | 1497 | 18994 | 440 | 11 | 45 | 11 | 0 | 0 | 178984 | 22928 | 247436 | 10 | 186 | 1213 | 7868 | 0 | 0 | 33 | 33 | 187 | 615 | 97 | 0 | 70 | 0 | 10 | 0 | 0 | 615 | 378 | 0 |
| SRR6246339 | 0 | 0 | 0 | 0 | 16 | 6741 | 1565 | 19058 | 482 | 22 | 71 | 14 | 0 | 0 | 174888 | 23684 | 246328 | 0 | 204 | 1130 | 7927 | 0 | 0 | 57 | 47 | 310 | 1020 | 160 | 0 | 114 | 0 | 0 | 0 | 0 | 577 | 401 | 0 |
| SRR6246340 | 0 | 0 | 0 | 0 | 12 | 6938 | 1516 | 17897 | 576 | 17 | 80 | 12 | 0 | 0 | 177832 | 24338 | 244927 | 0 | 240 | 1132 | 7869 | 0 | 0 | 47 | 58 | 328 | 980 | 152 | 0 | 93 | 0 | 0 | 0 | 0 | 578 | 357 | 0 |
| SRR6246341 | 0 | 0 | 0 | 0 | 14 | 17 | 0 | 44 | 0 | 16 | 74 | 0 | 0 | 0 | 359864 | 10477 | 270652 | 0 | 336 | 2573 | 48487 | 0 | 0 | 0 | 11 | 29 | 11 | 25 | 0 | 35 | 0 | 0 | 0 | 0 | 72 | 128 | 0 |

### Supp Table S8.3.Columns_BX_to_DH

| | | | | | | | | | | | | | | | | | | | | | | | | | | | | | | | | | | | | | |
|---|---|---|---|---|---|---|---|---|---|---|---|---|---|---|---|---|---|---|---|---|---|---|---|---|---|---|---|---|---|---|---|---|---|---|---|---|---|
| SRR6246342 | 0 | 0 | 0 | 0 | 22 | 31 | 0 | 54 | 0 | 16 | 77 | 0 | 0 | 0 | 364236 | 10344 | 268703 | 0 | 321 | 2613 | 48339 | 0 | 0 | 0 | 13 | 15 | 17 | 22 | 0 | 39 | 0 | 0 | 0 | 0 | 60 | 192 | 0 |
| SRR6246343 | 0 | 0 | 0 | 0 | 16 | 49 | 0 | 49 | 0 | 17 | 83 | 11 | 0 | 0 | 362313 | 10527 | 270714 | 0 | 287 | 2542 | 48085 | 0 | 0 | 0 | 16 | 20 | 14 | 19 | 0 | 34 | 0 | 0 | 0 | 0 | 57 | 213 | 0 |
| SRR6246344 | 0 | 0 | 0 | 0 | 0 | 15 | 15 | 63 | 0 | 52 | 316 | 0 | 0 | 0 | 363256 | 10355 | 268532 | 0 | 336 | 2590 | 48550 | 0 | 0 | 0 | 21 | 26 | 0 | 24 | 0 | 136 | 0 | 0 | 0 | 0 | 60 | 276 | 0 |
| SRR6246345 | 0 | 0 | 0 | 0 | 22 | 57 | 0 | 29 | 0 | 22 | 221 | 22 | 0 | 0 | 202618 | 15003 | 202816 | 0 | 229 | 1862 | 26577 | 0 | 0 | 22 | 22 | 0 | 0 | 66 | 0 | 199 | 0 | 0 | 0 | 0 | 53 | 301 | 0 |
| SRR6246346 | 0 | 0 | 0 | 0 | 46 | 0 | 0 | 61 | 0 | 46 | 70 | 0 | 0 | 0 | 203628 | 13367 | 206310 | 0 | 460 | 2118 | 28300 | 0 | 0 | 0 | 0 | 0 | 0 | 0 | 0 | 47 | 0 | 0 | 0 | 0 | 144 | 366 | 0 |
| SRR6246347 | 0 | 0 | 0 | 0 | 0 | 0 | 0 | 0 | 0 | 44 | 263 | 0 | 0 | 0 | 207003 | 14583 | 203657 | 0 | 231 | 1902 | 27781 | 0 | 0 | 22 | 0 | 44 | 22 | 22 | 0 | 285 | 0 | 0 | 0 | 0 | 87 | 300 | 0 |
| SRR6246348 | 0 | 0 | 0 | 0 | 0 | 0 | 0 | 0 | 0 | 0 | 137 | 23 | 0 | 0 | 204540 | 14300 | 201312 | 0 | 357 | 1545 | 30010 | 0 | 0 | 0 | 0 | 0 | 23 | 23 | 0 | 229 | 0 | 0 | 0 | 0 | 18 | 204 | 0 |
| SRR6246349 | 0 | 0 | 0 | 0 | 456 | 0 | 0 | 222 | 222 | 0 | 83 | 0 | 0 | 0 | 100511 | 55321 | 105821 | 0 | 0 | 3336 | 19569 | 0 | 41 | 0 | 0 | 0 | 0 | 0 | 0 | 415 | 0 | 0 | 0 | 0 | 362 | 368 | 41 |
| SRR6246350 | 0 | 0 | 0 | 0 | 137 | 0 | 0 | 226 | 226 | 43 | 49 | 43 | 0 | 0 | 107216 | 48556 | 108297 | 0 | 226 | 1583 | 22619 | 0 | 0 | 0 | 0 | 0 | 0 | 0 | 0 | 29 | 0 | 0 | 43 | 0 | 492 | 511 | 0 |
| SRR6246351 | 0 | 0 | 0 | 0 | 408 | 0 | 0 | 0 | 0 | 0 | 122 | 0 | 0 | 0 | 108495 | 51134 | 107581 | 0 | 421 | 841 | 17662 | 0 | 0 | 0 | 0 | 0 | 0 | 0 | 0 | 326 | 0 | 0 | 0 | 0 | 354 | 405 | 41 |
| SRR6246352 | 0 | 0 | 0 | 0 | 124 | 0 | 0 | 228 | 0 | 42 | 48 | 0 | 0 | 0 | 113164 | 53656 | 109989 | 0 | 228 | 3650 | 19165 | 0 | 0 | 0 | 0 | 0 | 0 | 0 | 0 | 57 | 0 | 0 | 0 | 0 | 487 | 333 | 42 |
| SRR6246353 | 17 | 0 | 0 | 0 | 16 | 377 | 166 | 453 | 56 | 25 | 63 | 0 | 0 | 0 | 366175 | 15318 | 414738 | 0 | 1893 | 3682 | 1906 | 0 | 0 | 87 | 39 | 58 | 164 | 12 | 0 | 119 | 0 | 0 | 0 | 0 | 583 | 631 | 0 |
| SRR6246354 | 17 | 0 | 0 | 0 | 31 | 362 | 146 | 432 | 56 | 49 | 151 | 0 | 0 | 0 | 365644 | 15105 | 415223 | 0 | 1915 | 3575 | 1974 | 0 | 0 | 189 | 101 | 125 | 392 | 20 | 0 | 258 | 0 | 0 | 0 | 0 | 541 | 696 | 0 |
| SRR6246355 | 10 | 0 | 0 | 0 | 14 | 363 | 175 | 479 | 60 | 22 | 59 | 0 | 0 | 0 | 365098 | 15347 | 414428 | 0 | 1933 | 3602 | 1994 | 0 | 0 | 84 | 43 | 55 | 160 | 21 | 0 | 109 | 0 | 0 | 0 | 0 | 513 | 589 | 0 |
| SRR6246356 | 13 | 0 | 0 | 0 | 38 | 360 | 156 | 423 | 61 | 56 | 138 | 0 | 0 | 0 | 366712 | 15104 | 415651 | 0 | 1924 | 3648 | 2017 | 0 | 0 | 195 | 97 | 137 | 368 | 22 | 0 | 258 | 0 | 0 | 0 | 0 | 562 | 526 | 0 |
| SRR6246358 | 0 | 0 | 0 | 0 | 22 | 15 | 0 | 30 | 0 | 33 | 33 | 0 | 0 | 0 | 813604 | 56841 | 823241 | 0 | 53 | 457 | 5433 | 0 | 0 | 0 | 0 | 11 | 22 | 0 | 0 | 0 | 0 | 0 | 0 | 0 | 18 | 0 | 0 |
| SRR6246359 | 0 | 0 | 0 | 0 | 21 | 0 | 0 | 22 | 0 | 21 | 32 | 0 | 0 | 0 | 811634 | 55722 | 826718 | 0 | 22 | 493 | 5237 | 0 | 0 | 0 | 0 | 11 | 11 | 0 | 0 | 32 | 0 | 0 | 0 | 0 | 24 | 0 | 0 |
| SRR6246360 | 0 | 0 | 0 | 0 | 33 | 23 | 0 | 15 | 0 | 22 | 54 | 0 | 0 | 0 | 815568 | 56498 | 825213 | 0 | 23 | 387 | 5484 | 0 | 0 | 11 | 33 | 44 | 33 | 0 | 0 | 33 | 0 | 0 | 0 | 0 | 36 | 0 | 0 |
| SRR6246361 | 0 | 0 | 0 | 0 | 24 | 42 | 0 | 42 | 0 | 24 | 48 | 0 | 0 | 0 | 299661 | 80081 | 337286 | 0 | 229 | 3579 | 20495 | 0 | 0 | 12 | 0 | 0 | 12 | 0 | 0 | 688 | 0 | 0 | 0 | 0 | 177 | 466 | 0 |
| SRR6246362 | 0 | 0 | 0 | 0 | 12 | 43 | 0 | 107 | 0 | 49 | 123 | 0 | 0 | 0 | 300109 | 80381 | 336687 | 0 | 342 | 3894 | 19768 | 0 | 0 | 12 | 0 | 12 | 12 | 0 | 0 | 615 | 0 | 0 | 0 | 0 | 108 | 106 | 12 |
| SRR6246363 | 0 | 0 | 12 | 0 | 18 | 21 | 0 | 84 | 0 | 11 | 21 | 0 | 0 | 0 | 299235 | 79906 | 335542 | 21 | 209 | 3930 | 19276 | 0 | 0 | 0 | 0 | 12 | 12 | 0 | 0 | 140 | 0 | 0 | 0 | 0 | 180 | 155 | 24 |
| SRR6246364 | 0 | 0 | 0 | 0 | 12 | 85 | 42 | 85 | 0 | 11 | 40 | 0 | 0 | 0 | 299668 | 79728 | 334802 | 21 | 338 | 4524 | 18965 | 0 | 0 | 11 | 0 | 0 | 15 | 0 | 0 | 145 | 0 | 0 | 0 | 0 | 305 | 369 | 25 |
| SRR6246365 | 0 | 0 | 0 | 0 | 59 | 0 | 0 | 0 | 0 | 29 | 0 | 0 | 0 | 0 | 194960 | 241911 | 179171 | 77 | 77 | 1391 | 5720 | 0 | 0 | 0 | 0 | 0 | 0 | 0 | 0 | 764 | 0 | 0 | 0 | 0 | 932 | 317 | 29 |
| SRR6246366 | 0 | 0 | 0 | 0 | 30 | 0 | 0 | 0 | 0 | 60 | 90 | 0 | 0 | 0 | 196459 | 244106 | 178583 | 0 | 241 | 1851 | 5553 | 0 | 0 | 0 | 0 | 0 | 0 | 0 | 0 | 571 | 0 | 0 | 0 | 0 | 332 | 642 | 0 |
| SRR6246367 | 0 | 0 | 0 | 0 | 116 | 0 | 0 | 0 | 0 | 29 | 29 | 0 | 0 | 0 | 194869 | 248105 | 178144 | 0 | 80 | 2151 | 4939 | 0 | 0 | 0 | 0 | 0 | 0 | 0 | 0 | 903 | 0 | 0 | 0 | 0 | 494 | 271 | 0 |
| SRR6246368 | 0 | 0 | 0 | 0 | 30 | 0 | 0 | 0 | 0 | 30 | 60 | 0 | 0 | 0 | 198333 | 249227 | 179613 | 0 | 0 | 2164 | 4808 | 0 | 0 | 0 | 0 | 0 | 0 | 0 | 0 | 629 | 0 | 0 | 0 | 0 | 442 | 320 | 0 |
| SRR6246369 | 13 | 0 | 0 | 0 | 41 | 248 | 101 | 522 | 15 | 259 | 125 | 0 | 0 | 0 | 556807 | 28220 | 537401 | 0 | 368 | 714 | 3907 | 0 | 0 | 132 | 57 | 48 | 54 | 10 | 0 | 281 | 0 | 0 | 0 | 0 | 564 | 228 | 0 |
| SRR6246370 | 13 | 18 | 0 | 0 | 54 | 254 | 99 | 504 | 20 | 273 | 118 | 0 | 0 | 0 | 558637 | 28510 | 535345 | 0 | 367 | 742 | 3842 | 0 | 0 | 91 | 62 | 49 | 61 | 0 | 0 | 270 | 0 | 0 | 0 | 0 | 538 | 88 | 0 |
| SRR6246371 | 0 | 13 | 0 | 0 | 11 | 227 | 101 | 498 | 18 | 62 | 36 | 0 | 0 | 0 | 556425 | 28186 | 536526 | 0 | 370 | 714 | 3890 | 0 | 0 | 28 | 16 | 34 | 13 | 11 | 0 | 84 | 0 | 0 | 0 | 0 | 601 | 290 | 0 |
| SRR6246372 | 11 | 0 | 0 | 0 | 32 | 232 | 106 | 499 | 16 | 229 | 102 | 0 | 0 | 0 | 558758 | 28506 | 536732 | 0 | 334 | 707 | 3737 | 0 | 0 | 120 | 64 | 43 | 46 | 0 | 0 | 269 | 0 | 0 | 0 | 0 | 573 | 350 | 0 |
| SRR6246373 | 0 | 0 | 0 | 0 | 12 | 0 | 13 | 13 | 0 | 15 | 14 | 0 | 0 | 0 | 589800 | 43872 | 432087 | 0 | 113 | 2564 | 28235 | 0 | 0 | 0 | 15 | 0 | 0 | 0 | 0 | 23 | 0 | 0 | 0 | 0 | 63 | 136 | 0 |
| SRR6246374 | 0 | 0 | 0 | 0 | 23 | 41 | 0 | 27 | 0 | 19 | 11 | 0 | 0 | 0 | 596992 | 43851 | 435117 | 0 | 75 | 2621 | 27568 | 0 | 0 | 0 | 16 | 16 | 0 | 0 | 0 | 14 | 0 | 0 | 0 | 0 | 90 | 421 | 0 |
| SRR6246375 | 0 | 0 | 0 | 0 | 15 | 27 | 0 | 27 | 0 | 14 | 15 | 0 | 0 | 0 | 593717 | 44327 | 432697 | 0 | 113 | 2453 | 26967 | 0 | 0 | 0 | 0 | 23 | 0 | 0 | 0 | 10 | 0 | 0 | 0 | 0 | 73 | 135 | 0 |
| SRR6246376 | 0 | 0 | 0 | 0 | 23 | 20 | 0 | 14 | 0 | 38 | 38 | 0 | 0 | 0 | 594420 | 45227 | 431706 | 0 | 54 | 2508 | 27164 | 0 | 0 | 0 | 15 | 0 | 0 | 0 | 0 | 46 | 0 | 0 | 0 | 0 | 58 | 142 | 0 |
| SRR6246379 | 0 | 0 | 0 | 0 | 322 | 0 | 0 | 520 | 0 | 0 | 322 | 0 | 0 | 0 | 417274 | 90289 | 445237 | 0 | 520 | 2601 | 62435 | 0 | 0 | 0 | 0 | 0 | 0 | 0 | 0 | 0 | 0 | 0 | 0 | 0 | 0 | 0 | 0 |
| SRR6246385 | 11 | 0 | 0 | 13 | 28 | 158 | 94 | 219 | 27 | 96 | 117 | 0 | 0 | 0 | 508646 | 18048 | 529872 | 0 | 1637 | 3223 | 3662 | 0 | 0 | 193 | 64 | 98 | 869 | 16 | 0 | 195 | 0 | 0 | 0 | 0 | 469 | 964 | 0 |
| SRR6246386 | 13 | 0 | 0 | 15 | 28 | 155 | 79 | 199 | 34 | 91 | 122 | 0 | 0 | 0 | 509995 | 18332 | 529746 | 0 | 1620 | 3197 | 3683 | 0 | 0 | 184 | 60 | 114 | 907 | 19 | 0 | 209 | 0 | 0 | 0 | 0 | 432 | 969 | 0 |
| SRR6246387 | 11 | 0 | 0 | 14 | 29 | 169 | 84 | 194 | 22 | 97 | 125 | 0 | 0 | 0 | 509020 | 18294 | 529260 | 0 | 1718 | 3165 | 3702 | 0 | 0 | 188 | 57 | 114 | 887 | 17 | 0 | 198 | 0 | 0 | 0 | 0 | 470 | 902 | 0 |
| SRR6246388 | 16 | 0 | 0 | 13 | 30 | 147 | 85 | 219 | 21 | 105 | 115 | 0 | 0 | 0 | 510581 | 18518 | 531426 | 0 | 1622 | 3130 | 3672 | 0 | 0 | 177 | 73 | 106 | 892 | 18 | 0 | 216 | 0 | 0 | 0 | 0 | 477 | 1021 | 0 |
| SRR6246389 | 0 | 0 | 0 | 0 | 60 | 165 | 0 | 62 | 0 | 17 | 27 | 0 | 0 | 0 | 552919 | 43484 | 430162 | 0 | 51 | 2429 | 29317 | 0 | 0 | 0 | 17 | 34 | 0 | 0 | 0 | 14 | 0 | 0 | 0 | 0 | 59 | 206 | 0 |
| SRR6246390 | 0 | 0 | 0 | 0 | 88 | 92 | 0 | 57 | 0 | 26 | 159 | 0 | 0 | 0 | 553360 | 43194 | 428122 | 0 | 46 | 2335 | 30169 | 0 | 0 | 0 | 0 | 44 | 0 | 0 | 0 | 88 | 0 | 0 | 0 | 0 | 93 | 71 | 0 |
| SRR6246391 | 0 | 0 | 0 | 0 | 51 | 165 | 0 | 68 | 0 | 26 | 14 | 0 | 0 | 0 | 550550 | 43351 | 429037 | 0 | 68 | 2365 | 30217 | 0 | 0 | 0 | 0 | 0 | 0 | 0 | 0 | 13 | 0 | 0 | 0 | 0 | 73 | 138 | 0 |
| SRR6246392 | 0 | 0 | 0 | 0 | 52 | 108 | 0 | 57 | 0 | 17 | 26 | 0 | 0 | 0 | 554170 | 43031 | 428661 | 0 | 85 | 2221 | 29639 | 0 | 0 | 0 | 0 | 0 | 0 | 0 | 0 | 14 | 0 | 0 | 0 | 0 | 103 | 35 | 0 |
| SRR6246393 | 0 | 0 | 14 | 0 | 0 | 0 | 0 | 0 | 0 | 31 | 45 | 14 | 0 | 0 | 342875 | 70755 | 506011 | 0 | 22 | 762 | 28734 | 0 | 0 | 0 | 0 | 0 | 14 | 14 | 0 | 211 | 0 | 0 | 0 | 0 | 357 | 243 | 0 |
| SRR6246394 | 0 | 0 | 0 | 0 | 0 | 0 | 0 | 22 | 0 | 90 | 254 | 0 | 0 | 0 | 347812 | 70691 | 506100 | 0 | 45 | 779 | 29394 | 0 | 0 | 0 | 0 | 0 | 0 | 0 | 0 | 761 | 0 | 0 | 0 | 0 | 251 | 251 | 0 |
| SRR6246395 | 0 | 0 | 0 | 0 | 0 | 0 | 0 | 0 | 0 | 18 | 45 | 0 | 0 | 0 | 343400 | 72523 | 503645 | 0 | 109 | 1064 | 30070 | 0 | 0 | 0 | 0 | 0 | 15 | 0 | 0 | 248 | 0 | 0 | 0 | 0 | 112 | 120 | 29 |
| SRR6246396 | 0 | 0 | 0 | 0 | 0 | 0 | 0 | 22 | 0 | 14 | 60 | 15 | 0 | 0 | 350366 | 71002 | 509502 | 0 | 0 | 981 | 27916 | 0 | 0 | 0 | 0 | 15 | 0 | 0 | 0 | 229 | 0 | 0 | 0 | 0 | 205 | 63 | 15 |
| SRR6246397 | 0 | 0 | 0 | 0 | 0 | 0 | 0 | 0 | 0 | 13 | 45 | 0 | 0 | 0 | 84620 | 27119 | 145029 | 0 | 32 | 1401 | 10474 | 0 | 0 | 0 | 0 | 0 | 0 | 0 | 0 | 92 | 0 | 0 | 0 | 0 | 758 | 624 | 0 |
| SRR6246398 | 0 | 0 | 0 | 0 | 0 | 0 | 0 | 0 | 0 | 0 | 20 | 0 | 0 | 0 | 84816 | 26570 | 145316 | 0 | 0 | 1981 | 9547 | 0 | 0 | 0 | 0 | 0 | 0 | 0 | 0 | 47 | 0 | 0 | 0 | 0 | 965 | 749 | 0 |
| SRR6246399 | 0 | 0 | 0 | 0 | 0 | 0 | 0 | 0 | 0 | 0 | 19 | 0 | 0 | 0 | 86188 | 27933 | 145880 | 0 | 96 | 1826 | 11342 | 0 | 0 | 0 | 0 | 0 | 0 | 0 | 0 | 74 | 0 | 0 | 0 | 0 | 852 | 750 | 0 |
| SRR6246400 | 0 | 0 | 0 | 0 | 0 | 0 | 33 | 0 | 0 | 0 | 26 | 0 | 0 | 0 | 87232 | 28625 | 144421 | 0 | 98 | 1334 | 10217 | 0 | 0 | 0 | 0 | 0 | 0 | 0 | 0 | 150 | 0 | 0 | 0 | 0 | 1463 | 657 | 0 |
| SRR6265480 | 29 | 12 | 10 | 27 | 41 | 686 | 243 | 1699 | 95 | 20 | 12 | 17 | 0 | 0 | 30957 | 7824 | 71435 | 21 | 63 | 8705 | 9485 | 0 | 0 | 14 | 42 | 17 | 13 | 0 | 0 | 135 | 0 | 11 | 0 | 0 | 501 | 0 | 0 |
| SRR6265481 | 0 | 0 | 25 | 48 | 25 | 340 | 118 | 783 | 49 | 21 | 80 | 0 | 0 | 0 | 45973 | 9366 | 130733 | 0 | 97 | 4388 | 11930 | 0 | 0 | 46 | 84 | 183 | 261 | 31 | 0 | 291 | 0 | 0 | 0 | 0 | 329 | 0 | 0 |
| SRR6265482 | 18 | 26 | 14 | 12 | 36 | 891 | 517 | 2672 | 71 | 22 | 30 | 24 | 0 | 0 | 18970 | 6217 | 74905 | 18 | 71 | 4792 | 7000 | 0 | 0 | 60 | 20 | 32 | 50 | 0 | 0 | 371 | 0 | 18 | 0 | 0 | 974 | 0 | 0 |
| SRR6265483 | 27 | 0 | 0 | 20 | 37 | 826 | 286 | 2097 | 127 | 25 | 40 | 17 | 0 | 0 | 34226 | 9238 | 82941 | 0 | 127 | 6960 | 12521 | 0 | 0 | 11 | 0 | 15 | 27 | 0 | 0 | 165 | 0 | 0 | 0 | 0 | 914 | 0 | 0 |
| SRR6265484 | 11 | 0 | 17 | 20 | 35 | 456 | 162 | 1293 | 57 | 18 | 65 | 33 | 0 | 0 | 18048 | 20198 | 81263 | 38 | 76 | 6466 | 12314 | 0 | 0 | 41 | 47 | 137 | 193 | 13 | 0 | 1054 | 0 | 0 | 0 | 0 | 790 | 0 | 0 |
| SRR6265485 | 24 | 0 | 16 | 10 | 25 | 385 | 192 | 685 | 60 | 13 | 25 | 25 | 0 | 0 | 16872 | 15483 | 75903 | 0 | 156 | 7943 | 8436 | 0 | 0 | 10 | 34 | 21 | 51 | 43 | 0 | 280 | 0 | 0 | 0 | 0 | 711 | 0 | 0 |
| SRR6265486 | 29 | 36 | 0 | 0 | 15 | 303 | 273 | 797 | 46 | 0 | 20 | 28 | 0 | 0 | 7799 | 16420 | 23386 | 23 | 46 | 5257 | 3702 | 0 | 0 | 15 | 0 | 0 | 18 | 0 | 0 | 466 | 0 | 0 | 0 | 0 | 1548 | 0 | 0 |
| SRR6265487 | 0 | 0 | 0 | 21 | 79 | 745 | 426 | 1703 | 106 | 0 | 33 | 32 | 0 | 0 | 11679 | 58550 | 54848 | 0 | 39 | 12154 | 14544 | 0 | 0 | 140 | 52 | 74 | 142 | 0 | 0 | 540 | 0 | 0 | 0 | 0 | 904 | 0 | 0 |
| SRR6265488 | 15 | 22 | 16 | 10 | 16 | 450 | 265 | 1164 | 38 | 14 | 11 | 14 | 0 | 0 | 27413 | 26655 | 75918 | 0 | 49 | 3856 | 6752 | 0 | 0 | 28 | 19 | 33 | 43 | 0 | 0 | 177 | 0 | 0 | 0 | 0 | 684 | 0 | 0 |
| SRR6265489 | 0 | 0 | 0 | 0 | 0 | 197 | 197 | 494 | 118 | 0 | 0 | 0 | 0 | 0 | 7660 | 28430 | 26960 | 59 | 237 | 5133 | 2646 | 0 | 0 | 0 | 0 | 0 | 0 | 0 | 0 | 50 | 0 | 178 | 0 | 0 | 54985 | 0 | 0 |
| SRR6265490 | 28 | 24 | 44 | 100 | 35 | 284 | 278 | 886 | 51 | 0 | 121 | 29 | 0 | 0 | 9387 | 44922 | 64148 | 10 | 137 | 9681 | 7053 | 0 | 0 | 44 | 25 | 90 | 374 | 115 | 0 | 1179 | 0 | 25 | 0 | 0 | 1144 | 0 | 0 |
| SRR6265491 | 12 | 0 | 0 | 14 | 27 | 607 | 258 | 1349 | 90 | 13 | 14 | 17 | 0 | 0 | 14890 | 28629 | 55710 | 11 | 34 | 7495 | 7518 | 0 | 0 | 12 | 28 | 12 | 19 | 0 | 0 | 183 | 0 | 11 | 0 | 0 | 1587 | 0 | 0 |
| SRR6306014 | 114 | 111 | 0 | 0 | 0 | 27 | 117 | 282 | 45 | 56 | 150 | 0 | 0 | 0 | 18414 | 19675 | 47489 | 0 | 78 | 10782 | 1838 | 0 | 0 | 27 | 0 | 0 | 0 | 0 | 0 | 417 | 0 | 13 | 0 | 0 | 3274 | 0 | 0 |
| SRR6306015 | 137 | 121 | 0 | 15 | 0 | 42 | 118 | 251 | 39 | 101 | 160 | 0 | 0 | 0 | 30730 | 22124 | 64626 | 0 | 80 | 24303 | 2449 | 0 | 0 | 27 | 0 | 0 | 0 | 0 | 0 | 366 | 0 | 21 | 0 | 0 | 2020 | 0 | 0 |
| SRR6313948 | 3262 | 1300 | 83 | 40 | 394 | 420 | 65 | 462 | 2451 | 1545 | 38 | 183 | 0 | 0 | 12965 | 12231 | 14412 | 11518 | 215 | 663 | 14235 | 119 | 189 | 287 | 13 | 20 | 12 | 0 | 325 | 51 | 162 | 19515 | 0 | 168 | 16312 | 1178 | 0 |
| SRR6313949 | 1310 | 397 | 33 | 0 | 288 | 1075 | 329 | 1650 | 7881 | 1847 | 0 | 166 | 0 | 0 | 6032 | 7774 | 6291 | 1643 | 425 | 1945 | 2677 | 171 | 249 | 288 | 12 | 0 | 0 | 0 | 704 | 20 | 180 | 3547 | 14 | 146 | 10549 | 586 | 0 |
| SRR6317622 | 14 | 42 | 21 | 127 | 70 | 65 | 97 | 357 | 0 | 14 | 92 | 0 | 0 | 0 | 26588 | 18908 | 118157 | 0 | 0 | 389 | 1005 | 0 | 0 | 15 | 0 | 0 | 0 | 0 | 0 | 22 | 0 | 0 | 0 | 0 | 4087 | 0 | 14 |
| SRR6317623 | 24 | 47 | 33 | 99 | 57 | 116 | 140 | 326 | 23 | 0 | 47 | 0 | 0 | 0 | 10107 | 7644 | 44736 | 47 | 0 | 349 | 885 | 0 | 0 | 132 | 0 | 0 | 0 | 0 | 0 | 16 | 0 | 0 | 0 | 0 | 4903 | 0 | 0 |
| SRR6317624 | 50 | 21 | 17 | 109 | 122 | 217 | 67 | 284 | 67 | 0 | 117 | 0 | 0 | 0 | 19549 | 13470 | 87151 | 0 | 0 | 400 | 1201 | 0 | 0 | 23 | 0 | 0 | 0 | 0 | 0 | 22 | 0 | 17 | 0 | 0 | 4628 | 0 | 0 |
| SRR6317625 | 50 | 28 | 50 | 107 | 64 | 166 | 119 | 71 | 0 | 0 | 15 | 0 | 0 | 0 | 8348 | 6068 | 36367 | 0 | 0 | 285 | 688 | 0 | 0 | 32 | 0 | 0 | 0 | 0 | 0 | 23 | 0 | 0 | 0 | 0 | 5914 | 0 | 0 |
| SRR6317626 | 0 | 0 | 11 | 35 | 60 | 109 | 181 | 181 | 91 | 14 | 68 | 11 | 0 | 0 | 61019 | 25981 | 172497 | 0 | 0 | 598 | 4439 | 0 | 0 | 68 | 0 | 0 | 0 | 0 | 0 | 23 | 0 | 54 | 0 | 0 | 3252 | 0 | 14 |
| SRR6317627 | 0 | 0 | 13 | 18 | 39 | 603 | 322 | 1367 | 67 | 78 | 455 | 33 | 0 | 0 | 115277 | 64519 | 281816 | 0 | 0 | 938 | 4503 | 0 | 0 | 52 | 0 | 0 | 0 | 0 | 0 | 1477 | 0 | 174 | 0 | 0 | 1797 | 0 | 15 |
| SRR6317628 | 0 | 0 | 27 | 43 | 50 | 209 | 235 | 287 | 130 | 0 | 50 | 0 | 0 | 0 | 4040 | 6972 | 22193 | 0 | 0 | 443 | 704 | 0 | 0 | 86 | 0 | 0 | 0 | 0 | 0 | 170 | 0 | 0 | 0 | 0 | 7320 | 0 | 0 |
| SRR6317629 | 0 | 0 | 12 | 63 | 56 | 237 | 281 | 296 | 74 | 0 | 56 | 13 | 0 | 0 | 13572 | 15523 | 59867 | 0 | 15 | 429 | 1774 | 0 | 0 | 90 | 0 | 0 | 0 | 0 | 0 | 14 | 0 | 30 | 0 | 0 | 6456 | 0 | 0 |
| SRR6317630 | 38 | 51 | 51 | 12 | 109 | 92 | 122 | 306 | 46 | 0 | 89 | 0 | 0 | 0 | 17923 | 14564 | 87452 | 0 | 15 | 444 | 2112 | 0 | 0 | 11 | 0 | 0 | 0 | 0 | 0 | 24 | 0 | 0 | 0 | 0 | 4680 | 0 | 0 |
| SRR6317631 | 16 | 28 | 34 | 146 | 84 | 81 | 147 | 309 | 98 | 0 | 156 | 0 | 0 | 0 | 5763 | 6889 | 35169 | 0 | 0 | 374 | 700 | 0 | 0 | 249 | 0 | 0 | 0 | 0 | 0 | 429 | 0 | 0 | 0 | 0 | 10035 | 0 | 0 |
| SRR6317632 | 25 | 25 | 33 | 123 | 152 | 112 | 160 | 255 | 0 | 0 | 148 | 0 | 0 | 0 | 10490 | 8421 | 49634 | 0 | 0 | 303 | 1182 | 0 | 0 | 288 | 0 | 0 | 0 | 0 | 0 | 358 | 0 | 0 | 0 | 0 | 5027 | 0 | 0 |
| SRR6317633 | 0 | 14 | 37 | 84 | 88 | 172 | 172 | 129 | 22 | 0 | 15 | 0 | 0 | 0 | 7561 | 5234 | 31508 | 0 | 0 | 452 | 1077 | 0 | 0 | 26 | 0 | 0 | 0 | 0 | 0 | 39 | 0 | 0 | 0 | 0 | 6037 | 0 | 0 |
| SRR6317634 | 0 | 12 | 15 | 54 | 54 | 151 | 252 | 353 | 50 | 0 | 79 | 0 | 0 | 0 | 65886 | 22027 | 174752 | 0 | 0 | 723 | 3717 | 0 | 0 | 131 | 0 | 0 | 0 | 0 | 0 | 264 | 0 | 118 | 0 | 0 | 2845 | 0 | 12 |
| SRR6317635 | 0 | 0 | 17 | 36 | 53 | 498 | 317 | 1404 | 91 | 51 | 338 | 58 | 0 | 0 | 99098 | 52638 | 252287 | 0 | 13 | 1203 | 4722 | 0 | 0 | 55 | 0 | 0 | 0 | 0 | 0 | 876 | 0 | 155 | 0 | 0 | 1570 | 0 | 12 |
| SRR6317636 | 11 | 14 | 20 | 105 | 62 | 244 | 244 | 401 | 52 | 0 | 54 | 0 | 0 | 0 | 5127 | 6505 | 28333 | 0 | 0 | 506 | 558 | 0 | 0 | 14 | 0 | 0 | 0 | 0 | 0 | 15 | 0 | 0 | 0 | 0 | 5968 | 0 | 0 |
| SRR6317637 | 0 | 0 | 0 | 28 | 30 | 250 | 367 | 500 | 50 | 0 | 67 | 13 | 0 | 0 | 10508 | 12381 | 52121 | 0 | 0 | 400 | 1418 | 0 | 0 | 92 | 0 | 0 | 0 | 0 | 0 | 283 | 0 | 83 | 0 | 0 | 9592 | 0 | 0 |
| SRR6317638 | 24 | 29 | 19 | 82 | 72 | 108 | 130 | 259 | 43 | 0 | 87 | 0 | 0 | 0 | 10658 | 8230 | 52947 | 0 | 0 | 346 | 1665 | 0 | 0 | 11 | 0 | 0 | 0 | 0 | 0 | 20 | 0 | 22 | 0 | 0 | 5151 | 0 | 0 |
| SRR6317639 | 34 | 41 | 15 | 128 | 45 | 252 | 213 | 369 | 58 | 0 | 41 | 0 | 0 | 0 | 10282 | 8154 | 50661 | 0 | 0 | 310 | 601 | 0 | 0 | 120 | 0 | 0 | 0 | 0 | 0 | 260 | 0 | 19 | 0 | 0 | 4239 | 0 | 0 |
| SRR6317640 | 24 | 29 | 38 | 59 | 115 | 153 | 102 | 166 | 127 | 0 | 11 | 0 | 0 | 0 | 13397 | 10557 | 63943 | 0 | 0 | 344 | 1108 | 0 | 0 | 24 | 0 | 0 | 0 | 0 | 0 | 36 | 0 | 0 | 0 | 0 | 5220 | 0 | 0 |
| SRR6317641 | 37 | 14 | 41 | 28 | 96 | 81 | 102 | 163 | 61 | 0 | 179 | 0 | 0 | 0 | 7132 | 5890 | 33886 | 0 | 0 | 264 | 914 | 0 | 0 | 371 | 0 | 0 | 0 | 0 | 0 | 385 | 0 | 0 | 0 | 0 | 5926 | 0 | 0 |
| SRR6317642 | 12 | 0 | 12 | 52 | 59 | 211 | 249 | 307 | 58 | 18 | 10 | 12 | 0 | 0 | 97947 | 17638 | 197546 | 0 | 0 | 767 | 7784 | 0 | 0 | 74 | 0 | 0 | 0 | 0 | 0 | 23 | 0 | 0 | 0 | 0 | 1780 | 0 | 0 |
| SRR6317643 | 12 | 0 | 28 | 46 | 20 | 386 | 367 | 1430 | 47 | 41 | 37 | 12 | 0 | 0 | 116773 | 49438 | 304330 | 0 | 0 | 611 | 3631 | 0 | 0 | 11 | 0 | 0 | 0 | 0 | 0 | 107 | 0 | 19 | 0 | 0 | 2507 | 0 | 0 |
| SRR6317644 | 24 | 0 | 19 | 84 | 79 | 138 | 241 | 241 | 34 | 0 | 49 | 0 | 0 | 0 | 6076 | 7358 | 35531 | 0 | 0 | 534 | 878 | 0 | 0 | 147 | 0 | 0 | 0 | 0 | 0 | 228 | 0 | 0 | 0 | 0 | 5394 | 0 | 0 |
| SRR6317645 | 13 | 11 | 0 | 68 | 45 | 228 | 456 | 309 | 81 | 11 | 70 | 0 | 0 | 0 | 14125 | 17653 | 64810 | 0 | 0 | 651 | 2799 | 0 | 0 | 55 | 0 | 0 | 0 | 0 | 0 | 30 | 0 | 65 | 0 | 0 | 6943 | 0 | 0 |
| SRR6324194 | 12 | 0 | 0 | 0 | 0 | 50 | 90 | 83 | 0 | 0 | 0 | 0 | 0 | 0 | 70199 | 45415 | 76544 | 83 | 0 | 1170 | 15653 | 0 | 0 | 16 | 0 | 0 | 0 | 0 | 0 | 0 | 0 | 0 | 0 | 0 | 8798 | 0 | 0 |
| SRR6324195 | 54 | 20 | 24 | 35 | 22 | 35 | 28 | 17 | 0 | 0 | 0 | 0 | 0 | 0 | 90597 | 54907 | 74236 | 100 | 52 | 2861 | 16489 | 0 | 0 | 42 | 0 | 39 | 24 | 0 | 0 | 20 | 0 | 0 | 0 | 0 | 9279 | 0 | 0 |
| SRR6363052 | 77 | 31 | 18 | 31 | 11 | 276 | 87 | 319 | 81 | 0 | 52 | 0 | 0 | 0 | 12995 | 26003 | 63605 | 135 | 124 | 2617 | 676 | 0 | 0 | 73 | 27 | 12 | 12 | 20 | 0 | 72 | 0 | 59 | 0 | 0 | 5632 | 601 | 0 |
| SRR6363053 | 875 | 386 | 157 | 290 | 122 | 203 | 98 | 166 | 43 | 0 | 33 | 16 | 0 | 0 | 25192 | 28918 | 99113 | 142 | 117 | 2966 | 2080 | 0 | 0 | 50 | 20 | 86 | 75 | 0 | 0 | 104 | 0 | 105 | 0 | 0 | 6812 | 489 | 0 |
| SRR6363054 | 47 | 16 | 65 | 13 | 105 | 271 | 154 | 154 | 37 | 0 | 21 | 0 | 0 | 0 | 57898 | 25931 | 192726 | 105 | 0 | 2866 | 1858 | 0 | 0 | 30 | 12 | 60 | 60 | 0 | 0 | 59 | 0 | 92 | 0 | 0 | 5639 | 326 | 0 |
| SRR6370846 | 0 | 0 | 0 | 0 | 0 | 23 | 0 | 92 | 69 | 40 | 72 | 0 | 0 | 0 | 0 | 2161 | 786 | 0 | 11 | 0 | 0 | 0 | 0 | 0 | 0 | 0 | 0 | 0 | 0 | 0 | 0 | 0 | 0 | 0 | 196 | 0 | 0 |
| SRR6370847 | 17 | 17 | 0 | 0 | 50 | 65 | 0 | 617 | 617 | 345 | 17 | 0 | 0 | 0 | 2663 | 56462 | 43299 | 0 | 0 | 227 | 130 | 0 | 0 | 84 | 0 | 0 | 0 | 0 | 0 | 0 | 0 | 0 | 0 | 0 | 4896 | 0 | 0 |
| SRR6370848 | 0 | 0 | 0 | 0 | 0 | 0 | 0 | 417 | 835 | 13 | 0 | 0 | 0 | 0 | 522 | 62180 | 50564 | 0 | 0 | 417 | 209 | 0 | 0 | 0 | 0 | 0 | 0 | 0 | 0 | 0 | 0 | 0 | 0 | 0 | 7516 | 0 | 0 |
| SRR6370849 | 27 | 0 | 0 | 0 | 20 | 314 | 200 | 429 | 2315 | 279 | 94 | 0 | 0 | 0 | 3515 | 26991 | 56059 | 0 | 0 | 229 | 457 | 0 | 0 | 24 | 27 | 0 | 0 | 0 | 0 | 10 | 0 | 0 | 0 | 0 | 2741 | 0 | 0 |
| SRR6370852 | 109 | 18 | 0 | 18 | 0 | 369 | 37 | 369 | 1514 | 173 | 36 | 0 | 0 | 0 | 923 | 15932 | 54752 | 0 | 74 | 369 | 74 | 0 | 0 | 91 | 0 | 0 | 45 | 0 | 0 | 0 | 0 | 0 | 0 | 0 | 1683 | 0 | 0 |
| SRR6370853 | 0 | 0 | 0 | 0 | 0 | 1595 | 0 | 5765 | 4661 | 277 | 0 | 0 | 0 | 0 | 368 | 31897 | 9914 | 0 | 245 | 0 | 0 | 0 | 0 | 69 | 13 | 0 | 0 | 0 | 0 | 0 | 0 | 0 | 0 | 0 | 10776 | 0 | 0 |
| SRR6370856 | 38 | 25 | 0 | 27 | 30 | 225 | 35 | 381 | 450 | 14 | 120 | 0 | 0 | 0 | 761 | 23237 | 25771 | 17 | 52 | 156 | 208 | 0 | 0 | 35 | 16 | 0 | 14 | 0 | 0 | 0 | 0 | 0 | 0 | 0 | 5624 | 0 | 0 |
| SRR6372849 | 142 | 41 | 0 | 13 | 349 | 35510 | 9953 | 39270 | 42773 | 148 | 168 | 17 | 60 | 30 | 65027 | 20872 | 33501 | 677 | 858 | 1838 | 2851 | 0 | 16 | 58 | 84 | 19 | 0 | 0 | 0 | 1243 | 0 | 19 | 0 | 26 | 6082 | 554 | 0 |
| SRR6372850 | 72 | 22 | 0 | 0 | 539 | 9952 | 4034 | 21786 | 18943 | 72 | 16 | 13 | 84 | 44 | 26827 | 16160 | 10361 | 776 | 576 | 853 | 1099 | 0 | 0 | 37 | 13 | 0 | 0 | 0 | 0 | 172 | 0 | 31 | 0 | 12 | 15511 | 513 | 0 |
| SRR6372851 | 13 | 35 | 0 | 0 | 162 | 10916 | 5526 | 18277 | 47085 | 22 | 11 | 28 | 77 | 49 | 18009 | 25581 | 8629 | 351 | 487 | 3885 | 1154 | 0 | 15 | 56 | 0 | 0 | 0 | 0 | 0 | 43 | 21 | 78 | 0 | 13 | 26118 | 403 | 0 |
| SRR6372852 | 53 | 10 | 11 | 21 | 343 | 12808 | 4542 | 14267 | 12235 | 60 | 56 | 11 | 13 | 14 | 3738 | 30238 | 4105 | 625 | 1334 | 762 | 1624 | 0 | 0 | 19 | 11 | 0 | 0 | 0 | 17 | 317 | 0 | 67 | 0 | 0 | 26134 | 71 | 0 |
| SRR6372853 | 18 | 31 | 26 | 21 | 256 | 12316 | 9934 | 27174 | 71928 | 19 | 63 | 15 | 12 | 16 | 6516 | 44757 | 10758 | 259 | 636 | 1018 | 494 | 0 | 0 | 10 | 38 | 17 | 0 | 0 | 14 | 246 | 0 | 25 | 0 | 0 | 8703 | 62 | 0 |
| SRR6372854 | 144 | 69 | 0 | 0 | 1382 | 21616 | 1763 | 17563 | 9399 | 203 | 27 | 35 | 217 | 57 | 4091 | 4330 | 1235 | 326 | 613 | 738 | 517 | 0 | 12 | 24 | 0 | 0 | 0 | 0 | 39 | 232 | 0 | 48 | 0 | 34 | 25885 | 344 | 0 |
| SRR6372855 | 0 | 0 | 0 | 0 | 30 | 54419 | 25776 | 94070 | 112701 | 21 | 0 | 0 | 0 | 0 | 228 | 37391 | 3461 | 312 | 104 | 83 | 145 | 0 | 0 | 39 | 11 | 0 | 0 | 0 | 0 | 11 | 0 | 21 | 0 | 0 | 31148 | 0 | 0 |
| SRR6372856 | 19 | 0 | 0 | 0 | 502 | 17890 | 4220 | 17594 | 21962 | 123 | 24 | 45 | 12 | 0 | 3617 | 41763 | 12414 | 424 | 613 | 1742 | 950 | 0 | 0 | 74 | 43 | 10 | 0 | 0 | 22 | 166 | 0 | 118 | 0 | 10 | 60990 | 0 | 0 |
| SRR6372857 | 32 | 14 | 0 | 0 | 119 | 40151 | 15672 | 37479 | 26755 | 12 | 24 | 11 | 12 | 0 | 771 | 88289 | 10734 | 734 | 755 | 977 | 290 | 0 | 0 | 85 | 27 | 14 | 0 | 0 | 0 | 221 | 0 | 32 | 0 | 14 | 20949 | 31 | 0 |
| SRR6389795 | 75 | 163 | 48 | 10 | 19 | 100 | 124 | 2469 | 242 | 0 | 250 | 75 | 0 | 0 | 13134 | 32336 | 82344 | 0 | 66 | 9016 | 2869 | 0 | 0 | 60 | 0 | 13 | 44 | 11 | 0 | 300 | 0 | 29 | 0 | 0 | 3004 | 0 | 23 |
| SRR6389796 | 46 | 124 | 48 | 63 | 86 | 73 | 61 | 1268 | 71 | 78 | 43 | 11 | 0 | 0 | 18129 | 27994 | 46519 | 0 | 441 | 74407 | 1345 | 0 | 0 | 15 | 0 | 0 | 61 | 0 | 0 | 79 | 0 | 11 | 0 | 0 | 536 | 0 | 0 |
| SRR6389797 | 116 | 321 | 20 | 23 | 22 | 25 | 43 | 478 | 56 | 32 | 49 | 11 | 0 | 0 | 38129 | 28991 | 71097 | 0 | 357 | 12135 | 2002 | 0 | 0 | 17 | 0 | 13 | 75 | 0 | 0 | 151 | 0 | 24 | 0 | 0 | 771 | 0 | 0 |
| SRR6389798 | 236 | 335 | 19 | 16 | 38 | 215 | 143 | 828 | 389 | 40 | 363 | 0 | 0 | 0 | 25219 | 53985 | 85311 | 13 | 761 | 24246 | 30324 | 0 | 0 | 269 | 0 | 13 | 38 | 0 | 0 | 465 | 0 | 16 | 0 | 0 | 2069 | 245 | 0 |
| SRR6389799 | 68 | 122 | 92 | 85 | 37 | 23 | 63 | 517 | 72 | 55 | 54 | 42 | 0 | 0 | 6843 | 38999 | 53763 | 12 | 549 | 63686 | 681 | 0 | 0 | 70 | 0 | 0 | 22 | 0 | 0 | 123 | 0 | 44 | 0 | 0 | 777 | 30 | 0 |
| SRR6389800 | 587 | 479 | 45 | 31 | 98 | 17 | 40 | 334 | 258 | 420 | 486 | 0 | 0 | 0 | 33203 | 20933 | 119794 | 0 | 452 | 10954 | 4069 | 0 | 0 | 398 | 42 | 63 | 207 | 0 | 0 | 1181 | 0 | 15 | 0 | 0 | 341 | 520 | 0 |
| SRR6389801 | 43 | 74 | 0 | 0 | 14 | 68 | 85 | 417 | 178 | 73 | 82 | 0 | 0 | 0 | 2923 | 6606 | 29886 | 0 | 27 | 30554 | 727 | 0 | 0 | 20 | 10 | 0 | 10 | 0 | 0 | 28 | 0 | 23 | 0 | 0 | 1841 | 747 | 0 |
| SRR6389802 | 2123 | 2715 | 15 | 0 | 558 | 416 | 487 | 3465 | 698 | 426 | 255 | 0 | 20 | 0 | 41272 | 18211 | 95987 | 15 | 185 | 6506 | 2326 | 0 | 0 | 400 | 0 | 0 | 24 | 0 | 0 | 660 | 0 | 42 | 0 | 0 | 2414 | 240 | 0 |
| SRR6389803 | 74 | 140 | 57 | 20 | 15 | 61 | 85 | 377 | 367 | 29 | 14 | 10 | 0 | 0 | 22312 | 104202 | 87144 | 15 | 752 | 7516 | 9043 | 0 | 0 | 19 | 0 | 0 | 43 | 0 | 0 | 28 | 0 | 27 | 0 | 0 | 2968 | 774 | 0 |
| SRR6389804 | 28 | 43 | 11 | 14 | 580 | 111 | 303 | 675 | 197 | 18 | 197 | 39 | 18 | 0 | 45921 | 5923 | 81863 | 47 | 238 | 6052 | 4980 | 0 | 0 | 16 | 0 | 0 | 11 | 0 | 0 | 17 | 0 | 21 | 0 | 0 | 12681 | 0 | 0 |
| SRR6389805 | 671 | 787 | 21 | 14 | 187 | 641 | 662 | 3652 | 1167 | 124 | 881 | 28 | 0 | 0 | 31816 | 73340 | 52434 | 68 | 481 | 5246 | 4347 | 0 | 0 | 118 | 0 | 0 | 0 | 0 | 0 | 1302 | 0 | 176 | 0 | 0 | 1882 | 1888 | 0 |
| SRR6389806 | 608 | 1236 | 94 | 63 | 109 | 175 | 181 | 744 | 199 | 181 | 238 | 0 | 0 | 0 | 22044 | 35156 | 79740 | 11 | 548 | 4848 | 4799 | 0 | 0 | 111 | 0 | 11 | 68 | 12 | 0 | 380 | 0 | 18 | 0 | 0 | 1338 | 812 | 0 |

### Supp Table S8.3.Columns_BX_to_DH

| | | | | | | | | | | | | | | | | | | | | | | | | | | | | | | | | | | | | | |
|---|---|---|---|---|---|---|---|---|---|---|---|---|---|---|---|---|---|---|---|---|---|---|---|---|---|---|---|---|---|---|---|---|---|---|---|---|---|
| SRR6389807 | 37 | 74 | 20 | 29 | 75 | 253 | 124 | 1417 | 232 | 41 | 44 | 62 | 0 | 0 | 43994 | 13655 | 78271 | 62 | 379 | 71322 | 570 | 0 | 0 | 86 | 14 | 47 | 119 | 0 | 0 | 101 | 0 | 0 | 0 | 0 | 232 | 722 | 0 |
| SRR6389808 | 154 | 460 | 14 | 19 | 24 | 38 | 76 | 685 | 139 | 28 | 52 | 68 | 0 | 0 | 47541 | 19946 | 53774 | 10 | 438 | 33010 | 1897 | 0 | 0 | 46 | 50 | 13 | 41 | 0 | 0 | 141 | 0 | 22 | 0 | 0 | 508 | 0 | 0 |
| SRR6389809 | 48 | 125 | 25 | 31 | 157 | 148 | 122 | 1060 | 162 | 154 | 36 | 136 | 0 | 0 | 59912 | 52666 | 112440 | 21 | 1140 | 28406 | 3309 | 0 | 0 | 19 | 0 | 0 | 101 | 0 | 0 | 74 | 0 | 0 | 0 | 0 | 936 | 1787 | 18 |
| SRR6389810 | 68 | 191 | 38 | 37 | 47 | 486 | 316 | 3302 | 228 | 16 | 18 | 18 | 0 | 0 | 42756 | 45050 | 64350 | 31 | 563 | 8359 | 1134 | 0 | 0 | 31 | 0 | 0 | 28 | 0 | 0 | 87 | 0 | 41 | 0 | 0 | 6039 | 116 | 0 |
| SRR6389811 | 349 | 1161 | 55 | 69 | 194 | 534 | 360 | 2567 | 108 | 16 | 904 | 53 | 0 | 0 | 26593 | 42965 | 77303 | 0 | 343 | 8917 | 735 | 0 | 0 | 206 | 0 | 22 | 160 | 0 | 0 | 1547 | 0 | 15 | 0 | 0 | 1416 | 0 | 0 |
| SRR6389812 | 11 | 29 | 0 | 0 | 97 | 217 | 203 | 2193 | 559 | 292 | 89 | 0 | 0 | 0 | 153721 | 61734 | 116468 | 42 | 629 | 4565 | 1000 | 0 | 0 | 12 | 0 | 0 | 24 | 0 | 0 | 21 | 0 | 46 | 0 | 0 | 2858 | 0 | 0 |
| SRR6389813 | 142 | 311 | 29 | 26 | 32 | 221 | 446 | 2343 | 139 | 0 | 74 | 86 | 0 | 0 | 2818 | 43591 | 71453 | 20 | 234 | 19338 | 1654 | 0 | 0 | 31 | 0 | 0 | 30 | 0 | 0 | 36 | 0 | 45 | 0 | 0 | 1347 | 0 | 14 |
| SRR6389814 | 204 | 293 | 11 | 0 | 97 | 360 | 244 | 2390 | 259 | 677 | 134 | 0 | 0 | 0 | 114815 | 10162 | 76349 | 34 | 198 | 1597 | 1754 | 0 | 0 | 170 | 0 | 17 | 25 | 0 | 0 | 241 | 0 | 17 | 0 | 0 | 1084 | 0 | 0 |
| SRR6389815 | 32 | 61 | 0 | 0 | 14 | 1745 | 1285 | 16769 | 626 | 17 | 14 | 127 | 0 | 0 | 23056 | 150958 | 71504 | 0 | 247 | 8714 | 1988 | 0 | 0 | 141 | 0 | 0 | 0 | 0 | 0 | 24 | 0 | 107 | 0 | 0 | 1724 | 589 | 15 |
| SRR6389816 | 1016 | 1323 | 59 | 26 | 44 | 50 | 34 | 255 | 54 | 1805 | 2824 | 20 | 0 | 0 | 99600 | 42390 | 209487 | 57 | 455 | 5129 | 4104 | 0 | 0 | 22 | 0 | 0 | 0 | 0 | 0 | 8442 | 0 | 106 | 0 | 0 | 284 | 2228 | 0 |
| SRR6389817 | 1326 | 1274 | 0 | 0 | 651 | 457 | 556 | 5242 | 1317 | 104 | 411 | 0 | 0 | 0 | 45326 | 42916 | 144743 | 33 | 154 | 5533 | 7333 | 0 | 0 | 42 | 0 | 0 | 0 | 0 | 0 | 397 | 0 | 24 | 0 | 0 | 1928 | 34 | 0 |
| SRR6389818 | 791 | 725 | 26 | 0 | 192 | 139 | 145 | 1059 | 364 | 1946 | 732 | 16 | 0 | 0 | 96377 | 7419 | 105067 | 18 | 244 | 5327 | 6686 | 0 | 0 | 435 | 0 | 22 | 35 | 0 | 0 | 801 | 0 | 26 | 0 | 0 | 319 | 0 | 0 |
| SRR6389819 | 67 | 81 | 29 | 13 | 101 | 56 | 136 | 685 | 176 | 75 | 20 | 0 | 0 | 0 | 86519 | 11038 | 89633 | 21 | 118 | 7929 | 2995 | 0 | 0 | 346 | 38 | 35 | 15 | 0 | 0 | 75 | 0 | 20 | 0 | 0 | 5285 | 380 | 0 |
| SRR6389820 | 45 | 61 | 0 | 0 | 40 | 214 | 454 | 4239 | 427 | 61 | 153 | 35 | 0 | 0 | 25526 | 122188 | 61296 | 40 | 1193 | 19474 | 2301 | 0 | 0 | 13 | 0 | 0 | 0 | 0 | 0 | 49 | 0 | 127 | 0 | 0 | 1423 | 1119 | 0 |
| SRR6409751 | 0 | 0 | 0 | 0 | 0 | 0 | 0 | 1728 | 0 | 0 | 0 | 0 | 0 | 0 | 0 | 149359 | 35904 | 0 | 0 | 0 | 9843 | 0 | 0 | 0 | 0 | 0 | 59 | 0 | 0 | 0 | 0 | 0 | 0 | 0 | 1174 | 0 | 0 |
| SRR6409760 | 0 | 0 | 0 | 0 | 0 | 0 | 0 | 0 | 0 | 0 | 0 | 0 | 0 | 0 | 8565 | 193248 | 43153 | 0 | 0 | 0 | 0 | 0 | 0 | 0 | 0 | 0 | 112 | 0 | 0 | 17 | 0 | 0 | 0 | 0 | 620 | 0 | 0 |
| SRR6409798 | 0 | 0 | 0 | 362 | 26 | 0 | 460 | 0 | 0 | 0 | 0 | 0 | 0 | 0 | 31085 | 50434 | 358710 | 0 | 0 | 37158 | 33807 | 0 | 0 | 0 | 0 | 199 | 114 | 0 | 0 | 15 | 0 | 0 | 0 | 0 | 0 | 0 | 0 |
| SRR6409800 | 0 | 0 | 0 | 0 | 0 | 0 | 0 | 0 | 0 | 0 | 0 | 0 | 0 | 0 | 146 | 78545 | 31027 | 0 | 0 | 0 | 9518 | 0 | 0 | 0 | 0 | 64 | 0 | 0 | 0 | 0 | 0 | 0 | 0 | 0 | 0 | 0 | 0 |
| SRR6409827 | 0 | 0 | 0 | 0 | 0 | 0 | 3160 | 1733 | 0 | 0 | 0 | 0 | 0 | 0 | 34465 | 51799 | 319821 | 0 | 19 | 54279 | 36088 | 0 | 0 | 87 | 0 | 0 | 425 | 23 | 0 | 0 | 0 | 0 | 0 | 0 | 0 | 0 | 0 |
| SRR6409832 | 0 | 0 | 0 | 0 | 0 | 0 | 0 | 0 | 0 | 0 | 0 | 0 | 0 | 0 | 10289 | 39421 | 33778 | 0 | 0 | 0 | 0 | 0 | 0 | 0 | 0 | 0 | 0 | 0 | 0 | 0 | 0 | 0 | 0 | 0 | 0 | 0 | 0 |
| SRR6409833 | 0 | 0 | 0 | 0 | 0 | 633 | 2620 | 0 | 0 | 0 | 0 | 0 | 0 | 0 | 23072 | 45660 | 315519 | 0 | 20 | 47690 | 29685 | 0 | 0 | 0 | 0 | 0 | 147 | 0 | 0 | 0 | 0 | 0 | 0 | 0 | 0 | 0 | 0 |
| SRR6409834 | 0 | 0 | 0 | 0 | 0 | 0 | 0 | 0 | 0 | 0 | 0 | 0 | 0 | 0 | 24680 | 67464 | 348481 | 0 | 0 | 29120 | 34613 | 0 | 0 | 7975 | 0 | 0 | 404 | 21 | 0 | 0 | 0 | 0 | 0 | 0 | 2254 | 0 | 0 |
| SRR6409845 | 0 | 0 | 0 | 0 | 0 | 0 | 0 | 16173 | 0 | 0 | 0 | 0 | 0 | 0 | 23201 | 125335 | 54503 | 0 | 0 | 0 | 13463 | 0 | 0 | 0 | 0 | 20 | 93 | 12 | 0 | 0 | 0 | 0 | 0 | 0 | 5453 | 0 | 0 |
| SRR6409851 | 0 | 0 | 0 | 27 | 0 | 0 | 182 | 0 | 0 | 0 | 0 | 0 | 0 | 0 | 24020 | 43037 | 368305 | 0 | 0 | 37670 | 30081 | 0 | 0 | 562 | 0 | 107 | 427 | 0 | 0 | 0 | 0 | 479 | 0 | 0 | 155 | 0 | 0 |
| SRR6409853 | 0 | 0 | 0 | 29 | 0 | 0 | 0 | 328 | 749 | 0 | 0 | 0 | 0 | 0 | 30642 | 37379 | 369202 | 0 | 0 | 34622 | 43572 | 0 | 0 | 0 | 0 | 224 | 460 | 0 | 0 | 0 | 0 | 0 | 0 | 0 | 0 | 0 | 0 |
| SRR6409859 | 0 | 0 | 0 | 187 | 0 | 0 | 852 | 0 | 0 | 0 | 0 | 0 | 0 | 0 | 23157 | 48062 | 333906 | 0 | 23 | 35402 | 37188 | 0 | 0 | 107 | 35 | 403 | 0 | 0 | 0 | 0 | 0 | 0 | 0 | 0 | 0 | 0 | 0 |
| SRR6409860 | 0 | 0 | 0 | 0 | 0 | 0 | 567 | 0 | 1275 | 0 | 0 | 0 | 0 | 0 | 26147 | 38092 | 344174 | 0 | 454 | 41752 | 30387 | 0 | 0 | 0 | 0 | 492 | 519 | 41 | 0 | 0 | 0 | 0 | 0 | 0 | 792 | 0 | 0 |
| SRR6409863 | 0 | 0 | 0 | 0 | 0 | 0 | 0 | 0 | 0 | 0 | 0 | 0 | 0 | 0 | 23708 | 37968 | 288435 | 0 | 0 | 33587 | 30816 | 0 | 0 | 0 | 0 | 714 | 398 | 0 | 0 | 0 | 0 | 0 | 0 | 0 | 0 | 0 | 0 |
| SRR6409866 | 0 | 0 | 0 | 0 | 0 | 0 | 0 | 0 | 0 | 0 | 0 | 0 | 0 | 0 | 32675 | 45645 | 348031 | 0 | 0 | 38531 | 24227 | 0 | 0 | 0 | 709 | 0 | 385 | 35 | 0 | 0 | 0 | 0 | 0 | 0 | 0 | 0 | 0 |
| SRR6409867 | 0 | 0 | 0 | 0 | 0 | 0 | 0 | 0 | 0 | 0 | 0 | 0 | 0 | 0 | 24650 | 43552 | 362201 | 0 | 0 | 46234 | 30636 | 0 | 0 | 0 | 0 | 249 | 1720 | 0 | 0 | 0 | 0 | 0 | 0 | 0 | 181 | 0 | 0 |
| SRR6409868 | 0 | 0 | 0 | 0 | 0 | 0 | 0 | 0 | 0 | 0 | 0 | 0 | 0 | 0 | 29867 | 45218 | 322997 | 0 | 12 | 23174 | 35723 | 0 | 0 | 0 | 0 | 0 | 539 | 0 | 0 | 0 | 0 | 12 | 0 | 0 | 0 | 0 | 0 |
| SRR6450851 | 38 | 0 | 0 | 19 | 95 | 0 | 0 | 424 | 1273 | 534 | 210 | 0 | 0 | 0 | 6367 | 386421 | 13454 | 0 | 0 | 1486 | 212 | 0 | 0 | 38 | 0 | 19 | 19 | 0 | 19 | 76 | 19 | 2122 | 0 | 95 | 2190 | 0 | 19 |
| SRR6450852 | 76 | 23 | 12 | 11 | 40 | 46 | 35 | 116 | 685 | 639 | 138 | 0 | 0 | 0 | 2928 | 251023 | 5754 | 186 | 58 | 186 | 209 | 12 | 0 | 15 | 47 | 47 | 21 | 12 | 12 | 23 | 24 | 4008 | 27 | 30 | 2193 | 0 | 29 |
| SRR6450853 | 363 | 17 | 66 | 33 | 264 | 367 | 0 | 407 | 1100 | 2231 | 661 | 0 | 0 | 0 | 4521 | 276720 | 6544 | 163 | 0 | 163 | 326 | 116 | 17 | 149 | 33 | 66 | 17 | 66 | 33 | 264 | 17 | 3625 | 66 | 132 | 1212 | 0 | 0 |
| SRR6450854 | 267 | 23 | 23 | 46 | 186 | 446 | 52 | 210 | 917 | 1743 | 407 | 23 | 0 | 12 | 2490 | 216230 | 4732 | 236 | 79 | 419 | 288 | 35 | 0 | 81 | 46 | 12 | 12 | 0 | 35 | 244 | 23 | 4770 | 46 | 198 | 1694 | 0 | 12 |
| SRR6450855 | 89 | 0 | 0 | 13 | 25 | 171 | 171 | 0 | 854 | 292 | 76 | 0 | 0 | 0 | 4951 | 275681 | 4695 | 0 | 0 | 171 | 0 | 13 | 0 | 13 | 0 | 25 | 13 | 13 | 0 | 25 | 0 | 3585 | 13 | 13 | 1690 | 0 | 0 |
| SRR6450856 | 44 | 24 | 41 | 11 | 68 | 109 | 87 | 240 | 1046 | 452 | 188 | 0 | 0 | 0 | 5492 | 334876 | 10319 | 240 | 131 | 349 | 305 | 48 | 12 | 13 | 15 | 48 | 15 | 11 | 12 | 61 | 36 | 4642 | 11 | 37 | 708 | 0 | 0 |
| SRR6450857 | 52 | 35 | 0 | 17 | 87 | 188 | 0 | 0 | 752 | 574 | 157 | 0 | 0 | 0 | 5261 | 375586 | 13795 | 188 | 0 | 564 | 0 | 17 | 0 | 35 | 17 | 0 | 52 | 0 | 0 | 35 | 0 | 3194 | 17 | 0 | 781 | 0 | 0 |
| SRR6450858 | 395 | 70 | 93 | 70 | 337 | 182 | 52 | 312 | 1094 | 2834 | 941 | 0 | 0 | 0 | 5755 | 350274 | 11497 | 78 | 130 | 312 | 495 | 0 | 12 | 93 | 81 | 35 | 35 | 23 | 46 | 267 | 12 | 4349 | 128 | 163 | 1795 | 0 | 0 |
| SRR6450859 | 553 | 111 | 277 | 111 | 166 | 332 | 0 | 221 | 886 | 2047 | 941 | 0 | 0 | 0 | 5756 | 388380 | 14841 | 332 | 553 | 664 | 443 | 166 | 0 | 166 | 0 | 111 | 111 | 55 | 0 | 498 | 55 | 3431 | 55 | 332 | 1263 | 0 | 0 |
| SRR6450860 | 71 | 35 | 27 | 35 | 98 | 325 | 0 | 325 | 974 | 275 | 160 | 0 | 0 | 0 | 4981 | 397581 | 10470 | 108 | 217 | 541 | 541 | 18 | 46 | 18 | 0 | 27 | 27 | 18 | 0 | 80 | 46 | 1732 | 27 | 27 | 2360 | 0 | 0 |
| SRR6450861 | 61 | 11 | 11 | 31 | 55 | 504 | 0 | 567 | 756 | 353 | 55 | 0 | 0 | 0 | 2141 | 182406 | 4050 | 252 | 63 | 126 | 315 | 11 | 0 | 11 | 0 | 0 | 31 | 0 | 0 | 17 | 31 | 4030 | 11 | 22 | 2061 | 0 | 0 |
| SRR6450862 | 320 | 101 | 34 | 22 | 251 | 456 | 94 | 559 | 798 | 1563 | 438 | 0 | 0 | 0 | 1689 | 128273 | 2784 | 462 | 66 | 190 | 145 | 52 | 17 | 71 | 27 | 30 | 20 | 0 | 27 | 116 | 15 | 5176 | 135 | 187 | 1172 | 0 | 32 |
| SRR6450863 | 384 | 70 | 47 | 24 | 274 | 165 | 41 | 206 | 741 | 2969 | 580 | 0 | 0 | 0 | 3313 | 281256 | 7108 | 206 | 62 | 206 | 226 | 24 | 24 | 16 | 16 | 16 | 39 | 0 | 39 | 157 | 39 | 4054 | 110 | 188 | 1631 | 0 | 16 |
| SRR6450867 | 322 | 212 | 387 | 1244 | 55 | 20 | 30 | 30 | 0 | 0 | 83 | 0 | 0 | 0 | 1963 | 2224 | 137 | 684 | 0 | 0 | 20 | 0 | 0 | 37 | 64 | 46 | 230 | 46 | 0 | 0 | 0 | 89 | 0 | 0 | 3941 | 0 | 0 |
| SRR6450868 | 20 | 10 | 13 | 0 | 0 | 65 | 32 | 0 | 32 | 10 | 20 | 0 | 0 | 0 | 7702 | 1752 | 182 | 324 | 0 | 0 | 32 | 0 | 0 | 0 | 0 | 0 | 0 | 0 | 0 | 0 | 0 | 0 | 151 | 0 | 4525 | 0 | 15 |
| SRR6450869 | 20 | 0 | 20 | 0 | 0 | 0 | 0 | 93 | 0 | 20 | 20 | 0 | 0 | 0 | 6163 | 2091 | 116 | 93 | 0 | 0 | 0 | 20 | 0 | 0 | 0 | 0 | 0 | 0 | 0 | 0 | 0 | 0 | 140 | 0 | 5344 | 0 | 0 |
| SRR6453341 | 19 | 11 | 0 | 0 | 174 | 246 | 123 | 931 | 2135 | 48 | 108 | 13 | 0 | 0 | 1027 | 48793 | 10541 | 0 | 0 | 110 | 82 | 0 | 0 | 11 | 26 | 12 | 0 | 0 | 0 | 123 | 0 | 27 | 0 | 0 | 96430 | 0 | 0 |
| SRR6453342 | 16 | 0 | 0 | 0 | 99 | 443 | 346 | 1525 | 5904 | 38 | 338 | 28 | 0 | 0 | 1125 | 69692 | 18560 | 32 | 0 | 303 | 76 | 0 | 0 | 0 | 17 | 15 | 10 | 0 | 0 | 138 | 0 | 11 | 0 | 0 | 22143 | 0 | 0 |
| SRR6453343 | 32 | 12 | 0 | 0 | 32 | 898 | 847 | 3908 | 11409 | 44 | 37 | 24 | 0 | 0 | 1379 | 103442 | 16729 | 0 | 0 | 164 | 101 | 0 | 0 | 0 | 20 | 15 | 10 | 0 | 0 | 31 | 0 | 25 | 0 | 0 | 62878 | 0 | 0 |
| SRR6453344 | 34 | 22 | 0 | 0 | 42 | 1049 | 1116 | 2898 | 10499 | 44 | 62 | 41 | 0 | 0 | 1331 | 117514 | 21788 | 23 | 34 | 192 | 90 | 0 | 0 | 14 | 24 | 17 | 29 | 16 | 0 | 37 | 0 | 11 | 0 | 0 | 31718 | 0 | 0 |
| SRR6453345 | 43 | 28 | 0 | 0 | 33 | 619 | 449 | 2197 | 7142 | 15 | 29 | 29 | 0 | 0 | 2507 | 99700 | 18227 | 20 | 30 | 160 | 80 | 0 | 0 | 12 | 21 | 19 | 18 | 15 | 0 | 19 | 0 | 20 | 0 | 0 | 28886 | 0 | 0 |
| SRR6453346 | 31 | 12 | 0 | 0 | 379 | 641 | 379 | 1780 | 4344 | 52 | 581 | 17 | 0 | 0 | 1662 | 108126 | 18502 | 0 | 13 | 222 | 92 | 0 | 0 | 11 | 21 | 11 | 14 | 0 | 0 | 280 | 0 | 39 | 0 | 0 | 21880 | 0 | 0 |
| SRR6453347 | 11 | 0 | 0 | 0 | 87 | 657 | 578 | 2076 | 6713 | 57 | 201 | 32 | 0 | 0 | 2943 | 57715 | 23747 | 53 | 0 | 420 | 66 | 0 | 0 | 0 | 11 | 0 | 0 | 0 | 0 | 118 | 0 | 0 | 0 | 0 | 27353 | 0 | 0 |
| SRR6453348 | 52 | 34 | 0 | 0 | 263 | 574 | 641 | 2182 | 6973 | 55 | 296 | 37 | 0 | 0 | 1507 | 99802 | 22912 | 34 | 22 | 247 | 225 | 0 | 0 | 16 | 17 | 23 | 29 | 11 | 0 | 200 | 0 | 22 | 0 | 0 | 25408 | 0 | 0 |
| SRR6453349 | 51 | 27 | 0 | 0 | 263 | 389 | 259 | 1138 | 3672 | 71 | 287 | 25 | 0 | 0 | 1295 | 86863 | 15815 | 19 | 19 | 130 | 93 | 0 | 0 | 0 | 17 | 17 | 20 | 0 | 0 | 145 | 0 | 19 | 0 | 0 | 32026 | 0 | 0 |
| SRR6453350 | 46 | 24 | 0 | 0 | 49 | 710 | 552 | 2338 | 7850 | 10 | 41 | 26 | 0 | 0 | 2683 | 125790 | 37675 | 14 | 21 | 331 | 138 | 0 | 0 | 16 | 22 | 10 | 21 | 0 | 0 | 22 | 0 | 14 | 0 | 0 | 21907 | 0 | 0 |
| SRR6453351 | 19 | 0 | 0 | 0 | 138 | 430 | 240 | 903 | 2956 | 89 | 433 | 29 | 0 | 0 | 1235 | 71043 | 18278 | 21 | 28 | 268 | 113 | 0 | 0 | 0 | 10 | 12 | 15 | 0 | 0 | 183 | 0 | 0 | 0 | 0 | 64991 | 0 | 0 |
| SRR6453352 | 32 | 12 | 0 | 0 | 37 | 1081 | 1295 | 4378 | 15415 | 29 | 67 | 26 | 0 | 0 | 1468 | 74698 | 18330 | 40 | 0 | 214 | 80 | 0 | 0 | 0 | 20 | 10 | 0 | 0 | 0 | 38 | 0 | 0 | 0 | 0 | 38556 | 0 | 0 |
| SRR6453353 | 17 | 0 | 0 | 0 | 132 | 566 | 370 | 1880 | 7268 | 68 | 239 | 34 | 0 | 0 | 968 | 76863 | 11981 | 31 | 16 | 275 | 79 | 0 | 0 | 0 | 15 | 0 | 0 | 0 | 0 | 230 | 0 | 39 | 0 | 0 | 27280 | 0 | 0 |
| SRR6453354 | 23 | 10 | 0 | 0 | 185 | 501 | 866 | 3598 | 16050 | 81 | 132 | 19 | 0 | 0 | 1439 | 103141 | 16646 | 31 | 52 | 302 | 83 | 0 | 0 | 16 | 36 | 24 | 27 | 0 | 0 | 189 | 0 | 10 | 0 | 0 | 14909 | 0 | 0 |
| SRR6453355 | 17 | 13 | 0 | 0 | 306 | 316 | 450 | 1283 | 5745 | 100 | 293 | 26 | 0 | 0 | 527 | 49508 | 11216 | 29 | 0 | 215 | 19 | 0 | 0 | 11 | 26 | 19 | 19 | 0 | 0 | 296 | 0 | 0 | 0 | 0 | 8633 | 0 | 0 |
| SRR6453356 | 32 | 23 | 0 | 0 | 32 | 363 | 462 | 1320 | 4844 | 21 | 28 | 37 | 0 | 0 | 1213 | 62644 | 17519 | 41 | 0 | 314 | 58 | 0 | 0 | 0 | 15 | 12 | 11 | 0 | 0 | 38 | 0 | 50 | 0 | 0 | 16627 | 0 | 0 |
| SRR6453357 | 30 | 12 | 0 | 0 | 31 | 815 | 1048 | 3947 | 17253 | 39 | 37 | 20 | 0 | 0 | 2119 | 90672 | 21982 | 35 | 35 | 233 | 70 | 0 | 0 | 19 | 46 | 27 | 19 | 10 | 0 | 35 | 0 | 23 | 0 | 0 | 47989 | 0 | 0 |
| SRR6453358 | 27 | 13 | 0 | 0 | 283 | 327 | 496 | 2269 | 10926 | 63 | 218 | 41 | 0 | 0 | 1822 | 69007 | 16919 | 27 | 16 | 115 | 38 | 0 | 0 | 11 | 33 | 20 | 18 | 0 | 0 | 332 | 0 | 0 | 0 | 0 | 9716 | 0 | 0 |
| SRR6453359 | 0 | 0 | 0 | 0 | 23 | 158 | 165 | 757 | 2382 | 15 | 37 | 25 | 0 | 0 | 1019 | 63937 | 17727 | 0 | 17 | 234 | 14 | 0 | 0 | 0 | 21 | 11 | 19 | 0 | 0 | 30 | 0 | 0 | 0 | 0 | 5511 | 0 | 0 |
| SRR6453360 | 20 | 0 | 0 | 0 | 122 | 795 | 515 | 2576 | 9348 | 32 | 196 | 24 | 0 | 0 | 1030 | 54158 | 14546 | 44 | 0 | 633 | 29 | 0 | 0 | 0 | 0 | 0 | 0 | 0 | 0 | 225 | 0 | 15 | 0 | 0 | 52698 | 0 | 0 |
| SRR6453361 | 15 | 0 | 0 | 0 | 30 | 323 | 421 | 1568 | 5655 | 13 | 36 | 22 | 0 | 0 | 763 | 56080 | 14396 | 43 | 12 | 342 | 55 | 0 | 0 | 16 | 27 | 20 | 30 | 10 | 0 | 41 | 0 | 37 | 0 | 0 | 16700 | 0 | 0 |
| SRR6453362 | 11 | 0 | 0 | 0 | 34 | 553 | 474 | 1772 | 6906 | 20 | 58 | 34 | 0 | 0 | 1357 | 47387 | 18766 | 69 | 21 | 394 | 59 | 0 | 0 | 0 | 22 | 15 | 19 | 0 | 0 | 83 | 0 | 0 | 0 | 0 | 22069 | 0 | 0 |
| SRR6453363 | 12 | 11 | 0 | 0 | 45 | 617 | 353 | 1658 | 6196 | 20 | 88 | 39 | 0 | 0 | 1546 | 62248 | 19627 | 35 | 41 | 306 | 188 | 0 | 0 | 0 | 13 | 16 | 0 | 0 | 0 | 88 | 0 | 24 | 0 | 0 | 37570 | 0 | 0 |
| SRR6453364 | 33 | 10 | 0 | 0 | 159 | 1050 | 613 | 3115 | 8593 | 43 | 187 | 30 | 0 | 0 | 1313 | 41332 | 11959 | 70 | 18 | 455 | 35 | 0 | 0 | 0 | 0 | 0 | 0 | 0 | 0 | 234 | 0 | 0 | 0 | 0 | 55421 | 0 | 0 |
| SRR6453365 | 23 | 18 | 0 | 0 | 63 | 570 | 841 | 2860 | 9654 | 14 | 35 | 23 | 0 | 0 | 3112 | 93230 | 18041 | 0 | 0 | 367 | 58 | 0 | 0 | 0 | 16 | 0 | 0 | 0 | 0 | 49 | 0 | 19 | 0 | 0 | 40446 | 0 | 0 |
| SRR6453366 | 14 | 0 | 0 | 0 | 31 | 559 | 584 | 2355 | 8201 | 17 | 53 | 34 | 0 | 0 | 1520 | 55505 | 21752 | 25 | 0 | 320 | 82 | 0 | 0 | 0 | 14 | 12 | 0 | 0 | 0 | 63 | 0 | 0 | 0 | 0 | 21552 | 0 | 0 |
| SRR6453367 | 0 | 0 | 0 | 0 | 20 | 548 | 317 | 891 | 2989 | 0 | 49 | 16 | 0 | 0 | 796 | 64908 | 22107 | 0 | 0 | 240 | 26 | 0 | 0 | 25 | 0 | 0 | 18 | 0 | 0 | 75 | 0 | 60 | 0 | 0 | 7975 | 0 | 0 |
| SRR6453368 | 0 | 0 | 0 | 0 | 233 | 980 | 468 | 1536 | 4325 | 0 | 438 | 32 | 0 | 0 | 828 | 59306 | 29408 | 33 | 44 | 425 | 33 | 0 | 0 | 19 | 24 | 14 | 28 | 0 | 0 | 791 | 0 | 120 | 0 | 0 | 5232 | 0 | 0 |
| SRR6453369 | 25 | 11 | 0 | 0 | 139 | 299 | 242 | 1290 | 4642 | 52 | 185 | 16 | 0 | 0 | 2246 | 58021 | 18096 | 12 | 12 | 92 | 104 | 0 | 0 | 0 | 20 | 0 | 0 | 0 | 0 | 118 | 0 | 35 | 0 | 0 | 43469 | 0 | 0 |
| SRR6453370 | 21 | 10 | 0 | 0 | 151 | 624 | 484 | 1936 | 9086 | 46 | 532 | 32 | 0 | 0 | 3060 | 103026 | 32378 | 16 | 16 | 250 | 203 | 0 | 0 | 0 | 0 | 0 | 0 | 0 | 0 | 177 | 0 | 16 | 0 | 0 | 17710 | 0 | 0 |
| SRR6453371 | 12 | 0 | 0 | 0 | 119 | 610 | 169 | 1376 | 4595 | 390 | 184 | 28 | 0 | 0 | 2051 | 44360 | 13362 | 13 | 13 | 195 | 130 | 0 | 0 | 0 | 14 | 0 | 0 | 0 | 0 | 251 | 0 | 104 | 0 | 0 | 81203 | 0 | 0 |
| SRR6453372 | 10 | 0 | 0 | 0 | 23 | 395 | 150 | 953 | 4504 | 129 | 41 | 10 | 0 | 0 | 1279 | 34732 | 13755 | 27 | 0 | 231 | 109 | 0 | 0 | 0 | 13 | 11 | 0 | 0 | 0 | 59 | 0 | 41 | 0 | 0 | 66251 | 0 | 0 |
| SRR6453373 | 36 | 14 | 0 | 0 | 183 | 1189 | 686 | 3114 | 11652 | 514 | 117 | 26 | 0 | 0 | 2277 | 65270 | 22193 | 33 | 17 | 151 | 167 | 0 | 0 | 12 | 31 | 20 | 0 | 0 | 0 | 216 | 0 | 84 | 0 | 0 | 60508 | 0 | 0 |
| SRR6453374 | 15 | 13 | 0 | 0 | 97 | 486 | 341 | 1261 | 4042 | 563 | 126 | 18 | 0 | 0 | 1251 | 52177 | 15156 | 31 | 21 | 207 | 300 | 0 | 0 | 0 | 15 | 0 | 0 | 0 | 0 | 248 | 0 | 93 | 0 | 0 | 86657 | 0 | 0 |
| SRR6453375 | 13 | 0 | 0 | 0 | 193 | 747 | 229 | 1072 | 4168 | 491 | 84 | 20 | 0 | 0 | 1060 | 42520 | 13493 | 12 | 12 | 253 | 157 | 0 | 0 | 0 | 17 | 0 | 0 | 0 | 0 | 124 | 0 | 72 | 0 | 0 | 54344 | 0 | 0 |
| SRR6453376 | 0 | 0 | 0 | 0 | 25 | 1102 | 513 | 3102 | 6332 | 55 | 15 | 25 | 0 | 0 | 1948 | 50735 | 17796 | 51 | 26 | 308 | 51 | 0 | 0 | 0 | 0 | 0 | 17 | 0 | 0 | 31 | 0 | 26 | 0 | 14 | 18055 | 0 | 0 |
| SRR6453377 | 18 | 0 | 0 | 0 | 95 | 1034 | 770 | 3018 | 9749 | 292 | 197 | 12 | 0 | 0 | 2954 | 66328 | 23216 | 21 | 21 | 222 | 53 | 0 | 0 | 11 | 34 | 18 | 0 | 0 | 0 | 192 | 0 | 74 | 0 | 0 | 45726 | 0 | 0 |
| SRR6453378 | 15 | 11 | 0 | 0 | 28 | 1033 | 1033 | 3202 | 12146 | 85 | 24 | 22 | 0 | 0 | 1270 | 69200 | 13368 | 17 | 0 | 152 | 102 | 0 | 0 | 0 | 20 | 0 | 0 | 0 | 0 | 43 | 0 | 152 | 0 | 0 | 50791 | 0 | 0 |
| SRR6453379 | 10 | 11 | 0 | 0 | 28 | 1188 | 422 | 2520 | 10702 | 116 | 26 | 24 | 0 | 0 | 3589 | 72158 | 20371 | 40 | 13 | 343 | 106 | 0 | 0 | 0 | 14 | 10 | 11 | 0 | 0 | 67 | 0 | 132 | 0 | 0 | 20887 | 0 | 0 |
| SRR6453380 | 0 | 0 | 0 | 0 | 59 | 1082 | 792 | 3509 | 11647 | 282 | 104 | 0 | 0 | 0 | 755 | 62288 | 10823 | 25 | 13 | 239 | 38 | 0 | 0 | 11 | 21 | 0 | 0 | 0 | 0 | 162 | 0 | 126 | 0 | 0 | 15140 | 0 | 0 |
| SRR6453381 | 15 | 0 | 0 | 0 | 24 | 416 | 244 | 1058 | 4466 | 29 | 44 | 34 | 0 | 0 | 1058 | 60431 | 16462 | 18 | 18 | 90 | 145 | 0 | 0 | 0 | 19 | 0 | 0 | 0 | 0 | 39 | 0 | 0 | 0 | 0 | 24934 | 0 | 0 |
| SRR6453382 | 19 | 0 | 0 | 0 | 103 | 625 | 731 | 3038 | 14938 | 52 | 205 | 42 | 0 | 0 | 1577 | 78929 | 24942 | 11 | 11 | 233 | 32 | 0 | 0 | 0 | 26 | 11 | 16 | 0 | 0 | 261 | 0 | 11 | 0 | 0 | 17154 | 0 | 0 |
| SRR6453383 | 24 | 11 | 0 | 0 | 25 | 461 | 226 | 726 | 3474 | 10 | 33 | 39 | 0 | 0 | 1453 | 98106 | 17568 | 0 | 10 | 206 | 157 | 0 | 0 | 0 | 10 | 0 | 0 | 0 | 0 | 47 | 0 | 10 | 0 | 0 | 18032 | 0 | 0 |
| SRR6453384 | 13 | 0 | 0 | 0 | 33 | 575 | 280 | 1044 | 3829 | 11 | 37 | 34 | 0 | 0 | 1362 | 51280 | 17814 | 23 | 0 | 280 | 98 | 0 | 0 | 0 | 13 | 0 | 0 | 0 | 0 | 40 | 0 | 0 | 0 | 0 | 48975 | 0 | 0 |
| SRR6453385 | 11 | 0 | 0 | 0 | 25 | 915 | 402 | 1563 | 5771 | 33 | 83 | 10 | 0 | 0 | 4174 | 66975 | 20158 | 11 | 11 | 123 | 190 | 0 | 0 | 0 | 12 | 0 | 0 | 0 | 0 | 78 | 0 | 33 | 0 | 0 | 114005 | 0 | 0 |
| SRR6453386 | 0 | 0 | 0 | 0 | 109 | 1185 | 0 | 456 | 1732 | 89 | 12 | 84 | 0 | 0 | 91 | 6298 | 2576 | 0 | 0 | 0 | 319 | 0 | 0 | 47 | 124 | 95 | 30 | 49 | 0 | 15 | 0 | 46 | 0 | 0 | 19395 | 0 | 0 |
| SRR6453387 | 13 | 0 | 0 | 0 | 45 | 1462 | 1180 | 5179 | 24434 | 43 | 0 | 10 | 0 | 0 | 53 | 14016 | 7888 | 18 | 0 | 0 | 53 | 0 | 0 | 26 | 134 | 80 | 46 | 58 | 0 | 24 | 0 | 18 | 0 | 0 | 5217 | 0 | 0 |
| SRR6453388 | 10 | 0 | 0 | 0 | 124 | 4287 | 4903 | 15273 | 91324 | 60 | 148 | 38 | 0 | 0 | 13 | 35277 | 14335 | 13 | 0 | 52 | 131 | 0 | 0 | 118 | 366 | 288 | 93 | 108 | 0 | 245 | 0 | 13 | 0 | 0 | 7097 | 0 | 0 |
| SRR6453389 | 0 | 0 | 0 | 0 | 67 | 5136 | 4394 | 15937 | 74637 | 135 | 48 | 21 | 0 | 0 | 225 | 26041 | 10771 | 11 | 0 | 67 | 202 | 0 | 0 | 68 | 189 | 123 | 51 | 52 | 0 | 130 | 0 | 34 | 0 | 0 | 19266 | 0 | 0 |
| SRR6453390 | 20 | 13 | 14 | 11 | 74 | 3126 | 3689 | 16489 | 71001 | 97 | 56 | 14 | 0 | 0 | 296 | 23092 | 15691 | 0 | 0 | 59 | 133 | 0 | 0 | 51 | 203 | 125 | 76 | 84 | 0 | 88 | 0 | 0 | 0 | 11 | 5511 | 0 | 0 |
| SRR6453391 | 16 | 0 | 0 | 0 | 99 | 2165 | 2447 | 9469 | 55813 | 91 | 75 | 61 | 0 | 0 | 269 | 31098 | 42463 | 13 | 38 | 51 | 205 | 0 | 0 | 152 | 408 | 302 | 151 | 137 | 0 | 143 | 0 | 0 | 0 | 0 | 6078 | 0 | 0 |
| SRR6453392 | 0 | 0 | 0 | 0 | 147 | 1910 | 2352 | 8416 | 47387 | 101 | 131 | 31 | 0 | 0 | 481 | 25500 | 18276 | 13 | 19 | 38 | 115 | 0 | 0 | 131 | 582 | 265 | 166 | 111 | 0 | 200 | 0 | 0 | 0 | 0 | 3337 | 0 | 0 |
| SRR6453393 | 15 | 0 | 11 | 17 | 173 | 2472 | 2389 | 9740 | 45088 | 206 | 146 | 38 | 0 | 0 | 213 | 20883 | 13434 | 0 | 0 | 37 | 74 | 0 | 0 | 182 | 670 | 416 | 306 | 192 | 0 | 352 | 0 | 19 | 0 | 0 | 5241 | 0 | 0 |
| SRR6453394 | 0 | 0 | 0 | 0 | 167 | 7633 | 7415 | 25994 | 96890 | 27 | 74 | 33 | 0 | 0 | 153 | 32460 | 9192 | 0 | 0 | 0 | 44 | 0 | 0 | 87 | 180 | 137 | 40 | 59 | 0 | 266 | 0 | 22 | 0 | 0 | 15074 | 0 | 0 |
| SRR6453395 | 0 | 0 | 0 | 0 | 138 | 3098 | 4052 | 16605 | 82843 | 84 | 88 | 41 | 0 | 0 | 254 | 26992 | 14612 | 0 | 0 | 51 | 91 | 0 | 0 | 147 | 543 | 336 | 188 | 161 | 0 | 221 | 0 | 0 | 0 | 0 | 6699 | 0 | 0 |
| SRR6453396 | 0 | 0 | 0 | 0 | 61 | 2895 | 3730 | 13544 | 62212 | 86 | 37 | 23 | 0 | 0 | 147 | 22850 | 10677 | 0 | 0 | 12 | 98 | 0 | 0 | 58 | 191 | 106 | 56 | 53 | 0 | 89 | 0 | 0 | 0 | 0 | 7118 | 0 | 0 |
| SRR6453397 | 0 | 0 | 0 | 0 | 103 | 3540 | 3346 | 13723 | 58139 | 67 | 44 | 17 | 0 | 0 | 242 | 17181 | 8854 | 0 | 0 | 73 | 48 | 0 | 0 | 92 | 206 | 162 | 70 | 70 | 0 | 92 | 0 | 0 | 0 | 0 | 8268 | 0 | 0 |
| SRR6453398 | 11 | 0 | 0 | 0 | 64 | 4217 | 3290 | 15671 | 58199 | 26 | 21 | 19 | 0 | 0 | 329 | 18445 | 8212 | 30 | 60 | 30 | 239 | 0 | 0 | 50 | 102 | 80 | 30 | 40 | 0 | 56 | 0 | 0 | 0 | 0 | 9451 | 0 | 0 |
| SRR6453399 | 12 | 0 | 0 | 0 | 51 | 4531 | 2796 | 9841 | 48346 | 22 | 64 | 21 | 0 | 0 | 108 | 28236 | 7674 | 0 | 0 | 27 | 67 | 0 | 0 | 22 | 61 | 65 | 17 | 19 | 0 | 111 | 0 | 81 | 0 | 0 | 14975 | 0 | 0 |
| SRR6453400 | 0 | 0 | 0 | 0 | 143 | 5544 | 6747 | 25514 | 108030 | 66 | 47 | 41 | 0 | 0 | 215 | 22946 | 9130 | 14 | 14 | 43 | 14 | 0 | 0 | 119 | 282 | 191 | 71 | 76 | 0 | 159 | 0 | 0 | 0 | 0 | 7809 | 0 | 0 |
| SRR6453401 | 20 | 0 | 0 | 0 | 56 | 3415 | 4635 | 19922 | 96686 | 23 | 47 | 28 | 0 | 0 | 257 | 26735 | 15478 | 11 | 0 | 54 | 118 | 0 | 0 | 41 | 176 | 110 | 71 | 51 | 0 | 101 | 0 | 0 | 0 | 0 | 3432 | 0 | 0 |
| SRR6453402 | 21 | 14 | 0 | 17 | 69 | 2351 | 1587 | 5321 | 32524 | 32 | 32 | 18 | 0 | 0 | 144 | 13170 | 7988 | 0 | 0 | 60 | 181 | 0 | 0 | 18 | 73 | 68 | 50 | 36 | 0 | 52 | 0 | 15 | 0 | 0 | 1341 | 0 | 0 |
| SRR6453403 | 0 | 0 | 0 | 0 | 174 | 1631 | 1244 | 5096 | 31861 | 120 | 161 | 42 | 0 | 0 | 515 | 32294 | 16671 | 11 | 21 | 86 | 54 | 0 | 0 | 79 | 399 | 302 | 185 | 108 | 0 | 225 | 0 | 0 | 0 | 0 | 3376 | 0 | 0 |
| SRR6453404 | 10 | 0 | 0 | 0 | 56 | 1854 | 1122 | 4262 | 21942 | 23 | 55 | 13 | 0 | 0 | 177 | 30639 | 9137 | 25 | 25 | 50 | 88 | 0 | 0 | 22 | 80 | 96 | 44 | 43 | 0 | 70 | 0 | 0 | 0 | 0 | 6909 | 0 | 0 |
| SRR6453405 | 22 | 0 | 0 | 10 | 44 | 2041 | 1843 | 9054 | 42497 | 20 | 24 | 15 | 0 | 0 | 198 | 14600 | 9859 | 20 | 20 | 79 | 277 | 0 | 0 | 28 | 88 | 72 | 28 | 30 | 0 | 46 | 0 | 0 | 0 | 0 | 5980 | 0 | 0 |
| SRR6453406 | 20 | 11 | 0 | 0 | 64 | 2191 | 2598 | 7526 | 47710 | 24 | 50 | 20 | 0 | 0 | 300 | 27592 | 16621 | 0 | 0 | 15 | 85 | 0 | 0 | 26 | 84 | 80 | 49 | 38 | 0 | 86 | 0 | 15 | 0 | 0 | 5573 | 0 | 0 |
| SRR6453407 | 0 | 0 | 0 | 0 | 158 | 3908 | 2880 | 11967 | 55218 | 43 | 118 | 34 | 0 | 0 | 206 | 32918 | 6669 | 37 | 0 | 0 | 37 | 0 | 0 | 60 | 228 | 223 | 103 | 103 | 0 | 211 | 0 | 37 | 0 | 0 | 6066 | 0 | 0 |
| SRR6453408 | 16 | 0 | 0 | 0 | 125 | 2453 | 1409 | 4833 | 26131 | 39 | 117 | 56 | 0 | 0 | 219 | 14094 | 6670 | 0 | 0 | 121 | 340 | 0 | 0 | 61 | 196 | 184 | 60 | 80 | 0 | 116 | 0 | 0 | 0 | 0 | 7308 | 0 | 0 |
| SRR6453409 | 13 | 0 | 0 | 0 | 76 | 918 | 615 | 2412 | 14100 | 19 | 45 | 28 | 0 | 0 | 303 | 10698 | 7668 | 20 | 20 | 49 | 127 | 0 | 0 | 22 | 103 | 82 | 51 | 59 | 0 | 72 | 0 | 0 | 0 | 0 | 2126 | 0 | 0 |
| SRR6453410 | 13 | 0 | 0 | 0 | 59 | 3513 | 2591 | 8457 | 45686 | 30 | 63 | 14 | 0 | 0 | 64 | 29444 | 9447 | 0 | 0 | 318 | 127 | 0 | 0 | 14 | 46 | 47 | 17 | 24 | 0 | 115 | 0 | 32 | 0 | 0 | 9688 | 0 | 0 |
| SRR6453411 | 19 | 12 | 0 | 0 | 60 | 3038 | 2669 | 10520 | 55130 | 15 | 33 | 16 | 0 | 0 | 326 | 31535 | 20916 | 0 | 0 | 60 | 403 | 0 | 0 | 28 | 120 | 75 | 42 | 39 | 0 | 69 | 0 | 0 | 0 | 0 | 9782 | 0 | 0 |
| SRR6453412 | 0 | 0 | 0 | 0 | 49 | 436 | 109 | 835 | 2578 | 63 | 18 | 15 | 0 | 0 | 218 | 7183 | 3990 | 0 | 0 | 36 | 218 | 0 | 0 | 81 | 124 | 105 | 59 | 69 | 0 | 12 | 0 | 0 | 0 | 0 | 8140 | 0 | 0 |
| SRR6453413 | 13 | 0 | 0 | 0 | 82 | 1597 | 1334 | 5243 | 22972 | 76 | 12 | 16 | 0 | 0 | 71 | 15265 | 7018 | 0 | 0 | 0 | 167 | 0 | 0 | 34 | 0 | 0 | 0 | 0 | 0 | 14 | 0 | 24 | 0 | 0 | 5379 | 0 | 0 |
| SRR6453414 | 0 | 0 | 0 | 0 | 158 | 3873 | 1238 | 6251 | 17393 | 0 | 140 | 18 | 0 | 0 | 233 | 13836 | 10275 | 25 | 0 | 25 | 37 | 0 | 0 | 68 | 165 | 90 | 196 | 55 | 0 | 451 | 0 | 25 | 0 | 0 | 1949 | 0 | 0 |
| SRR6453415 | 0 | 0 | 0 | 0 | 136 | 3878 | 1565 | 7685 | 23217 | 0 | 307 | 32 | 0 | 0 | 601 | 54190 | 15947 | 35 | 21 | 56 | 203 | 0 | 0 | 44 | 116 | 79 | 128 | 37 | 0 | 1073 | 0 | 84 | 0 | 15 | 5813 | 0 | 0 |
| SRR6453416 | 0 | 0 | 0 | 24 | 22 | 1958 | 932 | 4600 | 16999 | 0 | 39 | 19 | 0 | 0 | 691 | 21611 | 54973 | 0 | 74 | 262 | 952 | 0 | 0 | 32 | 106 | 30 | 49 | 27 | 0 | 135 | 0 | 20 | 0 | 12 | 5975 | 0 | 0 |
| SRR6453417 | 0 | 0 | 0 | 15 | 32 | 1807 | 484 | 2519 | 8105 | 0 | 27 | 29 | 0 | 0 | 73 | 8266 | 8001 | 0 | 0 | 37 | 265 | 0 | 0 | 13 | 38 | 21 | 42 | 16 | 0 | 79 | 0 | 91 | 0 | 0 | 2877 | 0 | 0 |

### Supp Table S8.3.Columns_BX_to_DH

| | | | | | | | | | | | | | | | | | | | | | | | | | | | | | | | | | | | | | |
|---|---|---|---|---|---|---|---|---|---|---|---|---|---|---|---|---|---|---|---|---|---|---|---|---|---|---|---|---|---|---|---|---|---|---|---|---|---|
| SRR6453418 | 0 | 10 | 0 | 14 | 258 | 2119 | 1851 | 7642 | 31731 | 232 | 96 | 65 | 0 | 0 | 239 | 24508 | 11539 | 15 | 0 | 45 | 179 | 0 | 0 | 79 | 375 | 299 | 345 | 302 | 0 | 339 | 0 | 15 | 0 | 17 | 6241 | 0 | 0 |
| SRR6453419 | 14 | 0 | 0 | 16 | 63 | 3070 | 2708 | 9554 | 46570 | 50 | 45 | 21 | 0 | 0 | 97 | 16233 | 8796 | 0 | 0 | 26 | 168 | 0 | 0 | 19 | 58 | 48 | 43 | 38 | 0 | 118 | 0 | 35 | 0 | 0 | 9207 | 0 | 0 |
| SRR6453420 | 0 | 0 | 0 | 0 | 58 | 2217 | 1953 | 7096 | 37266 | 67 | 42 | 27 | 0 | 0 | 190 | 15464 | 7490 | 0 | 0 | 99 | 173 | 0 | 0 | 18 | 69 | 51 | 32 | 19 | 0 | 102 | 0 | 0 | 0 | 11 | 6074 | 0 | 0 |
| SRR6453421 | 35 | 0 | 0 | 11 | 167 | 1273 | 656 | 2856 | 13696 | 204 | 88 | 73 | 0 | 0 | 409 | 21343 | 52734 | 0 | 19 | 96 | 5148 | 0 | 0 | 93 | 281 | 172 | 140 | 93 | 0 | 225 | 0 | 46 | 0 | 10 | 27609 | 0 | 0 |
| SRR6453422 | 13 | 15 | 0 | 0 | 48 | 649 | 410 | 1865 | 8176 | 75 | 12 | 28 | 0 | 0 | 7820 | 17749 | 75578 | 0 | 0 | 0 | 130 | 0 | 0 | 20 | 60 | 49 | 41 | 32 | 0 | 14 | 0 | 0 | 0 | 0 | 5119 | 0 | 0 |
| SRR6453423 | 0 | 0 | 0 | 0 | 79 | 942 | 603 | 2334 | 10803 | 407 | 11 | 51 | 0 | 0 | 1260 | 23161 | 25195 | 0 | 0 | 22 | 110 | 0 | 0 | 88 | 362 | 209 | 170 | 106 | 0 | 35 | 0 | 11 | 0 | 0 | 14783 | 0 | 0 |
| SRR6453424 | 0 | 0 | 0 | 12 | 182 | 1646 | 484 | 1997 | 6507 | 0 | 192 | 17 | 0 | 0 | 170 | 5510 | 9713 | 0 | 0 | 44 | 251 | 0 | 0 | 71 | 270 | 110 | 279 | 90 | 0 | 497 | 0 | 50 | 0 | 0 | 1624 | 0 | 0 |
| SRR6453425 | 0 | 0 | 0 | 0 | 30 | 4582 | 1723 | 9646 | 19088 | 0 | 92 | 0 | 0 | 0 | 1299 | 29964 | 27437 | 33 | 25 | 57 | 229 | 0 | 0 | 16 | 63 | 19 | 23 | 27 | 0 | 267 | 0 | 57 | 0 | 0 | 5181 | 0 | 0 |
| SRR6453426 | 0 | 0 | 0 | 13 | 91 | 1092 | 608 | 3407 | 10978 | 0 | 164 | 50 | 0 | 0 | 148 | 10153 | 14161 | 0 | 0 | 74 | 328 | 0 | 0 | 80 | 224 | 112 | 178 | 68 | 0 | 456 | 0 | 66 | 0 | 20 | 2121 | 0 | 0 |
| SRR6453427 | 0 | 0 | 0 | 0 | 89 | 2682 | 930 | 4949 | 14784 | 0 | 141 | 40 | 0 | 0 | 212 | 20621 | 12073 | 27 | 0 | 44 | 97 | 0 | 0 | 91 | 245 | 101 | 171 | 68 | 0 | 470 | 0 | 71 | 0 | 17 | 3104 | 0 | 0 |
| SRR6453428 | 0 | 0 | 0 | 11 | 28 | 3161 | 1365 | 8165 | 25007 | 0 | 31 | 36 | 0 | 0 | 148 | 16360 | 15163 | 0 | 34 | 102 | 159 | 0 | 0 | 16 | 44 | 21 | 42 | 21 | 0 | 113 | 0 | 80 | 0 | 16 | 1574 | 0 | 0 |
| SRR6454160 | 292 | 53 | 27 | 186 | 106 | 59 | 51 | 107 | 0 | 0 | 80 | 0 | 0 | 0 | 46852 | 64384 | 195435 | 33 | 434 | 167244 | 933 | 0 | 0 | 372 | 27 | 133 | 319 | 80 | 0 | 612 | 0 | 0 | 0 | 0 | 431 | 248 | 0 |
| SRR6454161 | 15 | 0 | 0 | 0 | 14 | 136 | 50 | 261 | 14 | 137 | 224 | 0 | 0 | 0 | 79487 | 106804 | 305862 | 79 | 152 | 53658 | 2036 | 0 | 0 | 106 | 0 | 0 | 18 | 0 | 0 | 1762 | 0 | 0 | 0 | 0 | 555 | 213 | 47 |
| SRR6454162 | 0 | 0 | 0 | 92 | 123 | 155 | 0 | 155 | 0 | 0 | 123 | 0 | 0 | 0 | 2628 | 34247 | 23116 | 0 | 0 | 618 | 618 | 0 | 0 | 61 | 31 | 92 | 184 | 0 | 0 | 92 | 0 | 0 | 0 | 0 | 856 | 0 | 0 |
| SRR6454163 | 114 | 62 | 21 | 10 | 31 | 64 | 36 | 145 | 0 | 21 | 114 | 10 | 21 | 0 | 56794 | 30932 | 248839 | 454 | 175 | 88239 | 832 | 0 | 0 | 72 | 0 | 0 | 41 | 0 | 0 | 733 | 0 | 0 | 10 | 0 | 395 | 223 | 0 |
| SRR6454164 | 0 | 31 | 0 | 0 | 153 | 0 | 0 | 0 | 0 | 0 | 31 | 0 | 0 | 0 | 4832 | 49353 | 43420 | 0 | 0 | 604 | 151 | 0 | 0 | 61 | 92 | 153 | 429 | 0 | 0 | 92 | 0 | 0 | 0 | 0 | 809 | 0 | 0 |
| SRR6454165 | 84 | 54 | 23 | 92 | 176 | 99 | 53 | 266 | 29 | 15 | 69 | 0 | 0 | 0 | 115545 | 72344 | 336011 | 446 | 147 | 68890 | 1843 | 0 | 0 | 123 | 0 | 0 | 69 | 0 | 0 | 667 | 0 | 0 | 0 | 0 | 533 | 314 | 15 |
| SRR6454166 | 0 | 0 | 0 | 32 | 0 | 0 | 179 | 0 | 0 | 0 | 95 | 0 | 0 | 0 | 5012 | 45323 | 42060 | 0 | 0 | 1611 | 179 | 0 | 0 | 221 | 95 | 316 | 633 | 0 | 0 | 95 | 0 | 0 | 0 | 0 | 725 | 0 | 0 |
| SRR6454167 | 131 | 108 | 0 | 12 | 24 | 62 | 42 | 177 | 13 | 0 | 239 | 0 | 0 | 0 | 76690 | 22985 | 233964 | 196 | 121 | 81261 | 473 | 0 | 0 | 96 | 0 | 0 | 72 | 0 | 0 | 1148 | 0 | 0 | 0 | 0 | 529 | 268 | 0 |
| SRR6454169 | 92 | 31 | 0 | 82 | 31 | 125 | 64 | 253 | 24 | 31 | 72 | 0 | 0 | 0 | 120128 | 44558 | 321829 | 383 | 79 | 52055 | 1513 | 0 | 0 | 51 | 0 | 10 | 31 | 0 | 0 | 492 | 0 | 0 | 0 | 0 | 569 | 245 | 10 |
| SRR6454170 | 0 | 20 | 20 | 20 | 41 | 0 | 0 | 0 | 0 | 0 | 265 | 20 | 0 | 0 | 5246 | 44455 | 67840 | 0 | 0 | 999 | 125 | 0 | 0 | 61 | 163 | 286 | 898 | 41 | 0 | 592 | 0 | 0 | 0 | 0 | 2084 | 0 | 0 |
| SRR6454171 | 18 | 18 | 44 | 80 | 27 | 52 | 19 | 73 | 0 | 0 | 80 | 0 | 0 | 0 | 70685 | 27252 | 221833 | 216 | 151 | 88916 | 450 | 0 | 0 | 142 | 0 | 35 | 98 | 0 | 0 | 577 | 0 | 0 | 0 | 0 | 310 | 370 | 0 |
| SRR6454172 | 0 | 0 | 0 | 33 | 66 | 0 | 0 | 0 | 0 | 0 | 312 | 0 | 0 | 0 | 4060 | 31769 | 50903 | 0 | 0 | 1059 | 177 | 0 | 0 | 33 | 164 | 279 | 623 | 0 | 0 | 558 | 0 | 0 | 0 | 0 | 1264 | 0 | 0 |
| SRR6454173 | 55 | 0 | 0 | 47 | 39 | 82 | 80 | 190 | 89 | 0 | 55 | 0 | 0 | 0 | 71909 | 53399 | 235359 | 276 | 97 | 53776 | 1884 | 0 | 0 | 70 | 0 | 39 | 31 | 16 | 0 | 421 | 0 | 17 | 0 | 0 | 232 | 638 | 0 |
| SRR6454174 | 0 | 0 | 32 | 0 | 32 | 0 | 0 | 182 | 0 | 0 | 429 | 0 | 0 | 0 | 9001 | 38003 | 48100 | 0 | 0 | 6637 | 182 | 0 | 0 | 16 | 0 | 79 | 556 | 32 | 0 | 302 | 0 | 0 | 0 | 0 | 4347 | 0 | 0 |
| SRR6454175 | 44 | 95 | 0 | 64 | 51 | 29 | 32 | 77 | 12 | 0 | 273 | 0 | 0 | 0 | 54068 | 84656 | 209316 | 36 | 180 | 102000 | 4052 | 0 | 0 | 171 | 0 | 13 | 114 | 13 | 0 | 1156 | 0 | 29 | 0 | 0 | 439 | 112 | 13 |
| SRR6454176 | 0 | 0 | 0 | 13 | 105 | 0 | 0 | 0 | 0 | 0 | 237 | 13 | 0 | 0 | 3445 | 27170 | 26056 | 0 | 0 | 842 | 77 | 0 | 0 | 66 | 26 | 79 | 670 | 26 | 0 | 11 | 0 | 0 | 0 | 0 | 557 | 0 | 13 |
| SRR6454177 | 182 | 91 | 26 | 143 | 91 | 104 | 97 | 276 | 31 | 0 | 130 | 0 | 0 | 0 | 126796 | 58445 | 321447 | 221 | 190 | 59457 | 529 | 0 | 0 | 1381 | 13 | 130 | 365 | 26 | 0 | 495 | 0 | 0 | 0 | 0 | 467 | 232 | 39 |
| SRR6454178 | 0 | 0 | 17 | 0 | 138 | 0 | 0 | 108 | 0 | 0 | 87 | 0 | 0 | 0 | 3991 | 43043 | 43798 | 0 | 0 | 755 | 216 | 0 | 0 | 156 | 87 | 277 | 796 | 17 | 0 | 398 | 0 | 0 | 0 | 0 | 2643 | 0 | 0 |
| SRR6454179 | 38 | 0 | 25 | 126 | 25 | 62 | 21 | 86 | 15 | 0 | 25 | 25 | 0 | 0 | 140532 | 57966 | 342564 | 517 | 66 | 41873 | 728 | 0 | 0 | 264 | 25 | 38 | 126 | 13 | 0 | 113 | 0 | 0 | 0 | 0 | 284 | 355 | 0 |
| SRR6454180 | 0 | 0 | 0 | 24 | 24 | 53 | 0 | 0 | 53 | 32 | 280 | 0 | 0 | 0 | 5388 | 30558 | 57678 | 0 | 0 | 2881 | 160 | 0 | 0 | 64 | 96 | 248 | 767 | 32 | 0 | 439 | 0 | 53 | 0 | 0 | 1273 | 0 | 16 |
| SRR6454181 | 10 | 20 | 0 | 50 | 20 | 22 | 16 | 44 | 11 | 30 | 273 | 40 | 0 | 0 | 83970 | 22280 | 260359 | 145 | 156 | 98173 | 621 | 0 | 0 | 40 | 0 | 0 | 0 | 0 | 0 | 15 | 0 | 0 | 0 | 0 | 287 | 281 | 20 |
| SRR6454182 | 0 | 0 | 0 | 27 | 20 | 0 | 0 | 37 | 0 | 0 | 208 | 20 | 0 | 0 | 4490 | 24098 | 40066 | 0 | 0 | 2320 | 112 | 0 | 0 | 54 | 13 | 61 | 350 | 0 | 0 | 276 | 0 | 0 | 0 | 0 | 1355 | 0 | 0 |
| SRR6454183 | 21 | 0 | 0 | 35 | 28 | 44 | 23 | 75 | 16 | 0 | 42 | 0 | 0 | 0 | 116704 | 51557 | 353594 | 224 | 75 | 52973 | 2281 | 0 | 0 | 78 | 0 | 14 | 28 | 0 | 0 | 275 | 0 | 14 | 0 | 0 | 341 | 293 | 0 |
| SRR6454184 | 0 | 0 | 0 | 16 | 48 | 104 | 0 | 0 | 0 | 0 | 97 | 0 | 0 | 0 | 10845 | 39154 | 28577 | 0 | 0 | 2086 | 0 | 0 | 0 | 113 | 65 | 161 | 242 | 0 | 0 | 597 | 0 | 0 | 0 | 0 | 2412 | 0 | 0 |
| SRR6454185 | 59 | 33 | 0 | 92 | 13 | 41 | 43 | 158 | 35 | 0 | 115 | 0 | 0 | 0 | 47671 | 19383 | 134075 | 69 | 153 | 56906 | 842 | 0 | 0 | 52 | 0 | 56 | 281 | 39 | 0 | 17 | 0 | 0 | 0 | 0 | 259 | 512 | 0 |
| SRR6454186 | 0 | 0 | 0 | 31 | 93 | 0 | 0 | 0 | 0 | 16 | 358 | 16 | 0 | 0 | 16468 | 47339 | 37871 | 0 | 0 | 962 | 120 | 0 | 0 | 93 | 62 | 155 | 653 | 31 | 0 | 404 | 0 | 0 | 0 | 0 | 2314 | 0 | 0 |
| SRR6454187 | 26 | 44 | 0 | 62 | 22 | 102 | 19 | 65 | 0 | 110 | 322 | 66 | 0 | 0 | 71708 | 28679 | 194031 | 183 | 136 | 77393 | 563 | 0 | 0 | 66 | 0 | 18 | 106 | 22 | 0 | 19 | 0 | 0 | 0 | 0 | 432 | 221 | 31 |
| SRR6454188 | 0 | 0 | 0 | 0 | 41 | 0 | 0 | 0 | 0 | 0 | 41 | 0 | 0 | 0 | 9624 | 46534 | 54019 | 0 | 0 | 3787 | 631 | 0 | 0 | 103 | 83 | 62 | 621 | 0 | 0 | 248 | 0 | 0 | 0 | 0 | 1627 | 0 | 0 |
| SRR6454189 | 0 | 0 | 0 | 22 | 89 | 88 | 69 | 170 | 0 | 22 | 22 | 0 | 0 | 0 | 124703 | 19430 | 291054 | 128 | 88 | 74567 | 96 | 0 | 0 | 469 | 0 | 45 | 112 | 0 | 0 | 313 | 0 | 0 | 0 | 0 | 531 | 428 | 0 |
| SRR6454190 | 0 | 0 | 0 | 0 | 75 | 0 | 0 | 0 | 0 | 0 | 50 | 25 | 0 | 0 | 9980 | 32309 | 38219 | 0 | 0 | 1174 | 587 | 0 | 0 | 125 | 100 | 299 | 499 | 25 | 0 | 125 | 0 | 0 | 0 | 0 | 788 | 0 | 0 |
| SRR6454191 | 15 | 15 | 23 | 53 | 76 | 89 | 28 | 172 | 31 | 0 | 38 | 0 | 0 | 0 | 135880 | 66393 | 340944 | 233 | 46 | 33891 | 1232 | 0 | 0 | 107 | 0 | 15 | 244 | 15 | 0 | 160 | 0 | 0 | 0 | 0 | 275 | 535 | 0 |
| SRR6454193 | 14 | 42 | 0 | 98 | 126 | 43 | 38 | 96 | 0 | 0 | 14 | 0 | 0 | 0 | 95228 | 20758 | 244983 | 77 | 81 | 62890 | 387 | 0 | 0 | 98 | 0 | 14 | 168 | 28 | 0 | 405 | 0 | 0 | 0 | 0 | 504 | 577 | 0 |
| SRR6454194 | 0 | 0 | 0 | 74 | 0 | 0 | 147 | 0 | 0 | 0 | 148 | 197 | 0 | 0 | 10147 | 30556 | 56713 | 0 | 0 | 1618 | 0 | 0 | 0 | 99 | 25 | 49 | 148 | 0 | 0 | 838 | 0 | 0 | 0 | 0 | 4167 | 0 | 25 |
| SRR6454195 | 53 | 71 | 0 | 53 | 195 | 60 | 30 | 148 | 0 | 0 | 18 | 0 | 0 | 0 | 145510 | 67871 | 345420 | 164 | 82 | 49352 | 676 | 0 | 0 | 515 | 18 | 71 | 160 | 0 | 0 | 213 | 0 | 0 | 0 | 0 | 434 | 452 | 18 |
| SRR6454196 | 0 | 25 | 0 | 0 | 0 | 0 | 0 | 0 | 0 | 0 | 74 | 0 | 0 | 0 | 19827 | 39885 | 87109 | 0 | 0 | 15953 | 684 | 0 | 0 | 74 | 99 | 74 | 966 | 50 | 0 | 25 | 0 | 0 | 0 | 0 | 638 | 0 | 0 |
| SRR6454197 | 148 | 282 | 28 | 247 | 14 | 32 | 24 | 83 | 0 | 28 | 353 | 0 | 0 | 0 | 67823 | 60246 | 224912 | 47 | 237 | 68988 | 1617 | 0 | 0 | 501 | 21 | 56 | 247 | 35 | 0 | 13 | 0 | 0 | 0 | 0 | 445 | 179 | 0 |
| SRR6454198 | 0 | 40 | 0 | 40 | 0 | 111 | 0 | 0 | 0 | 0 | 332 | 13 | 0 | 0 | 13909 | 45893 | 75868 | 0 | 111 | 10237 | 779 | 0 | 0 | 40 | 93 | 173 | 1169 | 93 | 0 | 319 | 0 | 0 | 0 | 0 | 3218 | 0 | 0 |
| SRR6454199 | 99 | 68 | 18 | 283 | 62 | 49 | 90 | 107 | 27 | 0 | 179 | 0 | 0 | 0 | 86578 | 111088 | 289933 | 26 | 167 | 56512 | 3117 | 0 | 0 | 209 | 12 | 31 | 154 | 12 | 0 | 690 | 0 | 0 | 0 | 0 | 397 | 158 | 0 |
| SRR6454201 | 37 | 37 | 80 | 299 | 270 | 49 | 17 | 78 | 0 | 22 | 109 | 0 | 0 | 0 | 35418 | 80468 | 264785 | 59 | 91 | 35123 | 1068 | 0 | 0 | 460 | 15 | 73 | 474 | 37 | 0 | 175 | 0 | 0 | 0 | 0 | 306 | 279 | 0 |
| SRR6454203 | 51 | 17 | 17 | 85 | 51 | 29 | 19 | 62 | 11 | 0 | 34 | 0 | 0 | 0 | 146932 | 65784 | 441229 | 265 | 56 | 39395 | 2061 | 0 | 0 | 51 | 0 | 0 | 119 | 0 | 0 | 322 | 0 | 0 | 0 | 0 | 149 | 326 | 0 |
| SRR6454205 | 86 | 74 | 0 | 74 | 37 | 51 | 54 | 124 | 31 | 12 | 258 | 12 | 0 | 0 | 109901 | 31879 | 307230 | 274 | 137 | 68803 | 662 | 0 | 0 | 86 | 0 | 25 | 12 | 0 | 0 | 799 | 0 | 0 | 0 | 0 | 369 | 261 | 12 |
| SRR6454206 | 0 | 0 | 28 | 0 | 83 | 0 | 0 | 0 | 0 | 0 | 194 | 0 | 0 | 0 | 6567 | 40346 | 36984 | 0 | 0 | 973 | 0 | 0 | 0 | 221 | 166 | 443 | 858 | 83 | 0 | 111 | 0 | 0 | 0 | 0 | 1201 | 0 | 0 |
| SRR6454207 | 91 | 57 | 11 | 137 | 46 | 137 | 110 | 377 | 130 | 11 | 148 | 11 | 0 | 0 | 101245 | 91205 | 251659 | 451 | 139 | 53225 | 1042 | 0 | 0 | 91 | 0 | 23 | 80 | 0 | 0 | 717 | 0 | 0 | 0 | 0 | 462 | 452 | 0 |
| SRR6454208 | 0 | 0 | 29 | 58 | 115 | 0 | 0 | 0 | 0 | 0 | 288 | 29 | 0 | 0 | 5306 | 50335 | 44384 | 0 | 0 | 2459 | 0 | 0 | 0 | 144 | 29 | 202 | 403 | 0 | 0 | 490 | 0 | 0 | 0 | 0 | 1736 | 0 | 0 |
| SRR6454209 | 34 | 0 | 0 | 42 | 34 | 38 | 43 | 190 | 23 | 101 | 361 | 0 | 0 | 0 | 124857 | 28031 | 313276 | 51 | 87 | 63535 | 597 | 0 | 0 | 176 | 0 | 17 | 59 | 0 | 0 | 1941 | 0 | 0 | 0 | 0 | 561 | 292 | 17 |
| SRR6454211 | 49 | 16 | 16 | 27 | 71 | 32 | 15 | 75 | 10 | 22 | 239 | 0 | 0 | 0 | 182328 | 44969 | 433343 | 107 | 53 | 46150 | 1418 | 0 | 0 | 76 | 0 | 0 | 65 | 0 | 0 | 1004 | 0 | 0 | 0 | 0 | 225 | 275 | 0 |
| SRR6454213 | 23 | 56 | 11 | 68 | 23 | 50 | 43 | 153 | 18 | 23 | 316 | 11 | 0 | 0 | 100140 | 29704 | 285783 | 73 | 103 | 87959 | 537 | 0 | 0 | 271 | 0 | 11 | 124 | 0 | 0 | 1513 | 0 | 0 | 0 | 0 | 502 | 467 | 0 |
| SRR6454214 | 0 | 0 | 0 | 54 | 0 | 0 | 0 | 0 | 0 | 0 | 54 | 0 | 0 | 0 | 1305 | 17077 | 13835 | 0 | 0 | 913 | 0 | 0 | 0 | 242 | 27 | 188 | 617 | 0 | 0 | 268 | 0 | 0 | 0 | 0 | 865 | 0 | 0 |
| SRR6454215 | 42 | 24 | 0 | 42 | 54 | 32 | 24 | 110 | 11 | 12 | 161 | 12 | 0 | 0 | 103911 | 52915 | 329689 | 117 | 87 | 74288 | 1151 | 0 | 0 | 113 | 0 | 0 | 77 | 0 | 0 | 786 | 0 | 0 | 0 | 0 | 425 | 276 | 12 |
| SRR6454216 | 49 | 0 | 0 | 0 | 0 | 0 | 0 | 0 | 0 | 0 | 49 | 0 | 0 | 0 | 16119 | 50753 | 105355 | 0 | 0 | 11514 | 0 | 0 | 0 | 49 | 0 | 0 | 0 | 0 | 0 | 0 | 0 | 0 | 0 | 0 | 1050 | 0 | 0 |
| SRR6454217 | 100 | 158 | 33 | 324 | 58 | 56 | 53 | 188 | 74 | 0 | 249 | 0 | 0 | 0 | 75460 | 128978 | 316737 | 36 | 144 | 50640 | 4347 | 0 | 0 | 83 | 25 | 66 | 382 | 17 | 0 | 972 | 0 | 29 | 0 | 0 | 330 | 225 | 17 |
| SRR6454218 | 0 | 0 | 0 | 0 | 0 | 0 | 0 | 0 | 0 | 0 | 81 | 0 | 0 | 0 | 7610 | 47257 | 73849 | 0 | 64 | 3645 | 192 | 0 | 0 | 81 | 54 | 108 | 325 | 0 | 0 | 189 | 0 | 0 | 0 | 0 | 2583 | 0 | 0 |
| SRR6454219 | 70 | 61 | 26 | 122 | 52 | 48 | 18 | 88 | 0 | 0 | 252 | 0 | 0 | 0 | 86546 | 122366 | 373207 | 36 | 201 | 83158 | 3395 | 0 | 0 | 408 | 0 | 17 | 209 | 17 | 0 | 1286 | 0 | 0 | 0 | 0 | 327 | 72 | 0 |
| SRR6454220 | 0 | 0 | 0 | 22 | 45 | 0 | 0 | 103 | 0 | 0 | 45 | 0 | 0 | 0 | 7301 | 35091 | 53472 | 0 | 0 | 8226 | 308 | 0 | 0 | 45 | 89 | 67 | 357 | 22 | 0 | 89 | 0 | 0 | 0 | 0 | 2599 | 0 | 0 |
| SRR6454221 | 158 | 113 | 34 | 147 | 79 | 30 | 29 | 87 | 0 | 0 | 226 | 0 | 11 | 0 | 66406 | 49142 | 222843 | 15 | 366 | 147219 | 2056 | 0 | 0 | 350 | 11 | 34 | 124 | 11 | 0 | 1038 | 0 | 0 | 0 | 0 | 626 | 107 | 11 |
| SRR6454222 | 0 | 0 | 25 | 0 | 25 | 0 | 0 | 0 | 0 | 0 | 0 | 0 | 0 | 0 | 14334 | 39130 | 103702 | 0 | 0 | 9673 | 117 | 0 | 0 | 50 | 25 | 50 | 124 | 0 | 0 | 25 | 0 | 0 | 0 | 0 | 2106 | 0 | 0 |
| SRR6454223 | 98 | 65 | 59 | 287 | 26 | 57 | 48 | 136 | 55 | 20 | 209 | 0 | 0 | 0 | 78976 | 104827 | 306371 | 43 | 228 | 78339 | 5079 | 0 | 0 | 150 | 13 | 20 | 157 | 0 | 0 | 876 | 0 | 0 | 0 | 0 | 515 | 106 | 0 |
| SRR6454224 | 0 | 0 | 0 | 0 | 0 | 0 | 0 | 141 | 0 | 0 | 26 | 0 | 0 | 0 | 8301 | 13508 | 37711 | 0 | 0 | 9989 | 141 | 0 | 0 | 52 | 26 | 52 | 78 | 0 | 0 | 52 | 0 | 0 | 0 | 0 | 750 | 0 | 0 |
| SRR6454225 | 140 | 98 | 42 | 154 | 14 | 45 | 36 | 142 | 20 | 0 | 363 | 0 | 0 | 0 | 40407 | 35108 | 148366 | 0 | 364 | 103352 | 632 | 0 | 0 | 237 | 0 | 14 | 168 | 28 | 0 | 1662 | 0 | 0 | 0 | 0 | 593 | 170 | 14 |
| SRR6454226 | 0 | 0 | 0 | 24 | 0 | 104 | 104 | 0 | 0 | 0 | 24 | 0 | 0 | 0 | 9966 | 19793 | 44385 | 0 | 0 | 8201 | 0 | 0 | 0 | 95 | 0 | 71 | 95 | 0 | 0 | 71 | 0 | 0 | 0 | 0 | 2549 | 0 | 0 |
| SRR6454227 | 114 | 93 | 45 | 247 | 126 | 36 | 37 | 125 | 17 | 0 | 264 | 0 | 0 | 0 | 67863 | 69651 | 282354 | 27 | 274 | 87579 | 2004 | 0 | 0 | 470 | 0 | 45 | 312 | 36 | 0 | 12 | 0 | 11 | 0 | 0 | 468 | 86 | 0 |
| SRR6454228 | 0 | 0 | 0 | 35 | 69 | 116 | 116 | 116 | 0 | 0 | 0 | 0 | 0 | 0 | 5468 | 21771 | 37554 | 0 | 0 | 7214 | 233 | 0 | 0 | 104 | 17 | 69 | 173 | 0 | 0 | 69 | 0 | 0 | 0 | 0 | 1814 | 0 | 17 |
| SRR6454229 | 67 | 167 | 83 | 233 | 33 | 44 | 55 | 147 | 21 | 0 | 583 | 0 | 0 | 0 | 32955 | 34003 | 139341 | 16 | 323 | 145309 | 1001 | 0 | 0 | 300 | 0 | 0 | 133 | 17 | 0 | 2300 | 0 | 0 | 0 | 0 | 600 | 140 | 0 |
| SRR6454230 | 0 | 0 | 0 | 0 | 19 | 0 | 0 | 0 | 0 | 0 | 0 | 19 | 0 | 0 | 3737 | 15800 | 36570 | 107 | 0 | 9395 | 0 | 0 | 0 | 267 | 95 | 38 | 133 | 19 | 0 | 133 | 0 | 0 | 0 | 0 | 888 | 0 | 0 |
| SRR6454231 | 52 | 104 | 26 | 155 | 52 | 18 | 33 | 94 | 15 | 26 | 233 | 13 | 0 | 0 | 81425 | 79178 | 269960 | 34 | 241 | 96643 | 2202 | 0 | 0 | 609 | 0 | 65 | 155 | 0 | 0 | 1321 | 0 | 0 | 0 | 0 | 609 | 102 | 0 |
| SRR6454232 | 16 | 0 | 0 | 0 | 16 | 0 | 0 | 0 | 0 | 16 | 32 | 49 | 0 | 0 | 9280 | 47344 | 85555 | 0 | 0 | 3269 | 0 | 0 | 0 | 0 | 32 | 49 | 65 | 0 | 0 | 97 | 0 | 0 | 0 | 0 | 1305 | 0 | 16 |
| SRR6454233 | 31 | 31 | 16 | 63 | 16 | 39 | 55 | 143 | 22 | 0 | 71 | 0 | 0 | 0 | 72413 | 93215 | 273677 | 33 | 166 | 69136 | 5439 | 0 | 0 | 39 | 0 | 0 | 31 | 0 | 0 | 817 | 0 | 37 | 0 | 0 | 453 | 127 | 0 |
| SRR6454234 | 38 | 19 | 0 | 0 | 19 | 59 | 37 | 202 | 44 | 19 | 75 | 0 | 0 | 0 | 44221 | 70128 | 268011 | 77 | 203 | 89051 | 2442 | 0 | 0 | 66 | 0 | 0 | 47 | 0 | 0 | 10 | 0 | 0 | 0 | 0 | 295 | 183 | 0 |
| SRR6454235 | 0 | 0 | 0 | 0 | 33 | 68 | 0 | 137 | 0 | 16 | 65 | 0 | 0 | 0 | 6355 | 19553 | 47217 | 68 | 0 | 2870 | 273 | 0 | 0 | 49 | 33 | 33 | 81 | 0 | 0 | 407 | 0 | 0 | 0 | 0 | 3222 | 0 | 0 |
| SRR6454236 | 62 | 33 | 0 | 86 | 19 | 56 | 29 | 104 | 11 | 0 | 71 | 0 | 0 | 0 | 170979 | 115890 | 449359 | 60 | 140 | 48809 | 6473 | 0 | 0 | 143 | 14 | 29 | 29 | 0 | 0 | 543 | 0 | 19 | 0 | 0 | 341 | 128 | 0 |
| SRR6454237 | 0 | 39 | 0 | 0 | 39 | 60 | 0 | 0 | 0 | 39 | 39 | 19 | 0 | 0 | 4424 | 17195 | 41319 | 0 | 0 | 1196 | 120 | 0 | 0 | 136 | 0 | 0 | 97 | 0 | 0 | 97 | 0 | 0 | 0 | 0 | 1412 | 0 | 0 |
| SRR6454238 | 48 | 0 | 0 | 24 | 0 | 19 | 29 | 85 | 0 | 0 | 48 | 0 | 0 | 0 | 73315 | 47608 | 296398 | 55 | 258 | 88078 | 1190 | 0 | 0 | 24 | 0 | 0 | 16 | 0 | 0 | 546 | 0 | 93 | 0 | 0 | 427 | 138 | 40 |
| SRR6454239 | 0 | 0 | 0 | 0 | 21 | 0 | 0 | 91 | 0 | 10 | 31 | 0 | 0 | 0 | 7364 | 29079 | 63769 | 0 | 0 | 2013 | 274 | 0 | 0 | 21 | 0 | 41 | 103 | 0 | 0 | 103 | 0 | 0 | 0 | 0 | 2143 | 0 | 0 |
| SRR6454240 | 43 | 16 | 21 | 124 | 48 | 33 | 37 | 94 | 26 | 0 | 59 | 0 | 0 | 0 | 78636 | 68435 | 321368 | 18 | 164 | 73195 | 3952 | 0 | 0 | 75 | 0 | 11 | 118 | 0 | 0 | 629 | 0 | 47 | 0 | 0 | 361 | 153 | 11 |
| SRR6454241 | 0 | 0 | 0 | 0 | 23 | 0 | 0 | 54 | 0 | 12 | 23 | 12 | 0 | 0 | 6137 | 17510 | 46476 | 0 | 0 | 2227 | 272 | 0 | 0 | 12 | 12 | 12 | 23 | 0 | 0 | 104 | 0 | 0 | 0 | 0 | 3999 | 0 | 0 |
| SRR6454242 | 20 | 30 | 10 | 55 | 0 | 60 | 41 | 148 | 12 | 20 | 75 | 0 | 0 | 0 | 150915 | 176123 | 440702 | 42 | 164 | 43063 | 5867 | 0 | 0 | 70 | 0 | 20 | 50 | 0 | 0 | 621 | 0 | 19 | 0 | 0 | 285 | 127 | 10 |
| SRR6454244 | 83 | 103 | 21 | 131 | 62 | 93 | 119 | 199 | 30 | 21 | 282 | 14 | 0 | 0 | 88742 | 84212 | 312322 | 32 | 220 | 86395 | 2433 | 0 | 0 | 269 | 14 | 48 | 165 | 0 | 0 | 12 | 0 | 27 | 0 | 0 | 837 | 125 | 0 |
| SRR6454245 | 0 | 0 | 0 | 0 | 0 | 0 | 0 | 141 | 0 | 27 | 0 | 0 | 0 | 0 | 11036 | 32583 | 52948 | 0 | 0 | 8630 | 424 | 0 | 0 | 110 | 0 | 27 | 192 | 27 | 0 | 82 | 0 | 0 | 0 | 0 | 1929 | 0 | 0 |
| SRR6454246 | 33 | 58 | 0 | 58 | 25 | 23 | 35 | 65 | 18 | 0 | 183 | 0 | 0 | 0 | 45986 | 64398 | 141127 | 16 | 134 | 100865 | 1577 | 0 | 0 | 25 | 0 | 17 | 66 | 0 | 0 | 979 | 0 | 28 | 0 | 0 | 264 | 205 | 0 |
| SRR6454247 | 75 | 34 | 21 | 48 | 62 | 74 | 79 | 163 | 60 | 0 | 55 | 0 | 0 | 0 | 64161 | 92995 | 302196 | 41 | 353 | 126258 | 3098 | 0 | 0 | 281 | 0 | 14 | 68 | 0 | 0 | 733 | 0 | 36 | 0 | 0 | 705 | 63 | 0 |
| SRR6454249 | 39 | 79 | 0 | 92 | 53 | 25 | 28 | 76 | 11 | 39 | 564 | 0 | 0 | 0 | 35944 | 31472 | 158894 | 25 | 344 | 126381 | 761 | 0 | 0 | 105 | 13 | 0 | 53 | 0 | 0 | 2048 | 0 | 0 | 0 | 0 | 501 | 131 | 13 |
| SRR6454251 | 17 | 33 | 0 | 42 | 42 | 38 | 73 | 126 | 66 | 0 | 150 | 0 | 0 | 0 | 59643 | 67075 | 218650 | 20 | 155 | 83483 | 2260 | 0 | 0 | 75 | 0 | 0 | 33 | 0 | 0 | 10 | 0 | 22 | 0 | 0 | 463 | 151 | 0 |
| SRR6454252 | 0 | 0 | 0 | 0 | 0 | 0 | 0 | 0 | 0 | 0 | 0 | 0 | 0 | 0 | 12377 | 38536 | 81435 | 0 | 0 | 14257 | 157 | 0 | 0 | 0 | 17 | 21 | 154 | 0 | 0 | 21 | 0 | 0 | 0 | 0 | 1697 | 0 | 0 |
| SRR6454253 | 56 | 28 | 14 | 105 | 35 | 77 | 57 | 196 | 25 | 0 | 154 | 0 | 0 | 0 | 90466 | 135641 | 362052 | 46 | 146 | 52049 | 6635 | 0 | 0 | 91 | 0 | 0 | 21 | 0 | 0 | 10 | 0 | 96 | 0 | 0 | 301 | 113 | 0 |
| SRR6454254 | 14 | 0 | 0 | 56 | 14 | 0 | 74 | 0 | 0 | 0 | 56 | 0 | 0 | 0 | 7788 | 48066 | 73895 | 0 | 0 | 7640 | 223 | 0 | 0 | 14 | 56 | 182 | 742 | 154 | 0 | 238 | 0 | 0 | 0 | 0 | 2624 | 0 | 0 |
| SRR6454255 | 34 | 20 | 13 | 47 | 20 | 48 | 38 | 113 | 16 | 0 | 154 | 0 | 0 | 0 | 81288 | 89979 | 312798 | 40 | 96 | 65024 | 3497 | 0 | 0 | 81 | 0 | 20 | 60 | 20 | 0 | 912 | 0 | 121 | 0 | 0 | 355 | 143 | 0 |
| SRR6454256 | 0 | 0 | 0 | 32 | 0 | 0 | 0 | 0 | 0 | 32 | 127 | 0 | 0 | 0 | 7355 | 29434 | 65409 | 0 | 0 | 9518 | 433 | 0 | 0 | 63 | 32 | 253 | 1108 | 32 | 0 | 127 | 0 | 0 | 0 | 0 | 755 | 0 | 0 |
| SRR6454257 | 65 | 122 | 0 | 86 | 72 | 24 | 29 | 70 | 30 | 22 | 108 | 0 | 0 | 0 | 73480 | 103801 | 292756 | 28 | 121 | 64262 | 6770 | 0 | 0 | 108 | 0 | 29 | 100 | 29 | 0 | 466 | 0 | 30 | 0 | 0 | 237 | 154 | 0 |
| SRR6454258 | 49 | 49 | 0 | 0 | 49 | 44 | 26 | 142 | 13 | 0 | 148 | 0 | 0 | 0 | 46424 | 40115 | 201113 | 86 | 266 | 129807 | 743 | 0 | 0 | 476 | 0 | 0 | 16 | 0 | 0 | 1051 | 0 | 0 | 0 | 0 | 883 | 92 | 0 |
| SRR6454259 | 78 | 59 | 0 | 0 | 49 | 52 | 63 | 164 | 64 | 0 | 107 | 0 | 0 | 0 | 59042 | 111841 | 224445 | 189 | 218 | 72213 | 3272 | 0 | 0 | 390 | 0 | 0 | 29 | 0 | 0 | 634 | 0 | 17 | 0 | 0 | 881 | 157 | 0 |
| SRR6454260 | 86 | 48 | 0 | 0 | 29 | 62 | 42 | 156 | 14 | 0 | 86 | 0 | 0 | 0 | 55402 | 40342 | 217050 | 162 | 182 | 104097 | 939 | 0 | 0 | 173 | 0 | 0 | 0 | 0 | 0 | 1150 | 0 | 0 | 0 | 0 | 806 | 130 | 0 |
| SRR6454261 | 38 | 51 | 13 | 26 | 0 | 34 | 23 | 83 | 0 | 0 | 83 | 0 | 0 | 0 | 93879 | 62217 | 308837 | 217 | 197 | 89086 | 2145 | 0 | 0 | 102 | 0 | 13 | 26 | 0 | 0 | 530 | 0 | 0 | 0 | 0 | 632 | 115 | 0 |
| SRR6454262 | 25 | 25 | 0 | 0 | 13 | 27 | 20 | 75 | 11 | 0 | 25 | 0 | 25 | 0 | 62921 | 90404 | 244045 | 206 | 196 | 94337 | 5369 | 0 | 0 | 76 | 0 | 13 | 13 | 0 | 0 | 418 | 0 | 11 | 0 | 0 | 477 | 82 | 0 |
| SRR6454263 | 32 | 13 | 13 | 101 | 57 | 122 | 94 | 311 | 176 | 13 | 13 | 0 | 0 | 0 | 49802 | 65171 | 188352 | 103 | 185 | 54777 | 2325 | 0 | 0 | 209 | 57 | 38 | 171 | 19 | 0 | 322 | 0 | 15 | 0 | 0 | 575 | 356 | 25 |
| SRR6454264 | 18 | 54 | 0 | 36 | 0 | 38 | 51 | 99 | 12 | 18 | 36 | 0 | 0 | 0 | 74792 | 102988 | 254302 | 192 | 190 | 71310 | 4152 | 0 | 0 | 268 | 0 | 54 | 0 | 18 | 0 | 499 | 0 | 0 | 0 | 0 | 677 | 90 | 0 |
| SRR6454265 | 30 | 45 | 0 | 30 | 30 | 80 | 44 | 151 | 12 | 0 | 15 | 0 | 0 | 0 | 87512 | 85507 | 290555 | 153 | 202 | 114452 | 4408 | 0 | 0 | 180 | 0 | 0 | 0 | 0 | 0 | 285 | 0 | 0 | 0 | 0 | 547 | 69 | 15 |
| SRR6454266 | 13 | 27 | 0 | 0 | 22 | 130 | 70 | 271 | 30 | 18 | 27 | 0 | 0 | 0 | 84068 | 103725 | 342880 | 193 | 128 | 49622 | 6630 | 0 | 0 | 121 | 0 | 0 | 0 | 0 | 0 | 407 | 0 | 26 | 0 | 0 | 553 | 124 | 0 |
| SRR6454267 | 55 | 16 | 0 | 32 | 21 | 59 | 40 | 137 | 36 | 0 | 37 | 0 | 0 | 0 | 87936 | 102049 | 320015 | 214 | 147 | 55685 | 6961 | 0 | 0 | 108 | 0 | 11 | 39 | 0 | 0 | 471 | 0 | 30 | 0 | 0 | 361 | 181 | 0 |
| SRR6454268 | 54 | 107 | 0 | 27 | 27 | 61 | 42 | 131 | 0 | 0 | 54 | 0 | 27 | 0 | 81362 | 38217 | 245610 | 81 | 364 | 145258 | 943 | 0 | 0 | 187 | 0 | 0 | 27 | 0 | 0 | 857 | 0 | 0 | 0 | 0 | 648 | 64 | 54 |
| SRR6454269 | 26 | 0 | 0 | 53 | 79 | 17 | 20 | 55 | 0 | 0 | 0 | 0 | 0 | 0 | 101314 | 70509 | 276292 | 135 | 194 | 80898 | 3169 | 0 | 0 | 132 | 0 | 26 | 106 | 0 | 0 | 846 | 0 | 0 | 0 | 0 | 396 | 74 | 0 |
| SRR6454270 | 28 | 0 | 0 | 28 | 28 | 60 | 110 | 181 | 48 | 0 | 42 | 0 | 14 | 0 | 95005 | 69809 | 293132 | 188 | 140 | 69336 | 3922 | 0 | 0 | 14 | 0 | 14 | 14 | 0 | 0 | 488 | 0 | 43 | 0 | 0 | 424 | 115 | 0 |
| SRR6454271 | 162 | 251 | 181 | 1267 | 60 | 50 | 93 | 196 | 20 | 0 | 474 | 0 | 0 | 0 | 42968 | 56317 | 141925 | 0 | 236 | 102702 | 1455 | 0 | 0 | 270 | 0 | 0 | 57 | 16 | 0 | 1359 | 0 | 0 | 0 | 0 | 897 | 73 | 16 |
| SRR6454272 | 103 | 206 | 87 | 865 | 103 | 44 | 95 | 188 | 12 | 56 | 619 | 0 | 0 | 0 | 74956 | 59599 | 155901 | 0 | 244 | 117587 | 6892 | 0 | 0 | 214 | 0 | 56 | 294 | 71 | 0 | 1666 | 0 | 0 | 0 | 0 | 404 | 138 | 16 |
| SRR6454273 | 89 | 35 | 97 | 248 | 80 | 83 | 114 | 1054 | 79 | 390 | 470 | 0 | 0 | 0 | 30775 | 9905 | 462143 | 0 | 56 | 22908 | 12473 | 0 | 0 | 470 | 62 | 106 | 602 | 97 | 0 | 13 | 0 | 13 | 0 | 0 | 76 | 0 | 0 |
| SRR6454274 | 38 | 29 | 57 | 219 | 124 | 102 | 151 | 1244 | 121 | 143 | 304 | 0 | 0 | 0 | 24649 | 14020 | 477184 | 0 | 76 | 18181 | 8980 | 0 | 0 | 618 | 48 | 171 | 837 | 162 | 0 | 533 | 0 | 13 | 0 | 0 | 95 | 0 | 0 |
| SRR6454275 | 37 | 85 | 37 | 117 | 117 | 0 | 31 | 66 | 0 | 16 | 123 | 43 | 0 | 0 | 95540 | 66890 | 280226 | 0 | 101 | 54768 | 828 | 0 | 0 | 266 | 0 | 16 | 85 | 16 | 0 | 357 | 0 | 0 | 0 | 0 | 237 | 147 | 32 |
| SRR6454276 | 80 | 60 | 25 | 166 | 45 | 11 | 184 | 237 | 146 | 10 | 297 | 20 | 0 | 0 | 35986 | 58739 | 147825 | 0 | 211 | 74618 | 1746 | 0 | 0 | 96 | 0 | 0 | 55 | 15 | 0 | 850 | 0 | 47 | 0 | 0 | 675 | 144 | 10 |
| SRR6454277 | 40 | 59 | 20 | 84 | 35 | 74 | 121 | 499 | 58 | 15 | 247 | 64 | 0 | 0 | 33954 | 27746 | 179902 | 0 | 229 | 107567 | 580 | 0 | 0 | 124 | 0 | 30 | 30 | 20 | 0 | 588 | 0 | 26 | 0 | 0 | 799 | 170 | 30 |
| SRR6454278 | 50 | 90 | 20 | 231 | 116 | 14 | 39 | 48 | 0 | 0 | 397 | 25 | 0 | 0 | 77836 | 39143 | 195155 | 0 | 122 | 138742 | 1587 | 0 | 0 | 201 | 0 | 0 | 45 | 25 | 0 | 1100 | 0 | 12 | 0 | 0 | 254 | 111 | 10 |
| SRR6454279 | 24 | 14 | 14 | 19 | 24 | 26 | 31 | 97 | 0 | 0 | 96 | 14 | 0 | 0 | 125563 | 57671 | 316880 | 14 | 154 | 69778 | 1189 | 0 | 0 | 719 | 0 | 29 | 48 | 0 | 0 | 376 | 0 | 0 | 0 | 0 | 705 | 170 | 24 |
| SRR6454280 | 11 | 27 | 0 | 69 | 69 | 45 | 134 | 286 | 29 | 0 | 34 | 19 | 0 | 0 | 54207 | 87109 | 258636 | 10 | 149 | 84600 | 1249 | 0 | 0 | 176 | 0 | 0 | 19 | 0 | 0 | 122 | 0 | 0 | 0 | 0 | 719 | 199 | 34 |
| SRR6454281 | 51 | 47 | 12 | 102 | 39 | 28 | 27 | 76 | 0 | 0 | 94 | 0 | 0 | 0 | 48337 | 52151 | 193635 | 0 | 146 | 110378 | 2206 | 0 | 0 | 177 | 0 | 0 | 20 | 0 | 0 | 443 | 0 | 0 | 0 | 0 | 752 | 131 | 0 |
| SRR6454282 | 37 | 28 | 0 | 51 | 89 | 25 | 59 | 87 | 0 | 0 | 103 | 0 | 0 | 0 | 93490 | 56046 | 271907 | 0 | 90 | 95384 | 3061 | 0 | 0 | 145 | 0 | 0 | 19 | 0 | 0 | 739 | 0 | 0 | 0 | 0 | 749 | 140 | 0 |

### Supp Table S8.3.Columns_BX_to_DH

| | | | | | | | | | | | | | | | | | | | | | | | | | | | | | | | | | | | | |
|---|---|---|---|---|---|---|---|---|---|---|---|---|---|---|---|---|---|---|---|---|---|---|---|---|---|---|---|---|---|---|---|---|---|---|---|---|---|
| SRR6454283 | 47 | 65 | 28 | 131 | 84 | 15 | 17 | 61 | 0 | 0 | 70 | 0 | 0 | 0 | 81954 | 56836 | 306026 | 24 | 115 | 64185 | 2249 | 0 | 0 | 252 | 0 | 0 | 0 | 0 | 0 | 471 | 0 | 0 | 0 | 0 | 912 | 134 | 33 |
| SRR6454284 | 70 | 38 | 22 | 162 | 65 | 61 | 76 | 163 | 20 | 0 | 16 | 0 | 0 | 0 | 53024 | 55584 | 199716 | 0 | 141 | 107859 | 1809 | 0 | 0 | 335 | 16 | 32 | 86 | 27 | 0 | 367 | 0 | 24 | 0 | 0 | 681 | 236 | 11 |
| SRR6454285 | 0 | 0 | 0 | 56 | 40 | 0 | 0 | 39 | 0 | 0 | 66 | 30 | 0 | 0 | 56543 | 75484 | 297696 | 0 | 54 | 36329 | 617 | 0 | 0 | 91 | 0 | 0 | 25 | 0 | 0 | 404 | 0 | 0 | 0 | 0 | 510 | 60 | 15 |
| SRR6454286 | 99 | 102 | 30 | 255 | 108 | 19 | 53 | 67 | 0 | 15 | 66 | 12 | 0 | 0 | 59697 | 48035 | 186149 | 0 | 128 | 195877 | 1927 | 0 | 0 | 336 | 0 | 39 | 150 | 33 | 0 | 336 | 0 | 26 | 0 | 0 | 441 | 89 | 0 |
| SRR6454287 | 27 | 27 | 0 | 243 | 81 | 76 | 378 | 832 | 0 | 0 | 324 | 0 | 0 | 0 | 1059 | 13965 | 84262 | 0 | 0 | 3403 | 1210 | 0 | 0 | 27 | 27 | 108 | 23 | 513 | 0 | 351 | 0 | 76 | 0 | 0 | 8473 | 0 | 0 |
| SRR6454288 | 0 | 0 | 83 | 124 | 165 | 360 | 854 | 2697 | 180 | 0 | 83 | 0 | 0 | 0 | 1168 | 15035 | 50981 | 0 | 0 | 1663 | 135 | 0 | 0 | 660 | 206 | 991 | 66 | 18 | 0 | 413 | 0 | 0 | 0 | 0 | 5078 | 0 | 41 |
| SRR6454289 | 67 | 17 | 34 | 84 | 118 | 63 | 347 | 2174 | 126 | 0 | 101 | 17 | 0 | 0 | 1575 | 14061 | 129244 | 0 | 0 | 3339 | 2520 | 0 | 0 | 555 | 488 | 1716 | 10058 | 3347 | 0 | 269 | 0 | 95 | 0 | 0 | 2499 | 0 | 0 |
| SRR6454290 | 112 | 33 | 11 | 56 | 190 | 313 | 417 | 9947 | 83 | 22 | 246 | 22 | 0 | 0 | 4734 | 8428 | 251422 | 0 | 42 | 7737 | 4525 | 0 | 0 | 67 | 0 | 33 | 89 | 56 | 0 | 22 | 0 | 21 | 0 | 0 | 3703 | 0 | 11 |
| SRR6454291 | 40 | 30 | 20 | 81 | 131 | 561 | 539 | 7347 | 97 | 0 | 10 | 0 | 0 | 0 | 23616 | 7181 | 349272 | 0 | 43 | 8944 | 2654 | 0 | 0 | 38 | 182 | 12 | 54 | 403 | 0 | 222 | 0 | 0 | 0 | 0 | 1773 | 0 | 0 |
| SRR6454292 | 29 | 39 | 49 | 39 | 98 | 442 | 385 | 5950 | 29 | 0 | 0 | 0 | 0 | 0 | 18606 | 8452 | 337592 | 0 | 29 | 12856 | 7334 | 0 | 0 | 50 | 128 | 481 | 34 | 108 | 0 | 157 | 0 | 0 | 0 | 0 | 2318 | 0 | 0 |
| SRR6454293 | 35 | 105 | 0 | 70 | 70 | 224 | 561 | 2916 | 0 | 0 | 35 | 0 | 0 | 0 | 1122 | 4585 | 50284 | 0 | 0 | 1626 | 280 | 0 | 0 | 12 | 70 | 420 | 34 | 210 | 0 | 105 | 0 | 0 | 0 | 0 | 3730 | 0 | 0 |
| SRR6454294 | 24 | 49 | 24 | 0 | 220 | 229 | 344 | 4124 | 69 | 0 | 49 | 0 | 0 | 0 | 848 | 3637 | 38918 | 0 | 0 | 1718 | 619 | 0 | 0 | 952 | 0 | 24 | 464 | 0 | 0 | 366 | 0 | 0 | 0 | 0 | 2467 | 0 | 0 |
| SRR6454295 | 13 | 13 | 39 | 66 | 105 | 652 | 426 | 9949 | 80 | 26 | 79 | 13 | 0 | 0 | 9936 | 4377 | 220995 | 0 | 0 | 6451 | 3698 | 0 | 0 | 1117 | 39 | 342 | 1090 | 144 | 0 | 394 | 0 | 0 | 13 | 0 | 1898 | 0 | 0 |
| SRR6454296 | 18 | 18 | 0 | 36 | 0 | 290 | 339 | 4474 | 0 | 0 | 53 | 0 | 0 | 0 | 3047 | 4276 | 104469 | 0 | 24 | 3483 | 2249 | 0 | 0 | 1834 | 142 | 534 | 2831 | 231 | 0 | 321 | 0 | 24 | 0 | 0 | 2725 | 0 | 0 |
| SRR6454298 | 85 | 47 | 0 | 0 | 39 | 1571 | 817 | 4678 | 208 | 47 | 0 | 0 | 0 | 0 | 240143 | 162839 | 510128 | 427 | 18 | 16457 | 9356 | 0 | 0 | 683 | 0 | 0 | 0 | 0 | 0 | 23 | 0 | 0 | 0 | 0 | 677 | 64 | 0 |
| SRR6454299 | 0 | 0 | 0 | 0 | 29 | 478 | 558 | 17896 | 100 | 21 | 0 | 0 | 0 | 0 | 2471 | 2854 | 53042 | 0 | 10 | 4344 | 1714 | 0 | 0 | 15 | 36 | 64 | 179 | 0 | 0 | 121 | 0 | 0 | 0 | 0 | 5276 | 0 | 0 |
| SRR6454300 | 0 | 0 | 0 | 0 | 27 | 646 | 704 | 17299 | 112 | 0 | 0 | 0 | 0 | 0 | 7927 | 6023 | 152057 | 0 | 0 | 9022 | 3795 | 0 | 0 | 798 | 0 | 27 | 41 | 0 | 0 | 131 | 0 | 21 | 0 | 0 | 5640 | 0 | 0 |
| SRR6454301 | 51 | 85 | 21 | 141 | 77 | 39 | 65 | 152 | 16 | 0 | 141 | 26 | 0 | 0 | 56195 | 41204 | 241475 | 0 | 168 | 124229 | 529 | 0 | 0 | 414 | 0 | 17 | 43 | 17 | 0 | 683 | 0 | 0 | 0 | 0 | 839 | 125 | 21 |
| SRR6454302 | 19 | 47 | 16 | 190 | 97 | 49 | 250 | 446 | 72 | 0 | 51 | 23 | 0 | 0 | 46616 | 78133 | 205094 | 14 | 118 | 70665 | 1728 | 0 | 0 | 175 | 16 | 62 | 264 | 70 | 0 | 385 | 0 | 127 | 0 | 0 | 1082 | 276 | 0 |
| SRR6454303 | 27 | 44 | 12 | 49 | 89 | 23 | 33 | 106 | 10 | 0 | 71 | 20 | 0 | 0 | 59380 | 34555 | 223877 | 12 | 224 | 87475 | 1525 | 0 | 0 | 158 | 0 | 52 | 98 | 42 | 0 | 458 | 0 | 0 | 0 | 0 | 362 | 225 | 20 |
| SRR6454304 | 21 | 13 | 13 | 59 | 113 | 41 | 103 | 208 | 45 | 0 | 62 | 35 | 0 | 0 | 78886 | 72875 | 285037 | 0 | 128 | 29968 | 1757 | 0 | 0 | 129 | 0 | 0 | 30 | 0 | 0 | 244 | 0 | 0 | 0 | 0 | 275 | 278 | 19 |
| SRR6454305 | 27 | 0 | 0 | 16 | 54 | 17 | 42 | 169 | 24 | 0 | 59 | 38 | 0 | 0 | 62983 | 76233 | 284447 | 0 | 135 | 55515 | 343 | 0 | 0 | 27 | 0 | 16 | 86 | 0 | 0 | 10 | 0 | 0 | 0 | 0 | 639 | 125 | 32 |
| SRR6454306 | 14 | 15 | 14 | 44 | 105 | 76 | 301 | 356 | 53 | 0 | 20 | 0 | 0 | 0 | 84764 | 134958 | 311055 | 0 | 86 | 41174 | 1191 | 0 | 0 | 255 | 0 | 0 | 31 | 0 | 0 | 141 | 0 | 0 | 0 | 0 | 497 | 248 | 20 |
| SRR6454307 | 17 | 21 | 0 | 31 | 40 | 34 | 84 | 137 | 16 | 0 | 54 | 17 | 0 | 0 | 46227 | 45565 | 196343 | 0 | 143 | 118924 | 1061 | 0 | 0 | 151 | 0 | 0 | 21 | 0 | 0 | 12 | 0 | 0 | 0 | 0 | 1239 | 128 | 19 |
| SRR6454308 | 0 | 17 | 0 | 39 | 124 | 53 | 143 | 279 | 44 | 0 | 28 | 25 | 0 | 0 | 43289 | 39569 | 195931 | 0 | 144 | 96520 | 1273 | 0 | 0 | 157 | 22 | 50 | 121 | 0 | 0 | 494 | 0 | 0 | 0 | 0 | 517 | 361 | 22 |
| SRR6454309 | 76 | 57 | 70 | 228 | 200 | 183 | 237 | 655 | 92 | 0 | 162 | 0 | 0 | 0 | 88189 | 58657 | 304498 | 0 | 174 | 58385 | 923 | 0 | 0 | 1043 | 0 | 54 | 143 | 38 | 0 | 501 | 0 | 0 | 0 | 0 | 1315 | 254 | 25 |
| SRR6454310 | 27 | 18 | 21 | 201 | 121 | 61 | 108 | 178 | 38 | 0 | 45 | 16 | 0 | 0 | 41818 | 46814 | 164345 | 17 | 159 | 84018 | 1336 | 0 | 0 | 129 | 51 | 133 | 435 | 139 | 0 | 451 | 0 | 181 | 0 | 0 | 413 | 575 | 0 |
| SRR6454311 | 11 | 11 | 11 | 119 | 65 | 246 | 225 | 615 | 20 | 0 | 108 | 0 | 0 | 0 | 1148 | 10583 | 129509 | 0 | 0 | 2644 | 1394 | 0 | 0 | 119 | 22 | 76 | 508 | 97 | 0 | 595 | 0 | 0 | 0 | 0 | 3323 | 0 | 11 |
| SRR6454312 | 33 | 33 | 18 | 90 | 51 | 202 | 157 | 546 | 30 | 13 | 62 | 48 | 0 | 0 | 2247 | 15610 | 224012 | 0 | 20 | 4738 | 5205 | 0 | 0 | 15 | 84 | 315 | 28 | 189 | 0 | 319 | 0 | 86 | 0 | 0 | 1891 | 0 | 0 |
| SRR6454313 | 23 | 46 | 0 | 35 | 104 | 355 | 591 | 2088 | 98 | 0 | 35 | 0 | 0 | 0 | 906 | 6267 | 51266 | 0 | 0 | 1871 | 630 | 0 | 0 | 406 | 128 | 395 | 3076 | 371 | 0 | 476 | 0 | 20 | 0 | 0 | 4058 | 0 | 12 |
| SRR6454314 | 31 | 54 | 0 | 39 | 31 | 156 | 168 | 360 | 12 | 0 | 46 | 31 | 0 | 0 | 1656 | 5631 | 88112 | 0 | 0 | 3913 | 4555 | 0 | 0 | 10 | 46 | 178 | 28 | 77 | 0 | 332 | 0 | 0 | 0 | 0 | 988 | 0 | 0 |
| SRR6454315 | 19 | 26 | 0 | 13 | 122 | 75 | 156 | 555 | 27 | 0 | 26 | 0 | 0 | 0 | 539 | 3231 | 16154 | 0 | 11 | 1935 | 588 | 0 | 0 | 608 | 13 | 102 | 999 | 51 | 0 | 416 | 0 | 11 | 0 | 0 | 3488 | 0 | 13 |
| SRR6454316 | 0 | 0 | 0 | 0 | 60 | 139 | 146 | 324 | 0 | 0 | 34 | 0 | 0 | 0 | 478 | 3615 | 18289 | 0 | 0 | 1796 | 524 | 0 | 0 | 299 | 26 | 128 | 14 | 0 | 0 | 307 | 0 | 0 | 0 | 0 | 3256 | 0 | 0 |
| SRR6454317 | 13 | 20 | 20 | 20 | 139 | 512 | 263 | 876 | 34 | 0 | 13 | 0 | 0 | 0 | 575 | 4617 | 26236 | 0 | 0 | 1341 | 417 | 0 | 0 | 727 | 535 | 13 | 39 | 126 | 0 | 211 | 0 | 0 | 0 | 0 | 4447 | 0 | 0 |
| SRR6454318 | 21 | 11 | 0 | 43 | 129 | 642 | 344 | 940 | 23 | 0 | 32 | 21 | 11 | 0 | 528 | 3661 | 18926 | 0 | 11 | 1594 | 390 | 0 | 0 | 557 | 386 | 12 | 30 | 129 | 0 | 343 | 0 | 11 | 0 | 0 | 4452 | 0 | 21 |
| SRR6454319 | 30 | 52 | 22 | 22 | 223 | 133 | 133 | 477 | 24 | 0 | 22 | 0 | 0 | 0 | 801 | 1983 | 18145 | 0 | 0 | 1949 | 467 | 0 | 0 | 1363 | 186 | 722 | 3247 | 127 | 0 | 246 | 0 | 0 | 0 | 0 | 2323 | 0 | 0 |
| SRR6454320 | 57 | 34 | 0 | 46 | 80 | 136 | 463 | 1281 | 0 | 0 | 80 | 0 | 0 | 0 | 363 | 3115 | 12917 | 0 | 0 | 1626 | 327 | 0 | 0 | 274 | 0 | 34 | 171 | 11 | 0 | 696 | 0 | 0 | 0 | 0 | 5963 | 0 | 0 |
| SRR6454321 | 0 | 30 | 15 | 76 | 45 | 101 | 355 | 710 | 25 | 0 | 76 | 15 | 0 | 0 | 1420 | 5274 | 31330 | 0 | 0 | 1851 | 279 | 0 | 0 | 332 | 106 | 665 | 4985 | 1209 | 0 | 106 | 0 | 0 | 0 | 0 | 3239 | 0 | 30 |
| SRR6454322 | 0 | 13 | 13 | 66 | 80 | 310 | 563 | 985 | 113 | 0 | 80 | 0 | 0 | 0 | 676 | 4765 | 34985 | 0 | 0 | 2506 | 141 | 0 | 0 | 319 | 53 | 439 | 31 | 651 | 0 | 359 | 0 | 28 | 0 | 0 | 3913 | 0 | 13 |
| SRR6454323 | 42 | 65 | 28 | 135 | 37 | 103 | 46 | 308 | 23 | 0 | 60 | 23 | 0 | 0 | 1905 | 5994 | 101660 | 0 | 0 | 3651 | 3297 | 0 | 0 | 172 | 0 | 107 | 28 | 130 | 0 | 139 | 0 | 46 | 0 | 0 | 1416 | 0 | 0 |
| SRR6454324 | 66 | 30 | 48 | 151 | 91 | 79 | 71 | 260 | 0 | 0 | 54 | 0 | 0 | 0 | 576 | 4907 | 21772 | 0 | 24 | 1523 | 276 | 0 | 0 | 604 | 54 | 537 | 3357 | 797 | 0 | 332 | 0 | 16 | 0 | 0 | 5301 | 0 | 0 |
| SRR6454325 | 72 | 85 | 55 | 358 | 176 | 31 | 43 | 102 | 0 | 0 | 140 | 0 | 0 | 0 | 84839 | 34619 | 229820 | 0 | 236 | 71100 | 599 | 0 | 0 | 325 | 0 | 13 | 104 | 16 | 0 | 674 | 0 | 0 | 0 | 0 | 440 | 202 | 23 |
| SRR6454326 | 38 | 35 | 23 | 234 | 61 | 266 | 577 | 927 | 355 | 12 | 42 | 18 | 0 | 0 | 40481 | 37573 | 161014 | 0 | 841 | 69315 | 621 | 0 | 0 | 295 | 39 | 131 | 507 | 229 | 0 | 282 | 0 | 0 | 0 | 0 | 947 | 688 | 39 |
| SRR6454331 | 21 | 11 | 32 | 182 | 150 | 252 | 582 | 1734 | 40 | 0 | 53 | 0 | 0 | 0 | 1244 | 5464 | 41096 | 0 | 26 | 2145 | 265 | 0 | 0 | 983 | 107 | 555 | 5074 | 1036 | 0 | 256 | 0 | 0 | 0 | 0 | 4522 | 0 | 11 |
| SRR6454332 | 0 | 79 | 34 | 181 | 90 | 206 | 511 | 3632 | 28 | 11 | 56 | 23 | 0 | 0 | 2412 | 3229 | 46024 | 0 | 21 | 2504 | 1007 | 0 | 0 | 15 | 147 | 12 | 88 | 17 | 0 | 271 | 0 | 0 | 0 | 0 | 1374 | 0 | 0 |
| SRR6454335 | 102 | 0 | 0 | 0 | 34 | 68 | 102 | 407 | 34 | 0 | 0 | 34 | 0 | 0 | 11103 | 8364 | 68991 | 68 | 170 | 53954 | 3803 | 0 | 0 | 614 | 0 | 34 | 34 | 0 | 0 | 0 | 0 | 34 | 0 | 0 | 2887 | 22 | 0 |
| SRR6454337 | 73 | 23 | 0 | 0 | 41 | 3305 | 1818 | 10029 | 898 | 14 | 0 | 0 | 0 | 0 | 104832 | 120009 | 339666 | 78 | 22 | 18182 | 2029 | 0 | 0 | 264 | 0 | 0 | 0 | 0 | 0 | 23 | 0 | 0 | 0 | 0 | 1330 | 126 | 23 |
| SRR6454338 | 82 | 52 | 0 | 0 | 15 | 5456 | 1959 | 16353 | 937 | 31 | 0 | 0 | 0 | 0 | 292757 | 152981 | 491170 | 270 | 16 | 2860 | 4756 | 0 | 0 | 675 | 0 | 0 | 0 | 0 | 0 | 0 | 0 | 0 | 0 | 0 | 776 | 89 | 15 |
| SRR6454339 | 55 | 25 | 0 | 0 | 0 | 7197 | 4576 | 23095 | 3087 | 31 | 0 | 0 | 0 | 0 | 86730 | 121381 | 217440 | 183 | 90 | 23771 | 4311 | 0 | 0 | 251 | 0 | 0 | 0 | 0 | 0 | 40 | 0 | 0 | 0 | 0 | 1668 | 133 | 12 |
| SRR6454340 | 17 | 22 | 0 | 0 | 39 | 247 | 356 | 901 | 126 | 11 | 0 | 0 | 0 | 0 | 49858 | 67988 | 165339 | 29 | 57 | 110959 | 826 | 0 | 0 | 853 | 28 | 17 | 22 | 0 | 0 | 17 | 0 | 23 | 0 | 0 | 1886 | 207 | 11 |
| SRR6454341 | 44 | 37 | 0 | 0 | 0 | 6188 | 3271 | 17075 | 1235 | 15 | 0 | 0 | 0 | 0 | 94780 | 178452 | 337012 | 132 | 26 | 8071 | 13055 | 0 | 0 | 314 | 15 | 26 | 15 | 0 | 0 | 26 | 0 | 19 | 0 | 0 | 1440 | 75 | 0 |
| SRR6454342 | 12 | 33 | 0 | 0 | 12 | 6673 | 2472 | 14394 | 953 | 50 | 0 | 0 | 0 | 0 | 95776 | 105052 | 363134 | 217 | 23 | 11580 | 11231 | 0 | 0 | 222 | 31 | 26 | 12 | 0 | 0 | 31 | 0 | 36 | 0 | 0 | 1181 | 158 | 0 |
| SRR6454343 | 66 | 30 | 0 | 0 | 30 | 4447 | 1781 | 11178 | 723 | 0 | 0 | 0 | 0 | 0 | 63242 | 80299 | 238351 | 129 | 60 | 23780 | 5005 | 0 | 0 | 224 | 17 | 31 | 0 | 0 | 0 | 46 | 0 | 32 | 0 | 0 | 506 | 122 | 0 |
| SRR6454345 | 0 | 0 | 0 | 0 | 0 | 317 | 221 | 745 | 69 | 0 | 0 | 17 | 0 | 0 | 483 | 1162 | 2957 | 14 | 14 | 1780 | 276 | 0 | 0 | 107 | 0 | 0 | 0 | 0 | 0 | 240 | 0 | 0 | 0 | 0 | 5288 | 0 | 0 |
| SRR6454347 | 34 | 41 | 0 | 0 | 88 | 80 | 95 | 598 | 43 | 0 | 0 | 15 | 0 | 0 | 728 | 8672 | 43356 | 40 | 0 | 4170 | 1963 | 0 | 0 | 610 | 0 | 0 | 0 | 0 | 0 | 0 | 0 | 115 | 0 | 0 | 10007 | 0 | 0 |
| SRR6454348 | 29 | 28 | 0 | 0 | 123 | 246 | 340 | 1201 | 86 | 0 | 0 | 23 | 0 | 0 | 1509 | 14347 | 60351 | 17 | 0 | 2141 | 2651 | 0 | 0 | 288 | 26 | 0 | 0 | 0 | 0 | 0 | 0 | 101 | 0 | 0 | 6952 | 0 | 0 |
| SRR6454349 | 14 | 50 | 0 | 0 | 116 | 213 | 420 | 1068 | 93 | 0 | 0 | 0 | 0 | 0 | 1032 | 16737 | 49467 | 16 | 0 | 3935 | 2250 | 0 | 0 | 306 | 105 | 0 | 0 | 0 | 0 | 0 | 0 | 67 | 0 | 0 | 8378 | 0 | 0 |
| SRR6454350 | 28 | 0 | 0 | 0 | 19 | 309 | 449 | 1431 | 70 | 0 | 0 | 0 | 0 | 0 | 589 | 9032 | 24523 | 28 | 14 | 2749 | 505 | 0 | 0 | 408 | 47 | 0 | 0 | 0 | 0 | 0 | 0 | 112 | 0 | 0 | 3920 | 0 | 28 |
| SRR6454351 | 29 | 0 | 0 | 0 | 39 | 2570 | 874 | 6006 | 601 | 0 | 0 | 0 | 0 | 0 | 88664 | 60011 | 411286 | 245 | 69 | 40729 | 2464 | 0 | 0 | 434 | 166 | 0 | 0 | 0 | 0 | 0 | 0 | 38 | 0 | 0 | 273 | 376 | 0 |
| SRR6454352 | 35 | 0 | 0 | 0 | 10 | 1966 | 953 | 5232 | 241 | 17 | 10 | 0 | 0 | 0 | 178712 | 97343 | 416845 | 435 | 22 | 16780 | 6091 | 0 | 0 | 207 | 10 | 0 | 0 | 0 | 0 | 0 | 0 | 14 | 0 | 0 | 1221 | 305 | 14 |
| SRR6454354 | 12 | 0 | 0 | 0 | 0 | 3492 | 1777 | 12957 | 672 | 31 | 0 | 19 | 0 | 0 | 81174 | 58365 | 351511 | 440 | 26 | 28922 | 9415 | 0 | 0 | 62 | 0 | 0 | 0 | 0 | 0 | 0 | 0 | 84 | 0 | 0 | 1007 | 198 | 12 |
| SRR6496456 | 40 | 18 | 14 | 14 | 12 | 5153 | 1342 | 2393 | 289 | 0 | 25 | 0 | 0 | 0 | 5960 | 4803 | 2163 | 36134 | 347 | 433 | 33 | 365 | 132 | 12 | 0 | 0 | 18 | 0 | 0 | 209 | 108 | 34033 | 149 | 0 | 1050 | 3826 | 15 |
| SRR6496457 | 15 | 0 | 0 | 0 | 0 | 4475 | 1810 | 3441 | 434 | 0 | 44 | 0 | 0 | 0 | 7593 | 5253 | 2285 | 63841 | 248 | 265 | 27 | 58 | 32 | 10 | 0 | 0 | 0 | 0 | 0 | 204 | 54 | 41721 | 161 | 0 | 1795 | 4827 | 0 |
| SRR6496458 | 60 | 14 | 12 | 0 | 20 | 8203 | 2284 | 4022 | 438 | 0 | 87 | 0 | 0 | 0 | 8091 | 5723 | 2951 | 50231 | 260 | 449 | 18 | 540 | 223 | 15 | 0 | 0 | 13 | 0 | 0 | 580 | 252 | 38468 | 301 | 0 | 1281 | 4282 | 0 |
| SRR6496459 | 18 | 0 | 0 | 0 | 0 | 13816 | 2816 | 9221 | 569 | 0 | 14 | 0 | 0 | 0 | 3193 | 3271 | 852 | 63805 | 136 | 341 | 14 | 171 | 47 | 0 | 0 | 0 | 0 | 0 | 0 | 131 | 138 | 21908 | 189 | 0 | 2393 | 4763 | 0 |
| SRR6496460 | 74 | 17 | 0 | 0 | 16 | 9680 | 2675 | 5149 | 773 | 0 | 27 | 0 | 0 | 0 | 7816 | 3799 | 2538 | 48281 | 284 | 527 | 29 | 1144 | 293 | 22 | 0 | 0 | 0 | 0 | 0 | 344 | 536 | 28612 | 349 | 0 | 1236 | 5547 | 0 |
| SRR6496461 | 0 | 0 | 15 | 58 | 63 | 155 | 184 | 325 | 116 | 0 | 147 | 0 | 0 | 0 | 21019 | 17372 | 49534 | 679 | 58 | 2327 | 1226 | 0 | 0 | 47 | 64 | 257 | 332 | 660 | 0 | 1009 | 0 | 1755 | 0 | 0 | 16911 | 390 | 10 |
| SRR6496462 | 0 | 0 | 16 | 63 | 64 | 170 | 151 | 316 | 102 | 0 | 154 | 0 | 0 | 0 | 21300 | 17361 | 48470 | 608 | 34 | 2243 | 1031 | 0 | 0 | 52 | 61 | 248 | 379 | 660 | 0 | 980 | 0 | 1800 | 0 | 0 | 16768 | 368 | 11 |
| SRR6496463 | 0 | 0 | 19 | 87 | 55 | 169 | 139 | 279 | 110 | 0 | 167 | 0 | 0 | 0 | 20869 | 16655 | 48355 | 603 | 75 | 2291 | 1021 | 0 | 0 | 50 | 61 | 283 | 366 | 695 | 0 | 962 | 0 | 1897 | 0 | 0 | 17430 | 375 | 15 |
| SRR6496464 | 0 | 0 | 14 | 22 | 21 | 166 | 205 | 292 | 54 | 0 | 60 | 0 | 0 | 0 | 21101 | 18269 | 48238 | 628 | 49 | 2445 | 999 | 0 | 0 | 19 | 20 | 94 | 133 | 210 | 0 | 355 | 0 | 1768 | 0 | 0 | 17128 | 364 | 13 |
| SRR6496465 | 0 | 0 | 21 | 86 | 66 | 172 | 172 | 415 | 115 | 10 | 166 | 0 | 0 | 0 | 20056 | 17227 | 47544 | 702 | 72 | 2736 | 1060 | 0 | 0 | 66 | 79 | 318 | 369 | 642 | 0 | 839 | 0 | 1619 | 0 | 0 | 17056 | 363 | 14 |
| SRR6498854 | 18 | 12 | 20 | 14 | 21 | 175 | 230 | 700 | 77 | 0 | 21 | 0 | 0 | 0 | 46343 | 197425 | 47075 | 0 | 88 | 5876 | 2462 | 0 | 0 | 37 | 19 | 56 | 109 | 121 | 0 | 25 | 0 | 0 | 0 | 0 | 1186 | 0 | 0 |
| SRR6498855 | 21 | 49 | 58 | 10 | 16 | 167 | 202 | 480 | 111 | 0 | 15 | 0 | 0 | 0 | 32647 | 204275 | 38679 | 0 | 90 | 4807 | 2386 | 0 | 0 | 39 | 11 | 42 | 103 | 101 | 0 | 32 | 0 | 0 | 0 | 0 | 1573 | 0 | 0 |
| SRR6498856 | 17 | 22 | 28 | 64 | 81 | 142 | 284 | 542 | 135 | 13 | 98 | 0 | 0 | 0 | 35237 | 194396 | 40510 | 0 | 68 | 5692 | 2035 | 0 | 0 | 148 | 121 | 348 | 755 | 642 | 0 | 161 | 0 | 0 | 0 | 0 | 1304 | 0 | 0 |
| SRR6498857 | 21 | 29 | 53 | 13 | 12 | 203 | 292 | 519 | 97 | 0 | 38 | 0 | 0 | 0 | 40735 | 231874 | 50066 | 0 | 122 | 5016 | 2581 | 0 | 0 | 50 | 18 | 61 | 94 | 122 | 0 | 116 | 0 | 0 | 0 | 0 | 1134 | 0 | 0 |
| SRR6498858 | 28 | 30 | 48 | 107 | 78 | 89 | 183 | 485 | 18 | 24 | 184 | 0 | 0 | 0 | 39118 | 166241 | 55701 | 12 | 71 | 4828 | 2609 | 0 | 11 | 276 | 374 | 1066 | 1153 | 933 | 0 | 529 | 0 | 0 | 0 | 0 | 1157 | 0 | 0 |
| SRR6498860 | 33 | 12 | 40 | 71 | 123 | 172 | 222 | 615 | 40 | 19 | 52 | 0 | 0 | 0 | 41684 | 193002 | 49339 | 10 | 101 | 5135 | 3652 | 0 | 15 | 231 | 221 | 544 | 1007 | 600 | 0 | 83 | 0 | 0 | 0 | 0 | 985 | 0 | 0 |
| SRR6498861 | 29 | 23 | 26 | 29 | 85 | 242 | 472 | 387 | 97 | 26 | 88 | 0 | 0 | 0 | 55388 | 151738 | 66062 | 24 | 85 | 5772 | 2239 | 0 | 0 | 193 | 69 | 258 | 366 | 164 | 0 | 222 | 0 | 0 | 0 | 0 | 1871 | 0 | 0 |
| SRR6498862 | 27 | 12 | 35 | 42 | 25 | 406 | 261 | 985 | 130 | 22 | 33 | 0 | 0 | 0 | 36705 | 187664 | 48216 | 29 | 72 | 4288 | 3476 | 0 | 0 | 56 | 85 | 272 | 261 | 188 | 0 | 97 | 0 | 0 | 0 | 0 | 809 | 0 | 0 |
| SRR6498863 | 32 | 18 | 41 | 48 | 83 | 151 | 194 | 463 | 237 | 32 | 131 | 0 | 0 | 0 | 42700 | 112529 | 57670 | 0 | 97 | 3830 | 3055 | 0 | 0 | 356 | 326 | 1117 | 563 | 278 | 0 | 182 | 0 | 0 | 0 | 0 | 860 | 0 | 0 |
| SRR6498864 | 13 | 11 | 11 | 24 | 32 | 218 | 180 | 526 | 58 | 18 | 39 | 0 | 0 | 0 | 41704 | 98832 | 67639 | 0 | 64 | 4115 | 3293 | 0 | 13 | 112 | 157 | 372 | 196 | 90 | 0 | 97 | 0 | 0 | 0 | 0 | 769 | 0 | 0 |
| SRR6498865 | 0 | 0 | 10 | 20 | 18 | 168 | 56 | 224 | 0 | 25 | 35 | 13 | 0 | 0 | 35996 | 89783 | 55548 | 0 | 56 | 3242 | 1956 | 0 | 0 | 96 | 89 | 281 | 175 | 73 | 0 | 122 | 0 | 0 | 0 | 0 | 1447 | 0 | 0 |
| SRR6498866 | 16 | 10 | 34 | 22 | 21 | 103 | 271 | 394 | 26 | 10 | 30 | 0 | 0 | 0 | 51133 | 121738 | 38115 | 0 | 77 | 2994 | 4052 | 0 | 12 | 51 | 22 | 65 | 124 | 91 | 0 | 30 | 0 | 0 | 0 | 0 | 815 | 0 | 0 |
| SRR6498867 | 0 | 0 | 20 | 10 | 52 | 56 | 80 | 215 | 0 | 0 | 32 | 0 | 0 | 0 | 28393 | 70798 | 36273 | 0 | 16 | 1506 | 3035 | 0 | 0 | 18 | 23 | 75 | 109 | 175 | 0 | 20 | 0 | 0 | 0 | 0 | 851 | 0 | 0 |
| SRR6498868 | 17 | 11 | 40 | 119 | 76 | 105 | 93 | 326 | 35 | 11 | 180 | 0 | 0 | 0 | 35553 | 98084 | 40467 | 0 | 35 | 1071 | 4226 | 0 | 0 | 226 | 254 | 782 | 762 | 705 | 0 | 544 | 0 | 0 | 0 | 0 | 2051 | 0 | 0 |
| SRR6498869 | 22 | 20 | 50 | 78 | 82 | 55 | 110 | 302 | 18 | 15 | 98 | 0 | 0 | 0 | 49699 | 93971 | 55853 | 0 | 0 | 2350 | 4644 | 0 | 0 | 305 | 240 | 660 | 637 | 690 | 0 | 127 | 0 | 0 | 0 | 0 | 970 | 0 | 0 |
| SRR6498870 | 18 | 11 | 18 | 62 | 94 | 44 | 114 | 180 | 26 | 12 | 199 | 0 | 0 | 0 | 62350 | 75573 | 64219 | 0 | 13 | 1642 | 3636 | 0 | 0 | 154 | 200 | 570 | 448 | 428 | 0 | 320 | 0 | 0 | 0 | 0 | 800 | 0 | 0 |
| SRR6498871 | 19 | 16 | 39 | 61 | 45 | 81 | 54 | 331 | 15 | 19 | 162 | 15 | 0 | 0 | 32155 | 94554 | 44724 | 0 | 39 | 1490 | 4482 | 0 | 0 | 207 | 172 | 507 | 311 | 215 | 0 | 304 | 0 | 12 | 0 | 0 | 1434 | 0 | 0 |
| SRR6705804 | 0 | 0 | 0 | 0 | 0 | 4732 | 2169 | 8948 | 502 | 0 | 0 | 0 | 0 | 0 | 28348 | 63035 | 323504 | 31 | 232 | 5867 | 232 | 0 | 0 | 20 | 0 | 0 | 0 | 0 | 0 | 0 | 0 | 15 | 0 | 0 | 217577 | 1130 | 0 |
| SRR6705805 | 0 | 0 | 0 | 0 | 0 | 16947 | 2036 | 21639 | 444 | 0 | 0 | 0 | 0 | 0 | 44153 | 74607 | 564466 | 20 | 287 | 12073 | 241 | 0 | 0 | 31 | 0 | 0 | 0 | 0 | 0 | 0 | 0 | 20 | 0 | 0 | 32697 | 749 | 0 |
| SRR6705806 | 0 | 0 | 0 | 0 | 0 | 4073 | 0 | 0 | 0 | 0 | 0 | 0 | 0 | 0 | 34623 | 51370 | 484589 | 0 | 0 | 0 | 0 | 0 | 0 | 0 | 0 | 0 | 0 | 0 | 0 | 0 | 0 | 0 | 0 | 0 | 1712 | 0 | 0 |
| SRR6705808 | 59 | 43 | 0 | 0 | 24 | 2155 | 967 | 4022 | 259 | 0 | 0 | 0 | 0 | 0 | 29433 | 113733 | 632489 | 53 | 87 | 1028 | 311 | 27 | 37 | 27 | 64 | 182 | 16 | 16 | 0 | 0 | 0 | 20 | 0 | 0 | 3383 | 2253 | 0 |
| SRR6705809 | 40 | 18 | 0 | 0 | 53 | 1894 | 1115 | 4062 | 137 | 0 | 0 | 0 | 0 | 0 | 99742 | 202527 | 471686 | 95 | 32 | 1463 | 347 | 0 | 0 | 73 | 10 | 0 | 0 | 0 | 0 | 0 | 0 | 95 | 0 | 0 | 1389 | 990 | 0 |
| SRR6705810 | 81 | 110 | 0 | 0 | 176 | 889 | 147 | 873 | 66 | 0 | 15 | 0 | 0 | 0 | 26890 | 33394 | 375117 | 13 | 395 | 1731 | 558 | 0 | 37 | 124 | 44 | 95 | 10 | 0 | 0 | 81 | 0 | 0 | 0 | 0 | 4743 | 3230 | 0 |
| SRR6705811 | 0 | 0 | 0 | 0 | 28 | 96 | 28 | 139 | 0 | 0 | 0 | 0 | 0 | 0 | 29685 | 50820 | 515468 | 0 | 351 | 4603 | 465 | 0 | 0 | 14 | 23 | 23 | 18 | 0 | 0 | 0 | 0 | 0 | 0 | 0 | 3064 | 67 | 0 |
| SRR6705812 | 0 | 0 | 0 | 0 | 0 | 30048 | 3771 | 29144 | 563 | 0 | 0 | 0 | 0 | 0 | 48673 | 51029 | 406966 | 26 | 486 | 2269 | 375 | 0 | 0 | 78 | 0 | 0 | 0 | 0 | 0 | 0 | 0 | 17 | 0 | 0 | 71668 | 1822 | 0 |
| SRR6705813 | 0 | 0 | 0 | 0 | 0 | 6201 | 2728 | 9603 | 501 | 0 | 0 | 0 | 0 | 0 | 23702 | 43494 | 247234 | 10 | 337 | 2074 | 337 | 0 | 0 | 43 | 0 | 0 | 0 | 0 | 0 | 21 | 0 | 0 | 0 | 0 | 157631 | 2302 | 0 |
| SRR6705814 | 20 | 0 | 0 | 0 | 14 | 1204 | 424 | 3747 | 53 | 27 | 80 | 0 | 0 | 0 | 77386 | 69992 | 543746 | 0 | 128 | 2464 | 602 | 0 | 0 | 36 | 90 | 18 | 38 | 67 | 0 | 22 | 0 | 14 | 0 | 0 | 1226 | 3356 | 0 |
| SRR6705815 | 52 | 17 | 0 | 0 | 187 | 3898 | 2193 | 9603 | 505 | 17 | 351 | 0 | 0 | 0 | 32808 | 74205 | 337125 | 0 | 356 | 6972 | 564 | 0 | 0 | 3159 | 139 | 317 | 1402 | 217 | 0 | 239 | 0 | 0 | 0 | 0 | 1231 | 6321 | 0 |
| SRR6705816 | 35 | 28 | 0 | 0 | 30 | 2169 | 1028 | 5257 | 180 | 0 | 25 | 0 | 0 | 0 | 30003 | 61915 | 339198 | 0 | 475 | 9587 | 319 | 0 | 0 | 628 | 69 | 163 | 329 | 0 | 0 | 83 | 0 | 10 | 0 | 0 | 3042 | 2444 | 0 |
| SRR6705817 | 43 | 56 | 0 | 0 | 19 | 1584 | 1065 | 2770 | 192 | 13 | 0 | 0 | 0 | 0 | 62178 | 73642 | 480578 | 0 | 533 | 3732 | 2564 | 0 | 14 | 168 | 32 | 39 | 51 | 0 | 0 | 83 | 0 | 28 | 0 | 0 | 4830 | 5868 | 0 |
| SRR6705818 | 20 | 37 | 0 | 15 | 24 | 1113 | 698 | 4996 | 198 | 0 | 28 | 0 | 0 | 0 | 20278 | 47265 | 314791 | 0 | 543 | 4224 | 652 | 0 | 0 | 137 | 13 | 35 | 80 | 108 | 0 | 42 | 0 | 0 | 0 | 0 | 2897 | 2680 | 0 |
| SRR6705819 | 67 | 64 | 15 | 59 | 88 | 1444 | 798 | 5375 | 286 | 0 | 35 | 0 | 0 | 0 | 40150 | 55713 | 480864 | 0 | 1039 | 5924 | 1411 | 0 | 49 | 75 | 17 | 44 | 65 | 75 | 0 | 35 | 0 | 11 | 0 | 0 | 1718 | 1782 | 0 |
| SRR6705820 | 0 | 0 | 0 | 0 | 16 | 924 | 629 | 3527 | 228 | 0 | 12 | 0 | 0 | 0 | 17128 | 87993 | 112111 | 0 | 266 | 7033 | 153 | 0 | 0 | 380 | 37 | 77 | 471 | 29 | 0 | 0 | 0 | 0 | 0 | 0 | 643 | 3439 | 0 |
| SRR6705821 | 60 | 35 | 0 | 17 | 204 | 821 | 635 | 3687 | 264 | 0 | 63 | 0 | 0 | 0 | 41151 | 63419 | 193584 | 18 | 650 | 14368 | 489 | 0 | 0 | 843 | 66 | 132 | 305 | 138 | 0 | 98 | 0 | 18 | 0 | 0 | 1105 | 3665 | 0 |
| SRR6705822 | 16 | 14 | 0 | 0 | 25 | 2349 | 929 | 4357 | 683 | 0 | 0 | 0 | 0 | 0 | 23111 | 100456 | 52228 | 0 | 270 | 12111 | 722 | 0 | 0 | 751 | 20 | 32 | 78 | 23 | 0 | 10 | 0 | 0 | 0 | 0 | 185 | 3534 | 0 |
| SRR6705823 | 53 | 53 | 0 | 0 | 159 | 898 | 1110 | 3572 | 402 | 0 | 74 | 0 | 0 | 0 | 46389 | 157950 | 242843 | 16 | 1136 | 2764 | 1876 | 22 | 133 | 1090 | 295 | 408 | 705 | 0 | 0 | 252 | 0 | 37 | 0 | 0 | 2228 | 4562 | 0 |
| SRR6705824 | 0 | 16 | 0 | 0 | 70 | 1030 | 536 | 4452 | 101 | 14 | 0 | 0 | 0 | 0 | 13969 | 100973 | 361728 | 0 | 31 | 4085 | 399 | 0 | 0 | 44 | 0 | 0 | 0 | 0 | 0 | 0 | 0 | 25 | 0 | 0 | 1619 | 1039 | 0 |
| SRR6705825 | 49 | 14 | 0 | 0 | 248 | 2253 | 1331 | 4919 | 360 | 20 | 0 | 0 | 0 | 0 | 61208 | 85661 | 586836 | 0 | 38 | 10219 | 126 | 37 | 196 | 268 | 14 | 0 | 0 | 0 | 0 | 0 | 17 | 34 | 0 | 0 | 237 | 344 | 0 |
| SRR6705826 | 21 | 20 | 0 | 0 | 31 | 752 | 359 | 1590 | 73 | 0 | 0 | 0 | 0 | 0 | 30883 | 28665 | 221733 | 0 | 29 | 1410 | 421 | 0 | 0 | 21 | 33 | 0 | 0 | 0 | 0 | 0 | 0 | 11 | 0 | 0 | 284 | 1212 | 0 |
| SRR6705827 | 109 | 42 | 0 | 0 | 395 | 387 | 218 | 899 | 35 | 25 | 42 | 0 | 0 | 0 | 28778 | 48024 | 574152 | 0 | 44 | 3210 | 1664 | 34 | 25 | 46 | 336 | 134 | 92 | 17 | 0 | 17 | 0 | 0 | 0 | 0 | 130 | 1198 | 0 |
| SRR6705828 | 0 | 0 | 0 | 0 | 33 | 24721 | 8994 | 68600 | 2895 | 0 | 11 | 0 | 0 | 0 | 9386 | 84862 | 227565 | 62 | 78 | 1243 | 409 | 35 | 42 | 33 | 0 | 0 | 0 | 0 | 0 | 15 | 0 | 56 | 0 | 0 | 2620 | 1018 | 0 |
| SRR6705829 | 79 | 75 | 0 | 0 | 99 | 451 | 276 | 968 | 50 | 12 | 18 | 0 | 0 | 0 | 79690 | 39398 | 361761 | 0 | 49 | 2781 | 1776 | 0 | 0 | 21 | 12 | 56 | 0 | 0 | 0 | 20 | 0 | 0 | 0 | 0 | 247 | 941 | 0 |
| SRR6705830 | 0 | 0 | 0 | 0 | 46 | 2077 | 362 | 1238 | 27 | 0 | 12 | 0 | 0 | 0 | 96306 | 84272 | 721009 | 0 | 129 | 534 | 1833 | 0 | 12 | 44 | 22 | 31 | 0 | 0 | 0 | 18 | 0 | 0 | 0 | 0 | 1388 | 902 | 0 |
| SRR6705831 | 0 | 0 | 0 | 0 | 41 | 3293 | 909 | 4021 | 211 | 0 | 19 | 0 | 0 | 0 | 30113 | 61766 | 275718 | 34 | 724 | 2143 | 1316 | 0 | 0 | 47 | 23 | 23 | 0 | 0 | 0 | 97 | 0 | 19 | 0 | 0 | 53322 | 585 | 0 |
| SRR6705832 | 0 | 0 | 0 | 0 | 12 | 1378 | 531 | 1584 | 69 | 0 | 0 | 0 | 0 | 0 | 33678 | 114982 | 516958 | 49 | 404 | 1880 | 443 | 0 | 0 | 20 | 0 | 0 | 0 | 0 | 0 | 17 | 0 | 30 | 0 | 0 | 15933 | 1391 | 0 |
| SRR6705833 | 0 | 0 | 0 | 0 | 31 | 6121 | 1301 | 3687 | 122 | 0 | 0 | 0 | 0 | 0 | 88795 | 85464 | 660337 | 26 | 226 | 978 | 1501 | 0 | 0 | 67 | 24 | 21 | 0 | 0 | 0 | 52 | 0 | 0 | 0 | 0 | 9602 | 709 | 0 |
| SRR6705834 | 16 | 15 | 0 | 0 | 45 | 817 | 733 | 3612 | 283 | 0 | 14 | 0 | 0 | 0 | 26034 | 46220 | 207098 | 0 | 819 | 6359 | 665 | 0 | 19 | 75 | 30 | 100 | 68 | 0 | 0 | 52 | 0 | 0 | 0 | 0 | 3496 | 3892 | 0 |
| SRR6705835 | 66 | 34 | 0 | 0 | 100 | 1115 | 763 | 3523 | 305 | 0 | 51 | 0 | 0 | 0 | 25400 | 71146 | 366469 | 10 | 717 | 8418 | 204 | 0 | 0 | 274 | 34 | 73 | 159 | 17 | 0 | 40 | 0 | 0 | 0 | 0 | 1290 | 3284 | 0 |
| SRR6705837 | 0 | 0 | 0 | 0 | 15 | 659 | 582 | 2547 | 80 | 0 | 12 | 0 | 0 | 0 | 43568 | 65908 | 442496 | 0 | 87 | 4683 | 415 | 0 | 0 | 277 | 23 | 27 | 102 | 0 | 0 | 40 | 0 | 0 | 0 | 0 | 1550 | 306 | 0 |
| SRR6705838 | 50 | 60 | 0 | 0 | 36 | 612 | 439 | 2725 | 114 | 0 | 26 | 0 | 0 | 0 | 43814 | 48926 | 213649 | 0 | 690 | 11945 | 1379 | 0 | 13 | 53 | 15 | 26 | 55 | 0 | 0 | 61 | 0 | 0 | 0 | 0 | 1657 | 569 | 0 |
| SRR6705839 | 25 | 11 | 0 | 0 | 10 | 3858 | 1906 | 11831 | 291 | 0 | 19 | 0 | 0 | 0 | 36631 | 65163 | 392383 | 0 | 112 | 1621 | 333 | 0 | 0 | 466 | 44 | 59 | 146 | 0 | 0 | 26 | 0 | 17 | 0 | 0 | 793 | 3765 | 0 |
| SRR6705840 | 32 | 38 | 0 | 0 | 178 | 1870 | 944 | 6289 | 276 | 0 | 141 | 0 | 0 | 0 | 27351 | 33889 | 228410 | 190 | 301 | 4041 | 1028 | 846 | 2389 | 457 | 273 | 470 | 281 | 0 | 0 | 227 | 38 | 126 | 0 | 0 | 4007 | 1395 | 0 |
| SRR6705841 | 27 | 31 | 0 | 0 | 27 | 170 | 129 | 950 | 0 | 0 | 0 | 0 | 0 | 0 | 49630 | 49751 | 468192 | 29 | 246 | 8295 | 1180 | 38 | 72 | 215 | 23 | 45 | 93 | 11 | 0 | 66 | 0 | 34 | 0 | 0 | 2217 | 3116 | 0 |

### Supp Table S8.3.Columns_BX_to_DH

| | | | | | | | | | | | | | | | | | | | | | | | | | | | | | | | | | | | | | |
|---|---|---|---|---|---|---|---|---|---|---|---|---|---|---|---|---|---|---|---|---|---|---|---|---|---|---|---|---|---|---|---|---|---|---|---|---|---|
| SRR6705842 | 44 | 34 | 0 | 0 | 16 | 471 | 262 | 1612 | 106 | 0 | 27 | 0 | 0 | 0 | 42482 | 36524 | 318282 | 0 | 1160 | 19382 | 2443 | 0 | 22 | 42 | 98 | 107 | 23 | 0 | 0 | 73 | 0 | 24 | 0 | 0 | 2949 | 260 | 0 |
| SRR6705843 | 32 | 41 | 0 | 0 | 64 | 639 | 393 | 2484 | 45 | 0 | 41 | 0 | 0 | 0 | 27864 | 86906 | 678309 | 0 | 64 | 1692 | 984 | 0 | 29 | 41 | 48 | 70 | 89 | 0 | 0 | 10 | 0 | 11 | 0 | 0 | 759 | 1015 | 0 |
| SRR6705844 | 0 | 0 | 0 | 0 | 45 | 300 | 105 | 1085 | 22 | 0 | 57 | 0 | 0 | 0 | 20643 | 87563 | 622133 | 25 | 87 | 1049 | 443 | 0 | 25 | 92 | 33 | 71 | 149 | 406 | 0 | 48 | 0 | 15 | 0 | 0 | 879 | 4964 | 0 |
| SRR6705845 | 58 | 61 | 17 | 22 | 68 | 667 | 325 | 2422 | 142 | 0 | 54 | 0 | 0 | 0 | 22043 | 58046 | 338797 | 49 | 413 | 2138 | 494 | 17 | 54 | 74 | 20 | 49 | 110 | 134 | 0 | 13 | 0 | 28 | 0 | 0 | 1824 | 8107 | 0 |
| SRR6705846 | 17 | 0 | 0 | 17 | 65 | 556 | 451 | 2557 | 174 | 0 | 50 | 0 | 0 | 0 | 13730 | 159610 | 349151 | 0 | 373 | 1740 | 359 | 0 | 17 | 580 | 55 | 158 | 370 | 29 | 0 | 76 | 0 | 21 | 0 | 0 | 1032 | 3375 | 0 |
| SRR6705847 | 0 | 22 | 0 | 0 | 238 | 479 | 358 | 1431 | 105 | 0 | 61 | 0 | 0 | 0 | 29164 | 107369 | 488938 | 11 | 269 | 1737 | 703 | 26 | 62 | 466 | 72 | 220 | 342 | 83 | 0 | 102 | 14 | 0 | 0 | 0 | 1103 | 1054 | 0 |
| SRR6705848 | 0 | 0 | 0 | 0 | 15 | 438 | 311 | 1221 | 85 | 0 | 16 | 0 | 0 | 0 | 9383 | 109175 | 284122 | 0 | 219 | 1370 | 297 | 0 | 0 | 44 | 0 | 13 | 22 | 24 | 0 | 18 | 0 | 28 | 0 | 0 | 638 | 2389 | 0 |
| SRR6705849 | 0 | 0 | 0 | 0 | 19 | 109 | 68 | 270 | 31 | 0 | 35 | 0 | 0 | 0 | 24278 | 113527 | 374753 | 157 | 362 | 2024 | 1693 | 0 | 35 | 17 | 24 | 11 | 20 | 17 | 0 | 16 | 0 | 321 | 0 | 0 | 1101 | 4160 | 0 |
| SRR6705850 | 0 | 16 | 0 | 0 | 51 | 750 | 359 | 2000 | 70 | 0 | 33 | 0 | 0 | 0 | 12993 | 60484 | 201660 | 16 | 344 | 2219 | 320 | 0 | 0 | 397 | 36 | 87 | 275 | 55 | 0 | 22 | 0 | 16 | 0 | 0 | 1980 | 4071 | 0 |
| SRR6705851 | 0 | 0 | 0 | 0 | 25 | 451 | 345 | 1867 | 47 | 0 | 27 | 0 | 0 | 0 | 17215 | 68748 | 289090 | 41 | 451 | 3078 | 550 | 0 | 11 | 214 | 20 | 25 | 50 | 25 | 0 | 36 | 0 | 23 | 0 | 0 | 1028 | 3296 | 0 |
| SRR6705852 | 0 | 11 | 0 | 11 | 63 | 831 | 230 | 2271 | 95 | 0 | 12 | 0 | 0 | 0 | 11389 | 73591 | 419718 | 28 | 367 | 2180 | 239 | 16 | 26 | 448 | 42 | 121 | 203 | 214 | 0 | 28 | 0 | 39 | 0 | 0 | 1855 | 2230 | 0 |
| SRR6705853 | 26 | 24 | 0 | 0 | 59 | 1697 | 435 | 3834 | 174 | 0 | 59 | 0 | 0 | 0 | 31270 | 54474 | 282357 | 123 | 644 | 1931 | 570 | 14 | 37 | 157 | 61 | 231 | 91 | 70 | 0 | 46 | 0 | 57 | 0 | 0 | 1370 | 3950 | 0 |
| SRR6705854 | 16 | 11 | 0 | 0 | 25 | 436 | 259 | 1143 | 58 | 13 | 10 | 0 | 0 | 0 | 48022 | 99634 | 471536 | 17 | 1175 | 2072 | 1261 | 21 | 42 | 30 | 29 | 50 | 50 | 0 | 0 | 13 | 0 | 11 | 0 | 0 | 1033 | 2125 | 0 |
| SRR6705855 | 12 | 0 | 0 | 0 | 71 | 417 | 216 | 1239 | 44 | 0 | 35 | 0 | 0 | 0 | 14191 | 82350 | 418405 | 19 | 941 | 2525 | 348 | 0 | 0 | 415 | 63 | 128 | 230 | 0 | 0 | 55 | 0 | 22 | 0 | 0 | 944 | 4763 | 0 |
| SRR6705856 | 210 | 155 | 12 | 0 | 158 | 298 | 202 | 1085 | 77 | 18 | 73 | 0 | 0 | 0 | 43075 | 54456 | 341987 | 0 | 600 | 6220 | 858 | 0 | 0 | 584 | 122 | 146 | 681 | 0 | 0 | 119 | 0 | 0 | 0 | 0 | 2442 | 1361 | 0 |
| SRR6705857 | 42 | 16 | 0 | 29 | 90 | 345 | 283 | 1181 | 87 | 0 | 11 | 0 | 0 | 0 | 67660 | 80092 | 552171 | 0 | 573 | 6593 | 922 | 0 | 0 | 131 | 13 | 26 | 66 | 14 | 0 | 93 | 0 | 0 | 0 | 0 | 768 | 2880 | 0 |
| SRR6705859 | 25 | 25 | 0 | 15 | 84 | 1617 | 601 | 2988 | 119 | 15 | 341 | 0 | 0 | 0 | 52454 | 94347 | 532890 | 0 | 52 | 1066 | 511 | 0 | 0 | 761 | 0 | 0 | 0 | 0 | 0 | 425 | 0 | 51 | 0 | 0 | 1451 | 2488 | 0 |
| SRR6705860 | 17 | 0 | 0 | 0 | 13 | 1669 | 1298 | 8374 | 274 | 10 | 41 | 0 | 0 | 0 | 16430 | 65467 | 298323 | 20 | 832 | 3843 | 515 | 10 | 0 | 55 | 13 | 30 | 19 | 0 | 0 | 33 | 0 | 91 | 0 | 0 | 19073 | 2163 | 0 |
| SRR6705861 | 24 | 0 | 0 | 0 | 17 | 130 | 114 | 761 | 12 | 0 | 22 | 0 | 0 | 0 | 37562 | 82884 | 541484 | 0 | 87 | 2199 | 1259 | 0 | 0 | 19 | 0 | 16 | 13 | 0 | 0 | 26 | 0 | 10 | 0 | 0 | 809 | 1457 | 0 |
| SRR6705862 | 24 | 22 | 0 | 0 | 31 | 269 | 197 | 834 | 35 | 0 | 39 | 0 | 0 | 0 | 64430 | 41696 | 357519 | 0 | 1174 | 16690 | 1687 | 0 | 0 | 26 | 34 | 47 | 20 | 0 | 0 | 48 | 0 | 0 | 0 | 0 | 1189 | 515 | 0 |
| SRR6705863 | 27 | 19 | 0 | 0 | 25 | 560 | 448 | 2624 | 85 | 33 | 113 | 0 | 0 | 0 | 83160 | 60428 | 532403 | 0 | 123 | 9186 | 1627 | 0 | 0 | 835 | 148 | 157 | 511 | 0 | 0 | 151 | 0 | 24 | 0 | 0 | 1200 | 3027 | 0 |
| SRR6705866 | 0 | 0 | 0 | 0 | 37 | 534 | 107 | 962 | 0 | 0 | 12 | 0 | 0 | 0 | 22123 | 65072 | 363775 | 0 | 267 | 2832 | 267 | 12 | 25 | 12 | 12 | 12 | 25 | 25 | 0 | 37 | 0 | 53 | 0 | 0 | 3243 | 2671 | 0 |
| SRR6705867 | 23 | 16 | 0 | 0 | 78 | 685 | 497 | 1182 | 137 | 0 | 116 | 0 | 0 | 0 | 34450 | 129941 | 422690 | 17 | 377 | 2929 | 480 | 0 | 47 | 116 | 0 | 70 | 70 | 31 | 0 | 171 | 0 | 34 | 0 | 0 | 2952 | 2967 | 0 |
| SRR6705868 | 0 | 11 | 0 | 0 | 11 | 523 | 296 | 979 | 159 | 0 | 17 | 0 | 0 | 0 | 18602 | 61841 | 321127 | 15 | 394 | 2898 | 197 | 0 | 19 | 22 | 12 | 19 | 26 | 0 | 0 | 40 | 0 | 0 | 0 | 0 | 9615 | 5524 | 0 |
| SRR6705869 | 38 | 56 | 0 | 0 | 28 | 302 | 192 | 715 | 52 | 0 | 27 | 0 | 0 | 0 | 36102 | 91460 | 475946 | 0 | 405 | 3737 | 265 | 0 | 56 | 22 | 13 | 44 | 21 | 15 | 0 | 50 | 0 | 15 | 0 | 0 | 1560 | 3579 | 13 |
| SRR6705870 | 0 | 11 | 0 | 0 | 13 | 626 | 525 | 3258 | 225 | 11 | 13 | 0 | 0 | 0 | 19427 | 138250 | 314282 | 18 | 561 | 3014 | 299 | 0 | 15 | 56 | 44 | 24 | 60 | 73 | 0 | 24 | 0 | 18 | 0 | 0 | 1598 | 4773 | 0 |
| SRR6705871 | 24 | 16 | 0 | 16 | 30 | 434 | 333 | 1113 | 63 | 14 | 12 | 0 | 0 | 0 | 27431 | 149053 | 271785 | 93 | 771 | 2946 | 813 | 0 | 26 | 87 | 22 | 39 | 120 | 86 | 0 | 12 | 0 | 291 | 0 | 0 | 1856 | 4185 | 0 |
| SRR6705874 | 0 | 0 | 0 | 0 | 0 | 537 | 701 | 1819 | 30 | 0 | 0 | 0 | 0 | 0 | 10301 | 80013 | 51751 | 0 | 45 | 1207 | 447 | 0 | 0 | 21 | 0 | 0 | 0 | 0 | 0 | 0 | 0 | 15 | 0 | 0 | 210 | 43 | 0 |
| SRR6705875 | 0 | 0 | 0 | 0 | 0 | 154 | 195 | 1111 | 15 | 0 | 0 | 0 | 0 | 0 | 20326 | 154794 | 126223 | 0 | 61 | 890 | 492 | 0 | 0 | 13 | 0 | 0 | 0 | 0 | 0 | 12 | 0 | 0 | 0 | 0 | 823 | 0 | 0 |
| SRR6705876 | 0 | 0 | 0 | 0 | 0 | 52 | 12 | 84 | 0 | 14 | 11 | 0 | 0 | 0 | 38804 | 291964 | 333582 | 0 | 0 | 128 | 1669 | 0 | 0 | 0 | 0 | 0 | 0 | 0 | 0 | 0 | 0 | 0 | 0 | 0 | 73 | 243 | 0 |
| SRR6705877 | 0 | 0 | 0 | 0 | 0 | 77 | 0 | 26 | 0 | 44 | 0 | 0 | 0 | 0 | 108930 | 14254 | 73904 | 0 | 0 | 115 | 1348 | 0 | 0 | 0 | 0 | 0 | 0 | 0 | 0 | 0 | 0 | 0 | 0 | 0 | 0 | 55 | 0 |
| SRR6705878 | 0 | 0 | 0 | 0 | 0 | 332 | 187 | 1810 | 15 | 11 | 10 | 0 | 0 | 0 | 17894 | 123638 | 205479 | 0 | 19 | 131 | 569 | 0 | 0 | 10 | 0 | 0 | 0 | 0 | 0 | 11 | 0 | 0 | 0 | 0 | 410 | 356 | 0 |
| SRR6705879 | 0 | 0 | 0 | 0 | 12 | 996 | 637 | 2851 | 239 | 0 | 17 | 0 | 0 | 0 | 9319 | 109563 | 75174 | 0 | 135 | 741 | 573 | 12 | 13 | 14 | 0 | 0 | 0 | 0 | 0 | 18 | 0 | 88 | 0 | 0 | 675 | 1042 | 0 |
| SRR6705880 | 0 | 0 | 0 | 0 | 0 | 2335 | 1511 | 10300 | 558 | 0 | 0 | 0 | 0 | 0 | 44049 | 58398 | 486757 | 0 | 77 | 3983 | 927 | 0 | 0 | 16 | 0 | 0 | 0 | 0 | 0 | 15 | 0 | 26 | 0 | 0 | 1820 | 613 | 0 |
| SRR6705881 | 0 | 0 | 0 | 0 | 0 | 976 | 1097 | 4333 | 428 | 0 | 0 | 0 | 0 | 0 | 21718 | 42080 | 419675 | 55 | 154 | 2676 | 263 | 0 | 0 | 22 | 0 | 0 | 0 | 0 | 0 | 0 | 0 | 22 | 0 | 0 | 9258 | 1179 | 0 |
| SRR6705882 | 0 | 0 | 0 | 0 | 0 | 3202 | 2780 | 14014 | 1035 | 0 | 0 | 0 | 0 | 0 | 67692 | 38357 | 525415 | 77 | 115 | 6729 | 997 | 0 | 0 | 14 | 0 | 0 | 0 | 0 | 0 | 0 | 0 | 0 | 0 | 0 | 6194 | 1876 | 0 |
| SRR6705883 | 0 | 0 | 0 | 0 | 13 | 351 | 188 | 2098 | 132 | 0 | 0 | 0 | 0 | 0 | 29197 | 63404 | 693732 | 0 | 65 | 4178 | 899 | 12 | 47 | 15 | 0 | 0 | 0 | 0 | 0 | 0 | 0 | 42 | 0 | 0 | 2378 | 1029 | 0 |
| SRR6705884 | 0 | 12 | 0 | 0 | 69 | 682 | 249 | 2303 | 100 | 0 | 19 | 0 | 0 | 0 | 19057 | 73210 | 316338 | 199 | 433 | 2405 | 913 | 15 | 41 | 86 | 35 | 116 | 88 | 135 | 0 | 26 | 0 | 233 | 0 | 0 | 1473 | 3132 | 0 |
| SRR6705885 | 0 | 13 | 0 | 0 | 14 | 518 | 141 | 2005 | 0 | 20 | 40 | 0 | 0 | 0 | 12926 | 97891 | 257227 | 151 | 254 | 1986 | 527 | 27 | 10 | 36 | 13 | 19 | 43 | 47 | 0 | 40 | 0 | 329 | 0 | 0 | 1460 | 2719 | 0 |
| SRR6705886 | 18 | 33 | 0 | 0 | 54 | 333 | 279 | 873 | 27 | 0 | 51 | 0 | 0 | 0 | 16055 | 85339 | 259287 | 45 | 315 | 2934 | 378 | 124 | 496 | 48 | 15 | 36 | 21 | 15 | 0 | 136 | 12 | 72 | 0 | 0 | 9274 | 2136 | 0 |
| SRR6705887 | 15 | 23 | 0 | 0 | 11 | 528 | 344 | 1133 | 34 | 0 | 12 | 0 | 0 | 0 | 16522 | 75919 | 274852 | 19 | 232 | 2575 | 286 | 27 | 109 | 21 | 22 | 23 | 37 | 15 | 0 | 18 | 0 | 19 | 0 | 0 | 6520 | 2326 | 0 |
| SRR6705888 | 21 | 21 | 0 | 0 | 14 | 386 | 259 | 1001 | 72 | 0 | 11 | 0 | 0 | 0 | 21749 | 103189 | 430523 | 48 | 350 | 2327 | 434 | 1429 | 6652 | 26 | 0 | 15 | 0 | 0 | 0 | 13 | 19 | 18 | 0 | 0 | 2553 | 3180 | 0 |
| SRR6705889 | 0 | 14 | 0 | 0 | 94 | 191 | 156 | 338 | 53 | 0 | 48 | 0 | 0 | 0 | 7575 | 79061 | 286528 | 13 | 289 | 2389 | 525 | 330 | 1076 | 53 | 0 | 21 | 41 | 20 | 0 | 118 | 0 | 27 | 0 | 0 | 3622 | 681 | 0 |
| SRR6705890 | 69 | 61 | 0 | 36 | 815 | 519 | 357 | 1983 | 100 | 0 | 79 | 0 | 0 | 0 | 32363 | 100208 | 411320 | 88 | 413 | 2177 | 2039 | 0 | 36 | 739 | 133 | 292 | 678 | 227 | 0 | 108 | 0 | 0 | 0 | 0 | 1387 | 2581 | 0 |
| SRR6705891 | 27 | 19 | 0 | 0 | 18 | 492 | 685 | 4263 | 479 | 0 | 31 | 0 | 0 | 0 | 12481 | 128943 | 329317 | 17 | 514 | 1935 | 278 | 0 | 0 | 74 | 10 | 37 | 65 | 24 | 0 | 41 | 0 | 39 | 0 | 0 | 1677 | 3919 | 0 |
| SRR6705892 | 13 | 28 | 0 | 0 | 223 | 102 | 86 | 371 | 39 | 0 | 38 | 0 | 0 | 0 | 30298 | 65458 | 550570 | 125 | 389 | 1651 | 1993 | 16 | 0 | 264 | 91 | 138 | 198 | 0 | 0 | 44 | 0 | 144 | 0 | 0 | 250 | 8211 | 0 |
| SRR6705893 | 10 | 0 | 0 | 0 | 28 | 342 | 444 | 2091 | 226 | 0 | 26 | 0 | 0 | 0 | 18440 | 86213 | 275724 | 55 | 444 | 2973 | 376 | 0 | 10 | 69 | 16 | 40 | 97 | 76 | 0 | 33 | 0 | 55 | 0 | 0 | 1404 | 4163 | 0 |
| SRR6705894 | 16 | 0 | 0 | 0 | 25 | 590 | 343 | 1555 | 15 | 0 | 0 | 0 | 0 | 0 | 45389 | 50244 | 270218 | 20 | 96 | 4680 | 730 | 12 | 62 | 14 | 10 | 0 | 0 | 0 | 0 | 0 | 0 | 17 | 0 | 0 | 980 | 115 | 0 |
| SRR6705895 | 104 | 0 | 0 | 0 | 91 | 1759 | 1351 | 3915 | 132 | 0 | 0 | 20 | 0 | 0 | 39225 | 44783 | 254466 | 40 | 40 | 3104 | 276 | 0 | 0 | 808 | 26 | 13 | 0 | 0 | 0 | 0 | 0 | 0 | 0 | 0 | 2311 | 1371 | 0 |
| SRR6705897 | 172 | 32 | 0 | 0 | 134 | 1770 | 1678 | 7576 | 68 | 0 | 0 | 0 | 0 | 0 | 44629 | 108052 | 297354 | 29 | 54 | 4392 | 112 | 0 | 0 | 433 | 19 | 0 | 0 | 0 | 0 | 51 | 0 | 0 | 0 | 0 | 1040 | 579 | 0 |
| SRR6705898 | 0 | 0 | 0 | 0 | 0 | 184 | 85 | 481 | 0 | 0 | 0 | 0 | 0 | 0 | 28759 | 56823 | 399600 | 0 | 105 | 3912 | 1119 | 0 | 0 | 0 | 0 | 0 | 0 | 0 | 0 | 12 | 0 | 0 | 0 | 0 | 255 | 0 | 0 |
| SRR6705899 | 0 | 0 | 0 | 0 | 14 | 618 | 1969 | 3764 | 154 | 0 | 0 | 0 | 0 | 0 | 10000 | 71204 | 227712 | 0 | 135 | 618 | 444 | 0 | 0 | 11 | 0 | 0 | 0 | 0 | 0 | 11 | 0 | 0 | 0 | 0 | 1520 | 0 | 0 |
| SRR6705900 | 0 | 0 | 0 | 0 | 0 | 796 | 256 | 497 | 0 | 0 | 0 | 0 | 0 | 0 | 24364 | 83617 | 254022 | 0 | 114 | 3182 | 582 | 0 | 0 | 0 | 0 | 0 | 0 | 0 | 0 | 0 | 0 | 0 | 0 | 0 | 246 | 33 | 0 |
| SRR6705901 | 28 | 18 | 0 | 0 | 0 | 201 | 227 | 649 | 21 | 0 | 0 | 0 | 0 | 0 | 53463 | 42046 | 225603 | 0 | 248 | 10250 | 1368 | 0 | 0 | 18 | 0 | 0 | 0 | 0 | 0 | 18 | 0 | 0 | 0 | 0 | 566 | 0 | 0 |
| SRR6705902 | 39 | 32 | 0 | 0 | 117 | 482 | 213 | 1365 | 23 | 0 | 95 | 0 | 0 | 0 | 46757 | 109258 | 631857 | 0 | 65 | 2780 | 862 | 0 | 0 | 374 | 138 | 300 | 791 | 18 | 0 | 321 | 0 | 20 | 0 | 0 | 296 | 3810 | 0 |
| SRR6705903 | 47 | 44 | 0 | 0 | 19 | 3046 | 1142 | 8006 | 435 | 12 | 13 | 0 | 0 | 0 | 31148 | 64044 | 387406 | 0 | 383 | 3457 | 7185 | 0 | 11 | 58 | 16 | 33 | 48 | 0 | 0 | 45 | 0 | 0 | 0 | 0 | 2117 | 1205 | 0 |
| SRR6726498 | 18 | 14 | 10 | 19 | 0 | 0 | 39 | 0 | 0 | 0 | 0 | 0 | 0 | 0 | 81087 | 11779 | 108027 | 0 | 0 | 1783 | 6018 | 0 | 0 | 0 | 0 | 55 | 78 | 0 | 0 | 0 | 0 | 0 | 0 | 0 | 6764 | 0 | 0 |
| SRR6726499 | 0 | 100 | 0 | 179 | 369 | 14 | 0 | 83 | 35 | 0 | 90 | 0 | 0 | 0 | 66503 | 23973 | 150841 | 14 | 14 | 2433 | 8373 | 0 | 0 | 249 | 130 | 458 | 319 | 110 | 0 | 0 | 0 | 0 | 0 | 0 | 5210 | 0 | 0 |
| SRR6757230 | 92 | 120 | 149 | 106 | 80 | 95 | 187 | 1275 | 95 | 0 | 56 | 0 | 0 | 0 | 28579 | 11160 | 100259 | 0 | 31 | 1860 | 25416 | 0 | 0 | 164 | 198 | 351 | 902 | 1034 | 0 | 120 | 0 | 14 | 0 | 0 | 2944 | 0 | 0 |
| SRR6757231 | 17 | 24 | 25 | 19 | 16 | 80 | 159 | 1261 | 86 | 0 | 12 | 0 | 0 | 0 | 29942 | 11525 | 102170 | 0 | 73 | 2104 | 26809 | 0 | 0 | 27 | 40 | 68 | 166 | 187 | 0 | 29 | 0 | 0 | 0 | 0 | 3267 | 0 | 0 |
| SRR6757232 | 10 | 19 | 16 | 13 | 14 | 241 | 328 | 2895 | 87 | 18 | 14 | 0 | 0 | 0 | 32516 | 10370 | 67863 | 0 | 53 | 1986 | 27869 | 0 | 0 | 25 | 21 | 40 | 97 | 24 | 0 | 44 | 0 | 0 | 0 | 0 | 1952 | 0 | 0 |
| SRR6757233 | 10 | 21 | 16 | 13 | 14 | 321 | 297 | 2756 | 92 | 10 | 15 | 0 | 0 | 0 | 32809 | 10610 | 71226 | 0 | 100 | 2299 | 28200 | 0 | 0 | 31 | 26 | 40 | 95 | 33 | 0 | 46 | 0 | 16 | 0 | 0 | 1865 | 0 | 0 |
| SRR6757234 | 47 | 48 | 14 | 16 | 49 | 147 | 215 | 2632 | 52 | 17 | 34 | 0 | 0 | 0 | 52463 | 4960 | 123429 | 0 | 50 | 1495 | 41460 | 0 | 0 | 15 | 18 | 39 | 102 | 112 | 0 | 25 | 0 | 0 | 0 | 0 | 826 | 0 | 0 |
| SRR6757235 | 30 | 40 | 16 | 16 | 39 | 125 | 206 | 2660 | 41 | 13 | 35 | 0 | 0 | 0 | 52676 | 5050 | 124307 | 0 | 50 | 1483 | 43677 | 0 | 0 | 15 | 17 | 38 | 110 | 117 | 0 | 22 | 0 | 0 | 0 | 0 | 824 | 0 | 0 |
| SRR6757236 | 24 | 21 | 14 | 10 | 15 | 158 | 222 | 3035 | 64 | 0 | 27 | 0 | 0 | 0 | 26916 | 5226 | 89819 | 0 | 77 | 1509 | 14066 | 0 | 0 | 20 | 21 | 44 | 122 | 11 | 0 | 34 | 0 | 0 | 0 | 0 | 1278 | 0 | 0 |
| SRR6757237 | 86 | 83 | 82 | 36 | 48 | 182 | 216 | 3004 | 72 | 0 | 29 | 0 | 0 | 0 | 27316 | 5573 | 92976 | 14 | 48 | 1509 | 14519 | 0 | 0 | 100 | 135 | 214 | 695 | 105 | 0 | 140 | 0 | 10 | 0 | 0 | 1254 | 0 | 0 |
| SRR6757238 | 29 | 38 | 57 | 56 | 35 | 0 | 25 | 25 | 37 | 0 | 28 | 0 | 0 | 0 | 35115 | 17926 | 117826 | 0 | 160 | 3544 | 45861 | 0 | 0 | 27 | 46 | 85 | 228 | 296 | 0 | 95 | 0 | 21 | 0 | 0 | 764 | 0 | 0 |
| SRR6757239 | 101 | 89 | 149 | 150 | 99 | 0 | 20 | 72 | 20 | 0 | 89 | 0 | 0 | 0 | 35002 | 17154 | 117321 | 0 | 92 | 3404 | 46906 | 0 | 0 | 25 | 132 | 216 | 615 | 828 | 0 | 248 | 0 | 10 | 0 | 0 | 854 | 0 | 0 |
| SRR6757240 | 136 | 165 | 293 | 258 | 175 | 0 | 37 | 452 | 51 | 12 | 69 | 0 | 0 | 0 | 56478 | 16909 | 154113 | 0 | 144 | 2878 | 36263 | 0 | 0 | 78 | 581 | 901 | 1737 | 2448 | 0 | 285 | 0 | 0 | 0 | 0 | 300 | 0 | 0 |
| SRR6757241 | 36 | 43 | 72 | 62 | 41 | 10 | 35 | 499 | 62 | 11 | 20 | 0 | 0 | 0 | 60272 | 16957 | 155110 | 0 | 158 | 3110 | 39892 | 0 | 0 | 19 | 145 | 231 | 425 | 588 | 0 | 74 | 0 | 0 | 0 | 0 | 333 | 0 | 0 |
| SRR6757242 | 82 | 141 | 145 | 185 | 105 | 12 | 96 | 313 | 80 | 0 | 53 | 0 | 0 | 0 | 39648 | 18375 | 84214 | 0 | 100 | 3231 | 33560 | 0 | 0 | 73 | 251 | 393 | 745 | 877 | 0 | 138 | 0 | 0 | 0 | 0 | 2984 | 0 | 0 |
| SRR6757243 | 20 | 33 | 34 | 47 | 27 | 28 | 84 | 321 | 84 | 0 | 16 | 0 | 0 | 0 | 40340 | 19194 | 86693 | 0 | 124 | 3220 | 36918 | 0 | 0 | 23 | 55 | 103 | 183 | 213 | 0 | 39 | 0 | 11 | 0 | 0 | 3010 | 0 | 0 |
| SRR6757244 | 90 | 124 | 152 | 166 | 92 | 22 | 103 | 603 | 124 | 0 | 102 | 0 | 0 | 0 | 32837 | 11180 | 135501 | 0 | 89 | 2615 | 67214 | 0 | 0 | 69 | 280 | 0 | 12 | 0 | 0 | 318 | 0 | 0 | 0 | 0 | 2065 | 0 | 0 |
| SRR6757245 | 77 | 113 | 158 | 152 | 84 | 21 | 69 | 577 | 135 | 0 | 87 | 0 | 0 | 0 | 30004 | 11109 | 137548 | 0 | 102 | 2490 | 65573 | 0 | 0 | 90 | 296 | 22 | 15 | 0 | 0 | 355 | 0 | 0 | 0 | 0 | 2154 | 0 | 0 |
| SRR6757246 | 51 | 77 | 73 | 80 | 52 | 320 | 435 | 5795 | 184 | 13 | 49 | 0 | 0 | 0 | 29366 | 5806 | 70659 | 0 | 101 | 1795 | 22953 | 0 | 0 | 128 | 155 | 279 | 705 | 747 | 0 | 133 | 0 | 0 | 0 | 0 | 1687 | 0 | 0 |
| SRR6757247 | 65 | 67 | 91 | 74 | 53 | 300 | 385 | 5561 | 169 | 17 | 61 | 0 | 0 | 0 | 28325 | 5731 | 71536 | 0 | 80 | 1779 | 22422 | 0 | 0 | 99 | 139 | 270 | 709 | 734 | 0 | 132 | 0 | 0 | 0 | 0 | 1607 | 0 | 0 |
| SRR6757248 | 56 | 62 | 109 | 95 | 84 | 229 | 294 | 3330 | 173 | 11 | 59 | 0 | 0 | 0 | 31217 | 12919 | 70223 | 0 | 67 | 2073 | 36318 | 0 | 0 | 101 | 176 | 334 | 586 | 826 | 0 | 113 | 0 | 0 | 0 | 0 | 2009 | 0 | 0 |
| SRR6757249 | 53 | 71 | 101 | 95 | 80 | 203 | 352 | 3589 | 126 | 0 | 58 | 0 | 0 | 0 | 31606 | 13305 | 72146 | 0 | 77 | 2100 | 39541 | 0 | 0 | 102 | 203 | 328 | 593 | 835 | 0 | 131 | 0 | 0 | 0 | 0 | 1975 | 0 | 0 |
| SRR6757250 | 11 | 16 | 17 | 23 | 13 | 117 | 220 | 1752 | 106 | 15 | 53 | 0 | 0 | 0 | 32415 | 11453 | 99563 | 0 | 53 | 1914 | 29625 | 0 | 0 | 25 | 37 | 58 | 121 | 136 | 0 | 22 | 0 | 0 | 0 | 0 | 2646 | 0 | 0 |
| SRR6757251 | 56 | 93 | 99 | 103 | 89 | 98 | 207 | 1702 | 98 | 23 | 60 | 0 | 0 | 0 | 31541 | 11306 | 102285 | 0 | 45 | 1962 | 30690 | 0 | 0 | 162 | 183 | 369 | 812 | 850 | 0 | 132 | 0 | 0 | 0 | 0 | 2678 | 0 | 0 |
| SRR6757252 | 70 | 50 | 101 | 130 | 82 | 32 | 144 | 918 | 59 | 0 | 67 | 0 | 0 | 0 | 39705 | 14760 | 137535 | 11 | 74 | 2274 | 47955 | 0 | 0 | 127 | 176 | 320 | 522 | 958 | 0 | 175 | 0 | 0 | 0 | 0 | 3472 | 0 | 0 |
| SRR6757253 | 16 | 14 | 18 | 26 | 18 | 26 | 89 | 1063 | 41 | 0 | 15 | 0 | 0 | 0 | 40557 | 15364 | 140084 | 0 | 37 | 2344 | 51848 | 0 | 0 | 23 | 36 | 63 | 114 | 209 | 0 | 40 | 0 | 0 | 0 | 0 | 3403 | 0 | 0 |
| SRR6757254 | 20 | 24 | 38 | 28 | 13 | 108 | 153 | 532 | 77 | 10 | 15 | 0 | 0 | 0 | 44787 | 13657 | 86622 | 0 | 61 | 2516 | 41811 | 0 | 0 | 17 | 41 | 58 | 151 | 138 | 0 | 43 | 0 | 0 | 0 | 0 | 1522 | 0 | 0 |
| SRR6757255 | 111 | 145 | 170 | 149 | 80 | 110 | 133 | 533 | 56 | 0 | 70 | 0 | 0 | 0 | 45095 | 13507 | 87484 | 10 | 71 | 2452 | 43421 | 0 | 0 | 95 | 210 | 346 | 819 | 723 | 0 | 236 | 0 | 0 | 0 | 0 | 1539 | 0 | 0 |
| SRR6757256 | 123 | 147 | 136 | 181 | 74 | 388 | 345 | 3406 | 174 | 12 | 47 | 0 | 0 | 0 | 38444 | 9870 | 54801 | 15 | 123 | 2521 | 34816 | 0 | 0 | 47 | 193 | 269 | 594 | 753 | 0 | 113 | 0 | 0 | 0 | 0 | 991 | 0 | 0 |
| SRR6757257 | 26 | 35 | 32 | 44 | 16 | 349 | 359 | 3170 | 173 | 13 | 13 | 0 | 0 | 0 | 38964 | 10497 | 56552 | 0 | 85 | 2406 | 36150 | 0 | 0 | 11 | 50 | 66 | 153 | 182 | 0 | 28 | 0 | 0 | 0 | 0 | 964 | 0 | 0 |
| SRR6757258 | 56 | 49 | 167 | 120 | 84 | 39 | 93 | 1269 | 30 | 0 | 99 | 0 | 0 | 0 | 58214 | 8824 | 212314 | 0 | 50 | 1896 | 16720 | 0 | 0 | 64 | 214 | 290 | 716 | 1305 | 0 | 222 | 0 | 0 | 0 | 0 | 1306 | 0 | 0 |
| SRR6757259 | 49 | 46 | 139 | 104 | 97 | 40 | 69 | 1250 | 38 | 0 | 89 | 0 | 0 | 0 | 60552 | 8903 | 211772 | 0 | 38 | 2016 | 18291 | 0 | 0 | 68 | 203 | 349 | 686 | 1183 | 0 | 203 | 0 | 0 | 0 | 0 | 1317 | 0 | 0 |
| SRR6757260 | 19 | 25 | 31 | 25 | 17 | 172 | 156 | 1432 | 84 | 0 | 14 | 0 | 0 | 0 | 27505 | 14149 | 76558 | 0 | 49 | 2061 | 45685 | 0 | 0 | 14 | 34 | 69 | 158 | 165 | 0 | 36 | 0 | 13 | 0 | 0 | 2226 | 0 | 0 |
| SRR6757261 | 22 | 20 | 30 | 28 | 16 | 134 | 158 | 1390 | 88 | 0 | 15 | 0 | 0 | 0 | 27897 | 14501 | 77670 | 15 | 111 | 2138 | 46881 | 0 | 0 | 12 | 32 | 61 | 165 | 178 | 0 | 34 | 0 | 0 | 0 | 0 | 2300 | 0 | 0 |
| SRR6757262 | 286 | 317 | 281 | 245 | 71 | 14 | 41 | 87 | 30 | 0 | 64 | 0 | 0 | 0 | 25939 | 14245 | 98809 | 14 | 98 | 2520 | 38104 | 0 | 0 | 59 | 367 | 702 | 1921 | 1833 | 0 | 300 | 0 | 11 | 0 | 0 | 1653 | 0 | 0 |
| SRR6757263 | 239 | 273 | 302 | 226 | 76 | 23 | 56 | 130 | 31 | 0 | 93 | 0 | 0 | 0 | 27000 | 14606 | 99751 | 14 | 62 | 2646 | 39351 | 0 | 0 | 74 | 372 | 648 | 1845 | 1778 | 0 | 289 | 0 | 0 | 0 | 0 | 1630 | 0 | 0 |
| SRR6757264 | 208 | 248 | 250 | 322 | 87 | 80 | 91 | 393 | 36 | 0 | 59 | 0 | 0 | 0 | 28943 | 14556 | 82928 | 0 | 120 | 2518 | 46903 | 0 | 0 | 69 | 281 | 380 | 817 | 874 | 0 | 97 | 0 | 0 | 0 | 0 | 671 | 0 | 0 |
| SRR6757265 | 212 | 241 | 246 | 279 | 99 | 102 | 67 | 418 | 45 | 0 | 58 | 0 | 0 | 0 | 31056 | 14985 | 84817 | 0 | 89 | 2838 | 52826 | 0 | 0 | 78 | 303 | 411 | 809 | 883 | 0 | 99 | 0 | 0 | 0 | 0 | 659 | 0 | 0 |
| SRR6757266 | 115 | 231 | 182 | 183 | 78 | 15 | 45 | 253 | 57 | 0 | 92 | 0 | 0 | 0 | 21228 | 11086 | 65979 | 0 | 57 | 2926 | 47428 | 0 | 0 | 49 | 608 | 1158 | 3695 | 2444 | 0 | 318 | 0 | 0 | 0 | 0 | 557 | 0 | 0 |
| SRR6757267 | 113 | 210 | 183 | 190 | 68 | 17 | 34 | 195 | 47 | 0 | 100 | 0 | 0 | 0 | 20654 | 10575 | 68369 | 10 | 94 | 2596 | 46360 | 0 | 0 | 57 | 578 | 1062 | 3483 | 2256 | 0 | 346 | 0 | 0 | 0 | 0 | 571 | 0 | 0 |
| SRR6757268 | 75 | 106 | 164 | 140 | 66 | 21 | 69 | 532 | 69 | 0 | 94 | 0 | 0 | 0 | 31658 | 11215 | 133510 | 0 | 30 | 1782 | 32534 | 0 | 0 | 51 | 217 | 518 | 1332 | 1856 | 0 | 299 | 0 | 0 | 0 | 0 | 2095 | 0 | 0 |
| SRR6757269 | 16 | 17 | 24 | 19 | 10 | 18 | 89 | 508 | 70 | 0 | 18 | 0 | 0 | 0 | 30236 | 11285 | 135594 | 0 | 37 | 1804 | 32900 | 0 | 0 | 43 | 33 | 86 | 220 | 315 | 0 | 51 | 0 | 0 | 0 | 0 | 2196 | 0 | 0 |
| SRR6757270 | 60 | 13 | 19 | 15 | 53 | 58 | 133 | 1132 | 58 | 0 | 11 | 0 | 0 | 0 | 31650 | 9019 | 126748 | 0 | 42 | 1963 | 23903 | 0 | 0 | 47 | 32 | 67 | 192 | 210 | 0 | 28 | 0 | 0 | 0 | 0 | 1336 | 0 | 0 |
| SRR6757271 | 11 | 15 | 17 | 16 | 57 | 85 | 99 | 1137 | 34 | 0 | 12 | 0 | 0 | 0 | 32309 | 9362 | 128747 | 0 | 44 | 2034 | 25826 | 0 | 0 | 49 | 36 | 66 | 198 | 226 | 0 | 34 | 0 | 0 | 0 | 0 | 1308 | 0 | 0 |
| SRR6757272 | 62 | 100 | 146 | 160 | 66 | 182 | 171 | 2187 | 82 | 0 | 79 | 0 | 0 | 0 | 31105 | 10196 | 95303 | 0 | 39 | 1562 | 26433 | 0 | 0 | 61 | 235 | 379 | 843 | 1122 | 0 | 195 | 0 | 0 | 0 | 0 | 1139 | 0 | 0 |
| SRR6757273 | 76 | 73 | 134 | 128 | 80 | 223 | 153 | 2374 | 103 | 0 | 89 | 0 | 0 | 0 | 32367 | 10294 | 97252 | 0 | 57 | 1717 | 29596 | 0 | 0 | 69 | 224 | 406 | 849 | 1132 | 0 | 204 | 0 | 0 | 0 | 0 | 1011 | 0 | 0 |
| SRR6757274 | 110 | 187 | 138 | 208 | 73 | 175 | 166 | 2024 | 90 | 0 | 72 | 0 | 0 | 0 | 26130 | 8532 | 64712 | 0 | 45 | 1451 | 28254 | 0 | 0 | 58 | 327 | 615 | 1366 | 985 | 0 | 232 | 0 | 0 | 0 | 0 | 611 | 0 | 0 |
| SRR6757275 | 18 | 30 | 22 | 31 | 12 | 221 | 218 | 2134 | 43 | 0 | 13 | 0 | 0 | 0 | 26554 | 8792 | 66016 | 0 | 56 | 1398 | 30541 | 0 | 0 | 10 | 56 | 92 | 216 | 163 | 0 | 36 | 0 | 0 | 0 | 0 | 658 | 0 | 0 |
| SRR6757276 | 72 | 11 | 17 | 18 | 11 | 35 | 87 | 720 | 41 | 0 | 12 | 0 | 0 | 0 | 28139 | 12818 | 153113 | 11 | 33 | 1891 | 26794 | 0 | 0 | 79 | 34 | 76 | 187 | 256 | 0 | 35 | 0 | 0 | 0 | 0 | 2372 | 0 | 0 |
| SRR6757277 | 76 | 102 | 158 | 145 | 93 | 55 | 44 | 854 | 51 | 0 | 110 | 0 | 0 | 0 | 29453 | 13021 | 156139 | 0 | 16 | 1835 | 29686 | 0 | 0 | 83 | 279 | 622 | 1611 | 2043 | 0 | 272 | 0 | 0 | 0 | 0 | 2478 | 0 | 0 |
| SRR6757373 | 0 | 0 | 0 | 0 | 0 | 1383 | 976 | 6126 | 197 | 30 | 0 | 0 | 0 | 0 | 27126 | 68904 | 299270 | 366 | 25 | 10478 | 4637 | 0 | 0 | 104 | 0 | 0 | 0 | 0 | 0 | 0 | 0 | 0 | 0 | 0 | 3593 | 72 | 0 |
| SRR6757374 | 0 | 0 | 0 | 0 | 30 | 1083 | 759 | 5211 | 183 | 0 | 0 | 0 | 0 | 0 | 57680 | 52835 | 199046 | 173 | 0 | 2170 | 1879 | 0 | 0 | 560 | 0 | 0 | 11 | 0 | 0 | 0 | 0 | 0 | 0 | 0 | 9502 | 348 | 0 |
| SRR6757375 | 0 | 0 | 0 | 0 | 0 | 2744 | 989 | 5376 | 59 | 0 | 0 | 0 | 0 | 0 | 1391 | 13476 | 31028 | 0 | 0 | 706 | 290 | 0 | 0 | 39 | 0 | 0 | 0 | 0 | 0 | 0 | 0 | 0 | 0 | 0 | 17100 | 95 | 0 |
| SRR6757376 | 0 | 0 | 0 | 0 | 0 | 2105 | 945 | 5271 | 89 | 29 | 0 | 0 | 0 | 0 | 49369 | 36935 | 281520 | 0 | 0 | 1857 | 3606 | 0 | 0 | 438 | 0 | 0 | 0 | 0 | 0 | 12 | 0 | 0 | 0 | 0 | 11324 | 472 | 0 |
| SRR6757377 | 0 | 0 | 0 | 0 | 0 | 1506 | 1041 | 5356 | 93 | 14 | 0 | 0 | 0 | 0 | 4603 | 34278 | 89759 | 0 | 0 | 4975 | 355 | 0 | 0 | 109 | 0 | 0 | 0 | 0 | 0 | 0 | 0 | 0 | 0 | 0 | 19105 | 83 | 0 |
| SRR6757378 | 0 | 0 | 0 | 0 | 59 | 36 | 61 | 190 | 27 | 39 | 0 | 0 | 0 | 0 | 78863 | 28164 | 351853 | 50 | 31 | 7655 | 4159 | 0 | 0 | 199 | 16 | 17 | 55 | 0 | 0 | 44 | 0 | 0 | 0 | 0 | 776 | 256 | 0 |
| SRR6757379 | 0 | 0 | 0 | 0 | 21 | 135 | 443 | 1622 | 174 | 21 | 0 | 0 | 0 | 0 | 17059 | 105332 | 261816 | 79 | 16 | 20873 | 1575 | 0 | 0 | 63 | 0 | 0 | 0 | 0 | 0 | 0 | 0 | 0 | 0 | 0 | 5101 | 121 | 0 |
| SRR6757380 | 0 | 0 | 0 | 0 | 41 | 189 | 273 | 965 | 142 | 0 | 0 | 0 | 0 | 0 | 41807 | 70679 | 281087 | 91 | 0 | 13624 | 3426 | 0 | 0 | 204 | 0 | 0 | 16 | 0 | 0 | 30 | 0 | 11 | 0 | 0 | 1450 | 870 | 0 |
| SRR6757381 | 0 | 0 | 0 | 0 | 25 | 111 | 616 | 4649 | 257 | 27 | 0 | 0 | 0 | 0 | 17919 | 155587 | 174430 | 15 | 15 | 14002 | 1282 | 0 | 0 | 15 | 0 | 0 | 0 | 0 | 0 | 18 | 0 | 15 | 0 | 0 | 4417 | 75 | 0 |
| SRR6757382 | 0 | 0 | 0 | 0 | 14 | 787 | 1614 | 3806 | 1025 | 22 | 0 | 0 | 0 | 0 | 67669 | 71320 | 217641 | 28 | 34 | 4327 | 1053 | 0 | 0 | 559 | 0 | 0 | 17 | 0 | 0 | 14 | 0 | 0 | 0 | 0 | 5791 | 275 | 0 |
| SRR6757383 | 0 | 0 | 0 | 0 | 0 | 1573 | 864 | 7911 | 183 | 22 | 0 | 0 | 0 | 0 | 5679 | 70433 | 160155 | 37 | 22 | 7552 | 512 | 0 | 0 | 148 | 0 | 0 | 0 | 0 | 0 | 0 | 0 | 0 | 0 | 0 | 25057 | 90 | 0 |
| SRR6757384 | 0 | 0 | 0 | 0 | 18 | 4354 | 3813 | 34008 | 3220 | 0 | 0 | 0 | 0 | 0 | 2831 | 14219 | 91385 | 0 | 0 | 1605 | 215 | 0 | 0 | 507 | 0 | 12 | 62 | 25 | 0 | 17 | 0 | 0 | 0 | 0 | 37422 | 899 | 0 |
| SRR6757385 | 0 | 0 | 0 | 0 | 0 | 4658 | 1201 | 8030 | 145 | 66 | 0 | 0 | 0 | 0 | 24127 | 81423 | 221534 | 54 | 0 | 6366 | 3199 | 0 | 0 | 305 | 0 | 0 | 0 | 0 | 0 | 11 | 0 | 0 | 0 | 0 | 5191 | 58 | 10 |
| SRR6757386 | 0 | 0 | 0 | 0 | 16 | 356 | 197 | 1135 | 56 | 18 | 0 | 0 | 0 | 0 | 84436 | 55227 | 324685 | 232 | 35 | 4652 | 6323 | 0 | 0 | 857 | 0 | 0 | 15 | 14 | 0 | 0 | 0 | 0 | 0 | 0 | 4995 | 460 | 0 |
| SRR6757387 | 0 | 0 | 0 | 0 | 0 | 2010 | 2590 | 31570 | 360 | 13 | 0 | 0 | 0 | 0 | 8806 | 34533 | 158595 | 28 | 0 | 2365 | 890 | 0 | 0 | 135 | 0 | 0 | 0 | 0 | 0 | 0 | 0 | 0 | 0 | 0 | 17505 | 67 | 0 |
| SRR6757388 | 0 | 0 | 0 | 0 | 0 | 761 | 557 | 3816 | 238 | 64 | 0 | 0 | 0 | 0 | 16486 | 35848 | 313478 | 238 | 54 | 5378 | 9071 | 0 | 0 | 119 | 19 | 13 | 26 | 0 | 0 | 35 | 0 | 18 | 0 | 0 | 2399 | 327 | 12 |
| SRR6757389 | 0 | 0 | 0 | 0 | 26 | 403 | 262 | 2192 | 94 | 11 | 0 | 0 | 0 | 0 | 16611 | 42603 | 129691 | 27 | 0 | 21492 | 1781 | 0 | 0 | 98 | 0 | 0 | 0 | 0 | 0 | 18 | 0 | 0 | 0 | 0 | 7164 | 57 | 0 |
| SRR6757390 | 19 | 18 | 0 | 23 | 174 | 42 | 23 | 141 | 0 | 55 | 24 | 0 | 0 | 0 | 242782 | 81539 | 372623 | 37 | 10 | 1460 | 5736 | 0 | 0 | 16 | 23 | 64 | 179 | 0 | 0 | 153 | 0 | 0 | 0 | 0 | 2158 | 185 | 0 |

### Supp Table S8.3.Columns_BX_to_DH

| | | | | | | | | | | | | | | | | | | | | | | | | | | | | | | | | | | | | | |
|---|---|---|---|---|---|---|---|---|---|---|---|---|---|---|---|---|---|---|---|---|---|---|---|---|---|---|---|---|---|---|---|---|---|---|---|---|---|
| SRR6757391 | 0 | 0 | 0 | 0 | 11 | 2230 | 1177 | 8007 | 242 | 0 | 0 | 0 | 0 | 0 | 15134 | 90785 | 207810 | 31 | 23 | 3704 | 1341 | 0 | 0 | 39 | 0 | 0 | 0 | 0 | 0 | 0 | 0 | 0 | 0 | 0 | 10645 | 131 | 0 |
| SRR6757392 | 0 | 0 | 0 | 0 | 14 | 438 | 306 | 1741 | 114 | 13 | 0 | 0 | 0 | 0 | 84923 | 50951 | 376620 | 163 | 73 | 7318 | 3470 | 0 | 0 | 251 | 0 | 27 | 66 | 25 | 0 | 16 | 0 | 0 | 0 | 0 | 1831 | 289 | 0 |
| SRR6757393 | 0 | 0 | 0 | 0 | 0 | 5280 | 1758 | 15761 | 213 | 0 | 0 | 0 | 0 | 0 | 9129 | 45212 | 157005 | 93 | 0 | 3629 | 1309 | 0 | 0 | 106 | 0 | 0 | 0 | 0 | 0 | 0 | 0 | 0 | 0 | 0 | 9569 | 91 | 0 |
| SRR6757394 | 0 | 0 | 0 | 0 | 0 | 1243 | 582 | 2186 | 56 | 20 | 0 | 0 | 0 | 0 | 25734 | 14006 | 303868 | 0 | 75 | 3546 | 629 | 0 | 0 | 115 | 0 | 0 | 18 | 0 | 0 | 0 | 0 | 0 | 0 | 0 | 3362 | 784 | 0 |
| SRR6757395 | 0 | 11 | 0 | 0 | 85 | 351 | 212 | 2023 | 102 | 24 | 0 | 0 | 0 | 0 | 27103 | 109488 | 286312 | 65 | 189 | 45333 | 3169 | 0 | 0 | 148 | 0 | 10 | 36 | 0 | 0 | 39 | 0 | 0 | 0 | 0 | 2403 | 151 | 0 |
| SRR6757396 | 0 | 0 | 0 | 0 | 75 | 199 | 203 | 1400 | 94 | 20 | 0 | 0 | 0 | 0 | 35813 | 81757 | 340367 | 58 | 36 | 20011 | 3511 | 0 | 0 | 580 | 25 | 41 | 116 | 20 | 0 | 36 | 0 | 0 | 0 | 0 | 1415 | 767 | 0 |
| SRR6757397 | 0 | 0 | 0 | 0 | 0 | 602 | 690 | 2430 | 103 | 0 | 0 | 0 | 0 | 0 | 6974 | 60412 | 116156 | 132 | 0 | 5733 | 440 | 0 | 0 | 22 | 0 | 0 | 0 | 0 | 0 | 0 | 0 | 29 | 0 | 0 | 14826 | 134 | 0 |
| SRR6757398 | 0 | 0 | 0 | 0 | 21 | 454 | 506 | 2034 | 103 | 17 | 0 | 0 | 0 | 0 | 26631 | 54908 | 115591 | 57 | 0 | 1390 | 460 | 0 | 0 | 342 | 0 | 0 | 24 | 0 | 0 | 19 | 0 | 0 | 0 | 0 | 14598 | 323 | 0 |
| SRR6757399 | 0 | 0 | 0 | 0 | 0 | 700 | 1215 | 3436 | 123 | 20 | 0 | 0 | 0 | 0 | 13414 | 67962 | 183499 | 0 | 0 | 2418 | 2467 | 0 | 0 | 29 | 0 | 0 | 0 | 0 | 0 | 0 | 0 | 0 | 0 | 0 | 10997 | 74 | 0 |
| SRR6757400 | 0 | 0 | 0 | 0 | 31 | 263 | 403 | 1962 | 107 | 22 | 0 | 0 | 0 | 0 | 204798 | 87140 | 380528 | 26 | 18 | 2529 | 2828 | 0 | 0 | 19 | 0 | 0 | 15 | 0 | 0 | 14 | 0 | 14 | 0 | 0 | 1776 | 286 | 0 |
| SRR6757401 | 0 | 0 | 0 | 0 | 24 | 2295 | 897 | 8519 | 211 | 20 | 0 | 0 | 0 | 0 | 20287 | 89186 | 220659 | 189 | 0 | 5719 | 1279 | 0 | 0 | 69 | 0 | 0 | 16 | 0 | 0 | 13 | 0 | 0 | 0 | 0 | 4707 | 115 | 0 |
| SRR6757402 | 0 | 0 | 0 | 0 | 11 | 409 | 368 | 1508 | 68 | 0 | 0 | 0 | 0 | 0 | 56290 | 96459 | 294147 | 351 | 27 | 10845 | 1479 | 0 | 0 | 145 | 0 | 0 | 0 | 0 | 0 | 0 | 0 | 0 | 0 | 0 | 5936 | 291 | 0 |
| SRR6757403 | 0 | 0 | 0 | 0 | 10 | 2065 | 885 | 10566 | 255 | 21 | 0 | 0 | 0 | 0 | 14099 | 91307 | 236358 | 134 | 27 | 11531 | 2977 | 0 | 0 | 11 | 0 | 0 | 0 | 0 | 0 | 0 | 0 | 0 | 0 | 0 | 6122 | 109 | 0 |
| SRR6757404 | 0 | 0 | 0 | 0 | 72 | 150 | 104 | 797 | 39 | 24 | 0 | 0 | 0 | 0 | 168514 | 77867 | 409762 | 117 | 70 | 2005 | 1796 | 0 | 0 | 18 | 0 | 17 | 34 | 0 | 0 | 16 | 0 | 0 | 0 | 0 | 738 | 250 | 0 |
| SRR6757405 | 0 | 0 | 0 | 0 | 10 | 345 | 564 | 3163 | 127 | 0 | 0 | 0 | 0 | 0 | 13588 | 106314 | 251026 | 45 | 0 | 16787 | 3054 | 0 | 0 | 38 | 0 | 0 | 0 | 0 | 0 | 0 | 0 | 0 | 0 | 0 | 7150 | 69 | 0 |
| SRR6757406 | 0 | 0 | 0 | 0 | 12 | 289 | 423 | 1514 | 141 | 0 | 0 | 0 | 0 | 0 | 23869 | 91482 | 390063 | 89 | 15 | 12046 | 6643 | 0 | 0 | 57 | 0 | 0 | 0 | 0 | 0 | 12 | 0 | 0 | 0 | 0 | 2412 | 168 | 0 |
| SRR6757407 | 0 | 0 | 0 | 0 | 0 | 1390 | 1066 | 10065 | 247 | 12 | 0 | 0 | 0 | 0 | 3582 | 72535 | 95052 | 43 | 26 | 5152 | 452 | 0 | 0 | 45 | 0 | 0 | 0 | 0 | 0 | 0 | 0 | 0 | 0 | 0 | 17044 | 100 | 0 |
| SRR6757408 | 0 | 0 | 0 | 0 | 0 | 522 | 370 | 2362 | 71 | 11 | 0 | 0 | 0 | 0 | 53432 | 50614 | 234919 | 412 | 39 | 5841 | 1762 | 0 | 0 | 56 | 0 | 0 | 0 | 0 | 0 | 0 | 0 | 0 | 0 | 0 | 10748 | 234 | 0 |
| SRR6757409 | 0 | 0 | 0 | 0 | 11 | 4504 | 1173 | 11069 | 56 | 0 | 0 | 0 | 0 | 0 | 29325 | 59591 | 334980 | 145 | 14 | 10566 | 3506 | 0 | 0 | 99 | 0 | 0 | 0 | 0 | 0 | 0 | 0 | 0 | 0 | 0 | 3606 | 87 | 0 |
| SRR6757410 | 0 | 0 | 0 | 0 | 64 | 1040 | 712 | 5780 | 153 | 0 | 0 | 0 | 0 | 0 | 41600 | 58727 | 311470 | 279 | 18 | 6280 | 2513 | 0 | 0 | 16 | 0 | 12 | 28 | 46 | 0 | 29 | 0 | 0 | 0 | 0 | 6073 | 354 | 0 |
| SRR6757411 | 0 | 0 | 0 | 0 | 12 | 384 | 491 | 3699 | 107 | 23 | 0 | 0 | 0 | 0 | 24622 | 85250 | 224182 | 15 | 0 | 12050 | 1596 | 0 | 0 | 84 | 0 | 0 | 0 | 0 | 0 | 0 | 0 | 15 | 0 | 0 | 4306 | 102 | 0 |
| SRR6757412 | 0 | 10 | 0 | 10 | 102 | 101 | 165 | 583 | 33 | 11 | 0 | 0 | 0 | 0 | 129127 | 94096 | 307748 | 72 | 0 | 4622 | 2603 | 0 | 0 | 22 | 15 | 34 | 137 | 40 | 0 | 13 | 0 | 20 | 0 | 0 | 1528 | 187 | 0 |
| SRR6757413 | 17 | 11 | 0 | 0 | 106 | 729 | 342 | 2840 | 198 | 55 | 0 | 0 | 0 | 0 | 47735 | 136051 | 399675 | 85 | 21 | 12062 | 6734 | 0 | 0 | 14 | 0 | 11 | 28 | 0 | 0 | 81 | 0 | 16 | 0 | 0 | 1586 | 131 | 0 |
| SRR6757414 | 0 | 0 | 0 | 0 | 12 | 567 | 562 | 2913 | 103 | 0 | 0 | 0 | 0 | 0 | 17774 | 50367 | 198508 | 87 | 30 | 7220 | 2055 | 0 | 0 | 330 | 0 | 0 | 11 | 13 | 0 | 0 | 0 | 0 | 0 | 0 | 6068 | 202 | 0 |
| SRR6757415 | 0 | 0 | 0 | 0 | 0 | 1179 | 1036 | 6135 | 191 | 20 | 0 | 0 | 0 | 0 | 2597 | 82081 | 89288 | 48 | 0 | 3288 | 214 | 0 | 0 | 61 | 0 | 0 | 0 | 0 | 0 | 0 | 0 | 12 | 0 | 0 | 28024 | 158 | 0 |
| SRR6757416 | 0 | 0 | 0 | 0 | 0 | 1491 | 909 | 4814 | 105 | 11 | 0 | 0 | 0 | 0 | 17935 | 44674 | 210579 | 197 | 31 | 4202 | 643 | 0 | 0 | 12 | 0 | 0 | 0 | 0 | 0 | 0 | 0 | 0 | 0 | 0 | 11062 | 264 | 0 |
| SRR6757417 | 0 | 0 | 0 | 0 | 20 | 1008 | 425 | 3831 | 119 | 0 | 0 | 0 | 0 | 0 | 33703 | 77177 | 299245 | 158 | 15 | 16976 | 3593 | 0 | 0 | 42 | 0 | 0 | 0 | 0 | 0 | 0 | 0 | 0 | 0 | 0 | 2791 | 86 | 0 |
| SRR6757418 | 0 | 13 | 0 | 0 | 78 | 26 | 42 | 264 | 30 | 23 | 0 | 0 | 0 | 0 | 39302 | 89008 | 336700 | 461 | 24 | 17546 | 3355 | 0 | 0 | 17 | 0 | 0 | 63 | 0 | 0 | 56 | 0 | 16 | 0 | 0 | 1219 | 260 | 12 |
| SRR6757419 | 0 | 0 | 0 | 0 | 33 | 988 | 335 | 2443 | 63 | 25 | 0 | 0 | 0 | 0 | 25303 | 63366 | 221531 | 46 | 30 | 33400 | 7168 | 0 | 0 | 63 | 0 | 0 | 28 | 0 | 0 | 15 | 0 | 0 | 0 | 0 | 1177 | 80 | 0 |
| SRR6757420 | 0 | 11 | 0 | 0 | 68 | 106 | 77 | 400 | 24 | 0 | 0 | 0 | 0 | 0 | 32907 | 46101 | 266888 | 351 | 43 | 18898 | 2516 | 0 | 0 | 145 | 0 | 15 | 72 | 0 | 0 | 33 | 0 | 0 | 0 | 0 | 860 | 236 | 0 |
| SRR6757421 | 0 | 0 | 0 | 0 | 0 | 1406 | 825 | 4347 | 136 | 10 | 0 | 0 | 0 | 0 | 35184 | 44831 | 305434 | 50 | 0 | 6614 | 3113 | 0 | 0 | 78 | 0 | 0 | 0 | 0 | 0 | 0 | 0 | 0 | 0 | 0 | 6899 | 121 | 0 |
| SRR6757422 | 0 | 0 | 0 | 0 | 11 | 370 | 261 | 1296 | 65 | 0 | 0 | 0 | 0 | 0 | 138123 | 80143 | 508060 | 73 | 15 | 5111 | 5666 | 0 | 0 | 159 | 0 | 0 | 0 | 0 | 0 | 0 | 0 | 0 | 0 | 0 | 3328 | 293 | 0 |
| SRR6757423 | 0 | 0 | 0 | 0 | 26 | 488 | 151 | 1197 | 19 | 0 | 43 | 0 | 0 | 0 | 95794 | 43357 | 482734 | 22 | 34 | 19790 | 3866 | 0 | 0 | 13 | 0 | 0 | 51 | 0 | 0 | 24 | 0 | 0 | 0 | 0 | 547 | 76 | 0 |
| SRR6757424 | 0 | 14 | 19 | 26 | 106 | 463 | 270 | 2551 | 42 | 21 | 13 | 0 | 0 | 0 | 48678 | 65100 | 283797 | 21 | 58 | 14232 | 1572 | 0 | 0 | 563 | 11 | 63 | 368 | 45 | 0 | 107 | 0 | 0 | 0 | 0 | 623 | 216 | 0 |
| SRR6757425 | 0 | 0 | 0 | 0 | 11 | 1318 | 670 | 3733 | 187 | 18 | 0 | 0 | 0 | 0 | 15241 | 55325 | 208869 | 0 | 11 | 14352 | 3294 | 0 | 0 | 68 | 0 | 0 | 0 | 11 | 0 | 10 | 0 | 0 | 0 | 0 | 10703 | 113 | 0 |
| SRR6757426 | 0 | 0 | 0 | 0 | 21 | 359 | 194 | 1314 | 98 | 31 | 0 | 0 | 0 | 0 | 53670 | 48869 | 242267 | 0 | 41 | 12431 | 5125 | 0 | 0 | 11 | 0 | 0 | 21 | 0 | 0 | 31 | 0 | 27 | 0 | 0 | 1326 | 285 | 12 |
| SRR6757427 | 0 | 0 | 0 | 23 | 39 | 341 | 228 | 1419 | 31 | 20 | 0 | 0 | 0 | 0 | 56703 | 64659 | 290406 | 18 | 64 | 31055 | 6973 | 0 | 0 | 154 | 0 | 15 | 99 | 17 | 0 | 37 | 0 | 0 | 0 | 0 | 851 | 79 | 0 |
| SRR6757428 | 0 | 0 | 0 | 0 | 101 | 39 | 63 | 280 | 28 | 36 | 0 | 0 | 0 | 0 | 170083 | 116059 | 362582 | 0 | 11 | 1451 | 957 | 0 | 0 | 729 | 23 | 18 | 19 | 0 | 0 | 42 | 0 | 0 | 0 | 0 | 816 | 354 | 31 |
| SRR6757429 | 0 | 0 | 0 | 0 | 0 | 911 | 948 | 4090 | 335 | 0 | 0 | 0 | 0 | 0 | 4109 | 62210 | 129853 | 0 | 0 | 2993 | 781 | 0 | 0 | 67 | 0 | 0 | 0 | 0 | 0 | 22 | 0 | 37 | 0 | 0 | 19865 | 113 | 0 |
| SRR6757430 | 0 | 0 | 0 | 0 | 11 | 327 | 256 | 1804 | 113 | 0 | 0 | 0 | 0 | 0 | 32699 | 78225 | 239355 | 54 | 0 | 4537 | 1137 | 0 | 0 | 112 | 0 | 0 | 0 | 0 | 0 | 42 | 0 | 18 | 0 | 0 | 7078 | 204 | 0 |
| SRR6757431 | 0 | 0 | 0 | 0 | 0 | 2043 | 1294 | 9016 | 92 | 0 | 0 | 0 | 0 | 0 | 3635 | 38392 | 81801 | 21 | 10 | 12785 | 390 | 0 | 0 | 111 | 0 | 0 | 0 | 0 | 0 | 0 | 0 | 0 | 0 | 0 | 12440 | 118 | 0 |
| SRR6757432 | 0 | 0 | 0 | 0 | 115 | 105 | 97 | 535 | 33 | 11 | 0 | 0 | 0 | 0 | 65594 | 69129 | 300501 | 86 | 20 | 10276 | 1472 | 0 | 0 | 341 | 20 | 20 | 32 | 0 | 0 | 27 | 0 | 0 | 0 | 0 | 1201 | 408 | 0 |
| SRR6757433 | 0 | 0 | 0 | 0 | 0 | 801 | 862 | 4051 | 99 | 0 | 0 | 0 | 0 | 0 | 4509 | 54127 | 128791 | 15 | 38 | 9971 | 565 | 0 | 0 | 40 | 0 | 0 | 0 | 0 | 0 | 0 | 0 | 0 | 0 | 0 | 21038 | 105 | 0 |
| SRR6757434 | 0 | 0 | 0 | 0 | 0 | 388 | 363 | 1436 | 88 | 27 | 0 | 13 | 0 | 0 | 52408 | 76135 | 326844 | 165 | 18 | 1507 | 6400 | 0 | 20 | 18 | 0 | 0 | 0 | 0 | 0 | 100 | 15 | 34 | 0 | 0 | 4983 | 1027 | 42 |
| SRR6757435 | 0 | 0 | 0 | 0 | 26 | 1933 | 720 | 6804 | 100 | 20 | 0 | 0 | 0 | 0 | 42905 | 78795 | 349870 | 73 | 0 | 7165 | 8573 | 0 | 0 | 126 | 0 | 0 | 0 | 0 | 0 | 0 | 0 | 12 | 0 | 0 | 2066 | 89 | 0 |
| SRR6757436 | 0 | 0 | 0 | 0 | 37 | 127 | 151 | 1079 | 55 | 23 | 0 | 0 | 0 | 0 | 117398 | 76703 | 348920 | 114 | 14 | 5290 | 5097 | 0 | 0 | 19 | 10 | 18 | 115 | 18 | 0 | 33 | 0 | 0 | 0 | 0 | 1803 | 334 | 0 |
| SRR6757437 | 0 | 0 | 0 | 0 | 41 | 218 | 180 | 2023 | 61 | 21 | 0 | 0 | 0 | 0 | 36571 | 71966 | 288837 | 49 | 48 | 40098 | 3676 | 0 | 0 | 82 | 0 | 0 | 11 | 0 | 0 | 31 | 0 | 0 | 0 | 0 | 1108 | 100 | 0 |
| SRR6757438 | 0 | 0 | 0 | 0 | 224 | 46 | 50 | 301 | 24 | 28 | 0 | 0 | 0 | 0 | 110978 | 82688 | 324488 | 68 | 18 | 5617 | 2469 | 0 | 0 | 26 | 11 | 12 | 72 | 0 | 0 | 129 | 0 | 10 | 0 | 0 | 2777 | 683 | 0 |
| SRR6757439 | 0 | 13 | 0 | 0 | 73 | 153 | 158 | 1338 | 101 | 18 | 0 | 0 | 0 | 0 | 21628 | 94575 | 242522 | 16 | 38 | 21564 | 5486 | 0 | 0 | 78 | 0 | 0 | 28 | 0 | 0 | 179 | 0 | 17 | 0 | 0 | 1708 | 159 | 0 |
| SRR6757440 | 0 | 12 | 0 | 0 | 152 | 21 | 20 | 263 | 48 | 42 | 24 | 37 | 0 | 0 | 210412 | 57866 | 315271 | 23 | 37 | 5936 | 6479 | 0 | 0 | 16 | 75 | 51 | 169 | 14 | 0 | 23 | 0 | 0 | 0 | 0 | 557 | 625 | 0 |
| SRR6757441 | 12 | 11 | 0 | 0 | 113 | 21 | 30 | 298 | 56 | 26 | 0 | 0 | 0 | 0 | 60595 | 97935 | 306927 | 33 | 0 | 24297 | 9334 | 0 | 0 | 91 | 0 | 0 | 22 | 0 | 0 | 155 | 0 | 0 | 0 | 0 | 892 | 77 | 0 |
| SRR6757442 | 0 | 0 | 0 | 0 | 37 | 60 | 44 | 234 | 24 | 31 | 0 | 0 | 0 | 0 | 214253 | 46193 | 388134 | 14 | 15 | 1692 | 2372 | 0 | 0 | 723 | 14 | 14 | 21 | 0 | 0 | 187 | 0 | 0 | 0 | 0 | 1850 | 364 | 0 |
| SRR6757443 | 0 | 0 | 0 | 0 | 15 | 232 | 387 | 1658 | 86 | 36 | 0 | 0 | 0 | 0 | 31264 | 125044 | 330114 | 0 | 17 | 9279 | 3548 | 0 | 0 | 57 | 0 | 0 | 0 | 0 | 0 | 0 | 0 | 0 | 0 | 0 | 7426 | 104 | 0 |
| SRR6757444 | 0 | 0 | 0 | 0 | 51 | 69 | 110 | 505 | 106 | 132 | 0 | 19 | 0 | 0 | 129064 | 70162 | 358974 | 23 | 74 | 7423 | 2515 | 0 | 0 | 12 | 15 | 16 | 67 | 0 | 0 | 40 | 0 | 0 | 0 | 0 | 1714 | 271 | 0 |
| SRR6757445 | 0 | 0 | 0 | 0 | 24 | 272 | 342 | 2481 | 113 | 16 | 0 | 0 | 0 | 0 | 31055 | 128986 | 291811 | 116 | 0 | 22994 | 3724 | 0 | 0 | 90 | 0 | 0 | 0 | 0 | 0 | 0 | 0 | 0 | 0 | 0 | 4974 | 98 | 0 |
| SRR6757446 | 0 | 0 | 0 | 0 | 0 | 468 | 663 | 2589 | 274 | 32 | 0 | 0 | 0 | 0 | 42223 | 78732 | 188112 | 106 | 0 | 9597 | 592 | 0 | 0 | 16 | 0 | 0 | 13 | 0 | 0 | 0 | 0 | 0 | 0 | 0 | 15516 | 201 | 0 |
| SRR6757447 | 0 | 0 | 0 | 0 | 0 | 311 | 458 | 3321 | 196 | 18 | 0 | 0 | 0 | 0 | 21298 | 129805 | 209503 | 25 | 0 | 24717 | 3141 | 0 | 0 | 10 | 0 | 0 | 0 | 0 | 0 | 0 | 0 | 0 | 0 | 0 | 13600 | 95 | 0 |
| SRR6757448 | 0 | 0 | 0 | 0 | 16 | 31 | 41 | 198 | 35 | 42 | 0 | 0 | 0 | 0 | 170690 | 53343 | 326105 | 225 | 18 | 6296 | 6298 | 0 | 0 | 13 | 16 | 18 | 78 | 20 | 0 | 27 | 0 | 0 | 0 | 0 | 1265 | 107 | 0 |
| SRR6757449 | 0 | 0 | 0 | 0 | 0 | 1609 | 953 | 5494 | 250 | 0 | 0 | 0 | 0 | 0 | 5846 | 66801 | 121213 | 0 | 28 | 16076 | 435 | 0 | 0 | 83 | 0 | 0 | 0 | 12 | 0 | 10 | 0 | 0 | 0 | 0 | 19934 | 38 | 0 |
| SRR6757450 | 0 | 0 | 0 | 0 | 0 | 1226 | 1385 | 4961 | 277 | 13 | 0 | 0 | 0 | 0 | 7286 | 22343 | 106152 | 25 | 0 | 13833 | 1309 | 0 | 0 | 280 | 11 | 11 | 52 | 18 | 0 | 0 | 0 | 0 | 0 | 0 | 6894 | 352 | 0 |
| SRR6757451 | 11 | 0 | 0 | 22 | 38 | 102 | 53 | 701 | 16 | 77 | 0 | 0 | 0 | 0 | 71335 | 51826 | 454920 | 78 | 0 | 7301 | 9229 | 0 | 0 | 15 | 0 | 0 | 11 | 22 | 0 | 18 | 0 | 0 | 0 | 0 | 407 | 104 | 0 |
| SRR6757452 | 24 | 31 | 0 | 0 | 120 | 32 | 22 | 172 | 26 | 21 | 0 | 11 | 0 | 0 | 86185 | 48033 | 270452 | 36 | 18 | 2544 | 4521 | 0 | 0 | 42 | 20 | 34 | 38 | 0 | 0 | 54 | 0 | 0 | 0 | 0 | 1546 | 1751 | 15 |
| SRR6757453 | 0 | 0 | 0 | 0 | 46 | 102 | 111 | 848 | 36 | 36 | 0 | 0 | 0 | 0 | 92564 | 77411 | 495803 | 127 | 32 | 22640 | 7925 | 0 | 0 | 168 | 0 | 0 | 14 | 0 | 0 | 14 | 0 | 0 | 0 | 0 | 536 | 113 | 0 |
| SRR6757454 | 0 | 0 | 0 | 0 | 132 | 129 | 182 | 1006 | 53 | 22 | 0 | 0 | 0 | 0 | 59763 | 65401 | 347931 | 46 | 89 | 32474 | 2138 | 0 | 0 | 30 | 15 | 20 | 56 | 0 | 0 | 67 | 0 | 0 | 0 | 0 | 1510 | 186 | 0 |
| SRR6757455 | 0 | 0 | 0 | 0 | 0 | 5890 | 1072 | 9031 | 140 | 45 | 0 | 0 | 0 | 0 | 12396 | 77347 | 201795 | 19 | 19 | 3215 | 2181 | 0 | 0 | 11 | 0 | 0 | 0 | 0 | 0 | 17 | 0 | 0 | 0 | 0 | 16917 | 101 | 0 |
| SRR6757456 | 0 | 0 | 0 | 0 | 21 | 423 | 275 | 1845 | 73 | 40 | 0 | 0 | 0 | 0 | 47493 | 63331 | 303554 | 153 | 0 | 1513 | 4765 | 0 | 0 | 665 | 0 | 0 | 23 | 0 | 0 | 51 | 0 | 0 | 0 | 0 | 2687 | 614 | 11 |
| SRR6757457 | 0 | 0 | 0 | 0 | 0 | 1465 | 839 | 5072 | 114 | 16 | 0 | 0 | 0 | 0 | 8947 | 54369 | 163933 | 49 | 0 | 9110 | 4136 | 0 | 0 | 136 | 0 | 0 | 0 | 0 | 0 | 14 | 0 | 0 | 0 | 0 | 13147 | 49 | 0 |
| SRR6757458 | 0 | 0 | 0 | 0 | 0 | 1342 | 744 | 3341 | 46 | 11 | 0 | 0 | 0 | 0 | 17094 | 37655 | 189040 | 42 | 0 | 2328 | 690 | 0 | 0 | 140 | 0 | 0 | 0 | 0 | 0 | 22 | 0 | 0 | 0 | 0 | 13804 | 404 | 0 |
| SRR6757459 | 0 | 0 | 0 | 0 | 18 | 293 | 172 | 1433 | 47 | 20 | 0 | 0 | 0 | 0 | 51403 | 70773 | 366000 | 155 | 20 | 18271 | 11876 | 0 | 0 | 76 | 0 | 0 | 0 | 0 | 0 | 27 | 0 | 0 | 0 | 0 | 1869 | 88 | 0 |
| SRR6757460 | 0 | 0 | 0 | 0 | 20 | 372 | 281 | 1786 | 99 | 67 | 0 | 0 | 0 | 0 | 120268 | 54446 | 286786 | 160 | 22 | 2222 | 2629 | 0 | 0 | 185 | 0 | 0 | 0 | 0 | 0 | 42 | 0 | 0 | 0 | 0 | 3522 | 518 | 0 |
| SRR6757461 | 0 | 0 | 0 | 0 | 0 | 12653 | 1728 | 11067 | 67 | 39 | 0 | 0 | 0 | 0 | 39619 | 38363 | 379013 | 75 | 0 | 1421 | 2397 | 0 | 0 | 210 | 0 | 0 | 0 | 0 | 0 | 0 | 0 | 0 | 0 | 0 | 4341 | 87 | 0 |
| SRR6757462 | 0 | 0 | 0 | 0 | 0 | 2945 | 456 | 3419 | 48 | 36 | 0 | 0 | 0 | 0 | 138916 | 33259 | 324390 | 37 | 11 | 2129 | 19003 | 0 | 0 | 95 | 0 | 0 | 0 | 0 | 0 | 17 | 0 | 0 | 0 | 0 | 1648 | 595 | 0 |
| SRR6757463 | 0 | 0 | 0 | 0 | 19 | 1404 | 552 | 3988 | 110 | 47 | 0 | 0 | 0 | 0 | 44074 | 99190 | 227808 | 55 | 0 | 4119 | 3592 | 0 | 0 | 11 | 0 | 0 | 0 | 0 | 0 | 17 | 0 | 0 | 0 | 0 | 2100 | 124 | 0 |
| SRR6757464 | 0 | 0 | 0 | 0 | 56 | 65 | 143 | 604 | 108 | 12 | 15 | 11 | 0 | 0 | 179085 | 21462 | 214564 | 475 | 33 | 2123 | 11431 | 0 | 0 | 110 | 10 | 0 | 22 | 0 | 0 | 10 | 0 | 25 | 0 | 0 | 774 | 598 | 17 |
| SRR6757465 | 0 | 0 | 0 | 0 | 16 | 2240 | 1431 | 19898 | 320 | 46 | 0 | 0 | 0 | 0 | 29514 | 88990 | 289008 | 256 | 24 | 8030 | 2088 | 0 | 0 | 133 | 0 | 0 | 0 | 0 | 0 | 25 | 0 | 0 | 0 | 0 | 3257 | 124 | 0 |
| SRR6757466 | 0 | 0 | 0 | 0 | 27 | 263 | 223 | 1173 | 71 | 90 | 0 | 0 | 0 | 0 | 119999 | 46745 | 225310 | 288 | 25 | 1691 | 7924 | 0 | 0 | 207 | 0 | 0 | 12 | 0 | 0 | 116 | 0 | 0 | 0 | 0 | 3753 | 1854 | 0 |
| SRR6757467 | 0 | 0 | 0 | 0 | 0 | 3179 | 1093 | 8280 | 27 | 0 | 0 | 0 | 0 | 0 | 14589 | 60814 | 182539 | 61 | 0 | 3515 | 1681 | 0 | 0 | 167 | 0 | 0 | 0 | 0 | 0 | 11 | 0 | 0 | 0 | 0 | 5652 | 101 | 0 |
| SRR6757468 | 0 | 0 | 0 | 0 | 0 | 2101 | 852 | 5633 | 87 | 0 | 0 | 0 | 0 | 0 | 19666 | 44337 | 171790 | 83 | 16 | 4250 | 1776 | 0 | 0 | 363 | 0 | 0 | 12 | 0 | 0 | 18 | 0 | 0 | 0 | 0 | 8066 | 274 | 0 |
| SRR6761159 | 0 | 0 | 0 | 0 | 0 | 0 | 0 | 1721 | 0 | 0 | 1193 | 0 | 0 | 0 | 7745 | 89693 | 299819 | 0 | 0 | 4303 | 18933 | 0 | 0 | 0 | 0 | 170 | 85 | 0 | 0 | 937 | 0 | 0 | 0 | 0 | 1808 | 0 | 0 |
| SRR6761162 | 0 | 0 | 0 | 0 | 0 | 0 | 0 | 0 | 0 | 0 | 0 | 0 | 0 | 0 | 4367 | 92676 | 263079 | 0 | 2183 | 4367 | 8734 | 0 | 0 | 0 | 0 | 0 | 0 | 0 | 0 | 0 | 0 | 0 | 0 | 0 | 2990 | 0 | 302 |
| SRR6761163 | 0 | 0 | 0 | 0 | 0 | 0 | 0 | 0 | 0 | 0 | 108 | 0 | 0 | 0 | 5837 | 47391 | 216557 | 0 | 0 | 0 | 10700 | 0 | 0 | 0 | 0 | 0 | 0 | 432 | 0 | 324 | 0 | 0 | 0 | 0 | 4199 | 0 | 324 |
| SRR6761165 | 0 | 0 | 0 | 0 | 0 | 0 | 0 | 2442 | 0 | 0 | 358 | 0 | 0 | 0 | 40293 | 61528 | 258133 | 0 | 0 | 3663 | 4884 | 0 | 0 | 0 | 0 | 0 | 0 | 0 | 0 | 716 | 0 | 0 | 0 | 0 | 6365 | 0 | 0 |
| SRR6761167 | 0 | 0 | 0 | 0 | 0 | 2857 | 2857 | 0 | 0 | 0 | 1153 | 0 | 0 | 0 | 17143 | 57312 | 383399 | 0 | 0 | 0 | 17143 | 0 | 0 | 0 | 0 | 0 | 0 | 0 | 0 | 0 | 0 | 0 | 0 | 0 | 5929 | 0 | 0 |
| SRR6761175 | 0 | 0 | 0 | 0 | 0 | 0 | 0 | 3867 | 0 | 0 | 430 | 0 | 0 | 0 | 12374 | 60090 | 343049 | 0 | 0 | 0 | 13148 | 0 | 0 | 0 | 0 | 86 | 0 | 0 | 0 | 1118 | 0 | 0 | 0 | 0 | 2242 | 0 | 0 |
| SRR6761179 | 0 | 0 | 0 | 0 | 0 | 0 | 0 | 1969 | 0 | 0 | 1714 | 0 | 0 | 0 | 9843 | 103529 | 271765 | 0 | 0 | 0 | 0 | 0 | 0 | 171 | 0 | 0 | 0 | 171 | 0 | 1371 | 0 | 0 | 0 | 0 | 0 | 0 | 0 |
| SRR6761181 | 0 | 0 | 0 | 0 | 0 | 585 | 585 | 1170 | 0 | 0 | 1003 | 0 | 0 | 0 | 10526 | 78277 | 360815 | 0 | 0 | 585 | 5848 | 0 | 0 | 0 | 0 | 0 | 0 | 111 | 0 | 1337 | 0 | 0 | 0 | 0 | 4169 | 0 | 0 |
| SRR6761182 | 0 | 0 | 0 | 0 | 0 | 1072 | 1072 | 536 | 0 | 0 | 377 | 0 | 0 | 0 | 12326 | 65100 | 310579 | 0 | 0 | 0 | 7503 | 0 | 0 | 0 | 0 | 0 | 0 | 0 | 0 | 965 | 0 | 0 | 0 | 0 | 1356 | 0 | 0 |
| SRR6761183 | 0 | 0 | 0 | 0 | 0 | 0 | 0 | 0 | 0 | 0 | 11 | 0 | 0 | 0 | 6028 | 88394 | 240584 | 0 | 0 | 1808 | 9042 | 0 | 0 | 0 | 0 | 0 | 0 | 0 | 0 | 17 | 0 | 0 | 0 | 0 | 2306 | 0 | 109 |
| SRR6761184 | 0 | 0 | 0 | 0 | 0 | 0 | 384 | 0 | 0 | 0 | 11 | 0 | 0 | 0 | 8452 | 53487 | 252888 | 0 | 0 | 768 | 6147 | 0 | 0 | 0 | 0 | 0 | 0 | 0 | 0 | 10 | 0 | 0 | 0 | 0 | 2140 | 0 | 0 |
| SRR6761188 | 0 | 0 | 0 | 0 | 0 | 1496 | 0 | 0 | 0 | 0 | 19 | 0 | 0 | 0 | 5484 | 53939 | 237757 | 0 | 0 | 1496 | 4985 | 0 | 0 | 0 | 0 | 0 | 38 | 0 | 0 | 15 | 0 | 0 | 0 | 0 | 2129 | 0 | 113 |
| SRR6761192 | 0 | 0 | 0 | 0 | 0 | 0 | 0 | 0 | 0 | 0 | 3404 | 0 | 0 | 0 | 19048 | 182353 | 338235 | 0 | 0 | 12698 | 0 | 0 | 0 | 0 | 0 | 0 | 0 | 0 | 0 | 681 | 0 | 0 | 0 | 0 | 2941 | 0 | 0 |
| SRR6761198 | 0 | 0 | 0 | 0 | 0 | 0 | 0 | 258 | 0 | 0 | 27 | 0 | 0 | 0 | 14179 | 67241 | 243602 | 0 | 0 | 0 | 9539 | 0 | 0 | 0 | 0 | 0 | 0 | 0 | 0 | 25 | 0 | 0 | 0 | 0 | 1396 | 0 | 0 |
| SRR6767691 | 0 | 0 | 0 | 0 | 0 | 0 | 0 | 0 | 0 | 0 | 1344 | 12 | 0 | 0 | 5014 | 10140 | 50598 | 2700 | 0 | 0 | 0 | 0 | 199 | 0 | 0 | 0 | 0 | 0 | 0 | 286 | 0 | 1929 | 0 | 0 | 993 | 0 | 0 |
| SRR6767701 | 0 | 0 | 0 | 0 | 0 | 0 | 0 | 0 | 0 | 0 | 323 | 24 | 0 | 0 | 551 | 17714 | 59845 | 0 | 0 | 787 | 0 | 58 | 15 | 0 | 0 | 0 | 0 | 0 | 0 | 185 | 48 | 0 | 0 | 0 | 410 | 0 | 0 |
| SRR6767712 | 0 | 0 | 0 | 0 | 0 | 0 | 0 | 0 | 0 | 0 | 1536 | 0 | 0 | 0 | 6218 | 13075 | 43617 | 1219 | 0 | 0 | 122 | 107 | 0 | 0 | 0 | 0 | 0 | 0 | 0 | 534 | 0 | 0 | 0 | 0 | 366 | 0 | 0 |
| SRR6767713 | 0 | 0 | 0 | 0 | 0 | 0 | 0 | 0 | 0 | 0 | 11 | 38 | 0 | 0 | 3489 | 6834 | 39718 | 1744 | 0 | 0 | 0 | 0 | 14 | 0 | 0 | 0 | 0 | 0 | 0 | 229 | 0 | 4579 | 0 | 0 | 2643 | 0 | 0 |
| SRR6821085 | 21 | 15 | 0 | 47 | 44 | 145 | 104 | 243 | 54 | 32 | 39 | 0 | 0 | 0 | 81690 | 76558 | 211401 | 68 | 186 | 19716 | 1559 | 0 | 0 | 175 | 124 | 16 | 31 | 12 | 0 | 18 | 0 | 0 | 0 | 0 | 1521 | 0 | 0 |
| SRR6821086 | 43 | 35 | 18 | 46 | 110 | 306 | 147 | 219 | 34 | 93 | 37 | 0 | 0 | 0 | 27807 | 171434 | 216923 | 61 | 956 | 28941 | 731 | 0 | 0 | 151 | 32 | 159 | 19 | 94 | 0 | 11 | 0 | 10 | 0 | 0 | 1327 | 0 | 16 |
| SRR6821087 | 13 | 0 | 0 | 0 | 25 | 1448 | 443 | 694 | 93 | 291 | 18 | 0 | 0 | 0 | 46304 | 174227 | 201685 | 21 | 144 | 10165 | 1182 | 0 | 0 | 126 | 38 | 88 | 129 | 0 | 0 | 225 | 0 | 69 | 0 | 0 | 2789 | 0 | 21 |
| SRR6821088 | 13 | 0 | 13 | 55 | 42 | 132 | 247 | 729 | 185 | 0 | 19 | 0 | 0 | 0 | 48544 | 42493 | 175315 | 0 | 353 | 6066 | 8326 | 0 | 0 | 308 | 101 | 334 | 1531 | 185 | 0 | 110 | 0 | 42 | 0 | 0 | 340 | 0 | 0 |
| SRR6821089 | 0 | 0 | 0 | 0 | 12 | 123 | 454 | 735 | 597 | 0 | 0 | 0 | 0 | 0 | 26636 | 20154 | 41712 | 0 | 196 | 587 | 1499 | 0 | 0 | 15 | 151 | 16 | 77 | 19 | 0 | 79 | 0 | 12 | 0 | 0 | 278 | 0 | 0 |
| SRR6821090 | 0 | 0 | 16 | 41 | 49 | 172 | 427 | 708 | 270 | 20 | 16 | 0 | 0 | 0 | 18555 | 41798 | 260532 | 11 | 1068 | 10775 | 54848 | 0 | 0 | 94 | 114 | 462 | 1269 | 282 | 0 | 186 | 0 | 29 | 0 | 0 | 258 | 0 | 0 |
| SRR6823984 | 3481 | 795 | 224 | 272 | 2702 | 2252 | 500 | 1736 | 3503 | 150 | 112 | 0 | 30 | 16 | 5677 | 52752 | 19506 | 328 | 3769 | 4660 | 1517 | 0 | 0 | 202 | 58 | 82 | 160 | 148 | 20 | 198 | 0 | 4066 | 0 | 0 | 9844 | 0 | 0 |
| SRR6823985 | 3304 | 733 | 59 | 53 | 2052 | 2412 | 594 | 1817 | 2813 | 499 | 137 | 28 | 56 | 18 | 1747 | 53854 | 9646 | 140 | 3775 | 2167 | 682 | 0 | 0 | 69 | 15 | 10 | 31 | 0 | 13 | 260 | 0 | 192 | 0 | 0 | 7880 | 0 | 0 |
| SRR6823986 | 5541 | 2737 | 450 | 312 | 192 | 2570 | 695 | 2594 | 3394 | 36 | 146 | 0 | 40 | 29 | 5252 | 6515 | 18978 | 1131 | 687 | 5204 | 8493 | 10 | 15 | 305 | 94 | 73 | 123 | 15 | 21 | 180 | 19 | 1616 | 0 | 19 | 5992 | 17 | 0 |
| SRR6823987 | 258 | 131 | 20 | 12 | 24 | 2292 | 760 | 3732 | 3824 | 35 | 12 | 0 | 19 | 0 | 4803 | 9292 | 21084 | 392 | 852 | 3271 | 4688 | 19 | 0 | 17 | 86 | 54 | 11 | 0 | 0 | 24 | 0 | 15389 | 11 | 13 | 3327 | 0 | 0 |
| SRR6823990 | 2957 | 728 | 574 | 217 | 1438 | 1755 | 348 | 1142 | 2433 | 427 | 141 | 13 | 16 | 0 | 4139 | 54237 | 47346 | 315 | 1705 | 1738 | 6870 | 0 | 0 | 122 | 85 | 50 | 107 | 107 | 13 | 78 | 0 | 315 | 0 | 0 | 8359 | 123 | 0 |
| SRR6823991 | 121 | 37 | 28 | 109 | 76 | 2766 | 1013 | 6079 | 4299 | 11 | 19 | 0 | 0 | 0 | 7558 | 24672 | 25091 | 958 | 3204 | 2793 | 5038 | 51 | 0 | 118 | 38 | 80 | 10 | 96 | 19 | 15 | 32 | 29848 | 0 | 22 | 3520 | 274 | 0 |
| SRR6829958 | 518 | 91 | 10 | 103 | 46 | 171 | 83 | 717 | 1620 | 129 | 107 | 130 | 10 | 15 | 53190 | 300329 | 240510 | 571 | 7001 | 9369 | 66858 | 0 | 0 | 28 | 34 | 21 | 27 | 0 | 0 | 126 | 0 | 31 | 0 | 15 | 4480 | 418 | 0 |
| SRR6829959 | 144 | 39 | 70 | 10 | 68 | 141 | 93 | 819 | 3907 | 90 | 16 | 20 | 0 | 0 | 31949 | 227827 | 439710 | 100 | 6711 | 21466 | 52807 | 12 | 19 | 34 | 64 | 30 | 31 | 0 | 11 | 12 | 0 | 26 | 0 | 74 | 2327 | 388 | 0 |
| SRR6829960 | 18193 | 2399 | 470 | 190 | 1169 | 208 | 43 | 424 | 999 | 3213 | 48 | 117 | 26 | 0 | 56939 | 370971 | 245352 | 494 | 6262 | 6713 | 48747 | 0 | 0 | 934 | 59 | 45 | 59 | 0 | 0 | 52 | 0 | 72 | 0 | 14 | 3351 | 612 | 0 |
| SRR6829961 | 84 | 16 | 87 | 109 | 34 | 288 | 134 | 770 | 3254 | 33 | 47 | 168 | 12 | 0 | 29569 | 211441 | 464621 | 90 | 8892 | 16891 | 43785 | 0 | 0 | 17 | 50 | 20 | 32 | 0 | 0 | 77 | 0 | 25 | 0 | 22 | 2743 | 329 | 0 |
| SRR6829962 | 321 | 48 | 160 | 78 | 43 | 91 | 48 | 320 | 1396 | 114 | 19 | 162 | 12 | 13 | 44097 | 325471 | 196850 | 574 | 5172 | 2911 | 40960 | 0 | 10 | 21 | 25 | 15 | 16 | 0 | 14 | 37 | 0 | 70 | 0 | 33 | 8458 | 497 | 0 |
| SRR6829963 | 4330 | 1007 | 38 | 48 | 1161 | 426 | 180 | 976 | 3877 | 1587 | 10 | 148 | 0 | 10 | 11925 | 229617 | 292031 | 88 | 3864 | 7380 | 48245 | 0 | 0 | 608 | 13 | 0 | 15 | 0 | 0 | 0 | 0 | 44 | 0 | 0 | 13971 | 170 | 0 |
| SRR6831609 | 0 | 0 | 0 | 0 | 22 | 20040 | 2138 | 9452 | 24015 | 346 | 22 | 22 | 0 | 0 | 2639 | 85875 | 57348 | 0 | 167 | 401 | 835 | 0 | 22 | 27 | 713 | 259 | 216 | 43 | 0 | 0 | 0 | 0 | 0 | 0 | 12207 | 23 | 0 |
| SRR6831611 | 0 | 0 | 0 | 0 | 20 | 10331 | 266 | 2178 | 1567 | 1573 | 39 | 0 | 0 | 0 | 3187 | 85504 | 45106 | 53 | 558 | 292 | 2310 | 0 | 0 | 3284 | 511 | 157 | 157 | 0 | 0 | 118 | 0 | 53 | 0 | 0 | 58981 | 60 | 0 |
| SRR6831613 | 0 | 0 | 0 | 0 | 115 | 17793 | 1230 | 8101 | 9403 | 344 | 0 | 0 | 0 | 0 | 3038 | 26587 | 3589 | 72 | 145 | 506 | 1157 | 29 | 0 | 2064 | 229 | 229 | 172 | 0 | 0 | 57 | 0 | 0 | 0 | 57 | 11457 | 25 | 0 |
| SRR6837863 | 0 | 0 | 0 | 0 | 0 | 330 | 177 | 457 | 115 | 0 | 0 | 0 | 0 | 0 | 88 | 879 | 224 | 319 | 0 | 169 | 0 | 14 | 11 | 56 | 0 | 0 | 0 | 0 | 0 | 0 | 49 | 4104 | 0 | 0 | 23135 | 0 | 69 |
| SRR6837867 | 0 | 0 | 0 | 0 | 0 | 377 | 263 | 500 | 100 | 0 | 0 | 0 | 0 | 0 | 195 | 963 | 420 | 209 | 0 | 154 | 32 | 0 | 0 | 24 | 0 | 0 | 0 | 0 | 0 | 0 | 97 | 3406 | 0 | 0 | 28358 | 0 | 62 |
| SRR6837868 | 0 | 0 | 0 | 0 | 0 | 244 | 237 | 361 | 127 | 0 | 0 | 0 | 0 | 0 | 317 | 1539 | 704 | 324 | 0 | 244 | 90 | 0 | 11 | 39 | 0 | 0 | 0 | 0 | 0 | 0 | 65 | 4975 | 0 | 0 | 26313 | 0 | 71 |
| SRR6880326 | 0 | 0 | 0 | 0 | 0 | 363 | 120 | 134 | 171 | 0 | 45 | 0 | 0 | 0 | 340 | 23969 | 24511 | 39 | 0 | 0 | 33 | 12 | 23 | 143 | 242 | 25 | 117 | 39 | 0 | 383 | 14 | 451 | 0 | 0 | 1223 | 0 | 0 |
| SRR6880328 | 0 | 0 | 0 | 0 | 0 | 219 | 98 | 98 | 44 | 0 | 21701 | 0 | 0 | 0 | 9146 | 59041 | 59944 | 1136 | 11 | 295 | 175 | 0 | 0 | 0 | 0 | 0 | 16 | 0 | 0 | 3130 | 0 | 153 | 0 | 0 | 469 | 0 | 0 |
| SRR6880329 | 0 | 0 | 0 | 0 | 0 | 203 | 45 | 45 | 15 | 0 | 24331 | 0 | 0 | 0 | 14573 | 59847 | 71770 | 946 | 0 | 225 | 90 | 0 | 0 | 0 | 0 | 0 | 23 | 0 | 0 | 2887 | 0 | 83 | 0 | 0 | 334 | 0 | 0 |
| SRR6880330 | 0 | 0 | 0 | 0 | 10 | 458 | 293 | 110 | 257 | 0 | 18095 | 0 | 0 | 0 | 5684 | 47636 | 47789 | 1045 | 0 | 165 | 165 | 0 | 12 | 0 | 0 | 13 | 34 | 16 | 0 | 1336 | 0 | 110 | 0 | 0 | 247 | 0 | 0 |

### Supp Table S8.3.Columns_BX_to_DH

| | | | | | | | | | | | | | | | | | | | | | | | | | | | | | | | | | | | | | |
|---|---|---|---|---|---|---|---|---|---|---|---|---|---|---|---|---|---|---|---|---|---|---|---|---|---|---|---|---|---|---|---|---|---|---|---|---|---|
| SRR6880331 | 0 | 0 | 0 | 0 | 0 | 452 | 151 | 215 | 194 | 0 | 23649 | 0 | 0 | 0 | 9440 | 46410 | 65608 | 645 | 0 | 215 | 344 | 0 | 15 | 0 | 0 | 0 | 38 | 0 | 0 | 1315 | 0 | 172 | 0 | 0 | 267 | 0 | 0 |
| SRR6880333 | 0 | 0 | 0 | 0 | 16 | 1439 | 441 | 544 | 606 | 0 | 2253 | 10 | 0 | 0 | 3029 | 28616 | 313136 | 1294 | 0 | 1742 | 413 | 0 | 15 | 25 | 0 | 15 | 29 | 12 | 0 | 210 | 0 | 516 | 0 | 0 | 1027 | 0 | 0 |
| SRR6880334 | 0 | 0 | 0 | 0 | 10 | 1534 | 901 | 797 | 685 | 0 | 1480 | 16 | 0 | 0 | 2816 | 35718 | 170224 | 2825 | 0 | 1057 | 667 | 0 | 0 | 11 | 0 | 12 | 28 | 14 | 0 | 158 | 0 | 312 | 0 | 0 | 773 | 0 | 0 |
| SRR6880335 | 0 | 0 | 0 | 0 | 19 | 1448 | 648 | 530 | 564 | 0 | 3374 | 19 | 0 | 0 | 2551 | 29945 | 332640 | 1895 | 0 | 2282 | 1372 | 0 | 0 | 15 | 0 | 14 | 29 | 16 | 0 | 316 | 0 | 724 | 0 | 0 | 893 | 0 | 0 |
| SRR6880336 | 0 | 0 | 0 | 0 | 11 | 3609 | 520 | 1041 | 1203 | 18 | 1937 | 0 | 0 | 0 | 168198 | 73326 | 464626 | 228 | 163 | 211 | 2390 | 57 | 105 | 0 | 0 | 0 | 0 | 0 | 0 | 2184 | 45 | 374 | 0 | 0 | 2353 | 219 | 0 |
| SRR6880337 | 0 | 0 | 0 | 0 | 25 | 2078 | 505 | 660 | 1046 | 0 | 745 | 0 | 0 | 0 | 116125 | 79490 | 490466 | 386 | 30 | 349 | 1016 | 30 | 96 | 52 | 18 | 74 | 86 | 39 | 0 | 164 | 18 | 334 | 0 | 0 | 3858 | 63 | 0 |
| SRR6880338 | 0 | 0 | 0 | 0 | 0 | 296 | 144 | 95 | 146 | 0 | 82 | 19 | 0 | 0 | 276 | 21568 | 20229 | 94 | 0 | 0 | 17 | 0 | 21 | 129 | 129 | 28 | 115 | 31 | 0 | 21 | 26 | 477 | 0 | 0 | 1749 | 0 | 0 |
| SRR6880339 | 0 | 0 | 0 | 0 | 0 | 166 | 73 | 54 | 111 | 0 | 2439 | 0 | 0 | 0 | 217 | 24691 | 35666 | 42 | 0 | 0 | 29 | 0 | 48 | 98 | 124 | 633 | 2222 | 901 | 0 | 715 | 21 | 405 | 0 | 0 | 4847 | 0 | 0 |
| SRR6880341 | 0 | 0 | 0 | 0 | 0 | 116 | 41 | 33 | 64 | 0 | 8050 | 0 | 0 | 0 | 10889 | 68649 | 529317 | 66 | 17 | 0 | 251 | 33 | 366 | 30 | 86 | 194 | 745 | 165 | 0 | 755 | 23 | 304 | 0 | 0 | 552 | 0 | 0 |
| SRR6880342 | 0 | 0 | 0 | 0 | 0 | 57 | 12 | 0 | 17 | 0 | 793 | 0 | 0 | 0 | 5119 | 128708 | 443398 | 61 | 0 | 0 | 36 | 11 | 98 | 0 | 13 | 57 | 28 | 48 | 0 | 171 | 12 | 144 | 0 | 0 | 1649 | 0 | 0 |
| SRR6880343 | 0 | 0 | 0 | 0 | 0 | 54 | 26 | 22 | 23 | 0 | 329 | 0 | 0 | 0 | 16624 | 131303 | 549975 | 58 | 0 | 20 | 88 | 44 | 25 | 19 | 22 | 47 | 17 | 60 | 0 | 65 | 38 | 77 | 0 | 0 | 1194 | 0 | 0 |
| SRR6880345 | 0 | 0 | 0 | 0 | 17 | 789 | 563 | 296 | 1045 | 0 | 7187 | 0 | 0 | 0 | 391082 | 55006 | 755061 | 122 | 182 | 430 | 4127 | 134 | 728 | 50 | 137 | 409 | 367 | 202 | 0 | 1866 | 136 | 174 | 0 | 0 | 1141 | 190 | 0 |
| SRR6880351 | 0 | 0 | 0 | 0 | 13 | 584 | 63 | 243 | 156 | 0 | 2287 | 0 | 0 | 0 | 132648 | 65273 | 529335 | 166 | 39 | 531 | 1539 | 38 | 424 | 0 | 0 | 33 | 19 | 15 | 0 | 717 | 52 | 1081 | 0 | 0 | 900 | 95 | 0 |
| SRR6880356 | 0 | 0 | 0 | 0 | 20 | 2251 | 287 | 519 | 482 | 0 | 2202 | 10 | 0 | 0 | 56922 | 65534 | 366095 | 482 | 0 | 500 | 648 | 241 | 457 | 20 | 37 | 107 | 128 | 57 | 0 | 393 | 109 | 1158 | 0 | 0 | 5595 | 60 | 0 |
| SRR6880357 | 0 | 0 | 0 | 0 | 20 | 1591 | 174 | 376 | 328 | 0 | 555 | 0 | 0 | 0 | 68787 | 108671 | 247883 | 473 | 29 | 212 | 1032 | 20 | 62 | 10 | 20 | 40 | 15 | 31 | 0 | 72 | 21 | 1032 | 0 | 0 | 7271 | 16 | 0 |
| SRR6898464 | 0 | 0 | 14 | 22 | 25 | 236 | 231 | 361 | 60 | 22 | 44 | 0 | 0 | 0 | 66745 | 18086 | 140035 | 0 | 52 | 1620 | 22282 | 0 | 12 | 90 | 168 | 339 | 940 | 631 | 0 | 142 | 0 | 63 | 0 | 0 | 555 | 1811 | 0 |
| SRR6914752 | 10834 | 462 | 210 | 168 | 273 | 374 | 87 | 125 | 312 | 0 | 0 | 0 | 0 | 0 | 36972 | 27843 | 71090 | 1409 | 200 | 7382 | 50 | 0 | 0 | 1890 | 168 | 588 | 1071 | 168 | 0 | 126 | 0 | 0 | 0 | 0 | 3771 | 3000 | 0 |
| SRR6914753 | 175 | 22 | 644 | 87 | 202 | 338 | 403 | 135 | 783 | 0 | 29 | 0 | 16 | 0 | 24909 | 25668 | 42427 | 1714 | 176 | 20716 | 58 | 0 | 0 | 18 | 125 | 15 | 28 | 551 | 0 | 179 | 0 | 0 | 0 | 0 | 3857 | 7659 | 0 |
| SRR6914754 | 42 | 226 | 319 | 186 | 120 | 1574 | 211 | 262 | 372 | 33 | 213 | 13 | 0 | 0 | 154971 | 124586 | 365257 | 771 | 1023 | 35289 | 1767 | 0 | 20 | 13 | 712 | 17 | 16 | 333 | 0 | 585 | 0 | 0 | 0 | 0 | 2687 | 4787 | 0 |
| SRR6914755 | 61 | 567 | 261 | 64 | 528 | 1355 | 472 | 508 | 1037 | 11 | 22 | 0 | 0 | 0 | 69715 | 106501 | 116274 | 1093 | 711 | 34689 | 597 | 0 | 0 | 21 | 361 | 26 | 36 | 300 | 0 | 295 | 0 | 0 | 0 | 0 | 2725 | 12032 | 14 |
| SRR6928219 | 3057 | 974 | 566 | 290 | 171 | 745 | 251 | 1024 | 5252 | 129 | 126 | 0 | 10 | 0 | 21877 | 70182 | 174987 | 243 | 2094 | 6293 | 5255 | 0 | 0 | 520 | 284 | 431 | 418 | 578 | 0 | 120 | 0 | 0 | 0 | 0 | 6856 | 0 | 0 |
| SRR6928220 | 407 | 208 | 71 | 42 | 16 | 444 | 147 | 706 | 3269 | 83 | 12 | 0 | 0 | 0 | 23244 | 55167 | 194101 | 210 | 3069 | 5057 | 6512 | 0 | 0 | 28 | 20 | 30 | 30 | 31 | 0 | 13 | 0 | 0 | 0 | 0 | 1454 | 0 | 0 |
| SRR6928221 | 1019 | 205 | 295 | 131 | 411 | 563 | 78 | 717 | 1576 | 926 | 269 | 0 | 0 | 0 | 12506 | 128679 | 160230 | 109 | 2774 | 4579 | 18267 | 0 | 0 | 849 | 324 | 310 | 326 | 567 | 0 | 158 | 0 | 17 | 0 | 0 | 1490 | 0 | 0 |
| SRR6928222 | 836 | 219 | 295 | 105 | 1334 | 170 | 38 | 188 | 1089 | 98 | 139 | 0 | 0 | 0 | 15424 | 195160 | 155805 | 97 | 1502 | 4489 | 6514 | 0 | 0 | 404 | 288 | 500 | 593 | 496 | 0 | 104 | 0 | 0 | 0 | 0 | 1249 | 0 | 0 |
| SRR6928223 | 7104 | 2138 | 1054 | 428 | 170 | 583 | 156 | 762 | 3952 | 67 | 143 | 0 | 26 | 19 | 8734 | 38715 | 101060 | 66 | 1609 | 5839 | 3510 | 0 | 0 | 471 | 241 | 399 | 481 | 398 | 0 | 122 | 0 | 12 | 0 | 0 | 5104 | 0 | 0 |
| SRR6928224 | 366 | 194 | 91 | 50 | 21 | 571 | 247 | 1231 | 3920 | 14 | 19 | 0 | 0 | 0 | 20045 | 38519 | 154449 | 303 | 3938 | 2952 | 4888 | 0 | 0 | 30 | 20 | 24 | 24 | 12 | 0 | 13 | 0 | 0 | 0 | 0 | 1090 | 0 | 0 |
| SRR6928225 | 1770 | 365 | 397 | 227 | 599 | 753 | 107 | 919 | 2969 | 947 | 215 | 0 | 21 | 0 | 5804 | 101481 | 79073 | 51 | 1468 | 4724 | 12919 | 0 | 0 | 682 | 342 | 378 | 374 | 695 | 0 | 131 | 0 | 10 | 0 | 0 | 1621 | 0 | 0 |
| SRR6928226 | 139 | 33 | 72 | 23 | 325 | 209 | 42 | 233 | 1092 | 25 | 34 | 0 | 0 | 0 | 18245 | 162008 | 210492 | 137 | 1464 | 4619 | 4921 | 0 | 0 | 77 | 58 | 96 | 103 | 87 | 0 | 28 | 0 | 0 | 0 | 0 | 1126 | 0 | 0 |
| SRR6930936 | 223 | 52 | 18 | 0 | 80 | 3680 | 2087 | 6790 | 25400 | 12 | 163 | 54 | 10 | 0 | 1244 | 42045 | 17607 | 349 | 924 | 3017 | 674 | 0 | 0 | 47 | 61 | 29 | 21 | 17 | 0 | 539 | 0 | 47 | 0 | 28 | 2611 | 0 | 0 |
| SRR6930937 | 116 | 24 | 0 | 0 | 85 | 7829 | 2366 | 19749 | 23598 | 24 | 155 | 32 | 12 | 0 | 2153 | 33121 | 33010 | 183 | 1411 | 1158 | 1005 | 0 | 0 | 122 | 96 | 33 | 11 | 10 | 0 | 224 | 0 | 0 | 0 | 47 | 2569 | 0 | 0 |
| SRR6930939 | 21 | 0 | 0 | 0 | 64 | 10134 | 1975 | 13096 | 13356 | 0 | 548 | 11 | 0 | 0 | 9718 | 31826 | 111435 | 260 | 1351 | 2391 | 1143 | 0 | 0 | 39 | 39 | 23 | 14 | 14 | 0 | 1728 | 0 | 0 | 0 | 23 | 1915 | 0 | 0 |
| SRR6930941 | 46 | 14 | 22 | 0 | 16 | 5890 | 1799 | 9869 | 17611 | 36 | 94 | 31 | 16 | 0 | 11197 | 50596 | 156845 | 357 | 1572 | 3675 | 1638 | 0 | 0 | 16 | 18 | 35 | 24 | 20 | 0 | 330 | 0 | 0 | 0 | 21 | 2612 | 0 | 0 |
| SRR6932941 | 0 | 0 | 0 | 0 | 0 | 440 | 233 | 781 | 66 | 0 | 0 | 0 | 0 | 0 | 1038 | 1452 | 2657 | 66 | 0 | 108 | 125 | 0 | 0 | 17 | 0 | 0 | 0 | 0 | 0 | 0 | 0 | 0 | 0 | 0 | 4820 | 283 | 0 |
| SRR6932942 | 0 | 0 | 0 | 0 | 0 | 90 | 145 | 361 | 72 | 0 | 0 | 0 | 0 | 0 | 1337 | 3084 | 4191 | 0 | 0 | 18 | 271 | 0 | 0 | 0 | 0 | 0 | 0 | 0 | 0 | 0 | 0 | 0 | 0 | 0 | 5496 | 122 | 0 |
| SRR6932943 | 90 | 15 | 0 | 0 | 52 | 1596 | 1145 | 2828 | 382 | 15 | 22 | 0 | 0 | 0 | 73359 | 115257 | 233061 | 104 | 0 | 87 | 260 | 0 | 0 | 13 | 0 | 0 | 0 | 0 | 0 | 0 | 0 | 0 | 0 | 0 | 8198 | 402 | 0 |
| SRR6932946 | 17 | 46 | 0 | 0 | 55 | 329 | 536 | 803 | 99 | 0 | 0 | 0 | 14 | 0 | 298646 | 107085 | 556366 | 0 | 0 | 77 | 1806 | 0 | 0 | 101 | 0 | 0 | 0 | 0 | 0 | 0 | 0 | 0 | 0 | 0 | 4194 | 113 | 0 |
| SRR6932949 | 0 | 0 | 0 | 0 | 0 | 147 | 379 | 325 | 418 | 0 | 0 | 0 | 0 | 0 | 31 | 47 | 47 | 0 | 0 | 54 | 0 | 0 | 0 | 36 | 0 | 0 | 0 | 0 | 0 | 0 | 0 | 0 | 0 | 0 | 15835 | 384 | 0 |
| SRR6932950 | 0 | 0 | 0 | 0 | 0 | 125 | 331 | 200 | 287 | 0 | 0 | 0 | 0 | 0 | 37 | 113 | 189 | 0 | 0 | 137 | 13 | 0 | 0 | 24 | 0 | 0 | 0 | 0 | 0 | 18 | 0 | 0 | 0 | 0 | 13877 | 378 | 0 |
| SRR6932951 | 0 | 0 | 0 | 0 | 0 | 155 | 369 | 201 | 110 | 0 | 0 | 0 | 0 | 0 | 19 | 38 | 152 | 0 | 0 | 45 | 0 | 0 | 0 | 21 | 0 | 0 | 0 | 0 | 0 | 24 | 0 | 0 | 0 | 0 | 12007 | 413 | 0 |
| SRR6932952 | 2749 | 732 | 0 | 0 | 678 | 117 | 309 | 667 | 315 | 0 | 0 | 0 | 27 | 38 | 16025 | 15601 | 43602 | 68 | 432 | 56619 | 7272 | 0 | 0 | 342 | 43 | 60 | 16 | 0 | 0 | 1871 | 0 | 0 | 0 | 0 | 4852 | 184 | 0 |
| SRR6932953 | 183 | 53 | 41 | 26 | 56 | 393 | 734 | 1157 | 620 | 15 | 0 | 0 | 26 | 41 | 38027 | 20750 | 78975 | 76 | 393 | 51572 | 14589 | 0 | 0 | 16 | 10 | 15 | 21 | 0 | 0 | 117 | 0 | 0 | 0 | 0 | 4846 | 339 | 0 |
| SRR6932954 | 176 | 61 | 0 | 11 | 22 | 535 | 1018 | 1508 | 498 | 0 | 0 | 0 | 11 | 27 | 43808 | 28731 | 85097 | 0 | 52 | 29381 | 12525 | 0 | 0 | 108 | 0 | 0 | 0 | 0 | 0 | 20 | 0 | 0 | 0 | 0 | 7207 | 168 | 11 |
| SRR6932955 | 41 | 0 | 0 | 0 | 15 | 1623 | 1214 | 2982 | 325 | 10 | 0 | 0 | 0 | 0 | 63179 | 151776 | 209739 | 12 | 0 | 84 | 204 | 0 | 0 | 46 | 0 | 0 | 0 | 0 | 0 | 0 | 0 | 0 | 0 | 0 | 9426 | 498 | 0 |
| SRR7062776 | 32 | 17 | 12 | 12 | 410 | 139 | 135 | 125 | 73 | 32 | 124 | 0 | 0 | 0 | 39821 | 32839 | 95639 | 0 | 1348 | 15752 | 576 | 0 | 0 | 97 | 247 | 280 | 514 | 57 | 0 | 309 | 0 | 0 | 0 | 0 | 522 | 0 | 0 |
| SRR7062777 | 31 | 19 | 0 | 14 | 361 | 141 | 102 | 149 | 65 | 37 | 154 | 0 | 0 | 0 | 47471 | 33422 | 108846 | 0 | 1308 | 18574 | 654 | 0 | 0 | 82 | 265 | 244 | 390 | 38 | 0 | 328 | 0 | 10 | 0 | 0 | 512 | 0 | 0 |
| SRR7062778 | 31 | 32 | 12 | 0 | 424 | 157 | 138 | 181 | 97 | 26 | 161 | 0 | 0 | 0 | 24711 | 49961 | 66459 | 16 | 2691 | 26860 | 1118 | 0 | 13 | 96 | 167 | 155 | 255 | 24 | 0 | 661 | 0 | 17 | 0 | 0 | 1101 | 0 | 13 |
| SRR7062779 | 37 | 30 | 16 | 16 | 422 | 137 | 101 | 137 | 68 | 25 | 167 | 0 | 0 | 0 | 42368 | 44230 | 100875 | 22 | 2046 | 23426 | 918 | 0 | 24 | 76 | 177 | 172 | 300 | 25 | 0 | 638 | 0 | 27 | 0 | 0 | 774 | 0 | 10 |
| SRR7062780 | 27 | 26 | 11 | 11 | 68 | 208 | 165 | 200 | 37 | 16 | 46 | 0 | 0 | 0 | 22785 | 86633 | 60933 | 45 | 5385 | 25223 | 509 | 0 | 17 | 18 | 30 | 27 | 63 | 29 | 0 | 632 | 18 | 75 | 14 | 0 | 1772 | 0 | 25 |
| SRR7062781 | 34 | 28 | 11 | 13 | 323 | 190 | 154 | 172 | 69 | 21 | 205 | 0 | 0 | 0 | 34333 | 85448 | 85207 | 25 | 5198 | 24850 | 605 | 0 | 14 | 91 | 113 | 92 | 278 | 33 | 0 | 2303 | 13 | 105 | 11 | 0 | 1672 | 0 | 31 |
| SRR7081766 | 0 | 0 | 0 | 0 | 0 | 0 | 0 | 0 | 0 | 0 | 60 | 0 | 0 | 0 | 2651 | 24732 | 68288 | 0 | 0 | 16021 | 4495 | 0 | 0 | 15 | 0 | 15 | 30 | 0 | 0 | 135 | 0 | 346 | 0 | 0 | 1924 | 0 | 45 |
| SRR7081767 | 0 | 0 | 0 | 0 | 20 | 0 | 0 | 0 | 0 | 0 | 162 | 61 | 0 | 0 | 1714 | 21488 | 44864 | 0 | 0 | 10853 | 2475 | 0 | 0 | 0 | 0 | 0 | 142 | 0 | 0 | 162 | 0 | 190 | 0 | 0 | 4250 | 0 | 182 |
| SRR7081768 | 0 | 0 | 0 | 0 | 36 | 67 | 0 | 0 | 0 | 0 | 24 | 0 | 0 | 0 | 2316 | 111494 | 60530 | 0 | 201 | 4935 | 806 | 0 | 0 | 12 | 0 | 20 | 55 | 0 | 0 | 40 | 0 | 0 | 0 | 0 | 1635 | 0 | 83 |
| SRR7081769 | 0 | 0 | 0 | 0 | 0 | 0 | 0 | 58 | 0 | 69 | 26 | 0 | 0 | 0 | 2245 | 57513 | 92572 | 0 | 0 | 1382 | 288 | 0 | 0 | 43 | 0 | 86 | 354 | 0 | 0 | 224 | 0 | 230 | 0 | 0 | 6801 | 0 | 17 |
| SRR7081770 | 0 | 0 | 0 | 0 | 0 | 0 | 106 | 0 | 0 | 0 | 78 | 0 | 0 | 0 | 2759 | 25686 | 154114 | 0 | 106 | 18145 | 1698 | 0 | 0 | 39 | 0 | 0 | 410 | 0 | 0 | 254 | 0 | 0 | 0 | 0 | 2047 | 0 | 293 |
| SRR7081771 | 0 | 0 | 0 | 0 | 0 | 0 | 0 | 0 | 0 | 0 | 73 | 31 | 0 | 0 | 1735 | 36331 | 34348 | 89 | 0 | 37322 | 801 | 0 | 0 | 135 | 0 | 31 | 323 | 31 | 0 | 62 | 0 | 89 | 0 | 0 | 2840 | 0 | 219 |
| SRR7081772 | 0 | 0 | 0 | 0 | 0 | 0 | 0 | 220 | 0 | 0 | 62 | 25 | 0 | 0 | 3305 | 43768 | 90988 | 0 | 551 | 14324 | 1212 | 0 | 0 | 0 | 0 | 0 | 50 | 0 | 0 | 112 | 0 | 220 | 0 | 0 | 2959 | 0 | 199 |
| SRR7081773 | 0 | 0 | 0 | 0 | 0 | 0 | 363 | 0 | 0 | 20 | 40 | 134 | 0 | 0 | 9010 | 60379 | 173209 | 0 | 73 | 9446 | 727 | 0 | 0 | 20 | 0 | 27 | 20 | 0 | 0 | 120 | 0 | 0 | 0 | 0 | 554 | 0 | 280 |
| SRR7081774 | 0 | 0 | 0 | 0 | 0 | 0 | 0 | 0 | 0 | 0 | 69 | 26 | 0 | 0 | 5125 | 49964 | 114719 | 0 | 0 | 8875 | 2125 | 0 | 0 | 60 | 0 | 0 | 103 | 0 | 0 | 172 | 0 | 625 | 0 | 0 | 1443 | 0 | 129 |
| SRR7081775 | 0 | 0 | 0 | 0 | 0 | 0 | 0 | 139 | 0 | 0 | 148 | 43 | 0 | 0 | 3607 | 57310 | 117070 | 69 | 0 | 7284 | 1387 | 0 | 0 | 0 | 0 | 35 | 17 | 0 | 0 | 321 | 0 | 0 | 0 | 0 | 3172 | 0 | 148 |
| SRR7081776 | 0 | 0 | 0 | 0 | 0 | 0 | 0 | 0 | 102 | 15 | 122 | 76 | 0 | 0 | 816 | 39183 | 48149 | 0 | 0 | 4184 | 510 | 0 | 0 | 0 | 0 | 0 | 0 | 0 | 0 | 46 | 0 | 0 | 0 | 0 | 664 | 0 | 244 |
| SRR7081777 | 0 | 0 | 0 | 0 | 0 | 48 | 95 | 0 | 0 | 0 | 165 | 0 | 0 | 0 | 381 | 23191 | 46290 | 0 | 0 | 6707 | 285 | 0 | 0 | 37 | 0 | 18 | 110 | 0 | 0 | 165 | 0 | 0 | 0 | 0 | 1359 | 0 | 73 |
| SRR7107416 | 162 | 90 | 134 | 229 | 297 | 0 | 621 | 0 | 2051 | 0 | 2088 | 0 | 0 | 0 | 108 | 25195 | 212 | 60 | 24 | 3408 | 0 | 0 | 0 | 0 | 0 | 0 | 20 | 0 | 0 | 412 | 0 | 30 | 0 | 0 | 17172 | 0 | 0 |
| SRR7107417 | 11 | 122 | 78 | 111 | 23 | 0 | 880 | 0 | 2572 | 0 | 119 | 0 | 0 | 0 | 164 | 36045 | 339 | 30 | 12 | 3887 | 0 | 0 | 0 | 0 | 0 | 0 | 0 | 0 | 0 | 26 | 0 | 13 | 0 | 0 | 10271 | 0 | 0 |
| SRR7107418 | 40 | 21 | 85 | 29 | 0 | 508 | 48 | 170 | 60 | 0 | 0 | 0 | 0 | 0 | 72720 | 4791 | 34605 | 93 | 864 | 19047 | 68405 | 0 | 0 | 12 | 0 | 0 | 0 | 0 | 0 | 141 | 0 | 0 | 0 | 0 | 3313 | 19 | 0 |
| SRR7107419 | 39 | 27 | 89 | 36 | 0 | 364 | 42 | 126 | 39 | 0 | 0 | 0 | 0 | 0 | 70103 | 3610 | 33132 | 69 | 884 | 18624 | 75641 | 0 | 0 | 142 | 0 | 11 | 30 | 0 | 0 | 597 | 0 | 0 | 0 | 0 | 1859 | 20 | 0 |
| SRR7107420 | 95 | 47 | 56 | 199 | 317 | 0 | 711 | 0 | 1614 | 0 | 747 | 0 | 0 | 0 | 0 | 26527 | 0 | 38 | 0 | 3661 | 0 | 0 | 0 | 0 | 0 | 0 | 0 | 0 | 0 | 0 | 0 | 19 | 0 | 0 | 10015 | 0 | 0 |
| SRR7107421 | 82 | 47 | 58 | 10 | 16 | 0 | 720 | 0 | 1569 | 0 | 42 | 0 | 0 | 0 | 0 | 25335 | 0 | 32 | 0 | 3345 | 0 | 0 | 0 | 0 | 0 | 0 | 0 | 0 | 0 | 0 | 0 | 0 | 0 | 0 | 6362 | 0 | 0 |
| SRR7126083 | 15 | 0 | 0 | 0 | 161 | 799 | 0 | 333 | 133 | 380 | 175 | 15 | 0 | 0 | 6988 | 348732 | 27296 | 0 | 266 | 67 | 1265 | 15 | 58 | 15 | 29 | 29 | 15 | 15 | 0 | 88 | 175 | 0 | 0 | 0 | 2394 | 0 | 0 |
| SRR7126084 | 481 | 99 | 72 | 27 | 171 | 709 | 225 | 1683 | 1400 | 353 | 22 | 0 | 0 | 0 | 58760 | 86996 | 160779 | 392 | 3978 | 5210 | 8870 | 0 | 0 | 18 | 73 | 75 | 12 | 71 | 0 | 25 | 0 | 14 | 0 | 0 | 15258 | 0 | 0 |
| SRR7126085 | 795 | 187 | 92 | 46 | 202 | 994 | 354 | 2190 | 2245 | 508 | 28 | 0 | 0 | 0 | 51596 | 91202 | 141253 | 316 | 4629 | 3736 | 16700 | 0 | 0 | 30 | 13 | 13 | 16 | 73 | 0 | 25 | 0 | 0 | 0 | 0 | 18471 | 0 | 0 |
| SRR7126086 | 5512 | 1522 | 545 | 408 | 2720 | 810 | 278 | 1745 | 1625 | 8211 | 278 | 0 | 15 | 0 | 38867 | 137289 | 107763 | 201 | 5332 | 1985 | 24651 | 0 | 0 | 143 | 64 | 49 | 58 | 32 | 0 | 267 | 0 | 0 | 0 | 0 | 14757 | 0 | 0 |
| SRR7126087 | 2190 | 363 | 297 | 223 | 1832 | 1044 | 248 | 2171 | 1316 | 2787 | 201 | 0 | 0 | 11 | 85357 | 150200 | 246797 | 246 | 4262 | 4951 | 18127 | 0 | 0 | 139 | 50 | 43 | 60 | 35 | 0 | 171 | 0 | 0 | 0 | 0 | 7683 | 0 | 0 |
| SRR7126088 | 3755 | 987 | 352 | 302 | 3312 | 524 | 155 | 925 | 1091 | 3852 | 217 | 0 | 0 | 0 | 30602 | 167088 | 110503 | 150 | 5315 | 2936 | 24036 | 0 | 0 | 112 | 46 | 44 | 57 | 25 | 0 | 255 | 0 | 16 | 0 | 0 | 11433 | 0 | 0 |
| SRR7126089 | 523 | 140 | 36 | 35 | 356 | 474 | 155 | 789 | 1178 | 707 | 49 | 10 | 0 | 0 | 37790 | 186370 | 112134 | 183 | 3795 | 4309 | 21780 | 0 | 0 | 11 | 53 | 76 | 58 | 44 | 0 | 39 | 0 | 0 | 0 | 0 | 14850 | 0 | 0 |
| SRR7126090 | 917 | 243 | 82 | 36 | 279 | 582 | 173 | 1454 | 1521 | 780 | 25 | 0 | 14 | 0 | 48635 | 113188 | 194914 | 123 | 4007 | 5644 | 23055 | 0 | 0 | 21 | 10 | 11 | 68 | 38 | 0 | 18 | 0 | 17 | 0 | 0 | 17095 | 0 | 0 |
| SRR7126091 | 602 | 129 | 63 | 26 | 168 | 653 | 145 | 1648 | 1409 | 256 | 19 | 0 | 16 | 0 | 67571 | 96155 | 245952 | 195 | 4237 | 6639 | 15575 | 0 | 0 | 17 | 80 | 80 | 102 | 54 | 0 | 12 | 0 | 11 | 0 | 0 | 9500 | 0 | 0 |
| SRR7126092 | 417 | 100 | 35 | 27 | 258 | 638 | 91 | 1229 | 916 | 542 | 41 | 0 | 13 | 0 | 53213 | 164025 | 175157 | 193 | 4267 | 4364 | 20999 | 0 | 0 | 13 | 62 | 39 | 55 | 27 | 0 | 39 | 0 | 0 | 0 | 0 | 13673 | 0 | 0 |
| SRR7126093 | 621 | 152 | 62 | 41 | 242 | 609 | 293 | 1590 | 1066 | 600 | 29 | 0 | 0 | 16 | 39507 | 130298 | 114996 | 259 | 7758 | 3310 | 24374 | 0 | 0 | 14 | 27 | 21 | 41 | 52 | 0 | 41 | 0 | 0 | 0 | 0 | 22009 | 0 | 0 |
| SRR7153253 | 0 | 11 | 14 | 84 | 29 | 33 | 15 | 109 | 0 | 91 | 67 | 0 | 0 | 0 | 44254 | 30749 | 275656 | 0 | 3178 | 29290 | 194364 | 0 | 0 | 128 | 72 | 15 | 79 | 25 | 0 | 13 | 0 | 0 | 0 | 0 | 185 | 0 | 0 |
| SRR7153254 | 12 | 19 | 0 | 77 | 15 | 51 | 16 | 147 | 0 | 0 | 100 | 0 | 0 | 0 | 50879 | 16573 | 230471 | 0 | 3785 | 25088 | 148108 | 0 | 0 | 110 | 87 | 29 | 144 | 11 | 0 | 24 | 0 | 0 | 0 | 0 | 183 | 0 | 0 |
| SRR7153255 | 0 | 25 | 13 | 120 | 18 | 29 | 23 | 195 | 22 | 0 | 43 | 0 | 0 | 0 | 30055 | 28615 | 227859 | 0 | 2956 | 21783 | 220014 | 0 | 0 | 47 | 62 | 356 | 1969 | 588 | 0 | 176 | 0 | 0 | 0 | 0 | 187 | 0 | 0 |
| SRR7153256 | 12 | 58 | 37 | 327 | 48 | 79 | 34 | 272 | 40 | 0 | 72 | 0 | 0 | 0 | 34674 | 20227 | 252276 | 0 | 2666 | 26283 | 173651 | 0 | 0 | 199 | 215 | 723 | 3368 | 1915 | 0 | 231 | 0 | 13 | 0 | 0 | 269 | 0 | 0 |
| SRR7153257 | 0 | 13 | 0 | 69 | 13 | 14 | 13 | 67 | 0 | 0 | 86 | 0 | 0 | 0 | 31759 | 19733 | 253006 | 0 | 4674 | 24817 | 141575 | 0 | 0 | 54 | 104 | 426 | 2629 | 1073 | 0 | 291 | 0 | 0 | 0 | 0 | 69 | 0 | 0 |
| SRR7153258 | 0 | 11 | 21 | 19 | 12 | 43 | 23 | 127 | 16 | 0 | 70 | 0 | 0 | 0 | 35259 | 36080 | 249232 | 0 | 3043 | 24299 | 199001 | 0 | 0 | 71 | 77 | 31 | 159 | 153 | 0 | 13 | 0 | 0 | 0 | 0 | 180 | 0 | 0 |
| SRR7153259 | 0 | 0 | 13 | 94 | 33 | 14 | 10 | 59 | 0 | 0 | 69 | 0 | 0 | 0 | 41728 | 24485 | 271644 | 0 | 4190 | 26157 | 138541 | 0 | 0 | 111 | 95 | 410 | 2164 | 926 | 0 | 256 | 0 | 0 | 0 | 0 | 134 | 0 | 0 |
| SRR7153260 | 0 | 14 | 17 | 155 | 36 | 26 | 13 | 76 | 0 | 0 | 59 | 0 | 0 | 0 | 66429 | 32223 | 409237 | 0 | 3881 | 25135 | 167285 | 0 | 0 | 199 | 132 | 496 | 1929 | 804 | 0 | 372 | 0 | 0 | 0 | 0 | 143 | 0 | 0 |
| SRR7153261 | 0 | 18 | 20 | 158 | 18 | 0 | 0 | 19 | 0 | 24 | 115 | 0 | 0 | 0 | 58021 | 25563 | 345212 | 0 | 4815 | 26927 | 130287 | 0 | 0 | 72 | 65 | 268 | 1586 | 1268 | 0 | 309 | 0 | 0 | 0 | 0 | 48 | 0 | 0 |
| SRR7153262 | 0 | 12 | 0 | 34 | 12 | 37 | 33 | 248 | 28 | 0 | 81 | 0 | 0 | 0 | 25861 | 23392 | 187557 | 0 | 2534 | 31049 | 98860 | 0 | 0 | 73 | 73 | 34 | 198 | 109 | 0 | 24 | 0 | 0 | 0 | 0 | 262 | 0 | 0 |
| SRR7153263 | 0 | 15 | 23 | 13 | 34 | 129 | 64 | 518 | 58 | 0 | 39 | 0 | 0 | 0 | 56839 | 30774 | 302357 | 0 | 3287 | 29968 | 109371 | 0 | 0 | 11 | 112 | 42 | 197 | 102 | 0 | 11 | 0 | 0 | 0 | 0 | 191 | 0 | 0 |
| SRR7153264 | 10 | 21 | 26 | 11 | 44 | 24 | 14 | 67 | 10 | 0 | 110 | 0 | 0 | 0 | 51272 | 30710 | 352788 | 0 | 3972 | 23115 | 177871 | 0 | 0 | 149 | 152 | 28 | 83 | 52 | 0 | 16 | 0 | 0 | 0 | 0 | 135 | 0 | 0 |
| SRR7153265 | 0 | 19 | 16 | 145 | 30 | 38 | 19 | 169 | 10 | 0 | 46 | 0 | 0 | 0 | 33042 | 29841 | 267975 | 0 | 2935 | 24018 | 213209 | 0 | 0 | 121 | 99 | 399 | 2059 | 1118 | 0 | 177 | 0 | 0 | 0 | 0 | 152 | 0 | 0 |
| SRR7153266 | 0 | 22 | 39 | 25 | 26 | 15 | 11 | 92 | 0 | 0 | 75 | 0 | 0 | 0 | 29058 | 22907 | 232998 | 0 | 3379 | 28469 | 192302 | 0 | 0 | 76 | 87 | 43 | 277 | 144 | 0 | 22 | 0 | 0 | 0 | 0 | 182 | 0 | 0 |
| SRR7153267 | 0 | 24 | 0 | 99 | 21 | 33 | 14 | 110 | 12 | 0 | 66 | 0 | 0 | 0 | 16991 | 33119 | 236170 | 0 | 3015 | 25508 | 248693 | 0 | 0 | 116 | 110 | 38 | 177 | 22 | 0 | 14 | 0 | 0 | 0 | 0 | 168 | 0 | 0 |
| SRR7153268 | 0 | 10 | 50 | 296 | 20 | 0 | 0 | 0 | 0 | 0 | 69 | 0 | 0 | 0 | 31307 | 59101 | 319832 | 0 | 3475 | 31537 | 164053 | 0 | 0 | 25 | 128 | 522 | 2177 | 1224 | 0 | 229 | 0 | 0 | 0 | 0 | 20 | 0 | 0 |
| SRR7153269 | 0 | 0 | 35 | 74 | 36 | 24 | 13 | 79 | 0 | 0 | 36 | 0 | 0 | 0 | 46211 | 43113 | 209009 | 0 | 3024 | 35155 | 163794 | 0 | 0 | 75 | 163 | 583 | 1878 | 143 | 0 | 180 | 0 | 19 | 0 | 0 | 168 | 0 | 0 |
| SRR7153270 | 0 | 14 | 12 | 121 | 22 | 11 | 0 | 56 | 0 | 0 | 77 | 0 | 0 | 0 | 40535 | 24753 | 268927 | 0 | 6491 | 25730 | 153469 | 0 | 0 | 60 | 0 | 0 | 0 | 0 | 0 | 310 | 0 | 0 | 0 | 0 | 48 | 0 | 0 |
| SRR7153271 | 0 | 0 | 0 | 18 | 12 | 45 | 24 | 169 | 14 | 0 | 15 | 0 | 0 | 0 | 24920 | 27456 | 211956 | 0 | 4222 | 17835 | 152711 | 0 | 0 | 20 | 11 | 61 | 362 | 191 | 0 | 46 | 0 | 0 | 0 | 0 | 157 | 0 | 0 |
| SRR7153272 | 0 | 0 | 0 | 59 | 16 | 45 | 22 | 158 | 22 | 0 | 54 | 0 | 0 | 0 | 32833 | 34433 | 235878 | 0 | 3997 | 26578 | 135102 | 0 | 0 | 63 | 107 | 32 | 170 | 88 | 0 | 16 | 0 | 0 | 0 | 0 | 164 | 0 | 0 |
| SRR7153273 | 0 | 0 | 14 | 70 | 15 | 64 | 54 | 195 | 18 | 0 | 90 | 0 | 0 | 0 | 90244 | 60586 | 446434 | 0 | 762 | 27779 | 21016 | 0 | 0 | 141 | 77 | 207 | 1092 | 879 | 0 | 247 | 0 | 41 | 0 | 0 | 635 | 0 | 0 |
| SRR7153274 | 0 | 31 | 0 | 72 | 0 | 67 | 22 | 334 | 21 | 0 | 73 | 0 | 0 | 0 | 19349 | 23382 | 168713 | 0 | 2757 | 24695 | 224749 | 0 | 0 | 58 | 79 | 435 | 2628 | 1065 | 0 | 239 | 0 | 0 | 0 | 0 | 197 | 0 | 0 |
| SRR7153275 | 10 | 26 | 30 | 252 | 17 | 40 | 12 | 139 | 20 | 0 | 121 | 0 | 0 | 0 | 30214 | 37526 | 199065 | 0 | 3871 | 28050 | 106082 | 0 | 0 | 79 | 123 | 602 | 2867 | 1407 | 0 | 473 | 0 | 12 | 0 | 0 | 116 | 0 | 0 |
| SRR7153276 | 0 | 12 | 0 | 92 | 17 | 144 | 113 | 857 | 51 | 0 | 28 | 0 | 0 | 0 | 21095 | 44438 | 180820 | 0 | 1595 | 11470 | 98798 | 0 | 0 | 38 | 51 | 306 | 1883 | 749 | 0 | 147 | 0 | 0 | 0 | 0 | 831 | 0 | 0 |
| SRR7153277 | 0 | 0 | 0 | 50 | 0 | 51 | 20 | 166 | 17 | 0 | 38 | 0 | 0 | 0 | 15747 | 26514 | 211087 | 0 | 2551 | 21380 | 276669 | 0 | 0 | 22 | 138 | 516 | 1686 | 472 | 0 | 141 | 0 | 0 | 0 | 0 | 125 | 0 | 0 |
| SRR7153278 | 0 | 0 | 13 | 57 | 11 | 83 | 49 | 231 | 30 | 58 | 180 | 17 | 0 | 0 | 31463 | 23372 | 166693 | 0 | 2252 | 16871 | 169258 | 0 | 0 | 17 | 353 | 486 | 923 | 743 | 0 | 415 | 0 | 0 | 0 | 0 | 161 | 0 | 0 |
| SRR7153279 | 0 | 0 | 0 | 45 | 17 | 92 | 53 | 295 | 37 | 14 | 67 | 0 | 0 | 0 | 112534 | 75666 | 358288 | 11 | 707 | 30403 | 13562 | 0 | 0 | 84 | 70 | 239 | 1439 | 800 | 0 | 220 | 0 | 18 | 0 | 0 | 719 | 0 | 0 |
| SRR7153280 | 0 | 16 | 0 | 10 | 11 | 92 | 28 | 296 | 17 | 0 | 60 | 0 | 0 | 0 | 25111 | 28287 | 181269 | 0 | 2525 | 27270 | 168485 | 0 | 0 | 49 | 81 | 46 | 250 | 115 | 0 | 25 | 0 | 0 | 0 | 0 | 206 | 0 | 0 |
| SRR7153281 | 0 | 13 | 28 | 113 | 31 | 37 | 16 | 98 | 0 | 0 | 98 | 0 | 0 | 0 | 23766 | 97106 | 188143 | 17 | 2895 | 27443 | 148614 | 0 | 0 | 103 | 128 | 22 | 105 | 61 | 0 | 23 | 0 | 17 | 0 | 0 | 172 | 0 | 0 |
| SRR7153282 | 0 | 0 | 0 | 60 | 11 | 64 | 27 | 263 | 19 | 0 | 50 | 0 | 0 | 0 | 25969 | 33784 | 189013 | 0 | 2994 | 19362 | 190496 | 0 | 0 | 48 | 79 | 274 | 1190 | 892 | 0 | 220 | 0 | 0 | 0 | 0 | 200 | 0 | 0 |
| SRR7153283 | 0 | 14 | 13 | 113 | 30 | 41 | 33 | 215 | 12 | 0 | 50 | 0 | 0 | 0 | 25434 | 54076 | 156735 | 13 | 2869 | 23377 | 122940 | 0 | 0 | 84 | 11 | 55 | 221 | 51 | 0 | 14 | 0 | 14 | 0 | 0 | 219 | 0 | 0 |
| SRR7153284 | 0 | 0 | 0 | 36 | 18 | 19 | 0 | 46 | 0 | 0 | 128 | 0 | 0 | 0 | 15871 | 56064 | 285974 | 0 | 1774 | 14675 | 136526 | 0 | 0 | 52 | 105 | 520 | 1402 | 473 | 0 | 704 | 0 | 0 | 0 | 0 | 94 | 0 | 0 |
| SRR7153285 | 0 | 19 | 12 | 53 | 24 | 34 | 17 | 137 | 16 | 0 | 66 | 0 | 0 | 0 | 23012 | 36817 | 195744 | 0 | 2384 | 23041 | 252246 | 0 | 0 | 56 | 121 | 542 | 2198 | 79 | 0 | 245 | 0 | 18 | 0 | 0 | 114 | 0 | 0 |
| SRR7153286 | 0 | 0 | 0 | 68 | 37 | 89 | 84 | 383 | 16 | 0 | 39 | 0 | 0 | 0 | 28316 | 54540 | 208846 | 0 | 2273 | 17643 | 191214 | 0 | 0 | 53 | 90 | 447 | 1569 | 377 | 0 | 164 | 0 | 0 | 0 | 0 | 266 | 0 | 0 |
| SRR7153287 | 0 | 0 | 22 | 193 | 17 | 0 | 0 | 19 | 0 | 0 | 143 | 0 | 0 | 0 | 40667 | 30557 | 263128 | 0 | 3477 | 26795 | 233347 | 0 | 0 | 33 | 112 | 644 | 2770 | 458 | 0 | 248 | 0 | 0 | 0 | 0 | 40 | 0 | 0 |
| SRR7153288 | 0 | 15 | 15 | 119 | 21 | 19 | 11 | 67 | 0 | 0 | 72 | 0 | 0 | 0 | 20529 | 52924 | 176720 | 0 | 3511 | 28225 | 162066 | 0 | 0 | 72 | 76 | 524 | 2618 | 305 | 0 | 252 | 0 | 0 | 0 | 0 | 96 | 0 | 0 |
| SRR7153289 | 0 | 17 | 0 | 59 | 0 | 30 | 11 | 121 | 13 | 0 | 55 | 0 | 0 | 0 | 19971 | 45107 | 193811 | 0 | 2610 | 21118 | 89386 | 0 | 0 | 70 | 90 | 29 | 124 | 44 | 0 | 22 | 0 | 0 | 0 | 0 | 217 | 0 | 0 |
| SRR7153290 | 0 | 19 | 0 | 82 | 25 | 44 | 18 | 232 | 11 | 0 | 35 | 0 | 0 | 0 | 18100 | 41637 | 184329 | 0 | 2405 | 23217 | 180084 | 0 | 0 | 43 | 15 | 55 | 229 | 61 | 0 | 16 | 0 | 16 | 0 | 0 | 131 | 0 | 0 |
| SRR7153291 | 0 | 0 | 0 | 28 | 19 | 166 | 73 | 424 | 13 | 0 | 98 | 0 | 0 | 0 | 94610 | 83351 | 351261 | 0 | 729 | 24185 | 21413 | 0 | 0 | 93 | 69 | 20 | 94 | 17 | 0 | 19 | 0 | 43 | 0 | 0 | 831 | 0 | 0 |
| SRR7153292 | 0 | 0 | 12 | 48 | 12 | 67 | 33 | 188 | 14 | 17 | 84 | 0 | 0 | 0 | 77097 | 97568 | 363044 | 21 | 795 | 28697 | 25127 | 0 | 0 | 117 | 65 | 18 | 79 | 24 | 0 | 20 | 0 | 98 | 0 | 0 | 647 | 0 | 0 |
| SRR7153293 | 0 | 0 | 10 | 68 | 17 | 32 | 40 | 114 | 16 | 12 | 148 | 0 | 0 | 0 | 49733 | 88056 | 260930 | 0 | 681 | 21524 | 19819 | 0 | 0 | 33 | 53 | 272 | 1570 | 935 | 0 | 367 | 0 | 23 | 0 | 0 | 346 | 0 | 0 |
| SRR7153294 | 0 | 0 | 0 | 57 | 11 | 40 | 18 | 111 | 0 | 0 | 83 | 0 | 0 | 0 | 17727 | 48885 | 203761 | 17 | 4317 | 25105 | 173431 | 0 | 0 | 44 | 81 | 402 | 1895 | 809 | 0 | 234 | 0 | 0 | 0 | 0 | 147 | 0 | 0 |
| SRR7153295 | 0 | 12 | 22 | 101 | 14 | 15 | 0 | 71 | 0 | 0 | 13 | 0 | 0 | 0 | 25045 | 70815 | 289505 | 19 | 5200 | 32380 | 123589 | 0 | 0 | 149 | 100 | 38 | 138 | 88 | 0 | 38 | 0 | 25 | 0 | 0 | 236 | 0 | 0 |
| SRR7153296 | 0 | 0 | 0 | 37 | 0 | 74 | 36 | 334 | 27 | 0 | 37 | 0 | 0 | 0 | 17634 | 32384 | 162312 | 0 | 2196 | 19081 | 163278 | 0 | 0 | 28 | 46 | 318 | 2126 | 707 | 0 | 160 | 0 | 0 | 0 | 0 | 390 | 0 | 0 |
| SRR7153297 | 0 | 0 | 0 | 11 | 0 | 82 | 48 | 282 | 27 | 37 | 281 | 67 | 0 | 0 | 22213 | 44079 | 176057 | 0 | 2457 | 21832 | 171750 | 0 | 0 | 13 | 58 | 148 | 520 | 127 | 0 | 855 | 0 | 12 | 0 | 0 | 458 | 0 | 26 |
| SRR7153298 | 0 | 0 | 10 | 89 | 12 | 48 | 18 | 137 | 0 | 0 | 42 | 0 | 0 | 0 | 25226 | 28304 | 181943 | 0 | 3024 | 26214 | 202705 | 0 | 0 | 32 | 59 | 367 | 1942 | 625 | 0 | 119 | 0 | 0 | 0 | 0 | 147 | 0 | 0 |

### Supp Table S8.3.Columns_BX_to_DH

| | | | | | | | | | | | | | | | | | | | | | | | | | | | | | | | | | | | | | |
|---|---|---|---|---|---|---|---|---|---|---|---|---|---|---|---|---|---|---|---|---|---|---|---|---|---|---|---|---|---|---|---|---|---|---|---|---|---|
| SRR7153299 | 10 | 23 | 0 | 85 | 23 | 22 | 13 | 126 | 11 | 0 | 81 | 0 | 0 | 0 | 16547 | 20509 | 214648 | 0 | 2975 | 20590 | 301136 | 0 | 0 | 83 | 491 | 2355 | 8123 | 170 | 0 | 426 | 0 | 0 | 0 | 0 | 98 | 0 | 0 |
| SRR7153300 | 0 | 0 | 13 | 91 | 21 | 16 | 0 | 84 | 0 | 0 | 92 | 0 | 0 | 0 | 30344 | 35828 | 191546 | 0 | 2578 | 14103 | 339113 | 0 | 0 | 44 | 75 | 33 | 186 | 43 | 0 | 31 | 0 | 0 | 0 | 0 | 112 | 0 | 0 |
| SRR7153301 | 0 | 0 | 0 | 32 | 14 | 339 | 424 | 900 | 142 | 66 | 72 | 0 | 0 | 0 | 30266 | 35747 | 175956 | 10 | 2676 | 35746 | 88648 | 0 | 0 | 22 | 204 | 319 | 658 | 531 | 0 | 219 | 0 | 0 | 0 | 0 | 328 | 0 | 0 |
| SRR7153302 | 0 | 0 | 0 | 34 | 0 | 47 | 23 | 136 | 0 | 0 | 40 | 0 | 0 | 0 | 31014 | 27562 | 226145 | 0 | 3205 | 18970 | 267509 | 0 | 0 | 53 | 157 | 583 | 2118 | 896 | 0 | 187 | 0 | 0 | 0 | 0 | 228 | 0 | 0 |
| SRR7153303 | 0 | 0 | 0 | 30 | 19 | 271 | 144 | 649 | 47 | 0 | 59 | 0 | 0 | 0 | 37350 | 81312 | 262386 | 0 | 327 | 20976 | 3401 | 0 | 0 | 31 | 50 | 139 | 793 | 123 | 0 | 277 | 0 | 17 | 0 | 0 | 1529 | 0 | 0 |
| SRR7153304 | 0 | 18 | 20 | 74 | 15 | 14 | 0 | 47 | 0 | 0 | 62 | 0 | 0 | 0 | 23386 | 42731 | 180332 | 28 | 4019 | 31215 | 135418 | 0 | 0 | 52 | 77 | 472 | 2561 | 458 | 0 | 244 | 0 | 18 | 0 | 0 | 113 | 0 | 0 |
| SRR7153305 | 0 | 0 | 17 | 83 | 0 | 39 | 19 | 231 | 19 | 0 | 46 | 0 | 0 | 0 | 20317 | 27933 | 163891 | 0 | 2967 | 20325 | 189397 | 0 | 0 | 39 | 53 | 40 | 255 | 96 | 0 | 17 | 0 | 0 | 0 | 0 | 180 | 0 | 0 |
| SRR7153306 | 0 | 0 | 18 | 47 | 17 | 37 | 22 | 136 | 26 | 0 | 11 | 0 | 0 | 0 | 29841 | 32027 | 202873 | 0 | 4205 | 21948 | 134515 | 0 | 0 | 98 | 46 | 138 | 470 | 183 | 0 | 34 | 0 | 13 | 0 | 0 | 196 | 0 | 0 |
| SRR7153307 | 0 | 22 | 10 | 129 | 21 | 27 | 18 | 136 | 0 | 0 | 64 | 0 | 0 | 0 | 34896 | 25027 | 199006 | 12 | 3484 | 33105 | 164622 | 0 | 0 | 55 | 108 | 555 | 2121 | 1176 | 0 | 230 | 0 | 0 | 0 | 0 | 141 | 0 | 0 |
| SRR7153308 | 0 | 14 | 13 | 57 | 18 | 105 | 32 | 298 | 21 | 0 | 43 | 0 | 0 | 0 | 19765 | 36231 | 179740 | 0 | 2792 | 18868 | 220863 | 0 | 0 | 72 | 132 | 566 | 2562 | 557 | 0 | 147 | 0 | 0 | 0 | 0 | 245 | 0 | 0 |
| SRR7153309 | 0 | 0 | 11 | 73 | 10 | 18 | 14 | 106 | 0 | 0 | 40 | 0 | 0 | 0 | 33093 | 37573 | 275332 | 0 | 3009 | 17916 | 314227 | 0 | 0 | 55 | 70 | 38 | 162 | 72 | 0 | 14 | 0 | 0 | 0 | 0 | 87 | 0 | 0 |
| SRR7153310 | 0 | 20 | 0 | 0 | 54 | 65 | 51 | 200 | 27 | 55 | 59 | 0 | 0 | 0 | 24260 | 36474 | 231437 | 0 | 2177 | 19810 | 237594 | 0 | 0 | 86 | 20 | 37 | 80 | 29 | 0 | 33 | 0 | 0 | 0 | 0 | 229 | 0 | 0 |
| SRR7153311 | 0 | 0 | 0 | 66 | 12 | 86 | 40 | 289 | 33 | 0 | 25 | 0 | 0 | 0 | 26751 | 28890 | 145035 | 0 | 2347 | 19431 | 160855 | 0 | 0 | 44 | 91 | 576 | 3377 | 485 | 0 | 164 | 0 | 10 | 0 | 0 | 284 | 0 | 0 |
| SRR7153312 | 0 | 0 | 0 | 74 | 20 | 120 | 85 | 411 | 44 | 0 | 84 | 0 | 0 | 0 | 46291 | 85793 | 271611 | 34 | 617 | 29843 | 16484 | 0 | 0 | 82 | 71 | 22 | 107 | 52 | 0 | 24 | 0 | 51 | 0 | 0 | 856 | 0 | 0 |
| SRR7153313 | 0 | 0 | 0 | 64 | 21 | 41 | 21 | 74 | 16 | 43 | 79 | 0 | 0 | 0 | 97248 | 167002 | 343618 | 36 | 489 | 27668 | 12912 | 0 | 0 | 79 | 58 | 152 | 24 | 21 | 0 | 17 | 0 | 150 | 0 | 0 | 290 | 0 | 0 |
| SRR7153314 | 0 | 0 | 28 | 62 | 0 | 38 | 11 | 93 | 0 | 0 | 60 | 0 | 0 | 0 | 25238 | 55633 | 182655 | 24 | 3601 | 28986 | 158196 | 0 | 0 | 81 | 14 | 54 | 182 | 23 | 0 | 17 | 0 | 14 | 0 | 0 | 151 | 0 | 0 |
| SRR7153315 | 0 | 13 | 11 | 10 | 12 | 27 | 17 | 122 | 22 | 22 | 46 | 0 | 0 | 0 | 19792 | 28084 | 165740 | 0 | 3014 | 22975 | 219694 | 0 | 0 | 32 | 51 | 41 | 230 | 133 | 0 | 29 | 0 | 0 | 0 | 0 | 240 | 0 | 0 |
| SRR7153316 | 0 | 13 | 17 | 21 | 10 | 21 | 17 | 78 | 14 | 0 | 18 | 0 | 0 | 0 | 18796 | 32570 | 134871 | 0 | 3503 | 31961 | 85845 | 0 | 0 | 10 | 14 | 106 | 563 | 151 | 0 | 48 | 0 | 0 | 0 | 0 | 93 | 0 | 0 |
| SRR7153317 | 0 | 11 | 0 | 16 | 18 | 54 | 48 | 179 | 23 | 123 | 83 | 0 | 0 | 0 | 91460 | 31574 | 473573 | 0 | 1039 | 23068 | 21367 | 0 | 0 | 64 | 83 | 206 | 869 | 34 | 0 | 423 | 0 | 0 | 0 | 0 | 497 | 0 | 0 |
| SRR7153318 | 0 | 0 | 11 | 28 | 17 | 214 | 32 | 326 | 35 | 49 | 524 | 30 | 0 | 0 | 47516 | 34088 | 278335 | 0 | 2314 | 24212 | 190876 | 0 | 0 | 37 | 425 | 467 | 494 | 109 | 0 | 1288 | 0 | 21 | 0 | 0 | 302 | 0 | 14 |
| SRR7153319 | 0 | 0 | 0 | 15 | 0 | 141 | 101 | 406 | 33 | 41 | 44 | 94 | 0 | 0 | 43120 | 102045 | 194272 | 21 | 581 | 36676 | 4597 | 0 | 0 | 34 | 26 | 127 | 764 | 144 | 0 | 178 | 0 | 169 | 0 | 0 | 881 | 0 | 23 |
| SRR7153320 | 0 | 0 | 0 | 75 | 18 | 107 | 71 | 418 | 31 | 0 | 85 | 0 | 0 | 0 | 69867 | 41150 | 331861 | 0 | 627 | 20238 | 37490 | 0 | 0 | 54 | 30 | 14 | 87 | 44 | 0 | 24 | 0 | 25 | 0 | 0 | 799 | 0 | 0 |
| SRR7153321 | 0 | 0 | 13 | 14 | 25 | 21 | 25 | 104 | 14 | 20 | 24 | 12 | 0 | 0 | 43759 | 67732 | 228199 | 0 | 519 | 46082 | 5356 | 0 | 0 | 18 | 12 | 56 | 313 | 160 | 0 | 60 | 0 | 17 | 0 | 0 | 244 | 0 | 0 |
| SRR7153322 | 15 | 11 | 0 | 12 | 33 | 40 | 24 | 115 | 0 | 20 | 45 | 0 | 0 | 0 | 51913 | 149855 | 293722 | 14 | 504 | 37376 | 16756 | 0 | 0 | 78 | 206 | 14 | 19 | 18 | 0 | 14 | 0 | 66 | 0 | 0 | 313 | 0 | 0 |
| SRR7153323 | 0 | 0 | 0 | 28 | 26 | 81 | 64 | 254 | 22 | 15 | 94 | 0 | 0 | 0 | 48743 | 72796 | 316010 | 0 | 604 | 20469 | 28759 | 0 | 0 | 49 | 57 | 202 | 752 | 515 | 0 | 331 | 0 | 74 | 0 | 0 | 878 | 0 | 0 |
| SRR7153324 | 0 | 23 | 23 | 68 | 23 | 40 | 12 | 82 | 0 | 29 | 103 | 0 | 0 | 0 | 32027 | 98824 | 206220 | 68 | 2561 | 24367 | 140675 | 0 | 0 | 121 | 10 | 26 | 74 | 168 | 0 | 17 | 0 | 79 | 0 | 0 | 728 | 0 | 0 |
| SRR7153325 | 0 | 0 | 0 | 12 | 0 | 122 | 73 | 380 | 25 | 16 | 82 | 0 | 0 | 0 | 63289 | 49033 | 366670 | 0 | 654 | 30358 | 43697 | 0 | 0 | 150 | 86 | 259 | 1249 | 111 | 0 | 380 | 0 | 29 | 0 | 0 | 1069 | 0 | 0 |
| SRR7153326 | 0 | 0 | 0 | 40 | 15 | 90 | 51 | 289 | 23 | 0 | 112 | 0 | 0 | 0 | 61595 | 90096 | 321158 | 18 | 629 | 25037 | 7585 | 0 | 0 | 59 | 46 | 134 | 805 | 342 | 0 | 324 | 0 | 35 | 0 | 0 | 908 | 0 | 0 |
| SRR7153327 | 0 | 0 | 0 | 45 | 27 | 180 | 77 | 562 | 30 | 0 | 97 | 0 | 0 | 0 | 64423 | 38981 | 336769 | 0 | 435 | 23175 | 27234 | 0 | 0 | 80 | 43 | 151 | 1159 | 150 | 0 | 290 | 0 | 30 | 0 | 0 | 1432 | 0 | 0 |
| SRR7153328 | 0 | 0 | 0 | 29 | 18 | 93 | 28 | 269 | 18 | 0 | 124 | 0 | 0 | 0 | 97737 | 61819 | 466482 | 0 | 997 | 21763 | 27594 | 0 | 0 | 12 | 19 | 54 | 135 | 61 | 0 | 25 | 0 | 19 | 0 | 0 | 679 | 0 | 0 |
| SRR7153329 | 0 | 0 | 0 | 38 | 24 | 149 | 101 | 501 | 44 | 29 | 70 | 0 | 0 | 0 | 57113 | 96622 | 314231 | 24 | 723 | 22043 | 13688 | 0 | 0 | 73 | 37 | 17 | 103 | 69 | 0 | 28 | 0 | 38 | 0 | 0 | 1055 | 0 | 0 |
| SRR7153330 | 0 | 0 | 0 | 29 | 12 | 57 | 54 | 205 | 19 | 88 | 95 | 24 | 0 | 0 | 94351 | 59877 | 306524 | 0 | 578 | 24069 | 19964 | 0 | 0 | 52 | 52 | 146 | 791 | 447 | 0 | 482 | 0 | 59 | 0 | 0 | 600 | 0 | 0 |
| SRR7153331 | 0 | 0 | 0 | 38 | 15 | 151 | 139 | 552 | 55 | 71 | 15 | 38 | 0 | 0 | 61520 | 44176 | 321332 | 0 | 505 | 19297 | 24644 | 0 | 0 | 49 | 55 | 25 | 161 | 59 | 0 | 56 | 0 | 26 | 0 | 0 | 1288 | 0 | 0 |
| SRR7162919 | 29 | 29 | 0 | 0 | 14 | 456 | 532 | 1003 | 1398 | 187 | 0 | 0 | 0 | 0 | 972 | 3733 | 2516 | 182 | 46 | 76 | 213 | 0 | 0 | 79 | 22 | 14 | 50 | 0 | 0 | 58 | 0 | 76 | 0 | 0 | 17693 | 6173 | 0 |
| SRR7162920 | 25 | 25 | 0 | 0 | 0 | 237 | 107 | 285 | 311 | 59 | 0 | 0 | 0 | 0 | 987 | 1562 | 1531 | 425 | 0 | 107 | 55 | 0 | 0 | 47 | 16 | 18 | 0 | 0 | 0 | 0 | 0 | 122 | 0 | 0 | 7921 | 6303 | 0 |
| SRR7162921 | 11 | 40 | 0 | 0 | 34 | 215 | 209 | 424 | 824 | 28 | 0 | 0 | 0 | 0 | 1320 | 2249 | 3240 | 167 | 10 | 66 | 36 | 0 | 0 | 91 | 40 | 57 | 11 | 0 | 0 | 28 | 0 | 12 | 0 | 0 | 4232 | 7315 | 0 |
| SRR7178585 | 14 | 0 | 14 | 71 | 71 | 51 | 42 | 267 | 27 | 0 | 124 | 0 | 0 | 0 | 29461 | 18167 | 179228 | 34 | 0 | 13045 | 1557 | 0 | 0 | 283 | 35 | 233 | 52 | 26 | 0 | 25 | 0 | 97 | 0 | 0 | 124 | 436 | 0 |
| SRR7178586 | 13 | 33 | 33 | 119 | 53 | 34 | 41 | 307 | 20 | 0 | 106 | 0 | 0 | 0 | 28628 | 17632 | 174332 | 31 | 14 | 13214 | 1442 | 0 | 0 | 12 | 46 | 237 | 63 | 33 | 0 | 24 | 0 | 97 | 0 | 0 | 107 | 405 | 0 |
| SRR7178587 | 32 | 0 | 16 | 32 | 111 | 48 | 32 | 281 | 21 | 16 | 207 | 0 | 0 | 0 | 39724 | 22993 | 233835 | 37 | 27 | 13896 | 1614 | 0 | 0 | 381 | 16 | 191 | 1255 | 779 | 0 | 651 | 0 | 80 | 0 | 0 | 106 | 494 | 0 |
| SRR7178588 | 18 | 0 | 0 | 106 | 53 | 34 | 15 | 240 | 0 | 0 | 195 | 0 | 0 | 0 | 43076 | 23253 | 240373 | 0 | 15 | 13746 | 1628 | 0 | 0 | 159 | 53 | 159 | 23 | 566 | 0 | 15 | 0 | 69 | 0 | 0 | 92 | 567 | 18 |
| SRR7178589 | 11 | 11 | 0 | 23 | 25 | 212 | 110 | 446 | 51 | 0 | 106 | 0 | 0 | 0 | 11121 | 7356 | 82935 | 0 | 120 | 183693 | 970 | 0 | 0 | 246 | 0 | 0 | 0 | 0 | 0 | 642 | 14 | 534 | 0 | 0 | 927 | 721 | 0 |
| SRR7178590 | 0 | 0 | 0 | 23 | 38 | 216 | 138 | 489 | 52 | 0 | 80 | 0 | 0 | 0 | 10638 | 6898 | 82350 | 13 | 156 | 184043 | 809 | 0 | 0 | 272 | 0 | 0 | 0 | 0 | 0 | 611 | 0 | 493 | 0 | 0 | 1076 | 711 | 0 |
| SRR7178591 | 0 | 0 | 19 | 0 | 19 | 207 | 144 | 495 | 32 | 0 | 94 | 0 | 0 | 0 | 14086 | 8536 | 94136 | 48 | 175 | 196704 | 830 | 0 | 19 | 244 | 0 | 0 | 0 | 0 | 0 | 23 | 0 | 303 | 0 | 0 | 836 | 770 | 0 |
| SRR7178592 | 0 | 0 | 11 | 11 | 44 | 95 | 110 | 473 | 16 | 0 | 44 | 0 | 0 | 0 | 16121 | 8324 | 97823 | 0 | 79 | 208716 | 820 | 0 | 0 | 142 | 0 | 0 | 11 | 0 | 0 | 25 | 0 | 379 | 0 | 0 | 722 | 831 | 0 |
| SRR7178593 | 0 | 0 | 22 | 186 | 75 | 34 | 33 | 167 | 0 | 0 | 106 | 0 | 0 | 0 | 30375 | 13279 | 265274 | 0 | 164 | 66699 | 3009 | 0 | 0 | 345 | 274 | 1162 | 5630 | 1220 | 0 | 208 | 0 | 0 | 0 | 0 | 102 | 1063 | 0 |
| SRR7178594 | 0 | 30 | 34 | 137 | 43 | 47 | 26 | 169 | 11 | 0 | 77 | 0 | 0 | 0 | 29415 | 13358 | 263240 | 0 | 191 | 67383 | 2998 | 0 | 0 | 11 | 244 | 45 | 182 | 42 | 0 | 231 | 0 | 15 | 0 | 0 | 97 | 1063 | 0 |
| SRR7178595 | 0 | 0 | 0 | 156 | 52 | 50 | 67 | 269 | 0 | 0 | 52 | 17 | 0 | 0 | 40273 | 15663 | 297754 | 0 | 252 | 73696 | 2934 | 0 | 0 | 329 | 346 | 1314 | 6190 | 1280 | 0 | 207 | 0 | 17 | 0 | 0 | 80 | 1156 | 0 |
| SRR7178596 | 0 | 0 | 24 | 48 | 96 | 43 | 43 | 195 | 0 | 0 | 96 | 0 | 0 | 0 | 43256 | 15810 | 305323 | 0 | 246 | 74371 | 3620 | 0 | 0 | 506 | 313 | 1519 | 6630 | 1326 | 0 | 555 | 0 | 14 | 0 | 0 | 69 | 1153 | 0 |
| SRR7178597 | 0 | 0 | 16 | 36 | 41 | 64 | 55 | 205 | 14 | 0 | 68 | 0 | 0 | 0 | 17870 | 10731 | 118806 | 0 | 168 | 93600 | 1594 | 0 | 0 | 132 | 98 | 21 | 76 | 127 | 0 | 127 | 0 | 0 | 0 | 0 | 158 | 478 | 0 |
| SRR7178598 | 0 | 0 | 0 | 28 | 42 | 94 | 52 | 308 | 25 | 0 | 125 | 0 | 0 | 0 | 21778 | 6747 | 140357 | 0 | 111 | 138467 | 1727 | 0 | 0 | 11 | 46 | 15 | 67 | 86 | 0 | 20 | 0 | 54 | 0 | 0 | 311 | 662 | 0 |
| SRR7178599 | 0 | 0 | 0 | 22 | 34 | 94 | 48 | 197 | 15 | 0 | 104 | 0 | 0 | 0 | 22220 | 6945 | 143518 | 0 | 132 | 139683 | 1774 | 0 | 0 | 128 | 41 | 15 | 61 | 99 | 0 | 20 | 0 | 46 | 0 | 0 | 271 | 647 | 0 |
| SRR7178600 | 0 | 0 | 0 | 0 | 35 | 126 | 42 | 218 | 50 | 0 | 22 | 0 | 0 | 0 | 25008 | 8237 | 141546 | 0 | 15 | 176865 | 1136 | 0 | 0 | 57 | 26 | 11 | 25 | 11 | 0 | 378 | 0 | 165 | 0 | 0 | 3762 | 495 | 0 |
| SRR7178601 | 0 | 0 | 0 | 0 | 41 | 130 | 72 | 203 | 36 | 0 | 189 | 0 | 0 | 0 | 24096 | 7830 | 137413 | 0 | 47 | 179533 | 1084 | 0 | 0 | 74 | 25 | 83 | 183 | 12 | 0 | 2314 | 0 | 170 | 0 | 0 | 3680 | 501 | 0 |
| SRR7178602 | 0 | 0 | 0 | 0 | 53 | 105 | 45 | 226 | 15 | 0 | 149 | 0 | 0 | 0 | 33965 | 8710 | 160186 | 0 | 15 | 200581 | 1280 | 0 | 0 | 75 | 16 | 85 | 165 | 0 | 0 | 3262 | 0 | 166 | 0 | 0 | 2772 | 554 | 11 |
| SRR7178603 | 0 | 0 | 0 | 0 | 38 | 105 | 15 | 135 | 30 | 0 | 216 | 0 | 0 | 0 | 34992 | 8775 | 165011 | 0 | 60 | 209980 | 1227 | 0 | 0 | 76 | 32 | 70 | 165 | 13 | 0 | 3666 | 0 | 135 | 0 | 0 | 2742 | 638 | 0 |
| SRR7178604 | 0 | 0 | 13 | 29 | 22 | 37 | 63 | 392 | 25 | 16 | 160 | 0 | 0 | 0 | 40226 | 22929 | 244064 | 88 | 0 | 11829 | 3609 | 0 | 0 | 106 | 0 | 12 | 41 | 34 | 0 | 35 | 0 | 582 | 0 | 0 | 172 | 459 | 0 |
| SRR7178605 | 15 | 0 | 12 | 37 | 40 | 46 | 58 | 416 | 31 | 0 | 14 | 0 | 0 | 0 | 39001 | 23076 | 241029 | 62 | 0 | 11723 | 3648 | 0 | 0 | 95 | 40 | 11 | 45 | 35 | 0 | 37 | 0 | 613 | 0 | 0 | 169 | 449 | 0 |
| SRR7178606 | 28 | 0 | 0 | 14 | 42 | 69 | 139 | 473 | 28 | 14 | 222 | 0 | 0 | 0 | 55351 | 28236 | 306603 | 42 | 14 | 12092 | 3259 | 14 | 14 | 180 | 28 | 194 | 846 | 818 | 0 | 680 | 0 | 459 | 0 | 0 | 155 | 506 | 0 |
| SRR7178607 | 0 | 0 | 0 | 19 | 97 | 71 | 71 | 496 | 27 | 19 | 292 | 0 | 0 | 0 | 58035 | 29764 | 324772 | 44 | 0 | 12453 | 4080 | 0 | 0 | 117 | 39 | 175 | 777 | 661 | 0 | 816 | 19 | 602 | 0 | 0 | 124 | 571 | 0 |
| SRR7178608 | 0 | 0 | 0 | 47 | 26 | 32 | 50 | 122 | 0 | 0 | 63 | 21 | 0 | 0 | 37161 | 10535 | 141998 | 0 | 135 | 145951 | 2283 | 0 | 0 | 324 | 0 | 0 | 0 | 0 | 0 | 282 | 0 | 142 | 0 | 0 | 143 | 978 | 14 |
| SRR7178609 | 0 | 0 | 17 | 63 | 17 | 53 | 31 | 139 | 12 | 15 | 72 | 37 | 0 | 0 | 35779 | 10428 | 140567 | 0 | 170 | 148301 | 2267 | 0 | 0 | 21 | 0 | 0 | 0 | 0 | 0 | 12 | 0 | 100 | 0 | 0 | 130 | 936 | 20 |
| SRR7178610 | 0 | 0 | 0 | 76 | 0 | 47 | 53 | 112 | 0 | 0 | 66 | 19 | 0 | 0 | 46711 | 11390 | 159389 | 0 | 136 | 149323 | 2494 | 0 | 0 | 332 | 0 | 0 | 0 | 0 | 0 | 380 | 0 | 88 | 0 | 0 | 129 | 1154 | 19 |
| SRR7178611 | 0 | 0 | 0 | 43 | 43 | 17 | 33 | 89 | 0 | 43 | 76 | 76 | 0 | 0 | 48385 | 11620 | 162266 | 0 | 162 | 153135 | 2456 | 0 | 0 | 206 | 0 | 0 | 0 | 0 | 0 | 358 | 0 | 72 | 0 | 0 | 100 | 1297 | 11 |
| SRR7178612 | 0 | 0 | 12 | 65 | 42 | 82 | 101 | 327 | 38 | 12 | 30 | 0 | 0 | 0 | 25604 | 29414 | 117212 | 19 | 151 | 129778 | 1668 | 0 | 0 | 11 | 97 | 29 | 171 | 39 | 0 | 111 | 0 | 327 | 0 | 0 | 535 | 466 | 0 |
| SRR7178613 | 13 | 0 | 11 | 97 | 58 | 56 | 119 | 364 | 19 | 0 | 34 | 0 | 0 | 0 | 24248 | 29610 | 116063 | 0 | 169 | 129813 | 1732 | 0 | 0 | 153 | 99 | 429 | 2033 | 492 | 0 | 79 | 0 | 276 | 0 | 0 | 427 | 431 | 0 |
| SRR7178614 | 0 | 0 | 0 | 67 | 67 | 21 | 103 | 309 | 41 | 29 | 57 | 0 | 0 | 0 | 33106 | 32447 | 134023 | 21 | 62 | 131846 | 2140 | 0 | 0 | 172 | 143 | 20 | 106 | 25 | 0 | 210 | 0 | 185 | 0 | 0 | 398 | 459 | 0 |
| SRR7178615 | 0 | 0 | 0 | 40 | 93 | 54 | 27 | 134 | 0 | 27 | 27 | 0 | 0 | 0 | 34001 | 33668 | 137020 | 0 | 54 | 137533 | 2599 | 0 | 0 | 212 | 106 | 624 | 3173 | 597 | 0 | 106 | 0 | 268 | 13 | 0 | 292 | 546 | 0 |
| SRR7178616 | 0 | 0 | 0 | 43 | 44 | 54 | 36 | 161 | 16 | 0 | 66 | 0 | 0 | 0 | 38569 | 10080 | 207318 | 47 | 70 | 105197 | 2317 | 0 | 0 | 11 | 23 | 78 | 70 | 14 | 0 | 39 | 0 | 778 | 0 | 0 | 660 | 681 | 0 |
| SRR7178617 | 0 | 0 | 0 | 46 | 30 | 42 | 65 | 191 | 25 | 0 | 10 | 0 | 0 | 0 | 37280 | 9931 | 203527 | 57 | 60 | 107011 | 2160 | 0 | 0 | 13 | 22 | 81 | 68 | 21 | 0 | 42 | 0 | 786 | 0 | 0 | 643 | 686 | 0 |
| SRR7178618 | 0 | 0 | 0 | 41 | 35 | 26 | 35 | 103 | 0 | 12 | 71 | 0 | 0 | 0 | 49193 | 11450 | 238703 | 78 | 103 | 106915 | 2406 | 0 | 0 | 59 | 18 | 124 | 654 | 94 | 0 | 495 | 0 | 586 | 0 | 0 | 444 | 851 | 0 |
| SRR7178619 | 0 | 0 | 0 | 34 | 61 | 33 | 33 | 91 | 0 | 14 | 68 | 0 | 0 | 0 | 51334 | 11751 | 246386 | 74 | 91 | 109987 | 2423 | 0 | 0 | 68 | 0 | 68 | 28 | 68 | 0 | 27 | 0 | 651 | 0 | 0 | 423 | 959 | 0 |
| SRR7178620 | 0 | 0 | 0 | 21 | 52 | 39 | 29 | 169 | 12 | 0 | 26 | 0 | 0 | 0 | 18975 | 7144 | 115658 | 0 | 104 | 159896 | 1870 | 0 | 0 | 78 | 71 | 17 | 68 | 107 | 0 | 21 | 0 | 18 | 0 | 0 | 351 | 706 | 0 |
| SRR7178621 | 0 | 0 | 0 | 16 | 32 | 43 | 19 | 162 | 0 | 0 | 48 | 0 | 0 | 0 | 18612 | 7192 | 114870 | 0 | 136 | 159247 | 1802 | 0 | 0 | 64 | 78 | 19 | 60 | 64 | 0 | 18 | 0 | 0 | 0 | 0 | 336 | 693 | 0 |
| SRR7178622 | 0 | 0 | 0 | 11 | 22 | 45 | 28 | 168 | 11 | 0 | 44 | 11 | 0 | 0 | 24956 | 8367 | 133666 | 0 | 140 | 171525 | 2022 | 0 | 0 | 44 | 132 | 14 | 37 | 121 | 0 | 13 | 0 | 0 | 0 | 0 | 339 | 783 | 0 |
| SRR7178623 | 0 | 0 | 0 | 0 | 106 | 43 | 50 | 179 | 0 | 0 | 61 | 0 | 0 | 0 | 24845 | 8393 | 137460 | 0 | 122 | 174850 | 2374 | 0 | 0 | 30 | 106 | 319 | 38 | 46 | 0 | 11 | 0 | 0 | 0 | 0 | 218 | 810 | 30 |
| SRR7178624 | 0 | 11 | 0 | 37 | 26 | 84 | 75 | 206 | 21 | 11 | 78 | 20 | 0 | 0 | 16912 | 7016 | 66182 | 30 | 164 | 115716 | 1126 | 0 | 0 | 69 | 57 | 210 | 853 | 155 | 0 | 336 | 0 | 329 | 0 | 0 | 473 | 828 | 0 |
| SRR7178625 | 0 | 16 | 0 | 40 | 40 | 71 | 80 | 242 | 17 | 11 | 95 | 0 | 0 | 0 | 15904 | 6722 | 64739 | 28 | 196 | 117011 | 1007 | 0 | 0 | 79 | 74 | 19 | 77 | 11 | 0 | 32 | 0 | 359 | 0 | 0 | 492 | 819 | 0 |
| SRR7178626 | 13 | 13 | 0 | 25 | 25 | 60 | 131 | 231 | 30 | 0 | 101 | 0 | 0 | 0 | 20575 | 7284 | 70539 | 10 | 201 | 121153 | 986 | 13 | 0 | 76 | 63 | 165 | 40 | 101 | 0 | 19 | 13 | 221 | 0 | 0 | 311 | 1004 | 0 |
| SRR7178627 | 0 | 14 | 0 | 28 | 56 | 47 | 0 | 244 | 0 | 14 | 56 | 28 | 0 | 0 | 22262 | 7146 | 71124 | 28 | 225 | 125124 | 1115 | 0 | 0 | 42 | 69 | 250 | 23 | 125 | 0 | 21 | 0 | 281 | 0 | 0 | 309 | 1030 | 14 |
| SRR7178628 | 0 | 0 | 0 | 21 | 20 | 36 | 21 | 144 | 12 | 0 | 54 | 0 | 0 | 0 | 15076 | 6247 | 117634 | 0 | 154 | 205988 | 2149 | 0 | 0 | 72 | 44 | 13 | 52 | 12 | 0 | 36 | 0 | 0 | 0 | 0 | 354 | 544 | 0 |
| SRR7178629 | 0 | 0 | 0 | 17 | 24 | 37 | 23 | 210 | 10 | 0 | 54 | 0 | 0 | 0 | 16377 | 6404 | 123934 | 0 | 154 | 214546 | 2389 | 0 | 0 | 58 | 45 | 12 | 43 | 14 | 0 | 22 | 0 | 0 | 0 | 0 | 364 | 610 | 0 |
| SRR7178630 | 0 | 0 | 0 | 16 | 25 | 36 | 17 | 163 | 17 | 0 | 54 | 0 | 0 | 0 | 16485 | 6484 | 126576 | 0 | 152 | 216011 | 2351 | 0 | 0 | 84 | 43 | 116 | 540 | 107 | 0 | 370 | 0 | 0 | 0 | 0 | 298 | 611 | 0 |
| SRR7178631 | 0 | 0 | 0 | 21 | 25 | 38 | 44 | 128 | 17 | 0 | 106 | 0 | 0 | 0 | 14344 | 7247 | 123419 | 0 | 134 | 155916 | 911 | 0 | 0 | 88 | 32 | 88 | 503 | 78 | 0 | 528 | 0 | 172 | 0 | 0 | 285 | 447 | 0 |
| SRR7178632 | 0 | 0 | 0 | 19 | 37 | 42 | 45 | 145 | 0 | 11 | 107 | 0 | 0 | 0 | 13160 | 6888 | 122222 | 0 | 106 | 157145 | 839 | 0 | 0 | 94 | 30 | 135 | 565 | 90 | 0 | 472 | 0 | 81 | 0 | 0 | 299 | 448 | 0 |
| SRR7178633 | 0 | 0 | 0 | 10 | 20 | 30 | 50 | 110 | 20 | 0 | 90 | 0 | 0 | 0 | 17649 | 7984 | 139772 | 0 | 80 | 165910 | 959 | 0 | 0 | 70 | 20 | 70 | 24 | 50 | 0 | 32 | 0 | 150 | 0 | 0 | 189 | 531 | 0 |
| SRR7178634 | 0 | 0 | 0 | 23 | 46 | 0 | 38 | 95 | 38 | 0 | 137 | 0 | 0 | 0 | 18565 | 7889 | 142576 | 0 | 143 | 167956 | 1008 | 0 | 0 | 68 | 0 | 103 | 536 | 34 | 0 | 605 | 0 | 152 | 0 | 0 | 191 | 606 | 0 |
| SRR7178635 | 0 | 14 | 0 | 43 | 35 | 146 | 75 | 278 | 25 | 0 | 58 | 0 | 0 | 0 | 14209 | 7235 | 62960 | 18 | 150 | 115370 | 1007 | 0 | 0 | 74 | 114 | 491 | 2321 | 182 | 0 | 209 | 0 | 68 | 0 | 0 | 379 | 794 | 0 |
| SRR7178636 | 0 | 11 | 0 | 38 | 40 | 99 | 83 | 279 | 38 | 0 | 60 | 0 | 0 | 0 | 13663 | 7264 | 62411 | 16 | 144 | 113771 | 948 | 0 | 0 | 124 | 124 | 28 | 144 | 13 | 0 | 16 | 0 | 83 | 0 | 0 | 379 | 790 | 0 |
| SRR7178637 | 0 | 0 | 13 | 40 | 47 | 137 | 160 | 259 | 15 | 0 | 33 | 0 | 0 | 0 | 19020 | 8619 | 77133 | 0 | 221 | 118585 | 1076 | 0 | 0 | 67 | 181 | 19 | 103 | 154 | 0 | 11 | 0 | 69 | 0 | 0 | 268 | 860 | 0 |
| SRR7178638 | 0 | 0 | 0 | 0 | 129 | 126 | 117 | 214 | 58 | 0 | 55 | 0 | 0 | 0 | 19342 | 8726 | 81175 | 19 | 185 | 119582 | 1059 | 0 | 0 | 111 | 111 | 629 | 2644 | 342 | 0 | 351 | 0 | 97 | 0 | 0 | 268 | 952 | 0 |
| SRR7178639 | 10 | 0 | 21 | 145 | 57 | 74 | 139 | 231 | 23 | 10 | 67 | 0 | 0 | 0 | 19592 | 8781 | 59854 | 0 | 157 | 116078 | 905 | 0 | 0 | 93 | 44 | 12 | 64 | 142 | 0 | 116 | 0 | 55 | 0 | 0 | 393 | 162 | 0 |
| SRR7178640 | 0 | 13 | 19 | 119 | 52 | 56 | 102 | 346 | 31 | 0 | 48 | 0 | 0 | 0 | 22030 | 8897 | 63557 | 0 | 122 | 117965 | 1058 | 0 | 0 | 97 | 55 | 14 | 60 | 193 | 0 | 110 | 0 | 25 | 0 | 0 | 393 | 167 | 0 |
| SRR7178641 | 10 | 17 | 13 | 104 | 67 | 67 | 109 | 197 | 31 | 10 | 50 | 0 | 0 | 0 | 22356 | 9062 | 64805 | 0 | 150 | 118559 | 996 | 0 | 0 | 70 | 54 | 221 | 1245 | 94 | 0 | 157 | 0 | 67 | 0 | 0 | 307 | 173 | 0 |
| SRR7178642 | 0 | 0 | 18 | 52 | 29 | 36 | 30 | 154 | 20 | 0 | 22 | 0 | 0 | 0 | 36516 | 7959 | 221181 | 0 | 174 | 116723 | 5522 | 0 | 0 | 77 | 11 | 87 | 45 | 22 | 0 | 66 | 0 | 0 | 0 | 0 | 169 | 415 | 0 |
| SRR7178643 | 0 | 0 | 0 | 40 | 33 | 49 | 29 | 190 | 0 | 0 | 12 | 0 | 0 | 0 | 39428 | 8469 | 234394 | 0 | 179 | 117839 | 5259 | 0 | 0 | 63 | 33 | 11 | 44 | 25 | 0 | 56 | 0 | 0 | 0 | 0 | 148 | 451 | 0 |
| SRR7178644 | 0 | 0 | 0 | 32 | 32 | 46 | 30 | 137 | 0 | 0 | 29 | 0 | 0 | 0 | 40519 | 8531 | 240314 | 0 | 182 | 119929 | 5418 | 0 | 0 | 80 | 29 | 156 | 36 | 18 | 0 | 71 | 0 | 0 | 0 | 0 | 148 | 448 | 0 |
| SRR7178645 | 0 | 0 | 30 | 56 | 66 | 0 | 16 | 61 | 0 | 0 | 56 | 0 | 0 | 0 | 64820 | 13192 | 350853 | 0 | 101 | 41373 | 6842 | 0 | 0 | 175 | 36 | 182 | 30 | 109 | 0 | 119 | 0 | 23 | 0 | 0 | 63 | 694 | 0 |
| SRR7178646 | 0 | 0 | 0 | 65 | 68 | 13 | 0 | 56 | 0 | 0 | 43 | 0 | 0 | 0 | 62496 | 13136 | 344001 | 0 | 79 | 42391 | 6585 | 0 | 0 | 188 | 28 | 198 | 34 | 142 | 0 | 90 | 0 | 26 | 0 | 0 | 68 | 671 | 0 |
| SRR7178647 | 0 | 13 | 27 | 67 | 27 | 12 | 0 | 54 | 0 | 0 | 67 | 0 | 0 | 0 | 86036 | 15092 | 412171 | 0 | 120 | 43862 | 7136 | 0 | 0 | 148 | 0 | 175 | 793 | 81 | 0 | 94 | 0 | 0 | 0 | 0 | 37 | 764 | 0 |
| SRR7178648 | 0 | 0 | 15 | 15 | 45 | 0 | 0 | 51 | 0 | 0 | 30 | 0 | 0 | 0 | 87502 | 15007 | 423150 | 0 | 108 | 43266 | 7388 | 0 | 0 | 106 | 15 | 76 | 11 | 61 | 0 | 227 | 0 | 32 | 0 | 0 | 46 | 966 | 0 |
| SRR7178649 | 0 | 0 | 0 | 130 | 87 | 38 | 53 | 287 | 31 | 0 | 163 | 0 | 0 | 0 | 35766 | 18830 | 222250 | 49 | 22 | 12581 | 5690 | 0 | 0 | 365 | 25 | 173 | 982 | 1210 | 0 | 968 | 0 | 69 | 0 | 0 | 280 | 577 | 0 |
| SRR7178650 | 0 | 14 | 14 | 48 | 57 | 68 | 68 | 293 | 33 | 0 | 158 | 0 | 0 | 0 | 38850 | 20129 | 243936 | 57 | 18 | 12378 | 5666 | 0 | 0 | 378 | 53 | 177 | 1278 | 1412 | 0 | 948 | 0 | 112 | 0 | 0 | 239 | 656 | 0 |
| SRR7178651 | 20 | 0 | 15 | 70 | 106 | 62 | 62 | 294 | 0 | 0 | 136 | 0 | 0 | 0 | 39729 | 21090 | 253585 | 62 | 25 | 12640 | 5655 | 0 | 0 | 13 | 30 | 181 | 53 | 52 | 0 | 33 | 0 | 103 | 0 | 0 | 253 | 647 | 0 |
| SRR7178652 | 0 | 0 | 0 | 22 | 52 | 32 | 30 | 124 | 0 | 0 | 47 | 0 | 0 | 0 | 38177 | 7444 | 198792 | 0 | 96 | 123476 | 2372 | 0 | 0 | 101 | 37 | 125 | 49 | 84 | 0 | 32 | 0 | 68 | 0 | 0 | 274 | 410 | 0 |
| SRR7178653 | 0 | 0 | 0 | 19 | 62 | 43 | 0 | 136 | 0 | 0 | 60 | 0 | 0 | 0 | 37413 | 7373 | 197868 | 0 | 111 | 122435 | 2175 | 0 | 0 | 115 | 31 | 150 | 602 | 55 | 0 | 406 | 0 | 47 | 0 | 0 | 259 | 393 | 0 |
| SRR7178654 | 0 | 0 | 0 | 10 | 62 | 38 | 13 | 102 | 19 | 0 | 73 | 0 | 0 | 0 | 52162 | 8877 | 244607 | 0 | 108 | 127534 | 2330 | 0 | 0 | 197 | 42 | 177 | 779 | 93 | 0 | 644 | 0 | 32 | 0 | 0 | 195 | 445 | 0 |
| SRR7178655 | 0 | 0 | 0 | 58 | 58 | 24 | 24 | 90 | 0 | 0 | 29 | 0 | 0 | 0 | 54920 | 8871 | 255035 | 16 | 98 | 130590 | 2800 | 0 | 0 | 58 | 44 | 233 | 596 | 29 | 0 | 596 | 0 | 49 | 0 | 0 | 184 | 437 | 0 |
| SRR7178656 | 0 | 0 | 13 | 76 | 79 | 42 | 24 | 197 | 22 | 0 | 71 | 0 | 0 | 0 | 13368 | 5809 | 79397 | 0 | 195 | 167633 | 652 | 0 | 0 | 119 | 53 | 198 | 39 | 102 | 0 | 14 | 0 | 10 | 0 | 0 | 373 | 150 | 0 |
| SRR7178657 | 0 | 0 | 0 | 63 | 68 | 56 | 28 | 191 | 0 | 0 | 56 | 0 | 0 | 0 | 13206 | 5788 | 78962 | 0 | 229 | 167154 | 665 | 0 | 0 | 122 | 41 | 124 | 741 | 97 | 0 | 307 | 0 | 0 | 0 | 0 | 377 | 136 | 0 |
| SRR7178658 | 0 | 0 | 0 | 77 | 55 | 33 | 33 | 339 | 0 | 11 | 44 | 0 | 0 | 0 | 17031 | 6235 | 84456 | 0 | 159 | 172937 | 751 | 0 | 0 | 208 | 33 | 131 | 1039 | 88 | 0 | 405 | 0 | 13 | 0 | 0 | 302 | 149 | 0 |
| SRR7178659 | 0 | 0 | 0 | 45 | 105 | 34 | 43 | 238 | 17 | 0 | 90 | 0 | 0 | 0 | 17406 | 6283 | 85827 | 0 | 179 | 173985 | 749 | 0 | 0 | 211 | 60 | 211 | 22 | 105 | 0 | 12 | 0 | 0 | 0 | 0 | 274 | 157 | 0 |
| SRR7178660 | 0 | 0 | 28 | 127 | 77 | 51 | 27 | 212 | 0 | 0 | 87 | 0 | 0 | 0 | 36477 | 10211 | 260253 | 0 | 125 | 142324 | 2211 | 0 | 0 | 12 | 55 | 22 | 84 | 32 | 0 | 32 | 0 | 21 | 0 | 0 | 409 | 1013 | 0 |
| SRR7178661 | 0 | 0 | 21 | 111 | 61 | 56 | 28 | 254 | 0 | 0 | 90 | 0 | 0 | 0 | 40299 | 10341 | 276563 | 0 | 129 | 144645 | 2362 | 0 | 0 | 252 | 66 | 422 | 1694 | 550 | 0 | 434 | 0 | 30 | 0 | 0 | 358 | 1104 | 0 |
| SRR7178662 | 17 | 0 | 20 | 110 | 73 | 77 | 17 | 199 | 0 | 0 | 98 | 0 | 0 | 0 | 41436 | 10597 | 283074 | 0 | 121 | 147488 | 2261 | 0 | 0 | 12 | 86 | 21 | 79 | 30 | 0 | 23 | 0 | 20 | 0 | 0 | 321 | 1137 | 0 |
| SRR7178663 | 0 | 0 | 0 | 41 | 77 | 55 | 44 | 269 | 19 | 0 | 86 | 0 | 0 | 0 | 36276 | 7679 | 175675 | 0 | 105 | 90584 | 4865 | 0 | 0 | 12 | 45 | 155 | 40 | 130 | 0 | 13 | 0 | 21 | 0 | 0 | 219 | 783 | 0 |
| SRR7178664 | 0 | 0 | 0 | 57 | 121 | 74 | 62 | 247 | 24 | 0 | 57 | 0 | 0 | 0 | 34497 | 7575 | 171395 | 0 | 66 | 91666 | 4644 | 0 | 0 | 234 | 49 | 170 | 820 | 136 | 0 | 253 | 0 | 42 | 0 | 0 | 222 | 751 | 0 |
| SRR7178665 | 0 | 0 | 0 | 0 | 75 | 97 | 82 | 171 | 15 | 19 | 56 | 19 | 0 | 0 | 46163 | 9034 | 215228 | 15 | 97 | 93412 | 4488 | 0 | 0 | 280 | 56 | 205 | 635 | 75 | 0 | 317 | 0 | 22 | 0 | 0 | 176 | 917 | 0 |
| SRR7178666 | 0 | 0 | 0 | 22 | 65 | 21 | 42 | 197 | 28 | 22 | 44 | 0 | 0 | 0 | 47665 | 9283 | 220653 | 0 | 56 | 95563 | 4385 | 0 | 0 | 174 | 44 | 130 | 17 | 65 | 0 | 435 | 0 | 42 | 0 | 0 | 174 | 1059 | 0 |
| SRR7178667 | 0 | 12 | 12 | 33 | 57 | 17 | 78 | 338 | 65 | 0 | 83 | 0 | 0 | 0 | 13196 | 6758 | 127400 | 0 | 65 | 179521 | 952 | 0 | 0 | 15 | 38 | 19 | 97 | 17 | 0 | 76 | 0 | 22 | 0 | 0 | 1141 | 320 | 0 |
| SRR7178668 | 0 | 0 | 0 | 42 | 60 | 45 | 90 | 366 | 41 | 0 | 93 | 0 | 0 | 0 | 13274 | 6404 | 124773 | 0 | 99 | 180762 | 891 | 0 | 0 | 16 | 33 | 19 | 117 | 19 | 0 | 91 | 0 | 33 | 0 | 0 | 1004 | 316 | 0 |
| SRR7178669 | 21 | 0 | 0 | 42 | 106 | 15 | 92 | 489 | 15 | 11 | 85 | 0 | 0 | 11 | 17376 | 8193 | 154343 | 0 | 92 | 190598 | 870 | 0 | 0 | 169 | 21 | 148 | 1311 | 264 | 0 | 1385 | 0 | 0 | 0 | 0 | 973 | 393 | 0 |
| SRR7178670 | 0 | 12 | 12 | 49 | 49 | 58 | 73 | 336 | 44 | 12 | 134 | 12 | 0 | 0 | 18330 | 8236 | 159370 | 0 | 102 | 194409 | 862 | 0 | 0 | 122 | 24 | 122 | 950 | 280 | 0 | 1680 | 0 | 15 | 0 | 0 | 835 | 409 | 0 |
| SRR7178671 | 11 | 0 | 16 | 76 | 33 | 39 | 23 | 148 | 0 | 0 | 60 | 0 | 0 | 0 | 75842 | 11637 | 322512 | 0 | 108 | 81624 | 3412 | 0 | 0 | 161 | 63 | 223 | 1094 | 362 | 0 | 283 | 0 | 0 | 0 | 0 | 153 | 497 | 0 |
| SRR7178672 | 0 | 11 | 21 | 92 | 58 | 43 | 44 | 95 | 0 | 0 | 50 | 0 | 0 | 0 | 74455 | 11527 | 320738 | 0 | 121 | 82265 | 3513 | 0 | 0 | 12 | 58 | 13 | 55 | 19 | 0 | 14 | 0 | 0 | 0 | 0 | 169 | 499 | 0 |

Supp Table S8.3.Columns_BX_to_DH

| | | | | | | | | | | | | | | | | | | | | | | | | | | | | | | | | | | | | | |
|---|---|---|---|---|---|---|---|---|---|---|---|---|---|---|---|---|---|---|---|---|---|---|---|---|---|---|---|---|---|---|---|---|---|---|---|---|---|
| SRR7178673 | 0 | 0 | 11 | 33 | 87 | 42 | 37 | 141 | 10 | 0 | 65 | 0 | 0 | 0 | 100333 | 12775 | 376004 | 0 | 136 | 81881 | 3490 | 0 | 0 | 272 | 54 | 381 | 1284 | 435 | 0 | 283 | 0 | 10 | 0 | 0 | 130 | 568 | 0 |
| SRR7178674 | 0 | 0 | 0 | 44 | 103 | 34 | 20 | 107 | 13 | 0 | 103 | 0 | 0 | 0 | 104111 | 13534 | 388171 | 0 | 121 | 83947 | 4136 | 0 | 0 | 279 | 73 | 279 | 30 | 381 | 0 | 12 | 0 | 0 | 0 | 0 | 103 | 565 | 15 |
| SRR7178675 | 0 | 0 | 0 | 96 | 28 | 11 | 11 | 74 | 0 | 14 | 60 | 14 | 0 | 0 | 97703 | 20093 | 368174 | 45 | 58 | 62854 | 5465 | 0 | 0 | 64 | 28 | 161 | 730 | 206 | 0 | 156 | 0 | 1082 | 0 | 0 | 86 | 268 | 18 |
| SRR7178676 | 0 | 0 | 0 | 98 | 64 | 20 | 28 | 64 | 0 | 17 | 90 | 30 | 0 | 0 | 95233 | 20134 | 361971 | 64 | 71 | 64115 | 5369 | 0 | 0 | 132 | 38 | 200 | 35 | 11 | 0 | 188 | 0 | 1194 | 0 | 0 | 105 | 228 | 34 |
| SRR7178677 | 0 | 0 | 19 | 0 | 58 | 17 | 0 | 60 | 0 | 0 | 19 | 19 | 0 | 0 | 124406 | 21609 | 408053 | 86 | 69 | 62585 | 5440 | 19 | 0 | 58 | 19 | 117 | 18 | 194 | 0 | 214 | 0 | 807 | 0 | 0 | 71 | 289 | 39 |
| SRR7178678 | 0 | 0 | 0 | 133 | 67 | 16 | 16 | 32 | 0 | 0 | 44 | 22 | 0 | 0 | 127092 | 21891 | 416205 | 63 | 71 | 62202 | 5637 | 0 | 0 | 133 | 0 | 133 | 732 | 155 | 0 | 266 | 0 | 836 | 0 | 0 | 55 | 333 | 89 |
| SRR7178679 | 22 | 0 | 38 | 12 | 73 | 32 | 24 | 110 | 0 | 0 | 73 | 0 | 0 | 0 | 35751 | 9243 | 200572 | 0 | 164 | 128207 | 2914 | 0 | 0 | 14 | 63 | 20 | 86 | 27 | 0 | 161 | 0 | 31 | 0 | 0 | 137 | 703 | 0 |
| SRR7178680 | 15 | 0 | 23 | 224 | 112 | 24 | 16 | 130 | 0 | 0 | 89 | 0 | 0 | 0 | 39674 | 9289 | 211917 | 0 | 198 | 128065 | 2882 | 0 | 0 | 13 | 70 | 16 | 81 | 24 | 0 | 182 | 0 | 18 | 0 | 0 | 110 | 775 | 0 |
| SRR7178681 | 0 | 12 | 36 | 228 | 84 | 13 | 27 | 130 | 0 | 0 | 124 | 0 | 0 | 0 | 40494 | 9497 | 216401 | 0 | 210 | 129337 | 3104 | 0 | 0 | 369 | 44 | 453 | 2204 | 641 | 0 | 168 | 0 | 33 | 0 | 0 | 102 | 793 | 0 |
| SRR7178682 | 0 | 0 | 26 | 83 | 105 | 97 | 66 | 219 | 19 | 0 | 67 | 0 | 0 | 0 | 36487 | 11857 | 240843 | 0 | 106 | 95172 | 3581 | 0 | 0 | 15 | 47 | 13 | 97 | 31 | 0 | 133 | 0 | 0 | 0 | 0 | 152 | 678 | 0 |
| SRR7178683 | 0 | 0 | 15 | 76 | 95 | 84 | 69 | 262 | 14 | 0 | 83 | 0 | 0 | 0 | 35155 | 11574 | 236100 | 0 | 99 | 96087 | 3451 | 0 | 0 | 17 | 28 | 15 | 110 | 31 | 0 | 132 | 0 | 0 | 0 | 0 | 152 | 658 | 0 |
| SRR7178684 | 0 | 0 | 0 | 109 | 124 | 91 | 76 | 228 | 0 | 0 | 102 | 0 | 0 | 0 | 44978 | 12784 | 256786 | 0 | 111 | 103248 | 3376 | 0 | 0 | 335 | 80 | 240 | 51 | 12 | 0 | 153 | 0 | 20 | 0 | 0 | 115 | 919 | 0 |
| SRR7178685 | 0 | 0 | 25 | 51 | 127 | 93 | 64 | 137 | 0 | 0 | 85 | 0 | 0 | 0 | 48510 | 12793 | 264480 | 0 | 156 | 104319 | 3332 | 0 | 0 | 187 | 17 | 170 | 40 | 374 | 0 | 221 | 0 | 0 | 0 | 0 | 106 | 1007 | 0 |
| SRR7178686 | 12 | 0 | 0 | 55 | 53 | 311 | 179 | 929 | 61 | 0 | 90 | 0 | 0 | 0 | 23656 | 13844 | 265227 | 10 | 15 | 33605 | 715 | 0 | 0 | 325 | 61 | 284 | 1035 | 190 | 0 | 1097 | 0 | 46 | 0 | 0 | 634 | 514 | 0 |
| SRR7178687 | 0 | 0 | 13 | 55 | 61 | 238 | 147 | 822 | 86 | 0 | 76 | 0 | 0 | 0 | 23781 | 13557 | 263888 | 0 | 15 | 33218 | 832 | 0 | 0 | 15 | 40 | 16 | 56 | 10 | 0 | 55 | 0 | 36 | 0 | 0 | 782 | 499 | 0 |
| SRR7178688 | 0 | 25 | 0 | 42 | 76 | 448 | 215 | 1272 | 197 | 0 | 127 | 17 | 0 | 0 | 31883 | 17099 | 331391 | 0 | 0 | 34749 | 770 | 0 | 0 | 347 | 34 | 356 | 33 | 296 | 0 | 33 | 0 | 54 | 0 | 0 | 640 | 545 | 0 |
| SRR7178689 | 0 | 12 | 0 | 23 | 94 | 160 | 91 | 1257 | 297 | 0 | 94 | 0 | 0 | 0 | 35438 | 17940 | 349555 | 0 | 0 | 36809 | 1006 | 0 | 0 | 375 | 35 | 270 | 29 | 282 | 0 | 30 | 0 | 0 | 0 | 0 | 516 | 633 | 12 |
| SRR7178690 | 0 | 0 | 15 | 46 | 31 | 69 | 39 | 222 | 0 | 0 | 100 | 0 | 0 | 0 | 58121 | 14700 | 377311 | 0 | 85 | 54567 | 5203 | 0 | 0 | 106 | 82 | 30 | 78 | 33 | 0 | 21 | 0 | 233 | 0 | 0 | 309 | 626 | 0 |
| SRR7178691 | 12 | 0 | 0 | 41 | 23 | 61 | 63 | 227 | 13 | 0 | 64 | 0 | 0 | 0 | 20022 | 11238 | 124321 | 0 | 177 | 95920 | 1586 | 0 | 0 | 197 | 73 | 299 | 1402 | 189 | 0 | 122 | 0 | 16 | 0 | 0 | 161 | 512 | 0 |
| SRR7178692 | 15 | 0 | 0 | 51 | 39 | 76 | 35 | 225 | 14 | 0 | 39 | 15 | 0 | 0 | 20637 | 11447 | 127767 | 0 | 187 | 97532 | 1663 | 0 | 0 | 204 | 75 | 304 | 1422 | 96 | 0 | 162 | 0 | 0 | 0 | 0 | 132 | 528 | 0 |
| SRR7178693 | 0 | 0 | 0 | 50 | 48 | 51 | 56 | 326 | 16 | 16 | 59 | 0 | 0 | 0 | 27439 | 11647 | 157785 | 0 | 125 | 140267 | 2252 | 0 | 0 | 12 | 66 | 15 | 57 | 165 | 0 | 133 | 0 | 366 | 0 | 0 | 213 | 742 | 0 |
| SRR7178694 | 0 | 0 | 14 | 55 | 36 | 48 | 46 | 310 | 20 | 10 | 60 | 0 | 0 | 0 | 26980 | 11525 | 155958 | 0 | 119 | 139895 | 2260 | 0 | 0 | 217 | 95 | 221 | 884 | 179 | 0 | 148 | 0 | 367 | 0 | 0 | 239 | 717 | 0 |
| SRR7178695 | 0 | 0 | 0 | 26 | 68 | 70 | 78 | 392 | 0 | 0 | 51 | 0 | 0 | 0 | 36158 | 12667 | 169151 | 0 | 148 | 147252 | 2018 | 0 | 0 | 264 | 26 | 13 | 37 | 221 | 0 | 204 | 0 | 183 | 0 | 0 | 184 | 872 | 0 |
| SRR7178696 | 0 | 24 | 24 | 24 | 83 | 34 | 134 | 347 | 45 | 12 | 59 | 12 | 0 | 0 | 37240 | 12685 | 174710 | 0 | 123 | 150035 | 2283 | 0 | 0 | 165 | 94 | 330 | 30 | 271 | 0 | 224 | 12 | 291 | 0 | 0 | 163 | 962 | 0 |
| SRR7178697 | 0 | 0 | 0 | 26 | 44 | 56 | 46 | 215 | 21 | 0 | 121 | 0 | 0 | 0 | 19870 | 6588 | 131668 | 0 | 163 | 135753 | 1811 | 0 | 0 | 114 | 47 | 175 | 770 | 79 | 0 | 319 | 0 | 48 | 0 | 0 | 313 | 608 | 0 |
| SRR7178698 | 0 | 0 | 0 | 35 | 29 | 67 | 29 | 223 | 31 | 0 | 82 | 0 | 0 | 0 | 64823 | 15193 | 397906 | 0 | 85 | 56008 | 5485 | 0 | 0 | 111 | 97 | 27 | 83 | 28 | 0 | 18 | 0 | 238 | 0 | 0 | 248 | 702 | 0 |
| SRR7178699 | 0 | 0 | 0 | 28 | 36 | 87 | 39 | 261 | 0 | 0 | 91 | 0 | 0 | 0 | 66476 | 15447 | 407925 | 0 | 128 | 57149 | 5561 | 0 | 0 | 104 | 104 | 24 | 76 | 26 | 0 | 18 | 0 | 234 | 0 | 0 | 279 | 695 | 0 |
| SRR7178700 | 0 | 0 | 0 | 65 | 53 | 62 | 23 | 249 | 16 | 0 | 72 | 0 | 0 | 0 | 48422 | 13159 | 293831 | 0 | 88 | 98001 | 3583 | 0 | 0 | 197 | 38 | 221 | 1044 | 115 | 0 | 362 | 0 | 60 | 0 | 0 | 495 | 978 | 0 |
| SRR7178701 | 0 | 12 | 0 | 85 | 48 | 66 | 28 | 210 | 0 | 0 | 71 | 0 | 0 | 0 | 47277 | 13045 | 292590 | 0 | 82 | 96789 | 3772 | 0 | 0 | 12 | 28 | 11 | 66 | 115 | 0 | 32 | 0 | 66 | 0 | 0 | 503 | 922 | 0 |
| SRR7178702 | 0 | 11 | 11 | 42 | 42 | 98 | 44 | 257 | 27 | 0 | 84 | 0 | 0 | 0 | 64061 | 15177 | 351380 | 0 | 142 | 100049 | 3292 | 0 | 0 | 264 | 32 | 391 | 1372 | 169 | 0 | 570 | 0 | 98 | 0 | 0 | 330 | 1004 | 0 |
| SRR7178703 | 15 | 0 | 15 | 29 | 147 | 91 | 34 | 205 | 23 | 0 | 88 | 15 | 0 | 0 | 66963 | 15708 | 363602 | 11 | 68 | 102647 | 3479 | 0 | 0 | 206 | 15 | 265 | 40 | 118 | 0 | 20 | 0 | 46 | 0 | 0 | 308 | 1111 | 0 |
| SRR7178704 | 0 | 0 | 0 | 34 | 62 | 62 | 58 | 539 | 29 | 0 | 50 | 0 | 0 | 0 | 29794 | 8755 | 155538 | 0 | 62 | 55725 | 1592 | 0 | 0 | 18 | 32 | 16 | 108 | 104 | 0 | 53 | 0 | 0 | 0 | 0 | 416 | 263 | 0 |
| SRR7178705 | 0 | 0 | 15 | 50 | 70 | 55 | 147 | 575 | 69 | 0 | 77 | 0 | 0 | 0 | 33795 | 9761 | 171581 | 0 | 18 | 56277 | 1627 | 0 | 0 | 14 | 42 | 16 | 99 | 104 | 0 | 40 | 0 | 0 | 0 | 0 | 396 | 284 | 0 |
| SRR7178706 | 0 | 16 | 10 | 36 | 54 | 43 | 96 | 449 | 48 | 0 | 57 | 0 | 0 | 0 | 33952 | 9911 | 180033 | 0 | 43 | 56652 | 1729 | 0 | 0 | 16 | 39 | 12 | 97 | 90 | 0 | 47 | 0 | 0 | 0 | 0 | 405 | 273 | 0 |
| SRR7178707 | 0 | 0 | 13 | 57 | 64 | 10 | 0 | 69 | 0 | 25 | 25 | 0 | 0 | 0 | 144774 | 24344 | 433631 | 0 | 53 | 35955 | 9168 | 0 | 0 | 153 | 38 | 210 | 19 | 89 | 0 | 38 | 0 | 59 | 0 | 0 | 79 | 427 | 0 |
| SRR7178708 | 0 | 0 | 12 | 18 | 61 | 10 | 10 | 48 | 0 | 0 | 25 | 0 | 0 | 0 | 141220 | 24475 | 429747 | 0 | 59 | 36122 | 9153 | 0 | 0 | 129 | 80 | 166 | 12 | 68 | 0 | 55 | 0 | 61 | 0 | 0 | 71 | 424 | 0 |
| SRR7178709 | 0 | 0 | 0 | 0 | 0 | 16 | 0 | 57 | 0 | 0 | 0 | 0 | 0 | 0 | 187066 | 28169 | 514161 | 0 | 25 | 35634 | 9164 | 0 | 0 | 176 | 101 | 101 | 12 | 25 | 0 | 25 | 0 | 74 | 0 | 0 | 64 | 498 | 0 |
| SRR7178710 | 0 | 0 | 0 | 34 | 34 | 0 | 10 | 21 | 0 | 0 | 0 | 0 | 0 | 0 | 194227 | 28724 | 533091 | 10 | 63 | 36375 | 11602 | 0 | 0 | 68 | 0 | 204 | 272 | 204 | 0 | 136 | 0 | 94 | 0 | 0 | 45 | 546 | 0 |
| SRR7178711 | 32 | 18 | 36 | 273 | 76 | 43 | 81 | 281 | 14 | 11 | 25 | 11 | 0 | 0 | 57087 | 12366 | 324055 | 0 | 62 | 68436 | 4603 | 0 | 0 | 500 | 72 | 342 | 2557 | 507 | 0 | 68 | 0 | 0 | 0 | 0 | 277 | 447 | 0 |
| SRR7178712 | 18 | 18 | 40 | 301 | 128 | 101 | 80 | 245 | 32 | 0 | 35 | 0 | 0 | 0 | 65238 | 12905 | 347412 | 0 | 96 | 69599 | 4951 | 0 | 0 | 549 | 89 | 434 | 2826 | 562 | 0 | 80 | 0 | 11 | 0 | 0 | 272 | 502 | 0 |
| SRR7178713 | 14 | 18 | 23 | 283 | 137 | 55 | 71 | 267 | 38 | 0 | 37 | 0 | 0 | 0 | 66494 | 13190 | 356585 | 0 | 49 | 70065 | 5173 | 0 | 0 | 457 | 59 | 434 | 2681 | 644 | 0 | 96 | 0 | 0 | 0 | 0 | 202 | 509 | 0 |
| SRR7178714 | 0 | 11 | 15 | 41 | 34 | 22 | 22 | 125 | 0 | 0 | 15 | 0 | 0 | 0 | 61535 | 11172 | 247294 | 0 | 164 | 110746 | 4200 | 0 | 0 | 134 | 45 | 187 | 38 | 119 | 0 | 63 | 0 | 11 | 0 | 0 | 100 | 331 | 0 |
| SRR7178715 | 18 | 0 | 14 | 70 | 49 | 26 | 37 | 141 | 13 | 0 | 0 | 0 | 0 | 0 | 59258 | 11094 | 244883 | 0 | 86 | 112769 | 4003 | 0 | 0 | 144 | 49 | 204 | 859 | 123 | 0 | 56 | 0 | 0 | 0 | 0 | 117 | 334 | 0 |
| SRR7178716 | 0 | 0 | 0 | 62 | 47 | 0 | 0 | 113 | 0 | 0 | 31 | 16 | 0 | 0 | 75451 | 11948 | 274872 | 0 | 121 | 114261 | 3825 | 0 | 16 | 109 | 31 | 156 | 24 | 31 | 0 | 93 | 0 | 0 | 16 | 0 | 75 | 394 | 0 |
| SRR7178717 | 0 | 0 | 35 | 70 | 70 | 16 | 40 | 72 | 0 | 0 | 0 | 0 | 0 | 0 | 78400 | 12147 | 280371 | 0 | 112 | 114591 | 3699 | 0 | 0 | 105 | 18 | 193 | 683 | 18 | 0 | 105 | 0 | 0 | 0 | 0 | 103 | 453 | 0 |
| SRR7178718 | 0 | 0 | 20 | 105 | 49 | 30 | 18 | 102 | 0 | 0 | 17 | 0 | 0 | 0 | 56548 | 12346 | 270192 | 0 | 98 | 115025 | 3264 | 0 | 0 | 147 | 56 | 253 | 1105 | 199 | 0 | 46 | 0 | 23 | 0 | 0 | 111 | 505 | 0 |
| SRR7178719 | 0 | 0 | 12 | 97 | 53 | 35 | 25 | 122 | 12 | 0 | 12 | 0 | 0 | 0 | 60860 | 12537 | 274809 | 0 | 122 | 118166 | 3110 | 0 | 0 | 10 | 63 | 17 | 75 | 10 | 0 | 69 | 0 | 21 | 0 | 0 | 118 | 623 | 0 |
| SRR7178720 | 0 | 12 | 12 | 118 | 58 | 24 | 26 | 88 | 0 | 0 | 26 | 0 | 0 | 0 | 62169 | 12800 | 281034 | 0 | 126 | 119003 | 3274 | 0 | 0 | 190 | 58 | 325 | 1333 | 209 | 0 | 51 | 0 | 29 | 0 | 0 | 105 | 607 | 0 |
| SRR7178721 | 0 | 16 | 44 | 11 | 76 | 25 | 37 | 138 | 0 | 52 | 40 | 0 | 0 | 0 | 157363 | 31409 | 469206 | 0 | 59 | 35171 | 10112 | 0 | 0 | 12 | 76 | 22 | 88 | 14 | 0 | 80 | 0 | 184 | 0 | 0 | 99 | 558 | 0 |
| SRR7178722 | 23 | 16 | 58 | 311 | 58 | 27 | 35 | 119 | 0 | 31 | 27 | 0 | 0 | 0 | 154218 | 31644 | 466447 | 0 | 54 | 35415 | 9975 | 12 | 27 | 272 | 66 | 598 | 1985 | 287 | 0 | 58 | 0 | 240 | 0 | 0 | 100 | 539 | 0 |
| SRR7178723 | 0 | 31 | 47 | 187 | 93 | 15 | 38 | 150 | 15 | 62 | 31 | 0 | 0 | 0 | 198990 | 34659 | 525161 | 15 | 83 | 35033 | 10184 | 16 | 0 | 327 | 187 | 14 | 56 | 264 | 0 | 109 | 0 | 128 | 0 | 0 | 59 | 654 | 16 |
| SRR7178724 | 22 | 0 | 22 | 215 | 194 | 0 | 28 | 171 | 0 | 22 | 43 | 22 | 0 | 0 | 207170 | 35503 | 538186 | 28 | 76 | 35279 | 10655 | 0 | 0 | 409 | 108 | 818 | 2841 | 366 | 0 | 86 | 0 | 114 | 0 | 0 | 64 | 696 | 0 |
| SRR7213733 | 0 | 0 | 0 | 0 | 95 | 2671 | 238 | 1777 | 2304 | 1022 | 122 | 36 | 0 | 0 | 3585 | 35745 | 25761 | 79 | 70 | 328 | 933 | 21 | 212 | 35 | 35 | 106 | 16 | 0 | 0 | 601 | 41 | 30 | 0 | 0 | 51946 | 10 | 0 |
| SRR7213734 | 0 | 0 | 0 | 0 | 143 | 1994 | 138 | 1451 | 1442 | 1031 | 122 | 59 | 40 | 0 | 3687 | 15370 | 8064 | 86 | 95 | 475 | 570 | 34 | 350 | 36 | 94 | 86 | 0 | 0 | 0 | 565 | 105 | 78 | 0 | 0 | 16828 | 11 | 0 |
| SRR7213735 | 0 | 0 | 0 | 0 | 66 | 4103 | 343 | 2972 | 2960 | 714 | 73 | 42 | 27 | 0 | 2732 | 26530 | 13221 | 80 | 46 | 274 | 971 | 33 | 237 | 80 | 141 | 245 | 44 | 13 | 0 | 445 | 68 | 34 | 0 | 0 | 20393 | 12 | 0 |
| SRR7226512 | 19 | 14 | 11 | 0 | 130 | 4618 | 5822 | 18696 | 6453 | 25 | 151 | 18 | 0 | 0 | 15422 | 87062 | 128495 | 0 | 180 | 14175 | 1135 | 0 | 0 | 27 | 20 | 40 | 24 | 30 | 0 | 229 | 0 | 51 | 0 | 0 | 1716 | 26 | 0 |
| SRR7226513 | 12 | 16 | 0 | 14 | 251 | 2544 | 3105 | 15174 | 6280 | 0 | 144 | 0 | 0 | 0 | 21159 | 65921 | 88256 | 15 | 194 | 5394 | 1913 | 0 | 0 | 35 | 46 | 40 | 29 | 27 | 0 | 127 | 0 | 0 | 0 | 0 | 7563 | 44 | 0 |
| SRR7226514 | 95 | 53 | 41 | 24 | 169 | 8008 | 14346 | 40468 | 19248 | 0 | 97 | 0 | 0 | 0 | 5406 | 40431 | 19180 | 0 | 412 | 6993 | 289 | 0 | 0 | 11 | 26 | 46 | 47 | 37 | 0 | 195 | 0 | 0 | 0 | 0 | 7084 | 5710 | 0 |
| SRR7226515 | 68 | 108 | 0 | 0 | 345 | 6658 | 15817 | 47253 | 23495 | 0 | 169 | 0 | 0 | 0 | 10101 | 39655 | 36965 | 0 | 180 | 1765 | 292 | 0 | 0 | 23 | 16 | 24 | 11 | 0 | 0 | 352 | 0 | 0 | 0 | 0 | 15956 | 3357 | 0 |
| SRR7226516 | 211 | 150 | 20 | 0 | 663 | 14408 | 15602 | 51177 | 27803 | 0 | 365 | 0 | 0 | 0 | 7931 | 29487 | 43453 | 0 | 700 | 8128 | 635 | 0 | 0 | 27 | 52 | 75 | 88 | 0 | 0 | 790 | 0 | 0 | 0 | 0 | 4744 | 9086 | 0 |
| SRR7226517 | 168 | 82 | 16 | 0 | 1642 | 9099 | 2829 | 12705 | 4502 | 34 | 553 | 19 | 0 | 0 | 70083 | 48082 | 150450 | 0 | 1476 | 59197 | 3170 | 11 | 16 | 33 | 156 | 271 | 594 | 0 | 0 | 1204 | 11 | 16 | 0 | 0 | 6242 | 490 | 0 |
| SRR7226518 | 38 | 21 | 12 | 0 | 536 | 4528 | 2777 | 9525 | 3220 | 12 | 248 | 29 | 0 | 0 | 89099 | 79809 | 272276 | 0 | 942 | 9959 | 3345 | 0 | 19 | 11 | 58 | 126 | 239 | 30 | 0 | 339 | 11 | 0 | 0 | 0 | 3001 | 294 | 13 |
| SRR7226519 | 23 | 14 | 15 | 0 | 1679 | 1408 | 1643 | 4528 | 1581 | 14 | 266 | 22 | 0 | 0 | 76238 | 95718 | 268366 | 0 | 600 | 5164 | 3347 | 16 | 16 | 16 | 33 | 76 | 116 | 11 | 0 | 327 | 0 | 0 | 0 | 0 | 2539 | 116 | 0 |
| SRR7226520 | 33 | 22 | 27 | 16 | 52 | 8891 | 10283 | 35555 | 19307 | 0 | 81 | 0 | 0 | 0 | 19609 | 76212 | 86356 | 0 | 613 | 8833 | 3120 | 0 | 0 | 33 | 20 | 30 | 37 | 28 | 0 | 303 | 0 | 0 | 0 | 0 | 5171 | 313 | 0 |
| SRR7226521 | 161 | 99 | 34 | 16 | 313 | 4219 | 4592 | 20118 | 5858 | 0 | 182 | 0 | 0 | 0 | 19167 | 48265 | 115564 | 0 | 290 | 5509 | 1616 | 0 | 0 | 21 | 64 | 137 | 99 | 13 | 0 | 1156 | 0 | 0 | 0 | 0 | 4206 | 1041 | 0 |
| SRR7226522 | 186 | 103 | 43 | 12 | 288 | 2841 | 8901 | 20690 | 11941 | 0 | 379 | 0 | 0 | 0 | 37688 | 98032 | 149052 | 0 | 391 | 12950 | 12495 | 0 | 0 | 11 | 36 | 64 | 56 | 0 | 0 | 1061 | 0 | 0 | 0 | 0 | 6812 | 212 | 0 |
| SRR7226523 | 108 | 51 | 42 | 16 | 188 | 10800 | 9683 | 42697 | 15044 | 0 | 104 | 0 | 0 | 0 | 9334 | 63607 | 18831 | 0 | 1109 | 10966 | 502 | 0 | 0 | 34 | 11 | 23 | 16 | 24 | 0 | 462 | 0 | 0 | 0 | 0 | 5093 | 4654 | 0 |
| SRR7226524 | 38 | 40 | 0 | 0 | 50 | 3170 | 8612 | 20622 | 5015 | 26 | 452 | 164 | 0 | 0 | 6481 | 56455 | 20856 | 0 | 604 | 4740 | 352 | 0 | 0 | 0 | 0 | 27 | 0 | 0 | 41 | 247 | 0 | 0 | 0 | 0 | 411 | 3555 | 27 |
| SRR7226525 | 105 | 79 | 17 | 46 | 121 | 8946 | 9506 | 35797 | 13095 | 0 | 72 | 0 | 0 | 0 | 22687 | 49168 | 72605 | 0 | 529 | 15642 | 641 | 0 | 11 | 12 | 20 | 26 | 23 | 24 | 0 | 252 | 0 | 10 | 0 | 0 | 3747 | 5986 | 0 |
| SRR7226526 | 232 | 204 | 45 | 16 | 233 | 9446 | 6840 | 33501 | 10171 | 0 | 105 | 0 | 0 | 0 | 26575 | 49754 | 40131 | 0 | 1079 | 11139 | 600 | 0 | 0 | 10 | 19 | 36 | 28 | 30 | 0 | 400 | 0 | 0 | 0 | 0 | 4404 | 5984 | 0 |
| SRR7233553 | 15 | 15 | 0 | 0 | 42 | 1073 | 1333 | 6067 | 25558 | 49 | 11 | 18 | 14 | 13 | 1175 | 98869 | 36204 | 0 | 11 | 68 | 102 | 0 | 0 | 21 | 0 | 0 | 0 | 0 | 0 | 44 | 0 | 23 | 0 | 0 | 69267 | 0 | 0 |
| SRR7233554 | 11 | 0 | 0 | 0 | 33 | 1866 | 2292 | 12166 | 42393 | 21 | 17 | 11 | 11 | 0 | 1279 | 105961 | 36592 | 32 | 21 | 139 | 139 | 0 | 0 | 19 | 15 | 0 | 0 | 0 | 0 | 54 | 0 | 0 | 0 | 0 | 61689 | 0 | 0 |
| SRR7233556 | 44 | 0 | 0 | 0 | 18 | 851 | 1526 | 7551 | 37510 | 13 | 11 | 11 | 0 | 0 | 796 | 99882 | 34640 | 55 | 0 | 133 | 55 | 0 | 0 | 14 | 11 | 0 | 0 | 0 | 0 | 46 | 0 | 11 | 0 | 0 | 44428 | 0 | 0 |
| SRR7233557 | 43 | 0 | 0 | 0 | 61 | 1256 | 1521 | 6911 | 32050 | 85 | 24 | 37 | 0 | 0 | 464 | 63718 | 21230 | 18 | 0 | 27 | 0 | 0 | 0 | 17 | 0 | 0 | 0 | 0 | 0 | 65 | 0 | 0 | 0 | 0 | 52715 | 0 | 0 |
| SRR7233558 | 54 | 0 | 0 | 0 | 111 | 1105 | 2008 | 11494 | 55238 | 136 | 81 | 82 | 0 | 0 | 1222 | 110590 | 43615 | 53 | 11 | 138 | 74 | 0 | 0 | 32 | 30 | 0 | 0 | 0 | 0 | 229 | 0 | 21 | 0 | 0 | 46195 | 0 | 0 |
| SRR7233559 | 11 | 0 | 0 | 0 | 24 | 1883 | 2453 | 10591 | 50191 | 20 | 12 | 46 | 14 | 15 | 632 | 53678 | 14868 | 25 | 0 | 12 | 74 | 0 | 0 | 29 | 0 | 0 | 0 | 0 | 0 | 11 | 0 | 0 | 0 | 0 | 54711 | 0 | 0 |
| SRR7233560 | 30 | 0 | 0 | 0 | 68 | 1828 | 2067 | 10652 | 43506 | 64 | 10 | 34 | 0 | 0 | 629 | 54663 | 13582 | 0 | 0 | 60 | 105 | 0 | 0 | 32 | 0 | 0 | 0 | 0 | 0 | 39 | 0 | 30 | 0 | 0 | 47419 | 0 | 0 |
| SRR7233561 | 22 | 22 | 0 | 0 | 72 | 896 | 1001 | 4384 | 20432 | 37 | 17 | 16 | 21 | 10 | 2576 | 58358 | 27876 | 15 | 15 | 75 | 239 | 0 | 0 | 22 | 20 | 0 | 0 | 0 | 0 | 49 | 0 | 15 | 0 | 0 | 76028 | 0 | 0 |
| SRR7233562 | 14 | 11 | 0 | 0 | 31 | 553 | 959 | 4174 | 20898 | 26 | 11 | 13 | 12 | 0 | 1408 | 78137 | 27380 | 31 | 0 | 86 | 61 | 0 | 0 | 28 | 0 | 0 | 0 | 0 | 0 | 27 | 0 | 12 | 0 | 0 | 32281 | 0 | 0 |
| SRR7233563 | 121 | 14 | 0 | 0 | 252 | 1731 | 2396 | 11824 | 57697 | 157 | 112 | 84 | 0 | 0 | 3462 | 160129 | 65600 | 46 | 0 | 556 | 185 | 0 | 0 | 17 | 18 | 0 | 0 | 0 | 0 | 443 | 0 | 31 | 0 | 0 | 45453 | 0 | 0 |
| SRR7233564 | 19 | 17 | 0 | 0 | 37 | 1979 | 2255 | 13751 | 59863 | 25 | 14 | 16 | 11 | 0 | 2881 | 127432 | 56594 | 58 | 44 | 422 | 218 | 0 | 0 | 21 | 23 | 0 | 0 | 0 | 0 | 60 | 0 | 58 | 0 | 0 | 48124 | 0 | 0 |
| SRR7233565 | 28 | 20 | 0 | 0 | 44 | 982 | 1926 | 9368 | 50468 | 26 | 13 | 15 | 12 | 0 | 4740 | 138930 | 60806 | 56 | 0 | 243 | 112 | 0 | 0 | 37 | 36 | 0 | 0 | 0 | 0 | 68 | 0 | 28 | 0 | 0 | 33272 | 0 | 0 |
| SRR7233566 | 192 | 17 | 0 | 0 | 284 | 1048 | 1781 | 8454 | 44042 | 145 | 118 | 75 | 0 | 0 | 5542 | 133176 | 61223 | 73 | 0 | 126 | 115 | 0 | 0 | 41 | 46 | 11 | 0 | 0 | 0 | 412 | 0 | 21 | 0 | 0 | 36599 | 0 | 0 |
| SRR7233568 | 55 | 11 | 0 | 0 | 20 | 1074 | 1348 | 6484 | 24995 | 15 | 59 | 46 | 17 | 0 | 1103 | 93794 | 39178 | 63 | 17 | 51 | 40 | 0 | 0 | 25 | 0 | 0 | 0 | 0 | 0 | 34 | 0 | 0 | 0 | 0 | 25043 | 0 | 0 |
| SRR7233570 | 46 | 0 | 0 | 0 | 38 | 2257 | 2536 | 11463 | 46397 | 44 | 22 | 46 | 14 | 0 | 1205 | 103939 | 36016 | 51 | 13 | 203 | 25 | 0 | 0 | 12 | 0 | 0 | 0 | 0 | 0 | 30 | 0 | 13 | 0 | 0 | 66767 | 0 | 0 |
| SRR7233571 | 19 | 0 | 0 | 0 | 37 | 1184 | 996 | 6023 | 28915 | 40 | 14 | 18 | 0 | 0 | 1400 | 110819 | 53349 | 75 | 19 | 348 | 85 | 0 | 0 | 19 | 0 | 0 | 0 | 0 | 0 | 88 | 0 | 28 | 0 | 0 | 64881 | 0 | 0 |
| SRR7233572 | 12 | 0 | 0 | 0 | 18 | 962 | 1420 | 6604 | 29609 | 31 | 32 | 42 | 0 | 0 | 352 | 67446 | 19819 | 73 | 0 | 80 | 13 | 0 | 0 | 21 | 0 | 0 | 0 | 0 | 0 | 25 | 0 | 13 | 0 | 0 | 44559 | 0 | 0 |
| SRR7262826 | 98 | 61 | 0 | 0 | 247 | 242 | 58 | 433 | 606 | 190 | 62 | 69 | 0 | 0 | 16985 | 513426 | 219400 | 143 | 627 | 8624 | 2032 | 0 | 18 | 161 | 16 | 0 | 0 | 0 | 0 | 392 | 0 | 0 | 0 | 131 | 6293 | 0 | 0 |
| SRR7262827 | 116 | 62 | 0 | 0 | 214 | 663 | 63 | 468 | 609 | 833 | 175 | 54 | 0 | 0 | 39400 | 378253 | 243632 | 251 | 1140 | 17476 | 4555 | 0 | 25 | 214 | 44 | 15 | 13 | 0 | 0 | 669 | 0 | 0 | 0 | 111 | 14680 | 0 | 0 |
| SRR7262828 | 27 | 13 | 15 | 16 | 35 | 209 | 29 | 218 | 485 | 198 | 27 | 69 | 0 | 0 | 31836 | 421655 | 208571 | 169 | 1123 | 24636 | 5421 | 11 | 30 | 26 | 40 | 15 | 16 | 0 | 0 | 84 | 0 | 16 | 0 | 17 | 7017 | 0 | 0 |
| SRR7262829 | 164 | 114 | 32 | 35 | 363 | 65 | 14 | 93 | 154 | 398 | 202 | 104 | 0 | 0 | 14310 | 560584 | 179447 | 39 | 567 | 7470 | 3153 | 0 | 23 | 92 | 23 | 33 | 61 | 0 | 0 | 576 | 0 | 28 | 0 | 63 | 4340 | 0 | 0 |
| SRR7262830 | 15 | 61 | 0 | 0 | 33 | 574 | 62 | 612 | 788 | 125 | 66 | 75 | 0 | 0 | 33157 | 403733 | 219380 | 347 | 932 | 16594 | 4418 | 16 | 21 | 34 | 22 | 0 | 0 | 0 | 0 | 55 | 0 | 23 | 0 | 19 | 18833 | 0 | 0 |
| SRR7288164 | 0 | 0 | 0 | 0 | 0 | 0 | 0 | 0 | 0 | 0 | 0 | 353 | 0 | 0 | 1065 | 93491 | 86391 | 0 | 0 | 2130 | 8520 | 0 | 0 | 0 | 0 | 0 | 0 | 0 | 0 | 0 | 0 | 0 | 0 | 0 | 3550 | 0 | 245 |
| SRR7288166 | 0 | 0 | 0 | 0 | 0 | 0 | 0 | 0 | 0 | 0 | 0 | 12 | 0 | 0 | 11066 | 87007 | 100928 | 0 | 0 | 4024 | 6036 | 0 | 0 | 0 | 0 | 0 | 0 | 0 | 0 | 0 | 0 | 0 | 0 | 0 | 0 | 0 | 229 |
| SRR7288167 | 0 | 0 | 0 | 0 | 0 | 0 | 0 | 0 | 0 | 0 | 0 | 11 | 0 | 0 | 5587 | 80000 | 98824 | 0 | 0 | 4469 | 1117 | 0 | 0 | 0 | 0 | 0 | 0 | 0 | 0 | 16 | 0 | 0 | 0 | 0 | 2353 | 0 | 94 |
| SRR7288236 | 0 | 0 | 0 | 0 | 0 | 0 | 0 | 0 | 0 | 0 | 0 | 63 | 0 | 0 | 2494 | 91041 | 28902 | 0 | 0 | 0 | 2494 | 0 | 0 | 0 | 0 | 0 | 0 | 0 | 0 | 0 | 0 | 0 | 0 | 0 | 1445 | 0 | 157 |
| SRR7288237 | 0 | 0 | 0 | 0 | 0 | 0 | 0 | 0 | 0 | 0 | 0 | 0 | 0 | 0 | 0 | 117820 | 23564 | 0 | 0 | 0 | 12887 | 0 | 0 | 0 | 0 | 0 | 32 | 0 | 0 | 0 | 0 | 0 | 0 | 0 | 4418 | 0 | 32 |
| SRR7288238 | 0 | 0 | 0 | 0 | 0 | 6881 | 0 | 0 | 0 | 0 | 0 | 15 | 0 | 0 | 6881 | 114710 | 29690 | 0 | 0 | 0 | 4587 | 0 | 0 | 0 | 0 | 0 | 15 | 0 | 0 | 0 | 0 | 0 | 0 | 0 | 1350 | 0 | 77 |
| SRR7288239 | 0 | 0 | 0 | 0 | 0 | 0 | 0 | 0 | 0 | 0 | 0 | 0 | 0 | 0 | 4651 | 97101 | 36232 | 0 | 0 | 0 | 9302 | 0 | 0 | 0 | 0 | 0 | 0 | 0 | 0 | 0 | 0 | 0 | 0 | 0 | 1449 | 0 | 62 |
| SRR7288244 | 0 | 0 | 0 | 0 | 0 | 15385 | 0 | 0 | 0 | 0 | 0 | 0 | 0 | 0 | 15385 | 182796 | 86022 | 0 | 0 | 15385 | 0 | 0 | 0 | 0 | 0 | 0 | 0 | 0 | 0 | 0 | 0 | 0 | 0 | 0 | 0 | 0 | 0 |
| SRR7288264 | 0 | 0 | 0 | 0 | 0 | 0 | 0 | 0 | 0 | 0 | 0 | 67 | 0 | 0 | 3836 | 80075 | 190875 | 0 | 0 | 2558 | 2558 | 0 | 0 | 0 | 0 | 0 | 0 | 0 | 0 | 22 | 0 | 0 | 0 | 0 | 931 | 0 | 135 |
| SRR7288265 | 0 | 0 | 0 | 0 | 0 | 0 | 0 | 0 | 0 | 0 | 0 | 0 | 0 | 0 | 13554 | 75000 | 202083 | 0 | 0 | 1506 | 3012 | 0 | 0 | 0 | 0 | 0 | 0 | 0 | 0 | 23 | 0 | 0 | 0 | 0 | 6250 | 0 | 91 |
| SRR7288266 | 0 | 0 | 0 | 0 | 0 | 0 | 0 | 0 | 0 | 0 | 0 | 88 | 0 | 0 | 11220 | 92885 | 196640 | 0 | 0 | 2805 | 4208 | 0 | 0 | 0 | 0 | 0 | 0 | 0 | 0 | 0 | 0 | 0 | 0 | 0 | 2964 | 0 | 110 |
| SRR7288267 | 0 | 0 | 0 | 0 | 0 | 0 | 0 | 0 | 0 | 0 | 23 | 23 | 0 | 0 | 14845 | 70448 | 219579 | 0 | 0 | 1350 | 5398 | 0 | 0 | 0 | 0 | 0 | 0 | 0 | 0 | 46 | 0 | 0 | 0 | 0 | 4575 | 0 | 182 |
| SRR7288277 | 0 | 0 | 0 | 0 | 0 | 0 | 0 | 0 | 0 | 0 | 0 | 0 | 0 | 0 | 30992 | 45499 | 469506 | 0 | 0 | 8264 | 10331 | 0 | 0 | 0 | 0 | 0 | 0 | 0 | 0 | 0 | 0 | 0 | 0 | 0 | 1936 | 0 | 70 |
| SRR7288300 | 0 | 0 | 0 | 0 | 0 | 0 | 0 | 0 | 0 | 0 | 0 | 50 | 0 | 0 | 116340 | 43467 | 190133 | 0 | 347 | 3299 | 8682 | 0 | 0 | 0 | 0 | 0 | 0 | 0 | 0 | 76 | 0 | 0 | 0 | 0 | 235 | 0 | 252 |
| SRR7288301 | 0 | 0 | 0 | 0 | 0 | 0 | 0 | 0 | 0 | 0 | 0 | 52 | 0 | 0 | 109935 | 43724 | 192241 | 0 | 0 | 2511 | 6277 | 0 | 0 | 0 | 0 | 0 | 0 | 0 | 0 | 52 | 0 | 0 | 0 | 0 | 241 | 0 | 412 |
| SRR7288302 | 0 | 0 | 0 | 0 | 0 | 0 | 0 | 0 | 0 | 0 | 50 | 25 | 0 | 0 | 116658 | 43109 | 194799 | 0 | 525 | 3854 | 10159 | 0 | 0 | 0 | 0 | 0 | 0 | 0 | 0 | 226 | 0 | 0 | 0 | 0 | 169 | 0 | 351 |
| SRR7288303 | 0 | 0 | 0 | 0 | 0 | 0 | 0 | 0 | 0 | 0 | 26 | 78 | 0 | 0 | 110282 | 43595 | 192949 | 0 | 0 | 2664 | 7814 | 0 | 0 | 0 | 0 | 0 | 0 | 0 | 0 | 130 | 0 | 0 | 0 | 0 | 137 | 0 | 389 |
| SRR7288308 | 0 | 0 | 0 | 0 | 0 | 0 | 0 | 0 | 0 | 0 | 0 | 0 | 0 | 0 | 5942 | 183719 | 128454 | 0 | 0 | 2377 | 9507 | 0 | 0 | 0 | 0 | 0 | 0 | 0 | 0 | 0 | 0 | 594 | 0 | 0 | 2240 | 0 | 394 |
| SRR7288309 | 0 | 0 | 0 | 0 | 0 | 0 | 0 | 0 | 0 | 0 | 0 | 0 | 0 | 0 | 9294 | 189755 | 116883 | 0 | 0 | 3717 | 620 | 0 | 0 | 0 | 0 | 0 | 0 | 37 | 0 | 0 | 0 | 0 | 0 | 0 | 2165 | 0 | 11 |
| SRR7288310 | 0 | 0 | 0 | 0 | 0 | 0 | 0 | 0 | 0 | 0 | 0 | 35 | 0 | 0 | 6086 | 198571 | 130714 | 0 | 0 | 2435 | 4261 | 0 | 0 | 0 | 0 | 0 | 0 | 0 | 0 | 0 | 0 | 0 | 0 | 0 | 2857 | 0 | 213 |
| SRR7288311 | 0 | 0 | 0 | 0 | 0 | 0 | 0 | 0 | 0 | 0 | 0 | 0 | 0 | 0 | 4748 | 180264 | 128982 | 0 | 0 | 2374 | 4154 | 0 | 0 | 0 | 0 | 0 | 0 | 0 | 0 | 0 | 0 | 0 | 0 | 0 | 777 | 0 | 186 |
| SRR7288320 | 0 | 0 | 0 | 0 | 0 | 0 | 0 | 0 | 0 | 0 | 0 | 25 | 0 | 0 | 8547 | 130693 | 310099 | 0 | 0 | 1603 | 534 | 0 | 0 | 0 | 0 | 0 | 0 | 0 | 0 | 0 | 0 | 0 | 0 | 0 | 1188 | 0 | 49 |
| SRR7288321 | 0 | 0 | 0 | 0 | 0 | 0 | 0 | 0 | 0 | 0 | 0 | 26 | 0 | 0 | 7391 | 137463 | 310564 | 0 | 0 | 1706 | 1137 | 0 | 0 | 0 | 0 | 0 | 0 | 0 | 0 | 0 | 0 | 0 | 0 | 0 | 1273 | 0 | 208 |
| SRR7288322 | 0 | 0 | 0 | 0 | 0 | 0 | 0 | 0 | 0 | 0 | 0 | 0 | 0 | 0 | 7403 | 137917 | 316375 | 0 | 0 | 1586 | 1586 | 0 | 0 | 0 | 0 | 0 | 0 | 0 | 0 | 0 | 0 | 0 | 0 | 0 | 1590 | 0 | 243 |
| SRR7288323 | 0 | 0 | 0 | 0 | 0 | 549 | 0 | 0 | 0 | 0 | 26 | 26 | 0 | 0 | 12624 | 137198 | 318182 | 0 | 0 | 1647 | 1098 | 0 | 0 | 0 | 0 | 0 | 0 | 0 | 0 | 0 | 0 | 0 | 0 | 0 | 1251 | 0 | 103 |
| SRR7288376 | 0 | 0 | 0 | 0 | 0 | 0 | 0 | 0 | 0 | 0 | 0 | 0 | 0 | 0 | 8403 | 59937 | 100946 | 0 | 0 | 5602 | 8403 | 0 | 0 | 0 | 0 | 0 | 0 | 0 | 0 | 0 | 0 | 0 | 0 | 0 | 3155 | 0 | 237 |
| SRR7288377 | 0 | 0 | 0 | 0 | 0 | 0 | 0 | 0 | 0 | 0 | 0 | 0 | 0 | 0 | 2857 | 63492 | 117460 | 0 | 0 | 0 | 2857 | 0 | 0 | 0 | 0 | 0 | 61 | 0 | 0 | 0 | 0 | 2857 | 0 | 0 | 0 | 0 | 61 |
| SRR7288378 | 0 | 0 | 0 | 0 | 0 | 0 | 0 | 0 | 0 | 0 | 0 | 0 | 0 | 0 | 5731 | 72848 | 102649 | 0 | 0 | 2865 | 0 | 0 | 0 | 0 | 0 | 0 | 0 | 0 | 0 | 0 | 0 | 0 | 0 | 0 | 3311 | 0 | 176 |
| SRR7288379 | 0 | 0 | 0 | 0 | 0 | 0 | 0 | 0 | 0 | 0 | 61 | 0 | 0 | 0 | 5988 | 88146 | 124620 | 0 | 0 | 0 | 0 | 0 | 0 | 0 | 0 | 0 | 0 | 0 | 0 | 0 | 0 | 0 | 0 | 0 | 6079 | 0 | 184 |
| SRR7288380 | 0 | 0 | 0 | 0 | 0 | 0 | 0 | 0 | 0 | 0 | 0 | 66 | 0 | 0 | 9363 | 124128 | 66946 | 0 | 0 | 0 | 5618 | 0 | 0 | 0 | 0 | 0 | 22 | 0 | 0 | 0 | 0 | 0 | 0 | 0 | 2789 | 0 | 88 |
| SRR7288381 | 0 | 0 | 0 | 0 | 0 | 0 | 0 | 0 | 0 | 0 | 23 | 0 | 0 | 0 | 9671 | 99567 | 92352 | 0 | 0 | 0 | 5803 | 0 | 0 | 0 | 0 | 0 | 0 | 0 | 0 | 0 | 0 | 0 | 0 | 0 | 1443 | 0 | 90 |
| SRR7288382 | 0 | 0 | 0 | 0 | 0 | 0 | 0 | 0 | 0 | 0 | 0 | 43 | 0 | 0 | 3540 | 133053 | 75630 | 0 | 0 | 0 | 1770 | 0 | 0 | 0 | 0 | 0 | 22 | 22 | 0 | 0 | 0 | 0 | 0 | 0 | 0 | 0 | 195 |

### Supp Table S8.3.Columns_BX_to_DH

| | | | | | | | | | | | | | | | | | | | | | | | | | | | | | | | | | | | | | |
|---|---|---|---|---|---|---|---|---|---|---|---|---|---|---|---|---|---|---|---|---|---|---|---|---|---|---|---|---|---|---|---|---|---|---|---|---|---|
| SRR7288383 | 0 | 0 | 0 | 0 | 0 | 0 | 0 | 0 | 0 | 0 | 0 | 21 | 0 | 0 | 3724 | 93391 | 89081 | 0 | 0 | 0 | 9311 | 0 | 0 | 0 | 0 | 0 | 0 | 0 | 0 | 0 | 0 | 0 | 0 | 0 | 1437 | 0 | 64 |
| SRR7288384 | 0 | 0 | 0 | 0 | 0 | 0 | 0 | 0 | 0 | 0 | 0 | 86 | 0 | 0 | 3749 | 129657 | 21116 | 937 | 0 | 937 | 4686 | 0 | 0 | 0 | 0 | 0 | 17 | 0 | 0 | 17 | 0 | 0 | 0 | 0 | 672 | 0 | 224 |
| SRR7288385 | 0 | 0 | 0 | 0 | 0 | 0 | 0 | 0 | 0 | 0 | 0 | 35 | 0 | 0 | 10377 | 127877 | 26044 | 0 | 0 | 3774 | 3774 | 0 | 0 | 0 | 0 | 0 | 0 | 0 | 0 | 0 | 0 | 0 | 0 | 0 | 827 | 0 | 315 |
| SRR7288386 | 0 | 0 | 0 | 0 | 0 | 0 | 0 | 0 | 0 | 0 | 17 | 84 | 0 | 0 | 6579 | 142711 | 21228 | 0 | 0 | 0 | 4699 | 0 | 0 | 0 | 0 | 0 | 17 | 0 | 0 | 0 | 0 | 0 | 0 | 0 | 767 | 0 | 134 |
| SRR7288387 | 0 | 0 | 0 | 0 | 0 | 0 | 0 | 0 | 0 | 0 | 0 | 17 | 0 | 0 | 5164 | 135512 | 22923 | 861 | 0 | 5164 | 3442 | 0 | 0 | 0 | 0 | 0 | 0 | 0 | 0 | 33 | 0 | 0 | 0 | 0 | 887 | 0 | 234 |
| SRR7288400 | 0 | 0 | 0 | 0 | 0 | 0 | 0 | 0 | 0 | 0 | 0 | 0 | 0 | 0 | 14925 | 40000 | 106667 | 0 | 0 | 14925 | 0 | 0 | 0 | 0 | 0 | 0 | 0 | 0 | 0 | 0 | 0 | 0 | 0 | 0 | 0 | 0 | 0 |
| SRR7288402 | 0 | 0 | 0 | 0 | 0 | 11628 | 0 | 0 | 0 | 0 | 0 | 0 | 0 | 0 | 11628 | 77922 | 298701 | 0 | 0 | 11628 | 0 | 0 | 0 | 0 | 0 | 488 | 0 | 0 | 0 | 0 | 0 | 0 | 0 | 0 | 0 | 0 | 488 |
| SRR7288472 | 0 | 0 | 0 | 0 | 0 | 0 | 2433 | 0 | 0 | 0 | 0 | 199 | 0 | 0 | 4866 | 105556 | 300000 | 0 | 0 | 0 | 0 | 0 | 0 | 0 | 0 | 0 | 0 | 0 | 0 | 0 | 0 | 0 | 0 | 0 | 0 | 0 | 199 |
| SRR7288474 | 0 | 0 | 0 | 0 | 0 | 0 | 0 | 0 | 0 | 0 | 0 | 291 | 0 | 0 | 6944 | 90909 | 315508 | 0 | 0 | 0 | 2315 | 0 | 0 | 0 | 0 | 0 | 0 | 0 | 0 | 0 | 0 | 0 | 0 | 0 | 2674 | 0 | 194 |
| SRR7288475 | 0 | 0 | 0 | 0 | 0 | 0 | 0 | 0 | 0 | 0 | 0 | 103 | 0 | 0 | 7317 | 150000 | 323529 | 0 | 0 | 0 | 2439 | 0 | 0 | 0 | 0 | 0 | 0 | 0 | 0 | 0 | 0 | 0 | 0 | 0 | 0 | 0 | 103 |
| SRR7288512 | 0 | 0 | 0 | 0 | 0 | 0 | 0 | 0 | 0 | 0 | 0 | 0 | 0 | 0 | 17272 | 307538 | 105474 | 0 | 0 | 2978 | 4169 | 0 | 0 | 0 | 0 | 0 | 50 | 0 | 0 | 0 | 0 | 0 | 0 | 0 | 1720 | 0 | 25 |
| SRR7288513 | 0 | 0 | 0 | 0 | 0 | 0 | 0 | 0 | 0 | 0 | 0 | 51 | 0 | 0 | 10903 | 335989 | 110384 | 0 | 0 | 1817 | 1817 | 0 | 0 | 0 | 0 | 0 | 0 | 0 | 0 | 0 | 0 | 0 | 0 | 0 | 1138 | 0 | 51 |
| SRR7288514 | 0 | 0 | 0 | 0 | 0 | 0 | 0 | 0 | 0 | 0 | 0 | 49 | 0 | 0 | 11786 | 331349 | 104592 | 0 | 0 | 1179 | 1768 | 0 | 0 | 0 | 0 | 0 | 24 | 0 | 0 | 0 | 0 | 0 | 0 | 0 | 1134 | 0 | 98 |
| SRR7288515 | 0 | 0 | 0 | 0 | 0 | 0 | 0 | 0 | 0 | 0 | 0 | 24 | 0 | 0 | 15734 | 339241 | 97480 | 0 | 0 | 1166 | 1748 | 0 | 0 | 0 | 0 | 0 | 48 | 0 | 0 | 0 | 0 | 0 | 0 | 0 | 1385 | 0 | 48 |
| SRR7293222 | 37 | 36 | 0 | 16 | 0 | 647 | 467 | 1817 | 156 | 20 | 130 | 11 | 0 | 0 | 87145 | 21488 | 240578 | 0 | 108 | 4135 | 8767 | 0 | 0 | 53 | 0 | 0 | 34 | 0 | 0 | 452 | 0 | 0 | 0 | 0 | 2052 | 28 | 0 |
| SRR7293223 | 25 | 59 | 0 | 17 | 0 | 673 | 365 | 1804 | 121 | 0 | 117 | 0 | 0 | 0 | 40506 | 11273 | 141776 | 0 | 98 | 3560 | 6564 | 0 | 0 | 34 | 0 | 0 | 16 | 0 | 0 | 451 | 0 | 13 | 0 | 0 | 3148 | 22 | 0 |
| SRR7293224 | 94 | 61 | 10 | 21 | 0 | 337 | 476 | 1107 | 207 | 0 | 127 | 0 | 0 | 0 | 17131 | 25384 | 91878 | 0 | 106 | 15533 | 4083 | 0 | 0 | 13 | 0 | 0 | 0 | 0 | 0 | 21 | 0 | 13 | 0 | 0 | 11447 | 89 | 0 |
| SRR7293225 | 51 | 33 | 0 | 85 | 0 | 190 | 170 | 389 | 53 | 0 | 88 | 0 | 0 | 0 | 64949 | 9253 | 194586 | 0 | 165 | 3767 | 11110 | 0 | 0 | 42 | 0 | 0 | 15 | 0 | 0 | 263 | 0 | 0 | 0 | 0 | 3087 | 119 | 18 |
| SRR7293226 | 34 | 48 | 0 | 14 | 0 | 287 | 134 | 664 | 67 | 31 | 174 | 0 | 0 | 0 | 57010 | 25634 | 161848 | 0 | 115 | 4377 | 8663 | 0 | 0 | 113 | 22 | 15 | 167 | 0 | 0 | 18 | 0 | 0 | 0 | 0 | 1515 | 136 | 12 |
| SRR7293227 | 19 | 26 | 0 | 13 | 0 | 366 | 156 | 656 | 31 | 18 | 75 | 0 | 0 | 0 | 58726 | 26511 | 154773 | 0 | 127 | 3498 | 11265 | 0 | 0 | 111 | 0 | 15 | 48 | 0 | 0 | 288 | 0 | 0 | 0 | 0 | 1266 | 116 | 0 |
| SRR7293228 | 69 | 57 | 0 | 0 | 0 | 217 | 903 | 630 | 315 | 0 | 55 | 0 | 0 | 0 | 18641 | 16834 | 92655 | 0 | 60 | 30768 | 2725 | 0 | 0 | 24 | 0 | 0 | 0 | 0 | 0 | 226 | 0 | 13 | 0 | 0 | 17808 | 36 | 0 |
| SRR7293229 | 67 | 53 | 0 | 0 | 0 | 1233 | 1058 | 1940 | 283 | 0 | 13 | 0 | 0 | 0 | 42565 | 11131 | 175967 | 17 | 75 | 2748 | 7096 | 0 | 0 | 36 | 0 | 0 | 0 | 0 | 0 | 182 | 0 | 0 | 0 | 0 | 8050 | 28 | 0 |
| SRR7293230 | 12 | 0 | 0 | 17 | 28 | 2914 | 352 | 1564 | 82 | 20 | 166 | 23 | 0 | 0 | 47742 | 26400 | 137478 | 25 | 199 | 4621 | 3312 | 0 | 0 | 101 | 0 | 0 | 0 | 0 | 0 | 378 | 0 | 82 | 0 | 0 | 746 | 412 | 15 |
| SRR7293231 | 0 | 14 | 0 | 0 | 21 | 1569 | 320 | 1123 | 56 | 42 | 116 | 26 | 0 | 0 | 53651 | 26650 | 121065 | 0 | 96 | 4568 | 2517 | 0 | 0 | 95 | 0 | 0 | 0 | 0 | 0 | 434 | 0 | 36 | 0 | 0 | 576 | 256 | 0 |
| SRR7293232 | 78 | 13 | 0 | 0 | 16 | 1760 | 1267 | 6727 | 424 | 0 | 91 | 0 | 0 | 0 | 11575 | 38501 | 50148 | 25 | 200 | 50873 | 605 | 0 | 0 | 15 | 0 | 0 | 0 | 0 | 0 | 31 | 0 | 112 | 0 | 0 | 11459 | 176 | 0 |
| SRR7293233 | 0 | 0 | 0 | 24 | 70 | 331 | 793 | 735 | 172 | 45 | 337 | 36 | 0 | 0 | 86063 | 27685 | 220028 | 0 | 290 | 8179 | 3072 | 0 | 0 | 33 | 0 | 0 | 19 | 0 | 0 | 552 | 0 | 20 | 0 | 0 | 825 | 301 | 12 |
| SRR7343626 | 0 | 0 | 0 | 21 | 0 | 0 | 76 | 228 | 76 | 0 | 0 | 0 | 0 | 0 | 531 | 15671 | 11865 | 0 | 0 | 0 | 0 | 0 | 0 | 41 | 0 | 21 | 0 | 0 | 0 | 62 | 0 | 0 | 0 | 0 | 11417 | 0 | 0 |
| SRR7343627 | 0 | 0 | 0 | 0 | 0 | 0 | 0 | 277 | 0 | 0 | 50 | 0 | 0 | 0 | 1385 | 59962 | 17469 | 0 | 0 | 0 | 0 | 0 | 0 | 100 | 0 | 0 | 0 | 0 | 0 | 149 | 0 | 0 | 0 | 0 | 4249 | 0 | 0 |
| SRR7343629 | 0 | 0 | 0 | 0 | 50 | 0 | 504 | 0 | 504 | 0 | 0 | 0 | 0 | 0 | 2017 | 104225 | 46009 | 0 | 0 | 0 | 504 | 0 | 0 | 149 | 0 | 0 | 0 | 0 | 0 | 298 | 0 | 0 | 0 | 0 | 16901 | 0 | 0 |
| SRR7343630 | 0 | 0 | 0 | 0 | 0 | 0 | 0 | 189 | 0 | 0 | 45 | 0 | 0 | 0 | 416 | 5640 | 4165 | 0 | 0 | 113 | 38 | 0 | 0 | 118 | 0 | 0 | 0 | 0 | 0 | 36 | 0 | 0 | 0 | 0 | 2343 | 0 | 0 |
| SRR7343631 | 0 | 0 | 0 | 0 | 21 | 0 | 347 | 173 | 0 | 0 | 21 | 0 | 0 | 0 | 1387 | 59166 | 25357 | 0 | 0 | 347 | 0 | 0 | 0 | 21 | 21 | 0 | 21 | 0 | 0 | 192 | 0 | 0 | 0 | 0 | 7578 | 0 | 0 |
| SRR7343632 | 0 | 0 | 0 | 0 | 124 | 0 | 321 | 321 | 0 | 0 | 41 | 0 | 0 | 0 | 6096 | 69113 | 39085 | 0 | 0 | 0 | 0 | 0 | 0 | 41 | 41 | 0 | 41 | 0 | 0 | 290 | 0 | 0 | 0 | 0 | 4766 | 0 | 0 |
| SRR7343633 | 0 | 0 | 0 | 0 | 10 | 427 | 570 | 712 | 0 | 0 | 134 | 31 | 0 | 0 | 2279 | 45273 | 24772 | 0 | 0 | 427 | 285 | 0 | 0 | 52 | 52 | 41 | 31 | 10 | 0 | 392 | 0 | 0 | 0 | 0 | 3132 | 0 | 0 |
| SRR7343635 | 0 | 0 | 0 | 0 | 39 | 249 | 249 | 1992 | 0 | 0 | 0 | 0 | 0 | 0 | 996 | 42237 | 14079 | 0 | 0 | 498 | 0 | 0 | 0 | 77 | 0 | 0 | 0 | 0 | 0 | 116 | 0 | 0 | 0 | 0 | 7431 | 0 | 0 |
| SRR7343636 | 0 | 0 | 0 | 0 | 61 | 0 | 156 | 78 | 78 | 0 | 85 | 12 | 0 | 0 | 938 | 43216 | 28516 | 0 | 0 | 391 | 235 | 0 | 0 | 73 | 12 | 12 | 37 | 0 | 0 | 341 | 0 | 0 | 0 | 0 | 6908 | 0 | 0 |
| SRR7343637 | 0 | 0 | 0 | 0 | 25 | 233 | 233 | 349 | 116 | 0 | 102 | 0 | 0 | 0 | 2209 | 24189 | 26750 | 0 | 0 | 581 | 116 | 0 | 0 | 102 | 102 | 203 | 51 | 0 | 0 | 102 | 0 | 0 | 0 | 0 | 3984 | 0 | 0 |
| SRR7343638 | 0 | 0 | 0 | 0 | 0 | 425 | 850 | 425 | 0 | 0 | 78 | 0 | 0 | 0 | 1699 | 34375 | 12500 | 0 | 0 | 850 | 0 | 0 | 0 | 156 | 0 | 0 | 156 | 0 | 0 | 78 | 0 | 0 | 0 | 0 | 3750 | 0 | 0 |
| SRR7343639 | 0 | 0 | 0 | 0 | 44 | 265 | 0 | 265 | 0 | 0 | 0 | 0 | 0 | 0 | 265 | 16192 | 3886 | 0 | 0 | 0 | 0 | 0 | 0 | 44 | 0 | 44 | 0 | 0 | 0 | 88 | 0 | 0 | 0 | 0 | 5181 | 0 | 0 |
| SRR7343640 | 0 | 0 | 0 | 19 | 0 | 0 | 439 | 526 | 0 | 0 | 77 | 0 | 0 | 0 | 2018 | 32728 | 16792 | 0 | 0 | 526 | 0 | 0 | 0 | 19 | 0 | 0 | 19 | 0 | 0 | 617 | 0 | 0 | 0 | 0 | 28101 | 0 | 0 |
| SRR7412243 | 14 | 0 | 28 | 43 | 78 | 59 | 59 | 119 | 30 | 0 | 0 | 0 | 0 | 0 | 238 | 26485 | 66617 | 0 | 0 | 1902 | 683 | 0 | 0 | 156 | 50 | 490 | 1357 | 931 | 0 | 43 | 0 | 0 | 0 | 0 | 2027 | 0 | 0 |
| SRR7412244 | 18 | 18 | 46 | 286 | 65 | 43 | 217 | 0 | 43 | 0 | 0 | 0 | 0 | 0 | 348 | 15406 | 22009 | 0 | 0 | 1435 | 1087 | 0 | 0 | 55 | 0 | 221 | 793 | 701 | 0 | 55 | 0 | 0 | 0 | 0 | 1853 | 0 | 0 |
| SRR7412245 | 0 | 23 | 31 | 19 | 13 | 53 | 0 | 0 | 0 | 0 | 0 | 0 | 0 | 0 | 106 | 20946 | 14684 | 106 | 0 | 2378 | 1110 | 0 | 0 | 10 | 31 | 22 | 122 | 89 | 0 | 16 | 0 | 0 | 0 | 0 | 2267 | 0 | 0 |
| SRR7412246 | 45 | 38 | 71 | 147 | 71 | 36 | 181 | 145 | 36 | 0 | 26 | 0 | 0 | 0 | 289 | 18717 | 47113 | 36 | 0 | 3146 | 723 | 0 | 0 | 90 | 26 | 289 | 750 | 526 | 0 | 64 | 0 | 0 | 0 | 0 | 1760 | 0 | 0 |
| SRR7412247 | 12 | 16 | 40 | 131 | 87 | 51 | 119 | 34 | 0 | 0 | 16 | 0 | 0 | 0 | 525 | 22835 | 54234 | 0 | 0 | 3048 | 2134 | 0 | 0 | 32 | 36 | 508 | 1338 | 885 | 0 | 24 | 0 | 0 | 0 | 0 | 1762 | 0 | 0 |
| SRR7412248 | 19 | 33 | 28 | 248 | 65 | 0 | 23 | 23 | 0 | 0 | 19 | 0 | 0 | 0 | 626 | 17954 | 84021 | 0 | 0 | 898 | 490 | 0 | 0 | 84 | 47 | 524 | 1496 | 1197 | 0 | 19 | 0 | 0 | 0 | 0 | 1553 | 0 | 0 |
| SRR7412249 | 14 | 23 | 70 | 25 | 12 | 20 | 121 | 20 | 40 | 0 | 0 | 0 | 0 | 0 | 20 | 11677 | 12405 | 0 | 0 | 1290 | 363 | 0 | 0 | 37 | 37 | 49 | 153 | 93 | 0 | 37 | 0 | 0 | 0 | 0 | 1899 | 0 | 0 |
| SRR7412250 | 0 | 14 | 21 | 178 | 96 | 0 | 116 | 87 | 29 | 0 | 0 | 14 | 0 | 0 | 29 | 14078 | 17910 | 0 | 0 | 349 | 58 | 0 | 0 | 165 | 21 | 432 | 1228 | 782 | 0 | 41 | 0 | 0 | 0 | 0 | 2317 | 0 | 0 |
| SRR7412251 | 17 | 33 | 0 | 149 | 99 | 64 | 0 | 0 | 0 | 0 | 17 | 0 | 0 | 0 | 890 | 17993 | 51232 | 0 | 0 | 1208 | 572 | 0 | 0 | 66 | 50 | 463 | 1784 | 1289 | 0 | 17 | 0 | 0 | 0 | 0 | 1241 | 0 | 0 |
| SRR7412252 | 0 | 12 | 20 | 160 | 152 | 54 | 67 | 13 | 13 | 0 | 0 | 0 | 0 | 0 | 364 | 19773 | 61227 | 0 | 13 | 620 | 431 | 0 | 0 | 66 | 53 | 410 | 1270 | 741 | 0 | 12 | 0 | 0 | 0 | 0 | 2041 | 0 | 0 |
| SRR7412253 | 12 | 0 | 43 | 223 | 168 | 179 | 209 | 179 | 60 | 0 | 25 | 0 | 0 | 0 | 1077 | 33411 | 108821 | 0 | 0 | 987 | 837 | 0 | 0 | 192 | 124 | 335 | 900 | 677 | 0 | 50 | 0 | 0 | 0 | 0 | 4896 | 0 | 0 |
| SRR7412254 | 0 | 0 | 26 | 158 | 198 | 86 | 86 | 86 | 0 | 0 | 26 | 0 | 0 | 0 | 429 | 22990 | 39258 | 0 | 0 | 771 | 1114 | 0 | 0 | 383 | 145 | 805 | 2296 | 1055 | 0 | 145 | 0 | 0 | 0 | 0 | 4706 | 0 | 0 |
| SRR7412255 | 10 | 62 | 21 | 243 | 124 | 33 | 148 | 33 | 33 | 0 | 16 | 0 | 0 | 0 | 346 | 17715 | 66222 | 0 | 0 | 825 | 1138 | 0 | 0 | 119 | 124 | 760 | 1587 | 372 | 0 | 36 | 0 | 0 | 0 | 0 | 1432 | 0 | 0 |
| SRR7412256 | 31 | 10 | 31 | 239 | 218 | 40 | 281 | 40 | 0 | 0 | 21 | 0 | 0 | 0 | 281 | 11544 | 20137 | 0 | 0 | 1488 | 80 | 0 | 10 | 364 | 31 | 270 | 727 | 457 | 0 | 62 | 0 | 0 | 0 | 0 | 2430 | 0 | 0 |
| SRR7412257 | 14 | 14 | 54 | 19 | 20 | 0 | 129 | 77 | 26 | 0 | 74 | 0 | 0 | 0 | 952 | 17720 | 40858 | 26 | 0 | 2135 | 566 | 0 | 0 | 16 | 0 | 26 | 85 | 52 | 0 | 11 | 0 | 26 | 0 | 0 | 6095 | 0 | 0 |
| SRR7412258 | 0 | 0 | 72 | 72 | 216 | 153 | 77 | 77 | 0 | 0 | 0 | 0 | 0 | 0 | 153 | 19939 | 21974 | 0 | 0 | 1225 | 383 | 0 | 0 | 18 | 72 | 24 | 93 | 80 | 0 | 0 | 0 | 0 | 0 | 0 | 2238 | 0 | 0 |
| SRR7412259 | 38 | 26 | 38 | 186 | 205 | 26 | 129 | 103 | 0 | 0 | 51 | 0 | 0 | 0 | 694 | 24751 | 108977 | 51 | 0 | 2082 | 1002 | 0 | 0 | 71 | 26 | 314 | 1046 | 526 | 0 | 45 | 0 | 77 | 0 | 0 | 3030 | 0 | 0 |
| SRR7412260 | 0 | 36 | 18 | 18 | 36 | 0 | 48 | 0 | 0 | 0 | 18 | 0 | 0 | 0 | 387 | 25442 | 65346 | 0 | 0 | 968 | 1549 | 0 | 0 | 252 | 18 | 504 | 1889 | 450 | 0 | 18 | 0 | 0 | 0 | 0 | 1473 | 0 | 0 |
| SRR7412261 | 21 | 21 | 69 | 236 | 340 | 0 | 51 | 25 | 0 | 0 | 21 | 0 | 0 | 0 | 203 | 27260 | 78786 | 0 | 0 | 1497 | 1725 | 0 | 0 | 104 | 0 | 340 | 1089 | 250 | 0 | 28 | 0 | 0 | 0 | 0 | 2521 | 0 | 0 |
| SRR7412262 | 0 | 17 | 26 | 111 | 12 | 87 | 0 | 58 | 0 | 0 | 0 | 0 | 0 | 0 | 58 | 11671 | 25699 | 0 | 0 | 899 | 435 | 0 | 0 | 77 | 60 | 38 | 111 | 37 | 0 | 26 | 0 | 0 | 0 | 0 | 1964 | 0 | 0 |
| SRR7412263 | 47 | 24 | 118 | 14 | 15 | 0 | 31 | 125 | 0 | 0 | 24 | 0 | 0 | 0 | 2345 | 29773 | 195424 | 0 | 0 | 750 | 782 | 0 | 0 | 21 | 47 | 38 | 138 | 114 | 0 | 47 | 0 | 0 | 0 | 0 | 1998 | 0 | 0 |
| SRR7412264 | 39 | 0 | 145 | 368 | 118 | 122 | 81 | 41 | 0 | 0 | 13 | 26 | 0 | 0 | 894 | 35794 | 139331 | 0 | 0 | 1382 | 1544 | 0 | 0 | 276 | 13 | 355 | 1210 | 855 | 0 | 0 | 0 | 0 | 0 | 0 | 5342 | 0 | 0 |
| SRR7412265 | 17 | 0 | 34 | 22 | 15 | 44 | 0 | 22 | 0 | 0 | 59 | 0 | 0 | 0 | 110 | 12675 | 19936 | 0 | 0 | 1585 | 286 | 0 | 0 | 101 | 67 | 44 | 133 | 119 | 0 | 59 | 0 | 44 | 0 | 0 | 5476 | 0 | 0 |
| SRR7412266 | 0 | 22 | 74 | 273 | 354 | 65 | 46 | 74 | 19 | 0 | 96 | 0 | 0 | 0 | 716 | 31193 | 73145 | 0 | 0 | 1162 | 390 | 0 | 0 | 118 | 81 | 679 | 1971 | 1462 | 0 | 406 | 0 | 0 | 0 | 0 | 4331 | 0 | 0 |
| SRR7412267 | 13 | 0 | 33 | 160 | 247 | 97 | 169 | 97 | 0 | 0 | 80 | 0 | 0 | 0 | 266 | 15966 | 33779 | 0 | 0 | 1281 | 242 | 0 | 0 | 73 | 20 | 280 | 1101 | 994 | 0 | 173 | 0 | 0 | 0 | 0 | 5470 | 0 | 0 |
| SRR7412268 | 19 | 0 | 44 | 16 | 26 | 17 | 86 | 52 | 17 | 0 | 56 | 0 | 0 | 0 | 362 | 18741 | 121785 | 0 | 0 | 483 | 362 | 0 | 0 | 44 | 75 | 29 | 50 | 70 | 0 | 21 | 0 | 0 | 0 | 0 | 2251 | 0 | 0 |
| SRR7412269 | 0 | 18 | 79 | 28 | 21 | 26 | 79 | 79 | 0 | 0 | 54 | 0 | 0 | 0 | 2909 | 23487 | 416425 | 0 | 0 | 1799 | 2698 | 0 | 0 | 67 | 24 | 37 | 77 | 67 | 0 | 48 | 0 | 0 | 0 | 0 | 2542 | 0 | 0 |
| SRR7412270 | 0 | 15 | 30 | 297 | 193 | 50 | 150 | 50 | 0 | 0 | 104 | 0 | 0 | 0 | 100 | 14353 | 31388 | 0 | 0 | 1102 | 300 | 0 | 0 | 104 | 45 | 387 | 1115 | 1294 | 0 | 149 | 0 | 50 | 0 | 0 | 2524 | 0 | 0 |
| SRR7412271 | 23 | 23 | 0 | 91 | 206 | 116 | 39 | 154 | 0 | 0 | 23 | 0 | 0 | 0 | 232 | 17379 | 25439 | 0 | 39 | 1506 | 425 | 0 | 0 | 57 | 126 | 719 | 2124 | 1541 | 0 | 91 | 0 | 0 | 0 | 0 | 2771 | 0 | 0 |
| SRR7412272 | 19 | 0 | 39 | 145 | 164 | 69 | 103 | 69 | 34 | 0 | 19 | 0 | 0 | 0 | 241 | 13115 | 25786 | 0 | 0 | 1170 | 344 | 0 | 0 | 154 | 48 | 270 | 1215 | 839 | 0 | 67 | 0 | 0 | 0 | 0 | 2924 | 0 | 0 |
| SRR7412273 | 49 | 16 | 57 | 130 | 146 | 70 | 187 | 94 | 0 | 0 | 69 | 0 | 0 | 0 | 1406 | 28250 | 230240 | 0 | 47 | 4029 | 1944 | 0 | 0 | 158 | 81 | 316 | 774 | 478 | 0 | 69 | 0 | 0 | 0 | 0 | 2765 | 0 | 0 |
| SRR7412274 | 0 | 0 | 39 | 186 | 264 | 75 | 112 | 0 | 37 | 0 | 137 | 0 | 0 | 0 | 972 | 33955 | 36797 | 0 | 0 | 2802 | 747 | 0 | 0 | 156 | 59 | 303 | 762 | 459 | 0 | 176 | 0 | 0 | 0 | 0 | 5449 | 0 | 0 |
| SRR7412275 | 0 | 39 | 32 | 181 | 97 | 44 | 66 | 110 | 0 | 0 | 78 | 0 | 0 | 0 | 1646 | 23615 | 151986 | 0 | 44 | 1822 | 2239 | 0 | 0 | 97 | 32 | 310 | 975 | 698 | 0 | 78 | 0 | 0 | 0 | 0 | 2301 | 0 | 0 |
| SRR7412276 | 18 | 0 | 36 | 11 | 27 | 0 | 63 | 63 | 0 | 0 | 18 | 0 | 0 | 0 | 317 | 28111 | 37878 | 0 | 0 | 1268 | 888 | 0 | 0 | 54 | 0 | 16 | 84 | 120 | 0 | 71 | 0 | 0 | 0 | 0 | 4751 | 0 | 0 |
| SRR7412277 | 32 | 16 | 0 | 121 | 10 | 33 | 33 | 130 | 0 | 0 | 32 | 0 | 0 | 0 | 293 | 23873 | 48068 | 0 | 0 | 1662 | 912 | 0 | 0 | 19 | 72 | 51 | 191 | 236 | 0 | 14 | 0 | 0 | 0 | 0 | 2662 | 0 | 0 |
| SRR7412278 | 56 | 0 | 28 | 12 | 16 | 117 | 0 | 0 | 0 | 0 | 14 | 0 | 0 | 0 | 176 | 28328 | 66259 | 0 | 0 | 1346 | 1521 | 0 | 0 | 0 | 42 | 53 | 58 | 38 | 0 | 0 | 0 | 0 | 0 | 0 | 1801 | 0 | 0 |
| SRR7412279 | 0 | 0 | 33 | 90 | 57 | 113 | 151 | 75 | 0 | 0 | 41 | 0 | 0 | 0 | 226 | 19789 | 60565 | 0 | 0 | 830 | 717 | 0 | 0 | 123 | 25 | 384 | 1177 | 842 | 0 | 65 | 0 | 38 | 0 | 0 | 2075 | 0 | 0 |
| SRR7412280 | 0 | 27 | 12 | 24 | 18 | 25 | 0 | 49 | 0 | 0 | 18 | 0 | 0 | 0 | 270 | 20328 | 58770 | 0 | 0 | 590 | 442 | 0 | 0 | 82 | 0 | 37 | 80 | 57 | 0 | 27 | 0 | 0 | 0 | 0 | 2918 | 0 | 0 |
| SRR7412281 | 20 | 0 | 0 | 20 | 16 | 41 | 82 | 0 | 0 | 0 | 0 | 0 | 0 | 0 | 778 | 12687 | 50274 | 41 | 0 | 1310 | 1146 | 0 | 0 | 13 | 49 | 17 | 61 | 83 | 0 | 98 | 0 | 0 | 0 | 0 | 2141 | 0 | 0 |
| SRR7412282 | 11 | 0 | 46 | 137 | 274 | 0 | 0 | 0 | 0 | 0 | 34 | 0 | 0 | 0 | 71 | 14734 | 10145 | 0 | 0 | 531 | 212 | 0 | 0 | 103 | 11 | 285 | 912 | 924 | 0 | 11 | 0 | 0 | 0 | 0 | 2778 | 0 | 0 |
| SRR7412283 | 0 | 0 | 70 | 21 | 140 | 33 | 67 | 33 | 0 | 0 | 44 | 0 | 0 | 0 | 167 | 11745 | 32580 | 0 | 0 | 1270 | 501 | 0 | 0 | 14 | 18 | 27 | 69 | 42 | 0 | 44 | 0 | 0 | 0 | 0 | 2333 | 0 | 0 |
| SRR7412284 | 45 | 0 | 0 | 13 | 11 | 0 | 86 | 0 | 0 | 0 | 0 | 0 | 0 | 0 | 173 | 5104 | 9641 | 0 | 0 | 864 | 605 | 0 | 0 | 45 | 36 | 54 | 248 | 96 | 0 | 45 | 0 | 0 | 0 | 0 | 780 | 0 | 0 |
| SRR7412285 | 47 | 41 | 123 | 450 | 210 | 0 | 93 | 19 | 0 | 0 | 35 | 0 | 0 | 0 | 837 | 32070 | 209159 | 0 | 0 | 1749 | 1730 | 0 | 0 | 245 | 0 | 111 | 765 | 631 | 0 | 41 | 0 | 0 | 0 | 0 | 1752 | 0 | 0 |
| SRR7412286 | 0 | 0 | 16 | 31 | 12 | 79 | 79 | 314 | 0 | 0 | 0 | 0 | 0 | 0 | 236 | 19104 | 31776 | 0 | 0 | 943 | 1022 | 0 | 0 | 15 | 0 | 64 | 55 | 47 | 0 | 24 | 0 | 0 | 0 | 0 | 1135 | 0 | 0 |
| SRR7412287 | 13 | 0 | 0 | 74 | 81 | 55 | 164 | 0 | 55 | 0 | 74 | 0 | 0 | 0 | 164 | 16878 | 19188 | 55 | 0 | 1747 | 710 | 0 | 0 | 34 | 0 | 128 | 571 | 383 | 0 | 114 | 0 | 109 | 0 | 0 | 2311 | 0 | 0 |
| SRR7412288 | 30 | 45 | 74 | 545 | 94 | 51 | 38 | 77 | 26 | 0 | 0 | 0 | 0 | 0 | 77 | 14620 | 15365 | 0 | 0 | 230 | 294 | 0 | 0 | 183 | 109 | 1230 | 3967 | 2098 | 0 | 15 | 0 | 13 | 0 | 0 | 2142 | 0 | 0 |
| SRR7412289 | 10 | 41 | 31 | 19 | 102 | 0 | 119 | 59 | 0 | 0 | 0 | 0 | 0 | 0 | 415 | 7201 | 16113 | 0 | 0 | 1661 | 356 | 0 | 0 | 92 | 61 | 40 | 125 | 47 | 0 | 92 | 0 | 0 | 0 | 0 | 540 | 0 | 10 |
| SRR7412290 | 14 | 28 | 28 | 42 | 83 | 0 | 113 | 113 | 0 | 0 | 0 | 0 | 0 | 0 | 113 | 19041 | 6720 | 0 | 0 | 2155 | 113 | 0 | 0 | 167 | 56 | 389 | 847 | 930 | 0 | 28 | 0 | 0 | 0 | 0 | 2688 | 0 | 0 |
| SRR7412291 | 16 | 0 | 16 | 104 | 40 | 135 | 135 | 135 | 67 | 0 | 0 | 0 | 0 | 0 | 944 | 23549 | 115926 | 0 | 0 | 2428 | 2428 | 0 | 0 | 40 | 0 | 0 | 0 | 0 | 0 | 80 | 0 | 0 | 0 | 0 | 1093 | 0 | 0 |
| SRR7412292 | 37 | 37 | 46 | 165 | 101 | 0 | 54 | 108 | 0 | 0 | 37 | 0 | 0 | 0 | 54 | 11023 | 6484 | 0 | 0 | 1720 | 108 | 0 | 0 | 192 | 0 | 393 | 878 | 430 | 0 | 18 | 0 | 0 | 0 | 0 | 5317 | 0 | 0 |
| SRR7412293 | 0 | 18 | 92 | 14 | 147 | 26 | 77 | 52 | 0 | 0 | 37 | 0 | 0 | 0 | 696 | 25863 | 87893 | 26 | 0 | 206 | 412 | 0 | 0 | 110 | 83 | 30 | 78 | 56 | 0 | 74 | 0 | 0 | 0 | 0 | 4225 | 0 | 0 |
| SRR7412294 | 21 | 43 | 100 | 285 | 86 | 117 | 821 | 234 | 0 | 0 | 21 | 0 | 0 | 0 | 59 | 21683 | 23706 | 0 | 0 | 2872 | 352 | 0 | 0 | 121 | 21 | 178 | 456 | 456 | 0 | 43 | 0 | 0 | 0 | 0 | 24188 | 0 | 0 |
| SRR7412295 | 18 | 18 | 13 | 50 | 30 | 29 | 29 | 29 | 0 | 0 | 52 | 0 | 0 | 0 | 143 | 14540 | 48435 | 0 | 0 | 1402 | 601 | 0 | 0 | 27 | 44 | 27 | 95 | 76 | 0 | 52 | 0 | 0 | 0 | 0 | 3982 | 0 | 0 |
| SRR7412296 | 12 | 49 | 61 | 73 | 24 | 105 | 0 | 209 | 0 | 0 | 12 | 0 | 0 | 0 | 784 | 14712 | 56983 | 0 | 0 | 1358 | 1358 | 0 | 0 | 73 | 37 | 475 | 2106 | 1083 | 0 | 37 | 0 | 0 | 0 | 0 | 1814 | 0 | 0 |
| SRR7412297 | 22 | 22 | 11 | 65 | 65 | 31 | 31 | 246 | 0 | 0 | 44 | 11 | 0 | 0 | 3934 | 14768 | 195770 | 0 | 0 | 2582 | 3043 | 0 | 0 | 27 | 33 | 42 | 289 | 103 | 0 | 16 | 0 | 0 | 0 | 0 | 1299 | 0 | 0 |
| SRR7412298 | 25 | 50 | 75 | 26 | 10 | 0 | 0 | 97 | 0 | 0 | 0 | 0 | 0 | 0 | 315 | 17172 | 40642 | 0 | 0 | 387 | 460 | 0 | 0 | 18 | 17 | 30 | 103 | 47 | 0 | 0 | 0 | 0 | 0 | 0 | 1381 | 0 | 0 |
| SRR7412299 | 0 | 18 | 53 | 125 | 107 | 0 | 92 | 46 | 46 | 0 | 0 | 0 | 0 | 0 | 322 | 11803 | 23751 | 0 | 0 | 1610 | 506 | 0 | 0 | 196 | 36 | 1265 | 4046 | 3208 | 0 | 0 | 0 | 0 | 0 | 0 | 507 | 0 | 0 |
| SRR7412300 | 45 | 0 | 45 | 189 | 123 | 70 | 56 | 139 | 0 | 0 | 22 | 0 | 0 | 0 | 920 | 10782 | 52766 | 0 | 0 | 1004 | 962 | 0 | 0 | 301 | 401 | 2618 | 9949 | 4011 | 0 | 33 | 0 | 0 | 0 | 0 | 1743 | 0 | 0 |
| SRR7412301 | 23 | 45 | 91 | 238 | 68 | 0 | 0 | 231 | 0 | 0 | 0 | 0 | 0 | 0 | 385 | 10778 | 20517 | 0 | 0 | 3461 | 1384 | 0 | 0 | 170 | 23 | 204 | 828 | 408 | 0 | 79 | 0 | 0 | 0 | 0 | 1169 | 0 | 0 |
| SRR7412302 | 13 | 21 | 68 | 295 | 73 | 14 | 42 | 14 | 0 | 0 | 0 | 0 | 0 | 0 | 209 | 23530 | 63903 | 0 | 0 | 1043 | 793 | 0 | 0 | 51 | 0 | 419 | 2392 | 1425 | 0 | 30 | 0 | 0 | 0 | 0 | 2024 | 0 | 0 |
| SRR7412303 | 18 | 18 | 13 | 13 | 86 | 98 | 25 | 25 | 0 | 0 | 61 | 0 | 0 | 0 | 615 | 19011 | 63613 | 25 | 0 | 2115 | 762 | 0 | 0 | 11 | 31 | 21 | 79 | 88 | 0 | 10 | 0 | 0 | 0 | 0 | 2515 | 0 | 0 |
| SRR7412304 | 0 | 17 | 42 | 18 | 67 | 0 | 100 | 67 | 67 | 0 | 17 | 0 | 0 | 0 | 268 | 13439 | 35167 | 0 | 0 | 1070 | 970 | 0 | 0 | 25 | 0 | 34 | 251 | 72 | 0 | 42 | 0 | 0 | 0 | 0 | 1633 | 0 | 0 |
| SRR7412305 | 34 | 11 | 69 | 15 | 12 | 0 | 18 | 18 | 0 | 0 | 0 | 0 | 0 | 0 | 319 | 17039 | 84479 | 0 | 0 | 938 | 584 | 0 | 0 | 21 | 46 | 25 | 78 | 78 | 0 | 46 | 0 | 18 | 0 | 0 | 2935 | 0 | 0 |
| SRR7412306 | 0 | 20 | 81 | 173 | 183 | 22 | 89 | 89 | 22 | 0 | 71 | 0 | 0 | 0 | 244 | 17798 | 7138 | 0 | 0 | 1086 | 310 | 0 | 0 | 31 | 0 | 0 | 20 | 0 | 0 | 71 | 0 | 0 | 0 | 0 | 1972 | 0 | 0 |
| SRR7412307 | 49 | 0 | 115 | 115 | 115 | 0 | 141 | 0 | 0 | 0 | 0 | 0 | 0 | 0 | 660 | 18920 | 78725 | 0 | 0 | 1179 | 802 | 0 | 0 | 230 | 82 | 853 | 1951 | 1164 | 0 | 0 | 0 | 0 | 0 | 0 | 1720 | 0 | 0 |
| SRR7412308 | 19 | 38 | 29 | 181 | 57 | 0 | 73 | 73 | 0 | 0 | 19 | 0 | 0 | 0 | 219 | 15630 | 30007 | 37 | 0 | 1424 | 1424 | 0 | 0 | 191 | 0 | 296 | 1098 | 1537 | 0 | 19 | 0 | 0 | 0 | 0 | 1649 | 0 | 0 |
| SRR7412309 | 25 | 33 | 75 | 27 | 12 | 0 | 0 | 0 | 0 | 0 | 25 | 0 | 0 | 0 | 400 | 13395 | 47492 | 0 | 0 | 1535 | 567 | 0 | 0 | 12 | 25 | 35 | 107 | 90 | 0 | 25 | 0 | 0 | 0 | 0 | 2816 | 0 | 0 |
| SRR7412310 | 19 | 23 | 38 | 222 | 76 | 0 | 39 | 19 | 39 | 0 | 0 | 0 | 0 | 0 | 289 | 17929 | 32751 | 0 | 0 | 1312 | 444 | 0 | 0 | 130 | 0 | 164 | 547 | 352 | 0 | 0 | 0 | 0 | 0 | 0 | 1885 | 0 | 0 |
| SRR7412311 | 41 | 0 | 58 | 181 | 123 | 76 | 38 | 38 | 0 | 0 | 0 | 0 | 0 | 0 | 0 | 13148 | 8024 | 0 | 0 | 1142 | 114 | 0 | 0 | 82 | 0 | 395 | 1151 | 904 | 0 | 33 | 0 | 0 | 0 | 0 | 3190 | 0 | 0 |
| SRR7412312 | 25 | 25 | 25 | 204 | 119 | 24 | 95 | 24 | 24 | 0 | 0 | 0 | 0 | 0 | 284 | 16776 | 29837 | 24 | 0 | 189 | 189 | 0 | 0 | 93 | 76 | 687 | 1808 | 1239 | 0 | 0 | 0 | 0 | 0 | 0 | 1153 | 0 | 0 |
| SRR7412313 | 0 | 0 | 19 | 76 | 105 | 33 | 131 | 99 | 164 | 0 | 19 | 0 | 0 | 0 | 394 | 19841 | 56116 | 0 | 0 | 1282 | 953 | 0 | 0 | 57 | 0 | 506 | 1920 | 955 | 0 | 115 | 0 | 0 | 0 | 0 | 2405 | 0 | 0 |
| SRR7412314 | 22 | 15 | 22 | 74 | 30 | 81 | 243 | 0 | 0 | 0 | 82 | 15 | 0 | 0 | 892 | 23124 | 32587 | 41 | 0 | 1419 | 1054 | 0 | 0 | 74 | 45 | 549 | 1596 | 950 | 0 | 97 | 0 | 0 | 0 | 0 | 2800 | 0 | 0 |
| SRR7412315 | 24 | 12 | 24 | 94 | 59 | 37 | 0 | 74 | 0 | 0 | 24 | 0 | 0 | 0 | 956 | 24956 | 95934 | 37 | 0 | 1508 | 1324 | 0 | 0 | 59 | 94 | 764 | 2516 | 1575 | 0 | 47 | 0 | 0 | 0 | 0 | 1427 | 0 | 0 |
| SRR7412316 | 0 | 18 | 109 | 41 | 29 | 80 | 80 | 0 | 27 | 0 | 55 | 0 | 0 | 0 | 664 | 22933 | 112197 | 0 | 0 | 930 | 1036 | 0 | 0 | 16 | 36 | 24 | 96 | 98 | 0 | 109 | 0 | 0 | 0 | 0 | 4020 | 0 | 0 |
| SRR7412317 | 12 | 17 | 40 | 121 | 75 | 97 | 129 | 64 | 0 | 0 | 0 | 0 | 0 | 0 | 1514 | 29220 | 174360 | 0 | 0 | 4542 | 3994 | 0 | 0 | 75 | 52 | 35 | 95 | 43 | 0 | 29 | 0 | 0 | 0 | 0 | 1735 | 0 | 0 |
| SRR7412318 | 0 | 25 | 38 | 115 | 140 | 76 | 76 | 0 | 0 | 0 | 0 | 0 | 0 | 0 | 76 | 11156 | 7387 | 0 | 0 | 1438 | 454 | 0 | 0 | 153 | 38 | 318 | 1209 | 713 | 0 | 38 | 0 | 0 | 0 | 0 | 1658 | 0 | 0 |
| SRR7412319 | 43 | 57 | 57 | 17 | 78 | 30 | 30 | 0 | 0 | 0 | 21 | 0 | 0 | 0 | 182 | 9457 | 8603 | 0 | 0 | 3146 | 272 | 0 | 0 | 11 | 21 | 32 | 92 | 70 | 0 | 57 | 0 | 0 | 0 | 0 | 3349 | 0 | 0 |
| SRR7412320 | 30 | 30 | 70 | 714 | 171 | 17 | 0 | 17 | 0 | 0 | 10 | 0 | 0 | 0 | 109 | 15264 | 38182 | 0 | 0 | 227 | 302 | 0 | 0 | 161 | 55 | 990 | 3754 | 2030 | 0 | 35 | 0 | 0 | 0 | 0 | 2849 | 0 | 0 |
| SRR7412321 | 19 | 19 | 47 | 123 | 142 | 23 | 46 | 23 | 46 | 0 | 19 | 0 | 0 | 0 | 228 | 11333 | 16170 | 23 | 0 | 615 | 91 | 0 | 0 | 66 | 0 | 0 | 0 | 0 | 0 | 76 | 0 | 0 | 0 | 0 | 1612 | 0 | 0 |
| SRR7412322 | 11 | 11 | 52 | 212 | 143 | 41 | 81 | 20 | 20 | 0 | 52 | 0 | 0 | 0 | 81 | 10245 | 34873 | 0 | 0 | 1765 | 690 | 0 | 0 | 155 | 0 | 298 | 3254 | 1999 | 0 | 109 | 0 | 20 | 0 | 0 | 3251 | 0 | 0 |
| SRR7412323 | 16 | 0 | 107 | 39 | 20 | 20 | 0 | 10 | 0 | 0 | 74 | 0 | 0 | 0 | 143 | 14549 | 64082 | 0 | 0 | 573 | 348 | 0 | 0 | 139 | 82 | 49 | 145 | 134 | 0 | 10 | 0 | 0 | 0 | 0 | 3736 | 0 | 0 |
| SRR7412324 | 36 | 55 | 45 | 400 | 237 | 20 | 0 | 0 | 0 | 0 | 36 | 0 | 0 | 0 | 0 | 7162 | 12028 | 0 | 0 | 642 | 181 | 0 | 0 | 127 | 27 | 418 | 1474 | 819 | 0 | 55 | 0 | 0 | 0 | 0 | 2387 | 0 | 0 |
| SRR7412325 | 0 | 43 | 98 | 314 | 228 | 0 | 31 | 62 | 0 | 0 | 0 | 11 | 0 | 0 | 342 | 8085 | 26119 | 0 | 0 | 934 | 124 | 0 | 0 | 238 | 65 | 683 | 1832 | 1583 | 0 | 54 | 0 | 0 | 0 | 0 | 977 | 0 | 0 |
| SRR7412326 | 0 | 20 | 56 | 16 | 105 | 33 | 66 | 44 | 0 | 0 | 0 | 0 | 0 | 0 | 685 | 25452 | 64696 | 11 | 0 | 1469 | 1005 | 0 | 0 | 117 | 105 | 37 | 93 | 37 | 0 | 52 | 0 | 0 | 0 | 0 | 1592 | 0 | 0 |
| SRR7412327 | 13 | 0 | 26 | 380 | 138 | 75 | 201 | 100 | 0 | 0 | 13 | 0 | 0 | 0 | 627 | 20497 | 56464 | 0 | 0 | 979 | 828 | 0 | 0 | 131 | 33 | 517 | 1317 | 439 | 0 | 59 | 0 | 0 | 0 | 0 | 1694 | 0 | 0 |

### Supp Table S8.3.Columns_BX_to_DH

| | | | | | | | | | | | | | | | | | | | | | | | | | | | | | | | | | | | | | |
|---|---|---|---|---|---|---|---|---|---|---|---|---|---|---|---|---|---|---|---|---|---|---|---|---|---|---|---|---|---|---|---|---|---|---|---|---|---|
| SRR7412328 | 23 | 0 | 46 | 179 | 242 | 52 | 79 | 210 | 52 | 0 | 81 | 0 | 0 | 0 | 1964 | 23883 | 265899 | 0 | 0 | 3746 | 1257 | 0 | 0 | 87 | 23 | 156 | 548 | 502 | 0 | 289 | 0 | 0 | 0 | 0 | 4053 | 0 | 12 |
| SRR7412329 | 19 | 0 | 47 | 21 | 44 | 0 | 86 | 173 | 86 | 0 | 18 | 0 | 0 | 0 | 1122 | 29928 | 275875 | 0 | 0 | 2200 | 2028 | 0 | 0 | 20 | 0 | 23 | 48 | 53 | 0 | 20 | 0 | 43 | 0 | 0 | 22734 | 0 | 0 |
| SRR7412330 | 0 | 16 | 88 | 239 | 271 | 250 | 584 | 751 | 0 | 0 | 80 | 0 | 0 | 0 | 834 | 21561 | 33290 | 0 | 83 | 2168 | 1501 | 0 | 0 | 72 | 0 | 64 | 199 | 287 | 0 | 152 | 0 | 0 | 0 | 0 | 11847 | 0 | 0 |
| SRR7412331 | 32 | 0 | 32 | 287 | 207 | 42 | 21 | 126 | 0 | 0 | 72 | 24 | 0 | 0 | 1033 | 22119 | 183583 | 0 | 0 | 1602 | 2087 | 0 | 0 | 0 | 0 | 0 | 0 | 0 | 0 | 96 | 0 | 0 | 0 | 0 | 3871 | 0 | 0 |
| SRR7412332 | 28 | 19 | 76 | 256 | 114 | 0 | 69 | 0 | 69 | 0 | 66 | 0 | 0 | 0 | 1583 | 31325 | 121857 | 0 | 0 | 1755 | 2718 | 0 | 0 | 57 | 28 | 407 | 918 | 1107 | 0 | 104 | 0 | 0 | 0 | 0 | 3371 | 0 | 0 |
| SRR7412333 | 0 | 0 | 19 | 117 | 132 | 225 | 75 | 200 | 0 | 0 | 44 | 0 | 0 | 0 | 626 | 23668 | 117935 | 50 | 25 | 2678 | 1827 | 0 | 0 | 83 | 29 | 239 | 804 | 468 | 0 | 29 | 0 | 0 | 0 | 0 | 3129 | 0 | 0 |
| SRR7412334 | 0 | 14 | 56 | 77 | 126 | 58 | 204 | 175 | 87 | 0 | 77 | 0 | 0 | 0 | 2763 | 22748 | 92932 | 29 | 29 | 1687 | 2094 | 0 | 0 | 28 | 49 | 328 | 2771 | 1222 | 0 | 91 | 0 | 0 | 0 | 0 | 1321 | 0 | 0 |
| SRR7412335 | 0 | 15 | 15 | 17 | 11 | 120 | 80 | 160 | 0 | 0 | 15 | 0 | 0 | 0 | 1604 | 26688 | 232328 | 40 | 0 | 1684 | 2726 | 0 | 0 | 60 | 30 | 33 | 58 | 58 | 0 | 15 | 0 | 0 | 0 | 0 | 680 | 0 | 0 |
| SRR7412336 | 19 | 0 | 0 | 169 | 225 | 25 | 127 | 127 | 51 | 0 | 38 | 0 | 0 | 0 | 1628 | 21105 | 71712 | 0 | 0 | 890 | 305 | 0 | 0 | 75 | 47 | 178 | 1107 | 900 | 0 | 169 | 0 | 0 | 0 | 0 | 3565 | 0 | 0 |
| SRR7412337 | 40 | 30 | 109 | 395 | 178 | 97 | 32 | 129 | 0 | 0 | 59 | 0 | 0 | 0 | 744 | 24645 | 32268 | 0 | 0 | 1327 | 162 | 0 | 0 | 612 | 158 | 563 | 2262 | 1896 | 0 | 49 | 0 | 0 | 0 | 0 | 2590 | 0 | 0 |
| SRR7412338 | 75 | 75 | 75 | 481 | 166 | 0 | 0 | 63 | 0 | 0 | 0 | 0 | 0 | 0 | 693 | 26056 | 139802 | 0 | 0 | 1512 | 2394 | 0 | 0 | 120 | 15 | 436 | 1520 | 1023 | 0 | 0 | 0 | 0 | 0 | 0 | 1438 | 0 | 0 |
| SRR7412339 | 13 | 0 | 13 | 11 | 108 | 34 | 34 | 34 | 0 | 0 | 135 | 0 | 0 | 0 | 203 | 15232 | 28465 | 0 | 0 | 1014 | 237 | 0 | 0 | 121 | 108 | 31 | 95 | 72 | 0 | 148 | 0 | 0 | 0 | 0 | 4094 | 0 | 0 |
| SRR7412340 | 48 | 0 | 84 | 180 | 204 | 57 | 0 | 0 | 0 | 0 | 301 | 0 | 0 | 0 | 509 | 17630 | 18079 | 0 | 0 | 848 | 396 | 0 | 0 | 132 | 60 | 361 | 974 | 1119 | 0 | 241 | 0 | 0 | 0 | 0 | 5827 | 0 | 12 |
| SRR7412341 | 37 | 0 | 19 | 141 | 141 | 55 | 110 | 0 | 55 | 0 | 262 | 0 | 0 | 0 | 166 | 7937 | 7513 | 0 | 0 | 1104 | 386 | 0 | 0 | 47 | 47 | 375 | 1124 | 1171 | 0 | 337 | 0 | 0 | 0 | 0 | 6243 | 0 | 0 |
| SRR7412342 | 0 | 15 | 15 | 60 | 11 | 0 | 139 | 70 | 0 | 0 | 0 | 0 | 0 | 0 | 975 | 28979 | 163512 | 0 | 0 | 4110 | 2438 | 0 | 0 | 53 | 0 | 35 | 86 | 71 | 0 | 0 | 0 | 0 | 0 | 0 | 1246 | 0 | 0 |
| SRR7412343 | 17 | 25 | 17 | 12 | 75 | 0 | 0 | 50 | 0 | 0 | 17 | 0 | 0 | 0 | 600 | 15737 | 27202 | 0 | 0 | 1600 | 650 | 0 | 0 | 17 | 17 | 26 | 159 | 52 | 0 | 42 | 0 | 0 | 0 | 0 | 4201 | 0 | 0 |
| SRR7412344 | 15 | 15 | 15 | 11 | 29 | 0 | 0 | 0 | 0 | 0 | 10 | 0 | 0 | 0 | 235 | 19541 | 36650 | 0 | 0 | 739 | 168 | 0 | 0 | 15 | 0 | 19 | 66 | 48 | 0 | 26 | 0 | 0 | 0 | 0 | 4683 | 0 | 0 |
| SRR7412345 | 39 | 98 | 20 | 17 | 59 | 78 | 78 | 156 | 0 | 0 | 0 | 0 | 0 | 0 | 156 | 17528 | 14542 | 0 | 0 | 2418 | 156 | 0 | 0 | 39 | 59 | 32 | 152 | 122 | 0 | 20 | 0 | 78 | 0 | 0 | 3246 | 0 | 0 |
| SRR7466734 | 23 | 68 | 20 | 16 | 41 | 133 | 95 | 304 | 38 | 0 | 74 | 0 | 20 | 0 | 29682 | 13595 | 21424 | 247 | 76 | 4015 | 875 | 0 | 0 | 10 | 31 | 29 | 138 | 66 | 0 | 62 | 0 | 0 | 0 | 0 | 7619 | 0 | 0 |
| SRR7466735 | 24 | 61 | 28 | 14 | 45 | 137 | 234 | 449 | 78 | 0 | 14 | 0 | 20 | 0 | 30115 | 13416 | 21086 | 312 | 78 | 4140 | 566 | 0 | 0 | 12 | 11 | 27 | 169 | 20 | 0 | 74 | 0 | 20 | 0 | 0 | 8026 | 0 | 0 |
| SRR7466736 | 81 | 194 | 0 | 47 | 121 | 444 | 1126 | 1195 | 222 | 0 | 179 | 0 | 0 | 0 | 65827 | 31710 | 47319 | 3123 | 34 | 4693 | 3977 | 0 | 0 | 100 | 37 | 96 | 433 | 19 | 0 | 488 | 0 | 17 | 0 | 0 | 5938 | 0 | 0 |
| SRR7466737 | 82 | 201 | 21 | 78 | 149 | 259 | 1035 | 1017 | 207 | 0 | 183 | 0 | 0 | 0 | 66105 | 30380 | 47024 | 3069 | 121 | 4810 | 3793 | 0 | 0 | 75 | 36 | 64 | 427 | 15 | 0 | 593 | 0 | 0 | 0 | 0 | 5583 | 0 | 0 |
| SRR7466738 | 24 | 70 | 14 | 15 | 42 | 459 | 968 | 1116 | 180 | 0 | 49 | 0 | 10 | 0 | 66576 | 30429 | 49993 | 3380 | 164 | 4413 | 3790 | 0 | 0 | 30 | 11 | 24 | 127 | 17 | 0 | 172 | 0 | 0 | 0 | 0 | 5508 | 0 | 0 |
| SRR7466739 | 85 | 200 | 0 | 60 | 114 | 319 | 1025 | 1042 | 202 | 0 | 195 | 0 | 0 | 0 | 66481 | 29136 | 50657 | 3259 | 134 | 4503 | 4217 | 0 | 0 | 82 | 42 | 75 | 443 | 19 | 0 | 552 | 0 | 0 | 0 | 0 | 6091 | 0 | 0 |
| SRR7466740 | 112 | 151 | 0 | 50 | 115 | 262 | 1070 | 1312 | 161 | 0 | 105 | 0 | 0 | 0 | 59560 | 28667 | 38114 | 3956 | 61 | 6438 | 2260 | 0 | 0 | 28 | 31 | 104 | 381 | 15 | 0 | 423 | 0 | 40 | 0 | 0 | 7818 | 0 | 0 |
| SRR7466741 | 117 | 155 | 0 | 51 | 124 | 205 | 1191 | 903 | 82 | 0 | 95 | 0 | 0 | 0 | 60937 | 28320 | 39257 | 3819 | 185 | 6693 | 2115 | 0 | 0 | 26 | 28 | 106 | 339 | 0 | 0 | 468 | 0 | 0 | 0 | 0 | 7182 | 0 | 0 |
| SRR7466742 | 24 | 40 | 0 | 14 | 30 | 148 | 853 | 837 | 213 | 0 | 20 | 0 | 0 | 0 | 50009 | 26700 | 38512 | 3004 | 164 | 5580 | 1625 | 0 | 0 | 37 | 36 | 29 | 94 | 20 | 0 | 117 | 0 | 0 | 0 | 0 | 7522 | 0 | 0 |
| SRR7466743 | 86 | 203 | 11 | 46 | 121 | 202 | 947 | 1008 | 242 | 0 | 74 | 0 | 0 | 0 | 61121 | 27244 | 36885 | 4111 | 242 | 6811 | 2136 | 0 | 0 | 45 | 24 | 105 | 393 | 0 | 0 | 428 | 0 | 20 | 0 | 0 | 7833 | 0 | 0 |
| SRR7466744 | 30 | 71 | 21 | 19 | 45 | 200 | 709 | 600 | 55 | 0 | 28 | 0 | 0 | 0 | 36784 | 24637 | 27721 | 2018 | 73 | 4873 | 2036 | 0 | 0 | 13 | 42 | 33 | 152 | 37 | 0 | 117 | 0 | 0 | 0 | 0 | 8087 | 0 | 0 |
| SRR7466745 | 212 | 424 | 16 | 94 | 207 | 308 | 726 | 923 | 44 | 0 | 178 | 10 | 0 | 0 | 42544 | 24010 | 28565 | 2682 | 110 | 5277 | 2814 | 0 | 0 | 52 | 47 | 152 | 740 | 24 | 0 | 633 | 0 | 0 | 0 | 0 | 7339 | 0 | 0 |
| SRR7466746 | 204 | 439 | 0 | 90 | 333 | 96 | 366 | 501 | 289 | 0 | 144 | 0 | 0 | 0 | 32946 | 17804 | 25132 | 944 | 58 | 4486 | 1232 | 0 | 0 | 105 | 66 | 249 | 1159 | 39 | 0 | 697 | 0 | 0 | 0 | 0 | 9901 | 0 | 0 |
| SRR7466747 | 247 | 530 | 27 | 104 | 390 | 153 | 401 | 630 | 191 | 0 | 134 | 12 | 0 | 0 | 31928 | 18017 | 24249 | 725 | 115 | 3969 | 973 | 0 | 0 | 122 | 36 | 292 | 1179 | 65 | 0 | 709 | 0 | 0 | 0 | 0 | 8607 | 0 | 0 |
| SRR7466748 | 250 | 526 | 24 | 97 | 285 | 218 | 501 | 850 | 327 | 0 | 100 | 0 | 0 | 0 | 35517 | 18611 | 23993 | 1198 | 44 | 4707 | 1089 | 0 | 0 | 97 | 88 | 247 | 1181 | 35 | 0 | 723 | 0 | 44 | 0 | 0 | 9260 | 0 | 0 |
| SRR7466749 | 28 | 72 | 11 | 14 | 46 | 150 | 514 | 793 | 300 | 0 | 17 | 0 | 11 | 0 | 37488 | 18347 | 26126 | 1007 | 64 | 5058 | 1436 | 0 | 0 | 14 | 63 | 35 | 159 | 52 | 0 | 96 | 0 | 43 | 0 | 0 | 8564 | 0 | 0 |
| SRR7466750 | 81 | 114 | 12 | 27 | 89 | 204 | 1045 | 908 | 159 | 0 | 133 | 0 | 0 | 0 | 43310 | 22683 | 35084 | 4224 | 91 | 4565 | 1771 | 0 | 0 | 22 | 24 | 82 | 301 | 0 | 0 | 604 | 0 | 23 | 0 | 0 | 9657 | 0 | 0 |
| SRR7466751 | 24 | 34 | 10 | 15 | 31 | 290 | 974 | 1553 | 132 | 0 | 41 | 0 | 0 | 0 | 51994 | 22299 | 35041 | 6582 | 158 | 6371 | 1948 | 0 | 0 | 25 | 30 | 21 | 79 | 0 | 0 | 184 | 0 | 0 | 0 | 0 | 8974 | 0 | 0 |
| SRR7466752 | 27 | 51 | 0 | 11 | 27 | 340 | 1073 | 1125 | 79 | 0 | 39 | 0 | 0 | 0 | 52440 | 21493 | 34816 | 5129 | 131 | 6019 | 2329 | 0 | 0 | 18 | 20 | 29 | 86 | 13 | 0 | 169 | 0 | 26 | 0 | 0 | 9424 | 0 | 0 |
| SRR7466753 | 21 | 39 | 0 | 11 | 26 | 127 | 1322 | 1144 | 152 | 0 | 41 | 0 | 0 | 0 | 48388 | 22074 | 35103 | 5362 | 127 | 5616 | 1931 | 0 | 0 | 26 | 11 | 21 | 78 | 11 | 0 | 196 | 0 | 0 | 0 | 0 | 9960 | 0 | 0 |
| SRR7466754 | 30 | 79 | 25 | 22 | 41 | 220 | 593 | 879 | 110 | 0 | 27 | 0 | 0 | 0 | 53972 | 23685 | 36125 | 1406 | 242 | 5052 | 2965 | 0 | 0 | 34 | 10 | 22 | 125 | 43 | 0 | 112 | 0 | 0 | 0 | 0 | 5637 | 0 | 0 |
| SRR7466755 | 34 | 78 | 0 | 24 | 39 | 89 | 557 | 446 | 89 | 0 | 24 | 0 | 0 | 0 | 54061 | 23196 | 34338 | 1739 | 201 | 4615 | 2876 | 0 | 0 | 12 | 37 | 18 | 115 | 31 | 0 | 101 | 0 | 22 | 0 | 0 | 6383 | 0 | 0 |
| SRR7466756 | 193 | 478 | 18 | 71 | 193 | 166 | 349 | 631 | 100 | 0 | 113 | 0 | 0 | 0 | 41614 | 23810 | 35580 | 947 | 150 | 4186 | 1993 | 0 | 0 | 65 | 39 | 113 | 653 | 27 | 0 | 679 | 0 | 17 | 0 | 0 | 6653 | 0 | 0 |
| SRR7466757 | 198 | 497 | 15 | 119 | 216 | 308 | 274 | 496 | 34 | 0 | 140 | 0 | 0 | 0 | 42362 | 24810 | 36557 | 993 | 154 | 4005 | 1968 | 0 | 0 | 73 | 43 | 119 | 643 | 43 | 0 | 628 | 0 | 34 | 0 | 0 | 6321 | 0 | 0 |
| SRR7466758 | 25 | 70 | 13 | 18 | 28 | 29 | 29 | 431 | 115 | 13 | 21 | 13 | 13 | 0 | 24252 | 18836 | 25778 | 0 | 115 | 3415 | 1607 | 0 | 0 | 10 | 54 | 10 | 63 | 13 | 0 | 103 | 0 | 29 | 0 | 0 | 6664 | 0 | 0 |
| SRR7466759 | 34 | 71 | 27 | 23 | 42 | 222 | 74 | 334 | 74 | 14 | 19 | 0 | 14 | 0 | 32406 | 20076 | 24183 | 0 | 111 | 5488 | 2076 | 0 | 0 | 11 | 54 | 12 | 74 | 0 | 0 | 136 | 0 | 37 | 0 | 0 | 6184 | 0 | 0 |
| SRR7466760 | 263 | 482 | 0 | 77 | 244 | 28 | 111 | 277 | 83 | 0 | 180 | 0 | 19 | 0 | 27453 | 18101 | 23430 | 0 | 138 | 4566 | 1522 | 0 | 0 | 32 | 39 | 103 | 501 | 0 | 0 | 835 | 0 | 0 | 0 | 0 | 7047 | 0 | 0 |
| SRR7466761 | 217 | 486 | 0 | 112 | 289 | 57 | 143 | 286 | 57 | 20 | 151 | 0 | 0 | 0 | 26094 | 19433 | 23110 | 0 | 86 | 3867 | 1747 | 0 | 0 | 59 | 20 | 85 | 638 | 20 | 0 | 848 | 0 | 0 | 0 | 0 | 6720 | 0 | 26 |
| SRR7466762 | 26 | 60 | 16 | 13 | 41 | 98 | 235 | 411 | 176 | 0 | 15 | 0 | 12 | 0 | 32015 | 12914 | 22262 | 529 | 39 | 4151 | 783 | 0 | 0 | 74 | 41 | 35 | 160 | 49 | 0 | 87 | 0 | 20 | 0 | 0 | 7410 | 0 | 0 |
| SRR7466763 | 31 | 61 | 21 | 10 | 50 | 72 | 241 | 386 | 145 | 0 | 62 | 0 | 0 | 0 | 37200 | 12849 | 23419 | 313 | 72 | 4364 | 964 | 0 | 0 | 15 | 62 | 23 | 151 | 54 | 0 | 78 | 0 | 48 | 0 | 0 | 7981 | 0 | 12 |
| SRR7466764 | 36 | 85 | 13 | 22 | 36 | 363 | 768 | 960 | 192 | 0 | 31 | 0 | 13 | 0 | 44557 | 23413 | 30394 | 2518 | 192 | 5570 | 2881 | 0 | 0 | 14 | 13 | 28 | 135 | 18 | 0 | 118 | 0 | 0 | 0 | 0 | 7835 | 0 | 0 |
| SRR7466765 | 34 | 79 | 26 | 18 | 31 | 152 | 910 | 910 | 152 | 0 | 27 | 0 | 0 | 0 | 44713 | 24134 | 30145 | 2318 | 87 | 5806 | 2795 | 0 | 0 | 15 | 44 | 28 | 153 | 28 | 0 | 112 | 0 | 0 | 0 | 0 | 7225 | 0 | 0 |
| SRR7541164 | 20 | 13 | 0 | 0 | 31 | 143 | 373 | 1239 | 104 | 0 | 11 | 0 | 0 | 0 | 40816 | 4415 | 176148 | 0 | 13 | 10200 | 4474 | 0 | 0 | 1741 | 31 | 126 | 824 | 19 | 0 | 111 | 0 | 0 | 0 | 0 | 306 | 0 | 0 |
| SRR7541165 | 0 | 0 | 0 | 0 | 0 | 96 | 198 | 492 | 88 | 0 | 25 | 0 | 0 | 0 | 42193 | 3446 | 104918 | 0 | 0 | 1007 | 2727 | 0 | 0 | 48 | 58 | 136 | 352 | 0 | 0 | 848 | 0 | 10 | 0 | 0 | 468 | 0 | 0 |
| SRR7541166 | 17 | 14 | 0 | 0 | 12 | 142 | 403 | 1621 | 122 | 0 | 17 | 0 | 0 | 0 | 8270 | 4787 | 147102 | 0 | 0 | 25134 | 1841 | 0 | 0 | 43 | 67 | 72 | 187 | 0 | 0 | 268 | 0 | 0 | 0 | 0 | 1132 | 0 | 0 |
| SRR7541167 | 0 | 0 | 0 | 0 | 18 | 325 | 365 | 1624 | 92 | 0 | 26 | 0 | 0 | 0 | 8192 | 5617 | 180401 | 13 | 0 | 8661 | 4081 | 0 | 0 | 43 | 20 | 24 | 31 | 0 | 0 | 82 | 0 | 0 | 0 | 0 | 462 | 0 | 0 |
| SRR7541168 | 225 | 83 | 26 | 15 | 97 | 583 | 713 | 2476 | 155 | 11 | 18 | 0 | 0 | 0 | 45059 | 4811 | 134406 | 58 | 19 | 24264 | 6657 | 0 | 0 | 1044 | 160 | 255 | 336 | 0 | 0 | 330 | 0 | 0 | 0 | 0 | 1240 | 0 | 0 |
| SRR7541169 | 38 | 21 | 0 | 0 | 43 | 3254 | 2879 | 17459 | 1488 | 0 | 30 | 43 | 0 | 0 | 886 | 12829 | 88520 | 0 | 0 | 1599 | 3197 | 0 | 0 | 12 | 55 | 53 | 24 | 0 | 0 | 57 | 0 | 216 | 0 | 0 | 1895 | 0 | 0 |
| SRR7541170 | 11 | 0 | 0 | 0 | 13 | 361 | 642 | 2082 | 223 | 0 | 33 | 0 | 0 | 0 | 14299 | 5131 | 150021 | 0 | 0 | 3627 | 1033 | 0 | 0 | 45 | 22 | 21 | 36 | 0 | 0 | 153 | 0 | 29 | 0 | 0 | 1853 | 0 | 0 |
| SRR7541171 | 12 | 45 | 10 | 12 | 71 | 1478 | 2745 | 7450 | 1186 | 0 | 0 | 0 | 0 | 0 | 3397 | 20736 | 98944 | 0 | 41 | 14060 | 1695 | 0 | 0 | 71 | 73 | 11 | 24 | 0 | 0 | 11 | 0 | 33 | 0 | 0 | 1400 | 17 | 0 |
| SRR7541172 | 23 | 10 | 0 | 0 | 57 | 314 | 490 | 1636 | 90 | 0 | 12 | 0 | 0 | 0 | 4483 | 7299 | 74793 | 0 | 0 | 2330 | 1412 | 0 | 0 | 30 | 67 | 11 | 28 | 0 | 0 | 27 | 0 | 12 | 0 | 0 | 4484 | 0 | 0 |
| SRR7541173 | 43 | 0 | 0 | 0 | 164 | 488 | 711 | 3093 | 91 | 0 | 0 | 0 | 0 | 0 | 23353 | 6297 | 127621 | 21 | 11 | 1657 | 3416 | 0 | 0 | 680 | 42 | 51 | 97 | 0 | 0 | 186 | 0 | 0 | 0 | 0 | 1738 | 0 | 0 |
| SRR7541174 | 33 | 10 | 0 | 0 | 14 | 80 | 83 | 349 | 21 | 0 | 24 | 0 | 0 | 0 | 12756 | 3384 | 104316 | 0 | 0 | 1832 | 2414 | 0 | 0 | 152 | 15 | 28 | 72 | 10 | 0 | 119 | 0 | 12 | 0 | 0 | 787 | 0 | 0 |
| SRR7541175 | 33 | 10 | 0 | 0 | 28 | 223 | 619 | 1430 | 131 | 0 | 16 | 0 | 0 | 0 | 3867 | 9581 | 81343 | 11 | 0 | 3995 | 6955 | 0 | 0 | 40 | 13 | 42 | 17 | 0 | 0 | 20 | 0 | 56 | 0 | 0 | 3623 | 0 | 0 |
| SRR7541176 | 78 | 47 | 0 | 0 | 42 | 306 | 1615 | 1295 | 126 | 0 | 13 | 0 | 0 | 0 | 6848 | 7672 | 137629 | 0 | 0 | 2153 | 2599 | 0 | 0 | 148 | 59 | 114 | 44 | 0 | 0 | 114 | 0 | 0 | 0 | 0 | 1463 | 0 | 0 |
| SRR7541177 | 114 | 37 | 0 | 15 | 85 | 151 | 737 | 1974 | 211 | 0 | 0 | 0 | 0 | 0 | 16245 | 5357 | 107781 | 31 | 0 | 1869 | 9938 | 0 | 0 | 366 | 185 | 230 | 194 | 0 | 0 | 72 | 0 | 0 | 0 | 0 | 922 | 0 | 0 |
| SRR7541178 | 24 | 0 | 0 | 10 | 24 | 116 | 227 | 750 | 68 | 11 | 13 | 21 | 0 | 0 | 14825 | 6373 | 130600 | 0 | 0 | 3863 | 5198 | 0 | 0 | 18 | 11 | 27 | 73 | 0 | 0 | 29 | 0 | 0 | 0 | 0 | 2411 | 0 | 0 |
| SRR7541179 | 12 | 0 | 0 | 0 | 45 | 1187 | 1369 | 6061 | 258 | 17 | 0 | 0 | 0 | 0 | 7481 | 7780 | 128299 | 0 | 13 | 9275 | 2690 | 0 | 0 | 157 | 59 | 59 | 117 | 0 | 0 | 118 | 0 | 11 | 0 | 0 | 2124 | 0 | 0 |
| SRR7541180 | 119 | 46 | 18 | 0 | 23 | 33 | 181 | 315 | 36 | 11 | 28 | 12 | 0 | 0 | 44509 | 9555 | 265600 | 35 | 44 | 14984 | 18801 | 0 | 0 | 20 | 24 | 59 | 104 | 0 | 0 | 30 | 0 | 0 | 0 | 0 | 425 | 0 | 0 |
| SRR7541181 | 72 | 49 | 32 | 35 | 83 | 808 | 1156 | 4032 | 144 | 0 | 0 | 0 | 0 | 0 | 16739 | 12947 | 146075 | 0 | 39 | 1596 | 3113 | 0 | 0 | 334 | 35 | 48 | 106 | 0 | 0 | 105 | 0 | 0 | 0 | 0 | 2592 | 0 | 0 |
| SRR7541182 | 97 | 24 | 0 | 19 | 55 | 75 | 103 | 298 | 31 | 0 | 12 | 0 | 0 | 0 | 10800 | 5880 | 150959 | 20 | 11 | 5666 | 5971 | 0 | 0 | 234 | 25 | 46 | 117 | 0 | 0 | 219 | 0 | 0 | 0 | 0 | 1326 | 0 | 0 |
| SRR7541183 | 46 | 0 | 0 | 0 | 95 | 534 | 476 | 1568 | 92 | 0 | 0 | 0 | 0 | 0 | 8076 | 10496 | 221771 | 0 | 0 | 4914 | 4087 | 0 | 0 | 145 | 44 | 39 | 116 | 0 | 0 | 142 | 0 | 12 | 0 | 0 | 1692 | 0 | 0 |
| SRR7541184 | 89 | 11 | 0 | 0 | 11 | 116 | 312 | 990 | 40 | 0 | 0 | 0 | 0 | 0 | 79173 | 6958 | 257268 | 13 | 0 | 1659 | 8532 | 0 | 0 | 40 | 87 | 74 | 14 | 0 | 0 | 99 | 0 | 0 | 0 | 0 | 430 | 0 | 0 |
| SRR7541185 | 52 | 10 | 0 | 0 | 12 | 189 | 313 | 532 | 42 | 0 | 10 | 0 | 0 | 0 | 28415 | 8369 | 232467 | 15 | 0 | 1534 | 2947 | 0 | 0 | 11 | 78 | 157 | 14 | 0 | 0 | 15 | 0 | 18 | 0 | 0 | 1307 | 0 | 0 |
| SRR7541186 | 31 | 10 | 0 | 0 | 51 | 142 | 154 | 489 | 19 | 0 | 0 | 0 | 0 | 0 | 2223 | 4962 | 118834 | 0 | 0 | 7520 | 2558 | 0 | 0 | 16 | 0 | 17 | 43 | 0 | 0 | 20 | 0 | 0 | 0 | 0 | 1212 | 0 | 0 |
| SRR7541187 | 0 | 0 | 0 | 0 | 0 | 5693 | 5657 | 14894 | 935 | 0 | 0 | 30 | 0 | 0 | 2051 | 9130 | 62524 | 0 | 0 | 3364 | 1882 | 0 | 0 | 16 | 12 | 0 | 11 | 0 | 0 | 23 | 0 | 36 | 0 | 0 | 3128 | 0 | 0 |
| SRR7541188 | 15 | 0 | 0 | 0 | 0 | 1306 | 3205 | 5058 | 1091 | 0 | 0 | 0 | 0 | 0 | 7434 | 9184 | 89302 | 0 | 13 | 3129 | 1591 | 0 | 0 | 51 | 0 | 19 | 27 | 0 | 0 | 84 | 0 | 0 | 0 | 0 | 3272 | 0 | 0 |
| SRR7541189 | 54 | 24 | 10 | 20 | 33 | 239 | 784 | 3125 | 160 | 0 | 0 | 0 | 0 | 0 | 1658 | 5541 | 72736 | 0 | 0 | 1278 | 2123 | 0 | 0 | 139 | 146 | 221 | 329 | 0 | 0 | 105 | 0 | 0 | 0 | 0 | 4075 | 0 | 0 |
| SRR7541190 | 44 | 18 | 0 | 19 | 55 | 180 | 571 | 1705 | 99 | 24 | 12 | 24 | 0 | 0 | 18994 | 4774 | 62393 | 28 | 0 | 1530 | 5062 | 0 | 0 | 226 | 202 | 150 | 251 | 0 | 0 | 108 | 0 | 73 | 0 | 0 | 1518 | 0 | 0 |
| SRR7541191 | 14 | 0 | 0 | 0 | 50 | 357 | 1134 | 1629 | 138 | 0 | 17 | 0 | 0 | 0 | 48135 | 11318 | 285387 | 0 | 0 | 1700 | 4911 | 0 | 0 | 16 | 39 | 97 | 11 | 0 | 0 | 16 | 0 | 12 | 0 | 0 | 719 | 0 | 0 |
| SRR7541192 | 41 | 11 | 0 | 12 | 26 | 439 | 762 | 2192 | 168 | 13 | 0 | 0 | 0 | 0 | 13040 | 14630 | 130492 | 0 | 10 | 6155 | 3158 | 0 | 0 | 225 | 16 | 24 | 59 | 0 | 0 | 133 | 0 | 25 | 0 | 0 | 2640 | 0 | 0 |
| SRR7541193 | 0 | 0 | 0 | 0 | 17 | 1463 | 1234 | 10569 | 253 | 0 | 0 | 12 | 0 | 0 | 4464 | 9986 | 135351 | 0 | 0 | 1957 | 3163 | 0 | 0 | 75 | 0 | 0 | 0 | 0 | 0 | 157 | 0 | 59 | 0 | 0 | 2454 | 0 | 0 |
| SRR7547853 | 42 | 11 | 16 | 35 | 43 | 1083 | 931 | 4231 | 11169 | 459 | 130 | 34 | 0 | 0 | 52296 | 78319 | 246988 | 4456 | 1949 | 4630 | 3416 | 0 | 0 | 12 | 30 | 19 | 24 | 15 | 0 | 355 | 0 | 112 | 0 | 21 | 14631 | 396 | 0 |
| SRR7547854 | 42 | 14 | 0 | 33 | 185 | 2149 | 1771 | 9164 | 35400 | 1147 | 156 | 19 | 0 | 0 | 10924 | 82212 | 330776 | 899 | 951 | 3153 | 7209 | 0 | 0 | 32 | 76 | 0 | 0 | 0 | 0 | 434 | 0 | 179 | 0 | 12 | 24592 | 11038 | 0 |
| SRR7547855 | 16 | 0 | 0 | 28 | 791 | 1000 | 819 | 2753 | 6715 | 3423 | 262 | 12 | 0 | 0 | 60408 | 156145 | 257528 | 1346 | 1295 | 11991 | 1154 | 0 | 0 | 58 | 195 | 209 | 185 | 241 | 0 | 1276 | 0 | 67 | 0 | 0 | 10176 | 276 | 0 |
| SRR7547856 | 32 | 10 | 0 | 19 | 181 | 1056 | 1135 | 5187 | 12086 | 4862 | 976 | 24 | 0 | 0 | 159250 | 76558 | 356271 | 1033 | 1346 | 2018 | 3791 | 0 | 0 | 52 | 169 | 105 | 52 | 69 | 0 | 2941 | 0 | 69 | 0 | 15 | 9405 | 5674 | 0 |
| SRR7547857 | 28 | 10 | 11 | 15 | 413 | 111 | 60 | 295 | 529 | 5279 | 620 | 11 | 0 | 0 | 333780 | 59218 | 519293 | 407 | 1041 | 660 | 11203 | 0 | 0 | 55 | 162 | 136 | 82 | 62 | 0 | 1557 | 0 | 17 | 0 | 0 | 3663 | 1857 | 0 |
| SRR7547858 | 34 | 0 | 0 | 31 | 519 | 630 | 426 | 2175 | 4469 | 4579 | 587 | 12 | 0 | 0 | 96983 | 154883 | 451007 | 1610 | 840 | 1637 | 3247 | 0 | 0 | 72 | 148 | 114 | 98 | 124 | 0 | 1499 | 0 | 37 | 0 | 10 | 6505 | 3897 | 0 |
| SRR7547859 | 24 | 0 | 12 | 16 | 22 | 102 | 52 | 226 | 654 | 1306 | 40 | 23 | 0 | 0 | 247992 | 259613 | 289438 | 773 | 580 | 6724 | 1749 | 0 | 17 | 21 | 12 | 13 | 10 | 10 | 0 | 156 | 0 | 23 | 0 | 0 | 3792 | 1563 | 0 |
| SRR7547860 | 16 | 0 | 0 | 10 | 498 | 2409 | 2847 | 8491 | 24254 | 3355 | 463 | 39 | 0 | 0 | 107846 | 95985 | 302584 | 1119 | 1179 | 2116 | 3663 | 0 | 0 | 107 | 274 | 335 | 290 | 287 | 0 | 1382 | 0 | 91 | 0 | 11 | 10572 | 6047 | 0 |
| SRR7547861 | 36 | 11 | 15 | 10 | 169 | 875 | 940 | 3226 | 9710 | 2005 | 203 | 11 | 0 | 0 | 59450 | 185828 | 119669 | 672 | 1059 | 3239 | 2900 | 0 | 13 | 18 | 47 | 53 | 42 | 96 | 0 | 348 | 0 | 123 | 0 | 11 | 12047 | 2989 | 0 |
| SRR7547862 | 42 | 0 | 0 | 31 | 34 | 1566 | 545 | 3829 | 7394 | 133 | 117 | 14 | 0 | 0 | 62159 | 133369 | 431706 | 620 | 3126 | 12239 | 15440 | 0 | 0 | 16 | 44 | 32 | 24 | 24 | 0 | 288 | 0 | 18 | 0 | 0 | 3042 | 1756 | 0 |
| SRR7547863 | 13 | 18 | 11 | 15 | 24 | 1197 | 730 | 3479 | 13043 | 356 | 88 | 18 | 0 | 0 | 54146 | 79404 | 327938 | 1932 | 5000 | 7884 | 7476 | 0 | 0 | 13 | 28 | 33 | 32 | 36 | 0 | 252 | 0 | 76 | 0 | 12 | 5719 | 9841 | 0 |
| SRR7547864 | 23 | 0 | 11 | 17 | 496 | 7768 | 4234 | 16687 | 14108 | 1015 | 159 | 11 | 0 | 0 | 40052 | 185810 | 362458 | 1248 | 1156 | 2785 | 2202 | 0 | 0 | 73 | 66 | 40 | 23 | 15 | 0 | 975 | 0 | 0 | 0 | 14 | 19881 | 2075 | 0 |
| SRR7547865 | 34 | 0 | 0 | 19 | 52 | 534 | 343 | 1736 | 3822 | 362 | 66 | 0 | 0 | 0 | 141478 | 131928 | 473962 | 1121 | 515 | 2358 | 10526 | 0 | 0 | 15 | 29 | 18 | 19 | 16 | 0 | 183 | 0 | 62 | 0 | 17 | 1335 | 149 | 0 |
| SRR7547866 | 26 | 28 | 38 | 59 | 33 | 1491 | 1400 | 6853 | 20447 | 123 | 108 | 0 | 0 | 0 | 109445 | 192888 | 223566 | 2909 | 556 | 5414 | 4441 | 0 | 0 | 24 | 36 | 23 | 31 | 28 | 0 | 312 | 0 | 0 | 0 | 0 | 6686 | 2132 | 0 |
| SRR7547867 | 57 | 0 | 0 | 10 | 204 | 617 | 544 | 2758 | 5371 | 2042 | 313 | 16 | 0 | 0 | 146735 | 79097 | 405314 | 1275 | 1798 | 1519 | 7483 | 0 | 0 | 45 | 120 | 84 | 48 | 75 | 0 | 1027 | 0 | 69 | 0 | 0 | 7728 | 4352 | 0 |
| SRR7547868 | 43 | 13 | 16 | 46 | 784 | 1225 | 931 | 4570 | 11535 | 2727 | 345 | 16 | 0 | 0 | 66268 | 202764 | 163892 | 2144 | 991 | 3518 | 3332 | 0 | 0 | 72 | 317 | 272 | 164 | 289 | 0 | 930 | 0 | 80 | 0 | 0 | 2580 | 3011 | 0 |
| SRR7547869 | 42 | 13 | 0 | 20 | 267 | 1719 | 1117 | 6856 | 17964 | 7964 | 695 | 25 | 0 | 0 | 67697 | 119187 | 258707 | 828 | 845 | 6157 | 7694 | 0 | 0 | 187 | 656 | 470 | 339 | 233 | 0 | 1570 | 0 | 102 | 0 | 10 | 9195 | 5954 | 0 |
| SRR7547870 | 21 | 0 | 10 | 19 | 76 | 586 | 401 | 1833 | 4669 | 3516 | 149 | 18 | 0 | 0 | 190260 | 107022 | 312619 | 1724 | 991 | 1955 | 12146 | 10 | 22 | 14 | 53 | 41 | 29 | 32 | 0 | 307 | 0 | 128 | 0 | 13 | 11889 | 6648 | 0 |
| SRR7547871 | 58 | 0 | 22 | 33 | 449 | 1660 | 604 | 3364 | 7163 | 3546 | 703 | 11 | 0 | 0 | 137261 | 164720 | 455251 | 1082 | 1097 | 5686 | 5394 | 0 | 0 | 311 | 647 | 559 | 654 | 629 | 0 | 1294 | 0 | 34 | 0 | 0 | 6187 | 3313 | 0 |
| SRR7547872 | 18 | 0 | 0 | 17 | 38 | 378 | 248 | 1312 | 2402 | 870 | 63 | 23 | 0 | 0 | 184652 | 63777 | 553944 | 840 | 991 | 1324 | 5161 | 0 | 0 | 14 | 29 | 19 | 13 | 13 | 0 | 209 | 0 | 14 | 0 | 13 | 7847 | 2224 | 0 |
| SRR7547873 | 28 | 25 | 28 | 40 | 25 | 517 | 315 | 1926 | 5530 | 1440 | 206 | 12 | 0 | 0 | 142693 | 84223 | 317283 | 4665 | 1130 | 3251 | 5254 | 0 | 21 | 21 | 50 | 31 | 22 | 33 | 0 | 449 | 0 | 130 | 0 | 17 | 3581 | 15550 | 0 |
| SRR7547874 | 57 | 15 | 13 | 51 | 502 | 435 | 306 | 1537 | 3281 | 4541 | 462 | 17 | 0 | 0 | 52236 | 111501 | 493156 | 948 | 1916 | 2114 | 2136 | 0 | 0 | 53 | 176 | 105 | 86 | 127 | 0 | 1193 | 0 | 29 | 0 | 0 | 8260 | 4568 | 0 |
| SRR7547875 | 56 | 19 | 0 | 14 | 283 | 578 | 282 | 1704 | 3450 | 3345 | 668 | 14 | 0 | 0 | 228423 | 111215 | 491728 | 736 | 1978 | 1796 | 5265 | 0 | 0 | 35 | 135 | 0 | 0 | 0 | 0 | 1603 | 0 | 21 | 0 | 0 | 9908 | 1269 | 0 |
| SRR7547876 | 16 | 0 | 0 | 0 | 34 | 7756 | 6546 | 31403 | 40655 | 173 | 47 | 32 | 0 | 0 | 35047 | 155562 | 185819 | 2648 | 1173 | 4858 | 1295 | 0 | 0 | 17 | 15 | 38 | 13 | 0 | 0 | 392 | 0 | 11 | 0 | 24 | 20508 | 2540 | 0 |
| SRR7547877 | 71 | 17 | 15 | 59 | 257 | 1257 | 826 | 4561 | 7145 | 2066 | 382 | 16 | 0 | 0 | 20926 | 151723 | 411680 | 925 | 2182 | 2923 | 6502 | 0 | 0 | 67 | 183 | 113 | 82 | 115 | 0 | 728 | 0 | 417 | 0 | 15 | 8994 | 5130 | 0 |
| SRR7547878 | 26 | 0 | 0 | 12 | 42 | 4434 | 2782 | 10228 | 15616 | 919 | 69 | 15 | 0 | 0 | 170583 | 78478 | 249209 | 2480 | 954 | 1842 | 5304 | 0 | 0 | 16 | 20 | 25 | 17 | 14 | 0 | 347 | 0 | 26 | 0 | 0 | 40042 | 11724 | 0 |
| SRR7547879 | 33 | 0 | 0 | 0 | 27 | 4168 | 2042 | 9612 | 10157 | 1580 | 41 | 16 | 0 | 0 | 89413 | 103106 | 354508 | 1247 | 658 | 3722 | 11731 | 0 | 0 | 41 | 13 | 48 | 48 | 16 | 0 | 216 | 0 | 21 | 0 | 14 | 24211 | 1930 | 0 |
| SRR7547880 | 0 | 0 | 0 | 0 | 161 | 5315 | 4969 | 16371 | 25271 | 1050 | 115 | 26 | 0 | 0 | 59864 | 133392 | 195013 | 5360 | 986 | 12440 | 1119 | 0 | 0 | 55 | 53 | 43 | 31 | 24 | 0 | 912 | 0 | 14 | 0 | 18 | 30658 | 1579 | 0 |
| SRR7547881 | 21 | 0 | 0 | 0 | 423 | 3096 | 2060 | 7631 | 12691 | 1163 | 235 | 20 | 0 | 0 | 69881 | 109588 | 181553 | 6725 | 977 | 2796 | 1822 | 0 | 0 | 50 | 48 | 41 | 24 | 29 | 0 | 1172 | 0 | 19 | 0 | 16 | 52889 | 3699 | 0 |
| SRR7547882 | 25 | 19 | 19 | 14 | 216 | 606 | 893 | 2835 | 9561 | 875 | 105 | 20 | 0 | 0 | 1901 | 102941 | 55327 | 311 | 541 | 959 | 1942 | 0 | 18 | 27 | 105 | 99 | 151 | 203 | 0 | 313 | 0 | 25 | 0 | 20 | 43876 | 367 | 0 |
| SRR7547883 | 19 | 0 | 0 | 0 | 225 | 246 | 309 | 1850 | 2362 | 251 | 91 | 29 | 0 | 0 | 11685 | 105509 | 83708 | 366 | 457 | 639 | 1615 | 0 | 0 | 52 | 204 | 156 | 43 | 187 | 0 | 385 | 0 | 0 | 0 | 0 | 35320 | 217 | 0 |
| SRR7547884 | 20 | 23 | 10 | 19 | 104 | 439 | 263 | 1547 | 4454 | 774 | 156 | 11 | 0 | 0 | 11236 | 212123 | 113537 | 238 | 652 | 3027 | 3292 | 0 | 0 | 32 | 117 | 52 | 62 | 156 | 0 | 465 | 0 | 22 | 0 | 15 | 6945 | 320 | 0 |
| SRR7547885 | 43 | 15 | 14 | 27 | 165 | 3046 | 1448 | 8143 | 10252 | 7119 | 348 | 23 | 0 | 0 | 266492 | 80627 | 421005 | 1568 | 1142 | 1524 | 8826 | 0 | 0 | 188 | 231 | 139 | 71 | 61 | 0 | 1871 | 0 | 15 | 0 | 0 | 26090 | 3385 | 0 |
| SRR7547886 | 31 | 13 | 10 | 24 | 60 | 12737 | 3464 | 23783 | 16946 | 547 | 82 | 17 | 14 | 0 | 49453 | 43612 | 124897 | 1825 | 508 | 2511 | 1199 | 0 | 0 | 21 | 28 | 20 | 49 | 45 | 0 | 328 | 0 | 42 | 0 | 15 | 45531 | 11877 | 0 |
| SRR7547887 | 24 | 0 | 0 | 0 | 34 | 3891 | 1876 | 7927 | 7664 | 490 | 48 | 15 | 0 | 0 | 200001 | 79365 | 249188 | 1861 | 1115 | 2207 | 12399 | 0 | 0 | 25 | 0 | 0 | 0 | 0 | 0 | 215 | 0 | 118 | 0 | 0 | 36684 | 9044 | 0 |
| SRR7547888 | 29 | 0 | 0 | 0 | 282 | 2519 | 1656 | 7066 | 8248 | 6182 | 358 | 25 | 0 | 0 | 39253 | 60161 | 158917 | 6603 | 1081 | 2252 | 5360 | 0 | 11 | 43 | 50 | 35 | 17 | 12 | 0 | 1810 | 0 | 53 | 0 | 15 | 76563 | 6248 | 0 |
| SRR7547889 | 22 | 0 | 0 | 0 | 170 | 6410 | 3978 | 17011 | 17388 | 11006 | 490 | 21 | 0 | 0 | 56167 | 81156 | 147732 | 1747 | 1133 | 2792 | 4875 | 28 | 34 | 62 | 97 | 65 | 42 | 35 | 0 | 1685 | 55 | 289 | 0 | 13 | 58495 | 17188 | 0 |
| SRR7547890 | 39 | 15 | 0 | 0 | 39 | 2397 | 1272 | 5450 | 5255 | 652 | 74 | 18 | 0 | 0 | 54358 | 85272 | 256381 | 1095 | 469 | 4917 | 3664 | 0 | 0 | 11 | 16 | 12 | 20 | 15 | 0 | 289 | 0 | 27 | 0 | 0 | 46137 | 9016 | 0 |
| SRR7547891 | 14 | 0 | 0 | 0 | 215 | 6195 | 4100 | 19283 | 18756 | 1090 | 203 | 26 | 0 | 0 | 42791 | 104003 | 200587 | 2348 | 878 | 5514 | 1170 | 0 | 0 | 113 | 164 | 86 | 37 | 41 | 0 | 1500 | 0 | 60 | 0 | 21 | 23381 | 6950 | 0 |
| SRR7547892 | 31 | 0 | 0 | 0 | 29 | 9059 | 6087 | 28163 | 33441 | 406 | 74 | 25 | 0 | 0 | 113311 | 122332 | 252156 | 2164 | 777 | 2203 | 20225 | 0 | 0 | 25 | 33 | 20 | 10 | 11 | 0 | 283 | 0 | 16 | 0 | 26 | 14437 | 1634 | 0 |
| SRR7547893 | 19 | 0 | 0 | 0 | 46 | 4293 | 2414 | 10151 | 11392 | 482 | 57 | 21 | 0 | 0 | 48453 | 81585 | 148161 | 1688 | 862 | 3112 | 6971 | 0 | 0 | 64 | 17 | 13 | 36 | 17 | 0 | 261 | 0 | 56 | 0 | 0 | 49451 | 2515 | 0 |
| SRR7547894 | 10 | 0 | 0 | 0 | 205 | 889 | 473 | 1907 | 2233 | 2254 | 366 | 21 | 0 | 0 | 141595 | 33273 | 338959 | 2211 | 888 | 4500 | 4004 | 0 | 0 | 38 | 70 | 57 | 34 | 22 | 0 | 1529 | 0 | 14 | 0 | 12 | 8751 | 12116 | 0 |
| SRR7547895 | 28 | 11 | 0 | 0 | 50 | 4417 | 2070 | 10957 | 11909 | 475 | 32 | 22 | 0 | 0 | 223381 | 77973 | 144355 | 1346 | 302 | 1729 | 5957 | 0 | 0 | 25 | 28 | 17 | 53 | 41 | 0 | 123 | 0 | 14 | 0 | 14 | 37927 | 2140 | 0 |
| SRR7547896 | 47 | 0 | 0 | 10 | 222 | 11183 | 9259 | 37502 | 45435 | 224 | 273 | 27 | 0 | 0 | 37165 | 74029 | 122768 | 2150 | 1399 | 7562 | 3632 | 0 | 0 | 116 | 106 | 62 | 36 | 24 | 0 | 2001 | 0 | 0 | 0 | 22 | 12130 | 6299 | 0 |

### Supp Table S8.3.Columns_BX_to_DH

| | | | | | | | | | | | | | | | | | | | | | | | | | | | | | | | | | | | | | |
|---|---|---|---|---|---|---|---|---|---|---|---|---|---|---|---|---|---|---|---|---|---|---|---|---|---|---|---|---|---|---|---|---|---|---|---|---|---|
| SRR7547897 | 27 | 0 | 0 | 15 | 107 | 15575 | 15045 | 45182 | 69603 | 165 | 39 | 16 | 0 | 0 | 4838 | 56425 | 72159 | 883 | 277 | 5256 | 441 | 0 | 0 | 21 | 20 | 24 | 22 | 11 | 0 | 152 | 0 | 18 | 0 | 0 | 24907 | 4919 | 0 |
| SRR7547898 | 80 | 15 | 11 | 15 | 247 | 6988 | 5698 | 28193 | 36595 | 385 | 334 | 29 | 0 | 0 | 34146 | 91370 | 182446 | 800 | 527 | 9307 | 3621 | 0 | 0 | 149 | 134 | 107 | 66 | 45 | 0 | 1300 | 0 | 12 | 0 | 27 | 17362 | 5308 | 0 |
| SRR7547899 | 17 | 0 | 0 | 0 | 411 | 11168 | 6866 | 30486 | 31704 | 542 | 455 | 20 | 0 | 0 | 17610 | 94415 | 111822 | 3473 | 567 | 3751 | 632 | 0 | 0 | 144 | 145 | 123 | 62 | 57 | 0 | 1803 | 0 | 15 | 0 | 0 | 40370 | 9944 | 0 |
| SRR7547900 | 13 | 0 | 0 | 0 | 257 | 8518 | 4386 | 21737 | 24924 | 2594 | 213 | 23 | 0 | 0 | 61984 | 72418 | 159623 | 2849 | 1790 | 2890 | 3190 | 0 | 0 | 93 | 130 | 0 | 0 | 0 | 0 | 1010 | 0 | 64 | 0 | 13 | 67547 | 2289 | 0 |
| SRR7547901 | 60 | 10 | 13 | 13 | 600 | 3902 | 4377 | 17243 | 38969 | 654 | 261 | 28 | 0 | 0 | 123539 | 174056 | 208259 | 3106 | 469 | 4089 | 4298 | 0 | 0 | 187 | 178 | 146 | 94 | 123 | 0 | 1331 | 0 | 18 | 0 | 22 | 10275 | 4366 | 0 |
| SRR7547902 | 31 | 0 | 0 | 0 | 369 | 8150 | 2865 | 17955 | 48806 | 510 | 284 | 21 | 0 | 0 | 85438 | 126799 | 301821 | 1551 | 6016 | 17641 | 2548 | 0 | 0 | 88 | 85 | 180 | 92 | 96 | 0 | 624 | 0 | 29 | 0 | 25 | 2599 | 7168 | 0 |
| SRR7547903 | 22 | 0 | 0 | 11 | 53 | 7015 | 5554 | 26551 | 29468 | 96 | 61 | 29 | 0 | 0 | 9137 | 70896 | 151449 | 2256 | 1936 | 7793 | 7676 | 0 | 0 | 20 | 22 | 17 | 11 | 35 | 0 | 208 | 0 | 82 | 0 | 21 | 31557 | 3075 | 0 |
| SRR7547904 | 0 | 0 | 0 | 0 | 205 | 7156 | 4971 | 23300 | 30519 | 563 | 285 | 35 | 0 | 0 | 61672 | 173243 | 320789 | 1476 | 784 | 5494 | 12197 | 0 | 0 | 230 | 201 | 123 | 71 | 49 | 0 | 1720 | 0 | 28 | 0 | 24 | 13857 | 704 | 0 |
| SRR7547905 | 15 | 0 | 0 | 0 | 375 | 5005 | 4972 | 17911 | 27918 | 867 | 190 | 25 | 0 | 0 | 64080 | 78399 | 294413 | 1359 | 1315 | 2452 | 6820 | 0 | 0 | 100 | 111 | 107 | 91 | 46 | 0 | 730 | 0 | 27 | 0 | 24 | 53347 | 4103 | 0 |
| SRR7547906 | 16 | 0 | 0 | 0 | 259 | 2369 | 1307 | 7567 | 7511 | 2138 | 88 | 34 | 0 | 0 | 10989 | 89784 | 90430 | 705 | 461 | 1166 | 1241 | 0 | 0 | 58 | 96 | 45 | 39 | 45 | 0 | 1069 | 0 | 0 | 0 | 0 | 64749 | 117 | 0 |
| SRR7547907 | 34 | 16 | 0 | 0 | 352 | 1059 | 1037 | 4022 | 6115 | 3233 | 55 | 34 | 0 | 0 | 10039 | 102862 | 55293 | 819 | 402 | 1565 | 1926 | 0 | 0 | 93 | 193 | 153 | 150 | 129 | 0 | 426 | 0 | 0 | 0 | 0 | 63499 | 118 | 0 |
| SRR7547908 | 113 | 35 | 40 | 41 | 526 | 4171 | 2425 | 9835 | 24766 | 388 | 212 | 24 | 0 | 0 | 6571 | 244545 | 64588 | 229 | 356 | 2391 | 4646 | 0 | 0 | 92 | 122 | 0 | 0 | 0 | 0 | 1049 | 0 | 17 | 0 | 28 | 9074 | 594 | 0 |
| SRR7547909 | 40 | 14 | 0 | 0 | 407 | 1414 | 918 | 4536 | 4122 | 2641 | 445 | 23 | 0 | 0 | 22071 | 54793 | 412244 | 4037 | 771 | 3962 | 33418 | 0 | 0 | 58 | 59 | 55 | 21 | 11 | 0 | 2090 | 0 | 27 | 0 | 17 | 12392 | 13705 | 0 |
| SRR7547910 | 36 | 13 | 0 | 22 | 379 | 1937 | 804 | 4583 | 5696 | 4883 | 332 | 19 | 0 | 0 | 179967 | 101359 | 422310 | 2541 | 447 | 4355 | 5121 | 0 | 0 | 121 | 125 | 107 | 97 | 50 | 0 | 1601 | 0 | 33 | 0 | 11 | 14847 | 7013 | 0 |
| SRR7547911 | 20 | 0 | 0 | 0 | 43 | 10121 | 5942 | 32622 | 33504 | 632 | 37 | 23 | 0 | 0 | 43742 | 131372 | 360886 | 3476 | 451 | 6173 | 2055 | 0 | 0 | 36 | 37 | 23 | 12 | 10 | 0 | 317 | 0 | 16 | 0 | 14 | 22538 | 10453 | 0 |
| SRR7547912 | 30 | 0 | 0 | 0 | 587 | 2468 | 2034 | 9971 | 12131 | 3137 | 276 | 20 | 0 | 0 | 73154 | 134353 | 227959 | 4769 | 374 | 2398 | 2566 | 0 | 0 | 124 | 148 | 122 | 47 | 44 | 0 | 1557 | 0 | 27 | 0 | 0 | 39128 | 2781 | 0 |
| SRR7547913 | 25 | 0 | 0 | 0 | 86 | 7520 | 6269 | 26958 | 28254 | 492 | 40 | 23 | 0 | 0 | 29258 | 136330 | 212521 | 1547 | 538 | 7807 | 1977 | 0 | 0 | 21 | 27 | 21 | 50 | 46 | 0 | 240 | 0 | 22 | 0 | 0 | 38217 | 5636 | 0 |
| SRR7547914 | 18 | 0 | 0 | 0 | 395 | 4917 | 3358 | 16341 | 18367 | 2293 | 156 | 26 | 0 | 0 | 89788 | 156334 | 271753 | 6686 | 505 | 2199 | 1349 | 0 | 0 | 83 | 67 | 45 | 24 | 17 | 0 | 1224 | 0 | 0 | 0 | 12 | 27411 | 2724 | 0 |
| SRR7547915 | 29 | 0 | 0 | 0 | 49 | 2475 | 1002 | 5623 | 6001 | 252 | 33 | 22 | 0 | 0 | 240706 | 70964 | 448172 | 2030 | 2088 | 2535 | 2028 | 0 | 0 | 22 | 25 | 18 | 13 | 10 | 0 | 219 | 0 | 0 | 0 | 18 | 8929 | 6383 | 0 |
| SRR7547916 | 0 | 0 | 0 | 0 | 160 | 12477 | 10123 | 55748 | 39995 | 205 | 228 | 37 | 0 | 0 | 48804 | 53077 | 249880 | 2531 | 1537 | 3279 | 1899 | 0 | 0 | 106 | 95 | 56 | 17 | 18 | 0 | 2570 | 0 | 17 | 0 | 28 | 31612 | 16559 | 0 |
| SRR7547917 | 107 | 15 | 34 | 22 | 418 | 1914 | 1593 | 8344 | 12116 | 1499 | 106 | 22 | 0 | 0 | 7085 | 197549 | 165697 | 660 | 1080 | 1604 | 2042 | 0 | 0 | 149 | 239 | 119 | 79 | 134 | 0 | 748 | 0 | 0 | 0 | 14 | 13946 | 331 | 0 |
| SRR7547918 | 17 | 0 | 0 | 0 | 259 | 3580 | 3720 | 19323 | 15575 | 1702 | 37 | 17 | 0 | 0 | 7166 | 94151 | 82196 | 972 | 137 | 846 | 1319 | 0 | 0 | 62 | 105 | 52 | 28 | 31 | 0 | 465 | 0 | 0 | 0 | 16 | 34042 | 262 | 0 |
| SRR7547919 | 15 | 0 | 0 | 0 | 262 | 4901 | 5161 | 24214 | 18694 | 2577 | 38 | 26 | 0 | 0 | 7312 | 35883 | 36739 | 993 | 353 | 1403 | 1045 | 0 | 0 | 118 | 242 | 118 | 80 | 78 | 0 | 312 | 0 | 0 | 0 | 0 | 76845 | 161 | 0 |
| SRR7547920 | 31 | 28 | 12 | 10 | 126 | 4474 | 2284 | 14368 | 19626 | 333 | 70 | 24 | 0 | 0 | 2872 | 133799 | 61674 | 818 | 507 | 838 | 2210 | 0 | 0 | 34 | 48 | 29 | 29 | 41 | 0 | 452 | 0 | 20 | 0 | 13 | 28299 | 394 | 0 |
| SRR7585373 | 14 | 12 | 15 | 66 | 32 | 12867 | 342 | 5661 | 345 | 15 | 40 | 0 | 0 | 0 | 27988 | 12379 | 153436 | 15 | 333 | 15162 | 2308 | 0 | 0 | 42 | 24 | 76 | 304 | 65 | 0 | 26 | 0 | 115 | 0 | 0 | 909 | 2297 | 0 |
| SRR7586110 | 103 | 23 | 25 | 14 | 550 | 1771 | 1308 | 6237 | 12539 | 1025 | 135 | 0 | 0 | 0 | 12496 | 44727 | 123823 | 52 | 279 | 1069 | 1143 | 0 | 0 | 41 | 171 | 114 | 59 | 79 | 0 | 241 | 0 | 0 | 0 | 0 | 5135 | 0 | 0 |
| SRR7586111 | 34 | 11 | 22 | 22 | 221 | 1726 | 1072 | 5760 | 11096 | 366 | 45 | 0 | 0 | 0 | 10407 | 44721 | 120623 | 61 | 281 | 894 | 853 | 0 | 0 | 14 | 59 | 40 | 21 | 28 | 0 | 84 | 0 | 15 | 0 | 0 | 4670 | 0 | 0 |
| SRR7586113 | 133 | 50 | 20 | 21 | 570 | 2689 | 1485 | 6391 | 11622 | 1041 | 367 | 0 | 0 | 0 | 10194 | 21427 | 43659 | 257 | 180 | 908 | 641 | 0 | 0 | 34 | 161 | 110 | 42 | 59 | 0 | 1566 | 0 | 0 | 0 | 0 | 3104 | 0 | 0 |
| SRR7586114 | 127 | 58 | 36 | 34 | 629 | 2318 | 1324 | 5607 | 10341 | 1033 | 371 | 15 | 0 | 0 | 8824 | 19288 | 42826 | 261 | 273 | 858 | 509 | 0 | 0 | 34 | 148 | 101 | 38 | 60 | 0 | 1563 | 0 | 0 | 0 | 0 | 3141 | 0 | 0 |
| SRR7586116 | 144 | 34 | 18 | 25 | 108 | 1327 | 1257 | 3951 | 12744 | 105 | 63 | 0 | 11 | 0 | 2688 | 21404 | 22655 | 316 | 862 | 2065 | 134 | 0 | 0 | 35 | 32 | 36 | 20 | 42 | 0 | 182 | 0 | 0 | 0 | 0 | 2156 | 0 | 0 |
| SRR7586117 | 545 | 170 | 151 | 163 | 470 | 1394 | 1402 | 4148 | 13102 | 408 | 233 | 19 | 14 | 0 | 2635 | 20715 | 21615 | 335 | 892 | 2185 | 142 | 0 | 0 | 34 | 117 | 141 | 69 | 150 | 0 | 676 | 0 | 11 | 0 | 0 | 2405 | 0 | 0 |
| SRR7586118 | 149 | 42 | 38 | 42 | 124 | 1645 | 1560 | 4876 | 15610 | 104 | 60 | 16 | 12 | 0 | 3181 | 20818 | 21704 | 349 | 1153 | 2776 | 158 | 0 | 0 | 36 | 31 | 38 | 17 | 40 | 0 | 179 | 0 | 0 | 0 | 0 | 2124 | 10 | 0 |
| SRR7586119 | 134 | 29 | 27 | 22 | 184 | 1686 | 1767 | 6842 | 23955 | 367 | 149 | 10 | 0 | 0 | 13636 | 48527 | 113477 | 303 | 643 | 5694 | 1433 | 14 | 25 | 24 | 68 | 68 | 43 | 42 | 0 | 618 | 0 | 0 | 0 | 0 | 2362 | 12 | 0 |
| SRR7586120 | 368 | 89 | 142 | 86 | 530 | 1685 | 1565 | 6565 | 23310 | 925 | 387 | 29 | 0 | 0 | 13615 | 46274 | 112522 | 311 | 649 | 5745 | 1374 | 13 | 21 | 66 | 168 | 177 | 113 | 105 | 0 | 1649 | 0 | 0 | 0 | 0 | 2212 | 12 | 0 |
| SRR7586122 | 265 | 76 | 46 | 37 | 733 | 815 | 587 | 2060 | 6631 | 744 | 399 | 0 | 0 | 0 | 3953 | 11581 | 20178 | 387 | 494 | 829 | 328 | 0 | 0 | 22 | 143 | 151 | 66 | 150 | 0 | 864 | 0 | 0 | 0 | 0 | 2149 | 0 | 0 |
| SRR7586123 | 288 | 106 | 78 | 61 | 796 | 956 | 841 | 2439 | 7827 | 727 | 400 | 21 | 0 | 0 | 4602 | 11026 | 19614 | 381 | 691 | 1077 | 351 | 0 | 0 | 20 | 139 | 156 | 65 | 138 | 0 | 860 | 0 | 0 | 0 | 0 | 2183 | 0 | 0 |
| SRR7586125 | 61 | 17 | 13 | 12 | 392 | 1046 | 692 | 3440 | 9747 | 918 | 198 | 13 | 0 | 0 | 8529 | 26068 | 53283 | 233 | 518 | 1280 | 1421 | 0 | 15 | 17 | 84 | 73 | 29 | 19 | 0 | 678 | 0 | 11 | 0 | 0 | 2530 | 0 | 0 |
| SRR7586126 | 171 | 59 | 19 | 19 | 1241 | 1135 | 705 | 3389 | 9987 | 2511 | 529 | 29 | 0 | 0 | 8464 | 24830 | 49727 | 327 | 525 | 1308 | 1221 | 0 | 14 | 51 | 217 | 174 | 70 | 38 | 0 | 1899 | 0 | 0 | 0 | 0 | 3085 | 0 | 0 |
| SRR7586128 | 145 | 51 | 21 | 16 | 285 | 1038 | 884 | 2813 | 10123 | 509 | 136 | 11 | 0 | 0 | 8137 | 37807 | 47793 | 373 | 999 | 2162 | 696 | 0 | 0 | 16 | 90 | 83 | 36 | 37 | 0 | 439 | 0 | 12 | 0 | 0 | 4236 | 12 | 0 |
| SRR7586129 | 146 | 59 | 39 | 23 | 309 | 1109 | 1037 | 3156 | 10808 | 480 | 134 | 11 | 0 | 0 | 8853 | 36097 | 44859 | 426 | 1152 | 2590 | 708 | 0 | 0 | 16 | 82 | 83 | 36 | 34 | 0 | 447 | 0 | 0 | 0 | 0 | 4100 | 0 | 0 |
| SRR7586131 | 118 | 35 | 32 | 10 | 256 | 1275 | 582 | 2367 | 6941 | 226 | 127 | 12 | 15 | 10 | 3906 | 20327 | 21044 | 289 | 380 | 664 | 457 | 0 | 0 | 28 | 44 | 42 | 16 | 36 | 0 | 163 | 0 | 0 | 0 | 0 | 5166 | 0 | 0 |
| SRR7586132 | 362 | 140 | 66 | 50 | 914 | 1441 | 634 | 2719 | 7798 | 671 | 387 | 25 | 15 | 0 | 4513 | 19409 | 20213 | 403 | 553 | 965 | 440 | 0 | 0 | 29 | 133 | 121 | 48 | 108 | 0 | 498 | 0 | 0 | 0 | 0 | 5006 | 0 | 0 |
| SRR7586134 | 145 | 40 | 20 | 17 | 330 | 1072 | 1030 | 3757 | 12959 | 421 | 148 | 11 | 0 | 0 | 9939 | 26268 | 65172 | 518 | 542 | 2282 | 910 | 0 | 0 | 15 | 87 | 89 | 39 | 32 | 0 | 402 | 0 | 19 | 0 | 0 | 3096 | 0 | 0 |
| SRR7586135 | 151 | 48 | 36 | 24 | 366 | 1089 | 1000 | 3714 | 13097 | 414 | 146 | 11 | 0 | 0 | 9548 | 25990 | 61196 | 420 | 567 | 2393 | 960 | 0 | 0 | 15 | 79 | 84 | 33 | 29 | 0 | 407 | 0 | 0 | 0 | 0 | 3320 | 11 | 0 |
| SRR7586137 | 138 | 40 | 30 | 18 | 447 | 1544 | 863 | 3717 | 7569 | 875 | 469 | 0 | 0 | 0 | 7564 | 21910 | 71231 | 224 | 192 | 1554 | 910 | 0 | 0 | 31 | 172 | 121 | 51 | 127 | 0 | 1423 | 0 | 16 | 0 | 0 | 3168 | 0 | 0 |
| SRR7586138 | 38 | 11 | 14 | 32 | 127 | 1299 | 761 | 3626 | 7474 | 220 | 119 | 13 | 0 | 0 | 6942 | 21392 | 70105 | 280 | 201 | 1623 | 991 | 0 | 0 | 35 | 44 | 36 | 14 | 29 | 0 | 389 | 0 | 0 | 0 | 0 | 2975 | 0 | 0 |
| SRR7586140 | 109 | 31 | 13 | 10 | 379 | 1125 | 847 | 3240 | 12889 | 554 | 239 | 12 | 0 | 0 | 16497 | 38278 | 108622 | 426 | 466 | 2214 | 1616 | 0 | 0 | 18 | 93 | 87 | 38 | 25 | 0 | 891 | 0 | 18 | 0 | 0 | 3969 | 0 | 0 |
| SRR7586141 | 115 | 37 | 27 | 17 | 415 | 1146 | 864 | 3376 | 13132 | 545 | 245 | 16 | 0 | 0 | 17654 | 35903 | 104128 | 437 | 465 | 2600 | 1742 | 0 | 0 | 16 | 88 | 90 | 34 | 24 | 0 | 901 | 0 | 16 | 0 | 0 | 4184 | 0 | 0 |
| SRR7586143 | 127 | 41 | 14 | 25 | 313 | 928 | 450 | 1856 | 5272 | 433 | 165 | 15 | 13 | 0 | 4750 | 18559 | 26978 | 275 | 189 | 1141 | 811 | 0 | 0 | 11 | 56 | 59 | 23 | 33 | 0 | 195 | 0 | 0 | 0 | 0 | 4410 | 0 | 0 |
| SRR7586144 | 335 | 133 | 56 | 46 | 870 | 938 | 451 | 1708 | 5422 | 1059 | 413 | 34 | 11 | 11 | 4745 | 17653 | 26262 | 340 | 231 | 1232 | 702 | 0 | 0 | 29 | 142 | 142 | 61 | 77 | 0 | 517 | 0 | 24 | 0 | 0 | 4806 | 0 | 0 |
| SRR7586146 | 75 | 13 | 0 | 0 | 426 | 2598 | 1294 | 6698 | 12405 | 291 | 53 | 0 | 13 | 0 | 11005 | 14308 | 51428 | 176 | 554 | 878 | 484 | 0 | 0 | 19 | 52 | 21 | 28 | 0 | 0 | 120 | 0 | 11 | 0 | 0 | 3688 | 0 | 0 |
| SRR7586147 | 234 | 52 | 0 | 0 | 1503 | 2628 | 1504 | 7025 | 12904 | 927 | 154 | 17 | 12 | 0 | 11757 | 13302 | 49887 | 172 | 560 | 989 | 614 | 0 | 0 | 71 | 156 | 65 | 29 | 0 | 0 | 379 | 0 | 0 | 0 | 0 | 3502 | 11 | 0 |
| SRR7586149 | 204 | 62 | 11 | 15 | 461 | 1588 | 688 | 3483 | 6481 | 1892 | 498 | 16 | 21 | 0 | 19006 | 16480 | 51315 | 191 | 210 | 652 | 2645 | 0 | 0 | 50 | 187 | 103 | 25 | 19 | 0 | 1697 | 0 | 11 | 0 | 0 | 2474 | 0 | 0 |
| SRR7586150 | 75 | 26 | 21 | 19 | 182 | 1641 | 666 | 3292 | 6074 | 655 | 182 | 28 | 19 | 0 | 17871 | 15067 | 48692 | 235 | 275 | 527 | 2855 | 0 | 0 | 21 | 63 | 33 | 29 | 14 | 0 | 587 | 0 | 11 | 0 | 0 | 2184 | 11 | 0 |
| SRR7586152 | 351 | 74 | 11 | 0 | 750 | 2411 | 1100 | 6810 | 14535 | 1602 | 483 | 11 | 19 | 0 | 31435 | 19449 | 85421 | 242 | 485 | 1146 | 1957 | 0 | 0 | 83 | 273 | 123 | 55 | 48 | 0 | 1822 | 0 | 15 | 0 | 0 | 3425 | 15 | 0 |
| SRR7586153 | 115 | 28 | 18 | 15 | 280 | 2745 | 1363 | 7447 | 16076 | 545 | 168 | 10 | 20 | 0 | 33478 | 19765 | 83832 | 286 | 569 | 1273 | 2319 | 0 | 0 | 30 | 88 | 44 | 18 | 13 | 0 | 631 | 0 | 15 | 0 | 0 | 3263 | 13 | 0 |
| SRR7586155 | 85 | 21 | 12 | 11 | 176 | 2851 | 1403 | 6963 | 10480 | 646 | 55 | 0 | 22 | 0 | 20873 | 13762 | 48100 | 257 | 520 | 1024 | 1322 | 0 | 13 | 26 | 54 | 20 | 22 | 0 | 0 | 141 | 0 | 0 | 0 | 0 | 2757 | 0 | 0 |
| SRR7586156 | 86 | 24 | 22 | 18 | 192 | 2675 | 1197 | 6010 | 9270 | 660 | 52 | 18 | 23 | 0 | 18877 | 13000 | 46691 | 333 | 594 | 922 | 1307 | 0 | 15 | 19 | 59 | 24 | 31 | 0 | 0 | 144 | 0 | 10 | 0 | 0 | 2678 | 0 | 0 |
| SRR7586158 | 710 | 141 | 39 | 35 | 707 | 1256 | 774 | 3975 | 8805 | 2177 | 465 | 0 | 16 | 0 | 26227 | 23723 | 95657 | 325 | 781 | 2492 | 5495 | 0 | 0 | 63 | 221 | 139 | 79 | 50 | 0 | 1064 | 0 | 14 | 0 | 0 | 2491 | 11 | 0 |
| SRR7586159 | 740 | 176 | 67 | 56 | 775 | 1144 | 707 | 3947 | 8490 | 2130 | 441 | 18 | 18 | 0 | 25988 | 22348 | 93052 | 384 | 813 | 2529 | 5348 | 0 | 0 | 59 | 212 | 147 | 74 | 50 | 0 | 1097 | 0 | 16 | 0 | 0 | 2709 | 0 | 0 |
| SRR7609258 | 71 | 0 | 0 | 0 | 62 | 12657 | 1046 | 14027 | 2355 | 53 | 11 | 0 | 0 | 0 | 2953 | 69141 | 39174 | 115 | 214 | 1807 | 2838 | 0 | 0 | 15 | 0 | 0 | 0 | 0 | 0 | 22 | 0 | 0 | 0 | 0 | 129682 | 65 | 0 |
| SRR7609259 | 20 | 106 | 0 | 0 | 43 | 945 | 88 | 566 | 503 | 50 | 0 | 0 | 0 | 0 | 3796 | 458680 | 50119 | 781 | 589 | 4108 | 7643 | 0 | 0 | 187 | 12 | 0 | 0 | 0 | 0 | 48 | 0 | 13 | 0 | 12 | 80366 | 479 | 0 |
| SRR7609260 | 147 | 40 | 0 | 0 | 66 | 711 | 60 | 523 | 498 | 154 | 10 | 15 | 0 | 0 | 3398 | 277022 | 54786 | 306 | 838 | 4866 | 474 | 0 | 10 | 200 | 0 | 0 | 0 | 0 | 13 | 90 | 0 | 0 | 0 | 0 | 143101 | 902 | 0 |
| SRR7609261 | 31 | 11 | 0 | 0 | 103 | 6887 | 810 | 5435 | 8168 | 592 | 18 | 0 | 0 | 0 | 32220 | 222619 | 96487 | 495 | 282 | 2304 | 1502 | 0 | 0 | 190 | 0 | 0 | 0 | 0 | 0 | 18 | 0 | 24 | 0 | 0 | 60620 | 2704 | 0 |
| SRR7609262 | 46 | 24 | 0 | 0 | 95 | 1274 | 229 | 1006 | 680 | 24 | 0 | 0 | 0 | 0 | 17149 | 327072 | 42515 | 809 | 171 | 1171 | 730 | 0 | 12 | 113 | 0 | 0 | 0 | 0 | 0 | 0 | 0 | 24 | 0 | 0 | 58952 | 62 | 0 |
| SRR7609263 | 40 | 12 | 0 | 0 | 88 | 3379 | 377 | 1633 | 394 | 12 | 12 | 0 | 0 | 0 | 6728 | 143374 | 55544 | 792 | 1070 | 5455 | 1190 | 0 | 0 | 196 | 0 | 0 | 0 | 0 | 0 | 0 | 0 | 0 | 0 | 0 | 145698 | 5603 | 0 |
| SRR7609264 | 25 | 0 | 0 | 0 | 119 | 1277 | 264 | 1205 | 984 | 24 | 11 | 0 | 0 | 0 | 8335 | 122028 | 57528 | 624 | 944 | 8104 | 912 | 0 | 0 | 46 | 0 | 0 | 0 | 0 | 0 | 22 | 0 | 0 | 0 | 0 | 111791 | 593 | 0 |
| SRR7609265 | 14 | 10 | 0 | 0 | 54 | 3483 | 447 | 2254 | 1089 | 93 | 0 | 0 | 0 | 0 | 2555 | 57702 | 29063 | 777 | 902 | 4140 | 783 | 0 | 0 | 158 | 0 | 0 | 0 | 0 | 0 | 13 | 0 | 0 | 0 | 0 | 207044 | 1012 | 0 |
| SRR7609266 | 34 | 0 | 0 | 0 | 53 | 2717 | 264 | 1846 | 1054 | 1017 | 0 | 0 | 0 | 0 | 7280 | 129802 | 61289 | 2431 | 471 | 3179 | 1496 | 0 | 0 | 88 | 0 | 0 | 0 | 0 | 74 | 11 | 0 | 0 | 0 | 22 | 80154 | 1494 | 0 |
| SRR7609267 | 33 | 14 | 0 | 0 | 81 | 744 | 194 | 808 | 489 | 18 | 0 | 0 | 0 | 0 | 13151 | 283790 | 64973 | 2016 | 366 | 1898 | 2167 | 0 | 0 | 65 | 0 | 0 | 0 | 0 | 25 | 0 | 0 | 15 | 0 | 0 | 49637 | 1193 | 0 |
| SRR7609268 | 34 | 14 | 0 | 0 | 11 | 1598 | 478 | 2687 | 3837 | 44 | 0 | 0 | 0 | 0 | 10477 | 165302 | 116497 | 2148 | 795 | 6867 | 1830 | 0 | 0 | 11 | 0 | 0 | 0 | 0 | 0 | 16 | 0 | 10 | 0 | 0 | 52391 | 415 | 0 |
| SRR7609269 | 33 | 0 | 0 | 0 | 39 | 7372 | 719 | 3156 | 1756 | 59 | 0 | 0 | 0 | 0 | 25939 | 156170 | 68442 | 4240 | 156 | 2253 | 4242 | 0 | 0 | 17 | 0 | 0 | 0 | 0 | 0 | 0 | 0 | 15 | 0 | 0 | 56990 | 666 | 0 |
| SRR7609270 | 62 | 12 | 0 | 0 | 74 | 1608 | 200 | 888 | 375 | 2261 | 0 | 11 | 0 | 0 | 18503 | 105707 | 43857 | 8366 | 89 | 792 | 1518 | 0 | 0 | 86 | 0 | 0 | 0 | 0 | 31 | 12 | 0 | 46 | 0 | 0 | 140032 | 0 | 0 |
| SRR7609271 | 33 | 0 | 0 | 0 | 41 | 1864 | 255 | 1214 | 926 | 17 | 0 | 0 | 0 | 0 | 9291 | 226350 | 67106 | 440 | 384 | 2546 | 7845 | 0 | 0 | 27 | 0 | 0 | 0 | 0 | 0 | 0 | 0 | 0 | 0 | 0 | 109977 | 659 | 0 |
| SRR7609272 | 39 | 0 | 0 | 0 | 11 | 3370 | 595 | 3362 | 3815 | 66 | 0 | 0 | 0 | 0 | 21706 | 169857 | 68890 | 3445 | 612 | 3727 | 2411 | 0 | 0 | 12 | 0 | 0 | 0 | 0 | 0 | 0 | 0 | 13 | 0 | 0 | 110487 | 266 | 0 |
| SRR7609273 | 16 | 0 | 0 | 0 | 43 | 10284 | 1153 | 6386 | 2825 | 7779 | 0 | 0 | 0 | 0 | 5570 | 112459 | 53938 | 188 | 49 | 158 | 1083 | 0 | 0 | 111 | 0 | 0 | 0 | 0 | 0 | 0 | 0 | 10 | 0 | 0 | 135406 | 1846 | 0 |
| SRR7609274 | 44 | 13 | 0 | 0 | 82 | 2567 | 332 | 1371 | 982 | 16 | 0 | 0 | 0 | 0 | 32553 | 173196 | 94624 | 1109 | 158 | 637 | 3723 | 0 | 0 | 157 | 0 | 0 | 0 | 0 | 0 | 17 | 0 | 10 | 0 | 0 | 80725 | 2085 | 0 |
| SRR7609275 | 103 | 24 | 0 | 0 | 563 | 2080 | 246 | 1092 | 1245 | 655 | 0 | 0 | 0 | 0 | 6426 | 316425 | 59694 | 1673 | 521 | 2314 | 17961 | 0 | 0 | 248 | 0 | 0 | 0 | 0 | 13 | 25 | 0 | 0 | 0 | 0 | 78639 | 1093 | 0 |
| SRR7609276 | 0 | 0 | 0 | 0 | 62 | 8941 | 1563 | 7942 | 9560 | 472 | 0 | 0 | 0 | 0 | 405 | 68360 | 12444 | 56 | 95 | 2182 | 111 | 0 | 0 | 103 | 0 | 0 | 0 | 0 | 0 | 0 | 0 | 0 | 0 | 0 | 219145 | 14 | 0 |
| SRR7609277 | 11 | 50 | 0 | 0 | 58 | 1342 | 171 | 703 | 625 | 59 | 0 | 0 | 0 | 0 | 23557 | 280136 | 73174 | 2399 | 958 | 4398 | 3520 | 0 | 0 | 13 | 0 | 0 | 0 | 0 | 0 | 22 | 0 | 11 | 0 | 0 | 66922 | 710 | 0 |
| SRR7609278 | 27 | 0 | 0 | 0 | 57 | 6005 | 681 | 3234 | 1694 | 20 | 0 | 0 | 0 | 0 | 232 | 74401 | 19997 | 94 | 1081 | 7945 | 459 | 0 | 0 | 15 | 0 | 11 | 12 | 0 | 0 | 0 | 0 | 0 | 0 | 0 | 234487 | 1407 | 0 |
| SRR7609279 | 72 | 17 | 0 | 0 | 70 | 3489 | 447 | 2100 | 1340 | 24 | 0 | 0 | 0 | 0 | 1834 | 225040 | 52025 | 830 | 169 | 701 | 740 | 0 | 0 | 165 | 0 | 0 | 0 | 0 | 0 | 0 | 0 | 0 | 0 | 0 | 113573 | 695 | 0 |
| SRR7609280 | 24 | 0 | 0 | 0 | 33 | 2815 | 520 | 2752 | 3023 | 20 | 0 | 0 | 0 | 0 | 5122 | 264110 | 53110 | 1734 | 220 | 1137 | 644 | 0 | 0 | 11 | 0 | 0 | 0 | 0 | 0 | 0 | 0 | 0 | 0 | 0 | 79339 | 1054 | 0 |
| SRR7609281 | 15 | 0 | 0 | 0 | 205 | 1825 | 151 | 908 | 567 | 781 | 0 | 0 | 0 | 0 | 3125 | 337394 | 57179 | 140 | 246 | 1435 | 776 | 0 | 0 | 149 | 0 | 0 | 0 | 0 | 0 | 0 | 0 | 0 | 0 | 0 | 127384 | 1746 | 0 |
| SRR7609282 | 142 | 49 | 0 | 0 | 365 | 1522 | 126 | 614 | 468 | 698 | 0 | 0 | 0 | 0 | 7331 | 207849 | 47766 | 1784 | 236 | 1553 | 1739 | 0 | 0 | 470 | 0 | 0 | 0 | 0 | 0 | 18 | 0 | 10 | 0 | 0 | 162129 | 854 | 0 |
| SRR7609283 | 129 | 51 | 0 | 0 | 593 | 1121 | 73 | 787 | 784 | 817 | 0 | 0 | 0 | 0 | 13200 | 243203 | 23521 | 468 | 76 | 322 | 714 | 0 | 0 | 176 | 0 | 0 | 0 | 0 | 11 | 13 | 0 | 0 | 0 | 0 | 198060 | 862 | 0 |
| SRR7609284 | 101 | 23 | 0 | 0 | 178 | 1308 | 300 | 1451 | 1827 | 27 | 0 | 0 | 0 | 0 | 19652 | 134533 | 55505 | 439 | 598 | 3274 | 27365 | 0 | 0 | 11 | 15 | 0 | 0 | 0 | 0 | 0 | 0 | 0 | 0 | 0 | 115207 | 1184 | 0 |
| SRR7609285 | 89 | 26 | 0 | 0 | 399 | 1118 | 93 | 738 | 638 | 613 | 0 | 15 | 0 | 0 | 11481 | 203969 | 31360 | 322 | 202 | 265 | 843 | 0 | 0 | 205 | 0 | 0 | 0 | 0 | 0 | 21 | 0 | 0 | 0 | 0 | 199727 | 1783 | 0 |
| SRR7609286 | 76 | 29 | 0 | 0 | 322 | 2322 | 243 | 1510 | 957 | 1090 | 0 | 0 | 0 | 0 | 4129 | 212368 | 26005 | 506 | 110 | 588 | 627 | 0 | 0 | 206 | 0 | 0 | 0 | 0 | 0 | 33 | 0 | 12 | 0 | 0 | 219022 | 200 | 0 |
| SRR7609287 | 216 | 82 | 0 | 0 | 313 | 1883 | 203 | 855 | 315 | 540 | 14 | 0 | 0 | 0 | 34231 | 486363 | 87016 | 838 | 99 | 1024 | 2220 | 0 | 0 | 247 | 0 | 0 | 0 | 0 | 0 | 65 | 0 | 0 | 0 | 0 | 40158 | 498 | 0 |
| SRR7609288 | 17 | 75 | 0 | 0 | 81 | 4348 | 304 | 1619 | 1436 | 58 | 10 | 0 | 0 | 0 | 41838 | 217948 | 180022 | 2193 | 433 | 6774 | 6704 | 0 | 0 | 34 | 15 | 0 | 18 | 13 | 0 | 39 | 0 | 0 | 0 | 0 | 135233 | 3783 | 0 |
| SRR7609289 | 28 | 15 | 0 | 0 | 80 | 4039 | 457 | 2244 | 762 | 64 | 0 | 17 | 10 | 10 | 29890 | 448733 | 56407 | 360 | 274 | 1922 | 668 | 0 | 0 | 52 | 0 | 0 | 0 | 0 | 19 | 16 | 0 | 16 | 0 | 13 | 46876 | 784 | 0 |
| SRR7609290 | 55 | 58 | 0 | 0 | 168 | 387 | 41 | 287 | 215 | 28 | 12 | 0 | 0 | 0 | 88658 | 151155 | 67387 | 2716 | 140 | 1639 | 4964 | 0 | 0 | 165 | 15 | 0 | 0 | 0 | 0 | 31 | 0 | 26 | 0 | 0 | 54155 | 2936 | 0 |
| SRR7609291 | 21 | 0 | 0 | 0 | 33 | 10097 | 799 | 4574 | 509 | 144 | 0 | 0 | 0 | 0 | 23366 | 596399 | 62307 | 285 | 140 | 8339 | 7635 | 0 | 0 | 37 | 0 | 0 | 0 | 0 | 0 | 0 | 0 | 0 | 0 | 0 | 17517 | 716 | 0 |
| SRR7609292 | 38 | 13 | 0 | 0 | 144 | 1828 | 371 | 2377 | 2432 | 53 | 0 | 0 | 0 | 0 | 11143 | 153864 | 106587 | 2962 | 557 | 6943 | 2504 | 0 | 0 | 16 | 0 | 0 | 0 | 0 | 0 | 0 | 0 | 15 | 0 | 0 | 43505 | 482 | 0 |
| SRR7609293 | 226 | 69 | 0 | 0 | 307 | 1777 | 255 | 1070 | 801 | 557 | 0 | 0 | 10 | 0 | 8853 | 241670 | 47952 | 548 | 191 | 769 | 987 | 0 | 0 | 159 | 0 | 0 | 0 | 0 | 0 | 15 | 0 | 17 | 0 | 0 | 140974 | 502 | 0 |
| SRR7609294 | 12 | 65 | 0 | 0 | 25 | 12937 | 950 | 6430 | 1400 | 27 | 0 | 17 | 0 | 0 | 9147 | 403739 | 91457 | 677 | 329 | 3012 | 2568 | 0 | 0 | 16 | 0 | 0 | 0 | 0 | 0 | 23 | 0 | 0 | 0 | 0 | 79875 | 1575 | 0 |
| SRR7609295 | 128 | 22 | 12 | 0 | 50 | 1934 | 113 | 561 | 174 | 1173 | 18 | 0 | 0 | 0 | 22964 | 163666 | 121525 | 838 | 2419 | 16638 | 2429 | 0 | 12 | 38 | 0 | 0 | 0 | 0 | 0 | 0 | 12 | 0 | 0 | 0 | 28220 | 1096 | 0 |
| SRR7609296 | 21 | 0 | 0 | 0 | 36 | 9872 | 654 | 3287 | 455 | 78 | 0 | 0 | 0 | 0 | 2376 | 146398 | 30401 | 119 | 139 | 257 | 683 | 0 | 0 | 23 | 0 | 0 | 0 | 0 | 0 | 0 | 0 | 0 | 0 | 0 | 132460 | 180 | 0 |
| SRR7609297 | 48 | 20 | 0 | 14 | 102 | 5055 | 398 | 1919 | 467 | 268 | 19 | 0 | 0 | 0 | 961 | 262438 | 92725 | 273 | 228 | 1418 | 2863 | 0 | 0 | 70 | 0 | 0 | 10 | 0 | 0 | 17 | 0 | 10 | 0 | 0 | 102959 | 947 | 0 |
| SRR7609298 | 31 | 19 | 0 | 0 | 121 | 7691 | 519 | 3353 | 1779 | 84 | 0 | 0 | 0 | 0 | 3964 | 95348 | 38296 | 267 | 512 | 1932 | 374 | 0 | 0 | 130 | 12 | 0 | 0 | 14 | 0 | 14 | 0 | 0 | 0 | 0 | 110047 | 582 | 0 |
| SRR7609299 | 58 | 14 | 0 | 0 | 46 | 1400 | 127 | 459 | 94 | 91 | 0 | 0 | 0 | 0 | 20123 | 134760 | 121566 | 845 | 295 | 2613 | 5071 | 0 | 0 | 38 | 0 | 0 | 22 | 0 | 0 | 18 | 0 | 0 | 0 | 0 | 50066 | 987 | 0 |
| SRR7609300 | 117 | 18 | 20 | 44 | 187 | 4112 | 329 | 1068 | 391 | 178 | 26 | 0 | 0 | 0 | 3839 | 321254 | 52177 | 273 | 311 | 1963 | 1131 | 0 | 0 | 50 | 12 | 0 | 18 | 20 | 0 | 88 | 0 | 0 | 0 | 0 | 88234 | 400 | 0 |
| SRR7609301 | 159 | 59 | 12 | 22 | 164 | 485 | 37 | 159 | 37 | 243 | 56 | 0 | 0 | 0 | 24201 | 215200 | 166599 | 2271 | 738 | 4180 | 11683 | 0 | 0 | 59 | 0 | 0 | 17 | 15 | 0 | 47 | 0 | 13 | 0 | 0 | 26865 | 1035 | 0 |
| SRR7609302 | 61 | 0 | 0 | 12 | 12 | 1500 | 168 | 823 | 368 | 169 | 21 | 0 | 0 | 0 | 1000 | 206463 | 89974 | 232 | 1668 | 7442 | 2196 | 0 | 0 | 70 | 15 | 31 | 46 | 12 | 0 | 28 | 0 | 0 | 0 | 0 | 57823 | 737 | 0 |
| SRR7644105 | 0 | 0 | 0 | 0 | 120 | 4818 | 4818 | 4380 | 0 | 0 | 30 | 0 | 0 | 0 | 11389 | 133209 | 218092 | 0 | 0 | 0 | 438 | 0 | 0 | 60 | 30 | 30 | 0 | 60 | 0 | 180 | 0 | 0 | 0 | 0 | 34696 | 0 | 210 |
| SRR7644106 | 0 | 0 | 0 | 0 | 130 | 1437 | 2875 | 5750 | 0 | 0 | 65 | 33 | 0 | 0 | 9583 | 146416 | 204739 | 0 | 0 | 0 | 958 | 0 | 0 | 65 | 98 | 33 | 0 | 65 | 0 | 260 | 0 | 0 | 0 | 0 | 34022 | 0 | 98 |
| SRR7644107 | 0 | 0 | 0 | 0 | 122 | 2207 | 5296 | 9709 | 0 | 0 | 61 | 0 | 0 | 0 | 8826 | 132241 | 208309 | 0 | 0 | 0 | 0 | 0 | 0 | 91 | 0 | 0 | 0 | 30 | 0 | 274 | 0 | 0 | 0 | 0 | 33938 | 0 | 91 |
| SRR7644108 | 0 | 0 | 0 | 0 | 96 | 3385 | 4352 | 6286 | 0 | 0 | 32 | 0 | 0 | 0 | 5319 | 150919 | 219404 | 0 | 0 | 0 | 967 | 0 | 0 | 64 | 64 | 128 | 0 | 96 | 0 | 160 | 0 | 0 | 0 | 0 | 32340 | 0 | 64 |
| SRR7644109 | 0 | 0 | 0 | 0 | 107 | 6846 | 2934 | 5379 | 978 | 0 | 53 | 27 | 0 | 0 | 6846 | 144676 | 163194 | 0 | 0 | 489 | 489 | 0 | 0 | 0 | 53 | 107 | 80 | 0 | 0 | 213 | 0 | 0 | 0 | 0 | 15046 | 0 | 107 |
| SRR7644110 | 0 | 0 | 0 | 0 | 29 | 7853 | 4712 | 3141 | 4188 | 0 | 87 | 0 | 0 | 0 | 5759 | 152752 | 179963 | 0 | 0 | 524 | 0 | 0 | 0 | 0 | 116 | 58 | 58 | 0 | 0 | 521 | 0 | 0 | 0 | 0 | 22882 | 0 | 58 |
| SRR7644111 | 0 | 0 | 0 | 0 | 54 | 4392 | 5368 | 5857 | 1464 | 0 | 54 | 27 | 0 | 0 | 9273 | 149263 | 189803 | 0 | 0 | 488 | 0 | 0 | 0 | 0 | 27 | 109 | 54 | 0 | 0 | 13 | 0 | 0 | 0 | 0 | 16585 | 0 | 54 |
| SRR7644112 | 0 | 0 | 0 | 0 | 142 | 5189 | 5708 | 7265 | 1038 | 0 | 29 | 29 | 0 | 0 | 7784 | 147321 | 186224 | 0 | 0 | 519 | 0 | 0 | 0 | 0 | 199 | 29 | 57 | 57 | 0 | 513 | 0 | 0 | 0 | 0 | 15306 | 12 | 85 |
| SRR7644113 | 0 | 0 | 0 | 0 | 89 | 4959 | 5543 | 16336 | 3209 | 0 | 59 | 0 | 0 | 0 | 5543 | 156509 | 241159 | 0 | 0 | 0 | 292 | 0 | 59 | 0 | 237 | 296 | 0 | 118 | 0 | 148 | 0 | 0 | 0 | 0 | 15425 | 0 | 118 |
| SRR7644114 | 0 | 0 | 0 | 0 | 126 | 4159 | 4798 | 15995 | 2559 | 0 | 31 | 0 | 0 | 0 | 4798 | 144793 | 249365 | 0 | 0 | 0 | 0 | 0 | 31 | 0 | 252 | 252 | 31 | 157 | 0 | 220 | 0 | 0 | 0 | 0 | 22015 | 0 | 157 |
| SRR7644115 | 0 | 0 | 0 | 0 | 149 | 4356 | 5601 | 14935 | 1556 | 0 | 59 | 0 | 0 | 0 | 6223 | 139012 | 253515 | 0 | 0 | 622 | 0 | 0 | 59 | 0 | 208 | 12 | 119 | 89 | 0 | 208 | 0 | 0 | 0 | 0 | 26115 | 0 | 59 |
| SRR7644116 | 0 | 0 | 0 | 0 | 156 | 3895 | 6816 | 12334 | 3246 | 0 | 0 | 0 | 0 | 0 | 5518 | 153465 | 257013 | 0 | 0 | 0 | 0 | 62 | 94 | 31 | 62 | 156 | 62 | 125 | 0 | 343 | 0 | 0 | 0 | 0 | 20215 | 0 | 250 |
| SRR7644117 | 0 | 0 | 0 | 0 | 27 | 4587 | 3670 | 13762 | 1376 | 0 | 53 | 53 | 0 | 0 | 5046 | 126271 | 173305 | 0 | 0 | 1835 | 459 | 0 | 0 | 53 | 80 | 106 | 0 | 80 | 0 | 212 | 0 | 0 | 0 | 0 | 16525 | 0 | 53 |
| SRR7644118 | 0 | 0 | 0 | 0 | 57 | 4482 | 4980 | 11454 | 3984 | 0 | 57 | 57 | 0 | 0 | 4482 | 110512 | 176101 | 0 | 0 | 1992 | 0 | 0 | 0 | 28 | 198 | 85 | 0 | 28 | 0 | 425 | 0 | 0 | 0 | 0 | 10782 | 0 | 113 |
| SRR7644119 | 0 | 0 | 0 | 0 | 27 | 4558 | 4558 | 13674 | 2279 | 0 | 53 | 53 | 0 | 0 | 10027 | 110522 | 169319 | 0 | 0 | 2735 | 0 | 0 | 0 | 80 | 133 | 159 | 27 | 106 | 0 | 451 | 0 | 0 | 0 | 0 | 10610 | 0 | 80 |
| SRR7644120 | 0 | 0 | 0 | 0 | 112 | 4006 | 4006 | 8513 | 3005 | 0 | 196 | 140 | 0 | 0 | 5508 | 115316 | 173642 | 0 | 0 | 1502 | 501 | 0 | 0 | 0 | 140 | 84 | 28 | 56 | 0 | 309 | 0 | 0 | 0 | 0 | 8459 | 0 | 196 |
| SRR7644121 | 0 | 0 | 0 | 0 | 52 | 3969 | 4820 | 14176 | 1701 | 0 | 52 | 0 | 0 | 0 | 146016 | 140467 | 338909 | 0 | 0 | 284 | 851 | 0 | 0 | 52 | 26 | 79 | 79 | 131 | 0 | 446 | 0 | 0 | 0 | 0 | 13749 | 0 | 184 |
| SRR7644122 | 0 | 0 | 0 | 0 | 57 | 4505 | 5405 | 21021 | 2402 | 0 | 0 | 0 | 0 | 0 | 157057 | 143667 | 337358 | 0 | 0 | 601 | 601 | 0 | 29 | 115 | 144 | 57 | 172 | 29 | 0 | 316 | 0 | 0 | 0 | 0 | 16757 | 0 | 201 |
| SRR7644123 | 0 | 0 | 0 | 0 | 107 | 4433 | 5819 | 19119 | 3879 | 0 | 134 | 0 | 0 | 0 | 141313 | 139524 | 328482 | 0 | 0 | 277 | 1108 | 0 | 27 | 80 | 188 | 161 | 161 | 107 | 0 | 375 | 0 | 0 | 0 | 0 | 19635 | 0 | 80 |

### Supp Table S8.3.Columns_BX_to_DH

| | | | | | | | | | | | | | | | | | | | | | | | | | | | | | | | | | | | | | |
|---|---|---|---|---|---|---|---|---|---|---|---|---|---|---|---|---|---|---|---|---|---|---|---|---|---|---|---|---|---|---|---|---|---|---|---|---|---|
| SRR7644124 | 0 | 0 | 0 | 0 | 0 | 3847 | 5327 | 13909 | 1776 | 0 | 85 | 0 | 0 | 0 | 155963 | 142610 | 337955 | 0 | 0 | 0 | 296 | 0 | 0 | 141 | 57 | 113 | 85 | 85 | 0 | 368 | 0 | 0 | 0 | 0 | 17083 | 0 | 198 |
| SRR7644125 | 0 | 0 | 0 | 0 | 92 | 7521 | 14523 | 20488 | 5446 | 0 | 23 | 0 | 0 | 0 | 4409 | 114981 | 120615 | 0 | 0 | 259 | 259 | 0 | 0 | 435 | 298 | 321 | 23 | 114 | 0 | 733 | 0 | 0 | 0 | 0 | 11268 | 0 | 0 |
| SRR7644126 | 0 | 0 | 0 | 0 | 99 | 9291 | 17737 | 19989 | 5068 | 0 | 223 | 25 | 0 | 0 | 2252 | 123233 | 138170 | 0 | 0 | 563 | 282 | 0 | 0 | 11 | 173 | 12 | 50 | 12 | 0 | 22 | 0 | 0 | 0 | 0 | 10670 | 0 | 0 |
| SRR7644128 | 0 | 0 | 0 | 0 | 184 | 9160 | 11515 | 24339 | 5496 | 0 | 230 | 46 | 0 | 0 | 2617 | 122781 | 115529 | 0 | 0 | 785 | 0 | 0 | 0 | 161 | 230 | 460 | 46 | 253 | 0 | 667 | 0 | 0 | 0 | 0 | 10253 | 0 | 0 |
| SRR7644129 | 0 | 0 | 0 | 0 | 50 | 11611 | 15293 | 19258 | 4248 | 0 | 173 | 50 | 0 | 0 | 3682 | 132492 | 134595 | 0 | 0 | 0 | 0 | 0 | 0 | 198 | 198 | 297 | 25 | 124 | 0 | 818 | 0 | 0 | 0 | 0 | 10515 | 0 | 0 |
| SRR7663662 | 0 | 0 | 0 | 0 | 0 | 0 | 0 | 0 | 0 | 0 | 30 | 0 | 0 | 0 | 65 | 510 | 255 | 33 | 0 | 141 | 22 | 0 | 0 | 0 | 0 | 0 | 0 | 0 | 0 | 19 | 0 | 1023 | 19 | 0 | 7827 | 0 | 175 |
| SRR7663663 | 0 | 0 | 0 | 0 | 13 | 0 | 0 | 0 | 0 | 0 | 82 | 14 | 0 | 0 | 670 | 4117 | 5846 | 31 | 16 | 218 | 327 | 0 | 0 | 0 | 0 | 0 | 0 | 0 | 0 | 275 | 0 | 1417 | 0 | 0 | 5495 | 0 | 329 |
| SRR7664346 | 16 | 0 | 0 | 0 | 145 | 717 | 775 | 2257 | 463 | 13 | 0 | 0 | 0 | 0 | 62525 | 130250 | 194739 | 21 | 102 | 27908 | 1401 | 0 | 0 | 365 | 19 | 30 | 31 | 0 | 0 | 34 | 0 | 0 | 0 | 0 | 13624 | 0 | 0 |
| SRR7664347 | 14 | 0 | 0 | 0 | 105 | 1443 | 1879 | 2782 | 502 | 19 | 0 | 0 | 0 | 0 | 29525 | 94428 | 110066 | 13 | 34 | 11708 | 721 | 0 | 0 | 419 | 10 | 21 | 20 | 0 | 0 | 38 | 0 | 0 | 0 | 0 | 15881 | 0 | 0 |
| SRR7690697 | 0 | 0 | 0 | 31 | 116 | 408 | 273 | 700 | 184 | 15 | 153 | 0 | 0 | 0 | 12510 | 22492 | 142467 | 0 | 91 | 60185 | 803 | 0 | 0 | 277 | 168 | 526 | 2265 | 1011 | 0 | 1174 | 0 | 45 | 0 | 0 | 2821 | 1200 | 0 |
| SRR7690698 | 0 | 0 | 0 | 24 | 22 | 290 | 183 | 467 | 113 | 0 | 37 | 0 | 0 | 0 | 4159 | 15099 | 66739 | 0 | 116 | 89107 | 238 | 0 | 0 | 40 | 53 | 268 | 1127 | 579 | 0 | 224 | 0 | 20 | 0 | 0 | 1922 | 1153 | 0 |
| SRR7690699 | 0 | 0 | 0 | 38 | 126 | 419 | 263 | 754 | 152 | 0 | 136 | 21 | 0 | 0 | 14853 | 29608 | 120132 | 0 | 105 | 79328 | 1704 | 0 | 0 | 429 | 202 | 697 | 2740 | 1246 | 0 | 917 | 0 | 38 | 0 | 0 | 2331 | 560 | 0 |
| SRR7690700 | 34 | 15 | 0 | 0 | 318 | 9440 | 6607 | 24844 | 33674 | 1045 | 141 | 22 | 0 | 0 | 20813 | 86891 | 194587 | 3601 | 935 | 21951 | 855 | 0 | 0 | 190 | 186 | 88 | 61 | 0 | 0 | 2854 | 0 | 20 | 0 | 0 | 21680 | 9372 | 0 |
| SRR7690701 | 24 | 13 | 0 | 0 | 154 | 10382 | 12765 | 37291 | 72443 | 723 | 25 | 24 | 0 | 0 | 10847 | 120434 | 124505 | 4148 | 696 | 18746 | 609 | 0 | 0 | 155 | 97 | 42 | 24 | 0 | 0 | 645 | 0 | 28 | 0 | 0 | 29637 | 7011 | 0 |
| SRR7690702 | 23 | 0 | 0 | 0 | 251 | 15638 | 12132 | 36876 | 71125 | 495 | 101 | 26 | 0 | 0 | 10030 | 102090 | 112704 | 2732 | 716 | 15362 | 566 | 0 | 0 | 138 | 317 | 151 | 53 | 0 | 0 | 2234 | 0 | 23 | 0 | 0 | 35080 | 8959 | 0 |
| SRR7690703 | 13 | 0 | 0 | 0 | 175 | 11762 | 11014 | 31117 | 73976 | 4032 | 22 | 36 | 0 | 0 | 31268 | 102268 | 247296 | 2112 | 958 | 5845 | 590 | 0 | 0 | 124 | 147 | 45 | 11 | 0 | 0 | 880 | 0 | 180 | 0 | 0 | 35418 | 6308 | 0 |
| SRR7690704 | 40 | 17 | 0 | 0 | 60 | 11350 | 6885 | 26698 | 34676 | 179 | 31 | 27 | 0 | 0 | 13794 | 90918 | 117666 | 2684 | 633 | 15919 | 504 | 0 | 0 | 27 | 22 | 23 | 22 | 0 | 0 | 184 | 0 | 49 | 0 | 18 | 43932 | 9454 | 0 |
| SRR7690705 | 15 | 0 | 0 | 0 | 226 | 17928 | 13741 | 46619 | 55953 | 974 | 0 | 23 | 0 | 0 | 5403 | 68167 | 63391 | 2714 | 251 | 3851 | 214 | 0 | 0 | 120 | 77 | 19 | 0 | 0 | 0 | 394 | 0 | 43 | 0 | 0 | 73038 | 8196 | 0 |
| SRR7690706 | 31 | 0 | 0 | 0 | 217 | 27963 | 14005 | 55715 | 65549 | 5135 | 56 | 29 | 0 | 0 | 4508 | 99126 | 99436 | 2149 | 300 | 9677 | 210 | 0 | 0 | 120 | 160 | 51 | 12 | 0 | 0 | 1924 | 0 | 48 | 0 | 18 | 55231 | 13305 | 0 |
| SRR7690707 | 0 | 0 | 0 | 46 | 88 | 369 | 213 | 426 | 203 | 0 | 54 | 0 | 0 | 0 | 5343 | 11882 | 55420 | 0 | 38 | 30702 | 397 | 0 | 0 | 205 | 129 | 736 | 6841 | 4378 | 0 | 419 | 0 | 203 | 0 | 0 | 2452 | 1421 | 0 |
| SRR7690708 | 33 | 16 | 0 | 0 | 263 | 21714 | 11231 | 41147 | 49168 | 1984 | 146 | 31 | 0 | 0 | 7766 | 74857 | 103003 | 1411 | 380 | 6950 | 353 | 0 | 0 | 123 | 175 | 66 | 45 | 0 | 0 | 2290 | 0 | 14 | 0 | 21 | 55893 | 12974 | 0 |
| SRR7690713 | 66 | 18 | 0 | 0 | 362 | 21962 | 8243 | 34959 | 44951 | 2071 | 168 | 28 | 0 | 0 | 12548 | 94816 | 272097 | 1520 | 708 | 13027 | 289 | 0 | 0 | 220 | 240 | 97 | 72 | 0 | 0 | 3879 | 0 | 15 | 0 | 17 | 29634 | 10856 | 0 |
| SRR7690714 | 35 | 17 | 0 | 0 | 608 | 23186 | 12954 | 49736 | 64626 | 2351 | 102 | 24 | 0 | 0 | 4887 | 79864 | 113936 | 933 | 256 | 3043 | 93 | 0 | 0 | 241 | 409 | 130 | 80 | 0 | 0 | 2924 | 0 | 0 | 0 | 14 | 61847 | 12158 | 0 |
| SRR7690715 | 0 | 0 | 0 | 15 | 30 | 592 | 412 | 1034 | 337 | 0 | 43 | 0 | 0 | 0 | 2374 | 12599 | 49092 | 15 | 0 | 20949 | 345 | 0 | 0 | 31 | 40 | 167 | 926 | 652 | 0 | 231 | 0 | 165 | 0 | 0 | 6806 | 1332 | 0 |
| SRR7690716 | 0 | 0 | 0 | 34 | 20 | 206 | 145 | 409 | 76 | 15 | 30 | 13 | 0 | 0 | 3057 | 9929 | 41732 | 0 | 90 | 47280 | 242 | 0 | 0 | 47 | 23 | 143 | 1112 | 781 | 0 | 141 | 0 | 25 | 0 | 0 | 2366 | 1027 | 0 |
| SRR7690717 | 25 | 0 | 0 | 0 | 452 | 12765 | 12976 | 56018 | 108305 | 3124 | 60 | 53 | 0 | 0 | 8006 | 140008 | 136345 | 3995 | 457 | 8513 | 573 | 0 | 20 | 280 | 347 | 63 | 36 | 0 | 0 | 1385 | 0 | 71 | 0 | 17 | 25980 | 8134 | 0 |
| SRR7690718 | 0 | 0 | 0 | 0 | 251 | 22043 | 16086 | 56710 | 91530 | 415 | 81 | 64 | 0 | 0 | 3083 | 96178 | 109841 | 2949 | 368 | 10267 | 275 | 0 | 0 | 176 | 449 | 165 | 63 | 0 | 0 | 1371 | 0 | 26 | 0 | 12 | 21331 | 5608 | 0 |
| SRR7690719 | 10 | 0 | 0 | 0 | 86 | 11870 | 8170 | 36439 | 56484 | 194 | 16 | 39 | 0 | 0 | 23424 | 98161 | 200320 | 3413 | 387 | 8020 | 579 | 0 | 0 | 40 | 52 | 12 | 27 | 0 | 0 | 588 | 0 | 71 | 0 | 15 | 21816 | 9533 | 0 |
| SRR7690720 | 11 | 18 | 0 | 0 | 43 | 18512 | 9003 | 38558 | 45945 | 414 | 18 | 32 | 0 | 0 | 13411 | 55862 | 169278 | 2301 | 886 | 9405 | 328 | 0 | 0 | 62 | 39 | 48 | 49 | 0 | 0 | 395 | 0 | 0 | 0 | 27 | 44689 | 17129 | 0 |
| SRR7690721 | 32 | 11 | 0 | 0 | 63 | 17813 | 10005 | 40082 | 45387 | 551 | 52 | 14 | 0 | 0 | 18493 | 50164 | 207873 | 2184 | 1673 | 9749 | 487 | 0 | 0 | 37 | 31 | 12 | 13 | 0 | 0 | 799 | 0 | 0 | 0 | 18 | 18706 | 14561 | 0 |
| SRR7690722 | 27 | 18 | 26 | 41 | 75 | 787 | 175 | 782 | 2271 | 91 | 129 | 124 | 0 | 0 | 13310 | 186016 | 289806 | 291 | 1634 | 12152 | 3645 | 0 | 0 | 26 | 109 | 86 | 170 | 15 | 34 | 16 | 0 | 15 | 0 | 47 | 29501 | 2418 | 0 |
| SRR7690723 | 243 | 161 | 0 | 0 | 802 | 2425 | 514 | 2682 | 7234 | 1357 | 52 | 80 | 0 | 0 | 3924 | 143925 | 133122 | 428 | 642 | 2640 | 685 | 0 | 12 | 445 | 37 | 27 | 55 | 0 | 20 | 176 | 0 | 29 | 0 | 57 | 80626 | 1937 | 0 |
| SRR7690724 | 28 | 16 | 25 | 20 | 27 | 993 | 107 | 1126 | 1158 | 576 | 84 | 164 | 0 | 0 | 46362 | 266854 | 442979 | 292 | 1025 | 6044 | 5307 | 84 | 119 | 25 | 47 | 25 | 102 | 0 | 17 | 16 | 17 | 32 | 0 | 82 | 8841 | 3657 | 0 |
| SRR7690726 | 474 | 229 | 12 | 22 | 943 | 695 | 203 | 893 | 2260 | 3321 | 95 | 79 | 0 | 0 | 34805 | 200126 | 459643 | 571 | 1603 | 11600 | 3593 | 0 | 0 | 546 | 198 | 136 | 132 | 0 | 0 | 338 | 0 | 15 | 10 | 53 | 13699 | 2766 | 0 |
| SRR7690727 | 38 | 0 | 0 | 0 | 291 | 16412 | 14914 | 56133 | 77953 | 921 | 34 | 35 | 0 | 0 | 8787 | 100330 | 173682 | 2612 | 590 | 13978 | 569 | 0 | 0 | 198 | 260 | 75 | 26 | 0 | 0 | 965 | 0 | 15 | 0 | 0 | 23367 | 3642 | 0 |
| SRR7690728 | 32 | 0 | 0 | 0 | 247 | 19319 | 22445 | 60123 | 111787 | 953 | 12 | 49 | 0 | 0 | 6198 | 88465 | 99185 | 2344 | 371 | 10669 | 566 | 0 | 0 | 174 | 126 | 44 | 18 | 0 | 0 | 455 | 0 | 29 | 0 | 0 | 29487 | 1785 | 0 |
| SRR7690729 | 18 | 37 | 0 | 0 | 115 | 19201 | 12151 | 49935 | 66073 | 220 | 14 | 27 | 0 | 0 | 6535 | 82188 | 128128 | 3317 | 618 | 17826 | 936 | 0 | 0 | 51 | 35 | 39 | 15 | 0 | 0 | 96 | 0 | 40 | 0 | 0 | 36383 | 8542 | 0 |
| SRR7690730 | 38 | 19 | 0 | 0 | 322 | 15011 | 9400 | 30865 | 38877 | 977 | 11 | 32 | 0 | 0 | 2595 | 31169 | 53662 | 876 | 182 | 8251 | 296 | 0 | 0 | 321 | 330 | 79 | 58 | 0 | 0 | 280 | 0 | 34 | 0 | 0 | 46460 | 10301 | 0 |
| SRR7690731 | 27 | 11 | 0 | 0 | 150 | 21616 | 11757 | 44318 | 65232 | 1909 | 23 | 39 | 0 | 0 | 5671 | 85637 | 91887 | 3693 | 549 | 18201 | 503 | 0 | 0 | 137 | 86 | 19 | 0 | 0 | 0 | 669 | 0 | 30 | 0 | 11 | 42470 | 11873 | 0 |
| SRR7690732 | 18 | 10 | 0 | 0 | 95 | 23770 | 10276 | 35173 | 39616 | 189 | 29 | 27 | 0 | 0 | 13369 | 52423 | 234803 | 1095 | 349 | 7085 | 670 | 0 | 0 | 45 | 39 | 13 | 15 | 0 | 0 | 684 | 0 | 11 | 0 | 13 | 28506 | 14779 | 0 |
| SRR7690733 | 40 | 15 | 0 | 0 | 59 | 31171 | 12185 | 45961 | 58002 | 172 | 18 | 15 | 0 | 0 | 7602 | 43850 | 88358 | 1662 | 494 | 3235 | 306 | 0 | 0 | 30 | 20 | 17 | 12 | 0 | 0 | 407 | 0 | 0 | 0 | 15 | 47709 | 12551 | 0 |
| SRR7690734 | 175 | 80 | 0 | 0 | 429 | 4902 | 647 | 4743 | 6527 | 1626 | 163 | 150 | 0 | 0 | 14428 | 220898 | 135500 | 1189 | 4175 | 7056 | 1995 | 0 | 0 | 462 | 47 | 19 | 32 | 0 | 43 | 404 | 0 | 330 | 0 | 37 | 16664 | 4010 | 0 |
| SRR7690736 | 657 | 347 | 18 | 20 | 2660 | 515 | 206 | 618 | 5605 | 1003 | 118 | 138 | 0 | 0 | 19345 | 282087 | 245804 | 559 | 1611 | 11570 | 1611 | 0 | 16 | 592 | 225 | 200 | 311 | 0 | 11 | 374 | 0 | 0 | 0 | 46 | 22138 | 2832 | 0 |
| SRR7690737 | 515 | 218 | 0 | 0 | 1777 | 916 | 283 | 946 | 2584 | 3316 | 14 | 11 | 0 | 0 | 23026 | 188797 | 307740 | 700 | 730 | 7358 | 4468 | 0 | 17 | 979 | 93 | 20 | 28 | 0 | 0 | 127 | 0 | 60 | 0 | 54 | 37471 | 3007 | 0 |
| SRR7690739 | 723 | 326 | 45 | 36 | 2014 | 2133 | 323 | 1234 | 3342 | 3895 | 51 | 18 | 0 | 0 | 13140 | 270664 | 302610 | 222 | 760 | 3323 | 2450 | 0 | 0 | 832 | 181 | 172 | 368 | 0 | 0 | 226 | 0 | 0 | 0 | 54 | 21520 | 2915 | 0 |
| SRR7690740 | 754 | 356 | 38 | 43 | 1927 | 1287 | 172 | 845 | 2319 | 3120 | 63 | 20 | 0 | 0 | 13666 | 279408 | 249164 | 187 | 565 | 2192 | 2565 | 10 | 33 | 593 | 199 | 154 | 262 | 23 | 0 | 272 | 10 | 0 | 0 | 13 | 19187 | 2841 | 0 |
| SRR7690741 | 0 | 0 | 0 | 43 | 18 | 539 | 522 | 1044 | 266 | 0 | 18 | 0 | 0 | 0 | 5820 | 13535 | 79979 | 0 | 90 | 71450 | 460 | 0 | 0 | 73 | 28 | 107 | 536 | 302 | 0 | 130 | 0 | 17 | 0 | 0 | 4113 | 1266 | 0 |
| SRR7690742 | 0 | 0 | 0 | 39 | 14 | 501 | 306 | 894 | 240 | 0 | 14 | 0 | 0 | 0 | 6339 | 10451 | 73458 | 0 | 108 | 66948 | 574 | 0 | 0 | 46 | 36 | 146 | 625 | 270 | 0 | 99 | 0 | 13 | 0 | 0 | 4492 | 1243 | 0 |
| SRR7690743 | 0 | 0 | 0 | 38 | 113 | 680 | 366 | 1161 | 370 | 0 | 40 | 0 | 0 | 0 | 3682 | 8328 | 54649 | 0 | 89 | 44839 | 689 | 0 | 0 | 292 | 77 | 393 | 2622 | 1055 | 0 | 330 | 0 | 38 | 0 | 0 | 6599 | 980 | 0 |
| SRR7690744 | 11 | 16 | 0 | 0 | 92 | 13358 | 8217 | 24696 | 40247 | 12 | 29 | 0 | 0 | 0 | 7246 | 56627 | 96615 | 2818 | 1585 | 53306 | 543 | 0 | 0 | 62 | 67 | 43 | 36 | 0 | 0 | 414 | 0 | 47 | 0 | 0 | 38050 | 11640 | 0 |
| SRR7690745 | 39 | 12 | 0 | 0 | 51 | 13653 | 7682 | 24325 | 45707 | 15 | 12 | 22 | 0 | 0 | 10496 | 68587 | 104931 | 3171 | 1230 | 21940 | 572 | 0 | 0 | 44 | 31 | 67 | 50 | 0 | 0 | 254 | 0 | 38 | 0 | 11 | 44422 | 14211 | 0 |
| SRR7690746 | 27 | 0 | 0 | 0 | 95 | 10460 | 6762 | 28012 | 69755 | 28 | 24 | 19 | 0 | 0 | 7048 | 82719 | 64160 | 1699 | 628 | 11307 | 1164 | 0 | 0 | 38 | 84 | 58 | 14 | 0 | 0 | 177 | 0 | 35 | 0 | 0 | 112690 | 9131 | 0 |
| SRR7690747 | 20 | 0 | 0 | 0 | 47 | 15259 | 8613 | 35575 | 41274 | 96 | 23 | 12 | 0 | 0 | 19809 | 72816 | 168039 | 4352 | 709 | 11982 | 633 | 0 | 0 | 54 | 31 | 24 | 12 | 0 | 0 | 120 | 0 | 35 | 0 | 0 | 67489 | 11537 | 0 |
| SRR7690748 | 68 | 11 | 0 | 0 | 298 | 15403 | 8854 | 32498 | 57473 | 2507 | 130 | 55 | 0 | 0 | 5234 | 98107 | 84003 | 2080 | 478 | 5184 | 389 | 0 | 0 | 105 | 210 | 89 | 26 | 0 | 0 | 2845 | 0 | 21 | 0 | 12 | 62369 | 13082 | 0 |
| SRR7690749 | 0 | 0 | 0 | 0 | 384 | 25044 | 11772 | 23824 | 39566 | 1556 | 107 | 16 | 0 | 0 | 6835 | 68982 | 92579 | 1172 | 154 | 2613 | 150 | 0 | 0 | 198 | 243 | 156 | 53 | 0 | 0 | 2329 | 0 | 12 | 0 | 0 | 43138 | 17527 | 0 |
| SRR7690750 | 37 | 0 | 0 | 0 | 300 | 9669 | 10507 | 32385 | 63522 | 1014 | 67 | 36 | 0 | 0 | 8726 | 104624 | 149103 | 4180 | 780 | 25326 | 582 | 0 | 0 | 175 | 220 | 120 | 46 | 0 | 0 | 1887 | 0 | 0 | 0 | 0 | 23611 | 3543 | 0 |
| SRR7690751 | 17 | 0 | 0 | 0 | 81 | 11076 | 11073 | 32318 | 59355 | 367 | 27 | 26 | 0 | 0 | 11719 | 105882 | 174883 | 4357 | 764 | 22328 | 862 | 0 | 0 | 46 | 64 | 47 | 24 | 0 | 0 | 573 | 0 | 19 | 0 | 0 | 29659 | 6879 | 0 |
| SRR7690752 | 26 | 14 | 0 | 0 | 43 | 15862 | 11096 | 43509 | 66761 | 492 | 21 | 14 | 0 | 0 | 11113 | 93956 | 150941 | 3718 | 486 | 13563 | 1106 | 0 | 0 | 53 | 16 | 20 | 14 | 0 | 0 | 138 | 0 | 17 | 0 | 0 | 55589 | 8879 | 0 |
| SRR7690753 | 12 | 17 | 0 | 0 | 68 | 15456 | 11786 | 34710 | 65683 | 539 | 68 | 23 | 0 | 0 | 13541 | 77852 | 149310 | 1906 | 953 | 12479 | 375 | 0 | 0 | 50 | 85 | 64 | 46 | 0 | 0 | 955 | 0 | 0 | 0 | 15 | 33693 | 15181 | 0 |
| SRR7690754 | 16 | 24 | 0 | 0 | 82 | 18444 | 12121 | 44552 | 61035 | 541 | 41 | 27 | 0 | 0 | 4170 | 57556 | 58966 | 1258 | 600 | 4055 | 185 | 0 | 0 | 56 | 79 | 25 | 23 | 0 | 0 | 600 | 0 | 32 | 0 | 11 | 58224 | 14294 | 0 |
| SRR7690755 | 57 | 18 | 10 | 14 | 210 | 17524 | 8698 | 40588 | 55687 | 1718 | 351 | 16 | 0 | 0 | 5882 | 38538 | 75652 | 1550 | 1017 | 6515 | 284 | 0 | 0 | 270 | 203 | 69 | 69 | 0 | 0 | 3007 | 0 | 0 | 0 | 27 | 22616 | 17086 | 0 |
| SRR7690756 | 36 | 11 | 0 | 0 | 38 | 14051 | 5641 | 26794 | 34284 | 342 | 33 | 21 | 0 | 0 | 15492 | 57664 | 197196 | 2238 | 685 | 12661 | 828 | 0 | 0 | 69 | 23 | 18 | 34 | 0 | 0 | 220 | 0 | 20 | 0 | 24 | 37741 | 14819 | 0 |
| SRR7690757 | 36 | 19 | 0 | 12 | 146 | 16307 | 6481 | 26341 | 34005 | 1661 | 236 | 19 | 0 | 0 | 21014 | 45985 | 115052 | 2331 | 963 | 14004 | 1284 | 0 | 0 | 201 | 203 | 128 | 79 | 0 | 0 | 3512 | 0 | 17 | 0 | 0 | 30048 | 14222 | 0 |
| SRR7690758 | 28 | 335 | 21 | 0 | 36 | 310 | 67 | 402 | 627 | 181 | 52 | 0 | 0 | 0 | 49485 | 289172 | 546242 | 76 | 383 | 2567 | 6867 | 49 | 21 | 20 | 122 | 136 | 192 | 0 | 0 | 164 | 28 | 0 | 0 | 17 | 3861 | 2160 | 0 |
| SRR7690759 | 64 | 38 | 25 | 29 | 179 | 415 | 183 | 590 | 4253 | 83 | 12 | 10 | 0 | 0 | 10700 | 225975 | 167381 | 706 | 1238 | 12959 | 3489 | 0 | 27 | 47 | 11 | 12 | 20 | 0 | 18 | 20 | 0 | 0 | 0 | 48 | 44907 | 1249 | 0 |
| SRR7690762 | 25 | 11 | 31 | 37 | 49 | 1516 | 197 | 1092 | 2957 | 130 | 88 | 43 | 0 | 0 | 14330 | 210075 | 295149 | 197 | 796 | 2752 | 1645 | 0 | 11 | 20 | 114 | 105 | 205 | 0 | 17 | 11 | 0 | 0 | 0 | 48 | 34130 | 2820 | 0 |
| SRR7690764 | 0 | 0 | 0 | 16 | 29 | 168 | 102 | 289 | 57 | 11 | 23 | 11 | 0 | 0 | 6728 | 10599 | 68380 | 0 | 99 | 54849 | 872 | 0 | 0 | 84 | 47 | 239 | 2291 | 1614 | 0 | 124 | 0 | 63 | 0 | 0 | 2292 | 1729 | 0 |
| SRR7690765 | 0 | 0 | 0 | 34 | 112 | 330 | 228 | 743 | 117 | 0 | 64 | 0 | 0 | 0 | 5249 | 8563 | 58570 | 0 | 121 | 56907 | 1034 | 0 | 0 | 182 | 124 | 502 | 4340 | 3018 | 0 | 422 | 0 | 15 | 0 | 0 | 3910 | 1088 | 0 |
| SRR7690766 | 23 | 0 | 0 | 0 | 45 | 16413 | 11825 | 47064 | 78701 | 443 | 27 | 32 | 0 | 0 | 14900 | 88823 | 308066 | 3716 | 844 | 11472 | 726 | 0 | 0 | 39 | 85 | 34 | 19 | 0 | 0 | 467 | 0 | 24 | 0 | 16 | 15412 | 8538 | 0 |
| SRR7690767 | 20 | 0 | 0 | 0 | 260 | 14413 | 16493 | 54140 | 103656 | 746 | 74 | 35 | 0 | 0 | 4351 | 122426 | 115032 | 4625 | 629 | 20226 | 351 | 0 | 0 | 196 | 215 | 113 | 54 | 0 | 0 | 1574 | 0 | 80 | 0 | 11 | 24671 | 7238 | 0 |
| SRR7690768 | 39 | 14 | 0 | 0 | 54 | 21241 | 10144 | 36623 | 47055 | 337 | 12 | 27 | 0 | 0 | 11117 | 66398 | 214628 | 1703 | 448 | 7224 | 615 | 0 | 0 | 40 | 17 | 47 | 42 | 0 | 0 | 370 | 0 | 13 | 0 | 23 | 37192 | 13610 | 0 |
| SRR7690769 | 19 | 180 | 16 | 18 | 58 | 2916 | 379 | 2712 | 8981 | 24 | 114 | 71 | 0 | 0 | 5671 | 155832 | 257053 | 190 | 1181 | 3951 | 802 | 0 | 0 | 33 | 107 | 119 | 13 | 0 | 0 | 17 | 0 | 29 | 0 | 52 | 43251 | 2247 | 0 |
| SRR7690770 | 648 | 351 | 27 | 20 | 1746 | 1096 | 184 | 958 | 2882 | 1340 | 138 | 78 | 0 | 0 | 12242 | 170469 | 338755 | 373 | 2127 | 24388 | 3748 | 0 | 13 | 954 | 191 | 126 | 222 | 0 | 13 | 566 | 0 | 14 | 0 | 95 | 15159 | 2806 | 0 |
| SRR7690773 | 23 | 10 | 0 | 11 | 25 | 2321 | 600 | 3332 | 7185 | 61 | 19 | 49 | 0 | 0 | 19983 | 155299 | 396295 | 418 | 963 | 3545 | 3245 | 0 | 0 | 27 | 76 | 57 | 72 | 0 | 15 | 140 | 0 | 39 | 11 | 11 | 51313 | 3571 | 0 |
| SRR7690775 | 19 | 0 | 0 | 0 | 393 | 30199 | 19123 | 95989 | 98108 | 61 | 215 | 12 | 0 | 0 | 5063 | 59050 | 111854 | 2195 | 880 | 9903 | 177 | 0 | 0 | 227 | 263 | 53 | 45 | 0 | 0 | 4189 | 0 | 10 | 0 | 12 | 27426 | 16221 | 0 |
| SRR7690777 | 45 | 0 | 0 | 0 | 334 | 12034 | 5245 | 21688 | 20610 | 50 | 69 | 0 | 0 | 0 | 49835 | 147505 | 324940 | 2994 | 1484 | 8379 | 1575 | 0 | 0 | 273 | 144 | 55 | 63 | 0 | 0 | 2067 | 0 | 27 | 0 | 0 | 32196 | 18901 | 0 |
| SRR7690778 | 0 | 0 | 0 | 0 | 214 | 10110 | 8059 | 26402 | 54527 | 518 | 113 | 18 | 0 | 0 | 2273 | 94783 | 33050 | 2020 | 364 | 9055 | 103 | 0 | 0 | 42 | 122 | 54 | 17 | 0 | 0 | 2841 | 0 | 16 | 0 | 0 | 19402 | 15913 | 0 |
| SRR7690779 | 36 | 13 | 0 | 13 | 165 | 9122 | 4918 | 21714 | 38837 | 2728 | 251 | 12 | 0 | 0 | 9776 | 31365 | 57250 | 1077 | 750 | 2563 | 438 | 0 | 0 | 86 | 122 | 69 | 18 | 0 | 0 | 1010 | 0 | 0 | 0 | 16 | 32693 | 11212 | 0 |
| SRR7690780 | 34 | 0 | 0 | 0 | 59 | 19714 | 10619 | 39067 | 54647 | 327 | 25 | 24 | 0 | 0 | 8478 | 55417 | 118371 | 2198 | 757 | 14744 | 435 | 0 | 0 | 100 | 74 | 19 | 16 | 0 | 0 | 601 | 0 | 0 | 0 | 19 | 46284 | 11336 | 0 |
| SRR7690781 | 13 | 32 | 0 | 0 | 93 | 5357 | 3403 | 13873 | 24502 | 660 | 32 | 39 | 0 | 0 | 30463 | 81948 | 310260 | 1856 | 1300 | 41720 | 742 | 0 | 0 | 36 | 50 | 24 | 19 | 0 | 0 | 631 | 0 | 18 | 0 | 27 | 20964 | 10345 | 0 |
| SRR7690783 | 680 | 390 | 21 | 38 | 1116 | 2545 | 290 | 1060 | 3976 | 1646 | 167 | 59 | 0 | 0 | 8903 | 198494 | 186945 | 365 | 736 | 9059 | 1688 | 0 | 0 | 578 | 161 | 161 | 362 | 15 | 0 | 483 | 0 | 0 | 0 | 19 | 37409 | 2704 | 0 |
| SRR7690788 | 34 | 26 | 40 | 35 | 87 | 1344 | 205 | 1000 | 3607 | 475 | 17 | 65 | 0 | 0 | 20258 | 149860 | 318199 | 545 | 1597 | 11840 | 3026 | 0 | 0 | 72 | 27 | 22 | 36 | 11 | 0 | 44 | 0 | 0 | 0 | 50 | 19793 | 3306 | 0 |
| SRR7690789 | 55 | 21 | 88 | 78 | 53 | 73 | 22 | 122 | 411 | 266 | 17 | 128 | 0 | 0 | 31776 | 270045 | 500004 | 292 | 455 | 2348 | 7601 | 0 | 0 | 19 | 174 | 151 | 19 | 15 | 0 | 34 | 0 | 0 | 0 | 46 | 2721 | 3141 | 0 |
| SRR7690790 | 66 | 0 | 0 | 0 | 686 | 10278 | 7186 | 33368 | 48011 | 44 | 581 | 14 | 0 | 0 | 15363 | 63752 | 209869 | 2282 | 2043 | 28418 | 372 | 0 | 0 | 172 | 591 | 203 | 171 | 0 | 0 | 10164 | 0 | 0 | 0 | 0 | 11381 | 18161 | 0 |
| SRR7690791 | 36 | 0 | 0 | 0 | 481 | 15006 | 10198 | 28605 | 39137 | 85 | 156 | 13 | 0 | 0 | 12957 | 199752 | 137896 | 2756 | 1044 | 7069 | 641 | 0 | 0 | 304 | 267 | 139 | 121 | 0 | 0 | 2883 | 0 | 0 | 0 | 0 | 50166 | 4554 | 0 |
| SRR7690792 | 26 | 12 | 0 | 0 | 236 | 4692 | 2453 | 8328 | 10716 | 79 | 65 | 19 | 0 | 0 | 112089 | 101392 | 573795 | 1778 | 1470 | 10931 | 6443 | 0 | 0 | 260 | 134 | 66 | 73 | 0 | 0 | 1360 | 0 | 26 | 0 | 39 | 13251 | 17850 | 0 |
| SRR7690793 | 19 | 0 | 0 | 0 | 420 | 13797 | 11732 | 40780 | 98844 | 1227 | 101 | 31 | 0 | 0 | 3382 | 95061 | 51906 | 2495 | 322 | 9458 | 228 | 0 | 0 | 162 | 209 | 94 | 63 | 0 | 0 | 2178 | 0 | 44 | 0 | 12 | 36118 | 15304 | 0 |
| SRR7690794 | 0 | 0 | 0 | 0 | 259 | 10742 | 5984 | 13185 | 24632 | 1682 | 12 | 14 | 0 | 0 | 8032 | 76864 | 77045 | 1336 | 101 | 4300 | 98 | 0 | 0 | 50 | 32 | 12 | 21 | 0 | 0 | 425 | 0 | 41 | 0 | 12 | 55063 | 8484 | 0 |
| SRR7690795 | 0 | 0 | 0 | 0 | 261 | 19779 | 10517 | 20618 | 35220 | 780 | 174 | 17 | 0 | 0 | 1494 | 110907 | 27616 | 1212 | 317 | 4525 | 88 | 0 | 0 | 33 | 113 | 99 | 94 | 12 | 0 | 2289 | 0 | 12 | 0 | 0 | 40295 | 8871 | 0 |
| SRR7690796 | 43 | 11 | 0 | 0 | 306 | 14001 | 6870 | 29437 | 44785 | 2636 | 152 | 36 | 0 | 0 | 12195 | 79513 | 208675 | 1866 | 903 | 27853 | 215 | 0 | 0 | 238 | 270 | 124 | 130 | 0 | 0 | 2864 | 0 | 18 | 0 | 35 | 37584 | 12667 | 0 |
| SRR7690797 | 38 | 18 | 89 | 85 | 128 | 2665 | 435 | 1451 | 6346 | 214 | 44 | 37 | 0 | 0 | 5567 | 168582 | 147097 | 515 | 620 | 3364 | 356 | 0 | 0 | 23 | 188 | 262 | 17 | 0 | 15 | 12 | 0 | 26 | 0 | 0 | 77314 | 1918 | 0 |
| SRR7690798 | 26 | 14 | 31 | 31 | 49 | 2620 | 334 | 2286 | 6508 | 55 | 61 | 64 | 0 | 0 | 10221 | 213866 | 340920 | 349 | 714 | 4116 | 592 | 0 | 14 | 45 | 10 | 184 | 15 | 11 | 0 | 281 | 0 | 38 | 11 | 45 | 24040 | 2129 | 0 |
| SRR7690800 | 29 | 19 | 0 | 0 | 48 | 1574 | 259 | 1498 | 3924 | 142 | 107 | 89 | 0 | 0 | 5530 | 173425 | 133813 | 711 | 1434 | 2350 | 1175 | 0 | 0 | 40 | 83 | 100 | 10 | 0 | 24 | 20 | 0 | 43 | 0 | 51 | 98841 | 1902 | 0 |
| SRR7690801 | 478 | 199 | 21 | 13 | 1218 | 806 | 117 | 789 | 2197 | 2013 | 13 | 38 | 0 | 0 | 49603 | 181710 | 470433 | 760 | 1566 | 10546 | 7473 | 0 | 0 | 863 | 123 | 47 | 55 | 0 | 17 | 182 | 0 | 0 | 0 | 30 | 19572 | 3578 | 0 |
| SRR7690802 | 12 | 144 | 0 | 0 | 35 | 2433 | 747 | 2668 | 12942 | 31 | 22 | 37 | 0 | 0 | 4236 | 159924 | 103959 | 1304 | 440 | 4998 | 645 | 0 | 13 | 29 | 61 | 41 | 31 | 0 | 19 | 122 | 0 | 29 | 0 | 31 | 63843 | 2303 | 0 |
| SRR7690803 | 385 | 192 | 17 | 23 | 1188 | 3216 | 498 | 2283 | 9530 | 848 | 83 | 92 | 0 | 0 | 8253 | 214144 | 218153 | 471 | 942 | 11921 | 815 | 0 | 21 | 737 | 184 | 212 | 466 | 0 | 11 | 297 | 0 | 0 | 0 | 36 | 36724 | 2342 | 0 |
| SRR7819267 | 0 | 0 | 0 | 0 | 0 | 7579 | 330 | 9424 | 395 | 87 | 0 | 0 | 0 | 0 | 0 | 961 | 824 | 0 | 0 | 66 | 66 | 0 | 0 | 10 | 0 | 0 | 0 | 0 | 15 | 0 | 36 | 0 | 0 | 0 | 74272 | 12 | 0 |
| SRR7819269 | 0 | 0 | 0 | 0 | 0 | 13032 | 649 | 8904 | 1179 | 143 | 0 | 0 | 0 | 0 | 59 | 2016 | 864 | 0 | 0 | 0 | 118 | 0 | 0 | 285 | 0 | 0 | 29 | 0 | 29 | 0 | 57 | 0 | 0 | 0 | 18142 | 0 | 0 |
| SRR7819272 | 0 | 0 | 0 | 0 | 33 | 10086 | 620 | 5002 | 785 | 164 | 0 | 0 | 33 | 0 | 289 | 8731 | 3358 | 0 | 0 | 0 | 455 | 20 | 0 | 0 | 0 | 0 | 0 | 0 | 0 | 0 | 26 | 0 | 0 | 13 | 70103 | 0 | 13 |
| SRR7851651 | 0 | 0 | 0 | 0 | 0 | 2837 | 2837 | 4309 | 7005 | 0 | 0 | 0 | 0 | 0 | 953 | 601 | 1363 | 5527 | 97 | 200 | 54 | 1505 | 316 | 189 | 0 | 0 | 0 | 0 | 118 | 0 | 148 | 1608 | 960 | 22 | 10691 | 21 | 246 |
| SRR7851652 | 0 | 0 | 0 | 0 | 0 | 2652 | 1882 | 2940 | 5314 | 0 | 0 | 0 | 0 | 0 | 566 | 583 | 1088 | 10794 | 70 | 246 | 0 | 380 | 113 | 46 | 0 | 0 | 0 | 0 | 52 | 0 | 22 | 4377 | 371 | 14 | 9151 | 0 | 106 |
| SRR7851653 | 0 | 0 | 0 | 0 | 0 | 5410 | 1777 | 3628 | 2445 | 0 | 0 | 0 | 0 | 0 | 290 | 222 | 509 | 1925 | 41 | 124 | 0 | 84 | 22 | 14 | 0 | 0 | 0 | 0 | 32 | 0 | 19 | 562 | 390 | 0 | 5901 | 13 | 53 |
| SRR7851655 | 0 | 0 | 0 | 0 | 0 | 2918 | 2388 | 4125 | 6623 | 0 | 0 | 0 | 0 | 0 | 703 | 319 | 1367 | 6544 | 79 | 152 | 0 | 1180 | 282 | 174 | 0 | 0 | 0 | 0 | 135 | 0 | 78 | 2865 | 991 | 19 | 11601 | 0 | 320 |
| SRR7851656 | 0 | 0 | 0 | 0 | 0 | 4758 | 2957 | 5019 | 7245 | 0 | 0 | 0 | 0 | 0 | 1168 | 983 | 2599 | 3048 | 59 | 405 | 0 | 279 | 57 | 65 | 0 | 0 | 0 | 0 | 27 | 0 | 25 | 731 | 355 | 17 | 8459 | 0 | 58 |
| SRR7901057 | 22 | 16 | 14 | 0 | 183 | 0 | 0 | 0 | 37 | 5686 | 37 | 20 | 0 | 0 | 1153 | 231 | 447 | 834 | 1176 | 2742 | 377 | 0 | 0 | 23 | 29 | 20 | 42 | 0 | 0 | 22 | 0 | 0 | 0 | 14 | 52594 | 0 | 0 |
| SRR7901058 | 71 | 51 | 0 | 0 | 330 | 0 | 0 | 17 | 55 | 6457 | 46 | 183 | 0 | 0 | 1199 | 1311 | 1971 | 1140 | 1275 | 2359 | 721 | 0 | 0 | 23 | 85 | 41 | 93 | 0 | 0 | 51 | 0 | 0 | 0 | 21 | 48278 | 0 | 0 |
| SRR7901059 | 23 | 23 | 0 | 0 | 164 | 20 | 0 | 28 | 57 | 1432 | 14 | 14 | 0 | 0 | 842 | 342 | 396 | 1212 | 2023 | 2680 | 143 | 0 | 0 | 11 | 33 | 19 | 52 | 0 | 0 | 15 | 0 | 0 | 0 | 0 | 54515 | 0 | 0 |
| SRR7903701 | 2138 | 839 | 621 | 166 | 1220 | 1135 | 225 | 1263 | 1729 | 175 | 89 | 17 | 13 | 0 | 13284 | 100235 | 61413 | 398 | 5318 | 1465 | 2268 | 0 | 0 | 1378 | 492 | 285 | 712 | 21 | 0 | 409 | 0 | 74 | 0 | 0 | 61679 | 2213 | 0 |
| SRR7903702 | 1716 | 681 | 475 | 140 | 1585 | 1199 | 252 | 1447 | 1791 | 192 | 69 | 0 | 12 | 0 | 18119 | 129220 | 86329 | 340 | 5412 | 1667 | 2616 | 0 | 0 | 1031 | 397 | 249 | 666 | 18 | 0 | 741 | 0 | 101 | 0 | 0 | 45275 | 2240 | 0 |
| SRR7903703 | 1708 | 609 | 474 | 153 | 1183 | 1237 | 211 | 1532 | 1903 | 132 | 63 | 11 | 0 | 0 | 16400 | 102615 | 78084 | 412 | 5495 | 1734 | 2364 | 0 | 0 | 1374 | 506 | 286 | 744 | 26 | 0 | 363 | 0 | 81 | 0 | 0 | 52897 | 2165 | 0 |
| SRR7903704 | 212 | 202 | 80 | 14 | 2346 | 0 | 0 | 11 | 16 | 1083 | 129 | 18 | 0 | 0 | 492 | 9886 | 2000 | 4687 | 37620 | 5600 | 270 | 0 | 0 | 245 | 532 | 386 | 1053 | 421 | 0 | 56 | 0 | 0 | 0 | 0 | 216603 | 233 | 0 |
| SRR7903705 | 231 | 166 | 78 | 21 | 1690 | 20 | 0 | 24 | 22 | 554 | 147 | 20 | 0 | 0 | 1076 | 22643 | 6382 | 4546 | 26941 | 2356 | 306 | 0 | 0 | 288 | 665 | 684 | 1647 | 627 | 0 | 85 | 0 | 0 | 0 | 14 | 170818 | 180 | 0 |
| SRR7903706 | 198 | 192 | 75 | 16 | 1411 | 15 | 0 | 0 | 17 | 964 | 131 | 24 | 0 | 0 | 720 | 10818 | 2812 | 5182 | 41450 | 3799 | 227 | 0 | 0 | 358 | 744 | 777 | 1673 | 810 | 0 | 88 | 0 | 0 | 0 | 16 | 210098 | 291 | 0 |
| SRR8063607 | 2431 | 1104 | 29 | 22 | 448 | 1085 | 205 | 1133 | 5857 | 816 | 62 | 11 | 15 | 0 | 4278 | 27452 | 19143 | 108 | 928 | 3073 | 976 | 0 | 0 | 160 | 91 | 22 | 11 | 15 | 0 | 91 | 0 | 0 | 0 | 11 | 1214 | 0 | 0 |
| SRR8063608 | 251 | 117 | 29 | 19 | 49 | 912 | 88 | 1265 | 5323 | 92 | 58 | 0 | 0 | 0 | 4323 | 33103 | 20963 | 59 | 647 | 2764 | 1412 | 0 | 0 | 11 | 88 | 0 | 0 | 0 | 0 | 78 | 0 | 0 | 0 | 0 | 1310 | 0 | 0 |
| SRR8063609 | 303 | 145 | 81 | 13 | 54 | 917 | 115 | 1197 | 6214 | 75 | 17 | 0 | 0 | 0 | 5183 | 32962 | 21588 | 127 | 1223 | 3820 | 1006 | 0 | 0 | 19 | 23 | 77 | 29 | 0 | 0 | 24 | 0 | 0 | 0 | 0 | 1400 | 0 | 0 |
| SRR8063610 | 89 | 37 | 0 | 0 | 43 | 199 | 21 | 192 | 2392 | 85 | 28 | 25 | 0 | 0 | 1892 | 196358 | 12702 | 37 | 5897 | 15603 | 1616 | 0 | 0 | 14 | 29 | 10 | 0 | 0 | 13 | 12 | 0 | 0 | 0 | 0 | 205 | 0 | 0 |
| SRR8063611 | 84 | 28 | 0 | 0 | 41 | 218 | 38 | 198 | 2159 | 74 | 46 | 32 | 0 | 0 | 2026 | 194755 | 13691 | 51 | 6863 | 16532 | 1920 | 0 | 0 | 12 | 31 | 0 | 0 | 0 | 0 | 18 | 0 | 0 | 0 | 0 | 260 | 0 | 0 |
| SRR8063612 | 77 | 24 | 0 | 0 | 43 | 178 | 23 | 161 | 2095 | 85 | 96 | 34 | 0 | 0 | 2373 | 204116 | 15903 | 51 | 7598 | 17626 | 1984 | 0 | 0 | 11 | 44 | 0 | 0 | 0 | 0 | 25 | 0 | 0 | 0 | 0 | 197 | 0 | 0 |
| SRR8063613 | 184 | 201 | 0 | 0 | 134 | 256 | 64 | 128 | 2110 | 503 | 251 | 50 | 0 | 0 | 767 | 107558 | 5814 | 64 | 2302 | 831 | 1343 | 0 | 0 | 17 | 0 | 17 | 0 | 0 | 67 | 101 | 0 | 64 | 0 | 17 | 8140 | 0 | 0 |
| SRR8063614 | 26 | 24 | 12 | 14 | 30 | 844 | 299 | 1319 | 5366 | 19 | 25 | 40 | 0 | 0 | 704 | 38364 | 8151 | 53 | 2498 | 2604 | 616 | 0 | 0 | 16 | 0 | 12 | 0 | 0 | 20 | 60 | 0 | 0 | 0 | 0 | 2108 | 0 | 0 |
| SRR8063615 | 57 | 25 | 0 | 0 | 49 | 1080 | 455 | 2047 | 7334 | 21 | 66 | 10 | 0 | 0 | 625 | 36202 | 3694 | 114 | 2103 | 3070 | 1194 | 0 | 0 | 33 | 0 | 0 | 0 | 0 | 33 | 16 | 0 | 0 | 0 | 0 | 2955 | 0 | 0 |
| SRR8074870 | 12 | 0 | 0 | 0 | 0 | 550 | 901 | 1842 | 4365 | 131 | 0 | 0 | 0 | 0 | 632 | 14087 | 1621 | 871 | 158 | 408 | 0 | 0 | 0 | 20 | 0 | 0 | 0 | 0 | 21 | 0 | 0 | 62 | 31 | 0 | 5128 | 0 | 21 |
| SRR8074871 | 0 | 0 | 0 | 0 | 0 | 553 | 857 | 1769 | 4265 | 148 | 0 | 0 | 0 | 12 | 584 | 14085 | 1602 | 834 | 158 | 408 | 0 | 0 | 0 | 11 | 0 | 0 | 0 | 0 | 37 | 0 | 0 | 68 | 37 | 0 | 5064 | 0 | 18 |

### Supp Table S8.3.Columns_BX_to_DH

| | | | | | | | | | | | | | | | | | | | | | | | | | | | | | | | | | | | | | |
|---|---|---|---|---|---|---|---|---|---|---|---|---|---|---|---|---|---|---|---|---|---|---|---|---|---|---|---|---|---|---|---|---|---|---|---|---|---|
| SRR8074873 | 22 | 0 | 0 | 0 | 0 | 447 | 987 | 1627 | 7272 | 59 | 0 | 0 | 0 | 0 | 1806 | 48228 | 6280 | 1545 | 216 | 1284 | 0 | 0 | 0 | 30 | 0 | 0 | 0 | 0 | 29 | 0 | 0 | 113 | 14 | 0 | 5069 | 0 | 18 |
| SRR8074874 | 0 | 0 | 0 | 0 | 0 | 299 | 890 | 1777 | 8223 | 25 | 0 | 0 | 0 | 0 | 6388 | 60311 | 12487 | 1436 | 1457 | 3749 | 14 | 0 | 0 | 19 | 0 | 0 | 0 | 0 | 15 | 23 | 0 | 153 | 0 | 0 | 4213 | 0 | 11 |
| SRR8074875 | 13 | 0 | 0 | 0 | 0 | 309 | 939 | 1780 | 8494 | 339 | 0 | 0 | 0 | 0 | 6064 | 58944 | 11992 | 1372 | 1260 | 3401 | 12 | 0 | 0 | 0 | 0 | 0 | 0 | 0 | 23 | 21 | 0 | 145 | 11 | 0 | 4504 | 0 | 13 |
| SRR8094756 | 162 | 57 | 14 | 39 | 75 | 240 | 269 | 654 | 1942 | 31 | 92 | 43 | 0 | 0 | 11316 | 117356 | 101036 | 1045 | 2356 | 3677 | 1612 | 42 | 14 | 77 | 64 | 42 | 136 | 0 | 37 | 91 | 156 | 2913 | 11 | 11 | 1865 | 0 | 0 |
| SRR8094757 | 1938 | 387 | 247 | 81 | 192 | 147 | 56 | 449 | 707 | 142 | 139 | 0 | 0 | 0 | 7217 | 58277 | 45765 | 9696 | 2200 | 876 | 795 | 15696 | 4504 | 275 | 40 | 75 | 219 | 24 | 45 | 31 | 2565 | 7204 | 67 | 117 | 687 | 0 | 0 |
| SRR8094758 | 371 | 105 | 14 | 35 | 85 | 104 | 75 | 213 | 778 | 31 | 60 | 91 | 0 | 0 | 24065 | 201376 | 155726 | 436 | 1471 | 2268 | 4944 | 33 | 17 | 26 | 17 | 20 | 73 | 10 | 42 | 14 | 10 | 5210 | 34 | 40 | 4313 | 0 | 12 |
| SRR8094759 | 65 | 37 | 0 | 0 | 50 | 359 | 97 | 960 | 1153 | 80 | 141 | 62 | 0 | 0 | 15802 | 174901 | 386444 | 96 | 2563 | 209390 | 19735 | 0 | 0 | 33 | 18 | 22 | 36 | 12 | 0 | 191 | 0 | 0 | 0 | 20 | 1222 | 71 | 0 |
| SRR8094760 | 106 | 35 | 20 | 13 | 42 | 1861 | 720 | 3122 | 5863 | 18 | 14 | 24 | 0 | 0 | 4933 | 603662 | 69093 | 530 | 690 | 11276 | 5443 | 0 | 0 | 25 | 0 | 0 | 27 | 0 | 0 | 49 | 0 | 30 | 0 | 70 | 13942 | 92 | 0 |
| SRR8094761 | 111 | 26 | 241 | 83 | 89 | 15011 | 2094 | 17245 | 10521 | 116 | 226 | 11 | 0 | 0 | 734 | 197436 | 85472 | 32 | 229 | 3357 | 2470 | 0 | 18 | 64 | 86 | 134 | 208 | 0 | 0 | 359 | 0 | 132 | 0 | 12 | 772 | 47 | 0 |
| SRR8094762 | 6150 | 1443 | 567 | 587 | 1410 | 6979 | 956 | 7821 | 6225 | 433 | 283 | 1089 | 0 | 0 | 8126 | 279465 | 306879 | 80 | 1734 | 16371 | 13149 | 0 | 12 | 1064 | 53 | 162 | 719 | 51 | 30 | 240 | 0 | 31 | 0 | 25 | 989 | 49 | 0 |
| SRR8094763 | 922 | 265 | 160 | 286 | 469 | 1390 | 125 | 867 | 2368 | 676 | 989 | 87 | 0 | 0 | 27431 | 236348 | 62816 | 1966 | 924 | 6483 | 30241 | 361 | 188 | 1055 | 258 | 379 | 767 | 107 | 16 | 289 | 302 | 15349 | 0 | 0 | 1586 | 14 | 0 |
| SRR8094764 | 146 | 38 | 16 | 17 | 89 | 437 | 203 | 1016 | 2843 | 39 | 141 | 162 | 0 | 0 | 3332 | 145528 | 11554 | 4739 | 766 | 2527 | 6326 | 0 | 0 | 156 | 162 | 174 | 25 | 31 | 167 | 19 | 0 | 26013 | 0 | 37 | 1471 | 0 | 0 |
| SRR8094765 | 43 | 12 | 12 | 14 | 18 | 584 | 90 | 586 | 1310 | 30 | 381 | 70 | 0 | 0 | 5492 | 86216 | 19724 | 6156 | 1944 | 3274 | 10364 | 128 | 31 | 50 | 78 | 148 | 27 | 15 | 41 | 51 | 0 | 27499 | 10 | 107 | 3084 | 0 | 0 |
| SRR8094766 | 73 | 18 | 197 | 189 | 110 | 506 | 116 | 854 | 1651 | 81 | 26 | 76 | 0 | 0 | 11842 | 231069 | 37805 | 1433 | 561 | 1876 | 16377 | 28 | 12 | 110 | 240 | 116 | 17 | 11 | 59 | 242 | 17 | 20540 | 0 | 80 | 1455 | 13 | 0 |
| SRR8094767 | 22 | 529 | 163 | 192 | 34 | 1485 | 300 | 2042 | 3976 | 20 | 174 | 17 | 0 | 0 | 3876 | 192998 | 30299 | 528 | 250 | 978 | 12392 | 58 | 52 | 41 | 140 | 157 | 477 | 17 | 0 | 122 | 12 | 9315 | 41 | 76 | 2214 | 0 | 0 |
| SRR8094768 | 149 | 43 | 27 | 32 | 205 | 352 | 209 | 929 | 3145 | 20 | 44 | 157 | 0 | 0 | 3762 | 153056 | 11480 | 4894 | 802 | 3461 | 2754 | 0 | 0 | 262 | 12 | 19 | 74 | 56 | 70 | 14 | 0 | 22525 | 0 | 11 | 2090 | 0 | 0 |
| SRR8094769 | 2804 | 682 | 410 | 400 | 1445 | 1459 | 293 | 1477 | 2860 | 536 | 185 | 12 | 0 | 0 | 4130 | 171258 | 17507 | 3979 | 568 | 1415 | 9737 | 0 | 0 | 4957 | 152 | 177 | 692 | 53 | 16 | 219 | 0 | 10314 | 0 | 0 | 6243 | 0 | 0 |
| SRR8094770 | 9517 | 2424 | 586 | 93 | 235 | 105 | 20 | 83 | 198 | 660 | 405 | 398 | 36 | 29 | 1352 | 4515 | 1551 | 23720 | 164 | 121 | 248 | 44 | 25 | 992 | 19 | 25 | 110 | 0 | 222 | 235 | 25 | 37707 | 12 | 52 | 4957 | 1713 | 0 |
| SRR8137033 | 167 | 76 | 51 | 16 | 22 | 372 | 160 | 744 | 1861 | 71 | 22 | 0 | 20 | 0 | 10793 | 13785 | 22360 | 160 | 266 | 1063 | 319 | 0 | 0 | 13 | 11 | 20 | 25 | 22 | 0 | 51 | 0 | 53 | 0 | 0 | 11831 | 0 | 0 |
| SRR8137034 | 374 | 209 | 175 | 85 | 167 | 473 | 165 | 514 | 1111 | 146 | 51 | 0 | 0 | 0 | 10861 | 31035 | 24330 | 309 | 1584 | 597 | 926 | 0 | 0 | 0 | 0 | 23 | 32 | 64 | 0 | 48 | 0 | 21 | 0 | 0 | 21711 | 0 | 0 |
| SRR8137035 | 247 | 143 | 29 | 17 | 64 | 87 | 0 | 260 | 476 | 35 | 21 | 21 | 12 | 0 | 9994 | 41213 | 23227 | 87 | 216 | 303 | 779 | 0 | 0 | 12 | 12 | 0 | 12 | 22 | 0 | 76 | 0 | 87 | 0 | 0 | 22405 | 0 | 0 |
| SRR8137036 | 1016 | 412 | 392 | 171 | 332 | 356 | 0 | 178 | 1212 | 181 | 151 | 10 | 20 | 10 | 5631 | 21398 | 19038 | 178 | 249 | 428 | 285 | 10 | 0 | 30 | 10 | 20 | 141 | 161 | 0 | 382 | 0 | 36 | 0 | 0 | 11346 | 0 | 0 |
| SRR8137037 | 406 | 333 | 104 | 61 | 35 | 83 | 0 | 413 | 413 | 0 | 33 | 0 | 0 | 0 | 69057 | 121915 | 250916 | 165 | 4130 | 578 | 13382 | 0 | 0 | 20 | 20 | 16 | 49 | 28 | 0 | 33 | 0 | 0 | 0 | 0 | 27608 | 21 | 0 |
| SRR8203922 | 0 | 0 | 0 | 0 | 71 | 0 | 0 | 0 | 0 | 16 | 0 | 0 | 0 | 0 | 26103 | 30611 | 45896 | 4869 | 1213 | 771 | 458 | 0 | 0 | 0 | 0 | 0 | 0 | 0 | 0 | 0 | 0 | 0 | 0 | 0 | 1687 | 0 | 0 |
| SRR8203923 | 0 | 0 | 0 | 0 | 46 | 0 | 0 | 0 | 0 | 11 | 0 | 0 | 0 | 0 | 24813 | 33675 | 47132 | 5103 | 1202 | 812 | 482 | 0 | 0 | 0 | 0 | 0 | 0 | 0 | 0 | 0 | 0 | 0 | 0 | 0 | 1739 | 0 | 0 |
| SRR8203924 | 0 | 0 | 0 | 0 | 36 | 0 | 0 | 0 | 0 | 14 | 0 | 0 | 0 | 0 | 24583 | 31922 | 43538 | 5213 | 1376 | 779 | 381 | 0 | 0 | 0 | 0 | 0 | 0 | 0 | 0 | 0 | 0 | 0 | 0 | 0 | 1157 | 0 | 0 |
| SRR8203925 | 0 | 0 | 11 | 0 | 43 | 0 | 0 | 0 | 0 | 14 | 0 | 0 | 0 | 0 | 24811 | 32543 | 46779 | 5195 | 1149 | 718 | 355 | 0 | 0 | 0 | 0 | 0 | 0 | 0 | 0 | 0 | 0 | 0 | 0 | 0 | 1546 | 0 | 0 |
| SRR8203926 | 0 | 0 | 0 | 0 | 28 | 13 | 22 | 0 | 0 | 13 | 0 | 0 | 0 | 0 | 26737 | 20653 | 35864 | 4048 | 1283 | 2556 | 315 | 0 | 0 | 0 | 0 | 0 | 13 | 0 | 0 | 0 | 0 | 80 | 0 | 0 | 1072 | 0 | 0 |
| SRR8203927 | 11 | 0 | 0 | 0 | 41 | 33 | 19 | 19 | 0 | 11 | 0 | 0 | 0 | 0 | 26580 | 21812 | 36703 | 4023 | 1357 | 2483 | 296 | 0 | 0 | 0 | 0 | 0 | 0 | 16 | 0 | 0 | 0 | 75 | 0 | 0 | 838 | 0 | 0 |
| SRR8203928 | 0 | 0 | 0 | 0 | 39 | 41 | 18 | 0 | 0 | 0 | 0 | 0 | 0 | 0 | 26908 | 21463 | 35901 | 4161 | 1378 | 2619 | 283 | 0 | 0 | 0 | 0 | 0 | 11 | 11 | 0 | 0 | 0 | 64 | 0 | 0 | 1100 | 0 | 0 |
| SRR8203929 | 0 | 0 | 0 | 0 | 27 | 28 | 0 | 0 | 0 | 0 | 0 | 0 | 0 | 0 | 26597 | 21727 | 36037 | 4061 | 1327 | 2748 | 322 | 0 | 0 | 0 | 0 | 0 | 0 | 13 | 0 | 11 | 11 | 107 | 0 | 0 | 1079 | 0 | 0 |
| SRR8203930 | 0 | 0 | 0 | 0 | 68 | 0 | 0 | 0 | 0 | 26 | 0 | 0 | 0 | 0 | 23561 | 17441 | 29617 | 2819 | 797 | 4189 | 685 | 0 | 0 | 0 | 0 | 0 | 0 | 0 | 0 | 11 | 0 | 0 | 0 | 0 | 2814 | 0 | 0 |
| SRR8203931 | 0 | 0 | 0 | 0 | 70 | 23 | 0 | 0 | 0 | 27 | 0 | 0 | 0 | 0 | 23120 | 18497 | 30249 | 2745 | 789 | 3950 | 649 | 0 | 0 | 0 | 0 | 0 | 0 | 0 | 0 | 0 | 0 | 12 | 0 | 0 | 2631 | 0 | 0 |
| SRR8203932 | 0 | 0 | 0 | 0 | 58 | 15 | 15 | 0 | 0 | 33 | 15 | 0 | 0 | 0 | 23074 | 18038 | 29221 | 2972 | 679 | 3788 | 668 | 0 | 0 | 0 | 0 | 0 | 0 | 0 | 0 | 0 | 0 | 0 | 0 | 0 | 2615 | 0 | 0 |
| SRR8203933 | 0 | 0 | 0 | 0 | 92 | 12 | 0 | 0 | 0 | 15 | 0 | 0 | 0 | 0 | 23742 | 17692 | 28661 | 2739 | 733 | 3973 | 603 | 0 | 0 | 0 | 0 | 0 | 0 | 0 | 0 | 0 | 0 | 0 | 0 | 0 | 2685 | 0 | 0 |
| SRR8203934 | 0 | 0 | 0 | 0 | 44 | 0 | 0 | 0 | 0 | 59 | 16 | 0 | 0 | 0 | 6496 | 7364 | 10713 | 3430 | 1018 | 3927 | 223 | 0 | 0 | 0 | 0 | 0 | 0 | 0 | 0 | 51 | 0 | 190 | 0 | 0 | 1791 | 0 | 0 |
| SRR8203935 | 0 | 0 | 0 | 0 | 45 | 0 | 0 | 0 | 0 | 34 | 0 | 0 | 0 | 0 | 6127 | 7326 | 11267 | 3460 | 1037 | 3922 | 225 | 0 | 0 | 0 | 0 | 0 | 0 | 0 | 0 | 57 | 0 | 201 | 0 | 0 | 1890 | 0 | 0 |
| SRR8203936 | 0 | 0 | 0 | 0 | 34 | 0 | 0 | 0 | 0 | 53 | 11 | 0 | 0 | 0 | 6308 | 7534 | 10877 | 3284 | 1030 | 3952 | 230 | 0 | 0 | 0 | 0 | 0 | 0 | 0 | 0 | 40 | 0 | 218 | 0 | 0 | 2038 | 0 | 0 |
| SRR8203937 | 0 | 0 | 0 | 0 | 56 | 0 | 0 | 0 | 0 | 56 | 21 | 0 | 0 | 0 | 6337 | 7262 | 10997 | 3361 | 1038 | 3928 | 233 | 0 | 0 | 0 | 0 | 0 | 0 | 0 | 0 | 45 | 0 | 212 | 0 | 0 | 1703 | 0 | 0 |
| SRR8203938 | 0 | 0 | 0 | 0 | 11 | 0 | 0 | 0 | 0 | 13 | 15 | 0 | 0 | 0 | 11630 | 12063 | 18378 | 2894 | 544 | 4130 | 458 | 0 | 0 | 0 | 0 | 0 | 0 | 0 | 0 | 41 | 0 | 99 | 0 | 0 | 2331 | 0 | 0 |
| SRR8203939 | 0 | 0 | 0 | 0 | 10 | 0 | 0 | 0 | 0 | 119 | 13 | 0 | 0 | 0 | 11901 | 12344 | 17735 | 2845 | 697 | 4119 | 366 | 0 | 0 | 0 | 0 | 0 | 0 | 0 | 0 | 38 | 0 | 118 | 0 | 0 | 2472 | 0 | 0 |
| SRR8203940 | 0 | 0 | 0 | 0 | 11 | 0 | 0 | 0 | 0 | 111 | 16 | 0 | 0 | 0 | 11949 | 12366 | 18343 | 2888 | 548 | 4146 | 437 | 0 | 0 | 0 | 0 | 0 | 0 | 0 | 0 | 33 | 0 | 100 | 0 | 0 | 2547 | 0 | 0 |
| SRR8203941 | 0 | 0 | 0 | 0 | 0 | 0 | 0 | 0 | 0 | 103 | 0 | 0 | 0 | 0 | 12103 | 12607 | 18525 | 2922 | 615 | 4188 | 398 | 0 | 0 | 0 | 0 | 0 | 0 | 0 | 0 | 40 | 0 | 140 | 0 | 0 | 2752 | 0 | 0 |
| SRR8203942 | 0 | 0 | 0 | 0 | 26 | 0 | 0 | 0 | 0 | 0 | 51 | 0 | 0 | 0 | 7347 | 4542 | 12407 | 1107 | 589 | 4604 | 412 | 0 | 0 | 0 | 0 | 0 | 0 | 10 | 0 | 57 | 15 | 324 | 0 | 0 | 2312 | 0 | 0 |
| SRR8203943 | 0 | 0 | 0 | 0 | 21 | 0 | 0 | 0 | 0 | 0 | 43 | 0 | 0 | 0 | 7419 | 4530 | 11887 | 1201 | 685 | 4575 | 417 | 0 | 0 | 0 | 0 | 0 | 0 | 0 | 0 | 649 | 21 | 311 | 0 | 0 | 2136 | 0 | 0 |
| SRR8203944 | 0 | 0 | 0 | 0 | 21 | 0 | 0 | 0 | 0 | 0 | 37 | 0 | 0 | 0 | 7115 | 4303 | 12530 | 1170 | 549 | 4311 | 332 | 0 | 0 | 0 | 0 | 0 | 0 | 0 | 0 | 581 | 26 | 374 | 0 | 0 | 2120 | 0 | 0 |
| SRR8203945 | 0 | 0 | 0 | 11 | 21 | 0 | 0 | 0 | 0 | 0 | 54 | 0 | 0 | 0 | 7153 | 4463 | 11880 | 1105 | 660 | 4758 | 451 | 0 | 0 | 0 | 0 | 0 | 0 | 0 | 0 | 45 | 16 | 352 | 0 | 0 | 2231 | 0 | 0 |
| SRR8203946 | 0 | 0 | 0 | 0 | 0 | 20 | 0 | 13 | 0 | 11 | 16 | 0 | 0 | 0 | 13675 | 6927 | 19084 | 836 | 212 | 4722 | 1085 | 0 | 0 | 0 | 0 | 0 | 0 | 0 | 0 | 67 | 18 | 104 | 0 | 0 | 5303 | 0 | 0 |
| SRR8203947 | 0 | 0 | 0 | 0 | 14 | 0 | 0 | 0 | 0 | 12 | 14 | 0 | 0 | 0 | 13751 | 6935 | 19659 | 736 | 205 | 4814 | 998 | 0 | 0 | 0 | 0 | 0 | 0 | 0 | 0 | 65 | 26 | 127 | 0 | 0 | 5579 | 0 | 0 |
| SRR8203948 | 0 | 0 | 0 | 0 | 10 | 10 | 0 | 0 | 0 | 86 | 75 | 0 | 0 | 0 | 13433 | 6877 | 19091 | 762 | 206 | 5016 | 1068 | 0 | 0 | 0 | 0 | 0 | 0 | 0 | 0 | 495 | 17 | 134 | 0 | 0 | 5488 | 0 | 0 |
| SRR8203949 | 0 | 0 | 0 | 0 | 26 | 0 | 0 | 0 | 0 | 15 | 12 | 0 | 0 | 0 | 13457 | 7391 | 18500 | 596 | 183 | 5126 | 984 | 0 | 0 | 0 | 0 | 0 | 0 | 0 | 0 | 63 | 15 | 116 | 0 | 0 | 5399 | 0 | 0 |
| SRR8248787 | 1988 | 1883 | 721 | 491 | 808 | 316 | 73 | 961 | 2229 | 2744 | 178 | 0 | 0 | 0 | 18283 | 49716 | 91491 | 335 | 2898 | 641 | 1239 | 0 | 0 | 166 | 84 | 79 | 180 | 33 | 0 | 40 | 0 | 12 | 0 | 0 | 5365 | 0 | 0 |
| SRR8248788 | 5019 | 4698 | 977 | 393 | 2018 | 341 | 79 | 764 | 1667 | 6656 | 166 | 10 | 14 | 0 | 14879 | 41216 | 66171 | 272 | 2995 | 605 | 1144 | 0 | 0 | 264 | 92 | 49 | 79 | 0 | 0 | 66 | 0 | 22 | 0 | 0 | 5952 | 0 | 0 |
| SRR8248789 | 4259 | 4185 | 999 | 406 | 1564 | 264 | 61 | 609 | 1561 | 6334 | 210 | 0 | 0 | 0 | 12656 | 40033 | 55021 | 349 | 2405 | 619 | 1200 | 0 | 0 | 141 | 68 | 53 | 97 | 0 | 0 | 74 | 0 | 0 | 0 | 0 | 5698 | 0 | 0 |
| SRR8248790 | 1152 | 1317 | 451 | 199 | 409 | 184 | 29 | 499 | 1258 | 1928 | 76 | 0 | 0 | 0 | 8609 | 39010 | 37200 | 163 | 1915 | 240 | 1351 | 0 | 0 | 28 | 20 | 28 | 73 | 56 | 0 | 35 | 0 | 0 | 0 | 0 | 6026 | 0 | 0 |
| SRR8248791 | 351 | 282 | 20 | 86 | 108 | 326 | 78 | 837 | 2046 | 600 | 52 | 10 | 14 | 0 | 6477 | 28890 | 26804 | 387 | 1671 | 318 | 1267 | 0 | 0 | 16 | 30 | 15 | 13 | 0 | 0 | 45 | 0 | 17 | 0 | 0 | 8156 | 0 | 0 |
| SRR8248792 | 8718 | 7294 | 472 | 150 | 1459 | 310 | 53 | 298 | 1655 | 5885 | 83 | 11 | 0 | 13 | 6260 | 23783 | 21496 | 204 | 1164 | 147 | 1515 | 0 | 0 | 146 | 32 | 13 | 19 | 0 | 0 | 138 | 0 | 0 | 0 | 0 | 3514 | 0 | 0 |
| SRR8272058 | 0 | 0 | 0 | 0 | 0 | 11 | 35 | 221 | 51 | 0 | 0 | 0 | 0 | 0 | 0 | 87 | 87 | 22 | 0 | 0 | 11 | 0 | 0 | 12 | 0 | 0 | 0 | 0 | 0 | 0 | 0 | 119 | 0 | 0 | 68289 | 0 | 23 |
| SRR8280405 | 0 | 0 | 0 | 0 | 0 | 0 | 73 | 0 | 0 | 131 | 87 | 0 | 0 | 0 | 25115 | 62533 | 133606 | 0 | 0 | 511 | 35117 | 0 | 0 | 0 | 0 | 0 | 0 | 0 | 0 | 44 | 0 | 0 | 0 | 0 | 155 | 0 | 0 |
| SRR8280408 | 0 | 0 | 0 | 0 | 0 | 0 | 0 | 92 | 0 | 0 | 110 | 0 | 0 | 0 | 19001 | 70175 | 119411 | 0 | 0 | 92 | 45438 | 0 | 0 | 166 | 0 | 0 | 0 | 0 | 0 | 0 | 0 | 0 | 0 | 0 | 142 | 0 | 0 |
| SRR8280414 | 0 | 0 | 0 | 0 | 0 | 0 | 146 | 146 | 0 | 0 | 246 | 0 | 0 | 0 | 38599 | 24404 | 138517 | 0 | 0 | 292 | 12647 | 0 | 0 | 0 | 22 | 0 | 0 | 0 | 0 | 67 | 0 | 0 | 0 | 0 | 169 | 0 | 134 |
| SRR8280415 | 0 | 0 | 0 | 0 | 0 | 0 | 0 | 0 | 0 | 0 | 324 | 0 | 0 | 0 | 40950 | 32024 | 125317 | 0 | 0 | 0 | 12147 | 0 | 0 | 36 | 36 | 0 | 0 | 0 | 0 | 0 | 0 | 0 | 0 | 0 | 329 | 0 | 288 |
| SRR8280416 | 0 | 0 | 0 | 0 | 0 | 0 | 0 | 0 | 0 | 0 | 16 | 0 | 0 | 0 | 28080 | 25518 | 72463 | 0 | 0 | 1228 | 24091 | 0 | 0 | 0 | 0 | 0 | 0 | 0 | 0 | 0 | 0 | 0 | 0 | 0 | 368 | 0 | 0 |
| SRR8280417 | 0 | 0 | 0 | 0 | 0 | 0 | 0 | 0 | 0 | 0 | 17 | 51 | 0 | 0 | 8235 | 50408 | 40382 | 0 | 0 | 341 | 8576 | 0 | 0 | 0 | 0 | 0 | 0 | 0 | 0 | 0 | 0 | 0 | 0 | 0 | 301 | 0 | 137 |
| SRR8280423 | 0 | 0 | 0 | 0 | 0 | 242 | 0 | 242 | 0 | 0 | 355 | 0 | 0 | 0 | 20194 | 75715 | 151441 | 0 | 0 | 0 | 42322 | 0 | 0 | 0 | 0 | 0 | 0 | 0 | 0 | 65 | 0 | 0 | 0 | 0 | 237 | 0 | 0 |
| SRR8280425 | 0 | 0 | 0 | 0 | 0 | 0 | 0 | 224 | 0 | 0 | 0 | 0 | 0 | 0 | 46679 | 35358 | 182093 | 0 | 0 | 449 | 16831 | 0 | 0 | 0 | 0 | 0 | 0 | 0 | 0 | 0 | 0 | 0 | 0 | 0 | 88 | 0 | 0 |
| SRR8280432 | 0 | 0 | 0 | 0 | 0 | 0 | 0 | 534 | 0 | 0 | 550 | 0 | 0 | 0 | 23241 | 59488 | 104076 | 0 | 0 | 178 | 19590 | 0 | 0 | 37 | 0 | 0 | 37 | 37 | 0 | 0 | 0 | 0 | 0 | 0 | 315 | 0 | 73 |
| SRR8280440 | 0 | 0 | 0 | 0 | 0 | 69 | 242 | 2036 | 0 | 0 | 1064 | 0 | 0 | 0 | 10316 | 80150 | 215694 | 0 | 0 | 242 | 9247 | 0 | 0 | 0 | 0 | 0 | 0 | 0 | 0 | 39 | 0 | 0 | 0 | 0 | 293 | 0 | 0 |
| SRR8280442 | 0 | 0 | 0 | 0 | 27 | 27 | 0 | 433 | 0 | 0 | 383 | 0 | 0 | 0 | 14923 | 46009 | 158256 | 0 | 0 | 433 | 14977 | 0 | 0 | 0 | 0 | 0 | 0 | 0 | 0 | 27 | 0 | 0 | 0 | 0 | 48 | 0 | 0 |
| SRR8280452 | 0 | 0 | 0 | 90 | 0 | 364 | 0 | 364 | 0 | 0 | 90 | 0 | 0 | 0 | 38937 | 36898 | 147199 | 0 | 0 | 0 | 6914 | 0 | 0 | 0 | 0 | 0 | 0 | 0 | 0 | 0 | 0 | 0 | 0 | 0 | 41 | 0 | 0 |
| SRR8280460 | 0 | 0 | 0 | 43 | 0 | 0 | 0 | 0 | 0 | 0 | 257 | 0 | 0 | 0 | 30750 | 32350 | 185805 | 0 | 44 | 530 | 21163 | 0 | 0 | 0 | 0 | 0 | 14 | 0 | 0 | 57 | 0 | 0 | 0 | 0 | 241 | 0 | 0 |
| SRR8280461 | 0 | 0 | 0 | 0 | 0 | 0 | 0 | 325 | 0 | 0 | 120 | 0 | 0 | 0 | 25520 | 53161 | 152436 | 0 | 0 | 0 | 49903 | 0 | 0 | 0 | 0 | 0 | 0 | 0 | 0 | 0 | 0 | 0 | 0 | 0 | 146 | 0 | 0 |
| SRR8280463 | 0 | 0 | 0 | 0 | 0 | 0 | 83 | 292 | 0 | 0 | 63 | 0 | 0 | 0 | 47544 | 23270 | 189357 | 0 | 0 | 42 | 11626 | 0 | 0 | 0 | 0 | 0 | 0 | 0 | 0 | 0 | 0 | 0 | 0 | 0 | 40 | 0 | 0 |
| SRR8280464 | 0 | 0 | 0 | 0 | 46 | 370 | 0 | 159 | 0 | 0 | 46 | 0 | 0 | 0 | 36169 | 40221 | 212124 | 0 | 0 | 159 | 36487 | 0 | 0 | 0 | 0 | 46 | 0 | 0 | 0 | 92 | 0 | 0 | 0 | 0 | 193 | 0 | 0 |
| SRR8280467 | 0 | 12 | 0 | 0 | 12 | 110 | 49 | 964 | 0 | 0 | 200 | 0 | 0 | 0 | 86488 | 25062 | 229753 | 0 | 37 | 268 | 6162 | 0 | 0 | 262 | 0 | 12 | 125 | 87 | 0 | 75 | 0 | 0 | 0 | 0 | 94 | 0 | 0 |
| SRR8280468 | 0 | 0 | 0 | 0 | 0 | 0 | 0 | 701 | 0 | 0 | 0 | 0 | 0 | 0 | 47335 | 45072 | 155270 | 0 | 0 | 701 | 9116 | 0 | 0 | 0 | 0 | 0 | 0 | 0 | 0 | 0 | 0 | 0 | 0 | 0 | 203 | 0 | 0 |
| SRR8280472 | 0 | 0 | 0 | 0 | 0 | 0 | 0 | 993 | 0 | 0 | 137 | 0 | 0 | 0 | 79087 | 24348 | 178896 | 0 | 0 | 331 | 10258 | 0 | 0 | 0 | 0 | 0 | 0 | 0 | 0 | 0 | 0 | 0 | 0 | 0 | 235 | 11 | 0 |
| SRR8280473 | 0 | 0 | 0 | 0 | 0 | 0 | 199 | 1791 | 0 | 0 | 96 | 0 | 0 | 0 | 39395 | 42821 | 148282 | 0 | 0 | 398 | 23677 | 0 | 0 | 0 | 0 | 0 | 0 | 0 | 0 | 48 | 0 | 0 | 0 | 0 | 411 | 0 | 0 |
| SRR8280476 | 0 | 0 | 0 | 0 | 0 | 0 | 0 | 0 | 0 | 0 | 147 | 0 | 0 | 0 | 50848 | 42005 | 126567 | 0 | 0 | 177 | 7062 | 0 | 0 | 221 | 0 | 0 | 0 | 0 | 0 | 0 | 0 | 0 | 0 | 0 | 405 | 0 | 0 |
| SRR8280479 | 0 | 0 | 0 | 0 | 0 | 0 | 0 | 0 | 0 | 0 | 110 | 0 | 0 | 0 | 71358 | 38609 | 183844 | 0 | 0 | 248 | 9167 | 0 | 0 | 0 | 0 | 0 | 0 | 0 | 0 | 0 | 0 | 0 | 0 | 0 | 207 | 0 | 442 |
| SRR8280481 | 0 | 0 | 0 | 0 | 0 | 0 | 0 | 0 | 0 | 0 | 258 | 129 | 0 | 0 | 62013 | 17880 | 161316 | 0 | 0 | 0 | 5802 | 0 | 0 | 0 | 0 | 0 | 0 | 0 | 0 | 0 | 0 | 0 | 0 | 0 | 104 | 0 | 0 |
| SRR8280483 | 0 | 0 | 0 | 0 | 0 | 0 | 274 | 0 | 0 | 0 | 68 | 0 | 0 | 0 | 16690 | 49017 | 19548 | 0 | 0 | 0 | 4651 | 0 | 0 | 0 | 0 | 0 | 0 | 0 | 0 | 0 | 0 | 0 | 0 | 0 | 248 | 0 | 0 |
| SRR8280485 | 0 | 0 | 0 | 0 | 0 | 0 | 0 | 214 | 0 | 0 | 299 | 0 | 0 | 0 | 13870 | 64214 | 26192 | 0 | 0 | 214 | 3427 | 0 | 0 | 0 | 0 | 54 | 0 | 0 | 0 | 68 | 0 | 0 | 0 | 0 | 672 | 0 | 81 |
| SRR8280487 | 0 | 0 | 0 | 0 | 0 | 0 | 0 | 0 | 0 | 0 | 119 | 0 | 0 | 0 | 26324 | 48670 | 201928 | 0 | 0 | 598 | 20640 | 0 | 0 | 0 | 0 | 0 | 0 | 0 | 0 | 119 | 0 | 0 | 0 | 0 | 133 | 0 | 0 |
| SRR8280490 | 0 | 0 | 0 | 0 | 0 | 0 | 0 | 571 | 0 | 0 | 0 | 0 | 0 | 0 | 18846 | 37871 | 217642 | 0 | 0 | 0 | 23415 | 0 | 0 | 0 | 0 | 142 | 142 | 0 | 0 | 0 | 0 | 0 | 0 | 0 | 197 | 0 | 0 |
| SRR8280494 | 0 | 0 | 0 | 0 | 0 | 0 | 504 | 0 | 0 | 0 | 11 | 0 | 0 | 0 | 16633 | 49693 | 128147 | 0 | 0 | 0 | 15121 | 0 | 0 | 0 | 0 | 0 | 0 | 0 | 0 | 0 | 0 | 0 | 0 | 0 | 358 | 0 | 0 |
| SRR8280497 | 0 | 0 | 0 | 0 | 0 | 0 | 45 | 270 | 0 | 54 | 151 | 0 | 0 | 0 | 189986 | 31075 | 198846 | 0 | 0 | 2474 | 23754 | 0 | 0 | 0 | 0 | 108 | 129 | 32 | 0 | 11 | 0 | 0 | 0 | 0 | 100 | 0 | 32 |
| SRR8280498 | 0 | 0 | 0 | 0 | 0 | 286 | 0 | 0 | 0 | 0 | 85 | 57 | 0 | 0 | 37122 | 23460 | 118878 | 0 | 0 | 143 | 21416 | 0 | 0 | 0 | 0 | 28 | 0 | 28 | 0 | 57 | 0 | 0 | 0 | 0 | 25 | 0 | 339 |
| SRR8280500 | 0 | 0 | 0 | 0 | 0 | 0 | 0 | 0 | 0 | 0 | 336 | 0 | 0 | 0 | 6996 | 71127 | 24190 | 0 | 0 | 328 | 3481 | 0 | 0 | 0 | 0 | 0 | 0 | 0 | 0 | 107 | 0 | 0 | 0 | 0 | 830 | 0 | 138 |
| SRR8280501 | 0 | 0 | 0 | 0 | 0 | 0 | 17 | 153 | 0 | 0 | 11 | 0 | 0 | 0 | 432091 | 9766 | 507850 | 0 | 44 | 1371 | 29948 | 0 | 0 | 195 | 429 | 2039 | 2190 | 584 | 0 | 31 | 0 | 0 | 0 | 0 | 236 | 484 | 0 |
| SRR8280502 | 0 | 0 | 0 | 0 | 0 | 0 | 0 | 0 | 0 | 0 | 311 | 0 | 0 | 0 | 69435 | 49106 | 183376 | 0 | 0 | 358 | 11453 | 0 | 0 | 0 | 0 | 0 | 0 | 0 | 0 | 0 | 0 | 0 | 0 | 0 | 99 | 0 | 104 |
| SRR8280503 | 0 | 0 | 0 | 0 | 0 | 0 | 27 | 167 | 0 | 0 | 23 | 0 | 0 | 0 | 534928 | 10356 | 550346 | 0 | 34 | 1089 | 26735 | 0 | 0 | 391 | 334 | 926 | 880 | 392 | 0 | 65 | 0 | 0 | 0 | 0 | 384 | 371 | 0 |
| SRR8280504 | 0 | 0 | 0 | 0 | 0 | 0 | 0 | 88 | 0 | 0 | 108 | 54 | 0 | 0 | 129904 | 44173 | 233600 | 0 | 0 | 88 | 17642 | 0 | 0 | 54 | 0 | 0 | 0 | 0 | 0 | 0 | 0 | 0 | 0 | 0 | 38 | 0 | 134 |
| SRR8280505 | 0 | 0 | 0 | 0 | 0 | 0 | 18 | 106 | 0 | 0 | 54 | 0 | 0 | 0 | 648967 | 13619 | 725023 | 0 | 69 | 1585 | 28177 | 0 | 0 | 169 | 285 | 743 | 1403 | 254 | 0 | 1095 | 0 | 0 | 0 | 0 | 2714 | 980 | 0 |
| SRR8280506 | 0 | 0 | 0 | 0 | 0 | 0 | 0 | 0 | 0 | 0 | 299 | 0 | 0 | 0 | 43781 | 62033 | 98805 | 0 | 0 | 923 | 15995 | 0 | 0 | 0 | 0 | 85 | 128 | 0 | 0 | 0 | 0 | 0 | 0 | 0 | 160 | 0 | 43 |
| SRR8280507 | 0 | 0 | 0 | 0 | 0 | 19 | 21 | 144 | 0 | 0 | 0 | 0 | 0 | 0 | 453436 | 14326 | 568118 | 0 | 87 | 2585 | 43249 | 0 | 0 | 190 | 562 | 1927 | 2054 | 708 | 0 | 52 | 0 | 0 | 0 | 0 | 401 | 720 | 0 |
| SRR8280509 | 0 | 0 | 0 | 0 | 0 | 26 | 22 | 145 | 0 | 0 | 0 | 0 | 0 | 0 | 542160 | 14008 | 610730 | 0 | 83 | 1945 | 38367 | 0 | 0 | 13 | 36 | 115 | 151 | 43 | 0 | 53 | 0 | 0 | 0 | 0 | 975 | 720 | 0 |
| SRR8280510 | 0 | 0 | 0 | 0 | 0 | 270 | 0 | 540 | 0 | 0 | 118 | 118 | 0 | 0 | 24028 | 50527 | 84968 | 0 | 0 | 810 | 13229 | 0 | 0 | 0 | 0 | 0 | 59 | 0 | 0 | 0 | 0 | 0 | 0 | 0 | 266 | 0 | 118 |
| SRR8280511 | 0 | 0 | 0 | 0 | 0 | 12 | 0 | 76 | 0 | 0 | 0 | 0 | 0 | 0 | 594789 | 5547 | 652301 | 0 | 97 | 1311 | 63250 | 0 | 0 | 13 | 33 | 185 | 196 | 49 | 0 | 18 | 0 | 0 | 0 | 0 | 1324 | 504 | 0 |
| SRR8280512 | 0 | 0 | 0 | 0 | 0 | 0 | 0 | 104 | 0 | 0 | 0 | 93 | 0 | 0 | 71013 | 26743 | 166219 | 0 | 0 | 416 | 21210 | 0 | 0 | 0 | 0 | 0 | 0 | 0 | 0 | 46 | 0 | 0 | 0 | 0 | 111 | 0 | 46 |
| SRR8280513 | 0 | 0 | 0 | 0 | 0 | 23 | 21 | 202 | 0 | 0 | 12 | 0 | 0 | 0 | 440559 | 15303 | 396862 | 0 | 55 | 736 | 49523 | 0 | 0 | 12 | 30 | 69 | 48 | 52 | 0 | 15 | 0 | 0 | 0 | 0 | 1585 | 363 | 0 |
| SRR8280515 | 0 | 0 | 0 | 0 | 0 | 0 | 0 | 94 | 0 | 0 | 184 | 61 | 0 | 0 | 52933 | 36530 | 160940 | 0 | 0 | 752 | 12693 | 0 | 0 | 0 | 61 | 0 | 0 | 0 | 0 | 0 | 0 | 0 | 0 | 0 | 178 | 0 | 0 |
| SRR8280519 | 0 | 0 | 0 | 0 | 0 | 316 | 0 | 789 | 0 | 0 | 219 | 0 | 0 | 0 | 47051 | 71341 | 282413 | 0 | 0 | 237 | 17052 | 0 | 0 | 0 | 0 | 0 | 37 | 0 | 0 | 0 | 0 | 0 | 0 | 0 | 92 | 0 | 0 |
| SRR8280521 | 0 | 0 | 0 | 0 | 0 | 94 | 56 | 394 | 0 | 0 | 14 | 0 | 0 | 0 | 38260 | 62626 | 310702 | 0 | 0 | 75 | 14193 | 0 | 0 | 0 | 34 | 0 | 0 | 25 | 0 | 204 | 0 | 0 | 0 | 0 | 86 | 0 | 0 |
| SRR8280525 | 0 | 0 | 0 | 0 | 0 | 0 | 0 | 0 | 0 | 0 | 94 | 0 | 0 | 0 | 40678 | 27632 | 230008 | 0 | 0 | 0 | 12403 | 0 | 0 | 0 | 94 | 0 | 0 | 94 | 0 | 0 | 0 | 0 | 0 | 0 | 35 | 0 | 0 |
| SRR8280531 | 0 | 0 | 0 | 0 | 0 | 0 | 0 | 197 | 0 | 0 | 271 | 0 | 0 | 0 | 48569 | 45625 | 150966 | 0 | 0 | 0 | 9279 | 0 | 0 | 0 | 0 | 0 | 54 | 0 | 0 | 54 | 0 | 0 | 0 | 0 | 182 | 0 | 0 |
| SRR8280533 | 0 | 0 | 0 | 0 | 0 | 0 | 0 | 82 | 0 | 120 | 120 | 0 | 0 | 0 | 21354 | 58434 | 124400 | 0 | 0 | 824 | 6431 | 0 | 0 | 0 | 0 | 0 | 0 | 0 | 0 | 120 | 0 | 0 | 0 | 0 | 71 | 0 | 60 |
| SRR8280536 | 0 | 0 | 0 | 0 | 0 | 98 | 0 | 197 | 0 | 0 | 467 | 0 | 0 | 0 | 30184 | 42783 | 191129 | 0 | 0 | 820 | 24278 | 0 | 0 | 0 | 0 | 0 | 16 | 0 | 0 | 16 | 0 | 0 | 0 | 0 | 131 | 0 | 0 |
| SRR8280539 | 0 | 0 | 0 | 0 | 0 | 61 | 20 | 344 | 0 | 0 | 72 | 0 | 0 | 0 | 24991 | 72088 | 173637 | 0 | 0 | 182 | 43007 | 0 | 0 | 0 | 27 | 0 | 0 | 0 | 0 | 0 | 0 | 0 | 0 | 0 | 91 | 0 | 45 |
| SRR8280541 | 0 | 0 | 0 | 0 | 0 | 0 | 0 | 34 | 0 | 0 | 329 | 110 | 0 | 0 | 37696 | 51329 | 182322 | 0 | 0 | 517 | 7718 | 0 | 0 | 0 | 0 | 73 | 0 | 146 | 0 | 73 | 0 | 0 | 0 | 0 | 212 | 0 | 73 |
| SRR8280543 | 0 | 0 | 0 | 0 | 0 | 2020 | 0 | 0 | 0 | 0 | 128 | 0 | 0 | 0 | 32323 | 34364 | 173114 | 0 | 0 | 0 | 34343 | 0 | 0 | 0 | 0 | 0 | 0 | 0 | 0 | 0 | 0 | 0 | 0 | 0 | 1297 | 0 | 0 |
| SRR8280544 | 0 | 0 | 0 | 0 | 0 | 540 | 270 | 270 | 0 | 0 | 0 | 0 | 0 | 0 | 26735 | 50929 | 151546 | 0 | 0 | 0 | 67243 | 0 | 0 | 0 | 0 | 0 | 0 | 0 | 0 | 0 | 0 | 0 | 0 | 0 | 404 | 0 | 0 |
| SRR8280546 | 0 | 0 | 0 | 0 | 0 | 0 | 0 | 0 | 0 | 0 | 118 | 0 | 0 | 0 | 105427 | 37430 | 176102 | 0 | 0 | 312 | 9357 | 0 | 0 | 0 | 0 | 0 | 0 | 0 | 0 | 118 | 0 | 0 | 0 | 0 | 246 | 0 | 0 |
| SRR8280548 | 0 | 0 | 0 | 0 | 0 | 0 | 2304 | 0 | 0 | 0 | 193 | 193 | 0 | 0 | 31106 | 31709 | 108922 | 0 | 0 | 0 | 19585 | 0 | 0 | 0 | 0 | 0 | 0 | 0 | 0 | 0 | 0 | 0 | 0 | 0 | 269 | 0 | 0 |
| SRR8280550 | 0 | 0 | 0 | 0 | 0 | 0 | 0 | 0 | 0 | 0 | 432 | 0 | 0 | 0 | 33650 | 30571 | 219371 | 0 | 0 | 107 | 12887 | 0 | 0 | 0 | 0 | 0 | 0 | 0 | 0 | 0 | 0 | 0 | 0 | 0 | 121 | 0 | 0 |
| SRR8280551 | 0 | 0 | 0 | 0 | 0 | 0 | 0 | 0 | 0 | 0 | 140 | 0 | 0 | 0 | 37789 | 33730 | 179018 | 0 | 0 | 0 | 11634 | 0 | 0 | 0 | 0 | 0 | 0 | 0 | 0 | 0 | 0 | 0 | 0 | 0 | 32 | 0 | 35 |
| SRR8280552 | 0 | 0 | 0 | 0 | 0 | 0 | 0 | 0 | 0 | 0 | 496 | 0 | 0 | 0 | 22101 | 41303 | 168403 | 0 | 0 | 0 | 57644 | 0 | 0 | 0 | 0 | 0 | 0 | 0 | 0 | 0 | 0 | 0 | 0 | 0 | 204 | 0 | 0 |
| SRR8280554 | 0 | 0 | 0 | 0 | 0 | 253 | 0 | 0 | 0 | 0 | 0 | 0 | 0 | 0 | 29613 | 65185 | 185845 | 0 | 0 | 0 | 24298 | 0 | 0 | 0 | 0 | 0 | 0 | 0 | 0 | 0 | 0 | 0 | 0 | 0 | 191 | 0 | 0 |
| SRR8280558 | 0 | 0 | 0 | 0 | 0 | 0 | 0 | 0 | 0 | 0 | 0 | 0 | 0 | 0 | 16675 | 32833 | 115909 | 0 | 0 | 1042 | 19020 | 0 | 0 | 0 | 0 | 0 | 0 | 0 | 0 | 219 | 0 | 0 | 0 | 0 | 297 | 0 | 0 |
| SRR8280560 | 0 | 0 | 0 | 0 | 0 | 0 | 0 | 299 | 0 | 0 | 294 | 0 | 0 | 0 | 40906 | 29305 | 254535 | 0 | 0 | 449 | 4639 | 0 | 0 | 0 | 0 | 0 | 0 | 0 | 0 | 0 | 0 | 0 | 0 | 0 | 104 | 0 | 0 |
| SRR8280561 | 0 | 0 | 0 | 0 | 0 | 73 | 0 | 0 | 0 | 0 | 125 | 0 | 0 | 0 | 51303 | 35955 | 221760 | 0 | 0 | 366 | 7977 | 0 | 0 | 0 | 0 | 0 | 0 | 0 | 0 | 31 | 0 | 0 | 0 | 0 | 113 | 0 | 0 |
| SRR8280562 | 0 | 0 | 0 | 0 | 0 | 100 | 0 | 200 | 0 | 0 | 106 | 0 | 0 | 0 | 44060 | 53279 | 175896 | 0 | 0 | 848 | 15568 | 0 | 0 | 0 | 35 | 0 | 35 | 0 | 0 | 106 | 0 | 0 | 0 | 0 | 56 | 0 | 0 |
| SRR8280564 | 0 | 0 | 0 | 0 | 0 | 51 | 0 | 256 | 0 | 0 | 340 | 0 | 0 | 0 | 28443 | 52423 | 153338 | 0 | 0 | 307 | 26290 | 0 | 0 | 0 | 24 | 49 | 0 | 24 | 0 | 24 | 0 | 0 | 0 | 0 | 111 | 0 | 0 |
| SRR8280566 | 0 | 0 | 0 | 0 | 0 | 0 | 0 | 75 | 0 | 0 | 182 | 0 | 0 | 0 | 61072 | 62310 | 148134 | 0 | 0 | 338 | 11854 | 0 | 0 | 0 | 0 | 0 | 0 | 26 | 0 | 52 | 0 | 0 | 0 | 0 | 91 | 0 | 0 |

### Supp Table S8.3.Columns_BX_to_DH

| | | | | | | | | | | | | | | | | | | | | | | | | | | | | | | | | | | | | | |
|---|---|---|---|---|---|---|---|---|---|---|---|---|---|---|---|---|---|---|---|---|---|---|---|---|---|---|---|---|---|---|---|---|---|---|---|---|---|
| SRR8280568 | 0 | 0 | 0 | 0 | 0 | 0 | 101 | 101 | 0 | 0 | 85 | 43 | 0 | 0 | 26869 | 44943 | 104683 | 0 | 0 | 404 | 12626 | 0 | 0 | 128 | 0 | 85 | 43 | 0 | 0 | 0 | 0 | 0 | 0 | 0 | 122 | 0 | 0 |
| SRR8280572 | 0 | 0 | 0 | 0 | 0 | 0 | 0 | 183 | 0 | 0 | 100 | 0 | 0 | 0 | 29078 | 25363 | 90353 | 0 | 0 | 1283 | 18387 | 0 | 0 | 0 | 0 | 0 | 0 | 14 | 0 | 29 | 0 | 0 | 0 | 0 | 161 | 0 | 0 |
| SRR8280573 | 0 | 0 | 0 | 0 | 0 | 0 | 0 | 45 | 0 | 0 | 115 | 19 | 0 | 0 | 48272 | 38065 | 153192 | 0 | 0 | 179 | 21211 | 0 | 0 | 96 | 77 | 58 | 77 | 211 | 0 | 77 | 0 | 0 | 0 | 0 | 161 | 0 | 173 |
| SRR8280574 | 0 | 0 | 0 | 0 | 0 | 0 | 0 | 0 | 0 | 0 | 305 | 76 | 0 | 0 | 8836 | 47722 | 36695 | 0 | 0 | 393 | 12566 | 0 | 0 | 0 | 0 | 0 | 0 | 0 | 0 | 0 | 0 | 0 | 0 | 0 | 602 | 0 | 191 |
| SRR8280576 | 0 | 0 | 0 | 0 | 0 | 655 | 0 | 0 | 0 | 0 | 898 | 0 | 0 | 0 | 17900 | 52595 | 17280 | 0 | 0 | 0 | 9823 | 0 | 0 | 0 | 0 | 0 | 0 | 0 | 0 | 0 | 0 | 0 | 0 | 0 | 72 | 0 | 0 |
| SRR8280580 | 0 | 0 | 0 | 0 | 0 | 0 | 0 | 0 | 0 | 0 | 443 | 40 | 0 | 0 | 66269 | 47291 | 195458 | 0 | 0 | 0 | 21357 | 0 | 0 | 0 | 40 | 242 | 81 | 0 | 0 | 0 | 0 | 0 | 0 | 0 | 153 | 0 | 40 |
| SRR8280582 | 0 | 0 | 0 | 0 | 0 | 0 | 0 | 0 | 0 | 0 | 827 | 0 | 0 | 0 | 42986 | 37236 | 161166 | 0 | 0 | 86 | 28142 | 0 | 0 | 0 | 0 | 0 | 0 | 0 | 0 | 0 | 0 | 0 | 0 | 0 | 98 | 0 | 0 |
| SRR8280583 | 0 | 0 | 0 | 0 | 0 | 0 | 0 | 0 | 0 | 30 | 113 | 0 | 0 | 0 | 616267 | 6308 | 383394 | 0 | 14 | 270 | 18939 | 0 | 12 | 142 | 178 | 723 | 996 | 326 | 0 | 356 | 0 | 0 | 0 | 0 | 69 | 406 | 0 |
| SRR8280584 | 0 | 0 | 0 | 0 | 0 | 0 | 0 | 0 | 0 | 0 | 245 | 0 | 0 | 0 | 39617 | 32414 | 145716 | 0 | 0 | 299 | 20657 | 0 | 0 | 0 | 0 | 0 | 147 | 0 | 0 | 0 | 0 | 0 | 0 | 0 | 193 | 0 | 49 |
| SRR8280589 | 0 | 0 | 0 | 0 | 0 | 0 | 0 | 0 | 0 | 0 | 529 | 0 | 0 | 0 | 17069 | 29398 | 71189 | 0 | 0 | 0 | 5061 | 0 | 0 | 0 | 0 | 0 | 0 | 0 | 0 | 0 | 0 | 0 | 0 | 0 | 62 | 0 | 66 |
| SRR8280590 | 0 | 0 | 0 | 0 | 0 | 0 | 0 | 507 | 0 | 0 | 110 | 74 | 0 | 0 | 38036 | 95982 | 213708 | 0 | 101 | 507 | 43311 | 0 | 0 | 0 | 0 | 184 | 0 | 0 | 0 | 0 | 0 | 0 | 0 | 0 | 26 | 0 | 74 |
| SRR8280592 | 0 | 0 | 0 | 0 | 0 | 0 | 0 | 0 | 0 | 0 | 78 | 0 | 0 | 0 | 29125 | 33080 | 156841 | 0 | 0 | 459 | 7683 | 0 | 0 | 39 | 0 | 0 | 0 | 0 | 0 | 0 | 0 | 0 | 0 | 0 | 15 | 0 | 0 |
| SRR8280593 | 0 | 0 | 0 | 0 | 0 | 0 | 0 | 163 | 0 | 0 | 50 | 66 | 0 | 0 | 18129 | 81867 | 143218 | 0 | 0 | 217 | 36528 | 0 | 0 | 0 | 0 | 17 | 17 | 0 | 0 | 0 | 0 | 0 | 0 | 0 | 69 | 0 | 33 |
| SRR8311061 | 0 | 0 | 0 | 0 | 61 | 0 | 0 | 0 | 0 | 0 | 31 | 0 | 0 | 0 | 46919 | 206908 | 391915 | 0 | 0 | 0 | 0 | 0 | 0 | 0 | 0 | 0 | 0 | 0 | 0 | 0 | 0 | 0 | 0 | 0 | 2833 | 0 | 77 |
| SRR8311062 | 0 | 0 | 0 | 0 | 104 | 1018 | 611 | 3596 | 0 | 0 | 122 | 0 | 0 | 0 | 53739 | 160967 | 524768 | 0 | 0 | 0 | 0 | 0 | 0 | 0 | 45 | 77 | 0 | 149 | 0 | 63 | 0 | 0 | 0 | 0 | 3203 | 0 | 63 |
| SRR8311063 | 0 | 0 | 0 | 0 | 346 | 0 | 0 | 0 | 0 | 0 | 154 | 0 | 0 | 0 | 61174 | 295490 | 389717 | 0 | 0 | 0 | 0 | 0 | 0 | 0 | 0 | 0 | 0 | 109 | 0 | 0 | 0 | 0 | 0 | 0 | 1644 | 0 | 0 |
| SRR8311064 | 0 | 0 | 0 | 0 | 14 | 0 | 0 | 2562 | 0 | 0 | 0 | 0 | 0 | 0 | 61746 | 189580 | 346935 | 0 | 0 | 0 | 0 | 0 | 0 | 0 | 0 | 0 | 0 | 0 | 0 | 0 | 0 | 0 | 0 | 0 | 3193 | 0 | 131 |
| SRR8311070 | 0 | 0 | 0 | 0 | 619 | 0 | 0 | 0 | 0 | 0 | 0 | 0 | 0 | 0 | 117552 | 205084 | 298806 | 0 | 0 | 0 | 0 | 0 | 0 | 0 | 0 | 0 | 0 | 0 | 0 | 0 | 0 | 0 | 0 | 0 | 1624 | 0 | 150 |
| SRR8311071 | 0 | 0 | 0 | 0 | 157 | 0 | 0 | 3274 | 0 | 0 | 0 | 0 | 0 | 0 | 38423 | 303260 | 310434 | 0 | 0 | 0 | 0 | 0 | 0 | 0 | 0 | 0 | 0 | 0 | 0 | 0 | 0 | 0 | 0 | 0 | 827 | 0 | 0 |
| SRR8311072 | 0 | 0 | 0 | 0 | 96 | 0 | 302 | 804 | 427 | 0 | 72 | 22 | 0 | 0 | 66161 | 271007 | 359177 | 0 | 0 | 0 | 0 | 0 | 0 | 0 | 14 | 43 | 0 | 12 | 0 | 34 | 0 | 0 | 0 | 0 | 1952 | 0 | 22 |
| SRR8311073 | 0 | 0 | 0 | 0 | 353 | 0 | 0 | 0 | 0 | 0 | 41 | 70 | 0 | 0 | 55890 | 200490 | 328161 | 0 | 0 | 0 | 1844 | 0 | 0 | 0 | 0 | 79 | 0 | 0 | 0 | 139 | 0 | 0 | 0 | 0 | 180 | 0 | 0 |
| SRR8311074 | 0 | 0 | 0 | 0 | 0 | 0 | 0 | 0 | 1596 | 0 | 0 | 0 | 0 | 0 | 76640 | 243862 | 321387 | 0 | 0 | 0 | 0 | 0 | 0 | 0 | 0 | 0 | 0 | 0 | 0 | 0 | 0 | 0 | 0 | 0 | 2862 | 0 | 0 |
| SRR8311076 | 0 | 0 | 0 | 0 | 197 | 0 | 1666 | 278 | 0 | 0 | 55 | 55 | 0 | 0 | 53927 | 172399 | 274601 | 0 | 0 | 0 | 0 | 0 | 0 | 0 | 0 | 48 | 77 | 26 | 0 | 0 | 0 | 0 | 0 | 0 | 1887 | 0 | 45 |
| SRR8311078 | 0 | 0 | 0 | 0 | 0 | 632 | 1328 | 948 | 1391 | 85 | 0 | 0 | 0 | 0 | 46532 | 287143 | 423584 | 0 | 0 | 0 | 0 | 0 | 67 | 0 | 0 | 71 | 0 | 0 | 0 | 0 | 0 | 0 | 0 | 0 | 1129 | 0 | 0 |
| SRR8311080 | 0 | 0 | 0 | 0 | 270 | 0 | 0 | 0 | 0 | 165 | 0 | 0 | 0 | 0 | 86416 | 212630 | 421958 | 0 | 0 | 1084 | 0 | 0 | 0 | 0 | 0 | 0 | 0 | 0 | 0 | 0 | 0 | 0 | 0 | 0 | 3274 | 0 | 0 |
| SRR8311081 | 0 | 0 | 0 | 0 | 0 | 0 | 0 | 0 | 0 | 0 | 0 | 0 | 0 | 0 | 70362 | 340568 | 332367 | 0 | 0 | 0 | 0 | 0 | 0 | 0 | 0 | 0 | 0 | 0 | 0 | 0 | 0 | 0 | 0 | 395 | 1068 | 0 | 229 |
| SRR8311082 | 0 | 0 | 0 | 0 | 0 | 0 | 0 | 0 | 0 | 0 | 0 | 0 | 0 | 0 | 39589 | 195928 | 498326 | 0 | 0 | 0 | 0 | 0 | 0 | 0 | 0 | 0 | 0 | 0 | 0 | 0 | 0 | 0 | 0 | 0 | 3393 | 0 | 0 |
| SRR8311085 | 0 | 0 | 0 | 0 | 71 | 0 | 540 | 848 | 0 | 0 | 0 | 0 | 0 | 0 | 87273 | 353943 | 286772 | 0 | 0 | 0 | 0 | 0 | 0 | 0 | 0 | 0 | 0 | 0 | 0 | 0 | 52 | 0 | 0 | 0 | 1025 | 0 | 257 |
| SRR8311087 | 0 | 0 | 0 | 0 | 0 | 528 | 0 | 791 | 0 | 0 | 0 | 0 | 0 | 0 | 85733 | 208299 | 372933 | 0 | 0 | 0 | 0 | 0 | 0 | 244 | 0 | 0 | 0 | 0 | 0 | 0 | 0 | 0 | 0 | 0 | 3296 | 12 | 0 |
| SRR8311089 | 0 | 0 | 0 | 0 | 0 | 0 | 0 | 2510 | 1159 | 0 | 0 | 116 | 0 | 0 | 53973 | 305301 | 259911 | 0 | 0 | 0 | 0 | 0 | 0 | 0 | 0 | 0 | 0 | 0 | 0 | 0 | 0 | 0 | 0 | 0 | 0 | 15 | 160 |
| SRR8311092 | 0 | 0 | 0 | 0 | 0 | 3479 | 0 | 6185 | 0 | 0 | 0 | 0 | 0 | 0 | 75384 | 268749 | 383383 | 0 | 0 | 0 | 0 | 0 | 0 | 0 | 0 | 146 | 0 | 479 | 0 | 0 | 0 | 0 | 0 | 0 | 1322 | 0 | 232 |
| SRR8311093 | 0 | 0 | 0 | 0 | 59 | 0 | 1397 | 898 | 0 | 0 | 76 | 296 | 0 | 0 | 39972 | 181344 | 377931 | 0 | 0 | 0 | 0 | 0 | 0 | 0 | 0 | 0 | 0 | 0 | 0 | 0 | 0 | 0 | 0 | 0 | 1641 | 0 | 194 |
| SRR8311095 | 0 | 0 | 0 | 0 | 0 | 0 | 0 | 2270 | 0 | 0 | 120 | 0 | 0 | 0 | 44046 | 245433 | 253117 | 0 | 0 | 0 | 0 | 0 | 0 | 0 | 0 | 103 | 0 | 0 | 0 | 0 | 0 | 0 | 0 | 0 | 2899 | 0 | 158 |
| SRR8311098 | 0 | 0 | 0 | 0 | 412 | 0 | 1838 | 0 | 0 | 0 | 0 | 0 | 0 | 0 | 46448 | 338673 | 278182 | 0 | 0 | 0 | 0 | 0 | 0 | 0 | 168 | 148 | 0 | 0 | 0 | 0 | 0 | 0 | 0 | 0 | 3742 | 0 | 0 |
| SRR8311105 | 0 | 0 | 0 | 0 | 0 | 238 | 0 | 0 | 1031 | 0 | 0 | 128 | 0 | 0 | 59388 | 291405 | 363501 | 0 | 0 | 0 | 0 | 0 | 0 | 0 | 39 | 83 | 0 | 0 | 0 | 0 | 108 | 0 | 0 | 0 | 2427 | 0 | 98 |
| SRR8311106 | 0 | 0 | 0 | 0 | 0 | 0 | 0 | 0 | 0 | 0 | 0 | 0 | 0 | 0 | 33219 | 184573 | 330816 | 0 | 0 | 0 | 0 | 0 | 0 | 0 | 0 | 0 | 0 | 0 | 0 | 0 | 0 | 0 | 0 | 0 | 2691 | 0 | 0 |
| SRR835324 | 18 | 19 | 0 | 0 | 23 | 25495 | 13150 | 85129 | 146242 | 310 | 38 | 0 | 11 | 0 | 4882 | 70585 | 45001 | 402 | 146 | 601 | 571 | 0 | 0 | 39 | 50 | 36 | 0 | 0 | 0 | 0 | 0 | 0 | 0 | 15 | 16869 | 0 | 0 |
| SRR835325 | 40 | 11 | 0 | 0 | 31 | 15071 | 16038 | 103465 | 203238 | 162 | 96 | 0 | 0 | 0 | 8524 | 118642 | 67620 | 505 | 136 | 441 | 50 | 0 | 0 | 41 | 84 | 80 | 11 | 35 | 0 | 0 | 0 | 0 | 0 | 0 | 19438 | 0 | 0 |
| SRR8354354 | 0 | 0 | 0 | 0 | 84 | 0 | 0 | 0 | 0 | 49 | 23 | 22 | 0 | 0 | 15036 | 222378 | 76750 | 0 | 0 | 284 | 0 | 0 | 0 | 0 | 0 | 0 | 0 | 0 | 0 | 15 | 53 | 0 | 0 | 0 | 8434 | 0 | 0 |
| SRR837775 | 112 | 69 | 26 | 23 | 35 | 194 | 352 | 1204 | 112 | 47 | 63 | 39 | 0 | 0 | 5560 | 1092 | 3989 | 21 | 21 | 1699 | 1609 | 0 | 0 | 192 | 20 | 88 | 181 | 15 | 0 | 703 | 0 | 17 | 0 | 0 | 13404 | 0 | 19 |
| SRR837776 | 84 | 41 | 13 | 15 | 30 | 426 | 534 | 1846 | 225 | 19 | 17 | 12 | 0 | 0 | 2935 | 584 | 3032 | 33 | 0 | 581 | 822 | 0 | 0 | 13 | 0 | 54 | 80 | 0 | 0 | 12 | 0 | 0 | 0 | 0 | 10688 | 0 | 0 |
| SRR837777 | 0 | 0 | 0 | 0 | 0 | 999 | 461 | 1021 | 150 | 0 | 0 | 0 | 0 | 0 | 453 | 84 | 1216 | 0 | 0 | 208 | 93 | 0 | 0 | 42 | 0 | 0 | 25 | 0 | 0 | 16 | 0 | 0 | 0 | 0 | 4081 | 0 | 0 |
| SRR837778 | 0 | 0 | 0 | 0 | 0 | 782 | 502 | 679 | 158 | 0 | 0 | 0 | 0 | 0 | 1440 | 354 | 1546 | 24 | 0 | 648 | 24 | 0 | 0 | 59 | 0 | 0 | 0 | 0 | 0 | 15 | 0 | 0 | 0 | 0 | 3782 | 0 | 0 |
| SRR837779 | 84 | 41 | 13 | 15 | 30 | 426 | 534 | 1846 | 225 | 19 | 17 | 12 | 0 | 0 | 2935 | 584 | 3032 | 33 | 0 | 581 | 822 | 0 | 0 | 13 | 0 | 54 | 80 | 0 | 0 | 12 | 0 | 0 | 0 | 0 | 10688 | 0 | 0 |
| SRR837780 | 112 | 69 | 26 | 23 | 35 | 189 | 343 | 1172 | 109 | 47 | 63 | 39 | 0 | 0 | 5412 | 1092 | 3989 | 20 | 20 | 1653 | 1566 | 0 | 0 | 192 | 20 | 88 | 181 | 15 | 0 | 703 | 0 | 17 | 0 | 0 | 13404 | 0 | 19 |
| SRR8380550 | 0 | 0 | 0 | 0 | 13 | 187 | 58 | 383 | 359 | 14 | 0 | 0 | 0 | 0 | 43082 | 77960 | 607352 | 61 | 2268 | 132278 | 2152 | 0 | 0 | 14 | 24 | 0 | 0 | 0 | 0 | 14 | 0 | 0 | 0 | 0 | 8028 | 101 | 0 |
| SRR8380552 | 125 | 64 | 0 | 0 | 158 | 822 | 403 | 1393 | 9709 | 3798 | 18 | 381 | 0 | 0 | 16409 | 283462 | 95575 | 122 | 1453 | 3693 | 3686 | 0 | 15 | 404 | 24 | 0 | 17 | 0 | 80 | 0 | 0 | 308 | 0 | 0 | 16451 | 13 | 0 |
| SRR8380553 | 94 | 77 | 0 | 0 | 71 | 475 | 460 | 505 | 3438 | 5419 | 23 | 251 | 0 | 0 | 9681 | 464560 | 46991 | 86 | 416 | 5218 | 12967 | 0 | 35 | 426 | 17 | 0 | 0 | 10 | 29 | 23 | 0 | 44 | 0 | 0 | 6022 | 208 | 0 |
| SRR8380554 | 0 | 0 | 0 | 0 | 65 | 58 | 73 | 183 | 1388 | 4164 | 16 | 41 | 0 | 0 | 3548 | 257211 | 114002 | 34 | 393 | 12708 | 1021 | 32 | 78 | 46 | 15 | 0 | 0 | 0 | 0 | 0 | 0 | 448 | 0 | 0 | 8529 | 0 | 0 |
| SRR8380555 | 0 | 0 | 0 | 0 | 54 | 3911 | 1737 | 4376 | 6817 | 359 | 0 | 0 | 0 | 0 | 31466 | 90855 | 300136 | 467 | 6863 | 54913 | 5692 | 0 | 0 | 283 | 21 | 0 | 0 | 0 | 0 | 15 | 0 | 49 | 0 | 0 | 6793 | 48 | 0 |
| SRR8380722 | 529 | 183 | 42 | 62 | 279 | 416 | 116 | 779 | 2524 | 1001 | 230 | 73 | 0 | 0 | 128133 | 262664 | 529095 | 29 | 1786 | 6571 | 4863 | 0 | 0 | 1060 | 750 | 871 | 2936 | 167 | 0 | 98 | 0 | 0 | 0 | 23 | 1996 | 0 | 0 |
| SRR8380723 | 193 | 61 | 0 | 0 | 177 | 2746 | 665 | 5291 | 15174 | 659 | 20 | 10 | 0 | 0 | 62433 | 118077 | 541315 | 262 | 3921 | 4733 | 1672 | 0 | 0 | 2291 | 182 | 66 | 91 | 0 | 0 | 41 | 0 | 0 | 0 | 0 | 6702 | 23 | 10 |
| SRR8380724 | 285 | 77 | 64 | 109 | 355 | 243 | 130 | 533 | 2677 | 80 | 74 | 0 | 0 | 0 | 74711 | 93991 | 809478 | 74 | 7569 | 5743 | 1192 | 0 | 0 | 1088 | 1034 | 1354 | 2710 | 509 | 0 | 90 | 0 | 0 | 0 | 0 | 572 | 29 | 0 |
| SRR8380725 | 147 | 57 | 0 | 0 | 94 | 3093 | 934 | 7110 | 12368 | 412 | 20 | 0 | 0 | 0 | 72755 | 224247 | 444455 | 178 | 3669 | 4959 | 1230 | 0 | 0 | 1328 | 102 | 57 | 77 | 0 | 0 | 40 | 0 | 0 | 0 | 0 | 2515 | 19 | 0 |
| SRR8380726 | 12 | 87 | 0 | 0 | 146 | 914 | 472 | 2944 | 7035 | 43 | 30 | 52 | 0 | 0 | 57995 | 334007 | 361174 | 86 | 2960 | 5008 | 4557 | 0 | 0 | 66 | 105 | 44 | 74 | 0 | 0 | 81 | 0 | 11 | 0 | 31 | 4050 | 0 | 0 |
| SRR8380727 | 112 | 33 | 0 | 0 | 155 | 1075 | 325 | 2776 | 6739 | 633 | 20 | 26 | 0 | 0 | 52854 | 109198 | 646060 | 114 | 4745 | 2598 | 1677 | 0 | 0 | 775 | 87 | 47 | 56 | 0 | 0 | 35 | 0 | 0 | 0 | 25 | 7877 | 10 | 0 |
| SRR8400006 | 0 | 0 | 0 | 0 | 15 | 54 | 15 | 104 | 0 | 0 | 12 | 0 | 0 | 0 | 7396 | 27953 | 76839 | 625 | 2146 | 3859 | 23974 | 0 | 0 | 18 | 11 | 0 | 27 | 0 | 0 | 15 | 0 | 0 | 13 | 16 | 7476 | 10451 | 53 |
| SRR8400007 | 0 | 0 | 0 | 0 | 0 | 3093 | 1046 | 6117 | 163 | 0 | 0 | 0 | 0 | 0 | 622 | 2919 | 4504 | 672 | 78 | 156 | 876 | 0 | 0 | 235 | 0 | 0 | 0 | 0 | 0 | 0 | 0 | 0 | 0 | 0 | 26327 | 3583 | 35 |
| SRR8400008 | 0 | 0 | 0 | 0 | 96 | 427 | 103 | 557 | 77 | 0 | 11 | 0 | 0 | 0 | 6200 | 13791 | 24785 | 2491 | 958 | 934 | 12526 | 0 | 0 | 397 | 0 | 0 | 15 | 11 | 0 | 0 | 0 | 57 | 16 | 18 | 11111 | 3393 | 37 |
| SRR8400009 | 0 | 0 | 0 | 0 | 163 | 1171 | 189 | 2777 | 195 | 0 | 0 | 0 | 0 | 0 | 2225 | 49854 | 106966 | 221 | 479 | 1376 | 6719 | 0 | 0 | 192 | 0 | 0 | 14 | 0 | 0 | 0 | 0 | 19 | 0 | 0 | 8228 | 16035 | 41 |
| SRR8400010 | 0 | 0 | 0 | 0 | 80 | 3845 | 1423 | 4363 | 459 | 0 | 0 | 0 | 0 | 0 | 2504 | 15403 | 13199 | 2230 | 850 | 567 | 5219 | 0 | 0 | 56 | 0 | 0 | 0 | 0 | 0 | 0 | 0 | 0 | 0 | 0 | 12579 | 3688 | 11 |
| SRR8417876 | 101 | 76 | 47 | 138 | 20 | 27 | 51 | 92 | 29 | 0 | 48 | 0 | 0 | 0 | 58811 | 61535 | 119514 | 354 | 56 | 5210 | 4894 | 0 | 0 | 45 | 29 | 135 | 378 | 190 | 0 | 82 | 0 | 7248 | 0 | 0 | 6284 | 0 | 0 |
| SRR8417877 | 47 | 37 | 34 | 14 | 32 | 46 | 104 | 124 | 41 | 0 | 57 | 0 | 0 | 0 | 64542 | 64643 | 108046 | 371 | 75 | 10174 | 7203 | 0 | 0 | 74 | 39 | 25 | 59 | 20 | 0 | 15 | 0 | 8552 | 0 | 0 | 6405 | 0 | 0 |
| SRR8417878 | 41 | 26 | 19 | 47 | 146 | 65 | 142 | 118 | 134 | 0 | 25 | 0 | 0 | 0 | 85772 | 30842 | 134958 | 274 | 100 | 14202 | 9577 | 0 | 0 | 91 | 21 | 87 | 159 | 15 | 0 | 83 | 0 | 8558 | 0 | 0 | 6312 | 0 | 0 |
| SRR8435171 | 30 | 0 | 0 | 0 | 54 | 7213 | 1232 | 8936 | 2612 | 1884 | 0 | 59 | 0 | 0 | 11698 | 30868 | 48292 | 928 | 309 | 6617 | 326 | 0 | 15 | 364 | 0 | 0 | 0 | 0 | 10 | 19 | 16 | 182 | 0 | 57 | 5794 | 11 | 18 |
| SRR8435172 | 15 | 0 | 0 | 0 | 0 | 6128 | 1375 | 7991 | 2030 | 485 | 0 | 131 | 0 | 0 | 6680 | 16460 | 22602 | 623 | 141 | 3025 | 276 | 10 | 16 | 91 | 0 | 0 | 0 | 0 | 35 | 0 | 0 | 662 | 17 | 103 | 14022 | 12 | 16 |
| SRR8481810 | 28 | 55 | 0 | 24 | 19 | 308 | 462 | 769 | 77 | 0 | 86 | 0 | 0 | 0 | 2201 | 4326 | 23368 | 15 | 0 | 3694 | 200 | 0 | 0 | 12 | 12 | 70 | 71 | 31 | 0 | 56 | 0 | 15 | 0 | 0 | 5401 | 0 | 0 |
| SRR8481811 | 0 | 22 | 0 | 16 | 10 | 440 | 378 | 794 | 174 | 0 | 32 | 24 | 0 | 0 | 3027 | 19198 | 26629 | 12 | 0 | 2854 | 502 | 0 | 0 | 59 | 27 | 15 | 66 | 11 | 0 | 39 | 0 | 174 | 0 | 0 | 14169 | 0 | 0 |
| SRR8481812 | 14 | 40 | 0 | 38 | 16 | 308 | 662 | 891 | 137 | 0 | 61 | 0 | 0 | 0 | 3449 | 22828 | 48239 | 11 | 11 | 3141 | 240 | 0 | 0 | 14 | 30 | 98 | 68 | 35 | 0 | 41 | 0 | 23 | 0 | 0 | 5489 | 0 | 0 |
| SRR8481813 | 0 | 0 | 0 | 0 | 12 | 529 | 246 | 601 | 184 | 0 | 40 | 27 | 0 | 0 | 11477 | 40237 | 67862 | 0 | 0 | 3734 | 1841 | 0 | 0 | 18 | 26 | 62 | 67 | 0 | 0 | 193 | 0 | 361 | 0 | 0 | 20762 | 0 | 0 |
| SRR8481814 | 21 | 49 | 0 | 0 | 25 | 317 | 346 | 480 | 67 | 0 | 51 | 0 | 0 | 0 | 4544 | 3338 | 52746 | 0 | 0 | 3411 | 538 | 0 | 0 | 13 | 29 | 10 | 83 | 12 | 0 | 34 | 0 | 0 | 0 | 0 | 3729 | 0 | 0 |
| SRR8481815 | 16 | 27 | 0 | 20 | 23 | 464 | 253 | 717 | 84 | 0 | 25 | 18 | 0 | 0 | 8070 | 10314 | 66933 | 0 | 11 | 2732 | 928 | 0 | 0 | 117 | 59 | 215 | 677 | 10 | 0 | 178 | 0 | 148 | 0 | 0 | 15008 | 0 | 0 |
| SRR8481816 | 28 | 12 | 19 | 63 | 44 | 217 | 248 | 348 | 117 | 10 | 100 | 0 | 0 | 0 | 4439 | 2452 | 50062 | 0 | 0 | 3491 | 417 | 0 | 0 | 156 | 60 | 249 | 1198 | 313 | 0 | 738 | 0 | 0 | 0 | 0 | 4022 | 0 | 0 |
| SRR8481817 | 0 | 0 | 0 | 0 | 0 | 391 | 197 | 619 | 137 | 0 | 27 | 26 | 0 | 0 | 12337 | 22324 | 64966 | 0 | 0 | 3028 | 1518 | 0 | 0 | 19 | 14 | 46 | 60 | 14 | 0 | 33 | 0 | 316 | 0 | 0 | 15642 | 0 | 0 |
| SRR8481818 | 11 | 14 | 14 | 73 | 43 | 58 | 137 | 195 | 94 | 0 | 38 | 0 | 0 | 0 | 2060 | 5542 | 37141 | 0 | 14 | 4069 | 665 | 0 | 0 | 23 | 39 | 28 | 224 | 36 | 0 | 196 | 0 | 0 | 0 | 0 | 4749 | 0 | 0 |
| SRR8481819 | 30 | 12 | 44 | 157 | 42 | 118 | 281 | 335 | 91 | 0 | 83 | 12 | 0 | 0 | 4120 | 17538 | 60960 | 0 | 54 | 4183 | 1123 | 0 | 0 | 16 | 133 | 44 | 208 | 38 | 0 | 37 | 0 | 54 | 0 | 0 | 4966 | 0 | 0 |
| SRR8481820 | 49 | 31 | 45 | 183 | 58 | 12 | 61 | 184 | 12 | 0 | 103 | 0 | 0 | 0 | 2610 | 11694 | 44580 | 0 | 25 | 4264 | 649 | 0 | 0 | 13 | 22 | 15 | 139 | 31 | 0 | 28 | 0 | 37 | 0 | 0 | 4975 | 0 | 0 |
| SRR8481821 | 0 | 0 | 0 | 11 | 0 | 353 | 132 | 406 | 82 | 0 | 24 | 29 | 0 | 0 | 7979 | 40861 | 57982 | 0 | 0 | 2054 | 3233 | 0 | 0 | 19 | 11 | 56 | 57 | 19 | 0 | 20 | 0 | 500 | 0 | 0 | 6971 | 0 | 0 |
| SRR8485055 | 35 | 35 | 14 | 17 | 147 | 1284 | 888 | 3167 | 1708 | 0 | 11 | 0 | 0 | 0 | 130120 | 261261 | 292330 | 0 | 2843 | 3637 | 819 | 0 | 0 | 11 | 20 | 29 | 49 | 30 | 0 | 38 | 0 | 0 | 0 | 0 | 979 | 0 | 0 |
| SRR8485056 | 91 | 32 | 26 | 18 | 1641 | 1143 | 885 | 3080 | 1012 | 41 | 106 | 0 | 0 | 0 | 56418 | 143439 | 300221 | 13 | 717 | 8000 | 1980 | 0 | 0 | 37 | 83 | 124 | 338 | 168 | 0 | 406 | 0 | 0 | 0 | 0 | 2555 | 12 | 0 |
| SRR8485057 | 34 | 26 | 18 | 21 | 85 | 833 | 648 | 2194 | 945 | 0 | 24 | 0 | 0 | 0 | 31854 | 239985 | 146842 | 0 | 541 | 3176 | 2952 | 0 | 0 | 24 | 51 | 90 | 168 | 79 | 0 | 80 | 0 | 0 | 0 | 0 | 1360 | 472 | 0 |
| SRR8485058 | 24 | 17 | 0 | 0 | 1206 | 560 | 447 | 1539 | 823 | 11 | 150 | 18 | 0 | 0 | 237015 | 95764 | 495445 | 0 | 908 | 1400 | 456 | 0 | 0 | 47 | 101 | 153 | 338 | 260 | 10 | 426 | 0 | 0 | 0 | 0 | 2066 | 81 | 0 |
| SRR8485059 | 60 | 54 | 54 | 59 | 44 | 203 | 161 | 633 | 337 | 0 | 18 | 0 | 0 | 0 | 130198 | 274157 | 171579 | 0 | 55 | 1013 | 1007 | 0 | 0 | 16 | 33 | 56 | 95 | 35 | 0 | 36 | 0 | 0 | 0 | 0 | 593 | 60 | 0 |
| SRR8485060 | 25 | 17 | 10 | 17 | 228 | 4152 | 3416 | 10888 | 7037 | 0 | 38 | 0 | 0 | 0 | 26061 | 76728 | 169335 | 0 | 767 | 1902 | 423 | 0 | 0 | 48 | 88 | 189 | 503 | 172 | 0 | 84 | 0 | 0 | 0 | 0 | 6048 | 88 | 0 |
| SRR8485061 | 55 | 34 | 16 | 19 | 1654 | 2004 | 1521 | 4951 | 2528 | 0 | 232 | 0 | 0 | 0 | 106632 | 208221 | 207568 | 0 | 1106 | 3982 | 471 | 0 | 0 | 131 | 126 | 240 | 548 | 282 | 0 | 956 | 0 | 0 | 0 | 0 | 1405 | 70 | 0 |
| SRR8485062 | 58 | 45 | 18 | 23 | 84 | 3266 | 2003 | 9915 | 3198 | 11 | 18 | 0 | 0 | 0 | 39926 | 100223 | 230510 | 0 | 699 | 9769 | 2861 | 0 | 0 | 75 | 12 | 19 | 26 | 34 | 0 | 46 | 0 | 0 | 0 | 0 | 4513 | 181 | 0 |
| SRR8485063 | 61 | 39 | 16 | 19 | 591 | 8944 | 6595 | 23582 | 8942 | 10 | 942 | 0 | 0 | 0 | 82380 | 182871 | 208949 | 0 | 790 | 2326 | 865 | 0 | 0 | 566 | 311 | 378 | 443 | 221 | 0 | 567 | 0 | 0 | 0 | 0 | 4064 | 218 | 0 |
| SRR8485064 | 68 | 52 | 10 | 23 | 1282 | 6246 | 4132 | 15770 | 7108 | 17 | 264 | 0 | 0 | 0 | 63167 | 54265 | 182390 | 0 | 775 | 1477 | 792 | 0 | 0 | 170 | 243 | 386 | 617 | 510 | 0 | 588 | 0 | 0 | 0 | 0 | 2943 | 344 | 0 |
| SRR8485065 | 23 | 16 | 0 | 19 | 24 | 1796 | 962 | 4437 | 1877 | 0 | 26 | 0 | 0 | 0 | 287546 | 103271 | 491113 | 0 | 1130 | 5938 | 6205 | 0 | 0 | 30 | 38 | 45 | 61 | 37 | 0 | 46 | 0 | 0 | 0 | 0 | 2567 | 19 | 0 |
| SRR8485066 | 33 | 38 | 0 | 26 | 335 | 3493 | 2288 | 8789 | 4198 | 0 | 434 | 0 | 0 | 0 | 26619 | 385039 | 79605 | 0 | 334 | 2439 | 480 | 0 | 0 | 216 | 136 | 166 | 150 | 210 | 0 | 423 | 0 | 0 | 0 | 0 | 2130 | 189 | 0 |
| SRR8485067 | 86 | 92 | 16 | 28 | 29 | 2509 | 1848 | 4753 | 2274 | 29 | 102 | 0 | 0 | 0 | 111518 | 471412 | 233031 | 0 | 1236 | 1041 | 198 | 61 | 96 | 48 | 11 | 17 | 21 | 11 | 0 | 73 | 11 | 0 | 0 | 0 | 409 | 162 | 0 |
| SRR8487217 | 59 | 30 | 50 | 12 | 135 | 2208 | 839 | 4088 | 1354 | 0 | 27 | 0 | 0 | 0 | 50025 | 263965 | 236427 | 0 | 692 | 4065 | 2416 | 0 | 0 | 51 | 94 | 125 | 221 | 80 | 0 | 86 | 0 | 0 | 0 | 0 | 2433 | 353 | 0 |
| SRR8487218 | 13 | 66 | 13 | 0 | 116 | 302 | 279 | 1009 | 373 | 10 | 83 | 0 | 0 | 0 | 151289 | 265516 | 278524 | 0 | 48 | 678 | 1810 | 0 | 0 | 23 | 58 | 84 | 78 | 92 | 0 | 32 | 0 | 0 | 0 | 0 | 2015 | 63 | 0 |
| SRR8487219 | 14 | 57 | 35 | 52 | 19 | 1141 | 651 | 2625 | 664 | 0 | 48 | 0 | 0 | 0 | 196971 | 72838 | 400916 | 20 | 563 | 11229 | 5683 | 0 | 0 | 15 | 24 | 32 | 50 | 42 | 0 | 12 | 0 | 0 | 0 | 0 | 494 | 406 | 0 |
| SRR8487220 | 95 | 66 | 19 | 31 | 14 | 457 | 236 | 997 | 390 | 0 | 103 | 0 | 0 | 0 | 161232 | 475985 | 183867 | 0 | 382 | 458 | 143 | 0 | 0 | 27 | 18 | 24 | 24 | 25 | 0 | 16 | 0 | 0 | 0 | 0 | 460 | 64 | 0 |
| SRR8487221 | 13 | 41 | 30 | 14 | 227 | 6186 | 4971 | 18035 | 11567 | 0 | 26 | 0 | 0 | 0 | 284466 | 70916 | 653634 | 0 | 600 | 3757 | 825 | 0 | 0 | 297 | 81 | 51 | 15 | 40 | 0 | 44 | 0 | 0 | 0 | 0 | 1031 | 17 | 0 |
| SRR8487222 | 81 | 51 | 38 | 52 | 906 | 17679 | 13444 | 47369 | 22117 | 0 | 51 | 0 | 0 | 0 | 41547 | 267294 | 295211 | 0 | 208 | 12029 | 1319 | 0 | 0 | 346 | 153 | 127 | 67 | 105 | 0 | 183 | 0 | 0 | 0 | 0 | 1632 | 80 | 0 |
| SRR8487223 | 27 | 17 | 40 | 11 | 70 | 7808 | 4681 | 16310 | 2679 | 45 | 33 | 0 | 0 | 0 | 98382 | 116898 | 403196 | 12 | 179 | 2431 | 3104 | 24 | 21 | 46 | 21 | 12 | 29 | 32 | 0 | 61 | 0 | 0 | 0 | 0 | 1213 | 90 | 0 |
| SRR8487224 | 20 | 0 | 0 | 0 | 506 | 6064 | 6037 | 16261 | 3440 | 14 | 577 | 0 | 0 | 0 | 124464 | 200106 | 276640 | 0 | 257 | 1445 | 1346 | 0 | 12 | 89 | 157 | 214 | 144 | 1668 | 0 | 3058 | 0 | 0 | 0 | 0 | 1833 | 131 | 0 |
| SRR8586304 | 0 | 0 | 0 | 0 | 0 | 28 | 0 | 45 | 0 | 47 | 43 | 0 | 0 | 0 | 326276 | 69732 | 766949 | 48 | 58 | 180 | 6240 | 0 | 0 | 0 | 0 | 0 | 0 | 0 | 0 | 66 | 0 | 0 | 0 | 0 | 58 | 22 | 0 |
| SRR871609 | 0 | 0 | 0 | 0 | 59 | 125 | 410 | 159 | 117 | 0 | 0 | 0 | 0 | 0 | 611 | 46322 | 19617 | 59 | 0 | 0 | 17 | 90 | 910 | 49 | 0 | 0 | 0 | 0 | 0 | 0 | 0 | 0 | 0 | 0 | 2920 | 938 | 0 |
| SRR871671 | 0 | 0 | 0 | 0 | 189 | 0 | 218 | 0 | 109 | 0 | 0 | 0 | 0 | 0 | 1579 | 159799 | 76923 | 0 | 0 | 0 | 0 | 23 | 53 | 83 | 0 | 0 | 0 | 0 | 0 | 0 | 0 | 0 | 0 | 0 | 0 | 90 | 0 |
| SRR873381 | 48 | 24 | 21 | 0 | 158 | 168 | 348 | 428 | 48 | 17 | 149 | 0 | 0 | 0 | 6104 | 39625 | 65197 | 12 | 30 | 9887 | 2351 | 0 | 0 | 123 | 12 | 13 | 31 | 0 | 0 | 25 | 0 | 0 | 0 | 0 | 4713 | 448 | 0 |
| SRR873401 | 81 | 55 | 0 | 0 | 29 | 109 | 314 | 596 | 114 | 0 | 104 | 29 | 0 | 0 | 5294 | 14963 | 56746 | 23 | 46 | 8125 | 9773 | 0 | 0 | 118 | 0 | 56 | 57 | 0 | 0 | 701 | 0 | 109 | 0 | 0 | 3583 | 82 | 0 |
| SRR873410 | 19 | 19 | 0 | 0 | 62 | 126 | 279 | 677 | 111 | 10 | 320 | 19 | 0 | 0 | 18100 | 31499 | 162904 | 19 | 77 | 13979 | 4535 | 0 | 0 | 308 | 57 | 323 | 478 | 0 | 0 | 1241 | 0 | 214 | 0 | 0 | 1541 | 252 | 0 |
| SRR8834226 | 0 | 0 | 0 | 0 | 25 | 0 | 0 | 0 | 0 | 0 | 100 | 0 | 0 | 0 | 0 | 357 | 0 | 0 | 0 | 0 | 0 | 0 | 0 | 0 | 0 | 0 | 0 | 0 | 0 | 141 | 0 | 0 | 0 | 0 | 83 | 0 | 0 |
| SRR8834227 | 0 | 0 | 0 | 0 | 0 | 0 | 16 | 52 | 24 | 0 | 0 | 0 | 0 | 0 | 92 | 511 | 3291 | 0 | 0 | 0 | 18 | 0 | 0 | 0 | 0 | 12 | 0 | 0 | 0 | 12 | 0 | 0 | 0 | 0 | 22 | 0 | 0 |
| SRR8839811 | 0 | 0 | 0 | 0 | 24 | 0 | 0 | 0 | 0 | 0 | 631 | 0 | 0 | 0 | 3275 | 256457 | 94585 | 0 | 653 | 100 | 4988 | 1270 | 2786 | 0 | 0 | 0 | 0 | 0 | 0 | 343 | 1275 | 31 | 0 | 12 | 4856 | 569 | 0 |
| SRR8839812 | 0 | 0 | 0 | 0 | 236 | 0 | 0 | 0 | 0 | 74 | 140 | 44 | 0 | 0 | 6078 | 864838 | 11617 | 0 | 0 | 0 | 2866 | 41 | 66 | 0 | 0 | 0 | 0 | 0 | 0 | 85 | 52 | 0 | 0 | 0 | 1306 | 142 | 0 |
| SRR8850891 | 465 | 174 | 286 | 100 | 313 | 697 | 616 | 4181 | 11695 | 856 | 233 | 0 | 0 | 0 | 77848 | 99957 | 214490 | 418 | 1219 | 708 | 1196 | 0 | 0 | 955 | 543 | 546 | 715 | 214 | 0 | 250 | 0 | 35 | 0 | 0 | 8599 | 0 | 0 |
| SRR8850892 | 63 | 23 | 43 | 17 | 36 | 710 | 563 | 3591 | 10469 | 113 | 27 | 16 | 0 | 0 | 80459 | 111219 | 244014 | 358 | 651 | 657 | 1133 | 0 | 0 | 117 | 83 | 64 | 92 | 10 | 0 | 35 | 0 | 59 | 0 | 0 | 10873 | 11 | 0 |
| SRR8850893 | 2512 | 597 | 1196 | 260 | 484 | 945 | 750 | 5051 | 12887 | 1599 | 142 | 18 | 0 | 0 | 19571 | 139891 | 61989 | 497 | 4956 | 1900 | 816 | 0 | 0 | 1253 | 954 | 671 | 720 | 89 | 0 | 183 | 0 | 33 | 0 | 0 | 8419 | 0 | 0 |
| SRR8850894 | 166 | 38 | 104 | 19 | 33 | 1061 | 1050 | 5686 | 13678 | 129 | 14 | 0 | 0 | 0 | 14501 | 101173 | 43347 | 380 | 5144 | 1000 | 525 | 0 | 0 | 115 | 77 | 56 | 62 | 13 | 0 | 19 | 0 | 11 | 0 | 0 | 5761 | 0 | 0 |
| SRR8850895 | 45 | 19 | 17 | 81 | 24 | 917 | 607 | 3928 | 7832 | 71 | 16 | 0 | 0 | 0 | 89599 | 93744 | 250051 | 417 | 773 | 880 | 1107 | 0 | 0 | 97 | 50 | 47 | 84 | 22 | 0 | 18 | 0 | 23 | 0 | 0 | 7542 | 0 | 0 |
| SRR8850896 | 306 | 127 | 81 | 66 | 85 | 784 | 499 | 2878 | 6553 | 808 | 113 | 0 | 0 | 0 | 94370 | 180877 | 224318 | 541 | 898 | 655 | 1752 | 0 | 0 | 745 | 406 | 373 | 637 | 401 | 0 | 89 | 0 | 57 | 0 | 0 | 7293 | 21 | 0 |
| SRR8850897 | 1409 | 436 | 543 | 149 | 233 | 693 | 558 | 2996 | 8552 | 1327 | 196 | 14 | 0 | 0 | 17020 | 140230 | 60680 | 475 | 2848 | 2143 | 975 | 0 | 0 | 330 | 214 | 256 | 437 | 262 | 0 | 352 | 0 | 64 | 0 | 0 | 10060 | 19 | 0 |
| SRR8850898 | 125 | 34 | 76 | 24 | 52 | 1397 | 834 | 5622 | 12518 | 150 | 19 | 15 | 0 | 0 | 29210 | 116358 | 187942 | 568 | 7152 | 4241 | 1806 | 0 | 0 | 80 | 69 | 58 | 56 | 13 | 0 | 26 | 0 | 276 | 0 | 0 | 9946 | 101 | 0 |
| SRR8850899 | 57 | 25 | 21 | 11 | 26 | 933 | 780 | 6013 | 11189 | 633 | 12 | 19 | 0 | 0 | 96358 | 107312 | 288807 | 990 | 1126 | 2345 | 2546 | 0 | 0 | 86 | 64 | 37 | 27 | 11 | 0 | 22 | 0 | 26 | 0 | 19 | 6973 | 0 | 0 |
| SRR8850900 | 404 | 156 | 214 | 77 | 158 | 1615 | 651 | 9791 | 8598 | 3291 | 138 | 14 | 0 | 0 | 118694 | 74640 | 320386 | 542 | 1314 | 1880 | 2808 | 0 | 0 | 1052 | 832 | 298 | 245 | 0 | 0 | 219 | 0 | 18 | 0 | 0 | 5023 | 0 | 0 |
| SRR8850901 | 1083 | 338 | 595 | 126 | 282 | 1248 | 980 | 6557 | 11750 | 4677 | 109 | 24 | 0 | 0 | 19952 | 106816 | 57611 | 735 | 3121 | 2777 | 2310 | 0 | 0 | 849 | 665 | 356 | 338 | 20 | 0 | 209 | 0 | 70 | 0 | 0 | 4769 | 0 | 0 |
| SRR8850902 | 88 | 32 | 38 | 61 | 24 | 878 | 583 | 4420 | 6697 | 376 | 78 | 20 | 0 | 0 | 12268 | 64994 | 38807 | 399 | 2719 | 1590 | 1673 | 0 | 0 | 71 | 46 | 15 | 92 | 0 | 0 | 27 | 0 | 157 | 0 | 0 | 13967 | 0 | 0 |
| SRR8858246 | 0 | 0 | 0 | 0 | 0 | 330 | 0 | 1666 | 0 | 0 | 0 | 0 | 0 | 0 | 507 | 152 | 742 | 0 | 0 | 1497 | 131 | 0 | 0 | 1025 | 0 | 0 | 0 | 0 | 0 | 0 | 0 | 0 | 0 | 0 | 23728 | 68 | 0 |
| SRR8858247 | 0 | 0 | 0 | 0 | 0 | 240 | 0 | 1835 | 155 | 70 | 0 | 0 | 0 | 0 | 682 | 115 | 2329 | 201 | 0 | 8232 | 0 | 0 | 0 | 75 | 0 | 0 | 0 | 0 | 0 | 0 | 0 | 240 | 0 | 0 | 21625 | 67 | 34 |
| SRR8858248 | 0 | 0 | 0 | 0 | 0 | 0 | 248 | 1356 | 0 | 0 | 0 | 29 | 0 | 0 | 580 | 0 | 518 | 0 | 0 | 673 | 0 | 0 | 0 | 73 | 0 | 0 | 0 | 0 | 0 | 0 | 0 | 0 | 0 | 0 | 42400 | 82 | 0 |

### Supp Table S8.3.Columns_BX_to_DH

| | | | | | | | | | | | | | | | | | | | | | | | | | | | | | | | | | | | | | |
|---|---|---|---|---|---|---|---|---|---|---|---|---|---|---|---|---|---|---|---|---|---|---|---|---|---|---|---|---|---|---|---|---|---|---|---|---|---|
| SRR8858254 | 0 | 0 | 0 | 0 | 0 | 0 | 0 | 1264 | 0 | 0 | 39 | 0 | 0 | 0 | 10323 | 7480 | 56245 | 0 | 0 | 10 | 1202 | 0 | 0 | 47 | 0 | 0 | 0 | 0 | 0 | 51 | 0 | 0 | 0 | 0 | 17092 | 79 | 0 |
| SRR8858258 | 0 | 0 | 0 | 0 | 0 | 34 | 0 | 0 | 475 | 0 | 0 | 0 | 0 | 0 | 0 | 316 | 0 | 0 | 0 | 68 | 0 | 0 | 0 | 40 | 0 | 0 | 0 | 0 | 0 | 0 | 0 | 0 | 0 | 0 | 20741 | 46 | 0 |
| SRR8907824 | 18 | 0 | 0 | 0 | 37 | 125 | 0 | 52 | 407 | 26 | 22 | 15 | 0 | 0 | 252 | 26394 | 6372 | 665 | 15 | 29 | 843 | 13 | 10 | 0 | 0 | 0 | 0 | 0 | 41 | 51 | 0 | 9491 | 0 | 0 | 2407 | 0 | 0 |
| SRR8932122 | 0 | 12 | 0 | 13 | 13 | 16 | 16 | 42 | 0 | 41 | 42 | 0 | 0 | 0 | 272141 | 43962 | 643054 | 14 | 62 | 6644 | 14379 | 0 | 0 | 26 | 0 | 0 | 18 | 0 | 0 | 45 | 0 | 187 | 0 | 0 | 244 | 67 | 0 |
| SRR8932123 | 0 | 0 | 0 | 0 | 0 | 44 | 158 | 311 | 20 | 16 | 32 | 0 | 0 | 0 | 80135 | 43871 | 422175 | 271 | 410 | 14860 | 4326 | 0 | 0 | 15 | 0 | 0 | 15 | 0 | 0 | 20 | 0 | 2520 | 0 | 0 | 1940 | 24 | 12 |
| SRR8932124 | 0 | 0 | 0 | 0 | 0 | 33 | 37 | 66 | 14 | 47 | 36 | 12 | 0 | 0 | 160361 | 28864 | 638574 | 22 | 82 | 13251 | 9765 | 0 | 0 | 28 | 0 | 11 | 22 | 0 | 0 | 59 | 0 | 25 | 0 | 0 | 943 | 485 | 16 |
| SRR8932125 | 0 | 0 | 0 | 0 | 0 | 137 | 188 | 351 | 57 | 99 | 56 | 15 | 0 | 0 | 41864 | 41548 | 372223 | 0 | 13 | 5319 | 2384 | 0 | 0 | 30 | 0 | 0 | 23 | 23 | 0 | 333 | 0 | 161 | 0 | 0 | 3430 | 504 | 12 |
| SRR8944248 | 0 | 0 | 0 | 0 | 0 | 0 | 0 | 2177 | 0 | 0 | 0 | 0 | 0 | 0 | 15602 | 527667 | 98565 | 0 | 0 | 0 | 726 | 0 | 0 | 2931 | 0 | 140 | 838 | 0 | 0 | 0 | 0 | 0 | 0 | 0 | 0 | 0 | 0 |
| SRR8944250 | 0 | 0 | 0 | 0 | 0 | 0 | 0 | 0 | 0 | 0 | 0 | 0 | 0 | 0 | 16813 | 225494 | 65649 | 0 | 0 | 0 | 1051 | 0 | 0 | 3734 | 0 | 747 | 6721 | 249 | 0 | 0 | 0 | 0 | 0 | 0 | 645 | 0 | 0 |
| SRR8944251 | 0 | 0 | 0 | 0 | 0 | 0 | 0 | 0 | 0 | 0 | 0 | 0 | 0 | 0 | 6478 | 376727 | 33690 | 0 | 0 | 0 | 0 | 0 | 0 | 1257 | 1048 | 419 | 629 | 629 | 0 | 0 | 0 | 0 | 0 | 0 | 0 | 0 | 0 |
| SRR8944252 | 0 | 0 | 0 | 0 | 0 | 0 | 0 | 0 | 0 | 0 | 0 | 0 | 0 | 0 | 12692 | 420459 | 186801 | 0 | 0 | 0 | 0 | 0 | 0 | 3423 | 590 | 4957 | 4367 | 0 | 0 | 0 | 0 | 0 | 0 | 0 | 439 | 0 | 0 |
| SRR8944257 | 0 | 0 | 0 | 0 | 0 | 0 | 0 | 3942 | 0 | 0 | 0 | 0 | 0 | 0 | 4730 | 240258 | 41262 | 0 | 0 | 0 | 0 | 0 | 0 | 0 | 0 | 3433 | 0 | 0 | 0 | 0 | 0 | 0 | 0 | 0 | 0 | 0 | 0 |
| SRR8944259 | 0 | 0 | 0 | 0 | 0 | 0 | 0 | 0 | 0 | 0 | 0 | 0 | 0 | 0 | 3719 | 310357 | 95277 | 0 | 0 | 0 | 2231 | 0 | 0 | 1506 | 0 | 502 | 1255 | 0 | 0 | 0 | 0 | 0 | 0 | 0 | 0 | 0 | 0 |
| SRR8956086 | 14 | 0 | 0 | 0 | 49 | 28 | 0 | 0 | 23 | 341 | 11 | 10 | 0 | 0 | 82249 | 138225 | 474981 | 11 | 684 | 181 | 5125 | 0 | 0 | 0 | 0 | 0 | 0 | 0 | 0 | 16 | 0 | 0 | 0 | 0 | 6382 | 312 | 0 |
| SRR8956087 | 14 | 0 | 0 | 0 | 121 | 19 | 0 | 0 | 16 | 691 | 11 | 0 | 0 | 0 | 122733 | 141630 | 479101 | 16 | 606 | 142 | 5699 | 0 | 0 | 0 | 0 | 0 | 0 | 0 | 0 | 0 | 0 | 15 | 0 | 0 | 5550 | 241 | 0 |
| SRR8956089 | 25 | 0 | 0 | 0 | 77 | 53 | 0 | 0 | 0 | 329 | 40 | 0 | 0 | 0 | 173051 | 187684 | 544152 | 24 | 1204 | 182 | 6763 | 0 | 0 | 12 | 12 | 0 | 0 | 0 | 0 | 30 | 0 | 0 | 0 | 0 | 4222 | 488 | 0 |
| SRR8956090 | 57 | 22 | 0 | 0 | 274 | 59 | 0 | 14 | 19 | 1229 | 67 | 14 | 0 | 0 | 233898 | 204253 | 566941 | 27 | 1387 | 383 | 11914 | 0 | 0 | 0 | 11 | 0 | 0 | 0 | 0 | 46 | 0 | 0 | 0 | 0 | 3016 | 306 | 0 |
| SRR8956091 | 43 | 13 | 0 | 0 | 354 | 38 | 0 | 11 | 14 | 1475 | 67 | 0 | 0 | 0 | 195957 | 185048 | 570345 | 23 | 1020 | 193 | 6246 | 0 | 0 | 10 | 12 | 0 | 0 | 0 | 0 | 35 | 0 | 29 | 0 | 0 | 3287 | 369 | 0 |
| SRR8992368 | 0 | 0 | 0 | 0 | 19 | 0 | 0 | 0 | 0 | 132 | 110 | 78 | 0 | 0 | 8274 | 439027 | 10336 | 19 | 172 | 121 | 674 | 0 | 0 | 0 | 0 | 0 | 0 | 0 | 0 | 29 | 0 | 83 | 0 | 16 | 1787 | 0 | 0 |
| SRR8992369 | 0 | 0 | 0 | 0 | 41 | 0 | 0 | 0 | 0 | 80 | 250 | 37 | 0 | 0 | 34877 | 534390 | 69867 | 94 | 552 | 571 | 10816 | 0 | 0 | 0 | 0 | 0 | 0 | 0 | 0 | 58 | 0 | 140 | 0 | 41 | 10474 | 0 | 0 |
| SRR8992370 | 0 | 0 | 0 | 0 | 271 | 0 | 0 | 0 | 0 | 968 | 2113 | 40 | 0 | 0 | 26750 | 594752 | 67553 | 93 | 465 | 623 | 11833 | 0 | 0 | 0 | 0 | 0 | 0 | 0 | 0 | 422 | 0 | 207 | 0 | 20 | 10287 | 0 | 0 |
| SRR8992371 | 0 | 0 | 0 | 0 | 410 | 0 | 0 | 0 | 0 | 2313 | 1935 | 37 | 0 | 0 | 36121 | 502451 | 78315 | 95 | 550 | 723 | 8151 | 0 | 0 | 0 | 0 | 0 | 0 | 0 | 0 | 443 | 0 | 132 | 0 | 39 | 10479 | 0 | 0 |
| SRR8992378 | 0 | 0 | 0 | 0 | 42 | 0 | 0 | 0 | 0 | 227 | 327 | 40 | 0 | 0 | 23430 | 557937 | 59698 | 83 | 482 | 485 | 11440 | 0 | 0 | 0 | 0 | 0 | 0 | 0 | 0 | 64 | 0 | 150 | 0 | 31 | 11803 | 0 | 0 |
| SRR8992379 | 0 | 0 | 0 | 0 | 13 | 0 | 0 | 0 | 0 | 71 | 120 | 88 | 0 | 0 | 7207 | 577849 | 9113 | 0 | 160 | 87 | 560 | 0 | 0 | 0 | 0 | 0 | 0 | 0 | 0 | 16 | 0 | 44 | 0 | 33 | 1073 | 0 | 0 |
| SRR8992380 | 0 | 0 | 0 | 0 | 444 | 0 | 0 | 0 | 0 | 2198 | 2733 | 22 | 0 | 0 | 36446 | 536289 | 81921 | 81 | 525 | 759 | 9018 | 0 | 0 | 0 | 0 | 0 | 0 | 0 | 0 | 511 | 0 | 153 | 0 | 51 | 8341 | 0 | 0 |
| SRR8992381 | 0 | 0 | 0 | 0 | 13 | 0 | 0 | 0 | 0 | 48 | 112 | 71 | 0 | 0 | 10296 | 777414 | 12079 | 15 | 177 | 46 | 353 | 0 | 0 | 0 | 0 | 0 | 0 | 0 | 0 | 16 | 0 | 30 | 0 | 24 | 613 | 0 | 0 |
| SRR8992382 | 0 | 0 | 0 | 0 | 27 | 0 | 0 | 0 | 0 | 82 | 222 | 78 | 0 | 0 | 8166 | 664737 | 13112 | 42 | 282 | 113 | 892 | 0 | 0 | 0 | 0 | 0 | 0 | 0 | 0 | 30 | 0 | 64 | 0 | 31 | 1588 | 0 | 0 |
| SRR8992383 | 0 | 0 | 0 | 0 | 25 | 0 | 0 | 0 | 0 | 66 | 248 | 55 | 0 | 0 | 12124 | 560769 | 18617 | 10 | 280 | 108 | 865 | 0 | 0 | 0 | 0 | 0 | 0 | 0 | 0 | 39 | 0 | 40 | 0 | 23 | 1775 | 0 | 0 |
| SRR8992384 | 0 | 0 | 0 | 0 | 36 | 0 | 0 | 0 | 0 | 119 | 314 | 41 | 0 | 0 | 24000 | 563110 | 53878 | 48 | 373 | 421 | 8852 | 0 | 0 | 0 | 0 | 0 | 0 | 0 | 0 | 55 | 0 | 192 | 0 | 45 | 10141 | 0 | 0 |
| SRR8992385 | 0 | 0 | 0 | 0 | 323 | 0 | 0 | 0 | 0 | 801 | 2498 | 32 | 0 | 0 | 21822 | 582324 | 49029 | 61 | 361 | 378 | 7989 | 0 | 0 | 0 | 0 | 0 | 0 | 0 | 0 | 474 | 0 | 289 | 0 | 33 | 10361 | 0 | 0 |
| SRR8992387 | 0 | 0 | 0 | 0 | 366 | 0 | 0 | 0 | 0 | 894 | 2298 | 37 | 0 | 0 | 23165 | 572006 | 49170 | 50 | 487 | 441 | 9370 | 0 | 0 | 0 | 0 | 0 | 0 | 0 | 0 | 442 | 0 | 274 | 0 | 35 | 12960 | 0 | 0 |
| SRR8992388 | 0 | 0 | 0 | 0 | 282 | 0 | 0 | 0 | 0 | 1087 | 3213 | 19 | 0 | 0 | 26407 | 526532 | 70706 | 76 | 473 | 468 | 9195 | 0 | 0 | 0 | 0 | 0 | 0 | 0 | 0 | 499 | 0 | 354 | 0 | 29 | 7370 | 0 | 0 |
| SRR8992389 | 0 | 0 | 0 | 0 | 30 | 0 | 0 | 0 | 0 | 62 | 183 | 23 | 0 | 0 | 39720 | 594213 | 83925 | 45 | 299 | 547 | 7239 | 0 | 0 | 0 | 0 | 0 | 0 | 0 | 0 | 25 | 0 | 118 | 0 | 28 | 4728 | 0 | 0 |
| SRR8992390 | 0 | 0 | 0 | 0 | 462 | 0 | 0 | 0 | 0 | 824 | 3060 | 35 | 0 | 0 | 27785 | 572531 | 61739 | 73 | 566 | 435 | 9083 | 0 | 0 | 0 | 0 | 0 | 0 | 0 | 0 | 555 | 0 | 193 | 0 | 28 | 11323 | 0 | 0 |
| SRR8992391 | 0 | 0 | 0 | 0 | 244 | 0 | 0 | 0 | 0 | 543 | 2274 | 28 | 0 | 0 | 20055 | 528210 | 32963 | 71 | 424 | 435 | 8507 | 0 | 0 | 0 | 0 | 0 | 0 | 0 | 0 | 479 | 0 | 170 | 0 | 24 | 12705 | 0 | 0 |
| SRR9003330 | 96 | 47 | 52 | 19 | 139 | 796 | 90 | 338 | 696 | 87 | 128 | 15 | 14 | 0 | 4893 | 97736 | 200981 | 25 | 1395 | 1372 | 6431 | 70 | 98 | 86 | 98 | 81 | 77 | 0 | 11 | 100 | 0 | 24 | 0 | 0 | 7469 | 0 | 0 |
| SRR9003332 | 0 | 0 | 0 | 0 | 0 | 3126 | 440 | 1701 | 1932 | 78 | 0 | 0 | 14 | 0 | 0 | 87 | 71 | 19 | 0 | 11 | 0 | 38 | 31 | 65 | 0 | 0 | 0 | 0 | 0 | 0 | 35 | 0 | 0 | 12 | 11322 | 61 | 0 |
| SRR9003333 | 2161 | 986 | 548 | 90 | 2027 | 1425 | 185 | 505 | 1136 | 97 | 70 | 17 | 18 | 13 | 5217 | 121959 | 208543 | 36 | 1190 | 1670 | 6631 | 23 | 64 | 76 | 73 | 80 | 131 | 0 | 13 | 79 | 0 | 13 | 0 | 16 | 9202 | 0 | 0 |
| SRR9003334 | 0 | 0 | 0 | 0 | 0 | 7992 | 1271 | 6025 | 4864 | 60 | 0 | 0 | 45 | 38 | 0 | 87 | 15 | 0 | 16 | 0 | 0 | 26 | 0 | 223 | 0 | 0 | 0 | 0 | 11 | 0 | 0 | 0 | 0 | 24 | 17952 | 18 | 0 |
| SRR9003336 | 2615 | 1273 | 623 | 70 | 2084 | 1121 | 131 | 417 | 1111 | 61 | 49 | 20 | 14 | 12 | 4715 | 115679 | 197455 | 44 | 1305 | 2031 | 6583 | 52 | 81 | 66 | 63 | 66 | 90 | 0 | 0 | 85 | 0 | 34 | 0 | 13 | 8525 | 0 | 0 |
| SRR9003338 | 0 | 0 | 0 | 0 | 0 | 1950 | 461 | 2110 | 3204 | 20 | 0 | 0 | 31 | 44 | 0 | 143 | 155 | 0 | 0 | 0 | 0 | 13 | 11 | 206 | 0 | 0 | 0 | 0 | 11 | 0 | 0 | 0 | 0 | 15 | 16731 | 26 | 15 |
| SRR9003339 | 140 | 63 | 57 | 11 | 104 | 956 | 130 | 441 | 1094 | 66 | 94 | 15 | 13 | 13 | 5330 | 128594 | 202901 | 30 | 1784 | 2073 | 7253 | 32 | 61 | 98 | 100 | 81 | 137 | 0 | 0 | 78 | 0 | 35 | 0 | 0 | 9766 | 0 | 0 |
| SRR9003341 | 46 | 12 | 0 | 0 | 58 | 2756 | 797 | 2719 | 3218 | 83 | 0 | 0 | 71 | 62 | 0 | 703 | 570 | 12 | 0 | 0 | 0 | 17 | 0 | 351 | 0 | 0 | 0 | 0 | 0 | 0 | 0 | 0 | 0 | 0 | 19520 | 17 | 17 |
| SRR9003342 | 126 | 60 | 32 | 45 | 141 | 1205 | 159 | 500 | 1258 | 98 | 39 | 13 | 10 | 13 | 4454 | 127705 | 191190 | 62 | 1341 | 1998 | 5598 | 44 | 87 | 147 | 126 | 113 | 137 | 0 | 0 | 84 | 0 | 14 | 0 | 0 | 9253 | 0 | 0 |
| SRR9018258 | 1875 | 863 | 87 | 143 | 137 | 1134 | 334 | 1445 | 5240 | 392 | 148 | 0 | 33 | 14 | 436 | 2198 | 451 | 947 | 208 | 957 | 676 | 32 | 0 | 83 | 80 | 58 | 22 | 0 | 0 | 92 | 11 | 2156 | 22 | 0 | 2872 | 0 | 0 |
| SRR9018259 | 1666 | 759 | 70 | 63 | 183 | 2479 | 627 | 3239 | 12960 | 636 | 61 | 0 | 27 | 13 | 4450 | 13482 | 8421 | 223 | 874 | 3011 | 760 | 82 | 0 | 74 | 95 | 49 | 33 | 0 | 0 | 64 | 0 | 978 | 0 | 0 | 2314 | 0 | 0 |
| SRR9018260 | 1977 | 787 | 247 | 113 | 66 | 127 | 24 | 167 | 779 | 1667 | 55 | 28 | 11 | 0 | 429 | 1820 | 658 | 191 | 103 | 40 | 32 | 31 | 0 | 0 | 0 | 0 | 0 | 0 | 23 | 0 | 16 | 199 | 0 | 0 | 2662 | 0 | 0 |
| SRR9018261 | 222 | 76 | 29 | 20 | 45 | 71 | 13 | 71 | 323 | 80 | 131 | 0 | 0 | 0 | 700 | 4114 | 899 | 314 | 84 | 182 | 403 | 30 | 0 | 43 | 12 | 27 | 55 | 14 | 12 | 16 | 11 | 11558 | 0 | 15 | 2990 | 0 | 0 |
| SRR952248 | 92 | 14 | 17 | 23 | 81 | 291 | 182 | 839 | 73 | 111 | 14 | 18 | 0 | 0 | 69123 | 9305 | 114408 | 14 | 180 | 3151 | 1400 | 0 | 0 | 10 | 10 | 51 | 259 | 187 | 0 | 118 | 0 | 12 | 0 | 0 | 876 | 0 | 0 |
| SRR952249 | 71 | 72 | 129 | 215 | 67 | 597 | 358 | 2069 | 124 | 178 | 56 | 11 | 0 | 0 | 54582 | 19343 | 97394 | 25 | 361 | 3628 | 1488 | 0 | 0 | 79 | 88 | 437 | 1336 | 919 | 0 | 98 | 0 | 36 | 0 | 0 | 1458 | 0 | 0 |
| SRR952288 | 70 | 73 | 18 | 31 | 66 | 553 | 337 | 2036 | 129 | 24 | 57 | 0 | 0 | 0 | 54538 | 19444 | 97715 | 31 | 363 | 3583 | 1420 | 0 | 0 | 10 | 11 | 57 | 169 | 121 | 0 | 12 | 0 | 27 | 0 | 0 | 1520 | 0 | 0 |
| SRR952289 | 20 | 25 | 90 | 94 | 79 | 513 | 326 | 1493 | 89 | 20 | 56 | 24 | 0 | 0 | 78406 | 11402 | 123326 | 26 | 252 | 3213 | 1409 | 0 | 0 | 13 | 108 | 33 | 109 | 12 | 0 | 11 | 0 | 22 | 0 | 0 | 867 | 0 | 12 |
| SRR952290 | 20 | 27 | 89 | 88 | 80 | 476 | 272 | 1359 | 82 | 20 | 54 | 24 | 0 | 0 | 75258 | 11281 | 123021 | 24 | 236 | 3109 | 1281 | 0 | 0 | 13 | 96 | 33 | 112 | 12 | 0 | 11 | 0 | 28 | 0 | 0 | 847 | 0 | 0 |
| SRR952309 | 125 | 27 | 17 | 61 | 100 | 196 | 140 | 235 | 39 | 0 | 28 | 0 | 0 | 0 | 8200 | 1970 | 15044 | 25 | 47 | 1996 | 269 | 0 | 0 | 18 | 54 | 11 | 72 | 15 | 0 | 13 | 0 | 16 | 0 | 0 | 1882 | 0 | 0 |
| SRR952310 | 105 | 280 | 23 | 82 | 88 | 208 | 142 | 258 | 24 | 0 | 79 | 0 | 0 | 0 | 17710 | 4850 | 36235 | 0 | 81 | 4413 | 574 | 0 | 0 | 166 | 71 | 276 | 1886 | 253 | 0 | 416 | 0 | 12 | 0 | 0 | 2177 | 0 | 0 |
| SRR952311 | 103 | 358 | 24 | 93 | 102 | 432 | 258 | 360 | 69 | 38 | 55 | 16 | 0 | 0 | 55394 | 9650 | 96919 | 14 | 195 | 5054 | 1118 | 0 | 0 | 255 | 115 | 253 | 1428 | 176 | 0 | 295 | 0 | 24 | 0 | 0 | 1271 | 0 | 0 |
| SRR952312 | 56 | 221 | 56 | 224 | 112 | 91 | 72 | 353 | 0 | 0 | 49 | 11 | 0 | 0 | 55602 | 5658 | 69220 | 0 | 118 | 2661 | 2643 | 0 | 0 | 77 | 46 | 375 | 970 | 291 | 0 | 105 | 0 | 36 | 0 | 0 | 599 | 0 | 0 |
| SRR952313 | 113 | 190 | 155 | 400 | 98 | 49 | 90 | 215 | 26 | 0 | 82 | 0 | 0 | 0 | 58371 | 5068 | 69119 | 0 | 36 | 3175 | 1876 | 0 | 0 | 61 | 63 | 490 | 1402 | 346 | 0 | 80 | 0 | 0 | 0 | 0 | 561 | 0 | 0 |
| SRR952314 | 191 | 312 | 186 | 422 | 188 | 39 | 10 | 110 | 10 | 0 | 48 | 0 | 0 | 0 | 5657 | 5925 | 20535 | 0 | 39 | 3805 | 865 | 0 | 0 | 75 | 43 | 335 | 1163 | 571 | 0 | 152 | 0 | 29 | 0 | 0 | 2694 | 0 | 0 |
| SRR952315 | 119 | 191 | 124 | 17 | 103 | 37 | 34 | 112 | 0 | 0 | 52 | 0 | 0 | 0 | 4654 | 6581 | 22485 | 30 | 49 | 3232 | 1186 | 0 | 0 | 58 | 54 | 16 | 64 | 81 | 0 | 150 | 0 | 34 | 0 | 0 | 2411 | 0 | 0 |
| SRR952316 | 0 | 0 | 0 | 0 | 245 | 0 | 0 | 96 | 0 | 0 | 123 | 0 | 0 | 0 | 13882 | 644 | 5342 | 0 | 100 | 5451 | 98 | 0 | 0 | 0 | 27 | 0 | 14 | 0 | 0 | 202 | 0 | 89 | 0 | 0 | 1226 | 0 | 0 |
| SRR952317 | 0 | 0 | 0 | 13 | 436 | 0 | 0 | 0 | 0 | 0 | 845 | 0 | 0 | 0 | 12018 | 540 | 7292 | 0 | 121 | 8232 | 39 | 0 | 0 | 0 | 0 | 0 | 26 | 0 | 0 | 556 | 0 | 0 | 0 | 0 | 428 | 0 | 0 |
| SRR952318 | 0 | 0 | 0 | 12 | 888 | 0 | 0 | 0 | 0 | 0 | 546 | 0 | 0 | 0 | 1899 | 499 | 1668 | 0 | 58 | 5686 | 11 | 0 | 0 | 0 | 14 | 0 | 13 | 16 | 0 | 1369 | 0 | 89 | 0 | 0 | 5832 | 0 | 0 |
| SRR952319 | 0 | 0 | 0 | 0 | 611 | 0 | 0 | 0 | 0 | 0 | 395 | 0 | 0 | 0 | 1092 | 175 | 975 | 64 | 107 | 5089 | 0 | 0 | 0 | 0 | 0 | 32 | 0 | 0 | 0 | 921 | 0 | 81 | 0 | 0 | 4953 | 0 | 0 |
| SRR9588764 | 0 | 0 | 0 | 0 | 49 | 2441 | 759 | 6795 | 8127 | 82 | 408 | 0 | 0 | 0 | 108298 | 23603 | 680533 | 226 | 292 | 5877 | 1210 | 0 | 0 | 96 | 264 | 58 | 30 | 42 | 0 | 912 | 0 | 0 | 0 | 34 | 3504 | 28 | 0 |
| SRR9588765 | 0 | 0 | 0 | 0 | 59 | 20943 | 3570 | 42926 | 35982 | 36 | 134 | 0 | 0 | 0 | 64452 | 31782 | 347860 | 342 | 284 | 2587 | 825 | 0 | 0 | 253 | 633 | 220 | 89 | 133 | 0 | 770 | 0 | 0 | 0 | 23 | 15880 | 21 | 0 |
| SRR9588766 | 0 | 0 | 0 | 0 | 13 | 8662 | 725 | 4528 | 6964 | 22 | 81 | 0 | 0 | 0 | 111113 | 36785 | 321769 | 849 | 1009 | 3608 | 5707 | 0 | 0 | 32 | 61 | 10 | 23 | 20 | 0 | 294 | 0 | 0 | 0 | 24 | 30628 | 23 | 0 |
| SRR9588767 | 12 | 0 | 0 | 0 | 110 | 3762 | 581 | 6939 | 5712 | 94 | 193 | 0 | 0 | 0 | 86347 | 87355 | 436194 | 1120 | 461 | 3483 | 631 | 0 | 0 | 270 | 0 | 0 | 0 | 0 | 0 | 738 | 0 | 0 | 0 | 19 | 11621 | 0 | 0 |
| SRR9618218 | 30 | 19 | 16 | 19 | 35 | 78 | 142 | 401 | 26 | 0 | 57 | 0 | 0 | 0 | 194 | 7485 | 24969 | 0 | 13 | 6402 | 776 | 0 | 0 | 45 | 0 | 65 | 71 | 11 | 0 | 19 | 0 | 0 | 0 | 0 | 1928 | 0 | 0 |
| SRR9618219 | 176 | 75 | 14 | 101 | 75 | 358 | 593 | 898 | 235 | 0 | 0 | 43 | 0 | 0 | 273 | 22345 | 35191 | 11 | 21 | 3917 | 417 | 0 | 0 | 213 | 0 | 37 | 39 | 0 | 0 | 108 | 0 | 21 | 0 | 0 | 8474 | 0 | 0 |
| SRR9618220 | 37 | 40 | 27 | 337 | 74 | 135 | 203 | 191 | 23 | 0 | 84 | 13 | 0 | 0 | 113 | 18487 | 27580 | 0 | 11 | 6504 | 1328 | 0 | 0 | 327 | 37 | 180 | 1012 | 261 | 0 | 251 | 0 | 23 | 0 | 0 | 3064 | 0 | 0 |
| SRR9618221 | 19 | 22 | 22 | 72 | 29 | 89 | 178 | 334 | 45 | 0 | 15 | 19 | 0 | 0 | 100 | 9442 | 27912 | 0 | 11 | 6058 | 367 | 0 | 0 | 34 | 17 | 45 | 28 | 12 | 0 | 79 | 0 | 11 | 0 | 0 | 2144 | 0 | 0 |
| SRR9618222 | 150 | 103 | 34 | 265 | 77 | 427 | 240 | 573 | 40 | 21 | 39 | 47 | 0 | 0 | 2507 | 68437 | 128486 | 13 | 27 | 6747 | 4667 | 0 | 0 | 94 | 0 | 21 | 94 | 13 | 0 | 197 | 0 | 0 | 0 | 0 | 5119 | 0 | 13 |
| SRR9627609 | 0 | 0 | 0 | 15 | 285 | 344 | 1575 | 2516 | 72 | 31 | 47 | 0 | 0 | 0 | 1014 | 28244 | 157246 | 54 | 0 | 253 | 0 | 0 | 0 | 14 | 14 | 46 | 35 | 11 | 0 | 426 | 0 | 0 | 0 | 0 | 16159 | 0 | 0 |
| SRR9627610 | 0 | 0 | 0 | 0 | 163 | 527 | 1527 | 3448 | 161 | 30 | 57 | 0 | 0 | 0 | 1063 | 19048 | 134133 | 18 | 0 | 107 | 0 | 0 | 0 | 14 | 0 | 16 | 0 | 0 | 0 | 162 | 0 | 0 | 0 | 0 | 18100 | 0 | 0 |
| SRR9627611 | 0 | 0 | 0 | 0 | 938 | 748 | 1794 | 2791 | 199 | 26 | 133 | 0 | 0 | 0 | 1146 | 20671 | 145146 | 33 | 0 | 399 | 0 | 0 | 0 | 57 | 0 | 17 | 15 | 0 | 0 | 1397 | 0 | 0 | 0 | 0 | 16174 | 0 | 0 |
| SRR9627612 | 0 | 0 | 0 | 17 | 222 | 309 | 1416 | 2506 | 139 | 0 | 23 | 0 | 0 | 0 | 944 | 20589 | 128326 | 46 | 0 | 255 | 0 | 0 | 0 | 15 | 19 | 11 | 11 | 0 | 0 | 392 | 0 | 0 | 0 | 0 | 12823 | 0 | 0 |
| SRR9627613 | 0 | 0 | 0 | 14 | 207 | 382 | 1208 | 2041 | 111 | 0 | 24 | 0 | 0 | 0 | 1284 | 19749 | 151836 | 14 | 28 | 222 | 0 | 0 | 0 | 15 | 0 | 41 | 34 | 11 | 0 | 337 | 0 | 0 | 0 | 0 | 11684 | 0 | 0 |
| SRR9627614 | 0 | 0 | 0 | 0 | 192 | 438 | 1363 | 2101 | 175 | 0 | 42 | 0 | 0 | 0 | 888 | 15966 | 88965 | 63 | 0 | 225 | 0 | 0 | 0 | 18 | 15 | 16 | 17 | 0 | 0 | 186 | 0 | 0 | 0 | 0 | 15917 | 0 | 0 |
| SRR9627651 | 21 | 19 | 0 | 0 | 416 | 149 | 2432 | 2620 | 2401 | 22 | 54 | 0 | 0 | 0 | 455 | 13907 | 41654 | 0 | 47 | 1993 | 0 | 0 | 0 | 45 | 0 | 0 | 23 | 0 | 0 | 142 | 0 | 0 | 0 | 0 | 15551 | 0 | 0 |
| SRR9627653 | 16 | 0 | 0 | 0 | 143 | 145 | 3266 | 2669 | 2960 | 17 | 10 | 0 | 0 | 0 | 145 | 4043 | 13541 | 0 | 0 | 421 | 0 | 0 | 0 | 44 | 0 | 0 | 0 | 0 | 0 | 29 | 0 | 0 | 0 | 0 | 19033 | 0 | 0 |
| SRR9627654 | 23 | 0 | 0 | 0 | 3563 | 204 | 3252 | 2151 | 2168 | 0 | 30 | 0 | 0 | 0 | 285 | 7507 | 22949 | 0 | 24 | 1614 | 0 | 0 | 0 | 43 | 0 | 0 | 17 | 0 | 0 | 907 | 0 | 0 | 0 | 0 | 13635 | 0 | 0 |
| SRR9627655 | 0 | 0 | 0 | 0 | 1970 | 153 | 2679 | 2166 | 2454 | 0 | 0 | 0 | 0 | 0 | 153 | 2998 | 9634 | 0 | 0 | 539 | 18 | 0 | 0 | 24 | 0 | 0 | 10 | 0 | 0 | 476 | 0 | 0 | 0 | 0 | 13393 | 0 | 0 |
| SRR9627656 | 13 | 0 | 0 | 0 | 513 | 60 | 2779 | 1959 | 2599 | 0 | 39 | 0 | 0 | 0 | 620 | 12618 | 36484 | 0 | 60 | 1766 | 13 | 0 | 0 | 23 | 0 | 0 | 36 | 0 | 0 | 170 | 0 | 0 | 0 | 0 | 14056 | 0 | 0 |
| SRR9627693 | 16 | 11 | 12 | 0 | 171 | 2537 | 2718 | 8903 | 1824 | 44 | 14 | 0 | 0 | 0 | 1652 | 31657 | 84175 | 0 | 244 | 686 | 172 | 0 | 0 | 0 | 16 | 32 | 32 | 63 | 0 | 173 | 0 | 0 | 0 | 0 | 15541 | 0 | 0 |
| SRR9627694 | 12 | 13 | 19 | 0 | 184 | 2590 | 2781 | 8926 | 2705 | 35 | 17 | 0 | 0 | 0 | 1290 | 31767 | 85742 | 0 | 248 | 1109 | 191 | 0 | 0 | 13 | 22 | 55 | 53 | 11 | 0 | 189 | 0 | 0 | 0 | 0 | 13943 | 0 | 0 |
| SRR9627695 | 24 | 0 | 11 | 0 | 181 | 2764 | 3107 | 9600 | 2470 | 39 | 18 | 0 | 0 | 0 | 1601 | 32400 | 85705 | 0 | 239 | 964 | 135 | 0 | 0 | 17 | 43 | 10 | 12 | 15 | 0 | 185 | 0 | 0 | 0 | 0 | 13809 | 0 | 0 |
| SRR9627696 | 0 | 0 | 11 | 15 | 103 | 1663 | 1684 | 5557 | 1010 | 0 | 12 | 0 | 0 | 0 | 2231 | 31614 | 99484 | 0 | 242 | 737 | 284 | 0 | 0 | 0 | 17 | 28 | 42 | 69 | 0 | 147 | 0 | 32 | 0 | 0 | 9753 | 0 | 0 |
| SRR9627697 | 0 | 0 | 0 | 16 | 790 | 1956 | 1919 | 5887 | 877 | 0 | 94 | 0 | 0 | 0 | 2152 | 28797 | 90693 | 18 | 258 | 687 | 258 | 0 | 0 | 0 | 0 | 23 | 33 | 61 | 0 | 1113 | 0 | 0 | 0 | 0 | 11787 | 0 | 0 |
| SRR9627698 | 0 | 0 | 10 | 22 | 134 | 1311 | 1682 | 6471 | 811 | 15 | 46 | 0 | 0 | 0 | 1826 | 33952 | 78595 | 0 | 83 | 114 | 91 | 0 | 0 | 0 | 13 | 34 | 43 | 12 | 0 | 76 | 0 | 0 | 0 | 0 | 13435 | 0 | 0 |
| SRR9648387 | 0 | 0 | 0 | 0 | 70 | 0 | 0 | 0 | 0 | 55 | 27 | 0 | 0 | 0 | 10588 | 856267 | 11532 | 22 | 0 | 0 | 2287 | 0 | 0 | 0 | 0 | 0 | 0 | 0 | 0 | 0 | 0 | 40 | 0 | 0 | 3825 | 242 | 0 |
| SRR9648388 | 0 | 0 | 0 | 0 | 403 | 0 | 0 | 0 | 0 | 0 | 8600 | 0 | 0 | 0 | 544 | 38395 | 1244 | 103 | 0 | 0 | 87 | 513 | 186 | 0 | 0 | 0 | 0 | 0 | 0 | 137 | 156 | 1886 | 0 | 0 | 102 | 0 | 0 |
| SRR9648389 | 0 | 0 | 0 | 0 | 15 | 0 | 0 | 0 | 0 | 14 | 252 | 0 | 0 | 0 | 1014 | 115595 | 1997 | 54 | 0 | 0 | 56 | 33 | 11 | 0 | 0 | 0 | 0 | 0 | 0 | 11 | 42 | 4021 | 0 | 0 | 295 | 0 | 0 |
| SRR965653 | 0 | 0 | 0 | 0 | 0 | 171 | 285 | 949 | 171 | 14 | 115 | 118 | 0 | 0 | 2277 | 53004 | 29194 | 0 | 417 | 4724 | 417 | 0 | 0 | 0 | 27 | 53 | 32 | 23 | 0 | 11 | 0 | 95 | 0 | 0 | 7259 | 0 | 0 |
| SRR965655 | 108 | 57 | 11 | 20 | 111 | 236 | 369 | 469 | 72 | 38 | 39 | 14 | 0 | 0 | 6921 | 123705 | 86209 | 0 | 83 | 5902 | 1103 | 0 | 0 | 150 | 0 | 36 | 38 | 0 | 0 | 73 | 0 | 14 | 0 | 0 | 2581 | 0 | 0 |
| SRR9833150 | 14 | 16 | 0 | 0 | 18 | 1008 | 317 | 6150 | 312 | 0 | 0 | 28 | 0 | 0 | 5510 | 26673 | 11666 | 26 | 41 | 19687 | 3254 | 0 | 0 | 21 | 0 | 0 | 0 | 0 | 0 | 40 | 0 | 153 | 0 | 0 | 3005 | 0 | 0 |
| SRR9833151 | 0 | 0 | 0 | 0 | 12 | 1002 | 406 | 5399 | 295 | 0 | 0 | 0 | 0 | 0 | 1681 | 6854 | 3033 | 39 | 11 | 3952 | 1503 | 0 | 0 | 15 | 0 | 0 | 0 | 0 | 0 | 0 | 0 | 89 | 0 | 0 | 2653 | 0 | 0 |
| SRR9833152 | 16 | 15 | 0 | 0 | 16 | 1109 | 505 | 5882 | 285 | 0 | 0 | 15 | 0 | 0 | 3943 | 21911 | 8354 | 58 | 58 | 9569 | 3507 | 0 | 0 | 15 | 0 | 0 | 0 | 0 | 0 | 17 | 0 | 232 | 0 | 0 | 3905 | 0 | 0 |
| SRR9833153 | 44 | 43 | 0 | 0 | 0 | 615 | 283 | 3349 | 135 | 0 | 0 | 19 | 0 | 0 | 4363 | 8093 | 6406 | 37 | 25 | 4517 | 4154 | 0 | 0 | 21 | 0 | 0 | 0 | 0 | 0 | 29 | 0 | 117 | 0 | 0 | 3961 | 0 | 0 |
| SRR9833154 | 20 | 12 | 0 | 0 | 11 | 680 | 267 | 2932 | 283 | 0 | 0 | 0 | 0 | 0 | 1491 | 7806 | 3151 | 27 | 22 | 2410 | 3177 | 0 | 0 | 19 | 0 | 0 | 0 | 0 | 0 | 11 | 0 | 38 | 0 | 0 | 3327 | 0 | 0 |
| SRR9833155 | 132 | 121 | 0 | 0 | 29 | 542 | 228 | 2687 | 171 | 0 | 0 | 24 | 0 | 0 | 11484 | 31041 | 19960 | 11 | 51 | 8175 | 9384 | 0 | 0 | 25 | 14 | 12 | 0 | 0 | 0 | 54 | 0 | 57 | 0 | 0 | 2836 | 0 | 0 |
| SRR9833156 | 40 | 56 | 0 | 0 | 31 | 404 | 241 | 2142 | 127 | 0 | 0 | 0 | 0 | 0 | 8519 | 55694 | 23106 | 23 | 95 | 6994 | 8610 | 0 | 0 | 17 | 16 | 22 | 20 | 0 | 0 | 120 | 0 | 73 | 0 | 0 | 2230 | 0 | 0 |
| SRR9833157 | 19 | 15 | 0 | 13 | 12 | 605 | 232 | 2567 | 242 | 0 | 0 | 0 | 0 | 0 | 4683 | 34548 | 13499 | 34 | 73 | 13348 | 10287 | 0 | 0 | 0 | 0 | 14 | 11 | 0 | 0 | 50 | 0 | 102 | 0 | 0 | 2691 | 0 | 0 |
| SRR9833158 | 17 | 22 | 0 | 19 | 30 | 1103 | 297 | 4676 | 269 | 0 | 38 | 32 | 0 | 0 | 4806 | 33533 | 14382 | 51 | 139 | 15492 | 3934 | 0 | 0 | 0 | 17 | 76 | 173 | 23 | 0 | 286 | 0 | 148 | 0 | 0 | 3284 | 0 | 0 |
| SRR9833159 | 0 | 0 | 0 | 13 | 22 | 5506 | 1374 | 19879 | 814 | 18 | 36 | 125 | 0 | 0 | 2839 | 33124 | 20303 | 36 | 80 | 4692 | 2238 | 0 | 0 | 17 | 0 | 30 | 67 | 36 | 0 | 170 | 0 | 338 | 0 | 0 | 5418 | 0 | 16 |
| SRR9833160 | 0 | 0 | 0 | 11 | 34 | 1986 | 724 | 10436 | 544 | 14 | 157 | 115 | 0 | 0 | 2056 | 40532 | 20934 | 27 | 74 | 8081 | 1300 | 0 | 0 | 11 | 0 | 45 | 142 | 66 | 0 | 893 | 0 | 425 | 0 | 0 | 6649 | 0 | 12 |
| SRR9833161 | 12 | 13 | 0 | 11 | 0 | 1403 | 813 | 6451 | 369 | 0 | 55 | 96 | 0 | 0 | 2121 | 40498 | 24753 | 30 | 30 | 4543 | 2209 | 0 | 0 | 0 | 0 | 16 | 23 | 17 | 0 | 212 | 0 | 38 | 0 | 0 | 10530 | 0 | 18 |
| SRR9833162 | 0 | 0 | 0 | 23 | 45 | 1786 | 907 | 11362 | 745 | 0 | 73 | 250 | 0 | 0 | 2064 | 25560 | 17049 | 19 | 44 | 6976 | 963 | 0 | 0 | 26 | 0 | 45 | 271 | 68 | 0 | 401 | 0 | 259 | 0 | 0 | 4391 | 0 | 21 |
| SRR9833163 | 0 | 0 | 0 | 0 | 34 | 3149 | 1430 | 17952 | 842 | 12 | 76 | 338 | 0 | 0 | 3629 | 34375 | 24592 | 23 | 49 | 8034 | 2419 | 0 | 0 | 15 | 11 | 60 | 146 | 54 | 0 | 437 | 0 | 155 | 0 | 0 | 3369 | 0 | 32 |
| SRR9833164 | 0 | 0 | 0 | 29 | 68 | 2338 | 880 | 12432 | 795 | 0 | 151 | 97 | 0 | 0 | 2466 | 23529 | 17789 | 14 | 83 | 4167 | 2152 | 0 | 0 | 40 | 23 | 164 | 388 | 131 | 0 | 613 | 0 | 394 | 0 | 0 | 4719 | 0 | 0 |
| SRR9856157 | 153 | 19 | 0 | 18 | 135 | 4514 | 9071 | 21005 | 103016 | 109 | 66 | 36 | 0 | 0 | 1285 | 34941 | 64785 | 265 | 3160 | 2242 | 897 | 0 | 0 | 77 | 23 | 13 | 21 | 29 | 0 | 13 | 0 | 0 | 0 | 0 | 1724 | 0 | 0 |
| SRR9856158 | 218 | 98 | 0 | 0 | 101 | 20192 | 7121 | 35592 | 35098 | 83 | 95 | 21 | 0 | 0 | 18872 | 70617 | 312746 | 195 | 8195 | 11358 | 2996 | 0 | 0 | 4737 | 193 | 92 | 211 | 18 | 0 | 533 | 0 | 0 | 0 | 34 | 470 | 0 | 0 |
| SRR9856159 | 12 | 60 | 19 | 24 | 204 | 5133 | 3872 | 12210 | 56206 | 100 | 229 | 87 | 0 | 0 | 11423 | 51766 | 232969 | 450 | 3622 | 7046 | 3827 | 0 | 0 | 99 | 12 | 179 | 20 | 59 | 0 | 20 | 0 | 0 | 0 | 39 | 720 | 0 | 0 |
| SRR9856160 | 191 | 26 | 0 | 0 | 170 | 3211 | 512 | 4166 | 18379 | 258 | 42 | 191 | 14 | 0 | 1274 | 7250 | 34958 | 23 | 799 | 3563 | 5251 | 0 | 0 | 76 | 251 | 119 | 133 | 0 | 0 | 12 | 0 | 0 | 0 | 30 | 2286 | 0 | 0 |
| SRR9856161 | 145 | 18 | 0 | 0 | 178 | 11445 | 4864 | 29656 | 45847 | 85 | 118 | 12 | 0 | 0 | 10122 | 31680 | 499274 | 399 | 9548 | 16090 | 3085 | 0 | 0 | 5522 | 601 | 121 | 242 | 0 | 0 | 764 | 0 | 0 | 0 | 30 | 488 | 0 | 0 |
| SRR9856162 | 150 | 42 | 19 | 0 | 197 | 13325 | 5478 | 25218 | 31796 | 61 | 98 | 0 | 0 | 0 | 3435 | 51683 | 263976 | 523 | 18525 | 11241 | 1514 | 0 | 0 | 8576 | 730 | 136 | 468 | 0 | 0 | 927 | 0 | 0 | 0 | 0 | 1002 | 0 | 0 |
| SRR9856163 | 101 | 15 | 0 | 21 | 211 | 26292 | 6315 | 40132 | 48667 | 70 | 486 | 24 | 0 | 0 | 7346 | 40060 | 346097 | 593 | 8791 | 9173 | 828 | 0 | 0 | 11424 | 1244 | 590 | 2195 | 31 | 0 | 1651 | 0 | 0 | 0 | 21 | 560 | 0 | 0 |
| SRR9856164 | 92 | 24 | 0 | 0 | 121 | 7090 | 6852 | 23119 | 52099 | 107 | 168 | 18 | 0 | 0 | 14366 | 29886 | 355628 | 345 | 7457 | 6269 | 2181 | 0 | 0 | 115 | 18 | 204 | 13 | 29 | 0 | 18 | 0 | 0 | 0 | 29 | 446 | 0 | 0 |
| SRR9856165 | 173 | 37 | 17 | 0 | 265 | 8627 | 8118 | 37454 | 77779 | 227 | 176 | 24 | 0 | 0 | 1756 | 20727 | 87646 | 229 | 3624 | 3402 | 1756 | 0 | 0 | 102 | 22 | 343 | 13 | 24 | 0 | 10 | 0 | 0 | 0 | 27 | 673 | 0 | 0 |
| SRR9856166 | 59 | 0 | 0 | 0 | 121 | 19750 | 10702 | 38427 | 64120 | 34 | 100 | 0 | 0 | 0 | 3680 | 25613 | 184363 | 317 | 5820 | 5881 | 540 | 0 | 0 | 79 | 10 | 221 | 436 | 12 | 0 | 19 | 0 | 0 | 0 | 0 | 596 | 0 | 0 |
| SRR9856167 | 93 | 25 | 0 | 0 | 74 | 3775 | 1926 | 14657 | 26742 | 263 | 119 | 33 | 0 | 0 | 1363 | 16824 | 75781 | 44 | 1942 | 9155 | 3184 | 0 | 0 | 62 | 27 | 236 | 14 | 49 | 0 | 14 | 0 | 0 | 0 | 11 | 995 | 0 | 0 |
| SRR9856168 | 196 | 28 | 19 | 14 | 97 | 10347 | 5981 | 20579 | 29668 | 52 | 416 | 21 | 0 | 0 | 14479 | 54160 | 248270 | 323 | 3074 | 11333 | 1777 | 0 | 0 | 3038 | 544 | 211 | 599 | 33 | 0 | 980 | 0 | 0 | 0 | 17 | 284 | 0 | 0 |
| SRR9856169 | 118 | 26 | 14 | 15 | 104 | 7352 | 4505 | 24753 | 45837 | 12 | 16 | 30 | 0 | 0 | 4821 | 42368 | 202107 | 350 | 4006 | 17287 | 1534 | 0 | 0 | 113 | 28 | 146 | 11 | 52 | 0 | 44 | 0 | 0 | 0 | 23 | 353 | 0 | 0 |
| SRR9856170 | 322 | 87 | 0 | 0 | 80 | 9849 | 6645 | 27125 | 43368 | 115 | 24 | 0 | 0 | 0 | 2680 | 8444 | 57344 | 204 | 2965 | 1557 | 800 | 0 | 0 | 2563 | 21 | 0 | 21 | 0 | 0 | 89 | 0 | 0 | 0 | 16 | 293 | 0 | 0 |

### Supp Table S8.3.Columns_BX_to_DH

| | | | | | | | | | | | | | | | | | | | | | | | | | | | | | | | | | | | | |
|---|---|---|---|---|---|---|---|---|---|---|---|---|---|---|---|---|---|---|---|---|---|---|---|---|---|---|---|---|---|---|---|---|---|---|---|---|---|
| SRR9856171 | 63 | 24 | 0 | 0 | 14 | 12409 | 4824 | 21060 | 30634 | 75 | 16 | 14 | 0 | 0 | 96 | 4568 | 3199 | 113 | 1195 | 647 | 87 | 0 | 0 | 25 | 0 | 0 | 11 | 0 | 0 | 48 | 0 | 0 | 0 | 25 | 268 | 0 | 0 |
| SRR9856172 | 184 | 26 | 0 | 14 | 62 | 14655 | 7805 | 27494 | 36115 | 53 | 81 | 0 | 0 | 0 | 1524 | 16838 | 88403 | 292 | 2147 | 8713 | 985 | 0 | 0 | 99 | 187 | 65 | 182 | 14 | 0 | 419 | 0 | 0 | 0 | 0 | 348 | 0 | 0 |
| SRR9856173 | 61 | 16 | 0 | 0 | 61 | 10232 | 4476 | 26649 | 69073 | 101 | 235 | 118 | 0 | 0 | 23290 | 56042 | 389972 | 508 | 5191 | 4492 | 2767 | 0 | 0 | 910 | 149 | 29 | 40 | 0 | 0 | 499 | 0 | 0 | 11 | 120 | 1090 | 0 | 0 |
| SRR9856174 | 13 | 51 | 0 | 17 | 54 | 1901 | 1298 | 5554 | 11140 | 82 | 35 | 30 | 0 | 0 | 13103 | 24254 | 193886 | 155 | 2149 | 3978 | 10104 | 0 | 0 | 73 | 99 | 29 | 81 | 10 | 0 | 126 | 0 | 0 | 0 | 15 | 1012 | 0 | 0 |
| SRR9856175 | 194 | 21 | 13 | 0 | 100 | 9700 | 3319 | 23625 | 20552 | 169 | 378 | 43 | 0 | 0 | 9613 | 55489 | 257852 | 159 | 3720 | 11152 | 4344 | 0 | 0 | 2270 | 600 | 217 | 507 | 46 | 0 | 567 | 0 | 0 | 0 | 13 | 256 | 0 | 0 |
| SRR9856176 | 11 | 44 | 14 | 18 | 110 | 4731 | 5952 | 14741 | 48453 | 145 | 142 | 40 | 0 | 0 | 5834 | 95152 | 265289 | 501 | 2146 | 13143 | 3924 | 0 | 0 | 91 | 12 | 129 | 19 | 29 | 0 | 22 | 0 | 0 | 0 | 27 | 1329 | 0 | 0 |
| SRR9856177 | 85 | 21 | 0 | 0 | 66 | 3963 | 1446 | 8062 | 16587 | 205 | 179 | 78 | 0 | 0 | 26541 | 65788 | 330955 | 934 | 2363 | 35206 | 4560 | 0 | 0 | 382 | 85 | 42 | 112 | 13 | 0 | 328 | 0 | 0 | 0 | 39 | 737 | 0 | 0 |
| SRR9856178 | 108 | 20 | 0 | 0 | 100 | 11786 | 2200 | 21812 | 74774 | 240 | 112 | 115 | 12 | 0 | 679 | 12424 | 56660 | 182 | 564 | 651 | 12284 | 0 | 0 | 46 | 28 | 321 | 10 | 23 | 0 | 24 | 0 | 0 | 0 | 41 | 9537 | 0 | 0 |
| SRR9856179 | 89 | 32 | 32 | 11 | 113 | 22074 | 11908 | 50729 | 62384 | 168 | 186 | 17 | 0 | 0 | 392 | 14373 | 80068 | 322 | 2794 | 2768 | 836 | 0 | 0 | 4937 | 639 | 271 | 1150 | 78 | 0 | 682 | 0 | 0 | 0 | 29 | 1230 | 0 | 0 |
| SRR9856180 | 123 | 48 | 26 | 19 | 71 | 16453 | 12200 | 43001 | 51012 | 81 | 200 | 0 | 0 | 0 | 1609 | 28197 | 138771 | 312 | 9655 | 7302 | 766 | 0 | 0 | 166 | 12 | 520 | 36 | 65 | 0 | 15 | 0 | 0 | 0 | 16 | 825 | 0 | 0 |
| SRR9856181 | 265 | 42 | 34 | 21 | 244 | 18336 | 8156 | 50142 | 79801 | 55 | 219 | 29 | 0 | 0 | 3589 | 31165 | 163887 | 290 | 6560 | 4362 | 2907 | 0 | 0 | 10802 | 1993 | 657 | 1626 | 21 | 0 | 1074 | 0 | 0 | 0 | 13 | 253 | 0 | 0 |
| SRR9856182 | 119 | 19 | 0 | 13 | 151 | 17281 | 9599 | 52277 | 118169 | 68 | 161 | 19 | 0 | 0 | 495 | 25842 | 67201 | 126 | 8623 | 2311 | 685 | 0 | 0 | 207 | 42 | 13 | 36 | 58 | 0 | 17 | 0 | 0 | 0 | 0 | 176 | 0 | 0 |
| SRR9856183 | 40 | 0 | 0 | 0 | 61 | 26729 | 7490 | 56809 | 82028 | 74 | 71 | 14 | 0 | 0 | 1871 | 23930 | 226772 | 280 | 4851 | 3919 | 3355 | 0 | 0 | 166 | 18 | 84 | 141 | 0 | 0 | 18 | 0 | 0 | 0 | 29 | 1328 | 0 | 0 |
| SRR9856184 | 148 | 36 | 45 | 41 | 86 | 6152 | 3881 | 16232 | 58372 | 47 | 205 | 34 | 0 | 0 | 554 | 23373 | 47534 | 48 | 6365 | 1070 | 864 | 0 | 0 | 108 | 27 | 16 | 64 | 183 | 0 | 246 | 0 | 0 | 0 | 19 | 582 | 0 | 0 |
| SRR9856185 | 210 | 22 | 15 | 0 | 102 | 18849 | 7634 | 23748 | 64858 | 36 | 44 | 58 | 0 | 0 | 507 | 13779 | 95006 | 101 | 574 | 676 | 7871 | 0 | 0 | 29 | 297 | 80 | 261 | 58 | 0 | 276 | 0 | 0 | 0 | 22 | 5957 | 0 | 0 |
| SRR9856186 | 482 | 85 | 22 | 0 | 70 | 4831 | 2554 | 11009 | 21121 | 288 | 51 | 87 | 0 | 0 | 3178 | 57674 | 120874 | 68 | 2625 | 3009 | 12382 | 0 | 0 | 4818 | 291 | 109 | 487 | 36 | 0 | 179 | 0 | 0 | 0 | 17 | 476 | 0 | 0 |
| SRR990360 | 0 | 0 | 0 | 0 | 0 | 6533 | 2456 | 48450 | 414 | 44 | 0 | 0 | 0 | 0 | 15 | 157 | 395 | 0 | 0 | 0 | 0 | 0 | 0 | 19 | 0 | 0 | 0 | 0 | 0 | 0 | 0 | 0 | 0 | 0 | 1111 | 310 | 0 |
| SRR9924500 | 21 | 21 | 16 | 61 | 86 | 75 | 140 | 275 | 68 | 0 | 241 | 17 | 0 | 0 | 5219 | 4619 | 9907 | 279 | 57 | 2304 | 544 | 0 | 0 | 86 | 59 | 290 | 934 | 150 | 0 | 1019 | 0 | 6736 | 0 | 0 | 5130 | 130 | 31 |
| SRR9924502 | 11 | 0 | 0 | 15 | 64 | 140 | 163 | 431 | 76 | 17 | 17 | 21 | 0 | 0 | 4498 | 4185 | 8752 | 163 | 20 | 2408 | 522 | 0 | 0 | 53 | 21 | 14 | 31 | 16 | 0 | 101 | 0 | 4458 | 0 | 0 | 11240 | 128 | 22 |
| SRR9924511 | 14 | 13 | 21 | 39 | 51 | 169 | 102 | 239 | 59 | 11 | 144 | 12 | 0 | 0 | 7402 | 5519 | 55883 | 153 | 31 | 2478 | 1153 | 11 | 0 | 56 | 69 | 309 | 817 | 244 | 0 | 510 | 0 | 3438 | 0 | 0 | 4397 | 170 | 14 |
| SRR9924512 | 11 | 0 | 10 | 33 | 48 | 120 | 96 | 297 | 29 | 0 | 145 | 0 | 0 | 0 | 9282 | 4801 | 62014 | 177 | 19 | 2306 | 1455 | 0 | 0 | 73 | 56 | 217 | 241 | 87 | 0 | 520 | 0 | 5311 | 0 | 0 | 4545 | 196 | 21 |
| SRR9924514 | 11 | 0 | 0 | 32 | 30 | 178 | 123 | 417 | 92 | 19 | 26 | 17 | 0 | 0 | 5169 | 3417 | 16714 | 49 | 43 | 2863 | 871 | 0 | 0 | 33 | 34 | 29 | 89 | 37 | 0 | 75 | 0 | 895 | 0 | 0 | 8656 | 155 | 22 |
| SRR9924515 | 0 | 0 | 12 | 22 | 25 | 87 | 92 | 249 | 39 | 0 | 14 | 27 | 0 | 0 | 6091 | 3995 | 36904 | 113 | 17 | 2278 | 1034 | 0 | 0 | 38 | 36 | 29 | 60 | 55 | 0 | 58 | 0 | 1972 | 0 | 0 | 4469 | 164 | 31 |
| SRR9924516 | 0 | 0 | 0 | 0 | 23 | 187 | 112 | 410 | 62 | 23 | 47 | 0 | 0 | 0 | 3526 | 2403 | 14230 | 50 | 25 | 1318 | 491 | 0 | 0 | 49 | 21 | 54 | 18 | 0 | 0 | 39 | 0 | 759 | 0 | 0 | 6780 | 167 | 19 |
| SRR9924517 | 12 | 16 | 0 | 22 | 29 | 195 | 179 | 379 | 114 | 15 | 29 | 20 | 0 | 0 | 5081 | 3235 | 16994 | 49 | 38 | 2727 | 693 | 0 | 0 | 52 | 34 | 23 | 72 | 36 | 0 | 83 | 0 | 514 | 0 | 0 | 9565 | 166 | 20 |
| SRR9924518 | 0 | 14 | 0 | 30 | 68 | 95 | 59 | 131 | 23 | 13 | 16 | 11 | 0 | 0 | 2701 | 1670 | 5805 | 325 | 13 | 1610 | 399 | 0 | 0 | 42 | 37 | 20 | 73 | 10 | 0 | 97 | 0 | 10343 | 0 | 0 | 3783 | 162 | 18 |
| SRR9924519 | 0 | 0 | 0 | 18 | 11 | 87 | 91 | 157 | 50 | 20 | 52 | 18 | 0 | 0 | 2673 | 2167 | 7314 | 416 | 31 | 2301 | 540 | 0 | 0 | 30 | 30 | 28 | 89 | 64 | 0 | 201 | 0 | 10347 | 0 | 0 | 5087 | 148 | 20 |
| SRR9924520 | 14 | 13 | 0 | 34 | 56 | 128 | 137 | 291 | 71 | 11 | 264 | 20 | 0 | 0 | 4724 | 3589 | 11919 | 79 | 53 | 3120 | 815 | 0 | 0 | 36 | 51 | 186 | 773 | 429 | 0 | 1130 | 0 | 1005 | 0 | 0 | 13222 | 126 | 19 |
| SRR9924521 | 11 | 0 | 0 | 24 | 68 | 101 | 84 | 155 | 32 | 10 | 22 | 15 | 0 | 0 | 2453 | 2264 | 6241 | 448 | 17 | 1790 | 439 | 0 | 0 | 36 | 29 | 23 | 89 | 20 | 0 | 117 | 0 | 9246 | 0 | 0 | 4517 | 146 | 25 |
| SRR9924522 | 0 | 10 | 0 | 0 | 37 | 155 | 208 | 529 | 83 | 31 | 117 | 21 | 0 | 0 | 3324 | 1902 | 6563 | 125 | 36 | 2349 | 381 | 0 | 0 | 25 | 25 | 60 | 240 | 23 | 0 | 820 | 0 | 1689 | 0 | 0 | 10378 | 139 | 25 |
| SRR9924523 | 12 | 0 | 0 | 15 | 37 | 176 | 150 | 326 | 75 | 16 | 36 | 14 | 0 | 0 | 3341 | 2006 | 6398 | 118 | 11 | 2138 | 631 | 0 | 0 | 30 | 20 | 16 | 65 | 17 | 0 | 165 | 0 | 2143 | 0 | 0 | 8618 | 147 | 16 |
| SRR9924524 | 0 | 0 | 0 | 16 | 25 | 153 | 112 | 288 | 72 | 15 | 27 | 26 | 0 | 0 | 21983 | 6164 | 40703 | 314 | 101 | 5085 | 3822 | 0 | 0 | 38 | 30 | 25 | 65 | 21 | 0 | 112 | 0 | 4581 | 0 | 0 | 6056 | 151 | 14 |
| SRR9924525 | 16 | 12 | 15 | 98 | 73 | 117 | 108 | 175 | 32 | 0 | 203 | 0 | 0 | 0 | 4516 | 3201 | 8245 | 737 | 29 | 2348 | 565 | 0 | 0 | 57 | 45 | 310 | 721 | 641 | 0 | 704 | 0 | 7289 | 0 | 0 | 4206 | 121 | 25 |
| SRR9924526 | 0 | 0 | 0 | 28 | 27 | 150 | 118 | 237 | 39 | 0 | 27 | 0 | 0 | 0 | 24507 | 6317 | 46325 | 279 | 61 | 4342 | 1561 | 0 | 0 | 39 | 33 | 31 | 85 | 39 | 0 | 101 | 0 | 6215 | 0 | 0 | 3366 | 131 | 40 |
| SRR9924527 | 0 | 0 | 0 | 17 | 138 | 159 | 157 | 198 | 41 | 12 | 177 | 14 | 0 | 0 | 12833 | 5373 | 24527 | 143 | 58 | 4106 | 1075 | 0 | 0 | 42 | 26 | 125 | 467 | 47 | 0 | 777 | 0 | 5824 | 0 | 0 | 6901 | 131 | 27 |
