## Supplementary material for "Single-nucleotide Differences and Cell Type Decide the Subcellular Localization of miRNA Isoforms (isomiRs), tRNA-derived Fragments (tRFs) and rRNA-derived Fragments (rRFs)": Supp. Table S8 (4 of 5)

### Supp Table S8.4.Columns_DI_to_ES

**Supplemental Table S8:** The abundances

#### 111 Short RNA (Figure 5)

| | >hg38_21_+_8433172_8446622_RNA45SN1_45S_50ntFlanks@12366.12383.18\|TAACTGGCTTGTGGCGGC | >hg38_21_+_8433172_8446622_RNA45SN1_45S_50ntFlanks@3705.3728.24\|TACCTGGTTGATCCTGCCAGTAGC | >hg38_21_+_8433172_8446622_RNA45SN1_45S_50ntFlanks@6649.6683.35\|TACGACTCTTAGCGGTGGATCACTCGGCTCGTGCG | >MI0000087\|hsa-mir-29a&WithFlank&7\|-\|130876741\|130876816@48.67.20(+1C)&[MIMAT0000086&hsa-miR-29a-3p&offsets\|0\|-2(+1C);m-26&7\|-\|130876749\|130876769&offsets\|0\|-1(+1C)]\|TAGCACCATCTGAAATCGGTC | >MI0000077\|hsa-mir-21&WithFlank&17\|+\|59841260\|59841343@14.34.21(+1U)&[MIMAT0000076&hsa-miR-21-5p&offsets\|0\|-1(+1U);m-1&17\|+\|59841273\|59841294&offsets\|0\|-1(+1U)]\|TAGCTTATCAGACTGATGTTGT | >hg38_21_+_8433172_8446622_RNA45SN1_45S_50ntFlanks@12654.12674.21\|TATGACTGAACGCCTCTAAGT | >hg38_21_+_8433172_8446622_RNA45SN1_45S_50ntFlanks@12605.12624.20\|TATGTGCTTGGCTGAGGAGC | >trna11_GluTTC_15_-_26327381_26327452@1.30.30,trna3_GluTTC_13_-_45492062_45492133@1.30.30\|TCCCACATGGTCTAGCGGTTAGGATTCCTG | >trna11_GluTTC_15_-_26327381_26327452@1.31.31,trna3_GluTTC_13_-_45492062_45492133@1.31.31\|TCCCACATGGTCTAGCGGTTAGGATTCCTGG | >trna11_GluTTC_15_-_26327381_26327452@1.33.33,trna3_GluTTC_13_-_45492062_45492133@1.33.33\|TCCCACATGGTCTAGCGGTTAGGATTCCTGGTT | >trna101_AlaAGC_6_-_28831462_28831533@58.75.18,trna102_AlaAGC_6_-_28806221_28806292@58.75.18,trna110_AlaTGC_6_-_28757547_28757618@58.75.18,trna113_AlaTGC_6_-_28726141_28726212@58.75.18,trna65_AlaAGC_6_+_28574933_28575004@58.75.18\|TCCCCGGCACCTCCACCA | >trna10_AlaCGC_6_+_26553731_26553802@58.75.18,trna119_AlaCGC_6_-_28641613_28641684@58.75.18,trna13_AlaCGC_2_+_157257281_157257352@58.75.18,trna13_AlaTGC_12_-_125406301_125406372@58.75.18,trna66_AlaTGC_6_+_28611222_28611293@58.75.18,trna8_AlaTGC_12_+_125424 | >hg38_21_+_8433172_8446622_RNA45SN1_45S_50ntFlanks@12992.13013.22\|TCCCTCGCTGCGATCTATTGAA | >trna134_GluTTC_1_-_16861774_16861845@1.33.33,trna5_GluTTC_1_+_17199078_17199149@1.33.33,trna84_GluTTC_1_-_161391883_161391954@1.33.33,trna94_GluTTC_1_-_149664355_149664427@1.33.33\|TCCCTGGTGGTCTAGTGGCTAGGATTCGGCGCT | >trnaMT_HisGTG_MT_+_12138_12206@32.70.39\|TGAATCTGACAACAGAGGCTTACGACCCCTTATTTACCC | >trnaMT_HisGTG_MT_+_12138_12206@32.71.40\|TGAATCTGACAACAGAGGCTTACGACCCCTTATTTACCCC | >hg38_21_+_8433172_8446622_RNA45SN1_45S_50ntFlanks@11785.11803.19\|TGAATGGATGAACGAGATT | >MI0000081\|hsa-mir-24-2&WithFlank&19\|-\|13836281\|13836365@56.77.22(+1U);MI0000080\|hsa-mir-24-1&WithFlank&9\|+\|95086015\|95086094@50.71.22(+1U)&[MIMAT0000080&hsa-miR-24-3p&offsets\|0\|0(+1U);m-36&19\|-\|13836288\|13836310&offsets\|0\|-1(+1U)];[MIMAT0000080_1&hsa- | >MI0003643\|hsa-mir-629&WithFlank&15\|-\|70079366\|70079474@28.49.22&[MIMAT0004810&hsa-miR-629-5p&offsets\|0\|+1;m-150&15\|-\|70079427\|70079447&offsets\|0\|+1]\|TGGGTTTACGTTGGGAGAACTT | >trnaMT_GluTTC_MT_-_14674_14742@41.72.32,trnalookalike8_GluTTC_5_-_93905172_93905240@41.72.32\|TGGTCGTGGTTGTAGTCCGTGCGAGAATACCA | >MI0000441\|hsa-mir-30b&WithFlank&8\|-\|134800514\|134800613@23.44.22&[MIMAT0000420&hsa-miR-30b-5p&offsets\|0\|0;m-76&8\|-\|134800570\|134800591&offsets\|0\|0]\|TGTAAACATCCTACACTCAGCT | >MI0000736\|hsa-mir-30c-1&WithFlank&1\|+\|40757278\|40757378@23.46.24;MI0000254\|hsa-mir-30c-2&WithFlank&6\|-\|71376954\|71377037@13.36.24&[MIMAT0000244&hsa-miR-30c-5p&offsets\|0\|+1;m-57&1\|+\|40757300\|40757323&offsets\|0\|0];[MIMAT0000244_1&hsa-miR-30c- | >MI0000088\|hsa-mir-30a&WithFlank&6\|-\|71403545\|71403627@12.33.22(+1A)&[MIMAT0000087&hsa-miR-30a-5p&offsets\|0\|0(+1A);m-15&6\|-\|71403595\|71403616&offsets\|0\|0(+1A)]\|TGTAAACATCCTCGACTGGAAGA | >MI0000088\|hsa-mir-30a&WithFlank&6\|-\|71403545\|71403627@12.35.24&[MIMAT0000087&hsa-miR-30a-5p&offsets\|0\|+2;m-15&6\|-\|71403595\|71403616&offsets\|0\|+2]\|TGTAAACATCCTCGACTGGAAGCT | >MI0000749\|hsa-mir-30e&WithFlank&1\|+\|40754349\|40754452@23.46.24&[MIMAT0000692&hsa-miR-30e-5p&offsets\|0\|+2;m-24&1\|+\|40754371\|40754394&offsets\|0\|0]\|TGTAAACATCCTTGACTGGAAGCT | >MI0003591\|hsa-mir-584&WithFlank&5\|-\|149062307\|149062415@22.42.21&[MIMAT0003249&hsa-miR-584-5p&offsets\|0\|-1;m-167&5\|-\|149062374\|149062394&offsets\|0\|0]\|TTATGGTTTGCCTGGGACTGA | >hg38_21_+_8433172_8446622_RNA45SN1_45S_50ntFlanks@12416.12433.18\|TTCGATGTCGGCTCTTCC | >hg38_21_+_8433172_8446622_RNA45SN1_45S_50ntFlanks@476.495.20\|TTCGTGATCGATGTGGTGAC | >hg38_21_+_8433172_8446622_RNA45SN1_45S_50ntFlanks@476.501.26\|TTCGTGATCGATGTGGTGACGTCGTG | >hg38_21_+_8433172_8446622_RNA45SN1_45S_50ntFlanks@476.503.28\|TTCGTGATCGATGTGGTGACGTCGTGCT | >hg38_21_+_8433172_8446622_RNA45SN1_45S_50ntFlanks@476.505.30\|TTCGTGATCGATGTGGTGACGTCGTGCTCT | >hg38_21_+_8433172_8446622_RNA45SN1_45S_50ntFlanks@476.507.32\|TTCGTGATCGATGTGGTGACGTCGTGCTCTCC | >hg38_21_+_8433172_8446622_RNA45SN1_45S_50ntFlanks@476.509.34\|TTCGTGATCGATGTGGTGACGTCGTGCTCTCCCG | >hg38_21_+_8433172_8446622_RNA45SN1_45S_50ntFlanks@476.510.35\|TTCGTGATCGATGTGGTGACGTCGTGCTCTCCCGG | >hg38_21_+_8433172_8446622_RNA45SN1_45S_50ntFlanks@476.518.43\|TTCGTGATCGATGTGGTGACGTCGTGCTCTCCCGGGCCGGGTC | >trnaMT_GluTTC_MT_-_14674_14742@40.72.33,trnalookalike8_GluTTC_5_-_93905172_93905240@40.72.33\|TTGGTCGTGGTTGTAGTCCGTGCGAGAATACCA | >MI0000098\|hsa-mir-96&WithFlank&7\|-\|129774686\|129774775@15.37.23&[MIMAT0000095&hsa-miR-96-5p&offsets\|0\|0;m-248&7\|-\|129774739\|129774761&offsets\|0\|0]\|TTTGGCACTAGCACATTTTTGCT |
|---|---|---|---|---|---|---|---|---|---|---|---|---|---|---|---|---|---|---|---|---|---|---|---|---|---|---|---|---|---|---|---|---|---|---|---|---|---|
| SRR10024150 | 10 | 572 | 291 | 0 | 0 | 10 | 0 | 244 | 441 | 1185 | 33225 | 85594 | 0 | 23 | 93 | 197 | 31 | 29 | 34 | 105 | 813 | 27 | 0 | 49 | 227 | 32 | 1375 | 15 | 31 | 368 | 419 | 388 | 128 | 82 | 41 | 221 | 0 |
| SRR10024151 | 0 | 210 | 14 | 0 | 0 | 0 | 0 | 291 | 979 | 6058 | 29883 | 74296 | 0 | 795 | 48 | 126 | 11 | 22 | 70 | 291 | 213 | 2492 | 0 | 11 | 146 | 72 | 14 | 0 | 47 | 21 | 19 | 18 | 174 | 146 | 28 | 572 | 19 |
| SRR10024152 | 10 | 129 | 221 | 0 | 0 | 26 | 0 | 151 | 407 | 814 | 2418 | 81225 | 31 | 140 | 23 | 47 | 21 | 18 | 41 | 175 | 181 | 1124 | 0 | 11 | 101 | 30 | 4633 | 0 | 36 | 211 | 350 | 345 | 134 | 93 | 10 | 186 | 30 |
| SRR10024153 | 0 | 622 | 353 | 0 | 0 | 15 | 0 | 200 | 478 | 1296 | 29719 | 65176 | 0 | 154 | 0 | 0 | 0 | 28 | 42 | 170 | 861 | 5039 | 0 | 22 | 262 | 28 | 861 | 15 | 65 | 1140 | 812 | 578 | 90 | 55 | 0 | 355 | 25 |
| SRR10024154 | 85 | 316 | 123 | 0 | 0 | 47 | 0 | 144 | 369 | 2070 | 1576 | 81731 | 0 | 48 | 32 | 80 | 0 | 19 | 30 | 160 | 502 | 1930 | 0 | 0 | 146 | 37 | 263 | 0 | 0 | 190 | 177 | 139 | 38 | 0 | 0 | 177 | 17 |
| SRR10024155 | 11 | 141 | 174 | 0 | 0 | 0 | 0 | 51 | 242 | 1122 | 27113 | 75994 | 18 | 26 | 13 | 13 | 0 | 17 | 44 | 1620 | 640 | 2278 | 0 | 14 | 160 | 56 | 733 | 11 | 24 | 342 | 275 | 208 | 46 | 27 | 0 | 10011 | 13 |
| SRR10024156 | 20 | 236 | 287 | 0 | 0 | 18 | 0 | 280 | 763 | 5014 | 1233 | 41702 | 0 | 249 | 47 | 156 | 0 | 21 | 62 | 498 | 336 | 1619 | 0 | 11 | 242 | 92 | 50 | 0 | 30 | 17 | 17 | 13 | 114 | 69 | 15 | 1324 | 31 |
| SRR10024158 | 58 | 312 | 128 | 0 | 0 | 41 | 0 | 201 | 660 | 2024 | 44520 | 85438 | 0 | 100 | 0 | 43 | 12 | 31 | 50 | 172 | 538 | 1908 | 0 | 36 | 194 | 55 | 3765 | 12 | 14 | 278 | 208 | 111 | 0 | 0 | 0 | 287 | 23 |
| SRR10024159 | 28 | 383 | 103 | 0 | 0 | 22 | 0 | 367 | 776 | 1976 | 35239 | 70491 | 17 | 85 | 0 | 14 | 17 | 28 | 64 | 325 | 403 | 2579 | 0 | 14 | 226 | 84 | 4083 | 11 | 48 | 557 | 509 | 355 | 70 | 56 | 0 | 494 | 15 |
| SRR10024161 | 0 | 565 | 179 | 0 | 0 | 0 | 0 | 86 | 343 | 660 | 42274 | 122330 | 11 | 69 | 0 | 0 | 0 | 23 | 41 | 137 | 688 | 16 | 0 | 14 | 221 | 42 | 20 | 0 | 19 | 447 | 210 | 198 | 65 | 19 | 0 | 360 | 0 |
| SRR10024162 | 12 | 508 | 11 | 0 | 0 | 0 | 0 | 369 | 1107 | 2361 | 2798 | 65322 | 0 | 179 | 74 | 74 | 0 | 25 | 56 | 306 | 388 | 13 | 0 | 19 | 271 | 59 | 24 | 25 | 75 | 25 | 21 | 16 | 237 | 146 | 22 | 411 | 20 |
| SRR10024163 | 0 | 210 | 625 | 0 | 0 | 0 | 0 | 70 | 167 | 914 | 5552 | 115411 | 14 | 167 | 21 | 35 | 23 | 15 | 26 | 119 | 321 | 1873 | 0 | 0 | 112 | 47 | 22 | 0 | 47 | 597 | 602 | 798 | 313 | 266 | 23 | 160 | 10 |
| SRR10024164 | 43 | 806 | 130 | 0 | 0 | 35 | 0 | 428 | 941 | 3537 | 317 | 44008 | 26 | 114 | 0 | 0 | 35 | 27 | 21 | 257 | 846 | 1839 | 0 | 0 | 199 | 37 | 6255 | 17 | 43 | 225 | 546 | 277 | 104 | 61 | 0 | 314 | 0 |
| SRR10024166 | 0 | 438 | 252 | 0 | 0 | 21 | 0 | 143 | 294 | 1134 | 4681 | 91207 | 0 | 42 | 0 | 25 | 17 | 16 | 34 | 151 | 440 | 1584 | 0 | 0 | 76 | 72 | 57 | 12 | 70 | 10 | 15 | 17 | 239 | 95 | 0 | 218 | 23 |
| SRR10024167 | 58 | 512 | 27 | 0 | 0 | 17 | 0 | 144 | 879 | 767 | 68030 | 137851 | 14 | 32 | 16 | 32 | 55 | 13 | 63 | 112 | 252 | 1496 | 0 | 36 | 121 | 144 | 2011 | 0 | 0 | 219 | 157 | 75 | 17 | 10 | 0 | 64 | 13 |
| SRR10024168 | 19 | 281 | 126 | 0 | 0 | 24 | 0 | 708 | 1416 | 2333 | 661 | 22751 | 0 | 193 | 0 | 32 | 0 | 52 | 63 | 386 | 528 | 5018 | 0 | 24 | 316 | 74 | 32 | 0 | 48 | 116 | 97 | 102 | 68 | 44 | 0 | 547 | 12 |
| SRR10024169 | 0 | 211 | 297 | 0 | 0 | 0 | 0 | 149 | 503 | 2423 | 3620 | 76935 | 13 | 251 | 0 | 34 | 0 | 22 | 59 | 297 | 337 | 11 | 0 | 13 | 129 | 45 | 57 | 0 | 27 | 31 | 20 | 13 | 91 | 35 | 0 | 263 | 19 |
| SRR10024170 | 17 | 408 | 299 | 0 | 0 | 0 | 0 | 103 | 308 | 1276 | 3425 | 107046 | 17 | 46 | 0 | 23 | 0 | 24 | 65 | 114 | 663 | 1801 | 0 | 0 | 157 | 106 | 51 | 0 | 17 | 239 | 477 | 421 | 143 | 48 | 0 | 387 | 30 |
| SRR10024171 | 32 | 271 | 116 | 0 | 0 | 0 | 0 | 246 | 1366 | 3641 | 39563 | 102585 | 0 | 172 | 74 | 86 | 0 | 17 | 53 | 197 | 194 | 820 | 0 | 0 | 164 | 55 | 65 | 12 | 54 | 244 | 242 | 180 | 72 | 22 | 25 | 135 | 17 |
| SRR10024173 | 0 | 16 | 95 | 0 | 13 | 18 | 0 | 359 | 1489 | 4073 | 14360 | 47495 | 0 | 137 | 0 | 17 | 23 | 37 | 60 | 736 | 698 | 1600 | 0 | 0 | 258 | 77 | 37 | 0 | 32 | 149 | 261 | 212 | 104 | 27 | 0 | 822 | 54 |
| SRR10024176 | 0 | 822 | 34 | 0 | 0 | 0 | 0 | 66 | 92 | 269 | 77709 | 148891 | 0 | 35 | 0 | 0 | 0 | 19 | 76 | 35 | 319 | 2185 | 0 | 27 | 100 | 54 | 363 | 0 | 0 | 120 | 72 | 39 | 0 | 0 | 0 | 31 | 11 |
| SRR10024178 | 0 | 173 | 256 | 0 | 0 | 0 | 0 | 99 | 291 | 957 | 6405 | 115822 | 0 | 43 | 14 | 0 | 19 | 18 | 56 | 170 | 243 | 1750 | 0 | 0 | 95 | 63 | 18 | 0 | 22 | 178 | 178 | 237 | 106 | 61 | 0 | 170 | 0 |
| SRR10024179 | 19 | 139 | 152 | 0 | 0 | 15 | 27 | 83 | 550 | 1083 | 31266 | 85336 | 13 | 100 | 0 | 0 | 40 | 21 | 89 | 383 | 186 | 1182 | 0 | 0 | 123 | 46 | 1466 | 0 | 23 | 125 | 103 | 57 | 19 | 21 | 0 | 1249 | 11 |
| SRR10024180 | 26 | 508 | 99 | 0 | 0 | 11 | 0 | 153 | 390 | 877 | 2002 | 69986 | 0 | 84 | 0 | 0 | 33 | 14 | 44 | 139 | 573 | 13 | 0 | 13 | 107 | 30 | 48 | 18 | 26 | 379 | 291 | 236 | 44 | 15 | 0 | 153 | 0 |
| SRR10024181 | 0 | 170 | 452 | 0 | 0 | 0 | 0 | 104 | 431 | 1788 | 3000 | 98862 | 0 | 183 | 91 | 235 | 0 | 19 | 83 | 131 | 154 | 956 | 0 | 24 | 135 | 58 | 650 | 16 | 28 | 452 | 427 | 295 | 75 | 38 | 35 | 457 | 24 |
| SRR10024182 | 0 | 309 | 463 | 0 | 0 | 0 | 0 | 161 | 644 | 2706 | 1040 | 57320 | 29 | 145 | 16 | 16 | 24 | 21 | 31 | 274 | 471 | 2474 | 0 | 0 | 143 | 36 | 12 | 0 | 34 | 599 | 14 | 11 | 237 | 130 | 14 | 451 | 0 |
| SRR10024183 | 0 | 565 | 400 | 0 | 0 | 15 | 12 | 70 | 218 | 1684 | 35903 | 152332 | 0 | 64 | 0 | 0 | 18 | 21 | 112 | 134 | 248 | 1001 | 0 | 0 | 118 | 150 | 1585 | 0 | 27 | 147 | 360 | 217 | 95 | 40 | 0 | 199 | 41 |
| SRR10024185 | 0 | 364 | 199 | 0 | 0 | 0 | 0 | 313 | 1201 | 2290 | 29736 | 68309 | 0 | 38 | 13 | 25 | 0 | 20 | 32 | 826 | 947 | 1910 | 0 | 0 | 131 | 90 | 1392 | 0 | 24 | 437 | 566 | 515 | 111 | 90 | 0 | 926 | 17 |
| SRR10024186 | 11 | 275 | 39 | 0 | 0 | 0 | 0 | 52 | 353 | 999 | 6995 | 121597 | 0 | 103 | 0 | 0 | 25 | 22 | 43 | 95 | 336 | 12 | 0 | 0 | 128 | 63 | 1097 | 11 | 17 | 253 | 176 | 176 | 47 | 22 | 0 | 207 | 14 |
| SRR10024187 | 0 | 15 | 132 | 0 | 10 | 0 | 0 | 88 | 272 | 754 | 2496 | 61024 | 13 | 18 | 0 | 35 | 31 | 55 | 28 | 79 | 3751 | 3780 | 0 | 10 | 149 | 196 | 43 | 0 | 50 | 120 | 535 | 309 | 170 | 69 | 0 | 79 | 47 |
| SRR10024188 | 0 | 407 | 173 | 0 | 0 | 12 | 0 | 111 | 194 | 1412 | 4226 | 79336 | 16 | 18 | 0 | 0 | 0 | 0 | 36 | 129 | 407 | 1126 | 0 | 21 | 89 | 72 | 9234 | 12 | 56 | 153 | 702 | 762 | 190 | 60 | 0 | 258 | 23 |
| SRR10024189 | 0 | 509 | 350 | 0 | 0 | 13 | 0 | 177 | 801 | 2026 | 27882 | 66588 | 0 | 47 | 0 | 0 | 13 | 19 | 41 | 212 | 710 | 1887 | 0 | 66 | 190 | 57 | 2710 | 13 | 54 | 467 | 792 | 630 | 171 | 54 | 0 | 188 | 26 |
| SRR10024190 | 0 | 209 | 72 | 0 | 0 | 28 | 0 | 337 | 654 | 3905 | 597 | 35896 | 36 | 139 | 0 | 0 | 0 | 0 | 19 | 1070 | 242 | 571 | 0 | 0 | 39 | 57 | 283 | 0 | 24 | 76 | 19 | 26 | 575 | 173 | 0 | 1804 | 18 |
| SRR10024191 | 11 | 1345 | 228 | 0 | 10 | 0 | 0 | 288 | 864 | 3088 | 643 | 50141 | 0 | 52 | 0 | 0 | 33 | 14 | 24 | 157 | 501 | 1230 | 0 | 0 | 225 | 32 | 1063 | 11 | 54 | 174 | 336 | 141 | 33 | 54 | 0 | 131 | 20 |
| SRR10024192 | 0 | 512 | 651 | 0 | 0 | 11 | 0 | 352 | 1478 | 2147 | 15168 | 33678 | 0 | 35 | 0 | 0 | 32 | 13 | 29 | 282 | 710 | 1878 | 0 | 21 | 143 | 31 | 1634 | 32 | 128 | 619 | 1623 | 897 | 406 | 149 | 0 | 422 | 25 |
| SRR10024193 | 0 | 16 | 449 | 0 | 0 | 0 | 0 | 454 | 1726 | 2217 | 37201 | 63897 | 0 | 345 | 73 | 91 | 0 | 24 | 57 | 200 | 631 | 4213 | 0 | 15 | 259 | 46 | 16 | 22 | 40 | 18 | 12 | 370 | 98 | 62 | 0 | 291 | 13 |
| SRR10024194 | 0 | 11 | 67 | 0 | 0 | 0 | 0 | 220 | 470 | 720 | 2673 | 89604 | 0 | 40 | 10 | 0 | 0 | 29 | 58 | 70 | 259 | 1293 | 0 | 31 | 136 | 68 | 36 | 12 | 15 | 103 | 152 | 85 | 50 | 21 | 0 | 120 | 13 |
| SRR10024195 | 0 | 71 | 303 | 0 | 0 | 20 | 0 | 231 | 381 | 1305 | 41914 | 89106 | 0 | 127 | 23 | 23 | 0 | 15 | 52 | 116 | 215 | 1706 | 0 | 11 | 157 | 87 | 2503 | 16 | 28 | 475 | 440 | 699 | 200 | 94 | 24 | 312 | 17 |
| SRR10024196 | 0 | 610 | 125 | 0 | 0 | 0 | 0 | 337 | 621 | 1578 | 36512 | 109056 | 0 | 71 | 0 | 0 | 31 | 12 | 16 | 284 | 653 | 1473 | 0 | 0 | 162 | 68 | 3052 | 0 | 23 | 391 | 438 | 454 | 125 | 63 | 0 | 177 | 12 |
| SRR10024197 | 30 | 441 | 91 | 0 | 0 | 17 | 0 | 216 | 293 | 1112 | 1833 | 57560 | 0 | 46 | 0 | 31 | 22 | 20 | 32 | 124 | 387 | 2279 | 0 | 0 | 184 | 37 | 24 | 0 | 30 | 164 | 112 | 121 | 69 | 30 | 13 | 46 | 15 |
| SRR10024198 | 13 | 210 | 130 | 0 | 0 | 16 | 0 | 157 | 668 | 2673 | 47475 | 92146 | 0 | 79 | 13 | 52 | 11 | 22 | 65 | 157 | 223 | 1514 | 0 | 0 | 178 | 70 | 21 | 0 | 11 | 157 | 117 | 101 | 72 | 27 | 0 | 301 | 20 |
| SRR10024199 | 13 | 368 | 232 | 0 | 0 | 13 | 0 | 196 | 908 | 3009 | 25268 | 67540 | 0 | 142 | 0 | 0 | 13 | 17 | 31 | 427 | 335 | 1021 | 0 | 0 | 144 | 53 | 4443 | 0 | 0 | 123 | 323 | 232 | 90 | 52 | 0 | 427 | 60 |
| SRR10024201 | 32 | 311 | 180 | 0 | 0 | 23 | 0 | 53 | 456 | 1070 | 1820 | 90408 | 0 | 18 | 18 | 18 | 18 | 13 | 48 | 193 | 322 | 2296 | 0 | 0 | 88 | 38 | 3239 | 0 | 27 | 812 | 550 | 406 | 72 | 72 | 0 | 298 | 0 |
| SRR10024202 | 0 | 391 | 263 | 0 | 0 | 0 | 0 | 205 | 375 | 3172 | 20496 | 75743 | 0 | 358 | 17 | 17 | 24 | 16 | 32 | 409 | 384 | 1250 | 0 | 0 | 168 | 53 | 1698 | 0 | 32 | 175 | 582 | 470 | 271 | 136 | 0 | 375 | 40 |
| SRR10024203 | 0 | 304 | 492 | 0 | 0 | 0 | 0 | 556 | 1768 | 3793 | 22382 | 43951 | 0 | 258 | 179 | 218 | 55 | 26 | 126 | 338 | 307 | 2570 | 0 | 11 | 247 | 99 | 465 | 37 | 46 | 19 | 12 | 584 | 106 | 60 | 23 | 497 | 13 |
| SRR10024204 | 29 | 221 | 111 | 0 | 0 | 36 | 0 | 70 | 279 | 823 | 1923 | 66553 | 0 | 125 | 0 | 0 | 16 | 23 | 62 | 98 | 321 | 1607 | 0 | 11 | 171 | 67 | 104 | 16 | 49 | 14 | 16 | 527 | 36 | 20 | 0 | 167 | 0 |
| SRR10024205 | 12 | 418 | 650 | 0 | 0 | 12 | 0 | 76 | 420 | 2482 | 12258 | 34061 | 46 | 76 | 0 | 76 | 35 | 13 | 12 | 993 | 554 | 1196 | 0 | 0 | 122 | 26 | 4495 | 0 | 128 | 1069 | 1464 | 1127 | 290 | 93 | 0 | 611 | 19 |
| SRR10024206 | 27 | 224 | 10 | 0 | 0 | 46 | 0 | 232 | 877 | 2609 | 30421 | 89888 | 0 | 254 | 0 | 32 | 19 | 26 | 44 | 243 | 610 | 2252 | 0 | 0 | 148 | 57 | 116 | 13 | 23 | 16 | 12 | 249 | 75 | 46 | 0 | 338 | 29 |
| SRR10024207 | 10 | 13 | 46 | 0 | 0 | 10 | 0 | 63 | 331 | 1197 | 38863 | 125342 | 10 | 16 | 0 | 0 | 15 | 16 | 42 | 110 | 454 | 1457 | 0 | 65 | 133 | 49 | 652 | 0 | 25 | 199 | 245 | 239 | 31 | 15 | 0 | 126 | 13 |
| SRR10024208 | 20 | 247 | 152 | 0 | 0 | 32 | 0 | 424 | 943 | 927 | 952 | 36605 | 16 | 79 | 126 | 79 | 16 | 32 | 27 | 283 | 720 | 2989 | 0 | 14 | 311 | 21 | 40 | 0 | 36 | 215 | 335 | 279 | 68 | 36 | 0 | 503 | 12 |
| SRR1003025 | 0 | 0 | 48 | 17 | 13 | 0 | 0 | 356 | 177 | 9553 | 2739 | 2228 | 0 | 1103 | 321 | 128 | 0 | 41 | 0 | 412 | 2251 | 3892 | 118 | 3838 | 2468 | 0 | 0 | 83 | 62 | 20 | 17 | 38 | 84 | 19 | 53 | 1597 | 0 |
| SRR1005875 | 0 | 0 | 0 | 0 | 0 | 0 | 0 | 3553 | 1093 | 5558 | 1184 | 1640 | 0 | 91 | 0 | 0 | 0 | 52 | 0 | 91 | 1510 | 608 | 0 | 689 | 13117 | 2384 | 19 | 0 | 0 | 0 | 0 | 0 | 0 | 0 | 0 | 456 | 0 |
| SRR10072377 | 0 | 2640 | 99 | 446 | 345 | 12 | 30 | 1049 | 7060 | 11742 | 61 | 201 | 0 | 707 | 280 | 500 | 20 | 3730 | 17 | 4835 | 2509 | 1610 | 157 | 3211 | 3015 | 14 | 0 | 0 | 40 | 32 | 35 | 0 | 0 | 0 | 0 | 7115 | 0 |
| SRR10072378 | 0 | 356 | 13 | 431 | 334 | 15 | 30 | 950 | 7096 | 11452 | 47 | 175 | 0 | 630 | 332 | 478 | 17 | 3710 | 15 | 4962 | 2440 | 1599 | 152 | 3248 | 3037 | 17 | 12 | 0 | 28 | 36 | 33 | 0 | 0 | 0 | 0 | 7108 | 0 |
| SRR10072379 | 0 | 3154 | 59 | 370 | 376 | 21 | 27 | 607 | 2495 | 5423 | 72 | 827 | 0 | 520 | 367 | 300 | 17 | 3880 | 0 | 5096 | 5093 | 3504 | 177 | 3074 | 2582 | 16 | 0 | 10 | 36 | 36 | 47 | 0 | 0 | 0 | 0 | 6824 | 12 |
| SRR10072380 | 0 | 3198 | 73 | 367 | 365 | 14 | 37 | 568 | 2409 | 5223 | 73 | 842 | 0 | 542 | 398 | 300 | 12 | 3845 | 0 | 4740 | 5094 | 3500 | 184 | 2994 | 2584 | 16 | 0 | 0 | 33 | 48 | 44 | 0 | 0 | 0 | 0 | 7182 | 20 |
| SRR10072381 | 0 | 412 | 52 | 385 | 256 | 26 | 23 | 808 | 3309 | 7787 | 666 | 1353 | 0 | 701 | 1049 | 1594 | 30 | 3362 | 0 | 2997 | 4753 | 3596 | 151 | 3330 | 3157 | 0 | 0 | 0 | 27 | 22 | 42 | 0 | 0 | 0 | 0 | 3018 | 0 |
| SRR10072382 | 0 | 406 | 64 | 371 | 253 | 34 | 23 | 659 | 2894 | 7584 | 687 | 1228 | 0 | 680 | 1117 | 1492 | 48 | 3317 | 0 | 3123 | 4681 | 3562 | 149 | 3244 | 3109 | 18 | 13 | 0 | 32 | 28 | 27 | 0 | 0 | 0 | 0 | 3067 | 0 |
| SRR10072383 | 0 | 238 | 59 | 361 | 266 | 0 | 19 | 108 | 1030 | 6828 | 488 | 780 | 0 | 217 | 1322 | 1301 | 0 | 3658 | 0 | 3858 | 4547 | 3193 | 237 | 3997 | 3774 | 20 | 0 | 16 | 108 | 87 | 108 | 19 | 0 | 12 | 12 | 4790 | 0 |
| SRR10072384 | 0 | 225 | 81 | 380 | 267 | 0 | 21 | 106 | 950 | 6028 | 369 | 918 | 0 | 433 | 1520 | 1246 | 0 | 3636 | 0 | 3473 | 4420 | 3143 | 234 | 3887 | 3670 | 22 | 0 | 0 | 76 | 95 | 62 | 30 | 16 | 0 | 0 | 4993 | 0 |
| SRR10072385 | 0 | 366 | 90 | 370 | 240 | 21 | 39 | 1084 | 8287 | 12340 | 617 | 1394 | 0 | 789 | 852 | 893 | 33 | 4117 | 0 | 2785 | 4227 | 3582 | 177 | 3289 | 3180 | 13 | 25 | 0 | 52 | 43 | 58 | 0 | 0 | 0 | 0 | 4142 | 0 |
| SRR10072386 | 0 | 3305 | 78 | 342 | 235 | 18 | 31 | 1048 | 7806 | 12132 | 166 | 1218 | 0 | 838 | 831 | 806 | 34 | 4094 | 0 | 2732 | 4204 | 3548 | 172 | 3194 | 3163 | 12 | 33 | 0 | 41 | 53 | 59 | 0 | 0 | 0 | 0 | 3907 | 0 |
| SRR10072387 | 0 | 2633 | 65 | 411 | 264 | 14 | 29 | 485 | 2251 | 4336 | 123 | 685 | 0 | 854 | 259 | 387 | 30 | 3538 | 0 | 1377 | 4114 | 3817 | 140 | 3073 | 3279 | 14 | 0 | 0 | 17 | 22 | 29 | 0 | 0 | 0 | 0 | 1592 | 0 |
| SRR10072388 | 0 | 366 | 71 | 395 | 254 | 18 | 47 | 448 | 2216 | 4036 | 132 | 622 | 0 | 829 | 274 | 356 | 33 | 3488 | 0 | 1290 | 3999 | 3814 | 135 | 3059 | 3196 | 17 | 0 | 0 | 13 | 24 | 26 | 0 | 0 | 0 | 0 | 1578 | 0 |
| SRR10072389 | 0 | 201 | 32 | 178 | 91 | 0 | 0 | 39 | 297 | 1616 | 142 | 272 | 0 | 103 | 1875 | 2108 | 0 | 4913 | 0 | 10785 | 3866 | 2324 | 318 | 7099 | 5804 | 30 | 0 | 0 | 13 | 31 | 37 | 11 | 37 | 37 | 0 | 14354 | 0 |
| SRR10072390 | 0 | 2634 | 39 | 163 | 88 | 0 | 0 | 37 | 261 | 1489 | 13 | 360 | 0 | 112 | 1414 | 1935 | 0 | 4935 | 0 | 10259 | 3814 | 2265 | 306 | 7041 | 5720 | 36 | 15 | 0 | 139 | 374 | 486 | 107 | 56 | 44 | 0 | 14576 | 0 |
| SRR10072391 | 0 | 454 | 69 | 358 | 350 | 0 | 17 | 337 | 1554 | 4644 | 53 | 855 | 0 | 329 | 1726 | 2113 | 0 | 3774 | 0 | 4932 | 5173 | 3703 | 281 | 4734 | 3606 | 15 | 0 | 0 | 40 | 50 | 62 | 10 | 12 | 0 | 0 | 6034 | 0 |
| SRR10072392 | 0 | 3629 | 65 | 329 | 347 | 0 | 14 | 250 | 1389 | 5320 | 291 | 646 | 0 | 283 | 1518 | 2123 | 0 | 3819 | 0 | 4998 | 5110 | 3646 | 262 | 4683 | 3542 | 11 | 11 | 11 | 64 | 65 | 60 | 0 | 0 | 0 | 0 | 6427 | 0 |
| SRR10072393 | 0 | 3536 | 67 | 385 | 424 | 0 | 23 | 648 | 4175 | 13481 | 324 | 491 | 0 | 536 | 471 | 663 | 16 | 3125 | 0 | 3451 | 3825 | 2567 | 218 | 4171 | 3107 | 18 | 11 | 0 | 59 | 57 | 44 | 0 | 16 | 0 | 0 | 9200 | 0 |
| SRR10072394 | 0 | 3372 | 95 | 382 | 404 | 0 | 21 | 585 | 3878 | 12963 | 41 | 531 | 0 | 570 | 438 | 687 | 12 | 3123 | 0 | 2981 | 3835 | 2623 | 209 | 4081 | 3030 | 23 | 21 | 0 | 46 | 49 | 53 | 0 | 0 | 0 | 0 | 8988 | 0 |
| SRR10072395 | 0 | 172 | 54 | 147 | 347 | 14 | 14 | 25 | 266 | 2812 | 773 | 1533 | 0 | 190 | 583 | 469 | 0 | 4321 | 0 | 3927 | 2383 | 2243 | 163 | 3614 | 2802 | 15 | 0 | 20 | 150 | 231 | 184 | 75 | 27 | 54 | 0 | 4345 | 12 |
| SRR10072396 | 0 | 7095 | 54 | 160 | 343 | 0 | 0 | 25 | 262 | 2961 | 787 | 1150 | 0 | 125 | 412 | 600 | 0 | 4321 | 0 | 4061 | 2396 | 2221 | 145 | 3577 | 2775 | 13 | 13 | 34 | 161 | 201 | 127 | 94 | 67 | 27 | 0 | 3973 | 12 |
| SRR10072397 | 0 | 3501 | 106 | 468 | 307 | 12 | 0 | 645 | 5675 | 20483 | 172 | 432 | 0 | 492 | 381 | 872 | 37 | 3932 | 11 | 3861 | 3674 | 2194 | 268 | 5155 | 3924 | 17 | 13 | 0 | 46 | 61 | 101 | 0 | 20 | 22 | 0 | 5875 | 11 |
| SRR10072398 | 0 | 423 | 11 | 462 | 328 | 0 | 15 | 606 | 5308 | 19960 | 152 | 343 | 0 | 383 | 486 | 753 | 48 | 3932 | 0 | 3833 | 3638 | 2201 | 231 | 4994 | 3853 | 15 | 0 | 0 | 38 | 61 | 14 | 13 | 12 | 15 | 0 | 6154 | 0 |
| SRR10072399 | 0 | 3969 | 87 | 239 | 175 | 17 | 30 | 590 | 2315 | 10002 | 125 | 244 | 0 | 1019 | 646 | 701 | 23 | 3835 | 12 | 3864 | 4531 | 4148 | 146 | 2945 | 2784 | 19 | 12 | 12 | 29 | 38 | 30 | 0 | 0 | 0 | 0 | 3380 | 0 |
| SRR10072400 | 0 | 3871 | 128 | 235 | 173 | 19 | 21 | 566 | 2256 | 9522 | 126 | 274 | 0 | 1007 | 683 | 737 | 28 | 3797 | 13 | 3743 | 4526 | 4103 | 159 | 2983 | 2719 | 15 | 0 | 0 | 27 | 21 | 35 | 0 | 0 | 0 | 0 | 3510 | 0 |
| SRR10077412 | 0 | 106 | 0 | 0 | 0 | 0 | 0 | 1109 | 268 | 765 | 23330 | 11149 | 0 | 115 | 0 | 19 | 0 | 368 | 0 | 440 | 1417 | 10332 | 159 | 92381 | 6256 | 336 | 0 | 48 | 58 | 237 | 126 | 0 | 19 | 24 | 0 | 1205 | 75 |
| SRR10077413 | 0 | 69 | 0 | 0 | 0 | 0 | 0 | 1130 | 273 | 1052 | 21378 | 10445 | 0 | 175 | 0 | 0 | 0 | 362 | 0 | 604 | 1251 | 9365 | 160 | 93898 | 6124 | 339 | 20 | 74 | 49 | 212 | 168 | 0 | 0 | 0 | 0 | 1325 | 72 |
| SRR10077414 | 0 | 137 | 0 | 0 | 0 | 0 | 0 | 707 | 323 | 849 | 1091 | 9818 | 0 | 121 | 0 | 0 | 0 | 362 | 0 | 424 | 1448 | 12 | 156 | 92269 | 6267 | 345 | 0 | 69 | 69 | 200 | 169 | 0 | 0 | 11 | 0 | 990 | 75 |
| SRR10077415 | 0 | 129 | 0 | 0 | 0 | 0 | 0 | 1099 | 157 | 1118 | 22134 | 9752 | 0 | 59 | 0 | 0 | 0 | 368 | 0 | 432 | 1291 | 9551 | 164 | 94609 | 6079 | 332 | 0 | 80 | 50 | 154 | 169 | 25 | 20 | 35 | 0 | 1452 | 77 |
| SRR10077416 | 0 | 154 | 0 | 0 | 0 | 0 | 0 | 153 | 22 | 44 | 35181 | 13762 | 0 | 0 | 0 | 0 | 0 | 423 | 0 | 88 | 1635 | 7752 | 179 | 100753 | 7064 | 354 | 0 | 67 | 14 | 111 | 0 | 0 | 0 | 14 | 0 | 350 | 116 |
| SRR10077417 | 0 | 132 | 0 | 0 | 0 | 0 | 0 | 132 | 88 | 88 | 1621 | 13802 | 0 | 44 | 0 | 0 | 0 | 428 | 0 | 220 | 1566 | 7370 | 179 | 101904 | 6869 | 357 | 0 | 49 | 24 | 108 | 15 | 0 | 0 | 0 | 0 | 220 | 116 |
| SRR10077418 | 0 | 84 | 0 | 0 | 0 | 0 | 0 | 72 | 48 | 120 | 35231 | 14682 | 0 | 24 | 0 | 0 | 0 | 423 | 0 | 144 | 1656 | 11 | 191 | 100494 | 7044 | 344 | 0 | 42 | 21 | 100 | 31 | 0 | 0 | 16 | 0 | 240 | 122 |
| SRR10077419 | 0 | 128 | 0 | 0 | 11 | 0 | 0 | 111 | 0 | 111 | 1611 | 14398 | 0 | 0 | 0 | 0 | 0 | 436 | 0 | 111 | 1584 | 7364 | 189 | 102690 | 6967 | 362 | 0 | 35 | 59 | 114 | 40 | 0 | 0 | 0 | 0 | 399 | 115 |
| SRR10077420 | 0 | 87 | 0 | 0 | 0 | 0 | 0 | 369 | 101 | 402 | 1401 | 12434 | 0 | 67 | 0 | 0 | 0 | 381 | 12 | 184 | 1426 | 10736 | 199 | 103136 | 9011 | 341 | 0 | 41 | 26 | 55 | 23 | 0 | 0 | 0 | 0 | 402 | 62 |
| SRR10077421 | 0 | 69 | 0 | 0 | 0 | 0 | 0 | 328 | 86 | 397 | 1449 | 13969 | 0 | 34 | 0 | 0 | 0 | 385 | 12 | 121 | 1242 | 10130 | 196 | 104865 | 8642 | 362 | 0 | 36 | 21 | 81 | 60 | 0 | 12 | 0 | 0 | 345 | 54 |
| SRR10077422 | 0 | 74 | 0 | 0 | 0 | 0 | 0 | 257 | 18 | 275 | 1397 | 13339 | 0 | 55 | 0 | 0 | 0 | 373 | 11 | 165 | 1364 | 15 | 187 | 102906 | 8849 | 353 | 0 | 38 | 38 | 61 | 32 | 0 | 0 | 0 | 0 | 312 | 55 |
| SRR10077423 | 0 | 87 | 0 | 0 | 0 | 0 | 0 | 244 | 122 | 383 | 1418 | 13286 | 0 | 52 | 0 | 0 | 0 | 384 | 0 | 122 | 1311 | 10141 | 200 | 105432 | 8820 | 332 | 0 | 51 | 48 | 72 | 33 | 0 | 0 | 0 | 0 | 383 | 49 |
| SRR10077424 | 0 | 206 | 0 | 0 | 0 | 0 | 0 | 400 | 128 | 608 | 3411 | 19130 | 0 | 304 | 0 | 0 | 0 | 617 | 0 | 184 | 1630 | 7827 | 191 | 89772 | 8868 | 316 | 0 | 51 | 58 | 130 | 65 | 11 | 0 | 0 | 0 | 384 | 111 |
| SRR10077425 | 0 | 199 | 0 | 0 | 0 | 0 | 0 | 433 | 188 | 678 | 3409 | 18169 | 0 | 368 | 0 | 0 | 0 | 598 | 0 | 106 | 1464 | 7298 | 188 | 90509 | 8628 | 308 | 0 | 74 | 37 | 144 | 63 | 0 | 0 | 18 | 0 | 400 | 119 |
| SRR10077426 | 0 | 181 | 0 | 0 | 0 | 0 | 0 | 377 | 228 | 780 | 3125 | 19141 | 0 | 351 | 0 | 0 | 0 | 613 | 10 | 167 | 1588 | 11 | 184 | 89084 | 8875 | 304 | 0 | 90 | 63 | 114 | 86 | 12 | 0 | 0 | 0 | 316 | 118 |
| SRR10077427 | 0 | 170 | 0 | 0 | 0 | 0 | 0 | 602 | 175 | 794 | 28825 | 19007 | 0 | 326 | 0 | 0 | 0 | 618 | 0 | 217 | 1523 | 7461 | 189 | 91564 | 8730 | 318 | 0 | 85 | 55 | 133 | 63 | 0 | 11 | 18 | 0 | 351 | 120 |

### Supp Table S8.4.Columns_DI_to_ES

| | | | | | | | | | | | | | | | | | | | | | | | | | | | | | | | | | | | | | |
|---|---|---|---|---|---|---|---|---|---|---|---|---|---|---|---|---|---|---|---|---|---|---|---|---|---|---|---|---|---|---|---|---|---|---|---|---|---|
| SRR10077428 | 0 | 184 | 28 | 0 | 13 | 0 | 0 | 939 | 476 | 3410 | 12955 | 8629 | 0 | 1972 | 0 | 0 | 0 | 640 | 0 | 835 | 1675 | 8528 | 236 | 118949 | 11552 | 236 | 14 | 42 | 99 | 198 | 212 | 42 | 50 | 78 | 0 | 1891 | 110 |
| SRR10077429 | 0 | 146 | 0 | 0 | 13 | 0 | 0 | 756 | 307 | 3320 | 1051 | 8223 | 0 | 1831 | 0 | 0 | 0 | 628 | 0 | 756 | 1607 | 8251 | 225 | 120543 | 11320 | 224 | 0 | 51 | 131 | 204 | 226 | 22 | 66 | 88 | 0 | 1914 | 102 |
| SRR10077430 | 0 | 266 | 0 | 0 | 11 | 0 | 0 | 750 | 375 | 3186 | 1002 | 7822 | 0 | 1924 | 0 | 0 | 0 | 630 | 0 | 612 | 1697 | 11 | 240 | 118434 | 11447 | 239 | 0 | 70 | 63 | 305 | 258 | 23 | 63 | 109 | 0 | 1924 | 111 |
| SRR10077431 | 0 | 240 | 36 | 0 | 0 | 0 | 0 | 788 | 365 | 3329 | 1095 | 8352 | 0 | 1706 | 0 | 0 | 0 | 669 | 0 | 788 | 1597 | 8406 | 246 | 121184 | 11237 | 210 | 0 | 58 | 36 | 10 | 247 | 15 | 44 | 80 | 0 | 1906 | 113 |
| SRR10077432 | 0 | 189 | 0 | 0 | 11 | 0 | 0 | 870 | 210 | 1148 | 2432 | 10381 | 0 | 376 | 0 | 0 | 0 | 648 | 0 | 272 | 1749 | 8500 | 237 | 111404 | 12392 | 194 | 0 | 88 | 63 | 200 | 179 | 21 | 25 | 21 | 0 | 481 | 87 |
| SRR10077433 | 0 | 218 | 0 | 0 | 14 | 0 | 0 | 614 | 269 | 1202 | 2405 | 10536 | 0 | 486 | 0 | 0 | 0 | 636 | 0 | 185 | 1606 | 8205 | 257 | 113195 | 12003 | 192 | 0 | 105 | 112 | 221 | 167 | 18 | 11 | 25 | 0 | 473 | 81 |
| SRR10077434 | 0 | 250 | 0 | 0 | 10 | 0 | 0 | 586 | 202 | 1276 | 15816 | 10732 | 0 | 530 | 0 | 0 | 0 | 653 | 14 | 195 | 10 | 13 | 250 | 110900 | 12296 | 197 | 0 | 108 | 54 | 247 | 181 | 0 | 0 | 23 | 0 | 481 | 80 |
| SRR10077435 | 0 | 212 | 0 | 0 | 12 | 0 | 0 | 670 | 322 | 1175 | 15355 | 10457 | 0 | 584 | 0 | 0 | 0 | 642 | 11 | 197 | 1664 | 8240 | 242 | 113952 | 12131 | 192 | 0 | 73 | 91 | 164 | 168 | 0 | 15 | 0 | 0 | 499 | 82 |
| SRR10116271 | 13 | 87 | 0 | 0 | 75 | 13 | 24 | 62 | 292 | 1237 | 40106 | 33430 | 0 | 329 | 12 | 0 | 18 | 1916 | 13 | 143 | 773 | 1867 | 110 | 2245 | 15 | 0 | 79 | 42 | 16 | 168 | 18 | 0 | 0 | 0 | 0 | 118 | 498 |
| SRR10116272 | 0 | 83 | 0 | 0 | 0 | 0 | 51 | 458 | 310 | 1164 | 5805 | 4870 | 0 | 301 | 0 | 0 | 60 | 1807 | 15 | 584 | 1883 | 2874 | 445 | 9863 | 8361 | 12 | 147 | 0 | 83 | 52 | 17 | 0 | 0 | 0 | 0 | 1474 | 98 |
| SRR10116273 | 12 | 23 | 13 | 0 | 28 | 0 | 29 | 101 | 140 | 289 | 52071 | 44081 | 0 | 188 | 0 | 13 | 53 | 3542 | 52 | 355 | 1787 | 4331 | 23 | 1389 | 25 | 14 | 68 | 32 | 185 | 32 | 56 | 0 | 0 | 0 | 0 | 534 | 1720 |
| SRR10116274 | 0 | 85 | 0 | 0 | 23 | 0 | 23 | 149 | 129 | 777 | 9859 | 7041 | 0 | 528 | 0 | 0 | 51 | 1696 | 21 | 269 | 2281 | 2010 | 342 | 5576 | 20 | 0 | 79 | 11 | 215 | 113 | 43 | 0 | 0 | 0 | 0 | 538 | 382 |
| SRR10116275 | 24 | 53 | 0 | 0 | 12 | 18 | 64 | 1528 | 12455 | 1928 | 27569 | 16466 | 0 | 339 | 29 | 49 | 79 | 1057 | 32 | 180 | 3402 | 3475 | 29 | 4781 | 3813 | 0 | 64 | 52 | 181 | 91 | 19 | 0 | 0 | 15 | 0 | 433 | 1004 |
| SRR10119011 | 0 | 12 | 0 | 0 | 0 | 0 | 0 | 230 | 124 | 2874 | 4419 | 2737 | 0 | 823 | 19 | 38 | 0 | 14 | 0 | 4808 | 134 | 512 | 0 | 534 | 76 | 31 | 14 | 0 | 0 | 0 | 0 | 0 | 0 | 20 | 0 | 22348 | 0 |
| SRR10119012 | 43 | 0 | 0 | 0 | 0 | 10 | 0 | 0 | 0 | 2348 | 0 | 0 | 0 | 0 | 0 | 0 | 24 | 0 | 0 | 3718 | 0 | 45 | 0 | 109 | 945 | 89 | 20 | 0 | 0 | 0 | 0 | 0 | 0 | 0 | 0 | 23875 | 0 |
| SRR10119014 | 0 | 0 | 0 | 0 | 0 | 0 | 0 | 4729 | 0 | 0 | 0 | 0 | 0 | 0 | 0 | 0 | 32 | 0 | 0 | 0 | 0 | 0 | 0 | 0 | 129 | 0 | 323 | 0 | 0 | 0 | 0 | 0 | 0 | 0 | 0 | 5255 | 0 |
| SRR10119022 | 0 | 12 | 0 | 0 | 0 | 0 | 0 | 86 | 81 | 1432 | 5348 | 1811 | 0 | 518 | 16 | 0 | 0 | 40 | 0 | 642 | 61 | 525 | 15 | 588 | 106 | 14 | 0 | 0 | 0 | 0 | 0 | 0 | 0 | 0 | 14 | 3434 | 0 |
| SRR10119033 | 61 | 0 | 0 | 0 | 0 | 0 | 0 | 0 | 617 | 1367 | 0 | 0 | 0 | 485 | 485 | 926 | 0 | 0 | 0 | 4189 | 30 | 114 | 13 | 175 | 776 | 86 | 511 | 0 | 0 | 0 | 0 | 0 | 0 | 0 | 0 | 17108 | 0 |
| SRR10119034 | 0 | 0 | 0 | 0 | 0 | 354 | 0 | 0 | 0 | 0 | 0 | 0 | 0 | 0 | 0 | 708 | 0 | 0 | 0 | 1719 | 0 | 18 | 0 | 51 | 429 | 56 | 399 | 0 | 0 | 0 | 0 | 0 | 0 | 0 | 0 | 22150 | 0 |
| SRR10119035 | 0 | 18 | 0 | 0 | 0 | 0 | 0 | 32 | 39 | 1755 | 6715 | 3008 | 0 | 2057 | 0 | 101 | 0 | 14 | 0 | 875 | 101 | 835 | 16 | 498 | 47 | 13 | 0 | 0 | 0 | 0 | 0 | 0 | 0 | 15 | 12 | 4166 | 0 |
| SRR10119037 | 56 | 0 | 0 | 0 | 0 | 796 | 0 | 3165 | 0 | 1934 | 0 | 0 | 0 | 0 | 0 | 0 | 0 | 0 | 0 | 0 | 13 | 0 | 13 | 195 | 526 | 37 | 903 | 0 | 0 | 0 | 0 | 0 | 0 | 0 | 0 | 12661 | 0 |
| SRR10119040 | 0 | 0 | 0 | 0 | 0 | 237 | 0 | 0 | 0 | 804 | 35 | 0 | 0 | 1433 | 874 | 245 | 0 | 0 | 0 | 1293 | 0 | 54 | 13 | 173 | 250 | 72 | 406 | 0 | 0 | 0 | 0 | 0 | 0 | 0 | 0 | 54532 | 0 |
| SRR10119044 | 0 | 0 | 0 | 0 | 0 | 41 | 0 | 2099 | 2624 | 1662 | 0 | 0 | 0 | 8922 | 350 | 0 | 54 | 0 | 0 | 3062 | 0 | 27 | 17 | 92 | 347 | 19 | 254 | 0 | 0 | 0 | 0 | 0 | 0 | 0 | 0 | 13209 | 0 |
| SRR10119046 | 0 | 0 | 0 | 0 | 0 | 0 | 0 | 46 | 38 | 1836 | 6131 | 2896 | 0 | 1806 | 30 | 96 | 0 | 24 | 0 | 923 | 47 | 727 | 13 | 443 | 67 | 34 | 0 | 0 | 0 | 0 | 0 | 0 | 0 | 15 | 13 | 4851 | 0 |
| SRR10119048 | 0 | 0 | 0 | 0 | 0 | 0 | 0 | 2263 | 0 | 4942 | 19 | 1132 | 0 | 1320 | 0 | 339 | 0 | 0 | 0 | 2942 | 26 | 114 | 0 | 184 | 753 | 195 | 212 | 0 | 0 | 0 | 0 | 0 | 0 | 0 | 0 | 12863 | 0 |
| SRR10119051 | 0 | 0 | 0 | 0 | 0 | 11 | 0 | 2770 | 0 | 0 | 0 | 0 | 0 | 0 | 0 | 0 | 0 | 0 | 0 | 3526 | 0 | 32 | 11 | 133 | 591 | 100 | 13 | 0 | 0 | 0 | 0 | 0 | 0 | 0 | 0 | 18887 | 0 |
| SRR10119052 | 69 | 0 | 0 | 0 | 0 | 471 | 0 | 0 | 0 | 2879 | 0 | 0 | 0 | 2629 | 501 | 125 | 0 | 0 | 0 | 2003 | 11 | 63 | 0 | 69 | 520 | 58 | 644 | 0 | 0 | 0 | 0 | 0 | 0 | 0 | 0 | 7260 | 0 |
| SRR10119056 | 209 | 0 | 0 | 0 | 0 | 225 | 0 | 2384 | 0 | 0 | 62 | 0 | 0 | 3774 | 0 | 0 | 0 | 0 | 0 | 4211 | 0 | 37 | 0 | 26 | 323 | 56 | 13 | 0 | 0 | 0 | 0 | 0 | 0 | 0 | 0 | 15176 | 0 |
| SRR10119057 | 0 | 11 | 0 | 0 | 0 | 0 | 0 | 49 | 0 | 485 | 7702 | 5433 | 0 | 127 | 0 | 86 | 0 | 32 | 0 | 11685 | 114 | 1282 | 19 | 515 | 67 | 17 | 0 | 0 | 0 | 0 | 0 | 0 | 0 | 0 | 10 | 65455 | 0 |
| SRR10119058 | 0 | 0 | 0 | 0 | 0 | 0 | 0 | 28 | 21 | 282 | 7731 | 6111 | 0 | 176 | 14 | 99 | 0 | 29 | 0 | 11371 | 128 | 1516 | 24 | 707 | 38 | 24 | 0 | 0 | 0 | 0 | 0 | 0 | 0 | 0 | 11 | 54503 | 0 |
| SRR10119059 | 0 | 10 | 0 | 0 | 0 | 0 | 11 | 44 | 0 | 1470 | 588 | 616 | 0 | 542 | 0 | 0 | 0 | 29 | 0 | 995 | 147 | 278 | 19 | 464 | 50 | 55 | 0 | 0 | 0 | 0 | 0 | 0 | 0 | 0 | 19 | 3527 | 0 |
| SRR10119063 | 0 | 0 | 0 | 0 | 0 | 0 | 0 | 706 | 0 | 1883 | 0 | 0 | 0 | 0 | 0 | 0 | 205 | 0 | 0 | 0 | 68 | 26 | 0 | 0 | 289 | 0 | 124 | 0 | 0 | 0 | 0 | 0 | 0 | 0 | 0 | 0 | 0 |
| SRR10119066 | 0 | 14 | 0 | 0 | 0 | 0 | 0 | 71 | 90 | 798 | 3240 | 2495 | 0 | 798 | 45 | 84 | 0 | 22 | 0 | 6003 | 159 | 1445 | 43 | 1640 | 94 | 21 | 0 | 0 | 0 | 0 | 0 | 0 | 0 | 11 | 17 | 25583 | 0 |
| SRR10119067 | 0 | 13 | 0 | 0 | 0 | 0 | 0 | 34 | 42 | 1013 | 3689 | 2414 | 0 | 646 | 50 | 98 | 0 | 28 | 0 | 7415 | 133 | 1219 | 27 | 1603 | 71 | 0 | 0 | 0 | 0 | 0 | 0 | 0 | 0 | 18 | 17 | 39394 | 0 |
| SRR10119068 | 0 | 13 | 0 | 0 | 0 | 0 | 0 | 41 | 35 | 787 | 2609 | 2451 | 0 | 2129 | 77 | 362 | 0 | 19 | 0 | 1730 | 72 | 577 | 0 | 658 | 71 | 16 | 0 | 0 | 0 | 0 | 0 | 0 | 0 | 11 | 11 | 9622 | 0 |
| SRR10119069 | 0 | 10 | 0 | 0 | 0 | 0 | 0 | 70 | 76 | 894 | 1634 | 1521 | 0 | 2044 | 106 | 513 | 0 | 39 | 0 | 1580 | 97 | 663 | 0 | 562 | 61 | 16 | 0 | 0 | 0 | 0 | 0 | 0 | 0 | 0 | 14 | 8067 | 0 |
| SRR10119070 | 0 | 13 | 0 | 0 | 0 | 0 | 0 | 0 | 76 | 2268 | 1287 | 1307 | 0 | 565 | 0 | 76 | 0 | 12 | 0 | 6175 | 25 | 368 | 19 | 628 | 79 | 0 | 0 | 0 | 0 | 0 | 0 | 0 | 0 | 0 | 0 | 35920 | 0 |
| SRR10119071 | 0 | 0 | 0 | 0 | 0 | 0 | 0 | 35 | 0 | 161 | 4789 | 3583 | 0 | 352 | 0 | 20 | 0 | 25 | 0 | 11793 | 47 | 561 | 19 | 932 | 89 | 25 | 0 | 0 | 0 | 0 | 0 | 0 | 0 | 0 | 0 | 60028 | 0 |
| SRR10119072 | 0 | 0 | 0 | 0 | 0 | 0 | 0 | 19 | 15 | 623 | 4103 | 1987 | 0 | 323 | 19 | 27 | 0 | 63 | 0 | 12768 | 160 | 465 | 30 | 903 | 69 | 83 | 0 | 0 | 0 | 0 | 0 | 0 | 0 | 0 | 0 | 72360 | 0 |
| SRR10119073 | 0 | 15 | 0 | 0 | 0 | 0 | 0 | 0 | 0 | 153 | 3062 | 3175 | 0 | 64 | 0 | 0 | 0 | 54 | 0 | 12185 | 142 | 808 | 11 | 855 | 140 | 17 | 0 | 0 | 0 | 0 | 0 | 0 | 0 | 0 | 0 | 57354 | 0 |
| SRR10119074 | 0 | 0 | 0 | 0 | 0 | 0 | 0 | 0 | 0 | 129 | 2501 | 2408 | 0 | 106 | 0 | 0 | 0 | 49 | 0 | 10049 | 105 | 429 | 0 | 456 | 134 | 60 | 0 | 0 | 0 | 0 | 0 | 0 | 0 | 0 | 0 | 59676 | 0 |
| SRR10119075 | 0 | 0 | 0 | 0 | 0 | 0 | 0 | 102 | 0 | 1790 | 13494 | 5057 | 0 | 1174 | 68 | 1192 | 0 | 16 | 0 | 2769 | 72 | 824 | 0 | 392 | 39 | 28 | 0 | 0 | 0 | 0 | 0 | 0 | 0 | 0 | 16 | 9037 | 0 |
| SRR10119076 | 0 | 0 | 0 | 0 | 0 | 0 | 0 | 95 | 48 | 3048 | 2112 | 3676 | 0 | 1065 | 64 | 878 | 0 | 15 | 0 | 2635 | 101 | 688 | 15 | 416 | 87 | 14 | 0 | 0 | 0 | 0 | 0 | 0 | 0 | 0 | 0 | 9657 | 0 |
| SRR10119077 | 0 | 14 | 0 | 0 | 0 | 0 | 0 | 83 | 28 | 710 | 3844 | 6460 | 0 | 1120 | 0 | 57 | 0 | 26 | 0 | 4447 | 110 | 1073 | 15 | 591 | 35 | 14 | 0 | 0 | 0 | 0 | 0 | 0 | 0 | 0 | 15 | 23428 | 0 |
| SRR10119078 | 0 | 22 | 0 | 0 | 0 | 0 | 0 | 41 | 27 | 793 | 6021 | 6135 | 0 | 590 | 0 | 63 | 0 | 18 | 0 | 6346 | 71 | 685 | 0 | 715 | 32 | 0 | 0 | 0 | 0 | 0 | 0 | 0 | 0 | 0 | 14 | 36240 | 0 |
| SRR10119079 | 0 | 0 | 0 | 0 | 0 | 0 | 0 | 102 | 45 | 2427 | 2764 | 929 | 0 | 1472 | 41 | 374 | 0 | 0 | 0 | 1202 | 80 | 407 | 0 | 272 | 26 | 25 | 0 | 0 | 0 | 0 | 0 | 0 | 11 | 47 | 19 | 4867 | 0 |
| SRR10119080 | 0 | 13 | 0 | 0 | 0 | 0 | 0 | 107 | 125 | 2648 | 3034 | 1380 | 0 | 1480 | 35 | 282 | 0 | 12 | 0 | 1567 | 68 | 297 | 19 | 254 | 34 | 46 | 0 | 0 | 0 | 0 | 0 | 0 | 0 | 18 | 11 | 6458 | 0 |
| SRR10119081 | 0 | 0 | 0 | 0 | 0 | 0 | 0 | 0 | 0 | 119 | 6470 | 2734 | 0 | 79 | 0 | 17 | 0 | 34 | 0 | 10741 | 33 | 367 | 22 | 679 | 71 | 14 | 0 | 0 | 0 | 0 | 0 | 0 | 0 | 0 | 0 | 52583 | 0 |
| SRR10119082 | 0 | 0 | 0 | 0 | 0 | 0 | 0 | 24 | 13 | 204 | 2198 | 2705 | 0 | 75 | 0 | 0 | 0 | 22 | 0 | 11770 | 31 | 312 | 0 | 542 | 76 | 17 | 0 | 0 | 0 | 0 | 0 | 0 | 0 | 0 | 0 | 57015 | 0 |
| SRR1014666 | 0 | 0 | 0 | 0 | 0 | 0 | 11 | 33 | 0 | 22 | 125 | 839 | 0 | 0 | 0 | 0 | 17 | 0 | 14 | 0 | 300 | 83 | 0 | 0 | 988 | 242 | 11 | 123 | 514 | 15 | 31 | 0 | 0 | 0 | 0 | 22 | 79 |
| SRR1014667 | 0 | 0 | 0 | 0 | 16 | 0 | 38 | 1472 | 129 | 1020 | 11 | 12 | 0 | 0 | 0 | 0 | 11 | 22 | 17 | 0 | 579 | 125 | 0 | 0 | 917 | 194 | 0 | 0 | 0 | 17 | 18 | 0 | 0 | 0 | 0 | 129 | 97 |
| SRR1014669 | 0 | 11 | 0 | 0 | 11 | 0 | 53 | 75 | 11 | 214 | 17 | 203 | 0 | 0 | 0 | 0 | 51 | 12 | 0 | 0 | 411 | 121 | 0 | 0 | 824 | 148 | 11 | 0 | 11 | 17 | 0 | 0 | 0 | 0 | 0 | 0 | 90 |
| SRR10197184 | 0 | 339 | 0 | 31 | 1659 | 14 | 16 | 89 | 211 | 3742 | 11164 | 15812 | 0 | 1524 | 0 | 0 | 11 | 780 | 188 | 134 | 142 | 584 | 171 | 13500 | 240 | 103 | 57 | 40 | 28 | 38 | 56 | 0 | 0 | 0 | 0 | 529 | 0 |
| SRR10197185 | 0 | 0 | 0 | 0 | 0 | 20 | 0 | 0 | 74 | 2738 | 1184 | 2664 | 28 | 3737 | 0 | 0 | 0 | 99 | 0 | 777 | 91 | 25 | 50 | 4633 | 33 | 66 | 1099 | 0 | 0 | 16 | 25 | 47 | 14 | 12 | 0 | 2146 | 0 |
| SRR10197186 | 0 | 180 | 0 | 0 | 141 | 0 | 0 | 305 | 265 | 2641 | 842 | 4600 | 0 | 2138 | 0 | 0 | 20 | 260 | 239 | 788 | 101 | 187 | 66 | 10964 | 79 | 97 | 35 | 21 | 12 | 13 | 24 | 0 | 0 | 0 | 0 | 1702 | 0 |
| SRR10197187 | 0 | 258 | 0 | 12 | 395 | 1352 | 62 | 977 | 333 | 1792 | 3723 | 5101 | 25 | 1687 | 0 | 0 | 249 | 316 | 290 | 154 | 122 | 381 | 82 | 13232 | 118 | 111 | 77 | 31 | 16 | 0 | 21 | 0 | 0 | 0 | 0 | 375 | 0 |
| SRR10197188 | 21 | 360 | 0 | 27 | 1043 | 1330 | 64 | 838 | 375 | 1576 | 3486 | 9111 | 32 | 1365 | 0 | 0 | 223 | 568 | 273 | 70 | 142 | 501 | 149 | 14281 | 159 | 87 | 70 | 25 | 15 | 12 | 18 | 0 | 0 | 0 | 0 | 278 | 0 |
| SRR10197189 | 0 | 157 | 0 | 0 | 123 | 1236 | 46 | 378 | 154 | 1438 | 2410 | 2863 | 13 | 1164 | 0 | 0 | 147 | 184 | 215 | 415 | 101 | 196 | 65 | 9956 | 61 | 52 | 168 | 22 | 12 | 0 | 33 | 0 | 0 | 0 | 0 | 827 | 0 |
| SRR10197190 | 16 | 31 | 0 | 29 | 1000 | 454 | 29 | 824 | 466 | 1293 | 4345 | 5830 | 0 | 812 | 0 | 0 | 68 | 541 | 185 | 135 | 151 | 357 | 119 | 11728 | 150 | 45 | 15 | 0 | 0 | 0 | 0 | 0 | 0 | 0 | 0 | 277 | 0 |
| SRR10197191 | 14 | 72 | 0 | 28 | 933 | 654 | 40 | 853 | 351 | 1888 | 5579 | 6563 | 13 | 1279 | 0 | 0 | 104 | 642 | 223 | 169 | 223 | 387 | 180 | 12706 | 129 | 71 | 23 | 11 | 0 | 0 | 0 | 0 | 0 | 0 | 0 | 267 | 0 |
| SRR10197192 | 13 | 21 | 0 | 19 | 988 | 626 | 34 | 697 | 315 | 1825 | 3883 | 5127 | 0 | 1095 | 0 | 0 | 77 | 639 | 214 | 527 | 177 | 376 | 146 | 12298 | 126 | 61 | 22 | 0 | 0 | 0 | 0 | 0 | 0 | 0 | 0 | 1713 | 0 |
| SRR10225092 | 0 | 788 | 88 | 0 | 0 | 0 | 0 | 333 | 0 | 2995 | 0 | 0 | 0 | 166 | 0 | 0 | 0 | 285 | 219 | 11980 | 613 | 876 | 0 | 876 | 1379 | 285 | 0 | 0 | 0 | 0 | 175 | 0 | 88 | 350 | 1664 | 28453 | 0 |
| SRR10225093 | 0 | 0 | 0 | 0 | 56 | 0 | 0 | 1600 | 559 | 27094 | 0 | 0 | 0 | 1148 | 0 | 0 | 0 | 155 | 0 | 36 | 724 | 1012 | 112 | 10243 | 2571 | 14 | 0 | 0 | 0 | 0 | 0 | 0 | 0 | 0 | 0 | 393 | 0 |
| SRR10225094 | 0 | 20 | 18 | 0 | 0 | 0 | 0 | 795 | 76 | 3482 | 0 | 0 | 0 | 265 | 38 | 38 | 0 | 244 | 0 | 5261 | 172 | 163 | 18 | 4774 | 235 | 0 | 0 | 0 | 11 | 33 | 22 | 0 | 13 | 40 | 76 | 46893 | 0 |
| SRR10225096 | 0 | 42 | 23 | 0 | 0 | 0 | 0 | 18969 | 923 | 2954 | 0 | 0 | 0 | 11031 | 415 | 554 | 0 | 478 | 0 | 969 | 281 | 84 | 0 | 1914 | 84 | 141 | 0 | 0 | 0 | 0 | 0 | 0 | 29 | 19 | 35 | 7246 | 28 |
| SRR10282881 | 0 | 0 | 0 | 0 | 0 | 0 | 0 | 233 | 89 | 19919 | 38 | 37 | 0 | 12279 | 0 | 16 | 16 | 59 | 17 | 30 | 652 | 1536 | 0 | 1884 | 10899 | 278 | 0 | 0 | 0 | 0 | 0 | 0 | 0 | 0 | 0 | 391 | 0 |
| SRR10282882 | 0 | 0 | 0 | 0 | 0 | 0 | 0 | 234 | 105 | 21650 | 26 | 25 | 0 | 12671 | 14 | 17 | 10 | 81 | 0 | 37 | 653 | 1597 | 0 | 2014 | 10072 | 258 | 0 | 0 | 0 | 0 | 0 | 0 | 0 | 0 | 0 | 339 | 0 |
| SRR10282883 | 0 | 0 | 0 | 0 | 0 | 0 | 0 | 108 | 46 | 14918 | 48 | 71 | 0 | 5786 | 58 | 23 | 0 | 64 | 18 | 40 | 999 | 2215 | 0 | 2592 | 13931 | 529 | 0 | 0 | 0 | 0 | 0 | 0 | 0 | 0 | 0 | 510 | 0 |
| SRR10282884 | 0 | 0 | 0 | 0 | 11 | 0 | 0 | 125 | 64 | 15933 | 47 | 50 | 0 | 5999 | 79 | 32 | 0 | 68 | 15 | 35 | 985 | 2288 | 0 | 2556 | 13230 | 510 | 0 | 0 | 0 | 0 | 0 | 0 | 0 | 0 | 0 | 468 | 0 |
| SRR10282885 | 0 | 0 | 0 | 0 | 0 | 0 | 14 | 196 | 61 | 10626 | 48 | 28 | 0 | 3977 | 0 | 0 | 22 | 71 | 11 | 20 | 682 | 1362 | 0 | 2358 | 10006 | 310 | 18 | 0 | 0 | 0 | 0 | 0 | 0 | 0 | 0 | 220 | 0 |
| SRR10282886 | 11 | 13 | 0 | 0 | 13 | 0 | 15 | 202 | 66 | 11106 | 23 | 24 | 0 | 3859 | 0 | 0 | 38 | 60 | 17 | 17 | 688 | 1463 | 0 | 2291 | 9293 | 269 | 21 | 0 | 0 | 0 | 0 | 0 | 0 | 0 | 0 | 168 | 0 |
| SRR10282890 | 0 | 0 | 0 | 0 | 17 | 0 | 0 | 275 | 62 | 6418 | 198 | 229 | 0 | 2799 | 0 | 14 | 24 | 77 | 20 | 56 | 729 | 1280 | 0 | 2426 | 12726 | 312 | 0 | 0 | 0 | 0 | 0 | 0 | 0 | 0 | 0 | 350 | 0 |
| SRR10282891 | 0 | 0 | 0 | 0 | 27 | 0 | 0 | 287 | 76 | 6948 | 124 | 186 | 0 | 2744 | 0 | 13 | 26 | 87 | 19 | 47 | 705 | 1418 | 0 | 2503 | 11943 | 287 | 0 | 0 | 0 | 0 | 0 | 0 | 0 | 0 | 0 | 291 | 0 |
| SRR10282892 | 0 | 0 | 0 | 0 | 0 | 0 | 0 | 209 | 26 | 3812 | 83 | 71 | 0 | 685 | 0 | 0 | 0 | 42 | 24 | 21 | 409 | 1023 | 0 | 1737 | 9832 | 425 | 0 | 0 | 0 | 0 | 0 | 0 | 0 | 0 | 0 | 82 | 0 |
| SRR10282893 | 0 | 0 | 0 | 0 | 0 | 0 | 0 | 226 | 31 | 4151 | 45 | 51 | 0 | 705 | 0 | 0 | 0 | 41 | 23 | 19 | 431 | 1101 | 0 | 1740 | 9248 | 399 | 0 | 0 | 0 | 0 | 0 | 0 | 0 | 0 | 0 | 73 | 0 |
| SRR10282894 | 0 | 0 | 0 | 0 | 0 | 0 | 0 | 238 | 16 | 972 | 157 | 68 | 0 | 888 | 0 | 0 | 0 | 81 | 24 | 26 | 117 | 355 | 0 | 693 | 2480 | 127 | 0 | 0 | 0 | 0 | 0 | 0 | 0 | 0 | 0 | 48 | 0 |
| SRR10282895 | 0 | 0 | 0 | 0 | 0 | 0 | 0 | 236 | 24 | 1081 | 44 | 50 | 0 | 879 | 0 | 0 | 0 | 86 | 23 | 21 | 100 | 352 | 0 | 699 | 2272 | 114 | 0 | 0 | 0 | 0 | 0 | 0 | 0 | 0 | 0 | 31 | 0 |
| SRR10323931 | 0 | 135 | 31 | 13 | 614 | 0 | 0 | 2657 | 6158 | 15834 | 23273 | 32928 | 0 | 159 | 0 | 0 | 29 | 628 | 289 | 69 | 983 | 2498 | 242 | 8917 | 1355 | 0 | 171 | 12 | 21 | 23 | 78 | 15 | 148 | 66 | 0 | 384 | 2202 |
| SRR10323932 | 0 | 1931 | 21 | 21 | 637 | 10 | 33 | 2144 | 7418 | 49116 | 8207 | 12334 | 0 | 856 | 0 | 0 | 43 | 460 | 83 | 75 | 992 | 1388 | 218 | 5002 | 924 | 0 | 215 | 370 | 475 | 561 | 2041 | 388 | 156 | 58 | 0 | 401 | 1787 |
| SRR10323933 | 0 | 143 | 23 | 13 | 316 | 0 | 0 | 7192 | 26257 | 64846 | 1355 | 1695 | 0 | 842 | 0 | 0 | 16 | 299 | 61 | 100 | 717 | 1758 | 170 | 5200 | 1486 | 0 | 39 | 96 | 23 | 28 | 115 | 35 | 27 | 16 | 0 | 460 | 2468 |
| SRR10323939 | 0 | 4505 | 0 | 17 | 644 | 0 | 19 | 477 | 918 | 8833 | 27629 | 35187 | 0 | 467 | 0 | 0 | 22 | 848 | 97 | 95 | 1917 | 3609 | 396 | 13626 | 972 | 0 | 277 | 248 | 425 | 508 | 973 | 114 | 47 | 18 | 0 | 306 | 911 |
| SRR10323940 | 11 | 2666 | 94 | 32 | 1338 | 11 | 0 | 3000 | 8850 | 29730 | 14899 | 21168 | 0 | 1869 | 0 | 0 | 58 | 1088 | 116 | 325 | 2238 | 2943 | 392 | 11140 | 1601 | 0 | 86 | 166 | 376 | 697 | 1814 | 200 | 94 | 64 | 0 | 1244 | 1951 |
| SRR10323941 | 0 | 1721 | 56 | 27 | 970 | 0 | 10 | 1879 | 4669 | 21770 | 1662 | 14497 | 0 | 1580 | 0 | 0 | 41 | 915 | 75 | 135 | 1880 | 2145 | 169 | 6121 | 1126 | 0 | 105 | 124 | 666 | 564 | 1124 | 111 | 139 | 84 | 0 | 463 | 1246 |
| SRR10385125 | 0 | 54 | 66 | 118 | 976 | 0 | 0 | 50 | 216 | 726 | 67613 | 70135 | 0 | 201 | 0 | 0 | 25 | 706 | 905 | 72 | 22 | 1040 | 0 | 25 | 2962 | 0 | 27 | 25 | 75 | 64 | 61 | 15 | 19 | 13 | 0 | 201 | 0 |
| SRR10385126 | 0 | 993 | 1274 | 109 | 680 | 0 | 0 | 103 | 189 | 863 | 57144 | 66360 | 0 | 189 | 0 | 0 | 23 | 603 | 928 | 342 | 13 | 1062 | 0 | 18 | 3798 | 0 | 14 | 299 | 818 | 1142 | 1049 | 326 | 406 | 339 | 18 | 641 | 0 |
| SRR10416439 | 0 | 126 | 727 | 809 | 116 | 0 | 0 | 246 | 159 | 332 | 83155 | 97452 | 0 | 113 | 93 | 126 | 0 | 367 | 81 | 379 | 275 | 809 | 0 | 0 | 3167 | 0 | 18 | 18 | 277 | 1270 | 1151 | 918 | 21 | 26 | 0 | 651 | 0 |
| SRR10416440 | 0 | 23 | 78 | 1054 | 377 | 0 | 0 | 156 | 102 | 258 | 10715 | 98809 | 0 | 39 | 0 | 39 | 0 | 1498 | 71 | 102 | 85 | 656 | 0 | 0 | 2796 | 0 | 28 | 0 | 49 | 172 | 128 | 91 | 0 | 27 | 0 | 328 | 0 |
| SRR10416441 | 0 | 37 | 140 | 1100 | 340 | 0 | 0 | 395 | 137 | 422 | 22898 | 113037 | 0 | 65 | 27 | 34 | 0 | 1541 | 43 | 110 | 181 | 820 | 0 | 0 | 2978 | 0 | 22 | 17 | 99 | 257 | 178 | 133 | 15 | 27 | 0 | 300 | 0 |
| SRR10416442 | 0 | 681 | 1185 | 904 | 271 | 0 | 0 | 461 | 434 | 310 | 13158 | 142374 | 0 | 55 | 48 | 28 | 10 | 683 | 41 | 324 | 1044 | 1275 | 0 | 11 | 3544 | 0 | 19 | 36 | 686 | 1484 | 1476 | 606 | 91 | 127 | 0 | 399 | 0 |
| SRR10416496 | 0 | 0 | 10 | 0 | 0 | 10 | 0 | 0 | 0 | 0 | 1587 | 3704 | 0 | 0 | 0 | 0 | 0 | 0 | 0 | 265 | 384 | 1425 | 0 | 0 | 348 | 10 | 0 | 0 | 0 | 0 | 0 | 0 | 0 | 0 | 0 | 529 | 0 |
| SRR10416497 | 0 | 0 | 0 | 0 | 0 | 0 | 0 | 0 | 0 | 0 | 1076 | 4661 | 0 | 0 | 0 | 0 | 0 | 0 | 0 | 0 | 422 | 1797 | 0 | 0 | 145 | 12 | 90 | 0 | 0 | 0 | 0 | 0 | 0 | 0 | 0 | 0 | 0 |
| SRR10416498 | 0 | 0 | 0 | 0 | 0 | 0 | 0 | 0 | 0 | 0 | 1807 | 5777 | 0 | 0 | 1005 | 1256 | 0 | 0 | 0 | 251 | 1058 | 4196 | 0 | 0 | 406 | 0 | 0 | 0 | 0 | 0 | 0 | 0 | 0 | 0 | 0 | 754 | 0 |
| SRR10416499 | 0 | 0 | 0 | 0 | 0 | 0 | 0 | 0 | 0 | 396 | 3566 | 4358 | 0 | 0 | 396 | 0 | 0 | 0 | 0 | 0 | 471 | 1718 | 0 | 0 | 427 | 0 | 45 | 0 | 0 | 0 | 0 | 0 | 0 | 0 | 0 | 396 | 0 |
| SRR10416500 | 15 | 0 | 15 | 0 | 0 | 0 | 0 | 575 | 0 | 287 | 11 | 6033 | 0 | 0 | 287 | 0 | 0 | 0 | 0 | 0 | 305 | 816 | 0 | 0 | 587 | 21 | 15 | 0 | 0 | 0 | 15 | 0 | 0 | 0 | 0 | 1436 | 0 |
| SRR10416501 | 0 | 0 | 0 | 0 | 0 | 0 | 0 | 313 | 0 | 313 | 1563 | 3126 | 0 | 0 | 313 | 0 | 0 | 0 | 0 | 0 | 577 | 1548 | 0 | 0 | 697 | 17 | 15 | 0 | 0 | 0 | 0 | 0 | 0 | 0 | 0 | 0 | 0 |
| SRR10416502 | 0 | 20 | 0 | 0 | 0 | 0 | 0 | 460 | 460 | 460 | 2761 | 10585 | 0 | 0 | 0 | 460 | 0 | 0 | 0 | 0 | 523 | 790 | 0 | 13 | 783 | 0 | 119 | 0 | 0 | 0 | 0 | 0 | 0 | 0 | 0 | 2301 | 0 |
| SRR10416503 | 0 | 22 | 22 | 0 | 0 | 0 | 0 | 0 | 0 | 416 | 864 | 3742 | 0 | 0 | 0 | 416 | 0 | 0 | 0 | 416 | 497 | 1281 | 0 | 21 | 767 | 26 | 109 | 0 | 0 | 0 | 0 | 0 | 0 | 0 | 0 | 832 | 0 |
| SRR10416504 | 0 | 0 | 21 | 0 | 0 | 0 | 0 | 0 | 0 | 0 | 3442 | 2295 | 0 | 0 | 1147 | 0 | 0 | 0 | 0 | 0 | 259 | 755 | 0 | 0 | 556 | 21 | 0 | 0 | 0 | 0 | 0 | 0 | 0 | 0 | 0 | 0 | 0 |
| SRR10416505 | 0 | 0 | 0 | 0 | 0 | 0 | 0 | 315 | 0 | 157 | 2520 | 12284 | 0 | 0 | 0 | 157 | 0 | 0 | 0 | 0 | 670 | 2401 | 0 | 0 | 556 | 0 | 10 | 0 | 0 | 0 | 0 | 0 | 0 | 0 | 0 | 472 | 0 |
| SRR10416506 | 0 | 0 | 0 | 0 | 0 | 0 | 18 | 386 | 0 | 772 | 3860 | 5017 | 0 | 0 | 0 | 386 | 0 | 0 | 0 | 386 | 550 | 1354 | 0 | 0 | 1021 | 14 | 53 | 0 | 0 | 0 | 0 | 0 | 0 | 0 | 35 | 386 | 0 |
| SRR10416507 | 0 | 0 | 30 | 0 | 0 | 0 | 0 | 0 | 0 | 0 | 461 | 3687 | 0 | 0 | 461 | 461 | 0 | 0 | 0 | 0 | 339 | 991 | 0 | 0 | 355 | 16 | 0 | 0 | 0 | 0 | 0 | 0 | 0 | 0 | 0 | 461 | 0 |
| SRR10416508 | 0 | 0 | 0 | 0 | 0 | 23 | 0 | 0 | 0 | 0 | 4545 | 5303 | 0 | 0 | 0 | 758 | 0 | 0 | 0 | 0 | 751 | 2043 | 0 | 0 | 545 | 32 | 45 | 0 | 0 | 23 | 0 | 0 | 0 | 0 | 0 | 379 | 0 |
| SRR10416509 | 0 | 0 | 0 | 0 | 0 | 0 | 0 | 0 | 0 | 0 | 4313 | 15527 | 0 | 0 | 647 | 216 | 0 | 0 | 0 | 216 | 544 | 2310 | 0 | 0 | 892 | 13 | 0 | 0 | 0 | 0 | 0 | 0 | 0 | 0 | 0 | 1294 | 0 |
| SRR10416510 | 0 | 0 | 63 | 0 | 0 | 0 | 0 | 1178 | 0 | 0 | 4819 | 7067 | 0 | 0 | 0 | 0 | 0 | 0 | 0 | 0 | 23 | 295 | 0 | 0 | 11 | 0 | 0 | 0 | 0 | 0 | 0 | 0 | 0 | 0 | 0 | 1178 | 0 |
| SRR10416513 | 0 | 0 | 0 | 0 | 0 | 0 | 0 | 133 | 0 | 133 | 674 | 3993 | 0 | 0 | 532 | 133 | 0 | 0 | 0 | 0 | 490 | 2161 | 0 | 0 | 623 | 13 | 0 | 0 | 0 | 0 | 0 | 0 | 0 | 0 | 0 | 399 | 0 |
| SRR10416514 | 0 | 0 | 0 | 0 | 0 | 0 | 0 | 0 | 0 | 1026 | 527 | 3079 | 0 | 0 | 0 | 2052 | 0 | 0 | 0 | 513 | 738 | 2263 | 0 | 25 | 641 | 27 | 47 | 0 | 0 | 0 | 0 | 0 | 0 | 23 | 0 | 2052 | 0 |
| SRR10416515 | 0 | 0 | 0 | 0 | 0 | 0 | 0 | 0 | 0 | 431 | 5603 | 3017 | 0 | 0 | 0 | 431 | 0 | 0 | 0 | 0 | 743 | 1190 | 0 | 0 | 769 | 16 | 23 | 0 | 0 | 0 | 0 | 0 | 0 | 0 | 23 | 862 | 0 |
| SRR10416516 | 0 | 0 | 0 | 0 | 0 | 0 | 0 | 0 | 0 | 0 | 1645 | 6803 | 0 | 0 | 400 | 0 | 0 | 0 | 0 | 400 | 770 | 2420 | 0 | 0 | 401 | 0 | 0 | 0 | 0 | 0 | 0 | 0 | 0 | 0 | 0 | 400 | 0 |
| SRR10416517 | 0 | 0 | 0 | 0 | 0 | 0 | 0 | 0 | 0 | 358 | 3224 | 6447 | 0 | 0 | 0 | 0 | 0 | 0 | 0 | 358 | 817 | 2368 | 0 | 0 | 576 | 13 | 0 | 0 | 0 | 0 | 0 | 0 | 0 | 0 | 0 | 358 | 0 |
| SRR10416518 | 0 | 0 | 55 | 0 | 0 | 0 | 0 | 0 | 0 | 968 | 1936 | 6131 | 0 | 0 | 323 | 323 | 0 | 0 | 0 | 323 | 1427 | 4153 | 0 | 0 | 330 | 10 | 139 | 0 | 0 | 0 | 0 | 0 | 0 | 0 | 28 | 0 | 0 |
| SRR10416519 | 16 | 48 | 16 | 0 | 0 | 0 | 0 | 0 | 0 | 521 | 1563 | 4690 | 0 | 521 | 0 | 0 | 0 | 0 | 0 | 3648 | 128 | 799 | 0 | 0 | 28 | 0 | 48 | 0 | 16 | 0 | 0 | 16 | 0 | 0 | 0 | 2606 | 0 |
| SRR10416520 | 0 | 0 | 0 | 0 | 0 | 0 | 0 | 649 | 0 | 0 | 1995 | 3245 | 0 | 0 | 0 | 649 | 0 | 0 | 0 | 649 | 129 | 994 | 0 | 0 | 49 | 0 | 39 | 0 | 0 | 0 | 0 | 0 | 0 | 0 | 0 | 1947 | 0 |
| SRR10416521 | 0 | 0 | 0 | 0 | 0 | 0 | 0 | 0 | 0 | 485 | 3880 | 8729 | 0 | 0 | 0 | 0 | 0 | 0 | 0 | 0 | 1367 | 3660 | 0 | 0 | 401 | 27 | 0 | 0 | 0 | 0 | 0 | 0 | 0 | 74 | 0 | 485 | 0 |
| SRR10416522 | 0 | 0 | 0 | 0 | 0 | 0 | 0 | 0 | 0 | 0 | 15287 | 18977 | 0 | 0 | 0 | 527 | 0 | 0 | 0 | 0 | 63 | 277 | 0 | 0 | 28 | 0 | 0 | 0 | 0 | 0 | 0 | 0 | 0 | 0 | 0 | 0 | 0 |
| SRR10416523 | 0 | 0 | 0 | 0 | 0 | 0 | 0 | 409 | 818 | 1635 | 7359 | 4906 | 0 | 0 | 818 | 1226 | 0 | 0 | 0 | 818 | 115 | 1533 | 0 | 0 | 35 | 0 | 0 | 0 | 0 | 0 | 0 | 0 | 0 | 0 | 0 | 3679 | 0 |
| SRR10416524 | 0 | 0 | 0 | 0 | 0 | 0 | 0 | 0 | 0 | 1325 | 8615 | 8615 | 0 | 663 | 0 | 0 | 0 | 0 | 0 | 0 | 60 | 494 | 0 | 0 | 52 | 0 | 30 | 0 | 0 | 0 | 0 | 0 | 0 | 0 | 0 | 1325 | 0 |
| SRR10416525 | 0 | 0 | 0 | 0 | 0 | 43 | 0 | 727 | 0 | 1453 | 6627 | 5814 | 0 | 0 | 0 | 0 | 0 | 0 | 0 | 727 | 36 | 277 | 0 | 0 | 0 | 0 | 0 | 0 | 0 | 0 | 0 | 0 | 0 | 43 | 0 | 1453 | 0 |
| SRR10416526 | 39 | 0 | 0 | 0 | 0 | 0 | 0 | 0 | 0 | 676 | 6281 | 11494 | 0 | 676 | 0 | 0 | 0 | 0 | 0 | 0 | 197 | 1430 | 0 | 0 | 76 | 0 | 0 | 0 | 0 | 0 | 0 | 0 | 0 | 39 | 0 | 0 | 0 |
| SRR10416527 | 0 | 0 | 0 | 0 | 0 | 0 | 0 | 736 | 0 | 736 | 2946 | 1473 | 0 | 0 | 0 | 736 | 0 | 0 | 0 | 736 | 107 | 500 | 0 | 0 | 12 | 0 | 0 | 0 | 0 | 0 | 0 | 0 | 0 | 0 | 0 | 1473 | 0 |
| SRR10416528 | 0 | 0 | 0 | 0 | 0 | 0 | 0 | 506 | 506 | 2024 | 9109 | 7591 | 0 | 0 | 0 | 2024 | 0 | 0 | 0 | 0 | 95 | 722 | 0 | 0 | 51 | 11 | 41 | 0 | 0 | 0 | 0 | 0 | 0 | 0 | 41 | 3543 | 0 |
| SRR10416529 | 0 | 0 | 55 | 0 | 0 | 0 | 0 | 903 | 0 | 0 | 5415 | 9928 | 0 | 0 | 903 | 0 | 0 | 0 | 0 | 0 | 54 | 246 | 0 | 0 | 54 | 0 | 0 | 0 | 0 | 0 | 0 | 0 | 0 | 0 | 0 | 903 | 0 |
| SRR10416530 | 0 | 0 | 0 | 0 | 0 | 0 | 0 | 749 | 0 | 0 | 3953 | 2994 | 0 | 0 | 749 | 1497 | 0 | 0 | 0 | 0 | 46 | 310 | 0 | 0 | 0 | 0 | 0 | 0 | 0 | 0 | 0 | 0 | 0 | 0 | 0 | 0 | 0 |
| SRR10416531 | 0 | 0 | 0 | 0 | 0 | 0 | 0 | 0 | 0 | 1105 | 6630 | 15470 | 0 | 0 | 0 | 0 | 0 | 0 | 0 | 0 | 42 | 314 | 0 | 0 | 30 | 0 | 99 | 0 | 0 | 0 | 0 | 0 | 0 | 0 | 0 | 1105 | 0 |
| SRR10416532 | 0 | 0 | 0 | 0 | 0 | 0 | 0 | 770 | 0 | 1540 | 16 | 20015 | 0 | 0 | 0 | 0 | 0 | 0 | 0 | 770 | 45 | 356 | 0 | 0 | 23 | 0 | 0 | 0 | 0 | 0 | 0 | 0 | 0 | 0 | 0 | 1540 | 0 |

Supp Table S8.4.Columns_DI_to_ES

| | | | | | | | | | | | | | | | | | | | | | | | | | | | | | | | | | | | | | |
|---|---|---|---|---|---|---|---|---|---|---|---|---|---|---|---|---|---|---|---|---|---|---|---|---|---|---|---|---|---|---|---|---|---|---|---|---|---|
| SRR10416533 | 0 | 0 | 0 | 0 | 0 | 0 | 0 | 0 | 0 | 2010 | 0 | 1005 | 0 | 0 | 0 | 0 | 0 | 0 | 0 | 0 | 421 | 2071 | 0 | 0 | 198 | 0 | 0 | 0 | 0 | 0 | 0 | 0 | 0 | 0 | 0 | 3015 | 0 |
| SRR10416534 | 0 | 0 | 0 | 0 | 0 | 0 | 0 | 1156 | 0 | 1156 | 2312 | 4624 | 0 | 0 | 0 | 1156 | 0 | 0 | 0 | 1156 | 93 | 471 | 0 | 0 | 42 | 0 | 0 | 0 | 0 | 0 | 0 | 0 | 0 | 0 | 0 | 1156 | 0 |
| SRR10416535 | 0 | 0 | 0 | 0 | 0 | 0 | 0 | 1679 | 0 | 0 | 10915 | 5877 | 0 | 0 | 0 | 840 | 0 | 0 | 0 | 840 | 17 | 152 | 0 | 0 | 0 | 0 | 0 | 0 | 0 | 0 | 0 | 0 | 0 | 0 | 0 | 840 | 0 |
| SRR10416536 | 0 | 0 | 44 | 0 | 0 | 0 | 0 | 0 | 0 | 721 | 10 | 7210 | 0 | 0 | 0 | 1442 | 0 | 0 | 0 | 0 | 0 | 186 | 0 | 0 | 0 | 0 | 0 | 0 | 0 | 0 | 0 | 0 | 0 | 44 | 0 | 721 | 0 |
| SRR10416537 | 0 | 0 | 0 | 0 | 0 | 0 | 0 | 0 | 0 | 0 | 30 | 19178 | 0 | 0 | 0 | 0 | 0 | 0 | 0 | 0 | 0 | 82 | 0 | 0 | 0 | 0 | 0 | 0 | 0 | 0 | 0 | 0 | 0 | 0 | 0 | 1096 | 0 |
| SRR10416538 | 0 | 0 | 0 | 0 | 0 | 0 | 0 | 1321 | 0 | 661 | 9247 | 13210 | 0 | 0 | 0 | 2642 | 0 | 0 | 0 | 0 | 18 | 191 | 0 | 0 | 11 | 0 | 0 | 0 | 0 | 0 | 0 | 0 | 0 | 0 | 0 | 1982 | 0 |
| SRR10416539 | 0 | 0 | 0 | 0 | 0 | 0 | 0 | 704 | 0 | 3519 | 4926 | 5630 | 0 | 0 | 0 | 0 | 0 | 0 | 0 | 704 | 134 | 457 | 0 | 0 | 475 | 23 | 67 | 0 | 0 | 0 | 0 | 0 | 0 | 0 | 0 | 704 | 0 |
| SRR10416540 | 0 | 0 | 0 | 0 | 0 | 0 | 0 | 0 | 0 | 806 | 15 | 12903 | 0 | 0 | 0 | 0 | 0 | 0 | 0 | 0 | 28 | 237 | 0 | 0 | 0 | 0 | 0 | 0 | 0 | 0 | 0 | 0 | 0 | 0 | 0 | 0 | 0 |
| SRR10416541 | 0 | 0 | 0 | 0 | 0 | 0 | 0 | 6438 | 0 | 2146 | 8584 | 6438 | 0 | 0 | 0 | 0 | 0 | 0 | 0 | 0 | 77 | 325 | 0 | 0 | 51 | 0 | 0 | 0 | 0 | 0 | 0 | 0 | 0 | 0 | 0 | 2146 | 0 |
| SRR10416542 | 0 | 0 | 0 | 0 | 0 | 0 | 0 | 1508 | 0 | 1508 | 1546 | 7541 | 0 | 0 | 1508 | 0 | 0 | 0 | 0 | 1508 | 73 | 715 | 0 | 0 | 16 | 0 | 0 | 0 | 0 | 0 | 0 | 0 | 0 | 90 | 0 | 0 | 0 |
| SRR10416543 | 0 | 0 | 0 | 0 | 0 | 0 | 0 | 526 | 526 | 1052 | 535 | 2631 | 0 | 0 | 0 | 0 | 0 | 0 | 0 | 0 | 1607 | 4692 | 0 | 0 | 1662 | 0 | 0 | 0 | 0 | 0 | 0 | 0 | 0 | 0 | 0 | 0 | 0 |
| SRR10416544 | 0 | 0 | 0 | 0 | 0 | 0 | 0 | 3467 | 0 | 693 | 4854 | 5548 | 0 | 0 | 0 | 2774 | 0 | 0 | 0 | 0 | 34 | 191 | 0 | 0 | 12 | 0 | 0 | 0 | 0 | 0 | 0 | 0 | 0 | 0 | 0 | 1387 | 0 |
| SRR10416545 | 0 | 0 | 19 | 0 | 0 | 19 | 0 | 1251 | 0 | 417 | 1289 | 2084 | 0 | 0 | 0 | 417 | 0 | 0 | 0 | 417 | 424 | 1779 | 0 | 0 | 110 | 0 | 0 | 0 | 0 | 0 | 0 | 0 | 0 | 0 | 0 | 1251 | 0 |
| SRR10416546 | 0 | 0 | 0 | 0 | 0 | 0 | 0 | 1224 | 0 | 2448 | 4361 | 7956 | 0 | 0 | 0 | 612 | 0 | 0 | 0 | 0 | 108 | 657 | 0 | 0 | 152 | 11 | 0 | 0 | 0 | 0 | 0 | 0 | 0 | 0 | 0 | 612 | 0 |
| SRR10416547 | 0 | 0 | 0 | 0 | 0 | 0 | 0 | 0 | 0 | 0 | 1577 | 3155 | 0 | 0 | 0 | 0 | 0 | 0 | 0 | 0 | 138 | 801 | 0 | 0 | 25 | 0 | 0 | 0 | 0 | 0 | 0 | 0 | 0 | 0 | 0 | 1577 | 0 |
| SRR10416549 | 0 | 0 | 23 | 0 | 0 | 0 | 0 | 561 | 280 | 1683 | 7852 | 7572 | 0 | 0 | 0 | 2243 | 0 | 0 | 0 | 280 | 109 | 1067 | 0 | 0 | 0 | 0 | 0 | 0 | 0 | 0 | 0 | 0 | 0 | 0 | 0 | 841 | 0 |
| SRR10416550 | 0 | 0 | 0 | 0 | 0 | 0 | 0 | 493 | 0 | 1971 | 15279 | 16757 | 0 | 0 | 0 | 493 | 0 | 0 | 0 | 0 | 169 | 1041 | 0 | 0 | 50 | 0 | 0 | 0 | 0 | 0 | 0 | 0 | 0 | 0 | 0 | 4929 | 0 |
| SRR10416551 | 0 | 0 | 0 | 0 | 0 | 0 | 0 | 941 | 471 | 471 | 1426 | 941 | 0 | 0 | 0 | 0 | 0 | 0 | 0 | 0 | 492 | 2545 | 0 | 0 | 97 | 0 | 0 | 0 | 0 | 0 | 0 | 0 | 0 | 0 | 0 | 1412 | 0 |
| SRR10416553 | 0 | 0 | 65 | 0 | 0 | 0 | 0 | 727 | 0 | 1455 | 749 | 4364 | 0 | 0 | 727 | 0 | 0 | 0 | 0 | 0 | 132 | 696 | 0 | 0 | 50 | 0 | 0 | 0 | 0 | 0 | 0 | 0 | 0 | 0 | 0 | 727 | 0 |
| SRR10416554 | 0 | 0 | 0 | 0 | 0 | 0 | 0 | 0 | 0 | 0 | 698 | 4190 | 0 | 0 | 698 | 0 | 0 | 0 | 0 | 0 | 651 | 2690 | 0 | 0 | 339 | 0 | 0 | 0 | 0 | 0 | 0 | 0 | 0 | 71 | 0 | 698 | 0 |
| SRR10416555 | 0 | 0 | 0 | 0 | 0 | 0 | 0 | 1168 | 0 | 0 | 7009 | 9346 | 0 | 0 | 0 | 0 | 0 | 0 | 0 | 1168 | 41 | 400 | 0 | 0 | 22 | 0 | 0 | 0 | 0 | 0 | 0 | 0 | 0 | 0 | 0 | 0 | 0 |
| SRR10416557 | 0 | 0 | 0 | 0 | 0 | 0 | 0 | 1862 | 0 | 1862 | 1862 | 1862 | 0 | 0 | 0 | 1862 | 0 | 0 | 0 | 1862 | 25 | 197 | 0 | 0 | 0 | 0 | 115 | 0 | 0 | 0 | 0 | 0 | 0 | 0 | 0 | 0 | 0 |
| SRR10416558 | 0 | 0 | 25 | 0 | 0 | 0 | 0 | 1347 | 0 | 673 | 4808 | 2020 | 0 | 0 | 673 | 2020 | 0 | 0 | 0 | 0 | 21 | 254 | 0 | 0 | 28 | 0 | 0 | 0 | 0 | 0 | 0 | 0 | 0 | 0 | 0 | 0 | 0 |
| SRR10416559 | 0 | 0 | 0 | 0 | 0 | 0 | 0 | 545 | 0 | 545 | 7088 | 5998 | 0 | 0 | 0 | 0 | 0 | 0 | 0 | 545 | 127 | 888 | 0 | 0 | 27 | 0 | 0 | 0 | 0 | 0 | 0 | 0 | 0 | 0 | 0 | 545 | 0 |
| SRR10416560 | 0 | 0 | 0 | 0 | 0 | 0 | 0 | 868 | 0 | 434 | 8681 | 4340 | 0 | 0 | 868 | 0 | 0 | 0 | 0 | 0 | 132 | 1050 | 0 | 0 | 0 | 0 | 23 | 0 | 0 | 0 | 0 | 0 | 0 | 0 | 0 | 868 | 0 |
| SRR10416562 | 0 | 0 | 0 | 0 | 0 | 0 | 0 | 0 | 0 | 0 | 722 | 722 | 0 | 0 | 0 | 0 | 0 | 0 | 0 | 722 | 96 | 625 | 0 | 0 | 23 | 0 | 0 | 0 | 0 | 0 | 0 | 0 | 0 | 0 | 0 | 2165 | 0 |
| SRR10416563 | 0 | 15 | 0 | 0 | 0 | 0 | 0 | 0 | 0 | 0 | 2737 | 2737 | 0 | 0 | 0 | 547 | 0 | 0 | 0 | 3831 | 19 | 284 | 0 | 0 | 0 | 0 | 15 | 0 | 0 | 0 | 0 | 0 | 0 | 0 | 30 | 2737 | 0 |
| SRR10416564 | 0 | 43 | 0 | 0 | 0 | 0 | 0 | 0 | 0 | 699 | 6988 | 6988 | 0 | 0 | 0 | 699 | 0 | 0 | 0 | 0 | 115 | 722 | 0 | 0 | 11 | 0 | 0 | 0 | 0 | 0 | 0 | 0 | 0 | 0 | 0 | 699 | 0 |
| SRR10416565 | 0 | 0 | 0 | 0 | 0 | 0 | 0 | 0 | 0 | 632 | 3793 | 1896 | 0 | 0 | 1264 | 1264 | 0 | 0 | 0 | 1264 | 135 | 1008 | 0 | 0 | 17 | 0 | 0 | 0 | 0 | 0 | 0 | 0 | 0 | 0 | 0 | 632 | 0 |
| SRR10416566 | 0 | 0 | 0 | 0 | 0 | 0 | 0 | 1672 | 0 | 1672 | 3344 | 3344 | 0 | 0 | 0 | 0 | 0 | 0 | 0 | 1672 | 12 | 67 | 0 | 0 | 0 | 0 | 0 | 0 | 0 | 0 | 0 | 0 | 0 | 0 | 0 | 1672 | 0 |
| SRR10416567 | 0 | 0 | 0 | 0 | 0 | 0 | 0 | 0 | 0 | 528 | 4749 | 9499 | 0 | 0 | 0 | 528 | 0 | 0 | 0 | 0 | 707 | 1926 | 0 | 0 | 58 | 0 | 321 | 0 | 0 | 0 | 0 | 0 | 0 | 0 | 0 | 528 | 0 |
| SRR10416568 | 0 | 0 | 0 | 0 | 0 | 0 | 0 | 1685 | 842 | 842 | 11 | 12637 | 0 | 0 | 0 | 2527 | 0 | 0 | 0 | 0 | 46 | 332 | 0 | 0 | 15 | 0 | 0 | 0 | 0 | 0 | 0 | 0 | 0 | 0 | 0 | 0 | 0 |
| SRR10416569 | 0 | 0 | 0 | 0 | 0 | 0 | 0 | 637 | 0 | 637 | 5242 | 10828 | 0 | 0 | 0 | 0 | 0 | 0 | 0 | 0 | 225 | 1754 | 0 | 0 | 38 | 0 | 48 | 0 | 0 | 0 | 0 | 0 | 0 | 48 | 0 | 637 | 0 |
| SRR10416570 | 0 | 0 | 0 | 0 | 0 | 0 | 0 | 0 | 0 | 1288 | 0 | 5151 | 0 | 0 | 644 | 0 | 0 | 0 | 0 | 0 | 2355 | 16 | 0 | 0 | 801 | 0 | 686 | 0 | 0 | 0 | 0 | 0 | 0 | 0 | 0 | 1932 | 0 |
| SRR10416571 | 0 | 0 | 0 | 0 | 0 | 0 | 0 | 0 | 0 | 770 | 6265 | 7704 | 0 | 0 | 0 | 770 | 0 | 0 | 0 | 0 | 56 | 241 | 0 | 0 | 0 | 0 | 0 | 0 | 0 | 0 | 0 | 0 | 0 | 0 | 0 | 2311 | 0 |
| SRR10416572 | 0 | 0 | 0 | 0 | 0 | 0 | 0 | 0 | 546 | 1092 | 3275 | 5459 | 0 | 0 | 0 | 546 | 0 | 0 | 0 | 546 | 1959 | 5147 | 0 | 0 | 365 | 13 | 131 | 0 | 0 | 0 | 0 | 0 | 0 | 0 | 0 | 546 | 0 |
| SRR10416573 | 0 | 0 | 0 | 0 | 0 | 0 | 0 | 1701 | 0 | 1701 | 1701 | 0 | 0 | 1701 | 0 | 0 | 0 | 0 | 0 | 0 | 114 | 735 | 0 | 0 | 17 | 0 | 0 | 0 | 0 | 0 | 0 | 0 | 0 | 0 | 0 | 0 | 0 |
| SRR10416574 | 0 | 0 | 0 | 0 | 0 | 0 | 0 | 0 | 0 | 0 | 16 | 21821 | 0 | 0 | 0 | 1505 | 0 | 0 | 0 | 0 | 90 | 759 | 0 | 0 | 10 | 0 | 0 | 0 | 0 | 0 | 0 | 0 | 0 | 0 | 0 | 0 | 0 |
| SRR10416575 | 0 | 42 | 0 | 0 | 0 | 0 | 0 | 0 | 0 | 0 | 0 | 0 | 0 | 0 | 0 | 0 | 0 | 0 | 0 | 0 | 34 | 456 | 0 | 0 | 0 | 0 | 0 | 0 | 0 | 0 | 0 | 0 | 0 | 0 | 0 | 3891 | 0 |
| SRR10416576 | 0 | 0 | 0 | 0 | 0 | 0 | 0 | 0 | 0 | 0 | 0 | 8840 | 0 | 0 | 1105 | 0 | 0 | 0 | 0 | 0 | 57 | 638 | 0 | 0 | 0 | 0 | 28 | 0 | 0 | 0 | 0 | 0 | 0 | 0 | 0 | 2210 | 0 |
| SRR10416577 | 0 | 0 | 0 | 0 | 0 | 0 | 0 | 682 | 0 | 0 | 5453 | 3408 | 0 | 0 | 0 | 0 | 0 | 0 | 0 | 682 | 173 | 866 | 0 | 0 | 35 | 0 | 71 | 0 | 0 | 0 | 0 | 0 | 0 | 0 | 0 | 1363 | 0 |
| SRR10416578 | 0 | 0 | 0 | 0 | 0 | 0 | 0 | 0 | 0 | 0 | 5966 | 3729 | 0 | 0 | 0 | 0 | 0 | 0 | 0 | 0 | 722 | 1736 | 0 | 0 | 191 | 0 | 292 | 0 | 0 | 0 | 0 | 0 | 0 | 0 | 0 | 0 | 0 |
| SRR10416579 | 0 | 0 | 0 | 0 | 0 | 0 | 0 | 1124 | 0 | 1124 | 7919 | 13483 | 0 | 0 | 0 | 0 | 0 | 0 | 0 | 0 | 10 | 185 | 0 | 0 | 28 | 0 | 0 | 0 | 0 | 0 | 0 | 0 | 0 | 0 | 0 | 1124 | 0 |
| SRR10416580 | 0 | 0 | 0 | 0 | 0 | 0 | 0 | 772 | 0 | 0 | 7053 | 6944 | 0 | 0 | 0 | 772 | 0 | 0 | 0 | 0 | 185 | 915 | 0 | 0 | 64 | 0 | 0 | 0 | 0 | 0 | 0 | 0 | 0 | 0 | 0 | 772 | 0 |
| SRR10416581 | 0 | 0 | 61 | 0 | 0 | 20 | 0 | 0 | 0 | 2239 | 767 | 2985 | 0 | 0 | 0 | 0 | 0 | 0 | 0 | 4478 | 63 | 540 | 0 | 0 | 0 | 0 | 20 | 0 | 0 | 0 | 0 | 0 | 0 | 0 | 20 | 3731 | 0 |
| SRR10416582 | 0 | 92 | 0 | 0 | 0 | 0 | 0 | 2804 | 0 | 0 | 6542 | 6542 | 0 | 0 | 0 | 935 | 0 | 0 | 0 | 0 | 92 | 626 | 0 | 0 | 11 | 0 | 0 | 0 | 0 | 0 | 0 | 0 | 0 | 0 | 0 | 935 | 0 |
| SRR10416583 | 0 | 0 | 0 | 0 | 0 | 0 | 0 | 0 | 0 | 1905 | 1905 | 1905 | 0 | 0 | 1905 | 0 | 0 | 0 | 0 | 0 | 936 | 3152 | 0 | 0 | 654 | 0 | 0 | 0 | 0 | 0 | 0 | 0 | 0 | 0 | 0 | 0 | 0 |
| SRR10416584 | 0 | 0 | 44 | 0 | 0 | 0 | 0 | 557 | 0 | 0 | 10 | 7804 | 0 | 0 | 0 | 557 | 0 | 0 | 0 | 0 | 147 | 736 | 0 | 0 | 19 | 0 | 44 | 0 | 0 | 0 | 0 | 0 | 0 | 0 | 0 | 1115 | 0 |
| SRR10416585 | 0 | 0 | 0 | 0 | 0 | 0 | 0 | 0 | 0 | 2028 | 2092 | 6085 | 0 | 0 | 0 | 0 | 0 | 0 | 0 | 0 | 27 | 375 | 0 | 0 | 16 | 0 | 0 | 0 | 0 | 0 | 0 | 0 | 0 | 0 | 0 | 0 | 0 |
| SRR10416586 | 0 | 0 | 0 | 0 | 0 | 0 | 0 | 1131 | 0 | 1131 | 12443 | 11312 | 0 | 0 | 1131 | 0 | 0 | 0 | 0 | 0 | 49 | 368 | 0 | 0 | 13 | 0 | 0 | 0 | 0 | 0 | 0 | 0 | 0 | 94 | 0 | 0 | 0 |
| SRR10416587 | 0 | 0 | 0 | 0 | 0 | 0 | 0 | 0 | 0 | 2004 | 2047 | 1002 | 0 | 0 | 0 | 1002 | 0 | 0 | 0 | 1002 | 175 | 756 | 0 | 0 | 71 | 0 | 0 | 0 | 0 | 0 | 0 | 0 | 0 | 65 | 0 | 2004 | 0 |
| SRR10416588 | 0 | 0 | 0 | 0 | 0 | 0 | 0 | 991 | 0 | 1982 | 4955 | 4955 | 0 | 0 | 0 | 1982 | 0 | 0 | 0 | 0 | 191 | 915 | 0 | 0 | 86 | 0 | 0 | 0 | 0 | 0 | 0 | 0 | 0 | 0 | 0 | 1982 | 0 |
| SRR10416589 | 0 | 0 | 0 | 0 | 0 | 49 | 0 | 0 | 0 | 0 | 4762 | 7143 | 0 | 0 | 0 | 0 | 0 | 0 | 0 | 3175 | 145 | 605 | 0 | 0 | 79 | 0 | 0 | 0 | 0 | 0 | 0 | 0 | 0 | 0 | 0 | 5556 | 0 |
| SRR10416590 | 0 | 0 | 0 | 0 | 0 | 0 | 0 | 794 | 0 | 2381 | 2381 | 3968 | 0 | 0 | 1587 | 0 | 0 | 0 | 0 | 1587 | 104 | 435 | 0 | 0 | 54 | 0 | 0 | 0 | 0 | 0 | 0 | 0 | 0 | 0 | 0 | 1587 | 0 |
| SRR10416591 | 0 | 0 | 0 | 0 | 0 | 0 | 0 | 3021 | 0 | 0 | 0 | 0 | 0 | 0 | 0 | 0 | 0 | 0 | 0 | 3021 | 57 | 263 | 0 | 0 | 25 | 0 | 0 | 0 | 0 | 0 | 0 | 0 | 0 | 0 | 0 | 3021 | 0 |
| SRR10416592 | 0 | 0 | 45 | 0 | 0 | 0 | 0 | 371 | 0 | 0 | 2703 | 7058 | 0 | 0 | 0 | 371 | 0 | 0 | 0 | 0 | 1627 | 4697 | 0 | 0 | 254 | 10 | 91 | 0 | 0 | 0 | 0 | 45 | 0 | 0 | 45 | 371 | 0 |
| SRR10416593 | 0 | 0 | 0 | 0 | 0 | 0 | 0 | 935 | 0 | 0 | 3742 | 5613 | 0 | 0 | 0 | 935 | 0 | 0 | 0 | 0 | 23 | 146 | 0 | 0 | 0 | 0 | 0 | 0 | 0 | 0 | 0 | 0 | 0 | 0 | 0 | 1871 | 0 |
| SRR10416594 | 0 | 0 | 0 | 0 | 0 | 0 | 0 | 1862 | 1862 | 931 | 4713 | 3724 | 0 | 931 | 931 | 931 | 0 | 0 | 0 | 0 | 74 | 288 | 0 | 0 | 16 | 0 | 0 | 0 | 0 | 0 | 0 | 0 | 0 | 0 | 0 | 0 | 0 |
| SRR10416595 | 0 | 0 | 0 | 0 | 0 | 42 | 0 | 0 | 0 | 492 | 499 | 985 | 0 | 0 | 492 | 0 | 0 | 0 | 0 | 0 | 129 | 552 | 0 | 0 | 10 | 0 | 0 | 0 | 0 | 0 | 0 | 0 | 0 | 0 | 42 | 492 | 0 |
| SRR10416596 | 0 | 0 | 0 | 0 | 0 | 0 | 0 | 0 | 0 | 0 | 1067 | 6289 | 0 | 0 | 0 | 1048 | 0 | 0 | 0 | 6289 | 31 | 220 | 0 | 0 | 0 | 0 | 0 | 0 | 0 | 0 | 0 | 0 | 0 | 0 | 0 | 6289 | 0 |
| SRR10416597 | 0 | 27 | 0 | 0 | 0 | 0 | 0 | 0 | 0 | 1644 | 1096 | 548 | 0 | 0 | 0 | 0 | 0 | 0 | 0 | 1096 | 109 | 523 | 0 | 0 | 32 | 0 | 0 | 0 | 0 | 0 | 0 | 0 | 0 | 0 | 0 | 2192 | 0 |
| SRR10416598 | 0 | 28 | 0 | 0 | 0 | 0 | 0 | 1182 | 0 | 1182 | 0 | 0 | 0 | 0 | 0 | 0 | 0 | 0 | 0 | 2364 | 107 | 460 | 0 | 0 | 0 | 0 | 0 | 0 | 0 | 0 | 0 | 0 | 0 | 0 | 0 | 3546 | 0 |
| SRR10416599 | 0 | 0 | 0 | 0 | 0 | 0 | 0 | 0 | 444 | 0 | 2797 | 7552 | 0 | 0 | 0 | 0 | 0 | 0 | 0 | 0 | 1671 | 4454 | 0 | 0 | 331 | 0 | 504 | 0 | 0 | 0 | 0 | 0 | 0 | 0 | 0 | 0 | 0 |
| SRR10416600 | 0 | 0 | 0 | 0 | 0 | 0 | 0 | 419 | 0 | 0 | 1675 | 2931 | 0 | 0 | 0 | 0 | 0 | 0 | 0 | 419 | 111 | 1066 | 0 | 0 | 0 | 0 | 0 | 0 | 0 | 0 | 0 | 0 | 0 | 0 | 0 | 419 | 0 |
| SRR10428553 | 15 | 0 | 0 | 0 | 0 | 10 | 0 | 7648 | 4006 | 7223 | 127 | 108 | 0 | 1083 | 0 | 0 | 0 | 231 | 0 | 18 | 989 | 910 | 156 | 3847 | 2369 | 54 | 16 | 0 | 0 | 0 | 0 | 0 | 0 | 0 | 0 | 310 | 0 |
| SRR10428554 | 43 | 0 | 0 | 0 | 0 | 18 | 12 | 643 | 319 | 5037 | 596 | 1563 | 0 | 987 | 11 | 0 | 22 | 116 | 0 | 426 | 732 | 554 | 23 | 906 | 2578 | 44 | 23 | 0 | 0 | 11 | 12 | 0 | 0 | 12 | 0 | 3359 | 0 |
| SRR10428555 | 54 | 0 | 0 | 0 | 0 | 26 | 27 | 3565 | 2991 | 11395 | 335 | 358 | 0 | 236 | 0 | 0 | 18 | 407 | 0 | 92 | 834 | 402 | 59 | 1097 | 1537 | 0 | 25 | 0 | 0 | 0 | 0 | 0 | 0 | 0 | 0 | 578 | 0 |
| SRR10428556 | 21 | 0 | 0 | 0 | 12 | 11 | 15 | 4323 | 1677 | 3679 | 1335 | 1392 | 0 | 2246 | 0 | 0 | 24 | 146 | 0 | 320 | 778 | 843 | 192 | 3072 | 2558 | 29 | 14 | 0 | 0 | 0 | 14 | 0 | 0 | 0 | 0 | 1021 | 0 |
| SRR10428557 | 76 | 0 | 0 | 0 | 0 | 36 | 46 | 378 | 173 | 2978 | 785 | 914 | 0 | 891 | 17 | 47 | 17 | 169 | 0 | 351 | 1174 | 2017 | 291 | 6435 | 3216 | 28 | 39 | 0 | 0 | 0 | 0 | 0 | 0 | 0 | 0 | 3099 | 0 |
| SRR10428558 | 43 | 0 | 0 | 0 | 0 | 27 | 12 | 606 | 326 | 3047 | 152 | 165 | 0 | 1017 | 0 | 0 | 17 | 126 | 0 | 145 | 992 | 1025 | 359 | 5048 | 2179 | 18 | 43 | 0 | 0 | 0 | 0 | 0 | 0 | 0 | 0 | 999 | 0 |
| SRR10428559 | 42 | 0 | 0 | 0 | 16 | 19 | 13 | 1506 | 770 | 5641 | 283 | 326 | 0 | 672 | 0 | 0 | 25 | 170 | 0 | 189 | 866 | 923 | 211 | 3925 | 3352 | 15 | 28 | 0 | 0 | 0 | 0 | 0 | 0 | 11 | 0 | 1885 | 0 |
| SRR10428560 | 0 | 0 | 0 | 0 | 11 | 0 | 0 | 2586 | 925 | 3217 | 202 | 196 | 0 | 515 | 0 | 0 | 70 | 126 | 0 | 25 | 930 | 680 | 194 | 3986 | 6833 | 36 | 16 | 0 | 0 | 0 | 0 | 0 | 0 | 0 | 0 | 185 | 0 |
| SRR10446207 | 0 | 707 | 134 | 0 | 148 | 0 | 0 | 161 | 2816 | 4335 | 948 | 1475 | 0 | 98 | 536 | 1332 | 0 | 1547 | 0 | 125 | 1139 | 2642 | 186 | 932 | 1470 | 0 | 0 | 23 | 241 | 77 | 11 | 17 | 54 | 82 | 0 | 787 | 0 |
| SRR10446208 | 0 | 70 | 106 | 0 | 158 | 0 | 0 | 361 | 3978 | 9207 | 563 | 1364 | 0 | 101 | 1420 | 4260 | 0 | 2334 | 0 | 0 | 1332 | 3534 | 225 | 1424 | 1624 | 0 | 0 | 32 | 13 | 32 | 16 | 16 | 48 | 54 | 0 | 11 | 0 |
| SRR10446209 | 0 | 599 | 251 | 0 | 171 | 0 | 0 | 219 | 2335 | 8245 | 2748 | 5059 | 0 | 24 | 462 | 1143 | 0 | 1546 | 0 | 122 | 720 | 3494 | 157 | 945 | 847 | 0 | 20 | 59 | 310 | 49 | 93 | 39 | 69 | 49 | 0 | 1265 | 0 |
| SRR10446210 | 0 | 69 | 11 | 0 | 361 | 0 | 0 | 450 | 8679 | 23402 | 16 | 1136 | 0 | 21 | 31031 | 47511 | 0 | 932 | 0 | 64 | 1115 | 4132 | 116 | 1601 | 1941 | 0 | 0 | 157 | 32 | 171 | 39 | 39 | 52 | 66 | 0 | 664 | 0 |
| SRR10446211 | 0 | 22 | 49 | 0 | 168 | 12 | 0 | 180 | 1112 | 7296 | 69 | 115 | 0 | 725 | 4979 | 11638 | 0 | 1648 | 25 | 162 | 391 | 1970 | 169 | 983 | 329 | 19 | 23 | 17 | 18 | 0 | 0 | 0 | 0 | 24 | 0 | 1024 | 0 |
| SRR10446212 | 0 | 379 | 251 | 0 | 292 | 0 | 0 | 238 | 1534 | 19983 | 529 | 1018 | 0 | 450 | 3293 | 11929 | 0 | 462 | 0 | 489 | 307 | 1165 | 103 | 487 | 303 | 11 | 0 | 140 | 461 | 70 | 32 | 32 | 45 | 124 | 16 | 1706 | 0 |
| SRR10531363 | 0 | 0 | 0 | 0 | 0 | 0 | 82 | 68 | 0 | 0 | 0 | 0 | 0 | 2684 | 0 | 0 | 0 | 0 | 47 | 818 | 0 | 0 | 45 | 60 | 110 | 185 | 1799 | 0 | 0 | 0 | 0 | 0 | 0 | 0 | 0 | 1381 | 0 |
| SRR10531364 | 0 | 0 | 0 | 0 | 0 | 22 | 0 | 0 | 0 | 725 | 0 | 0 | 0 | 3977 | 0 | 0 | 0 | 0 | 49 | 1775 | 0 | 0 | 63 | 107 | 20 | 414 | 89 | 0 | 0 | 0 | 0 | 0 | 0 | 0 | 0 | 3787 | 0 |
| SRR10531365 | 0 | 0 | 0 | 0 | 0 | 0 | 0 | 0 | 4237 | 0 | 0 | 0 | 0 | 3074 | 0 | 1035 | 0 | 0 | 10 | 1720 | 0 | 0 | 53 | 35 | 0 | 167 | 1154 | 0 | 0 | 0 | 0 | 0 | 0 | 0 | 0 | 1656 | 0 |
| SRR10531367 | 0 | 0 | 0 | 0 | 0 | 0 | 0 | 0 | 2366 | 0 | 0 | 0 | 0 | 3512 | 0 | 0 | 143 | 0 | 34 | 18772 | 0 | 0 | 68 | 36 | 59 | 501 | 35 | 0 | 0 | 0 | 0 | 0 | 0 | 0 | 0 | 20211 | 12 |
| SRR10531368 | 0 | 0 | 0 | 0 | 0 | 0 | 150 | 0 | 708 | 2157 | 0 | 0 | 0 | 9296 | 0 | 0 | 151 | 0 | 46 | 7194 | 0 | 0 | 63 | 74 | 28 | 464 | 24 | 0 | 0 | 0 | 0 | 0 | 0 | 0 | 0 | 9358 | 0 |
| SRR10531369 | 0 | 0 | 0 | 0 | 0 | 0 | 0 | 0 | 3497 | 0 | 0 | 0 | 0 | 2543 | 0 | 0 | 0 | 0 | 19 | 6135 | 0 | 0 | 60 | 72 | 49 | 857 | 2132 | 0 | 0 | 0 | 0 | 0 | 0 | 0 | 0 | 18681 | 0 |
| SRR10531370 | 0 | 0 | 0 | 0 | 0 | 0 | 0 | 0 | 607 | 0 | 0 | 0 | 104 | 2896 | 0 | 0 | 0 | 0 | 104 | 9887 | 0 | 0 | 16 | 40 | 100 | 420 | 4663 | 0 | 0 | 0 | 0 | 60 | 0 | 0 | 0 | 22457 | 0 |
| SRR10544939 | 0 | 63 | 0 | 0 | 0 | 0 | 0 | 0 | 0 | 36 | 4979 | 2053 | 0 | 39 | 0 | 0 | 0 | 595 | 182 | 0 | 611 | 1140 | 0 | 0 | 449 | 0 | 0 | 0 | 0 | 0 | 0 | 0 | 0 | 0 | 0 | 0 | 301 |
| SRR10544940 | 0 | 41 | 0 | 0 | 0 | 0 | 0 | 0 | 0 | 16 | 4493 | 780 | 0 | 19 | 0 | 0 | 0 | 441 | 122 | 0 | 420 | 992 | 0 | 0 | 193 | 0 | 0 | 0 | 0 | 0 | 0 | 0 | 0 | 0 | 0 | 0 | 225 |
| SRR10544941 | 0 | 143 | 0 | 0 | 0 | 0 | 0 | 0 | 0 | 46 | 10875 | 2946 | 0 | 48 | 0 | 0 | 0 | 864 | 147 | 0 | 666 | 1720 | 0 | 0 | 305 | 0 | 0 | 0 | 0 | 0 | 0 | 0 | 0 | 0 | 0 | 0 | 335 |
| SRR10544942 | 0 | 86 | 0 | 0 | 0 | 0 | 0 | 0 | 0 | 0 | 2739 | 1092 | 0 | 21 | 0 | 0 | 0 | 726 | 163 | 0 | 838 | 1148 | 0 | 0 | 341 | 0 | 0 | 0 | 0 | 0 | 0 | 0 | 0 | 0 | 0 | 0 | 186 |
| SRR10544943 | 0 | 17 | 0 | 0 | 0 | 0 | 0 | 0 | 0 | 25 | 3380 | 1589 | 0 | 31 | 0 | 0 | 0 | 584 | 267 | 0 | 581 | 1623 | 0 | 0 | 211 | 0 | 0 | 0 | 0 | 0 | 0 | 0 | 0 | 0 | 0 | 0 | 210 |
| SRR10544944 | 0 | 53 | 0 | 0 | 0 | 0 | 0 | 0 | 0 | 73 | 10886 | 5293 | 0 | 55 | 16 | 16 | 0 | 705 | 297 | 0 | 1629 | 2516 | 0 | 0 | 547 | 0 | 0 | 0 | 0 | 0 | 0 | 0 | 0 | 0 | 0 | 0 | 876 |
| SRR10544945 | 0 | 0 | 0 | 0 | 0 | 0 | 0 | 0 | 0 | 0 | 88 | 185 | 0 | 12 | 28 | 23 | 0 | 167 | 105 | 83 | 732 | 635 | 0 | 160 | 321 | 0 | 0 | 0 | 0 | 0 | 0 | 0 | 0 | 0 | 0 | 111 | 119 |
| SRR10544946 | 0 | 0 | 0 | 0 | 0 | 0 | 0 | 0 | 0 | 10 | 402 | 280 | 0 | 14 | 0 | 10 | 0 | 64 | 67 | 88 | 546 | 567 | 0 | 106 | 152 | 0 | 0 | 0 | 0 | 0 | 0 | 0 | 0 | 0 | 0 | 119 | 79 |
| SRR10544947 | 0 | 0 | 0 | 0 | 0 | 0 | 0 | 0 | 0 | 17 | 1255 | 403 | 0 | 23 | 0 | 17 | 0 | 109 | 42 | 252 | 603 | 790 | 0 | 148 | 328 | 0 | 0 | 0 | 0 | 0 | 0 | 0 | 0 | 0 | 0 | 550 | 127 |
| SRR10544948 | 0 | 0 | 0 | 0 | 0 | 0 | 0 | 0 | 0 | 11 | 259 | 264 | 0 | 17 | 0 | 0 | 0 | 103 | 161 | 68 | 546 | 461 | 0 | 121 | 116 | 0 | 0 | 0 | 0 | 0 | 0 | 0 | 0 | 0 | 0 | 63 | 85 |
| SRR10544949 | 0 | 0 | 0 | 0 | 0 | 0 | 0 | 0 | 0 | 11 | 1202 | 523 | 0 | 17 | 11 | 17 | 0 | 105 | 132 | 200 | 636 | 425 | 0 | 219 | 149 | 0 | 0 | 0 | 0 | 0 | 0 | 0 | 0 | 0 | 0 | 195 | 75 |
| SRR10544950 | 0 | 0 | 0 | 0 | 0 | 0 | 0 | 0 | 10 | 0 | 206 | 378 | 0 | 26 | 0 | 10 | 0 | 77 | 189 | 109 | 728 | 490 | 0 | 217 | 175 | 0 | 0 | 0 | 0 | 0 | 0 | 0 | 0 | 0 | 0 | 124 | 133 |
| SRR10756575 | 0 | 100 | 70 | 165 | 125 | 14 | 27 | 3031 | 4388 | 24323 | 6040 | 32165 | 0 | 1145 | 0 | 0 | 41 | 423 | 97 | 1542 | 245 | 1245 | 0 | 43 | 1141 | 146 | 24 | 31 | 34 | 56 | 62 | 10 | 19 | 18 | 210 | 6255 | 0 |
| SRR10756576 | 0 | 67 | 50 | 186 | 162 | 15 | 38 | 1085 | 1274 | 4001 | 4597 | 28883 | 11 | 487 | 0 | 17 | 55 | 530 | 56 | 5118 | 219 | 1145 | 0 | 26 | 1365 | 36 | 116 | 46 | 40 | 84 | 80 | 15 | 16 | 14 | 109 | 23707 | 0 |
| SRR10756577 | 0 | 174 | 435 | 136 | 151 | 0 | 70 | 1940 | 1981 | 7265 | 6503 | 44947 | 23 | 361 | 0 | 0 | 55 | 534 | 87 | 1038 | 303 | 1121 | 0 | 41 | 1205 | 114 | 700 | 78 | 1198 | 2031 | 1488 | 219 | 252 | 191 | 53 | 2963 | 0 |
| SRR10756578 | 0 | 356 | 422 | 227 | 200 | 0 | 24 | 868 | 1450 | 5829 | 3489 | 6734 | 10 | 1062 | 0 | 0 | 103 | 806 | 183 | 1033 | 466 | 1448 | 0 | 42 | 1192 | 49 | 938 | 63 | 813 | 1784 | 1151 | 148 | 203 | 208 | 179 | 4148 | 0 |
| SRR10756579 | 0 | 122 | 26 | 189 | 186 | 12 | 17 | 1411 | 1594 | 6843 | 2046 | 18531 | 29 | 911 | 0 | 0 | 49 | 640 | 108 | 1348 | 391 | 1329 | 0 | 39 | 1202 | 109 | 16 | 58 | 41 | 76 | 71 | 11 | 12 | 13 | 171 | 3747 | 0 |
| SRR10756581 | 0 | 171 | 34 | 280 | 178 | 12 | 17 | 1405 | 1965 | 6389 | 4904 | 8180 | 12 | 1000 | 0 | 13 | 0 | 1043 | 200 | 2748 | 1005 | 1501 | 0 | 55 | 2040 | 39 | 218 | 33 | 44 | 114 | 157 | 18 | 25 | 25 | 294 | 10059 | 0 |
| SRR10756582 | 0 | 59 | 1627 | 85 | 124 | 15 | 0 | 1243 | 1362 | 13472 | 3843 | 5974 | 30 | 1612 | 0 | 15 | 15 | 540 | 52 | 1273 | 272 | 1264 | 0 | 53 | 1383 | 264 | 185 | 30 | 1116 | 2151 | 4052 | 495 | 991 | 798 | 798 | 4213 | 0 |
| SRR10756583 | 0 | 84 | 1089 | 186 | 202 | 50 | 25 | 1321 | 1146 | 7960 | 20601 | 23579 | 17 | 987 | 0 | 16 | 50 | 567 | 142 | 1178 | 512 | 1350 | 0 | 75 | 1413 | 175 | 360 | 42 | 1357 | 2496 | 2019 | 134 | 184 | 209 | 109 | 4386 | 0 |
| SRR10875785 | 0 | 37 | 101 | 369 | 198 | 0 | 29 | 9943 | 7974 | 8092 | 781 | 15844 | 57 | 267 | 196 | 339 | 245 | 622 | 24 | 1030 | 627 | 429 | 0 | 17 | 1307 | 21 | 1540 | 11 | 1030 | 924 | 2576 | 728 | 491 | 303 | 0 | 1930 | 2825 |
| SRR10875786 | 33 | 0 | 0 | 133 | 220 | 17 | 72 | 12743 | 10318 | 3899 | 228 | 4412 | 21 | 19 | 177 | 224 | 98 | 515 | 15 | 513 | 728 | 516 | 0 | 68 | 1024 | 34 | 156 | 0 | 15 | 21 | 66 | 25 | 19 | 15 | 0 | 1091 | 2023 |
| SRR10953234 | 0 | 727 | 351 | 667 | 2520 | 0 | 0 | 1204 | 3473 | 14574 | 6359 | 10433 | 0 | 2168 | 187 | 357 | 0 | 275 | 105 | 5209 | 527 | 790 | 0 | 32 | 2085 | 0 | 0 | 0 | 91 | 164 | 345 | 54 | 102 | 117 | 196 | 2204 | 0 |
| SRR10953235 | 0 | 636 | 322 | 553 | 2023 | 0 | 0 | 881 | 2406 | 10863 | 3958 | 8765 | 0 | 1588 | 170 | 273 | 0 | 335 | 118 | 4056 | 459 | 899 | 0 | 33 | 2190 | 0 | 0 | 0 | 69 | 173 | 339 | 55 | 73 | 90 | 189 | 1628 | 0 |
| SRR10953236 | 0 | 651 | 304 | 632 | 2461 | 0 | 0 | 904 | 2354 | 12764 | 3540 | 11131 | 0 | 1590 | 334 | 631 | 0 | 246 | 82 | 4834 | 475 | 796 | 0 | 37 | 2331 | 0 | 0 | 0 | 115 | 221 | 373 | 48 | 73 | 94 | 167 | 2020 | 0 |
| SRR10953237 | 0 | 219 | 112 | 701 | 2467 | 0 | 0 | 619 | 1857 | 9894 | 3978 | 8745 | 0 | 936 | 246 | 576 | 0 | 395 | 138 | 5577 | 723 | 840 | 0 | 34 | 2472 | 0 | 0 | 0 | 35 | 75 | 164 | 20 | 30 | 33 | 67 | 2377 | 0 |
| SRR10953238 | 0 | 271 | 68 | 1020 | 1861 | 0 | 0 | 919 | 2815 | 9965 | 3139 | 7576 | 0 | 2324 | 100 | 152 | 0 | 345 | 56 | 3905 | 735 | 727 | 0 | 26 | 1866 | 0 | 0 | 0 | 30 | 63 | 113 | 17 | 27 | 34 | 63 | 1723 | 0 |
| SRR10953239 | 0 | 823 | 202 | 530 | 1413 | 0 | 0 | 1215 | 2972 | 12481 | 4573 | 9497 | 0 | 3077 | 113 | 159 | 0 | 348 | 56 | 4822 | 450 | 575 | 0 | 26 | 1633 | 0 | 0 | 0 | 92 | 220 | 436 | 60 | 90 | 100 | 156 | 2431 | 0 |
| SRR1107492 | 0 | 0 | 0 | 0 | 57 | 0 | 0 | 3268 | 1021 | 5571 | 5273 | 5329 | 0 | 0 | 167 | 130 | 0 | 236 | 0 | 390 | 2823 | 2679 | 0 | 61 | 11081 | 52 | 0 | 17 | 131 | 177 | 142 | 21 | 137 | 175 | 0 | 2098 | 0 |
| SRR1107493 | 0 | 0 | 0 | 0 | 54 | 0 | 0 | 6458 | 3085 | 6998 | 4164 | 5167 | 0 | 19 | 77 | 77 | 0 | 189 | 0 | 617 | 2564 | 2608 | 0 | 69 | 11675 | 76 | 0 | 0 | 61 | 21 | 12 | 28 | 22 | 28 | 0 | 2506 | 0 |
| SRR1107494 | 0 | 0 | 42 | 0 | 38 | 0 | 0 | 4229 | 1318 | 7228 | 332 | 3325 | 0 | 276 | 88 | 38 | 0 | 244 | 0 | 376 | 2104 | 1848 | 0 | 48 | 8306 | 51 | 0 | 0 | 53 | 13 | 12 | 18 | 49 | 15 | 68 | 1920 | 0 |
| SRR1107495 | 0 | 0 | 33 | 0 | 39 | 0 | 0 | 4010 | 1555 | 8061 | 291 | 3682 | 0 | 327 | 41 | 0 | 0 | 227 | 0 | 273 | 2001 | 1847 | 0 | 48 | 8796 | 53 | 0 | 15 | 58 | 11 | 54 | 15 | 32 | 18 | 71 | 1596 | 0 |
| SRR1107496 | 0 | 11 | 44 | 0 | 40 | 0 | 0 | 3641 | 865 | 5180 | 2435 | 1785 | 0 | 246 | 71 | 32 | 0 | 330 | 0 | 460 | 2303 | 2056 | 0 | 44 | 10515 | 40 | 0 | 0 | 63 | 98 | 53 | 15 | 51 | 111 | 86 | 2348 | 0 |
| SRR1107497 | 0 | 0 | 55 | 0 | 44 | 0 | 0 | 3188 | 1033 | 5567 | 2380 | 1931 | 0 | 314 | 45 | 90 | 0 | 281 | 0 | 629 | 2284 | 1915 | 0 | 46 | 11315 | 31 | 0 | 13 | 59 | 10 | 30 | 0 | 47 | 10 | 55 | 2559 | 0 |
| SRR1159512 | 0 | 64 | 68 | 0 | 13 | 0 | 0 | 4058 | 3149 | 1578 | 31063 | 27640 | 0 | 2365 | 0 | 0 | 0 | 1034 | 17 | 443 | 2052 | 6001 | 130 | 49217 | 6496 | 277 | 0 | 69 | 25 | 47 | 37 | 11 | 31 | 68 | 11 | 1358 | 69 |
| SRR1207050 | 0 | 0 | 65 | 0 | 0 | 0 | 0 | 2562 | 3630 | 832 | 269 | 233 | 0 | 0 | 126 | 114 | 0 | 622 | 0 | 55 | 24 | 82 | 29 | 417 | 22 | 0 | 0 | 32 | 0 | 0 | 0 | 0 | 0 | 0 | 0 | 83 | 0 |
| SRR1207051 | 0 | 31 | 111 | 0 | 0 | 0 | 0 | 8313 | 4226 | 31813 | 704 | 823 | 15 | 12 | 0 | 16 | 52 | 901 | 0 | 95 | 51 | 133 | 0 | 370 | 36 | 14 | 0 | 27 | 25 | 49 | 0 | 0 | 0 | 34 | 0 | 467 | 0 |
| SRR1207053 | 0 | 0 | 40 | 0 | 0 | 0 | 0 | 8184 | 3208 | 10874 | 85 | 80 | 0 | 0 | 0 | 0 | 23 | 776 | 0 | 40 | 14 | 31 | 0 | 115 | 35 | 0 | 0 | 728 | 20 | 11 | 0 | 0 | 0 | 11 | 0 | 194 | 0 |
| SRR1207054 | 0 | 0 | 19 | 0 | 0 | 0 | 0 | 3564 | 3342 | 4429 | 90 | 100 | 0 | 0 | 0 | 0 | 0 | 732 | 0 | 34 | 0 | 32 | 14 | 371 | 18 | 0 | 0 | 0 | 0 | 0 | 0 | 0 | 0 | 0 | 0 | 95 | 0 |

### Supp Table S8.4.Columns_DI_to_ES

| | | | | | | | | | | | | | | | | | | | | | | | | | | | | | | | | | | | | |
|---|---|---|---|---|---|---|---|---|---|---|---|---|---|---|---|---|---|---|---|---|---|---|---|---|---|---|---|---|---|---|---|---|---|---|---|---|---|
| SRR1207055 | 0 | 104 | 0 | 0 | 0 | 0 | 0 | 3612 | 1662 | 1375 | 41 | 28 | 0 | 0 | 0 | 0 | 23 | 884 | 0 | 12 | 0 | 22 | 0 | 283 | 28 | 0 | 0 | 165 | 20 | 0 | 0 | 0 | 0 | 0 | 0 | 19 | 0 |
| SRR1232072 | 0 | 0 | 0 | 0 | 22 | 12 | 0 | 9200 | 389 | 18686 | 97 | 60 | 0 | 404 | 685 | 519 | 16 | 62 | 0 | 93 | 1344 | 439 | 2237 | 8233 | 4211 | 0 | 12 | 0 | 0 | 0 | 0 | 0 | 0 | 0 | 0 | 600 | 124 |
| SRR1232073 | 0 | 0 | 0 | 0 | 24 | 29 | 0 | 7634 | 471 | 22361 | 1526 | 298 | 0 | 1349 | 261 | 219 | 29 | 314 | 0 | 79 | 1229 | 1100 | 754 | 2442 | 4516 | 0 | 27 | 0 | 0 | 0 | 0 | 0 | 0 | 0 | 0 | 566 | 33 |
| SRR1240812 | 0 | 88 | 13 | 0 | 0 | 0 | 0 | 47 | 29 | 260 | 4073 | 6663 | 0 | 883 | 23 | 26 | 0 | 3691 | 73 | 4993 | 4323 | 13959 | 116 | 131928 | 7933 | 277 | 0 | 21 | 40 | 46 | 31 | 12 | 17 | 28 | 22 | 2077 | 61 |
| SRR1240813 | 0 | 111 | 12 | 0 | 0 | 0 | 0 | 97 | 17 | 258 | 12752 | 6774 | 0 | 414 | 17 | 38 | 0 | 1884 | 30 | 4431 | 3107 | 10144 | 103 | 121796 | 5778 | 121 | 0 | 11 | 20 | 32 | 28 | 13 | 18 | 24 | 34 | 4232 | 62 |
| SRR1240814 | 0 | 269 | 48 | 0 | 13 | 0 | 0 | 146 | 86 | 1574 | 224 | 2309 | 0 | 886 | 0 | 0 | 0 | 1411 | 23 | 8599 | 2003 | 6500 | 75 | 69924 | 4737 | 0 | 0 | 20 | 39 | 116 | 202 | 39 | 78 | 133 | 104 | 11814 | 39 |
| SRR1240815 | 0 | 328 | 44 | 0 | 14 | 0 | 0 | 101 | 70 | 1099 | 1950 | 2244 | 0 | 1168 | 0 | 0 | 0 | 1429 | 22 | 9107 | 2257 | 6787 | 61 | 78801 | 4240 | 0 | 0 | 22 | 47 | 153 | 224 | 43 | 81 | 132 | 106 | 14948 | 42 |
| SRR1240816 | 0 | 162 | 48 | 0 | 0 | 0 | 0 | 20 | 0 | 92 | 18312 | 11783 | 0 | 195 | 38 | 88 | 0 | 128 | 158 | 2871 | 2359 | 3351 | 0 | 9223 | 8639 | 0 | 0 | 12 | 16 | 46 | 41 | 11 | 14 | 22 | 19 | 1926 | 40 |
| SRR1240817 | 0 | 1564 | 52 | 0 | 0 | 0 | 0 | 16 | 0 | 89 | 7845 | 10270 | 0 | 210 | 31 | 80 | 0 | 107 | 134 | 2648 | 2244 | 2914 | 0 | 8668 | 7843 | 0 | 0 | 111 | 139 | 465 | 373 | 129 | 123 | 185 | 187 | 2191 | 41 |
| SRR1240818 | 0 | 180 | 0 | 0 | 0 | 0 | 47 | 79 | 25 | 1203 | 2607 | 2064 | 0 | 3110 | 0 | 25 | 10 | 525 | 0 | 5210 | 5819 | 5125 | 128 | 142283 | 1978 | 43 | 70 | 0 | 13 | 19 | 39 | 0 | 13 | 18 | 13 | 22194 | 115 |
| SRR1241600 | 0 | 29 | 19 | 0 | 0 | 0 | 11 | 12 | 0 | 0 | 9971 | 5202 | 0 | 129 | 0 | 0 | 0 | 47 | 114 | 48 | 203 | 291 | 0 | 455 | 564 | 0 | 0 | 14 | 0 | 0 | 0 | 11 | 31 | 10 | 0 | 175 | 101 |
| SRR1241601 | 0 | 45 | 40 | 14 | 0 | 0 | 0 | 28 | 27 | 24 | 21211 | 5676 | 0 | 297 | 0 | 19 | 0 | 29 | 58 | 111 | 72 | 188 | 0 | 1370 | 938 | 0 | 0 | 81 | 14 | 0 | 0 | 0 | 132 | 156 | 0 | 403 | 43 |
| SRR1241602 | 0 | 27 | 12 | 0 | 13 | 0 | 0 | 16 | 17 | 15 | 8332 | 2937 | 0 | 214 | 0 | 0 | 0 | 27 | 40 | 66 | 173 | 693 | 0 | 2105 | 1093 | 0 | 11 | 179 | 11 | 0 | 0 | 0 | 38 | 26 | 0 | 178 | 0 |
| SRR1241603 | 0 | 16 | 195 | 0 | 0 | 0 | 0 | 139 | 54 | 1225 | 1961 | 1476 | 0 | 184 | 52 | 304 | 0 | 0 | 57 | 2632 | 1612 | 771 | 11 | 4296 | 14228 | 0 | 0 | 1292 | 1010 | 1883 | 1308 | 300 | 980 | 1252 | 53 | 5096 | 28 |
| SRR1241604 | 0 | 0 | 193 | 0 | 0 | 0 | 0 | 1412 | 1852 | 2212 | 4892 | 2492 | 0 | 559 | 62 | 257 | 0 | 0 | 44 | 1393 | 1715 | 1309 | 0 | 6875 | 13168 | 0 | 0 | 824 | 607 | 1087 | 714 | 185 | 680 | 919 | 62 | 2817 | 83 |
| SRR1241605 | 0 | 0 | 180 | 0 | 0 | 0 | 0 | 127 | 64 | 1635 | 962 | 1822 | 0 | 347 | 75 | 425 | 0 | 0 | 47 | 3107 | 1845 | 1000 | 0 | 5659 | 8898 | 0 | 0 | 1097 | 621 | 1038 | 910 | 327 | 865 | 1192 | 62 | 5984 | 61 |
| SRR1264353 | 0 | 13 | 0 | 0 | 86 | 0 | 0 | 320 | 1090 | 11376 | 350 | 10992 | 0 | 3140 | 0 | 0 | 0 | 3585 | 213 | 609 | 3395 | 13902 | 192 | 23618 | 3151 | 28 | 0 | 0 | 11 | 0 | 15 | 0 | 0 | 0 | 0 | 3108 | 1842 |
| SRR1264354 | 0 | 15 | 0 | 0 | 0 | 0 | 0 | 106 | 1727 | 26791 | 38247 | 21715 | 0 | 740 | 0 | 71 | 0 | 329 | 1352 | 388 | 6144 | 21994 | 35 | 9554 | 26000 | 0 | 0 | 0 | 16 | 19 | 15 | 16 | 12 | 0 | 0 | 1093 | 3417 |
| SRR1270153 | 0 | 144 | 0 | 0 | 0 | 0 | 0 | 1213 | 4080 | 1407 | 628 | 357 | 0 | 0 | 90 | 225 | 0 | 12916 | 0 | 17 | 1780 | 2326 | 87 | 382 | 62 | 235 | 17 | 0 | 0 | 0 | 0 | 0 | 0 | 0 | 0 | 14 | 0 |
| SRR1290784 | 0 | 16 | 113 | 0 | 0 | 0 | 0 | 253 | 40 | 1827 | 3306 | 11113 | 0 | 16 | 24 | 32 | 0 | 12 | 55 | 3629 | 1246 | 4663 | 0 | 14 | 19254 | 0 | 0 | 144 | 50 | 18 | 98 | 49 | 111 | 19 | 47 | 4410 | 40 |
| SRR1290785 | 0 | 33 | 76 | 0 | 0 | 0 | 0 | 181 | 18 | 1142 | 17349 | 14214 | 0 | 12 | 72 | 145 | 0 | 17 | 44 | 3274 | 1613 | 4233 | 0 | 20 | 18101 | 0 | 0 | 99 | 35 | 20 | 92 | 14 | 37 | 12 | 67 | 4398 | 41 |
| SRR1290786 | 0 | 300 | 71 | 0 | 0 | 0 | 0 | 167 | 18 | 1517 | 3031 | 10745 | 0 | 0 | 145 | 246 | 0 | 15 | 49 | 3333 | 1733 | 5488 | 0 | 60 | 19567 | 0 | 0 | 77 | 28 | 206 | 940 | 28 | 41 | 118 | 39 | 3675 | 36 |
| SRR1290787 | 0 | 230 | 108 | 0 | 0 | 0 | 0 | 415 | 41 | 3155 | 12803 | 9453 | 0 | 30 | 11 | 28 | 0 | 0 | 39 | 6555 | 1608 | 5094 | 0 | 14 | 18434 | 0 | 0 | 36 | 29 | 140 | 980 | 31 | 71 | 246 | 16 | 21977 | 31 |
| SRR1290788 | 0 | 225 | 97 | 0 | 0 | 0 | 0 | 124 | 19 | 1716 | 14304 | 11614 | 0 | 27 | 31 | 27 | 0 | 0 | 49 | 4561 | 1320 | 4419 | 0 | 13 | 17307 | 0 | 0 | 61 | 37 | 194 | 1011 | 18 | 65 | 185 | 11 | 16548 | 19 |
| SRR1290789 | 0 | 236 | 50 | 0 | 0 | 0 | 0 | 117 | 48 | 1251 | 2973 | 13136 | 0 | 0 | 11 | 0 | 0 | 0 | 52 | 2492 | 1112 | 4244 | 0 | 13 | 17143 | 0 | 0 | 90 | 32 | 273 | 1550 | 22 | 38 | 104 | 0 | 4217 | 30 |
| SRR1290790 | 0 | 273 | 87 | 0 | 0 | 0 | 0 | 170 | 30 | 1286 | 3719 | 13254 | 0 | 0 | 0 | 34 | 0 | 12 | 57 | 2186 | 1345 | 4150 | 0 | 13 | 16192 | 0 | 18 | 84 | 36 | 167 | 822 | 17 | 54 | 190 | 27 | 3268 | 13 |
| SRR1290791 | 0 | 14 | 159 | 0 | 0 | 0 | 0 | 269 | 31 | 1422 | 19423 | 18792 | 0 | 22 | 0 | 0 | 0 | 21 | 50 | 3819 | 1164 | 3227 | 0 | 11 | 15393 | 0 | 16 | 113 | 56 | 22 | 134 | 70 | 113 | 18 | 38 | 4195 | 28 |
| SRR1290792 | 0 | 28 | 93 | 0 | 0 | 0 | 0 | 215 | 49 | 1751 | 17881 | 13339 | 0 | 0 | 24 | 73 | 0 | 14 | 39 | 3200 | 1456 | 5077 | 0 | 14 | 20989 | 0 | 23 | 67 | 40 | 21 | 106 | 51 | 68 | 15 | 33 | 3416 | 32 |
| SRR1290793 | 0 | 487 | 70 | 0 | 0 | 0 | 0 | 174 | 16 | 666 | 7209 | 14932 | 0 | 0 | 13 | 0 | 0 | 18 | 40 | 4826 | 1594 | 4813 | 0 | 14 | 22093 | 0 | 0 | 62 | 42 | 342 | 1364 | 29 | 64 | 175 | 13 | 4070 | 43 |
| SRR1290794 | 0 | 280 | 46 | 0 | 0 | 0 | 0 | 54 | 16 | 337 | 18424 | 17130 | 0 | 0 | 0 | 27 | 0 | 12 | 71 | 3491 | 1408 | 3766 | 0 | 12 | 18657 | 0 | 0 | 22 | 31 | 163 | 908 | 23 | 29 | 56 | 14 | 4443 | 24 |
| SRR1290795 | 0 | 21 | 52 | 0 | 0 | 0 | 0 | 110 | 15 | 727 | 13563 | 13491 | 0 | 0 | 0 | 0 | 0 | 11 | 64 | 4955 | 963 | 2961 | 0 | 12 | 16132 | 0 | 0 | 28 | 18 | 24 | 124 | 20 | 43 | 10 | 0 | 7048 | 20 |
| SRR1290796 | 0 | 152 | 50 | 0 | 0 | 0 | 0 | 167 | 26 | 1398 | 4944 | 13500 | 0 | 13 | 180 | 132 | 0 | 13 | 49 | 3143 | 1140 | 3538 | 0 | 0 | 14702 | 0 | 0 | 20 | 14 | 231 | 725 | 16 | 25 | 103 | 15 | 6755 | 20 |
| SRR1290797 | 0 | 20 | 12 | 0 | 0 | 0 | 0 | 193 | 49 | 1579 | 12609 | 10092 | 0 | 0 | 27 | 37 | 0 | 0 | 37 | 3843 | 1157 | 3374 | 0 | 11 | 12821 | 0 | 0 | 33 | 13 | 31 | 86 | 10 | 46 | 22 | 41 | 10439 | 18 |
| SRR1290798 | 0 | 96 | 101 | 0 | 0 | 0 | 0 | 182 | 34 | 1082 | 3248 | 7824 | 0 | 30 | 121 | 111 | 0 | 0 | 29 | 2569 | 1075 | 3011 | 0 | 0 | 9772 | 0 | 0 | 72 | 53 | 630 | 1847 | 41 | 117 | 361 | 94 | 6597 | 29 |
| SRR1290799 | 0 | 57 | 62 | 0 | 0 | 0 | 0 | 171 | 39 | 1203 | 2071 | 3258 | 0 | 58 | 45 | 86 | 0 | 0 | 11 | 998 | 1028 | 2721 | 0 | 0 | 5329 | 0 | 12 | 0 | 31 | 23 | 63 | 24 | 10 | 50 | 40 | 8885 | 27 |
| SRR1290800 | 0 | 35 | 53 | 0 | 0 | 0 | 0 | 141 | 34 | 1066 | 2386 | 2981 | 0 | 56 | 245 | 463 | 0 | 0 | 14 | 979 | 1067 | 2909 | 0 | 0 | 7347 | 0 | 58 | 0 | 18 | 83 | 211 | 13 | 35 | 194 | 64 | 7336 | 26 |
| SRR1290801 | 0 | 12 | 10 | 0 | 0 | 0 | 0 | 298 | 36 | 2122 | 1348 | 2088 | 0 | 22 | 383 | 529 | 0 | 11 | 45 | 1824 | 1186 | 5015 | 0 | 18 | 21848 | 0 | 31 | 0 | 22 | 11 | 39 | 18 | 43 | 26 | 40 | 6941 | 37 |
| SRR1290802 | 0 | 18 | 126 | 0 | 0 | 0 | 0 | 111 | 15 | 1067 | 3097 | 11382 | 0 | 12 | 119 | 265 | 0 | 12 | 55 | 3470 | 1474 | 5370 | 0 | 23 | 24653 | 0 | 0 | 73 | 42 | 33 | 64 | 39 | 114 | 26 | 72 | 5816 | 24 |
| SRR1290803 | 0 | 261 | 71 | 0 | 0 | 0 | 0 | 124 | 0 | 1266 | 2330 | 8677 | 0 | 0 | 183 | 501 | 0 | 12 | 49 | 2971 | 1633 | 5681 | 0 | 16 | 22457 | 0 | 0 | 63 | 19 | 300 | 729 | 16 | 73 | 238 | 82 | 4401 | 19 |
| SRR1290804 | 0 | 54 | 77 | 0 | 11 | 0 | 0 | 127 | 10 | 509 | 5772 | 12967 | 0 | 0 | 134 | 209 | 0 | 23 | 44 | 9919 | 1968 | 4285 | 0 | 18 | 22343 | 0 | 0 | 105 | 53 | 37 | 106 | 36 | 52 | 17 | 82 | 40454 | 28 |
| SRR1290805 | 0 | 38 | 74 | 0 | 0 | 0 | 0 | 357 | 36 | 1529 | 13892 | 9844 | 0 | 12 | 103 | 115 | 0 | 22 | 44 | 11439 | 1573 | 4739 | 0 | 18 | 21925 | 0 | 0 | 54 | 26 | 43 | 152 | 46 | 85 | 35 | 63 | 47387 | 21 |
| SRR1290806 | 0 | 32 | 46 | 0 | 0 | 0 | 0 | 350 | 57 | 1710 | 13745 | 10700 | 0 | 0 | 24 | 33 | 0 | 12 | 65 | 5936 | 1744 | 5337 | 0 | 25 | 22302 | 0 | 0 | 104 | 21 | 51 | 125 | 31 | 80 | 23 | 40 | 23581 | 23 |
| SRR1290807 | 0 | 322 | 115 | 0 | 0 | 0 | 0 | 297 | 41 | 1927 | 13507 | 13237 | 0 | 0 | 14 | 35 | 0 | 20 | 60 | 5358 | 1365 | 4150 | 0 | 16 | 19926 | 0 | 0 | 81 | 26 | 516 | 1437 | 34 | 110 | 325 | 60 | 6429 | 26 |
| SRR1290808 | 0 | 407 | 109 | 0 | 0 | 0 | 0 | 304 | 89 | 1655 | 2076 | 9723 | 0 | 19 | 0 | 13 | 0 | 20 | 62 | 4808 | 1691 | 4387 | 0 | 15 | 20553 | 0 | 11 | 114 | 25 | 521 | 1232 | 67 | 164 | 404 | 47 | 6387 | 30 |
| SRR1290809 | 0 | 15 | 111 | 0 | 0 | 0 | 0 | 211 | 55 | 1496 | 4540 | 8448 | 0 | 21 | 45 | 87 | 0 | 14 | 62 | 3171 | 1484 | 4664 | 0 | 38 | 22408 | 0 | 0 | 99 | 59 | 30 | 90 | 59 | 11 | 28 | 104 | 5800 | 35 |
| SRR1292461 | 0 | 99 | 91 | 0 | 0 | 0 | 0 | 0 | 0 | 0 | 38120 | 64812 | 0 | 0 | 0 | 0 | 0 | 83 | 30 | 13 | 39 | 223 | 0 | 0 | 877 | 0 | 26 | 362 | 289 | 318 | 11 | 15 | 0 | 0 | 0 | 27 | 19 |
| SRR1304304 | 0 | 0 | 0 | 0 | 0 | 0 | 0 | 0 | 0 | 19 | 1284 | 2742 | 0 | 16 | 0 | 0 | 0 | 10 | 97 | 0 | 873 | 2593 | 23 | 20218 | 6376 | 0 | 53 | 109 | 352 | 277 | 267 | 25 | 30 | 42 | 0 | 41 | 65 |
| SRR1304307 | 0 | 0 | 0 | 0 | 0 | 0 | 0 | 29 | 12 | 40 | 455 | 3092 | 0 | 0 | 0 | 0 | 15 | 31 | 11 | 23 | 356 | 1720 | 97 | 11844 | 4807 | 0 | 35 | 127 | 180 | 253 | 30 | 0 | 0 | 0 | 0 | 29 | 92 |
| SRR1304309 | 0 | 0 | 0 | 0 | 0 | 0 | 0 | 0 | 15 | 27 | 281 | 2118 | 0 | 64 | 0 | 0 | 0 | 37 | 0 | 686 | 124 | 1419 | 94 | 20524 | 516 | 25 | 55 | 2581 | 315 | 865 | 91 | 0 | 15 | 28 | 0 | 727 | 0 |
| SRR1304313 | 0 | 0 | 48 | 63 | 26 | 0 | 0 | 589 | 666 | 5483 | 921 | 979 | 0 | 349 | 837 | 520 | 0 | 17 | 0 | 1324 | 1278 | 2461 | 0 | 826 | 6868 | 0 | 0 | 358 | 228 | 422 | 294 | 79 | 261 | 714 | 115 | 4569 | 0 |
| SRR1393733 | 0 | 52 | 0 | 0 | 0 | 125 | 0 | 198 | 143 | 250 | 71 | 107 | 0 | 425 | 0 | 0 | 0 | 4487 | 581 | 324 | 2230 | 5007 | 67 | 37463 | 16 | 59 | 59 | 854 | 8710 | 5861 | 1948 | 192 | 24 | 0 | 0 | 242 | 127 |
| SRR1510077 | 0 | 0 | 0 | 0 | 0 | 0 | 0 | 202 | 70 | 3352 | 74 | 245 | 0 | 3026 | 450 | 1312 | 0 | 117 | 12 | 699 | 1169 | 2878 | 444 | 11147 | 6586 | 0 | 0 | 0 | 0 | 0 | 0 | 0 | 13 | 0 | 0 | 2282 | 0 |
| SRR1510078 | 0 | 0 | 0 | 0 | 0 | 0 | 0 | 412 | 152 | 3881 | 81 | 220 | 0 | 2278 | 861 | 1207 | 0 | 126 | 12 | 565 | 1584 | 2929 | 414 | 11027 | 6910 | 35 | 16 | 0 | 0 | 0 | 0 | 0 | 0 | 0 | 0 | 2536 | 0 |
| SRR1510079 | 0 | 18 | 36 | 0 | 0 | 0 | 0 | 94 | 67 | 2296 | 59 | 38 | 0 | 1856 | 930 | 1949 | 0 | 97 | 13 | 293 | 1359 | 2329 | 514 | 9113 | 3238 | 0 | 0 | 0 | 0 | 0 | 0 | 0 | 0 | 0 | 0 | 991 | 0 |
| SRR1510080 | 0 | 0 | 0 | 0 | 0 | 0 | 0 | 327 | 104 | 3221 | 51 | 72 | 0 | 1050 | 27 | 51 | 0 | 121 | 0 | 856 | 1721 | 2445 | 382 | 10322 | 4048 | 186 | 0 | 0 | 0 | 0 | 0 | 0 | 12 | 17 | 0 | 3043 | 0 |
| SRR1510081 | 0 | 0 | 0 | 0 | 0 | 0 | 0 | 600 | 228 | 4583 | 79 | 146 | 0 | 1086 | 2767 | 3685 | 0 | 109 | 0 | 449 | 771 | 1042 | 594 | 6815 | 2796 | 23 | 15 | 0 | 0 | 0 | 0 | 13 | 0 | 0 | 15 | 3260 | 0 |
| SRR1510082 | 0 | 0 | 20 | 0 | 0 | 0 | 0 | 199 | 257 | 4927 | 103 | 147 | 0 | 635 | 5616 | 7976 | 0 | 107 | 18 | 761 | 1506 | 3546 | 414 | 8118 | 2902 | 22 | 0 | 0 | 0 | 10 | 0 | 0 | 12 | 19 | 13 | 3979 | 0 |
| SRR1510083 | 0 | 12 | 0 | 0 | 0 | 0 | 0 | 97 | 40 | 2096 | 114 | 120 | 0 | 1641 | 5760 | 14083 | 0 | 76 | 18 | 1039 | 818 | 3566 | 369 | 9152 | 5765 | 80 | 0 | 0 | 10 | 0 | 0 | 0 | 0 | 20 | 17 | 3122 | 0 |
| SRR1510084 | 0 | 31 | 0 | 0 | 0 | 0 | 0 | 227 | 72 | 5001 | 49 | 92 | 0 | 1704 | 53 | 35 | 0 | 106 | 16 | 202 | 2247 | 3818 | 250 | 7007 | 2695 | 0 | 0 | 0 | 11 | 0 | 0 | 0 | 0 | 28 | 0 | 658 | 0 |
| SRR1510085 | 0 | 24 | 0 | 0 | 0 | 0 | 14 | 99 | 59 | 5439 | 128 | 60 | 0 | 1835 | 1373 | 2615 | 0 | 103 | 16 | 651 | 1831 | 2979 | 305 | 8781 | 4323 | 35 | 0 | 0 | 0 | 0 | 0 | 0 | 0 | 0 | 0 | 1919 | 0 |
| SRR1510086 | 0 | 0 | 0 | 0 | 0 | 0 | 0 | 3288 | 1340 | 1678 | 20 | 16 | 0 | 629 | 274 | 318 | 0 | 105 | 14 | 100 | 2253 | 1603 | 568 | 11580 | 5849 | 36 | 33 | 0 | 0 | 0 | 0 | 0 | 0 | 10 | 0 | 353 | 0 |
| SRR1510087 | 0 | 13 | 28 | 0 | 0 | 0 | 0 | 80 | 0 | 1284 | 127 | 136 | 0 | 3197 | 1279 | 4396 | 0 | 103 | 11 | 2091 | 1399 | 4505 | 99 | 2460 | 1092 | 104 | 0 | 0 | 0 | 0 | 0 | 0 | 24 | 56 | 35 | 7882 | 0 |
| SRR1510088 | 0 | 11 | 0 | 0 | 0 | 0 | 0 | 338 | 47 | 748 | 76 | 66 | 0 | 1355 | 2838 | 1252 | 0 | 97 | 0 | 304 | 2221 | 4197 | 432 | 13613 | 7929 | 34 | 0 | 0 | 14 | 22 | 0 | 0 | 0 | 12 | 0 | 1201 | 0 |
| SRR1510089 | 0 | 15 | 11 | 0 | 0 | 0 | 0 | 70 | 35 | 1812 | 89 | 80 | 0 | 1996 | 1688 | 6003 | 0 | 71 | 22 | 1217 | 1231 | 4422 | 174 | 6424 | 2693 | 36 | 0 | 0 | 0 | 0 | 0 | 0 | 21 | 15 | 15 | 3726 | 0 |
| SRR1510090 | 0 | 39 | 29 | 0 | 0 | 0 | 0 | 118 | 118 | 10773 | 41 | 19 | 0 | 3648 | 2756 | 4519 | 0 | 242 | 0 | 362 | 2031 | 3252 | 455 | 9506 | 4227 | 39 | 10 | 0 | 0 | 0 | 21 | 0 | 0 | 0 | 26 | 856 | 0 |
| SRR1510091 | 0 | 0 | 16 | 0 | 0 | 0 | 0 | 168 | 56 | 1236 | 82 | 94 | 0 | 3433 | 1474 | 2683 | 0 | 132 | 0 | 184 | 1223 | 2023 | 227 | 5049 | 3038 | 156 | 22 | 0 | 0 | 0 | 0 | 0 | 0 | 22 | 16 | 682 | 0 |
| SRR1510092 | 0 | 10 | 13 | 0 | 0 | 0 | 0 | 22 | 40 | 6194 | 64 | 55 | 0 | 4347 | 352 | 1529 | 0 | 119 | 0 | 443 | 950 | 3609 | 45 | 1188 | 736 | 406 | 0 | 0 | 0 | 0 | 0 | 0 | 0 | 0 | 10 | 1499 | 0 |
| SRR1510093 | 0 | 0 | 18 | 0 | 0 | 0 | 0 | 292 | 137 | 6528 | 61 | 48 | 0 | 2523 | 2537 | 9386 | 0 | 100 | 13 | 18 | 1297 | 4386 | 358 | 9722 | 5283 | 102 | 0 | 0 | 0 | 0 | 0 | 0 | 0 | 12 | 10 | 58 | 0 |
| SRR1510094 | 0 | 0 | 0 | 0 | 0 | 0 | 0 | 360 | 164 | 5410 | 31 | 83 | 0 | 893 | 1112 | 1494 | 0 | 73 | 0 | 1003 | 723 | 1311 | 294 | 6207 | 3163 | 489 | 0 | 0 | 0 | 0 | 0 | 0 | 0 | 20 | 0 | 3706 | 0 |
| SRR1510095 | 0 | 22 | 19 | 0 | 0 | 0 | 0 | 41 | 28 | 1728 | 68 | 76 | 0 | 1875 | 2368 | 2255 | 0 | 132 | 0 | 113 | 1683 | 2820 | 235 | 7945 | 5612 | 233 | 0 | 0 | 0 | 0 | 0 | 0 | 19 | 19 | 11 | 458 | 0 |
| SRR1510096 | 0 | 18 | 0 | 0 | 0 | 0 | 0 | 97 | 0 | 3103 | 127 | 111 | 0 | 701 | 1079 | 3569 | 0 | 179 | 10 | 1083 | 2205 | 3296 | 337 | 7025 | 4530 | 27 | 0 | 0 | 0 | 0 | 0 | 0 | 0 | 0 | 0 | 4706 | 0 |
| SRR1510097 | 0 | 14 | 32 | 0 | 0 | 0 | 0 | 776 | 166 | 9768 | 487 | 795 | 0 | 1512 | 16 | 52 | 0 | 34 | 34 | 519 | 706 | 1317 | 88 | 2561 | 2882 | 0 | 18 | 0 | 0 | 0 | 14 | 0 | 0 | 45 | 18 | 1613 | 0 |
| SRR1510098 | 0 | 13 | 15 | 0 | 0 | 0 | 0 | 896 | 80 | 4861 | 127 | 578 | 0 | 1272 | 716 | 684 | 0 | 71 | 25 | 636 | 841 | 2793 | 113 | 3851 | 1981 | 0 | 0 | 0 | 0 | 0 | 13 | 0 | 13 | 38 | 25 | 2295 | 0 |
| SRR1510099 | 0 | 0 | 0 | 0 | 0 | 11 | 0 | 375 | 86 | 5452 | 334 | 429 | 0 | 2250 | 822 | 1387 | 0 | 19 | 39 | 867 | 224 | 2497 | 38 | 1224 | 770 | 0 | 0 | 0 | 0 | 0 | 0 | 0 | 27 | 52 | 0 | 2923 | 0 |
| SRR1510100 | 15 | 19 | 10 | 0 | 0 | 0 | 0 | 117 | 51 | 2270 | 756 | 3124 | 0 | 1496 | 1182 | 1919 | 16 | 24 | 0 | 394 | 617 | 4999 | 33 | 1421 | 729 | 0 | 0 | 0 | 0 | 0 | 0 | 0 | 0 | 24 | 10 | 1598 | 0 |
| SRR1510101 | 0 | 33 | 0 | 0 | 0 | 0 | 11 | 176 | 64 | 6717 | 124 | 59 | 0 | 6654 | 1030 | 995 | 0 | 145 | 23 | 280 | 1822 | 3219 | 112 | 3248 | 1618 | 25 | 0 | 0 | 0 | 0 | 19 | 0 | 22 | 24 | 14 | 1172 | 0 |
| SRR1510102 | 0 | 19 | 0 | 0 | 0 | 0 | 0 | 257 | 63 | 4089 | 233 | 216 | 0 | 1137 | 1459 | 1780 | 0 | 123 | 14 | 714 | 2154 | 3237 | 259 | 7643 | 5155 | 42 | 10 | 0 | 11 | 0 | 0 | 0 | 0 | 17 | 10 | 1964 | 0 |
| SRR1510103 | 0 | 16 | 0 | 0 | 0 | 0 | 0 | 168 | 58 | 1504 | 207 | 153 | 0 | 1772 | 837 | 779 | 0 | 173 | 14 | 142 | 2572 | 3761 | 272 | 8300 | 5661 | 0 | 0 | 0 | 12 | 20 | 12 | 15 | 0 | 20 | 13 | 588 | 0 |
| SRR1510104 | 0 | 16 | 0 | 0 | 0 | 0 | 0 | 66 | 96 | 3089 | 208 | 201 | 0 | 3262 | 507 | 768 | 0 | 67 | 17 | 404 | 1534 | 4572 | 74 | 2277 | 959 | 103 | 0 | 0 | 0 | 0 | 0 | 0 | 0 | 0 | 10 | 1565 | 0 |
| SRR1510105 | 0 | 44 | 0 | 0 | 0 | 0 | 0 | 30 | 12 | 625 | 95 | 216 | 0 | 428 | 4484 | 10049 | 0 | 218 | 18 | 1896 | 2194 | 3594 | 354 | 8926 | 5936 | 147 | 0 | 0 | 20 | 0 | 16 | 0 | 0 | 24 | 12 | 5290 | 0 |
| SRR1510106 | 0 | 0 | 0 | 0 | 0 | 0 | 0 | 299 | 73 | 4997 | 64 | 96 | 0 | 1651 | 1087 | 1120 | 0 | 133 | 0 | 540 | 1464 | 2776 | 150 | 4326 | 1847 | 389 | 0 | 0 | 0 | 0 | 10 | 0 | 17 | 19 | 0 | 2281 | 0 |
| SRR1510107 | 0 | 14 | 0 | 0 | 0 | 14 | 0 | 112 | 92 | 3444 | 34 | 29 | 0 | 1818 | 670 | 963 | 0 | 172 | 13 | 383 | 1886 | 2516 | 345 | 9007 | 4683 | 45 | 0 | 0 | 14 | 22 | 0 | 0 | 0 | 12 | 17 | 1155 | 0 |
| SRR1510108 | 0 | 17 | 0 | 0 | 0 | 0 | 0 | 147 | 20 | 1798 | 35 | 180 | 0 | 3290 | 7617 | 12147 | 0 | 55 | 14 | 539 | 856 | 4410 | 100 | 2750 | 1018 | 20 | 0 | 0 | 0 | 0 | 0 | 0 | 0 | 0 | 23 | 1704 | 0 |
| SRR1510109 | 0 | 19 | 0 | 0 | 0 | 0 | 0 | 270 | 43 | 3089 | 90 | 54 | 0 | 1091 | 2703 | 4466 | 0 | 87 | 16 | 528 | 1369 | 3281 | 428 | 12270 | 5830 | 18 | 0 | 0 | 0 | 13 | 14 | 0 | 18 | 31 | 12 | 2404 | 0 |
| SRR1510110 | 0 | 15 | 0 | 0 | 0 | 0 | 0 | 90 | 25 | 804 | 128 | 339 | 0 | 1334 | 3325 | 7767 | 0 | 42 | 13 | 1029 | 1161 | 5084 | 84 | 2902 | 894 | 13 | 0 | 0 | 0 | 0 | 0 | 0 | 0 | 0 | 0 | 4336 | 0 |
| SRR1510111 | 0 | 26 | 21 | 0 | 0 | 0 | 0 | 261 | 62 | 1971 | 121 | 129 | 0 | 2014 | 1333 | 1438 | 0 | 102 | 11 | 358 | 1838 | 1906 | 277 | 5896 | 3678 | 65 | 17 | 0 | 17 | 21 | 17 | 0 | 17 | 13 | 26 | 1634 | 0 |
| SRR1510112 | 0 | 0 | 35 | 0 | 0 | 0 | 0 | 224 | 105 | 1464 | 30 | 22 | 0 | 1046 | 1777 | 1577 | 0 | 172 | 0 | 692 | 1564 | 2076 | 453 | 15277 | 3324 | 0 | 13 | 0 | 11 | 0 | 16 | 0 | 0 | 24 | 16 | 3758 | 0 |
| SRR1510113 | 0 | 22 | 24 | 0 | 0 | 0 | 0 | 88 | 39 | 3126 | 139 | 112 | 0 | 1165 | 3069 | 9328 | 0 | 116 | 11 | 986 | 1173 | 2797 | 503 | 10865 | 5460 | 170 | 0 | 0 | 13 | 20 | 17 | 15 | 13 | 33 | 24 | 2989 | 0 |
| SRR1510114 | 0 | 0 | 12 | 0 | 0 | 0 | 0 | 426 | 215 | 2633 | 76 | 187 | 0 | 1507 | 2862 | 2223 | 0 | 148 | 11 | 311 | 1696 | 2336 | 453 | 7653 | 3554 | 23 | 20 | 0 | 0 | 0 | 0 | 0 | 0 | 12 | 0 | 1217 | 0 |
| SRR1510115 | 0 | 0 | 0 | 0 | 0 | 0 | 0 | 520 | 81 | 1518 | 34 | 36 | 0 | 1819 | 7516 | 6288 | 0 | 126 | 0 | 324 | 1677 | 2978 | 390 | 8620 | 5062 | 36 | 0 | 0 | 0 | 0 | 11 | 0 | 0 | 0 | 0 | 1437 | 0 |
| SRR1510116 | 0 | 0 | 19 | 0 | 0 | 0 | 0 | 180 | 34 | 1976 | 39 | 118 | 0 | 1921 | 2570 | 4732 | 0 | 31 | 11 | 1022 | 587 | 2882 | 124 | 3029 | 962 | 64 | 0 | 0 | 0 | 0 | 0 | 0 | 19 | 19 | 12 | 3488 | 0 |
| SRR1510117 | 0 | 0 | 13 | 0 | 0 | 0 | 0 | 87 | 16 | 678 | 94 | 101 | 0 | 1965 | 2423 | 6569 | 0 | 64 | 10 | 1827 | 631 | 3161 | 132 | 2032 | 619 | 16 | 0 | 0 | 0 | 0 | 0 | 0 | 21 | 20 | 17 | 8306 | 0 |
| SRR1510118 | 0 | 0 | 0 | 0 | 0 | 0 | 0 | 103 | 82 | 1839 | 88 | 95 | 0 | 679 | 4481 | 4193 | 0 | 99 | 21 | 875 | 1909 | 4359 | 177 | 3603 | 1491 | 71 | 0 | 0 | 0 | 0 | 11 | 0 | 16 | 32 | 25 | 4330 | 0 |
| SRR1510119 | 0 | 19 | 16 | 0 | 0 | 0 | 0 | 142 | 61 | 2855 | 81 | 123 | 0 | 415 | 3294 | 6292 | 0 | 165 | 11 | 594 | 2470 | 4310 | 363 | 10335 | 5344 | 334 | 0 | 0 | 10 | 17 | 16 | 0 | 19 | 0 | 16 | 2322 | 0 |
| SRR1510120 | 0 | 19 | 15 | 0 | 0 | 0 | 0 | 147 | 70 | 2062 | 149 | 195 | 0 | 472 | 2238 | 2686 | 0 | 115 | 0 | 819 | 2048 | 3830 | 375 | 9831 | 5641 | 259 | 12 | 0 | 0 | 12 | 15 | 0 | 26 | 29 | 25 | 3258 | 0 |
| SRR1510121 | 0 | 16 | 28 | 0 | 0 | 0 | 0 | 393 | 72 | 3678 | 326 | 362 | 0 | 1069 | 3442 | 3280 | 0 | 106 | 0 | 351 | 1478 | 1658 | 325 | 5756 | 2815 | 43 | 12 | 0 | 12 | 26 | 21 | 0 | 32 | 35 | 37 | 2152 | 0 |
| SRR1519906 | 87 | 0 | 0 | 0 | 24 | 0 | 0 | 915 | 1177 | 7322 | 0 | 262 | 0 | 0 | 0 | 0 | 79 | 0 | 0 | 0 | 16 | 16 | 231 | 820 | 1704 | 96 | 51 | 0 | 0 | 0 | 0 | 0 | 0 | 0 | 0 | 915 | 0 |
| SRR1519919 | 107 | 0 | 0 | 0 | 0 | 11 | 0 | 352 | 387 | 9773 | 12 | 1582 | 0 | 668 | 0 | 70 | 32 | 11 | 19 | 352 | 174 | 674 | 123 | 963 | 4108 | 1284 | 53 | 0 | 0 | 0 | 0 | 0 | 0 | 0 | 0 | 1512 | 0 |
| SRR1525198 | 0 | 0 | 74 | 0 | 0 | 0 | 0 | 579 | 219 | 2161 | 1176 | 1689 | 0 | 123 | 137 | 710 | 0 | 0 | 76 | 30 | 329 | 1414 | 16 | 10914 | 17401 | 0 | 11 | 2885 | 1095 | 2366 | 1095 | 109 | 844 | 1179 | 60 | 88 | 287 |
| SRR1525199 | 0 | 0 | 59 | 0 | 0 | 0 | 0 | 676 | 114 | 4322 | 1843 | 1911 | 0 | 114 | 142 | 1093 | 11 | 0 | 57 | 17 | 384 | 1429 | 20 | 11840 | 19085 | 0 | 0 | 2765 | 1219 | 3103 | 1186 | 84 | 723 | 979 | 78 | 72 | 372 |
| SRR1525200 | 0 | 0 | 61 | 0 | 0 | 0 | 0 | 621 | 106 | 4668 | 1344 | 1315 | 0 | 245 | 55 | 310 | 27 | 0 | 85 | 11 | 388 | 1659 | 22 | 16345 | 22394 | 0 | 34 | 2247 | 867 | 2766 | 1680 | 96 | 895 | 922 | 89 | 55 | 337 |
| SRR1525201 | 0 | 0 | 88 | 0 | 0 | 0 | 0 | 610 | 170 | 7272 | 777 | 1560 | 0 | 220 | 73 | 565 | 0 | 0 | 63 | 36 | 387 | 1436 | 33 | 12384 | 15551 | 0 | 20 | 1383 | 779 | 2390 | 1122 | 100 | 814 | 949 | 100 | 122 | 314 |
| SRR1525202 | 0 | 0 | 0 | 0 | 0 | 0 | 0 | 419 | 375 | 2033 | 271 | 6360 | 0 | 214 | 70 | 515 | 0 | 0 | 100 | 92 | 460 | 2564 | 17 | 15414 | 22931 | 0 | 0 | 68 | 24 | 72 | 48 | 11 | 28 | 42 | 13 | 144 | 264 |
| SRR1525203 | 0 | 0 | 64 | 0 | 0 | 0 | 0 | 450 | 177 | 3344 | 620 | 1496 | 0 | 175 | 147 | 712 | 16 | 0 | 92 | 38 | 330 | 1585 | 30 | 9822 | 14304 | 0 | 16 | 54 | 24 | 86 | 40 | 131 | 31 | 43 | 70 | 194 | 342 |
| SRR1563015 | 0 | 0 | 40 | 0 | 0 | 0 | 0 | 577 | 82 | 3411 | 2149 | 9702 | 0 | 82 | 223 | 232 | 0 | 0 | 10 | 680 | 1570 | 5495 | 0 | 0 | 23578 | 0 | 12 | 26 | 155 | 586 | 682 | 50 | 216 | 409 | 0 | 3009 | 39 |
| SRR1563018 | 0 | 0 | 23 | 0 | 0 | 0 | 0 | 285 | 44 | 479 | 239 | 5527 | 0 | 0 | 884 | 416 | 0 | 0 | 23 | 186 | 2293 | 2894 | 0 | 22 | 14659 | 0 | 20 | 131 | 20 | 65 | 113 | 135 | 27 | 97 | 49 | 1391 | 43 |
| SRR1563057 | 0 | 0 | 31 | 0 | 0 | 0 | 0 | 357 | 50 | 1155 | 1055 | 6947 | 10 | 75 | 440 | 357 | 10 | 0 | 25 | 258 | 2118 | 4916 | 0 | 0 | 15544 | 0 | 10 | 12 | 13 | 80 | 92 | 207 | 23 | 79 | 41 | 1213 | 71 |
| SRR1563058 | 12 | 48 | 0 | 0 | 0 | 19 | 0 | 297 | 66 | 3103 | 430 | 1774 | 0 | 464 | 142 | 41 | 0 | 0 | 20 | 25 | 5528 | 1819 | 47 | 709 | 14762 | 0 | 70 | 0 | 0 | 0 | 11 | 0 | 0 | 0 | 0 | 104 | 0 |
| SRR1563059 | 0 | 0 | 287 | 0 | 28 | 0 | 0 | 623 | 277 | 952 | 992 | 8724 | 0 | 75 | 159 | 225 | 10 | 25 | 0 | 1299 | 2509 | 3492 | 0 | 1016 | 15003 | 0 | 24 | 253 | 509 | 1884 | 913 | 104 | 421 | 1237 | 115 | 4599 | 0 |
| SRR1563060 | 0 | 30 | 0 | 0 | 78 | 12 | 17 | 294 | 93 | 5658 | 92 | 502 | 0 | 549 | 23 | 0 | 15 | 28 | 0 | 1244 | 1739 | 1387 | 18 | 1719 | 13080 | 0 | 12 | 0 | 0 | 0 | 0 | 0 | 0 | 0 | 0 | 13541 | 0 |
| SRR1563061 | 0 | 0 | 16 | 0 | 0 | 0 | 0 | 488 | 84 | 1167 | 1197 | 2714 | 0 | 320 | 480 | 1014 | 0 | 54 | 39 | 663 | 399 | 700 | 0 | 542 | 4840 | 0 | 0 | 18 | 29 | 48 | 36 | 0 | 16 | 39 | 0 | 3073 | 0 |
| SRR1563062 | 18 | 0 | 0 | 0 | 30 | 22 | 32 | 907 | 89 | 8164 | 15 | 253 | 0 | 535 | 15 | 30 | 0 | 40 | 15 | 149 | 240 | 510 | 60 | 2304 | 8203 | 0 | 27 | 0 | 0 | 0 | 0 | 0 | 0 | 0 | 0 | 788 | 0 |
| SRR1563063 | 0 | 0 | 121 | 0 | 34 | 0 | 0 | 815 | 81 | 2720 | 735 | 4908 | 0 | 135 | 31 | 23 | 0 | 148 | 13 | 1105 | 561 | 1598 | 0 | 24 | 12645 | 0 | 16 | 213 | 756 | 1762 | 1381 | 333 | 507 | 906 | 53 | 4672 | 0 |
| SRR1563064 | 0 | 33 | 0 | 0 | 42 | 15 | 10 | 528 | 86 | 6197 | 749 | 614 | 0 | 381 | 129 | 98 | 0 | 38 | 0 | 283 | 1340 | 1039 | 27 | 431 | 11283 | 0 | 99 | 0 | 0 | 14 | 0 | 0 | 0 | 0 | 11 | 1965 | 0 |
| SRR1575598 | 20 | 0 | 0 | 0 | 0 | 0 | 14 | 1281 | 43 | 0 | 406 | 790 | 0 | 0 | 0 | 0 | 20 | 690 | 193 | 0 | 1791 | 9288 | 219 | 67285 | 10994 | 64 | 0 | 44 | 0 | 0 | 0 | 0 | 0 | 0 | 0 | 0 | 118 |
| SRR1575599 | 43 | 0 | 0 | 0 | 0 | 0 | 20 | 52583 | 1398 | 0 | 56 | 225 | 0 | 0 | 0 | 0 | 0 | 85 | 70 | 0 | 1066 | 8366 | 151 | 62234 | 15632 | 78 | 0 | 16 | 17 | 19 | 17 | 0 | 0 | 0 | 0 | 0 | 97 |
| SRR1575601 | 10 | 0 | 0 | 0 | 16 | 0 | 20 | 2446 | 77 | 0 | 15 | 103 | 0 | 0 | 0 | 0 | 19 | 214 | 75 | 0 | 1071 | 3523 | 155 | 16177 | 2857 | 36 | 0 | 18 | 0 | 0 | 0 | 0 | 0 | 0 | 0 | 0 | 179 |
| SRR1575602 | 37 | 0 | 0 | 0 | 0 | 0 | 14 | 34968 | 1271 | 40 | 40 | 104 | 0 | 67 | 0 | 0 | 11 | 61 | 73 | 94 | 1278 | 5465 | 226 | 67834 | 16291 | 40 | 0 | 19 | 16 | 29 | 32 | 33 | 0 | 0 | 0 | 399 | 85 |
| SRR1592244 | 0 | 0 | 0 | 0 | 0 | 0 | 0 | 132 | 22 | 84 | 19419 | 31508 | 0 | 77 | 26 | 128 | 0 | 167 | 26 | 4202 | 202 | 331 | 0 | 51 | 256 | 1370 | 0 | 0 | 0 | 0 | 0 | 0 | 0 | 0 | 12 | 10601 | 0 |
| SRR1592245 | 0 | 0 | 0 | 0 | 0 | 0 | 0 | 39 | 0 | 802 | 6699 | 10600 | 0 | 317 | 81 | 421 | 0 | 100 | 29 | 3438 | 225 | 341 | 0 | 37 | 240 | 1716 | 0 | 0 | 0 | 0 | 0 | 0 | 0 | 0 | 24 | 8941 | 0 |
| SRR1592246 | 0 | 0 | 0 | 0 | 0 | 0 | 0 | 20 | 0 | 1376 | 1541 | 1978 | 0 | 529 | 47 | 171 | 0 | 99 | 30 | 1638 | 262 | 251 | 0 | 31 | 136 | 1136 | 0 | 0 | 0 | 0 | 0 | 0 | 0 | 0 | 28 | 4259 | 0 |
| SRR1592247 | 0 | 0 | 0 | 0 | 0 | 0 | 0 | 40 | 0 | 795 | 306 | 604 | 0 | 620 | 909 | 2642 | 0 | 27 | 23 | 4172 | 176 | 760 | 0 | 42 | 216 | 1904 | 0 | 0 | 0 | 0 | 0 | 0 | 0 | 0 | 22 | 10951 | 0 |
| SRR1592248 | 0 | 0 | 0 | 0 | 0 | 0 | 0 | 16 | 13 | 2293 | 202 | 232 | 0 | 670 | 374 | 881 | 0 | 79 | 21 | 2931 | 222 | 430 | 0 | 72 | 410 | 964 | 0 | 0 | 0 | 0 | 0 | 0 | 0 | 0 | 22 | 6905 | 0 |
| SRR1596226 | 0 | 15 | 0 | 0 | 0 | 16 | 0 | 208 | 26 | 355 | 157 | 1837 | 0 | 0 | 4393 | 2608 | 0 | 524 | 0 | 841 | 2099 | 3034 | 299 | 5946 | 4215 | 16 | 0 | 0 | 0 | 0 | 16 | 0 | 0 | 0 | 0 | 6516 | 0 |

### Supp Table S8.4.Columns_DI_to_ES

| | | | | | | | | | | | | | | | | | | | | | | | | | | | | | | | | | | | | | |
|---|---|---|---|---|---|---|---|---|---|---|---|---|---|---|---|---|---|---|---|---|---|---|---|---|---|---|---|---|---|---|---|---|---|---|---|---|---|
| SRR1596227 | 0 | 32 | 11 | 0 | 17 | 11 | 0 | 264 | 88 | 364 | 108 | 1720 | 0 | 0 | 7007 | 4495 | 0 | 666 | 0 | 703 | 1835 | 2526 | 410 | 3823 | 2731 | 0 | 0 | 0 | 0 | 0 | 0 | 0 | 0 | 14 | 0 | 6228 | 26 |
| SRR1596228 | 0 | 124 | 10 | 0 | 36 | 0 | 0 | 1725 | 1462 | 642 | 157 | 3355 | 0 | 0 | 11002 | 10507 | 0 | 577 | 0 | 841 | 1686 | 13 | 493 | 3724 | 33 | 12 | 0 | 0 | 0 | 0 | 0 | 0 | 10 | 13 | 0 | 8235 | 0 |
| SRR1596229 | 0 | 34 | 0 | 0 | 30 | 14 | 0 | 1851 | 1437 | 849 | 1894 | 3288 | 0 | 109 | 7991 | 6859 | 0 | 596 | 0 | 1023 | 1449 | 1444 | 557 | 3894 | 5659 | 15 | 0 | 0 | 0 | 0 | 0 | 0 | 0 | 14 | 0 | 8644 | 0 |
| SRR1596230 | 0 | 494 | 74 | 0 | 33 | 0 | 0 | 395 | 189 | 503 | 125 | 1770 | 0 | 63 | 13152 | 7250 | 0 | 766 | 0 | 764 | 1635 | 15 | 336 | 3234 | 43 | 0 | 0 | 0 | 17 | 24 | 17 | 10 | 10 | 17 | 0 | 5561 | 24 |
| SRR1596231 | 0 | 415 | 81 | 0 | 0 | 0 | 0 | 381 | 148 | 391 | 117 | 1565 | 0 | 85 | 18839 | 10815 | 0 | 714 | 0 | 592 | 1504 | 2492 | 319 | 2903 | 7091 | 0 | 0 | 0 | 41 | 49 | 16 | 20 | 0 | 12 | 0 | 5444 | 30 |
| SRR1608128 | 0 | 70 | 0 | 0 | 80 | 0 | 0 | 2184 | 3438 | 310 | 759 | 836 | 0 | 0 | 0 | 0 | 0 | 64 | 18 | 232 | 22 | 10 | 33 | 29 | 26 | 11 | 13 | 42 | 0 | 0 | 0 | 0 | 0 | 0 | 0 | 929 | 0 |
| SRR1608138 | 0 | 28 | 0 | 21 | 147 | 0 | 0 | 1349 | 1002 | 62 | 20 | 285 | 0 | 0 | 0 | 0 | 0 | 388 | 19 | 149 | 82 | 0 | 25 | 16 | 19 | 0 | 10 | 54 | 0 | 0 | 0 | 0 | 0 | 0 | 0 | 903 | 0 |
| SRR1608143 | 0 | 14 | 0 | 0 | 0 | 15 | 0 | 8827 | 7031 | 871 | 492 | 1491 | 0 | 0 | 0 | 0 | 0 | 193 | 0 | 113 | 29 | 0 | 0 | 0 | 25 | 0 | 28 | 12 | 0 | 0 | 0 | 0 | 0 | 0 | 0 | 450 | 18 |
| SRR1608150 | 0 | 51 | 0 | 0 | 140 | 0 | 0 | 1863 | 2516 | 408 | 37 | 966 | 0 | 0 | 0 | 0 | 0 | 213 | 0 | 27 | 38 | 0 | 40 | 38 | 23 | 0 | 24 | 27 | 0 | 0 | 0 | 0 | 0 | 0 | 0 | 354 | 0 |
| SRR1608158 | 0 | 15 | 0 | 0 | 74 | 0 | 0 | 1824 | 2699 | 133 | 59 | 193 | 0 | 0 | 0 | 0 | 0 | 125 | 0 | 237 | 30 | 0 | 231 | 102 | 14 | 25 | 0 | 18 | 0 | 0 | 0 | 0 | 0 | 0 | 0 | 1038 | 0 |
| SRR1608160 | 0 | 55 | 0 | 0 | 82 | 0 | 0 | 958 | 1563 | 134 | 50 | 202 | 0 | 0 | 0 | 34 | 0 | 74 | 0 | 319 | 37 | 0 | 78 | 39 | 13 | 0 | 0 | 19 | 0 | 0 | 0 | 0 | 0 | 0 | 0 | 1647 | 0 |
| SRR1608163 | 0 | 97 | 0 | 0 | 115 | 0 | 0 | 1379 | 1020 | 147 | 138 | 1709 | 0 | 0 | 0 | 0 | 0 | 170 | 0 | 37 | 57 | 12 | 100 | 85 | 31 | 0 | 0 | 45 | 0 | 0 | 0 | 0 | 0 | 0 | 0 | 138 | 0 |
| SRR1608165 | 0 | 11 | 0 | 0 | 91 | 0 | 0 | 1329 | 1938 | 181 | 214 | 415 | 0 | 0 | 0 | 0 | 0 | 64 | 0 | 32 | 23 | 0 | 33 | 20 | 0 | 0 | 0 | 73 | 17 | 0 | 0 | 0 | 0 | 0 | 0 | 246 | 0 |
| SRR1608166 | 0 | 82 | 0 | 0 | 128 | 0 | 13 | 1276 | 2352 | 125 | 375 | 600 | 0 | 0 | 0 | 0 | 0 | 84 | 17 | 250 | 37 | 0 | 41 | 23 | 16 | 0 | 0 | 26 | 0 | 0 | 0 | 0 | 0 | 0 | 0 | 1076 | 0 |
| SRR1608167 | 0 | 56 | 0 | 0 | 161 | 0 | 0 | 880 | 1123 | 137 | 243 | 395 | 0 | 0 | 0 | 0 | 0 | 45 | 12 | 288 | 29 | 0 | 18 | 10 | 14 | 0 | 0 | 17 | 0 | 0 | 0 | 0 | 0 | 0 | 0 | 774 | 0 |
| SRR1608168 | 0 | 183 | 0 | 0 | 29 | 0 | 12 | 1354 | 2610 | 297 | 17 | 162 | 0 | 0 | 0 | 0 | 0 | 20 | 0 | 43 | 163 | 0 | 36 | 31 | 16 | 0 | 0 | 28 | 18 | 0 | 0 | 0 | 0 | 0 | 0 | 259 | 0 |
| SRR1608169 | 0 | 101 | 0 | 0 | 173 | 0 | 0 | 543 | 696 | 164 | 297 | 481 | 0 | 0 | 10 | 20 | 0 | 78 | 0 | 338 | 70 | 0 | 35 | 18 | 12 | 0 | 13 | 40 | 0 | 0 | 0 | 0 | 0 | 0 | 0 | 2366 | 0 |
| SRR1608170 | 0 | 48 | 0 | 0 | 137 | 0 | 0 | 567 | 840 | 65 | 1055 | 1571 | 0 | 0 | 0 | 0 | 0 | 181 | 0 | 93 | 70 | 0 | 35 | 25 | 16 | 0 | 0 | 22 | 0 | 0 | 0 | 0 | 0 | 0 | 0 | 258 | 0 |
| SRR1608171 | 0 | 156 | 0 | 0 | 80 | 0 | 0 | 1577 | 2369 | 281 | 213 | 332 | 0 | 0 | 0 | 0 | 0 | 57 | 0 | 128 | 29 | 0 | 24 | 13 | 0 | 0 | 0 | 66 | 0 | 0 | 0 | 0 | 0 | 0 | 0 | 486 | 0 |
| SRR1608172 | 0 | 74 | 0 | 0 | 164 | 0 | 12 | 765 | 1138 | 169 | 27 | 187 | 0 | 0 | 0 | 0 | 0 | 166 | 18 | 71 | 43 | 0 | 20 | 15 | 20 | 14 | 11 | 18 | 0 | 0 | 0 | 0 | 0 | 0 | 0 | 1005 | 0 |
| SRR1608174 | 0 | 37 | 0 | 0 | 213 | 0 | 0 | 1835 | 3071 | 26 | 481 | 403 | 0 | 0 | 0 | 0 | 0 | 69 | 0 | 260 | 75 | 12 | 36 | 44 | 25 | 0 | 14 | 10 | 25 | 0 | 0 | 0 | 0 | 0 | 0 | 1431 | 0 |
| SRR1608175 | 0 | 208 | 0 | 0 | 138 | 0 | 0 | 1204 | 845 | 949 | 60 | 97 | 0 | 0 | 0 | 0 | 0 | 161 | 0 | 45 | 99 | 0 | 24 | 12 | 15 | 0 | 0 | 31 | 0 | 0 | 0 | 0 | 0 | 0 | 0 | 449 | 0 |
| SRR1608177 | 0 | 36 | 0 | 0 | 177 | 0 | 0 | 1011 | 1455 | 90 | 159 | 139 | 0 | 0 | 0 | 0 | 0 | 121 | 0 | 69 | 15 | 0 | 47 | 56 | 22 | 0 | 0 | 28 | 0 | 0 | 0 | 0 | 0 | 0 | 0 | 499 | 0 |
| SRR1608178 | 0 | 326 | 0 | 0 | 118 | 0 | 0 | 1089 | 2141 | 181 | 28 | 454 | 0 | 0 | 0 | 0 | 0 | 139 | 19 | 327 | 31 | 0 | 20 | 16 | 35 | 15 | 0 | 89 | 47 | 0 | 0 | 0 | 0 | 0 | 0 | 817 | 0 |
| SRR1608179 | 0 | 16 | 0 | 0 | 197 | 0 | 0 | 842 | 1358 | 285 | 23 | 421 | 0 | 0 | 27 | 14 | 0 | 92 | 0 | 68 | 23 | 0 | 14 | 0 | 17 | 0 | 0 | 119 | 0 | 0 | 0 | 0 | 0 | 11 | 0 | 761 | 0 |
| SRR1608180 | 0 | 331 | 0 | 0 | 218 | 0 | 0 | 1195 | 2011 | 189 | 37 | 423 | 0 | 0 | 0 | 29 | 0 | 127 | 0 | 131 | 17 | 0 | 36 | 24 | 18 | 0 | 11 | 39 | 32 | 11 | 0 | 0 | 0 | 0 | 0 | 597 | 0 |
| SRR1608181 | 0 | 102 | 0 | 0 | 156 | 0 | 0 | 923 | 1603 | 97 | 29 | 369 | 0 | 0 | 0 | 0 | 0 | 135 | 0 | 224 | 128 | 15 | 55 | 39 | 23 | 0 | 11 | 20 | 0 | 0 | 0 | 0 | 0 | 0 | 0 | 1050 | 0 |
| SRR1608182 | 0 | 72 | 0 | 0 | 121 | 0 | 0 | 917 | 1220 | 318 | 15 | 163 | 0 | 0 | 0 | 0 | 0 | 104 | 15 | 104 | 21 | 0 | 18 | 0 | 11 | 0 | 12 | 51 | 0 | 0 | 0 | 0 | 0 | 0 | 0 | 296 | 0 |
| SRR1608184 | 0 | 16 | 0 | 0 | 154 | 0 | 0 | 438 | 776 | 40 | 577 | 736 | 0 | 0 | 0 | 0 | 0 | 90 | 0 | 119 | 34 | 0 | 33 | 17 | 17 | 0 | 0 | 73 | 45 | 0 | 0 | 0 | 0 | 0 | 0 | 856 | 0 |
| SRR1608185 | 0 | 94 | 0 | 16 | 84 | 0 | 0 | 1639 | 1666 | 66 | 344 | 555 | 0 | 0 | 0 | 0 | 0 | 130 | 0 | 185 | 107 | 0 | 73 | 25 | 21 | 0 | 0 | 29 | 0 | 0 | 0 | 0 | 0 | 0 | 0 | 436 | 0 |
| SRR1608186 | 0 | 143 | 0 | 0 | 110 | 0 | 0 | 1708 | 2719 | 238 | 23 | 297 | 0 | 0 | 0 | 0 | 0 | 51 | 12 | 74 | 31 | 0 | 14 | 11 | 14 | 0 | 0 | 71 | 0 | 0 | 0 | 0 | 0 | 0 | 0 | 446 | 0 |
| SRR1608189 | 0 | 75 | 0 | 0 | 143 | 0 | 0 | 675 | 1476 | 76 | 52 | 751 | 0 | 0 | 0 | 0 | 0 | 115 | 14 | 59 | 40 | 0 | 20 | 10 | 19 | 0 | 11 | 36 | 0 | 0 | 0 | 0 | 0 | 0 | 0 | 261 | 0 |
| SRR1608190 | 0 | 11 | 0 | 0 | 126 | 0 | 0 | 973 | 778 | 22 | 126 | 1860 | 0 | 0 | 0 | 11 | 0 | 147 | 0 | 0 | 246 | 17 | 117 | 60 | 16 | 0 | 14 | 55 | 32 | 0 | 0 | 0 | 0 | 0 | 0 | 0 | 0 |
| SRR1608191 | 0 | 13 | 0 | 0 | 131 | 0 | 0 | 642 | 465 | 44 | 77 | 1959 | 0 | 0 | 0 | 0 | 0 | 220 | 13 | 122 | 112 | 20 | 57 | 31 | 46 | 0 | 0 | 70 | 0 | 0 | 0 | 0 | 0 | 0 | 0 | 742 | 0 |
| SRR1608192 | 0 | 125 | 0 | 0 | 111 | 0 | 0 | 1989 | 3271 | 241 | 144 | 433 | 0 | 0 | 0 | 0 | 0 | 101 | 14 | 176 | 56 | 0 | 46 | 24 | 13 | 0 | 15 | 26 | 0 | 0 | 0 | 0 | 0 | 0 | 0 | 1459 | 0 |
| SRR1608193 | 0 | 237 | 0 | 0 | 154 | 0 | 0 | 989 | 1931 | 177 | 233 | 243 | 0 | 0 | 0 | 0 | 0 | 100 | 13 | 56 | 54 | 0 | 34 | 28 | 17 | 0 | 0 | 37 | 10 | 0 | 0 | 0 | 0 | 0 | 0 | 373 | 0 |
| SRR1608194 | 0 | 245 | 0 | 0 | 170 | 0 | 0 | 1115 | 1904 | 268 | 10 | 297 | 0 | 0 | 15 | 0 | 0 | 120 | 26 | 74 | 63 | 0 | 23 | 21 | 14 | 0 | 0 | 78 | 15 | 0 | 0 | 0 | 0 | 11 | 0 | 402 | 0 |
| SRR1608196 | 0 | 41 | 0 | 0 | 86 | 0 | 0 | 952 | 1736 | 159 | 30 | 476 | 0 | 0 | 0 | 0 | 0 | 163 | 0 | 85 | 24 | 0 | 69 | 27 | 15 | 0 | 13 | 0 | 0 | 0 | 0 | 0 | 0 | 0 | 0 | 487 | 0 |
| SRR1608201 | 0 | 72 | 0 | 0 | 96 | 0 | 0 | 1307 | 1466 | 159 | 41 | 565 | 0 | 0 | 0 | 0 | 0 | 173 | 14 | 88 | 160 | 0 | 41 | 15 | 26 | 0 | 0 | 98 | 34 | 0 | 0 | 0 | 0 | 0 | 0 | 300 | 0 |
| SRR1608208 | 0 | 19 | 0 | 0 | 135 | 0 | 0 | 505 | 974 | 94 | 1726 | 2465 | 0 | 0 | 0 | 0 | 0 | 219 | 0 | 47 | 76 | 10 | 42 | 37 | 34 | 0 | 17 | 50 | 0 | 0 | 0 | 0 | 0 | 0 | 0 | 610 | 0 |
| SRR1608213 | 0 | 178 | 0 | 0 | 135 | 0 | 0 | 763 | 1697 | 57 | 438 | 496 | 0 | 0 | 0 | 0 | 0 | 117 | 15 | 172 | 29 | 0 | 27 | 30 | 34 | 0 | 0 | 267 | 24 | 0 | 0 | 0 | 0 | 0 | 0 | 934 | 0 |
| SRR1608214 | 0 | 16 | 0 | 0 | 108 | 0 | 0 | 1165 | 1804 | 153 | 264 | 319 | 0 | 0 | 0 | 0 | 0 | 166 | 15 | 42 | 29 | 0 | 23 | 11 | 14 | 0 | 0 | 59 | 0 | 0 | 0 | 0 | 0 | 0 | 0 | 236 | 0 |
| SRR1608222 | 0 | 0 | 0 | 0 | 20 | 0 | 0 | 2048 | 3104 | 32 | 96 | 576 | 0 | 0 | 0 | 0 | 0 | 43 | 0 | 352 | 20 | 0 | 411 | 113 | 0 | 14 | 0 | 19 | 0 | 0 | 0 | 0 | 0 | 0 | 0 | 640 | 0 |
| SRR1608232 | 0 | 45 | 0 | 0 | 89 | 0 | 0 | 4718 | 6150 | 618 | 228 | 586 | 0 | 0 | 0 | 0 | 0 | 74 | 11 | 358 | 26 | 0 | 251 | 133 | 15 | 21 | 0 | 10 | 0 | 0 | 0 | 0 | 0 | 0 | 0 | 716 | 0 |
| SRR1608237 | 0 | 48 | 0 | 0 | 44 | 12 | 0 | 9951 | 18496 | 1472 | 18 | 319 | 0 | 11 | 0 | 0 | 0 | 166 | 0 | 99 | 50 | 0 | 505 | 148 | 0 | 16 | 0 | 15 | 0 | 0 | 0 | 0 | 0 | 0 | 0 | 220 | 0 |
| SRR1608254 | 0 | 23 | 0 | 0 | 40 | 0 | 0 | 1186 | 2155 | 140 | 242 | 281 | 0 | 0 | 0 | 0 | 0 | 93 | 0 | 268 | 64 | 0 | 359 | 117 | 0 | 24 | 18 | 10 | 0 | 0 | 0 | 0 | 0 | 0 | 0 | 612 | 0 |
| SRR1608255 | 0 | 37 | 0 | 0 | 38 | 0 | 12 | 1520 | 2584 | 91 | 20 | 426 | 0 | 0 | 0 | 0 | 0 | 79 | 0 | 426 | 46 | 0 | 471 | 185 | 0 | 18 | 0 | 10 | 0 | 0 | 0 | 0 | 0 | 0 | 0 | 1074 | 0 |
| SRR1608256 | 0 | 12 | 0 | 0 | 27 | 13 | 0 | 528 | 959 | 173 | 14 | 441 | 0 | 0 | 0 | 0 | 0 | 154 | 10 | 221 | 95 | 10 | 644 | 175 | 0 | 25 | 15 | 0 | 0 | 0 | 0 | 0 | 0 | 0 | 0 | 787 | 0 |
| SRR1608257 | 0 | 45 | 0 | 0 | 88 | 0 | 0 | 1101 | 2251 | 220 | 17 | 294 | 0 | 0 | 0 | 0 | 0 | 194 | 0 | 171 | 54 | 0 | 171 | 77 | 16 | 12 | 0 | 11 | 0 | 0 | 0 | 0 | 0 | 0 | 0 | 465 | 0 |
| SRR1608260 | 0 | 43 | 0 | 0 | 42 | 0 | 0 | 1408 | 2569 | 152 | 11 | 400 | 0 | 0 | 0 | 0 | 0 | 150 | 11 | 190 | 97 | 14 | 431 | 184 | 0 | 10 | 15 | 15 | 0 | 0 | 0 | 0 | 0 | 0 | 0 | 628 | 0 |
| SRR1608263 | 0 | 56 | 0 | 0 | 29 | 0 | 0 | 982 | 1767 | 229 | 753 | 1243 | 0 | 0 | 0 | 0 | 0 | 75 | 0 | 327 | 86 | 13 | 547 | 218 | 0 | 14 | 0 | 27 | 0 | 0 | 0 | 0 | 0 | 0 | 0 | 720 | 0 |
| SRR1608264 | 0 | 22 | 0 | 0 | 34 | 0 | 0 | 2531 | 5036 | 723 | 181 | 491 | 0 | 0 | 0 | 0 | 0 | 62 | 10 | 362 | 30 | 0 | 402 | 175 | 10 | 20 | 0 | 20 | 0 | 0 | 0 | 0 | 0 | 0 | 0 | 517 | 0 |
| SRR1608266 | 0 | 56 | 0 | 0 | 67 | 0 | 0 | 1373 | 1601 | 92 | 595 | 1144 | 0 | 0 | 0 | 0 | 0 | 42 | 0 | 503 | 36 | 0 | 296 | 84 | 0 | 13 | 0 | 60 | 0 | 0 | 0 | 0 | 0 | 0 | 0 | 595 | 0 |
| SRR1608267 | 0 | 27 | 0 | 0 | 30 | 0 | 0 | 893 | 1839 | 189 | 26 | 595 | 0 | 0 | 0 | 0 | 0 | 42 | 0 | 298 | 43 | 0 | 535 | 173 | 0 | 39 | 0 | 17 | 12 | 0 | 0 | 0 | 0 | 0 | 0 | 487 | 0 |
| SRR1608269 | 0 | 71 | 0 | 0 | 37 | 0 | 0 | 4982 | 7373 | 707 | 306 | 404 | 0 | 0 | 0 | 0 | 0 | 87 | 0 | 337 | 55 | 0 | 515 | 160 | 0 | 13 | 13 | 57 | 22 | 0 | 0 | 0 | 0 | 0 | 0 | 1010 | 0 |
| SRR1608270 | 0 | 71 | 0 | 0 | 123 | 0 | 0 | 4369 | 8524 | 796 | 118 | 388 | 0 | 0 | 0 | 0 | 0 | 77 | 0 | 214 | 31 | 0 | 175 | 70 | 0 | 18 | 0 | 96 | 0 | 12 | 0 | 0 | 0 | 0 | 0 | 660 | 0 |
| SRR1608271 | 0 | 28 | 0 | 0 | 68 | 0 | 0 | 9800 | 6693 | 478 | 199 | 398 | 0 | 0 | 0 | 0 | 0 | 52 | 12 | 159 | 12 | 0 | 115 | 53 | 0 | 56 | 0 | 0 | 0 | 0 | 0 | 0 | 0 | 0 | 0 | 797 | 0 |
| SRR1608274 | 0 | 142 | 0 | 0 | 90 | 0 | 0 | 5064 | 6787 | 672 | 20 | 588 | 0 | 0 | 0 | 0 | 0 | 103 | 14 | 168 | 56 | 0 | 350 | 106 | 12 | 26 | 0 | 53 | 0 | 0 | 0 | 0 | 0 | 0 | 0 | 273 | 0 |
| SRR1608276 | 0 | 89 | 0 | 0 | 86 | 0 | 0 | 5699 | 9342 | 1439 | 264 | 676 | 0 | 0 | 0 | 0 | 0 | 75 | 13 | 176 | 49 | 0 | 346 | 123 | 11 | 26 | 0 | 14 | 14 | 0 | 14 | 0 | 0 | 0 | 0 | 969 | 0 |
| SRR1608277 | 0 | 99 | 0 | 0 | 88 | 0 | 0 | 5973 | 10366 | 546 | 345 | 603 | 0 | 0 | 0 | 0 | 0 | 81 | 12 | 0 | 64 | 12 | 321 | 157 | 16 | 17 | 0 | 18 | 0 | 0 | 0 | 0 | 0 | 0 | 0 | 0 | 0 |
| SRR1608278 | 0 | 29 | 0 | 0 | 31 | 0 | 0 | 2924 | 3688 | 133 | 565 | 1063 | 0 | 0 | 0 | 0 | 0 | 49 | 0 | 233 | 34 | 0 | 514 | 110 | 0 | 11 | 0 | 26 | 10 | 0 | 0 | 0 | 0 | 0 | 0 | 199 | 0 |
| SRR1608281 | 0 | 65 | 0 | 0 | 50 | 0 | 0 | 2873 | 3757 | 276 | 227 | 552 | 0 | 0 | 0 | 0 | 0 | 60 | 10 | 304 | 43 | 0 | 405 | 197 | 12 | 19 | 0 | 23 | 0 | 0 | 0 | 0 | 0 | 0 | 0 | 829 | 0 |
| SRR1608282 | 0 | 61 | 0 | 0 | 40 | 0 | 0 | 3272 | 5448 | 242 | 15 | 662 | 0 | 0 | 0 | 0 | 0 | 102 | 0 | 204 | 52 | 0 | 505 | 180 | 0 | 23 | 0 | 21 | 0 | 0 | 0 | 0 | 0 | 0 | 0 | 675 | 0 |
| SRR1608283 | 0 | 16 | 0 | 0 | 36 | 0 | 0 | 12169 | 20412 | 2094 | 15 | 218 | 0 | 0 | 0 | 0 | 0 | 72 | 0 | 262 | 30 | 0 | 358 | 140 | 0 | 25 | 0 | 19 | 0 | 0 | 0 | 0 | 0 | 0 | 0 | 523 | 0 |
| SRR1608286 | 0 | 21 | 0 | 0 | 41 | 0 | 0 | 2402 | 2953 | 236 | 433 | 1260 | 0 | 0 | 0 | 0 | 0 | 60 | 0 | 197 | 21 | 0 | 491 | 160 | 0 | 17 | 0 | 25 | 0 | 0 | 0 | 0 | 0 | 0 | 0 | 709 | 0 |
| SRR1608287 | 0 | 57 | 0 | 0 | 43 | 0 | 0 | 5092 | 7152 | 529 | 362 | 696 | 0 | 0 | 0 | 0 | 0 | 112 | 0 | 195 | 61 | 0 | 436 | 115 | 0 | 26 | 0 | 60 | 0 | 0 | 0 | 0 | 0 | 0 | 0 | 417 | 0 |
| SRR1608289 | 0 | 30 | 0 | 0 | 27 | 0 | 0 | 1901 | 3530 | 39 | 310 | 737 | 0 | 0 | 0 | 0 | 0 | 64 | 0 | 427 | 44 | 0 | 460 | 134 | 0 | 17 | 0 | 47 | 25 | 0 | 0 | 0 | 0 | 0 | 0 | 388 | 0 |
| SRR1608290 | 0 | 46 | 0 | 0 | 40 | 0 | 0 | 2415 | 4695 | 168 | 238 | 570 | 0 | 0 | 0 | 0 | 0 | 55 | 0 | 268 | 26 | 0 | 370 | 158 | 0 | 21 | 0 | 32 | 14 | 0 | 0 | 0 | 0 | 0 | 0 | 570 | 0 |
| SRR1608302 | 0 | 78 | 0 | 0 | 74 | 0 | 0 | 1061 | 2593 | 59 | 538 | 943 | 0 | 0 | 0 | 0 | 0 | 49 | 0 | 472 | 24 | 0 | 247 | 94 | 0 | 12 | 0 | 67 | 0 | 0 | 0 | 0 | 0 | 0 | 0 | 589 | 0 |
| SRR1608307 | 0 | 32 | 0 | 0 | 35 | 0 | 0 | 1807 | 3763 | 99 | 322 | 396 | 0 | 0 | 0 | 0 | 0 | 43 | 0 | 223 | 31 | 0 | 434 | 144 | 0 | 18 | 0 | 34 | 0 | 0 | 0 | 0 | 0 | 0 | 0 | 347 | 0 |
| SRR1608308 | 0 | 149 | 0 | 0 | 37 | 0 | 0 | 3772 | 6872 | 336 | 13 | 672 | 0 | 0 | 0 | 0 | 0 | 59 | 0 | 299 | 43 | 0 | 348 | 166 | 12 | 14 | 0 | 28 | 11 | 0 | 0 | 0 | 0 | 0 | 0 | 822 | 0 |
| SRR1634379 | 0 | 572 | 24 | 0 | 0 | 0 | 0 | 1228 | 69 | 788 | 303 | 536 | 0 | 179 | 23 | 325 | 0 | 1364 | 17 | 518 | 961 | 1467 | 395 | 3174 | 2822 | 0 | 0 | 0 | 0 | 0 | 0 | 0 | 0 | 0 | 0 | 3300 | 0 |
| SRR1634381 | 0 | 324 | 46 | 0 | 17 | 0 | 0 | 836 | 70 | 444 | 448 | 548 | 0 | 413 | 113 | 813 | 0 | 3031 | 0 | 331 | 2128 | 2382 | 437 | 6797 | 5302 | 40 | 0 | 0 | 0 | 0 | 0 | 13 | 0 | 0 | 0 | 1843 | 29 |
| SRR1634384 | 0 | 107 | 40 | 0 | 0 | 0 | 0 | 289 | 37 | 264 | 702 | 648 | 0 | 422 | 103 | 903 | 0 | 2696 | 0 | 943 | 1993 | 2669 | 312 | 8128 | 3322 | 0 | 0 | 0 | 0 | 0 | 0 | 0 | 0 | 0 | 0 | 3281 | 0 |
| SRR1634388 | 0 | 70 | 17 | 0 | 0 | 0 | 0 | 291 | 28 | 189 | 427 | 472 | 0 | 286 | 42 | 247 | 0 | 2439 | 0 | 319 | 1627 | 1755 | 201 | 3352 | 2546 | 24 | 0 | 0 | 0 | 0 | 0 | 0 | 0 | 0 | 0 | 1398 | 15 |
| SRR1634390 | 0 | 521 | 33 | 0 | 0 | 0 | 0 | 773 | 51 | 545 | 264 | 252 | 0 | 391 | 0 | 42 | 0 | 2814 | 0 | 187 | 2116 | 1808 | 360 | 7066 | 10733 | 27 | 0 | 0 | 0 | 19 | 14 | 12 | 0 | 0 | 0 | 995 | 31 |
| SRR1634392 | 0 | 315 | 31 | 0 | 0 | 0 | 0 | 393 | 29 | 250 | 195 | 539 | 0 | 240 | 61 | 306 | 0 | 2285 | 0 | 281 | 1864 | 2499 | 281 | 4464 | 4678 | 17 | 0 | 0 | 0 | 0 | 0 | 12 | 0 | 0 | 0 | 1211 | 28 |
| SRR1634394 | 0 | 396 | 30 | 0 | 0 | 0 | 0 | 617 | 48 | 290 | 288 | 392 | 0 | 551 | 0 | 46 | 0 | 2574 | 0 | 189 | 1440 | 1417 | 164 | 2828 | 3094 | 17 | 0 | 0 | 0 | 0 | 0 | 0 | 0 | 0 | 0 | 952 | 0 |
| SRR1634395 | 0 | 188 | 12 | 0 | 60 | 0 | 0 | 1041 | 97 | 616 | 240 | 350 | 0 | 171 | 101 | 461 | 0 | 2627 | 12 | 391 | 1645 | 1413 | 249 | 4120 | 15 | 14 | 0 | 0 | 0 | 23 | 21 | 17 | 0 | 0 | 0 | 1820 | 42 |
| SRR1634398 | 0 | 693 | 36 | 0 | 62 | 0 | 0 | 808 | 73 | 597 | 474 | 251 | 0 | 601 | 34 | 50 | 0 | 2287 | 15 | 152 | 1432 | 1253 | 107 | 1698 | 13 | 36 | 0 | 0 | 0 | 15 | 11 | 0 | 0 | 0 | 0 | 861 | 72 |
| SRR1634399 | 0 | 148 | 11 | 0 | 24 | 0 | 0 | 1018 | 92 | 588 | 580 | 675 | 0 | 219 | 17 | 107 | 0 | 2979 | 21 | 187 | 1990 | 2270 | 314 | 5179 | 9121 | 0 | 0 | 0 | 0 | 17 | 22 | 21 | 0 | 0 | 0 | 1246 | 27 |
| SRR1634400 | 0 | 450 | 24 | 0 | 13 | 0 | 0 | 1124 | 95 | 658 | 686 | 665 | 0 | 195 | 18 | 103 | 0 | 2933 | 15 | 191 | 2048 | 2191 | 315 | 5297 | 8937 | 20 | 0 | 0 | 0 | 26 | 21 | 21 | 0 | 0 | 0 | 1117 | 29 |
| SRR1634406 | 0 | 294 | 29 | 0 | 11 | 0 | 0 | 592 | 60 | 351 | 417 | 1361 | 0 | 387 | 15 | 104 | 0 | 2179 | 0 | 446 | 2132 | 2186 | 387 | 4173 | 4017 | 15 | 0 | 0 | 0 | 0 | 0 | 0 | 0 | 0 | 11 | 2535 | 15 |
| SRR1634409 | 0 | 266 | 11 | 0 | 83 | 0 | 0 | 861 | 56 | 464 | 143 | 279 | 0 | 238 | 13 | 35 | 0 | 1587 | 13 | 258 | 1734 | 1119 | 183 | 3414 | 16 | 29 | 11 | 11 | 0 | 23 | 17 | 13 | 0 | 0 | 0 | 1112 | 53 |
| SRR1634411 | 0 | 71 | 20 | 0 | 25 | 0 | 0 | 283 | 56 | 303 | 310 | 450 | 0 | 301 | 86 | 371 | 0 | 1896 | 0 | 936 | 2404 | 1622 | 378 | 8508 | 10 | 21 | 0 | 0 | 0 | 13 | 0 | 12 | 0 | 0 | 0 | 3755 | 13 |
| SRR1634412 | 0 | 71 | 20 | 0 | 19 | 0 | 0 | 254 | 25 | 309 | 185 | 482 | 0 | 286 | 91 | 382 | 0 | 1887 | 0 | 931 | 2355 | 1627 | 394 | 8533 | 15 | 19 | 0 | 0 | 0 | 12 | 0 | 18 | 0 | 0 | 0 | 3577 | 15 |
| SRR1634414 | 0 | 239 | 15 | 0 | 27 | 0 | 0 | 836 | 72 | 504 | 249 | 365 | 0 | 252 | 0 | 29 | 0 | 2527 | 13 | 213 | 1877 | 1426 | 271 | 4792 | 11 | 24 | 0 | 0 | 0 | 11 | 12 | 0 | 0 | 0 | 0 | 1096 | 64 |
| SRR1634415 | 0 | 176 | 15 | 0 | 26 | 0 | 0 | 917 | 61 | 468 | 238 | 611 | 0 | 117 | 0 | 22 | 0 | 1649 | 11 | 295 | 1433 | 1901 | 73 | 3243 | 6581 | 26 | 0 | 0 | 0 | 0 | 0 | 17 | 0 | 0 | 0 | 1727 | 15 |
| SRR1634417 | 0 | 619 | 23 | 0 | 18 | 0 | 0 | 886 | 49 | 429 | 235 | 368 | 0 | 31 | 23 | 115 | 0 | 1005 | 0 | 346 | 2649 | 2182 | 1027 | 15653 | 19 | 10 | 10 | 0 | 0 | 15 | 18 | 11 | 0 | 0 | 0 | 1983 | 28 |
| SRR1636959 | 0 | 134 | 32 | 0 | 0 | 0 | 14 | 446 | 24 | 450 | 498 | 2113 | 0 | 34 | 2007 | 5289 | 0 | 187 | 0 | 1234 | 2365 | 3120 | 205 | 5859 | 4761 | 301 | 55 | 0 | 12 | 0 | 22 | 0 | 34 | 71 | 40 | 22349 | 0 |
| SRR1636960 | 0 | 14 | 12 | 0 | 0 | 0 | 33 | 176 | 18 | 191 | 1312 | 1755 | 0 | 18 | 1722 | 6349 | 0 | 306 | 0 | 789 | 2674 | 2203 | 158 | 6238 | 7697 | 175 | 40 | 0 | 28 | 59 | 54 | 0 | 35 | 99 | 212 | 22686 | 0 |
| SRR1636961 | 0 | 11 | 11 | 0 | 0 | 0 | 22 | 203 | 17 | 185 | 406 | 2317 | 0 | 14 | 1782 | 5860 | 0 | 302 | 0 | 700 | 2790 | 2369 | 157 | 6583 | 8782 | 169 | 19 | 0 | 19 | 64 | 72 | 0 | 11 | 55 | 91 | 22803 | 0 |
| SRR1636962 | 0 | 22 | 34 | 0 | 0 | 0 | 0 | 342 | 14 | 392 | 665 | 2235 | 0 | 23 | 1279 | 1154 | 0 | 99 | 0 | 402 | 1764 | 4474 | 68 | 8586 | 21600 | 328 | 0 | 22 | 127 | 102 | 112 | 34 | 75 | 174 | 93 | 4072 | 0 |
| SRR1636963 | 0 | 11 | 26 | 0 | 0 | 0 | 0 | 793 | 39 | 460 | 2448 | 3467 | 0 | 25 | 520 | 575 | 0 | 115 | 0 | 505 | 1371 | 3806 | 63 | 6905 | 15423 | 402 | 0 | 22 | 88 | 70 | 86 | 36 | 65 | 172 | 120 | 5628 | 0 |
| SRR1636964 | 0 | 12 | 42 | 0 | 0 | 0 | 0 | 918 | 28 | 473 | 1419 | 4124 | 0 | 0 | 501 | 566 | 0 | 125 | 11 | 353 | 1257 | 4333 | 64 | 5953 | 13471 | 411 | 0 | 0 | 48 | 90 | 72 | 30 | 54 | 11 | 138 | 3414 | 0 |
| SRR1636965 | 0 | 28 | 0 | 0 | 0 | 0 | 156 | 595 | 57 | 924 | 2278 | 2927 | 0 | 1666 | 176 | 431 | 65 | 119 | 0 | 155 | 1314 | 2895 | 106 | 7088 | 13348 | 372 | 367 | 0 | 0 | 14 | 19 | 0 | 11 | 28 | 44 | 2372 | 0 |
| SRR1636966 | 0 | 0 | 0 | 0 | 0 | 0 | 131 | 1866 | 117 | 1533 | 27 | 850 | 0 | 766 | 50 | 50 | 51 | 62 | 11 | 17 | 826 | 3342 | 66 | 2989 | 10470 | 158 | 148 | 0 | 0 | 0 | 0 | 0 | 0 | 0 | 0 | 250 | 11 |
| SRR1636967 | 0 | 0 | 32 | 0 | 0 | 0 | 231 | 840 | 43 | 926 | 47 | 1508 | 0 | 496 | 151 | 86 | 55 | 86 | 14 | 2305 | 1012 | 4964 | 34 | 3212 | 8448 | 171 | 171 | 0 | 18 | 23 | 23 | 0 | 0 | 0 | 14 | 46254 | 0 |
| SRR1636968 | 0 | 13 | 27 | 0 | 0 | 0 | 12 | 480 | 27 | 520 | 3587 | 6677 | 0 | 106 | 1244 | 3381 | 0 | 227 | 11 | 203 | 2251 | 2658 | 49 | 12282 | 26983 | 35 | 33 | 12 | 63 | 280 | 370 | 27 | 64 | 191 | 128 | 4630 | 15 |
| SRR1636969 | 0 | 13 | 12 | 0 | 0 | 0 | 23 | 1067 | 21 | 479 | 2631 | 4305 | 0 | 195 | 448 | 1571 | 0 | 295 | 17 | 243 | 1854 | 2886 | 51 | 14339 | 23661 | 40 | 41 | 0 | 41 | 278 | 298 | 31 | 61 | 187 | 133 | 6660 | 16 |
| SRR1636970 | 0 | 16 | 14 | 0 | 0 | 0 | 11 | 979 | 36 | 309 | 1784 | 3040 | 0 | 235 | 488 | 1537 | 0 | 393 | 11 | 161 | 2001 | 3281 | 62 | 13615 | 22501 | 66 | 36 | 16 | 67 | 350 | 364 | 49 | 110 | 268 | 243 | 2671 | 36 |
| SRR1642941 | 0 | 1167 | 26 | 225 | 369 | 0 | 45 | 738 | 2213 | 3804 | 621 | 5957 | 0 | 873 | 3026 | 3073 | 43 | 936 | 32 | 1090 | 600 | 840 | 80 | 1981 | 1503 | 0 | 73 | 0 | 0 | 0 | 11 | 0 | 0 | 0 | 0 | 1807 | 13 |
| SRR1642942 | 0 | 1332 | 32 | 97 | 220 | 0 | 40 | 151 | 647 | 716 | 745 | 2444 | 0 | 3531 | 2130 | 7320 | 62 | 852 | 209 | 2080 | 1299 | 1306 | 0 | 129 | 1352 | 0 | 142 | 18 | 51 | 28 | 42 | 0 | 0 | 0 | 0 | 5410 | 26 |
| SRR1642943 | 0 | 509 | 20 | 247 | 386 | 12 | 19 | 2565 | 9490 | 12131 | 76 | 60 | 0 | 469 | 140 | 301 | 24 | 685 | 0 | 713 | 702 | 512 | 74 | 1506 | 1713 | 0 | 25 | 0 | 0 | 0 | 0 | 0 | 0 | 0 | 0 | 2140 | 0 |
| SRR1642944 | 0 | 1345 | 33 | 189 | 546 | 0 | 47 | 706 | 1504 | 4967 | 7223 | 12031 | 0 | 524 | 934 | 1572 | 54 | 775 | 30 | 1139 | 756 | 765 | 0 | 371 | 653 | 0 | 95 | 0 | 15 | 11 | 18 | 0 | 0 | 0 | 0 | 2256 | 41 |
| SRR1642945 | 0 | 1032 | 25 | 187 | 313 | 0 | 47 | 1098 | 3659 | 6656 | 117 | 258 | 0 | 748 | 46 | 94 | 75 | 650 | 12 | 380 | 1046 | 641 | 46 | 943 | 1431 | 0 | 38 | 0 | 0 | 0 | 0 | 0 | 0 | 0 | 0 | 927 | 0 |
| SRR1642946 | 0 | 545 | 10 | 180 | 618 | 0 | 13 | 1730 | 6400 | 6684 | 475 | 4219 | 0 | 1501 | 615 | 1874 | 23 | 328 | 44 | 368 | 1101 | 763 | 0 | 286 | 2398 | 0 | 28 | 0 | 0 | 0 | 0 | 0 | 0 | 0 | 0 | 970 | 18 |
| SRR1642947 | 0 | 475 | 23 | 86 | 246 | 0 | 17 | 514 | 2036 | 3087 | 985 | 1765 | 0 | 355 | 973 | 1680 | 28 | 513 | 22 | 840 | 745 | 992 | 57 | 1407 | 1480 | 0 | 21 | 0 | 0 | 0 | 0 | 0 | 0 | 0 | 0 | 1862 | 0 |
| SRR1642948 | 0 | 471 | 16 | 243 | 737 | 17 | 21 | 1307 | 3739 | 6097 | 312 | 1829 | 0 | 292 | 1939 | 4005 | 19 | 804 | 18 | 792 | 751 | 748 | 53 | 1212 | 1538 | 0 | 34 | 0 | 18 | 12 | 17 | 0 | 0 | 0 | 0 | 1365 | 54 |
| SRR1642949 | 0 | 1009 | 19 | 120 | 469 | 0 | 39 | 521 | 1271 | 2218 | 444 | 5098 | 0 | 527 | 865 | 1717 | 71 | 883 | 39 | 913 | 644 | 859 | 68 | 1086 | 936 | 0 | 40 | 0 | 0 | 0 | 0 | 0 | 0 | 0 | 0 | 1684 | 0 |
| SRR1642950 | 0 | 1224 | 23 | 195 | 584 | 0 | 41 | 328 | 1047 | 1215 | 1983 | 3778 | 0 | 77 | 2193 | 2737 | 37 | 717 | 0 | 663 | 903 | 665 | 30 | 967 | 1296 | 0 | 55 | 0 | 0 | 0 | 0 | 0 | 0 | 0 | 0 | 1257 | 0 |
| SRR1642951 | 0 | 844 | 23 | 268 | 472 | 0 | 13 | 912 | 2764 | 4398 | 530 | 5901 | 0 | 681 | 1284 | 2255 | 25 | 1072 | 25 | 1101 | 751 | 1052 | 61 | 1067 | 1231 | 11 | 27 | 0 | 0 | 0 | 12 | 0 | 0 | 0 | 0 | 2231 | 0 |
| SRR1642952 | 0 | 492 | 15 | 134 | 691 | 12 | 19 | 926 | 2652 | 3115 | 666 | 4145 | 0 | 202 | 791 | 1457 | 29 | 373 | 45 | 1133 | 428 | 364 | 24 | 1134 | 2537 | 0 | 39 | 0 | 16 | 0 | 11 | 0 | 0 | 0 | 0 | 2549 | 184 |
| SRR1642953 | 0 | 786 | 15 | 208 | 299 | 11 | 13 | 1614 | 6476 | 9798 | 1467 | 2408 | 0 | 980 | 87 | 220 | 11 | 821 | 11 | 820 | 1116 | 1236 | 100 | 2085 | 3037 | 0 | 19 | 0 | 17 | 15 | 16 | 0 | 0 | 0 | 0 | 2281 | 0 |
| SRR1642954 | 0 | 776 | 26 | 722 | 1481 | 14 | 34 | 1988 | 9207 | 11416 | 3576 | 3781 | 0 | 1608 | 1305 | 950 | 10 | 772 | 24 | 909 | 1083 | 946 | 56 | 2286 | 3316 | 0 | 17 | 20 | 12 | 22 | 32 | 16 | 0 | 0 | 0 | 1582 | 28 |
| SRR1642955 | 0 | 1290 | 23 | 98 | 251 | 0 | 45 | 1255 | 7608 | 11441 | 13 | 17 | 0 | 661 | 108 | 135 | 100 | 692 | 17 | 288 | 384 | 456 | 49 | 1451 | 1728 | 12 | 27 | 0 | 0 | 0 | 0 | 0 | 0 | 0 | 0 | 879 | 0 |
| SRR1642956 | 0 | 1965 | 48 | 154 | 703 | 0 | 71 | 813 | 2752 | 4548 | 7103 | 8329 | 0 | 2492 | 216 | 296 | 63 | 483 | 243 | 516 | 817 | 899 | 11 | 357 | 2379 | 0 | 90 | 14 | 66 | 26 | 46 | 0 | 0 | 0 | 0 | 1109 | 28 |
| SRR1642957 | 0 | 2475 | 157 | 148 | 925 | 0 | 27 | 9590 | 27883 | 51795 | 2236 | 1635 | 0 | 519 | 536 | 653 | 28 | 612 | 28 | 2171 | 1480 | 1023 | 92 | 1292 | 1623 | 0 | 58 | 0 | 31 | 19 | 32 | 0 | 0 | 0 | 0 | 3729 | 12 |
| SRR1642958 | 0 | 701 | 14 | 280 | 495 | 0 | 26 | 1995 | 5900 | 10712 | 3554 | 4612 | 0 | 3189 | 143 | 215 | 27 | 1127 | 34 | 1480 | 1373 | 899 | 90 | 1289 | 1113 | 0 | 21 | 0 | 19 | 0 | 22 | 0 | 0 | 0 | 0 | 3390 | 0 |
| SRR1642959 | 0 | 1966 | 28 | 188 | 544 | 18 | 42 | 1164 | 3229 | 4997 | 394 | 5385 | 0 | 893 | 316 | 451 | 62 | 842 | 26 | 1867 | 964 | 1191 | 63 | 1017 | 1128 | 0 | 98 | 0 | 12 | 0 | 12 | 0 | 0 | 0 | 0 | 3491 | 0 |
| SRR1642960 | 0 | 697 | 11 | 185 | 614 | 18 | 18 | 934 | 2403 | 4033 | 2273 | 2977 | 0 | 918 | 344 | 459 | 35 | 731 | 27 | 765 | 660 | 1034 | 49 | 1099 | 1323 | 0 | 57 | 0 | 12 | 0 | 12 | 0 | 0 | 0 | 0 | 1492 | 31 |
| SRR1642961 | 0 | 501 | 11 | 174 | 218 | 0 | 12 | 1164 | 4841 | 7961 | 224 | 715 | 0 | 563 | 14 | 63 | 17 | 774 | 11 | 400 | 880 | 856 | 68 | 1849 | 2087 | 0 | 32 | 0 | 11 | 0 | 14 | 0 | 0 | 0 | 0 | 960 | 0 |
| SRR1642962 | 0 | 898 | 11 | 239 | 659 | 0 | 13 | 480 | 1096 | 3622 | 9227 | 11126 | 0 | 376 | 449 | 1367 | 23 | 700 | 0 | 689 | 936 | 1122 | 27 | 1024 | 1908 | 0 | 25 | 0 | 19 | 12 | 16 | 0 | 0 | 0 | 0 | 1576 | 99 |
| SRR1642963 | 0 | 1443 | 58 | 179 | 452 | 19 | 34 | 2203 | 7730 | 9051 | 513 | 3546 | 0 | 543 | 209 | 213 | 53 | 588 | 18 | 1434 | 1799 | 1549 | 160 | 4618 | 5977 | 0 | 65 | 0 | 17 | 15 | 18 | 0 | 0 | 0 | 0 | 2451 | 0 |

### Supp Table S8.4.Columns_DI_to_ES

| | | | | | | | | | | | | | | | | | | | | | | | | | | | | | | | | | | | | | |
|---|---|---|---|---|---|---|---|---|---|---|---|---|---|---|---|---|---|---|---|---|---|---|---|---|---|---|---|---|---|---|---|---|---|---|---|---|---|
| SRR1642964 | 0 | 828 | 39 | 143 | 211 | 0 | 31 | 389 | 1691 | 2304 | 380 | 4676 | 0 | 243 | 525 | 554 | 30 | 988 | 0 | 982 | 2963 | 942 | 22 | 492 | 710 | 0 | 24 | 19 | 15 | 10 | 17 | 21 | 0 | 0 | 0 | 2770 | 0 |
| SRR1642965 | 0 | 873 | 0 | 211 | 541 | 14 | 20 | 2156 | 4371 | 3308 | 3527 | 5117 | 0 | 298 | 40 | 60 | 29 | 1160 | 31 | 536 | 701 | 699 | 101 | 1710 | 1326 | 0 | 50 | 0 | 22 | 0 | 14 | 0 | 0 | 0 | 0 | 527 | 11 |
| SRR1642966 | 0 | 2253 | 0 | 249 | 582 | 0 | 59 | 1417 | 3276 | 3870 | 461 | 1231 | 0 | 1203 | 46 | 74 | 84 | 644 | 52 | 404 | 562 | 1284 | 127 | 5765 | 7290 | 0 | 115 | 0 | 37 | 0 | 14 | 0 | 0 | 0 | 0 | 771 | 498 |
| SRR1642967 | 0 | 893 | 0 | 292 | 529 | 15 | 15 | 844 | 2120 | 2393 | 424 | 3112 | 0 | 733 | 0 | 0 | 28 | 907 | 29 | 357 | 939 | 1015 | 96 | 1427 | 1612 | 0 | 47 | 0 | 16 | 0 | 10 | 0 | 0 | 0 | 0 | 422 | 13 |
| SRR1642968 | 0 | 2989 | 0 | 302 | 705 | 11 | 46 | 689 | 2383 | 1572 | 385 | 5714 | 0 | 365 | 32 | 65 | 86 | 1093 | 34 | 697 | 1003 | 1018 | 87 | 1430 | 1116 | 0 | 115 | 12 | 27 | 0 | 13 | 0 | 0 | 0 | 0 | 721 | 0 |
| SRR1642969 | 0 | 836 | 0 | 313 | 375 | 11 | 11 | 1439 | 7359 | 4414 | 932 | 723 | 0 | 235 | 19 | 60 | 14 | 727 | 13 | 578 | 1072 | 1154 | 120 | 2665 | 3069 | 0 | 30 | 0 | 16 | 0 | 0 | 0 | 0 | 0 | 0 | 1100 | 0 |
| SRR1642970 | 0 | 896 | 14 | 212 | 1029 | 0 | 12 | 1014 | 2449 | 3075 | 2039 | 1025 | 0 | 426 | 345 | 583 | 19 | 883 | 19 | 345 | 970 | 1103 | 52 | 1728 | 2142 | 0 | 23 | 0 | 37 | 19 | 28 | 0 | 0 | 0 | 0 | 480 | 0 |
| SRR1642971 | 0 | 864 | 0 | 302 | 714 | 18 | 18 | 1299 | 6455 | 2758 | 2822 | 4575 | 0 | 246 | 25 | 13 | 20 | 727 | 24 | 229 | 959 | 1264 | 111 | 3396 | 3440 | 0 | 29 | 0 | 15 | 0 | 13 | 0 | 0 | 0 | 0 | 441 | 0 |
| SRR1642972 | 0 | 935 | 0 | 196 | 945 | 16 | 11 | 2936 | 6676 | 6027 | 744 | 8222 | 0 | 175 | 124 | 72 | 26 | 667 | 25 | 495 | 1600 | 954 | 52 | 1386 | 3187 | 0 | 43 | 12 | 18 | 17 | 26 | 0 | 0 | 0 | 0 | 608 | 70 |
| SRR1642973 | 0 | 1305 | 0 | 428 | 685 | 14 | 15 | 640 | 1384 | 1279 | 8420 | 11572 | 0 | 324 | 23 | 23 | 22 | 737 | 24 | 527 | 1182 | 1154 | 90 | 1959 | 2348 | 0 | 47 | 0 | 23 | 14 | 14 | 0 | 0 | 0 | 0 | 1031 | 0 |
| SRR1642974 | 0 | 4172 | 0 | 195 | 557 | 0 | 42 | 1432 | 4396 | 3120 | 626 | 11204 | 0 | 879 | 14 | 28 | 65 | 1213 | 30 | 411 | 908 | 1497 | 34 | 577 | 516 | 10 | 109 | 0 | 22 | 13 | 17 | 0 | 0 | 0 | 0 | 482 | 15 |
| SRR1642975 | 0 | 963 | 0 | 134 | 686 | 11 | 27 | 310 | 826 | 766 | 1531 | 17541 | 0 | 99 | 33 | 46 | 26 | 981 | 29 | 991 | 1273 | 1543 | 98 | 1387 | 1365 | 0 | 14 | 12 | 21 | 21 | 19 | 0 | 0 | 0 | 0 | 1327 | 0 |
| SRR1642976 | 0 | 1026 | 23 | 392 | 340 | 12 | 21 | 752 | 2448 | 2716 | 3866 | 5101 | 0 | 91 | 19 | 105 | 17 | 926 | 20 | 240 | 564 | 840 | 85 | 2566 | 1774 | 0 | 33 | 11 | 14 | 13 | 15 | 0 | 0 | 0 | 0 | 244 | 0 |
| SRR1642977 | 0 | 990 | 25 | 254 | 521 | 0 | 17 | 1476 | 5483 | 12330 | 663 | 941 | 0 | 1343 | 81 | 86 | 26 | 1023 | 37 | 391 | 775 | 1380 | 105 | 1788 | 933 | 0 | 17 | 0 | 19 | 11 | 26 | 0 | 0 | 0 | 0 | 618 | 0 |
| SRR1642978 | 0 | 2535 | 41 | 335 | 727 | 17 | 44 | 644 | 2249 | 5816 | 275 | 1211 | 0 | 594 | 706 | 690 | 31 | 510 | 0 | 1284 | 895 | 1067 | 12 | 511 | 1486 | 0 | 124 | 12 | 23 | 11 | 22 | 0 | 0 | 0 | 0 | 2596 | 0 |
| SRR1642979 | 0 | 2136 | 32 | 250 | 1167 | 0 | 22 | 2381 | 5808 | 8833 | 762 | 9057 | 0 | 403 | 439 | 260 | 36 | 1567 | 40 | 1638 | 1022 | 1355 | 113 | 1086 | 790 | 14 | 61 | 0 | 31 | 19 | 26 | 0 | 0 | 0 | 0 | 2058 | 51 |
| SRR1642980 | 0 | 1995 | 31 | 126 | 764 | 12 | 58 | 1213 | 3431 | 6188 | 2605 | 2053 | 0 | 413 | 910 | 1185 | 79 | 352 | 29 | 1640 | 372 | 499 | 0 | 553 | 806 | 0 | 128 | 15 | 38 | 38 | 53 | 21 | 0 | 0 | 0 | 3238 | 19 |
| SRR1642981 | 0 | 2146 | 22 | 134 | 393 | 14 | 47 | 1880 | 6188 | 7516 | 437 | 2515 | 0 | 2239 | 40 | 69 | 40 | 1058 | 27 | 356 | 860 | 1411 | 78 | 978 | 842 | 0 | 106 | 0 | 11 | 0 | 0 | 0 | 0 | 0 | 0 | 867 | 0 |
| SRR1642982 | 0 | 3239 | 36 | 183 | 794 | 0 | 53 | 1464 | 4779 | 15166 | 604 | 966 | 0 | 813 | 155 | 205 | 59 | 964 | 16 | 218 | 954 | 1712 | 39 | 1514 | 2046 | 0 | 146 | 0 | 23 | 17 | 23 | 0 | 0 | 0 | 0 | 451 | 41 |
| SRR1642983 | 0 | 2942 | 28 | 231 | 305 | 0 | 43 | 2074 | 8573 | 12179 | 453 | 407 | 0 | 1116 | 73 | 104 | 68 | 1041 | 0 | 459 | 1391 | 1576 | 50 | 911 | 1291 | 0 | 99 | 0 | 20 | 12 | 22 | 0 | 0 | 0 | 0 | 884 | 0 |
| SRR1642984 | 0 | 982 | 21 | 266 | 1618 | 0 | 14 | 2100 | 6821 | 11432 | 2154 | 2294 | 0 | 731 | 1517 | 848 | 21 | 1122 | 19 | 995 | 1723 | 927 | 37 | 670 | 1056 | 0 | 28 | 14 | 11 | 24 | 12 | 0 | 0 | 0 | 0 | 1851 | 33 |
| SRR1642985 | 0 | 1199 | 22 | 183 | 257 | 13 | 19 | 1035 | 2882 | 4321 | 1125 | 5375 | 0 | 1804 | 78 | 74 | 24 | 918 | 0 | 788 | 781 | 961 | 50 | 849 | 798 | 11 | 53 | 10 | 11 | 0 | 16 | 0 | 0 | 0 | 0 | 1662 | 0 |
| SRR1642986 | 0 | 1476 | 20 | 198 | 339 | 0 | 21 | 1934 | 6071 | 7433 | 3824 | 4335 | 0 | 2060 | 99 | 176 | 29 | 842 | 12 | 626 | 855 | 1504 | 39 | 831 | 739 | 0 | 73 | 0 | 15 | 0 | 17 | 0 | 0 | 0 | 0 | 1890 | 0 |
| SRR1642987 | 0 | 906 | 12 | 322 | 202 | 12 | 22 | 1485 | 5349 | 4348 | 5650 | 8621 | 0 | 409 | 200 | 334 | 35 | 853 | 20 | 643 | 546 | 950 | 37 | 890 | 1181 | 0 | 52 | 0 | 0 | 0 | 12 | 0 | 0 | 0 | 0 | 1636 | 0 |
| SRR1642988 | 0 | 3294 | 25 | 427 | 1127 | 10 | 40 | 3564 | 10916 | 19100 | 6558 | 6740 | 0 | 356 | 2226 | 1888 | 66 | 1048 | 43 | 356 | 1832 | 1448 | 13 | 238 | 1138 | 0 | 81 | 0 | 16 | 18 | 35 | 0 | 0 | 0 | 0 | 588 | 146 |
| SRR1657194 | 0 | 53 | 82 | 0 | 0 | 0 | 0 | 404 | 54 | 1223 | 13670 | 13121 | 0 | 408 | 27 | 50 | 0 | 49 | 61 | 747 | 3798 | 9206 | 12 | 20478 | 18173 | 0 | 0 | 18 | 23 | 54 | 27 | 25 | 106 | 18 | 33 | 2124 | 137 |
| SRR1657195 | 0 | 77 | 12 | 0 | 0 | 0 | 0 | 329 | 99 | 1838 | 15007 | 8767 | 0 | 609 | 17 | 42 | 0 | 78 | 60 | 1627 | 4998 | 9145 | 24 | 25004 | 21264 | 0 | 0 | 56 | 226 | 314 | 137 | 20 | 50 | 79 | 19 | 8124 | 124 |
| SRR1657196 | 0 | 607 | 41 | 0 | 0 | 0 | 0 | 367 | 69 | 1345 | 18748 | 11980 | 0 | 503 | 18 | 51 | 0 | 91 | 77 | 641 | 4681 | 8033 | 16 | 21311 | 20556 | 0 | 14 | 211 | 569 | 1409 | 630 | 66 | 249 | 354 | 30 | 2336 | 104 |
| SRR1657197 | 0 | 663 | 111 | 0 | 0 | 0 | 0 | 472 | 74 | 2520 | 1016 | 9165 | 0 | 766 | 236 | 368 | 0 | 33 | 12 | 707 | 5710 | 6402 | 20 | 25602 | 9829 | 0 | 0 | 152 | 166 | 345 | 221 | 14 | 97 | 401 | 180 | 1636 | 131 |
| SRR1657198 | 0 | 60 | 0 | 0 | 0 | 0 | 0 | 291 | 71 | 1305 | 27124 | 16592 | 0 | 569 | 0 | 0 | 0 | 33 | 17 | 303 | 4647 | 5447 | 40 | 28153 | 9236 | 0 | 21 | 25 | 30 | 61 | 27 | 19 | 40 | 51 | 0 | 402 | 78 |
| SRR1686806 | 0 | 0 | 0 | 0 | 25 | 0 | 73 | 0 | 0 | 7238 | 220 | 251 | 0 | 2490 | 268 | 390 | 30 | 437 | 20 | 1117 | 571 | 2398 | 822 | 32282 | 5336 | 15 | 51 | 0 | 0 | 0 | 0 | 0 | 0 | 0 | 0 | 19447 | 157 |
| SRR1707574 | 0 | 35 | 0 | 0 | 0 | 0 | 0 | 18612 | 24 | 0 | 3645 | 1206 | 0 | 26 | 0 | 0 | 18 | 0 | 0 | 462 | 262 | 525 | 13 | 3437 | 2826 | 21 | 0 | 92 | 0 | 11 | 11 | 0 | 37 | 98 | 0 | 673 | 0 |
| SRR1707575 | 0 | 44 | 12 | 0 | 0 | 0 | 0 | 194 | 35 | 146 | 17587 | 2996 | 0 | 2657 | 0 | 0 | 0 | 0 | 0 | 521 | 0 | 25 | 0 | 128 | 102 | 23 | 12 | 33 | 0 | 0 | 0 | 21 | 0 | 0 | 0 | 639 | 0 |
| SRR1708906 | 0 | 157 | 55 | 0 | 40 | 0 | 0 | 1305 | 423 | 3005 | 24266 | 15565 | 0 | 0 | 441 | 324 | 0 | 410 | 27 | 999 | 1088 | 3783 | 0 | 37 | 15619 | 0 | 18 | 58 | 47 | 421 | 774 | 37 | 186 | 573 | 42 | 2096 | 0 |
| SRR1708907 | 0 | 126 | 17 | 0 | 16 | 0 | 0 | 5732 | 440 | 2190 | 23128 | 15694 | 0 | 16 | 16 | 16 | 0 | 558 | 23 | 1014 | 894 | 4217 | 0 | 119 | 6889 | 0 | 19 | 17 | 36 | 16 | 37 | 31 | 51 | 102 | 0 | 939 | 0 |
| SRR1708908 | 0 | 120 | 0 | 0 | 14 | 12 | 0 | 3053 | 224 | 931 | 16677 | 12203 | 0 | 14 | 0 | 0 | 0 | 503 | 32 | 541 | 870 | 3547 | 0 | 35 | 7462 | 0 | 0 | 36 | 29 | 10 | 23 | 14 | 0 | 19 | 0 | 931 | 0 |
| SRR1708909 | 0 | 16 | 32 | 0 | 12 | 0 | 0 | 2414 | 260 | 2263 | 6150 | 4131 | 0 | 17 | 101 | 40 | 15 | 467 | 31 | 1865 | 790 | 3863 | 0 | 131 | 4986 | 0 | 0 | 0 | 0 | 14 | 23 | 16 | 33 | 13 | 10 | 2441 | 0 |
| SRR1708910 | 0 | 146 | 13 | 0 | 24 | 0 | 0 | 7255 | 420 | 1124 | 2829 | 4613 | 0 | 0 | 0 | 0 | 0 | 365 | 26 | 361 | 962 | 4065 | 0 | 12 | 5620 | 0 | 0 | 22 | 41 | 128 | 19 | 20 | 0 | 26 | 0 | 903 | 0 |
| SRR1708911 | 0 | 236 | 20 | 0 | 17 | 12 | 0 | 3460 | 228 | 558 | 1752 | 12348 | 0 | 0 | 0 | 11 | 0 | 549 | 25 | 353 | 1099 | 5034 | 0 | 15 | 10361 | 0 | 0 | 24 | 24 | 199 | 212 | 0 | 12 | 20 | 0 | 467 | 0 |
| SRR1708912 | 0 | 124 | 26 | 0 | 33 | 0 | 0 | 2249 | 1402 | 1845 | 3522 | 21006 | 0 | 50 | 61 | 30 | 0 | 432 | 18 | 1059 | 1055 | 3551 | 0 | 50 | 13828 | 0 | 14 | 58 | 35 | 118 | 323 | 12 | 35 | 69 | 0 | 1361 | 0 |
| SRR1708913 | 0 | 14 | 0 | 0 | 31 | 0 | 0 | 4739 | 1418 | 738 | 3757 | 10887 | 0 | 24 | 15 | 0 | 0 | 604 | 23 | 471 | 855 | 3114 | 0 | 167 | 9047 | 0 | 0 | 59 | 23 | 82 | 11 | 0 | 0 | 12 | 0 | 379 | 0 |
| SRR1708914 | 0 | 159 | 0 | 0 | 45 | 0 | 0 | 1751 | 897 | 1230 | 1211 | 11284 | 0 | 29 | 43 | 14 | 0 | 462 | 19 | 1562 | 1225 | 2641 | 0 | 71 | 13647 | 0 | 0 | 65 | 50 | 189 | 11 | 20 | 79 | 99 | 0 | 4384 | 0 |
| SRR1708915 | 0 | 128 | 39 | 0 | 63 | 0 | 0 | 2193 | 1648 | 1362 | 3246 | 9525 | 0 | 0 | 175 | 56 | 0 | 406 | 30 | 1215 | 897 | 3018 | 0 | 254 | 11673 | 0 | 0 | 50 | 26 | 109 | 205 | 28 | 76 | 185 | 0 | 2409 | 0 |
| SRR1708916 | 0 | 138 | 0 | 0 | 35 | 0 | 0 | 1400 | 350 | 637 | 2657 | 12294 | 0 | 0 | 0 | 0 | 0 | 524 | 31 | 870 | 1202 | 3626 | 0 | 21 | 13720 | 0 | 0 | 41 | 70 | 192 | 496 | 0 | 43 | 60 | 0 | 1346 | 0 |
| SRR1708917 | 0 | 108 | 24 | 0 | 33 | 0 | 0 | 1906 | 777 | 1616 | 2425 | 12111 | 0 | 0 | 39 | 0 | 0 | 414 | 30 | 1310 | 1226 | 3524 | 0 | 22 | 14856 | 0 | 0 | 42 | 42 | 148 | 280 | 28 | 54 | 144 | 0 | 2032 | 0 |
| SRR1708918 | 0 | 216 | 0 | 0 | 42 | 0 | 0 | 1603 | 508 | 1171 | 34393 | 19540 | 0 | 22 | 0 | 11 | 13 | 427 | 22 | 597 | 873 | 2872 | 0 | 111 | 11804 | 0 | 13 | 78 | 39 | 88 | 274 | 0 | 0 | 26 | 0 | 796 | 0 |
| SRR1708919 | 0 | 138 | 30 | 0 | 56 | 0 | 0 | 1222 | 1101 | 2556 | 13993 | 9543 | 0 | 56 | 47 | 19 | 14 | 546 | 27 | 1763 | 1069 | 2459 | 0 | 495 | 13457 | 0 | 0 | 66 | 66 | 136 | 11 | 0 | 30 | 102 | 0 | 1726 | 0 |
| SRR1708920 | 0 | 128 | 25 | 0 | 64 | 0 | 0 | 3086 | 1291 | 1710 | 17599 | 9100 | 0 | 24 | 42 | 12 | 0 | 419 | 25 | 1031 | 1266 | 3169 | 0 | 163 | 12248 | 0 | 0 | 35 | 40 | 97 | 173 | 31 | 25 | 83 | 0 | 2825 | 0 |
| SRR1708921 | 14 | 115 | 30 | 0 | 46 | 0 | 0 | 2006 | 1164 | 2544 | 1835 | 7382 | 0 | 78 | 78 | 17 | 12 | 395 | 15 | 1615 | 1080 | 3179 | 0 | 501 | 11983 | 0 | 0 | 38 | 71 | 85 | 206 | 16 | 51 | 131 | 0 | 2892 | 0 |
| SRR1708922 | 0 | 125 | 49 | 0 | 42 | 0 | 0 | 1415 | 413 | 1791 | 16380 | 8068 | 0 | 0 | 44 | 29 | 0 | 461 | 32 | 1068 | 1276 | 3133 | 0 | 47 | 14034 | 0 | 0 | 44 | 39 | 148 | 14 | 28 | 65 | 86 | 0 | 1488 | 0 |
| SRR1708923 | 0 | 186 | 49 | 0 | 30 | 0 | 0 | 2217 | 739 | 4053 | 15841 | 11395 | 0 | 48 | 72 | 12 | 0 | 476 | 23 | 1871 | 1212 | 3612 | 0 | 44 | 14197 | 0 | 0 | 24 | 30 | 97 | 216 | 12 | 33 | 73 | 0 | 2086 | 0 |
| SRR1708924 | 0 | 126 | 24 | 0 | 30 | 0 | 0 | 2250 | 195 | 4189 | 5387 | 11047 | 0 | 35 | 91 | 44 | 0 | 486 | 20 | 809 | 1285 | 4116 | 0 | 35 | 10498 | 0 | 22 | 28 | 17 | 18 | 39 | 28 | 78 | 15 | 0 | 1599 | 0 |
| SRR1708925 | 0 | 161 | 41 | 0 | 30 | 0 | 0 | 4156 | 458 | 5764 | 2942 | 7735 | 0 | 69 | 41 | 32 | 0 | 463 | 19 | 1517 | 961 | 3475 | 0 | 167 | 8730 | 0 | 14 | 23 | 30 | 24 | 30 | 32 | 79 | 11 | 0 | 2520 | 0 |
| SRR1708926 | 0 | 144 | 36 | 0 | 19 | 0 | 0 | 7587 | 434 | 4449 | 5134 | 3864 | 0 | 83 | 251 | 151 | 0 | 426 | 29 | 709 | 711 | 3615 | 0 | 31 | 7118 | 0 | 0 | 17 | 39 | 157 | 38 | 41 | 88 | 14 | 58 | 2219 | 0 |
| SRR1708927 | 0 | 134 | 25 | 0 | 22 | 0 | 0 | 1668 | 341 | 2820 | 3028 | 12919 | 0 | 45 | 61 | 30 | 0 | 511 | 25 | 1365 | 992 | 3534 | 0 | 166 | 11735 | 0 | 0 | 41 | 59 | 410 | 562 | 16 | 88 | 177 | 14 | 1706 | 0 |
| SRR1708928 | 0 | 132 | 19 | 0 | 35 | 0 | 0 | 1175 | 301 | 5371 | 9778 | 9107 | 0 | 45 | 45 | 0 | 0 | 537 | 18 | 1130 | 1289 | 3447 | 0 | 13 | 13712 | 0 | 0 | 38 | 60 | 545 | 1170 | 31 | 50 | 136 | 0 | 2644 | 0 |
| SRR1708929 | 0 | 172 | 0 | 0 | 16 | 0 | 0 | 1856 | 209 | 4037 | 2552 | 13866 | 0 | 44 | 11 | 0 | 0 | 474 | 20 | 870 | 1644 | 5126 | 0 | 16 | 12802 | 0 | 0 | 14 | 20 | 129 | 19 | 18 | 35 | 62 | 0 | 1828 | 0 |
| SRR1708930 | 0 | 155 | 15 | 0 | 33 | 0 | 0 | 3699 | 1892 | 4260 | 3660 | 7112 | 0 | 70 | 286 | 147 | 0 | 371 | 24 | 667 | 1032 | 3492 | 0 | 48 | 13026 | 0 | 13 | 59 | 42 | 10 | 11 | 0 | 45 | 185 | 0 | 1899 | 0 |
| SRR1708931 | 0 | 154 | 23 | 0 | 37 | 0 | 0 | 3600 | 1204 | 3114 | 20431 | 10558 | 0 | 83 | 276 | 133 | 0 | 438 | 20 | 928 | 1059 | 3483 | 0 | 184 | 11780 | 0 | 0 | 38 | 41 | 210 | 402 | 0 | 49 | 128 | 0 | 2115 | 0 |
| SRR1708932 | 0 | 206 | 0 | 0 | 42 | 0 | 0 | 2324 | 1370 | 831 | 30604 | 23075 | 0 | 34 | 0 | 11 | 0 | 454 | 18 | 494 | 1173 | 3065 | 0 | 44 | 11491 | 0 | 0 | 62 | 51 | 11 | 11 | 0 | 14 | 23 | 0 | 1707 | 0 |
| SRR1708933 | 0 | 172 | 12 | 0 | 35 | 0 | 0 | 1306 | 1084 | 547 | 4871 | 19161 | 0 | 23 | 29 | 0 | 0 | 808 | 26 | 656 | 1078 | 3009 | 0 | 188 | 11740 | 0 | 0 | 112 | 121 | 357 | 417 | 0 | 42 | 88 | 0 | 1300 | 0 |
| SRR1708934 | 0 | 297 | 37 | 0 | 29 | 0 | 0 | 2339 | 1359 | 1978 | 5225 | 10827 | 0 | 41 | 187 | 136 | 0 | 617 | 22 | 1308 | 1883 | 4137 | 0 | 18 | 14525 | 0 | 0 | 44 | 61 | 604 | 784 | 20 | 143 | 222 | 13 | 1612 | 0 |
| SRR1708935 | 0 | 85 | 10 | 0 | 51 | 0 | 0 | 2158 | 2118 | 818 | 23680 | 13670 | 0 | 0 | 0 | 0 | 0 | 387 | 21 | 389 | 1491 | 3683 | 0 | 26 | 13392 | 0 | 0 | 64 | 39 | 162 | 193 | 0 | 21 | 35 | 0 | 609 | 0 |
| SRR1708936 | 0 | 161 | 66 | 0 | 46 | 0 | 0 | 2079 | 728 | 3238 | 13657 | 8924 | 0 | 274 | 341 | 211 | 0 | 391 | 20 | 633 | 962 | 2721 | 0 | 35 | 11664 | 0 | 11 | 43 | 62 | 103 | 184 | 0 | 34 | 96 | 0 | 3252 | 0 |
| SRR1708937 | 0 | 153 | 0 | 0 | 46 | 0 | 0 | 696 | 118 | 342 | 6530 | 19796 | 0 | 12 | 0 | 0 | 0 | 355 | 19 | 136 | 1123 | 2496 | 0 | 131 | 10785 | 0 | 0 | 56 | 64 | 80 | 54 | 0 | 0 | 0 | 0 | 330 | 0 |
| SRR1708938 | 0 | 173 | 0 | 0 | 40 | 0 | 0 | 1466 | 324 | 571 | 3902 | 15414 | 0 | 42 | 0 | 0 | 0 | 378 | 17 | 457 | 1245 | 2363 | 0 | 60 | 11023 | 0 | 0 | 62 | 136 | 198 | 170 | 0 | 15 | 19 | 0 | 1129 | 0 |
| SRR1708939 | 0 | 172 | 13 | 0 | 45 | 0 | 0 | 1156 | 493 | 1266 | 17923 | 12727 | 0 | 155 | 0 | 0 | 0 | 569 | 24 | 832 | 1145 | 2684 | 0 | 293 | 10908 | 0 | 0 | 42 | 137 | 221 | 198 | 0 | 20 | 52 | 0 | 2333 | 0 |
| SRR1708940 | 0 | 141 | 21 | 0 | 48 | 0 | 0 | 2152 | 1093 | 2253 | 4225 | 10600 | 0 | 87 | 0 | 19 | 0 | 381 | 27 | 684 | 1488 | 3101 | 0 | 31 | 11844 | 0 | 0 | 62 | 116 | 307 | 456 | 0 | 27 | 71 | 0 | 857 | 0 |
| SRR1708941 | 0 | 183 | 13 | 0 | 32 | 0 | 0 | 2685 | 906 | 2256 | 19197 | 13256 | 0 | 122 | 0 | 0 | 0 | 483 | 17 | 682 | 1534 | 3900 | 0 | 26 | 13455 | 0 | 0 | 32 | 73 | 105 | 228 | 17 | 26 | 49 | 0 | 1165 | 0 |
| SRR1708942 | 17 | 141 | 17 | 0 | 30 | 0 | 0 | 1775 | 238 | 2506 | 3925 | 23758 | 0 | 25 | 74 | 90 | 0 | 579 | 16 | 879 | 1099 | 4432 | 0 | 30 | 10317 | 0 | 26 | 66 | 26 | 23 | 42 | 0 | 49 | 89 | 0 | 1307 | 0 |
| SRR1708943 | 0 | 15 | 0 | 0 | 16 | 0 | 0 | 3401 | 199 | 703 | 19170 | 11543 | 0 | 20 | 0 | 0 | 0 | 470 | 28 | 488 | 706 | 4052 | 0 | 77 | 3531 | 0 | 23 | 0 | 13 | 10 | 13 | 0 | 0 | 0 | 0 | 528 | 0 |
| SRR1708944 | 0 | 110 | 11 | 0 | 19 | 0 | 0 | 4024 | 365 | 2770 | 5901 | 4730 | 0 | 46 | 19 | 11 | 0 | 577 | 30 | 623 | 799 | 3343 | 0 | 30 | 5287 | 0 | 0 | 13 | 46 | 175 | 460 | 27 | 22 | 94 | 0 | 1391 | 0 |
| SRR1708945 | 0 | 28 | 53 | 0 | 28 | 0 | 0 | 3417 | 498 | 3031 | 5489 | 3788 | 0 | 66 | 127 | 66 | 0 | 578 | 22 | 1300 | 1005 | 5322 | 0 | 105 | 5835 | 0 | 16 | 29 | 16 | 14 | 18 | 21 | 56 | 15 | 11 | 1919 | 0 |
| SRR1708946 | 0 | 69 | 0 | 0 | 24 | 0 | 0 | 2046 | 179 | 2803 | 8789 | 7029 | 0 | 33 | 27 | 40 | 0 | 576 | 28 | 505 | 1236 | 3559 | 0 | 12 | 12686 | 0 | 20 | 11 | 34 | 13 | 26 | 37 | 37 | 66 | 0 | 2445 | 0 |
| SRR1708947 | 0 | 95 | 36 | 0 | 13 | 0 | 0 | 6084 | 391 | 2287 | 5445 | 4256 | 0 | 92 | 98 | 44 | 0 | 561 | 24 | 498 | 922 | 4138 | 0 | 11 | 6193 | 0 | 18 | 11 | 0 | 84 | 296 | 26 | 47 | 126 | 0 | 2082 | 0 |
| SRR1708948 | 19 | 139 | 0 | 0 | 28 | 0 | 0 | 2404 | 1113 | 2340 | 4337 | 11560 | 0 | 30 | 49 | 15 | 11 | 552 | 18 | 628 | 1101 | 3458 | 0 | 49 | 11338 | 0 | 24 | 30 | 51 | 193 | 261 | 15 | 0 | 38 | 0 | 1622 | 0 |
| SRR1708949 | 0 | 13 | 0 | 0 | 19 | 0 | 0 | 4876 | 705 | 2786 | 2852 | 7320 | 0 | 67 | 67 | 27 | 0 | 655 | 21 | 1136 | 938 | 3134 | 0 | 130 | 8044 | 0 | 0 | 48 | 31 | 139 | 18 | 0 | 20 | 102 | 0 | 2125 | 0 |
| SRR1708950 | 0 | 74 | 0 | 0 | 49 | 0 | 0 | 3709 | 891 | 3991 | 16457 | 7933 | 0 | 93 | 18 | 35 | 0 | 406 | 26 | 507 | 1346 | 3525 | 0 | 65 | 11143 | 0 | 0 | 27 | 38 | 15 | 15 | 11 | 25 | 41 | 0 | 3603 | 0 |
| SRR1708951 | 0 | 70 | 14 | 0 | 32 | 0 | 0 | 1817 | 699 | 2314 | 15070 | 7815 | 0 | 178 | 333 | 82 | 0 | 476 | 19 | 733 | 1002 | 3076 | 0 | 167 | 11750 | 0 | 0 | 59 | 54 | 240 | 240 | 0 | 72 | 176 | 0 | 3548 | 0 |
| SRR1708952 | 0 | 83 | 17 | 0 | 37 | 0 | 0 | 1850 | 600 | 3031 | 14002 | 9217 | 0 | 65 | 27 | 12 | 22 | 463 | 24 | 442 | 1266 | 3192 | 0 | 31 | 12570 | 0 | 0 | 69 | 81 | 19 | 22 | 0 | 47 | 91 | 0 | 2954 | 0 |
| SRR1708953 | 0 | 88 | 18 | 0 | 37 | 0 | 0 | 1619 | 790 | 2219 | 4593 | 14234 | 0 | 79 | 53 | 0 | 0 | 483 | 22 | 728 | 1248 | 3501 | 0 | 29 | 12779 | 0 | 0 | 33 | 47 | 171 | 312 | 12 | 25 | 71 | 0 | 2749 | 0 |
| SRR1708954 | 24 | 137 | 13 | 0 | 51 | 0 | 0 | 938 | 253 | 1738 | 20660 | 12121 | 0 | 147 | 28 | 18 | 0 | 461 | 17 | 722 | 1133 | 2461 | 0 | 41 | 11630 | 0 | 13 | 55 | 48 | 102 | 119 | 0 | 15 | 31 | 0 | 1977 | 0 |
| SRR1708955 | 0 | 76 | 28 | 0 | 84 | 0 | 0 | 1430 | 639 | 3486 | 21358 | 8367 | 0 | 141 | 13 | 0 | 0 | 332 | 16 | 835 | 1186 | 2820 | 0 | 169 | 11098 | 0 | 10 | 49 | 74 | 133 | 142 | 0 | 27 | 43 | 0 | 2596 | 0 |
| SRR1708956 | 0 | 67 | 29 | 0 | 45 | 0 | 0 | 1891 | 461 | 3539 | 14628 | 7060 | 0 | 349 | 247 | 46 | 0 | 320 | 18 | 764 | 1354 | 2557 | 0 | 50 | 11102 | 0 | 0 | 51 | 96 | 232 | 195 | 0 | 29 | 67 | 0 | 5448 | 0 |
| SRR1708957 | 0 | 131 | 80 | 0 | 49 | 0 | 0 | 953 | 278 | 1661 | 3027 | 6938 | 0 | 251 | 577 | 98 | 0 | 433 | 21 | 913 | 1324 | 2406 | 0 | 116 | 11426 | 0 | 0 | 41 | 168 | 10 | 149 | 12 | 41 | 101 | 0 | 3814 | 0 |
| SRR1708958 | 0 | 84 | 0 | 0 | 46 | 0 | 0 | 985 | 272 | 3361 | 20063 | 12402 | 0 | 367 | 52 | 21 | 0 | 432 | 18 | 559 | 1425 | 2737 | 0 | 16 | 9739 | 0 | 0 | 52 | 87 | 230 | 211 | 0 | 15 | 21 | 0 | 2288 | 0 |
| SRR1708959 | 0 | 147 | 37 | 0 | 47 | 0 | 0 | 1340 | 427 | 3839 | 6004 | 7153 | 0 | 229 | 49 | 19 | 0 | 388 | 13 | 940 | 1787 | 2975 | 0 | 28 | 11659 | 0 | 0 | 29 | 82 | 95 | 176 | 0 | 32 | 65 | 0 | 3900 | 0 |
| SRR1708960 | 0 | 104 | 45 | 0 | 39 | 0 | 0 | 1323 | 296 | 6049 | 3111 | 14635 | 0 | 41 | 448 | 152 | 0 | 412 | 11 | 1075 | 1048 | 3061 | 0 | 34 | 11933 | 0 | 55 | 22 | 16 | 312 | 550 | 22 | 116 | 222 | 27 | 1881 | 0 |
| SRR1708961 | 0 | 227 | 21 | 0 | 37 | 0 | 0 | 1426 | 235 | 6070 | 3445 | 9797 | 0 | 57 | 181 | 60 | 0 | 635 | 19 | 1604 | 1573 | 4185 | 0 | 223 | 14480 | 0 | 12 | 0 | 59 | 400 | 784 | 32 | 94 | 221 | 0 | 3144 | 0 |
| SRR1708962 | 0 | 67 | 10 | 0 | 30 | 0 | 0 | 1982 | 218 | 4011 | 3503 | 15102 | 0 | 71 | 24 | 30 | 0 | 425 | 22 | 849 | 1086 | 3364 | 0 | 36 | 11000 | 0 | 12 | 29 | 39 | 12 | 32 | 33 | 29 | 45 | 0 | 1941 | 0 |
| SRR1708963 | 0 | 111 | 0 | 0 | 14 | 0 | 0 | 2499 | 274 | 1691 | 12272 | 9447 | 0 | 53 | 14 | 19 | 10 | 477 | 30 | 759 | 702 | 3370 | 0 | 113 | 6929 | 0 | 14 | 19 | 25 | 16 | 24 | 14 | 54 | 87 | 0 | 1326 | 0 |
| SRR1708964 | 0 | 75 | 19 | 0 | 24 | 0 | 0 | 2061 | 348 | 4916 | 1158 | 6013 | 0 | 15 | 44 | 37 | 0 | 652 | 25 | 452 | 1041 | 2549 | 0 | 0 | 9839 | 0 | 11 | 23 | 87 | 17 | 26 | 53 | 68 | 151 | 11 | 2395 | 0 |
| SRR1708965 | 0 | 98 | 0 | 0 | 28 | 0 | 0 | 1321 | 248 | 3876 | 2051 | 10386 | 0 | 36 | 15 | 0 | 0 | 439 | 20 | 474 | 1455 | 3992 | 0 | 16 | 14206 | 0 | 20 | 20 | 20 | 171 | 307 | 0 | 18 | 30 | 0 | 1277 | 0 |
| SRR1708966 | 0 | 80 | 11 | 0 | 48 | 0 | 0 | 2189 | 668 | 5279 | 3005 | 7793 | 0 | 176 | 1011 | 558 | 0 | 426 | 19 | 696 | 980 | 3449 | 0 | 50 | 13765 | 0 | 15 | 23 | 40 | 169 | 239 | 0 | 38 | 176 | 0 | 2304 | 0 |
| SRR1708967 | 0 | 63 | 0 | 0 | 40 | 0 | 0 | 1729 | 343 | 1263 | 5119 | 17924 | 0 | 43 | 25 | 0 | 0 | 412 | 21 | 337 | 1028 | 3143 | 0 | 157 | 11699 | 0 | 13 | 33 | 21 | 146 | 11 | 0 | 0 | 0 | 0 | 442 | 0 |
| SRR1708968 | 0 | 122 | 0 | 0 | 41 | 0 | 0 | 941 | 176 | 1041 | 22206 | 15540 | 0 | 50 | 25 | 13 | 0 | 433 | 23 | 395 | 1200 | 2669 | 0 | 62 | 12994 | 0 | 0 | 41 | 91 | 447 | 284 | 13 | 22 | 31 | 0 | 1329 | 0 |
| SRR1708969 | 0 | 93 | 0 | 0 | 44 | 0 | 0 | 925 | 518 | 1837 | 13720 | 9674 | 0 | 83 | 35 | 21 | 12 | 536 | 28 | 898 | 983 | 2743 | 0 | 176 | 12320 | 0 | 12 | 59 | 87 | 331 | 238 | 22 | 49 | 96 | 0 | 2237 | 0 |
| SRR1708970 | 0 | 95 | 0 | 0 | 48 | 0 | 0 | 1286 | 511 | 1267 | 24803 | 11503 | 0 | 31 | 0 | 0 | 0 | 558 | 30 | 328 | 1539 | 3210 | 0 | 37 | 12080 | 0 | 0 | 56 | 67 | 18 | 19 | 13 | 16 | 19 | 0 | 1458 | 0 |
| SRR1708971 | 0 | 318 | 0 | 0 | 47 | 0 | 0 | 1302 | 849 | 1176 | 2507 | 10720 | 0 | 57 | 0 | 0 | 0 | 602 | 13 | 535 | 1989 | 4735 | 0 | 40 | 13262 | 0 | 14 | 53 | 91 | 265 | 252 | 11 | 11 | 25 | 0 | 925 | 0 |
| SRR1708972 | 0 | 120 | 0 | 0 | 42 | 0 | 0 | 1130 | 351 | 2169 | 28606 | 15322 | 0 | 196 | 14 | 0 | 0 | 413 | 17 | 405 | 1325 | 3024 | 0 | 33 | 10441 | 0 | 15 | 30 | 51 | 10 | 11 | 0 | 13 | 0 | 0 | 1134 | 0 |
| SRR1708973 | 0 | 101 | 0 | 0 | 51 | 0 | 0 | 308 | 62 | 448 | 8994 | 16349 | 0 | 37 | 0 | 0 | 18 | 368 | 18 | 70 | 1094 | 2154 | 0 | 98 | 9144 | 0 | 15 | 57 | 67 | 93 | 29 | 0 | 0 | 0 | 0 | 263 | 0 |
| SRR1708974 | 0 | 98 | 11 | 0 | 41 | 0 | 0 | 1051 | 250 | 4505 | 9698 | 13295 | 0 | 321 | 16 | 12 | 0 | 330 | 15 | 595 | 1756 | 2713 | 0 | 47 | 11087 | 0 | 0 | 37 | 94 | 12 | 110 | 0 | 21 | 24 | 0 | 2815 | 0 |
| SRR1708975 | 0 | 184 | 19 | 0 | 66 | 0 | 0 | 1123 | 422 | 2650 | 21838 | 10188 | 0 | 220 | 37 | 0 | 14 | 287 | 17 | 857 | 1158 | 2508 | 0 | 103 | 10235 | 0 | 0 | 28 | 80 | 10 | 138 | 0 | 16 | 25 | 0 | 2320 | 0 |
| SRR1708976 | 0 | 63 | 18 | 0 | 41 | 0 | 0 | 1529 | 412 | 3689 | 17286 | 8895 | 0 | 327 | 48 | 0 | 0 | 396 | 19 | 517 | 1468 | 2873 | 0 | 24 | 12249 | 0 | 0 | 66 | 113 | 290 | 280 | 0 | 25 | 37 | 0 | 1799 | 0 |
| SRR1708977 | 0 | 46 | 48 | 0 | 42 | 0 | 0 | 1554 | 615 | 4564 | 10453 | 7287 | 0 | 453 | 239 | 74 | 0 | 349 | 19 | 535 | 1369 | 2703 | 0 | 22 | 11146 | 0 | 0 | 40 | 68 | 108 | 133 | 0 | 31 | 72 | 0 | 2460 | 0 |
| SRR1708978 | 0 | 124 | 0 | 0 | 14 | 0 | 0 | 3940 | 312 | 2020 | 16213 | 8854 | 0 | 17 | 30 | 26 | 0 | 498 | 28 | 702 | 783 | 3942 | 0 | 24 | 7308 | 0 | 11 | 19 | 28 | 18 | 32 | 11 | 56 | 103 | 0 | 1456 | 0 |
| SRR1708979 | 0 | 123 | 0 | 0 | 18 | 0 | 0 | 6378 | 355 | 2073 | 2410 | 6914 | 0 | 31 | 66 | 10 | 0 | 454 | 28 | 696 | 557 | 3097 | 0 | 81 | 4291 | 0 | 34 | 13 | 15 | 265 | 420 | 28 | 43 | 65 | 0 | 1332 | 0 |
| SRR1708980 | 0 | 102 | 0 | 0 | 51 | 0 | 0 | 2371 | 426 | 3038 | 20540 | 11400 | 0 | 19 | 28 | 0 | 0 | 361 | 23 | 871 | 1334 | 3525 | 0 | 51 | 13783 | 0 | 0 | 46 | 36 | 194 | 436 | 12 | 48 | 106 | 0 | 1963 | 0 |
| SRR1708981 | 0 | 12 | 0 | 0 | 12 | 0 | 0 | 2684 | 237 | 1293 | 12518 | 9502 | 0 | 19 | 14 | 14 | 0 | 618 | 35 | 924 | 1039 | 5609 | 0 | 116 | 6769 | 0 | 0 | 18 | 29 | 187 | 21 | 14 | 28 | 72 | 0 | 1020 | 0 |
| SRR1708982 | 0 | 110 | 0 | 0 | 27 | 0 | 0 | 1825 | 235 | 3028 | 15794 | 8898 | 0 | 76 | 0 | 0 | 0 | 528 | 22 | 984 | 1308 | 3389 | 0 | 12 | 12142 | 0 | 0 | 39 | 49 | 406 | 919 | 12 | 81 | 169 | 0 | 3002 | 0 |
| SRR1708983 | 0 | 102 | 24 | 0 | 11 | 0 | 0 | 7787 | 436 | 2356 | 7898 | 5408 | 0 | 78 | 17 | 19 | 0 | 513 | 33 | 569 | 841 | 4524 | 0 | 0 | 5222 | 0 | 0 | 0 | 0 | 63 | 245 | 17 | 26 | 102 | 0 | 1079 | 0 |
| SRR1708984 | 0 | 129 | 19 | 0 | 43 | 0 | 0 | 3840 | 3165 | 2466 | 18592 | 14153 | 0 | 30 | 167 | 53 | 0 | 410 | 21 | 751 | 828 | 2719 | 0 | 35 | 11598 | 0 | 16 | 49 | 0 | 121 | 15 | 11 | 44 | 80 | 0 | 1313 | 0 |
| SRR1708985 | 0 | 68 | 36 | 0 | 63 | 0 | 0 | 4666 | 3563 | 2487 | 4182 | 7301 | 0 | 63 | 497 | 242 | 0 | 484 | 21 | 1189 | 1331 | 3767 | 0 | 225 | 10816 | 0 | 11 | 43 | 42 | 134 | 314 | 15 | 54 | 158 | 17 | 2698 | 0 |
| SRR1708986 | 0 | 11 | 40 | 0 | 44 | 0 | 0 | 3087 | 3241 | 1713 | 1897 | 10094 | 0 | 62 | 154 | 23 | 0 | 407 | 14 | 864 | 1707 | 4162 | 0 | 57 | 11427 | 0 | 10 | 91 | 54 | 12 | 28 | 17 | 94 | 12 | 0 | 3758 | 0 |
| SRR1708987 | 11 | 109 | 26 | 0 | 47 | 0 | 0 | 1799 | 801 | 731 | 3493 | 12535 | 0 | 121 | 57 | 51 | 0 | 378 | 22 | 604 | 926 | 3086 | 0 | 179 | 13291 | 0 | 0 | 101 | 83 | 266 | 10 | 23 | 60 | 176 | 0 | 1309 | 0 |
| SRR1708988 | 49 | 168 | 0 | 0 | 30 | 0 | 0 | 1778 | 646 | 1169 | 12110 | 15936 | 0 | 85 | 32 | 17 | 32 | 777 | 23 | 720 | 1829 | 3986 | 0 | 15 | 12548 | 0 | 10 | 60 | 87 | 365 | 329 | 15 | 32 | 65 | 0 | 1440 | 0 |
| SRR1708989 | 0 | 171 | 11 | 0 | 29 | 0 | 0 | 2272 | 669 | 977 | 14810 | 13190 | 0 | 111 | 17 | 13 | 15 | 469 | 11 | 767 | 1263 | 3283 | 0 | 16 | 12239 | 0 | 0 | 103 | 97 | 14 | 15 | 0 | 27 | 42 | 0 | 1016 | 0 |
| SRR1708990 | 0 | 97 | 49 | 0 | 60 | 0 | 0 | 3055 | 1476 | 3454 | 3747 | 10038 | 0 | 237 | 116 | 60 | 0 | 499 | 19 | 566 | 1142 | 2747 | 0 | 93 | 10568 | 0 | 14 | 92 | 43 | 84 | 12 | 25 | 70 | 10 | 0 | 4216 | 0 |
| SRR1708991 | 0 | 100 | 24 | 0 | 69 | 0 | 0 | 2661 | 1143 | 4012 | 14916 | 7633 | 0 | 396 | 52 | 0 | 0 | 296 | 18 | 850 | 1238 | 2783 | 0 | 263 | 12158 | 0 | 14 | 31 | 77 | 126 | 10 | 0 | 22 | 61 | 0 | 3501 | 0 |
| SRR1708992 | 0 | 97 | 39 | 0 | 47 | 0 | 0 | 1983 | 823 | 1983 | 2243 | 9460 | 0 | 130 | 207 | 78 | 0 | 411 | 28 | 1050 | 1347 | 2482 | 0 | 106 | 13084 | 0 | 0 | 71 | 58 | 220 | 236 | 0 | 94 | 152 | 29 | 6000 | 0 |
| SRR1708993 | 0 | 100 | 0 | 0 | 44 | 0 | 0 | 1809 | 596 | 1463 | 16862 | 12804 | 0 | 278 | 0 | 0 | 0 | 345 | 21 | 644 | 1198 | 2944 | 0 | 163 | 9637 | 0 | 0 | 31 | 103 | 131 | 212 | 0 | 12 | 56 | 0 | 2581 | 0 |
| SRR1708994 | 14 | 30 | 0 | 0 | 43 | 17 | 0 | 3395 | 637 | 2429 | 12509 | 11428 | 0 | 110 | 0 | 0 | 11 | 196 | 15 | 602 | 1190 | 2872 | 0 | 15 | 10294 | 0 | 0 | 29 | 86 | 17 | 20 | 0 | 14 | 16 | 0 | 747 | 0 |
| SRR1708995 | 0 | 166 | 30 | 0 | 43 | 0 | 0 | 2184 | 689 | 2214 | 14986 | 14276 | 0 | 179 | 23 | 18 | 0 | 253 | 0 | 707 | 1786 | 2406 | 0 | 10 | 9481 | 0 | 0 | 48 | 75 | 109 | 177 | 0 | 23 | 40 | 0 | 941 | 0 |
| SRR1708996 | 15 | 90 | 34 | 0 | 42 | 0 | 0 | 2090 | 450 | 4219 | 15004 | 8821 | 0 | 13 | 86 | 20 | 0 | 379 | 19 | 1065 | 917 | 3153 | 0 | 38 | 12902 | 0 | 68 | 43 | 25 | 402 | 534 | 21 | 76 | 193 | 0 | 2466 | 0 |

### Supp Table S8.4.Columns_DI_to_ES

| | | | | | | | | | | | | | | | | | |
|---|---|---|---|---|---|---|---|---|---|---|---|---|---|---|---|---|---|
| SRR1708997 | 17 | 31 | 0 | 0 | 19 | 0 | 0 | 6217 | 455 | 4322 | 11256 | 7550 | 0 | 46 | 162 | 162 | 0 |
| SRR1708998 | 0 | 106 | 28 | 0 | 30 | 0 | 0 | 2307 | 354 | 4199 | 2279 | 11853 | 0 | 17 | 43 | 43 | 0 |
| SRR1708999 | 0 | 190 | 0 | 0 | 14 | 0 | 0 | 2708 | 183 | 744 | 1945 | 8801 | 0 | 0 | 0 | 0 | 0 |
| SRR1709000 | 16 | 195 | 24 | 0 | 31 | 0 | 0 | 4295 | 1733 | 3149 | 6988 | 9200 | 0 | 56 | 199 | 59 | 0 |
| SRR1709001 | 0 | 154 | 0 | 0 | 38 | 0 | 0 | 3704 | 2351 | 1373 | 3918 | 15336 | 0 | 21 | 14 | 28 | 0 |
| SRR1709002 | 0 | 132 | 21 | 0 | 34 | 0 | 0 | 1940 | 1360 | 2052 | 4344 | 10938 | 0 | 69 | 416 | 249 | 0 |
| SRR1709003 | 0 | 128 | 18 | 0 | 34 | 12 | 0 | 1142 | 641 | 661 | 3353 | 13888 | 0 | 27 | 13 | 20 | 18 |
| SRR1709004 | 0 | 12 | 0 | 0 | 16 | 0 | 0 | 4132 | 379 | 1236 | 15682 | 12190 | 0 | 25 | 12 | 12 | 0 |
| SRR1709005 | 0 | 144 | 23 | 0 | 38 | 0 | 0 | 5552 | 717 | 3652 | 2801 | 8426 | 0 | 16 | 55 | 66 | 0 |
| SRR1709006 | 0 | 171 | 24 | 0 | 30 | 0 | 0 | 4654 | 607 | 5386 | 1564 | 4948 | 0 | 38 | 175 | 50 | 0 |
| SRR1709007 | 0 | 136 | 29 | 0 | 25 | 13 | 0 | 2624 | 356 | 3182 | 16754 | 8911 | 0 | 19 | 154 | 106 | 0 |
| SRR1709008 | 0 | 184 | 11 | 0 | 17 | 0 | 0 | 3903 | 224 | 1801 | 15278 | 8950 | 0 | 27 | 27 | 0 | 0 |
| SRR1709009 | 0 | 134 | 11 | 0 | 20 | 0 | 0 | 2323 | 280 | 3397 | 1357 | 12589 | 0 | 68 | 0 | 30 | 0 |
| SRR1709010 | 0 | 21 | 23 | 0 | 86 | 0 | 0 | 1891 | 1425 | 1224 | 17205 | 10579 | 0 | 17 | 23 | 29 | 0 |
| SRR1709011 | 0 | 267 | 14 | 0 | 33 | 0 | 0 | 1947 | 564 | 2127 | 9352 | 11662 | 0 | 76 | 168 | 86 | 0 |
| SRR1709012 | 0 | 140 | 27 | 0 | 40 | 0 | 0 | 2286 | 532 | 816 | 18318 | 10568 | 0 | 57 | 192 | 144 | 17 |
| SRR1709013 | 0 | 100 | 12 | 0 | 39 | 0 | 0 | 1621 | 371 | 576 | 25872 | 14129 | 0 | 76 | 15 | 0 | 0 |
| SRR1709014 | 0 | 207 | 24 | 0 | 45 | 0 | 0 | 2561 | 807 | 1100 | 2590 | 11380 | 0 | 51 | 79 | 39 | 0 |
| SRR1709015 | 0 | 140 | 14 | 0 | 30 | 0 | 0 | 1544 | 624 | 931 | 20171 | 15154 | 0 | 33 | 13 | 17 | 0 |
| SRR1709016 | 0 | 126 | 49 | 0 | 57 | 0 | 0 | 2573 | 1300 | 2048 | 20113 | 10538 | 0 | 151 | 34 | 49 | 0 |
| SRR1709017 | 0 | 18 | 77 | 0 | 95 | 0 | 0 | 1565 | 1702 | 2947 | 2684 | 9205 | 0 | 182 | 50 | 33 | 0 |
| SRR1709018 | 0 | 202 | 24 | 0 | 46 | 0 | 0 | 1769 | 543 | 972 | 10112 | 22791 | 0 | 88 | 0 | 0 | 0 |
| SRR1709019 | 0 | 144 | 31 | 0 | 57 | 0 | 0 | 932 | 450 | 1101 | 20424 | 15569 | 0 | 209 | 0 | 13 | 10 |
| SRR1709020 | 0 | 118 | 45 | 0 | 41 | 0 | 0 | 1175 | 433 | 2425 | 17642 | 12138 | 0 | 123 | 160 | 93 | 0 |
| SRR1709021 | 0 | 15 | 49 | 0 | 93 | 0 | 0 | 1552 | 1134 | 2365 | 11185 | 9301 | 0 | 156 | 0 | 0 | 0 |
| SRR1732015 | 0 | 0 | 0 | 0 | 37 | 39 | 13 | 5543 | 506 | 4694 | 1009 | 5162 | 0 | 478 | 0 | 43 | 20 |
| SRR1732016 | 0 | 0 | 0 | 0 | 136 | 45 | 54 | 28201 | 1211 | 3311 | 298 | 423 | 0 | 719 | 0 | 0 | 18 |
| SRR1748132 | 0 | 25 | 0 | 0 | 0 | 24 | 0 | 775 | 986 | 3426 | 3174 | 1554 | 0 | 1201 | 0 | 0 | 0 |
| SRR1748133 | 0 | 24 | 0 | 0 | 21 | 25 | 0 | 822 | 608 | 3221 | 422 | 636 | 0 | 1258 | 0 | 0 | 0 |
| SRR1759212 | 0 | 155 | 16 | 0 | 0 | 0 | 0 | 216 | 53 | 11982 | 307 | 98 | 0 | 2081 | 726 | 1898 | 0 |
| SRR1759213 | 0 | 11 | 0 | 0 | 0 | 0 | 0 | 607 | 54 | 11454 | 525 | 121 | 0 | 2624 | 870 | 2455 | 0 |
| SRR1759214 | 0 | 104 | 18 | 0 | 0 | 0 | 0 | 181 | 33 | 3699 | 714 | 301 | 0 | 497 | 1437 | 3335 | 0 |
| SRR1759215 | 0 | 144 | 0 | 0 | 0 | 0 | 0 | 186 | 63 | 2912 | 783 | 452 | 0 | 585 | 616 | 1095 | 0 |
| SRR1759216 | 0 | 120 | 0 | 0 | 0 | 0 | 0 | 224 | 48 | 7936 | 573 | 135 | 0 | 710 | 3866 | 7622 | 0 |
| SRR1759217 | 0 | 67 | 0 | 0 | 0 | 0 | 0 | 70 | 37 | 2127 | 478 | 362 | 0 | 399 | 467 | 2229 | 0 |
| SRR1759218 | 0 | 170 | 19 | 0 | 0 | 0 | 0 | 163 | 37 | 2275 | 1185 | 625 | 0 | 392 | 322 | 821 | 0 |
| SRR1759219 | 0 | 90 | 16 | 0 | 0 | 0 | 0 | 325 | 33 | 3338 | 352 | 163 | 0 | 747 | 1174 | 3898 | 0 |
| SRR1759220 | 0 | 83 | 0 | 0 | 0 | 0 | 0 | 118 | 56 | 3501 | 646 | 314 | 0 | 506 | 175 | 440 | 0 |
| SRR1759221 | 0 | 0 | 0 | 0 | 0 | 0 | 0 | 57 | 84 | 4135 | 200 | 53 | 0 | 1648 | 770 | 1144 | 0 |
| SRR1759222 | 0 | 23 | 0 | 0 | 0 | 0 | 0 | 228 | 32 | 2565 | 624 | 1414 | 0 | 1958 | 13 | 64 | 0 |
| SRR1759223 | 0 | 46 | 11 | 0 | 0 | 0 | 0 | 143 | 62 | 3233 | 51 | 30 | 0 | 605 | 2137 | 5393 | 0 |
| SRR1759224 | 0 | 11 | 0 | 0 | 0 | 0 | 0 | 152 | 52 | 7834 | 149 | 49 | 0 | 584 | 112 | 163 | 0 |
| SRR1759225 | 0 | 0 | 0 | 0 | 0 | 0 | 0 | 345 | 69 | 3649 | 102 | 67 | 0 | 919 | 149 | 271 | 0 |
| SRR1759226 | 0 | 0 | 0 | 0 | 0 | 0 | 0 | 93 | 21 | 2149 | 261 | 152 | 0 | 740 | 43 | 82 | 0 |
| SRR1759227 | 0 | 15 | 0 | 0 | 0 | 0 | 0 | 179 | 143 | 5856 | 138 | 126 | 0 | 1699 | 153 | 359 | 0 |
| SRR1759228 | 0 | 0 | 12 | 0 | 0 | 0 | 0 | 431 | 262 | 6962 | 239 | 195 | 0 | 1381 | 72 | 66 | 0 |
| SRR1759229 | 0 | 15 | 16 | 0 | 0 | 0 | 0 | 149 | 75 | 4068 | 84 | 46 | 0 | 633 | 530 | 1021 | 0 |
| SRR1759230 | 0 | 0 | 0 | 0 | 0 | 0 | 0 | 360 | 94 | 2695 | 105 | 77 | 0 | 1958 | 207 | 125 | 0 |
| SRR1759231 | 0 | 0 | 31 | 0 | 0 | 0 | 0 | 86 | 83 | 6761 | 126 | 79 | 0 | 1797 | 156 | 375 | 0 |
| SRR1759232 | 0 | 15 | 26 | 0 | 0 | 0 | 0 | 287 | 85 | 3345 | 467 | 385 | 0 | 684 | 100 | 263 | 0 |
| SRR1759233 | 0 | 0 | 0 | 0 | 0 | 0 | 0 | 193 | 211 | 8302 | 268 | 143 | 0 | 1023 | 95 | 103 | 0 |
| SRR1759234 | 0 | 45 | 24 | 0 | 0 | 0 | 0 | 124 | 83 | 6208 | 143 | 20 | 0 | 1989 | 436 | 768 | 0 |
| SRR1759235 | 0 | 55 | 23 | 0 | 0 | 0 | 0 | 301 | 50 | 5091 | 597 | 346 | 0 | 391 | 618 | 885 | 0 |
| SRR1759236 | 0 | 28 | 17 | 0 | 0 | 0 | 0 | 114 | 53 | 5976 | 245 | 72 | 0 | 740 | 3835 | 8310 | 0 |
| SRR1759237 | 0 | 50 | 11 | 0 | 0 | 0 | 0 | 152 | 55 | 3399 | 308 | 164 | 0 | 600 | 5800 | 11312 | 0 |
| SRR1759238 | 0 | 45 | 25 | 0 | 0 | 0 | 0 | 150 | 35 | 3407 | 135 | 53 | 0 | 1161 | 1042 | 2709 | 0 |
| SRR1759239 | 0 | 13 | 0 | 0 | 0 | 0 | 0 | 186 | 37 | 443 | 685 | 236 | 0 | 994 | 317 | 185 | 13 |
| SRR1759240 | 0 | 29 | 19 | 0 | 0 | 0 | 0 | 172 | 44 | 4541 | 180 | 93 | 0 | 547 | 1754 | 4614 | 0 |
| SRR1759241 | 0 | 60 | 18 | 0 | 0 | 0 | 0 | 175 | 28 | 1065 | 120 | 78 | 0 | 502 | 1138 | 1687 | 0 |
| SRR1759242 | 0 | 65 | 51 | 0 | 0 | 0 | 0 | 220 | 669 | 15949 | 56 | 48 | 0 | 930 | 511 | 609 | 0 |
| SRR1759243 | 0 | 126 | 13 | 0 | 0 | 0 | 0 | 143 | 102 | 6891 | 188 | 36 | 0 | 2086 | 344 | 639 | 0 |
| SRR1759244 | 0 | 171 | 12 | 0 | 0 | 0 | 0 | 195 | 42 | 6171 | 493 | 227 | 0 | 955 | 963 | 3557 | 0 |
| SRR1759245 | 0 | 154 | 15 | 0 | 0 | 0 | 0 | 132 | 70 | 7454 | 467 | 83 | 0 | 493 | 2320 | 2646 | 0 |
| SRR1759246 | 0 | 31 | 47 | 0 | 0 | 0 | 0 | 138 | 63 | 8077 | 92 | 51 | 0 | 3097 | 536 | 1826 | 0 |
| SRR1759247 | 0 | 19 | 0 | 0 | 0 | 0 | 0 | 274 | 232 | 9910 | 164 | 109 | 0 | 1057 | 266 | 295 | 0 |
| SRR1759248 | 0 | 67 | 37 | 0 | 0 | 0 | 0 | 91 | 28 | 1270 | 203 | 133 | 0 | 470 | 1527 | 4047 | 0 |
| SRR1759249 | 0 | 37 | 14 | 0 | 0 | 0 | 0 | 280 | 132 | 21811 | 120 | 56 | 0 | 3978 | 466 | 1010 | 0 |
| SRR1759250 | 0 | 80 | 25 | 0 | 0 | 0 | 0 | 125 | 63 | 10727 | 487 | 393 | 0 | 5167 | 135 | 342 | 0 |
| SRR1759251 | 0 | 26 | 11 | 0 | 0 | 0 | 0 | 76 | 82 | 18338 | 39 | 24 | 0 | 833 | 1236 | 3656 | 0 |
| SRR1759252 | 0 | 48 | 23 | 0 | 0 | 0 | 0 | 175 | 248 | 7270 | 315 | 179 | 0 | 722 | 2663 | 6023 | 12 |
| SRR1759253 | 0 | 15 | 0 | 0 | 0 | 0 | 0 | 102 | 40 | 3952 | 408 | 280 | 0 | 1388 | 4183 | 9236 | 0 |
| SRR1759254 | 0 | 66 | 12 | 0 | 0 | 0 | 0 | 231 | 204 | 6916 | 205 | 75 | 0 | 1380 | 1524 | 4186 | 0 |
| SRR1759255 | 0 | 12 | 0 | 0 | 0 | 0 | 0 | 230 | 93 | 31659 | 148 | 96 | 0 | 4204 | 1024 | 2722 | 0 |
| SRR1759256 | 0 | 74 | 11 | 0 | 0 | 0 | 0 | 242 | 197 | 4914 | 452 | 195 | 0 | 464 | 1465 | 3049 | 0 |
| SRR1759257 | 0 | 166 | 0 | 0 | 0 | 0 | 0 | 131 | 319 | 30182 | 59 | 0 | 0 | 5733 | 284 | 880 | 0 |
| SRR1759258 | 0 | 35 | 132 | 0 | 0 | 0 | 0 | 833 | 419 | 3938 | 986 | 636 | 0 | 578 | 1805 | 2626 | 31 |
| SRR1759259 | 0 | 51 | 86 | 0 | 0 | 0 | 0 | 133 | 85 | 3056 | 1049 | 1072 | 0 | 571 | 441 | 680 | 0 |
| SRR1759260 | 0 | 69 | 0 | 0 | 0 | 0 | 0 | 220 | 58 | 4275 | 276 | 42 | 0 | 586 | 1445 | 1875 | 0 |
| SRR1759261 | 0 | 19 | 22 | 0 | 0 | 0 | 0 | 203 | 115 | 3417 | 296 | 142 | 0 | 1227 | 2799 | 2019 | 0 |
| SRR1759262 | 0 | 58 | 0 | 0 | 0 | 0 | 0 | 200 | 368 | 6042 | 269 | 61 | 0 | 562 | 464 | 945 | 0 |
| SRR1759263 | 0 | 55 | 16 | 0 | 0 | 0 | 0 | 161 | 50 | 6171 | 543 | 336 | 0 | 1462 | 435 | 1335 | 0 |
| SRR1759264 | 0 | 12 | 13 | 0 | 0 | 0 | 0 | 113 | 31 | 5149 | 197 | 142 | 0 | 1435 | 123 | 402 | 0 |
| SRR1759265 | 0 | 47 | 0 | 0 | 0 | 0 | 0 | 215 | 634 | 31505 | 41 | 29 | 0 | 4340 | 95 | 412 | 0 |
| SRR1759266 | 0 | 184 | 13 | 0 | 0 | 0 | 0 | 133 | 220 | 16452 | 184 | 25 | 0 | 992 | 130 | 501 | 0 |
| SRR1759267 | 0 | 90 | 16 | 0 | 0 | 0 | 0 | 299 | 67 | 4651 | 299 | 151 | 0 | 1447 | 22 | 24 | 0 |
| SRR1759268 | 0 | 22 | 0 | 0 | 0 | 0 | 0 | 163 | 159 | 7364 | 630 | 360 | 0 | 858 | 79 | 157 | 0 |
| SRR1759269 | 0 | 31 | 13 | 0 | 0 | 0 | 0 | 128 | 172 | 16162 | 182 | 70 | 0 | 4930 | 0 | 0 | 0 |
| SRR1759270 | 0 | 141 | 0 | 0 | 0 | 0 | 0 | 172 | 36 | 11735 | 365 | 88 | 0 | 1198 | 1246 | 4476 | 0 |
| SRR1759271 | 0 | 16 | 10 | 0 | 0 | 0 | 0 | 237 | 64 | 5747 | 263 | 222 | 0 | 1451 | 193 | 460 | 0 |
| SRR1759272 | 0 | 21 | 0 | 0 | 0 | 0 | 0 | 180 | 68 | 16726 | 129 | 73 | 0 | 4142 | 30 | 82 | 0 |
| SRR1759273 | 0 | 79 | 0 | 0 | 0 | 0 | 0 | 183 | 274 | 3552 | 280 | 131 | 0 | 766 | 2669 | 3656 | 0 |
| SRR1759274 | 0 | 133 | 0 | 0 | 0 | 0 | 0 | 208 | 64 | 2324 | 239 | 80 | 0 | 969 | 1986 | 1121 | 0 |
| SRR1759275 | 0 | 58 | 0 | 0 | 0 | 0 | 0 | 137 | 36 | 1684 | 694 | 351 | 0 | 767 | 1170 | 2895 | 0 |
| SRR1777167 | 0 | 22 | 33 | 0 | 0 | 0 | 11 | 920 | 544 | 4827 | 993 | 14256 | 0 | 335 | 75 | 17 | 15 |
| SRR1820679 | 0 | 499 | 79 | 41 | 12 | 0 | 15 | 961 | 3563 | 3703 | 45 | 1481 | 0 | 1181 | 1561 | 2102 | 27 |
| SRR1820680 | 0 | 1938 | 10 | 30 | 0 | 0 | 50 | 534 | 1133 | 898 | 898 | 4682 | 0 | 21 | 64 | 21 | 47 |
| SRR1820681 | 0 | 1163 | 77 | 41 | 0 | 0 | 13 | 647 | 1410 | 2637 | 15 | 846 | 0 | 1360 | 4129 | 4561 | 23 |
| SRR1820682 | 0 | 2518 | 14 | 56 | 13 | 0 | 46 | 757 | 1964 | 1944 | 1105 | 6792 | 0 | 450 | 61 | 41 | 52 |
| SRR1820683 | 0 | 825 | 61 | 66 | 0 | 0 | 0 | 541 | 1721 | 3655 | 36 | 1605 | 0 | 1160 | 3152 | 4428 | 23 |
| SRR1820684 | 0 | 156 | 13 | 46 | 11 | 0 | 39 | 1251 | 2348 | 1583 | 4033 | 9394 | 0 | 817 | 0 | 26 | 70 |
| SRR1820685 | 0 | 231 | 26 | 41 | 17 | 0 | 21 | 577 | 1570 | 3526 | 20 | 1635 | 0 | 1218 | 4487 | 6218 | 22 |
| SRR1820686 | 0 | 198 | 0 | 37 | 0 | 10 | 31 | 796 | 1906 | 1422 | 2816 | 9642 | 0 | 455 | 28 | 0 | 74 |
| SRR1820687 | 0 | 301 | 27 | 23 | 18 | 0 | 12 | 3154 | 6093 | 8061 | 400 | 2027 | 0 | 1887 | 76 | 81 | 52 |
| SRR1820688 | 0 | 244 | 26 | 22 | 0 | 0 | 11 | 3598 | 7172 | 10191 | 731 | 2484 | 0 | 1538 | 29 | 18 | 20 |
| SRR1820689 | 0 | 911 | 0 | 58 | 0 | 0 | 46 | 4320 | 6168 | 4488 | 348 | 2297 | 0 | 1090 | 39 | 14 | 140 |
| SRR1820690 | 0 | 1280 | 0 | 28 | 0 | 11 | 79 | 2384 | 4320 | 3334 | 194 | 2548 | 0 | 488 | 75 | 40 | 151 |
| SRR1853804 | 0 | 0 | 0 | 0 | 0 | 0 | 0 | 8106 | 376 | 308 | 12758 | 20283 | 0 | 103 | 0 | 68 | 0 |
| SRR1853805 | 0 | 0 | 13 | 0 | 0 | 0 | 0 | 8281 | 0 | 414 | 16354 | 22306 | 0 | 104 | 0 | 104 | 0 |
| SRR1853806 | 0 | 0 | 20 | 0 | 0 | 0 | 0 | 5661 | 300 | 2847 | 11555 | 16517 | 0 | 133 | 33 | 83 | 0 |
| SRR1853807 | 0 | 0 | 40 | 0 | 0 | 0 | 0 | 2227 | 66 | 1375 | 333 | 16833 | 0 | 131 | 0 | 0 | 0 |
| SRR1853808 | 0 | 0 | 103 | 0 | 0 | 0 | 0 | 3232 | 75 | 1879 | 10673 | 20143 | 0 | 225 | 0 | 75 | 0 |
| SRR1853809 | 0 | 0 | 22 | 0 | 0 | 0 | 15 | 3477 | 343 | 1288 | 7040 | 11032 | 0 | 215 | 86 | 0 | 0 |
| SRR1917323 | 0 | 17 | 59 | 81 | 128 | 0 | 18 | 97 | 211 | 2434 | 4734 | 13721 | 0 | 422 | 828 | 1034 | 0 |
| SRR1917324 | 0 | 409 | 1309 | 46 | 74 | 0 | 15 | 88 | 132 | 2202 | 3112 | 7338 | 0 | 326 | 578 | 652 | 0 |
| SRR1917325 | 0 | 39 | 65 | 131 | 234 | 0 | 10 | 101 | 306 | 4026 | 3214 | 13811 | 0 | 877 | 648 | 735 | 0 |
| SRR1917326 | 0 | 332 | 0 | 1501 | 100 | 0 | 0 | 1507 | 5041 | 5552 | 35 | 12 | 0 | 1061 | 0 | 0 | 0 |
| SRR1917327 | 0 | 46 | 0 | 540 | 65 | 0 | 0 | 536 | 2757 | 4179 | 101 | 36 | 0 | 3466 | 0 | 0 | 0 |
| SRR1917328 | 0 | 0 | 0 | 39 | 0 | 0 | 0 | 1080 | 6409 | 3398 | 10 | 203 | 0 | 1145 | 157 | 55 | 0 |
| SRR1917329 | 0 | 65 | 81 | 107 | 170 | 0 | 0 | 384 | 1453 | 6172 | 12980 | 18948 | 0 | 686 | 1129 | 1139 | 0 |
| SRR1917330 | 0 | 30 | 83 | 131 | 153 | 0 | 11 | 452 | 1373 | 6590 | 4902 | 11047 | 0 | 730 | 1421 | 1195 | 10 |
| SRR1917331 | 0 | 415 | 711 | 174 | 183 | 0 | 0 | 339 | 1137 | 6411 | 4515 | 13512 | 0 | 765 | 1377 | 2779 | 0 |
| SRR1917332 | 0 | 0 | 11 | 103 | 64 | 0 | 0 | 827 | 1347 | 6713 | 71 | 0 | 0 | 1655 | 95 | 95 | 0 |
| SRR1917333 | 0 | 0 | 0 | 74 | 34 | 0 | 0 | 401 | 763 | 6785 | 20 | 0 | 0 | 3232 | 100 | 40 | 0 |
| SRR1917334 | 0 | 0 | 0 | 89 | 29 | 0 | 12 | 446 | 1522 | 6331 | 83 | 13 | 0 | 5344 | 115 | 57 | 16 |

| | | | | | | | | | | | | | | | | | | | |
|---|---|---|---|---|---|---|---|---|---|---|---|---|---|---|---|---|---|---|---|
| SRR1708997 | 410 | 32 | 1441 | 620 | 2658 | 0 | 132 | 8233 | 0 | 17 | 31 | 31 | 195 | 392 | 0 | 52 | 159 | 27 | 3166 | 0 |
| SRR1708998 | 457 | 28 | 1175 | 1153 | 3937 | 0 | 45 | 12676 | 0 | 0 | 34 | 52 | 338 | 682 | 44 | 98 | 186 | 0 | 2687 | 0 |
| SRR1708999 | 512 | 24 | 466 | 619 | 3485 | 0 | 125 | 5733 | 0 | 0 | 23 | 12 | 232 | 399 | 12 | 21 | 19 | 0 | 650 | 0 |
| SRR1709000 | 556 | 19 | 1272 | 1203 | 3670 | 0 | 41 | 11418 | 0 | 16 | 50 | 38 | 140 | 425 | 17 | 86 | 300 | 0 | 2062 | 0 |
| SRR1709001 | 509 | 24 | 624 | 1087 | 3510 | 0 | 188 | 12755 | 0 | 15 | 55 | 38 | 148 | 311 | 0 | 23 | 35 | 0 | 1262 | 0 |
| SRR1709002 | 472 | 21 | 1514 | 1626 | 3655 | 0 | 58 | 13888 | 0 | 0 | 38 | 60 | 12 | 20 | 18 | 105 | 15 | 41 | 9044 | 0 |
| SRR1709003 | 452 | 20 | 861 | 982 | 2885 | 0 | 140 | 11578 | 0 | 0 | 74 | 77 | 255 | 220 | 24 | 30 | 110 | 0 | 1743 | 0 |
| SRR1709004 | 695 | 22 | 895 | 780 | 3736 | 0 | 24 | 6737 | 0 | 33 | 29 | 20 | 15 | 38 | 16 | 49 | 114 | 0 | 795 | 0 |
| SRR1709005 | 490 | 23 | 1697 | 992 | 3742 | 0 | 170 | 7951 | 0 | 17 | 17 | 30 | 18 | 36 | 32 | 73 | 13 | 0 | 2634 | 0 |
| SRR1709006 | 576 | 26 | 2133 | 1235 | 4119 | 0 | 41 | 9647 | 0 | 0 | 41 | 37 | 18 | 37 | 35 | 100 | 15 | 22 | 3860 | 0 |
| SRR1709007 | 448 | 30 | 1798 | 826 | 3569 | 0 | 149 | 10700 | 0 | 0 | 40 | 13 | 350 | 713 | 37 | 80 | 195 | 0 | 2672 | 0 |
| SRR1709008 | 612 | 29 | 586 | 989 | 3601 | 0 | 0 | 7747 | 0 | 0 | 39 | 11 | 12 | 35 | 37 | 24 | 52 | 0 | 1111 | 0 |
| SRR1709009 | 512 | 20 | 1150 | 941 | 3660 | 0 | 14 | 10866 | 0 | 0 | 33 | 13 | 140 | 430 | 33 | 53 | 90 | 0 | 3253 | 0 |
| SRR1709010 | 382 | 12 | 1040 | 1551 | 4356 | 0 | 41 | 13154 | 0 | 18 | 125 | 53 | 15 | 16 | 10 | 30 | 112 | 0 | 787 | 0 |
| SRR1709011 | 445 | 16 | 1339 | 1618 | 4509 | 0 | 161 | 13160 | 0 | 0 | 80 | 69 | 303 | 484 | 17 | 56 | 186 | 0 | 1513 | 0 |
| SRR1709012 | 385 | 21 | 799 | 1205 | 3168 | 0 | 48 | 13146 | 0 | 0 | 81 | 81 | 16 | 18 | 15 | 91 | 226 | 12 | 3840 | 0 |
| SRR1709013 | 427 | 15 | 629 | 999 | 3249 | 0 | 146 | 10597 | 0 | 0 | 66 | 63 | 19 | 12 | 0 | 27 | 76 | 0 | 833 | 0 |
| SRR1709014 | 467 | 24 | 773 | 1148 | 3167 | 0 | 17 | 13130 | 0 | 0 | 80 | 118 | 1069 | 984 | 18 | 71 | 174 | 0 | 1523 | 0 |
| SRR1709015 | 404 | 13 | 664 | 1816 | 3792 | 0 | 22 | 15191 | 0 | 0 | 70 | 57 | 215 | 15 | 19 | 57 | 121 | 0 | 1061 | 0 |
| SRR1709016 | 286 | 17 | 631 | 1085 | 2372 | 0 | 33 | 11242 | 0 | 12 | 71 | 83 | 242 | 234 | 0 | 47 | 145 | 0 | 2641 | 0 |
| SRR1709017 | 265 | 0 | 2165 | 1305 | 2296 | 0 | 94 | 9943 | 0 | 0 | 85 | 12 | 17 | 13 | 0 | 34 | 107 | 0 | 1498 | 0 |
| SRR1709018 | 247 | 15 | 789 | 1253 | 2088 | 0 | 18 | 8913 | 0 | 0 | 84 | 159 | 318 | 179 | 0 | 27 | 24 | 0 | 1471 | 0 |
| SRR1709019 | 240 | 14 | 730 | 1045 | 1795 | 0 | 73 | 9766 | 0 | 0 | 86 | 180 | 12 | 201 | 0 | 16 | 50 | 0 | 749 | 0 |
| SRR1709020 | 258 | 20 | 1067 | 1831 | 2885 | 0 | 17 | 12782 | 0 | 0 | 36 | 69 | 15 | 24 | 15 | 40 | 97 | 0 | 1478 | 0 |
| SRR1709021 | 267 | 0 | 1092 | 1826 | 2850 | 0 | 13 | 9769 | 0 | 0 | 89 | 143 | 12 | 13 | 11 | 18 | 47 | 0 | 1231 | 0 |
| SRR1732015 | 42 | 154 | 2165 | 1977 | 2208 | 22 | 8887 | 5123 | 62 | 15 | 11 | 24 | 11 | 26 | 20 | 15 | 23 | 0 | 3280 | 20 |
| SRR1732016 | 281 | 49 | 317 | 1720 | 1888 | 341 | 15687 | 3175 | 127 | 29 | 0 | 0 | 0 | 0 | 0 | 0 | 0 | 0 | 604 | 43 |
| SRR1748132 | 429 | 0 | 1062 | 671 | 1209 | 550 | 3842 | 970 | 13 | 22 | 0 | 0 | 28 | 11 | 0 | 12 | 14 | 0 | 9022 | 0 |
| SRR1748133 | 783 | 13 | 1472 | 1073 | 1682 | 179 | 3357 | 7180 | 19 | 0 | 0 | 0 | 0 | 0 | 0 | 0 | 0 | 0 | 5994 | 0 |
| SRR1759212 | 207 | 0 | 0 | 3290 | 5079 | 247 | 9377 | 11102 | 105 | 0 | 0 | 0 | 13 | 13 | 0 | 0 | 0 | 0 | 29 | 0 |
| SRR1759213 | 266 | 0 | 1303 | 3222 | 5832 | 341 | 12853 | 12171 | 181 | 0 | 0 | 0 | 0 | 0 | 0 | 0 | 0 | 0 | 4171 | 0 |
| SRR1759214 | 164 | 0 | 1455 | 2825 | 5816 | 296 | 12088 | 12570 | 196 | 0 | 0 | 0 | 0 | 12 | 0 | 0 | 0 | 0 | 5865 | 0 |
| SRR1759215 | 257 | 0 | 1575 | 3196 | 5577 | 280 | 12869 | 12316 | 204 | 0 | 0 | 0 | 10 | 0 | 0 | 0 | 0 | 0 | 5565 | 0 |
| SRR1759216 | 142 | 0 | 611 | 2934 | 6399 | 315 | 12140 | 12312 | 199 | 0 | 0 | 0 | 0 | 0 | 0 | 0 | 0 | 0 | 2287 | 0 |
| SRR1759217 | 552 | 0 | 1065 | 3042 | 3942 | 256 | 10366 | 8661 | 149 | 0 | 0 | 0 | 0 | 0 | 0 | 0 | 0 | 0 | 4186 | 0 |
| SRR1759218 | 340 | 0 | 1506 | 2373 | 3980 | 207 | 8758 | 9990 | 296 | 15 | 0 | 0 | 0 | 16 | 0 | 0 | 11 | 0 | 6410 | 0 |
| SRR1759219 | 234 | 0 | 1027 | 3162 | 13 | 316 | 10969 | 11782 | 153 | 24 | 0 | 0 | 14 | 12 | 0 | 0 | 0 | 10 | 4942 | 0 |
| SRR1759220 | 180 | 0 | 2630 | 2969 | 4889 | 304 | 10801 | 10733 | 283 | 0 | 0 | 0 | 0 | 0 | 0 | 0 | 16 | 0 | 12393 | 0 |
| SRR1759221 | 78 | 0 | 312 | 3594 | 5044 | 362 | 12486 | 7852 | 63 | 16 | 0 | 17 | 13 | 12 | 0 | 17 | 26 | 12 | 2247 | 0 |
| SRR1759222 | 85 | 0 | 1606 | 1540 | 3140 | 271 | 8919 | 5935 | 1364 | 17 | 0 | 23 | 28 | 17 | 0 | 20 | 46 | 0 | 4246 | 0 |
| SRR1759223 | 145 | 0 | 1097 | 2696 | 6063 | 583 | 16842 | 11781 | 183 | 0 | 0 | 17 | 18 | 28 | 0 | 24 | 34 | 13 | 6175 | 0 |
| SRR1759224 | 42 | 0 | 1445 | 3205 | 3320 | 579 | 23660 | 5553 | 68 | 0 | 0 | 0 | 33 | 33 | 0 | 22 | 46 | 0 | 10525 | 0 |
| SRR1759225 | 62 | 0 | 368 | 2443 | 4641 | 396 | 9795 | 7419 | 468 | 0 | 0 | 17 | 13 | 14 | 0 | 13 | 27 | 0 | 1735 | 0 |
| SRR1759226 | 55 | 0 | 1328 | 2047 | 2220 | 400 | 21895 | 3090 | 223 | 11 | 0 | 0 | 20 | 25 | 0 | 23 | 26 | 0 | 8050 | 0 |
| SRR1759227 | 81 | 0 | 397 | 3080 | 4434 | 423 | 11340 | 7662 | 268 | 19 | 0 | 0 | 17 | 12 | 0 | 13 | 43 | 15 | 2088 | 0 |
| SRR1759228 | 79 | 0 | 122 | 1824 | 2527 | 320 | 7624 | 5808 | 931 | 25 | 0 | 0 | 0 | 0 | 12 | 23 | 27 | 0 | 857 | 0 |
| SRR1759229 | 117 | 0 | 628 | 2459 | 5044 | 507 | 13070 | 10183 | 229 | 12 | 0 | 0 | 10 | 14 | 0 | 10 | 22 | 20 | 3455 | 0 |
| SRR1759230 | 61 | 0 | 296 | 2302 | 3263 | 574 | 13282 | 6412 | 122 | 17 | 0 | 0 | 16 | 0 | 0 | 11 | 16 | 0 | 2351 | 0 |
| SRR1759231 | 105 | 0 | 216 | 2401 | 3017 | 258 | 8447 | 6742 | 562 | 0 | 0 | 0 | 22 | 20 | 0 | 16 | 31 | 0 | 765 | 0 |
| SRR1759232 | 53 | 0 | 527 | 2563 | 4122 | 492 | 12286 | 6900 | 148 | 17 | 0 | 17 | 50 | 20 | 0 | 43 | 76 | 13 | 3874 | 0 |
| SRR1759233 | 56 | 0 | 164 | 2766 | 3532 | 463 | 12648 | 6578 | 230 | 35 | 0 | 11 | 27 | 16 | 0 | 19 | 21 | 0 | 997 | 0 |
| SRR1759234 | 190 | 0 | 498 | 2064 | 4650 | 304 | 9820 | 10440 | 297 | 0 | 0 | 15 | 10 | 10 | 0 | 0 | 37 | 0 | 2195 | 0 |
| SRR1759235 | 149 | 0 | 467 | 2332 | 4735 | 192 | 7837 | 9169 | 552 | 0 | 0 | 0 | 18 | 0 | 0 | 12 | 17 | 10 | 2277 | 0 |
| SRR1759236 | 99 | 0 | 822 | 2514 | 5517 | 554 | 14002 | 10058 | 270 | 0 | 0 | 11 | 18 | 23 | 0 | 0 | 59 | 24 | 3322 | 0 |
| SRR1759237 | 171 | 0 | 1007 | 1977 | 3610 | 364 | 11044 | 8124 | 677 | 0 | 0 | 12 | 21 | 23 | 12 | 14 | 37 | 35 | 4232 | 0 |
| SRR1759238 | 164 | 0 | 832 | 2729 | 4375 | 467 | 13861 | 9732 | 157 | 0 | 0 | 21 | 40 | 15 | 11 | 21 | 34 | 49 | 4011 | 0 |
| SRR1759239 | 297 | 14 | 154 | 1524 | 2569 | 454 | 11743 | 12419 | 133 | 21 | 0 | 31 | 30 | 16 | 0 | 0 | 0 | 0 | 798 | 0 |
| SRR1759240 | 154 | 0 | 1364 | 1954 | 5120 | 414 | 12878 | 8518 | 396 | 0 | 0 | 15 | 28 | 20 | 0 | 22 | 52 | 22 | 6140 | 0 |
| SRR1759241 | 189 | 0 | 1233 | 2623 | 5315 | 483 | 13338 | 9943 | 324 | 0 | 0 | 23 | 34 | 25 | 0 | 24 | 59 | 14 | 6427 | 0 |
| SRR1759242 | 200 | 0 | 151 | 1727 | 2695 | 237 | 7407 | 5932 | 808 | 0 | 0 | 16 | 27 | 37 | 11 | 14 | 63 | 30 | 767 | 0 |
| SRR1759243 | 228 | 0 | 365 | 2055 | 3645 | 330 | 10352 | 7991 | 562 | 0 | 0 | 28 | 27 | 23 | 0 | 10 | 40 | 15 | 1548 | 0 |
| SRR1759244 | 435 | 0 | 752 | 3529 | 4670 | 283 | 11445 | 10216 | 140 | 0 | 0 | 0 | 0 | 0 | 0 | 0 | 0 | 0 | 2038 | 0 |
| SRR1759245 | 172 | 0 | 356 | 3565 | 5760 | 291 | 12812 | 13323 | 129 | 0 | 0 | 0 | 0 | 0 | 0 | 0 | 0 | 0 | 1076 | 0 |
| SRR1759246 | 187 | 0 | 473 | 3090 | 3850 | 477 | 9908 | 6454 | 58 | 0 | 0 | 0 | 11 | 0 | 0 | 16 | 55 | 19 | 3316 | 0 |
| SRR1759247 | 65 | 0 | 215 | 2788 | 3342 | 460 | 11279 | 6879 | 377 | 0 | 0 | 13 | 19 | 0 | 0 | 30 | 28 | 0 | 1267 | 0 |
| SRR1759248 | 136 | 0 | 969 | 3004 | 5510 | 498 | 16554 | 11617 | 135 | 0 | 0 | 13 | 32 | 16 | 0 | 15 | 69 | 14 | 3863 | 0 |
| SRR1759249 | 170 | 0 | 898 | 3234 | 6704 | 786 | 19385 | 11446 | 148 | 0 | 0 | 0 | 0 | 0 | 0 | 0 | 21 | 11 | 4612 | 0 |
| SRR1759250 | 190 | 0 | 1968 | 1799 | 4343 | 360 | 13912 | 10566 | 619 | 0 | 0 | 11 | 0 | 18 | 0 | 15 | 18 | 0 | 7709 | 0 |
| SRR1759251 | 144 | 0 | 523 | 2594 | 5488 | 657 | 18256 | 11172 | 181 | 11 | 0 | 0 | 20 | 11 | 0 | 0 | 26 | 11 | 2640 | 0 |
| SRR1759252 | 175 | 0 | 534 | 2126 | 4545 | 567 | 14165 | 10196 | 75 | 21 | 0 | 23 | 14 | 25 | 0 | 18 | 64 | 18 | 2295 | 13 |
| SRR1759253 | 87 | 0 | 1460 | 1881 | 6367 | 390 | 12288 | 9091 | 391 | 0 | 0 | 0 | 20 | 17 | 0 | 22 | 66 | 20 | 5160 | 0 |
| SRR1759254 | 243 | 0 | 1134 | 2774 | 5129 | 688 | 18690 | 11112 | 185 | 0 | 0 | 18 | 16 | 14 | 0 | 12 | 15 | 12 | 3941 | 0 |
| SRR1759255 | 60 | 0 | 641 | 2043 | 6913 | 493 | 15577 | 9427 | 164 | 0 | 0 | 0 | 11 | 0 | 0 | 11 | 15 | 0 | 2836 | 0 |
| SRR1759256 | 154 | 0 | 1479 | 2574 | 5521 | 537 | 14321 | 10153 | 132 | 0 | 0 | 31 | 47 | 36 | 0 | 17 | 43 | 14 | 5296 | 0 |
| SRR1759257 | 212 | 0 | 223 | 2161 | 3448 | 475 | 12571 | 9740 | 256 | 0 | 0 | 14 | 0 | 0 | 0 | 0 | 0 | 0 | 493 | 0 |
| SRR1759258 | 150 | 0 | 169 | 2498 | 4763 | 326 | 8137 | 8264 | 71 | 155 | 0 | 58 | 84 | 62 | 24 | 53 | 113 | 103 | 1415 | 0 |
| SRR1759259 | 167 | 0 | 1083 | 2424 | 3826 | 440 | 12022 | 7658 | 257 | 0 | 0 | 46 | 69 | 76 | 23 | 66 | 126 | 35 | 4459 | 0 |
| SRR1759260 | 134 | 0 | 218 | 2484 | 4711 | 266 | 11131 | 12624 | 152 | 0 | 0 | 14 | 14 | 0 | 0 | 0 | 0 | 0 | 991 | 0 |
| SRR1759261 | 78 | 0 | 383 | 3442 | 4503 | 363 | 11042 | 7496 | 170 | 36 | 0 | 19 | 36 | 19 | 0 | 12 | 35 | 22 | 2315 | 0 |
| SRR1759262 | 196 | 0 | 257 | 2902 | 3894 | 325 | 10986 | 11206 | 172 | 0 | 0 | 0 | 10 | 0 | 0 | 0 | 0 | 0 | 2354 | 0 |
| SRR1759263 | 81 | 0 | 695 | 2590 | 5197 | 321 | 9185 | 11 | 509 | 11 | 0 | 35 | 19 | 14 | 0 | 11 | 25 | 0 | 2119 | 0 |
| SRR1759264 | 62 | 0 | 812 | 3304 | 4557 | 609 | 18375 | 8055 | 139 | 12 | 0 | 0 | 17 | 23 | 0 | 12 | 25 | 0 | 3489 | 0 |
| SRR1759265 | 106 | 0 | 146 | 1876 | 2356 | 371 | 9427 | 6701 | 961 | 12 | 0 | 0 | 31 | 0 | 0 | 0 | 24 | 0 | 488 | 0 |
| SRR1759266 | 247 | 0 | 227 | 3212 | 4484 | 413 | 14197 | 13173 | 65 | 13 | 0 | 0 | 0 | 0 | 0 | 0 | 0 | 0 | 904 | 0 |
| SRR1759267 | 199 | 0 | 718 | 3396 | 3455 | 777 | 15375 | 7306 | 57 | 25 | 0 | 11 | 23 | 25 | 0 | 14 | 0 | 0 | 7840 | 0 |
| SRR1759268 | 85 | 0 | 566 | 3005 | 3321 | 725 | 15424 | 6266 | 67 | 44 | 0 | 12 | 24 | 22 | 15 | 0 | 44 | 0 | 2770 | 0 |
| SRR1759269 | 88 | 0 | 498 | 4084 | 5133 | 387 | 11130 | 8113 | 416 | 13 | 0 | 14 | 14 | 0 | 0 | 0 | 16 | 0 | 1439 | 0 |
| SRR1759270 | 198 | 0 | 406 | 3187 | 4501 | 261 | 10312 | 12256 | 98 | 0 | 0 | 15 | 17 | 0 | 0 | 0 | 0 | 13 | 985 | 0 |
| SRR1759271 | 45 | 0 | 673 | 3073 | 4656 | 412 | 12662 | 7132 | 297 | 12 | 0 | 14 | 14 | 12 | 0 | 18 | 26 | 0 | 2679 | 0 |
| SRR1759272 | 101 | 0 | 627 | 2254 | 2840 | 477 | 14694 | 6020 | 178 | 17 | 0 | 15 | 19 | 32 | 0 | 32 | 32 | 0 | 3539 | 0 |
| SRR1759273 | 513 | 0 | 511 | 3201 | 4009 | 1169 | 19330 | 5230 | 51 | 23 | 0 | 17 | 23 | 17 | 0 | 25 | 12 | 0 | 2516 | 0 |
| SRR1759274 | 176 | 0 | 250 | 2962 | 4725 | 339 | 12810 | 14311 | 66 | 10 | 0 | 35 | 24 | 0 | 0 | 0 | 0 | 0 | 1016 | 0 |
| SRR1759275 | 310 | 0 | 285 | 2327 | 3706 | 206 | 7848 | 8398 | 335 | 0 | 0 | 17 | 22 | 0 | 0 | 12 | 24 | 13 | 1195 | 0 |
| SRR1777167 | 167 | 190 | 351 | 43 | 602 | 0 | 2006 | 647 | 52 | 45 | 0 | 12 | 51 | 27 | 19 | 50 | 11 | 0 | 1021 | 35 |
| SRR1820679 | 1588 | 0 | 3423 | 488 | 1402 | 29 | 3799 | 1587 | 0 | 0 | 0 | 15 | 12 | 19 | 0 | 0 | 0 | 10 | 4824 | 0 |
| SRR1820680 | 640 | 0 | 1390 | 612 | 1213 | 27 | 1821 | 880 | 0 | 14 | 0 | 18 | 0 | 0 | 0 | 0 | 0 | 0 | 1154 | 0 |
| SRR1820681 | 979 | 0 | 4163 | 562 | 1488 | 28 | 4026 | 1671 | 0 | 0 | 0 | 21 | 0 | 15 | 0 | 0 | 0 | 0 | 4527 | 0 |
| SRR1820682 | 742 | 0 | 2209 | 591 | 1354 | 25 | 2333 | 846 | 0 | 39 | 0 | 21 | 0 | 19 | 0 | 0 | 0 | 0 | 3151 | 0 |
| SRR1820683 | 1567 | 0 | 3017 | 472 | 1488 | 28 | 3924 | 1636 | 0 | 0 | 0 | 0 | 0 | 0 | 0 | 0 | 0 | 0 | 4525 | 0 |
| SRR1820684 | 982 | 0 | 2374 | 320 | 996 | 16 | 1872 | 757 | 14 | 33 | 0 | 31 | 18 | 29 | 0 | 0 | 0 | 0 | 1966 | 0 |
| SRR1820685 | 1121 | 0 | 3878 | 394 | 1176 | 23 | 3674 | 1266 | 0 | 0 | 0 | 13 | 12 | 12 | 10 | 0 | 0 | 10 | 4487 | 0 |
| SRR1820686 | 730 | 0 | 1223 | 424 | 1388 | 19 | 1936 | 733 | 0 | 10 | 0 | 22 | 16 | 23 | 0 | 0 | 0 | 0 | 1109 | 0 |
| SRR1820687 | 1004 | 0 | 7206 | 561 | 1273 | 23 | 3932 | 1015 | 0 | 12 | 0 | 19 | 16 | 24 | 0 | 0 | 0 | 0 | 5262 | 0 |
| SRR1820688 | 846 | 0 | 3747 | 619 | 1015 | 25 | 3527 | 1086 | 0 | 32 | 0 | 17 | 14 | 26 | 0 | 0 | 0 | 0 | 3868 | 0 |
| SRR1820689 | 1194 | 13 | 3008 | 380 | 712 | 33 | 5035 | 1036 | 0 | 26 | 0 | 16 | 12 | 24 | 0 | 0 | 0 | 0 | 2202 | 0 |
| SRR1820690 | 745 | 0 | 2899 | 560 | 732 | 29 | 3224 | 987 | 0 | 41 | 0 | 17 | 0 | 13 | 0 | 0 | 0 | 0 | 1860 | 0 |
| SRR1853804 | 45 | 0 | 376 | 460 | 1066 | 0 | 0 | 1021 | 0 | 0 | 0 | 0 | 39 | 21 | 0 | 15 | 48 | 18 | 1573 | 0 |
| SRR1853805 | 32 | 0 | 52 | 424 | 1152 | 0 | 0 | 1399 | 0 | 0 | 0 | 0 | 61 | 18 | 0 | 18 | 53 | 53 | 932 | 0 |
| SRR1853806 | 142 | 0 | 300 | 570 | 1303 | 0 | 0 | 1578 | 0 | 12 | 0 | 20 | 60 | 28 | 11 | 28 | 54 | 19 | 1282 | 0 |
| SRR1853807 | 67 | 10 | 524 | 306 | 841 | 0 | 0 | 932 | 0 | 32 | 0 | 21 | 53 | 71 | 13 | 17 | 61 | 32 | 7532 | 0 |
| SRR1853808 | 146 | 11 | 601 | 675 | 995 | 0 | 0 | 1362 | 0 | 35 | 0 | 16 | 51 | 119 | 13 | 51 | 93 | 51 | 8042 | 0 |
| SRR1853809 | 66 | 0 | 773 | 399 | 1027 | 0 | 0 | 1994 | 0 | 55 | 0 | 15 | 57 | 127 | 20 | 70 | 70 | 135 | 3821 | 0 |
| SRR1917323 | 428 | 113 | 2546 | 170 | 285 | 0 | 59 | 2173 | 44 | 52 | 99 | 34 | 35 | 192 | 76 | 152 | 131 | 0 | 9210 | 1521 |
| SRR1917324 | 252 | 124 | 8115 | 89 | 185 | 0 | 53 | 1306 | 48 | 611 | 82 | 594 | 517 | 4051 | 1574 | 392 | 381 | 13 | 37369 | 836 |
| SRR1917325 | 503 | 150 | 4087 | 184 | 350 | 0 | 65 | 2132 | 39 | 13 | 95 | 28 | 50 | 265 | 87 | 28 | 19 | 0 | 12477 | 1196 |
| SRR1917326 | 3721 | 950 | 0 | 284 | 366 | 0 | 181 | 678 | 92 | 0 | 0 | 0 | 0 | 0 | 0 | 0 | 0 | 0 | 12 | 47 |
| SRR1917327 | 1079 | 620 | 0 | 138 | 391 | 0 | 136 | 556 | 143 | 0 | 0 | 0 | 0 | 0 | 0 | 0 | 0 | 0 | 12 | 94 |
| SRR1917328 | 397 | 413 | 92 | 24 | 42 | 0 | 20 | 83 | 56 | 0 | 0 | 0 | 0 | 0 | 0 | 0 | 0 | 0 | 55 | 11 |
| SRR1917329 | 547 | 171 | 2304 | 235 | 494 | 0 | 84 | 1746 | 37 | 116 | 63 | 27 | 34 | 135 | 48 | 16 | 13 | 0 | 8114 | 1214 |
| SRR1917330 | 466 | 164 | 4157 | 208 | 399 | 0 | 96 | 1620 | 37 | 36 | 45 | 25 | 36 | 193 | 78 | 25 | 16 | 14 | 16100 | 1090 |
| SRR1917331 | 687 | 187 | 2466 | 241 | 579 | 0 | 82 | 1866 | 35 | 474 | 38 | 226 | 330 | 1512 | 678 | 315 | 217 | 26 | 8254 | 1228 |
| SRR1917332 | 281 | 316 | 95 | 99 | 92 | 0 | 21 | 146 | 14 | 21 | 0 | 0 | 0 | 0 | 0 | 0 | 0 | 0 | 47 | 25 |
| SRR1917333 | 280 | 366 | 20 | 51 | 74 | 0 | 74 | 154 | 34 | 0 | 0 | 0 | 0 | 0 | 0 | 0 | 0 | 0 | 80 | 11 |
| SRR1917334 | 266 | 400 | 0 | 43 | 62 | 0 | 66 | 159 | 23 | 36 | 0 | 0 | 0 | 0 | 0 | 0 | 0 | 0 | 19 | 27 |

### Supp Table S8.4.Columns_DI_to_ES

| | | | | | | | | | | | | | | | | | | | | | | | | | | | | | | | | | | | | | |
|---|---|---|---|---|---|---|---|---|---|---|---|---|---|---|---|---|---|---|---|---|---|---|---|---|---|---|---|---|---|---|---|---|---|---|---|---|---|
| SRR1917335 | 0 | 28 | 69 | 93 | 162 | 0 | 0 | 140 | 343 | 3241 | 5826 | 14249 | 0 | 476 | 883 | 864 | 0 | 390 | 130 | 3665 | 177 | 314 | 0 | 63 | 2168 | 39 | 20 | 93 | 31 | 35 | 183 | 89 | 37 | 23 | 0 | 11244 | 1283 |
| SRR1917336 | 0 | 326 | 965 | 133 | 174 | 0 | 24 | 132 | 341 | 3169 | 8098 | 13443 | 0 | 368 | 661 | 533 | 11 | 682 | 113 | 5579 | 191 | 344 | 0 | 58 | 2073 | 39 | 698 | 134 | 733 | 1030 | 4156 | 1660 | 452 | 281 | 0 | 16027 | 1893 |
| SRR1917337 | 0 | 41 | 51 | 239 | 258 | 0 | 32 | 133 | 335 | 2883 | 7896 | 13231 | 0 | 566 | 491 | 560 | 14 | 1007 | 139 | 5226 | 290 | 550 | 0 | 54 | 2208 | 39 | 82 | 71 | 38 | 53 | 171 | 63 | 26 | 18 | 0 | 17161 | 1724 |
| SRR1917338 | 0 | 0 | 0 | 29 | 0 | 0 | 0 | 365 | 990 | 2198 | 19 | 0 | 0 | 4353 | 0 | 0 | 0 | 55 | 274 | 14 | 16 | 84 | 0 | 52 | 106 | 61 | 15 | 0 | 0 | 0 | 0 | 0 | 0 | 0 | 0 | 14 | 0 |
| SRR1917339 | 0 | 0 | 0 | 24 | 0 | 0 | 0 | 463 | 2203 | 3129 | 41 | 0 | 0 | 3149 | 31 | 10 | 0 | 95 | 380 | 0 | 0 | 128 | 0 | 185 | 147 | 86 | 15 | 0 | 0 | 0 | 18 | 0 | 0 | 0 | 0 | 10 | 0 |
| SRR1917340 | 0 | 0 | 0 | 24 | 17 | 0 | 0 | 431 | 1948 | 3526 | 90 | 17 | 0 | 3901 | 34 | 0 | 0 | 145 | 416 | 0 | 36 | 95 | 0 | 155 | 90 | 88 | 45 | 0 | 0 | 0 | 12 | 0 | 0 | 0 | 0 | 50 | 0 |
| SRR1922516 | 0 | 0 | 0 | 0 | 0 | 0 | 0 | 323 | 176 | 4058 | 8176 | 14778 | 0 | 1162 | 637 | 1067 | 0 | 54 | 22 | 2169 | 462 | 2430 | 12 | 3315 | 2227 | 130 | 0 | 0 | 0 | 0 | 0 | 0 | 0 | 0 | 0 | 7273 | 15 |
| SRR1922517 | 0 | 0 | 0 | 0 | 0 | 0 | 0 | 349 | 137 | 4248 | 10018 | 20310 | 0 | 896 | 543 | 914 | 0 | 49 | 15 | 2144 | 305 | 1807 | 0 | 1970 | 1281 | 139 | 0 | 0 | 0 | 0 | 0 | 0 | 0 | 0 | 0 | 3914 | 16 |
| SRR1922518 | 0 | 0 | 0 | 0 | 0 | 0 | 0 | 292 | 135 | 3432 | 6393 | 10270 | 0 | 841 | 559 | 960 | 0 | 45 | 29 | 2682 | 529 | 2645 | 26 | 3387 | 2214 | 116 | 0 | 0 | 0 | 0 | 0 | 0 | 0 | 0 | 0 | 7875 | 14 |
| SRR1922519 | 0 | 0 | 0 | 0 | 0 | 0 | 0 | 264 | 60 | 2403 | 3330 | 16280 | 0 | 578 | 468 | 842 | 0 | 51 | 15 | 1866 | 374 | 1797 | 13 | 2177 | 1382 | 109 | 0 | 0 | 0 | 0 | 0 | 0 | 0 | 0 | 0 | 2545 | 14 |
| SRR1922520 | 0 | 0 | 0 | 0 | 0 | 0 | 0 | 279 | 74 | 2356 | 1367 | 11856 | 0 | 581 | 404 | 702 | 0 | 47 | 13 | 1431 | 335 | 1188 | 11 | 2240 | 1418 | 65 | 0 | 0 | 0 | 0 | 0 | 0 | 0 | 0 | 0 | 3272 | 18 |
| SRR1922521 | 0 | 0 | 0 | 0 | 0 | 0 | 0 | 267 | 79 | 3195 | 2822 | 17171 | 0 | 861 | 551 | 929 | 0 | 25 | 24 | 1174 | 336 | 2302 | 21 | 3572 | 2291 | 160 | 0 | 0 | 0 | 0 | 0 | 0 | 0 | 0 | 0 | 4279 | 0 |
| SRR1922522 | 0 | 0 | 0 | 0 | 0 | 0 | 0 | 287 | 100 | 3028 | 1342 | 13790 | 0 | 368 | 474 | 627 | 0 | 49 | 22 | 1358 | 423 | 1565 | 0 | 2836 | 1988 | 47 | 0 | 0 | 0 | 0 | 0 | 0 | 0 | 0 | 0 | 3396 | 14 |
| SRR1922523 | 0 | 0 | 0 | 0 | 0 | 0 | 0 | 225 | 72 | 2588 | 2254 | 21290 | 0 | 656 | 354 | 492 | 0 | 25 | 24 | 728 | 392 | 2015 | 36 | 3716 | 2356 | 140 | 0 | 0 | 0 | 0 | 0 | 0 | 0 | 0 | 0 | 2741 | 0 |
| SRR1922524 | 0 | 0 | 0 | 0 | 0 | 0 | 0 | 249 | 104 | 3632 | 3060 | 15270 | 0 | 488 | 511 | 794 | 0 | 36 | 22 | 1795 | 329 | 1165 | 11 | 2673 | 1754 | 84 | 0 | 0 | 0 | 0 | 0 | 0 | 0 | 12 | 0 | 4735 | 0 |
| SRR1922525 | 0 | 0 | 0 | 0 | 0 | 0 | 0 | 299 | 83 | 2981 | 1514 | 13998 | 0 | 909 | 488 | 698 | 0 | 38 | 22 | 1751 | 425 | 2276 | 27 | 4094 | 2554 | 144 | 0 | 0 | 0 | 0 | 0 | 0 | 0 | 0 | 0 | 4101 | 12 |
| SRR1922526 | 0 | 0 | 0 | 0 | 0 | 0 | 0 | 261 | 138 | 2500 | 1064 | 11498 | 0 | 840 | 391 | 500 | 0 | 325 | 18 | 2050 | 415 | 1469 | 0 | 2804 | 1819 | 172 | 0 | 0 | 0 | 0 | 0 | 0 | 0 | 10 | 0 | 3514 | 0 |
| SRR1922527 | 0 | 0 | 0 | 0 | 0 | 0 | 0 | 201 | 71 | 2717 | 9204 | 15337 | 0 | 420 | 524 | 844 | 0 | 28 | 18 | 1100 | 459 | 2353 | 15 | 4939 | 2920 | 82 | 0 | 0 | 0 | 0 | 0 | 0 | 0 | 0 | 0 | 4323 | 0 |
| SRR1973407 | 0 | 32 | 117 | 0 | 11 | 21 | 0 | 2036 | 814 | 23680 | 0 | 0 | 0 | 475 | 1018 | 1493 | 0 | 71 | 14 | 746 | 1251 | 4080 | 39 | 6151 | 5385 | 37 | 0 | 21 | 0 | 53 | 42 | 64 | 21 | 64 | 42 | 5564 | 93 |
| SRR1973408 | 0 | 35 | 59 | 0 | 15 | 12 | 0 | 1317 | 852 | 22859 | 0 | 0 | 0 | 775 | 930 | 852 | 0 | 107 | 14 | 1162 | 1082 | 3495 | 57 | 6437 | 4285 | 40 | 0 | 0 | 35 | 59 | 59 | 47 | 12 | 47 | 130 | 5114 | 31 |
| SRR1973409 | 0 | 20 | 89 | 0 | 16 | 0 | 0 | 1702 | 1513 | 21839 | 0 | 0 | 0 | 1229 | 804 | 520 | 0 | 183 | 0 | 520 | 1876 | 4792 | 61 | 5384 | 7203 | 19 | 0 | 0 | 30 | 30 | 20 | 0 | 89 | 128 | 49 | 2931 | 37 |
| SRR1973410 | 0 | 30 | 60 | 0 | 0 | 0 | 20 | 460 | 115 | 7069 | 0 | 0 | 0 | 1667 | 1379 | 1035 | 0 | 286 | 17 | 1379 | 2614 | 6011 | 61 | 6037 | 10163 | 23 | 0 | 0 | 70 | 30 | 79 | 40 | 20 | 119 | 30 | 6552 | 76 |
| SRR1973411 | 0 | 0 | 124 | 0 | 11 | 0 | 0 | 724 | 0 | 9413 | 0 | 0 | 0 | 1207 | 2896 | 724 | 0 | 150 | 11 | 724 | 2437 | 5855 | 60 | 5555 | 10519 | 37 | 0 | 31 | 31 | 31 | 0 | 31 | 0 | 93 | 31 | 3621 | 67 |
| SRR1973412 | 0 | 0 | 11 | 0 | 21 | 0 | 15 | 645 | 369 | 5622 | 0 | 0 | 0 | 1475 | 645 | 461 | 0 | 217 | 0 | 645 | 2591 | 6162 | 59 | 5149 | 9556 | 26 | 0 | 45 | 60 | 75 | 15 | 15 | 60 | 195 | 60 | 5806 | 35 |
| SRR1973413 | 0 | 45 | 225 | 0 | 14 | 0 | 11 | 1162 | 155 | 8987 | 0 | 0 | 0 | 1550 | 310 | 775 | 0 | 150 | 0 | 1782 | 2367 | 5501 | 71 | 6773 | 8973 | 31 | 0 | 11 | 90 | 90 | 45 | 11 | 112 | 270 | 11 | 5888 | 17 |
| SRR1973414 | 0 | 0 | 191 | 0 | 0 | 0 | 0 | 1036 | 188 | 4710 | 0 | 0 | 0 | 1225 | 659 | 471 | 0 | 156 | 0 | 848 | 2681 | 5602 | 82 | 7928 | 7703 | 49 | 0 | 15 | 73 | 88 | 15 | 15 | 147 | 117 | 59 | 7160 | 39 |
| SRR1973415 | 0 | 0 | 81 | 0 | 21 | 0 | 0 | 121 | 242 | 2421 | 0 | 0 | 0 | 2179 | 484 | 363 | 0 | 222 | 0 | 726 | 2566 | 6530 | 88 | 7931 | 7143 | 31 | 0 | 16 | 97 | 16 | 32 | 32 | 64 | 145 | 0 | 6658 | 19 |
| SRR1973952 | 0 | 2799 | 26 | 0 | 821 | 0 | 13 | 146 | 49 | 2092 | 10606 | 35515 | 64 | 195 | 49 | 97 | 13 | 263 | 143 | 1557 | 25 | 307 | 25 | 1201 | 1038 | 0 | 18842 | 129 | 993 | 2205 | 20002 | 4114 | 2760 | 1290 | 310 | 1995 | 3428 |
| SRR1973953 | 0 | 1040 | 14 | 16 | 750 | 0 | 10 | 487 | 1680 | 2048 | 6134 | 10488 | 0 | 925 | 80 | 408 | 13 | 566 | 550 | 1233 | 1804 | 241 | 0 | 272 | 2172 | 0 | 92 | 15 | 404 | 153 | 279 | 52 | 48 | 61 | 118 | 2923 | 1741 |
| SRR1973956 | 0 | 17 | 0 | 0 | 0 | 0 | 14 | 604 | 439 | 1152 | 0 | 219 | 0 | 1427 | 0 | 55 | 31 | 1159 | 193 | 110 | 0 | 0 | 0 | 579 | 193 | 0 | 24 | 0 | 0 | 0 | 0 | 0 | 0 | 0 | 0 | 0 | 0 |
| SRR1973958 | 0 | 0 | 0 | 0 | 88 | 0 | 0 | 1734 | 3287 | 3154 | 0 | 0 | 0 | 261 | 0 | 0 | 22 | 175 | 0 | 0 | 350 | 175 | 0 | 1402 | 0 | 0 | 0 | 0 | 0 | 0 | 0 | 0 | 0 | 0 | 0 | 0 | 0 |
| SRR1973964 | 0 | 0 | 0 | 0 | 0 | 0 | 0 | 5646 | 18863 | 1294 | 0 | 0 | 0 | 76 | 0 | 0 | 0 | 0 | 162 | 0 | 0 | 0 | 0 | 284 | 122 | 0 | 0 | 0 | 0 | 0 | 0 | 0 | 0 | 0 | 0 | 0 | 41 |
| SRR1973966 | 0 | 752 | 58 | 18 | 69 | 0 | 0 | 1125 | 2177 | 2696 | 5454 | 16188 | 0 | 3042 | 74 | 470 | 18 | 2613 | 89 | 1719 | 386 | 563 | 13 | 7381 | 3332 | 23 | 18 | 0 | 51 | 23 | 50 | 19 | 16 | 12 | 0 | 3376 | 137 |
| SRR1973967 | 0 | 108 | 0 | 61 | 91 | 0 | 13 | 1176 | 4334 | 3722 | 215 | 4624 | 0 | 2208 | 32 | 177 | 16 | 3189 | 390 | 467 | 444 | 85 | 0 | 4364 | 3913 | 0 | 33 | 13 | 59 | 46 | 60 | 14 | 10 | 12 | 42 | 870 | 353 |
| SRR1979367 | 0 | 373 | 0 | 0 | 15 | 33 | 0 | 0 | 29 | 181 | 534 | 11416 | 33 | 17 | 111 | 93 | 0 | 2856 | 48 | 0 | 2443 | 7822 | 13 | 435 | 1462 | 0 | 0 | 381 | 91 | 17 | 17 | 0 | 0 | 17 | 0 | 0 | 6953 |
| SRR1988280 | 0 | 25 | 10 | 0 | 0 | 0 | 0 | 737 | 292 | 24681 | 4975 | 10716 | 0 | 1142 | 0 | 0 | 0 | 27 | 22 | 366 | 639 | 4265 | 0 | 3064 | 2570 | 0 | 0 | 0 | 0 | 25 | 25 | 15 | 27 | 17 | 0 | 5054 | 0 |
| SRR1988281 | 0 | 31 | 10 | 0 | 0 | 0 | 0 | 784 | 423 | 22213 | 1251 | 12191 | 0 | 1228 | 0 | 0 | 0 | 33 | 27 | 371 | 702 | 4935 | 12 | 3427 | 2977 | 0 | 12 | 0 | 10 | 109 | 113 | 0 | 38 | 65 | 16 | 6255 | 0 |
| SRR1988282 | 0 | 26 | 0 | 0 | 0 | 0 | 0 | 712 | 391 | 25560 | 1135 | 10273 | 0 | 992 | 0 | 0 | 0 | 47 | 22 | 436 | 771 | 4778 | 0 | 2883 | 3110 | 0 | 0 | 0 | 0 | 28 | 27 | 12 | 23 | 32 | 12 | 6923 | 0 |
| SRR1988283 | 0 | 10 | 0 | 0 | 0 | 0 | 0 | 980 | 213 | 6428 | 2173 | 5955 | 0 | 9009 | 0 | 0 | 0 | 370 | 0 | 1852 | 663 | 1771 | 0 | 10316 | 3305 | 242 | 35 | 16 | 18 | 15 | 15 | 22 | 48 | 47 | 23 | 17161 | 0 |
| SRR1988284 | 0 | 23 | 0 | 0 | 10 | 0 | 0 | 1214 | 236 | 11007 | 8301 | 7186 | 0 | 12797 | 0 | 0 | 0 | 311 | 0 | 1643 | 520 | 2027 | 0 | 9140 | 2746 | 252 | 28 | 93 | 13 | 60 | 64 | 15 | 53 | 56 | 15 | 13687 | 0 |
| SRR1988285 | 0 | 24 | 0 | 0 | 0 | 0 | 0 | 1158 | 368 | 10180 | 1425 | 6344 | 0 | 10137 | 0 | 0 | 0 | 313 | 0 | 2203 | 548 | 2013 | 0 | 8703 | 2670 | 259 | 32 | 69 | 11 | 71 | 54 | 17 | 42 | 65 | 13 | 19207 | 0 |
| SRR1988286 | 0 | 46 | 0 | 0 | 0 | 0 | 0 | 768 | 184 | 6851 | 2496 | 6086 | 0 | 14233 | 0 | 0 | 0 | 503 | 0 | 1770 | 682 | 1708 | 0 | 9406 | 3849 | 239 | 20 | 10 | 12 | 13 | 14 | 19 | 30 | 47 | 17 | 13010 | 0 |
| SRR1988287 | 0 | 11 | 0 | 0 | 0 | 0 | 0 | 275 | 212 | 17799 | 1026 | 3030 | 0 | 1562 | 0 | 0 | 0 | 35 | 24 | 267 | 482 | 3363 | 0 | 2882 | 5193 | 11 | 0 | 16 | 18 | 15 | 21 | 26 | 17 | 24 | 14 | 1446 | 61 |
| SRR1988288 | 0 | 52 | 0 | 0 | 0 | 0 | 0 | 310 | 207 | 15858 | 2278 | 2525 | 0 | 1732 | 0 | 0 | 0 | 32 | 21 | 261 | 476 | 3551 | 0 | 2849 | 5764 | 0 | 0 | 23 | 20 | 41 | 70 | 38 | 55 | 100 | 17 | 1802 | 82 |
| SRR1988289 | 0 | 22 | 0 | 0 | 0 | 0 | 0 | 358 | 290 | 15244 | 1247 | 2523 | 0 | 2071 | 0 | 0 | 0 | 90 | 37 | 400 | 491 | 3020 | 0 | 3980 | 4900 | 44 | 19 | 15 | 14 | 39 | 56 | 18 | 35 | 67 | 0 | 5098 | 54 |
| SRR1988290 | 0 | 24 | 0 | 0 | 0 | 0 | 0 | 328 | 299 | 15437 | 985 | 2089 | 0 | 1352 | 0 | 0 | 0 | 28 | 34 | 343 | 523 | 3557 | 0 | 2839 | 5052 | 0 | 26 | 0 | 14 | 14 | 18 | 17 | 14 | 26 | 13 | 3502 | 64 |
| SRR1988291 | 0 | 40 | 20 | 0 | 0 | 0 | 0 | 400 | 96 | 5075 | 1104 | 17722 | 0 | 400 | 0 | 0 | 0 | 22 | 11 | 1237 | 434 | 4229 | 0 | 2081 | 2155 | 0 | 19 | 0 | 11 | 128 | 89 | 14 | 35 | 62 | 13 | 18577 | 0 |
| SRR1988292 | 0 | 39 | 20 | 0 | 0 | 0 | 0 | 500 | 189 | 5325 | 6483 | 14266 | 0 | 464 | 0 | 12 | 0 | 32 | 19 | 1726 | 574 | 4319 | 0 | 2559 | 2414 | 0 | 25 | 0 | 10 | 167 | 107 | 0 | 40 | 64 | 15 | 26415 | 0 |
| SRR1988293 | 0 | 18 | 12 | 0 | 0 | 0 | 0 | 473 | 124 | 4672 | 6632 | 16518 | 0 | 387 | 0 | 11 | 0 | 28 | 15 | 1595 | 394 | 3802 | 0 | 2097 | 1775 | 0 | 26 | 0 | 15 | 129 | 107 | 13 | 40 | 77 | 20 | 27762 | 0 |
| SRR1988294 | 0 | 30 | 28 | 0 | 0 | 0 | 0 | 434 | 113 | 5195 | 6705 | 16453 | 0 | 398 | 0 | 12 | 0 | 27 | 16 | 1254 | 393 | 3969 | 0 | 1979 | 1832 | 0 | 19 | 0 | 10 | 23 | 19 | 11 | 46 | 14 | 12 | 21060 | 0 |
| SRR1988295 | 0 | 14 | 0 | 0 | 10 | 0 | 0 | 939 | 288 | 14179 | 692 | 4594 | 0 | 6918 | 0 | 0 | 0 | 196 | 0 | 1147 | 291 | 2662 | 60 | 7248 | 1607 | 13 | 14 | 0 | 13 | 35 | 60 | 12 | 49 | 94 | 0 | 10127 | 34 |
| SRR1988296 | 0 | 17 | 0 | 0 | 16 | 0 | 0 | 1089 | 280 | 10438 | 641 | 3534 | 0 | 4688 | 0 | 0 | 0 | 268 | 0 | 699 | 335 | 2829 | 66 | 9124 | 1852 | 11 | 30 | 0 | 14 | 28 | 58 | 12 | 42 | 88 | 19 | 9822 | 55 |
| SRR1988297 | 0 | 24 | 0 | 0 | 0 | 0 | 0 | 1050 | 266 | 12899 | 662 | 3499 | 0 | 5885 | 0 | 0 | 0 | 205 | 0 | 756 | 325 | 3068 | 70 | 9151 | 2353 | 14 | 40 | 12 | 18 | 27 | 60 | 17 | 43 | 116 | 23 | 8111 | 41 |
| SRR1988298 | 0 | 16 | 0 | 0 | 12 | 0 | 0 | 780 | 240 | 11671 | 3362 | 4266 | 0 | 8092 | 0 | 0 | 0 | 210 | 10 | 556 | 354 | 2820 | 66 | 8450 | 2032 | 11 | 41 | 0 | 12 | 26 | 40 | 10 | 25 | 10 | 12 | 5335 | 45 |
| SRR1988299 | 0 | 29 | 18 | 0 | 0 | 0 | 0 | 529 | 210 | 4203 | 9265 | 14965 | 0 | 409 | 30 | 90 | 0 | 14 | 17 | 2985 | 383 | 5177 | 98 | 10184 | 2026 | 28 | 0 | 0 | 0 | 157 | 127 | 0 | 58 | 139 | 43 | 18010 | 0 |
| SRR1988300 | 0 | 40 | 27 | 0 | 0 | 0 | 0 | 998 | 151 | 5233 | 6087 | 12200 | 0 | 243 | 20 | 59 | 0 | 21 | 15 | 3080 | 626 | 7329 | 116 | 14154 | 3335 | 25 | 0 | 0 | 14 | 22 | 18 | 27 | 42 | 14 | 32 | 16494 | 0 |
| SRR1988301 | 0 | 39 | 35 | 0 | 0 | 0 | 0 | 841 | 132 | 4366 | 806 | 12766 | 0 | 278 | 13 | 40 | 0 | 14 | 12 | 2822 | 366 | 4756 | 102 | 9984 | 2158 | 24 | 13 | 0 | 15 | 22 | 14 | 12 | 65 | 14 | 32 | 18569 | 0 |
| SRR1988302 | 0 | 37 | 19 | 0 | 0 | 0 | 0 | 541 | 97 | 4661 | 1214 | 14326 | 0 | 262 | 23 | 68 | 0 | 18 | 14 | 3789 | 418 | 6239 | 92 | 11506 | 2600 | 19 | 12 | 0 | 0 | 197 | 107 | 12 | 52 | 118 | 27 | 23620 | 0 |
| SRR1988303 | 0 | 51 | 16 | 0 | 0 | 0 | 0 | 1080 | 139 | 6163 | 8098 | 14102 | 0 | 292 | 33 | 80 | 0 | 14 | 12 | 2664 | 283 | 3474 | 27 | 5164 | 1606 | 12 | 11 | 0 | 14 | 131 | 86 | 0 | 38 | 78 | 23 | 14526 | 0 |
| SRR1988304 | 0 | 0 | 16 | 0 | 0 | 0 | 0 | 564 | 135 | 5936 | 456 | 10180 | 0 | 485 | 34 | 23 | 0 | 0 | 10 | 3397 | 257 | 2815 | 30 | 5661 | 1356 | 11 | 15 | 0 | 11 | 153 | 135 | 16 | 50 | 105 | 32 | 17448 | 0 |
| SRR1988305 | 0 | 34 | 15 | 0 | 0 | 0 | 0 | 727 | 161 | 4837 | 1038 | 13429 | 0 | 336 | 20 | 40 | 0 | 14 | 11 | 2597 | 219 | 3106 | 29 | 4613 | 1230 | 0 | 10 | 0 | 17 | 125 | 104 | 14 | 49 | 101 | 31 | 16187 | 0 |
| SRR1988306 | 0 | 29 | 22 | 0 | 0 | 0 | 0 | 715 | 55 | 4542 | 492 | 12363 | 0 | 363 | 22 | 44 | 0 | 0 | 12 | 3102 | 259 | 3932 | 26 | 6714 | 1978 | 0 | 16 | 0 | 11 | 18 | 13 | 16 | 62 | 15 | 36 | 13561 | 0 |
| SRR1988307 | 0 | 54 | 22 | 0 | 0 | 0 | 0 | 536 | 125 | 2571 | 9713 | 14313 | 0 | 553 | 0 | 17 | 0 | 86 | 23 | 3386 | 695 | 8112 | 72 | 29789 | 2652 | 0 | 36 | 10 | 23 | 23 | 23 | 22 | 77 | 17 | 31 | 14405 | 0 |
| SRR1988308 | 0 | 25 | 16 | 0 | 0 | 0 | 0 | 770 | 203 | 5580 | 1692 | 13029 | 0 | 461 | 0 | 0 | 0 | 48 | 25 | 1600 | 882 | 8484 | 48 | 19201 | 3411 | 0 | 39 | 0 | 21 | 18 | 18 | 12 | 54 | 15 | 25 | 16827 | 0 |
| SRR1988309 | 0 | 63 | 0 | 0 | 0 | 0 | 0 | 486 | 46 | 3276 | 8195 | 15905 | 0 | 459 | 0 | 0 | 0 | 47 | 21 | 4203 | 709 | 7349 | 45 | 21454 | 2550 | 0 | 0 | 13 | 13 | 19 | 22 | 15 | 12 | 32 | 37 | 13252 | 0 |
| SRR1988310 | 0 | 77 | 19 | 0 | 0 | 0 | 0 | 646 | 109 | 1950 | 12391 | 22423 | 0 | 327 | 0 | 0 | 0 | 44 | 30 | 1365 | 570 | 6327 | 22 | 9761 | 2982 | 0 | 0 | 0 | 13 | 280 | 216 | 20 | 59 | 115 | 41 | 6394 | 0 |
| SRR1988311 | 0 | 65 | 35 | 0 | 0 | 0 | 0 | 3127 | 432 | 9540 | 10315 | 24013 | 0 | 1223 | 0 | 0 | 0 | 41 | 12 | 1295 | 516 | 3793 | 15 | 4310 | 1811 | 0 | 0 | 10 | 33 | 272 | 241 | 10 | 57 | 145 | 29 | 9196 | 0 |
| SRR1988312 | 0 | 36 | 40 | 0 | 0 | 0 | 0 | 2803 | 490 | 8962 | 1000 | 18792 | 0 | 1229 | 16 | 0 | 0 | 29 | 14 | 1341 | 419 | 3383 | 11 | 4880 | 2297 | 10 | 0 | 0 | 46 | 16 | 14 | 15 | 86 | 12 | 40 | 4923 | 0 |
| SRR2052873 | 0 | 15 | 197 | 0 | 0 | 0 | 0 | 189 | 49 | 1327 | 771 | 1916 | 0 | 558 | 28 | 58 | 0 | 309 | 11 | 7639 | 1324 | 6256 | 0 | 7974 | 9495 | 24 | 54 | 192 | 523 | 1203 | 702 | 88 | 383 | 1040 | 35 | 90912 | 0 |
| SRR2052874 | 0 | 30 | 275 | 0 | 0 | 0 | 0 | 242 | 49 | 1190 | 899 | 1731 | 0 | 651 | 18 | 47 | 10 | 738 | 12 | 7417 | 1359 | 6150 | 0 | 7545 | 9697 | 32 | 60 | 164 | 721 | 1068 | 721 | 93 | 341 | 961 | 35 | 80265 | 0 |
| SRR2052875 | 0 | 18 | 292 | 0 | 0 | 0 | 0 | 284 | 68 | 1298 | 285 | 1477 | 0 | 1143 | 16 | 22 | 0 | 708 | 14 | 9932 | 1287 | 6789 | 0 | 8939 | 9953 | 27 | 73 | 187 | 665 | 936 | 554 | 79 | 442 | 1161 | 31 | 78108 | 0 |
| SRR2052877 | 0 | 23 | 142 | 0 | 0 | 0 | 0 | 299 | 62 | 1326 | 670 | 1483 | 0 | 327 | 0 | 15 | 0 | 123 | 17 | 11937 | 736 | 4605 | 0 | 6638 | 7697 | 24 | 48 | 100 | 209 | 539 | 504 | 50 | 252 | 797 | 26 | 115060 | 0 |
| SRR2052878 | 0 | 19 | 22 | 0 | 0 | 0 | 0 | 311 | 14 | 1584 | 180 | 1144 | 0 | 452 | 0 | 17 | 12 | 142 | 10 | 8549 | 1002 | 5011 | 0 | 10394 | 8600 | 28 | 11 | 17 | 32 | 111 | 114 | 13 | 55 | 131 | 39 | 107073 | 0 |
| SRR2053027 | 0 | 49 | 279 | 0 | 0 | 0 | 0 | 406 | 161 | 1171 | 3248 | 5238 | 0 | 656 | 0 | 0 | 0 | 583 | 13 | 11654 | 1451 | 9079 | 0 | 8685 | 10301 | 0 | 14 | 37 | 229 | 639 | 353 | 29 | 208 | 568 | 22 | 87654 | 0 |
| SRR2053028 | 0 | 58 | 225 | 0 | 0 | 11 | 0 | 462 | 92 | 1475 | 1991 | 3391 | 0 | 277 | 0 | 0 | 0 | 240 | 15 | 13091 | 1302 | 7172 | 0 | 11175 | 11175 | 12 | 27 | 63 | 234 | 759 | 518 | 70 | 182 | 623 | 17 | 103848 | 0 |
| SRR2053029 | 0 | 39 | 423 | 0 | 0 | 0 | 0 | 486 | 103 | 788 | 1164 | 3300 | 0 | 299 | 0 | 0 | 0 | 553 | 15 | 16468 | 1271 | 8814 | 0 | 9247 | 9727 | 11 | 21 | 65 | 343 | 644 | 495 | 45 | 303 | 679 | 42 | 104201 | 0 |
| SRR2061931 | 0 | 17 | 50 | 0 | 63 | 0 | 0 | 699 | 325 | 2410 | 197 | 1313 | 0 | 337 | 36 | 72 | 0 | 79 | 0 | 639 | 1339 | 6347 | 83 | 3653 | 5431 | 30 | 17 | 0 | 17 | 0 | 0 | 0 | 0 | 13 | 17 | 1602 | 18 |
| SRR2061932 | 14 | 0 | 54 | 0 | 10 | 14 | 0 | 1158 | 356 | 1113 | 2805 | 1647 | 0 | 223 | 45 | 134 | 0 | 61 | 0 | 401 | 580 | 3845 | 51 | 1719 | 3611 | 10 | 0 | 0 | 0 | 0 | 14 | 0 | 0 | 0 | 27 | 935 | 0 |
| SRR2061933 | 0 | 13 | 18 | 0 | 52 | 0 | 0 | 580 | 120 | 1721 | 2101 | 1120 | 0 | 460 | 20 | 20 | 0 | 108 | 0 | 1301 | 1527 | 6588 | 201 | 5392 | 3944 | 30 | 0 | 0 | 0 | 0 | 0 | 0 | 0 | 0 | 26 | 2701 | 26 |
| SRR2061934 | 0 | 10 | 0 | 0 | 28 | 0 | 23 | 1962 | 561 | 2430 | 5887 | 2803 | 0 | 374 | 93 | 93 | 23 | 126 | 0 | 467 | 687 | 3871 | 84 | 3268 | 3576 | 14 | 0 | 0 | 0 | 0 | 0 | 0 | 23 | 0 | 46 | 934 | 14 |
| SRR2061935 | 0 | 51 | 76 | 0 | 0 | 0 | 0 | 1061 | 0 | 2654 | 20 | 1380 | 0 | 955 | 106 | 318 | 25 | 63 | 21 | 318 | 655 | 2685 | 127 | 1670 | 2262 | 21 | 0 | 0 | 0 | 0 | 0 | 0 | 0 | 0 | 25 | 531 | 0 |
| SRR2061936 | 0 | 16 | 128 | 0 | 93 | 16 | 0 | 850 | 170 | 1938 | 1394 | 1224 | 0 | 238 | 0 | 0 | 0 | 225 | 0 | 1394 | 2191 | 7090 | 639 | 7429 | 3493 | 18 | 0 | 0 | 0 | 0 | 0 | 0 | 0 | 27 | 21 | 2652 | 21 |
| SRR2061937 | 0 | 43 | 101 | 0 | 99 | 14 | 0 | 281 | 241 | 2613 | 36 | 603 | 0 | 322 | 0 | 0 | 36 | 208 | 0 | 1085 | 2307 | 8137 | 597 | 11074 | 3589 | 0 | 0 | 0 | 0 | 14 | 0 | 0 | 0 | 0 | 22 | 2291 | 0 |
| SRR2061938 | 0 | 0 | 29 | 0 | 85 | 0 | 0 | 849 | 356 | 1041 | 109 | 1671 | 0 | 301 | 55 | 301 | 0 | 171 | 0 | 1342 | 1619 | 7625 | 304 | 8668 | 4353 | 19 | 0 | 0 | 0 | 0 | 0 | 0 | 0 | 0 | 29 | 2465 | 16 |
| SRR2061940 | 0 | 94 | 24 | 0 | 22 | 24 | 0 | 875 | 146 | 2186 | 1749 | 583 | 24 | 0 | 146 | 292 | 0 | 133 | 0 | 1020 | 877 | 3663 | 322 | 2653 | 2053 | 22 | 0 | 24 | 24 | 0 | 0 | 0 | 0 | 0 | 0 | 1458 | 22 |
| SRR2061941 | 0 | 61 | 619 | 0 | 30 | 0 | 0 | 114 | 392 | 10424 | 97 | 1316 | 0 | 13498 | 63 | 38 | 0 | 2872 | 52 | 228 | 913 | 5003 | 71 | 22435 | 10739 | 1117 | 12 | 49 | 85 | 97 | 97 | 36 | 24 | 243 | 61 | 557 | 11 |
| SRR2061942 | 0 | 33 | 37 | 0 | 16 | 0 | 0 | 256 | 223 | 19429 | 435 | 1895 | 0 | 10248 | 33 | 54 | 0 | 1022 | 19 | 338 | 970 | 5563 | 28 | 16010 | 8140 | 892 | 0 | 22 | 37 | 52 | 67 | 0 | 33 | 15 | 48 | 1666 | 31 |
| SRR2061943 | 0 | 36 | 18 | 0 | 48 | 0 | 0 | 837 | 267 | 18805 | 461 | 1642 | 0 | 6838 | 14 | 0 | 0 | 535 | 0 | 253 | 1264 | 7243 | 92 | 21322 | 7654 | 708 | 0 | 33 | 62 | 49 | 41 | 15 | 54 | 21 | 51 | 2682 | 102 |
| SRR2061944 | 0 | 54 | 168 | 0 | 14 | 0 | 0 | 182 | 207 | 23046 | 269 | 2600 | 0 | 7301 | 49 | 73 | 0 | 459 | 19 | 583 | 543 | 3935 | 43 | 20064 | 6959 | 545 | 0 | 0 | 0 | 54 | 35 | 15 | 54 | 124 | 40 | 3365 | 24 |
| SRR2061945 | 0 | 17 | 201 | 0 | 23 | 0 | 0 | 331 | 171 | 19196 | 2909 | 1856 | 0 | 8903 | 89 | 18 | 0 | 839 | 0 | 337 | 1063 | 5374 | 38 | 17356 | 8986 | 887 | 0 | 28 | 35 | 42 | 62 | 0 | 38 | 104 | 73 | 2122 | 26 |
| SRR2061946 | 0 | 45 | 85 | 0 | 35 | 0 | 0 | 461 | 196 | 15593 | 693 | 930 | 0 | 10672 | 26 | 11 | 0 | 647 | 0 | 164 | 1550 | 7077 | 94 | 20697 | 8382 | 774 | 0 | 14 | 30 | 34 | 11 | 0 | 16 | 41 | 37 | 1036 | 88 |
| SRR2061947 | 0 | 37 | 214 | 0 | 21 | 0 | 0 | 227 | 277 | 15499 | 2602 | 2007 | 0 | 7189 | 78 | 57 | 0 | 947 | 21 | 312 | 1009 | 5200 | 44 | 20355 | 10942 | 755 | 0 | 47 | 107 | 65 | 102 | 19 | 61 | 140 | 75 | 1205 | 36 |
| SRR2061948 | 0 | 24 | 173 | 0 | 20 | 0 | 0 | 321 | 228 | 18949 | 2618 | 1527 | 0 | 8518 | 57 | 43 | 0 | 643 | 0 | 371 | 1156 | 5681 | 39 | 18130 | 9438 | 873 | 0 | 27 | 27 | 24 | 47 | 20 | 20 | 98 | 71 | 2932 | 47 |
| SRR2061949 | 0 | 45 | 15 | 0 | 39 | 0 | 0 | 774 | 177 | 20692 | 1885 | 1134 | 0 | 8042 | 12 | 0 | 0 | 440 | 0 | 236 | 1572 | 7165 | 77 | 19019 | 7684 | 600 | 0 | 17 | 20 | 36 | 20 | 0 | 47 | 53 | 45 | 2157 | 102 |
| SRR2061950 | 0 | 50 | 168 | 0 | 12 | 0 | 0 | 206 | 124 | 5214 | 1933 | 13188 | 0 | 1046 | 198 | 140 | 0 | 327 | 25 | 362 | 1000 | 7452 | 295 | 49895 | 2399 | 32 | 0 | 0 | 50 | 20 | 17 | 45 | 18 | 49 | 15 | 1400 | 31 |
| SRR2061951 | 0 | 28 | 1550 | 0 | 16 | 0 | 0 | 189 | 116 | 6205 | 2530 | 11224 | 0 | 1351 | 152 | 128 | 0 | 406 | 24 | 347 | 1466 | 9125 | 263 | 52406 | 2731 | 49 | 0 | 45 | 52 | 104 | 149 | 28 | 121 | 377 | 204 | 535 | 11 |
| SRR2061952 | 0 | 43 | 133 | 0 | 12 | 0 | 0 | 160 | 63 | 3572 | 4524 | 7443 | 0 | 1159 | 59 | 52 | 0 | 424 | 12 | 376 | 2026 | 11386 | 292 | 58536 | 3725 | 46 | 0 | 46 | 12 | 18 | 15 | 28 | 17 | 37 | 17 | 1100 | 40 |
| SRR2061953 | 0 | 19 | 293 | 0 | 13 | 0 | 0 | 52 | 148 | 2810 | 2510 | 12986 | 0 | 2030 | 129 | 180 | 0 | 1098 | 46 | 245 | 1744 | 10575 | 344 | 64410 | 4022 | 88 | 0 | 28 | 16 | 21 | 23 | 23 | 18 | 38 | 16 | 290 | 35 |
| SRR2061954 | 0 | 32 | 189 | 0 | 15 | 0 | 0 | 162 | 77 | 4921 | 16574 | 10421 | 0 | 1140 | 223 | 223 | 0 | 353 | 15 | 223 | 1969 | 9311 | 348 | 56572 | 3355 | 55 | 0 | 44 | 11 | 21 | 18 | 28 | 15 | 48 | 26 | 693 | 32 |
| SRR2061955 | 0 | 25 | 113 | 0 | 13 | 0 | 0 | 142 | 57 | 3796 | 3385 | 7932 | 0 | 1481 | 96 | 45 | 0 | 333 | 0 | 159 | 2537 | 11754 | 386 | 54702 | 3655 | 47 | 0 | 56 | 16 | 18 | 11 | 49 | 13 | 24 | 13 | 528 | 71 |
| SRR2061956 | 0 | 23 | 2238 | 0 | 27 | 0 | 0 | 115 | 173 | 4379 | 2869 | 13131 | 0 | 1423 | 178 | 235 | 0 | 585 | 44 | 319 | 1678 | 9670 | 345 | 56983 | 3633 | 95 | 0 | 46 | 186 | 199 | 196 | 39 | 212 | 493 | 186 | 581 | 27 |
| SRR2061957 | 0 | 45 | 166 | 0 | 11 | 0 | 0 | 176 | 98 | 5463 | 15415 | 9972 | 0 | 1235 | 111 | 118 | 0 | 350 | 0 | 261 | 2032 | 9492 | 307 | 55895 | 3404 | 60 | 0 | 22 | 16 | 21 | 20 | 26 | 14 | 46 | 22 | 810 | 28 |
| SRR2061958 | 0 | 65 | 1204 | 0 | 14 | 0 | 0 | 140 | 80 | 4508 | 2680 | 5963 | 0 | 1559 | 85 | 45 | 0 | 447 | 0 | 375 | 2045 | 10047 | 308 | 49945 | 3116 | 88 | 0 | 36 | 130 | 114 | 120 | 23 | 111 | 273 | 127 | 1614 | 59 |
| SRR2069629 | 0 | 76 | 42 | 40 | 23 | 0 | 0 | 1415 | 2056 | 3405 | 437 | 2031 | 0 | 1944 | 66 | 72 | 0 | 250 | 2749 | 488 | 983 | 4690 | 48 | 6165 | 6707 | 0 | 0 | 10 | 49 | 17 | 12 | 0 | 0 | 0 | 0 | 470 | 727 |
| SRR2069630 | 0 | 81 | 15 | 82 | 55 | 0 | 0 | 743 | 838 | 244 | 576 | 11655 | 0 | 42 | 42 | 90 | 17 | 331 | 173 | 69 | 2393 | 1490 | 172 | 6393 | 4587 | 0 | 42 | 66 | 39 | 0 | 0 | 0 | 0 | 0 | 0 | 42 | 1104 |
| SRR2084358 | 0 | 147 | 0 | 0 | 51 | 0 | 0 | 8681 | 2085 | 18798 | 699 | 6938 | 0 | 854 | 0 | 34 | 98 | 32 | 120 | 1435 | 698 | 929 | 0 | 255 | 3735 | 12 | 0 | 147 | 82 | 32 | 47 | 392 | 27 | 44 | 49 | 1401 | 631 |
| SRR2084359 | 0 | 12 | 70 | 0 | 18 | 0 | 0 | 7867 | 2051 | 27873 | 512 | 5285 | 0 | 941 | 72 | 0 | 0 | 68 | 104 | 2365 | 633 | 1403 | 0 | 241 | 4491 | 0 | 35 | 106 | 109 | 67 | 92 | 19 | 38 | 112 | 70 | 8350 | 815 |
| SRR2085173 | 0 | 0 | 0 | 0 | 0 | 0 | 22 | 1538 | 173 | 0 | 45 | 35 | 0 | 0 | 0 | 0 | 0 | 2891 | 119 | 47 | 52 | 70 | 11 | 1442 | 1332 | 0 | 0 | 3064 | 305 | 57 | 33 | 0 | 0 | 0 | 0 | 80 | 112 |
| SRR2103379 | 0 | 15 | 41 | 0 | 15 | 0 | 0 | 292 | 56 | 919 | 39775 | 20876 | 0 | 199 | 222 | 156 | 33 | 70 | 63 | 0 | 1268 | 2781 | 0 | 34 | 8970 | 66 | 17 | 110 | 74 | 169 | 139 | 19 | 38 | 94 | 33 | 0 | 545 |
| SRR2103380 | 0 | 13 | 39 | 0 | 0 | 0 | 0 | 265 | 48 | 858 | 54740 | 20520 | 0 | 288 | 140 | 114 | 26 | 70 | 48 | 0 | 1119 | 2619 | 0 | 30 | 7352 | 64 | 15 | 94 | 72 | 147 | 136 | 19 | 50 | 116 | 33 | 0 | 523 |
| SRR2103381 | 0 | 15 | 10 | 0 | 0 | 0 | 0 | 244 | 34 | 755 | 44415 | 19457 | 0 | 240 | 153 | 111 | 19 | 76 | 47 | 0 | 1184 | 2479 | 0 | 27 | 7522 | 55 | 0 | 22 | 14 | 39 | 33 | 24 | 44 | 22 | 38 | 0 | 481 |
| SRR2103382 | 0 | 51 | 43 | 0 | 11 | 0 | 11 | 568 | 59 | 646 | 18506 | 15015 | 0 | 161 | 93 | 252 | 33 | 277 | 19 | 0 | 2239 | 4079 | 0 | 20 | 13196 | 0 | 310 | 40 | 146 | 427 | 176 | 27 | 50 | 124 | 87 | 0 | 288 |
| SRR2103383 | 0 | 58 | 47 | 0 | 15 | 0 | 0 | 474 | 44 | 579 | 13762 | 11401 | 0 | 146 | 99 | 226 | 18 | 285 | 19 | 0 | 2542 | 4616 | 0 | 46 | 16434 | 10 | 213 | 33 | 134 | 426 | 197 | 24 | 49 | 105 | 60 | 25 | 278 |
| SRR2103384 | 0 | 40 | 92 | 0 | 0 | 0 | 0 | 501 | 66 | 791 | 11676 | 10159 | 0 | 227 | 67 | 138 | 16 | 191 | 19 | 0 | 2043 | 3354 | 0 | 29 | 10011 | 14 | 114 | 214 | 164 | 625 | 240 | 27 | 53 | 163 | 64 | 23 | 192 |
| SRR2105940 | 0 | 0 | 0 | 0 | 16 | 0 | 0 | 0 | 0 | 157 | 0 | 88 | 0 | 0 | 0 | 0 | 10 | 37 | 17 | 160 | 53 | 478 | 0 | 287 | 705 | 0 | 23 | 11 | 0 | 0 | 0 | 0 | 0 | 0 | 0 | 723 | 115 |
| SRR2105943 | 0 | 0 | 0 | 0 | 11 | 0 | 0 | 0 | 0 | 0 | 0 | 0 | 0 | 0 | 0 | 0 | 0 | 30 | 0 | 186 | 162 | 448 | 11 | 462 | 449 | 0 | 13 | 0 | 0 | 13 | 0 | 0 | 0 | 0 | 0 | 656 | 89 |
| SRR2105944 | 0 | 0 | 0 | 0 | 0 | 0 | 0 | 99 | 0 | 0 | 0 | 0 | 0 | 0 | 0 | 0 | 0 | 54 | 0 | 51 | 252 | 474 | 85 | 17868 | 687 | 0 | 31 | 0 | 0 | 19 | 0 | 0 | 0 | 0 | 0 | 88 | 88 |
| SRR2105949 | 0 | 0 | 0 | 0 | 18 | 0 | 0 | 0 | 0 | 0 | 272 | 89 | 0 | 0 | 0 | 0 | 0 | 29 | 0 | 0 | 46 | 380 | 53 | 4342 | 427 | 0 | 39 | 0 | 0 | 0 | 0 | 0 | 0 | 0 | 0 | 432 | 42 |
| SRR2105951 | 0 | 0 | 0 | 0 | 25 | 0 | 0 | 0 | 0 | 0 | 0 | 0 | 0 | 0 | 0 | 0 | 0 | 23 | 0 | 377 | 38 | 185 | 0 | 4092 | 277 | 103 | 64 | 0 | 0 | 0 | 0 | 0 | 0 | 0 | 0 | 1577 | 16 |
| SRR2106873 | 0 | 438 | 0 | 0 | 20 | 0 | 0 | 507 | 198 | 51976 | 2369 | 17486 | 0 | 804 | 0 | 0 | 0 | 515 | 12 | 629 | 2107 | 5567 | 17 | 11738 | 10935 | 45 | 0 | 118 | 258 | 823 | 1116 | 23 | 168 | 390 | 23 | 1066 | 142 |
| SRR2106874 | 0 | 16 | 11 | 0 | 20 | 0 | 0 | 917 | 324 | 51751 | 9983 | 7695 | 0 | 1068 | 0 | 0 | 0 | 559 | 25 | 1371 | 1331 | 3490 | 27 | 12818 | 11137 | 99 | 0 | 108 | 39 | 105 | 105 | 46 | 13 | 29 | 11 | 2256 | 119 |
| SRR2106875 | 0 | 18 | 73 | 0 | 18 | 0 | 0 | 882 | 323 | 91352 | 8710 | 7620 | 0 | 918 | 0 | 0 | 0 | 633 | 30 | 1462 | 1900 | 5770 | 21 | 12337 | 11925 | 88 | 0 | 75 | 48 | 67 | 72 | 50 | 29 | 78 | 31 | 2789 | 193 |
| SRR2106876 | 0 | 143 | 112 | 0 | 20 | 0 | 0 | 465 | 190 | 47970 | 287 | 5215 | 0 | 1858 | 63 | 169 | 0 | 992 | 28 | 3906 | 1691 | 5006 | 0 | 16929 | 17876 | 156 | 0 | 19 | 81 | 116 | 136 | 0 | 43 | 174 | 39 | 4603 | 106 |
| SRR2106877 | 0 | 10 | 98 | 0 | 37 | 0 | 0 | 768 | 697 | 38481 | 4305 | 3609 | 0 | 1340 | 214 | 125 | 0 | 928 | 21 | 3251 | 1359 | 3048 | 24 | 8860 | 11130 | 35 | 0 | 67 | 10 | 23 | 35 | 24 | 13 | 26 | 15 | 8772 | 121 |

### Supp Table S8.4.Columns_DI_to_ES

| | | | | | | | | | | | | | | | | | | | | | | | | | | | | | | | | | | | | | |
|---|---|---|---|---|---|---|---|---|---|---|---|---|---|---|---|---|---|---|---|---|---|---|---|---|---|---|---|---|---|---|---|---|---|---|---|---|---|
| SRR2106878 | 0 | 129 | 117 | 0 | 26 | 0 | 0 | 1259 | 846 | 88258 | 165 | 3302 | 0 | 1176 | 165 | 144 | 0 | 936 | 25 | 2312 | 881 | 3338 | 15 | 10894 | 13165 | 52 | 0 | 49 | 135 | 461 | 485 | 37 | 215 | 799 | 448 | 3550 | 81 |
| SRR2106879 | 0 | 16 | 80 | 0 | 30 | 0 | 0 | 970 | 657 | 93924 | 459 | 5259 | 0 | 1035 | 43 | 86 | 0 | 544 | 27 | 2899 | 1391 | 4492 | 32 | 8561 | 15186 | 45 | 0 | 42 | 10 | 20 | 16 | 36 | 86 | 13 | 0 | 8610 | 109 |
| SRR2106880 | 0 | 43 | 0 | 0 | 17 | 0 | 0 | 637 | 176 | 43483 | 10927 | 10038 | 0 | 1317 | 0 | 0 | 0 | 943 | 29 | 3245 | 1747 | 4336 | 25 | 10207 | 16238 | 55 | 0 | 10 | 42 | 37 | 25 | 17 | 44 | 20 | 0 | 5392 | 128 |
| SRR2106881 | 0 | 29 | 48 | 0 | 22 | 0 | 0 | 463 | 246 | 52901 | 7125 | 7067 | 0 | 883 | 14 | 0 | 0 | 1388 | 26 | 4475 | 2353 | 5234 | 0 | 9914 | 16769 | 32 | 0 | 36 | 15 | 23 | 22 | 27 | 63 | 12 | 12 | 5213 | 147 |
| SRR2106882 | 0 | 400 | 0 | 0 | 14 | 0 | 0 | 413 | 75 | 9781 | 16259 | 23323 | 0 | 910 | 0 | 0 | 0 | 853 | 15 | 1341 | 2408 | 4937 | 0 | 5057 | 11327 | 60 | 0 | 22 | 53 | 241 | 279 | 26 | 93 | 184 | 15 | 2881 | 54 |
| SRR2106883 | 0 | 674 | 23 | 0 | 14 | 0 | 0 | 6428 | 488 | 8828 | 593 | 4941 | 0 | 1331 | 149 | 163 | 0 | 938 | 32 | 3844 | 2175 | 5541 | 0 | 7992 | 17203 | 69 | 0 | 25 | 111 | 226 | 247 | 55 | 173 | 326 | 111 | 4984 | 49 |
| SRR2106884 | 0 | 513 | 11 | 0 | 16 | 0 | 0 | 3425 | 355 | 12044 | 2229 | 15222 | 0 | 1069 | 61 | 76 | 0 | 907 | 22 | 2128 | 2145 | 5063 | 0 | 6596 | 14245 | 56 | 0 | 31 | 85 | 215 | 270 | 33 | 147 | 272 | 44 | 3481 | 51 |
| SRR2124760 | 0 | 0 | 0 | 0 | 12 | 0 | 0 | 509 | 106 | 1274 | 97 | 481 | 0 | 197 | 77 | 827 | 0 | 78 | 0 | 3085 | 855 | 2297 | 257 | 4534 | 3063 | 36 | 0 | 0 | 0 | 0 | 0 | 0 | 0 | 0 | 14 | 7685 | 0 |
| SRR2124761 | 0 | 0 | 10 | 0 | 0 | 0 | 0 | 218 | 69 | 1294 | 73 | 151 | 0 | 264 | 37 | 76 | 0 | 212 | 0 | 946 | 1375 | 2471 | 314 | 4788 | 3348 | 27 | 0 | 0 | 0 | 0 | 0 | 0 | 0 | 0 | 23 | 2297 | 0 |
| SRR2124762 | 0 | 0 | 13 | 0 | 0 | 0 | 0 | 141 | 50 | 700 | 148 | 152 | 0 | 195 | 174 | 310 | 0 | 280 | 0 | 494 | 1252 | 2871 | 497 | 6683 | 4131 | 0 | 0 | 0 | 0 | 0 | 0 | 0 | 0 | 0 | 0 | 1323 | 0 |
| SRR2124763 | 0 | 0 | 27 | 0 | 0 | 0 | 0 | 271 | 138 | 1599 | 74 | 122 | 0 | 189 | 160 | 229 | 0 | 372 | 0 | 531 | 1519 | 2163 | 519 | 6658 | 3945 | 14 | 0 | 0 | 0 | 0 | 0 | 0 | 0 | 14 | 13 | 1456 | 0 |
| SRR2124764 | 0 | 0 | 22 | 0 | 14 | 0 | 0 | 271 | 75 | 2514 | 617 | 372 | 0 | 312 | 583 | 612 | 0 | 116 | 0 | 2439 | 1027 | 2573 | 274 | 5293 | 4539 | 24 | 0 | 0 | 0 | 0 | 0 | 0 | 0 | 10 | 16 | 6618 | 0 |
| SRR2124765 | 0 | 0 | 14 | 0 | 12 | 0 | 0 | 378 | 85 | 2055 | 114 | 333 | 0 | 119 | 793 | 600 | 0 | 149 | 0 | 3868 | 1359 | 4048 | 327 | 6418 | 6438 | 19 | 0 | 0 | 0 | 0 | 0 | 0 | 0 | 0 | 11 | 10325 | 0 |
| SRR2124766 | 0 | 0 | 17 | 0 | 0 | 0 | 0 | 585 | 44 | 1058 | 161 | 209 | 0 | 63 | 539 | 759 | 0 | 87 | 0 | 2468 | 958 | 3586 | 250 | 4348 | 5372 | 25 | 0 | 0 | 0 | 0 | 0 | 0 | 0 | 0 | 16 | 5122 | 0 |
| SRR2124767 | 0 | 0 | 0 | 0 | 0 | 0 | 0 | 143 | 43 | 1118 | 99 | 104 | 0 | 67 | 795 | 853 | 0 | 91 | 0 | 1218 | 1536 | 5032 | 333 | 6642 | 8578 | 0 | 0 | 0 | 0 | 0 | 0 | 0 | 0 | 0 | 12 | 3067 | 0 |
| SRR2124768 | 0 | 0 | 23 | 0 | 17 | 0 | 0 | 140 | 55 | 514 | 77 | 96 | 0 | 78 | 374 | 390 | 0 | 104 | 0 | 502 | 1625 | 3821 | 460 | 6537 | 7384 | 0 | 0 | 0 | 0 | 0 | 0 | 0 | 0 | 10 | 12 | 1278 | 0 |
| SRR2124769 | 0 | 0 | 15 | 0 | 0 | 0 | 0 | 303 | 128 | 1019 | 274 | 309 | 0 | 82 | 1957 | 2015 | 0 | 107 | 0 | 2469 | 1679 | 4211 | 381 | 6594 | 7516 | 13 | 0 | 0 | 0 | 0 | 0 | 0 | 0 | 0 | 0 | 7164 | 0 |
| SRR2124770 | 0 | 0 | 0 | 0 | 21 | 0 | 0 | 92 | 19 | 435 | 143 | 144 | 0 | 113 | 171 | 208 | 0 | 90 | 0 | 831 | 1952 | 4295 | 415 | 9660 | 7883 | 0 | 0 | 0 | 0 | 0 | 0 | 0 | 0 | 0 | 0 | 2789 | 0 |
| SRR2124771 | 0 | 0 | 10 | 0 | 0 | 0 | 0 | 271 | 134 | 1246 | 214 | 244 | 0 | 107 | 214 | 241 | 0 | 51 | 0 | 2289 | 2032 | 4574 | 388 | 5929 | 7148 | 0 | 0 | 0 | 0 | 0 | 0 | 0 | 0 | 0 | 13 | 6462 | 0 |
| SRR2124772 | 0 | 0 | 14 | 0 | 0 | 0 | 0 | 149 | 24 | 537 | 214 | 202 | 0 | 53 | 192 | 178 | 0 | 64 | 0 | 1047 | 1255 | 4309 | 220 | 5795 | 8732 | 11 | 0 | 0 | 0 | 0 | 0 | 0 | 0 | 0 | 13 | 3199 | 0 |
| SRR2124773 | 0 | 0 | 11 | 0 | 12 | 0 | 0 | 702 | 183 | 1460 | 447 | 622 | 0 | 48 | 846 | 878 | 0 | 72 | 0 | 6510 | 1840 | 5156 | 409 | 5859 | 7830 | 12 | 0 | 0 | 0 | 0 | 0 | 0 | 0 | 0 | 11 | 17120 | 0 |
| SRR2124774 | 0 | 0 | 24 | 0 | 21 | 0 | 12 | 109 | 42 | 486 | 54 | 119 | 0 | 37 | 118 | 104 | 0 | 103 | 0 | 951 | 1624 | 3039 | 450 | 5783 | 6679 | 0 | 0 | 0 | 0 | 0 | 0 | 0 | 0 | 0 | 0 | 2298 | 0 |
| SRR2124775 | 0 | 0 | 29 | 0 | 20 | 0 | 0 | 137 | 26 | 574 | 139 | 209 | 0 | 63 | 117 | 113 | 0 | 132 | 0 | 1582 | 2052 | 2766 | 458 | 6032 | 5849 | 12 | 0 | 0 | 0 | 0 | 0 | 0 | 0 | 11 | 17 | 3531 | 0 |
| SRR2124776 | 0 | 0 | 21 | 0 | 0 | 0 | 0 | 143 | 28 | 1035 | 154 | 203 | 0 | 63 | 356 | 403 | 0 | 94 | 0 | 1407 | 1200 | 3217 | 230 | 4585 | 6180 | 29 | 0 | 0 | 0 | 0 | 0 | 0 | 0 | 0 | 17 | 3393 | 0 |
| SRR2124777 | 0 | 0 | 36 | 0 | 0 | 0 | 0 | 311 | 171 | 1473 | 125 | 218 | 0 | 57 | 547 | 648 | 0 | 120 | 0 | 1141 | 1449 | 3649 | 243 | 5069 | 6098 | 15 | 0 | 0 | 0 | 0 | 0 | 0 | 0 | 18 | 25 | 2931 | 0 |
| SRR2124778 | 0 | 0 | 14 | 0 | 0 | 0 | 0 | 136 | 21 | 524 | 252 | 201 | 0 | 41 | 127 | 113 | 0 | 74 | 0 | 802 | 1159 | 3518 | 383 | 6950 | 5424 | 0 | 0 | 0 | 0 | 0 | 0 | 0 | 0 | 0 | 0 | 2033 | 0 |
| SRR2124779 | 0 | 0 | 14 | 0 | 12 | 0 | 0 | 189 | 38 | 936 | 97 | 121 | 0 | 153 | 122 | 124 | 0 | 119 | 0 | 863 | 1144 | 2464 | 383 | 5381 | 4222 | 24 | 0 | 0 | 0 | 0 | 0 | 0 | 0 | 0 | 0 | 2219 | 0 |
| SRR2126527 | 0 | 26 | 15 | 0 | 13 | 0 | 0 | 413 | 57 | 1174 | 663 | 526 | 0 | 919 | 327 | 245 | 0 | 56 | 37 | 2034 | 1553 | 2406 | 20 | 5498 | 11123 | 0 | 0 | 0 | 43 | 80 | 89 | 44 | 178 | 266 | 171 | 8851 | 263 |
| SRR2126528 | 0 | 30 | 82 | 0 | 14 | 0 | 0 | 407 | 61 | 1163 | 1050 | 580 | 0 | 924 | 340 | 168 | 0 | 72 | 38 | 2323 | 1561 | 2305 | 21 | 5575 | 11116 | 0 | 0 | 0 | 318 | 511 | 565 | 302 | 1028 | 1466 | 1132 | 10035 | 272 |
| SRR2126529 | 0 | 23 | 99 | 0 | 11 | 0 | 0 | 388 | 47 | 931 | 711 | 573 | 0 | 921 | 301 | 161 | 0 | 67 | 29 | 2194 | 1644 | 2568 | 23 | 5497 | 10716 | 0 | 0 | 0 | 298 | 547 | 531 | 282 | 1031 | 1454 | 985 | 8501 | 276 |
| SRR2126530 | 0 | 16 | 15 | 0 | 11 | 0 | 0 | 469 | 108 | 1061 | 571 | 578 | 0 | 889 | 469 | 241 | 0 | 64 | 33 | 1665 | 2394 | 2222 | 24 | 4778 | 10064 | 14 | 0 | 0 | 34 | 57 | 64 | 42 | 161 | 271 | 198 | 7125 | 251 |
| SRR2126531 | 0 | 25 | 72 | 0 | 12 | 0 | 0 | 474 | 94 | 848 | 620 | 507 | 0 | 727 | 397 | 258 | 0 | 69 | 30 | 1884 | 2580 | 2211 | 19 | 4695 | 9350 | 0 | 0 | 10 | 189 | 289 | 310 | 172 | 813 | 1186 | 904 | 7990 | 213 |
| SRR2126532 | 0 | 22 | 17 | 0 | 11 | 0 | 0 | 547 | 62 | 779 | 640 | 508 | 0 | 853 | 228 | 197 | 0 | 91 | 44 | 1954 | 1945 | 2410 | 23 | 5180 | 10143 | 0 | 0 | 0 | 55 | 93 | 109 | 71 | 226 | 373 | 233 | 8313 | 230 |
| SRR2135530 | 0 | 143 | 33 | 44 | 163 | 0 | 11 | 629 | 1781 | 35023 | 4054 | 5667 | 0 | 795 | 123 | 177 | 31 | 810 | 68 | 5022 | 612 | 975 | 0 | 51 | 6012 | 0 | 18 | 23 | 34 | 62 | 129 | 42 | 16 | 14 | 0 | 12485 | 0 |
| SRR2135531 | 0 | 1201 | 287 | 34 | 181 | 0 | 0 | 415 | 1740 | 24337 | 6167 | 8632 | 0 | 446 | 518 | 1178 | 0 | 1524 | 80 | 2795 | 1489 | 1894 | 0 | 47 | 7164 | 0 | 79 | 22 | 173 | 155 | 349 | 159 | 110 | 116 | 72 | 7102 | 0 |
| SRR2135532 | 0 | 251 | 35 | 51 | 286 | 0 | 0 | 341 | 1439 | 15237 | 16757 | 14448 | 0 | 376 | 94 | 173 | 0 | 1619 | 138 | 2430 | 1699 | 2115 | 0 | 36 | 6054 | 0 | 11 | 33 | 27 | 26 | 47 | 18 | 61 | 11 | 0 | 9174 | 0 |
| SRR2229584 | 0 | 0 | 0 | 0 | 0 | 0 | 0 | 0 | 0 | 368 | 40 | 404 | 0 | 18 | 263 | 561 | 0 | 0 | 638 | 0 | 28 | 59 | 0 | 0 | 814 | 0 | 0 | 74 | 11 | 19 | 43 | 12 | 12 | 57 | 32 | 18 | 0 |
| SRR2229585 | 0 | 0 | 21 | 0 | 0 | 0 | 0 | 43 | 0 | 1109 | 52 | 512 | 0 | 128 | 213 | 341 | 0 | 21 | 337 | 0 | 80 | 365 | 0 | 31 | 2289 | 0 | 0 | 27 | 0 | 24 | 14 | 0 | 14 | 21 | 16 | 0 | 0 |
| SRR2229586 | 0 | 0 | 25 | 0 | 0 | 0 | 0 | 0 | 0 | 0 | 404 | 118 | 0 | 0 | 0 | 0 | 0 | 0 | 0 | 0 | 0 | 0 | 0 | 0 | 0 | 0 | 0 | 136 | 0 | 237 | 326 | 68 | 337 | 1008 | 721 | 17 | 0 |
| SRR2229587 | 0 | 0 | 0 | 0 | 0 | 0 | 0 | 0 | 0 | 0 | 16 | 27 | 0 | 0 | 0 | 0 | 0 | 0 | 0 | 27 | 0 | 0 | 0 | 0 | 0 | 0 | 0 | 48 | 0 | 74 | 171 | 26 | 171 | 485 | 512 | 0 | 0 |
| SRR2229588 | 20 | 0 | 0 | 0 | 0 | 0 | 0 | 16098 | 130 | 1402 | 441 | 675 | 0 | 182 | 208 | 156 | 0 | 0 | 59 | 0 | 103 | 336 | 0 | 0 | 1068 | 0 | 0 | 11 | 0 | 0 | 0 | 20 | 24 | 20 | 14 | 0 | 49 |
| SRR2229589 | 0 | 0 | 0 | 0 | 0 | 12 | 0 | 15135 | 120 | 933 | 765 | 1387 | 0 | 359 | 72 | 502 | 0 | 0 | 65 | 0 | 73 | 384 | 0 | 0 | 850 | 0 | 0 | 49 | 0 | 0 | 16 | 0 | 0 | 16 | 12 | 0 | 44 |
| SRR2287868 | 0 | 18 | 16 | 0 | 0 | 0 | 0 | 21 | 20 | 295 | 15 | 0 | 0 | 66 | 115 | 250 | 0 | 350 | 0 | 449 | 2712 | 3083 | 734 | 11560 | 7500 | 59 | 79 | 0 | 14 | 25 | 21 | 0 | 20 | 18 | 31 | 3603 | 0 |
| SRR2289624 | 0 | 0 | 0 | 0 | 0 | 0 | 40 | 486 | 55 | 2462 | 262 | 19 | 0 | 595 | 230 | 601 | 0 | 259 | 10 | 442 | 3082 | 2521 | 673 | 10838 | 7216 | 93 | 17 | 0 | 0 | 0 | 0 | 0 | 0 | 0 | 0 | 2413 | 0 |
| SRR2289625 | 0 | 18 | 0 | 0 | 0 | 0 | 16 | 456 | 281 | 1397 | 352 | 103 | 0 | 184 | 110 | 172 | 0 | 124 | 0 | 1117 | 1745 | 2003 | 423 | 8860 | 4407 | 162 | 42 | 0 | 38 | 36 | 26 | 12 | 20 | 30 | 12 | 5196 | 0 |
| SRR2289626 | 0 | 27 | 0 | 0 | 0 | 0 | 0 | 104 | 124 | 8579 | 229 | 19 | 0 | 313 | 163 | 177 | 0 | 201 | 0 | 185 | 1581 | 1620 | 645 | 7899 | 4122 | 76 | 17 | 0 | 0 | 0 | 0 | 0 | 0 | 0 | 0 | 539 | 0 |
| SRR2294377 | 0 | 17 | 25 | 0 | 0 | 0 | 0 | 212 | 114 | 5552 | 218 | 123 | 0 | 731 | 196 | 380 | 0 | 278 | 0 | 366 | 1865 | 2111 | 476 | 6696 | 5092 | 488 | 14 | 0 | 15 | 14 | 0 | 15 | 0 | 12 | 11 | 1469 | 0 |
| SRR2294380 | 0 | 32 | 23 | 0 | 0 | 0 | 0 | 198 | 398 | 6669 | 750 | 162 | 0 | 1303 | 54 | 109 | 0 | 441 | 0 | 173 | 779 | 940 | 266 | 4672 | 4166 | 923 | 52 | 0 | 52 | 37 | 42 | 32 | 33 | 55 | 20 | 1179 | 0 |
| SRR2294739 | 0 | 17 | 61 | 0 | 0 | 0 | 0 | 23 | 51 | 613 | 276 | 21 | 0 | 228 | 14 | 45 | 0 | 315 | 0 | 155 | 608 | 848 | 277 | 4705 | 3506 | 714 | 171 | 0 | 20 | 28 | 34 | 20 | 32 | 24 | 35 | 1354 | 0 |
| SRR2294742 | 0 | 18 | 37 | 0 | 0 | 0 | 0 | 588 | 945 | 9088 | 230 | 175 | 0 | 547 | 119 | 248 | 0 | 390 | 0 | 238 | 635 | 872 | 205 | 3196 | 3037 | 857 | 0 | 0 | 28 | 36 | 37 | 25 | 56 | 48 | 39 | 1012 | 0 |
| SRR2294746 | 0 | 0 | 11 | 0 | 0 | 0 | 0 | 694 | 617 | 5970 | 110 | 70 | 0 | 93 | 49 | 104 | 0 | 133 | 0 | 249 | 552 | 685 | 158 | 3120 | 2550 | 727 | 15 | 0 | 22 | 26 | 0 | 15 | 19 | 22 | 22 | 1314 | 0 |
| SRR2294747 | 0 | 14 | 0 | 0 | 0 | 0 | 0 | 23 | 112 | 20872 | 177 | 0 | 0 | 1059 | 105 | 174 | 0 | 199 | 0 | 38 | 768 | 1012 | 446 | 6785 | 3344 | 452 | 10 | 0 | 0 | 17 | 35 | 0 | 0 | 0 | 0 | 121 | 0 |
| SRR2294748 | 0 | 14 | 32 | 0 | 0 | 0 | 0 | 629 | 505 | 7751 | 262 | 184 | 0 | 303 | 466 | 704 | 0 | 212 | 0 | 240 | 773 | 994 | 181 | 2461 | 2853 | 706 | 0 | 0 | 15 | 19 | 19 | 33 | 27 | 30 | 24 | 774 | 0 |
| SRR2294820 | 0 | 31 | 36 | 0 | 0 | 0 | 0 | 140 | 426 | 5463 | 480 | 673 | 0 | 536 | 52 | 137 | 0 | 415 | 0 | 214 | 628 | 804 | 311 | 4603 | 3611 | 791 | 46 | 14 | 95 | 66 | 92 | 59 | 66 | 120 | 42 | 960 | 0 |
| SRR2296788 | 0 | 12 | 77 | 0 | 0 | 0 | 0 | 634 | 330 | 1876 | 5678 | 9024 | 0 | 51 | 482 | 406 | 0 | 388 | 13 | 1014 | 723 | 12 | 0 | 1056 | 63112 | 57 | 0 | 30 | 12 | 25 | 25 | 54 | 34 | 70 | 72 | 3168 | 0 |
| SRR2296789 | 0 | 22 | 91 | 0 | 0 | 0 | 0 | 378 | 151 | 1020 | 256 | 13110 | 0 | 151 | 264 | 718 | 0 | 314 | 0 | 1700 | 478 | 34992 | 0 | 1479 | 35995 | 51 | 0 | 15 | 11 | 57 | 48 | 107 | 42 | 121 | 148 | 2191 | 0 |
| SRR2296790 | 0 | 50 | 66 | 0 | 0 | 0 | 0 | 480 | 240 | 1761 | 104 | 14328 | 0 | 320 | 160 | 320 | 0 | 468 | 0 | 1041 | 445 | 36950 | 0 | 1045 | 67413 | 69 | 0 | 17 | 83 | 116 | 166 | 100 | 249 | 432 | 913 | 2001 | 0 |
| SRR2296791 | 0 | 66 | 66 | 0 | 0 | 0 | 0 | 376 | 250 | 1670 | 178 | 9809 | 0 | 250 | 501 | 626 | 13 | 1276 | 21 | 918 | 585 | 38824 | 0 | 991 | 65875 | 97 | 0 | 13 | 19 | 39 | 30 | 11 | 19 | 81 | 116 | 1002 | 0 |
| SRR2296792 | 0 | 19 | 12 | 0 | 0 | 0 | 0 | 689 | 215 | 1593 | 5897 | 6844 | 0 | 43 | 517 | 818 | 0 | 409 | 0 | 2497 | 760 | 45521 | 0 | 2175 | 55850 | 60 | 0 | 12 | 17 | 58 | 54 | 15 | 44 | 125 | 114 | 3401 | 0 |
| SRR2296793 | 0 | 76 | 41 | 0 | 0 | 0 | 0 | 651 | 204 | 1466 | 8223 | 10666 | 0 | 163 | 692 | 285 | 0 | 404 | 15 | 1425 | 509 | 44068 | 0 | 967 | 81856 | 59 | 0 | 21 | 19 | 25 | 17 | 69 | 26 | 50 | 40 | 1954 | 0 |
| SRR2296794 | 0 | 66 | 66 | 0 | 0 | 0 | 0 | 213 | 305 | 1889 | 5422 | 5971 | 0 | 0 | 548 | 579 | 0 | 749 | 35 | 1767 | 496 | 33430 | 0 | 1371 | 64817 | 88 | 0 | 0 | 146 | 219 | 113 | 27 | 192 | 742 | 517 | 3655 | 0 |
| SRR2296795 | 0 | 90 | 27 | 0 | 0 | 0 | 0 | 822 | 286 | 1714 | 109 | 7644 | 0 | 107 | 214 | 429 | 0 | 374 | 0 | 1036 | 756 | 57066 | 0 | 1820 | 127796 | 109 | 0 | 0 | 13 | 37 | 30 | 63 | 26 | 51 | 42 | 2107 | 0 |
| SRR2296797 | 0 | 52 | 47 | 0 | 0 | 19 | 0 | 374 | 156 | 702 | 478 | 10016 | 0 | 47 | 265 | 125 | 0 | 797 | 25 | 1014 | 491 | 36945 | 0 | 2442 | 63729 | 69 | 0 | 14 | 15 | 18 | 22 | 28 | 28 | 82 | 59 | 1623 | 0 |
| SRR2296798 | 0 | 13 | 38 | 0 | 0 | 0 | 0 | 397 | 132 | 2119 | 97 | 9601 | 0 | 132 | 199 | 199 | 0 | 379 | 24 | 728 | 358 | 11 | 0 | 1066 | 69590 | 56 | 0 | 51 | 38 | 215 | 126 | 38 | 253 | 341 | 873 | 1986 | 19 |
| SRR2296799 | 0 | 104 | 52 | 0 | 0 | 0 | 0 | 652 | 223 | 1509 | 17952 | 17627 | 0 | 51 | 206 | 154 | 0 | 438 | 0 | 840 | 709 | 41429 | 0 | 1196 | 46571 | 64 | 0 | 62 | 104 | 338 | 182 | 89 | 167 | 552 | 547 | 2778 | 0 |
| SRR2296800 | 0 | 18 | 12 | 0 | 0 | 0 | 0 | 542 | 271 | 993 | 120 | 6637 | 0 | 45 | 542 | 632 | 0 | 430 | 14 | 2438 | 711 | 43850 | 0 | 2643 | 56032 | 42 | 0 | 14 | 84 | 47 | 38 | 70 | 44 | 119 | 104 | 5102 | 0 |
| SRR2296801 | 0 | 251 | 33 | 0 | 0 | 19 | 0 | 526 | 195 | 1326 | 14855 | 18442 | 0 | 78 | 175 | 273 | 0 | 404 | 0 | 1501 | 1080 | 22 | 0 | 2278 | 77641 | 66 | 0 | 0 | 104 | 109 | 166 | 14 | 128 | 384 | 356 | 2866 | 0 |
| SRR2296802 | 0 | 266 | 48 | 0 | 16 | 0 | 0 | 650 | 325 | 974 | 21954 | 21629 | 0 | 130 | 130 | 65 | 0 | 514 | 16 | 2208 | 1044 | 44791 | 12 | 3228 | 50135 | 69 | 0 | 48 | 72 | 459 | 241 | 97 | 386 | 700 | 241 | 4092 | 0 |
| SRR2296805 | 0 | 19 | 52 | 0 | 0 | 0 | 0 | 597 | 223 | 1908 | 16619 | 17064 | 0 | 116 | 62 | 98 | 0 | 344 | 11 | 909 | 646 | 39401 | 0 | 971 | 46334 | 51 | 0 | 55 | 92 | 26 | 28 | 64 | 31 | 85 | 59 | 2015 | 0 |
| SRR2296806 | 0 | 145 | 40 | 0 | 0 | 0 | 0 | 343 | 86 | 729 | 10892 | 12864 | 0 | 129 | 429 | 472 | 0 | 337 | 13 | 1544 | 666 | 32107 | 0 | 2083 | 50843 | 40 | 0 | 158 | 119 | 606 | 567 | 79 | 395 | 870 | 975 | 2444 | 0 |
| SRR2296807 | 0 | 25 | 56 | 0 | 0 | 0 | 0 | 506 | 217 | 1646 | 11164 | 11352 | 0 | 58 | 650 | 578 | 0 | 364 | 0 | 737 | 803 | 45110 | 0 | 1180 | 62186 | 52 | 0 | 36 | 10 | 26 | 26 | 72 | 27 | 88 | 61 | 3033 | 0 |
| SRR2296808 | 0 | 69 | 52 | 0 | 0 | 0 | 0 | 605 | 55 | 1265 | 168 | 10667 | 0 | 55 | 715 | 275 | 0 | 362 | 0 | 1265 | 652 | 33054 | 0 | 2126 | 48413 | 60 | 0 | 17 | 16 | 36 | 50 | 20 | 36 | 70 | 82 | 4729 | 0 |
| SRR2296809 | 0 | 35 | 35 | 0 | 0 | 0 | 0 | 323 | 172 | 818 | 12306 | 13846 | 0 | 54 | 280 | 506 | 0 | 347 | 16 | 732 | 1141 | 51330 | 0 | 2844 | 79043 | 31 | 0 | 0 | 12 | 21 | 25 | 20 | 17 | 59 | 41 | 3025 | 0 |
| SRR2296810 | 0 | 26 | 61 | 0 | 0 | 0 | 0 | 340 | 136 | 975 | 8186 | 8300 | 0 | 23 | 386 | 295 | 0 | 492 | 15 | 1633 | 747 | 38323 | 12 | 3735 | 55485 | 73 | 0 | 0 | 16 | 53 | 35 | 35 | 24 | 95 | 66 | 3197 | 0 |
| SRR2353417 | 0 | 11 | 20 | 0 | 0 | 0 | 0 | 138 | 27 | 4769 | 469 | 130 | 0 | 526 | 351 | 1210 | 0 | 205 | 0 | 1139 | 3576 | 6067 | 282 | 12655 | 12582 | 92 | 0 | 0 | 0 | 11 | 0 | 0 | 0 | 0 | 0 | 5003 | 0 |
| SRR2353418 | 0 | 134 | 0 | 0 | 0 | 0 | 0 | 314 | 134 | 18801 | 369 | 105 | 0 | 3560 | 689 | 1543 | 0 | 181 | 0 | 0 | 3012 | 5626 | 278 | 11842 | 12754 | 196 | 0 | 0 | 0 | 0 | 0 | 0 | 0 | 0 | 0 | 0 | 0 |
| SRR2353419 | 0 | 56 | 14 | 0 | 0 | 0 | 0 | 108 | 28 | 7402 | 661 | 287 | 0 | 383 | 505 | 1454 | 0 | 112 | 0 | 1458 | 3158 | 6681 | 217 | 11326 | 12175 | 139 | 0 | 0 | 0 | 0 | 0 | 0 | 0 | 0 | 0 | 6207 | 0 |
| SRR2353420 | 0 | 130 | 18 | 0 | 0 | 0 | 0 | 201 | 15 | 1736 | 690 | 241 | 0 | 182 | 853 | 1942 | 0 | 121 | 0 | 1047 | 3253 | 7623 | 247 | 12934 | 13029 | 114 | 11 | 0 | 0 | 0 | 0 | 0 | 0 | 0 | 0 | 5913 | 0 |
| SRR2353421 | 0 | 59 | 13 | 0 | 0 | 0 | 0 | 187 | 52 | 8265 | 604 | 433 | 0 | 608 | 676 | 1933 | 0 | 140 | 0 | 952 | 2469 | 5789 | 232 | 12083 | 11947 | 244 | 25 | 0 | 0 | 13 | 0 | 0 | 0 | 13 | 0 | 4417 | 0 |
| SRR2353422 | 0 | 10 | 18 | 0 | 0 | 0 | 0 | 340 | 40 | 9243 | 474 | 143 | 0 | 1901 | 555 | 1586 | 0 | 163 | 0 | 845 | 3049 | 6741 | 275 | 13312 | 13862 | 182 | 0 | 0 | 12 | 0 | 0 | 0 | 0 | 0 | 0 | 3019 | 0 |
| SRR2353423 | 0 | 149 | 12 | 0 | 0 | 0 | 0 | 108 | 26 | 4755 | 608 | 298 | 0 | 320 | 483 | 1332 | 0 | 197 | 0 | 1509 | 2669 | 5736 | 298 | 12793 | 13578 | 267 | 0 | 0 | 0 | 0 | 15 | 0 | 0 | 0 | 0 | 6589 | 0 |
| SRR2353424 | 0 | 134 | 25 | 0 | 0 | 0 | 0 | 270 | 59 | 12731 | 384 | 222 | 0 | 2858 | 736 | 2595 | 0 | 269 | 0 | 617 | 2065 | 4410 | 228 | 9769 | 11013 | 302 | 0 | 0 | 0 | 0 | 10 | 0 | 0 | 0 | 0 | 2036 | 0 |
| SRR2353425 | 0 | 88 | 12 | 0 | 0 | 0 | 0 | 327 | 70 | 5106 | 430 | 160 | 0 | 1198 | 165 | 343 | 0 | 136 | 0 | 537 | 2837 | 6051 | 304 | 13113 | 12962 | 61 | 11 | 0 | 0 | 14 | 15 | 11 | 0 | 16 | 0 | 2467 | 0 |
| SRR2353426 | 0 | 85 | 13 | 0 | 0 | 0 | 0 | 313 | 35 | 5145 | 514 | 182 | 0 | 638 | 864 | 2525 | 0 | 112 | 0 | 840 | 3144 | 7780 | 277 | 12996 | 13466 | 172 | 0 | 0 | 0 | 0 | 0 | 0 | 0 | 0 | 0 | 4139 | 0 |
| SRR2353427 | 0 | 110 | 0 | 0 | 0 | 0 | 0 | 313 | 35 | 4926 | 519 | 75 | 0 | 1136 | 198 | 349 | 0 | 123 | 0 | 625 | 2875 | 5847 | 269 | 13123 | 13598 | 59 | 0 | 0 | 13 | 12 | 11 | 0 | 0 | 0 | 0 | 2781 | 0 |
| SRR2353428 | 0 | 12 | 0 | 0 | 0 | 0 | 0 | 194 | 46 | 5443 | 377 | 115 | 0 | 1174 | 131 | 287 | 0 | 241 | 0 | 568 | 2362 | 4538 | 243 | 11251 | 12073 | 183 | 0 | 0 | 12 | 15 | 0 | 0 | 0 | 0 | 0 | 2299 | 0 |
| SRR2353429 | 0 | 51 | 13 | 0 | 0 | 0 | 0 | 291 | 62 | 3462 | 388 | 108 | 0 | 369 | 855 | 1267 | 0 | 154 | 0 | 445 | 2660 | 6061 | 277 | 12077 | 12487 | 114 | 0 | 0 | 0 | 0 | 0 | 0 | 0 | 0 | 0 | 2238 | 0 |
| SRR2353430 | 0 | 124 | 0 | 0 | 0 | 0 | 0 | 185 | 48 | 6635 | 157 | 65 | 0 | 2375 | 337 | 1050 | 0 | 253 | 0 | 293 | 2560 | 5326 | 234 | 10231 | 9048 | 187 | 0 | 0 | 0 | 10 | 0 | 0 | 0 | 0 | 0 | 1632 | 0 |
| SRR2353431 | 0 | 12 | 17 | 0 | 0 | 0 | 0 | 206 | 53 | 3630 | 446 | 155 | 0 | 240 | 1735 | 1898 | 0 | 159 | 0 | 610 | 3003 | 5917 | 296 | 11721 | 11599 | 56 | 0 | 0 | 17 | 19 | 11 | 0 | 0 | 11 | 0 | 3051 | 0 |
| SRR2353432 | 0 | 121 | 0 | 0 | 0 | 0 | 0 | 181 | 34 | 4882 | 342 | 87 | 0 | 832 | 293 | 713 | 0 | 248 | 0 | 513 | 3216 | 5841 | 264 | 12299 | 11632 | 93 | 0 | 0 | 22 | 26 | 0 | 0 | 0 | 0 | 0 | 1991 | 0 |
| SRR2353433 | 0 | 210 | 0 | 0 | 0 | 0 | 0 | 125 | 73 | 13076 | 128 | 45 | 0 | 1961 | 217 | 570 | 0 | 207 | 0 | 245 | 3040 | 5086 | 269 | 10751 | 11882 | 155 | 0 | 0 | 0 | 0 | 0 | 0 | 0 | 0 | 0 | 805 | 0 |
| SRR2353434 | 0 | 108 | 17 | 0 | 0 | 0 | 0 | 262 | 41 | 4660 | 575 | 268 | 0 | 549 | 657 | 1362 | 0 | 177 | 0 | 593 | 2698 | 5188 | 271 | 11087 | 10775 | 209 | 0 | 0 | 0 | 16 | 0 | 0 | 0 | 0 | 0 | 2894 | 0 |
| SRR2353435 | 0 | 14 | 20 | 0 | 0 | 0 | 0 | 217 | 64 | 3463 | 974 | 429 | 0 | 537 | 2605 | 3086 | 0 | 213 | 0 | 728 | 2848 | 4898 | 241 | 10286 | 10641 | 327 | 0 | 0 | 0 | 10 | 0 | 0 | 0 | 0 | 0 | 2599 | 0 |
| SRR2353436 | 0 | 123 | 0 | 0 | 0 | 0 | 0 | 121 | 94 | 22761 | 76 | 30 | 0 | 4964 | 404 | 1226 | 0 | 295 | 0 | 190 | 2130 | 3317 | 251 | 9347 | 10101 | 282 | 0 | 0 | 0 | 0 | 0 | 0 | 0 | 0 | 0 | 527 | 0 |
| SRR2353437 | 0 | 91 | 19 | 0 | 0 | 0 | 0 | 551 | 39 | 11627 | 584 | 209 | 0 | 1852 | 486 | 2187 | 0 | 186 | 0 | 875 | 3040 | 6446 | 272 | 12994 | 12344 | 124 | 0 | 0 | 0 | 13 | 0 | 0 | 0 | 0 | 0 | 3066 | 0 |
| SRR2353438 | 0 | 138 | 13 | 0 | 0 | 0 | 0 | 307 | 212 | 31335 | 260 | 88 | 0 | 4278 | 516 | 1035 | 0 | 208 | 0 | 711 | 2388 | 4148 | 217 | 8135 | 9910 | 286 | 0 | 0 | 0 | 13 | 15 | 0 | 0 | 12 | 0 | 2972 | 0 |
| SRR2353439 | 0 | 148 | 11 | 0 | 0 | 0 | 0 | 134 | 233 | 25684 | 252 | 79 | 0 | 4391 | 96 | 347 | 0 | 222 | 0 | 283 | 1864 | 3398 | 184 | 8495 | 10080 | 453 | 0 | 0 | 0 | 0 | 0 | 0 | 0 | 0 | 0 | 1112 | 0 |
| SRR2353440 | 0 | 106 | 25 | 0 | 0 | 0 | 0 | 220 | 39 | 11292 | 408 | 147 | 0 | 1461 | 168 | 487 | 0 | 187 | 0 | 366 | 2796 | 5336 | 243 | 11751 | 12759 | 288 | 0 | 0 | 0 | 0 | 13 | 0 | 0 | 0 | 0 | 1753 | 0 |
| SRR2353441 | 0 | 120 | 25 | 0 | 0 | 0 | 0 | 102 | 51 | 9519 | 344 | 73 | 0 | 1290 | 296 | 1008 | 0 | 225 | 0 | 298 | 2069 | 3839 | 201 | 9074 | 10294 | 318 | 0 | 0 | 0 | 0 | 0 | 0 | 0 | 0 | 0 | 1107 | 0 |
| SRR2353442 | 0 | 195 | 16 | 0 | 0 | 0 | 0 | 158 | 91 | 14777 | 201 | 49 | 0 | 2913 | 276 | 883 | 0 | 240 | 0 | 330 | 2558 | 4095 | 297 | 10865 | 11350 | 189 | 0 | 0 | 0 | 0 | 0 | 0 | 0 | 0 | 0 | 1565 | 0 |
| SRR2353443 | 0 | 112 | 15 | 0 | 0 | 0 | 0 | 113 | 171 | 12848 | 390 | 114 | 0 | 604 | 752 | 1344 | 0 | 179 | 0 | 196 | 3293 | 4869 | 228 | 10337 | 10953 | 167 | 0 | 0 | 0 | 0 | 0 | 0 | 0 | 0 | 0 | 935 | 0 |
| SRR2353444 | 0 | 151 | 0 | 0 | 0 | 0 | 0 | 146 | 80 | 15749 | 171 | 71 | 0 | 3499 | 231 | 1022 | 0 | 270 | 0 | 456 | 2782 | 4545 | 249 | 11644 | 12076 | 213 | 0 | 0 | 0 | 0 | 0 | 0 | 0 | 0 | 0 | 1762 | 0 |
| SRR2353445 | 0 | 145 | 13 | 0 | 0 | 0 | 0 | 137 | 30 | 14622 | 356 | 149 | 0 | 4148 | 865 | 2391 | 0 | 230 | 0 | 286 | 2813 | 5061 | 284 | 11429 | 10802 | 157 | 0 | 0 | 0 | 0 | 0 | 0 | 0 | 0 | 0 | 1074 | 0 |
| SRR2422969 | 0 | 96 | 127 | 0 | 0 | 0 | 0 | 0 | 438 | 1241 | 1679 | 2118 | 0 | 2483 | 365 | 438 | 0 | 4271 | 61 | 1460 | 805 | 2061 | 26 | 4635 | 1037 | 0 | 96 | 159 | 96 | 64 | 64 | 32 | 64 | 127 | 64 | 1314 | 53 |
| SRR2422970 | 0 | 100 | 33 | 0 | 0 | 0 | 0 | 0 | 158 | 1736 | 1263 | 1815 | 0 | 2525 | 395 | 316 | 33 | 4299 | 41 | 947 | 762 | 2214 | 0 | 4855 | 937 | 0 | 0 | 67 | 133 | 133 | 67 | 33 | 33 | 133 | 0 | 631 | 64 |
| SRR2422971 | 0 | 181 | 0 | 0 | 0 | 0 | 0 | 70 | 139 | 1323 | 1253 | 2228 | 0 | 2994 | 278 | 418 | 0 | 4431 | 46 | 835 | 718 | 1978 | 0 | 4561 | 1088 | 0 | 0 | 60 | 212 | 272 | 60 | 151 | 151 | 121 | 60 | 2019 | 88 |
| SRR2422972 | 0 | 186 | 31 | 0 | 0 | 31 | 0 | 0 | 0 | 1162 | 1234 | 2468 | 0 | 3267 | 363 | 290 | 0 | 4218 | 43 | 1234 | 828 | 2124 | 17 | 4737 | 940 | 0 | 0 | 10 | 217 | 217 | 155 | 31 | 62 | 248 | 124 | 1742 | 60 |
| SRR2454328 | 0 | 67 | 26 | 0 | 19 | 0 | 0 | 344 | 115 | 1097 | 2965 | 4189 | 0 | 1653 | 0 | 0 | 0 | 665 | 78 | 119 | 2156 | 4226 | 610 | 43282 | 9032 | 99 | 0 | 180 | 125 | 175 | 141 | 25 | 29 | 45 | 0 | 675 | 55 |
| SRR2454329 | 0 | 62 | 20 | 0 | 14 | 0 | 0 | 289 | 93 | 966 | 3384 | 4045 | 0 | 1665 | 0 | 0 | 0 | 612 | 84 | 152 | 2340 | 4404 | 629 | 43382 | 8986 | 92 | 0 | 16 | 14 | 24 | 16 | 21 | 28 | 38 | 0 | 700 | 56 |
| SRR2454330 | 0 | 63 | 25 | 0 | 15 | 0 | 0 | 201 | 73 | 636 | 4608 | 4422 | 0 | 1279 | 0 | 0 | 0 | 601 | 79 | 115 | 2372 | 4939 | 769 | 48483 | 9182 | 103 | 0 | 142 | 140 | 276 | 151 | 29 | 25 | 28 | 0 | 452 | 55 |
| SRR2454331 | 0 | 76 | 27 | 0 | 18 | 0 | 0 | 254 | 80 | 816 | 4468 | 4492 | 0 | 1203 | 0 | 0 | 0 | 540 | 75 | 111 | 2448 | 4989 | 830 | 50983 | 9579 | 84 | 0 | 141 | 127 | 248 | 175 | 30 | 30 | 44 | 0 | 566 | 52 |
| SRR2454332 | 0 | 174 | 133 | 0 | 0 | 0 | 0 | 284 | 182 | 2098 | 1557 | 1700 | 0 | 1735 | 0 | 0 | 0 | 729 | 108 | 224 | 2055 | 6700 | 420 | 41756 | 11028 | 25 | 0 | 137 | 13 | 24 | 24 | 135 | 155 | 152 | 0 | 1113 | 132 |
| SRR2454333 | 0 | 10 | 116 | 0 | 12 | 0 | 0 | 230 | 169 | 1641 | 879 | 1759 | 0 | 2014 | 0 | 0 | 0 | 734 | 101 | 232 | 2615 | 7986 | 423 | 41486 | 11545 | 34 | 0 | 150 | 15 | 32 | 24 | 106 | 160 | 122 | 0 | 979 | 157 |
| SRR2454334 | 0 | 254 | 131 | 0 | 12 | 0 | 0 | 217 | 148 | 1571 | 455 | 1478 | 0 | 1409 | 0 | 0 | 0 | 901 | 107 | 250 | 2595 | 8762 | 284 | 46868 | 12665 | 60 | 0 | 208 | 363 | 702 | 492 | 90 | 117 | 172 | 0 | 1571 | 178 |
| SRR2454335 | 0 | 306 | 41 | 0 | 11 | 0 | 0 | 111 | 123 | 368 | 3047 | 3321 | 0 | 654 | 0 | 0 | 0 | 707 | 146 | 111 | 3014 | 8929 | 324 | 45574 | 13274 | 76 | 0 | 303 | 441 | 743 | 344 | 34 | 34 | 41 | 0 | 630 | 159 |
| SRR2454336 | 0 | 290 | 17 | 0 | 12 | 0 | 10 | 107 | 59 | 487 | 3078 | 3179 | 0 | 626 | 0 | 0 | 10 | 711 | 127 | 102 | 2878 | 8655 | 395 | 43699 | 12259 | 69 | 0 | 168 | 378 | 584 | 378 | 31 | 10 | 35 | 0 | 391 | 130 |
| SRR2454337 | 0 | 343 | 10 | 0 | 16 | 0 | 0 | 71 | 76 | 503 | 3489 | 3651 | 0 | 533 | 0 | 0 | 0 | 599 | 133 | 46 | 3295 | 8554 | 435 | 46588 | 12912 | 76 | 0 | 267 | 364 | 651 | 388 | 31 | 21 | 21 | 0 | 432 | 147 |
| SRR2487412 | 0 | 11 | 32 | 0 | 29 | 0 | 0 | 328 | 573 | 3987 | 328 | 765 | 0 | 1311 | 27 | 109 | 11 | 117 | 0 | 0 | 1096 | 3603 | 1595 | 11556 | 3137 | 17 | 11 | 0 | 16 | 32 | 21 | 0 | 0 | 32 | 16 | 0 | 0 |
| SRR2487413 | 0 | 12 | 49 | 0 | 27 | 0 | 0 | 399 | 307 | 3403 | 21 | 583 | 0 | 950 | 31 | 61 | 0 | 119 | 0 | 0 | 890 | 3037 | 1616 | 11556 | 20 | 17 | 12 | 0 | 24 | 42 | 18 | 18 | 24 | 0 | 0 | 0 | 0 |
| SRR2487414 | 0 | 26 | 26 | 0 | 28 | 0 | 0 | 300 | 464 | 4531 | 328 | 682 | 0 | 1174 | 82 | 82 | 0 | 136 | 0 | 0 | 1127 | 3602 | 1599 | 11399 | 3278 | 17 | 0 | 0 | 31 | 36 | 41 | 0 | 31 | 26 | 0 | 0 | 0 |
| SRR2487415 | 14 | 28 | 111 | 0 | 0 | 14 | 0 | 665 | 385 | 2450 | 770 | 1540 | 0 | 175 | 1225 | 2485 | 0 | 114 | 0 | 0 | 1978 | 20 | 4592 | 39661 | 26 | 0 | 0 | 42 | 139 | 152 | 222 | 14 | 69 | 97 | 0 | 0 | 0 |
| SRR2487416 | 0 | 16 | 64 | 0 | 0 | 0 | 0 | 638 | 399 | 1994 | 11 | 1515 | 0 | 199 | 1874 | 1994 | 32 | 105 | 0 | 0 | 1626 | 8843 | 4603 | 40039 | 39 | 0 | 0 | 16 | 129 | 241 | 161 | 0 | 112 | 129 | 0 | 0 | 0 |
| SRR2487417 | 0 | 41 | 68 | 0 | 0 | 41 | 0 | 492 | 105 | 2671 | 773 | 1757 | 14 | 316 | 1195 | 2530 | 14 | 107 | 0 | 0 | 1982 | 10176 | 4543 | 39459 | 29 | 0 | 0 | 14 | 68 | 178 | 123 | 0 | 0 | 41 | 0 | 0 | 0 |
| SRR2487418 | 0 | 121 | 80 | 0 | 55 | 0 | 0 | 290 | 207 | 1779 | 1117 | 910 | 0 | 1283 | 41 | 0 | 0 | 125 | 0 | 538 | 1691 | 4163 | 1955 | 13919 | 20 | 12 | 0 | 20 | 20 | 161 | 80 | 0 | 0 | 40 | 0 | 786 | 0 |

### Supp Table S8.4.Columns_DI_to_ES

| | | | | | | | | | | | | | | | | | | | | | | | | | | | | | | | | | | | | | |
|---|---|---|---|---|---|---|---|---|---|---|---|---|---|---|---|---|---|---|---|---|---|---|---|---|---|---|---|---|---|---|---|---|---|---|---|---|---|
| SRR2487419 | 0 | 46 | 138 | 0 | 64 | 0 | 0 | 279 | 186 | 2324 | 883 | 930 | 0 | 1162 | 139 | 46 | 0 | 128 | 0 | 279 | 1196 | 3416 | 1944 | 14141 | 3594 | 17 | 0 | 23 | 69 | 161 | 69 | 0 | 0 | 69 | 0 | 697 | 0 |
| SRR2487420 | 0 | 40 | 60 | 0 | 53 | 0 | 0 | 412 | 41 | 2553 | 20 | 700 | 0 | 1029 | 41 | 0 | 0 | 134 | 0 | 453 | 1694 | 4142 | 1939 | 13853 | 21 | 22 | 0 | 0 | 60 | 239 | 99 | 20 | 0 | 20 | 0 | 741 | 0 |
| SRR2487422 | 0 | 25 | 0 | 0 | 12 | 0 | 0 | 247 | 49 | 2270 | 25 | 1924 | 0 | 345 | 197 | 49 | 0 | 97 | 0 | 3454 | 1446 | 8304 | 3471 | 41268 | 4083 | 0 | 0 | 0 | 98 | 74 | 49 | 0 | 25 | 49 | 25 | 8487 | 28 |
| SRR2487423 | 0 | 83 | 28 | 0 | 14 | 28 | 28 | 282 | 56 | 2426 | 23 | 2200 | 0 | 508 | 338 | 226 | 0 | 107 | 0 | 4344 | 1168 | 7100 | 3491 | 41930 | 4065 | 0 | 0 | 0 | 28 | 110 | 28 | 0 | 55 | 55 | 0 | 9195 | 26 |
| SRR2487424 | 0 | 120 | 24 | 0 | 16 | 0 | 0 | 151 | 301 | 2711 | 1757 | 1657 | 0 | 552 | 351 | 251 | 24 | 106 | 0 | 3414 | 1452 | 8318 | 3391 | 40988 | 4188 | 0 | 0 | 0 | 143 | 215 | 191 | 24 | 0 | 0 | 0 | 7782 | 29 |
| SRR2487426 | 0 | 35 | 189 | 0 | 77 | 12 | 0 | 562 | 458 | 6431 | 31 | 937 | 0 | 1894 | 21 | 83 | 0 | 101 | 0 | 3392 | 2000 | 5641 | 1036 | 17718 | 23 | 15 | 12 | 0 | 106 | 106 | 24 | 35 | 83 | 106 | 0 | 11468 | 19 |
| SRR2487427 | 0 | 27 | 188 | 0 | 85 | 0 | 0 | 525 | 621 | 6397 | 501 | 716 | 0 | 2220 | 48 | 0 | 13 | 95 | 0 | 3771 | 1422 | 4681 | 1106 | 17885 | 30 | 0 | 13 | 13 | 134 | 175 | 108 | 0 | 54 | 0 | 0 | 11410 | 17 |
| SRR2487428 | 0 | 12 | 187 | 0 | 78 | 23 | 0 | 505 | 484 | 7113 | 631 | 547 | 0 | 1936 | 63 | 21 | 12 | 94 | 0 | 3809 | 1998 | 5773 | 977 | 17674 | 15 | 17 | 47 | 12 | 58 | 82 | 94 | 12 | 35 | 70 | 0 | 11237 | 15 |
| SRR2487430 | 0 | 0 | 66 | 0 | 30 | 16 | 0 | 402 | 522 | 7059 | 272 | 587 | 0 | 827 | 22 | 22 | 16 | 89 | 0 | 337 | 1351 | 5470 | 3144 | 28909 | 15 | 0 | 0 | 0 | 82 | 132 | 115 | 0 | 33 | 49 | 0 | 1327 | 30 |
| SRR2487431 | 19 | 38 | 94 | 0 | 29 | 38 | 0 | 432 | 469 | 6979 | 284 | 518 | 0 | 654 | 62 | 12 | 0 | 76 | 0 | 382 | 1134 | 4963 | 3225 | 29332 | 20 | 0 | 0 | 0 | 19 | 56 | 113 | 0 | 75 | 131 | 0 | 1443 | 32 |
| SRR2487432 | 0 | 0 | 146 | 0 | 23 | 0 | 16 | 608 | 369 | 6634 | 43 | 554 | 0 | 836 | 43 | 33 | 16 | 88 | 0 | 369 | 1380 | 5526 | 3118 | 28980 | 3265 | 0 | 0 | 16 | 32 | 49 | 130 | 0 | 0 | 81 | 0 | 1162 | 36 |
| SRR2487434 | 0 | 54 | 27 | 0 | 111 | 0 | 0 | 291 | 373 | 3525 | 134 | 172 | 0 | 1337 | 0 | 0 | 0 | 55 | 0 | 158 | 1604 | 3399 | 2141 | 9020 | 18 | 15 | 0 | 14 | 41 | 68 | 47 | 0 | 14 | 47 | 0 | 263 | 0 |
| SRR2487435 | 0 | 23 | 0 | 0 | 116 | 0 | 0 | 328 | 306 | 3430 | 18 | 169 | 0 | 1568 | 11 | 0 | 0 | 56 | 0 | 104 | 1177 | 2881 | 2194 | 9268 | 4417 | 14 | 0 | 16 | 62 | 54 | 31 | 0 | 16 | 23 | 0 | 257 | 0 |
| SRR2487436 | 0 | 40 | 27 | 0 | 103 | 13 | 0 | 319 | 314 | 3630 | 86 | 186 | 0 | 1410 | 0 | 0 | 13 | 64 | 0 | 181 | 1617 | 3517 | 2137 | 9013 | 17 | 13 | 0 | 0 | 66 | 53 | 27 | 0 | 0 | 40 | 0 | 300 | 0 |
| SRR2487438 | 0 | 45 | 33 | 0 | 14 | 0 | 0 | 630 | 661 | 3910 | 28 | 285 | 0 | 1057 | 142 | 275 | 0 | 127 | 0 | 783 | 2444 | 7424 | 4063 | 38377 | 5455 | 0 | 0 | 11 | 67 | 67 | 33 | 11 | 78 | 45 | 11 | 1352 | 0 |
| SRR2487439 | 0 | 51 | 25 | 0 | 11 | 13 | 0 | 699 | 665 | 3727 | 24 | 329 | 0 | 1098 | 208 | 324 | 0 | 109 | 0 | 913 | 2088 | 6647 | 4138 | 39056 | 5278 | 0 | 13 | 0 | 64 | 76 | 76 | 38 | 13 | 76 | 0 | 1387 | 0 |
| SRR2487441 | 0 | 39 | 78 | 0 | 28 | 0 | 0 | 571 | 245 | 5876 | 13 | 938 | 0 | 367 | 1020 | 979 | 0 | 57 | 0 | 1387 | 1627 | 7356 | 2000 | 20984 | 17 | 0 | 0 | 13 | 0 | 26 | 105 | 0 | 105 | 144 | 13 | 8161 | 0 |
| SRR2487443 | 0 | 54 | 40 | 0 | 22 | 0 | 0 | 791 | 208 | 6455 | 17 | 1083 | 0 | 375 | 750 | 958 | 0 | 73 | 0 | 1458 | 1429 | 6567 | 2049 | 21417 | 19 | 12 | 0 | 0 | 40 | 27 | 54 | 0 | 67 | 134 | 13 | 8287 | 0 |
| SRR2487445 | 0 | 17 | 34 | 0 | 37 | 0 | 0 | 847 | 265 | 7620 | 688 | 1217 | 0 | 423 | 900 | 794 | 0 | 60 | 0 | 1164 | 1619 | 7265 | 2013 | 20409 | 16 | 15 | 0 | 17 | 17 | 34 | 67 | 0 | 51 | 168 | 0 | 7937 | 0 |
| SRR2487447 | 0 | 50 | 83 | 0 | 31 | 17 | 0 | 360 | 154 | 6952 | 772 | 978 | 0 | 412 | 824 | 566 | 0 | 79 | 0 | 1699 | 1511 | 6550 | 2038 | 21091 | 2541 | 0 | 0 | 17 | 0 | 116 | 116 | 0 | 83 | 200 | 17 | 7930 | 0 |
| SRR2487449 | 0 | 34 | 246 | 0 | 0 | 0 | 0 | 267 | 285 | 2028 | 20 | 109 | 0 | 103 | 759 | 656 | 0 | 139 | 0 | 1190 | 3302 | 13 | 3796 | 59461 | 32 | 0 | 0 | 11 | 0 | 90 | 123 | 0 | 56 | 190 | 22 | 4001 | 0 |
| SRR2487451 | 0 | 11 | 218 | 0 | 0 | 0 | 0 | 403 | 248 | 1644 | 137 | 118 | 0 | 186 | 757 | 652 | 0 | 132 | 0 | 1396 | 2838 | 9677 | 3890 | 60877 | 29 | 0 | 0 | 0 | 34 | 115 | 69 | 23 | 103 | 160 | 34 | 4741 | 0 |
| SRR2487453 | 0 | 14 | 158 | 0 | 0 | 14 | 0 | 359 | 187 | 1811 | 133 | 148 | 0 | 226 | 585 | 539 | 0 | 120 | 0 | 1327 | 3244 | 10588 | 3708 | 58991 | 28 | 0 | 0 | 0 | 14 | 57 | 101 | 29 | 129 | 201 | 14 | 4144 | 0 |
| SRR2487455 | 0 | 14 | 213 | 0 | 0 | 14 | 0 | 246 | 269 | 1853 | 20 | 131 | 0 | 131 | 715 | 561 | 0 | 135 | 0 | 1261 | 2863 | 9876 | 3698 | 59936 | 30 | 0 | 0 | 0 | 14 | 156 | 57 | 0 | 99 | 340 | 0 | 4644 | 0 |
| SRR2487457 | 0 | 54 | 244 | 0 | 40 | 0 | 0 | 497 | 455 | 8297 | 24 | 414 | 0 | 641 | 517 | 372 | 0 | 84 | 0 | 1366 | 1504 | 9648 | 4004 | 31866 | 4064 | 0 | 0 | 0 | 54 | 41 | 190 | 14 | 54 | 68 | 14 | 8401 | 0 |
| SRR2487459 | 0 | 0 | 185 | 0 | 51 | 0 | 0 | 986 | 263 | 7737 | 18 | 263 | 0 | 767 | 351 | 285 | 0 | 84 | 0 | 1271 | 1223 | 8675 | 4123 | 32566 | 4103 | 0 | 0 | 0 | 43 | 100 | 14 | 0 | 85 | 100 | 0 | 8066 | 0 |
| SRR2487461 | 0 | 18 | 216 | 0 | 47 | 0 | 0 | 702 | 322 | 7844 | 17 | 293 | 0 | 498 | 468 | 234 | 0 | 60 | 0 | 849 | 1404 | 9440 | 4031 | 31372 | 36 | 0 | 0 | 18 | 36 | 54 | 126 | 0 | 36 | 18 | 18 | 8166 | 0 |
| SRR2487463 | 0 | 18 | 141 | 0 | 57 | 0 | 0 | 743 | 468 | 9406 | 275 | 275 | 0 | 633 | 385 | 303 | 0 | 75 | 0 | 1705 | 1251 | 8670 | 4091 | 31727 | 34 | 0 | 0 | 0 | 106 | 53 | 230 | 18 | 53 | 106 | 18 | 8746 | 0 |
| SRR2487465 | 0 | 33 | 100 | 0 | 0 | 17 | 0 | 5919 | 4998 | 3573 | 58 | 44 | 0 | 88 | 2733 | 2236 | 0 | 117 | 0 | 1995 | 3393 | 14 | 4886 | 58655 | 55 | 0 | 0 | 0 | 17 | 166 | 83 | 0 | 66 | 415 | 17 | 9873 | 0 |
| SRR2487467 | 0 | 69 | 223 | 0 | 0 | 0 | 0 | 5494 | 4945 | 3460 | 22 | 45 | 0 | 186 | 2755 | 1878 | 0 | 110 | 0 | 2057 | 2908 | 12002 | 4998 | 60008 | 7562 | 0 | 0 | 34 | 0 | 120 | 120 | 0 | 154 | 429 | 34 | 9778 | 0 |
| SRR2487469 | 0 | 21 | 107 | 0 | 0 | 0 | 0 | 6502 | 5342 | 3692 | 86 | 38 | 0 | 230 | 2417 | 1467 | 21 | 116 | 0 | 1956 | 3325 | 13260 | 4825 | 57402 | 7751 | 0 | 0 | 0 | 86 | 107 | 86 | 43 | 86 | 322 | 0 | 10175 | 0 |
| SRR2487471 | 0 | 43 | 107 | 0 | 0 | 0 | 0 | 6523 | 5340 | 3986 | 85 | 66 | 0 | 95 | 2490 | 1581 | 0 | 114 | 0 | 2073 | 3006 | 12264 | 4924 | 58354 | 49 | 0 | 0 | 21 | 21 | 21 | 64 | 0 | 214 | 278 | 0 | 9951 | 0 |
| SRR2487473 | 0 | 85 | 509 | 0 | 51 | 0 | 0 | 3147 | 1905 | 6542 | 414 | 1946 | 0 | 166 | 455 | 497 | 0 | 30 | 0 | 3271 | 2441 | 9988 | 5137 | 39215 | 4626 | 0 | 0 | 0 | 153 | 271 | 424 | 34 | 153 | 525 | 17 | 16687 | 0 |
| SRR2487475 | 0 | 104 | 383 | 0 | 49 | 0 | 0 | 2974 | 2555 | 6953 | 17 | 1801 | 0 | 209 | 754 | 628 | 0 | 45 | 0 | 2806 | 2017 | 8714 | 5161 | 40254 | 39 | 0 | 0 | 52 | 87 | 400 | 296 | 35 | 174 | 418 | 52 | 16838 | 0 |
| SRR2487477 | 0 | 109 | 438 | 0 | 66 | 22 | 0 | 3646 | 2520 | 7398 | 483 | 1608 | 0 | 107 | 268 | 268 | 0 | 45 | 0 | 2788 | 2312 | 9786 | 5088 | 38844 | 4429 | 0 | 0 | 0 | 109 | 416 | 241 | 88 | 175 | 525 | 0 | 19139 | 0 |
| SRR2487479 | 0 | 22 | 522 | 0 | 61 | 0 | 0 | 2971 | 2397 | 7296 | 678 | 1668 | 0 | 208 | 625 | 573 | 0 | 34 | 0 | 3127 | 1998 | 8946 | 5216 | 39421 | 4415 | 0 | 0 | 43 | 152 | 413 | 239 | 43 | 217 | 522 | 43 | 18658 | 0 |
| SRR2487481 | 0 | 28 | 308 | 0 | 0 | 0 | 0 | 128 | 244 | 1567 | 13 | 193 | 0 | 154 | 2196 | 2312 | 0 | 131 | 0 | 2890 | 2993 | 12 | 3677 | 67067 | 29 | 0 | 0 | 0 | 70 | 28 | 196 | 42 | 42 | 280 | 0 | 8015 | 0 |
| SRR2487483 | 0 | 29 | 10 | 0 | 11 | 0 | 0 | 167 | 167 | 1377 | 13 | 129 | 0 | 39 | 1957 | 2291 | 0 | 152 | 0 | 2871 | 2766 | 12145 | 3842 | 69041 | 26 | 0 | 0 | 0 | 14 | 115 | 158 | 29 | 101 | 259 | 29 | 8033 | 0 |
| SRR2487485 | 0 | 0 | 343 | 0 | 11 | 0 | 0 | 290 | 205 | 1537 | 14 | 324 | 0 | 68 | 1861 | 1912 | 0 | 135 | 0 | 3124 | 3021 | 13210 | 3688 | 66970 | 3304 | 0 | 0 | 0 | 54 | 36 | 54 | 72 | 54 | 234 | 0 | 7990 | 0 |
| SRR2487487 | 0 | 36 | 341 | 0 | 18 | 18 | 0 | 215 | 182 | 1488 | 182 | 232 | 0 | 149 | 2117 | 1786 | 0 | 127 | 0 | 2811 | 2743 | 12517 | 3688 | 67766 | 28 | 0 | 0 | 0 | 54 | 90 | 54 | 54 | 143 | 376 | 0 | 8880 | 0 |
| SRR2487489 | 0 | 0 | 34 | 0 | 52 | 0 | 0 | 517 | 1046 | 36407 | 35 | 12 | 0 | 2824 | 70 | 46 | 0 | 156 | 0 | 99 | 2318 | 7848 | 488 | 25014 | 6927 | 16 | 0 | 0 | 0 | 16 | 13 | 0 | 16 | 66 | 0 | 720 | 14 |
| SRR2487491 | 0 | 16 | 45 | 0 | 28 | 0 | 0 | 450 | 1116 | 36278 | 42 | 24 | 0 | 2988 | 114 | 54 | 0 | 133 | 11 | 66 | 1960 | 6785 | 511 | 25850 | 14 | 11 | 0 | 0 | 23 | 0 | 23 | 0 | 0 | 45 | 0 | 630 | 0 |
| SRR2487493 | 0 | 16 | 63 | 0 | 45 | 0 | 0 | 596 | 1162 | 36725 | 59 | 22 | 0 | 2927 | 81 | 44 | 0 | 131 | 0 | 74 | 2315 | 7698 | 478 | 24559 | 6812 | 0 | 0 | 0 | 43 | 16 | 55 | 0 | 16 | 63 | 0 | 662 | 10 |
| SRR2487495 | 0 | 12 | 36 | 0 | 73 | 0 | 0 | 480 | 1167 | 37611 | 14 | 14 | 0 | 3078 | 50 | 43 | 0 | 140 | 0 | 86 | 2051 | 7165 | 480 | 25658 | 6932 | 0 | 0 | 0 | 36 | 0 | 16 | 0 | 0 | 68 | 0 | 515 | 0 |
| SRR2487497 | 0 | 0 | 215 | 0 | 0 | 0 | 0 | 433 | 213 | 3361 | 13 | 202 | 0 | 551 | 504 | 427 | 17 | 186 | 0 | 1316 | 3615 | 10 | 3689 | 55348 | 27 | 0 | 0 | 0 | 26 | 34 | 17 | 0 | 69 | 181 | 0 | 5015 | 0 |
| SRR2487499 | 0 | 0 | 11 | 0 | 12 | 0 | 0 | 362 | 172 | 3453 | 19 | 215 | 0 | 449 | 590 | 584 | 0 | 193 | 0 | 1413 | 3040 | 10504 | 3863 | 57164 | 3933 | 0 | 0 | 0 | 0 | 0 | 44 | 0 | 27 | 169 | 0 | 4866 | 0 |
| SRR2487501 | 0 | 44 | 288 | 0 | 0 | 11 | 0 | 428 | 306 | 3384 | 14 | 206 | 0 | 359 | 420 | 443 | 11 | 187 | 0 | 1451 | 3397 | 11770 | 3664 | 55042 | 3993 | 0 | 0 | 0 | 11 | 66 | 55 | 11 | 22 | 144 | 0 | 4904 | 0 |
| SRR2487503 | 0 | 22 | 286 | 0 | 0 | 0 | 0 | 425 | 186 | 3518 | 13 | 171 | 0 | 574 | 455 | 447 | 0 | 180 | 0 | 1498 | 3124 | 10699 | 3693 | 55851 | 21 | 12 | 0 | 0 | 44 | 55 | 55 | 0 | 33 | 132 | 22 | 5345 | 0 |
| SRR2487504 | 0 | 120 | 300 | 0 | 13 | 0 | 0 | 2109 | 1768 | 8842 | 136 | 340 | 0 | 680 | 612 | 408 | 0 | 42 | 0 | 816 | 4222 | 7480 | 2068 | 17451 | 5509 | 0 | 0 | 60 | 180 | 120 | 120 | 0 | 60 | 60 | 0 | 1905 | 0 |
| SRR2487505 | 0 | 41 | 0 | 0 | 35 | 0 | 0 | 2646 | 1694 | 11909 | 371 | 370 | 0 | 582 | 370 | 212 | 0 | 53 | 0 | 741 | 3708 | 7402 | 2544 | 20370 | 33 | 0 | 0 | 0 | 0 | 165 | 165 | 41 | 165 | 165 | 0 | 2117 | 0 |
| SRR2487507 | 0 | 0 | 455 | 0 | 60 | 0 | 0 | 2245 | 2551 | 11379 | 255 | 408 | 0 | 255 | 357 | 918 | 41 | 35 | 0 | 918 | 3216 | 6652 | 2399 | 20451 | 5830 | 0 | 41 | 0 | 0 | 41 | 83 | 83 | 41 | 124 | 0 | 2755 | 0 |
| SRR2487508 | 0 | 63 | 569 | 0 | 36 | 0 | 0 | 1283 | 1354 | 7628 | 143 | 143 | 0 | 570 | 143 | 356 | 0 | 49 | 0 | 784 | 3200 | 6169 | 2115 | 17998 | 27 | 0 | 0 | 63 | 0 | 126 | 126 | 0 | 63 | 126 | 0 | 1711 | 0 |
| SRR2487509 | 0 | 66 | 12 | 0 | 38 | 0 | 0 | 2252 | 1051 | 9310 | 377 | 450 | 0 | 676 | 75 | 225 | 0 | 55 | 0 | 601 | 3997 | 7329 | 2156 | 18132 | 17 | 0 | 0 | 0 | 0 | 66 | 66 | 0 | 132 | 198 | 0 | 1502 | 0 |
| SRR2487510 | 0 | 68 | 270 | 0 | 40 | 0 | 0 | 2685 | 1119 | 7758 | 224 | 448 | 0 | 224 | 149 | 298 | 0 | 50 | 0 | 970 | 2987 | 6021 | 2033 | 18572 | 16 | 0 | 0 | 0 | 0 | 68 | 0 | 68 | 135 | 135 | 0 | 2014 | 0 |
| SRR2487512 | 0 | 32 | 11 | 0 | 32 | 0 | 0 | 1916 | 2229 | 11221 | 235 | 274 | 0 | 704 | 352 | 313 | 0 | 44 | 0 | 665 | 3798 | 7613 | 2320 | 20041 | 5963 | 0 | 0 | 0 | 32 | 96 | 224 | 32 | 32 | 224 | 32 | 2737 | 0 |
| SRR2487514 | 0 | 67 | 13 | 0 | 44 | 0 | 0 | 2083 | 2328 | 11642 | 287 | 204 | 0 | 572 | 449 | 572 | 0 | 50 | 0 | 940 | 3293 | 6808 | 2308 | 20783 | 37 | 0 | 0 | 0 | 33 | 200 | 133 | 0 | 133 | 100 | 0 | 2533 | 0 |
| SRR2487516 | 0 | 0 | 50 | 0 | 13 | 0 | 0 | 87 | 87 | 1914 | 218 | 305 | 0 | 0 | 870 | 522 | 0 | 69 | 0 | 1436 | 3276 | 15 | 3711 | 46704 | 22 | 0 | 0 | 0 | 0 | 101 | 0 | 0 | 101 | 251 | 0 | 3828 | 0 |
| SRR2487517 | 0 | 0 | 131 | 0 | 11 | 0 | 0 | 204 | 116 | 1951 | 175 | 582 | 0 | 87 | 1106 | 1194 | 0 | 86 | 0 | 2096 | 3100 | 12 | 3925 | 54313 | 38 | 0 | 0 | 0 | 33 | 33 | 98 | 33 | 33 | 196 | 0 | 5648 | 16 |
| SRR2487519 | 0 | 0 | 32 | 0 | 17 | 0 | 0 | 287 | 115 | 2615 | 201 | 345 | 0 | 86 | 891 | 718 | 0 | 83 | 0 | 1839 | 2845 | 12 | 4210 | 54779 | 46 | 0 | 0 | 0 | 65 | 32 | 65 | 0 | 0 | 129 | 32 | 5344 | 0 |
| SRR2487520 | 0 | 51 | 51 | 0 | 22 | 0 | 0 | 181 | 45 | 1628 | 136 | 452 | 0 | 45 | 1086 | 678 | 0 | 74 | 0 | 1583 | 2608 | 10478 | 3737 | 48365 | 4384 | 0 | 0 | 0 | 51 | 103 | 0 | 0 | 0 | 205 | 0 | 4071 | 0 |
| SRR2487521 | 0 | 0 | 111 | 0 | 0 | 0 | 0 | 301 | 100 | 1657 | 151 | 452 | 0 | 50 | 602 | 1205 | 0 | 78 | 0 | 1657 | 3290 | 13 | 3490 | 46673 | 4420 | 0 | 0 | 0 | 111 | 167 | 111 | 0 | 56 | 167 | 0 | 3816 | 0 |
| SRR2487522 | 0 | 56 | 169 | 0 | 0 | 0 | 0 | 48 | 0 | 2276 | 97 | 436 | 0 | 0 | 387 | 484 | 0 | 65 | 0 | 1162 | 2570 | 13 | 3465 | 47678 | 11 | 0 | 0 | 0 | 0 | 169 | 225 | 0 | 0 | 281 | 0 | 4601 | 0 |
| SRR2487524 | 0 | 0 | 127 | 0 | 10 | 0 | 0 | 217 | 43 | 2104 | 174 | 325 | 0 | 108 | 1150 | 1323 | 0 | 69 | 0 | 1583 | 3196 | 13001 | 4171 | 54452 | 49 | 0 | 0 | 25 | 51 | 76 | 51 | 25 | 76 | 51 | 25 | 4816 | 0 |
| SRR2487526 | 0 | 52 | 104 | 0 | 18 | 0 | 0 | 89 | 134 | 2165 | 157 | 223 | 0 | 45 | 1116 | 1339 | 0 | 83 | 0 | 1830 | 2595 | 11617 | 4075 | 56150 | 5182 | 0 | 0 | 0 | 26 | 182 | 78 | 26 | 104 | 287 | 26 | 5201 | 0 |
| SRR2487529 | 0 | 35 | 80 | 0 | 31 | 0 | 0 | 502 | 244 | 7105 | 16 | 651 | 0 | 610 | 1925 | 1817 | 0 | 72 | 0 | 786 | 2475 | 8714 | 3196 | 29598 | 40 | 0 | 0 | 18 | 160 | 186 | 231 | 62 | 115 | 328 | 0 | 5681 | 0 |
| SRR2487531 | 0 | 64 | 156 | 0 | 38 | 0 | 0 | 601 | 279 | 7532 | 335 | 531 | 0 | 307 | 2040 | 1593 | 0 | 58 | 0 | 699 | 2086 | 7902 | 3314 | 30501 | 35 | 0 | 0 | 0 | 137 | 238 | 247 | 27 | 128 | 348 | 18 | 6036 | 0 |
| SRR2487533 | 0 | 45 | 136 | 0 | 40 | 0 | 0 | 371 | 230 | 7527 | 477 | 583 | 0 | 512 | 1714 | 1361 | 0 | 72 | 0 | 901 | 2422 | 8695 | 3226 | 29753 | 36 | 0 | 0 | 11 | 147 | 170 | 215 | 79 | 102 | 249 | 11 | 5654 | 0 |
| SRR2487535 | 0 | 56 | 101 | 0 | 40 | 0 | 0 | 364 | 260 | 6796 | 295 | 555 | 0 | 537 | 1612 | 1543 | 0 | 64 | 0 | 694 | 2157 | 8068 | 3350 | 29740 | 5116 | 0 | 22 | 11 | 112 | 235 | 235 | 56 | 146 | 403 | 22 | 5496 | 0 |
| SRR2487537 | 0 | 76 | 137 | 0 | 11 | 0 | 0 | 201 | 132 | 4445 | 14 | 170 | 0 | 271 | 1015 | 1007 | 0 | 124 | 0 | 1200 | 4021 | 11863 | 3926 | 51759 | 6463 | 0 | 0 | 30 | 30 | 121 | 182 | 0 | 106 | 212 | 0 | 4283 | 0 |
| SRR2487539 | 0 | 63 | 189 | 0 | 12 | 0 | 0 | 134 | 126 | 4343 | 11 | 110 | 0 | 189 | 867 | 1253 | 0 | 126 | 0 | 1434 | 3587 | 10658 | 4000 | 53329 | 6422 | 0 | 0 | 0 | 32 | 63 | 110 | 32 | 110 | 284 | 0 | 4162 | 0 |
| SRR2487541 | 0 | 39 | 137 | 0 | 15 | 20 | 0 | 129 | 119 | 4461 | 109 | 129 | 0 | 169 | 835 | 1053 | 20 | 117 | 0 | 1013 | 3919 | 11669 | 3950 | 51941 | 55 | 0 | 0 | 39 | 78 | 117 | 176 | 20 | 59 | 176 | 0 | 4252 | 0 |
| SRR2487543 | 0 | 19 | 136 | 0 | 0 | 0 | 0 | 146 | 175 | 4937 | 12 | 156 | 0 | 165 | 826 | 982 | 0 | 111 | 0 | 1302 | 3648 | 10797 | 3921 | 52031 | 6353 | 0 | 0 | 0 | 77 | 155 | 58 | 0 | 58 | 174 | 0 | 4529 | 0 |
| SRR2487545 | 0 | 92 | 183 | 0 | 45 | 0 | 0 | 814 | 571 | 8598 | 41 | 74 | 0 | 1581 | 0 | 10 | 0 | 106 | 0 | 34 | 2940 | 9753 | 3227 | 28657 | 5772 | 13 | 0 | 31 | 61 | 336 | 245 | 61 | 92 | 275 | 61 | 111 | 0 |
| SRR2487547 | 0 | 62 | 218 | 0 | 59 | 0 | 0 | 699 | 623 | 8358 | 14 | 62 | 0 | 1659 | 10 | 21 | 0 | 85 | 0 | 42 | 2452 | 8405 | 3286 | 29317 | 22 | 21 | 0 | 0 | 125 | 218 | 125 | 31 | 125 | 498 | 31 | 93 | 0 |
| SRR2487549 | 0 | 78 | 78 | 0 | 54 | 0 | 0 | 886 | 532 | 8647 | 10 | 64 | 0 | 1729 | 30 | 21 | 0 | 132 | 0 | 43 | 3115 | 9504 | 3174 | 28936 | 5693 | 13 | 0 | 0 | 39 | 392 | 157 | 39 | 78 | 392 | 0 | 106 | 0 |
| SRR2487551 | 0 | 195 | 78 | 0 | 47 | 0 | 0 | 690 | 581 | 8635 | 63 | 46 | 0 | 1722 | 25 | 13 | 0 | 117 | 0 | 21 | 2609 | 8581 | 3069 | 28637 | 25 | 16 | 0 | 0 | 0 | 234 | 234 | 78 | 156 | 273 | 0 | 84 | 0 |
| SRR2487553 | 0 | 20 | 40 | 0 | 0 | 0 | 0 | 392 | 677 | 8863 | 0 | 0 | 0 | 379 | 3289 | 5896 | 0 | 160 | 0 | 341 | 3839 | 24553 | 4223 | 99576 | 10002 | 0 | 0 | 0 | 20 | 61 | 61 | 20 | 101 | 243 | 121 | 890 | 0 |
| SRR2487555 | 0 | 41 | 124 | 0 | 0 | 0 | 0 | 382 | 675 | 8939 | 0 | 0 | 0 | 446 | 3364 | 6215 | 0 | 137 | 0 | 350 | 3153 | 20371 | 4350 | 102576 | 9792 | 0 | 0 | 0 | 0 | 21 | 124 | 21 | 166 | 166 | 0 | 844 | 0 |
| SRR2487560 | 0 | 208 | 346 | 0 | 71 | 69 | 0 | 196 | 124 | 2738 | 71 | 124 | 0 | 267 | 89 | 18 | 0 | 64 | 0 | 89 | 3193 | 9441 | 3343 | 29983 | 4580 | 0 | 0 | 0 | 0 | 69 | 138 | 0 | 69 | 277 | 0 | 391 | 0 |
| SRR2487561 | 0 | 0 | 18 | 0 | 110 | 0 | 0 | 327 | 192 | 3139 | 0 | 0 | 0 | 359 | 135 | 125 | 0 | 187 | 0 | 135 | 2666 | 15911 | 3931 | 52190 | 7539 | 0 | 0 | 0 | 19 | 57 | 132 | 0 | 76 | 114 | 0 | 489 | 0 |
| SRR2487563 | 0 | 19 | 27 | 0 | 99 | 0 | 0 | 361 | 139 | 3222 | 0 | 0 | 0 | 444 | 145 | 165 | 0 | 140 | 0 | 103 | 2196 | 13743 | 4185 | 53132 | 7990 | 0 | 0 | 0 | 78 | 0 | 39 | 0 | 19 | 97 | 19 | 506 | 0 |
| SRR2487564 | 0 | 0 | 744 | 0 | 86 | 0 | 0 | 356 | 225 | 2735 | 169 | 112 | 0 | 412 | 37 | 19 | 0 | 86 | 0 | 150 | 2495 | 7919 | 3215 | 30991 | 4132 | 0 | 0 | 0 | 0 | 74 | 74 | 0 | 74 | 74 | 0 | 300 | 0 |
| SRR2487565 | 0 | 77 | 153 | 0 | 85 | 0 | 77 | 314 | 235 | 2608 | 78 | 137 | 0 | 392 | 39 | 39 | 0 | 97 | 12 | 59 | 2832 | 9232 | 2965 | 30390 | 4198 | 0 | 77 | 0 | 77 | 384 | 153 | 0 | 0 | 77 | 0 | 216 | 0 |
| SRR2487566 | 0 | 0 | 314 | 0 | 59 | 0 | 0 | 414 | 217 | 2998 | 158 | 138 | 0 | 414 | 59 | 20 | 0 | 59 | 0 | 79 | 2476 | 7782 | 3094 | 30949 | 4176 | 0 | 0 | 0 | 0 | 314 | 0 | 0 | 79 | 314 | 0 | 414 | 0 |
| SRR2487568 | 0 | 29 | 632 | 0 | 55 | 0 | 0 | 310 | 182 | 3207 | 0 | 0 | 0 | 401 | 165 | 236 | 0 | 117 | 0 | 91 | 2654 | 15990 | 4265 | 53070 | 7625 | 0 | 0 | 0 | 15 | 29 | 15 | 15 | 59 | 147 | 29 | 517 | 0 |
| SRR2487570 | 0 | 30 | 22 | 0 | 92 | 0 | 0 | 299 | 149 | 3118 | 0 | 13 | 0 | 350 | 171 | 183 | 0 | 155 | 0 | 111 | 2221 | 13848 | 3730 | 54447 | 7890 | 0 | 0 | 0 | 0 | 0 | 46 | 15 | 61 | 152 | 30 | 503 | 0 |
| SRR2487572 | 0 | 65 | 422 | 0 | 0 | 0 | 0 | 589 | 499 | 3076 | 19 | 123 | 0 | 74 | 196 | 524 | 0 | 77 | 0 | 352 | 3506 | 12265 | 4089 | 57651 | 5055 | 0 | 0 | 0 | 0 | 130 | 162 | 0 | 65 | 260 | 0 | 941 | 0 |
| SRR2487573 | 0 | 0 | 567 | 0 | 19 | 0 | 0 | 582 | 589 | 4347 | 0 | 15 | 0 | 191 | 1224 | 2441 | 0 | 141 | 0 | 429 | 2912 | 18274 | 5174 | 93497 | 8491 | 19 | 0 | 0 | 0 | 30 | 30 | 30 | 179 | 298 | 60 | 1706 | 0 |
| SRR2487575 | 0 | 60 | 210 | 0 | 0 | 0 | 0 | 536 | 528 | 4255 | 0 | 23 | 0 | 219 | 1086 | 2512 | 0 | 119 | 0 | 483 | 2523 | 18026 | 4708 | 96088 | 8528 | 0 | 0 | 0 | 0 | 60 | 150 | 30 | 90 | 300 | 30 | 1509 | 0 |
| SRR2487576 | 0 | 34 | 335 | 0 | 12 | 0 | 0 | 610 | 559 | 3345 | 93 | 169 | 0 | 229 | 263 | 500 | 0 | 88 | 0 | 347 | 2884 | 10348 | 4174 | 59074 | 4869 | 0 | 0 | 0 | 134 | 101 | 67 | 67 | 0 | 235 | 0 | 1220 | 0 |
| SRR2487577 | 0 | 36 | 254 | 0 | 0 | 0 | 0 | 604 | 531 | 3311 | 64 | 146 | 0 | 183 | 238 | 457 | 0 | 77 | 0 | 329 | 3525 | 15 | 4105 | 56927 | 4894 | 0 | 0 | 0 | 36 | 72 | 72 | 0 | 72 | 217 | 0 | 1098 | 0 |
| SRR2487578 | 0 | 0 | 328 | 0 | 14 | 36 | 0 | 522 | 421 | 3390 | 92 | 101 | 0 | 183 | 211 | 495 | 0 | 105 | 0 | 247 | 2848 | 10296 | 4285 | 59029 | 4854 | 0 | 0 | 0 | 0 | 73 | 146 | 73 | 73 | 219 | 0 | 962 | 0 |
| SRR2487580 | 0 | 0 | 14 | 0 | 15 | 0 | 0 | 567 | 549 | 4338 | 18 | 18 | 0 | 136 | 1210 | 2703 | 0 | 81 | 0 | 549 | 2834 | 18962 | 4724 | 96943 | 10 | 0 | 0 | 0 | 0 | 91 | 23 | 23 | 45 | 295 | 45 | 1493 | 15 |
| SRR2487582 | 0 | 23 | 162 | 0 | 0 | 0 | 0 | 519 | 495 | 4433 | 0 | 18 | 0 | 199 | 1176 | 3407 | 0 | 137 | 0 | 603 | 2468 | 17154 | 5081 | 97662 | 13 | 0 | 0 | 0 | 0 | 23 | 93 | 23 | 93 | 232 | 46 | 1417 | 0 |
| SRR2487584 | 0 | 46 | 29 | 0 | 47 | 0 | 0 | 1296 | 1404 | 5020 | 29 | 1835 | 0 | 270 | 108 | 162 | 0 | 94 | 0 | 486 | 1426 | 6270 | 1124 | 18401 | 2401 | 0 | 0 | 0 | 15 | 18 | 0 | 0 | 0 | 0 | 0 | 1404 | 0 |
| SRR2487585 | 0 | 12 | 27 | 0 | 67 | 0 | 0 | 691 | 691 | 7913 | 0 | 154 | 0 | 307 | 691 | 615 | 0 | 90 | 0 | 1076 | 1275 | 12060 | 1790 | 34882 | 4095 | 0 | 0 | 0 | 0 | 0 | 0 | 0 | 0 | 14 | 14 | 2151 | 0 |
| SRR2487587 | 0 | 65 | 60 | 0 | 43 | 0 | 0 | 1060 | 757 | 7646 | 151 | 76 | 0 | 227 | 984 | 1060 | 0 | 128 | 0 | 606 | 1215 | 10021 | 1684 | 35224 | 4328 | 0 | 0 | 0 | 0 | 0 | 0 | 0 | 0 | 37 | 0 | 2725 | 0 |
| SRR2487588 | 0 | 252 | 27 | 0 | 44 | 12 | 0 | 617 | 729 | 5386 | 3254 | 2581 | 0 | 449 | 168 | 56 | 0 | 79 | 0 | 617 | 1081 | 5420 | 1095 | 18247 | 2221 | 0 | 0 | 0 | 12 | 0 | 23 | 0 | 16 | 19 | 0 | 1627 | 0 |
| SRR2487589 | 0 | 260 | 24 | 0 | 27 | 12 | 0 | 1381 | 1020 | 4742 | 3181 | 2701 | 0 | 660 | 60 | 120 | 0 | 78 | 0 | 480 | 1379 | 6221 | 1127 | 17804 | 2342 | 0 | 0 | 0 | 0 | 20 | 12 | 0 | 0 | 12 | 0 | 2041 | 0 |
| SRR2487590 | 0 | 290 | 46 | 0 | 33 | 17 | 0 | 530 | 1002 | 6128 | 2062 | 2829 | 0 | 471 | 59 | 354 | 0 | 61 | 0 | 707 | 1007 | 5268 | 1181 | 18628 | 2240 | 0 | 0 | 0 | 21 | 0 | 17 | 0 | 0 | 21 | 0 | 2239 | 12 |
| SRR2487592 | 0 | 18 | 39 | 0 | 71 | 0 | 0 | 543 | 965 | 7361 | 181 | 60 | 0 | 543 | 845 | 845 | 0 | 125 | 0 | 483 | 1104 | 11808 | 1781 | 35388 | 17 | 0 | 0 | 0 | 0 | 0 | 14 | 0 | 0 | 11 | 0 | 3379 | 0 |
| SRR2487594 | 0 | 40 | 73 | 0 | 18 | 0 | 0 | 735 | 796 | 7163 | 122 | 245 | 0 | 490 | 979 | 1041 | 0 | 92 | 0 | 612 | 1165 | 10502 | 1683 | 35702 | 4068 | 0 | 0 | 0 | 0 | 0 | 11 | 0 | 11 | 18 | 15 | 3183 | 0 |
| SRR2487597 | 0 | 53 | 148 | 0 | 12 | 0 | 0 | 193 | 180 | 2136 | 360 | 425 | 0 | 116 | 4967 | 3449 | 0 | 182 | 0 | 2522 | 2279 | 17 | 3021 | 51928 | 3129 | 0 | 0 | 11 | 53 | 106 | 42 | 11 | 63 | 159 | 21 | 5623 | 0 |
| SRR2487598 | 0 | 23 | 135 | 0 | 0 | 0 | 0 | 269 | 226 | 2248 | 22 | 636 | 0 | 85 | 4156 | 3563 | 0 | 180 | 0 | 2531 | 2801 | 17 | 2975 | 50815 | 3208 | 0 | 0 | 0 | 90 | 79 | 102 | 11 | 34 | 248 | 0 | 6574 | 0 |
| SRR2487599 | 0 | 23 | 194 | 0 | 0 | 0 | 0 | 237 | 139 | 1936 | 26 | 696 | 0 | 167 | 4749 | 3565 | 11 | 167 | 0 | 2326 | 2152 | 12 | 2960 | 51906 | 3229 | 0 | 0 | 0 | 34 | 159 | 80 | 11 | 34 | 273 | 11 | 5822 | 0 |
| SRR2487600 | 0 | 0 | 14 | 0 | 0 | 0 | 0 | 237 | 237 | 3022 | 65 | 108 | 0 | 108 | 23122 | 22949 | 0 | 152 | 0 | 3217 | 2509 | 20683 | 3507 | 84344 | 16 | 14 | 0 | 0 | 16 | 16 | 47 | 16 | 11 | 16 | 63 | 9046 | 0 |
| SRR2487602 | 0 | 32 | 28 | 0 | 15 | 0 | 0 | 309 | 331 | 2997 | 44 | 44 | 0 | 66 | 22326 | 22612 | 0 | 217 | 0 | 2777 | 1887 | 18401 | 3301 | 84219 | 15 | 0 | 0 | 0 | 32 | 32 | 0 | 0 | 16 | 29 | 15 | 9080 | 15 |
| SRR2487604 | 0 | 0 | 27 | 0 | 0 | 0 | 0 | 175 | 175 | 3387 | 117 | 88 | 0 | 146 | 21459 | 19590 | 0 | 319 | 0 | 3095 | 2499 | 20789 | 3667 | 81977 | 5145 | 0 | 0 | 0 | 0 | 0 | 20 | 20 | 20 | 18 | 0 | 8583 | 0 |
| SRR2487606 | 0 | 21 | 18 | 0 | 18 | 0 | 0 | 343 | 200 | 3317 | 86 | 57 | 0 | 172 | 19843 | 19157 | 0 | 222 | 0 | 3002 | 2147 | 19079 | 3353 | 82827 | 18 | 0 | 0 | 0 | 0 | 41 | 104 | 0 | 62 | 16 | 41 | 8520 | 0 |
| SRR2487607 | 0 | 20 | 121 | 0 | 0 | 0 | 0 | 138 | 138 | 1989 | 313 | 575 | 0 | 88 | 4590 | 3589 | 0 | 175 | 0 | 2401 | 2929 | 15 | 2859 | 51217 | 3382 | 0 | 0 | 10 | 20 | 70 | 80 | 10 | 70 | 221 | 10 | 6216 | 0 |
| SRR2487609 | 0 | 23 | 81 | 0 | 97 | 0 | 0 | 407 | 343 | 4577 | 60 | 582 | 0 | 1212 | 32 | 32 | 0 | 38 | 0 | 96 | 1674 | 5626 | 2336 | 19193 | 4059 | 0 | 23 | 12 | 35 | 115 | 81 | 23 | 12 | 173 | 0 | 287 | 0 |
| SRR2487610 | 23 | 35 | 46 | 0 | 96 | 0 | 0 | 440 | 290 | 3883 | 89 | 689 | 0 | 1319 | 33 | 25 | 12 | 34 | 0 | 141 | 2095 | 6629 | 2241 | 18950 | 4250 | 0 | 0 | 35 | 23 | 208 | 58 | 35 | 69 | 173 | 12 | 365 | 0 |
| SRR2487611 | 0 | 12 | 47 | 0 | 93 | 0 | 0 | 350 | 242 | 4553 | 80 | 609 | 0 | 1234 | 33 | 67 | 0 | 30 | 0 | 108 | 1618 | 5456 | 2266 | 19470 | 4171 | 0 | 35 | 23 | 94 | 164 | 106 | 23 | 59 | 129 | 0 | 300 | 0 |
| SRR2487612 | 0 | 0 | 67 | 0 | 0 | 0 | 0 | 331 | 442 | 5080 | 0 | 0 | 0 | 1126 | 110 | 243 | 0 | 39 | 0 | 110 | 2005 | 12834 | 3577 | 38679 | 16 | 0 | 0 | 0 | 0 | 168 | 67 | 0 | 101 | 168 | 168 | 442 | 0 |
| SRR2487614 | 0 | 0 | 140 | 0 | 185 | 0 | 0 | 387 | 205 | 5377 | 114 | 0 | 0 | 1253 | 137 | 182 | 0 | 21 | 0 | 114 | 1457 | 10606 | 3221 | 37379 | 9129 | 0 | 0 | 0 | 0 | 0 | 35 | 35 | 70 | 15 | 105 | 684 | 0 |
| SRR2487616 | 0 | 0 | 45 | 0 | 99 | 0 | 0 | 309 | 281 | 6152 | 0 | 56 | 0 | 1264 | 140 | 225 | 0 | 50 | 0 | 0 | 1889 | 12503 | 3256 | 38007 | 8352 | 0 | 0 | 0 | 0 | 45 | 45 | 0 | 45 | 0 | 11 | 562 | 0 |
| SRR2487618 | 0 | 46 | 91 | 0 | 73 | 0 | 0 | 281 | 281 | 5824 | 56 | 56 | 0 | 1350 | 141 | 338 | 0 | 48 | 0 | 141 | 1985 | 10941 | 3171 | 37907 | 13 | 0 | 0 | 0 | 46 | 0 | 0 | 91 | 91 | 11 | 46 | 309 | 0 |
| SRR2487619 | 0 | 0 | 32 | 0 | 74 | 0 | 0 | 340 | 287 | 4429 | 679 | 551 | 0 | 1237 | 15 | 0 | 11 | 26 | 0 | 75 | 2225 | 6837 | 2248 | 19318 | 4392 | 0 | 11 | 0 | 117 | 191 | 74 | 11 | 42 | 117 | 21 | 415 | 0 |
| SRR2487621 | 0 | 26 | 26 | 0 | 11 | 0 | 0 | 1226 | 704 | 12052 | 287 | 235 | 0 | 287 | 652 | 939 | 26 | 84 | 0 | 887 | 2724 | 10 | 2742 | 42382 | 15 | 0 | 0 | 0 | 0 | 51 | 77 | 0 | 128 | 77 | 0 | 2139 | 28 |
| SRR2487622 | 0 | 27 | 191 | 0 | 16 | 0 | 0 | 1143 | 976 | 11068 | 335 | 307 | 0 | 335 | 641 | 613 | 0 | 92 | 0 | 697 | 3110 | 11395 | 2816 | 41343 | 5419 | 0 | 0 | 27 | 55 | 0 | 55 | 0 | 0 | 191 | 0 | 1227 | 27 |
| SRR2487623 | 0 | 28 | 83 | 0 | 21 | 0 | 0 | 1009 | 981 | 11662 | 10 | 308 | 0 | 392 | 645 | 533 | 0 | 77 | 0 | 645 | 2539 | 9752 | 2766 | 42357 | 5509 | 0 | 0 | 0 | 55 | 55 | 28 | 0 | 28 | 28 | 0 | 1822 | 22 |
| SRR2487624 | 0 | 0 | 92 | 0 | 19 | 0 | 0 | 1045 | 1216 | 13271 | 34 | 17 | 0 | 257 | 3528 | 3459 | 0 | 93 | 0 | 788 | 2590 | 17871 | 3335 | 67988 | 32 | 0 | 0 | 15 | 0 | 31 | 15 | 0 | 46 | 11 | 0 | 2123 | 37 |
| SRR2487626 | 0 | 0 | 126 | 0 | 0 | 0 | 0 | 1180 | 1092 | 13825 | 18 | 35 | 0 | 493 | 3804 | 3769 | 0 | 107 | 0 | 810 | 2299 | 16216 | 3542 | 69437 | 31 | 0 | 0 | 0 | 16 | 47 | 47 | 0 | 47 | 79 | 31 | 2078 | 42 |
| SRR2487628 | 0 | 40 | 139 | 0 | 0 | 0 | 0 | 1128 | 998 | 13948 | 65 | 22 | 0 | 369 | 3167 | 2733 | 0 | 140 | 0 | 564 | 2793 | 17622 | 3078 | 67037 | 27 | 0 | 0 | 0 | 0 | 20 | 60 | 20 | 0 | 20 | 0 | 2061 | 14 |

### Supp Table S8.4.Columns_DI_to_ES

| | | | | | | | | | | | | | | | | | | | | | | | | | | | | | | | | | | | | | |
|---|---|---|---|---|---|---|---|---|---|---|---|---|---|---|---|---|---|---|---|---|---|---|---|---|---|---|---|---|---|---|---|---|---|---|---|---|---|
| SRR2487630 | 0 | 0 | 179 | 0 | 23 | 0 | 0 | 1157 | 1026 | 13749 | 44 | 87 | 0 | 240 | 2924 | 2553 | 0 | 73 | 0 | 742 | 2558 | 16762 | 3531 | 68898 | 39 | 0 | 0 | 0 | 0 | 0 | 20 | 20 | 40 | 159 | 0 | 2531 | 37 |
| SRR2487631 | 0 | 24 | 97 | 0 | 12 | 0 | 0 | 1241 | 918 | 11068 | 199 | 323 | 0 | 248 | 943 | 571 | 0 | 84 | 0 | 869 | 3283 | 11649 | 2671 | 41996 | 5638 | 0 | 0 | 0 | 24 | 0 | 48 | 24 | 0 | 145 | 24 | 1886 | 23 |
| SRR2487633 | 0 | 0 | 37 | 0 | 97 | 0 | 0 | 420 | 315 | 6771 | 0 | 0 | 0 | 242 | 54 | 70 | 0 | 65 | 0 | 57 | 1921 | 9306 | 1531 | 33764 | 8501 | 0 | 0 | 0 | 0 | 74 | 18 | 0 | 74 | 74 | 92 | 382 | 13 |
| SRR2487635 | 0 | 19 | 19 | 0 | 100 | 0 | 0 | 507 | 293 | 6893 | 0 | 0 | 0 | 244 | 69 | 99 | 0 | 127 | 0 | 56 | 1387 | 8110 | 1747 | 35800 | 8370 | 0 | 0 | 0 | 39 | 58 | 77 | 0 | 58 | 232 | 39 | 379 | 0 |
| SRR2487637 | 0 | 24 | 95 | 0 | 90 | 0 | 0 | 487 | 331 | 6812 | 0 | 12 | 0 | 259 | 72 | 76 | 0 | 90 | 0 | 64 | 1962 | 9591 | 1726 | 34105 | 8361 | 0 | 0 | 0 | 48 | 48 | 119 | 24 | 24 | 143 | 48 | 339 | 0 |
| SRR2487639 | 0 | 24 | 97 | 0 | 39 | 0 | 0 | 497 | 326 | 7150 | 0 | 12 | 0 | 267 | 52 | 103 | 0 | 110 | 0 | 60 | 1648 | 8387 | 1655 | 35383 | 8575 | 0 | 0 | 0 | 0 | 48 | 24 | 24 | 72 | 169 | 72 | 322 | 0 |
| SRR2487641 | 0 | 0 | 206 | 0 | 23 | 0 | 0 | 843 | 465 | 7448 | 0 | 0 | 0 | 187 | 948 | 1391 | 23 | 75 | 0 | 509 | 2886 | 16775 | 5022 | 97341 | 8890 | 0 | 0 | 0 | 69 | 114 | 69 | 23 | 114 | 183 | 0 | 1730 | 0 |
| SRR2487643 | 0 | 0 | 214 | 0 | 24 | 0 | 0 | 789 | 417 | 7504 | 0 | 18 | 0 | 179 | 856 | 1269 | 0 | 163 | 0 | 637 | 2536 | 15135 | 5330 | 98569 | 13 | 12 | 0 | 0 | 24 | 95 | 24 | 24 | 142 | 190 | 71 | 1838 | 0 |
| SRR2487645 | 0 | 0 | 91 | 0 | 15 | 0 | 0 | 869 | 498 | 7448 | 0 | 11 | 0 | 205 | 991 | 1245 | 0 | 95 | 0 | 526 | 2851 | 15967 | 5125 | 96144 | 8510 | 0 | 0 | 0 | 30 | 182 | 30 | 0 | 30 | 212 | 0 | 1787 | 0 |
| SRR2487647 | 0 | 0 | 92 | 0 | 21 | 0 | 0 | 827 | 546 | 7949 | 17 | 0 | 0 | 149 | 827 | 1202 | 0 | 114 | 0 | 518 | 2616 | 15164 | 4984 | 97718 | 9093 | 0 | 0 | 0 | 0 | 31 | 123 | 0 | 123 | 338 | 61 | 1825 | 0 |
| SRR2487648 | 0 | 84 | 63 | 0 | 69 | 0 | 0 | 342 | 241 | 4607 | 316 | 304 | 0 | 532 | 266 | 177 | 0 | 69 | 0 | 152 | 2441 | 7852 | 1877 | 27129 | 4925 | 0 | 0 | 0 | 42 | 251 | 105 | 0 | 21 | 84 | 42 | 329 | 13 |
| SRR2487649 | 0 | 0 | 12 | 0 | 119 | 0 | 0 | 305 | 247 | 6344 | 29 | 15 | 0 | 784 | 1002 | 799 | 0 | 77 | 0 | 160 | 2180 | 14321 | 2571 | 53351 | 15 | 26 | 0 | 0 | 0 | 0 | 72 | 24 | 143 | 167 | 24 | 523 | 26 |
| SRR2487651 | 0 | 24 | 143 | 0 | 147 | 0 | 0 | 412 | 99 | 6096 | 43 | 28 | 0 | 583 | 966 | 909 | 0 | 139 | 0 | 171 | 1907 | 11948 | 2774 | 53382 | 22 | 0 | 0 | 0 | 0 | 166 | 143 | 0 | 24 | 143 | 24 | 384 | 16 |
| SRR2487652 | 0 | 63 | 63 | 0 | 81 | 0 | 0 | 470 | 287 | 4669 | 326 | 313 | 0 | 352 | 222 | 235 | 0 | 66 | 0 | 78 | 1855 | 6637 | 1876 | 27868 | 4818 | 0 | 0 | 21 | 126 | 148 | 211 | 42 | 0 | 126 | 21 | 261 | 11 |
| SRR2487653 | 0 | 92 | 252 | 0 | 78 | 0 | 0 | 332 | 332 | 4535 | 19 | 263 | 0 | 415 | 194 | 207 | 0 | 60 | 0 | 124 | 2291 | 7917 | 1805 | 27189 | 11 | 0 | 0 | 23 | 160 | 114 | 114 | 0 | 69 | 137 | 0 | 249 | 0 |
| SRR2487654 | 0 | 69 | 115 | 0 | 78 | 0 | 0 | 461 | 224 | 4990 | 16 | 266 | 0 | 419 | 168 | 70 | 0 | 53 | 0 | 196 | 1743 | 6154 | 1871 | 27636 | 4694 | 0 | 0 | 23 | 138 | 229 | 252 | 46 | 161 | 92 | 23 | 294 | 11 |
| SRR2487656 | 0 | 37 | 130 | 0 | 130 | 0 | 0 | 358 | 197 | 5713 | 58 | 46 | 0 | 486 | 1018 | 1099 | 0 | 82 | 0 | 127 | 2162 | 14468 | 2435 | 50988 | 9050 | 0 | 0 | 0 | 56 | 130 | 11 | 19 | 56 | 14 | 74 | 428 | 0 |
| SRR2487658 | 0 | 0 | 12 | 0 | 78 | 0 | 0 | 424 | 212 | 5931 | 0 | 35 | 0 | 600 | 1024 | 1188 | 0 | 42 | 0 | 153 | 1739 | 12333 | 2792 | 53000 | 22 | 0 | 0 | 0 | 38 | 76 | 172 | 0 | 76 | 172 | 96 | 424 | 14 |
| SRR2487661 | 0 | 28 | 57 | 0 | 0 | 0 | 0 | 251 | 266 | 7940 | 11 | 0 | 0 | 1069 | 2322 | 2056 | 0 | 199 | 0 | 359 | 2512 | 22933 | 4541 | 82935 | 13 | 0 | 0 | 0 | 0 | 0 | 57 | 0 | 85 | 198 | 0 | 2470 | 13 |
| SRR2487663 | 0 | 29 | 58 | 0 | 22 | 0 | 0 | 190 | 301 | 7915 | 0 | 0 | 0 | 1123 | 2231 | 1984 | 0 | 217 | 0 | 373 | 2054 | 19932 | 4829 | 85922 | 18 | 0 | 0 | 0 | 29 | 117 | 175 | 58 | 58 | 204 | 0 | 2696 | 43 |
| SRR2487665 | 0 | 37 | 222 | 0 | 0 | 0 | 0 | 230 | 183 | 7922 | 0 | 0 | 0 | 1179 | 1894 | 1659 | 0 | 211 | 0 | 338 | 2346 | 21136 | 4460 | 83253 | 9337 | 0 | 0 | 37 | 0 | 0 | 148 | 37 | 37 | 297 | 0 | 2467 | 37 |
| SRR2487667 | 0 | 0 | 75 | 0 | 15 | 0 | 0 | 200 | 232 | 8029 | 0 | 0 | 0 | 1055 | 1793 | 1752 | 0 | 240 | 0 | 362 | 2136 | 20027 | 4598 | 83848 | 12 | 0 | 0 | 0 | 0 | 0 | 75 | 0 | 75 | 75 | 37 | 2927 | 15 |
| SRR2487668 | 0 | 0 | 182 | 0 | 74 | 0 | 0 | 250 | 264 | 3429 | 14 | 125 | 0 | 722 | 0 | 0 | 0 | 167 | 0 | 847 | 1903 | 3853 | 1876 | 14372 | 5021 | 20 | 0 | 0 | 73 | 73 | 109 | 36 | 219 | 291 | 0 | 3845 | 0 |
| SRR2487669 | 0 | 0 | 165 | 0 | 48 | 0 | 0 | 300 | 114 | 4141 | 31 | 52 | 0 | 849 | 0 | 10 | 0 | 206 | 0 | 787 | 2099 | 5695 | 2170 | 22961 | 15 | 32 | 0 | 0 | 0 | 66 | 66 | 0 | 132 | 430 | 33 | 4607 | 0 |
| SRR2487671 | 0 | 33 | 234 | 0 | 85 | 0 | 0 | 247 | 226 | 4471 | 0 | 21 | 0 | 833 | 10 | 10 | 0 | 254 | 0 | 822 | 1634 | 5072 | 2289 | 23600 | 23 | 15 | 0 | 0 | 0 | 0 | 67 | 33 | 133 | 334 | 0 | 5160 | 15 |
| SRR2487672 | 0 | 38 | 225 | 0 | 64 | 0 | 0 | 358 | 201 | 4053 | 72 | 86 | 0 | 630 | 0 | 0 | 0 | 195 | 0 | 1017 | 1591 | 3269 | 1869 | 14881 | 5096 | 29 | 0 | 0 | 150 | 188 | 263 | 0 | 113 | 113 | 0 | 3709 | 0 |
| SRR2487673 | 0 | 0 | 276 | 0 | 67 | 0 | 0 | 260 | 214 | 3887 | 31 | 138 | 0 | 719 | 0 | 15 | 0 | 172 | 0 | 765 | 1888 | 3848 | 1798 | 14622 | 4910 | 12 | 0 | 0 | 40 | 197 | 197 | 0 | 0 | 158 | 0 | 3566 | 0 |
| SRR2487674 | 0 | 0 | 319 | 0 | 62 | 0 | 0 | 339 | 200 | 3852 | 46 | 62 | 0 | 740 | 15 | 0 | 0 | 163 | 0 | 678 | 1541 | 3225 | 1800 | 15042 | 4824 | 28 | 0 | 0 | 40 | 160 | 80 | 0 | 80 | 200 | 40 | 3297 | 0 |
| SRR2487676 | 0 | 0 | 11 | 0 | 83 | 0 | 0 | 282 | 191 | 4302 | 0 | 50 | 0 | 698 | 25 | 0 | 0 | 268 | 0 | 872 | 1794 | 6358 | 2260 | 24572 | 27 | 26 | 0 | 0 | 0 | 0 | 78 | 0 | 157 | 15 | 78 | 4841 | 0 |
| SRR2487678 | 0 | 0 | 161 | 0 | 53 | 0 | 0 | 212 | 203 | 4421 | 0 | 25 | 0 | 830 | 0 | 17 | 0 | 204 | 0 | 838 | 1472 | 5436 | 2353 | 24204 | 8204 | 0 | 0 | 0 | 27 | 27 | 215 | 27 | 81 | 511 | 54 | 5166 | 0 |
| SRR2487688 | 0 | 45 | 295 | 0 | 0 | 0 | 0 | 219 | 197 | 2055 | 154 | 503 | 0 | 393 | 525 | 437 | 0 | 214 | 0 | 2273 | 4019 | 12225 | 3353 | 46242 | 5212 | 0 | 0 | 45 | 45 | 181 | 159 | 0 | 91 | 23 | 0 | 5748 | 0 |
| SRR2487689 | 0 | 0 | 1119 | 0 | 0 | 0 | 0 | 447 | 503 | 2681 | 0 | 56 | 0 | 447 | 2457 | 3351 | 0 | 336 | 0 | 2513 | 3336 | 11 | 3943 | 68845 | 38 | 0 | 0 | 0 | 0 | 236 | 118 | 0 | 59 | 177 | 0 | 8600 | 0 |
| SRR2487691 | 0 | 0 | 896 | 0 | 0 | 0 | 0 | 444 | 333 | 2500 | 0 | 56 | 0 | 500 | 1944 | 3055 | 0 | 282 | 0 | 2667 | 3198 | 16061 | 3887 | 68415 | 7764 | 0 | 0 | 0 | 60 | 60 | 119 | 0 | 60 | 299 | 0 | 9000 | 0 |
| SRR2487692 | 0 | 0 | 415 | 0 | 0 | 0 | 0 | 205 | 182 | 2070 | 228 | 432 | 0 | 523 | 341 | 796 | 0 | 219 | 0 | 2207 | 3277 | 10545 | 3466 | 47655 | 4965 | 0 | 0 | 23 | 69 | 207 | 138 | 0 | 23 | 207 | 0 | 5005 | 0 |
| SRR2487693 | 0 | 51 | 616 | 0 | 0 | 0 | 0 | 198 | 322 | 2133 | 248 | 372 | 0 | 496 | 422 | 595 | 0 | 213 | 0 | 2207 | 3870 | 16 | 3510 | 46035 | 12 | 0 | 0 | 0 | 128 | 154 | 231 | 26 | 103 | 77 | 0 | 4960 | 0 |
| SRR2487694 | 0 | 25 | 561 | 0 | 0 | 0 | 0 | 174 | 174 | 2267 | 224 | 349 | 0 | 424 | 473 | 424 | 25 | 193 | 0 | 2242 | 3176 | 10189 | 3417 | 47473 | 14 | 0 | 0 | 0 | 102 | 255 | 102 | 76 | 102 | 280 | 0 | 5008 | 0 |
| SRR2487696 | 0 | 46 | 911 | 0 | 0 | 0 | 0 | 128 | 213 | 3320 | 0 | 85 | 0 | 511 | 2554 | 3065 | 0 | 204 | 0 | 2511 | 3571 | 17135 | 3827 | 70333 | 7398 | 0 | 0 | 0 | 0 | 137 | 0 | 0 | 182 | 91 | 46 | 7279 | 26 |
| SRR2487698 | 0 | 0 | 67 | 0 | 17 | 0 | 0 | 218 | 261 | 2785 | 44 | 0 | 0 | 435 | 2263 | 3220 | 0 | 210 | 0 | 3133 | 3373 | 14857 | 3740 | 72160 | 7586 | 0 | 0 | 0 | 0 | 12 | 92 | 0 | 92 | 19 | 92 | 8136 | 0 |
| SRR2487700 | 0 | 13 | 156 | 0 | 108 | 13 | 0 | 237 | 219 | 5270 | 28 | 748 | 0 | 456 | 73 | 73 | 0 | 89 | 0 | 310 | 2789 | 6481 | 1460 | 17039 | 19 | 0 | 0 | 26 | 52 | 143 | 195 | 13 | 52 | 235 | 26 | 1313 | 0 |
| SRR2487701 | 0 | 0 | 21 | 0 | 94 | 0 | 0 | 197 | 230 | 6542 | 165 | 33 | 0 | 526 | 132 | 164 | 0 | 105 | 0 | 329 | 2325 | 11114 | 2315 | 33080 | 9155 | 0 | 0 | 0 | 24 | 122 | 11 | 0 | 73 | 19 | 24 | 1907 | 0 |
| SRR2487703 | 0 | 24 | 316 | 0 | 101 | 0 | 0 | 511 | 191 | 7434 | 64 | 64 | 0 | 638 | 383 | 191 | 0 | 81 | 0 | 160 | 1975 | 9987 | 1854 | 32572 | 9141 | 20 | 0 | 0 | 0 | 73 | 49 | 0 | 49 | 146 | 97 | 2393 | 0 |
| SRR2487704 | 0 | 26 | 170 | 0 | 92 | 0 | 13 | 281 | 131 | 5416 | 25 | 543 | 0 | 356 | 19 | 112 | 0 | 97 | 0 | 244 | 2112 | 5260 | 1471 | 17578 | 4889 | 15 | 0 | 13 | 39 | 39 | 105 | 0 | 13 | 131 | 0 | 1462 | 0 |
| SRR2487705 | 0 | 57 | 157 | 0 | 104 | 14 | 0 | 400 | 220 | 5678 | 400 | 520 | 0 | 320 | 0 | 40 | 0 | 93 | 0 | 320 | 2566 | 6319 | 1412 | 16934 | 5006 | 11 | 0 | 0 | 86 | 72 | 100 | 0 | 14 | 243 | 0 | 1499 | 0 |
| SRR2487706 | 0 | 57 | 86 | 0 | 92 | 14 | 0 | 303 | 303 | 5577 | 525 | 586 | 0 | 485 | 40 | 20 | 0 | 94 | 0 | 364 | 1995 | 5353 | 1504 | 17388 | 4888 | 14 | 0 | 0 | 57 | 172 | 115 | 0 | 29 | 172 | 14 | 1475 | 0 |
| SRR2487708 | 0 | 0 | 30 | 0 | 117 | 0 | 0 | 104 | 234 | 6722 | 0 | 26 | 0 | 599 | 182 | 417 | 0 | 84 | 0 | 313 | 2523 | 11818 | 2079 | 32881 | 18 | 17 | 0 | 0 | 19 | 75 | 15 | 0 | 19 | 18 | 38 | 2215 | 0 |
| SRR2487710 | 0 | 39 | 271 | 0 | 94 | 0 | 0 | 316 | 132 | 6658 | 26 | 26 | 0 | 395 | 211 | 132 | 0 | 153 | 0 | 474 | 1756 | 9937 | 2207 | 34703 | 25 | 0 | 0 | 19 | 58 | 19 | 58 | 0 | 39 | 329 | 58 | 2579 | 0 |
| SRR2529523 | 0 | 19 | 34 | 0 | 33 | 0 | 0 | 1908 | 751 | 25571 | 10817 | 12740 | 0 | 210 | 0 | 0 | 0 | 502 | 26 | 376 | 1666 | 6491 | 0 | 1042 | 8488 | 0 | 0 | 13 | 0 | 13 | 0 | 0 | 0 | 19 | 0 | 1217 | 0 |
| SRR2529524 | 0 | 22 | 37 | 0 | 54 | 0 | 0 | 1837 | 710 | 23516 | 12430 | 12955 | 0 | 170 | 0 | 0 | 0 | 488 | 21 | 340 | 1635 | 6581 | 0 | 1053 | 14 | 0 | 0 | 0 | 17 | 0 | 17 | 0 | 12 | 20 | 0 | 1173 | 0 |
| SRR2529525 | 0 | 25 | 39 | 0 | 47 | 0 | 0 | 600 | 146 | 8019 | 327 | 6366 | 0 | 92 | 0 | 0 | 0 | 165 | 32 | 569 | 1160 | 6234 | 0 | 750 | 16946 | 0 | 0 | 0 | 10 | 27 | 31 | 0 | 41 | 12 | 19 | 2360 | 0 |
| SRR2529526 | 0 | 17 | 21 | 0 | 78 | 0 | 0 | 685 | 212 | 7928 | 2439 | 6239 | 0 | 65 | 0 | 0 | 0 | 182 | 28 | 759 | 1222 | 6252 | 0 | 748 | 17188 | 0 | 0 | 0 | 19 | 45 | 30 | 0 | 31 | 66 | 17 | 2561 | 0 |
| SRR2529527 | 0 | 12 | 13 | 0 | 70 | 0 | 0 | 1507 | 627 | 44987 | 3750 | 5582 | 0 | 729 | 11 | 14 | 0 | 286 | 20 | 239 | 1168 | 4914 | 0 | 1022 | 17118 | 0 | 0 | 0 | 15 | 44 | 51 | 0 | 44 | 13 | 11 | 919 | 0 |
| SRR2529528 | 0 | 79 | 0 | 0 | 86 | 0 | 0 | 1422 | 624 | 44920 | 3609 | 5529 | 0 | 689 | 14 | 0 | 0 | 297 | 17 | 368 | 1180 | 4853 | 0 | 1019 | 17142 | 0 | 0 | 0 | 17 | 49 | 45 | 0 | 44 | 75 | 0 | 899 | 0 |
| SRR2529529 | 0 | 53 | 43 | 0 | 38 | 0 | 0 | 2059 | 1009 | 44174 | 269 | 8641 | 0 | 222 | 0 | 20 | 0 | 332 | 18 | 747 | 1966 | 7891 | 0 | 430 | 7652 | 0 | 0 | 0 | 11 | 23 | 14 | 0 | 0 | 30 | 11 | 1736 | 0 |
| SRR2529530 | 0 | 23 | 77 | 0 | 45 | 0 | 0 | 2216 | 1036 | 41385 | 238 | 8845 | 0 | 394 | 21 | 0 | 0 | 319 | 15 | 932 | 2034 | 7875 | 0 | 415 | 7663 | 0 | 0 | 0 | 14 | 28 | 19 | 0 | 0 | 40 | 0 | 2154 | 0 |
| SRR2529531 | 0 | 109 | 23 | 0 | 43 | 0 | 0 | 1696 | 806 | 23869 | 948 | 14573 | 0 | 237 | 0 | 12 | 0 | 556 | 27 | 498 | 2088 | 7115 | 0 | 1292 | 9989 | 0 | 0 | 0 | 12 | 43 | 49 | 0 | 31 | 55 | 0 | 1885 | 0 |
| SRR2529532 | 0 | 80 | 46 | 0 | 58 | 0 | 0 | 1232 | 549 | 22926 | 937 | 14361 | 0 | 146 | 0 | 0 | 0 | 557 | 22 | 573 | 2098 | 7021 | 0 | 1308 | 9976 | 0 | 0 | 0 | 16 | 53 | 53 | 0 | 38 | 61 | 13 | 2294 | 0 |
| SRR2529533 | 0 | 10 | 53 | 0 | 50 | 0 | 0 | 2435 | 773 | 29559 | 655 | 11199 | 0 | 57 | 14 | 14 | 0 | 479 | 33 | 702 | 1931 | 6762 | 0 | 974 | 8182 | 0 | 0 | 0 | 15 | 26 | 27 | 0 | 16 | 36 | 0 | 1919 | 0 |
| SRR2529534 | 0 | 58 | 51 | 0 | 67 | 0 | 0 | 2356 | 790 | 29546 | 632 | 10990 | 0 | 220 | 15 | 15 | 0 | 453 | 28 | 966 | 1927 | 6737 | 0 | 963 | 8190 | 0 | 0 | 0 | 0 | 26 | 25 | 0 | 15 | 33 | 0 | 2049 | 0 |
| SRR2529535 | 0 | 46 | 11 | 0 | 78 | 0 | 0 | 6299 | 1854 | 99447 | 381 | 1505 | 0 | 528 | 0 | 0 | 23 | 155 | 15 | 103 | 2171 | 4605 | 0 | 865 | 16462 | 0 | 0 | 0 | 0 | 0 | 0 | 0 | 0 | 0 | 0 | 1219 | 0 |
| SRR2529536 | 0 | 53 | 13 | 0 | 107 | 0 | 0 | 6050 | 1852 | 102065 | 1128 | 1453 | 0 | 540 | 0 | 0 | 20 | 160 | 16 | 101 | 2197 | 4581 | 0 | 886 | 17 | 0 | 0 | 0 | 0 | 0 | 0 | 0 | 0 | 0 | 0 | 1282 | 0 |
| SRR2529537 | 0 | 46 | 11 | 0 | 69 | 0 | 0 | 2601 | 803 | 28117 | 583 | 6051 | 0 | 1473 | 0 | 0 | 11 | 153 | 11 | 196 | 818 | 3030 | 0 | 346 | 10080 | 0 | 0 | 0 | 27 | 32 | 22 | 0 | 16 | 49 | 22 | 1209 | 0 |
| SRR2529538 | 0 | 35 | 12 | 0 | 99 | 0 | 0 | 2778 | 806 | 27548 | 2671 | 5920 | 0 | 1549 | 0 | 0 | 14 | 127 | 16 | 240 | 855 | 3067 | 0 | 344 | 10174 | 0 | 0 | 0 | 37 | 37 | 32 | 0 | 0 | 29 | 12 | 1024 | 0 |
| SRR2529539 | 0 | 50 | 12 | 0 | 74 | 0 | 0 | 3166 | 1203 | 21347 | 1049 | 8336 | 0 | 757 | 0 | 13 | 0 | 159 | 0 | 254 | 893 | 3187 | 0 | 514 | 11534 | 0 | 0 | 11 | 32 | 46 | 24 | 0 | 16 | 28 | 16 | 730 | 0 |
| SRR2529540 | 0 | 25 | 0 | 0 | 108 | 0 | 0 | 3201 | 1216 | 21994 | 4343 | 8625 | 0 | 820 | 18 | 23 | 0 | 147 | 0 | 189 | 885 | 3186 | 0 | 521 | 19 | 0 | 0 | 0 | 36 | 41 | 30 | 0 | 10 | 31 | 14 | 852 | 0 |
| SRR2529541 | 0 | 34 | 0 | 0 | 41 | 0 | 0 | 3652 | 1583 | 109480 | 2870 | 3828 | 0 | 675 | 0 | 0 | 0 | 214 | 13 | 189 | 1762 | 6244 | 0 | 303 | 6799 | 0 | 0 | 0 | 19 | 0 | 0 | 0 | 0 | 15 | 0 | 1829 | 0 |
| SRR2529542 | 0 | 35 | 29 | 0 | 42 | 0 | 0 | 3325 | 1538 | 106858 | 435 | 3899 | 0 | 715 | 13 | 0 | 0 | 221 | 12 | 223 | 1738 | 6263 | 0 | 293 | 6943 | 0 | 0 | 0 | 23 | 17 | 0 | 0 | 0 | 16 | 0 | 1806 | 0 |
| SRR2529543 | 0 | 59 | 15 | 0 | 44 | 0 | 0 | 4208 | 1201 | 75770 | 3278 | 4629 | 0 | 930 | 0 | 0 | 19 | 189 | 0 | 136 | 1468 | 5009 | 0 | 787 | 6763 | 0 | 0 | 0 | 17 | 19 | 0 | 0 | 17 | 23 | 0 | 1004 | 0 |
| SRR2529544 | 0 | 11 | 12 | 0 | 69 | 0 | 0 | 3953 | 1171 | 75448 | 2845 | 4142 | 0 | 941 | 21 | 0 | 27 | 199 | 10 | 98 | 1450 | 4916 | 0 | 730 | 6776 | 0 | 0 | 0 | 16 | 21 | 0 | 0 | 16 | 0 | 0 | 1164 | 0 |
| SRR2529545 | 0 | 43 | 34 | 0 | 52 | 0 | 0 | 1487 | 812 | 79526 | 303 | 3973 | 0 | 337 | 0 | 0 | 0 | 204 | 17 | 362 | 1532 | 6023 | 0 | 666 | 6309 | 0 | 11 | 0 | 0 | 18 | 0 | 0 | 0 | 20 | 11 | 2411 | 0 |
| SRR2529546 | 0 | 46 | 25 | 0 | 59 | 0 | 0 | 1604 | 916 | 78037 | 277 | 3754 | 0 | 573 | 25 | 0 | 0 | 213 | 18 | 356 | 1543 | 5993 | 0 | 667 | 6355 | 0 | 0 | 0 | 14 | 28 | 0 | 0 | 0 | 25 | 0 | 2214 | 0 |
| SRR2529547 | 0 | 136 | 43 | 0 | 62 | 0 | 0 | 2399 | 600 | 10576 | 206 | 7469 | 0 | 109 | 27 | 0 | 0 | 285 | 17 | 436 | 2487 | 6452 | 0 | 1248 | 21325 | 0 | 0 | 49 | 29 | 35 | 29 | 0 | 14 | 23 | 0 | 1227 | 0 |
| SRR2529548 | 0 | 152 | 39 | 0 | 87 | 0 | 0 | 2369 | 948 | 11038 | 229 | 8947 | 0 | 0 | 28 | 0 | 0 | 280 | 19 | 725 | 2557 | 16 | 0 | 1296 | 39 | 0 | 12 | 27 | 18 | 21 | 27 | 0 | 24 | 33 | 0 | 1003 | 0 |
| SRR2529549 | 0 | 33 | 18 | 0 | 69 | 0 | 0 | 924 | 337 | 4058 | 2847 | 8266 | 0 | 0 | 0 | 0 | 0 | 124 | 23 | 549 | 1367 | 11 | 0 | 741 | 17485 | 0 | 0 | 0 | 28 | 24 | 50 | 18 | 28 | 74 | 31 | 2273 | 0 |
| SRR2529550 | 0 | 27 | 18 | 0 | 78 | 0 | 0 | 1241 | 294 | 3300 | 3121 | 7343 | 0 | 26 | 0 | 0 | 0 | 129 | 21 | 614 | 1345 | 6994 | 0 | 735 | 17483 | 0 | 0 | 0 | 0 | 27 | 52 | 0 | 18 | 11 | 0 | 2264 | 0 |
| SRR2529551 | 0 | 22 | 0 | 0 | 53 | 0 | 0 | 1387 | 812 | 5769 | 791 | 11885 | 0 | 91 | 0 | 0 | 0 | 273 | 15 | 584 | 1273 | 6011 | 0 | 1028 | 18432 | 11 | 0 | 11 | 12 | 48 | 45 | 0 | 54 | 17 | 22 | 1479 | 0 |
| SRR2529552 | 0 | 113 | 13 | 0 | 79 | 0 | 0 | 1586 | 711 | 5882 | 7439 | 12869 | 0 | 115 | 19 | 19 | 0 | 278 | 21 | 596 | 1258 | 6018 | 0 | 1035 | 32 | 0 | 0 | 14 | 0 | 49 | 58 | 17 | 44 | 54 | 16 | 1538 | 0 |
| SRR2529553 | 0 | 44 | 57 | 0 | 35 | 0 | 0 | 3240 | 1112 | 16263 | 252 | 12325 | 0 | 64 | 32 | 64 | 0 | 264 | 22 | 699 | 2007 | 8752 | 0 | 423 | 8485 | 0 | 0 | 0 | 13 | 29 | 0 | 0 | 10 | 41 | 0 | 2033 | 0 |
| SRR2529554 | 0 | 40 | 53 | 0 | 49 | 0 | 0 | 2419 | 1452 | 16484 | 257 | 12904 | 0 | 0 | 0 | 0 | 0 | 280 | 17 | 968 | 1954 | 8752 | 0 | 429 | 17 | 12 | 13 | 0 | 16 | 11 | 13 | 0 | 19 | 43 | 0 | 2452 | 0 |
| SRR2529555 | 0 | 67 | 41 | 0 | 44 | 0 | 0 | 2196 | 1078 | 9135 | 18172 | 21230 | 0 | 0 | 20 | 20 | 0 | 609 | 33 | 804 | 2135 | 8315 | 0 | 1635 | 9754 | 10 | 0 | 0 | 13 | 46 | 46 | 0 | 28 | 98 | 15 | 3215 | 0 |
| SRR2529556 | 0 | 52 | 30 | 0 | 44 | 0 | 0 | 1807 | 1024 | 8530 | 927 | 18662 | 0 | 56 | 0 | 37 | 0 | 618 | 32 | 633 | 2149 | 8529 | 0 | 1627 | 9798 | 10 | 0 | 0 | 10 | 40 | 40 | 0 | 27 | 70 | 17 | 3259 | 0 |
| SRR2529557 | 0 | 56 | 56 | 0 | 54 | 0 | 0 | 2374 | 1213 | 8097 | 390 | 12422 | 0 | 26 | 0 | 0 | 0 | 360 | 23 | 686 | 1888 | 8086 | 0 | 997 | 8241 | 0 | 0 | 16 | 28 | 39 | 23 | 0 | 26 | 37 | 21 | 2796 | 0 |
| SRR2529558 | 0 | 62 | 62 | 0 | 71 | 0 | 0 | 2542 | 1001 | 8655 | 373 | 13605 | 0 | 54 | 0 | 0 | 0 | 365 | 32 | 703 | 1867 | 8055 | 0 | 1022 | 8349 | 0 | 0 | 22 | 19 | 41 | 24 | 0 | 33 | 10 | 14 | 2083 | 0 |
| SRR2529559 | 0 | 168 | 23 | 0 | 74 | 0 | 0 | 2290 | 511 | 27157 | 5992 | 7628 | 0 | 20 | 0 | 41 | 0 | 272 | 15 | 818 | 2403 | 5397 | 0 | 1233 | 21475 | 0 | 0 | 0 | 17 | 21 | 15 | 0 | 0 | 15 | 0 | 2065 | 0 |
| SRR2529560 | 0 | 183 | 24 | 0 | 86 | 0 | 0 | 2083 | 804 | 27864 | 269 | 7342 | 0 | 62 | 0 | 21 | 0 | 302 | 22 | 866 | 2414 | 5488 | 0 | 1212 | 39 | 0 | 0 | 0 | 13 | 11 | 11 | 0 | 0 | 0 | 0 | 2331 | 0 |
| SRR2529561 | 0 | 66 | 16 | 0 | 68 | 0 | 0 | 991 | 631 | 3040 | 3660 | 8625 | 0 | 11 | 0 | 0 | 0 | 235 | 33 | 912 | 1241 | 5823 | 0 | 813 | 17310 | 0 | 0 | 0 | 0 | 17 | 28 | 0 | 24 | 44 | 0 | 2522 | 0 |
| SRR2529562 | 0 | 55 | 17 | 0 | 83 | 0 | 0 | 1166 | 577 | 3072 | 3580 | 8810 | 0 | 0 | 0 | 0 | 0 | 242 | 36 | 912 | 1269 | 5766 | 0 | 762 | 17275 | 0 | 0 | 0 | 10 | 10 | 19 | 0 | 25 | 41 | 0 | 2483 | 0 |
| SRR2529563 | 0 | 75 | 18 | 0 | 62 | 0 | 0 | 1530 | 1070 | 5295 | 492 | 9852 | 0 | 11 | 21 | 32 | 0 | 225 | 15 | 524 | 1313 | 5909 | 0 | 1010 | 20781 | 11 | 0 | 0 | 0 | 28 | 37 | 0 | 18 | 49 | 11 | 1498 | 0 |
| SRR2529564 | 0 | 71 | 14 | 0 | 87 | 0 | 0 | 1374 | 1000 | 5231 | 513 | 9418 | 0 | 44 | 11 | 22 | 0 | 209 | 21 | 571 | 1313 | 5919 | 0 | 985 | 35 | 10 | 0 | 0 | 11 | 25 | 27 | 0 | 11 | 30 | 12 | 1385 | 0 |
| SRR2529565 | 0 | 56 | 34 | 0 | 36 | 0 | 0 | 1992 | 956 | 18049 | 173 | 11275 | 0 | 80 | 80 | 80 | 0 | 250 | 17 | 1195 | 2204 | 8981 | 0 | 426 | 8685 | 0 | 0 | 0 | 0 | 0 | 0 | 0 | 0 | 22 | 0 | 3347 | 0 |
| SRR2529566 | 0 | 121 | 26 | 0 | 51 | 0 | 0 | 2238 | 870 | 18731 | 187 | 10899 | 0 | 0 | 0 | 0 | 0 | 255 | 16 | 1077 | 2215 | 8937 | 0 | 449 | 17 | 0 | 0 | 0 | 13 | 13 | 16 | 0 | 0 | 22 | 13 | 2901 | 0 |
| SRR2529567 | 0 | 68 | 40 | 0 | 28 | 0 | 0 | 2240 | 392 | 15876 | 15904 | 16772 | 0 | 28 | 0 | 0 | 0 | 658 | 26 | 728 | 1855 | 7799 | 0 | 1199 | 9050 | 0 | 13 | 0 | 13 | 26 | 13 | 0 | 11 | 38 | 11 | 3108 | 0 |
| SRR2529568 | 0 | 16 | 31 | 0 | 53 | 0 | 0 | 2010 | 566 | 14696 | 528 | 18575 | 0 | 85 | 0 | 0 | 0 | 644 | 30 | 481 | 1891 | 7648 | 0 | 1181 | 14 | 13 | 0 | 0 | 0 | 0 | 20 | 0 | 0 | 48 | 0 | 2747 | 0 |
| SRR2529569 | 0 | 25 | 63 | 0 | 60 | 0 | 0 | 2364 | 1229 | 42689 | 122 | 8557 | 0 | 0 | 0 | 0 | 0 | 247 | 22 | 993 | 1871 | 8442 | 0 | 868 | 8774 | 0 | 0 | 0 | 28 | 13 | 16 | 0 | 16 | 0 | 0 | 4822 | 0 |
| SRR2529570 | 0 | 23 | 45 | 0 | 80 | 0 | 0 | 1850 | 1200 | 41958 | 6601 | 7752 | 0 | 0 | 0 | 0 | 0 | 255 | 30 | 1300 | 1917 | 8402 | 0 | 904 | 14 | 0 | 0 | 0 | 0 | 0 | 13 | 0 | 0 | 0 | 0 | 3751 | 0 |
| SRR2529571 | 0 | 13 | 26 | 0 | 41 | 0 | 0 | 1708 | 687 | 16589 | 519 | 12997 | 0 | 59 | 0 | 20 | 0 | 506 | 32 | 432 | 1615 | 29 | 0 | 972 | 7999 | 0 | 0 | 16 | 12 | 16 | 12 | 0 | 0 | 20 | 12 | 1610 | 0 |
| SRR2529572 | 0 | 16 | 28 | 0 | 48 | 0 | 0 | 1683 | 461 | 16547 | 484 | 13862 | 0 | 40 | 0 | 0 | 0 | 478 | 24 | 461 | 1623 | 30 | 0 | 991 | 14 | 0 | 0 | 10 | 16 | 0 | 14 | 0 | 16 | 20 | 0 | 1402 | 0 |
| SRR2529573 | 0 | 40 | 25 | 0 | 55 | 0 | 0 | 947 | 348 | 9030 | 254 | 6406 | 0 | 70 | 0 | 0 | 0 | 140 | 26 | 765 | 1449 | 6693 | 0 | 730 | 17322 | 0 | 0 | 0 | 14 | 45 | 60 | 0 | 42 | 67 | 20 | 2408 | 0 |
| SRR2529574 | 0 | 35 | 24 | 0 | 81 | 0 | 0 | 895 | 425 | 9518 | 258 | 6337 | 0 | 115 | 18 | 0 | 0 | 159 | 31 | 620 | 1429 | 6742 | 0 | 728 | 21 | 0 | 0 | 0 | 11 | 37 | 52 | 0 | 26 | 13 | 17 | 2331 | 0 |
| SRR2529575 | 0 | 82 | 15 | 0 | 44 | 0 | 0 | 1815 | 499 | 5726 | 534 | 9184 | 0 | 109 | 27 | 0 | 0 | 247 | 13 | 535 | 1199 | 5254 | 0 | 904 | 16447 | 0 | 0 | 15 | 15 | 59 | 30 | 13 | 44 | 68 | 0 | 2196 | 0 |
| SRR2529576 | 0 | 78 | 12 | 0 | 67 | 0 | 0 | 1749 | 682 | 5486 | 4909 | 9588 | 0 | 183 | 29 | 0 | 0 | 257 | 20 | 528 | 1163 | 5267 | 0 | 863 | 16 | 0 | 0 | 0 | 17 | 49 | 64 | 0 | 53 | 87 | 18 | 2901 | 0 |
| SRR2529577 | 0 | 46 | 52 | 0 | 37 | 0 | 0 | 2812 | 1146 | 40050 | 5547 | 10520 | 0 | 208 | 26 | 104 | 0 | 344 | 13 | 1120 | 2046 | 7848 | 0 | 428 | 7963 | 0 | 0 | 13 | 0 | 20 | 15 | 0 | 17 | 15 | 11 | 3333 | 0 |
| SRR2529578 | 0 | 40 | 55 | 0 | 41 | 0 | 0 | 2660 | 1207 | 35120 | 228 | 9236 | 0 | 148 | 74 | 49 | 0 | 346 | 11 | 887 | 2012 | 7900 | 0 | 419 | 12 | 0 | 0 | 0 | 11 | 13 | 17 | 0 | 0 | 21 | 0 | 3522 | 0 |
| SRR2529579 | 0 | 11 | 16 | 0 | 42 | 0 | 0 | 1728 | 1136 | 5777 | 1102 | 21268 | 0 | 16 | 16 | 0 | 0 | 636 | 21 | 608 | 2051 | 6428 | 0 | 1330 | 10386 | 0 | 0 | 0 | 19 | 81 | 56 | 0 | 0 | 53 | 18 | 3073 | 0 |
| SRR2529580 | 0 | 100 | 20 | 0 | 59 | 0 | 0 | 1889 | 928 | 6400 | 1103 | 21643 | 0 | 33 | 0 | 16 | 0 | 641 | 23 | 586 | 2117 | 6515 | 0 | 1325 | 10467 | 0 | 0 | 0 | 15 | 74 | 44 | 0 | 20 | 53 | 21 | 2948 | 0 |
| SRR2529581 | 0 | 46 | 54 | 0 | 50 | 0 | 0 | 2215 | 711 | 22552 | 430 | 12901 | 0 | 102 | 20 | 0 | 0 | 423 | 33 | 955 | 1810 | 6753 | 0 | 1004 | 7749 | 0 | 0 | 12 | 0 | 35 | 18 | 0 | 18 | 30 | 0 | 2885 | 0 |
| SRR2529582 | 0 | 57 | 39 | 0 | 69 | 0 | 0 | 2201 | 976 | 23027 | 459 | 12188 | 0 | 42 | 21 | 0 | 0 | 439 | 26 | 934 | 1762 | 6788 | 0 | 957 | 7735 | 0 | 0 | 0 | 25 | 32 | 35 | 0 | 0 | 34 | 15 | 3073 | 0 |
| SRR2529583 | 0 | 16 | 14 | 0 | 77 | 0 | 0 | 2013 | 798 | 17342 | 6051 | 7401 | 0 | 25 | 25 | 0 | 0 | 170 | 12 | 442 | 2074 | 10 | 0 | 814 | 15679 | 0 | 0 | 0 | 19 | 28 | 0 | 0 | 0 | 16 | 0 | 2209 | 0 |
| SRR2529584 | 0 | 131 | 16 | 0 | 104 | 0 | 0 | 1721 | 1047 | 16672 | 497 | 7367 | 0 | 72 | 12 | 12 | 0 | 173 | 15 | 770 | 2055 | 4511 | 0 | 830 | 15749 | 0 | 0 | 0 | 24 | 22 | 11 | 0 | 0 | 13 | 0 | 2347 | 0 |
| SRR2529585 | 0 | 53 | 21 | 0 | 101 | 0 | 0 | 2054 | 682 | 3612 | 4015 | 9348 | 0 | 54 | 0 | 0 | 0 | 79 | 10 | 434 | 852 | 2807 | 0 | 314 | 8887 | 0 | 0 | 17 | 23 | 32 | 11 | 0 | 20 | 34 | 20 | 1829 | 0 |
| SRR2529586 | 0 | 44 | 13 | 0 | 138 | 0 | 0 | 1654 | 677 | 3408 | 496 | 8870 | 0 | 38 | 0 | 0 | 0 | 103 | 0 | 338 | 837 | 2794 | 0 | 340 | 9035 | 0 | 0 | 14 | 25 | 36 | 19 | 0 | 16 | 31 | 19 | 1477 | 0 |
| SRR2529587 | 0 | 53 | 11 | 0 | 92 | 0 | 0 | 2515 | 959 | 4331 | 4817 | 9189 | 0 | 103 | 0 | 0 | 0 | 65 | 0 | 384 | 738 | 2232 | 11 | 359 | 8211 | 0 | 0 | 24 | 33 | 56 | 16 | 0 | 21 | 48 | 18 | 932 | 0 |
| SRR2529588 | 0 | 46 | 0 | 0 | 103 | 0 | 0 | 2253 | 1196 | 4422 | 5380 | 9823 | 0 | 91 | 0 | 14 | 0 | 73 | 0 | 350 | 734 | 2219 | 0 | 362 | 8251 | 0 | 0 | 31 | 49 | 60 | 18 | 0 | 13 | 55 | 20 | 958 | 0 |
| SRR2529589 | 0 | 49 | 22 | 0 | 37 | 0 | 0 | 4244 | 3627 | 17884 | 7678 | 11402 | 0 | 135 | 19 | 0 | 0 | 154 | 0 | 675 | 1824 | 6456 | 0 | 302 | 5974 | 0 | 0 | 0 | 27 | 22 | 13 | 0 | 0 | 11 | 0 | 2817 | 0 |
| SRR2529590 | 0 | 41 | 20 | 0 | 49 | 0 | 0 | 3643 | 3348 | 18451 | 7601 | 11146 | 0 | 138 | 0 | 0 | 0 | 176 | 12 | 492 | 1825 | 6445 | 0 | 303 | 5890 | 0 | 0 | 0 | 18 | 25 | 0 | 0 | 14 | 25 | 0 | 2993 | 0 |
| SRR2529591 | 0 | 61 | 24 | 0 | 34 | 0 | 0 | 4884 | 6541 | 9536 | 7791 | 11309 | 0 | 0 | 0 | 0 | 0 | 122 | 14 | 407 | 1311 | 5385 | 0 | 674 | 5997 | 0 | 0 | 0 | 13 | 24 | 13 | 0 | 11 | 37 | 19 | 2326 | 0 |
| SRR2529592 | 0 | 66 | 25 | 0 | 59 | 0 | 0 | 4661 | 6196 | 8356 | 282 | 10431 | 0 | 57 | 0 | 0 | 0 | 119 | 0 | 625 | 1341 | 5417 | 0 | 691 | 5880 | 0 | 0 | 0 | 49 | 27 | 11 | 0 | 16 | 38 | 22 | 1961 | 0 |
| SRR2529593 | 0 | 41 | 33 | 0 | 61 | 0 | 0 | 4245 | 5487 | 20620 | 255 | 9646 | 0 | 58 | 0 | 29 | 0 | 175 | 17 | 838 | 1482 | 21 | 0 | 553 | 5603 | 0 | 0 | 0 | 15 | 12 | 18 | 0 | 15 | 38 | 0 | 2195 | 0 |
| SRR2529594 | 0 | 49 | 30 | 0 | 65 | 0 | 0 | 4408 | 5572 | 21276 | 7623 | 9000 | 0 | 31 | 31 | 0 | 0 | 164 | 16 | 429 | 1415 | 27 | 0 | 578 | 13 | 0 | 0 | 0 | 27 | 30 | 18 | 0 | 12 | 30 | 18 | 2541 | 0 |
| SRR2648184 | 0 | 46 | 0 | 0 | 0 | 0 | 0 | 170 | 149 | 955 | 1709 | 6415 | 0 | 493 | 21 | 14 | 19 | 3599 | 118 | 561 | 3958 | 8764 | 18 | 3361 | 17801 | 15 | 14 | 19 | 189 | 46 | 18 | 0 | 0 | 0 | 0 | 2084 | 1249 |
| SRR2648185 | 11 | 46 | 0 | 0 | 10 | 0 | 0 | 183 | 208 | 922 | 1746 | 6209 | 0 | 468 | 28 | 11 | 25 | 3293 | 113 | 521 | 3737 | 8287 | 17 | 3339 | 18522 | 13 | 12 | 26 | 24 | 40 | 24 | 0 | 0 | 0 | 0 | 1868 | 1449 |
| SRR2648186 | 0 | 47 | 0 | 0 | 0 | 0 | 0 | 322 | 259 | 851 | 1394 | 7121 | 0 | 592 | 37 | 16 | 12 | 3354 | 107 | 698 | 3597 | 9522 | 20 | 3619 | 16815 | 11 | 0 | 15 | 107 | 29 | 19 | 0 | 0 | 0 | 0 | 2363 | 998 |
| SRR2648187 | 0 | 141 | 65 | 0 | 0 | 0 | 0 | 218 | 297 | 4794 | 5653 | 6393 | 0 | 544 | 0 | 15 | 0 | 2847 | 139 | 225 | 3613 | 10757 | 15 | 2387 | 23595 | 0 | 0 | 43 | 52 | 18 | 141 | 12 | 19 | 25 | 0 | 613 | 732 |
| SRR2648188 | 0 | 151 | 42 | 0 | 0 | 0 | 0 | 342 | 301 | 5851 | 1471 | 6744 | 0 | 555 | 0 | 10 | 0 | 3401 | 144 | 237 | 3828 | 11597 | 14 | 2526 | 24503 | 0 | 0 | 31 | 1405 | 466 | 140 | 14 | 19 | 28 | 0 | 511 | 691 |
| SRR2648189 | 0 | 111 | 55 | 0 | 0 | 0 | 0 | 315 | 306 | 4828 | 1764 | 7201 | 0 | 812 | 12 | 12 | 0 | 3299 | 143 | 309 | 3849 | 12804 | 11 | 2356 | 22274 | 0 | 0 | 35 | 1090 | 280 | 109 | 0 | 15 | 27 | 0 | 596 | 605 |

### Supp Table S8.4.Columns_DI_to_ES

| | | | | | | | | | | | | | | | | | | | | | | | | | | | | | | | | | | | | | |
|---|---|---|---|---|---|---|---|---|---|---|---|---|---|---|---|---|---|---|---|---|---|---|---|---|---|---|---|---|---|---|---|---|---|---|---|---|---|
| SRR2728221 | 0 | 0 | 0 | 0 | 0 | 0 | 0 | 268 | 0 | 0 | 43 | 4550 | 0 | 0 | 0 | 0 | 0 | 0 | 0 | 0 | 0 | 28 | 0 | 0 | 31 | 34 | 0 | 0 | 0 | 0 | 0 | 0 | 0 | 0 | 0 | 0 | 0 |
| SRR2731278 | 0 | 362 | 136 | 946 | 5209 | 0 | 0 | 1143 | 6027 | 74624 | 11 | 18 | 0 | 2407 | 2857 | 5863 | 0 | 3221 | 27 | 661 | 1206 | 1220 | 768 | 10199 | 2975 | 0 | 0 | 0 | 16 | 16 | 22 | 11 | 0 | 0 | 0 | 1447 | 206 |
| SRR2731279 | 0 | 78 | 49 | 223 | 618 | 0 | 0 | 2024 | 5866 | 9329 | 18 | 23 | 0 | 901 | 1309 | 2244 | 0 | 2268 | 86 | 948 | 1033 | 2096 | 157 | 6931 | 3650 | 30 | 0 | 0 | 19 | 24 | 14 | 17 | 0 | 0 | 14 | 2276 | 15 |
| SRR2731280 | 0 | 147 | 137 | 107 | 171 | 13 | 0 | 1559 | 1678 | 4279 | 155 | 2285 | 11 | 196 | 155 | 54 | 0 | 2517 | 72 | 4356 | 2974 | 17 | 282 | 8500 | 938 | 0 | 16 | 0 | 24 | 15 | 46 | 30 | 12 | 12 | 0 | 9367 | 0 |
| SRR2731281 | 0 | 1521 | 405 | 380 | 2006 | 0 | 0 | 2768 | 7707 | 28681 | 87 | 159 | 0 | 671 | 5379 | 9655 | 0 | 2848 | 48 | 883 | 2570 | 2604 | 636 | 9888 | 2324 | 0 | 0 | 0 | 92 | 67 | 92 | 29 | 16 | 14 | 0 | 2281 | 109 |
| SRR2731282 | 0 | 297 | 249 | 735 | 1503 | 0 | 0 | 1508 | 3951 | 42347 | 22 | 50 | 0 | 1614 | 4015 | 4300 | 0 | 2874 | 42 | 1537 | 2163 | 2945 | 516 | 10832 | 3372 | 0 | 0 | 0 | 47 | 59 | 115 | 34 | 27 | 24 | 14 | 3357 | 45 |
| SRR2731283 | 0 | 196 | 81 | 452 | 881 | 0 | 0 | 1512 | 4169 | 6794 | 35 | 52 | 0 | 1251 | 1887 | 2858 | 0 | 2070 | 106 | 1800 | 1111 | 4320 | 258 | 21686 | 6805 | 0 | 0 | 0 | 18 | 15 | 31 | 17 | 18 | 16 | 21 | 4119 | 378 |
| SRR2731284 | 0 | 219 | 54 | 517 | 2148 | 0 | 0 | 2165 | 6353 | 8813 | 25 | 57 | 0 | 336 | 14187 | 13821 | 0 | 3883 | 38 | 1128 | 1780 | 2030 | 157 | 4223 | 3574 | 0 | 0 | 0 | 12 | 12 | 16 | 0 | 0 | 0 | 0 | 2336 | 272 |
| SRR2731285 | 0 | 334 | 54 | 1310 | 3233 | 0 | 0 | 2669 | 7216 | 12988 | 70 | 142 | 0 | 1019 | 1315 | 1673 | 0 | 2089 | 38 | 2117 | 1944 | 3918 | 608 | 12954 | 4703 | 0 | 0 | 0 | 12 | 14 | 19 | 17 | 0 | 0 | 0 | 4141 | 173 |
| SRR2731286 | 0 | 290 | 37 | 751 | 1611 | 11 | 0 | 1443 | 5446 | 40918 | 16 | 27 | 0 | 3666 | 707 | 1142 | 0 | 3023 | 292 | 600 | 1633 | 1313 | 323 | 12683 | 5497 | 0 | 0 | 0 | 12 | 13 | 17 | 0 | 0 | 0 | 0 | 1587 | 47 |
| SRR2731287 | 0 | 135 | 66 | 442 | 759 | 0 | 0 | 1663 | 5684 | 95132 | 10 | 20 | 0 | 1639 | 655 | 739 | 0 | 1901 | 24 | 2715 | 958 | 3728 | 922 | 41087 | 3072 | 0 | 0 | 0 | 26 | 27 | 64 | 14 | 10 | 13 | 22 | 5607 | 214 |
| SRR2731288 | 0 | 933 | 182 | 849 | 1137 | 0 | 0 | 684 | 2182 | 5371 | 23 | 27 | 0 | 1454 | 15297 | 17732 | 0 | 2684 | 31 | 1194 | 3159 | 4787 | 1265 | 28139 | 5792 | 0 | 0 | 0 | 75 | 53 | 73 | 29 | 13 | 0 | 11 | 3423 | 92 |
| SRR2731289 | 0 | 102 | 92 | 163 | 139 | 0 | 0 | 2376 | 4550 | 34660 | 0 | 24 | 0 | 1703 | 7189 | 8099 | 0 | 837 | 44 | 4163 | 384 | 843 | 163 | 12009 | 1224 | 0 | 0 | 0 | 37 | 33 | 145 | 36 | 23 | 13 | 22 | 10627 | 92 |
| SRR2731290 | 0 | 110 | 132 | 122 | 109 | 0 | 0 | 3152 | 7186 | 36838 | 17 | 35 | 0 | 585 | 1761 | 2199 | 0 | 1954 | 57 | 6427 | 1386 | 2247 | 595 | 24542 | 1631 | 0 | 0 | 0 | 21 | 17 | 75 | 24 | 23 | 11 | 18 | 12649 | 0 |
| SRR2731291 | 0 | 250 | 165 | 207 | 600 | 0 | 0 | 3487 | 7548 | 46797 | 0 | 0 | 0 | 3278 | 1883 | 1819 | 0 | 1460 | 32 | 1417 | 382 | 839 | 188 | 10850 | 1104 | 0 | 0 | 0 | 97 | 86 | 292 | 37 | 14 | 13 | 15 | 3412 | 93 |
| SRR2731292 | 0 | 111 | 213 | 101 | 101 | 0 | 0 | 3168 | 6225 | 18884 | 11 | 35 | 0 | 879 | 23446 | 18025 | 0 | 772 | 75 | 20080 | 665 | 1789 | 431 | 24874 | 1469 | 0 | 0 | 0 | 34 | 54 | 446 | 129 | 38 | 36 | 21 | 23137 | 53 |
| SRR2731293 | 0 | 166 | 62 | 1342 | 2493 | 0 | 0 | 1160 | 4220 | 22900 | 16 | 40 | 0 | 942 | 3858 | 6894 | 0 | 2596 | 88 | 1180 | 2861 | 3628 | 383 | 11430 | 4838 | 0 | 0 | 0 | 11 | 14 | 11 | 0 | 0 | 0 | 0 | 2654 | 206 |
| SRR2731294 | 0 | 1291 | 277 | 328 | 747 | 0 | 0 | 1685 | 4851 | 5532 | 102 | 204 | 0 | 245 | 5293 | 5997 | 0 | 3687 | 35 | 2349 | 3211 | 6506 | 903 | 20125 | 4016 | 0 | 0 | 0 | 54 | 38 | 58 | 25 | 0 | 12 | 11 | 4419 | 19 |
| SRR2731295 | 0 | 229 | 57 | 867 | 1451 | 0 | 0 | 791 | 1728 | 2867 | 288 | 93 | 0 | 217 | 3204 | 9027 | 0 | 3645 | 45 | 3683 | 1694 | 3103 | 244 | 7898 | 5527 | 0 | 0 | 10 | 46 | 22 | 36 | 13 | 45 | 44 | 22 | 8854 | 98 |
| SRR2731296 | 0 | 237 | 195 | 767 | 1691 | 0 | 0 | 1755 | 4654 | 19625 | 31 | 31 | 0 | 591 | 2454 | 2983 | 0 | 3836 | 31 | 1190 | 2481 | 4329 | 744 | 18012 | 5018 | 0 | 0 | 0 | 18 | 29 | 52 | 11 | 0 | 0 | 0 | 3625 | 23 |
| SRR2731297 | 0 | 268 | 110 | 874 | 2304 | 0 | 0 | 2111 | 6345 | 23772 | 31 | 57 | 0 | 406 | 814 | 1755 | 0 | 4180 | 18 | 1369 | 2989 | 6271 | 1259 | 22913 | 6484 | 0 | 0 | 0 | 0 | 15 | 32 | 0 | 0 | 0 | 0 | 2724 | 128 |
| SRR2731298 | 0 | 19 | 24 | 155 | 499 | 0 | 0 | 1028 | 1418 | 34285 | 25 | 216 | 0 | 1915 | 2461 | 720 | 0 | 1159 | 80 | 578 | 351 | 538 | 38 | 2501 | 1177 | 0 | 16 | 26 | 62 | 18 | 55 | 24 | 21 | 23 | 23 | 6495 | 29 |
| SRR2731299 | 0 | 85 | 62 | 395 | 1841 | 0 | 0 | 916 | 3380 | 27878 | 11 | 23 | 0 | 1304 | 5366 | 8837 | 0 | 3393 | 44 | 1904 | 2534 | 3157 | 579 | 13442 | 2718 | 0 | 0 | 0 | 15 | 11 | 22 | 28 | 13 | 13 | 19 | 3417 | 61 |
| SRR2731300 | 0 | 355 | 55 | 665 | 849 | 0 | 0 | 3203 | 8084 | 9144 | 80 | 162 | 0 | 255 | 4206 | 4440 | 0 | 4469 | 42 | 3337 | 3760 | 5251 | 953 | 10778 | 2868 | 0 | 0 | 0 | 12 | 16 | 25 | 25 | 11 | 0 | 0 | 5172 | 29 |
| SRR2731301 | 0 | 1284 | 258 | 731 | 864 | 0 | 0 | 1971 | 5629 | 58742 | 41 | 53 | 0 | 1526 | 634 | 1366 | 0 | 4808 | 58 | 1273 | 2611 | 3756 | 775 | 20017 | 3809 | 0 | 0 | 0 | 23 | 18 | 31 | 10 | 0 | 0 | 0 | 2310 | 56 |
| SRR2731302 | 0 | 746 | 278 | 508 | 641 | 0 | 0 | 1134 | 4427 | 49022 | 15 | 27 | 0 | 1296 | 8549 | 9455 | 0 | 5154 | 73 | 788 | 2786 | 7083 | 1105 | 34152 | 5519 | 0 | 0 | 0 | 33 | 44 | 83 | 19 | 0 | 11 | 17 | 1449 | 0 |
| SRR2731303 | 0 | 706 | 307 | 140 | 616 | 26 | 0 | 833 | 1604 | 20537 | 426 | 866 | 0 | 819 | 5877 | 9600 | 0 | 1754 | 48 | 2174 | 1655 | 1429 | 189 | 5341 | 1219 | 12 | 15 | 0 | 90 | 35 | 131 | 58 | 26 | 26 | 14 | 4237 | 20 |
| SRR2731304 | 0 | 1403 | 346 | 256 | 1834 | 0 | 0 | 2102 | 6123 | 11578 | 94 | 155 | 0 | 635 | 1642 | 1912 | 0 | 2030 | 52 | 1478 | 1583 | 2364 | 511 | 11516 | 2555 | 0 | 0 | 0 | 73 | 44 | 62 | 18 | 13 | 11 | 0 | 2997 | 33 |
| SRR2731305 | 0 | 482 | 210 | 738 | 1438 | 0 | 0 | 1453 | 5212 | 12371 | 34 | 54 | 0 | 934 | 1037 | 4902 | 0 | 3907 | 31 | 1486 | 2765 | 6558 | 1048 | 18433 | 4691 | 0 | 0 | 0 | 15 | 30 | 48 | 17 | 0 | 0 | 0 | 2887 | 31 |
| SRR2731306 | 0 | 359 | 229 | 529 | 1511 | 0 | 0 | 2034 | 6343 | 28271 | 47 | 87 | 0 | 720 | 3133 | 4094 | 0 | 2671 | 46 | 3632 | 1989 | 3998 | 480 | 17206 | 4205 | 0 | 0 | 0 | 16 | 26 | 63 | 15 | 0 | 0 | 13 | 5497 | 67 |
| SRR2731307 | 0 | 108 | 64 | 895 | 675 | 0 | 0 | 1354 | 4845 | 7726 | 53 | 46 | 0 | 118 | 9835 | 16366 | 0 | 3644 | 37 | 3488 | 3487 | 9609 | 854 | 25483 | 13721 | 0 | 0 | 0 | 13 | 18 | 36 | 31 | 16 | 13 | 17 | 9247 | 23 |
| SRR2731308 | 0 | 125 | 105 | 864 | 2032 | 0 | 0 | 2563 | 7810 | 23774 | 40 | 70 | 0 | 1050 | 2519 | 3012 | 0 | 2305 | 64 | 1351 | 1013 | 2467 | 143 | 5397 | 3788 | 0 | 0 | 0 | 13 | 23 | 13 | 13 | 0 | 0 | 0 | 2997 | 56 |
| SRR2731309 | 0 | 396 | 173 | 711 | 1116 | 0 | 0 | 2181 | 6055 | 15640 | 39 | 64 | 0 | 455 | 19288 | 18944 | 0 | 4346 | 58 | 2783 | 2957 | 6914 | 1042 | 23485 | 5149 | 0 | 0 | 0 | 17 | 27 | 58 | 13 | 0 | 0 | 0 | 4604 | 10 |
| SRR2731310 | 0 | 321 | 559 | 518 | 1774 | 0 | 0 | 2166 | 5520 | 32836 | 29 | 72 | 0 | 1343 | 3843 | 5639 | 0 | 3015 | 20 | 2141 | 1755 | 5969 | 848 | 24027 | 3348 | 0 | 0 | 0 | 47 | 83 | 165 | 43 | 21 | 30 | 36 | 5975 | 31 |
| SRR2731311 | 0 | 120 | 46 | 1241 | 1075 | 0 | 0 | 1128 | 5052 | 53340 | 27 | 33 | 0 | 1821 | 1330 | 1760 | 0 | 3456 | 57 | 839 | 4105 | 4685 | 836 | 23401 | 6421 | 0 | 0 | 0 | 11 | 19 | 12 | 0 | 0 | 0 | 0 | 1422 | 235 |
| SRR2731312 | 0 | 102 | 92 | 687 | 1408 | 0 | 0 | 2262 | 5058 | 27671 | 23 | 103 | 0 | 1679 | 1084 | 1247 | 0 | 3668 | 48 | 1230 | 2693 | 3483 | 337 | 10970 | 3235 | 0 | 0 | 0 | 0 | 20 | 14 | 14 | 0 | 0 | 0 | 2742 | 34 |
| SRR2731313 | 0 | 788 | 221 | 810 | 1871 | 0 | 0 | 1809 | 5796 | 24432 | 21 | 50 | 0 | 1177 | 1206 | 1491 | 0 | 3972 | 41 | 1216 | 2062 | 2246 | 331 | 7351 | 3165 | 0 | 0 | 0 | 15 | 17 | 26 | 12 | 0 | 0 | 0 | 1934 | 0 |
| SRR2731314 | 0 | 293 | 43 | 615 | 2009 | 0 | 0 | 2042 | 6329 | 8925 | 65 | 140 | 0 | 249 | 1337 | 1351 | 0 | 4572 | 54 | 1569 | 3459 | 5115 | 1601 | 15749 | 3984 | 0 | 0 | 0 | 28 | 11 | 15 | 0 | 0 | 0 | 0 | 3408 | 13 |
| SRR2731315 | 0 | 160 | 112 | 243 | 838 | 0 | 0 | 1890 | 6714 | 28767 | 34 | 67 | 0 | 1653 | 4208 | 3987 | 0 | 2836 | 29 | 1375 | 2339 | 4459 | 618 | 13436 | 2223 | 0 | 0 | 0 | 29 | 12 | 20 | 20 | 14 | 15 | 13 | 3131 | 49 |
| SRR2731316 | 0 | 487 | 289 | 655 | 1589 | 0 | 0 | 2451 | 7454 | 11781 | 40 | 172 | 0 | 471 | 479 | 742 | 0 | 3497 | 58 | 2263 | 3650 | 4382 | 880 | 12472 | 2869 | 0 | 0 | 0 | 16 | 23 | 48 | 15 | 0 | 0 | 0 | 5886 | 18 |
| SRR2731317 | 0 | 556 | 302 | 1155 | 2092 | 0 | 0 | 2446 | 7825 | 14024 | 54 | 94 | 0 | 926 | 1187 | 1291 | 0 | 3803 | 33 | 1791 | 3632 | 3404 | 593 | 13024 | 3061 | 0 | 0 | 0 | 23 | 41 | 65 | 13 | 0 | 0 | 0 | 3817 | 0 |
| SRR2731318 | 0 | 83 | 33 | 393 | 730 | 0 | 15 | 3656 | 8036 | 26334 | 104 | 329 | 0 | 1606 | 239 | 272 | 0 | 1547 | 94 | 645 | 1416 | 1816 | 487 | 14020 | 3195 | 0 | 0 | 0 | 12 | 14 | 30 | 23 | 10 | 23 | 0 | 4817 | 12 |
| SRR2731319 | 0 | 191 | 201 | 534 | 887 | 0 | 0 | 2898 | 7475 | 15280 | 102 | 185 | 0 | 331 | 3988 | 5371 | 0 | 3192 | 29 | 2717 | 1688 | 3638 | 468 | 11184 | 3028 | 0 | 0 | 0 | 22 | 25 | 16 | 15 | 0 | 11 | 0 | 5311 | 31 |
| SRR2731320 | 0 | 126 | 241 | 515 | 1575 | 0 | 0 | 2795 | 7812 | 18525 | 57 | 158 | 0 | 1358 | 3459 | 2517 | 0 | 2282 | 60 | 2009 | 1488 | 3206 | 1019 | 23747 | 4819 | 0 | 0 | 0 | 33 | 78 | 189 | 67 | 37 | 22 | 22 | 11136 | 70 |
| SRR2731321 | 0 | 145 | 21 | 477 | 545 | 0 | 18 | 5934 | 12016 | 17023 | 142 | 441 | 0 | 663 | 234 | 323 | 0 | 2089 | 32 | 1358 | 4009 | 4663 | 1217 | 17789 | 2724 | 0 | 0 | 0 | 18 | 24 | 50 | 28 | 13 | 33 | 11 | 9537 | 0 |
| SRR2731322 | 0 | 304 | 249 | 326 | 388 | 0 | 0 | 1576 | 2523 | 2895 | 129 | 343 | 0 | 132 | 1716 | 2013 | 0 | 4118 | 61 | 2786 | 2107 | 4027 | 996 | 11516 | 1868 | 0 | 0 | 0 | 12 | 0 | 25 | 28 | 10 | 0 | 0 | 8283 | 19 |
| SRR2731323 | 0 | 157 | 63 | 469 | 1622 | 11 | 0 | 1452 | 6031 | 7889 | 87 | 119 | 0 | 4216 | 956 | 1499 | 0 | 1069 | 162 | 2241 | 2853 | 1460 | 1309 | 27288 | 6043 | 0 | 0 | 0 | 15 | 25 | 44 | 15 | 0 | 0 | 0 | 3396 | 96 |
| SRR2731324 | 0 | 772 | 160 | 1074 | 3806 | 0 | 0 | 1777 | 5127 | 11777 | 34 | 85 | 0 | 426 | 1319 | 1653 | 0 | 2887 | 47 | 1347 | 1798 | 2243 | 287 | 7575 | 5427 | 0 | 0 | 0 | 19 | 12 | 23 | 0 | 0 | 0 | 0 | 1983 | 50 |
| SRR2731325 | 0 | 272 | 46 | 1315 | 2628 | 0 | 0 | 1792 | 5057 | 11600 | 46 | 93 | 0 | 606 | 692 | 2070 | 0 | 3892 | 45 | 1417 | 6162 | 5505 | 1335 | 21452 | 3772 | 0 | 0 | 0 | 22 | 29 | 17 | 11 | 0 | 0 | 0 | 2938 | 141 |
| SRR2731326 | 0 | 466 | 260 | 384 | 1776 | 0 | 0 | 3589 | 10125 | 16039 | 45 | 116 | 0 | 2147 | 806 | 873 | 0 | 3738 | 37 | 2637 | 1495 | 2079 | 218 | 6387 | 2328 | 0 | 0 | 0 | 0 | 19 | 41 | 13 | 0 | 0 | 0 | 4716 | 15 |
| SRR2731327 | 0 | 403 | 59 | 745 | 1262 | 0 | 0 | 2395 | 6424 | 8885 | 59 | 135 | 0 | 867 | 5556 | 6054 | 0 | 4730 | 31 | 1334 | 3007 | 3985 | 613 | 10228 | 3006 | 0 | 0 | 0 | 16 | 18 | 10 | 0 | 0 | 0 | 0 | 2416 | 29 |
| SRR2731328 | 0 | 57 | 65 | 427 | 1639 | 0 | 0 | 1746 | 5887 | 21396 | 38 | 52 | 0 | 1288 | 1779 | 1495 | 0 | 2577 | 38 | 838 | 1228 | 1533 | 160 | 4133 | 2108 | 0 | 0 | 0 | 0 | 0 | 23 | 0 | 0 | 0 | 0 | 1783 | 46 |
| SRR2731329 | 0 | 723 | 272 | 507 | 1316 | 0 | 0 | 1399 | 5128 | 14165 | 97 | 85 | 0 | 2857 | 4144 | 2152 | 0 | 2390 | 83 | 862 | 664 | 1725 | 80 | 11709 | 3357 | 0 | 0 | 0 | 51 | 78 | 121 | 35 | 27 | 33 | 15 | 2050 | 0 |
| SRR2731330 | 0 | 84 | 56 | 233 | 360 | 34 | 12 | 277 | 1758 | 30231 | 0 | 0 | 0 | 7099 | 163 | 231 | 0 | 719 | 214 | 1046 | 642 | 418 | 369 | 15060 | 3072 | 39 | 12 | 0 | 92 | 112 | 214 | 66 | 38 | 53 | 22 | 3506 | 0 |
| SRR2731331 | 0 | 96 | 58 | 120 | 314 | 0 | 13 | 1370 | 3362 | 19683 | 90 | 264 | 0 | 811 | 8404 | 6763 | 0 | 2496 | 47 | 3561 | 2334 | 17 | 2099 | 32350 | 2909 | 0 | 0 | 0 | 24 | 36 | 36 | 30 | 14 | 10 | 0 | 26720 | 13 |
| SRR2731332 | 0 | 260 | 393 | 371 | 553 | 0 | 0 | 2155 | 7019 | 24086 | 52 | 153 | 0 | 727 | 2025 | 1899 | 0 | 2723 | 18 | 4040 | 2304 | 3473 | 403 | 13286 | 1875 | 0 | 0 | 0 | 13 | 33 | 53 | 20 | 0 | 0 | 0 | 8408 | 63 |
| SRR2731333 | 0 | 485 | 172 | 319 | 1525 | 0 | 0 | 819 | 3394 | 26824 | 78 | 54 | 0 | 6655 | 747 | 984 | 0 | 1726 | 168 | 654 | 1578 | 2442 | 135 | 6919 | 5839 | 0 | 0 | 0 | 251 | 167 | 309 | 97 | 30 | 31 | 0 | 2088 | 118 |
| SRR2731335 | 0 | 93 | 57 | 801 | 2706 | 0 | 0 | 1380 | 3535 | 24575 | 11 | 27 | 0 | 966 | 3898 | 5772 | 0 | 1555 | 36 | 1185 | 854 | 2545 | 751 | 16784 | 6788 | 0 | 0 | 0 | 17 | 22 | 12 | 14 | 0 | 0 | 10 | 2887 | 422 |
| SRR2731336 | 0 | 73 | 72 | 214 | 280 | 14 | 0 | 461 | 2623 | 16268 | 24 | 62 | 0 | 4145 | 747 | 1928 | 0 | 1328 | 223 | 7997 | 2689 | 1353 | 437 | 22435 | 4726 | 0 | 0 | 0 | 39 | 55 | 58 | 37 | 18 | 19 | 0 | 20258 | 107 |
| SRR2731337 | 0 | 225 | 94 | 158 | 496 | 0 | 0 | 1301 | 4325 | 48318 | 75 | 132 | 0 | 891 | 4155 | 3503 | 0 | 3183 | 80 | 2506 | 2343 | 3762 | 823 | 13180 | 2633 | 0 | 0 | 0 | 30 | 40 | 32 | 30 | 13 | 16 | 0 | 6505 | 10 |
| SRR2731338 | 0 | 29 | 33 | 260 | 472 | 0 | 21 | 1196 | 4008 | 25550 | 67 | 201 | 0 | 449 | 2846 | 2351 | 0 | 1563 | 35 | 1664 | 2244 | 2405 | 1241 | 26245 | 3234 | 0 | 0 | 0 | 17 | 32 | 63 | 57 | 27 | 19 | 12 | 27703 | 25 |
| SRR2731339 | 0 | 187 | 30 | 405 | 697 | 0 | 15 | 1854 | 3325 | 4963 | 23 | 1005 | 0 | 945 | 191 | 287 | 19 | 6163 | 90 | 395 | 1485 | 3441 | 360 | 9425 | 1957 | 0 | 0 | 0 | 20 | 27 | 24 | 11 | 0 | 0 | 0 | 5872 | 0 |
| SRR2731340 | 0 | 201 | 404 | 317 | 958 | 0 | 0 | 1561 | 7369 | 23634 | 60 | 148 | 0 | 1301 | 4150 | 2393 | 0 | 1355 | 53 | 1269 | 827 | 1863 | 147 | 12533 | 2156 | 0 | 0 | 0 | 75 | 101 | 190 | 68 | 32 | 31 | 23 | 3535 | 24 |
| SRR2731341 | 0 | 98 | 59 | 574 | 1163 | 0 | 0 | 2441 | 6231 | 12220 | 22 | 174 | 0 | 608 | 1206 | 1541 | 0 | 2999 | 37 | 2383 | 2316 | 4736 | 1121 | 17034 | 2910 | 0 | 0 | 0 | 11 | 30 | 18 | 18 | 0 | 0 | 0 | 5367 | 37 |
| SRR2731342 | 0 | 702 | 175 | 733 | 1147 | 0 | 0 | 4697 | 12398 | 17159 | 42 | 111 | 0 | 1054 | 596 | 1042 | 0 | 3805 | 110 | 1260 | 3325 | 5992 | 1008 | 14450 | 3505 | 0 | 0 | 0 | 15 | 20 | 25 | 0 | 0 | 0 | 0 | 2719 | 14 |
| SRR2731343 | 0 | 250 | 42 | 441 | 1629 | 0 | 0 | 2769 | 8166 | 11160 | 75 | 155 | 0 | 1641 | 350 | 763 | 0 | 2214 | 99 | 1380 | 1985 | 3453 | 362 | 14929 | 6520 | 14 | 0 | 0 | 13 | 11 | 19 | 0 | 0 | 0 | 0 | 2562 | 151 |
| SRR2731344 | 0 | 409 | 96 | 691 | 1722 | 0 | 0 | 1789 | 4923 | 7858 | 122 | 248 | 0 | 546 | 319 | 997 | 0 | 5473 | 64 | 999 | 3305 | 7017 | 1026 | 17795 | 2250 | 0 | 0 | 0 | 22 | 21 | 30 | 21 | 13 | 10 | 0 | 2105 | 39 |
| SRR2731345 | 0 | 319 | 290 | 208 | 614 | 13 | 0 | 1417 | 1450 | 8267 | 218 | 3521 | 0 | 1033 | 1939 | 2059 | 0 | 2249 | 131 | 1820 | 1410 | 3224 | 851 | 12869 | 1152 | 0 | 21 | 17 | 79 | 48 | 169 | 52 | 15 | 27 | 0 | 4799 | 0 |
| SRR2731346 | 0 | 143 | 40 | 415 | 1458 | 0 | 0 | 2048 | 6371 | 53616 | 31 | 35 | 0 | 4333 | 1268 | 1471 | 0 | 3954 | 50 | 604 | 1688 | 2173 | 285 | 6670 | 2312 | 0 | 0 | 0 | 30 | 33 | 13 | 0 | 0 | 0 | 0 | 1636 | 0 |
| SRR2731347 | 0 | 670 | 155 | 443 | 1281 | 12 | 0 | 1488 | 4004 | 12789 | 89 | 242 | 0 | 453 | 15525 | 14240 | 0 | 2329 | 92 | 3300 | 2128 | 5405 | 900 | 18515 | 3094 | 13 | 0 | 0 | 53 | 70 | 100 | 30 | 13 | 10 | 14 | 6098 | 81 |
| SRR2731348 | 0 | 195 | 146 | 447 | 1710 | 0 | 0 | 2726 | 8924 | 28612 | 13 | 39 | 0 | 1185 | 4250 | 6714 | 0 | 2668 | 29 | 913 | 1570 | 2132 | 539 | 8743 | 2613 | 0 | 0 | 0 | 26 | 19 | 10 | 0 | 0 | 0 | 0 | 1908 | 41 |
| SRR2731349 | 0 | 96 | 23 | 115 | 649 | 0 | 25 | 609 | 1257 | 19171 | 737 | 1263 | 0 | 2519 | 2124 | 870 | 0 | 1722 | 100 | 1370 | 2401 | 2125 | 610 | 11001 | 1370 | 0 | 29 | 16 | 21 | 47 | 19 | 64 | 24 | 27 | 0 | 3693 | 0 |
| SRR2731350 | 0 | 845 | 373 | 723 | 1812 | 0 | 0 | 1549 | 4964 | 22076 | 132 | 165 | 0 | 1343 | 11907 | 13001 | 0 | 2062 | 57 | 1605 | 1429 | 5446 | 562 | 19588 | 2641 | 0 | 0 | 0 | 37 | 45 | 56 | 13 | 10 | 0 | 0 | 3370 | 23 |
| SRR2910706 | 0 | 0 | 39 | 0 | 39 | 0 | 0 | 3766 | 1391 | 6022 | 917 | 4282 | 0 | 305 | 57 | 28 | 0 | 322 | 0 | 476 | 2582 | 2233 | 0 | 48 | 7622 | 48 | 0 | 0 | 54 | 129 | 72 | 11 | 48 | 136 | 73 | 2314 | 0 |
| SRR2910737 | 0 | 0 | 37 | 0 | 47 | 0 | 0 | 3742 | 1228 | 7423 | 1041 | 3436 | 0 | 156 | 43 | 28 | 0 | 252 | 0 | 472 | 2133 | 1877 | 0 | 46 | 6859 | 48 | 0 | 12 | 58 | 108 | 60 | 13 | 46 | 106 | 37 | 1342 | 0 |
| SRR2910738 | 0 | 12 | 42 | 0 | 40 | 0 | 0 | 3852 | 1121 | 8235 | 2625 | 1893 | 0 | 303 | 50 | 30 | 0 | 256 | 0 | 535 | 2374 | 2073 | 0 | 57 | 10938 | 52 | 0 | 12 | 58 | 13 | 60 | 18 | 44 | 18 | 61 | 1893 | 0 |
| SRR2912507 | 0 | 90 | 0 | 0 | 10 | 0 | 0 | 219 | 707 | 52 | 0 | 0 | 0 | 298 | 0 | 160 | 0 | 41 | 41 | 0 | 41 | 190 | 0 | 10 | 226 | 0 | 0 | 645 | 931 | 241 | 75 | 18 | 0 | 0 | 0 | 0 | 36 |
| SRR2912508 | 0 | 226 | 0 | 0 | 0 | 18 | 0 | 302 | 289 | 47 | 0 | 0 | 0 | 560 | 0 | 161 | 0 | 59 | 21 | 0 | 16 | 193 | 0 | 27 | 193 | 0 | 0 | 1341 | 2111 | 575 | 192 | 64 | 0 | 0 | 0 | 0 | 75 |
| SRR2912510 | 0 | 15 | 0 | 0 | 0 | 0 | 0 | 474 | 547 | 69 | 10 | 0 | 0 | 194 | 0 | 124 | 0 | 107 | 25 | 0 | 25 | 87 | 0 | 0 | 371 | 0 | 0 | 253 | 335 | 62 | 0 | 0 | 0 | 0 | 0 | 0 | 51 |
| SRR2912511 | 0 | 13 | 0 | 0 | 12 | 0 | 0 | 361 | 513 | 73 | 0 | 0 | 0 | 163 | 0 | 62 | 0 | 81 | 29 | 0 | 0 | 140 | 0 | 12 | 302 | 0 | 0 | 169 | 284 | 71 | 26 | 0 | 0 | 0 | 0 | 0 | 52 |
| SRR2912513 | 0 | 12 | 0 | 0 | 25 | 0 | 0 | 339 | 1788 | 45 | 10 | 0 | 0 | 116 | 34 | 404 | 0 | 450 | 0 | 0 | 22 | 133 | 0 | 883 | 448 | 0 | 0 | 26 | 28 | 11 | 0 | 0 | 0 | 0 | 0 | 0 | 61 |
| SRR2912514 | 0 | 86 | 0 | 0 | 0 | 0 | 0 | 402 | 534 | 62 | 0 | 0 | 0 | 442 | 19 | 791 | 0 | 269 | 0 | 0 | 19 | 253 | 0 | 438 | 240 | 0 | 0 | 404 | 467 | 159 | 62 | 12 | 0 | 0 | 0 | 0 | 53 |
| SRR2912515 | 0 | 13 | 0 | 0 | 0 | 0 | 0 | 391 | 1443 | 44 | 0 | 0 | 0 | 152 | 15 | 420 | 0 | 416 | 10 | 0 | 0 | 148 | 0 | 849 | 313 | 0 | 0 | 130 | 170 | 48 | 15 | 0 | 0 | 0 | 0 | 0 | 48 |
| SRR2913202 | 0 | 0 | 0 | 0 | 13 | 0 | 0 | 339 | 61 | 585 | 84 | 269 | 0 | 179 | 207 | 179 | 0 | 103 | 0 | 3320 | 1084 | 2285 | 232 | 4517 | 3659 | 39 | 0 | 0 | 0 | 0 | 0 | 0 | 0 | 0 | 0 | 8039 | 0 |
| SRR2913203 | 0 | 0 | 37 | 0 | 0 | 0 | 0 | 214 | 74 | 1187 | 317 | 171 | 0 | 157 | 203 | 417 | 0 | 82 | 0 | 1219 | 405 | 1083 | 79 | 1658 | 2148 | 25 | 0 | 0 | 0 | 0 | 0 | 0 | 0 | 13 | 30 | 3391 | 0 |
| SRR2913204 | 0 | 0 | 40 | 0 | 20 | 0 | 0 | 124 | 48 | 844 | 299 | 292 | 0 | 269 | 69 | 162 | 0 | 143 | 0 | 974 | 875 | 1768 | 283 | 3553 | 2657 | 15 | 0 | 0 | 0 | 0 | 0 | 0 | 0 | 0 | 14 | 2061 | 0 |
| SRR2913205 | 0 | 0 | 15 | 0 | 0 | 0 | 0 | 114 | 20 | 277 | 68 | 166 | 0 | 171 | 43 | 84 | 0 | 116 | 0 | 1569 | 1362 | 2578 | 372 | 5016 | 4048 | 0 | 0 | 0 | 0 | 0 | 0 | 0 | 0 | 0 | 0 | 3809 | 0 |
| SRR2913206 | 0 | 0 | 0 | 0 | 24 | 0 | 0 | 401 | 37 | 424 | 364 | 280 | 0 | 47 | 205 | 247 | 0 | 39 | 16 | 2629 | 917 | 2595 | 243 | 4069 | 4939 | 0 | 0 | 0 | 0 | 0 | 0 | 0 | 0 | 0 | 0 | 6905 | 0 |
| SRR2913207 | 0 | 0 | 20 | 0 | 12 | 0 | 0 | 22 | 10 | 216 | 115 | 67 | 0 | 263 | 27 | 28 | 0 | 78 | 0 | 95 | 1423 | 3928 | 405 | 4139 | 4088 | 36 | 0 | 0 | 0 | 0 | 0 | 0 | 22 | 26 | 27 | 184 | 0 |
| SRR2913208 | 0 | 0 | 11 | 0 | 0 | 0 | 0 | 303 | 79 | 708 | 700 | 830 | 0 | 149 | 1137 | 1566 | 0 | 117 | 10 | 3434 | 1072 | 2709 | 176 | 5077 | 4553 | 0 | 0 | 0 | 0 | 0 | 0 | 0 | 0 | 0 | 0 | 7458 | 0 |
| SRR2913209 | 0 | 0 | 11 | 0 | 0 | 0 | 0 | 495 | 82 | 1703 | 155 | 540 | 0 | 860 | 385 | 717 | 0 | 145 | 0 | 3087 | 892 | 1658 | 309 | 3302 | 1981 | 16 | 0 | 0 | 0 | 0 | 0 | 0 | 0 | 0 | 0 | 7239 | 0 |
| SRR2913210 | 0 | 0 | 0 | 0 | 0 | 0 | 0 | 310 | 55 | 675 | 166 | 348 | 0 | 210 | 513 | 798 | 0 | 155 | 0 | 1948 | 692 | 1690 | 269 | 4327 | 3388 | 32 | 0 | 0 | 0 | 0 | 0 | 0 | 0 | 0 | 0 | 5448 | 0 |
| SRR2913211 | 0 | 0 | 22 | 0 | 0 | 0 | 0 | 495 | 99 | 836 | 125 | 1078 | 0 | 165 | 1990 | 2958 | 0 | 112 | 0 | 2276 | 1030 | 2033 | 410 | 4812 | 3198 | 19 | 0 | 0 | 0 | 0 | 0 | 0 | 0 | 0 | 0 | 5707 | 0 |
| SRR2913212 | 0 | 0 | 0 | 0 | 0 | 0 | 0 | 412 | 188 | 1763 | 199 | 171 | 0 | 188 | 230 | 485 | 0 | 79 | 0 | 22 | 578 | 1194 | 155 | 2283 | 2256 | 49 | 0 | 0 | 0 | 0 | 0 | 0 | 0 | 0 | 11 | 53 | 0 |
| SRR2913213 | 0 | 0 | 0 | 0 | 0 | 0 | 0 | 646 | 45 | 455 | 63 | 564 | 0 | 264 | 155 | 127 | 0 | 226 | 0 | 18 | 760 | 1503 | 365 | 3264 | 2496 | 0 | 0 | 0 | 0 | 0 | 0 | 0 | 0 | 0 | 11 | 36 | 0 |
| SRR2913214 | 0 | 0 | 0 | 0 | 0 | 0 | 0 | 467 | 39 | 576 | 45 | 483 | 0 | 54 | 350 | 467 | 0 | 133 | 0 | 4461 | 1038 | 3329 | 216 | 4932 | 5380 | 0 | 0 | 0 | 0 | 0 | 0 | 0 | 0 | 0 | 11 | 10782 | 0 |
| SRR2913215 | 0 | 0 | 13 | 0 | 0 | 0 | 10 | 210 | 59 | 830 | 397 | 465 | 0 | 91 | 812 | 994 | 0 | 143 | 0 | 1419 | 1575 | 3444 | 327 | 5512 | 4730 | 23 | 0 | 0 | 0 | 0 | 0 | 0 | 0 | 0 | 10 | 3216 | 0 |
| SRR2913216 | 0 | 0 | 0 | 0 | 0 | 0 | 0 | 156 | 31 | 366 | 47 | 159 | 0 | 370 | 35 | 90 | 0 | 112 | 0 | 850 | 893 | 1842 | 321 | 4110 | 2079 | 14 | 0 | 0 | 0 | 0 | 0 | 0 | 0 | 0 | 13 | 2008 | 0 |
| SRR2913217 | 0 | 0 | 0 | 0 | 0 | 0 | 0 | 443 | 56 | 865 | 112 | 527 | 0 | 225 | 583 | 858 | 0 | 119 | 0 | 2256 | 789 | 1745 | 244 | 3131 | 2773 | 21 | 0 | 0 | 0 | 0 | 0 | 0 | 0 | 0 | 14 | 5089 | 0 |
| SRR2913218 | 0 | 0 | 0 | 0 | 0 | 0 | 0 | 96 | 13 | 199 | 78 | 103 | 0 | 53 | 80 | 99 | 0 | 131 | 0 | 653 | 1410 | 3362 | 224 | 4380 | 4633 | 0 | 0 | 0 | 0 | 0 | 0 | 0 | 0 | 0 | 0 | 1607 | 0 |
| SRR2913219 | 0 | 0 | 10 | 0 | 0 | 0 | 0 | 309 | 0 | 653 | 619 | 667 | 0 | 89 | 763 | 894 | 0 | 220 | 0 | 2722 | 1143 | 3230 | 269 | 5688 | 4370 | 16 | 0 | 0 | 0 | 0 | 0 | 0 | 0 | 0 | 0 | 6490 | 0 |
| SRR2913220 | 0 | 0 | 0 | 0 | 24 | 0 | 0 | 132 | 47 | 679 | 236 | 194 | 0 | 230 | 194 | 275 | 0 | 107 | 0 | 646 | 1231 | 3182 | 379 | 5794 | 4787 | 39 | 0 | 0 | 0 | 0 | 0 | 0 | 0 | 0 | 13 | 1877 | 0 |
| SRR2913221 | 0 | 0 | 0 | 0 | 11 | 0 | 0 | 59 | 16 | 316 | 290 | 117 | 0 | 134 | 136 | 150 | 0 | 110 | 0 | 251 | 1273 | 2970 | 449 | 6212 | 5166 | 0 | 0 | 0 | 0 | 0 | 0 | 0 | 0 | 0 | 0 | 577 | 0 |
| SRR2913222 | 0 | 0 | 13 | 0 | 0 | 0 | 0 | 133 | 74 | 1186 | 393 | 265 | 0 | 233 | 335 | 389 | 0 | 119 | 0 | 507 | 1215 | 3352 | 242 | 5287 | 5614 | 0 | 0 | 0 | 0 | 0 | 0 | 0 | 0 | 0 | 0 | 1514 | 0 |
| SRR2913223 | 0 | 0 | 13 | 0 | 0 | 0 | 0 | 164 | 33 | 342 | 120 | 72 | 0 | 138 | 67 | 109 | 0 | 44 | 0 | 332 | 413 | 983 | 66 | 1320 | 1874 | 17 | 0 | 0 | 0 | 0 | 0 | 0 | 0 | 0 | 17 | 860 | 0 |
| SRR2913224 | 0 | 0 | 16 | 0 | 0 | 0 | 0 | 124 | 31 | 302 | 158 | 213 | 0 | 119 | 236 | 320 | 0 | 177 | 0 | 566 | 1163 | 2447 | 228 | 4365 | 3671 | 17 | 0 | 0 | 0 | 0 | 0 | 0 | 0 | 0 | 14 | 1452 | 0 |
| SRR2913225 | 0 | 0 | 12 | 0 | 0 | 0 | 0 | 230 | 131 | 1205 | 344 | 487 | 0 | 137 | 534 | 748 | 0 | 175 | 0 | 748 | 1242 | 2561 | 285 | 4830 | 3950 | 0 | 0 | 0 | 0 | 0 | 0 | 0 | 0 | 0 | 0 | 2025 | 0 |
| SRR2915342 | 0 | 752 | 30 | 0 | 0 | 0 | 15 | 1399 | 2339 | 6543 | 4906 | 13194 | 0 | 5463 | 13 | 16 | 66 | 896 | 82 | 3116 | 51 | 390 | 12 | 13596 | 1540 | 0 | 15 | 26 | 26 | 112 | 72 | 44 | 26 | 21 | 11 | 3534 | 0 |
| SRR2915343 | 0 | 714 | 34 | 0 | 0 | 0 | 14 | 1454 | 2930 | 6873 | 3353 | 10503 | 0 | 5113 | 24 | 19 | 52 | 1135 | 71 | 2212 | 59 | 491 | 10 | 12416 | 2082 | 17 | 14 | 24 | 30 | 109 | 82 | 48 | 15 | 15 | 15 | 3378 | 18 |
| SRR2915344 | 0 | 1712 | 78 | 0 | 0 | 0 | 16 | 1361 | 3172 | 7510 | 6467 | 11093 | 0 | 5937 | 14 | 30 | 95 | 1215 | 71 | 3625 | 51 | 659 | 15 | 15137 | 2052 | 0 | 16 | 21 | 64 | 389 | 244 | 206 | 44 | 39 | 37 | 4471 | 11 |
| SRR2915345 | 0 | 3297 | 280 | 0 | 14 | 0 | 48 | 2130 | 4638 | 15840 | 20642 | 51309 | 0 | 1365 | 22 | 57 | 67 | 1318 | 40 | 3263 | 412 | 1321 | 0 | 8713 | 16 | 42 | 46 | 95 | 189 | 747 | 625 | 117 | 127 | 88 | 50 | 3992 | 320 |
| SRR2915346 | 0 | 983 | 84 | 0 | 14 | 0 | 18 | 2389 | 4621 | 14434 | 17623 | 44638 | 0 | 1920 | 55 | 124 | 21 | 1525 | 47 | 2334 | 418 | 1773 | 0 | 8708 | 7965 | 50 | 16 | 25 | 37 | 168 | 137 | 28 | 30 | 22 | 24 | 2930 | 304 |
| SRR2915347 | 0 | 1022 | 93 | 10 | 16 | 0 | 14 | 2181 | 4891 | 12505 | 12749 | 38465 | 0 | 1698 | 44 | 169 | 12 | 1872 | 41 | 2072 | 648 | 2104 | 0 | 8187 | 8337 | 56 | 10 | 24 | 46 | 190 | 136 | 28 | 28 | 23 | 28 | 4204 | 395 |
| SRR2915348 | 0 | 480 | 74 | 0 | 16 | 0 | 25 | 2579 | 4811 | 11683 | 3059 | 8807 | 0 | 717 | 11 | 0 | 10 | 974 | 91 | 4888 | 197 | 716 | 22 | 10779 | 1783 | 10 | 38 | 13 | 25 | 74 | 67 | 34 | 34 | 29 | 23 | 6015 | 209 |
| SRR2915349 | 0 | 671 | 66 | 0 | 19 | 0 | 29 | 2960 | 6164 | 11568 | 15144 | 13233 | 0 | 798 | 0 | 0 | 13 | 1341 | 94 | 4782 | 184 | 1016 | 23 | 10811 | 2139 | 13 | 42 | 12 | 26 | 72 | 46 | 21 | 18 | 14 | 28 | 5347 | 248 |
| SRR2915350 | 0 | 867 | 80 | 0 | 15 | 0 | 30 | 2914 | 5262 | 11802 | 2571 | 13013 | 0 | 886 | 10 | 10 | 19 | 1234 | 90 | 4748 | 163 | 1120 | 23 | 10888 | 2015 | 15 | 17 | 13 | 28 | 84 | 67 | 32 | 29 | 24 | 11 | 5880 | 251 |
| SRR2915905 | 0 | 88 | 64 | 0 | 0 | 0 | 0 | 1513 | 268 | 5234 | 85 | 46 | 0 | 313 | 141 | 61 | 0 | 368 | 0 | 162 | 1099 | 1372 | 281 | 6306 | 3072 | 26 | 0 | 0 | 21 | 34 | 38 | 0 | 24 | 31 | 18 | 886 | 13 |
| SRR2915915 | 0 | 29 | 20 | 0 | 0 | 0 | 0 | 480 | 740 | 4943 | 130 | 54 | 0 | 401 | 21 | 0 | 0 | 268 | 0 | 116 | 1055 | 524 | 186 | 1255 | 1374 | 0 | 0 | 0 | 33 | 80 | 87 | 21 | 40 | 47 | 0 | 1121 | 0 |
| SRR2915918 | 0 | 44 | 32 | 0 | 33 | 0 | 0 | 227 | 151 | 3011 | 119 | 170 | 0 | 820 | 18 | 51 | 0 | 705 | 0 | 153 | 1214 | 2400 | 252 | 9034 | 5300 | 37 | 0 | 0 | 20 | 30 | 31 | 0 | 28 | 50 | 28 | 447 | 0 |
| SRR2915921 | 0 | 0 | 0 | 0 | 54 | 0 | 0 | 348 | 180 | 4563 | 836 | 622 | 0 | 1103 | 18 | 0 | 17 | 319 | 0 | 142 | 853 | 1946 | 255 | 8957 | 4271 | 30 | 26 | 0 | 0 | 0 | 0 | 0 | 0 | 0 | 0 | 558 | 0 |
| SRR2915923 | 0 | 169 | 18 | 0 | 29 | 0 | 12 | 56 | 26 | 2736 | 1018 | 67 | 0 | 718 | 0 | 0 | 11 | 224 | 0 | 54 | 1154 | 3316 | 318 | 11914 | 5208 | 15 | 36 | 0 | 0 | 20 | 12 | 0 | 13 | 11 | 0 | 225 | 19 |
| SRR2915926 | 0 | 12 | 15 | 0 | 15 | 0 | 0 | 113 | 159 | 4791 | 85 | 78 | 0 | 1324 | 0 | 0 | 0 | 253 | 0 | 41 | 825 | 1890 | 189 | 14150 | 5631 | 20 | 19 | 0 | 11 | 12 | 12 | 0 | 11 | 23 | 11 | 331 | 169 |
| SRR2915928 | 0 | 22 | 46 | 0 | 58 | 0 | 0 | 132 | 59 | 5495 | 437 | 351 | 0 | 477 | 18 | 12 | 24 | 460 | 0 | 361 | 1325 | 2069 | 353 | 11515 | 4969 | 66 | 32 | 0 | 0 | 14 | 0 | 0 | 0 | 0 | 0 | 1712 | 0 |
| SRR2960039 | 0 | 45 | 0 | 296 | 55 | 0 | 45 | 220 | 0 | 0 | 25 | 32 | 0 | 0 | 0 | 0 | 50 | 1220 | 16 | 0 | 108 | 1169 | 0 | 28 | 501 | 0 | 0 | 313 | 134 | 196 | 17 | 0 | 0 | 0 | 0 | 0 | 0 |

### Supp Table S8.4.Columns_DI_to_ES

| | | | | | | | | | | | | | | | | | | | | | | | | | | | | | | | | | | | | | |
|---|---|---|---|---|---|---|---|---|---|---|---|---|---|---|---|---|---|---|---|---|---|---|---|---|---|---|---|---|---|---|---|---|---|---|---|---|---|
| SRR2976280 | 0 | 0 | 0 | 0 | 11 | 0 | 0 | 4293 | 156 | 52 | 719 | 5679 | 0 | 0 | 0 | 0 | 35 | 0 | 0 | 0 | 1564 | 705 | 880 | 29325 | 1550 | 0 | 0 | 0 | 0 | 0 | 0 | 0 | 0 | 0 | 0 | 0 | 50 |
| SRR2976305 | 0 | 0 | 0 | 0 | 0 | 0 | 0 | 130 | 0 | 208 | 832 | 3161 | 0 | 0 | 0 | 0 | 0 | 0 | 0 | 0 | 1476 | 984 | 681 | 22911 | 1146 | 0 | 232 | 0 | 0 | 24 | 0 | 0 | 0 | 0 | 0 | 0 | 34 |
| SRR3092114 | 0 | 222 | 0 | 0 | 0 | 0 | 0 | 0 | 18 | 73 | 617 | 20049 | 0 | 0 | 0 | 0 | 0 | 93 | 1228 | 476 | 818 | 1472 | 0 | 9663 | 9465 | 0 | 0 | 59 | 58 | 197 | 478 | 49 | 0 | 0 | 0 | 531 | 19 |
| SRR3092115 | 0 | 148 | 0 | 0 | 0 | 0 | 0 | 0 | 13 | 40 | 12964 | 17280 | 0 | 0 | 0 | 0 | 0 | 122 | 1104 | 824 | 887 | 1643 | 0 | 8757 | 7728 | 0 | 0 | 11 | 16 | 63 | 161 | 14 | 0 | 0 | 0 | 1089 | 33 |
| SRR3103222 | 0 | 29 | 16 | 0 | 37 | 10 | 0 | 3857 | 2170 | 3388 | 1043 | 6926 | 0 | 304 | 0 | 0 | 16 | 343 | 0 | 157 | 1208 | 1305 | 401 | 4640 | 3224 | 0 | 0 | 21 | 34 | 78 | 50 | 0 | 0 | 21 | 0 | 416 | 12 |
| SRR3103223 | 0 | 28 | 99 | 0 | 69 | 0 | 0 | 2626 | 1868 | 13772 | 0 | 231 | 0 | 64 | 0 | 20 | 0 | 299 | 0 | 128 | 109 | 129 | 66 | 1392 | 654 | 0 | 14 | 14 | 14 | 57 | 14 | 0 | 28 | 57 | 14 | 1170 | 0 |
| SRR3103224 | 0 | 26 | 116 | 0 | 61 | 0 | 0 | 7952 | 3305 | 6283 | 327 | 2978 | 0 | 0 | 33 | 65 | 0 | 253 | 0 | 164 | 123 | 158 | 63 | 692 | 534 | 0 | 0 | 13 | 32 | 39 | 39 | 0 | 19 | 26 | 13 | 1964 | 0 |
| SRR3103225 | 0 | 0 | 52 | 0 | 50 | 0 | 0 | 631 | 276 | 2484 | 64 | 56 | 0 | 158 | 50 | 54 | 0 | 272 | 0 | 24 | 1277 | 1520 | 422 | 6307 | 3467 | 0 | 0 | 0 | 0 | 19 | 33 | 14 | 28 | 76 | 28 | 150 | 0 |
| SRR3103226 | 0 | 21 | 42 | 0 | 113 | 0 | 0 | 7684 | 5809 | 11397 | 148 | 735 | 0 | 110 | 110 | 184 | 0 | 279 | 0 | 404 | 113 | 187 | 42 | 1059 | 433 | 13 | 21 | 0 | 42 | 0 | 21 | 21 | 0 | 63 | 0 | 4559 | 10 |
| SRR3103227 | 0 | 0 | 131 | 0 | 41 | 0 | 0 | 1492 | 1249 | 4605 | 29 | 94 | 0 | 98 | 0 | 0 | 0 | 245 | 0 | 156 | 1188 | 1063 | 421 | 4194 | 4039 | 0 | 10 | 30 | 10 | 51 | 91 | 20 | 40 | 61 | 20 | 314 | 0 |
| SRR3103228 | 0 | 24 | 27 | 0 | 15 | 0 | 0 | 2809 | 1772 | 3095 | 101 | 519 | 0 | 251 | 31 | 40 | 30 | 107 | 0 | 145 | 610 | 834 | 324 | 2125 | 2326 | 0 | 0 | 0 | 0 | 20 | 20 | 0 | 13 | 60 | 20 | 549 | 0 |
| SRR3103229 | 0 | 0 | 53 | 0 | 53 | 0 | 0 | 1603 | 954 | 4953 | 881 | 929 | 0 | 263 | 167 | 103 | 0 | 160 | 0 | 141 | 737 | 718 | 355 | 4019 | 2228 | 0 | 0 | 0 | 14 | 32 | 25 | 0 | 32 | 84 | 53 | 911 | 0 |
| SRR3103230 | 0 | 0 | 76 | 0 | 39 | 0 | 0 | 401 | 265 | 2527 | 88 | 79 | 0 | 118 | 0 | 0 | 0 | 232 | 0 | 53 | 1070 | 941 | 436 | 3306 | 4061 | 0 | 0 | 0 | 34 | 17 | 42 | 0 | 51 | 25 | 42 | 98 | 0 |
| SRR3103231 | 0 | 0 | 0 | 0 | 27 | 0 | 0 | 5883 | 4042 | 10603 | 100 | 238 | 0 | 150 | 28 | 37 | 0 | 138 | 0 | 72 | 825 | 861 | 473 | 4787 | 3409 | 0 | 0 | 0 | 23 | 0 | 41 | 0 | 18 | 55 | 55 | 341 | 0 |
| SRR3103232 | 0 | 18 | 89 | 0 | 31 | 0 | 0 | 2902 | 1319 | 2950 | 1511 | 1199 | 0 | 96 | 0 | 0 | 0 | 186 | 0 | 480 | 731 | 837 | 454 | 3029 | 2832 | 0 | 0 | 0 | 18 | 36 | 18 | 0 | 18 | 18 | 18 | 1631 | 0 |
| SRR3103233 | 0 | 59 | 36 | 0 | 34 | 0 | 0 | 2696 | 1470 | 1805 | 163 | 825 | 0 | 108 | 0 | 12 | 0 | 174 | 0 | 114 | 649 | 1148 | 339 | 3283 | 2772 | 0 | 0 | 24 | 12 | 12 | 24 | 0 | 12 | 18 | 12 | 329 | 0 |
| SRR3103234 | 0 | 11 | 67 | 0 | 28 | 0 | 0 | 1087 | 419 | 6333 | 50 | 80 | 0 | 156 | 14 | 26 | 0 | 98 | 0 | 53 | 1008 | 1275 | 355 | 3402 | 3914 | 0 | 0 | 0 | 0 | 48 | 41 | 0 | 30 | 41 | 26 | 129 | 0 |
| SRR3103235 | 0 | 0 | 55 | 0 | 22 | 0 | 0 | 2581 | 1504 | 4677 | 118 | 132 | 0 | 294 | 0 | 0 | 0 | 91 | 0 | 124 | 1001 | 1211 | 769 | 4562 | 3287 | 0 | 0 | 13 | 0 | 0 | 32 | 0 | 19 | 48 | 23 | 303 | 0 |
| SRR3103236 | 0 | 0 | 17 | 0 | 30 | 0 | 0 | 6652 | 6790 | 16607 | 306 | 487 | 0 | 68 | 11 | 15 | 0 | 117 | 0 | 50 | 827 | 1489 | 494 | 4076 | 3035 | 0 | 10 | 10 | 10 | 17 | 27 | 20 | 20 | 54 | 17 | 199 | 0 |
| SRR3103237 | 0 | 0 | 128 | 0 | 26 | 0 | 0 | 469 | 291 | 5167 | 52 | 77 | 0 | 153 | 0 | 0 | 0 | 148 | 0 | 31 | 1421 | 1323 | 546 | 4781 | 4283 | 0 | 19 | 0 | 0 | 48 | 19 | 0 | 19 | 76 | 38 | 108 | 0 |
| SRR3103238 | 0 | 0 | 50 | 0 | 37 | 0 | 0 | 660 | 431 | 3559 | 18 | 39 | 0 | 89 | 0 | 0 | 0 | 236 | 0 | 66 | 1296 | 1176 | 583 | 4691 | 3663 | 0 | 0 | 90 | 32 | 23 | 23 | 12 | 26 | 29 | 12 | 429 | 0 |
| SRR3103239 | 0 | 0 | 0 | 0 | 20 | 0 | 0 | 13704 | 8178 | 9401 | 200 | 2255 | 0 | 88 | 1975 | 1636 | 0 | 124 | 0 | 280 | 617 | 900 | 509 | 4772 | 2735 | 0 | 0 | 0 | 0 | 33 | 14 | 0 | 0 | 37 | 44 | 1709 | 0 |
| SRR3103240 | 0 | 0 | 0 | 0 | 101 | 0 | 0 | 3162 | 1442 | 13674 | 376 | 205 | 0 | 282 | 585 | 378 | 0 | 207 | 0 | 119 | 474 | 1499 | 232 | 14634 | 2994 | 0 | 0 | 0 | 21 | 30 | 30 | 0 | 48 | 88 | 30 | 538 | 0 |
| SRR3103241 | 0 | 0 | 0 | 0 | 94 | 0 | 0 | 4432 | 1927 | 9533 | 112 | 690 | 0 | 100 | 3764 | 2328 | 0 | 164 | 0 | 167 | 354 | 1797 | 175 | 18713 | 4145 | 0 | 0 | 0 | 25 | 44 | 38 | 19 | 32 | 82 | 25 | 1013 | 18 |
| SRR3103242 | 0 | 32 | 46 | 0 | 62 | 19 | 0 | 709 | 347 | 4246 | 993 | 1158 | 0 | 591 | 71 | 102 | 0 | 175 | 0 | 291 | 782 | 1039 | 80 | 4437 | 3027 | 0 | 0 | 42 | 190 | 121 | 93 | 0 | 162 | 15 | 51 | 2552 | 58 |
| SRR3103243 | 0 | 17 | 0 | 0 | 98 | 35 | 64 | 553 | 125 | 3530 | 1373 | 8557 | 0 | 18 | 36 | 36 | 35 | 87 | 12 | 71 | 159 | 233 | 0 | 150 | 862 | 0 | 31 | 119 | 127 | 54 | 37 | 0 | 0 | 27 | 0 | 285 | 83 |
| SRR3103244 | 0 | 0 | 131 | 0 | 126 | 0 | 0 | 6103 | 3444 | 13947 | 18 | 367 | 0 | 123 | 25 | 15 | 0 | 303 | 0 | 320 | 124 | 116 | 21 | 809 | 464 | 15 | 30 | 0 | 59 | 125 | 53 | 12 | 53 | 101 | 18 | 6368 | 0 |
| SRR3103245 | 0 | 51 | 143 | 0 | 128 | 0 | 0 | 1301 | 814 | 22192 | 1670 | 5746 | 0 | 49 | 42 | 14 | 0 | 224 | 0 | 570 | 259 | 304 | 34 | 919 | 518 | 10 | 27 | 18 | 54 | 89 | 80 | 0 | 68 | 128 | 12 | 4216 | 19 |
| SRR3103246 | 0 | 0 | 17 | 0 | 17 | 21 | 23 | 436 | 339 | 6904 | 320 | 1599 | 0 | 365 | 45 | 47 | 0 | 104 | 22 | 215 | 393 | 481 | 0 | 324 | 1138 | 0 | 10 | 13 | 52 | 62 | 66 | 15 | 55 | 111 | 39 | 1765 | 57 |
| SRR3103247 | 0 | 34 | 64 | 0 | 53 | 0 | 0 | 604 | 406 | 4654 | 279 | 1470 | 0 | 496 | 63 | 18 | 0 | 124 | 14 | 622 | 717 | 1444 | 111 | 3220 | 4275 | 0 | 24 | 64 | 157 | 73 | 83 | 44 | 88 | 196 | 29 | 1119 | 0 |
| SRR3103248 | 0 | 0 | 118 | 0 | 30 | 0 | 11 | 305 | 275 | 6407 | 162 | 210 | 0 | 662 | 54 | 57 | 11 | 87 | 26 | 66 | 464 | 1179 | 64 | 1754 | 6068 | 0 | 0 | 0 | 13 | 42 | 29 | 0 | 27 | 72 | 15 | 291 | 25 |
| SRR3103249 | 0 | 0 | 24 | 0 | 48 | 0 | 0 | 242 | 229 | 3580 | 236 | 189 | 0 | 939 | 32 | 30 | 18 | 136 | 69 | 61 | 591 | 481 | 43 | 3807 | 3250 | 26 | 17 | 27 | 34 | 32 | 35 | 19 | 37 | 61 | 18 | 829 | 16 |
| SRR3103250 | 0 | 0 | 55 | 0 | 25 | 0 | 0 | 1999 | 1091 | 12452 | 2721 | 2147 | 0 | 196 | 0 | 15 | 0 | 202 | 13 | 246 | 464 | 1075 | 96 | 2319 | 3789 | 14 | 0 | 47 | 30 | 74 | 55 | 0 | 36 | 71 | 11 | 1587 | 50 |
| SRR3103251 | 0 | 0 | 108 | 0 | 78 | 0 | 0 | 1184 | 821 | 5025 | 11197 | 9035 | 0 | 145 | 181 | 266 | 0 | 103 | 0 | 205 | 1110 | 737 | 95 | 3722 | 3142 | 0 | 0 | 43 | 129 | 90 | 61 | 0 | 32 | 72 | 14 | 713 | 0 |
| SRR3103252 | 0 | 0 | 0 | 0 | 29 | 0 | 0 | 1055 | 747 | 4757 | 373 | 3475 | 0 | 260 | 65 | 16 | 15 | 95 | 0 | 244 | 1008 | 552 | 23 | 538 | 2040 | 0 | 0 | 0 | 15 | 0 | 15 | 0 | 31 | 31 | 0 | 779 | 37 |
| SRR3103253 | 0 | 13 | 0 | 0 | 45 | 0 | 0 | 701 | 382 | 3431 | 546 | 489 | 0 | 329 | 0 | 0 | 0 | 295 | 0 | 107 | 434 | 772 | 286 | 3194 | 1639 | 12 | 0 | 22 | 43 | 43 | 39 | 0 | 30 | 22 | 0 | 359 | 23 |
| SRR3103254 | 0 | 19 | 0 | 0 | 42 | 0 | 0 | 859 | 295 | 3183 | 580 | 480 | 0 | 63 | 37 | 47 | 0 | 119 | 0 | 56 | 598 | 848 | 62 | 996 | 4996 | 0 | 0 | 187 | 90 | 71 | 84 | 23 | 126 | 236 | 68 | 243 | 46 |
| SRR3103255 | 0 | 64 | 24 | 0 | 29 | 0 | 0 | 758 | 434 | 3149 | 4553 | 5236 | 0 | 2343 | 74 | 37 | 0 | 35 | 13 | 579 | 2054 | 875 | 37 | 1284 | 3266 | 0 | 0 | 64 | 54 | 68 | 63 | 11 | 77 | 12 | 13 | 2377 | 0 |
| SRR3103256 | 0 | 19 | 30 | 0 | 13 | 0 | 0 | 273 | 115 | 2081 | 220 | 299 | 0 | 656 | 0 | 0 | 0 | 161 | 0 | 77 | 498 | 2231 | 91 | 9425 | 5403 | 0 | 11 | 0 | 49 | 27 | 30 | 0 | 49 | 93 | 22 | 637 | 26 |
| SRR3103257 | 0 | 26 | 63 | 0 | 37 | 0 | 0 | 732 | 305 | 8077 | 813 | 2536 | 0 | 272 | 140 | 223 | 0 | 86 | 0 | 134 | 1302 | 772 | 103 | 2444 | 2562 | 0 | 0 | 15 | 26 | 20 | 27 | 0 | 26 | 43 | 0 | 656 | 39 |
| SRR3103258 | 0 | 0 | 55 | 0 | 42 | 10 | 0 | 374 | 260 | 2486 | 828 | 474 | 0 | 568 | 139 | 222 | 0 | 129 | 16 | 87 | 634 | 1891 | 148 | 4874 | 4869 | 0 | 0 | 67 | 19 | 15 | 21 | 116 | 17 | 26 | 191 | 467 | 0 |
| SRR3103259 | 0 | 0 | 0 | 0 | 27 | 0 | 0 | 6187 | 5212 | 11627 | 532 | 622 | 0 | 249 | 92 | 62 | 0 | 103 | 0 | 96 | 800 | 977 | 325 | 4021 | 2729 | 0 | 0 | 0 | 0 | 18 | 15 | 0 | 23 | 44 | 13 | 615 | 0 |
| SRR3103260 | 0 | 0 | 0 | 0 | 17 | 0 | 0 | 1738 | 811 | 10202 | 2217 | 1485 | 0 | 1642 | 334 | 281 | 0 | 59 | 36 | 117 | 296 | 1172 | 42 | 886 | 4691 | 0 | 0 | 0 | 29 | 27 | 50 | 16 | 51 | 108 | 34 | 760 | 0 |
| SRR3103261 | 0 | 0 | 13 | 0 | 10 | 0 | 11 | 3466 | 2222 | 7308 | 359 | 1140 | 0 | 802 | 230 | 188 | 11 | 54 | 39 | 88 | 305 | 931 | 21 | 635 | 4877 | 16 | 0 | 0 | 40 | 76 | 110 | 38 | 89 | 244 | 69 | 814 | 0 |
| SRR3103262 | 0 | 15 | 62 | 0 | 60 | 0 | 0 | 267 | 122 | 1436 | 741 | 625 | 0 | 507 | 15 | 30 | 12 | 121 | 14 | 99 | 617 | 1039 | 39 | 4251 | 3097 | 0 | 0 | 66 | 97 | 100 | 50 | 0 | 100 | 147 | 62 | 518 | 54 |
| SRR3103263 | 0 | 14 | 19 | 0 | 249 | 23 | 19 | 988 | 437 | 7657 | 795 | 4825 | 0 | 114 | 0 | 0 | 19 | 141 | 0 | 166 | 94 | 85 | 13 | 409 | 386 | 57 | 28 | 65 | 125 | 185 | 74 | 0 | 28 | 42 | 0 | 944 | 44 |
| SRR3103264 | 0 | 0 | 40 | 0 | 24 | 0 | 24 | 876 | 802 | 11484 | 0 | 53 | 0 | 329 | 57 | 70 | 0 | 167 | 0 | 125 | 279 | 141 | 20 | 846 | 520 | 0 | 18 | 0 | 0 | 0 | 0 | 0 | 0 | 0 | 14 | 1905 | 0 |
| SRR3103265 | 0 | 0 | 109 | 0 | 137 | 0 | 11 | 476 | 263 | 2579 | 923 | 683 | 0 | 64 | 28 | 44 | 0 | 131 | 0 | 25 | 1816 | 905 | 92 | 2367 | 1640 | 0 | 20 | 0 | 14 | 23 | 34 | 0 | 43 | 100 | 46 | 224 | 16 |
| SRR3103266 | 0 | 0 | 86 | 0 | 0 | 0 | 0 | 640 | 466 | 6302 | 21 | 157 | 0 | 90 | 0 | 0 | 0 | 138 | 15 | 67 | 69 | 166 | 0 | 1032 | 576 | 0 | 0 | 19 | 67 | 19 | 77 | 29 | 105 | 211 | 96 | 1083 | 101 |
| SRR3103267 | 0 | 0 | 33 | 0 | 81 | 0 | 0 | 179 | 197 | 3209 | 431 | 242 | 0 | 553 | 0 | 0 | 0 | 114 | 0 | 27 | 601 | 556 | 106 | 2030 | 1922 | 0 | 0 | 70 | 149 | 100 | 100 | 33 | 70 | 133 | 21 | 156 | 0 |
| SRR3103268 | 0 | 23 | 127 | 0 | 24 | 0 | 0 | 328 | 300 | 2359 | 70 | 81 | 0 | 497 | 11 | 14 | 0 | 165 | 35 | 14 | 576 | 1028 | 34 | 1382 | 6904 | 0 | 13 | 0 | 39 | 49 | 88 | 39 | 65 | 95 | 26 | 89 | 20 |
| SRR3103269 | 0 | 0 | 84 | 0 | 74 | 0 | 0 | 442 | 243 | 7357 | 483 | 296 | 0 | 269 | 107 | 59 | 23 | 36 | 0 | 16 | 932 | 845 | 78 | 2604 | 3329 | 0 | 14 | 28 | 17 | 353 | 274 | 79 | 316 | 511 | 84 | 88 | 35 |
| SRR3103270 | 0 | 0 | 120 | 0 | 20 | 0 | 0 | 644 | 348 | 1415 | 95 | 65 | 0 | 179 | 0 | 0 | 0 | 267 | 0 | 45 | 917 | 897 | 425 | 3218 | 3431 | 0 | 0 | 12 | 24 | 132 | 36 | 24 | 36 | 96 | 24 | 90 | 0 |
| SRR3103271 | 0 | 0 | 110 | 0 | 63 | 0 | 0 | 713 | 538 | 4135 | 430 | 445 | 0 | 342 | 33 | 82 | 0 | 135 | 0 | 24 | 1385 | 1012 | 92 | 5071 | 3541 | 0 | 0 | 39 | 91 | 58 | 97 | 13 | 103 | 78 | 52 | 89 | 0 |
| SRR3103272 | 0 | 0 | 38 | 0 | 23 | 0 | 0 | 248 | 176 | 1945 | 176 | 174 | 0 | 256 | 0 | 0 | 19 | 120 | 12 | 41 | 712 | 667 | 63 | 895 | 2585 | 0 | 0 | 0 | 38 | 57 | 19 | 0 | 19 | 19 | 19 | 145 | 33 |
| SRR3103273 | 0 | 0 | 35 | 0 | 56 | 0 | 0 | 392 | 297 | 4468 | 91 | 65 | 0 | 454 | 62 | 59 | 0 | 223 | 0 | 19 | 367 | 1234 | 95 | 4056 | 3900 | 12 | 0 | 35 | 78 | 61 | 43 | 17 | 121 | 121 | 61 | 98 | 84 |
| SRR3103274 | 0 | 0 | 0 | 0 | 41 | 0 | 0 | 521 | 186 | 2608 | 750 | 561 | 0 | 27 | 0 | 0 | 0 | 114 | 0 | 43 | 669 | 1020 | 32 | 813 | 3851 | 0 | 0 | 100 | 54 | 73 | 87 | 19 | 122 | 179 | 60 | 186 | 54 |
| SRR3103275 | 0 | 0 | 17 | 0 | 25 | 0 | 14 | 376 | 162 | 2582 | 269 | 139 | 0 | 999 | 0 | 0 | 0 | 73 | 12 | 46 | 2717 | 1081 | 108 | 4308 | 5337 | 0 | 0 | 31 | 51 | 50 | 67 | 15 | 55 | 109 | 24 | 271 | 0 |
| SRR3103276 | 0 | 0 | 16 | 0 | 13 | 0 | 21 | 478 | 208 | 2987 | 562 | 471 | 0 | 613 | 0 | 15 | 16 | 130 | 0 | 54 | 450 | 1863 | 136 | 10189 | 5272 | 14 | 14 | 23 | 51 | 42 | 66 | 14 | 68 | 122 | 14 | 205 | 30 |
| SRR3103277 | 0 | 0 | 60 | 0 | 33 | 0 | 0 | 267 | 109 | 2224 | 189 | 118 | 0 | 213 | 0 | 0 | 0 | 156 | 0 | 18 | 1835 | 667 | 68 | 2741 | 2929 | 10 | 0 | 30 | 18 | 36 | 71 | 0 | 18 | 137 | 24 | 99 | 32 |
| SRR3103278 | 0 | 0 | 17 | 0 | 34 | 0 | 17 | 1558 | 799 | 4234 | 79 | 77 | 0 | 342 | 0 | 0 | 0 | 175 | 0 | 14 | 1119 | 1206 | 217 | 4196 | 3058 | 0 | 17 | 30 | 604 | 276 | 371 | 155 | 526 | 716 | 181 | 64 | 0 |
| SRR3103279 | 0 | 0 | 25 | 0 | 30 | 0 | 0 | 1260 | 827 | 4868 | 79 | 144 | 0 | 132 | 0 | 0 | 0 | 138 | 0 | 25 | 892 | 759 | 486 | 3351 | 2333 | 0 | 0 | 0 | 0 | 13 | 30 | 18 | 20 | 44 | 24 | 143 | 0 |
| SRR3103280 | 0 | 0 | 31 | 0 | 68 | 0 | 0 | 1138 | 739 | 13968 | 747 | 963 | 0 | 121 | 102 | 99 | 0 | 119 | 0 | 50 | 775 | 686 | 189 | 3665 | 1893 | 0 | 0 | 48 | 26 | 12 | 118 | 38 | 19 | 29 | 87 | 285 | 52 |
| SRR3103281 | 0 | 0 | 516 | 0 | 100 | 0 | 0 | 1028 | 551 | 8719 | 324 | 946 | 0 | 137 | 105 | 74 | 0 | 150 | 0 | 37 | 625 | 593 | 108 | 2599 | 1591 | 17 | 0 | 951 | 534 | 344 | 215 | 80 | 413 | 757 | 136 | 198 | 26 |
| SRR3103386 | 0 | 0 | 130 | 0 | 29 | 0 | 0 | 1128 | 98 | 196 | 0 | 0 | 0 | 98 | 0 | 49 | 0 | 381 | 0 | 147 | 1654 | 1877 | 0 | 132 | 4256 | 399 | 0 | 0 | 65 | 162 | 0 | 0 | 0 | 0 | 32 | 1569 | 0 |
| SRR3103387 | 0 | 0 | 0 | 0 | 23 | 0 | 0 | 365 | 0 | 182 | 0 | 0 | 0 | 91 | 91 | 91 | 0 | 310 | 0 | 274 | 1443 | 1528 | 0 | 190 | 5836 | 327 | 0 | 0 | 0 | 95 | 63 | 0 | 63 | 126 | 32 | 6932 | 0 |
| SRR3103388 | 0 | 0 | 0 | 0 | 16 | 0 | 44 | 543 | 0 | 102 | 0 | 0 | 0 | 170 | 0 | 0 | 0 | 414 | 0 | 238 | 1177 | 1428 | 0 | 78 | 2954 | 626 | 0 | 0 | 0 | 175 | 88 | 0 | 0 | 44 | 131 | 3021 | 0 |
| SRR3103389 | 0 | 0 | 0 | 0 | 25 | 0 | 0 | 328 | 123 | 738 | 0 | 0 | 0 | 41 | 82 | 0 | 0 | 217 | 0 | 287 | 2209 | 2831 | 0 | 170 | 8826 | 408 | 0 | 32 | 0 | 0 | 0 | 0 | 0 | 0 | 63 | 3238 | 0 |
| SRR3103390 | 0 | 0 | 56 | 0 | 0 | 0 | 0 | 961 | 148 | 443 | 0 | 0 | 0 | 0 | 0 | 74 | 0 | 211 | 0 | 296 | 2065 | 2664 | 0 | 260 | 8190 | 684 | 0 | 0 | 28 | 84 | 84 | 0 | 0 | 56 | 0 | 4138 | 0 |
| SRR3103391 | 0 | 0 | 0 | 0 | 30 | 0 | 0 | 421 | 94 | 328 | 0 | 0 | 0 | 0 | 47 | 47 | 0 | 505 | 0 | 328 | 3184 | 2377 | 0 | 163 | 8712 | 464 | 0 | 0 | 0 | 62 | 42 | 21 | 0 | 62 | 145 | 4402 | 0 |
| SRR3103392 | 0 | 77 | 19 | 0 | 21 | 0 | 0 | 52 | 52 | 302 | 0 | 0 | 0 | 26 | 0 | 0 | 0 | 684 | 0 | 131 | 2350 | 2365 | 0 | 255 | 11011 | 1025 | 0 | 0 | 19 | 77 | 39 | 39 | 19 | 58 | 213 | 1365 | 0 |
| SRR3103393 | 0 | 0 | 0 | 0 | 0 | 0 | 0 | 0 | 315 | 0 | 0 | 0 | 0 | 0 | 0 | 0 | 0 | 563 | 0 | 315 | 1185 | 1195 | 0 | 88 | 2903 | 494 | 0 | 0 | 0 | 0 | 0 | 0 | 0 | 86 | 0 | 945 | 0 |
| SRR3103394 | 0 | 17 | 59 | 0 | 55 | 0 | 0 | 226 | 34 | 75 | 0 | 0 | 0 | 67 | 25 | 42 | 0 | 528 | 0 | 1156 | 3319 | 2221 | 0 | 171 | 7099 | 358 | 0 | 0 | 42 | 184 | 50 | 17 | 67 | 50 | 100 | 15151 | 0 |
| SRR3103395 | 0 | 0 | 0 | 0 | 32 | 0 | 0 | 422 | 0 | 241 | 0 | 0 | 0 | 0 | 241 | 241 | 0 | 990 | 0 | 241 | 2226 | 2027 | 0 | 173 | 6194 | 306 | 0 | 0 | 0 | 21 | 21 | 21 | 0 | 21 | 21 | 4647 | 0 |
| SRR3103396 | 0 | 0 | 0 | 0 | 31 | 0 | 0 | 221 | 0 | 258 | 0 | 0 | 0 | 37 | 37 | 0 | 0 | 719 | 0 | 626 | 1794 | 1853 | 0 | 82 | 7005 | 526 | 0 | 0 | 0 | 24 | 24 | 0 | 0 | 24 | 0 | 11483 | 0 |
| SRR3103397 | 0 | 0 | 23 | 0 | 20 | 0 | 0 | 314 | 70 | 244 | 0 | 0 | 0 | 0 | 0 | 0 | 0 | 491 | 0 | 140 | 1673 | 1924 | 0 | 161 | 4861 | 593 | 0 | 0 | 23 | 23 | 23 | 47 | 0 | 70 | 23 | 3350 | 0 |
| SRR3103400 | 0 | 41 | 41 | 0 | 30 | 0 | 0 | 144 | 41 | 658 | 0 | 0 | 0 | 51 | 21 | 51 | 0 | 426 | 0 | 165 | 2610 | 2956 | 0 | 131 | 10834 | 583 | 0 | 0 | 41 | 0 | 41 | 0 | 0 | 124 | 41 | 1471 | 0 |
| SRR3103401 | 0 | 0 | 0 | 0 | 75 | 0 | 0 | 472 | 34 | 640 | 0 | 0 | 0 | 67 | 0 | 135 | 0 | 433 | 0 | 67 | 2685 | 2706 | 0 | 48 | 8658 | 176 | 0 | 109 | 0 | 328 | 109 | 0 | 219 | 0 | 109 | 1685 | 0 |
| SRR3103402 | 0 | 81 | 0 | 0 | 46 | 0 | 0 | 223 | 28 | 1141 | 0 | 0 | 0 | 84 | 0 | 0 | 0 | 381 | 0 | 84 | 2636 | 2759 | 0 | 67 | 8788 | 275 | 0 | 0 | 0 | 325 | 0 | 0 | 244 | 81 | 0 | 2561 | 0 |
| SRR3103403 | 0 | 0 | 162 | 0 | 36 | 0 | 0 | 320 | 20 | 740 | 0 | 0 | 0 | 40 | 100 | 120 | 0 | 330 | 0 | 160 | 2600 | 3778 | 0 | 138 | 10951 | 174 | 0 | 0 | 0 | 323 | 108 | 0 | 54 | 108 | 54 | 1699 | 0 |
| SRR3103404 | 0 | 0 | 57 | 0 | 19 | 0 | 0 | 85 | 0 | 717 | 0 | 0 | 0 | 73 | 49 | 85 | 0 | 514 | 0 | 36 | 2590 | 3646 | 0 | 206 | 11403 | 666 | 0 | 0 | 0 | 114 | 0 | 0 | 0 | 57 | 0 | 1228 | 0 |
| SRR3103405 | 0 | 54 | 161 | 0 | 86 | 0 | 0 | 111 | 28 | 474 | 0 | 0 | 54 | 84 | 14 | 42 | 0 | 633 | 19 | 56 | 2660 | 2226 | 0 | 103 | 11646 | 189 | 0 | 0 | 0 | 161 | 268 | 0 | 107 | 0 | 0 | 683 | 0 |
| SRR3103406 | 0 | 33 | 33 | 0 | 16 | 0 | 0 | 292 | 0 | 526 | 0 | 0 | 0 | 0 | 234 | 263 | 0 | 223 | 0 | 438 | 2210 | 5388 | 0 | 211 | 9187 | 928 | 0 | 0 | 0 | 134 | 100 | 33 | 0 | 33 | 33 | 5114 | 0 |
| SRR3103407 | 0 | 58 | 0 | 0 | 31 | 0 | 0 | 90 | 70 | 1235 | 0 | 0 | 0 | 80 | 40 | 70 | 0 | 229 | 0 | 40 | 2186 | 2891 | 0 | 158 | 7203 | 390 | 0 | 0 | 0 | 58 | 0 | 0 | 0 | 0 | 117 | 603 | 0 |
| SRR3103408 | 0 | 0 | 0 | 0 | 37 | 0 | 0 | 1081 | 60 | 600 | 0 | 0 | 0 | 120 | 240 | 480 | 0 | 202 | 0 | 180 | 1848 | 3412 | 0 | 190 | 5322 | 367 | 0 | 0 | 114 | 228 | 114 | 0 | 0 | 0 | 0 | 3722 | 0 |
| SRR3103409 | 0 | 62 | 62 | 0 | 22 | 0 | 0 | 655 | 0 | 179 | 0 | 0 | 0 | 60 | 119 | 60 | 0 | 178 | 0 | 357 | 1575 | 3627 | 0 | 130 | 5051 | 417 | 0 | 0 | 0 | 62 | 0 | 0 | 124 | 0 | 0 | 3454 | 0 |
| SRR3138807 | 0 | 0 | 0 | 16 | 293 | 32 | 0 | 16501 | 4802 | 1262 | 0 | 0 | 0 | 222 | 65 | 24 | 17 | 34 | 15 | 0 | 26 | 12 | 0 | 170 | 36 | 0 | 13 | 0 | 15 | 28 | 39 | 0 | 0 | 15 | 0 | 0 | 0 |
| SRR3138808 | 0 | 0 | 0 | 0 | 214 | 15 | 0 | 15782 | 7566 | 1514 | 0 | 0 | 0 | 229 | 0 | 0 | 19 | 15 | 13 | 0 | 20 | 0 | 0 | 104 | 27 | 17 | 19 | 0 | 0 | 13 | 22 | 0 | 0 | 0 | 0 | 0 | 0 |
| SRR3138809 | 0 | 0 | 0 | 0 | 211 | 19 | 0 | 14646 | 7389 | 1480 | 0 | 0 | 0 | 252 | 0 | 0 | 20 | 23 | 11 | 0 | 15 | 0 | 0 | 79 | 26 | 17 | 20 | 0 | 11 | 0 | 19 | 0 | 0 | 0 | 0 | 0 | 0 |
| SRR3138810 | 0 | 0 | 0 | 12 | 279 | 150 | 0 | 13448 | 3636 | 1354 | 0 | 0 | 0 | 267 | 93 | 41 | 29 | 29 | 11 | 0 | 17 | 0 | 0 | 341 | 55 | 16 | 32 | 0 | 12 | 0 | 21 | 0 | 0 | 0 | 0 | 0 | 0 |
| SRR3199254 | 0 | 211 | 63 | 158 | 18 | 0 | 0 | 256 | 240 | 2706 | 8458 | 10551 | 0 | 297 | 202 | 420 | 11 | 432 | 54 | 308 | 545 | 1446 | 0 | 1819 | 8058 | 16 | 0 | 113 | 45 | 252 | 338 | 86 | 73 | 36 | 0 | 316 | 5411 |
| SRR3199255 | 0 | 75 | 18 | 0 | 0 | 0 | 55 | 279 | 239 | 2173 | 14860 | 24611 | 0 | 231 | 15 | 53 | 42 | 117 | 108 | 86 | 542 | 3329 | 19 | 11711 | 8883 | 24 | 11 | 80 | 73 | 27 | 23 | 53 | 69 | 55 | 0 | 193 | 4792 |
| SRR3199256 | 0 | 187 | 41 | 49 | 0 | 0 | 12 | 45 | 110 | 897 | 3252 | 1885 | 0 | 270 | 180 | 70 | 17 | 92 | 175 | 68 | 582 | 1620 | 29 | 5450 | 6150 | 0 | 0 | 1829 | 340 | 319 | 286 | 37 | 37 | 70 | 0 | 98 | 2993 |
| SRR3199257 | 10 | 30 | 50 | 0 | 0 | 0 | 70 | 457 | 123 | 334 | 359 | 7000 | 0 | 33 | 473 | 501 | 0 | 10 | 29 | 22 | 212 | 1473 | 18 | 5418 | 3965 | 0 | 50 | 341 | 311 | 1334 | 251 | 150 | 241 | 191 | 0 | 11 | 109 |
| SRR3199258 | 0 | 31 | 25 | 0 | 0 | 0 | 84 | 103 | 114 | 754 | 302 | 560 | 0 | 62 | 65 | 32 | 42 | 0 | 45 | 26 | 382 | 1054 | 20 | 4162 | 4771 | 0 | 45 | 109 | 84 | 89 | 103 | 187 | 53 | 45 | 0 | 21 | 46 |
| SRR3199259 | 0 | 16 | 17 | 0 | 0 | 0 | 160 | 1035 | 965 | 1102 | 174 | 748 | 0 | 112 | 20 | 0 | 33 | 24 | 45 | 53 | 370 | 1118 | 25 | 3380 | 4219 | 14 | 32 | 11 | 25 | 44 | 46 | 56 | 49 | 44 | 0 | 22 | 68 |
| SRR3199260 | 0 | 18 | 15 | 0 | 0 | 0 | 48 | 527 | 327 | 1715 | 947 | 2216 | 0 | 86 | 20 | 12 | 24 | 27 | 41 | 0 | 504 | 895 | 34 | 4347 | 6378 | 0 | 19 | 0 | 84 | 126 | 214 | 113 | 92 | 87 | 0 | 0 | 79 |
| SRR3199261 | 0 | 37 | 0 | 0 | 0 | 0 | 104 | 705 | 586 | 3577 | 168 | 667 | 0 | 321 | 101 | 25 | 31 | 111 | 18 | 41 | 349 | 570 | 40 | 2006 | 2791 | 0 | 24 | 34 | 24 | 37 | 31 | 58 | 37 | 18 | 0 | 0 | 152 |
| SRR3199262 | 0 | 0 | 0 | 0 | 0 | 0 | 144 | 1508 | 809 | 3371 | 1423 | 1488 | 0 | 240 | 85 | 25 | 27 | 72 | 16 | 30 | 275 | 505 | 50 | 2394 | 3280 | 0 | 78 | 16 | 47 | 43 | 86 | 183 | 70 | 62 | 0 | 0 | 86 |
| SRR3199263 | 0 | 178 | 44 | 351 | 0 | 0 | 44 | 75 | 107 | 144 | 1673 | 9026 | 0 | 48 | 0 | 48 | 89 | 81 | 106 | 27 | 419 | 348 | 15 | 2882 | 2843 | 16 | 89 | 1201 | 712 | 12 | 1379 | 178 | 178 | 178 | 0 | 48 | 0 |
| SRR3235777 | 0 | 23 | 278 | 193 | 222 | 0 | 0 | 527 | 2957 | 2422 | 5396 | 11820 | 0 | 1666 | 153 | 467 | 0 | 2717 | 361 | 6297 | 192 | 1411 | 81 | 15783 | 4202 | 65 | 100 | 402 | 660 | 1375 | 1641 | 344 | 266 | 320 | 35 | 7894 | 201 |
| SRR3235778 | 0 | 42 | 1184 | 153 | 256 | 0 | 0 | 1072 | 3121 | 6146 | 138 | 8401 | 15 | 729 | 261 | 701 | 0 | 1858 | 500 | 8951 | 200 | 776 | 42 | 9534 | 2834 | 129 | 15 | 218 | 1406 | 1495 | 2181 | 471 | 598 | 732 | 15 | 9322 | 270 |
| SRR3235779 | 0 | 12 | 404 | 137 | 255 | 0 | 0 | 737 | 3006 | 3667 | 1278 | 6833 | 0 | 280 | 172 | 737 | 0 | 2338 | 500 | 22977 | 923 | 609 | 188 | 23436 | 5026 | 73 | 65 | 818 | 2003 | 2165 | 5573 | 1030 | 847 | 1016 | 92 | 26670 | 930 |
| SRR3235780 | 0 | 19 | 537 | 228 | 437 | 13 | 0 | 1590 | 5501 | 6632 | 168 | 7421 | 16 | 200 | 895 | 2921 | 0 | 3677 | 24 | 11178 | 334 | 747 | 184 | 26279 | 5275 | 62 | 35 | 154 | 1549 | 757 | 1692 | 391 | 383 | 407 | 24 | 13275 | 256 |
| SRR3235781 | 0 | 38 | 20 | 121 | 352 | 0 | 0 | 768 | 3737 | 3130 | 1954 | 7900 | 0 | 85 | 370 | 1517 | 0 | 2229 | 115 | 48784 | 683 | 891 | 242 | 29473 | 4424 | 13 | 31 | 18 | 89 | 52 | 160 | 30 | 25 | 32 | 10 | 32168 | 312 |
| SRR3279553 | 0 | 0 | 0 | 0 | 237 | 0 | 0 | 300 | 0 | 900 | 0 | 0 | 0 | 0 | 0 | 0 | 0 | 237 | 0 | 300 | 0 | 473 | 0 | 947 | 710 | 0 | 0 | 0 | 0 | 0 | 0 | 0 | 0 | 0 | 0 | 1200 | 0 |
| SRR3279554 | 0 | 59 | 0 | 0 | 88 | 0 | 29 | 1220 | 390 | 2390 | 49 | 0 | 0 | 49 | 195 | 0 | 0 | 176 | 0 | 98 | 29 | 147 | 59 | 411 | 793 | 0 | 29 | 29 | 0 | 59 | 0 | 29 | 0 | 0 | 0 | 49 | 0 |
| SRR3279555 | 0 | 0 | 0 | 0 | 0 | 0 | 0 | 174 | 0 | 174 | 350 | 58 | 0 | 0 | 0 | 58 | 0 | 1345 | 0 | 116 | 336 | 168 | 0 | 1345 | 1682 | 0 | 0 | 0 | 0 | 0 | 0 | 0 | 0 | 0 | 0 | 174 | 252 |
| SRR3279556 | 0 | 68 | 0 | 0 | 52 | 0 | 0 | 210 | 0 | 93 | 47 | 47 | 0 | 23 | 93 | 70 | 10 | 1015 | 0 | 140 | 338 | 143 | 26 | 677 | 2291 | 26 | 14 | 20 | 0 | 0 | 0 | 0 | 0 | 0 | 0 | 583 | 117 |
| SRR3279557 | 0 | 103 | 0 | 0 | 21 | 0 | 11 | 713 | 127 | 949 | 12 | 41 | 0 | 22 | 46 | 52 | 0 | 348 | 21 | 106 | 391 | 242 | 18 | 1703 | 3739 | 78 | 55 | 17 | 20 | 37 | 22 | 17 | 0 | 0 | 0 | 415 | 103 |
| SRR3279558 | 0 | 96 | 0 | 0 | 0 | 0 | 0 | 91 | 0 | 272 | 91 | 0 | 0 | 0 | 363 | 0 | 0 | 151 | 75 | 181 | 414 | 226 | 0 | 1770 | 3427 | 75 | 19 | 0 | 0 | 39 | 19 | 0 | 0 | 0 | 0 | 363 | 113 |
| SRR3279559 | 0 | 24 | 0 | 0 | 0 | 0 | 0 | 2097 | 445 | 1723 | 158 | 57 | 0 | 0 | 115 | 14 | 0 | 184 | 0 | 0 | 105 | 112 | 26 | 1122 | 1435 | 56 | 0 | 18 | 12 | 36 | 0 | 0 | 0 | 0 | 0 | 0 | 49 |
| SRR3279560 | 0 | 0 | 0 | 0 | 19 | 0 | 0 | 1068 | 114 | 667 | 57 | 114 | 0 | 114 | 76 | 38 | 0 | 343 | 0 | 210 | 139 | 195 | 56 | 1179 | 1540 | 19 | 12 | 12 | 0 | 37 | 0 | 0 | 25 | 0 | 0 | 496 | 102 |
| SRR3279561 | 0 | 0 | 0 | 0 | 0 | 0 | 0 | 241 | 29 | 300 | 0 | 0 | 0 | 34 | 221 | 197 | 12 | 242 | 25 | 128 | 346 | 267 | 49 | 3387 | 2715 | 0 | 12 | 20 | 0 | 0 | 0 | 16 | 0 | 0 | 0 | 275 | 173 |
| SRR3279562 | 0 | 0 | 0 | 0 | 20 | 0 | 0 | 1039 | 168 | 503 | 151 | 302 | 0 | 17 | 1425 | 821 | 0 | 174 | 20 | 218 | 204 | 163 | 0 | 347 | 2123 | 51 | 0 | 0 | 12 | 19 | 0 | 0 | 12 | 0 | 0 | 570 | 184 |
| SRR3279563 | 0 | 0 | 0 | 0 | 0 | 0 | 0 | 1533 | 307 | 690 | 78 | 307 | 0 | 153 | 383 | 153 | 0 | 370 | 0 | 153 | 154 | 216 | 62 | 1571 | 1386 | 92 | 0 | 12 | 0 | 0 | 0 | 0 | 0 | 0 | 0 | 613 | 277 |
| SRR3279564 | 0 | 14 | 0 | 0 | 30 | 0 | 0 | 403 | 68 | 447 | 0 | 10 | 0 | 19 | 61 | 34 | 0 | 124 | 25 | 51 | 249 | 154 | 35 | 1056 | 1922 | 45 | 0 | 0 | 0 | 0 | 0 | 0 | 0 | 0 | 0 | 126 | 95 |
| SRR3279565 | 0 | 48 | 0 | 0 | 12 | 0 | 0 | 783 | 69 | 634 | 81 | 173 | 0 | 0 | 1820 | 1751 | 0 | 517 | 24 | 369 | 409 | 204 | 84 | 806 | 2044 | 48 | 0 | 0 | 0 | 0 | 0 | 0 | 0 | 11 | 0 | 2247 | 385 |
| SRR3279566 | 0 | 18 | 0 | 0 | 41 | 0 | 0 | 251 | 0 | 293 | 209 | 84 | 0 | 0 | 126 | 0 | 0 | 902 | 0 | 126 | 451 | 164 | 82 | 328 | 1968 | 41 | 0 | 0 | 0 | 0 | 11 | 0 | 0 | 0 | 0 | 293 | 41 |
| SRR3279567 | 0 | 11 | 0 | 0 | 17 | 0 | 0 | 45 | 0 | 352 | 99 | 0 | 0 | 0 | 18 | 0 | 0 | 29 | 55 | 108 | 170 | 129 | 48 | 1690 | 8000 | 24 | 18 | 17 | 48 | 40 | 22 | 27 | 0 | 0 | 0 | 199 | 184 |
| SRR3279568 | 0 | 28 | 0 | 0 | 0 | 0 | 14 | 1034 | 49 | 1206 | 0 | 123 | 0 | 0 | 443 | 197 | 14 | 188 | 0 | 197 | 102 | 273 | 17 | 154 | 11 | 0 | 0 | 0 | 0 | 0 | 0 | 0 | 0 | 0 | 0 | 418 | 51 |
| SRR3279569 | 0 | 0 | 0 | 0 | 19 | 0 | 0 | 1172 | 146 | 934 | 0 | 37 | 0 | 128 | 165 | 37 | 16 | 192 | 19 | 92 | 154 | 249 | 19 | 557 | 1305 | 19 | 0 | 0 | 0 | 0 | 0 | 0 | 0 | 0 | 0 | 238 | 96 |
| SRR3279570 | 0 | 0 | 0 | 0 | 19 | 0 | 0 | 302 | 0 | 201 | 103 | 352 | 0 | 50 | 201 | 453 | 0 | 103 | 19 | 453 | 75 | 282 | 19 | 338 | 1173 | 0 | 0 | 0 | 0 | 0 | 0 | 0 | 0 | 0 | 0 | 905 | 66 |
| SRR3279571 | 0 | 14 | 0 | 0 | 24 | 0 | 0 | 3162 | 435 | 1506 | 50 | 17 | 0 | 84 | 67 | 50 | 0 | 163 | 0 | 117 | 54 | 97 | 18 | 368 | 1074 | 24 | 0 | 0 | 0 | 14 | 0 | 27 | 0 | 0 | 0 | 318 | 24 |

Supp Table S8.4.Columns_DI_to_ES

| | | | | | | | | | | | | | | | | | | | | | | | | | | | | | | | | | | | | | |
|---|---|---|---|---|---|---|---|---|---|---|---|---|---|---|---|---|---|---|---|---|---|---|---|---|---|---|---|---|---|---|---|---|---|---|---|---|---|
| SRR3279572 | 0 | 0 | 0 | 0 | 13 | 0 | 0 | 1370 | 202 | 810 | 22 | 40 | 0 | 0 | 1213 | 421 | 0 | 119 | 13 | 90 | 297 | 271 | 52 | 2132 | 1778 | 0 | 0 | 39 | 22 | 45 | 0 | 0 | 11 | 0 | 0 | 269 | 119 |
| SRR3279573 | 0 | 11 | 0 | 0 | 39 | 0 | 0 | 220 | 18 | 257 | 111 | 92 | 0 | 18 | 18 | 55 | 0 | 283 | 47 | 147 | 157 | 189 | 55 | 495 | 1974 | 39 | 0 | 0 | 0 | 0 | 0 | 0 | 0 | 0 | 0 | 1138 | 165 |
| SRR3279574 | 0 | 0 | 0 | 0 | 23 | 0 | 0 | 235 | 23 | 258 | 71 | 446 | 0 | 94 | 2396 | 1832 | 0 | 749 | 0 | 235 | 328 | 140 | 0 | 1358 | 1639 | 0 | 0 | 0 | 0 | 0 | 0 | 0 | 0 | 0 | 0 | 963 | 164 |
| SRR3279575 | 0 | 12 | 0 | 0 | 50 | 0 | 0 | 336 | 36 | 509 | 10 | 15 | 0 | 15 | 143 | 199 | 0 | 580 | 14 | 178 | 231 | 195 | 54 | 780 | 2104 | 0 | 13 | 0 | 0 | 32 | 13 | 21 | 0 | 0 | 0 | 825 | 163 |
| SRR3279576 | 0 | 71 | 0 | 0 | 42 | 0 | 12 | 147 | 13 | 281 | 26 | 38 | 0 | 19 | 89 | 121 | 12 | 482 | 17 | 128 | 274 | 200 | 42 | 624 | 2012 | 33 | 0 | 0 | 0 | 0 | 0 | 0 | 0 | 0 | 0 | 574 | 125 |
| SRR3279577 | 0 | 0 | 0 | 0 | 21 | 0 | 12 | 1345 | 227 | 1142 | 41 | 122 | 0 | 65 | 356 | 105 | 0 | 210 | 0 | 275 | 242 | 273 | 105 | 1336 | 2525 | 11 | 20 | 0 | 0 | 0 | 12 | 0 | 0 | 0 | 0 | 794 | 95 |
| SRR3279578 | 0 | 13 | 0 | 0 | 36 | 0 | 0 | 1149 | 151 | 233 | 164 | 383 | 0 | 14 | 109 | 27 | 0 | 575 | 23 | 192 | 210 | 524 | 23 | 492 | 5271 | 0 | 0 | 10 | 0 | 0 | 0 | 0 | 0 | 0 | 0 | 547 | 68 |
| SRR3279579 | 0 | 0 | 0 | 0 | 52 | 0 | 0 | 862 | 164 | 821 | 125 | 82 | 0 | 123 | 2300 | 1807 | 0 | 732 | 0 | 123 | 157 | 366 | 0 | 366 | 2457 | 105 | 0 | 0 | 0 | 15 | 0 | 0 | 0 | 0 | 0 | 1273 | 52 |
| SRR3279580 | 0 | 14 | 0 | 0 | 37 | 0 | 12 | 279 | 31 | 417 | 32 | 24 | 0 | 0 | 51 | 12 | 0 | 189 | 0 | 67 | 213 | 158 | 58 | 935 | 2219 | 0 | 21 | 12 | 0 | 0 | 0 | 0 | 0 | 0 | 0 | 181 | 58 |
| SRR3279581 | 0 | 17 | 0 | 0 | 0 | 0 | 0 | 422 | 60 | 572 | 25 | 60 | 0 | 0 | 899 | 445 | 0 | 742 | 0 | 122 | 247 | 1171 | 52 | 651 | 13077 | 0 | 17 | 30 | 0 | 17 | 30 | 0 | 0 | 0 | 0 | 353 | 833 |
| SRR3279582 | 0 | 0 | 0 | 0 | 23 | 0 | 0 | 1463 | 206 | 1365 | 17 | 49 | 0 | 90 | 551 | 206 | 0 | 462 | 0 | 25 | 158 | 146 | 11 | 2713 | 2026 | 45 | 0 | 0 | 0 | 0 | 0 | 11 | 0 | 0 | 0 | 148 | 23 |
| SRR3279583 | 0 | 23 | 0 | 0 | 0 | 0 | 14 | 321 | 64 | 305 | 81 | 129 | 0 | 0 | 1671 | 803 | 0 | 443 | 0 | 289 | 353 | 166 | 14 | 443 | 11 | 48 | 0 | 0 | 0 | 18 | 23 | 0 | 0 | 0 | 0 | 739 | 249 |
| SRR3279584 | 0 | 16 | 0 | 0 | 34 | 0 | 0 | 250 | 51 | 511 | 193 | 163 | 0 | 20 | 32114 | 19768 | 0 | 845 | 21 | 604 | 1034 | 671 | 74 | 1691 | 24 | 29 | 0 | 0 | 13 | 19 | 13 | 0 | 0 | 0 | 0 | 1088 | 194 |
| SRR3279585 | 0 | 11 | 0 | 0 | 0 | 0 | 0 | 162 | 32 | 356 | 65 | 162 | 0 | 32 | 4267 | 3976 | 0 | 352 | 32 | 65 | 96 | 128 | 32 | 416 | 1151 | 0 | 0 | 0 | 0 | 0 | 0 | 0 | 0 | 0 | 0 | 582 | 64 |
| SRR3279586 | 0 | 0 | 0 | 0 | 64 | 0 | 0 | 1178 | 41 | 528 | 82 | 122 | 0 | 41 | 325 | 284 | 0 | 544 | 0 | 122 | 224 | 320 | 32 | 1537 | 2146 | 0 | 0 | 0 | 0 | 0 | 0 | 0 | 0 | 0 | 0 | 731 | 64 |
| SRR3317221 | 0 | 41 | 0 | 0 | 854 | 0 | 0 | 408 | 1222 | 2445 | 8557 | 13040 | 0 | 0 | 0 | 0 | 0 | 113 | 89 | 0 | 175 | 136 | 0 | 111 | 631 | 0 | 0 | 143 | 68 | 11 | 22 | 179 | 36 | 36 | 0 | 1630 | 806 |
| SRR3477269 | 0 | 0 | 0 | 0 | 0 | 0 | 0 | 505 | 94 | 115 | 0 | 0 | 0 | 185 | 0 | 0 | 0 | 0 | 0 | 0 | 0 | 262 | 0 | 1837 | 1575 | 0 | 0 | 0 | 0 | 0 | 0 | 0 | 0 | 0 | 0 | 0 | 0 |
| SRR3477271 | 0 | 0 | 0 | 0 | 0 | 0 | 0 | 280 | 49 | 51 | 0 | 0 | 0 | 151 | 0 | 0 | 0 | 0 | 0 | 0 | 140 | 280 | 0 | 1260 | 1120 | 0 | 0 | 0 | 0 | 0 | 0 | 0 | 0 | 0 | 0 | 0 | 0 |
| SRR3490374 | 0 | 0 | 63 | 0 | 0 | 0 | 0 | 0 | 148 | 4883 | 0 | 0 | 0 | 2220 | 3551 | 1924 | 0 | 136 | 0 | 3107 | 3723 | 2154 | 692 | 4515 | 1569 | 78 | 0 | 0 | 0 | 0 | 0 | 0 | 63 | 0 | 0 | 5327 | 642 |
| SRR3490375 | 0 | 0 | 0 | 0 | 0 | 0 | 0 | 0 | 0 | 296 | 0 | 0 | 0 | 593 | 2074 | 1185 | 0 | 96 | 0 | 2074 | 1248 | 514 | 209 | 7082 | 696 | 0 | 0 | 0 | 0 | 0 | 0 | 0 | 0 | 0 | 0 | 3852 | 0 |
| SRR3490376 | 0 | 0 | 59 | 0 | 0 | 0 | 0 | 156 | 52 | 3070 | 0 | 0 | 0 | 1457 | 468 | 312 | 0 | 199 | 0 | 5516 | 1178 | 1594 | 738 | 8316 | 1515 | 0 | 0 | 0 | 0 | 15 | 0 | 0 | 0 | 0 | 0 | 13635 | 0 |
| SRR3490377 | 0 | 0 | 0 | 0 | 0 | 0 | 0 | 199 | 0 | 943 | 0 | 0 | 0 | 298 | 496 | 447 | 0 | 132 | 0 | 7097 | 616 | 1794 | 1106 | 8460 | 1985 | 0 | 0 | 0 | 0 | 0 | 0 | 0 | 0 | 0 | 0 | 12557 | 0 |
| SRR3490378 | 0 | 0 | 0 | 0 | 0 | 48 | 24 | 172 | 0 | 430 | 0 | 0 | 0 | 86 | 344 | 172 | 0 | 46 | 0 | 6106 | 758 | 2157 | 1936 | 15199 | 2740 | 0 | 0 | 0 | 0 | 0 | 24 | 0 | 24 | 0 | 24 | 13932 | 18 |
| SRR3490379 | 0 | 0 | 0 | 0 | 0 | 0 | 0 | 392 | 0 | 5096 | 0 | 0 | 0 | 392 | 3332 | 1176 | 0 | 88 | 0 | 2548 | 506 | 1732 | 1462 | 19798 | 2872 | 0 | 0 | 0 | 0 | 0 | 92 | 0 | 0 | 0 | 92 | 7448 | 0 |
| SRR3490380 | 0 | 0 | 0 | 0 | 0 | 43 | 0 | 156 | 0 | 311 | 0 | 0 | 0 | 156 | 467 | 156 | 0 | 71 | 0 | 5761 | 740 | 1714 | 1514 | 12692 | 2014 | 0 | 0 | 0 | 0 | 0 | 0 | 0 | 0 | 0 | 0 | 10587 | 0 |
| SRR3490382 | 0 | 0 | 0 | 12 | 0 | 0 | 0 | 0 | 0 | 6264 | 0 | 0 | 0 | 1139 | 0 | 569 | 0 | 386 | 0 | 2563 | 982 | 596 | 140 | 1123 | 784 | 0 | 0 | 0 | 0 | 0 | 0 | 0 | 0 | 202 | 403 | 3132 | 94 |
| SRR3490383 | 0 | 0 | 13 | 0 | 15 | 13 | 0 | 215 | 215 | 1075 | 0 | 0 | 0 | 215 | 430 | 215 | 0 | 201 | 0 | 6662 | 943 | 1449 | 308 | 2317 | 2374 | 0 | 0 | 13 | 0 | 13 | 25 | 0 | 0 | 0 | 13 | 30518 | 30 |
| SRR3490384 | 0 | 0 | 186 | 0 | 21 | 0 | 0 | 454 | 181 | 6259 | 0 | 0 | 0 | 1361 | 0 | 91 | 0 | 351 | 0 | 998 | 1759 | 1223 | 333 | 2308 | 3401 | 0 | 0 | 47 | 0 | 0 | 0 | 0 | 0 | 0 | 47 | 8617 | 18 |
| SRR3490385 | 0 | 51 | 51 | 0 | 0 | 0 | 0 | 658 | 658 | 2303 | 0 | 0 | 0 | 0 | 329 | 658 | 0 | 286 | 0 | 8226 | 570 | 1301 | 382 | 3548 | 2814 | 0 | 0 | 0 | 0 | 101 | 101 | 0 | 0 | 304 | 51 | 34551 | 34 |
| SRR3490386 | 0 | 50 | 0 | 0 | 0 | 0 | 0 | 264 | 396 | 3697 | 0 | 0 | 0 | 2377 | 0 | 132 | 0 | 148 | 0 | 3829 | 928 | 1530 | 271 | 3017 | 3040 | 0 | 0 | 0 | 50 | 50 | 0 | 0 | 0 | 0 | 0 | 21653 | 38 |
| SRR3490387 | 0 | 73 | 109 | 0 | 0 | 0 | 0 | 529 | 0 | 6267 | 0 | 0 | 0 | 831 | 151 | 76 | 0 | 430 | 0 | 2114 | 1556 | 1859 | 811 | 5330 | 2463 | 19 | 0 | 0 | 0 | 36 | 73 | 0 | 109 | 73 | 109 | 9968 | 37 |
| SRR3490388 | 0 | 0 | 200 | 15 | 0 | 0 | 0 | 454 | 454 | 454 | 0 | 0 | 0 | 1362 | 0 | 0 | 0 | 380 | 0 | 1362 | 2063 | 1112 | 263 | 3248 | 366 | 0 | 0 | 0 | 0 | 0 | 0 | 0 | 0 | 0 | 200 | 4541 | 0 |
| SRR3490389 | 0 | 0 | 153 | 0 | 13 | 0 | 0 | 165 | 494 | 1153 | 0 | 0 | 0 | 823 | 165 | 0 | 0 | 160 | 13 | 8564 | 1185 | 529 | 125 | 7820 | 595 | 0 | 0 | 77 | 0 | 77 | 0 | 0 | 230 | 0 | 0 | 20916 | 10 |
| SRR3490390 | 0 | 0 | 12 | 22 | 0 | 65 | 0 | 0 | 274 | 3150 | 0 | 0 | 0 | 3561 | 411 | 274 | 0 | 196 | 0 | 2876 | 1462 | 1571 | 87 | 1506 | 109 | 22 | 0 | 0 | 0 | 0 | 0 | 0 | 0 | 0 | 0 | 11231 | 22 |
| SRR3490391 | 0 | 0 | 0 | 0 | 0 | 0 | 0 | 0 | 0 | 3176 | 0 | 0 | 0 | 3176 | 1361 | 3630 | 0 | 143 | 12 | 907 | 940 | 440 | 285 | 3414 | 856 | 0 | 0 | 0 | 0 | 0 | 0 | 0 | 0 | 0 | 0 | 2722 | 12 |
| SRR3490392 | 0 | 34 | 17 | 0 | 0 | 0 | 0 | 262 | 52 | 1782 | 0 | 0 | 0 | 2201 | 52 | 52 | 0 | 275 | 0 | 3564 | 779 | 1237 | 607 | 6311 | 987 | 0 | 0 | 0 | 0 | 17 | 0 | 0 | 0 | 17 | 17 | 7756 | 0 |
| SRR3490393 | 0 | 0 | 117 | 0 | 16 | 0 | 0 | 348 | 174 | 8516 | 0 | 0 | 0 | 1738 | 58 | 116 | 0 | 375 | 0 | 1217 | 2394 | 1717 | 619 | 5850 | 2643 | 11 | 0 | 0 | 0 | 23 | 23 | 0 | 0 | 23 | 70 | 5967 | 11 |
| SRR3490394 | 0 | 33 | 133 | 0 | 0 | 0 | 0 | 41 | 0 | 11738 | 0 | 0 | 0 | 2157 | 415 | 456 | 0 | 396 | 0 | 995 | 2325 | 1685 | 773 | 5641 | 2153 | 0 | 0 | 0 | 0 | 100 | 0 | 33 | 100 | 66 | 0 | 3526 | 14 |
| SRR3493372 | 0 | 0 | 0 | 253 | 423 | 0 | 0 | 204 | 505 | 4118 | 0 | 0 | 0 | 1332 | 16 | 19 | 11 | 1608 | 123 | 0 | 374 | 252 | 84 | 5540 | 762 | 19 | 13 | 0 | 0 | 0 | 0 | 0 | 0 | 0 | 0 | 14 | 1333 |
| SRR3493373 | 0 | 0 | 0 | 294 | 427 | 13 | 0 | 1628 | 3280 | 8701 | 36 | 30 | 0 | 1821 | 0 | 0 | 15 | 1347 | 289 | 27 | 166 | 84 | 86 | 7793 | 981 | 38 | 23 | 0 | 0 | 0 | 0 | 0 | 0 | 0 | 0 | 33 | 723 |
| SRR3534933 | 27 | 37 | 0 | 0 | 25 | 0 | 27 | 7257 | 540 | 1790 | 12563 | 9477 | 0 | 236 | 0 | 21 | 0 | 191 | 52 | 876 | 976 | 2241 | 0 | 3250 | 6543 | 0 | 0 | 12 | 126 | 22 | 17 | 62 | 16 | 33 | 71 | 4118 | 108 |
| SRR3534934 | 0 | 54 | 12 | 0 | 30 | 0 | 12 | 4055 | 1110 | 3232 | 12622 | 9081 | 0 | 612 | 16 | 20 | 0 | 180 | 52 | 1165 | 967 | 2101 | 0 | 2630 | 6238 | 0 | 23 | 17 | 168 | 25 | 19 | 89 | 17 | 38 | 51 | 6318 | 99 |
| SRR3534935 | 0 | 30 | 25 | 0 | 19 | 0 | 0 | 1918 | 247 | 2485 | 24932 | 18747 | 0 | 486 | 0 | 0 | 0 | 293 | 12 | 2454 | 1198 | 3477 | 16 | 8940 | 7863 | 215 | 0 | 281 | 76 | 216 | 217 | 47 | 197 | 496 | 63 | 7828 | 131 |
| SRR3534936 | 18 | 54 | 31 | 0 | 18 | 0 | 0 | 1206 | 209 | 2711 | 28992 | 13312 | 0 | 441 | 0 | 0 | 0 | 292 | 0 | 4243 | 1267 | 3828 | 13 | 9851 | 8168 | 301 | 0 | 25 | 78 | 12 | 17 | 65 | 13 | 36 | 78 | 9214 | 145 |
| SRR3534937 | 16 | 31 | 54 | 0 | 27 | 0 | 0 | 9250 | 595 | 2747 | 34464 | 16038 | 0 | 354 | 11 | 19 | 0 | 346 | 13 | 1860 | 1454 | 3896 | 25 | 12166 | 7242 | 171 | 0 | 36 | 81 | 10 | 12 | 25 | 98 | 21 | 10 | 6085 | 245 |
| SRR3534938 | 33 | 43 | 69 | 0 | 25 | 13 | 10 | 3269 | 383 | 2361 | 23251 | 14170 | 0 | 286 | 0 | 0 | 0 | 343 | 12 | 4510 | 1440 | 3964 | 23 | 12837 | 7123 | 183 | 0 | 36 | 114 | 189 | 298 | 35 | 158 | 455 | 98 | 9166 | 179 |
| SRR3566024 | 0 | 42 | 0 | 0 | 13 | 0 | 0 | 191 | 764 | 9121 | 78271 | 60888 | 0 | 716 | 0 | 0 | 0 | 168 | 714 | 573 | 2926 | 12114 | 0 | 3534 | 8274 | 0 | 17 | 25 | 12 | 23 | 34 | 11 | 16 | 18 | 0 | 907 | 2780 |
| SRR3566026 | 0 | 23 | 0 | 0 | 0 | 0 | 19 | 406 | 343 | 6022 | 3921 | 52357 | 0 | 343 | 0 | 94 | 0 | 175 | 597 | 437 | 2537 | 12142 | 14 | 3543 | 8030 | 0 | 19 | 19 | 13 | 29 | 23 | 18 | 15 | 28 | 38 | 1092 | 2681 |
| SRR3566150 | 0 | 146 | 21 | 0 | 0 | 16 | 0 | 506 | 384 | 3237 | 5413 | 67676 | 0 | 344 | 0 | 20 | 42 | 123 | 465 | 587 | 2857 | 12610 | 0 | 3510 | 8697 | 0 | 0 | 10 | 94 | 10 | 10 | 31 | 0 | 31 | 0 | 850 | 2177 |
| SRR3566151 | 0 | 109 | 0 | 0 | 0 | 27 | 16 | 297 | 297 | 5100 | 3956 | 70068 | 0 | 480 | 0 | 46 | 16 | 95 | 432 | 892 | 2502 | 12944 | 0 | 3332 | 8530 | 0 | 11 | 0 | 11 | 27 | 33 | 33 | 44 | 33 | 0 | 572 | 1874 |
| SRR3566152 | 0 | 62 | 0 | 0 | 0 | 39 | 16 | 223 | 255 | 4178 | 101997 | 82509 | 0 | 32 | 0 | 32 | 39 | 174 | 435 | 191 | 3099 | 13693 | 0 | 3875 | 8118 | 0 | 0 | 0 | 23 | 54 | 16 | 0 | 0 | 0 | 0 | 319 | 2342 |
| SRR3566153 | 0 | 39 | 0 | 0 | 0 | 0 | 0 | 83 | 111 | 2558 | 128155 | 84316 | 0 | 278 | 0 | 56 | 20 | 198 | 140 | 334 | 2796 | 12259 | 23 | 3057 | 7522 | 0 | 13 | 13 | 0 | 0 | 40 | 0 | 13 | 13 | 20 | 306 | 2406 |
| SRR3566154 | 0 | 30 | 0 | 0 | 0 | 0 | 0 | 301 | 0 | 4216 | 168 | 68654 | 0 | 301 | 0 | 0 | 0 | 0 | 78 | 2108 | 2733 | 11478 | 0 | 5466 | 8277 | 0 | 0 | 0 | 0 | 0 | 0 | 0 | 0 | 0 | 0 | 903 | 1562 |
| SRR3566155 | 0 | 318 | 0 | 0 | 0 | 212 | 0 | 701 | 234 | 2103 | 309 | 77355 | 0 | 0 | 0 | 0 | 0 | 0 | 0 | 0 | 2546 | 13667 | 67 | 5427 | 8442 | 0 | 0 | 0 | 0 | 0 | 0 | 0 | 0 | 0 | 0 | 701 | 2546 |
| SRR3582392 | 0 | 106 | 0 | 0 | 0 | 0 | 0 | 141 | 352 | 14347 | 515 | 38470 | 0 | 281 | 70 | 0 | 0 | 267 | 845 | 492 | 2956 | 13480 | 0 | 4119 | 10621 | 0 | 0 | 0 | 88 | 212 | 194 | 88 | 106 | 88 | 53 | 914 | 3194 |
| SRR3582393 | 0 | 13 | 0 | 0 | 0 | 0 | 0 | 396 | 1266 | 15270 | 43279 | 42488 | 0 | 554 | 0 | 0 | 19 | 184 | 752 | 396 | 2863 | 12932 | 15 | 3974 | 10093 | 0 | 0 | 39 | 16 | 32 | 24 | 16 | 39 | 26 | 19 | 1029 | 3261 |
| SRR3623374 | 10 | 1410 | 0 | 0 | 0 | 10 | 10 | 0 | 17 | 0 | 8414 | 13991 | 10 | 0 | 26 | 43 | 10 | 619 | 15 | 43 | 333 | 143 | 0 | 19 | 833 | 286 | 62 | 0 | 0 | 0 | 0 | 0 | 0 | 0 | 0 | 69 | 0 |
| SRR3624069 | 0 | 1034 | 20 | 478 | 1399 | 24 | 18 | 1224 | 1086 | 2066 | 152 | 1051 | 0 | 520 | 874 | 1984 | 68 | 1620 | 22 | 1826 | 562 | 633 | 31 | 986 | 1795 | 74 | 278 | 70 | 118 | 64 | 386 | 22 | 18 | 28 | 11 | 1787 | 274 |
| SRR3624070 | 0 | 273 | 10 | 301 | 867 | 0 | 13 | 875 | 2228 | 5146 | 722 | 1074 | 0 | 747 | 1364 | 2385 | 23 | 1209 | 17 | 2111 | 532 | 591 | 33 | 1002 | 1622 | 56 | 27 | 35 | 60 | 38 | 354 | 23 | 15 | 20 | 14 | 1502 | 379 |
| SRR3624071 | 0 | 271 | 15 | 331 | 665 | 0 | 20 | 183 | 507 | 3111 | 500 | 2221 | 0 | 668 | 1158 | 1807 | 19 | 1397 | 20 | 2644 | 439 | 734 | 29 | 936 | 1752 | 56 | 35 | 35 | 66 | 47 | 432 | 25 | 21 | 19 | 15 | 2692 | 454 |
| SRR3624072 | 0 | 138 | 11 | 206 | 655 | 0 | 0 | 230 | 330 | 5202 | 185 | 990 | 0 | 660 | 2330 | 3634 | 19 | 996 | 20 | 2985 | 298 | 408 | 19 | 645 | 1347 | 36 | 15 | 17 | 51 | 52 | 588 | 40 | 29 | 38 | 23 | 2503 | 494 |
| SRR3624073 | 0 | 534 | 10 | 383 | 1319 | 0 | 18 | 139 | 535 | 4191 | 854 | 1293 | 0 | 550 | 894 | 799 | 40 | 1596 | 20 | 2107 | 484 | 509 | 24 | 592 | 1401 | 33 | 29 | 21 | 56 | 43 | 239 | 17 | 32 | 12 | 0 | 1301 | 451 |
| SRR3624074 | 0 | 1143 | 54 | 335 | 1215 | 0 | 16 | 89 | 389 | 3805 | 1144 | 1931 | 0 | 401 | 1072 | 1306 | 103 | 1315 | 25 | 1960 | 538 | 639 | 36 | 787 | 1654 | 34 | 19 | 87 | 240 | 227 | 2323 | 130 | 91 | 114 | 14 | 1683 | 624 |
| SRR3675003 | 0 | 90 | 0 | 0 | 0 | 0 | 0 | 126 | 76 | 2457 | 189 | 37 | 0 | 2434 | 75 | 160 | 21 | 0 | 61 | 238 | 61 | 121 | 0 | 667 | 3215 | 0 | 46 | 0 | 0 | 18 | 13 | 0 | 0 | 0 | 0 | 3651 | 0 |
| SRR3675004 | 0 | 13 | 0 | 0 | 0 | 0 | 0 | 121 | 77 | 3243 | 87 | 14 | 0 | 2551 | 53 | 112 | 11 | 0 | 25 | 141 | 764 | 560 | 0 | 866 | 3312 | 0 | 36 | 0 | 11 | 19 | 15 | 0 | 0 | 0 | 0 | 2143 | 0 |
| SRR3675005 | 0 | 16 | 42 | 0 | 0 | 0 | 0 | 450 | 232 | 4276 | 2373 | 1327 | 0 | 121 | 673 | 892 | 0 | 0 | 39 | 233 | 1300 | 501 | 0 | 3090 | 12372 | 0 | 45 | 462 | 336 | 1195 | 904 | 206 | 373 | 484 | 62 | 811 | 19 |
| SRR3675006 | 0 | 62 | 0 | 0 | 0 | 0 | 0 | 179 | 94 | 3056 | 22 | 14 | 0 | 2409 | 60 | 125 | 17 | 0 | 0 | 192 | 152 | 0 | 0 | 456 | 1317 | 0 | 55 | 12 | 0 | 17 | 17 | 0 | 11 | 0 | 13 | 2549 | 0 |
| SRR3675007 | 0 | 16 | 0 | 0 | 0 | 0 | 0 | 97 | 99 | 3613 | 33 | 0 | 0 | 3590 | 33 | 91 | 0 | 0 | 0 | 146 | 191 | 96 | 0 | 478 | 2009 | 0 | 11 | 0 | 0 | 18 | 17 | 0 | 0 | 0 | 0 | 2204 | 0 |
| SRR3675008 | 0 | 12 | 0 | 0 | 0 | 0 | 0 | 129 | 76 | 3527 | 97 | 14 | 0 | 2945 | 71 | 173 | 0 | 0 | 0 | 211 | 58 | 116 | 0 | 347 | 1676 | 0 | 27 | 0 | 0 | 11 | 0 | 0 | 0 | 0 | 0 | 3114 | 0 |
| SRR3713941 | 63 | 356 | 11 | 60 | 470 | 34 | 31 | 10394 | 4410 | 81 | 407 | 686 | 15 | 122 | 0 | 0 | 67 | 2497 | 113 | 174 | 397 | 974 | 20 | 4378 | 2166 | 159 | 19 | 0 | 27 | 23 | 29 | 34 | 21 | 15 | 0 | 139 | 225 |
| SRR3713942 | 0 | 61 | 0 | 22 | 195 | 0 | 0 | 1046 | 897 | 3737 | 17 | 797 | 0 | 11310 | 149 | 149 | 0 | 865 | 43 | 249 | 65 | 195 | 22 | 1687 | 346 | 303 | 0 | 0 | 0 | 0 | 0 | 0 | 0 | 0 | 0 | 299 | 43 |
| SRR3713943 | 142 | 354 | 0 | 17 | 604 | 42 | 219 | 13289 | 5945 | 366 | 43 | 1232 | 25 | 133 | 0 | 0 | 430 | 177 | 205 | 616 | 211 | 359 | 0 | 63 | 1277 | 0 | 44 | 0 | 66 | 44 | 49 | 0 | 0 | 0 | 0 | 266 | 616 |
| SRR3713945 | 41 | 1048 | 106 | 119 | 378 | 13 | 67 | 21045 | 12225 | 11679 | 3666 | 2868 | 33 | 233 | 323 | 332 | 70 | 834 | 140 | 1622 | 374 | 1458 | 108 | 20921 | 739 | 255 | 35 | 98 | 90 | 21 | 53 | 12 | 17 | 16 | 0 | 1362 | 45 |
| SRR3713946 | 19 | 220 | 26 | 503 | 688 | 0 | 22 | 13710 | 14530 | 15132 | 15323 | 10101 | 0 | 396 | 301 | 478 | 28 | 1917 | 217 | 2720 | 1923 | 2453 | 149 | 21930 | 1051 | 176 | 0 | 31 | 43 | 10 | 20 | 16 | 17 | 13 | 0 | 2242 | 119 |
| SRR3879751 | 0 | 17 | 0 | 21 | 26 | 0 | 25 | 491 | 1096 | 12618 | 192 | 302 | 0 | 4510 | 0 | 0 | 0 | 4653 | 272 | 44 | 695 | 1818 | 13 | 2103 | 894 | 364 | 27 | 0 | 11 | 0 | 0 | 0 | 0 | 0 | 0 | 265 | 314 |
| SRR3879752 | 0 | 40 | 0 | 18 | 20 | 0 | 11 | 435 | 1244 | 12152 | 774 | 378 | 0 | 4758 | 0 | 0 | 14 | 4591 | 275 | 61 | 619 | 2561 | 10 | 2007 | 1021 | 302 | 126 | 0 | 11 | 0 | 0 | 0 | 0 | 0 | 0 | 122 | 359 |
| SRR3879754 | 0 | 79 | 0 | 26 | 19 | 0 | 0 | 589 | 2850 | 28920 | 147 | 1151 | 0 | 7747 | 0 | 0 | 11 | 1908 | 147 | 14 | 928 | 2817 | 41 | 6419 | 1021 | 106 | 34 | 0 | 12 | 0 | 0 | 0 | 0 | 0 | 0 | 35 | 123 |
| SRR3879755 | 0 | 21 | 0 | 0 | 26 | 0 | 0 | 530 | 2678 | 24792 | 1474 | 1294 | 0 | 7045 | 0 | 0 | 0 | 2028 | 207 | 36 | 1272 | 3071 | 10 | 7360 | 1191 | 134 | 13 | 0 | 16 | 0 | 0 | 0 | 0 | 0 | 0 | 36 | 132 |
| SRR3879756 | 0 | 60 | 0 | 35 | 12 | 0 | 0 | 935 | 3507 | 32551 | 1365 | 1235 | 0 | 9071 | 17 | 0 | 0 | 1787 | 231 | 11 | 1055 | 2977 | 12 | 5544 | 873 | 91 | 35 | 0 | 36 | 0 | 0 | 0 | 0 | 0 | 0 | 45 | 156 |
| SRR3879757 | 0 | 18 | 0 | 0 | 24 | 0 | 0 | 265 | 707 | 4505 | 2473 | 1104 | 0 | 4063 | 44 | 0 | 0 | 2543 | 758 | 221 | 2201 | 2763 | 0 | 1565 | 2029 | 147 | 14 | 0 | 0 | 0 | 0 | 0 | 0 | 0 | 0 | 442 | 318 |
| SRR3879758 | 0 | 0 | 0 | 0 | 25 | 0 | 0 | 187 | 710 | 5902 | 63 | 1793 | 0 | 5080 | 37 | 37 | 0 | 1718 | 622 | 336 | 2166 | 2738 | 25 | 1469 | 1568 | 25 | 16 | 0 | 11 | 0 | 0 | 0 | 0 | 0 | 0 | 448 | 124 |
| SRR3879759 | 0 | 14 | 0 | 0 | 0 | 0 | 0 | 304 | 895 | 5064 | 2836 | 1891 | 0 | 5351 | 51 | 17 | 0 | 1944 | 404 | 152 | 1671 | 2779 | 0 | 1108 | 1915 | 38 | 43 | 0 | 16 | 0 | 0 | 0 | 0 | 0 | 0 | 675 | 197 |
| SRR3879760 | 0 | 27 | 0 | 14 | 0 | 0 | 0 | 774 | 2130 | 6905 | 2323 | 2646 | 0 | 7034 | 65 | 0 | 0 | 4079 | 154 | 452 | 768 | 3534 | 0 | 6706 | 964 | 140 | 45 | 0 | 21 | 0 | 0 | 0 | 0 | 0 | 0 | 258 | 335 |
| SRR3879761 | 0 | 0 | 0 | 0 | 0 | 0 | 0 | 578 | 1576 | 5096 | 4150 | 3152 | 0 | 8564 | 53 | 0 | 0 | 1868 | 152 | 53 | 543 | 2400 | 11 | 7666 | 966 | 239 | 21 | 0 | 0 | 0 | 0 | 0 | 0 | 0 | 0 | 263 | 347 |
| SRR3879766 | 0 | 987 | 125 | 0 | 20 | 0 | 0 | 2091 | 5146 | 343 | 66070 | 65611 | 0 | 73 | 0 | 0 | 14 | 3442 | 161 | 86 | 348 | 1190 | 0 | 2772 | 1866 | 191 | 71 | 160 | 1841 | 507 | 897 | 166 | 56 | 56 | 0 | 328 | 801 |
| SRR3879767 | 0 | 13 | 0 | 0 | 0 | 0 | 0 | 457 | 1715 | 1029 | 37 | 2059 | 0 | 114 | 0 | 0 | 0 | 37 | 595 | 3202 | 0 | 15 | 0 | 9847 | 164 | 0 | 1358 | 0 | 0 | 0 | 283 | 18 | 0 | 16 | 0 | 8120 | 104 |
| SRR3879768 | 0 | 59 | 22 | 0 | 0 | 0 | 0 | 107 | 82 | 365 | 40156 | 35991 | 0 | 150 | 0 | 0 | 10 | 401 | 497 | 200 | 137 | 248 | 0 | 1071 | 1555 | 24 | 17 | 37 | 239 | 66 | 225 | 51 | 12 | 17 | 0 | 412 | 283 |
| SRR3879769 | 0 | 34 | 11 | 0 | 0 | 0 | 0 | 396 | 422 | 330 | 1262 | 20422 | 0 | 330 | 0 | 13 | 21 | 259 | 101 | 304 | 15 | 139 | 0 | 12222 | 284 | 74 | 52 | 36 | 83 | 40 | 394 | 50 | 27 | 15 | 0 | 1135 | 175 |
| SRR3879770 | 0 | 0 | 0 | 0 | 0 | 0 | 0 | 464 | 121 | 767 | 1896 | 4115 | 0 | 726 | 0 | 20 | 0 | 160 | 354 | 726 | 0 | 16 | 0 | 1860 | 80 | 35 | 1387 | 0 | 0 | 12 | 855 | 36 | 16 | 56 | 0 | 2118 | 16 |
| SRR3923813 | 0 | 0 | 0 | 0 | 16 | 0 | 0 | 24 | 0 | 207 | 275 | 1787 | 0 | 0 | 0 | 0 | 0 | 0 | 0 | 0 | 145 | 242 | 0 | 0 | 272 | 50 | 0 | 0 | 0 | 10 | 0 | 0 | 0 | 0 | 0 | 0 | 37 |
| SRR3931357 | 0 | 224 | 77 | 0 | 11 | 0 | 0 | 197 | 39 | 529 | 4998 | 7299 | 0 | 121 | 23 | 20 | 11 | 112 | 31 | 1978 | 3589 | 7978 | 26 | 45834 | 14505 | 0 | 15 | 81 | 293 | 478 | 461 | 37 | 120 | 185 | 31 | 4460 | 152 |
| SRR3931358 | 0 | 246 | 72 | 0 | 14 | 0 | 0 | 187 | 17 | 180 | 16850 | 7359 | 0 | 69 | 17 | 17 | 0 | 133 | 22 | 961 | 4780 | 6108 | 27 | 37079 | 10791 | 13 | 35 | 146 | 396 | 553 | 478 | 65 | 107 | 183 | 32 | 2976 | 129 |
| SRR3931359 | 0 | 12 | 12 | 0 | 15 | 0 | 34 | 832 | 14 | 46 | 1163 | 570 | 0 | 78 | 28 | 28 | 11 | 219 | 21 | 5379 | 2266 | 4211 | 34 | 30445 | 11065 | 0 | 55 | 12 | 35 | 69 | 113 | 13 | 61 | 119 | 23 | 43825 | 130 |
| SRR3931360 | 0 | 10 | 46 | 0 | 13 | 0 | 29 | 761 | 0 | 0 | 894 | 463 | 0 | 73 | 33 | 13 | 24 | 218 | 22 | 5806 | 2343 | 4011 | 29 | 37000 | 11089 | 0 | 188 | 48 | 120 | 276 | 411 | 49 | 206 | 379 | 94 | 53476 | 96 |
| SRR3931984 | 10 | 30 | 0 | 0 | 0 | 30 | 22 | 455 | 111 | 70 | 1509 | 2667 | 0 | 0 | 0 | 0 | 0 | 1294 | 0 | 15 | 1765 | 2736 | 108 | 6713 | 29380 | 0 | 66 | 145 | 356 | 498 | 174 | 0 | 0 | 0 | 0 | 22 | 0 |
| SRR3931987 | 11 | 23 | 0 | 0 | 0 | 35 | 32 | 300 | 90 | 15 | 297 | 4121 | 0 | 0 | 0 | 0 | 12 | 1502 | 0 | 0 | 2138 | 2957 | 295 | 7463 | 26149 | 0 | 143 | 82 | 103 | 108 | 29 | 0 | 0 | 0 | 0 | 15 | 0 |
| SRR3931988 | 0 | 20 | 0 | 0 | 0 | 13 | 0 | 481 | 175 | 109 | 1152 | 1439 | 0 | 0 | 0 | 0 | 0 | 1561 | 0 | 12 | 1967 | 19 | 44 | 6246 | 12 | 0 | 21 | 147 | 36 | 44 | 16 | 11 | 0 | 0 | 0 | 39 | 0 |
| SRR3992213 | 0 | 239 | 203 | 307 | 33 | 0 | 0 | 1448 | 2298 | 82264 | 36462 | 25473 | 0 | 12087 | 0 | 0 | 0 | 3661 | 102 | 2847 | 23 | 636 | 0 | 6216 | 88 | 42 | 35 | 0 | 83 | 56 | 215 | 80 | 53 | 127 | 0 | 14035 | 0 |
| SRR3992215 | 0 | 314 | 32 | 194 | 124 | 0 | 12 | 283 | 905 | 7240 | 1012 | 90786 | 0 | 1301 | 0 | 0 | 0 | 3500 | 33 | 509 | 39 | 202 | 47 | 3741 | 331 | 21 | 21 | 0 | 23 | 24 | 42 | 24 | 0 | 12 | 0 | 1640 | 0 |
| SRR3992474 | 0 | 88 | 113 | 0 | 30 | 0 | 0 | 4599 | 412 | 265 | 7113 | 23267 | 0 | 0 | 0 | 0 | 0 | 978 | 0 | 0 | 982 | 1907 | 26 | 7543 | 8360 | 0 | 0 | 0 | 0 | 0 | 0 | 0 | 0 | 0 | 0 | 0 | 0 |
| SRR3992475 | 0 | 120 | 59 | 0 | 49 | 0 | 0 | 3531 | 509 | 185 | 2176 | 22721 | 0 | 0 | 0 | 0 | 12 | 1215 | 0 | 0 | 949 | 1778 | 31 | 6329 | 7666 | 0 | 0 | 0 | 0 | 0 | 0 | 0 | 0 | 0 | 0 | 0 | 0 |
| SRR3992476 | 0 | 17 | 19 | 0 | 33 | 17 | 13 | 6681 | 790 | 642 | 1223 | 2644 | 0 | 70 | 21 | 0 | 27 | 1198 | 45 | 1146 | 1437 | 2599 | 93 | 13151 | 3911 | 29 | 16 | 49 | 15 | 53 | 54 | 21 | 40 | 10 | 61 | 5230 | 171 |
| SRR3992477 | 0 | 31 | 34 | 0 | 38 | 0 | 0 | 2125 | 385 | 1529 | 7831 | 6631 | 0 | 155 | 81 | 25 | 0 | 1169 | 37 | 1187 | 1648 | 2444 | 83 | 14244 | 4375 | 34 | 0 | 93 | 64 | 202 | 109 | 57 | 124 | 136 | 158 | 6227 | 155 |
| SRR3996364 | 0 | 0 | 0 | 0 | 0 | 0 | 0 | 8146 | 1381 | 33809 | 0 | 0 | 0 | 2249 | 39 | 39 | 0 | 263 | 20 | 79 | 716 | 2997 | 234 | 37864 | 8569 | 0 | 0 | 114 | 27 | 94 | 47 | 0 | 27 | 34 | 80 | 1006 | 72 |
| SRR3996365 | 0 | 0 | 0 | 0 | 0 | 0 | 0 | 3741 | 914 | 27510 | 0 | 0 | 0 | 1168 | 34 | 17 | 0 | 387 | 22 | 220 | 697 | 2805 | 245 | 56001 | 10778 | 10 | 0 | 54 | 15 | 147 | 54 | 0 | 31 | 23 | 77 | 1185 | 87 |
| SRR3996366 | 0 | 0 | 0 | 0 | 0 | 0 | 0 | 1658 | 254 | 37948 | 0 | 0 | 0 | 1980 | 39 | 10 | 0 | 377 | 34 | 361 | 625 | 2427 | 203 | 42521 | 9777 | 14 | 0 | 10 | 67 | 156 | 76 | 17 | 25 | 51 | 25 | 1805 | 56 |
| SRR3996367 | 0 | 0 | 0 | 0 | 0 | 0 | 0 | 2521 | 142 | 50143 | 0 | 0 | 0 | 2151 | 0 | 0 | 0 | 415 | 23 | 399 | 696 | 2725 | 282 | 53061 | 11082 | 12 | 0 | 10 | 35 | 180 | 83 | 35 | 48 | 62 | 48 | 2379 | 97 |
| SRR3996368 | 0 | 0 | 0 | 0 | 15 | 0 | 0 | 5959 | 1431 | 55417 | 0 | 0 | 0 | 520 | 26 | 0 | 0 | 650 | 23 | 91 | 584 | 2031 | 353 | 71255 | 14010 | 0 | 0 | 17 | 12 | 37 | 21 | 0 | 0 | 0 | 0 | 612 | 67 |
| SRR3996369 | 0 | 0 | 0 | 0 | 18 | 0 | 0 | 3957 | 921 | 44138 | 0 | 0 | 0 | 607 | 0 | 0 | 0 | 387 | 26 | 133 | 419 | 1307 | 430 | 88297 | 15128 | 0 | 0 | 0 | 11 | 22 | 33 | 0 | 0 | 15 | 0 | 1044 | 54 |
| SRR4000625 | 0 | 2831 | 0 | 38 | 85 | 13 | 35 | 412 | 648 | 2891 | 1178 | 7234 | 0 | 916 | 10 | 41 | 100 | 3351 | 164 | 9765 | 122 | 1328 | 0 | 1910 | 324 | 24 | 83 | 0 | 0 | 0 | 22 | 0 | 0 | 0 | 0 | 6565 | 127 |
| SRR4000626 | 0 | 2761 | 10 | 45 | 69 | 11 | 41 | 363 | 736 | 2933 | 1233 | 7307 | 0 | 860 | 0 | 0 | 100 | 3407 | 202 | 9981 | 109 | 1351 | 19 | 1883 | 328 | 40 | 107 | 0 | 0 | 0 | 24 | 0 | 0 | 0 | 0 | 6830 | 109 |
| SRR4000627 | 0 | 2318 | 0 | 0 | 12 | 10 | 40 | 39 | 77 | 406 | 18061 | 11937 | 0 | 155 | 0 | 0 | 104 | 2728 | 181 | 1507 | 78 | 1848 | 0 | 1403 | 243 | 49 | 59 | 0 | 0 | 0 | 0 | 0 | 0 | 0 | 0 | 1256 | 86 |
| SRR4000628 | 0 | 2233 | 0 | 0 | 0 | 14 | 41 | 39 | 39 | 232 | 17352 | 12220 | 0 | 252 | 0 | 19 | 109 | 2695 | 124 | 1704 | 70 | 1850 | 0 | 1508 | 288 | 49 | 61 | 0 | 0 | 0 | 0 | 0 | 0 | 0 | 0 | 1104 | 128 |
| SRR4000629 | 0 | 3087 | 0 | 0 | 33 | 0 | 40 | 312 | 439 | 1722 | 2582 | 18578 | 0 | 982 | 12 | 46 | 145 | 1930 | 188 | 12617 | 100 | 2039 | 0 | 1772 | 167 | 91 | 60 | 0 | 0 | 0 | 41 | 0 | 0 | 0 | 0 | 7325 | 70 |
| SRR4000630 | 0 | 1933 | 0 | 15 | 24 | 13 | 25 | 279 | 535 | 1827 | 2647 | 17749 | 0 | 1187 | 12 | 23 | 92 | 1994 | 237 | 12279 | 107 | 2089 | 0 | 1742 | 128 | 82 | 33 | 0 | 0 | 0 | 30 | 0 | 0 | 0 | 0 | 7553 | 73 |
| SRR4000631 | 0 | 2635 | 0 | 18 | 22 | 15 | 38 | 299 | 610 | 2917 | 1704 | 12624 | 0 | 1423 | 0 | 12 | 150 | 2373 | 408 | 36353 | 148 | 2364 | 0 | 1691 | 319 | 49 | 50 | 0 | 0 | 0 | 13 | 0 | 0 | 0 | 0 | 25104 | 135 |
| SRR4000632 | 0 | 1716 | 0 | 23 | 32 | 12 | 29 | 274 | 448 | 3224 | 20799 | 13094 | 0 | 1568 | 0 | 0 | 90 | 2210 | 275 | 37765 | 140 | 2440 | 0 | 1728 | 325 | 68 | 36 | 0 | 0 | 0 | 15 | 0 | 0 | 0 | 0 | 26151 | 77 |
| SRR4000633 | 0 | 1667 | 0 | 0 | 38 | 14 | 36 | 186 | 338 | 804 | 3265 | 19221 | 0 | 746 | 12 | 82 | 72 | 1561 | 188 | 47 | 89 | 1468 | 0 | 1597 | 203 | 32 | 30 | 0 | 0 | 0 | 21 | 0 | 0 | 0 | 0 | 0 | 104 |
| SRR4000634 | 0 | 2837 | 0 | 17 | 36 | 16 | 62 | 277 | 458 | 711 | 39721 | 20078 | 0 | 639 | 12 | 36 | 120 | 1534 | 176 | 36 | 94 | 1560 | 0 | 1566 | 198 | 62 | 49 | 0 | 0 | 0 | 36 | 0 | 0 | 0 | 0 | 12 | 77 |
| SRR4000635 | 0 | 2140 | 0 | 0 | 60 | 13 | 44 | 82 | 227 | 216 | 21953 | 12460 | 0 | 391 | 10 | 21 | 78 | 2456 | 189 | 7733 | 124 | 2329 | 0 | 2262 | 241 | 22 | 50 | 0 | 0 | 0 | 13 | 0 | 0 | 0 | 0 | 3264 | 137 |
| SRR4000636 | 0 | 2099 | 0 | 15 | 45 | 13 | 38 | 101 | 161 | 302 | 2091 | 12510 | 0 | 503 | 0 | 0 | 75 | 2393 | 170 | 6838 | 115 | 2363 | 13 | 2238 | 225 | 10 | 43 | 0 | 0 | 0 | 15 | 0 | 0 | 0 | 0 | 3570 | 123 |
| SRR4000638 | 0 | 1442 | 0 | 49 | 451 | 12 | 27 | 0 | 23 | 698 | 512 | 7858 | 0 | 608 | 0 | 0 | 137 | 2288 | 218 | 1846 | 157 | 2364 | 22 | 2360 | 228 | 67 | 26 | 0 | 0 | 0 | 0 | 0 | 0 | 0 | 0 | 1463 | 94 |
| SRR4000639 | 0 | 2108 | 0 | 68 | 168 | 24 | 52 | 0 | 24 | 71 | 802 | 13803 | 0 | 236 | 0 | 0 | 249 | 2042 | 232 | 1272 | 172 | 2262 | 20 | 1658 | 232 | 52 | 49 | 0 | 0 | 0 | 0 | 0 | 0 | 0 | 0 | 1107 | 68 |
| SRR4000640 | 0 | 2018 | 0 | 112 | 215 | 22 | 48 | 0 | 24 | 143 | 835 | 15039 | 0 | 430 | 0 | 48 | 241 | 1961 | 251 | 1194 | 191 | 2264 | 44 | 1766 | 243 | 48 | 48 | 0 | 0 | 0 | 0 | 0 | 0 | 0 | 0 | 1146 | 36 |

### Supp Table S8.4.Columns_DI_to_ES

| | | | | | | | | | | | | | | | | | | | | | | | | | | | | | | | | | | | | | |
|---|---|---|---|---|---|---|---|---|---|---|---|---|---|---|---|---|---|---|---|---|---|---|---|---|---|---|---|---|---|---|---|---|---|---|---|---|---|
| SRR4000641 | 0 | 2402 | 0 | 85 | 332 | 15 | 49 | 0 | 20 | 341 | 1545 | 20056 | 0 | 341 | 20 | 80 | 270 | 2274 | 268 | 1203 | 151 | 2511 | 28 | 2111 | 242 | 68 | 47 | 0 | 0 | 0 | 0 | 0 | 0 | 0 | 0 | 1003 | 52 |
| SRR4000642 | 0 | 2459 | 0 | 82 | 284 | 24 | 38 | 0 | 44 | 377 | 35137 | 21348 | 0 | 266 | 0 | 67 | 281 | 2217 | 223 | 1507 | 162 | 2417 | 26 | 2295 | 244 | 61 | 40 | 0 | 0 | 0 | 0 | 0 | 0 | 0 | 0 | 1618 | 49 |
| SRR4000643 | 0 | 843 | 0 | 108 | 490 | 15 | 35 | 0 | 87 | 729 | 268 | 5015 | 0 | 846 | 29 | 233 | 124 | 2905 | 166 | 2741 | 94 | 1903 | 36 | 1262 | 216 | 72 | 24 | 0 | 0 | 0 | 0 | 0 | 0 | 0 | 0 | 2624 | 94 |
| SRR4000644 | 0 | 1517 | 0 | 94 | 382 | 31 | 48 | 66 | 33 | 1022 | 8836 | 5011 | 0 | 890 | 0 | 264 | 231 | 2874 | 252 | 2802 | 187 | 2096 | 43 | 1044 | 259 | 65 | 46 | 0 | 0 | 0 | 0 | 0 | 0 | 0 | 0 | 2736 | 180 |
| SRR4000645 | 0 | 1157 | 0 | 69 | 230 | 16 | 20 | 61 | 61 | 184 | 48225 | 25967 | 0 | 123 | 0 | 0 | 124 | 1775 | 128 | 1472 | 164 | 2115 | 15 | 1647 | 321 | 51 | 20 | 0 | 0 | 0 | 0 | 0 | 0 | 0 | 0 | 797 | 77 |
| SRR4000649 | 0 | 1761 | 0 | 130 | 514 | 17 | 39 | 481 | 766 | 1536 | 816 | 4144 | 0 | 481 | 0 | 0 | 169 | 2696 | 282 | 3502 | 136 | 1103 | 76 | 5293 | 932 | 23 | 36 | 0 | 0 | 0 | 36 | 0 | 0 | 0 | 0 | 2045 | 147 |
| SRR4000650 | 0 | 1781 | 0 | 131 | 502 | 17 | 44 | 458 | 930 | 1626 | 851 | 4122 | 0 | 518 | 0 | 0 | 188 | 2643 | 291 | 3637 | 132 | 1114 | 82 | 5296 | 923 | 22 | 37 | 0 | 0 | 0 | 37 | 0 | 0 | 0 | 0 | 2267 | 145 |
| SRR4011144 | 0 | 0 | 0 | 0 | 0 | 0 | 0 | 0 | 0 | 0 | 238 | 8086 | 0 | 0 | 238 | 0 | 0 | 349 | 0 | 0 | 42 | 735 | 0 | 139 | 1582 | 589 | 0 | 0 | 0 | 0 | 250 | 0 | 322 | 393 | 0 | 238 | 0 |
| SRR4044175 | 78 | 0 | 0 | 0 | 0 | 15 | 52 | 88379 | 4367 | 174 | 270 | 68 | 0 | 61 | 0 | 0 | 31 | 541 | 53 | 23 | 628 | 1825 | 20 | 12154 | 3745 | 320 | 0 | 0 | 0 | 16 | 13 | 0 | 0 | 16 | 0 | 200 | 86 |
| SRR4044176 | 200 | 0 | 0 | 0 | 0 | 0 | 121 | 101990 | 4727 | 346 | 93 | 70 | 0 | 64 | 0 | 0 | 79 | 462 | 31 | 19 | 639 | 1810 | 23 | 10766 | 3048 | 349 | 0 | 0 | 0 | 16 | 10 | 0 | 0 | 21 | 0 | 158 | 142 |
| SRR4044177 | 28 | 0 | 0 | 0 | 0 | 0 | 19 | 91846 | 4532 | 1303 | 129 | 28 | 0 | 705 | 0 | 0 | 15 | 541 | 57 | 113 | 614 | 1478 | 52 | 8971 | 2557 | 220 | 0 | 0 | 13 | 10 | 10 | 12 | 14 | 34 | 0 | 1619 | 111 |
| SRR4044178 | 64 | 0 | 0 | 0 | 0 | 24 | 48 | 84744 | 4553 | 233 | 104 | 82 | 0 | 78 | 0 | 0 | 25 | 641 | 65 | 71 | 722 | 1949 | 27 | 12902 | 4072 | 440 | 0 | 0 | 13 | 23 | 17 | 0 | 16 | 11 | 0 | 976 | 112 |
| SRR4044179 | 106 | 0 | 0 | 0 | 0 | 0 | 64 | 107706 | 4836 | 361 | 52 | 38 | 0 | 70 | 0 | 0 | 54 | 408 | 53 | 30 | 642 | 1998 | 30 | 11794 | 3684 | 266 | 0 | 0 | 0 | 14 | 15 | 0 | 12 | 31 | 0 | 230 | 189 |
| SRR4044180 | 62 | 0 | 0 | 0 | 0 | 0 | 45 | 76393 | 3649 | 1020 | 49 | 39 | 0 | 410 | 0 | 0 | 42 | 662 | 84 | 73 | 643 | 1321 | 48 | 10040 | 3519 | 165 | 0 | 0 | 14 | 27 | 23 | 15 | 24 | 45 | 0 | 924 | 91 |
| SRR4044206 | 0 | 144 | 0 | 216 | 263 | 0 | 24 | 172 | 1130 | 11718 | 26 | 89 | 0 | 250 | 30 | 30 | 10 | 275 | 444 | 208 | 1396 | 263 | 216 | 5684 | 2161 | 0 | 11 | 0 | 0 | 0 | 0 | 0 | 0 | 0 | 0 | 345 | 339 |
| SRR4044210 | 0 | 32 | 0 | 67 | 391 | 0 | 0 | 3734 | 8049 | 9128 | 42 | 41 | 0 | 1162 | 1784 | 1286 | 24 | 508 | 374 | 913 | 33 | 200 | 1207 | 26630 | 2447 | 33 | 14 | 0 | 0 | 0 | 0 | 0 | 0 | 0 | 0 | 1992 | 25 |
| SRR4044211 | 0 | 0 | 0 | 16 | 439 | 0 | 15 | 445 | 714 | 6224 | 80 | 143 | 0 | 1397 | 12876 | 12908 | 35 | 298 | 94 | 5589 | 78 | 705 | 548 | 19751 | 1488 | 110 | 259 | 0 | 0 | 0 | 17 | 0 | 0 | 0 | 0 | 5208 | 31 |
| SRR4270461 | 0 | 3389 | 306 | 92 | 1733 | 0 | 17 | 612 | 2246 | 3748 | 12191 | 11862 | 0 | 4153 | 121 | 718 | 89 | 2074 | 90 | 2818 | 771 | 4047 | 443 | 27748 | 1782 | 18 | 14 | 57 | 455 | 272 | 343 | 109 | 90 | 66 | 115 | 1356 | 363 |
| SRR4270462 | 0 | 2911 | 368 | 85 | 1191 | 0 | 34 | 510 | 1719 | 2012 | 16676 | 14744 | 0 | 3091 | 117 | 334 | 170 | 1644 | 110 | 3163 | 581 | 2881 | 338 | 27402 | 1542 | 14 | 17 | 114 | 588 | 391 | 562 | 162 | 122 | 102 | 110 | 1123 | 291 |
| SRR4292754 | 0 | 59 | 0 | 0 | 10 | 0 | 0 | 8136 | 1676 | 321 | 1446 | 2029 | 0 | 94 | 0 | 0 | 0 | 1194 | 12 | 531 | 1478 | 16283 | 144 | 222263 | 18050 | 78 | 0 | 60 | 55 | 17 | 57 | 13 | 18 | 19 | 10 | 962 | 394 |
| SRR4292755 | 0 | 654 | 0 | 0 | 0 | 0 | 0 | 7872 | 3155 | 305 | 3010 | 1998 | 0 | 151 | 0 | 0 | 0 | 2357 | 26 | 675 | 1491 | 15417 | 147 | 228369 | 17130 | 125 | 0 | 77 | 74 | 157 | 60 | 0 | 22 | 27 | 0 | 838 | 101 |
| SRR4292756 | 0 | 121 | 0 | 0 | 16 | 0 | 0 | 181 | 347 | 453 | 1400 | 1859 | 0 | 164 | 0 | 0 | 0 | 2083 | 27 | 921 | 2251 | 13775 | 137 | 178543 | 15793 | 131 | 0 | 14 | 99 | 11 | 58 | 12 | 12 | 14 | 0 | 1159 | 229 |
| SRR4292757 | 0 | 144 | 0 | 0 | 12 | 0 | 0 | 176 | 306 | 450 | 1639 | 1462 | 0 | 202 | 0 | 0 | 0 | 1625 | 22 | 1088 | 2040 | 13607 | 132 | 185700 | 17526 | 133 | 0 | 14 | 10 | 18 | 69 | 11 | 17 | 15 | 0 | 1207 | 288 |
| SRR4296311 | 0 | 990 | 433 | 434 | 2239 | 0 | 18 | 3506 | 917 | 17583 | 3721 | 8845 | 0 | 6688 | 270 | 647 | 0 | 2480 | 72 | 593 | 292 | 1360 | 351 | 11295 | 646 | 10 | 79 | 101 | 293 | 670 | 1800 | 425 | 197 | 201 | 88 | 5016 | 1035 |
| SRR4296312 | 23 | 1183 | 349 | 436 | 2280 | 11 | 11 | 4201 | 1961 | 16524 | 4761 | 7702 | 0 | 4901 | 280 | 140 | 11 | 2493 | 70 | 1680 | 294 | 1417 | 329 | 11567 | 714 | 0 | 113 | 79 | 484 | 608 | 1600 | 349 | 237 | 135 | 90 | 5601 | 976 |
| SRR4296313 | 0 | 132 | 37 | 990 | 2554 | 0 | 61 | 2890 | 1020 | 15754 | 51 | 7707 | 0 | 4817 | 227 | 510 | 24 | 4855 | 110 | 510 | 345 | 1486 | 381 | 11218 | 801 | 0 | 81 | 101 | 38 | 61 | 91 | 20 | 12 | 17 | 97 | 3457 | 866 |
| SRR4296314 | 0 | 1552 | 450 | 885 | 2478 | 0 | 54 | 3206 | 1566 | 13272 | 49 | 7531 | 0 | 5592 | 298 | 1118 | 11 | 4803 | 98 | 895 | 278 | 1229 | 385 | 10236 | 767 | 0 | 48 | 64 | 439 | 589 | 958 | 225 | 86 | 166 | 112 | 4846 | 899 |
| SRR4296315 | 0 | 1453 | 345 | 672 | 2500 | 0 | 11 | 2598 | 585 | 21628 | 4936 | 7664 | 0 | 4611 | 260 | 520 | 0 | 4787 | 42 | 974 | 305 | 1228 | 384 | 9612 | 574 | 0 | 33 | 111 | 401 | 629 | 1765 | 456 | 306 | 195 | 84 | 2403 | 968 |
| SRR4296316 | 11 | 1950 | 222 | 691 | 2284 | 0 | 0 | 3107 | 1195 | 20672 | 34 | 8723 | 0 | 4780 | 0 | 358 | 0 | 4836 | 31 | 119 | 233 | 1249 | 396 | 9757 | 524 | 0 | 44 | 133 | 521 | 521 | 1718 | 410 | 366 | 211 | 89 | 2151 | 907 |
| SRR4296317 | 0 | 1032 | 287 | 617 | 2693 | 0 | 0 | 2870 | 1380 | 29637 | 95 | 7285 | 0 | 5519 | 276 | 331 | 0 | 3001 | 36 | 0 | 265 | 1098 | 478 | 10693 | 463 | 0 | 61 | 99 | 381 | 629 | 2523 | 546 | 381 | 287 | 287 | 2263 | 950 |
| SRR4296318 | 0 | 52 | 20 | 594 | 2631 | 0 | 0 | 2857 | 1111 | 29367 | 68 | 8096 | 0 | 6826 | 238 | 635 | 0 | 2970 | 32 | 397 | 273 | 917 | 439 | 10040 | 440 | 0 | 16 | 158 | 24 | 42 | 152 | 32 | 29 | 13 | 15 | 1270 | 870 |
| SRR4296319 | 0 | 23 | 100 | 347 | 3338 | 0 | 0 | 1891 | 1121 | 80328 | 2521 | 2521 | 0 | 11906 | 140 | 280 | 0 | 1666 | 46 | 210 | 237 | 652 | 438 | 8223 | 311 | 0 | 48 | 53 | 14 | 28 | 64 | 16 | 14 | 62 | 96 | 3642 | 801 |
| SRR4296320 | 0 | 328 | 117 | 323 | 3305 | 0 | 0 | 1668 | 1289 | 76455 | 2602 | 2552 | 0 | 11269 | 303 | 303 | 0 | 1602 | 45 | 505 | 255 | 715 | 449 | 8244 | 298 | 0 | 26 | 31 | 163 | 246 | 724 | 150 | 107 | 88 | 81 | 3108 | 843 |
| SRR4296321 | 0 | 363 | 136 | 429 | 3234 | 0 | 0 | 2838 | 1497 | 65330 | 60 | 3555 | 0 | 9324 | 405 | 592 | 0 | 2044 | 39 | 281 | 225 | 702 | 440 | 8872 | 318 | 0 | 34 | 34 | 132 | 184 | 643 | 156 | 108 | 100 | 78 | 2807 | 659 |
| SRR4296322 | 0 | 301 | 117 | 519 | 3163 | 0 | 0 | 2277 | 1822 | 62092 | 29 | 3264 | 0 | 10475 | 531 | 759 | 0 | 2076 | 34 | 455 | 257 | 735 | 404 | 8932 | 327 | 0 | 15 | 29 | 146 | 126 | 738 | 102 | 107 | 44 | 78 | 2505 | 635 |
| SRR4296979 | 0 | 84 | 0 | 0 | 0 | 0 | 0 | 1026 | 867 | 14771 | 160 | 2759 | 0 | 1013 | 307 | 1093 | 0 | 24 | 44 | 1853 | 1872 | 8234 | 0 | 11 | 13274 | 0 | 14 | 0 | 0 | 40 | 55 | 0 | 32 | 64 | 55 | 22876 | 105 |
| SRR4296985 | 0 | 0 | 0 | 0 | 15 | 0 | 0 | 421 | 961 | 3244 | 51 | 2163 | 0 | 3004 | 0 | 0 | 0 | 188 | 60 | 60 | 413 | 930 | 45 | 953 | 15036 | 0 | 15 | 0 | 0 | 0 | 0 | 0 | 0 | 0 | 0 | 60 | 15 |
| SRR4296986 | 0 | 0 | 0 | 0 | 17 | 0 | 0 | 363 | 363 | 8407 | 69 | 1179 | 0 | 3326 | 30 | 0 | 0 | 73 | 82 | 91 | 514 | 1503 | 65 | 980 | 12954 | 0 | 50 | 0 | 0 | 0 | 0 | 0 | 0 | 0 | 0 | 514 | 13 |
| SRR4296987 | 0 | 0 | 0 | 0 | 23 | 0 | 0 | 843 | 589 | 58786 | 63 | 61 | 0 | 5779 | 111 | 39 | 0 | 23 | 0 | 19 | 3893 | 11039 | 0 | 248 | 20340 | 0 | 0 | 0 | 0 | 0 | 0 | 0 | 0 | 0 | 0 | 202 | 134 |
| SRR4296988 | 0 | 0 | 0 | 0 | 17 | 0 | 0 | 843 | 613 | 51563 | 66 | 77 | 0 | 6231 | 112 | 40 | 0 | 18 | 14 | 21 | 3654 | 10482 | 0 | 349 | 18404 | 0 | 0 | 0 | 0 | 0 | 0 | 0 | 0 | 0 | 0 | 295 | 157 |
| SRR4328127 | 0 | 0 | 0 | 40 | 20 | 0 | 0 | 0 | 50 | 586 | 34 | 50 | 0 | 67 | 84 | 0 | 0 | 613 | 119 | 837 | 89 | 425 | 0 | 267 | 2065 | 731 | 100 | 0 | 0 | 0 | 0 | 0 | 0 | 0 | 0 | 1121 | 49 |
| SRR4328130 | 0 | 0 | 0 | 32 | 64 | 0 | 0 | 0 | 0 | 1484 | 742 | 495 | 0 | 1484 | 0 | 247 | 0 | 321 | 80 | 1237 | 80 | 450 | 16 | 321 | 1993 | 321 | 27 | 0 | 0 | 0 | 0 | 0 | 0 | 0 | 0 | 1237 | 32 |
| SRR4328135 | 0 | 0 | 0 | 0 | 0 | 0 | 0 | 0 | 0 | 3699 | 0 | 0 | 0 | 231 | 0 | 0 | 0 | 0 | 143 | 1773 | 0 | 57 | 0 | 0 | 918 | 315 | 75 | 0 | 0 | 0 | 0 | 0 | 0 | 0 | 0 | 1464 | 0 |
| SRR4328136 | 0 | 0 | 0 | 38 | 19 | 0 | 0 | 0 | 0 | 60 | 0 | 0 | 0 | 40 | 20 | 0 | 0 | 917 | 191 | 3421 | 19 | 172 | 0 | 287 | 2714 | 707 | 42 | 0 | 0 | 0 | 0 | 0 | 0 | 0 | 0 | 1301 | 0 |
| SRR4328137 | 0 | 0 | 0 | 187 | 0 | 0 | 0 | 0 | 0 | 115 | 0 | 0 | 0 | 69 | 0 | 46 | 0 | 998 | 249 | 2123 | 62 | 686 | 0 | 624 | 2931 | 811 | 14 | 0 | 0 | 0 | 0 | 0 | 0 | 0 | 0 | 1108 | 0 |
| SRR4409201 | 0 | 0 | 10 | 0 | 0 | 0 | 0 | 53 | 102 | 1073 | 146 | 171 | 0 | 168 | 0 | 11 | 0 | 694 | 0 | 0 | 1104 | 768 | 443 | 3486 | 1018 | 0 | 0 | 0 | 0 | 0 | 0 | 0 | 0 | 0 | 19 | 0 | 0 |
| SRR4409202 | 0 | 0 | 33 | 0 | 0 | 0 | 0 | 206 | 36 | 2287 | 4274 | 1897 | 0 | 2134 | 473 | 1281 | 0 | 83 | 0 | 0 | 81 | 296 | 0 | 15 | 656 | 0 | 0 | 0 | 0 | 0 | 0 | 0 | 15 | 21 | 201 | 31 | 14 |
| SRR4409203 | 0 | 0 | 0 | 0 | 0 | 0 | 0 | 125 | 200 | 2327 | 738 | 338 | 0 | 228 | 0 | 0 | 0 | 803 | 0 | 168 | 803 | 812 | 290 | 2942 | 485 | 0 | 0 | 0 | 0 | 0 | 0 | 0 | 0 | 0 | 18 | 990 | 0 |
| SRR4409204 | 0 | 0 | 0 | 0 | 0 | 0 | 0 | 271 | 122 | 3981 | 297 | 1633 | 0 | 402 | 114 | 166 | 0 | 87 | 62 | 1187 | 149 | 416 | 0 | 43 | 1055 | 180 | 0 | 0 | 0 | 0 | 0 | 0 | 0 | 0 | 11 | 20805 | 25 |
| SRR4417245 | 0 | 3589 | 0 | 0 | 0 | 34 | 0 | 0 | 33 | 33 | 347 | 955 | 0 | 0 | 0 | 0 | 0 | 5139 | 33 | 178 | 2435 | 2201 | 0 | 85 | 1613 | 436 | 13 | 0 | 0 | 0 | 0 | 0 | 0 | 0 | 0 | 147 | 0 |
| SRR4417246 | 0 | 40 | 0 | 0 | 0 | 24 | 0 | 0 | 0 | 0 | 769 | 1844 | 0 | 0 | 0 | 0 | 0 | 3896 | 25 | 29 | 3589 | 1416 | 0 | 26 | 944 | 554 | 0 | 0 | 0 | 0 | 0 | 0 | 0 | 0 | 0 | 36 | 0 |
| SRR4417247 | 0 | 139 | 0 | 0 | 0 | 29 | 0 | 18 | 97 | 42 | 74 | 210 | 0 | 0 | 0 | 0 | 0 | 6426 | 0 | 729 | 2929 | 2537 | 0 | 95 | 1608 | 225 | 0 | 0 | 0 | 0 | 0 | 0 | 0 | 0 | 0 | 639 | 0 |
| SRR4417248 | 0 | 127 | 0 | 0 | 0 | 36 | 0 | 45 | 252 | 54 | 474 | 1877 | 0 | 0 | 0 | 0 | 0 | 5509 | 15 | 241 | 2497 | 2492 | 0 | 35 | 1073 | 217 | 0 | 0 | 0 | 0 | 0 | 0 | 0 | 0 | 0 | 194 | 0 |
| SRR4417249 | 0 | 2091 | 0 | 0 | 11 | 63 | 0 | 0 | 0 | 0 | 788 | 4021 | 0 | 0 | 0 | 0 | 0 | 3679 | 0 | 0 | 3831 | 2489 | 0 | 26 | 803 | 202 | 0 | 0 | 0 | 0 | 0 | 0 | 0 | 0 | 0 | 0 | 0 |
| SRR4417250 | 0 | 5972 | 0 | 0 | 11 | 41 | 0 | 30 | 108 | 136 | 296 | 1826 | 0 | 0 | 0 | 0 | 0 | 6332 | 0 | 591 | 3149 | 4341 | 0 | 34 | 1440 | 114 | 0 | 0 | 0 | 0 | 0 | 0 | 0 | 0 | 0 | 572 | 0 |
| SRR4423247 | 0 | 89 | 78 | 0 | 16 | 0 | 0 | 279 | 128 | 663 | 1728 | 6367 | 0 | 189 | 0 | 25 | 0 | 137 | 12 | 978 | 1432 | 3280 | 313 | 21273 | 6898 | 203 | 0 | 120 | 203 | 608 | 402 | 83 | 392 | 846 | 415 | 2616 | 37 |
| SRR4423248 | 0 | 30 | 25 | 0 | 35 | 0 | 24 | 393 | 494 | 978 | 613 | 5116 | 0 | 228 | 11 | 0 | 0 | 272 | 23 | 531 | 2248 | 4002 | 423 | 19411 | 7217 | 227 | 103 | 23 | 40 | 75 | 141 | 52 | 156 | 707 | 155 | 1923 | 79 |
| SRR4423250 | 0 | 66 | 84 | 0 | 12 | 0 | 0 | 382 | 108 | 1151 | 14824 | 10406 | 0 | 487 | 100 | 228 | 0 | 508 | 20 | 2355 | 898 | 5721 | 468 | 63166 | 4198 | 1363 | 26 | 27 | 157 | 1423 | 1404 | 80 | 176 | 479 | 262 | 21705 | 165 |
| SRR4423251 | 0 | 120 | 68 | 0 | 11 | 0 | 0 | 188 | 94 | 2048 | 8228 | 5408 | 0 | 380 | 33 | 135 | 0 | 534 | 71 | 9422 | 581 | 2467 | 95 | 14240 | 6022 | 0 | 0 | 36 | 169 | 1294 | 3473 | 171 | 460 | 1003 | 712 | 39133 | 61 |
| SRR4473670 | 0 | 0 | 0 | 0 | 0 | 0 | 0 | 647 | 123 | 2217 | 1990 | 8449 | 0 | 859 | 162 | 437 | 0 | 163 | 10 | 1666 | 729 | 2802 | 0 | 2707 | 4642 | 274 | 0 | 0 | 0 | 17 | 46 | 26 | 35 | 13 | 41 | 47476 | 20 |
| SRR4734676 | 0 | 43 | 29 | 0 | 34 | 0 | 0 | 481 | 204 | 257 | 5899 | 38148 | 0 | 0 | 0 | 0 | 0 | 413 | 0 | 228 | 2106 | 5209 | 0 | 108 | 12 | 177 | 18 | 0 | 90 | 294 | 68 | 10 | 0 | 29 | 0 | 544 | 0 |
| SRR4734677 | 0 | 101 | 34 | 0 | 23 | 17 | 0 | 635 | 276 | 470 | 23789 | 27684 | 0 | 28 | 28 | 0 | 0 | 400 | 0 | 166 | 2023 | 5066 | 0 | 159 | 16 | 279 | 17 | 0 | 134 | 905 | 117 | 17 | 17 | 218 | 17 | 497 | 0 |
| SRR4734678 | 0 | 66 | 27 | 0 | 0 | 27 | 13 | 173 | 87 | 0 | 974 | 36605 | 0 | 0 | 0 | 0 | 0 | 100 | 0 | 650 | 387 | 2579 | 0 | 15 | 1088 | 114 | 0 | 0 | 0 | 93 | 40 | 13 | 27 | 13 | 0 | 1170 | 0 |
| SRR4734679 | 0 | 82 | 68 | 0 | 0 | 0 | 0 | 142 | 95 | 0 | 720 | 58704 | 0 | 0 | 0 | 47 | 0 | 150 | 0 | 332 | 1154 | 5355 | 0 | 40 | 1067 | 229 | 14 | 27 | 0 | 14 | 41 | 0 | 14 | 55 | 14 | 427 | 0 |
| SRR4734680 | 0 | 67 | 51 | 0 | 0 | 0 | 0 | 203 | 68 | 0 | 571 | 46199 | 0 | 0 | 0 | 0 | 0 | 165 | 0 | 338 | 1120 | 4710 | 0 | 31 | 1270 | 214 | 0 | 0 | 17 | 135 | 17 | 17 | 17 | 17 | 17 | 473 | 0 |
| SRR4734681 | 0 | 69 | 54 | 0 | 18 | 0 | 0 | 567 | 118 | 47 | 999 | 29226 | 0 | 24 | 0 | 0 | 0 | 293 | 0 | 519 | 2250 | 4548 | 0 | 100 | 4186 | 352 | 0 | 38 | 130 | 353 | 77 | 15 | 38 | 38 | 0 | 897 | 0 |
| SRR4734682 | 0 | 57 | 32 | 0 | 11 | 16 | 0 | 382 | 42 | 85 | 42298 | 61389 | 0 | 0 | 0 | 0 | 0 | 201 | 0 | 424 | 1899 | 5355 | 0 | 32 | 1969 | 265 | 0 | 0 | 16 | 73 | 24 | 0 | 24 | 0 | 0 | 1061 | 0 |
| SRR4734683 | 0 | 47 | 20 | 0 | 0 | 0 | 0 | 141 | 20 | 54 | 55079 | 60929 | 0 | 40 | 0 | 0 | 0 | 111 | 0 | 357 | 854 | 5636 | 0 | 22 | 947 | 160 | 0 | 11 | 0 | 77 | 16 | 0 | 15 | 18 | 0 | 990 | 0 |
| SRR4734684 | 0 | 97 | 52 | 0 | 10 | 0 | 0 | 216 | 127 | 140 | 32823 | 48189 | 0 | 13 | 0 | 0 | 0 | 269 | 0 | 289 | 2091 | 6533 | 0 | 36 | 2597 | 137 | 36 | 24 | 93 | 12 | 135 | 30 | 45 | 79 | 0 | 977 | 0 |
| SRR4734685 | 0 | 17 | 26 | 0 | 0 | 10 | 0 | 100 | 84 | 42 | 6243 | 54868 | 0 | 33 | 33 | 25 | 0 | 88 | 0 | 652 | 1227 | 6104 | 0 | 30 | 910 | 208 | 0 | 0 | 17 | 62 | 31 | 14 | 19 | 33 | 0 | 1522 | 0 |
| SRR4734686 | 0 | 67 | 42 | 0 | 17 | 0 | 0 | 411 | 123 | 108 | 2933 | 30266 | 0 | 22 | 0 | 0 | 0 | 226 | 0 | 159 | 1335 | 4353 | 0 | 63 | 5528 | 180 | 13 | 25 | 93 | 15 | 249 | 13 | 34 | 67 | 0 | 404 | 0 |
| SRR4787084 | 0 | 2625 | 0 | 0 | 0 | 0 | 0 | 988 | 1729 | 11359 | 0 | 0 | 0 | 247 | 864 | 864 | 0 | 787 | 0 | 123 | 902 | 4114 | 192 | 5042 | 245 | 192 | 69 | 0 | 0 | 0 | 0 | 0 | 0 | 0 | 0 | 864 | 0 |
| SRR4787085 | 0 | 71 | 0 | 0 | 0 | 0 | 0 | 2766 | 178 | 7405 | 359 | 535 | 0 | 3122 | 3122 | 3212 | 0 | 109 | 0 | 803 | 1846 | 3085 | 339 | 9876 | 187 | 415 | 0 | 0 | 0 | 36 | 0 | 36 | 0 | 0 | 0 | 4282 | 0 |
| SRR4787086 | 0 | 207 | 0 | 0 | 0 | 0 | 0 | 666 | 932 | 12385 | 400 | 266 | 0 | 666 | 1332 | 2131 | 64 | 843 | 15 | 133 | 1274 | 5641 | 169 | 4977 | 3336 | 243 | 0 | 0 | 64 | 0 | 0 | 0 | 0 | 0 | 0 | 1065 | 0 |
| SRR4787087 | 0 | 7254 | 45 | 0 | 0 | 0 | 0 | 1239 | 1594 | 9119 | 89 | 177 | 0 | 531 | 797 | 2390 | 45 | 726 | 10 | 177 | 1404 | 5229 | 266 | 5369 | 3419 | 228 | 0 | 0 | 0 | 45 | 0 | 0 | 0 | 0 | 0 | 797 | 0 |
| SRR4787088 | 0 | 24694 | 0 | 0 | 0 | 0 | 0 | 1507 | 879 | 7159 | 379 | 377 | 0 | 502 | 879 | 4145 | 0 | 1807 | 0 | 126 | 6701 | 10 | 208 | 4257 | 221 | 129 | 66 | 0 | 0 | 0 | 0 | 0 | 0 | 0 | 0 | 502 | 0 |
| SRR4787089 | 0 | 1008 | 0 | 0 | 16 | 0 | 0 | 1096 | 548 | 7491 | 737 | 548 | 0 | 731 | 1644 | 5116 | 0 | 1699 | 0 | 365 | 6979 | 10594 | 201 | 4097 | 255 | 145 | 0 | 0 | 0 | 0 | 0 | 0 | 0 | 0 | 0 | 365 | 0 |
| SRR4787090 | 0 | 2225 | 49 | 80 | 15 | 13 | 0 | 1729 | 3911 | 16185 | 303 | 367 | 0 | 4300 | 1772 | 4516 | 36 | 311 | 17 | 1707 | 1386 | 1467 | 161 | 6632 | 93 | 213 | 22 | 0 | 31 | 13 | 27 | 22 | 0 | 0 | 0 | 3220 | 0 |
| SRR4787091 | 0 | 433 | 0 | 10 | 0 | 0 | 0 | 1390 | 1112 | 8065 | 556 | 834 | 0 | 278 | 556 | 5006 | 0 | 1703 | 10 | 278 | 7417 | 14 | 94 | 4137 | 196 | 219 | 0 | 0 | 0 | 0 | 157 | 0 | 0 | 0 | 0 | 278 | 0 |
| SRR4787092 | 0 | 2096 | 96 | 99 | 17 | 19 | 24 | 1395 | 4255 | 16698 | 389 | 366 | 0 | 4003 | 1510 | 3477 | 43 | 296 | 10 | 1647 | 1362 | 1486 | 124 | 6328 | 78 | 203 | 29 | 0 | 63 | 19 | 24 | 0 | 0 | 0 | 0 | 2905 | 0 |
| SRR4787093 | 0 | 1807 | 51 | 116 | 0 | 28 | 28 | 1622 | 3749 | 15719 | 18 | 241 | 0 | 4363 | 1293 | 4538 | 28 | 350 | 13 | 1907 | 1416 | 1516 | 162 | 6602 | 59 | 229 | 41 | 0 | 32 | 18 | 51 | 0 | 0 | 0 | 0 | 3157 | 0 |
| SRR4787095 | 0 | 374 | 39 | 0 | 14 | 0 | 0 | 14 | 339 | 11985 | 49 | 94 | 0 | 1007 | 654 | 2598 | 0 | 566 | 16 | 210 | 530 | 1384 | 570 | 7967 | 2038 | 413 | 0 | 21 | 249 | 30 | 56 | 17 | 17 | 30 | 0 | 983 | 0 |
| SRR4787096 | 0 | 251 | 10 | 0 | 19 | 0 | 0 | 45 | 295 | 13583 | 80 | 83 | 0 | 754 | 869 | 3430 | 10 | 611 | 16 | 212 | 495 | 1391 | 487 | 7474 | 1911 | 395 | 10 | 10 | 348 | 31 | 51 | 31 | 41 | 26 | 0 | 911 | 0 |
| SRR4787097 | 0 | 31 | 0 | 0 | 0 | 0 | 0 | 151 | 103 | 4070 | 86 | 22 | 0 | 5739 | 202 | 942 | 35 | 482 | 0 | 159 | 91 | 1353 | 89 | 412 | 307 | 368 | 0 | 0 | 35 | 0 | 0 | 0 | 0 | 0 | 0 | 1316 | 72 |
| SRR4787098 | 0 | 0 | 16 | 0 | 0 | 0 | 0 | 21 | 103 | 4075 | 53 | 18 | 0 | 6791 | 195 | 678 | 0 | 334 | 0 | 13 | 112 | 1045 | 73 | 345 | 300 | 347 | 0 | 0 | 22 | 0 | 0 | 0 | 0 | 10 | 0 | 1148 | 66 |
| SRR4787099 | 0 | 13 | 24 | 0 | 0 | 0 | 0 | 67 | 64 | 4374 | 40 | 40 | 0 | 6011 | 187 | 468 | 0 | 385 | 0 | 70 | 73 | 1004 | 70 | 456 | 325 | 371 | 0 | 0 | 46 | 0 | 0 | 0 | 33 | 19 | 11 | 1251 | 52 |
| SRR4787100 | 0 | 117 | 34 | 0 | 0 | 18 | 0 | 148 | 539 | 15742 | 26 | 175 | 0 | 1253 | 768 | 2647 | 0 | 1956 | 0 | 195 | 717 | 5417 | 98 | 757 | 749 | 428 | 22 | 0 | 58 | 0 | 31 | 25 | 28 | 52 | 0 | 1448 | 103 |
| SRR4787101 | 0 | 109 | 21 | 0 | 0 | 0 | 0 | 106 | 529 | 13909 | 27 | 139 | 0 | 1625 | 668 | 1842 | 0 | 1703 | 0 | 150 | 558 | 4761 | 108 | 687 | 691 | 430 | 16 | 0 | 59 | 16 | 0 | 11 | 0 | 13 | 0 | 1447 | 141 |
| SRR4787102 | 0 | 12 | 34 | 0 | 0 | 0 | 0 | 195 | 452 | 15424 | 269 | 229 | 0 | 1529 | 796 | 2314 | 0 | 1704 | 0 | 183 | 568 | 4553 | 103 | 873 | 807 | 405 | 22 | 0 | 56 | 17 | 19 | 15 | 17 | 34 | 0 | 1564 | 94 |
| SRR4787103 | 0 | 33 | 25 | 0 | 20 | 0 | 0 | 146 | 671 | 22285 | 243 | 251 | 0 | 210 | 785 | 2200 | 15 | 3165 | 0 | 154 | 1652 | 6152 | 116 | 1660 | 2039 | 316 | 40 | 0 | 137 | 30 | 46 | 51 | 40 | 15 | 10 | 1383 | 155 |
| SRR4787104 | 0 | 42 | 49 | 0 | 19 | 0 | 0 | 187 | 813 | 23907 | 54 | 289 | 0 | 295 | 657 | 2156 | 0 | 3149 | 0 | 181 | 1420 | 6177 | 125 | 1720 | 1900 | 321 | 35 | 0 | 116 | 14 | 60 | 39 | 32 | 49 | 0 | 1554 | 125 |
| SRR4787105 | 0 | 0 | 21 | 0 | 0 | 25 | 0 | 188 | 129 | 24748 | 69 | 20 | 0 | 1664 | 248 | 540 | 0 | 28 | 0 | 441 | 3130 | 8055 | 75 | 5549 | 3594 | 584 | 12 | 0 | 0 | 41 | 29 | 12 | 37 | 104 | 0 | 4491 | 71 |
| SRR4787106 | 0 | 19 | 38 | 0 | 0 | 0 | 0 | 275 | 153 | 30047 | 69 | 0 | 0 | 1452 | 420 | 665 | 32 | 44 | 0 | 420 | 3420 | 8786 | 87 | 5614 | 3747 | 587 | 57 | 0 | 0 | 45 | 25 | 0 | 38 | 89 | 0 | 4554 | 93 |
| SRR4787107 | 0 | 15 | 21 | 0 | 0 | 0 | 0 | 298 | 91 | 33971 | 246 | 111 | 0 | 822 | 723 | 1056 | 0 | 109 | 0 | 766 | 2721 | 6200 | 92 | 6244 | 4959 | 290 | 15 | 0 | 0 | 21 | 42 | 12 | 12 | 91 | 0 | 5262 | 181 |
| SRR4787108 | 0 | 16 | 31 | 0 | 0 | 25 | 0 | 164 | 68 | 20134 | 209 | 113 | 0 | 419 | 679 | 1182 | 0 | 60 | 0 | 1024 | 1274 | 2982 | 108 | 10172 | 3246 | 131 | 41 | 0 | 0 | 16 | 44 | 0 | 13 | 56 | 0 | 8003 | 89 |
| SRR4787109 | 0 | 22 | 14 | 0 | 0 | 18 | 0 | 299 | 130 | 32394 | 22 | 26 | 0 | 1197 | 360 | 681 | 14 | 46 | 0 | 590 | 3319 | 8752 | 70 | 5396 | 3988 | 553 | 25 | 0 | 0 | 40 | 11 | 0 | 14 | 97 | 11 | 4628 | 89 |
| SRR4787110 | 0 | 42 | 21 | 0 | 0 | 12 | 0 | 346 | 124 | 33501 | 51 | 85 | 0 | 905 | 509 | 760 | 15 | 82 | 0 | 477 | 3681 | 7529 | 95 | 5681 | 5144 | 447 | 24 | 0 | 0 | 36 | 39 | 0 | 24 | 63 | 0 | 3991 | 150 |
| SRR4787118 | 0 | 0 | 0 | 0 | 0 | 0 | 0 | 0 | 0 | 0 | 0 | 0 | 0 | 908 | 0 | 1735 | 0 | 0 | 0 | 0 | 0 | 0 | 0 | 3144 | 0 | 0 | 0 | 0 | 0 | 0 | 0 | 0 | 0 | 0 | 0 | 0 | 0 |
| SRR4787119 | 0 | 0 | 0 | 0 | 0 | 0 | 0 | 0 | 0 | 0 | 0 | 0 | 0 | 1735 | 0 | 0 | 0 | 0 | 0 | 0 | 0 | 0 | 6992 | 8114 | 4113 | 0 | 0 | 0 | 0 | 0 | 0 | 0 | 0 | 0 | 0 | 0 | 0 |
| SRR4787120 | 0 | 45 | 51 | 0 | 18 | 0 | 0 | 56 | 433 | 13654 | 30 | 111 | 0 | 433 | 1492 | 6175 | 0 | 901 | 17 | 232 | 585 | 2201 | 409 | 7588 | 2577 | 447 | 14 | 46 | 11 | 55 | 23 | 28 | 28 | 32 | 0 | 1063 | 0 |
| SRR4787121 | 0 | 89 | 0 | 0 | 0 | 0 | 0 | 19 | 47 | 1709 | 0 | 0 | 0 | 2642 | 748 | 2851 | 0 | 139 | 24 | 159 | 28 | 511 | 292 | 2566 | 430 | 457 | 0 | 12 | 45 | 0 | 0 | 14 | 12 | 16 | 18 | 850 | 0 |
| SRR4787122 | 0 | 26 | 24 | 0 | 0 | 0 | 0 | 27 | 53 | 1503 | 0 | 43 | 15 | 2554 | 837 | 3150 | 0 | 110 | 22 | 249 | 33 | 493 | 211 | 2005 | 321 | 460 | 24 | 18 | 46 | 0 | 0 | 0 | 22 | 31 | 0 | 804 | 0 |
| SRR4787123 | 0 | 73 | 39 | 0 | 0 | 0 | 0 | 0 | 15 | 1194 | 0 | 0 | 0 | 2750 | 462 | 2050 | 0 | 113 | 23 | 97 | 23 | 416 | 280 | 2135 | 323 | 387 | 18 | 0 | 91 | 0 | 14 | 0 | 0 | 0 | 20 | 660 | 0 |
| SRR4787124 | 0 | 0 | 0 | 0 | 0 | 0 | 0 | 0 | 0 | 636 | 0 | 0 | 0 | 3844 | 302 | 297 | 0 | 0 | 0 | 0 | 0 | 336 | 467 | 4054 | 0 | 385 | 0 | 0 | 0 | 0 | 0 | 0 | 0 | 0 | 0 | 299 | 0 |
| SRR4787125 | 0 | 0 | 0 | 0 | 0 | 0 | 0 | 0 | 0 | 238 | 0 | 0 | 0 | 3108 | 159 | 603 | 0 | 102 | 0 | 0 | 0 | 168 | 394 | 6187 | 711 | 733 | 0 | 0 | 200 | 0 | 0 | 0 | 0 | 0 | 0 | 120 | 0 |
| SRR4787126 | 0 | 0 | 0 | 0 | 0 | 0 | 0 | 0 | 0 | 156 | 0 | 0 | 0 | 3230 | 77 | 264 | 0 | 0 | 0 | 89 | 132 | 147 | 586 | 5099 | 491 | 978 | 0 | 0 | 29 | 0 | 0 | 0 | 0 | 0 | 0 | 545 | 0 |
| SRR4787127 | 0 | 0 | 0 | 0 | 0 | 0 | 0 | 0 | 0 | 0 | 0 | 0 | 0 | 3865 | 0 | 0 | 0 | 0 | 0 | 0 | 0 | 0 | 0 | 11518 | 0 | 473 | 0 | 0 | 0 | 0 | 0 | 0 | 0 | 0 | 0 | 2300 | 0 |
| SRR4787132 | 0 | 47 | 0 | 0 | 0 | 24 | 0 | 3830 | 225 | 5970 | 338 | 282 | 0 | 3267 | 1577 | 3098 | 0 | 104 | 0 | 1295 | 1910 | 3513 | 282 | 9678 | 5510 | 394 | 0 | 0 | 0 | 0 | 0 | 0 | 0 | 0 | 0 | 4675 | 0 |
| SRR4787133 | 33 | 67 | 0 | 0 | 11 | 0 | 0 | 3887 | 793 | 7695 | 239 | 555 | 0 | 3094 | 2459 | 3491 | 0 | 122 | 0 | 952 | 1612 | 2936 | 343 | 9803 | 195 | 343 | 33 | 0 | 0 | 0 | 33 | 0 | 0 | 0 | 0 | 4522 | 0 |
| SRR4787134 | 0 | 0 | 22 | 0 | 0 | 0 | 0 | 50 | 40 | 2121 | 0 | 0 | 0 | 9593 | 12 | 12 | 0 | 165 | 0 | 59 | 61 | 325 | 112 | 613 | 356 | 355 | 0 | 0 | 0 | 0 | 0 | 0 | 0 | 0 | 0 | 1424 | 38 |
| SRR4787135 | 0 | 564 | 18 | 0 | 24 | 0 | 0 | 193 | 635 | 22393 | 30 | 335 | 0 | 157 | 820 | 1855 | 14 | 3644 | 0 | 114 | 1921 | 6971 | 113 | 1562 | 1768 | 325 | 42 | 14 | 194 | 55 | 42 | 32 | 18 | 28 | 0 | 1149 | 194 |
| SRR4787136 | 0 | 115 | 34 | 0 | 0 | 0 | 0 | 214 | 489 | 17636 | 95 | 90 | 0 | 1665 | 670 | 2081 | 0 | 1387 | 14 | 296 | 476 | 3442 | 102 | 1409 | 1483 | 304 | 16 | 29 | 13 | 16 | 0 | 32 | 21 | 56 | 11 | 1151 | 169 |
| SRR4787137 | 0 | 39 | 0 | 0 | 0 | 0 | 0 | 92 | 68 | 2374 | 0 | 0 | 0 | 7557 | 36 | 16 | 0 | 166 | 0 | 12 | 16 | 346 | 120 | 944 | 529 | 431 | 0 | 22 | 43 | 0 | 0 | 0 | 17 | 47 | 0 | 1153 | 48 |
| SRR4787138 | 0 | 27 | 15 | 0 | 0 | 0 | 0 | 210 | 265 | 6842 | 18 | 15 | 0 | 5758 | 51 | 161 | 11 | 709 | 0 | 33 | 217 | 1592 | 116 | 1027 | 1173 | 231 | 0 | 18 | 75 | 0 | 0 | 0 | 0 | 28 | 0 | 918 | 87 |
| SRR4787139 | 0 | 408 | 27 | 0 | 0 | 0 | 0 | 236 | 372 | 10237 | 19 | 36 | 0 | 5107 | 130 | 328 | 0 | 954 | 0 | 92 | 301 | 2541 | 123 | 1122 | 1326 | 261 | 0 | 12 | 69 | 0 | 14 | 10 | 0 | 25 | 0 | 1021 | 78 |
| SRR4787140 | 0 | 1090 | 49 | 0 | 0 | 0 | 0 | 382 | 1155 | 25336 | 69 | 34 | 0 | 2433 | 249 | 677 | 0 | 1866 | 0 | 77 | 702 | 4530 | 142 | 1690 | 2081 | 198 | 15 | 15 | 107 | 26 | 34 | 17 | 17 | 43 | 0 | 952 | 110 |
| SRR4787141 | 0 | 61 | 23 | 0 | 0 | 0 | 0 | 260 | 424 | 9402 | 51 | 0 | 0 | 5126 | 131 | 284 | 13 | 922 | 0 | 53 | 268 | 2389 | 124 | 1039 | 1084 | 250 | 10 | 0 | 11 | 0 | 15 | 0 | 0 | 19 | 0 | 999 | 79 |
| SRR4787142 | 0 | 1096 | 44 | 0 | 13 | 0 | 0 | 401 | 1274 | 30841 | 68 | 100 | 0 | 1763 | 321 | 645 | 0 | 2061 | 0 | 120 | 775 | 4944 | 142 | 1658 | 2054 | 193 | 28 | 23 | 158 | 30 | 26 | 30 | 30 | 23 | 0 | 1082 | 79 |
| SRR4787143 | 0 | 56 | 53 | 0 | 12 | 0 | 0 | 266 | 1064 | 22720 | 60 | 82 | 0 | 3191 | 236 | 528 | 0 | 1736 | 0 | 60 | 557 | 3916 | 126 | 1502 | 1720 | 176 | 24 | 33 | 12 | 15 | 22 | 26 | 24 | 48 | 0 | 1120 | 91 |
| SRR4836291 | 0 | 0 | 18 | 0 | 16 | 0 | 0 | 148 | 27 | 1140 | 10422 | 6814 | 0 | 215 | 684 | 1583 | 0 | 262 | 28 | 2133 | 3250 | 2224 | 22 | 6731 | 7429 | 1219 | 14 | 0 | 10 | 28 | 15 | 13 | 21 | 16 | 10 | 18135 | 19 |
| SRR4836293 | 0 | 0 | 13 | 0 | 19 | 0 | 0 | 148 | 16 | 1004 | 11377 | 6190 | 0 | 263 | 840 | 2074 | 0 | 270 | 23 | 2009 | 2989 | 2022 | 31 | 6252 | 6652 | 1310 | 13 | 0 | 19 | 22 | 57 | 0 | 22 | 57 | 35 | 19592 | 12 |
| SRR4836294 | 0 | 0 | 0 | 0 | 11 | 0 | 0 | 128 | 0 | 1134 | 10621 | 6548 | 0 | 306 | 572 | 1706 | 0 | 284 | 32 | 2169 | 3165 | 2026 | 16 | 6554 | 6888 | 1262 | 10 | 0 | 0 | 12 | 28 | 0 | 12 | 16 | 14 | 19131 | 18 |
| SRR4836295 | 0 | 36 | 0 | 0 | 16 | 26 | 0 | 541 | 123 | 418 | 17050 | 11129 | 0 | 369 | 0 | 0 | 0 | 585 | 29 | 1155 | 903 | 435 | 89 | 4855 | 1745 | 845 | 52 | 62 | 21 | 36 | 82 | 26 | 21 | 52 | 118 | 10859 | 0 |
| SRR4836297 | 0 | 38 | 0 | 0 | 19 | 0 | 0 | 519 | 183 | 427 | 19341 | 11318 | 0 | 275 | 0 | 0 | 19 | 564 | 34 | 1098 | 718 | 390 | 99 | 4751 | 1479 | 963 | 44 | 44 | 19 | 50 | 57 | 38 | 57 | 76 | 113 | 11837 | 0 |

### Supp Table S8.4.Columns_DI_to_ES

| | | | | | | | | | | | | | | | | | | | | | | | | | | | | | | | | | | | | | |
|---|---|---|---|---|---|---|---|---|---|---|---|---|---|---|---|---|---|---|---|---|---|---|---|---|---|---|---|---|---|---|---|---|---|---|---|---|---|
| SRR4836299 | 0 | 30 | 11 | 0 | 16 | 11 | 11 | 463 | 392 | 606 | 952 | 10477 | 0 | 267 | 0 | 0 | 19 | 561 | 33 | 1105 | 820 | 395 | 98 | 4698 | 1493 | 833 | 10 | 19 | 19 | 11 | 18 | 26 | 37 | 45 | 23 | 10923 | 0 |
| SRR4836301 | 0 | 11 | 21 | 0 | 19 | 0 | 21 | 148 | 17 | 513 | 3470 | 2261 | 0 | 52 | 52 | 61 | 21 | 270 | 37 | 3861 | 3683 | 2369 | 16 | 11787 | 7764 | 1285 | 57 | 32 | 17 | 22 | 23 | 27 | 32 | 19 | 21 | 40283 | 0 |
| SRR4836302 | 0 | 0 | 24 | 0 | 26 | 0 | 24 | 126 | 21 | 546 | 339 | 2112 | 0 | 74 | 21 | 84 | 20 | 182 | 23 | 3446 | 3330 | 2192 | 42 | 11498 | 7127 | 1249 | 154 | 98 | 61 | 56 | 70 | 11 | 65 | 61 | 48 | 39616 | 16 |
| SRR4836303 | 0 | 10 | 26 | 0 | 19 | 0 | 35 | 70 | 19 | 518 | 3314 | 2399 | 0 | 45 | 38 | 83 | 13 | 184 | 33 | 3871 | 3574 | 2057 | 31 | 10995 | 7556 | 1107 | 182 | 93 | 34 | 67 | 60 | 17 | 76 | 71 | 54 | 41504 | 19 |
| SRR4836305 | 0 | 0 | 44 | 0 | 19 | 0 | 44 | 136 | 24 | 1002 | 553 | 2004 | 0 | 2650 | 184 | 581 | 25 | 265 | 12 | 1203 | 3234 | 2572 | 33 | 7988 | 7079 | 949 | 32 | 44 | 54 | 44 | 54 | 0 | 59 | 11 | 11 | 12277 | 0 |
| SRR4836307 | 0 | 0 | 36 | 0 | 0 | 0 | 30 | 111 | 22 | 782 | 3292 | 1934 | 0 | 2584 | 162 | 650 | 12 | 322 | 0 | 1181 | 2934 | 2403 | 30 | 7342 | 6595 | 981 | 163 | 66 | 54 | 66 | 60 | 42 | 78 | 90 | 72 | 12460 | 0 |
| SRR4836308 | 0 | 15 | 22 | 0 | 0 | 0 | 10 | 133 | 18 | 946 | 789 | 2126 | 0 | 2598 | 164 | 610 | 29 | 270 | 0 | 1330 | 3252 | 2246 | 20 | 7453 | 6627 | 920 | 29 | 13 | 54 | 11 | 11 | 11 | 58 | 17 | 12 | 12691 | 0 |
| SRR4836310 | 0 | 19 | 12 | 0 | 0 | 0 | 0 | 87 | 16 | 442 | 731 | 2887 | 0 | 1341 | 229 | 371 | 12 | 240 | 26 | 1625 | 1266 | 1273 | 15 | 8108 | 2754 | 2718 | 90 | 43 | 82 | 140 | 101 | 12 | 66 | 101 | 97 | 12501 | 0 |
| SRR4836311 | 0 | 24 | 24 | 0 | 14 | 0 | 14 | 101 | 0 | 434 | 5820 | 3137 | 0 | 1695 | 161 | 242 | 0 | 236 | 39 | 1463 | 1125 | 1219 | 29 | 7499 | 2552 | 2617 | 19 | 11 | 18 | 31 | 15 | 28 | 12 | 16 | 18 | 13718 | 0 |
| SRR4836312 | 0 | 11 | 43 | 0 | 0 | 0 | 11 | 59 | 12 | 381 | 949 | 2863 | 0 | 1426 | 217 | 422 | 0 | 225 | 33 | 1625 | 1149 | 1155 | 26 | 7591 | 2569 | 2408 | 17 | 11 | 11 | 28 | 18 | 45 | 19 | 15 | 19 | 13161 | 0 |
| SRR4836313 | 0 | 10 | 20 | 0 | 16 | 0 | 36 | 363 | 73 | 1275 | 22395 | 20964 | 0 | 290 | 0 | 0 | 11 | 373 | 14 | 2851 | 7292 | 1701 | 23 | 6192 | 4993 | 1958 | 67 | 182 | 110 | 258 | 176 | 37 | 199 | 235 | 49 | 15842 | 0 |
| SRR4836314 | 0 | 0 | 19 | 0 | 0 | 0 | 29 | 396 | 109 | 1415 | 21875 | 20172 | 0 | 307 | 0 | 20 | 14 | 363 | 27 | 3207 | 7731 | 1772 | 26 | 6732 | 5385 | 1891 | 21 | 63 | 28 | 70 | 57 | 36 | 49 | 70 | 34 | 16342 | 0 |
| SRR4836315 | 0 | 0 | 27 | 0 | 13 | 0 | 23 | 338 | 107 | 1188 | 23115 | 21209 | 0 | 214 | 0 | 17 | 17 | 369 | 23 | 2590 | 7817 | 1618 | 26 | 5868 | 4755 | 1672 | 58 | 219 | 105 | 293 | 175 | 53 | 184 | 281 | 43 | 17250 | 0 |
| SRR4836316 | 0 | 14 | 0 | 0 | 0 | 0 | 0 | 88 | 10 | 801 | 2227 | 3608 | 0 | 973 | 0 | 26 | 12 | 251 | 109 | 2115 | 1761 | 2721 | 50 | 15732 | 7711 | 1935 | 17 | 20 | 28 | 23 | 18 | 14 | 17 | 23 | 10 | 13954 | 0 |
| SRR4836317 | 0 | 11 | 0 | 0 | 0 | 0 | 0 | 109 | 0 | 767 | 2357 | 3524 | 0 | 963 | 0 | 22 | 0 | 261 | 96 | 2279 | 1931 | 2816 | 46 | 16765 | 7794 | 1941 | 92 | 103 | 56 | 117 | 97 | 11 | 72 | 81 | 49 | 13455 | 0 |
| SRR4836318 | 0 | 0 | 0 | 0 | 10 | 0 | 0 | 98 | 13 | 701 | 2922 | 3835 | 0 | 810 | 0 | 32 | 0 | 214 | 101 | 2229 | 1976 | 2542 | 53 | 14825 | 6561 | 1651 | 12 | 20 | 32 | 20 | 17 | 23 | 17 | 19 | 36 | 14354 | 0 |
| SRR4836320 | 0 | 17 | 50 | 0 | 0 | 0 | 21 | 35 | 17 | 1067 | 1137 | 1249 | 0 | 972 | 0 | 17 | 0 | 554 | 21 | 704 | 5738 | 2001 | 33 | 5466 | 9081 | 1006 | 22 | 41 | 27 | 34 | 19 | 17 | 16 | 27 | 13 | 2524 | 86 |
| SRR4836321 | 0 | 20 | 32 | 0 | 0 | 0 | 12 | 38 | 31 | 1062 | 1753 | 1338 | 0 | 997 | 0 | 14 | 16 | 520 | 24 | 766 | 6298 | 2063 | 38 | 5890 | 9810 | 952 | 25 | 39 | 24 | 41 | 19 | 52 | 12 | 22 | 12 | 2674 | 80 |
| SRR4836323 | 0 | 17 | 34 | 17 | 13 | 0 | 14 | 33 | 22 | 1004 | 1923 | 1405 | 0 | 920 | 0 | 14 | 14 | 490 | 30 | 800 | 6540 | 1902 | 31 | 5328 | 8477 | 807 | 23 | 45 | 29 | 43 | 19 | 27 | 17 | 22 | 11 | 2693 | 112 |
| SRR4836325 | 0 | 0 | 11 | 0 | 12 | 0 | 13 | 264 | 36 | 1176 | 1297 | 12155 | 0 | 300 | 12 | 24 | 0 | 241 | 12 | 3156 | 1005 | 1453 | 29 | 10387 | 2701 | 689 | 40 | 68 | 35 | 39 | 48 | 35 | 20 | 40 | 38 | 40918 | 18 |
| SRR4836326 | 0 | 0 | 0 | 0 | 12 | 0 | 21 | 241 | 23 | 1080 | 1329 | 11724 | 0 | 287 | 0 | 92 | 0 | 215 | 0 | 2954 | 1076 | 1575 | 28 | 11267 | 2872 | 675 | 47 | 53 | 26 | 41 | 37 | 11 | 23 | 32 | 54 | 39894 | 18 |
| SRR4836327 | 0 | 0 | 0 | 0 | 12 | 0 | 0 | 219 | 30 | 745 | 16637 | 13299 | 0 | 288 | 30 | 60 | 11 | 176 | 0 | 3208 | 1080 | 1364 | 29 | 9500 | 2362 | 582 | 42 | 397 | 37 | 300 | 289 | 51 | 137 | 229 | 37 | 42004 | 13 |
| SRR4836328 | 0 | 0 | 15 | 0 | 11 | 0 | 12 | 235 | 67 | 1140 | 6035 | 4291 | 0 | 570 | 22 | 34 | 0 | 261 | 19 | 2369 | 2692 | 1528 | 26 | 3063 | 5314 | 184 | 19 | 10 | 10 | 10 | 25 | 0 | 22 | 29 | 15 | 24720 | 0 |
| SRR4836330 | 0 | 0 | 13 | 0 | 15 | 0 | 14 | 182 | 32 | 1303 | 5670 | 3759 | 0 | 513 | 21 | 32 | 11 | 248 | 22 | 2381 | 2910 | 1512 | 21 | 3349 | 5805 | 196 | 62 | 15 | 0 | 23 | 27 | 0 | 20 | 30 | 23 | 23995 | 0 |
| SRR4836332 | 0 | 0 | 19 | 0 | 13 | 0 | 18 | 212 | 0 | 1169 | 6424 | 4169 | 0 | 405 | 46 | 74 | 0 | 181 | 18 | 2338 | 3014 | 1467 | 29 | 2971 | 4700 | 160 | 51 | 11 | 15 | 27 | 27 | 0 | 0 | 39 | 27 | 24554 | 14 |
| SRR4836334 | 0 | 0 | 13 | 0 | 22 | 0 | 11 | 393 | 137 | 2065 | 778 | 11999 | 0 | 1229 | 34 | 85 | 0 | 136 | 67 | 2748 | 1081 | 2487 | 101 | 11401 | 6697 | 904 | 14 | 304 | 91 | 89 | 46 | 16 | 78 | 115 | 23 | 20107 | 0 |
| SRR4836335 | 0 | 0 | 18 | 0 | 17 | 0 | 14 | 546 | 132 | 2217 | 13796 | 11795 | 0 | 1009 | 0 | 33 | 12 | 121 | 68 | 3523 | 1159 | 2632 | 114 | 12660 | 6932 | 851 | 22 | 303 | 94 | 85 | 37 | 11 | 73 | 108 | 14 | 19536 | 0 |
| SRR4836336 | 0 | 0 | 16 | 0 | 18 | 0 | 10 | 358 | 124 | 1763 | 995 | 12699 | 0 | 1102 | 28 | 69 | 10 | 120 | 57 | 3278 | 1097 | 2278 | 118 | 10871 | 6027 | 837 | 71 | 1135 | 267 | 304 | 169 | 74 | 269 | 343 | 40 | 19819 | 0 |
| SRR4836337 | 0 | 18 | 23 | 0 | 0 | 0 | 30 | 161 | 17 | 914 | 8183 | 3832 | 0 | 752 | 391 | 1075 | 32 | 306 | 21 | 3076 | 3478 | 2280 | 28 | 18195 | 3595 | 2475 | 32 | 40 | 13 | 12 | 10 | 0 | 27 | 10 | 10 | 32457 | 17 |
| SRR4836339 | 0 | 0 | 12 | 0 | 14 | 0 | 35 | 218 | 14 | 787 | 8427 | 3764 | 0 | 773 | 296 | 1005 | 18 | 281 | 20 | 3190 | 3565 | 1947 | 40 | 14907 | 3031 | 2315 | 110 | 46 | 16 | 49 | 37 | 11 | 39 | 47 | 53 | 34851 | 13 |
| SRR4836341 | 0 | 11 | 26 | 0 | 0 | 0 | 32 | 265 | 28 | 885 | 2256 | 3299 | 0 | 872 | 275 | 970 | 21 | 295 | 22 | 2787 | 3778 | 2352 | 34 | 19498 | 3948 | 2722 | 117 | 37 | 29 | 56 | 39 | 14 | 49 | 56 | 56 | 30491 | 15 |
| SRR4836343 | 0 | 15 | 25 | 0 | 16 | 0 | 41 | 283 | 14 | 889 | 8256 | 3620 | 0 | 688 | 342 | 1013 | 32 | 295 | 19 | 2952 | 3877 | 2168 | 34 | 16971 | 3384 | 2301 | 32 | 33 | 29 | 45 | 11 | 11 | 36 | 41 | 38 | 33359 | 11 |
| SRR4836344 | 0 | 0 | 24 | 0 | 0 | 0 | 0 | 42 | 0 | 346 | 1729 | 5676 | 0 | 1158 | 187 | 631 | 0 | 103 | 63 | 2821 | 2752 | 3361 | 0 | 11502 | 10279 | 4374 | 54 | 54 | 39 | 39 | 28 | 39 | 21 | 42 | 23 | 21315 | 113 |
| SRR4836345 | 0 | 46 | 31 | 0 | 0 | 0 | 0 | 48 | 0 | 267 | 8822 | 6774 | 0 | 1184 | 299 | 715 | 20 | 69 | 65 | 3547 | 2726 | 2823 | 0 | 9353 | 8548 | 3892 | 56 | 76 | 36 | 224 | 239 | 41 | 193 | 356 | 229 | 24678 | 106 |
| SRR4836346 | 0 | 38 | 23 | 0 | 0 | 0 | 0 | 70 | 0 | 329 | 2119 | 5527 | 0 | 1233 | 209 | 575 | 0 | 106 | 70 | 3100 | 2843 | 3606 | 19 | 12499 | 10996 | 4355 | 65 | 65 | 34 | 298 | 161 | 42 | 161 | 306 | 180 | 21002 | 92 |
| SRR4836347 | 0 | 27 | 23 | 0 | 11 | 0 | 0 | 44 | 0 | 294 | 8668 | 6436 | 0 | 1124 | 274 | 624 | 0 | 86 | 75 | 3351 | 2946 | 3274 | 0 | 10685 | 9528 | 3884 | 43 | 73 | 43 | 224 | 174 | 27 | 166 | 364 | 205 | 23877 | 71 |
| SRR4836348 | 0 | 0 | 37 | 0 | 13 | 0 | 0 | 332 | 72 | 1889 | 29643 | 27004 | 0 | 260 | 43 | 721 | 0 | 116 | 49 | 3273 | 1167 | 1983 | 23 | 7306 | 5004 | 4081 | 0 | 17 | 16 | 14 | 15 | 0 | 17 | 33 | 10 | 26904 | 28 |
| SRR4836349 | 0 | 0 | 10 | 0 | 25 | 0 | 0 | 202 | 47 | 1785 | 31441 | 29051 | 0 | 171 | 93 | 885 | 0 | 117 | 40 | 2871 | 1205 | 1642 | 28 | 6039 | 4162 | 3600 | 30 | 12 | 17 | 10 | 24 | 0 | 18 | 37 | 21 | 29284 | 12 |
| SRR4836350 | 0 | 0 | 29 | 0 | 13 | 0 | 0 | 238 | 43 | 1968 | 2506 | 26063 | 0 | 260 | 130 | 584 | 0 | 76 | 32 | 2855 | 1244 | 2232 | 23 | 7835 | 5083 | 4187 | 39 | 12 | 13 | 51 | 14 | 39 | 17 | 36 | 14 | 25089 | 34 |
| SRR4836351 | 0 | 0 | 40 | 0 | 13 | 0 | 13 | 198 | 44 | 1862 | 2603 | 27462 | 0 | 143 | 55 | 628 | 0 | 107 | 32 | 2678 | 1243 | 1967 | 20 | 6942 | 4555 | 3640 | 30 | 53 | 0 | 10 | 20 | 13 | 16 | 30 | 21 | 26360 | 23 |
| SRR4836352 | 0 | 0 | 27 | 0 | 0 | 0 | 0 | 142 | 24 | 1250 | 12298 | 12354 | 0 | 546 | 0 | 87 | 0 | 260 | 24 | 5829 | 1804 | 2933 | 27 | 5283 | 9493 | 1464 | 13 | 118 | 25 | 52 | 20 | 30 | 41 | 65 | 43 | 31778 | 0 |
| SRR4836353 | 0 | 0 | 28 | 0 | 0 | 0 | 10 | 144 | 17 | 1129 | 12833 | 13461 | 0 | 560 | 34 | 76 | 0 | 272 | 22 | 6060 | 1887 | 2625 | 21 | 4031 | 7448 | 1228 | 11 | 102 | 25 | 57 | 27 | 29 | 34 | 51 | 35 | 34350 | 0 |
| SRR4836354 | 0 | 0 | 22 | 0 | 0 | 0 | 0 | 246 | 31 | 1431 | 11341 | 12176 | 0 | 639 | 0 | 49 | 10 | 316 | 19 | 5858 | 1866 | 3073 | 23 | 5542 | 9775 | 1424 | 29 | 105 | 19 | 47 | 34 | 21 | 38 | 53 | 43 | 31315 | 0 |
| SRR4836355 | 0 | 0 | 25 | 0 | 0 | 0 | 0 | 183 | 12 | 1126 | 1955 | 12053 | 0 | 448 | 12 | 94 | 11 | 299 | 26 | 5808 | 1778 | 2607 | 16 | 4680 | 8042 | 1132 | 48 | 390 | 86 | 175 | 102 | 20 | 118 | 188 | 139 | 31347 | 0 |
| SRR4836356 | 0 | 0 | 17 | 0 | 25 | 0 | 14 | 171 | 45 | 356 | 4443 | 4109 | 11 | 289 | 45 | 185 | 0 | 214 | 17 | 4109 | 3782 | 1472 | 12 | 1156 | 6737 | 1155 | 75 | 0 | 11 | 36 | 34 | 0 | 35 | 47 | 35 | 48237 | 11 |
| SRR4836357 | 0 | 0 | 27 | 0 | 25 | 0 | 14 | 164 | 47 | 344 | 576 | 4391 | 0 | 242 | 55 | 234 | 0 | 161 | 13 | 4274 | 3815 | 1284 | 15 | 920 | 5541 | 931 | 27 | 0 | 0 | 15 | 21 | 0 | 28 | 15 | 16 | 50131 | 0 |
| SRR4836359 | 0 | 0 | 14 | 0 | 18 | 0 | 14 | 133 | 28 | 489 | 768 | 4210 | 0 | 256 | 28 | 234 | 10 | 211 | 10 | 3887 | 3965 | 1549 | 14 | 1184 | 7072 | 1125 | 70 | 0 | 15 | 39 | 40 | 14 | 24 | 61 | 45 | 44364 | 0 |
| SRR4836361 | 0 | 0 | 20 | 0 | 25 | 0 | 0 | 127 | 24 | 446 | 4665 | 4213 | 0 | 247 | 60 | 241 | 13 | 170 | 16 | 4038 | 4141 | 1396 | 22 | 1084 | 6145 | 945 | 64 | 14 | 12 | 43 | 52 | 0 | 22 | 35 | 38 | 49365 | 16 |
| SRR4836362 | 0 | 0 | 63 | 0 | 16 | 0 | 83 | 850 | 435 | 1791 | 4180 | 15930 | 0 | 1060 | 639 | 2608 | 18 | 344 | 12 | 1541 | 513 | 2548 | 139 | 29885 | 4836 | 814 | 169 | 197 | 71 | 115 | 91 | 37 | 168 | 253 | 119 | 14560 | 132 |
| SRR4836364 | 0 | 0 | 15 | 0 | 17 | 0 | 21 | 908 | 296 | 1725 | 4126 | 17034 | 0 | 887 | 619 | 2865 | 33 | 324 | 14 | 1661 | 503 | 2305 | 99 | 23983 | 3963 | 673 | 36 | 38 | 13 | 27 | 23 | 31 | 35 | 59 | 25 | 16168 | 142 |
| SRR4836366 | 0 | 0 | 51 | 0 | 12 | 0 | 80 | 979 | 434 | 1757 | 28070 | 15398 | 0 | 1297 | 588 | 2837 | 21 | 340 | 13 | 1604 | 535 | 2754 | 129 | 32786 | 5174 | 745 | 173 | 167 | 78 | 134 | 75 | 32 | 159 | 269 | 135 | 14731 | 133 |
| SRR4836368 | 0 | 0 | 56 | 0 | 21 | 0 | 91 | 797 | 380 | 1369 | 5227 | 16244 | 0 | 925 | 561 | 3026 | 24 | 317 | 19 | 1449 | 521 | 2597 | 117 | 27993 | 4445 | 665 | 165 | 175 | 59 | 146 | 84 | 35 | 162 | 288 | 128 | 15228 | 130 |
| SRR4836369 | 0 | 0 | 24 | 0 | 0 | 0 | 11 | 566 | 123 | 1764 | 24715 | 28376 | 0 | 57 | 75 | 160 | 0 | 159 | 23 | 1623 | 1543 | 1573 | 32 | 7706 | 2600 | 1476 | 37 | 87 | 19 | 70 | 118 | 16 | 86 | 171 | 57 | 14197 | 0 |
| SRR4836370 | 0 | 0 | 37 | 0 | 11 | 0 | 14 | 677 | 69 | 1263 | 2038 | 28822 | 0 | 69 | 80 | 195 | 0 | 160 | 34 | 1377 | 1388 | 1207 | 25 | 6388 | 2111 | 1468 | 50 | 82 | 43 | 70 | 96 | 34 | 70 | 146 | 60 | 15576 | 0 |
| SRR4836371 | 0 | 0 | 30 | 0 | 18 | 0 | 0 | 724 | 121 | 1283 | 2168 | 27543 | 0 | 77 | 77 | 208 | 0 | 196 | 18 | 1217 | 1503 | 1388 | 32 | 7349 | 2404 | 1580 | 43 | 85 | 21 | 69 | 87 | 25 | 105 | 123 | 64 | 15098 | 0 |
| SRR4836373 | 0 | 0 | 36 | 0 | 16 | 0 | 29 | 621 | 131 | 1478 | 2680 | 26180 | 0 | 122 | 70 | 254 | 0 | 157 | 32 | 1295 | 1550 | 1372 | 48 | 7453 | 2448 | 1379 | 56 | 86 | 27 | 53 | 102 | 21 | 91 | 125 | 56 | 15587 | 0 |
| SRR4836375 | 0 | 0 | 10 | 0 | 18 | 0 | 15 | 133 | 27 | 773 | 3047 | 9407 | 0 | 312 | 2165 | 4538 | 17 | 124 | 38 | 3792 | 7681 | 2441 | 77 | 4927 | 5500 | 719 | 11 | 18 | 23 | 48 | 58 | 47 | 64 | 115 | 48 | 42968 | 48 |
| SRR4836376 | 0 | 0 | 10 | 0 | 25 | 0 | 17 | 149 | 15 | 651 | 2454 | 9931 | 0 | 269 | 2278 | 4272 | 21 | 87 | 42 | 3765 | 7078 | 2036 | 61 | 4230 | 4862 | 756 | 50 | 94 | 121 | 236 | 232 | 56 | 320 | 585 | 201 | 44619 | 58 |
| SRR4836378 | 0 | 0 | 16 | 0 | 0 | 0 | 18 | 120 | 14 | 688 | 12369 | 9684 | 0 | 294 | 2118 | 4505 | 14 | 111 | 57 | 3817 | 6975 | 2235 | 64 | 4886 | 5431 | 763 | 72 | 83 | 133 | 264 | 260 | 72 | 286 | 638 | 189 | 45025 | 46 |
| SRR4836380 | 0 | 0 | 0 | 0 | 19 | 0 | 13 | 155 | 0 | 718 | 3078 | 9472 | 0 | 279 | 2209 | 4532 | 18 | 104 | 49 | 3833 | 7404 | 2122 | 77 | 4827 | 5194 | 676 | 12 | 17 | 30 | 54 | 52 | 15 | 53 | 123 | 34 | 43867 | 42 |
| SRR4836382 | 0 | 0 | 25 | 0 | 11 | 0 | 0 | 188 | 32 | 1351 | 8117 | 3131 | 0 | 879 | 172 | 611 | 0 | 172 | 0 | 4724 | 1429 | 1630 | 57 | 7192 | 3788 | 881 | 20 | 120 | 28 | 36 | 24 | 0 | 28 | 35 | 30 | 18315 | 0 |
| SRR4836383 | 0 | 0 | 25 | 0 | 14 | 0 | 0 | 174 | 40 | 1195 | 1307 | 3005 | 0 | 761 | 134 | 594 | 0 | 135 | 12 | 4882 | 1364 | 1409 | 40 | 6120 | 3291 | 727 | 18 | 118 | 25 | 33 | 30 | 0 | 28 | 45 | 13 | 19434 | 0 |
| SRR4836384 | 0 | 0 | 27 | 0 | 13 | 0 | 0 | 272 | 39 | 1483 | 8281 | 3218 | 0 | 978 | 175 | 751 | 0 | 144 | 26 | 4377 | 1365 | 1532 | 65 | 7005 | 3711 | 834 | 16 | 110 | 29 | 34 | 27 | 0 | 29 | 56 | 27 | 18452 | 13 |
| SRR4836386 | 0 | 0 | 22 | 0 | 13 | 0 | 0 | 166 | 57 | 1324 | 8462 | 3162 | 0 | 903 | 234 | 639 | 0 | 110 | 25 | 5051 | 1556 | 1527 | 52 | 6912 | 3636 | 747 | 14 | 114 | 23 | 32 | 17 | 0 | 28 | 39 | 28 | 18502 | 0 |
| SRR4836388 | 0 | 0 | 24 | 0 | 12 | 0 | 0 | 351 | 54 | 1614 | 12974 | 7781 | 0 | 581 | 736 | 2371 | 0 | 139 | 13 | 2323 | 2298 | 1314 | 18 | 1141 | 4319 | 1626 | 11 | 72 | 32 | 39 | 37 | 21 | 30 | 48 | 21 | 32392 | 0 |
| SRR4836390 | 0 | 0 | 22 | 0 | 14 | 0 | 12 | 341 | 26 | 1457 | 13140 | 7967 | 0 | 579 | 767 | 2420 | 0 | 134 | 17 | 2471 | 2137 | 1147 | 18 | 915 | 3734 | 1522 | 40 | 325 | 121 | 129 | 159 | 32 | 113 | 214 | 109 | 33744 | 0 |
| SRR4836391 | 0 | 0 | 12 | 0 | 21 | 0 | 0 | 343 | 57 | 1420 | 13785 | 7917 | 0 | 531 | 735 | 2563 | 0 | 165 | 13 | 2497 | 2172 | 1349 | 0 | 1148 | 4120 | 1676 | 39 | 274 | 154 | 160 | 145 | 25 | 127 | 195 | 98 | 32091 | 0 |
| SRR4836392 | 0 | 0 | 25 | 0 | 18 | 0 | 13 | 458 | 65 | 1471 | 12876 | 7651 | 0 | 600 | 703 | 2232 | 0 | 149 | 12 | 2297 | 2209 | 1208 | 0 | 1078 | 4054 | 1555 | 35 | 63 | 36 | 36 | 37 | 11 | 37 | 55 | 28 | 31390 | 0 |
| SRR4836394 | 0 | 0 | 40 | 0 | 20 | 0 | 0 | 417 | 54 | 1321 | 14584 | 11121 | 0 | 143 | 184 | 536 | 0 | 498 | 21 | 1505 | 3341 | 1185 | 34 | 6091 | 3055 | 3177 | 215 | 0 | 0 | 31 | 29 | 0 | 18 | 48 | 43 | 16982 | 0 |
| SRR4836396 | 0 | 0 | 13 | 0 | 23 | 0 | 0 | 436 | 23 | 1249 | 15808 | 11323 | 0 | 83 | 158 | 572 | 0 | 463 | 17 | 1693 | 3138 | 951 | 23 | 5166 | 2521 | 2878 | 62 | 12 | 0 | 28 | 25 | 0 | 26 | 10 | 11 | 17779 | 0 |
| SRR4836398 | 0 | 0 | 51 | 0 | 20 | 0 | 0 | 374 | 36 | 1338 | 15413 | 11134 | 0 | 115 | 122 | 633 | 0 | 501 | 14 | 1259 | 3125 | 1122 | 37 | 5688 | 2831 | 3093 | 218 | 0 | 0 | 38 | 37 | 0 | 27 | 41 | 40 | 17161 | 0 |
| SRR4836399 | 0 | 0 | 12 | 0 | 31 | 0 | 11 | 456 | 60 | 1177 | 2692 | 10490 | 0 | 93 | 115 | 528 | 0 | 465 | 22 | 1352 | 3223 | 1039 | 30 | 5994 | 2882 | 2924 | 62 | 0 | 0 | 30 | 32 | 0 | 22 | 15 | 13 | 16708 | 0 |
| SRR4836400 | 0 | 0 | 75 | 0 | 12 | 0 | 0 | 274 | 31 | 1065 | 18237 | 8210 | 0 | 1018 | 204 | 313 | 0 | 386 | 37 | 901 | 2637 | 3374 | 22 | 6470 | 10648 | 1455 | 49 | 588 | 323 | 366 | 199 | 52 | 222 | 280 | 251 | 5045 | 0 |
| SRR4836402 | 0 | 11 | 12 | 0 | 10 | 0 | 0 | 301 | 30 | 883 | 19203 | 8097 | 0 | 873 | 271 | 321 | 11 | 355 | 28 | 873 | 2543 | 2796 | 24 | 5597 | 9000 | 1372 | 47 | 103 | 60 | 67 | 45 | 54 | 31 | 59 | 44 | 5117 | 0 |
| SRR4836404 | 0 | 10 | 73 | 0 | 0 | 0 | 0 | 365 | 48 | 1000 | 19175 | 8059 | 0 | 933 | 221 | 317 | 0 | 342 | 24 | 654 | 2491 | 3009 | 26 | 6240 | 10191 | 1471 | 66 | 598 | 339 | 345 | 200 | 24 | 228 | 252 | 266 | 5251 | 0 |
| SRR4836406 | 0 | 20 | 12 | 0 | 0 | 0 | 0 | 268 | 31 | 895 | 19197 | 7483 | 0 | 995 | 230 | 329 | 0 | 335 | 28 | 788 | 2711 | 2878 | 18 | 6366 | 10181 | 1332 | 42 | 99 | 63 | 78 | 36 | 17 | 43 | 50 | 41 | 5310 | 0 |
| SRR4836407 | 0 | 0 | 28 | 0 | 14 | 0 | 0 | 215 | 59 | 1301 | 29121 | 16761 | 0 | 127 | 49 | 108 | 0 | 287 | 49 | 1535 | 1932 | 1799 | 42 | 12848 | 3515 | 2625 | 21 | 10 | 0 | 21 | 25 | 24 | 18 | 21 | 13 | 12321 | 0 |
| SRR4836408 | 0 | 0 | 30 | 0 | 14 | 0 | 0 | 485 | 86 | 1178 | 28042 | 16383 | 0 | 133 | 57 | 190 | 0 | 278 | 50 | 1539 | 1960 | 1938 | 57 | 15206 | 4239 | 2655 | 0 | 25 | 25 | 15 | 24 | 22 | 17 | 26 | 13 | 11907 | 0 |
| SRR4836410 | 0 | 0 | 28 | 0 | 17 | 0 | 0 | 339 | 95 | 1136 | 30102 | 16845 | 0 | 126 | 47 | 150 | 0 | 245 | 55 | 1625 | 1965 | 1661 | 45 | 13261 | 3425 | 2299 | 13 | 31 | 21 | 63 | 101 | 28 | 59 | 82 | 56 | 12042 | 0 |
| SRR4836412 | 0 | 0 | 31 | 0 | 0 | 0 | 0 | 105 | 0 | 708 | 1913 | 886 | 0 | 223 | 0 | 52 | 0 | 229 | 16 | 874 | 2758 | 1867 | 38 | 8768 | 6681 | 1253 | 61 | 31 | 25 | 52 | 46 | 17 | 44 | 59 | 61 | 6248 | 0 |
| SRR4836414 | 0 | 0 | 17 | 0 | 0 | 0 | 0 | 71 | 0 | 657 | 787 | 893 | 0 | 263 | 16 | 80 | 10 | 227 | 13 | 796 | 2769 | 2005 | 29 | 10032 | 7553 | 1260 | 56 | 35 | 29 | 41 | 54 | 21 | 43 | 43 | 64 | 5884 | 0 |
| SRR4836415 | 0 | 0 | 20 | 0 | 0 | 0 | 0 | 70 | 14 | 683 | 1788 | 960 | 0 | 206 | 23 | 70 | 0 | 205 | 36 | 840 | 2907 | 1821 | 28 | 9288 | 6330 | 1240 | 21 | 14 | 17 | 13 | 14 | 15 | 34 | 16 | 18 | 6305 | 0 |
| SRR4836416 | 0 | 0 | 16 | 0 | 11 | 0 | 0 | 209 | 87 | 1701 | 2877 | 11236 | 0 | 462 | 366 | 1387 | 0 | 326 | 67 | 1692 | 4752 | 2690 | 35 | 11830 | 6901 | 2071 | 17 | 137 | 15 | 34 | 22 | 13 | 31 | 40 | 29 | 8793 | 0 |
| SRR4836417 | 0 | 0 | 16 | 0 | 0 | 0 | 0 | 275 | 86 | 1798 | 2771 | 10524 | 0 | 404 | 508 | 1265 | 0 | 421 | 67 | 1532 | 4817 | 2901 | 44 | 13703 | 8270 | 2032 | 12 | 139 | 25 | 34 | 22 | 12 | 27 | 36 | 34 | 8553 | 0 |
| SRR4836419 | 0 | 0 | 11 | 0 | 14 | 0 | 0 | 284 | 99 | 1660 | 3449 | 10696 | 0 | 340 | 411 | 1397 | 0 | 319 | 78 | 1419 | 4980 | 2709 | 34 | 12239 | 7097 | 1878 | 12 | 347 | 53 | 82 | 60 | 25 | 73 | 75 | 69 | 8199 | 0 |
| SRR4836421 | 0 | 18 | 27 | 0 | 16 | 0 | 13 | 442 | 75 | 809 | 7730 | 4122 | 0 | 360 | 749 | 3245 | 35 | 245 | 38 | 3140 | 2889 | 1778 | 46 | 5307 | 4756 | 3354 | 23 | 31 | 43 | 11 | 41 | 14 | 11 | 21 | 14 | 53746 | 44 |
| SRR4836422 | 0 | 13 | 34 | 0 | 18 | 0 | 72 | 521 | 57 | 1049 | 7046 | 4090 | 0 | 330 | 700 | 2873 | 19 | 218 | 40 | 3214 | 2889 | 1745 | 62 | 6526 | 5345 | 3402 | 135 | 19 | 53 | 70 | 44 | 32 | 67 | 122 | 67 | 51232 | 26 |
| SRR4836423 | 0 | 0 | 0 | 0 | 24 | 0 | 14 | 409 | 44 | 971 | 2425 | 4158 | 0 | 278 | 795 | 3278 | 38 | 235 | 39 | 3351 | 2963 | 1579 | 48 | 5678 | 4635 | 3180 | 29 | 38 | 38 | 12 | 13 | 13 | 15 | 20 | 14 | 51333 | 38 |
| SRR4836425 | 0 | 0 | 0 | 0 | 14 | 0 | 27 | 572 | 79 | 1425 | 1049 | 7949 | 0 | 1038 | 97 | 299 | 20 | 555 | 18 | 2146 | 1628 | 2499 | 64 | 15557 | 4249 | 735 | 164 | 31 | 43 | 270 | 106 | 20 | 254 | 368 | 188 | 40248 | 45 |
| SRR4836427 | 0 | 0 | 34 | 0 | 12 | 0 | 19 | 570 | 97 | 1544 | 9097 | 7720 | 0 | 1184 | 70 | 307 | 0 | 533 | 16 | 2518 | 1789 | 2623 | 94 | 18339 | 5004 | 743 | 132 | 19 | 42 | 215 | 117 | 15 | 177 | 355 | 159 | 39196 | 44 |
| SRR4836428 | 0 | 16 | 25 | 0 | 16 | 0 | 31 | 511 | 112 | 1204 | 1390 | 7404 | 0 | 1218 | 105 | 294 | 28 | 476 | 14 | 2330 | 1701 | 2385 | 69 | 15636 | 4098 | 663 | 156 | 13 | 28 | 256 | 147 | 25 | 134 | 372 | 150 | 38925 | 46 |
| SRR4836429 | 0 | 12 | 37 | 0 | 28 | 0 | 0 | 346 | 32 | 841 | 3814 | 6549 | 0 | 422 | 177 | 551 | 0 | 377 | 19 | 7084 | 5532 | 3436 | 373 | 20873 | 6870 | 1283 | 107 | 107 | 35 | 81 | 66 | 23 | 61 | 135 | 144 | 73619 | 0 |
| SRR4836431 | 0 | 11 | 31 | 0 | 27 | 0 | 0 | 313 | 39 | 913 | 3687 | 6256 | 0 | 499 | 224 | 472 | 0 | 367 | 16 | 7312 | 5604 | 3700 | 418 | 24584 | 7992 | 1250 | 13 | 11 | 25 | 68 | 15 | 0 | 11 | 21 | 20 | 69483 | 0 |
| SRR4836433 | 0 | 12 | 31 | 0 | 39 | 0 | 0 | 363 | 33 | 742 | 4525 | 6360 | 0 | 405 | 177 | 628 | 0 | 341 | 0 | 7354 | 5688 | 3133 | 413 | 21077 | 6729 | 1117 | 50 | 11 | 17 | 13 | 12 | 19 | 62 | 19 | 21 | 72372 | 0 |
| SRR4836434 | 0 | 0 | 0 | 0 | 16 | 0 | 0 | 483 | 69 | 4344 | 11411 | 9825 | 0 | 896 | 86 | 69 | 0 | 236 | 0 | 1948 | 1920 | 1644 | 32 | 9425 | 4897 | 1569 | 32 | 29 | 0 | 45 | 16 | 0 | 32 | 45 | 11 | 23443 | 0 |
| SRR4836435 | 0 | 10 | 0 | 0 | 13 | 0 | 16 | 515 | 83 | 4151 | 11440 | 9099 | 0 | 1146 | 17 | 166 | 0 | 212 | 13 | 2441 | 1950 | 1942 | 29 | 10161 | 5143 | 1594 | 26 | 21 | 0 | 18 | 16 | 0 | 10 | 12 | 13 | 22747 | 0 |
| SRR4836437 | 0 | 0 | 13 | 0 | 13 | 0 | 0 | 470 | 201 | 4256 | 10634 | 9224 | 0 | 873 | 27 | 81 | 0 | 192 | 19 | 2189 | 1942 | 1912 | 42 | 10064 | 4955 | 1514 | 30 | 19 | 11 | 25 | 12 | 0 | 30 | 30 | 0 | 22987 | 0 |
| SRR4836439 | 0 | 0 | 11 | 0 | 20 | 0 | 0 | 586 | 160 | 3782 | 10733 | 9095 | 0 | 1012 | 27 | 67 | 0 | 199 | 0 | 1997 | 2000 | 1959 | 31 | 10181 | 5062 | 1521 | 14 | 15 | 17 | 15 | 19 | 0 | 26 | 14 | 0 | 22864 | 0 |
| SRR4836440 | 0 | 19 | 38 | 0 | 11 | 0 | 19 | 251 | 58 | 2741 | 35512 | 25167 | 0 | 637 | 19 | 0 | 0 | 203 | 51 | 618 | 394 | 1856 | 32 | 9421 | 4451 | 1209 | 48 | 77 | 77 | 21 | 30 | 19 | 22 | 36 | 38 | 4613 | 17 |
| SRR4836441 | 0 | 0 | 56 | 0 | 0 | 0 | 0 | 263 | 94 | 2971 | 33642 | 24484 | 0 | 470 | 0 | 56 | 0 | 191 | 34 | 715 | 428 | 2019 | 17 | 10272 | 4800 | 1315 | 47 | 66 | 66 | 281 | 216 | 28 | 178 | 347 | 38 | 4382 | 34 |
| SRR4836443 | 0 | 15 | 0 | 0 | 11 | 0 | 0 | 416 | 119 | 2988 | 2207 | 23715 | 0 | 684 | 0 | 45 | 15 | 186 | 64 | 476 | 423 | 2072 | 29 | 10192 | 4754 | 1237 | 31 | 61 | 77 | 15 | 21 | 61 | 20 | 33 | 54 | 4505 | 19 |
| SRR4836445 | 0 | 31 | 46 | 0 | 12 | 0 | 0 | 256 | 121 | 2941 | 34247 | 24762 | 0 | 558 | 15 | 15 | 15 | 179 | 56 | 633 | 407 | 2006 | 32 | 10392 | 4631 | 1250 | 84 | 76 | 53 | 206 | 252 | 23 | 199 | 367 | 76 | 4630 | 32 |
| SRR4836446 | 0 | 0 | 35 | 0 | 0 | 0 | 0 | 528 | 33 | 2059 | 989 | 7861 | 0 | 584 | 33 | 22 | 11 | 174 | 89 | 4679 | 4808 | 2711 | 156 | 12064 | 9690 | 2388 | 46 | 145 | 67 | 137 | 143 | 11 | 97 | 191 | 30 | 19036 | 11 |
| SRR4836447 | 0 | 13 | 24 | 0 | 0 | 0 | 16 | 494 | 32 | 2203 | 988 | 7661 | 0 | 774 | 32 | 54 | 0 | 143 | 63 | 4717 | 4875 | 2828 | 94 | 12874 | 10091 | 2409 | 61 | 177 | 66 | 140 | 122 | 21 | 90 | 148 | 53 | 17719 | 0 |
| SRR4836448 | 0 | 0 | 36 | 0 | 0 | 0 | 0 | 461 | 133 | 2334 | 10958 | 8261 | 0 | 674 | 18 | 44 | 11 | 166 | 45 | 5164 | 4833 | 3005 | 138 | 13403 | 10154 | 2463 | 58 | 135 | 73 | 154 | 133 | 17 | 88 | 161 | 47 | 18553 | 0 |
| SRR4836450 | 0 | 0 | 29 | 0 | 14 | 0 | 0 | 536 | 53 | 2205 | 1191 | 7650 | 0 | 782 | 0 | 35 | 11 | 162 | 82 | 4682 | 4798 | 2978 | 145 | 13413 | 10140 | 2243 | 21 | 56 | 26 | 42 | 42 | 13 | 33 | 56 | 15 | 18200 | 0 |
| SRR4836452 | 0 | 0 | 0 | 0 | 14 | 0 | 12 | 316 | 105 | 3003 | 939 | 11274 | 0 | 790 | 0 | 0 | 0 | 436 | 16 | 3661 | 496 | 2354 | 14 | 18213 | 5035 | 348 | 27 | 11 | 20 | 36 | 11 | 0 | 17 | 25 | 12 | 27473 | 206 |
| SRR4836453 | 0 | 0 | 0 | 0 | 16 | 0 | 0 | 438 | 129 | 2754 | 1013 | 12020 | 0 | 618 | 0 | 0 | 0 | 418 | 16 | 3732 | 576 | 2391 | 21 | 19822 | 5216 | 324 | 20 | 14 | 0 | 12 | 12 | 12 | 16 | 23 | 16 | 27257 | 187 |
| SRR4836454 | 0 | 0 | 0 | 0 | 0 | 0 | 0 | 310 | 124 | 2830 | 1195 | 11508 | 0 | 640 | 0 | 0 | 0 | 479 | 0 | 3099 | 589 | 2500 | 22 | 20167 | 5432 | 311 | 70 | 60 | 22 | 51 | 60 | 0 | 41 | 83 | 35 | 27581 | 168 |
| SRR4836456 | 0 | 0 | 0 | 0 | 13 | 0 | 0 | 438 | 146 | 3500 | 26606 | 12209 | 0 | 813 | 0 | 0 | 0 | 401 | 15 | 3604 | 527 | 2456 | 27 | 20018 | 5129 | 305 | 10 | 11 | 13 | 14 | 15 | 0 | 32 | 21 | 32 | 28586 | 179 |
| SRR4836458 | 0 | 0 | 0 | 0 | 14 | 0 | 0 | 789 | 136 | 3972 | 22772 | 26146 | 0 | 354 | 0 | 0 | 0 | 155 | 34 | 2204 | 918 | 2190 | 26 | 8886 | 4010 | 1302 | 17 | 29 | 23 | 31 | 34 | 17 | 20 | 44 | 11 | 17358 | 0 |
| SRR4836459 | 0 | 0 | 0 | 0 | 12 | 0 | 0 | 608 | 185 | 3434 | 21399 | 27105 | 0 | 476 | 0 | 0 | 0 | 171 | 42 | 1981 | 1043 | 2288 | 34 | 9822 | 4323 | 1268 | 10 | 34 | 23 | 31 | 21 | 0 | 22 | 29 | 17 | 17938 | 0 |
| SRR4836461 | 0 | 0 | 14 | 0 | 0 | 0 | 0 | 580 | 0 | 3790 | 1052 | 26801 | 0 | 311 | 0 | 0 | 0 | 138 | 27 | 1843 | 905 | 2295 | 34 | 9489 | 4396 | 1290 | 23 | 80 | 23 | 100 | 103 | 14 | 91 | 137 | 0 | 18372 | 0 |
| SRR4836463 | 0 | 0 | 12 | 0 | 0 | 0 | 0 | 512 | 143 | 3950 | 1082 | 25134 | 0 | 430 | 0 | 0 | 0 | 178 | 28 | 2006 | 944 | 2219 | 23 | 9614 | 4398 | 1131 | 12 | 95 | 25 | 104 | 109 | 12 | 65 | 136 | 12 | 17540 | 11 |
| SRR4836464 | 0 | 11 | 221 | 0 | 0 | 0 | 14 | 250 | 92 | 1346 | 1901 | 5595 | 0 | 1254 | 0 | 13 | 0 | 1521 | 0 | 5562 | 10568 | 3257 | 298 | 9892 | 4580 | 3469 | 86 | 54 | 276 | 119 | 54 | 0 | 32 | 62 | 29 | 22280 | 81 |
| SRR4836466 | 0 | 22 | 179 | 0 | 0 | 0 | 0 | 385 | 128 | 1218 | 1933 | 5199 | 0 | 1250 | 0 | 0 | 0 | 1579 | 0 | 4795 | 10991 | 3375 | 308 | 10515 | 4856 | 3406 | 65 | 47 | 302 | 121 | 44 | 0 | 31 | 50 | 35 | 21450 | 72 |
| SRR4836468 | 0 | 0 | 47 | 0 | 0 | 0 | 0 | 334 | 104 | 1444 | 12650 | 5282 | 0 | 1231 | 0 | 10 | 0 | 1584 | 0 | 4828 | 10967 | 3353 | 350 | 10613 | 4875 | 3392 | 20 | 10 | 60 | 29 | 13 | 12 | 34 | 13 | 40 | 22614 | 72 |
| SRR4836469 | 0 | 13 | 212 | 0 | 10 | 0 | 0 | 345 | 103 | 1435 | 2403 | 5014 | 0 | 1106 | 0 | 0 | 0 | 1532 | 0 | 4813 | 10989 | 3322 | 331 | 10881 | 4909 | 3283 | 68 | 46 | 279 | 127 | 54 | 11 | 41 | 57 | 24 | 21823 | 88 |
| SRR4836471 | 11 | 0 | 18 | 0 | 25 | 0 | 0 | 507 | 123 | 2283 | 14515 | 16871 | 0 | 859 | 0 | 25 | 0 | 52 | 21 | 1031 | 1373 | 2568 | 110 | 8209 | 4431 | 864 | 41 | 22 | 19 | 23 | 21 | 33 | 16 | 31 | 43 | 10424 | 0 |
| SRR4836473 | 0 | 0 | 105 | 0 | 19 | 0 | 0 | 746 | 110 | 2331 | 2257 | 16337 | 0 | 1029 | 12 | 58 | 0 | 85 | 12 | 972 | 1432 | 3110 | 139 | 10891 | 5482 | 964 | 42 | 153 | 125 | 197 | 131 | 36 | 155 | 223 | 24 | 10022 | 0 |
| SRR4836474 | 0 | 0 | 14 | 0 | 18 | 0 | 17 | 594 | 151 | 2503 | 2258 | 15685 | 0 | 937 | 0 | 29 | 10 | 65 | 15 | 1059 | 1479 | 3119 | 155 | 11080 | 5620 | 914 | 39 | 20 | 18 | 21 | 18 | 35 | 25 | 34 | 37 | 9615 | 0 |
| SRR4836475 | 0 | 0 | 0 | 0 | 0 | 0 | 21 | 554 | 35 | 1431 | 24238 | 16366 | 0 | 312 | 35 | 69 | 0 | 147 | 14 | 1500 | 1993 | 2927 | 35 | 11031 | 5338 | 1488 | 84 | 113 | 46 | 104 | 154 | 38 | 113 | 154 | 25 | 11715 | 0 |
| SRR4836477 | 0 | 0 | 40 | 0 | 0 | 0 | 12 | 434 | 16 | 1448 | 2697 | 14256 | 0 | 434 | 16 | 33 | 0 | 121 | 16 | 1334 | 2084 | 3773 | 44 | 15101 | 6871 | 1673 | 13 | 12 | 18 | 18 | 18 | 0 | 23 | 30 | 15 | 11285 | 0 |
| SRR4836479 | 0 | 0 | 16 | 0 | 0 | 0 | 0 | 673 | 33 | 1596 | 21274 | 15181 | 0 | 466 | 42 | 33 | 13 | 165 | 18 | 1480 | 2109 | 3848 | 59 | 15493 | 7166 | 1562 | 103 | 81 | 38 | 75 | 119 | 22 | 106 | 156 | 34 | 10583 | 0 |

### Supp Table S8.4.Columns_DI_to_ES

| | | | | | | | | | | | | | | | | | | | | | | | | | | | | | | | | | | | | | |
|---|---|---|---|---|---|---|---|---|---|---|---|---|---|---|---|---|---|---|---|---|---|---|---|---|---|---|---|---|---|---|---|---|---|---|---|---|---|
| SRR4836481 | 0 | 0 | 33 | 0 | 0 | 0 | 0 | 175 | 39 | 526 | 11846 | 6468 | 0 | 1111 | 19 | 39 | 12 | 66 | 14 | 1266 | 801 | 1113 | 26 | 3150 | 2802 | 548 | 29 | 28 | 27 | 53 | 26 | 37 | 32 | 27 | 16 | 7579 | 0 |
| SRR4836482 | 0 | 0 | 21 | 0 | 0 | 0 | 0 | 306 | 56 | 641 | 10666 | 4909 | 0 | 1295 | 0 | 28 | 0 | 54 | 13 | 933 | 834 | 1333 | 30 | 4605 | 3643 | 758 | 15 | 125 | 89 | 186 | 89 | 34 | 107 | 43 | 21 | 6698 | 15 |
| SRR4836483 | 0 | 0 | 11 | 0 | 0 | 0 | 0 | 279 | 84 | 474 | 701 | 5842 | 0 | 1339 | 0 | 42 | 0 | 90 | 0 | 865 | 862 | 1432 | 43 | 4389 | 3791 | 611 | 16 | 42 | 30 | 39 | 25 | 28 | 25 | 25 | 19 | 6777 | 11 |
| SRR4836484 | 0 | 11 | 11 | 0 | 11 | 0 | 0 | 884 | 48 | 788 | 15023 | 17244 | 0 | 406 | 48 | 119 | 0 | 66 | 0 | 9673 | 1899 | 3540 | 77 | 6844 | 9202 | 723 | 34 | 274 | 354 | 663 | 560 | 160 | 628 | 469 | 183 | 12993 | 0 |
| SRR4836485 | 0 | 0 | 77 | 0 | 12 | 0 | 0 | 682 | 68 | 990 | 851 | 17574 | 0 | 461 | 17 | 68 | 0 | 98 | 15 | 9503 | 1909 | 4381 | 86 | 9187 | 12327 | 861 | 0 | 22 | 26 | 44 | 33 | 14 | 39 | 47 | 86 | 12916 | 0 |
| SRR4836487 | 0 | 17 | 95 | 0 | 0 | 0 | 0 | 606 | 52 | 1385 | 14355 | 17922 | 0 | 519 | 52 | 87 | 0 | 104 | 15 | 8865 | 1959 | 4365 | 73 | 9429 | 12299 | 764 | 0 | 24 | 25 | 54 | 35 | 52 | 31 | 49 | 10 | 12311 | 0 |
| SRR4836489 | 0 | 0 | 14 | 0 | 0 | 0 | 24 | 279 | 40 | 877 | 12934 | 6308 | 0 | 473 | 0 | 35 | 13 | 689 | 33 | 2591 | 1006 | 2050 | 28 | 11345 | 4781 | 349 | 44 | 11 | 22 | 11 | 15 | 13 | 11 | 13 | 19 | 26870 | 87 |
| SRR4836491 | 0 | 0 | 0 | 0 | 0 | 0 | 30 | 297 | 55 | 986 | 3405 | 5779 | 0 | 583 | 0 | 17 | 16 | 744 | 38 | 2517 | 1039 | 2550 | 28 | 15221 | 6333 | 394 | 45 | 11 | 14 | 11 | 13 | 12 | 12 | 16 | 14 | 23782 | 86 |
| SRR4836492 | 0 | 0 | 0 | 0 | 0 | 0 | 21 | 323 | 32 | 911 | 11814 | 5567 | 0 | 511 | 0 | 0 | 0 | 783 | 38 | 2496 | 1081 | 2565 | 29 | 15534 | 6305 | 356 | 46 | 34 | 18 | 15 | 16 | 11 | 12 | 15 | 11 | 24877 | 96 |
| SRR4836493 | 0 | 0 | 37 | 0 | 31 | 0 | 0 | 595 | 181 | 6297 | 25207 | 18146 | 0 | 499 | 11 | 53 | 0 | 426 | 25 | 1083 | 611 | 2434 | 57 | 8664 | 5969 | 398 | 21 | 41 | 34 | 103 | 123 | 15 | 53 | 85 | 14 | 7804 | 0 |
| SRR4836494 | 0 | 0 | 42 | 0 | 35 | 0 | 0 | 716 | 201 | 7420 | 3168 | 16473 | 0 | 380 | 0 | 22 | 0 | 504 | 31 | 1014 | 635 | 3030 | 62 | 12346 | 8112 | 481 | 21 | 33 | 42 | 103 | 105 | 26 | 64 | 105 | 12 | 7248 | 10 |
| SRR4836495 | 0 | 0 | 47 | 0 | 27 | 0 | 0 | 796 | 218 | 7363 | 3020 | 15905 | 0 | 450 | 0 | 23 | 0 | 472 | 25 | 856 | 653 | 3084 | 69 | 12603 | 7976 | 459 | 20 | 38 | 40 | 105 | 123 | 27 | 65 | 91 | 18 | 7453 | 0 |
| SRR4836496 | 0 | 0 | 0 | 0 | 13 | 0 | 23 | 429 | 92 | 1620 | 4719 | 16008 | 0 | 412 | 132 | 383 | 0 | 248 | 30 | 1179 | 1019 | 1866 | 41 | 15036 | 4008 | 404 | 15 | 46 | 15 | 23 | 23 | 16 | 20 | 26 | 12 | 15785 | 29 |
| SRR4836498 | 0 | 0 | 17 | 0 | 0 | 0 | 18 | 351 | 61 | 1619 | 29180 | 17875 | 0 | 428 | 183 | 504 | 0 | 250 | 18 | 1459 | 926 | 1539 | 20 | 12827 | 3266 | 402 | 54 | 197 | 29 | 92 | 73 | 14 | 68 | 95 | 63 | 16080 | 38 |
| SRR4836500 | 0 | 0 | 17 | 0 | 15 | 0 | 18 | 345 | 93 | 1460 | 28875 | 17709 | 0 | 395 | 223 | 360 | 0 | 266 | 19 | 1316 | 931 | 1753 | 33 | 14547 | 3649 | 441 | 18 | 44 | 24 | 23 | 19 | 16 | 18 | 24 | 15 | 16321 | 35 |
| SRR4836502 | 0 | 0 | 12 | 0 | 13 | 0 | 20 | 275 | 74 | 1846 | 4732 | 15122 | 0 | 386 | 143 | 476 | 11 | 263 | 20 | 1280 | 932 | 1814 | 48 | 17175 | 4452 | 396 | 14 | 47 | 14 | 26 | 23 | 16 | 13 | 26 | 14 | 15132 | 29 |
| SRR4836503 | 0 | 0 | 0 | 0 | 12 | 0 | 11 | 111 | 23 | 2785 | 2948 | 1662 | 0 | 1668 | 178 | 403 | 0 | 517 | 37 | 1331 | 4764 | 2751 | 59 | 8718 | 10021 | 1032 | 112 | 18 | 0 | 44 | 39 | 0 | 36 | 41 | 48 | 6662 | 16 |
| SRR4836504 | 0 | 0 | 0 | 0 | 27 | 0 | 0 | 76 | 11 | 2599 | 3160 | 1771 | 0 | 1557 | 191 | 401 | 0 | 494 | 41 | 1226 | 4326 | 2257 | 60 | 7170 | 8737 | 981 | 21 | 11 | 18 | 43 | 45 | 11 | 25 | 34 | 49 | 6951 | 11 |
| SRR4836505 | 0 | 0 | 13 | 0 | 11 | 0 | 15 | 81 | 13 | 2594 | 3083 | 1601 | 0 | 1567 | 196 | 386 | 0 | 478 | 40 | 1218 | 4421 | 2569 | 62 | 8069 | 9517 | 1057 | 20 | 17 | 13 | 49 | 43 | 11 | 21 | 54 | 34 | 6707 | 16 |
| SRR4836507 | 0 | 0 | 12 | 0 | 13 | 0 | 17 | 104 | 11 | 2865 | 2451 | 1466 | 0 | 1555 | 189 | 403 | 0 | 575 | 30 | 1228 | 4698 | 2746 | 63 | 9679 | 10901 | 1038 | 91 | 0 | 22 | 39 | 24 | 10 | 32 | 44 | 44 | 6301 | 19 |
| SRR4836509 | 0 | 14 | 0 | 0 | 16 | 0 | 21 | 35 | 0 | 867 | 4376 | 1182 | 0 | 1454 | 72 | 302 | 14 | 654 | 57 | 270 | 3732 | 2777 | 72 | 10591 | 7214 | 698 | 37 | 142 | 23 | 45 | 27 | 14 | 31 | 39 | 23 | 3158 | 0 |
| SRR4836511 | 0 | 0 | 16 | 0 | 14 | 0 | 22 | 47 | 11 | 749 | 3526 | 1268 | 0 | 1361 | 78 | 340 | 0 | 511 | 66 | 292 | 3282 | 2152 | 70 | 9165 | 6090 | 629 | 163 | 511 | 85 | 159 | 113 | 48 | 116 | 157 | 94 | 3522 | 0 |
| SRR4836512 | 0 | 23 | 13 | 0 | 18 | 0 | 20 | 50 | 0 | 837 | 3591 | 1235 | 0 | 1360 | 76 | 310 | 10 | 686 | 49 | 279 | 3549 | 2469 | 70 | 10274 | 6914 | 747 | 45 | 143 | 24 | 44 | 22 | 17 | 26 | 38 | 27 | 3311 | 0 |
| SRR4836513 | 0 | 17 | 12 | 0 | 17 | 0 | 14 | 52 | 0 | 868 | 3408 | 1204 | 0 | 1503 | 72 | 307 | 0 | 636 | 60 | 297 | 3542 | 2704 | 81 | 12452 | 8002 | 725 | 39 | 143 | 24 | 37 | 26 | 13 | 32 | 43 | 23 | 3124 | 0 |
| SRR4836514 | 0 | 0 | 26 | 0 | 0 | 0 | 0 | 403 | 37 | 955 | 5158 | 16829 | 0 | 242 | 1184 | 3268 | 0 | 165 | 13 | 725 | 1890 | 2022 | 67 | 7573 | 7672 | 616 | 0 | 336 | 50 | 124 | 162 | 23 | 104 | 186 | 103 | 4310 | 0 |
| SRR4836515 | 0 | 0 | 30 | 0 | 0 | 0 | 0 | 383 | 83 | 925 | 35074 | 17516 | 0 | 292 | 1183 | 3308 | 0 | 162 | 16 | 775 | 1763 | 1686 | 58 | 6293 | 6575 | 579 | 0 | 107 | 16 | 39 | 49 | 14 | 34 | 58 | 32 | 4733 | 0 |
| SRR4836517 | 0 | 0 | 23 | 0 | 0 | 0 | 0 | 322 | 47 | 959 | 34197 | 17747 | 0 | 197 | 1093 | 3389 | 0 | 154 | 18 | 786 | 1745 | 1880 | 56 | 7132 | 7365 | 607 | 0 | 289 | 51 | 117 | 156 | 22 | 104 | 142 | 101 | 4584 | 0 |
| SRR4836519 | 0 | 0 | 25 | 0 | 11 | 0 | 0 | 404 | 140 | 1059 | 5160 | 15547 | 0 | 263 | 1187 | 3129 | 0 | 162 | 15 | 819 | 1769 | 2022 | 56 | 9113 | 8562 | 662 | 0 | 313 | 59 | 111 | 173 | 13 | 95 | 158 | 99 | 4176 | 0 |
| SRR4836521 | 0 | 13 | 54 | 0 | 13 | 0 | 0 | 88 | 53 | 1612 | 842 | 3762 | 0 | 768 | 322 | 733 | 0 | 171 | 12 | 1653 | 1129 | 1189 | 45 | 3033 | 1815 | 355 | 42 | 20 | 38 | 67 | 40 | 0 | 74 | 76 | 51 | 13210 | 0 |
| SRR4836522 | 0 | 0 | 62 | 0 | 13 | 0 | 0 | 56 | 64 | 1680 | 5362 | 4478 | 0 | 643 | 257 | 748 | 0 | 149 | 0 | 1881 | 1003 | 897 | 31 | 2532 | 1421 | 271 | 45 | 26 | 37 | 74 | 57 | 14 | 37 | 91 | 40 | 14019 | 0 |
| SRR4836523 | 0 | 0 | 57 | 0 | 0 | 0 | 0 | 192 | 52 | 1776 | 5216 | 4217 | 0 | 607 | 370 | 866 | 0 | 187 | 10 | 1635 | 1037 | 1054 | 56 | 2848 | 1659 | 344 | 92 | 24 | 40 | 51 | 54 | 16 | 65 | 75 | 73 | 13405 | 0 |
| SRR4836524 | 0 | 0 | 39 | 0 | 0 | 0 | 0 | 82 | 44 | 1759 | 870 | 3661 | 0 | 760 | 382 | 787 | 0 | 188 | 0 | 1257 | 1124 | 1223 | 50 | 3603 | 1973 | 325 | 35 | 26 | 35 | 59 | 59 | 11 | 48 | 56 | 82 | 12731 | 0 |
| SRR4836525 | 0 | 0 | 30 | 0 | 13 | 0 | 0 | 416 | 74 | 678 | 30338 | 13864 | 0 | 274 | 439 | 1453 | 0 | 181 | 19 | 644 | 1856 | 1656 | 73 | 9591 | 6159 | 659 | 0 | 561 | 71 | 138 | 135 | 13 | 109 | 185 | 115 | 4069 | 0 |
| SRR4836526 | 0 | 0 | 21 | 0 | 14 | 0 | 0 | 376 | 88 | 759 | 30770 | 14836 | 0 | 162 | 450 | 1718 | 0 | 151 | 16 | 597 | 1732 | 1397 | 68 | 8176 | 5163 | 642 | 0 | 650 | 84 | 129 | 141 | 17 | 132 | 196 | 129 | 4734 | 0 |
| SRR4836528 | 0 | 0 | 12 | 0 | 19 | 0 | 0 | 427 | 40 | 501 | 4422 | 14305 | 0 | 214 | 394 | 1509 | 0 | 206 | 10 | 474 | 1759 | 1513 | 72 | 9222 | 6048 | 668 | 0 | 177 | 18 | 35 | 40 | 15 | 28 | 55 | 34 | 4472 | 0 |
| SRR4836529 | 0 | 14 | 36 | 0 | 11 | 0 | 0 | 348 | 104 | 733 | 5279 | 12842 | 0 | 234 | 385 | 1571 | 0 | 208 | 15 | 598 | 1837 | 1628 | 82 | 10985 | 6800 | 621 | 0 | 607 | 62 | 141 | 138 | 13 | 103 | 186 | 119 | 4114 | 0 |
| SRR4836531 | 0 | 11 | 26 | 0 | 0 | 0 | 57 | 203 | 64 | 709 | 21178 | 13772 | 0 | 843 | 70 | 47 | 28 | 60 | 12 | 657 | 3155 | 2007 | 0 | 7089 | 4777 | 498 | 170 | 105 | 62 | 153 | 105 | 23 | 159 | 269 | 28 | 5947 | 0 |
| SRR4836533 | 0 | 0 | 19 | 0 | 0 | 0 | 58 | 192 | 34 | 593 | 18948 | 13283 | 0 | 1028 | 56 | 62 | 28 | 61 | 10 | 751 | 3289 | 2064 | 22 | 8160 | 5444 | 977 | 202 | 80 | 61 | 147 | 116 | 22 | 169 | 346 | 14 | 5326 | 0 |
| SRR4836534 | 0 | 10 | 14 | 0 | 0 | 0 | 46 | 138 | 52 | 707 | 5748 | 13541 | 0 | 832 | 62 | 59 | 32 | 81 | 13 | 625 | 3362 | 1836 | 15 | 6914 | 4881 | 831 | 34 | 17 | 12 | 22 | 15 | 36 | 34 | 53 | 17 | 5763 | 0 |
| SRR4836535 | 0 | 0 | 12 | 0 | 24 | 0 | 0 | 203 | 92 | 1723 | 3365 | 2186 | 0 | 170 | 15 | 30 | 0 | 376 | 11 | 2115 | 4160 | 2657 | 104 | 17679 | 5318 | 1930 | 194 | 20 | 33 | 53 | 51 | 16 | 78 | 88 | 31 | 16594 | 11 |
| SRR4836536 | 0 | 0 | 24 | 0 | 0 | 0 | 20 | 214 | 96 | 1782 | 3243 | 1957 | 0 | 118 | 18 | 39 | 0 | 364 | 0 | 2238 | 4054 | 2756 | 152 | 20951 | 6276 | 1940 | 173 | 16 | 24 | 75 | 60 | 10 | 60 | 75 | 15 | 15915 | 0 |
| SRR4836537 | 0 | 0 | 12 | 0 | 16 | 0 | 0 | 197 | 84 | 1746 | 1640 | 2028 | 0 | 122 | 0 | 29 | 0 | 359 | 0 | 2349 | 4164 | 2497 | 103 | 18109 | 5357 | 1606 | 40 | 21 | 26 | 16 | 12 | 23 | 17 | 22 | 0 | 16597 | 16 |
| SRR4836539 | 0 | 0 | 0 | 0 | 14 | 0 | 59 | 241 | 14 | 255 | 722 | 1270 | 0 | 1070 | 0 | 0 | 41 | 942 | 32 | 3186 | 5014 | 5242 | 88 | 17601 | 17300 | 294 | 252 | 24 | 0 | 13 | 12 | 0 | 0 | 0 | 0 | 40393 | 18 |
| SRR4836541 | 0 | 0 | 0 | 0 | 0 | 0 | 41 | 313 | 20 | 280 | 715 | 1140 | 0 | 1034 | 0 | 0 | 41 | 1046 | 35 | 3679 | 4938 | 5668 | 67 | 20361 | 18856 | 253 | 239 | 12 | 0 | 16 | 16 | 0 | 0 | 16 | 0 | 37913 | 0 |
| SRR4836543 | 0 | 14 | 0 | 0 | 13 | 0 | 54 | 317 | 17 | 266 | 2447 | 1166 | 0 | 1017 | 0 | 10 | 36 | 871 | 15 | 3678 | 5411 | 5083 | 57 | 17530 | 16764 | 259 | 261 | 15 | 0 | 14 | 0 | 0 | 0 | 22 | 0 | 40481 | 0 |
| SRR4836544 | 0 | 0 | 88 | 0 | 38 | 0 | 0 | 302 | 0 | 5862 | 14322 | 18674 | 0 | 91 | 0 | 0 | 15 | 142 | 0 | 5137 | 777 | 1507 | 62 | 5925 | 2739 | 498 | 71 | 159 | 103 | 244 | 177 | 59 | 159 | 168 | 32 | 29128 | 0 |
| SRR4836545 | 0 | 0 | 61 | 0 | 44 | 0 | 0 | 591 | 148 | 7654 | 15337 | 17760 | 0 | 59 | 0 | 0 | 0 | 118 | 0 | 4078 | 756 | 1595 | 70 | 7398 | 3192 | 487 | 89 | 182 | 84 | 265 | 170 | 32 | 136 | 228 | 23 | 29374 | 0 |
| SRR4836546 | 0 | 0 | 13 | 0 | 47 | 0 | 0 | 437 | 91 | 7320 | 14367 | 17500 | 0 | 36 | 0 | 0 | 0 | 132 | 0 | 4097 | 768 | 1475 | 52 | 6108 | 2604 | 462 | 18 | 36 | 13 | 42 | 34 | 34 | 33 | 39 | 28 | 30374 | 0 |
| SRR4836548 | 0 | 0 | 13 | 0 | 0 | 0 | 13 | 161 | 21 | 636 | 23929 | 14891 | 0 | 1257 | 14 | 21 | 0 | 71 | 17 | 1460 | 4813 | 1604 | 20 | 6274 | 3557 | 3536 | 15 | 20 | 67 | 33 | 15 | 13 | 13 | 16 | 18 | 12258 | 18 |
| SRR4836550 | 0 | 0 | 13 | 0 | 0 | 0 | 0 | 190 | 0 | 524 | 22538 | 13908 | 18 | 1360 | 14 | 14 | 0 | 56 | 19 | 1483 | 4924 | 1741 | 31 | 7530 | 4016 | 3670 | 17 | 12 | 75 | 33 | 19 | 22 | 15 | 18 | 27 | 11194 | 12 |
| SRR4836552 | 0 | 11 | 150 | 0 | 0 | 0 | 0 | 126 | 0 | 534 | 23480 | 14077 | 0 | 1290 | 25 | 0 | 0 | 55 | 29 | 1618 | 5099 | 1524 | 27 | 6538 | 3393 | 3263 | 115 | 109 | 67 | 298 | 171 | 22 | 96 | 150 | 27 | 11673 | 15 |
| SRR4836553 | 0 | 0 | 10 | 0 | 13 | 0 | 48 | 1105 | 833 | 1880 | 585 | 6646 | 0 | 919 | 14 | 86 | 35 | 244 | 0 | 4206 | 2278 | 1953 | 64 | 6311 | 2057 | 550 | 41 | 51 | 36 | 56 | 26 | 11 | 57 | 58 | 22 | 47169 | 0 |
| SRR4836554 | 0 | 0 | 55 | 0 | 13 | 0 | 51 | 1751 | 740 | 1850 | 7743 | 6007 | 0 | 740 | 0 | 14 | 25 | 209 | 0 | 4654 | 2304 | 2308 | 91 | 7095 | 2605 | 535 | 220 | 178 | 161 | 305 | 186 | 38 | 178 | 292 | 42 | 44551 | 0 |
| SRR4836555 | 0 | 13 | 47 | 0 | 20 | 0 | 47 | 1309 | 846 | 1457 | 973 | 6429 | 0 | 890 | 0 | 52 | 16 | 189 | 0 | 4833 | 2367 | 2019 | 95 | 6032 | 2061 | 449 | 43 | 46 | 34 | 61 | 26 | 32 | 47 | 69 | 32 | 46906 | 0 |
| SRR4836556 | 0 | 0 | 29 | 0 | 16 | 0 | 29 | 151 | 43 | 949 | 1799 | 23262 | 19 | 43 | 0 | 43 | 19 | 86 | 35 | 1833 | 4135 | 3488 | 145 | 21128 | 8808 | 2387 | 160 | 398 | 117 | 451 | 267 | 49 | 218 | 199 | 24 | 11707 | 18 |
| SRR4836560 | 12 | 0 | 12 | 0 | 0 | 0 | 19 | 86 | 34 | 1344 | 38944 | 24797 | 27 | 86 | 0 | 17 | 39 | 61 | 38 | 1590 | 4189 | 3591 | 132 | 22817 | 9120 | 2444 | 159 | 422 | 70 | 376 | 256 | 46 | 170 | 136 | 0 | 12080 | 0 |
| SRR4836561 | 0 | 0 | 23 | 0 | 0 | 0 | 20 | 139 | 104 | 1320 | 39531 | 21225 | 0 | 69 | 0 | 0 | 27 | 72 | 36 | 1841 | 4187 | 3480 | 132 | 22801 | 9530 | 2169 | 153 | 368 | 102 | 399 | 231 | 90 | 204 | 172 | 0 | 11290 | 12 |
| SRR4836562 | 0 | 12 | 0 | 13 | 0 | 0 | 0 | 130 | 20 | 1093 | 2997 | 10005 | 0 | 3832 | 10 | 15 | 0 | 76 | 15 | 624 | 1054 | 3042 | 196 | 11222 | 8700 | 4055 | 14 | 66 | 32 | 33 | 22 | 12 | 14 | 20 | 24 | 2929 | 22 |
| SRR4836563 | 0 | 0 | 0 | 0 | 0 | 0 | 0 | 170 | 35 | 1101 | 15501 | 10322 | 0 | 3733 | 0 | 10 | 0 | 62 | 10 | 670 | 1067 | 3234 | 189 | 11862 | 9197 | 3899 | 10 | 72 | 37 | 33 | 19 | 0 | 14 | 24 | 20 | 2502 | 32 |
| SRR4836565 | 0 | 0 | 0 | 0 | 0 | 0 | 0 | 154 | 20 | 1428 | 15051 | 10262 | 0 | 3960 | 0 | 32 | 0 | 53 | 13 | 574 | 1179 | 3329 | 201 | 11906 | 9227 | 4190 | 74 | 495 | 215 | 293 | 151 | 26 | 109 | 196 | 23 | 2900 | 30 |
| SRR4836567 | 0 | 0 | 0 | 12 | 0 | 0 | 0 | 135 | 25 | 1148 | 14910 | 10313 | 0 | 3539 | 0 | 21 | 0 | 56 | 13 | 603 | 1156 | 3320 | 187 | 12355 | 9377 | 3943 | 91 | 477 | 258 | 268 | 186 | 26 | 114 | 176 | 13 | 2628 | 18 |
| SRR4836569 | 0 | 18 | 62 | 0 | 18 | 0 | 51 | 662 | 87 | 835 | 24960 | 13527 | 0 | 477 | 11 | 0 | 29 | 198 | 17 | 456 | 520 | 1295 | 29 | 11757 | 2496 | 190 | 135 | 29 | 153 | 491 | 899 | 51 | 106 | 113 | 11 | 2701 | 181 |
| SRR4836570 | 0 | 36 | 61 | 0 | 24 | 0 | 47 | 597 | 64 | 843 | 25174 | 13056 | 0 | 544 | 0 | 0 | 18 | 219 | 16 | 608 | 540 | 1369 | 33 | 12774 | 2639 | 203 | 143 | 36 | 179 | 29 | 54 | 54 | 111 | 143 | 0 | 2624 | 183 |
| SRR4836571 | 0 | 38 | 46 | 0 | 24 | 0 | 49 | 636 | 86 | 851 | 24729 | 13770 | 0 | 550 | 0 | 0 | 29 | 197 | 18 | 370 | 548 | 1389 | 34 | 12798 | 2639 | 187 | 78 | 26 | 159 | 25 | 47 | 40 | 72 | 139 | 0 | 2647 | 180 |
| SRR4836572 | 0 | 15 | 64 | 0 | 24 | 0 | 41 | 683 | 51 | 862 | 24787 | 13041 | 0 | 509 | 0 | 0 | 26 | 202 | 15 | 495 | 548 | 1388 | 36 | 13059 | 2636 | 187 | 149 | 44 | 152 | 27 | 50 | 23 | 120 | 152 | 0 | 2782 | 168 |
| SRR4896694 | 0 | 1249 | 22 | 0 | 13 | 0 | 14 | 98 | 272 | 1182 | 38 | 56 | 0 | 482 | 0 | 0 | 19 | 509 | 10 | 217 | 835 | 573 | 105 | 2695 | 2991 | 39 | 122 | 19 | 22 | 46 | 67 | 31 | 24 | 21 | 24 | 2817 | 0 |
| SRR4896695 | 0 | 96 | 104 | 0 | 11 | 12 | 0 | 1016 | 517 | 3022 | 123 | 1331 | 0 | 360 | 351 | 185 | 0 | 312 | 0 | 1063 | 681 | 489 | 93 | 2049 | 2644 | 0 | 0 | 84 | 56 | 68 | 100 | 40 | 20 | 32 | 0 | 4500 | 26 |
| SRR4896698 | 0 | 332 | 64 | 0 | 0 | 0 | 0 | 1034 | 142 | 15611 | 566 | 652 | 0 | 1459 | 95 | 52 | 0 | 558 | 0 | 600 | 1400 | 985 | 87 | 3500 | 2928 | 31 | 0 | 59 | 21 | 75 | 59 | 38 | 32 | 11 | 0 | 1748 | 0 |
| SRR4896701 | 0 | 411 | 95 | 0 | 44 | 0 | 0 | 128 | 593 | 7911 | 116 | 87 | 0 | 2748 | 31 | 15 | 0 | 287 | 100 | 149 | 1514 | 1400 | 56 | 3123 | 9569 | 0 | 0 | 408 | 138 | 120 | 73 | 43 | 85 | 33 | 30 | 431 | 844 |
| SRR4896708 | 0 | 298 | 28 | 0 | 55 | 0 | 0 | 18 | 61 | 4333 | 273 | 163 | 0 | 3327 | 43 | 23 | 10 | 567 | 68 | 171 | 1198 | 955 | 105 | 2639 | 3021 | 0 | 29 | 59 | 106 | 137 | 133 | 81 | 77 | 31 | 0 | 611 | 187 |
| SRR5022482 | 0 | 887 | 39 | 0 | 0 | 0 | 0 | 235 | 113 | 762 | 13365 | 13928 | 0 | 716 | 23 | 33 | 0 | 71 | 43 | 339 | 5780 | 4133 | 0 | 12 | 14422 | 0 | 15 | 49 | 106 | 227 | 208 | 193 | 91 | 148 | 47 | 1033 | 59 |
| SRR5022483 | 0 | 55 | 0 | 0 | 0 | 0 | 0 | 94 | 146 | 2935 | 3183 | 1416 | 0 | 3082 | 0 | 0 | 0 | 29 | 19 | 114 | 3254 | 5318 | 0 | 65 | 10750 | 0 | 0 | 0 | 0 | 0 | 0 | 0 | 0 | 25 | 0 | 1722 | 0 |
| SRR5022484 | 0 | 67 | 15 | 0 | 0 | 0 | 0 | 823 | 234 | 3857 | 5682 | 5476 | 0 | 789 | 79 | 64 | 0 | 14 | 39 | 289 | 3801 | 7003 | 0 | 18 | 21623 | 0 | 145 | 12 | 20 | 15 | 89 | 52 | 18 | 18 | 0 | 1039 | 33 |
| SRR5022485 | 16 | 12 | 0 | 0 | 0 | 0 | 0 | 381 | 286 | 8681 | 1756 | 1092 | 0 | 3706 | 0 | 0 | 0 | 12 | 31 | 181 | 1502 | 4324 | 14 | 190 | 6847 | 0 | 0 | 0 | 0 | 27 | 23 | 35 | 121 | 0 | 0 | 2066 | 0 |
| SRR5029397 | 12 | 111 | 0 | 45 | 36 | 24 | 15 | 362 | 608 | 547 | 155 | 2234 | 0 | 123 | 0 | 75 | 19 | 1539 | 87 | 2070 | 2383 | 663 | 58 | 3030 | 606 | 45 | 37 | 25 | 33 | 33 | 68 | 21 | 0 | 0 | 0 | 2049 | 21 |
| SRR5029398 | 0 | 247 | 0 | 46 | 71 | 25 | 0 | 173 | 66 | 25 | 319 | 3697 | 0 | 0 | 41 | 156 | 0 | 1765 | 65 | 1317 | 1766 | 610 | 12 | 1092 | 1153 | 30 | 22 | 0 | 32 | 32 | 68 | 14 | 0 | 0 | 0 | 1153 | 89 |
| SRR5029399 | 0 | 81 | 55 | 39 | 45 | 28 | 0 | 475 | 797 | 3548 | 821 | 1841 | 0 | 193 | 1875 | 2588 | 0 | 1164 | 21 | 4419 | 2141 | 760 | 37 | 1301 | 562 | 0 | 96 | 0 | 13 | 25 | 49 | 12 | 33 | 0 | 0 | 11124 | 10 |
| SRR5029400 | 0 | 674 | 0 | 66 | 133 | 44 | 16 | 201 | 187 | 716 | 100 | 1017 | 14 | 94 | 676 | 388 | 0 | 1337 | 172 | 930 | 1536 | 339 | 20 | 1295 | 1118 | 34 | 115 | 19 | 206 | 333 | 384 | 35 | 0 | 0 | 0 | 1519 | 101 |
| SRR5029401 | 0 | 271 | 0 | 65 | 74 | 55 | 15 | 141 | 128 | 192 | 1623 | 1891 | 0 | 26 | 0 | 26 | 0 | 1064 | 208 | 1597 | 2654 | 686 | 15 | 1294 | 1139 | 17 | 139 | 0 | 82 | 91 | 167 | 19 | 0 | 0 | 0 | 1290 | 39 |
| SRR5029402 | 0 | 216 | 0 | 43 | 95 | 35 | 16 | 304 | 101 | 203 | 118 | 1178 | 0 | 25 | 25 | 51 | 15 | 986 | 144 | 3420 | 3829 | 999 | 35 | 1584 | 2809 | 50 | 143 | 15 | 106 | 104 | 293 | 41 | 0 | 0 | 0 | 3382 | 76 |
| SRR5029403 | 0 | 106 | 15 | 59 | 170 | 22 | 0 | 416 | 808 | 2234 | 10654 | 10603 | 0 | 154 | 559 | 594 | 0 | 1068 | 39 | 806 | 1250 | 516 | 18 | 1098 | 838 | 13 | 35 | 11 | 20 | 80 | 13 | 17 | 0 | 0 | 0 | 713 | 31 |
| SRR5029404 | 0 | 268 | 0 | 61 | 59 | 183 | 20 | 0 | 77 | 103 | 1670 | 1824 | 0 | 0 | 51 | 51 | 0 | 931 | 92 | 1772 | 2153 | 543 | 50 | 1985 | 1108 | 54 | 180 | 17 | 64 | 72 | 38 | 0 | 0 | 0 | 0 | 1156 | 51 |
| SRR5029405 | 0 | 241 | 0 | 37 | 50 | 172 | 10 | 121 | 311 | 190 | 79 | 1227 | 11 | 69 | 52 | 363 | 15 | 1099 | 178 | 4043 | 3684 | 1080 | 50 | 1785 | 1958 | 0 | 94 | 0 | 184 | 133 | 224 | 33 | 0 | 0 | 0 | 4285 | 51 |
| SRR5029406 | 0 | 208 | 13 | 75 | 79 | 60 | 0 | 250 | 238 | 413 | 62 | 901 | 0 | 50 | 13 | 300 | 14 | 1900 | 114 | 7198 | 3231 | 979 | 0 | 879 | 2339 | 35 | 92 | 39 | 220 | 196 | 888 | 202 | 15 | 0 | 0 | 7811 | 253 |
| SRR5029407 | 0 | 107 | 0 | 40 | 103 | 10 | 0 | 96 | 66 | 96 | 3918 | 9010 | 0 | 22 | 30 | 89 | 0 | 1464 | 50 | 214 | 3094 | 869 | 73 | 2372 | 904 | 0 | 40 | 11 | 11 | 14 | 13 | 10 | 0 | 0 | 0 | 236 | 0 |
| SRR5029408 | 0 | 128 | 0 | 55 | 65 | 17 | 0 | 23 | 34 | 23 | 5072 | 6562 | 0 | 0 | 11 | 85 | 0 | 1348 | 137 | 625 | 2627 | 918 | 27 | 1248 | 1189 | 30 | 43 | 0 | 13 | 16 | 15 | 0 | 0 | 0 | 0 | 409 | 57 |
| SRR5029409 | 0 | 74 | 0 | 32 | 174 | 50 | 34 | 362 | 444 | 411 | 6416 | 10398 | 0 | 49 | 247 | 428 | 0 | 917 | 56 | 987 | 1432 | 449 | 54 | 2018 | 691 | 12 | 0 | 22 | 18 | 33 | 11 | 0 | 0 | 0 | 0 | 1563 | 12 |
| SRR5029410 | 0 | 63 | 12 | 27 | 36 | 0 | 0 | 416 | 1125 | 6582 | 1362 | 2349 | 0 | 383 | 351 | 946 | 0 | 1030 | 66 | 6940 | 1684 | 644 | 60 | 3643 | 1072 | 0 | 56 | 0 | 28 | 27 | 84 | 18 | 56 | 16 | 0 | 8873 | 17 |
| SRR5029411 | 0 | 308 | 0 | 71 | 189 | 99 | 0 | 24 | 94 | 94 | 147 | 4512 | 0 | 0 | 0 | 94 | 0 | 2671 | 55 | 5339 | 2220 | 798 | 0 | 430 | 1017 | 61 | 141 | 15 | 120 | 231 | 302 | 27 | 0 | 0 | 0 | 3945 | 203 |
| SRR5029412 | 0 | 907 | 18 | 51 | 68 | 45 | 14 | 460 | 1004 | 4847 | 120 | 1818 | 14 | 478 | 177 | 236 | 0 | 1262 | 36 | 2243 | 1973 | 598 | 47 | 1491 | 615 | 0 | 235 | 25 | 115 | 296 | 50 | 0 | 0 | 0 | 0 | 3938 | 10 |
| SRR5029414 | 0 | 94 | 0 | 61 | 79 | 43 | 12 | 16 | 109 | 31 | 1899 | 1774 | 0 | 16 | 31 | 78 | 20 | 1879 | 0 | 3979 | 3591 | 998 | 66 | 4956 | 1590 | 23 | 0 | 19 | 25 | 23 | 0 | 0 | 0 | 0 | 0 | 825 | 47 |
| SRR5029415 | 0 | 106 | 19 | 66 | 160 | 19 | 0 | 133 | 0 | 664 | 4778 | 3318 | 0 | 0 | 796 | 3451 | 0 | 1259 | 47 | 929 | 2227 | 489 | 0 | 1475 | 893 | 38 | 55 | 0 | 35 | 19 | 29 | 0 | 0 | 0 | 0 | 1195 | 113 |
| SRR5029416 | 0 | 166 | 36 | 38 | 88 | 21 | 13 | 828 | 1224 | 3387 | 502 | 6893 | 10 | 291 | 1186 | 2290 | 0 | 1300 | 56 | 6200 | 2229 | 1222 | 42 | 1279 | 1243 | 0 | 18 | 0 | 36 | 33 | 85 | 15 | 34 | 0 | 0 | 7908 | 34 |
| SRR5029417 | 0 | 821 | 0 | 79 | 164 | 33 | 0 | 71 | 106 | 88 | 209 | 2743 | 0 | 0 | 0 | 0 | 11 | 1295 | 59 | 389 | 3032 | 638 | 44 | 1695 | 1135 | 35 | 174 | 22 | 93 | 77 | 86 | 0 | 0 | 0 | 0 | 425 | 60 |
| SRR5029418 | 0 | 136 | 0 | 45 | 41 | 32 | 0 | 159 | 167 | 351 | 1132 | 1403 | 0 | 56 | 0 | 80 | 0 | 1395 | 180 | 5262 | 3901 | 969 | 75 | 3647 | 1674 | 51 | 58 | 0 | 85 | 124 | 335 | 72 | 0 | 0 | 0 | 4457 | 24 |
| SRR5029419 | 0 | 182 | 0 | 69 | 72 | 49 | 13 | 60 | 133 | 99 | 266 | 3297 | 0 | 0 | 13 | 53 | 0 | 1321 | 90 | 3283 | 1776 | 869 | 14 | 1012 | 873 | 81 | 131 | 0 | 18 | 18 | 461 | 59 | 0 | 0 | 0 | 2461 | 96 |
| SRR5029420 | 0 | 1307 | 0 | 34 | 304 | 16 | 27 | 651 | 848 | 3699 | 337 | 5555 | 0 | 184 | 74 | 86 | 0 | 1180 | 186 | 1389 | 983 | 498 | 16 | 1211 | 1194 | 0 | 62 | 0 | 259 | 178 | 405 | 89 | 0 | 0 | 0 | 2151 | 0 |
| SRR5029422 | 0 | 89 | 53 | 105 | 315 | 20 | 0 | 594 | 1187 | 1682 | 5539 | 9125 | 0 | 25 | 1014 | 1929 | 0 | 1497 | 69 | 4031 | 1416 | 692 | 36 | 1036 | 1346 | 23 | 12 | 87 | 33 | 31 | 45 | 147 | 20 | 0 | 0 | 4229 | 146 |
| SRR5029424 | 0 | 135 | 0 | 0 | 19 | 0 | 0 | 453 | 1415 | 1132 | 10073 | 20769 | 117 | 57 | 0 | 0 | 14 | 69 | 45 | 1415 | 760 | 483 | 0 | 24 | 3972 | 82 | 864 | 0 | 45 | 41 | 54 | 0 | 0 | 0 | 0 | 1415 | 161 |
| SRR5029425 | 0 | 336 | 0 | 56 | 95 | 22 | 26 | 124 | 48 | 163 | 1234 | 2028 | 0 | 0 | 0 | 144 | 0 | 1372 | 87 | 517 | 2870 | 683 | 43 | 2011 | 1266 | 27 | 243 | 14 | 46 | 49 | 101 | 20 | 0 | 0 | 0 | 498 | 58 |
| SRR5029426 | 0 | 897 | 0 | 61 | 91 | 19 | 13 | 264 | 351 | 472 | 791 | 16653 | 0 | 22 | 154 | 286 | 0 | 1665 | 48 | 1384 | 2303 | 888 | 60 | 2475 | 680 | 171 | 143 | 32 | 123 | 118 | 171 | 18 | 0 | 0 | 0 | 1318 | 48 |
| SRR5029427 | 0 | 341 | 0 | 64 | 83 | 67 | 19 | 295 | 402 | 496 | 146 | 958 | 0 | 33 | 87 | 382 | 10 | 1615 | 42 | 1929 | 3753 | 861 | 63 | 2828 | 1464 | 49 | 155 | 0 | 89 | 187 | 442 | 124 | 0 | 0 | 0 | 3055 | 26 |
| SRR5029428 | 0 | 155 | 0 | 0 | 24 | 0 | 0 | 270 | 622 | 974 | 6842 | 17335 | 49 | 406 | 0 | 0 | 0 | 52 | 41 | 3218 | 324 | 218 | 0 | 11 | 2680 | 83 | 439 | 0 | 89 | 40 | 40 | 191 | 13 | 13 | 0 | 7815 | 230 |
| SRR5029429 | 0 | 1178 | 21 | 27 | 55 | 26 | 0 | 597 | 1377 | 2173 | 1559 | 4014 | 0 | 116 | 83 | 382 | 0 | 850 | 25 | 2737 | 1595 | 488 | 30 | 1144 | 455 | 0 | 212 | 16 | 227 | 160 | 362 | 83 | 0 | 0 | 0 | 5606 | 0 |
| SRR5029430 | 0 | 69 | 37 | 37 | 223 | 37 | 0 | 539 | 971 | 2715 | 529 | 15948 | 0 | 180 | 162 | 647 | 0 | 1389 | 28 | 1546 | 1277 | 512 | 27 | 1228 | 421 | 17 | 14 | 0 | 143 | 17 | 29 | 79 | 0 | 0 | 0 | 2571 | 49 |
| SRR5029431 | 0 | 55 | 0 | 52 | 44 | 28 | 12 | 46 | 0 | 15 | 1300 | 2033 | 0 | 0 | 15 | 61 | 0 | 1220 | 107 | 871 | 3056 | 872 | 61 | 2482 | 1096 | 61 | 36 | 0 | 16 | 15 | 18 | 0 | 0 | 0 | 0 | 780 | 21 |
| SRR5029432 | 0 | 76 | 22 | 74 | 104 | 41 | 13 | 1569 | 3288 | 11370 | 1275 | 3040 | 0 | 599 | 640 | 537 | 0 | 1594 | 42 | 1368 | 1649 | 706 | 58 | 1643 | 860 | 11 | 98 | 0 | 14 | 27 | 38 | 85 | 0 | 0 | 0 | 2916 | 10 |
| SRR5029433 | 0 | 206 | 0 | 68 | 114 | 40 | 39 | 74 | 44 | 15 | 72 | 1565 | 0 | 0 | 0 | 15 | 50 | 1750 | 118 | 768 | 4748 | 776 | 14 | 1355 | 1648 | 33 | 813 | 0 | 15 | 31 | 66 | 0 | 0 | 0 | 0 | 473 | 153 |
| SRR5029434 | 0 | 47 | 0 | 36 | 93 | 12 | 11 | 15 | 44 | 671 | 53 | 2189 | 0 | 0 | 15 | 117 | 16 | 899 | 141 | 3648 | 4702 | 909 | 38 | 1311 | 2142 | 104 | 34 | 0 | 86 | 72 | 168 | 29 | 0 | 0 | 0 | 2816 | 45 |
| SRR5029435 | 0 | 1293 | 0 | 62 | 80 | 78 | 28 | 493 | 987 | 3406 | 224 | 6797 | 0 | 318 | 159 | 239 | 0 | 1282 | 53 | 2022 | 1797 | 556 | 44 | 1761 | 622 | 0 | 106 | 0 | 131 | 197 | 437 | 72 | 0 | 0 | 0 | 3215 | 0 |
| SRR5029436 | 0 | 141 | 43 | 72 | 178 | 14 | 0 | 490 | 853 | 1816 | 6902 | 13227 | 0 | 182 | 648 | 995 | 0 | 1477 | 113 | 2827 | 2207 | 473 | 15 | 1290 | 872 | 62 | 18 | 14 | 59 | 41 | 103 | 16 | 17 | 26 | 0 | 3782 | 95 |
| SRR5029437 | 0 | 0 | 0 | 12 | 39 | 0 | 0 | 375 | 1009 | 3026 | 23 | 117 | 217 | 188 | 0 | 0 | 12 | 167 | 74 | 94 | 37 | 0 | 0 | 88 | 7477 | 65 | 34 | 0 | 133 | 72 | 259 | 0 | 0 | 0 | 0 | 563 | 186 |
| SRR5029438 | 0 | 53 | 0 | 107 | 271 | 0 | 14 | 1062 | 1142 | 3754 | 6454 | 11589 | 0 | 152 | 942 | 1206 | 0 | 1248 | 65 | 2133 | 1755 | 668 | 32 | 1197 | 1181 | 32 | 64 | 29 | 20 | 21 | 39 | 50 | 0 | 0 | 0 | 1893 | 182 |
| SRR5029439 | 0 | 91 | 0 | 145 | 307 | 15 | 10 | 457 | 753 | 1520 | 3799 | 7939 | 12 | 109 | 464 | 451 | 15 | 1259 | 84 | 2215 | 1484 | 521 | 14 | 734 | 967 | 28 | 68 | 32 | 26 | 23 | 44 | 24 | 0 | 0 | 0 | 5316 | 209 |
| SRR5029440 | 0 | 21 | 0 | 0 | 38 | 0 | 0 | 808 | 2208 | 2488 | 15 | 466 | 151 | 31 | 0 | 0 | 10 | 125 | 49 | 529 | 1453 | 103 | 0 | 47 | 4869 | 56 | 356 | 0 | 182 | 83 | 17 | 16 | 0 | 0 | 0 | 746 | 250 |
| SRR5029441 | 0 | 181 | 0 | 104 | 106 | 14 | 15 | 268 | 105 | 58 | 779 | 1280 | 0 | 0 | 0 | 12 | 12 | 1882 | 108 | 605 | 3439 | 913 | 48 | 2351 | 1326 | 76 | 107 | 17 | 57 | 48 | 136 | 25 | 0 | 0 | 0 | 535 | 134 |
| SRR5029442 | 0 | 1065 | 0 | 40 | 271 | 44 | 11 | 226 | 251 | 193 | 4014 | 6385 | 0 | 17 | 109 | 277 | 0 | 1627 | 61 | 1533 | 2750 | 710 | 11 | 755 | 1562 | 60 | 177 | 40 | 237 | 252 | 368 | 74 | 0 | 0 | 0 | 1936 | 226 |
| SRR5029444 | 0 | 773 | 15 | 90 | 216 | 62 | 77 | 413 | 578 | 3467 | 12 | 2476 | 0 | 330 | 83 | 0 | 0 | 1000 | 99 | 83 | 829 | 297 | 21 | 1271 | 1802 | 27 | 46 | 0 | 77 | 46 | 46 | 0 | 0 | 0 | 0 | 495 | 0 |
| SRR5029446 | 0 | 236 | 0 | 0 | 17 | 0 | 0 | 541 | 270 | 595 | 16216 | 31622 | 174 | 0 | 0 | 0 | 0 | 71 | 49 | 3189 | 582 | 421 | 0 | 28 | 3047 | 96 | 1694 | 0 | 62 | 22 | 44 | 0 | 0 | 0 | 0 | 3405 | 144 |
| SRR5029447 | 0 | 165 | 0 | 60 | 127 | 37 | 0 | 145 | 116 | 116 | 266 | 3921 | 0 | 19 | 19 | 145 | 0 | 1495 | 96 | 540 | 3276 | 1004 | 53 | 1849 | 1545 | 26 | 31 | 16 | 28 | 22 | 33 | 20 | 0 | 0 | 0 | 434 | 100 |
| SRR5029448 | 0 | 673 | 13 | 37 | 73 | 24 | 0 | 639 | 769 | 1226 | 14268 | 28106 | 0 | 26 | 417 | 639 | 0 | 1099 | 41 | 1513 | 1384 | 732 | 45 | 1601 | 583 | 0 | 71 | 0 | 103 | 203 | 340 | 42 | 0 | 0 | 0 | 1252 | 11 |
| SRR5029452 | 0 | 11 | 19 | 0 | 25 | 0 | 0 | 1112 | 1939 | 6343 | 15 | 15 | 124 | 601 | 30 | 15 | 0 | 135 | 86 | 406 | 0 | 0 | 0 | 39 | 5963 | 78 | 170 | 0 | 60 | 23 | 124 | 15 | 0 | 0 | 0 | 481 | 209 |
| SRR5029455 | 0 | 62 | 0 | 90 | 306 | 34 | 0 | 240 | 470 | 1291 | 5918 | 10961 | 0 | 37 | 1005 | 1586 | 25 | 1076 | 100 | 5015 | 1083 | 720 | 11 | 686 | 1288 | 0 | 48 | 31 | 37 | 42 | 77 | 84 | 13 | 0 | 0 | 7301 | 0 |
| SRR5029456 | 0 | 207 | 0 | 99 | 181 | 0 | 12 | 344 | 356 | 310 | 459 | 7333 | 0 | 23 | 230 | 69 | 0 | 1421 | 107 | 230 | 2537 | 824 | 36 | 2406 | 1641 | 13 | 48 | 26 | 51 | 31 | 62 | 42 | 0 | 0 | 0 | 448 | 92 |

### Supp Table S8.4.Columns_DI_to_ES

| | | | | | | | | | | | | | | | | | | | | | | | | | | | | | | | | | | | | | |
|---|---|---|---|---|---|---|---|---|---|---|---|---|---|---|---|---|---|---|---|---|---|---|---|---|---|---|---|---|---|---|---|---|---|---|---|---|---|
| SRR5029457 | 0 | 1085 | 0 | 80 | 299 | 16 | 16 | 576 | 559 | 699 | 12146 | 20485 | 0 | 26 | 26 | 61 | 0 | 1437 | 57 | 707 | 1274 | 696 | 39 | 1582 | 605 | 11 | 86 | 0 | 56 | 61 | 105 | 19 | 0 | 0 | 0 | 943 | 0 |
| SRR5029459 | 0 | 1065 | 35 | 46 | 268 | 13 | 0 | 584 | 981 | 3333 | 774 | 6729 | 0 | 210 | 334 | 879 | 13 | 1158 | 103 | 1610 | 1420 | 421 | 27 | 1561 | 944 | 13 | 113 | 32 | 262 | 262 | 795 | 163 | 19 | 0 | 0 | 4138 | 42 |
| SRR5029460 | 0 | 196 | 0 | 47 | 185 | 44 | 17 | 360 | 455 | 654 | 3280 | 7167 | 0 | 85 | 95 | 265 | 0 | 1100 | 68 | 512 | 1831 | 816 | 37 | 2120 | 1411 | 28 | 27 | 0 | 29 | 15 | 23 | 0 | 0 | 0 | 0 | 683 | 21 |
| SRR5029462 | 0 | 129 | 83 | 58 | 135 | 11 | 0 | 840 | 1464 | 3198 | 3451 | 7236 | 0 | 99 | 379 | 1319 | 14 | 1638 | 56 | 4535 | 1810 | 799 | 43 | 1825 | 811 | 0 | 63 | 0 | 13 | 24 | 48 | 11 | 16 | 16 | 0 | 8320 | 12 |
| SRR5029463 | 0 | 1156 | 0 | 64 | 292 | 41 | 29 | 677 | 968 | 1500 | 167 | 11177 | 0 | 97 | 0 | 0 | 0 | 1278 | 13 | 387 | 1108 | 585 | 44 | 1889 | 662 | 18 | 117 | 0 | 117 | 158 | 194 | 12 | 0 | 0 | 0 | 339 | 0 |
| SRR5029465 | 0 | 79 | 0 | 0 | 15 | 0 | 0 | 370 | 329 | 643 | 145 | 6202 | 89 | 246 | 14 | 0 | 24 | 40 | 50 | 246 | 433 | 217 | 0 | 21 | 2739 | 89 | 2057 | 0 | 12 | 0 | 107 | 18 | 0 | 0 | 0 | 452 | 193 |
| SRR5029466 | 0 | 883 | 29 | 58 | 130 | 33 | 39 | 693 | 1340 | 2372 | 45 | 1802 | 0 | 77 | 354 | 1032 | 12 | 1446 | 88 | 3249 | 2065 | 783 | 36 | 1959 | 1647 | 18 | 179 | 10 | 230 | 214 | 453 | 35 | 0 | 0 | 0 | 3588 | 153 |
| SRR5029467 | 0 | 107 | 0 | 52 | 150 | 15 | 0 | 44 | 98 | 77 | 221 | 2657 | 0 | 0 | 0 | 77 | 0 | 1188 | 147 | 339 | 2864 | 850 | 28 | 1743 | 1921 | 82 | 23 | 17 | 39 | 14 | 20 | 0 | 0 | 0 | 0 | 405 | 75 |
| SRR5029469 | 0 | 836 | 0 | 36 | 135 | 15 | 24 | 161 | 184 | 368 | 6456 | 6168 | 0 | 23 | 242 | 472 | 0 | 778 | 95 | 437 | 2513 | 624 | 0 | 1189 | 1487 | 28 | 94 | 24 | 162 | 79 | 170 | 20 | 0 | 0 | 0 | 667 | 127 |
| SRR5029471 | 0 | 144 | 0 | 41 | 191 | 0 | 18 | 663 | 647 | 5409 | 6199 | 10474 | 0 | 136 | 72 | 176 | 11 | 1793 | 146 | 1358 | 2932 | 391 | 21 | 1388 | 712 | 22 | 47 | 15 | 23 | 24 | 38 | 35 | 0 | 0 | 0 | 1861 | 124 |
| SRR5029472 | 0 | 1387 | 18 | 27 | 60 | 0 | 16 | 392 | 724 | 1720 | 687 | 7114 | 0 | 219 | 1094 | 1169 | 0 | 946 | 106 | 2278 | 1974 | 594 | 56 | 2724 | 1608 | 16 | 40 | 13 | 364 | 257 | 578 | 96 | 19 | 0 | 0 | 5115 | 24 |
| SRR5029473 | 0 | 1043 | 16 | 72 | 161 | 11 | 24 | 164 | 215 | 339 | 888 | 1159 | 0 | 17 | 633 | 1091 | 13 | 1829 | 178 | 2414 | 2927 | 804 | 0 | 720 | 1806 | 115 | 175 | 29 | 191 | 263 | 927 | 212 | 10 | 0 | 0 | 3324 | 114 |
| SRR5029474 | 0 | 962 | 30 | 61 | 231 | 0 | 15 | 711 | 940 | 2296 | 4115 | 6929 | 0 | 102 | 609 | 813 | 0 | 1443 | 104 | 1934 | 1595 | 606 | 20 | 1110 | 1284 | 0 | 75 | 15 | 223 | 194 | 507 | 154 | 12 | 0 | 0 | 4953 | 36 |
| SRR5029476 | 0 | 1219 | 29 | 29 | 144 | 0 | 0 | 718 | 1122 | 3086 | 336 | 2053 | 0 | 441 | 192 | 396 | 0 | 1143 | 60 | 1143 | 1610 | 500 | 26 | 1401 | 794 | 0 | 107 | 0 | 134 | 232 | 442 | 80 | 0 | 0 | 0 | 2498 | 36 |
| SRR5029477 | 0 | 755 | 0 | 127 | 99 | 84 | 48 | 126 | 151 | 417 | 181 | 2600 | 0 | 63 | 38 | 151 | 0 | 1554 | 156 | 656 | 2454 | 691 | 10 | 1115 | 1562 | 108 | 146 | 11 | 192 | 261 | 181 | 0 | 0 | 0 | 0 | 997 | 129 |
| SRR5029478 | 0 | 120 | 0 | 61 | 109 | 17 | 35 | 401 | 413 | 522 | 220 | 3035 | 0 | 36 | 73 | 413 | 21 | 1459 | 158 | 2113 | 2215 | 547 | 11 | 1115 | 1299 | 173 | 90 | 27 | 33 | 35 | 70 | 12 | 0 | 0 | 0 | 2428 | 207 |
| SRR5029480 | 0 | 83 | 11 | 40 | 83 | 22 | 0 | 762 | 772 | 1306 | 9350 | 19581 | 0 | 40 | 99 | 505 | 0 | 958 | 43 | 2968 | 1193 | 671 | 36 | 1094 | 690 | 0 | 42 | 0 | 10 | 121 | 30 | 46 | 0 | 0 | 0 | 2434 | 14 |
| SRR5029481 | 0 | 723 | 0 | 47 | 120 | 16 | 0 | 184 | 184 | 184 | 232 | 4812 | 0 | 0 | 172 | 551 | 0 | 1819 | 89 | 1229 | 3816 | 1275 | 12 | 858 | 3116 | 66 | 65 | 11 | 227 | 151 | 298 | 52 | 0 | 0 | 0 | 1574 | 281 |
| SRR5029482 | 0 | 245 | 0 | 70 | 118 | 26 | 26 | 101 | 166 | 650 | 319 | 680 | 0 | 95 | 248 | 189 | 14 | 1073 | 96 | 615 | 727 | 124 | 31 | 2105 | 1800 | 54 | 18 | 13 | 34 | 17 | 29 | 0 | 0 | 0 | 0 | 804 | 67 |
| SRR5029483 | 0 | 443 | 0 | 67 | 71 | 99 | 28 | 190 | 202 | 320 | 1804 | 2623 | 0 | 154 | 36 | 95 | 0 | 1581 | 158 | 807 | 3690 | 1170 | 106 | 4365 | 2196 | 24 | 46 | 14 | 140 | 107 | 156 | 12 | 0 | 0 | 0 | 297 | 37 |
| SRR5029484 | 0 | 236 | 20 | 199 | 504 | 0 | 13 | 922 | 863 | 2369 | 2155 | 3924 | 0 | 60 | 3284 | 1459 | 13 | 1415 | 84 | 3133 | 3256 | 658 | 10 | 550 | 1264 | 70 | 33 | 20 | 104 | 48 | 145 | 42 | 16 | 0 | 0 | 7486 | 85 |
| SRR5029486 | 0 | 183 | 18 | 65 | 427 | 16 | 12 | 464 | 436 | 982 | 699 | 4187 | 0 | 141 | 391 | 200 | 19 | 1179 | 198 | 491 | 1379 | 423 | 10 | 714 | 1060 | 44 | 91 | 26 | 77 | 42 | 84 | 15 | 0 | 0 | 0 | 1141 | 113 |
| SRR5029487 | 0 | 545 | 0 | 83 | 133 | 38 | 26 | 81 | 56 | 30 | 370 | 3440 | 0 | 0 | 0 | 45 | 12 | 1597 | 85 | 303 | 4200 | 1060 | 35 | 1400 | 1841 | 29 | 154 | 19 | 119 | 80 | 100 | 13 | 0 | 0 | 0 | 338 | 70 |
| SRR5029488 | 0 | 839 | 0 | 56 | 196 | 0 | 15 | 307 | 460 | 1303 | 70 | 9274 | 0 | 0 | 690 | 2683 | 0 | 1323 | 90 | 843 | 2623 | 701 | 17 | 1513 | 1205 | 56 | 118 | 0 | 235 | 368 | 250 | 44 | 0 | 0 | 0 | 1150 | 101 |
| SRR5029489 | 0 | 149 | 26 | 60 | 252 | 13 | 17 | 474 | 433 | 527 | 972 | 1558 | 0 | 23 | 41 | 334 | 0 | 1323 | 60 | 3027 | 2451 | 819 | 19 | 973 | 2060 | 59 | 37 | 25 | 30 | 38 | 99 | 22 | 12 | 0 | 0 | 3156 | 131 |
| SRR5029490 | 0 | 786 | 22 | 31 | 241 | 0 | 15 | 547 | 716 | 1804 | 5548 | 8739 | 0 | 147 | 2295 | 868 | 13 | 1299 | 83 | 1359 | 1638 | 510 | 11 | 928 | 1321 | 74 | 188 | 0 | 120 | 107 | 268 | 70 | 0 | 0 | 0 | 2475 | 297 |
| SRR5029491 | 0 | 1172 | 0 | 80 | 301 | 12 | 0 | 755 | 989 | 1340 | 5449 | 7301 | 0 | 45 | 1250 | 2068 | 0 | 1333 | 109 | 2311 | 1780 | 491 | 19 | 1460 | 1257 | 25 | 107 | 44 | 401 | 235 | 359 | 44 | 0 | 0 | 0 | 2895 | 76 |
| SRR5029492 | 0 | 1546 | 0 | 59 | 245 | 37 | 0 | 435 | 837 | 3067 | 4798 | 6883 | 0 | 459 | 250 | 258 | 0 | 1212 | 111 | 797 | 863 | 450 | 12 | 1423 | 1156 | 21 | 105 | 23 | 191 | 99 | 179 | 21 | 0 | 0 | 0 | 749 | 12 |
| SRR5029493 | 0 | 1037 | 0 | 34 | 148 | 15 | 0 | 472 | 405 | 512 | 6984 | 10107 | 0 | 73 | 259 | 299 | 0 | 1316 | 100 | 631 | 1548 | 720 | 20 | 1236 | 1319 | 17 | 198 | 28 | 284 | 276 | 402 | 52 | 0 | 0 | 0 | 1010 | 135 |
| SRR5029494 | 0 | 232 | 0 | 44 | 183 | 12 | 0 | 432 | 611 | 1418 | 137 | 1883 | 0 | 229 | 1571 | 509 | 0 | 1944 | 72 | 909 | 1133 | 506 | 0 | 915 | 1256 | 34 | 34 | 0 | 42 | 50 | 72 | 13 | 0 | 0 | 0 | 1609 | 267 |
| SRR5029495 | 0 | 947 | 25 | 77 | 128 | 0 | 28 | 329 | 423 | 924 | 129 | 1214 | 0 | 86 | 47 | 392 | 14 | 1555 | 99 | 4228 | 2481 | 809 | 16 | 1254 | 1839 | 54 | 55 | 19 | 240 | 321 | 540 | 72 | 12 | 0 | 0 | 4862 | 285 |
| SRR5029496 | 0 | 175 | 0 | 54 | 130 | 11 | 17 | 198 | 198 | 252 | 422 | 4530 | 0 | 34 | 1825 | 1383 | 0 | 1629 | 110 | 1274 | 2140 | 849 | 14 | 853 | 1932 | 97 | 29 | 32 | 29 | 30 | 48 | 24 | 0 | 0 | 0 | 1383 | 191 |
| SRR5029497 | 0 | 1064 | 0 | 72 | 247 | 34 | 23 | 216 | 593 | 782 | 266 | 4051 | 0 | 81 | 979 | 817 | 0 | 1191 | 66 | 683 | 1753 | 517 | 39 | 2119 | 1372 | 0 | 100 | 29 | 271 | 210 | 286 | 28 | 0 | 0 | 0 | 862 | 64 |
| SRR5029498 | 0 | 150 | 0 | 62 | 200 | 11 | 29 | 99 | 59 | 316 | 2369 | 2211 | 0 | 79 | 59 | 415 | 38 | 1146 | 113 | 2369 | 2673 | 517 | 0 | 571 | 2069 | 62 | 21 | 12 | 117 | 103 | 145 | 48 | 0 | 0 | 0 | 3257 | 151 |
| SRR5029499 | 0 | 199 | 0 | 37 | 159 | 43 | 20 | 620 | 1247 | 2957 | 1576 | 3002 | 0 | 224 | 246 | 231 | 0 | 1180 | 106 | 523 | 1744 | 498 | 26 | 1429 | 1657 | 32 | 15 | 0 | 30 | 29 | 44 | 29 | 0 | 0 | 0 | 463 | 77 |
| SRR5029500 | 0 | 593 | 11 | 10 | 136 | 53 | 21 | 2179 | 319 | 531 | 109 | 9830 | 0 | 266 | 159 | 744 | 11 | 1147 | 86 | 372 | 975 | 500 | 0 | 1834 | 1051 | 0 | 191 | 0 | 233 | 201 | 222 | 11 | 0 | 11 | 0 | 531 | 91 |
| SRR5029501 | 0 | 145 | 0 | 24 | 113 | 0 | 15 | 124 | 171 | 311 | 790 | 11226 | 0 | 0 | 630 | 366 | 15 | 900 | 66 | 327 | 1784 | 466 | 25 | 1124 | 910 | 51 | 66 | 0 | 17 | 54 | 17 | 0 | 0 | 0 | 0 | 591 | 297 |
| SRR5029502 | 0 | 121 | 0 | 64 | 174 | 0 | 0 | 1726 | 1931 | 7292 | 4134 | 4748 | 11 | 363 | 681 | 1454 | 22 | 966 | 67 | 681 | 927 | 366 | 0 | 1076 | 696 | 89 | 67 | 44 | 79 | 88 | 124 | 11 | 44 | 33 | 0 | 818 | 274 |
| SRR5029503 | 0 | 183 | 0 | 54 | 233 | 16 | 15 | 330 | 453 | 501 | 7431 | 10022 | 0 | 18 | 1337 | 601 | 0 | 989 | 76 | 542 | 1528 | 400 | 0 | 383 | 798 | 19 | 60 | 27 | 30 | 29 | 37 | 31 | 0 | 0 | 0 | 1195 | 187 |
| SRR5029504 | 0 | 793 | 0 | 46 | 109 | 18 | 23 | 142 | 119 | 230 | 154 | 2058 | 0 | 16 | 127 | 404 | 0 | 1271 | 136 | 783 | 3397 | 955 | 18 | 1734 | 2206 | 62 | 178 | 0 | 241 | 188 | 252 | 22 | 0 | 0 | 0 | 831 | 119 |
| SRR5029505 | 0 | 977 | 0 | 59 | 108 | 31 | 20 | 187 | 182 | 105 | 1588 | 2472 | 0 | 23 | 18 | 146 | 16 | 1275 | 97 | 937 | 3082 | 879 | 16 | 931 | 1962 | 40 | 320 | 27 | 209 | 126 | 209 | 15 | 0 | 0 | 0 | 914 | 164 |
| SRR5029506 | 0 | 1423 | 67 | 30 | 158 | 0 | 0 | 679 | 858 | 1641 | 3228 | 4917 | 10 | 76 | 13750 | 3277 | 17 | 1033 | 95 | 1775 | 2062 | 526 | 0 | 356 | 1453 | 0 | 127 | 16 | 288 | 252 | 607 | 152 | 16 | 10 | 0 | 3232 | 170 |
| SRR5029507 | 0 | 1104 | 12 | 59 | 236 | 21 | 0 | 1017 | 754 | 1029 | 5646 | 9246 | 0 | 69 | 1326 | 1417 | 0 | 1087 | 81 | 3383 | 974 | 345 | 0 | 391 | 1298 | 39 | 69 | 12 | 230 | 306 | 686 | 52 | 12 | 0 | 0 | 4469 | 140 |
| SRR5029508 | 0 | 134 | 0 | 31 | 132 | 11 | 40 | 291 | 262 | 291 | 2591 | 3377 | 0 | 146 | 728 | 582 | 27 | 1125 | 193 | 1485 | 1465 | 407 | 15 | 1352 | 1912 | 51 | 31 | 53 | 82 | 50 | 83 | 33 | 0 | 0 | 0 | 1135 | 112 |
| SRR5029509 | 0 | 528 | 0 | 49 | 195 | 12 | 0 | 343 | 658 | 829 | 371 | 4384 | 0 | 210 | 1201 | 1010 | 42 | 898 | 146 | 867 | 2012 | 439 | 0 | 430 | 1262 | 26 | 503 | 0 | 164 | 113 | 246 | 24 | 0 | 0 | 0 | 1286 | 244 |
| SRR5029510 | 0 | 868 | 0 | 92 | 236 | 0 | 0 | 622 | 334 | 968 | 739 | 6097 | 0 | 23 | 1769 | 1158 | 0 | 991 | 75 | 974 | 1782 | 513 | 15 | 896 | 1035 | 65 | 110 | 12 | 252 | 204 | 412 | 75 | 0 | 0 | 0 | 1429 | 242 |
| SRR5029511 | 0 | 826 | 25 | 66 | 343 | 0 | 18 | 160 | 243 | 347 | 1543 | 2085 | 0 | 21 | 1390 | 862 | 0 | 1526 | 72 | 2175 | 2792 | 906 | 21 | 817 | 2101 | 52 | 220 | 22 | 288 | 279 | 505 | 76 | 0 | 0 | 0 | 3148 | 148 |
| SRR5029512 | 0 | 1114 | 27 | 98 | 201 | 14 | 23 | 280 | 536 | 762 | 2148 | 3205 | 0 | 0 | 3471 | 2630 | 0 | 1409 | 52 | 1883 | 2277 | 731 | 32 | 1464 | 1505 | 48 | 98 | 21 | 219 | 227 | 389 | 66 | 11 | 0 | 0 | 2497 | 168 |
| SRR5029513 | 0 | 944 | 0 | 52 | 188 | 10 | 24 | 111 | 146 | 257 | 1502 | 3544 | 0 | 39 | 338 | 449 | 14 | 1152 | 128 | 646 | 2718 | 379 | 11 | 985 | 2002 | 53 | 148 | 83 | 649 | 528 | 966 | 62 | 17 | 0 | 0 | 1922 | 91 |
| SRR5029514 | 0 | 129 | 0 | 24 | 246 | 18 | 25 | 296 | 212 | 360 | 838 | 13135 | 0 | 21 | 53 | 212 | 18 | 982 | 52 | 265 | 1146 | 428 | 16 | 970 | 648 | 18 | 44 | 25 | 27 | 13 | 18 | 20 | 0 | 0 | 0 | 233 | 74 |
| SRR5029515 | 0 | 164 | 0 | 32 | 164 | 19 | 21 | 96 | 122 | 226 | 1532 | 20406 | 0 | 17 | 530 | 486 | 25 | 903 | 136 | 278 | 1177 | 333 | 0 | 765 | 884 | 88 | 30 | 37 | 33 | 20 | 20 | 0 | 0 | 0 | 0 | 486 | 144 |
| SRR5029516 | 0 | 286 | 17 | 168 | 363 | 28 | 0 | 536 | 798 | 1809 | 123 | 3206 | 0 | 62 | 5726 | 5114 | 0 | 2407 | 88 | 7160 | 2175 | 854 | 12 | 490 | 1480 | 36 | 43 | 0 | 37 | 23 | 66 | 11 | 0 | 0 | 0 | 9667 | 438 |
| SRR5029870 | 0 | 28 | 55 | 0 | 0 | 0 | 0 | 2661 | 939 | 3600 | 77 | 1670 | 0 | 104 | 70 | 243 | 0 | 32 | 0 | 2400 | 1725 | 7688 | 0 | 0 | 18944 | 0 | 0 | 0 | 180 | 50 | 41 | 111 | 11 | 31 | 28 | 19410 | 0 |
| SRR5029871 | 0 | 57 | 38 | 0 | 0 | 0 | 0 | 2103 | 851 | 3380 | 146 | 6310 | 0 | 75 | 75 | 0 | 0 | 28 | 0 | 2354 | 1800 | 6907 | 0 | 0 | 17628 | 0 | 38 | 0 | 283 | 34 | 33 | 19 | 302 | 13 | 38 | 15349 | 0 |
| SRR5029872 | 0 | 15 | 61 | 0 | 0 | 0 | 0 | 2218 | 953 | 3930 | 1031 | 1557 | 0 | 156 | 58 | 136 | 0 | 26 | 0 | 2627 | 1400 | 6927 | 0 | 0 | 18614 | 0 | 0 | 0 | 138 | 49 | 44 | 107 | 12 | 24 | 15 | 19495 | 0 |
| SRR5029873 | 0 | 118 | 98 | 0 | 0 | 0 | 0 | 2471 | 618 | 2847 | 144 | 5721 | 0 | 107 | 27 | 81 | 20 | 27 | 0 | 2659 | 1459 | 6248 | 0 | 0 | 16586 | 0 | 98 | 0 | 236 | 36 | 27 | 20 | 118 | 295 | 0 | 15954 | 0 |
| SRR5029874 | 0 | 57 | 114 | 0 | 0 | 0 | 0 | 2600 | 879 | 4102 | 1318 | 1886 | 0 | 128 | 220 | 92 | 0 | 29 | 0 | 2930 | 1727 | 7936 | 0 | 0 | 15 | 0 | 14 | 0 | 213 | 52 | 38 | 57 | 128 | 21 | 28 | 19208 | 0 |
| SRR5029875 | 0 | 19 | 58 | 0 | 0 | 0 | 0 | 2106 | 624 | 3379 | 142 | 5147 | 0 | 52 | 26 | 26 | 0 | 27 | 0 | 2002 | 1815 | 7001 | 0 | 0 | 17986 | 0 | 58 | 0 | 193 | 35 | 26 | 58 | 173 | 15 | 39 | 14869 | 0 |
| SRR5029876 | 0 | 30 | 105 | 0 | 0 | 0 | 0 | 2434 | 974 | 4100 | 54 | 1779 | 0 | 206 | 112 | 94 | 0 | 29 | 0 | 2808 | 1484 | 7172 | 0 | 0 | 18479 | 0 | 15 | 0 | 285 | 1124 | 959 | 75 | 165 | 794 | 60 | 20557 | 0 |
| SRR5029877 | 0 | 40 | 99 | 0 | 0 | 0 | 79 | 2513 | 838 | 3729 | 3026 | 5026 | 0 | 81 | 0 | 162 | 20 | 27 | 0 | 2216 | 1465 | 6134 | 0 | 0 | 16712 | 0 | 60 | 20 | 218 | 1131 | 873 | 40 | 159 | 536 | 0 | 16753 | 0 |
| SRR5029878 | 0 | 0 | 252 | 0 | 0 | 0 | 0 | 1049 | 766 | 4290 | 556 | 1259 | 0 | 619 | 273 | 336 | 0 | 64 | 0 | 304 | 1191 | 4050 | 35 | 1404 | 9002 | 0 | 0 | 10 | 151 | 1362 | 999 | 161 | 474 | 969 | 40 | 787 | 0 |
| SRR5029879 | 0 | 0 | 101 | 0 | 0 | 0 | 0 | 1495 | 664 | 4485 | 2104 | 2879 | 0 | 720 | 0 | 221 | 0 | 52 | 0 | 166 | 1278 | 3553 | 29 | 1082 | 8167 | 0 | 0 | 0 | 505 | 1515 | 656 | 50 | 454 | 858 | 0 | 720 | 0 |
| SRR5029880 | 0 | 32 | 281 | 0 | 10 | 0 | 0 | 943 | 721 | 4150 | 64 | 1298 | 0 | 655 | 244 | 233 | 0 | 82 | 0 | 300 | 1062 | 3822 | 42 | 1298 | 8736 | 0 | 0 | 22 | 184 | 1328 | 961 | 108 | 508 | 1047 | 22 | 688 | 0 |
| SRR5029882 | 0 | 21 | 226 | 0 | 11 | 0 | 0 | 973 | 757 | 4478 | 854 | 1352 | 0 | 606 | 238 | 357 | 0 | 71 | 0 | 346 | 1166 | 3965 | 30 | 1315 | 9188 | 0 | 0 | 10 | 216 | 54 | 39 | 134 | 18 | 37 | 51 | 790 | 0 |
| SRR5029883 | 0 | 0 | 11 | 0 | 15 | 0 | 0 | 999 | 555 | 4441 | 37 | 3053 | 0 | 278 | 222 | 278 | 0 | 79 | 0 | 56 | 1339 | 3545 | 26 | 1356 | 8215 | 0 | 0 | 51 | 254 | 50 | 44 | 204 | 11 | 29 | 51 | 611 | 0 |
| SRR5029884 | 0 | 0 | 178 | 0 | 0 | 0 | 0 | 1225 | 650 | 4335 | 64 | 1138 | 0 | 856 | 228 | 260 | 0 | 72 | 0 | 152 | 1086 | 3833 | 36 | 1396 | 8945 | 0 | 0 | 10 | 261 | 1475 | 920 | 167 | 471 | 837 | 31 | 683 | 0 |
| SRR5029885 | 0 | 0 | 52 | 0 | 0 | 0 | 0 | 1483 | 742 | 3479 | 32 | 2852 | 0 | 456 | 57 | 57 | 0 | 57 | 0 | 285 | 1150 | 3170 | 36 | 1105 | 7604 | 0 | 0 | 0 | 105 | 1467 | 786 | 157 | 262 | 1100 | 0 | 913 | 0 |
| SRR5029886 | 0 | 10 | 19 | 0 | 0 | 21 | 0 | 2177 | 629 | 952 | 75 | 4194 | 0 | 16 | 16 | 97 | 0 | 45 | 0 | 661 | 1724 | 7364 | 0 | 20 | 10695 | 0 | 0 | 21 | 62 | 135 | 175 | 11 | 41 | 71 | 10 | 903 | 0 |
| SRR5029887 | 0 | 55 | 120 | 0 | 0 | 0 | 0 | 2069 | 739 | 905 | 3306 | 11174 | 0 | 37 | 0 | 18 | 0 | 40 | 0 | 443 | 1760 | 6852 | 0 | 19 | 10194 | 14 | 0 | 0 | 46 | 126 | 108 | 138 | 19 | 32 | 0 | 905 | 0 |
| SRR5029888 | 0 | 11 | 15 | 0 | 0 | 0 | 0 | 1952 | 610 | 837 | 63 | 4061 | 0 | 105 | 105 | 139 | 11 | 55 | 0 | 540 | 1312 | 6423 | 0 | 21 | 10214 | 13 | 0 | 33 | 133 | 159 | 173 | 67 | 39 | 68 | 56 | 1133 | 0 |
| SRR5029889 | 0 | 29 | 134 | 0 | 0 | 0 | 0 | 2269 | 942 | 962 | 3500 | 9655 | 0 | 96 | 0 | 38 | 0 | 53 | 0 | 481 | 1341 | 5967 | 0 | 17 | 9386 | 11 | 0 | 0 | 48 | 112 | 121 | 115 | 20 | 38 | 0 | 981 | 0 |
| SRR5029890 | 0 | 32 | 255 | 0 | 0 | 0 | 0 | 1763 | 643 | 1005 | 74 | 4185 | 0 | 16 | 0 | 148 | 0 | 44 | 0 | 593 | 1700 | 7561 | 0 | 21 | 10628 | 17 | 0 | 0 | 74 | 2484 | 2548 | 159 | 637 | 1062 | 53 | 840 | 0 |
| SRR5029891 | 0 | 0 | 132 | 0 | 0 | 0 | 0 | 1719 | 673 | 598 | 3644 | 11213 | 0 | 75 | 19 | 37 | 0 | 43 | 0 | 449 | 1747 | 7050 | 0 | 15 | 10287 | 14 | 0 | 19 | 76 | 2531 | 2323 | 132 | 416 | 756 | 0 | 953 | 0 |
| SRR5029892 | 0 | 22 | 239 | 0 | 0 | 0 | 0 | 2066 | 665 | 840 | 1541 | 4096 | 0 | 70 | 18 | 158 | 0 | 42 | 0 | 508 | 1330 | 6561 | 0 | 19 | 10422 | 13 | 0 | 0 | 87 | 2730 | 2458 | 174 | 555 | 1022 | 22 | 823 | 0 |
| SRR5029893 | 0 | 0 | 154 | 0 | 10 | 0 | 0 | 2315 | 720 | 856 | 3773 | 11048 | 0 | 58 | 0 | 97 | 0 | 46 | 0 | 506 | 1417 | 5881 | 0 | 13 | 9703 | 19 | 0 | 0 | 58 | 117 | 121 | 77 | 22 | 34 | 0 | 797 | 0 |
| SRR5029894 | 0 | 25 | 80 | 0 | 0 | 12 | 0 | 1537 | 367 | 1927 | 780 | 1790 | 0 | 11 | 184 | 298 | 0 | 21 | 0 | 505 | 1399 | 8710 | 0 | 227 | 16670 | 0 | 18 | 0 | 31 | 44 | 32 | 49 | 24 | 84 | 12 | 2168 | 13 |
| SRR5029895 | 0 | 0 | 23 | 0 | 0 | 15 | 0 | 1979 | 199 | 1580 | 142 | 4908 | 0 | 61 | 92 | 107 | 0 | 30 | 0 | 552 | 1371 | 8069 | 0 | 184 | 15784 | 0 | 61 | 0 | 84 | 581 | 559 | 31 | 214 | 650 | 0 | 1963 | 15 |
| SRR5029896 | 0 | 13 | 79 | 0 | 0 | 0 | 0 | 1826 | 270 | 2084 | 79 | 1716 | 0 | 37 | 294 | 257 | 0 | 15 | 0 | 662 | 1098 | 7485 | 0 | 226 | 13 | 0 | 20 | 0 | 73 | 457 | 536 | 53 | 179 | 999 | 20 | 2010 | 15 |
| SRR5029897 | 0 | 32 | 0 | 0 | 0 | 0 | 0 | 1774 | 431 | 1870 | 130 | 4970 | 0 | 16 | 16 | 64 | 0 | 22 | 0 | 719 | 1078 | 6798 | 0 | 208 | 14636 | 0 | 48 | 0 | 88 | 568 | 512 | 48 | 184 | 648 | 0 | 1758 | 17 |
| SRR5029898 | 0 | 19 | 38 | 0 | 0 | 0 | 0 | 1711 | 342 | 1758 | 55 | 1664 | 0 | 59 | 212 | 295 | 13 | 18 | 0 | 720 | 1336 | 8581 | 0 | 216 | 13 | 0 | 13 | 0 | 44 | 41 | 40 | 44 | 27 | 81 | 0 | 2230 | 10 |
| SRR5029899 | 0 | 31 | 47 | 0 | 0 | 0 | 0 | 2105 | 220 | 1586 | 147 | 5513 | 0 | 110 | 63 | 94 | 0 | 18 | 0 | 518 | 1345 | 7931 | 0 | 182 | 15830 | 0 | 39 | 0 | 94 | 648 | 453 | 47 | 187 | 616 | 0 | 1948 | 0 |
| SRR5029900 | 0 | 0 | 51 | 0 | 0 | 0 | 0 | 1682 | 348 | 1970 | 62 | 1874 | 0 | 96 | 180 | 300 | 0 | 27 | 0 | 396 | 1114 | 7754 | 0 | 196 | 16040 | 0 | 13 | 0 | 45 | 515 | 540 | 32 | 270 | 927 | 19 | 2115 | 0 |
| SRR5029901 | 0 | 32 | 40 | 0 | 0 | 0 | 0 | 1844 | 340 | 1569 | 2070 | 4982 | 0 | 49 | 81 | 226 | 0 | 19 | 0 | 534 | 1109 | 6893 | 0 | 211 | 14765 | 0 | 40 | 0 | 72 | 495 | 590 | 40 | 207 | 606 | 0 | 1828 | 0 |
| SRR5029902 | 0 | 48 | 157 | 0 | 0 | 0 | 0 | 2984 | 709 | 1736 | 88 | 5502 | 0 | 49 | 24 | 122 | 0 | 32 | 0 | 1859 | 1711 | 4392 | 0 | 0 | 9243 | 10 | 0 | 0 | 102 | 491 | 484 | 34 | 273 | 593 | 14 | 3864 | 0 |
| SRR5029904 | 0 | 22 | 238 | 0 | 0 | 0 | 0 | 3003 | 670 | 1636 | 90 | 5122 | 0 | 27 | 80 | 295 | 0 | 40 | 0 | 2333 | 1380 | 3870 | 0 | 0 | 10 | 0 | 0 | 0 | 119 | 581 | 596 | 22 | 194 | 626 | 22 | 4961 | 0 |
| SRR5029905 | 0 | 0 | 223 | 0 | 0 | 0 | 0 | 3128 | 417 | 1251 | 5422 | 9385 | 0 | 0 | 0 | 209 | 0 | 30 | 17 | 2920 | 1366 | 3495 | 0 | 0 | 7727 | 22 | 0 | 0 | 112 | 725 | 279 | 56 | 167 | 279 | 0 | 4380 | 0 |
| SRR5029906 | 0 | 0 | 20 | 0 | 0 | 14 | 0 | 2673 | 807 | 1412 | 106 | 4539 | 0 | 0 | 126 | 151 | 0 | 25 | 0 | 2295 | 1739 | 4454 | 0 | 0 | 9043 | 12 | 0 | 0 | 10 | 29 | 32 | 0 | 12 | 32 | 0 | 4413 | 0 |
| SRR5029908 | 0 | 14 | 14 | 0 | 0 | 14 | 0 | 2418 | 762 | 1472 | 94 | 4941 | 0 | 0 | 79 | 158 | 0 | 39 | 0 | 2365 | 1386 | 3972 | 0 | 0 | 9030 | 11 | 0 | 0 | 80 | 34 | 36 | 14 | 12 | 41 | 29 | 4809 | 0 |
| SRR5029910 | 0 | 0 | 246 | 0 | 0 | 0 | 0 | 962 | 707 | 15367 | 67 | 1483 | 0 | 116 | 0 | 23 | 0 | 0 | 0 | 313 | 759 | 3705 | 0 | 0 | 10974 | 0 | 0 | 0 | 225 | 1659 | 676 | 61 | 164 | 328 | 20 | 1831 | 0 |
| SRR5029911 | 0 | 0 | 276 | 0 | 0 | 0 | 0 | 999 | 999 | 13662 | 73 | 2301 | 0 | 66 | 0 | 26 | 0 | 13 | 0 | 302 | 724 | 3775 | 0 | 0 | 11746 | 0 | 0 | 0 | 360 | 50 | 21 | 21 | 64 | 339 | 0 | 1486 | 0 |
| SRR5029912 | 0 | 0 | 245 | 0 | 0 | 0 | 0 | 928 | 979 | 15714 | 903 | 1627 | 0 | 140 | 0 | 25 | 0 | 29 | 0 | 280 | 740 | 3570 | 0 | 0 | 11337 | 0 | 0 | 0 | 223 | 50 | 18 | 156 | 200 | 13 | 45 | 1907 | 0 |
| SRR5029913 | 0 | 0 | 288 | 0 | 0 | 0 | 0 | 1068 | 944 | 14001 | 1341 | 2067 | 0 | 109 | 0 | 0 | 0 | 20 | 0 | 233 | 686 | 3556 | 0 | 0 | 11181 | 0 | 0 | 0 | 177 | 55 | 24 | 0 | 177 | 266 | 22 | 1725 | 0 |
| SRR5029914 | 0 | 0 | 299 | 0 | 0 | 0 | 0 | 915 | 818 | 15072 | 976 | 1574 | 0 | 110 | 12 | 0 | 0 | 13 | 0 | 342 | 697 | 3717 | 0 | 0 | 12 | 0 | 0 | 0 | 11 | 49 | 23 | 149 | 320 | 320 | 64 | 1648 | 0 |
| SRR5029915 | 0 | 0 | 196 | 0 | 0 | 0 | 0 | 1248 | 895 | 13686 | 1017 | 2197 | 0 | 95 | 41 | 0 | 0 | 0 | 0 | 217 | 725 | 3785 | 0 | 0 | 11614 | 0 | 22 | 22 | 283 | 42 | 21 | 174 | 22 | 174 | 0 | 1736 | 0 |
| SRR5029916 | 0 | 0 | 129 | 0 | 0 | 0 | 0 | 839 | 888 | 15640 | 851 | 1677 | 0 | 111 | 12 | 0 | 0 | 20 | 0 | 321 | 700 | 3565 | 0 | 0 | 11074 | 0 | 0 | 0 | 193 | 55 | 27 | 0 | 215 | 279 | 0 | 1875 | 0 |
| SRR5029917 | 0 | 0 | 284 | 0 | 0 | 0 | 0 | 1238 | 867 | 13879 | 1142 | 2160 | 0 | 151 | 14 | 0 | 0 | 18 | 0 | 385 | 699 | 3555 | 0 | 15 | 11033 | 0 | 0 | 22 | 240 | 1793 | 721 | 66 | 153 | 175 | 44 | 1637 | 0 |
| SRR5029918 | 0 | 0 | 382 | 0 | 0 | 0 | 0 | 2389 | 3256 | 1733 | 30 | 3021 | 0 | 47 | 94 | 117 | 0 | 16 | 0 | 1639 | 5040 | 7712 | 0 | 10 | 14572 | 0 | 0 | 0 | 183 | 1196 | 482 | 50 | 399 | 1479 | 83 | 2670 | 0 |
| SRR5029919 | 0 | 16 | 202 | 0 | 0 | 0 | 0 | 2815 | 3363 | 2070 | 3044 | 9496 | 0 | 46 | 15 | 61 | 0 | 28 | 0 | 2070 | 5058 | 7166 | 0 | 0 | 14043 | 0 | 0 | 0 | 234 | 1097 | 654 | 48 | 387 | 968 | 24 | 3226 | 0 |
| SRR5029920 | 0 | 0 | 354 | 0 | 0 | 0 | 0 | 1992 | 2845 | 2250 | 1061 | 3233 | 0 | 26 | 155 | 78 | 0 | 22 | 0 | 1604 | 4025 | 6573 | 0 | 0 | 14342 | 0 | 18 | 0 | 177 | 939 | 638 | 89 | 478 | 1417 | 35 | 3957 | 0 |
| SRR5029921 | 0 | 0 | 118 | 0 | 0 | 0 | 0 | 2821 | 3597 | 2187 | 198 | 10412 | 0 | 48 | 48 | 63 | 17 | 25 | 0 | 1807 | 4122 | 6232 | 0 | 0 | 13253 | 0 | 0 | 0 | 168 | 1083 | 638 | 34 | 285 | 957 | 0 | 3550 | 0 |
| SRR5029922 | 0 | 0 | 356 | 0 | 0 | 17 | 0 | 2105 | 3158 | 1885 | 38 | 3084 | 0 | 49 | 24 | 147 | 0 | 17 | 0 | 1713 | 4888 | 7799 | 0 | 0 | 11 | 0 | 0 | 0 | 186 | 1102 | 644 | 51 | 509 | 1271 | 17 | 2962 | 0 |
| SRR5029923 | 0 | 0 | 229 | 0 | 0 | 0 | 0 | 3008 | 3515 | 2072 | 191 | 9316 | 0 | 92 | 31 | 61 | 0 | 20 | 0 | 1658 | 5183 | 7069 | 0 | 0 | 14156 | 0 | 0 | 0 | 163 | 1136 | 605 | 41 | 392 | 834 | 0 | 2747 | 0 |
| SRR5029924 | 0 | 0 | 18 | 0 | 0 | 0 | 0 | 2676 | 3307 | 1994 | 884 | 2903 | 0 | 76 | 50 | 126 | 0 | 20 | 0 | 1994 | 4139 | 6946 | 0 | 11 | 14898 | 0 | 0 | 0 | 156 | 50 | 35 | 87 | 27 | 55 | 35 | 3610 | 0 |
| SRR5029925 | 0 | 0 | 268 | 0 | 12 | 0 | 0 | 2932 | 3474 | 1514 | 2518 | 9562 | 0 | 48 | 16 | 64 | 0 | 17 | 0 | 2120 | 4106 | 6231 | 0 | 10 | 13388 | 0 | 0 | 17 | 226 | 1104 | 702 | 75 | 234 | 861 | 0 | 2901 | 0 |
| SRR5029926 | 0 | 0 | 24 | 0 | 0 | 0 | 0 | 1370 | 474 | 3266 | 87 | 8877 | 0 | 105 | 79 | 53 | 0 | 14 | 0 | 869 | 1510 | 9801 | 0 | 27 | 17818 | 0 | 14 | 0 | 16 | 26 | 22 | 24 | 103 | 31 | 24 | 2081 | 0 |
| SRR5029927 | 0 | 0 | 0 | 0 | 0 | 0 | 0 | 1247 | 410 | 1904 | 478 | 24600 | 0 | 98 | 0 | 16 | 0 | 19 | 0 | 755 | 1598 | 8614 | 0 | 33 | 16594 | 0 | 305 | 0 | 67 | 455 | 278 | 22 | 56 | 205 | 0 | 1428 | 0 |
| SRR5029928 | 0 | 0 | 17 | 0 | 0 | 0 | 0 | 981 | 462 | 3751 | 68 | 8742 | 0 | 87 | 115 | 29 | 0 | 16 | 0 | 1298 | 1163 | 8399 | 0 | 22 | 21 | 0 | 95 | 0 | 26 | 215 | 189 | 43 | 60 | 336 | 0 | 2741 | 0 |
| SRR5029929 | 0 | 0 | 17 | 0 | 0 | 23 | 0 | 1120 | 414 | 2050 | 7065 | 25123 | 0 | 52 | 17 | 17 | 0 | 17 | 0 | 793 | 1325 | 7563 | 0 | 29 | 15402 | 0 | 24 | 12 | 17 | 23 | 14 | 46 | 58 | 11 | 0 | 1826 | 0 |
| SRR5029930 | 0 | 0 | 16 | 0 | 0 | 0 | 0 | 1172 | 307 | 3601 | 3238 | 10245 | 0 | 56 | 56 | 140 | 16 | 29 | 0 | 810 | 1632 | 9691 | 0 | 39 | 14 | 0 | 178 | 0 | 32 | 252 | 178 | 0 | 138 | 333 | 0 | 2205 | 0 |
| SRR5029931 | 0 | 0 | 0 | 0 | 0 | 11 | 0 | 1207 | 347 | 2199 | 405 | 26623 | 0 | 66 | 17 | 33 | 0 | 27 | 0 | 695 | 1657 | 8826 | 0 | 30 | 16729 | 0 | 22 | 0 | 62 | 21 | 18 | 17 | 90 | 11 | 0 | 1670 | 0 |
| SRR5029932 | 0 | 0 | 0 | 0 | 0 | 0 | 0 | 1347 | 253 | 3423 | 107 | 9007 | 0 | 112 | 28 | 28 | 0 | 0 | 0 | 1010 | 1346 | 8367 | 0 | 40 | 17080 | 0 | 75 | 0 | 17 | 309 | 134 | 33 | 100 | 326 | 0 | 2806 | 0 |
| SRR5029933 | 0 | 23 | 0 | 0 | 0 | 0 | 0 | 1305 | 373 | 2119 | 422 | 24227 | 0 | 136 | 0 | 34 | 0 | 18 | 0 | 780 | 1277 | 7589 | 0 | 28 | 15674 | 0 | 25 | 0 | 35 | 25 | 17 | 0 | 98 | 16 | 0 | 1577 | 0 |
| SRR5029934 | 0 | 0 | 93 | 0 | 0 | 0 | 0 | 637 | 361 | 1801 | 57 | 1047 | 0 | 18 | 0 | 18 | 0 | 41 | 0 | 1782 | 2314 | 5384 | 0 | 12 | 12752 | 0 | 0 | 26 | 106 | 914 | 768 | 40 | 344 | 980 | 26 | 6455 | 0 |
| SRR5029935 | 0 | 0 | 45 | 0 | 14 | 0 | 0 | 602 | 367 | 1835 | 91 | 2549 | 0 | 61 | 20 | 0 | 0 | 66 | 14 | 2202 | 2399 | 5011 | 0 | 0 | 12066 | 12 | 15 | 30 | 135 | 823 | 838 | 30 | 150 | 344 | 0 | 6168 | 0 |
| SRR5029936 | 0 | 0 | 115 | 0 | 0 | 0 | 0 | 592 | 233 | 1784 | 61 | 1058 | 0 | 33 | 0 | 27 | 14 | 59 | 0 | 1598 | 1825 | 4744 | 0 | 0 | 12238 | 15 | 0 | 0 | 72 | 949 | 762 | 0 | 431 | 676 | 29 | 6550 | 0 |
| SRR5029937 | 0 | 31 | 47 | 0 | 15 | 0 | 0 | 731 | 408 | 1849 | 1107 | 2751 | 0 | 32 | 0 | 21 | 0 | 45 | 0 | 2117 | 2002 | 4317 | 0 | 11 | 11111 | 13 | 0 | 0 | 78 | 767 | 626 | 0 | 235 | 360 | 16 | 6266 | 0 |
| SRR5029938 | 0 | 14 | 191 | 0 | 0 | 0 | 0 | 535 | 338 | 1573 | 382 | 1032 | 0 | 51 | 13 | 38 | 0 | 46 | 0 | 2012 | 2294 | 5512 | 0 | 0 | 12825 | 13 | 0 | 14 | 137 | 30 | 41 | 41 | 11 | 28 | 27 | 6578 | 0 |
| SRR5029939 | 0 | 15 | 61 | 0 | 0 | 0 | 0 | 652 | 420 | 1640 | 1030 | 2639 | 0 | 53 | 11 | 0 | 0 | 63 | 0 | 2081 | 2478 | 5102 | 0 | 0 | 12165 | 0 | 0 | 0 | 107 | 28 | 28 | 46 | 168 | 13 | 0 | 6171 | 0 |
| SRR5029940 | 0 | 0 | 56 | 0 | 11 | 0 | 0 | 651 | 361 | 1799 | 51 | 942 | 0 | 39 | 0 | 26 | 0 | 41 | 0 | 1812 | 1947 | 4783 | 0 | 0 | 12400 | 0 | 0 | 0 | 84 | 829 | 885 | 56 | 393 | 773 | 42 | 6559 | 0 |
| SRR5029941 | 0 | 0 | 78 | 0 | 11 | 0 | 0 | 587 | 246 | 1676 | 1174 | 2808 | 0 | 64 | 0 | 0 | 0 | 30 | 0 | 2274 | 2022 | 4167 | 0 | 0 | 11488 | 13 | 16 | 0 | 109 | 876 | 735 | 63 | 203 | 313 | 16 | 6374 | 0 |
| SRR5029942 | 0 | 0 | 112 | 0 | 0 | 0 | 0 | 918 | 184 | 4335 | 29 | 2865 | 0 | 257 | 514 | 735 | 14 | 14 | 0 | 735 | 3902 | 10693 | 0 | 29 | 19554 | 0 | 0 | 0 | 98 | 252 | 252 | 42 | 182 | 518 | 0 | 8817 | 0 |
| SRR5029943 | 0 | 0 | 66 | 0 | 0 | 0 | 0 | 1525 | 210 | 3412 | 4079 | 10788 | 0 | 267 | 152 | 191 | 0 | 0 | 0 | 1201 | 4366 | 9529 | 0 | 13 | 17533 | 0 | 11 | 0 | 116 | 319 | 215 | 33 | 138 | 369 | 0 | 6137 | 0 |
| SRR5029944 | 0 | 0 | 12 | 0 | 0 | 0 | 0 | 1240 | 240 | 4120 | 25 | 2920 | 0 | 400 | 200 | 720 | 0 | 0 | 0 | 1160 | 3203 | 9417 | 0 | 0 | 15 | 0 | 0 | 0 | 108 | 24 | 15 | 31 | 20 | 48 | 31 | 9199 | 0 |
| SRR5029945 | 0 | 23 | 63 | 0 | 0 | 0 | 0 | 1317 | 216 | 3399 | 206 | 11574 | 0 | 255 | 79 | 118 | 0 | 14 | 0 | 1395 | 3591 | 8285 | 0 | 12 | 16358 | 0 | 17 | 12 | 126 | 18 | 138 | 29 | 109 | 18 | 0 | 6032 | 0 |

### Supp Table S8.4.Columns_DI_to_ES

| | | | | | | | | | | | | | | | | | | | | | | | | | | | | | | | | | | | | | |
|---|---|---|---|---|---|---|---|---|---|---|---|---|---|---|---|---|---|---|---|---|---|---|---|---|---|---|---|---|---|---|---|---|---|---|---|---|---|
| SRR5029946 | 0 | 0 | 14 | 0 | 0 | 0 | 0 | 996 | 153 | 3906 | 1149 | 2719 | 0 | 383 | 383 | 651 | 0 | 0 | 0 | 1225 | 4286 | 10917 | 0 | 0 | 19130 | 0 | 0 | 0 | 102 | 27 | 11 | 73 | 21 | 62 | 29 | 7927 | 0 |
| SRR5029947 | 0 | 11 | 89 | 0 | 0 | 0 | 0 | 1337 | 172 | 3075 | 211 | 11728 | 0 | 344 | 38 | 191 | 0 | 11 | 0 | 1012 | 4502 | 9620 | 0 | 11 | 17926 | 0 | 11 | 0 | 95 | 334 | 239 | 22 | 78 | 317 | 0 | 6074 | 0 |
| SRR5029948 | 0 | 30 | 136 | 0 | 0 | 0 | 0 | 702 | 273 | 4293 | 859 | 2498 | 0 | 312 | 390 | 585 | 0 | 22 | 0 | 1444 | 3498 | 9879 | 0 | 0 | 15 | 0 | 0 | 0 | 121 | 318 | 273 | 61 | 152 | 652 | 0 | 8976 | 0 |
| SRR5029949 | 0 | 12 | 104 | 0 | 0 | 0 | 0 | 1304 | 261 | 4033 | 4374 | 12179 | 0 | 301 | 201 | 161 | 0 | 14 | 12 | 843 | 3490 | 8474 | 0 | 12 | 16744 | 0 | 0 | 23 | 86 | 19 | 16 | 58 | 115 | 16 | 0 | 6120 | 0 |
| SRR5029950 | 0 | 0 | 158 | 0 | 0 | 0 | 0 | 1444 | 769 | 1537 | 3458 | 9540 | 0 | 159 | 742 | 610 | 0 | 90 | 0 | 1047 | 942 | 5114 | 0 | 0 | 12355 | 0 | 0 | 0 | 275 | 825 | 531 | 28 | 247 | 541 | 65 | 3313 | 0 |
| SRR5029951 | 0 | 0 | 128 | 0 | 0 | 0 | 0 | 2015 | 756 | 3149 | 4283 | 10077 | 0 | 0 | 126 | 1008 | 0 | 84 | 0 | 1134 | 1187 | 5361 | 0 | 0 | 12475 | 0 | 0 | 43 | 256 | 812 | 684 | 0 | 214 | 214 | 0 | 3401 | 0 |
| SRR5029952 | 0 | 0 | 14 | 0 | 0 | 0 | 0 | 1509 | 747 | 1624 | 3535 | 8393 | 0 | 201 | 719 | 805 | 0 | 80 | 0 | 805 | 895 | 4772 | 0 | 0 | 12371 | 0 | 0 | 0 | 17 | 63 | 44 | 55 | 21 | 46 | 85 | 3392 | 0 |
| SRR5029953 | 0 | 0 | 44 | 0 | 0 | 0 | 0 | 1548 | 774 | 1935 | 23 | 8255 | 0 | 387 | 129 | 258 | 0 | 88 | 0 | 774 | 759 | 4862 | 0 | 0 | 12330 | 0 | 44 | 44 | 220 | 617 | 617 | 44 | 352 | 396 | 44 | 3869 | 0 |
| SRR5029954 | 0 | 0 | 158 | 0 | 0 | 0 | 0 | 1428 | 824 | 1373 | 3212 | 8937 | 0 | 206 | 673 | 728 | 0 | 80 | 0 | 686 | 946 | 5063 | 0 | 0 | 12372 | 0 | 24 | 0 | 230 | 814 | 508 | 34 | 225 | 532 | 48 | 3226 | 0 |
| SRR5029955 | 0 | 0 | 44 | 0 | 0 | 0 | 0 | 1274 | 637 | 1147 | 4587 | 9939 | 0 | 127 | 255 | 0 | 0 | 107 | 21 | 1019 | 980 | 5346 | 0 | 14 | 12809 | 0 | 0 | 0 | 480 | 916 | 872 | 44 | 305 | 436 | 44 | 3186 | 0 |
| SRR5029956 | 0 | 0 | 15 | 0 | 0 | 0 | 0 | 1446 | 723 | 1473 | 3739 | 9314 | 0 | 125 | 626 | 751 | 0 | 73 | 0 | 820 | 847 | 4820 | 0 | 0 | 12300 | 0 | 0 | 0 | 17 | 61 | 35 | 29 | 20 | 57 | 63 | 3503 | 0 |
| SRR5029957 | 0 | 0 | 177 | 0 | 0 | 0 | 0 | 1425 | 907 | 1425 | 3758 | 8941 | 0 | 389 | 259 | 389 | 0 | 43 | 0 | 518 | 1046 | 4784 | 0 | 0 | 12131 | 0 | 0 | 44 | 221 | 1060 | 707 | 88 | 132 | 662 | 44 | 3499 | 0 |
| SRR5029958 | 0 | 0 | 0 | 0 | 0 | 0 | 0 | 1341 | 473 | 2426 | 38 | 2130 | 0 | 79 | 79 | 138 | 15 | 0 | 0 | 1795 | 764 | 6512 | 0 | 0 | 15945 | 0 | 0 | 0 | 118 | 75 | 43 | 88 | 11 | 30 | 29 | 3747 | 0 |
| SRR5029959 | 0 | 0 | 0 | 0 | 0 | 0 | 0 | 1570 | 432 | 1874 | 246 | 10267 | 0 | 32 | 32 | 64 | 0 | 0 | 15 | 1233 | 902 | 5745 | 0 | 0 | 14185 | 0 | 0 | 0 | 147 | 50 | 41 | 82 | 114 | 13 | 0 | 2819 | 0 |
| SRR5029960 | 0 | 16 | 63 | 0 | 0 | 0 | 0 | 1355 | 525 | 2164 | 1071 | 2361 | 0 | 22 | 44 | 219 | 0 | 10 | 19 | 1705 | 792 | 6280 | 0 | 0 | 15980 | 0 | 0 | 0 | 32 | 1515 | 1010 | 111 | 379 | 631 | 47 | 3454 | 0 |
| SRR5029961 | 0 | 0 | 34 | 0 | 0 | 25 | 0 | 1674 | 452 | 1808 | 4201 | 11197 | 0 | 50 | 0 | 0 | 0 | 16 | 11 | 1456 | 788 | 5434 | 0 | 0 | 13556 | 0 | 0 | 0 | 110 | 1169 | 754 | 68 | 178 | 246 | 0 | 3163 | 0 |
| SRR5029962 | 0 | 15 | 30 | 0 | 0 | 0 | 0 | 1301 | 567 | 2078 | 1028 | 2203 | 0 | 105 | 84 | 105 | 0 | 14 | 12 | 1469 | 890 | 6481 | 0 | 0 | 15806 | 0 | 0 | 0 | 90 | 68 | 60 | 165 | 17 | 26 | 30 | 3316 | 0 |
| SRR5029963 | 0 | 0 | 42 | 0 | 0 | 0 | 0 | 1516 | 521 | 1597 | 230 | 10691 | 0 | 81 | 0 | 33 | 0 | 12 | 10 | 1255 | 894 | 5776 | 0 | 0 | 14319 | 0 | 0 | 0 | 166 | 64 | 39 | 91 | 83 | 11 | 0 | 2852 | 0 |
| SRR5029964 | 0 | 0 | 108 | 0 | 0 | 0 | 0 | 1698 | 503 | 2431 | 964 | 2159 | 0 | 42 | 42 | 168 | 0 | 14 | 12 | 1551 | 850 | 6447 | 0 | 12 | 15777 | 0 | 0 | 0 | 108 | 72 | 56 | 62 | 16 | 29 | 31 | 3814 | 0 |
| SRR5029965 | 0 | 0 | 0 | 0 | 0 | 0 | 0 | 1515 | 539 | 1684 | 4394 | 11330 | 0 | 101 | 0 | 118 | 0 | 10 | 0 | 1448 | 815 | 5372 | 0 | 0 | 13824 | 0 | 17 | 0 | 110 | 1212 | 737 | 102 | 136 | 331 | 0 | 2643 | 0 |
| SRR5029966 | 0 | 11 | 95 | 0 | 0 | 0 | 0 | 1563 | 568 | 3931 | 74 | 2668 | 0 | 47 | 79 | 95 | 0 | 17 | 16 | 600 | 1607 | 9507 | 31 | 1990 | 20853 | 0 | 0 | 0 | 149 | 1708 | 1125 | 53 | 329 | 859 | 127 | 1942 | 0 |
| SRR5029967 | 0 | 0 | 50 | 0 | 0 | 0 | 0 | 1215 | 380 | 2969 | 569 | 11885 | 0 | 21 | 32 | 0 | 0 | 19 | 11 | 486 | 1604 | 8159 | 23 | 1701 | 18494 | 0 | 0 | 22 | 188 | 126 | 60 | 72 | 165 | 22 | 22 | 1870 | 0 |
| SRR5029968 | 0 | 12 | 82 | 0 | 0 | 23 | 0 | 1515 | 383 | 3935 | 73 | 2159 | 0 | 52 | 35 | 17 | 0 | 25 | 14 | 401 | 1251 | 8367 | 38 | 2114 | 20292 | 0 | 0 | 0 | 128 | 142 | 86 | 105 | 30 | 65 | 70 | 2124 | 0 |
| SRR5029969 | 0 | 0 | 41 | 0 | 0 | 0 | 0 | 1609 | 641 | 3657 | 6267 | 12332 | 0 | 101 | 0 | 0 | 0 | 21 | 0 | 484 | 1353 | 7294 | 26 | 1745 | 17374 | 0 | 0 | 17 | 220 | 2164 | 1224 | 58 | 180 | 342 | 23 | 1587 | 0 |
| SRR5029970 | 0 | 0 | 121 | 0 | 0 | 0 | 0 | 1228 | 647 | 3918 | 1179 | 2357 | 0 | 50 | 33 | 66 | 0 | 24 | 0 | 598 | 1607 | 9583 | 23 | 1964 | 20540 | 11 | 0 | 0 | 11 | 127 | 88 | 77 | 14 | 57 | 66 | 1926 | 12 |
| SRR5029971 | 0 | 0 | 45 | 0 | 0 | 0 | 0 | 1501 | 427 | 2838 | 6104 | 12406 | 0 | 99 | 33 | 11 | 0 | 20 | 13 | 471 | 1612 | 8320 | 29 | 1725 | 18712 | 0 | 0 | 0 | 153 | 1898 | 1181 | 45 | 175 | 378 | 0 | 1611 | 0 |
| SRR5029972 | 0 | 11 | 79 | 0 | 0 | 0 | 0 | 1208 | 569 | 4106 | 1173 | 2501 | 0 | 17 | 52 | 52 | 0 | 22 | 15 | 500 | 1324 | 8333 | 36 | 2027 | 20274 | 11 | 0 | 0 | 136 | 1781 | 1304 | 57 | 284 | 851 | 79 | 1777 | 0 |
| SRR5029973 | 0 | 0 | 47 | 0 | 0 | 0 | 0 | 1548 | 417 | 3293 | 516 | 11799 | 0 | 88 | 33 | 0 | 0 | 23 | 12 | 582 | 1366 | 7216 | 21 | 1781 | 17474 | 0 | 0 | 0 | 187 | 108 | 64 | 47 | 12 | 19 | 18 | 1624 | 0 |
| SRR5029974 | 0 | 16 | 74 | 0 | 0 | 0 | 0 | 3294 | 1193 | 5824 | 2267 | 9738 | 0 | 0 | 286 | 191 | 0 | 52 | 0 | 1647 | 1612 | 10048 | 0 | 15 | 20520 | 0 | 0 | 0 | 66 | 54 | 49 | 41 | 21 | 46 | 41 | 2315 | 0 |
| SRR5029975 | 0 | 12 | 18 | 0 | 0 | 0 | 0 | 2740 | 802 | 2852 | 5182 | 21102 | 0 | 19 | 19 | 56 | 0 | 58 | 12 | 839 | 1660 | 8997 | 0 | 23 | 19268 | 0 | 0 | 0 | 89 | 713 | 541 | 48 | 125 | 178 | 30 | 1230 | 0 |
| SRR5029976 | 0 | 0 | 90 | 0 | 0 | 0 | 0 | 3387 | 821 | 4721 | 83 | 8749 | 0 | 0 | 282 | 308 | 0 | 49 | 22 | 1514 | 1388 | 9168 | 0 | 22 | 20603 | 0 | 0 | 0 | 63 | 51 | 49 | 81 | 13 | 47 | 18 | 2129 | 0 |
| SRR5029977 | 0 | 0 | 25 | 0 | 0 | 0 | 0 | 2814 | 512 | 2814 | 5687 | 22158 | 0 | 0 | 20 | 20 | 0 | 49 | 0 | 767 | 1321 | 8103 | 0 | 18 | 18010 | 0 | 0 | 0 | 37 | 47 | 32 | 43 | 74 | 15 | 0 | 1259 | 0 |
| SRR5029978 | 0 | 0 | 60 | 0 | 0 | 0 | 0 | 3099 | 659 | 5294 | 100 | 8808 | 0 | 0 | 146 | 342 | 0 | 54 | 10 | 1025 | 1672 | 10242 | 0 | 20 | 15 | 0 | 0 | 0 | 95 | 629 | 551 | 78 | 189 | 577 | 26 | 2074 | 0 |
| SRR5029979 | 0 | 31 | 24 | 0 | 0 | 0 | 0 | 2086 | 801 | 2943 | 320 | 21329 | 0 | 0 | 56 | 93 | 0 | 59 | 11 | 633 | 1656 | 9132 | 0 | 20 | 19351 | 0 | 0 | 0 | 122 | 43 | 30 | 43 | 116 | 17 | 0 | 1472 | 0 |
| SRR5029980 | 0 | 0 | 78 | 0 | 0 | 0 | 0 | 3567 | 1090 | 5549 | 102 | 9512 | 0 | 0 | 372 | 272 | 0 | 41 | 12 | 1139 | 1327 | 9198 | 0 | 15 | 18 | 0 | 0 | 0 | 44 | 48 | 41 | 78 | 20 | 47 | 17 | 2353 | 0 |
| SRR5029981 | 0 | 0 | 50 | 0 | 0 | 0 | 0 | 2897 | 610 | 2763 | 277 | 21650 | 0 | 0 | 76 | 0 | 0 | 54 | 18 | 915 | 1423 | 8172 | 0 | 19 | 18196 | 0 | 0 | 0 | 87 | 615 | 534 | 31 | 161 | 230 | 12 | 1391 | 0 |
| SRR5029982 | 0 | 0 | 69 | 0 | 0 | 0 | 0 | 643 | 877 | 785 | 933 | 2899 | 0 | 28 | 0 | 0 | 0 | 34 | 0 | 1980 | 1125 | 4264 | 0 | 11 | 12622 | 0 | 0 | 14 | 152 | 635 | 829 | 83 | 304 | 649 | 14 | 9877 | 0 |
| SRR5029983 | 0 | 0 | 0 | 0 | 0 | 0 | 0 | 869 | 1073 | 971 | 14 | 2351 | 0 | 0 | 0 | 0 | 0 | 29 | 0 | 2760 | 1148 | 4370 | 0 | 0 | 13241 | 0 | 0 | 0 | 94 | 281 | 25 | 0 | 281 | 13 | 0 | 9609 | 0 |
| SRR5029984 | 0 | 15 | 60 | 0 | 0 | 0 | 0 | 809 | 908 | 847 | 106 | 2328 | 0 | 15 | 15 | 15 | 0 | 33 | 0 | 2320 | 1016 | 3998 | 0 | 0 | 11 | 0 | 0 | 0 | 105 | 719 | 824 | 90 | 255 | 554 | 30 | 10531 | 0 |
| SRR5029986 | 0 | 0 | 28 | 0 | 0 | 0 | 0 | 886 | 974 | 879 | 123 | 2461 | 0 | 59 | 0 | 0 | 0 | 36 | 0 | 2205 | 1153 | 4168 | 0 | 0 | 12 | 0 | 0 | 14 | 57 | 19 | 18 | 142 | 255 | 13 | 42 | 10020 | 0 |
| SRR5029988 | 15 | 0 | 88 | 0 | 0 | 0 | 0 | 833 | 1116 | 781 | 123 | 2626 | 0 | 15 | 0 | 0 | 15 | 30 | 0 | 2068 | 982 | 4063 | 0 | 11 | 12718 | 0 | 0 | 29 | 88 | 876 | 862 | 73 | 102 | 613 | 15 | 10252 | 0 |
| SRR5029990 | 0 | 0 | 102 | 0 | 0 | 0 | 0 | 568 | 142 | 3141 | 12 | 621 | 0 | 337 | 160 | 195 | 0 | 116 | 0 | 532 | 1772 | 4717 | 0 | 26 | 14374 | 12 | 0 | 0 | 62 | 252 | 285 | 22 | 132 | 296 | 29 | 1686 | 0 |
| SRR5029991 | 0 | 0 | 45 | 0 | 0 | 0 | 0 | 448 | 149 | 2418 | 28 | 2000 | 0 | 269 | 179 | 30 | 0 | 120 | 14 | 448 | 1732 | 4136 | 0 | 23 | 12857 | 0 | 0 | 15 | 60 | 40 | 31 | 25 | 55 | 31 | 0 | 1403 | 18 |
| SRR5029992 | 0 | 0 | 76 | 0 | 0 | 0 | 0 | 761 | 247 | 3425 | 14 | 438 | 0 | 304 | 304 | 152 | 0 | 136 | 0 | 666 | 1489 | 4193 | 0 | 13 | 13948 | 0 | 0 | 0 | 52 | 268 | 288 | 20 | 80 | 280 | 24 | 1865 | 0 |
| SRR5029993 | 0 | 0 | 26 | 0 | 0 | 0 | 0 | 855 | 127 | 3358 | 887 | 1774 | 0 | 253 | 63 | 32 | 0 | 114 | 12 | 444 | 1439 | 3695 | 0 | 24 | 12045 | 0 | 0 | 0 | 12 | 45 | 35 | 26 | 53 | 20 | 16 | 1647 | 0 |
| SRR5029994 | 0 | 0 | 14 | 0 | 0 | 0 | 0 | 893 | 130 | 3218 | 205 | 800 | 0 | 298 | 242 | 130 | 0 | 104 | 0 | 539 | 1712 | 4816 | 0 | 27 | 14310 | 0 | 0 | 0 | 45 | 39 | 31 | 34 | 12 | 43 | 10 | 1786 | 15 |
| SRR5029995 | 0 | 0 | 46 | 0 | 0 | 0 | 0 | 763 | 244 | 3021 | 25 | 1678 | 0 | 336 | 61 | 92 | 0 | 113 | 12 | 427 | 1728 | 4507 | 0 | 14 | 13279 | 14 | 0 | 20 | 61 | 42 | 28 | 10 | 61 | 23 | 10 | 1373 | 0 |
| SRR5029996 | 0 | 0 | 14 | 0 | 0 | 0 | 0 | 835 | 241 | 3339 | 170 | 408 | 0 | 241 | 315 | 204 | 0 | 102 | 0 | 612 | 1501 | 4391 | 0 | 41 | 14101 | 15 | 0 | 0 | 51 | 40 | 34 | 19 | 14 | 37 | 43 | 1929 | 10 |
| SRR5029997 | 0 | 0 | 31 | 0 | 0 | 0 | 0 | 902 | 64 | 2770 | 1031 | 1900 | 0 | 64 | 97 | 161 | 0 | 119 | 0 | 741 | 1653 | 3735 | 0 | 28 | 12176 | 17 | 0 | 0 | 47 | 303 | 272 | 10 | 63 | 199 | 0 | 1836 | 0 |
| SRR5029998 | 0 | 0 | 177 | 0 | 0 | 0 | 0 | 842 | 232 | 1117 | 47 | 661 | 0 | 335 | 618 | 953 | 0 | 42 | 0 | 7937 | 1017 | 4723 | 0 | 20 | 12214 | 0 | 0 | 0 | 0 | 372 | 514 | 80 | 257 | 1073 | 27 | 83353 | 0 |
| SRR5029999 | 0 | 11 | 91 | 0 | 0 | 11 | 0 | 878 | 150 | 830 | 1148 | 2357 | 0 | 301 | 307 | 319 | 0 | 31 | 0 | 7227 | 1170 | 4256 | 0 | 14 | 11149 | 0 | 37 | 16 | 69 | 486 | 433 | 107 | 203 | 748 | 48 | 68161 | 0 |
| SRR5030000 | 0 | 0 | 66 | 0 | 0 | 0 | 0 | 839 | 255 | 1223 | 310 | 620 | 0 | 265 | 657 | 903 | 0 | 28 | 0 | 7929 | 887 | 4432 | 0 | 0 | 11673 | 0 | 0 | 0 | 66 | 339 | 528 | 75 | 245 | 1094 | 19 | 85927 | 0 |
| SRR5030001 | 0 | 0 | 61 | 0 | 0 | 0 | 0 | 840 | 241 | 865 | 177 | 2496 | 0 | 303 | 278 | 365 | 11 | 33 | 0 | 7681 | 966 | 4086 | 0 | 11 | 10462 | 0 | 22 | 11 | 61 | 412 | 507 | 89 | 167 | 708 | 0 | 68440 | 0 |
| SRR5030002 | 0 | 0 | 12 | 0 | 0 | 0 | 0 | 960 | 326 | 1277 | 37 | 775 | 0 | 361 | 652 | 854 | 0 | 43 | 0 | 7997 | 1032 | 4915 | 0 | 10 | 12122 | 0 | 18 | 0 | 36 | 36 | 33 | 81 | 18 | 98 | 0 | 83944 | 0 |
| SRR5030003 | 0 | 16 | 125 | 0 | 0 | 0 | 11 | 831 | 222 | 887 | 1170 | 2371 | 0 | 240 | 252 | 382 | 0 | 39 | 0 | 7253 | 1073 | 4210 | 0 | 0 | 11222 | 0 | 44 | 0 | 65 | 582 | 376 | 93 | 234 | 691 | 0 | 66973 | 0 |
| SRR5030004 | 0 | 0 | 110 | 0 | 0 | 0 | 0 | 781 | 222 | 1189 | 28 | 568 | 0 | 364 | 772 | 825 | 0 | 26 | 0 | 7913 | 961 | 4540 | 0 | 21 | 12013 | 0 | 18 | 0 | 46 | 35 | 40 | 101 | 26 | 85 | 0 | 85048 | 0 |
| SRR5030005 | 0 | 11 | 67 | 0 | 0 | 0 | 0 | 947 | 229 | 971 | 152 | 2413 | 0 | 346 | 334 | 359 | 0 | 35 | 0 | 7648 | 994 | 3929 | 0 | 14 | 10662 | 0 | 11 | 28 | 44 | 32 | 27 | 72 | 10 | 47 | 11 | 68116 | 0 |
| SRR5030006 | 0 | 25 | 24 | 0 | 0 | 0 | 0 | 2580 | 1564 | 2526 | 45 | 4157 | 0 | 134 | 67 | 94 | 0 | 16 | 0 | 775 | 1374 | 5597 | 0 | 12 | 10 | 0 | 0 | 17 | 15 | 101 | 79 | 59 | 14 | 31 | 42 | 2379 | 0 |
| SRR5030008 | 0 | 0 | 27 | 0 | 0 | 0 | 0 | 2571 | 1936 | 2586 | 526 | 4073 | 0 | 186 | 77 | 108 | 0 | 21 | 0 | 650 | 1046 | 4818 | 0 | 16 | 10 | 10 | 0 | 18 | 234 | 93 | 73 | 36 | 10 | 27 | 45 | 2803 | 0 |
| SRR5030009 | 0 | 0 | 14 | 0 | 13 | 0 | 0 | 3085 | 2244 | 2805 | 982 | 8414 | 0 | 421 | 0 | 140 | 0 | 34 | 0 | 280 | 1165 | 4258 | 0 | 13 | 9087 | 13 | 0 | 0 | 11 | 86 | 77 | 0 | 157 | 37 | 0 | 2945 | 0 |
| SRR5030010 | 0 | 0 | 619 | 0 | 0 | 17 | 0 | 2711 | 1656 | 3136 | 469 | 4617 | 0 | 117 | 44 | 59 | 0 | 17 | 0 | 689 | 1385 | 5677 | 0 | 18 | 10160 | 0 | 0 | 0 | 284 | 2346 | 1822 | 34 | 292 | 653 | 26 | 2491 | 0 |
| SRR5030011 | 0 | 0 | 25 | 0 | 0 | 0 | 0 | 3165 | 1646 | 2026 | 1195 | 10888 | 0 | 0 | 0 | 0 | 0 | 21 | 0 | 633 | 1512 | 5313 | 0 | 21 | 9408 | 0 | 0 | 0 | 14 | 66 | 61 | 0 | 155 | 17 | 0 | 2026 | 0 |
| SRR5030012 | 0 | 0 | 596 | 0 | 0 | 0 | 0 | 2660 | 1828 | 3007 | 589 | 4684 | 0 | 212 | 0 | 45 | 0 | 15 | 0 | 1103 | 1136 | 4940 | 0 | 14 | 9902 | 0 | 0 | 18 | 333 | 2473 | 1692 | 88 | 228 | 631 | 53 | 3068 | 0 |
| SRR5030014 | 0 | 0 | 372 | 0 | 0 | 0 | 0 | 838 | 327 | 4702 | 94 | 2373 | 0 | 337 | 120 | 207 | 0 | 58 | 25 | 1883 | 2224 | 8138 | 0 | 0 | 15 | 0 | 0 | 0 | 28 | 537 | 744 | 165 | 165 | 1433 | 55 | 4659 | 0 |
| SRR5030015 | 0 | 14 | 301 | 0 | 0 | 0 | 0 | 779 | 348 | 3226 | 4143 | 9733 | 0 | 264 | 42 | 83 | 0 | 55 | 18 | 1780 | 2308 | 7095 | 0 | 12 | 17750 | 0 | 0 | 14 | 137 | 710 | 683 | 55 | 191 | 710 | 0 | 4255 | 0 |
| SRR5030016 | 0 | 0 | 395 | 0 | 0 | 0 | 0 | 717 | 323 | 4293 | 81 | 2523 | 0 | 311 | 72 | 120 | 0 | 66 | 17 | 2128 | 1843 | 7431 | 0 | 12 | 19413 | 0 | 0 | 0 | 46 | 593 | 881 | 137 | 441 | 1459 | 15 | 5094 | 0 |
| SRR5030017 | 0 | 14 | 172 | 0 | 0 | 0 | 0 | 826 | 391 | 3405 | 297 | 9693 | 0 | 362 | 29 | 43 | 0 | 51 | 23 | 1565 | 1943 | 6223 | 0 | 0 | 16774 | 0 | 0 | 14 | 129 | 19 | 28 | 57 | 172 | 18 | 0 | 4129 | 0 |
| SRR5030018 | 0 | 0 | 26 | 0 | 0 | 0 | 0 | 788 | 259 | 4424 | 104 | 2420 | 0 | 360 | 191 | 214 | 0 | 73 | 21 | 2071 | 2215 | 8368 | 0 | 0 | 18 | 0 | 0 | 0 | 129 | 30 | 36 | 129 | 18 | 58 | 29 | 4874 | 0 |
| SRR5030019 | 14 | 14 | 223 | 0 | 0 | 0 | 0 | 826 | 356 | 3602 | 4185 | 8827 | 0 | 299 | 28 | 57 | 0 | 67 | 22 | 1708 | 2343 | 7331 | 0 | 0 | 17914 | 0 | 0 | 14 | 56 | 696 | 710 | 70 | 153 | 543 | 14 | 3417 | 0 |
| SRR5030020 | 0 | 29 | 24 | 0 | 0 | 0 | 0 | 800 | 394 | 4535 | 97 | 2320 | 0 | 325 | 128 | 139 | 0 | 56 | 26 | 2482 | 1851 | 7472 | 0 | 0 | 19457 | 0 | 0 | 0 | 88 | 25 | 35 | 177 | 13 | 65 | 29 | 5115 | 0 |
| SRR5030021 | 0 | 0 | 216 | 0 | 0 | 0 | 0 | 718 | 352 | 3090 | 4628 | 9564 | 0 | 410 | 0 | 0 | 0 | 51 | 15 | 1655 | 1896 | 6234 | 0 | 0 | 16867 | 0 | 0 | 0 | 101 | 16 | 30 | 57 | 230 | 23 | 0 | 3837 | 0 |
| SRR5030022 | 0 | 12 | 30 | 0 | 0 | 0 | 0 | 2328 | 924 | 3320 | 67 | 6744 | 0 | 34 | 548 | 171 | 0 | 15 | 0 | 7326 | 1354 | 4347 | 0 | 78 | 13 | 0 | 0 | 0 | 48 | 182 | 145 | 24 | 206 | 218 | 36 | 16020 | 0 |
| SRR5030023 | 0 | 0 | 19 | 0 | 0 | 22 | 0 | 3103 | 742 | 2294 | 2699 | 10052 | 0 | 0 | 270 | 135 | 0 | 0 | 0 | 7556 | 1335 | 4372 | 0 | 89 | 10508 | 0 | 0 | 22 | 90 | 179 | 179 | 45 | 157 | 224 | 22 | 12346 | 0 |
| SRR5030024 | 0 | 0 | 29 | 0 | 0 | 0 | 0 | 2590 | 1036 | 3478 | 2109 | 6364 | 0 | 0 | 222 | 481 | 0 | 12 | 0 | 6772 | 1176 | 3995 | 0 | 84 | 9578 | 0 | 13 | 0 | 39 | 220 | 12 | 78 | 194 | 19 | 39 | 15911 | 0 |
| SRR5030025 | 0 | 0 | 488 | 0 | 0 | 0 | 0 | 2035 | 814 | 2577 | 47 | 8275 | 0 | 68 | 271 | 271 | 0 | 13 | 0 | 6376 | 1186 | 4083 | 0 | 84 | 9919 | 0 | 0 | 0 | 93 | 326 | 349 | 47 | 163 | 140 | 23 | 13294 | 0 |
| SRR5030026 | 0 | 12 | 684 | 0 | 0 | 0 | 0 | 1690 | 1267 | 2781 | 59 | 7075 | 0 | 70 | 563 | 387 | 0 | 13 | 0 | 6864 | 1345 | 4299 | 0 | 66 | 11 | 0 | 0 | 0 | 137 | 249 | 298 | 87 | 75 | 448 | 37 | 14115 | 0 |
| SRR5030027 | 0 | 0 | 653 | 0 | 0 | 23 | 0 | 2229 | 743 | 2432 | 44 | 11484 | 0 | 0 | 0 | 68 | 0 | 0 | 0 | 5202 | 1354 | 4467 | 0 | 74 | 10501 | 0 | 0 | 0 | 68 | 180 | 360 | 23 | 23 | 315 | 0 | 11890 | 0 |
| SRR5030028 | 0 | 0 | 43 | 0 | 0 | 13 | 0 | 2057 | 1249 | 3196 | 2498 | 7128 | 0 | 37 | 588 | 257 | 0 | 11 | 0 | 6797 | 1220 | 4089 | 0 | 72 | 9520 | 0 | 13 | 0 | 13 | 202 | 12 | 38 | 177 | 14 | 25 | 15394 | 0 |
| SRR5030030 | 0 | 0 | 79 | 0 | 0 | 0 | 0 | 1199 | 264 | 949 | 72 | 3031 | 0 | 26 | 26 | 40 | 0 | 19 | 0 | 1502 | 829 | 7557 | 0 | 0 | 16691 | 0 | 0 | 0 | 79 | 787 | 1024 | 45 | 360 | 607 | 34 | 3057 | 0 |
| SRR5030031 | 0 | 0 | 23 | 0 | 0 | 23 | 0 | 1705 | 316 | 758 | 260 | 15052 | 0 | 0 | 21 | 0 | 0 | 18 | 12 | 1663 | 852 | 6384 | 0 | 0 | 14849 | 0 | 0 | 12 | 116 | 741 | 961 | 58 | 185 | 289 | 12 | 2358 | 0 |
| SRR5030032 | 0 | 0 | 36 | 0 | 0 | 12 | 0 | 1129 | 226 | 818 | 89 | 3047 | 0 | 28 | 0 | 0 | 0 | 22 | 13 | 1580 | 743 | 6845 | 0 | 0 | 16181 | 0 | 0 | 12 | 97 | 860 | 1066 | 24 | 363 | 593 | 12 | 2906 | 0 |
| SRR5030033 | 0 | 0 | 72 | 0 | 0 | 24 | 0 | 1568 | 269 | 560 | 5064 | 14721 | 0 | 22 | 0 | 0 | 0 | 18 | 11 | 1232 | 723 | 5974 | 0 | 0 | 13941 | 0 | 0 | 36 | 108 | 880 | 1036 | 36 | 120 | 229 | 0 | 2420 | 0 |
| SRR5030034 | 0 | 0 | 92 | 0 | 0 | 12 | 0 | 1298 | 219 | 943 | 1011 | 3251 | 0 | 0 | 55 | 14 | 0 | 18 | 0 | 1434 | 821 | 7451 | 0 | 0 | 14 | 0 | 0 | 0 | 69 | 49 | 54 | 58 | 17 | 25 | 12 | 3074 | 0 |
| SRR5030035 | 0 | 0 | 59 | 0 | 0 | 24 | 0 | 1716 | 551 | 636 | 262 | 13858 | 0 | 42 | 0 | 0 | 0 | 16 | 13 | 1314 | 910 | 6630 | 0 | 0 | 14706 | 0 | 24 | 0 | 71 | 791 | 814 | 94 | 153 | 271 | 0 | 2034 | 0 |
| SRR5030036 | 0 | 0 | 118 | 0 | 0 | 0 | 0 | 1299 | 377 | 1160 | 964 | 3367 | 0 | 0 | 14 | 56 | 0 | 15 | 10 | 1355 | 677 | 7144 | 0 | 0 | 11 | 0 | 0 | 0 | 71 | 976 | 941 | 24 | 447 | 647 | 12 | 3241 | 0 |
| SRR5030038 | 0 | 13 | 76 | 0 | 0 | 0 | 0 | 2164 | 953 | 2872 | 220 | 8399 | 0 | 14 | 68 | 95 | 25 | 27 | 0 | 2301 | 1331 | 4938 | 0 | 0 | 12 | 35 | 0 | 38 | 229 | 1829 | 889 | 63 | 254 | 775 | 13 | 17043 | 0 |
| SRR5030039 | 0 | 0 | 47 | 0 | 0 | 0 | 0 | 2064 | 756 | 2355 | 4274 | 11832 | 0 | 0 | 58 | 58 | 24 | 28 | 0 | 1919 | 1355 | 5193 | 0 | 0 | 14689 | 25 | 0 | 47 | 15 | 88 | 32 | 47 | 118 | 18 | 0 | 15467 | 0 |
| SRR5030040 | 0 | 27 | 54 | 0 | 0 | 0 | 0 | 2337 | 697 | 2715 | 228 | 7680 | 0 | 15 | 15 | 58 | 13 | 19 | 0 | 2279 | 1180 | 4707 | 0 | 0 | 11 | 34 | 13 | 13 | 94 | 64 | 33 | 27 | 175 | 26 | 27 | 17741 | 0 |
| SRR5030041 | 0 | 24 | 24 | 0 | 10 | 0 | 0 | 2816 | 851 | 2552 | 142 | 11851 | 0 | 0 | 0 | 88 | 24 | 28 | 0 | 2141 | 1219 | 4690 | 0 | 0 | 13925 | 30 | 0 | 0 | 268 | 2215 | 876 | 24 | 170 | 487 | 0 | 14227 | 0 |
| SRR5030042 | 0 | 0 | 26 | 0 | 0 | 0 | 0 | 2221 | 985 | 3178 | 231 | 8383 | 0 | 0 | 111 | 56 | 0 | 23 | 0 | 1985 | 1298 | 4947 | 0 | 0 | 14 | 44 | 0 | 13 | 181 | 1875 | 931 | 65 | 207 | 634 | 0 | 16905 | 0 |
| SRR5030043 | 0 | 0 | 48 | 0 | 0 | 0 | 0 | 2594 | 1066 | 2882 | 161 | 11095 | 0 | 0 | 0 | 29 | 0 | 27 | 0 | 1643 | 1326 | 5008 | 0 | 0 | 14887 | 30 | 0 | 0 | 191 | 1811 | 977 | 24 | 238 | 477 | 0 | 14293 | 0 |
| SRR5030044 | 0 | 13 | 26 | 0 | 0 | 0 | 0 | 2156 | 1165 | 2986 | 3409 | 8391 | 0 | 44 | 29 | 44 | 26 | 24 | 0 | 2229 | 1221 | 4664 | 0 | 0 | 13842 | 32 | 13 | 0 | 250 | 1618 | 908 | 13 | 289 | 605 | 39 | 17568 | 0 |
| SRR5030045 | 0 | 0 | 24 | 0 | 0 | 0 | 0 | 2392 | 1063 | 2421 | 133 | 11457 | 0 | 0 | 30 | 0 | 0 | 26 | 0 | 1654 | 1248 | 4769 | 0 | 0 | 14069 | 26 | 0 | 49 | 219 | 1951 | 902 | 24 | 146 | 366 | 0 | 15236 | 0 |
| SRR5030046 | 0 | 22 | 29 | 0 | 12 | 15 | 0 | 6499 | 2401 | 12652 | 924 | 2415 | 0 | 0 | 152 | 290 | 0 | 42 | 0 | 1159 | 1548 | 4471 | 0 | 0 | 7941 | 11 | 15 | 0 | 149 | 51 | 54 | 59 | 20 | 42 | 30 | 9631 | 0 |
| SRR5030048 | 0 | 16 | 491 | 0 | 13 | 0 | 0 | 6206 | 3051 | 13227 | 64 | 2355 | 0 | 44 | 222 | 326 | 0 | 54 | 0 | 1422 | 1330 | 3846 | 0 | 0 | 7700 | 0 | 16 | 0 | 233 | 933 | 901 | 97 | 330 | 740 | 56 | 11257 | 0 |
| SRR5030050 | 0 | 0 | 549 | 0 | 15 | 0 | 0 | 6640 | 2448 | 13139 | 75 | 2405 | 0 | 14 | 253 | 295 | 0 | 42 | 0 | 1280 | 1528 | 4347 | 0 | 0 | 8040 | 0 | 0 | 0 | 259 | 1006 | 678 | 122 | 358 | 800 | 69 | 10550 | 0 |
| SRR5030052 | 0 | 0 | 520 | 0 | 13 | 0 | 0 | 6080 | 2418 | 13777 | 69 | 2661 | 0 | 29 | 329 | 286 | 0 | 39 | 0 | 1431 | 1347 | 3886 | 0 | 0 | 7681 | 0 | 0 | 0 | 205 | 961 | 709 | 47 | 394 | 685 | 55 | 10415 | 0 |
| SRR5030053 | 0 | 0 | 320 | 0 | 11 | 0 | 0 | 6488 | 2102 | 11239 | 21 | 6305 | 0 | 0 | 91 | 91 | 0 | 34 | 0 | 1279 | 1417 | 3437 | 0 | 11 | 7232 | 15 | 0 | 0 | 91 | 502 | 1598 | 0 | 91 | 502 | 46 | 9046 | 0 |
| SRR5030054 | 0 | 0 | 102 | 0 | 0 | 0 | 0 | 1895 | 1701 | 6660 | 102 | 1607 | 0 | 382 | 78 | 161 | 15 | 0 | 0 | 537 | 1496 | 7542 | 0 | 0 | 15438 | 0 | 0 | 0 | 117 | 1180 | 1165 | 15 | 451 | 1296 | 58 | 676 | 12 |
| SRR5030055 | 0 | 0 | 0 | 0 | 0 | 0 | 0 | 1837 | 1816 | 6579 | 1645 | 4016 | 0 | 320 | 85 | 21 | 48 | 16 | 0 | 577 | 1543 | 6656 | 0 | 0 | 14441 | 0 | 0 | 0 | 96 | 31 | 23 | 48 | 13 | 17 | 0 | 534 | 0 |
| SRR5030056 | 0 | 16 | 172 | 0 | 0 | 0 | 0 | 1976 | 1583 | 6453 | 92 | 1637 | 0 | 458 | 77 | 185 | 0 | 0 | 0 | 482 | 1189 | 6791 | 0 | 0 | 13 | 0 | 0 | 16 | 220 | 1207 | 1160 | 78 | 517 | 1333 | 0 | 708 | 0 |
| SRR5030057 | 0 | 0 | 153 | 0 | 0 | 0 | 0 | 2447 | 1713 | 5739 | 1935 | 3715 | 0 | 356 | 67 | 89 | 0 | 15 | 0 | 690 | 1299 | 5908 | 0 | 0 | 13232 | 0 | 0 | 0 | 255 | 1429 | 1224 | 0 | 357 | 969 | 0 | 623 | 0 |
| SRR5030058 | 0 | 0 | 135 | 0 | 0 | 0 | 0 | 1774 | 1466 | 7083 | 662 | 1580 | 0 | 325 | 46 | 86 | 0 | 0 | 0 | 553 | 1534 | 7655 | 0 | 0 | 15260 | 0 | 0 | 0 | 225 | 30 | 33 | 60 | 16 | 38 | 30 | 719 | 0 |
| SRR5030059 | 0 | 0 | 148 | 0 | 0 | 0 | 0 | 1914 | 1675 | 5808 | 1523 | 4177 | 0 | 239 | 65 | 131 | 0 | 19 | 0 | 674 | 1578 | 6914 | 0 | 0 | 14090 | 0 | 49 | 98 | 197 | 31 | 31 | 49 | 246 | 18 | 0 | 1088 | 12 |
| SRR5030060 | 0 | 15 | 108 | 0 | 0 | 0 | 0 | 1770 | 1834 | 6960 | 105 | 1539 | 0 | 359 | 93 | 98 | 15 | 13 | 0 | 411 | 1240 | 6719 | 0 | 0 | 14990 | 0 | 0 | 0 | 169 | 28 | 37 | 31 | 16 | 37 | 61 | 735 | 11 |
| SRR5030061 | 0 | 0 | 151 | 0 | 0 | 0 | 0 | 1972 | 1748 | 5984 | 1793 | 4056 | 0 | 583 | 45 | 67 | 50 | 15 | 0 | 740 | 1314 | 5911 | 0 | 0 | 13500 | 0 | 0 | 101 | 101 | 29 | 29 | 0 | 352 | 13 | 0 | 762 | 15 |
| SRR5030062 | 0 | 0 | 104 | 0 | 0 | 13 | 0 | 1839 | 658 | 8890 | 2530 | 5769 | 0 | 34 | 51 | 67 | 0 | 11 | 14 | 1063 | 921 | 5467 | 0 | 0 | 14829 | 0 | 0 | 0 | 130 | 14 | 11 | 26 | 78 | 13 | 13 | 3289 | 0 |
| SRR5030064 | 0 | 14 | 153 | 0 | 0 | 0 | 0 | 1974 | 471 | 9253 | 2952 | 6537 | 0 | 0 | 91 | 72 | 0 | 11 | 15 | 1213 | 789 | 5100 | 0 | 0 | 17 | 0 | 0 | 0 | 84 | 516 | 209 | 98 | 181 | 600 | 14 | 3133 | 0 |
| SRR5030066 | 0 | 0 | 147 | 0 | 0 | 0 | 0 | 2123 | 621 | 8611 | 2813 | 6713 | 0 | 52 | 17 | 52 | 0 | 15 | 12 | 1104 | 878 | 5550 | 0 | 0 | 14843 | 0 | 0 | 0 | 67 | 495 | 267 | 13 | 134 | 361 | 0 | 3486 | 0 |
| SRR5030068 | 0 | 13 | 121 | 0 | 0 | 13 | 0 | 1744 | 541 | 8058 | 144 | 6122 | 0 | 17 | 122 | 192 | 0 | 12 | 14 | 1256 | 801 | 5246 | 0 | 0 | 14415 | 0 | 0 | 13 | 40 | 430 | 283 | 27 | 121 | 457 | 13 | 3436 | 0 |
| SRR5030069 | 0 | 0 | 125 | 0 | 0 | 0 | 0 | 2737 | 821 | 8850 | 41 | 8120 | 0 | 0 | 182 | 0 | 0 | 0 | 15 | 1186 | 829 | 5098 | 0 | 0 | 14983 | 0 | 0 | 0 | 63 | 375 | 125 | 0 | 125 | 0 | 0 | 4015 | 0 |
| SRR5030070 | 0 | 0 | 189 | 0 | 11 | 0 | 0 | 1408 | 548 | 7484 | 40 | 2138 | 0 | 639 | 13 | 13 | 0 | 30 | 0 | 2921 | 3497 | 9378 | 0 | 14 | 12 | 0 | 0 | 0 | 70 | 348 | 348 | 0 | 169 | 756 | 30 | 4225 | 0 |
| SRR5030071 | 0 | 21 | 185 | 0 | 0 | 0 | 0 | 1606 | 364 | 7728 | 2152 | 6031 | 0 | 697 | 30 | 30 | 0 | 44 | 0 | 2303 | 3600 | 8441 | 0 | 0 | 17933 | 0 | 0 | 0 | 41 | 288 | 452 | 62 | 164 | 329 | 21 | 3515 | 0 |
| SRR5030072 | 0 | 32 | 290 | 0 | 12 | 0 | 0 | 1331 | 637 | 8060 | 977 | 2181 | 0 | 637 | 0 | 71 | 0 | 34 | 0 | 3060 | 2853 | 8150 | 0 | 0 | 18379 | 0 | 11 | 0 | 107 | 290 | 494 | 54 | 183 | 558 | 0 | 4844 | 0 |
| SRR5030073 | 0 | 43 | 149 | 0 | 17 | 0 | 0 | 1026 | 673 | 6154 | 2051 | 5225 | 0 | 385 | 0 | 0 | 0 | 32 | 0 | 2660 | 2920 | 7558 | 0 | 0 | 16954 | 0 | 0 | 21 | 106 | 18 | 20 | 43 | 149 | 14 | 0 | 3077 | 0 |
| SRR5030074 | 0 | 10 | 14 | 0 | 15 | 0 | 0 | 953 | 681 | 7404 | 858 | 2069 | 0 | 558 | 27 | 14 | 0 | 33 | 12 | 2845 | 3471 | 9418 | 0 | 0 | 13 | 0 | 0 | 0 | 72 | 17 | 17 | 20 | 12 | 35 | 10 | 4328 | 0 |
| SRR5030076 | 0 | 0 | 302 | 0 | 23 | 0 | 0 | 1358 | 590 | 7696 | 56 | 1893 | 0 | 398 | 14 | 41 | 0 | 33 | 0 | 2840 | 2939 | 8342 | 0 | 12 | 18513 | 0 | 0 | 0 | 63 | 385 | 469 | 10 | 208 | 448 | 31 | 4527 | 0 |
| SRR5030077 | 0 | 0 | 12 | 0 | 13 | 0 | 0 | 1480 | 598 | 7024 | 56 | 5040 | 0 | 472 | 0 | 0 | 0 | 32 | 0 | 3024 | 2880 | 7527 | 0 | 11 | 16795 | 0 | 0 | 0 | 85 | 17 | 14 | 64 | 212 | 191 | 21 | 3937 | 0 |
| SRR5030078 | 0 | 0 | 12 | 0 | 0 | 0 | 0 | 1100 | 180 | 3099 | 580 | 1440 | 0 | 140 | 11358 | 10018 | 0 | 45 | 0 | 0 | 1564 | 4228 | 0 | 0 | 7010 | 108 | 0 | 0 | 14 | 24 | 24 | 85 | 19 | 44 | 74 | 0 | 0 |
| SRR5030079 | 0 | 0 | 189 | 0 | 0 | 0 | 17 | 1464 | 237 | 2967 | 1464 | 3561 | 0 | 277 | 4668 | 3640 | 0 | 47 | 0 | 0 | 1699 | 3983 | 0 | 0 | 6709 | 116 | 0 | 0 | 378 | 549 | 429 | 69 | 258 | 567 | 0 | 0 | 0 |
| SRR5030080 | 0 | 11 | 206 | 0 | 0 | 0 | 0 | 1074 | 215 | 3739 | 236 | 1397 | 0 | 215 | 11132 | 10981 | 0 | 49 | 0 | 0 | 1305 | 3495 | 0 | 0 | 6955 | 105 | 0 | 11 | 263 | 481 | 492 | 160 | 469 | 996 | 11 | 21 | 0 |
| SRR5030081 | 0 | 0 | 143 | 0 | 0 | 0 | 0 | 1350 | 245 | 2986 | 20 | 3845 | 0 | 327 | 4458 | 3845 | 0 | 53 | 0 | 0 | 1366 | 3419 | 0 | 0 | 6276 | 106 | 0 | 0 | 12 | 22 | 16 | 36 | 89 | 27 | 18 | 0 | 0 |
| SRR5030082 | 0 | 0 | 12 | 0 | 0 | 0 | 0 | 940 | 286 | 2799 | 18 | 1348 | 0 | 163 | 10502 | 11135 | 0 | 48 | 0 | 0 | 1596 | 4170 | 0 | 0 | 7019 | 108 | 0 | 11 | 18 | 24 | 26 | 131 | 15 | 46 | 33 | 0 | 0 |
| SRR5030083 | 0 | 0 | 123 | 0 | 0 | 0 | 0 | 921 | 280 | 2162 | 29 | 4044 | 0 | 400 | 5085 | 3684 | 0 | 51 | 0 | 0 | 1749 | 3945 | 0 | 0 | 6782 | 100 | 0 | 0 | 11 | 24 | 20 | 35 | 229 | 21 | 0 | 0 | 0 |

### Supp Table S8.4.Columns_DI_to_ES

| | | | | | | | | | | | | | | | | | | | | | | | | | | | | | | | | | | | | |
|---|---|---|---|---|---|---|---|---|---|---|---|---|---|---|---|---|---|---|---|---|---|---|---|---|---|---|---|---|---|---|---|---|---|---|---|---|
| SRR5030084 | 0 | 0 | 13 | 0 | 0 | 0 | 0 | 989 | 350 | 3173 | 19 | 1690 | 0 | 103 | 10282 | 11003 | 0 | 48 | 0 | 0 | 1252 | 3708 | 0 | 0 | 6859 | 129 | 11 | 0 | 15 | 16 | 18 | 90 | 17 | 50 | 34 | 0 | 0 |
| SRR5030085 | 0 | 0 | 107 | 0 | 13 | 0 | 18 | 1203 | 281 | 2486 | 28 | 3729 | 0 | 160 | 4571 | 3649 | 0 | 50 | 0 | 0 | 1279 | 3405 | 0 | 0 | 6597 | 133 | 36 | 0 | 10 | 21 | 15 | 107 | 160 | 24 | 0 | 0 | 0 |
| SRR5030086 | 0 | 0 | 79 | 0 | 0 | 0 | 0 | 1900 | 649 | 7625 | 36 | 3295 | 0 | 289 | 192 | 168 | 0 | 59 | 0 | 1924 | 1748 | 4170 | 0 | 96 | 9340 | 14 | 139 | 0 | 59 | 347 | 297 | 20 | 99 | 307 | 0 | 18810 | 0 |
| SRR5030087 | 0 | 0 | 68 | 0 | 0 | 0 | 0 | 2491 | 531 | 7390 | 1184 | 4450 | 0 | 122 | 122 | 122 | 0 | 62 | 0 | 2082 | 1630 | 4396 | 0 | 84 | 9574 | 17 | 145 | 0 | 77 | 230 | 324 | 26 | 94 | 247 | 26 | 16128 | 0 |
| SRR5030088 | 0 | 0 | 47 | 0 | 0 | 0 | 0 | 1799 | 437 | 7299 | 822 | 3315 | 0 | 129 | 103 | 129 | 0 | 46 | 0 | 2159 | 1437 | 3969 | 0 | 82 | 13 | 13 | 11 | 0 | 58 | 34 | 24 | 10 | 13 | 30 | 26 | 18838 | 0 |
| SRR5030090 | 0 | 0 | 60 | 0 | 0 | 0 | 0 | 2001 | 395 | 6646 | 1013 | 3064 | 0 | 222 | 173 | 173 | 10 | 48 | 0 | 2372 | 1709 | 4134 | 0 | 89 | 9182 | 25 | 125 | 0 | 40 | 281 | 276 | 10 | 115 | 281 | 0 | 19246 | 0 |
| SRR5030091 | 0 | 0 | 34 | 0 | 0 | 0 | 0 | 2419 | 287 | 7217 | 29 | 5290 | 0 | 369 | 123 | 123 | 0 | 69 | 0 | 2050 | 1722 | 4251 | 0 | 89 | 9517 | 15 | 20 | 0 | 60 | 38 | 28 | 26 | 14 | 19 | 0 | 15090 | 0 |
| SRR5030092 | 0 | 0 | 12 | 0 | 0 | 0 | 0 | 1933 | 678 | 7708 | 904 | 3917 | 0 | 151 | 326 | 176 | 0 | 66 | 0 | 2461 | 1422 | 3980 | 0 | 89 | 9062 | 21 | 13 | 0 | 61 | 37 | 25 | 15 | 14 | 38 | 20 | 16948 | 0 |
| SRR5030093 | 0 | 0 | 62 | 0 | 0 | 0 | 0 | 2302 | 293 | 6027 | 28 | 4102 | 0 | 209 | 42 | 126 | 0 | 76 | 0 | 2135 | 1475 | 4035 | 0 | 95 | 9316 | 24 | 168 | 18 | 62 | 265 | 336 | 0 | 168 | 212 | 0 | 15822 | 0 |
| SRR5030094 | 0 | 0 | 0 | 0 | 0 | 0 | 0 | 1502 | 108 | 3449 | 541 | 1226 | 0 | 204 | 336 | 192 | 0 | 123 | 0 | 1610 | 1701 | 6163 | 0 | 38 | 11503 | 34 | 88 | 0 | 25 | 208 | 113 | 0 | 29 | 140 | 0 | 12209 | 0 |
| SRR5030095 | 0 | 0 | 0 | 0 | 0 | 0 | 0 | 1040 | 0 | 2674 | 1200 | 3566 | 0 | 149 | 149 | 149 | 0 | 114 | 0 | 1486 | 1804 | 5334 | 0 | 0 | 10563 | 35 | 75 | 0 | 21 | 42 | 54 | 21 | 0 | 41 | 21 | 8914 | 0 |
| SRR5030096 | 0 | 0 | 15 | 0 | 0 | 0 | 0 | 1230 | 298 | 3872 | 298 | 1114 | 0 | 181 | 246 | 272 | 0 | 123 | 0 | 1658 | 1338 | 5423 | 0 | 40 | 11352 | 27 | 37 | 0 | 32 | 71 | 26 | 0 | 28 | 47 | 0 | 12329 | 0 |
| SRR5030097 | 0 | 0 | 21 | 0 | 0 | 0 | 0 | 1812 | 0 | 2718 | 1224 | 5134 | 0 | 0 | 0 | 0 | 0 | 119 | 0 | 1510 | 1443 | 4825 | 0 | 55 | 9255 | 46 | 511 | 0 | 21 | 255 | 85 | 0 | 64 | 192 | 21 | 8153 | 0 |
| SRR5030098 | 0 | 0 | 17 | 0 | 0 | 0 | 0 | 1457 | 291 | 3970 | 23 | 1396 | 0 | 194 | 304 | 364 | 0 | 130 | 0 | 1809 | 1737 | 6249 | 0 | 25 | 11735 | 39 | 123 | 0 | 33 | 200 | 115 | 0 | 33 | 170 | 0 | 11971 | 0 |
| SRR5030100 | 0 | 0 | 18 | 0 | 0 | 0 | 0 | 1539 | 250 | 3467 | 33 | 1239 | 0 | 200 | 263 | 275 | 0 | 114 | 0 | 1715 | 1410 | 5623 | 0 | 52 | 11466 | 42 | 23 | 0 | 26 | 56 | 24 | 0 | 32 | 43 | 0 | 11502 | 0 |
| SRR5030101 | 0 | 0 | 0 | 0 | 18 | 0 | 0 | 1401 | 0 | 4046 | 1261 | 2646 | 0 | 311 | 156 | 467 | 21 | 100 | 0 | 2646 | 1516 | 4612 | 0 | 27 | 9950 | 54 | 213 | 0 | 43 | 256 | 149 | 0 | 43 | 85 | 0 | 11360 | 0 |
| SRR5030102 | 0 | 15 | 231 | 0 | 0 | 0 | 0 | 2035 | 383 | 2059 | 2179 | 7662 | 0 | 24 | 48 | 168 | 0 | 29 | 0 | 2275 | 2881 | 9153 | 0 | 12 | 17912 | 0 | 0 | 0 | 46 | 1002 | 570 | 39 | 247 | 616 | 39 | 4621 | 0 |
| SRR5030103 | 0 | 0 | 122 | 0 | 0 | 0 | 0 | 2167 | 298 | 1463 | 4714 | 13059 | 0 | 0 | 0 | 54 | 0 | 25 | 0 | 1382 | 2942 | 8364 | 0 | 0 | 16785 | 0 | 19 | 19 | 122 | 873 | 676 | 47 | 206 | 469 | 19 | 2818 | 0 |
| SRR5030104 | 0 | 0 | 193 | 0 | 0 | 0 | 0 | 2323 | 297 | 2002 | 98 | 6797 | 0 | 25 | 49 | 173 | 0 | 37 | 0 | 1458 | 2324 | 8148 | 0 | 0 | 11 | 0 | 0 | 0 | 117 | 956 | 889 | 59 | 226 | 730 | 0 | 4276 | 0 |
| SRR5030105 | 0 | 0 | 117 | 0 | 0 | 0 | 0 | 1997 | 485 | 1484 | 4252 | 14867 | 0 | 0 | 29 | 114 | 0 | 31 | 0 | 1541 | 2406 | 7504 | 0 | 0 | 15912 | 0 | 20 | 0 | 107 | 59 | 38 | 29 | 156 | 22 | 29 | 3253 | 0 |
| SRR5030106 | 0 | 0 | 21 | 0 | 0 | 0 | 0 | 2618 | 374 | 1940 | 100 | 6382 | 0 | 23 | 117 | 374 | 0 | 28 | 11 | 2291 | 2900 | 9190 | 0 | 0 | 17734 | 0 | 16 | 0 | 79 | 68 | 51 | 79 | 14 | 55 | 63 | 4418 | 0 |
| SRR5030107 | 0 | 29 | 105 | 0 | 0 | 0 | 0 | 2253 | 318 | 1140 | 185 | 14046 | 0 | 27 | 0 | 27 | 0 | 19 | 0 | 1484 | 3038 | 8706 | 0 | 0 | 17175 | 0 | 29 | 19 | 95 | 912 | 665 | 57 | 171 | 447 | 0 | 2518 | 0 |
| SRR5030108 | 0 | 25 | 205 | 0 | 0 | 0 | 0 | 2469 | 312 | 2349 | 90 | 5945 | 0 | 48 | 72 | 192 | 0 | 32 | 0 | 2349 | 2375 | 8095 | 0 | 0 | 11 | 0 | 16 | 0 | 90 | 893 | 631 | 49 | 270 | 574 | 49 | 4459 | 0 |
| SRR5030109 | 0 | 29 | 98 | 0 | 0 | 0 | 0 | 1954 | 440 | 1349 | 172 | 14534 | 0 | 28 | 55 | 83 | 0 | 36 | 11 | 1982 | 2373 | 7470 | 0 | 0 | 16060 | 10 | 0 | 20 | 39 | 1039 | 637 | 39 | 167 | 461 | 29 | 3193 | 0 |
| SRR5030110 | 0 | 0 | 60 | 0 | 0 | 0 | 0 | 1408 | 340 | 2670 | 850 | 3228 | 0 | 267 | 49 | 0 | 0 | 30 | 0 | 534 | 795 | 5379 | 0 | 13 | 12121 | 0 | 0 | 0 | 99 | 58 | 35 | 60 | 16 | 29 | 60 | 1966 | 0 |
| SRR5030111 | 0 | 0 | 33 | 0 | 0 | 0 | 0 | 1653 | 611 | 1833 | 190 | 11516 | 0 | 323 | 0 | 0 | 0 | 29 | 0 | 593 | 825 | 4737 | 0 | 16 | 10901 | 0 | 0 | 0 | 106 | 763 | 357 | 45 | 84 | 189 | 11 | 1186 | 0 |
| SRR5030112 | 0 | 0 | 117 | 0 | 0 | 11 | 0 | 1336 | 210 | 2515 | 707 | 2567 | 0 | 105 | 26 | 157 | 0 | 36 | 0 | 576 | 658 | 5059 | 0 | 11 | 12164 | 0 | 0 | 0 | 96 | 55 | 33 | 21 | 117 | 27 | 64 | 1991 | 0 |
| SRR5030113 | 0 | 0 | 52 | 0 | 0 | 0 | 0 | 1772 | 343 | 1962 | 3296 | 12077 | 0 | 190 | 38 | 0 | 0 | 24 | 0 | 571 | 739 | 4311 | 0 | 0 | 10355 | 0 | 0 | 0 | 115 | 55 | 28 | 23 | 81 | 10 | 12 | 1505 | 0 |
| SRR5030114 | 0 | 0 | 41 | 0 | 0 | 0 | 0 | 1560 | 446 | 2749 | 743 | 3318 | 0 | 297 | 124 | 25 | 0 | 27 | 0 | 768 | 825 | 5416 | 0 | 0 | 12355 | 0 | 0 | 10 | 71 | 754 | 397 | 61 | 163 | 306 | 20 | 2031 | 0 |
| SRR5030115 | 0 | 0 | 40 | 0 | 0 | 0 | 0 | 1638 | 331 | 1932 | 162 | 11408 | 0 | 239 | 0 | 0 | 0 | 34 | 0 | 570 | 800 | 4674 | 0 | 11 | 10894 | 0 | 0 | 0 | 136 | 47 | 30 | 40 | 113 | 12 | 0 | 1362 | 0 |
| SRR5030116 | 0 | 0 | 52 | 0 | 0 | 0 | 0 | 1818 | 227 | 2449 | 783 | 2701 | 0 | 303 | 25 | 0 | 0 | 31 | 0 | 732 | 715 | 5143 | 0 | 11 | 11814 | 0 | 0 | 10 | 197 | 778 | 280 | 31 | 197 | 425 | 41 | 1565 | 0 |
| SRR5030117 | 0 | 0 | 17 | 0 | 0 | 0 | 0 | 1537 | 426 | 2297 | 189 | 11224 | 0 | 222 | 37 | 19 | 0 | 31 | 0 | 574 | 687 | 4389 | 0 | 12 | 10186 | 0 | 0 | 23 | 86 | 53 | 24 | 46 | 86 | 12 | 0 | 1537 | 0 |
| SRR5030118 | 0 | 10 | 31 | 0 | 0 | 0 | 28 | 1584 | 116 | 2151 | 786 | 1584 | 0 | 150 | 35 | 46 | 21 | 38 | 13 | 3747 | 1361 | 6660 | 0 | 11 | 13 | 0 | 25 | 0 | 10 | 72 | 38 | 17 | 12 | 48 | 24 | 41781 | 24 |
| SRR5030119 | 0 | 22 | 41 | 0 | 0 | 0 | 22 | 1703 | 184 | 1726 | 106 | 2244 | 0 | 115 | 12 | 23 | 13 | 23 | 0 | 3694 | 1360 | 6561 | 0 | 0 | 16843 | 0 | 143 | 0 | 38 | 460 | 225 | 16 | 92 | 178 | 16 | 38875 | 29 |
| SRR5030120 | 0 | 11 | 30 | 0 | 0 | 11 | 22 | 1757 | 99 | 2103 | 656 | 1398 | 0 | 148 | 12 | 49 | 22 | 20 | 0 | 3068 | 1262 | 6372 | 0 | 0 | 16024 | 0 | 31 | 0 | 19 | 67 | 43 | 19 | 18 | 43 | 15 | 42542 | 22 |
| SRR5030121 | 0 | 13 | 23 | 0 | 0 | 13 | 23 | 1751 | 95 | 1632 | 100 | 2370 | 0 | 95 | 0 | 0 | 13 | 27 | 13 | 3550 | 1282 | 6329 | 0 | 0 | 16298 | 0 | 152 | 0 | 33 | 447 | 228 | 23 | 63 | 189 | 0 | 38273 | 23 |
| SRR5030122 | 0 | 11 | 21 | 0 | 0 | 0 | 0 | 1338 | 213 | 1894 | 59 | 1291 | 0 | 178 | 36 | 71 | 11 | 32 | 0 | 3434 | 1371 | 6660 | 0 | 0 | 16242 | 0 | 21 | 0 | 50 | 78 | 41 | 14 | 19 | 45 | 11 | 42552 | 24 |
| SRR5030123 | 0 | 0 | 0 | 0 | 0 | 13 | 29 | 1651 | 177 | 1828 | 94 | 2229 | 0 | 118 | 0 | 35 | 0 | 25 | 15 | 3844 | 1348 | 6712 | 0 | 0 | 17156 | 0 | 158 | 0 | 61 | 470 | 183 | 23 | 97 | 122 | 0 | 38135 | 28 |
| SRR5030124 | 0 | 18 | 58 | 0 | 0 | 0 | 11 | 1448 | 109 | 1990 | 772 | 1713 | 0 | 97 | 36 | 36 | 0 | 26 | 0 | 3739 | 1315 | 6270 | 0 | 0 | 16180 | 0 | 127 | 0 | 29 | 312 | 254 | 11 | 116 | 181 | 0 | 43161 | 21 |
| SRR5030125 | 0 | 0 | 16 | 0 | 0 | 0 | 20 | 1470 | 215 | 1948 | 91 | 2091 | 0 | 131 | 48 | 12 | 16 | 32 | 12 | 3644 | 1224 | 6294 | 0 | 0 | 16612 | 0 | 125 | 0 | 36 | 525 | 292 | 20 | 98 | 184 | 0 | 38928 | 24 |
| SRR5030126 | 0 | 0 | 0 | 0 | 0 | 0 | 0 | 2278 | 742 | 1695 | 3655 | 8528 | 0 | 0 | 212 | 318 | 0 | 30 | 0 | 900 | 663 | 3502 | 0 | 67 | 9865 | 0 | 15 | 0 | 59 | 39 | 35 | 37 | 12 | 30 | 0 | 1112 | 13 |
| SRR5030127 | 0 | 0 | 0 | 0 | 0 | 0 | 0 | 2571 | 741 | 1133 | 7016 | 12376 | 0 | 0 | 174 | 44 | 21 | 25 | 0 | 915 | 609 | 3488 | 0 | 64 | 10317 | 0 | 21 | 14 | 34 | 558 | 489 | 41 | 48 | 124 | 0 | 1046 | 14 |
| SRR5030128 | 0 | 0 | 16 | 0 | 0 | 0 | 0 | 2081 | 636 | 1272 | 3757 | 7399 | 0 | 116 | 173 | 405 | 0 | 29 | 0 | 578 | 501 | 3359 | 0 | 65 | 9691 | 0 | 47 | 0 | 31 | 431 | 329 | 31 | 78 | 282 | 16 | 1850 | 18 |
| SRR5030130 | 0 | 0 | 22 | 0 | 0 | 0 | 0 | 2436 | 498 | 1827 | 3875 | 7585 | 0 | 55 | 388 | 221 | 15 | 31 | 0 | 1052 | 573 | 3598 | 0 | 71 | 10034 | 0 | 0 | 0 | 60 | 47 | 34 | 37 | 82 | 20 | 0 | 1605 | 12 |
| SRR5030131 | 0 | 0 | 0 | 0 | 0 | 14 | 0 | 2169 | 753 | 1018 | 6729 | 12218 | 0 | 0 | 44 | 44 | 28 | 26 | 0 | 753 | 643 | 3653 | 0 | 74 | 10306 | 0 | 56 | 28 | 77 | 43 | 26 | 35 | 70 | 14 | 0 | 841 | 17 |
| SRR5030132 | 0 | 0 | 15 | 0 | 0 | 0 | 0 | 2403 | 335 | 1789 | 4080 | 9670 | 0 | 112 | 224 | 559 | 0 | 31 | 0 | 1006 | 547 | 3398 | 0 | 67 | 9965 | 0 | 15 | 0 | 54 | 322 | 292 | 23 | 115 | 238 | 0 | 783 | 24 |
| SRR5030133 | 0 | 0 | 0 | 0 | 0 | 0 | 0 | 2672 | 223 | 980 | 6413 | 11892 | 0 | 45 | 45 | 178 | 0 | 39 | 0 | 757 | 521 | 3355 | 0 | 68 | 9858 | 0 | 58 | 36 | 115 | 42 | 22 | 14 | 93 | 14 | 0 | 891 | 16 |
| SRR5030134 | 0 | 0 | 17 | 0 | 0 | 0 | 0 | 2452 | 240 | 799 | 666 | 1759 | 0 | 147 | 306 | 413 | 0 | 28 | 0 | 3931 | 931 | 3847 | 0 | 0 | 9472 | 0 | 36 | 0 | 29 | 55 | 49 | 17 | 29 | 95 | 0 | 34362 | 0 |
| SRR5030135 | 0 | 0 | 27 | 0 | 10 | 0 | 21 | 2385 | 118 | 736 | 97 | 5242 | 0 | 29 | 177 | 206 | 0 | 19 | 0 | 3710 | 982 | 3565 | 0 | 0 | 8809 | 0 | 74 | 0 | 24 | 66 | 38 | 0 | 21 | 47 | 0 | 23970 | 0 |
| SRR5030136 | 0 | 0 | 21 | 0 | 0 | 0 | 0 | 2639 | 258 | 846 | 875 | 1807 | 0 | 258 | 359 | 373 | 0 | 26 | 0 | 4647 | 897 | 3735 | 0 | 0 | 9110 | 0 | 26 | 0 | 11 | 68 | 47 | 21 | 22 | 82 | 0 | 36398 | 0 |
| SRR5030137 | 0 | 0 | 11 | 0 | 13 | 0 | 0 | 2222 | 185 | 617 | 2654 | 5833 | 0 | 123 | 154 | 185 | 18 | 24 | 0 | 3518 | 894 | 3361 | 0 | 0 | 8255 | 0 | 301 | 0 | 28 | 262 | 106 | 11 | 64 | 152 | 0 | 25645 | 0 |
| SRR5030138 | 0 | 0 | 17 | 0 | 11 | 0 | 0 | 2283 | 135 | 716 | 57 | 1608 | 0 | 216 | 378 | 324 | 0 | 27 | 0 | 3864 | 940 | 3964 | 0 | 0 | 9602 | 0 | 112 | 0 | 25 | 191 | 140 | 0 | 88 | 280 | 11 | 34586 | 0 |
| SRR5030139 | 0 | 0 | 10 | 0 | 0 | 0 | 14 | 2459 | 270 | 630 | 87 | 5367 | 0 | 90 | 210 | 210 | 0 | 23 | 0 | 3058 | 1018 | 3494 | 0 | 0 | 9039 | 0 | 329 | 0 | 28 | 267 | 156 | 28 | 73 | 184 | 0 | 25366 | 0 |
| SRR5030140 | 0 | 0 | 30 | 0 | 11 | 0 | 0 | 2531 | 320 | 946 | 918 | 1572 | 11 | 167 | 376 | 515 | 0 | 19 | 0 | 3630 | 951 | 3740 | 0 | 0 | 9493 | 0 | 39 | 0 | 11 | 64 | 50 | 13 | 32 | 94 | 0 | 36217 | 0 |
| SRR5030141 | 0 | 0 | 11 | 0 | 0 | 0 | 14 | 1893 | 186 | 496 | 2544 | 5181 | 0 | 31 | 93 | 248 | 11 | 28 | 0 | 3227 | 922 | 3342 | 0 | 0 | 8330 | 0 | 95 | 0 | 12 | 66 | 38 | 18 | 27 | 38 | 0 | 27210 | 0 |
| SRR5030142 | 0 | 0 | 35 | 0 | 0 | 0 | 14 | 1149 | 119 | 331 | 116 | 368 | 0 | 23 | 286 | 300 | 0 | 47 | 14 | 11789 | 2478 | 11270 | 0 | 15 | 19152 | 0 | 14 | 0 | 21 | 72 | 42 | 35 | 17 | 84 | 17 | 133688 | 0 |
| SRR5030143 | 0 | 0 | 30 | 0 | 0 | 0 | 30 | 1042 | 80 | 321 | 12 | 1416 | 0 | 0 | 160 | 134 | 0 | 76 | 38 | 12153 | 2828 | 10201 | 0 | 0 | 17551 | 0 | 90 | 0 | 30 | 780 | 330 | 0 | 210 | 570 | 0 | 121231 | 0 |
| SRR5030144 | 0 | 0 | 38 | 0 | 0 | 0 | 15 | 1130 | 134 | 310 | 34 | 474 | 0 | 33 | 292 | 289 | 0 | 43 | 16 | 11872 | 2153 | 10048 | 0 | 13 | 18264 | 0 | 38 | 0 | 42 | 77 | 44 | 54 | 18 | 70 | 19 | 136610 | 0 |
| SRR5030145 | 0 | 0 | 0 | 0 | 16 | 0 | 0 | 1286 | 196 | 307 | 196 | 1258 | 0 | 28 | 224 | 224 | 0 | 56 | 24 | 11570 | 2130 | 9227 | 0 | 16 | 16451 | 0 | 159 | 0 | 64 | 893 | 510 | 0 | 191 | 638 | 0 | 128361 | 0 |
| SRR5030146 | 0 | 14 | 25 | 0 | 0 | 0 | 25 | 992 | 131 | 271 | 44 | 412 | 0 | 23 | 280 | 315 | 0 | 32 | 14 | 11619 | 2571 | 11382 | 0 | 11 | 18732 | 0 | 25 | 0 | 29 | 72 | 50 | 47 | 18 | 74 | 14 | 132320 | 0 |
| SRR5030147 | 0 | 0 | 0 | 0 | 0 | 0 | 31 | 988 | 192 | 192 | 12 | 1400 | 0 | 27 | 192 | 82 | 0 | 31 | 23 | 11612 | 2521 | 9990 | 0 | 0 | 17815 | 0 | 93 | 0 | 31 | 60 | 60 | 0 | 93 | 45 | 0 | 121174 | 0 |
| SRR5030148 | 0 | 19 | 19 | 0 | 0 | 0 | 0 | 1085 | 136 | 308 | 40 | 403 | 0 | 44 | 347 | 338 | 0 | 31 | 23 | 11640 | 2110 | 10118 | 0 | 13 | 18455 | 0 | 48 | 0 | 48 | 690 | 338 | 41 | 171 | 731 | 15 | 134827 | 0 |
| SRR5030149 | 0 | 94 | 0 | 0 | 0 | 0 | 31 | 866 | 84 | 335 | 10 | 1202 | 0 | 28 | 252 | 140 | 0 | 32 | 0 | 11122 | 2294 | 9304 | 0 | 0 | 16400 | 0 | 94 | 0 | 0 | 786 | 346 | 0 | 189 | 974 | 0 | 124050 | 0 |
| SRR5030150 | 0 | 0 | 0 | 0 | 0 | 0 | 27 | 817 | 228 | 839 | 145 | 2231 | 0 | 74 | 44 | 44 | 30 | 51 | 0 | 2503 | 904 | 4723 | 0 | 0 | 12435 | 0 | 119 | 0 | 27 | 242 | 165 | 11 | 64 | 139 | 0 | 21719 | 0 |
| SRR5030151 | 0 | 0 | 0 | 0 | 0 | 0 | 79 | 1274 | 469 | 536 | 1274 | 2816 | 0 | 134 | 0 | 67 | 0 | 56 | 0 | 2883 | 668 | 4584 | 0 | 0 | 12801 | 0 | 237 | 0 | 0 | 394 | 197 | 0 | 59 | 99 | 0 | 21792 | 19 |
| SRR5030152 | 0 | 0 | 0 | 0 | 0 | 0 | 30 | 786 | 214 | 651 | 1184 | 2415 | 0 | 103 | 48 | 56 | 20 | 26 | 0 | 2622 | 854 | 4451 | 0 | 0 | 11850 | 0 | 17 | 0 | 30 | 43 | 33 | 0 | 11 | 26 | 0 | 22720 | 0 |
| SRR5030153 | 0 | 0 | 0 | 0 | 0 | 0 | 0 | 1385 | 346 | 692 | 21 | 3393 | 0 | 208 | 0 | 0 | 20 | 39 | 29 | 2354 | 680 | 4516 | 0 | 0 | 11888 | 0 | 143 | 0 | 61 | 348 | 184 | 20 | 123 | 82 | 0 | 17517 | 0 |
| SRR5030154 | 0 | 0 | 14 | 0 | 0 | 19 | 12 | 843 | 273 | 767 | 1086 | 2817 | 0 | 182 | 38 | 38 | 19 | 46 | 11 | 2574 | 927 | 4783 | 0 | 0 | 12191 | 0 | 21 | 0 | 38 | 43 | 27 | 12 | 14 | 27 | 0 | 22445 | 0 |
| SRR5030155 | 20 | 20 | 0 | 0 | 0 | 0 | 20 | 951 | 272 | 747 | 1087 | 3193 | 0 | 0 | 68 | 0 | 11 | 38 | 0 | 2582 | 942 | 4573 | 0 | 0 | 12384 | 0 | 40 | 0 | 0 | 51 | 29 | 0 | 41 | 22 | 0 | 17326 | 0 |
| SRR5030156 | 0 | 0 | 0 | 0 | 0 | 14 | 39 | 924 | 193 | 948 | 131 | 2565 | 0 | 177 | 15 | 62 | 41 | 32 | 0 | 2558 | 834 | 4606 | 0 | 0 | 11870 | 0 | 20 | 0 | 24 | 43 | 30 | 0 | 53 | 28 | 0 | 23042 | 0 |
| SRR5030157 | 0 | 0 | 0 | 0 | 0 | 0 | 41 | 833 | 69 | 1111 | 32 | 3959 | 0 | 69 | 69 | 0 | 62 | 39 | 0 | 2431 | 814 | 4286 | 0 | 0 | 12290 | 0 | 62 | 0 | 62 | 145 | 145 | 0 | 21 | 186 | 0 | 19863 | 0 |
| SRR5030158 | 0 | 0 | 37 | 0 | 0 | 0 | 21 | 1314 | 154 | 532 | 23 | 801 | 0 | 212 | 154 | 128 | 0 | 11 | 0 | 1250 | 1396 | 6809 | 0 | 0 | 15851 | 0 | 41 | 0 | 49 | 59 | 54 | 41 | 17 | 54 | 21 | 11236 | 0 |
| SRR5030159 | 0 | 18 | 29 | 0 | 0 | 0 | 35 | 1480 | 144 | 578 | 373 | 2203 | 12 | 301 | 84 | 36 | 0 | 15 | 0 | 1396 | 1445 | 6319 | 0 | 0 | 15251 | 0 | 12 | 0 | 14 | 76 | 43 | 29 | 11 | 28 | 12 | 10736 | 0 |
| SRR5030160 | 0 | 13 | 53 | 0 | 0 | 0 | 31 | 1329 | 152 | 637 | 31 | 866 | 0 | 145 | 138 | 132 | 31 | 16 | 0 | 1509 | 1189 | 6157 | 0 | 0 | 15629 | 0 | 40 | 0 | 12 | 62 | 52 | 62 | 20 | 54 | 27 | 12166 | 0 |
| SRR5030161 | 0 | 0 | 31 | 0 | 0 | 0 | 31 | 1504 | 149 | 584 | 28 | 2126 | 0 | 274 | 37 | 50 | 24 | 21 | 0 | 1442 | 1198 | 5653 | 0 | 0 | 14127 | 0 | 153 | 0 | 73 | 594 | 447 | 37 | 110 | 251 | 0 | 10929 | 0 |
| SRR5030162 | 0 | 25 | 34 | 0 | 0 | 0 | 29 | 1195 | 166 | 684 | 159 | 876 | 0 | 166 | 186 | 93 | 25 | 17 | 0 | 1388 | 1378 | 6905 | 0 | 0 | 11 | 0 | 55 | 0 | 105 | 639 | 425 | 38 | 135 | 467 | 21 | 12072 | 0 |
| SRR5030163 | 0 | 0 | 59 | 0 | 0 | 0 | 53 | 1506 | 85 | 498 | 50 | 2235 | 0 | 231 | 97 | 24 | 12 | 14 | 0 | 1433 | 1467 | 6513 | 0 | 0 | 15550 | 0 | 172 | 0 | 47 | 622 | 338 | 36 | 124 | 249 | 0 | 10142 | 0 |
| SRR5030164 | 0 | 26 | 34 | 0 | 0 | 0 | 22 | 1266 | 150 | 619 | 157 | 755 | 0 | 245 | 170 | 102 | 22 | 16 | 0 | 1320 | 1210 | 6180 | 0 | 0 | 10 | 0 | 52 | 0 | 116 | 533 | 486 | 39 | 185 | 495 | 22 | 12857 | 0 |
| SRR5030165 | 0 | 0 | 31 | 0 | 0 | 12 | 67 | 1613 | 88 | 466 | 554 | 1928 | 0 | 227 | 50 | 25 | 18 | 18 | 0 | 1046 | 1203 | 5721 | 0 | 0 | 14390 | 0 | 11 | 0 | 11 | 65 | 46 | 24 | 16 | 35 | 0 | 10205 | 0 |
| SRR5030166 | 0 | 26 | 16 | 0 | 0 | 13 | 0 | 3057 | 975 | 1689 | 59 | 1951 | 0 | 116 | 233 | 277 | 0 | 0 | 0 | 1660 | 636 | 4352 | 0 | 0 | 11194 | 0 | 0 | 0 | 182 | 69 | 78 | 78 | 27 | 66 | 13 | 3829 | 0 |
| SRR5030167 | 0 | 19 | 229 | 0 | 0 | 0 | 0 | 3060 | 998 | 1685 | 167 | 5147 | 0 | 169 | 91 | 156 | 19 | 0 | 0 | 1271 | 669 | 4051 | 0 | 0 | 10634 | 12 | 0 | 0 | 10 | 60 | 57 | 95 | 13 | 31 | 0 | 3228 | 0 |
| SRR5030168 | 0 | 14 | 396 | 0 | 11 | 0 | 0 | 3072 | 1061 | 1694 | 52 | 2043 | 0 | 95 | 301 | 206 | 14 | 10 | 0 | 1900 | 602 | 3825 | 0 | 0 | 13 | 0 | 0 | 0 | 339 | 1429 | 1811 | 156 | 523 | 1259 | 28 | 3467 | 0 |
| SRR5030169 | 0 | 40 | 201 | 0 | 0 | 10 | 0 | 3132 | 1062 | 1380 | 1987 | 4967 | 0 | 166 | 179 | 166 | 0 | 0 | 0 | 1228 | 559 | 3663 | 0 | 0 | 9815 | 0 | 0 | 0 | 241 | 1574 | 1414 | 70 | 441 | 752 | 20 | 2953 | 0 |
| SRR5030170 | 0 | 40 | 21 | 0 | 12 | 0 | 0 | 3152 | 918 | 1622 | 826 | 1790 | 0 | 199 | 199 | 214 | 0 | 0 | 0 | 1423 | 669 | 4275 | 0 | 0 | 11115 | 0 | 0 | 0 | 148 | 67 | 109 | 81 | 25 | 69 | 27 | 3213 | 0 |
| SRR5030171 | 0 | 58 | 283 | 0 | 11 | 19 | 0 | 2843 | 1207 | 1475 | 2240 | 5593 | 0 | 94 | 80 | 188 | 0 | 0 | 0 | 1475 | 708 | 4116 | 0 | 0 | 10516 | 12 | 0 | 0 | 156 | 1461 | 1481 | 88 | 409 | 838 | 0 | 2951 | 0 |
| SRR5030172 | 0 | 28 | 13 | 0 | 0 | 14 | 0 | 3052 | 1038 | 1720 | 36 | 1875 | 14 | 201 | 279 | 341 | 0 | 11 | 0 | 1828 | 534 | 3900 | 0 | 0 | 11 | 0 | 0 | 0 | 194 | 69 | 74 | 111 | 29 | 59 | 42 | 3300 | 0 |
| SRR5030173 | 0 | 20 | 178 | 0 | 0 | 0 | 0 | 3492 | 955 | 1651 | 2346 | 5511 | 0 | 164 | 68 | 82 | 0 | 0 | 0 | 1351 | 547 | 3700 | 0 | 0 | 9970 | 11 | 0 | 0 | 286 | 1450 | 1500 | 79 | 474 | 839 | 20 | 3042 | 11 |
| SRR5030174 | 0 | 0 | 0 | 0 | 0 | 0 | 17 | 1266 | 127 | 439 | 22 | 201 | 0 | 134 | 134 | 134 | 0 | 20 | 0 | 6190 | 768 | 4174 | 0 | 0 | 10889 | 0 | 21 | 0 | 0 | 137 | 137 | 18 | 59 | 174 | 0 | 70369 | 0 |
| SRR5030175 | 0 | 0 | 0 | 0 | 0 | 0 | 26 | 1226 | 121 | 534 | 44 | 692 | 0 | 134 | 85 | 24 | 0 | 20 | 0 | 5838 | 764 | 3759 | 0 | 0 | 9612 | 0 | 30 | 0 | 0 | 45 | 38 | 17 | 15 | 31 | 0 | 55295 | 0 |
| SRR5030176 | 0 | 0 | 0 | 0 | 0 | 13 | 0 | 1185 | 103 | 469 | 135 | 151 | 0 | 175 | 127 | 159 | 0 | 16 | 0 | 6530 | 734 | 4104 | 0 | 0 | 10875 | 0 | 10 | 0 | 0 | 47 | 47 | 0 | 21 | 51 | 0 | 71468 | 0 |
| SRR5030177 | 0 | 0 | 0 | 0 | 0 | 0 | 22 | 1419 | 101 | 494 | 431 | 811 | 0 | 177 | 51 | 13 | 12 | 14 | 0 | 5701 | 637 | 3455 | 0 | 0 | 9301 | 0 | 101 | 0 | 0 | 169 | 142 | 0 | 81 | 110 | 0 | 57141 | 0 |
| SRR5030178 | 0 | 0 | 0 | 0 | 0 | 0 | 0 | 1186 | 62 | 470 | 20 | 300 | 0 | 77 | 92 | 62 | 0 | 13 | 0 | 6182 | 735 | 4136 | 0 | 0 | 10788 | 0 | 11 | 0 | 0 | 49 | 36 | 20 | 21 | 53 | 0 | 70648 | 0 |
| SRR5030179 | 0 | 0 | 0 | 0 | 0 | 19 | 0 | 1537 | 74 | 508 | 53 | 1078 | 0 | 161 | 25 | 87 | 19 | 16 | 0 | 5874 | 788 | 3804 | 0 | 0 | 9827 | 0 | 27 | 0 | 12 | 44 | 33 | 14 | 12 | 33 | 0 | 57121 | 0 |
| SRR5030180 | 0 | 0 | 0 | 0 | 0 | 0 | 0 | 1272 | 147 | 527 | 93 | 209 | 0 | 124 | 140 | 101 | 0 | 0 | 0 | 6475 | 656 | 4171 | 0 | 13 | 10664 | 0 | 22 | 0 | 0 | 49 | 30 | 11 | 25 | 63 | 0 | 70830 | 0 |
| SRR5030181 | 0 | 0 | 0 | 0 | 0 | 0 | 22 | 1243 | 63 | 477 | 565 | 829 | 0 | 138 | 25 | 75 | 0 | 12 | 0 | 5939 | 710 | 3580 | 0 | 0 | 9461 | 0 | 19 | 0 | 0 | 40 | 39 | 0 | 13 | 28 | 0 | 56112 | 0 |
| SRR5030182 | 0 | 0 | 165 | 0 | 0 | 0 | 0 | 1586 | 659 | 2040 | 1751 | 3791 | 0 | 62 | 41 | 21 | 0 | 16 | 0 | 1133 | 1420 | 7332 | 0 | 0 | 12 | 0 | 0 | 0 | 96 | 851 | 920 | 27 | 316 | 851 | 27 | 2596 | 0 |
| SRR5030183 | 12 | 23 | 127 | 0 | 0 | 23 | 0 | 1824 | 471 | 1667 | 218 | 11687 | 0 | 39 | 20 | 39 | 0 | 0 | 0 | 1176 | 1393 | 6662 | 0 | 0 | 11700 | 0 | 12 | 0 | 104 | 1025 | 1359 | 58 | 299 | 507 | 0 | 2275 | 0 |
| SRR5030184 | 0 | 15 | 224 | 0 | 0 | 0 | 0 | 1371 | 348 | 1937 | 93 | 4004 | 0 | 65 | 22 | 65 | 0 | 21 | 12 | 1175 | 1145 | 6303 | 0 | 0 | 11851 | 0 | 0 | 0 | 104 | 760 | 1013 | 134 | 283 | 864 | 30 | 2633 | 0 |
| SRR5030185 | 0 | 0 | 131 | 0 | 0 | 36 | 0 | 2141 | 572 | 1971 | 4960 | 12570 | 0 | 42 | 0 | 64 | 0 | 19 | 0 | 784 | 1121 | 5816 | 0 | 0 | 10732 | 0 | 0 | 24 | 143 | 968 | 1123 | 60 | 323 | 609 | 36 | 2586 | 0 |
| SRR5030186 | 0 | 28 | 198 | 0 | 0 | 0 | 0 | 1835 | 536 | 2165 | 78 | 4556 | 0 | 62 | 21 | 41 | 0 | 16 | 0 | 1051 | 1397 | 7369 | 0 | 0 | 15 | 0 | 0 | 0 | 128 | 879 | 964 | 85 | 340 | 638 | 14 | 2660 | 0 |
| SRR5030187 | 0 | 35 | 221 | 0 | 0 | 0 | 12 | 2117 | 699 | 2017 | 201 | 12062 | 0 | 60 | 0 | 0 | 12 | 19 | 0 | 1178 | 1434 | 6668 | 0 | 0 | 11647 | 0 | 0 | 0 | 152 | 921 | 1002 | 58 | 291 | 489 | 0 | 2556 | 0 |
| SRR5030188 | 0 | 29 | 189 | 0 | 0 | 0 | 0 | 2020 | 587 | 2151 | 1499 | 4432 | 0 | 65 | 43 | 0 | 0 | 23 | 0 | 1086 | 1121 | 6599 | 0 | 13 | 11 | 0 | 0 | 0 | 116 | 29 | 51 | 87 | 16 | 43 | 58 | 2911 | 0 |
| SRR5030189 | 0 | 24 | 84 | 0 | 0 | 0 | 0 | 1910 | 651 | 1826 | 4638 | 11270 | 0 | 21 | 21 | 0 | 0 | 17 | 0 | 1343 | 1088 | 5711 | 0 | 0 | 10792 | 0 | 0 | 36 | 167 | 27 | 24 | 60 | 13 | 17 | 12 | 2414 | 0 |
| SRR5030190 | 0 | 0 | 19 | 0 | 0 | 0 | 0 | 1232 | 188 | 1241 | 103 | 1542 | 0 | 66 | 113 | 85 | 0 | 16 | 19 | 1589 | 1240 | 6351 | 0 | 0 | 11 | 0 | 19 | 0 | 0 | 56 | 91 | 0 | 54 | 133 | 0 | 7756 | 0 |
| SRR5030191 | 0 | 18 | 0 | 0 | 13 | 0 | 0 | 1395 | 184 | 1065 | 38 | 2313 | 0 | 110 | 73 | 37 | 35 | 22 | 18 | 1469 | 1282 | 6616 | 0 | 0 | 19998 | 0 | 27 | 0 | 0 | 21 | 11 | 0 | 18 | 53 | 0 | 6022 | 0 |
| SRR5030192 | 0 | 0 | 20 | 0 | 0 | 0 | 0 | 1324 | 232 | 1283 | 89 | 1819 | 0 | 101 | 182 | 131 | 0 | 0 | 20 | 2092 | 1122 | 6014 | 0 | 0 | 19083 | 0 | 45 | 0 | 0 | 77 | 77 | 15 | 27 | 161 | 0 | 7437 | 0 |
| SRR5030193 | 0 | 18 | 0 | 0 | 0 | 0 | 0 | 1427 | 113 | 1352 | 38 | 1840 | 0 | 75 | 38 | 38 | 0 | 23 | 18 | 1803 | 1133 | 5950 | 0 | 14 | 19537 | 0 | 28 | 0 | 0 | 64 | 183 | 18 | 28 | 128 | 0 | 7249 | 0 |
| SRR5030194 | 0 | 0 | 12 | 0 | 0 | 0 | 0 | 1231 | 106 | 1135 | 91 | 1712 | 0 | 154 | 125 | 115 | 0 | 16 | 21 | 1769 | 1283 | 6274 | 0 | 0 | 19034 | 0 | 17 | 0 | 0 | 41 | 98 | 0 | 45 | 122 | 0 | 7799 | 0 |
| SRR5030196 | 0 | 0 | 17 | 0 | 0 | 0 | 0 | 1202 | 187 | 1350 | 109 | 1676 | 0 | 108 | 138 | 138 | 19 | 22 | 22 | 1754 | 1144 | 6044 | 0 | 0 | 18820 | 0 | 32 | 0 | 0 | 16 | 14 | 0 | 36 | 31 | 0 | 7500 | 0 |
| SRR5030197 | 18 | 0 | 0 | 0 | 0 | 0 | 0 | 1500 | 262 | 1312 | 1237 | 2587 | 0 | 37 | 37 | 75 | 0 | 33 | 19 | 1762 | 1209 | 6073 | 0 | 0 | 19894 | 0 | 55 | 0 | 0 | 92 | 147 | 0 | 46 | 119 | 0 | 7498 | 0 |
| SRR5030198 | 0 | 0 | 14 | 0 | 0 | 0 | 0 | 2248 | 348 | 2502 | 2168 | 4068 | 0 | 80 | 94 | 134 | 0 | 24 | 14 | 3064 | 1098 | 6316 | 0 | 20 | 11 | 19 | 200 | 0 | 0 | 291 | 301 | 64 | 132 | 455 | 0 | 25825 | 0 |
| SRR5030199 | 0 | 0 | 22 | 0 | 0 | 0 | 0 | 2738 | 418 | 1717 | 132 | 5569 | 0 | 209 | 70 | 93 | 0 | 33 | 13 | 2831 | 1048 | 6696 | 0 | 15 | 18076 | 31 | 225 | 0 | 0 | 502 | 385 | 29 | 102 | 196 | 0 | 20628 | 0 |
| SRR5030200 | 0 | 0 | 15 | 0 | 0 | 0 | 0 | 2203 | 358 | 2132 | 2518 | 4664 | 0 | 286 | 72 | 72 | 0 | 30 | 16 | 3462 | 920 | 6234 | 0 | 14 | 17157 | 20 | 27 | 0 | 34 | 49 | 38 | 39 | 13 | 39 | 34 | 27740 | 0 |
| SRR5030201 | 0 | 0 | 0 | 0 | 0 | 0 | 0 | 2878 | 411 | 2346 | 120 | 5466 | 0 | 266 | 73 | 97 | 0 | 17 | 0 | 2854 | 999 | 6306 | 0 | 25 | 17169 | 21 | 26 | 0 | 23 | 48 | 31 | 46 | 16 | 29 | 0 | 21500 | 0 |
| SRR5030202 | 0 | 0 | 42 | 0 | 0 | 0 | 0 | 2339 | 316 | 2394 | 2325 | 4127 | 0 | 234 | 96 | 124 | 0 | 19 | 11 | 2930 | 1043 | 6564 | 0 | 24 | 16 | 17 | 19 | 0 | 14 | 47 | 40 | 47 | 15 | 44 | 33 | 24638 | 0 |
| SRR5030203 | 0 | 0 | 15 | 0 | 11 | 0 | 0 | 2615 | 315 | 1816 | 3777 | 6173 | 0 | 242 | 0 | 97 | 0 | 17 | 13 | 3147 | 994 | 6841 | 0 | 21 | 18469 | 21 | 26 | 0 | 52 | 44 | 38 | 30 | 17 | 33 | 0 | 21546 | 0 |
| SRR5030204 | 0 | 0 | 14 | 0 | 0 | 0 | 19 | 2455 | 435 | 1992 | 2665 | 4377 | 0 | 140 | 126 | 112 | 0 | 18 | 13 | 3170 | 1026 | 6229 | 0 | 14 | 17192 | 25 | 12 | 0 | 24 | 52 | 40 | 47 | 15 | 43 | 0 | 26119 | 0 |
| SRR5030205 | 0 | 0 | 38 | 0 | 0 | 0 | 15 | 2379 | 294 | 1913 | 3237 | 5665 | 0 | 172 | 25 | 98 | 0 | 34 | 13 | 3188 | 927 | 6276 | 0 | 17 | 17397 | 17 | 197 | 23 | 45 | 371 | 258 | 23 | 76 | 212 | 0 | 23517 | 0 |
| SRR5030206 | 0 | 12 | 62 | 0 | 0 | 0 | 0 | 1811 | 539 | 1175 | 211 | 4866 | 0 | 41 | 55 | 235 | 0 | 18 | 0 | 4825 | 1405 | 4628 | 0 | 0 | 14 | 0 | 125 | 0 | 87 | 811 | 225 | 44 | 156 | 480 | 0 | 36953 | 0 |
| SRR5030207 | 0 | 0 | 79 | 0 | 12 | 0 | 0 | 3444 | 574 | 1148 | 50 | 6410 | 0 | 96 | 0 | 287 | 0 | 25 | 21 | 5453 | 1454 | 4667 | 0 | 0 | 16997 | 0 | 79 | 0 | 40 | 75 | 18 | 79 | 10 | 38 | 0 | 36545 | 0 |
| SRR5030208 | 0 | 0 | 47 | 0 | 0 | 0 | 0 | 1387 | 693 | 1146 | 2743 | 4597 | 0 | 15 | 181 | 181 | 0 | 11 | 14 | 5080 | 1301 | 4355 | 0 | 0 | 16243 | 0 | 10 | 0 | 113 | 59 | 17 | 47 | 126 | 20 | 13 | 40562 | 0 |
| SRR5030209 | 0 | 0 | 0 | 0 | 13 | 0 | 0 | 2133 | 508 | 508 | 3656 | 7109 | 0 | 0 | 102 | 305 | 0 | 17 | 26 | 5484 | 1403 | 4534 | 0 | 13 | 16267 | 0 | 82 | 0 | 16 | 37 | 24 | 0 | 21 | 21 | 0 | 36966 | 0 |
| SRR5030210 | 0 | 13 | 45 | 0 | 0 | 0 | 0 | 1627 | 538 | 1104 | 190 | 4557 | 13 | 0 | 42 | 325 | 0 | 22 | 0 | 4882 | 1393 | 4530 | 0 | 0 | 12 | 0 | 134 | 0 | 77 | 807 | 288 | 32 | 173 | 403 | 0 | 37247 | 0 |
| SRR5030211 | 0 | 0 | 41 | 0 | 0 | 0 | 0 | 1254 | 386 | 1061 | 38 | 5885 | 0 | 0 | 96 | 96 | 0 | 25 | 17 | 4438 | 1429 | 4431 | 0 | 0 | 16930 | 0 | 10 | 0 | 13 | 48 | 15 | 0 | 10 | 20 | 0 | 38877 | 0 |

### Supp Table S8.4.Columns_DI_to_ES

| | | | | | | | | | | | | | | | | | | | | | | | | | | | | | | | | | | | | | |
|---|---|---|---|---|---|---|---|---|---|---|---|---|---|---|---|---|---|---|---|---|---|---|---|---|---|---|---|---|---|---|---|---|---|---|---|---|---|
| SRR5030212 | 0 | 13 | 39 | 0 | 13 | 0 | 0 | 1647 | 659 | 859 | 185 | 4396 | 13 | 29 | 57 | 358 | 0 | 16 | 0 | 4983 | 1254 | 4373 | 0 | 0 | 16161 | 0 | 97 | 0 | 65 | 55 | 20 | 32 | 148 | 26 | 13 | 38807 | 0 |
| SRR5030213 | 0 | 0 | 83 | 0 | 0 | 0 | 0 | 1902 | 801 | 801 | 3505 | 6909 | 42 | 100 | 0 | 100 | 0 | 17 | 0 | 4506 | 1423 | 4520 | 0 | 0 | 15832 | 0 | 42 | 0 | 125 | 499 | 457 | 0 | 208 | 291 | 0 | 36347 | 0 |
| SRR5030214 | 0 | 0 | 47 | 0 | 13 | 0 | 0 | 883 | 612 | 1936 | 106 | 4598 | 0 | 147 | 170 | 260 | 0 | 89 | 14 | 3873 | 1321 | 5635 | 0 | 0 | 11462 | 0 | 29 | 0 | 83 | 33 | 20 | 14 | 58 | 20 | 0 | 22003 | 0 |
| SRR5030216 | 0 | 0 | 81 | 0 | 13 | 0 | 0 | 968 | 380 | 1838 | 85 | 4264 | 0 | 184 | 196 | 282 | 0 | 85 | 18 | 4215 | 1148 | 5388 | 0 | 0 | 11360 | 0 | 42 | 0 | 85 | 39 | 18 | 35 | 54 | 21 | 0 | 21861 | 0 |
| SRR5030217 | 0 | 0 | 12 | 0 | 0 | 74 | 0 | 236 | 354 | 2358 | 1081 | 5542 | 0 | 118 | 0 | 118 | 0 | 100 | 31 | 4599 | 1266 | 5449 | 0 | 23 | 11755 | 0 | 37 | 0 | 16 | 40 | 16 | 0 | 12 | 16 | 0 | 18514 | 0 |
| SRR5030218 | 0 | 0 | 52 | 0 | 12 | 0 | 0 | 799 | 465 | 1943 | 1228 | 4482 | 0 | 155 | 119 | 238 | 0 | 94 | 15 | 3707 | 1372 | 5659 | 0 | 0 | 11424 | 0 | 22 | 0 | 77 | 33 | 21 | 22 | 66 | 21 | 11 | 21812 | 0 |
| SRR5030220 | 0 | 0 | 67 | 0 | 13 | 0 | 0 | 778 | 474 | 2213 | 1155 | 4292 | 0 | 146 | 61 | 316 | 0 | 93 | 10 | 3793 | 1239 | 5316 | 0 | 0 | 11452 | 0 | 15 | 0 | 52 | 303 | 198 | 34 | 67 | 153 | 15 | 21277 | 0 |
| SRR5030221 | 37 | 0 | 16 | 0 | 0 | 0 | 0 | 1294 | 353 | 1764 | 1073 | 4233 | 0 | 353 | 0 | 118 | 0 | 123 | 0 | 3645 | 1235 | 5671 | 0 | 0 | 11019 | 0 | 37 | 0 | 20 | 32 | 12 | 0 | 74 | 74 | 0 | 19050 | 0 |
| SRR5030222 | 0 | 0 | 0 | 0 | 0 | 0 | 0 | 1025 | 150 | 812 | 70 | 2819 | 0 | 0 | 43 | 0 | 0 | 18 | 0 | 983 | 1109 | 5384 | 0 | 0 | 13425 | 0 | 0 | 0 | 197 | 737 | 426 | 33 | 328 | 573 | 98 | 2542 | 15 |
| SRR5030223 | 0 | 0 | 13 | 0 | 0 | 0 | 0 | 1078 | 136 | 581 | 2481 | 5014 | 0 | 15 | 0 | 0 | 0 | 14 | 17 | 1093 | 1250 | 5612 | 0 | 0 | 13783 | 0 | 0 | 0 | 11 | 66 | 44 | 17 | 15 | 26 | 0 | 2367 | 0 |
| SRR5030224 | 0 | 0 | 18 | 0 | 0 | 0 | 18 | 663 | 160 | 846 | 61 | 2971 | 0 | 0 | 23 | 0 | 0 | 0 | 23 | 709 | 1013 | 5524 | 0 | 0 | 13287 | 0 | 0 | 18 | 105 | 66 | 44 | 35 | 28 | 64 | 0 | 2857 | 0 |
| SRR5030225 | 0 | 0 | 0 | 0 | 0 | 13 | 0 | 946 | 100 | 569 | 335 | 4224 | 0 | 15 | 0 | 0 | 0 | 21 | 16 | 862 | 1087 | 5277 | 0 | 0 | 13239 | 0 | 13 | 0 | 153 | 776 | 495 | 53 | 189 | 298 | 0 | 2716 | 0 |
| SRR5030226 | 0 | 0 | 17 | 0 | 0 | 0 | 0 | 892 | 178 | 691 | 1561 | 2676 | 0 | 0 | 0 | 0 | 0 | 13 | 16 | 1026 | 1145 | 5373 | 0 | 0 | 18 | 0 | 0 | 0 | 67 | 917 | 483 | 33 | 317 | 450 | 0 | 2631 | 0 |
| SRR5030227 | 0 | 0 | 0 | 0 | 0 | 17 | 0 | 1101 | 177 | 593 | 2570 | 4948 | 0 | 0 | 0 | 0 | 0 | 20 | 16 | 862 | 1165 | 5518 | 0 | 0 | 13816 | 0 | 0 | 22 | 95 | 68 | 40 | 35 | 13 | 23 | 0 | 2617 | 0 |
| SRR5030229 | 0 | 0 | 0 | 0 | 0 | 18 | 0 | 1104 | 149 | 556 | 2639 | 4839 | 0 | 23 | 0 | 0 | 0 | 16 | 14 | 955 | 1043 | 5307 | 0 | 0 | 13238 | 0 | 0 | 0 | 101 | 873 | 491 | 18 | 149 | 268 | 0 | 2725 | 0 |
| SRR5030230 | 0 | 0 | 0 | 0 | 0 | 0 | 0 | 1898 | 500 | 3746 | 749 | 899 | 0 | 0 | 150 | 300 | 0 | 0 | 11 | 7593 | 765 | 5859 | 0 | 15 | 13 | 0 | 34 | 0 | 34 | 1761 | 1355 | 34 | 373 | 1457 | 102 | 87417 | 95 |
| SRR5030231 | 0 | 17 | 25 | 0 | 0 | 21 | 0 | 1815 | 344 | 2488 | 3145 | 4445 | 0 | 75 | 0 | 67 | 0 | 14 | 0 | 6522 | 771 | 4986 | 0 | 0 | 10432 | 0 | 38 | 0 | 131 | 80 | 36 | 38 | 220 | 25 | 17 | 60465 | 79 |
| SRR5030232 | 36 | 0 | 0 | 0 | 0 | 0 | 0 | 1579 | 105 | 3369 | 16 | 1211 | 0 | 53 | 211 | 263 | 0 | 0 | 0 | 8843 | 588 | 5357 | 0 | 0 | 12 | 0 | 0 | 0 | 73 | 129 | 61 | 109 | 21 | 91 | 0 | 84167 | 89 |
| SRR5030233 | 0 | 13 | 31 | 0 | 0 | 0 | 0 | 1572 | 410 | 2400 | 386 | 4481 | 0 | 53 | 23 | 46 | 0 | 19 | 0 | 6714 | 703 | 4695 | 0 | 0 | 9839 | 0 | 31 | 0 | 113 | 1831 | 907 | 52 | 292 | 645 | 22 | 60629 | 70 |
| SRR5030234 | 0 | 0 | 35 | 0 | 0 | 0 | 0 | 2165 | 619 | 2938 | 619 | 1237 | 0 | 0 | 52 | 258 | 0 | 22 | 0 | 7836 | 710 | 5804 | 0 | 11 | 12144 | 0 | 0 | 35 | 0 | 1381 | 932 | 104 | 380 | 1346 | 69 | 87380 | 78 |
| SRR5030235 | 0 | 17 | 13 | 0 | 0 | 13 | 0 | 1729 | 442 | 2430 | 3055 | 4762 | 0 | 61 | 15 | 61 | 0 | 13 | 0 | 7055 | 755 | 4967 | 0 | 0 | 10443 | 0 | 17 | 0 | 133 | 83 | 44 | 86 | 12 | 31 | 13 | 60818 | 73 |
| SRR5030236 | 0 | 0 | 0 | 0 | 11 | 0 | 0 | 1682 | 210 | 3416 | 578 | 1419 | 0 | 0 | 210 | 105 | 0 | 0 | 0 | 7779 | 595 | 5597 | 0 | 0 | 11 | 0 | 0 | 0 | 35 | 138 | 55 | 70 | 30 | 83 | 70 | 87354 | 105 |
| SRR5030237 | 0 | 17 | 35 | 0 | 0 | 0 | 0 | 1582 | 390 | 2522 | 404 | 4892 | 0 | 0 | 15 | 46 | 0 | 15 | 0 | 6765 | 695 | 4709 | 0 | 0 | 9944 | 0 | 31 | 0 | 109 | 1873 | 974 | 70 | 327 | 572 | 0 | 60224 | 86 |
| SRR5030238 | 0 | 0 | 0 | 0 | 0 | 0 | 0 | 1911 | 221 | 598 | 129 | 4264 | 0 | 65 | 78 | 91 | 0 | 47 | 21 | 22114 | 1748 | 9491 | 0 | 11 | 16 | 0 | 36 | 0 | 20 | 115 | 49 | 100 | 23 | 65 | 36 | 88131 | 0 |
| SRR5030239 | 0 | 0 | 25 | 0 | 0 | 0 | 12 | 2030 | 257 | 327 | 83 | 5692 | 0 | 23 | 23 | 23 | 0 | 33 | 23 | 22348 | 1723 | 9525 | 0 | 15 | 23629 | 0 | 74 | 12 | 15 | 110 | 51 | 74 | 20 | 42 | 12 | 75513 | 0 |
| SRR5030240 | 0 | 15 | 15 | 0 | 0 | 0 | 31 | 1680 | 168 | 714 | 130 | 4283 | 0 | 56 | 84 | 70 | 0 | 34 | 21 | 23991 | 1526 | 8836 | 0 | 0 | 11 | 0 | 100 | 0 | 200 | 1779 | 570 | 62 | 377 | 832 | 23 | 90492 | 0 |
| SRR5030241 | 0 | 26 | 0 | 0 | 0 | 0 | 0 | 1178 | 221 | 638 | 109 | 5718 | 0 | 49 | 98 | 25 | 0 | 53 | 16 | 22306 | 1467 | 8923 | 0 | 0 | 22874 | 0 | 52 | 0 | 361 | 2124 | 708 | 39 | 245 | 451 | 13 | 78892 | 0 |
| SRR5030242 | 0 | 22 | 29 | 0 | 0 | 0 | 0 | 1599 | 203 | 542 | 1626 | 4648 | 0 | 54 | 54 | 68 | 0 | 41 | 17 | 22753 | 1688 | 9523 | 0 | 0 | 22692 | 0 | 44 | 0 | 220 | 1589 | 784 | 29 | 344 | 857 | 22 | 90308 | 0 |
| SRR5030243 | 0 | 13 | 13 | 0 | 0 | 13 | 0 | 1948 | 192 | 625 | 109 | 6374 | 0 | 48 | 48 | 0 | 0 | 49 | 20 | 21911 | 1711 | 9896 | 0 | 0 | 23934 | 0 | 88 | 0 | 16 | 121 | 59 | 51 | 17 | 32 | 51 | 78457 | 0 |
| SRR5030244 | 0 | 0 | 15 | 0 | 0 | 0 | 0 | 1930 | 96 | 725 | 1670 | 4941 | 0 | 55 | 82 | 109 | 0 | 36 | 18 | 22597 | 1517 | 9038 | 0 | 0 | 22189 | 0 | 98 | 23 | 128 | 1650 | 578 | 30 | 360 | 765 | 45 | 90099 | 0 |
| SRR5030245 | 0 | 0 | 0 | 0 | 0 | 13 | 26 | 1966 | 170 | 461 | 117 | 5752 | 0 | 49 | 0 | 24 | 0 | 50 | 14 | 20944 | 1539 | 8827 | 0 | 0 | 22597 | 0 | 52 | 0 | 258 | 2035 | 683 | 52 | 296 | 644 | 13 | 76035 | 0 |
| SRR5030246 | 11 | 0 | 0 | 0 | 20 | 0 | 0 | 1519 | 250 | 3806 | 1068 | 1920 | 0 | 117 | 234 | 401 | 0 | 133 | 0 | 2220 | 1888 | 6216 | 0 | 23 | 11587 | 0 | 0 | 0 | 86 | 706 | 428 | 139 | 214 | 984 | 32 | 20231 | 0 |
| SRR5030247 | 0 | 35 | 0 | 0 | 28 | 35 | 0 | 2104 | 537 | 3250 | 194 | 5977 | 0 | 232 | 58 | 131 | 0 | 120 | 13 | 1915 | 1905 | 5542 | 0 | 0 | 11182 | 0 | 0 | 0 | 124 | 52 | 28 | 62 | 117 | 26 | 21 | 17396 | 0 |
| SRR5030248 | 0 | 0 | 0 | 0 | 21 | 11 | 11 | 2020 | 372 | 4200 | 868 | 2003 | 0 | 124 | 337 | 549 | 0 | 127 | 0 | 2144 | 1390 | 5155 | 0 | 0 | 11196 | 0 | 0 | 0 | 114 | 45 | 31 | 46 | 125 | 72 | 46 | 20274 | 0 |
| SRR5030249 | 0 | 0 | 22 | 0 | 31 | 22 | 0 | 2225 | 686 | 3489 | 153 | 6004 | 0 | 137 | 30 | 46 | 0 | 114 | 0 | 1966 | 1503 | 4906 | 0 | 12 | 10376 | 0 | 0 | 0 | 129 | 884 | 374 | 50 | 101 | 374 | 0 | 17706 | 0 |
| SRR5030250 | 0 | 11 | 22 | 0 | 39 | 11 | 0 | 1969 | 252 | 3870 | 42 | 1985 | 0 | 135 | 202 | 353 | 0 | 109 | 0 | 1901 | 1771 | 6267 | 0 | 0 | 13 | 0 | 0 | 0 | 108 | 530 | 660 | 54 | 249 | 887 | 43 | 20359 | 0 |
| SRR5030251 | 14 | 14 | 35 | 0 | 41 | 49 | 0 | 2286 | 371 | 3088 | 2657 | 6057 | 0 | 148 | 59 | 45 | 0 | 126 | 12 | 2153 | 2014 | 5640 | 0 | 0 | 11247 | 0 | 0 | 0 | 168 | 925 | 400 | 35 | 154 | 477 | 0 | 16167 | 0 |
| SRR5030252 | 0 | 22 | 11 | 0 | 37 | 33 | 0 | 2301 | 279 | 3451 | 767 | 1900 | 0 | 192 | 279 | 523 | 0 | 108 | 16 | 2144 | 1466 | 5557 | 0 | 0 | 11335 | 0 | 0 | 0 | 56 | 56 | 38 | 78 | 15 | 65 | 33 | 21246 | 0 |
| SRR5030253 | 0 | 22 | 0 | 0 | 32 | 0 | 0 | 2488 | 427 | 3159 | 2900 | 6029 | 0 | 107 | 92 | 46 | 0 | 134 | 15 | 1832 | 1468 | 4860 | 0 | 0 | 10596 | 0 | 0 | 0 | 72 | 695 | 351 | 57 | 93 | 487 | 14 | 16926 | 0 |
| SRR5030254 | 0 | 0 | 0 | 0 | 0 | 0 | 0 | 1432 | 168 | 2667 | 9826 | 13601 | 0 | 14 | 0 | 14 | 0 | 0 | 0 | 814 | 1102 | 4504 | 0 | 0 | 8394 | 0 | 11 | 12 | 16 | 25 | 14 | 0 | 0 | 0 | 0 | 1165 | 116 |
| SRR5030255 | 0 | 0 | 0 | 0 | 0 | 0 | 0 | 1691 | 244 | 2694 | 618 | 13297 | 0 | 0 | 43 | 43 | 0 | 0 | 0 | 845 | 873 | 3753 | 0 | 0 | 8169 | 0 | 16 | 0 | 0 | 18 | 19 | 0 | 0 | 10 | 0 | 1404 | 105 |
| SRR5030256 | 0 | 0 | 0 | 0 | 0 | 0 | 11 | 1492 | 335 | 2510 | 653 | 14055 | 0 | 14 | 98 | 70 | 0 | 0 | 0 | 823 | 1095 | 4423 | 0 | 0 | 8269 | 0 | 0 | 0 | 24 | 20 | 11 | 0 | 0 | 11 | 0 | 1241 | 102 |
| SRR5030257 | 0 | 0 | 0 | 0 | 0 | 0 | 13 | 1334 | 170 | 2767 | 9537 | 13212 | 0 | 43 | 85 | 0 | 0 | 0 | 0 | 1022 | 906 | 3928 | 0 | 0 | 8131 | 0 | 15 | 0 | 10 | 14 | 21 | 0 | 0 | 0 | 0 | 1405 | 109 |
| SRR5030258 | 0 | 14 | 0 | 0 | 0 | 0 | 0 | 230 | 148 | 11471 | 301 | 7861 | 0 | 0 | 181 | 0 | 0 | 30 | 11 | 443 | 1256 | 7758 | 0 | 0 | 16020 | 59 | 0 | 48 | 24 | 48 | 45 | 0 | 17 | 41 | 0 | 919 | 0 |
| SRR5030259 | 0 | 0 | 0 | 0 | 0 | 0 | 0 | 155 | 86 | 11004 | 267 | 8748 | 0 | 17 | 69 | 121 | 0 | 25 | 0 | 654 | 956 | 6609 | 0 | 0 | 15675 | 62 | 0 | 40 | 18 | 43 | 18 | 0 | 0 | 29 | 0 | 999 | 0 |
| SRR5030260 | 0 | 17 | 0 | 0 | 0 | 0 | 0 | 153 | 119 | 11037 | 5127 | 7750 | 0 | 0 | 136 | 34 | 0 | 26 | 0 | 681 | 1173 | 7656 | 0 | 0 | 14 | 65 | 0 | 59 | 14 | 21 | 24 | 0 | 31 | 21 | 0 | 1039 | 0 |
| SRR5030261 | 0 | 17 | 0 | 0 | 0 | 0 | 0 | 239 | 120 | 11991 | 4800 | 8489 | 0 | 17 | 68 | 85 | 0 | 24 | 0 | 393 | 986 | 6783 | 0 | 0 | 11 | 60 | 0 | 35 | 21 | 28 | 52 | 0 | 14 | 45 | 0 | 1025 | 0 |
| SRR5030262 | 0 | 0 | 36 | 0 | 0 | 10 | 0 | 955 | 159 | 4975 | 10847 | 11512 | 10 | 58 | 0 | 14 | 0 | 0 | 14 | 116 | 3044 | 9165 | 0 | 0 | 13 | 0 | 0 | 102 | 51 | 13 | 12 | 31 | 76 | 10 | 0 | 651 | 0 |
| SRR5030263 | 0 | 0 | 37 | 0 | 0 | 11 | 0 | 1028 | 224 | 4664 | 10923 | 11877 | 26 | 89 | 0 | 0 | 0 | 0 | 27 | 134 | 2367 | 7532 | 0 | 0 | 13716 | 0 | 11 | 79 | 32 | 185 | 185 | 42 | 85 | 143 | 0 | 522 | 0 |
| SRR5030265 | 10 | 10 | 36 | 0 | 0 | 10 | 0 | 883 | 157 | 4515 | 752 | 11410 | 0 | 57 | 0 | 14 | 0 | 0 | 25 | 199 | 2392 | 7734 | 0 | 0 | 13656 | 0 | 15 | 108 | 46 | 17 | 11 | 26 | 57 | 12 | 0 | 769 | 0 |
| SRR5030266 | 0 | 38 | 38 | 0 | 0 | 0 | 0 | 54 | 54 | 968 | 12607 | 13226 | 0 | 376 | 54 | 108 | 15 | 0 | 38 | 484 | 1471 | 5825 | 0 | 0 | 11 | 0 | 0 | 91 | 30 | 83 | 83 | 15 | 0 | 38 | 0 | 860 | 87 |
| SRR5030267 | 16 | 55 | 0 | 0 | 0 | 0 | 0 | 28 | 112 | 811 | 13066 | 13122 | 0 | 392 | 56 | 84 | 0 | 0 | 53 | 476 | 1132 | 4804 | 0 | 0 | 10 | 0 | 0 | 118 | 24 | 63 | 110 | 0 | 55 | 31 | 0 | 811 | 81 |
| SRR5030268 | 0 | 31 | 23 | 0 | 0 | 0 | 0 | 160 | 27 | 453 | 454 | 13011 | 0 | 267 | 133 | 53 | 23 | 0 | 37 | 347 | 1378 | 5586 | 0 | 0 | 10008 | 0 | 0 | 69 | 39 | 85 | 85 | 0 | 15 | 46 | 0 | 933 | 91 |
| SRR5030270 | 0 | 0 | 46 | 0 | 0 | 20 | 0 | 373 | 11 | 9836 | 1419 | 13147 | 0 | 22 | 22 | 0 | 26 | 0 | 48 | 307 | 2527 | 9096 | 0 | 0 | 25 | 0 | 0 | 66 | 125 | 11 | 191 | 20 | 59 | 131 | 0 | 680 | 0 |
| SRR5030271 | 0 | 28 | 76 | 0 | 0 | 0 | 0 | 401 | 35 | 10415 | 16443 | 13164 | 0 | 12 | 24 | 0 | 0 | 0 | 45 | 330 | 2105 | 7866 | 0 | 0 | 20897 | 0 | 14 | 90 | 187 | 11 | 194 | 0 | 62 | 139 | 0 | 649 | 0 |
| SRR5030272 | 0 | 0 | 80 | 0 | 0 | 13 | 0 | 289 | 33 | 9944 | 1467 | 13169 | 0 | 33 | 0 | 0 | 13 | 0 | 45 | 323 | 2590 | 9109 | 0 | 0 | 14 | 0 | 0 | 27 | 154 | 13 | 201 | 0 | 74 | 167 | 0 | 690 | 0 |
| SRR5030273 | 0 | 20 | 27 | 0 | 0 | 13 | 0 | 348 | 112 | 9875 | 1417 | 12481 | 0 | 11 | 11 | 0 | 33 | 11 | 45 | 393 | 2143 | 8087 | 0 | 0 | 20774 | 0 | 0 | 47 | 234 | 214 | 234 | 0 | 87 | 147 | 0 | 472 | 0 |
| SRR5030274 | 0 | 0 | 33 | 0 | 0 | 0 | 13 | 490 | 70 | 4919 | 566 | 16001 | 0 | 0 | 105 | 70 | 13 | 0 | 10 | 228 | 1125 | 4004 | 0 | 0 | 6775 | 0 | 0 | 26 | 13 | 66 | 113 | 13 | 40 | 33 | 0 | 910 | 0 |
| SRR5030275 | 0 | 35 | 14 | 0 | 0 | 0 | 0 | 579 | 149 | 4894 | 10031 | 16121 | 0 | 0 | 131 | 37 | 0 | 0 | 13 | 187 | 900 | 3427 | 0 | 0 | 6609 | 0 | 21 | 28 | 0 | 28 | 118 | 0 | 14 | 21 | 0 | 841 | 0 |
| SRR5030276 | 0 | 14 | 0 | 0 | 0 | 0 | 0 | 473 | 146 | 5080 | 10761 | 16442 | 0 | 0 | 182 | 36 | 0 | 0 | 12 | 291 | 1142 | 3953 | 0 | 0 | 6730 | 0 | 14 | 0 | 27 | 47 | 95 | 14 | 27 | 41 | 0 | 838 | 0 |
| SRR5030277 | 0 | 0 | 20 | 0 | 11 | 0 | 0 | 480 | 107 | 4497 | 569 | 15854 | 0 | 0 | 231 | 36 | 0 | 0 | 11 | 444 | 919 | 3461 | 0 | 0 | 6694 | 0 | 27 | 34 | 14 | 68 | 81 | 0 | 14 | 41 | 0 | 835 | 0 |
| SRR5030278 | 0 | 0 | 17 | 0 | 21 | 0 | 0 | 627 | 74 | 2710 | 494 | 4019 | 0 | 0 | 83 | 28 | 0 | 0 | 42 | 230 | 1986 | 6723 | 0 | 0 | 14544 | 0 | 0 | 12 | 37 | 49 | 59 | 0 | 15 | 66 | 0 | 857 | 0 |
| SRR5030279 | 0 | 0 | 41 | 0 | 17 | 0 | 0 | 563 | 49 | 2686 | 4869 | 4049 | 0 | 0 | 49 | 0 | 0 | 0 | 53 | 237 | 1623 | 5801 | 0 | 0 | 18 | 0 | 0 | 15 | 38 | 59 | 54 | 31 | 38 | 92 | 0 | 1116 | 0 |
| SRR5030280 | 0 | 0 | 27 | 0 | 21 | 0 | 0 | 553 | 38 | 2583 | 465 | 4403 | 0 | 0 | 29 | 0 | 0 | 0 | 39 | 391 | 2048 | 6511 | 0 | 0 | 10 | 0 | 0 | 12 | 40 | 57 | 65 | 17 | 40 | 67 | 0 | 886 | 0 |
| SRR5030281 | 0 | 0 | 15 | 0 | 15 | 0 | 0 | 408 | 57 | 2706 | 484 | 4216 | 0 | 0 | 47 | 19 | 0 | 0 | 39 | 275 | 1630 | 5732 | 0 | 0 | 14067 | 0 | 0 | 12 | 47 | 52 | 42 | 22 | 20 | 75 | 0 | 912 | 0 |
| SRR5030282 | 0 | 21 | 10 | 0 | 0 | 0 | 0 | 587 | 129 | 348 | 2404 | 43955 | 0 | 10 | 0 | 0 | 0 | 0 | 11 | 149 | 876 | 5027 | 0 | 0 | 2450 | 0 | 0 | 0 | 42 | 452 | 192 | 16 | 0 | 36 | 0 | 697 | 0 |
| SRR5030283 | 0 | 16 | 0 | 0 | 0 | 0 | 0 | 561 | 97 | 281 | 24788 | 45227 | 0 | 32 | 0 | 0 | 0 | 0 | 18 | 151 | 710 | 4436 | 0 | 0 | 2377 | 0 | 0 | 0 | 87 | 451 | 103 | 0 | 27 | 33 | 0 | 615 | 0 |
| SRR5030284 | 11 | 11 | 0 | 0 | 0 | 0 | 0 | 595 | 136 | 334 | 24091 | 45876 | 0 | 21 | 0 | 0 | 0 | 0 | 12 | 136 | 851 | 4875 | 0 | 0 | 2367 | 0 | 16 | 0 | 32 | 12 | 127 | 0 | 21 | 42 | 0 | 699 | 0 |
| SRR5030285 | 11 | 21 | 0 | 0 | 0 | 0 | 0 | 523 | 147 | 314 | 24070 | 45094 | 0 | 10 | 0 | 0 | 0 | 0 | 14 | 178 | 726 | 4446 | 0 | 0 | 2397 | 0 | 11 | 0 | 69 | 290 | 132 | 0 | 11 | 53 | 0 | 816 | 0 |
| SRR5030287 | 0 | 35 | 0 | 0 | 0 | 0 | 0 | 1405 | 107 | 160 | 837 | 29815 | 0 | 0 | 0 | 0 | 0 | 0 | 18 | 213 | 1325 | 5562 | 0 | 0 | 10873 | 0 | 272 | 0 | 114 | 552 | 219 | 18 | 26 | 70 | 0 | 356 | 0 |
| SRR5030289 | 17 | 26 | 0 | 0 | 0 | 0 | 0 | 1351 | 191 | 208 | 14583 | 29704 | 0 | 0 | 0 | 17 | 0 | 0 | 18 | 191 | 1354 | 5531 | 0 | 0 | 10690 | 0 | 180 | 0 | 197 | 650 | 162 | 17 | 0 | 85 | 0 | 433 | 0 |
| SRR5030290 | 0 | 0 | 16 | 0 | 0 | 21 | 0 | 524 | 69 | 1294 | 15082 | 39422 | 0 | 62 | 0 | 0 | 0 | 15 | 19 | 678 | 1281 | 6027 | 0 | 43 | 4171 | 0 | 42 | 0 | 78 | 32 | 17 | 0 | 26 | 84 | 10 | 12424 | 0 |
| SRR5030291 | 0 | 22 | 27 | 0 | 0 | 0 | 0 | 673 | 63 | 1393 | 1979 | 38379 | 0 | 87 | 0 | 24 | 0 | 17 | 0 | 981 | 1030 | 5243 | 0 | 56 | 4099 | 0 | 22 | 0 | 87 | 32 | 20 | 11 | 43 | 87 | 0 | 12827 | 0 |
| SRR5030292 | 0 | 21 | 32 | 0 | 0 | 0 | 0 | 557 | 110 | 1294 | 16260 | 39737 | 0 | 47 | 0 | 0 | 0 | 10 | 0 | 1043 | 1239 | 5890 | 0 | 41 | 4058 | 0 | 16 | 0 | 59 | 458 | 277 | 16 | 48 | 101 | 0 | 12833 | 0 |
| SRR5030293 | 0 | 0 | 32 | 0 | 0 | 11 | 0 | 494 | 78 | 1254 | 16126 | 38881 | 0 | 63 | 0 | 0 | 0 | 0 | 11 | 901 | 1032 | 5350 | 0 | 77 | 4024 | 0 | 16 | 11 | 80 | 573 | 313 | 11 | 53 | 90 | 0 | 13250 | 0 |
| SRR5030294 | 0 | 27 | 33 | 0 | 0 | 0 | 0 | 331 | 88 | 1128 | 24885 | 51968 | 0 | 68 | 0 | 0 | 0 | 0 | 12 | 506 | 997 | 5085 | 0 | 56 | 3361 | 0 | 13 | 13 | 73 | 15 | 12 | 13 | 20 | 93 | 0 | 5282 | 0 |
| SRR5030295 | 14 | 42 | 49 | 0 | 0 | 0 | 0 | 300 | 100 | 1308 | 2457 | 50747 | 0 | 40 | 0 | 0 | 0 | 13 | 0 | 489 | 856 | 4462 | 0 | 41 | 3183 | 0 | 21 | 14 | 97 | 14 | 12 | 35 | 49 | 132 | 0 | 5573 | 0 |
| SRR5030296 | 0 | 27 | 47 | 0 | 0 | 0 | 0 | 305 | 138 | 1405 | 25533 | 52137 | 0 | 69 | 0 | 0 | 0 | 10 | 13 | 511 | 986 | 5011 | 0 | 50 | 3312 | 0 | 0 | 0 | 74 | 15 | 13 | 40 | 61 | 61 | 0 | 5661 | 0 |
| SRR5030297 | 0 | 20 | 34 | 0 | 0 | 0 | 0 | 367 | 60 | 1112 | 26006 | 50940 | 0 | 70 | 0 | 0 | 0 | 0 | 0 | 725 | 881 | 4451 | 0 | 50 | 3236 | 0 | 0 | 14 | 95 | 405 | 236 | 27 | 74 | 95 | 0 | 5978 | 0 |
| SRR5034616 | 12 | 0 | 0 | 0 | 0 | 0 | 37 | 352 | 176 | 1986 | 45 | 5229 | 0 | 251 | 0 | 50 | 0 | 0 | 0 | 1182 | 397 | 1949 | 0 | 206 | 6161 | 327 | 37 | 0 | 0 | 0 | 0 | 0 | 0 | 0 | 0 | 21569 | 0 |
| SRR5034617 | 0 | 0 | 0 | 0 | 0 | 0 | 0 | 709 | 150 | 1608 | 2388 | 12790 | 0 | 103 | 75 | 168 | 0 | 0 | 0 | 610 | 382 | 1630 | 0 | 195 | 5002 | 512 | 79 | 0 | 0 | 0 | 12 | 0 | 0 | 0 | 0 | 6641 | 0 |
| SRR5034618 | 0 | 0 | 0 | 0 | 0 | 0 | 0 | 292 | 99 | 1905 | 2955 | 15250 | 0 | 116 | 163 | 671 | 0 | 12 | 0 | 741 | 998 | 3769 | 0 | 339 | 8271 | 905 | 43 | 0 | 0 | 0 | 0 | 0 | 0 | 0 | 19 | 5776 | 0 |
| SRR5034619 | 226 | 15 | 0 | 0 | 0 | 0 | 0 | 506 | 80 | 2102 | 1064 | 5908 | 0 | 506 | 0 | 53 | 0 | 26 | 0 | 1543 | 622 | 2728 | 0 | 245 | 8427 | 495 | 36 | 0 | 0 | 0 | 0 | 0 | 0 | 0 | 0 | 18894 | 0 |
| SRR5034620 | 0 | 0 | 0 | 0 | 0 | 0 | 0 | 471 | 163 | 4452 | 105 | 13827 | 0 | 181 | 163 | 290 | 0 | 0 | 0 | 398 | 128 | 710 | 0 | 116 | 3326 | 196 | 25 | 0 | 0 | 0 | 18 | 0 | 0 | 0 | 12 | 6316 | 0 |
| SRR5034621 | 0 | 0 | 0 | 0 | 0 | 0 | 0 | 561 | 187 | 2294 | 406 | 12296 | 0 | 137 | 230 | 453 | 0 | 0 | 0 | 482 | 246 | 953 | 0 | 58 | 3576 | 312 | 66 | 0 | 0 | 16 | 13 | 0 | 0 | 0 | 41 | 4559 | 0 |
| SRR5034622 | 0 | 0 | 0 | 0 | 0 | 0 | 0 | 642 | 130 | 2641 | 1289 | 5327 | 0 | 162 | 110 | 368 | 0 | 26 | 0 | 1101 | 1140 | 3744 | 0 | 561 | 11047 | 875 | 29 | 0 | 0 | 0 | 11 | 0 | 0 | 0 | 17 | 6470 | 0 |
| SRR5034623 | 0 | 0 | 0 | 0 | 0 | 0 | 0 | 300 | 718 | 11201 | 46 | 111 | 0 | 30 | 20 | 29 | 0 | 0 | 0 | 14 | 117 | 244 | 0 | 60 | 6262 | 204 | 0 | 0 | 0 | 0 | 0 | 0 | 0 | 0 | 0 | 260 | 0 |
| SRR5034625 | 0 | 0 | 0 | 0 | 0 | 0 | 0 | 405 | 1015 | 11779 | 43 | 157 | 0 | 205 | 771 | 767 | 0 | 0 | 0 | 90 | 429 | 1356 | 21 | 368 | 8151 | 630 | 26 | 0 | 0 | 0 | 0 | 0 | 0 | 0 | 0 | 948 | 0 |
| SRR5034626 | 0 | 0 | 0 | 0 | 11 | 0 | 0 | 5284 | 9478 | 23704 | 17 | 38 | 0 | 44 | 80 | 89 | 0 | 0 | 0 | 54 | 253 | 433 | 35 | 174 | 4693 | 380 | 0 | 0 | 0 | 0 | 0 | 0 | 0 | 0 | 44 | 1195 | 0 |
| SRR5034627 | 0 | 0 | 0 | 0 | 0 | 0 | 0 | 154 | 406 | 2623 | 12 | 55 | 0 | 160 | 62 | 86 | 10 | 0 | 0 | 135 | 29 | 148 | 0 | 66 | 775 | 147 | 0 | 0 | 0 | 0 | 0 | 0 | 0 | 0 | 0 | 449 | 0 |
| SRR5034628 | 48 | 0 | 0 | 0 | 0 | 0 | 0 | 604 | 411 | 10342 | 110 | 357 | 0 | 384 | 1728 | 2880 | 0 | 0 | 0 | 165 | 89 | 369 | 0 | 216 | 2762 | 310 | 16 | 0 | 0 | 0 | 0 | 0 | 0 | 0 | 0 | 3347 | 0 |
| SRR5034629 | 0 | 0 | 0 | 0 | 0 | 22 | 0 | 876 | 794 | 13382 | 46 | 137 | 0 | 465 | 265 | 164 | 22 | 0 | 0 | 0 | 105 | 231 | 13 | 257 | 5862 | 298 | 44 | 0 | 0 | 0 | 0 | 0 | 0 | 0 | 11 | 0 | 0 |
| SRR5034630 | 12 | 0 | 0 | 0 | 0 | 12 | 24 | 1103 | 874 | 14634 | 125 | 271 | 0 | 62 | 125 | 291 | 24 | 0 | 0 | 0 | 303 | 1273 | 60 | 711 | 7794 | 520 | 47 | 0 | 0 | 0 | 35 | 0 | 0 | 0 | 12 | 104 | 0 |
| SRR5034632 | 11 | 0 | 0 | 0 | 0 | 0 | 0 | 612 | 3503 | 11752 | 102 | 340 | 0 | 476 | 85 | 238 | 0 | 0 | 0 | 221 | 270 | 1024 | 60 | 873 | 6811 | 352 | 42 | 0 | 0 | 0 | 0 | 0 | 0 | 0 | 0 | 1412 | 0 |
| SRR5034634 | 0 | 0 | 0 | 0 | 0 | 0 | 0 | 1644 | 4632 | 6541 | 50 | 149 | 0 | 232 | 17 | 50 | 0 | 0 | 0 | 66 | 317 | 1396 | 55 | 739 | 8905 | 384 | 37 | 0 | 0 | 0 | 0 | 0 | 0 | 0 | 0 | 797 | 0 |
| SRR5034635 | 0 | 0 | 0 | 0 | 0 | 0 | 0 | 906 | 1600 | 8484 | 170 | 142 | 0 | 85 | 85 | 297 | 0 | 0 | 0 | 127 | 491 | 1789 | 42 | 522 | 6946 | 524 | 48 | 0 | 0 | 0 | 0 | 0 | 0 | 0 | 0 | 1841 | 0 |
| SRR5034636 | 126 | 0 | 0 | 0 | 0 | 0 | 0 | 969 | 4018 | 11721 | 10 | 206 | 0 | 302 | 95 | 127 | 0 | 0 | 0 | 79 | 394 | 1934 | 26 | 480 | 8409 | 455 | 0 | 0 | 0 | 0 | 0 | 0 | 0 | 0 | 0 | 1032 | 0 |
| SRR5034637 | 51 | 0 | 0 | 0 | 0 | 0 | 0 | 1912 | 8002 | 13084 | 0 | 403 | 0 | 252 | 2114 | 4781 | 0 | 0 | 0 | 101 | 164 | 737 | 35 | 491 | 4485 | 177 | 26 | 0 | 0 | 0 | 0 | 0 | 0 | 0 | 0 | 1963 | 0 |
| SRR5034639 | 0 | 0 | 0 | 0 | 0 | 25 | 0 | 1083 | 2728 | 19335 | 40 | 562 | 0 | 1966 | 120 | 201 | 0 | 0 | 0 | 281 | 808 | 3193 | 121 | 1240 | 10154 | 882 | 0 | 0 | 0 | 0 | 0 | 0 | 0 | 0 | 0 | 2046 | 0 |
| SRR5034641 | 0 | 0 | 0 | 0 | 0 | 0 | 0 | 138 | 355 | 5951 | 39 | 217 | 0 | 493 | 99 | 493 | 0 | 0 | 0 | 20 | 67 | 249 | 11 | 263 | 3690 | 769 | 0 | 0 | 0 | 0 | 24 | 0 | 0 | 0 | 0 | 788 | 0 |
| SRR5034642 | 27 | 0 | 0 | 0 | 0 | 16 | 0 | 351 | 2313 | 3150 | 0 | 0 | 0 | 24 | 16 | 29 | 0 | 0 | 0 | 14 | 65 | 292 | 0 | 81 | 1248 | 170 | 22 | 0 | 0 | 0 | 16 | 0 | 0 | 0 | 0 | 167 | 0 |
| SRR5034643 | 0 | 0 | 0 | 0 | 0 | 26 | 0 | 574 | 906 | 15645 | 26 | 133 | 0 | 42 | 56 | 84 | 0 | 0 | 0 | 75 | 300 | 1305 | 0 | 197 | 6056 | 695 | 11 | 0 | 0 | 0 | 0 | 0 | 0 | 0 | 0 | 808 | 0 |
| SRR5034644 | 35 | 0 | 0 | 0 | 0 | 0 | 0 | 2142 | 2269 | 14697 | 59 | 109 | 0 | 254 | 186 | 172 | 0 | 0 | 0 | 59 | 629 | 2380 | 109 | 2123 | 6523 | 753 | 29 | 0 | 0 | 0 | 0 | 0 | 0 | 0 | 0 | 607 | 0 |
| SRR5034645 | 0 | 0 | 0 | 0 | 0 | 48 | 0 | 406 | 950 | 6030 | 61 | 169 | 0 | 307 | 973 | 1203 | 0 | 0 | 0 | 61 | 34 | 127 | 0 | 81 | 2433 | 159 | 133 | 0 | 0 | 0 | 0 | 0 | 0 | 0 | 0 | 199 | 0 |
| SRR5034646 | 0 | 0 | 0 | 0 | 0 | 24 | 0 | 994 | 1154 | 10774 | 46 | 57 | 0 | 229 | 1188 | 1314 | 0 | 0 | 0 | 183 | 273 | 1029 | 29 | 333 | 5453 | 396 | 0 | 0 | 0 | 0 | 0 | 0 | 0 | 0 | 0 | 1234 | 0 |
| SRR5034647 | 0 | 0 | 0 | 0 | 0 | 0 | 0 | 2170 | 2348 | 21299 | 24 | 0 | 0 | 0 | 49 | 97 | 0 | 0 | 0 | 0 | 151 | 739 | 22 | 249 | 7403 | 274 | 0 | 0 | 0 | 0 | 0 | 0 | 0 | 0 | 0 | 91 | 0 |
| SRR5034648 | 0 | 0 | 15 | 0 | 0 | 0 | 15 | 536 | 1266 | 11006 | 24 | 243 | 0 | 24 | 1656 | 2118 | 30 | 10 | 0 | 195 | 416 | 1816 | 39 | 752 | 7422 | 346 | 0 | 0 | 0 | 0 | 0 | 0 | 0 | 0 | 0 | 1534 | 0 |
| SRR5034649 | 28 | 0 | 0 | 0 | 0 | 19 | 0 | 1615 | 7516 | 6933 | 25 | 74 | 0 | 90 | 11 | 60 | 0 | 0 | 0 | 19 | 279 | 1473 | 18 | 648 | 10732 | 501 | 19 | 0 | 0 | 0 | 0 | 0 | 0 | 0 | 0 | 238 | 0 |
| SRR5034650 | 0 | 0 | 0 | 0 | 0 | 0 | 0 | 1401 | 1326 | 10486 | 16 | 294 | 0 | 316 | 1650 | 1552 | 0 | 0 | 0 | 68 | 208 | 578 | 29 | 221 | 2788 | 205 | 0 | 0 | 0 | 0 | 0 | 0 | 15 | 25 | 60 | 1808 | 0 |
| SRR5034651 | 0 | 0 | 0 | 0 | 14 | 0 | 0 | 366 | 1085 | 5043 | 68 | 230 | 0 | 108 | 3566 | 3850 | 0 | 0 | 0 | 95 | 163 | 711 | 0 | 147 | 2372 | 197 | 28 | 0 | 0 | 0 | 0 | 0 | 0 | 0 | 14 | 976 | 0 |
| SRR5034652 | 0 | 0 | 0 | 0 | 0 | 0 | 0 | 3334 | 2407 | 16862 | 13 | 189 | 0 | 80 | 219 | 434 | 0 | 0 | 0 | 45 | 179 | 550 | 0 | 175 | 2748 | 166 | 0 | 0 | 0 | 0 | 0 | 0 | 0 | 0 | 0 | 498 | 0 |
| SRR5034653 | 0 | 0 | 0 | 0 | 0 | 0 | 0 | 2639 | 2531 | 15595 | 44 | 84 | 0 | 76 | 426 | 482 | 0 | 0 | 0 | 117 | 252 | 702 | 13 | 188 | 3509 | 228 | 17 | 0 | 0 | 0 | 0 | 0 | 0 | 0 | 12 | 723 | 0 |
| SRR5034654 | 0 | 0 | 0 | 0 | 0 | 0 | 0 | 1909 | 2411 | 18497 | 28 | 172 | 0 | 42 | 94 | 78 | 0 | 0 | 0 | 71 | 561 | 1491 | 17 | 315 | 5068 | 551 | 0 | 0 | 0 | 0 | 0 | 0 | 0 | 0 | 0 | 687 | 0 |
| SRR5034655 | 0 | 0 | 0 | 0 | 0 | 0 | 0 | 1526 | 2415 | 9635 | 21 | 106 | 0 | 90 | 422 | 677 | 0 | 0 | 0 | 40 | 204 | 622 | 0 | 155 | 4259 | 297 | 26 | 0 | 0 | 0 | 0 | 0 | 0 | 0 | 0 | 172 | 0 |
| SRR5034656 | 20 | 0 | 0 | 0 | 0 | 0 | 0 | 1100 | 3870 | 4922 | 13 | 10 | 0 | 156 | 274 | 392 | 0 | 0 | 0 | 38 | 493 | 1395 | 15 | 330 | 5346 | 472 | 0 | 0 | 0 | 0 | 0 | 0 | 0 | 0 | 0 | 169 | 0 |
| SRR5034657 | 0 | 0 | 0 | 0 | 0 | 0 | 0 | 1291 | 6351 | 11620 | 11 | 89 | 0 | 89 | 12 | 19 | 0 | 0 | 0 | 35 | 178 | 604 | 32 | 447 | 8121 | 146 | 13 | 0 | 0 | 0 | 0 | 0 | 0 | 0 | 0 | 406 | 0 |
| SRR5034658 | 0 | 0 | 17 | 0 | 0 | 0 | 0 | 993 | 2878 | 27470 | 47 | 113 | 0 | 89 | 295 | 423 | 0 | 0 | 0 | 62 | 216 | 783 | 26 | 399 | 4041 | 281 | 0 | 0 | 0 | 0 | 0 | 0 | 0 | 0 | 0 | 795 | 0 |
| SRR5034659 | 0 | 0 | 0 | 0 | 0 | 0 | 0 | 1710 | 2980 | 31891 | 15 | 26 | 0 | 28 | 13 | 29 | 0 | 0 | 0 | 12 | 270 | 828 | 25 | 469 | 3931 | 236 | 0 | 0 | 0 | 0 | 0 | 0 | 0 | 0 | 0 | 146 | 0 |
| SRR5034660 | 0 | 0 | 0 | 0 | 0 | 0 | 0 | 1765 | 4475 | 31584 | 33 | 29 | 0 | 0 | 60 | 75 | 0 | 0 | 0 | 11 | 185 | 500 | 16 | 300 | 3890 | 466 | 0 | 0 | 0 | 0 | 0 | 0 | 0 | 0 | 0 | 94 | 0 |
| SRR5034661 | 18 | 0 | 0 | 0 | 0 | 0 | 0 | 393 | 1654 | 3137 | 53 | 43 | 0 | 21 | 66 | 91 | 0 | 0 | 0 | 33 | 571 | 1035 | 13 | 1401 | 1727 | 72 | 0 | 0 | 0 | 0 | 0 | 0 | 0 | 0 | 0 | 646 | 0 |
| SRR5034662 | 0 | 0 | 0 | 0 | 0 | 0 | 0 | 1214 | 1826 | 19065 | 34 | 116 | 0 | 38 | 202 | 203 | 0 | 0 | 0 | 48 | 410 | 1489 | 11 | 187 | 5744 | 488 | 0 | 0 | 0 | 0 | 0 | 0 | 0 | 0 | 0 | 383 | 0 |
| SRR5034663 | 0 | 0 | 0 | 0 | 0 | 0 | 0 | 1817 | 6132 | 14084 | 26 | 26 | 0 | 190 | 23 | 13 | 0 | 0 | 0 | 87 | 458 | 1820 | 63 | 944 | 5698 | 204 | 19 | 0 | 0 | 0 | 0 | 0 | 0 | 14 | 0 | 2136 | 0 |
| SRR5054304 | 0 | 568 | 83 | 36 | 1059 | 0 | 14 | 393 | 150 | 433 | 32878 | 68781 | 0 | 98 | 0 | 0 | 59 | 1398 | 150 | 40 | 1075 | 1557 | 601 | 13959 | 472 | 38 | 214 | 628 | 1808 | 4777 | 15656 | 2110 | 757 | 349 | 23 | 35 | 689 |
| SRR5054305 | 0 | 114 | 1116 | 36 | 1321 | 26 | 13 | 935 | 1111 | 4703 | 19651 | 37867 | 0 | 1499 | 0 | 0 | 86 | 1215 | 166 | 59 | 692 | 966 | 521 | 15042 | 603 | 32 | 114 | 479 | 1845 | 8189 | 25162 | 2839 | 2722 | 1595 | 83 | 206 | 956 |

### Supp Table S8.4.Columns_DI_to_ES

| | | | | | | | | | | | | | | | | | | | | | | | | | | | | | | | | | | | | | |
|---|---|---|---|---|---|---|---|---|---|---|---|---|---|---|---|---|---|---|---|---|---|---|---|---|---|---|---|---|---|---|---|---|---|---|---|---|---|
| SRR5087508 | 0 | 56 | 22 | 0 | 60 | 0 | 0 | 7778 | 8898 | 6465 | 2175 | 1340 | 0 | 1344 | 0 | 0 | 22 | 243 | 0 | 326 | 2441 | 2002 | 256 | 12018 | 6677 | 11 | 0 | 0 | 27 | 44 | 27 | 0 | 0 | 14 | 0 | 1184 | 280 |
| SRR5087510 | 0 | 91 | 25 | 0 | 72 | 0 | 0 | 7955 | 9091 | 6025 | 2848 | 1623 | 0 | 1283 | 0 | 0 | 26 | 280 | 0 | 345 | 2689 | 2335 | 256 | 11178 | 6781 | 16 | 0 | 0 | 29 | 39 | 29 | 0 | 13 | 23 | 0 | 1020 | 292 |
| SRR5087512 | 0 | 86 | 36 | 0 | 55 | 0 | 0 | 7568 | 7373 | 5176 | 2457 | 1718 | 0 | 1436 | 0 | 0 | 27 | 256 | 0 | 360 | 2567 | 2433 | 216 | 11109 | 6857 | 14 | 0 | 0 | 36 | 37 | 21 | 0 | 15 | 21 | 0 | 1064 | 332 |
| SRR5087516 | 0 | 45 | 27 | 0 | 86 | 0 | 0 | 4653 | 5014 | 2762 | 2538 | 776 | 0 | 1425 | 0 | 0 | 17 | 339 | 0 | 188 | 2598 | 2030 | 262 | 14227 | 7841 | 20 | 13 | 0 | 21 | 28 | 14 | 0 | 0 | 13 | 0 | 680 | 386 |
| SRR5087517 | 0 | 54 | 23 | 0 | 83 | 0 | 0 | 5538 | 5575 | 2663 | 1576 | 616 | 0 | 1396 | 0 | 0 | 13 | 310 | 0 | 205 | 2658 | 1919 | 355 | 13299 | 7519 | 20 | 0 | 0 | 14 | 22 | 20 | 0 | 12 | 13 | 0 | 670 | 332 |
| SRR5087519 | 0 | 44 | 31 | 0 | 76 | 0 | 0 | 4684 | 4388 | 2040 | 1613 | 612 | 0 | 1594 | 0 | 0 | 17 | 334 | 0 | 172 | 2688 | 1987 | 326 | 12903 | 7786 | 17 | 12 | 0 | 21 | 26 | 14 | 0 | 0 | 12 | 0 | 605 | 337 |
| SRR5087522 | 0 | 142 | 37 | 0 | 56 | 0 | 0 | 7915 | 5226 | 22093 | 1014 | 4110 | 0 | 1511 | 0 | 0 | 11 | 216 | 0 | 846 | 2945 | 3143 | 241 | 10946 | 7728 | 0 | 0 | 30 | 73 | 99 | 85 | 23 | 36 | 82 | 35 | 2210 | 222 |
| SRR5087527 | 0 | 117 | 38 | 0 | 56 | 0 | 0 | 6714 | 4886 | 20849 | 1015 | 4333 | 0 | 1690 | 0 | 0 | 13 | 269 | 0 | 1411 | 3165 | 3257 | 260 | 11390 | 7818 | 10 | 0 | 22 | 82 | 100 | 80 | 20 | 28 | 55 | 33 | 2428 | 210 |
| SRR5087554 | 0 | 13 | 43 | 0 | 62 | 0 | 0 | 7673 | 5383 | 23012 | 872 | 3289 | 0 | 1358 | 0 | 0 | 19 | 210 | 0 | 1075 | 2910 | 2938 | 278 | 10718 | 7598 | 11 | 0 | 23 | 80 | 78 | 74 | 14 | 19 | 61 | 37 | 2251 | 219 |
| SRR5090283 | 0 | 765 | 18 | 41 | 632 | 0 | 57 | 1856 | 2226 | 8015 | 16678 | 16719 | 0 | 673 | 467 | 525 | 142 | 457 | 129 | 2661 | 58 | 515 | 0 | 24 | 976 | 0 | 53 | 0 | 18 | 15 | 42 | 12 | 0 | 0 | 0 | 1292 | 0 |
| SRR5092303 | 0 | 138 | 0 | 0 | 252 | 0 | 0 | 257 | 428 | 599 | 300 | 1498 | 0 | 86 | 0 | 0 | 20 | 525 | 1229 | 3338 | 56 | 198 | 0 | 88 | 222 | 1508 | 236 | 0 | 0 | 0 | 39 | 0 | 0 | 0 | 0 | 1926 | 0 |
| SRR5092304 | 0 | 47 | 0 | 14 | 293 | 0 | 0 | 0 | 384 | 432 | 12 | 2784 | 23 | 48 | 0 | 0 | 47 | 367 | 751 | 1200 | 81 | 247 | 0 | 73 | 463 | 1481 | 187 | 0 | 0 | 47 | 70 | 0 | 0 | 0 | 0 | 912 | 0 |
| SRR5092305 | 0 | 110 | 22 | 0 | 176 | 0 | 0 | 209 | 209 | 1149 | 104 | 1775 | 22 | 104 | 0 | 0 | 44 | 203 | 467 | 313 | 28 | 100 | 0 | 61 | 502 | 801 | 198 | 0 | 0 | 0 | 0 | 0 | 0 | 0 | 0 | 418 | 0 |
| SRR5092836 | 0 | 38 | 0 | 0 | 30 | 38 | 0 | 1802 | 150 | 11363 | 400 | 400 | 0 | 4405 | 0 | 0 | 0 | 80 | 0 | 0 | 384 | 1862 | 968 | 66754 | 4478 | 20 | 38 | 76 | 38 | 0 | 0 | 0 | 38 | 0 | 0 | 50 | 80 |
| SRR5100734 | 0 | 0 | 0 | 0 | 559 | 0 | 112 | 7488 | 2825 | 12 | 0 | 0 | 15 | 0 | 0 | 0 | 0 | 559 | 70 | 77 | 350 | 1818 | 210 | 18111 | 350 | 0 | 21 | 0 | 0 | 0 | 0 | 0 | 0 | 0 | 0 | 234 | 140 |
| SRR5100735 | 0 | 0 | 0 | 17 | 52 | 0 | 35 | 3470 | 1678 | 69 | 22 | 13 | 0 | 560 | 49 | 17 | 0 | 121 | 243 | 174 | 121 | 1490 | 52 | 10481 | 208 | 260 | 212 | 0 | 24 | 16 | 151 | 97 | 106 | 101 | 28 | 475 | 35 |
| SRR5104206 | 0 | 13 | 0 | 0 | 0 | 0 | 0 | 500 | 73 | 2754 | 4425 | 26614 | 0 | 1950 | 0 | 0 | 0 | 112 | 0 | 0 | 978 | 9861 | 17 | 13569 | 1429 | 55 | 0 | 0 | 0 | 37 | 14 | 0 | 17 | 39 | 0 | 0 | 0 |
| SRR5104207 | 0 | 16 | 14 | 0 | 0 | 0 | 0 | 479 | 109 | 2796 | 45944 | 28779 | 0 | 2444 | 0 | 0 | 13 | 124 | 0 | 0 | 1147 | 11982 | 0 | 7806 | 961 | 56 | 0 | 0 | 12 | 21 | 11 | 0 | 12 | 19 | 0 | 0 | 0 |
| SRR5104208 | 0 | 16 | 12 | 0 | 0 | 0 | 0 | 458 | 122 | 3168 | 5151 | 26615 | 0 | 2397 | 0 | 0 | 0 | 120 | 0 | 0 | 1093 | 13499 | 0 | 7260 | 941 | 60 | 0 | 0 | 11 | 26 | 24 | 0 | 11 | 16 | 0 | 0 | 0 |
| SRR5104209 | 10 | 69 | 10 | 0 | 0 | 0 | 0 | 355 | 91 | 2888 | 4629 | 25722 | 0 | 3954 | 0 | 0 | 17 | 100 | 0 | 0 | 793 | 9609 | 0 | 6207 | 659 | 45 | 0 | 0 | 0 | 0 | 13 | 0 | 11 | 18 | 0 | 0 | 0 |
| SRR5104210 | 0 | 37 | 15 | 0 | 0 | 0 | 20 | 507 | 100 | 3394 | 34047 | 25063 | 0 | 3527 | 0 | 0 | 13 | 78 | 0 | 0 | 791 | 6169 | 52 | 1378 | 551 | 82 | 0 | 0 | 17 | 15 | 15 | 0 | 19 | 25 | 0 | 0 | 0 |
| SRR5104211 | 15 | 99 | 19 | 0 | 0 | 0 | 0 | 441 | 66 | 2273 | 44880 | 26810 | 0 | 3223 | 0 | 0 | 0 | 56 | 0 | 0 | 813 | 6069 | 37 | 906 | 335 | 49 | 0 | 0 | 0 | 0 | 0 | 0 | 11 | 15 | 0 | 0 | 0 |
| SRR5104212 | 0 | 92 | 13 | 0 | 0 | 0 | 0 | 208 | 54 | 2712 | 5835 | 35297 | 0 | 2419 | 0 | 0 | 0 | 73 | 0 | 0 | 668 | 5781 | 37 | 1231 | 820 | 52 | 0 | 0 | 0 | 12 | 12 | 0 | 0 | 14 | 0 | 0 | 0 |
| SRR5104213 | 0 | 54 | 12 | 0 | 0 | 0 | 0 | 482 | 82 | 2173 | 3452 | 31386 | 0 | 2854 | 0 | 0 | 0 | 54 | 0 | 0 | 780 | 6397 | 51 | 1229 | 742 | 61 | 0 | 0 | 0 | 13 | 0 | 0 | 0 | 12 | 0 | 0 | 0 |
| SRR5104214 | 0 | 81 | 12 | 0 | 0 | 0 | 0 | 257 | 105 | 1250 | 55055 | 29946 | 0 | 2384 | 0 | 0 | 0 | 42 | 0 | 0 | 517 | 3855 | 27 | 558 | 290 | 42 | 0 | 0 | 0 | 0 | 0 | 0 | 0 | 0 | 0 | 0 | 0 |
| SRR5104215 | 11 | 78 | 31 | 0 | 0 | 0 | 0 | 368 | 54 | 2632 | 5114 | 36380 | 0 | 3715 | 0 | 0 | 0 | 37 | 0 | 0 | 422 | 3837 | 26 | 527 | 352 | 50 | 0 | 0 | 0 | 0 | 0 | 0 | 0 | 11 | 0 | 0 | 0 |
| SRR5104216 | 13 | 100 | 18 | 0 | 0 | 0 | 0 | 497 | 83 | 2372 | 5240 | 26057 | 0 | 3532 | 0 | 0 | 0 | 76 | 0 | 0 | 709 | 4946 | 40 | 731 | 483 | 78 | 0 | 0 | 13 | 0 | 16 | 0 | 0 | 21 | 0 | 0 | 0 |
| SRR5104217 | 0 | 82 | 20 | 0 | 0 | 0 | 0 | 168 | 13 | 1032 | 4097 | 19795 | 0 | 2051 | 0 | 0 | 0 | 70 | 0 | 0 | 665 | 3659 | 26 | 503 | 298 | 57 | 0 | 0 | 0 | 12 | 0 | 0 | 0 | 22 | 0 | 0 | 0 |
| SRR5104218 | 14 | 78 | 19 | 0 | 0 | 0 | 0 | 444 | 30 | 1470 | 4879 | 32825 | 0 | 3137 | 0 | 0 | 13 | 47 | 0 | 0 | 540 | 3386 | 20 | 271 | 205 | 74 | 0 | 0 | 0 | 12 | 0 | 0 | 0 | 12 | 0 | 0 | 0 |
| SRR5104219 | 0 | 62 | 34 | 0 | 0 | 0 | 0 | 544 | 111 | 2084 | 4964 | 34255 | 0 | 3877 | 10 | 0 | 0 | 40 | 0 | 0 | 487 | 3456 | 26 | 424 | 300 | 67 | 0 | 0 | 0 | 11 | 11 | 0 | 0 | 20 | 0 | 0 | 0 |
| SRR5104220 | 11 | 35 | 16 | 0 | 0 | 0 | 0 | 407 | 54 | 1580 | 37706 | 32008 | 0 | 4142 | 0 | 0 | 0 | 46 | 0 | 0 | 359 | 2620 | 30 | 567 | 372 | 98 | 0 | 0 | 0 | 13 | 13 | 0 | 14 | 12 | 0 | 0 | 0 |
| SRR5104221 | 0 | 47 | 10 | 0 | 0 | 0 | 0 | 309 | 18 | 679 | 42355 | 36400 | 0 | 2003 | 0 | 0 | 0 | 44 | 0 | 0 | 513 | 2691 | 31 | 446 | 244 | 51 | 0 | 0 | 0 | 0 | 0 | 0 | 0 | 0 | 0 | 0 | 0 |
| SRR5104222 | 0 | 37 | 12 | 0 | 0 | 0 | 0 | 79 | 0 | 863 | 4384 | 20410 | 0 | 3646 | 0 | 0 | 0 | 46 | 0 | 0 | 459 | 2212 | 15 | 349 | 161 | 61 | 0 | 0 | 0 | 12 | 0 | 0 | 0 | 0 | 0 | 0 | 0 |
| SRR5104223 | 0 | 11 | 0 | 0 | 0 | 0 | 0 | 443 | 21 | 1634 | 4998 | 33053 | 0 | 6948 | 0 | 0 | 12 | 28 | 0 | 0 | 545 | 3215 | 11 | 132 | 119 | 62 | 0 | 0 | 0 | 0 | 0 | 0 | 0 | 19 | 0 | 0 | 0 |
| SRR5104224 | 0 | 36 | 0 | 0 | 0 | 0 | 0 | 63 | 0 | 457 | 2824 | 15690 | 0 | 2505 | 0 | 0 | 0 | 39 | 0 | 0 | 462 | 2370 | 21 | 372 | 189 | 48 | 0 | 0 | 0 | 0 | 0 | 0 | 0 | 13 | 0 | 0 | 0 |
| SRR5104225 | 0 | 83 | 13 | 0 | 0 | 0 | 0 | 182 | 16 | 1343 | 40022 | 32884 | 0 | 7186 | 0 | 0 | 12 | 70 | 0 | 0 | 543 | 2570 | 25 | 387 | 202 | 73 | 0 | 0 | 0 | 0 | 0 | 0 | 13 | 15 | 0 | 0 | 0 |
| SRR5116489 | 0 | 37 | 0 | 0 | 13 | 0 | 0 | 588 | 0 | 4926 | 0 | 0 | 0 | 809 | 0 | 0 | 0 | 111 | 0 | 441 | 1564 | 2241 | 19 | 14664 | 3125 | 630 | 0 | 73 | 37 | 440 | 110 | 0 | 0 | 110 | 0 | 1323 | 76 |
| SRR5116490 | 0 | 13 | 92 | 0 | 0 | 0 | 13 | 424 | 49 | 5555 | 0 | 0 | 0 | 3144 | 16 | 0 | 0 | 327 | 0 | 1743 | 1628 | 5688 | 36 | 21974 | 4640 | 815 | 0 | 66 | 26 | 263 | 132 | 53 | 171 | 369 | 26 | 2932 | 121 |
| SRR5116491 | 0 | 21 | 43 | 0 | 22 | 0 | 0 | 650 | 78 | 4317 | 0 | 0 | 0 | 676 | 13 | 26 | 0 | 141 | 0 | 1157 | 1387 | 1884 | 18 | 9820 | 4773 | 676 | 0 | 107 | 149 | 448 | 213 | 107 | 256 | 362 | 21 | 2626 | 244 |
| SRR5116492 | 0 | 0 | 46 | 0 | 13 | 0 | 0 | 784 | 228 | 7894 | 0 | 0 | 0 | 2416 | 13 | 0 | 0 | 134 | 0 | 3681 | 1260 | 4578 | 17 | 16047 | 5411 | 893 | 0 | 16 | 12 | 18 | 16 | 73 | 19 | 51 | 32 | 10791 | 72 |
| SRR5116493 | 0 | 0 | 0 | 0 | 0 | 0 | 0 | 962 | 124 | 14186 | 0 | 0 | 0 | 1211 | 62 | 0 | 0 | 193 | 0 | 1800 | 1813 | 3294 | 57 | 18246 | 5742 | 898 | 0 | 173 | 252 | 268 | 189 | 63 | 79 | 425 | 0 | 3880 | 200 |
| SRR5116494 | 0 | 0 | 11 | 0 | 0 | 0 | 0 | 437 | 238 | 8262 | 0 | 0 | 0 | 3972 | 0 | 0 | 0 | 354 | 0 | 2463 | 1109 | 6112 | 44 | 27882 | 4768 | 1252 | 0 | 236 | 224 | 325 | 258 | 22 | 112 | 269 | 11 | 2860 | 64 |
| SRR5116495 | 0 | 0 | 0 | 0 | 14 | 0 | 0 | 1041 | 256 | 14409 | 0 | 0 | 0 | 1844 | 0 | 0 | 0 | 157 | 0 | 4780 | 1318 | 3725 | 17 | 12399 | 5859 | 1198 | 0 | 138 | 76 | 277 | 177 | 24 | 157 | 339 | 0 | 8365 | 114 |
| SRR5116496 | 0 | 0 | 29 | 0 | 0 | 0 | 0 | 1463 | 310 | 6825 | 0 | 0 | 0 | 1507 | 0 | 0 | 0 | 161 | 0 | 3368 | 1161 | 3258 | 30 | 17545 | 6769 | 955 | 0 | 147 | 98 | 314 | 265 | 59 | 147 | 157 | 0 | 5850 | 163 |
| SRR5121477 | 0 | 36 | 0 | 0 | 31 | 0 | 0 | 263 | 71 | 5230 | 580 | 2442 | 0 | 0 | 609 | 615 | 0 | 188 | 0 | 2109 | 2656 | 7939 | 0 | 59 | 28549 | 0 | 0 | 19 | 12 | 20 | 15 | 0 | 12 | 24 | 12 | 6724 | 92 |
| SRR5121478 | 0 | 290 | 0 | 0 | 50 | 0 | 0 | 448 | 200 | 3520 | 112 | 2118 | 0 | 0 | 2642 | 3539 | 0 | 114 | 10 | 4789 | 3990 | 8577 | 0 | 116 | 31256 | 0 | 0 | 0 | 55 | 104 | 243 | 27 | 100 | 208 | 166 | 5351 | 163 |
| SRR5121479 | 0 | 22 | 41 | 0 | 0 | 0 | 0 | 252 | 75 | 1107 | 6123 | 5936 | 0 | 262 | 23 | 0 | 0 | 237 | 219 | 621 | 206 | 2485 | 0 | 0 | 14994 | 0 | 0 | 1436 | 1092 | 999 | 816 | 214 | 753 | 850 | 123 | 2583 | 49 |
| SRR5121480 | 0 | 13 | 14 | 0 | 53 | 0 | 0 | 57 | 0 | 726 | 1703 | 6111 | 0 | 1219 | 63 | 73 | 0 | 196 | 89 | 163 | 0 | 1644 | 0 | 0 | 8657 | 0 | 0 | 118 | 111 | 183 | 517 | 72 | 174 | 468 | 390 | 983 | 0 |
| SRR5121481 | 0 | 0 | 13 | 0 | 0 | 0 | 0 | 204 | 34 | 366 | 6672 | 8802 | 0 | 47 | 115 | 25 | 0 | 0 | 14 | 423 | 937 | 1785 | 0 | 0 | 11585 | 0 | 0 | 91 | 149 | 392 | 269 | 45 | 223 | 383 | 153 | 1469 | 99 |
| SRR5121482 | 0 | 0 | 107 | 0 | 0 | 0 | 0 | 212 | 42 | 309 | 2126 | 5580 | 0 | 23 | 31 | 21 | 0 | 0 | 42 | 369 | 806 | 1734 | 0 | 12 | 9433 | 0 | 12 | 79 | 13 | 28 | 60 | 22 | 49 | 77 | 17 | 1419 | 462 |
| SRR5121483 | 0 | 46 | 0 | 0 | 0 | 0 | 0 | 278 | 45 | 1850 | 2336 | 2927 | 0 | 0 | 1751 | 3502 | 0 | 0 | 0 | 6746 | 1821 | 4535 | 0 | 32 | 12968 | 0 | 0 | 0 | 0 | 12 | 15 | 10 | 26 | 47 | 0 | 28730 | 87 |
| SRR5121484 | 0 | 17 | 81 | 0 | 0 | 0 | 0 | 229 | 35 | 1017 | 388 | 2619 | 0 | 533 | 22 | 18 | 0 | 28 | 129 | 924 | 806 | 2045 | 0 | 22 | 6231 | 0 | 0 | 17 | 16 | 17 | 32 | 167 | 21 | 39 | 72 | 3552 | 102 |
| SRR5121485 | 0 | 0 | 84 | 0 | 24 | 14 | 0 | 193 | 104 | 339 | 1205 | 5790 | 0 | 99 | 0 | 0 | 0 | 17 | 16 | 274 | 1261 | 1389 | 21 | 3576 | 7627 | 0 | 32 | 17 | 14 | 87 | 89 | 41 | 31 | 60 | 52 | 3305 | 0 |
| SRR5121486 | 0 | 12 | 0 | 0 | 25 | 0 | 0 | 51 | 0 | 60 | 2318 | 4643 | 0 | 35 | 0 | 0 | 0 | 18 | 13 | 3550 | 1194 | 2840 | 55 | 3243 | 14911 | 0 | 21 | 0 | 0 | 17 | 16 | 0 | 0 | 10 | 0 | 36534 | 12 |
| SRR5121487 | 22 | 0 | 155 | 0 | 28 | 0 | 0 | 685 | 249 | 280 | 784 | 4614 | 0 | 194 | 0 | 10 | 0 | 25 | 30 | 424 | 1479 | 1905 | 17 | 2661 | 7899 | 0 | 58 | 206 | 177 | 421 | 716 | 62 | 279 | 662 | 742 | 2724 | 0 |
| SRR5127200 | 0 | 0 | 30 | 0 | 31 | 0 | 0 | 2738 | 463 | 7812 | 2176 | 3347 | 0 | 537 | 13 | 21 | 0 | 41 | 0 | 518 | 1067 | 1621 | 0 | 32 | 6625 | 0 | 0 | 11 | 25 | 165 | 115 | 13 | 65 | 227 | 88 | 1092 | 0 |
| SRR5127201 | 0 | 121 | 16 | 0 | 10 | 0 | 0 | 752 | 386 | 27865 | 3706 | 5675 | 0 | 2015 | 42 | 37 | 0 | 558 | 0 | 811 | 503 | 3231 | 32 | 38823 | 5445 | 83 | 18 | 11 | 11 | 136 | 136 | 13 | 34 | 81 | 0 | 2942 | 0 |
| SRR5127202 | 0 | 93 | 76 | 0 | 0 | 0 | 0 | 643 | 445 | 38854 | 4231 | 3042 | 0 | 6683 | 668 | 445 | 0 | 287 | 11 | 797 | 636 | 2694 | 0 | 15837 | 7471 | 33 | 0 | 11 | 86 | 23 | 29 | 37 | 12 | 26 | 36 | 1627 | 0 |
| SRR5127203 | 0 | 283 | 96 | 0 | 12 | 0 | 0 | 280 | 169 | 25640 | 3866 | 2739 | 0 | 2378 | 180 | 115 | 0 | 1374 | 0 | 993 | 1597 | 4194 | 11 | 19481 | 11798 | 198 | 0 | 35 | 108 | 436 | 738 | 70 | 204 | 467 | 72 | 2052 | 160 |
| SRR5127204 | 0 | 257 | 62 | 0 | 28 | 0 | 0 | 318 | 70 | 3220 | 1575 | 3602 | 0 | 1022 | 1528 | 1337 | 0 | 141 | 0 | 4781 | 1863 | 12374 | 562 | 99107 | 5907 | 60 | 0 | 21 | 28 | 81 | 101 | 16 | 57 | 104 | 15 | 9338 | 82 |
| SRR5127205 | 0 | 211 | 15 | 0 | 0 | 0 | 0 | 1121 | 335 | 13747 | 4696 | 2656 | 0 | 725 | 518 | 524 | 0 | 378 | 0 | 1730 | 752 | 2242 | 11 | 13009 | 7268 | 16 | 0 | 18 | 10 | 63 | 223 | 10 | 29 | 84 | 15 | 3508 | 0 |
| SRR5127206 | 0 | 13 | 66 | 0 | 16 | 0 | 0 | 792 | 580 | 55107 | 4783 | 3164 | 0 | 2434 | 0 | 0 | 0 | 366 | 0 | 796 | 683 | 2568 | 18 | 17401 | 5624 | 25 | 0 | 17 | 23 | 106 | 16 | 16 | 50 | 109 | 30 | 1638 | 0 |
| SRR5127207 | 0 | 0 | 0 | 0 | 0 | 0 | 0 | 3148 | 381 | 31071 | 436 | 3168 | 0 | 1745 | 90 | 60 | 0 | 210 | 0 | 421 | 584 | 1126 | 20 | 3679 | 3146 | 64 | 0 | 0 | 0 | 0 | 0 | 0 | 0 | 0 | 0 | 1354 | 0 |
| SRR5127208 | 0 | 15 | 43 | 0 | 16 | 0 | 0 | 319 | 122 | 21672 | 2672 | 2540 | 0 | 2804 | 404 | 252 | 0 | 849 | 10 | 831 | 2006 | 4458 | 0 | 11974 | 10979 | 197 | 0 | 113 | 55 | 14 | 41 | 35 | 93 | 14 | 23 | 1167 | 389 |
| SRR5127209 | 0 | 64 | 98 | 0 | 0 | 0 | 0 | 1985 | 188 | 18505 | 8645 | 7214 | 0 | 1987 | 431 | 268 | 0 | 54 | 0 | 1094 | 1472 | 18 | 143 | 10706 | 11128 | 0 | 0 | 16 | 64 | 122 | 182 | 37 | 73 | 140 | 15 | 6491 | 0 |
| SRR5127210 | 0 | 15 | 72 | 0 | 13 | 0 | 0 | 464 | 35 | 15275 | 587 | 2327 | 0 | 2484 | 23 | 64 | 0 | 212 | 13 | 3116 | 1493 | 8271 | 19 | 57270 | 9725 | 74 | 0 | 57 | 81 | 21 | 43 | 75 | 17 | 34 | 52 | 6935 | 0 |
| SRR5127211 | 0 | 0 | 12 | 0 | 16 | 0 | 0 | 1432 | 443 | 124247 | 738 | 728 | 0 | 6996 | 30 | 38 | 0 | 89 | 10 | 569 | 719 | 976 | 53 | 13119 | 4114 | 48 | 0 | 0 | 0 | 0 | 13 | 0 | 0 | 22 | 18 | 7964 | 0 |
| SRR5127212 | 0 | 229 | 47 | 0 | 28 | 0 | 0 | 190 | 104 | 17145 | 3122 | 1939 | 0 | 1969 | 99 | 69 | 0 | 888 | 0 | 1432 | 1076 | 4643 | 105 | 36449 | 7544 | 165 | 12 | 29 | 36 | 162 | 147 | 22 | 68 | 94 | 18 | 4056 | 330 |
| SRR5127213 | 0 | 11 | 45 | 0 | 16 | 0 | 0 | 803 | 360 | 51569 | 3633 | 3450 | 0 | 4769 | 33 | 22 | 0 | 320 | 0 | 1357 | 500 | 4616 | 17 | 38597 | 8072 | 173 | 0 | 31 | 102 | 26 | 27 | 62 | 11 | 26 | 70 | 2038 | 0 |
| SRR5127214 | 0 | 165 | 81 | 0 | 25 | 0 | 0 | 1998 | 754 | 68519 | 3306 | 2206 | 0 | 2947 | 35 | 61 | 0 | 385 | 0 | 1001 | 531 | 6759 | 63 | 53985 | 10344 | 0 | 0 | 18 | 39 | 165 | 255 | 23 | 71 | 162 | 13 | 2570 | 0 |
| SRR5127215 | 0 | 132 | 88 | 0 | 28 | 0 | 0 | 1053 | 957 | 55264 | 2198 | 1197 | 0 | 2104 | 18 | 36 | 0 | 285 | 0 | 475 | 931 | 2146 | 30 | 8169 | 5621 | 0 | 0 | 15 | 22 | 137 | 229 | 24 | 57 | 140 | 46 | 961 | 0 |
| SRR5127216 | 0 | 261 | 64 | 0 | 11 | 0 | 0 | 412 | 72 | 5718 | 16660 | 8025 | 0 | 1962 | 336 | 109 | 0 | 626 | 0 | 1211 | 772 | 4973 | 25 | 32819 | 6813 | 143 | 23 | 30 | 40 | 70 | 131 | 11 | 47 | 77 | 0 | 3042 | 291 |
| SRR5127217 | 0 | 149 | 58 | 0 | 26 | 0 | 0 | 1258 | 1733 | 60588 | 1298 | 824 | 0 | 2201 | 0 | 0 | 0 | 380 | 0 | 459 | 646 | 2033 | 20 | 15383 | 5988 | 55 | 0 | 10 | 21 | 57 | 109 | 17 | 60 | 118 | 24 | 714 | 0 |
| SRR5127218 | 0 | 12 | 99 | 0 | 0 | 0 | 0 | 848 | 300 | 13529 | 12137 | 11487 | 0 | 1586 | 56 | 44 | 0 | 460 | 0 | 1392 | 620 | 4271 | 19 | 33949 | 5564 | 23 | 0 | 18 | 34 | 100 | 14 | 28 | 69 | 10 | 16 | 4426 | 0 |
| SRR5127219 | 0 | 0 | 0 | 0 | 0 | 0 | 0 | 2854 | 664 | 8633 | 256 | 272 | 0 | 235 | 246 | 407 | 0 | 27 | 0 | 149 | 1171 | 724 | 0 | 26 | 4381 | 0 | 28 | 0 | 0 | 20 | 12 | 0 | 11 | 29 | 17 | 1522 | 10 |
| SRR5127220 | 0 | 120 | 30 | 0 | 0 | 0 | 0 | 564 | 177 | 17743 | 10637 | 7530 | 0 | 4580 | 14 | 46 | 0 | 193 | 19 | 1055 | 921 | 3158 | 0 | 12590 | 6068 | 88 | 0 | 99 | 75 | 206 | 278 | 38 | 109 | 167 | 19 | 1736 | 0 |
| SRR5127221 | 0 | 166 | 50 | 0 | 0 | 0 | 0 | 1112 | 377 | 17121 | 7858 | 5638 | 0 | 10517 | 489 | 362 | 0 | 207 | 0 | 546 | 439 | 1995 | 0 | 14592 | 4410 | 48 | 0 | 25 | 61 | 15 | 25 | 57 | 11 | 17 | 49 | 1170 | 0 |
| SRR5127222 | 0 | 153 | 97 | 0 | 19 | 0 | 0 | 913 | 638 | 19655 | 3006 | 4386 | 0 | 3631 | 0 | 0 | 0 | 297 | 0 | 1469 | 589 | 1174 | 37 | 8301 | 3206 | 16 | 0 | 40 | 42 | 111 | 226 | 31 | 59 | 152 | 17 | 7225 | 0 |
| SRR5127223 | 0 | 132 | 45 | 0 | 12 | 0 | 0 | 347 | 185 | 14046 | 3362 | 6021 | 0 | 6531 | 22 | 41 | 0 | 278 | 0 | 1088 | 522 | 1916 | 11 | 12608 | 4575 | 80 | 0 | 148 | 68 | 222 | 374 | 65 | 235 | 374 | 49 | 1785 | 0 |
| SRR5127224 | 0 | 43 | 78 | 0 | 14 | 0 | 0 | 234 | 91 | 7818 | 2020 | 6703 | 0 | 764 | 767 | 741 | 0 | 546 | 0 | 2011 | 1810 | 5323 | 130 | 29341 | 7004 | 567 | 0 | 59 | 83 | 26 | 26 | 40 | 15 | 21 | 17 | 13171 | 289 |
| SRR5127225 | 0 | 3057 | 74 | 0 | 0 | 0 | 0 | 1124 | 293 | 9186 | 1000 | 4585 | 0 | 1161 | 373 | 618 | 0 | 617 | 0 | 1874 | 760 | 1988 | 0 | 14415 | 7011 | 33 | 0 | 75 | 158 | 658 | 683 | 126 | 343 | 672 | 129 | 2748 | 0 |
| SRR5127226 | 0 | 11 | 57 | 0 | 0 | 0 | 0 | 597 | 285 | 16074 | 4400 | 9200 | 0 | 3002 | 22 | 37 | 0 | 300 | 0 | 1584 | 716 | 2740 | 11 | 17511 | 4886 | 52 | 0 | 23 | 35 | 13 | 21 | 20 | 78 | 14 | 27 | 2219 | 0 |
| SRR5127227 | 0 | 184 | 48 | 0 | 0 | 0 | 0 | 1552 | 392 | 14277 | 6871 | 3600 | 0 | 2658 | 109 | 136 | 0 | 519 | 0 | 1191 | 718 | 3047 | 12 | 20858 | 6621 | 37 | 0 | 43 | 36 | 150 | 234 | 18 | 104 | 178 | 28 | 3298 | 0 |
| SRR5127228 | 0 | 0 | 24 | 0 | 13 | 0 | 0 | 1042 | 280 | 25763 | 856 | 973 | 0 | 8615 | 29 | 14 | 0 | 114 | 0 | 1297 | 642 | 1110 | 85 | 24784 | 2610 | 143 | 25 | 0 | 43 | 175 | 253 | 37 | 119 | 287 | 189 | 9018 | 0 |
| SRR5127229 | 0 | 10 | 60 | 0 | 0 | 0 | 0 | 849 | 489 | 9944 | 7952 | 12684 | 0 | 2399 | 10 | 22 | 0 | 285 | 0 | 957 | 472 | 1851 | 0 | 4117 | 5397 | 79 | 0 | 56 | 39 | 105 | 14 | 31 | 104 | 13 | 28 | 1109 | 0 |
| SRR5127230 | 0 | 11 | 75 | 0 | 14 | 0 | 0 | 395 | 86 | 3790 | 8761 | 6321 | 0 | 1314 | 0 | 0 | 0 | 157 | 0 | 3224 | 2038 | 14933 | 152 | 141544 | 7393 | 52 | 0 | 38 | 43 | 12 | 17 | 27 | 81 | 14 | 24 | 4601 | 57 |
| SRR5127231 | 0 | 22 | 0 | 0 | 0 | 0 | 0 | 16652 | 756 | 48507 | 1618 | 5638 | 0 | 3888 | 13 | 22 | 0 | 176 | 0 | 1268 | 330 | 971 | 99 | 11629 | 2893 | 35 | 0 | 0 | 0 | 27 | 30 | 0 | 23 | 48 | 23 | 10758 | 0 |
| SRR5127232 | 0 | 0 | 0 | 0 | 0 | 0 | 0 | 159 | 71 | 1258 | 1542 | 2623 | 0 | 53 | 35 | 18 | 0 | 0 | 0 | 18 | 73 | 76 | 0 | 0 | 5675 | 156 | 0 | 0 | 0 | 0 | 0 | 0 | 0 | 0 | 0 | 106 | 19 |
| SRR5127233 | 0 | 13 | 50 | 0 | 12 | 0 | 0 | 1143 | 754 | 26055 | 3670 | 1904 | 0 | 6584 | 19 | 17 | 0 | 338 | 0 | 424 | 575 | 2023 | 15 | 14680 | 4599 | 0 | 0 | 0 | 19 | 51 | 115 | 13 | 37 | 50 | 13 | 968 | 0 |
| SRR5127234 | 0 | 42 | 39 | 0 | 16 | 0 | 0 | 867 | 316 | 20880 | 7129 | 3433 | 0 | 3576 | 259 | 97 | 0 | 140 | 0 | 462 | 2384 | 7878 | 212 | 21997 | 11233 | 0 | 0 | 0 | 29 | 53 | 74 | 0 | 32 | 50 | 0 | 1340 | 244 |
| SRR5127235 | 0 | 197 | 71 | 0 | 20 | 0 | 0 | 475 | 64 | 5797 | 3745 | 2995 | 0 | 3129 | 160 | 51 | 0 | 211 | 0 | 2482 | 1568 | 14559 | 82 | 86436 | 8239 | 25 | 0 | 65 | 88 | 475 | 814 | 82 | 318 | 605 | 74 | 6932 | 117 |
| SRR5127236 | 0 | 12 | 0 | 0 | 0 | 0 | 0 | 292 | 73 | 2796 | 8139 | 6778 | 0 | 1595 | 0 | 10 | 0 | 383 | 0 | 2460 | 456 | 2487 | 26 | 25549 | 5255 | 94 | 0 | 0 | 0 | 0 | 16 | 0 | 0 | 0 | 0 | 5313 | 0 |
| SRR5136285 | 0 | 0 | 0 | 0 | 0 | 0 | 0 | 263 | 394 | 8532 | 0 | 0 | 0 | 1444 | 0 | 0 | 47 | 0 | 0 | 131 | 859 | 1797 | 78 | 6015 | 2344 | 78 | 43 | 0 | 0 | 0 | 0 | 0 | 0 | 0 | 0 | 0 | 0 |
| SRR5136286 | 0 | 0 | 0 | 0 | 0 | 0 | 0 | 257 | 0 | 8863 | 0 | 128 | 0 | 1413 | 0 | 0 | 22 | 0 | 0 | 0 | 733 | 570 | 0 | 4643 | 896 | 0 | 12 | 0 | 0 | 0 | 0 | 0 | 0 | 0 | 0 | 128 | 81 |
| SRR5136287 | 0 | 0 | 0 | 0 | 0 | 0 | 0 | 122 | 243 | 7666 | 261 | 122 | 0 | 1582 | 122 | 122 | 54 | 0 | 0 | 0 | 987 | 1063 | 0 | 6456 | 2051 | 0 | 36 | 0 | 0 | 0 | 0 | 0 | 0 | 0 | 0 | 365 | 0 |
| SRR5136288 | 0 | 0 | 0 | 0 | 0 | 0 | 0 | 0 | 373 | 7331 | 0 | 0 | 0 | 870 | 0 | 124 | 44 | 0 | 0 | 0 | 704 | 1509 | 0 | 9257 | 2314 | 0 | 44 | 0 | 0 | 0 | 0 | 0 | 0 | 0 | 0 | 124 | 101 |
| SRR5136289 | 0 | 0 | 0 | 0 | 0 | 0 | 0 | 122 | 244 | 7918 | 0 | 122 | 0 | 731 | 0 | 0 | 37 | 0 | 0 | 0 | 855 | 986 | 197 | 5655 | 2104 | 66 | 34 | 0 | 0 | 0 | 0 | 0 | 0 | 0 | 0 | 0 | 0 |
| SRR5136290 | 0 | 0 | 0 | 0 | 0 | 0 | 0 | 268 | 268 | 8031 | 134 | 134 | 0 | 1071 | 0 | 0 | 13 | 0 | 0 | 0 | 800 | 655 | 0 | 5674 | 1673 | 0 | 10 | 0 | 0 | 0 | 0 | 0 | 0 | 0 | 0 | 0 | 73 |
| SRR5136291 | 0 | 0 | 0 | 0 | 0 | 0 | 0 | 0 | 507 | 9500 | 127 | 0 | 0 | 1773 | 0 | 0 | 71 | 148 | 0 | 127 | 742 | 1187 | 297 | 6902 | 1855 | 148 | 28 | 0 | 0 | 0 | 0 | 0 | 0 | 0 | 0 | 380 | 74 |
| SRR5136292 | 0 | 0 | 0 | 0 | 0 | 0 | 0 | 0 | 371 | 8030 | 132 | 124 | 0 | 1359 | 0 | 0 | 21 | 0 | 0 | 124 | 555 | 1572 | 185 | 7492 | 1757 | 0 | 14 | 0 | 0 | 0 | 0 | 0 | 0 | 0 | 0 | 247 | 0 |
| SRR5136293 | 0 | 0 | 0 | 0 | 0 | 0 | 0 | 32 | 64 | 2023 | 0 | 0 | 0 | 1136 | 0 | 0 | 14 | 26 | 0 | 0 | 409 | 281 | 77 | 4446 | 1175 | 77 | 33 | 0 | 0 | 0 | 0 | 0 | 0 | 0 | 0 | 0 | 26 |
| SRR5136294 | 0 | 0 | 0 | 0 | 0 | 0 | 0 | 82 | 107 | 1687 | 17 | 16 | 0 | 1012 | 0 | 0 | 19 | 28 | 0 | 0 | 498 | 304 | 28 | 4232 | 1272 | 55 | 26 | 0 | 0 | 0 | 0 | 0 | 0 | 0 | 0 | 0 | 28 |
| SRR5136295 | 0 | 0 | 0 | 0 | 0 | 0 | 0 | 70 | 117 | 1632 | 16 | 0 | 0 | 1054 | 0 | 0 | 24 | 0 | 51 | 0 | 563 | 409 | 51 | 4376 | 1049 | 77 | 22 | 0 | 0 | 0 | 0 | 0 | 0 | 0 | 0 | 0 | 26 |
| SRR5136296 | 0 | 0 | 0 | 0 | 27 | 0 | 0 | 57 | 106 | 1779 | 0 | 16 | 0 | 1281 | 0 | 0 | 25 | 27 | 27 | 0 | 440 | 495 | 0 | 4013 | 1127 | 82 | 40 | 0 | 0 | 0 | 0 | 0 | 0 | 0 | 0 | 24 | 27 |
| SRR5136297 | 0 | 0 | 0 | 0 | 0 | 0 | 0 | 57 | 106 | 1974 | 0 | 14 | 0 | 1061 | 0 | 0 | 28 | 0 | 0 | 0 | 553 | 448 | 53 | 4133 | 1027 | 26 | 21 | 0 | 0 | 0 | 0 | 0 | 0 | 0 | 0 | 0 | 0 |
| SRR5136298 | 0 | 0 | 0 | 0 | 27 | 0 | 0 | 29 | 94 | 1889 | 0 | 0 | 0 | 1068 | 0 | 0 | 18 | 0 | 0 | 0 | 568 | 541 | 27 | 4922 | 1271 | 81 | 35 | 0 | 0 | 0 | 0 | 0 | 0 | 0 | 0 | 0 | 27 |
| SRR5136299 | 0 | 0 | 0 | 0 | 52 | 0 | 0 | 0 | 111 | 1859 | 0 | 0 | 0 | 1177 | 0 | 0 | 11 | 26 | 0 | 0 | 618 | 489 | 26 | 4584 | 1159 | 26 | 26 | 0 | 0 | 0 | 0 | 0 | 0 | 0 | 0 | 21 | 52 |
| SRR5136300 | 0 | 0 | 0 | 0 | 83 | 0 | 0 | 36 | 72 | 1654 | 0 | 0 | 0 | 1163 | 0 | 0 | 0 | 55 | 28 | 0 | 553 | 304 | 28 | 4785 | 1079 | 28 | 39 | 0 | 0 | 0 | 0 | 0 | 0 | 0 | 0 | 0 | 0 |
| SRR5139121 | 0 | 87 | 61 | 0 | 19 | 0 | 0 | 996 | 226 | 14859 | 490 | 2303 | 0 | 741 | 51 | 57 | 0 | 365 | 0 | 2026 | 665 | 3362 | 63 | 41192 | 8463 | 63 | 0 | 36 | 61 | 16 | 17 | 31 | 88 | 16 | 43 | 12720 | 0 |
| SRR5145238 | 0 | 850 | 31 | 51 | 130 | 0 | 0 | 353 | 931 | 2347 | 5091 | 15536 | 0 | 1067 | 122 | 172 | 37 | 518 | 1757 | 3406 | 2354 | 5036 | 125 | 24240 | 25414 | 0 | 10 | 36 | 108 | 92 | 101 | 20 | 15 | 17 | 37 | 1461 | 3838 |
| SRR5145239 | 0 | 3085 | 210 | 34 | 80 | 0 | 10 | 586 | 1070 | 4429 | 2914 | 3724 | 0 | 2310 | 30 | 153 | 22 | 278 | 1079 | 2635 | 950 | 3779 | 186 | 20179 | 14650 | 0 | 0 | 54 | 690 | 943 | 1482 | 413 | 480 | 375 | 54 | 1080 | 2155 |
| SRR5145240 | 0 | 1348 | 20 | 48 | 260 | 0 | 0 | 351 | 742 | 2393 | 655 | 475 | 0 | 1348 | 162 | 126 | 55 | 347 | 3236 | 2160 | 793 | 3035 | 294 | 18653 | 13481 | 0 | 0 | 16 | 411 | 171 | 253 | 61 | 60 | 55 | 0 | 2325 | 1444 |
| SRR5145241 | 0 | 2991 | 50 | 57 | 266 | 0 | 0 | 96 | 496 | 1575 | 2240 | 1256 | 0 | 2399 | 78 | 31 | 88 | 292 | 3276 | 503 | 797 | 2138 | 496 | 19006 | 14183 | 0 | 15 | 21 | 919 | 576 | 1632 | 369 | 267 | 245 | 11 | 308 | 1712 |
| SRR5145242 | 0 | 1027 | 52 | 50 | 169 | 10 | 0 | 605 | 859 | 1858 | 96 | 175 | 0 | 1675 | 251 | 135 | 52 | 273 | 5520 | 1162 | 774 | 2754 | 356 | 13557 | 12887 | 0 | 0 | 11 | 165 | 45 | 67 | 18 | 20 | 16 | 0 | 1428 | 1315 |
| SRR5145243 | 0 | 1545 | 13 | 38 | 175 | 0 | 24 | 300 | 560 | 1167 | 7358 | 34656 | 0 | 931 | 0 | 25 | 73 | 192 | 3647 | 2732 | 587 | 2340 | 359 | 12878 | 14790 | 0 | 21 | 33 | 170 | 62 | 67 | 11 | 26 | 18 | 0 | 1041 | 1470 |
| SRR5145244 | 0 | 1503 | 75 | 67 | 184 | 0 | 0 | 505 | 860 | 2081 | 6886 | 3521 | 0 | 1408 | 424 | 131 | 88 | 337 | 3914 | 2006 | 1514 | 4572 | 192 | 13903 | 21267 | 0 | 0 | 17 | 172 | 85 | 79 | 16 | 14 | 16 | 0 | 1090 | 2465 |
| SRR5145245 | 0 | 1471 | 132 | 44 | 127 | 0 | 0 | 738 | 2706 | 16197 | 1489 | 5591 | 0 | 3124 | 208 | 584 | 29 | 210 | 1678 | 6740 | 682 | 3297 | 252 | 14193 | 14776 | 0 | 11 | 35 | 325 | 414 | 504 | 123 | 67 | 69 | 39 | 3068 | 1550 |
| SRR5163971 | 0 | 149 | 51 | 172 | 1508 | 32 | 22 | 1321 | 9543 | 33901 | 1022 | 2234 | 0 | 1212 | 72 | 244 | 34 | 540 | 17 | 190 | 2226 | 497 | 437 | 3441 | 629 | 0 | 32 | 0 | 0 | 103 | 174 | 14 | 18 | 14 | 14 | 570 | 13 |
| SRR5163972 | 0 | 114 | 43 | 165 | 1469 | 35 | 12 | 1060 | 9557 | 33758 | 64 | 2473 | 12 | 1625 | 71 | 141 | 12 | 545 | 19 | 265 | 2197 | 469 | 446 | 3476 | 589 | 0 | 43 | 0 | 27 | 55 | 173 | 31 | 0 | 24 | 24 | 406 | 17 |
| SRR5163973 | 0 | 108 | 83 | 277 | 7590 | 64 | 17 | 906 | 4803 | 87807 | 134 | 1005 | 0 | 1210 | 0 | 66 | 45 | 1678 | 23 | 86 | 956 | 394 | 504 | 4239 | 584 | 15 | 45 | 13 | 59 | 15 | 47 | 102 | 55 | 53 | 74 | 148 | 656 |
| SRR5163974 | 0 | 63 | 82 | 255 | 7372 | 39 | 44 | 808 | 4830 | 85622 | 647 | 932 | 0 | 1027 | 19 | 38 | 53 | 1642 | 30 | 86 | 931 | 395 | 473 | 4294 | 531 | 21 | 77 | 24 | 58 | 228 | 663 | 58 | 92 | 82 | 73 | 133 | 646 |
| SRR5163975 | 12 | 105 | 25 | 112 | 4565 | 39 | 21 | 1020 | 6092 | 43568 | 450 | 5071 | 0 | 1449 | 31 | 55 | 34 | 534 | 99 | 514 | 481 | 422 | 320 | 6294 | 615 | 0 | 52 | 31 | 78 | 172 | 636 | 148 | 154 | 143 | 29 | 826 | 20 |
| SRR5163976 | 0 | 120 | 32 | 111 | 4498 | 55 | 39 | 1222 | 5971 | 43640 | 3667 | 5009 | 0 | 1763 | 0 | 40 | 42 | 529 | 92 | 822 | 500 | 391 | 304 | 6135 | 581 | 0 | 68 | 39 | 78 | 185 | 560 | 113 | 152 | 120 | 23 | 842 | 26 |
| SRR5163977 | 0 | 19 | 15 | 161 | 7493 | 45 | 0 | 843 | 4311 | 26154 | 399 | 14835 | 0 | 1276 | 1204 | 2794 | 47 | 240 | 91 | 1927 | 1477 | 317 | 117 | 1302 | 373 | 15 | 50 | 34 | 54 | 19 | 58 | 47 | 11 | 54 | 26 | 3950 | 57 |

### Supp Table S8.4.Columns_DI_to_ES

| | | | | | | | | | | | | | | | | | |
|---|---|---|---|---|---|---|---|---|---|---|---|---|---|---|---|---|---|
| SRR5163978 | 22 | 134 | 79 | 174 | 7407 | 41 | 18 | 852 | 4501 | 27128 | 11526 | 16180 | 0 | 1338 | 821 | 2524 | 36 |
| SRR5163979 | 0 | 93 | 63 | 181 | 4193 | 76 | 15 | 1211 | 3675 | 42343 | 1605 | 3354 | 0 | 1895 | 31 | 41 | 30 |
| SRR5163980 | 0 | 11 | 41 | 163 | 4095 | 78 | 24 | 1142 | 3727 | 42577 | 115 | 3324 | 0 | 2149 | 0 | 84 | 24 |
| SRR5163981 | 0 | 188 | 74 | 127 | 956 | 40 | 15 | 3794 | 42691 | 38134 | 200 | 307 | 0 | 636 | 89 | 151 | 61 |
| SRR5163982 | 0 | 137 | 48 | 205 | 4731 | 37 | 13 | 609 | 3852 | 30569 | 436 | 3515 | 11 | 1336 | 83 | 61 | 47 |
| SRR5163983 | 0 | 149 | 39 | 126 | 1148 | 34 | 0 | 552 | 3343 | 31562 | 205 | 3319 | 0 | 1277 | 158 | 308 | 20 |
| SRR5163984 | 0 | 13 | 50 | 106 | 1105 | 34 | 0 | 333 | 3466 | 32195 | 94 | 3251 | 0 | 1469 | 118 | 255 | 46 |
| SRR5163985 | 0 | 140 | 0 | 13 | 192 | 53 | 29 | 1306 | 8492 | 36799 | 55 | 11432 | 0 | 653 | 435 | 327 | 23 |
| SRR5163986 | 0 | 88 | 28 | 13 | 173 | 14 | 0 | 1427 | 9903 | 35250 | 68 | 11918 | 0 | 587 | 0 | 504 | 37 |
| SRR5163987 | 0 | 151 | 64 | 121 | 2364 | 63 | 15 | 1909 | 13357 | 28990 | 1268 | 2138 | 0 | 1192 | 48 | 133 | 32 |
| SRR5168444 | 0 | 0 | 23 | 428 | 116 | 0 | 0 | 10271 | 46813 | 24378 | 24 | 20 | 0 | 180 | 256 | 642 | 25 |
| SRR5168445 | 0 | 148 | 57 | 461 | 183 | 0 | 12 | 2089 | 11981 | 8316 | 0 | 0 | 0 | 536 | 93 | 97 | 0 |
| SRR5168447 | 0 | 546 | 140 | 517 | 64 | 0 | 0 | 719 | 2152 | 5317 | 253 | 308 | 0 | 268 | 510 | 569 | 40 |
| SRR5168448 | 0 | 85 | 0 | 212 | 134 | 0 | 14 | 2654 | 22554 | 35273 | 23 | 11 | 0 | 718 | 125 | 194 | 140 |
| SRR5168451 | 0 | 99 | 11 | 445 | 132 | 0 | 0 | 1045 | 3013 | 5637 | 11 | 0 | 0 | 423 | 357 | 560 | 22 |
| SRR5168454 | 0 | 106 | 18 | 431 | 69 | 0 | 0 | 9940 | 32386 | 13052 | 0 | 0 | 0 | 130 | 341 | 75 | 0 |
| SRR5168458 | 0 | 31 | 44 | 394 | 87 | 0 | 11 | 293 | 1333 | 3473 | 0 | 0 | 0 | 268 | 82 | 23 | 0 |
| SRR5168460 | 0 | 48 | 0 | 954 | 121 | 0 | 0 | 2652 | 19407 | 18586 | 0 | 0 | 0 | 222 | 76 | 24 | 29 |
| SRR5168465 | 0 | 39 | 14 | 441 | 398 | 0 | 0 | 2243 | 6833 | 58444 | 37 | 14 | 0 | 290 | 1376 | 2035 | 0 |
| SRR5168467 | 0 | 18 | 0 | 2320 | 176 | 0 | 0 | 698 | 4422 | 52026 | 0 | 0 | 0 | 533 | 10 | 11 | 20 |
| SRR5168471 | 0 | 130 | 131 | 1371 | 500 | 0 | 0 | 1875 | 8424 | 268768 | 0 | 0 | 0 | 255 | 261 | 483 | 0 |
| SRR5168475 | 0 | 40 | 12 | 233 | 23 | 0 | 0 | 6990 | 34760 | 22460 | 0 | 18 | 0 | 115 | 87 | 154 | 36 |
| SRR5171728 | 0 | 27 | 15 | 0 | 0 | 0 | 0 | 269 | 77 | 1306 | 2535 | 2803 | 0 | 230 | 0 | 77 | 0 |
| SRR5171729 | 0 | 36 | 0 | 0 | 0 | 0 | 0 | 195 | 65 | 3122 | 69 | 2895 | 0 | 2016 | 0 | 163 | 0 |
| SRR5171730 | 0 | 16 | 11 | 0 | 0 | 0 | 0 | 416 | 59 | 2616 | 1486 | 2438 | 0 | 297 | 59 | 0 | 0 |
| SRR5171731 | 0 | 12 | 15 | 0 | 0 | 0 | 0 | 199 | 0 | 1870 | 46 | 3620 | 0 | 438 | 40 | 40 | 0 |
| SRR5171732 | 0 | 0 | 0 | 0 | 0 | 0 | 0 | 476 | 0 | 1811 | 1048 | 858 | 0 | 95 | 0 | 95 | 0 |
| SRR5171733 | 0 | 19 | 19 | 0 | 11 | 0 | 0 | 21 | 21 | 2022 | 834 | 938 | 0 | 855 | 0 | 0 | 0 |
| SRR5195030 | 0 | 0 | 0 | 0 | 0 | 0 | 0 | 2602 | 1827 | 41717 | 1235 | 936 | 0 | 13569 | 0 | 0 | 0 |
| SRR5195032 | 0 | 11 | 0 | 0 | 0 | 0 | 0 | 957 | 630 | 67448 | 1925 | 3547 | 0 | 4985 | 0 | 0 | 0 |
| SRR5195033 | 0 | 31 | 0 | 0 | 0 | 0 | 0 | 1937 | 1080 | 38755 | 583 | 611 | 0 | 18397 | 11 | 0 | 0 |
| SRR5195034 | 0 | 0 | 22 | 0 | 0 | 0 | 0 | 725 | 388 | 41631 | 616 | 668 | 0 | 8191 | 0 | 11 | 0 |
| SRR5224010 | 0 | 492 | 175 | 100 | 180 | 0 | 12 | 1834 | 5200 | 21731 | 1507 | 1699 | 0 | 1563 | 72 | 56 | 46 |
| SRR5224011 | 0 | 77 | 36 | 54 | 166 | 0 | 0 | 2234 | 4586 | 36112 | 1290 | 1187 | 0 | 3015 | 15 | 22 | 27 |
| SRR5224012 | 0 | 81 | 35 | 64 | 205 | 0 | 0 | 1154 | 2341 | 17756 | 1355 | 1665 | 0 | 4375 | 26 | 11 | 46 |
| SRR5224013 | 0 | 170 | 27 | 49 | 192 | 0 | 16 | 1492 | 3259 | 38106 | 3083 | 3198 | 0 | 4847 | 619 | 998 | 16 |
| SRR5224014 | 0 | 106 | 23 | 39 | 108 | 0 | 16 | 1677 | 2917 | 12197 | 457 | 1640 | 0 | 2169 | 136 | 191 | 19 |
| SRR5224015 | 0 | 308 | 37 | 14 | 87 | 0 | 16 | 1964 | 3652 | 26519 | 2358 | 3092 | 0 | 2905 | 614 | 926 | 27 |
| SRR5224016 | 0 | 365 | 38 | 17 | 112 | 0 | 29 | 1270 | 2170 | 6558 | 1249 | 5861 | 0 | 3494 | 786 | 944 | 37 |
| SRR5224017 | 0 | 275 | 43 | 20 | 59 | 0 | 23 | 644 | 1400 | 10026 | 1364 | 1288 | 0 | 3147 | 365 | 446 | 116 |
| SRR5224018 | 0 | 1253 | 107 | 18 | 152 | 0 | 33 | 1091 | 1755 | 11893 | 1025 | 7173 | 0 | 3629 | 734 | 1510 | 118 |
| SRR5224019 | 0 | 235 | 25 | 60 | 163 | 0 | 19 | 1290 | 2820 | 21512 | 5280 | 4840 | 0 | 6514 | 541 | 711 | 19 |
| SRR5224020 | 0 | 253 | 26 | 0 | 57 | 0 | 27 | 1614 | 2932 | 13419 | 3667 | 4357 | 0 | 4618 | 845 | 1008 | 27 |
| SRR5224021 | 0 | 273 | 21 | 40 | 140 | 0 | 25 | 682 | 2058 | 26896 | 1756 | 5594 | 0 | 3698 | 497 | 986 | 21 |
| SRR5224022 | 0 | 314 | 35 | 10 | 98 | 0 | 23 | 1293 | 2327 | 13381 | 3733 | 3645 | 0 | 4314 | 703 | 1099 | 15 |
| SRR5224023 | 0 | 953 | 66 | 43 | 218 | 0 | 21 | 763 | 1530 | 12144 | 846 | 2743 | 0 | 3710 | 223 | 299 | 121 |
| SRR5224024 | 0 | 1126 | 102 | 13 | 113 | 0 | 31 | 1624 | 2402 | 10583 | 7802 | 7902 | 0 | 7012 | 583 | 998 | 108 |
| SRR5230550 | 0 | 13 | 22 | 170 | 82 | 0 | 11 | 3707 | 5780 | 3530 | 0 | 28 | 0 | 444 | 42 | 173 | 0 |
| SRR5230551 | 0 | 37 | 15 | 17 | 46 | 0 | 0 | 1148 | 1928 | 1220 | 0 | 0 | 0 | 233 | 0 | 18 | 0 |
| SRR5230552 | 13 | 36 | 18 | 63 | 59 | 13 | 31 | 7931 | 7644 | 2983 | 28 | 251 | 0 | 461 | 0 | 36 | 38 |
| SRR5230553 | 0 | 26 | 0 | 70 | 247 | 0 | 0 | 2819 | 3379 | 1647 | 33 | 46 | 0 | 560 | 119 | 244 | 22 |
| SRR5230554 | 0 | 11 | 12 | 74 | 97 | 0 | 0 | 951 | 1656 | 4734 | 20 | 42 | 0 | 749 | 35 | 76 | 24 |
| SRR5230555 | 0 | 17 | 11 | 206 | 140 | 0 | 0 | 3500 | 5846 | 3445 | 39 | 39 | 0 | 951 | 39 | 179 | 12 |
| SRR5230556 | 0 | 0 | 15 | 25 | 18 | 20 | 0 | 2519 | 7669 | 1965 | 13 | 86 | 0 | 2671 | 55 | 106 | 12 |
| SRR5230557 | 0 | 13 | 11 | 90 | 117 | 0 | 0 | 2391 | 4003 | 2428 | 29 | 70 | 0 | 856 | 33 | 149 | 20 |
| SRR5230558 | 0 | 18 | 10 | 110 | 69 | 0 | 13 | 749 | 1188 | 2030 | 16 | 125 | 0 | 699 | 34 | 44 | 41 |
| SRR5230559 | 0 | 14 | 16 | 40 | 40 | 0 | 16 | 1368 | 2305 | 3049 | 0 | 35 | 0 | 1435 | 84 | 320 | 0 |
| SRR5230560 | 0 | 0 | 12 | 31 | 40 | 0 | 0 | 3435 | 3852 | 3007 | 22 | 81 | 0 | 531 | 43 | 141 | 19 |
| SRR5230561 | 0 | 0 | 0 | 29 | 48 | 0 | 14 | 3425 | 4297 | 1852 | 17 | 51 | 0 | 217 | 11 | 28 | 0 |
| SRR5230562 | 0 | 0 | 0 | 26 | 35 | 0 | 0 | 1813 | 1556 | 1590 | 11 | 40 | 0 | 1249 | 134 | 183 | 0 |
| SRR5230563 | 0 | 14 | 16 | 42 | 28 | 0 | 12 | 5667 | 4303 | 1250 | 29 | 160 | 0 | 431 | 23 | 67 | 11 |
| SRR5230564 | 0 | 16 | 0 | 73 | 43 | 0 | 0 | 1563 | 1765 | 1974 | 13 | 42 | 0 | 751 | 151 | 316 | 16 |
| SRR5230565 | 0 | 25 | 0 | 36 | 29 | 15 | 0 | 213 | 502 | 901 | 60 | 119 | 0 | 298 | 0 | 17 | 35 |
| SRR5230566 | 0 | 13 | 0 | 189 | 39 | 0 | 0 | 1674 | 2839 | 1339 | 0 | 37 | 0 | 423 | 31 | 173 | 0 |
| SRR5230567 | 0 | 0 | 0 | 28 | 22 | 0 | 19 | 3819 | 3553 | 1583 | 29 | 23 | 0 | 394 | 35 | 47 | 28 |
| SRR5230568 | 0 | 0 | 0 | 34 | 38 | 0 | 10 | 2551 | 2361 | 1241 | 11 | 16 | 0 | 2719 | 42 | 95 | 12 |
| SRR5230569 | 0 | 17 | 11 | 75 | 133 | 42 | 0 | 2412 | 3237 | 2526 | 11 | 38 | 0 | 669 | 95 | 285 | 0 |
| SRR5230570 | 0 | 23 | 20 | 38 | 49 | 0 | 0 | 2407 | 2888 | 1595 | 21 | 77 | 0 | 795 | 17 | 43 | 15 |
| SRR5230571 | 0 | 57 | 15 | 46 | 84 | 0 | 51 | 2250 | 3388 | 2155 | 97 | 144 | 0 | 340 | 34 | 44 | 29 |
| SRR5230572 | 0 | 28 | 20 | 106 | 95 | 0 | 0 | 2371 | 5304 | 4657 | 0 | 39 | 0 | 673 | 116 | 154 | 0 |
| SRR5230573 | 0 | 41 | 25 | 413 | 102 | 0 | 0 | 3413 | 6721 | 2752 | 20 | 20 | 0 | 326 | 75 | 271 | 0 |
| SRR5230574 | 0 | 25 | 14 | 56 | 56 | 0 | 21 | 2707 | 4430 | 1265 | 90 | 297 | 0 | 723 | 25 | 51 | 17 |
| SRR5230575 | 0 | 26 | 28 | 65 | 28 | 0 | 0 | 2890 | 4624 | 1041 | 58 | 246 | 0 | 780 | 29 | 72 | 14 |
| SRR5230576 | 0 | 32 | 15 | 205 | 240 | 0 | 0 | 1275 | 2140 | 1071 | 37 | 113 | 0 | 392 | 23 | 66 | 19 |
| SRR5230577 | 0 | 15 | 20 | 191 | 226 | 0 | 0 | 1342 | 2128 | 1168 | 10 | 131 | 0 | 338 | 33 | 44 | 20 |
| SRR5230578 | 0 | 17 | 16 | 34 | 67 | 0 | 0 | 186 | 638 | 5469 | 15 | 70 | 0 | 718 | 221 | 384 | 45 |
| SRR5230579 | 0 | 37 | 27 | 54 | 77 | 0 | 16 | 313 | 560 | 5456 | 0 | 78 | 0 | 677 | 286 | 573 | 27 |
| SRR5230580 | 0 | 0 | 0 | 120 | 136 | 0 | 10 | 1730 | 3990 | 9819 | 0 | 21 | 0 | 2811 | 255 | 1654 | 25 |
| SRR5230581 | 0 | 0 | 22 | 79 | 110 | 0 | 0 | 1471 | 2898 | 8954 | 0 | 0 | 0 | 3244 | 389 | 2379 | 22 |
| SRR5230582 | 0 | 21 | 50 | 42 | 94 | 0 | 0 | 985 | 2041 | 2692 | 0 | 46 | 0 | 1374 | 176 | 602 | 11 |
| SRR5230583 | 0 | 0 | 86 | 40 | 80 | 0 | 0 | 981 | 2491 | 2589 | 0 | 59 | 0 | 1432 | 255 | 451 | 0 |
| SRR5230584 | 0 | 0 | 0 | 45 | 83 | 0 | 0 | 596 | 851 | 1516 | 19 | 148 | 16 | 1064 | 41 | 164 | 35 |
| SRR5230585 | 0 | 0 | 17 | 48 | 101 | 0 | 0 | 639 | 881 | 1141 | 69 | 207 | 22 | 985 | 35 | 190 | 43 |
| SRR5230586 | 0 | 0 | 12 | 116 | 36 | 0 | 0 | 8707 | 8322 | 3469 | 13 | 31 | 0 | 222 | 0 | 36 | 12 |
| SRR5230587 | 0 | 0 | 0 | 104 | 22 | 0 | 15 | 8248 | 8607 | 3847 | 0 | 16 | 0 | 456 | 65 | 33 | 0 |
| SRR5230588 | 0 | 0 | 12 | 60 | 92 | 0 | 0 | 1935 | 3275 | 3815 | 17 | 83 | 0 | 1125 | 193 | 309 | 0 |
| SRR5230589 | 0 | 0 | 0 | 44 | 74 | 0 | 15 | 1889 | 2954 | 3469 | 0 | 69 | 0 | 1133 | 34 | 172 | 0 |
| SRR5230590 | 0 | 28 | 0 | 13 | 27 | 0 | 28 | 1959 | 2612 | 1959 | 163 | 490 | 14 | 0 | 0 | 163 | 0 |
| SRR5230591 | 0 | 37 | 0 | 18 | 33 | 0 | 32 | 2486 | 2435 | 1000 | 369 | 715 | 0 | 229 | 0 | 0 | 25 |
| SRR5230592 | 0 | 0 | 0 | 10 | 58 | 48 | 48 | 1364 | 1636 | 1364 | 0 | 273 | 0 | 0 | 0 | 0 | 48 |
| SRR5230593 | 0 | 76 | 0 | 34 | 59 | 12 | 35 | 1532 | 1705 | 1647 | 105 | 211 | 0 | 86 | 0 | 0 | 10 |
| SRR5230594 | 0 | 27 | 25 | 103 | 105 | 0 | 0 | 5136 | 8962 | 2484 | 0 | 26 | 0 | 645 | 21 | 142 | 0 |
| SRR5230595 | 0 | 23 | 28 | 81 | 81 | 0 | 0 | 5021 | 9370 | 2577 | 0 | 0 | 0 | 644 | 27 | 242 | 0 |
| SRR5230596 | 0 | 22 | 15 | 46 | 50 | 0 | 14 | 2338 | 2812 | 1824 | 84 | 179 | 0 | 913 | 17 | 76 | 25 |
| SRR5230597 | 0 | 15 | 17 | 50 | 64 | 0 | 20 | 2475 | 2974 | 1767 | 53 | 236 | 0 | 883 | 17 | 79 | 26 |
| SRR5230600 | 0 | 16 | 0 | 52 | 74 | 0 | 0 | 2964 | 3594 | 483 | 35 | 35 | 0 | 116 | 0 | 20 | 0 |
| SRR5230601 | 0 | 27 | 10 | 50 | 84 | 0 | 10 | 2465 | 3009 | 512 | 32 | 48 | 0 | 128 | 16 | 16 | 0 |
| SRR5230602 | 0 | 12 | 16 | 50 | 62 | 11 | 0 | 2363 | 2076 | 2967 | 14 | 38 | 0 | 1067 | 304 | 615 | 19 |
| SRR5230603 | 0 | 0 | 22 | 20 | 85 | 0 | 0 | 2353 | 2180 | 3087 | 0 | 86 | 0 | 842 | 302 | 561 | 37 |
| SRR5230604 | 0 | 16 | 14 | 68 | 89 | 19 | 17 | 2933 | 2622 | 1732 | 82 | 203 | 0 | 651 | 16 | 65 | 14 |
| SRR5230605 | 0 | 24 | 12 | 70 | 74 | 23 | 18 | 2701 | 2481 | 1945 | 101 | 220 | 0 | 510 | 0 | 44 | 24 |
| SRR5230606 | 0 | 0 | 0 | 60 | 139 | 0 | 25 | 1042 | 1153 | 3014 | 18 | 113 | 0 | 1091 | 101 | 152 | 22 |
| SRR5230607 | 0 | 0 | 17 | 49 | 137 | 0 | 35 | 825 | 985 | 3229 | 29 | 87 | 0 | 1144 | 58 | 116 | 0 |
| SRR5230608 | 0 | 0 | 0 | 35 | 73 | 0 | 0 | 3822 | 4440 | 2413 | 0 | 58 | 0 | 1313 | 63 | 227 | 0 |
| SRR5230609 | 0 | 12 | 12 | 27 | 57 | 0 | 0 | 3326 | 4525 | 2337 | 0 | 30 | 0 | 1378 | 90 | 150 | 0 |
| SRR5230610 | 0 | 29 | 0 | 65 | 126 | 0 | 0 | 2710 | 2830 | 1144 | 60 | 241 | 0 | 482 | 0 | 60 | 20 |
| SRR5230611 | 0 | 29 | 0 | 68 | 99 | 0 | 17 | 2681 | 2734 | 910 | 34 | 174 | 0 | 331 | 0 | 0 | 25 |
| SRR5230613 | 0 | 25 | 0 | 77 | 100 | 0 | 31 | 5392 | 8710 | 1091 | 22 | 58 | 0 | 87 | 0 | 0 | 29 |
| SRR5230614 | 13 | 23 | 0 | 49 | 64 | 23 | 43 | 1777 | 2081 | 2549 | 94 | 47 | 10 | 585 | 0 | 0 | 150 |
| SRR5230615 | 0 | 17 | 0 | 55 | 75 | 27 | 59 | 1655 | 1831 | 2715 | 25 | 41 | 0 | 482 | 0 | 0 | 153 |
| SRR5230617 | 0 | 23 | 0 | 48 | 156 | 0 | 19 | 4500 | 11364 | 1287 | 41 | 105 | 0 | 264 | 0 | 23 | 35 |
| SRR5230618 | 0 | 33 | 0 | 40 | 98 | 0 | 55 | 2083 | 3067 | 1215 | 58 | 116 | 0 | 521 | 0 | 0 | 22 |
| SRR5230619 | 0 | 44 | 0 | 36 | 118 | 0 | 41 | 2107 | 2329 | 1181 | 44 | 182 | 0 | 562 | 12 | 16 | 48 |
| SRR5230620 | 0 | 14 | 0 | 38 | 55 | 0 | 26 | 2545 | 2472 | 2034 | 62 | 235 | 0 | 609 | 22 | 64 | 22 |
| SRR5230621 | 0 | 11 | 0 | 39 | 70 | 0 | 0 | 2692 | 2892 | 1907 | 108 | 154 | 0 | 769 | 15 | 62 | 23 |
| SRR5230622 | 0 | 10 | 0 | 14 | 50 | 0 | 0 | 881 | 2337 | 4241 | 34 | 49 | 0 | 803 | 63 | 175 | 0 |
| SRR5230623 | 0 | 0 | 0 | 17 | 30 | 0 | 0 | 2063 | 3803 | 4980 | 11 | 27 | 0 | 941 | 139 | 220 | 0 |
| SRR5230624 | 0 | 18 | 39 | 14 | 33 | 14 | 11 | 3519 | 4659 | 2141 | 28 | 79 | 0 | 426 | 82 | 464 | 0 |
| SRR5230625 | 0 | 19 | 18 | 51 | 123 | 0 | 0 | 2727 | 3420 | 2395 | 14 | 37 | 0 | 2026 | 95 | 335 | 0 |
| SRR5230626 | 0 | 25 | 43 | 17 | 23 | 0 | 16 | 3563 | 3781 | 1681 | 25 | 36 | 0 | 432 | 35 | 99 | 0 |
| SRR5230627 | 0 | 13 | 14 | 71 | 57 | 0 | 0 | 1126 | 1988 | 1723 | 15 | 61 | 0 | 855 | 297 | 591 | 0 |
| SRR5230628 | 0 | 13 | 23 | 0 | 19 | 0 | 11 | 1902 | 2884 | 1220 | 13 | 80 | 0 | 256 | 83 | 161 | 0 |
| SRR5230629 | 0 | 30 | 12 | 23 | 34 | 0 | 24 | 2981 | 5110 | 4290 | 14 | 25 | 0 | 197 | 37 | 101 | 11 |
| SRR5230630 | 0 | 0 | 10 | 41 | 39 | 0 | 10 | 1353 | 4453 | 2312 | 16 | 39 | 0 | 561 | 56 | 321 | 0 |

| | | | | | | | | | | | | | | | | | | | |
|---|---|---|---|---|---|---|---|---|---|---|---|---|---|---|---|---|---|---|---|
| SRR5163978 | 246 | 99 | 2038 | 1397 | 300 | 110 | 1302 | 388 | 0 | 30 | 36 | 43 | 165 | 424 | 46 | 73 | 46 | 28 | 3710 | 63 |
| SRR5163979 | 648 | 27 | 238 | 277 | 274 | 158 | 1049 | 414 | 0 | 36 | 0 | 13 | 76 | 124 | 20 | 15 | 13 | 0 | 238 | 10 |
| SRR5163980 | 649 | 14 | 101 | 271 | 252 | 163 | 1087 | 411 | 0 | 27 | 11 | 19 | 35 | 12 | 0 | 16 | 0 | 11 | 302 | 12 |
| SRR5163981 | 550 | 20 | 292 | 1486 | 372 | 483 | 3078 | 751 | 0 | 86 | 0 | 23 | 64 | 202 | 43 | 37 | 42 | 42 | 799 | 0 |
| SRR5163982 | 1128 | 16 | 352 | 47 | 1233 | 457 | 5078 | 1203 | 0 | 69 | 13 | 43 | 127 | 21 | 60 | 84 | 118 | 31 | 711 | 364 |
| SRR5163983 | 460 | 16 | 260 | 2571 | 517 | 445 | 3631 | 533 | 0 | 20 | 0 | 24 | 66 | 278 | 17 | 24 | 19 | 31 | 544 | 0 |
| SRR5163984 | 493 | 20 | 196 | 2614 | 502 | 448 | 3524 | 508 | 0 | 34 | 0 | 0 | 50 | 23 | 25 | 25 | 25 | 21 | 529 | 0 |
| SRR5163985 | 68 | 0 | 218 | 221 | 52 | 219 | 670 | 72 | 0 | 35 | 0 | 0 | 41 | 199 | 18 | 29 | 0 | 0 | 1633 | 53 |
| SRR5163986 | 62 | 0 | 839 | 201 | 39 | 230 | 667 | 71 | 0 | 14 | 0 | 14 | 56 | 205 | 19 | 0 | 19 | 0 | 1930 | 54 |
| SRR5163987 | 571 | 12 | 298 | 1345 | 445 | 486 | 3472 | 586 | 0 | 46 | 0 | 28 | 143 | 460 | 46 | 61 | 51 | 30 | 668 | 0 |
| SRR5168444 | 747 | 74 | 5958 | 177 | 149 | 912 | 5594 | 1229 | 11 | 101 | 0 | 78 | 71 | 103 | 13 | 16 | 12 | 10 | 26790 | 0 |
| SRR5168445 | 6848 | 87 | 367 | 631 | 2649 | 87 | 7566 | 5457 | 0 | 23 | 0 | 13 | 18 | 31 | 21 | 21 | 27 | 45 | 1280 | 0 |
| SRR5168447 | 3210 | 80 | 2875 | 682 | 1279 | 238 | 9071 | 7398 | 19 | 31 | 21 | 101 | 110 | 97 | 15 | 13 | 12 | 0 | 9048 | 0 |
| SRR5168448 | 994 | 111 | 861 | 1784 | 924 | 1210 | 11177 | 3235 | 0 | 35 | 0 | 14 | 91 | 85 | 22 | 0 | 0 | 0 | 3298 | 83 |
| SRR5168451 | 4458 | 174 | 1519 | 1905 | 2649 | 351 | 21475 | 9156 | 0 | 0 | 0 | 0 | 13 | 16 | 0 | 0 | 0 | 0 | 5714 | 0 |
| SRR5168454 | 3031 | 29 | 340 | 381 | 459 | 169 | 3185 | 3216 | 12 | 0 | 0 | 39 | 22 | 24 | 13 | 0 | 0 | 0 | 1103 | 0 |
| SRR5168458 | 5799 | 46 | 661 | 867 | 1171 | 444 | 8303 | 3910 | 14 | 40 | 0 | 30 | 20 | 11 | 0 | 0 | 0 | 0 | 2186 | 0 |
| SRR5168460 | 3024 | 79 | 52 | 571 | 624 | 252 | 7500 | 4895 | 0 | 24 | 0 | 18 | 25 | 29 | 12 | 0 | 0 | 0 | 150 | 0 |
| SRR5168465 | 2263 | 47 | 765 | 386 | 917 | 659 | 18356 | 4721 | 30 | 0 | 22 | 13 | 123 | 11 | 24 | 30 | 34 | 0 | 1466 | 0 |
| SRR5168467 | 4612 | 102 | 14 | 597 | 515 | 482 | 14080 | 8634 | 34 | 0 | 0 | 0 | 0 | 0 | 0 | 0 | 0 | 0 | 31 | 0 |
| SRR5168471 | 1540 | 20 | 183 | 1095 | 549 | 392 | 10053 | 4361 | 0 | 0 | 0 | 23 | 26 | 30 | 0 | 0 | 0 | 0 | 527 | 0 |
| SRR5168475 | 1947 | 139 | 108 | 1252 | 973 | 157 | 9648 | 5305 | 20 | 20 | 0 | 0 | 0 | 0 | 0 | 0 | 0 | 0 | 378 | 0 |
| SRR5171728 | 198 | 198 | 1843 | 624 | 1030 | 0 | 347 | 20703 | 178 | 0 | 21 | 0 | 13 | 50 | 42 | 50 | 48 | 199 | 5185 | 0 |
| SRR5171729 | 297 | 190 | 943 | 428 | 999 | 0 | 321 | 22174 | 107 | 0 | 63 | 11 | 20 | 34 | 38 | 51 | 74 | 257 | 2862 | 0 |
| SRR5171730 | 160 | 175 | 2319 | 859 | 2432 | 0 | 422 | 21464 | 73 | 0 | 16 | 0 | 16 | 16 | 10 | 16 | 30 | 31 | 5411 | 0 |
| SRR5171731 | 128 | 279 | 1432 | 512 | 1536 | 0 | 582 | 22490 | 105 | 0 | 15 | 0 | 10 | 16 | 13 | 18 | 26 | 72 | 4773 | 0 |
| SRR5171732 | 120 | 120 | 2859 | 361 | 922 | 0 | 882 | 23897 | 160 | 0 | 0 | 0 | 0 | 22 | 12 | 14 | 27 | 58 | 8576 | 0 |
| SRR5171733 | 98 | 109 | 1689 | 337 | 1154 | 0 | 686 | 26492 | 76 | 0 | 17 | 0 | 13 | 26 | 28 | 33 | 45 | 158 | 8026 | 0 |
| SRR5195030 | 170 | 0 | 278 | 200 | 1409 | 0 | 6870 | 1615 | 95 | 0 | 19 | 16 | 34 | 36 | 0 | 22 | 59 | 0 | 2329 | 0 |
| SRR5195032 | 401 | 15 | 803 | 311 | 3051 | 10 | 5773 | 1122 | 62 | 0 | 0 | 0 | 0 | 0 | 0 | 0 | 22 | 0 | 5733 | 0 |
| SRR5195033 | 232 | 0 | 1914 | 216 | 3692 | 31 | 19822 | 1949 | 150 | 0 | 0 | 0 | 0 | 0 | 0 | 0 | 17 | 0 | 14333 | 0 |
| SRR5195034 | 198 | 0 | 1448 | 246 | 4469 | 0 | 11664 | 1157 | 39 | 0 | 0 | 0 | 14 | 11 | 0 | 33 | 51 | 18 | 14587 | 0 |
| SRR5224010 | 2069 | 26 | 5901 | 153 | 915 | 216 | 7091 | 855 | 0 | 0 | 11 | 114 | 154 | 301 | 81 | 40 | 41 | 40 | 2871 | 0 |
| SRR5224011 | 2146 | 26 | 4549 | 102 | 973 | 232 | 7295 | 737 | 0 | 0 | 0 | 14 | 17 | 34 | 12 | 30 | 28 | 29 | 1976 | 0 |
| SRR5224012 | 2082 | 27 | 3550 | 99 | 895 | 288 | 7518 | 686 | 0 | 0 | 12 | 20 | 23 | 60 | 18 | 15 | 13 | 20 | 2586 | 0 |
| SRR5224013 | 1100 | 154 | 3358 | 36 | 836 | 49 | 12735 | 1450 | 37 | 17 | 13 | 32 | 64 | 93 | 38 | 33 | 22 | 16 | 1601 | 0 |
| SRR5224014 | 1358 | 48 | 4242 | 24 | 272 | 120 | 7301 | 470 | 15 | 19 | 0 | 36 | 36 | 97 | 33 | 21 | 12 | 30 | 2282 | 0 |
| SRR5224015 | 1200 | 201 | 3692 | 12 | 520 | 37 | 10716 | 1056 | 84 | 21 | 15 | 38 | 52 | 67 | 36 | 12 | 23 | 15 | 1871 | 0 |
| SRR5224016 | 1175 | 164 | 5110 | 19 | 351 | 59 | 8547 | 643 | 46 | 14 | 26 | 47 | 55 | 71 | 41 | 13 | 27 | 26 | 2822 | 0 |
| SRR5224017 | 719 | 230 | 3448 | 0 | 207 | 28 | 12436 | 603 | 23 | 41 | 12 | 129 | 166 | 483 | 141 | 37 | 24 | 21 | 2467 | 0 |
| SRR5224018 | 1437 | 158 | 5502 | 16 | 331 | 42 | 8232 | 662 | 46 | 0 | 22 | 147 | 191 | 220 | 124 | 44 | 30 | 15 | 2964 | 0 |
| SRR5224019 | 1540 | 271 | 2675 | 21 | 690 | 32 | 13942 | 1362 | 61 | 19 | 15 | 34 | 68 | 83 | 31 | 25 | 17 | 14 | 1422 | 0 |
| SRR5224020 | 919 | 158 | 4419 | 0 | 345 | 23 | 11166 | 831 | 33 | 14 | 17 | 39 | 52 | 90 | 51 | 17 | 11 | 25 | 2844 | 0 |
| SRR5224021 | 1467 | 278 | 3123 | 14 | 685 | 28 | 13491 | 1209 | 49 | 21 | 20 | 34 | 61 | 84 | 27 | 25 | 20 | 26 | 1585 | 0 |
| SRR5224022 | 1237 | 185 | 4480 | 10 | 426 | 23 | 12287 | 854 | 40 | 23 | 19 | 53 | 77 | 117 | 65 | 23 | 15 | 28 | 2497 | 0 |
| SRR5224023 | 1908 | 337 | 2957 | 33 | 538 | 34 | 16127 | 1446 | 45 | 22 | 19 | 118 | 266 | 319 | 108 | 26 | 16 | 17 | 1450 | 0 |
| SRR5224024 | 959 | 180 | 3504 | 12 | 450 | 20 | 10250 | 927 | 62 | 16 | 19 | 118 | 152 | 260 | 110 | 61 | 45 | 48 | 1660 | 0 |
| SRR5230550 | 374 | 434 | 467 | 100 | 345 | 0 | 147 | 3214 | 151 | 0 | 0 | 0 | 0 | 0 | 0 | 0 | 0 | 0 | 1475 | 0 |
| SRR5230551 | 53 | 150 | 108 | 27 | 42 | 0 | 282 | 3780 | 89 | 0 | 0 | 0 | 0 | 0 | 0 | 0 | 0 | 0 | 179 | 0 |
| SRR5230552 | 424 | 224 | 214 | 157 | 540 | 0 | 180 | 3423 | 231 | 57 | 0 | 0 | 0 | 0 | 0 | 0 | 0 | 0 | 481 | 31 |
| SRR5230553 | 521 | 319 | 72 | 39 | 300 | 0 | 314 | 3259 | 458 | 23 | 0 | 0 | 0 | 0 | 0 | 0 | 0 | 0 | 191 | 0 |
| SRR5230554 | 409 | 43 | 113 | 80 | 359 | 0 | 153 | 4472 | 437 | 0 | 0 | 0 | 0 | 0 | 0 | 0 | 0 | 0 | 335 | 26 |
| SRR5230555 | 596 | 442 | 218 | 96 | 446 | 0 | 342 | 5273 | 309 | 0 | 0 | 0 | 0 | 0 | 0 | 0 | 0 | 0 | 756 | 12 |
| SRR5230556 | 171 | 477 | 464 | 26 | 387 | 17 | 320 | 1008 | 171 | 37 | 0 | 0 | 0 | 0 | 0 | 0 | 0 | 0 | 655 | 22 |
| SRR5230557 | 320 | 412 | 211 | 90 | 446 | 10 | 490 | 3708 | 246 | 11 | 0 | 0 | 0 | 0 | 0 | 0 | 0 | 0 | 356 | 29 |
| SRR5230558 | 541 | 1044 | 68 | 101 | 496 | 0 | 210 | 3764 | 397 | 19 | 0 | 0 | 0 | 0 | 0 | 0 | 0 | 0 | 289 | 22 |
| SRR5230559 | 502 | 643 | 547 | 99 | 884 | 0 | 337 | 2567 | 617 | 13 | 0 | 0 | 0 | 0 | 0 | 0 | 0 | 0 | 808 | 11 |
| SRR5230560 | 198 | 347 | 320 | 57 | 331 | 0 | 179 | 3377 | 239 | 0 | 0 | 0 | 0 | 0 | 0 | 0 | 0 | 0 | 433 | 12 |
| SRR5230561 | 78 | 367 | 131 | 18 | 57 | 0 | 149 | 3082 | 146 | 0 | 0 | 0 | 0 | 0 | 0 | 0 | 0 | 0 | 239 | 0 |
| SRR5230562 | 317 | 386 | 525 | 35 | 433 | 0 | 392 | 2385 | 235 | 16 | 0 | 0 | 0 | 0 | 0 | 0 | 0 | 0 | 956 | 26 |
| SRR5230563 | 384 | 143 | 164 | 32 | 381 | 13 | 381 | 2496 | 175 | 0 | 0 | 0 | 0 | 0 | 0 | 0 | 0 | 0 | 362 | 24 |
| SRR5230564 | 530 | 433 | 187 | 46 | 610 | 0 | 398 | 3846 | 393 | 13 | 0 | 0 | 0 | 0 | 0 | 0 | 0 | 0 | 520 | 14 |
| SRR5230565 | 88 | 283 | 77 | 124 | 217 | 0 | 200 | 2811 | 60 | 67 | 0 | 0 | 0 | 0 | 0 | 0 | 0 | 0 | 111 | 68 |
| SRR5230566 | 931 | 730 | 189 | 43 | 438 | 0 | 276 | 3302 | 211 | 0 | 0 | 0 | 0 | 0 | 0 | 0 | 0 | 0 | 359 | 0 |
| SRR5230567 | 121 | 168 | 210 | 22 | 97 | 0 | 109 | 1913 | 107 | 20 | 0 | 0 | 0 | 0 | 0 | 0 | 0 | 0 | 417 | 0 |
| SRR5230568 | 175 | 238 | 226 | 16 | 141 | 13 | 811 | 1777 | 191 | 24 | 0 | 0 | 0 | 0 | 0 | 0 | 0 | 0 | 784 | 0 |
| SRR5230569 | 517 | 253 | 399 | 45 | 289 | 23 | 430 | 5176 | 324 | 11 | 0 | 0 | 0 | 0 | 0 | 0 | 0 | 0 | 563 | 0 |
| SRR5230570 | 211 | 148 | 77 | 30 | 151 | 0 | 173 | 2679 | 151 | 12 | 0 | 0 | 0 | 0 | 0 | 0 | 0 | 0 | 279 | 12 |
| SRR5230571 | 265 | 133 | 128 | 156 | 138 | 0 | 83 | 4442 | 86 | 13 | 0 | 0 | 0 | 0 | 0 | 0 | 0 | 0 | 426 | 19 |
| SRR5230572 | 566 | 393 | 206 | 171 | 467 | 0 | 131 | 4528 | 175 | 0 | 0 | 0 | 0 | 0 | 0 | 0 | 0 | 0 | 442 | 19 |
| SRR5230573 | 729 | 106 | 321 | 126 | 349 | 0 | 270 | 3645 | 144 | 0 | 0 | 0 | 0 | 0 | 0 | 0 | 0 | 0 | 2155 | 18 |
| SRR5230574 | 466 | 175 | 267 | 45 | 535 | 16 | 434 | 2279 | 366 | 30 | 0 | 0 | 0 | 0 | 0 | 0 | 0 | 0 | 1007 | 11 |
| SRR5230575 | 490 | 250 | 173 | 40 | 485 | 12 | 458 | 2478 | 372 | 33 | 0 | 0 | 0 | 0 | 0 | 0 | 0 | 0 | 867 | 13 |
| SRR5230576 | 1085 | 1259 | 478 | 77 | 366 | 0 | 253 | 6540 | 422 | 18 | 0 | 0 | 0 | 0 | 0 | 0 | 0 | 0 | 1251 | 17 |
| SRR5230577 | 1086 | 1337 | 415 | 77 | 378 | 0 | 251 | 6805 | 430 | 24 | 0 | 0 | 0 | 0 | 0 | 0 | 0 | 0 | 1015 | 25 |
| SRR5230578 | 172 | 297 | 56 | 131 | 211 | 0 | 60 | 3801 | 78 | 17 | 0 | 0 | 0 | 0 | 0 | 0 | 0 | 0 | 124 | 74 |
| SRR5230579 | 163 | 352 | 52 | 149 | 206 | 0 | 68 | 4232 | 91 | 0 | 0 | 0 | 0 | 11 | 0 | 0 | 0 | 0 | 117 | 107 |
| SRR5230580 | 277 | 481 | 413 | 18 | 118 | 19 | 397 | 6753 | 161 | 19 | 0 | 0 | 0 | 0 | 0 | 0 | 0 | 0 | 627 | 12 |
| SRR5230581 | 283 | 468 | 433 | 35 | 102 | 35 | 380 | 7236 | 155 | 15 | 0 | 0 | 0 | 0 | 0 | 0 | 0 | 0 | 173 | 13 |
| SRR5230582 | 402 | 527 | 445 | 37 | 333 | 0 | 301 | 3407 | 438 | 29 | 0 | 0 | 0 | 15 | 0 | 0 | 0 | 0 | 565 | 0 |
| SRR5230583 | 442 | 572 | 392 | 40 | 424 | 0 | 334 | 3536 | 480 | 31 | 0 | 0 | 0 | 24 | 0 | 0 | 0 | 0 | 745 | 0 |
| SRR5230584 | 518 | 1710 | 583 | 46 | 338 | 20 | 547 | 3768 | 412 | 60 | 0 | 0 | 0 | 0 | 0 | 0 | 0 | 0 | 1215 | 38 |
| SRR5230585 | 498 | 1699 | 691 | 36 | 330 | 24 | 590 | 3908 | 380 | 77 | 0 | 0 | 0 | 0 | 0 | 0 | 0 | 0 | 1020 | 31 |
| SRR5230586 | 739 | 243 | 33 | 71 | 628 | 0 | 386 | 3710 | 252 | 0 | 0 | 0 | 0 | 0 | 0 | 0 | 0 | 0 | 156 | 39 |
| SRR5230587 | 715 | 217 | 33 | 78 | 542 | 0 | 390 | 3864 | 316 | 0 | 0 | 0 | 0 | 0 | 0 | 0 | 0 | 0 | 98 | 22 |
| SRR5230588 | 302 | 211 | 259 | 47 | 408 | 0 | 338 | 3610 | 403 | 0 | 0 | 0 | 0 | 0 | 0 | 0 | 0 | 0 | 651 | 0 |
| SRR5230589 | 356 | 257 | 275 | 40 | 430 | 20 | 375 | 3789 | 425 | 0 | 0 | 0 | 0 | 0 | 0 | 0 | 0 | 0 | 859 | 0 |
| SRR5230590 | 217 | 85 | 0 | 71 | 204 | 0 | 113 | 2015 | 131 | 28 | 0 | 0 | 0 | 0 | 0 | 0 | 0 | 0 | 327 | 12 |
| SRR5230591 | 230 | 82 | 229 | 72 | 179 | 0 | 111 | 2192 | 169 | 26 | 0 | 0 | 0 | 0 | 0 | 0 | 0 | 0 | 486 | 17 |
| SRR5230592 | 103 | 92 | 0 | 127 | 267 | 0 | 89 | 3877 | 79 | 0 | 0 | 0 | 0 | 0 | 0 | 0 | 0 | 0 | 273 | 31 |
| SRR5230593 | 106 | 89 | 38 | 166 | 246 | 0 | 95 | 4058 | 109 | 24 | 0 | 0 | 0 | 0 | 0 | 0 | 0 | 0 | 29 | 47 |
| SRR5230594 | 414 | 132 | 202 | 45 | 164 | 21 | 509 | 7481 | 113 | 11 | 0 | 0 | 0 | 0 | 0 | 0 | 0 | 0 | 438 | 0 |
| SRR5230595 | 390 | 153 | 54 | 35 | 191 | 22 | 617 | 7890 | 137 | 11 | 0 | 0 | 0 | 0 | 0 | 17 | 0 | 0 | 430 | 0 |
| SRR5230596 | 908 | 331 | 224 | 93 | 978 | 0 | 295 | 4333 | 532 | 17 | 0 | 0 | 0 | 0 | 0 | 0 | 0 | 0 | 544 | 0 |
| SRR5230597 | 974 | 342 | 297 | 89 | 1034 | 0 | 317 | 4355 | 551 | 17 | 0 | 0 | 0 | 0 | 0 | 0 | 0 | 0 | 542 | 0 |
| SRR5230600 | 338 | 172 | 38 | 16 | 210 | 0 | 326 | 4440 | 214 | 25 | 0 | 0 | 0 | 0 | 0 | 0 | 0 | 0 | 106 | 12 |
| SRR5230601 | 337 | 170 | 96 | 16 | 207 | 0 | 340 | 4590 | 248 | 20 | 0 | 0 | 0 | 0 | 0 | 0 | 0 | 0 | 32 | 10 |
| SRR5230602 | 312 | 109 | 304 | 68 | 539 | 0 | 335 | 4330 | 212 | 15 | 0 | 0 | 0 | 14 | 0 | 0 | 0 | 0 | 1064 | 23 |
| SRR5230603 | 324 | 109 | 345 | 72 | 741 | 0 | 311 | 4755 | 253 | 0 | 0 | 0 | 0 | 0 | 0 | 0 | 0 | 0 | 669 | 38 |
| SRR5230604 | 783 | 334 | 127 | 68 | 820 | 14 | 422 | 4991 | 591 | 42 | 0 | 0 | 0 | 0 | 0 | 0 | 0 | 0 | 253 | 13 |
| SRR5230605 | 734 | 352 | 145 | 48 | 894 | 0 | 405 | 5177 | 656 | 50 | 0 | 0 | 0 | 0 | 0 | 0 | 0 | 0 | 271 | 21 |
| SRR5230606 | 793 | 302 | 134 | 67 | 1134 | 0 | 396 | 4997 | 736 | 26 | 0 | 0 | 0 | 0 | 0 | 0 | 0 | 0 | 263 | 0 |
| SRR5230607 | 868 | 314 | 174 | 81 | 1135 | 17 | 390 | 5310 | 802 | 29 | 0 | 0 | 0 | 0 | 0 | 0 | 0 | 0 | 362 | 0 |
| SRR5230608 | 169 | 182 | 82 | 46 | 313 | 0 | 207 | 4541 | 115 | 15 | 0 | 0 | 0 | 0 | 0 | 0 | 0 | 0 | 217 | 30 |
| SRR5230609 | 168 | 215 | 240 | 54 | 345 | 13 | 221 | 4816 | 148 | 36 | 0 | 0 | 0 | 0 | 0 | 0 | 0 | 0 | 479 | 20 |
| SRR5230610 | 411 | 209 | 60 | 53 | 651 | 11 | 373 | 5442 | 190 | 0 | 0 | 0 | 0 | 0 | 0 | 0 | 0 | 0 | 120 | 15 |
| SRR5230611 | 462 | 211 | 127 | 72 | 597 | 11 | 403 | 5801 | 183 | 16 | 0 | 0 | 0 | 0 | 0 | 0 | 0 | 0 | 202 | 13 |
| SRR5230613 | 804 | 213 | 50 | 53 | 267 | 25 | 980 | 4995 | 153 | 0 | 0 | 0 | 0 | 0 | 0 | 0 | 0 | 0 | 74 | 0 |
| SRR5230614 | 523 | 158 | 23 | 35 | 247 | 20 | 597 | 5423 | 79 | 293 | 0 | 0 | 0 | 0 | 0 | 0 | 0 | 0 | 0 | 15 |
| SRR5230615 | 526 | 207 | 14 | 58 | 238 | 17 | 636 | 5438 | 91 | 360 | 0 | 0 | 0 | 0 | 0 | 0 | 0 | 0 | 20 | 17 |
| SRR5230617 | 857 | 199 | 79 | 42 | 327 | 0 | 597 | 5234 | 516 | 23 | 0 | 0 | 0 | 0 | 0 | 0 | 0 | 0 | 73 | 0 |
| SRR5230618 | 400 | 243 | 116 | 184 | 443 | 0 | 195 | 4369 | 253 | 38 | 0 | 0 | 0 | 0 | 0 | 0 | 0 | 0 | 174 | 34 |
| SRR5230619 | 453 | 222 | 200 | 200 | 448 | 0 | 201 | 4599 | 256 | 35 | 0 | 0 | 0 | 0 | 0 | 0 | 0 | 0 | 224 | 50 |
| SRR5230620 | 932 | 145 | 538 | 54 | 791 | 0 | 295 | 4789 | 1044 | 27 | 0 | 0 | 0 | 0 | 0 | 0 | 0 | 0 | 1296 | 0 |
| SRR5230621 | 1071 | 129 | 492 | 63 | 763 | 0 | 296 | 5026 | 1164 | 27 | 0 | 0 | 0 | 0 | 0 | 0 | 0 | 0 | 1461 | 0 |
| SRR5230622 | 72 | 190 | 331 | 39 | 47 | 0 | 76 | 4523 | 85 | 0 | 0 | 0 | 0 | 0 | 0 | 0 | 0 | 0 | 550 | 11 |
| SRR5230623 | 134 | 172 | 97 | 45 | 307 | 0 | 617 | 3924 | 168 | 0 | 0 | 0 | 0 | 0 | 0 | 0 | 0 | 0 | 346 | 12 |
| SRR5230624 | 736 | 150 | 975 | 98 | 352 | 0 | 345 | 6751 | 544 | 18 | 0 | 0 | 0 | 0 | 0 | 0 | 0 | 0 | 2475 | 10 |
| SRR5230625 | 573 | 142 | 426 | 100 | 721 | 25 | 719 | 5462 | 286 | 0 | 0 | 0 | 0 | 0 | 0 | 0 | 0 | 0 | 475 | 24 |
| SRR5230626 | 162 | 134 | 98 | 59 | 268 | 0 | 143 | 4671 | 96 | 0 | 0 | 0 | 0 | 0 | 0 | 0 | 0 | 0 | 247 | 26 |
| SRR5230627 | 513 | 428 | 120 | 77 | 698 | 0 | 531 | 5220 | 217 | 0 | 0 | 0 | 0 | 0 | 0 | 0 | 0 | 0 | 421 | 43 |
| SRR5230628 | 348 | 125 | 319 | 45 | 246 | 0 | 223 | 5096 | 288 | 13 | 0 | 0 | 0 | 0 | 0 | 0 | 0 | 0 | 694 | 16 |
| SRR5230629 | 573 | 250 | 121 | 162 | 345 | 0 | 78 | 5186 | 114 | 0 | 0 | 0 | 0 | 0 | 0 | 0 | 0 | 0 | 236 | 58 |
| SRR5230630 | 1243 | 1140 | 111 | 92 | 590 | 18 | 461 | 10394 | 669 | 11 | 0 | 0 | 0 | 0 | 0 | 0 | 0 | 0 | 417 | 26 |

### Supp Table S8.4.Columns_DI_to_ES

| | | | | | | | | | | | | | | | | | | | | | | | | | | | | | | | | | | | | | |
|---|---|---|---|---|---|---|---|---|---|---|---|---|---|---|---|---|---|---|---|---|---|---|---|---|---|---|---|---|---|---|---|---|---|---|---|---|---|
| SRR5230631 | 0 | 0 | 13 | 56 | 103 | 0 | 0 | 1331 | 2073 | 3710 | 12 | 21 | 0 | 1107 | 65 | 189 | 0 | 363 | 123 | 227 | 42 | 247 | 0 | 290 | 3463 | 426 | 11 | 0 | 0 | 0 | 0 | 0 | 0 | 0 | 0 | 535 | 0 |
| SRR5230632 | 0 | 10 | 28 | 31 | 45 | 0 | 0 | 4744 | 5316 | 3021 | 19 | 56 | 0 | 424 | 71 | 338 | 0 | 415 | 295 | 280 | 74 | 232 | 0 | 240 | 7069 | 257 | 0 | 0 | 0 | 0 | 0 | 0 | 0 | 0 | 0 | 726 | 13 |
| SRR5230633 | 0 | 12 | 17 | 18 | 24 | 25 | 0 | 3511 | 3491 | 1982 | 0 | 72 | 0 | 630 | 283 | 906 | 0 | 324 | 356 | 456 | 62 | 425 | 0 | 303 | 5488 | 509 | 22 | 0 | 0 | 0 | 0 | 0 | 0 | 0 | 0 | 1063 | 17 |
| SRR5230634 | 0 | 0 | 0 | 45 | 49 | 0 | 0 | 608 | 1691 | 23430 | 0 | 13 | 0 | 2832 | 13 | 19 | 73 | 61 | 397 | 351 | 14 | 96 | 50 | 211 | 790 | 1043 | 87 | 0 | 0 | 0 | 0 | 0 | 0 | 0 | 0 | 564 | 26 |
| SRR5230635 | 0 | 0 | 0 | 0 | 19 | 0 | 37 | 3518 | 7159 | 26926 | 0 | 0 | 23 | 2609 | 818 | 705 | 28 | 43 | 422 | 1259 | 0 | 58 | 19 | 91 | 592 | 470 | 139 | 0 | 0 | 0 | 0 | 0 | 0 | 0 | 0 | 2854 | 13 |
| SRR5230636 | 0 | 0 | 0 | 0 | 23 | 0 | 43 | 655 | 1564 | 17952 | 0 | 19 | 0 | 2865 | 330 | 392 | 46 | 31 | 507 | 311 | 0 | 90 | 27 | 118 | 298 | 838 | 73 | 0 | 0 | 0 | 0 | 0 | 0 | 0 | 0 | 1545 | 0 |
| SRR5230637 | 0 | 16 | 0 | 0 | 24 | 20 | 32 | 8437 | 23701 | 15142 | 0 | 0 | 0 | 1332 | 188 | 403 | 12 | 56 | 562 | 278 | 0 | 96 | 23 | 73 | 329 | 341 | 119 | 0 | 0 | 0 | 0 | 0 | 0 | 0 | 0 | 585 | 34 |
| SRR5230638 | 0 | 0 | 0 | 0 | 38 | 0 | 16 | 767 | 2423 | 6569 | 0 | 14 | 0 | 4311 | 764 | 2119 | 24 | 160 | 765 | 2159 | 0 | 406 | 36 | 165 | 643 | 2343 | 77 | 0 | 0 | 0 | 0 | 0 | 0 | 0 | 0 | 3951 | 0 |
| SRR5230639 | 0 | 0 | 0 | 0 | 29 | 0 | 15 | 847 | 3398 | 5848 | 0 | 0 | 22 | 2502 | 1088 | 1221 | 0 | 162 | 463 | 1176 | 0 | 245 | 27 | 225 | 859 | 1713 | 95 | 0 | 0 | 0 | 0 | 0 | 0 | 0 | 0 | 2401 | 11 |
| SRR5230640 | 0 | 0 | 0 | 0 | 28 | 0 | 0 | 1472 | 5351 | 4463 | 0 | 21 | 0 | 1346 | 255 | 780 | 0 | 76 | 220 | 979 | 0 | 107 | 33 | 80 | 374 | 611 | 39 | 0 | 0 | 0 | 0 | 0 | 0 | 0 | 0 | 2006 | 0 |
| SRR5230641 | 0 | 0 | 0 | 0 | 12 | 0 | 53 | 180 | 322 | 8960 | 0 | 0 | 0 | 8803 | 785 | 1790 | 33 | 87 | 508 | 530 | 0 | 213 | 37 | 213 | 254 | 633 | 55 | 0 | 0 | 0 | 0 | 0 | 0 | 0 | 0 | 1913 | 21 |
| SRR5230642 | 0 | 0 | 0 | 25 | 58 | 0 | 0 | 519 | 2604 | 59060 | 0 | 0 | 0 | 1562 | 0 | 32 | 53 | 28 | 549 | 278 | 18 | 81 | 13 | 69 | 494 | 309 | 13 | 0 | 0 | 0 | 0 | 0 | 0 | 0 | 0 | 428 | 56 |
| SRR5230643 | 0 | 0 | 0 | 0 | 30 | 0 | 0 | 1384 | 4362 | 8217 | 0 | 0 | 0 | 2151 | 1066 | 1646 | 54 | 127 | 628 | 1360 | 16 | 196 | 62 | 215 | 389 | 2168 | 44 | 0 | 0 | 0 | 0 | 0 | 0 | 0 | 0 | 2831 | 0 |
| SRR5230644 | 0 | 0 | 0 | 0 | 45 | 0 | 0 | 1230 | 3800 | 7873 | 0 | 19 | 0 | 2555 | 717 | 1529 | 0 | 124 | 459 | 2377 | 0 | 252 | 46 | 185 | 458 | 1351 | 74 | 0 | 0 | 0 | 0 | 0 | 0 | 0 | 0 | 4755 | 0 |
| SRR5230645 | 0 | 10 | 0 | 0 | 40 | 0 | 0 | 207 | 807 | 10386 | 30 | 52 | 0 | 3125 | 1654 | 2498 | 0 | 162 | 35 | 657 | 12 | 298 | 64 | 276 | 626 | 2761 | 44 | 0 | 0 | 0 | 0 | 0 | 0 | 0 | 0 | 1891 | 29 |
| SRR5230646 | 0 | 0 | 0 | 0 | 42 | 0 | 0 | 1880 | 8408 | 6700 | 0 | 0 | 0 | 2606 | 538 | 659 | 22 | 100 | 1233 | 1090 | 0 | 186 | 47 | 186 | 432 | 1010 | 20 | 0 | 0 | 0 | 0 | 0 | 0 | 0 | 0 | 2819 | 0 |
| SRR5230647 | 0 | 0 | 0 | 0 | 10 | 0 | 0 | 52 | 671 | 8289 | 0 | 0 | 0 | 3370 | 262 | 521 | 0 | 0 | 222 | 495 | 0 | 18 | 0 | 29 | 180 | 330 | 24 | 0 | 0 | 0 | 0 | 0 | 0 | 0 | 0 | 1522 | 0 |
| SRR5230648 | 0 | 0 | 0 | 0 | 32 | 15 | 10 | 288 | 755 | 16578 | 16 | 12 | 0 | 2606 | 1219 | 1690 | 43 | 110 | 637 | 624 | 13 | 271 | 35 | 191 | 924 | 773 | 189 | 0 | 0 | 0 | 0 | 0 | 0 | 0 | 0 | 1927 | 21 |
| SRR5230649 | 0 | 0 | 0 | 0 | 64 | 19 | 57 | 427 | 925 | 13583 | 0 | 0 | 0 | 4050 | 212 | 634 | 123 | 102 | 480 | 1097 | 0 | 238 | 58 | 288 | 445 | 988 | 198 | 0 | 0 | 0 | 0 | 0 | 0 | 0 | 0 | 4774 | 0 |
| SRR5230650 | 0 | 0 | 0 | 0 | 16 | 0 | 0 | 116 | 142 | 5518 | 0 | 0 | 0 | 3707 | 673 | 860 | 0 | 13 | 110 | 375 | 0 | 39 | 0 | 22 | 308 | 393 | 32 | 0 | 0 | 0 | 0 | 0 | 0 | 0 | 0 | 1643 | 0 |
| SRR5230651 | 0 | 0 | 0 | 15 | 75 | 14 | 0 | 697 | 2640 | 9734 | 0 | 30 | 0 | 3400 | 847 | 1833 | 21 | 176 | 1296 | 447 | 0 | 324 | 49 | 242 | 719 | 2152 | 63 | 0 | 0 | 0 | 0 | 0 | 0 | 0 | 0 | 747 | 19 |
| SRR5230652 | 0 | 0 | 0 | 20 | 57 | 12 | 170 | 252 | 729 | 32973 | 0 | 0 | 0 | 7365 | 371 | 890 | 280 | 98 | 1160 | 362 | 0 | 164 | 45 | 201 | 388 | 670 | 80 | 0 | 0 | 0 | 0 | 0 | 0 | 0 | 0 | 845 | 65 |
| SRR5230653 | 0 | 0 | 0 | 0 | 37 | 14 | 11 | 162 | 436 | 11916 | 0 | 0 | 0 | 3692 | 137 | 343 | 12 | 72 | 585 | 596 | 0 | 198 | 59 | 231 | 658 | 481 | 72 | 0 | 0 | 0 | 0 | 0 | 0 | 0 | 0 | 1814 | 21 |
| SRR5230654 | 0 | 0 | 0 | 0 | 31 | 16 | 20 | 518 | 846 | 26015 | 0 | 0 | 0 | 2354 | 998 | 3063 | 115 | 86 | 1913 | 841 | 13 | 154 | 66 | 278 | 408 | 511 | 76 | 0 | 0 | 0 | 0 | 0 | 0 | 0 | 0 | 1920 | 37 |
| SRR5230655 | 0 | 0 | 0 | 0 | 30 | 22 | 0 | 1522 | 6275 | 5440 | 0 | 15 | 0 | 1824 | 140 | 395 | 0 | 124 | 491 | 1143 | 0 | 288 | 42 | 261 | 641 | 1036 | 46 | 0 | 0 | 0 | 0 | 0 | 0 | 0 | 0 | 2900 | 0 |
| SRR5230656 | 0 | 0 | 0 | 0 | 59 | 0 | 0 | 3439 | 12543 | 5723 | 0 | 0 | 0 | 2205 | 101 | 437 | 33 | 88 | 204 | 311 | 0 | 81 | 67 | 203 | 841 | 510 | 112 | 0 | 0 | 0 | 0 | 0 | 0 | 0 | 0 | 655 | 0 |
| SRR5230657 | 0 | 0 | 0 | 10 | 39 | 0 | 0 | 643 | 3719 | 6733 | 0 | 10 | 0 | 2541 | 148 | 335 | 28 | 117 | 1097 | 578 | 0 | 390 | 66 | 282 | 689 | 430 | 37 | 0 | 0 | 0 | 0 | 0 | 0 | 0 | 0 | 1337 | 63 |
| SRR5230658 | 0 | 0 | 0 | 15 | 62 | 0 | 18 | 2871 | 9660 | 6230 | 0 | 0 | 0 | 2096 | 251 | 392 | 28 | 230 | 524 | 439 | 0 | 324 | 42 | 240 | 1047 | 775 | 56 | 0 | 0 | 0 | 0 | 0 | 0 | 0 | 0 | 748 | 0 |
| SRR5230659 | 0 | 0 | 0 | 13 | 51 | 0 | 14 | 805 | 2999 | 6823 | 0 | 0 | 0 | 3794 | 96 | 400 | 0 | 68 | 392 | 268 | 13 | 268 | 42 | 191 | 291 | 1434 | 36 | 0 | 0 | 0 | 0 | 0 | 0 | 0 | 0 | 550 | 13 |
| SRR5230660 | 0 | 0 | 0 | 0 | 42 | 0 | 0 | 839 | 4096 | 5912 | 0 | 0 | 0 | 1823 | 636 | 563 | 33 | 108 | 253 | 481 | 0 | 138 | 47 | 144 | 1004 | 802 | 66 | 0 | 0 | 0 | 0 | 0 | 0 | 0 | 0 | 1168 | 16 |
| SRR5230661 | 0 | 0 | 0 | 0 | 17 | 0 | 50 | 504 | 1660 | 2623 | 0 | 0 | 0 | 1764 | 53 | 96 | 0 | 68 | 189 | 163 | 0 | 204 | 26 | 102 | 790 | 395 | 97 | 0 | 0 | 0 | 0 | 0 | 0 | 0 | 0 | 937 | 12 |
| SRR5230662 | 0 | 0 | 0 | 0 | 19 | 0 | 0 | 968 | 3652 | 4738 | 0 | 16 | 0 | 2637 | 150 | 71 | 10 | 28 | 182 | 598 | 0 | 40 | 31 | 86 | 903 | 304 | 32 | 0 | 0 | 0 | 0 | 0 | 0 | 0 | 0 | 1440 | 0 |
| SRR5230663 | 0 | 0 | 0 | 0 | 39 | 0 | 0 | 192 | 578 | 4987 | 0 | 0 | 0 | 1985 | 532 | 1278 | 17 | 162 | 2000 | 430 | 11 | 383 | 22 | 166 | 616 | 1107 | 13 | 0 | 0 | 0 | 0 | 0 | 0 | 0 | 0 | 951 | 15 |
| SRR5230664 | 0 | 0 | 0 | 0 | 19 | 0 | 0 | 396 | 1459 | 1388 | 0 | 0 | 0 | 675 | 281 | 505 | 0 | 39 | 746 | 580 | 0 | 104 | 16 | 109 | 267 | 1501 | 0 | 0 | 0 | 0 | 0 | 0 | 0 | 0 | 0 | 1505 | 0 |
| SRR5230665 | 0 | 0 | 0 | 11 | 51 | 16 | 54 | 136 | 714 | 8769 | 0 | 0 | 0 | 2869 | 3282 | 3603 | 148 | 213 | 1306 | 640 | 16 | 348 | 57 | 438 | 992 | 1748 | 81 | 0 | 0 | 0 | 0 | 0 | 0 | 0 | 0 | 1639 | 45 |
| SRR5230666 | 0 | 0 | 0 | 11 | 50 | 0 | 13 | 2556 | 9456 | 14124 | 0 | 0 | 0 | 977 | 935 | 4701 | 43 | 100 | 1432 | 408 | 14 | 77 | 37 | 229 | 281 | 507 | 0 | 0 | 0 | 0 | 0 | 0 | 0 | 0 | 0 | 1101 | 0 |
| SRR5230667 | 0 | 0 | 0 | 0 | 11 | 0 | 0 | 1259 | 3515 | 3987 | 0 | 13 | 0 | 1928 | 105 | 157 | 0 | 33 | 146 | 498 | 0 | 33 | 16 | 47 | 714 | 357 | 12 | 0 | 0 | 0 | 0 | 0 | 0 | 0 | 0 | 1207 | 0 |
| SRR5230668 | 0 | 0 | 0 | 0 | 35 | 0 | 0 | 235 | 960 | 6176 | 0 | 10 | 0 | 2477 | 1511 | 2682 | 18 | 113 | 908 | 854 | 0 | 228 | 22 | 181 | 442 | 896 | 20 | 0 | 0 | 0 | 0 | 0 | 0 | 0 | 0 | 1838 | 0 |
| SRR5230669 | 0 | 0 | 0 | 0 | 13 | 0 | 35 | 1839 | 10049 | 3151 | 0 | 0 | 0 | 1561 | 344 | 1096 | 0 | 158 | 749 | 3538 | 12 | 365 | 42 | 144 | 430 | 565 | 45 | 0 | 0 | 0 | 0 | 0 | 0 | 0 | 0 | 7220 | 0 |
| SRR5230670 | 0 | 0 | 0 | 0 | 36 | 0 | 0 | 2697 | 12158 | 2139 | 0 | 0 | 0 | 677 | 79 | 145 | 0 | 102 | 230 | 132 | 0 | 126 | 63 | 208 | 1105 | 503 | 54 | 0 | 0 | 0 | 0 | 0 | 0 | 0 | 0 | 557 | 16 |
| SRR5230671 | 0 | 0 | 0 | 0 | 17 | 0 | 0 | 859 | 3992 | 4517 | 0 | 0 | 0 | 2179 | 72 | 40 | 0 | 26 | 77 | 914 | 37 | 220 | 21 | 87 | 657 | 291 | 37 | 0 | 0 | 0 | 0 | 0 | 0 | 0 | 0 | 1193 | 38 |
| SRR5230673 | 0 | 0 | 0 | 0 | 21 | 0 | 0 | 909 | 4099 | 4279 | 0 | 0 | 0 | 2761 | 558 | 1001 | 11 | 60 | 251 | 1934 | 0 | 186 | 41 | 106 | 217 | 832 | 34 | 0 | 0 | 0 | 0 | 0 | 0 | 0 | 0 | 4943 | 0 |
| SRR5230674 | 0 | 0 | 0 | 0 | 51 | 20 | 22 | 487 | 1623 | 4095 | 0 | 0 | 11 | 7895 | 317 | 544 | 66 | 132 | 268 | 350 | 18 | 293 | 50 | 341 | 856 | 596 | 44 | 0 | 0 | 0 | 0 | 0 | 0 | 0 | 0 | 1287 | 0 |
| SRR5230675 | 0 | 0 | 0 | 14 | 48 | 0 | 0 | 465 | 3028 | 6612 | 0 | 0 | 0 | 1348 | 0 | 42 | 43 | 102 | 730 | 212 | 11 | 50 | 0 | 43 | 884 | 154 | 15 | 0 | 0 | 0 | 0 | 0 | 0 | 0 | 0 | 774 | 0 |
| SRR5230676 | 0 | 0 | 0 | 0 | 20 | 13 | 23 | 671 | 2340 | 6208 | 0 | 0 | 0 | 2945 | 81 | 325 | 66 | 108 | 431 | 1381 | 0 | 228 | 48 | 181 | 655 | 659 | 89 | 0 | 0 | 0 | 0 | 0 | 0 | 0 | 0 | 3871 | 0 |
| SRR5230677 | 0 | 0 | 0 | 0 | 0 | 0 | 0 | 1498 | 6171 | 1423 | 0 | 0 | 0 | 708 | 65 | 133 | 0 | 58 | 141 | 369 | 0 | 91 | 17 | 177 | 622 | 612 | 0 | 0 | 0 | 0 | 0 | 0 | 0 | 0 | 0 | 660 | 0 |
| SRR5230678 | 0 | 0 | 0 | 0 | 22 | 0 | 0 | 1897 | 7952 | 2847 | 0 | 0 | 0 | 1004 | 80 | 125 | 0 | 128 | 234 | 498 | 0 | 229 | 43 | 189 | 553 | 1083 | 37 | 0 | 0 | 0 | 0 | 0 | 0 | 0 | 0 | 1361 | 0 |
| SRR5230679 | 0 | 0 | 0 | 0 | 25 | 0 | 0 | 1855 | 7139 | 3537 | 0 | 0 | 0 | 2788 | 76 | 76 | 0 | 73 | 175 | 694 | 0 | 145 | 113 | 486 | 657 | 412 | 36 | 0 | 0 | 0 | 0 | 0 | 0 | 0 | 0 | 1744 | 0 |
| SRR5230680 | 0 | 0 | 0 | 0 | 38 | 0 | 0 | 1118 | 4428 | 4101 | 0 | 11 | 0 | 1428 | 166 | 258 | 25 | 20 | 238 | 390 | 0 | 25 | 0 | 62 | 986 | 330 | 17 | 0 | 0 | 0 | 0 | 0 | 0 | 0 | 0 | 1187 | 0 |
| SRR5230681 | 0 | 0 | 0 | 0 | 12 | 0 | 0 | 684 | 2504 | 7576 | 0 | 0 | 0 | 2409 | 60 | 127 | 0 | 52 | 397 | 383 | 0 | 135 | 31 | 193 | 208 | 543 | 18 | 0 | 0 | 0 | 0 | 0 | 0 | 0 | 0 | 922 | 15 |
| SRR5230682 | 0 | 0 | 0 | 0 | 0 | 0 | 0 | 365 | 2008 | 5559 | 0 | 0 | 0 | 1748 | 2044 | 1220 | 14 | 231 | 796 | 426 | 13 | 393 | 21 | 142 | 342 | 3037 | 25 | 0 | 0 | 0 | 0 | 0 | 0 | 0 | 0 | 714 | 0 |
| SRR5230683 | 0 | 0 | 0 | 0 | 31 | 24 | 0 | 1136 | 4173 | 4936 | 0 | 11 | 0 | 1516 | 0 | 0 | 20 | 69 | 175 | 1448 | 0 | 105 | 21 | 106 | 777 | 524 | 45 | 0 | 0 | 0 | 0 | 0 | 0 | 0 | 0 | 3807 | 0 |
| SRR5230684 | 0 | 0 | 18 | 0 | 48 | 18 | 14 | 208 | 840 | 3858 | 13 | 0 | 13 | 6949 | 271 | 538 | 0 | 192 | 239 | 133 | 0 | 506 | 131 | 959 | 769 | 761 | 36 | 0 | 0 | 0 | 0 | 0 | 0 | 0 | 0 | 241 | 0 |
| SRR5230685 | 0 | 0 | 0 | 0 | 23 | 0 | 0 | 1111 | 3716 | 7197 | 0 | 31 | 0 | 1674 | 237 | 564 | 0 | 153 | 213 | 510 | 0 | 191 | 33 | 166 | 993 | 880 | 73 | 0 | 0 | 0 | 0 | 0 | 0 | 0 | 0 | 1746 | 0 |
| SRR5230686 | 0 | 0 | 0 | 0 | 19 | 0 | 0 | 1686 | 6312 | 4947 | 0 | 18 | 11 | 2161 | 906 | 1343 | 0 | 201 | 299 | 955 | 34 | 490 | 78 | 236 | 814 | 1034 | 80 | 0 | 0 | 0 | 0 | 0 | 11 | 0 | 0 | 2263 | 13 |
| SRR5230687 | 0 | 0 | 0 | 0 | 39 | 0 | 0 | 540 | 2408 | 6256 | 12 | 25 | 0 | 1604 | 716 | 908 | 0 | 297 | 304 | 1111 | 15 | 695 | 38 | 305 | 983 | 1982 | 62 | 0 | 0 | 0 | 0 | 0 | 0 | 0 | 0 | 2613 | 0 |
| SRR5230688 | 0 | 0 | 0 | 0 | 31 | 0 | 0 | 397 | 1521 | 8361 | 0 | 0 | 0 | 1936 | 388 | 866 | 15 | 44 | 180 | 736 | 0 | 93 | 0 | 47 | 965 | 423 | 11 | 0 | 0 | 0 | 0 | 0 | 0 | 0 | 0 | 2882 | 17 |
| SRR5230689 | 0 | 0 | 0 | 0 | 0 | 0 | 0 | 1249 | 3986 | 6773 | 0 | 0 | 12 | 2586 | 698 | 2146 | 0 | 46 | 104 | 1441 | 0 | 53 | 62 | 251 | 243 | 156 | 0 | 0 | 0 | 0 | 0 | 0 | 0 | 0 | 0 | 3476 | 0 |
| SRR5230690 | 0 | 0 | 0 | 0 | 45 | 0 | 0 | 1054 | 3726 | 6437 | 11 | 24 | 0 | 2169 | 681 | 1709 | 0 | 266 | 401 | 1483 | 16 | 632 | 50 | 320 | 1099 | 1320 | 35 | 0 | 0 | 0 | 0 | 0 | 0 | 0 | 0 | 4006 | 0 |
| SRR5230691 | 0 | 0 | 0 | 0 | 0 | 0 | 0 | 400 | 1063 | 3786 | 0 | 14 | 0 | 2896 | 477 | 774 | 0 | 30 | 103 | 1076 | 0 | 101 | 20 | 62 | 83 | 1119 | 39 | 0 | 0 | 0 | 0 | 0 | 0 | 0 | 0 | 2864 | 0 |
| SRR5230692 | 0 | 0 | 0 | 0 | 133 | 0 | 0 | 551 | 1571 | 13760 | 0 | 0 | 0 | 1863 | 1803 | 4643 | 28 | 147 | 636 | 1009 | 14 | 323 | 43 | 272 | 604 | 939 | 29 | 0 | 0 | 0 | 0 | 0 | 0 | 0 | 0 | 2254 | 0 |
| SRR5230693 | 0 | 18 | 0 | 0 | 19 | 29 | 25 | 1406 | 5129 | 2788 | 0 | 12 | 0 | 1910 | 398 | 1030 | 25 | 315 | 314 | 414 | 0 | 702 | 42 | 235 | 714 | 1258 | 93 | 0 | 0 | 0 | 0 | 0 | 0 | 0 | 0 | 839 | 0 |
| SRR5230694 | 0 | 0 | 0 | 0 | 36 | 0 | 31 | 2527 | 12720 | 3303 | 0 | 0 | 0 | 551 | 10 | 21 | 17 | 99 | 299 | 222 | 0 | 73 | 0 | 32 | 341 | 322 | 62 | 0 | 0 | 0 | 0 | 0 | 0 | 0 | 0 | 1095 | 0 |
| SRR5230695 | 0 | 0 | 0 | 0 | 15 | 0 | 22 | 1596 | 5477 | 4015 | 0 | 0 | 0 | 1751 | 471 | 1033 | 0 | 118 | 250 | 590 | 0 | 204 | 33 | 264 | 693 | 840 | 35 | 0 | 0 | 0 | 0 | 0 | 0 | 0 | 10 | 1061 | 0 |
| SRR5230696 | 0 | 0 | 0 | 0 | 36 | 0 | 19 | 1000 | 5034 | 3727 | 0 | 0 | 0 | 1098 | 624 | 877 | 13 | 78 | 770 | 455 | 0 | 80 | 21 | 77 | 172 | 664 | 37 | 0 | 0 | 0 | 0 | 0 | 0 | 0 | 0 | 1185 | 0 |
| SRR5230697 | 0 | 0 | 0 | 0 | 0 | 0 | 0 | 1140 | 3315 | 3351 | 0 | 18 | 0 | 1944 | 550 | 1018 | 0 | 99 | 165 | 830 | 10 | 343 | 0 | 170 | 399 | 776 | 47 | 0 | 0 | 0 | 0 | 0 | 0 | 0 | 0 | 1913 | 0 |
| SRR5230698 | 0 | 0 | 0 | 10 | 18 | 0 | 0 | 455 | 1832 | 5651 | 0 | 0 | 0 | 2101 | 1235 | 1559 | 0 | 160 | 371 | 192 | 0 | 322 | 27 | 188 | 449 | 2203 | 16 | 0 | 0 | 0 | 0 | 0 | 0 | 0 | 0 | 595 | 10 |
| SRR5230699 | 0 | 0 | 0 | 12 | 42 | 0 | 0 | 452 | 2044 | 8476 | 0 | 0 | 0 | 1908 | 1174 | 3182 | 29 | 176 | 1544 | 821 | 24 | 259 | 24 | 117 | 702 | 654 | 16 | 0 | 0 | 0 | 0 | 0 | 0 | 0 | 0 | 1710 | 16 |
| SRR5230700 | 0 | 0 | 0 | 0 | 22 | 0 | 20 | 539 | 2015 | 5491 | 0 | 0 | 0 | 2261 | 515 | 1069 | 12 | 71 | 162 | 742 | 0 | 127 | 25 | 102 | 612 | 1009 | 37 | 0 | 0 | 0 | 0 | 0 | 0 | 0 | 0 | 2497 | 0 |
| SRR5230701 | 0 | 0 | 0 | 0 | 19 | 0 | 0 | 972 | 3958 | 5272 | 0 | 0 | 0 | 1875 | 129 | 335 | 22 | 42 | 324 | 272 | 0 | 219 | 29 | 116 | 413 | 637 | 19 | 0 | 0 | 0 | 0 | 0 | 0 | 0 | 0 | 838 | 16 |
| SRR5230702 | 0 | 0 | 0 | 0 | 56 | 0 | 25 | 413 | 1118 | 8819 | 0 | 0 | 0 | 2713 | 911 | 2311 | 21 | 135 | 629 | 0 | 16 | 239 | 49 | 242 | 507 | 1379 | 40 | 0 | 0 | 0 | 0 | 0 | 0 | 0 | 0 | 20 | 11 |
| SRR5230703 | 0 | 0 | 0 | 0 | 12 | 0 | 15 | 182 | 652 | 5562 | 0 | 38 | 15 | 2053 | 1782 | 2046 | 18 | 72 | 304 | 395 | 0 | 181 | 11 | 72 | 633 | 787 | 53 | 0 | 0 | 0 | 0 | 0 | 0 | 0 | 0 | 1224 | 17 |
| SRR5230704 | 0 | 0 | 0 | 0 | 0 | 0 | 0 | 118 | 677 | 5705 | 0 | 0 | 0 | 3637 | 255 | 196 | 0 | 82 | 387 | 288 | 14 | 167 | 80 | 185 | 195 | 1143 | 47 | 0 | 0 | 0 | 0 | 0 | 0 | 0 | 0 | 526 | 21 |
| SRR5230705 | 0 | 0 | 0 | 0 | 0 | 0 | 0 | 480 | 1361 | 3642 | 0 | 0 | 0 | 1679 | 65 | 179 | 0 | 12 | 159 | 196 | 0 | 25 | 0 | 22 | 340 | 359 | 38 | 0 | 0 | 0 | 0 | 0 | 0 | 0 | 0 | 1082 | 0 |
| SRR5230706 | 0 | 0 | 0 | 0 | 26 | 0 | 0 | 306 | 923 | 5802 | 0 | 0 | 0 | 2695 | 600 | 1271 | 22 | 121 | 608 | 225 | 20 | 297 | 14 | 116 | 647 | 531 | 0 | 0 | 0 | 0 | 0 | 0 | 0 | 0 | 0 | 1097 | 33 |
| SRR5230707 | 0 | 0 | 0 | 0 | 45 | 0 | 0 | 286 | 1313 | 5559 | 0 | 0 | 0 | 1639 | 303 | 463 | 13 | 230 | 831 | 192 | 13 | 446 | 10 | 153 | 1609 | 1221 | 0 | 0 | 0 | 0 | 0 | 0 | 0 | 0 | 0 | 867 | 0 |
| SRR5230708 | 0 | 0 | 0 | 13 | 50 | 0 | 0 | 834 | 3500 | 6919 | 0 | 0 | 36 | 3202 | 337 | 518 | 29 | 514 | 559 | 966 | 36 | 715 | 45 | 258 | 1896 | 1702 | 58 | 0 | 0 | 0 | 0 | 0 | 0 | 0 | 0 | 2155 | 0 |
| SRR5230709 | 0 | 0 | 0 | 0 | 47 | 0 | 21 | 657 | 2664 | 4518 | 0 | 0 | 0 | 2534 | 375 | 399 | 30 | 203 | 667 | 422 | 29 | 521 | 59 | 266 | 1587 | 944 | 119 | 0 | 0 | 0 | 0 | 0 | 0 | 0 | 0 | 1337 | 0 |
| SRR5230710 | 0 | 0 | 0 | 0 | 14 | 0 | 0 | 303 | 843 | 4915 | 0 | 29 | 0 | 3305 | 250 | 1117 | 0 | 118 | 412 | 149 | 0 | 327 | 14 | 192 | 381 | 1859 | 0 | 0 | 0 | 0 | 0 | 0 | 0 | 0 | 0 | 688 | 0 |
| SRR5230711 | 0 | 0 | 0 | 0 | 42 | 0 | 0 | 891 | 3893 | 5220 | 0 | 0 | 0 | 3639 | 0 | 80 | 0 | 94 | 137 | 284 | 0 | 172 | 74 | 374 | 946 | 511 | 97 | 0 | 0 | 0 | 0 | 0 | 0 | 0 | 0 | 672 | 0 |
| SRR5230712 | 13 | 0 | 0 | 0 | 62 | 0 | 0 | 595 | 2204 | 5852 | 10 | 35 | 11 | 2694 | 800 | 1344 | 19 | 116 | 251 | 1544 | 0 | 185 | 27 | 114 | 1672 | 455 | 38 | 0 | 0 | 0 | 0 | 0 | 0 | 0 | 0 | 3048 | 0 |
| SRR5230713 | 0 | 0 | 0 | 11 | 57 | 0 | 0 | 387 | 1035 | 8898 | 0 | 11 | 0 | 3977 | 340 | 907 | 38 | 219 | 482 | 280 | 29 | 351 | 59 | 394 | 736 | 1773 | 20 | 0 | 0 | 0 | 0 | 0 | 0 | 0 | 0 | 627 | 40 |
| SRR5230714 | 0 | 0 | 0 | 0 | 52 | 0 | 0 | 424 | 2176 | 6957 | 0 | 0 | 0 | 2146 | 1350 | 2007 | 0 | 101 | 654 | 472 | 0 | 102 | 13 | 89 | 428 | 858 | 32 | 0 | 0 | 0 | 0 | 0 | 0 | 0 | 0 | 1657 | 10 |
| SRR5230716 | 0 | 0 | 0 | 0 | 38 | 0 | 0 | 1020 | 2783 | 4852 | 0 | 0 | 0 | 2081 | 24 | 117 | 21 | 65 | 233 | 161 | 0 | 118 | 24 | 188 | 2253 | 762 | 0 | 0 | 0 | 0 | 0 | 0 | 0 | 0 | 0 | 621 | 0 |
| SRR5230717 | 10 | 0 | 0 | 0 | 18 | 0 | 0 | 611 | 2556 | 4124 | 0 | 0 | 0 | 3033 | 115 | 154 | 0 | 203 | 356 | 361 | 12 | 432 | 40 | 259 | 1323 | 697 | 71 | 0 | 0 | 0 | 0 | 0 | 0 | 0 | 0 | 785 | 13 |
| SRR5230718 | 0 | 0 | 0 | 0 | 32 | 0 | 0 | 1133 | 4254 | 3766 | 0 | 14 | 0 | 2421 | 200 | 269 | 0 | 294 | 244 | 273 | 31 | 443 | 68 | 396 | 1163 | 991 | 37 | 0 | 0 | 0 | 0 | 0 | 0 | 0 | 0 | 636 | 0 |
| SRR5230719 | 0 | 0 | 0 | 0 | 16 | 0 | 0 | 475 | 2100 | 4626 | 0 | 0 | 0 | 3117 | 461 | 808 | 0 | 155 | 229 | 323 | 15 | 463 | 27 | 235 | 370 | 1013 | 25 | 0 | 0 | 0 | 0 | 0 | 0 | 0 | 0 | 618 | 0 |
| SRR5230720 | 0 | 0 | 0 | 0 | 11 | 0 | 0 | 243 | 1453 | 2946 | 0 | 0 | 0 | 2590 | 822 | 1146 | 11 | 155 | 445 | 1751 | 16 | 491 | 19 | 115 | 562 | 2113 | 35 | 0 | 0 | 0 | 0 | 0 | 0 | 0 | 0 | 4424 | 0 |
| SRR5230721 | 0 | 0 | 0 | 0 | 28 | 0 | 0 | 670 | 2455 | 3829 | 0 | 0 | 0 | 4471 | 267 | 358 | 0 | 68 | 440 | 304 | 0 | 132 | 47 | 236 | 549 | 357 | 19 | 0 | 0 | 0 | 0 | 0 | 0 | 0 | 0 | 444 | 0 |
| SRR5230722 | 0 | 0 | 0 | 0 | 34 | 0 | 19 | 464 | 1553 | 3452 | 0 | 16 | 0 | 3992 | 850 | 2268 | 0 | 199 | 209 | 679 | 0 | 441 | 46 | 357 | 425 | 1099 | 29 | 0 | 0 | 0 | 0 | 0 | 0 | 0 | 0 | 1814 | 0 |
| SRR5230723 | 0 | 0 | 0 | 0 | 41 | 12 | 0 | 3757 | 17921 | 1291 | 0 | 0 | 0 | 644 | 45 | 111 | 20 | 455 | 607 | 270 | 16 | 349 | 57 | 382 | 2174 | 663 | 12 | 0 | 0 | 0 | 0 | 0 | 0 | 0 | 0 | 764 | 0 |
| SRR5230724 | 0 | 0 | 0 | 0 | 21 | 0 | 0 | 2355 | 8160 | 1488 | 0 | 0 | 0 | 1081 | 56 | 172 | 0 | 75 | 130 | 78 | 0 | 43 | 18 | 120 | 1278 | 396 | 27 | 0 | 0 | 0 | 0 | 0 | 0 | 0 | 0 | 314 | 0 |
| SRR5230725 | 0 | 0 | 0 | 0 | 56 | 0 | 16 | 499 | 2419 | 7552 | 0 | 11 | 0 | 2509 | 394 | 838 | 28 | 141 | 238 | 490 | 0 | 193 | 26 | 180 | 842 | 1025 | 39 | 0 | 0 | 0 | 0 | 0 | 0 | 0 | 0 | 970 | 0 |
| SRR5230726 | 0 | 0 | 0 | 0 | 16 | 0 | 0 | 969 | 4474 | 2679 | 0 | 20 | 0 | 1002 | 287 | 336 | 21 | 123 | 348 | 747 | 0 | 148 | 20 | 131 | 894 | 878 | 34 | 0 | 0 | 0 | 0 | 0 | 0 | 0 | 0 | 1886 | 0 |
| SRR5230727 | 0 | 0 | 0 | 0 | 0 | 0 | 0 | 1110 | 4935 | 3392 | 0 | 0 | 0 | 1886 | 585 | 1034 | 12 | 36 | 272 | 515 | 0 | 75 | 12 | 73 | 664 | 370 | 26 | 0 | 0 | 0 | 0 | 0 | 0 | 0 | 0 | 1027 | 0 |
| SRR5230728 | 0 | 0 | 0 | 0 | 52 | 0 | 0 | 383 | 879 | 13086 | 0 | 0 | 0 | 2329 | 404 | 1179 | 15 | 125 | 1058 | 271 | 22 | 237 | 30 | 260 | 521 | 803 | 43 | 0 | 0 | 0 | 0 | 0 | 0 | 0 | 0 | 578 | 16 |
| SRR5230729 | 0 | 0 | 0 | 11 | 54 | 11 | 20 | 443 | 1541 | 6192 | 0 | 0 | 0 | 2342 | 325 | 659 | 14 | 205 | 528 | 768 | 0 | 337 | 25 | 253 | 977 | 1855 | 11 | 0 | 0 | 0 | 0 | 0 | 0 | 0 | 0 | 1817 | 0 |
| SRR5230732 | 0 | 0 | 12 | 0 | 54 | 0 | 29 | 399 | 1579 | 7820 | 0 | 10 | 0 | 3105 | 451 | 392 | 49 | 262 | 1177 | 219 | 19 | 507 | 37 | 228 | 1524 | 1175 | 78 | 0 | 0 | 0 | 0 | 0 | 0 | 0 | 0 | 663 | 0 |
| SRR5230733 | 0 | 0 | 0 | 0 | 21 | 0 | 23 | 577 | 1820 | 6503 | 11 | 11 | 0 | 3728 | 55 | 36 | 11 | 63 | 155 | 69 | 0 | 135 | 31 | 257 | 960 | 691 | 62 | 0 | 0 | 0 | 0 | 0 | 0 | 0 | 0 | 420 | 0 |
| SRR5230734 | 0 | 0 | 0 | 0 | 47 | 0 | 154 | 627 | 1396 | 20169 | 0 | 0 | 0 | 5267 | 158 | 137 | 119 | 122 | 321 | 145 | 0 | 328 | 140 | 1141 | 635 | 603 | 161 | 0 | 0 | 0 | 0 | 0 | 0 | 0 | 0 | 461 | 10 |
| SRR5230735 | 0 | 11 | 0 | 0 | 55 | 0 | 13 | 387 | 915 | 8550 | 0 | 31 | 17 | 2703 | 318 | 553 | 15 | 145 | 303 | 250 | 0 | 363 | 78 | 542 | 867 | 2115 | 63 | 0 | 0 | 0 | 0 | 0 | 0 | 0 | 0 | 718 | 0 |
| SRR5230736 | 0 | 0 | 0 | 0 | 19 | 17 | 0 | 850 | 3041 | 4845 | 0 | 0 | 0 | 2895 | 50 | 27 | 0 | 50 | 174 | 355 | 0 | 129 | 16 | 126 | 1297 | 507 | 21 | 0 | 0 | 0 | 0 | 0 | 0 | 0 | 0 | 1268 | 10 |
| SRR5230737 | 0 | 30 | 0 | 20 | 76 | 25 | 10 | 1385 | 5519 | 6125 | 0 | 0 | 0 | 1158 | 84 | 206 | 50 | 278 | 31 | 606 | 32 | 239 | 48 | 394 | 1339 | 1081 | 10 | 0 | 0 | 0 | 0 | 0 | 0 | 0 | 0 | 1101 | 0 |
| SRR5230738 | 0 | 0 | 0 | 0 | 60 | 17 | 0 | 975 | 3304 | 4947 | 18 | 28 | 20 | 1812 | 153 | 265 | 0 | 151 | 169 | 186 | 0 | 205 | 43 | 188 | 1439 | 618 | 38 | 0 | 0 | 0 | 0 | 0 | 0 | 0 | 0 | 656 | 0 |
| SRR5230739 | 0 | 0 | 0 | 0 | 55 | 0 | 22 | 4096 | 13419 | 1080 | 0 | 0 | 0 | 433 | 36 | 63 | 52 | 110 | 318 | 226 | 0 | 319 | 70 | 297 | 354 | 1158 | 49 | 0 | 0 | 0 | 0 | 0 | 0 | 0 | 0 | 373 | 0 |
| SRR5230740 | 0 | 0 | 0 | 0 | 33 | 0 | 0 | 613 | 1486 | 7819 | 0 | 0 | 0 | 2664 | 54 | 64 | 52 | 120 | 683 | 88 | 10 | 271 | 13 | 147 | 1016 | 433 | 40 | 0 | 0 | 0 | 0 | 0 | 0 | 0 | 0 | 201 | 16 |
| SRR5230741 | 0 | 0 | 0 | 0 | 44 | 0 | 18 | 465 | 1138 | 6542 | 0 | 0 | 0 | 4440 | 881 | 1477 | 35 | 174 | 602 | 310 | 0 | 489 | 180 | 1783 | 483 | 691 | 35 | 0 | 0 | 0 | 0 | 0 | 0 | 0 | 0 | 852 | 18 |
| SRR5230742 | 0 | 0 | 0 | 18 | 120 | 0 | 0 | 4409 | 13340 | 14076 | 0 | 0 | 0 | 1689 | 50 | 50 | 0 | 416 | 376 | 519 | 26 | 229 | 45 | 496 | 2273 | 1561 | 63 | 0 | 0 | 0 | 0 | 0 | 0 | 0 | 0 | 2014 | 0 |
| SRR5230743 | 0 | 0 | 0 | 0 | 39 | 0 | 0 | 820 | 3142 | 6823 | 0 | 0 | 0 | 1837 | 143 | 225 | 27 | 213 | 1113 | 425 | 10 | 512 | 32 | 288 | 1012 | 884 | 15 | 0 | 0 | 0 | 0 | 0 | 0 | 0 | 0 | 913 | 25 |
| SRR5230744 | 0 | 0 | 0 | 0 | 21 | 0 | 11 | 445 | 1265 | 6320 | 16 | 0 | 0 | 6023 | 306 | 587 | 0 | 151 | 174 | 190 | 0 | 207 | 86 | 493 | 369 | 571 | 34 | 0 | 0 | 0 | 0 | 0 | 0 | 0 | 0 | 481 | 0 |
| SRR5230745 | 0 | 0 | 0 | 23 | 130 | 13 | 35 | 1478 | 4137 | 19127 | 0 | 31 | 0 | 2138 | 428 | 1046 | 123 | 217 | 1286 | 623 | 16 | 380 | 46 | 327 | 1475 | 654 | 57 | 0 | 0 | 0 | 0 | 0 | 0 | 0 | 0 | 1834 | 13 |
| SRR5230746 | 0 | 0 | 18 | 15 | 51 | 27 | 0 | 750 | 1634 | 6191 | 18 | 26 | 23 | 3386 | 132 | 431 | 23 | 168 | 265 | 506 | 12 | 550 | 84 | 394 | 1596 | 1324 | 141 | 0 | 0 | 0 | 0 | 0 | 0 | 0 | 0 | 1979 | 0 |
| SRR5230747 | 0 | 0 | 0 | 0 | 49 | 0 | 0 | 1991 | 6217 | 3707 | 0 | 14 | 0 | 929 | 148 | 322 | 0 | 255 | 141 | 869 | 17 | 420 | 59 | 356 | 1729 | 1048 | 54 | 0 | 0 | 0 | 0 | 0 | 0 | 0 | 0 | 1882 | 0 |
| SRR5230748 | 0 | 0 | 0 | 0 | 50 | 16 | 0 | 834 | 3518 | 5597 | 0 | 35 | 0 | 2287 | 401 | 545 | 32 | 247 | 528 | 1066 | 16 | 474 | 40 | 272 | 1848 | 1702 | 38 | 0 | 0 | 0 | 0 | 0 | 0 | 0 | 0 | 2649 | 0 |
| SRR5230749 | 0 | 0 | 14 | 0 | 41 | 0 | 0 | 1104 | 3395 | 7809 | 0 | 0 | 0 | 2014 | 183 | 249 | 14 | 141 | 257 | 420 | 10 | 292 | 28 | 143 | 1498 | 746 | 42 | 0 | 0 | 0 | 0 | 0 | 0 | 0 | 0 | 1367 | 13 |
| SRR5230750 | 0 | 0 | 0 | 0 | 23 | 0 | 0 | 2312 | 7497 | 3830 | 0 | 0 | 0 | 809 | 49 | 42 | 0 | 140 | 155 | 266 | 0 | 105 | 44 | 242 | 1535 | 437 | 38 | 0 | 0 | 0 | 0 | 0 | 0 | 0 | 0 | 474 | 0 |
| SRR5230751 | 0 | 0 | 0 | 0 | 26 | 0 | 0 | 1415 | 4048 | 2465 | 0 | 0 | 0 | 577 | 40 | 104 | 0 | 53 | 255 | 152 | 0 | 41 | 20 | 92 | 1617 | 386 | 0 | 0 | 0 | 0 | 0 | 0 | 0 | 0 | 0 | 517 | 0 |
| SRR5230752 | 0 | 0 | 0 | 0 | 37 | 28 | 55 | 766 | 2415 | 4996 | 0 | 0 | 17 | 2509 | 120 | 154 | 17 | 262 | 318 | 172 | 0 | 330 | 57 | 440 | 1403 | 1661 | 133 | 0 | 0 | 0 | 0 | 0 | 0 | 0 | 0 | 672 | 0 |
| SRR5230753 | 11 | 0 | 0 | 0 | 82 | 0 | 14 | 1113 | 4384 | 5838 | 0 | 0 | 0 | 5194 | 129 | 202 | 20 | 461 | 422 | 594 | 13 | 748 | 65 | 519 | 2075 | 914 | 84 | 0 | 0 | 0 | 0 | 0 | 0 | 0 | 0 | 1409 | 0 |
| SRR5231309 | 117 | 0 | 0 | 0 | 20 | 0 | 16 | 96888 | 5863 | 2478 | 16 | 0 | 0 | 399 | 29 | 64 | 13 | 490 | 0 | 38 | 320 | 2545 | 473 | 187588 | 14750 | 0 | 0 | 0 | 0 | 0 | 0 | 0 | 12 | 19 | 11 | 131 | 251 |
| SRR5231310 | 104 | 0 | 0 | 0 | 20 | 0 | 42 | 112609 | 5766 | 2297 | 15 | 22 | 0 | 385 | 0 | 11 | 30 | 594 | 0 | 43 | 357 | 2130 | 827 | 140974 | 11797 | 20 | 0 | 0 | 0 | 0 | 0 | 0 | 0 | 14 | 0 | 166 | 209 |
| SRR5231311 | 48 | 0 | 0 | 0 | 24 | 0 | 11 | 100386 | 5029 | 3680 | 10 | 12 | 0 | 404 | 31 | 43 | 21 | 495 | 0 | 53 | 469 | 2322 | 824 | 148985 | 12164 | 14 | 0 | 0 | 0 | 0 | 0 | 0 | 0 | 0 | 0 | 213 | 189 |
| SRR5231312 | 414 | 0 | 0 | 0 | 18 | 0 | 159 | 96405 | 5310 | 1113 | 28 | 13 | 0 | 209 | 0 | 13 | 49 | 650 | 0 | 47 | 267 | 1830 | 371 | 164556 | 12662 | 12 | 17 | 0 | 0 | 0 | 0 | 0 | 0 | 0 | 0 | 100 | 158 |
| SRR5231313 | 15 | 0 | 0 | 0 | 18 | 0 | 15 | 85359 | 5497 | 1088 | 0 | 0 | 0 | 273 | 0 | 17 | 14 | 611 | 0 | 82 | 340 | 1965 | 542 | 193973 | 12843 | 0 | 0 | 0 | 0 | 0 | 0 | 0 | 0 | 14 | 0 | 188 | 146 |

### Supp Table S8.4.Columns_DI_to_ES

| | | | | | | | | | | | | | | | | | | | | | | | | | | | | | | | | | | | | | |
|---|---|---|---|---|---|---|---|---|---|---|---|---|---|---|---|---|---|---|---|---|---|---|---|---|---|---|---|---|---|---|---|---|---|---|---|---|---|
| SRR5231314 | 29 | 0 | 0 | 0 | 31 | 0 | 0 | 64066 | 3963 | 3297 | 29 | 10 | 0 | 915 | 20 | 22 | 13 | 445 | 0 | 81 | 440 | 1957 | 696 | 178040 | 13311 | 16 | 0 | 0 | 0 | 0 | 0 | 0 | 0 | 0 | 0 | 330 | 237 |
| SRR5239182 | 0 | 0 | 0 | 0 | 0 | 0 | 0 | 1474 | 140 | 14379 | 113 | 552 | 0 | 502 | 268 | 397 | 0 | 0 | 60 | 5840 | 57 | 1796 | 11 | 4744 | 10834 | 0 | 20 | 0 | 0 | 20 | 18 | 0 | 15 | 39 | 0 | 25508 | 81 |
| SRR5239183 | 0 | 0 | 0 | 0 | 0 | 0 | 0 | 7196 | 198 | 1537 | 286 | 205 | 0 | 4751 | 132 | 190 | 0 | 28 | 42 | 1625 | 42 | 582 | 0 | 1482 | 1108 | 14 | 0 | 0 | 0 | 0 | 14 | 0 | 13 | 23 | 24 | 3638 | 125 |
| SRR5239184 | 0 | 0 | 0 | 0 | 0 | 0 | 12 | 3535 | 142 | 6496 | 258 | 216 | 0 | 1420 | 563 | 710 | 0 | 0 | 67 | 5386 | 17 | 724 | 0 | 837 | 1936 | 0 | 16 | 0 | 0 | 10 | 13 | 0 | 32 | 136 | 12 | 22013 | 34 |
| SRR5239185 | 0 | 0 | 0 | 0 | 0 | 0 | 0 | 2200 | 200 | 5240 | 40 | 287 | 0 | 713 | 33 | 27 | 0 | 14 | 108 | 6221 | 101 | 1717 | 0 | 5240 | 10591 | 0 | 20 | 0 | 0 | 20 | 17 | 0 | 0 | 20 | 0 | 45370 | 157 |
| SRR5239186 | 0 | 0 | 0 | 0 | 0 | 0 | 0 | 1547 | 315 | 15000 | 402 | 343 | 0 | 291 | 249 | 388 | 0 | 0 | 66 | 4660 | 76 | 2168 | 0 | 1636 | 10933 | 0 | 20 | 0 | 0 | 25 | 34 | 0 | 13 | 31 | 0 | 40732 | 76 |
| SRR5239187 | 0 | 0 | 0 | 0 | 0 | 0 | 10 | 2334 | 114 | 10667 | 167 | 548 | 0 | 1277 | 267 | 292 | 0 | 0 | 52 | 4184 | 64 | 1059 | 0 | 602 | 2910 | 25 | 15 | 0 | 0 | 13 | 12 | 10 | 14 | 69 | 16 | 29675 | 25 |
| SRR5239188 | 0 | 0 | 0 | 0 | 0 | 0 | 0 | 1434 | 131 | 8461 | 82 | 276 | 0 | 608 | 340 | 332 | 0 | 15 | 50 | 5631 | 86 | 1591 | 0 | 2653 | 5687 | 0 | 13 | 0 | 0 | 20 | 22 | 0 | 23 | 73 | 0 | 52726 | 39 |
| SRR5275577 | 20 | 0 | 0 | 0 | 0 | 11 | 29 | 23869 | 454 | 295 | 365 | 44 | 0 | 61 | 0 | 0 | 99 | 15 | 137 | 617 | 666 | 3236 | 11 | 13224 | 6718 | 0 | 0 | 0 | 12 | 39 | 110 | 10 | 23 | 36 | 0 | 10317 | 26 |
| SRR5275578 | 27 | 0 | 0 | 0 | 0 | 14 | 46 | 22909 | 411 | 308 | 655 | 86 | 0 | 64 | 0 | 0 | 154 | 19 | 153 | 590 | 716 | 3136 | 16 | 12082 | 6082 | 0 | 0 | 0 | 14 | 35 | 93 | 0 | 16 | 29 | 0 | 10409 | 32 |
| SRR5275579 | 20 | 0 | 0 | 0 | 0 | 11 | 34 | 23643 | 351 | 276 | 79 | 50 | 0 | 43 | 0 | 0 | 105 | 14 | 101 | 608 | 663 | 3225 | 13 | 13152 | 6576 | 0 | 0 | 0 | 12 | 43 | 117 | 0 | 22 | 33 | 0 | 12730 | 30 |
| SRR5275580 | 18 | 0 | 0 | 0 | 0 | 23 | 25 | 48518 | 965 | 713 | 436 | 260 | 0 | 157 | 0 | 0 | 71 | 15 | 92 | 765 | 750 | 2539 | 0 | 9187 | 5529 | 0 | 23 | 14 | 21 | 59 | 108 | 10 | 20 | 42 | 0 | 12031 | 48 |
| SRR5275581 | 42 | 0 | 0 | 0 | 0 | 23 | 71 | 47847 | 824 | 517 | 426 | 244 | 0 | 110 | 0 | 0 | 183 | 18 | 93 | 678 | 724 | 2456 | 0 | 8460 | 4685 | 0 | 31 | 13 | 62 | 154 | 239 | 24 | 43 | 75 | 0 | 11376 | 37 |
| SRR5275582 | 21 | 0 | 0 | 0 | 0 | 14 | 35 | 42743 | 853 | 842 | 490 | 62 | 0 | 131 | 0 | 0 | 63 | 20 | 80 | 1002 | 705 | 2615 | 0 | 9697 | 5578 | 0 | 17 | 0 | 42 | 144 | 318 | 42 | 102 | 215 | 18 | 21635 | 46 |
| SRR5284195 | 0 | 166 | 24 | 196 | 552 | 0 | 24 | 3053 | 7674 | 16107 | 21366 | 21703 | 0 | 3017 | 0 | 0 | 11 | 1534 | 31 | 1524 | 186 | 347 | 136 | 6747 | 575 | 10 | 10 | 16 | 42 | 12 | 48 | 20 | 20 | 21 | 0 | 2446 | 0 |
| SRR5284196 | 0 | 147 | 43 | 134 | 706 | 0 | 21 | 2883 | 6710 | 25704 | 8338 | 10698 | 0 | 4458 | 0 | 0 | 33 | 1411 | 20 | 1937 | 193 | 502 | 138 | 5887 | 483 | 0 | 16 | 0 | 24 | 14 | 28 | 13 | 14 | 22 | 16 | 4218 | 0 |
| SRR5284197 | 0 | 142 | 27 | 557 | 433 | 0 | 28 | 1647 | 6758 | 25870 | 2072 | 10467 | 0 | 5702 | 0 | 0 | 32 | 2415 | 17 | 877 | 169 | 554 | 135 | 12339 | 1028 | 18 | 36 | 0 | 26 | 15 | 72 | 24 | 30 | 35 | 12 | 1690 | 0 |
| SRR5284198 | 0 | 638 | 89 | 303 | 589 | 0 | 35 | 2902 | 9238 | 23276 | 994 | 6295 | 0 | 7225 | 0 | 0 | 27 | 1818 | 18 | 1554 | 148 | 531 | 184 | 10477 | 652 | 20 | 52 | 0 | 39 | 32 | 92 | 37 | 42 | 50 | 0 | 2600 | 0 |
| SRR5284199 | 0 | 541 | 129 | 86 | 1235 | 0 | 27 | 2157 | 5577 | 15896 | 1337 | 11047 | 0 | 3050 | 0 | 0 | 22 | 1051 | 17 | 787 | 131 | 321 | 134 | 4186 | 509 | 0 | 12 | 0 | 34 | 42 | 32 | 17 | 19 | 27 | 0 | 1700 | 0 |
| SRR5284200 | 0 | 301 | 131 | 55 | 1133 | 0 | 17 | 2298 | 6133 | 20644 | 1514 | 6847 | 0 | 3202 | 0 | 0 | 15 | 805 | 12 | 911 | 138 | 239 | 177 | 3268 | 323 | 0 | 16 | 0 | 27 | 28 | 59 | 23 | 27 | 25 | 0 | 1700 | 0 |
| SRR5284201 | 0 | 227 | 20 | 532 | 840 | 0 | 25 | 2029 | 9030 | 22361 | 3716 | 4961 | 0 | 6917 | 0 | 0 | 22 | 2151 | 16 | 950 | 178 | 539 | 196 | 14207 | 1143 | 20 | 21 | 0 | 10 | 15 | 30 | 27 | 13 | 14 | 0 | 1700 | 0 |
| SRR5284202 | 0 | 443 | 69 | 410 | 827 | 0 | 45 | 2687 | 8989 | 22812 | 4275 | 4250 | 0 | 14231 | 0 | 0 | 76 | 1475 | 17 | 866 | 98 | 537 | 198 | 10821 | 541 | 23 | 32 | 0 | 23 | 37 | 122 | 10 | 0 | 11 | 0 | 1488 | 0 |
| SRR5284203 | 0 | 42 | 19 | 117 | 1279 | 0 | 12 | 6255 | 16780 | 56303 | 4259 | 5017 | 0 | 6086 | 0 | 0 | 29 | 635 | 37 | 837 | 92 | 188 | 75 | 3407 | 351 | 0 | 10 | 0 | 15 | 38 | 20 | 12 | 17 | 20 | 0 | 2687 | 0 |
| SRR5284204 | 0 | 567 | 143 | 43 | 1060 | 0 | 30 | 3983 | 8633 | 24454 | 1545 | 7689 | 0 | 2671 | 0 | 0 | 37 | 658 | 22 | 2341 | 103 | 212 | 99 | 2072 | 293 | 0 | 30 | 11 | 38 | 44 | 60 | 35 | 33 | 43 | 0 | 6521 | 0 |
| SRR5284205 | 0 | 148 | 13 | 484 | 603 | 0 | 11 | 1435 | 3444 | 15334 | 1040 | 5533 | 0 | 6055 | 0 | 0 | 10 | 1631 | 48 | 1198 | 145 | 436 | 150 | 10248 | 828 | 0 | 15 | 11 | 39 | 21 | 62 | 18 | 18 | 20 | 0 | 4221 | 0 |
| SRR5284206 | 0 | 166 | 24 | 487 | 718 | 0 | 35 | 1665 | 6868 | 18143 | 5451 | 4872 | 0 | 7551 | 0 | 0 | 27 | 1457 | 34 | 1655 | 172 | 572 | 198 | 8908 | 684 | 0 | 12 | 0 | 13 | 13 | 65 | 14 | 13 | 15 | 0 | 4241 | 0 |
| SRR5317818 | 0 | 10 | 0 | 0 | 37 | 0 | 0 | 1735 | 1487 | 6164 | 772 | 828 | 0 | 547 | 0 | 0 | 0 | 305 | 0 | 4795 | 3282 | 853 | 1219 | 8536 | 2138 | 0 | 0 | 0 | 0 | 24 | 32 | 11 | 0 | 12 | 0 | 15793 | 0 |
| SRR5317819 | 0 | 11 | 13 | 0 | 24 | 0 | 0 | 8398 | 10462 | 6826 | 371 | 584 | 0 | 336 | 25 | 19 | 0 | 489 | 13 | 2012 | 1904 | 1812 | 2118 | 18843 | 5660 | 34 | 0 | 0 | 0 | 12 | 13 | 0 | 0 | 16 | 0 | 8774 | 0 |
| SRR5317820 | 0 | 0 | 0 | 0 | 22 | 10 | 0 | 662 | 343 | 589 | 485 | 1274 | 0 | 154 | 12 | 19 | 0 | 164 | 0 | 5829 | 1425 | 1099 | 759 | 8774 | 2099 | 0 | 0 | 0 | 0 | 0 | 11 | 0 | 0 | 14 | 0 | 17058 | 0 |
| SRR5317821 | 0 | 0 | 0 | 0 | 0 | 0 | 0 | 1938 | 10543 | 16037 | 352 | 303 | 0 | 576 | 0 | 0 | 14 | 123 | 0 | 278 | 1610 | 1071 | 8940 | 11061 | 3747 | 29 | 35 | 0 | 0 | 0 | 0 | 0 | 0 | 0 | 0 | 1861 | 0 |
| SRR5317822 | 0 | 12 | 0 | 0 | 40 | 0 | 0 | 3744 | 3617 | 3792 | 201 | 255 | 0 | 163 | 0 | 0 | 0 | 380 | 0 | 1705 | 4438 | 1310 | 949 | 9591 | 3133 | 0 | 0 | 0 | 0 | 20 | 29 | 0 | 0 | 20 | 0 | 20028 | 0 |
| SRR5317823 | 0 | 0 | 0 | 0 | 0 | 0 | 0 | 5657 | 6890 | 4058 | 60 | 129 | 0 | 459 | 15 | 18 | 0 | 481 | 0 | 2157 | 2186 | 2487 | 4443 | 23169 | 4443 | 36 | 0 | 0 | 0 | 0 | 0 | 0 | 0 | 0 | 0 | 20238 | 0 |
| SRR5317824 | 0 | 0 | 0 | 0 | 30 | 23 | 0 | 838 | 769 | 4410 | 86 | 129 | 0 | 444 | 0 | 0 | 0 | 201 | 0 | 6361 | 1088 | 624 | 663 | 13318 | 2232 | 0 | 0 | 0 | 0 | 0 | 0 | 0 | 0 | 0 | 0 | 26040 | 10 |
| SRR5317825 | 0 | 14 | 11 | 0 | 0 | 0 | 0 | 534 | 419 | 1067 | 0 | 0 | 0 | 321 | 0 | 0 | 0 | 476 | 0 | 1088 | 1121 | 1338 | 5011 | 28823 | 9764 | 14 | 0 | 0 | 0 | 0 | 0 | 0 | 0 | 0 | 0 | 15919 | 38 |
| SRR5317826 | 0 | 0 | 0 | 0 | 61 | 0 | 0 | 2062 | 2217 | 1858 | 152 | 125 | 0 | 600 | 0 | 0 | 11 | 809 | 0 | 1510 | 1267 | 1270 | 1486 | 12748 | 2538 | 28 | 0 | 0 | 0 | 0 | 12 | 0 | 0 | 0 | 0 | 14877 | 0 |
| SRR5317827 | 0 | 0 | 0 | 0 | 0 | 0 | 0 | 1524 | 1397 | 1145 | 82 | 136 | 0 | 1187 | 20 | 0 | 0 | 475 | 0 | 4111 | 1257 | 1840 | 2251 | 18900 | 1935 | 28 | 0 | 0 | 0 | 0 | 0 | 0 | 0 | 19 | 0 | 55479 | 0 |
| SRR5387779 | 0 | 0 | 41 | 0 | 0 | 0 | 0 | 119 | 0 | 830 | 5810 | 9130 | 0 | 0 | 474 | 237 | 0 | 0 | 0 | 0 | 630 | 1456 | 0 | 14 | 156 | 17 | 41 | 0 | 0 | 0 | 0 | 0 | 0 | 14 | 27 | 237 | 0 |
| SRR5387780 | 0 | 0 | 30 | 0 | 0 | 0 | 0 | 494 | 71 | 776 | 5852 | 9378 | 0 | 0 | 141 | 71 | 0 | 0 | 0 | 71 | 580 | 2361 | 0 | 25 | 250 | 24 | 38 | 0 | 0 | 0 | 0 | 0 | 0 | 0 | 0 | 705 | 0 |
| SRR5387781 | 0 | 0 | 18 | 0 | 0 | 0 | 0 | 405 | 135 | 1483 | 15573 | 29933 | 0 | 0 | 67 | 270 | 12 | 0 | 0 | 0 | 320 | 1213 | 0 | 0 | 473 | 37 | 62 | 0 | 0 | 0 | 0 | 0 | 12 | 12 | 12 | 135 | 0 |
| SRR5387782 | 0 | 0 | 22 | 0 | 0 | 0 | 0 | 65 | 65 | 1687 | 92 | 11227 | 0 | 65 | 0 | 130 | 0 | 0 | 0 | 324 | 483 | 1529 | 0 | 0 | 998 | 38 | 0 | 0 | 0 | 0 | 0 | 0 | 0 | 0 | 0 | 260 | 0 |
| SRR5387783 | 17 | 0 | 42 | 0 | 0 | 0 | 0 | 82 | 82 | 164 | 147 | 18967 | 0 | 0 | 328 | 657 | 17 | 0 | 0 | 164 | 469 | 1369 | 0 | 0 | 258 | 23 | 134 | 0 | 0 | 0 | 0 | 0 | 0 | 0 | 0 | 164 | 0 |
| SRR5387784 | 31 | 0 | 16 | 0 | 0 | 0 | 0 | 116 | 58 | 465 | 70 | 9181 | 0 | 0 | 465 | 639 | 0 | 0 | 0 | 58 | 532 | 2157 | 0 | 0 | 597 | 35 | 110 | 0 | 0 | 0 | 0 | 0 | 0 | 0 | 0 | 174 | 0 |
| SRR5387785 | 13 | 0 | 0 | 0 | 0 | 0 | 0 | 45 | 91 | 453 | 7973 | 9513 | 0 | 45 | 181 | 91 | 0 | 0 | 0 | 45 | 477 | 1306 | 0 | 0 | 610 | 43 | 40 | 0 | 0 | 0 | 0 | 0 | 0 | 13 | 0 | 136 | 0 |
| SRR5387786 | 0 | 0 | 0 | 0 | 0 | 0 | 0 | 156 | 0 | 467 | 300 | 17814 | 0 | 0 | 233 | 78 | 0 | 0 | 0 | 78 | 481 | 1661 | 0 | 0 | 120 | 15 | 81 | 0 | 0 | 0 | 0 | 0 | 0 | 0 | 0 | 233 | 0 |
| SRR5387787 | 27 | 0 | 18 | 0 | 0 | 0 | 0 | 0 | 599 | 1198 | 8026 | 12458 | 0 | 120 | 0 | 359 | 18 | 0 | 0 | 0 | 399 | 2036 | 0 | 0 | 127 | 26 | 10 | 0 | 0 | 0 | 0 | 0 | 0 | 0 | 0 | 599 | 0 |
| SRR5387788 | 41 | 0 | 0 | 0 | 0 | 0 | 0 | 149 | 0 | 1188 | 5272 | 12994 | 0 | 0 | 223 | 149 | 0 | 0 | 0 | 0 | 615 | 2501 | 0 | 0 | 670 | 33 | 387 | 0 | 0 | 0 | 0 | 0 | 0 | 0 | 0 | 371 | 0 |
| SRR5387789 | 0 | 0 | 0 | 0 | 0 | 0 | 0 | 75 | 0 | 900 | 247 | 18823 | 0 | 0 | 75 | 75 | 0 | 0 | 0 | 0 | 422 | 1516 | 0 | 0 | 246 | 30 | 39 | 0 | 0 | 0 | 0 | 0 | 0 | 0 | 0 | 75 | 0 |
| SRR5387790 | 14 | 0 | 14 | 0 | 0 | 0 | 0 | 130 | 0 | 649 | 13818 | 20045 | 0 | 0 | 130 | 130 | 0 | 0 | 0 | 65 | 363 | 1633 | 0 | 0 | 329 | 36 | 29 | 0 | 0 | 0 | 0 | 0 | 0 | 0 | 0 | 130 | 0 |
| SRR5387791 | 0 | 0 | 35 | 0 | 0 | 0 | 0 | 196 | 196 | 489 | 9790 | 8419 | 0 | 0 | 294 | 294 | 0 | 0 | 0 | 0 | 747 | 11 | 0 | 0 | 159 | 20 | 0 | 0 | 0 | 0 | 0 | 0 | 0 | 18 | 35 | 98 | 0 |
| SRR5387792 | 0 | 0 | 20 | 0 | 0 | 0 | 0 | 179 | 0 | 626 | 50 | 5812 | 0 | 0 | 358 | 268 | 0 | 0 | 0 | 89 | 709 | 2305 | 0 | 0 | 462 | 18 | 20 | 0 | 0 | 0 | 10 | 0 | 10 | 31 | 0 | 715 | 0 |
| SRR5387793 | 0 | 0 | 29 | 0 | 0 | 0 | 0 | 125 | 0 | 748 | 16214 | 14592 | 0 | 125 | 0 | 374 | 0 | 0 | 0 | 0 | 301 | 1270 | 0 | 0 | 182 | 28 | 144 | 0 | 0 | 0 | 0 | 14 | 0 | 14 | 0 | 125 | 0 |
| SRR5387794 | 0 | 0 | 0 | 0 | 0 | 0 | 0 | 63 | 126 | 1260 | 5608 | 9641 | 0 | 0 | 63 | 189 | 0 | 0 | 0 | 63 | 428 | 1748 | 0 | 0 | 234 | 36 | 31 | 0 | 0 | 0 | 0 | 0 | 0 | 0 | 0 | 630 | 0 |
| SRR5387795 | 11 | 11 | 32 | 0 | 0 | 0 | 0 | 341 | 0 | 683 | 8531 | 12172 | 0 | 114 | 455 | 0 | 0 | 0 | 0 | 341 | 120 | 320 | 0 | 0 | 96 | 13 | 194 | 0 | 0 | 0 | 0 | 11 | 22 | 22 | 0 | 114 | 0 |
| SRR5387796 | 0 | 0 | 0 | 0 | 0 | 0 | 0 | 0 | 207 | 828 | 6625 | 9731 | 0 | 0 | 1035 | 207 | 0 | 0 | 0 | 207 | 289 | 882 | 0 | 0 | 167 | 19 | 134 | 0 | 0 | 0 | 0 | 0 | 0 | 27 | 13 | 828 | 0 |
| SRR5387797 | 0 | 0 | 16 | 0 | 0 | 0 | 0 | 112 | 224 | 1121 | 101 | 9472 | 0 | 56 | 1009 | 785 | 0 | 0 | 0 | 112 | 213 | 747 | 0 | 0 | 272 | 47 | 24 | 0 | 0 | 0 | 0 | 0 | 0 | 0 | 0 | 392 | 0 |
| SRR5387798 | 0 | 0 | 0 | 0 | 0 | 36 | 0 | 314 | 0 | 1258 | 41 | 9040 | 0 | 0 | 943 | 1494 | 0 | 0 | 0 | 157 | 420 | 1395 | 0 | 0 | 694 | 35 | 15 | 0 | 0 | 0 | 0 | 0 | 0 | 0 | 0 | 472 | 0 |
| SRR5387799 | 16 | 0 | 0 | 0 | 0 | 0 | 0 | 438 | 164 | 602 | 197 | 12586 | 0 | 55 | 164 | 219 | 0 | 0 | 0 | 55 | 265 | 934 | 0 | 0 | 139 | 17 | 62 | 0 | 0 | 0 | 16 | 0 | 0 | 16 | 0 | 164 | 0 |
| SRR5387800 | 0 | 0 | 58 | 0 | 0 | 0 | 0 | 421 | 105 | 1262 | 125 | 22197 | 0 | 0 | 736 | 316 | 0 | 0 | 0 | 316 | 189 | 773 | 0 | 0 | 94 | 19 | 17 | 0 | 0 | 0 | 0 | 0 | 0 | 0 | 17 | 421 | 0 |
| SRR5387801 | 0 | 0 | 38 | 0 | 0 | 0 | 0 | 206 | 0 | 1238 | 10829 | 22587 | 0 | 103 | 103 | 309 | 0 | 0 | 0 | 0 | 214 | 1256 | 0 | 0 | 185 | 25 | 113 | 0 | 0 | 0 | 0 | 0 | 0 | 0 | 28 | 206 | 0 |
| SRR5387802 | 0 | 0 | 19 | 0 | 0 | 0 | 0 | 164 | 164 | 1150 | 102 | 11833 | 0 | 0 | 164 | 247 | 0 | 0 | 0 | 82 | 242 | 1158 | 0 | 0 | 411 | 27 | 89 | 0 | 0 | 0 | 0 | 0 | 0 | 0 | 0 | 575 | 0 |
| SRR5387803 | 13 | 0 | 13 | 0 | 0 | 0 | 0 | 162 | 0 | 810 | 6885 | 12150 | 0 | 0 | 405 | 324 | 0 | 0 | 0 | 162 | 384 | 1763 | 0 | 0 | 375 | 24 | 0 | 0 | 0 | 0 | 0 | 0 | 0 | 0 | 13 | 567 | 0 |
| SRR5387804 | 0 | 0 | 0 | 0 | 0 | 0 | 0 | 163 | 0 | 980 | 6287 | 13962 | 0 | 82 | 82 | 572 | 16 | 0 | 0 | 82 | 578 | 1957 | 0 | 0 | 616 | 19 | 31 | 0 | 0 | 0 | 0 | 0 | 0 | 0 | 0 | 490 | 0 |
| SRR5387805 | 0 | 0 | 0 | 0 | 0 | 0 | 0 | 614 | 102 | 1637 | 7469 | 10231 | 0 | 102 | 307 | 307 | 19 | 0 | 0 | 512 | 87 | 248 | 0 | 0 | 45 | 31 | 28 | 0 | 0 | 0 | 0 | 0 | 0 | 0 | 0 | 1535 | 0 |
| SRR5387806 | 0 | 0 | 16 | 0 | 0 | 0 | 0 | 0 | 0 | 1727 | 93 | 13904 | 0 | 0 | 545 | 727 | 0 | 0 | 0 | 273 | 336 | 754 | 0 | 0 | 494 | 23 | 0 | 0 | 0 | 0 | 0 | 0 | 0 | 16 | 0 | 727 | 0 |
| SRR5387807 | 12 | 0 | 36 | 0 | 0 | 36 | 0 | 255 | 191 | 509 | 9297 | 14964 | 0 | 0 | 64 | 191 | 12 | 0 | 0 | 64 | 423 | 1472 | 0 | 20 | 391 | 35 | 47 | 0 | 0 | 0 | 0 | 0 | 0 | 36 | 0 | 127 | 0 |
| SRR5387808 | 0 | 0 | 20 | 0 | 0 | 20 | 0 | 486 | 122 | 669 | 115 | 15385 | 0 | 0 | 426 | 182 | 0 | 0 | 0 | 122 | 778 | 10 | 0 | 12 | 171 | 21 | 79 | 0 | 0 | 0 | 0 | 0 | 0 | 0 | 0 | 365 | 0 |
| SRR5387809 | 0 | 0 | 35 | 0 | 0 | 0 | 0 | 305 | 152 | 609 | 6245 | 12491 | 0 | 0 | 609 | 152 | 0 | 0 | 0 | 305 | 344 | 1767 | 0 | 0 | 158 | 12 | 71 | 0 | 0 | 0 | 0 | 0 | 0 | 35 | 0 | 1066 | 0 |
| SRR5387810 | 0 | 0 | 0 | 0 | 0 | 0 | 0 | 626 | 104 | 939 | 54 | 9707 | 0 | 104 | 313 | 522 | 0 | 0 | 0 | 104 | 355 | 904 | 0 | 0 | 952 | 51 | 21 | 0 | 0 | 0 | 0 | 0 | 0 | 0 | 0 | 1566 | 0 |
| SRR5387811 | 101 | 0 | 0 | 0 | 0 | 0 | 0 | 370 | 0 | 1201 | 6375 | 9424 | 0 | 185 | 277 | 92 | 0 | 0 | 0 | 92 | 510 | 1546 | 0 | 0 | 346 | 25 | 366 | 0 | 0 | 0 | 0 | 0 | 16 | 0 | 0 | 462 | 0 |
| SRR5387812 | 42 | 0 | 14 | 0 | 0 | 0 | 0 | 202 | 67 | 539 | 228 | 14832 | 0 | 0 | 0 | 67 | 0 | 0 | 0 | 135 | 527 | 1723 | 0 | 0 | 509 | 39 | 285 | 0 | 0 | 0 | 0 | 0 | 0 | 0 | 0 | 202 | 0 |
| SRR5387813 | 0 | 0 | 0 | 0 | 0 | 0 | 0 | 391 | 0 | 880 | 117 | 17110 | 0 | 98 | 196 | 98 | 30 | 0 | 0 | 196 | 154 | 726 | 0 | 0 | 131 | 23 | 106 | 0 | 0 | 0 | 0 | 0 | 0 | 0 | 15 | 196 | 0 |
| SRR5387814 | 0 | 0 | 53 | 0 | 0 | 0 | 0 | 85 | 85 | 940 | 51 | 12216 | 0 | 0 | 0 | 85 | 0 | 0 | 0 | 342 | 509 | 1900 | 0 | 0 | 353 | 29 | 123 | 0 | 0 | 0 | 0 | 0 | 0 | 0 | 0 | 427 | 0 |
| SRR5387815 | 0 | 0 | 0 | 0 | 0 | 0 | 0 | 170 | 0 | 933 | 99 | 15265 | 0 | 0 | 85 | 85 | 0 | 0 | 0 | 594 | 594 | 1870 | 0 | 0 | 653 | 26 | 56 | 0 | 0 | 0 | 0 | 0 | 0 | 0 | 0 | 933 | 0 |
| SRR5387816 | 19 | 19 | 28 | 0 | 0 | 0 | 0 | 313 | 104 | 730 | 157 | 16065 | 0 | 0 | 104 | 104 | 0 | 0 | 0 | 417 | 566 | 1954 | 0 | 0 | 210 | 21 | 103 | 0 | 0 | 0 | 0 | 0 | 0 | 0 | 0 | 209 | 0 |
| SRR5387817 | 0 | 26 | 65 | 0 | 0 | 0 | 0 | 78 | 78 | 314 | 10279 | 14359 | 0 | 0 | 314 | 157 | 0 | 0 | 0 | 78 | 586 | 2150 | 0 | 0 | 216 | 27 | 39 | 0 | 0 | 0 | 0 | 0 | 0 | 0 | 0 | 314 | 0 |
| SRR5387818 | 0 | 0 | 0 | 0 | 0 | 0 | 0 | 316 | 0 | 126 | 159 | 13259 | 0 | 0 | 126 | 316 | 0 | 0 | 0 | 316 | 485 | 2693 | 0 | 13 | 263 | 36 | 26 | 0 | 0 | 0 | 0 | 0 | 0 | 0 | 0 | 442 | 0 |
| SRR5387819 | 0 | 0 | 34 | 0 | 0 | 0 | 0 | 313 | 45 | 625 | 137 | 11838 | 0 | 0 | 357 | 447 | 25 | 0 | 0 | 45 | 569 | 1165 | 0 | 19 | 218 | 19 | 118 | 0 | 0 | 0 | 0 | 0 | 0 | 17 | 25 | 134 | 0 |
| SRR5387820 | 0 | 0 | 0 | 0 | 0 | 0 | 0 | 188 | 0 | 471 | 164 | 28448 | 0 | 0 | 188 | 377 | 19 | 0 | 0 | 377 | 714 | 1712 | 0 | 21 | 287 | 13 | 58 | 0 | 0 | 0 | 0 | 0 | 0 | 0 | 29 | 377 | 0 |
| SRR5387821 | 17 | 0 | 26 | 0 | 0 | 35 | 0 | 307 | 102 | 2455 | 7057 | 9512 | 0 | 0 | 1125 | 921 | 0 | 0 | 0 | 409 | 458 | 1695 | 0 | 17 | 504 | 53 | 78 | 0 | 0 | 0 | 0 | 0 | 0 | 0 | 17 | 614 | 0 |
| SRR5387822 | 0 | 0 | 13 | 0 | 0 | 0 | 0 | 243 | 0 | 1213 | 95 | 12296 | 0 | 0 | 890 | 1861 | 0 | 0 | 0 | 81 | 179 | 913 | 0 | 0 | 247 | 27 | 67 | 0 | 0 | 0 | 0 | 0 | 0 | 0 | 0 | 566 | 0 |
| SRR5387823 | 0 | 0 | 0 | 0 | 0 | 0 | 0 | 284 | 71 | 711 | 14573 | 17488 | 0 | 71 | 71 | 142 | 0 | 0 | 0 | 0 | 366 | 1543 | 0 | 0 | 347 | 30 | 24 | 0 | 0 | 0 | 0 | 0 | 0 | 0 | 0 | 213 | 0 |
| SRR5387824 | 0 | 0 | 0 | 0 | 0 | 0 | 0 | 222 | 0 | 1035 | 10419 | 13079 | 0 | 74 | 148 | 369 | 0 | 0 | 0 | 74 | 489 | 1795 | 0 | 0 | 724 | 30 | 18 | 0 | 0 | 0 | 0 | 0 | 0 | 18 | 0 | 222 | 0 |
| SRR5387825 | 0 | 0 | 18 | 0 | 0 | 28 | 0 | 327 | 0 | 327 | 7444 | 10062 | 0 | 82 | 409 | 82 | 0 | 0 | 0 | 409 | 592 | 1557 | 0 | 0 | 248 | 27 | 110 | 0 | 0 | 0 | 0 | 0 | 0 | 0 | 0 | 409 | 0 |
| SRR5387826 | 12 | 12 | 12 | 0 | 0 | 0 | 0 | 0 | 118 | 589 | 6949 | 12366 | 0 | 0 | 471 | 353 | 0 | 0 | 0 | 118 | 866 | 2839 | 0 | 22 | 418 | 17 | 195 | 0 | 0 | 0 | 0 | 0 | 0 | 24 | 24 | 236 | 0 |
| SRR5387827 | 16 | 0 | 47 | 0 | 0 | 0 | 0 | 327 | 163 | 2288 | 6211 | 10788 | 0 | 0 | 490 | 981 | 16 | 0 | 0 | 327 | 218 | 761 | 0 | 0 | 181 | 27 | 95 | 0 | 16 | 0 | 0 | 0 | 0 | 16 | 0 | 327 | 0 |
| SRR5387828 | 0 | 0 | 37 | 0 | 0 | 0 | 0 | 84 | 0 | 670 | 6702 | 13152 | 0 | 0 | 419 | 838 | 0 | 0 | 0 | 84 | 388 | 1434 | 0 | 0 | 304 | 24 | 82 | 0 | 0 | 0 | 0 | 0 | 0 | 37 | 0 | 251 | 0 |
| SRR5387829 | 0 | 0 | 15 | 0 | 0 | 25 | 0 | 117 | 0 | 391 | 279 | 18985 | 0 | 39 | 196 | 196 | 0 | 0 | 0 | 39 | 285 | 1234 | 0 | 0 | 101 | 13 | 241 | 0 | 0 | 0 | 0 | 0 | 0 | 25 | 0 | 274 | 0 |
| SRR5387830 | 19 | 0 | 13 | 0 | 0 | 13 | 0 | 502 | 143 | 1362 | 81 | 9680 | 0 | 143 | 215 | 287 | 13 | 0 | 0 | 72 | 384 | 1337 | 0 | 0 | 553 | 40 | 129 | 0 | 0 | 0 | 0 | 0 | 0 | 0 | 0 | 287 | 0 |
| SRR5387831 | 19 | 0 | 28 | 0 | 0 | 0 | 0 | 344 | 115 | 344 | 17775 | 17202 | 0 | 0 | 0 | 115 | 0 | 0 | 0 | 115 | 223 | 1387 | 0 | 11 | 178 | 17 | 19 | 0 | 0 | 0 | 0 | 0 | 0 | 0 | 0 | 115 | 0 |
| SRR5387832 | 0 | 0 | 31 | 0 | 0 | 0 | 0 | 357 | 0 | 1191 | 10599 | 17149 | 0 | 0 | 0 | 119 | 0 | 0 | 0 | 0 | 400 | 1623 | 0 | 12 | 252 | 21 | 114 | 0 | 0 | 0 | 0 | 10 | 10 | 31 | 0 | 119 | 0 |
| SRR5387833 | 0 | 12 | 35 | 0 | 0 | 0 | 0 | 437 | 0 | 984 | 10062 | 11484 | 0 | 0 | 109 | 219 | 12 | 0 | 0 | 109 | 442 | 1590 | 0 | 0 | 131 | 18 | 81 | 0 | 0 | 0 | 0 | 0 | 0 | 23 | 0 | 219 | 0 |
| SRR5387834 | 0 | 0 | 36 | 0 | 0 | 0 | 0 | 0 | 83 | 581 | 4315 | 8963 | 0 | 0 | 332 | 166 | 0 | 0 | 0 | 83 | 397 | 1640 | 0 | 0 | 302 | 24 | 73 | 0 | 0 | 0 | 0 | 0 | 0 | 27 | 0 | 0 | 0 |
| SRR5387835 | 21 | 0 | 14 | 0 | 0 | 0 | 0 | 77 | 77 | 307 | 8748 | 10360 | 0 | 0 | 77 | 307 | 0 | 0 | 0 | 77 | 345 | 1547 | 0 | 0 | 496 | 63 | 124 | 0 | 0 | 0 | 0 | 0 | 0 | 21 | 0 | 384 | 0 |
| SRR5387836 | 0 | 0 | 22 | 0 | 0 | 15 | 0 | 262 | 0 | 350 | 67 | 9178 | 0 | 0 | 612 | 350 | 0 | 0 | 0 | 437 | 257 | 1345 | 0 | 0 | 312 | 27 | 126 | 0 | 0 | 0 | 0 | 0 | 0 | 0 | 0 | 175 | 0 |
| SRR5387837 | 0 | 0 | 0 | 0 | 0 | 0 | 0 | 482 | 96 | 868 | 52 | 9158 | 0 | 0 | 289 | 193 | 0 | 0 | 0 | 193 | 487 | 1365 | 0 | 0 | 592 | 24 | 80 | 0 | 0 | 0 | 0 | 0 | 0 | 0 | 0 | 578 | 0 |
| SRR5387838 | 0 | 0 | 38 | 0 | 0 | 0 | 0 | 194 | 0 | 1361 | 7291 | 18180 | 0 | 0 | 681 | 875 | 0 | 0 | 0 | 389 | 480 | 1666 | 0 | 0 | 505 | 27 | 82 | 0 | 0 | 0 | 0 | 0 | 0 | 13 | 13 | 486 | 0 |
| SRR5387839 | 11 | 11 | 64 | 0 | 0 | 21 | 0 | 347 | 116 | 694 | 72 | 10992 | 0 | 0 | 0 | 116 | 0 | 0 | 0 | 0 | 379 | 1542 | 0 | 0 | 266 | 33 | 193 | 0 | 0 | 0 | 0 | 0 | 11 | 0 | 11 | 347 | 0 |
| SRR5387840 | 0 | 0 | 29 | 0 | 0 | 43 | 0 | 177 | 0 | 707 | 2474 | 8481 | 0 | 0 | 0 | 530 | 0 | 0 | 0 | 0 | 516 | 2255 | 0 | 0 | 142 | 27 | 358 | 0 | 0 | 0 | 0 | 0 | 14 | 29 | 0 | 177 | 0 |
| SRR5387841 | 0 | 0 | 33 | 0 | 0 | 0 | 0 | 337 | 84 | 590 | 8853 | 11551 | 0 | 84 | 422 | 169 | 0 | 0 | 0 | 337 | 88 | 438 | 0 | 0 | 57 | 21 | 25 | 0 | 0 | 0 | 0 | 0 | 0 | 0 | 0 | 84 | 0 |
| SRR5387842 | 0 | 0 | 44 | 0 | 0 | 0 | 0 | 186 | 93 | 929 | 6133 | 13566 | 0 | 0 | 279 | 186 | 26 | 0 | 0 | 93 | 300 | 1067 | 0 | 0 | 330 | 18 | 18 | 0 | 0 | 0 | 0 | 0 | 0 | 0 | 0 | 279 | 0 |
| SRR5387843 | 0 | 0 | 39 | 0 | 0 | 0 | 0 | 217 | 72 | 1158 | 8830 | 10929 | 0 | 72 | 145 | 72 | 0 | 0 | 0 | 145 | 410 | 1258 | 0 | 0 | 390 | 48 | 52 | 0 | 0 | 0 | 0 | 0 | 0 | 0 | 0 | 362 | 0 |
| SRR5387844 | 0 | 18 | 0 | 0 | 0 | 18 | 0 | 78 | 0 | 1095 | 96 | 9858 | 0 | 78 | 78 | 156 | 0 | 0 | 0 | 0 | 799 | 2485 | 0 | 0 | 425 | 38 | 110 | 0 | 0 | 0 | 0 | 0 | 0 | 0 | 18 | 78 | 0 |
| SRR5387845 | 0 | 0 | 10 | 0 | 0 | 0 | 0 | 560 | 187 | 871 | 48 | 11141 | 0 | 0 | 373 | 124 | 0 | 0 | 0 | 0 | 592 | 1361 | 0 | 12 | 441 | 26 | 163 | 0 | 0 | 0 | 0 | 0 | 10 | 10 | 0 | 187 | 0 |
| SRR5387846 | 25 | 0 | 51 | 0 | 0 | 0 | 0 | 247 | 82 | 1151 | 11595 | 16859 | 0 | 0 | 164 | 493 | 0 | 0 | 0 | 82 | 788 | 2477 | 0 | 13 | 312 | 11 | 169 | 0 | 0 | 0 | 0 | 0 | 0 | 17 | 0 | 329 | 0 |
| SRR5387847 | 0 | 11 | 11 | 0 | 0 | 0 | 0 | 383 | 64 | 638 | 326 | 23621 | 0 | 64 | 192 | 64 | 0 | 0 | 0 | 128 | 570 | 2076 | 0 | 0 | 183 | 39 | 100 | 0 | 0 | 0 | 0 | 0 | 0 | 11 | 0 | 64 | 0 |
| SRR5387848 | 14 | 0 | 14 | 0 | 0 | 0 | 0 | 429 | 0 | 1165 | 15764 | 18708 | 0 | 0 | 184 | 245 | 14 | 0 | 0 | 245 | 482 | 1573 | 0 | 0 | 416 | 36 | 75 | 0 | 0 | 0 | 0 | 0 | 0 | 0 | 0 | 613 | 0 |
| SRR5387849 | 0 | 0 | 15 | 0 | 0 | 0 | 0 | 273 | 0 | 491 | 20446 | 24590 | 0 | 0 | 55 | 164 | 0 | 0 | 0 | 55 | 473 | 2086 | 0 | 0 | 331 | 25 | 185 | 0 | 0 | 0 | 0 | 0 | 0 | 15 | 0 | 109 | 0 |
| SRR5387850 | 0 | 0 | 46 | 0 | 0 | 0 | 0 | 399 | 80 | 718 | 14994 | 17706 | 0 | 0 | 160 | 0 | 0 | 0 | 0 | 0 | 315 | 1328 | 0 | 0 | 275 | 16 | 174 | 0 | 0 | 0 | 0 | 0 | 0 | 0 | 0 | 80 | 0 |
| SRR5387851 | 0 | 0 | 28 | 0 | 0 | 0 | 0 | 501 | 0 | 1837 | 9521 | 13028 | 0 | 167 | 1002 | 334 | 0 | 0 | 0 | 167 | 446 | 1781 | 0 | 0 | 214 | 16 | 55 | 0 | 0 | 0 | 0 | 0 | 0 | 14 | 14 | 668 | 0 |
| SRR5387852 | 0 | 0 | 27 | 0 | 0 | 0 | 0 | 395 | 0 | 633 | 7039 | 11152 | 0 | 0 | 316 | 316 | 0 | 0 | 0 | 0 | 331 | 1245 | 0 | 0 | 530 | 48 | 55 | 0 | 0 | 0 | 0 | 0 | 0 | 0 | 0 | 158 | 0 |
| SRR5387853 | 0 | 0 | 0 | 0 | 0 | 0 | 0 | 0 | 0 | 1263 | 7579 | 7452 | 0 | 0 | 379 | 253 | 0 | 0 | 0 | 126 | 439 | 1475 | 0 | 23 | 342 | 27 | 177 | 0 | 0 | 0 | 0 | 0 | 10 | 10 | 0 | 1010 | 0 |
| SRR5387854 | 21 | 0 | 0 | 0 | 0 | 0 | 0 | 132 | 0 | 662 | 12051 | 12780 | 0 | 0 | 464 | 265 | 0 | 0 | 0 | 199 | 255 | 1239 | 0 | 21 | 384 | 31 | 106 | 0 | 0 | 0 | 0 | 0 | 0 | 0 | 0 | 66 | 0 |
| SRR5387855 | 0 | 0 | 18 | 0 | 0 | 0 | 0 | 582 | 0 | 582 | 118 | 15131 | 0 | 73 | 145 | 145 | 0 | 0 | 0 | 73 | 526 | 1870 | 0 | 0 | 150 | 23 | 0 | 0 | 0 | 0 | 0 | 0 | 0 | 0 | 0 | 73 | 0 |
| SRR5387856 | 0 | 0 | 14 | 0 | 0 | 0 | 0 | 138 | 0 | 276 | 12412 | 18273 | 0 | 0 | 138 | 138 | 0 | 0 | 0 | 0 | 400 | 1493 | 0 | 0 | 372 | 24 | 20 | 0 | 0 | 0 | 0 | 0 | 0 | 0 | 0 | 276 | 0 |
| SRR5387857 | 11 | 0 | 33 | 0 | 0 | 0 | 0 | 466 | 93 | 745 | 114 | 14813 | 0 | 0 | 186 | 93 | 0 | 0 | 0 | 93 | 598 | 2133 | 0 | 0 | 560 | 21 | 111 | 0 | 0 | 0 | 11 | 0 | 0 | 0 | 0 | 186 | 0 |
| SRR5387858 | 0 | 0 | 16 | 0 | 0 | 0 | 0 | 205 | 0 | 547 | 9843 | 21396 | 0 | 0 | 137 | 273 | 0 | 0 | 0 | 68 | 556 | 2555 | 0 | 0 | 291 | 28 | 90 | 0 | 0 | 0 | 0 | 0 | 0 | 33 | 0 | 273 | 0 |
| SRR5387859 | 0 | 0 | 13 | 0 | 0 | 0 | 0 | 0 | 0 | 2103 | 44 | 9178 | 0 | 0 | 191 | 0 | 0 | 0 | 0 | 191 | 296 | 878 | 0 | 0 | 146 | 18 | 26 | 0 | 0 | 0 | 0 | 0 | 0 | 26 | 0 | 382 | 0 |
| SRR5387860 | 11 | 0 | 32 | 0 | 0 | 0 | 0 | 277 | 0 | 739 | 79 | 16070 | 0 | 92 | 0 | 277 | 11 | 0 | 0 | 185 | 656 | 2411 | 0 | 18 | 413 | 18 | 85 | 0 | 0 | 0 | 0 | 0 | 0 | 32 | 11 | 277 | 0 |
| SRR5387861 | 0 | 20 | 10 | 0 | 0 | 0 | 0 | 185 | 0 | 277 | 8876 | 15718 | 0 | 0 | 185 | 185 | 0 | 0 | 0 | 185 | 337 | 1376 | 0 | 0 | 215 | 17 | 71 | 0 | 0 | 0 | 0 | 0 | 0 | 10 | 20 | 92 | 0 |
| SRR5387862 | 25 | 0 | 0 | 0 | 0 | 0 | 0 | 273 | 0 | 1749 | 7322 | 13770 | 0 | 0 | 55 | 219 | 0 | 0 | 0 | 0 | 279 | 1303 | 0 | 0 | 546 | 43 | 67 | 0 | 0 | 0 | 0 | 0 | 0 | 0 | 0 | 219 | 0 |
| SRR5387863 | 0 | 0 | 29 | 0 | 0 | 0 | 0 | 236 | 157 | 1810 | 80 | 7396 | 0 | 0 | 79 | 0 | 0 | 0 | 0 | 315 | 569 | 1328 | 0 | 17 | 936 | 53 | 76 | 0 | 0 | 0 | 0 | 0 | 0 | 0 | 19 | 393 | 0 |
| SRR5387864 | 0 | 0 | 23 | 0 | 0 | 0 | 0 | 221 | 147 | 2138 | 80 | 8700 | 0 | 0 | 147 | 221 | 0 | 0 | 0 | 74 | 479 | 1879 | 0 | 20 | 734 | 31 | 76 | 0 | 0 | 0 | 0 | 0 | 0 | 15 | 0 | 811 | 0 |
| SRR5387865 | 0 | 13 | 13 | 0 | 0 | 0 | 0 | 205 | 205 | 1435 | 123 | 14756 | 0 | 102 | 102 | 0 | 0 | 0 | 0 | 102 | 324 | 1743 | 0 | 0 | 228 | 81 | 46 | 0 | 0 | 0 | 0 | 0 | 0 | 0 | 0 | 512 | 0 |
| SRR5387866 | 19 | 0 | 0 | 0 | 0 | 0 | 0 | 644 | 59 | 819 | 79 | 8074 | 0 | 59 | 59 | 0 | 18 | 0 | 0 | 176 | 435 | 1401 | 0 | 0 | 584 | 37 | 0 | 0 | 0 | 0 | 0 | 0 | 0 | 0 | 0 | 1638 | 0 |

### Supp Table S8.4.Columns_DI_to_ES

| | | | | | | | | | | | | | | | | | | | | | | | | | | | | | | | | | | | | | |
|---|---|---|---|---|---|---|---|---|---|---|---|---|---|---|---|---|---|---|---|---|---|---|---|---|---|---|---|---|---|---|---|---|---|---|---|---|---|
| SRR5387867 | 0 | 0 | 55 | 0 | 0 | 0 | 0 | 250 | 0 | 125 | 164 | 19672 | 0 | 0 | 187 | 312 | 0 | 0 | 0 | 250 | 432 | 1373 | 0 | 0 | 303 | 14 | 79 | 0 | 0 | 0 | 0 | 0 | 0 | 0 | 0 | 250 | 0 |
| SRR5387868 | 0 | 0 | 21 | 0 | 0 | 0 | 0 | 62 | 0 | 435 | 6653 | 16976 | 0 | 0 | 124 | 124 | 0 | 0 | 0 | 124 | 346 | 1888 | 0 | 0 | 223 | 21 | 31 | 0 | 0 | 0 | 0 | 10 | 0 | 21 | 0 | 124 | 0 |
| SRR5387869 | 0 | 0 | 32 | 0 | 0 | 0 | 0 | 676 | 338 | 789 | 9468 | 10144 | 0 | 0 | 113 | 451 | 0 | 0 | 0 | 113 | 548 | 11 | 0 | 0 | 310 | 14 | 22 | 0 | 0 | 0 | 0 | 11 | 11 | 22 | 0 | 564 | 0 |
| SRR5387870 | 0 | 0 | 23 | 0 | 0 | 0 | 0 | 260 | 195 | 845 | 89 | 12281 | 0 | 0 | 260 | 325 | 0 | 0 | 0 | 195 | 306 | 1842 | 0 | 0 | 377 | 34 | 15 | 0 | 0 | 0 | 0 | 0 | 0 | 0 | 0 | 65 | 0 |
| SRR5387871 | 0 | 0 | 12 | 0 | 0 | 12 | 0 | 0 | 0 | 582 | 37 | 6633 | 0 | 0 | 116 | 233 | 0 | 0 | 0 | 116 | 371 | 1436 | 0 | 0 | 225 | 11 | 71 | 0 | 0 | 0 | 0 | 0 | 0 | 12 | 0 | 349 | 0 |
| SRR5387872 | 20 | 0 | 0 | 0 | 0 | 0 | 0 | 208 | 104 | 625 | 3753 | 8756 | 0 | 0 | 104 | 104 | 0 | 0 | 0 | 0 | 522 | 2328 | 0 | 0 | 650 | 27 | 119 | 0 | 0 | 0 | 0 | 0 | 0 | 0 | 0 | 730 | 0 |
| SRR5387873 | 0 | 36 | 73 | 0 | 0 | 0 | 0 | 268 | 0 | 535 | 17800 | 21146 | 0 | 0 | 268 | 0 | 0 | 0 | 0 | 0 | 417 | 1923 | 0 | 0 | 188 | 18 | 73 | 0 | 0 | 0 | 0 | 0 | 0 | 0 | 0 | 669 | 0 |
| SRR5387874 | 0 | 0 | 34 | 0 | 0 | 0 | 0 | 174 | 58 | 811 | 184 | 17961 | 0 | 0 | 232 | 174 | 0 | 0 | 0 | 116 | 648 | 3063 | 0 | 0 | 457 | 26 | 127 | 0 | 0 | 0 | 0 | 0 | 0 | 0 | 0 | 232 | 0 |
| SRR5387875 | 0 | 0 | 24 | 0 | 0 | 0 | 0 | 241 | 361 | 1805 | 7462 | 11433 | 0 | 0 | 120 | 241 | 0 | 0 | 0 | 0 | 406 | 1649 | 0 | 12 | 377 | 34 | 73 | 0 | 0 | 0 | 0 | 0 | 0 | 0 | 0 | 1083 | 0 |
| SRR5387876 | 0 | 0 | 0 | 0 | 0 | 0 | 0 | 0 | 119 | 833 | 4402 | 10350 | 0 | 0 | 357 | 238 | 0 | 0 | 0 | 0 | 273 | 1261 | 0 | 0 | 194 | 18 | 54 | 0 | 0 | 0 | 0 | 0 | 0 | 0 | 0 | 357 | 0 |
| SRR5387877 | 11 | 23 | 23 | 0 | 0 | 34 | 0 | 140 | 140 | 561 | 6036 | 8282 | 0 | 0 | 1123 | 561 | 11 | 0 | 0 | 0 | 586 | 1993 | 0 | 0 | 313 | 20 | 80 | 0 | 0 | 0 | 0 | 0 | 0 | 0 | 0 | 0 | 0 |
| SRR5387878 | 0 | 0 | 46 | 0 | 0 | 0 | 0 | 79 | 79 | 393 | 129 | 13040 | 0 | 0 | 628 | 864 | 0 | 0 | 0 | 236 | 556 | 2108 | 0 | 0 | 405 | 26 | 0 | 0 | 0 | 0 | 0 | 0 | 0 | 0 | 15 | 393 | 0 |
| SRR5387879 | 12 | 31 | 25 | 0 | 0 | 0 | 0 | 429 | 86 | 686 | 10292 | 9005 | 0 | 0 | 600 | 686 | 0 | 0 | 0 | 257 | 278 | 1626 | 0 | 0 | 244 | 55 | 93 | 0 | 0 | 0 | 0 | 0 | 0 | 0 | 0 | 172 | 0 |
| SRR5387880 | 0 | 0 | 25 | 0 | 0 | 0 | 0 | 419 | 210 | 1257 | 80 | 9535 | 0 | 105 | 314 | 314 | 0 | 0 | 0 | 0 | 269 | 1271 | 0 | 0 | 228 | 31 | 41 | 0 | 0 | 0 | 0 | 0 | 0 | 0 | 0 | 105 | 0 |
| SRR5387881 | 0 | 0 | 0 | 0 | 0 | 15 | 0 | 208 | 208 | 834 | 232 | 22256 | 0 | 156 | 156 | 156 | 0 | 0 | 0 | 52 | 99 | 646 | 0 | 0 | 172 | 59 | 108 | 0 | 0 | 0 | 0 | 0 | 0 | 0 | 0 | 469 | 0 |
| SRR5387882 | 0 | 0 | 0 | 0 | 0 | 12 | 0 | 162 | 108 | 1082 | 230 | 18119 | 0 | 54 | 108 | 108 | 0 | 0 | 0 | 0 | 289 | 1602 | 0 | 20 | 316 | 51 | 67 | 0 | 0 | 0 | 0 | 0 | 0 | 0 | 0 | 108 | 0 |
| SRR5387883 | 0 | 0 | 0 | 0 | 0 | 0 | 0 | 476 | 0 | 2220 | 3409 | 6104 | 0 | 79 | 0 | 396 | 0 | 0 | 0 | 79 | 292 | 1087 | 0 | 0 | 502 | 31 | 38 | 0 | 0 | 0 | 0 | 0 | 0 | 0 | 0 | 713 | 0 |
| SRR5387884 | 0 | 0 | 17 | 0 | 0 | 0 | 0 | 390 | 98 | 1560 | 88 | 11115 | 0 | 0 | 488 | 390 | 0 | 0 | 0 | 293 | 481 | 1968 | 0 | 0 | 294 | 24 | 61 | 0 | 0 | 0 | 0 | 0 | 0 | 0 | 0 | 390 | 0 |
| SRR5387885 | 10 | 10 | 41 | 0 | 0 | 10 | 0 | 94 | 188 | 1035 | 5455 | 8934 | 0 | 0 | 282 | 376 | 21 | 0 | 0 | 0 | 646 | 2106 | 0 | 0 | 257 | 30 | 103 | 0 | 0 | 0 | 0 | 0 | 0 | 0 | 0 | 376 | 0 |
| SRR5387886 | 0 | 0 | 37 | 0 | 0 | 18 | 0 | 0 | 0 | 1435 | 81 | 16067 | 0 | 96 | 96 | 669 | 0 | 0 | 0 | 96 | 256 | 2321 | 0 | 0 | 90 | 28 | 229 | 0 | 0 | 0 | 0 | 0 | 0 | 0 | 0 | 383 | 0 |
| SRR5387887 | 11 | 11 | 11 | 0 | 0 | 0 | 0 | 79 | 79 | 395 | 11704 | 10439 | 0 | 0 | 79 | 79 | 0 | 0 | 0 | 158 | 412 | 1790 | 0 | 0 | 221 | 32 | 11 | 11 | 0 | 0 | 0 | 0 | 0 | 0 | 0 | 237 | 0 |
| SRR5387888 | 0 | 0 | 15 | 0 | 0 | 0 | 0 | 283 | 57 | 622 | 155 | 11881 | 0 | 0 | 113 | 0 | 0 | 0 | 0 | 0 | 522 | 1957 | 0 | 0 | 939 | 37 | 0 | 0 | 0 | 0 | 0 | 0 | 0 | 0 | 0 | 0 | 0 |
| SRR5387889 | 0 | 0 | 0 | 0 | 0 | 19 | 0 | 153 | 77 | 613 | 9883 | 16242 | 0 | 77 | 460 | 153 | 0 | 0 | 0 | 77 | 105 | 573 | 0 | 0 | 57 | 26 | 202 | 0 | 0 | 0 | 0 | 0 | 29 | 0 | 0 | 460 | 0 |
| SRR5387890 | 0 | 0 | 38 | 0 | 0 | 19 | 0 | 346 | 0 | 1175 | 210 | 20527 | 0 | 0 | 0 | 138 | 0 | 0 | 0 | 138 | 366 | 1268 | 0 | 0 | 447 | 49 | 85 | 0 | 0 | 0 | 0 | 0 | 0 | 0 | 28 | 346 | 0 |
| SRR5387891 | 0 | 0 | 67 | 0 | 0 | 0 | 0 | 55 | 0 | 775 | 9804 | 13404 | 0 | 55 | 111 | 111 | 0 | 0 | 0 | 55 | 993 | 2949 | 0 | 0 | 472 | 40 | 96 | 0 | 0 | 0 | 0 | 0 | 0 | 0 | 0 | 222 | 0 |
| SRR5387892 | 20 | 0 | 54 | 0 | 0 | 0 | 0 | 108 | 108 | 1456 | 219 | 15849 | 0 | 0 | 485 | 216 | 0 | 0 | 0 | 54 | 749 | 11 | 0 | 0 | 811 | 39 | 68 | 0 | 0 | 0 | 0 | 0 | 0 | 0 | 0 | 108 | 0 |
| SRR5387893 | 0 | 12 | 12 | 0 | 0 | 0 | 0 | 0 | 60 | 838 | 8500 | 14606 | 0 | 0 | 419 | 419 | 0 | 0 | 0 | 0 | 529 | 2629 | 0 | 0 | 664 | 43 | 12 | 0 | 0 | 0 | 0 | 0 | 0 | 12 | 0 | 239 | 0 |
| SRR5387894 | 0 | 0 | 0 | 0 | 0 | 0 | 0 | 131 | 131 | 589 | 115 | 10471 | 0 | 0 | 262 | 196 | 0 | 0 | 0 | 65 | 357 | 1450 | 0 | 0 | 419 | 30 | 0 | 0 | 0 | 0 | 0 | 0 | 0 | 0 | 0 | 524 | 0 |
| SRR5387895 | 29 | 0 | 35 | 0 | 0 | 0 | 0 | 407 | 0 | 1425 | 8751 | 13839 | 0 | 0 | 305 | 407 | 0 | 0 | 0 | 0 | 284 | 1086 | 0 | 0 | 569 | 65 | 82 | 0 | 0 | 0 | 0 | 0 | 0 | 12 | 12 | 1425 | 0 |
| SRR5387896 | 20 | 0 | 20 | 0 | 0 | 0 | 0 | 599 | 120 | 1557 | 42 | 6828 | 0 | 0 | 120 | 0 | 0 | 0 | 0 | 0 | 405 | 1434 | 0 | 0 | 296 | 37 | 132 | 0 | 0 | 0 | 0 | 0 | 0 | 13 | 0 | 958 | 0 |
| SRR5387897 | 11 | 11 | 32 | 0 | 0 | 0 | 0 | 444 | 148 | 740 | 13544 | 20574 | 0 | 0 | 74 | 148 | 0 | 0 | 0 | 0 | 519 | 2226 | 0 | 0 | 358 | 27 | 32 | 0 | 0 | 0 | 0 | 11 | 0 | 11 | 0 | 222 | 0 |
| SRR5387898 | 0 | 0 | 0 | 0 | 0 | 0 | 0 | 208 | 138 | 900 | 184 | 18557 | 0 | 0 | 208 | 138 | 0 | 0 | 0 | 69 | 581 | 1763 | 0 | 0 | 371 | 16 | 17 | 0 | 0 | 0 | 0 | 0 | 0 | 0 | 0 | 277 | 0 |
| SRR5387899 | 28 | 0 | 28 | 0 | 0 | 18 | 0 | 186 | 0 | 929 | 8267 | 14583 | 0 | 0 | 279 | 93 | 0 | 0 | 0 | 0 | 421 | 1163 | 0 | 0 | 507 | 31 | 138 | 0 | 0 | 0 | 0 | 0 | 0 | 18 | 0 | 279 | 0 |
| SRR5387900 | 0 | 17 | 35 | 0 | 0 | 26 | 0 | 408 | 0 | 1089 | 100 | 15793 | 0 | 0 | 0 | 272 | 0 | 0 | 0 | 136 | 208 | 684 | 0 | 0 | 192 | 40 | 35 | 0 | 0 | 0 | 0 | 0 | 0 | 0 | 0 | 1361 | 0 |
| SRR5387901 | 0 | 0 | 0 | 0 | 0 | 0 | 0 | 347 | 208 | 2013 | 4304 | 6040 | 0 | 69 | 208 | 278 | 0 | 0 | 0 | 555 | 313 | 969 | 0 | 31 | 930 | 50 | 25 | 0 | 0 | 0 | 0 | 0 | 0 | 0 | 0 | 1666 | 0 |
| SRR5387902 | 0 | 0 | 0 | 0 | 0 | 0 | 0 | 650 | 72 | 3176 | 147 | 9890 | 0 | 72 | 361 | 650 | 0 | 0 | 0 | 1155 | 257 | 905 | 0 | 11 | 316 | 28 | 18 | 0 | 0 | 0 | 0 | 0 | 0 | 0 | 0 | 4331 | 0 |
| SRR5387903 | 15 | 0 | 0 | 0 | 0 | 15 | 0 | 246 | 0 | 1084 | 15169 | 19356 | 0 | 49 | 99 | 49 | 0 | 0 | 0 | 49 | 288 | 1387 | 0 | 0 | 438 | 41 | 31 | 0 | 0 | 0 | 0 | 0 | 0 | 0 | 0 | 197 | 0 |
| SRR5387904 | 18 | 0 | 0 | 0 | 0 | 0 | 0 | 0 | 0 | 955 | 7759 | 11341 | 0 | 0 | 119 | 119 | 0 | 0 | 0 | 239 | 175 | 765 | 0 | 0 | 82 | 10 | 100 | 0 | 0 | 0 | 0 | 0 | 0 | 0 | 0 | 597 | 0 |
| SRR5387905 | 0 | 0 | 12 | 0 | 0 | 0 | 0 | 240 | 60 | 421 | 160 | 18094 | 0 | 60 | 60 | 180 | 0 | 0 | 0 | 120 | 431 | 2464 | 0 | 0 | 177 | 27 | 89 | 0 | 0 | 0 | 0 | 0 | 0 | 0 | 0 | 180 | 0 |
| SRR5387906 | 0 | 0 | 11 | 0 | 0 | 23 | 0 | 139 | 0 | 418 | 7663 | 14282 | 0 | 0 | 139 | 209 | 0 | 0 | 0 | 70 | 472 | 2231 | 0 | 0 | 271 | 30 | 107 | 0 | 0 | 0 | 0 | 0 | 0 | 0 | 0 | 557 | 0 |
| SRR5387907 | 0 | 0 | 19 | 0 | 0 | 126 | 0 | 239 | 60 | 479 | 8855 | 18189 | 0 | 0 | 60 | 239 | 30 | 0 | 0 | 0 | 371 | 1852 | 0 | 0 | 287 | 24 | 12 | 0 | 0 | 0 | 0 | 0 | 0 | 0 | 0 | 180 | 0 |
| SRR5387908 | 0 | 0 | 15 | 0 | 0 | 0 | 0 | 567 | 71 | 1063 | 9425 | 12898 | 0 | 0 | 71 | 213 | 0 | 0 | 0 | 213 | 661 | 2869 | 0 | 0 | 500 | 48 | 37 | 0 | 0 | 0 | 0 | 0 | 0 | 0 | 0 | 354 | 0 |
| SRR5387909 | 0 | 0 | 27 | 0 | 0 | 36 | 0 | 382 | 0 | 1529 | 6498 | 9684 | 0 | 127 | 637 | 1019 | 0 | 0 | 0 | 382 | 327 | 1259 | 0 | 0 | 90 | 0 | 108 | 0 | 0 | 0 | 0 | 0 | 0 | 0 | 0 | 510 | 0 |
| SRR5387910 | 26 | 0 | 17 | 0 | 0 | 0 | 0 | 0 | 0 | 1088 | 6893 | 12456 | 0 | 0 | 363 | 0 | 0 | 0 | 0 | 484 | 220 | 1199 | 0 | 0 | 223 | 28 | 69 | 0 | 0 | 0 | 0 | 0 | 0 | 17 | 17 | 0 | 0 |
| SRR5387911 | 25 | 0 | 34 | 0 | 0 | 17 | 0 | 524 | 0 | 974 | 59 | 7938 | 0 | 0 | 674 | 524 | 17 | 0 | 0 | 75 | 250 | 757 | 0 | 0 | 253 | 31 | 126 | 0 | 0 | 0 | 0 | 0 | 0 | 0 | 0 | 225 | 0 |
| SRR5387912 | 0 | 0 | 23 | 0 | 0 | 0 | 0 | 100 | 0 | 300 | 7892 | 11888 | 0 | 0 | 100 | 599 | 0 | 0 | 0 | 100 | 170 | 923 | 0 | 0 | 177 | 29 | 139 | 0 | 0 | 0 | 0 | 0 | 0 | 23 | 0 | 699 | 0 |
| SRR5387913 | 0 | 35 | 58 | 0 | 0 | 23 | 0 | 98 | 98 | 1473 | 14629 | 12862 | 0 | 98 | 491 | 98 | 0 | 0 | 0 | 0 | 308 | 1195 | 0 | 0 | 363 | 33 | 58 | 0 | 0 | 0 | 12 | 0 | 12 | 12 | 23 | 295 | 0 |
| SRR5387914 | 29 | 0 | 59 | 0 | 0 | 0 | 0 | 81 | 81 | 1141 | 109 | 11162 | 0 | 0 | 81 | 244 | 0 | 0 | 0 | 81 | 584 | 2221 | 0 | 0 | 293 | 18 | 234 | 0 | 0 | 0 | 0 | 0 | 0 | 0 | 39 | 163 | 0 |
| SRR5387915 | 0 | 15 | 15 | 0 | 0 | 0 | 0 | 442 | 221 | 1990 | 82 | 5140 | 0 | 0 | 332 | 111 | 15 | 0 | 0 | 0 | 534 | 1216 | 0 | 0 | 611 | 33 | 46 | 0 | 0 | 0 | 0 | 0 | 0 | 0 | 15 | 111 | 0 |
| SRR5387916 | 0 | 0 | 27 | 0 | 0 | 0 | 0 | 82 | 329 | 1234 | 6253 | 9708 | 0 | 82 | 329 | 494 | 0 | 0 | 0 | 165 | 250 | 1234 | 0 | 0 | 303 | 41 | 44 | 0 | 0 | 0 | 0 | 0 | 0 | 0 | 18 | 576 | 0 |
| SRR5387917 | 32 | 0 | 24 | 0 | 0 | 0 | 0 | 87 | 0 | 1303 | 64 | 12595 | 0 | 0 | 87 | 434 | 0 | 0 | 0 | 174 | 206 | 1146 | 0 | 0 | 326 | 33 | 113 | 0 | 0 | 0 | 0 | 16 | 0 | 16 | 0 | 174 | 0 |
| SRR5387918 | 33 | 0 | 41 | 0 | 0 | 0 | 0 | 275 | 92 | 918 | 6057 | 9912 | 0 | 92 | 0 | 184 | 0 | 0 | 0 | 92 | 261 | 1649 | 0 | 0 | 148 | 16 | 261 | 0 | 0 | 0 | 0 | 0 | 0 | 16 | 0 | 459 | 0 |
| SRR5387919 | 0 | 0 | 30 | 0 | 0 | 0 | 0 | 185 | 62 | 617 | 5365 | 13074 | 0 | 123 | 62 | 62 | 0 | 0 | 0 | 185 | 274 | 739 | 0 | 0 | 475 | 43 | 120 | 0 | 0 | 0 | 0 | 0 | 0 | 0 | 0 | 123 | 0 |
| SRR5387920 | 0 | 12 | 37 | 0 | 0 | 0 | 0 | 185 | 0 | 1478 | 136 | 18746 | 0 | 0 | 0 | 92 | 0 | 0 | 0 | 0 | 512 | 1995 | 0 | 0 | 229 | 36 | 134 | 0 | 0 | 12 | 0 | 0 | 0 | 12 | 0 | 277 | 0 |
| SRR5387921 | 22 | 0 | 0 | 0 | 0 | 0 | 0 | 355 | 0 | 533 | 21 | 7106 | 0 | 0 | 0 | 178 | 44 | 0 | 0 | 178 | 254 | 1132 | 0 | 0 | 108 | 25 | 22 | 0 | 0 | 0 | 0 | 0 | 0 | 22 | 0 | 178 | 0 |
| SRR5387922 | 20 | 0 | 0 | 0 | 0 | 40 | 0 | 445 | 148 | 148 | 74 | 9501 | 0 | 0 | 891 | 148 | 0 | 0 | 0 | 148 | 231 | 1392 | 0 | 0 | 110 | 20 | 0 | 0 | 0 | 0 | 0 | 0 | 0 | 20 | 20 | 742 | 0 |
| SRR5387923 | 0 | 0 | 68 | 0 | 0 | 0 | 0 | 389 | 97 | 584 | 6910 | 12847 | 0 | 97 | 584 | 487 | 0 | 0 | 0 | 195 | 263 | 688 | 0 | 0 | 207 | 17 | 215 | 0 | 0 | 0 | 0 | 0 | 11 | 23 | 0 | 779 | 0 |
| SRR5387924 | 0 | 0 | 16 | 0 | 0 | 0 | 0 | 333 | 166 | 832 | 37 | 7901 | 0 | 0 | 582 | 166 | 0 | 0 | 0 | 0 | 598 | 2296 | 0 | 0 | 414 | 24 | 288 | 0 | 0 | 0 | 0 | 0 | 0 | 0 | 0 | 333 | 0 |
| SRR5387925 | 17 | 0 | 0 | 0 | 0 | 0 | 0 | 104 | 70 | 800 | 270 | 18046 | 0 | 0 | 35 | 243 | 0 | 0 | 0 | 104 | 303 | 1546 | 0 | 23 | 528 | 47 | 61 | 0 | 0 | 0 | 0 | 0 | 0 | 0 | 0 | 35 | 0 |
| SRR5387926 | 10 | 0 | 26 | 0 | 0 | 0 | 0 | 214 | 54 | 857 | 266 | 22502 | 0 | 0 | 54 | 0 | 0 | 0 | 0 | 107 | 371 | 1631 | 0 | 19 | 443 | 34 | 84 | 0 | 0 | 0 | 0 | 0 | 0 | 0 | 0 | 214 | 0 |
| SRR5387927 | 0 | 28 | 57 | 0 | 0 | 0 | 0 | 268 | 89 | 626 | 188 | 12601 | 0 | 0 | 179 | 0 | 0 | 0 | 0 | 447 | 314 | 1055 | 0 | 0 | 220 | 22 | 57 | 0 | 0 | 0 | 0 | 0 | 0 | 0 | 0 | 894 | 0 |
| SRR5387928 | 0 | 0 | 0 | 0 | 0 | 0 | 0 | 168 | 421 | 1179 | 10194 | 11205 | 0 | 168 | 0 | 168 | 0 | 0 | 0 | 421 | 317 | 1332 | 0 | 11 | 326 | 37 | 65 | 0 | 0 | 0 | 0 | 0 | 0 | 0 | 0 | 1348 | 0 |
| SRR5387929 | 0 | 0 | 15 | 0 | 0 | 0 | 0 | 352 | 201 | 755 | 12132 | 16512 | 0 | 0 | 50 | 50 | 0 | 0 | 0 | 0 | 284 | 897 | 0 | 0 | 163 | 38 | 46 | 0 | 0 | 0 | 0 | 0 | 15 | 0 | 0 | 352 | 0 |
| SRR5387930 | 0 | 0 | 12 | 0 | 0 | 0 | 0 | 648 | 0 | 1189 | 6159 | 15451 | 0 | 0 | 216 | 432 | 0 | 0 | 0 | 216 | 794 | 2812 | 0 | 0 | 225 | 0 | 111 | 0 | 0 | 0 | 0 | 12 | 0 | 0 | 0 | 216 | 0 |
| SRR5387931 | 0 | 0 | 43 | 0 | 0 | 17 | 0 | 207 | 207 | 1243 | 192 | 15848 | 0 | 0 | 0 | 0 | 0 | 0 | 0 | 104 | 285 | 1117 | 0 | 0 | 218 | 43 | 204 | 0 | 0 | 0 | 0 | 0 | 0 | 0 | 0 | 518 | 0 |
| SRR5387932 | 0 | 0 | 24 | 0 | 0 | 0 | 0 | 76 | 76 | 765 | 8718 | 15906 | 0 | 0 | 76 | 76 | 0 | 0 | 0 | 76 | 360 | 1457 | 0 | 0 | 377 | 39 | 213 | 0 | 0 | 0 | 0 | 0 | 16 | 0 | 0 | 76 | 0 |
| SRR5387933 | 14 | 0 | 0 | 0 | 0 | 14 | 0 | 229 | 0 | 1068 | 63 | 10909 | 0 | 0 | 0 | 76 | 0 | 0 | 0 | 0 | 243 | 787 | 0 | 0 | 462 | 54 | 203 | 0 | 0 | 0 | 0 | 0 | 0 | 0 | 20 | 229 | 0 |
| SRR5387934 | 0 | 0 | 12 | 0 | 0 | 12 | 0 | 229 | 57 | 860 | 172 | 16687 | 0 | 0 | 115 | 115 | 12 | 0 | 0 | 0 | 467 | 1635 | 0 | 0 | 750 | 29 | 56 | 0 | 0 | 0 | 0 | 0 | 0 | 0 | 0 | 57 | 0 |
| SRR5387935 | 0 | 0 | 21 | 0 | 0 | 0 | 0 | 338 | 113 | 677 | 100 | 11277 | 0 | 226 | 226 | 226 | 0 | 0 | 0 | 338 | 293 | 916 | 0 | 0 | 700 | 54 | 115 | 0 | 0 | 0 | 0 | 0 | 21 | 0 | 21 | 113 | 0 |
| SRR5387936 | 0 | 0 | 17 | 0 | 0 | 0 | 0 | 83 | 0 | 908 | 12716 | 15110 | 0 | 0 | 165 | 248 | 0 | 0 | 0 | 0 | 537 | 2206 | 0 | 0 | 304 | 25 | 66 | 0 | 0 | 0 | 0 | 0 | 0 | 0 | 0 | 413 | 0 |
| SRR5387937 | 21 | 0 | 27 | 0 | 0 | 0 | 0 | 219 | 219 | 947 | 7941 | 16027 | 0 | 0 | 146 | 437 | 0 | 0 | 0 | 291 | 230 | 1093 | 0 | 0 | 293 | 38 | 144 | 0 | 0 | 0 | 0 | 0 | 0 | 0 | 14 | 583 | 0 |
| SRR5387938 | 14 | 0 | 0 | 0 | 0 | 0 | 0 | 86 | 172 | 1547 | 9195 | 16327 | 0 | 0 | 258 | 172 | 0 | 0 | 0 | 344 | 232 | 1009 | 0 | 0 | 288 | 27 | 86 | 0 | 0 | 0 | 0 | 0 | 0 | 14 | 0 | 430 | 0 |
| SRR5387939 | 0 | 0 | 42 | 0 | 0 | 0 | 0 | 379 | 0 | 1212 | 11594 | 15913 | 0 | 0 | 76 | 303 | 0 | 0 | 0 | 76 | 546 | 2086 | 0 | 0 | 347 | 46 | 84 | 0 | 0 | 0 | 0 | 0 | 17 | 17 | 0 | 227 | 0 |
| SRR5387940 | 19 | 0 | 19 | 0 | 0 | 15 | 0 | 575 | 72 | 575 | 4099 | 8845 | 0 | 0 | 216 | 360 | 27 | 0 | 0 | 144 | 600 | 1860 | 0 | 0 | 466 | 27 | 38 | 0 | 0 | 0 | 0 | 0 | 0 | 0 | 0 | 431 | 0 |
| SRR5387941 | 0 | 15 | 29 | 0 | 0 | 0 | 0 | 399 | 67 | 932 | 207 | 15770 | 0 | 67 | 67 | 333 | 0 | 0 | 0 | 67 | 727 | 2949 | 0 | 0 | 68 | 0 | 154 | 0 | 0 | 0 | 0 | 0 | 0 | 15 | 15 | 200 | 0 |
| SRR5387942 | 0 | 0 | 0 | 0 | 0 | 133 | 0 | 371 | 124 | 1113 | 36 | 7547 | 0 | 124 | 371 | 0 | 28 | 0 | 0 | 247 | 605 | 1747 | 0 | 0 | 339 | 18 | 15 | 0 | 0 | 0 | 0 | 0 | 0 | 0 | 0 | 866 | 0 |
| SRR5387943 | 0 | 32 | 64 | 0 | 0 | 0 | 0 | 0 | 0 | 216 | 20261 | 23602 | 0 | 108 | 108 | 431 | 0 | 0 | 0 | 0 | 471 | 2569 | 0 | 0 | 257 | 56 | 16 | 0 | 0 | 16 | 0 | 0 | 16 | 16 | 32 | 0 | 0 |
| SRR5387944 | 0 | 0 | 0 | 0 | 0 | 0 | 0 | 347 | 0 | 782 | 203 | 18242 | 0 | 87 | 87 | 174 | 0 | 0 | 0 | 174 | 480 | 2131 | 0 | 0 | 223 | 21 | 96 | 0 | 0 | 0 | 0 | 0 | 0 | 0 | 0 | 521 | 0 |
| SRR5387945 | 0 | 19 | 29 | 0 | 0 | 0 | 0 | 0 | 0 | 486 | 9718 | 12633 | 0 | 0 | 69 | 139 | 0 | 0 | 0 | 69 | 504 | 1632 | 0 | 0 | 344 | 42 | 48 | 0 | 0 | 0 | 0 | 0 | 0 | 0 | 0 | 69 | 0 |
| SRR5387946 | 0 | 0 | 0 | 0 | 0 | 0 | 0 | 272 | 0 | 1019 | 12232 | 14611 | 0 | 0 | 136 | 476 | 0 | 0 | 0 | 136 | 373 | 2061 | 0 | 0 | 142 | 27 | 90 | 0 | 0 | 0 | 0 | 0 | 0 | 0 | 0 | 476 | 0 |
| SRR5387947 | 11 | 0 | 45 | 0 | 0 | 0 | 0 | 330 | 330 | 989 | 5716 | 11103 | 0 | 0 | 220 | 220 | 11 | 0 | 0 | 220 | 524 | 2312 | 0 | 0 | 151 | 12 | 34 | 0 | 0 | 0 | 0 | 11 | 0 | 34 | 11 | 0 | 0 |
| SRR5387948 | 0 | 0 | 18 | 0 | 0 | 18 | 0 | 568 | 63 | 1452 | 75 | 8209 | 0 | 0 | 126 | 189 | 0 | 0 | 0 | 0 | 415 | 1332 | 0 | 0 | 707 | 29 | 18 | 0 | 0 | 0 | 0 | 0 | 0 | 0 | 0 | 505 | 0 |
| SRR5387949 | 13 | 26 | 26 | 0 | 0 | 13 | 0 | 202 | 0 | 806 | 11492 | 11895 | 0 | 0 | 403 | 302 | 0 | 0 | 0 | 0 | 843 | 2404 | 0 | 0 | 333 | 30 | 258 | 0 | 0 | 0 | 0 | 0 | 13 | 0 | 0 | 302 | 0 |
| SRR5387950 | 0 | 0 | 23 | 0 | 0 | 0 | 0 | 796 | 0 | 1094 | 7560 | 12534 | 0 | 0 | 99 | 298 | 0 | 0 | 0 | 497 | 254 | 1445 | 0 | 0 | 209 | 29 | 77 | 0 | 0 | 0 | 0 | 0 | 0 | 0 | 15 | 199 | 0 |
| SRR5387951 | 0 | 0 | 22 | 0 | 0 | 0 | 0 | 113 | 0 | 1020 | 87 | 11330 | 0 | 0 | 453 | 340 | 0 | 0 | 0 | 340 | 169 | 844 | 0 | 0 | 138 | 22 | 66 | 0 | 11 | 0 | 0 | 0 | 0 | 44 | 0 | 793 | 0 |
| SRR5387952 | 0 | 10 | 0 | 0 | 0 | 20 | 0 | 0 | 168 | 504 | 3445 | 7310 | 0 | 0 | 252 | 252 | 10 | 0 | 0 | 252 | 487 | 2183 | 0 | 0 | 303 | 26 | 152 | 0 | 0 | 0 | 0 | 0 | 0 | 10 | 10 | 420 | 0 |
| SRR5387953 | 0 | 0 | 0 | 0 | 0 | 0 | 0 | 78 | 78 | 702 | 14894 | 17311 | 0 | 78 | 78 | 78 | 0 | 0 | 0 | 390 | 234 | 1004 | 0 | 0 | 164 | 19 | 115 | 0 | 0 | 0 | 0 | 0 | 0 | 0 | 0 | 468 | 0 |
| SRR5387954 | 30 | 0 | 0 | 0 | 0 | 0 | 0 | 376 | 47 | 941 | 8984 | 12840 | 0 | 141 | 235 | 94 | 0 | 0 | 0 | 141 | 237 | 1091 | 0 | 0 | 279 | 34 | 85 | 0 | 0 | 0 | 0 | 0 | 0 | 0 | 0 | 94 | 0 |
| SRR5387955 | 11 | 0 | 44 | 0 | 0 | 11 | 0 | 296 | 99 | 690 | 10443 | 21576 | 0 | 0 | 197 | 394 | 0 | 0 | 0 | 493 | 463 | 1434 | 0 | 0 | 102 | 17 | 55 | 0 | 0 | 0 | 0 | 0 | 0 | 11 | 0 | 296 | 0 |
| SRR5387956 | 0 | 0 | 23 | 0 | 0 | 0 | 0 | 0 | 0 | 1267 | 8536 | 19495 | 0 | 0 | 55 | 330 | 0 | 0 | 0 | 55 | 307 | 1228 | 0 | 0 | 314 | 35 | 38 | 0 | 0 | 0 | 0 | 0 | 0 | 0 | 0 | 330 | 0 |
| SRR5387957 | 0 | 0 | 0 | 0 | 0 | 0 | 0 | 213 | 0 | 1280 | 181 | 15898 | 0 | 53 | 0 | 107 | 0 | 0 | 0 | 213 | 393 | 1501 | 0 | 0 | 398 | 48 | 40 | 0 | 0 | 0 | 0 | 0 | 0 | 16 | 0 | 267 | 0 |
| SRR5387958 | 10 | 0 | 21 | 0 | 0 | 0 | 0 | 96 | 48 | 240 | 12659 | 24887 | 0 | 0 | 144 | 96 | 0 | 0 | 0 | 48 | 92 | 557 | 0 | 0 | 108 | 20 | 16 | 0 | 0 | 0 | 0 | 0 | 0 | 0 | 10 | 671 | 0 |
| SRR5387959 | 25 | 0 | 0 | 0 | 0 | 0 | 0 | 180 | 60 | 1214 | 5414 | 10526 | 0 | 0 | 421 | 241 | 0 | 0 | 0 | 0 | 129 | 425 | 0 | 0 | 240 | 41 | 74 | 0 | 0 | 0 | 0 | 0 | 0 | 12 | 12 | 541 | 0 |
| SRR5387960 | 15 | 0 | 0 | 0 | 0 | 0 | 0 | 105 | 0 | 735 | 147 | 16486 | 0 | 0 | 315 | 525 | 0 | 0 | 0 | 315 | 239 | 1068 | 0 | 0 | 315 | 29 | 30 | 0 | 0 | 0 | 0 | 0 | 0 | 0 | 22 | 945 | 0 |
| SRR5387961 | 27 | 0 | 0 | 0 | 0 | 18 | 0 | 335 | 112 | 1228 | 8598 | 10719 | 0 | 112 | 782 | 893 | 0 | 0 | 0 | 112 | 371 | 1599 | 0 | 0 | 414 | 32 | 281 | 0 | 0 | 0 | 0 | 0 | 0 | 0 | 18 | 223 | 0 |
| SRR5387962 | 31 | 0 | 12 | 0 | 0 | 0 | 0 | 493 | 246 | 1724 | 67 | 7143 | 0 | 82 | 1149 | 575 | 0 | 0 | 0 | 328 | 613 | 2080 | 0 | 0 | 1212 | 53 | 149 | 0 | 0 | 0 | 0 | 0 | 0 | 0 | 0 | 328 | 0 |
| SRR5387963 | 0 | 0 | 17 | 0 | 0 | 0 | 0 | 316 | 189 | 694 | 221 | 22915 | 0 | 63 | 0 | 253 | 0 | 0 | 0 | 189 | 264 | 1238 | 0 | 0 | 331 | 34 | 75 | 0 | 0 | 0 | 0 | 0 | 0 | 0 | 0 | 63 | 0 |
| SRR5387964 | 0 | 0 | 18 | 0 | 0 | 0 | 0 | 0 | 0 | 883 | 165 | 20824 | 0 | 74 | 74 | 294 | 0 | 0 | 0 | 74 | 317 | 1228 | 0 | 0 | 373 | 20 | 102 | 0 | 0 | 0 | 0 | 0 | 0 | 0 | 0 | 147 | 0 |
| SRR5387965 | 13 | 0 | 20 | 0 | 0 | 13 | 0 | 375 | 75 | 1126 | 213 | 14866 | 0 | 0 | 150 | 0 | 0 | 0 | 0 | 0 | 229 | 908 | 0 | 0 | 215 | 23 | 119 | 0 | 0 | 0 | 0 | 0 | 0 | 0 | 0 | 225 | 0 |
| SRR5387966 | 39 | 0 | 0 | 0 | 0 | 0 | 0 | 170 | 0 | 1361 | 120 | 9610 | 0 | 0 | 255 | 255 | 0 | 0 | 0 | 85 | 446 | 1800 | 0 | 0 | 421 | 22 | 378 | 0 | 0 | 0 | 0 | 0 | 0 | 0 | 0 | 510 | 0 |
| SRR5387967 | 14 | 0 | 22 | 0 | 0 | 0 | 0 | 115 | 0 | 516 | 182 | 13521 | 0 | 0 | 344 | 401 | 0 | 0 | 0 | 115 | 454 | 2741 | 0 | 17 | 193 | 34 | 122 | 0 | 0 | 0 | 0 | 0 | 0 | 0 | 0 | 172 | 0 |
| SRR5387968 | 0 | 0 | 31 | 0 | 0 | 0 | 0 | 192 | 0 | 480 | 5851 | 9304 | 0 | 0 | 192 | 288 | 0 | 0 | 0 | 96 | 373 | 1363 | 0 | 18 | 311 | 21 | 117 | 0 | 0 | 0 | 0 | 0 | 0 | 0 | 0 | 96 | 0 |
| SRR5387969 | 0 | 0 | 16 | 0 | 0 | 0 | 0 | 259 | 52 | 1347 | 10257 | 19841 | 0 | 104 | 570 | 518 | 0 | 0 | 0 | 104 | 341 | 1321 | 0 | 0 | 259 | 34 | 56 | 0 | 0 | 0 | 0 | 0 | 0 | 0 | 16 | 207 | 0 |
| SRR5387970 | 0 | 0 | 26 | 0 | 0 | 0 | 0 | 161 | 0 | 1130 | 137 | 18167 | 0 | 0 | 323 | 242 | 17 | 0 | 0 | 242 | 532 | 872 | 0 | 0 | 712 | 25 | 51 | 0 | 0 | 0 | 0 | 0 | 0 | 0 | 0 | 161 | 0 |
| SRR5387971 | 21 | 10 | 0 | 0 | 0 | 10 | 0 | 59 | 0 | 413 | 12986 | 13931 | 0 | 0 | 236 | 0 | 0 | 0 | 0 | 59 | 757 | 10 | 0 | 14 | 282 | 33 | 463 | 0 | 0 | 0 | 0 | 10 | 0 | 10 | 0 | 177 | 0 |
| SRR5387972 | 23 | 0 | 15 | 0 | 0 | 0 | 0 | 194 | 0 | 1549 | 161 | 15687 | 0 | 0 | 97 | 194 | 0 | 0 | 0 | 97 | 269 | 1263 | 0 | 13 | 303 | 28 | 115 | 0 | 0 | 0 | 0 | 0 | 0 | 0 | 0 | 194 | 0 |
| SRR5387973 | 10 | 0 | 21 | 0 | 0 | 21 | 0 | 235 | 118 | 588 | 11987 | 17158 | 0 | 0 | 235 | 235 | 0 | 0 | 0 | 235 | 382 | 810 | 0 | 0 | 316 | 19 | 164 | 0 | 0 | 0 | 10 | 10 | 0 | 21 | 10 | 235 | 0 |
| SRR5387974 | 20 | 20 | 30 | 0 | 0 | 20 | 0 | 488 | 244 | 1463 | 4756 | 8171 | 0 | 122 | 488 | 366 | 0 | 0 | 0 | 122 | 423 | 988 | 0 | 0 | 473 | 28 | 60 | 0 | 0 | 0 | 0 | 0 | 20 | 0 | 30 | 732 | 0 |
| SRR5387975 | 0 | 18 | 18 | 0 | 0 | 0 | 0 | 477 | 318 | 1112 | 4608 | 7151 | 0 | 0 | 318 | 0 | 0 | 0 | 0 | 159 | 333 | 852 | 0 | 0 | 137 | 27 | 109 | 0 | 0 | 0 | 0 | 0 | 0 | 55 | 0 | 159 | 0 |
| SRR5387976 | 0 | 0 | 23 | 0 | 0 | 0 | 0 | 81 | 0 | 1543 | 62 | 7469 | 0 | 0 | 325 | 244 | 0 | 0 | 0 | 731 | 191 | 1046 | 0 | 0 | 247 | 56 | 17 | 0 | 0 | 0 | 0 | 0 | 0 | 12 | 0 | 1218 | 0 |
| SRR5387977 | 0 | 0 | 0 | 0 | 0 | 0 | 0 | 150 | 0 | 751 | 14862 | 17564 | 0 | 0 | 225 | 225 | 0 | 0 | 0 | 75 | 595 | 2099 | 0 | 0 | 494 | 35 | 67 | 0 | 0 | 0 | 0 | 0 | 0 | 0 | 19 | 600 | 0 |
| SRR5387978 | 0 | 0 | 19 | 0 | 0 | 0 | 0 | 460 | 0 | 1380 | 9279 | 17485 | 0 | 0 | 77 | 460 | 0 | 0 | 0 | 0 | 989 | 16 | 0 | 0 | 294 | 38 | 57 | 0 | 0 | 0 | 0 | 0 | 0 | 0 | 0 | 307 | 0 |
| SRR5387979 | 0 | 0 | 24 | 0 | 0 | 12 | 0 | 108 | 0 | 864 | 86 | 9395 | 0 | 0 | 108 | 0 | 0 | 0 | 0 | 432 | 605 | 1982 | 0 | 0 | 184 | 15 | 147 | 0 | 0 | 0 | 0 | 0 | 0 | 12 | 0 | 0 | 0 |
| SRR5387980 | 0 | 0 | 0 | 0 | 0 | 0 | 0 | 227 | 76 | 529 | 8308 | 21677 | 0 | 0 | 0 | 378 | 0 | 0 | 0 | 227 | 353 | 2798 | 0 | 0 | 141 | 34 | 54 | 0 | 0 | 0 | 0 | 0 | 0 | 27 | 0 | 302 | 0 |
| SRR5387981 | 0 | 14 | 14 | 0 | 0 | 0 | 0 | 404 | 404 | 674 | 184 | 16842 | 0 | 135 | 0 | 269 | 0 | 0 | 0 | 135 | 281 | 1149 | 0 | 0 | 122 | 27 | 83 | 0 | 0 | 0 | 0 | 0 | 14 | 0 | 0 | 539 | 0 |
| SRR5387982 | 0 | 0 | 11 | 0 | 0 | 11 | 0 | 342 | 114 | 1710 | 91 | 15507 | 0 | 0 | 0 | 114 | 0 | 0 | 0 | 114 | 352 | 1531 | 0 | 0 | 401 | 24 | 79 | 0 | 0 | 0 | 0 | 0 | 0 | 0 | 0 | 114 | 0 |
| SRR5387983 | 0 | 0 | 24 | 0 | 0 | 39 | 0 | 532 | 0 | 532 | 152 | 16431 | 0 | 0 | 473 | 532 | 0 | 0 | 0 | 0 | 927 | 2545 | 0 | 0 | 241 | 20 | 474 | 0 | 0 | 0 | 0 | 0 | 0 | 24 | 16 | 118 | 0 |
| SRR5387984 | 0 | 0 | 22 | 0 | 0 | 11 | 0 | 162 | 54 | 703 | 10281 | 15314 | 0 | 0 | 325 | 325 | 0 | 0 | 0 | 0 | 586 | 2369 | 0 | 0 | 320 | 26 | 332 | 0 | 0 | 0 | 0 | 0 | 0 | 0 | 0 | 216 | 0 |
| SRR5387985 | 0 | 0 | 13 | 0 | 0 | 0 | 0 | 223 | 49 | 569 | 5862 | 15287 | 0 | 49 | 173 | 49 | 13 | 0 | 0 | 49 | 621 | 1066 | 0 | 0 | 769 | 34 | 31 | 0 | 0 | 0 | 0 | 0 | 0 | 13 | 13 | 99 | 0 |
| SRR5387986 | 0 | 0 | 20 | 0 | 0 | 0 | 0 | 144 | 202 | 1038 | 177 | 13322 | 0 | 0 | 115 | 115 | 0 | 0 | 0 | 58 | 334 | 1169 | 0 | 0 | 759 | 40 | 15 | 0 | 0 | 0 | 0 | 0 | 0 | 15 | 0 | 58 | 0 |
| SRR5387987 | 0 | 0 | 111 | 0 | 0 | 0 | 0 | 103 | 207 | 827 | 162 | 25859 | 0 | 0 | 310 | 310 | 0 | 0 | 0 | 103 | 501 | 1898 | 0 | 0 | 229 | 25 | 50 | 0 | 0 | 0 | 0 | 0 | 0 | 20 | 20 | 207 | 0 |
| SRR5387988 | 0 | 0 | 26 | 0 | 0 | 0 | 0 | 0 | 87 | 871 | 14197 | 33447 | 0 | 0 | 174 | 174 | 0 | 0 | 0 | 0 | 260 | 1435 | 0 | 0 | 136 | 27 | 26 | 0 | 0 | 0 | 0 | 0 | 0 | 0 | 0 | 610 | 0 |
| SRR5387989 | 11 | 0 | 83 | 0 | 0 | 0 | 0 | 162 | 81 | 405 | 111 | 12238 | 0 | 0 | 486 | 324 | 0 | 0 | 0 | 81 | 156 | 931 | 0 | 0 | 186 | 30 | 78 | 0 | 0 | 0 | 0 | 0 | 0 | 22 | 17 | 567 | 0 |
| SRR5387990 | 0 | 0 | 23 | 0 | 0 | 38 | 0 | 362 | 121 | 604 | 8933 | 11227 | 0 | 121 | 0 | 0 | 0 | 0 | 0 | 241 | 334 | 2041 | 0 | 0 | 137 | 18 | 121 | 0 | 0 | 0 | 0 | 0 | 0 | 0 | 0 | 604 | 0 |

### Supp Table S8.4.Columns_DI_to_ES

| | | | | | | | | | | | | | | | | | | | | | | | | | | | | | | | | | | | | | |
|---|---|---|---|---|---|---|---|---|---|---|---|---|---|---|---|---|---|---|---|---|---|---|---|---|---|---|---|---|---|---|---|---|---|---|---|---|---|
| SRR5387991 | 0 | 0 | 31 | 0 | 0 | 15 | 0 | 113 | 113 | 1639 | 110 | 11246 | 0 | 0 | 113 | 396 | 0 | 0 | 0 | 57 | 783 | 2169 | 0 | 15 | 574 | 33 | 100 | 0 | 0 | 0 | 0 | 0 | 0 | 23 | 0 | 57 | 0 |
| SRR5387992 | 35 | 0 | 35 | 0 | 0 | 17 | 0 | 327 | 0 | 654 | 8254 | 18879 | 0 | 0 | 82 | 82 | 0 | 0 | 0 | 0 | 519 | 1811 | 0 | 21 | 518 | 19 | 105 | 0 | 0 | 0 | 0 | 0 | 0 | 0 | 0 | 0 | 0 |
| SRR5387993 | 0 | 0 | 34 | 0 | 0 | 0 | 0 | 355 | 89 | 444 | 7898 | 22185 | 0 | 0 | 89 | 177 | 0 | 0 | 0 | 89 | 529 | 2516 | 0 | 10 | 255 | 27 | 275 | 0 | 0 | 0 | 0 | 0 | 26 | 17 | 34 | 621 | 0 |
| SRR5387994 | 20 | 0 | 34 | 0 | 0 | 0 | 0 | 187 | 62 | 933 | 7343 | 21469 | 0 | 0 | 187 | 560 | 0 | 0 | 0 | 0 | 479 | 1978 | 0 | 11 | 443 | 29 | 69 | 0 | 0 | 0 | 0 | 0 | 0 | 0 | 0 | 187 | 0 |
| SRR5387995 | 0 | 0 | 26 | 0 | 0 | 0 | 0 | 141 | 0 | 1054 | 5553 | 14058 | 0 | 0 | 422 | 351 | 0 | 0 | 0 | 141 | 279 | 1956 | 0 | 0 | 288 | 28 | 113 | 0 | 0 | 0 | 0 | 0 | 0 | 0 | 0 | 70 | 0 |
| SRR5387996 | 0 | 0 | 42 | 0 | 0 | 0 | 0 | 510 | 113 | 851 | 9299 | 13495 | 0 | 0 | 170 | 113 | 0 | 0 | 0 | 340 | 494 | 1890 | 0 | 0 | 360 | 16 | 49 | 0 | 0 | 0 | 0 | 0 | 0 | 14 | 0 | 340 | 0 |
| SRR5387997 | 0 | 0 | 15 | 0 | 0 | 22 | 0 | 328 | 328 | 1771 | 7281 | 9839 | 0 | 66 | 590 | 197 | 0 | 0 | 0 | 131 | 649 | 1746 | 0 | 0 | 426 | 36 | 59 | 0 | 0 | 0 | 0 | 0 | 0 | 0 | 0 | 131 | 0 |
| SRR5387998 | 0 | 0 | 11 | 0 | 0 | 0 | 0 | 210 | 140 | 701 | 174 | 16892 | 0 | 210 | 210 | 140 | 0 | 0 | 0 | 210 | 456 | 1447 | 0 | 0 | 577 | 27 | 28 | 0 | 0 | 0 | 0 | 0 | 0 | 11 | 0 | 421 | 0 |
| SRR5387999 | 0 | 0 | 0 | 0 | 0 | 0 | 0 | 250 | 0 | 1001 | 7590 | 10175 | 0 | 167 | 0 | 167 | 0 | 0 | 0 | 250 | 47 | 198 | 0 | 0 | 93 | 22 | 47 | 0 | 0 | 0 | 0 | 0 | 0 | 12 | 0 | 917 | 0 |
| SRR5388000 | 0 | 0 | 17 | 0 | 0 | 0 | 0 | 722 | 0 | 963 | 9268 | 11435 | 0 | 120 | 120 | 120 | 0 | 0 | 0 | 361 | 270 | 918 | 0 | 0 | 154 | 20 | 34 | 0 | 0 | 0 | 0 | 0 | 0 | 0 | 0 | 1083 | 0 |
| SRR5388001 | 45 | 0 | 0 | 0 | 0 | 0 | 0 | 475 | 119 | 1187 | 5462 | 6412 | 0 | 0 | 712 | 712 | 0 | 0 | 0 | 0 | 657 | 2212 | 0 | 0 | 334 | 34 | 67 | 0 | 0 | 0 | 0 | 0 | 0 | 0 | 0 | 712 | 0 |
| SRR5388002 | 0 | 0 | 0 | 0 | 0 | 14 | 0 | 849 | 283 | 1887 | 110 | 10188 | 0 | 0 | 472 | 849 | 0 | 0 | 0 | 0 | 281 | 1294 | 0 | 0 | 624 | 45 | 56 | 0 | 0 | 0 | 0 | 0 | 0 | 0 | 0 | 0 | 0 |
| SRR5388003 | 0 | 0 | 39 | 0 | 0 | 26 | 0 | 367 | 183 | 1466 | 3299 | 11180 | 0 | 0 | 1466 | 916 | 0 | 0 | 0 | 0 | 532 | 2510 | 0 | 0 | 77 | 18 | 92 | 0 | 0 | 0 | 0 | 13 | 13 | 0 | 26 | 916 | 0 |
| SRR5388004 | 0 | 13 | 0 | 0 | 0 | 26 | 0 | 294 | 98 | 392 | 16841 | 18408 | 0 | 98 | 587 | 196 | 0 | 0 | 0 | 0 | 283 | 1084 | 0 | 0 | 389 | 59 | 45 | 0 | 0 | 0 | 0 | 0 | 13 | 0 | 0 | 98 | 0 |
| SRR5388005 | 14 | 0 | 0 | 0 | 0 | 0 | 0 | 293 | 49 | 976 | 6049 | 13123 | 0 | 49 | 293 | 244 | 0 | 0 | 0 | 146 | 172 | 670 | 0 | 0 | 270 | 42 | 83 | 0 | 0 | 0 | 0 | 0 | 0 | 0 | 0 | 439 | 0 |
| SRR5388006 | 0 | 0 | 24 | 0 | 0 | 0 | 0 | 337 | 67 | 1011 | 92 | 11658 | 0 | 0 | 202 | 67 | 0 | 0 | 0 | 0 | 348 | 949 | 0 | 0 | 710 | 45 | 98 | 0 | 0 | 0 | 0 | 0 | 0 | 0 | 0 | 270 | 0 |
| SRR5388007 | 62 | 0 | 0 | 0 | 0 | 31 | 0 | 297 | 297 | 2131 | 5749 | 12837 | 0 | 50 | 0 | 99 | 0 | 0 | 0 | 297 | 245 | 729 | 0 | 0 | 436 | 45 | 142 | 0 | 0 | 0 | 0 | 0 | 0 | 0 | 0 | 496 | 0 |
| SRR5388008 | 24 | 0 | 10 | 0 | 0 | 10 | 0 | 82 | 122 | 1673 | 156 | 14528 | 0 | 41 | 163 | 41 | 0 | 0 | 0 | 163 | 192 | 1173 | 0 | 0 | 459 | 39 | 165 | 0 | 0 | 0 | 0 | 0 | 0 | 0 | 0 | 816 | 0 |
| SRR5388009 | 0 | 0 | 10 | 0 | 0 | 0 | 0 | 414 | 0 | 621 | 63 | 14802 | 0 | 0 | 104 | 518 | 0 | 0 | 0 | 207 | 543 | 16 | 0 | 0 | 375 | 26 | 339 | 0 | 0 | 0 | 0 | 0 | 21 | 10 | 10 | 104 | 0 |
| SRR5388010 | 0 | 0 | 23 | 0 | 0 | 0 | 0 | 222 | 0 | 997 | 3657 | 7756 | 0 | 111 | 554 | 443 | 0 | 0 | 0 | 0 | 393 | 2089 | 0 | 0 | 534 | 29 | 166 | 0 | 0 | 0 | 0 | 0 | 0 | 15 | 15 | 554 | 0 |
| SRR5388011 | 23 | 0 | 11 | 0 | 0 | 23 | 0 | 328 | 82 | 820 | 13864 | 18130 | 0 | 0 | 1066 | 246 | 0 | 0 | 0 | 0 | 316 | 1127 | 0 | 11 | 403 | 31 | 124 | 0 | 0 | 0 | 0 | 23 | 11 | 34 | 11 | 82 | 0 |
| SRR5388012 | 21 | 10 | 10 | 0 | 0 | 10 | 0 | 432 | 72 | 1512 | 102 | 9213 | 0 | 0 | 504 | 936 | 0 | 0 | 0 | 0 | 425 | 1374 | 0 | 15 | 532 | 47 | 199 | 0 | 0 | 0 | 0 | 0 | 0 | 0 | 10 | 216 | 0 |
| SRR5388013 | 0 | 0 | 14 | 0 | 0 | 0 | 0 | 113 | 38 | 719 | 11802 | 14942 | 0 | 0 | 265 | 227 | 0 | 0 | 0 | 76 | 589 | 1951 | 0 | 0 | 1065 | 37 | 92 | 0 | 0 | 0 | 0 | 0 | 0 | 0 | 0 | 265 | 0 |
| SRR5388014 | 0 | 0 | 0 | 0 | 0 | 0 | 0 | 52 | 52 | 940 | 95 | 10446 | 0 | 0 | 470 | 418 | 0 | 0 | 0 | 104 | 671 | 2319 | 0 | 0 | 969 | 63 | 94 | 0 | 0 | 0 | 0 | 0 | 0 | 0 | 23 | 575 | 0 |
| SRR5388015 | 35 | 0 | 43 | 0 | 0 | 0 | 0 | 203 | 68 | 608 | 24389 | 24591 | 0 | 0 | 68 | 68 | 0 | 0 | 0 | 135 | 259 | 1130 | 0 | 0 | 299 | 23 | 157 | 0 | 0 | 0 | 0 | 0 | 0 | 0 | 0 | 0 | 0 |
| SRR5388016 | 18 | 0 | 23 | 0 | 0 | 18 | 0 | 93 | 140 | 979 | 4660 | 14679 | 0 | 0 | 559 | 746 | 0 | 0 | 0 | 93 | 532 | 2390 | 0 | 0 | 1391 | 55 | 123 | 0 | 0 | 0 | 0 | 0 | 0 | 23 | 0 | 513 | 0 |
| SRR5388017 | 0 | 19 | 19 | 0 | 0 | 19 | 0 | 236 | 0 | 118 | 290 | 19236 | 0 | 0 | 0 | 236 | 0 | 0 | 0 | 59 | 353 | 2252 | 0 | 0 | 163 | 26 | 201 | 0 | 0 | 0 | 0 | 0 | 0 | 0 | 0 | 236 | 0 |
| SRR5388018 | 0 | 0 | 0 | 0 | 0 | 37 | 0 | 511 | 0 | 937 | 187 | 20104 | 0 | 0 | 170 | 0 | 0 | 0 | 0 | 256 | 463 | 10 | 0 | 0 | 287 | 21 | 125 | 0 | 0 | 0 | 0 | 0 | 0 | 0 | 0 | 170 | 0 |
| SRR5388019 | 11 | 0 | 56 | 0 | 0 | 11 | 0 | 90 | 180 | 270 | 18647 | 11531 | 0 | 0 | 90 | 180 | 0 | 0 | 0 | 0 | 530 | 2028 | 0 | 0 | 335 | 44 | 146 | 0 | 0 | 0 | 0 | 0 | 11 | 11 | 0 | 270 | 0 |
| SRR5388020 | 0 | 0 | 28 | 0 | 0 | 0 | 0 | 432 | 216 | 432 | 7780 | 14371 | 0 | 0 | 216 | 324 | 0 | 0 | 0 | 432 | 558 | 2291 | 0 | 0 | 233 | 21 | 121 | 0 | 0 | 0 | 0 | 0 | 0 | 28 | 0 | 108 | 0 |
| SRR5388021 | 0 | 17 | 50 | 0 | 0 | 17 | 0 | 188 | 0 | 1191 | 145 | 14419 | 0 | 125 | 0 | 125 | 0 | 0 | 0 | 0 | 428 | 1813 | 0 | 0 | 273 | 64 | 125 | 0 | 0 | 0 | 0 | 0 | 0 | 0 | 0 | 125 | 0 |
| SRR5388022 | 0 | 0 | 25 | 0 | 0 | 0 | 0 | 198 | 66 | 1586 | 4032 | 8328 | 0 | 198 | 264 | 66 | 0 | 0 | 0 | 198 | 339 | 1395 | 0 | 0 | 575 | 31 | 69 | 0 | 0 | 0 | 0 | 0 | 0 | 25 | 13 | 330 | 0 |
| SRR5388023 | 42 | 0 | 31 | 0 | 0 | 0 | 0 | 771 | 386 | 1832 | 4821 | 7906 | 10 | 96 | 482 | 675 | 0 | 0 | 0 | 193 | 391 | 1492 | 0 | 0 | 460 | 44 | 84 | 0 | 0 | 0 | 0 | 21 | 0 | 10 | 0 | 193 | 0 |
| SRR5388024 | 0 | 0 | 0 | 0 | 0 | 0 | 0 | 234 | 0 | 1053 | 10994 | 18012 | 0 | 0 | 468 | 234 | 0 | 0 | 0 | 936 | 297 | 1137 | 0 | 0 | 350 | 22 | 41 | 0 | 0 | 0 | 0 | 0 | 0 | 16 | 16 | 702 | 0 |
| SRR5388025 | 0 | 0 | 49 | 0 | 0 | 0 | 0 | 0 | 0 | 473 | 195 | 17798 | 0 | 95 | 473 | 189 | 0 | 0 | 0 | 0 | 392 | 2067 | 0 | 0 | 296 | 40 | 16 | 0 | 0 | 16 | 0 | 0 | 0 | 0 | 0 | 0 | 0 |
| SRR5388026 | 31 | 0 | 61 | 0 | 0 | 0 | 0 | 98 | 0 | 787 | 8366 | 14370 | 0 | 0 | 98 | 492 | 15 | 0 | 0 | 295 | 499 | 12 | 0 | 0 | 348 | 41 | 31 | 0 | 0 | 0 | 0 | 0 | 0 | 15 | 0 | 295 | 0 |
| SRR5388027 | 0 | 0 | 12 | 0 | 0 | 0 | 0 | 286 | 0 | 10675 | 9436 | 10008 | 0 | 191 | 477 | 286 | 12 | 0 | 0 | 191 | 179 | 715 | 0 | 0 | 245 | 22 | 24 | 0 | 0 | 0 | 0 | 0 | 0 | 0 | 0 | 381 | 0 |
| SRR5388028 | 0 | 0 | 0 | 0 | 0 | 0 | 0 | 333 | 0 | 22768 | 11217 | 10107 | 0 | 111 | 444 | 666 | 0 | 0 | 0 | 333 | 266 | 1184 | 0 | 0 | 564 | 38 | 0 | 0 | 0 | 0 | 0 | 0 | 19 | 0 | 0 | 444 | 0 |
| SRR5388029 | 0 | 0 | 0 | 0 | 0 | 25 | 0 | 195 | 0 | 585 | 219 | 13918 | 0 | 65 | 195 | 325 | 0 | 0 | 0 | 0 | 209 | 1087 | 0 | 0 | 301 | 27 | 33 | 0 | 0 | 0 | 0 | 0 | 0 | 17 | 0 | 130 | 0 |
| SRR5388030 | 0 | 17 | 0 | 0 | 0 | 0 | 0 | 242 | 0 | 805 | 111 | 12483 | 0 | 81 | 403 | 483 | 0 | 0 | 0 | 161 | 136 | 710 | 0 | 0 | 246 | 28 | 17 | 0 | 0 | 0 | 0 | 0 | 0 | 0 | 0 | 81 | 0 |
| SRR5388031 | 0 | 0 | 0 | 0 | 0 | 0 | 0 | 322 | 59 | 614 | 297 | 11290 | 0 | 59 | 88 | 117 | 0 | 0 | 0 | 59 | 199 | 1372 | 0 | 0 | 238 | 40 | 112 | 0 | 0 | 0 | 0 | 0 | 0 | 0 | 0 | 234 | 0 |
| SRR5388032 | 17 | 0 | 17 | 0 | 0 | 66 | 0 | 188 | 0 | 1125 | 80 | 15565 | 0 | 0 | 94 | 0 | 17 | 0 | 0 | 0 | 649 | 2635 | 0 | 0 | 608 | 28 | 116 | 0 | 0 | 0 | 0 | 0 | 0 | 0 | 0 | 281 | 0 |
| SRR5388033 | 20 | 0 | 29 | 0 | 0 | 0 | 0 | 129 | 129 | 645 | 156 | 13729 | 0 | 0 | 193 | 129 | 0 | 0 | 0 | 129 | 431 | 2004 | 0 | 0 | 150 | 28 | 0 | 0 | 0 | 0 | 0 | 0 | 0 | 0 | 0 | 129 | 0 |
| SRR5388034 | 0 | 0 | 42 | 0 | 0 | 0 | 0 | 307 | 384 | 922 | 11983 | 21046 | 0 | 0 | 77 | 154 | 0 | 0 | 0 | 154 | 441 | 2506 | 0 | 0 | 567 | 47 | 42 | 0 | 0 | 0 | 0 | 0 | 0 | 17 | 0 | 0 | 0 |
| SRR5388035 | 0 | 0 | 59 | 0 | 0 | 0 | 0 | 92 | 184 | 829 | 17786 | 15114 | 0 | 0 | 184 | 184 | 0 | 0 | 0 | 276 | 351 | 1213 | 0 | 0 | 335 | 42 | 201 | 0 | 0 | 0 | 0 | 0 | 0 | 59 | 12 | 184 | 0 |
| SRR5388036 | 0 | 19 | 48 | 0 | 0 | 0 | 0 | 170 | 85 | 680 | 74 | 11212 | 0 | 0 | 255 | 85 | 0 | 0 | 0 | 170 | 451 | 1409 | 0 | 0 | 406 | 28 | 154 | 0 | 0 | 0 | 0 | 0 | 19 | 0 | 0 | 255 | 0 |
| SRR5388037 | 16 | 0 | 0 | 0 | 0 | 0 | 0 | 163 | 327 | 531 | 170 | 9847 | 0 | 0 | 776 | 531 | 0 | 0 | 0 | 0 | 265 | 1015 | 0 | 0 | 161 | 44 | 165 | 0 | 0 | 0 | 0 | 0 | 16 | 25 | 0 | 327 | 0 |
| SRR5388038 | 0 | 11 | 33 | 0 | 0 | 0 | 0 | 193 | 64 | 322 | 8571 | 12051 | 0 | 0 | 516 | 322 | 0 | 0 | 0 | 64 | 523 | 1434 | 0 | 0 | 472 | 45 | 142 | 0 | 0 | 0 | 0 | 0 | 0 | 0 | 0 | 193 | 0 |
| SRR5388039 | 0 | 0 | 0 | 0 | 0 | 0 | 0 | 349 | 0 | 1395 | 4359 | 9851 | 0 | 0 | 0 | 872 | 0 | 0 | 0 | 349 | 543 | 2510 | 0 | 0 | 116 | 19 | 207 | 0 | 0 | 0 | 0 | 0 | 0 | 14 | 28 | 436 | 0 |
| SRR5388040 | 0 | 0 | 22 | 0 | 0 | 15 | 0 | 419 | 0 | 1886 | 58 | 9638 | 0 | 140 | 279 | 559 | 0 | 0 | 0 | 70 | 340 | 1290 | 0 | 0 | 462 | 32 | 75 | 0 | 0 | 0 | 0 | 0 | 15 | 0 | 0 | 210 | 0 |
| SRR5388041 | 0 | 18 | 27 | 0 | 0 | 0 | 0 | 96 | 96 | 861 | 74 | 14640 | 0 | 0 | 191 | 191 | 0 | 0 | 0 | 0 | 541 | 1507 | 0 | 0 | 189 | 25 | 118 | 0 | 0 | 0 | 0 | 0 | 0 | 0 | 0 | 287 | 0 |
| SRR5388042 | 0 | 17 | 0 | 0 | 0 | 17 | 0 | 179 | 269 | 808 | 70 | 13102 | 0 | 0 | 179 | 90 | 0 | 0 | 0 | 90 | 386 | 1790 | 0 | 0 | 176 | 24 | 69 | 0 | 0 | 0 | 0 | 0 | 17 | 0 | 17 | 179 | 0 |
| SRR5388043 | 0 | 0 | 0 | 0 | 0 | 137 | 0 | 290 | 0 | 232 | 188 | 15639 | 0 | 0 | 0 | 58 | 32 | 0 | 0 | 174 | 513 | 1473 | 0 | 0 | 526 | 23 | 18 | 0 | 0 | 0 | 0 | 0 | 0 | 0 | 0 | 174 | 0 |
| SRR5388044 | 0 | 0 | 15 | 0 | 0 | 10 | 0 | 285 | 0 | 1059 | 363 | 20293 | 0 | 0 | 82 | 204 | 0 | 0 | 0 | 163 | 645 | 2657 | 0 | 0 | 755 | 30 | 182 | 0 | 0 | 0 | 0 | 0 | 0 | 0 | 25 | 326 | 0 |
| SRR5388045 | 0 | 0 | 58 | 0 | 0 | 0 | 0 | 401 | 134 | 1335 | 4674 | 8145 | 0 | 0 | 267 | 267 | 0 | 0 | 0 | 801 | 93 | 409 | 0 | 0 | 78 | 21 | 115 | 0 | 0 | 0 | 0 | 0 | 0 | 25 | 0 | 2270 | 0 |
| SRR5388046 | 25 | 0 | 0 | 0 | 0 | 0 | 0 | 300 | 0 | 801 | 5406 | 7809 | 0 | 0 | 0 | 100 | 0 | 0 | 0 | 0 | 402 | 1205 | 0 | 0 | 447 | 19 | 101 | 0 | 0 | 0 | 0 | 0 | 0 | 0 | 0 | 400 | 0 |
| SRR5388047 | 0 | 0 | 0 | 0 | 0 | 0 | 0 | 92 | 92 | 878 | 492 | 17646 | 0 | 0 | 92 | 231 | 0 | 0 | 0 | 185 | 361 | 1305 | 0 | 0 | 736 | 42 | 52 | 0 | 0 | 0 | 0 | 0 | 0 | 0 | 0 | 231 | 0 |
| SRR5388048 | 0 | 0 | 33 | 0 | 0 | 0 | 0 | 173 | 173 | 1443 | 22276 | 17832 | 0 | 0 | 173 | 231 | 0 | 0 | 0 | 58 | 328 | 1237 | 0 | 0 | 715 | 38 | 0 | 0 | 0 | 0 | 0 | 0 | 0 | 0 | 0 | 58 | 0 |
| SRR5388049 | 0 | 0 | 22 | 0 | 0 | 0 | 0 | 139 | 0 | 1252 | 12524 | 17325 | 0 | 0 | 348 | 765 | 0 | 0 | 0 | 70 | 646 | 1811 | 0 | 13 | 523 | 21 | 104 | 0 | 0 | 0 | 0 | 0 | 15 | 0 | 0 | 626 | 0 |
| SRR5388050 | 12 | 0 | 19 | 0 | 0 | 0 | 0 | 130 | 65 | 1172 | 8723 | 14127 | 0 | 65 | 391 | 716 | 0 | 0 | 0 | 0 | 692 | 1976 | 0 | 12 | 669 | 22 | 137 | 0 | 0 | 0 | 0 | 0 | 0 | 0 | 0 | 456 | 0 |
| SRR5388051 | 0 | 0 | 0 | 0 | 0 | 0 | 0 | 595 | 0 | 1289 | 6643 | 11997 | 0 | 0 | 496 | 991 | 0 | 0 | 0 | 297 | 274 | 1466 | 0 | 0 | 668 | 57 | 41 | 0 | 0 | 0 | 0 | 0 | 0 | 0 | 0 | 198 | 0 |
| SRR5388052 | 0 | 0 | 0 | 0 | 0 | 0 | 0 | 432 | 62 | 494 | 106 | 15815 | 0 | 62 | 247 | 432 | 0 | 0 | 0 | 0 | 278 | 1562 | 0 | 0 | 412 | 46 | 34 | 0 | 0 | 0 | 0 | 0 | 11 | 0 | 0 | 247 | 0 |
| SRR5388053 | 49 | 0 | 37 | 0 | 0 | 0 | 0 | 246 | 197 | 690 | 152 | 9653 | 0 | 0 | 148 | 197 | 0 | 0 | 0 | 197 | 327 | 1061 | 0 | 0 | 344 | 38 | 457 | 0 | 0 | 0 | 0 | 0 | 0 | 0 | 19 | 443 | 0 |
| SRR5388054 | 38 | 0 | 15 | 0 | 0 | 0 | 0 | 520 | 0 | 832 | 9252 | 15800 | 0 | 208 | 0 | 728 | 0 | 0 | 0 | 104 | 411 | 1682 | 0 | 0 | 396 | 27 | 241 | 0 | 0 | 0 | 0 | 0 | 0 | 0 | 0 | 832 | 0 |
| SRR5388055 | 19 | 0 | 0 | 0 | 0 | 0 | 0 | 328 | 0 | 2052 | 7223 | 6895 | 0 | 0 | 328 | 246 | 0 | 0 | 0 | 0 | 760 | 2770 | 0 | 0 | 383 | 33 | 76 | 0 | 0 | 0 | 0 | 0 | 19 | 19 | 0 | 328 | 0 |
| SRR5388056 | 0 | 0 | 15 | 0 | 0 | 0 | 0 | 245 | 82 | 818 | 139 | 13327 | 0 | 0 | 491 | 899 | 0 | 0 | 0 | 82 | 641 | 1856 | 0 | 0 | 587 | 34 | 0 | 0 | 0 | 0 | 0 | 0 | 0 | 0 | 0 | 245 | 0 |
| SRR5388057 | 0 | 0 | 0 | 0 | 0 | 0 | 0 | 225 | 112 | 900 | 29 | 6072 | 0 | 0 | 337 | 450 | 0 | 0 | 0 | 112 | 498 | 1721 | 0 | 0 | 362 | 37 | 88 | 0 | 0 | 0 | 0 | 0 | 0 | 18 | 18 | 675 | 0 |
| SRR5388058 | 0 | 0 | 24 | 0 | 0 | 0 | 0 | 443 | 177 | 1684 | 29 | 4873 | 0 | 89 | 709 | 620 | 0 | 0 | 0 | 177 | 283 | 1241 | 0 | 0 | 623 | 59 | 18 | 0 | 0 | 0 | 0 | 0 | 12 | 0 | 0 | 532 | 0 |
| SRR5388059 | 29 | 0 | 14 | 0 | 0 | 14 | 0 | 359 | 90 | 719 | 5571 | 10513 | 0 | 90 | 180 | 90 | 0 | 0 | 0 | 270 | 490 | 1686 | 0 | 0 | 392 | 37 | 187 | 0 | 0 | 0 | 0 | 0 | 0 | 0 | 0 | 270 | 0 |
| SRR5388060 | 0 | 13 | 27 | 0 | 0 | 20 | 0 | 0 | 0 | 1047 | 44 | 12826 | 0 | 0 | 0 | 87 | 0 | 0 | 0 | 175 | 425 | 1984 | 0 | 0 | 229 | 19 | 182 | 0 | 0 | 0 | 0 | 0 | 0 | 0 | 0 | 262 | 0 |
| SRR5388061 | 0 | 0 | 34 | 0 | 0 | 0 | 0 | 250 | 0 | 873 | 56 | 11106 | 0 | 0 | 624 | 873 | 0 | 0 | 0 | 0 | 716 | 2499 | 0 | 0 | 512 | 29 | 114 | 0 | 0 | 11 | 0 | 0 | 23 | 0 | 0 | 374 | 0 |
| SRR5388062 | 0 | 0 | 0 | 0 | 0 | 0 | 0 | 300 | 300 | 1202 | 102 | 13921 | 0 | 100 | 300 | 300 | 0 | 0 | 0 | 300 | 383 | 1414 | 0 | 0 | 656 | 50 | 55 | 0 | 0 | 0 | 0 | 0 | 0 | 31 | 0 | 501 | 0 |
| SRR5388063 | 11 | 11 | 43 | 0 | 0 | 0 | 0 | 140 | 0 | 351 | 179 | 11510 | 0 | 0 | 70 | 140 | 0 | 0 | 0 | 0 | 570 | 2039 | 0 | 0 | 192 | 24 | 64 | 0 | 0 | 0 | 0 | 0 | 11 | 0 | 0 | 0 | 0 |
| SRR5388064 | 0 | 0 | 18 | 0 | 0 | 0 | 0 | 319 | 80 | 239 | 120 | 11566 | 0 | 80 | 239 | 319 | 0 | 0 | 0 | 239 | 575 | 2516 | 0 | 0 | 230 | 27 | 88 | 0 | 0 | 0 | 0 | 0 | 0 | 0 | 0 | 399 | 0 |
| SRR5388065 | 0 | 14 | 42 | 0 | 0 | 0 | 0 | 217 | 72 | 433 | 13147 | 14592 | 0 | 0 | 72 | 0 | 0 | 0 | 0 | 144 | 419 | 1432 | 0 | 0 | 292 | 26 | 69 | 0 | 0 | 0 | 0 | 0 | 0 | 0 | 0 | 144 | 0 |
| SRR5388066 | 0 | 0 | 0 | 0 | 0 | 0 | 0 | 391 | 56 | 1283 | 8650 | 19308 | 0 | 0 | 112 | 56 | 0 | 0 | 0 | 56 | 239 | 1180 | 0 | 0 | 427 | 31 | 0 | 0 | 0 | 0 | 0 | 0 | 0 | 0 | 0 | 56 | 0 |
| SRR5388067 | 0 | 15 | 0 | 0 | 0 | 0 | 0 | 456 | 0 | 1293 | 73 | 8292 | 0 | 0 | 609 | 456 | 15 | 0 | 0 | 304 | 368 | 1601 | 0 | 0 | 626 | 28 | 45 | 0 | 0 | 0 | 0 | 0 | 0 | 0 | 0 | 989 | 0 |
| SRR5388068 | 17 | 0 | 26 | 0 | 0 | 0 | 0 | 358 | 215 | 1147 | 7884 | 15195 | 0 | 0 | 215 | 358 | 0 | 0 | 0 | 72 | 605 | 2938 | 0 | 0 | 810 | 39 | 120 | 0 | 0 | 0 | 0 | 0 | 0 | 0 | 0 | 645 | 0 |
| SRR5388069 | 24 | 0 | 0 | 0 | 0 | 0 | 0 | 139 | 0 | 905 | 214 | 15451 | 0 | 209 | 209 | 209 | 0 | 0 | 0 | 70 | 439 | 1593 | 0 | 0 | 265 | 20 | 158 | 0 | 0 | 0 | 0 | 0 | 0 | 0 | 16 | 139 | 0 |
| SRR5388070 | 16 | 0 | 24 | 0 | 0 | 0 | 0 | 256 | 0 | 852 | 7411 | 15929 | 0 | 0 | 0 | 511 | 0 | 0 | 0 | 511 | 256 | 1396 | 0 | 0 | 208 | 24 | 211 | 0 | 0 | 0 | 0 | 0 | 0 | 32 | 16 | 596 | 0 |
| SRR5388071 | 0 | 0 | 12 | 0 | 0 | 12 | 0 | 271 | 181 | 543 | 71 | 10761 | 0 | 90 | 181 | 181 | 12 | 0 | 0 | 0 | 119 | 526 | 0 | 0 | 151 | 33 | 204 | 0 | 0 | 0 | 0 | 0 | 12 | 0 | 0 | 362 | 0 |
| SRR5388072 | 20 | 0 | 20 | 0 | 0 | 0 | 0 | 413 | 275 | 895 | 140 | 11906 | 0 | 69 | 138 | 69 | 0 | 0 | 0 | 206 | 488 | 1408 | 0 | 0 | 534 | 39 | 247 | 0 | 0 | 0 | 0 | 0 | 0 | 20 | 0 | 275 | 0 |
| SRR5388073 | 0 | 18 | 18 | 0 | 0 | 0 | 0 | 523 | 65 | 523 | 11434 | 13133 | 0 | 65 | 588 | 327 | 0 | 0 | 0 | 196 | 265 | 1148 | 0 | 0 | 226 | 42 | 18 | 0 | 0 | 0 | 0 | 0 | 0 | 0 | 0 | 196 | 0 |
| SRR5388074 | 0 | 0 | 39 | 0 | 0 | 0 | 0 | 304 | 0 | 1366 | 6602 | 10244 | 0 | 152 | 379 | 531 | 0 | 0 | 0 | 76 | 640 | 2115 | 0 | 0 | 821 | 23 | 67 | 0 | 0 | 0 | 0 | 0 | 0 | 0 | 0 | 228 | 0 |
| SRR5388075 | 0 | 0 | 0 | 0 | 0 | 0 | 0 | 104 | 0 | 782 | 8970 | 17315 | 0 | 104 | 522 | 991 | 0 | 0 | 0 | 52 | 335 | 1982 | 0 | 0 | 282 | 39 | 139 | 0 | 0 | 0 | 0 | 0 | 22 | 0 | 0 | 417 | 0 |
| SRR5388076 | 0 | 0 | 25 | 0 | 0 | 0 | 0 | 231 | 0 | 925 | 4473 | 13186 | 0 | 77 | 231 | 308 | 0 | 0 | 0 | 231 | 251 | 1231 | 0 | 0 | 398 | 28 | 68 | 0 | 0 | 0 | 0 | 0 | 0 | 34 | 25 | 231 | 0 |
| SRR5388077 | 0 | 10 | 21 | 0 | 0 | 0 | 0 | 283 | 94 | 754 | 13281 | 11585 | 0 | 0 | 0 | 94 | 10 | 0 | 0 | 0 | 381 | 1027 | 0 | 0 | 241 | 26 | 73 | 0 | 0 | 0 | 0 | 0 | 0 | 0 | 10 | 94 | 0 |
| SRR5388078 | 13 | 0 | 0 | 0 | 0 | 16 | 0 | 917 | 108 | 647 | 4478 | 9334 | 0 | 0 | 108 | 108 | 0 | 0 | 0 | 378 | 517 | 1835 | 0 | 0 | 1010 | 37 | 0 | 0 | 0 | 0 | 0 | 0 | 0 | 0 | 0 | 2482 | 0 |
| SRR5388079 | 0 | 0 | 38 | 0 | 0 | 0 | 0 | 164 | 0 | 573 | 117 | 18088 | 0 | 0 | 327 | 327 | 0 | 0 | 0 | 82 | 500 | 1555 | 0 | 0 | 113 | 24 | 87 | 0 | 0 | 0 | 0 | 0 | 0 | 0 | 0 | 82 | 0 |
| SRR5388080 | 0 | 0 | 17 | 0 | 0 | 17 | 0 | 196 | 0 | 1669 | 7559 | 15805 | 0 | 0 | 196 | 196 | 0 | 0 | 0 | 393 | 328 | 825 | 0 | 0 | 401 | 29 | 60 | 0 | 0 | 0 | 0 | 0 | 0 | 0 | 0 | 491 | 0 |
| SRR5388081 | 0 | 17 | 17 | 0 | 0 | 0 | 0 | 331 | 0 | 663 | 114 | 10998 | 0 | 0 | 66 | 265 | 0 | 0 | 0 | 0 | 773 | 2284 | 0 | 0 | 387 | 25 | 111 | 0 | 0 | 0 | 0 | 0 | 0 | 0 | 0 | 265 | 0 |
| SRR5388082 | 0 | 0 | 0 | 0 | 0 | 0 | 0 | 126 | 0 | 1325 | 7758 | 15326 | 0 | 0 | 63 | 252 | 0 | 0 | 0 | 63 | 343 | 1612 | 0 | 0 | 216 | 15 | 27 | 0 | 0 | 0 | 0 | 0 | 0 | 20 | 0 | 252 | 0 |
| SRR5388083 | 0 | 0 | 0 | 0 | 0 | 0 | 0 | 54 | 163 | 599 | 257 | 18140 | 0 | 54 | 218 | 272 | 15 | 0 | 0 | 54 | 595 | 2114 | 0 | 0 | 588 | 42 | 23 | 0 | 0 | 0 | 0 | 0 | 0 | 15 | 0 | 109 | 0 |
| SRR5388084 | 19 | 0 | 28 | 0 | 0 | 0 | 0 | 146 | 73 | 1164 | 7785 | 12806 | 0 | 73 | 364 | 73 | 0 | 0 | 0 | 0 | 451 | 11 | 0 | 0 | 404 | 29 | 19 | 0 | 0 | 0 | 0 | 0 | 0 | 0 | 0 | 218 | 0 |
| SRR5388085 | 0 | 0 | 21 | 0 | 0 | 21 | 0 | 383 | 110 | 383 | 6901 | 10462 | 0 | 110 | 55 | 55 | 10 | 0 | 0 | 164 | 287 | 888 | 0 | 0 | 214 | 27 | 73 | 0 | 0 | 10 | 0 | 0 | 10 | 10 | 21 | 164 | 0 |
| SRR5388086 | 0 | 0 | 0 | 0 | 0 | 0 | 0 | 217 | 72 | 1012 | 245 | 16124 | 0 | 0 | 0 | 145 | 0 | 0 | 0 | 72 | 625 | 2077 | 0 | 0 | 451 | 17 | 70 | 0 | 0 | 0 | 0 | 0 | 0 | 0 | 0 | 72 | 0 |
| SRR5388087 | 0 | 0 | 23 | 0 | 0 | 0 | 0 | 148 | 118 | 502 | 9126 | 16156 | 0 | 0 | 89 | 30 | 0 | 0 | 0 | 0 | 199 | 596 | 0 | 0 | 388 | 56 | 30 | 0 | 0 | 0 | 0 | 0 | 0 | 23 | 0 | 177 | 0 |
| SRR5388088 | 0 | 0 | 39 | 0 | 0 | 20 | 0 | 135 | 135 | 471 | 130 | 21013 | 0 | 0 | 67 | 202 | 13 | 0 | 0 | 202 | 187 | 876 | 0 | 0 | 276 | 27 | 26 | 0 | 0 | 0 | 0 | 0 | 0 | 0 | 0 | 67 | 0 |
| SRR5388089 | 0 | 23 | 38 | 0 | 0 | 0 | 0 | 190 | 0 | 822 | 181 | 11944 | 0 | 0 | 316 | 190 | 0 | 0 | 0 | 0 | 414 | 1802 | 0 | 0 | 419 | 49 | 215 | 0 | 0 | 0 | 0 | 0 | 0 | 23 | 0 | 379 | 0 |
| SRR5388090 | 12 | 0 | 12 | 0 | 0 | 0 | 0 | 608 | 0 | 1215 | 157 | 12423 | 0 | 0 | 68 | 68 | 0 | 0 | 0 | 203 | 429 | 1958 | 0 | 0 | 707 | 33 | 178 | 0 | 0 | 0 | 0 | 0 | 0 | 0 | 0 | 68 | 0 |
| SRR5388091 | 0 | 11 | 0 | 0 | 0 | 0 | 0 | 209 | 0 | 838 | 104 | 13406 | 0 | 0 | 279 | 628 | 0 | 0 | 0 | 140 | 366 | 1438 | 0 | 0 | 415 | 32 | 102 | 0 | 0 | 0 | 0 | 0 | 0 | 0 | 0 | 70 | 0 |
| SRR5388092 | 0 | 13 | 0 | 0 | 0 | 0 | 0 | 267 | 0 | 1268 | 67 | 13283 | 0 | 0 | 534 | 467 | 0 | 0 | 0 | 67 | 453 | 1802 | 0 | 0 | 375 | 29 | 74 | 0 | 0 | 0 | 0 | 13 | 0 | 0 | 0 | 133 | 0 |
| SRR5388093 | 0 | 0 | 18 | 0 | 0 | 0 | 0 | 410 | 164 | 1968 | 14758 | 11724 | 0 | 0 | 246 | 656 | 0 | 0 | 0 | 492 | 184 | 576 | 0 | 0 | 86 | 16 | 20 | 0 | 0 | 0 | 0 | 0 | 0 | 11 | 0 | 2542 | 0 |
| SRR5388094 | 13 | 0 | 0 | 0 | 0 | 19 | 0 | 447 | 0 | 1660 | 116 | 7278 | 0 | 0 | 319 | 511 | 0 | 0 | 0 | 255 | 590 | 1819 | 0 | 0 | 666 | 31 | 44 | 0 | 0 | 0 | 0 | 0 | 0 | 0 | 0 | 766 | 0 |
| SRR5388095 | 0 | 0 | 0 | 0 | 0 | 0 | 0 | 418 | 0 | 627 | 8366 | 12341 | 0 | 0 | 209 | 314 | 12 | 0 | 0 | 105 | 311 | 975 | 0 | 0 | 207 | 26 | 0 | 0 | 0 | 0 | 0 | 0 | 0 | 12 | 12 | 105 | 0 |
| SRR5388096 | 0 | 0 | 22 | 0 | 0 | 0 | 0 | 222 | 148 | 1187 | 87 | 10976 | 0 | 0 | 297 | 297 | 0 | 0 | 0 | 148 | 292 | 1544 | 0 | 0 | 356 | 32 | 0 | 0 | 0 | 0 | 0 | 0 | 0 | 14 | 14 | 297 | 0 |
| SRR5388097 | 0 | 0 | 19 | 0 | 0 | 0 | 0 | 295 | 0 | 1475 | 7078 | 10224 | 0 | 0 | 0 | 98 | 0 | 0 | 0 | 0 | 498 | 1654 | 0 | 0 | 262 | 29 | 85 | 0 | 0 | 0 | 0 | 0 | 0 | 28 | 19 | 492 | 0 |
| SRR5388098 | 12 | 0 | 35 | 0 | 0 | 0 | 0 | 232 | 116 | 579 | 8576 | 14254 | 0 | 0 | 0 | 0 | 12 | 0 | 0 | 116 | 606 | 2402 | 0 | 0 | 154 | 21 | 117 | 0 | 0 | 0 | 0 | 0 | 0 | 0 | 12 | 811 | 0 |
| SRR5388099 | 0 | 16 | 55 | 0 | 0 | 24 | 0 | 354 | 89 | 1152 | 137 | 17185 | 0 | 0 | 0 | 89 | 0 | 0 | 0 | 177 | 370 | 1185 | 0 | 0 | 227 | 27 | 39 | 0 | 0 | 0 | 0 | 0 | 0 | 0 | 0 | 266 | 0 |
| SRR5388100 | 0 | 0 | 65 | 0 | 0 | 11 | 0 | 447 | 0 | 1901 | 9280 | 14647 | 0 | 112 | 335 | 447 | 0 | 0 | 0 | 112 | 288 | 1158 | 0 | 0 | 437 | 45 | 65 | 0 | 0 | 0 | 0 | 0 | 0 | 11 | 11 | 447 | 0 |
| SRR5388101 | 18 | 0 | 0 | 0 | 0 | 0 | 0 | 150 | 0 | 826 | 11784 | 12385 | 0 | 0 | 375 | 150 | 0 | 0 | 0 | 75 | 348 | 1863 | 0 | 0 | 219 | 20 | 92 | 0 | 0 | 0 | 0 | 0 | 0 | 0 | 0 | 225 | 0 |
| SRR5388102 | 0 | 0 | 0 | 0 | 0 | 0 | 0 | 599 | 75 | 823 | 133 | 14448 | 0 | 0 | 299 | 374 | 0 | 0 | 0 | 0 | 365 | 2083 | 0 | 0 | 159 | 19 | 69 | 0 | 0 | 0 | 0 | 0 | 0 | 0 | 35 | 150 | 0 |
| SRR5388103 | 21 | 0 | 11 | 0 | 0 | 0 | 0 | 157 | 79 | 315 | 178 | 15496 | 0 | 0 | 157 | 157 | 0 | 0 | 0 | 79 | 589 | 1248 | 0 | 0 | 225 | 30 | 64 | 0 | 0 | 0 | 0 | 0 | 0 | 11 | 11 | 157 | 0 |
| SRR5388104 | 0 | 0 | 22 | 0 | 0 | 0 | 0 | 192 | 0 | 1153 | 6535 | 10090 | 0 | 0 | 96 | 288 | 0 | 0 | 0 | 192 | 448 | 1837 | 0 | 0 | 235 | 23 | 59 | 0 | 0 | 0 | 0 | 0 | 0 | 0 | 0 | 480 | 0 |
| SRR5388105 | 0 | 12 | 25 | 0 | 0 | 12 | 0 | 0 | 52 | 518 | 261 | 16634 | 0 | 0 | 0 | 0 | 0 | 0 | 0 | 52 | 375 | 1813 | 0 | 0 | 385 | 45 | 100 | 0 | 0 | 0 | 0 | 0 | 0 | 37 | 25 | 155 | 0 |
| SRR5388106 | 0 | 0 | 0 | 0 | 0 | 0 | 0 | 90 | 0 | 403 | 222 | 21273 | 0 | 0 | 0 | 0 | 0 | 0 | 0 | 134 | 395 | 2091 | 0 | 0 | 338 | 35 | 108 | 0 | 0 | 0 | 0 | 0 | 0 | 0 | 0 | 134 | 0 |
| SRR5388107 | 12 | 0 | 24 | 0 | 0 | 0 | 0 | 0 | 97 | 0 | 161 | 16739 | 0 | 0 | 0 | 194 | 0 | 0 | 0 | 0 | 226 | 785 | 0 | 0 | 168 | 14 | 145 | 0 | 0 | 0 | 0 | 0 | 0 | 12 | 12 | 0 | 0 |
| SRR5388108 | 21 | 0 | 0 | 0 | 0 | 0 | 0 | 0 | 0 | 528 | 5193 | 10386 | 0 | 0 | 176 | 176 | 0 | 0 | 0 | 88 | 381 | 1416 | 0 | 0 | 267 | 16 | 150 | 0 | 0 | 0 | 0 | 0 | 0 | 0 | 0 | 792 | 0 |
| SRR5388109 | 0 | 0 | 11 | 0 | 0 | 0 | 0 | 354 | 59 | 413 | 84 | 12397 | 0 | 0 | 236 | 177 | 0 | 0 | 0 | 472 | 245 | 961 | 0 | 11 | 437 | 45 | 86 | 0 | 0 | 0 | 0 | 0 | 0 | 0 | 0 | 1122 | 0 |
| SRR5388110 | 0 | 0 | 19 | 0 | 0 | 0 | 0 | 420 | 158 | 1523 | 11080 | 15701 | 0 | 0 | 263 | 158 | 0 | 0 | 0 | 263 | 267 | 1068 | 0 | 15 | 350 | 46 | 63 | 0 | 0 | 0 | 0 | 0 | 0 | 0 | 0 | 1155 | 0 |
| SRR5388111 | 25 | 0 | 0 | 0 | 0 | 12 | 0 | 219 | 0 | 951 | 2853 | 6658 | 0 | 0 | 219 | 439 | 0 | 0 | 0 | 219 | 672 | 2228 | 0 | 0 | 1066 | 54 | 161 | 0 | 0 | 0 | 0 | 0 | 0 | 12 | 19 | 366 | 0 |
| SRR5388112 | 11 | 0 | 11 | 0 | 0 | 22 | 0 | 215 | 215 | 503 | 7541 | 14293 | 0 | 0 | 359 | 72 | 0 | 0 | 0 | 144 | 526 | 1838 | 0 | 0 | 684 | 41 | 230 | 0 | 0 | 0 | 0 | 0 | 0 | 0 | 27 | 503 | 0 |
| SRR5388113 | 23 | 0 | 15 | 0 | 0 | 23 | 0 | 81 | 0 | 894 | 87 | 13661 | 0 | 81 | 163 | 0 | 0 | 0 | 0 | 163 | 318 | 1287 | 0 | 0 | 313 | 30 | 186 | 0 | 0 | 0 | 0 | 0 | 0 | 0 | 0 | 244 | 0 |
| SRR5388114 | 40 | 0 | 0 | 0 | 0 | 35 | 0 | 209 | 0 | 1515 | 10345 | 15517 | 0 | 0 | 104 | 261 | 0 | 0 | 0 | 52 | 246 | 1755 | 0 | 0 | 354 | 77 | 230 | 0 | 0 | 0 | 0 | 0 | 0 | 0 | 0 | 261 | 0 |

### Supp Table S8.4.Columns_DI_to_ES

| | | | | | | | | | | | | | | | | | | | | | | | | | | | | | | | | | | | | | |
|---|---|---|---|---|---|---|---|---|---|---|---|---|---|---|---|---|---|---|---|---|---|---|---|---|---|---|---|---|---|---|---|---|---|---|---|---|---|
| SRR5388115 | 16 | 0 | 33 | 0 | 0 | 0 | 0 | 200 | 0 | 1300 | 245 | 17899 | 0 | 0 | 300 | 200 | 0 | 0 | 0 | 100 | 471 | 1923 | 0 | 0 | 980 | 32 | 25 | 0 | 0 | 0 | 0 | 0 | 25 | 0 | 0 | 150 | 0 |
| SRR5388116 | 0 | 0 | 19 | 0 | 0 | 0 | 0 | 243 | 0 | 971 | 74 | 10922 | 0 | 0 | 162 | 485 | 0 | 0 | 0 | 81 | 306 | 1344 | 0 | 0 | 500 | 24 | 67 | 0 | 0 | 0 | 0 | 0 | 0 | 0 | 0 | 324 | 0 |
| SRR5388117 | 12 | 0 | 12 | 0 | 0 | 0 | 0 | 70 | 70 | 525 | 4793 | 16792 | 0 | 0 | 35 | 70 | 0 | 0 | 0 | 35 | 348 | 1447 | 0 | 0 | 335 | 29 | 29 | 0 | 0 | 0 | 0 | 0 | 0 | 0 | 0 | 245 | 0 |
| SRR5388118 | 0 | 0 | 12 | 0 | 0 | 0 | 0 | 181 | 121 | 1266 | 4882 | 16515 | 0 | 0 | 60 | 121 | 0 | 0 | 0 | 181 | 373 | 1641 | 0 | 0 | 578 | 39 | 53 | 0 | 0 | 0 | 0 | 0 | 0 | 0 | 0 | 121 | 0 |
| SRR5388119 | 0 | 0 | 54 | 0 | 0 | 0 | 0 | 435 | 87 | 1826 | 7042 | 12693 | 0 | 0 | 609 | 956 | 0 | 0 | 0 | 87 | 104 | 603 | 0 | 0 | 158 | 41 | 0 | 0 | 0 | 0 | 0 | 0 | 0 | 18 | 18 | 1130 | 0 |
| SRR5388120 | 0 | 0 | 13 | 0 | 0 | 0 | 0 | 278 | 46 | 2499 | 8468 | 12309 | 0 | 93 | 833 | 879 | 0 | 0 | 0 | 1018 | 130 | 421 | 0 | 0 | 85 | 32 | 26 | 0 | 0 | 0 | 0 | 0 | 0 | 13 | 0 | 3702 | 0 |
| SRR5388121 | 0 | 0 | 15 | 0 | 0 | 0 | 0 | 57 | 114 | 515 | 14528 | 19446 | 0 | 0 | 114 | 172 | 0 | 0 | 0 | 114 | 302 | 1376 | 0 | 0 | 38 | 10 | 30 | 0 | 0 | 0 | 0 | 0 | 0 | 0 | 0 | 515 | 0 |
| SRR5388122 | 0 | 0 | 51 | 0 | 0 | 0 | 0 | 0 | 0 | 829 | 185 | 17407 | 0 | 64 | 0 | 64 | 0 | 0 | 0 | 0 | 299 | 1140 | 0 | 11 | 159 | 19 | 25 | 0 | 0 | 0 | 0 | 0 | 0 | 0 | 0 | 383 | 0 |
| SRR5388123 | 0 | 0 | 0 | 0 | 0 | 12 | 0 | 314 | 0 | 419 | 84 | 7379 | 0 | 0 | 366 | 105 | 0 | 0 | 0 | 105 | 563 | 1186 | 0 | 0 | 562 | 62 | 0 | 0 | 0 | 0 | 12 | 0 | 0 | 0 | 12 | 157 | 0 |
| SRR5388124 | 0 | 0 | 0 | 0 | 0 | 0 | 0 | 122 | 61 | 488 | 8236 | 11286 | 0 | 0 | 305 | 305 | 0 | 0 | 0 | 61 | 311 | 1608 | 0 | 0 | 406 | 39 | 0 | 0 | 0 | 0 | 0 | 0 | 0 | 0 | 0 | 549 | 0 |
| SRR5388125 | 16 | 0 | 0 | 0 | 0 | 0 | 0 | 223 | 0 | 816 | 132 | 13136 | 0 | 0 | 223 | 297 | 22 | 0 | 0 | 223 | 304 | 1421 | 0 | 0 | 257 | 16 | 11 | 0 | 0 | 0 | 0 | 0 | 0 | 0 | 0 | 891 | 0 |
| SRR5388126 | 0 | 0 | 12 | 0 | 0 | 0 | 0 | 111 | 0 | 996 | 51 | 8411 | 0 | 0 | 664 | 221 | 0 | 0 | 0 | 111 | 536 | 1783 | 0 | 0 | 414 | 14 | 25 | 0 | 0 | 0 | 0 | 0 | 0 | 0 | 25 | 0 | 0 |
| SRR5388127 | 11 | 11 | 54 | 0 | 0 | 22 | 11 | 475 | 0 | 665 | 77 | 14064 | 0 | 0 | 0 | 0 | 11 | 0 | 0 | 0 | 327 | 1445 | 0 | 0 | 271 | 24 | 258 | 0 | 0 | 0 | 11 | 0 | 11 | 11 | 0 | 285 | 0 |
| SRR5388128 | 0 | 0 | 39 | 0 | 0 | 0 | 0 | 325 | 65 | 1170 | 10790 | 14885 | 0 | 0 | 195 | 195 | 0 | 0 | 0 | 195 | 520 | 1972 | 0 | 0 | 587 | 37 | 140 | 0 | 0 | 0 | 0 | 0 | 0 | 0 | 0 | 390 | 0 |
| SRR5388129 | 0 | 0 | 20 | 0 | 0 | 10 | 0 | 530 | 132 | 1456 | 79 | 10062 | 0 | 0 | 530 | 265 | 0 | 0 | 0 | 132 | 309 | 960 | 0 | 0 | 253 | 36 | 102 | 0 | 0 | 0 | 0 | 0 | 0 | 10 | 0 | 265 | 0 |
| SRR5388130 | 23 | 0 | 23 | 0 | 0 | 0 | 0 | 641 | 92 | 1191 | 7970 | 13558 | 0 | 0 | 183 | 641 | 0 | 0 | 0 | 92 | 304 | 1341 | 0 | 0 | 149 | 44 | 199 | 0 | 0 | 0 | 0 | 0 | 0 | 0 | 0 | 275 | 0 |
| SRR5388131 | 0 | 0 | 43 | 0 | 0 | 0 | 0 | 197 | 0 | 591 | 95 | 12336 | 0 | 0 | 197 | 66 | 0 | 0 | 0 | 66 | 619 | 1044 | 0 | 0 | 370 | 28 | 115 | 0 | 14 | 0 | 0 | 0 | 0 | 0 | 0 | 131 | 0 |
| SRR5388132 | 0 | 0 | 25 | 0 | 0 | 0 | 0 | 228 | 65 | 1044 | 167 | 11358 | 0 | 0 | 65 | 326 | 0 | 0 | 0 | 65 | 589 | 2057 | 0 | 0 | 761 | 30 | 54 | 0 | 0 | 0 | 0 | 0 | 0 | 0 | 0 | 294 | 0 |
| SRR5388133 | 0 | 0 | 40 | 0 | 0 | 0 | 0 | 293 | 42 | 629 | 139 | 12364 | 0 | 42 | 84 | 126 | 0 | 0 | 0 | 0 | 675 | 1714 | 0 | 0 | 368 | 36 | 20 | 0 | 0 | 0 | 0 | 0 | 0 | 0 | 0 | 84 | 0 |
| SRR5388134 | 0 | 0 | 0 | 0 | 0 | 0 | 0 | 65 | 0 | 1505 | 10928 | 16752 | 0 | 65 | 65 | 131 | 0 | 0 | 0 | 131 | 374 | 1617 | 0 | 0 | 427 | 42 | 0 | 0 | 0 | 0 | 0 | 0 | 0 | 0 | 0 | 654 | 0 |
| SRR5388135 | 16 | 0 | 16 | 0 | 0 | 0 | 0 | 212 | 106 | 318 | 80 | 9114 | 0 | 0 | 106 | 212 | 0 | 0 | 0 | 212 | 334 | 1595 | 0 | 0 | 317 | 40 | 47 | 0 | 0 | 0 | 0 | 0 | 16 | 0 | 0 | 636 | 0 |
| SRR5388136 | 0 | 0 | 48 | 0 | 0 | 19 | 0 | 422 | 0 | 1548 | 51 | 8446 | 0 | 0 | 0 | 282 | 0 | 0 | 0 | 422 | 505 | 1763 | 0 | 0 | 240 | 24 | 68 | 0 | 0 | 0 | 0 | 0 | 0 | 0 | 0 | 141 | 0 |
| SRR5388137 | 0 | 0 | 19 | 0 | 0 | 25 | 0 | 0 | 84 | 674 | 8598 | 17449 | 0 | 0 | 84 | 506 | 12 | 0 | 0 | 0 | 595 | 2098 | 0 | 0 | 358 | 23 | 50 | 0 | 0 | 0 | 0 | 0 | 0 | 0 | 12 | 421 | 0 |
| SRR5388138 | 12 | 0 | 0 | 0 | 0 | 0 | 0 | 150 | 75 | 2252 | 6756 | 14488 | 0 | 0 | 150 | 150 | 0 | 0 | 0 | 150 | 228 | 1108 | 0 | 0 | 489 | 50 | 70 | 0 | 0 | 0 | 0 | 0 | 0 | 0 | 0 | 225 | 0 |
| SRR5388139 | 0 | 0 | 37 | 0 | 0 | 0 | 0 | 0 | 91 | 817 | 120 | 18871 | 0 | 0 | 454 | 544 | 0 | 0 | 0 | 0 | 575 | 2302 | 0 | 0 | 184 | 15 | 46 | 0 | 0 | 0 | 0 | 0 | 0 | 18 | 0 | 363 | 0 |
| SRR5388140 | 0 | 0 | 13 | 0 | 0 | 13 | 0 | 504 | 78 | 737 | 21216 | 23505 | 0 | 0 | 78 | 116 | 0 | 0 | 0 | 116 | 730 | 1604 | 0 | 11 | 429 | 25 | 20 | 0 | 0 | 0 | 0 | 0 | 0 | 0 | 0 | 272 | 0 |
| SRR5388141 | 0 | 0 | 32 | 0 | 0 | 11 | 0 | 484 | 0 | 363 | 5203 | 10647 | 0 | 0 | 726 | 363 | 21 | 0 | 0 | 121 | 471 | 1641 | 0 | 0 | 339 | 25 | 75 | 0 | 0 | 0 | 0 | 0 | 11 | 0 | 0 | 363 | 0 |
| SRR5388142 | 37 | 0 | 22 | 0 | 0 | 74 | 0 | 509 | 0 | 917 | 27 | 6826 | 0 | 0 | 306 | 306 | 0 | 0 | 0 | 0 | 444 | 1727 | 0 | 0 | 973 | 44 | 52 | 0 | 0 | 0 | 0 | 0 | 0 | 0 | 0 | 408 | 0 |
| SRR5388143 | 0 | 10 | 31 | 0 | 0 | 0 | 0 | 175 | 0 | 874 | 28756 | 31378 | 0 | 0 | 0 | 175 | 0 | 0 | 0 | 175 | 269 | 1509 | 0 | 0 | 252 | 10 | 41 | 0 | 0 | 0 | 0 | 0 | 20 | 0 | 20 | 175 | 0 |
| SRR5388144 | 0 | 12 | 48 | 0 | 0 | 0 | 0 | 0 | 0 | 1484 | 100 | 15234 | 0 | 0 | 198 | 198 | 12 | 0 | 0 | 99 | 559 | 10 | 0 | 0 | 188 | 18 | 96 | 0 | 0 | 0 | 0 | 0 | 0 | 12 | 12 | 396 | 0 |
| SRR5388145 | 11 | 0 | 0 | 0 | 0 | 11 | 0 | 401 | 80 | 1364 | 10595 | 10675 | 0 | 0 | 80 | 401 | 0 | 0 | 0 | 80 | 190 | 631 | 0 | 0 | 273 | 45 | 98 | 0 | 0 | 0 | 0 | 0 | 22 | 33 | 0 | 321 | 0 |
| SRR5388146 | 0 | 0 | 43 | 0 | 0 | 11 | 0 | 618 | 0 | 1340 | 4638 | 6390 | 0 | 0 | 722 | 412 | 0 | 0 | 0 | 103 | 695 | 1860 | 0 | 0 | 428 | 30 | 203 | 0 | 0 | 0 | 21 | 0 | 0 | 11 | 0 | 206 | 0 |
| SRR5388147 | 25 | 25 | 50 | 0 | 0 | 25 | 0 | 513 | 599 | 2739 | 10013 | 14377 | 0 | 86 | 428 | 513 | 0 | 0 | 0 | 342 | 200 | 866 | 0 | 25 | 596 | 73 | 150 | 0 | 0 | 0 | 0 | 0 | 0 | 17 | 0 | 513 | 0 |
| SRR5388148 | 0 | 0 | 15 | 0 | 0 | 0 | 0 | 402 | 67 | 1810 | 102 | 11396 | 0 | 67 | 134 | 402 | 0 | 0 | 0 | 201 | 344 | 1291 | 0 | 27 | 703 | 51 | 126 | 0 | 0 | 0 | 0 | 0 | 0 | 0 | 0 | 268 | 0 |
| SRR5388149 | 0 | 0 | 35 | 0 | 0 | 0 | 0 | 143 | 72 | 1505 | 11757 | 13908 | 0 | 0 | 215 | 358 | 0 | 0 | 0 | 0 | 506 | 2057 | 0 | 0 | 665 | 45 | 114 | 0 | 0 | 0 | 0 | 0 | 0 | 0 | 0 | 215 | 0 |
| SRR5388150 | 0 | 0 | 32 | 0 | 0 | 0 | 0 | 123 | 0 | 615 | 175 | 19114 | 0 | 0 | 123 | 369 | 0 | 0 | 0 | 0 | 483 | 2163 | 0 | 0 | 439 | 32 | 151 | 0 | 0 | 0 | 0 | 0 | 0 | 24 | 16 | 184 | 0 |
| SRR5388151 | 0 | 20 | 0 | 0 | 0 | 0 | 0 | 247 | 165 | 495 | 11629 | 15918 | 0 | 0 | 0 | 165 | 0 | 0 | 0 | 0 | 468 | 1502 | 0 | 0 | 303 | 33 | 59 | 0 | 0 | 0 | 0 | 0 | 0 | 0 | 0 | 165 | 0 |
| SRR5388152 | 0 | 0 | 35 | 0 | 0 | 0 | 0 | 0 | 0 | 343 | 9737 | 12480 | 0 | 69 | 69 | 69 | 0 | 0 | 0 | 0 | 363 | 2092 | 0 | 0 | 233 | 52 | 35 | 0 | 0 | 0 | 0 | 0 | 0 | 0 | 0 | 206 | 0 |
| SRR5388153 | 17 | 0 | 17 | 0 | 0 | 0 | 0 | 355 | 118 | 946 | 9341 | 10051 | 0 | 0 | 355 | 355 | 0 | 0 | 0 | 236 | 344 | 1284 | 0 | 0 | 219 | 27 | 50 | 0 | 0 | 0 | 0 | 0 | 0 | 67 | 0 | 473 | 0 |
| SRR5388154 | 0 | 23 | 0 | 0 | 0 | 0 | 0 | 386 | 129 | 1350 | 12926 | 16270 | 0 | 0 | 64 | 257 | 0 | 0 | 0 | 0 | 189 | 1455 | 0 | 20 | 328 | 53 | 46 | 0 | 0 | 0 | 0 | 0 | 0 | 23 | 0 | 257 | 0 |
| SRR5388155 | 35 | 0 | 0 | 0 | 0 | 0 | 0 | 253 | 84 | 1179 | 9769 | 12127 | 0 | 0 | 0 | 337 | 14 | 0 | 0 | 168 | 126 | 556 | 0 | 0 | 128 | 28 | 35 | 0 | 0 | 0 | 0 | 0 | 0 | 0 | 0 | 674 | 0 |
| SRR5388156 | 11 | 0 | 62 | 0 | 0 | 0 | 0 | 300 | 120 | 1018 | 115 | 11862 | 0 | 0 | 60 | 120 | 0 | 0 | 0 | 180 | 273 | 1143 | 0 | 0 | 326 | 28 | 28 | 0 | 0 | 0 | 0 | 0 | 0 | 0 | 0 | 659 | 0 |
| SRR5388157 | 12 | 0 | 12 | 0 | 0 | 0 | 0 | 625 | 89 | 536 | 116 | 12145 | 0 | 179 | 268 | 179 | 12 | 0 | 0 | 0 | 91 | 518 | 0 | 0 | 90 | 17 | 117 | 0 | 0 | 0 | 0 | 0 | 0 | 12 | 0 | 357 | 0 |
| SRR5388158 | 0 | 0 | 12 | 0 | 0 | 0 | 0 | 139 | 139 | 1387 | 34 | 11926 | 0 | 0 | 139 | 139 | 0 | 0 | 0 | 0 | 420 | 1608 | 0 | 0 | 176 | 0 | 59 | 0 | 0 | 0 | 0 | 0 | 0 | 0 | 0 | 139 | 0 |
| SRR5388159 | 0 | 0 | 63 | 0 | 0 | 0 | 0 | 174 | 87 | 1568 | 7752 | 10801 | 0 | 0 | 523 | 523 | 0 | 0 | 0 | 0 | 402 | 1345 | 0 | 0 | 133 | 22 | 94 | 0 | 0 | 0 | 0 | 0 | 21 | 10 | 21 | 348 | 0 |
| SRR5388160 | 0 | 0 | 24 | 0 | 0 | 0 | 0 | 190 | 95 | 1807 | 9700 | 11982 | 0 | 0 | 0 | 190 | 0 | 0 | 0 | 285 | 160 | 1197 | 0 | 0 | 224 | 48 | 40 | 0 | 0 | 0 | 0 | 0 | 0 | 16 | 0 | 571 | 0 |
| SRR5388161 | 0 | 0 | 0 | 0 | 0 | 0 | 0 | 329 | 55 | 877 | 202 | 14141 | 0 | 0 | 493 | 164 | 0 | 0 | 0 | 0 | 176 | 835 | 0 | 0 | 176 | 21 | 100 | 0 | 0 | 0 | 0 | 0 | 0 | 44 | 0 | 110 | 0 |
| SRR5388162 | 0 | 0 | 0 | 0 | 0 | 19 | 0 | 341 | 85 | 426 | 22681 | 21828 | 0 | 0 | 256 | 256 | 0 | 0 | 0 | 171 | 435 | 1671 | 0 | 0 | 201 | 30 | 139 | 0 | 0 | 0 | 0 | 0 | 0 | 0 | 0 | 256 | 0 |
| SRR5388163 | 0 | 0 | 51 | 0 | 0 | 13 | 0 | 128 | 51 | 332 | 11435 | 21109 | 0 | 0 | 0 | 153 | 0 | 0 | 0 | 51 | 701 | 2765 | 0 | 0 | 276 | 24 | 199 | 0 | 0 | 0 | 0 | 0 | 0 | 0 | 0 | 51 | 0 |
| SRR5388164 | 12 | 0 | 37 | 0 | 0 | 0 | 0 | 60 | 0 | 1147 | 86 | 18711 | 0 | 0 | 181 | 60 | 0 | 0 | 0 | 0 | 396 | 1857 | 0 | 0 | 161 | 18 | 175 | 0 | 0 | 0 | 12 | 0 | 12 | 0 | 0 | 181 | 0 |
| SRR5388165 | 0 | 0 | 16 | 0 | 0 | 0 | 0 | 205 | 0 | 410 | 12019 | 16527 | 0 | 68 | 137 | 68 | 0 | 0 | 0 | 546 | 327 | 591 | 0 | 0 | 189 | 18 | 170 | 0 | 0 | 0 | 0 | 0 | 0 | 11 | 16 | 1502 | 0 |
| SRR5388166 | 22 | 0 | 33 | 0 | 0 | 0 | 0 | 581 | 0 | 332 | 62 | 11957 | 0 | 0 | 83 | 83 | 0 | 0 | 0 | 83 | 297 | 1002 | 0 | 0 | 112 | 0 | 417 | 0 | 0 | 0 | 0 | 11 | 0 | 11 | 11 | 166 | 0 |
| SRR5388167 | 0 | 0 | 0 | 0 | 0 | 0 | 0 | 184 | 0 | 184 | 4469 | 5944 | 0 | 0 | 230 | 92 | 0 | 0 | 0 | 2027 | 193 | 938 | 0 | 0 | 161 | 25 | 292 | 0 | 0 | 0 | 0 | 0 | 0 | 0 | 0 | 7326 | 0 |
| SRR5388168 | 31 | 0 | 0 | 0 | 0 | 20 | 0 | 305 | 204 | 1425 | 77 | 7789 | 0 | 0 | 764 | 255 | 0 | 0 | 0 | 305 | 569 | 2274 | 0 | 0 | 508 | 28 | 13 | 0 | 0 | 16 | 0 | 0 | 0 | 0 | 0 | 407 | 0 |
| SRR5388169 | 0 | 0 | 0 | 0 | 0 | 0 | 0 | 249 | 62 | 529 | 4944 | 15579 | 0 | 0 | 187 | 280 | 0 | 0 | 0 | 0 | 249 | 1680 | 0 | 0 | 113 | 43 | 45 | 0 | 0 | 0 | 0 | 0 | 0 | 0 | 0 | 124 | 0 |
| SRR5388170 | 0 | 0 | 0 | 0 | 0 | 0 | 0 | 204 | 0 | 245 | 4649 | 10277 | 0 | 0 | 82 | 245 | 0 | 0 | 0 | 0 | 573 | 2984 | 0 | 13 | 199 | 19 | 109 | 0 | 0 | 0 | 0 | 0 | 0 | 0 | 0 | 0 | 0 |
| SRR5388171 | 0 | 0 | 0 | 0 | 0 | 0 | 0 | 100 | 60 | 220 | 204 | 15253 | 0 | 0 | 80 | 140 | 0 | 0 | 0 | 0 | 394 | 3652 | 0 | 0 | 99 | 16 | 220 | 0 | 0 | 0 | 0 | 0 | 0 | 20 | 0 | 0 | 0 |
| SRR5388172 | 0 | 0 | 0 | 0 | 0 | 0 | 0 | 462 | 0 | 336 | 3235 | 12604 | 0 | 0 | 42 | 210 | 0 | 0 | 0 | 0 | 291 | 2678 | 0 | 0 | 66 | 11 | 194 | 0 | 0 | 0 | 0 | 0 | 18 | 18 | 0 | 0 | 0 |
| SRR5388173 | 0 | 0 | 20 | 0 | 0 | 0 | 0 | 177 | 59 | 883 | 4770 | 20375 | 0 | 59 | 177 | 177 | 0 | 0 | 0 | 177 | 238 | 1517 | 0 | 0 | 105 | 20 | 69 | 0 | 0 | 0 | 0 | 0 | 0 | 0 | 0 | 294 | 0 |
| SRR5388174 | 0 | 12 | 37 | 0 | 0 | 24 | 12 | 63 | 0 | 756 | 58 | 16066 | 0 | 0 | 63 | 126 | 0 | 0 | 0 | 0 | 301 | 1409 | 0 | 0 | 119 | 21 | 61 | 12 | 0 | 0 | 0 | 0 | 12 | 0 | 0 | 126 | 0 |
| SRR5388175 | 0 | 0 | 25 | 0 | 0 | 12 | 0 | 177 | 59 | 1064 | 105 | 16262 | 0 | 0 | 59 | 0 | 0 | 0 | 0 | 237 | 213 | 910 | 0 | 0 | 122 | 19 | 106 | 0 | 0 | 0 | 0 | 0 | 0 | 0 | 0 | 0 | 0 |
| SRR5388176 | 10 | 10 | 10 | 0 | 0 | 0 | 0 | 316 | 0 | 947 | 71 | 15950 | 0 | 0 | 0 | 79 | 10 | 0 | 0 | 158 | 215 | 928 | 0 | 0 | 169 | 29 | 62 | 0 | 0 | 0 | 0 | 0 | 0 | 0 | 21 | 158 | 0 |
| SRR5388177 | 0 | 0 | 40 | 0 | 0 | 0 | 0 | 49 | 49 | 395 | 289 | 17768 | 0 | 99 | 247 | 543 | 10 | 0 | 0 | 49 | 392 | 1997 | 0 | 0 | 159 | 33 | 10 | 0 | 0 | 0 | 0 | 0 | 0 | 0 | 0 | 99 | 0 |
| SRR5388178 | 0 | 0 | 33 | 0 | 0 | 0 | 0 | 226 | 0 | 316 | 5013 | 16710 | 0 | 0 | 316 | 135 | 0 | 0 | 0 | 135 | 265 | 1974 | 0 | 0 | 68 | 13 | 11 | 0 | 0 | 0 | 0 | 0 | 11 | 11 | 0 | 135 | 0 |
| SRR5388179 | 0 | 0 | 0 | 0 | 0 | 0 | 0 | 98 | 49 | 636 | 101 | 18253 | 0 | 0 | 49 | 391 | 15 | 0 | 0 | 49 | 479 | 1820 | 0 | 0 | 150 | 24 | 69 | 0 | 0 | 0 | 0 | 0 | 0 | 0 | 0 | 98 | 0 |
| SRR5388180 | 0 | 0 | 0 | 0 | 0 | 0 | 0 | 648 | 65 | 1231 | 3627 | 16970 | 0 | 0 | 194 | 583 | 0 | 0 | 0 | 130 | 418 | 2137 | 0 | 0 | 83 | 38 | 72 | 0 | 0 | 0 | 0 | 0 | 0 | 0 | 0 | 518 | 0 |
| SRR5388181 | 0 | 0 | 10 | 0 | 0 | 21 | 0 | 147 | 0 | 809 | 3605 | 12212 | 0 | 0 | 147 | 368 | 0 | 0 | 0 | 0 | 723 | 2287 | 0 | 0 | 408 | 17 | 92 | 0 | 0 | 0 | 0 | 0 | 10 | 10 | 0 | 147 | 0 |
| SRR5388182 | 0 | 0 | 21 | 0 | 0 | 0 | 0 | 206 | 69 | 721 | 103 | 12261 | 0 | 0 | 34 | 172 | 0 | 0 | 0 | 103 | 664 | 2604 | 0 | 0 | 416 | 27 | 151 | 0 | 0 | 0 | 0 | 0 | 0 | 34 | 0 | 206 | 0 |
| SRR5388183 | 34 | 0 | 20 | 0 | 0 | 14 | 0 | 585 | 73 | 512 | 10827 | 13606 | 0 | 0 | 439 | 805 | 0 | 0 | 0 | 146 | 538 | 1820 | 0 | 0 | 343 | 36 | 224 | 0 | 0 | 0 | 0 | 0 | 0 | 0 | 0 | 293 | 0 |
| SRR5388184 | 0 | 0 | 11 | 0 | 0 | 22 | 0 | 254 | 0 | 592 | 4568 | 7867 | 0 | 0 | 508 | 254 | 22 | 0 | 0 | 0 | 372 | 2146 | 0 | 0 | 219 | 21 | 306 | 0 | 0 | 0 | 0 | 0 | 0 | 22 | 11 | 254 | 0 |
| SRR5388185 | 0 | 0 | 31 | 0 | 0 | 0 | 0 | 0 | 0 | 328 | 59 | 10538 | 0 | 0 | 218 | 328 | 0 | 0 | 0 | 55 | 941 | 4019 | 0 | 0 | 300 | 22 | 93 | 0 | 0 | 0 | 0 | 0 | 0 | 0 | 0 | 109 | 0 |
| SRR5388186 | 0 | 0 | 17 | 0 | 0 | 0 | 0 | 173 | 58 | 404 | 112 | 8369 | 0 | 0 | 115 | 115 | 0 | 0 | 0 | 58 | 213 | 1079 | 0 | 0 | 59 | 18 | 119 | 0 | 0 | 0 | 0 | 0 | 17 | 0 | 0 | 58 | 0 |
| SRR5388187 | 0 | 0 | 36 | 0 | 0 | 0 | 0 | 286 | 0 | 954 | 2195 | 6204 | 0 | 0 | 382 | 954 | 0 | 0 | 0 | 382 | 626 | 2463 | 0 | 0 | 514 | 40 | 85 | 0 | 12 | 0 | 0 | 0 | 0 | 24 | 12 | 286 | 0 |
| SRR5388188 | 14 | 0 | 14 | 0 | 0 | 0 | 0 | 118 | 177 | 591 | 3723 | 6973 | 0 | 0 | 414 | 1241 | 0 | 0 | 0 | 118 | 607 | 2576 | 0 | 0 | 337 | 36 | 85 | 0 | 0 | 0 | 0 | 0 | 28 | 35 | 0 | 236 | 0 |
| SRR5388189 | 0 | 0 | 0 | 0 | 0 | 0 | 0 | 160 | 0 | 266 | 188 | 9292 | 0 | 0 | 346 | 266 | 0 | 0 | 0 | 80 | 443 | 2167 | 0 | 0 | 194 | 18 | 271 | 0 | 0 | 0 | 0 | 0 | 0 | 18 | 0 | 213 | 0 |
| SRR5388190 | 0 | 0 | 0 | 0 | 0 | 0 | 0 | 258 | 52 | 568 | 9352 | 11573 | 0 | 0 | 155 | 258 | 0 | 0 | 0 | 52 | 561 | 2096 | 0 | 0 | 313 | 29 | 136 | 0 | 0 | 0 | 0 | 0 | 0 | 0 | 0 | 155 | 0 |
| SRR5388191 | 0 | 0 | 53 | 0 | 0 | 0 | 0 | 197 | 0 | 592 | 83 | 13807 | 0 | 0 | 394 | 789 | 18 | 0 | 0 | 0 | 416 | 1493 | 0 | 0 | 275 | 20 | 53 | 0 | 0 | 0 | 0 | 0 | 18 | 18 | 18 | 197 | 0 |
| SRR5388192 | 11 | 0 | 17 | 0 | 0 | 11 | 0 | 301 | 50 | 1255 | 101 | 12098 | 0 | 0 | 1205 | 1104 | 0 | 0 | 0 | 151 | 843 | 2102 | 0 | 0 | 663 | 27 | 63 | 0 | 0 | 0 | 0 | 0 | 0 | 0 | 11 | 502 | 0 |
| SRR5388193 | 12 | 0 | 0 | 0 | 0 | 0 | 0 | 76 | 0 | 865 | 191 | 14201 | 0 | 51 | 280 | 560 | 0 | 0 | 0 | 76 | 326 | 1837 | 0 | 13 | 194 | 32 | 120 | 0 | 0 | 0 | 0 | 0 | 0 | 0 | 0 | 229 | 0 |
| SRR5388194 | 19 | 0 | 0 | 0 | 0 | 0 | 0 | 364 | 156 | 937 | 64 | 11296 | 0 | 0 | 208 | 104 | 0 | 0 | 0 | 104 | 307 | 1767 | 0 | 12 | 169 | 44 | 112 | 0 | 0 | 0 | 0 | 0 | 0 | 19 | 0 | 364 | 0 |
| SRR5388195 | 0 | 0 | 40 | 0 | 0 | 0 | 0 | 79 | 79 | 556 | 7427 | 22958 | 0 | 40 | 159 | 40 | 0 | 0 | 0 | 79 | 461 | 2157 | 0 | 0 | 275 | 30 | 179 | 0 | 0 | 0 | 0 | 0 | 0 | 20 | 0 | 159 | 0 |
| SRR5388196 | 0 | 0 | 0 | 0 | 0 | 0 | 0 | 91 | 91 | 1149 | 6680 | 19738 | 0 | 60 | 60 | 151 | 0 | 0 | 0 | 91 | 493 | 2202 | 0 | 0 | 421 | 34 | 77 | 0 | 0 | 0 | 0 | 0 | 0 | 0 | 0 | 212 | 0 |
| SRR5388197 | 0 | 0 | 36 | 0 | 0 | 0 | 0 | 113 | 28 | 310 | 154 | 8906 | 0 | 0 | 169 | 395 | 0 | 0 | 0 | 56 | 614 | 2312 | 0 | 10 | 174 | 24 | 14 | 0 | 0 | 0 | 0 | 0 | 14 | 0 | 0 | 197 | 0 |
| SRR5388198 | 0 | 0 | 37 | 0 | 0 | 12 | 0 | 115 | 0 | 519 | 60 | 8368 | 0 | 0 | 404 | 462 | 12 | 0 | 0 | 115 | 486 | 1748 | 0 | 11 | 158 | 27 | 49 | 0 | 0 | 0 | 0 | 0 | 0 | 25 | 0 | 173 | 0 |
| SRR5388199 | 0 | 0 | 0 | 0 | 0 | 0 | 0 | 128 | 85 | 85 | 7809 | 9260 | 0 | 0 | 171 | 256 | 0 | 0 | 0 | 1323 | 289 | 1015 | 0 | 0 | 357 | 85 | 14 | 0 | 0 | 0 | 0 | 0 | 0 | 0 | 0 | 3926 | 0 |
| SRR5388200 | 0 | 0 | 33 | 0 | 0 | 13 | 0 | 77 | 0 | 540 | 6325 | 12804 | 0 | 0 | 309 | 617 | 13 | 0 | 0 | 77 | 264 | 1475 | 0 | 0 | 148 | 42 | 60 | 0 | 0 | 0 | 0 | 0 | 0 | 20 | 0 | 231 | 0 |
| SRR5388201 | 0 | 0 | 0 | 0 | 0 | 0 | 0 | 94 | 47 | 566 | 76 | 11235 | 0 | 0 | 330 | 661 | 0 | 0 | 0 | 94 | 218 | 1785 | 0 | 0 | 94 | 12 | 40 | 0 | 0 | 0 | 0 | 0 | 0 | 16 | 0 | 94 | 0 |
| SRR5388202 | 0 | 0 | 11 | 0 | 0 | 0 | 0 | 287 | 72 | 502 | 40 | 9317 | 0 | 0 | 215 | 430 | 0 | 0 | 0 | 215 | 464 | 1927 | 0 | 0 | 384 | 36 | 54 | 0 | 0 | 11 | 0 | 0 | 0 | 0 | 11 | 143 | 0 |
| SRR5388203 | 59 | 0 | 25 | 0 | 0 | 0 | 0 | 209 | 0 | 471 | 123 | 10771 | 0 | 0 | 157 | 366 | 0 | 0 | 0 | 52 | 477 | 1379 | 0 | 0 | 295 | 25 | 279 | 0 | 0 | 0 | 0 | 0 | 0 | 0 | 0 | 157 | 0 |
| SRR5388204 | 24 | 0 | 12 | 0 | 0 | 0 | 0 | 0 | 83 | 498 | 1992 | 12948 | 0 | 83 | 166 | 249 | 0 | 0 | 0 | 83 | 501 | 1567 | 0 | 0 | 186 | 11 | 424 | 0 | 0 | 0 | 0 | 0 | 0 | 12 | 24 | 249 | 0 |
| SRR5388205 | 0 | 0 | 18 | 0 | 0 | 0 | 0 | 241 | 0 | 241 | 210 | 17678 | 0 | 0 | 160 | 80 | 0 | 0 | 0 | 80 | 179 | 1124 | 0 | 0 | 55 | 13 | 18 | 0 | 0 | 0 | 0 | 0 | 18 | 0 | 0 | 40 | 0 |
| SRR5388206 | 0 | 16 | 0 | 0 | 0 | 0 | 0 | 438 | 27 | 219 | 5881 | 11982 | 0 | 0 | 219 | 164 | 0 | 0 | 0 | 27 | 579 | 2599 | 0 | 13 | 112 | 0 | 40 | 0 | 0 | 0 | 0 | 0 | 0 | 16 | 0 | 109 | 0 |
| SRR5388207 | 0 | 0 | 0 | 0 | 0 | 0 | 0 | 248 | 413 | 578 | 8344 | 9831 | 0 | 0 | 165 | 330 | 0 | 0 | 0 | 83 | 531 | 2049 | 0 | 0 | 162 | 28 | 154 | 0 | 0 | 0 | 0 | 0 | 0 | 0 | 0 | 248 | 0 |
| SRR5388208 | 0 | 0 | 28 | 0 | 0 | 28 | 0 | 217 | 72 | 435 | 75 | 16887 | 0 | 0 | 290 | 0 | 0 | 0 | 0 | 145 | 191 | 1136 | 0 | 0 | 78 | 19 | 195 | 0 | 0 | 0 | 0 | 0 | 0 | 0 | 0 | 0 | 0 |
| SRR5388209 | 0 | 0 | 0 | 0 | 0 | 0 | 0 | 0 | 0 | 72 | 7040 | 16486 | 0 | 36 | 144 | 180 | 0 | 0 | 0 | 72 | 231 | 1220 | 0 | 0 | 85 | 10 | 28 | 0 | 0 | 0 | 0 | 0 | 0 | 0 | 14 | 72 | 0 |
| SRR5388210 | 0 | 0 | 22 | 0 | 0 | 0 | 0 | 89 | 0 | 89 | 2406 | 10562 | 0 | 0 | 134 | 134 | 0 | 0 | 0 | 0 | 297 | 1705 | 0 | 0 | 88 | 0 | 0 | 0 | 0 | 0 | 0 | 0 | 0 | 0 | 0 | 0 | 0 |
| SRR5388211 | 0 | 0 | 0 | 0 | 0 | 101 | 0 | 51 | 51 | 510 | 114 | 12552 | 0 | 0 | 153 | 0 | 12 | 0 | 0 | 204 | 585 | 2567 | 0 | 0 | 335 | 22 | 80 | 0 | 0 | 0 | 0 | 0 | 0 | 0 | 0 | 306 | 0 |
| SRR5388212 | 0 | 18 | 0 | 0 | 0 | 0 | 0 | 339 | 85 | 847 | 171 | 14900 | 0 | 0 | 85 | 169 | 0 | 0 | 0 | 85 | 478 | 2421 | 0 | 0 | 299 | 15 | 100 | 0 | 0 | 0 | 0 | 0 | 0 | 18 | 0 | 339 | 0 |
| SRR5388213 | 0 | 0 | 0 | 0 | 0 | 0 | 0 | 77 | 116 | 39 | 5848 | 10069 | 0 | 0 | 0 | 426 | 0 | 0 | 0 | 0 | 464 | 2666 | 0 | 0 | 103 | 15 | 0 | 0 | 0 | 0 | 0 | 0 | 0 | 0 | 0 | 116 | 0 |
| SRR5388214 | 0 | 0 | 11 | 0 | 0 | 0 | 0 | 74 | 0 | 110 | 13095 | 20525 | 0 | 0 | 257 | 331 | 32 | 0 | 0 | 37 | 313 | 2050 | 0 | 0 | 83 | 0 | 11 | 0 | 0 | 0 | 0 | 0 | 0 | 0 | 11 | 110 | 0 |
| SRR5388215 | 0 | 0 | 10 | 0 | 0 | 10 | 0 | 556 | 278 | 649 | 4633 | 11491 | 0 | 0 | 93 | 93 | 0 | 0 | 0 | 0 | 255 | 1188 | 0 | 0 | 119 | 24 | 31 | 0 | 0 | 0 | 0 | 0 | 0 | 0 | 0 | 185 | 0 |
| SRR5388216 | 0 | 0 | 0 | 0 | 0 | 20 | 0 | 103 | 34 | 515 | 16096 | 18807 | 0 | 34 | 69 | 137 | 0 | 0 | 0 | 34 | 670 | 2320 | 0 | 0 | 344 | 49 | 90 | 0 | 0 | 0 | 0 | 0 | 0 | 0 | 0 | 103 | 0 |
| SRR5388217 | 0 | 36 | 18 | 0 | 0 | 0 | 0 | 132 | 132 | 527 | 16212 | 26756 | 0 | 0 | 0 | 264 | 0 | 0 | 0 | 264 | 234 | 672 | 0 | 0 | 98 | 21 | 18 | 0 | 0 | 0 | 0 | 0 | 0 | 0 | 0 | 264 | 0 |
| SRR5388218 | 0 | 0 | 0 | 0 | 0 | 0 | 0 | 442 | 63 | 695 | 13766 | 17239 | 0 | 0 | 316 | 316 | 0 | 0 | 0 | 63 | 860 | 3402 | 0 | 0 | 256 | 13 | 58 | 0 | 0 | 0 | 0 | 0 | 0 | 0 | 0 | 253 | 0 |
| SRR5388219 | 18 | 0 | 18 | 0 | 0 | 0 | 0 | 626 | 0 | 876 | 10759 | 11385 | 0 | 0 | 125 | 500 | 0 | 0 | 0 | 626 | 652 | 3174 | 0 | 0 | 303 | 24 | 98 | 0 | 0 | 0 | 0 | 0 | 18 | 0 | 0 | 626 | 0 |
| SRR5388220 | 0 | 0 | 13 | 0 | 0 | 0 | 0 | 552 | 0 | 1380 | 58 | 8555 | 0 | 0 | 414 | 414 | 13 | 0 | 0 | 276 | 319 | 1884 | 0 | 0 | 149 | 17 | 60 | 0 | 0 | 0 | 0 | 0 | 0 | 0 | 0 | 690 | 0 |
| SRR5388221 | 17 | 0 | 17 | 0 | 0 | 17 | 0 | 143 | 36 | 784 | 414 | 14643 | 0 | 0 | 214 | 321 | 0 | 0 | 0 | 107 | 852 | 2885 | 0 | 0 | 334 | 40 | 212 | 0 | 0 | 0 | 0 | 0 | 0 | 0 | 13 | 178 | 0 |
| SRR5388222 | 0 | 0 | 31 | 0 | 0 | 0 | 0 | 115 | 0 | 459 | 414 | 19066 | 0 | 0 | 115 | 57 | 10 | 0 | 0 | 0 | 880 | 3328 | 0 | 0 | 351 | 20 | 51 | 0 | 0 | 0 | 0 | 10 | 10 | 10 | 0 | 0 | 0 |
| SRR5388223 | 11 | 11 | 11 | 0 | 0 | 0 | 0 | 141 | 0 | 352 | 10340 | 13857 | 0 | 0 | 141 | 492 | 0 | 0 | 0 | 211 | 382 | 1297 | 0 | 0 | 209 | 18 | 142 | 0 | 0 | 0 | 0 | 0 | 0 | 0 | 22 | 0 | 0 |
| SRR5388224 | 0 | 10 | 10 | 0 | 0 | 20 | 0 | 161 | 81 | 1211 | 201 | 16229 | 0 | 0 | 161 | 242 | 0 | 0 | 0 | 0 | 848 | 4005 | 0 | 0 | 388 | 21 | 82 | 0 | 0 | 0 | 0 | 0 | 0 | 0 | 0 | 161 | 0 |
| SRR5388225 | 0 | 18 | 27 | 0 | 0 | 18 | 0 | 240 | 0 | 360 | 23645 | 23705 | 0 | 0 | 120 | 120 | 0 | 0 | 0 | 60 | 300 | 2592 | 0 | 0 | 114 | 13 | 108 | 0 | 0 | 0 | 0 | 0 | 0 | 18 | 0 | 360 | 0 |
| SRR5388226 | 0 | 0 | 0 | 0 | 0 | 0 | 0 | 668 | 0 | 223 | 303 | 19764 | 0 | 0 | 223 | 501 | 0 | 0 | 0 | 56 | 322 | 11 | 0 | 15 | 170 | 18 | 123 | 0 | 0 | 0 | 0 | 0 | 0 | 19 | 19 | 278 | 0 |
| SRR5388227 | 27 | 27 | 27 | 0 | 0 | 0 | 0 | 106 | 0 | 2124 | 26 | 12211 | 0 | 212 | 106 | 425 | 13 | 0 | 0 | 319 | 127 | 651 | 0 | 0 | 91 | 22 | 80 | 0 | 0 | 0 | 0 | 0 | 13 | 13 | 13 | 637 | 0 |
| SRR5388228 | 0 | 0 | 0 | 0 | 0 | 0 | 0 | 106 | 53 | 1275 | 14871 | 18164 | 0 | 159 | 372 | 372 | 0 | 0 | 0 | 53 | 594 | 3390 | 0 | 0 | 477 | 27 | 66 | 0 | 0 | 0 | 0 | 0 | 0 | 11 | 0 | 425 | 0 |
| SRR5388229 | 0 | 0 | 11 | 0 | 0 | 0 | 0 | 864 | 0 | 756 | 74 | 10803 | 0 | 0 | 108 | 432 | 0 | 0 | 0 | 324 | 301 | 1117 | 0 | 0 | 93 | 12 | 22 | 0 | 0 | 0 | 0 | 0 | 0 | 0 | 0 | 432 | 0 |
| SRR5388230 | 0 | 0 | 14 | 0 | 0 | 0 | 0 | 202 | 135 | 944 | 251 | 16188 | 0 | 0 | 202 | 337 | 0 | 0 | 0 | 0 | 450 | 2477 | 0 | 0 | 136 | 13 | 36 | 0 | 0 | 0 | 0 | 0 | 0 | 0 | 0 | 607 | 0 |
| SRR5388231 | 0 | 0 | 41 | 0 | 0 | 0 | 0 | 207 | 103 | 465 | 11837 | 16644 | 0 | 52 | 103 | 258 | 0 | 0 | 0 | 52 | 412 | 1409 | 0 | 0 | 178 | 16 | 91 | 0 | 0 | 0 | 0 | 0 | 0 | 16 | 16 | 362 | 0 |
| SRR5388232 | 0 | 0 | 36 | 0 | 0 | 0 | 0 | 1002 | 200 | 701 | 17031 | 15328 | 0 | 0 | 401 | 601 | 0 | 0 | 0 | 100 | 449 | 2298 | 0 | 0 | 222 | 20 | 134 | 0 | 0 | 0 | 0 | 0 | 0 | 18 | 0 | 501 | 0 |
| SRR5388233 | 0 | 0 | 0 | 0 | 0 | 0 | 0 | 306 | 102 | 883 | 324 | 16504 | 0 | 0 | 34 | 272 | 0 | 0 | 0 | 136 | 638 | 2097 | 0 | 23 | 470 | 59 | 37 | 0 | 0 | 0 | 0 | 0 | 0 | 0 | 0 | 340 | 0 |
| SRR5388234 | 0 | 0 | 16 | 0 | 0 | 0 | 0 | 484 | 161 | 807 | 16137 | 19364 | 0 | 0 | 242 | 484 | 0 | 0 | 0 | 242 | 831 | 13 | 0 | 11 | 201 | 18 | 80 | 0 | 0 | 0 | 0 | 0 | 0 | 32 | 0 | 161 | 0 |
| SRR5388235 | 0 | 0 | 0 | 0 | 0 | 0 | 0 | 362 | 72 | 1737 | 7817 | 10785 | 0 | 217 | 0 | 217 | 0 | 0 | 0 | 434 | 494 | 1729 | 0 | 0 | 535 | 29 | 46 | 0 | 0 | 0 | 0 | 0 | 0 | 0 | 0 | 1013 | 0 |
| SRR5388236 | 0 | 0 | 12 | 0 | 0 | 0 | 0 | 0 | 343 | 1030 | 164 | 16907 | 0 | 172 | 86 | 343 | 0 | 0 | 0 | 86 | 468 | 2232 | 0 | 0 | 268 | 17 | 70 | 0 | 0 | 0 | 0 | 0 | 0 | 18 | 0 | 86 | 0 |
| SRR5388237 | 0 | 0 | 0 | 0 | 0 | 0 | 0 | 464 | 199 | 1325 | 198 | 13719 | 0 | 0 | 199 | 199 | 0 | 0 | 0 | 265 | 526 | 2202 | 0 | 0 | 504 | 47 | 45 | 0 | 0 | 0 | 0 | 0 | 0 | 0 | 0 | 663 | 0 |
| SRR5388238 | 0 | 0 | 0 | 0 | 0 | 0 | 0 | 460 | 46 | 552 | 341 | 18230 | 0 | 0 | 92 | 46 | 0 | 0 | 0 | 276 | 816 | 11 | 0 | 0 | 453 | 19 | 168 | 0 | 0 | 0 | 0 | 0 | 0 | 17 | 0 | 92 | 0 |

### Supp Table S8.4.Columns_DI_to_ES

| | | | | | | | | | | | | | | | | | | | | | | | | | | | | | | | | | | | | | |
|---|---|---|---|---|---|---|---|---|---|---|---|---|---|---|---|---|---|---|---|---|---|---|---|---|---|---|---|---|---|---|---|---|---|---|---|---|---|
| SRR5388239 | 0 | 0 | 0 | 0 | 0 | 0 | 0 | 483 | 0 | 2011 | 132 | 13194 | 0 | 0 | 161 | 322 | 0 | 0 | 0 | 483 | 566 | 2496 | 0 | 0 | 358 | 23 | 57 | 0 | 0 | 0 | 0 | 0 | 0 | 0 | 0 | 805 | 0 |
| SRR5388240 | 0 | 0 | 18 | 0 | 0 | 0 | 0 | 704 | 0 | 1143 | 12224 | 15478 | 0 | 0 | 264 | 176 | 18 | 0 | 0 | 176 | 612 | 2677 | 0 | 0 | 347 | 18 | 97 | 0 | 0 | 0 | 0 | 0 | 0 | 0 | 0 | 791 | 0 |
| SRR5388241 | 0 | 0 | 0 | 0 | 0 | 11 | 0 | 474 | 0 | 1658 | 21077 | 24471 | 0 | 0 | 316 | 474 | 0 | 0 | 0 | 158 | 669 | 2990 | 0 | 0 | 550 | 45 | 81 | 0 | 0 | 0 | 0 | 0 | 0 | 0 | 0 | 474 | 0 |
| SRR5388242 | 0 | 0 | 0 | 0 | 0 | 0 | 0 | 426 | 61 | 2254 | 17790 | 22115 | 0 | 61 | 61 | 305 | 0 | 0 | 0 | 366 | 654 | 2017 | 0 | 0 | 986 | 48 | 105 | 0 | 0 | 0 | 0 | 0 | 0 | 0 | 0 | 731 | 0 |
| SRR5388243 | 0 | 0 | 14 | 0 | 0 | 0 | 0 | 691 | 63 | 1696 | 7475 | 14637 | 0 | 63 | 377 | 251 | 0 | 0 | 0 | 63 | 557 | 1983 | 0 | 0 | 360 | 24 | 27 | 0 | 0 | 0 | 0 | 0 | 0 | 0 | 0 | 314 | 0 |
| SRR5388244 | 0 | 0 | 20 | 0 | 0 | 20 | 0 | 189 | 94 | 1417 | 152 | 19645 | 0 | 0 | 0 | 189 | 0 | 0 | 0 | 94 | 524 | 11 | 0 | 0 | 189 | 10 | 51 | 0 | 0 | 0 | 0 | 0 | 0 | 31 | 20 | 189 | 0 |
| SRR5388245 | 0 | 0 | 0 | 0 | 0 | 0 | 0 | 160 | 213 | 587 | 10617 | 11364 | 0 | 0 | 0 | 160 | 0 | 0 | 0 | 0 | 397 | 2195 | 0 | 0 | 180 | 28 | 53 | 0 | 0 | 0 | 0 | 0 | 0 | 0 | 0 | 213 | 0 |
| SRR5388246 | 0 | 0 | 0 | 0 | 0 | 0 | 0 | 871 | 73 | 508 | 199 | 14806 | 0 | 0 | 145 | 218 | 0 | 0 | 0 | 73 | 431 | 2839 | 0 | 0 | 282 | 12 | 23 | 0 | 0 | 0 | 0 | 0 | 0 | 0 | 0 | 363 | 0 |
| SRR5388247 | 0 | 0 | 15 | 0 | 0 | 0 | 0 | 286 | 0 | 524 | 557 | 23837 | 0 | 0 | 191 | 286 | 0 | 0 | 0 | 0 | 755 | 3940 | 0 | 11 | 143 | 22 | 37 | 0 | 0 | 0 | 0 | 0 | 0 | 0 | 0 | 95 | 0 |
| SRR5388248 | 0 | 0 | 13 | 0 | 0 | 0 | 0 | 438 | 49 | 779 | 345 | 29555 | 0 | 49 | 146 | 146 | 0 | 0 | 0 | 49 | 717 | 4215 | 0 | 0 | 194 | 12 | 116 | 0 | 0 | 0 | 0 | 0 | 0 | 0 | 19 | 292 | 0 |
| SRR5388249 | 38 | 0 | 95 | 0 | 0 | 0 | 0 | 129 | 0 | 904 | 36 | 12264 | 0 | 0 | 387 | 645 | 19 | 0 | 0 | 0 | 214 | 1013 | 0 | 0 | 49 | 12 | 38 | 0 | 0 | 0 | 0 | 0 | 0 | 0 | 0 | 387 | 0 |
| SRR5388250 | 0 | 0 | 12 | 0 | 0 | 24 | 0 | 418 | 0 | 1150 | 14955 | 18197 | 0 | 0 | 627 | 1360 | 0 | 0 | 0 | 105 | 601 | 2900 | 0 | 0 | 189 | 16 | 18 | 0 | 0 | 0 | 0 | 0 | 0 | 0 | 0 | 418 | 0 |
| SRR5388251 | 0 | 0 | 0 | 0 | 0 | 0 | 0 | 0 | 0 | 1041 | 45 | 7852 | 0 | 0 | 0 | 0 | 0 | 0 | 0 | 95 | 716 | 2010 | 0 | 0 | 304 | 12 | 131 | 0 | 0 | 0 | 0 | 0 | 0 | 0 | 0 | 189 | 0 |
| SRR5388252 | 0 | 0 | 0 | 0 | 0 | 0 | 0 | 415 | 166 | 746 | 158 | 12603 | 0 | 0 | 166 | 332 | 0 | 0 | 0 | 332 | 593 | 2512 | 0 | 0 | 417 | 24 | 219 | 0 | 0 | 0 | 0 | 0 | 0 | 25 | 0 | 83 | 0 |
| SRR5388253 | 20 | 0 | 0 | 0 | 0 | 0 | 0 | 285 | 285 | 1140 | 5698 | 7407 | 0 | 0 | 427 | 427 | 20 | 0 | 0 | 142 | 305 | 1124 | 0 | 0 | 303 | 43 | 228 | 0 | 0 | 0 | 0 | 0 | 0 | 0 | 0 | 285 | 0 |
| SRR5388254 | 0 | 0 | 27 | 0 | 0 | 0 | 0 | 382 | 95 | 573 | 148 | 18618 | 0 | 0 | 477 | 764 | 0 | 0 | 0 | 191 | 754 | 2402 | 0 | 0 | 403 | 24 | 186 | 0 | 0 | 0 | 0 | 0 | 0 | 0 | 0 | 0 | 0 |
| SRR5388255 | 0 | 0 | 10 | 0 | 0 | 0 | 0 | 177 | 0 | 1326 | 10346 | 18216 | 0 | 0 | 265 | 442 | 10 | 0 | 0 | 0 | 64 | 668 | 0 | 0 | 40 | 25 | 104 | 0 | 0 | 0 | 0 | 0 | 0 | 0 | 0 | 354 | 0 |
| SRR5388256 | 0 | 0 | 0 | 0 | 0 | 0 | 0 | 48 | 0 | 571 | 6095 | 16903 | 0 | 0 | 333 | 381 | 0 | 0 | 0 | 48 | 267 | 2222 | 0 | 0 | 161 | 20 | 166 | 0 | 0 | 0 | 0 | 0 | 0 | 0 | 0 | 333 | 0 |
| SRR5388257 | 0 | 0 | 11 | 0 | 0 | 0 | 0 | 254 | 85 | 423 | 13198 | 21235 | 0 | 85 | 169 | 169 | 0 | 0 | 0 | 85 | 69 | 582 | 0 | 0 | 22 | 20 | 11 | 0 | 0 | 0 | 0 | 0 | 0 | 0 | 0 | 169 | 0 |
| SRR5388258 | 0 | 23 | 58 | 0 | 0 | 0 | 0 | 582 | 194 | 517 | 368 | 18290 | 0 | 0 | 452 | 129 | 12 | 0 | 0 | 65 | 612 | 13 | 0 | 22 | 172 | 28 | 81 | 0 | 0 | 0 | 0 | 12 | 0 | 0 | 12 | 259 | 0 |
| SRR5388259 | 0 | 0 | 0 | 0 | 0 | 0 | 0 | 247 | 0 | 544 | 19469 | 25992 | 0 | 49 | 198 | 346 | 0 | 0 | 0 | 0 | 703 | 11 | 0 | 0 | 183 | 13 | 191 | 0 | 0 | 0 | 0 | 0 | 0 | 14 | 20 | 395 | 0 |
| SRR5388260 | 0 | 0 | 28 | 0 | 0 | 0 | 0 | 83 | 83 | 998 | 302 | 20216 | 0 | 83 | 499 | 499 | 0 | 0 | 0 | 83 | 290 | 2019 | 0 | 0 | 208 | 27 | 170 | 0 | 0 | 0 | 0 | 0 | 0 | 0 | 28 | 83 | 0 |
| SRR5388261 | 0 | 0 | 27 | 0 | 0 | 14 | 0 | 68 | 68 | 473 | 78 | 7702 | 0 | 68 | 68 | 203 | 0 | 0 | 0 | 68 | 664 | 2112 | 0 | 15 | 314 | 26 | 122 | 0 | 0 | 0 | 0 | 0 | 0 | 0 | 20 | 878 | 0 |
| SRR5388262 | 0 | 0 | 19 | 0 | 0 | 0 | 0 | 145 | 0 | 723 | 154 | 9682 | 0 | 0 | 434 | 289 | 0 | 0 | 0 | 145 | 823 | 2471 | 0 | 23 | 578 | 42 | 89 | 0 | 0 | 0 | 0 | 0 | 0 | 0 | 0 | 361 | 0 |
| SRR5388263 | 0 | 0 | 88 | 0 | 0 | 0 | 0 | 635 | 53 | 582 | 111 | 19841 | 0 | 0 | 159 | 53 | 0 | 0 | 0 | 0 | 800 | 2912 | 0 | 0 | 142 | 16 | 29 | 0 | 0 | 0 | 0 | 0 | 0 | 29 | 20 | 265 | 0 |
| SRR5388264 | 0 | 0 | 11 | 0 | 0 | 0 | 0 | 287 | 72 | 934 | 19898 | 30027 | 0 | 144 | 0 | 216 | 0 | 0 | 0 | 72 | 620 | 2544 | 0 | 0 | 226 | 11 | 34 | 0 | 0 | 0 | 0 | 0 | 0 | 0 | 0 | 72 | 0 |
| SRR5388265 | 0 | 0 | 0 | 0 | 0 | 0 | 0 | 211 | 84 | 253 | 203 | 12725 | 0 | 0 | 42 | 337 | 0 | 0 | 0 | 126 | 541 | 4047 | 0 | 16 | 173 | 16 | 48 | 0 | 0 | 0 | 0 | 0 | 0 | 19 | 0 | 169 | 0 |
| SRR5388266 | 0 | 0 | 0 | 0 | 0 | 0 | 0 | 465 | 0 | 388 | 212 | 14814 | 0 | 0 | 465 | 310 | 0 | 0 | 0 | 0 | 307 | 2134 | 0 | 0 | 81 | 0 | 15 | 0 | 0 | 0 | 0 | 0 | 0 | 0 | 15 | 388 | 0 |
| SRR5388267 | 11 | 0 | 0 | 0 | 0 | 0 | 0 | 170 | 85 | 765 | 5952 | 12924 | 0 | 0 | 170 | 510 | 0 | 0 | 0 | 340 | 571 | 2244 | 0 | 28 | 376 | 0 | 66 | 0 | 0 | 0 | 0 | 0 | 0 | 11 | 11 | 340 | 0 |
| SRR5388268 | 0 | 0 | 25 | 0 | 0 | 25 | 0 | 137 | 0 | 957 | 335 | 20958 | 0 | 0 | 91 | 273 | 0 | 0 | 0 | 46 | 405 | 1908 | 0 | 0 | 227 | 24 | 133 | 0 | 0 | 0 | 0 | 0 | 0 | 0 | 13 | 364 | 0 |
| SRR5388269 | 0 | 0 | 28 | 0 | 0 | 0 | 0 | 371 | 93 | 557 | 20137 | 21437 | 0 | 0 | 278 | 278 | 0 | 0 | 0 | 0 | 488 | 2371 | 0 | 23 | 292 | 29 | 83 | 0 | 0 | 0 | 0 | 0 | 0 | 18 | 0 | 742 | 0 |
| SRR5388270 | 0 | 0 | 0 | 0 | 0 | 0 | 0 | 479 | 213 | 213 | 246 | 16972 | 0 | 0 | 106 | 160 | 0 | 0 | 0 | 106 | 586 | 2956 | 0 | 19 | 162 | 0 | 74 | 0 | 0 | 0 | 0 | 0 | 19 | 0 | 0 | 213 | 0 |
| SRR5388271 | 0 | 0 | 30 | 0 | 0 | 30 | 0 | 873 | 0 | 873 | 3274 | 6985 | 0 | 0 | 0 | 873 | 0 | 0 | 0 | 437 | 355 | 11 | 0 | 0 | 171 | 19 | 89 | 0 | 0 | 0 | 0 | 0 | 0 | 59 | 0 | 218 | 0 |
| SRR5388272 | 0 | 0 | 0 | 0 | 0 | 0 | 0 | 300 | 400 | 1201 | 8209 | 12213 | 0 | 0 | 200 | 1001 | 0 | 0 | 0 | 501 | 662 | 4267 | 0 | 0 | 301 | 24 | 39 | 0 | 0 | 0 | 0 | 0 | 0 | 0 | 0 | 100 | 0 |
| SRR5388273 | 0 | 0 | 16 | 0 | 0 | 0 | 0 | 204 | 0 | 680 | 84 | 9116 | 0 | 68 | 136 | 204 | 0 | 0 | 0 | 340 | 564 | 2389 | 0 | 0 | 224 | 20 | 56 | 0 | 0 | 0 | 0 | 0 | 0 | 0 | 0 | 612 | 0 |
| SRR5388274 | 0 | 0 | 0 | 0 | 0 | 12 | 0 | 683 | 152 | 3261 | 14486 | 16610 | 0 | 0 | 228 | 455 | 0 | 0 | 0 | 76 | 334 | 1818 | 0 | 0 | 281 | 20 | 30 | 0 | 0 | 0 | 0 | 0 | 12 | 0 | 0 | 607 | 0 |
| SRR5388275 | 0 | 0 | 0 | 0 | 0 | 0 | 0 | 189 | 0 | 1229 | 11435 | 17294 | 0 | 0 | 189 | 95 | 0 | 0 | 0 | 284 | 98 | 771 | 0 | 0 | 29 | 32 | 19 | 0 | 0 | 0 | 0 | 0 | 0 | 0 | 0 | 473 | 0 |
| SRR5388276 | 0 | 0 | 0 | 0 | 0 | 0 | 0 | 125 | 42 | 583 | 372 | 19618 | 0 | 0 | 292 | 125 | 0 | 0 | 0 | 83 | 599 | 2734 | 0 | 0 | 445 | 38 | 28 | 0 | 0 | 0 | 0 | 0 | 11 | 11 | 0 | 125 | 0 |
| SRR5388277 | 10 | 0 | 51 | 0 | 0 | 0 | 0 | 186 | 93 | 836 | 240 | 28800 | 0 | 93 | 0 | 186 | 0 | 0 | 0 | 93 | 408 | 1712 | 0 | 0 | 198 | 14 | 10 | 0 | 0 | 10 | 0 | 0 | 0 | 0 | 0 | 0 | 0 |
| SRR5388278 | 0 | 0 | 0 | 0 | 0 | 0 | 0 | 364 | 0 | 607 | 214 | 18891 | 0 | 61 | 243 | 607 | 0 | 0 | 0 | 121 | 554 | 2759 | 0 | 0 | 178 | 29 | 50 | 0 | 0 | 0 | 0 | 0 | 0 | 0 | 0 | 304 | 0 |
| SRR5388279 | 0 | 0 | 18 | 0 | 0 | 0 | 0 | 219 | 55 | 329 | 83 | 9804 | 0 | 0 | 110 | 329 | 0 | 0 | 0 | 110 | 693 | 3012 | 0 | 0 | 134 | 11 | 62 | 0 | 0 | 0 | 0 | 0 | 0 | 18 | 0 | 0 | 0 |
| SRR5388280 | 0 | 0 | 0 | 0 | 0 | 0 | 0 | 120 | 60 | 420 | 14577 | 19196 | 0 | 60 | 300 | 300 | 0 | 0 | 0 | 120 | 636 | 3873 | 0 | 0 | 111 | 10 | 27 | 0 | 0 | 0 | 0 | 0 | 0 | 0 | 13 | 300 | 0 |
| SRR5388281 | 0 | 0 | 24 | 0 | 0 | 0 | 0 | 393 | 262 | 1310 | 4062 | 6421 | 0 | 0 | 1048 | 655 | 0 | 0 | 0 | 0 | 260 | 1009 | 0 | 0 | 155 | 50 | 24 | 0 | 0 | 0 | 0 | 0 | 0 | 24 | 0 | 786 | 0 |
| SRR5388282 | 0 | 0 | 14 | 0 | 0 | 0 | 0 | 108 | 108 | 541 | 84 | 7899 | 0 | 0 | 757 | 433 | 0 | 0 | 0 | 0 | 344 | 2223 | 0 | 0 | 104 | 16 | 0 | 0 | 0 | 0 | 0 | 0 | 0 | 28 | 28 | 325 | 0 |
| SRR5388283 | 0 | 0 | 26 | 0 | 0 | 13 | 0 | 482 | 132 | 745 | 5524 | 12802 | 0 | 0 | 175 | 219 | 0 | 0 | 0 | 175 | 253 | 1150 | 0 | 0 | 72 | 23 | 46 | 0 | 0 | 0 | 0 | 0 | 0 | 13 | 0 | 44 | 0 |
| SRR5388284 | 0 | 0 | 33 | 0 | 0 | 0 | 0 | 171 | 34 | 137 | 467 | 15881 | 0 | 0 | 137 | 171 | 0 | 0 | 0 | 34 | 610 | 3447 | 0 | 0 | 227 | 19 | 43 | 0 | 0 | 0 | 0 | 0 | 0 | 0 | 0 | 205 | 0 |
| SRR5388285 | 0 | 0 | 28 | 0 | 0 | 0 | 0 | 222 | 111 | 1443 | 64 | 15759 | 0 | 111 | 111 | 444 | 0 | 0 | 0 | 0 | 262 | 1281 | 0 | 0 | 227 | 59 | 42 | 0 | 0 | 0 | 0 | 0 | 0 | 0 | 0 | 444 | 0 |
| SRR5388286 | 0 | 0 | 30 | 0 | 0 | 0 | 0 | 240 | 0 | 480 | 156 | 17826 | 0 | 0 | 0 | 320 | 0 | 0 | 0 | 80 | 318 | 2081 | 0 | 0 | 219 | 53 | 44 | 0 | 0 | 0 | 0 | 0 | 0 | 0 | 0 | 160 | 0 |
| SRR5388287 | 0 | 0 | 0 | 0 | 0 | 0 | 0 | 202 | 50 | 302 | 616 | 14911 | 0 | 50 | 302 | 151 | 0 | 0 | 0 | 101 | 285 | 1578 | 0 | 13 | 141 | 24 | 0 | 0 | 0 | 0 | 0 | 0 | 0 | 17 | 11 | 76 | 0 |
| SRR5388288 | 0 | 0 | 0 | 0 | 0 | 0 | 0 | 318 | 91 | 500 | 273 | 14310 | 0 | 0 | 136 | 273 | 0 | 0 | 0 | 91 | 583 | 3312 | 0 | 21 | 289 | 20 | 40 | 0 | 0 | 0 | 0 | 0 | 0 | 0 | 0 | 318 | 0 |
| SRR5388289 | 0 | 0 | 12 | 0 | 0 | 0 | 0 | 37 | 0 | 37 | 126 | 12512 | 0 | 0 | 112 | 112 | 23 | 0 | 0 | 37 | 599 | 2744 | 0 | 0 | 316 | 40 | 0 | 0 | 0 | 0 | 0 | 0 | 0 | 12 | 0 | 186 | 0 |
| SRR5388290 | 0 | 0 | 19 | 0 | 0 | 0 | 0 | 378 | 126 | 1007 | 18508 | 21026 | 0 | 63 | 0 | 126 | 0 | 0 | 0 | 189 | 482 | 2374 | 0 | 0 | 269 | 40 | 28 | 0 | 0 | 0 | 0 | 0 | 0 | 0 | 0 | 63 | 0 |
| SRR5388291 | 12 | 0 | 12 | 0 | 0 | 0 | 0 | 184 | 0 | 629 | 233 | 15552 | 0 | 0 | 236 | 210 | 0 | 0 | 0 | 79 | 636 | 2857 | 0 | 0 | 365 | 37 | 152 | 0 | 0 | 0 | 0 | 0 | 0 | 12 | 0 | 105 | 0 |
| SRR5388292 | 0 | 0 | 23 | 0 | 0 | 0 | 0 | 0 | 118 | 2071 | 312 | 22014 | 0 | 0 | 59 | 651 | 0 | 0 | 0 | 118 | 537 | 2360 | 0 | 0 | 366 | 40 | 173 | 0 | 0 | 0 | 0 | 0 | 0 | 0 | 15 | 237 | 0 |
| SRR5388293 | 0 | 0 | 0 | 0 | 0 | 21 | 0 | 352 | 117 | 880 | 5572 | 9737 | 0 | 0 | 117 | 528 | 0 | 0 | 0 | 117 | 830 | 2334 | 0 | 0 | 295 | 24 | 145 | 0 | 0 | 0 | 0 | 0 | 0 | 0 | 0 | 117 | 0 |
| SRR5388294 | 0 | 0 | 0 | 0 | 0 | 0 | 0 | 485 | 138 | 761 | 156 | 15782 | 0 | 0 | 277 | 208 | 0 | 0 | 0 | 69 | 705 | 3290 | 0 | 0 | 267 | 23 | 230 | 0 | 0 | 0 | 0 | 0 | 0 | 0 | 0 | 138 | 0 |
| SRR5476966 | 0 | 0 | 0 | 0 | 0 | 0 | 0 | 8902 | 0 | 0 | 38576 | 14837 | 0 | 0 | 0 | 0 | 0 | 32 | 0 | 0 | 352 | 544 | 0 | 1535 | 1087 | 0 | 0 | 0 | 0 | 0 | 0 | 0 | 0 | 0 | 0 | 0 | 64 |
| SRR5476967 | 0 | 0 | 0 | 0 | 0 | 0 | 0 | 13255 | 0 | 2946 | 24 | 8837 | 0 | 0 | 0 | 0 | 0 | 0 | 0 | 0 | 374 | 747 | 0 | 1075 | 1261 | 0 | 0 | 0 | 0 | 0 | 0 | 0 | 0 | 0 | 0 | 0 | 140 |
| SRR5476969 | 0 | 0 | 933 | 0 | 0 | 0 | 0 | 0 | 0 | 0 | 12 | 11662 | 0 | 0 | 0 | 0 | 0 | 0 | 0 | 0 | 0 | 0 | 0 | 0 | 0 | 0 | 0 | 0 | 933 | 0 | 0 | 0 | 0 | 933 | 0 | 0 | 0 |
| SRR5517463 | 0 | 13 | 15 | 127 | 165 | 0 | 95 | 193 | 435 | 2650 | 59 | 271 | 25 | 245 | 171 | 528 | 36 | 480 | 227 | 8610 | 131 | 2412 | 153 | 45721 | 2654 | 105 | 55 | 33 | 140 | 114 | 125 | 16 | 19 | 12 | 0 | 26343 | 414 |
| SRR5517464 | 0 | 17 | 67 | 211 | 421 | 0 | 93 | 205 | 631 | 5810 | 42 | 294 | 15 | 334 | 302 | 1492 | 38 | 677 | 216 | 18766 | 387 | 2984 | 283 | 53915 | 3811 | 108 | 47 | 38 | 152 | 211 | 246 | 28 | 40 | 27 | 17 | 52417 | 459 |
| SRR5517465 | 0 | 0 | 84 | 128 | 290 | 0 | 94 | 209 | 640 | 7106 | 54 | 212 | 15 | 453 | 833 | 3357 | 36 | 528 | 244 | 19506 | 186 | 2416 | 196 | 48950 | 2990 | 111 | 461 | 236 | 1202 | 1649 | 2223 | 338 | 719 | 509 | 20 | 67331 | 398 |
| SRR5517466 | 0 | 12 | 38 | 156 | 311 | 0 | 115 | 230 | 624 | 3999 | 226 | 384 | 20 | 254 | 144 | 811 | 53 | 505 | 201 | 12530 | 173 | 2256 | 210 | 51345 | 3559 | 99 | 619 | 304 | 1328 | 1607 | 1517 | 205 | 262 | 191 | 0 | 37572 | 506 |
| SRR5517467 | 0 | 11 | 363 | 318 | 327 | 0 | 25 | 406 | 1029 | 9700 | 495 | 476 | 13 | 623 | 1168 | 2553 | 25 | 713 | 240 | 7623 | 151 | 3204 | 243 | 46438 | 3480 | 126 | 159 | 798 | 2309 | 2457 | 2894 | 404 | 634 | 529 | 0 | 17388 | 567 |
| SRR5517468 | 0 | 0 | 446 | 235 | 312 | 0 | 39 | 396 | 1148 | 10008 | 142 | 474 | 18 | 525 | 1128 | 2518 | 18 | 667 | 236 | 10238 | 249 | 2794 | 244 | 47574 | 3553 | 137 | 210 | 1058 | 3001 | 3413 | 3908 | 447 | 792 | 705 | 23 | 22721 | 515 |
| SRR5517469 | 0 | 0 | 14 | 255 | 336 | 0 | 29 | 589 | 1188 | 10483 | 411 | 513 | 18 | 546 | 444 | 990 | 13 | 484 | 217 | 8167 | 97 | 2206 | 222 | 44506 | 2953 | 111 | 23 | 56 | 156 | 321 | 289 | 27 | 35 | 29 | 0 | 20709 | 475 |
| SRR5517470 | 0 | 0 | 28 | 208 | 404 | 0 | 28 | 434 | 1120 | 11708 | 438 | 501 | 18 | 676 | 1215 | 2723 | 26 | 482 | 217 | 9643 | 134 | 2005 | 276 | 47790 | 2905 | 132 | 21 | 57 | 202 | 312 | 364 | 38 | 77 | 62 | 21 | 24547 | 510 |
| SRR5575011 | 0 | 28 | 30 | 0 | 19 | 0 | 0 | 197 | 976 | 1288 | 1059 | 6606 | 0 | 10 | 654 | 83 | 0 | 2956 | 16 | 83 | 422 | 298 | 0 | 16 | 1911 | 0 | 0 | 0 | 20 | 22 | 0 | 0 | 40 | 11 | 47 | 436 | 13 |
| SRR5575012 | 0 | 52 | 27 | 0 | 14 | 0 | 0 | 170 | 887 | 1057 | 556 | 3611 | 0 | 0 | 529 | 143 | 0 | 2739 | 57 | 72 | 161 | 138 | 0 | 23 | 2215 | 0 | 0 | 0 | 16 | 18 | 13 | 0 | 53 | 73 | 40 | 358 | 20 |
| SRR5575013 | 0 | 592 | 59 | 0 | 36 | 0 | 0 | 170 | 682 | 903 | 100 | 5299 | 0 | 0 | 767 | 187 | 0 | 3484 | 19 | 119 | 342 | 213 | 0 | 14 | 1842 | 0 | 0 | 0 | 121 | 31 | 0 | 0 | 28 | 104 | 76 | 630 | 19 |
| SRR5575014 | 0 | 563 | 41 | 0 | 26 | 0 | 0 | 206 | 946 | 1629 | 121 | 4448 | 0 | 0 | 590 | 47 | 0 | 3642 | 45 | 56 | 464 | 351 | 0 | 19 | 2157 | 0 | 0 | 0 | 109 | 17 | 22 | 15 | 54 | 96 | 109 | 459 | 29 |
| SRR5575015 | 0 | 78 | 41 | 0 | 33 | 0 | 0 | 205 | 804 | 1587 | 1258 | 5206 | 0 | 0 | 366 | 80 | 0 | 3075 | 50 | 58 | 530 | 408 | 0 | 15 | 2415 | 0 | 0 | 0 | 11 | 28 | 13 | 0 | 41 | 65 | 32 | 219 | 19 |
| SRR5575016 | 0 | 530 | 42 | 0 | 25 | 0 | 0 | 118 | 690 | 1058 | 117 | 4598 | 0 | 0 | 823 | 176 | 0 | 3315 | 54 | 29 | 308 | 241 | 0 | 18 | 1905 | 0 | 0 | 0 | 130 | 27 | 15 | 15 | 66 | 121 | 42 | 558 | 18 |
| SRR5575017 | 0 | 67 | 48 | 0 | 31 | 0 | 0 | 156 | 912 | 1381 | 1068 | 4914 | 0 | 0 | 501 | 123 | 0 | 2639 | 18 | 99 | 494 | 326 | 0 | 25 | 2297 | 0 | 0 | 0 | 11 | 13 | 0 | 11 | 35 | 75 | 53 | 378 | 33 |
| SRR5575018 | 0 | 83 | 20 | 0 | 47 | 0 | 0 | 268 | 702 | 1812 | 1410 | 5992 | 0 | 0 | 574 | 134 | 0 | 3536 | 14 | 51 | 837 | 587 | 0 | 23 | 2851 | 0 | 0 | 0 | 11 | 57 | 24 | 0 | 31 | 73 | 49 | 223 | 21 |
| SRR5575019 | 0 | 799 | 34 | 0 | 28 | 0 | 0 | 195 | 667 | 793 | 222 | 8079 | 0 | 14 | 584 | 209 | 0 | 3369 | 45 | 222 | 390 | 236 | 0 | 16 | 1666 | 0 | 34 | 0 | 63 | 0 | 13 | 0 | 24 | 91 | 52 | 459 | 20 |
| SRR5644067 | 0 | 41 | 127 | 0 | 93 | 0 | 0 | 630 | 324 | 2676 | 96 | 169 | 0 | 163 | 1447 | 1095 | 0 | 183 | 0 | 2759 | 4522 | 1989 | 110 | 3317 | 8810 | 0 | 15 | 37 | 82 | 71 | 63 | 15 | 63 | 74 | 60 | 31235 | 0 |
| SRR5644068 | 0 | 122 | 76 | 0 | 40 | 0 | 0 | 201 | 196 | 1131 | 15 | 17 | 0 | 278 | 308 | 55 | 15 | 748 | 0 | 56 | 7742 | 2905 | 90 | 3344 | 17240 | 80 | 0 | 0 | 61 | 122 | 91 | 0 | 137 | 243 | 91 | 802 | 23 |
| SRR5644069 | 0 | 118 | 55 | 0 | 71 | 0 | 0 | 444 | 579 | 2893 | 100 | 103 | 0 | 28 | 1319 | 1542 | 16 | 450 | 0 | 289 | 3421 | 1012 | 229 | 5064 | 3664 | 0 | 39 | 0 | 16 | 39 | 39 | 16 | 39 | 79 | 71 | 2318 | 0 |
| SRR5644070 | 0 | 0 | 13 | 0 | 43 | 0 | 0 | 1755 | 536 | 3003 | 135 | 220 | 0 | 21 | 955 | 1556 | 13 | 184 | 0 | 1654 | 2395 | 1596 | 267 | 5048 | 7387 | 0 | 13 | 0 | 0 | 30 | 79 | 10 | 43 | 119 | 23 | 29322 | 0 |
| SRR5644071 | 0 | 250 | 116 | 0 | 0 | 18 | 0 | 494 | 425 | 2440 | 226 | 136 | 0 | 98 | 1660 | 1128 | 18 | 244 | 0 | 215 | 4753 | 4628 | 148 | 8906 | 12533 | 0 | 0 | 0 | 22 | 40 | 40 | 0 | 27 | 58 | 0 | 3494 | 11 |
| SRR5644072 | 0 | 38 | 0 | 0 | 80 | 0 | 0 | 1060 | 716 | 4471 | 18 | 153 | 0 | 294 | 12 | 12 | 26 | 528 | 0 | 121 | 1020 | 1041 | 172 | 3669 | 9879 | 0 | 0 | 0 | 11 | 18 | 0 | 0 | 11 | 22 | 0 | 1896 | 0 |
| SRR5644073 | 0 | 51 | 140 | 0 | 0 | 0 | 0 | 934 | 516 | 3104 | 22 | 40 | 0 | 184 | 40 | 72 | 0 | 508 | 0 | 575 | 2205 | 2044 | 534 | 6673 | 8179 | 0 | 16 | 0 | 0 | 16 | 19 | 0 | 29 | 29 | 19 | 11045 | 0 |
| SRR5644074 | 0 | 166 | 33 | 0 | 21 | 16 | 16 | 4588 | 4636 | 7659 | 72 | 334 | 0 | 112 | 112 | 107 | 19 | 247 | 0 | 773 | 2505 | 2465 | 336 | 6771 | 11169 | 0 | 0 | 0 | 0 | 47 | 33 | 0 | 11 | 42 | 0 | 2605 | 0 |
| SRR5644075 | 0 | 42 | 42 | 0 | 54 | 0 | 0 | 446 | 240 | 7464 | 46 | 58 | 0 | 282 | 27 | 21 | 12 | 336 | 0 | 200 | 1790 | 1172 | 624 | 7125 | 6477 | 0 | 12 | 0 | 0 | 30 | 12 | 12 | 30 | 24 | 60 | 3211 | 10 |
| SRR5644076 | 0 | 255 | 31 | 0 | 30 | 15 | 10 | 1170 | 603 | 1829 | 243 | 418 | 0 | 134 | 14 | 20 | 23 | 576 | 0 | 911 | 2388 | 1825 | 369 | 4073 | 9737 | 0 | 34 | 0 | 34 | 44 | 44 | 0 | 0 | 23 | 0 | 12775 | 0 |
| SRR5644077 | 0 | 24 | 30 | 0 | 49 | 28 | 0 | 765 | 543 | 3447 | 62 | 130 | 0 | 166 | 10 | 0 | 25 | 378 | 0 | 338 | 2795 | 1931 | 682 | 8051 | 12262 | 0 | 15 | 0 | 0 | 32 | 25 | 18 | 0 | 12 | 0 | 1715 | 0 |
| SRR5644078 | 0 | 104 | 31 | 0 | 34 | 26 | 20 | 1111 | 838 | 2976 | 275 | 472 | 0 | 147 | 112 | 89 | 37 | 471 | 0 | 478 | 1719 | 1665 | 456 | 6012 | 12438 | 0 | 16 | 0 | 0 | 16 | 11 | 0 | 0 | 13 | 0 | 1068 | 0 |
| SRR5644085 | 0 | 16 | 18 | 0 | 43 | 11 | 43 | 572 | 282 | 1409 | 871 | 1272 | 0 | 111 | 2032 | 896 | 29 | 135 | 0 | 359 | 3602 | 1607 | 41 | 985 | 7936 | 0 | 18 | 0 | 25 | 61 | 79 | 0 | 25 | 61 | 18 | 1127 | 125 |
| SRR5644086 | 0 | 59 | 0 | 0 | 50 | 0 | 17 | 681 | 341 | 933 | 147 | 116 | 0 | 307 | 294 | 388 | 25 | 611 | 0 | 1031 | 10040 | 1313 | 259 | 5740 | 4904 | 0 | 27 | 0 | 0 | 22 | 13 | 0 | 20 | 22 | 18 | 18615 | 10 |
| SRR5644087 | 0 | 33 | 49 | 0 | 32 | 0 | 0 | 667 | 290 | 1938 | 23 | 33 | 0 | 149 | 98 | 215 | 0 | 347 | 0 | 372 | 1882 | 925 | 688 | 7309 | 9184 | 0 | 0 | 0 | 0 | 33 | 33 | 0 | 16 | 77 | 49 | 7041 | 0 |
| SRR5644088 | 0 | 85 | 48 | 0 | 36 | 0 | 0 | 560 | 704 | 2165 | 43 | 62 | 0 | 72 | 190 | 259 | 0 | 569 | 0 | 529 | 2108 | 1389 | 654 | 4961 | 8568 | 0 | 14 | 0 | 10 | 17 | 17 | 27 | 17 | 88 | 34 | 7606 | 0 |
| SRR5657537 | 0 | 0 | 0 | 0 | 13 | 0 | 0 | 9210 | 5301 | 5261 | 874 | 1070 | 0 | 665 | 14 | 15 | 0 | 246 | 0 | 4904 | 1908 | 2710 | 3343 | 19412 | 4166 | 31 | 0 | 0 | 18 | 20 | 30 | 0 | 13 | 27 | 0 | 47093 | 0 |
| SRR5657538 | 0 | 0 | 0 | 0 | 56 | 17 | 0 | 699 | 463 | 1694 | 475 | 743 | 0 | 139 | 0 | 0 | 12 | 418 | 0 | 3089 | 2050 | 1709 | 1187 | 14175 | 5231 | 10 | 0 | 0 | 0 | 0 | 13 | 0 | 0 | 0 | 0 | 21741 | 24 |
| SRR5657539 | 0 | 0 | 0 | 0 | 11 | 12 | 0 | 23380 | 19870 | 6144 | 883 | 1150 | 0 | 210 | 14 | 0 | 0 | 444 | 0 | 4692 | 1666 | 1766 | 3475 | 21546 | 5498 | 21 | 0 | 0 | 0 | 17 | 14 | 0 | 0 | 0 | 0 | 19324 | 0 |
| SRR5657540 | 0 | 32 | 0 | 0 | 22 | 21 | 0 | 1329 | 849 | 2336 | 557 | 2122 | 0 | 279 | 0 | 0 | 0 | 175 | 0 | 2559 | 2231 | 867 | 3816 | 30143 | 3202 | 0 | 21 | 0 | 0 | 46 | 29 | 0 | 0 | 0 | 0 | 27227 | 0 |
| SRR5657541 | 0 | 23 | 0 | 0 | 0 | 0 | 0 | 2823 | 1975 | 1676 | 864 | 1635 | 0 | 499 | 0 | 0 | 11 | 336 | 0 | 2134 | 2430 | 2180 | 4044 | 12094 | 4172 | 16 | 0 | 0 | 13 | 35 | 35 | 0 | 12 | 11 | 0 | 8595 | 16 |
| SRR5657542 | 0 | 22 | 0 | 0 | 49 | 18 | 0 | 314 | 384 | 5013 | 634 | 1446 | 0 | 380 | 0 | 0 | 0 | 356 | 0 | 1617 | 1200 | 637 | 213 | 2206 | 2073 | 0 | 15 | 0 | 10 | 32 | 15 | 0 | 0 | 0 | 0 | 11846 | 20 |
| SRR5657543 | 0 | 27 | 0 | 0 | 18 | 22 | 0 | 1534 | 1960 | 6192 | 1150 | 2005 | 0 | 329 | 0 | 0 | 20 | 333 | 0 | 2234 | 2910 | 1881 | 4551 | 16503 | 4893 | 0 | 0 | 0 | 0 | 14 | 12 | 0 | 0 | 0 | 0 | 6324 | 11 |
| SRR5657544 | 0 | 0 | 0 | 0 | 31 | 0 | 0 | 12595 | 4518 | 4534 | 95 | 90 | 0 | 704 | 0 | 0 | 0 | 627 | 18 | 327 | 1973 | 645 | 1087 | 8526 | 818 | 50 | 0 | 0 | 0 | 0 | 0 | 0 | 0 | 10 | 0 | 5034 | 0 |
| SRR5657545 | 0 | 550 | 0 | 0 | 34 | 0 | 0 | 827 | 1499 | 402 | 392 | 640 | 0 | 0 | 30 | 33 | 22 | 607 | 0 | 386 | 2830 | 27 | 4222 | 16807 | 43 | 12 | 20 | 16 | 11 | 0 | 0 | 0 | 0 | 0 | 0 | 2752 | 50 |
| SRR5657546 | 0 | 68 | 0 | 0 | 95 | 0 | 0 | 984 | 169 | 1237 | 241 | 822 | 0 | 94 | 0 | 0 | 0 | 428 | 0 | 427 | 1226 | 700 | 528 | 4945 | 1649 | 0 | 13 | 20 | 11 | 33 | 24 | 26 | 0 | 0 | 0 | 6497 | 64 |
| SRR5658693 | 0 | 0 | 29 | 0 | 17 | 0 | 11 | 5793 | 849 | 3753 | 472 | 2247 | 0 | 275 | 0 | 0 | 0 | 121 | 0 | 388 | 329 | 893 | 0 | 20 | 2368 | 33 | 11 | 0 | 0 | 15 | 28 | 0 | 21 | 58 | 21 | 1123 | 0 |
| SRR5658694 | 0 | 0 | 44 | 0 | 15 | 0 | 0 | 3849 | 1233 | 3494 | 2127 | 1766 | 0 | 311 | 0 | 0 | 0 | 136 | 22 | 528 | 401 | 1109 | 0 | 21 | 2763 | 49 | 0 | 0 | 32 | 38 | 47 | 0 | 42 | 148 | 52 | 1572 | 0 |
| SRR5658695 | 0 | 0 | 32 | 0 | 16 | 0 | 0 | 4923 | 746 | 2401 | 2768 | 3093 | 0 | 259 | 0 | 0 | 0 | 168 | 11 | 259 | 423 | 991 | 0 | 26 | 2415 | 36 | 0 | 0 | 34 | 38 | 115 | 16 | 66 | 145 | 21 | 1276 | 0 |
| SRR5658696 | 0 | 0 | 31 | 0 | 25 | 0 | 0 | 4248 | 864 | 2924 | 3041 | 2265 | 0 | 238 | 0 | 0 | 0 | 124 | 0 | 354 | 569 | 1381 | 0 | 30 | 3496 | 53 | 0 | 0 | 32 | 46 | 78 | 11 | 51 | 167 | 28 | 1401 | 0 |
| SRR5658697 | 0 | 0 | 42 | 0 | 24 | 0 | 0 | 3659 | 893 | 2992 | 4436 | 3250 | 0 | 352 | 0 | 0 | 0 | 155 | 11 | 378 | 699 | 1721 | 0 | 37 | 3661 | 55 | 0 | 13 | 85 | 11 | 16 | 14 | 10 | 25 | 37 | 1958 | 0 |
| SRR5687455 | 0 | 0 | 0 | 0 | 22 | 0 | 0 | 2201 | 313 | 28760 | 224 | 251 | 0 | 264 | 25 | 159 | 0 | 32 | 0 | 7956 | 654 | 4032 | 69 | 51172 | 13917 | 319 | 0 | 80 | 52 | 122 | 378 | 27 | 70 | 132 | 0 | 26035 | 46 |
| SRR5687456 | 0 | 0 | 0 | 0 | 25 | 0 | 14 | 3126 | 339 | 31757 | 212 | 177 | 0 | 243 | 31 | 243 | 0 | 32 | 0 | 6172 | 664 | 4000 | 75 | 57375 | 15194 | 320 | 20 | 71 | 54 | 241 | 528 | 37 | 120 | 198 | 0 | 25075 | 51 |
| SRR5687457 | 0 | 0 | 0 | 0 | 18 | 0 | 12 | 2578 | 232 | 26886 | 43 | 178 | 0 | 383 | 39 | 85 | 0 | 37 | 0 | 8811 | 516 | 3054 | 62 | 50759 | 12009 | 283 | 18 | 68 | 55 | 39 | 76 | 36 | 17 | 34 | 0 | 42975 | 32 |
| SRR5687458 | 0 | 0 | 0 | 0 | 21 | 0 | 0 | 1868 | 321 | 25697 | 181 | 284 | 0 | 160 | 53 | 177 | 0 | 24 | 0 | 7040 | 591 | 3222 | 70 | 47922 | 12855 | 316 | 19 | 107 | 56 | 232 | 485 | 21 | 70 | 127 | 0 | 30429 | 40 |
| SRR5687459 | 0 | 0 | 0 | 0 | 21 | 0 | 0 | 1923 | 416 | 24615 | 174 | 241 | 0 | 286 | 27 | 72 | 0 | 30 | 0 | 7382 | 659 | 3618 | 69 | 49861 | 13218 | 337 | 0 | 14 | 53 | 25 | 64 | 22 | 53 | 15 | 0 | 25317 | 43 |
| SRR5687461 | 0 | 0 | 0 | 0 | 37 | 0 | 0 | 2261 | 291 | 29769 | 390 | 268 | 0 | 480 | 0 | 42 | 15 | 29 | 0 | 1392 | 979 | 1978 | 87 | 55602 | 14295 | 128 | 11 | 32 | 73 | 161 | 294 | 10 | 22 | 44 | 0 | 4080 | 73 |
| SRR5687462 | 0 | 0 | 0 | 0 | 28 | 0 | 0 | 2082 | 337 | 34641 | 352 | 207 | 0 | 779 | 12 | 35 | 0 | 37 | 0 | 2974 | 874 | 2451 | 70 | 45249 | 11030 | 151 | 0 | 24 | 13 | 39 | 68 | 14 | 35 | 13 | 0 | 11830 | 60 |
| SRR5687463 | 0 | 0 | 0 | 0 | 27 | 0 | 0 | 2109 | 384 | 30740 | 139 | 249 | 0 | 794 | 13 | 13 | 12 | 36 | 0 | 3440 | 857 | 2371 | 66 | 43954 | 10637 | 159 | 0 | 30 | 15 | 58 | 85 | 12 | 32 | 11 | 0 | 10937 | 54 |
| SRR5687464 | 0 | 0 | 0 | 0 | 34 | 0 | 0 | 1535 | 120 | 14371 | 75 | 314 | 0 | 246 | 10 | 10 | 0 | 35 | 0 | 1933 | 764 | 1708 | 65 | 46527 | 10006 | 137 | 21 | 42 | 64 | 121 | 167 | 0 | 12 | 25 | 0 | 5892 | 40 |
| SRR5687465 | 0 | 0 | 0 | 0 | 26 | 0 | 0 | 15730 | 3656 | 48889 | 67 | 50 | 0 | 744 | 0 | 0 | 0 | 44 | 0 | 2402 | 597 | 3271 | 75 | 49637 | 9694 | 265 | 14 | 37 | 201 | 418 | 745 | 43 | 168 | 285 | 0 | 19922 | 36 |
| SRR5687466 | 0 | 0 | 0 | 0 | 24 | 0 | 0 | 22113 | 4032 | 48033 | 58 | 54 | 0 | 454 | 0 | 18 | 14 | 46 | 0 | 1993 | 565 | 2827 | 58 | 56244 | 10162 | 242 | 11 | 40 | 40 | 72 | 114 | 36 | 24 | 38 | 0 | 17817 | 50 |
| SRR5687467 | 0 | 0 | 0 | 0 | 21 | 0 | 12 | 21250 | 4047 | 51731 | 44 | 35 | 0 | 484 | 0 | 0 | 0 | 38 | 0 | 2377 | 613 | 3087 | 54 | 51362 | 9135 | 234 | 14 | 42 | 220 | 417 | 683 | 44 | 186 | 301 | 0 | 22275 | 41 |
| SRR5687468 | 0 | 0 | 0 | 0 | 32 | 0 | 0 | 21366 | 3101 | 47079 | 51 | 62 | 0 | 220 | 0 | 0 | 0 | 51 | 0 | 1230 | 475 | 1646 | 79 | 74508 | 11933 | 167 | 14 | 29 | 188 | 548 | 636 | 20 | 115 | 105 | 0 | 11136 | 83 |
| SRR5687469 | 0 | 0 | 0 | 0 | 29 | 0 | 0 | 17684 | 2870 | 46298 | 74 | 61 | 0 | 394 | 0 | 0 | 0 | 42 | 0 | 2882 | 676 | 2730 | 65 | 53517 | 9724 | 224 | 13 | 11 | 43 | 86 | 117 | 43 | 18 | 37 | 0 | 23371 | 61 |
| SRR5753916 | 0 | 195 | 25 | 209 | 926 | 0 | 20 | 495 | 1904 | 35162 | 98743 | 27304 | 0 | 2509 | 25 | 48 | 0 | 1759 | 786 | 160 | 856 | 4390 | 684 | 34699 | 2250 | 24 | 12 | 108 | 227 | 745 | 393 | 43 | 46 | 45 | 54 | 304 | 1295 |
| SRR5755813 | 0 | 30 | 136 | 0 | 65 | 0 | 0 | 513 | 54 | 324 | 9959 | 17341 | 0 | 13 | 0 | 13 | 0 | 114 | 0 | 3617 | 1940 | 2467 | 0 | 0 | 8743 | 0 | 156 | 10 | 101 | 212 | 96 | 10 | 60 | 181 | 76 | 47406 | 0 |
| SRR5755814 | 0 | 26 | 104 | 0 | 33 | 0 | 0 | 146 | 0 | 513 | 224 | 28923 | 0 | 0 | 0 | 0 | 0 | 245 | 0 | 952 | 2641 | 3284 | 0 | 17 | 9554 | 0 | 26 | 26 | 11 | 14 | 130 | 0 | 78 | 156 | 26 | 44812 | 0 |

### Supp Table S8.4.Columns_DI_to_ES

| | | | | | | | | | | | | | | | | | | | | | | | | | | | | | | | | | | | | | |
|---|---|---|---|---|---|---|---|---|---|---|---|---|---|---|---|---|---|---|---|---|---|---|---|---|---|---|---|---|---|---|---|---|---|---|---|---|---|
| SRR5755815 | 35 | 105 | 0 | 0 | 18 | 0 | 0 | 116 | 0 | 290 | 11445 | 20392 | 0 | 0 | 0 | 0 | 0 | 828 | 0 | 116 | 1949 | 6897 | 0 | 39 | 20152 | 53 | 15 | 0 | 35 | 16 | 10 | 0 | 175 | 209 | 70 | 4183 | 0 |
| SRR5755818 | 0 | 12 | 12 | 0 | 17 | 0 | 0 | 260 | 71 | 2413 | 569 | 9546 | 0 | 47 | 0 | 12 | 0 | 289 | 0 | 2922 | 1058 | 3119 | 0 | 0 | 8874 | 0 | 81 | 0 | 50 | 112 | 54 | 0 | 27 | 19 | 19 | 75315 | 0 |
| SRR5755819 | 0 | 0 | 67 | 0 | 35 | 0 | 0 | 439 | 165 | 0 | 600 | 17156 | 0 | 0 | 27 | 27 | 0 | 137 | 0 | 3459 | 1593 | 2112 | 0 | 0 | 7363 | 0 | 11 | 11 | 90 | 10 | 123 | 0 | 90 | 190 | 67 | 240976 | 0 |
| SRR5755820 | 0 | 0 | 177 | 0 | 0 | 29 | 0 | 144 | 144 | 288 | 437 | 34301 | 0 | 0 | 0 | 0 | 0 | 178 | 0 | 2306 | 2251 | 2460 | 0 | 0 | 8099 | 0 | 29 | 0 | 177 | 471 | 59 | 0 | 147 | 236 | 29 | 173092 | 0 |
| SRR5755821 | 0 | 60 | 20 | 0 | 21 | 0 | 0 | 401 | 60 | 220 | 7073 | 10960 | 0 | 40 | 20 | 80 | 30 | 909 | 13 | 120 | 2833 | 5127 | 0 | 41 | 23104 | 23 | 139 | 0 | 90 | 21 | 149 | 0 | 50 | 100 | 110 | 2064 | 0 |
| SRR5755824 | 0 | 0 | 56 | 0 | 19 | 0 | 0 | 780 | 98 | 2146 | 450 | 25461 | 0 | 98 | 0 | 0 | 0 | 305 | 0 | 1171 | 1053 | 3345 | 0 | 11 | 8406 | 0 | 0 | 0 | 56 | 197 | 28 | 0 | 14 | 85 | 28 | 50288 | 0 |
| SRR5755828 | 0 | 19 | 23 | 0 | 19 | 0 | 0 | 683 | 163 | 601 | 6826 | 17912 | 0 | 61 | 61 | 82 | 0 | 1749 | 15 | 71 | 3660 | 4729 | 0 | 21 | 22516 | 19 | 84 | 0 | 136 | 262 | 90 | 0 | 29 | 90 | 48 | 846 | 0 |
| SRR5755830 | 0 | 11 | 109 | 0 | 24 | 0 | 0 | 340 | 0 | 534 | 10640 | 34495 | 0 | 243 | 146 | 146 | 0 | 88 | 0 | 146 | 2727 | 4673 | 0 | 156 | 14076 | 0 | 11 | 0 | 434 | 727 | 206 | 11 | 98 | 282 | 109 | 1263 | 0 |
| SRR5755831 | 0 | 0 | 28 | 0 | 16 | 0 | 0 | 1114 | 134 | 2651 | 57 | 1894 | 0 | 290 | 22 | 45 | 0 | 544 | 0 | 67 | 803 | 304 | 0 | 0 | 11205 | 0 | 45 | 0 | 38 | 11 | 31 | 17 | 10 | 38 | 55 | 1225 | 0 |
| SRR5755833 | 0 | 18 | 10 | 0 | 67 | 0 | 0 | 495 | 71 | 310 | 761 | 14503 | 0 | 18 | 0 | 0 | 0 | 93 | 0 | 982 | 1740 | 2181 | 0 | 0 | 8384 | 0 | 18 | 12 | 94 | 12 | 12 | 0 | 118 | 16 | 153 | 6349 | 0 |
| SRR5755835 | 0 | 25 | 19 | 0 | 26 | 0 | 0 | 79 | 11 | 192 | 1014 | 10586 | 0 | 23 | 19 | 26 | 13 | 505 | 13 | 165 | 1729 | 4991 | 0 | 25 | 20676 | 10 | 181 | 13 | 75 | 11 | 244 | 31 | 50 | 219 | 50 | 1646 | 0 |
| SRR5755837 | 0 | 23 | 12 | 0 | 29 | 0 | 0 | 305 | 54 | 305 | 8508 | 21889 | 0 | 161 | 36 | 0 | 0 | 34 | 0 | 2436 | 2270 | 4109 | 0 | 140 | 13586 | 0 | 15 | 0 | 13 | 19 | 70 | 14 | 47 | 22 | 80 | 11733 | 0 |
| SRR5755838 | 0 | 42 | 21 | 0 | 16 | 0 | 0 | 263 | 84 | 675 | 3720 | 9011 | 0 | 74 | 0 | 16 | 0 | 337 | 0 | 395 | 1487 | 3338 | 0 | 0 | 9579 | 0 | 54 | 0 | 67 | 121 | 96 | 0 | 59 | 100 | 33 | 3348 | 0 |
| SRR5755842 | 0 | 30 | 60 | 0 | 18 | 0 | 0 | 168 | 149 | 130 | 169 | 8589 | 15 | 19 | 298 | 540 | 0 | 725 | 0 | 429 | 1713 | 6710 | 0 | 32 | 19418 | 41 | 105 | 0 | 90 | 18 | 135 | 0 | 90 | 15 | 10 | 1602 | 0 |
| SRR5755844 | 0 | 17 | 67 | 0 | 35 | 0 | 0 | 82 | 0 | 247 | 4533 | 12198 | 0 | 0 | 247 | 247 | 0 | 14 | 0 | 1401 | 1641 | 4312 | 0 | 126 | 12248 | 0 | 33 | 17 | 33 | 133 | 133 | 0 | 67 | 183 | 83 | 8407 | 0 |
| SRR5755847 | 0 | 0 | 75 | 0 | 60 | 0 | 0 | 451 | 90 | 541 | 5319 | 12170 | 0 | 90 | 0 | 90 | 0 | 60 | 0 | 180 | 1554 | 3014 | 0 | 0 | 8072 | 0 | 0 | 0 | 37 | 224 | 149 | 37 | 75 | 299 | 149 | 1893 | 0 |
| SRR5755849 | 0 | 31 | 16 | 0 | 29 | 0 | 0 | 239 | 22 | 261 | 1511 | 4447 | 0 | 22 | 76 | 163 | 0 | 769 | 21 | 141 | 1835 | 4675 | 0 | 45 | 22892 | 37 | 16 | 16 | 93 | 14 | 109 | 31 | 78 | 124 | 10 | 609 | 0 |
| SRR5755851 | 0 | 36 | 83 | 0 | 16 | 0 | 12 | 506 | 0 | 276 | 68 | 9932 | 0 | 322 | 0 | 92 | 12 | 48 | 0 | 2253 | 1763 | 4590 | 11 | 32 | 13796 | 0 | 12 | 0 | 83 | 130 | 130 | 0 | 12 | 18 | 11 | 7311 | 0 |
| SRR5755854 | 0 | 34 | 103 | 0 | 41 | 0 | 0 | 184 | 31 | 490 | 2053 | 3569 | 0 | 92 | 0 | 0 | 0 | 280 | 0 | 5514 | 1885 | 2776 | 0 | 0 | 10867 | 0 | 68 | 0 | 188 | 325 | 188 | 17 | 85 | 188 | 103 | 70092 | 0 |
| SRR5755855 | 0 | 41 | 142 | 0 | 31 | 0 | 0 | 301 | 65 | 840 | 3122 | 6439 | 20 | 108 | 0 | 0 | 0 | 276 | 0 | 3402 | 2721 | 2962 | 0 | 0 | 12823 | 0 | 20 | 0 | 142 | 12 | 81 | 20 | 101 | 142 | 203 | 37749 | 0 |
| SRR5755856 | 0 | 109 | 95 | 0 | 21 | 0 | 0 | 113 | 0 | 366 | 8040 | 14170 | 14 | 0 | 122 | 113 | 0 | 838 | 20 | 166 | 2562 | 6057 | 0 | 28 | 23180 | 17 | 136 | 0 | 122 | 380 | 177 | 27 | 149 | 136 | 109 | 1325 | 0 |
| SRR5755858 | 0 | 19 | 99 | 0 | 14 | 0 | 0 | 253 | 14 | 380 | 398 | 14695 | 14 | 98 | 14 | 42 | 0 | 43 | 0 | 5273 | 2389 | 3715 | 35 | 123 | 13265 | 0 | 202 | 0 | 113 | 169 | 61 | 0 | 33 | 80 | 56 | 33355 | 0 |
| SRR5755861 | 0 | 33 | 154 | 0 | 28 | 0 | 0 | 163 | 25 | 733 | 2482 | 4312 | 0 | 107 | 13 | 13 | 0 | 253 | 0 | 9733 | 2759 | 3382 | 0 | 0 | 8884 | 0 | 110 | 0 | 44 | 143 | 93 | 11 | 49 | 115 | 99 | 94813 | 0 |
| SRR5755862 | 0 | 0 | 118 | 0 | 23 | 0 | 0 | 227 | 32 | 982 | 308 | 5488 | 0 | 260 | 0 | 32 | 0 | 163 | 0 | 6665 | 3722 | 2190 | 0 | 12 | 10317 | 0 | 135 | 17 | 93 | 15 | 59 | 0 | 42 | 118 | 25 | 58190 | 0 |
| SRR5755863 | 0 | 76 | 118 | 0 | 0 | 0 | 0 | 211 | 22 | 592 | 1725 | 3894 | 17 | 15 | 48 | 133 | 0 | 814 | 19 | 592 | 2274 | 6891 | 0 | 34 | 25476 | 13 | 13 | 17 | 109 | 13 | 252 | 67 | 101 | 227 | 67 | 3639 | 0 |
| SRR5755865 | 0 | 13 | 52 | 0 | 25 | 0 | 0 | 881 | 39 | 1077 | 108 | 7286 | 0 | 98 | 98 | 98 | 0 | 33 | 10 | 5406 | 2187 | 4103 | 0 | 75 | 15729 | 0 | 12 | 0 | 45 | 109 | 32 | 0 | 58 | 11 | 64 | 29339 | 0 |
| SRR5755866 | 0 | 24 | 71 | 0 | 0 | 0 | 0 | 493 | 81 | 4691 | 644 | 4990 | 0 | 104 | 0 | 52 | 0 | 409 | 0 | 2632 | 1161 | 3994 | 0 | 0 | 8990 | 0 | 59 | 0 | 71 | 153 | 94 | 0 | 35 | 141 | 94 | 28424 | 0 |
| SRR5755868 | 0 | 26 | 157 | 0 | 13 | 0 | 0 | 214 | 43 | 363 | 401 | 10089 | 0 | 64 | 0 | 21 | 0 | 182 | 0 | 9362 | 5015 | 8065 | 0 | 0 | 4589 | 0 | 118 | 0 | 0 | 26 | 131 | 39 | 26 | 144 | 105 | 174304 | 0 |
| SRR5755870 | 0 | 39 | 59 | 0 | 11 | 0 | 0 | 204 | 35 | 915 | 769 | 4798 | 0 | 22 | 22 | 48 | 0 | 960 | 0 | 335 | 3676 | 10129 | 0 | 22 | 16248 | 15 | 379 | 0 | 65 | 190 | 216 | 26 | 105 | 275 | 124 | 3019 | 0 |
| SRR5755875 | 0 | 67 | 172 | 0 | 63 | 0 | 0 | 631 | 47 | 1722 | 332 | 2931 | 0 | 118 | 0 | 0 | 0 | 491 | 0 | 1215 | 2280 | 1702 | 0 | 83 | 10398 | 0 | 22 | 0 | 45 | 90 | 75 | 15 | 67 | 97 | 104 | 13223 | 0 |
| SRR5755876 | 0 | 22 | 88 | 0 | 28 | 0 | 0 | 712 | 135 | 1703 | 1251 | 1357 | 0 | 298 | 0 | 0 | 0 | 360 | 0 | 375 | 2507 | 1473 | 0 | 55 | 14 | 0 | 0 | 0 | 29 | 133 | 125 | 15 | 74 | 133 | 88 | 2502 | 0 |
| SRR5755877 | 0 | 45 | 51 | 0 | 16 | 0 | 0 | 122 | 26 | 372 | 2576 | 3926 | 0 | 0 | 75 | 142 | 12 | 1249 | 0 | 201 | 3005 | 8153 | 24 | 3653 | 23203 | 0 | 391 | 0 | 149 | 335 | 221 | 54 | 78 | 212 | 123 | 1524 | 0 |
| SRR5755879 | 0 | 19 | 96 | 0 | 15 | 0 | 0 | 134 | 38 | 414 | 4154 | 11130 | 0 | 210 | 57 | 89 | 0 | 71 | 0 | 4715 | 3149 | 6356 | 102 | 995 | 21 | 0 | 13 | 0 | 111 | 12 | 55 | 0 | 19 | 10 | 60 | 36022 | 0 |
| SRR5755880 | 0 | 25 | 25 | 0 | 17 | 0 | 0 | 286 | 61 | 1735 | 998 | 1936 | 0 | 109 | 0 | 21 | 0 | 446 | 0 | 551 | 1338 | 2091 | 62 | 1269 | 11937 | 0 | 47 | 0 | 76 | 136 | 110 | 0 | 38 | 144 | 72 | 6833 | 0 |
| SRR5755882 | 0 | 38 | 76 | 0 | 91 | 0 | 0 | 350 | 27 | 364 | 191 | 4649 | 0 | 27 | 0 | 0 | 0 | 207 | 0 | 94 | 2015 | 1421 | 0 | 0 | 7306 | 0 | 13 | 0 | 76 | 166 | 89 | 13 | 38 | 127 | 38 | 1172 | 0 |
| SRR5755884 | 0 | 150 | 20 | 0 | 14 | 0 | 0 | 159 | 49 | 55 | 5248 | 12727 | 30 | 0 | 46 | 108 | 0 | 819 | 0 | 108 | 2230 | 5045 | 0 | 14 | 48 | 0 | 90 | 0 | 125 | 399 | 198 | 23 | 68 | 201 | 128 | 480 | 0 |
| SRR5755887 | 0 | 32 | 16 | 0 | 0 | 0 | 0 | 221 | 70 | 1265 | 217 | 3390 | 0 | 12 | 0 | 12 | 0 | 230 | 0 | 128 | 872 | 1270 | 0 | 0 | 8944 | 0 | 0 | 0 | 65 | 65 | 48 | 32 | 0 | 113 | 97 | 952 | 0 |
| SRR5755889 | 0 | 27 | 62 | 0 | 62 | 0 | 0 | 133 | 25 | 630 | 3749 | 7306 | 0 | 17 | 0 | 17 | 0 | 67 | 0 | 581 | 2169 | 3691 | 0 | 0 | 10613 | 0 | 96 | 21 | 82 | 274 | 246 | 0 | 62 | 205 | 75 | 2695 | 0 |
| SRR5755890 | 0 | 62 | 132 | 0 | 13 | 0 | 0 | 37 | 31 | 566 | 3730 | 7789 | 0 | 31 | 0 | 0 | 0 | 284 | 0 | 410 | 3161 | 4509 | 0 | 0 | 8407 | 0 | 155 | 0 | 108 | 333 | 147 | 31 | 116 | 302 | 46 | 1324 | 0 |
| SRR5755891 | 0 | 49 | 74 | 0 | 10 | 0 | 0 | 92 | 15 | 144 | 2049 | 4613 | 12 | 10 | 13 | 22 | 0 | 1174 | 14 | 113 | 4219 | 13759 | 0 | 10 | 11612 | 16 | 129 | 0 | 61 | 92 | 178 | 55 | 105 | 320 | 111 | 459 | 0 |
| SRR5755894 | 0 | 23 | 46 | 0 | 0 | 0 | 0 | 479 | 319 | 4252 | 6632 | 12828 | 0 | 36 | 29 | 44 | 0 | 424 | 0 | 1241 | 3041 | 10965 | 0 | 0 | 7347 | 0 | 83 | 0 | 32 | 46 | 120 | 32 | 46 | 120 | 69 | 6777 | 0 |
| SRR5755896 | 0 | 111 | 258 | 0 | 64 | 0 | 0 | 128 | 25 | 324 | 4810 | 9871 | 0 | 103 | 0 | 0 | 0 | 404 | 0 | 118 | 1993 | 1288 | 0 | 0 | 12 | 0 | 41 | 35 | 276 | 16 | 252 | 29 | 106 | 211 | 59 | 545 | 0 |
| SRR5755897 | 0 | 40 | 96 | 0 | 22 | 0 | 0 | 161 | 37 | 784 | 966 | 1758 | 0 | 128 | 0 | 0 | 0 | 485 | 0 | 33 | 1764 | 750 | 0 | 12 | 8770 | 0 | 0 | 0 | 128 | 295 | 128 | 40 | 80 | 80 | 104 | 376 | 0 |
| SRR5755898 | 0 | 72 | 101 | 0 | 16 | 0 | 0 | 330 | 86 | 115 | 538 | 20189 | 14 | 14 | 72 | 115 | 0 | 474 | 0 | 373 | 1632 | 11 | 0 | 20 | 23383 | 12 | 12 | 14 | 10 | 27 | 12 | 14 | 115 | 13 | 101 | 1934 | 0 |
| SRR5755901 | 0 | 35 | 31 | 0 | 11 | 0 | 0 | 170 | 38 | 1006 | 1631 | 2658 | 0 | 79 | 0 | 0 | 0 | 520 | 0 | 53 | 1021 | 1399 | 0 | 0 | 7767 | 0 | 40 | 0 | 106 | 278 | 84 | 13 | 84 | 119 | 93 | 339 | 0 |
| SRR5755903 | 0 | 89 | 129 | 0 | 72 | 0 | 0 | 66 | 13 | 296 | 2558 | 5256 | 0 | 90 | 0 | 0 | 0 | 259 | 0 | 117 | 2147 | 1381 | 0 | 0 | 7430 | 0 | 94 | 0 | 124 | 11 | 174 | 0 | 47 | 155 | 80 | 1032 | 0 |
| SRR5755905 | 0 | 140 | 42 | 0 | 11 | 0 | 0 | 94 | 16 | 158 | 1902 | 5394 | 0 | 21 | 0 | 26 | 0 | 1056 | 13 | 99 | 2430 | 4502 | 0 | 17 | 34 | 0 | 155 | 0 | 159 | 390 | 254 | 76 | 61 | 174 | 87 | 647 | 0 |
| SRR5755908 | 0 | 40 | 0 | 0 | 13 | 0 | 0 | 142 | 29 | 808 | 492 | 575 | 0 | 174 | 0 | 0 | 16 | 416 | 0 | 0 | 1171 | 818 | 0 | 0 | 8526 | 0 | 119 | 0 | 48 | 119 | 40 | 16 | 0 | 119 | 48 | 262 | 0 |
| SRR5755910 | 0 | 89 | 163 | 0 | 92 | 0 | 0 | 64 | 39 | 762 | 911 | 1091 | 0 | 207 | 0 | 0 | 0 | 370 | 0 | 26 | 1493 | 586 | 0 | 0 | 9433 | 0 | 16 | 0 | 71 | 160 | 114 | 0 | 54 | 73 | 68 | 266 | 0 |
| SRR5755911 | 0 | 16 | 149 | 0 | 39 | 0 | 0 | 140 | 63 | 2636 | 149 | 339 | 0 | 595 | 0 | 0 | 0 | 421 | 0 | 14 | 379 | 106 | 0 | 0 | 11985 | 0 | 0 | 0 | 86 | 133 | 39 | 0 | 63 | 102 | 117 | 218 | 0 |
| SRR5755912 | 0 | 131 | 52 | 0 | 15 | 0 | 0 | 103 | 17 | 163 | 3184 | 5717 | 0 | 15 | 0 | 0 | 0 | 963 | 16 | 235 | 1847 | 4641 | 0 | 13 | 40 | 0 | 194 | 14 | 111 | 11 | 207 | 31 | 83 | 152 | 104 | 1347 | 0 |
| SRR5755914 | 0 | 50 | 54 | 0 | 13 | 0 | 0 | 178 | 20 | 217 | 7245 | 19444 | 0 | 39 | 39 | 20 | 0 | 56 | 0 | 1579 | 2256 | 4607 | 0 | 44 | 13092 | 0 | 105 | 0 | 84 | 184 | 42 | 0 | 42 | 176 | 29 | 16127 | 0 |
| SRR5755915 | 0 | 40 | 28 | 0 | 21 | 0 | 0 | 98 | 68 | 2425 | 199 | 153 | 0 | 311 | 0 | 0 | 0 | 732 | 0 | 0 | 203 | 150 | 0 | 0 | 11765 | 0 | 17 | 17 | 57 | 137 | 74 | 17 | 23 | 85 | 114 | 104 | 0 |
| SRR5755917 | 0 | 86 | 151 | 0 | 50 | 0 | 0 | 457 | 102 | 173 | 12879 | 29244 | 0 | 0 | 102 | 152 | 14 | 418 | 12 | 254 | 1780 | 3713 | 0 | 0 | 15570 | 0 | 65 | 22 | 165 | 19 | 11 | 36 | 86 | 201 | 179 | 1921 | 0 |
| SRR5755918 | 0 | 27 | 11 | 0 | 26 | 14 | 0 | 641 | 51 | 987 | 5509 | 9788 | 0 | 26 | 13 | 51 | 0 | 118 | 0 | 333 | 2386 | 3070 | 0 | 0 | 9739 | 0 | 0 | 0 | 129 | 12 | 88 | 14 | 81 | 217 | 142 | 1102 | 0 |
| SRR5755919 | 0 | 49 | 97 | 0 | 16 | 19 | 0 | 363 | 25 | 88 | 4649 | 13378 | 0 | 0 | 94 | 250 | 0 | 435 | 15 | 175 | 1905 | 8622 | 0 | 20 | 25278 | 11 | 87 | 0 | 97 | 369 | 282 | 0 | 117 | 369 | 78 | 763 | 0 |
| SRR5755922 | 0 | 23 | 63 | 0 | 18 | 0 | 0 | 868 | 283 | 3703 | 6650 | 12624 | 0 | 50 | 61 | 172 | 0 | 310 | 0 | 252 | 1112 | 3669 | 0 | 0 | 10902 | 0 | 17 | 0 | 40 | 126 | 69 | 0 | 69 | 109 | 91 | 1625 | 0 |
| SRR5755924 | 0 | 18 | 243 | 0 | 43 | 12 | 0 | 1371 | 225 | 281 | 900 | 12916 | 0 | 0 | 11 | 22 | 0 | 71 | 0 | 753 | 1884 | 2756 | 0 | 0 | 9864 | 0 | 0 | 0 | 107 | 228 | 125 | 0 | 86 | 228 | 104 | 3676 | 0 |
| SRR5755925 | 0 | 51 | 250 | 0 | 15 | 0 | 0 | 1316 | 114 | 878 | 9596 | 13354 | 0 | 0 | 0 | 19 | 0 | 100 | 0 | 687 | 2268 | 3462 | 0 | 0 | 9752 | 0 | 11 | 0 | 114 | 290 | 125 | 0 | 46 | 171 | 80 | 4712 | 0 |
| SRR5755926 | 0 | 47 | 58 | 0 | 19 | 16 | 0 | 479 | 134 | 51 | 682 | 9058 | 0 | 0 | 0 | 65 | 0 | 430 | 11 | 221 | 1735 | 6590 | 0 | 21 | 21349 | 0 | 32 | 0 | 110 | 12 | 147 | 26 | 100 | 11 | 105 | 1056 | 0 |
| SRR5755928 | 0 | 21 | 124 | 0 | 25 | 0 | 0 | 1188 | 160 | 297 | 5801 | 14686 | 0 | 46 | 0 | 46 | 0 | 87 | 0 | 1096 | 2394 | 5807 | 0 | 85 | 14751 | 0 | 32 | 0 | 86 | 140 | 86 | 0 | 11 | 107 | 59 | 7400 | 0 |
| SRR5755929 | 0 | 0 | 74 | 0 | 14 | 0 | 0 | 803 | 349 | 4819 | 10022 | 15504 | 0 | 35 | 0 | 70 | 0 | 115 | 0 | 454 | 1005 | 4774 | 0 | 0 | 10602 | 0 | 0 | 0 | 25 | 136 | 37 | 0 | 50 | 99 | 25 | 2933 | 0 |
| SRR5755931 | 0 | 13 | 159 | 0 | 56 | 0 | 0 | 219 | 68 | 506 | 475 | 3801 | 0 | 46 | 0 | 0 | 0 | 200 | 0 | 615 | 1941 | 3104 | 0 | 0 | 8902 | 0 | 35 | 13 | 88 | 203 | 97 | 18 | 88 | 229 | 128 | 3254 | 0 |
| SRR5755933 | 0 | 50 | 141 | 0 | 29 | 0 | 30 | 135 | 15 | 176 | 2127 | 4018 | 10 | 15 | 0 | 30 | 0 | 646 | 16 | 331 | 2216 | 8304 | 0 | 30 | 23167 | 0 | 71 | 0 | 101 | 141 | 161 | 10 | 91 | 282 | 81 | 1525 | 0 |
| SRR5755936 | 0 | 70 | 44 | 0 | 14 | 0 | 0 | 178 | 36 | 718 | 1060 | 1018 | 0 | 82 | 0 | 11 | 0 | 481 | 0 | 42 | 1589 | 3094 | 0 | 0 | 12494 | 0 | 36 | 0 | 15 | 62 | 44 | 15 | 44 | 131 | 95 | 440 | 0 |
| SRR5755938 | 0 | 59 | 238 | 0 | 51 | 0 | 0 | 323 | 41 | 249 | 4210 | 6597 | 0 | 33 | 0 | 0 | 0 | 123 | 0 | 489 | 1921 | 2266 | 0 | 0 | 10758 | 0 | 20 | 20 | 172 | 495 | 145 | 20 | 106 | 271 | 92 | 3290 | 0 |
| SRR5755939 | 0 | 103 | 194 | 0 | 31 | 0 | 0 | 210 | 31 | 462 | 1191 | 3500 | 0 | 43 | 0 | 0 | 0 | 157 | 0 | 223 | 2837 | 2027 | 0 | 0 | 11913 | 0 | 36 | 0 | 94 | 326 | 183 | 0 | 49 | 234 | 149 | 1349 | 0 |
| SRR5755940 | 0 | 78 | 102 | 0 | 13 | 0 | 0 | 241 | 42 | 120 | 9631 | 22736 | 24 | 0 | 12 | 90 | 0 | 736 | 0 | 229 | 1880 | 5990 | 0 | 26 | 21982 | 10 | 141 | 0 | 12 | 30 | 17 | 63 | 71 | 15 | 165 | 1379 | 0 |
| SRR5755943 | 0 | 60 | 60 | 0 | 23 | 0 | 0 | 261 | 123 | 629 | 1973 | 5570 | 0 | 25 | 11 | 57 | 0 | 457 | 0 | 64 | 1593 | 3338 | 0 | 0 | 10141 | 0 | 0 | 0 | 80 | 292 | 136 | 20 | 93 | 169 | 139 | 457 | 0 |
| SRR5755946 | 0 | 15 | 12 | 0 | 26 | 0 | 0 | 696 | 105 | 717 | 318 | 12574 | 0 | 42 | 84 | 84 | 0 | 148 | 0 | 0 | 2470 | 2977 | 0 | 0 | 8555 | 0 | 15 | 0 | 206 | 14 | 175 | 0 | 84 | 221 | 183 | 0 | 0 |
| SRR5755947 | 16 | 91 | 58 | 0 | 0 | 0 | 0 | 295 | 88 | 88 | 3949 | 10689 | 0 | 0 | 104 | 651 | 0 | 802 | 12 | 0 | 2120 | 7940 | 0 | 27 | 22773 | 0 | 91 | 0 | 124 | 774 | 297 | 0 | 140 | 371 | 272 | 0 | 0 |
| SRR5755950 | 0 | 41 | 52 | 0 | 0 | 0 | 0 | 735 | 233 | 1668 | 5524 | 9003 | 0 | 0 | 0 | 143 | 0 | 473 | 0 | 0 | 1262 | 4005 | 0 | 17 | 8929 | 0 | 41 | 0 | 52 | 11 | 124 | 0 | 31 | 145 | 155 | 0 | 0 |
| SRR5755952 | 0 | 38 | 113 | 0 | 75 | 0 | 0 | 238 | 11 | 329 | 7219 | 11464 | 0 | 34 | 23 | 11 | 0 | 132 | 0 | 306 | 1592 | 2363 | 0 | 0 | 8487 | 0 | 28 | 19 | 179 | 19 | 169 | 0 | 104 | 11 | 10 | 2100 | 0 |
| SRR5755953 | 0 | 52 | 104 | 0 | 39 | 0 | 0 | 94 | 0 | 408 | 1939 | 3633 | 0 | 86 | 0 | 17 | 0 | 312 | 0 | 169 | 2389 | 2314 | 0 | 0 | 9968 | 0 | 36 | 10 | 52 | 249 | 83 | 16 | 47 | 223 | 176 | 780 | 0 |
| SRR5755954 | 0 | 83 | 0 | 0 | 19 | 0 | 17 | 86 | 25 | 123 | 4378 | 10821 | 0 | 12 | 0 | 172 | 0 | 740 | 19 | 344 | 1563 | 5961 | 0 | 44 | 24381 | 0 | 299 | 17 | 83 | 316 | 233 | 17 | 133 | 199 | 199 | 1808 | 0 |
| SRR5755956 | 0 | 48 | 85 | 0 | 16 | 0 | 0 | 271 | 71 | 343 | 4170 | 12054 | 0 | 43 | 71 | 428 | 0 | 99 | 0 | 2585 | 2473 | 5603 | 0 | 79 | 14592 | 0 | 115 | 0 | 71 | 182 | 59 | 0 | 41 | 160 | 85 | 11411 | 0 |
| SRR5755957 | 0 | 49 | 36 | 0 | 0 | 12 | 0 | 203 | 28 | 780 | 1011 | 1461 | 0 | 99 | 0 | 47 | 0 | 580 | 0 | 118 | 1099 | 1906 | 0 | 0 | 9441 | 0 | 12 | 0 | 85 | 170 | 73 | 0 | 36 | 170 | 109 | 383 | 0 |
| SRR5755959 | 0 | 38 | 236 | 0 | 58 | 0 | 0 | 529 | 81 | 278 | 1293 | 20059 | 0 | 27 | 63 | 45 | 0 | 93 | 0 | 377 | 2124 | 1837 | 0 | 0 | 8755 | 0 | 16 | 11 | 181 | 472 | 230 | 11 | 121 | 219 | 252 | 2313 | 0 |
| SRR5755960 | 0 | 35 | 11 | 0 | 25 | 0 | 0 | 178 | 32 | 356 | 2693 | 4084 | 0 | 172 | 13 | 38 | 0 | 264 | 0 | 114 | 3057 | 1913 | 0 | 0 | 8106 | 0 | 0 | 0 | 229 | 13 | 132 | 18 | 53 | 10 | 150 | 819 | 0 |
| SRR5755961 | 0 | 130 | 100 | 0 | 0 | 0 | 0 | 191 | 38 | 63 | 2064 | 13041 | 0 | 0 | 156 | 428 | 0 | 1305 | 14 | 325 | 2219 | 6860 | 0 | 35 | 24688 | 14 | 50 | 0 | 138 | 37 | 15 | 38 | 125 | 18 | 11 | 1645 | 0 |
| SRR5755963 | 0 | 0 | 134 | 0 | 25 | 0 | 0 | 832 | 200 | 233 | 202 | 15405 | 0 | 200 | 67 | 499 | 0 | 66 | 0 | 1264 | 2242 | 5782 | 0 | 127 | 12836 | 0 | 34 | 0 | 141 | 429 | 94 | 27 | 80 | 228 | 201 | 6555 | 0 |
| SRR5755964 | 0 | 0 | 80 | 0 | 11 | 0 | 0 | 293 | 90 | 712 | 598 | 3777 | 0 | 86 | 0 | 50 | 0 | 638 | 0 | 50 | 1517 | 2167 | 0 | 30 | 10099 | 30 | 45 | 0 | 98 | 12 | 134 | 0 | 98 | 214 | 276 | 446 | 0 |
| SRR5755966 | 0 | 15 | 221 | 0 | 57 | 0 | 0 | 269 | 35 | 467 | 495 | 6247 | 0 | 92 | 35 | 50 | 0 | 294 | 0 | 602 | 2041 | 2878 | 0 | 0 | 8362 | 0 | 15 | 0 | 80 | 321 | 151 | 0 | 65 | 216 | 201 | 1771 | 0 |
| SRR5755967 | 0 | 103 | 127 | 0 | 14 | 0 | 0 | 58 | 0 | 553 | 1629 | 3059 | 16 | 116 | 0 | 39 | 0 | 287 | 0 | 286 | 2603 | 3048 | 0 | 0 | 8650 | 0 | 32 | 40 | 40 | 206 | 135 | 0 | 71 | 254 | 190 | 645 | 0 |
| SRR5755968 | 0 | 40 | 29 | 0 | 25 | 0 | 0 | 183 | 26 | 144 | 8970 | 23785 | 0 | 13 | 13 | 118 | 12 | 508 | 0 | 400 | 2248 | 7201 | 0 | 16 | 18370 | 15 | 138 | 0 | 63 | 259 | 127 | 12 | 58 | 190 | 63 | 1520 | 0 |
| SRR5755971 | 0 | 12 | 27 | 0 | 14 | 0 | 0 | 324 | 143 | 1872 | 1732 | 15522 | 0 | 50 | 56 | 69 | 0 | 189 | 0 | 418 | 1143 | 2793 | 0 | 0 | 8291 | 0 | 24 | 0 | 42 | 118 | 54 | 0 | 27 | 63 | 33 | 1697 | 0 |
| SRR5755973 | 0 | 91 | 114 | 0 | 94 | 0 | 0 | 311 | 13 | 298 | 5848 | 10875 | 0 | 60 | 26 | 26 | 0 | 309 | 0 | 99 | 2008 | 1584 | 0 | 0 | 10013 | 0 | 15 | 0 | 106 | 227 | 129 | 15 | 91 | 205 | 99 | 549 | 0 |
| SRR5755974 | 0 | 16 | 62 | 0 | 57 | 0 | 0 | 174 | 38 | 480 | 2507 | 3598 | 0 | 52 | 0 | 52 | 0 | 326 | 0 | 42 | 1875 | 882 | 0 | 0 | 10736 | 0 | 29 | 0 | 41 | 94 | 53 | 0 | 33 | 53 | 57 | 447 | 0 |
| SRR5755975 | 0 | 136 | 54 | 0 | 23 | 0 | 0 | 149 | 47 | 91 | 13182 | 25409 | 0 | 15 | 55 | 138 | 0 | 1034 | 13 | 135 | 2757 | 6043 | 0 | 19 | 21121 | 12 | 165 | 0 | 165 | 22 | 12 | 33 | 99 | 144 | 120 | 757 | 0 |
| SRR5755978 | 0 | 34 | 17 | 0 | 15 | 0 | 0 | 146 | 31 | 608 | 381 | 5090 | 0 | 100 | 39 | 39 | 0 | 260 | 0 | 54 | 1070 | 786 | 0 | 11 | 9700 | 15 | 0 | 0 | 17 | 151 | 101 | 17 | 0 | 75 | 50 | 470 | 0 |
| SRR5755980 | 0 | 94 | 130 | 0 | 56 | 0 | 0 | 317 | 54 | 244 | 3063 | 18375 | 0 | 18 | 0 | 0 | 12 | 337 | 0 | 3843 | 3117 | 2076 | 0 | 0 | 7497 | 0 | 21 | 30 | 109 | 252 | 115 | 12 | 55 | 79 | 82 | 105287 | 0 |
| SRR5755981 | 0 | 40 | 223 | 0 | 29 | 0 | 0 | 520 | 87 | 395 | 1749 | 26689 | 0 | 58 | 0 | 0 | 0 | 319 | 0 | 1666 | 3580 | 2128 | 0 | 0 | 7399 | 0 | 0 | 25 | 184 | 323 | 144 | 0 | 65 | 79 | 124 | 42297 | 0 |
| SRR5755982 | 0 | 47 | 50 | 0 | 13 | 0 | 0 | 202 | 21 | 305 | 3222 | 20726 | 0 | 14 | 14 | 60 | 12 | 820 | 11 | 496 | 2675 | 6537 | 0 | 26 | 19192 | 12 | 24 | 0 | 122 | 14 | 151 | 15 | 95 | 163 | 95 | 5783 | 0 |
| SRR5755984 | 0 | 18 | 99 | 0 | 27 | 0 | 0 | 351 | 50 | 226 | 816 | 20392 | 0 | 75 | 0 | 17 | 0 | 61 | 0 | 2785 | 2873 | 4981 | 0 | 24 | 12743 | 0 | 139 | 0 | 68 | 10 | 58 | 0 | 22 | 65 | 71 | 46505 | 0 |
| SRR5755985 | 0 | 12 | 73 | 0 | 19 | 0 | 0 | 929 | 206 | 2858 | 1766 | 17010 | 0 | 57 | 0 | 0 | 0 | 361 | 0 | 312 | 1749 | 2209 | 0 | 0 | 8476 | 0 | 24 | 12 | 57 | 166 | 97 | 0 | 16 | 73 | 73 | 4027 | 0 |
| SRR5755987 | 0 | 62 | 84 | 0 | 93 | 0 | 0 | 287 | 29 | 504 | 6751 | 9971 | 0 | 74 | 0 | 0 | 0 | 346 | 0 | 139 | 2200 | 1715 | 0 | 12 | 7975 | 0 | 35 | 22 | 93 | 177 | 80 | 0 | 31 | 35 | 88 | 1591 | 0 |
| SRR5755988 | 15 | 39 | 112 | 0 | 34 | 0 | 0 | 366 | 54 | 941 | 2536 | 3370 | 0 | 150 | 0 | 0 | 0 | 530 | 0 | 54 | 1970 | 1058 | 0 | 0 | 10237 | 0 | 0 | 0 | 97 | 146 | 73 | 0 | 20 | 54 | 63 | 348 | 0 |
| SRR5755989 | 0 | 102 | 74 | 0 | 25 | 0 | 0 | 176 | 29 | 129 | 7343 | 16001 | 0 | 0 | 35 | 158 | 0 | 843 | 0 | 188 | 1920 | 5401 | 0 | 23 | 21832 | 15 | 157 | 37 | 157 | 19 | 17 | 74 | 55 | 11 | 185 | 1355 | 0 |
| SRR5755991 | 0 | 15 | 139 | 0 | 14 | 0 | 0 | 498 | 51 | 249 | 8886 | 21413 | 0 | 51 | 26 | 0 | 0 | 50 | 0 | 1310 | 2596 | 4548 | 22 | 127 | 11226 | 0 | 119 | 0 | 119 | 15 | 81 | 0 | 29 | 98 | 71 | 8720 | 0 |
| SRR5755992 | 0 | 30 | 17 | 0 | 16 | 0 | 0 | 518 | 75 | 1040 | 7308 | 8673 | 0 | 35 | 0 | 0 | 0 | 347 | 0 | 101 | 1265 | 1761 | 0 | 0 | 8144 | 0 | 0 | 0 | 30 | 124 | 30 | 0 | 13 | 34 | 34 | 1330 | 0 |
| SRR5755994 | 0 | 117 | 106 | 0 | 46 | 0 | 0 | 323 | 102 | 63 | 4012 | 19543 | 0 | 0 | 0 | 30 | 0 | 644 | 0 | 140 | 2566 | 3542 | 0 | 14 | 18942 | 0 | 17 | 22 | 263 | 687 | 196 | 11 | 61 | 218 | 296 | 714 | 0 |
| SRR5755995 | 0 | 33 | 209 | 0 | 24 | 0 | 0 | 445 | 133 | 491 | 2115 | 19172 | 0 | 25 | 0 | 0 | 19 | 408 | 0 | 175 | 2479 | 2778 | 0 | 0 | 11200 | 0 | 0 | 23 | 11 | 15 | 88 | 0 | 79 | 168 | 144 | 3552 | 0 |
| SRR5755996 | 0 | 189 | 61 | 0 | 20 | 0 | 0 | 127 | 43 | 14 | 8589 | 8571 | 0 | 0 | 0 | 12 | 0 | 1127 | 14 | 39 | 2659 | 5353 | 0 | 29 | 24628 | 12 | 58 | 32 | 324 | 937 | 313 | 16 | 104 | 218 | 210 | 335 | 0 |
| SRR5755997 | 0 | 0 | 0 | 0 | 0 | 0 | 0 | 551 | 193 | 2478 | 52 | 4846 | 0 | 83 | 0 | 28 | 37 | 1532 | 41 | 138 | 1054 | 3961 | 0 | 37 | 38111 | 0 | 205 | 0 | 28 | 0 | 19 | 0 | 0 | 0 | 0 | 220 | 0 |
| SRR5755998 | 0 | 19 | 100 | 0 | 17 | 0 | 0 | 620 | 26 | 149 | 5975 | 19218 | 0 | 122 | 0 | 26 | 0 | 29 | 0 | 341 | 2207 | 4571 | 18 | 129 | 13251 | 0 | 15 | 0 | 97 | 356 | 67 | 0 | 48 | 156 | 100 | 6124 | 0 |
| SRR5755999 | 0 | 28 | 56 | 0 | 14 | 0 | 0 | 419 | 285 | 662 | 2472 | 11223 | 0 | 0 | 0 | 11 | 11 | 367 | 0 | 195 | 1161 | 2778 | 0 | 0 | 9779 | 0 | 0 | 0 | 156 | 10 | 139 | 11 | 28 | 228 | 122 | 6717 | 0 |
| SRR5756000 | 163 | 0 | 0 | 0 | 0 | 0 | 0 | 0 | 0 | 1461 | 3704 | 16070 | 0 | 0 | 0 | 0 | 0 | 0 | 0 | 0 | 550 | 1520 | 0 | 0 | 4552 | 45 | 0 | 0 | 0 | 0 | 0 | 0 | 0 | 0 | 0 | 0 | 35 |
| SRR5756001 | 0 | 48 | 12 | 0 | 80 | 0 | 0 | 544 | 71 | 444 | 2665 | 24364 | 0 | 15 | 0 | 0 | 0 | 416 | 0 | 1195 | 2543 | 2087 | 0 | 10 | 11272 | 0 | 20 | 24 | 117 | 12 | 133 | 0 | 81 | 109 | 113 | 49131 | 0 |
| SRR5756002 | 0 | 24 | 16 | 0 | 40 | 0 | 0 | 802 | 98 | 785 | 1092 | 33709 | 0 | 49 | 0 | 0 | 0 | 464 | 0 | 376 | 2625 | 2214 | 0 | 0 | 11079 | 0 | 0 | 24 | 235 | 21 | 186 | 0 | 89 | 219 | 89 | 7773 | 0 |
| SRR5756003 | 0 | 111 | 146 | 0 | 17 | 0 | 0 | 315 | 58 | 111 | 2715 | 17209 | 15 | 0 | 18 | 67 | 0 | 1082 | 13 | 184 | 2386 | 6036 | 0 | 33 | 23643 | 10 | 211 | 15 | 222 | 705 | 341 | 19 | 123 | 330 | 172 | 2047 | 0 |
| SRR5756005 | 0 | 0 | 18 | 0 | 24 | 0 | 0 | 668 | 171 | 342 | 495 | 25231 | 0 | 0 | 31 | 62 | 0 | 91 | 0 | 714 | 2935 | 5354 | 0 | 57 | 14895 | 0 | 25 | 0 | 163 | 16 | 89 | 15 | 64 | 139 | 59 | 7872 | 0 |
| SRR5756006 | 0 | 46 | 166 | 0 | 16 | 0 | 0 | 1043 | 210 | 2530 | 2607 | 31581 | 0 | 53 | 0 | 15 | 0 | 491 | 0 | 323 | 1538 | 2618 | 0 | 12 | 10572 | 0 | 0 | 0 | 179 | 15 | 206 | 13 | 66 | 266 | 133 | 14120 | 0 |
| SRR5756007 | 0 | 0 | 0 | 0 | 0 | 0 | 0 | 0 | 0 | 0 | 4545 | 10000 | 0 | 0 | 0 | 0 | 0 | 0 | 0 | 0 | 414 | 1067 | 0 | 0 | 4005 | 70 | 0 | 0 | 0 | 0 | 0 | 0 | 0 | 0 | 0 | 0 | 0 |
| SRR5756008 | 0 | 12 | 152 | 0 | 59 | 0 | 0 | 1210 | 104 | 2644 | 1590 | 1642 | 0 | 570 | 0 | 17 | 0 | 493 | 0 | 69 | 1036 | 417 | 0 | 0 | 14679 | 0 | 0 | 0 | 49 | 116 | 67 | 24 | 24 | 79 | 79 | 380 | 0 |
| SRR5756009 | 0 | 124 | 162 | 0 | 0 | 0 | 0 | 157 | 27 | 144 | 6740 | 13998 | 0 | 0 | 54 | 175 | 0 | 1100 | 11 | 634 | 2824 | 8628 | 0 | 28 | 22890 | 13 | 119 | 0 | 92 | 11 | 222 | 11 | 76 | 11 | 108 | 4414 | 0 |
| SRR5756011 | 0 | 21 | 110 | 0 | 20 | 0 | 0 | 253 | 59 | 177 | 856 | 17930 | 0 | 84 | 34 | 67 | 0 | 81 | 17 | 2705 | 2720 | 4094 | 19 | 122 | 12773 | 0 | 85 | 0 | 73 | 11 | 66 | 0 | 23 | 120 | 56 | 33568 | 0 |
| SRR5756012 | 0 | 22 | 62 | 0 | 21 | 0 | 0 | 478 | 96 | 2311 | 319 | 1531 | 0 | 213 | 0 | 22 | 0 | 575 | 0 | 81 | 836 | 682 | 0 | 0 | 13285 | 0 | 0 | 0 | 53 | 84 | 65 | 12 | 31 | 50 | 31 | 754 | 0 |
| SRR5756014 | 0 | 22 | 175 | 0 | 136 | 0 | 0 | 443 | 124 | 3162 | 118 | 628 | 0 | 294 | 15 | 0 | 0 | 518 | 0 | 113 | 943 | 402 | 0 | 20 | 11011 | 0 | 0 | 0 | 85 | 166 | 45 | 0 | 18 | 72 | 58 | 752 | 0 |
| SRR5756016 | 0 | 103 | 41 | 0 | 13 | 0 | 0 | 110 | 14 | 312 | 4622 | 11011 | 0 | 0 | 60 | 185 | 0 | 1011 | 17 | 396 | 2152 | 4974 | 0 | 40 | 23495 | 27 | 180 | 0 | 103 | 327 | 203 | 37 | 69 | 126 | 37 | 3833 | 0 |
| SRR5756017 | 0 | 0 | 20 | 0 | 0 | 0 | 0 | 558 | 0 | 2699 | 468 | 1675 | 0 | 0 | 372 | 838 | 0 | 1143 | 30 | 93 | 352 | 1626 | 0 | 40 | 40916 | 0 | 41 | 0 | 81 | 20 | 20 | 0 | 20 | 0 | 0 | 745 | 0 |
| SRR5756018 | 0 | 0 | 117 | 0 | 30 | 0 | 0 | 377 | 47 | 346 | 433 | 20065 | 0 | 94 | 47 | 47 | 0 | 139 | 0 | 3520 | 2369 | 4674 | 0 | 47 | 13740 | 17 | 52 | 0 | 104 | 191 | 39 | 0 | 17 | 65 | 48 | 42157 | 0 |
| SRR5756019 | 0 | 0 | 52 | 0 | 21 | 0 | 0 | 1172 | 98 | 10332 | 1343 | 1539 | 0 | 195 | 0 | 49 | 0 | 548 | 0 | 49 | 734 | 808 | 0 | 45 | 13303 | 60 | 0 | 0 | 13 | 143 | 78 | 39 | 39 | 52 | 52 | 562 | 0 |
| SRR5756021 | 0 | 67 | 210 | 0 | 70 | 0 | 0 | 235 | 45 | 437 | 4965 | 8261 | 0 | 117 | 0 | 0 | 0 | 409 | 0 | 1643 | 2324 | 1414 | 0 | 0 | 7678 | 0 | 19 | 22 | 232 | 221 | 101 | 11 | 60 | 154 | 135 | 31663 | 0 |
| SRR5756022 | 0 | 0 | 129 | 0 | 29 | 0 | 0 | 887 | 53 | 1191 | 5056 | 6155 | 0 | 265 | 0 | 40 | 18 | 333 | 0 | 238 | 2352 | 1059 | 0 | 0 | 9216 | 0 | 0 | 0 | 92 | 178 | 92 | 12 | 18 | 104 | 111 | 2634 | 0 |

### Supp Table S8.4.Columns_DI_to_ES

| | | | | | | | | | | | | | | | | | | | | | | | | | | | | | | | | | | | | | |
|---|---|---|---|---|---|---|---|---|---|---|---|---|---|---|---|---|---|---|---|---|---|---|---|---|---|---|---|---|---|---|---|---|---|---|---|---|---|
| SRR5756023 | 0 | 98 | 15 | 0 | 12 | 0 | 0 | 127 | 29 | 179 | 8168 | 19066 | 0 | 29 | 23 | 40 | 15 | 1277 | 17 | 341 | 2281 | 6996 | 0 | 25 | 21851 | 17 | 204 | 0 | 129 | 356 | 121 | 30 | 76 | 204 | 83 | 3365 | 0 |
| SRR5756025 | 0 | 18 | 94 | 0 | 20 | 0 | 0 | 552 | 30 | 268 | 493 | 17303 | 0 | 179 | 15 | 15 | 0 | 83 | 0 | 2879 | 2390 | 5040 | 17 | 106 | 11762 | 0 | 138 | 0 | 165 | 179 | 71 | 0 | 31 | 107 | 58 | 44092 | 0 |
| SRR5756026 | 0 | 0 | 69 | 0 | 25 | 0 | 0 | 963 | 178 | 2468 | 745 | 6548 | 0 | 235 | 0 | 0 | 0 | 576 | 0 | 157 | 1365 | 1620 | 0 | 0 | 10465 | 0 | 38 | 0 | 63 | 107 | 63 | 0 | 69 | 57 | 63 | 2033 | 0 |
| SRR5756027 | 0 | 44 | 81 | 0 | 74 | 0 | 0 | 284 | 51 | 795 | 3631 | 5736 | 0 | 116 | 0 | 0 | 0 | 386 | 0 | 858 | 2307 | 1324 | 0 | 0 | 8718 | 0 | 47 | 14 | 47 | 145 | 61 | 0 | 27 | 78 | 37 | 14480 | 0 |
| SRR5756028 | 0 | 15 | 62 | 0 | 29 | 0 | 0 | 1655 | 41 | 1241 | 8397 | 8521 | 0 | 41 | 0 | 0 | 15 | 297 | 0 | 331 | 2305 | 1163 | 0 | 0 | 9221 | 0 | 0 | 0 | 31 | 93 | 62 | 0 | 46 | 77 | 39 | 1779 | 0 |
| SRR5756029 | 0 | 87 | 25 | 0 | 15 | 0 | 0 | 172 | 24 | 244 | 8790 | 15953 | 25 | 12 | 24 | 109 | 14 | 1007 | 0 | 247 | 2234 | 6117 | 0 | 20 | 21862 | 20 | 201 | 0 | 31 | 213 | 131 | 0 | 56 | 143 | 53 | 3069 | 0 |
| SRR5756031 | 0 | 14 | 70 | 0 | 15 | 0 | 0 | 193 | 19 | 256 | 1159 | 17444 | 0 | 56 | 12 | 81 | 0 | 89 | 0 | 3914 | 2355 | 4577 | 14 | 77 | 12158 | 0 | 13 | 0 | 86 | 11 | 53 | 0 | 20 | 61 | 25 | 65059 | 0 |
| SRR5756032 | 0 | 33 | 13 | 0 | 17 | 0 | 0 | 1015 | 83 | 2729 | 4010 | 5300 | 0 | 166 | 0 | 0 | 0 | 477 | 0 | 191 | 1425 | 1415 | 0 | 0 | 11037 | 0 | 40 | 0 | 20 | 93 | 93 | 0 | 40 | 46 | 60 | 1622 | 0 |
| SRR5756033 | 0 | 20 | 11 | 0 | 54 | 0 | 0 | 274 | 89 | 218 | 5883 | 8180 | 0 | 0 | 0 | 16 | 0 | 110 | 0 | 5528 | 1911 | 2861 | 0 | 0 | 9442 | 0 | 13 | 27 | 147 | 11 | 147 | 13 | 60 | 10 | 74 | 126604 | 0 |
| SRR5756034 | 0 | 22 | 279 | 0 | 19 | 0 | 0 | 287 | 107 | 343 | 1925 | 11628 | 0 | 42 | 0 | 10 | 0 | 256 | 0 | 3757 | 2738 | 2890 | 0 | 0 | 8435 | 0 | 18 | 0 | 146 | 288 | 79 | 0 | 47 | 142 | 34 | 106060 | 0 |
| SRR5756035 | 0 | 80 | 60 | 0 | 0 | 0 | 0 | 83 | 25 | 130 | 1720 | 11005 | 0 | 0 | 32 | 105 | 16 | 1050 | 15 | 783 | 2075 | 7693 | 0 | 21 | 21986 | 18 | 333 | 0 | 112 | 305 | 152 | 24 | 72 | 192 | 52 | 10345 | 0 |
| SRR5756037 | 0 | 20 | 13 | 0 | 22 | 0 | 0 | 666 | 101 | 295 | 7474 | 15563 | 0 | 67 | 51 | 25 | 0 | 81 | 0 | 1104 | 2707 | 5330 | 0 | 53 | 13224 | 0 | 62 | 0 | 10 | 11 | 62 | 0 | 42 | 99 | 32 | 20956 | 0 |
| SRR5756038 | 0 | 11 | 21 | 0 | 12 | 0 | 0 | 397 | 169 | 1121 | 9153 | 11563 | 0 | 20 | 0 | 20 | 0 | 425 | 0 | 3094 | 1071 | 3851 | 0 | 0 | 9448 | 0 | 16 | 0 | 53 | 90 | 32 | 0 | 27 | 64 | 21 | 79690 | 0 |
| SRR5756040 | 0 | 26 | 115 | 0 | 74 | 0 | 0 | 169 | 34 | 135 | 1222 | 7846 | 0 | 21 | 0 | 0 | 0 | 286 | 0 | 4871 | 2053 | 2504 | 0 | 0 | 7857 | 0 | 30 | 21 | 137 | 248 | 64 | 0 | 68 | 175 | 43 | 151016 | 0 |
| SRR5756041 | 0 | 11 | 85 | 0 | 26 | 0 | 0 | 505 | 32 | 688 | 6118 | 10687 | 0 | 54 | 0 | 22 | 0 | 339 | 0 | 1150 | 2272 | 3169 | 0 | 0 | 10780 | 0 | 17 | 23 | 40 | 91 | 85 | 11 | 45 | 80 | 11 | 31136 | 0 |
| SRR5756042 | 0 | 156 | 94 | 0 | 13 | 0 | 0 | 162 | 96 | 118 | 798 | 11281 | 0 | 96 | 22 | 125 | 0 | 1523 | 15 | 463 | 4373 | 10455 | 0 | 29 | 20606 | 23 | 211 | 0 | 94 | 17 | 164 | 16 | 62 | 156 | 39 | 5674 | 0 |
| SRR5756044 | 0 | 55 | 220 | 0 | 47 | 0 | 0 | 173 | 0 | 217 | 215 | 14739 | 0 | 22 | 22 | 43 | 0 | 147 | 0 | 2579 | 5483 | 7035 | 0 | 59 | 14985 | 0 | 98 | 18 | 104 | 250 | 37 | 0 | 12 | 73 | 43 | 56008 | 0 |
| SRR5756045 | 0 | 14 | 49 | 0 | 13 | 0 | 0 | 685 | 86 | 3189 | 353 | 12284 | 0 | 43 | 0 | 21 | 0 | 267 | 0 | 642 | 1618 | 2593 | 0 | 13 | 9240 | 0 | 0 | 0 | 49 | 98 | 56 | 0 | 56 | 56 | 35 | 19645 | 0 |
| SRR5756047 | 0 | 23 | 11 | 0 | 51 | 0 | 0 | 350 | 76 | 419 | 13949 | 24563 | 0 | 103 | 21 | 27 | 0 | 102 | 0 | 913 | 2026 | 1945 | 0 | 0 | 8293 | 0 | 14 | 14 | 147 | 12 | 128 | 23 | 101 | 10 | 151 | 7201 | 0 |
| SRR5756048 | 0 | 74 | 22 | 0 | 14 | 0 | 0 | 355 | 70 | 527 | 1575 | 12972 | 0 | 536 | 0 | 28 | 0 | 538 | 0 | 145 | 4318 | 2348 | 0 | 0 | 7386 | 0 | 32 | 0 | 199 | 14 | 106 | 14 | 79 | 148 | 171 | 1922 | 0 |
| SRR5756049 | 0 | 122 | 101 | 0 | 10 | 0 | 0 | 198 | 40 | 115 | 3231 | 23972 | 0 | 84 | 96 | 313 | 0 | 872 | 16 | 183 | 2973 | 7669 | 0 | 18 | 22366 | 11 | 115 | 0 | 118 | 515 | 265 | 17 | 91 | 240 | 94 | 1456 | 0 |
| SRR5756051 | 0 | 39 | 206 | 0 | 27 | 0 | 0 | 313 | 157 | 438 | 6564 | 17240 | 0 | 1430 | 104 | 83 | 0 | 163 | 0 | 877 | 4058 | 6429 | 12 | 114 | 13084 | 0 | 70 | 19 | 260 | 388 | 101 | 12 | 39 | 163 | 93 | 7660 | 0 |
| SRR5756052 | 0 | 47 | 53 | 0 | 13 | 0 | 0 | 568 | 129 | 1749 | 9921 | 15691 | 0 | 220 | 15 | 53 | 0 | 229 | 0 | 98 | 1984 | 3091 | 0 | 0 | 9365 | 0 | 18 | 0 | 29 | 233 | 105 | 0 | 18 | 111 | 70 | 1757 | 0 |
| SRR5756054 | 0 | 39 | 245 | 0 | 72 | 0 | 0 | 546 | 94 | 528 | 11653 | 18945 | 0 | 28 | 0 | 0 | 0 | 122 | 0 | 1149 | 1818 | 2153 | 0 | 0 | 7581 | 0 | 39 | 13 | 148 | 290 | 174 | 0 | 161 | 232 | 77 | 10287 | 0 |
| SRR5756055 | 0 | 56 | 325 | 0 | 31 | 0 | 0 | 289 | 54 | 596 | 9259 | 15324 | 0 | 569 | 0 | 0 | 0 | 235 | 0 | 271 | 4210 | 2591 | 0 | 0 | 6725 | 0 | 21 | 14 | 169 | 473 | 155 | 14 | 56 | 162 | 28 | 3276 | 0 |
| SRR5756056 | 0 | 182 | 55 | 0 | 0 | 0 | 0 | 224 | 50 | 146 | 2154 | 26229 | 11 | 207 | 50 | 112 | 0 | 1032 | 25 | 140 | 3478 | 8444 | 0 | 12 | 19716 | 13 | 111 | 0 | 133 | 25 | 11 | 28 | 66 | 12 | 44 | 987 | 0 |
| SRR5756058 | 0 | 24 | 213 | 0 | 32 | 0 | 0 | 407 | 81 | 419 | 8658 | 19528 | 0 | 221 | 58 | 47 | 0 | 127 | 0 | 1210 | 4885 | 5665 | 0 | 77 | 12135 | 0 | 82 | 0 | 125 | 313 | 67 | 0 | 27 | 116 | 15 | 9915 | 0 |
| SRR5756059 | 0 | 0 | 56 | 0 | 20 | 0 | 0 | 663 | 90 | 1733 | 16874 | 27360 | 0 | 60 | 15 | 30 | 0 | 226 | 0 | 121 | 2201 | 4094 | 0 | 0 | 8626 | 0 | 19 | 0 | 37 | 289 | 205 | 0 | 65 | 149 | 0 | 844 | 0 |
| SRR5756061 | 0 | 27 | 160 | 0 | 57 | 0 | 0 | 241 | 31 | 415 | 4600 | 7572 | 0 | 51 | 0 | 0 | 0 | 171 | 0 | 6694 | 2080 | 1643 | 0 | 0 | 8439 | 0 | 66 | 0 | 160 | 164 | 84 | 0 | 71 | 84 | 97 | 99247 | 0 |
| SRR5756062 | 0 | 20 | 89 | 0 | 35 | 0 | 0 | 530 | 84 | 1009 | 852 | 10419 | 0 | 488 | 17 | 0 | 0 | 256 | 0 | 673 | 2924 | 1947 | 0 | 0 | 9688 | 0 | 38 | 0 | 38 | 11 | 31 | 0 | 24 | 44 | 55 | 11521 | 0 |
| SRR5756063 | 0 | 126 | 32 | 0 | 13 | 0 | 0 | 158 | 90 | 174 | 2847 | 12085 | 0 | 142 | 52 | 97 | 13 | 1011 | 14 | 479 | 3765 | 7470 | 0 | 13 | 20804 | 0 | 151 | 0 | 139 | 24 | 158 | 39 | 58 | 177 | 55 | 3255 | 0 |
| SRR5756065 | 0 | 34 | 92 | 0 | 35 | 0 | 0 | 194 | 61 | 233 | 607 | 9116 | 0 | 587 | 0 | 28 | 0 | 111 | 0 | 4287 | 3517 | 6107 | 0 | 81 | 14802 | 0 | 121 | 0 | 95 | 244 | 49 | 0 | 34 | 69 | 32 | 92258 | 0 |
| SRR5756066 | 0 | 46 | 39 | 0 | 15 | 0 | 0 | 376 | 157 | 1758 | 5902 | 9912 | 0 | 678 | 0 | 16 | 0 | 556 | 0 | 919 | 1929 | 2638 | 0 | 0 | 9514 | 0 | 25 | 0 | 93 | 14 | 72 | 0 | 21 | 90 | 38 | 18868 | 0 |
| SRR5756068 | 0 | 39 | 248 | 0 | 45 | 0 | 0 | 155 | 25 | 362 | 1355 | 8979 | 0 | 151 | 11 | 0 | 16 | 147 | 0 | 4481 | 3911 | 2777 | 0 | 0 | 7678 | 0 | 48 | 0 | 103 | 261 | 132 | 0 | 58 | 148 | 48 | 149959 | 0 |
| SRR5756069 | 0 | 20 | 22 | 0 | 23 | 0 | 0 | 274 | 41 | 539 | 9593 | 18140 | 0 | 257 | 0 | 33 | 0 | 190 | 0 | 597 | 4948 | 2723 | 0 | 0 | 7073 | 0 | 0 | 0 | 114 | 14 | 118 | 0 | 65 | 155 | 98 | 12165 | 0 |
| SRR5756070 | 0 | 82 | 139 | 0 | 22 | 0 | 0 | 246 | 0 | 232 | 13598 | 33307 | 16 | 0 | 49 | 105 | 16 | 638 | 18 | 309 | 2088 | 5591 | 0 | 27 | 22790 | 11 | 131 | 0 | 163 | 645 | 318 | 57 | 139 | 327 | 155 | 1551 | 0 |
| SRR5756072 | 0 | 42 | 12 | 0 | 27 | 0 | 0 | 460 | 12 | 621 | 615 | 17580 | 0 | 37 | 12 | 12 | 0 | 90 | 0 | 2572 | 4051 | 7062 | 11 | 57 | 13177 | 0 | 117 | 0 | 108 | 12 | 78 | 0 | 24 | 60 | 18 | 43632 | 0 |
| SRR5756073 | 0 | 44 | 95 | 0 | 0 | 0 | 0 | 977 | 244 | 3891 | 1451 | 36759 | 0 | 0 | 15 | 46 | 0 | 659 | 0 | 183 | 2636 | 4647 | 0 | 0 | 9355 | 0 | 0 | 0 | 153 | 190 | 88 | 0 | 22 | 139 | 58 | 4303 | 0 |
| SRR5756075 | 0 | 25 | 150 | 0 | 55 | 0 | 0 | 193 | 29 | 236 | 5512 | 7520 | 0 | 50 | 0 | 0 | 0 | 138 | 0 | 6841 | 1904 | 2340 | 0 | 0 | 8970 | 0 | 13 | 31 | 50 | 188 | 157 | 13 | 63 | 106 | 119 | 161038 | 0 |
| SRR5756076 | 0 | 57 | 127 | 0 | 22 | 0 | 0 | 332 | 36 | 668 | 6602 | 11256 | 0 | 36 | 0 | 20 | 0 | 124 | 0 | 3111 | 3674 | 3346 | 0 | 0 | 9002 | 0 | 0 | 0 | 51 | 96 | 70 | 0 | 32 | 93 | 42 | 106379 | 0 |
| SRR5756077 | 0 | 28 | 179 | 0 | 15 | 0 | 0 | 277 | 71 | 223 | 7937 | 21480 | 0 | 0 | 71 | 89 | 14 | 738 | 11 | 429 | 1794 | 6316 | 0 | 19 | 20100 | 16 | 166 | 28 | 152 | 359 | 317 | 69 | 193 | 331 | 248 | 2759 | 0 |
| SRR5756079 | 0 | 76 | 274 | 0 | 26 | 0 | 0 | 485 | 67 | 568 | 8170 | 23207 | 0 | 33 | 17 | 33 | 0 | 106 | 0 | 1788 | 4629 | 6227 | 0 | 27 | 11639 | 0 | 72 | 13 | 166 | 233 | 72 | 0 | 36 | 76 | 49 | 26532 | 0 |
| SRR5756080 | 0 | 39 | 57 | 0 | 14 | 0 | 0 | 557 | 218 | 3688 | 9511 | 16322 | 0 | 71 | 13 | 31 | 0 | 341 | 0 | 325 | 2193 | 4340 | 0 | 0 | 9420 | 0 | 14 | 0 | 25 | 68 | 54 | 0 | 38 | 80 | 46 | 5622 | 0 |
| SRR5756082 | 0 | 69 | 107 | 0 | 89 | 0 | 0 | 178 | 18 | 361 | 1620 | 13319 | 0 | 18 | 0 | 0 | 0 | 139 | 0 | 2473 | 3192 | 3436 | 0 | 0 | 8352 | 0 | 33 | 0 | 102 | 10 | 79 | 0 | 51 | 79 | 26 | 76284 | 0 |
| SRR5756083 | 0 | 28 | 114 | 0 | 44 | 0 | 0 | 145 | 36 | 1261 | 1036 | 10937 | 0 | 29 | 0 | 15 | 0 | 251 | 0 | 3363 | 3318 | 5127 | 0 | 0 | 11011 | 0 | 19 | 11 | 58 | 111 | 69 | 0 | 36 | 78 | 36 | 74025 | 0 |
| SRR5756084 | 0 | 108 | 93 | 0 | 19 | 0 | 0 | 193 | 45 | 193 | 873 | 13222 | 0 | 0 | 30 | 112 | 0 | 543 | 17 | 1175 | 1685 | 5755 | 0 | 37 | 23099 | 11 | 162 | 23 | 100 | 185 | 11 | 23 | 69 | 12 | 39 | 6879 | 0 |
| SRR5756086 | 0 | 62 | 12 | 0 | 50 | 0 | 0 | 371 | 39 | 302 | 626 | 16487 | 0 | 59 | 0 | 39 | 0 | 125 | 0 | 2966 | 4678 | 6498 | 0 | 71 | 14437 | 0 | 15 | 13 | 58 | 16 | 58 | 0 | 19 | 67 | 43 | 36193 | 0 |
| SRR5756087 | 0 | 17 | 52 | 0 | 17 | 0 | 0 | 364 | 81 | 4981 | 449 | 13546 | 0 | 40 | 61 | 81 | 0 | 134 | 0 | 1053 | 2450 | 4937 | 0 | 0 | 10117 | 0 | 29 | 0 | 40 | 63 | 35 | 0 | 23 | 69 | 12 | 20936 | 0 |
| SRR5756089 | 0 | 12 | 181 | 0 | 59 | 0 | 0 | 536 | 94 | 308 | 10983 | 18416 | 0 | 27 | 0 | 13 | 0 | 91 | 0 | 1339 | 1860 | 2384 | 0 | 0 | 8737 | 0 | 0 | 29 | 146 | 16 | 105 | 18 | 41 | 12 | 10 | 19515 | 0 |
| SRR5756090 | 0 | 66 | 203 | 0 | 36 | 0 | 0 | 248 | 58 | 1196 | 2045 | 19827 | 0 | 58 | 0 | 29 | 0 | 282 | 0 | 838 | 4003 | 3924 | 0 | 0 | 9574 | 0 | 0 | 0 | 148 | 221 | 92 | 0 | 34 | 131 | 120 | 10821 | 0 |
| SRR5756091 | 0 | 102 | 72 | 0 | 12 | 0 | 0 | 139 | 36 | 51 | 2213 | 19532 | 0 | 0 | 10 | 72 | 0 | 1190 | 17 | 236 | 2795 | 8278 | 0 | 19 | 21713 | 21 | 38 | 13 | 102 | 352 | 110 | 0 | 47 | 186 | 59 | 1742 | 0 |
| SRR5756093 | 0 | 81 | 14 | 0 | 30 | 0 | 0 | 347 | 29 | 337 | 851 | 21845 | 11 | 19 | 0 | 19 | 0 | 110 | 0 | 1214 | 3855 | 7341 | 11 | 50 | 16281 | 0 | 51 | 11 | 103 | 12 | 86 | 0 | 22 | 103 | 76 | 10470 | 0 |
| SRR5756094 | 0 | 48 | 48 | 0 | 21 | 0 | 0 | 353 | 83 | 4257 | 14514 | 22593 | 0 | 93 | 0 | 10 | 0 | 309 | 0 | 415 | 1891 | 4913 | 0 | 0 | 10400 | 0 | 0 | 0 | 82 | 120 | 77 | 0 | 36 | 108 | 70 | 5538 | 0 |
| SRR5756096 | 0 | 64 | 282 | 0 | 90 | 0 | 0 | 341 | 77 | 557 | 3696 | 6283 | 0 | 85 | 0 | 0 | 0 | 522 | 0 | 765 | 2684 | 1720 | 0 | 0 | 9714 | 0 | 30 | 0 | 126 | 204 | 82 | 0 | 25 | 78 | 87 | 19483 | 0 |
| SRR5756097 | 0 | 36 | 20 | 0 | 50 | 0 | 0 | 3110 | 648 | 1598 | 209 | 8251 | 0 | 216 | 43 | 0 | 0 | 726 | 0 | 173 | 2927 | 1143 | 0 | 0 | 13838 | 0 | 0 | 0 | 118 | 145 | 54 | 0 | 36 | 99 | 81 | 2074 | 0 |
| SRR5756098 | 0 | 94 | 46 | 0 | 13 | 0 | 0 | 231 | 68 | 171 | 4971 | 18673 | 0 | 0 | 33 | 111 | 0 | 558 | 0 | 577 | 2379 | 6323 | 0 | 39 | 19601 | 15 | 369 | 11 | 120 | 357 | 187 | 17 | 82 | 118 | 111 | 6357 | 0 |
| SRR5756100 | 0 | 0 | 115 | 0 | 22 | 0 | 0 | 918 | 55 | 356 | 11069 | 25947 | 0 | 55 | 48 | 32 | 0 | 20 | 0 | 1204 | 2012 | 4253 | 24 | 229 | 12238 | 0 | 47 | 18 | 71 | 236 | 68 | 0 | 17 | 95 | 48 | 37428 | 0 |
| SRR5756101 | 0 | 60 | 48 | 0 | 21 | 0 | 0 | 2250 | 479 | 3854 | 1000 | 23346 | 0 | 74 | 0 | 111 | 0 | 474 | 0 | 240 | 2164 | 3370 | 0 | 0 | 11414 | 0 | 0 | 0 | 84 | 107 | 48 | 0 | 30 | 42 | 48 | 2416 | 0 |
| SRR5756103 | 0 | 15 | 12 | 0 | 65 | 0 | 0 | 983 | 176 | 328 | 681 | 21504 | 0 | 25 | 25 | 0 | 0 | 104 | 0 | 277 | 1934 | 2385 | 0 | 0 | 8895 | 0 | 0 | 37 | 171 | 14 | 13 | 0 | 97 | 164 | 82 | 454 | 0 |
| SRR5756104 | 0 | 17 | 245 | 0 | 26 | 0 | 0 | 651 | 55 | 669 | 3292 | 28318 | 0 | 31 | 31 | 0 | 0 | 117 | 0 | 172 | 2631 | 3201 | 0 | 0 | 8270 | 0 | 0 | 29 | 134 | 279 | 122 | 0 | 65 | 238 | 98 | 209 | 0 |
| SRR5756105 | 0 | 40 | 117 | 0 | 11 | 0 | 0 | 167 | 26 | 36 | 5852 | 21389 | 0 | 0 | 148 | 295 | 0 | 350 | 12 | 71 | 2114 | 7839 | 0 | 38 | 20737 | 0 | 17 | 23 | 263 | 19 | 343 | 33 | 183 | 437 | 137 | 200 | 0 |
| SRR5756106 | 0 | 18 | 0 | 0 | 0 | 0 | 0 | 486 | 22 | 1325 | 3534 | 13673 | 36 | 133 | 133 | 110 | 0 | 399 | 14 | 66 | 1074 | 10794 | 0 | 86 | 26188 | 0 | 425 | 0 | 18 | 109 | 0 | 0 | 0 | 45 | 36 | 177 | 0 |
| SRR5756107 | 0 | 20 | 201 | 0 | 26 | 0 | 0 | 359 | 63 | 225 | 1477 | 32978 | 0 | 63 | 36 | 54 | 0 | 26 | 0 | 198 | 2474 | 4436 | 0 | 60 | 13684 | 0 | 29 | 27 | 96 | 14 | 61 | 0 | 34 | 142 | 108 | 494 | 0 |
| SRR5756108 | 0 | 16 | 77 | 0 | 15 | 0 | 0 | 600 | 215 | 2764 | 2172 | 27496 | 0 | 0 | 0 | 0 | 0 | 65 | 0 | 0 | 1371 | 3654 | 0 | 16 | 8274 | 0 | 16 | 0 | 88 | 10 | 243 | 0 | 77 | 170 | 134 | 422 | 0 |
| SRR5756110 | 0 | 106 | 0 | 0 | 0 | 0 | 0 | 0 | 0 | 0 | 12410 | 14755 | 0 | 138 | 0 | 0 | 0 | 0 | 0 | 276 | 675 | 249 | 0 | 44 | 778 | 19 | 354 | 0 | 0 | 35 | 0 | 0 | 0 | 35 | 0 | 0 | 0 |
| SRR5756114 | 0 | 71 | 0 | 0 | 0 | 0 | 0 | 0 | 0 | 340 | 16403 | 25242 | 0 | 0 | 0 | 0 | 0 | 0 | 0 | 255 | 499 | 224 | 0 | 0 | 1131 | 27 | 47 | 0 | 0 | 0 | 0 | 0 | 24 | 0 | 0 | 85 | 0 |
| SRR5756115 | 35 | 70 | 35 | 0 | 0 | 0 | 0 | 110 | 0 | 221 | 14794 | 29587 | 0 | 0 | 0 | 0 | 0 | 0 | 0 | 110 | 686 | 221 | 0 | 29 | 822 | 25 | 35 | 0 | 0 | 70 | 0 | 0 | 0 | 0 | 0 | 0 | 0 |
| SRR5756116 | 0 | 0 | 135 | 0 | 0 | 0 | 0 | 0 | 0 | 252 | 8579 | 8832 | 0 | 0 | 0 | 0 | 0 | 0 | 0 | 0 | 879 | 245 | 0 | 21 | 744 | 21 | 0 | 68 | 0 | 0 | 0 | 0 | 0 | 0 | 0 | 505 | 0 |
| SRR5756118 | 0 | 85 | 0 | 0 | 0 | 43 | 0 | 0 | 0 | 0 | 56 | 17646 | 0 | 0 | 0 | 0 | 0 | 0 | 0 | 223 | 695 | 258 | 0 | 0 | 592 | 13 | 171 | 0 | 0 | 0 | 0 | 0 | 0 | 0 | 0 | 447 | 0 |
| SRR5756120 | 0 | 108 | 27 | 0 | 0 | 0 | 0 | 221 | 0 | 221 | 123 | 24167 | 0 | 0 | 0 | 0 | 54 | 0 | 0 | 662 | 312 | 208 | 0 | 0 | 898 | 22 | 244 | 0 | 27 | 27 | 0 | 0 | 0 | 27 | 0 | 331 | 0 |
| SRR5756121 | 0 | 176 | 0 | 0 | 0 | 0 | 0 | 188 | 0 | 0 | 8828 | 15965 | 0 | 0 | 0 | 0 | 0 | 0 | 0 | 563 | 701 | 326 | 0 | 0 | 664 | 0 | 527 | 0 | 0 | 0 | 0 | 0 | 0 | 0 | 0 | 376 | 0 |
| SRR5756122 | 0 | 27 | 0 | 0 | 0 | 27 | 0 | 119 | 0 | 475 | 89 | 24457 | 0 | 0 | 0 | 0 | 0 | 0 | 0 | 237 | 413 | 231 | 0 | 0 | 1692 | 24 | 106 | 0 | 0 | 0 | 0 | 0 | 27 | 0 | 0 | 237 | 12 |
| SRR5756123 | 0 | 143 | 29 | 0 | 0 | 0 | 0 | 105 | 0 | 0 | 15136 | 19866 | 0 | 0 | 0 | 0 | 0 | 0 | 0 | 420 | 344 | 230 | 0 | 0 | 976 | 38 | 0 | 0 | 0 | 0 | 0 | 0 | 0 | 0 | 0 | 420 | 0 |
| SRR5756125 | 0 | 32 | 0 | 0 | 0 | 0 | 0 | 99 | 0 | 198 | 177 | 26188 | 0 | 0 | 0 | 0 | 0 | 0 | 0 | 99 | 858 | 307 | 0 | 0 | 2434 | 65 | 95 | 0 | 0 | 0 | 0 | 0 | 0 | 0 | 0 | 496 | 14 |
| SRR5756128 | 0 | 46 | 93 | 0 | 0 | 46 | 0 | 245 | 0 | 734 | 4890 | 8802 | 0 | 0 | 0 | 0 | 0 | 0 | 0 | 245 | 825 | 215 | 0 | 0 | 686 | 0 | 371 | 0 | 0 | 0 | 0 | 0 | 0 | 46 | 0 | 0 | 0 |
| SRR5756130 | 0 | 45 | 0 | 0 | 0 | 0 | 0 | 0 | 0 | 659 | 80 | 20153 | 0 | 0 | 0 | 0 | 23 | 0 | 0 | 132 | 430 | 205 | 0 | 0 | 1752 | 39 | 45 | 0 | 0 | 0 | 0 | 0 | 0 | 23 | 0 | 527 | 0 |
| SRR5756131 | 0 | 204 | 26 | 0 | 0 | 0 | 0 | 0 | 0 | 0 | 165 | 22652 | 0 | 85 | 0 | 0 | 0 | 0 | 0 | 85 | 645 | 321 | 0 | 0 | 653 | 26 | 102 | 0 | 0 | 0 | 0 | 0 | 26 | 26 | 0 | 85 | 0 |
| SRR5756132 | 0 | 0 | 0 | 0 | 0 | 0 | 0 | 367 | 122 | 856 | 101 | 18843 | 0 | 0 | 0 | 0 | 0 | 0 | 0 | 245 | 680 | 267 | 0 | 0 | 1270 | 54 | 357 | 0 | 0 | 0 | 0 | 0 | 0 | 0 | 0 | 0 | 0 |
| SRR5756133 | 0 | 109 | 0 | 0 | 0 | 36 | 0 | 121 | 243 | 121 | 109 | 19679 | 0 | 243 | 0 | 0 | 0 | 0 | 0 | 243 | 441 | 186 | 0 | 0 | 578 | 27 | 36 | 0 | 0 | 0 | 0 | 0 | 36 | 36 | 0 | 607 | 0 |
| SRR5756135 | 37 | 184 | 74 | 0 | 0 | 37 | 0 | 0 | 199 | 199 | 50 | 16132 | 0 | 0 | 0 | 0 | 0 | 0 | 0 | 597 | 781 | 368 | 0 | 0 | 635 | 19 | 809 | 0 | 0 | 0 | 0 | 0 | 0 | 37 | 37 | 797 | 0 |
| SRR5756136 | 0 | 72 | 72 | 0 | 0 | 0 | 0 | 0 | 160 | 160 | 42 | 8982 | 0 | 0 | 160 | 0 | 0 | 0 | 0 | 0 | 1191 | 513 | 0 | 59 | 915 | 0 | 506 | 0 | 0 | 0 | 0 | 0 | 72 | 0 | 36 | 481 | 0 |
| SRR5756137 | 18 | 89 | 53 | 0 | 0 | 0 | 0 | 261 | 0 | 522 | 8873 | 13309 | 0 | 0 | 522 | 0 | 18 | 0 | 0 | 1044 | 301 | 228 | 0 | 0 | 1306 | 38 | 89 | 0 | 0 | 0 | 0 | 0 | 0 | 18 | 0 | 1044 | 11 |
| SRR5756138 | 0 | 57 | 0 | 0 | 0 | 0 | 57 | 834 | 208 | 417 | 40 | 11461 | 0 | 0 | 0 | 0 | 0 | 0 | 0 | 417 | 446 | 264 | 0 | 0 | 1016 | 16 | 114 | 0 | 0 | 0 | 0 | 0 | 29 | 29 | 29 | 0 | 0 |
| SRR5756139 | 0 | 69 | 35 | 0 | 0 | 0 | 0 | 0 | 162 | 162 | 54 | 10989 | 0 | 0 | 162 | 0 | 0 | 0 | 0 | 0 | 822 | 386 | 0 | 0 | 957 | 15 | 104 | 0 | 0 | 0 | 0 | 0 | 0 | 35 | 0 | 162 | 0 |
| SRR5756140 | 29 | 145 | 87 | 0 | 0 | 0 | 0 | 0 | 0 | 213 | 43 | 7041 | 0 | 0 | 0 | 0 | 0 | 0 | 0 | 1280 | 480 | 268 | 0 | 38 | 1164 | 22 | 87 | 0 | 0 | 0 | 0 | 0 | 0 | 29 | 0 | 640 | 0 |
| SRR5756141 | 16 | 48 | 64 | 0 | 0 | 0 | 0 | 0 | 0 | 851 | 12526 | 21039 | 0 | 0 | 0 | 243 | 32 | 0 | 0 | 851 | 429 | 259 | 0 | 56 | 1781 | 21 | 64 | 0 | 0 | 0 | 0 | 0 | 0 | 16 | 0 | 730 | 15 |
| SRR5756142 | 18 | 36 | 36 | 0 | 0 | 0 | 0 | 266 | 0 | 1465 | 75 | 19039 | 0 | 0 | 0 | 133 | 0 | 0 | 0 | 0 | 454 | 191 | 0 | 54 | 1405 | 26 | 55 | 0 | 0 | 0 | 0 | 0 | 18 | 18 | 0 | 266 | 0 |
| SRR5756143 | 36 | 0 | 18 | 0 | 0 | 0 | 0 | 137 | 0 | 822 | 61 | 11094 | 0 | 0 | 137 | 137 | 0 | 0 | 0 | 411 | 512 | 216 | 0 | 0 | 1777 | 37 | 126 | 0 | 0 | 0 | 0 | 0 | 0 | 0 | 0 | 685 | 0 |
| SRR5756144 | 0 | 20 | 41 | 0 | 0 | 0 | 0 | 146 | 291 | 437 | 89 | 16162 | 0 | 0 | 0 | 0 | 20 | 0 | 0 | 582 | 511 | 201 | 0 | 0 | 1200 | 36 | 102 | 0 | 0 | 0 | 0 | 0 | 0 | 0 | 0 | 582 | 0 |
| SRR5756145 | 0 | 115 | 76 | 0 | 0 | 0 | 0 | 0 | 0 | 458 | 7100 | 9849 | 0 | 0 | 0 | 0 | 76 | 0 | 0 | 229 | 407 | 203 | 0 | 0 | 643 | 13 | 191 | 0 | 0 | 0 | 0 | 0 | 0 | 76 | 38 | 458 | 0 |
| SRR5756146 | 0 | 59 | 40 | 0 | 0 | 20 | 0 | 439 | 219 | 439 | 20 | 7682 | 0 | 0 | 0 | 219 | 20 | 0 | 0 | 658 | 686 | 388 | 0 | 0 | 1005 | 20 | 712 | 20 | 20 | 0 | 0 | 0 | 0 | 0 | 0 | 1317 | 0 |
| SRR5756147 | 0 | 156 | 39 | 0 | 0 | 0 | 0 | 184 | 0 | 736 | 11222 | 17476 | 0 | 0 | 0 | 0 | 0 | 0 | 0 | 552 | 689 | 306 | 0 | 0 | 702 | 13 | 156 | 0 | 0 | 39 | 0 | 0 | 0 | 39 | 0 | 184 | 0 |
| SRR5756148 | 27 | 81 | 0 | 0 | 0 | 0 | 0 | 384 | 192 | 575 | 39 | 7670 | 0 | 0 | 0 | 0 | 0 | 0 | 0 | 192 | 431 | 211 | 0 | 0 | 951 | 0 | 54 | 0 | 0 | 0 | 0 | 0 | 0 | 0 | 0 | 0 | 0 |
| SRR5756149 | 0 | 38 | 0 | 0 | 0 | 0 | 0 | 0 | 195 | 1173 | 12314 | 13683 | 0 | 0 | 195 | 0 | 0 | 0 | 0 | 782 | 1213 | 563 | 0 | 0 | 1197 | 22 | 113 | 0 | 0 | 0 | 0 | 0 | 0 | 0 | 0 | 977 | 0 |
| SRR5756150 | 0 | 116 | 39 | 0 | 0 | 0 | 0 | 2447 | 0 | 1030 | 18290 | 17903 | 0 | 0 | 0 | 129 | 0 | 0 | 0 | 0 | 441 | 216 | 0 | 0 | 1220 | 31 | 58 | 0 | 0 | 19 | 0 | 0 | 0 | 19 | 0 | 386 | 0 |
| SRR5756151 | 38 | 115 | 268 | 0 | 0 | 0 | 0 | 0 | 0 | 2580 | 11963 | 29557 | 0 | 0 | 0 | 0 | 0 | 0 | 0 | 469 | 646 | 305 | 0 | 0 | 858 | 19 | 344 | 0 | 0 | 0 | 0 | 0 | 0 | 0 | 0 | 235 | 0 |
| SRR5756152 | 0 | 87 | 35 | 0 | 0 | 0 | 0 | 460 | 115 | 805 | 133 | 17130 | 0 | 0 | 0 | 0 | 0 | 0 | 0 | 115 | 735 | 287 | 0 | 0 | 1151 | 37 | 52 | 0 | 0 | 0 | 0 | 0 | 0 | 0 | 17 | 115 | 0 |
| SRR5756153 | 47 | 24 | 47 | 0 | 0 | 0 | 0 | 158 | 0 | 316 | 5690 | 8219 | 0 | 0 | 0 | 0 | 0 | 0 | 0 | 790 | 467 | 222 | 0 | 0 | 977 | 25 | 591 | 0 | 0 | 0 | 0 | 0 | 47 | 24 | 0 | 632 | 0 |
| SRR5756154 | 0 | 45 | 45 | 0 | 0 | 0 | 0 | 0 | 0 | 403 | 6240 | 7850 | 0 | 0 | 0 | 0 | 0 | 0 | 0 | 805 | 893 | 344 | 0 | 0 | 797 | 17 | 317 | 0 | 0 | 0 | 0 | 0 | 0 | 45 | 0 | 805 | 0 |
| SRR5756155 | 0 | 28 | 0 | 0 | 0 | 0 | 0 | 0 | 0 | 545 | 112 | 23417 | 0 | 0 | 136 | 0 | 0 | 0 | 0 | 681 | 327 | 180 | 0 | 0 | 698 | 11 | 126 | 0 | 0 | 0 | 0 | 0 | 0 | 14 | 0 | 272 | 0 |
| SRR5756156 | 0 | 123 | 123 | 0 | 0 | 0 | 0 | 0 | 137 | 684 | 73 | 12859 | 0 | 0 | 0 | 137 | 0 | 0 | 0 | 410 | 493 | 303 | 0 | 0 | 1470 | 29 | 41 | 0 | 0 | 21 | 21 | 0 | 21 | 62 | 0 | 684 | 0 |
| SRR5756157 | 38 | 38 | 76 | 0 | 0 | 0 | 0 | 0 | 0 | 721 | 69 | 8510 | 0 | 0 | 0 | 0 | 0 | 0 | 0 | 577 | 394 | 191 | 0 | 0 | 1377 | 26 | 152 | 0 | 0 | 19 | 0 | 0 | 38 | 38 | 0 | 433 | 0 |
| SRR5756158 | 0 | 45 | 23 | 0 | 0 | 0 | 0 | 194 | 0 | 774 | 9293 | 9681 | 0 | 0 | 0 | 0 | 0 | 0 | 0 | 387 | 745 | 258 | 0 | 0 | 1429 | 31 | 23 | 0 | 0 | 0 | 0 | 0 | 45 | 68 | 0 | 387 | 0 |
| SRR5756159 | 63 | 63 | 0 | 0 | 0 | 0 | 0 | 180 | 0 | 1082 | 54 | 11727 | 0 | 0 | 0 | 0 | 42 | 0 | 0 | 541 | 358 | 256 | 0 | 0 | 2408 | 25 | 84 | 0 | 0 | 21 | 0 | 0 | 0 | 0 | 0 | 1624 | 11 |
| SRR5756160 | 53 | 53 | 0 | 0 | 0 | 0 | 0 | 0 | 0 | 463 | 75 | 23606 | 0 | 0 | 0 | 0 | 0 | 0 | 0 | 231 | 462 | 232 | 0 | 0 | 1317 | 38 | 0 | 0 | 0 | 0 | 0 | 0 | 53 | 0 | 0 | 926 | 0 |
| SRR5756162 | 58 | 58 | 0 | 0 | 0 | 0 | 0 | 0 | 0 | 286 | 12879 | 17172 | 0 | 0 | 0 | 0 | 0 | 0 | 0 | 859 | 494 | 158 | 0 | 65 | 1483 | 41 | 349 | 0 | 0 | 0 | 0 | 0 | 0 | 116 | 0 | 0 | 0 |
| SRR5756164 | 0 | 45 | 0 | 0 | 0 | 0 | 0 | 0 | 0 | 525 | 9552 | 24772 | 0 | 0 | 0 | 0 | 0 | 0 | 0 | 420 | 214 | 152 | 0 | 0 | 2808 | 43 | 245 | 0 | 0 | 22 | 0 | 0 | 0 | 0 | 0 | 210 | 0 |
| SRR5756166 | 25 | 50 | 0 | 0 | 0 | 0 | 0 | 192 | 0 | 96 | 14910 | 30396 | 0 | 0 | 0 | 0 | 0 | 0 | 0 | 289 | 367 | 189 | 0 | 45 | 1500 | 42 | 0 | 0 | 0 | 0 | 0 | 0 | 0 | 0 | 0 | 481 | 0 |
| SRR5756167 | 30 | 30 | 0 | 0 | 0 | 30 | 0 | 0 | 0 | 864 | 48 | 15335 | 0 | 0 | 0 | 0 | 30 | 0 | 0 | 432 | 300 | 133 | 0 | 0 | 2112 | 52 | 295 | 0 | 0 | 0 | 30 | 0 | 0 | 0 | 0 | 432 | 11 |
| SRR5756168 | 36 | 71 | 0 | 0 | 0 | 0 | 0 | 220 | 0 | 220 | 10984 | 13840 | 0 | 0 | 0 | 0 | 36 | 0 | 0 | 220 | 325 | 150 | 0 | 0 | 1416 | 24 | 284 | 0 | 36 | 0 | 0 | 0 | 36 | 36 | 0 | 0 | 13 |
| SRR5756177 | 0 | 0 | 219 | 0 | 0 | 110 | 0 | 0 | 0 | 793 | 14 | 8323 | 0 | 0 | 0 | 0 | 0 | 0 | 0 | 0 | 877 | 307 | 0 | 0 | 656 | 11 | 439 | 0 | 0 | 0 | 0 | 0 | 0 | 0 | 0 | 0 | 0 |
| SRR5756178 | 0 | 89 | 0 | 0 | 0 | 0 | 0 | 0 | 0 | 0 | 7056 | 10976 | 0 | 0 | 0 | 0 | 0 | 0 | 0 | 392 | 1100 | 448 | 0 | 0 | 844 | 13 | 714 | 0 | 0 | 0 | 0 | 0 | 0 | 0 | 0 | 0 | 0 |
| SRR5756179 | 0 | 83 | 165 | 0 | 0 | 0 | 0 | 0 | 0 | 1046 | 10115 | 6976 | 0 | 0 | 0 | 0 | 0 | 0 | 0 | 0 | 524 | 158 | 0 | 0 | 1075 | 18 | 0 | 0 | 0 | 0 | 0 | 0 | 0 | 83 | 0 | 0 | 0 |
| SRR5756180 | 28 | 0 | 0 | 0 | 0 | 28 | 0 | 115 | 0 | 345 | 115 | 20488 | 0 | 0 | 0 | 0 | 0 | 0 | 0 | 345 | 251 | 143 | 0 | 0 | 1347 | 33 | 56 | 0 | 28 | 0 | 0 | 0 | 0 | 56 | 0 | 115 | 0 |
| SRR5756182 | 0 | 0 | 0 | 0 | 0 | 0 | 0 | 417 | 0 | 0 | 19 | 8750 | 0 | 0 | 0 | 0 | 0 | 0 | 0 | 0 | 425 | 206 | 0 | 0 | 563 | 13 | 666 | 0 | 0 | 0 | 0 | 0 | 0 | 0 | 0 | 417 | 0 |
| SRR5756183 | 0 | 125 | 34 | 0 | 0 | 0 | 0 | 135 | 90 | 135 | 8038 | 15444 | 0 | 45 | 90 | 0 | 0 | 0 | 0 | 452 | 399 | 226 | 0 | 0 | 597 | 12 | 114 | 0 | 0 | 0 | 11 | 0 | 23 | 34 | 23 | 452 | 0 |
| SRR5756184 | 0 | 30 | 0 | 0 | 0 | 30 | 0 | 127 | 0 | 253 | 5190 | 10506 | 0 | 0 | 127 | 0 | 0 | 0 | 0 | 127 | 441 | 242 | 0 | 0 | 556 | 15 | 836 | 0 | 0 | 0 | 0 | 0 | 0 | 30 | 0 | 253 | 0 |
| SRR5756185 | 17 | 35 | 17 | 0 | 0 | 0 | 0 | 67 | 133 | 266 | 15234 | 32863 | 0 | 67 | 0 | 0 | 0 | 0 | 0 | 399 | 435 | 185 | 0 | 0 | 404 | 24 | 226 | 0 | 0 | 0 | 0 | 0 | 0 | 0 | 17 | 200 | 0 |
| SRR5756186 | 0 | 98 | 16 | 0 | 0 | 16 | 0 | 126 | 0 | 315 | 252 | 30768 | 0 | 63 | 0 | 0 | 16 | 0 | 0 | 315 | 470 | 187 | 0 | 0 | 404 | 19 | 164 | 0 | 16 | 0 | 16 | 0 | 16 | 33 | 16 | 315 | 0 |
| SRR5804909 | 0 | 0 | 0 | 0 | 0 | 0 | 0 | 437 | 219 | 36 | 109 | 36 | 0 | 219 | 0 | 0 | 0 | 4471 | 124 | 0 | 0 | 497 | 0 | 5713 | 124 | 0 | 103 | 0 | 0 | 0 | 0 | 0 | 0 | 0 | 0 | 0 | 124 |
| SRR5804912 | 0 | 0 | 0 | 0 | 142 | 0 | 0 | 818 | 220 | 315 | 315 | 252 | 0 | 94 | 0 | 31 | 0 | 2747 | 0 | 0 | 0 | 663 | 47 | 33348 | 237 | 47 | 98 | 0 | 0 | 0 | 0 | 0 | 0 | 0 | 0 | 0 | 0 |
| SRR5804914 | 0 | 0 | 0 | 0 | 135 | 0 | 0 | 339 | 339 | 0 | 0 | 0 | 0 | 452 | 0 | 0 | 0 | 13819 | 135 | 0 | 406 | 271 | 0 | 10839 | 0 | 0 | 769 | 0 | 0 | 0 | 0 | 0 | 0 | 0 | 0 | 0 | 135 |
| SRR5817623 | 0 | 123 | 18 | 0 | 43 | 0 | 0 | 82 | 1318 | 11451 | 10480 | 7654 | 0 | 220 | 20 | 40 | 15 | 1307 | 141 | 772 | 468 | 1013 | 0 | 944 | 6293 | 186 | 0 | 113 | 16 | 177 | 171 | 91 | 14 | 149 | 31 | 2505 | 1737 |
| SRR5817624 | 0 | 140 | 20 | 0 | 55 | 0 | 0 | 198 | 5229 | 14804 | 7151 | 6299 | 0 | 546 | 11 | 26 | 24 | 1867 | 111 | 985 | 577 | 1323 | 0 | 611 | 5376 | 242 | 0 | 124 | 13 | 117 | 133 | 74 | 151 | 127 | 18 | 5009 | 1808 |

### Supp Table S8.4.Columns_DI_to_ES

| | | | | | | | | | | | | | | | | | | | | | | | | | | | | | | | | | | | | | |
|---|---|---|---|---|---|---|---|---|---|---|---|---|---|---|---|---|---|---|---|---|---|---|---|---|---|---|---|---|---|---|---|---|---|---|---|---|---|
| SRR5817625 | 0 | 1927 | 351 | 0 | 42 | 0 | 0 | 102 | 1155 | 10870 | 6248 | 7932 | 0 | 198 | 42 | 132 | 15 | 1238 | 139 | 534 | 465 | 1056 | 0 | 875 | 5717 | 170 | 0 | 112 | 238 | 176 | 164 | 83 | 202 | 113 | 37 | 2131 | 1602 |
| SRR5817626 | 0 | 1620 | 334 | 0 | 51 | 0 | 0 | 152 | 2921 | 13262 | 10709 | 6419 | 0 | 438 | 25 | 93 | 15 | 1536 | 119 | 847 | 547 | 1193 | 0 | 616 | 5742 | 209 | 0 | 122 | 213 | 116 | 150 | 73 | 146 | 127 | 25 | 3473 | 1734 |
| SRR5821803 | 0 | 19 | 0 | 0 | 42 | 0 | 0 | 325 | 125 | 1761 | 55485 | 20915 | 0 | 27 | 0 | 0 | 0 | 2772 | 11 | 860 | 1345 | 1554 | 107 | 43231 | 4075 | 713 | 0 | 30 | 27 | 27 | 12 | 0 | 16 | 27 | 0 | 3941 | 20 |
| SRR5821804 | 0 | 12 | 0 | 0 | 21 | 0 | 0 | 3225 | 978 | 8399 | 4027 | 6340 | 0 | 16 | 0 | 0 | 34 | 632 | 11 | 45 | 993 | 5386 | 297 | 38856 | 4713 | 683 | 0 | 92 | 23 | 33 | 17 | 0 | 0 | 12 | 0 | 94 | 53 |
| SRR5821805 | 0 | 14 | 59 | 0 | 0 | 0 | 0 | 675 | 244 | 3020 | 10847 | 16857 | 0 | 17 | 10 | 0 | 18 | 483 | 22 | 143 | 840 | 8192 | 118 | 60250 | 6708 | 645 | 0 | 857 | 185 | 283 | 212 | 71 | 216 | 220 | 0 | 696 | 52 |
| SRR5832003 | 0 | 0 | 0 | 0 | 0 | 0 | 0 | 366 | 81 | 3012 | 0 | 0 | 0 | 651 | 2239 | 3460 | 0 | 204 | 26 | 1343 | 1629 | 4286 | 230 | 9082 | 7211 | 128 | 0 | 0 | 0 | 0 | 0 | 0 | 0 | 0 | 0 | 6269 | 0 |
| SRR5832004 | 0 | 0 | 0 | 0 | 0 | 0 | 0 | 82 | 16 | 2189 | 0 | 0 | 0 | 1251 | 1152 | 1350 | 0 | 127 | 0 | 214 | 1612 | 4450 | 408 | 8393 | 6435 | 35 | 0 | 0 | 0 | 0 | 0 | 0 | 0 | 62 | 0 | 1416 | 0 |
| SRR5832005 | 0 | 0 | 0 | 0 | 0 | 0 | 0 | 1051 | 150 | 1437 | 0 | 0 | 0 | 1780 | 772 | 236 | 0 | 183 | 18 | 107 | 1225 | 3447 | 320 | 7607 | 5623 | 82 | 0 | 0 | 0 | 14 | 0 | 0 | 0 | 14 | 14 | 858 | 0 |
| SRR5832006 | 0 | 0 | 0 | 0 | 0 | 0 | 0 | 190 | 0 | 3520 | 0 | 0 | 0 | 999 | 143 | 714 | 0 | 130 | 0 | 1332 | 1490 | 4456 | 391 | 9722 | 6829 | 87 | 0 | 0 | 0 | 0 | 0 | 0 | 0 | 86 | 0 | 8753 | 0 |
| SRR5832007 | 0 | 0 | 0 | 0 | 0 | 0 | 0 | 3196 | 1960 | 4705 | 0 | 0 | 0 | 4620 | 70 | 23 | 0 | 238 | 0 | 0 | 1690 | 1848 | 396 | 8502 | 6918 | 0 | 0 | 0 | 0 | 42 | 0 | 0 | 42 | 0 | 0 | 86 | 0 |
| SRR5832008 | 0 | 0 | 0 | 0 | 0 | 0 | 0 | 715 | 54 | 3216 | 0 | 0 | 0 | 3555 | 411 | 232 | 0 | 78 | 0 | 54 | 1306 | 3373 | 370 | 9516 | 6220 | 39 | 0 | 0 | 12 | 61 | 0 | 0 | 0 | 25 | 12 | 322 | 0 |
| SRR5832009 | 0 | 36 | 0 | 11 | 0 | 0 | 0 | 549 | 72 | 4416 | 0 | 0 | 0 | 1026 | 286 | 740 | 0 | 239 | 11 | 597 | 1566 | 3270 | 325 | 8448 | 5896 | 51 | 0 | 0 | 0 | 0 | 36 | 0 | 0 | 0 | 0 | 3509 | 0 |
| SRR5832010 | 0 | 56 | 0 | 0 | 0 | 0 | 0 | 150 | 37 | 1200 | 0 | 0 | 0 | 1275 | 975 | 1275 | 0 | 149 | 11 | 262 | 1110 | 2905 | 345 | 8626 | 5593 | 122 | 0 | 0 | 0 | 0 | 0 | 0 | 0 | 28 | 0 | 1612 | 0 |
| SRR5832011 | 0 | 116 | 0 | 0 | 0 | 0 | 0 | 336 | 70 | 6569 | 0 | 0 | 0 | 2201 | 278 | 927 | 0 | 195 | 13 | 220 | 1701 | 3146 | 356 | 9539 | 6225 | 54 | 0 | 0 | 0 | 0 | 0 | 0 | 58 | 0 | 0 | 811 | 0 |
| SRR5832012 | 0 | 0 | 0 | 0 | 0 | 0 | 0 | 281 | 14 | 3721 | 0 | 0 | 0 | 1306 | 211 | 913 | 0 | 160 | 0 | 449 | 1674 | 4405 | 525 | 10643 | 6565 | 30 | 0 | 0 | 0 | 0 | 0 | 0 | 0 | 0 | 65 | 2106 | 0 |
| SRR5832013 | 0 | 0 | 42 | 0 | 0 | 0 | 0 | 385 | 0 | 1874 | 0 | 0 | 0 | 769 | 1442 | 1298 | 0 | 209 | 0 | 433 | 1337 | 3169 | 428 | 8753 | 5832 | 19 | 0 | 0 | 0 | 0 | 42 | 0 | 0 | 0 | 84 | 1682 | 0 |
| SRR5832014 | 0 | 0 | 0 | 0 | 13 | 0 | 0 | 209 | 38 | 3807 | 0 | 0 | 0 | 1332 | 152 | 685 | 0 | 286 | 0 | 1142 | 1234 | 3032 | 312 | 8240 | 5883 | 169 | 0 | 0 | 0 | 122 | 0 | 0 | 0 | 0 | 0 | 5405 | 0 |
| SRR5832015 | 0 | 0 | 0 | 0 | 0 | 0 | 0 | 248 | 124 | 4541 | 0 | 0 | 0 | 1836 | 3871 | 2332 | 0 | 84 | 0 | 199 | 989 | 3148 | 341 | 9145 | 6505 | 125 | 0 | 0 | 0 | 0 | 37 | 0 | 37 | 0 | 0 | 546 | 0 |
| SRR5832016 | 0 | 0 | 0 | 0 | 0 | 0 | 29 | 143 | 48 | 4220 | 0 | 0 | 0 | 2665 | 2094 | 968 | 0 | 75 | 0 | 79 | 1705 | 3828 | 343 | 9787 | 7029 | 33 | 0 | 0 | 0 | 29 | 0 | 0 | 0 | 0 | 0 | 286 | 0 |
| SRR5832017 | 0 | 151 | 0 | 0 | 0 | 0 | 0 | 215 | 481 | 23782 | 0 | 0 | 0 | 6110 | 131 | 360 | 0 | 108 | 0 | 51 | 1010 | 1893 | 240 | 7616 | 6037 | 207 | 0 | 0 | 0 | 50 | 0 | 0 | 50 | 50 | 0 | 234 | 0 |
| SRR5832018 | 0 | 25 | 25 | 0 | 0 | 0 | 0 | 364 | 77 | 8238 | 0 | 0 | 0 | 1732 | 1196 | 2861 | 0 | 123 | 0 | 316 | 1950 | 4962 | 409 | 13789 | 7962 | 72 | 0 | 0 | 25 | 0 | 0 | 0 | 0 | 0 | 0 | 1493 | 0 |
| SRR5832019 | 0 | 109 | 0 | 0 | 0 | 0 | 0 | 0 | 63 | 12821 | 0 | 0 | 0 | 3631 | 125 | 388 | 0 | 155 | 0 | 200 | 2024 | 4020 | 407 | 14882 | 8232 | 53 | 0 | 0 | 0 | 0 | 36 | 0 | 0 | 0 | 0 | 338 | 0 |
| SRR5832020 | 0 | 63 | 63 | 0 | 0 | 0 | 0 | 165 | 61 | 7310 | 0 | 0 | 0 | 2441 | 918 | 1548 | 0 | 98 | 0 | 86 | 1836 | 5096 | 523 | 12252 | 7758 | 35 | 0 | 0 | 0 | 0 | 63 | 0 | 0 | 31 | 0 | 557 | 0 |
| SRR5832021 | 0 | 0 | 0 | 0 | 0 | 19 | 0 | 649 | 45 | 2505 | 0 | 0 | 0 | 1677 | 1834 | 1700 | 0 | 73 | 0 | 268 | 1588 | 3842 | 312 | 10668 | 6437 | 179 | 0 | 0 | 0 | 19 | 19 | 0 | 0 | 19 | 0 | 1006 | 0 |
| SRR5832022 | 0 | 26 | 26 | 0 | 0 | 0 | 0 | 485 | 1355 | 22563 | 0 | 0 | 0 | 3845 | 1765 | 2017 | 0 | 114 | 0 | 32 | 2070 | 2449 | 584 | 12816 | 9608 | 68 | 0 | 0 | 0 | 26 | 0 | 0 | 26 | 52 | 0 | 353 | 0 |
| SRR5832023 | 0 | 65 | 0 | 0 | 0 | 0 | 0 | 292 | 172 | 2855 | 0 | 0 | 0 | 894 | 4609 | 3766 | 0 | 72 | 0 | 224 | 1855 | 3971 | 557 | 11187 | 6882 | 96 | 0 | 0 | 22 | 22 | 0 | 0 | 0 | 0 | 22 | 1444 | 0 |
| SRR5832024 | 0 | 0 | 0 | 0 | 0 | 0 | 0 | 146 | 55 | 11937 | 0 | 0 | 0 | 2099 | 548 | 2026 | 0 | 138 | 0 | 164 | 2277 | 4021 | 435 | 13395 | 8361 | 77 | 0 | 0 | 0 | 0 | 56 | 0 | 0 | 0 | 0 | 511 | 0 |
| SRR5832025 | 0 | 0 | 0 | 0 | 0 | 0 | 0 | 707 | 128 | 1199 | 0 | 0 | 0 | 1798 | 6444 | 2462 | 0 | 71 | 0 | 171 | 1914 | 5635 | 397 | 14294 | 8721 | 71 | 0 | 0 | 0 | 0 | 0 | 0 | 0 | 0 | 0 | 664 | 0 |
| SRR5832026 | 0 | 0 | 0 | 0 | 0 | 0 | 0 | 194 | 111 | 3515 | 0 | 0 | 0 | 1875 | 2612 | 1500 | 0 | 80 | 0 | 97 | 1531 | 3827 | 299 | 11583 | 7471 | 109 | 0 | 0 | 16 | 16 | 0 | 16 | 0 | 0 | 16 | 389 | 0 |
| SRR5832027 | 0 | 33 | 0 | 0 | 0 | 0 | 0 | 245 | 20 | 2759 | 0 | 0 | 0 | 899 | 1186 | 2800 | 0 | 68 | 0 | 327 | 2092 | 4629 | 366 | 12272 | 8267 | 62 | 0 | 0 | 0 | 33 | 0 | 0 | 0 | 0 | 0 | 1778 | 0 |
| SRR5832028 | 0 | 27 | 0 | 0 | 0 | 0 | 0 | 276 | 110 | 2207 | 0 | 0 | 0 | 1343 | 1067 | 1159 | 0 | 71 | 0 | 166 | 1520 | 4375 | 292 | 10409 | 6914 | 94 | 0 | 0 | 0 | 14 | 14 | 0 | 0 | 0 | 0 | 920 | 0 |
| SRR5832029 | 0 | 0 | 0 | 0 | 0 | 0 | 0 | 216 | 60 | 12213 | 0 | 0 | 0 | 1860 | 120 | 408 | 0 | 109 | 0 | 348 | 1786 | 2832 | 376 | 11404 | 6756 | 118 | 0 | 0 | 0 | 0 | 0 | 0 | 0 | 0 | 0 | 852 | 0 |
| SRR5868085 | 0 | 1497 | 481 | 70 | 876 | 0 | 0 | 989 | 2310 | 21876 | 13236 | 15705 | 0 | 2777 | 55 | 16 | 0 | 323 | 62 | 1709 | 703 | 3344 | 745 | 82524 | 996 | 0 | 20 | 61 | 89 | 1770 | 7586 | 887 | 736 | 460 | 15 | 7730 | 726 |
| SRR5901584 | 0 | 21 | 10 | 0 | 16 | 0 | 0 | 428 | 100 | 4266 | 6815 | 36642 | 0 | 37 | 687 | 1080 | 0 | 282 | 16 | 218 | 1751 | 3007 | 0 | 3510 | 4356 | 0 | 55 | 16 | 85 | 19 | 16 | 0 | 68 | 35 | 0 | 824 | 211 |
| SRR5901585 | 0 | 13 | 81 | 0 | 23 | 0 | 0 | 805 | 130 | 6831 | 4457 | 25768 | 0 | 48 | 633 | 943 | 0 | 163 | 0 | 208 | 1587 | 2722 | 0 | 4129 | 4461 | 0 | 12 | 14 | 70 | 17 | 15 | 0 | 52 | 28 | 0 | 840 | 163 |
| SRR5901586 | 0 | 10 | 11 | 0 | 22 | 0 | 0 | 884 | 192 | 7538 | 3525 | 22678 | 0 | 49 | 618 | 981 | 0 | 205 | 0 | 217 | 1668 | 3356 | 15 | 4105 | 4018 | 0 | 16 | 27 | 66 | 22 | 18 | 36 | 71 | 41 | 0 | 933 | 152 |
| SRR5901587 | 0 | 154 | 44 | 0 | 12 | 0 | 0 | 681 | 90 | 3416 | 6392 | 33452 | 0 | 37 | 363 | 437 | 0 | 274 | 31 | 330 | 2122 | 4039 | 0 | 4837 | 6939 | 0 | 65 | 15 | 102 | 207 | 178 | 17 | 54 | 229 | 0 | 885 | 334 |
| SRR5901588 | 0 | 235 | 45 | 0 | 13 | 0 | 0 | 596 | 156 | 3403 | 18978 | 45957 | 0 | 28 | 434 | 636 | 0 | 275 | 33 | 278 | 2070 | 3927 | 0 | 3538 | 6116 | 0 | 78 | 22 | 135 | 286 | 189 | 22 | 50 | 289 | 0 | 676 | 370 |
| SRR5901589 | 0 | 19 | 45 | 0 | 14 | 0 | 0 | 793 | 104 | 3575 | 4598 | 26655 | 0 | 24 | 357 | 405 | 0 | 193 | 20 | 277 | 2346 | 3987 | 0 | 4679 | 6341 | 0 | 73 | 28 | 11 | 32 | 25 | 14 | 57 | 27 | 0 | 853 | 350 |
| SRR5901590 | 0 | 57 | 0 | 0 | 0 | 0 | 0 | 176 | 15 | 272 | 4987 | 13565 | 0 | 0 | 11 | 65 | 0 | 63 | 32 | 14 | 4844 | 9839 | 0 | 5708 | 8728 | 0 | 86 | 72 | 33 | 63 | 36 | 50 | 56 | 11 | 0 | 144 | 1216 |
| SRR5901591 | 0 | 101 | 0 | 0 | 0 | 0 | 0 | 115 | 10 | 245 | 8080 | 11323 | 0 | 0 | 0 | 19 | 13 | 62 | 30 | 13 | 4577 | 8090 | 0 | 5254 | 8243 | 0 | 54 | 64 | 358 | 848 | 403 | 32 | 60 | 112 | 0 | 101 | 1446 |
| SRR5901592 | 0 | 0 | 0 | 0 | 0 | 0 | 0 | 95 | 0 | 179 | 6599 | 8863 | 0 | 0 | 0 | 24 | 0 | 48 | 39 | 11 | 4462 | 9113 | 0 | 5689 | 8158 | 0 | 13 | 49 | 30 | 60 | 37 | 24 | 33 | 78 | 0 | 78 | 1330 |
| SRR5901593 | 0 | 108 | 0 | 0 | 0 | 0 | 0 | 83 | 13 | 120 | 7560 | 14757 | 0 | 0 | 13 | 25 | 0 | 55 | 58 | 126 | 3458 | 15421 | 0 | 11146 | 16322 | 0 | 45 | 56 | 18 | 30 | 27 | 24 | 94 | 14 | 0 | 278 | 668 |
| SRR5901594 | 0 | 64 | 0 | 0 | 0 | 0 | 0 | 40 | 0 | 154 | 8453 | 20139 | 0 | 0 | 13 | 22 | 0 | 29 | 45 | 92 | 2910 | 14491 | 0 | 11264 | 15077 | 0 | 40 | 60 | 22 | 34 | 29 | 20 | 91 | 17 | 0 | 306 | 753 |
| SRR5901595 | 0 | 96 | 0 | 0 | 0 | 0 | 0 | 44 | 0 | 145 | 6264 | 16087 | 0 | 0 | 0 | 16 | 0 | 54 | 62 | 106 | 3856 | 16372 | 0 | 13731 | 17562 | 0 | 56 | 58 | 16 | 40 | 29 | 25 | 65 | 15 | 0 | 282 | 809 |
| SRR5922753 | 0 | 32064 | 132 | 10 | 108 | 0 | 0 | 524 | 6725 | 70992 | 2423 | 6059 | 0 | 717 | 21 | 17 | 18 | 5474 | 25 | 201 | 5435 | 4375 | 53 | 3790 | 3746 | 0 | 0 | 100 | 366 | 244 | 130 | 67 | 27 | 20 | 0 | 448 | 0 |
| SRR5922754 | 0 | 753 | 14 | 10 | 96 | 0 | 0 | 288 | 2952 | 24785 | 46735 | 21576 | 0 | 164 | 0 | 0 | 61 | 4047 | 22 | 55 | 5370 | 3717 | 54 | 3129 | 3090 | 0 | 14 | 129 | 11 | 98 | 47 | 11 | 0 | 0 | 0 | 101 | 0 |
| SRR5922756 | 0 | 800 | 0 | 18 | 115 | 20 | 0 | 365 | 575 | 4907 | 8946 | 20726 | 0 | 47 | 0 | 0 | 32 | 5668 | 24 | 0 | 5163 | 3777 | 55 | 5026 | 2542 | 0 | 0 | 21 | 10 | 17 | 0 | 0 | 0 | 0 | 0 | 23 | 0 |
| SRR5925650 | 34 | 0 | 0 | 0 | 11 | 0 | 25 | 85792 | 2069 | 389 | 819 | 498 | 0 | 55 | 147 | 81 | 28 | 890 | 0 | 37 | 234 | 1162 | 130 | 19187 | 2310 | 85 | 0 | 0 | 0 | 0 | 0 | 0 | 0 | 12 | 0 | 387 | 312 |
| SRR5925651 | 37 | 10 | 0 | 0 | 34 | 0 | 17 | 59442 | 2337 | 686 | 3510 | 1688 | 0 | 316 | 175 | 95 | 34 | 943 | 0 | 228 | 1189 | 3897 | 48 | 32773 | 7386 | 398 | 0 | 0 | 0 | 11 | 0 | 0 | 0 | 0 | 0 | 381 | 117 |
| SRR5925652 | 0 | 57 | 17 | 0 | 35 | 0 | 0 | 1086 | 87 | 1841 | 17063 | 11596 | 0 | 1014 | 23 | 13 | 0 | 288 | 49 | 2622 | 2173 | 5424 | 45 | 27657 | 10316 | 26 | 12 | 65 | 35 | 63 | 65 | 26 | 61 | 93 | 15 | 7374 | 42 |
| SRR5925665 | 0 | 0 | 20 | 0 | 36 | 0 | 0 | 1963 | 84 | 1337 | 4433 | 3867 | 0 | 361 | 12 | 36 | 0 | 268 | 50 | 96 | 1903 | 2321 | 17 | 128 | 9181 | 43 | 0 | 69 | 40 | 188 | 253 | 59 | 30 | 54 | 406 | 2277 | 212 |
| SRR5925666 | 0 | 0 | 16 | 0 | 36 | 0 | 0 | 1883 | 51 | 1052 | 392 | 3333 | 0 | 399 | 25 | 51 | 0 | 235 | 42 | 68 | 1957 | 2236 | 18 | 140 | 9591 | 47 | 0 | 78 | 39 | 151 | 171 | 35 | 39 | 74 | 18 | 2035 | 199 |
| SRR5925667 | 0 | 12 | 32 | 0 | 46 | 0 | 0 | 1235 | 56 | 1695 | 369 | 3986 | 0 | 595 | 45 | 22 | 12 | 322 | 50 | 101 | 2387 | 2722 | 0 | 231 | 8308 | 68 | 0 | 154 | 57 | 17 | 18 | 69 | 53 | 81 | 25 | 2403 | 177 |
| SRR5925668 | 0 | 11 | 24 | 0 | 45 | 0 | 0 | 1259 | 40 | 1444 | 575 | 4101 | 0 | 508 | 40 | 13 | 0 | 262 | 49 | 92 | 1921 | 2519 | 0 | 199 | 12314 | 40 | 0 | 121 | 26 | 197 | 273 | 34 | 32 | 76 | 350 | 1609 | 254 |
| SRR5925669 | 0 | 16 | 29 | 0 | 49 | 0 | 0 | 1446 | 85 | 1446 | 3621 | 4180 | 0 | 535 | 85 | 36 | 0 | 283 | 39 | 36 | 2263 | 2609 | 0 | 243 | 11561 | 56 | 0 | 135 | 57 | 14 | 23 | 37 | 45 | 65 | 19 | 1920 | 207 |
| SRR5925670 | 0 | 0 | 13 | 0 | 57 | 0 | 0 | 1401 | 69 | 1355 | 347 | 3894 | 0 | 414 | 69 | 46 | 0 | 277 | 43 | 80 | 2407 | 2439 | 0 | 202 | 9862 | 52 | 13 | 105 | 48 | 302 | 280 | 83 | 44 | 105 | 412 | 2090 | 208 |
| SRR5925671 | 0 | 0 | 57 | 0 | 45 | 0 | 15 | 1891 | 16 | 1157 | 553 | 5011 | 0 | 223 | 32 | 16 | 0 | 321 | 37 | 64 | 3608 | 2183 | 61 | 264 | 9255 | 46 | 0 | 130 | 50 | 271 | 214 | 46 | 38 | 76 | 298 | 2155 | 235 |
| SRR5925672 | 0 | 0 | 37 | 0 | 54 | 0 | 0 | 2187 | 48 | 1521 | 488 | 5173 | 0 | 304 | 0 | 19 | 0 | 297 | 39 | 38 | 3070 | 2354 | 53 | 315 | 8874 | 52 | 0 | 139 | 45 | 234 | 250 | 66 | 70 | 90 | 504 | 1959 | 231 |
| SRR5925673 | 0 | 18 | 29 | 0 | 48 | 0 | 0 | 2527 | 39 | 1678 | 3662 | 3119 | 0 | 207 | 49 | 109 | 0 | 423 | 30 | 188 | 2691 | 3222 | 11 | 117 | 12181 | 27 | 12 | 94 | 29 | 103 | 177 | 50 | 0 | 77 | 18 | 4817 | 172 |
| SRR5925674 | 0 | 15 | 15 | 0 | 41 | 0 | 0 | 1904 | 88 | 1313 | 4115 | 3546 | 0 | 372 | 44 | 153 | 0 | 489 | 39 | 66 | 2365 | 3468 | 11 | 102 | 13463 | 34 | 0 | 74 | 22 | 140 | 133 | 22 | 22 | 59 | 21 | 2714 | 230 |
| SRR5925675 | 12 | 0 | 20 | 0 | 40 | 0 | 12 | 1763 | 67 | 2723 | 459 | 3917 | 0 | 826 | 78 | 78 | 0 | 370 | 43 | 179 | 1810 | 2548 | 10 | 210 | 7671 | 49 | 20 | 18 | 15 | 67 | 35 | 0 | 20 | 12 | 18 | 3024 | 197 |
| SRR5925676 | 0 | 0 | 27 | 0 | 55 | 0 | 0 | 1923 | 113 | 2317 | 549 | 4128 | 0 | 603 | 72 | 16 | 0 | 415 | 33 | 32 | 2901 | 2580 | 14 | 242 | 9180 | 55 | 0 | 42 | 27 | 83 | 86 | 44 | 18 | 39 | 22 | 2156 | 194 |
| SRR5925677 | 0 | 0 | 0 | 0 | 55 | 0 | 0 | 1494 | 76 | 1580 | 385 | 4449 | 0 | 433 | 22 | 65 | 0 | 332 | 51 | 54 | 2258 | 2300 | 28 | 475 | 9219 | 67 | 0 | 76 | 45 | 112 | 85 | 36 | 27 | 18 | 18 | 1905 | 251 |
| SRR5925678 | 0 | 0 | 10 | 0 | 39 | 0 | 0 | 1368 | 36 | 1670 | 479 | 4914 | 0 | 480 | 27 | 44 | 0 | 371 | 53 | 89 | 2496 | 2743 | 22 | 346 | 4931 | 83 | 10 | 74 | 30 | 151 | 104 | 17 | 10 | 44 | 26 | 2008 | 167 |
| SRR5925679 | 0 | 0 | 19 | 0 | 42 | 0 | 0 | 1190 | 10 | 1741 | 436 | 5143 | 0 | 334 | 49 | 79 | 0 | 417 | 45 | 98 | 3457 | 3251 | 14 | 221 | 12854 | 44 | 0 | 107 | 31 | 157 | 153 | 50 | 42 | 31 | 18 | 3068 | 188 |
| SRR5925680 | 0 | 17 | 14 | 0 | 38 | 0 | 0 | 1622 | 56 | 1541 | 693 | 5460 | 0 | 248 | 43 | 19 | 0 | 359 | 38 | 74 | 3027 | 3323 | 12 | 185 | 13552 | 37 | 0 | 74 | 36 | 119 | 112 | 48 | 14 | 45 | 295 | 2284 | 183 |
| SRR5925681 | 0 | 17 | 54 | 0 | 43 | 0 | 0 | 1193 | 16 | 984 | 992 | 5188 | 0 | 319 | 88 | 126 | 0 | 385 | 75 | 33 | 3233 | 2496 | 0 | 64 | 8741 | 27 | 0 | 155 | 42 | 172 | 147 | 88 | 42 | 38 | 289 | 1028 | 138 |
| SRR5925682 | 0 | 19 | 15 | 0 | 45 | 0 | 0 | 1190 | 21 | 1001 | 5514 | 6060 | 0 | 252 | 35 | 70 | 0 | 322 | 66 | 0 | 2640 | 2787 | 0 | 103 | 11573 | 33 | 0 | 115 | 50 | 12 | 169 | 27 | 15 | 54 | 19 | 602 | 183 |
| SRR5925683 | 0 | 0 | 22 | 0 | 55 | 0 | 0 | 1238 | 76 | 1162 | 332 | 2891 | 0 | 586 | 0 | 0 | 0 | 309 | 45 | 57 | 2795 | 2813 | 22 | 250 | 12871 | 38 | 0 | 103 | 49 | 139 | 121 | 27 | 31 | 18 | 22 | 1606 | 206 |
| SRR5925684 | 0 | 0 | 26 | 0 | 37 | 0 | 0 | 1198 | 78 | 1187 | 4021 | 4603 | 0 | 235 | 11 | 45 | 0 | 328 | 51 | 78 | 3636 | 2823 | 21 | 185 | 13218 | 26 | 0 | 124 | 62 | 176 | 191 | 88 | 52 | 57 | 22 | 1568 | 176 |
| SRR5925685 | 0 | 0 | 28 | 0 | 0 | 0 | 0 | 920 | 42 | 2341 | 1678 | 1296 | 0 | 1152 | 328 | 305 | 0 | 116 | 109 | 0 | 792 | 751 | 0 | 0 | 5139 | 0 | 0 | 11 | 53 | 131 | 128 | 50 | 18 | 53 | 15 | 0 | 106 |
| SRR5925686 | 0 | 0 | 23 | 0 | 0 | 0 | 0 | 862 | 45 | 2202 | 1846 | 1218 | 0 | 1315 | 350 | 259 | 0 | 103 | 106 | 0 | 774 | 748 | 0 | 0 | 4742 | 0 | 0 | 235 | 68 | 174 | 140 | 34 | 45 | 61 | 246 | 13 | 89 |
| SRR5925687 | 0 | 0 | 31 | 0 | 0 | 0 | 0 | 847 | 73 | 1939 | 322 | 2143 | 0 | 798 | 440 | 473 | 0 | 71 | 91 | 0 | 805 | 922 | 0 | 0 | 6260 | 0 | 0 | 166 | 66 | 175 | 197 | 26 | 35 | 44 | 485 | 0 | 130 |
| SRR5925688 | 0 | 0 | 12 | 0 | 0 | 0 | 0 | 943 | 33 | 2048 | 2146 | 1561 | 0 | 1065 | 236 | 203 | 0 | 72 | 88 | 0 | 771 | 684 | 0 | 0 | 5235 | 0 | 0 | 10 | 90 | 114 | 138 | 37 | 0 | 16 | 19 | 0 | 97 |
| SRR5925689 | 0 | 0 | 32 | 0 | 0 | 0 | 0 | 625 | 51 | 2224 | 2828 | 1926 | 0 | 1345 | 378 | 327 | 0 | 97 | 84 | 0 | 700 | 890 | 0 | 0 | 6224 | 0 | 0 | 15 | 61 | 148 | 94 | 36 | 50 | 40 | 21 | 0 | 150 |
| SRR5925690 | 0 | 0 | 27 | 0 | 0 | 0 | 0 | 874 | 53 | 2336 | 2528 | 1926 | 0 | 966 | 357 | 390 | 0 | 88 | 81 | 0 | 814 | 835 | 0 | 0 | 5686 | 0 | 0 | 12 | 104 | 156 | 104 | 40 | 34 | 24 | 22 | 0 | 111 |
| SRR5925691 | 11 | 0 | 21 | 0 | 0 | 0 | 0 | 828 | 0 | 2403 | 1838 | 1299 | 0 | 983 | 478 | 404 | 0 | 91 | 97 | 0 | 774 | 1039 | 0 | 0 | 6129 | 0 | 0 | 165 | 74 | 130 | 151 | 56 | 32 | 53 | 291 | 13 | 75 |
| SRR5925692 | 0 | 0 | 36 | 0 | 0 | 0 | 0 | 724 | 44 | 2008 | 2310 | 1668 | 0 | 988 | 359 | 365 | 0 | 124 | 103 | 0 | 829 | 966 | 0 | 0 | 6024 | 0 | 0 | 10 | 46 | 154 | 121 | 79 | 39 | 52 | 18 | 0 | 71 |
| SRR5925693 | 0 | 0 | 32 | 0 | 0 | 0 | 0 | 805 | 25 | 2345 | 263 | 1327 | 0 | 799 | 428 | 409 | 0 | 79 | 110 | 0 | 695 | 851 | 0 | 0 | 6432 | 0 | 0 | 14 | 61 | 14 | 120 | 48 | 16 | 68 | 20 | 0 | 78 |
| SRR5925694 | 0 | 0 | 21 | 0 | 18 | 0 | 0 | 755 | 19 | 2441 | 1706 | 1744 | 0 | 888 | 469 | 368 | 0 | 90 | 109 | 0 | 613 | 773 | 0 | 0 | 5814 | 0 | 0 | 15 | 90 | 10 | 10 | 63 | 36 | 54 | 24 | 13 | 95 |
| SRR5925695 | 0 | 0 | 42 | 0 | 0 | 0 | 0 | 436 | 26 | 1008 | 358 | 1815 | 0 | 560 | 221 | 156 | 0 | 100 | 77 | 0 | 888 | 768 | 0 | 0 | 5284 | 0 | 0 | 175 | 42 | 156 | 104 | 28 | 0 | 28 | 316 | 0 | 105 |
| SRR5925696 | 0 | 0 | 33 | 0 | 10 | 0 | 0 | 398 | 23 | 1318 | 406 | 2336 | 0 | 567 | 222 | 192 | 0 | 94 | 75 | 0 | 803 | 778 | 0 | 0 | 5231 | 0 | 0 | 113 | 80 | 155 | 160 | 52 | 0 | 38 | 13 | 15 | 108 |
| SRR5925697 | 0 | 0 | 33 | 0 | 24 | 0 | 0 | 258 | 46 | 1335 | 3060 | 2526 | 0 | 1009 | 201 | 155 | 0 | 86 | 149 | 0 | 746 | 816 | 0 | 0 | 6662 | 0 | 0 | 310 | 105 | 172 | 133 | 41 | 21 | 43 | 380 | 27 | 182 |
| SRR5925698 | 0 | 0 | 44 | 0 | 0 | 0 | 0 | 532 | 43 | 848 | 316 | 1812 | 0 | 582 | 208 | 187 | 0 | 108 | 146 | 0 | 932 | 823 | 0 | 0 | 5862 | 0 | 0 | 331 | 94 | 178 | 202 | 35 | 25 | 59 | 365 | 14 | 158 |
| SRR5925699 | 0 | 0 | 34 | 0 | 0 | 0 | 0 | 714 | 36 | 1561 | 2261 | 1781 | 0 | 858 | 342 | 358 | 0 | 91 | 85 | 0 | 864 | 836 | 0 | 0 | 5997 | 0 | 0 | 11 | 66 | 151 | 98 | 43 | 33 | 38 | 21 | 0 | 104 |
| SRR5925700 | 0 | 0 | 34 | 0 | 18 | 0 | 0 | 580 | 14 | 1188 | 2133 | 1783 | 0 | 723 | 351 | 329 | 0 | 98 | 89 | 0 | 947 | 851 | 0 | 0 | 6101 | 0 | 0 | 194 | 97 | 147 | 126 | 34 | 29 | 51 | 412 | 0 | 122 |
| SRR5925701 | 0 | 0 | 32 | 0 | 0 | 0 | 0 | 667 | 36 | 1832 | 1760 | 1244 | 0 | 823 | 443 | 361 | 0 | 116 | 98 | 0 | 801 | 847 | 0 | 0 | 6359 | 0 | 0 | 226 | 92 | 162 | 131 | 42 | 29 | 62 | 300 | 0 | 97 |
| SRR5925702 | 12 | 0 | 23 | 0 | 11 | 0 | 0 | 654 | 28 | 1655 | 2181 | 1648 | 0 | 781 | 419 | 362 | 0 | 96 | 94 | 0 | 756 | 961 | 0 | 0 | 6246 | 0 | 0 | 183 | 51 | 97 | 82 | 43 | 47 | 39 | 327 | 0 | 100 |
| SRR5925703 | 0 | 0 | 27 | 0 | 17 | 0 | 0 | 610 | 33 | 1687 | 267 | 1647 | 0 | 735 | 433 | 263 | 11 | 102 | 96 | 0 | 796 | 839 | 0 | 0 | 5123 | 0 | 0 | 12 | 23 | 11 | 126 | 73 | 31 | 50 | 19 | 0 | 90 |
| SRR5925704 | 11 | 0 | 37 | 0 | 10 | 0 | 0 | 694 | 35 | 1468 | 1278 | 1408 | 0 | 622 | 279 | 303 | 0 | 98 | 87 | 0 | 839 | 683 | 0 | 0 | 4844 | 0 | 0 | 204 | 85 | 169 | 176 | 54 | 29 | 83 | 16 | 0 | 83 |
| SRR5925705 | 0 | 0 | 0 | 0 | 20 | 0 | 0 | 1659 | 29 | 2396 | 246 | 5738 | 0 | 698 | 16 | 29 | 0 | 823 | 82 | 7255 | 4874 | 1346 | 0 | 167 | 7215 | 40 | 0 | 33 | 0 | 68 | 120 | 32 | 33 | 69 | 40 | 40907 | 96 |
| SRR5925706 | 0 | 0 | 0 | 0 | 23 | 0 | 0 | 1027 | 25 | 2546 | 176 | 4680 | 0 | 1183 | 59 | 84 | 0 | 134 | 48 | 6359 | 1188 | 2419 | 0 | 197 | 9504 | 47 | 0 | 46 | 0 | 15 | 23 | 32 | 37 | 45 | 14 | 32341 | 52 |
| SRR5925707 | 0 | 0 | 0 | 0 | 24 | 0 | 0 | 1849 | 31 | 2450 | 87 | 2710 | 0 | 563 | 0 | 43 | 0 | 446 | 78 | 7609 | 4079 | 1239 | 0 | 151 | 7731 | 38 | 0 | 60 | 12 | 11 | 23 | 36 | 62 | 89 | 39 | 42289 | 108 |
| SRR5925708 | 0 | 0 | 0 | 0 | 23 | 0 | 0 | 1154 | 47 | 2161 | 176 | 3925 | 0 | 1239 | 25 | 38 | 0 | 159 | 45 | 7868 | 1155 | 2336 | 0 | 187 | 9801 | 51 | 0 | 38 | 0 | 102 | 127 | 19 | 26 | 40 | 13 | 37457 | 70 |
| SRR5925709 | 21 | 0 | 122 | 0 | 27 | 0 | 0 | 2043 | 51 | 2472 | 461 | 3823 | 0 | 288 | 67 | 176 | 0 | 592 | 71 | 368 | 2971 | 962 | 0 | 0 | 5296 | 0 | 16 | 106 | 58 | 388 | 303 | 133 | 144 | 218 | 308 | 2456 | 29 |
| SRR5925710 | 0 | 0 | 0 | 0 | 58 | 0 | 0 | 984 | 0 | 984 | 984 | 3937 | 0 | 0 | 0 | 0 | 0 | 232 | 0 | 984 | 2725 | 1276 | 0 | 0 | 6725 | 0 | 0 | 0 | 0 | 0 | 0 | 0 | 0 | 0 | 27 | 7874 | 0 |
| SRR5925711 | 0 | 0 | 0 | 0 | 20 | 0 | 0 | 2332 | 142 | 4307 | 1053 | 12412 | 0 | 2045 | 92 | 95 | 0 | 222 | 82 | 2050 | 1733 | 2259 | 0 | 47 | 9310 | 21 | 0 | 53 | 11 | 12 | 11 | 26 | 33 | 45 | 20 | 10295 | 40 |
| SRR5925712 | 0 | 0 | 0 | 0 | 59 | 0 | 0 | 3194 | 130 | 432 | 880 | 5532 | 0 | 135 | 0 | 0 | 0 | 605 | 635 | 790 | 1455 | 347 | 0 | 28 | 7445 | 27 | 0 | 31 | 11 | 48 | 70 | 31 | 36 | 82 | 30 | 21568 | 64 |
| SRR5925713 | 0 | 0 | 13 | 0 | 27 | 0 | 0 | 2941 | 47 | 4952 | 227 | 3116 | 0 | 495 | 28 | 39 | 0 | 657 | 81 | 4721 | 7043 | 1707 | 0 | 76 | 11333 | 16 | 0 | 22 | 0 | 82 | 23 | 39 | 50 | 13 | 81 | 38641 | 123 |
| SRR5925714 | 0 | 0 | 0 | 0 | 26 | 0 | 0 | 2303 | 72 | 4950 | 111 | 3854 | 0 | 519 | 19 | 72 | 0 | 992 | 86 | 4059 | 9661 | 1999 | 0 | 80 | 11213 | 23 | 0 | 22 | 0 | 12 | 27 | 39 | 45 | 15 | 10 | 36884 | 106 |
| SRR5925715 | 0 | 0 | 0 | 0 | 30 | 0 | 0 | 1673 | 92 | 7738 | 656 | 2872 | 0 | 590 | 128 | 122 | 0 | 118 | 44 | 4297 | 1462 | 2565 | 12 | 148 | 9711 | 47 | 0 | 27 | 0 | 85 | 11 | 22 | 31 | 42 | 14 | 33890 | 55 |
| SRR5925716 | 0 | 0 | 0 | 0 | 19 | 0 | 0 | 1427 | 96 | 6649 | 722 | 3280 | 0 | 779 | 164 | 142 | 0 | 127 | 40 | 3454 | 1601 | 2732 | 10 | 157 | 10126 | 51 | 0 | 12 | 0 | 53 | 79 | 0 | 11 | 22 | 14 | 32861 | 51 |
| SRR5925717 | 0 | 0 | 0 | 0 | 24 | 0 | 0 | 1608 | 58 | 2908 | 293 | 4152 | 0 | 1274 | 229 | 213 | 0 | 129 | 50 | 4800 | 1018 | 2266 | 0 | 77 | 11911 | 0 | 0 | 0 | 0 | 85 | 144 | 10 | 18 | 33 | 18 | 34009 | 27 |
| SRR5925718 | 0 | 0 | 0 | 0 | 24 | 0 | 0 | 1358 | 66 | 2665 | 226 | 4285 | 0 | 1343 | 248 | 189 | 0 | 146 | 45 | 5217 | 984 | 2193 | 0 | 70 | 12654 | 0 | 0 | 0 | 0 | 83 | 159 | 14 | 24 | 26 | 26 | 35916 | 28 |
| SRR5925719 | 0 | 0 | 13 | 0 | 0 | 0 | 0 | 782 | 34 | 2248 | 418 | 1974 | 0 | 748 | 271 | 209 | 14 | 62 | 96 | 21 | 401 | 757 | 0 | 0 | 5027 | 0 | 0 | 15 | 35 | 110 | 95 | 41 | 33 | 39 | 11 | 101 | 46 |
| SRR5925720 | 0 | 0 | 22 | 0 | 0 | 0 | 0 | 850 | 43 | 2267 | 387 | 1790 | 0 | 888 | 283 | 241 | 14 | 74 | 87 | 14 | 421 | 680 | 0 | 0 | 4972 | 0 | 0 | 14 | 52 | 98 | 86 | 40 | 38 | 52 | 12 | 52 | 46 |
| SRR5925721 | 10 | 0 | 11 | 0 | 0 | 0 | 0 | 549 | 92 | 3221 | 424 | 1445 | 0 | 1048 | 516 | 358 | 0 | 65 | 125 | 16 | 520 | 832 | 0 | 0 | 5539 | 0 | 0 | 92 | 15 | 51 | 44 | 19 | 24 | 28 | 20 | 98 | 47 |
| SRR5925722 | 0 | 0 | 14 | 0 | 0 | 0 | 0 | 619 | 31 | 2965 | 84 | 1391 | 0 | 1002 | 455 | 302 | 10 | 75 | 131 | 41 | 480 | 824 | 0 | 0 | 5463 | 0 | 0 | 123 | 12 | 46 | 59 | 22 | 24 | 34 | 12 | 72 | 42 |
| SRR5925723 | 0 | 0 | 0 | 0 | 0 | 13 | 0 | 870 | 49 | 2820 | 114 | 2820 | 0 | 629 | 74 | 74 | 0 | 54 | 122 | 10 | 519 | 585 | 0 | 0 | 4438 | 0 | 0 | 72 | 0 | 40 | 50 | 0 | 14 | 23 | 0 | 49 | 21 |
| SRR5925724 | 0 | 0 | 0 | 0 | 0 | 0 | 0 | 836 | 79 | 3003 | 539 | 2533 | 0 | 594 | 64 | 59 | 0 | 53 | 152 | 15 | 427 | 570 | 0 | 0 | 4712 | 0 | 0 | 13 | 0 | 25 | 43 | 17 | 13 | 17 | 0 | 94 | 35 |
| SRR5925725 | 14 | 0 | 0 | 0 | 0 | 0 | 0 | 623 | 29 | 2203 | 146 | 2697 | 0 | 900 | 115 | 81 | 0 | 55 | 190 | 0 | 468 | 718 | 0 | 0 | 6044 | 0 | 0 | 77 | 0 | 25 | 43 | 0 | 0 | 20 | 0 | 72 | 39 |
| SRR5925726 | 12 | 0 | 0 | 0 | 0 | 10 | 0 | 696 | 41 | 2449 | 170 | 1769 | 0 | 902 | 92 | 67 | 12 | 66 | 207 | 0 | 437 | 647 | 0 | 0 | 6553 | 0 | 0 | 14 | 0 | 17 | 33 | 0 | 0 | 16 | 0 | 57 | 51 |
| SRR5925727 | 0 | 0 | 0 | 0 | 0 | 0 | 0 | 0 | 0 | 0 | 0 | 0 | 0 | 0 | 0 | 0 | 0 | 447 | 0 | 0 | 2234 | 447 | 0 | 0 | 7149 | 0 | 0 | 0 | 0 | 0 | 0 | 0 | 0 | 0 | 0 | 8130 | 0 |
| SRR5925728 | 0 | 0 | 0 | 0 | 0 | 0 | 0 | 1417 | 62 | 4019 | 90 | 1126 | 0 | 2236 | 179 | 299 | 0 | 52 | 133 | 12 | 373 | 877 | 0 | 0 | 6298 | 0 | 0 | 31 | 15 | 46 | 87 | 46 | 39 | 57 | 40 | 105 | 40 |
| SRR5925729 | 0 | 0 | 0 | 0 | 0 | 0 | 0 | 742 | 21 | 2917 | 597 | 2547 | 0 | 639 | 103 | 58 | 0 | 56 | 80 | 0 | 412 | 636 | 0 | 0 | 4693 | 0 | 0 | 314 | 25 | 80 | 53 | 21 | 25 | 17 | 0 | 87 | 31 |
| SRR5925730 | 0 | 0 | 0 | 0 | 0 | 0 | 0 | 675 | 30 | 2814 | 162 | 2272 | 0 | 764 | 77 | 64 | 0 | 56 | 86 | 0 | 404 | 593 | 0 | 0 | 4517 | 0 | 0 | 286 | 24 | 58 | 55 | 15 | 21 | 31 | 0 | 64 | 28 |
| SRR5925731 | 0 | 0 | 0 | 0 | 0 | 0 | 0 | 1078 | 44 | 4030 | 307 | 1699 | 0 | 1567 | 320 | 270 | 0 | 23 | 122 | 31 | 239 | 499 | 0 | 0 | 4424 | 0 | 0 | 80 | 27 | 102 | 112 | 57 | 65 | 47 | 36 | 176 | 24 |
| SRR5925732 | 0 | 0 | 13 | 0 | 0 | 0 | 0 | 1131 | 16 | 4048 | 27 | 792 | 0 | 1026 | 226 | 170 | 0 | 30 | 147 | 0 | 270 | 553 | 0 | 0 | 5294 | 0 | 0 | 124 | 41 | 129 | 170 | 41 | 41 | 73 | 20 | 113 | 38 |
| SRR5930976 | 0 | 120 | 24 | 0 | 26 | 0 | 0 | 321 | 78 | 1523 | 18528 | 8258 | 0 | 1604 | 17 | 15 | 0 | 303 | 34 | 2449 | 2018 | 5687 | 45 | 49946 | 9515 | 30 | 0 | 22 | 116 | 31 | 31 | 86 | 19 | 31 | 50 | 7563 | 23 |
| SRR5930977 | 19 | 0 | 0 | 0 | 0 | 13 | 47 | 31232 | 1392 | 1006 | 1752 | 2607 | 0 | 30 | 0 | 0 | 46 | 280 | 18 | 1723 | 360 | 1778 | 71 | 7613 | 1290 | 150 | 0 | 0 | 0 | 0 | 0 | 0 | 11 | 16 | 26 | 4632 | 93 |
| SRR5930978 | 11 | 22 | 0 | 0 | 33 | 13 | 18 | 77242 | 2702 | 251 | 2928 | 1558 | 10 | 89 | 104 | 46 | 41 | 1480 | 0 | 150 | 1053 | 3500 | 117 | 27420 | 5037 | 203 | 15 | 18 | 34 | 33 | 24 | 0 | 12 | 13 | 0 | 412 | 116 |
| SRR5936591 | 0 | 0 | 0 | 0 | 11 | 0 | 0 | 247 | 57 | 1427 | 14564 | 9053 | 0 | 564 | 0 | 0 | 0 | 180 | 78 | 476 | 470 | 1645 | 41 | 17150 | 4676 | 0 | 0 | 51 | 13 | 68 | 65 | 37 | 73 | 73 | 17 | 2890 | 61 |
| SRR5936592 | 0 | 0 | 0 | 0 | 14 | 13 | 141 | 74803 | 1583 | 64 | 350 | 470 | 0 | 0 | 0 | 0 | 149 | 126 | 51 | 20 | 674 | 1376 | 222 | 21999 | 3197 | 48 | 0 | 0 | 0 | 11 | 0 | 0 | 0 | 0 | 0 | 225 | 168 |

### Supp Table S8.4.Columns_DI_to_ES

| | | | | | | | | | | | | | | | | | | | | | | | | | | | | | | | | | | | | | |
|---|---|---|---|---|---|---|---|---|---|---|---|---|---|---|---|---|---|---|---|---|---|---|---|---|---|---|---|---|---|---|---|---|---|---|---|---|---|
| SRR5951878 | 0 | 109 | 85 | 244 | 1517 | 0 | 0 | 48 | 235 | 139 | 2666 | 8879 | 0 | 29 | 41 | 542 | 0 | 373 | 115 | 8027 | 520 | 299 | 0 | 74 | 4387 | 62 | 95 | 24 | 35 | 23 | 49 | 183 | 67 | 26 | 0 | 8838 | 1618 |
| SRR5951879 | 0 | 101 | 65 | 886 | 1121 | 0 | 0 | 55 | 231 | 644 | 1262 | 6598 | 0 | 57 | 330 | 1553 | 0 | 1153 | 80 | 2237 | 607 | 968 | 0 | 55 | 4629 | 59 | 77 | 99 | 12 | 12 | 11 | 37 | 13 | 16 | 0 | 3512 | 1870 |
| SRR5951880 | 0 | 0 | 0 | 0 | 10 | 0 | 0 | 25 | 0 | 0 | 76 | 4934 | 0 | 0 | 0 | 0 | 0 | 0 | 0 | 401 | 0 | 0 | 0 | 0 | 0 | 36 | 0 | 0 | 0 | 0 | 0 | 0 | 0 | 0 | 0 | 776 | 0 |
| SRR5951881 | 0 | 164 | 219 | 188 | 1104 | 0 | 0 | 145 | 461 | 253 | 2002 | 9234 | 0 | 158 | 1705 | 5166 | 0 | 998 | 118 | 8419 | 330 | 193 | 0 | 304 | 2947 | 264 | 200 | 17 | 286 | 13 | 34 | 173 | 55 | 32 | 0 | 11274 | 2083 |
| SRR5951882 | 0 | 209 | 479 | 711 | 1138 | 0 | 0 | 118 | 371 | 1446 | 1400 | 13694 | 0 | 348 | 499 | 2657 | 0 | 1951 | 99 | 4593 | 386 | 439 | 10 | 288 | 3048 | 349 | 17 | 265 | 193 | 233 | 404 | 60 | 29 | 29 | 12 | 7163 | 2240 |
| SRR5951883 | 0 | 0 | 0 | 0 | 10 | 0 | 0 | 0 | 0 | 0 | 100 | 5227 | 48 | 0 | 18 | 18 | 0 | 0 | 0 | 512 | 0 | 0 | 30 | 0 | 0 | 200 | 0 | 0 | 0 | 0 | 32 | 0 | 0 | 0 | 0 | 2313 | 0 |
| SRR5951884 | 0 | 13 | 412 | 92 | 229 | 0 | 0 | 172 | 67 | 689 | 7089 | 28550 | 0 | 105 | 1235 | 5382 | 13 | 484 | 177 | 2994 | 141 | 256 | 0 | 111 | 2441 | 154 | 103 | 179 | 627 | 1720 | 3677 | 633 | 320 | 194 | 43 | 28430 | 1051 |
| SRR5951885 | 0 | 36 | 494 | 79 | 177 | 0 | 0 | 221 | 55 | 669 | 848 | 23042 | 0 | 179 | 711 | 3692 | 0 | 472 | 186 | 3492 | 139 | 245 | 0 | 101 | 2364 | 158 | 61 | 147 | 536 | 1751 | 3573 | 702 | 246 | 258 | 25 | 27714 | 1061 |
| SRR5951886 | 0 | 20 | 419 | 81 | 147 | 0 | 0 | 155 | 87 | 601 | 5820 | 25365 | 0 | 242 | 1731 | 7401 | 0 | 431 | 220 | 2008 | 117 | 251 | 0 | 99 | 2001 | 189 | 131 | 124 | 660 | 1492 | 3504 | 750 | 242 | 261 | 30 | 20103 | 1025 |
| SRR5951887 | 0 | 0 | 449 | 102 | 287 | 0 | 0 | 143 | 60 | 645 | 4622 | 24575 | 0 | 260 | 215 | 2132 | 0 | 338 | 106 | 2399 | 68 | 150 | 0 | 12 | 1629 | 92 | 209 | 1025 | 898 | 4749 | 7407 | 1052 | 327 | 435 | 71 | 6583 | 804 |
| SRR5951888 | 0 | 0 | 487 | 93 | 301 | 0 | 0 | 124 | 72 | 534 | 1181 | 21675 | 0 | 255 | 265 | 1870 | 0 | 340 | 117 | 2138 | 63 | 142 | 0 | 12 | 1462 | 99 | 458 | 781 | 786 | 4664 | 8623 | 1093 | 317 | 403 | 71 | 6263 | 789 |
| SRR5951889 | 0 | 0 | 21 | 91 | 297 | 0 | 0 | 135 | 50 | 745 | 4635 | 26962 | 0 | 405 | 212 | 2412 | 0 | 382 | 116 | 2539 | 81 | 186 | 0 | 11 | 1699 | 104 | 15 | 30 | 33 | 180 | 347 | 52 | 15 | 19 | 75 | 8043 | 851 |
| SRR5951890 | 0 | 27 | 603 | 76 | 201 | 0 | 0 | 114 | 55 | 498 | 7644 | 38119 | 0 | 216 | 158 | 1983 | 0 | 235 | 90 | 2587 | 92 | 229 | 0 | 162 | 2320 | 75 | 467 | 458 | 640 | 3313 | 5732 | 907 | 271 | 304 | 65 | 6395 | 1548 |
| SRR5951891 | 0 | 18 | 30 | 66 | 171 | 0 | 0 | 90 | 45 | 459 | 1335 | 27023 | 0 | 254 | 234 | 1522 | 0 | 319 | 104 | 2457 | 75 | 232 | 0 | 153 | 2054 | 82 | 36 | 19 | 33 | 201 | 340 | 55 | 16 | 18 | 75 | 7306 | 1488 |
| SRR5951892 | 0 | 22 | 25 | 66 | 212 | 0 | 0 | 69 | 50 | 557 | 5634 | 27916 | 0 | 274 | 99 | 1269 | 0 | 280 | 89 | 3709 | 91 | 182 | 0 | 188 | 2127 | 93 | 19 | 18 | 37 | 247 | 468 | 68 | 18 | 17 | 62 | 10281 | 1628 |
| SRR5951893 | 0 | 14 | 494 | 71 | 219 | 0 | 0 | 156 | 45 | 516 | 1172 | 14717 | 0 | 475 | 720 | 6139 | 0 | 496 | 121 | 4134 | 87 | 254 | 28 | 2012 | 1823 | 189 | 1501 | 141 | 479 | 889 | 2419 | 571 | 159 | 198 | 41 | 20656 | 955 |
| SRR5951894 | 0 | 22 | 286 | 90 | 252 | 0 | 0 | 142 | 71 | 653 | 10346 | 30285 | 0 | 192 | 369 | 1966 | 0 | 695 | 209 | 2718 | 101 | 197 | 68 | 4680 | 2011 | 305 | 26 | 386 | 1195 | 2160 | 4121 | 1234 | 462 | 406 | 122 | 17421 | 983 |
| SRR5951895 | 0 | 17 | 14 | 89 | 255 | 0 | 0 | 97 | 75 | 570 | 1037 | 14041 | 18 | 364 | 895 | 5799 | 0 | 490 | 106 | 4540 | 114 | 266 | 33 | 2196 | 1981 | 165 | 89 | 148 | 18 | 35 | 91 | 19 | 177 | 183 | 38 | 22305 | 1058 |
| SRR5951896 | 0 | 89 | 79 | 133 | 252 | 0 | 0 | 673 | 1991 | 2850 | 1915 | 9315 | 0 | 522 | 12 | 75 | 0 | 1828 | 211 | 7696 | 128 | 819 | 175 | 24197 | 2429 | 27 | 82 | 116 | 116 | 315 | 374 | 52 | 23 | 35 | 18 | 11074 | 98 |
| SRR5951897 | 0 | 99 | 42 | 307 | 418 | 0 | 0 | 637 | 2385 | 9866 | 815 | 5711 | 0 | 206 | 26 | 160 | 0 | 2134 | 96 | 4337 | 345 | 952 | 354 | 18314 | 3364 | 15 | 30 | 87 | 197 | 144 | 14 | 47 | 25 | 24 | 10 | 5046 | 84 |
| SRR5951898 | 0 | 0 | 0 | 0 | 0 | 0 | 0 | 28 | 0 | 28 | 1816 | 5587 | 0 | 0 | 0 | 0 | 0 | 0 | 0 | 1341 | 0 | 0 | 435 | 0 | 0 | 11 | 12 | 0 | 0 | 0 | 0 | 0 | 0 | 0 | 0 | 8381 | 0 |
| SRR5951899 | 0 | 83 | 86 | 180 | 347 | 0 | 0 | 1090 | 3299 | 24788 | 1937 | 11426 | 0 | 179 | 101 | 578 | 0 | 985 | 91 | 8377 | 118 | 271 | 280 | 13906 | 1962 | 24 | 97 | 164 | 353 | 379 | 451 | 93 | 31 | 23 | 56 | 8552 | 61 |
| SRR5951900 | 0 | 123 | 94 | 733 | 537 | 0 | 0 | 864 | 2302 | 21777 | 3378 | 12747 | 0 | 295 | 79 | 424 | 0 | 1737 | 102 | 4030 | 200 | 472 | 766 | 18812 | 3473 | 35 | 68 | 202 | 14 | 11 | 18 | 74 | 26 | 30 | 13 | 3912 | 52 |
| SRR5951901 | 0 | 0 | 0 | 0 | 0 | 0 | 0 | 0 | 0 | 0 | 1860 | 8225 | 0 | 0 | 0 | 0 | 0 | 0 | 0 | 816 | 0 | 0 | 449 | 0 | 0 | 0 | 0 | 0 | 0 | 0 | 0 | 0 | 0 | 0 | 0 | 4635 | 0 |
| SRR5951902 | 0 | 107 | 89 | 58 | 246 | 0 | 0 | 585 | 2070 | 2222 | 2115 | 10571 | 0 | 392 | 49 | 219 | 0 | 1617 | 118 | 8703 | 141 | 575 | 199 | 20211 | 1759 | 20 | 211 | 109 | 150 | 257 | 222 | 36 | 19 | 21 | 13 | 13420 | 101 |
| SRR5951903 | 0 | 90 | 92 | 140 | 542 | 0 | 0 | 567 | 2645 | 2202 | 3372 | 14176 | 0 | 140 | 252 | 707 | 0 | 2296 | 46 | 4319 | 336 | 1556 | 527 | 25061 | 2308 | 12 | 17 | 75 | 158 | 10 | 205 | 18 | 12 | 0 | 0 | 4275 | 74 |
| SRR5951904 | 0 | 0 | 0 | 0 | 0 | 0 | 0 | 0 | 0 | 0 | 2363 | 8710 | 11 | 0 | 0 | 0 | 0 | 0 | 0 | 1993 | 0 | 0 | 625 | 0 | 0 | 0 | 0 | 0 | 0 | 0 | 0 | 0 | 0 | 11 | 0 | 10817 | 0 |
| SRR5951905 | 0 | 13 | 126 | 106 | 241 | 0 | 13 | 581 | 1548 | 3062 | 774 | 13249 | 0 | 525 | 0 | 39 | 0 | 2161 | 150 | 11258 | 163 | 730 | 229 | 24836 | 2433 | 36 | 21 | 133 | 19 | 16 | 17 | 39 | 19 | 32 | 36 | 15795 | 87 |
| SRR5951906 | 0 | 96 | 192 | 250 | 381 | 0 | 0 | 578 | 1866 | 3255 | 9264 | 41078 | 0 | 370 | 42 | 196 | 0 | 2605 | 72 | 4645 | 138 | 2047 | 319 | 28697 | 2919 | 21 | 61 | 45 | 125 | 11 | 16 | 25 | 26 | 25 | 0 | 6994 | 59 |
| SRR5951907 | 0 | 0 | 0 | 0 | 0 | 0 | 0 | 0 | 0 | 0 | 87 | 5775 | 13 | 0 | 0 | 0 | 0 | 0 | 0 | 2635 | 0 | 0 | 432 | 0 | 0 | 12 | 0 | 0 | 0 | 0 | 0 | 0 | 0 | 0 | 0 | 11314 | 0 |
| SRR5951908 | 0 | 0 | 250 | 60 | 67 | 0 | 0 | 594 | 325 | 5277 | 7659 | 31974 | 0 | 671 | 167 | 1548 | 0 | 607 | 163 | 4978 | 38 | 452 | 131 | 14996 | 924 | 47 | 338 | 212 | 451 | 2439 | 4117 | 563 | 409 | 324 | 234 | 9575 | 83 |
| SRR5951909 | 0 | 0 | 274 | 72 | 45 | 0 | 0 | 408 | 243 | 4851 | 2703 | 29402 | 0 | 523 | 109 | 1105 | 0 | 536 | 173 | 3845 | 41 | 453 | 112 | 17259 | 1075 | 42 | 329 | 54 | 364 | 1936 | 3551 | 551 | 240 | 359 | 135 | 8877 | 87 |
| SRR5951910 | 0 | 0 | 250 | 56 | 46 | 0 | 0 | 750 | 363 | 7072 | 1416 | 25955 | 0 | 671 | 120 | 1425 | 0 | 453 | 145 | 4825 | 41 | 355 | 115 | 16834 | 967 | 37 | 457 | 45 | 367 | 1990 | 4820 | 749 | 295 | 346 | 155 | 11131 | 78 |
| SRR5951911 | 0 | 0 | 184 | 105 | 44 | 0 | 0 | 782 | 433 | 10083 | 5792 | 24392 | 0 | 994 | 49 | 487 | 0 | 714 | 119 | 4507 | 47 | 350 | 112 | 17627 | 1079 | 90 | 1706 | 47 | 322 | 751 | 1899 | 280 | 125 | 166 | 64 | 15402 | 70 |
| SRR5951912 | 0 | 0 | 158 | 109 | 66 | 0 | 0 | 754 | 515 | 15037 | 5271 | 20263 | 0 | 687 | 67 | 931 | 0 | 719 | 107 | 5049 | 52 | 405 | 131 | 16662 | 1084 | 88 | 836 | 51 | 289 | 628 | 1530 | 234 | 127 | 154 | 77 | 15635 | 87 |
| SRR5951913 | 0 | 0 | 10 | 102 | 60 | 0 | 0 | 938 | 469 | 23432 | 917 | 17680 | 0 | 637 | 108 | 1244 | 0 | 599 | 93 | 4285 | 52 | 385 | 154 | 18022 | 1057 | 76 | 43 | 73 | 16 | 29 | 91 | 17 | 151 | 181 | 88 | 15331 | 95 |
| SRR6020460 | 0 | 0 | 0 | 0 | 0 | 27 | 51 | 73 | 36 | 140 | 4776 | 5559 | 0 | 24 | 0 | 0 | 16 | 957 | 0 | 3884 | 3803 | 2024 | 160 | 14941 | 1529 | 0 | 43 | 0 | 0 | 13 | 0 | 0 | 0 | 0 | 0 | 14377 | 0 |
| SRR6020461 | 0 | 26 | 0 | 0 | 0 | 10 | 18 | 60 | 47 | 188 | 461 | 2425 | 0 | 43 | 0 | 0 | 0 | 2541 | 0 | 2809 | 4155 | 2019 | 266 | 18044 | 4222 | 0 | 14 | 0 | 0 | 11 | 0 | 0 | 0 | 0 | 0 | 9611 | 0 |
| SRR6020462 | 0 | 14 | 0 | 0 | 0 | 26 | 68 | 290 | 41 | 407 | 612 | 2107 | 0 | 107 | 0 | 0 | 17 | 1404 | 0 | 5622 | 2503 | 2752 | 152 | 13158 | 1967 | 10 | 11 | 0 | 0 | 0 | 0 | 0 | 0 | 0 | 0 | 24689 | 0 |
| SRR6020463 | 0 | 19 | 0 | 0 | 0 | 17 | 16 | 245 | 56 | 623 | 473 | 1868 | 0 | 88 | 0 | 0 | 0 | 1529 | 0 | 6581 | 3175 | 3270 | 135 | 14958 | 2072 | 0 | 10 | 0 | 0 | 19 | 0 | 0 | 0 | 0 | 0 | 16711 | 0 |
| SRR6020464 | 0 | 22 | 0 | 0 | 0 | 22 | 11 | 240 | 47 | 300 | 3731 | 4897 | 0 | 0 | 0 | 0 | 14 | 1890 | 0 | 4418 | 3858 | 2465 | 235 | 13719 | 2180 | 0 | 18 | 0 | 0 | 0 | 0 | 0 | 0 | 0 | 0 | 12320 | 0 |
| SRR6020465 | 0 | 18 | 0 | 0 | 0 | 19 | 48 | 155 | 16 | 149 | 654 | 7394 | 0 | 0 | 0 | 0 | 14 | 1345 | 0 | 2996 | 3731 | 2195 | 208 | 14016 | 2290 | 0 | 31 | 0 | 0 | 0 | 0 | 0 | 0 | 0 | 0 | 8882 | 0 |
| SRR6020466 | 0 | 18 | 0 | 0 | 0 | 27 | 73 | 503 | 78 | 461 | 1288 | 2229 | 0 | 24 | 0 | 0 | 0 | 2029 | 0 | 8635 | 3306 | 2633 | 279 | 22613 | 2767 | 0 | 20 | 0 | 0 | 0 | 0 | 0 | 0 | 0 | 0 | 17684 | 0 |
| SRR6020467 | 12 | 21 | 0 | 0 | 0 | 20 | 68 | 146 | 19 | 161 | 1085 | 5554 | 0 | 15 | 0 | 0 | 17 | 1321 | 0 | 4916 | 4877 | 2817 | 233 | 15187 | 2644 | 11 | 15 | 0 | 0 | 0 | 0 | 0 | 0 | 0 | 0 | 10231 | 0 |
| SRR6020468 | 0 | 25 | 0 | 0 | 0 | 22 | 115 | 96 | 22 | 401 | 858 | 6442 | 0 | 37 | 0 | 0 | 14 | 1076 | 0 | 3955 | 2862 | 1751 | 197 | 11280 | 1754 | 0 | 18 | 0 | 0 | 10 | 0 | 0 | 0 | 0 | 0 | 7881 | 0 |
| SRR6020469 | 0 | 17 | 0 | 0 | 0 | 19 | 49 | 150 | 30 | 261 | 497 | 1272 | 0 | 76 | 0 | 0 | 0 | 831 | 0 | 6583 | 4876 | 3441 | 224 | 21408 | 3066 | 19 | 0 | 0 | 0 | 0 | 0 | 0 | 0 | 0 | 0 | 25895 | 0 |
| SRR6020470 | 15 | 28 | 0 | 0 | 0 | 0 | 16 | 93 | 23 | 209 | 438 | 9166 | 0 | 0 | 0 | 0 | 12 | 1456 | 0 | 1787 | 3490 | 2092 | 135 | 12370 | 1555 | 0 | 43 | 0 | 12 | 0 | 0 | 0 | 0 | 0 | 0 | 5941 | 0 |
| SRR6020471 | 0 | 12 | 0 | 0 | 0 | 34 | 17 | 201 | 18 | 492 | 5261 | 6829 | 0 | 82 | 0 | 0 | 17 | 2212 | 0 | 3793 | 2827 | 2692 | 141 | 12052 | 1790 | 0 | 12 | 0 | 0 | 0 | 0 | 0 | 0 | 0 | 0 | 18555 | 0 |
| SRR6020472 | 19 | 15 | 0 | 0 | 0 | 23 | 24 | 82 | 82 | 124 | 298 | 7868 | 0 | 27 | 0 | 0 | 29 | 836 | 0 | 2019 | 2579 | 1231 | 229 | 10177 | 1573 | 0 | 11 | 0 | 0 | 0 | 0 | 0 | 0 | 0 | 0 | 6426 | 0 |
| SRR6020473 | 11 | 18 | 0 | 0 | 0 | 24 | 96 | 107 | 0 | 210 | 3958 | 5168 | 0 | 33 | 0 | 0 | 19 | 1385 | 0 | 3101 | 3122 | 2548 | 144 | 14149 | 1863 | 13 | 23 | 0 | 0 | 0 | 0 | 0 | 0 | 0 | 0 | 12392 | 0 |
| SRR6020474 | 0 | 18 | 0 | 0 | 0 | 21 | 115 | 174 | 14 | 150 | 755 | 2816 | 0 | 55 | 0 | 0 | 13 | 656 | 0 | 5301 | 3144 | 2490 | 128 | 12485 | 2026 | 0 | 26 | 0 | 0 | 0 | 0 | 0 | 0 | 0 | 0 | 11511 | 14 |
| SRR6020475 | 0 | 10 | 0 | 0 | 0 | 22 | 14 | 248 | 64 | 248 | 6954 | 7000 | 0 | 28 | 0 | 0 | 10 | 1236 | 0 | 3459 | 2210 | 1673 | 111 | 8982 | 1549 | 0 | 28 | 0 | 0 | 0 | 0 | 0 | 0 | 0 | 0 | 8899 | 0 |
| SRR6020476 | 0 | 20 | 0 | 0 | 0 | 13 | 13 | 238 | 54 | 506 | 168 | 2026 | 0 | 38 | 0 | 0 | 0 | 1387 | 0 | 9853 | 1952 | 1995 | 107 | 10813 | 1491 | 0 | 15 | 0 | 0 | 0 | 10 | 0 | 0 | 12 | 0 | 31609 | 0 |
| SRR6020477 | 13 | 19 | 0 | 0 | 0 | 19 | 11 | 235 | 36 | 295 | 4618 | 4546 | 0 | 30 | 0 | 0 | 12 | 938 | 0 | 4154 | 3166 | 2509 | 136 | 13847 | 1855 | 0 | 19 | 0 | 0 | 14 | 0 | 0 | 0 | 0 | 0 | 14024 | 0 |
| SRR6020478 | 0 | 15 | 0 | 0 | 0 | 22 | 72 | 200 | 65 | 352 | 2290 | 3617 | 0 | 18 | 0 | 0 | 17 | 1153 | 0 | 4533 | 4002 | 2233 | 235 | 16977 | 2619 | 0 | 19 | 0 | 0 | 0 | 0 | 0 | 0 | 0 | 0 | 13030 | 0 |
| SRR6020479 | 0 | 16 | 0 | 0 | 0 | 24 | 16 | 103 | 11 | 77 | 1036 | 1340 | 0 | 30 | 0 | 0 | 15 | 1227 | 0 | 5677 | 3823 | 1916 | 234 | 17655 | 1953 | 0 | 28 | 0 | 0 | 0 | 0 | 0 | 0 | 0 | 0 | 25996 | 10 |
| SRR6020480 | 0 | 30 | 0 | 0 | 0 | 23 | 41 | 120 | 16 | 129 | 510 | 860 | 0 | 36 | 0 | 0 | 0 | 1954 | 0 | 6270 | 4039 | 2977 | 281 | 23902 | 3873 | 0 | 22 | 0 | 0 | 12 | 0 | 0 | 0 | 0 | 0 | 26374 | 10 |
| SRR6020481 | 12 | 16 | 0 | 0 | 0 | 27 | 22 | 227 | 19 | 149 | 2763 | 3561 | 0 | 0 | 0 | 0 | 14 | 1030 | 0 | 2867 | 3149 | 2177 | 205 | 12129 | 2077 | 0 | 46 | 0 | 0 | 0 | 0 | 0 | 0 | 0 | 0 | 8289 | 13 |
| SRR6020482 | 11 | 24 | 0 | 0 | 0 | 11 | 56 | 314 | 44 | 721 | 938 | 1240 | 0 | 68 | 0 | 0 | 10 | 1442 | 0 | 6454 | 4233 | 3540 | 220 | 21423 | 3343 | 0 | 14 | 0 | 0 | 0 | 0 | 0 | 0 | 11 | 0 | 18521 | 0 |
| SRR6020483 | 12 | 14 | 0 | 0 | 0 | 28 | 90 | 185 | 40 | 298 | 836 | 8516 | 0 | 32 | 0 | 0 | 23 | 1745 | 0 | 2847 | 2987 | 2198 | 155 | 12469 | 1790 | 0 | 23 | 0 | 0 | 0 | 0 | 0 | 0 | 0 | 0 | 5283 | 0 |
| SRR6020484 | 11 | 36 | 0 | 0 | 0 | 19 | 67 | 159 | 20 | 188 | 170 | 2538 | 0 | 25 | 0 | 0 | 11 | 1240 | 0 | 6171 | 4043 | 2546 | 334 | 18269 | 2997 | 0 | 25 | 0 | 0 | 0 | 0 | 0 | 0 | 0 | 0 | 17462 | 0 |
| SRR6020485 | 0 | 13 | 0 | 0 | 0 | 21 | 62 | 268 | 78 | 437 | 218 | 3562 | 0 | 42 | 0 | 0 | 12 | 1201 | 0 | 5283 | 4224 | 2428 | 225 | 19809 | 2575 | 0 | 22 | 0 | 0 | 0 | 0 | 0 | 0 | 0 | 0 | 22656 | 0 |
| SRR6020486 | 0 | 14 | 0 | 0 | 0 | 22 | 50 | 91 | 46 | 152 | 2445 | 4198 | 0 | 20 | 0 | 0 | 0 | 1174 | 0 | 3365 | 3195 | 1702 | 163 | 12358 | 1422 | 0 | 43 | 0 | 0 | 0 | 0 | 0 | 0 | 0 | 0 | 21026 | 0 |
| SRR6020487 | 0 | 11 | 0 | 0 | 0 | 37 | 24 | 107 | 49 | 410 | 555 | 6537 | 0 | 41 | 0 | 0 | 12 | 1703 | 0 | 3068 | 2637 | 1675 | 112 | 12451 | 1310 | 0 | 44 | 0 | 0 | 0 | 0 | 0 | 0 | 0 | 0 | 14469 | 0 |
| SRR6020488 | 14 | 15 | 0 | 0 | 0 | 27 | 74 | 215 | 22 | 281 | 716 | 2089 | 0 | 85 | 0 | 0 | 0 | 1001 | 0 | 4933 | 4588 | 2620 | 145 | 14430 | 2216 | 11 | 18 | 0 | 0 | 0 | 0 | 0 | 0 | 0 | 0 | 14410 | 0 |
| SRR6020489 | 0 | 13 | 0 | 0 | 0 | 14 | 14 | 16 | 0 | 23 | 1782 | 3329 | 0 | 0 | 0 | 0 | 19 | 1768 | 0 | 1313 | 3470 | 1079 | 157 | 10200 | 2009 | 0 | 48 | 0 | 0 | 0 | 0 | 0 | 0 | 0 | 0 | 2610 | 0 |
| SRR6020490 | 0 | 12 | 0 | 0 | 0 | 25 | 107 | 67 | 0 | 113 | 807 | 5676 | 0 | 46 | 0 | 0 | 24 | 1147 | 0 | 3403 | 3065 | 1702 | 182 | 13915 | 2069 | 16 | 36 | 0 | 0 | 0 | 0 | 0 | 0 | 0 | 0 | 7257 | 15 |
| SRR6020491 | 0 | 0 | 0 | 0 | 0 | 32 | 139 | 261 | 24 | 103 | 826 | 6447 | 0 | 16 | 0 | 0 | 17 | 1132 | 0 | 2634 | 3081 | 2026 | 156 | 12333 | 1761 | 10 | 35 | 0 | 0 | 0 | 0 | 0 | 0 | 0 | 0 | 5703 | 0 |
| SRR6020492 | 0 | 30 | 0 | 0 | 0 | 18 | 12 | 213 | 58 | 599 | 717 | 957 | 0 | 37 | 17 | 0 | 0 | 1136 | 0 | 13738 | 4074 | 5140 | 382 | 17469 | 3520 | 0 | 12 | 0 | 0 | 0 | 0 | 0 | 0 | 14 | 0 | 31048 | 0 |
| SRR6020493 | 0 | 16 | 0 | 0 | 0 | 22 | 89 | 355 | 42 | 96 | 2609 | 4563 | 0 | 12 | 0 | 0 | 0 | 1649 | 0 | 3228 | 3417 | 1972 | 179 | 14433 | 1455 | 0 | 22 | 0 | 0 | 0 | 0 | 0 | 0 | 0 | 0 | 10136 | 0 |
| SRR6020494 | 11 | 0 | 0 | 0 | 0 | 23 | 15 | 36 | 0 | 135 | 8596 | 12177 | 0 | 45 | 0 | 0 | 17 | 1427 | 0 | 1445 | 3436 | 1623 | 197 | 13072 | 1901 | 0 | 44 | 0 | 0 | 0 | 0 | 0 | 0 | 0 | 0 | 3437 | 0 |
| SRR6020495 | 0 | 0 | 0 | 0 | 0 | 10 | 27 | 246 | 70 | 308 | 5752 | 5066 | 0 | 35 | 0 | 0 | 16 | 751 | 0 | 5260 | 833 | 761 | 88 | 10731 | 868 | 0 | 19 | 0 | 0 | 0 | 0 | 0 | 0 | 0 | 0 | 16931 | 0 |
| SRR6020496 | 13 | 17 | 0 | 0 | 0 | 11 | 37 | 160 | 30 | 190 | 2313 | 3370 | 0 | 15 | 0 | 0 | 11 | 1613 | 0 | 4573 | 2260 | 1806 | 107 | 13781 | 1195 | 0 | 35 | 0 | 0 | 0 | 0 | 0 | 0 | 0 | 0 | 12478 | 0 |
| SRR6020497 | 12 | 17 | 0 | 0 | 0 | 16 | 11 | 48 | 0 | 34 | 773 | 2440 | 0 | 13 | 0 | 0 | 21 | 874 | 0 | 1352 | 4480 | 2354 | 271 | 17753 | 2669 | 0 | 15 | 0 | 0 | 0 | 0 | 0 | 0 | 0 | 0 | 10119 | 0 |
| SRR6020498 | 0 | 21 | 0 | 0 | 0 | 11 | 0 | 166 | 83 | 533 | 518 | 640 | 0 | 93 | 0 | 0 | 0 | 2373 | 0 | 10578 | 2927 | 1844 | 164 | 15590 | 2559 | 0 | 16 | 0 | 0 | 23 | 19 | 0 | 18 | 18 | 11 | 26885 | 15 |
| SRR6020499 | 10 | 29 | 0 | 0 | 0 | 19 | 16 | 413 | 39 | 443 | 478 | 6701 | 0 | 20 | 0 | 0 | 16 | 1659 | 0 | 4074 | 3295 | 2306 | 226 | 13018 | 2481 | 0 | 23 | 0 | 0 | 0 | 0 | 0 | 0 | 0 | 0 | 9722 | 0 |
| SRR6020500 | 0 | 0 | 0 | 0 | 0 | 0 | 19 | 979 | 45 | 377 | 233 | 653 | 0 | 57 | 0 | 0 | 0 | 1610 | 0 | 10753 | 2547 | 2773 | 140 | 15234 | 2370 | 0 | 0 | 0 | 0 | 0 | 0 | 0 | 0 | 0 | 0 | 58654 | 25 |
| SRR6020501 | 0 | 16 | 0 | 0 | 0 | 19 | 12 | 131 | 11 | 98 | 408 | 2036 | 0 | 15 | 0 | 0 | 12 | 960 | 0 | 2820 | 2442 | 1978 | 99 | 10168 | 1328 | 0 | 26 | 0 | 13 | 13 | 0 | 0 | 0 | 0 | 0 | 20540 | 0 |
| SRR6020502 | 0 | 18 | 0 | 0 | 0 | 19 | 17 | 43 | 14 | 101 | 1727 | 1554 | 0 | 19 | 0 | 0 | 15 | 1880 | 0 | 3248 | 3380 | 1688 | 198 | 14341 | 2661 | 0 | 51 | 0 | 0 | 0 | 0 | 0 | 0 | 0 | 0 | 15822 | 0 |
| SRR6020503 | 0 | 10 | 0 | 0 | 0 | 18 | 74 | 197 | 32 | 117 | 2431 | 3348 | 0 | 43 | 0 | 0 | 17 | 957 | 0 | 5299 | 3711 | 2414 | 210 | 14523 | 2447 | 19 | 24 | 0 | 0 | 0 | 0 | 0 | 0 | 0 | 0 | 13696 | 0 |
| SRR6020504 | 12 | 20 | 0 | 0 | 0 | 21 | 21 | 97 | 39 | 280 | 1573 | 9177 | 0 | 0 | 0 | 0 | 15 | 1727 | 0 | 3534 | 3561 | 2298 | 169 | 15834 | 1973 | 0 | 32 | 0 | 0 | 10 | 0 | 0 | 0 | 0 | 0 | 15642 | 0 |
| SRR6020505 | 0 | 23 | 0 | 0 | 0 | 19 | 64 | 119 | 26 | 180 | 2402 | 5590 | 0 | 37 | 0 | 0 | 0 | 883 | 0 | 3572 | 3897 | 2600 | 146 | 12840 | 2096 | 0 | 20 | 0 | 13 | 15 | 0 | 0 | 0 | 0 | 0 | 15272 | 0 |
| SRR6020506 | 12 | 11 | 0 | 0 | 0 | 18 | 14 | 78 | 13 | 152 | 302 | 1920 | 0 | 17 | 0 | 0 | 12 | 951 | 0 | 4486 | 3384 | 1552 | 144 | 13340 | 1707 | 12 | 20 | 0 | 0 | 0 | 0 | 0 | 0 | 0 | 0 | 27343 | 0 |
| SRR6020507 | 0 | 15 | 0 | 0 | 0 | 11 | 19 | 63 | 14 | 134 | 8894 | 4796 | 0 | 176 | 0 | 0 | 0 | 1518 | 0 | 2627 | 2643 | 2044 | 110 | 11817 | 1808 | 0 | 45 | 0 | 0 | 0 | 0 | 0 | 0 | 0 | 0 | 6359 | 0 |
| SRR6020508 | 0 | 31 | 0 | 0 | 0 | 17 | 11 | 547 | 63 | 667 | 221 | 3032 | 0 | 82 | 0 | 0 | 10 | 2249 | 0 | 6057 | 3586 | 2820 | 241 | 18743 | 2258 | 0 | 15 | 0 | 0 | 0 | 0 | 0 | 0 | 10 | 0 | 14705 | 0 |
| SRR6020509 | 0 | 18 | 0 | 0 | 0 | 0 | 131 | 353 | 45 | 281 | 596 | 7310 | 0 | 27 | 0 | 0 | 14 | 1560 | 0 | 2989 | 3114 | 1905 | 155 | 9941 | 1282 | 0 | 37 | 0 | 0 | 0 | 0 | 0 | 0 | 0 | 0 | 8778 | 11 |
| SRR6020510 | 17 | 0 | 0 | 0 | 0 | 22 | 88 | 122 | 14 | 266 | 615 | 5694 | 0 | 14 | 0 | 0 | 20 | 743 | 0 | 1807 | 2888 | 1705 | 172 | 11496 | 1374 | 0 | 24 | 0 | 0 | 0 | 0 | 0 | 0 | 0 | 0 | 6442 | 11 |
| SRR6020511 | 11 | 16 | 0 | 0 | 0 | 20 | 40 | 217 | 0 | 202 | 836 | 5740 | 0 | 15 | 0 | 0 | 12 | 1728 | 0 | 5592 | 3883 | 2402 | 529 | 16519 | 4434 | 0 | 27 | 0 | 0 | 0 | 0 | 0 | 0 | 0 | 0 | 7816 | 0 |
| SRR6020512 | 12 | 18 | 0 | 0 | 0 | 20 | 80 | 156 | 29 | 181 | 6521 | 8980 | 0 | 53 | 0 | 0 | 17 | 1407 | 0 | 2903 | 4209 | 2336 | 186 | 12328 | 2052 | 0 | 25 | 0 | 0 | 0 | 0 | 0 | 0 | 0 | 0 | 5641 | 0 |
| SRR6020513 | 0 | 0 | 0 | 0 | 0 | 36 | 22 | 145 | 36 | 227 | 649 | 6924 | 0 | 0 | 0 | 0 | 20 | 1510 | 0 | 5139 | 2940 | 1714 | 119 | 11020 | 1397 | 16 | 19 | 0 | 0 | 0 | 0 | 0 | 0 | 0 | 0 | 12399 | 0 |
| SRR6020514 | 0 | 13 | 0 | 0 | 0 | 12 | 15 | 94 | 28 | 283 | 1055 | 10099 | 0 | 19 | 0 | 0 | 11 | 1270 | 0 | 2371 | 2697 | 2180 | 115 | 10820 | 1475 | 0 | 42 | 0 | 0 | 0 | 0 | 0 | 0 | 0 | 0 | 5574 | 0 |
| SRR6020515 | 0 | 0 | 0 | 0 | 0 | 40 | 77 | 80 | 0 | 80 | 5137 | 4492 | 0 | 0 | 13 | 0 | 16 | 725 | 0 | 2602 | 4301 | 2743 | 180 | 15989 | 2202 | 0 | 21 | 0 | 0 | 0 | 0 | 0 | 0 | 0 | 0 | 11399 | 0 |
| SRR6020516 | 0 | 26 | 0 | 0 | 0 | 25 | 34 | 425 | 86 | 425 | 296 | 3330 | 0 | 18 | 0 | 0 | 0 | 826 | 0 | 6709 | 3079 | 2514 | 143 | 12326 | 2359 | 0 | 0 | 0 | 0 | 0 | 0 | 0 | 0 | 0 | 0 | 19105 | 0 |
| SRR6020517 | 0 | 12 | 0 | 0 | 0 | 18 | 40 | 287 | 26 | 228 | 533 | 842 | 0 | 65 | 0 | 0 | 0 | 1419 | 0 | 9105 | 3580 | 2616 | 302 | 21374 | 3513 | 12 | 14 | 0 | 0 | 0 | 0 | 0 | 0 | 0 | 0 | 38884 | 38 |
| SRR6020518 | 0 | 0 | 0 | 0 | 0 | 22 | 72 | 415 | 37 | 323 | 7041 | 6644 | 0 | 83 | 0 | 0 | 0 | 737 | 0 | 4180 | 962 | 1083 | 29 | 4988 | 499 | 0 | 0 | 0 | 0 | 0 | 0 | 0 | 0 | 0 | 0 | 11406 | 0 |
| SRR6020519 | 0 | 18 | 0 | 0 | 0 | 19 | 62 | 434 | 53 | 413 | 627 | 3110 | 0 | 26 | 0 | 0 | 12 | 1338 | 0 | 4556 | 3664 | 2985 | 180 | 15244 | 2409 | 0 | 0 | 0 | 0 | 0 | 0 | 0 | 0 | 0 | 0 | 14846 | 0 |
| SRR6020520 | 14 | 36 | 0 | 0 | 0 | 16 | 65 | 227 | 21 | 185 | 1285 | 2068 | 0 | 37 | 11 | 0 | 14 | 1349 | 0 | 3929 | 4323 | 2775 | 291 | 15141 | 4055 | 0 | 16 | 0 | 10 | 0 | 0 | 0 | 0 | 0 | 0 | 12221 | 0 |
| SRR6020521 | 0 | 16 | 0 | 0 | 0 | 20 | 91 | 208 | 39 | 358 | 3988 | 3675 | 0 | 69 | 0 | 0 | 16 | 1377 | 0 | 5301 | 4044 | 2742 | 233 | 13718 | 2582 | 16 | 22 | 0 | 0 | 0 | 0 | 0 | 0 | 0 | 0 | 17769 | 0 |
| SRR6020522 | 0 | 0 | 0 | 0 | 0 | 18 | 24 | 396 | 36 | 234 | 442 | 6418 | 0 | 0 | 0 | 0 | 18 | 1694 | 0 | 4177 | 4321 | 2276 | 228 | 13688 | 2389 | 0 | 15 | 0 | 0 | 0 | 0 | 0 | 0 | 0 | 0 | 9254 | 0 |
| SRR6020523 | 0 | 20 | 0 | 0 | 0 | 23 | 13 | 399 | 102 | 352 | 1642 | 3116 | 0 | 46 | 0 | 0 | 0 | 1123 | 0 | 7661 | 3502 | 1528 | 230 | 15398 | 2447 | 16 | 27 | 0 | 0 | 15 | 0 | 0 | 0 | 0 | 0 | 19199 | 17 |
| SRR6020524 | 17 | 0 | 0 | 0 | 0 | 24 | 17 | 89 | 18 | 243 | 927 | 9485 | 0 | 0 | 0 | 0 | 15 | 1010 | 0 | 2158 | 3663 | 1937 | 142 | 13370 | 1576 | 0 | 36 | 0 | 0 | 0 | 0 | 0 | 0 | 0 | 0 | 6343 | 0 |
| SRR6020525 | 0 | 26 | 0 | 0 | 0 | 13 | 31 | 88 | 0 | 120 | 618 | 5335 | 0 | 57 | 0 | 0 | 0 | 1877 | 0 | 3099 | 3719 | 2233 | 497 | 15115 | 3434 | 0 | 44 | 0 | 0 | 0 | 0 | 0 | 0 | 0 | 0 | 4661 | 11 |
| SRR6020526 | 0 | 21 | 0 | 0 | 0 | 19 | 43 | 260 | 14 | 137 | 220 | 3413 | 0 | 27 | 0 | 0 | 15 | 956 | 0 | 6580 | 3342 | 2687 | 212 | 16611 | 2587 | 11 | 14 | 0 | 0 | 0 | 0 | 0 | 0 | 0 | 0 | 13173 | 0 |
| SRR6020527 | 0 | 27 | 0 | 0 | 0 | 16 | 12 | 71 | 16 | 110 | 432 | 2171 | 0 | 0 | 0 | 0 | 11 | 2466 | 0 | 3826 | 3866 | 1460 | 283 | 24006 | 4272 | 0 | 44 | 0 | 0 | 10 | 10 | 0 | 0 | 0 | 0 | 14561 | 0 |
| SRR6020528 | 0 | 16 | 0 | 0 | 0 | 17 | 0 | 305 | 103 | 384 | 1191 | 1142 | 0 | 20 | 0 | 0 | 0 | 1535 | 0 | 9269 | 2126 | 10 | 111 | 13799 | 1955 | 0 | 13 | 0 | 0 | 0 | 0 | 0 | 24 | 17 | 19 | 35660 | 13 |
| SRR6020529 | 13 | 13 | 0 | 0 | 0 | 20 | 15 | 140 | 30 | 118 | 368 | 3063 | 0 | 15 | 0 | 0 | 28 | 803 | 0 | 2163 | 2783 | 2043 | 122 | 11726 | 1580 | 0 | 36 | 0 | 0 | 0 | 0 | 0 | 0 | 0 | 0 | 9183 | 0 |
| SRR6020530 | 14 | 0 | 0 | 0 | 0 | 25 | 23 | 139 | 0 | 241 | 6039 | 6826 | 0 | 19 | 0 | 0 | 30 | 1155 | 0 | 1547 | 3169 | 2259 | 174 | 13008 | 1903 | 0 | 38 | 0 | 0 | 0 | 0 | 0 | 0 | 0 | 0 | 6197 | 0 |
| SRR6020531 | 0 | 11 | 0 | 0 | 0 | 19 | 25 | 153 | 18 | 141 | 813 | 6596 | 0 | 12 | 0 | 0 | 13 | 1538 | 0 | 2863 | 2861 | 1689 | 155 | 12856 | 1566 | 0 | 26 | 0 | 0 | 0 | 0 | 0 | 0 | 0 | 0 | 8456 | 0 |
| SRR6020532 | 18 | 30 | 0 | 0 | 0 | 21 | 253 | 70 | 15 | 448 | 2951 | 4108 | 0 | 10 | 0 | 0 | 0 | 2197 | 0 | 5269 | 5294 | 1889 | 243 | 13142 | 1876 | 0 | 53 | 0 | 0 | 0 | 0 | 0 | 0 | 0 | 0 | 10282 | 0 |
| SRR6025030 | 0 | 605 | 64 | 138 | 550 | 0 | 0 | 499 | 2577 | 25247 | 198 | 231 | 0 | 621 | 6904 | 4446 | 0 | 1142 | 18 | 155 | 806 | 489 | 84 | 1595 | 1253 | 0 | 59 | 0 | 233 | 50 | 174 | 51 | 25 | 29 | 0 | 477 | 16 |
| SRR6025031 | 0 | 0 | 32 | 0 | 0 | 0 | 0 | 0 | 0 | 1450 | 2348 | 2655 | 0 | 77 | 1128 | 7596 | 0 | 1864 | 85 | 1097 | 2029 | 3860 | 0 | 648 | 8900 | 628 | 0 | 0 | 0 | 0 | 0 | 0 | 0 | 0 | 0 | 0 | 0 |
| SRR6025033 | 0 | 0 | 0 | 0 | 0 | 0 | 0 | 0 | 0 | 1408 | 1280 | 1771 | 0 | 0 | 43 | 8598 | 0 | 1371 | 0 | 0 | 2508 | 3533 | 0 | 677 | 5720 | 177 | 0 | 11 | 0 | 11 | 0 | 0 | 0 | 0 | 11 | 981 | 0 |
| SRR6025035 | 0 | 335 | 16 | 0 | 32 | 0 | 0 | 229 | 372 | 6575 | 144 | 62 | 0 | 300 | 3285 | 1520 | 0 | 335 | 0 | 126 | 3394 | 2210 | 318 | 7295 | 5256 | 0 | 0 | 0 | 11 | 28 | 15 | 0 | 0 | 12 | 0 | 605 | 0 |
| SRR6025036 | 0 | 77 | 0 | 0 | 20 | 0 | 0 | 861 | 420 | 2373 | 161 | 427 | 0 | 1304 | 0 | 0 | 0 | 2127 | 0 | 941 | 16748 | 2998 | 1028 | 14277 | 3941 | 115 | 0 | 0 | 0 | 12 | 12 | 0 | 0 | 0 | 0 | 1755 | 0 |
| SRR6025037 | 0 | 329 | 11 | 0 | 31 | 0 | 0 | 290 | 412 | 7322 | 37 | 21 | 0 | 287 | 8424 | 4431 | 0 | 414 | 0 | 127 | 3571 | 2532 | 394 | 9130 | 6189 | 0 | 0 | 0 | 0 | 17 | 13 | 0 | 0 | 0 | 0 | 676 | 0 |
| SRR6025038 | 0 | 0 | 0 | 0 | 44 | 0 | 0 | 0 | 0 | 549 | 0 | 4653 | 0 | 135 | 18 | 18 | 0 | 2384 | 30 | 0 | 3103 | 5109 | 25 | 1220 | 7749 | 395 | 0 | 0 | 23 | 0 | 0 | 0 | 0 | 0 | 0 | 0 | 0 |
| SRR6025041 | 0 | 149 | 0 | 22 | 0 | 0 | 12 | 494 | 431 | 2026 | 174 | 134 | 0 | 2244 | 353 | 564 | 22 | 188 | 22 | 1676 | 26 | 376 | 23 | 3038 | 491 | 72 | 44 | 0 | 33 | 0 | 36 | 0 | 0 | 0 | 0 | 1783 | 0 |
| SRR6025042 | 0 | 35 | 13 | 0 | 0 | 0 | 0 | 36 | 155 | 9943 | 409 | 2148 | 0 | 663 | 9971 | 9999 | 0 | 1366 | 58 | 0 | 2771 | 3739 | 0 | 447 | 8963 | 481 | 0 | 0 | 13 | 13 | 0 | 0 | 0 | 0 | 0 | 0 | 0 |
| SRR6025043 | 0 | 59 | 0 | 0 | 38 | 0 | 0 | 328 | 498 | 7877 | 123 | 39 | 0 | 359 | 1119 | 462 | 0 | 412 | 0 | 141 | 3796 | 2549 | 404 | 7916 | 6099 | 0 | 0 | 0 | 0 | 29 | 16 | 0 | 0 | 13 | 0 | 608 | 0 |
| SRR6025044 | 0 | 400 | 29 | 0 | 0 | 0 | 0 | 221 | 155 | 7001 | 138 | 50 | 0 | 2323 | 136 | 202 | 0 | 270 | 0 | 758 | 4430 | 7399 | 319 | 19595 | 7686 | 270 | 0 | 0 | 17 | 27 | 21 | 0 | 0 | 15 | 0 | 2080 | 0 |
| SRR6025047 | 0 | 19 | 0 | 73 | 107 | 18 | 42 | 980 | 1096 | 3594 | 2042 | 3497 | 0 | 2443 | 0 | 0 | 13 | 2905 | 116 | 358 | 222 | 767 | 129 | 3080 | 721 | 147 | 42 | 0 | 45 | 24 | 10 | 14 | 0 | 0 | 0 | 543 | 16 |
| SRR6025048 | 0 | 40 | 16 | 0 | 22 | 12 | 11 | 1704 | 1332 | 2956 | 234 | 413 | 0 | 3051 | 64 | 145 | 0 | 1164 | 95 | 1225 | 125 | 376 | 92 | 4801 | 383 | 100 | 15 | 0 | 26 | 10 | 26 | 17 | 0 | 0 | 12 | 1439 | 0 |
| SRR6025050 | 0 | 21 | 86 | 46 | 124 | 0 | 15 | 470 | 1448 | 25541 | 75 | 174 | 0 | 637 | 9083 | 4635 | 0 | 434 | 27 | 263 | 306 | 200 | 39 | 1653 | 715 | 0 | 11 | 0 | 14 | 45 | 27 | 52 | 34 | 50 | 0 | 1223 | 0 |
| SRR6025051 | 0 | 94 | 0 | 0 | 23 | 0 | 0 | 1278 | 469 | 2744 | 184 | 701 | 0 | 1623 | 11 | 0 | 0 | 2179 | 0 | 1164 | 11862 | 2423 | 1248 | 12594 | 3618 | 139 | 0 | 0 | 10 | 28 | 28 | 0 | 0 | 0 | 0 | 1687 | 17 |

### Supp Table S8.4.Columns_DI_to_ES

| | | | | | | | | | | | | | | | | | | | | | | | | | | | | | | | | | | | | | |
|---|---|---|---|---|---|---|---|---|---|---|---|---|---|---|---|---|---|---|---|---|---|---|---|---|---|---|---|---|---|---|---|---|---|---|---|---|---|
| SRR6025053 | 0 | 40 | 13 | 0 | 0 | 0 | 0 | 221 | 176 | 6291 | 49 | 68 | 0 | 1575 | 2138 | 2828 | 0 | 179 | 0 | 622 | 3883 | 6475 | 380 | 18298 | 7269 | 153 | 0 | 0 | 0 | 15 | 10 | 0 | 0 | 0 | 0 | 2285 | 0 |
| SRR6025055 | 0 | 61 | 10 | 120 | 37 | 19 | 0 | 395 | 1320 | 22859 | 212 | 1089 | 11 | 2842 | 2661 | 3653 | 0 | 683 | 11 | 1114 | 1134 | 1563 | 115 | 3818 | 1690 | 229 | 97 | 0 | 10 | 41 | 97 | 44 | 61 | 25 | 11 | 1484 | 0 |
| SRR6025056 | 0 | 614 | 29 | 133 | 416 | 0 | 12 | 1165 | 4770 | 7551 | 163 | 5407 | 0 | 1740 | 31 | 62 | 0 | 5963 | 52 | 373 | 4944 | 882 | 285 | 2909 | 977 | 137 | 15 | 0 | 217 | 82 | 206 | 35 | 41 | 21 | 0 | 482 | 43 |
| SRR6025058 | 0 | 276 | 21 | 0 | 0 | 0 | 0 | 276 | 176 | 6895 | 75 | 32 | 0 | 1855 | 1351 | 1717 | 0 | 203 | 0 | 627 | 4021 | 8299 | 392 | 21081 | 8261 | 160 | 0 | 0 | 0 | 14 | 0 | 0 | 0 | 19 | 11 | 2623 | 0 |
| SRR6025061 | 0 | 28 | 13 | 0 | 0 | 0 | 0 | 138 | 142 | 4911 | 378 | 241 | 0 | 3021 | 1913 | 2639 | 0 | 180 | 0 | 836 | 1908 | 3303 | 455 | 14341 | 3594 | 478 | 10 | 0 | 12 | 12 | 0 | 0 | 13 | 23 | 12 | 2573 | 0 |
| SRR6025062 | 0 | 50 | 16 | 0 | 0 | 0 | 0 | 273 | 191 | 9689 | 111 | 82 | 0 | 2633 | 471 | 556 | 0 | 153 | 0 | 795 | 1889 | 3440 | 479 | 16065 | 4978 | 262 | 0 | 0 | 0 | 17 | 14 | 0 | 14 | 21 | 11 | 3796 | 0 |
| SRR6025065 | 0 | 70 | 11 | 0 | 40 | 0 | 0 | 151 | 346 | 5126 | 159 | 115 | 0 | 378 | 7305 | 3744 | 0 | 326 | 0 | 77 | 1364 | 1202 | 548 | 5914 | 3148 | 0 | 0 | 0 | 19 | 12 | 11 | 0 | 0 | 23 | 27 | 654 | 0 |
| SRR6025066 | 0 | 231 | 132 | 0 | 0 | 0 | 0 | 97 | 511 | 9813 | 25 | 123 | 0 | 2609 | 4240 | 8196 | 0 | 508 | 0 | 818 | 827 | 3545 | 61 | 1410 | 469 | 83 | 0 | 0 | 47 | 19 | 0 | 17 | 12 | 37 | 31 | 693 | 0 |
| SRR6025067 | 0 | 53 | 0 | 0 | 41 | 0 | 0 | 253 | 529 | 9272 | 26 | 15 | 0 | 322 | 4381 | 2097 | 0 | 417 | 0 | 73 | 1298 | 1398 | 548 | 7837 | 5072 | 0 | 0 | 0 | 0 | 20 | 18 | 0 | 16 | 21 | 14 | 796 | 0 |
| SRR6025068 | 0 | 74 | 0 | 0 | 20 | 0 | 0 | 398 | 152 | 3558 | 1092 | 1420 | 0 | 1939 | 17 | 17 | 0 | 1663 | 10 | 209 | 15 | 1244 | 1431 | 14314 | 2995 | 164 | 0 | 0 | 0 | 21 | 17 | 0 | 0 | 0 | 15 | 696 | 0 |
| SRR6025069 | 0 | 59 | 0 | 0 | 17 | 0 | 0 | 314 | 143 | 2048 | 2675 | 2842 | 0 | 2575 | 15 | 54 | 0 | 1417 | 0 | 147 | 5438 | 1145 | 1636 | 10605 | 2053 | 296 | 0 | 0 | 15 | 27 | 32 | 11 | 17 | 12 | 20 | 538 | 0 |
| SRR6025070 | 0 | 67 | 45 | 37 | 68 | 17 | 40 | 515 | 878 | 7986 | 96 | 82 | 0 | 1363 | 1900 | 1099 | 65 | 374 | 18 | 110 | 20 | 276 | 16 | 742 | 501 | 0 | 41 | 0 | 29 | 35 | 52 | 16 | 0 | 0 | 0 | 266 | 0 |
| SRR6025071 | 0 | 62 | 31 | 0 | 0 | 10 | 13 | 785 | 778 | 3485 | 49 | 27 | 0 | 1299 | 16633 | 13743 | 19 | 100 | 13 | 378 | 0 | 88 | 10 | 688 | 143 | 0 | 19 | 0 | 26 | 0 | 67 | 20 | 0 | 0 | 13 | 830 | 0 |
| SRR6025072 | 0 | 0 | 0 | 0 | 0 | 15 | 0 | 190 | 93 | 18689 | 64 | 79 | 0 | 3085 | 1175 | 1848 | 0 | 22 | 0 | 559 | 1568 | 7972 | 235 | 9397 | 4400 | 413 | 0 | 0 | 0 | 0 | 16 | 0 | 0 | 30 | 0 | 2547 | 0 |
| SRR6025073 | 0 | 0 | 0 | 0 | 25 | 0 | 0 | 909 | 214 | 15672 | 145 | 258 | 0 | 3496 | 36 | 46 | 0 | 208 | 0 | 166 | 7320 | 5923 | 804 | 12303 | 3888 | 232 | 0 | 0 | 0 | 0 | 0 | 0 | 14 | 14 | 0 | 1007 | 0 |
| SRR6025075 | 0 | 10 | 0 | 0 | 29 | 14 | 0 | 1243 | 278 | 20680 | 302 | 626 | 0 | 4139 | 0 | 19 | 0 | 188 | 0 | 95 | 3434 | 2747 | 953 | 10781 | 2378 | 312 | 0 | 0 | 0 | 0 | 0 | 0 | 0 | 18 | 0 | 474 | 0 |
| SRR6025079 | 0 | 27 | 37 | 0 | 12 | 0 | 0 | 740 | 956 | 5052 | 0 | 0 | 0 | 1253 | 14522 | 11724 | 0 | 121 | 0 | 398 | 10 | 180 | 15 | 1370 | 320 | 0 | 0 | 0 | 11 | 11 | 76 | 32 | 23 | 20 | 25 | 839 | 0 |
| SRR6025080 | 0 | 638 | 33 | 117 | 396 | 13 | 19 | 766 | 2146 | 10508 | 1386 | 2638 | 0 | 1622 | 21 | 57 | 25 | 3231 | 43 | 853 | 13 | 631 | 346 | 5489 | 1235 | 102 | 32 | 0 | 96 | 48 | 115 | 26 | 21 | 13 | 25 | 804 | 33 |
| SRR6025082 | 0 | 2396 | 56 | 123 | 19 | 0 | 20 | 1226 | 3358 | 21614 | 462 | 282 | 0 | 2756 | 2662 | 4812 | 59 | 377 | 15 | 1684 | 830 | 1367 | 0 | 8141 | 1469 | 141 | 22 | 11 | 40 | 23 | 65 | 13 | 0 | 0 | 0 | 1731 | 0 |
| SRR6025083 | 0 | 650 | 81 | 115 | 488 | 11 | 20 | 631 | 2527 | 26132 | 77 | 72 | 0 | 565 | 7902 | 4478 | 22 | 857 | 21 | 459 | 567 | 393 | 95 | 1785 | 1368 | 0 | 33 | 0 | 169 | 59 | 124 | 37 | 16 | 17 | 25 | 732 | 0 |
| SRR6025084 | 0 | 64 | 95 | 261 | 32 | 13 | 23 | 368 | 1259 | 21977 | 566 | 384 | 0 | 3587 | 899 | 1044 | 46 | 588 | 15 | 2624 | 782 | 1515 | 164 | 6641 | 2396 | 149 | 56 | 0 | 80 | 55 | 95 | 38 | 27 | 21 | 0 | 2519 | 0 |
| SRR6025085 | 0 | 521 | 12 | 52 | 221 | 17 | 29 | 1979 | 6554 | 20675 | 115 | 122 | 0 | 677 | 8750 | 7050 | 13 | 639 | 21 | 385 | 540 | 422 | 0 | 2768 | 1084 | 0 | 15 | 0 | 47 | 24 | 11 | 15 | 0 | 0 | 11 | 527 | 0 |
| SRR6025086 | 0 | 518 | 82 | 0 | 84 | 0 | 0 | 118 | 502 | 3267 | 189 | 863 | 0 | 2812 | 93 | 204 | 0 | 5759 | 42 | 106 | 6091 | 2874 | 435 | 3807 | 739 | 82 | 0 | 0 | 99 | 28 | 31 | 17 | 28 | 64 | 47 | 152 | 0 |
| SRR6025087 | 0 | 33 | 136 | 0 | 121 | 0 | 0 | 95 | 1225 | 9369 | 17 | 22 | 0 | 437 | 22092 | 14721 | 0 | 1647 | 10 | 60 | 1922 | 2628 | 121 | 1112 | 672 | 0 | 0 | 0 | 84 | 20 | 20 | 22 | 19 | 40 | 35 | 116 | 0 |
| SRR6025088 | 0 | 695 | 37 | 48 | 257 | 13 | 22 | 3197 | 9008 | 17249 | 889 | 2119 | 0 | 2091 | 24 | 101 | 33 | 2311 | 43 | 923 | 4528 | 714 | 0 | 7767 | 809 | 98 | 15 | 0 | 40 | 23 | 11 | 11 | 0 | 0 | 12 | 1080 | 16 |
| SRR6025091 | 0 | 10 | 43 | 138 | 0 | 0 | 46 | 207 | 291 | 5032 | 328 | 205 | 14 | 2117 | 501 | 910 | 51 | 348 | 27 | 586 | 21 | 675 | 44 | 2328 | 795 | 149 | 41 | 0 | 17 | 29 | 43 | 14 | 0 | 0 | 0 | 823 | 0 |
| SRR6025092 | 0 | 0 | 0 | 0 | 28 | 0 | 0 | 246 | 501 | 27142 | 11 | 0 | 0 | 473 | 6948 | 4432 | 0 | 71 | 0 | 95 | 1924 | 5131 | 348 | 6765 | 6450 | 0 | 0 | 0 | 0 | 0 | 0 | 0 | 12 | 31 | 0 | 732 | 0 |
| SRR6025093 | 0 | 52 | 28 | 32 | 65 | 18 | 18 | 1537 | 2669 | 12096 | 117 | 28 | 0 | 1344 | 8404 | 8730 | 13 | 300 | 21 | 110 | 25 | 234 | 0 | 1274 | 556 | 0 | 12 | 0 | 18 | 26 | 32 | 0 | 0 | 0 | 0 | 230 | 0 |
| SRR6025094 | 0 | 0 | 0 | 0 | 44 | 0 | 0 | 512 | 899 | 38545 | 20 | 0 | 0 | 615 | 2006 | 1356 | 0 | 36 | 0 | 62 | 1045 | 2951 | 390 | 7061 | 4643 | 62 | 0 | 0 | 0 | 12 | 0 | 0 | 0 | 32 | 0 | 624 | 0 |
| SRR6025095 | 0 | 61 | 30 | 0 | 11 | 16 | 12 | 598 | 1021 | 5161 | 48 | 44 | 13 | 1516 | 5611 | 2571 | 0 | 180 | 14 | 296 | 16 | 133 | 11 | 532 | 172 | 0 | 39 | 0 | 57 | 32 | 91 | 34 | 28 | 29 | 0 | 763 | 0 |
| SRR6025096 | 0 | 68 | 58 | 118 | 0 | 0 | 31 | 181 | 232 | 5022 | 370 | 109 | 0 | 2567 | 404 | 705 | 58 | 335 | 27 | 550 | 25 | 768 | 44 | 2678 | 862 | 145 | 32 | 0 | 33 | 42 | 73 | 33 | 0 | 16 | 0 | 847 | 0 |
| SRR6025097 | 0 | 245 | 28 | 49 | 0 | 0 | 25 | 596 | 786 | 6702 | 70 | 47 | 0 | 2706 | 1195 | 2620 | 70 | 157 | 31 | 668 | 17 | 519 | 0 | 4629 | 661 | 124 | 17 | 0 | 14 | 29 | 50 | 18 | 0 | 0 | 0 | 931 | 0 |
| SRR6025099 | 0 | 0 | 0 | 0 | 0 | 0 | 0 | 0 | 0 | 6132 | 0 | 0 | 0 | 1003 | 2072 | 6447 | 0 | 1051 | 59 | 478 | 2171 | 4262 | 0 | 928 | 6406 | 475 | 0 | 0 | 0 | 0 | 0 | 0 | 0 | 0 | 0 | 54 | 0 |
| SRR6025101 | 0 | 40 | 13 | 0 | 0 | 0 | 0 | 312 | 251 | 2624 | 10 | 0 | 0 | 2114 | 3447 | 8327 | 0 | 151 | 31 | 1740 | 26 | 607 | 39 | 7439 | 753 | 92 | 0 | 0 | 14 | 14 | 19 | 26 | 21 | 22 | 16 | 2430 | 0 |
| SRR6025103 | 0 | 31 | 15 | 0 | 17 | 13 | 23 | 1609 | 917 | 1796 | 718 | 1114 | 0 | 2698 | 32 | 60 | 24 | 996 | 92 | 1290 | 155 | 301 | 88 | 3842 | 315 | 131 | 27 | 0 | 26 | 11 | 19 | 18 | 0 | 0 | 0 | 1457 | 0 |
| SRR6025104 | 0 | 130 | 0 | 0 | 13 | 22 | 49 | 1168 | 387 | 486 | 979 | 2053 | 10 | 937 | 57 | 45 | 51 | 539 | 60 | 1220 | 103 | 196 | 89 | 3177 | 134 | 205 | 152 | 0 | 14 | 0 | 29 | 0 | 0 | 0 | 0 | 1292 | 0 |
| SRR6025105 | 0 | 0 | 0 | 0 | 0 | 0 | 0 | 274 | 320 | 22199 | 28 | 62 | 0 | 3679 | 1048 | 1795 | 0 | 23 | 0 | 494 | 722 | 3859 | 319 | 9897 | 2381 | 650 | 0 | 0 | 0 | 0 | 0 | 0 | 0 | 16 | 20 | 1371 | 0 |
| SRR6025106 | 0 | 88 | 27 | 0 | 0 | 0 | 16 | 308 | 357 | 2428 | 101 | 59 | 0 | 2104 | 2399 | 4635 | 22 | 124 | 21 | 1462 | 12 | 311 | 27 | 3494 | 404 | 118 | 33 | 0 | 22 | 11 | 52 | 16 | 13 | 14 | 10 | 1954 | 0 |
| SRR6025108 | 0 | 0 | 0 | 0 | 0 | 0 | 0 | 12 | 548 | 4935 | 0 | 0 | 0 | 71 | 19894 | 25265 | 0 | 1291 | 0 | 0 | 2457 | 4037 | 0 | 301 | 8138 | 279 | 0 | 0 | 0 | 0 | 0 | 0 | 0 | 0 | 0 | 0 | 0 |
| SRR6025111 | 0 | 349 | 101 | 244 | 37 | 0 | 0 | 354 | 1257 | 26049 | 138 | 866 | 13 | 3222 | 1721 | 2294 | 17 | 726 | 20 | 1068 | 1111 | 1856 | 142 | 4273 | 1683 | 215 | 105 | 0 | 63 | 46 | 139 | 38 | 42 | 38 | 13 | 1355 | 0 |
| SRR6025113 | 0 | 26 | 22 | 70 | 180 | 0 | 0 | 1640 | 3851 | 10562 | 51 | 4080 | 0 | 1906 | 267 | 153 | 0 | 4106 | 57 | 686 | 2893 | 537 | 151 | 3893 | 986 | 153 | 22 | 0 | 84 | 77 | 16 | 22 | 11 | 11 | 0 | 1182 | 29 |
| SRR6025114 | 0 | 37 | 86 | 30 | 113 | 0 | 0 | 1387 | 4569 | 9138 | 87 | 5136 | 0 | 1260 | 63 | 32 | 0 | 3140 | 65 | 1071 | 2476 | 468 | 147 | 3590 | 949 | 196 | 36 | 0 | 14 | 50 | 14 | 43 | 32 | 32 | 14 | 1607 | 33 |
| SRR6025116 | 0 | 14 | 10 | 0 | 17 | 16 | 0 | 674 | 959 | 24978 | 37 | 83 | 0 | 491 | 8435 | 4321 | 0 | 135 | 25 | 467 | 92 | 113 | 15 | 1417 | 216 | 0 | 34 | 0 | 33 | 11 | 36 | 11 | 33 | 61 | 0 | 2530 | 0 |
| SRR6025119 | 0 | 60 | 20 | 0 | 0 | 13 | 0 | 792 | 675 | 2783 | 90 | 29 | 0 | 1185 | 15650 | 9635 | 20 | 112 | 14 | 446 | 10 | 91 | 10 | 705 | 166 | 0 | 27 | 0 | 25 | 0 | 74 | 20 | 11 | 0 | 12 | 788 | 0 |
| SRR6025120 | 0 | 0 | 0 | 0 | 0 | 17 | 0 | 319 | 168 | 20694 | 51 | 37 | 0 | 2342 | 471 | 794 | 0 | 18 | 0 | 523 | 760 | 4258 | 260 | 10658 | 2851 | 379 | 0 | 0 | 0 | 12 | 0 | 0 | 0 | 18 | 0 | 1833 | 0 |
| SRR6025121 | 0 | 0 | 0 | 0 | 25 | 0 | 0 | 924 | 473 | 15500 | 643 | 951 | 0 | 5700 | 32 | 60 | 0 | 182 | 0 | 108 | 2663 | 1958 | 1218 | 8943 | 1603 | 492 | 0 | 0 | 10 | 0 | 0 | 0 | 0 | 0 | 0 | 351 | 0 |
| SRR6025123 | 0 | 15 | 18 | 84 | 97 | 13 | 41 | 754 | 685 | 3898 | 904 | 1908 | 0 | 2679 | 23 | 65 | 27 | 2975 | 126 | 212 | 223 | 791 | 128 | 3214 | 780 | 159 | 21 | 0 | 23 | 18 | 38 | 0 | 0 | 0 | 13 | 407 | 18 |
| SRR6025125 | 0 | 72 | 32 | 0 | 0 | 0 | 13 | 245 | 237 | 2121 | 171 | 66 | 0 | 2258 | 3096 | 7797 | 25 | 136 | 25 | 1499 | 16 | 361 | 25 | 3789 | 443 | 169 | 21 | 0 | 21 | 0 | 51 | 20 | 12 | 0 | 14 | 1886 | 0 |
| SRR6025126 | 0 | 87 | 10 | 0 | 17 | 0 | 17 | 1632 | 1079 | 2380 | 648 | 838 | 0 | 2875 | 81 | 156 | 25 | 1028 | 104 | 1179 | 151 | 292 | 97 | 4034 | 329 | 125 | 19 | 0 | 23 | 0 | 50 | 0 | 0 | 0 | 15 | 1362 | 0 |
| SRR6025128 | 0 | 11 | 49 | 46 | 66 | 0 | 38 | 467 | 777 | 7160 | 68 | 48 | 0 | 1259 | 6836 | 5664 | 31 | 416 | 22 | 88 | 12 | 236 | 22 | 684 | 472 | 0 | 31 | 0 | 24 | 26 | 46 | 14 | 0 | 0 | 11 | 241 | 0 |
| SRR6040044 | 0 | 194 | 65 | 14 | 463 | 0 | 0 | 1117 | 8115 | 20543 | 5518 | 6231 | 0 | 3997 | 0 | 0 | 0 | 255 | 1792 | 834 | 975 | 3214 | 409 | 14550 | 2463 | 0 | 0 | 78 | 128 | 227 | 373 | 67 | 52 | 46 | 12 | 2125 | 623 |
| SRR6040045 | 0 | 2895 | 52 | 13 | 418 | 0 | 0 | 1105 | 8505 | 15814 | 5264 | 6718 | 0 | 3709 | 0 | 0 | 0 | 223 | 1641 | 799 | 1062 | 3316 | 392 | 13524 | 2150 | 0 | 0 | 1152 | 2067 | 3435 | 6012 | 1183 | 1035 | 686 | 10 | 2107 | 656 |
| SRR6076852 | 0 | 0 | 0 | 0 | 110 | 0 | 0 | 4630 | 8044 | 2668 | 510 | 1648 | 16 | 3178 | 0 | 0 | 39 | 76 | 148 | 0 | 17 | 232 | 140 | 224 | 310 | 308 | 610 | 0 | 23 | 0 | 0 | 0 | 0 | 0 | 0 | 314 | 48 |
| SRR6076853 | 0 | 21 | 0 | 0 | 82 | 0 | 28 | 7073 | 7857 | 4861 | 346 | 760 | 0 | 5483 | 0 | 0 | 152 | 52 | 120 | 138 | 13 | 186 | 97 | 175 | 314 | 292 | 561 | 0 | 0 | 0 | 0 | 0 | 0 | 0 | 0 | 115 | 42 |
| SRR6076854 | 0 | 96 | 0 | 0 | 27 | 11 | 21 | 1845 | 3436 | 6004 | 434 | 958 | 0 | 6040 | 54 | 36 | 53 | 23 | 26 | 145 | 12 | 154 | 57 | 133 | 113 | 75 | 277 | 0 | 43 | 53 | 43 | 11 | 11 | 0 | 0 | 253 | 0 |
| SRR6076855 | 0 | 96 | 0 | 0 | 73 | 15 | 0 | 2711 | 4490 | 3897 | 762 | 988 | 0 | 3868 | 0 | 0 | 0 | 77 | 28 | 85 | 22 | 372 | 90 | 185 | 288 | 266 | 355 | 0 | 0 | 0 | 15 | 0 | 0 | 0 | 0 | 141 | 51 |
| SRR6076856 | 0 | 50 | 0 | 0 | 55 | 22 | 112 | 2051 | 2162 | 5975 | 29 | 1338 | 112 | 2876 | 0 | 0 | 17 | 80 | 24 | 111 | 26 | 230 | 85 | 172 | 210 | 316 | 324 | 0 | 0 | 0 | 17 | 0 | 0 | 0 | 0 | 178 | 22 |
| SRR6076858 | 0 | 31 | 0 | 0 | 85 | 166 | 96 | 2968 | 4186 | 1674 | 799 | 1370 | 0 | 3120 | 0 | 0 | 57 | 49 | 97 | 0 | 24 | 211 | 102 | 255 | 269 | 453 | 642 | 0 | 0 | 0 | 31 | 0 | 0 | 0 | 0 | 152 | 31 |
| SRR6076861 | 0 | 38 | 0 | 0 | 104 | 0 | 0 | 1710 | 2603 | 4113 | 21 | 167 | 32 | 5221 | 16 | 0 | 0 | 94 | 104 | 172 | 23 | 342 | 103 | 238 | 341 | 416 | 195 | 0 | 11 | 0 | 27 | 0 | 16 | 0 | 0 | 285 | 41 |
| SRR6153965 | 0 | 17 | 0 | 0 | 0 | 0 | 0 | 219 | 0 | 9786 | 124 | 144 | 0 | 1665 | 7782 | 4476 | 0 | 147 | 35 | 444 | 1009 | 3390 | 20 | 206 | 48118 | 24 | 0 | 0 | 0 | 0 | 0 | 0 | 0 | 0 | 0 | 2448 | 0 |
| SRR6153968 | 0 | 0 | 0 | 0 | 0 | 0 | 0 | 221 | 18 | 1205 | 2468 | 325 | 0 | 2331 | 4649 | 4133 | 0 | 364 | 125 | 440 | 1285 | 2501 | 18 | 233 | 6808 | 21 | 0 | 0 | 0 | 0 | 0 | 0 | 0 | 0 | 0 | 2463 | 21 |
| SRR6153976 | 0 | 0 | 0 | 0 | 0 | 0 | 0 | 108 | 68 | 20529 | 17 | 0 | 0 | 592 | 562 | 739 | 0 | 49 | 97 | 946 | 195 | 225 | 0 | 116 | 609 | 30 | 0 | 0 | 0 | 0 | 0 | 0 | 0 | 0 | 0 | 10682 | 0 |
| SRR6153984 | 0 | 0 | 0 | 0 | 0 | 0 | 0 | 238 | 34 | 559 | 4257 | 975 | 0 | 913 | 221 | 656 | 0 | 378 | 81 | 1085 | 504 | 1376 | 0 | 45 | 441 | 81 | 12 | 0 | 0 | 0 | 0 | 0 | 0 | 0 | 0 | 2738 | 0 |
| SRR6153998 | 0 | 16 | 0 | 0 | 0 | 0 | 0 | 16 | 11 | 2363 | 326 | 75 | 0 | 1401 | 79 | 106 | 0 | 425 | 44 | 449 | 1637 | 1142 | 16 | 1247 | 10453 | 115 | 17 | 0 | 0 | 0 | 0 | 0 | 0 | 0 | 0 | 4069 | 11 |
| SRR6154002 | 0 | 23 | 16 | 0 | 28 | 0 | 0 | 616 | 227 | 12211 | 386 | 265 | 0 | 133 | 298 | 422 | 0 | 427 | 71 | 2764 | 1554 | 2347 | 21 | 285 | 8627 | 28 | 0 | 0 | 0 | 0 | 0 | 0 | 0 | 0 | 0 | 13940 | 40 |
| SRR6154003 | 0 | 0 | 0 | 0 | 35 | 0 | 0 | 793 | 182 | 6574 | 91 | 96 | 0 | 885 | 4680 | 671 | 11 | 377 | 29 | 48 | 598 | 946 | 0 | 85 | 7873 | 15 | 11 | 0 | 0 | 0 | 0 | 0 | 0 | 0 | 0 | 301 | 15 |
| SRR6154004 | 0 | 48 | 0 | 0 | 0 | 0 | 0 | 149 | 19 | 2148 | 550 | 171 | 0 | 1107 | 1392 | 1466 | 0 | 281 | 49 | 613 | 1570 | 2430 | 0 | 136 | 13039 | 23 | 0 | 0 | 0 | 0 | 0 | 0 | 0 | 0 | 0 | 3991 | 0 |
| SRR6154005 | 0 | 48 | 0 | 0 | 11 | 0 | 0 | 71 | 13 | 4337 | 43 | 13 | 0 | 860 | 633 | 987 | 0 | 263 | 35 | 558 | 570 | 1147 | 11 | 155 | 8989 | 56 | 0 | 0 | 0 | 0 | 0 | 0 | 0 | 0 | 0 | 9584 | 22 |
| SRR6154006 | 0 | 0 | 0 | 0 | 0 | 0 | 0 | 1403 | 402 | 7332 | 68 | 13 | 0 | 358 | 8547 | 3402 | 0 | 309 | 50 | 220 | 312 | 592 | 0 | 69 | 4825 | 15 | 0 | 0 | 0 | 0 | 0 | 0 | 0 | 0 | 0 | 1412 | 25 |
| SRR6203722 | 0 | 37 | 0 | 0 | 0 | 12 | 0 | 44 | 163 | 5962 | 178 | 252 | 0 | 2388 | 341 | 489 | 0 | 105 | 0 | 0 | 1892 | 1457 | 427 | 11217 | 3418 | 529 | 12 | 0 | 12 | 25 | 37 | 0 | 12 | 25 | 0 | 371 | 0 |
| SRR6203757 | 0 | 23 | 0 | 0 | 22 | 0 | 0 | 188 | 345 | 5984 | 110 | 78 | 0 | 2585 | 78 | 282 | 0 | 156 | 0 | 0 | 1756 | 1515 | 851 | 14989 | 4288 | 325 | 0 | 0 | 0 | 0 | 23 | 0 | 0 | 0 | 0 | 329 | 0 |
| SRR6203760 | 0 | 0 | 0 | 0 | 0 | 0 | 0 | 1483 | 2966 | 4152 | 0 | 0 | 0 | 593 | 0 | 297 | 0 | 0 | 0 | 593 | 12 | 37 | 92 | 2128 | 1006 | 40 | 0 | 0 | 0 | 120 | 0 | 0 | 0 | 0 | 0 | 6524 | 0 |
| SRR6203761 | 0 | 0 | 0 | 0 | 0 | 0 | 0 | 1563 | 782 | 4689 | 0 | 0 | 0 | 2345 | 0 | 0 | 0 | 0 | 0 | 0 | 17 | 17 | 75 | 1942 | 977 | 17 | 0 | 0 | 0 | 0 | 0 | 0 | 0 | 0 | 0 | 4689 | 0 |
| SRR6203772 | 0 | 57 | 19 | 0 | 0 | 0 | 0 | 184 | 240 | 5431 | 184 | 127 | 0 | 1921 | 212 | 240 | 0 | 138 | 0 | 64 | 1712 | 1001 | 600 | 9419 | 2947 | 378 | 0 | 0 | 19 | 0 | 19 | 0 | 0 | 0 | 19 | 367 | 0 |
| SRR6203807 | 0 | 0 | 0 | 0 | 67 | 0 | 0 | 281 | 197 | 5194 | 337 | 309 | 0 | 1348 | 955 | 983 | 0 | 76 | 0 | 112 | 1018 | 485 | 185 | 3941 | 2799 | 142 | 0 | 0 | 0 | 0 | 0 | 0 | 0 | 0 | 0 | 758 | 0 |
| SRR6224558 | 48 | 178 | 0 | 0 | 162 | 0 | 29 | 155 | 135 | 232 | 118 | 77 | 0 | 0 | 19 | 0 | 99 | 620 | 211 | 135 | 103 | 85 | 0 | 47 | 404 | 1414 | 97 | 0 | 0 | 0 | 0 | 0 | 0 | 0 | 0 | 232 | 0 |
| SRR6226650 | 0 | 24 | 0 | 92 | 67 | 91 | 24 | 2455 | 13676 | 3558 | 984 | 3452 | 0 | 107 | 0 | 0 | 417 | 998 | 48 | 593 | 484 | 135 | 459 | 2965 | 320 | 13 | 1241 | 10 | 68 | 180 | 350 | 18 | 0 | 0 | 0 | 1993 | 14 |
| SRR6226651 | 0 | 58 | 0 | 89 | 93 | 139 | 39 | 4125 | 21949 | 9975 | 123 | 1636 | 0 | 146 | 0 | 0 | 572 | 742 | 64 | 598 | 320 | 136 | 351 | 2895 | 617 | 15 | 379 | 0 | 119 | 660 | 1564 | 68 | 21 | 13 | 0 | 2381 | 31 |
| SRR6226652 | 0 | 58 | 0 | 143 | 28 | 48 | 68 | 10945 | 67336 | 14848 | 50 | 774 | 0 | 565 | 0 | 0 | 253 | 862 | 55 | 1252 | 275 | 159 | 381 | 2088 | 387 | 13 | 288 | 0 | 51 | 228 | 370 | 54 | 16 | 13 | 0 | 4382 | 27 |
| SRR6226653 | 19 | 39 | 0 | 142 | 371 | 45 | 55 | 8895 | 47947 | 8091 | 246 | 329 | 0 | 109 | 0 | 0 | 34 | 862 | 53 | 167 | 375 | 285 | 164 | 1685 | 300 | 11 | 64 | 0 | 20 | 38 | 10 | 0 | 0 | 0 | 0 | 574 | 0 |
| SRR6226654 | 13 | 381 | 17 | 195 | 208 | 54 | 24 | 6100 | 52315 | 38115 | 146 | 1880 | 0 | 1756 | 0 | 0 | 822 | 1798 | 27 | 136 | 729 | 289 | 271 | 2110 | 395 | 11 | 734 | 15 | 59 | 175 | 433 | 71 | 32 | 47 | 0 | 335 | 19 |
| SRR6226655 | 0 | 99 | 37 | 61 | 117 | 171 | 26 | 6185 | 38753 | 46503 | 705 | 2016 | 0 | 814 | 0 | 0 | 382 | 1266 | 19 | 145 | 699 | 248 | 256 | 2027 | 235 | 14 | 1220 | 13 | 91 | 127 | 509 | 201 | 58 | 26 | 0 | 227 | 39 |
| SRR6226656 | 0 | 36 | 0 | 37 | 28 | 15 | 0 | 4646 | 32264 | 4996 | 443 | 2518 | 0 | 95 | 0 | 0 | 46 | 1080 | 17 | 85 | 688 | 171 | 292 | 2320 | 186 | 18 | 545 | 0 | 14 | 11 | 44 | 19 | 41 | 19 | 0 | 124 | 14 |
| SRR6226657 | 0 | 15 | 12 | 63 | 48 | 108 | 32 | 3811 | 35799 | 24951 | 161 | 350 | 0 | 250 | 0 | 0 | 577 | 1387 | 35 | 92 | 566 | 209 | 238 | 2054 | 106 | 13 | 7922 | 0 | 0 | 78 | 388 | 208 | 104 | 63 | 0 | 369 | 0 |
| SRR6230411 | 0 | 118 | 93 | 52 | 1127 | 12 | 54 | 208 | 460 | 6223 | 6519 | 29475 | 35 | 1479 | 10 | 0 | 23 | 266 | 347 | 1561 | 2180 | 717 | 71 | 5782 | 1034 | 37 | 368 | 788 | 4782 | 4480 | 9437 | 652 | 449 | 331 | 50 | 4703 | 293 |
| SRR6230412 | 11 | 44 | 51 | 67 | 2529 | 21 | 15 | 217 | 591 | 6586 | 25867 | 60288 | 61 | 3062 | 38 | 24 | 29 | 318 | 264 | 178 | 4817 | 867 | 94 | 5930 | 1434 | 33 | 46 | 32 | 162 | 374 | 1024 | 165 | 188 | 152 | 33 | 849 | 239 |
| SRR6230413 | 0 | 1258 | 65 | 59 | 2733 | 0 | 0 | 606 | 2776 | 7298 | 27488 | 46461 | 0 | 1952 | 0 | 0 | 10 | 358 | 425 | 195 | 3360 | 880 | 79 | 5463 | 1813 | 24 | 43 | 672 | 1773 | 5055 | 8859 | 1618 | 1465 | 891 | 12 | 412 | 223 |
| SRR6246305 | 0 | 105 | 30 | 0 | 0 | 0 | 0 | 42 | 31 | 9474 | 32 | 45 | 11 | 4160 | 0 | 0 | 10 | 1631 | 15 | 132 | 1724 | 1922 | 150 | 6184 | 3594 | 152 | 67 | 0 | 22 | 20 | 21 | 19 | 34 | 28 | 38 | 1366 | 143 |
| SRR6246306 | 0 | 425 | 22 | 0 | 0 | 0 | 0 | 47 | 42 | 9762 | 30 | 30 | 17 | 4183 | 0 | 0 | 0 | 1673 | 16 | 152 | 1603 | 1797 | 161 | 6130 | 3292 | 147 | 245 | 0 | 67 | 80 | 94 | 26 | 31 | 25 | 47 | 1330 | 114 |
| SRR6246307 | 0 | 104 | 21 | 0 | 11 | 0 | 0 | 40 | 25 | 9364 | 33 | 22 | 12 | 4218 | 0 | 0 | 0 | 1676 | 19 | 137 | 1642 | 1850 | 147 | 6099 | 3551 | 158 | 66 | 0 | 17 | 20 | 26 | 27 | 32 | 20 | 39 | 1305 | 132 |
| SRR6246308 | 0 | 408 | 26 | 0 | 0 | 0 | 0 | 46 | 47 | 9407 | 27 | 29 | 12 | 4277 | 0 | 0 | 0 | 1634 | 17 | 136 | 1629 | 1802 | 141 | 5962 | 3317 | 152 | 270 | 0 | 87 | 81 | 76 | 21 | 31 | 23 | 41 | 1306 | 146 |
| SRR6246309 | 0 | 11 | 0 | 0 | 21 | 0 | 0 | 144 | 43 | 6423 | 17 | 0 | 0 | 1145 | 0 | 0 | 0 | 1794 | 0 | 17 | 2484 | 2394 | 198 | 12521 | 6858 | 189 | 31 | 0 | 0 | 0 | 0 | 0 | 0 | 0 | 10 | 71 | 225 |
| SRR6246310 | 0 | 13 | 0 | 0 | 23 | 0 | 0 | 176 | 51 | 6361 | 13 | 12 | 0 | 1101 | 10 | 0 | 10 | 1773 | 0 | 16 | 2208 | 2292 | 230 | 12194 | 6572 | 203 | 20 | 0 | 0 | 0 | 0 | 0 | 0 | 0 | 0 | 83 | 235 |
| SRR6246311 | 0 | 15 | 17 | 0 | 17 | 0 | 0 | 165 | 48 | 6348 | 16 | 0 | 0 | 1145 | 0 | 0 | 0 | 1806 | 12 | 17 | 2504 | 2507 | 197 | 12449 | 6898 | 208 | 24 | 0 | 0 | 10 | 0 | 0 | 0 | 12 | 15 | 76 | 216 |
| SRR6246312 | 0 | 87 | 0 | 0 | 20 | 0 | 0 | 172 | 52 | 6412 | 16 | 13 | 0 | 1138 | 13 | 0 | 0 | 1797 | 0 | 27 | 2278 | 2303 | 233 | 12544 | 6709 | 200 | 29 | 0 | 0 | 0 | 0 | 0 | 0 | 0 | 15 | 78 | 237 |
| SRR6246313 | 0 | 72 | 0 | 0 | 20 | 12 | 0 | 189 | 42 | 8444 | 25 | 0 | 0 | 1726 | 0 | 0 | 0 | 1059 | 11 | 0 | 1976 | 2574 | 291 | 14332 | 6705 | 201 | 20 | 0 | 0 | 0 | 0 | 0 | 0 | 0 | 0 | 24 | 291 |
| SRR6246314 | 0 | 69 | 0 | 0 | 0 | 0 | 0 | 189 | 29 | 8448 | 33 | 0 | 0 | 1757 | 0 | 0 | 16 | 927 | 20 | 0 | 1799 | 2320 | 360 | 14673 | 6282 | 202 | 33 | 0 | 0 | 0 | 0 | 0 | 0 | 0 | 0 | 20 | 302 |
| SRR6246315 | 10 | 74 | 0 | 0 | 0 | 0 | 0 | 150 | 29 | 8600 | 19 | 0 | 0 | 1769 | 0 | 0 | 15 | 945 | 11 | 0 | 2025 | 2673 | 308 | 14530 | 7117 | 199 | 17 | 0 | 0 | 0 | 0 | 0 | 0 | 0 | 0 | 27 | 330 |
| SRR6246316 | 10 | 66 | 15 | 0 | 26 | 0 | 0 | 158 | 38 | 8519 | 34 | 0 | 0 | 1707 | 0 | 0 | 17 | 982 | 0 | 0 | 2013 | 2461 | 307 | 14722 | 6677 | 210 | 32 | 0 | 0 | 0 | 0 | 0 | 0 | 0 | 0 | 14 | 346 |
| SRR6246317 | 24 | 0 | 0 | 0 | 19 | 14 | 0 | 623 | 390 | 8922 | 779 | 545 | 0 | 5494 | 0 | 0 | 0 | 77 | 0 | 39 | 1328 | 5516 | 260 | 20320 | 8307 | 404 | 0 | 0 | 0 | 0 | 0 | 0 | 0 | 0 | 0 | 0 | 67 |
| SRR6246318 | 15 | 0 | 0 | 0 | 0 | 15 | 0 | 589 | 236 | 7694 | 589 | 471 | 0 | 5142 | 0 | 0 | 0 | 79 | 0 | 0 | 988 | 4643 | 198 | 19876 | 7370 | 346 | 24 | 0 | 0 | 0 | 0 | 0 | 0 | 0 | 0 | 39 | 119 |
| SRR6246319 | 37 | 0 | 0 | 0 | 0 | 14 | 0 | 617 | 347 | 7750 | 15 | 501 | 0 | 5128 | 39 | 0 | 0 | 116 | 19 | 0 | 1375 | 5056 | 203 | 19845 | 7884 | 310 | 0 | 0 | 0 | 0 | 0 | 0 | 0 | 0 | 0 | 0 | 136 |
| SRR6246320 | 19 | 0 | 0 | 0 | 0 | 19 | 0 | 859 | 468 | 8508 | 20 | 546 | 0 | 4605 | 78 | 0 | 0 | 108 | 0 | 78 | 1218 | 4951 | 206 | 20640 | 8360 | 314 | 0 | 0 | 0 | 0 | 0 | 0 | 0 | 0 | 0 | 39 | 88 |
| SRR6246321 | 0 | 18 | 0 | 0 | 0 | 0 | 0 | 170 | 96 | 39391 | 107 | 1052 | 0 | 775 | 1795 | 2422 | 0 | 284 | 0 | 1179 | 769 | 1673 | 105 | 4511 | 3525 | 60 | 56 | 0 | 11 | 14 | 0 | 0 | 11 | 18 | 12 | 19037 | 119 |
| SRR6246322 | 0 | 17 | 0 | 0 | 0 | 0 | 0 | 178 | 122 | 40339 | 888 | 1010 | 0 | 555 | 1564 | 2385 | 0 | 504 | 16 | 1032 | 504 | 1527 | 155 | 4588 | 3410 | 124 | 37 | 0 | 13 | 15 | 11 | 0 | 12 | 12 | 12 | 20303 | 109 |
| SRR6246323 | 0 | 26 | 0 | 0 | 0 | 0 | 0 | 169 | 169 | 39939 | 1140 | 1066 | 0 | 707 | 1953 | 2501 | 0 | 435 | 15 | 1340 | 730 | 1577 | 118 | 4297 | 3840 | 96 | 74 | 0 | 21 | 17 | 0 | 0 | 16 | 26 | 10 | 20582 | 140 |
| SRR6246324 | 0 | 27 | 11 | 0 | 0 | 0 | 13 | 164 | 109 | 40923 | 1071 | 1180 | 0 | 623 | 1682 | 2655 | 0 | 352 | 0 | 1191 | 712 | 1392 | 84 | 4284 | 3565 | 77 | 63 | 0 | 14 | 18 | 17 | 0 | 14 | 37 | 11 | 20429 | 115 |
| SRR6246325 | 0 | 25 | 0 | 0 | 12 | 0 | 0 | 164 | 80 | 7443 | 0 | 0 | 0 | 755 | 104 | 39 | 0 | 899 | 12 | 0 | 2272 | 1970 | 177 | 9379 | 7417 | 78 | 20 | 0 | 0 | 0 | 0 | 0 | 0 | 0 | 0 | 49 | 357 |
| SRR6246326 | 0 | 29 | 14 | 0 | 0 | 0 | 0 | 166 | 94 | 7338 | 0 | 0 | 0 | 697 | 89 | 46 | 0 | 836 | 0 | 0 | 1878 | 1652 | 218 | 9688 | 6966 | 129 | 26 | 0 | 0 | 0 | 0 | 0 | 0 | 0 | 19 | 42 | 279 |
| SRR6246327 | 0 | 20 | 0 | 0 | 19 | 0 | 0 | 154 | 88 | 7482 | 0 | 0 | 0 | 704 | 104 | 50 | 0 | 858 | 19 | 0 | 2216 | 1785 | 182 | 9227 | 7159 | 113 | 0 | 0 | 0 | 0 | 0 | 0 | 0 | 0 | 0 | 38 | 369 |
| SRR6246328 | 0 | 27 | 0 | 0 | 28 | 0 | 0 | 163 | 75 | 7249 | 0 | 0 | 0 | 693 | 104 | 46 | 0 | 904 | 20 | 0 | 1888 | 1624 | 164 | 9386 | 6806 | 116 | 13 | 0 | 0 | 0 | 0 | 0 | 0 | 0 | 0 | 50 | 288 |
| SRR6246329 | 0 | 11 | 0 | 0 | 0 | 0 | 0 | 382 | 84 | 17125 | 0 | 13 | 0 | 2878 | 40 | 25 | 0 | 272 | 14 | 0 | 1882 | 2370 | 186 | 10009 | 6975 | 115 | 0 | 0 | 0 | 0 | 0 | 0 | 0 | 0 | 0 | 31 | 306 |
| SRR6246330 | 0 | 11 | 0 | 0 | 0 | 0 | 0 | 372 | 103 | 17193 | 0 | 0 | 0 | 2758 | 26 | 26 | 0 | 264 | 0 | 0 | 1767 | 2160 | 194 | 10441 | 6699 | 164 | 0 | 0 | 0 | 0 | 0 | 0 | 0 | 0 | 0 | 22 | 343 |
| SRR6246331 | 0 | 0 | 19 | 0 | 0 | 0 | 0 | 425 | 86 | 17081 | 13 | 0 | 0 | 2935 | 42 | 23 | 0 | 227 | 0 | 11 | 1863 | 2492 | 156 | 10030 | 7249 | 142 | 17 | 0 | 0 | 0 | 0 | 0 | 0 | 0 | 0 | 29 | 208 |
| SRR6246332 | 0 | 11 | 0 | 0 | 15 | 0 | 0 | 427 | 72 | 16807 | 0 | 10 | 0 | 2796 | 35 | 30 | 0 | 279 | 0 | 0 | 1539 | 2353 | 137 | 10683 | 7040 | 181 | 13 | 0 | 0 | 0 | 0 | 0 | 0 | 0 | 0 | 37 | 314 |
| SRR6246333 | 19 | 0 | 0 | 0 | 59 | 19 | 0 | 671 | 214 | 11774 | 244 | 275 | 0 | 1373 | 702 | 0 | 0 | 44 | 0 | 0 | 1664 | 5612 | 221 | 15186 | 7203 | 147 | 0 | 0 | 0 | 0 | 0 | 0 | 0 | 0 | 0 | 61 | 191 |
| SRR6246334 | 20 | 0 | 0 | 0 | 15 | 0 | 0 | 512 | 352 | 10631 | 128 | 160 | 0 | 1249 | 768 | 32 | 0 | 92 | 15 | 32 | 1384 | 4522 | 185 | 15287 | 7336 | 200 | 0 | 0 | 0 | 0 | 0 | 0 | 0 | 0 | 0 | 96 | 292 |
| SRR6246335 | 0 | 0 | 0 | 0 | 0 | 0 | 0 | 601 | 330 | 12376 | 150 | 421 | 0 | 1442 | 841 | 0 | 19 | 58 | 0 | 0 | 1941 | 5085 | 188 | 14329 | 7635 | 174 | 0 | 0 | 0 | 0 | 0 | 0 | 0 | 0 | 0 | 60 | 232 |
| SRR6246336 | 0 | 0 | 0 | 0 | 0 | 0 | 20 | 434 | 248 | 11430 | 124 | 186 | 0 | 1797 | 681 | 62 | 0 | 60 | 0 | 0 | 1388 | 4692 | 166 | 14438 | 7000 | 106 | 0 | 0 | 0 | 0 | 0 | 0 | 0 | 0 | 0 | 31 | 196 |
| SRR6246337 | 0 | 30 | 0 | 0 | 18 | 0 | 15 | 77 | 106 | 5213 | 1299 | 1029 | 0 | 4954 | 366 | 568 | 12 | 1342 | 0 | 904 | 1410 | 2262 | 96 | 7211 | 3727 | 124 | 156 | 0 | 0 | 0 | 0 | 0 | 0 | 0 | 0 | 16044 | 234 |
| SRR6246338 | 0 | 13 | 0 | 0 | 38 | 0 | 0 | 137 | 127 | 5098 | 125 | 744 | 0 | 4805 | 225 | 626 | 0 | 1334 | 14 | 842 | 1300 | 2228 | 81 | 7028 | 4016 | 91 | 93 | 0 | 0 | 0 | 0 | 0 | 0 | 0 | 0 | 15364 | 253 |
| SRR6246339 | 0 | 22 | 0 | 0 | 23 | 0 | 14 | 139 | 83 | 4806 | 158 | 796 | 0 | 5102 | 370 | 787 | 14 | 1400 | 14 | 963 | 1395 | 2286 | 90 | 7021 | 3974 | 108 | 148 | 0 | 0 | 0 | 0 | 0 | 0 | 0 | 0 | 15955 | 149 |
| SRR6246340 | 0 | 32 | 0 | 0 | 23 | 0 | 10 | 192 | 134 | 4942 | 125 | 691 | 0 | 4549 | 509 | 557 | 0 | 1217 | 14 | 1008 | 1189 | 2142 | 99 | 7465 | 3500 | 150 | 164 | 0 | 0 | 0 | 0 | 0 | 0 | 0 | 0 | 15518 | 258 |
| SRR6246341 | 0 | 0 | 0 | 0 | 100 | 0 | 0 | 59 | 42 | 12039 | 0 | 0 | 0 | 7599 | 15 | 0 | 13 | 214 | 14 | 17 | 813 | 870 | 71 | 6178 | 2311 | 128 | 18 | 0 | 0 | 0 | 0 | 0 | 0 | 0 | 0 | 119 | 57 |

### Supp Table S8.4.Columns_DI_to_ES

| | | | | | | | | | | | | | | | | | | | | | | | | | | | | | | | | | | | | | |
|---|---|---|---|---|---|---|---|---|---|---|---|---|---|---|---|---|---|---|---|---|---|---|---|---|---|---|---|---|---|---|---|---|---|---|---|---|---|
| SRR6246342 | 0 | 12 | 0 | 0 | 29 | 0 | 0 | 62 | 26 | 12169 | 15 | 10 | 0 | 7131 | 28 | 15 | 13 | 133 | 15 | 10 | 678 | 796 | 133 | 6454 | 2269 | 162 | 0 | 0 | 0 | 0 | 0 | 0 | 0 | 0 | 0 | 100 | 118 |
| SRR6246343 | 0 | 0 | 0 | 0 | 28 | 0 | 0 | 75 | 41 | 11840 | 0 | 0 | 0 | 7509 | 27 | 24 | 0 | 99 | 0 | 12 | 880 | 1008 | 57 | 6972 | 2215 | 114 | 0 | 0 | 0 | 0 | 0 | 0 | 0 | 0 | 0 | 122 | 142 |
| SRR6246344 | 0 | 11 | 0 | 0 | 29 | 0 | 0 | 73 | 65 | 11917 | 15 | 0 | 0 | 7346 | 20 | 0 | 0 | 131 | 15 | 0 | 784 | 711 | 73 | 6665 | 2657 | 145 | 0 | 0 | 0 | 0 | 0 | 0 | 0 | 0 | 0 | 100 | 174 |
| SRR6246345 | 0 | 0 | 0 | 0 | 0 | 0 | 22 | 344 | 86 | 19789 | 57 | 29 | 0 | 25861 | 57 | 29 | 0 | 0 | 0 | 0 | 1256 | 2060 | 50 | 8090 | 3517 | 50 | 0 | 0 | 0 | 0 | 0 | 0 | 0 | 0 | 0 | 57 | 50 |
| SRR6246346 | 0 | 0 | 0 | 0 | 52 | 0 | 0 | 307 | 61 | 19829 | 184 | 61 | 0 | 23635 | 0 | 31 | 0 | 0 | 0 | 92 | 837 | 2040 | 105 | 8579 | 3348 | 52 | 0 | 0 | 0 | 0 | 0 | 0 | 0 | 0 | 0 | 31 | 157 |
| SRR6246347 | 22 | 0 | 0 | 0 | 100 | 22 | 22 | 202 | 86 | 18300 | 58 | 29 | 0 | 25850 | 29 | 29 | 0 | 0 | 0 | 29 | 1348 | 1547 | 150 | 8686 | 4243 | 200 | 22 | 0 | 0 | 0 | 0 | 0 | 0 | 0 | 0 | 86 | 100 |
| SRR6246348 | 0 | 0 | 0 | 0 | 102 | 0 | 0 | 238 | 89 | 20947 | 89 | 59 | 0 | 26355 | 0 | 0 | 23 | 51 | 0 | 30 | 714 | 1835 | 408 | 6882 | 3518 | 306 | 0 | 0 | 0 | 0 | 0 | 0 | 0 | 0 | 23 | 149 | 102 |
| SRR6246349 | 41 | 0 | 0 | 0 | 41 | 0 | 0 | 2224 | 889 | 6893 | 889 | 1112 | 0 | 6671 | 0 | 0 | 0 | 0 | 0 | 0 | 1268 | 1922 | 164 | 6501 | 2944 | 123 | 0 | 0 | 0 | 0 | 0 | 0 | 0 | 0 | 0 | 0 | 41 |
| SRR6246350 | 0 | 0 | 0 | 0 | 43 | 0 | 0 | 679 | 0 | 8595 | 906 | 0 | 0 | 10857 | 226 | 0 | 43 | 0 | 0 | 0 | 1021 | 1659 | 0 | 7531 | 2893 | 298 | 0 | 0 | 0 | 0 | 0 | 0 | 0 | 0 | 0 | 0 | 85 |
| SRR6246351 | 41 | 0 | 0 | 0 | 40 | 41 | 0 | 1472 | 1051 | 9041 | 1053 | 631 | 0 | 9672 | 0 | 0 | 0 | 40 | 0 | 0 | 1092 | 2549 | 162 | 7688 | 3601 | 243 | 0 | 0 | 0 | 0 | 0 | 0 | 0 | 0 | 0 | 0 | 81 |
| SRR6246352 | 0 | 0 | 0 | 0 | 83 | 0 | 0 | 1141 | 228 | 9582 | 229 | 1369 | 0 | 8898 | 0 | 0 | 0 | 0 | 0 | 0 | 873 | 1830 | 125 | 6488 | 3702 | 208 | 0 | 0 | 0 | 0 | 0 | 0 | 0 | 0 | 0 | 0 | 125 |
| SRR6246353 | 0 | 231 | 0 | 0 | 0 | 0 | 0 | 13 | 32 | 2316 | 52 | 31 | 0 | 8924 | 0 | 0 | 22 | 4074 | 19 | 86 | 1042 | 999 | 0 | 3754 | 3529 | 186 | 100 | 0 | 0 | 0 | 0 | 0 | 0 | 0 | 0 | 264 | 421 |
| SRR6246354 | 0 | 515 | 0 | 0 | 0 | 0 | 0 | 16 | 35 | 2235 | 65 | 25 | 0 | 8973 | 0 | 0 | 25 | 4154 | 24 | 89 | 939 | 1060 | 0 | 4023 | 3463 | 229 | 247 | 0 | 0 | 0 | 0 | 0 | 0 | 0 | 0 | 276 | 365 |
| SRR6246355 | 0 | 215 | 0 | 0 | 0 | 0 | 0 | 15 | 29 | 2213 | 82 | 36 | 0 | 8913 | 0 | 0 | 19 | 4101 | 19 | 92 | 910 | 1135 | 29 | 3722 | 3473 | 263 | 100 | 0 | 0 | 0 | 0 | 0 | 0 | 0 | 0 | 259 | 431 |
| SRR6246356 | 0 | 529 | 0 | 0 | 0 | 0 | 0 | 20 | 25 | 2300 | 46 | 43 | 0 | 8979 | 0 | 0 | 27 | 4293 | 0 | 101 | 1114 | 1163 | 24 | 3845 | 3410 | 222 | 241 | 0 | 0 | 0 | 0 | 0 | 0 | 0 | 0 | 268 | 429 |
| SRR6246358 | 0 | 11 | 0 | 0 | 221 | 0 | 11 | 69 | 0 | 1996 | 0 | 0 | 11 | 7681 | 0 | 0 | 0 | 1106 | 0 | 0 | 221 | 885 | 221 | 5307 | 3317 | 0 | 33 | 0 | 0 | 0 | 0 | 0 | 0 | 0 | 0 | 0 | 221 |
| SRR6246359 | 0 | 21 | 0 | 0 | 0 | 0 | 0 | 60 | 0 | 2040 | 0 | 0 | 0 | 8136 | 0 | 0 | 0 | 423 | 0 | 0 | 845 | 423 | 211 | 2958 | 3169 | 423 | 21 | 0 | 0 | 0 | 0 | 0 | 0 | 0 | 0 | 52 | 423 |
| SRR6246360 | 0 | 11 | 0 | 0 | 0 | 0 | 0 | 53 | 0 | 1820 | 0 | 0 | 0 | 7775 | 0 | 0 | 0 | 437 | 0 | 0 | 1312 | 219 | 0 | 3717 | 2186 | 0 | 22 | 0 | 0 | 0 | 0 | 0 | 0 | 0 | 0 | 106 | 219 |
| SRR6246361 | 12 | 12 | 0 | 0 | 0 | 0 | 0 | 187 | 21 | 9093 | 21 | 0 | 0 | 34810 | 0 | 0 | 12 | 104 | 0 | 0 | 621 | 2020 | 207 | 7872 | 4972 | 104 | 48 | 0 | 0 | 0 | 0 | 0 | 0 | 0 | 0 | 62 | 207 |
| SRR6246362 | 12 | 25 | 0 | 0 | 0 | 0 | 0 | 214 | 21 | 8365 | 64 | 86 | 0 | 36070 | 0 | 0 | 37 | 106 | 0 | 0 | 638 | 1913 | 159 | 7068 | 6005 | 53 | 49 | 0 | 0 | 0 | 0 | 0 | 0 | 0 | 0 | 171 | 213 |
| SRR6246363 | 12 | 0 | 0 | 0 | 52 | 12 | 0 | 188 | 21 | 8676 | 146 | 63 | 0 | 35583 | 0 | 0 | 0 | 206 | 0 | 21 | 1445 | 1961 | 103 | 8517 | 6710 | 310 | 18 | 0 | 0 | 0 | 0 | 0 | 0 | 0 | 0 | 42 | 413 |
| SRR6246364 | 15 | 0 | 0 | 0 | 0 | 0 | 0 | 148 | 42 | 9239 | 85 | 63 | 0 | 34779 | 0 | 0 | 25 | 422 | 53 | 21 | 1581 | 1687 | 158 | 7748 | 6536 | 53 | 25 | 0 | 0 | 0 | 0 | 0 | 0 | 0 | 0 | 42 | 422 |
| SRR6246365 | 0 | 0 | 0 | 0 | 0 | 0 | 0 | 464 | 77 | 1391 | 232 | 77 | 0 | 2860 | 0 | 0 | 0 | 136 | 0 | 77 | 1448 | 2625 | 136 | 11179 | 8644 | 272 | 0 | 0 | 0 | 0 | 0 | 0 | 0 | 0 | 0 | 77 | 317 |
| SRR6246366 | 30 | 0 | 0 | 0 | 0 | 0 | 0 | 402 | 80 | 1288 | 322 | 80 | 0 | 2495 | 0 | 0 | 30 | 92 | 0 | 0 | 1009 | 2432 | 46 | 11196 | 7663 | 367 | 30 | 0 | 0 | 0 | 0 | 0 | 0 | 0 | 0 | 0 | 229 |
| SRR6246367 | 29 | 0 | 0 | 0 | 0 | 0 | 0 | 398 | 80 | 2470 | 80 | 80 | 0 | 3027 | 0 | 0 | 58 | 45 | 0 | 0 | 1222 | 2353 | 45 | 11402 | 8777 | 226 | 58 | 0 | 0 | 0 | 0 | 0 | 0 | 0 | 0 | 0 | 90 |
| SRR6246368 | 30 | 0 | 0 | 0 | 46 | 0 | 30 | 240 | 80 | 2484 | 160 | 80 | 0 | 3045 | 0 | 0 | 30 | 137 | 0 | 0 | 867 | 2785 | 91 | 11824 | 8856 | 274 | 0 | 0 | 0 | 0 | 0 | 0 | 0 | 0 | 0 | 0 | 228 |
| SRR6246369 | 0 | 33 | 0 | 0 | 0 | 0 | 0 | 19 | 28 | 4823 | 0 | 0 | 0 | 6318 | 0 | 0 | 16 | 1797 | 114 | 57 | 998 | 570 | 29 | 11977 | 9924 | 171 | 77 | 0 | 0 | 0 | 0 | 0 | 0 | 0 | 0 | 141 | 314 |
| SRR6246370 | 0 | 32 | 0 | 0 | 0 | 0 | 0 | 26 | 18 | 4806 | 0 | 0 | 0 | 6366 | 0 | 0 | 15 | 1673 | 59 | 53 | 1233 | 587 | 59 | 12151 | 9421 | 205 | 83 | 0 | 0 | 0 | 0 | 0 | 0 | 0 | 0 | 182 | 411 |
| SRR6246371 | 0 | 33 | 0 | 0 | 0 | 0 | 0 | 16 | 22 | 4862 | 0 | 0 | 0 | 6455 | 0 | 0 | 19 | 1537 | 29 | 50 | 899 | 870 | 29 | 10759 | 10092 | 145 | 22 | 0 | 0 | 0 | 0 | 0 | 0 | 0 | 0 | 135 | 551 |
| SRR6246372 | 0 | 28 | 0 | 0 | 0 | 0 | 0 | 15 | 25 | 4863 | 10 | 0 | 0 | 6355 | 0 | 0 | 19 | 1749 | 29 | 42 | 875 | 758 | 29 | 10962 | 9359 | 379 | 65 | 0 | 0 | 0 | 0 | 10 | 0 | 0 | 0 | 135 | 408 |
| SRR6246373 | 0 | 0 | 0 | 0 | 136 | 0 | 0 | 120 | 33 | 5966 | 0 | 0 | 0 | 6869 | 0 | 0 | 0 | 0 | 0 | 0 | 1091 | 955 | 0 | 8184 | 3819 | 273 | 23 | 0 | 0 | 0 | 0 | 0 | 0 | 0 | 0 | 60 | 0 |
| SRR6246374 | 0 | 0 | 16 | 0 | 0 | 0 | 0 | 88 | 34 | 5806 | 0 | 0 | 0 | 6512 | 0 | 0 | 0 | 281 | 0 | 0 | 983 | 1264 | 0 | 6879 | 3369 | 140 | 31 | 0 | 0 | 0 | 0 | 0 | 0 | 0 | 0 | 34 | 281 |
| SRR6246375 | 0 | 0 | 0 | 0 | 0 | 0 | 15 | 80 | 53 | 6053 | 0 | 0 | 0 | 6840 | 0 | 0 | 23 | 405 | 0 | 0 | 811 | 1487 | 135 | 8785 | 4190 | 270 | 15 | 0 | 0 | 0 | 0 | 0 | 0 | 0 | 0 | 20 | 405 |
| SRR6246376 | 0 | 0 | 0 | 0 | 0 | 0 | 0 | 61 | 41 | 5887 | 0 | 0 | 0 | 6732 | 0 | 0 | 0 | 142 | 0 | 0 | 1282 | 1852 | 0 | 10687 | 3705 | 142 | 46 | 0 | 0 | 0 | 0 | 0 | 0 | 0 | 0 | 0 | 285 |
| SRR6246379 | 0 | 322 | 0 | 0 | 0 | 0 | 0 | 0 | 0 | 520 | 0 | 0 | 0 | 27575 | 0 | 0 | 0 | 0 | 0 | 0 | 0 | 0 | 0 | 17196 | 2646 | 0 | 0 | 0 | 0 | 0 | 0 | 0 | 0 | 0 | 0 | 0 | 0 |
| SRR6246385 | 0 | 382 | 0 | 0 | 17 | 0 | 0 | 56 | 138 | 2089 | 166 | 105 | 0 | 7570 | 0 | 0 | 46 | 3603 | 25 | 85 | 1625 | 1802 | 53 | 3870 | 5228 | 212 | 446 | 0 | 0 | 0 | 0 | 0 | 0 | 0 | 10 | 202 | 413 |
| SRR6246386 | 0 | 390 | 0 | 0 | 16 | 0 | 0 | 52 | 144 | 2088 | 148 | 89 | 0 | 7490 | 0 | 0 | 48 | 3731 | 23 | 94 | 1708 | 1816 | 43 | 3905 | 5177 | 230 | 443 | 0 | 0 | 11 | 0 | 0 | 0 | 0 | 11 | 221 | 379 |
| SRR6246387 | 0 | 353 | 10 | 0 | 0 | 0 | 0 | 68 | 161 | 2088 | 165 | 83 | 0 | 7499 | 0 | 0 | 46 | 3628 | 37 | 91 | 1674 | 1993 | 39 | 4050 | 5231 | 224 | 444 | 0 | 0 | 0 | 0 | 0 | 0 | 0 | 0 | 201 | 350 |
| SRR6246388 | 0 | 380 | 0 | 0 | 14 | 0 | 0 | 66 | 144 | 2052 | 158 | 95 | 0 | 7597 | 0 | 0 | 39 | 3698 | 38 | 87 | 1643 | 1915 | 24 | 3936 | 5069 | 267 | 439 | 0 | 0 | 0 | 0 | 0 | 0 | 0 | 12 | 203 | 456 |
| SRR6246389 | 0 | 0 | 0 | 0 | 137 | 0 | 17 | 85 | 11 | 3297 | 11 | 17 | 0 | 11134 | 0 | 0 | 26 | 722 | 34 | 11 | 481 | 515 | 69 | 3815 | 2543 | 69 | 26 | 0 | 0 | 0 | 0 | 0 | 0 | 0 | 0 | 250 | 241 |
| SRR6246390 | 0 | 18 | 0 | 0 | 0 | 0 | 0 | 92 | 29 | 3608 | 23 | 17 | 0 | 10946 | 0 | 0 | 18 | 747 | 36 | 23 | 569 | 498 | 142 | 3804 | 2382 | 142 | 18 | 0 | 0 | 0 | 0 | 0 | 0 | 0 | 0 | 132 | 178 |
| SRR6246391 | 0 | 0 | 0 | 0 | 69 | 0 | 26 | 68 | 28 | 3605 | 11 | 0 | 0 | 11173 | 0 | 0 | 0 | 414 | 0 | 23 | 518 | 518 | 276 | 4386 | 1830 | 138 | 17 | 0 | 0 | 0 | 0 | 0 | 0 | 0 | 0 | 182 | 276 |
| SRR6246392 | 0 | 0 | 0 | 0 | 105 | 0 | 26 | 85 | 28 | 3283 | 11 | 0 | 0 | 10932 | 0 | 0 | 0 | 803 | 0 | 11 | 454 | 733 | 244 | 4222 | 2477 | 70 | 35 | 0 | 0 | 0 | 0 | 0 | 0 | 0 | 0 | 142 | 384 |
| SRR6246393 | 14 | 0 | 0 | 0 | 122 | 0 | 0 | 327 | 65 | 4375 | 87 | 109 | 0 | 31934 | 0 | 0 | 14 | 61 | 61 | 22 | 425 | 1337 | 0 | 6319 | 3646 | 182 | 13 | 0 | 0 | 0 | 0 | 0 | 0 | 0 | 0 | 22 | 61 |
| SRR6246394 | 0 | 0 | 0 | 0 | 0 | 0 | 0 | 312 | 67 | 5051 | 22 | 0 | 15 | 30440 | 0 | 0 | 0 | 376 | 0 | 22 | 627 | 1882 | 251 | 5959 | 3513 | 314 | 0 | 0 | 0 | 0 | 0 | 0 | 0 | 0 | 0 | 45 | 251 |
| SRR6246395 | 15 | 0 | 0 | 0 | 60 | 0 | 15 | 347 | 65 | 4472 | 87 | 22 | 0 | 32631 | 0 | 0 | 15 | 299 | 0 | 22 | 478 | 1314 | 299 | 6214 | 3764 | 120 | 14 | 0 | 0 | 0 | 0 | 0 | 0 | 0 | 0 | 43 | 60 |
| SRR6246396 | 0 | 15 | 0 | 0 | 125 | 0 | 0 | 327 | 44 | 5191 | 44 | 65 | 0 | 31929 | 0 | 0 | 15 | 63 | 0 | 44 | 0 | 938 | 438 | 6438 | 3875 | 250 | 15 | 0 | 0 | 0 | 0 | 0 | 0 | 0 | 0 | 22 | 125 |
| SRR6246397 | 19 | 0 | 0 | 0 | 19 | 0 | 0 | 127 | 127 | 2133 | 350 | 255 | 0 | 6113 | 0 | 0 | 0 | 57 | 0 | 32 | 1475 | 3990 | 113 | 11366 | 8605 | 322 | 0 | 0 | 0 | 0 | 0 | 0 | 0 | 0 | 0 | 159 | 151 |
| SRR6246398 | 11 | 0 | 0 | 0 | 79 | 0 | 0 | 130 | 65 | 1851 | 260 | 487 | 0 | 5553 | 0 | 0 | 0 | 0 | 0 | 0 | 887 | 3429 | 118 | 11372 | 7706 | 315 | 0 | 0 | 0 | 0 | 0 | 0 | 0 | 0 | 0 | 65 | 256 |
| SRR6246399 | 19 | 0 | 0 | 0 | 19 | 0 | 0 | 128 | 64 | 1602 | 352 | 513 | 0 | 5255 | 0 | 0 | 0 | 115 | 19 | 0 | 1365 | 4172 | 154 | 11169 | 8113 | 308 | 0 | 0 | 0 | 0 | 0 | 0 | 0 | 0 | 0 | 96 | 173 |
| SRR6246400 | 20 | 0 | 0 | 0 | 19 | 0 | 13 | 228 | 98 | 2245 | 293 | 293 | 0 | 5792 | 0 | 0 | 0 | 77 | 19 | 0 | 889 | 3674 | 213 | 11853 | 7947 | 387 | 0 | 0 | 0 | 0 | 0 | 0 | 0 | 0 | 0 | 33 | 213 |
| SRR6265480 | 0 | 10 | 12 | 0 | 53 | 0 | 0 | 401 | 116 | 1773 | 2912 | 3936 | 0 | 148 | 42 | 11 | 0 | 196 | 19 | 1055 | 386 | 1899 | 0 | 0 | 12789 | 0 | 14 | 13 | 12 | 43 | 11 | 0 | 23 | 13 | 13 | 6985 | 0 |
| SRR6265481 | 0 | 0 | 0 | 0 | 75 | 0 | 0 | 229 | 90 | 2766 | 124 | 1033 | 0 | 166 | 769 | 478 | 0 | 398 | 15 | 714 | 340 | 1749 | 0 | 0 | 14445 | 0 | 0 | 0 | 0 | 21 | 26 | 0 | 0 | 32 | 43 | 7258 | 0 |
| SRR6265482 | 20 | 12 | 0 | 0 | 71 | 0 | 0 | 338 | 143 | 1461 | 227 | 5255 | 0 | 71 | 18 | 0 | 0 | 248 | 30 | 1229 | 258 | 1799 | 0 | 0 | 12357 | 0 | 12 | 38 | 16 | 56 | 77 | 14 | 28 | 36 | 0 | 4987 | 0 |
| SRR6265483 | 20 | 15 | 0 | 0 | 55 | 0 | 0 | 508 | 159 | 2383 | 162 | 8263 | 0 | 95 | 0 | 0 | 0 | 296 | 21 | 953 | 102 | 1593 | 0 | 0 | 10749 | 0 | 20 | 0 | 17 | 32 | 32 | 0 | 15 | 25 | 0 | 6006 | 0 |
| SRR6265484 | 16 | 0 | 0 | 0 | 46 | 0 | 0 | 761 | 190 | 4745 | 3414 | 4155 | 0 | 48 | 181 | 162 | 0 | 317 | 11 | 1626 | 127 | 1455 | 0 | 0 | 9251 | 0 | 18 | 0 | 10 | 16 | 23 | 11 | 20 | 37 | 22 | 5544 | 0 |
| SRR6265485 | 16 | 0 | 0 | 0 | 51 | 0 | 0 | 288 | 336 | 1142 | 270 | 3737 | 0 | 60 | 312 | 156 | 0 | 273 | 16 | 1586 | 119 | 1566 | 0 | 0 | 9913 | 0 | 0 | 21 | 0 | 10 | 25 | 10 | 16 | 12 | 12 | 3978 | 0 |
| SRR6265486 | 20 | 0 | 0 | 0 | 45 | 0 | 0 | 584 | 182 | 1639 | 9172 | 7738 | 0 | 121 | 0 | 23 | 0 | 221 | 15 | 1032 | 100 | 1541 | 0 | 0 | 10002 | 0 | 10 | 72 | 42 | 113 | 122 | 14 | 32 | 41 | 0 | 3315 | 0 |
| SRR6265487 | 0 | 0 | 16 | 0 | 48 | 0 | 0 | 542 | 155 | 13286 | 2787 | 2593 | 0 | 387 | 0 | 0 | 0 | 243 | 11 | 1839 | 344 | 1242 | 0 | 0 | 7918 | 0 | 19 | 0 | 30 | 51 | 34 | 0 | 34 | 49 | 14 | 8989 | 0 |
| SRR6265488 | 0 | 13 | 16 | 0 | 50 | 0 | 0 | 189 | 26 | 1750 | 573 | 2348 | 0 | 355 | 11 | 15 | 0 | 228 | 21 | 1656 | 434 | 1380 | 0 | 0 | 9325 | 0 | 17 | 0 | 13 | 12 | 11 | 17 | 29 | 10 | 25 | 9966 | 29 |
| SRR6265489 | 0 | 0 | 0 | 0 | 51 | 0 | 0 | 2468 | 178 | 2586 | 289 | 6871 | 0 | 197 | 39 | 0 | 0 | 171 | 21 | 1007 | 125 | 1182 | 0 | 0 | 7649 | 0 | 0 | 0 | 0 | 0 | 0 | 0 | 0 | 0 | 0 | 5410 | 0 |
| SRR6265490 | 0 | 12 | 13 | 0 | 47 | 0 | 0 | 597 | 162 | 3408 | 845 | 4835 | 0 | 147 | 344 | 542 | 0 | 334 | 18 | 2192 | 83 | 1236 | 0 | 0 | 8900 | 0 | 0 | 17 | 0 | 46 | 51 | 0 | 36 | 95 | 85 | 7625 | 0 |
| SRR6265491 | 0 | 0 | 0 | 0 | 50 | 0 | 0 | 809 | 112 | 1933 | 3293 | 4630 | 0 | 225 | 45 | 56 | 0 | 282 | 24 | 2101 | 167 | 1148 | 0 | 0 | 8814 | 0 | 0 | 0 | 0 | 29 | 33 | 0 | 18 | 22 | 0 | 6866 | 0 |
| SRR6306014 | 0 | 0 | 0 | 0 | 14 | 0 | 0 | 1166 | 833 | 20609 | 439 | 1054 | 0 | 349 | 0 | 12 | 0 | 58 | 0 | 1440 | 167 | 2834 | 21 | 27295 | 7218 | 108 | 24 | 232 | 117 | 515 | 584 | 90 | 419 | 916 | 16 | 4328 | 67 |
| SRR6306015 | 0 | 0 | 0 | 0 | 19 | 0 | 0 | 1109 | 987 | 21580 | 233 | 566 | 0 | 420 | 18 | 23 | 0 | 79 | 0 | 1205 | 195 | 2918 | 20 | 29651 | 8162 | 100 | 0 | 242 | 172 | 587 | 699 | 114 | 501 | 1028 | 0 | 4205 | 61 |
| SRR6313948 | 28 | 733 | 117 | 57 | 532 | 81 | 170 | 16826 | 1111 | 18745 | 5332 | 10579 | 11 | 2605 | 0 | 0 | 185 | 478 | 76 | 373 | 883 | 1912 | 757 | 45234 | 1002 | 55 | 114 | 95 | 168 | 758 | 2162 | 914 | 1214 | 555 | 0 | 2414 | 825 |
| SRR6313949 | 16 | 1557 | 59 | 38 | 993 | 179 | 330 | 2431 | 452 | 5553 | 17563 | 33696 | 20 | 1924 | 0 | 0 | 462 | 664 | 66 | 185 | 1094 | 3483 | 360 | 14031 | 302 | 21 | 110 | 90 | 94 | 486 | 882 | 269 | 256 | 144 | 0 | 637 | 177 |
| SRR6317622 | 0 | 49 | 63 | 0 | 13 | 0 | 0 | 584 | 324 | 2886 | 367 | 8917 | 0 | 746 | 292 | 259 | 0 | 852 | 0 | 2010 | 274 | 552 | 24 | 7765 | 1831 | 74 | 0 | 63 | 35 | 16 | 11 | 35 | 12 | 17 | 92 | 7944 | 11 |
| SRR6317623 | 0 | 76 | 61 | 0 | 0 | 0 | 0 | 303 | 186 | 2026 | 14066 | 7219 | 0 | 233 | 116 | 70 | 0 | 712 | 0 | 1444 | 320 | 684 | 32 | 9848 | 1964 | 94 | 0 | 47 | 76 | 14 | 109 | 28 | 80 | 132 | 52 | 12925 | 0 |
| SRR6317624 | 0 | 15 | 63 | 0 | 18 | 0 | 0 | 400 | 200 | 1835 | 1344 | 12627 | 0 | 634 | 67 | 67 | 0 | 1060 | 0 | 1902 | 406 | 640 | 36 | 9032 | 2168 | 72 | 21 | 55 | 59 | 13 | 14 | 25 | 96 | 117 | 80 | 5688 | 0 |
| SRR6317625 | 0 | 92 | 21 | 0 | 14 | 0 | 0 | 332 | 166 | 1707 | 1262 | 16197 | 0 | 308 | 71 | 24 | 0 | 1096 | 0 | 759 | 400 | 514 | 27 | 8303 | 2169 | 61 | 21 | 43 | 50 | 13 | 15 | 0 | 107 | 71 | 71 | 3557 | 0 |
| SRR6317626 | 0 | 57 | 30 | 0 | 22 | 0 | 0 | 707 | 362 | 41452 | 8859 | 5816 | 0 | 996 | 272 | 290 | 0 | 379 | 0 | 1250 | 200 | 287 | 38 | 5548 | 1144 | 0 | 0 | 14 | 14 | 51 | 41 | 0 | 32 | 84 | 22 | 2881 | 0 |
| SRR6317627 | 0 | 14 | 0 | 0 | 28 | 0 | 26 | 1106 | 958 | 23787 | 3397 | 1876 | 0 | 998 | 643 | 549 | 13 | 497 | 0 | 677 | 216 | 208 | 51 | 6399 | 1085 | 0 | 67 | 0 | 11 | 23 | 19 | 0 | 0 | 23 | 22 | 6057 | 0 |
| SRR6317628 | 0 | 50 | 33 | 0 | 20 | 0 | 0 | 391 | 78 | 5682 | 11572 | 6750 | 0 | 547 | 209 | 130 | 0 | 427 | 0 | 1460 | 234 | 482 | 21 | 5406 | 893 | 42 | 0 | 23 | 30 | 96 | 86 | 20 | 43 | 60 | 43 | 7167 | 0 |
| SRR6317629 | 0 | 67 | 21 | 0 | 22 | 0 | 0 | 680 | 384 | 24113 | 7333 | 5308 | 0 | 325 | 237 | 281 | 0 | 367 | 0 | 1419 | 176 | 326 | 33 | 6795 | 1243 | 15 | 0 | 12 | 19 | 44 | 46 | 0 | 17 | 31 | 27 | 5825 | 0 |
| SRR6317630 | 0 | 99 | 38 | 0 | 15 | 0 | 0 | 566 | 367 | 4454 | 12168 | 6887 | 0 | 490 | 306 | 107 | 0 | 792 | 0 | 1867 | 415 | 741 | 29 | 8732 | 2130 | 75 | 0 | 58 | 113 | 15 | 12 | 38 | 120 | 12 | 133 | 8433 | 0 |
| SRR6317631 | 0 | 149 | 28 | 0 | 21 | 0 | 0 | 228 | 114 | 3044 | 935 | 8563 | 0 | 358 | 358 | 212 | 0 | 925 | 0 | 1400 | 395 | 762 | 31 | 9187 | 2373 | 63 | 0 | 34 | 22 | 96 | 131 | 12 | 59 | 90 | 87 | 8759 | 0 |
| SRR6317632 | 0 | 189 | 37 | 0 | 17 | 0 | 0 | 415 | 224 | 2395 | 1601 | 14051 | 0 | 351 | 48 | 80 | 0 | 993 | 0 | 1469 | 476 | 712 | 27 | 8880 | 2470 | 62 | 0 | 45 | 49 | 238 | 201 | 0 | 95 | 136 | 103 | 4487 | 0 |
| SRR6317633 | 0 | 12 | 28 | 0 | 16 | 0 | 0 | 345 | 172 | 2025 | 32378 | 16889 | 0 | 259 | 65 | 65 | 0 | 1168 | 0 | 1163 | 436 | 612 | 24 | 9007 | 2446 | 53 | 0 | 65 | 47 | 11 | 14 | 14 | 47 | 93 | 56 | 3727 | 0 |
| SRR6317634 | 0 | 74 | 44 | 0 | 21 | 0 | 0 | 521 | 387 | 33221 | 10883 | 7637 | 0 | 858 | 320 | 185 | 0 | 477 | 0 | 1009 | 249 | 381 | 35 | 6383 | 1351 | 11 | 15 | 0 | 30 | 54 | 30 | 0 | 0 | 37 | 37 | 2708 | 0 |
| SRR6317635 | 0 | 34 | 16 | 0 | 29 | 0 | 20 | 1714 | 1533 | 36472 | 4127 | 2749 | 0 | 1003 | 2109 | 1850 | 12 | 429 | 0 | 1281 | 271 | 320 | 67 | 8092 | 1328 | 0 | 39 | 0 | 0 | 30 | 21 | 0 | 11 | 30 | 28 | 12789 | 0 |
| SRR6317636 | 0 | 65 | 37 | 0 | 20 | 0 | 0 | 558 | 157 | 7447 | 11824 | 7081 | 0 | 401 | 279 | 262 | 0 | 402 | 0 | 1308 | 287 | 545 | 27 | 8528 | 1971 | 28 | 0 | 17 | 31 | 85 | 10 | 0 | 40 | 62 | 76 | 7325 | 0 |
| SRR6317637 | 0 | 50 | 0 | 0 | 27 | 0 | 20 | 467 | 300 | 19881 | 602 | 4954 | 0 | 500 | 183 | 150 | 0 | 328 | 0 | 1084 | 219 | 405 | 36 | 7926 | 1457 | 18 | 13 | 0 | 12 | 52 | 30 | 0 | 12 | 23 | 17 | 5120 | 0 |
| SRR6317638 | 0 | 91 | 43 | 0 | 11 | 0 | 0 | 303 | 238 | 3048 | 678 | 10096 | 0 | 432 | 108 | 65 | 0 | 652 | 0 | 1470 | 298 | 616 | 35 | 8750 | 2172 | 76 | 0 | 63 | 67 | 13 | 13 | 0 | 101 | 135 | 48 | 6399 | 0 |
| SRR6317639 | 0 | 158 | 30 | 0 | 14 | 0 | 0 | 466 | 213 | 2929 | 575 | 7256 | 0 | 427 | 349 | 175 | 0 | 665 | 0 | 1416 | 364 | 844 | 32 | 10522 | 2327 | 106 | 0 | 45 | 45 | 188 | 147 | 19 | 64 | 135 | 79 | 11931 | 0 |
| SRR6317640 | 0 | 12 | 32 | 0 | 13 | 0 | 0 | 293 | 178 | 2140 | 1708 | 13270 | 0 | 382 | 102 | 127 | 0 | 966 | 0 | 1401 | 468 | 689 | 27 | 8864 | 2481 | 72 | 0 | 32 | 44 | 16 | 12 | 18 | 94 | 12 | 85 | 5858 | 0 |
| SRR6317641 | 0 | 161 | 32 | 0 | 11 | 0 | 0 | 284 | 325 | 1626 | 1805 | 17881 | 0 | 244 | 20 | 61 | 0 | 1585 | 0 | 792 | 436 | 586 | 24 | 7667 | 2199 | 59 | 0 | 18 | 14 | 73 | 110 | 23 | 50 | 60 | 55 | 3210 | 0 |
| SRR6317642 | 0 | 52 | 37 | 0 | 18 | 0 | 0 | 441 | 288 | 28680 | 7841 | 5310 | 0 | 997 | 268 | 268 | 12 | 377 | 0 | 824 | 207 | 317 | 34 | 5980 | 1236 | 13 | 15 | 12 | 28 | 46 | 46 | 0 | 25 | 46 | 46 | 3087 | 0 |
| SRR6317643 | 0 | 51 | 32 | 0 | 37 | 0 | 18 | 696 | 583 | 22687 | 4073 | 2351 | 0 | 1016 | 1872 | 1129 | 0 | 494 | 0 | 865 | 262 | 258 | 79 | 5914 | 1347 | 0 | 23 | 13 | 17 | 53 | 38 | 0 | 17 | 48 | 10 | 7525 | 0 |
| SRR6317644 | 0 | 60 | 43 | 0 | 17 | 0 | 0 | 740 | 172 | 6523 | 10086 | 6369 | 0 | 327 | 482 | 413 | 0 | 519 | 0 | 1497 | 248 | 490 | 27 | 9130 | 1910 | 29 | 0 | 16 | 27 | 125 | 125 | 19 | 30 | 84 | 81 | 8193 | 0 |
| SRR6317645 | 0 | 52 | 23 | 0 | 16 | 0 | 0 | 488 | 374 | 23482 | 584 | 4215 | 0 | 504 | 293 | 244 | 0 | 389 | 0 | 1286 | 231 | 411 | 33 | 7807 | 1539 | 15 | 0 | 0 | 27 | 34 | 29 | 0 | 29 | 36 | 45 | 7567 | 0 |
| SRR6324194 | 0 | 0 | 0 | 0 | 0 | 0 | 0 | 1277 | 349 | 2026 | 455 | 878 | 0 | 3606 | 0 | 0 | 0 | 108 | 0 | 0 | 36 | 200 | 0 | 6295 | 774 | 13 | 0 | 82 | 12 | 34 | 24 | 12 | 0 | 13 | 0 | 0 | 0 |
| SRR6324195 | 0 | 0 | 0 | 0 | 0 | 0 | 0 | 1831 | 557 | 4161 | 1101 | 393 | 0 | 7120 | 0 | 0 | 0 | 167 | 15 | 25 | 55 | 196 | 0 | 6190 | 790 | 12 | 0 | 49 | 17 | 24 | 46 | 22 | 42 | 42 | 12 | 241 | 0 |
| SRR6363052 | 0 | 0 | 94 | 0 | 10 | 33 | 0 | 1660 | 276 | 2477 | 1565 | 5008 | 0 | 627 | 0 | 0 | 19 | 354 | 0 | 2536 | 618 | 1512 | 39 | 37216 | 3637 | 11 | 0 | 19 | 13 | 34 | 46 | 20 | 30 | 57 | 18 | 23751 | 160 |
| SRR6363053 | 0 | 0 | 61 | 0 | 11 | 38 | 0 | 1643 | 338 | 2861 | 8658 | 4701 | 0 | 1188 | 0 | 0 | 0 | 285 | 0 | 1212 | 589 | 1429 | 41 | 35098 | 2972 | 0 | 11 | 328 | 215 | 538 | 701 | 317 | 474 | 916 | 188 | 6720 | 135 |
| SRR6363054 | 0 | 0 | 51 | 0 | 13 | 61 | 0 | 1021 | 141 | 1439 | 987 | 3666 | 0 | 1052 | 0 | 0 | 0 | 359 | 0 | 1864 | 543 | 1345 | 55 | 35173 | 3464 | 13 | 0 | 26 | 18 | 39 | 39 | 14 | 27 | 40 | 79 | 17020 | 123 |
| SRR6370846 | 0 | 0 | 0 | 0 | 24 | 0 | 0 | 0 | 11 | 0 | 19299 | 17556 | 0 | 0 | 0 | 0 | 12 | 10 | 3554 | 0 | 106 | 89 | 0 | 0 | 143 | 56 | 11 | 0 | 0 | 0 | 0 | 0 | 0 | 0 | 0 | 0 | 60 |
| SRR6370847 | 0 | 193 | 25 | 0 | 102 | 0 | 0 | 65 | 292 | 1299 | 1432 | 45760 | 0 | 32 | 0 | 0 | 0 | 24 | 96 | 292 | 222 | 221 | 0 | 0 | 652 | 46 | 353 | 0 | 17 | 17 | 17 | 0 | 0 | 0 | 0 | 260 | 427 |
| SRR6370848 | 0 | 85 | 0 | 0 | 86 | 0 | 0 | 0 | 104 | 835 | 205 | 15340 | 0 | 104 | 0 | 0 | 18 | 30 | 269 | 104 | 169 | 257 | 0 | 275 | 1020 | 117 | 0 | 0 | 24 | 0 | 12 | 0 | 0 | 0 | 0 | 313 | 190 |
| SRR6370849 | 10 | 77 | 0 | 0 | 55 | 0 | 0 | 314 | 715 | 3430 | 15863 | 34212 | 0 | 286 | 0 | 0 | 10 | 56 | 31 | 0 | 302 | 224 | 0 | 70 | 656 | 53 | 37 | 0 | 0 | 17 | 24 | 0 | 0 | 30 | 0 | 0 | 82 |
| SRR6370852 | 0 | 73 | 0 | 10 | 101 | 0 | 0 | 111 | 443 | 3066 | 499 | 36713 | 0 | 111 | 0 | 0 | 27 | 89 | 449 | 74 | 495 | 360 | 0 | 104 | 1040 | 53 | 27 | 0 | 27 | 27 | 64 | 0 | 0 | 0 | 0 | 37 | 162 |
| SRR6370853 | 0 | 43 | 0 | 10 | 88 | 0 | 0 | 123 | 245 | 859 | 89 | 10916 | 0 | 123 | 0 | 0 | 0 | 30 | 233 | 0 | 170 | 198 | 0 | 0 | 763 | 98 | 277 | 0 | 22 | 0 | 0 | 0 | 0 | 0 | 0 | 368 | 212 |
| SRR6370856 | 0 | 30 | 0 | 0 | 57 | 0 | 0 | 52 | 225 | 1661 | 35106 | 33791 | 0 | 138 | 0 | 0 | 11 | 40 | 191 | 0 | 240 | 187 | 0 | 94 | 516 | 58 | 125 | 0 | 19 | 11 | 46 | 0 | 16 | 19 | 0 | 35 | 235 |
| SRR6372849 | 0 | 262 | 116 | 0 | 53 | 0 | 30 | 393 | 1071 | 18423 | 684 | 755 | 0 | 1000 | 103 | 110 | 43 | 121 | 176 | 1890 | 87 | 1060 | 0 | 1019 | 2802 | 0 | 19 | 47 | 211 | 211 | 445 | 89 | 60 | 50 | 19 | 4367 | 701 |
| SRR6372850 | 0 | 357 | 99 | 12 | 47 | 0 | 22 | 315 | 922 | 10520 | 308 | 6324 | 0 | 607 | 0 | 0 | 49 | 222 | 285 | 1383 | 55 | 1230 | 0 | 918 | 3095 | 0 | 17 | 48 | 148 | 416 | 578 | 72 | 44 | 56 | 0 | 1829 | 468 |
| SRR6372851 | 0 | 37 | 30 | 0 | 14 | 0 | 15 | 463 | 1052 | 7498 | 164 | 1855 | 0 | 545 | 0 | 29 | 42 | 154 | 165 | 8613 | 60 | 829 | 0 | 803 | 2561 | 0 | 13 | 12 | 41 | 32 | 61 | 19 | 87 | 79 | 0 | 7761 | 319 |
| SRR6372852 | 0 | 74 | 60 | 0 | 0 | 0 | 20 | 625 | 698 | 4411 | 12476 | 28467 | 0 | 2897 | 116 | 104 | 0 | 558 | 359 | 4384 | 44 | 798 | 0 | 780 | 1549 | 0 | 21 | 22 | 49 | 191 | 140 | 20 | 29 | 13 | 14 | 2977 | 56 |
| SRR6372853 | 0 | 91 | 64 | 0 | 19 | 0 | 12 | 358 | 487 | 4838 | 372 | 9940 | 0 | 926 | 1135 | 1450 | 28 | 369 | 350 | 1691 | 160 | 1597 | 0 | 1396 | 2804 | 0 | 58 | 12 | 69 | 173 | 67 | 15 | 17 | 21 | 13 | 2875 | 99 |
| SRR6372854 | 0 | 28 | 99 | 0 | 22 | 0 | 63 | 287 | 287 | 3785 | 17755 | 43567 | 16 | 1102 | 48 | 163 | 20 | 849 | 1205 | 1418 | 40 | 1749 | 12 | 2331 | 4982 | 0 | 321 | 37 | 318 | 1469 | 713 | 70 | 46 | 53 | 0 | 1102 | 125 |
| SRR6372855 | 0 | 78 | 0 | 0 | 180 | 0 | 14 | 291 | 1101 | 935 | 9970 | 10967 | 0 | 810 | 21 | 0 | 31 | 947 | 62 | 561 | 279 | 4309 | 0 | 1625 | 6461 | 10 | 15 | 0 | 0 | 0 | 18 | 0 | 0 | 0 | 0 | 789 | 0 |
| SRR6372856 | 0 | 191 | 43 | 11 | 99 | 0 | 28 | 1175 | 1655 | 2503 | 5742 | 6636 | 0 | 1911 | 36 | 82 | 28 | 1249 | 89 | 3034 | 233 | 6834 | 11 | 2895 | 9611 | 18 | 78 | 0 | 40 | 36 | 42 | 0 | 0 | 0 | 0 | 4317 | 0 |
| SRR6372857 | 0 | 88 | 16 | 0 | 66 | 0 | 19 | 797 | 2355 | 2223 | 1052 | 6484 | 0 | 1119 | 0 | 16 | 17 | 2155 | 83 | 1283 | 736 | 14951 | 0 | 4383 | 10505 | 0 | 33 | 0 | 21 | 34 | 10 | 0 | 0 | 0 | 0 | 1827 | 22 |
| SRR6389795 | 0 | 0 | 21 | 0 | 0 | 0 | 0 | 343 | 51 | 1340 | 0 | 30 | 0 | 579 | 3602 | 7479 | 0 | 28 | 16 | 8849 | 998 | 9301 | 101 | 27869 | 12990 | 295 | 0 | 0 | 10 | 61 | 90 | 31 | 139 | 652 | 13 | 67087 | 248 |
| SRR6389796 | 0 | 0 | 95 | 0 | 0 | 0 | 0 | 191 | 47 | 3394 | 11 | 29 | 0 | 2164 | 582 | 1539 | 0 | 62 | 0 | 3504 | 814 | 12959 | 64 | 99236 | 9307 | 11 | 0 | 14 | 94 | 67 | 75 | 29 | 99 | 476 | 24 | 26027 | 166 |
| SRR6389797 | 0 | 0 | 15 | 0 | 0 | 0 | 0 | 301 | 68 | 12176 | 10 | 11 | 0 | 3247 | 914 | 2777 | 0 | 67 | 0 | 6464 | 277 | 11632 | 164 | 70562 | 10596 | 413 | 0 | 26 | 53 | 178 | 221 | 93 | 378 | 1418 | 62 | 45591 | 83 |
| SRR6389798 | 0 | 0 | 24 | 0 | 0 | 0 | 0 | 298 | 252 | 9976 | 27 | 65 | 0 | 189 | 263 | 269 | 0 | 0 | 0 | 1265 | 519 | 2165 | 47 | 5921 | 4266 | 0 | 0 | 0 | 402 | 836 | 667 | 131 | 553 | 1056 | 14 | 9919 | 258 |
| SRR6389799 | 0 | 0 | 31 | 0 | 0 | 0 | 0 | 147 | 42 | 3274 | 19 | 41 | 0 | 1494 | 259 | 536 | 0 | 11 | 14 | 5924 | 427 | 25217 | 148 | 163283 | 11996 | 144 | 0 | 12 | 85 | 85 | 106 | 29 | 116 | 468 | 22 | 28138 | 79 |
| SRR6389800 | 0 | 0 | 70 | 0 | 13 | 0 | 0 | 2279 | 2404 | 10888 | 24 | 60 | 0 | 3102 | 27 | 59 | 0 | 28 | 0 | 1751 | 750 | 17882 | 871 | 97927 | 7550 | 352 | 0 | 14 | 282 | 1536 | 1660 | 150 | 832 | 1905 | 0 | 13727 | 356 |
| SRR6389801 | 0 | 0 | 40 | 0 | 0 | 0 | 0 | 47 | 12 | 313 | 80 | 458 | 11 | 29 | 69 | 25 | 0 | 46 | 17 | 3569 | 1714 | 1789 | 0 | 97 | 5668 | 0 | 0 | 26 | 23 | 164 | 119 | 20 | 74 | 106 | 0 | 22857 | 158 |
| SRR6389802 | 0 | 0 | 215 | 0 | 20 | 0 | 0 | 565 | 430 | 10952 | 114 | 686 | 0 | 1222 | 0 | 14 | 0 | 35 | 0 | 21648 | 849 | 5530 | 133 | 65536 | 5385 | 237 | 0 | 238 | 387 | 5061 | 4113 | 466 | 1963 | 2803 | 0 | 71051 | 66 |
| SRR6389803 | 0 | 19 | 213 | 0 | 0 | 0 | 0 | 213 | 253 | 1843 | 326 | 337 | 0 | 746 | 61 | 158 | 0 | 36 | 16 | 8682 | 767 | 5025 | 0 | 1133 | 17997 | 15 | 14 | 24 | 16 | 126 | 128 | 20 | 99 | 225 | 0 | 36828 | 745 |
| SRR6389804 | 0 | 32 | 351 | 0 | 0 | 11 | 0 | 1340 | 1460 | 5318 | 220 | 1148 | 0 | 92 | 82 | 103 | 0 | 64 | 0 | 2224 | 1330 | 11895 | 376 | 85448 | 9634 | 62 | 0 | 64 | 17 | 78 | 96 | 10 | 48 | 88 | 0 | 4214 | 274 |
| SRR6389805 | 0 | 0 | 72 | 0 | 0 | 0 | 0 | 1056 | 404 | 4680 | 105 | 408 | 0 | 1317 | 393 | 381 | 0 | 89 | 17 | 5074 | 2163 | 3979 | 0 | 496 | 18170 | 13 | 10 | 17 | 738 | 1282 | 1158 | 194 | 667 | 1169 | 0 | 35567 | 357 |
| SRR6389806 | 0 | 0 | 90 | 0 | 0 | 0 | 0 | 565 | 427 | 9390 | 625 | 2321 | 0 | 2558 | 199 | 457 | 0 | 57 | 0 | 6042 | 1555 | 8212 | 43 | 28558 | 11204 | 503 | 13 | 129 | 202 | 2512 | 2603 | 218 | 1389 | 3490 | 80 | 23164 | 146 |

### Supp Table S8.4.Columns_DI_to_ES

| | | | | | | | | | | | | | | | | | | | | | | | | | | | | | | | | | | | | | |
|---|---|---|---|---|---|---|---|---|---|---|---|---|---|---|---|---|---|---|---|---|---|---|---|---|---|---|---|---|---|---|---|---|---|---|---|---|---|
| SRR6389807 | 0 | 0 | 57 | 0 | 0 | 0 | 0 | 262 | 211 | 929 | 70 | 75 | 0 | 505 | 192 | 262 | 0 | 270 | 0 | 1508 | 1454 | 11515 | 363 | 63820 | 8337 | 899 | 14 | 11 | 25 | 109 | 77 | 19 | 97 | 192 | 0 | 16345 | 179 |
| SRR6389808 | 0 | 0 | 24 | 0 | 0 | 0 | 0 | 343 | 58 | 4985 | 25 | 24 | 0 | 6305 | 53 | 246 | 0 | 71 | 11 | 6649 | 183 | 17814 | 57 | 85909 | 11527 | 0 | 0 | 12 | 30 | 269 | 239 | 75 | 276 | 1257 | 30 | 33756 | 46 |
| SRR6389809 | 0 | 0 | 117 | 0 | 0 | 0 | 0 | 859 | 263 | 3529 | 130 | 270 | 0 | 404 | 3505 | 9195 | 0 | 70 | 0 | 3114 | 1342 | 7839 | 174 | 23474 | 16091 | 385 | 17 | 52 | 12 | 66 | 53 | 16 | 83 | 244 | 0 | 16696 | 531 |
| SRR6389810 | 0 | 0 | 26 | 0 | 0 | 0 | 0 | 931 | 416 | 9424 | 293 | 1268 | 0 | 3757 | 78 | 185 | 0 | 142 | 0 | 8207 | 1049 | 10790 | 329 | 47853 | 4122 | 413 | 0 | 57 | 38 | 132 | 94 | 33 | 119 | 402 | 0 | 32020 | 76 |
| SRR6389811 | 0 | 0 | 124 | 0 | 0 | 0 | 0 | 184 | 74 | 9284 | 161 | 600 | 0 | 4371 | 1316 | 3672 | 0 | 128 | 19 | 7677 | 883 | 4626 | 23 | 19263 | 6693 | 317 | 15 | 48 | 129 | 949 | 1220 | 400 | 2053 | 7242 | 58 | 53988 | 86 |
| SRR6389812 | 0 | 0 | 68 | 0 | 0 | 0 | 0 | 658 | 374 | 8369 | 51 | 218 | 0 | 2015 | 0 | 0 | 0 | 36 | 0 | 3168 | 802 | 8400 | 39 | 24923 | 10472 | 67 | 0 | 25 | 15 | 32 | 33 | 195 | 27 | 49 | 0 | 19602 | 191 |
| SRR6389813 | 0 | 0 | 12 | 0 | 0 | 0 | 0 | 1054 | 357 | 10813 | 165 | 333 | 0 | 109 | 654 | 581 | 0 | 124 | 0 | 5194 | 835 | 13504 | 116 | 95063 | 9715 | 518 | 0 | 21 | 67 | 351 | 301 | 72 | 219 | 700 | 0 | 23322 | 51 |
| SRR6389814 | 0 | 0 | 45 | 0 | 0 | 0 | 0 | 303 | 184 | 14311 | 110 | 813 | 0 | 3886 | 0 | 0 | 0 | 77 | 0 | 4479 | 1135 | 7359 | 60 | 22249 | 14830 | 117 | 0 | 25 | 129 | 478 | 626 | 76 | 315 | 778 | 0 | 32119 | 94 |
| SRR6389815 | 0 | 0 | 0 | 0 | 0 | 0 | 0 | 1854 | 199 | 7979 | 74 | 298 | 0 | 500 | 1803 | 2059 | 0 | 0 | 25 | 6756 | 732 | 3108 | 27 | 6540 | 11900 | 0 | 20 | 0 | 117 | 39 | 50 | 13 | 36 | 144 | 0 | 76034 | 224 |
| SRR6389816 | 0 | 0 | 27 | 0 | 0 | 0 | 0 | 297 | 303 | 4450 | 93 | 55 | 0 | 677 | 61 | 144 | 0 | 102 | 0 | 1297 | 540 | 12343 | 134 | 71681 | 45643 | 376 | 46 | 578 | 750 | 2749 | 2857 | 626 | 4263 | 5596 | 0 | 7807 | 459 |
| SRR6389817 | 0 | 0 | 467 | 0 | 0 | 0 | 0 | 1082 | 589 | 7198 | 191 | 1364 | 0 | 203 | 28 | 20 | 0 | 45 | 0 | 11523 | 592 | 5462 | 137 | 25925 | 8887 | 0 | 0 | 34 | 518 | 2363 | 3407 | 167 | 1164 | 3043 | 0 | 41275 | 285 |
| SRR6389818 | 0 | 0 | 111 | 0 | 0 | 0 | 0 | 215 | 132 | 5952 | 238 | 256 | 0 | 2128 | 105 | 185 | 0 | 100 | 17 | 2816 | 1125 | 3645 | 10 | 18125 | 8949 | 13 | 0 | 51 | 259 | 2520 | 3404 | 263 | 1282 | 3424 | 0 | 9935 | 126 |
| SRR6389819 | 0 | 0 | 68 | 0 | 0 | 0 | 0 | 1597 | 313 | 13632 | 183 | 740 | 0 | 485 | 13 | 86 | 0 | 0 | 73 | 1644 | 1631 | 4086 | 0 | 10667 | 14624 | 38 | 0 | 27 | 34 | 105 | 159 | 15 | 77 | 167 | 0 | 11359 | 573 |
| SRR6389820 | 0 | 0 | 27 | 0 | 0 | 0 | 0 | 2142 | 307 | 22364 | 67 | 480 | 0 | 566 | 283 | 644 | 0 | 75 | 29 | 4243 | 741 | 4142 | 0 | 969 | 10341 | 270 | 0 | 18 | 14 | 103 | 88 | 12 | 43 | 92 | 0 | 38630 | 194 |
| SRR6409751 | 0 | 0 | 0 | 0 | 0 | 0 | 0 | 0 | 1602 | 3162 | 0 | 0 | 0 | 3393 | 0 | 0 | 0 | 0 | 0 | 0 | 0 | 16 | 182 | 7129 | 449 | 118 | 0 | 0 | 0 | 0 | 0 | 0 | 0 | 0 | 0 | 1096 | 0 |
| SRR6409760 | 0 | 0 | 0 | 0 | 0 | 0 | 0 | 0 | 0 | 3340 | 0 | 0 | 0 | 8102 | 0 | 0 | 0 | 0 | 62 | 0 | 0 | 64 | 0 | 5433 | 140 | 0 | 0 | 0 | 0 | 0 | 0 | 0 | 0 | 0 | 0 | 9194 | 0 |
| SRR6409798 | 0 | 0 | 0 | 0 | 0 | 0 | 0 | 0 | 0 | 2941 | 60 | 155 | 0 | 3194 | 15 | 13 | 0 | 49 | 0 | 0 | 1374 | 1845 | 528 | 22018 | 1340 | 76 | 0 | 0 | 0 | 0 | 0 | 0 | 0 | 0 | 0 | 5676 | 0 |
| SRR6409800 | 0 | 0 | 0 | 0 | 0 | 0 | 0 | 0 | 0 | 0 | 0 | 0 | 0 | 0 | 0 | 293 | 0 | 0 | 0 | 0 | 0 | 0 | 0 | 1210 | 0 | 0 | 0 | 0 | 0 | 0 | 0 | 0 | 0 | 0 | 0 | 0 | 0 |
| SRR6409827 | 0 | 0 | 0 | 0 | 0 | 0 | 0 | 0 | 0 | 3795 | 291 | 263 | 0 | 33 | 0 | 0 | 0 | 235 | 0 | 1843 | 663 | 1786 | 422 | 24016 | 1457 | 48 | 0 | 0 | 0 | 0 | 0 | 0 | 0 | 0 | 0 | 7217 | 0 |
| SRR6409832 | 0 | 0 | 0 | 0 | 0 | 0 | 0 | 25271 | 0 | 0 | 0 | 0 | 0 | 47 | 0 | 0 | 0 | 0 | 0 | 0 | 0 | 0 | 90 | 358 | 0 | 0 | 0 | 0 | 0 | 0 | 0 | 0 | 0 | 0 | 0 | 0 | 0 |
| SRR6409833 | 0 | 0 | 0 | 0 | 0 | 0 | 0 | 0 | 0 | 3936 | 0 | 145 | 0 | 728 | 28 | 0 | 0 | 112 | 0 | 1704 | 689 | 2092 | 803 | 28447 | 2264 | 0 | 0 | 0 | 0 | 0 | 13 | 0 | 0 | 0 | 0 | 4401 | 0 |
| SRR6409834 | 0 | 0 | 0 | 0 | 0 | 0 | 0 | 0 | 0 | 8867 | 501 | 0 | 0 | 19 | 0 | 13 | 0 | 187 | 0 | 1317 | 892 | 1674 | 447 | 16735 | 730 | 60 | 0 | 0 | 0 | 0 | 0 | 0 | 0 | 2517 | 0 | 2300 | 0 |
| SRR6409845 | 0 | 0 | 0 | 0 | 0 | 0 | 0 | 0 | 0 | 0 | 0 | 0 | 0 | 0 | 0 | 0 | 0 | 0 | 0 | 0 | 0 | 0 | 0 | 157 | 52 | 21 | 0 | 0 | 0 | 0 | 0 | 0 | 0 | 0 | 0 | 15411 | 0 |
| SRR6409851 | 0 | 0 | 0 | 0 | 0 | 0 | 0 | 809 | 0 | 2072 | 90 | 188 | 0 | 1744 | 0 | 12 | 0 | 75 | 0 | 1711 | 931 | 1910 | 510 | 23721 | 1793 | 179 | 0 | 0 | 0 | 0 | 0 | 0 | 0 | 0 | 0 | 5310 | 0 |
| SRR6409853 | 0 | 0 | 0 | 0 | 0 | 0 | 0 | 0 | 0 | 3744 | 0 | 96 | 0 | 2851 | 0 | 0 | 0 | 115 | 0 | 3033 | 971 | 1732 | 661 | 23111 | 1425 | 100 | 0 | 0 | 0 | 0 | 0 | 0 | 0 | 0 | 0 | 3829 | 0 |
| SRR6409859 | 0 | 0 | 0 | 0 | 0 | 0 | 0 | 0 | 0 | 2337 | 0 | 0 | 0 | 1782 | 39 | 31 | 0 | 0 | 0 | 771 | 885 | 1668 | 450 | 23013 | 2729 | 146 | 0 | 0 | 0 | 0 | 0 | 0 | 0 | 0 | 0 | 7868 | 0 |
| SRR6409860 | 0 | 0 | 0 | 0 | 0 | 0 | 0 | 0 | 0 | 3316 | 69 | 0 | 0 | 536 | 0 | 21 | 0 | 60 | 0 | 561 | 869 | 2675 | 283 | 22226 | 1763 | 132 | 0 | 0 | 0 | 0 | 0 | 0 | 0 | 0 | 0 | 5019 | 0 |
| SRR6409863 | 0 | 0 | 0 | 0 | 0 | 0 | 0 | 2936 | 0 | 9564 | 0 | 0 | 0 | 4998 | 0 | 63 | 0 | 247 | 0 | 0 | 701 | 1780 | 470 | 14792 | 1451 | 260 | 0 | 0 | 0 | 0 | 0 | 0 | 0 | 0 | 0 | 1023 | 0 |
| SRR6409866 | 0 | 0 | 0 | 0 | 0 | 0 | 0 | 0 | 0 | 1915 | 0 | 0 | 0 | 1242 | 19 | 0 | 0 | 0 | 0 | 0 | 766 | 959 | 313 | 13911 | 583 | 71 | 0 | 0 | 0 | 0 | 0 | 0 | 0 | 0 | 0 | 6720 | 0 |
| SRR6409867 | 0 | 0 | 463 | 0 | 0 | 0 | 0 | 0 | 0 | 3867 | 0 | 0 | 0 | 182 | 0 | 19 | 0 | 134 | 0 | 2155 | 1227 | 1738 | 605 | 23042 | 1707 | 224 | 0 | 0 | 0 | 0 | 0 | 0 | 0 | 0 | 0 | 7817 | 0 |
| SRR6409868 | 0 | 0 | 0 | 0 | 0 | 0 | 0 | 0 | 0 | 2255 | 319 | 0 | 0 | 24 | 0 | 24 | 0 | 131 | 0 | 4037 | 0 | 1207 | 166 | 19799 | 706 | 162 | 0 | 0 | 0 | 0 | 0 | 0 | 0 | 0 | 0 | 7721 | 0 |
| SRR6450851 | 0 | 229 | 0 | 321 | 1732 | 0 | 19 | 4032 | 3820 | 1698 | 1273 | 3183 | 19 | 637 | 0 | 212 | 95 | 2053 | 770 | 0 | 1026 | 2822 | 385 | 35407 | 1732 | 192 | 0 | 76 | 38 | 57 | 57 | 0 | 0 | 19 | 0 | 0 | 321 |
| SRR6450852 | 18 | 112 | 11 | 58 | 647 | 18 | 29 | 8447 | 5472 | 1173 | 111 | 1824 | 15 | 418 | 46 | 128 | 47 | 1375 | 700 | 23 | 887 | 2516 | 379 | 33185 | 1136 | 168 | 0 | 62 | 56 | 92 | 93 | 17 | 47 | 17 | 0 | 70 | 235 |
| SRR6450853 | 116 | 727 | 33 | 72 | 650 | 0 | 50 | 7658 | 4807 | 1589 | 2281 | 2281 | 99 | 570 | 81 | 41 | 248 | 1276 | 879 | 41 | 939 | 2588 | 361 | 32476 | 1107 | 144 | 0 | 413 | 330 | 248 | 314 | 33 | 149 | 33 | 0 | 41 | 349 |
| SRR6450854 | 139 | 744 | 23 | 60 | 997 | 0 | 35 | 8806 | 5635 | 1022 | 80 | 2673 | 105 | 393 | 26 | 79 | 256 | 1375 | 952 | 52 | 1050 | 2379 | 544 | 33237 | 1246 | 68 | 23 | 511 | 325 | 244 | 384 | 35 | 35 | 12 | 12 | 210 | 227 |
| SRR6450855 | 0 | 165 | 0 | 0 | 1022 | 0 | 13 | 7340 | 4097 | 1366 | 2561 | 3756 | 13 | 512 | 0 | 0 | 64 | 851 | 766 | 0 | 639 | 2554 | 341 | 31760 | 1149 | 85 | 25 | 114 | 51 | 38 | 51 | 0 | 25 | 25 | 0 | 0 | 255 |
| SRR6450856 | 48 | 223 | 31 | 62 | 1077 | 0 | 24 | 6494 | 6363 | 1722 | 75 | 1831 | 12 | 436 | 174 | 327 | 22 | 1454 | 592 | 44 | 738 | 2554 | 508 | 29774 | 1477 | 162 | 24 | 63 | 98 | 96 | 96 | 11 | 36 | 20 | 12 | 87 | 308 |
| SRR6450857 | 17 | 209 | 17 | 60 | 1559 | 0 | 0 | 7328 | 5073 | 1315 | 14 | 1691 | 0 | 564 | 0 | 188 | 52 | 1439 | 959 | 0 | 1379 | 2459 | 360 | 36220 | 1499 | 60 | 35 | 17 | 35 | 104 | 87 | 17 | 17 | 52 | 0 | 0 | 60 |
| SRR6450858 | 35 | 848 | 81 | 92 | 874 | 0 | 35 | 7421 | 6015 | 2109 | 51 | 2005 | 58 | 547 | 52 | 130 | 244 | 1143 | 782 | 0 | 782 | 2286 | 376 | 29400 | 1066 | 69 | 12 | 395 | 290 | 348 | 406 | 46 | 81 | 46 | 23 | 26 | 299 |
| SRR6450859 | 55 | 553 | 55 | 137 | 1197 | 0 | 0 | 5756 | 4649 | 2103 | 14 | 2546 | 166 | 1218 | 0 | 111 | 277 | 1539 | 787 | 0 | 855 | 2462 | 308 | 32489 | 1642 | 103 | 0 | 332 | 332 | 166 | 664 | 55 | 221 | 0 | 0 | 0 | 376 |
| SRR6450860 | 0 | 213 | 27 | 55 | 878 | 0 | 0 | 5089 | 4439 | 1624 | 17 | 2057 | 18 | 541 | 217 | 325 | 18 | 1536 | 549 | 108 | 1070 | 2634 | 466 | 28342 | 1180 | 137 | 0 | 89 | 106 | 89 | 124 | 18 | 18 | 18 | 27 | 108 | 329 |
| SRR6450861 | 31 | 94 | 0 | 79 | 806 | 0 | 11 | 7305 | 5668 | 630 | 29 | 2015 | 31 | 441 | 63 | 126 | 28 | 1297 | 826 | 0 | 1042 | 1887 | 295 | 29600 | 1199 | 157 | 31 | 110 | 33 | 28 | 77 | 11 | 11 | 31 | 0 | 378 | 216 |
| SRR6450862 | 126 | 930 | 47 | 53 | 1081 | 15 | 54 | 7212 | 6073 | 562 | 1018 | 1314 | 76 | 160 | 15 | 30 | 254 | 1529 | 838 | 124 | 872 | 2690 | 383 | 31703 | 1270 | 150 | 15 | 502 | 364 | 359 | 384 | 62 | 86 | 34 | 0 | 650 | 273 |
| SRR6450863 | 102 | 399 | 24 | 60 | 701 | 16 | 39 | 9199 | 5701 | 1852 | 1729 | 2120 | 78 | 370 | 62 | 226 | 337 | 1379 | 994 | 21 | 1070 | 2667 | 211 | 34950 | 1153 | 181 | 16 | 321 | 290 | 258 | 399 | 78 | 102 | 86 | 0 | 165 | 264 |
| SRR6450867 | 0 | 175 | 18 | 0 | 0 | 0 | 0 | 1358 | 1259 | 79 | 1279 | 724 | 0 | 40 | 20 | 0 | 0 | 221 | 251 | 50 | 562 | 847 | 25 | 1003 | 3098 | 0 | 0 | 221 | 4385 | 571 | 101 | 129 | 230 | 415 | 129 | 1061 | 226 |
| SRR6450868 | 28 | 18 | 0 | 0 | 0 | 0 | 0 | 97 | 906 | 162 | 16 | 485 | 0 | 32 | 0 | 0 | 20 | 2219 | 255 | 32 | 302 | 536 | 872 | 3929 | 1180 | 0 | 0 | 724 | 1548 | 84 | 0 | 10 | 0 | 0 | 0 | 97 | 80 |
| SRR6450869 | 30 | 0 | 0 | 0 | 21 | 0 | 20 | 0 | 1027 | 0 | 560 | 560 | 0 | 280 | 0 | 0 | 0 | 2490 | 322 | 0 | 150 | 730 | 708 | 4143 | 1073 | 0 | 0 | 908 | 1727 | 120 | 30 | 20 | 0 | 0 | 0 | 0 | 150 |
| SRR6453341 | 0 | 481 | 24 | 0 | 0 | 0 | 0 | 315 | 657 | 602 | 48811 | 51138 | 0 | 123 | 561 | 1013 | 34 | 27 | 49 | 4134 | 90 | 941 | 0 | 46 | 201 | 80 | 240 | 0 | 14 | 0 | 29 | 0 | 0 | 0 | 0 | 1341 | 17 |
| SRR6453342 | 0 | 701 | 14 | 0 | 0 | 0 | 20 | 368 | 552 | 1276 | 7447 | 110779 | 0 | 108 | 151 | 292 | 54 | 22 | 34 | 898 | 133 | 976 | 0 | 0 | 203 | 87 | 490 | 0 | 0 | 0 | 14 | 0 | 0 | 0 | 0 | 616 | 26 |
| SRR6453343 | 0 | 108 | 24 | 0 | 0 | 0 | 19 | 379 | 683 | 1290 | 49228 | 82721 | 0 | 152 | 228 | 278 | 40 | 23 | 36 | 1670 | 165 | 1157 | 0 | 0 | 293 | 29 | 50 | 0 | 32 | 12 | 45 | 19 | 0 | 0 | 0 | 620 | 22 |
| SRR6453344 | 0 | 118 | 42 | 0 | 0 | 0 | 17 | 124 | 462 | 992 | 68090 | 103646 | 0 | 101 | 203 | 214 | 31 | 22 | 29 | 823 | 215 | 1742 | 0 | 0 | 259 | 47 | 28 | 0 | 17 | 0 | 36 | 14 | 0 | 0 | 0 | 451 | 18 |
| SRR6453345 | 0 | 102 | 43 | 0 | 0 | 0 | 14 | 420 | 849 | 689 | 54288 | 102532 | 0 | 110 | 539 | 649 | 35 | 41 | 41 | 1588 | 227 | 2019 | 0 | 13 | 394 | 56 | 52 | 0 | 21 | 11 | 49 | 16 | 0 | 0 | 0 | 819 | 47 |
| SRR6453346 | 0 | 454 | 59 | 0 | 0 | 0 | 14 | 275 | 523 | 576 | 51950 | 74654 | 0 | 65 | 183 | 288 | 18 | 27 | 46 | 1269 | 211 | 1942 | 0 | 0 | 438 | 52 | 384 | 0 | 23 | 0 | 27 | 16 | 0 | 0 | 0 | 641 | 83 |
| SRR6453347 | 0 | 1024 | 15 | 0 | 0 | 0 | 23 | 355 | 683 | 1314 | 5002 | 87057 | 0 | 92 | 197 | 184 | 78 | 20 | 29 | 1261 | 164 | 977 | 0 | 17 | 248 | 34 | 96 | 0 | 0 | 0 | 13 | 0 | 0 | 0 | 0 | 617 | 24 |
| SRR6453348 | 0 | 746 | 38 | 0 | 0 | 0 | 16 | 225 | 675 | 776 | 4868 | 87536 | 0 | 67 | 686 | 754 | 33 | 26 | 37 | 2114 | 190 | 1142 | 0 | 0 | 257 | 95 | 175 | 0 | 20 | 12 | 51 | 18 | 13 | 0 | 14 | 979 | 27 |
| SRR6453349 | 0 | 596 | 30 | 0 | 0 | 0 | 11 | 268 | 389 | 897 | 4995 | 85984 | 0 | 74 | 194 | 370 | 32 | 18 | 49 | 953 | 158 | 1350 | 0 | 0 | 284 | 64 | 269 | 0 | 16 | 0 | 43 | 25 | 0 | 0 | 0 | 592 | 20 |
| SRR6453350 | 0 | 99 | 52 | 0 | 0 | 0 | 13 | 255 | 504 | 938 | 6613 | 69593 | 0 | 62 | 366 | 372 | 28 | 28 | 37 | 1221 | 259 | 2920 | 0 | 11 | 271 | 52 | 38 | 0 | 22 | 15 | 52 | 28 | 0 | 0 | 10 | 524 | 11 |
| SRR6453351 | 0 | 942 | 19 | 0 | 0 | 0 | 14 | 205 | 430 | 903 | 10496 | 112289 | 0 | 134 | 219 | 282 | 39 | 31 | 49 | 656 | 157 | 1036 | 0 | 0 | 393 | 59 | 72 | 0 | 19 | 0 | 26 | 12 | 0 | 0 | 0 | 332 | 37 |
| SRR6453352 | 0 | 250 | 24 | 0 | 0 | 0 | 15 | 280 | 507 | 868 | 87698 | 104607 | 0 | 133 | 187 | 360 | 15 | 31 | 43 | 1388 | 158 | 1642 | 0 | 11 | 281 | 47 | 23 | 0 | 17 | 0 | 39 | 11 | 0 | 0 | 0 | 561 | 27 |
| SRR6453353 | 0 | 531 | 19 | 0 | 0 | 0 | 23 | 433 | 944 | 1329 | 9142 | 113114 | 0 | 71 | 228 | 315 | 52 | 22 | 45 | 1447 | 140 | 938 | 0 | 0 | 336 | 44 | 93 | 0 | 12 | 0 | 27 | 13 | 0 | 0 | 0 | 590 | 16 |
| SRR6453354 | 0 | 896 | 24 | 0 | 0 | 0 | 15 | 302 | 740 | 1533 | 63480 | 89292 | 0 | 42 | 344 | 469 | 48 | 21 | 22 | 1523 | 203 | 1636 | 0 | 0 | 203 | 80 | 238 | 0 | 17 | 0 | 22 | 17 | 0 | 0 | 0 | 490 | 18 |
| SRR6453355 | 0 | 1038 | 44 | 0 | 0 | 0 | 17 | 297 | 603 | 704 | 16542 | 158414 | 0 | 48 | 34 | 96 | 45 | 25 | 44 | 503 | 303 | 2264 | 0 | 43 | 335 | 56 | 108 | 0 | 15 | 0 | 29 | 12 | 0 | 0 | 0 | 182 | 20 |
| SRR6453356 | 0 | 88 | 25 | 0 | 0 | 0 | 14 | 429 | 726 | 701 | 8058 | 98424 | 0 | 25 | 165 | 297 | 46 | 27 | 31 | 834 | 255 | 2056 | 0 | 12 | 289 | 73 | 155 | 0 | 14 | 12 | 23 | 21 | 0 | 0 | 0 | 338 | 0 |
| SRR6453357 | 0 | 187 | 23 | 0 | 0 | 0 | 15 | 477 | 757 | 792 | 4998 | 81736 | 0 | 105 | 198 | 314 | 11 | 43 | 56 | 1653 | 165 | 1813 | 0 | 36 | 217 | 54 | 17 | 0 | 17 | 0 | 45 | 12 | 0 | 0 | 0 | 629 | 12 |
| SRR6453358 | 0 | 1388 | 37 | 0 | 0 | 12 | 47 | 147 | 262 | 475 | 54682 | 89586 | 0 | 76 | 109 | 153 | 90 | 20 | 10 | 600 | 216 | 1880 | 0 | 0 | 172 | 70 | 131 | 0 | 0 | 0 | 26 | 11 | 0 | 0 | 0 | 202 | 13 |
| SRR6453359 | 0 | 156 | 24 | 0 | 0 | 0 | 23 | 117 | 282 | 458 | 25218 | 161660 | 0 | 17 | 79 | 121 | 43 | 24 | 14 | 241 | 358 | 1544 | 0 | 48 | 312 | 54 | 50 | 0 | 0 | 0 | 12 | 10 | 0 | 0 | 0 | 134 | 31 |
| SRR6453360 | 0 | 835 | 16 | 0 | 0 | 0 | 18 | 427 | 471 | 1089 | 38275 | 60784 | 0 | 265 | 442 | 486 | 77 | 25 | 42 | 1840 | 131 | 1174 | 0 | 0 | 264 | 39 | 228 | 0 | 22 | 11 | 35 | 0 | 0 | 0 | 0 | 677 | 21 |
| SRR6453361 | 0 | 130 | 38 | 0 | 0 | 0 | 17 | 238 | 561 | 939 | 69058 | 124763 | 0 | 55 | 134 | 268 | 25 | 32 | 50 | 1013 | 293 | 1686 | 0 | 12 | 372 | 88 | 11 | 0 | 12 | 0 | 23 | 18 | 0 | 0 | 0 | 348 | 15 |
| SRR6453362 | 0 | 290 | 21 | 0 | 0 | 0 | 29 | 234 | 388 | 803 | 17595 | 137051 | 0 | 53 | 106 | 117 | 21 | 21 | 41 | 383 | 226 | 1176 | 0 | 32 | 318 | 33 | 16 | 0 | 16 | 0 | 19 | 0 | 0 | 0 | 0 | 213 | 27 |
| SRR6453363 | 0 | 375 | 30 | 0 | 0 | 0 | 22 | 306 | 594 | 1370 | 13155 | 131796 | 0 | 71 | 129 | 170 | 20 | 24 | 75 | 1399 | 216 | 1393 | 0 | 16 | 506 | 85 | 13 | 0 | 15 | 12 | 33 | 15 | 0 | 0 | 0 | 470 | 22 |
| SRR6453364 | 0 | 681 | 25 | 0 | 0 | 0 | 13 | 280 | 648 | 788 | 41056 | 59886 | 0 | 210 | 683 | 753 | 80 | 26 | 44 | 2853 | 81 | 1007 | 0 | 0 | 200 | 59 | 113 | 0 | 17 | 0 | 40 | 0 | 0 | 0 | 0 | 945 | 28 |
| SRR6453365 | 0 | 167 | 11 | 0 | 0 | 0 | 16 | 416 | 1111 | 1749 | 69858 | 84208 | 0 | 68 | 251 | 232 | 33 | 28 | 48 | 3440 | 211 | 1711 | 0 | 10 | 315 | 61 | 26 | 0 | 18 | 13 | 31 | 19 | 0 | 0 | 0 | 1131 | 21 |
| SRR6453366 | 0 | 312 | 23 | 0 | 0 | 0 | 24 | 251 | 703 | 1068 | 13117 | 133941 | 0 | 94 | 113 | 245 | 21 | 26 | 19 | 1099 | 183 | 1094 | 0 | 0 | 294 | 88 | 24 | 0 | 11 | 0 | 17 | 0 | 0 | 0 | 0 | 333 | 29 |
| SRR6453367 | 0 | 31 | 27 | 0 | 0 | 14 | 34 | 736 | 1670 | 2423 | 67335 | 117722 | 0 | 34 | 146 | 188 | 104 | 28 | 55 | 265 | 203 | 1669 | 0 | 15 | 243 | 76 | 11 | 0 | 0 | 0 | 0 | 0 | 0 | 0 | 0 | 223 | 0 |
| SRR6453368 | 0 | 321 | 41 | 0 | 11 | 0 | 28 | 458 | 1013 | 2560 | 56515 | 126537 | 0 | 33 | 142 | 458 | 164 | 29 | 67 | 305 | 301 | 2003 | 0 | 56 | 287 | 68 | 64 | 0 | 0 | 12 | 0 | 0 | 0 | 0 | 0 | 251 | 0 |
| SRR6453369 | 0 | 968 | 19 | 0 | 0 | 0 | 0 | 219 | 426 | 841 | 5700 | 86945 | 0 | 150 | 230 | 323 | 47 | 20 | 42 | 2419 | 85 | 853 | 0 | 0 | 167 | 60 | 169 | 0 | 22 | 0 | 33 | 14 | 0 | 0 | 0 | 933 | 17 |
| SRR6453370 | 0 | 609 | 20 | 0 | 0 | 0 | 16 | 453 | 796 | 1592 | 3318 | 79665 | 0 | 78 | 375 | 437 | 52 | 19 | 39 | 1155 | 158 | 1362 | 0 | 10 | 186 | 58 | 61 | 0 | 0 | 0 | 37 | 0 | 0 | 0 | 0 | 578 | 13 |
| SRR6453371 | 0 | 535 | 13 | 0 | 0 | 0 | 21 | 1233 | 2739 | 4920 | 3616 | 83523 | 0 | 78 | 312 | 389 | 59 | 47 | 72 | 2116 | 103 | 1003 | 0 | 0 | 304 | 40 | 51 | 0 | 11 | 0 | 22 | 0 | 0 | 0 | 0 | 805 | 39 |
| SRR6453372 | 0 | 230 | 11 | 0 | 0 | 0 | 33 | 803 | 1851 | 4287 | 58909 | 70884 | 0 | 122 | 218 | 286 | 35 | 23 | 39 | 1157 | 94 | 866 | 0 | 63 | 158 | 61 | 15 | 0 | 0 | 0 | 17 | 0 | 0 | 0 | 0 | 463 | 21 |
| SRR6453373 | 0 | 1150 | 24 | 0 | 0 | 0 | 20 | 2344 | 6362 | 6814 | 2588 | 49255 | 0 | 117 | 536 | 837 | 57 | 40 | 53 | 3114 | 162 | 1643 | 0 | 26 | 196 | 56 | 75 | 0 | 22 | 15 | 44 | 18 | 0 | 0 | 0 | 1323 | 28 |
| SRR6453374 | 0 | 756 | 14 | 0 | 0 | 0 | 13 | 961 | 2998 | 4228 | 5617 | 91247 | 0 | 165 | 403 | 486 | 45 | 32 | 48 | 2729 | 94 | 1066 | 0 | 27 | 204 | 45 | 61 | 0 | 12 | 10 | 29 | 13 | 0 | 0 | 0 | 982 | 70 |
| SRR6453375 | 0 | 676 | 23 | 0 | 0 | 0 | 15 | 1301 | 2674 | 4469 | 5469 | 79076 | 0 | 120 | 205 | 301 | 65 | 28 | 51 | 1566 | 92 | 806 | 0 | 137 | 343 | 45 | 174 | 0 | 0 | 0 | 19 | 0 | 0 | 0 | 0 | 614 | 42 |
| SRR6453376 | 0 | 53 | 57 | 0 | 0 | 0 | 26 | 743 | 1410 | 2282 | 69449 | 102546 | 0 | 77 | 308 | 410 | 60 | 21 | 14 | 385 | 296 | 1705 | 0 | 0 | 258 | 16 | 118 | 0 | 17 | 0 | 0 | 0 | 0 | 0 | 0 | 128 | 22 |
| SRR6453377 | 0 | 653 | 17 | 0 | 0 | 0 | 16 | 2437 | 8051 | 6236 | 4973 | 63613 | 0 | 95 | 338 | 570 | 53 | 28 | 25 | 1678 | 238 | 2182 | 0 | 11 | 266 | 27 | 59 | 0 | 17 | 0 | 38 | 10 | 0 | 0 | 0 | 654 | 12 |
| SRR6453378 | 0 | 191 | 16 | 0 | 0 | 0 | 16 | 1592 | 3269 | 5692 | 41180 | 72722 | 0 | 169 | 525 | 711 | 14 | 25 | 29 | 3235 | 130 | 981 | 0 | 43 | 213 | 32 | 30 | 0 | 18 | 0 | 36 | 11 | 0 | 0 | 0 | 1101 | 23 |
| SRR6453379 | 0 | 340 | 20 | 0 | 0 | 0 | 18 | 2190 | 6400 | 4618 | 3073 | 61769 | 0 | 92 | 818 | 726 | 14 | 41 | 72 | 3550 | 246 | 1521 | 0 | 15 | 306 | 52 | 12 | 0 | 18 | 0 | 13 | 22 | 0 | 0 | 0 | 1320 | 22 |
| SRR6453380 | 0 | 719 | 0 | 0 | 0 | 0 | 11 | 1283 | 3182 | 2553 | 42876 | 68383 | 0 | 50 | 641 | 704 | 49 | 51 | 61 | 2239 | 244 | 2577 | 0 | 16 | 376 | 48 | 289 | 0 | 0 | 0 | 12 | 0 | 0 | 0 | 0 | 654 | 0 |
| SRR6453381 | 0 | 143 | 13 | 0 | 0 | 0 | 18 | 108 | 398 | 823 | 8102 | 105477 | 0 | 54 | 172 | 190 | 13 | 19 | 52 | 1627 | 115 | 1068 | 0 | 11 | 264 | 64 | 17 | 0 | 0 | 0 | 25 | 0 | 0 | 0 | 0 | 768 | 52 |
| SRR6453382 | 0 | 1582 | 15 | 0 | 0 | 0 | 34 | 233 | 773 | 1165 | 7404 | 129076 | 0 | 64 | 222 | 275 | 96 | 37 | 57 | 762 | 184 | 1034 | 0 | 0 | 214 | 64 | 83 | 0 | 0 | 0 | 19 | 0 | 0 | 0 | 0 | 360 | 18 |
| SRR6453383 | 0 | 118 | 17 | 0 | 0 | 0 | 14 | 1168 | 2866 | 1139 | 7219 | 99835 | 0 | 88 | 98 | 118 | 43 | 24 | 46 | 1354 | 175 | 1759 | 0 | 12 | 199 | 45 | 57 | 0 | 11 | 0 | 32 | 20 | 0 | 0 | 0 | 569 | 12 |
| SRR6453384 | 0 | 187 | 20 | 0 | 0 | 0 | 20 | 310 | 704 | 1355 | 72527 | 98862 | 0 | 129 | 129 | 212 | 22 | 28 | 52 | 1612 | 108 | 860 | 0 | 10 | 252 | 60 | 19 | 0 | 0 | 0 | 33 | 0 | 0 | 0 | 0 | 598 | 13 |
| SRR6453385 | 0 | 101 | 12 | 0 | 0 | 0 | 19 | 1473 | 4041 | 2690 | 58376 | 103759 | 0 | 257 | 469 | 670 | 13 | 51 | 103 | 3226 | 63 | 912 | 0 | 0 | 125 | 112 | 31 | 0 | 0 | 0 | 16 | 0 | 0 | 0 | 0 | 1328 | 43 |
| SRR6453386 | 0 | 193 | 0 | 0 | 0 | 14 | 0 | 1595 | 1413 | 2734 | 44974 | 52675 | 0 | 547 | 1549 | 4511 | 59 | 196 | 180 | 5969 | 13 | 218 | 0 | 73 | 152 | 1343 | 67 | 0 | 0 | 0 | 11 | 0 | 0 | 0 | 0 | 4557 | 0 |
| SRR6453387 | 0 | 187 | 0 | 0 | 15 | 0 | 14 | 846 | 1039 | 1709 | 93401 | 129778 | 0 | 810 | 528 | 1321 | 38 | 212 | 121 | 1762 | 23 | 409 | 0 | 70 | 246 | 1246 | 23 | 0 | 0 | 0 | 19 | 0 | 0 | 0 | 12 | 828 | 0 |
| SRR6453388 | 0 | 3332 | 27 | 0 | 59 | 14 | 37 | 577 | 944 | 1953 | 3207 | 64553 | 0 | 328 | 275 | 459 | 154 | 286 | 399 | 12009 | 75 | 669 | 0 | 154 | 393 | 1449 | 36 | 0 | 0 | 0 | 28 | 0 | 0 | 0 | 13 | 6621 | 0 |
| SRR6453389 | 0 | 493 | 10 | 0 | 56 | 13 | 15 | 1551 | 1551 | 9115 | 22219 | 30075 | 0 | 495 | 337 | 899 | 47 | 385 | 289 | 16195 | 52 | 686 | 0 | 153 | 857 | 853 | 24 | 0 | 0 | 0 | 16 | 0 | 0 | 0 | 11 | 11509 | 0 |
| SRR6453390 | 0 | 2042 | 15 | 0 | 86 | 0 | 23 | 741 | 1452 | 10548 | 14098 | 112230 | 0 | 126 | 185 | 504 | 44 | 413 | 300 | 3852 | 111 | 698 | 0 | 139 | 498 | 1137 | 11 | 0 | 13 | 0 | 10 | 0 | 0 | 0 | 18 | 1993 | 0 |
| SRR6453391 | 0 | 2490 | 20 | 0 | 38 | 0 | 20 | 602 | 1153 | 8072 | 5442 | 80773 | 0 | 256 | 115 | 282 | 143 | 270 | 159 | 19719 | 72 | 404 | 0 | 152 | 391 | 1575 | 36 | 0 | 0 | 0 | 26 | 0 | 0 | 0 | 14 | 8803 | 0 |
| SRR6453392 | 0 | 4881 | 24 | 0 | 67 | 0 | 26 | 596 | 1135 | 7692 | 19489 | 93812 | 0 | 160 | 147 | 500 | 153 | 273 | 218 | 6371 | 87 | 535 | 0 | 99 | 178 | 2136 | 18 | 0 | 0 | 0 | 16 | 0 | 0 | 0 | 0 | 3487 | 0 |
| SRR6453393 | 0 | 10726 | 38 | 0 | 75 | 0 | 27 | 787 | 1444 | 9055 | 107813 | 110739 | 0 | 296 | 120 | 491 | 99 | 327 | 261 | 7342 | 80 | 494 | 0 | 139 | 468 | 1723 | 28 | 0 | 0 | 0 | 24 | 0 | 0 | 0 | 14 | 3481 | 0 |
| SRR6453394 | 0 | 2016 | 32 | 0 | 94 | 17 | 37 | 523 | 589 | 1832 | 2467 | 49503 | 0 | 174 | 240 | 850 | 140 | 421 | 350 | 6150 | 52 | 801 | 0 | 45 | 768 | 600 | 37 | 0 | 0 | 0 | 13 | 0 | 0 | 0 | 0 | 4274 | 0 |
| SRR6453395 | 0 | 4654 | 25 | 0 | 53 | 12 | 34 | 335 | 1056 | 1869 | 7618 | 81361 | 0 | 335 | 142 | 650 | 143 | 297 | 225 | 9394 | 70 | 558 | 0 | 84 | 363 | 1442 | 40 | 0 | 0 | 0 | 19 | 0 | 0 | 0 | 0 | 4976 | 0 |
| SRR6453396 | 0 | 2233 | 11 | 0 | 44 | 0 | 17 | 356 | 503 | 2098 | 9615 | 85779 | 0 | 196 | 135 | 589 | 72 | 286 | 282 | 6220 | 70 | 433 | 0 | 77 | 264 | 1545 | 16 | 0 | 0 | 0 | 13 | 0 | 0 | 0 | 0 | 3803 | 0 |
| SRR6453397 | 0 | 728 | 0 | 0 | 36 | 0 | 18 | 412 | 412 | 655 | 104107 | 78408 | 0 | 218 | 121 | 388 | 142 | 234 | 182 | 8922 | 42 | 359 | 0 | 66 | 443 | 1376 | 19 | 0 | 0 | 0 | 0 | 0 | 0 | 0 | 11 | 5528 | 0 |
| SRR6453398 | 0 | 601 | 13 | 0 | 58 | 15 | 16 | 269 | 778 | 2213 | 2792 | 59515 | 0 | 449 | 120 | 1196 | 77 | 312 | 174 | 5294 | 32 | 637 | 0 | 136 | 819 | 1152 | 22 | 0 | 0 | 0 | 12 | 0 | 0 | 0 | 0 | 3469 | 0 |
| SRR6453399 | 0 | 274 | 24 | 0 | 65 | 10 | 13 | 296 | 350 | 202 | 1458 | 26271 | 0 | 148 | 376 | 1156 | 32 | 295 | 210 | 20328 | 41 | 561 | 0 | 189 | 810 | 829 | 15 | 0 | 0 | 0 | 21 | 0 | 0 | 0 | 0 | 13700 | 0 |
| SRR6453400 | 0 | 2057 | 16 | 0 | 69 | 16 | 31 | 172 | 387 | 1031 | 63448 | 65425 | 0 | 301 | 86 | 358 | 199 | 337 | 316 | 8409 | 40 | 555 | 0 | 57 | 378 | 1108 | 46 | 0 | 0 | 0 | 0 | 0 | 0 | 0 | 0 | 4957 | 0 |
| SRR6453401 | 0 | 1151 | 27 | 0 | 67 | 0 | 29 | 353 | 728 | 2880 | 7834 | 95519 | 0 | 128 | 171 | 482 | 67 | 324 | 304 | 11080 | 98 | 524 | 0 | 106 | 299 | 2641 | 26 | 0 | 0 | 0 | 23 | 0 | 0 | 0 | 0 | 5545 | 0 |
| SRR6453402 | 0 | 542 | 11 | 0 | 30 | 0 | 14 | 431 | 809 | 1292 | 10281 | 116134 | 0 | 159 | 60 | 325 | 63 | 173 | 234 | 2812 | 64 | 526 | 0 | 111 | 239 | 1639 | 21 | 0 | 0 | 0 | 29 | 0 | 0 | 0 | 12 | 1822 | 0 |
| SRR6453403 | 0 | 1447 | 24 | 0 | 21 | 0 | 26 | 933 | 1480 | 13635 | 6189 | 68538 | 0 | 375 | 247 | 858 | 136 | 142 | 103 | 9000 | 25 | 226 | 0 | 159 | 202 | 2678 | 40 | 0 | 0 | 0 | 20 | 0 | 0 | 0 | 11 | 4945 | 0 |
| SRR6453404 | 0 | 160 | 23 | 0 | 40 | 15 | 27 | 504 | 479 | 416 | 38184 | 32005 | 0 | 315 | 328 | 1538 | 31 | 175 | 178 | 14994 | 18 | 294 | 0 | 73 | 358 | 1080 | 17 | 0 | 0 | 0 | 20 | 0 | 0 | 0 | 13 | 10971 | 0 |
| SRR6453405 | 0 | 701 | 15 | 0 | 17 | 0 | 22 | 911 | 971 | 11709 | 3337 | 67758 | 0 | 357 | 277 | 1090 | 40 | 232 | 306 | 20327 | 22 | 265 | 0 | 53 | 239 | 1182 | 16 | 0 | 11 | 0 | 30 | 0 | 0 | 0 | 20 | 12383 | 0 |
| SRR6453406 | 0 | 586 | 19 | 0 | 41 | 14 | 18 | 392 | 753 | 1714 | 10556 | 95128 | 0 | 369 | 208 | 546 | 65 | 159 | 252 | 6873 | 40 | 349 | 0 | 97 | 169 | 1184 | 17 | 0 | 0 | 0 | 24 | 0 | 0 | 0 | 0 | 4313 | 0 |
| SRR6453407 | 0 | 418 | 15 | 0 | 21 | 12 | 23 | 393 | 617 | 748 | 80069 | 59313 | 0 | 337 | 617 | 2076 | 120 | 211 | 242 | 7835 | 24 | 272 | 0 | 117 | 478 | 1324 | 33 | 0 | 0 | 0 | 11 | 0 | 0 | 0 | 0 | 7087 | 0 |
| SRR6453408 | 0 | 854 | 13 | 0 | 12 | 19 | 32 | 243 | 486 | 1603 | 38273 | 46797 | 0 | 340 | 389 | 1360 | 150 | 179 | 387 | 22342 | 30 | 284 | 0 | 198 | 503 | 2210 | 82 | 0 | 0 | 0 | 17 | 0 | 0 | 0 | 0 | 18165 | 0 |
| SRR6453409 | 0 | 196 | 21 | 0 | 54 | 15 | 27 | 586 | 859 | 3222 | 75539 | 90488 | 0 | 566 | 146 | 937 | 39 | 164 | 206 | 8046 | 17 | 302 | 0 | 67 | 274 | 1252 | 10 | 0 | 0 | 0 | 18 | 0 | 0 | 0 | 14 | 5488 | 0 |
| SRR6453410 | 0 | 214 | 18 | 0 | 36 | 14 | 13 | 715 | 1145 | 2559 | 37976 | 46385 | 0 | 302 | 382 | 1097 | 47 | 214 | 164 | 11016 | 28 | 342 | 0 | 164 | 500 | 1054 | 17 | 0 | 0 | 0 | 13 | 0 | 0 | 0 | 0 | 8981 | 0 |
| SRR6453411 | 0 | 641 | 22 | 0 | 18 | 14 | 12 | 1450 | 2197 | 6598 | 5980 | 70841 | 0 | 772 | 215 | 755 | 44 | 262 | 216 | 9233 | 77 | 516 | 0 | 122 | 256 | 2063 | 15 | 0 | 0 | 0 | 21 | 0 | 0 | 0 | 0 | 5775 | 0 |
| SRR6453412 | 0 | 338 | 10 | 0 | 0 | 0 | 12 | 762 | 617 | 1779 | 96431 | 68221 | 0 | 581 | 436 | 1307 | 42 | 245 | 165 | 3485 | 19 | 320 | 0 | 31 | 112 | 1267 | 16 | 0 | 0 | 0 | 18 | 0 | 0 | 0 | 13 | 3122 | 0 |
| SRR6453413 | 0 | 294 | 13 | 0 | 21 | 0 | 14 | 763 | 2192 | 3384 | 98322 | 109737 | 0 | 310 | 214 | 1430 | 54 | 245 | 153 | 2717 | 16 | 430 | 0 | 50 | 178 | 698 | 12 | 0 | 0 | 0 | 13 | 0 | 0 | 0 | 11 | 2407 | 0 |
| SRR6453414 | 0 | 802 | 17 | 0 | 107 | 12 | 59 | 993 | 2353 | 10321 | 75946 | 97764 | 0 | 172 | 147 | 772 | 200 | 237 | 343 | 2831 | 68 | 468 | 0 | 186 | 446 | 3488 | 0 | 0 | 0 | 0 | 0 | 0 | 0 | 0 | 10 | 2304 | 0 |
| SRR6453415 | 0 | 804 | 19 | 0 | 161 | 16 | 38 | 929 | 2480 | 3773 | 66318 | 74374 | 0 | 140 | 140 | 748 | 189 | 320 | 292 | 3549 | 129 | 667 | 0 | 202 | 714 | 2079 | 13 | 0 | 0 | 0 | 0 | 0 | 0 | 0 | 11 | 3025 | 0 |
| SRR6453416 | 0 | 415 | 15 | 13 | 178 | 10 | 12 | 3816 | 11668 | 30390 | 71335 | 82118 | 0 | 74 | 168 | 550 | 106 | 557 | 250 | 1361 | 237 | 969 | 0 | 397 | 872 | 2435 | 18 | 0 | 0 | 0 | 0 | 0 | 0 | 0 | 20 | 1287 | 0 |
| SRR6453417 | 0 | 101 | 22 | 16 | 112 | 17 | 11 | 1196 | 2565 | 4482 | 77093 | 98105 | 0 | 110 | 91 | 812 | 35 | 272 | 263 | 2401 | 73 | 555 | 0 | 150 | 342 | 2526 | 21 | 0 | 0 | 0 | 0 | 0 | 0 | 0 | 13 | 2163 | 0 |

Supp Table S8.4.Columns_DI_to_ES

| | | | | | | | | | | | | | | | | | | | | | | | | | | | | | | | | | | | | | |
|---|---|---|---|---|---|---|---|---|---|---|---|---|---|---|---|---|---|---|---|---|---|---|---|---|---|---|---|---|---|---|---|---|---|---|---|---|---|
| SRR6453418 | 0 | 897 | 32 | 0 | 53 | 0 | 27 | 284 | 970 | 1761 | 90836 | 67716 | 0 | 209 | 209 | 373 | 105 | 295 | 267 | 6716 | 88 | 808 | 0 | 139 | 472 | 1295 | 65 | 0 | 0 | 0 | 16 | 0 | 0 | 0 | 29 | 5642 | 0 |
| SRR6453419 | 0 | 560 | 23 | 0 | 77 | 11 | 15 | 697 | 891 | 3185 | 70962 | 107678 | 0 | 168 | 159 | 653 | 42 | 239 | 416 | 6960 | 69 | 599 | 0 | 100 | 289 | 1221 | 36 | 0 | 11 | 0 | 10 | 0 | 0 | 0 | 18 | 3935 | 0 |
| SRR6453420 | 0 | 489 | 12 | 0 | 26 | 11 | 23 | 330 | 1195 | 651 | 89860 | 104364 | 0 | 140 | 223 | 799 | 61 | 229 | 425 | 8521 | 51 | 436 | 0 | 95 | 229 | 1274 | 43 | 0 | 0 | 0 | 13 | 0 | 0 | 0 | 11 | 5085 | 0 |
| SRR6453421 | 0 | 796 | 18 | 0 | 25 | 14 | 35 | 2408 | 2450 | 6614 | 10181 | 69666 | 0 | 463 | 170 | 486 | 99 | 830 | 305 | 6352 | 176 | 1782 | 0 | 356 | 607 | 1739 | 357 | 0 | 0 | 0 | 58 | 18 | 10 | 0 | 44 | 3604 | 0 |
| SRR6453422 | 0 | 470 | 13 | 0 | 0 | 0 | 16 | 608 | 1503 | 3511 | 22074 | 125085 | 0 | 266 | 451 | 1154 | 110 | 314 | 255 | 2377 | 27 | 393 | 0 | 157 | 232 | 1174 | 45 | 0 | 0 | 0 | 16 | 10 | 0 | 10 | 13 | 970 | 0 |
| SRR6453423 | 0 | 683 | 0 | 0 | 22 | 17 | 36 | 2323 | 3462 | 9422 | 9123 | 98419 | 0 | 1096 | 373 | 1128 | 110 | 364 | 244 | 2071 | 40 | 503 | 0 | 65 | 242 | 1379 | 71 | 0 | 0 | 0 | 0 | 0 | 0 | 0 | 0 | 1096 | 0 |
| SRR6453424 | 0 | 549 | 19 | 23 | 100 | 15 | 42 | 691 | 1633 | 5269 | 16883 | 115024 | 0 | 82 | 144 | 747 | 165 | 282 | 209 | 1922 | 71 | 439 | 0 | 147 | 461 | 1942 | 16 | 0 | 0 | 0 | 0 | 0 | 0 | 0 | 63 | 1489 | 0 |
| SRR6453425 | 0 | 102 | 17 | 14 | 188 | 19 | 40 | 1715 | 4084 | 10202 | 5054 | 52119 | 0 | 433 | 155 | 841 | 41 | 460 | 512 | 1968 | 175 | 1196 | 0 | 205 | 1154 | 1144 | 0 | 0 | 0 | 0 | 0 | 0 | 0 | 0 | 0 | 1650 | 0 |
| SRR6453426 | 0 | 890 | 13 | 14 | 90 | 19 | 30 | 1626 | 2586 | 17456 | 8550 | 101295 | 0 | 49 | 0 | 33 | 229 | 272 | 264 | 3260 | 105 | 515 | 0 | 143 | 536 | 2493 | 19 | 0 | 0 | 0 | 0 | 0 | 0 | 0 | 33 | 2767 | 0 |
| SRR6453427 | 0 | 840 | 17 | 0 | 161 | 19 | 48 | 2134 | 5409 | 35385 | 11423 | 67414 | 0 | 35 | 89 | 151 | 223 | 379 | 277 | 2886 | 160 | 576 | 0 | 172 | 516 | 1501 | 10 | 0 | 0 | 0 | 0 | 0 | 0 | 0 | 0 | 2373 | 0 |
| SRR6453428 | 0 | 329 | 14 | 16 | 186 | 13 | 30 | 1615 | 4412 | 13817 | 80262 | 101686 | 0 | 136 | 57 | 296 | 61 | 279 | 347 | 2843 | 112 | 626 | 0 | 165 | 478 | 1929 | 13 | 0 | 0 | 0 | 0 | 0 | 0 | 0 | 0 | 2308 | 0 |
| SRR6454160 | 0 | 160 | 27 | 0 | 29 | 0 | 0 | 175 | 27 | 487 | 115 | 214 | 0 | 21 | 36 | 18 | 27 | 329 | 0 | 74 | 1383 | 506 | 356 | 8952 | 4343 | 0 | 0 | 0 | 399 | 425 | 213 | 53 | 292 | 266 | 27 | 279 | 84 |
| SRR6454161 | 0 | 0 | 0 | 0 | 33 | 18 | 21 | 343 | 307 | 5191 | 377 | 113 | 0 | 34 | 0 | 0 | 79 | 81 | 0 | 0 | 2288 | 1006 | 463 | 10920 | 5226 | 0 | 92 | 11 | 46 | 31 | 52 | 0 | 30 | 46 | 0 | 37 | 30 |
| SRR6454162 | 0 | 31 | 61 | 0 | 0 | 0 | 0 | 155 | 0 | 0 | 247 | 26747 | 0 | 0 | 0 | 0 | 0 | 0 | 0 | 0 | 94 | 554 | 0 | 0 | 150 | 37 | 31 | 0 | 0 | 0 | 0 | 0 | 0 | 92 | 0 | 928 | 0 |
| SRR6454163 | 0 | 10 | 21 | 0 | 74 | 0 | 0 | 1468 | 479 | 4194 | 164 | 145 | 0 | 43 | 30 | 46 | 10 | 191 | 0 | 20 | 729 | 1270 | 224 | 8558 | 4324 | 0 | 31 | 10 | 330 | 1115 | 475 | 41 | 413 | 723 | 0 | 92 | 81 |
| SRR6454164 | 0 | 0 | 0 | 0 | 0 | 0 | 0 | 151 | 0 | 604 | 165 | 22195 | 0 | 151 | 0 | 0 | 0 | 0 | 0 | 302 | 85 | 567 | 0 | 0 | 154 | 38 | 31 | 0 | 0 | 0 | 0 | 31 | 61 | 0 | 0 | 1057 | 0 |
| SRR6454165 | 15 | 31 | 31 | 0 | 136 | 0 | 0 | 746 | 401 | 4301 | 238 | 191 | 0 | 85 | 20 | 22 | 69 | 219 | 0 | 16 | 2961 | 1357 | 441 | 13180 | 3936 | 0 | 445 | 61 | 330 | 1005 | 583 | 77 | 192 | 215 | 0 | 120 | 44 |
| SRR6454166 | 32 | 0 | 0 | 0 | 0 | 0 | 0 | 358 | 0 | 358 | 143 | 23447 | 0 | 0 | 179 | 0 | 0 | 0 | 0 | 537 | 105 | 577 | 0 | 0 | 169 | 23 | 158 | 0 | 0 | 0 | 0 | 0 | 0 | 0 | 0 | 895 | 0 |
| SRR6454167 | 0 | 72 | 0 | 0 | 89 | 12 | 12 | 220 | 104 | 1386 | 198 | 139 | 12 | 15 | 39 | 109 | 0 | 199 | 0 | 14 | 825 | 1327 | 215 | 10666 | 6438 | 0 | 12 | 120 | 526 | 1052 | 514 | 96 | 430 | 574 | 0 | 81 | 93 |
| SRR6454169 | 21 | 31 | 51 | 0 | 83 | 0 | 10 | 912 | 431 | 2550 | 272 | 180 | 0 | 42 | 40 | 18 | 21 | 283 | 0 | 0 | 1696 | 981 | 389 | 12903 | 3782 | 0 | 62 | 62 | 431 | 1097 | 462 | 31 | 267 | 421 | 0 | 44 | 50 |
| SRR6454170 | 20 | 20 | 20 | 0 | 0 | 0 | 0 | 250 | 0 | 749 | 213 | 25981 | 0 | 0 | 0 | 0 | 0 | 0 | 0 | 999 | 129 | 656 | 0 | 0 | 173 | 38 | 41 | 0 | 0 | 0 | 0 | 0 | 20 | 82 | 0 | 2123 | 0 |
| SRR6454171 | 18 | 18 | 0 | 0 | 88 | 0 | 18 | 152 | 103 | 1563 | 150 | 80 | 0 | 35 | 24 | 16 | 35 | 255 | 0 | 0 | 879 | 1658 | 141 | 11522 | 5884 | 0 | 18 | 0 | 98 | 284 | 257 | 27 | 160 | 231 | 18 | 31 | 95 |
| SRR6454172 | 0 | 0 | 66 | 0 | 0 | 0 | 0 | 265 | 88 | 353 | 362 | 30097 | 0 | 0 | 0 | 88 | 0 | 0 | 0 | 441 | 200 | 810 | 0 | 0 | 172 | 23 | 16 | 0 | 0 | 0 | 0 | 0 | 16 | 49 | 0 | 530 | 0 |
| SRR6454173 | 0 | 0 | 0 | 0 | 122 | 0 | 0 | 661 | 263 | 1917 | 241 | 157 | 0 | 97 | 24 | 13 | 31 | 266 | 0 | 30 | 5609 | 1525 | 429 | 12496 | 4102 | 0 | 218 | 23 | 156 | 491 | 148 | 16 | 125 | 172 | 0 | 75 | 77 |
| SRR6454174 | 0 | 16 | 0 | 0 | 0 | 16 | 0 | 546 | 0 | 91 | 126 | 13092 | 0 | 182 | 0 | 0 | 0 | 0 | 0 | 182 | 105 | 690 | 0 | 0 | 63 | 13 | 127 | 0 | 0 | 0 | 0 | 0 | 16 | 32 | 16 | 182 | 0 |
| SRR6454175 | 0 | 0 | 25 | 0 | 29 | 0 | 0 | 1920 | 498 | 3171 | 59 | 60 | 0 | 53 | 166 | 115 | 38 | 144 | 0 | 23 | 1191 | 279 | 514 | 8321 | 3252 | 0 | 140 | 13 | 146 | 343 | 343 | 83 | 191 | 279 | 57 | 61 | 18 |
| SRR6454176 | 0 | 13 | 39 | 0 | 0 | 0 | 0 | 230 | 77 | 383 | 23117 | 26102 | 0 | 0 | 77 | 0 | 0 | 0 | 0 | 77 | 309 | 1380 | 0 | 0 | 184 | 26 | 92 | 0 | 0 | 0 | 0 | 0 | 13 | 0 | 0 | 383 | 0 |
| SRR6454177 | 0 | 0 | 65 | 0 | 124 | 0 | 0 | 573 | 299 | 4372 | 617 | 486 | 0 | 37 | 0 | 18 | 0 | 265 | 0 | 16 | 880 | 1588 | 213 | 12244 | 5298 | 0 | 13 | 39 | 456 | 677 | 378 | 117 | 208 | 782 | 0 | 97 | 69 |
| SRR6454178 | 17 | 17 | 87 | 0 | 0 | 17 | 0 | 108 | 0 | 216 | 22436 | 24809 | 0 | 0 | 0 | 216 | 0 | 0 | 0 | 108 | 201 | 890 | 0 | 0 | 151 | 24 | 35 | 0 | 0 | 0 | 0 | 17 | 35 | 17 | 0 | 539 | 0 |
| SRR6454179 | 0 | 0 | 38 | 0 | 123 | 0 | 0 | 425 | 277 | 2217 | 265 | 117 | 0 | 71 | 100 | 82 | 25 | 309 | 0 | 0 | 2412 | 1144 | 441 | 13155 | 4079 | 0 | 201 | 101 | 327 | 528 | 365 | 88 | 138 | 365 | 25 | 33 | 60 |
| SRR6454180 | 0 | 16 | 0 | 0 | 0 | 0 | 0 | 53 | 0 | 0 | 471 | 22031 | 0 | 0 | 0 | 0 | 0 | 0 | 0 | 267 | 214 | 599 | 0 | 0 | 245 | 42 | 80 | 0 | 0 | 0 | 0 | 0 | 0 | 0 | 0 | 213 | 0 |
| SRR6454181 | 10 | 0 | 20 | 0 | 65 | 0 | 10 | 253 | 108 | 1782 | 72 | 74 | 0 | 43 | 11 | 0 | 10 | 424 | 0 | 14 | 583 | 1016 | 171 | 10821 | 5080 | 0 | 50 | 20 | 172 | 353 | 343 | 50 | 192 | 394 | 20 | 42 | 66 |
| SRR6454182 | 0 | 0 | 13 | 0 | 0 | 0 | 0 | 112 | 0 | 299 | 29820 | 27575 | 0 | 75 | 0 | 0 | 0 | 0 | 0 | 112 | 203 | 1026 | 0 | 0 | 189 | 46 | 94 | 0 | 0 | 0 | 0 | 0 | 0 | 13 | 0 | 75 | 0 |
| SRR6454183 | 0 | 0 | 14 | 0 | 88 | 0 | 0 | 909 | 505 | 3687 | 62 | 60 | 0 | 68 | 149 | 324 | 71 | 205 | 0 | 0 | 2068 | 895 | 373 | 11175 | 3425 | 0 | 317 | 42 | 183 | 360 | 219 | 0 | 92 | 247 | 14 | 34 | 35 |
| SRR6454184 | 16 | 65 | 65 | 0 | 0 | 0 | 0 | 104 | 0 | 104 | 234 | 19395 | 0 | 0 | 0 | 0 | 0 | 0 | 0 | 0 | 161 | 590 | 0 | 0 | 123 | 12 | 81 | 0 | 16 | 0 | 0 | 0 | 16 | 32 | 0 | 313 | 0 |
| SRR6454185 | 0 | 0 | 29 | 0 | 83 | 0 | 0 | 493 | 169 | 2313 | 174 | 195 | 0 | 163 | 106 | 91 | 0 | 218 | 0 | 43 | 2088 | 3288 | 292 | 13507 | 6383 | 0 | 0 | 13 | 92 | 190 | 173 | 0 | 88 | 255 | 36 | 455 | 74 |
| SRR6454186 | 0 | 16 | 31 | 0 | 0 | 0 | 0 | 0 | 120 | 240 | 155 | 16228 | 0 | 0 | 120 | 120 | 31 | 0 | 0 | 240 | 110 | 496 | 0 | 0 | 102 | 15 | 155 | 0 | 0 | 0 | 0 | 0 | 0 | 0 | 16 | 1202 | 0 |
| SRR6454187 | 31 | 0 | 18 | 0 | 110 | 18 | 13 | 158 | 84 | 1329 | 127 | 96 | 0 | 51 | 105 | 62 | 110 | 122 | 0 | 0 | 1362 | 1252 | 396 | 12582 | 4561 | 0 | 159 | 35 | 97 | 265 | 185 | 31 | 97 | 168 | 18 | 15 | 30 |
| SRR6454188 | 0 | 41 | 21 | 0 | 0 | 0 | 0 | 158 | 0 | 158 | 20038 | 16882 | 0 | 158 | 0 | 0 | 0 | 0 | 0 | 473 | 243 | 635 | 0 | 0 | 329 | 42 | 186 | 0 | 0 | 21 | 0 | 0 | 0 | 21 | 21 | 473 | 0 |
| SRR6454189 | 0 | 0 | 0 | 0 | 154 | 0 | 0 | 224 | 88 | 1767 | 205 | 248 | 0 | 22 | 37 | 54 | 0 | 277 | 0 | 0 | 720 | 1103 | 234 | 13899 | 8559 | 0 | 67 | 0 | 380 | 1229 | 357 | 45 | 313 | 335 | 0 | 22 | 108 |
| SRR6454190 | 50 | 25 | 25 | 0 | 0 | 25 | 0 | 587 | 0 | 196 | 19765 | 20548 | 0 | 0 | 0 | 0 | 0 | 0 | 0 | 196 | 240 | 625 | 0 | 0 | 247 | 32 | 274 | 0 | 0 | 0 | 0 | 0 | 0 | 25 | 0 | 1761 | 0 |
| SRR6454191 | 0 | 0 | 0 | 0 | 110 | 0 | 0 | 623 | 415 | 4078 | 235 | 153 | 0 | 80 | 25 | 20 | 122 | 359 | 0 | 0 | 4256 | 1115 | 425 | 13127 | 4110 | 0 | 228 | 53 | 221 | 442 | 122 | 30 | 91 | 228 | 23 | 84 | 50 |
| SRR6454193 | 0 | 0 | 14 | 0 | 125 | 0 | 0 | 244 | 120 | 1258 | 145 | 196 | 0 | 77 | 98 | 65 | 0 | 315 | 0 | 29 | 1087 | 1616 | 274 | 14734 | 7620 | 0 | 42 | 56 | 363 | 754 | 573 | 28 | 126 | 363 | 14 | 112 | 116 |
| SRR6454194 | 0 | 25 | 49 | 0 | 0 | 0 | 0 | 294 | 0 | 294 | 24706 | 18677 | 0 | 0 | 0 | 147 | 0 | 0 | 0 | 147 | 172 | 516 | 0 | 0 | 174 | 35 | 49 | 0 | 25 | 0 | 0 | 0 | 0 | 0 | 0 | 441 | 0 |
| SRR6454195 | 0 | 0 | 89 | 0 | 107 | 0 | 0 | 349 | 200 | 3217 | 327 | 194 | 0 | 84 | 0 | 13 | 18 | 288 | 0 | 17 | 1828 | 953 | 453 | 16290 | 4358 | 0 | 107 | 124 | 391 | 1297 | 462 | 89 | 53 | 391 | 0 | 26 | 49 |
| SRR6454196 | 0 | 0 | 25 | 0 | 0 | 0 | 0 | 228 | 456 | 228 | 43 | 14813 | 0 | 0 | 0 | 0 | 0 | 0 | 0 | 0 | 48 | 301 | 0 | 0 | 26 | 0 | 0 | 0 | 0 | 0 | 0 | 0 | 25 | 0 | 0 | 228 | 0 |
| SRR6454197 | 0 | 14 | 56 | 0 | 34 | 0 | 0 | 540 | 139 | 2047 | 97 | 92 | 0 | 25 | 159 | 251 | 42 | 234 | 0 | 37 | 831 | 812 | 223 | 7725 | 4585 | 0 | 14 | 21 | 78 | 254 | 261 | 134 | 240 | 522 | 21 | 214 | 41 |
| SRR6454198 | 0 | 13 | 27 | 0 | 0 | 0 | 0 | 334 | 111 | 556 | 8568 | 16691 | 0 | 0 | 111 | 0 | 0 | 0 | 0 | 111 | 133 | 580 | 0 | 0 | 97 | 19 | 66 | 0 | 0 | 0 | 0 | 0 | 13 | 40 | 27 | 779 | 0 |
| SRR6454199 | 0 | 0 | 43 | 0 | 35 | 0 | 0 | 803 | 237 | 2468 | 91 | 89 | 0 | 56 | 262 | 287 | 43 | 200 | 0 | 52 | 11 | 511 | 469 | 10514 | 2523 | 0 | 228 | 49 | 111 | 179 | 265 | 111 | 234 | 246 | 68 | 135 | 27 |
| SRR6454201 | 0 | 15 | 80 | 0 | 64 | 0 | 0 | 720 | 290 | 5875 | 88 | 114 | 0 | 306 | 30 | 47 | 15 | 168 | 0 | 23 | 1307 | 1280 | 349 | 5643 | 3544 | 24 | 0 | 0 | 66 | 190 | 139 | 58 | 124 | 204 | 15 | 62 | 44 |
| SRR6454203 | 0 | 17 | 85 | 0 | 116 | 0 | 0 | 485 | 218 | 3693 | 51 | 55 | 0 | 103 | 140 | 63 | 0 | 365 | 0 | 0 | 2517 | 1015 | 533 | 13662 | 3970 | 0 | 135 | 17 | 457 | 948 | 508 | 51 | 322 | 728 | 51 | 33 | 52 |
| SRR6454205 | 0 | 12 | 12 | 0 | 111 | 0 | 0 | 337 | 126 | 1661 | 84 | 121 | 0 | 33 | 190 | 157 | 25 | 250 | 0 | 26 | 437 | 534 | 299 | 12557 | 5005 | 0 | 61 | 61 | 418 | 688 | 492 | 61 | 270 | 602 | 12 | 106 | 94 |
| SRR6454206 | 0 | 0 | 0 | 0 | 0 | 0 | 0 | 122 | 0 | 122 | 265 | 31375 | 0 | 0 | 0 | 0 | 0 | 0 | 0 | 0 | 154 | 1077 | 0 | 0 | 94 | 16 | 28 | 0 | 0 | 0 | 0 | 0 | 83 | 28 | 0 | 122 | 0 |
| SRR6454207 | 0 | 0 | 46 | 0 | 92 | 0 | 0 | 505 | 449 | 2730 | 339 | 357 | 0 | 79 | 366 | 202 | 68 | 270 | 0 | 58 | 2611 | 1336 | 743 | 17279 | 3300 | 0 | 262 | 137 | 524 | 820 | 478 | 46 | 342 | 660 | 11 | 204 | 49 |
| SRR6454208 | 0 | 0 | 115 | 0 | 0 | 0 | 0 | 0 | 129 | 0 | 25495 | 18765 | 0 | 0 | 0 | 0 | 0 | 0 | 0 | 259 | 222 | 984 | 0 | 0 | 135 | 18 | 29 | 0 | 0 | 0 | 0 | 0 | 0 | 0 | 0 | 777 | 0 |
| SRR6454209 | 17 | 0 | 0 | 0 | 94 | 0 | 0 | 473 | 203 | 2769 | 81 | 92 | 0 | 25 | 82 | 66 | 0 | 429 | 0 | 23 | 709 | 726 | 208 | 12515 | 5963 | 0 | 59 | 50 | 370 | 824 | 353 | 25 | 294 | 462 | 0 | 94 | 58 |
| SRR6454211 | 0 | 0 | 81 | 0 | 89 | 0 | 0 | 446 | 211 | 2127 | 23 | 27 | 0 | 45 | 39 | 25 | 60 | 278 | 0 | 0 | 2054 | 808 | 399 | 13799 | 3510 | 0 | 608 | 49 | 54 | 423 | 222 | 27 | 201 | 499 | 22 | 27 | 32 |
| SRR6454213 | 0 | 11 | 23 | 0 | 102 | 0 | 0 | 426 | 182 | 2765 | 158 | 135 | 0 | 16 | 78 | 88 | 90 | 239 | 0 | 16 | 886 | 1197 | 303 | 14065 | 7437 | 0 | 56 | 45 | 395 | 722 | 350 | 34 | 226 | 531 | 0 | 114 | 94 |
| SRR6454214 | 0 | 0 | 27 | 0 | 0 | 0 | 0 | 0 | 0 | 0 | 179 | 17874 | 0 | 0 | 0 | 0 | 0 | 0 | 0 | 130 | 158 | 680 | 0 | 0 | 129 | 24 | 107 | 0 | 0 | 0 | 0 | 0 | 0 | 27 | 0 | 391 | 0 |
| SRR6454215 | 0 | 12 | 18 | 0 | 105 | 0 | 0 | 483 | 198 | 2863 | 90 | 56 | 0 | 52 | 54 | 33 | 71 | 112 | 0 | 0 | 1449 | 840 | 433 | 14802 | 4676 | 0 | 577 | 30 | 101 | 577 | 244 | 24 | 119 | 286 | 18 | 27 | 38 |
| SRR6454216 | 0 | 0 | 0 | 0 | 0 | 0 | 0 | 628 | 0 | 628 | 12142 | 23655 | 0 | 0 | 0 | 0 | 147 | 0 | 0 | 209 | 36 | 142 | 0 | 0 | 82 | 10 | 49 | 0 | 0 | 0 | 0 | 0 | 0 | 0 | 0 | 628 | 0 |
| SRR6454217 | 0 | 33 | 33 | 0 | 38 | 0 | 0 | 2315 | 834 | 3890 | 68 | 85 | 0 | 67 | 342 | 323 | 50 | 242 | 0 | 61 | 3603 | 881 | 319 | 9890 | 3537 | 0 | 66 | 0 | 249 | 515 | 357 | 91 | 249 | 374 | 58 | 449 | 32 |
| SRR6454218 | 0 | 27 | 27 | 0 | 0 | 0 | 0 | 128 | 0 | 128 | 15476 | 28138 | 0 | 0 | 0 | 64 | 0 | 0 | 0 | 0 | 172 | 707 | 0 | 0 | 133 | 17 | 0 | 0 | 0 | 0 | 0 | 0 | 0 | 0 | 0 | 0 | 0 |
| SRR6454219 | 0 | 0 | 17 | 0 | 34 | 0 | 0 | 1379 | 382 | 4342 | 77 | 67 | 0 | 27 | 23 | 22 | 122 | 108 | 0 | 20 | 1058 | 381 | 509 | 8527 | 2404 | 0 | 130 | 17 | 104 | 322 | 269 | 52 | 209 | 408 | 26 | 74 | 22 |
| SRR6454220 | 0 | 0 | 0 | 0 | 0 | 0 | 0 | 0 | 103 | 308 | 8432 | 11825 | 0 | 0 | 0 | 0 | 0 | 0 | 0 | 103 | 127 | 612 | 0 | 0 | 71 | 13 | 424 | 0 | 0 | 0 | 0 | 0 | 0 | 22 | 0 | 103 | 0 |
| SRR6454221 | 0 | 11 | 45 | 0 | 49 | 0 | 0 | 584 | 133 | 2159 | 94 | 93 | 0 | 32 | 64 | 80 | 0 | 167 | 0 | 26 | 447 | 350 | 253 | 5698 | 3463 | 0 | 45 | 11 | 203 | 440 | 372 | 113 | 282 | 406 | 11 | 92 | 32 |
| SRR6454222 | 25 | 0 | 0 | 0 | 0 | 0 | 0 | 233 | 0 | 350 | 9323 | 10954 | 0 | 0 | 0 | 0 | 0 | 0 | 0 | 117 | 70 | 363 | 0 | 0 | 103 | 22 | 471 | 0 | 0 | 0 | 0 | 0 | 25 | 25 | 0 | 233 | 0 |
| SRR6454223 | 0 | 13 | 33 | 0 | 40 | 0 | 0 | 802 | 278 | 2425 | 71 | 65 | 0 | 48 | 166 | 155 | 78 | 146 | 0 | 44 | 1275 | 377 | 338 | 6143 | 2672 | 0 | 137 | 20 | 150 | 327 | 425 | 98 | 248 | 287 | 52 | 101 | 19 |
| SRR6454224 | 0 | 26 | 0 | 0 | 0 | 0 | 0 | 0 | 141 | 281 | 19837 | 36860 | 0 | 0 | 0 | 0 | 0 | 0 | 0 | 0 | 26 | 174 | 0 | 0 | 22 | 0 | 52 | 0 | 0 | 0 | 0 | 0 | 0 | 0 | 0 | 141 | 0 |
| SRR6454225 | 0 | 0 | 84 | 0 | 30 | 0 | 0 | 427 | 98 | 1541 | 69 | 124 | 0 | 18 | 225 | 164 | 28 | 123 | 0 | 54 | 547 | 519 | 178 | 8197 | 5626 | 0 | 42 | 42 | 140 | 489 | 377 | 56 | 405 | 237 | 28 | 183 | 45 |
| SRR6454226 | 24 | 0 | 0 | 0 | 0 | 0 | 0 | 104 | 104 | 311 | 134 | 23461 | 0 | 0 | 0 | 0 | 24 | 0 | 0 | 0 | 51 | 219 | 0 | 0 | 44 | 0 | 47 | 0 | 0 | 0 | 0 | 0 | 47 | 0 | 0 | 0 | 0 |
| SRR6454227 | 0 | 0 | 77 | 0 | 38 | 0 | 0 | 1276 | 318 | 3132 | 134 | 101 | 0 | 34 | 81 | 67 | 89 | 212 | 0 | 47 | 1257 | 492 | 344 | 8529 | 2492 | 0 | 182 | 32 | 146 | 308 | 255 | 101 | 178 | 223 | 53 | 60 | 18 |
| SRR6454228 | 0 | 17 | 17 | 0 | 0 | 0 | 0 | 0 | 0 | 116 | 13845 | 21408 | 0 | 0 | 0 | 0 | 0 | 0 | 0 | 0 | 77 | 365 | 0 | 0 | 49 | 14 | 69 | 0 | 0 | 0 | 0 | 17 | 17 | 0 | 0 | 116 | 0 |
| SRR6454229 | 0 | 17 | 100 | 0 | 39 | 0 | 0 | 223 | 69 | 1243 | 69 | 85 | 0 | 10 | 175 | 136 | 17 | 161 | 0 | 27 | 425 | 383 | 255 | 10923 | 5706 | 0 | 67 | 33 | 133 | 683 | 600 | 150 | 200 | 417 | 17 | 115 | 36 |
| SRR6454230 | 0 | 38 | 0 | 0 | 0 | 0 | 19 | 0 | 0 | 107 | 17081 | 23060 | 0 | 107 | 0 | 0 | 0 | 0 | 0 | 0 | 110 | 388 | 0 | 0 | 90 | 21 | 95 | 0 | 0 | 0 | 0 | 0 | 0 | 0 | 0 | 107 | 0 |
| SRR6454231 | 0 | 0 | 78 | 0 | 37 | 0 | 0 | 736 | 276 | 3037 | 71 | 81 | 0 | 40 | 92 | 52 | 13 | 127 | 0 | 47 | 1032 | 358 | 389 | 7170 | 3181 | 0 | 246 | 52 | 155 | 376 | 324 | 91 | 168 | 427 | 65 | 152 | 16 |
| SRR6454232 | 0 | 0 | 0 | 0 | 0 | 32 | 0 | 316 | 0 | 316 | 12549 | 17611 | 0 | 0 | 105 | 0 | 0 | 0 | 0 | 0 | 109 | 619 | 0 | 0 | 63 | 0 | 453 | 0 | 0 | 0 | 0 | 0 | 0 | 0 | 0 | 211 | 0 |
| SRR6454233 | 0 | 31 | 16 | 0 | 50 | 0 | 24 | 1072 | 430 | 2824 | 36 | 42 | 0 | 26 | 87 | 64 | 0 | 114 | 0 | 15 | 1903 | 294 | 421 | 7817 | 3169 | 0 | 39 | 47 | 149 | 385 | 298 | 63 | 181 | 212 | 31 | 94 | 22 |
| SRR6454234 | 0 | 47 | 28 | 0 | 38 | 0 | 38 | 629 | 226 | 2888 | 55 | 61 | 0 | 60 | 50 | 55 | 19 | 95 | 0 | 0 | 3622 | 809 | 490 | 10841 | 3400 | 0 | 47 | 19 | 38 | 113 | 94 | 38 | 85 | 150 | 19 | 120 | 20 |
| SRR6454235 | 0 | 16 | 33 | 0 | 0 | 0 | 0 | 0 | 0 | 205 | 12574 | 24600 | 0 | 0 | 0 | 0 | 0 | 0 | 0 | 68 | 172 | 805 | 0 | 0 | 193 | 21 | 114 | 0 | 0 | 0 | 0 | 0 | 16 | 0 | 0 | 342 | 0 |
| SRR6454236 | 0 | 14 | 14 | 0 | 35 | 0 | 19 | 1521 | 425 | 4216 | 69 | 85 | 0 | 43 | 57 | 35 | 162 | 200 | 0 | 22 | 2001 | 424 | 479 | 7370 | 3085 | 0 | 424 | 33 | 71 | 190 | 171 | 48 | 119 | 167 | 0 | 114 | 19 |
| SRR6454237 | 0 | 0 | 0 | 0 | 0 | 0 | 0 | 0 | 0 | 239 | 225 | 22419 | 0 | 0 | 0 | 0 | 0 | 0 | 0 | 60 | 181 | 760 | 0 | 0 | 175 | 37 | 58 | 0 | 0 | 0 | 0 | 0 | 0 | 0 | 0 | 0 | 0 |
| SRR6454238 | 0 | 80 | 0 | 0 | 58 | 0 | 32 | 1137 | 287 | 1249 | 67 | 46 | 0 | 13 | 21 | 0 | 80 | 90 | 0 | 0 | 2036 | 303 | 648 | 7173 | 4665 | 0 | 24 | 16 | 209 | 321 | 185 | 16 | 112 | 136 | 0 | 17 | 27 |
| SRR6454239 | 0 | 31 | 10 | 0 | 0 | 0 | 10 | 46 | 46 | 183 | 343 | 35587 | 0 | 46 | 0 | 0 | 0 | 0 | 0 | 137 | 110 | 397 | 0 | 58 | 500 | 42 | 103 | 0 | 0 | 0 | 0 | 0 | 0 | 0 | 0 | 229 | 0 |
| SRR6454240 | 0 | 21 | 21 | 0 | 47 | 0 | 32 | 938 | 426 | 2408 | 85 | 91 | 0 | 94 | 18 | 17 | 48 | 425 | 0 | 39 | 2402 | 525 | 379 | 8006 | 3340 | 0 | 253 | 11 | 167 | 290 | 269 | 27 | 64 | 145 | 21 | 161 | 11 |
| SRR6454241 | 0 | 0 | 0 | 0 | 0 | 0 | 0 | 163 | 0 | 272 | 16239 | 28513 | 0 | 0 | 0 | 0 | 23 | 0 | 0 | 0 | 139 | 505 | 0 | 0 | 303 | 28 | 70 | 0 | 0 | 0 | 0 | 0 | 12 | 0 | 0 | 109 | 0 |
| SRR6454242 | 0 | 50 | 15 | 0 | 47 | 0 | 0 | 1395 | 631 | 4772 | 66 | 117 | 0 | 52 | 110 | 96 | 35 | 240 | 0 | 15 | 1351 | 414 | 470 | 7390 | 3617 | 0 | 115 | 55 | 80 | 250 | 195 | 35 | 90 | 150 | 30 | 89 | 21 |
| SRR6454244 | 0 | 21 | 34 | 0 | 46 | 0 | 0 | 754 | 273 | 2619 | 114 | 84 | 0 | 23 | 17 | 0 | 90 | 180 | 0 | 48 | 1877 | 346 | 670 | 8264 | 3565 | 0 | 83 | 76 | 213 | 454 | 344 | 103 | 255 | 330 | 21 | 102 | 17 |
| SRR6454245 | 0 | 27 | 0 | 0 | 0 | 0 | 27 | 283 | 0 | 566 | 6084 | 11602 | 0 | 141 | 0 | 0 | 0 | 0 | 0 | 0 | 202 | 532 | 0 | 0 | 131 | 0 | 27 | 0 | 0 | 0 | 0 | 0 | 0 | 0 | 0 | 141 | 0 |
| SRR6454246 | 0 | 0 | 42 | 0 | 41 | 0 | 0 | 465 | 328 | 1641 | 26 | 28 | 0 | 57 | 55 | 36 | 33 | 178 | 0 | 14 | 1324 | 421 | 496 | 10813 | 3860 | 0 | 75 | 33 | 125 | 365 | 340 | 75 | 183 | 266 | 33 | 44 | 12 |
| SRR6454247 | 0 | 27 | 21 | 0 | 45 | 0 | 0 | 1204 | 358 | 2301 | 154 | 185 | 0 | 18 | 0 | 0 | 171 | 159 | 0 | 53 | 2104 | 299 | 474 | 5169 | 2210 | 0 | 192 | 62 | 96 | 219 | 205 | 48 | 144 | 185 | 0 | 93 | 17 |
| SRR6454249 | 0 | 26 | 66 | 0 | 40 | 0 | 0 | 242 | 59 | 887 | 36 | 92 | 0 | 23 | 34 | 30 | 13 | 153 | 0 | 18 | 321 | 350 | 166 | 7694 | 5549 | 0 | 13 | 26 | 144 | 223 | 263 | 39 | 184 | 315 | 26 | 48 | 48 |
| SRR6454251 | 0 | 0 | 17 | 0 | 40 | 0 | 0 | 597 | 256 | 1923 | 18 | 29 | 0 | 64 | 30 | 16 | 92 | 153 | 0 | 19 | 2167 | 354 | 429 | 14812 | 3847 | 0 | 125 | 25 | 109 | 359 | 242 | 42 | 150 | 150 | 0 | 98 | 22 |
| SRR6454252 | 0 | 0 | 0 | 0 | 0 | 0 | 0 | 157 | 157 | 313 | 13160 | 13787 | 0 | 0 | 0 | 0 | 0 | 0 | 0 | 157 | 70 | 295 | 0 | 0 | 31 | 0 | 42 | 0 | 0 | 0 | 0 | 0 | 0 | 0 | 0 | 313 | 0 |
| SRR6454253 | 0 | 14 | 70 | 0 | 37 | 0 | 0 | 2957 | 1047 | 6858 | 45 | 66 | 0 | 86 | 175 | 164 | 91 | 116 | 0 | 27 | 604 | 190 | 587 | 6371 | 2350 | 0 | 245 | 0 | 231 | 384 | 335 | 77 | 210 | 308 | 28 | 91 | 17 |
| SRR6454254 | 0 | 14 | 14 | 0 | 0 | 0 | 0 | 519 | 0 | 148 | 115 | 10458 | 0 | 0 | 74 | 148 | 42 | 0 | 0 | 0 | 184 | 558 | 0 | 0 | 174 | 13 | 168 | 0 | 0 | 14 | 0 | 14 | 28 | 14 | 42 | 0 | 0 |
| SRR6454255 | 0 | 0 | 34 | 0 | 37 | 0 | 0 | 1953 | 668 | 3341 | 36 | 54 | 0 | 82 | 96 | 70 | 67 | 202 | 0 | 0 | 2219 | 369 | 388 | 8013 | 3342 | 0 | 114 | 34 | 114 | 208 | 315 | 60 | 94 | 174 | 20 | 0 | 15 |
| SRR6454256 | 0 | 0 | 32 | 0 | 0 | 0 | 0 | 0 | 0 | 288 | 43 | 7499 | 0 | 0 | 144 | 0 | 0 | 0 | 0 | 144 | 82 | 427 | 0 | 10 | 31 | 0 | 190 | 0 | 0 | 0 | 0 | 0 | 0 | 0 | 0 | 144 | 0 |
| SRR6454257 | 0 | 22 | 72 | 0 | 33 | 0 | 0 | 752 | 362 | 1972 | 34 | 52 | 0 | 112 | 111 | 94 | 22 | 186 | 0 | 42 | 4076 | 552 | 543 | 9349 | 2425 | 0 | 373 | 14 | 129 | 194 | 179 | 29 | 100 | 330 | 57 | 102 | 15 |
| SRR6454258 | 0 | 0 | 33 | 0 | 37 | 0 | 0 | 329 | 81 | 1425 | 141 | 158 | 0 | 17 | 0 | 11 | 0 | 125 | 0 | 25 | 587 | 670 | 208 | 7274 | 5470 | 0 | 66 | 16 | 213 | 148 | 181 | 82 | 115 | 181 | 0 | 56 | 32 |
| SRR6454259 | 0 | 29 | 49 | 0 | 26 | 0 | 0 | 674 | 259 | 2876 | 183 | 143 | 0 | 94 | 12 | 0 | 156 | 69 | 0 | 23 | 1591 | 702 | 440 | 11685 | 3877 | 0 | 332 | 59 | 98 | 176 | 117 | 29 | 156 | 215 | 0 | 47 | 22 |
| SRR6454260 | 0 | 29 | 0 | 0 | 36 | 0 | 0 | 494 | 92 | 1442 | 188 | 162 | 0 | 22 | 0 | 19 | 48 | 154 | 0 | 22 | 598 | 696 | 211 | 10111 | 6016 | 0 | 19 | 19 | 144 | 259 | 163 | 134 | 96 | 153 | 0 | 42 | 64 |
| SRR6454261 | 0 | 13 | 0 | 0 | 32 | 0 | 0 | 738 | 189 | 2356 | 86 | 47 | 0 | 43 | 0 | 17 | 13 | 90 | 0 | 12 | 1998 | 988 | 566 | 14960 | 3758 | 17 | 77 | 38 | 64 | 57 | 147 | 57 | 89 | 185 | 13 | 35 | 26 |
| SRR6454262 | 0 | 0 | 0 | 0 | 25 | 13 | 0 | 827 | 224 | 1815 | 71 | 55 | 0 | 60 | 11 | 0 | 25 | 77 | 0 | 0 | 1213 | 300 | 629 | 8480 | 3248 | 0 | 63 | 25 | 114 | 139 | 139 | 13 | 89 | 114 | 0 | 20 | 20 |
| SRR6454263 | 0 | 89 | 32 | 0 | 29 | 0 | 0 | 542 | 191 | 1142 | 388 | 445 | 0 | 114 | 24 | 32 | 76 | 294 | 0 | 58 | 5197 | 1576 | 345 | 10391 | 5087 | 0 | 44 | 13 | 95 | 76 | 152 | 13 | 38 | 44 | 0 | 266 | 38 |
| SRR6454264 | 0 | 36 | 36 | 0 | 28 | 0 | 0 | 940 | 313 | 2847 | 38 | 70 | 0 | 28 | 10 | 12 | 0 | 86 | 0 | 24 | 2084 | 497 | 550 | 7909 | 2896 | 0 | 107 | 54 | 143 | 196 | 178 | 107 | 89 | 178 | 18 | 57 | 21 |
| SRR6454265 | 0 | 0 | 15 | 0 | 29 | 0 | 0 | 801 | 195 | 2661 | 57 | 83 | 15 | 21 | 12 | 0 | 60 | 80 | 0 | 23 | 1083 | 273 | 497 | 5875 | 2388 | 0 | 75 | 90 | 315 | 315 | 315 | 135 | 75 | 105 | 15 | 32 | 23 |
| SRR6454266 | 0 | 31 | 13 | 0 | 31 | 0 | 0 | 2947 | 531 | 3187 | 198 | 130 | 0 | 62 | 32 | 20 | 81 | 86 | 0 | 15 | 1879 | 522 | 466 | 8862 | 3660 | 0 | 170 | 40 | 224 | 116 | 143 | 36 | 116 | 148 | 0 | 94 | 29 |
| SRR6454267 | 0 | 18 | 24 | 0 | 39 | 0 | 0 | 1950 | 479 | 2831 | 268 | 87 | 0 | 71 | 36 | 31 | 95 | 201 | 0 | 34 | 3850 | 633 | 400 | 10073 | 4257 | 0 | 226 | 26 | 166 | 173 | 208 | 50 | 89 | 179 | 0 | 194 | 36 |
| SRR6454268 | 0 | 0 | 0 | 0 | 41 | 0 | 0 | 537 | 103 | 1933 | 189 | 131 | 0 | 11 | 0 | 11 | 80 | 96 | 0 | 17 | 247 | 293 | 230 | 6375 | 4884 | 0 | 54 | 80 | 241 | 161 | 294 | 80 | 107 | 80 | 0 | 86 | 42 |
| SRR6454269 | 0 | 26 | 0 | 0 | 31 | 0 | 0 | 389 | 98 | 1554 | 36 | 30 | 0 | 18 | 0 | 0 | 79 | 88 | 0 | 0 | 620 | 256 | 342 | 5784 | 2869 | 0 | 132 | 26 | 212 | 317 | 185 | 53 | 185 | 159 | 0 | 21 | 29 |
| SRR6454270 | 0 | 0 | 14 | 0 | 43 | 0 | 0 | 1329 | 506 | 1906 | 85 | 51 | 0 | 29 | 69 | 47 | 42 | 68 | 0 | 11 | 2002 | 267 | 557 | 7149 | 3979 | 0 | 84 | 56 | 112 | 139 | 139 | 42 | 125 | 125 | 14 | 30 | 35 |
| SRR6454271 | 0 | 0 | 19 | 0 | 127 | 0 | 0 | 247 | 148 | 1803 | 33 | 53 | 0 | 43 | 246 | 209 | 0 | 345 | 0 | 409 | 302 | 303 | 198 | 6458 | 2898 | 0 | 22 | 32 | 216 | 687 | 481 | 229 | 356 | 356 | 48 | 814 | 27 |
| SRR6454272 | 0 | 24 | 127 | 0 | 268 | 0 | 0 | 145 | 66 | 1200 | 20 | 21 | 0 | 104 | 279 | 333 | 16 | 447 | 0 | 133 | 948 | 576 | 443 | 8820 | 2693 | 14 | 159 | 48 | 294 | 595 | 420 | 238 | 286 | 151 | 103 | 190 | 21 |
| SRR6454273 | 0 | 0 | 53 | 0 | 113 | 0 | 0 | 117 | 40 | 1610 | 35 | 164 | 0 | 196 | 160 | 327 | 0 | 199 | 0 | 541 | 1837 | 3371 | 0 | 515 | 12388 | 1573 | 18 | 0 | 35 | 204 | 177 | 44 | 151 | 328 | 80 | 3082 | 0 |
| SRR6454274 | 0 | 19 | 48 | 0 | 111 | 0 | 0 | 161 | 39 | 3155 | 125 | 315 | 0 | 171 | 108 | 269 | 0 | 268 | 0 | 1258 | 2165 | 3782 | 0 | 484 | 10558 | 2006 | 19 | 0 | 76 | 247 | 67 | 67 | 76 | 219 | 105 | 8031 | 0 |
| SRR6454275 | 0 | 0 | 48 | 0 | 281 | 0 | 16 | 304 | 233 | 3308 | 11 | 22 | 0 | 145 | 45 | 99 | 0 | 502 | 0 | 47 | 795 | 965 | 304 | 9002 | 3302 | 0 | 11 | 0 | 101 | 346 | 133 | 128 | 272 | 410 | 43 | 132 | 16 |
| SRR6454276 | 0 | 0 | 20 | 0 | 246 | 10 | 0 | 1102 | 437 | 4279 | 18 | 70 | 0 | 157 | 725 | 466 | 0 | 211 | 0 | 192 | 1861 | 933 | 343 | 12217 | 2576 | 0 | 55 | 25 | 196 | 593 | 493 | 146 | 377 | 558 | 25 | 660 | 0 |
| SRR6454277 | 0 | 0 | 30 | 0 | 190 | 44 | 20 | 422 | 100 | 2556 | 16 | 92 | 0 | 63 | 417 | 440 | 49 | 240 | 0 | 185 | 573 | 1259 | 289 | 12919 | 5216 | 0 | 15 | 0 | 109 | 138 | 183 | 25 | 183 | 376 | 35 | 485 | 23 |
| SRR6454278 | 0 | 0 | 50 | 0 | 248 | 10 | 0 | 269 | 191 | 1665 | 13 | 15 | 0 | 54 | 141 | 110 | 10 | 373 | 0 | 37 | 456 | 283 | 200 | 7807 | 2902 | 0 | 45 | 25 | 206 | 869 | 603 | 146 | 442 | 477 | 50 | 117 | 0 |
| SRR6454279 | 24 | 0 | 19 | 0 | 43 | 29 | 0 | 136 | 100 | 2917 | 123 | 181 | 0 | 38 | 0 | 0 | 14 | 301 | 0 | 12 | 576 | 725 | 226 | 9973 | 4679 | 0 | 140 | 48 | 111 | 275 | 203 | 63 | 174 | 121 | 19 | 45 | 18 |
| SRR6454280 | 23 | 0 | 19 | 0 | 43 | 19 | 11 | 1220 | 479 | 4088 | 68 | 127 | 0 | 155 | 0 | 10 | 27 | 248 | 0 | 118 | 892 | 670 | 418 | 7136 | 2848 | 17 | 191 | 0 | 73 | 187 | 88 | 42 | 54 | 92 | 0 | 299 | 12 |
| SRR6454281 | 0 | 0 | 12 | 0 | 49 | 0 | 0 | 567 | 215 | 1377 | 118 | 58 | 0 | 48 | 0 | 0 | 149 | 409 | 0 | 19 | 584 | 469 | 350 | 5111 | 2528 | 0 | 78 | 20 | 255 | 640 | 310 | 137 | 192 | 149 | 24 | 46 | 13 |
| SRR6454282 | 0 | 14 | 14 | 0 | 54 | 0 | 0 | 100 | 107 | 761 | 94 | 110 | 0 | 73 | 0 | 12 | 0 | 539 | 0 | 33 | 494 | 340 | 266 | 5607 | 2460 | 0 | 267 | 23 | 431 | 646 | 332 | 89 | 154 | 140 | 0 | 108 | 10 |

### Supp Table S8.4.Columns_DI_to_ES

| | | | | | | | | | | | | | | | | | | | | | | | | | | | | | | | | | | | | | |
|---|---|---|---|---|---|---|---|---|---|---|---|---|---|---|---|---|---|---|---|---|---|---|---|---|---|---|---|---|---|---|---|---|---|---|---|---|---|
| SRR6454283 | 14 | 0 | 28 | 0 | 54 | 0 | 0 | 458 | 260 | 3108 | 92 | 67 | 0 | 67 | 0 | 0 | 19 | 392 | 0 | 0 | 639 | 574 | 252 | 4865 | 2727 | 0 | 51 | 23 | 201 | 588 | 327 | 149 | 168 | 154 | 0 | 28 | 10 |
| SRR6454284 | 0 | 43 | 16 | 0 | 44 | 0 | 0 | 608 | 272 | 980 | 186 | 222 | 0 | 66 | 16 | 11 | 0 | 415 | 0 | 51 | 3301 | 771 | 417 | 7653 | 2614 | 0 | 108 | 27 | 627 | 962 | 438 | 124 | 200 | 184 | 22 | 232 | 18 |
| SRR6454285 | 25 | 0 | 20 | 0 | 33 | 10 | 0 | 59 | 75 | 2305 | 34 | 94 | 0 | 258 | 0 | 10 | 15 | 341 | 0 | 18 | 705 | 1139 | 59 | 3876 | 4204 | 45 | 10 | 0 | 15 | 25 | 10 | 10 | 20 | 30 | 15 | 61 | 0 |
| SRR6454286 | 0 | 33 | 15 | 0 | 44 | 0 | 0 | 887 | 360 | 1191 | 58 | 76 | 0 | 58 | 41 | 29 | 12 | 398 | 0 | 81 | 2925 | 532 | 463 | 9983 | 1645 | 0 | 105 | 33 | 180 | 678 | 510 | 132 | 243 | 249 | 33 | 244 | 0 |
| SRR6454287 | 0 | 27 | 162 | 0 | 121 | 0 | 0 | 1286 | 0 | 681 | 10 | 3706 | 0 | 0 | 605 | 378 | 0 | 107 | 0 | 681 | 1405 | 5040 | 0 | 353 | 11164 | 1431 | 27 | 0 | 81 | 243 | 54 | 27 | 162 | 324 | 54 | 1966 | 0 |
| SRR6454288 | 0 | 0 | 41 | 0 | 111 | 0 | 0 | 764 | 45 | 180 | 989 | 3371 | 0 | 45 | 764 | 944 | 0 | 144 | 0 | 944 | 1609 | 5733 | 0 | 405 | 12768 | 1854 | 0 | 0 | 83 | 165 | 206 | 41 | 124 | 702 | 0 | 1753 | 0 |
| SRR6454289 | 0 | 17 | 34 | 0 | 80 | 0 | 0 | 977 | 126 | 662 | 73 | 6931 | 0 | 0 | 851 | 1418 | 17 | 222 | 0 | 693 | 2080 | 5610 | 0 | 492 | 13170 | 1818 | 17 | 0 | 34 | 336 | 34 | 0 | 84 | 235 | 151 | 2804 | 0 |
| SRR6454290 | 0 | 0 | 100 | 0 | 121 | 0 | 0 | 730 | 209 | 709 | 36 | 3045 | 0 | 63 | 730 | 667 | 0 | 371 | 0 | 897 | 1738 | 4405 | 0 | 287 | 15300 | 968 | 0 | 22 | 112 | 11 | 145 | 22 | 179 | 16 | 45 | 5026 | 0 |
| SRR6454291 | 0 | 30 | 10 | 0 | 11 | 10 | 0 | 626 | 54 | 227 | 8145 | 16647 | 0 | 11 | 0 | 43 | 10 | 479 | 0 | 227 | 2706 | 3549 | 0 | 375 | 12033 | 2021 | 40 | 0 | 111 | 575 | 161 | 40 | 40 | 141 | 0 | 842 | 0 |
| SRR6454292 | 0 | 59 | 20 | 0 | 18 | 0 | 0 | 357 | 43 | 100 | 818 | 21131 | 0 | 29 | 0 | 29 | 20 | 443 | 0 | 143 | 2618 | 3055 | 0 | 421 | 11978 | 1804 | 167 | 0 | 98 | 451 | 118 | 0 | 69 | 128 | 98 | 1013 | 0 |
| SRR6454293 | 0 | 0 | 35 | 0 | 11 | 35 | 0 | 393 | 112 | 168 | 9141 | 19179 | 0 | 0 | 0 | 56 | 0 | 168 | 0 | 168 | 1375 | 3547 | 0 | 240 | 9582 | 1625 | 0 | 70 | 140 | 280 | 35 | 0 | 140 | 210 | 35 | 673 | 0 |
| SRR6454294 | 0 | 24 | 49 | 0 | 0 | 0 | 0 | 137 | 92 | 115 | 3620 | 13310 | 0 | 23 | 46 | 69 | 24 | 196 | 0 | 206 | 1459 | 3867 | 0 | 349 | 10037 | 2896 | 24 | 24 | 293 | 854 | 220 | 49 | 146 | 171 | 24 | 916 | 0 |
| SRR6454295 | 0 | 13 | 26 | 0 | 16 | 13 | 0 | 412 | 120 | 306 | 626 | 18741 | 0 | 13 | 0 | 13 | 13 | 393 | 0 | 186 | 2047 | 3740 | 0 | 206 | 11934 | 1120 | 39 | 53 | 118 | 499 | 144 | 13 | 79 | 131 | 66 | 865 | 0 |
| SRR6454296 | 0 | 36 | 53 | 0 | 13 | 0 | 0 | 411 | 73 | 508 | 408 | 21574 | 0 | 0 | 24 | 0 | 18 | 333 | 0 | 145 | 2148 | 3710 | 0 | 212 | 12852 | 1034 | 142 | 0 | 71 | 338 | 125 | 18 | 53 | 107 | 18 | 532 | 0 |
| SRR6454298 | 0 | 0 | 23 | 0 | 41 | 0 | 0 | 5369 | 1182 | 3728 | 130 | 276 | 0 | 771 | 0 | 0 | 0 | 241 | 0 | 94 | 1629 | 305 | 611 | 9777 | 2202 | 0 | 16 | 23 | 272 | 279 | 155 | 47 | 85 | 62 | 0 | 1277 | 0 |
| SRR6454299 | 0 | 29 | 0 | 0 | 13 | 0 | 0 | 329 | 60 | 259 | 587 | 24363 | 0 | 20 | 0 | 10 | 0 | 236 | 17 | 468 | 1574 | 2931 | 0 | 1109 | 18418 | 716 | 43 | 0 | 14 | 200 | 64 | 0 | 14 | 29 | 0 | 1126 | 0 |
| SRR6454300 | 0 | 22 | 0 | 0 | 25 | 0 | 0 | 208 | 50 | 108 | 1834 | 20539 | 0 | 25 | 17 | 13 | 27 | 402 | 0 | 333 | 2211 | 2360 | 0 | 836 | 20945 | 532 | 65 | 0 | 51 | 266 | 75 | 0 | 0 | 0 | 0 | 950 | 0 |
| SRR6454301 | 0 | 0 | 21 | 0 | 49 | 13 | 0 | 572 | 233 | 3328 | 110 | 202 | 0 | 50 | 24 | 20 | 26 | 380 | 0 | 58 | 536 | 679 | 207 | 8460 | 3428 | 0 | 111 | 21 | 183 | 474 | 269 | 115 | 277 | 273 | 17 | 140 | 15 |
| SRR6454302 | 16 | 210 | 0 | 0 | 55 | 0 | 0 | 4410 | 1179 | 3746 | 173 | 520 | 0 | 210 | 138 | 95 | 19 | 375 | 0 | 198 | 3213 | 1365 | 343 | 12274 | 3331 | 0 | 35 | 0 | 140 | 323 | 210 | 23 | 113 | 140 | 31 | 753 | 16 |
| SRR6454303 | 22 | 0 | 20 | 0 | 46 | 0 | 0 | 586 | 257 | 1430 | 173 | 124 | 0 | 79 | 21 | 27 | 22 | 386 | 0 | 34 | 1072 | 1115 | 304 | 7386 | 3642 | 0 | 20 | 0 | 148 | 256 | 165 | 39 | 66 | 106 | 0 | 197 | 18 |
| SRR6454304 | 24 | 0 | 38 | 0 | 35 | 0 | 0 | 185 | 211 | 2237 | 63 | 168 | 0 | 330 | 24 | 36 | 94 | 349 | 0 | 58 | 2319 | 1066 | 363 | 12429 | 3012 | 0 | 132 | 24 | 64 | 86 | 67 | 19 | 32 | 99 | 0 | 191 | 0 |
| SRR6454305 | 16 | 0 | 16 | 0 | 36 | 27 | 11 | 106 | 120 | 4060 | 133 | 357 | 0 | 157 | 0 | 0 | 38 | 383 | 0 | 29 | 907 | 1800 | 87 | 6257 | 5390 | 0 | 215 | 11 | 59 | 75 | 65 | 38 | 65 | 113 | 11 | 85 | 0 |
| SRR6454306 | 20 | 0 | 20 | 0 | 25 | 10 | 0 | 257 | 155 | 3283 | 112 | 179 | 0 | 163 | 0 | 0 | 97 | 463 | 0 | 38 | 1776 | 1065 | 317 | 10831 | 3401 | 10 | 4880 | 0 | 71 | 110 | 48 | 24 | 27 | 54 | 10 | 77 | 12 |
| SRR6454307 | 12 | 0 | 17 | 0 | 34 | 17 | 0 | 359 | 137 | 2130 | 123 | 249 | 0 | 58 | 13 | 0 | 14 | 351 | 0 | 22 | 516 | 651 | 158 | 5736 | 3879 | 0 | 59 | 21 | 71 | 217 | 161 | 57 | 106 | 106 | 12 | 97 | 14 |
| SRR6454308 | 11 | 41 | 30 | 0 | 31 | 0 | 11 | 239 | 144 | 568 | 160 | 200 | 0 | 78 | 0 | 16 | 25 | 485 | 0 | 69 | 2743 | 1113 | 263 | 9289 | 4023 | 0 | 41 | 28 | 138 | 251 | 144 | 19 | 55 | 97 | 14 | 525 | 20 |
| SRR6454309 | 13 | 0 | 70 | 0 | 44 | 16 | 0 | 221 | 150 | 5392 | 100 | 369 | 0 | 65 | 54 | 56 | 25 | 778 | 0 | 599 | 1310 | 1461 | 215 | 7588 | 5257 | 0 | 193 | 22 | 162 | 422 | 282 | 76 | 152 | 216 | 44 | 2011 | 18 |
| SRR6454310 | 0 | 88 | 29 | 0 | 28 | 0 | 0 | 2593 | 965 | 1313 | 368 | 333 | 0 | 119 | 191 | 151 | 18 | 776 | 0 | 143 | 6901 | 1484 | 372 | 12340 | 4040 | 0 | 25 | 23 | 168 | 263 | 129 | 29 | 62 | 109 | 64 | 758 | 18 |
| SRR6454311 | 0 | 11 | 43 | 0 | 17 | 0 | 0 | 1066 | 164 | 2439 | 108 | 7090 | 0 | 61 | 41 | 20 | 0 | 162 | 0 | 369 | 1619 | 4149 | 0 | 229 | 10738 | 1279 | 22 | 11 | 87 | 433 | 205 | 22 | 108 | 249 | 11 | 861 | 0 |
| SRR6454312 | 0 | 18 | 13 | 0 | 18 | 0 | 0 | 752 | 101 | 932 | 1408 | 13342 | 0 | 22 | 0 | 39 | 57 | 378 | 0 | 219 | 2176 | 3960 | 0 | 297 | 13904 | 1701 | 299 | 0 | 106 | 213 | 112 | 0 | 66 | 130 | 79 | 848 | 0 |
| SRR6454313 | 0 | 0 | 70 | 0 | 17 | 0 | 0 | 945 | 138 | 355 | 129 | 9000 | 0 | 39 | 177 | 98 | 0 | 174 | 0 | 571 | 1567 | 4264 | 0 | 267 | 12127 | 1635 | 0 | 23 | 58 | 197 | 70 | 12 | 35 | 116 | 23 | 1024 | 0 |
| SRR6454314 | 0 | 0 | 23 | 0 | 13 | 0 | 0 | 396 | 126 | 696 | 2929 | 12729 | 0 | 36 | 18 | 48 | 54 | 187 | 0 | 96 | 1475 | 4938 | 0 | 291 | 13662 | 2206 | 208 | 0 | 46 | 178 | 39 | 15 | 39 | 54 | 77 | 504 | 0 |
| SRR6454315 | 0 | 13 | 45 | 0 | 0 | 0 | 0 | 819 | 248 | 124 | 3418 | 12749 | 0 | 0 | 0 | 0 | 13 | 237 | 0 | 113 | 2087 | 4183 | 0 | 458 | 11623 | 2299 | 147 | 19 | 250 | 749 | 166 | 19 | 122 | 237 | 58 | 501 | 0 |
| SRR6454316 | 0 | 0 | 68 | 0 | 0 | 0 | 0 | 1295 | 116 | 146 | 2651 | 8925 | 0 | 15 | 0 | 23 | 17 | 164 | 0 | 200 | 1728 | 4108 | 0 | 380 | 10809 | 1614 | 77 | 0 | 188 | 11 | 154 | 17 | 43 | 145 | 34 | 370 | 0 |
| SRR6454317 | 0 | 20 | 0 | 0 | 0 | 0 | 0 | 1101 | 120 | 235 | 2907 | 6178 | 0 | 53 | 19 | 96 | 0 | 192 | 0 | 110 | 2274 | 4466 | 0 | 199 | 10045 | 1702 | 0 | 13 | 145 | 482 | 211 | 20 | 66 | 152 | 13 | 254 | 0 |
| SRR6454318 | 0 | 32 | 54 | 0 | 0 | 0 | 0 | 780 | 57 | 436 | 450 | 9416 | 0 | 80 | 11 | 23 | 21 | 188 | 0 | 183 | 2024 | 5049 | 0 | 236 | 11382 | 1974 | 21 | 21 | 75 | 418 | 161 | 21 | 11 | 129 | 21 | 310 | 0 |
| SRR6454319 | 0 | 45 | 67 | 0 | 0 | 0 | 15 | 701 | 62 | 124 | 1128 | 14306 | 0 | 0 | 0 | 0 | 0 | 200 | 0 | 176 | 2359 | 3613 | 0 | 519 | 9678 | 1476 | 156 | 0 | 350 | 1139 | 246 | 0 | 97 | 223 | 67 | 667 | 0 |
| SRR6454320 | 11 | 23 | 46 | 0 | 0 | 0 | 11 | 872 | 109 | 336 | 220 | 9968 | 0 | 0 | 0 | 55 | 0 | 156 | 0 | 182 | 1708 | 4431 | 0 | 299 | 12760 | 1689 | 23 | 34 | 194 | 12 | 274 | 57 | 148 | 308 | 0 | 327 | 0 |
| SRR6454321 | 0 | 30 | 60 | 0 | 0 | 0 | 0 | 862 | 177 | 583 | 144 | 7986 | 0 | 51 | 177 | 279 | 15 | 108 | 0 | 583 | 1670 | 4767 | 0 | 395 | 11682 | 1899 | 15 | 15 | 45 | 272 | 106 | 0 | 91 | 227 | 121 | 811 | 0 |
| SRR6454322 | 0 | 0 | 53 | 0 | 0 | 0 | 0 | 648 | 169 | 563 | 155 | 10164 | 0 | 28 | 84 | 113 | 0 | 114 | 0 | 1014 | 1588 | 4502 | 0 | 388 | 11055 | 1674 | 40 | 0 | 40 | 186 | 106 | 13 | 53 | 199 | 93 | 1520 | 0 |
| SRR6454323 | 0 | 14 | 0 | 0 | 0 | 0 | 0 | 924 | 240 | 1289 | 3571 | 10325 | 0 | 80 | 46 | 91 | 19 | 169 | 0 | 194 | 1701 | 5097 | 0 | 437 | 12058 | 1290 | 162 | 0 | 70 | 241 | 56 | 0 | 28 | 139 | 139 | 1141 | 0 |
| SRR6454324 | 0 | 30 | 72 | 0 | 0 | 0 | 0 | 1570 | 316 | 552 | 2438 | 6122 | 0 | 32 | 55 | 142 | 0 | 157 | 0 | 260 | 2041 | 4186 | 0 | 517 | 10781 | 1525 | 12 | 0 | 60 | 332 | 199 | 18 | 97 | 235 | 18 | 465 | 0 |
| SRR6454325 | 16 | 0 | 75 | 0 | 44 | 0 | 0 | 295 | 115 | 1787 | 87 | 173 | 0 | 115 | 41 | 47 | 13 | 464 | 0 | 44 | 609 | 535 | 248 | 6388 | 3915 | 0 | 55 | 42 | 111 | 251 | 146 | 85 | 169 | 267 | 23 | 87 | 18 |
| SRR6454326 | 19 | 32 | 20 | 0 | 34 | 12 | 0 | 217 | 110 | 864 | 939 | 1768 | 0 | 208 | 397 | 245 | 18 | 1207 | 0 | 322 | 1986 | 1831 | 312 | 10330 | 5041 | 0 | 37 | 0 | 69 | 105 | 80 | 19 | 34 | 73 | 34 | 1745 | 26 |
| SRR6454331 | 0 | 32 | 64 | 0 | 0 | 0 | 0 | 635 | 159 | 278 | 370 | 10802 | 0 | 26 | 53 | 159 | 11 | 203 | 0 | 675 | 1992 | 4346 | 0 | 280 | 12366 | 1157 | 0 | 0 | 107 | 246 | 96 | 0 | 96 | 214 | 53 | 1231 | 0 |
| SRR6454332 | 0 | 11 | 56 | 0 | 0 | 0 | 0 | 667 | 92 | 192 | 5909 | 19671 | 0 | 28 | 28 | 64 | 11 | 310 | 0 | 270 | 2542 | 4468 | 0 | 305 | 12919 | 1426 | 113 | 11 | 79 | 316 | 68 | 11 | 11 | 147 | 147 | 951 | 0 |
| SRR6454335 | 0 | 0 | 0 | 0 | 0 | 0 | 0 | 679 | 136 | 645 | 151 | 10594 | 0 | 102 | 0 | 0 | 0 | 263 | 24 | 68 | 1016 | 4867 | 22 | 824 | 14099 | 1111 | 34 | 0 | 239 | 649 | 341 | 34 | 171 | 0 | 0 | 1121 | 0 |
| SRR6454337 | 0 | 64 | 23 | 0 | 59 | 0 | 0 | 2241 | 1137 | 3848 | 121 | 227 | 0 | 795 | 0 | 0 | 0 | 166 | 0 | 213 | 1873 | 666 | 466 | 12745 | 2992 | 0 | 55 | 0 | 164 | 241 | 155 | 27 | 27 | 27 | 0 | 2121 | 0 |
| SRR6454338 | 0 | 31 | 0 | 0 | 54 | 0 | 15 | 2559 | 538 | 1577 | 97 | 235 | 0 | 287 | 0 | 0 | 10 | 281 | 0 | 192 | 1648 | 446 | 385 | 8074 | 2106 | 0 | 113 | 26 | 510 | 659 | 309 | 72 | 52 | 31 | 0 | 3099 | 0 |
| SRR6454339 | 0 | 15 | 0 | 0 | 51 | 0 | 0 | 3337 | 912 | 3233 | 53 | 156 | 0 | 131 | 16 | 11 | 0 | 168 | 0 | 274 | 1107 | 366 | 305 | 7860 | 3030 | 0 | 21 | 0 | 175 | 285 | 166 | 34 | 40 | 34 | 0 | 3695 | 0 |
| SRR6454340 | 0 | 342 | 0 | 0 | 32 | 0 | 0 | 746 | 172 | 453 | 1860 | 1796 | 0 | 706 | 0 | 11 | 0 | 379 | 0 | 178 | 5055 | 1370 | 473 | 12822 | 3086 | 0 | 39 | 39 | 56 | 95 | 50 | 0 | 0 | 11 | 0 | 821 | 0 |
| SRR6454341 | 0 | 55 | 0 | 0 | 22 | 0 | 0 | 2822 | 737 | 1015 | 83 | 190 | 0 | 198 | 18 | 11 | 0 | 321 | 0 | 310 | 4018 | 503 | 462 | 7989 | 1504 | 0 | 33 | 0 | 118 | 229 | 137 | 18 | 22 | 33 | 0 | 8769 | 0 |
| SRR6454342 | 0 | 101 | 0 | 0 | 21 | 0 | 0 | 1826 | 385 | 532 | 186 | 225 | 0 | 342 | 0 | 0 | 17 | 434 | 0 | 205 | 3169 | 693 | 365 | 8202 | 2379 | 0 | 17 | 0 | 52 | 101 | 68 | 0 | 0 | 0 | 0 | 5942 | 0 |
| SRR6454343 | 0 | 90 | 0 | 0 | 36 | 0 | 0 | 2009 | 578 | 871 | 556 | 417 | 0 | 53 | 0 | 0 | 0 | 495 | 0 | 574 | 5874 | 726 | 451 | 10759 | 2655 | 0 | 43 | 14 | 158 | 314 | 220 | 20 | 64 | 80 | 0 | 17045 | 0 |
| SRR6454345 | 0 | 17 | 0 | 0 | 11 | 0 | 0 | 221 | 14 | 262 | 3118 | 10734 | 0 | 0 | 0 | 0 | 33 | 100 | 11 | 276 | 979 | 3704 | 0 | 673 | 22176 | 1015 | 107 | 0 | 25 | 355 | 182 | 0 | 17 | 41 | 0 | 428 | 0 |
| SRR6454347 | 0 | 0 | 0 | 0 | 0 | 0 | 0 | 2573 | 288 | 828 | 1037 | 10550 | 0 | 13 | 0 | 0 | 83 | 179 | 0 | 85 | 1398 | 4315 | 0 | 338 | 21045 | 636 | 26 | 21 | 263 | 1233 | 456 | 13 | 128 | 0 | 0 | 270 | 0 |
| SRR6454348 | 0 | 19 | 0 | 0 | 0 | 0 | 0 | 1873 | 624 | 1117 | 2486 | 11512 | 0 | 155 | 0 | 0 | 57 | 706 | 14 | 337 | 2303 | 5816 | 0 | 330 | 17723 | 1062 | 48 | 0 | 304 | 1074 | 313 | 16 | 98 | 0 | 0 | 875 | 0 |
| SRR6454349 | 0 | 0 | 0 | 0 | 0 | 0 | 0 | 1763 | 410 | 1141 | 830 | 9627 | 0 | 275 | 0 | 0 | 0 | 268 | 0 | 550 | 1826 | 5960 | 0 | 227 | 10825 | 1800 | 0 | 0 | 188 | 781 | 309 | 33 | 97 | 0 | 0 | 1674 | 0 |
| SRR6454350 | 0 | 0 | 0 | 0 | 0 | 0 | 0 | 1108 | 533 | 645 | 5471 | 14210 | 0 | 267 | 0 | 0 | 0 | 524 | 0 | 533 | 1786 | 7025 | 0 | 425 | 18192 | 1434 | 114 | 0 | 85 | 256 | 57 | 0 | 28 | 0 | 0 | 758 | 0 |
| SRR6454351 | 0 | 49 | 0 | 0 | 48 | 0 | 0 | 222 | 145 | 292 | 82 | 69 | 0 | 101 | 0 | 0 | 0 | 716 | 0 | 612 | 3136 | 785 | 442 | 13444 | 5359 | 0 | 0 | 0 | 98 | 195 | 122 | 24 | 24 | 0 | 0 | 11181 | 38 |
| SRR6454352 | 0 | 14 | 0 | 0 | 75 | 0 | 0 | 1375 | 529 | 1508 | 16 | 20 | 0 | 63 | 0 | 0 | 0 | 447 | 0 | 387 | 1163 | 485 | 468 | 18931 | 7979 | 0 | 10 | 0 | 118 | 463 | 539 | 31 | 107 | 0 | 0 | 6356 | 27 |
| SRR6454354 | 0 | 0 | 0 | 0 | 58 | 0 | 0 | 1408 | 381 | 1822 | 22 | 18 | 0 | 49 | 0 | 0 | 0 | 300 | 0 | 681 | 1431 | 431 | 410 | 14075 | 7038 | 0 | 12 | 0 | 74 | 263 | 263 | 0 | 70 | 0 | 0 | 10966 | 19 |
| SRR6496456 | 0 | 53 | 0 | 0 | 63 | 0 | 20 | 31599 | 4502 | 522 | 1192 | 4494 | 0 | 61 | 0 | 13 | 101 | 4512 | 97 | 71 | 957 | 4148 | 108 | 28098 | 8282 | 0 | 11 | 0 | 15 | 14 | 16 | 0 | 10 | 16 | 0 | 344 | 5049 |
| SRR6496457 | 0 | 48 | 0 | 0 | 79 | 0 | 15 | 16163 | 4241 | 704 | 925 | 2945 | 0 | 43 | 0 | 12 | 113 | 5059 | 94 | 140 | 713 | 4281 | 120 | 27192 | 8178 | 0 | 0 | 0 | 0 | 13 | 14 | 0 | 0 | 0 | 0 | 1296 | 4895 |
| SRR6496458 | 0 | 129 | 0 | 0 | 50 | 0 | 27 | 26009 | 3919 | 868 | 1307 | 4141 | 0 | 48 | 0 | 0 | 151 | 3975 | 66 | 139 | 563 | 3654 | 114 | 25974 | 9081 | 0 | 0 | 0 | 18 | 36 | 36 | 0 | 0 | 22 | 0 | 712 | 5724 |
| SRR6496459 | 0 | 72 | 0 | 0 | 51 | 0 | 15 | 11559 | 3043 | 300 | 1100 | 9098 | 0 | 18 | 0 | 0 | 188 | 4362 | 86 | 109 | 858 | 3651 | 119 | 28964 | 9050 | 0 | 13 | 0 | 0 | 13 | 18 | 0 | 0 | 0 | 0 | 582 | 4360 |
| SRR6496460 | 0 | 121 | 0 | 0 | 61 | 0 | 30 | 18820 | 2779 | 639 | 5997 | 9688 | 0 | 42 | 0 | 0 | 191 | 3653 | 74 | 196 | 628 | 3469 | 129 | 31489 | 12419 | 0 | 11 | 0 | 13 | 33 | 31 | 0 | 0 | 0 | 0 | 439 | 4515 |
| SRR6496461 | 0 | 36 | 17 | 0 | 26 | 0 | 0 | 2623 | 48 | 3398 | 398 | 281 | 0 | 528 | 790 | 766 | 24 | 962 | 16 | 1682 | 1044 | 4675 | 352 | 106731 | 11707 | 68 | 0 | 0 | 0 | 10 | 11 | 0 | 17 | 36 | 41 | 10558 | 248 |
| SRR6496462 | 0 | 44 | 11 | 0 | 26 | 14 | 0 | 2399 | 78 | 3333 | 516 | 272 | 0 | 491 | 740 | 885 | 14 | 965 | 12 | 1683 | 981 | 4596 | 352 | 104967 | 12107 | 76 | 0 | 0 | 0 | 16 | 10 | 0 | 20 | 52 | 53 | 10669 | 257 |
| SRR6496463 | 0 | 57 | 23 | 0 | 25 | 0 | 0 | 2475 | 70 | 3466 | 458 | 368 | 0 | 503 | 722 | 871 | 21 | 882 | 14 | 1688 | 1049 | 4466 | 377 | 107176 | 12487 | 57 | 0 | 0 | 0 | 13 | 11 | 0 | 14 | 46 | 32 | 10218 | 258 |
| SRR6496464 | 0 | 16 | 17 | 0 | 29 | 11 | 0 | 2660 | 97 | 3493 | 521 | 268 | 0 | 531 | 755 | 872 | 10 | 954 | 16 | 1710 | 1044 | 4655 | 350 | 105781 | 11984 | 62 | 14 | 0 | 0 | 16 | 13 | 0 | 17 | 18 | 19 | 10346 | 258 |
| SRR6496465 | 0 | 41 | 21 | 0 | 10 | 10 | 0 | 2607 | 43 | 3710 | 387 | 201 | 0 | 759 | 759 | 974 | 24 | 913 | 10 | 1547 | 1069 | 4643 | 368 | 107170 | 12805 | 76 | 0 | 0 | 0 | 14 | 21 | 0 | 28 | 38 | 73 | 10486 | 237 |
| SRR6498854 | 0 | 41 | 18 | 0 | 0 | 0 | 0 | 1313 | 1762 | 8262 | 4377 | 3513 | 0 | 7069 | 131 | 55 | 0 | 203 | 0 | 306 | 390 | 3871 | 19 | 32247 | 3828 | 164 | 10 | 22 | 16 | 19 | 43 | 14 | 30 | 43 | 26 | 2407 | 0 |
| SRR6498855 | 0 | 19 | 12 | 0 | 0 | 0 | 0 | 682 | 960 | 6456 | 10706 | 7270 | 0 | 9857 | 132 | 111 | 0 | 181 | 0 | 334 | 402 | 5254 | 23 | 36063 | 2047 | 94 | 0 | 61 | 35 | 19 | 11 | 15 | 12 | 23 | 10 | 2045 | 0 |
| SRR6498856 | 0 | 43 | 33 | 0 | 0 | 0 | 0 | 777 | 1226 | 9398 | 3028 | 3409 | 0 | 8372 | 213 | 45 | 0 | 108 | 0 | 268 | 439 | 4660 | 27 | 33614 | 4573 | 236 | 0 | 35 | 26 | 81 | 34 | 0 | 42 | 83 | 46 | 2116 | 0 |
| SRR6498857 | 0 | 12 | 23 | 0 | 0 | 0 | 0 | 706 | 1055 | 4448 | 9009 | 6939 | 0 | 8400 | 187 | 24 | 0 | 316 | 0 | 414 | 425 | 5376 | 23 | 31332 | 2801 | 96 | 0 | 51 | 12 | 16 | 14 | 14 | 11 | 24 | 55 | 1859 | 0 |
| SRR6498858 | 0 | 73 | 17 | 0 | 0 | 0 | 0 | 846 | 1018 | 4012 | 827 | 4059 | 0 | 6237 | 195 | 41 | 0 | 245 | 11 | 633 | 465 | 4651 | 24 | 27108 | 2962 | 202 | 10 | 36 | 25 | 109 | 61 | 13 | 79 | 136 | 64 | 3586 | 0 |
| SRR6498860 | 0 | 65 | 27 | 0 | 0 | 0 | 0 | 958 | 1100 | 7808 | 7899 | 5972 | 0 | 3571 | 151 | 30 | 0 | 160 | 0 | 444 | 580 | 4072 | 13 | 29679 | 2757 | 173 | 13 | 19 | 19 | 100 | 86 | 0 | 25 | 83 | 21 | 2663 | 0 |
| SRR6498861 | 0 | 23 | 23 | 0 | 0 | 0 | 0 | 1029 | 859 | 4526 | 412 | 4078 | 0 | 3171 | 97 | 36 | 0 | 128 | 0 | 218 | 249 | 1846 | 22 | 73345 | 4448 | 28 | 20 | 52 | 39 | 200 | 173 | 13 | 52 | 167 | 0 | 2989 | 0 |
| SRR6498862 | 0 | 44 | 17 | 0 | 0 | 0 | 0 | 1173 | 1043 | 5012 | 4896 | 4186 | 0 | 2926 | 232 | 29 | 0 | 215 | 0 | 348 | 658 | 3811 | 15 | 23903 | 4510 | 142 | 0 | 30 | 25 | 15 | 35 | 12 | 12 | 18 | 35 | 2462 | 0 |
| SRR6498863 | 0 | 25 | 25 | 0 | 0 | 0 | 0 | 1517 | 968 | 4088 | 6670 | 4873 | 0 | 2894 | 183 | 22 | 0 | 246 | 0 | 269 | 366 | 3840 | 26 | 39674 | 2874 | 39 | 11 | 11 | 34 | 110 | 97 | 16 | 53 | 94 | 39 | 3303 | 0 |
| SRR6498864 | 0 | 14 | 23 | 0 | 0 | 0 | 0 | 1361 | 1284 | 2568 | 3127 | 2555 | 0 | 1907 | 141 | 19 | 0 | 370 | 0 | 437 | 614 | 3820 | 12 | 24125 | 2880 | 128 | 26 | 36 | 24 | 36 | 25 | 23 | 35 | 34 | 14 | 4577 | 0 |
| SRR6498865 | 0 | 18 | 18 | 0 | 0 | 0 | 0 | 1789 | 2124 | 2906 | 76 | 4527 | 0 | 2906 | 56 | 0 | 0 | 281 | 0 | 559 | 452 | 4562 | 0 | 33370 | 2500 | 33 | 10 | 10 | 18 | 33 | 30 | 0 | 25 | 71 | 0 | 4919 | 0 |
| SRR6498866 | 0 | 11 | 25 | 0 | 12 | 0 | 0 | 845 | 1271 | 7672 | 4162 | 2962 | 0 | 7382 | 97 | 90 | 0 | 190 | 20 | 542 | 325 | 3518 | 24 | 25341 | 1964 | 230 | 17 | 21 | 31 | 13 | 11 | 10 | 31 | 16 | 14 | 2497 | 0 |
| SRR6498867 | 0 | 20 | 0 | 0 | 0 | 0 | 0 | 534 | 422 | 3744 | 19957 | 11974 | 0 | 8015 | 191 | 112 | 0 | 86 | 13 | 207 | 274 | 3050 | 29 | 28159 | 1157 | 91 | 0 | 18 | 13 | 35 | 25 | 0 | 41 | 14 | 47 | 1769 | 0 |
| SRR6498868 | 0 | 61 | 11 | 0 | 0 | 0 | 0 | 885 | 966 | 3388 | 5844 | 4168 | 0 | 7206 | 186 | 35 | 0 | 165 | 0 | 768 | 334 | 4049 | 0 | 18887 | 1287 | 273 | 0 | 18 | 26 | 60 | 24 | 0 | 44 | 97 | 47 | 5134 | 0 |
| SRR6498869 | 0 | 48 | 23 | 0 | 0 | 0 | 0 | 1024 | 878 | 3556 | 10468 | 7808 | 0 | 6756 | 229 | 18 | 0 | 171 | 0 | 238 | 327 | 4361 | 21 | 28741 | 1398 | 84 | 0 | 20 | 32 | 83 | 45 | 17 | 55 | 113 | 48 | 2030 | 0 |
| SRR6498870 | 0 | 24 | 14 | 0 | 0 | 0 | 0 | 423 | 484 | 2659 | 3297 | 2513 | 0 | 6853 | 158 | 48 | 0 | 294 | 13 | 484 | 356 | 2790 | 17 | 33151 | 1842 | 215 | 0 | 12 | 25 | 141 | 53 | 0 | 46 | 113 | 35 | 3856 | 0 |
| SRR6498871 | 0 | 39 | 0 | 0 | 0 | 0 | 0 | 1255 | 674 | 2430 | 11218 | 7382 | 0 | 11137 | 196 | 42 | 0 | 150 | 0 | 466 | 261 | 3936 | 17 | 17443 | 1135 | 63 | 0 | 27 | 26 | 81 | 43 | 0 | 89 | 151 | 24 | 2642 | 0 |
| SRR6705804 | 0 | 44 | 0 | 0 | 0 | 0 | 0 | 208 | 247 | 1397 | 37 | 232 | 0 | 170 | 1668 | 1166 | 0 | 1634 | 0 | 23 | 935 | 1596 | 56 | 1650 | 3091 | 11 | 0 | 0 | 0 | 0 | 0 | 0 | 0 | 0 | 0 | 69 | 39 |
| SRR6705805 | 0 | 49 | 0 | 0 | 14 | 0 | 0 | 104 | 124 | 1481 | 78 | 137 | 0 | 385 | 2056 | 1964 | 0 | 918 | 0 | 26 | 556 | 579 | 102 | 1605 | 2415 | 0 | 0 | 0 | 0 | 0 | 0 | 0 | 0 | 0 | 0 | 33 | 0 |
| SRR6705806 | 0 | 0 | 0 | 0 | 0 | 0 | 0 | 0 | 0 | 2037 | 0 | 0 | 0 | 0 | 0 | 2037 | 0 | 0 | 0 | 0 | 2451 | 1225 | 0 | 0 | 3676 | 0 | 0 | 0 | 0 | 0 | 0 | 0 | 0 | 0 | 0 | 0 | 0 |
| SRR6705808 | 0 | 96 | 0 | 0 | 13 | 0 | 0 | 375 | 447 | 3544 | 108 | 289 | 0 | 433 | 33 | 27 | 0 | 3845 | 24 | 137 | 2491 | 4860 | 173 | 1355 | 4049 | 0 | 27 | 0 | 0 | 27 | 32 | 21 | 0 | 0 | 0 | 408 | 16 |
| SRR6705809 | 0 | 0 | 0 | 0 | 16 | 0 | 0 | 3977 | 694 | 726 | 810 | 800 | 0 | 168 | 53 | 84 | 0 | 1888 | 23 | 53 | 708 | 1555 | 40 | 1005 | 2581 | 28 | 0 | 10 | 28 | 53 | 38 | 0 | 0 | 0 | 0 | 21 | 22 |
| SRR6705810 | 0 | 59 | 0 | 0 | 11 | 0 | 0 | 92 | 35 | 317 | 36 | 29 | 0 | 468 | 47 | 23 | 0 | 2142 | 13 | 73 | 986 | 1246 | 128 | 2636 | 7751 | 0 | 0 | 0 | 44 | 44 | 22 | 29 | 102 | 22 | 0 | 398 | 0 |
| SRR6705811 | 0 | 18 | 0 | 0 | 19 | 0 | 0 | 72 | 70 | 327 | 14 | 16 | 0 | 531 | 0 | 0 | 0 | 1028 | 0 | 0 | 258 | 257 | 58 | 829 | 738 | 0 | 0 | 0 | 0 | 0 | 0 | 0 | 0 | 0 | 0 | 38 | 0 |
| SRR6705812 | 0 | 224 | 0 | 0 | 20 | 0 | 0 | 68 | 171 | 2124 | 56 | 316 | 0 | 392 | 4829 | 3720 | 0 | 1454 | 30 | 26 | 1693 | 1112 | 166 | 2267 | 4219 | 21 | 0 | 0 | 0 | 0 | 0 | 0 | 0 | 0 | 0 | 111 | 36 |
| SRR6705813 | 0 | 242 | 0 | 0 | 15 | 0 | 0 | 102 | 133 | 1645 | 736 | 572 | 0 | 204 | 3361 | 2411 | 0 | 3280 | 33 | 61 | 2812 | 2033 | 127 | 1771 | 3877 | 0 | 0 | 0 | 0 | 0 | 0 | 0 | 0 | 0 | 0 | 102 | 58 |
| SRR6705814 | 0 | 41 | 13 | 0 | 67 | 0 | 0 | 150 | 214 | 2500 | 13 | 0 | 0 | 326 | 25 | 61 | 0 | 3834 | 29 | 69 | 638 | 168 | 344 | 2763 | 5628 | 13 | 0 | 0 | 10 | 0 | 0 | 0 | 0 | 0 | 0 | 313 | 43 |
| SRR6705815 | 0 | 1015 | 52 | 0 | 146 | 0 | 0 | 82 | 139 | 1547 | 17 | 12 | 0 | 163 | 389 | 1225 | 0 | 4182 | 12 | 497 | 2827 | 885 | 127 | 1014 | 3500 | 0 | 0 | 22 | 56 | 48 | 35 | 17 | 0 | 13 | 0 | 1784 | 175 |
| SRR6705816 | 0 | 340 | 0 | 0 | 27 | 0 | 0 | 115 | 102 | 1320 | 24 | 14 | 0 | 305 | 81 | 37 | 0 | 2965 | 25 | 312 | 2022 | 682 | 182 | 1771 | 3396 | 0 | 0 | 28 | 14 | 0 | 0 | 16 | 0 | 0 | 0 | 1337 | 40 |
| SRR6705817 | 0 | 48 | 0 | 0 | 54 | 0 | 0 | 84 | 234 | 3373 | 182 | 149 | 0 | 617 | 47 | 19 | 0 | 2000 | 36 | 79 | 1970 | 1855 | 32 | 612 | 5228 | 25 | 0 | 20 | 31 | 24 | 16 | 23 | 0 | 20 | 0 | 523 | 243 |
| SRR6705818 | 0 | 63 | 22 | 0 | 48 | 0 | 0 | 128 | 116 | 1497 | 101 | 74 | 0 | 303 | 209 | 489 | 0 | 3417 | 29 | 182 | 1909 | 1774 | 371 | 2337 | 4529 | 0 | 0 | 17 | 27 | 15 | 0 | 0 | 0 | 0 | 32 | 392 | 17 |
| SRR6705819 | 0 | 11 | 13 | 0 | 88 | 0 | 0 | 121 | 209 | 2685 | 253 | 211 | 0 | 196 | 1217 | 1365 | 0 | 3376 | 18 | 409 | 1468 | 1513 | 77 | 948 | 2899 | 40 | 36 | 44 | 92 | 49 | 31 | 36 | 51 | 36 | 64 | 804 | 36 |
| SRR6705820 | 0 | 311 | 0 | 0 | 62 | 0 | 0 | 40 | 66 | 1576 | 12 | 26 | 0 | 136 | 87 | 349 | 0 | 4312 | 32 | 147 | 4491 | 1980 | 164 | 1706 | 2751 | 0 | 0 | 0 | 0 | 0 | 0 | 0 | 0 | 0 | 0 | 470 | 30 |
| SRR6705821 | 0 | 365 | 66 | 0 | 150 | 0 | 0 | 193 | 335 | 3615 | 11 | 25 | 0 | 236 | 264 | 1217 | 0 | 7107 | 14 | 128 | 2128 | 787 | 249 | 2511 | 3238 | 0 | 0 | 0 | 43 | 35 | 12 | 0 | 12 | 0 | 12 | 425 | 59 |
| SRR6705822 | 0 | 247 | 0 | 0 | 55 | 0 | 0 | 24 | 95 | 2000 | 222 | 151 | 0 | 63 | 659 | 1603 | 0 | 5103 | 39 | 905 | 8362 | 3772 | 83 | 975 | 1947 | 0 | 0 | 14 | 11 | 20 | 42 | 42 | 17 | 12 | 0 | 1897 | 262 |
| SRR6705823 | 0 | 83 | 40 | 0 | 69 | 0 | 0 | 497 | 1157 | 15541 | 27 | 137 | 0 | 122 | 2124 | 1591 | 0 | 6580 | 26 | 851 | 2400 | 1607 | 51 | 1023 | 2861 | 39 | 0 | 21 | 18 | 21 | 32 | 27 | 35 | 18 | 0 | 2151 | 364 |
| SRR6705824 | 0 | 0 | 0 | 0 | 0 | 0 | 0 | 296 | 105 | 627 | 0 | 35 | 0 | 521 | 21 | 14 | 0 | 951 | 0 | 106 | 181 | 352 | 170 | 1483 | 2543 | 12 | 0 | 0 | 0 | 0 | 0 | 0 | 0 | 0 | 0 | 445 | 13 |
| SRR6705825 | 0 | 0 | 0 | 0 | 24 | 0 | 0 | 79 | 68 | 128 | 29 | 55 | 0 | 488 | 34 | 23 | 0 | 1347 | 0 | 134 | 300 | 209 | 75 | 1087 | 930 | 10 | 26 | 12 | 49 | 49 | 61 | 55 | 0 | 0 | 0 | 194 | 12 |
| SRR6705826 | 0 | 0 | 0 | 0 | 15 | 0 | 0 | 93 | 25 | 173 | 14 | 19 | 0 | 494 | 13 | 0 | 0 | 779 | 26 | 135 | 250 | 114 | 248 | 2382 | 2525 | 0 | 0 | 0 | 15 | 14 | 21 | 0 | 0 | 0 | 0 | 697 | 26 |
| SRR6705827 | 0 | 0 | 0 | 0 | 25 | 0 | 0 | 64 | 24 | 88 | 0 | 0 | 0 | 194 | 0 | 0 | 0 | 1971 | 12 | 267 | 667 | 294 | 105 | 1255 | 1636 | 22 | 34 | 0 | 126 | 76 | 0 | 25 | 0 | 0 | 0 | 1256 | 228 |
| SRR6705828 | 0 | 0 | 0 | 0 | 10 | 0 | 0 | 515 | 207 | 414 | 851 | 997 | 0 | 168 | 398 | 274 | 0 | 1663 | 0 | 84 | 879 | 1285 | 160 | 1266 | 2355 | 0 | 0 | 0 | 14 | 17 | 0 | 0 | 0 | 0 | 0 | 308 | 28 |
| SRR6705829 | 0 | 0 | 0 | 0 | 13 | 0 | 0 | 57 | 34 | 230 | 22 | 0 | 0 | 629 | 0 | 0 | 0 | 677 | 31 | 229 | 775 | 181 | 45 | 1044 | 1697 | 31 | 20 | 0 | 16 | 34 | 24 | 28 | 30 | 0 | 0 | 1984 | 27 |
| SRR6705830 | 0 | 34 | 0 | 0 | 94 | 0 | 0 | 1105 | 4392 | 19799 | 16 | 0 | 0 | 824 | 1640 | 469 | 0 | 3693 | 31 | 36 | 1378 | 1413 | 254 | 4911 | 6707 | 0 | 0 | 0 | 0 | 0 | 0 | 0 | 0 | 0 | 0 | 174 | 105 |
| SRR6705831 | 0 | 171 | 0 | 0 | 98 | 0 | 0 | 298 | 724 | 2769 | 392 | 351 | 0 | 358 | 11256 | 12407 | 0 | 2493 | 25 | 283 | 1191 | 1535 | 105 | 1937 | 2869 | 0 | 0 | 0 | 0 | 0 | 0 | 0 | 0 | 0 | 0 | 664 | 83 |
| SRR6705832 | 0 | 15 | 0 | 0 | 20 | 0 | 0 | 344 | 482 | 4025 | 236 | 197 | 0 | 699 | 2145 | 1545 | 0 | 2732 | 16 | 59 | 1173 | 2806 | 90 | 1704 | 2939 | 0 | 0 | 0 | 0 | 0 | 0 | 0 | 0 | 0 | 0 | 295 | 35 |
| SRR6705833 | 0 | 113 | 0 | 0 | 78 | 0 | 0 | 1247 | 2999 | 13533 | 60 | 65 | 0 | 578 | 3349 | 1670 | 0 | 2169 | 11 | 50 | 1687 | 1800 | 143 | 3172 | 3970 | 0 | 0 | 0 | 0 | 0 | 0 | 0 | 0 | 0 | 0 | 308 | 95 |
| SRR6705834 | 0 | 14 | 18 | 0 | 69 | 0 | 0 | 66 | 247 | 1692 | 71 | 88 | 0 | 124 | 165 | 149 | 0 | 4651 | 48 | 506 | 2332 | 1038 | 47 | 658 | 1677 | 52 | 0 | 19 | 42 | 22 | 18 | 25 | 23 | 22 | 0 | 1967 | 82 |
| SRR6705835 | 0 | 92 | 29 | 0 | 50 | 0 | 0 | 58 | 82 | 1295 | 0 | 10 | 0 | 236 | 22 | 70 | 0 | 3985 | 31 | 168 | 3532 | 1278 | 201 | 1813 | 3109 | 0 | 0 | 11 | 57 | 54 | 11 | 26 | 17 | 20 | 0 | 620 | 28 |
| SRR6705837 | 0 | 112 | 0 | 0 | 18 | 0 | 0 | 50 | 56 | 576 | 0 | 13 | 0 | 299 | 0 | 0 | 0 | 1252 | 0 | 133 | 543 | 213 | 247 | 2151 | 1792 | 0 | 0 | 0 | 0 | 0 | 0 | 0 | 0 | 0 | 0 | 480 | 0 |
| SRR6705838 | 0 | 18 | 12 | 0 | 75 | 0 | 0 | 81 | 139 | 802 | 48 | 67 | 0 | 180 | 70 | 34 | 0 | 2857 | 16 | 206 | 790 | 426 | 74 | 814 | 1670 | 18 | 0 | 0 | 11 | 13 | 11 | 24 | 0 | 12 | 0 | 880 | 58 |
| SRR6705839 | 0 | 232 | 0 | 0 | 65 | 0 | 0 | 65 | 86 | 867 | 12 | 15 | 0 | 294 | 0 | 0 | 0 | 3286 | 22 | 199 | 3505 | 931 | 215 | 1795 | 4366 | 0 | 0 | 0 | 11 | 0 | 0 | 0 | 0 | 0 | 0 | 805 | 19 |
| SRR6705840 | 0 | 22 | 41 | 0 | 70 | 0 | 0 | 600 | 533 | 1005 | 72 | 47 | 0 | 119 | 71 | 29 | 0 | 3872 | 15 | 425 | 1908 | 928 | 101 | 1073 | 2557 | 21 | 27 | 0 | 0 | 14 | 0 | 19 | 11 | 0 | 0 | 2340 | 58 |
| SRR6705841 | 0 | 151 | 0 | 0 | 56 | 0 | 0 | 146 | 133 | 756 | 0 | 0 | 0 | 222 | 0 | 0 | 0 | 3084 | 41 | 53 | 482 | 280 | 224 | 2497 | 6017 | 12 | 0 | 0 | 0 | 0 | 11 | 0 | 0 | 0 | 0 | 191 | 39 |

### Supp Table S8.4.Columns_DI_to_ES

| | | | | | | | | | | | | | | | | | | | | | | | | | | | | | | | | | | | | |
|---|---|---|---|---|---|---|---|---|---|---|---|---|---|---|---|---|---|---|---|---|---|---|---|---|---|---|---|---|---|---|---|---|---|---|---|---|---|
| SRR6705842 | 0 | 12 | 41 | 0 | 91 | 0 | 0 | 63 | 85 | 670 | 0 | 0 | 0 | 279 | 22 | 12 | 0 | 2939 | 0 | 50 | 746 | 402 | 105 | 1114 | 1262 | 0 | 0 | 0 | 17 | 24 | 0 | 24 | 12 | 0 | 0 | 273 | 17 |
| SRR6705843 | 0 | 11 | 0 | 0 | 55 | 0 | 0 | 57 | 80 | 756 | 0 | 0 | 0 | 425 | 23 | 0 | 0 | 3595 | 37 | 87 | 1593 | 611 | 320 | 3723 | 3826 | 11 | 0 | 0 | 0 | 0 | 0 | 0 | 0 | 0 | 0 | 374 | 26 |
| SRR6705844 | 0 | 148 | 13 | 0 | 21 | 0 | 0 | 156 | 242 | 1763 | 36 | 36 | 0 | 445 | 48 | 91 | 0 | 9044 | 21 | 15 | 2339 | 823 | 104 | 1040 | 3138 | 0 | 0 | 0 | 0 | 0 | 0 | 0 | 0 | 0 | 0 | 159 | 180 |
| SRR6705845 | 0 | 27 | 10 | 0 | 42 | 0 | 0 | 156 | 242 | 2036 | 1105 | 1061 | 0 | 518 | 642 | 1973 | 0 | 10634 | 0 | 189 | 4861 | 3827 | 40 | 393 | 2007 | 27 | 0 | 0 | 30 | 13 | 0 | 23 | 32 | 42 | 15 | 957 | 475 |
| SRR6705846 | 0 | 188 | 20 | 0 | 28 | 0 | 0 | 108 | 204 | 2348 | 35 | 108 | 0 | 239 | 70 | 129 | 0 | 7208 | 0 | 85 | 6722 | 5050 | 72 | 667 | 1930 | 12 | 0 | 0 | 0 | 0 | 0 | 0 | 0 | 12 | 0 | 585 | 274 |
| SRR6705847 | 0 | 93 | 37 | 0 | 41 | 0 | 0 | 114 | 244 | 3721 | 201 | 170 | 0 | 412 | 356 | 611 | 0 | 7941 | 0 | 49 | 5617 | 3531 | 50 | 497 | 1286 | 10 | 13 | 0 | 0 | 0 | 0 | 0 | 0 | 14 | 0 | 459 | 202 |
| SRR6705848 | 0 | 40 | 0 | 0 | 14 | 0 | 0 | 42 | 99 | 1066 | 39 | 233 | 0 | 78 | 56 | 134 | 0 | 6187 | 19 | 35 | 7915 | 6052 | 77 | 594 | 1887 | 14 | 0 | 0 | 0 | 0 | 0 | 0 | 0 | 0 | 0 | 325 | 217 |
| SRR6705849 | 0 | 20 | 0 | 0 | 0 | 0 | 0 | 638 | 689 | 4198 | 31 | 34 | 0 | 102 | 72 | 20 | 0 | 8872 | 16 | 14 | 6183 | 2868 | 24 | 585 | 2185 | 41 | 0 | 0 | 0 | 0 | 0 | 0 | 0 | 0 | 0 | 61 | 2575 |
| SRR6705850 | 0 | 293 | 0 | 0 | 24 | 0 | 0 | 180 | 367 | 2469 | 53 | 578 | 0 | 227 | 86 | 180 | 0 | 7376 | 12 | 188 | 6202 | 5356 | 83 | 677 | 2310 | 0 | 0 | 0 | 0 | 0 | 0 | 0 | 0 | 0 | 15 | 641 | 262 |
| SRR6705851 | 0 | 127 | 0 | 0 | 89 | 0 | 0 | 193 | 304 | 1785 | 304 | 369 | 0 | 217 | 41 | 164 | 0 | 5814 | 0 | 82 | 4464 | 4610 | 53 | 692 | 2446 | 0 | 0 | 0 | 0 | 0 | 0 | 0 | 0 | 0 | 0 | 550 | 500 |
| SRR6705852 | 0 | 70 | 12 | 0 | 18 | 0 | 0 | 113 | 163 | 1347 | 178 | 206 | 0 | 297 | 65 | 158 | 0 | 8563 | 14 | 46 | 4898 | 3187 | 96 | 585 | 2462 | 0 | 11 | 0 | 0 | 0 | 0 | 0 | 0 | 0 | 0 | 260 | 172 |
| SRR6705853 | 0 | 181 | 0 | 0 | 60 | 0 | 0 | 179 | 253 | 1616 | 345 | 518 | 0 | 314 | 76 | 145 | 0 | 10483 | 17 | 187 | 4206 | 1959 | 65 | 647 | 2228 | 29 | 16 | 0 | 0 | 0 | 0 | 0 | 19 | 14 | 0 | 1410 | 858 |
| SRR6705854 | 0 | 14 | 21 | 0 | 48 | 0 | 0 | 58 | 95 | 4379 | 220 | 162 | 0 | 819 | 168 | 274 | 14 | 6656 | 0 | 65 | 6118 | 2792 | 66 | 495 | 1719 | 0 | 16 | 0 | 11 | 0 | 0 | 0 | 14 | 11 | 0 | 632 | 902 |
| SRR6705855 | 0 | 268 | 0 | 0 | 45 | 0 | 0 | 88 | 157 | 3118 | 16 | 53 | 0 | 191 | 19 | 16 | 0 | 9074 | 14 | 75 | 6073 | 4163 | 60 | 641 | 2675 | 0 | 0 | 0 | 0 | 10 | 0 | 0 | 0 | 0 | 0 | 267 | 210 |
| SRR6705856 | 0 | 228 | 36 | 0 | 47 | 0 | 0 | 73 | 121 | 1705 | 32 | 16 | 0 | 333 | 0 | 0 | 0 | 1945 | 23 | 197 | 1007 | 744 | 107 | 1244 | 1231 | 0 | 0 | 15 | 52 | 49 | 43 | 55 | 61 | 61 | 0 | 482 | 30 |
| SRR6705857 | 0 | 24 | 0 | 0 | 51 | 0 | 0 | 95 | 133 | 3972 | 14 | 26 | 0 | 234 | 20 | 30 | 0 | 2848 | 23 | 79 | 1673 | 929 | 254 | 1847 | 3769 | 0 | 0 | 13 | 29 | 26 | 32 | 0 | 13 | 0 | 0 | 274 | 22 |
| SRR6705859 | 0 | 143 | 15 | 0 | 30 | 0 | 0 | 150 | 115 | 1776 | 0 | 0 | 0 | 172 | 44 | 266 | 0 | 2160 | 15 | 204 | 3372 | 486 | 196 | 2187 | 4956 | 0 | 0 | 0 | 20 | 0 | 0 | 30 | 0 | 0 | 0 | 800 | 33 |
| SRR6705860 | 0 | 48 | 0 | 0 | 49 | 0 | 0 | 297 | 115 | 642 | 11 | 0 | 0 | 250 | 0 | 0 | 0 | 4168 | 29 | 472 | 2179 | 877 | 125 | 1095 | 3216 | 57 | 41 | 0 | 10 | 0 | 0 | 0 | 14 | 0 | 0 | 2833 | 127 |
| SRR6705861 | 0 | 18 | 0 | 0 | 15 | 0 | 0 | 87 | 81 | 1310 | 0 | 0 | 0 | 360 | 0 | 10 | 0 | 1885 | 15 | 99 | 638 | 285 | 353 | 2741 | 3905 | 0 | 0 | 0 | 0 | 0 | 0 | 0 | 0 | 0 | 0 | 519 | 23 |
| SRR6705862 | 0 | 28 | 0 | 0 | 112 | 0 | 0 | 72 | 77 | 1135 | 0 | 0 | 0 | 522 | 34 | 31 | 0 | 2544 | 0 | 107 | 537 | 244 | 142 | 1369 | 1472 | 0 | 0 | 0 | 11 | 0 | 0 | 0 | 0 | 0 | 0 | 512 | 37 |
| SRR6705863 | 0 | 1635 | 0 | 0 | 61 | 0 | 0 | 64 | 80 | 1381 | 11 | 0 | 0 | 243 | 144 | 128 | 0 | 3271 | 35 | 120 | 1047 | 468 | 222 | 2826 | 4411 | 0 | 0 | 0 | 0 | 0 | 0 | 0 | 0 | 0 | 0 | 360 | 75 |
| SRR6705866 | 0 | 17 | 0 | 0 | 0 | 0 | 0 | 53 | 53 | 802 | 381 | 855 | 0 | 214 | 1176 | 2084 | 0 | 4452 | 0 | 214 | 2999 | 2390 | 141 | 890 | 2062 | 0 | 0 | 12 | 12 | 0 | 25 | 0 | 0 | 12 | 0 | 321 | 141 |
| SRR6705867 | 0 | 225 | 0 | 0 | 45 | 0 | 16 | 137 | 497 | 1199 | 308 | 274 | 0 | 377 | 6561 | 3221 | 0 | 6039 | 0 | 120 | 2199 | 1913 | 151 | 798 | 2154 | 0 | 0 | 0 | 23 | 0 | 16 | 0 | 0 | 0 | 0 | 874 | 166 |
| SRR6705868 | 0 | 153 | 0 | 0 | 61 | 0 | 0 | 68 | 61 | 637 | 1017 | 1183 | 0 | 159 | 11592 | 13306 | 0 | 5456 | 23 | 106 | 3532 | 2804 | 84 | 996 | 2651 | 0 | 0 | 0 | 12 | 0 | 0 | 0 | 0 | 0 | 0 | 660 | 169 |
| SRR6705869 | 0 | 43 | 38 | 0 | 39 | 0 | 0 | 140 | 155 | 811 | 265 | 310 | 0 | 243 | 89320 | 47748 | 0 | 6166 | 26 | 103 | 2407 | 1789 | 77 | 708 | 1648 | 0 | 13 | 0 | 25 | 25 | 19 | 0 | 19 | 31 | 0 | 627 | 142 |
| SRR6705870 | 0 | 36 | 13 | 0 | 28 | 0 | 0 | 184 | 336 | 2186 | 120 | 106 | 0 | 248 | 345 | 778 | 0 | 8197 | 13 | 106 | 4305 | 2596 | 114 | 971 | 2715 | 10 | 0 | 0 | 0 | 0 | 0 | 0 | 0 | 0 | 0 | 796 | 287 |
| SRR6705871 | 0 | 15 | 33 | 0 | 61 | 0 | 0 | 270 | 270 | 2035 | 125 | 493 | 0 | 489 | 6296 | 4943 | 0 | 7732 | 0 | 76 | 5290 | 2878 | 58 | 474 | 1196 | 33 | 0 | 0 | 0 | 12 | 0 | 0 | 14 | 0 | 0 | 556 | 2378 |
| SRR6705874 | 0 | 0 | 0 | 0 | 0 | 0 | 0 | 45 | 253 | 1535 | 0 | 0 | 0 | 134 | 0 | 15 | 0 | 232 | 0 | 60 | 164 | 188 | 77 | 569 | 430 | 0 | 0 | 0 | 0 | 0 | 0 | 0 | 0 | 0 | 0 | 209 | 0 |
| SRR6705875 | 0 | 0 | 0 | 0 | 49 | 0 | 0 | 102 | 208 | 1995 | 0 | 0 | 0 | 338 | 0 | 0 | 0 | 563 | 21 | 39 | 295 | 545 | 112 | 2493 | 1845 | 0 | 0 | 0 | 0 | 0 | 0 | 0 | 0 | 0 | 0 | 381 | 0 |
| SRR6705876 | 0 | 0 | 0 | 0 | 0 | 0 | 0 | 564 | 3098 | 20887 | 0 | 0 | 0 | 232 | 0 | 0 | 0 | 798 | 0 | 0 | 555 | 278 | 104 | 5589 | 5520 | 0 | 0 | 0 | 0 | 0 | 0 | 0 | 0 | 0 | 0 | 96 | 0 |
| SRR6705877 | 0 | 36 | 0 | 0 | 576 | 0 | 0 | 652 | 4779 | 11967 | 0 | 0 | 0 | 173 | 0 | 0 | 0 | 686 | 55 | 0 | 274 | 192 | 165 | 4447 | 1976 | 0 | 0 | 0 | 0 | 0 | 0 | 0 | 0 | 0 | 0 | 26 | 0 |
| SRR6705878 | 0 | 0 | 0 | 0 | 20 | 0 | 0 | 467 | 1370 | 4972 | 0 | 0 | 0 | 208 | 0 | 0 | 0 | 798 | 0 | 0 | 814 | 245 | 308 | 2940 | 3576 | 0 | 0 | 0 | 0 | 0 | 0 | 0 | 0 | 0 | 0 | 84 | 0 |
| SRR6705879 | 0 | 0 | 0 | 0 | 23 | 0 | 0 | 1035 | 494 | 1593 | 56 | 24 | 0 | 104 | 24 | 16 | 0 | 1055 | 0 | 64 | 855 | 898 | 93 | 955 | 2354 | 38 | 0 | 0 | 0 | 0 | 0 | 0 | 0 | 0 | 0 | 470 | 64 |
| SRR6705880 | 0 | 0 | 0 | 0 | 19 | 0 | 0 | 541 | 215 | 1459 | 240 | 137 | 0 | 197 | 412 | 232 | 0 | 930 | 0 | 309 | 299 | 1071 | 203 | 1525 | 1577 | 0 | 0 | 0 | 0 | 0 | 0 | 0 | 0 | 0 | 0 | 506 | 0 |
| SRR6705881 | 0 | 0 | 0 | 0 | 65 | 0 | 0 | 636 | 494 | 724 | 2238 | 1634 | 0 | 132 | 801 | 570 | 0 | 2023 | 32 | 22 | 795 | 2640 | 60 | 809 | 924 | 42 | 0 | 0 | 0 | 0 | 0 | 0 | 0 | 0 | 0 | 263 | 41 |
| SRR6705882 | 0 | 0 | 0 | 0 | 41 | 0 | 0 | 403 | 288 | 1150 | 153 | 38 | 0 | 230 | 1419 | 1169 | 0 | 1176 | 0 | 58 | 523 | 841 | 195 | 2519 | 2570 | 0 | 0 | 0 | 0 | 0 | 0 | 0 | 0 | 0 | 0 | 690 | 17 |
| SRR6705883 | 0 | 0 | 0 | 0 | 24 | 0 | 0 | 345 | 144 | 426 | 439 | 385 | 0 | 257 | 161 | 130 | 0 | 974 | 16 | 196 | 412 | 1622 | 38 | 612 | 833 | 32 | 0 | 0 | 0 | 0 | 0 | 0 | 0 | 0 | 0 | 888 | 36 |
| SRR6705884 | 0 | 47 | 0 | 0 | 20 | 0 | 0 | 349 | 241 | 1839 | 103 | 336 | 0 | 194 | 139 | 210 | 0 | 7125 | 0 | 58 | 3283 | 3395 | 36 | 584 | 2082 | 14 | 0 | 0 | 0 | 0 | 0 | 0 | 0 | 0 | 0 | 464 | 682 |
| SRR6705885 | 0 | 23 | 13 | 0 | 36 | 0 | 0 | 348 | 292 | 1732 | 292 | 518 | 0 | 188 | 226 | 311 | 0 | 8391 | 0 | 85 | 5995 | 3751 | 108 | 709 | 2351 | 0 | 20 | 0 | 0 | 0 | 0 | 0 | 0 | 0 | 27 | 574 | 314 |
| SRR6705886 | 0 | 118 | 18 | 0 | 65 | 0 | 0 | 189 | 108 | 2016 | 954 | 846 | 0 | 540 | 6624 | 6876 | 0 | 7132 | 13 | 99 | 1722 | 1993 | 39 | 621 | 2097 | 13 | 0 | 0 | 0 | 0 | 0 | 0 | 0 | 0 | 0 | 450 | 246 |
| SRR6705887 | 0 | 62 | 0 | 0 | 64 | 0 | 0 | 53 | 102 | 620 | 201 | 1457 | 0 | 184 | 20477 | 13816 | 0 | 5681 | 20 | 73 | 3855 | 2731 | 94 | 792 | 2197 | 0 | 0 | 0 | 0 | 10 | 0 | 0 | 0 | 10 | 0 | 586 | 109 |
| SRR6705888 | 0 | 26 | 0 | 0 | 64 | 0 | 0 | 36 | 205 | 2291 | 193 | 145 | 0 | 555 | 22647 | 1941 | 0 | 5585 | 13 | 60 | 3928 | 1951 | 64 | 601 | 1292 | 0 | 0 | 0 | 0 | 10 | 0 | 14 | 0 | 0 | 0 | 313 | 1030 |
| SRR6705889 | 0 | 279 | 15 | 15 | 69 | 0 | 0 | 102 | 111 | 1076 | 471 | 681 | 0 | 534 | 3202 | 2402 | 0 | 4363 | 0 | 80 | 5247 | 3608 | 74 | 548 | 1836 | 0 | 0 | 0 | 0 | 0 | 0 | 0 | 0 | 0 | 0 | 218 | 74 |
| SRR6705890 | 0 | 115 | 184 | 0 | 21 | 0 | 0 | 288 | 325 | 4060 | 407 | 513 | 0 | 200 | 2853 | 4717 | 0 | 6362 | 0 | 175 | 6069 | 3475 | 42 | 442 | 2340 | 19 | 32 | 11 | 18 | 14 | 22 | 25 | 14 | 18 | 0 | 945 | 1239 |
| SRR6705891 | 0 | 19 | 0 | 0 | 17 | 0 | 0 | 124 | 402 | 3814 | 240 | 218 | 0 | 334 | 64 | 146 | 0 | 9223 | 11 | 158 | 7563 | 4947 | 56 | 541 | 2388 | 0 | 0 | 0 | 12 | 0 | 0 | 0 | 17 | 12 | 0 | 899 | 199 |
| SRR6705892 | 0 | 91 | 69 | 0 | 63 | 0 | 0 | 321 | 1168 | 7359 | 84 | 68 | 0 | 269 | 24 | 37 | 0 | 6969 | 17 | 29 | 6351 | 2804 | 29 | 272 | 938 | 26 | 0 | 0 | 13 | 0 | 0 | 0 | 19 | 28 | 0 | 123 | 1471 |
| SRR6705893 | 0 | 38 | 0 | 0 | 26 | 0 | 0 | 164 | 219 | 1750 | 458 | 301 | 0 | 239 | 451 | 943 | 0 | 9373 | 0 | 253 | 6777 | 5252 | 102 | 797 | 2319 | 10 | 10 | 0 | 0 | 10 | 0 | 10 | 10 | 0 | 0 | 930 | 230 |
| SRR6705894 | 55 | 68 | 0 | 0 | 106 | 0 | 0 | 189 | 93 | 1141 | 267 | 201 | 0 | 407 | 0 | 0 | 0 | 2160 | 25 | 34 | 1366 | 1242 | 41 | 1110 | 4856 | 0 | 62 | 0 | 0 | 0 | 0 | 0 | 0 | 0 | 0 | 47 | 34 |
| SRR6705895 | 33 | 78 | 0 | 0 | 20 | 0 | 0 | 86 | 161 | 1023 | 276 | 218 | 0 | 230 | 23 | 29 | 0 | 1076 | 26 | 57 | 1549 | 1087 | 130 | 1590 | 3878 | 11 | 208 | 0 | 26 | 26 | 46 | 0 | 0 | 0 | 0 | 126 | 37 |
| SRR6705897 | 38 | 51 | 0 | 0 | 0 | 0 | 0 | 137 | 49 | 484 | 27 | 112 | 0 | 171 | 0 | 39 | 0 | 835 | 13 | 98 | 970 | 455 | 141 | 1206 | 1819 | 0 | 45 | 0 | 0 | 38 | 76 | 0 | 0 | 0 | 0 | 157 | 18 |
| SRR6705898 | 0 | 0 | 0 | 0 | 12 | 0 | 0 | 0 | 52 | 382 | 0 | 0 | 0 | 367 | 0 | 0 | 0 | 164 | 0 | 47 | 126 | 68 | 71 | 890 | 371 | 0 | 0 | 0 | 0 | 0 | 0 | 0 | 0 | 0 | 0 | 423 | 0 |
| SRR6705899 | 0 | 0 | 0 | 0 | 0 | 0 | 0 | 77 | 58 | 830 | 58 | 0 | 0 | 425 | 0 | 0 | 0 | 111 | 0 | 347 | 114 | 129 | 51 | 482 | 380 | 0 | 0 | 0 | 0 | 0 | 0 | 0 | 0 | 0 | 0 | 1023 | 0 |
| SRR6705900 | 0 | 0 | 0 | 0 | 0 | 0 | 0 | 14 | 71 | 739 | 14 | 14 | 0 | 369 | 0 | 14 | 0 | 132 | 0 | 14 | 189 | 95 | 41 | 748 | 580 | 0 | 0 | 0 | 0 | 0 | 0 | 0 | 0 | 0 | 0 | 313 | 0 |
| SRR6705901 | 0 | 0 | 0 | 0 | 0 | 0 | 0 | 23 | 0 | 70 | 0 | 0 | 0 | 506 | 0 | 0 | 0 | 149 | 0 | 133 | 63 | 39 | 48 | 548 | 180 | 0 | 0 | 0 | 0 | 0 | 0 | 0 | 0 | 0 | 0 | 652 | 0 |
| SRR6705902 | 0 | 279 | 49 | 0 | 96 | 0 | 0 | 118 | 150 | 1399 | 0 | 0 | 0 | 170 | 13 | 19 | 0 | 2668 | 11 | 208 | 1494 | 293 | 123 | 1836 | 5539 | 0 | 0 | 0 | 0 | 11 | 0 | 18 | 0 | 0 | 0 | 768 | 43 |
| SRR6705903 | 0 | 26 | 25 | 0 | 120 | 0 | 0 | 208 | 382 | 2331 | 15 | 0 | 0 | 114 | 233 | 79 | 0 | 2533 | 16 | 659 | 1659 | 640 | 84 | 653 | 1702 | 53 | 0 | 0 | 28 | 0 | 12 | 12 | 12 | 0 | 0 | 3241 | 52 |
| SRR6726498 | 0 | 0 | 0 | 0 | 0 | 0 | 0 | 1269 | 467 | 3480 | 431 | 1791 | 0 | 5130 | 0 | 0 | 0 | 186 | 0 | 218 | 40 | 78 | 0 | 607 | 373 | 45 | 0 | 21 | 0 | 0 | 0 | 0 | 0 | 18 | 0 | 599 | 0 |
| SRR6726499 | 0 | 0 | 130 | 0 | 0 | 0 | 0 | 908 | 395 | 2814 | 344 | 1823 | 0 | 10154 | 0 | 0 | 0 | 183 | 0 | 0 | 42 | 131 | 0 | 1261 | 416 | 52 | 0 | 0 | 0 | 0 | 229 | 0 | 30 | 189 | 120 | 187 | 0 |
| SRR6757230 | 0 | 21 | 26 | 0 | 0 | 0 | 0 | 566 | 103 | 6105 | 3316 | 5182 | 0 | 879 | 1461 | 1593 | 0 | 21 | 39 | 1300 | 378 | 2060 | 0 | 33 | 3701 | 837 | 0 | 29 | 28 | 155 | 186 | 53 | 151 | 381 | 228 | 7940 | 0 |
| SRR6757231 | 0 | 16 | 27 | 0 | 0 | 0 | 0 | 570 | 111 | 6043 | 3617 | 5393 | 0 | 926 | 1359 | 1480 | 0 | 15 | 37 | 1194 | 349 | 1985 | 0 | 31 | 3283 | 784 | 0 | 25 | 17 | 30 | 34 | 42 | 29 | 72 | 39 | 8239 | 0 |
| SRR6757232 | 0 | 0 | 26 | 0 | 0 | 0 | 0 | 348 | 67 | 5844 | 5998 | 11033 | 0 | 983 | 2280 | 2848 | 0 | 28 | 45 | 983 | 431 | 2270 | 0 | 37 | 3500 | 702 | 0 | 0 | 0 | 17 | 24 | 19 | 22 | 40 | 20 | 7496 | 0 |
| SRR6757233 | 0 | 0 | 23 | 0 | 0 | 0 | 0 | 409 | 84 | 6122 | 6363 | 10511 | 0 | 1095 | 1797 | 2548 | 0 | 26 | 45 | 1204 | 388 | 2089 | 0 | 34 | 3165 | 686 | 0 | 16 | 0 | 25 | 24 | 21 | 21 | 46 | 14 | 8036 | 0 |
| SRR6757234 | 0 | 15 | 16 | 0 | 0 | 0 | 0 | 209 | 43 | 2072 | 2781 | 6841 | 0 | 482 | 745 | 1217 | 0 | 39 | 35 | 1623 | 428 | 1695 | 0 | 45 | 3274 | 595 | 0 | 11 | 11 | 16 | 12 | 20 | 12 | 32 | 30 | 8031 | 0 |
| SRR6757235 | 0 | 11 | 12 | 0 | 0 | 0 | 0 | 194 | 55 | 2220 | 2799 | 6785 | 0 | 548 | 729 | 1190 | 0 | 45 | 28 | 1503 | 358 | 1610 | 0 | 33 | 2933 | 594 | 0 | 17 | 11 | 16 | 13 | 15 | 12 | 34 | 33 | 8281 | 0 |
| SRR6757236 | 0 | 11 | 16 | 0 | 0 | 0 | 0 | 243 | 98 | 3585 | 6189 | 13644 | 0 | 520 | 750 | 921 | 0 | 46 | 61 | 2221 | 444 | 1697 | 0 | 33 | 3197 | 637 | 0 | 23 | 22 | 31 | 30 | 45 | 20 | 48 | 46 | 9739 | 0 |
| SRR6757237 | 0 | 11 | 14 | 0 | 0 | 0 | 0 | 250 | 86 | 3549 | 1805 | 13051 | 0 | 628 | 710 | 888 | 0 | 35 | 49 | 1958 | 420 | 1590 | 0 | 23 | 2738 | 610 | 0 | 24 | 17 | 184 | 157 | 23 | 118 | 252 | 58 | 9684 | 0 |
| SRR6757238 | 0 | 0 | 12 | 0 | 0 | 0 | 0 | 673 | 107 | 9046 | 88 | 690 | 0 | 1757 | 2726 | 2743 | 0 | 37 | 22 | 858 | 236 | 2694 | 0 | 69 | 4597 | 179 | 0 | 0 | 12 | 34 | 48 | 21 | 37 | 91 | 82 | 3330 | 0 |
| SRR6757239 | 0 | 0 | 27 | 0 | 0 | 0 | 0 | 692 | 89 | 8721 | 151 | 740 | 0 | 1534 | 2617 | 2437 | 0 | 41 | 23 | 859 | 197 | 2421 | 0 | 62 | 3943 | 183 | 0 | 0 | 0 | 110 | 140 | 29 | 91 | 256 | 220 | 3490 | 0 |
| SRR6757240 | 0 | 0 | 42 | 0 | 0 | 0 | 0 | 430 | 93 | 4826 | 102 | 389 | 0 | 793 | 651 | 884 | 0 | 34 | 45 | 2055 | 300 | 2525 | 0 | 60 | 5967 | 314 | 0 | 0 | 15 | 127 | 205 | 81 | 182 | 518 | 322 | 12896 | 0 |
| SRR6757241 | 0 | 0 | 14 | 0 | 0 | 0 | 0 | 429 | 106 | 5088 | 250 | 424 | 0 | 880 | 564 | 913 | 0 | 28 | 33 | 2070 | 307 | 2314 | 0 | 38 | 5390 | 354 | 0 | 0 | 15 | 34 | 46 | 16 | 54 | 128 | 91 | 14104 | 0 |
| SRR6757242 | 0 | 22 | 28 | 0 | 0 | 0 | 0 | 1146 | 176 | 20209 | 1034 | 1194 | 0 | 1748 | 5531 | 3663 | 0 | 25 | 36 | 770 | 352 | 2478 | 0 | 23 | 4686 | 569 | 0 | 0 | 16 | 80 | 129 | 57 | 139 | 390 | 173 | 4577 | 0 |
| SRR6757243 | 0 | 22 | 32 | 0 | 0 | 0 | 0 | 1013 | 169 | 19793 | 957 | 1086 | 0 | 1582 | 4937 | 3592 | 0 | 26 | 35 | 687 | 366 | 2473 | 0 | 27 | 4909 | 554 | 0 | 0 | 15 | 22 | 31 | 12 | 37 | 109 | 46 | 4301 | 0 |
| SRR6757244 | 0 | 21 | 27 | 0 | 0 | 0 | 0 | 651 | 105 | 6910 | 761 | 1093 | 0 | 1278 | 2438 | 2682 | 0 | 28 | 29 | 1323 | 234 | 2559 | 0 | 26 | 4970 | 203 | 0 | 0 | 23 | 106 | 169 | 41 | 154 | 382 | 239 | 5144 | 0 |
| SRR6757245 | 0 | 28 | 40 | 0 | 0 | 0 | 0 | 526 | 108 | 5971 | 244 | 919 | 0 | 1192 | 2189 | 2448 | 0 | 26 | 31 | 1126 | 257 | 2629 | 0 | 20 | 5227 | 188 | 0 | 0 | 19 | 94 | 171 | 48 | 155 | 419 | 226 | 4766 | 0 |
| SRR6757246 | 0 | 20 | 14 | 0 | 0 | 0 | 0 | 311 | 111 | 7568 | 3733 | 6086 | 0 | 1059 | 454 | 679 | 0 | 15 | 35 | 0 | 366 | 2069 | 0 | 10 | 3221 | 1161 | 0 | 29 | 17 | 36 | 84 | 29 | 113 | 352 | 114 | 0 | 0 |
| SRR6757247 | 0 | 23 | 15 | 0 | 0 | 0 | 0 | 376 | 106 | 6837 | 815 | 5561 | 0 | 921 | 368 | 613 | 0 | 15 | 38 | 0 | 353 | 2113 | 0 | 14 | 3268 | 1192 | 0 | 28 | 12 | 48 | 99 | 31 | 99 | 324 | 146 | 0 | 0 |
| SRR6757248 | 0 | 18 | 19 | 0 | 0 | 0 | 0 | 620 | 121 | 8350 | 2729 | 3486 | 0 | 1265 | 1717 | 2392 | 0 | 25 | 0 | 1615 | 341 | 1905 | 0 | 22 | 2776 | 806 | 0 | 0 | 0 | 81 | 71 | 29 | 102 | 283 | 140 | 12953 | 0 |
| SRR6757249 | 0 | 22 | 16 | 0 | 0 | 0 | 0 | 570 | 130 | 7537 | 2441 | 3482 | 0 | 1106 | 1645 | 2406 | 0 | 22 | 0 | 1680 | 366 | 1906 | 0 | 20 | 2920 | 820 | 0 | 0 | 10 | 76 | 84 | 35 | 101 | 280 | 154 | 13398 | 0 |
| SRR6757250 | 0 | 19 | 24 | 0 | 0 | 0 | 0 | 455 | 103 | 5476 | 3811 | 5591 | 0 | 764 | 1339 | 1367 | 0 | 17 | 33 | 1537 | 394 | 1774 | 0 | 18 | 3035 | 706 | 0 | 17 | 15 | 13 | 18 | 47 | 18 | 52 | 29 | 8654 | 0 |
| SRR6757251 | 0 | 26 | 35 | 0 | 0 | 0 | 0 | 399 | 64 | 5524 | 986 | 5362 | 0 | 715 | 1325 | 1258 | 0 | 14 | 35 | 1374 | 377 | 1895 | 0 | 16 | 3090 | 695 | 0 | 19 | 22 | 87 | 113 | 44 | 109 | 314 | 174 | 8551 | 0 |
| SRR6757252 | 0 | 13 | 23 | 0 | 0 | 0 | 0 | 649 | 141 | 12330 | 1221 | 1309 | 0 | 1394 | 2013 | 2638 | 0 | 25 | 27 | 1024 | 328 | 2072 | 0 | 35 | 3643 | 348 | 0 | 0 | 12 | 98 | 112 | 35 | 91 | 229 | 171 | 9880 | 0 |
| SRR6757253 | 0 | 19 | 19 | 0 | 0 | 0 | 0 | 620 | 111 | 11756 | 1118 | 1152 | 0 | 1230 | 1743 | 2504 | 0 | 25 | 32 | 1144 | 347 | 2011 | 0 | 40 | 3776 | 352 | 0 | 0 | 0 | 17 | 27 | 36 | 16 | 51 | 36 | 9838 | 0 |
| SRR6757254 | 0 | 13 | 23 | 0 | 0 | 0 | 0 | 1050 | 196 | 10809 | 3818 | 6686 | 0 | 1209 | 1500 | 1437 | 0 | 43 | 26 | 640 | 514 | 1757 | 0 | 54 | 3621 | 394 | 0 | 14 | 11 | 55 | 62 | 11 | 51 | 130 | 46 | 1902 | 0 |
| SRR6757255 | 0 | 15 | 27 | 0 | 0 | 0 | 0 | 1097 | 184 | 10719 | 3924 | 7057 | 0 | 1156 | 1539 | 1452 | 0 | 43 | 23 | 597 | 475 | 1591 | 0 | 55 | 3173 | 341 | 0 | 12 | 62 | 324 | 346 | 84 | 263 | 749 | 280 | 1883 | 0 |
| SRR6757256 | 0 | 12 | 17 | 0 | 0 | 0 | 0 | 812 | 102 | 5945 | 3438 | 5789 | 0 | 820 | 1864 | 2807 | 0 | 41 | 21 | 845 | 380 | 1634 | 0 | 37 | 3222 | 420 | 0 | 0 | 41 | 173 | 248 | 64 | 230 | 685 | 298 | 8955 | 0 |
| SRR6757257 | 0 | 0 | 16 | 0 | 0 | 0 | 0 | 690 | 102 | 6034 | 3593 | 5946 | 0 | 1043 | 1846 | 2568 | 0 | 36 | 26 | 856 | 347 | 1535 | 0 | 45 | 2849 | 397 | 0 | 0 | 37 | 44 | 72 | 23 | 60 | 175 | 76 | 8961 | 0 |
| SRR6757258 | 0 | 14 | 16 | 0 | 0 | 0 | 0 | 675 | 121 | 4486 | 838 | 3419 | 0 | 1564 | 1523 | 1874 | 0 | 45 | 23 | 373 | 433 | 1190 | 0 | 48 | 2378 | 254 | 0 | 0 | 13 | 118 | 141 | 34 | 113 | 324 | 205 | 1911 | 0 |
| SRR6757259 | 0 | 12 | 17 | 0 | 0 | 0 | 0 | 617 | 140 | 4521 | 1939 | 3706 | 0 | 1676 | 1436 | 1713 | 0 | 40 | 20 | 372 | 389 | 1136 | 0 | 39 | 2042 | 228 | 0 | 0 | 17 | 119 | 149 | 26 | 99 | 337 | 206 | 2145 | 13 |
| SRR6757260 | 0 | 45 | 32 | 0 | 0 | 0 | 0 | 810 | 130 | 5077 | 4442 | 7747 | 0 | 684 | 1286 | 1490 | 0 | 33 | 31 | 991 | 291 | 1566 | 0 | 0 | 3010 | 638 | 0 | 30 | 39 | 32 | 38 | 10 | 35 | 87 | 44 | 5213 | 0 |
| SRR6757261 | 0 | 42 | 32 | 0 | 0 | 0 | 0 | 771 | 143 | 4899 | 4271 | 8124 | 0 | 739 | 1171 | 1472 | 0 | 26 | 31 | 1034 | 261 | 1457 | 0 | 0 | 2651 | 592 | 0 | 23 | 49 | 33 | 44 | 13 | 33 | 89 | 49 | 5322 | 0 |
| SRR6757262 | 0 | 19 | 33 | 0 | 0 | 0 | 0 | 1044 | 123 | 6112 | 1285 | 2269 | 0 | 1211 | 2449 | 1886 | 0 | 36 | 30 | 566 | 331 | 2212 | 0 | 23 | 4263 | 438 | 0 | 0 | 47 | 364 | 541 | 92 | 405 | 1163 | 517 | 2397 | 0 |
| SRR6757263 | 0 | 20 | 39 | 0 | 0 | 0 | 0 | 903 | 133 | 6443 | 1363 | 2183 | 0 | 1233 | 2364 | 1873 | 0 | 29 | 26 | 598 | 301 | 2056 | 0 | 19 | 3702 | 405 | 0 | 12 | 40 | 376 | 574 | 116 | 357 | 1106 | 479 | 2468 | 0 |
| SRR6757264 | 0 | 18 | 26 | 0 | 0 | 0 | 0 | 1525 | 324 | 13118 | 885 | 3299 | 0 | 1689 | 1217 | 1241 | 0 | 11 | 0 | 278 | 346 | 1478 | 0 | 35 | 2074 | 459 | 0 | 15 | 103 | 348 | 628 | 154 | 550 | 2016 | 527 | 1663 | 0 |
| SRR6757265 | 0 | 20 | 33 | 0 | 0 | 0 | 0 | 1470 | 303 | 12896 | 1910 | 3342 | 0 | 1716 | 1164 | 1186 | 0 | 11 | 12 | 290 | 359 | 1517 | 0 | 38 | 2185 | 454 | 0 | 18 | 97 | 339 | 655 | 157 | 589 | 2064 | 500 | 1700 | 0 |
| SRR6757266 | 0 | 11 | 32 | 0 | 0 | 0 | 0 | 664 | 72 | 3141 | 747 | 970 | 0 | 1050 | 1510 | 1672 | 0 | 18 | 17 | 1027 | 267 | 2466 | 0 | 0 | 4215 | 324 | 0 | 0 | 38 | 147 | 291 | 92 | 387 | 1342 | 419 | 4443 | 0 |
| SRR6757267 | 0 | 15 | 35 | 0 | 0 | 0 | 0 | 575 | 74 | 3100 | 189 | 1015 | 0 | 1005 | 1345 | 1345 | 0 | 17 | 22 | 975 | 244 | 2267 | 0 | 0 | 3621 | 325 | 0 | 0 | 29 | 160 | 285 | 85 | 380 | 1371 | 461 | 4367 | 0 |
| SRR6757268 | 0 | 28 | 25 | 0 | 0 | 0 | 0 | 1016 | 143 | 5329 | 807 | 1035 | 0 | 1122 | 2309 | 2014 | 0 | 30 | 23 | 1606 | 238 | 1924 | 0 | 50 | 3275 | 337 | 0 | 0 | 35 | 119 | 189 | 49 | 194 | 584 | 296 | 4941 | 0 |
| SRR6757269 | 0 | 11 | 34 | 0 | 0 | 0 | 0 | 1063 | 156 | 4833 | 261 | 965 | 0 | 962 | 1969 | 1743 | 0 | 32 | 35 | 1427 | 240 | 1973 | 0 | 56 | 3436 | 303 | 0 | 0 | 23 | 24 | 31 | 49 | 29 | 99 | 48 | 4490 | 0 |
| SRR6757270 | 0 | 22 | 16 | 0 | 0 | 0 | 0 | 682 | 89 | 4790 | 1423 | 2299 | 0 | 764 | 1597 | 1999 | 0 | 43 | 29 | 2116 | 342 | 1397 | 0 | 15 | 2853 | 534 | 0 | 12 | 21 | 13 | 25 | 39 | 24 | 60 | 35 | 8946 | 0 |
| SRR6757271 | 0 | 29 | 12 | 0 | 0 | 0 | 0 | 648 | 106 | 4737 | 1460 | 2305 | 0 | 778 | 1497 | 1782 | 0 | 50 | 27 | 2260 | 326 | 1391 | 0 | 16 | 2900 | 555 | 0 | 19 | 24 | 15 | 26 | 41 | 24 | 69 | 40 | 9392 | 0 |
| SRR6757272 | 0 | 26 | 21 | 0 | 0 | 0 | 0 | 560 | 79 | 3636 | 1373 | 3303 | 0 | 879 | 1307 | 2011 | 0 | 19 | 25 | 1727 | 335 | 1479 | 0 | 54 | 2727 | 425 | 0 | 0 | 22 | 83 | 155 | 48 | 171 | 539 | 268 | 11435 | 0 |
| SRR6757273 | 0 | 18 | 25 | 0 | 0 | 0 | 0 | 574 | 124 | 3551 | 2119 | 3339 | 0 | 937 | 1363 | 1992 | 0 | 20 | 19 | 1887 | 323 | 1468 | 0 | 56 | 2814 | 408 | 0 | 0 | 27 | 92 | 161 | 53 | 189 | 521 | 281 | 12110 | 0 |
| SRR6757274 | 0 | 13 | 27 | 0 | 0 | 0 | 0 | 710 | 72 | 2586 | 2802 | 4873 | 0 | 523 | 982 | 1254 | 0 | 22 | 24 | 1528 | 282 | 1267 | 0 | 15 | 2558 | 722 | 0 | 17 | 33 | 171 | 277 | 88 | 339 | 1067 | 296 | 9086 | 0 |
| SRR6757275 | 0 | 0 | 23 | 0 | 0 | 0 | 0 | 678 | 94 | 2688 | 2840 | 4756 | 0 | 528 | 916 | 1218 | 0 | 20 | 27 | 1442 | 293 | 1363 | 0 | 11 | 2654 | 741 | 0 | 19 | 38 | 25 | 49 | 14 | 57 | 175 | 49 | 9560 | 0 |
| SRR6757276 | 0 | 20 | 25 | 0 | 0 | 0 | 0 | 791 | 79 | 3073 | 484 | 1722 | 0 | 916 | 2744 | 2948 | 0 | 18 | 21 | 1117 | 220 | 1392 | 0 | 57 | 2483 | 299 | 0 | 0 | 23 | 13 | 16 | 47 | 17 | 64 | 36 | 5939 | 0 |
| SRR6757277 | 0 | 28 | 25 | 0 | 0 | 0 | 0 | 803 | 103 | 3287 | 1448 | 1772 | 0 | 906 | 2844 | 2856 | 0 | 14 | 22 | 1155 | 229 | 1358 | 0 | 56 | 2555 | 301 | 0 | 13 | 37 | 105 | 197 | 50 | 130 | 588 | 276 | 5854 | 0 |
| SRR6757373 | 0 | 24 | 0 | 0 | 49 | 0 | 0 | 3789 | 1195 | 2559 | 4452 | 8723 | 0 | 116 | 0 | 0 | 0 | 345 | 0 | 100 | 3624 | 2092 | 1702 | 11537 | 4577 | 11 | 12 | 0 | 10 | 0 | 0 | 0 | 0 | 0 | 0 | 372 | 0 |
| SRR6757374 | 0 | 52 | 0 | 0 | 126 | 0 | 0 | 3121 | 1078 | 3543 | 1653 | 10357 | 0 | 47 | 0 | 0 | 0 | 243 | 0 | 19 | 3925 | 1398 | 362 | 3520 | 4592 | 0 | 11 | 0 | 23 | 15 | 0 | 0 | 0 | 0 | 0 | 258 | 45 |
| SRR6757375 | 0 | 0 | 0 | 0 | 24 | 0 | 0 | 89 | 52 | 164 | 1448 | 20330 | 0 | 0 | 0 | 0 | 0 | 164 | 0 | 0 | 3986 | 1410 | 2538 | 12365 | 2664 | 0 | 11 | 0 | 0 | 0 | 0 | 0 | 0 | 0 | 0 | 59 | 0 |
| SRR6757376 | 0 | 39 | 0 | 0 | 134 | 0 | 0 | 968 | 486 | 2928 | 14023 | 13770 | 0 | 61 | 14 | 0 | 20 | 225 | 0 | 19 | 3096 | 678 | 62 | 847 | 2618 | 0 | 18 | 0 | 15 | 0 | 0 | 0 | 0 | 0 | 0 | 117 | 204 |
| SRR6757377 | 0 | 0 | 0 | 0 | 14 | 0 | 0 | 288 | 144 | 406 | 956 | 14992 | 0 | 17 | 0 | 0 | 0 | 253 | 0 | 59 | 4510 | 1594 | 2563 | 16719 | 3280 | 18 | 17 | 0 | 0 | 0 | 0 | 0 | 0 | 0 | 0 | 322 | 0 |
| SRR6757378 | 0 | 36 | 10 | 0 | 105 | 13 | 0 | 2200 | 1251 | 5890 | 443 | 425 | 0 | 323 | 0 | 0 | 0 | 287 | 0 | 128 | 3020 | 1594 | 210 | 2579 | 4368 | 0 | 0 | 0 | 45 | 54 | 26 | 0 | 0 | 0 | 0 | 220 | 55 |
| SRR6757379 | 0 | 15 | 0 | 0 | 18 | 0 | 0 | 1155 | 491 | 1867 | 101 | 894 | 0 | 253 | 24 | 87 | 0 | 365 | 0 | 823 | 4525 | 3774 | 4216 | 38047 | 5289 | 0 | 0 | 0 | 11 | 34 | 15 | 0 | 0 | 0 | 0 | 1527 | 0 |
| SRR6757380 | 0 | 19 | 0 | 0 | 49 | 0 | 0 | 1660 | 845 | 3076 | 362 | 1358 | 0 | 80 | 178 | 309 | 0 | 273 | 0 | 1114 | 3089 | 2685 | 787 | 11980 | 7313 | 0 | 0 | 0 | 18 | 41 | 19 | 0 | 0 | 0 | 0 | 1675 | 22 |
| SRR6757381 | 0 | 0 | 0 | 0 | 21 | 0 | 0 | 2352 | 1454 | 3715 | 742 | 969 | 0 | 146 | 10 | 20 | 0 | 383 | 0 | 2912 | 2594 | 2307 | 2056 | 19429 | 4847 | 12 | 0 | 0 | 12 | 20 | 26 | 0 | 0 | 0 | 0 | 8318 | 0 |
| SRR6757382 | 0 | 32 | 0 | 0 | 59 | 0 | 0 | 759 | 379 | 1948 | 964 | 6813 | 0 | 345 | 57 | 96 | 0 | 357 | 0 | 357 | 1889 | 1439 | 322 | 3648 | 3284 | 0 | 13 | 0 | 21 | 54 | 29 | 0 | 0 | 0 | 0 | 470 | 55 |
| SRR6757383 | 0 | 18 | 0 | 0 | 17 | 0 | 0 | 1266 | 329 | 922 | 605 | 8833 | 0 | 51 | 15 | 0 | 0 | 283 | 0 | 95 | 3607 | 1907 | 1746 | 11661 | 12 | 0 | 0 | 0 | 11 | 12 | 0 | 0 | 0 | 0 | 0 | 432 | 0 |
| SRR6757384 | 0 | 40 | 0 | 0 | 43 | 0 | 0 | 123 | 51 | 368 | 16927 | 15741 | 0 | 31 | 82 | 82 | 10 | 216 | 0 | 133 | 4551 | 1329 | 110 | 1092 | 12 | 0 | 0 | 69 | 28 | 20 | 0 | 0 | 0 | 0 | 0 | 450 | 136 |
| SRR6757385 | 0 | 31 | 0 | 0 | 19 | 0 | 0 | 996 | 576 | 2310 | 3188 | 8154 | 0 | 183 | 75 | 65 | 34 | 224 | 0 | 38 | 4306 | 1309 | 1736 | 10422 | 2798 | 13 | 32 | 0 | 0 | 0 | 0 | 0 | 0 | 0 | 0 | 264 | 0 |
| SRR6757386 | 0 | 32 | 0 | 0 | 60 | 0 | 0 | 2953 | 1371 | 5116 | 1129 | 2272 | 0 | 454 | 0 | 0 | 16 | 314 | 0 | 15 | 2549 | 1253 | 804 | 5309 | 4497 | 0 | 0 | 17 | 0 | 12 | 0 | 0 | 0 | 0 | 0 | 42 | 41 |
| SRR6757387 | 0 | 29 | 0 | 0 | 104 | 0 | 0 | 450 | 248 | 636 | 20968 | 36148 | 0 | 23 | 11 | 0 | 0 | 418 | 0 | 56 | 2548 | 1093 | 763 | 5364 | 4374 | 0 | 30 | 23 | 0 | 0 | 0 | 0 | 0 | 0 | 0 | 146 | 0 |
| SRR6757388 | 0 | 36 | 0 | 0 | 59 | 13 | 14 | 1100 | 415 | 2746 | 5075 | 6710 | 0 | 98 | 42 | 30 | 33 | 219 | 0 | 42 | 4649 | 1280 | 183 | 2288 | 5121 | 0 | 25 | 38 | 117 | 49 | 19 | 0 | 0 | 0 | 0 | 266 | 40 |
| SRR6757389 | 0 | 0 | 0 | 0 | 11 | 0 | 0 | 860 | 524 | 1533 | 534 | 11731 | 0 | 54 | 0 | 0 | 0 | 218 | 0 | 276 | 3599 | 1263 | 3635 | 19134 | 2306 | 0 | 12 | 0 | 11 | 22 | 29 | 0 | 0 | 0 | 0 | 1223 | 0 |
| SRR6757390 | 0 | 35 | 26 | 0 | 72 | 0 | 0 | 1974 | 1270 | 7829 | 1126 | 1187 | 0 | 199 | 0 | 0 | 0 | 191 | 0 | 55 | 5377 | 738 | 235 | 1775 | 3159 | 0 | 23 | 0 | 37 | 32 | 45 | 16 | 14 | 20 | 0 | 279 | 69 |

### Supp Table S8.4.Columns_DI_to_ES

| | | | | | | | | | | | | | | | | | | | | | | | | | | | | | | | | | | | | | |
|---|---|---|---|---|---|---|---|---|---|---|---|---|---|---|---|---|---|---|---|---|---|---|---|---|---|---|---|---|---|---|---|---|---|---|---|---|---|
| SRR6757391 | 0 | 19 | 0 | 0 | 20 | 0 | 0 | 647 | 226 | 1692 | 1490 | 23952 | 0 | 101 | 179 | 140 | 0 | 278 | 0 | 211 | 5133 | 2485 | 1783 | 14124 | 3772 | 11 | 0 | 10 | 13 | 10 | 0 | 0 | 0 | 0 | 0 | 717 | 0 |
| SRR6757392 | 0 | 46 | 0 | 0 | 104 | 0 | 0 | 1387 | 430 | 2617 | 4882 | 7940 | 0 | 214 | 621 | 292 | 11 | 291 | 0 | 37 | 2786 | 1531 | 362 | 3637 | 4719 | 0 | 16 | 32 | 37 | 30 | 0 | 0 | 0 | 0 | 0 | 121 | 67 |
| SRR6757393 | 0 | 17 | 0 | 0 | 17 | 0 | 0 | 939 | 235 | 527 | 1084 | 16338 | 0 | 36 | 36 | 36 | 0 | 249 | 0 | 135 | 5264 | 1772 | 2283 | 12326 | 2813 | 0 | 34 | 0 | 0 | 0 | 0 | 0 | 0 | 0 | 0 | 704 | 0 |
| SRR6757394 | 0 | 16 | 0 | 0 | 52 | 0 | 0 | 108 | 70 | 2045 | 11601 | 13289 | 0 | 122 | 1032 | 896 | 29 | 167 | 0 | 52 | 1504 | 606 | 222 | 3037 | 1552 | 15 | 53 | 50 | 0 | 13 | 0 | 0 | 0 | 0 | 0 | 117 | 27 |
| SRR6757395 | 0 | 13 | 13 | 0 | 11 | 0 | 0 | 1562 | 663 | 2805 | 1308 | 2434 | 0 | 157 | 167 | 194 | 0 | 326 | 0 | 1164 | 5888 | 2884 | 2911 | 18425 | 5500 | 0 | 0 | 0 | 0 | 29 | 28 | 0 | 0 | 0 | 0 | 3339 | 11 |
| SRR6757396 | 0 | 31 | 0 | 0 | 47 | 0 | 0 | 2060 | 766 | 3468 | 1743 | 3487 | 0 | 119 | 960 | 1278 | 0 | 204 | 0 | 442 | 19259 | 2027 | 950 | 8188 | 5630 | 0 | 0 | 11 | 23 | 26 | 30 | 0 | 0 | 0 | 0 | 781 | 82 |
| SRR6757397 | 0 | 0 | 0 | 0 | 15 | 0 | 0 | 1637 | 389 | 1072 | 584 | 6967 | 0 | 73 | 15 | 66 | 0 | 332 | 0 | 198 | 3131 | 2257 | 1885 | 13164 | 3793 | 11 | 0 | 0 | 0 | 11 | 0 | 0 | 0 | 0 | 0 | 448 | 0 |
| SRR6757398 | 0 | 58 | 0 | 0 | 147 | 0 | 0 | 971 | 385 | 1385 | 18294 | 21196 | 0 | 75 | 103 | 155 | 19 | 357 | 0 | 138 | 1396 | 969 | 88 | 817 | 2882 | 0 | 24 | 17 | 21 | 58 | 21 | 0 | 0 | 0 | 0 | 241 | 42 |
| SRR6757399 | 0 | 0 | 0 | 0 | 18 | 0 | 0 | 810 | 258 | 1387 | 4381 | 9450 | 0 | 221 | 110 | 61 | 0 | 207 | 0 | 466 | 2781 | 1462 | 2759 | 19934 | 2982 | 15 | 24 | 0 | 0 | 0 | 0 | 0 | 0 | 0 | 0 | 1227 | 0 |
| SRR6757400 | 0 | 27 | 0 | 0 | 90 | 0 | 0 | 2197 | 1145 | 4192 | 680 | 1095 | 0 | 93 | 165 | 135 | 0 | 278 | 0 | 200 | 5185 | 1356 | 556 | 5217 | 4739 | 0 | 11 | 0 | 45 | 65 | 32 | 0 | 0 | 0 | 0 | 900 | 52 |
| SRR6757401 | 0 | 27 | 0 | 0 | 13 | 0 | 0 | 3499 | 822 | 1626 | 1322 | 11196 | 0 | 57 | 70 | 189 | 0 | 317 | 0 | 0 | 6200 | 2618 | 2379 | 17002 | 4292 | 0 | 11 | 0 | 14 | 29 | 13 | 0 | 0 | 0 | 0 | 0 | 0 |
| SRR6757402 | 0 | 22 | 0 | 0 | 51 | 0 | 0 | 4237 | 1200 | 2417 | 2762 | 6420 | 0 | 87 | 50 | 186 | 0 | 367 | 0 | 0 | 4578 | 1625 | 862 | 5640 | 4370 | 0 | 20 | 10 | 17 | 21 | 0 | 0 | 0 | 0 | 0 | 0 | 104 |
| SRR6757403 | 0 | 15 | 0 | 0 | 19 | 0 | 0 | 2353 | 597 | 1100 | 587 | 5960 | 0 | 114 | 0 | 13 | 0 | 310 | 0 | 241 | 4546 | 1996 | 2046 | 11959 | 3816 | 0 | 25 | 0 | 0 | 12 | 0 | 0 | 0 | 0 | 0 | 1609 | 14 |
| SRR6757404 | 0 | 46 | 12 | 0 | 111 | 0 | 0 | 1732 | 844 | 3449 | 658 | 633 | 0 | 604 | 0 | 0 | 0 | 345 | 0 | 61 | 1746 | 1711 | 385 | 3171 | 5460 | 0 | 0 | 12 | 22 | 64 | 51 | 0 | 0 | 0 | 0 | 116 | 65 |
| SRR6757405 | 0 | 0 | 0 | 0 | 15 | 0 | 0 | 1427 | 609 | 1454 | 1636 | 3308 | 0 | 155 | 27 | 36 | 0 | 213 | 0 | 891 | 3085 | 1750 | 2507 | 20199 | 3168 | 14 | 0 | 0 | 0 | 12 | 10 | 0 | 0 | 0 | 0 | 2381 | 0 |
| SRR6757406 | 0 | 0 | 0 | 0 | 34 | 0 | 0 | 2828 | 905 | 2123 | 2657 | 4060 | 0 | 260 | 59 | 67 | 0 | 212 | 0 | 557 | 2584 | 1296 | 1774 | 12755 | 2915 | 0 | 0 | 0 | 0 | 15 | 0 | 0 | 0 | 0 | 0 | 980 | 23 |
| SRR6757407 | 0 | 0 | 0 | 0 | 19 | 0 | 0 | 1296 | 324 | 896 | 5314 | 7429 | 0 | 26 | 0 | 17 | 0 | 280 | 0 | 418 | 3954 | 2169 | 2661 | 13681 | 4457 | 0 | 0 | 0 | 0 | 16 | 0 | 0 | 0 | 0 | 0 | 1552 | 15 |
| SRR6757408 | 0 | 11 | 0 | 0 | 90 | 0 | 0 | 5114 | 1213 | 3645 | 6926 | 7258 | 0 | 42 | 14 | 0 | 0 | 234 | 0 | 42 | 2208 | 1014 | 513 | 3167 | 3409 | 0 | 12 | 0 | 0 | 0 | 0 | 0 | 0 | 0 | 0 | 116 | 51 |
| SRR6757409 | 0 | 15 | 0 | 0 | 26 | 0 | 0 | 1983 | 735 | 2811 | 2526 | 5187 | 0 | 171 | 14 | 16 | 0 | 282 | 0 | 54 | 5597 | 1343 | 1867 | 12513 | 2874 | 0 | 10 | 0 | 0 | 0 | 0 | 0 | 0 | 0 | 0 | 520 | 0 |
| SRR6757410 | 0 | 36 | 0 | 0 | 55 | 0 | 0 | 5403 | 1377 | 3786 | 5166 | 6927 | 0 | 138 | 71 | 196 | 11 | 281 | 0 | 86 | 6584 | 2051 | 1184 | 7805 | 6100 | 0 | 0 | 19 | 30 | 26 | 16 | 0 | 0 | 0 | 0 | 377 | 58 |
| SRR6757411 | 0 | 0 | 0 | 0 | 14 | 0 | 0 | 2732 | 1028 | 1619 | 128 | 1374 | 0 | 207 | 15 | 23 | 0 | 322 | 0 | 192 | 4918 | 2115 | 2559 | 20383 | 3824 | 0 | 10 | 0 | 0 | 23 | 13 | 0 | 0 | 0 | 0 | 1458 | 0 |
| SRR6757412 | 0 | 17 | 0 | 0 | 73 | 0 | 0 | 2205 | 863 | 2531 | 1342 | 2524 | 0 | 137 | 38 | 74 | 0 | 162 | 0 | 210 | 2415 | 1339 | 218 | 2908 | 3636 | 0 | 21 | 15 | 49 | 93 | 45 | 0 | 0 | 0 | 0 | 445 | 39 |
| SRR6757413 | 0 | 29 | 0 | 0 | 16 | 0 | 0 | 1193 | 408 | 2197 | 873 | 1553 | 0 | 432 | 107 | 179 | 0 | 338 | 0 | 883 | 6040 | 3028 | 3899 | 23818 | 4935 | 0 | 0 | 0 | 24 | 50 | 43 | 11 | 11 | 0 | 0 | 2197 | 11 |
| SRR6757414 | 0 | 22 | 0 | 0 | 111 | 0 | 0 | 1499 | 744 | 2349 | 8382 | 12181 | 0 | 261 | 269 | 201 | 0 | 275 | 0 | 49 | 2456 | 1103 | 532 | 3567 | 3109 | 0 | 14 | 15 | 14 | 0 | 0 | 0 | 0 | 0 | 0 | 128 | 41 |
| SRR6757415 | 0 | 0 | 0 | 0 | 24 | 0 | 0 | 560 | 262 | 691 | 532 | 10329 | 0 | 24 | 12 | 0 | 0 | 289 | 0 | 131 | 3867 | 17 | 1811 | 11199 | 4141 | 11 | 12 | 0 | 0 | 0 | 0 | 0 | 0 | 0 | 0 | 203 | 0 |
| SRR6757416 | 0 | 16 | 0 | 0 | 54 | 0 | 0 | 1154 | 354 | 1277 | 2797 | 11547 | 0 | 105 | 0 | 0 | 11 | 214 | 0 | 22 | 3156 | 1881 | 957 | 5277 | 4631 | 0 | 18 | 15 | 0 | 0 | 0 | 0 | 0 | 0 | 0 | 61 | 51 |
| SRR6757417 | 0 | 16 | 0 | 0 | 15 | 0 | 0 | 2117 | 750 | 2013 | 899 | 4188 | 0 | 279 | 17 | 56 | 0 | 280 | 0 | 410 | 5838 | 2441 | 2941 | 20133 | 3713 | 12 | 0 | 0 | 12 | 18 | 12 | 0 | 0 | 0 | 0 | 2860 | 0 |
| SRR6757418 | 0 | 49 | 0 | 0 | 59 | 0 | 0 | 4158 | 1470 | 3102 | 399 | 275 | 0 | 196 | 0 | 10 | 0 | 319 | 0 | 110 | 6275 | 3731 | 1905 | 18488 | 6777 | 0 | 0 | 0 | 26 | 57 | 50 | 0 | 0 | 0 | 0 | 195 | 55 |
| SRR6757419 | 0 | 14 | 0 | 0 | 23 | 0 | 0 | 1367 | 680 | 3686 | 2332 | 4653 | 0 | 352 | 55 | 67 | 0 | 172 | 0 | 407 | 4258 | 2451 | 1913 | 15714 | 4837 | 20 | 0 | 0 | 0 | 21 | 21 | 0 | 0 | 0 | 0 | 2783 | 0 |
| SRR6757420 | 0 | 27 | 0 | 0 | 105 | 0 | 0 | 2629 | 921 | 4542 | 999 | 1563 | 0 | 290 | 24 | 46 | 0 | 271 | 0 | 123 | 4032 | 2334 | 486 | 6058 | 5766 | 0 | 14 | 16 | 20 | 26 | 36 | 0 | 0 | 15 | 0 | 257 | 48 |
| SRR6757421 | 0 | 18 | 0 | 0 | 27 | 0 | 0 | 1872 | 610 | 1937 | 5200 | 10336 | 0 | 115 | 29 | 29 | 0 | 246 | 0 | 0 | 3799 | 2039 | 3028 | 20268 | 3743 | 0 | 0 | 0 | 0 | 0 | 0 | 0 | 0 | 0 | 0 | 0 | 0 |
| SRR6757422 | 0 | 16 | 0 | 0 | 62 | 0 | 0 | 2236 | 690 | 2759 | 2813 | 3397 | 0 | 737 | 0 | 22 | 0 | 255 | 0 | 0 | 2647 | 2124 | 1829 | 10775 | 5553 | 0 | 19 | 0 | 12 | 11 | 0 | 0 | 0 | 0 | 0 | 11 | 52 |
| SRR6757423 | 0 | 0 | 0 | 0 | 17 | 0 | 0 | 1288 | 706 | 3710 | 305 | 1554 | 0 | 292 | 357 | 314 | 0 | 300 | 0 | 160 | 4747 | 1598 | 2760 | 15037 | 2503 | 0 | 11 | 0 | 0 | 0 | 0 | 0 | 0 | 0 | 0 | 1082 | 0 |
| SRR6757424 | 0 | 26 | 20 | 0 | 52 | 0 | 0 | 992 | 684 | 5521 | 1270 | 2372 | 0 | 139 | 730 | 870 | 0 | 247 | 0 | 456 | 3160 | 1947 | 958 | 9232 | 6188 | 0 | 14 | 0 | 16 | 66 | 24 | 0 | 0 | 17 | 0 | 5617 | 13 |
| SRR6757425 | 0 | 15 | 0 | 0 | 13 | 0 | 0 | 549 | 264 | 1636 | 3821 | 6951 | 0 | 176 | 176 | 88 | 0 | 211 | 0 | 110 | 4287 | 2434 | 2820 | 19356 | 4392 | 16 | 0 | 0 | 0 | 0 | 0 | 0 | 0 | 0 | 0 | 329 | 0 |
| SRR6757426 | 0 | 36 | 0 | 0 | 67 | 0 | 0 | 563 | 302 | 2565 | 1704 | 2522 | 0 | 172 | 73 | 47 | 0 | 302 | 0 | 122 | 1640 | 1862 | 435 | 4381 | 4890 | 0 | 18 | 0 | 16 | 37 | 21 | 0 | 0 | 0 | 0 | 392 | 63 |
| SRR6757427 | 0 | 12 | 13 | 0 | 14 | 0 | 0 | 1238 | 723 | 5315 | 783 | 2048 | 0 | 246 | 209 | 285 | 0 | 283 | 0 | 123 | 4728 | 1698 | 2608 | 18541 | 3945 | 0 | 15 | 0 | 0 | 28 | 13 | 0 | 0 | 11 | 0 | 750 | 0 |
| SRR6757428 | 0 | 71 | 0 | 0 | 78 | 0 | 0 | 366 | 222 | 2580 | 505 | 409 | 0 | 353 | 20 | 17 | 0 | 433 | 0 | 52 | 1280 | 1513 | 239 | 4170 | 5200 | 0 | 64 | 0 | 41 | 96 | 60 | 0 | 13 | 0 | 0 | 145 | 131 |
| SRR6757429 | 0 | 13 | 0 | 0 | 10 | 0 | 0 | 353 | 93 | 706 | 3774 | 7102 | 0 | 0 | 19 | 56 | 0 | 301 | 0 | 502 | 4877 | 2375 | 3341 | 20791 | 3764 | 11 | 0 | 0 | 0 | 11 | 0 | 0 | 0 | 0 | 0 | 874 | 0 |
| SRR6757430 | 0 | 0 | 0 | 0 | 58 | 0 | 0 | 1584 | 601 | 1512 | 681 | 5710 | 0 | 71 | 286 | 202 | 0 | 320 | 0 | 476 | 1514 | 1660 | 773 | 7285 | 4229 | 0 | 0 | 0 | 0 | 0 | 0 | 0 | 0 | 0 | 0 | 595 | 45 |
| SRR6757431 | 0 | 0 | 0 | 0 | 14 | 0 | 0 | 277 | 154 | 452 | 6500 | 12969 | 0 | 10 | 21 | 0 | 0 | 272 | 0 | 82 | 4153 | 1256 | 2079 | 9373 | 2275 | 13 | 15 | 0 | 0 | 0 | 0 | 0 | 0 | 0 | 0 | 380 | 0 |
| SRR6757432 | 0 | 67 | 0 | 0 | 40 | 0 | 0 | 1021 | 531 | 1927 | 547 | 633 | 0 | 92 | 38 | 23 | 0 | 227 | 0 | 77 | 5109 | 2301 | 1275 | 12564 | 5278 | 0 | 0 | 0 | 32 | 90 | 45 | 16 | 0 | 0 | 0 | 201 | 38 |
| SRR6757433 | 0 | 0 | 0 | 0 | 24 | 0 | 0 | 511 | 198 | 816 | 4844 | 7499 | 0 | 46 | 0 | 38 | 0 | 294 | 0 | 107 | 3062 | 15 | 1970 | 10802 | 3159 | 0 | 25 | 0 | 0 | 0 | 0 | 0 | 0 | 0 | 0 | 145 | 0 |
| SRR6757434 | 0 | 29 | 0 | 0 | 66 | 0 | 23 | 2076 | 794 | 4098 | 6342 | 7048 | 0 | 47 | 91 | 29 | 27 | 377 | 10 | 56 | 2308 | 1064 | 193 | 3668 | 6166 | 0 | 29 | 0 | 17 | 21 | 0 | 0 | 0 | 0 | 0 | 124 | 411 |
| SRR6757435 | 0 | 18 | 0 | 0 | 18 | 0 | 0 | 2712 | 999 | 2200 | 255 | 1713 | 0 | 244 | 53 | 56 | 0 | 287 | 0 | 470 | 3791 | 2526 | 3305 | 24710 | 5238 | 12 | 0 | 0 | 13 | 43 | 26 | 0 | 0 | 0 | 0 | 1748 | 0 |
| SRR6757436 | 0 | 60 | 0 | 0 | 92 | 0 | 0 | 3663 | 1762 | 4278 | 949 | 804 | 0 | 100 | 859 | 525 | 0 | 227 | 0 | 185 | 1373 | 2341 | 281 | 4672 | 7972 | 0 | 0 | 0 | 28 | 48 | 28 | 0 | 0 | 0 | 0 | 347 | 115 |
| SRR6757437 | 0 | 13 | 0 | 0 | 21 | 0 | 0 | 1969 | 953 | 3803 | 248 | 560 | 0 | 250 | 40 | 43 | 0 | 241 | 0 | 435 | 4602 | 2373 | 2854 | 22071 | 3762 | 0 | 0 | 0 | 10 | 40 | 21 | 0 | 0 | 0 | 0 | 3695 | 0 |
| SRR6757438 | 0 | 44 | 11 | 0 | 82 | 0 | 0 | 4837 | 2006 | 7191 | 593 | 867 | 0 | 106 | 0 | 18 | 0 | 336 | 0 | 222 | 6009 | 1889 | 923 | 13267 | 5848 | 0 | 0 | 0 | 29 | 81 | 59 | 10 | 0 | 0 | 0 | 484 | 68 |
| SRR6757439 | 0 | 28 | 0 | 0 | 23 | 0 | 0 | 1931 | 1318 | 5344 | 197 | 573 | 0 | 275 | 363 | 369 | 0 | 291 | 0 | 1501 | 4866 | 3998 | 3201 | 31378 | 6697 | 0 | 0 | 0 | 15 | 57 | 35 | 0 | 0 | 13 | 0 | 6006 | 0 |
| SRR6757440 | 0 | 49 | 23 | 0 | 71 | 0 | 0 | 1076 | 562 | 4774 | 157 | 161 | 0 | 510 | 204 | 142 | 0 | 274 | 0 | 312 | 2688 | 2876 | 172 | 2706 | 7197 | 0 | 0 | 0 | 33 | 81 | 56 | 0 | 15 | 15 | 0 | 885 | 52 |
| SRR6757441 | 0 | 26 | 0 | 0 | 15 | 0 | 0 | 2468 | 1512 | 7318 | 244 | 365 | 0 | 368 | 78 | 70 | 0 | 357 | 0 | 0 | 5545 | 5496 | 4135 | 65501 | 4714 | 0 | 0 | 0 | 0 | 50 | 41 | 0 | 13 | 24 | 0 | 0 | 0 |
| SRR6757442 | 0 | 30 | 0 | 0 | 122 | 0 | 0 | 980 | 823 | 10276 | 1250 | 1000 | 0 | 305 | 245 | 99 | 0 | 258 | 0 | 0 | 5080 | 1283 | 290 | 4292 | 3296 | 0 | 0 | 14 | 19 | 31 | 17 | 0 | 0 | 0 | 0 | 0 | 55 |
| SRR6757443 | 0 | 0 | 0 | 0 | 27 | 0 | 0 | 3187 | 1658 | 3007 | 1658 | 3067 | 0 | 69 | 69 | 95 | 0 | 381 | 0 | 670 | 3554 | 2976 | 3509 | 36841 | 5582 | 0 | 0 | 0 | 0 | 17 | 0 | 0 | 0 | 0 | 0 | 1160 | 0 |
| SRR6757444 | 0 | 28 | 0 | 0 | 96 | 0 | 0 | 1104 | 757 | 4284 | 453 | 640 | 0 | 215 | 22 | 16 | 0 | 315 | 0 | 513 | 2365 | 2662 | 220 | 4917 | 7582 | 0 | 0 | 0 | 53 | 46 | 43 | 0 | 0 | 0 | 0 | 883 | 31 |
| SRR6757445 | 0 | 0 | 0 | 0 | 22 | 0 | 0 | 3304 | 1013 | 1790 | 1037 | 1825 | 0 | 159 | 136 | 163 | 0 | 369 | 0 | 590 | 3632 | 2296 | 2999 | 25835 | 4267 | 0 | 0 | 0 | 11 | 20 | 19 | 0 | 0 | 0 | 0 | 2900 | 0 |
| SRR6757446 | 0 | 19 | 0 | 0 | 44 | 0 | 0 | 1317 | 300 | 1334 | 879 | 5081 | 0 | 80 | 71 | 62 | 0 | 305 | 0 | 345 | 2188 | 1879 | 1016 | 10159 | 4335 | 0 | 20 | 0 | 0 | 0 | 0 | 0 | 0 | 0 | 0 | 601 | 0 |
| SRR6757447 | 0 | 0 | 0 | 0 | 18 | 0 | 0 | 2037 | 597 | 1742 | 1309 | 2781 | 0 | 131 | 0 | 82 | 0 | 306 | 0 | 761 | 3516 | 2176 | 2602 | 24597 | 4718 | 0 | 0 | 0 | 0 | 17 | 10 | 0 | 0 | 0 | 0 | 2691 | 0 |
| SRR6757448 | 0 | 20 | 0 | 0 | 107 | 0 | 0 | 1128 | 440 | 1746 | 328 | 413 | 0 | 186 | 189 | 69 | 0 | 458 | 0 | 62 | 1111 | 1033 | 112 | 2627 | 4574 | 0 | 0 | 22 | 54 | 83 | 45 | 0 | 0 | 0 | 0 | 152 | 33 |
| SRR6757449 | 0 | 0 | 0 | 0 | 23 | 0 | 0 | 305 | 139 | 703 | 4930 | 8140 | 0 | 28 | 102 | 370 | 0 | 275 | 0 | 213 | 1505 | 977 | 862 | 5797 | 2590 | 14 | 11 | 0 | 0 | 0 | 0 | 0 | 0 | 0 | 0 | 740 | 0 |
| SRR6757450 | 0 | 25 | 0 | 0 | 32 | 0 | 0 | 378 | 118 | 789 | 14757 | 13456 | 0 | 92 | 2015 | 3257 | 16 | 203 | 0 | 336 | 4882 | 2258 | 658 | 6956 | 6260 | 12 | 0 | 33 | 16 | 23 | 13 | 0 | 0 | 0 | 0 | 739 | 102 |
| SRR6757451 | 0 | 18 | 0 | 0 | 14 | 0 | 0 | 2928 | 785 | 1668 | 105 | 343 | 0 | 225 | 0 | 0 | 0 | 360 | 0 | 14 | 4939 | 2057 | 2886 | 18338 | 4748 | 0 | 0 | 0 | 11 | 53 | 29 | 0 | 0 | 0 | 0 | 176 | 0 |
| SRR6757452 | 0 | 46 | 25 | 0 | 33 | 0 | 0 | 668 | 291 | 2564 | 1301 | 854 | 0 | 54 | 0 | 0 | 0 | 146 | 0 | 20 | 1514 | 2258 | 268 | 9790 | 6833 | 54 | 0 | 12 | 194 | 10 | 89 | 24 | 16 | 20 | 0 | 43 | 165 |
| SRR6757453 | 0 | 12 | 0 | 0 | 0 | 0 | 0 | 2971 | 827 | 2199 | 215 | 437 | 0 | 383 | 24 | 29 | 0 | 360 | 0 | 104 | 4917 | 2012 | 4231 | 20493 | 3773 | 0 | 0 | 0 | 12 | 25 | 20 | 0 | 0 | 0 | 0 | 511 | 0 |
| SRR6757454 | 0 | 17 | 27 | 0 | 52 | 0 | 0 | 3349 | 1384 | 6156 | 1455 | 2217 | 0 | 153 | 27 | 58 | 0 | 360 | 0 | 342 | 3389 | 1738 | 943 | 8047 | 5689 | 0 | 0 | 13 | 0 | 42 | 22 | 0 | 0 | 0 | 0 | 1541 | 19 |
| SRR6757455 | 0 | 26 | 0 | 0 | 13 | 0 | 0 | 662 | 158 | 1258 | 394 | 6375 | 0 | 130 | 28 | 56 | 0 | 303 | 0 | 158 | 3819 | 1713 | 2675 | 14722 | 3720 | 0 | 14 | 0 | 0 | 11 | 0 | 0 | 0 | 0 | 0 | 531 | 0 |
| SRR6757456 | 0 | 56 | 0 | 0 | 107 | 0 | 0 | 4358 | 1590 | 6933 | 5295 | 4912 | 0 | 49 | 0 | 0 | 0 | 271 | 0 | 37 | 1646 | 834 | 81 | 1996 | 3270 | 0 | 0 | 12 | 27 | 35 | 15 | 0 | 0 | 0 | 0 | 147 | 160 |
| SRR6757457 | 0 | 20 | 0 | 0 | 13 | 0 | 0 | 953 | 309 | 643 | 3973 | 8931 | 0 | 41 | 0 | 0 | 0 | 221 | 0 | 98 | 3943 | 1484 | 2733 | 14082 | 3612 | 12 | 0 | 0 | 0 | 0 | 0 | 0 | 0 | 0 | 0 | 342 | 0 |
| SRR6757458 | 0 | 11 | 0 | 0 | 60 | 15 | 0 | 874 | 265 | 1358 | 4336 | 20128 | 0 | 12 | 15 | 12 | 11 | 235 | 0 | 42 | 3737 | 1221 | 583 | 3196 | 3427 | 0 | 0 | 0 | 0 | 0 | 0 | 0 | 0 | 0 | 0 | 169 | 44 |
| SRR6757459 | 0 | 11 | 0 | 0 | 17 | 0 | 0 | 3377 | 1112 | 3919 | 524 | 1531 | 0 | 237 | 0 | 0 | 0 | 404 | 0 | 125 | 4546 | 2436 | 2967 | 20718 | 4766 | 0 | 0 | 0 | 0 | 12 | 10 | 0 | 0 | 0 | 0 | 369 | 0 |
| SRR6757460 | 0 | 44 | 0 | 0 | 63 | 0 | 0 | 3272 | 1260 | 3515 | 1678 | 3992 | 0 | 51 | 46 | 0 | 13 | 243 | 0 | 71 | 4176 | 883 | 177 | 2052 | 4399 | 0 | 13 | 12 | 18 | 19 | 0 | 0 | 0 | 0 | 0 | 138 | 80 |
| SRR6757461 | 0 | 17 | 0 | 0 | 15 | 0 | 0 | 6848 | 1775 | 1897 | 8174 | 21657 | 0 | 55 | 0 | 0 | 0 | 322 | 0 | 83 | 5772 | 12 | 3500 | 16605 | 3173 | 0 | 16 | 0 | 0 | 0 | 0 | 0 | 0 | 0 | 0 | 582 | 0 |
| SRR6757462 | 0 | 40 | 0 | 0 | 168 | 13 | 0 | 1124 | 582 | 2421 | 3251 | 5335 | 0 | 1114 | 0 | 0 | 17 | 232 | 0 | 20 | 2734 | 747 | 291 | 3242 | 4054 | 0 | 11 | 0 | 19 | 0 | 0 | 0 | 0 | 0 | 0 | 76 | 38 |
| SRR6757463 | 0 | 14 | 0 | 0 | 41 | 0 | 0 | 3693 | 1682 | 4818 | 5501 | 10316 | 0 | 89 | 76 | 30 | 0 | 296 | 0 | 194 | 3944 | 2541 | 2620 | 19307 | 4600 | 0 | 0 | 0 | 0 | 14 | 0 | 0 | 0 | 0 | 0 | 620 | 0 |
| SRR6757464 | 0 | 12 | 19 | 0 | 144 | 0 | 0 | 824 | 206 | 911 | 1323 | 1350 | 0 | 1651 | 0 | 0 | 11 | 241 | 0 | 35 | 1744 | 2553 | 232 | 3779 | 11373 | 0 | 0 | 34 | 11 | 111 | 51 | 0 | 0 | 14 | 0 | 141 | 58 |
| SRR6757465 | 0 | 22 | 0 | 0 | 41 | 0 | 0 | 3711 | 971 | 1747 | 485 | 2365 | 0 | 143 | 0 | 21 | 0 | 357 | 0 | 514 | 4605 | 3183 | 2574 | 18975 | 4834 | 0 | 0 | 0 | 12 | 22 | 0 | 0 | 0 | 0 | 0 | 4560 | 13 |
| SRR6757466 | 0 | 61 | 0 | 0 | 153 | 0 | 0 | 1315 | 443 | 1752 | 2259 | 2260 | 0 | 101 | 23 | 22 | 0 | 268 | 0 | 109 | 3031 | 2351 | 134 | 2490 | 9274 | 0 | 0 | 0 | 103 | 95 | 53 | 0 | 0 | 14 | 0 | 201 | 94 |
| SRR6757467 | 0 | 15 | 0 | 0 | 35 | 0 | 0 | 1643 | 546 | 1154 | 2012 | 14360 | 0 | 107 | 0 | 23 | 22 | 236 | 0 | 99 | 3580 | 877 | 1536 | 8313 | 2390 | 14 | 12 | 19 | 0 | 0 | 0 | 0 | 0 | 0 | 0 | 619 | 0 |
| SRR6757468 | 0 | 25 | 0 | 0 | 81 | 12 | 0 | 829 | 325 | 979 | 2986 | 15135 | 0 | 115 | 56 | 67 | 0 | 239 | 0 | 36 | 6603 | 1390 | 866 | 6256 | 3367 | 0 | 0 | 21 | 0 | 0 | 0 | 0 | 0 | 0 | 0 | 123 | 34 |
| SRR6761159 | 0 | 0 | 0 | 0 | 0 | 0 | 0 | 6885 | 0 | 4303 | 0 | 0 | 0 | 6885 | 0 | 0 | 0 | 12 | 0 | 3442 | 44 | 227 | 391 | 634 | 1880 | 29 | 0 | 0 | 0 | 0 | 0 | 0 | 0 | 0 | 0 | 861 | 0 |
| SRR6761162 | 0 | 0 | 0 | 0 | 0 | 0 | 0 | 0 | 0 | 0 | 0 | 0 | 0 | 2183 | 0 | 0 | 0 | 0 | 0 | 0 | 0 | 26 | 31 | 48 | 1921 | 17 | 0 | 0 | 0 | 0 | 0 | 0 | 0 | 0 | 0 | 0 | 0 |
| SRR6761163 | 0 | 0 | 0 | 0 | 0 | 0 | 0 | 1946 | 0 | 1946 | 973 | 0 | 0 | 5837 | 973 | 0 | 0 | 0 | 0 | 0 | 38 | 130 | 179 | 216 | 1169 | 30 | 108 | 0 | 0 | 0 | 0 | 0 | 0 | 0 | 0 | 2918 | 0 |
| SRR6761165 | 0 | 0 | 0 | 0 | 0 | 0 | 0 | 4884 | 0 | 3663 | 0 | 0 | 0 | 2442 | 0 | 0 | 0 | 0 | 0 | 0 | 34 | 211 | 252 | 302 | 1570 | 29 | 90 | 0 | 0 | 0 | 0 | 0 | 0 | 0 | 0 | 0 | 0 |
| SRR6761167 | 0 | 0 | 0 | 0 | 0 | 0 | 0 | 8571 | 0 | 8571 | 0 | 0 | 0 | 14286 | 0 | 0 | 384 | 0 | 0 | 0 | 0 | 88 | 213 | 264 | 1983 | 65 | 0 | 0 | 0 | 0 | 0 | 0 | 0 | 0 | 0 | 8571 | 0 |
| SRR6761175 | 0 | 0 | 0 | 0 | 0 | 0 | 0 | 773 | 2320 | 1547 | 0 | 0 | 0 | 4640 | 8507 | 1547 | 0 | 0 | 0 | 0 | 24 | 174 | 169 | 211 | 1927 | 18 | 0 | 0 | 0 | 0 | 0 | 0 | 0 | 0 | 0 | 3867 | 0 |
| SRR6761179 | 0 | 0 | 0 | 0 | 0 | 0 | 0 | 0 | 0 | 0 | 0 | 0 | 0 | 0 | 0 | 0 | 0 | 0 | 0 | 0 | 40 | 127 | 115 | 140 | 2264 | 41 | 0 | 0 | 0 | 0 | 0 | 0 | 0 | 0 | 0 | 5906 | 0 |
| SRR6761181 | 111 | 0 | 0 | 0 | 0 | 56 | 0 | 585 | 0 | 1754 | 0 | 0 | 0 | 3509 | 0 | 1170 | 0 | 0 | 0 | 3509 | 52 | 93 | 120 | 212 | 3553 | 16 | 111 | 0 | 0 | 0 | 0 | 0 | 0 | 0 | 0 | 8772 | 0 |
| SRR6761182 | 0 | 0 | 0 | 0 | 0 | 0 | 0 | 536 | 0 | 4823 | 544 | 1072 | 0 | 3215 | 0 | 0 | 0 | 0 | 0 | 536 | 34 | 105 | 108 | 128 | 2265 | 17 | 126 | 0 | 0 | 0 | 0 | 0 | 0 | 0 | 0 | 3751 | 0 |
| SRR6761183 | 0 | 0 | 0 | 0 | 0 | 0 | 0 | 0 | 0 | 5425 | 0 | 603 | 0 | 6028 | 1206 | 0 | 0 | 0 | 0 | 0 | 56 | 227 | 72 | 268 | 3051 | 71 | 55 | 0 | 0 | 0 | 0 | 0 | 0 | 0 | 0 | 3014 | 0 |
| SRR6761184 | 0 | 0 | 0 | 0 | 0 | 32 | 0 | 384 | 0 | 7683 | 0 | 1153 | 0 | 1153 | 384 | 384 | 32 | 0 | 0 | 768 | 19 | 90 | 51 | 150 | 4055 | 53 | 32 | 0 | 0 | 0 | 0 | 0 | 0 | 0 | 0 | 2305 | 0 |
| SRR6761188 | 0 | 0 | 0 | 0 | 0 | 0 | 0 | 499 | 0 | 1496 | 502 | 0 | 0 | 2991 | 0 | 0 | 0 | 24 | 0 | 499 | 46 | 423 | 170 | 377 | 3568 | 87 | 75 | 0 | 0 | 0 | 0 | 0 | 0 | 0 | 0 | 1994 | 0 |
| SRR6761192 | 0 | 0 | 0 | 0 | 0 | 0 | 0 | 0 | 0 | 3175 | 0 | 0 | 0 | 0 | 0 | 0 | 0 | 120 | 0 | 0 | 86 | 395 | 189 | 429 | 2213 | 103 | 0 | 0 | 0 | 0 | 0 | 0 | 0 | 0 | 0 | 6349 | 0 |
| SRR6761198 | 0 | 0 | 0 | 0 | 0 | 71 | 0 | 0 | 0 | 4125 | 1289 | 1031 | 0 | 2578 | 516 | 0 | 0 | 19 | 0 | 0 | 110 | 596 | 193 | 691 | 3661 | 55 | 18 | 18 | 0 | 0 | 0 | 0 | 0 | 0 | 0 | 5156 | 0 |
| SRR6767691 | 0 | 0 | 0 | 0 | 0 | 0 | 62 | 7714 | 0 | 0 | 654 | 1286 | 0 | 771 | 0 | 0 | 50 | 0 | 0 | 0 | 65 | 92 | 44 | 483 | 3417 | 669 | 386 | 0 | 0 | 0 | 0 | 0 | 0 | 0 | 0 | 643 | 0 |
| SRR6767701 | 0 | 0 | 0 | 0 | 0 | 0 | 0 | 5038 | 0 | 1338 | 27 | 0 | 0 | 0 | 0 | 0 | 261 | 0 | 0 | 1338 | 0 | 11 | 0 | 20 | 2567 | 208 | 0 | 0 | 0 | 0 | 0 | 0 | 0 | 0 | 0 | 787 | 0 |
| SRR6767712 | 0 | 0 | 0 | 0 | 0 | 0 | 13 | 11583 | 0 | 0 | 0 | 0 | 0 | 0 | 0 | 0 | 13 | 0 | 0 | 0 | 24 | 20 | 15 | 359 | 2990 | 162 | 107 | 0 | 0 | 0 | 0 | 0 | 0 | 0 | 0 | 0 | 0 |
| SRR6767713 | 0 | 0 | 0 | 0 | 0 | 0 | 0 | 9049 | 0 | 0 | 0 | 0 | 0 | 0 | 0 | 0 | 95 | 12 | 17 | 0 | 20 | 0 | 15 | 407 | 3283 | 308 | 258 | 0 | 0 | 0 | 0 | 0 | 0 | 0 | 0 | 218 | 0 |
| SRR6821085 | 0 | 173 | 11 | 0 | 37 | 0 | 0 | 998 | 715 | 9659 | 1697 | 1498 | 0 | 1667 | 227 | 348 | 28 | 228 | 0 | 246 | 780 | 1142 | 103 | 3412 | 3429 | 77 | 11 | 0 | 11 | 17 | 15 | 11 | 0 | 22 | 0 | 471 | 12 |
| SRR6821086 | 0 | 116 | 35 | 0 | 86 | 0 | 19 | 1338 | 992 | 10256 | 303 | 356 | 0 | 191 | 110 | 226 | 10 | 476 | 0 | 145 | 691 | 838 | 698 | 2068 | 2577 | 15 | 40 | 0 | 13 | 0 | 0 | 0 | 22 | 23 | 0 | 560 | 0 |
| SRR6821087 | 0 | 210 | 0 | 0 | 94 | 11 | 28 | 838 | 652 | 10380 | 1460 | 1822 | 0 | 144 | 63 | 105 | 156 | 396 | 0 | 446 | 438 | 327 | 227 | 1205 | 2559 | 19 | 78 | 0 | 0 | 0 | 0 | 0 | 0 | 0 | 0 | 1499 | 0 |
| SRR6821088 | 0 | 101 | 23 | 0 | 0 | 0 | 0 | 838 | 614 | 5666 | 528 | 469 | 0 | 1046 | 880 | 901 | 0 | 145 | 0 | 252 | 4905 | 6311 | 393 | 11865 | 4850 | 114 | 0 | 0 | 36 | 39 | 32 | 13 | 16 | 55 | 0 | 723 | 0 |
| SRR6821089 | 0 | 66 | 0 | 0 | 0 | 0 | 0 | 143 | 222 | 8486 | 953 | 892 | 0 | 2776 | 1061 | 2752 | 0 | 157 | 0 | 822 | 3326 | 8151 | 85 | 438 | 162 | 461 | 0 | 0 | 0 | 0 | 0 | 0 | 0 | 0 | 0 | 3905 | 0 |
| SRR6821090 | 0 | 63 | 0 | 0 | 0 | 0 | 0 | 369 | 356 | 7746 | 175 | 95 | 0 | 2528 | 2082 | 1356 | 0 | 153 | 0 | 262 | 2885 | 4274 | 446 | 8681 | 3388 | 29 | 0 | 0 | 0 | 14 | 22 | 0 | 0 | 29 | 0 | 1104 | 0 |
| SRR6823984 | 0 | 2385 | 1190 | 58 | 1572 | 0 | 0 | 438 | 328 | 10040 | 29198 | 24475 | 0 | 2440 | 16 | 94 | 22 | 1705 | 339 | 469 | 925 | 4813 | 1648 | 39308 | 804 | 24 | 12 | 425 | 1440 | 1045 | 3909 | 497 | 366 | 324 | 118 | 2455 | 418 |
| SRR6823985 | 0 | 2861 | 585 | 37 | 596 | 18 | 10 | 35 | 175 | 3530 | 45155 | 37606 | 0 | 2272 | 17 | 0 | 10 | 1191 | 357 | 891 | 591 | 3939 | 1060 | 34639 | 458 | 39 | 122 | 703 | 845 | 797 | 3042 | 448 | 275 | 176 | 15 | 4701 | 345 |
| SRR6823986 | 0 | 4002 | 67 | 302 | 2571 | 21 | 0 | 751 | 558 | 2998 | 18432 | 22335 | 0 | 485 | 40 | 48 | 13 | 2396 | 204 | 259 | 734 | 2519 | 703 | 17613 | 1421 | 74 | 13 | 1466 | 5219 | 2967 | 4916 | 1290 | 1087 | 866 | 169 | 1083 | 1038 |
| SRR6823987 | 0 | 277 | 72 | 698 | 2060 | 0 | 16 | 1497 | 1520 | 3329 | 1536 | 19317 | 0 | 668 | 35 | 35 | 43 | 2765 | 252 | 426 | 390 | 2395 | 715 | 16456 | 1022 | 107 | 0 | 81 | 247 | 180 | 269 | 68 | 41 | 32 | 11 | 1751 | 569 |
| SRR6823990 | 0 | 3362 | 838 | 177 | 1869 | 0 | 0 | 513 | 1871 | 9105 | 8294 | 16885 | 0 | 4288 | 298 | 166 | 13 | 589 | 162 | 1175 | 335 | 210 | 21 | 845 | 437 | 75 | 38 | 138 | 1061 | 1026 | 2122 | 367 | 267 | 254 | 232 | 3046 | 977 |
| SRR6823991 | 0 | 688 | 47 | 861 | 1513 | 0 | 16 | 3669 | 6079 | 6928 | 413 | 19223 | 0 | 904 | 137 | 55 | 16 | 2376 | 173 | 630 | 640 | 327 | 23 | 889 | 504 | 57 | 86 | 64 | 76 | 55 | 119 | 20 | 12 | 10 | 54 | 1177 | 1244 |
| SRR6829958 | 0 | 63 | 187 | 21 | 82 | 10 | 0 | 1039 | 2215 | 13006 | 901 | 2289 | 0 | 2293 | 0 | 0 | 0 | 1136 | 20 | 1945 | 781 | 630 | 183 | 1529 | 338 | 0 | 179 | 15 | 186 | 97 | 412 | 57 | 38 | 26 | 65 | 9449 | 374 |
| SRR6829959 | 0 | 81 | 129 | 31 | 139 | 26 | 15 | 1855 | 3120 | 21393 | 487 | 730 | 12 | 1293 | 0 | 0 | 11 | 1111 | 10 | 757 | 827 | 902 | 265 | 2598 | 339 | 0 | 45 | 0 | 137 | 36 | 167 | 28 | 20 | 13 | 23 | 3737 | 208 |
| SRR6829960 | 0 | 2374 | 250 | 26 | 115 | 0 | 19 | 1555 | 3876 | 22373 | 4641 | 8963 | 0 | 2655 | 0 | 0 | 0 | 1113 | 102 | 1142 | 758 | 550 | 172 | 1307 | 450 | 0 | 393 | 56 | 322 | 4077 | 15032 | 1500 | 1179 | 781 | 51 | 3014 | 390 |
| SRR6829961 | 0 | 76 | 131 | 27 | 146 | 22 | 20 | 1448 | 2331 | 19511 | 1321 | 2680 | 17 | 1243 | 0 | 0 | 45 | 1119 | 72 | 392 | 882 | 591 | 301 | 2416 | 274 | 0 | 354 | 22 | 262 | 29 | 116 | 16 | 376 | 191 | 30 | 1847 | 181 |
| SRR6829962 | 0 | 63 | 170 | 21 | 105 | 22 | 16 | 1549 | 3367 | 15547 | 2211 | 5263 | 15 | 3974 | 0 | 0 | 24 | 828 | 30 | 817 | 499 | 535 | 136 | 1055 | 333 | 0 | 225 | 35 | 206 | 64 | 290 | 31 | 23 | 16 | 23 | 1523 | 272 |
| SRR6829963 | 0 | 2218 | 113 | 17 | 64 | 31 | 10 | 1591 | 2127 | 18444 | 481 | 4264 | 23 | 1257 | 0 | 0 | 15 | 734 | 30 | 536 | 656 | 318 | 268 | 1654 | 122 | 0 | 767 | 17 | 253 | 810 | 5936 | 925 | 474 | 244 | 10 | 2211 | 181 |
| SRR6831609 | 0 | 324 | 0 | 24 | 255 | 0 | 0 | 1202 | 1570 | 2639 | 1904 | 7615 | 0 | 835 | 0 | 0 | 0 | 531 | 44 | 4910 | 396 | 922 | 0 | 259 | 3595 | 434 | 22 | 22 | 0 | 22 | 22 | 0 | 0 | 0 | 0 | 7949 | 0 |
| SRR6831611 | 0 | 157 | 20 | 19 | 173 | 20 | 0 | 1593 | 2045 | 2284 | 717 | 2709 | 0 | 2231 | 27 | 0 | 0 | 506 | 130 | 11632 | 210 | 903 | 21 | 335 | 2647 | 382 | 138 | 0 | 0 | 0 | 0 | 0 | 0 | 0 | 0 | 18696 | 0 |
| SRR6831613 | 0 | 29 | 0 | 24 | 262 | 57 | 0 | 4195 | 8028 | 6582 | 13 | 1085 | 0 | 1881 | 0 | 0 | 0 | 502 | 84 | 7160 | 167 | 1122 | 11 | 225 | 3884 | 242 | 29 | 0 | 29 | 0 | 0 | 0 | 0 | 0 | 0 | 9620 | 12 |
| SRR6837863 | 0 | 0 | 0 | 0 | 0 | 12 | 0 | 4239 | 2058 | 188 | 273 | 2430 | 0 | 0 | 0 | 0 | 63 | 574 | 0 | 19 | 2096 | 3228 | 57 | 30334 | 25809 | 0 | 14 | 68 | 105 | 217 | 76 | 0 | 0 | 0 | 0 | 12 | 783 |
| SRR6837867 | 12 | 13 | 0 | 0 | 0 | 0 | 0 | 3997 | 3452 | 759 | 1367 | 2830 | 0 | 18 | 0 | 0 | 95 | 702 | 0 | 27 | 2179 | 3324 | 55 | 27913 | 22540 | 0 | 12 | 54 | 68 | 65 | 22 | 0 | 0 | 0 | 0 | 23 | 717 |
| SRR6837868 | 0 | 0 | 0 | 0 | 0 | 0 | 0 | 6404 | 4775 | 1656 | 208 | 1990 | 0 | 37 | 0 | 0 | 87 | 578 | 0 | 37 | 1953 | 3299 | 50 | 29679 | 10 | 14 | 11 | 74 | 72 | 123 | 61 | 0 | 0 | 0 | 0 | 60 | 745 |
| SRR6880326 | 0 | 0 | 0 | 0 | 0 | 25 | 58 | 1721 | 0 | 33 | 345 | 519 | 0 | 62 | 97 | 155 | 62 | 87 | 44 | 2169 | 348 | 4373 | 0 | 968 | 9717 | 2176 | 0 | 0 | 0 | 0 | 0 | 0 | 0 | 0 | 0 | 1495 | 0 |
| SRR6880328 | 0 | 23 | 0 | 0 | 0 | 18 | 30 | 3071 | 33 | 109 | 10600 | 7125 | 0 | 11 | 0 | 33 | 28 | 0 | 13 | 1377 | 75 | 133 | 0 | 14 | 3637 | 164 | 0 | 0 | 0 | 0 | 0 | 0 | 0 | 0 | 0 | 951 | 31 |
| SRR6880329 | 0 | 13 | 0 | 0 | 0 | 22 | 46 | 2733 | 38 | 150 | 1526 | 8807 | 0 | 0 | 0 | 0 | 33 | 0 | 11 | 1126 | 82 | 129 | 0 | 14 | 3425 | 142 | 11 | 0 | 0 | 0 | 0 | 0 | 0 | 0 | 0 | 721 | 35 |
| SRR6880330 | 0 | 15 | 0 | 0 | 0 | 16 | 42 | 3410 | 128 | 293 | 5152 | 6197 | 0 | 37 | 37 | 55 | 20 | 0 | 14 | 1613 | 64 | 120 | 0 | 17 | 3305 | 135 | 0 | 0 | 0 | 0 | 0 | 0 | 0 | 0 | 0 | 1852 | 38 |

### Supp Table S8.4.Columns_DI_to_ES

| | | | | | | | | | | | | | | | | | | | | | | | | | | | | | | | | | | | | | |
|---|---|---|---|---|---|---|---|---|---|---|---|---|---|---|---|---|---|---|---|---|---|---|---|---|---|---|---|---|---|---|---|---|---|---|---|---|---|
| SRR6880331 | 0 | 20 | 0 | 0 | 0 | 15 | 48 | 1634 | 0 | 323 | 440 | 7677 | 0 | 65 | 65 | 22 | 23 | 0 | 15 | 1032 | 48 | 97 | 0 | 12 | 2301 | 112 | 0 | 0 | 0 | 0 | 0 | 0 | 0 | 0 | 0 | 860 | 32 |
| SRR6880333 | 10 | 10 | 0 | 0 | 0 | 90 | 202 | 13206 | 151 | 434 | 2286 | 17158 | 0 | 117 | 41 | 262 | 196 | 223 | 49 | 1535 | 2598 | 2397 | 0 | 214 | 10945 | 290 | 62 | 0 | 0 | 0 | 0 | 0 | 0 | 0 | 0 | 2885 | 0 |
| SRR6880334 | 12 | 10 | 0 | 0 | 0 | 45 | 78 | 15597 | 191 | 225 | 26429 | 37962 | 0 | 52 | 69 | 711 | 110 | 196 | 36 | 2227 | 2652 | 1782 | 0 | 247 | 9317 | 391 | 31 | 0 | 0 | 0 | 0 | 0 | 0 | 0 | 0 | 3345 | 0 |
| SRR6880335 | 0 | 25 | 0 | 0 | 0 | 94 | 180 | 17042 | 244 | 581 | 18549 | 21126 | 0 | 59 | 101 | 429 | 164 | 322 | 62 | 1499 | 2098 | 2176 | 0 | 164 | 10818 | 234 | 52 | 0 | 0 | 0 | 0 | 0 | 0 | 0 | 0 | 2577 | 0 |
| SRR6880336 | 0 | 0 | 0 | 0 | 0 | 11 | 20 | 23836 | 130 | 309 | 49 | 49 | 0 | 276 | 33 | 16 | 11 | 482 | 1886 | 3902 | 88 | 285 | 110 | 1359 | 7586 | 285 | 0 | 0 | 0 | 0 | 0 | 0 | 0 | 0 | 0 | 5561 | 0 |
| SRR6880337 | 0 | 0 | 0 | 0 | 0 | 35 | 33 | 21346 | 193 | 735 | 15 | 52 | 0 | 267 | 364 | 319 | 27 | 444 | 322 | 2679 | 318 | 1140 | 67 | 912 | 12160 | 110 | 0 | 0 | 0 | 0 | 0 | 0 | 0 | 0 | 0 | 11723 | 20 |
| SRR6880338 | 0 | 0 | 0 | 0 | 0 | 19 | 65 | 2826 | 0 | 17 | 385 | 560 | 0 | 20 | 27 | 58 | 96 | 87 | 21 | 2747 | 379 | 4138 | 0 | 926 | 8748 | 2034 | 0 | 0 | 0 | 0 | 0 | 0 | 0 | 0 | 0 | 1554 | 0 |
| SRR6880339 | 0 | 0 | 0 | 0 | 0 | 21 | 34 | 2174 | 0 | 13 | 244 | 303 | 0 | 20 | 13 | 38 | 37 | 82 | 34 | 2682 | 425 | 3507 | 0 | 878 | 9543 | 1303 | 0 | 0 | 0 | 0 | 0 | 0 | 0 | 0 | 0 | 1681 | 0 |
| SRR6880341 | 0 | 0 | 0 | 0 | 0 | 16 | 56 | 1389 | 36 | 193 | 135 | 246 | 0 | 213 | 983 | 1041 | 33 | 20 | 23 | 549 | 137 | 1564 | 0 | 358 | 5521 | 1254 | 16 | 0 | 0 | 0 | 0 | 0 | 0 | 0 | 0 | 558 | 12 |
| SRR6880342 | 0 | 0 | 0 | 0 | 0 | 11 | 11 | 1629 | 182 | 126 | 80 | 79 | 0 | 127 | 93 | 165 | 55 | 19 | 16 | 570 | 115 | 935 | 13 | 634 | 4881 | 1254 | 0 | 0 | 0 | 0 | 0 | 0 | 0 | 0 | 0 | 565 | 0 |
| SRR6880343 | 0 | 0 | 0 | 0 | 0 | 0 | 40 | 2149 | 120 | 189 | 55 | 70 | 0 | 203 | 106 | 104 | 35 | 24 | 31 | 334 | 116 | 977 | 0 | 391 | 5661 | 794 | 0 | 0 | 0 | 0 | 0 | 0 | 0 | 0 | 0 | 299 | 0 |
| SRR6880345 | 0 | 31 | 0 | 0 | 0 | 57 | 151 | 6161 | 133 | 1234 | 14 | 85 | 0 | 322 | 5728 | 4209 | 57 | 1163 | 366 | 1945 | 581 | 1012 | 60 | 536 | 6424 | 40 | 0 | 0 | 0 | 0 | 0 | 0 | 0 | 0 | 0 | 7084 | 45 |
| SRR6880351 | 0 | 0 | 0 | 0 | 0 | 24 | 133 | 3647 | 54 | 3959 | 24 | 34 | 0 | 1997 | 521 | 614 | 72 | 47 | 12 | 643 | 964 | 1186 | 2913 | 39258 | 3220 | 59 | 0 | 0 | 0 | 0 | 0 | 0 | 0 | 0 | 0 | 2342 | 33 |
| SRR6880356 | 0 | 0 | 0 | 0 | 0 | 35 | 212 | 22963 | 167 | 482 | 74 | 37 | 0 | 528 | 204 | 195 | 62 | 349 | 206 | 1686 | 390 | 1489 | 124 | 1999 | 18009 | 57 | 0 | 0 | 0 | 0 | 0 | 0 | 0 | 0 | 0 | 7355 | 0 |
| SRR6880357 | 0 | 0 | 0 | 0 | 0 | 11 | 68 | 29866 | 241 | 231 | 21 | 154 | 0 | 800 | 135 | 68 | 18 | 329 | 237 | 617 | 440 | 885 | 66 | 1046 | 19230 | 42 | 0 | 0 | 0 | 0 | 0 | 0 | 0 | 0 | 0 | 2507 | 21 |
| SRR6898464 | 0 | 30 | 11 | 0 | 0 | 0 | 12 | 440 | 81 | 3776 | 11173 | 7880 | 0 | 1343 | 2435 | 2920 | 0 | 328 | 12 | 578 | 837 | 1135 | 0 | 17 | 3476 | 167 | 0 | 0 | 0 | 10 | 10 | 0 | 0 | 18 | 26 | 1472 | 32 |
| SRR6914752 | 0 | 210 | 42 | 0 | 110 | 0 | 0 | 287 | 162 | 549 | 21048 | 9614 | 0 | 62 | 0 | 0 | 42 | 54 | 49 | 262 | 934 | 278 | 0 | 130 | 5535 | 156 | 0 | 2729 | 1386 | 2834 | 7769 | 504 | 462 | 441 | 273 | 998 | 912 |
| SRR6914753 | 0 | 65 | 240 | 54 | 68 | 0 | 13 | 51 | 84 | 341 | 14187 | 10493 | 0 | 40 | 0 | 0 | 0 | 86 | 28 | 596 | 1463 | 556 | 0 | 153 | 7410 | 304 | 0 | 44 | 17 | 27 | 49 | 14 | 641 | 628 | 109 | 1253 | 1360 |
| SRR6914754 | 0 | 339 | 86 | 0 | 104 | 0 | 13 | 330 | 422 | 2593 | 5099 | 4989 | 0 | 344 | 0 | 0 | 0 | 167 | 50 | 64 | 1794 | 346 | 0 | 327 | 5864 | 121 | 0 | 785 | 812 | 12 | 23 | 206 | 153 | 253 | 166 | 229 | 726 |
| SRR6914755 | 0 | 63 | 267 | 74 | 56 | 0 | 11 | 47 | 171 | 2007 | 6421 | 6354 | 0 | 187 | 15 | 15 | 0 | 276 | 35 | 330 | 2544 | 741 | 0 | 248 | 5990 | 171 | 0 | 15 | 12 | 13 | 27 | 372 | 250 | 328 | 42 | 806 | 1753 |
| SRR6928219 | 0 | 3754 | 86 | 53 | 1986 | 0 | 0 | 1251 | 5061 | 20730 | 8828 | 19186 | 0 | 2634 | 76 | 184 | 0 | 653 | 111 | 220 | 441 | 211 | 141 | 8950 | 602 | 82 | 10 | 195 | 473 | 620 | 1838 | 550 | 562 | 339 | 80 | 751 | 347 |
| SRR6928220 | 0 | 417 | 65 | 68 | 1947 | 0 | 0 | 954 | 2923 | 19616 | 44152 | 51297 | 0 | 2051 | 53 | 63 | 0 | 416 | 446 | 635 | 799 | 307 | 403 | 13688 | 586 | 59 | 0 | 105 | 65 | 95 | 234 | 121 | 120 | 66 | 14 | 1604 | 160 |
| SRR6928221 | 0 | 1559 | 116 | 29 | 3422 | 0 | 0 | 1604 | 8934 | 65618 | 10006 | 53000 | 0 | 2082 | 31 | 70 | 16 | 358 | 203 | 397 | 635 | 249 | 358 | 9709 | 421 | 17 | 35 | 73 | 201 | 602 | 1511 | 109 | 155 | 129 | 22 | 1260 | 274 |
| SRR6928222 | 0 | 3444 | 447 | 38 | 2208 | 0 | 0 | 3393 | 24415 | 24937 | 23121 | 33758 | 0 | 1540 | 86 | 116 | 0 | 242 | 249 | 1459 | 596 | 246 | 314 | 6141 | 246 | 53 | 35 | 270 | 124 | 204 | 493 | 138 | 106 | 76 | 20 | 3374 | 70 |
| SRR6928223 | 0 | 3643 | 88 | 59 | 2491 | 0 | 0 | 654 | 3251 | 15013 | 20726 | 35475 | 0 | 990 | 46 | 98 | 0 | 537 | 173 | 226 | 667 | 215 | 154 | 9014 | 459 | 80 | 22 | 349 | 556 | 1251 | 4368 | 1180 | 1108 | 625 | 78 | 652 | 403 |
| SRR6928224 | 0 | 522 | 58 | 97 | 1554 | 0 | 0 | 527 | 2037 | 11761 | 50684 | 52827 | 0 | 2256 | 29 | 45 | 30 | 513 | 919 | 644 | 797 | 329 | 475 | 15962 | 751 | 59 | 16 | 157 | 86 | 103 | 228 | 97 | 116 | 73 | 13 | 1609 | 200 |
| SRR6928225 | 0 | 1761 | 134 | 38 | 2059 | 0 | 0 | 1071 | 6627 | 45823 | 78393 | 64651 | 0 | 1065 | 58 | 151 | 0 | 400 | 319 | 556 | 828 | 204 | 344 | 9379 | 462 | 43 | 19 | 87 | 210 | 740 | 2024 | 203 | 193 | 146 | 35 | 1815 | 201 |
| SRR6928226 | 0 | 673 | 86 | 48 | 3109 | 0 | 0 | 4020 | 27674 | 28749 | 22787 | 29002 | 0 | 1685 | 54 | 87 | 0 | 246 | 260 | 1055 | 538 | 253 | 386 | 7154 | 310 | 48 | 16 | 65 | 30 | 48 | 101 | 21 | 16 | 16 | 24 | 2492 | 64 |
| SRR6930936 | 0 | 1682 | 30 | 37 | 2340 | 0 | 24 | 1023 | 2006 | 4506 | 10639 | 16842 | 0 | 599 | 93 | 279 | 68 | 793 | 31 | 6976 | 683 | 459 | 59 | 4069 | 1059 | 28 | 78 | 26 | 76 | 53 | 201 | 51 | 28 | 19 | 0 | 6674 | 101 |
| SRR6930937 | 0 | 1897 | 38 | 131 | 5030 | 19 | 21 | 325 | 1290 | 4539 | 9859 | 15424 | 0 | 1076 | 71 | 223 | 82 | 1477 | 56 | 5960 | 1021 | 797 | 83 | 5678 | 1813 | 25 | 39 | 29 | 110 | 146 | 290 | 30 | 16 | 0 | 0 | 5036 | 177 |
| SRR6930939 | 0 | 1356 | 30 | 124 | 9220 | 0 | 16 | 208 | 1143 | 5457 | 12005 | 14188 | 0 | 780 | 156 | 520 | 92 | 2019 | 68 | 1819 | 490 | 638 | 137 | 5864 | 1626 | 27 | 16 | 30 | 73 | 62 | 39 | 0 | 0 | 0 | 0 | 1767 | 133 |
| SRR6930941 | 0 | 343 | 47 | 79 | 5201 | 0 | 14 | 554 | 1120 | 6539 | 1930 | 18701 | 0 | 947 | 185 | 500 | 16 | 1015 | 51 | 3293 | 709 | 772 | 150 | 7875 | 1787 | 26 | 38 | 26 | 16 | 31 | 111 | 48 | 17 | 14 | 0 | 2984 | 132 |
| SRR6932941 | 0 | 0 | 0 | 0 | 10 | 0 | 0 | 108 | 17 | 66 | 177 | 1586 | 0 | 17 | 0 | 0 | 0 | 45 | 96 | 58 | 568 | 3469 | 0 | 265 | 7229 | 553 | 0 | 25 | 11 | 18 | 12 | 0 | 0 | 12 | 0 | 174 | 137 |
| SRR6932942 | 0 | 0 | 0 | 0 | 11 | 0 | 0 | 560 | 199 | 18 | 141 | 2385 | 0 | 0 | 18 | 0 | 0 | 0 | 55 | 90 | 521 | 1097 | 0 | 620 | 2870 | 299 | 0 | 53 | 12 | 70 | 48 | 0 | 0 | 0 | 0 | 289 | 100 |
| SRR6932943 | 0 | 60 | 0 | 0 | 42 | 0 | 0 | 798 | 659 | 69 | 203 | 1353 | 0 | 0 | 0 | 0 | 0 | 87 | 54 | 243 | 424 | 1679 | 0 | 2774 | 12554 | 768 | 37 | 22 | 120 | 22 | 25 | 127 | 0 | 0 | 0 | 104 | 124 |
| SRR6932946 | 0 | 0 | 0 | 0 | 23 | 0 | 0 | 643 | 750 | 742 | 505 | 275 | 0 | 689 | 0 | 0 | 0 | 14 | 40 | 880 | 139 | 327 | 0 | 690 | 3685 | 388 | 14 | 43 | 10 | 20 | 27 | 34 | 0 | 0 | 0 | 1186 | 114 |
| SRR6932949 | 0 | 24 | 0 | 0 | 32 | 0 | 0 | 0 | 0 | 0 | 376 | 3630 | 0 | 0 | 0 | 0 | 0 | 37 | 91 | 46 | 133 | 362 | 0 | 425 | 4416 | 342 | 0 | 154 | 0 | 0 | 0 | 0 | 0 | 0 | 0 | 23 | 30 |
| SRR6932950 | 0 | 12 | 0 | 0 | 33 | 0 | 0 | 0 | 0 | 13 | 448 | 3412 | 0 | 0 | 0 | 0 | 0 | 40 | 128 | 19 | 99 | 264 | 0 | 357 | 4746 | 363 | 0 | 16 | 0 | 0 | 0 | 0 | 0 | 0 | 0 | 62 | 28 |
| SRR6932951 | 0 | 21 | 0 | 0 | 32 | 0 | 0 | 0 | 0 | 19 | 296 | 2248 | 0 | 0 | 0 | 0 | 0 | 43 | 125 | 45 | 140 | 272 | 0 | 326 | 4139 | 300 | 0 | 104 | 0 | 0 | 0 | 0 | 0 | 0 | 0 | 13 | 30 |
| SRR6932952 | 0 | 190 | 694 | 0 | 14 | 0 | 0 | 296 | 179 | 8105 | 527 | 3580 | 0 | 1019 | 0 | 0 | 0 | 18 | 0 | 389 | 1086 | 1115 | 0 | 1364 | 4209 | 381 | 0 | 277 | 352 | 1166 | 1524 | 553 | 358 | 320 | 0 | 4309 | 266 |
| SRR6932953 | 0 | 13 | 35 | 0 | 0 | 0 | 0 | 166 | 98 | 6890 | 2556 | 2269 | 0 | 1550 | 30 | 0 | 0 | 25 | 0 | 333 | 859 | 802 | 0 | 2485 | 6101 | 369 | 21 | 23 | 20 | 76 | 107 | 31 | 17 | 67 | 0 | 3933 | 263 |
| SRR6932954 | 0 | 54 | 157 | 0 | 13 | 0 | 0 | 104 | 67 | 5542 | 646 | 914 | 0 | 1062 | 0 | 0 | 0 | 36 | 13 | 379 | 205 | 208 | 0 | 3169 | 6849 | 543 | 0 | 34 | 17 | 80 | 64 | 27 | 60 | 22 | 16 | 3284 | 37 |
| SRR6932955 | 10 | 76 | 0 | 0 | 44 | 0 | 0 | 998 | 625 | 60 | 53 | 457 | 0 | 36 | 0 | 0 | 0 | 101 | 47 | 108 | 407 | 298 | 0 | 439 | 10291 | 537 | 20 | 31 | 19 | 26 | 31 | 20 | 0 | 0 | 0 | 12 | 187 |
| SRR7062776 | 0 | 1304 | 44 | 0 | 21 | 0 | 0 | 50 | 267 | 1594 | 670 | 1249 | 0 | 1308 | 125 | 102 | 49 | 1192 | 32 | 113 | 2673 | 22464 | 99 | 41928 | 12361 | 0 | 11 | 0 | 28 | 23 | 17 | 15 | 11 | 24 | 0 | 71 | 0 |
| SRR7062777 | 0 | 1349 | 29 | 0 | 21 | 0 | 0 | 55 | 219 | 1506 | 1262 | 1042 | 0 | 1065 | 114 | 71 | 56 | 1101 | 29 | 119 | 2584 | 19791 | 113 | 39621 | 12415 | 0 | 18 | 0 | 56 | 23 | 18 | 0 | 14 | 21 | 0 | 62 | 0 |
| SRR7062778 | 0 | 1046 | 48 | 0 | 12 | 0 | 0 | 47 | 177 | 2386 | 1325 | 1268 | 0 | 1785 | 99 | 84 | 52 | 1180 | 28 | 99 | 2559 | 22642 | 105 | 41710 | 12370 | 0 | 30 | 0 | 54 | 27 | 23 | 19 | 26 | 51 | 0 | 100 | 0 |
| SRR7062779 | 0 | 994 | 44 | 0 | 14 | 0 | 0 | 32 | 248 | 2809 | 1167 | 978 | 0 | 1787 | 117 | 84 | 63 | 1176 | 32 | 107 | 2500 | 20922 | 113 | 43594 | 12950 | 0 | 26 | 0 | 61 | 26 | 23 | 22 | 21 | 43 | 0 | 82 | 0 |
| SRR7062780 | 27 | 194 | 26 | 0 | 10 | 32 | 12 | 397 | 275 | 3153 | 1622 | 1667 | 0 | 2713 | 131 | 101 | 11 | 1177 | 44 | 165 | 2539 | 24736 | 108 | 44783 | 12934 | 0 | 24 | 0 | 11 | 25 | 28 | 12 | 17 | 43 | 0 | 117 | 0 |
| SRR7062781 | 17 | 904 | 35 | 0 | 12 | 27 | 0 | 421 | 282 | 3169 | 1016 | 1582 | 0 | 2896 | 92 | 98 | 61 | 1125 | 35 | 145 | 2549 | 23678 | 109 | 46013 | 13269 | 0 | 23 | 0 | 66 | 31 | 26 | 17 | 21 | 33 | 0 | 105 | 0 |
| SRR7081766 | 30 | 0 | 0 | 0 | 0 | 90 | 0 | 807 | 922 | 3919 | 942 | 1037 | 0 | 576 | 0 | 0 | 30 | 0 | 0 | 807 | 144 | 152 | 13 | 193 | 5823 | 287 | 120 | 0 | 0 | 0 | 0 | 0 | 0 | 0 | 0 | 5533 | 0 |
| SRR7081767 | 20 | 0 | 0 | 0 | 0 | 20 | 0 | 2285 | 762 | 4189 | 952 | 952 | 0 | 1142 | 0 | 0 | 61 | 0 | 0 | 952 | 263 | 238 | 32 | 641 | 6330 | 273 | 61 | 0 | 0 | 0 | 0 | 0 | 0 | 0 | 0 | 4760 | 0 |
| SRR7081768 | 12 | 0 | 0 | 0 | 0 | 0 | 0 | 4196 | 10037 | 32998 | 28 | 1309 | 0 | 201 | 0 | 0 | 0 | 16 | 0 | 67 | 415 | 330 | 54 | 893 | 10163 | 215 | 36 | 0 | 0 | 0 | 0 | 0 | 0 | 0 | 0 | 369 | 0 |
| SRR7081769 | 26 | 0 | 0 | 0 | 0 | 35 | 0 | 3167 | 3973 | 27635 | 230 | 691 | 0 | 288 | 0 | 0 | 0 | 32 | 12 | 345 | 411 | 393 | 21 | 397 | 33520 | 147 | 26 | 0 | 0 | 0 | 0 | 0 | 0 | 0 | 0 | 173 | 0 |
| SRR7081770 | 59 | 0 | 0 | 0 | 0 | 0 | 0 | 2653 | 531 | 7322 | 1273 | 1698 | 0 | 743 | 0 | 0 | 0 | 0 | 0 | 106 | 53 | 161 | 0 | 311 | 4059 | 119 | 156 | 0 | 0 | 20 | 0 | 0 | 0 | 0 | 0 | 3077 | 0 |
| SRR7081771 | 104 | 0 | 0 | 0 | 0 | 10 | 0 | 534 | 178 | 2802 | 2358 | 979 | 0 | 133 | 0 | 0 | 0 | 0 | 0 | 0 | 147 | 97 | 0 | 241 | 5508 | 255 | 42 | 0 | 0 | 0 | 0 | 0 | 0 | 0 | 0 | 623 | 0 |
| SRR7081772 | 25 | 0 | 0 | 0 | 0 | 12 | 0 | 1322 | 331 | 18841 | 25 | 3526 | 0 | 1102 | 0 | 0 | 0 | 15 | 0 | 331 | 209 | 279 | 21 | 498 | 2649 | 315 | 75 | 0 | 0 | 0 | 0 | 0 | 0 | 0 | 0 | 2204 | 0 |
| SRR7081773 | 0 | 0 | 0 | 0 | 0 | 0 | 0 | 2034 | 2180 | 31098 | 799 | 1017 | 0 | 1598 | 363 | 291 | 0 | 0 | 0 | 1381 | 44 | 267 | 12 | 261 | 409 | 146 | 73 | 0 | 0 | 0 | 0 | 0 | 0 | 0 | 0 | 3342 | 0 |
| SRR7081774 | 34 | 0 | 0 | 0 | 0 | 0 | 0 | 2625 | 500 | 12750 | 625 | 1250 | 0 | 625 | 0 | 125 | 0 | 0 | 0 | 1000 | 177 | 259 | 22 | 399 | 3600 | 300 | 60 | 0 | 0 | 0 | 0 | 0 | 0 | 0 | 0 | 2625 | 0 |
| SRR7081775 | 35 | 0 | 0 | 0 | 0 | 0 | 0 | 1387 | 902 | 8255 | 1387 | 2706 | 17 | 416 | 0 | 0 | 0 | 0 | 0 | 347 | 101 | 130 | 0 | 112 | 5152 | 298 | 87 | 0 | 0 | 0 | 0 | 0 | 0 | 0 | 0 | 3052 | 0 |
| SRR7081776 | 0 | 0 | 0 | 0 | 0 | 0 | 0 | 510 | 408 | 2959 | 1633 | 4796 | 0 | 510 | 0 | 0 | 0 | 0 | 0 | 204 | 44 | 83 | 0 | 124 | 3224 | 149 | 107 | 0 | 0 | 0 | 0 | 0 | 0 | 0 | 0 | 1429 | 0 |
| SRR7081777 | 0 | 0 | 0 | 0 | 0 | 0 | 0 | 714 | 48 | 1379 | 5946 | 13033 | 0 | 190 | 0 | 0 | 0 | 0 | 0 | 238 | 95 | 247 | 0 | 332 | 5603 | 951 | 0 | 0 | 0 | 0 | 0 | 0 | 0 | 0 | 0 | 2093 | 0 |
| SRR7107416 | 0 | 0 | 457 | 0 | 0 | 0 | 0 | 1441 | 943 | 461 | 951 | 2521 | 0 | 2344 | 393 | 39 | 0 | 0 | 0 | 1525 | 0 | 95 | 89 | 664 | 163 | 0 | 0 | 111 | 108 | 0 | 0 | 121 | 525 | 1943 | 0 | 6400 | 0 |
| SRR7107417 | 0 | 0 | 25 | 0 | 0 | 0 | 0 | 587 | 1119 | 512 | 846 | 3401 | 0 | 2749 | 152 | 25 | 0 | 0 | 0 | 1598 | 0 | 87 | 123 | 729 | 182 | 0 | 0 | 10 | 134 | 0 | 0 | 163 | 40 | 124 | 0 | 7236 | 0 |
| SRR7107418 | 0 | 0 | 0 | 0 | 45 | 0 | 0 | 0 | 0 | 12873 | 1295 | 498 | 0 | 16288 | 57 | 23 | 0 | 41 | 0 | 313 | 1212 | 4630 | 113 | 28508 | 4785 | 0 | 0 | 44 | 0 | 18 | 27 | 30 | 160 | 0 | 0 | 1290 | 104 |
| SRR7107419 | 0 | 0 | 0 | 0 | 58 | 0 | 0 | 0 | 0 | 13823 | 2036 | 750 | 0 | 19551 | 36 | 20 | 0 | 46 | 0 | 177 | 1206 | 4951 | 107 | 28847 | 4922 | 0 | 0 | 389 | 0 | 262 | 380 | 36 | 1943 | 0 | 0 | 842 | 85 |
| SRR7107420 | 0 | 0 | 562 | 0 | 0 | 0 | 0 | 1325 | 7824 | 334 | 891 | 3344 | 0 | 469 | 523 | 185 | 0 | 0 | 0 | 647 | 0 | 0 | 261 | 0 | 0 | 0 | 0 | 219 | 214 | 0 | 0 | 44 | 449 | 1738 | 0 | 3413 | 0 |
| SRR7107421 | 0 | 0 | 28 | 0 | 0 | 0 | 0 | 1134 | 7323 | 228 | 1135 | 4398 | 0 | 286 | 384 | 164 | 0 | 0 | 0 | 738 | 0 | 0 | 270 | 0 | 0 | 0 | 0 | 11 | 192 | 0 | 0 | 46 | 23 | 80 | 0 | 3242 | 0 |
| SRR7126083 | 0 | 0 | 0 | 0 | 17 | 15 | 0 | 67 | 1065 | 3727 | 67 | 67 | 0 | 2063 | 466 | 932 | 0 | 100 | 452 | 599 | 17 | 151 | 0 | 109 | 1139 | 502 | 102 | 0 | 0 | 15 | 0 | 0 | 0 | 0 | 0 | 2063 | 17 |
| SRR7126084 | 0 | 23 | 55 | 39 | 305 | 0 | 0 | 234 | 622 | 3862 | 3467 | 14846 | 0 | 12880 | 0 | 50 | 0 | 1314 | 568 | 2535 | 316 | 1607 | 228 | 13391 | 1056 | 0 | 62 | 19 | 35 | 259 | 493 | 54 | 43 | 42 | 33 | 2908 | 199 |
| SRR7126085 | 0 | 30 | 73 | 46 | 307 | 0 | 0 | 307 | 956 | 4477 | 1732 | 14085 | 0 | 11254 | 13 | 55 | 0 | 1380 | 665 | 4262 | 433 | 1410 | 288 | 16173 | 1204 | 0 | 58 | 18 | 62 | 412 | 876 | 103 | 84 | 93 | 56 | 4722 | 337 |
| SRR7126086 | 0 | 205 | 671 | 44 | 260 | 0 | 0 | 182 | 446 | 2335 | 1672 | 15809 | 0 | 11412 | 10 | 53 | 0 | 1440 | 762 | 3630 | 332 | 1085 | 318 | 14055 | 1650 | 0 | 81 | 143 | 671 | 3590 | 6611 | 823 | 907 | 883 | 346 | 2752 | 457 |
| SRR7126087 | 0 | 104 | 555 | 66 | 619 | 0 | 0 | 130 | 423 | 3535 | 4563 | 8549 | 0 | 12923 | 0 | 58 | 0 | 1422 | 741 | 1642 | 369 | 1302 | 291 | 14312 | 1448 | 0 | 56 | 73 | 393 | 2723 | 2566 | 206 | 167 | 217 | 276 | 2398 | 256 |
| SRR7126088 | 0 | 93 | 939 | 26 | 269 | 0 | 0 | 75 | 310 | 2909 | 1486 | 15421 | 0 | 10374 | 0 | 86 | 0 | 1066 | 584 | 2481 | 193 | 1157 | 192 | 12592 | 1268 | 0 | 72 | 97 | 454 | 3133 | 5350 | 627 | 582 | 485 | 338 | 2112 | 318 |
| SRR7126089 | 0 | 23 | 101 | 19 | 146 | 0 | 0 | 186 | 480 | 3053 | 8151 | 16590 | 0 | 11823 | 19 | 120 | 0 | 990 | 457 | 6150 | 210 | 1371 | 181 | 10250 | 982 | 0 | 56 | 13 | 48 | 283 | 594 | 103 | 84 | 83 | 49 | 4382 | 218 |
| SRR7126090 | 0 | 25 | 89 | 38 | 218 | 11 | 0 | 90 | 403 | 4263 | 1840 | 14186 | 10 | 13105 | 0 | 67 | 0 | 896 | 515 | 2622 | 286 | 1349 | 183 | 10391 | 941 | 0 | 65 | 69 | 43 | 342 | 898 | 147 | 118 | 111 | 81 | 2935 | 326 |
| SRR7126091 | 0 | 27 | 59 | 52 | 488 | 0 | 0 | 127 | 486 | 4174 | 3257 | 12439 | 0 | 12814 | 14 | 34 | 0 | 986 | 505 | 1164 | 391 | 1385 | 277 | 14839 | 1037 | 0 | 12 | 14 | 40 | 328 | 635 | 71 | 50 | 56 | 31 | 1867 | 249 |
| SRR7126092 | 0 | 15 | 86 | 31 | 294 | 0 | 0 | 148 | 409 | 3493 | 1916 | 11224 | 10 | 11931 | 20 | 82 | 0 | 1172 | 540 | 1665 | 271 | 1386 | 248 | 11882 | 1173 | 0 | 18 | 12 | 53 | 344 | 506 | 63 | 49 | 50 | 36 | 1779 | 273 |
| SRR7126093 | 0 | 18 | 76 | 39 | 303 | 0 | 0 | 197 | 423 | 2915 | 7758 | 15956 | 0 | 10183 | 0 | 73 | 0 | 1280 | 516 | 2509 | 289 | 1234 | 282 | 14135 | 1535 | 0 | 82 | 13 | 63 | 502 | 867 | 78 | 83 | 94 | 47 | 2278 | 441 |
| SRR7153253 | 0 | 0 | 11 | 0 | 0 | 0 | 16 | 210 | 22 | 1903 | 426 | 564 | 0 | 4921 | 13 | 23 | 33 | 724 | 11 | 21 | 7012 | 1714 | 1023 | 9376 | 3512 | 703 | 36 | 0 | 16 | 16 | 18 | 0 | 25 | 48 | 25 | 67 | 0 |
| SRR7153254 | 0 | 0 | 0 | 0 | 0 | 0 | 0 | 224 | 29 | 2953 | 1246 | 1439 | 0 | 7293 | 77 | 77 | 0 | 509 | 0 | 44 | 5114 | 1888 | 1052 | 7563 | 3419 | 389 | 0 | 0 | 21 | 24 | 20 | 0 | 32 | 57 | 15 | 119 | 0 |
| SRR7153255 | 0 | 0 | 16 | 0 | 0 | 0 | 0 | 271 | 40 | 3945 | 666 | 668 | 0 | 6121 | 158 | 88 | 0 | 543 | 0 | 53 | 5325 | 2352 | 1143 | 9559 | 3857 | 619 | 0 | 0 | 17 | 17 | 20 | 0 | 27 | 40 | 49 | 99 | 0 |
| SRR7153256 | 0 | 13 | 35 | 0 | 0 | 0 | 0 | 356 | 36 | 2541 | 1156 | 1701 | 0 | 5166 | 42 | 33 | 0 | 714 | 0 | 44 | 5856 | 1867 | 1262 | 11491 | 3141 | 377 | 0 | 0 | 30 | 55 | 41 | 27 | 50 | 106 | 85 | 83 | 10 |
| SRR7153257 | 0 | 0 | 13 | 0 | 0 | 0 | 0 | 189 | 21 | 3337 | 917 | 378 | 0 | 5308 | 31 | 40 | 0 | 611 | 10 | 21 | 5933 | 2386 | 1216 | 7897 | 4345 | 539 | 0 | 0 | 0 | 16 | 11 | 0 | 24 | 34 | 49 | 68 | 0 |
| SRR7153258 | 0 | 0 | 10 | 0 | 0 | 0 | 0 | 178 | 34 | 2136 | 857 | 984 | 0 | 4053 | 118 | 83 | 0 | 450 | 0 | 25 | 5003 | 1818 | 1125 | 8078 | 2993 | 534 | 0 | 0 | 17 | 18 | 22 | 0 | 48 | 68 | 93 | 80 | 0 |
| SRR7153259 | 0 | 11 | 13 | 0 | 10 | 0 | 0 | 234 | 29 | 2424 | 1196 | 472 | 0 | 6620 | 61 | 48 | 0 | 598 | 0 | 18 | 5518 | 1890 | 1147 | 9233 | 3264 | 268 | 0 | 0 | 0 | 11 | 12 | 0 | 13 | 53 | 33 | 36 | 0 |
| SRR7153260 | 0 | 12 | 16 | 0 | 12 | 0 | 0 | 255 | 27 | 2453 | 377 | 493 | 0 | 4931 | 12 | 0 | 0 | 1122 | 0 | 18 | 6556 | 1606 | 1203 | 10715 | 2765 | 596 | 0 | 0 | 25 | 42 | 29 | 14 | 44 | 62 | 49 | 42 | 0 |
| SRR7153261 | 0 | 0 | 26 | 0 | 0 | 0 | 0 | 131 | 13 | 1778 | 111 | 136 | 0 | 5971 | 25 | 22 | 0 | 671 | 0 | 24 | 6192 | 1654 | 1206 | 14620 | 4304 | 442 | 0 | 0 | 0 | 0 | 11 | 0 | 23 | 39 | 75 | 55 | 13 |
| SRR7153262 | 0 | 12 | 0 | 0 | 11 | 0 | 0 | 536 | 35 | 3118 | 1080 | 1060 | 0 | 5408 | 35 | 31 | 0 | 623 | 0 | 50 | 5998 | 2360 | 1089 | 10803 | 3651 | 514 | 0 | 0 | 0 | 0 | 11 | 0 | 15 | 29 | 39 | 308 | 0 |
| SRR7153263 | 0 | 0 | 16 | 0 | 0 | 0 | 0 | 311 | 58 | 2162 | 741 | 1242 | 0 | 7780 | 26 | 52 | 0 | 1609 | 0 | 38 | 7032 | 1616 | 1243 | 11493 | 2782 | 500 | 0 | 0 | 24 | 37 | 24 | 16 | 34 | 59 | 79 | 95 | 0 |
| SRR7153264 | 0 | 0 | 12 | 0 | 0 | 0 | 0 | 113 | 21 | 1880 | 419 | 580 | 0 | 7017 | 21 | 19 | 0 | 665 | 0 | 23 | 6142 | 1330 | 1020 | 9908 | 2394 | 606 | 0 | 0 | 23 | 32 | 22 | 0 | 79 | 88 | 97 | 75 | 0 |
| SRR7153265 | 0 | 0 | 16 | 0 | 11 | 0 | 0 | 686 | 58 | 3095 | 638 | 761 | 0 | 8178 | 21 | 24 | 0 | 1030 | 0 | 33 | 7055 | 2533 | 1248 | 11379 | 3160 | 577 | 0 | 0 | 26 | 21 | 20 | 0 | 43 | 59 | 51 | 92 | 0 |
| SRR7153266 | 0 | 0 | 14 | 0 | 0 | 0 | 0 | 363 | 25 | 2429 | 582 | 356 | 0 | 6784 | 58 | 63 | 0 | 670 | 11 | 18 | 5644 | 2115 | 1592 | 11812 | 3964 | 572 | 0 | 0 | 19 | 29 | 17 | 0 | 40 | 67 | 14 | 67 | 0 |
| SRR7153267 | 0 | 0 | 16 | 0 | 0 | 0 | 0 | 678 | 39 | 3111 | 488 | 639 | 0 | 4898 | 42 | 30 | 0 | 669 | 0 | 11 | 6169 | 1850 | 1388 | 8348 | 2734 | 286 | 0 | 0 | 0 | 22 | 11 | 0 | 28 | 58 | 23 | 49 | 0 |
| SRR7153268 | 0 | 0 | 22 | 0 | 0 | 0 | 0 | 139 | 30 | 1745 | 40 | 51 | 0 | 5397 | 24 | 25 | 0 | 755 | 0 | 11 | 5166 | 2180 | 1711 | 14643 | 4750 | 264 | 0 | 0 | 0 | 0 | 12 | 0 | 20 | 31 | 81 | 31 | 0 |
| SRR7153269 | 0 | 20 | 24 | 0 | 15 | 0 | 0 | 485 | 43 | 2540 | 1117 | 620 | 0 | 3832 | 27 | 26 | 0 | 611 | 0 | 25 | 7380 | 1757 | 1515 | 8487 | 3141 | 248 | 0 | 0 | 11 | 11 | 14 | 0 | 0 | 44 | 35 | 56 | 0 |
| SRR7153270 | 0 | 0 | 14 | 0 | 0 | 0 | 0 | 111 | 19 | 2657 | 607 | 306 | 0 | 5485 | 669 | 489 | 0 | 660 | 11 | 29 | 5068 | 2097 | 1286 | 10437 | 3536 | 411 | 0 | 0 | 11 | 11 | 0 | 11 | 25 | 59 | 53 | 77 | 0 |
| SRR7153271 | 0 | 0 | 0 | 0 | 10 | 0 | 0 | 201 | 37 | 2686 | 561 | 660 | 0 | 4264 | 188 | 169 | 0 | 729 | 11 | 85 | 6375 | 2077 | 1271 | 8789 | 3471 | 534 | 0 | 0 | 0 | 0 | 0 | 0 | 14 | 34 | 49 | 189 | 0 |
| SRR7153272 | 0 | 14 | 11 | 0 | 0 | 0 | 0 | 191 | 32 | 2791 | 624 | 678 | 0 | 5253 | 36 | 36 | 0 | 574 | 0 | 35 | 5843 | 1836 | 1172 | 8249 | 2972 | 457 | 0 | 0 | 15 | 0 | 12 | 11 | 19 | 32 | 27 | 90 | 0 |
| SRR7153273 | 0 | 40 | 0 | 0 | 12 | 0 | 0 | 920 | 92 | 2218 | 1435 | 1344 | 0 | 2194 | 26 | 18 | 0 | 987 | 0 | 131 | 9748 | 2165 | 1220 | 15420 | 3795 | 462 | 0 | 0 | 31 | 59 | 31 | 0 | 17 | 35 | 31 | 193 | 0 |
| SRR7153274 | 0 | 0 | 0 | 0 | 0 | 0 | 0 | 388 | 49 | 3764 | 698 | 1795 | 0 | 3550 | 174 | 232 | 0 | 679 | 12 | 58 | 5772 | 2486 | 1093 | 8351 | 3722 | 468 | 0 | 0 | 0 | 0 | 13 | 0 | 18 | 55 | 27 | 124 | 0 |
| SRR7153275 | 0 | 0 | 0 | 0 | 0 | 0 | 0 | 370 | 64 | 3325 | 1311 | 719 | 0 | 6683 | 183 | 222 | 0 | 670 | 0 | 31 | 6202 | 2140 | 1192 | 9887 | 3859 | 214 | 0 | 0 | 23 | 43 | 23 | 0 | 38 | 96 | 111 | 102 | 0 |
| SRR7153276 | 0 | 0 | 0 | 0 | 0 | 0 | 0 | 458 | 158 | 4093 | 1052 | 2267 | 0 | 8201 | 214 | 218 | 0 | 925 | 0 | 220 | 6229 | 2635 | 1178 | 9831 | 2982 | 518 | 0 | 0 | 11 | 16 | 0 | 0 | 17 | 34 | 34 | 434 | 11 |
| SRR7153277 | 0 | 0 | 0 | 0 | 0 | 0 | 0 | 334 | 34 | 2689 | 841 | 856 | 0 | 3483 | 173 | 217 | 0 | 587 | 0 | 0 | 4494 | 1741 | 1052 | 6390 | 2973 | 209 | 0 | 0 | 0 | 10 | 0 | 0 | 12 | 36 | 28 | 0 | 0 |
| SRR7153278 | 0 | 0 | 0 | 0 | 0 | 0 | 0 | 288 | 28 | 3979 | 993 | 1010 | 0 | 8945 | 94 | 58 | 0 | 464 | 0 | 45 | 5490 | 2142 | 1560 | 8788 | 4351 | 154 | 0 | 0 | 0 | 0 | 0 | 0 | 14 | 25 | 19 | 130 | 0 |
| SRR7153279 | 0 | 31 | 11 | 0 | 11 | 0 | 0 | 797 | 122 | 2599 | 1156 | 1190 | 0 | 4027 | 20 | 20 | 0 | 939 | 0 | 166 | 9589 | 2772 | 1231 | 15294 | 4001 | 447 | 13 | 0 | 20 | 35 | 18 | 0 | 12 | 35 | 21 | 227 | 0 |
| SRR7153280 | 0 | 0 | 0 | 0 | 0 | 0 | 0 | 586 | 49 | 3608 | 997 | 1383 | 0 | 7311 | 85 | 85 | 0 | 520 | 0 | 0 | 5609 | 2262 | 1061 | 8393 | 3762 | 232 | 0 | 0 | 13 | 17 | 0 | 0 | 19 | 32 | 33 | 0 | 0 |
| SRR7153281 | 0 | 13 | 20 | 0 | 11 | 0 | 0 | 418 | 48 | 2370 | 850 | 402 | 0 | 4252 | 61 | 90 | 0 | 630 | 0 | 28 | 4746 | 1454 | 1400 | 8183 | 4129 | 127 | 0 | 0 | 0 | 19 | 21 | 0 | 23 | 52 | 67 | 75 | 0 |
| SRR7153282 | 0 | 0 | 0 | 0 | 10 | 0 | 0 | 271 | 56 | 2943 | 733 | 1508 | 0 | 6935 | 82 | 79 | 0 | 809 | 0 | 61 | 6098 | 2300 | 1244 | 8354 | 3908 | 452 | 0 | 0 | 13 | 0 | 0 | 0 | 11 | 24 | 30 | 137 | 0 |
| SRR7153283 | 0 | 0 | 17 | 0 | 0 | 0 | 0 | 294 | 56 | 2235 | 849 | 661 | 0 | 5904 | 26 | 32 | 0 | 698 | 0 | 29 | 5613 | 1807 | 1147 | 6530 | 3282 | 209 | 0 | 0 | 16 | 35 | 21 | 0 | 38 | 54 | 44 | 83 | 0 |
| SRR7153284 | 0 | 17 | 0 | 0 | 0 | 0 | 0 | 183 | 19 | 1218 | 1019 | 285 | 0 | 5170 | 37 | 50 | 0 | 783 | 0 | 0 | 7264 | 1566 | 1166 | 5722 | 2332 | 148 | 0 | 0 | 0 | 0 | 0 | 0 | 0 | 0 | 23 | 0 | 0 |
| SRR7153285 | 0 | 14 | 15 | 0 | 0 | 0 | 0 | 584 | 54 | 2737 | 608 | 830 | 0 | 7644 | 55 | 68 | 0 | 585 | 0 | 16 | 6042 | 1761 | 1154 | 6837 | 2535 | 214 | 0 | 0 | 23 | 21 | 17 | 0 | 22 | 44 | 0 | 59 | 0 |
| SRR7153286 | 0 | 0 | 28 | 0 | 10 | 0 | 0 | 386 | 65 | 4597 | 1164 | 1736 | 0 | 8131 | 107 | 103 | 0 | 1080 | 0 | 63 | 5768 | 2066 | 1319 | 6633 | 3100 | 380 | 0 | 0 | 21 | 18 | 14 | 0 | 18 | 43 | 24 | 151 | 0 |
| SRR7153287 | 0 | 0 | 18 | 0 | 17 | 0 | 0 | 129 | 31 | 2362 | 99 | 130 | 0 | 4260 | 112 | 128 | 0 | 777 | 0 | 0 | 6002 | 1605 | 1219 | 8041 | 3428 | 219 | 0 | 0 | 0 | 0 | 0 | 0 | 18 | 30 | 73 | 42 | 0 |
| SRR7153288 | 0 | 0 | 13 | 0 | 0 | 0 | 0 | 336 | 76 | 2619 | 309 | 337 | 0 | 5117 | 27 | 38 | 0 | 643 | 0 | 17 | 5653 | 1756 | 1025 | 5908 | 3397 | 136 | 0 | 0 | 17 | 0 | 12 | 0 | 40 | 81 | 44 | 56 | 0 |
| SRR7153289 | 0 | 0 | 0 | 0 | 0 | 0 | 0 | 267 | 28 | 1755 | 407 | 511 | 0 | 4138 | 145 | 177 | 0 | 791 | 0 | 27 | 5740 | 1688 | 1154 | 7466 | 3690 | 164 | 0 | 0 | 11 | 0 | 0 | 0 | 0 | 21 | 15 | 65 | 0 |
| SRR7153290 | 0 | 0 | 0 | 0 | 0 | 0 | 0 | 658 | 96 | 3948 | 437 | 591 | 0 | 5488 | 36 | 49 | 0 | 666 | 0 | 27 | 5655 | 2353 | 1389 | 8142 | 3957 | 127 | 0 | 0 | 11 | 14 | 12 | 0 | 19 | 35 | 27 | 55 | 0 |
| SRR7153291 | 0 | 33 | 11 | 0 | 18 | 0 | 0 | 892 | 108 | 3267 | 1088 | 1690 | 0 | 3274 | 61 | 59 | 0 | 960 | 0 | 169 | 9123 | 2287 | 1213 | 13012 | 3315 | 511 | 0 | 0 | 26 | 49 | 24 | 0 | 15 | 29 | 0 | 277 | 0 |
| SRR7153292 | 0 | 36 | 0 | 0 | 16 | 0 | 0 | 931 | 150 | 2769 | 1081 | 1215 | 0 | 3344 | 82 | 68 | 0 | 1142 | 0 | 81 | 10361 | 2080 | 1102 | 13421 | 4469 | 541 | 11 | 0 | 0 | 23 | 23 | 0 | 0 | 15 | 13 | 186 | 0 |
| SRR7153293 | 0 | 14 | 0 | 0 | 16 | 0 | 0 | 710 | 232 | 3504 | 705 | 486 | 0 | 4280 | 145 | 162 | 0 | 1018 | 0 | 339 | 11338 | 3869 | 1149 | 16890 | 5206 | 403 | 0 | 0 | 11 | 24 | 24 | 0 | 19 | 48 | 34 | 541 | 0 |
| SRR7153294 | 0 | 0 | 0 | 0 | 0 | 0 | 0 | 315 | 40 | 1843 | 705 | 652 | 0 | 3413 | 44 | 77 | 0 | 723 | 0 | 18 | 6299 | 1706 | 1014 | 5549 | 3096 | 268 | 0 | 0 | 0 | 0 | 0 | 0 | 14 | 25 | 42 | 68 | 0 |
| SRR7153295 | 0 | 0 | 0 | 0 | 0 | 0 | 0 | 388 | 50 | 2377 | 524 | 192 | 0 | 6723 | 0 | 14 | 0 | 1123 | 0 | 31 | 9152 | 1633 | 1207 | 8600 | 4137 | 569 | 0 | 0 | 0 | 0 | 0 | 12 | 12 | 42 | 83 | 133 | 0 |
| SRR7153296 | 0 | 0 | 0 | 0 | 0 | 0 | 0 | 505 | 58 | 2763 | 949 | 1832 | 0 | 9467 | 104 | 152 | 0 | 613 | 0 | 77 | 5740 | 1953 | 873 | 5313 | 3286 | 526 | 0 | 0 | 0 | 0 | 0 | 0 | 0 | 14 | 19 | 192 | 0 |
| SRR7153297 | 0 | 0 | 0 | 0 | 0 | 0 | 28 | 353 | 81 | 3341 | 999 | 1247 | 0 | 6916 | 94 | 92 | 27 | 398 | 0 | 82 | 5205 | 1889 | 1134 | 5699 | 3207 | 326 | 34 | 0 | 0 | 0 | 0 | 0 | 0 | 0 | 0 | 311 | 0 |
| SRR7153298 | 0 | 0 | 0 | 0 | 0 | 0 | 0 | 273 | 33 | 2223 | 781 | 752 | 0 | 4109 | 23 | 32 | 0 | 620 | 0 | 32 | 5386 | 2066 | 1131 | 7810 | 3901 | 361 | 0 | 0 | 0 | 11 | 13 | 0 | 17 | 15 | 30 | 53 | 0 |

### Supp Table S8.4.Columns_DI_to_ES

| | | | | | | | | | | | | | | | | | | | | | | | | | | | | | | | | | | | | | |
|---|---|---|---|---|---|---|---|---|---|---|---|---|---|---|---|---|---|---|---|---|---|---|---|---|---|---|---|---|---|---|---|---|---|---|---|---|---|
| SRR7153299 | 0 | 0 | 14 | 0 | 0 | 0 | 0 | 452 | 32 | 2540 | 272 | 322 | 0 | 5804 | 24 | 18 | 0 | 762 | 10 | 29 | 6717 | 2819 | 1178 | 9553 | 4425 | 590 | 0 | 0 | 22 | 16 | 23 | 0 | 26 | 79 | 29 | 84 | 0 |
| SRR7153300 | 0 | 0 | 15 | 0 | 0 | 0 | 0 | 210 | 21 | 3437 | 481 | 651 | 0 | 4058 | 36 | 33 | 0 | 716 | 0 | 27 | 5384 | 2079 | 1157 | 6299 | 3750 | 320 | 0 | 0 | 11 | 14 | 14 | 11 | 27 | 41 | 44 | 71 | 0 |
| SRR7153301 | 0 | 0 | 14 | 0 | 13 | 0 | 0 | 196 | 57 | 3059 | 1141 | 1474 | 0 | 13347 | 77 | 72 | 0 | 616 | 0 | 30 | 5397 | 1512 | 1279 | 7165 | 3189 | 148 | 0 | 0 | 0 | 11 | 0 | 0 | 11 | 24 | 27 | 141 | 11 |
| SRR7153302 | 0 | 0 | 0 | 0 | 0 | 0 | 0 | 342 | 32 | 1983 | 546 | 656 | 0 | 6934 | 29 | 40 | 0 | 755 | 11 | 27 | 6769 | 1876 | 1124 | 6594 | 3624 | 336 | 0 | 0 | 0 | 0 | 0 | 0 | 17 | 29 | 13 | 58 | 0 |
| SRR7153303 | 0 | 32 | 14 | 0 | 19 | 0 | 0 | 581 | 199 | 4076 | 689 | 1964 | 0 | 4979 | 81 | 51 | 0 | 784 | 0 | 492 | 11471 | 2768 | 1034 | 10244 | 3662 | 167 | 0 | 0 | 21 | 54 | 37 | 0 | 12 | 45 | 11 | 895 | 0 |
| SRR7153304 | 0 | 0 | 0 | 0 | 0 | 0 | 0 | 576 | 64 | 3049 | 251 | 313 | 0 | 5495 | 16 | 19 | 0 | 719 | 0 | 27 | 5449 | 2273 | 954 | 9027 | 4359 | 340 | 0 | 0 | 0 | 14 | 11 | 0 | 29 | 50 | 44 | 74 | 15 |
| SRR7153305 | 0 | 0 | 0 | 0 | 0 | 0 | 0 | 388 | 71 | 2993 | 522 | 957 | 0 | 11551 | 446 | 503 | 0 | 659 | 0 | 30 | 4979 | 2108 | 1231 | 7968 | 3343 | 257 | 0 | 0 | 0 | 16 | 13 | 0 | 15 | 37 | 69 | 68 | 0 |
| SRR7153306 | 0 | 23 | 12 | 0 | 0 | 0 | 0 | 581 | 90 | 3480 | 442 | 637 | 0 | 6572 | 77 | 124 | 0 | 714 | 0 | 33 | 6234 | 2100 | 1436 | 7608 | 3944 | 122 | 0 | 0 | 0 | 12 | 0 | 0 | 13 | 18 | 30 | 81 | 0 |
| SRR7153307 | 0 | 0 | 10 | 0 | 0 | 0 | 0 | 385 | 19 | 2762 | 844 | 638 | 0 | 4335 | 99 | 76 | 0 | 646 | 0 | 23 | 6438 | 1910 | 1080 | 8286 | 3205 | 398 | 0 | 0 | 12 | 15 | 15 | 0 | 26 | 36 | 57 | 52 | 0 |
| SRR7153308 | 0 | 0 | 15 | 0 | 0 | 0 | 0 | 488 | 57 | 3036 | 937 | 1109 | 0 | 6348 | 52 | 60 | 0 | 800 | 0 | 42 | 6292 | 2321 | 1046 | 7471 | 3209 | 310 | 0 | 0 | 10 | 14 | 16 | 0 | 21 | 22 | 29 | 102 | 0 |
| SRR7153309 | 0 | 0 | 10 | 0 | 0 | 0 | 0 | 247 | 30 | 2821 | 1438 | 621 | 0 | 6961 | 37 | 43 | 0 | 1054 | 0 | 21 | 6393 | 2096 | 1386 | 8795 | 3077 | 223 | 0 | 0 | 0 | 16 | 10 | 0 | 16 | 42 | 49 | 66 | 0 |
| SRR7153310 | 0 | 10 | 32 | 0 | 11 | 0 | 0 | 372 | 54 | 3160 | 1010 | 957 | 0 | 8789 | 19 | 11 | 0 | 810 | 0 | 32 | 6802 | 1804 | 1005 | 6940 | 3110 | 415 | 0 | 0 | 14 | 25 | 25 | 0 | 20 | 61 | 0 | 71 | 0 |
| SRR7153311 | 0 | 0 | 11 | 0 | 0 | 0 | 0 | 449 | 90 | 2484 | 1779 | 1774 | 0 | 8054 | 70 | 143 | 0 | 711 | 13 | 94 | 5776 | 2362 | 1200 | 7671 | 3810 | 423 | 0 | 0 | 13 | 13 | 0 | 0 | 18 | 19 | 32 | 291 | 0 |
| SRR7153312 | 0 | 55 | 12 | 0 | 11 | 0 | 0 | 1033 | 224 | 4071 | 1111 | 1377 | 0 | 2736 | 15 | 39 | 0 | 1138 | 0 | 260 | 12327 | 2879 | 1021 | 13340 | 4687 | 574 | 0 | 0 | 27 | 58 | 50 | 0 | 16 | 43 | 35 | 416 | 0 |
| SRR7153313 | 0 | 38 | 13 | 0 | 20 | 0 | 0 | 799 | 141 | 2618 | 501 | 319 | 0 | 2611 | 16 | 13 | 0 | 961 | 0 | 31 | 7034 | 1560 | 966 | 8884 | 4033 | 326 | 26 | 0 | 45 | 43 | 24 | 0 | 11 | 46 | 26 | 104 | 0 |
| SRR7153314 | 0 | 0 | 0 | 0 | 0 | 0 | 0 | 414 | 39 | 1649 | 336 | 421 | 0 | 8197 | 0 | 10 | 0 | 759 | 0 | 10 | 5965 | 1558 | 1042 | 6995 | 3603 | 217 | 0 | 0 | 0 | 11 | 0 | 0 | 19 | 29 | 27 | 47 | 0 |
| SRR7153315 | 0 | 0 | 16 | 0 | 0 | 0 | 0 | 479 | 47 | 2268 | 848 | 779 | 0 | 6538 | 50 | 46 | 0 | 611 | 0 | 112 | 5910 | 2269 | 1276 | 6839 | 2895 | 275 | 0 | 0 | 12 | 10 | 11 | 0 | 17 | 26 | 28 | 383 | 0 |
| SRR7153316 | 0 | 0 | 0 | 0 | 0 | 0 | 0 | 317 | 50 | 3519 | 410 | 649 | 0 | 7036 | 71 | 97 | 0 | 782 | 0 | 51 | 4900 | 2421 | 1059 | 8122 | 3630 | 213 | 0 | 0 | 0 | 0 | 0 | 0 | 21 | 12 | 14 | 171 | 0 |
| SRR7153317 | 0 | 60 | 0 | 0 | 14 | 0 | 0 | 162 | 35 | 2754 | 462 | 759 | 0 | 1917 | 40 | 18 | 43 | 899 | 0 | 196 | 9025 | 2444 | 1267 | 18612 | 4249 | 675 | 62 | 0 | 30 | 81 | 42 | 0 | 26 | 79 | 0 | 386 | 0 |
| SRR7153318 | 0 | 0 | 0 | 0 | 13 | 0 | 22 | 143 | 29 | 1296 | 767 | 892 | 0 | 5888 | 72 | 45 | 25 | 414 | 0 | 33 | 5139 | 1535 | 1265 | 10932 | 2721 | 269 | 17 | 0 | 0 | 0 | 0 | 0 | 13 | 23 | 16 | 202 | 0 |
| SRR7153319 | 0 | 21 | 0 | 0 | 17 | 0 | 22 | 973 | 361 | 2754 | 1369 | 924 | 0 | 3755 | 19 | 19 | 46 | 1069 | 0 | 284 | 12720 | 3224 | 1053 | 13255 | 4165 | 473 | 145 | 0 | 11 | 25 | 21 | 0 | 0 | 18 | 0 | 471 | 0 |
| SRR7153320 | 0 | 38 | 0 | 0 | 12 | 0 | 0 | 789 | 59 | 3808 | 2954 | 3120 | 0 | 4110 | 61 | 61 | 0 | 846 | 0 | 281 | 10899 | 3034 | 1083 | 13725 | 4294 | 677 | 0 | 0 | 14 | 53 | 28 | 0 | 14 | 50 | 24 | 327 | 0 |
| SRR7153321 | 0 | 15 | 22 | 0 | 25 | 0 | 0 | 597 | 208 | 4313 | 307 | 250 | 0 | 1312 | 23 | 26 | 0 | 1220 | 0 | 188 | 10959 | 3524 | 1117 | 12982 | 5701 | 828 | 13 | 0 | 15 | 36 | 33 | 0 | 18 | 12 | 41 | 355 | 12 |
| SRR7153322 | 0 | 47 | 12 | 0 | 24 | 0 | 10 | 910 | 167 | 2297 | 1205 | 588 | 0 | 2222 | 0 | 0 | 14 | 1325 | 0 | 86 | 9486 | 1818 | 881 | 11738 | 4275 | 612 | 23 | 0 | 33 | 62 | 52 | 12 | 38 | 51 | 0 | 201 | 11 |
| SRR7153323 | 0 | 34 | 15 | 0 | 17 | 0 | 0 | 1051 | 88 | 4180 | 2311 | 2356 | 0 | 3980 | 190 | 173 | 0 | 858 | 0 | 161 | 10438 | 2518 | 1358 | 13166 | 4461 | 221 | 14 | 0 | 18 | 33 | 27 | 0 | 13 | 27 | 27 | 281 | 0 |
| SRR7153324 | 10 | 0 | 21 | 0 | 11 | 0 | 0 | 1099 | 199 | 2731 | 1129 | 562 | 0 | 6373 | 0 | 0 | 0 | 574 | 0 | 22 | 5488 | 1251 | 939 | 7744 | 2736 | 223 | 0 | 0 | 0 | 14 | 17 | 0 | 21 | 33 | 0 | 112 | 11 |
| SRR7153325 | 0 | 49 | 0 | 0 | 19 | 0 | 0 | 722 | 71 | 2847 | 783 | 1737 | 0 | 5176 | 191 | 183 | 0 | 1027 | 0 | 212 | 11504 | 2961 | 1155 | 17488 | 4869 | 604 | 14 | 0 | 0 | 13 | 14 | 0 | 13 | 27 | 0 | 411 | 0 |
| SRR7153326 | 0 | 49 | 11 | 0 | 28 | 0 | 0 | 807 | 202 | 3603 | 826 | 933 | 0 | 2822 | 64 | 95 | 0 | 1045 | 0 | 184 | 9407 | 2443 | 831 | 8136 | 4858 | 655 | 0 | 0 | 24 | 40 | 23 | 0 | 16 | 40 | 19 | 325 | 15 |
| SRR7153327 | 0 | 55 | 20 | 0 | 15 | 0 | 0 | 902 | 109 | 5644 | 781 | 2161 | 0 | 3761 | 41 | 44 | 0 | 1013 | 0 | 346 | 14279 | 2675 | 1145 | 14544 | 3460 | 830 | 0 | 0 | 23 | 74 | 46 | 0 | 19 | 57 | 0 | 452 | 0 |
| SRR7153328 | 0 | 51 | 0 | 0 | 12 | 0 | 0 | 369 | 56 | 1939 | 1875 | 1726 | 0 | 4295 | 51 | 57 | 0 | 1197 | 0 | 106 | 12521 | 2037 | 1080 | 12619 | 4206 | 318 | 0 | 0 | 14 | 29 | 18 | 0 | 14 | 26 | 25 | 190 | 0 |
| SRR7153329 | 0 | 25 | 11 | 0 | 19 | 0 | 0 | 991 | 131 | 3461 | 813 | 1728 | 0 | 3251 | 16 | 26 | 0 | 932 | 0 | 242 | 10894 | 3053 | 1219 | 14128 | 4661 | 929 | 47 | 0 | 12 | 34 | 20 | 0 | 21 | 31 | 0 | 305 | 0 |
| SRR7153330 | 0 | 14 | 0 | 0 | 16 | 0 | 0 | 875 | 115 | 3280 | 1295 | 1354 | 0 | 6177 | 108 | 126 | 21 | 993 | 0 | 205 | 10648 | 2660 | 1155 | 14534 | 4590 | 622 | 80 | 0 | 11 | 32 | 19 | 0 | 0 | 20 | 20 | 234 | 0 |
| SRR7153331 | 0 | 28 | 0 | 0 | 18 | 0 | 13 | 605 | 175 | 5012 | 2432 | 2229 | 0 | 2174 | 722 | 679 | 17 | 1025 | 0 | 512 | 10604 | 3194 | 960 | 14177 | 4948 | 631 | 58 | 0 | 11 | 30 | 20 | 0 | 11 | 35 | 16 | 717 | 0 |
| SRR7162919 | 0 | 4686 | 0 | 0 | 35 | 0 | 0 | 46 | 167 | 441 | 4360 | 4482 | 0 | 122 | 0 | 0 | 14 | 842 | 100 | 668 | 1989 | 2149 | 0 | 85 | 4487 | 18 | 0 | 14 | 0 | 79 | 0 | 0 | 0 | 22 | 0 | 4223 | 596 |
| SRR7162920 | 0 | 14146 | 0 | 0 | 14 | 0 | 0 | 30 | 33 | 148 | 1819 | 3549 | 0 | 26 | 0 | 0 | 0 | 2009 | 33 | 447 | 1938 | 1533 | 0 | 46 | 3369 | 12 | 0 | 0 | 0 | 41 | 0 | 0 | 0 | 11 | 0 | 1893 | 3462 |
| SRR7162921 | 0 | 21 | 0 | 0 | 28 | 0 | 0 | 66 | 42 | 114 | 381 | 2402 | 17 | 36 | 0 | 0 | 0 | 1321 | 68 | 502 | 2389 | 2149 | 0 | 112 | 7155 | 18 | 0 | 23 | 11 | 85 | 11 | 0 | 11 | 0 | 0 | 1535 | 1089 |
| SRR7178585 | 0 | 18 | 25 | 0 | 0 | 0 | 0 | 532 | 164 | 1892 | 153 | 439 | 0 | 334 | 47 | 42 | 0 | 284 | 0 | 137 | 723 | 588 | 89 | 1424 | 1160 | 0 | 0 | 0 | 0 | 39 | 28 | 11 | 21 | 49 | 25 | 237 | 0 |
| SRR7178586 | 0 | 23 | 33 | 0 | 10 | 0 | 0 | 505 | 176 | 1874 | 131 | 404 | 0 | 377 | 39 | 43 | 16 | 262 | 0 | 108 | 761 | 642 | 89 | 1534 | 1246 | 0 | 0 | 0 | 0 | 23 | 30 | 0 | 20 | 36 | 33 | 239 | 0 |
| SRR7178587 | 0 | 48 | 64 | 0 | 0 | 0 | 0 | 642 | 234 | 2229 | 196 | 382 | 0 | 340 | 32 | 42 | 16 | 294 | 0 | 159 | 388 | 346 | 65 | 881 | 674 | 0 | 0 | 0 | 0 | 32 | 48 | 0 | 16 | 16 | 32 | 265 | 0 |
| SRR7178588 | 0 | 0 | 18 | 0 | 18 | 0 | 0 | 652 | 226 | 2295 | 191 | 417 | 0 | 383 | 15 | 15 | 0 | 277 | 0 | 103 | 304 | 268 | 35 | 622 | 522 | 0 | 0 | 0 | 0 | 18 | 35 | 0 | 53 | 53 | 88 | 177 | 0 |
| SRR7178589 | 0 | 41 | 14 | 0 | 31 | 0 | 0 | 897 | 97 | 386 | 2277 | 3233 | 0 | 60 | 133 | 110 | 0 | 444 | 0 | 423 | 668 | 828 | 85 | 2062 | 1963 | 0 | 0 | 0 | 16 | 45 | 23 | 14 | 16 | 32 | 18 | 506 | 12 |
| SRR7178590 | 0 | 34 | 11 | 0 | 34 | 0 | 0 | 783 | 56 | 432 | 503 | 3157 | 0 | 35 | 147 | 173 | 0 | 426 | 0 | 428 | 730 | 907 | 88 | 2248 | 2146 | 0 | 0 | 0 | 0 | 42 | 23 | 0 | 13 | 23 | 0 | 506 | 11 |
| SRR7178591 | 0 | 75 | 0 | 0 | 45 | 0 | 0 | 877 | 175 | 431 | 2169 | 3302 | 0 | 16 | 64 | 112 | 0 | 427 | 0 | 542 | 360 | 458 | 44 | 1166 | 1096 | 11 | 0 | 0 | 0 | 0 | 56 | 0 | 37 | 28 | 0 | 750 | 0 |
| SRR7178592 | 0 | 11 | 22 | 0 | 53 | 0 | 0 | 962 | 47 | 584 | 2366 | 3312 | 0 | 79 | 79 | 95 | 0 | 377 | 0 | 505 | 256 | 327 | 34 | 817 | 815 | 0 | 0 | 0 | 11 | 55 | 76 | 0 | 44 | 22 | 0 | 457 | 0 |
| SRR7178593 | 0 | 93 | 35 | 0 | 30 | 0 | 0 | 105 | 44 | 361 | 242 | 571 | 0 | 251 | 202 | 423 | 0 | 435 | 0 | 85 | 1023 | 1253 | 91 | 1425 | 2719 | 28 | 0 | 0 | 22 | 66 | 13 | 13 | 13 | 49 | 75 | 207 | 18 |
| SRR7178594 | 0 | 77 | 17 | 0 | 26 | 0 | 0 | 88 | 44 | 344 | 245 | 606 | 0 | 296 | 185 | 327 | 0 | 423 | 0 | 87 | 969 | 1133 | 80 | 1356 | 2650 | 33 | 0 | 17 | 13 | 64 | 13 | 0 | 0 | 51 | 86 | 142 | 15 |
| SRR7178595 | 0 | 121 | 17 | 0 | 28 | 0 | 0 | 135 | 56 | 443 | 443 | 600 | 0 | 191 | 219 | 466 | 17 | 336 | 0 | 67 | 731 | 949 | 75 | 1096 | 2184 | 26 | 17 | 0 | 52 | 69 | 35 | 35 | 0 | 52 | 86 | 213 | 11 |
| SRR7178596 | 0 | 217 | 24 | 0 | 27 | 0 | 0 | 137 | 43 | 405 | 58 | 578 | 0 | 145 | 304 | 513 | 0 | 352 | 0 | 36 | 885 | 995 | 61 | 1085 | 2379 | 23 | 0 | 0 | 0 | 24 | 24 | 24 | 0 | 0 | 72 | 152 | 0 |
| SRR7178597 | 0 | 0 | 0 | 0 | 30 | 0 | 0 | 177 | 100 | 330 | 700 | 944 | 0 | 239 | 39 | 68 | 0 | 500 | 0 | 84 | 766 | 1187 | 110 | 1778 | 2176 | 47 | 11 | 16 | 32 | 25 | 18 | 0 | 18 | 30 | 30 | 221 | 18 |
| SRR7178598 | 0 | 14 | 23 | 0 | 52 | 0 | 0 | 158 | 39 | 336 | 587 | 903 | 0 | 170 | 30 | 109 | 0 | 366 | 0 | 123 | 989 | 1057 | 72 | 1700 | 2904 | 29 | 0 | 0 | 12 | 26 | 0 | 0 | 16 | 0 | 21 | 222 | 19 |
| SRR7178599 | 0 | 0 | 26 | 0 | 128 | 0 | 0 | 177 | 51 | 428 | 259 | 863 | 0 | 96 | 28 | 86 | 0 | 397 | 0 | 162 | 918 | 940 | 68 | 1530 | 2644 | 27 | 0 | 0 | 0 | 17 | 14 | 0 | 14 | 0 | 22 | 190 | 13 |
| SRR7178600 | 0 | 11 | 0 | 0 | 23 | 0 | 0 | 911 | 80 | 524 | 1236 | 1423 | 0 | 115 | 27 | 15 | 0 | 375 | 0 | 604 | 830 | 697 | 41 | 3805 | 1445 | 12 | 0 | 0 | 0 | 22 | 0 | 0 | 0 | 0 | 0 | 1393 | 0 |
| SRR7178601 | 0 | 48 | 0 | 0 | 22 | 0 | 0 | 971 | 29 | 507 | 1442 | 1355 | 0 | 83 | 0 | 22 | 0 | 340 | 0 | 620 | 907 | 800 | 51 | 4149 | 1626 | 14 | 0 | 0 | 0 | 19 | 13 | 0 | 0 | 0 | 0 | 1402 | 11 |
| SRR7178602 | 0 | 69 | 11 | 0 | 16 | 0 | 0 | 1340 | 15 | 632 | 964 | 1445 | 0 | 166 | 0 | 30 | 0 | 398 | 0 | 693 | 475 | 454 | 24 | 2139 | 827 | 0 | 11 | 0 | 0 | 80 | 16 | 0 | 0 | 11 | 0 | 1731 | 0 |
| SRR7178603 | 0 | 133 | 13 | 0 | 32 | 0 | 0 | 1167 | 75 | 748 | 63 | 1452 | 0 | 180 | 15 | 0 | 0 | 337 | 0 | 584 | 354 | 297 | 13 | 1573 | 575 | 15 | 19 | 0 | 0 | 0 | 0 | 0 | 0 | 13 | 0 | 1332 | 0 |
| SRR7178604 | 0 | 13 | 0 | 0 | 0 | 0 | 0 | 3041 | 685 | 4339 | 125 | 349 | 0 | 206 | 39 | 84 | 13 | 182 | 0 | 110 | 589 | 510 | 53 | 1349 | 1560 | 0 | 0 | 0 | 0 | 16 | 35 | 0 | 0 | 16 | 19 | 227 | 0 |
| SRR7178605 | 0 | 0 | 0 | 0 | 0 | 0 | 0 | 3128 | 675 | 4316 | 45 | 371 | 0 | 168 | 21 | 83 | 0 | 193 | 0 | 70 | 564 | 493 | 41 | 1252 | 1494 | 0 | 0 | 0 | 0 | 28 | 12 | 0 | 15 | 18 | 46 | 215 | 0 |
| SRR7178606 | 0 | 28 | 0 | 0 | 11 | 0 | 0 | 3669 | 695 | 4878 | 69 | 292 | 0 | 146 | 42 | 97 | 14 | 170 | 0 | 83 | 445 | 492 | 51 | 1045 | 1188 | 0 | 14 | 0 | 0 | 14 | 14 | 0 | 14 | 28 | 14 | 292 | 0 |
| SRR7178607 | 0 | 19 | 19 | 0 | 0 | 0 | 0 | 3815 | 788 | 4806 | 142 | 319 | 0 | 212 | 44 | 97 | 0 | 146 | 0 | 124 | 478 | 463 | 78 | 1195 | 1273 | 0 | 0 | 0 | 0 | 0 | 0 | 0 | 0 | 39 | 19 | 212 | 0 |
| SRR7178608 | 0 | 0 | 0 | 0 | 41 | 0 | 0 | 284 | 48 | 244 | 368 | 875 | 0 | 93 | 61 | 147 | 19 | 408 | 0 | 117 | 679 | 809 | 57 | 1199 | 1931 | 25 | 56 | 0 | 0 | 28 | 14 | 0 | 23 | 23 | 56 | 374 | 14 |
| SRR7178609 | 0 | 11 | 13 | 0 | 30 | 0 | 0 | 304 | 53 | 224 | 752 | 809 | 0 | 107 | 96 | 157 | 0 | 373 | 0 | 119 | 714 | 868 | 62 | 1267 | 2064 | 21 | 20 | 13 | 11 | 24 | 11 | 0 | 17 | 17 | 52 | 389 | 17 |
| SRR7178610 | 0 | 38 | 28 | 0 | 45 | 0 | 19 | 324 | 53 | 360 | 100 | 761 | 0 | 106 | 94 | 88 | 0 | 387 | 0 | 100 | 384 | 480 | 43 | 706 | 1179 | 23 | 0 | 0 | 19 | 19 | 0 | 0 | 28 | 19 | 57 | 312 | 12 |
| SRR7178611 | 0 | 22 | 22 | 0 | 53 | 0 | 22 | 368 | 67 | 362 | 440 | 585 | 0 | 162 | 33 | 84 | 22 | 385 | 0 | 106 | 271 | 351 | 19 | 513 | 844 | 20 | 54 | 11 | 0 | 43 | 11 | 0 | 22 | 11 | 65 | 312 | 0 |
| SRR7178612 | 0 | 16 | 25 | 0 | 32 | 0 | 0 | 1586 | 283 | 428 | 199 | 1662 | 0 | 76 | 245 | 384 | 0 | 390 | 0 | 245 | 783 | 908 | 95 | 1496 | 2006 | 31 | 16 | 0 | 25 | 21 | 14 | 12 | 19 | 35 | 46 | 629 | 0 |
| SRR7178613 | 0 | 22 | 38 | 0 | 33 | 0 | 0 | 1575 | 339 | 577 | 220 | 1782 | 0 | 100 | 345 | 377 | 0 | 348 | 0 | 213 | 712 | 890 | 97 | 1425 | 1909 | 33 | 16 | 0 | 25 | 20 | 0 | 11 | 20 | 31 | 61 | 728 | 0 |
| SRR7178614 | 0 | 0 | 48 | 0 | 26 | 0 | 0 | 1811 | 473 | 556 | 55 | 1728 | 0 | 21 | 617 | 370 | 0 | 323 | 0 | 206 | 595 | 754 | 86 | 1131 | 1541 | 32 | 19 | 0 | 29 | 19 | 19 | 19 | 0 | 29 | 76 | 514 | 0 |
| SRR7178615 | 0 | 0 | 53 | 0 | 27 | 0 | 0 | 1929 | 295 | 402 | 45 | 1661 | 0 | 54 | 456 | 589 | 40 | 343 | 0 | 134 | 693 | 784 | 79 | 1231 | 1655 | 19 | 13 | 0 | 53 | 27 | 40 | 0 | 27 | 27 | 53 | 750 | 0 |
| SRR7178616 | 0 | 18 | 11 | 0 | 31 | 0 | 0 | 6096 | 241 | 353 | 825 | 1035 | 0 | 150 | 34 | 187 | 70 | 519 | 0 | 226 | 948 | 874 | 58 | 1842 | 2370 | 23 | 15 | 0 | 0 | 18 | 11 | 0 | 0 | 0 | 25 | 301 | 14 |
| SRR7178617 | 0 | 15 | 14 | 0 | 34 | 0 | 0 | 6105 | 303 | 323 | 833 | 1032 | 0 | 191 | 52 | 214 | 71 | 515 | 0 | 214 | 972 | 945 | 70 | 1998 | 2544 | 18 | 11 | 0 | 0 | 20 | 0 | 0 | 0 | 23 | 22 | 360 | 0 |
| SRR7178618 | 0 | 35 | 18 | 0 | 32 | 0 | 0 | 6839 | 302 | 336 | 716 | 983 | 0 | 233 | 26 | 95 | 106 | 585 | 0 | 259 | 521 | 501 | 43 | 1060 | 1369 | 23 | 41 | 12 | 0 | 18 | 12 | 0 | 0 | 0 | 41 | 379 | 0 |
| SRR7178619 | 0 | 34 | 14 | 0 | 30 | 0 | 0 | 7187 | 264 | 338 | 684 | 841 | 0 | 223 | 41 | 91 | 34 | 496 | 0 | 206 | 372 | 398 | 31 | 734 | 1049 | 19 | 14 | 0 | 20 | 34 | 0 | 0 | 0 | 0 | 14 | 256 | 0 |
| SRR7178620 | 0 | 33 | 17 | 0 | 30 | 0 | 0 | 71 | 20 | 379 | 415 | 665 | 0 | 74 | 44 | 95 | 0 | 378 | 0 | 108 | 873 | 974 | 98 | 2693 | 2937 | 13 | 0 | 0 | 0 | 14 | 0 | 0 | 0 | 0 | 0 | 155 | 16 |
| SRR7178621 | 0 | 32 | 0 | 0 | 28 | 0 | 0 | 93 | 31 | 359 | 409 | 747 | 0 | 67 | 28 | 105 | 0 | 368 | 0 | 103 | 798 | 944 | 83 | 2548 | 2775 | 15 | 11 | 0 | 0 | 16 | 21 | 0 | 0 | 0 | 30 | 171 | 15 |
| SRR7178622 | 0 | 22 | 11 | 0 | 32 | 0 | 0 | 73 | 39 | 291 | 319 | 605 | 0 | 56 | 34 | 179 | 0 | 290 | 0 | 56 | 729 | 820 | 62 | 2119 | 2335 | 15 | 0 | 0 | 22 | 22 | 22 | 0 | 0 | 0 | 33 | 134 | 11 |
| SRR7178623 | 0 | 30 | 15 | 0 | 35 | 0 | 0 | 115 | 29 | 481 | 49 | 509 | 0 | 50 | 43 | 136 | 15 | 339 | 0 | 115 | 716 | 886 | 69 | 2242 | 2361 | 0 | 15 | 0 | 30 | 46 | 15 | 0 | 0 | 0 | 61 | 122 | 10 |
| SRR7178624 | 0 | 20 | 23 | 0 | 37 | 0 | 0 | 1703 | 164 | 603 | 792 | 1114 | 0 | 72 | 182 | 320 | 0 | 449 | 0 | 131 | 913 | 1333 | 51 | 1915 | 2940 | 35 | 32 | 11 | 14 | 40 | 14 | 0 | 11 | 23 | 20 | 349 | 19 |
| SRR7178625 | 0 | 11 | 21 | 0 | 29 | 0 | 0 | 1710 | 134 | 518 | 865 | 1050 | 0 | 60 | 165 | 384 | 16 | 434 | 0 | 174 | 978 | 1429 | 57 | 2057 | 3200 | 46 | 21 | 0 | 19 | 37 | 21 | 0 | 0 | 19 | 37 | 390 | 17 |
| SRR7178626 | 0 | 51 | 25 | 0 | 43 | 0 | 0 | 1851 | 141 | 493 | 865 | 1137 | 0 | 60 | 121 | 272 | 25 | 475 | 0 | 201 | 546 | 780 | 35 | 1145 | 1718 | 43 | 0 | 25 | 0 | 51 | 13 | 0 | 0 | 63 | 38 | 332 | 0 |
| SRR7178627 | 0 | 14 | 14 | 0 | 49 | 0 | 14 | 2117 | 187 | 656 | 70 | 993 | 0 | 94 | 122 | 122 | 14 | 445 | 0 | 131 | 414 | 645 | 23 | 834 | 1333 | 42 | 14 | 28 | 28 | 83 | 0 | 0 | 14 | 42 | 28 | 347 | 0 |
| SRR7178628 | 0 | 33 | 0 | 0 | 35 | 0 | 0 | 55 | 34 | 266 | 210 | 613 | 0 | 50 | 88 | 154 | 0 | 460 | 0 | 115 | 874 | 947 | 102 | 2065 | 2957 | 19 | 13 | 0 | 0 | 15 | 0 | 0 | 0 | 0 | 0 | 251 | 19 |
| SRR7178629 | 0 | 22 | 0 | 0 | 47 | 0 | 0 | 76 | 27 | 284 | 352 | 498 | 0 | 146 | 111 | 183 | 0 | 460 | 0 | 95 | 956 | 1022 | 109 | 2129 | 3045 | 18 | 17 | 0 | 0 | 0 | 0 | 0 | 0 | 0 | 0 | 247 | 12 |
| SRR7178630 | 0 | 32 | 14 | 0 | 113 | 0 | 0 | 78 | 30 | 253 | 149 | 469 | 0 | 42 | 99 | 156 | 0 | 497 | 0 | 101 | 847 | 915 | 100 | 1990 | 2804 | 15 | 14 | 0 | 0 | 20 | 0 | 0 | 0 | 0 | 16 | 277 | 13 |
| SRR7178631 | 0 | 23 | 14 | 0 | 53 | 0 | 0 | 218 | 35 | 358 | 196 | 876 | 0 | 111 | 52 | 119 | 0 | 641 | 0 | 125 | 798 | 824 | 88 | 2096 | 2069 | 16 | 12 | 0 | 0 | 18 | 0 | 0 | 0 | 0 | 18 | 242 | 0 |
| SRR7178632 | 0 | 24 | 0 | 0 | 37 | 0 | 0 | 187 | 42 | 350 | 562 | 889 | 0 | 78 | 42 | 92 | 0 | 590 | 0 | 140 | 826 | 897 | 104 | 2298 | 2334 | 12 | 0 | 0 | 0 | 19 | 11 | 0 | 0 | 13 | 0 | 255 | 12 |
| SRR7178633 | 0 | 60 | 0 | 0 | 64 | 0 | 0 | 250 | 10 | 330 | 69 | 740 | 0 | 80 | 40 | 60 | 20 | 685 | 0 | 110 | 476 | 459 | 66 | 1148 | 1258 | 13 | 10 | 0 | 10 | 0 | 10 | 0 | 0 | 40 | 0 | 410 | 0 |
| SRR7178634 | 0 | 68 | 0 | 0 | 68 | 0 | 0 | 219 | 0 | 304 | 45 | 827 | 0 | 152 | 0 | 67 | 0 | 614 | 0 | 133 | 304 | 363 | 37 | 847 | 917 | 0 | 11 | 11 | 23 | 0 | 0 | 0 | 11 | 0 | 11 | 219 | 11 |
| SRR7178635 | 0 | 33 | 27 | 0 | 26 | 0 | 0 | 528 | 130 | 469 | 1259 | 1817 | 0 | 91 | 232 | 276 | 0 | 272 | 0 | 169 | 782 | 1365 | 78 | 1748 | 2841 | 26 | 11 | 0 | 16 | 28 | 11 | 0 | 11 | 16 | 30 | 276 | 13 |
| SRR7178636 | 0 | 24 | 32 | 0 | 24 | 0 | 0 | 552 | 88 | 471 | 590 | 1817 | 0 | 101 | 167 | 225 | 0 | 246 | 0 | 180 | 727 | 1274 | 65 | 1650 | 2668 | 28 | 0 | 0 | 11 | 12 | 17 | 0 | 0 | 21 | 20 | 266 | 15 |
| SRR7178637 | 0 | 33 | 47 | 0 | 31 | 0 | 0 | 557 | 114 | 649 | 1144 | 1518 | 0 | 107 | 259 | 237 | 0 | 240 | 0 | 160 | 620 | 1057 | 61 | 1387 | 2284 | 20 | 0 | 0 | 0 | 33 | 20 | 0 | 20 | 40 | 54 | 229 | 0 |
| SRR7178638 | 0 | 55 | 46 | 0 | 14 | 0 | 0 | 496 | 126 | 642 | 933 | 1516 | 0 | 68 | 301 | 292 | 0 | 249 | 0 | 165 | 627 | 1088 | 63 | 1478 | 2396 | 18 | 0 | 18 | 0 | 18 | 28 | 0 | 18 | 55 | 74 | 136 | 10 |
| SRR7178639 | 0 | 16 | 47 | 0 | 27 | 0 | 0 | 633 | 231 | 540 | 762 | 1173 | 0 | 171 | 42 | 92 | 0 | 404 | 0 | 148 | 527 | 723 | 76 | 1825 | 1681 | 36 | 10 | 13 | 26 | 41 | 16 | 18 | 13 | 34 | 54 | 600 | 0 |
| SRR7178640 | 0 | 19 | 45 | 0 | 38 | 0 | 0 | 626 | 173 | 656 | 722 | 1088 | 0 | 351 | 81 | 107 | 0 | 403 | 0 | 188 | 533 | 762 | 89 | 1876 | 1704 | 29 | 13 | 0 | 26 | 26 | 19 | 0 | 0 | 32 | 39 | 676 | 0 |
| SRR7178641 | 0 | 13 | 40 | 0 | 110 | 0 | 0 | 757 | 239 | 742 | 127 | 1162 | 0 | 156 | 41 | 21 | 0 | 444 | 0 | 135 | 494 | 675 | 82 | 1760 | 1566 | 35 | 13 | 13 | 27 | 17 | 0 | 10 | 23 | 37 | 40 | 581 | 0 |
| SRR7178642 | 0 | 11 | 0 | 0 | 25 | 0 | 0 | 109 | 57 | 306 | 367 | 687 | 0 | 128 | 89 | 186 | 0 | 537 | 0 | 55 | 860 | 975 | 116 | 1428 | 2739 | 24 | 18 | 0 | 15 | 17 | 0 | 0 | 0 | 15 | 13 | 209 | 14 |
| SRR7178643 | 0 | 19 | 14 | 0 | 46 | 0 | 0 | 82 | 29 | 302 | 173 | 574 | 0 | 188 | 97 | 190 | 0 | 551 | 0 | 64 | 891 | 990 | 119 | 1481 | 2951 | 22 | 23 | 0 | 0 | 16 | 0 | 0 | 0 | 0 | 0 | 230 | 10 |
| SRR7178644 | 0 | 19 | 12 | 0 | 112 | 0 | 0 | 121 | 52 | 332 | 150 | 624 | 0 | 159 | 87 | 166 | 0 | 535 | 0 | 71 | 824 | 930 | 97 | 1402 | 2621 | 19 | 34 | 0 | 0 | 15 | 12 | 0 | 0 | 0 | 44 | 182 | 12 |
| SRR7178645 | 0 | 23 | 20 | 0 | 39 | 0 | 0 | 357 | 50 | 454 | 205 | 348 | 0 | 259 | 14 | 35 | 0 | 448 | 0 | 22 | 720 | 762 | 69 | 1133 | 2027 | 38 | 13 | 0 | 0 | 23 | 13 | 0 | 23 | 20 | 43 | 83 | 16 |
| SRR7178646 | 0 | 25 | 31 | 0 | 39 | 0 | 0 | 388 | 39 | 435 | 186 | 375 | 0 | 244 | 11 | 35 | 0 | 452 | 0 | 24 | 717 | 831 | 83 | 1243 | 2213 | 36 | 0 | 12 | 12 | 19 | 22 | 0 | 0 | 22 | 40 | 80 | 13 |
| SRR7178647 | 0 | 27 | 81 | 0 | 58 | 0 | 0 | 338 | 43 | 504 | 167 | 314 | 0 | 260 | 12 | 43 | 0 | 533 | 0 | 54 | 440 | 482 | 43 | 671 | 1289 | 34 | 0 | 0 | 40 | 27 | 13 | 0 | 13 | 27 | 27 | 74 | 11 |
| SRR7178648 | 0 | 30 | 61 | 0 | 61 | 0 | 0 | 516 | 79 | 592 | 202 | 217 | 0 | 267 | 0 | 25 | 0 | 477 | 0 | 22 | 269 | 354 | 40 | 508 | 869 | 34 | 0 | 15 | 30 | 45 | 30 | 15 | 15 | 30 | 15 | 72 | 15 |
| SRR7178649 | 0 | 25 | 25 | 0 | 0 | 0 | 0 | 1438 | 314 | 3448 | 296 | 475 | 0 | 307 | 195 | 427 | 0 | 255 | 0 | 141 | 728 | 662 | 77 | 1623 | 2046 | 0 | 0 | 0 | 18 | 47 | 51 | 11 | 25 | 51 | 54 | 473 | 0 |
| SRR7178650 | 0 | 34 | 24 | 0 | 13 | 0 | 0 | 1426 | 295 | 3774 | 287 | 413 | 0 | 453 | 236 | 418 | 0 | 229 | 0 | 158 | 762 | 726 | 81 | 1795 | 2117 | 0 | 0 | 0 | 19 | 53 | 38 | 0 | 29 | 24 | 57 | 525 | 0 |
| SRR7178651 | 0 | 96 | 40 | 0 | 24 | 0 | 0 | 1423 | 308 | 3871 | 255 | 460 | 0 | 262 | 246 | 441 | 0 | 247 | 0 | 110 | 697 | 698 | 77 | 1567 | 1908 | 0 | 0 | 0 | 20 | 20 | 15 | 0 | 10 | 10 | 30 | 474 | 0 |
| SRR7178652 | 0 | 0 | 15 | 0 | 34 | 0 | 0 | 303 | 17 | 386 | 365 | 603 | 0 | 122 | 11 | 0 | 0 | 373 | 0 | 51 | 701 | 799 | 108 | 2456 | 2119 | 13 | 0 | 12 | 15 | 22 | 12 | 0 | 15 | 15 | 0 | 104 | 0 |
| SRR7178653 | 0 | 31 | 0 | 0 | 39 | 0 | 0 | 318 | 28 | 446 | 195 | 606 | 0 | 164 | 0 | 13 | 0 | 376 | 0 | 72 | 681 | 712 | 104 | 2318 | 2047 | 23 | 0 | 0 | 0 | 17 | 19 | 0 | 12 | 0 | 21 | 119 | 0 |
| SRR7178654 | 0 | 21 | 42 | 0 | 44 | 0 | 0 | 414 | 57 | 420 | 267 | 637 | 0 | 127 | 0 | 19 | 10 | 277 | 0 | 76 | 546 | 553 | 86 | 1965 | 1638 | 18 | 0 | 10 | 21 | 31 | 21 | 10 | 10 | 21 | 31 | 83 | 0 |
| SRR7178655 | 0 | 29 | 87 | 0 | 27 | 0 | 0 | 359 | 16 | 531 | 32 | 473 | 0 | 131 | 0 | 33 | 0 | 359 | 0 | 82 | 609 | 630 | 78 | 2215 | 1698 | 15 | 0 | 0 | 29 | 15 | 58 | 0 | 0 | 15 | 15 | 90 | 0 |
| SRR7178656 | 0 | 23 | 28 | 0 | 35 | 0 | 0 | 117 | 24 | 153 | 431 | 755 | 0 | 64 | 32 | 30 | 0 | 454 | 0 | 107 | 925 | 967 | 122 | 2245 | 1967 | 16 | 18 | 20 | 23 | 46 | 15 | 0 | 25 | 51 | 48 | 201 | 0 |
| SRR7178657 | 0 | 24 | 51 | 0 | 48 | 0 | 0 | 117 | 32 | 235 | 544 | 729 | 0 | 54 | 26 | 48 | 0 | 463 | 0 | 98 | 853 | 929 | 119 | 2147 | 1907 | 18 | 32 | 0 | 32 | 63 | 27 | 0 | 19 | 49 | 37 | 233 | 0 |
| SRR7178658 | 0 | 33 | 11 | 0 | 49 | 0 | 0 | 100 | 80 | 345 | 48 | 531 | 0 | 53 | 27 | 60 | 0 | 390 | 0 | 120 | 680 | 725 | 89 | 1744 | 1485 | 14 | 11 | 11 | 33 | 55 | 11 | 0 | 0 | 22 | 22 | 239 | 0 |
| SRR7178659 | 0 | 60 | 90 | 0 | 43 | 0 | 0 | 119 | 26 | 306 | 511 | 579 | 0 | 51 | 34 | 51 | 0 | 395 | 0 | 94 | 711 | 785 | 90 | 1738 | 1496 | 11 | 45 | 30 | 45 | 196 | 15 | 0 | 45 | 75 | 60 | 179 | 0 |
| SRR7178660 | 0 | 29 | 26 | 0 | 16 | 0 | 0 | 323 | 41 | 336 | 441 | 640 | 0 | 125 | 128 | 205 | 0 | 345 | 0 | 107 | 771 | 709 | 74 | 1774 | 3686 | 10 | 0 | 0 | 26 | 81 | 68 | 26 | 22 | 28 | 81 | 172 | 15 |
| SRR7178661 | 0 | 45 | 35 | 0 | 21 | 0 | 0 | 330 | 46 | 340 | 440 | 576 | 0 | 347 | 118 | 211 | 0 | 333 | 0 | 129 | 767 | 739 | 82 | 1843 | 3837 | 0 | 0 | 0 | 17 | 59 | 19 | 0 | 26 | 19 | 68 | 161 | 13 |
| SRR7178662 | 0 | 59 | 24 | 0 | 84 | 0 | 0 | 344 | 31 | 332 | 426 | 553 | 0 | 172 | 82 | 211 | 0 | 328 | 0 | 148 | 735 | 694 | 75 | 1705 | 3479 | 0 | 0 | 0 | 27 | 64 | 39 | 15 | 0 | 15 | 54 | 157 | 13 |
| SRR7178663 | 0 | 12 | 29 | 0 | 18 | 0 | 0 | 473 | 38 | 496 | 360 | 559 | 0 | 105 | 40 | 40 | 0 | 250 | 0 | 88 | 646 | 701 | 37 | 1188 | 1923 | 0 | 0 | 0 | 24 | 20 | 24 | 0 | 12 | 12 | 24 | 252 | 0 |
| SRR7178664 | 0 | 15 | 34 | 0 | 26 | 0 | 0 | 450 | 52 | 478 | 396 | 549 | 0 | 98 | 42 | 68 | 0 | 250 | 0 | 88 | 691 | 731 | 33 | 1306 | 2049 | 0 | 0 | 0 | 11 | 30 | 38 | 0 | 23 | 34 | 34 | 221 | 0 |
| SRR7178665 | 0 | 19 | 0 | 0 | 25 | 0 | 0 | 476 | 52 | 609 | 334 | 520 | 0 | 178 | 22 | 67 | 0 | 246 | 0 | 67 | 342 | 446 | 15 | 684 | 1121 | 10 | 0 | 0 | 19 | 93 | 19 | 19 | 19 | 19 | 0 | 141 | 0 |
| SRR7178666 | 0 | 44 | 22 | 0 | 41 | 0 | 0 | 515 | 14 | 592 | 275 | 472 | 0 | 212 | 21 | 28 | 0 | 257 | 0 | 78 | 280 | 335 | 11 | 570 | 925 | 0 | 0 | 0 | 0 | 44 | 0 | 0 | 22 | 0 | 22 | 169 | 0 |
| SRR7178667 | 0 | 50 | 12 | 0 | 11 | 0 | 0 | 151 | 39 | 199 | 744 | 1684 | 0 | 117 | 0 | 0 | 0 | 342 | 0 | 995 | 594 | 528 | 71 | 1683 | 880 | 0 | 17 | 0 | 0 | 43 | 36 | 0 | 24 | 14 | 0 | 2038 | 0 |
| SRR7178668 | 0 | 53 | 22 | 0 | 15 | 0 | 0 | 107 | 41 | 251 | 797 | 1845 | 0 | 107 | 0 | 0 | 0 | 319 | 0 | 1031 | 652 | 563 | 68 | 1764 | 943 | 0 | 18 | 0 | 0 | 35 | 18 | 0 | 22 | 29 | 20 | 2116 | 0 |
| SRR7178669 | 0 | 63 | 74 | 0 | 15 | 0 | 0 | 153 | 92 | 382 | 870 | 1527 | 0 | 122 | 15 | 0 | 0 | 324 | 0 | 1252 | 333 | 288 | 41 | 1000 | 544 | 0 | 53 | 11 | 11 | 0 | 21 | 11 | 42 | 11 | 53 | 2168 | 0 |
| SRR7178670 | 0 | 73 | 24 | 0 | 18 | 12 | 12 | 219 | 44 | 248 | 33 | 1577 | 0 | 190 | 0 | 0 | 12 | 351 | 0 | 1008 | 264 | 230 | 21 | 694 | 347 | 0 | 12 | 0 | 0 | 61 | 49 | 12 | 12 | 0 | 24 | 1738 | 0 |
| SRR7178671 | 0 | 27 | 24 | 0 | 47 | 0 | 0 | 123 | 28 | 524 | 284 | 500 | 0 | 150 | 48 | 245 | 0 | 503 | 0 | 89 | 1007 | 865 | 121 | 1893 | 3092 | 45 | 0 | 11 | 22 | 14 | 24 | 14 | 30 | 27 | 57 | 225 | 13 |
| SRR7178672 | 0 | 26 | 16 | 0 | 45 | 0 | 0 | 138 | 46 | 485 | 177 | 500 | 0 | 197 | 44 | 303 | 0 | 503 | 0 | 97 | 917 | 818 | 97 | 1756 | 2824 | 44 | 0 | 11 | 24 | 26 | 0 | 0 | 0 | 18 | 39 | 255 | 10 |

### Supp Table S8.4.Columns_DI_to_ES

| | | | | | | | | | | | | | | | | | | | | | | | | | | | | | | | | | | | | | |
|---|---|---|---|---|---|---|---|---|---|---|---|---|---|---|---|---|---|---|---|---|---|---|---|---|---|---|---|---|---|---|---|---|---|---|---|---|---|
| SRR7178673 | 0 | 54 | 22 | 0 | 70 | 0 | 0 | 162 | 73 | 576 | 204 | 392 | 0 | 162 | 47 | 330 | 11 | 402 | 0 | 73 | 795 | 643 | 79 | 1532 | 2289 | 38 | 11 | 11 | 33 | 22 | 22 | 0 | 11 | 11 | 98 | 272 | 0 |
| SRR7178674 | 0 | 15 | 29 | 0 | 63 | 0 | 0 | 147 | 34 | 597 | 39 | 429 | 0 | 141 | 74 | 315 | 0 | 450 | 0 | 107 | 755 | 709 | 96 | 1597 | 2458 | 28 | 0 | 29 | 59 | 0 | 29 | 0 | 0 | 15 | 29 | 221 | 0 |
| SRR7178675 | 0 | 0 | 32 | 0 | 47 | 0 | 0 | 6579 | 202 | 439 | 62 | 305 | 0 | 213 | 18 | 63 | 0 | 366 | 0 | 66 | 609 | 585 | 122 | 2453 | 1664 | 44 | 60 | 14 | 23 | 0 | 23 | 0 | 32 | 28 | 64 | 428 | 0 |
| SRR7178676 | 0 | 0 | 34 | 0 | 40 | 0 | 0 | 6711 | 246 | 419 | 236 | 305 | 0 | 234 | 30 | 69 | 0 | 337 | 0 | 102 | 615 | 634 | 129 | 2624 | 1759 | 52 | 34 | 0 | 21 | 21 | 13 | 0 | 0 | 17 | 60 | 414 | 0 |
| SRR7178677 | 0 | 19 | 0 | 0 | 74 | 0 | 0 | 6710 | 266 | 498 | 17 | 223 | 0 | 146 | 34 | 34 | 19 | 305 | 0 | 69 | 341 | 345 | 60 | 1444 | 886 | 29 | 97 | 0 | 0 | 19 | 0 | 0 | 39 | 19 | 0 | 403 | 0 |
| SRR7178678 | 0 | 22 | 22 | 0 | 59 | 0 | 0 | 7348 | 268 | 536 | 181 | 221 | 0 | 229 | 16 | 63 | 0 | 351 | 0 | 71 | 266 | 233 | 59 | 904 | 575 | 53 | 89 | 0 | 0 | 22 | 67 | 0 | 67 | 44 | 89 | 244 | 0 |
| SRR7178679 | 0 | 0 | 25 | 0 | 23 | 0 | 0 | 424 | 43 | 209 | 268 | 455 | 0 | 131 | 66 | 193 | 0 | 395 | 0 | 52 | 886 | 949 | 66 | 1265 | 3197 | 19 | 0 | 0 | 35 | 54 | 28 | 25 | 25 | 44 | 136 | 131 | 17 |
| SRR7178680 | 0 | 31 | 54 | 0 | 40 | 0 | 0 | 451 | 37 | 252 | 169 | 398 | 0 | 305 | 52 | 174 | 0 | 414 | 0 | 55 | 940 | 1037 | 73 | 1328 | 3169 | 14 | 0 | 15 | 31 | 85 | 27 | 19 | 12 | 35 | 132 | 94 | 12 |
| SRR7178681 | 0 | 28 | 64 | 0 | 113 | 0 | 0 | 492 | 42 | 219 | 134 | 354 | 0 | 142 | 57 | 215 | 0 | 413 | 0 | 52 | 874 | 980 | 63 | 1182 | 2950 | 20 | 0 | 0 | 28 | 64 | 32 | 12 | 32 | 32 | 112 | 102 | 12 |
| SRR7178682 | 0 | 11 | 33 | 0 | 21 | 0 | 0 | 171 | 34 | 430 | 383 | 680 | 0 | 214 | 182 | 315 | 0 | 296 | 0 | 108 | 947 | 1032 | 57 | 988 | 2024 | 0 | 0 | 0 | 36 | 15 | 21 | 0 | 12 | 27 | 41 | 289 | 10 |
| SRR7178683 | 0 | 20 | 41 | 0 | 21 | 0 | 0 | 161 | 47 | 402 | 390 | 644 | 0 | 184 | 161 | 364 | 0 | 275 | 0 | 104 | 1030 | 1083 | 69 | 1069 | 2144 | 13 | 0 | 0 | 25 | 29 | 14 | 0 | 18 | 27 | 81 | 242 | 0 |
| SRR7178684 | 0 | 29 | 44 | 0 | 28 | 0 | 0 | 238 | 51 | 572 | 86 | 678 | 0 | 121 | 157 | 344 | 0 | 349 | 0 | 147 | 574 | 582 | 33 | 634 | 1314 | 11 | 15 | 15 | 36 | 36 | 73 | 0 | 29 | 65 | 51 | 283 | 0 |
| SRR7178685 | 0 | 34 | 51 | 0 | 36 | 0 | 0 | 234 | 68 | 586 | 303 | 606 | 0 | 176 | 93 | 239 | 0 | 316 | 0 | 83 | 425 | 468 | 31 | 441 | 983 | 0 | 0 | 17 | 68 | 34 | 25 | 17 | 25 | 76 | 85 | 249 | 0 |
| SRR7178686 | 0 | 39 | 18 | 0 | 12 | 0 | 0 | 184 | 77 | 1251 | 210 | 1588 | 0 | 133 | 31 | 71 | 0 | 194 | 0 | 1128 | 693 | 681 | 75 | 2082 | 1543 | 0 | 0 | 0 | 16 | 71 | 39 | 12 | 27 | 22 | 47 | 1751 | 0 |
| SRR7178687 | 0 | 23 | 17 | 0 | 10 | 0 | 0 | 173 | 61 | 1182 | 258 | 1471 | 0 | 157 | 76 | 117 | 0 | 194 | 0 | 923 | 645 | 622 | 68 | 1968 | 1450 | 0 | 0 | 0 | 0 | 63 | 38 | 0 | 27 | 28 | 27 | 1918 | 0 |
| SRR7178688 | 0 | 34 | 25 | 0 | 10 | 0 | 0 | 161 | 72 | 1469 | 1021 | 1755 | 0 | 143 | 90 | 107 | 25 | 169 | 0 | 1146 | 539 | 530 | 59 | 1599 | 1248 | 0 | 0 | 0 | 34 | 85 | 0 | 17 | 42 | 68 | 34 | 1791 | 0 |
| SRR7178689 | 0 | 47 | 35 | 0 | 0 | 12 | 0 | 229 | 69 | 1486 | 50 | 1555 | 0 | 183 | 114 | 91 | 0 | 164 | 0 | 869 | 570 | 581 | 66 | 1874 | 1359 | 0 | 0 | 0 | 12 | 94 | 0 | 0 | 35 | 35 | 35 | 2012 | 0 |
| SRR7178690 | 0 | 16 | 20 | 0 | 33 | 0 | 0 | 938 | 98 | 720 | 249 | 754 | 0 | 218 | 336 | 1237 | 0 | 498 | 0 | 112 | 869 | 797 | 117 | 2012 | 3070 | 25 | 0 | 0 | 0 | 18 | 0 | 0 | 0 | 11 | 29 | 204 | 22 |
| SRR7178691 | 0 | 26 | 15 | 0 | 39 | 0 | 0 | 158 | 79 | 332 | 277 | 921 | 0 | 369 | 16 | 100 | 0 | 449 | 0 | 79 | 787 | 1227 | 112 | 1916 | 2285 | 45 | 12 | 0 | 26 | 17 | 0 | 0 | 0 | 32 | 26 | 190 | 12 |
| SRR7178692 | 0 | 24 | 21 | 0 | 143 | 0 | 0 | 168 | 111 | 393 | 243 | 884 | 0 | 241 | 30 | 71 | 0 | 476 | 0 | 90 | 680 | 1126 | 97 | 1714 | 2073 | 42 | 12 | 12 | 15 | 18 | 0 | 0 | 0 | 33 | 18 | 203 | 12 |
| SRR7178693 | 0 | 0 | 30 | 0 | 32 | 0 | 0 | 2069 | 118 | 397 | 931 | 1163 | 0 | 83 | 60 | 78 | 0 | 432 | 0 | 0 | 886 | 1177 | 76 | 1335 | 2812 | 38 | 14 | 0 | 0 | 25 | 0 | 0 | 0 | 16 | 29 | 0 | 13 |
| SRR7178694 | 0 | 16 | 34 | 0 | 32 | 0 | 0 | 2036 | 136 | 367 | 404 | 1050 | 0 | 79 | 77 | 101 | 0 | 411 | 0 | 0 | 832 | 1080 | 63 | 1214 | 2648 | 46 | 10 | 0 | 12 | 10 | 21 | 0 | 28 | 29 | 19 | 0 | 14 |
| SRR7178695 | 0 | 26 | 17 | 0 | 37 | 0 | 0 | 2471 | 191 | 479 | 853 | 792 | 0 | 61 | 78 | 87 | 0 | 400 | 0 | 0 | 662 | 869 | 67 | 1046 | 2363 | 31 | 17 | 0 | 34 | 26 | 17 | 0 | 26 | 26 | 17 | 0 | 0 |
| SRR7178696 | 0 | 0 | 12 | 0 | 23 | 0 | 0 | 2608 | 168 | 560 | 828 | 1019 | 0 | 78 | 56 | 112 | 12 | 410 | 0 | 0 | 745 | 1029 | 48 | 1172 | 2562 | 35 | 0 | 12 | 35 | 35 | 35 | 0 | 12 | 12 | 12 | 0 | 12 |
| SRR7178697 | 0 | 16 | 14 | 0 | 38 | 0 | 0 | 197 | 31 | 355 | 283 | 945 | 0 | 86 | 38 | 115 | 0 | 385 | 0 | 121 | 954 | 1002 | 60 | 1599 | 2835 | 35 | 0 | 0 | 18 | 23 | 12 | 0 | 0 | 0 | 25 | 259 | 15 |
| SRR7178698 | 0 | 19 | 0 | 0 | 37 | 0 | 0 | 953 | 107 | 947 | 450 | 606 | 0 | 461 | 321 | 1276 | 0 | 504 | 0 | 116 | 926 | 833 | 118 | 2113 | 3090 | 20 | 10 | 0 | 0 | 16 | 10 | 0 | 0 | 12 | 31 | 174 | 20 |
| SRR7178699 | 0 | 17 | 30 | 0 | 120 | 0 | 0 | 1024 | 73 | 841 | 192 | 671 | 0 | 222 | 328 | 1235 | 0 | 494 | 0 | 115 | 814 | 766 | 112 | 1984 | 2879 | 18 | 0 | 0 | 0 | 15 | 11 | 0 | 13 | 0 | 25 | 208 | 16 |
| SRR7178700 | 0 | 38 | 38 | 0 | 24 | 0 | 0 | 492 | 75 | 694 | 554 | 901 | 0 | 135 | 39 | 67 | 0 | 273 | 0 | 199 | 756 | 611 | 45 | 1621 | 2430 | 15 | 0 | 0 | 0 | 50 | 22 | 0 | 0 | 19 | 26 | 352 | 14 |
| SRR7178701 | 0 | 25 | 21 | 0 | 20 | 0 | 0 | 480 | 49 | 746 | 245 | 951 | 0 | 95 | 41 | 105 | 0 | 253 | 0 | 227 | 696 | 538 | 48 | 1464 | 2320 | 0 | 0 | 0 | 16 | 35 | 28 | 0 | 18 | 21 | 28 | 388 | 14 |
| SRR7178702 | 0 | 21 | 32 | 0 | 27 | 0 | 0 | 497 | 71 | 772 | 559 | 728 | 0 | 124 | 35 | 124 | 0 | 234 | 0 | 248 | 562 | 443 | 37 | 1257 | 1934 | 19 | 0 | 11 | 21 | 42 | 21 | 11 | 0 | 32 | 21 | 302 | 10 |
| SRR7178703 | 0 | 74 | 44 | 0 | 10 | 0 | 0 | 605 | 114 | 890 | 525 | 856 | 0 | 114 | 68 | 125 | 15 | 235 | 0 | 160 | 623 | 533 | 48 | 1332 | 2092 | 15 | 0 | 0 | 15 | 44 | 59 | 29 | 15 | 29 | 44 | 262 | 15 |
| SRR7178704 | 0 | 0 | 22 | 0 | 0 | 0 | 0 | 149 | 91 | 1061 | 156 | 837 | 0 | 145 | 676 | 796 | 0 | 169 | 0 | 547 | 464 | 604 | 55 | 1763 | 895 | 0 | 0 | 0 | 0 | 36 | 20 | 0 | 20 | 20 | 60 | 1409 | 0 |
| SRR7178705 | 0 | 0 | 30 | 0 | 0 | 0 | 0 | 120 | 124 | 1076 | 538 | 763 | 0 | 386 | 708 | 878 | 0 | 161 | 0 | 616 | 489 | 668 | 58 | 1878 | 991 | 0 | 0 | 0 | 0 | 0 | 22 | 0 | 22 | 20 | 17 | 1448 | 0 |
| SRR7178706 | 0 | 0 | 31 | 0 | 22 | 0 | 0 | 172 | 96 | 1098 | 104 | 740 | 0 | 129 | 654 | 802 | 0 | 160 | 0 | 640 | 465 | 607 | 49 | 1704 | 875 | 0 | 0 | 0 | 0 | 0 | 13 | 0 | 34 | 26 | 34 | 1461 | 0 |
| SRR7178707 | 0 | 19 | 19 | 0 | 30 | 0 | 0 | 660 | 214 | 947 | 204 | 278 | 0 | 247 | 0 | 18 | 0 | 413 | 0 | 33 | 604 | 421 | 142 | 2104 | 1726 | 24 | 0 | 0 | 13 | 0 | 0 | 0 | 0 | 32 | 0 | 120 | 0 |
| SRR7178708 | 0 | 18 | 0 | 0 | 32 | 0 | 0 | 602 | 201 | 1028 | 212 | 291 | 0 | 260 | 0 | 20 | 0 | 427 | 0 | 59 | 555 | 380 | 142 | 1961 | 1624 | 29 | 0 | 18 | 0 | 0 | 0 | 0 | 0 | 25 | 12 | 110 | 0 |
| SRR7178709 | 0 | 101 | 0 | 0 | 36 | 0 | 0 | 687 | 196 | 1038 | 131 | 204 | 0 | 221 | 0 | 16 | 0 | 374 | 0 | 65 | 457 | 313 | 104 | 1584 | 1273 | 18 | 0 | 0 | 25 | 25 | 0 | 25 | 25 | 25 | 0 | 106 | 14 |
| SRR7178710 | 0 | 0 | 34 | 0 | 25 | 0 | 0 | 680 | 209 | 994 | 14 | 272 | 0 | 157 | 0 | 10 | 0 | 371 | 0 | 42 | 488 | 423 | 104 | 1905 | 1438 | 21 | 0 | 0 | 0 | 68 | 0 | 0 | 0 | 0 | 0 | 136 | 0 |
| SRR7178711 | 0 | 18 | 32 | 0 | 22 | 0 | 0 | 214 | 90 | 309 | 191 | 1089 | 0 | 181 | 105 | 147 | 0 | 305 | 0 | 128 | 783 | 731 | 76 | 1092 | 1916 | 22 | 0 | 14 | 14 | 65 | 32 | 14 | 22 | 32 | 122 | 566 | 0 |
| SRR7178712 | 0 | 18 | 40 | 0 | 29 | 0 | 0 | 181 | 106 | 378 | 135 | 963 | 0 | 441 | 96 | 175 | 0 | 288 | 0 | 191 | 846 | 810 | 84 | 1146 | 1958 | 15 | 0 | 0 | 40 | 44 | 44 | 0 | 40 | 35 | 75 | 526 | 0 |
| SRR7178713 | 0 | 23 | 64 | 0 | 79 | 0 | 0 | 180 | 87 | 382 | 126 | 965 | 0 | 153 | 93 | 185 | 0 | 302 | 0 | 185 | 781 | 709 | 74 | 1064 | 1753 | 17 | 0 | 0 | 50 | 50 | 37 | 0 | 32 | 50 | 82 | 447 | 0 |
| SRR7178714 | 0 | 15 | 11 | 0 | 60 | 0 | 0 | 120 | 52 | 324 | 164 | 477 | 0 | 164 | 25 | 27 | 0 | 374 | 0 | 35 | 721 | 646 | 84 | 1401 | 1614 | 48 | 19 | 11 | 45 | 34 | 11 | 0 | 0 | 41 | 37 | 185 | 0 |
| SRR7178715 | 0 | 14 | 18 | 0 | 53 | 0 | 0 | 160 | 58 | 353 | 166 | 555 | 0 | 157 | 26 | 26 | 0 | 404 | 0 | 42 | 800 | 661 | 83 | 1528 | 1721 | 48 | 14 | 11 | 49 | 42 | 18 | 11 | 28 | 32 | 49 | 194 | 0 |
| SRR7178716 | 0 | 16 | 0 | 0 | 66 | 0 | 0 | 156 | 26 | 408 | 399 | 356 | 0 | 35 | 17 | 17 | 0 | 395 | 0 | 35 | 419 | 364 | 54 | 841 | 912 | 48 | 16 | 16 | 31 | 31 | 16 | 0 | 31 | 62 | 31 | 173 | 14 |
| SRR7178717 | 0 | 18 | 70 | 0 | 86 | 0 | 0 | 209 | 56 | 353 | 42 | 425 | 0 | 120 | 0 | 16 | 0 | 402 | 0 | 16 | 318 | 269 | 23 | 566 | 688 | 53 | 35 | 35 | 35 | 35 | 0 | 0 | 18 | 70 | 35 | 160 | 0 |
| SRR7178718 | 0 | 0 | 30 | 0 | 31 | 0 | 0 | 274 | 63 | 355 | 281 | 501 | 0 | 175 | 17 | 28 | 0 | 394 | 0 | 44 | 764 | 670 | 103 | 1332 | 1953 | 35 | 17 | 0 | 12 | 20 | 0 | 0 | 10 | 29 | 57 | 166 | 11 |
| SRR7178719 | 0 | 12 | 25 | 0 | 42 | 0 | 0 | 276 | 42 | 364 | 184 | 465 | 0 | 378 | 21 | 28 | 0 | 432 | 0 | 39 | 797 | 696 | 98 | 1414 | 2060 | 27 | 14 | 0 | 21 | 30 | 12 | 0 | 25 | 21 | 49 | 170 | 12 |
| SRR7178720 | 0 | 12 | 36 | 0 | 144 | 0 | 0 | 257 | 44 | 346 | 315 | 491 | 0 | 167 | 26 | 22 | 0 | 405 | 0 | 46 | 755 | 622 | 90 | 1282 | 1911 | 33 | 14 | 0 | 12 | 17 | 0 | 0 | 12 | 19 | 55 | 150 | 11 |
| SRR7178721 | 0 | 28 | 28 | 0 | 15 | 0 | 0 | 1312 | 93 | 482 | 186 | 467 | 0 | 307 | 120 | 172 | 0 | 347 | 0 | 64 | 832 | 579 | 154 | 2073 | 2070 | 19 | 0 | 20 | 20 | 48 | 36 | 20 | 20 | 64 | 96 | 233 | 13 |
| SRR7178722 | 0 | 12 | 39 | 0 | 0 | 0 | 0 | 1245 | 114 | 497 | 497 | 452 | 0 | 269 | 86 | 198 | 0 | 338 | 0 | 69 | 748 | 537 | 144 | 1954 | 1978 | 22 | 0 | 16 | 39 | 43 | 35 | 12 | 35 | 51 | 113 | 282 | 14 |
| SRR7178723 | 0 | 0 | 62 | 0 | 19 | 0 | 16 | 1353 | 128 | 594 | 376 | 413 | 0 | 271 | 113 | 203 | 0 | 291 | 0 | 83 | 619 | 485 | 140 | 1539 | 1551 | 17 | 0 | 16 | 62 | 62 | 47 | 16 | 16 | 31 | 124 | 180 | 15 |
| SRR7178724 | 0 | 65 | 0 | 0 | 18 | 0 | 0 | 1519 | 142 | 408 | 38 | 456 | 0 | 228 | 123 | 237 | 0 | 332 | 0 | 47 | 703 | 533 | 131 | 1649 | 1800 | 13 | 0 | 43 | 65 | 43 | 22 | 22 | 0 | 108 | 65 | 190 | 0 |
| SRR7213733 | 0 | 78 | 0 | 15 | 31 | 0 | 17 | 5759 | 5978 | 1281 | 50 | 258 | 0 | 616 | 50 | 119 | 43 | 1443 | 275 | 5859 | 101 | 177 | 0 | 720 | 8182 | 1401 | 0 | 0 | 0 | 0 | 0 | 0 | 0 | 0 | 0 | 10506 | 0 |
| SRR7213734 | 0 | 50 | 0 | 30 | 77 | 0 | 17 | 11078 | 8254 | 1140 | 130 | 432 | 0 | 699 | 52 | 43 | 34 | 1526 | 322 | 3100 | 81 | 110 | 0 | 724 | 9748 | 1418 | 40 | 0 | 0 | 0 | 0 | 0 | 0 | 0 | 0 | 5595 | 0 |
| SRR7213735 | 0 | 69 | 0 | 21 | 81 | 11 | 11 | 7600 | 5406 | 1051 | 160 | 560 | 0 | 400 | 57 | 229 | 42 | 1591 | 287 | 7852 | 62 | 163 | 0 | 772 | 9842 | 1545 | 31 | 0 | 0 | 0 | 0 | 0 | 0 | 0 | 0 | 9315 | 0 |
| SRR7226512 | 0 | 112 | 29 | 0 | 31 | 0 | 0 | 294 | 1340 | 2475 | 89 | 224 | 0 | 32 | 1267 | 2946 | 0 | 760 | 0 | 851 | 150 | 744 | 98 | 1172 | 1078 | 0 | 13 | 37 | 40 | 46 | 12 | 30 | 13 | 23 | 0 | 19173 | 0 |
| SRR7226513 | 0 | 138 | 25 | 0 | 18 | 0 | 0 | 688 | 2110 | 8939 | 112 | 646 | 0 | 776 | 1574 | 7017 | 0 | 1298 | 18 | 734 | 535 | 1549 | 114 | 1734 | 924 | 0 | 88 | 0 | 24 | 11 | 12 | 0 | 11 | 20 | 16 | 10419 | 0 |
| SRR7226514 | 0 | 114 | 27 | 0 | 127 | 0 | 0 | 128 | 342 | 12263 | 113 | 680 | 0 | 383 | 2169 | 7204 | 0 | 3026 | 0 | 1113 | 154 | 3252 | 275 | 12252 | 1583 | 154 | 33 | 31 | 102 | 41 | 40 | 13 | 30 | 33 | 33 | 7888 | 18 |
| SRR7226515 | 0 | 124 | 19 | 0 | 66 | 0 | 0 | 58 | 167 | 2523 | 38 | 307 | 0 | 1145 | 509 | 716 | 0 | 1773 | 16 | 188 | 85 | 970 | 500 | 8240 | 1036 | 182 | 25 | 110 | 424 | 59 | 54 | 53 | 50 | 56 | 11 | 3644 | 0 |
| SRR7226516 | 0 | 618 | 67 | 0 | 62 | 0 | 0 | 147 | 513 | 9991 | 35 | 294 | 0 | 406 | 0 | 0 | 0 | 2779 | 11 | 1973 | 193 | 1017 | 264 | 3707 | 759 | 102 | 31 | 143 | 394 | 83 | 97 | 46 | 141 | 149 | 0 | 7387 | 12 |
| SRR7226517 | 0 | 473 | 110 | 0 | 177 | 0 | 0 | 308 | 1521 | 10680 | 137 | 403 | 0 | 201 | 0 | 13 | 0 | 4031 | 22 | 1094 | 293 | 1849 | 175 | 6060 | 1158 | 0 | 81 | 27 | 304 | 169 | 109 | 23 | 58 | 69 | 0 | 9414 | 0 |
| SRR7226518 | 0 | 49 | 34 | 0 | 198 | 10 | 0 | 228 | 814 | 7234 | 98 | 372 | 0 | 170 | 48 | 193 | 0 | 3565 | 16 | 1365 | 197 | 1274 | 210 | 5859 | 825 | 0 | 28 | 15 | 88 | 36 | 23 | 23 | 17 | 28 | 0 | 15426 | 0 |
| SRR7226519 | 0 | 40 | 68 | 0 | 229 | 0 | 0 | 119 | 454 | 4413 | 106 | 176 | 0 | 192 | 27 | 57 | 0 | 2269 | 14 | 1379 | 127 | 647 | 197 | 5700 | 665 | 0 | 22 | 21 | 64 | 16 | 16 | 15 | 14 | 26 | 0 | 18476 | 0 |
| SRR7226520 | 0 | 183 | 29 | 0 | 0 | 0 | 0 | 89 | 421 | 2718 | 339 | 1308 | 0 | 142 | 81 | 222 | 0 | 650 | 0 | 1777 | 91 | 774 | 39 | 464 | 611 | 0 | 12 | 27 | 53 | 22 | 17 | 31 | 13 | 21 | 26 | 30520 | 0 |
| SRR7226521 | 0 | 388 | 44 | 0 | 16 | 0 | 0 | 80 | 456 | 2145 | 190 | 665 | 0 | 359 | 29 | 82 | 0 | 719 | 19 | 2411 | 164 | 1396 | 167 | 1118 | 1029 | 37 | 16 | 61 | 314 | 100 | 104 | 43 | 82 | 101 | 21 | 47403 | 0 |
| SRR7226522 | 0 | 340 | 60 | 0 | 0 | 0 | 0 | 137 | 610 | 4276 | 243 | 1216 | 0 | 271 | 61 | 181 | 0 | 342 | 0 | 720 | 72 | 528 | 56 | 536 | 612 | 0 | 26 | 46 | 226 | 78 | 121 | 56 | 79 | 129 | 14 | 10339 | 0 |
| SRR7226523 | 0 | 151 | 18 | 0 | 133 | 0 | 0 | 85 | 484 | 8497 | 370 | 579 | 0 | 225 | 403 | 2149 | 0 | 2796 | 26 | 812 | 213 | 2358 | 218 | 7795 | 1540 | 91 | 10 | 26 | 89 | 20 | 41 | 14 | 23 | 30 | 22 | 12192 | 0 |
| SRR7226524 | 0 | 31 | 0 | 0 | 118 | 0 | 13 | 13 | 30 | 388 | 11 | 18 | 0 | 250 | 110 | 257 | 0 | 1090 | 13 | 991 | 121 | 1660 | 64 | 10294 | 1033 | 57 | 278 | 0 | 23 | 13 | 15 | 16 | 0 | 16 | 0 | 21991 | 0 |
| SRR7226525 | 0 | 90 | 13 | 0 | 114 | 0 | 0 | 100 | 335 | 7751 | 74 | 621 | 0 | 273 | 2488 | 3012 | 0 | 1963 | 14 | 769 | 275 | 2839 | 141 | 8606 | 1622 | 23 | 31 | 32 | 104 | 53 | 49 | 27 | 51 | 71 | 17 | 6971 | 0 |
| SRR7226526 | 0 | 162 | 27 | 0 | 62 | 0 | 0 | 61 | 267 | 3836 | 378 | 946 | 0 | 274 | 847 | 1513 | 0 | 2328 | 24 | 644 | 251 | 3091 | 36 | 9313 | 1867 | 73 | 11 | 36 | 175 | 93 | 73 | 62 | 113 | 175 | 31 | 7682 | 0 |
| SRR7233553 | 0 | 267 | 20 | 0 | 0 | 0 | 29 | 508 | 418 | 655 | 44743 | 116231 | 0 | 79 | 11 | 23 | 20 | 23 | 48 | 599 | 351 | 2013 | 0 | 0 | 89 | 55 | 79 | 17 | 14 | 44 | 35 | 20 | 0 | 0 | 0 | 373 | 0 |
| SRR7233554 | 0 | 303 | 18 | 0 | 0 | 0 | 21 | 384 | 554 | 810 | 38224 | 102719 | 0 | 139 | 11 | 0 | 15 | 23 | 49 | 1098 | 382 | 2137 | 0 | 0 | 146 | 64 | 49 | 0 | 11 | 27 | 25 | 13 | 0 | 0 | 0 | 800 | 0 |
| SRR7233556 | 0 | 251 | 0 | 0 | 0 | 0 | 16 | 354 | 453 | 431 | 56502 | 126955 | 0 | 88 | 11 | 11 | 13 | 20 | 60 | 608 | 259 | 1577 | 0 | 0 | 72 | 72 | 39 | 0 | 10 | 17 | 18 | 0 | 0 | 0 | 0 | 531 | 0 |
| SRR7233557 | 0 | 714 | 0 | 0 | 0 | 0 | 13 | 246 | 364 | 237 | 54075 | 106692 | 0 | 73 | 0 | 18 | 72 | 13 | 36 | 473 | 279 | 2139 | 0 | 0 | 22 | 56 | 226 | 11 | 47 | 23 | 81 | 0 | 0 | 0 | 0 | 228 | 0 |
| SRR7233558 | 0 | 1459 | 12 | 0 | 0 | 0 | 15 | 382 | 552 | 521 | 34545 | 70237 | 0 | 53 | 11 | 21 | 87 | 13 | 35 | 722 | 369 | 2250 | 0 | 0 | 57 | 54 | 153 | 0 | 47 | 20 | 98 | 11 | 0 | 0 | 0 | 637 | 0 |
| SRR7233559 | 0 | 198 | 0 | 0 | 0 | 0 | 24 | 260 | 384 | 260 | 21603 | 77579 | 0 | 37 | 0 | 0 | 18 | 18 | 70 | 508 | 449 | 2938 | 0 | 0 | 42 | 74 | 59 | 17 | 16 | 35 | 31 | 22 | 0 | 0 | 0 | 396 | 0 |
| SRR7233560 | 0 | 836 | 0 | 0 | 0 | 0 | 22 | 255 | 165 | 255 | 1307 | 58713 | 0 | 45 | 0 | 0 | 75 | 23 | 66 | 345 | 385 | 2832 | 0 | 0 | 97 | 68 | 184 | 12 | 57 | 21 | 87 | 0 | 0 | 0 | 0 | 330 | 0 |
| SRR7233561 | 0 | 143 | 33 | 0 | 0 | 0 | 31 | 276 | 388 | 351 | 49795 | 131500 | 0 | 164 | 15 | 0 | 17 | 22 | 41 | 680 | 262 | 2611 | 0 | 25 | 72 | 42 | 81 | 14 | 17 | 49 | 45 | 32 | 0 | 0 | 0 | 582 | 0 |
| SRR7233562 | 0 | 215 | 17 | 0 | 0 | 0 | 16 | 227 | 166 | 203 | 59007 | 173905 | 0 | 12 | 0 | 0 | 18 | 16 | 48 | 381 | 408 | 2478 | 0 | 23 | 47 | 47 | 100 | 10 | 17 | 38 | 33 | 18 | 0 | 0 | 0 | 246 | 0 |
| SRR7233563 | 0 | 1931 | 21 | 0 | 0 | 0 | 17 | 711 | 804 | 1066 | 1407 | 56244 | 0 | 201 | 15 | 15 | 62 | 27 | 43 | 2040 | 551 | 2290 | 0 | 0 | 155 | 57 | 193 | 11 | 66 | 23 | 171 | 26 | 0 | 0 | 0 | 1561 | 0 |
| SRR7233564 | 0 | 259 | 24 | 0 | 0 | 0 | 23 | 480 | 873 | 931 | 1470 | 52078 | 0 | 102 | 0 | 15 | 14 | 27 | 41 | 1717 | 676 | 2803 | 0 | 0 | 189 | 41 | 38 | 0 | 12 | 40 | 33 | 22 | 0 | 0 | 0 | 1528 | 0 |
| SRR7233565 | 0 | 374 | 35 | 0 | 0 | 0 | 16 | 879 | 1477 | 2085 | 40071 | 114277 | 0 | 196 | 0 | 0 | 11 | 13 | 27 | 645 | 542 | 3934 | 0 | 0 | 37 | 19 | 53 | 13 | 16 | 54 | 38 | 24 | 0 | 0 | 0 | 683 | 0 |
| SRR7233566 | 0 | 2209 | 36 | 0 | 0 | 0 | 11 | 681 | 1540 | 1781 | 38961 | 122321 | 0 | 157 | 0 | 21 | 66 | 12 | 22 | 1236 | 442 | 2799 | 0 | 0 | 66 | 28 | 362 | 13 | 107 | 41 | 243 | 36 | 0 | 0 | 0 | 744 | 0 |
| SRR7233568 | 0 | 342 | 14 | 0 | 0 | 0 | 18 | 120 | 171 | 171 | 42677 | 190217 | 0 | 29 | 0 | 0 | 12 | 26 | 58 | 183 | 768 | 3841 | 0 | 11 | 107 | 77 | 51 | 12 | 46 | 21 | 13 | 0 | 0 | 0 | 0 | 160 | 0 |
| SRR7233570 | 0 | 170 | 17 | 0 | 0 | 0 | 17 | 330 | 520 | 621 | 1034 | 39004 | 0 | 76 | 0 | 25 | 13 | 31 | 68 | 1243 | 672 | 3451 | 0 | 45 | 136 | 44 | 127 | 0 | 11 | 20 | 21 | 13 | 0 | 0 | 0 | 1103 | 0 |
| SRR7233571 | 0 | 254 | 21 | 0 | 0 | 0 | 25 | 536 | 583 | 639 | 39580 | 110029 | 0 | 207 | 0 | 0 | 14 | 22 | 46 | 836 | 497 | 2838 | 0 | 10 | 72 | 40 | 151 | 0 | 13 | 23 | 27 | 18 | 0 | 0 | 0 | 573 | 0 |
| SRR7233572 | 0 | 329 | 0 | 0 | 0 | 0 | 14 | 199 | 179 | 192 | 7752 | 126400 | 0 | 60 | 0 | 0 | 14 | 23 | 30 | 358 | 651 | 3571 | 0 | 0 | 64 | 46 | 177 | 0 | 12 | 21 | 19 | 0 | 0 | 0 | 0 | 285 | 0 |
| SRR7262826 | 0 | 322 | 61 | 179 | 268 | 61 | 19 | 1243 | 8210 | 75685 | 91 | 537 | 0 | 11927 | 0 | 0 | 18 | 2771 | 28 | 688 | 675 | 595 | 42 | 568 | 471 | 0 | 31 | 0 | 28 | 34 | 92 | 28 | 29 | 23 | 0 | 654 | 35 |
| SRR7262827 | 0 | 434 | 79 | 284 | 408 | 52 | 20 | 1632 | 7172 | 42545 | 253 | 309 | 0 | 7530 | 0 | 0 | 20 | 2780 | 34 | 794 | 1415 | 985 | 61 | 765 | 855 | 0 | 34 | 0 | 36 | 47 | 90 | 38 | 34 | 30 | 0 | 806 | 38 |
| SRR7262828 | 0 | 46 | 14 | 166 | 274 | 25 | 12 | 1895 | 9037 | 43499 | 101 | 166 | 0 | 5949 | 0 | 0 | 15 | 2560 | 39 | 905 | 1128 | 854 | 76 | 969 | 797 | 0 | 16 | 0 | 45 | 48 | 15 | 50 | 39 | 38 | 0 | 1089 | 27 |
| SRR7262829 | 0 | 101 | 146 | 80 | 111 | 44 | 15 | 1848 | 9034 | 136378 | 131 | 259 | 0 | 12358 | 0 | 0 | 0 | 2158 | 23 | 715 | 535 | 481 | 45 | 413 | 302 | 0 | 38 | 0 | 16 | 24 | 69 | 35 | 55 | 55 | 0 | 616 | 36 |
| SRR7262830 | 0 | 65 | 60 | 290 | 363 | 44 | 13 | 2068 | 8575 | 39394 | 160 | 251 | 0 | 6744 | 0 | 0 | 14 | 2923 | 32 | 811 | 1510 | 1125 | 75 | 887 | 941 | 0 | 16 | 0 | 60 | 41 | 13 | 37 | 26 | 24 | 0 | 809 | 43 |
| SRR7288164 | 77 | 0 | 0 | 0 | 0 | 31 | 0 | 1065 | 0 | 0 | 0 | 0 | 0 | 0 | 0 | 1065 | 0 | 0 | 0 | 1065 | 18 | 23 | 0 | 97 | 991 | 397 | 322 | 0 | 0 | 0 | 0 | 0 | 0 | 0 | 0 | 1065 | 0 |
| SRR7288166 | 107 | 0 | 0 | 0 | 0 | 31 | 0 | 0 | 0 | 1006 | 1006 | 0 | 0 | 0 | 0 | 0 | 0 | 0 | 0 | 0 | 12 | 27 | 0 | 90 | 916 | 401 | 168 | 0 | 0 | 0 | 0 | 0 | 0 | 0 | 0 | 1006 | 0 |
| SRR7288167 | 79 | 0 | 0 | 0 | 0 | 47 | 0 | 2235 | 0 | 1117 | 0 | 0 | 0 | 0 | 0 | 0 | 0 | 0 | 0 | 0 | 0 | 25 | 11 | 74 | 939 | 406 | 283 | 0 | 0 | 0 | 0 | 0 | 0 | 0 | 0 | 1117 | 0 |
| SRR7288236 | 31 | 0 | 0 | 0 | 0 | 79 | 0 | 0 | 0 | 0 | 0 | 2494 | 0 | 0 | 0 | 0 | 0 | 0 | 0 | 0 | 0 | 0 | 0 | 0 | 205 | 96 | 79 | 0 | 0 | 0 | 0 | 0 | 0 | 0 | 0 | 0 | 0 |
| SRR7288237 | 64 | 0 | 0 | 0 | 0 | 32 | 0 | 0 | 0 | 0 | 0 | 0 | 0 | 0 | 0 | 0 | 0 | 0 | 0 | 0 | 0 | 0 | 0 | 0 | 191 | 116 | 80 | 0 | 0 | 0 | 0 | 0 | 0 | 0 | 0 | 0 | 0 |
| SRR7288238 | 31 | 0 | 0 | 0 | 0 | 62 | 0 | 0 | 0 | 0 | 0 | 0 | 0 | 0 | 0 | 0 | 0 | 0 | 0 | 0 | 0 | 0 | 0 | 0 | 194 | 100 | 62 | 0 | 0 | 0 | 0 | 0 | 0 | 0 | 0 | 0 | 0 |
| SRR7288239 | 62 | 0 | 0 | 0 | 0 | 62 | 0 | 0 | 0 | 0 | 0 | 0 | 0 | 0 | 0 | 0 | 0 | 0 | 0 | 0 | 0 | 0 | 0 | 0 | 191 | 114 | 77 | 0 | 0 | 0 | 0 | 0 | 0 | 0 | 0 | 0 | 0 |
| SRR7288244 | 0 | 0 | 0 | 0 | 0 | 0 | 0 | 0 | 0 | 0 | 0 | 0 | 0 | 0 | 0 | 0 | 0 | 110 | 55 | 0 | 0 | 0 | 0 | 276 | 1212 | 331 | 1705 | 0 | 0 | 0 | 0 | 0 | 0 | 0 | 0 | 0 | 0 |
| SRR7288264 | 22 | 0 | 0 | 0 | 0 | 112 | 0 | 0 | 0 | 0 | 2594 | 1279 | 0 | 0 | 0 | 0 | 0 | 0 | 0 | 0 | 65 | 18 | 24 | 151 | 1229 | 421 | 180 | 0 | 0 | 0 | 0 | 0 | 0 | 0 | 0 | 1279 | 0 |
| SRR7288265 | 69 | 0 | 0 | 0 | 0 | 69 | 0 | 1506 | 0 | 0 | 1527 | 1506 | 0 | 0 | 0 | 0 | 0 | 12 | 0 | 0 | 57 | 24 | 18 | 175 | 1157 | 365 | 69 | 0 | 0 | 0 | 0 | 0 | 0 | 0 | 0 | 0 | 0 |
| SRR7288266 | 44 | 0 | 0 | 0 | 0 | 199 | 0 | 0 | 0 | 0 | 0 | 1403 | 0 | 0 | 0 | 0 | 0 | 0 | 0 | 0 | 80 | 18 | 24 | 151 | 1314 | 437 | 287 | 0 | 0 | 0 | 0 | 0 | 0 | 0 | 0 | 1403 | 0 |
| SRR7288267 | 0 | 0 | 0 | 0 | 0 | 68 | 0 | 2699 | 0 | 1350 | 1350 | 1350 | 0 | 0 | 0 | 0 | 0 | 0 | 0 | 0 | 87 | 36 | 21 | 180 | 1221 | 441 | 91 | 0 | 0 | 0 | 0 | 0 | 0 | 0 | 0 | 0 | 12 |
| SRR7288277 | 140 | 0 | 0 | 0 | 0 | 0 | 0 | 0 | 0 | 6198 | 0 | 2066 | 0 | 2066 | 0 | 0 | 0 | 11 | 0 | 0 | 34 | 92 | 11 | 264 | 803 | 505 | 140 | 0 | 0 | 0 | 0 | 0 | 0 | 0 | 0 | 2066 | 0 |
| SRR7288300 | 101 | 0 | 0 | 0 | 0 | 25 | 0 | 3473 | 0 | 16496 | 695 | 0 | 0 | 1563 | 0 | 0 | 0 | 0 | 0 | 0 | 108 | 38 | 25 | 92 | 1438 | 705 | 227 | 0 | 0 | 0 | 0 | 0 | 0 | 0 | 0 | 0 | 0 |
| SRR7288301 | 103 | 0 | 0 | 0 | 0 | 26 | 0 | 4304 | 179 | 16499 | 359 | 0 | 0 | 1435 | 0 | 0 | 0 | 29 | 0 | 0 | 94 | 23 | 23 | 94 | 1243 | 779 | 258 | 0 | 0 | 0 | 0 | 0 | 0 | 0 | 0 | 179 | 0 |
| SRR7288302 | 75 | 0 | 0 | 0 | 0 | 100 | 0 | 2803 | 0 | 14714 | 351 | 175 | 0 | 1226 | 0 | 0 | 0 | 0 | 0 | 0 | 92 | 19 | 13 | 79 | 1387 | 779 | 176 | 0 | 0 | 0 | 0 | 0 | 0 | 0 | 0 | 0 | 0 |
| SRR7288303 | 0 | 0 | 0 | 0 | 0 | 78 | 0 | 2131 | 0 | 15805 | 355 | 355 | 0 | 1066 | 178 | 0 | 0 | 0 | 0 | 0 | 82 | 23 | 20 | 75 | 1303 | 705 | 337 | 0 | 0 | 0 | 0 | 0 | 0 | 0 | 0 | 355 | 0 |
| SRR7288308 | 36 | 0 | 0 | 0 | 0 | 107 | 0 | 594 | 0 | 0 | 594 | 594 | 0 | 594 | 0 | 0 | 0 | 0 | 0 | 0 | 60 | 174 | 57 | 389 | 2479 | 617 | 215 | 0 | 0 | 0 | 0 | 0 | 0 | 0 | 0 | 0 | 0 |
| SRR7288309 | 75 | 0 | 0 | 0 | 0 | 112 | 0 | 0 | 0 | 0 | 620 | 1859 | 0 | 1239 | 0 | 0 | 0 | 23 | 0 | 0 | 72 | 125 | 26 | 473 | 2189 | 601 | 224 | 0 | 0 | 0 | 0 | 0 | 0 | 0 | 0 | 0 | 0 |
| SRR7288310 | 106 | 0 | 0 | 0 | 0 | 106 | 0 | 0 | 0 | 1217 | 3043 | 609 | 0 | 609 | 609 | 0 | 0 | 0 | 0 | 0 | 78 | 205 | 53 | 392 | 2499 | 585 | 213 | 0 | 0 | 0 | 0 | 0 | 0 | 0 | 0 | 0 | 0 |
| SRR7288311 | 37 | 0 | 0 | 0 | 0 | 74 | 0 | 593 | 0 | 593 | 1187 | 593 | 0 | 1187 | 593 | 0 | 0 | 19 | 0 | 0 | 45 | 165 | 29 | 407 | 2190 | 659 | 260 | 0 | 0 | 0 | 0 | 0 | 0 | 0 | 0 | 0 | 0 |
| SRR7288320 | 0 | 0 | 0 | 0 | 0 | 49 | 0 | 0 | 0 | 2137 | 2151 | 534 | 0 | 534 | 0 | 0 | 25 | 40 | 0 | 1603 | 185 | 76 | 0 | 266 | 3904 | 1365 | 172 | 0 | 0 | 0 | 0 | 0 | 0 | 0 | 0 | 534 | 0 |
| SRR7288321 | 104 | 0 | 0 | 0 | 0 | 26 | 0 | 569 | 0 | 1706 | 1149 | 1706 | 0 | 569 | 0 | 0 | 0 | 58 | 0 | 0 | 116 | 83 | 0 | 296 | 3567 | 1178 | 156 | 0 | 0 | 0 | 0 | 0 | 0 | 0 | 0 | 569 | 0 |
| SRR7288322 | 24 | 24 | 0 | 0 | 0 | 49 | 0 | 529 | 0 | 1058 | 1598 | 1058 | 0 | 1058 | 0 | 0 | 0 | 26 | 0 | 0 | 143 | 88 | 0 | 359 | 3846 | 1200 | 97 | 0 | 0 | 0 | 0 | 0 | 0 | 0 | 0 | 1586 | 0 |
| SRR7288323 | 26 | 0 | 0 | 0 | 0 | 51 | 0 | 0 | 0 | 3293 | 1667 | 3293 | 0 | 1098 | 0 | 0 | 0 | 54 | 0 | 0 | 144 | 98 | 0 | 288 | 3633 | 1339 | 154 | 0 | 0 | 0 | 0 | 0 | 0 | 0 | 0 | 0 | 0 |
| SRR7288376 | 0 | 0 | 0 | 0 | 0 | 59 | 0 | 0 | 0 | 0 | 0 | 0 | 0 | 0 | 0 | 0 | 0 | 0 | 0 | 0 | 18 | 33 | 0 | 96 | 2787 | 546 | 178 | 0 | 0 | 0 | 0 | 0 | 0 | 0 | 0 | 2801 | 0 |
| SRR7288377 | 0 | 0 | 0 | 0 | 0 | 122 | 0 | 0 | 0 | 0 | 2857 | 5714 | 0 | 0 | 0 | 0 | 0 | 0 | 0 | 0 | 19 | 42 | 0 | 154 | 2595 | 509 | 245 | 0 | 0 | 0 | 0 | 0 | 0 | 0 | 0 | 0 | 0 |
| SRR7288378 | 294 | 0 | 0 | 0 | 0 | 0 | 0 | 0 | 0 | 2865 | 2941 | 0 | 0 | 0 | 2865 | 0 | 0 | 0 | 0 | 0 | 26 | 33 | 18 | 172 | 2943 | 562 | 588 | 0 | 0 | 0 | 0 | 0 | 0 | 0 | 0 | 0 | 0 |
| SRR7288379 | 0 | 0 | 0 | 0 | 0 | 184 | 0 | 0 | 0 | 0 | 2994 | 2994 | 0 | 0 | 2994 | 0 | 0 | 0 | 0 | 0 | 15 | 19 | 0 | 144 | 2630 | 501 | 490 | 0 | 0 | 0 | 0 | 0 | 0 | 0 | 0 | 0 | 0 |
| SRR7288380 | 22 | 0 | 0 | 0 | 0 | 66 | 0 | 0 | 0 | 0 | 0 | 0 | 0 | 0 | 0 | 0 | 0 | 0 | 0 | 0 | 26 | 29 | 11 | 57 | 620 | 227 | 44 | 0 | 0 | 0 | 0 | 0 | 0 | 0 | 0 | 0 | 0 |
| SRR7288381 | 23 | 0 | 0 | 0 | 0 | 23 | 0 | 0 | 0 | 1934 | 0 | 0 | 0 | 0 | 0 | 1934 | 0 | 0 | 0 | 0 | 35 | 29 | 0 | 72 | 638 | 283 | 136 | 0 | 0 | 0 | 0 | 0 | 0 | 0 | 0 | 0 | 0 |
| SRR7288382 | 22 | 0 | 0 | 0 | 0 | 108 | 0 | 0 | 0 | 1770 | 0 | 0 | 0 | 0 | 0 | 0 | 0 | 0 | 0 | 0 | 17 | 28 | 11 | 96 | 607 | 270 | 303 | 0 | 0 | 0 | 0 | 0 | 0 | 0 | 0 | 0 | 0 |

### Supp Table S8.4.Columns_DI_to_ES

| | | | | | | | | | | | | | | | | | | | | | | | | | | | | | | | | | | | | | |
|---|---|---|---|---|---|---|---|---|---|---|---|---|---|---|---|---|---|---|---|---|---|---|---|---|---|---|---|---|---|---|---|---|---|---|---|---|---|
| SRR7288383 | 107 | 0 | 0 | 0 | 0 | 43 | 0 | 0 | 0 | 0 | 0 | 0 | 0 | 0 | 0 | 1862 | 0 | 0 | 0 | 0 | 0 | 19 | 0 | 94 | 597 | 261 | 129 | 0 | 0 | 0 | 0 | 0 | 0 | 0 | 0 | 0 | 0 |
| SRR7288384 | 34 | 0 | 0 | 0 | 0 | 206 | 0 | 0 | 0 | 2812 | 937 | 0 | 0 | 0 | 1874 | 937 | 17 | 21 | 0 | 0 | 124 | 139 | 54 | 1138 | 4597 | 708 | 327 | 0 | 0 | 0 | 0 | 0 | 0 | 0 | 0 | 0 | 0 |
| SRR7288385 | 35 | 0 | 0 | 0 | 0 | 105 | 0 | 0 | 943 | 1887 | 3774 | 0 | 0 | 1887 | 0 | 0 | 0 | 24 | 0 | 0 | 137 | 92 | 64 | 1301 | 4349 | 666 | 210 | 0 | 0 | 0 | 0 | 0 | 0 | 0 | 0 | 0 | 0 |
| SRR7288386 | 17 | 0 | 0 | 0 | 15 | 269 | 0 | 0 | 0 | 3759 | 940 | 4699 | 0 | 0 | 940 | 940 | 0 | 38 | 0 | 0 | 124 | 121 | 53 | 1193 | 4793 | 607 | 252 | 0 | 0 | 0 | 0 | 0 | 0 | 0 | 0 | 0 | 0 |
| SRR7288387 | 50 | 0 | 0 | 0 | 0 | 200 | 0 | 0 | 861 | 861 | 1748 | 0 | 0 | 861 | 1721 | 0 | 0 | 23 | 0 | 0 | 117 | 132 | 70 | 1279 | 4137 | 655 | 13 | 0 | 0 | 0 | 0 | 0 | 0 | 0 | 0 | 0 | 12 |
| SRR7288400 | 0 | 0 | 0 | 0 | 0 | 0 | 0 | 0 | 0 | 0 | 0 | 0 | 0 | 0 | 0 | 0 | 0 | 23 | 0 | 0 | 0 | 69 | 0 | 277 | 3621 | 530 | 971 | 0 | 0 | 0 | 0 | 0 | 0 | 0 | 0 | 14925 | 0 |
| SRR7288402 | 0 | 0 | 0 | 0 | 0 | 0 | 0 | 0 | 0 | 0 | 0 | 0 | 0 | 0 | 0 | 0 | 0 | 0 | 0 | 0 | 22 | 22 | 22 | 201 | 3769 | 557 | 488 | 0 | 0 | 0 | 0 | 0 | 0 | 0 | 0 | 0 | 0 |
| SRR7288472 | 0 | 0 | 0 | 0 | 0 | 199 | 0 | 0 | 0 | 7299 | 0 | 2433 | 0 | 2433 | 0 | 0 | 0 | 33 | 0 | 0 | 41 | 25 | 0 | 41 | 1189 | 336 | 0 | 0 | 0 | 0 | 0 | 0 | 0 | 0 | 0 | 4866 | 0 |
| SRR7288474 | 0 | 0 | 0 | 0 | 0 | 0 | 0 | 0 | 0 | 2315 | 0 | 2315 | 0 | 0 | 0 | 0 | 0 | 32 | 0 | 0 | 57 | 24 | 0 | 122 | 1142 | 429 | 291 | 0 | 0 | 0 | 0 | 0 | 0 | 0 | 0 | 2315 | 0 |
| SRR7288475 | 103 | 0 | 0 | 0 | 0 | 0 | 0 | 4878 | 0 | 0 | 0 | 2439 | 0 | 0 | 0 | 0 | 0 | 17 | 0 | 0 | 17 | 34 | 0 | 60 | 1088 | 442 | 103 | 0 | 0 | 0 | 0 | 0 | 0 | 0 | 0 | 2439 | 0 |
| SRR7288512 | 25 | 0 | 0 | 0 | 0 | 74 | 0 | 0 | 0 | 0 | 607 | 0 | 0 | 0 | 0 | 0 | 0 | 14 | 0 | 0 | 37 | 0 | 0 | 63 | 1178 | 328 | 74 | 0 | 0 | 0 | 0 | 0 | 0 | 0 | 0 | 1787 | 0 |
| SRR7288513 | 51 | 0 | 0 | 0 | 0 | 76 | 0 | 0 | 0 | 0 | 0 | 2423 | 0 | 0 | 0 | 0 | 0 | 18 | 0 | 606 | 48 | 0 | 0 | 79 | 1095 | 339 | 178 | 0 | 0 | 0 | 0 | 0 | 0 | 0 | 0 | 3634 | 0 |
| SRR7288514 | 0 | 0 | 0 | 0 | 0 | 49 | 0 | 589 | 0 | 589 | 0 | 589 | 0 | 589 | 0 | 0 | 0 | 12 | 0 | 1768 | 22 | 18 | 0 | 69 | 1167 | 338 | 73 | 0 | 0 | 0 | 0 | 0 | 0 | 0 | 0 | 3536 | 0 |
| SRR7288515 | 0 | 0 | 0 | 0 | 0 | 48 | 0 | 0 | 0 | 0 | 0 | 3497 | 0 | 583 | 0 | 583 | 0 | 15 | 0 | 1166 | 26 | 0 | 0 | 69 | 1152 | 359 | 121 | 0 | 0 | 0 | 0 | 0 | 0 | 0 | 0 | 5245 | 0 |
| SRR7293222 | 0 | 0 | 0 | 0 | 0 | 0 | 0 | 1716 | 747 | 9271 | 78 | 568 | 0 | 224 | 26 | 34 | 0 | 0 | 0 | 2086 | 2145 | 2910 | 0 | 25 | 24 | 0 | 0 | 69 | 249 | 435 | 276 | 45 | 258 | 684 | 0 | 7473 | 333 |
| SRR7293223 | 0 | 0 | 0 | 0 | 0 | 0 | 0 | 1867 | 603 | 10441 | 73 | 886 | 0 | 245 | 51 | 54 | 0 | 0 | 0 | 1578 | 1816 | 2676 | 0 | 25 | 23 | 0 | 0 | 34 | 141 | 273 | 187 | 37 | 200 | 407 | 0 | 6160 | 304 |
| SRR7293224 | 0 | 0 | 0 | 0 | 0 | 0 | 0 | 681 | 198 | 8439 | 71 | 871 | 0 | 821 | 139 | 329 | 0 | 0 | 0 | 4361 | 768 | 1670 | 0 | 0 | 2861 | 0 | 0 | 20 | 66 | 13 | 13 | 25 | 102 | 21 | 12 | 15836 | 53 |
| SRR7293225 | 12 | 12 | 0 | 0 | 0 | 0 | 0 | 1059 | 301 | 19537 | 22 | 583 | 0 | 1492 | 131 | 68 | 15 | 0 | 0 | 131 | 1391 | 2109 | 0 | 14 | 19 | 0 | 12 | 0 | 214 | 393 | 269 | 27 | 230 | 13 | 12 | 1599 | 158 |
| SRR7293226 | 0 | 0 | 0 | 0 | 0 | 0 | 0 | 1773 | 734 | 11465 | 262 | 861 | 0 | 639 | 38 | 23 | 0 | 0 | 0 | 558 | 1843 | 2861 | 0 | 20 | 9420 | 0 | 0 | 84 | 212 | 15 | 225 | 29 | 10 | 23 | 21 | 3020 | 198 |
| SRR7293227 | 0 | 0 | 0 | 0 | 0 | 0 | 0 | 6287 | 3635 | 12124 | 272 | 1522 | 0 | 488 | 256 | 155 | 0 | 0 | 0 | 369 | 1672 | 2498 | 0 | 18 | 17 | 0 | 0 | 81 | 106 | 204 | 121 | 36 | 118 | 261 | 0 | 2227 | 159 |
| SRR7293228 | 0 | 0 | 0 | 0 | 0 | 0 | 0 | 1069 | 260 | 5374 | 99 | 1056 | 0 | 68 | 0 | 26 | 0 | 0 | 0 | 996 | 1330 | 2047 | 0 | 10 | 6029 | 0 | 19 | 134 | 184 | 24 | 13 | 31 | 188 | 12 | 0 | 2282 | 112 |
| SRR7293229 | 0 | 0 | 0 | 0 | 0 | 0 | 0 | 1066 | 267 | 10452 | 92 | 616 | 0 | 366 | 17 | 0 | 18 | 0 | 0 | 283 | 1787 | 2399 | 0 | 17 | 26 | 0 | 0 | 13 | 293 | 528 | 408 | 49 | 249 | 306 | 0 | 1199 | 228 |
| SRR7293230 | 17 | 0 | 13 | 0 | 0 | 0 | 0 | 17994 | 13970 | 18672 | 22 | 739 | 0 | 1131 | 163 | 102 | 15 | 0 | 0 | 224 | 1373 | 2885 | 0 | 27 | 13 | 0 | 0 | 0 | 35 | 28 | 35 | 0 | 28 | 80 | 0 | 2211 | 229 |
| SRR7293231 | 0 | 0 | 0 | 0 | 0 | 0 | 0 | 17839 | 12442 | 13737 | 251 | 968 | 0 | 1615 | 17 | 0 | 10 | 0 | 0 | 277 | 957 | 2030 | 0 | 22 | 11 | 0 | 0 | 0 | 15 | 26 | 24 | 0 | 15 | 28 | 0 | 2130 | 125 |
| SRR7293232 | 0 | 0 | 14 | 0 | 0 | 0 | 0 | 1635 | 736 | 11899 | 28 | 867 | 0 | 605 | 37 | 156 | 0 | 0 | 0 | 986 | 658 | 1391 | 0 | 13 | 2855 | 0 | 0 | 0 | 18 | 79 | 15 | 0 | 45 | 12 | 0 | 8642 | 72 |
| SRR7293233 | 13 | 0 | 19 | 0 | 0 | 0 | 0 | 22053 | 8914 | 13376 | 49 | 286 | 0 | 2243 | 172 | 143 | 15 | 0 | 0 | 229 | 1040 | 1645 | 0 | 39 | 12 | 0 | 20 | 0 | 17 | 28 | 25 | 0 | 19 | 38 | 0 | 3759 | 184 |
| SRR7343626 | 0 | 0 | 0 | 0 | 0 | 21 | 0 | 0 | 0 | 76 | 9485 | 22839 | 0 | 0 | 0 | 0 | 0 | 0 | 0 | 0 | 259 | 964 | 0 | 19 | 154 | 0 | 144 | 0 | 0 | 0 | 0 | 0 | 0 | 0 | 0 | 0 | 0 |
| SRR7343627 | 0 | 0 | 0 | 0 | 0 | 0 | 0 | 554 | 0 | 554 | 55 | 11638 | 0 | 0 | 0 | 0 | 0 | 0 | 0 | 0 | 213 | 1903 | 0 | 0 | 57 | 0 | 100 | 0 | 0 | 0 | 0 | 0 | 0 | 0 | 0 | 0 | 0 |
| SRR7343629 | 0 | 0 | 0 | 0 | 0 | 0 | 0 | 504 | 0 | 2017 | 31 | 9581 | 0 | 0 | 0 | 0 | 0 | 0 | 0 | 0 | 148 | 970 | 0 | 32 | 85 | 0 | 50 | 0 | 0 | 0 | 0 | 0 | 0 | 0 | 0 | 504 | 0 |
| SRR7343630 | 27 | 0 | 0 | 0 | 0 | 0 | 0 | 0 | 0 | 38 | 542 | 27235 | 0 | 0 | 0 | 0 | 27 | 0 | 0 | 0 | 766 | 2671 | 0 | 0 | 504 | 0 | 27 | 0 | 0 | 0 | 0 | 0 | 0 | 0 | 0 | 0 | 0 |
| SRR7343631 | 0 | 0 | 0 | 0 | 0 | 0 | 0 | 347 | 0 | 0 | 25481 | 15601 | 0 | 0 | 0 | 0 | 0 | 0 | 0 | 0 | 121 | 1255 | 0 | 0 | 63 | 0 | 0 | 0 | 0 | 0 | 0 | 0 | 0 | 0 | 0 | 0 | 0 |
| SRR7343632 | 0 | 0 | 0 | 0 | 0 | 0 | 0 | 962 | 0 | 0 | 20 | 13475 | 0 | 0 | 0 | 0 | 41 | 0 | 0 | 0 | 207 | 941 | 0 | 0 | 106 | 0 | 41 | 0 | 0 | 0 | 0 | 0 | 0 | 0 | 0 | 0 | 0 |
| SRR7343633 | 10 | 10 | 0 | 0 | 0 | 0 | 0 | 570 | 0 | 427 | 77 | 14245 | 0 | 0 | 0 | 0 | 10 | 0 | 0 | 0 | 422 | 2085 | 0 | 30 | 312 | 0 | 21 | 0 | 0 | 0 | 0 | 0 | 0 | 0 | 0 | 427 | 0 |
| SRR7343635 | 0 | 0 | 0 | 0 | 0 | 0 | 0 | 0 | 0 | 249 | 25 | 7221 | 0 | 249 | 0 | 0 | 0 | 0 | 0 | 0 | 321 | 890 | 0 | 0 | 203 | 0 | 0 | 0 | 0 | 0 | 0 | 39 | 0 | 0 | 0 | 747 | 0 |
| SRR7343636 | 0 | 12 | 0 | 0 | 0 | 0 | 0 | 704 | 0 | 235 | 112 | 11023 | 0 | 0 | 0 | 0 | 49 | 0 | 0 | 0 | 279 | 1074 | 0 | 41 | 210 | 0 | 49 | 0 | 0 | 0 | 0 | 0 | 0 | 0 | 0 | 78 | 0 |
| SRR7343637 | 0 | 0 | 0 | 0 | 0 | 0 | 0 | 116 | 0 | 698 | 98 | 12791 | 0 | 116 | 0 | 0 | 0 | 0 | 0 | 0 | 445 | 2317 | 0 | 0 | 496 | 0 | 76 | 0 | 0 | 0 | 0 | 0 | 25 | 0 | 0 | 0 | 0 |
| SRR7343638 | 0 | 0 | 0 | 0 | 0 | 0 | 0 | 0 | 0 | 0 | 18 | 6797 | 0 | 425 | 0 | 0 | 0 | 0 | 0 | 0 | 288 | 1108 | 0 | 0 | 130 | 0 | 156 | 0 | 0 | 0 | 0 | 0 | 0 | 0 | 0 | 850 | 0 |
| SRR7343639 | 44 | 0 | 0 | 0 | 0 | 0 | 0 | 265 | 0 | 0 | 65 | 10348 | 0 | 0 | 0 | 0 | 0 | 0 | 0 | 0 | 177 | 1087 | 0 | 18 | 45 | 0 | 354 | 0 | 0 | 44 | 0 | 0 | 0 | 0 | 0 | 0 | 0 |
| SRR7343640 | 0 | 0 | 0 | 0 | 0 | 0 | 0 | 614 | 0 | 175 | 15618 | 28955 | 0 | 0 | 0 | 0 | 0 | 0 | 0 | 0 | 284 | 1624 | 0 | 0 | 183 | 0 | 58 | 0 | 0 | 0 | 0 | 0 | 0 | 0 | 0 | 0 | 0 |
| SRR7412243 | 0 | 128 | 28 | 0 | 46 | 0 | 0 | 565 | 238 | 89 | 30548 | 24308 | 0 | 0 | 30 | 89 | 0 | 440 | 0 | 59 | 916 | 1342 | 0 | 11 | 2135 | 405 | 0 | 0 | 28 | 14 | 0 | 0 | 28 | 43 | 121 | 624 | 0 |
| SRR7412244 | 0 | 101 | 0 | 0 | 46 | 0 | 0 | 217 | 174 | 174 | 648 | 27784 | 0 | 0 | 87 | 43 | 0 | 398 | 15 | 87 | 804 | 1462 | 0 | 17 | 2320 | 185 | 0 | 18 | 92 | 18 | 37 | 0 | 55 | 65 | 277 | 435 | 0 |
| SRR7412245 | 0 | 11 | 0 | 0 | 61 | 0 | 0 | 476 | 317 | 106 | 32770 | 23256 | 0 | 53 | 211 | 317 | 0 | 289 | 0 | 106 | 947 | 1479 | 0 | 23 | 947 | 426 | 0 | 0 | 47 | 0 | 0 | 0 | 31 | 17 | 86 | 1903 | 0 |
| SRR7412246 | 0 | 90 | 26 | 0 | 72 | 0 | 0 | 398 | 72 | 217 | 41261 | 25061 | 0 | 0 | 36 | 325 | 0 | 434 | 10 | 253 | 1077 | 1480 | 0 | 12 | 1205 | 250 | 0 | 19 | 13 | 26 | 26 | 13 | 32 | 83 | 173 | 832 | 0 |
| SRR7412247 | 0 | 103 | 0 | 0 | 33 | 0 | 0 | 356 | 119 | 102 | 1724 | 25251 | 0 | 0 | 152 | 203 | 0 | 261 | 0 | 119 | 624 | 1027 | 0 | 10 | 1864 | 159 | 0 | 0 | 0 | 40 | 12 | 0 | 40 | 95 | 119 | 2608 | 0 |
| SRR7412248 | 0 | 136 | 0 | 0 | 21 | 0 | 0 | 264 | 204 | 38 | 2225 | 14451 | 0 | 0 | 30 | 30 | 0 | 204 | 0 | 68 | 595 | 1137 | 0 | 0 | 1987 | 227 | 0 | 14 | 51 | 23 | 19 | 0 | 47 | 168 | 168 | 196 | 0 |
| SRR7412249 | 0 | 10 | 19 | 0 | 44 | 0 | 0 | 806 | 262 | 161 | 34894 | 26871 | 0 | 0 | 161 | 161 | 0 | 235 | 0 | 181 | 645 | 1079 | 0 | 0 | 2343 | 102 | 0 | 19 | 10 | 28 | 19 | 0 | 37 | 11 | 24 | 927 | 0 |
| SRR7412250 | 0 | 89 | 14 | 0 | 53 | 0 | 0 | 494 | 204 | 145 | 35253 | 19255 | 0 | 0 | 29 | 0 | 0 | 372 | 0 | 87 | 881 | 1412 | 0 | 22 | 3118 | 247 | 0 | 0 | 55 | 14 | 0 | 21 | 21 | 14 | 103 | 436 | 0 |
| SRR7412251 | 0 | 0 | 50 | 0 | 28 | 0 | 0 | 382 | 64 | 64 | 515 | 26137 | 0 | 0 | 64 | 127 | 17 | 249 | 0 | 191 | 942 | 822 | 0 | 74 | 2707 | 593 | 0 | 17 | 50 | 33 | 17 | 0 | 33 | 116 | 99 | 445 | 0 |
| SRR7412252 | 0 | 102 | 25 | 0 | 39 | 0 | 0 | 499 | 351 | 94 | 35260 | 25431 | 0 | 13 | 67 | 135 | 0 | 310 | 0 | 13 | 910 | 1286 | 0 | 28 | 2965 | 253 | 0 | 0 | 37 | 12 | 0 | 16 | 45 | 53 | 123 | 310 | 0 |
| SRR7412253 | 0 | 62 | 56 | 0 | 51 | 0 | 0 | 30 | 179 | 150 | 1026 | 25331 | 0 | 0 | 0 | 30 | 0 | 398 | 0 | 419 | 1301 | 1591 | 0 | 58 | 2293 | 771 | 0 | 25 | 74 | 93 | 19 | 0 | 31 | 81 | 155 | 1256 | 0 |
| SRR7412254 | 0 | 40 | 40 | 0 | 39 | 0 | 0 | 257 | 214 | 171 | 785 | 27128 | 0 | 0 | 86 | 471 | 0 | 401 | 0 | 471 | 913 | 1289 | 0 | 234 | 3374 | 887 | 0 | 26 | 66 | 40 | 0 | 0 | 26 | 66 | 79 | 1629 | 0 |
| SRR7412255 | 0 | 140 | 41 | 0 | 51 | 0 | 0 | 445 | 214 | 82 | 31331 | 25230 | 0 | 16 | 33 | 33 | 0 | 251 | 0 | 82 | 650 | 1044 | 0 | 15 | 2733 | 446 | 0 | 10 | 83 | 31 | 21 | 0 | 36 | 93 | 134 | 264 | 0 |
| SRR7412256 | 0 | 42 | 31 | 0 | 30 | 0 | 0 | 322 | 80 | 40 | 26494 | 24001 | 0 | 0 | 40 | 0 | 0 | 600 | 11 | 201 | 830 | 1303 | 0 | 0 | 623 | 262 | 0 | 10 | 42 | 31 | 31 | 10 | 10 | 73 | 94 | 201 | 0 |
| SRR7412257 | 0 | 88 | 95 | 0 | 61 | 0 | 0 | 695 | 232 | 77 | 1063 | 16181 | 0 | 0 | 154 | 129 | 14 | 448 | 0 | 386 | 616 | 1087 | 0 | 24 | 3393 | 183 | 0 | 0 | 41 | 20 | 0 | 0 | 61 | 47 | 14 | 566 | 0 |
| SRR7412258 | 0 | 36 | 0 | 0 | 13 | 0 | 0 | 536 | 153 | 0 | 30772 | 29777 | 0 | 77 | 0 | 0 | 0 | 573 | 0 | 383 | 785 | 1367 | 0 | 81 | 2035 | 385 | 0 | 72 | 36 | 0 | 0 | 0 | 0 | 0 | 108 | 77 | 0 |
| SRR7412259 | 0 | 45 | 19 | 0 | 47 | 0 | 0 | 797 | 283 | 308 | 19456 | 13468 | 0 | 51 | 26 | 26 | 0 | 398 | 0 | 206 | 792 | 1014 | 0 | 23 | 2873 | 507 | 0 | 0 | 51 | 38 | 0 | 0 | 45 | 115 | 180 | 386 | 0 |
| SRR7412260 | 0 | 0 | 0 | 0 | 13 | 0 | 0 | 532 | 97 | 97 | 713 | 28412 | 0 | 0 | 48 | 0 | 18 | 193 | 0 | 0 | 469 | 971 | 0 | 36 | 1367 | 658 | 36 | 0 | 54 | 0 | 36 | 18 | 0 | 18 | 90 | 339 | 0 |
| SRR7412261 | 0 | 97 | 83 | 0 | 49 | 0 | 0 | 583 | 228 | 228 | 29175 | 24050 | 0 | 0 | 0 | 51 | 14 | 259 | 0 | 228 | 894 | 1071 | 0 | 0 | 1698 | 76 | 0 | 0 | 104 | 21 | 28 | 0 | 42 | 90 | 250 | 533 | 0 |
| SRR7412262 | 0 | 12 | 0 | 0 | 68 | 0 | 0 | 406 | 87 | 29 | 1557 | 31371 | 0 | 29 | 0 | 29 | 0 | 230 | 0 | 87 | 746 | 1221 | 0 | 15 | 2412 | 155 | 0 | 26 | 34 | 0 | 17 | 0 | 43 | 43 | 11 | 406 | 0 |
| SRR7412263 | 0 | 24 | 24 | 0 | 20 | 0 | 0 | 500 | 125 | 63 | 670 | 14760 | 0 | 31 | 31 | 63 | 0 | 259 | 0 | 125 | 621 | 1062 | 0 | 46 | 1818 | 1019 | 0 | 24 | 17 | 71 | 24 | 0 | 47 | 47 | 15 | 344 | 0 |
| SRR7412264 | 0 | 66 | 13 | 0 | 54 | 0 | 13 | 569 | 122 | 41 | 21375 | 12313 | 0 | 0 | 41 | 41 | 0 | 384 | 0 | 0 | 1282 | 1759 | 0 | 35 | 1973 | 571 | 26 | 39 | 79 | 79 | 13 | 26 | 66 | 171 | 184 | 0 | 0 |
| SRR7412265 | 0 | 126 | 67 | 0 | 48 | 0 | 0 | 330 | 132 | 154 | 25301 | 23562 | 0 | 44 | 0 | 66 | 0 | 302 | 0 | 264 | 702 | 1074 | 0 | 36 | 2912 | 358 | 0 | 17 | 76 | 76 | 34 | 17 | 51 | 15 | 24 | 705 | 0 |
| SRR7412266 | 0 | 177 | 59 | 0 | 59 | 0 | 0 | 177 | 102 | 121 | 2603 | 14469 | 0 | 0 | 19 | 195 | 0 | 252 | 14 | 734 | 915 | 1378 | 0 | 108 | 2084 | 564 | 30 | 52 | 244 | 162 | 30 | 0 | 44 | 118 | 81 | 1747 | 0 |
| SRR7412267 | 0 | 73 | 53 | 0 | 58 | 0 | 0 | 266 | 97 | 48 | 21584 | 12979 | 0 | 0 | 0 | 48 | 0 | 498 | 0 | 773 | 747 | 1003 | 0 | 66 | 1801 | 414 | 0 | 20 | 107 | 53 | 33 | 13 | 40 | 67 | 107 | 628 | 0 |
| SRR7412268 | 0 | 107 | 56 | 0 | 58 | 0 | 0 | 190 | 86 | 121 | 23276 | 19396 | 0 | 0 | 69 | 103 | 0 | 529 | 15 | 414 | 668 | 950 | 0 | 70 | 3601 | 503 | 0 | 0 | 19 | 107 | 50 | 0 | 81 | 12 | 16 | 431 | 0 |
| SRR7412269 | 0 | 60 | 30 | 0 | 71 | 0 | 0 | 212 | 159 | 317 | 35046 | 23487 | 0 | 26 | 0 | 26 | 0 | 340 | 0 | 476 | 1430 | 1811 | 0 | 75 | 3330 | 325 | 12 | 30 | 48 | 42 | 18 | 0 | 30 | 60 | 14 | 635 | 0 |
| SRR7412270 | 0 | 30 | 74 | 0 | 38 | 0 | 0 | 200 | 50 | 300 | 22332 | 14671 | 0 | 50 | 50 | 0 | 0 | 201 | 0 | 401 | 903 | 1157 | 0 | 23 | 1658 | 309 | 15 | 59 | 119 | 30 | 15 | 0 | 104 | 149 | 149 | 2053 | 0 |
| SRR7412271 | 0 | 23 | 23 | 0 | 29 | 0 | 0 | 656 | 154 | 154 | 764 | 21470 | 0 | 0 | 193 | 270 | 11 | 444 | 15 | 232 | 664 | 1302 | 0 | 55 | 2809 | 590 | 0 | 11 | 23 | 23 | 11 | 0 | 23 | 46 | 263 | 425 | 0 |
| SRR7412272 | 0 | 58 | 19 | 0 | 29 | 0 | 0 | 895 | 138 | 172 | 29667 | 21717 | 0 | 0 | 0 | 34 | 0 | 275 | 0 | 103 | 667 | 1139 | 0 | 41 | 2856 | 581 | 0 | 0 | 19 | 0 | 19 | 0 | 48 | 67 | 154 | 482 | 0 |
| SRR7412273 | 0 | 61 | 81 | 0 | 73 | 0 | 0 | 492 | 281 | 328 | 814 | 17640 | 0 | 23 | 0 | 70 | 0 | 525 | 0 | 211 | 1159 | 1305 | 0 | 14 | 2179 | 142 | 0 | 0 | 57 | 69 | 28 | 12 | 12 | 53 | 97 | 2132 | 0 |
| SRR7412274 | 0 | 29 | 20 | 0 | 50 | 0 | 0 | 299 | 187 | 112 | 467 | 16553 | 0 | 75 | 149 | 37 | 0 | 384 | 0 | 561 | 724 | 1059 | 0 | 17 | 2989 | 196 | 0 | 20 | 78 | 39 | 29 | 20 | 49 | 127 | 137 | 4484 | 0 |
| SRR7412275 | 0 | 45 | 19 | 0 | 45 | 0 | 0 | 1119 | 307 | 373 | 1036 | 20676 | 0 | 22 | 22 | 44 | 0 | 201 | 0 | 307 | 644 | 841 | 0 | 41 | 2116 | 367 | 0 | 0 | 26 | 26 | 78 | 13 | 52 | 58 | 271 | 615 | 0 |
| SRR7412276 | 18 | 71 | 0 | 0 | 34 | 0 | 0 | 507 | 254 | 63 | 23397 | 17247 | 0 | 0 | 0 | 0 | 0 | 320 | 0 | 1015 | 492 | 1156 | 0 | 15 | 2829 | 185 | 0 | 0 | 0 | 18 | 0 | 0 | 45 | 10 | 71 | 2600 | 0 |
| SRR7412277 | 0 | 32 | 32 | 0 | 18 | 0 | 0 | 424 | 196 | 98 | 17531 | 14533 | 0 | 33 | 228 | 391 | 0 | 497 | 13 | 293 | 893 | 1501 | 0 | 168 | 2522 | 1013 | 0 | 0 | 48 | 56 | 48 | 0 | 48 | 72 | 19 | 815 | 0 |
| SRR7412278 | 0 | 14 | 42 | 0 | 27 | 0 | 0 | 585 | 59 | 176 | 32824 | 29782 | 0 | 59 | 59 | 0 | 0 | 305 | 0 | 0 | 566 | 797 | 0 | 21 | 2058 | 486 | 0 | 0 | 14 | 28 | 0 | 14 | 85 | 56 | 85 | 410 | 0 |
| SRR7412279 | 0 | 57 | 33 | 0 | 27 | 0 | 0 | 528 | 226 | 226 | 544 | 17651 | 0 | 0 | 226 | 415 | 0 | 453 | 0 | 339 | 538 | 1097 | 0 | 18 | 2281 | 1178 | 0 | 0 | 33 | 0 | 0 | 0 | 16 | 74 | 172 | 302 | 0 |
| SRR7412280 | 0 | 18 | 0 | 0 | 27 | 0 | 0 | 393 | 221 | 123 | 614 | 11966 | 0 | 0 | 0 | 123 | 0 | 145 | 0 | 123 | 676 | 1005 | 0 | 40 | 2116 | 412 | 0 | 0 | 64 | 0 | 0 | 0 | 45 | 19 | 11 | 614 | 0 |
| SRR7412281 | 0 | 39 | 30 | 0 | 24 | 0 | 0 | 491 | 41 | 164 | 34024 | 32345 | 0 | 0 | 123 | 164 | 0 | 290 | 0 | 164 | 483 | 1089 | 0 | 29 | 2661 | 459 | 0 | 0 | 59 | 0 | 0 | 30 | 39 | 59 | 14 | 491 | 0 |
| SRR7412282 | 0 | 46 | 68 | 0 | 18 | 0 | 0 | 389 | 35 | 71 | 995 | 22221 | 0 | 0 | 35 | 35 | 0 | 175 | 0 | 106 | 645 | 1132 | 0 | 23 | 2591 | 549 | 0 | 23 | 34 | 34 | 23 | 0 | 23 | 137 | 160 | 708 | 0 |
| SRR7412283 | 0 | 61 | 0 | 0 | 12 | 0 | 0 | 368 | 33 | 67 | 977 | 25769 | 0 | 33 | 33 | 100 | 0 | 350 | 0 | 201 | 573 | 1121 | 0 | 35 | 2937 | 604 | 0 | 0 | 53 | 18 | 18 | 0 | 61 | 105 | 10 | 1003 | 0 |
| SRR7412284 | 0 | 0 | 0 | 0 | 33 | 0 | 0 | 475 | 0 | 86 | 656 | 28779 | 0 | 0 | 0 | 43 | 0 | 188 | 0 | 346 | 513 | 988 | 0 | 17 | 2533 | 157 | 0 | 0 | 36 | 36 | 0 | 0 | 36 | 89 | 10 | 1945 | 0 |
| SRR7412285 | 0 | 12 | 53 | 0 | 63 | 0 | 0 | 335 | 74 | 298 | 1363 | 21658 | 0 | 0 | 19 | 74 | 0 | 480 | 0 | 223 | 1114 | 1442 | 0 | 11 | 2514 | 334 | 0 | 0 | 53 | 29 | 18 | 0 | 53 | 117 | 351 | 856 | 0 |
| SRR7412286 | 0 | 24 | 0 | 0 | 48 | 0 | 0 | 550 | 0 | 236 | 37421 | 30818 | 0 | 0 | 236 | 314 | 0 | 423 | 13 | 157 | 824 | 1357 | 0 | 28 | 2855 | 478 | 0 | 16 | 24 | 40 | 24 | 0 | 32 | 24 | 45 | 2594 | 0 |
| SRR7412287 | 0 | 0 | 27 | 0 | 81 | 0 | 0 | 928 | 437 | 273 | 34936 | 22982 | 0 | 0 | 0 | 0 | 0 | 319 | 0 | 382 | 595 | 1421 | 0 | 0 | 1361 | 271 | 0 | 0 | 13 | 20 | 13 | 20 | 40 | 60 | 128 | 1638 | 0 |
| SRR7412288 | 0 | 50 | 0 | 0 | 35 | 0 | 0 | 192 | 154 | 38 | 24251 | 16611 | 0 | 13 | 13 | 26 | 0 | 267 | 0 | 77 | 874 | 1329 | 0 | 52 | 2158 | 1538 | 0 | 30 | 134 | 45 | 40 | 0 | 129 | 387 | 278 | 435 | 0 |
| SRR7412289 | 0 | 10 | 20 | 0 | 34 | 0 | 10 | 297 | 0 | 59 | 31081 | 34700 | 0 | 0 | 0 | 178 | 0 | 155 | 0 | 297 | 594 | 875 | 0 | 70 | 2419 | 312 | 0 | 0 | 20 | 20 | 61 | 10 | 41 | 13 | 17 | 830 | 0 |
| SRR7412290 | 0 | 0 | 42 | 0 | 35 | 0 | 0 | 340 | 0 | 113 | 19737 | 18149 | 0 | 0 | 0 | 113 | 0 | 528 | 14 | 0 | 1102 | 1224 | 0 | 0 | 283 | 1266 | 0 | 0 | 42 | 14 | 14 | 0 | 14 | 97 | 111 | 1021 | 0 |
| SRR7412291 | 0 | 40 | 16 | 0 | 21 | 0 | 0 | 742 | 202 | 405 | 411 | 20165 | 0 | 0 | 0 | 135 | 0 | 323 | 0 | 405 | 683 | 990 | 0 | 0 | 2736 | 197 | 0 | 0 | 40 | 16 | 16 | 16 | 0 | 56 | 143 | 809 | 0 |
| SRR7412292 | 0 | 55 | 27 | 0 | 35 | 0 | 0 | 1021 | 323 | 54 | 512 | 21340 | 0 | 0 | 54 | 54 | 0 | 428 | 0 | 108 | 845 | 1443 | 0 | 0 | 1869 | 512 | 0 | 0 | 37 | 0 | 37 | 0 | 46 | 119 | 119 | 1021 | 0 |
| SRR7412293 | 0 | 0 | 0 | 0 | 35 | 0 | 0 | 258 | 206 | 180 | 553 | 11569 | 0 | 26 | 0 | 103 | 0 | 204 | 0 | 77 | 738 | 1213 | 0 | 59 | 2111 | 1323 | 0 | 28 | 83 | 55 | 28 | 18 | 46 | 120 | 13 | 1056 | 0 |
| SRR7412294 | 0 | 43 | 43 | 0 | 61 | 0 | 0 | 2110 | 528 | 176 | 223 | 10961 | 0 | 59 | 117 | 59 | 0 | 340 | 0 | 352 | 561 | 968 | 0 | 44 | 3662 | 311 | 0 | 0 | 43 | 0 | 14 | 0 | 64 | 114 | 150 | 762 | 0 |
| SRR7412295 | 0 | 70 | 35 | 0 | 70 | 0 | 0 | 200 | 29 | 286 | 1135 | 35403 | 0 | 0 | 0 | 29 | 0 | 442 | 0 | 315 | 1146 | 1460 | 0 | 29 | 928 | 605 | 18 | 70 | 10 | 61 | 35 | 0 | 70 | 15 | 67 | 1059 | 0 |
| SRR7412296 | 0 | 0 | 24 | 0 | 29 | 0 | 0 | 784 | 261 | 157 | 38038 | 35007 | 0 | 0 | 0 | 366 | 0 | 269 | 0 | 261 | 714 | 886 | 0 | 0 | 1738 | 214 | 0 | 24 | 49 | 73 | 61 | 0 | 85 | 170 | 24 | 1097 | 0 |
| SRR7412297 | 0 | 11 | 0 | 0 | 30 | 0 | 0 | 830 | 277 | 461 | 15306 | 18103 | 0 | 61 | 31 | 154 | 0 | 123 | 0 | 184 | 589 | 984 | 0 | 69 | 2713 | 385 | 0 | 0 | 27 | 22 | 0 | 0 | 38 | 33 | 76 | 1260 | 0 |
| SRR7412298 | 0 | 17 | 0 | 0 | 22 | 0 | 0 | 436 | 24 | 24 | 33105 | 24369 | 0 | 0 | 24 | 48 | 0 | 239 | 0 | 0 | 711 | 1114 | 0 | 44 | 1899 | 794 | 0 | 0 | 50 | 42 | 33 | 0 | 75 | 14 | 23 | 0 | 0 |
| SRR7412299 | 0 | 0 | 0 | 0 | 14 | 0 | 0 | 276 | 138 | 46 | 396 | 23271 | 0 | 0 | 184 | 184 | 0 | 242 | 15 | 138 | 415 | 847 | 0 | 61 | 1673 | 738 | 18 | 0 | 18 | 53 | 0 | 0 | 125 | 267 | 232 | 690 | 0 |
| SRR7412300 | 0 | 45 | 45 | 0 | 13 | 0 | 0 | 390 | 70 | 84 | 1312 | 21923 | 0 | 14 | 293 | 307 | 0 | 280 | 13 | 56 | 453 | 737 | 0 | 45 | 1848 | 966 | 0 | 22 | 56 | 33 | 22 | 22 | 22 | 156 | 167 | 1144 | 0 |
| SRR7412301 | 0 | 23 | 45 | 0 | 36 | 0 | 0 | 231 | 0 | 0 | 23379 | 26686 | 0 | 0 | 77 | 0 | 0 | 197 | 15 | 154 | 835 | 1352 | 0 | 40 | 3105 | 168 | 0 | 0 | 68 | 11 | 45 | 34 | 57 | 250 | 204 | 1769 | 0 |
| SRR7412302 | 0 | 47 | 0 | 0 | 23 | 0 | 0 | 459 | 320 | 181 | 2276 | 19190 | 0 | 14 | 28 | 70 | 0 | 179 | 0 | 125 | 747 | 1098 | 0 | 15 | 1569 | 652 | 0 | 17 | 68 | 17 | 21 | 13 | 120 | 184 | 227 | 807 | 0 |
| SRR7412303 | 0 | 18 | 31 | 0 | 28 | 0 | 0 | 738 | 394 | 246 | 19332 | 18889 | 0 | 0 | 25 | 74 | 0 | 99 | 0 | 221 | 531 | 629 | 0 | 0 | 1886 | 105 | 0 | 49 | 61 | 18 | 24 | 0 | 55 | 24 | 14 | 1500 | 0 |
| SRR7412304 | 0 | 59 | 0 | 0 | 35 | 0 | 0 | 234 | 167 | 401 | 22679 | 16992 | 0 | 33 | 0 | 0 | 0 | 57 | 0 | 167 | 490 | 911 | 0 | 20 | 1906 | 426 | 0 | 33 | 50 | 25 | 17 | 42 | 42 | 32 | 14 | 1606 | 0 |
| SRR7412305 | 0 | 34 | 69 | 0 | 24 | 0 | 0 | 584 | 195 | 71 | 24409 | 20302 | 0 | 18 | 18 | 35 | 0 | 324 | 0 | 53 | 927 | 1227 | 0 | 85 | 2654 | 625 | 0 | 34 | 138 | 69 | 23 | 0 | 57 | 10 | 126 | 478 | 0 |
| SRR7412306 | 0 | 153 | 61 | 0 | 36 | 0 | 0 | 443 | 200 | 155 | 1310 | 19024 | 0 | 0 | 0 | 0 | 0 | 223 | 0 | 222 | 526 | 884 | 0 | 19 | 2564 | 244 | 0 | 31 | 51 | 51 | 31 | 0 | 31 | 112 | 143 | 1330 | 0 |
| SRR7412307 | 0 | 16 | 33 | 0 | 42 | 0 | 0 | 377 | 94 | 47 | 41773 | 28194 | 0 | 0 | 47 | 47 | 16 | 376 | 0 | 0 | 868 | 1318 | 0 | 27 | 2271 | 437 | 0 | 16 | 33 | 33 | 33 | 33 | 82 | 82 | 148 | 189 | 0 |
| SRR7412308 | 0 | 86 | 38 | 0 | 55 | 0 | 0 | 548 | 37 | 146 | 42232 | 35151 | 0 | 37 | 256 | 37 | 0 | 242 | 0 | 146 | 923 | 1165 | 0 | 18 | 2974 | 390 | 0 | 0 | 57 | 0 | 0 | 0 | 19 | 162 | 220 | 730 | 0 |
| SRR7412309 | 0 | 83 | 0 | 0 | 40 | 0 | 0 | 634 | 167 | 200 | 1079 | 20619 | 0 | 0 | 33 | 0 | 0 | 334 | 11 | 200 | 625 | 1172 | 0 | 11 | 1512 | 72 | 0 | 17 | 75 | 17 | 0 | 0 | 75 | 14 | 26 | 467 | 0 |
| SRR7412310 | 0 | 57 | 23 | 0 | 35 | 0 | 0 | 444 | 309 | 135 | 1562 | 20881 | 0 | 0 | 39 | 96 | 0 | 538 | 0 | 116 | 1225 | 1634 | 12 | 13 | 3472 | 178 | 0 | 0 | 69 | 0 | 11 | 0 | 31 | 57 | 191 | 521 | 0 |
| SRR7412311 | 0 | 123 | 25 | 0 | 48 | 0 | 0 | 495 | 152 | 152 | 28702 | 18500 | 0 | 0 | 0 | 76 | 0 | 341 | 0 | 0 | 901 | 1387 | 0 | 10 | 2228 | 124 | 0 | 0 | 58 | 0 | 16 | 0 | 25 | 132 | 99 | 381 | 0 |
| SRR7412312 | 0 | 102 | 25 | 0 | 31 | 0 | 0 | 260 | 166 | 189 | 38020 | 26883 | 0 | 0 | 47 | 142 | 17 | 292 | 0 | 71 | 753 | 1478 | 0 | 18 | 3053 | 246 | 0 | 0 | 25 | 0 | 17 | 0 | 34 | 153 | 68 | 166 | 0 |
| SRR7412313 | 0 | 57 | 0 | 0 | 21 | 0 | 0 | 99 | 99 | 493 | 27972 | 27742 | 0 | 33 | 164 | 131 | 0 | 198 | 0 | 197 | 529 | 767 | 0 | 66 | 2960 | 870 | 0 | 29 | 0 | 0 | 0 | 0 | 0 | 105 | 38 | 559 | 0 |
| SRR7412314 | 0 | 0 | 22 | 0 | 11 | 0 | 0 | 5313 | 324 | 41 | 576 | 21617 | 0 | 0 | 203 | 81 | 0 | 297 | 11 | 446 | 575 | 918 | 0 | 33 | 1813 | 1028 | 0 | 0 | 15 | 0 | 15 | 0 | 0 | 22 | 15 | 1176 | 0 |
| SRR7412315 | 0 | 59 | 35 | 0 | 18 | 0 | 0 | 184 | 147 | 74 | 43426 | 36586 | 0 | 37 | 221 | 331 | 0 | 385 | 0 | 221 | 746 | 992 | 0 | 16 | 1035 | 773 | 0 | 24 | 118 | 24 | 12 | 12 | 47 | 141 | 129 | 368 | 0 |
| SRR7412316 | 0 | 11 | 36 | 0 | 39 | 0 | 0 | 186 | 27 | 213 | 28767 | 21277 | 0 | 53 | 0 | 53 | 0 | 225 | 0 | 292 | 1001 | 1338 | 0 | 94 | 2736 | 431 | 0 | 27 | 82 | 100 | 18 | 27 | 36 | 27 | 41 | 1089 | 0 |
| SRR7412317 | 0 | 0 | 12 | 0 | 23 | 0 | 0 | 1256 | 64 | 290 | 516 | 13915 | 0 | 0 | 0 | 32 | 0 | 221 | 0 | 225 | 624 | 920 | 0 | 17 | 1466 | 446 | 0 | 35 | 35 | 0 | 12 | 0 | 46 | 58 | 98 | 1127 | 0 |
| SRR7412318 | 0 | 13 | 13 | 0 | 14 | 0 | 0 | 606 | 0 | 76 | 257 | 21798 | 0 | 0 | 0 | 0 | 13 | 168 | 11 | 151 | 505 | 927 | 0 | 48 | 2035 | 515 | 0 | 0 | 64 | 13 | 0 | 0 | 38 | 64 | 178 | 1060 | 0 |
| SRR7412319 | 0 | 21 | 36 | 0 | 19 | 0 | 0 | 1271 | 484 | 0 | 498 | 15581 | 0 | 30 | 61 | 0 | 0 | 266 | 17 | 151 | 547 | 1042 | 0 | 26 | 1646 | 271 | 0 | 21 | 85 | 43 | 71 | 14 | 36 | 16 | 12 | 1331 | 0 |
| SRR7412320 | 0 | 271 | 30 | 0 | 49 | 0 | 0 | 134 | 59 | 0 | 46531 | 34594 | 0 | 0 | 0 | 17 | 0 | 445 | 0 | 92 | 1377 | 1577 | 0 | 31 | 1891 | 1183 | 0 | 20 | 196 | 55 | 60 | 0 | 101 | 221 | 332 | 411 | 0 |
| SRR7412321 | 0 | 85 | 38 | 0 | 24 | 0 | 0 | 251 | 68 | 91 | 1345 | 29659 | 0 | 0 | 0 | 68 | 0 | 302 | 0 | 228 | 528 | 812 | 0 | 80 | 3242 | 778 | 0 | 0 | 133 | 57 | 19 | 28 | 114 | 180 | 142 | 319 | 0 |
| SRR7412322 | 0 | 40 | 17 | 0 | 37 | 0 | 0 | 325 | 122 | 101 | 924 | 19167 | 0 | 41 | 41 | 122 | 0 | 271 | 0 | 467 | 620 | 1079 | 0 | 0 | 2041 | 204 | 0 | 11 | 46 | 57 | 29 | 0 | 63 | 86 | 149 | 2434 | 0 |
| SRR7412323 | 0 | 156 | 41 | 0 | 23 | 0 | 0 | 10 | 72 | 51 | 2215 | 13973 | 0 | 10 | 20 | 82 | 0 | 375 | 12 | 184 | 696 | 1242 | 0 | 92 | 1891 | 1352 | 0 | 0 | 13 | 82 | 49 | 16 | 82 | 15 | 26 | 522 | 0 |
| SRR7412324 | 0 | 36 | 27 | 0 | 31 | 0 | 0 | 0 | 161 | 80 | 1568 | 21066 | 0 | 0 | 0 | 20 | 0 | 282 | 0 | 181 | 555 | 1015 | 0 | 35 | 1987 | 417 | 0 | 45 | 91 | 18 | 36 | 0 | 55 | 155 | 273 | 2127 | 0 |
| SRR7412325 | 0 | 65 | 76 | 0 | 47 | 0 | 0 | 871 | 31 | 124 | 1125 | 30837 | 0 | 0 | 31 | 31 | 0 | 378 | 0 | 62 | 812 | 1310 | 0 | 36 | 3024 | 232 | 0 | 43 | 76 | 11 | 11 | 0 | 22 | 54 | 379 | 373 | 0 |
| SRR7412326 | 0 | 105 | 0 | 0 | 46 | 0 | 0 | 420 | 133 | 144 | 2951 | 25190 | 0 | 0 | 22 | 88 | 0 | 328 | 0 | 121 | 708 | 1195 | 0 | 24 | 3142 | 506 | 0 | 12 | 60 | 12 | 36 | 16 | 64 | 113 | 10 | 342 | 0 |
| SRR7412327 | 0 | 92 | 39 | 0 | 37 | 0 | 0 | 326 | 201 | 125 | 45622 | 29185 | 0 | 25 | 0 | 0 | 0 | 245 | 11 | 75 | 670 | 1155 | 0 | 16 | 2534 | 267 | 0 | 26 | 33 | 26 | 20 | 13 | 66 | 118 | 354 | 226 | 0 |

### Supp Table S8.4.Columns_DI_to_ES

| | | | | | | | | | | | | | | | | | | | | | | | | | | | | | | | | | | | | | |
|---|---|---|---|---|---|---|---|---|---|---|---|---|---|---|---|---|---|---|---|---|---|---|---|---|---|---|---|---|---|---|---|---|---|---|---|---|---|
| SRR7412328 | 0 | 92 | 17 | 0 | 29 | 0 | 0 | 288 | 183 | 288 | 948 | 18047 | 0 | 0 | 0 | 52 | 0 | 276 | 0 | 655 | 447 | 619 | 0 | 51 | 2345 | 205 | 0 | 0 | 144 | 63 | 52 | 29 | 69 | 115 | 150 | 1703 | 13 |
| SRR7412329 | 0 | 16 | 28 | 0 | 66 | 0 | 0 | 345 | 0 | 129 | 18421 | 14193 | 0 | 43 | 43 | 43 | 0 | 331 | 0 | 820 | 1241 | 1144 | 0 | 140 | 3742 | 288 | 0 | 0 | 13 | 12 | 0 | 0 | 0 | 56 | 16 | 1251 | 0 |
| SRR7412330 | 0 | 80 | 72 | 0 | 51 | 0 | 0 | 0 | 0 | 250 | 355 | 19181 | 0 | 167 | 0 | 0 | 0 | 207 | 0 | 917 | 1261 | 1358 | 0 | 0 | 232 | 177 | 0 | 24 | 16 | 16 | 24 | 0 | 24 | 16 | 112 | 4420 | 0 |
| SRR7412331 | 0 | 80 | 32 | 0 | 76 | 0 | 0 | 422 | 148 | 358 | 1232 | 19962 | 0 | 0 | 21 | 42 | 0 | 221 | 0 | 169 | 1006 | 1305 | 0 | 92 | 2372 | 956 | 0 | 40 | 72 | 80 | 56 | 0 | 56 | 183 | 271 | 780 | 0 |
| SRR7412332 | 0 | 85 | 57 | 0 | 32 | 0 | 0 | 619 | 103 | 138 | 570 | 15415 | 0 | 0 | 69 | 0 | 0 | 211 | 0 | 172 | 767 | 806 | 0 | 17 | 2633 | 223 | 0 | 38 | 66 | 38 | 47 | 0 | 38 | 123 | 170 | 1101 | 0 |
| SRR7412333 | 0 | 19 | 0 | 0 | 35 | 0 | 0 | 576 | 175 | 175 | 21073 | 16293 | 0 | 0 | 50 | 100 | 0 | 317 | 0 | 425 | 831 | 1122 | 0 | 40 | 2908 | 369 | 0 | 15 | 34 | 19 | 0 | 0 | 24 | 93 | 83 | 2252 | 0 |
| SRR7412334 | 0 | 56 | 0 | 0 | 33 | 0 | 0 | 407 | 262 | 145 | 596 | 25276 | 0 | 29 | 87 | 87 | 0 | 228 | 0 | 349 | 507 | 874 | 0 | 24 | 1488 | 214 | 0 | 21 | 35 | 35 | 0 | 0 | 28 | 84 | 70 | 2821 | 0 |
| SRR7412335 | 0 | 0 | 60 | 0 | 12 | 0 | 0 | 842 | 120 | 241 | 999 | 28262 | 0 | 40 | 0 | 0 | 0 | 213 | 0 | 120 | 679 | 1030 | 0 | 66 | 1920 | 351 | 0 | 15 | 45 | 30 | 30 | 15 | 60 | 75 | 28 | 722 | 0 |
| SRR7412336 | 0 | 56 | 28 | 0 | 20 | 0 | 0 | 153 | 382 | 356 | 825 | 19941 | 0 | 0 | 153 | 153 | 0 | 326 | 13 | 483 | 527 | 972 | 0 | 88 | 3740 | 961 | 0 | 38 | 113 | 38 | 28 | 0 | 28 | 131 | 178 | 661 | 0 |
| SRR7412337 | 0 | 69 | 49 | 0 | 42 | 0 | 0 | 129 | 32 | 129 | 609 | 20357 | 0 | 0 | 194 | 453 | 0 | 762 | 0 | 97 | 1603 | 1644 | 0 | 14 | 781 | 613 | 0 | 0 | 69 | 109 | 40 | 0 | 0 | 79 | 257 | 1974 | 0 |
| SRR7412338 | 0 | 60 | 30 | 0 | 32 | 0 | 0 | 756 | 126 | 126 | 25261 | 14993 | 0 | 0 | 0 | 0 | 0 | 245 | 0 | 0 | 1638 | 2023 | 0 | 35 | 1845 | 393 | 0 | 60 | 60 | 105 | 150 | 15 | 90 | 271 | 421 | 630 | 0 |
| SRR7412339 | 0 | 81 | 67 | 0 | 34 | 0 | 0 | 101 | 135 | 34 | 947 | 23429 | 0 | 0 | 270 | 270 | 13 | 583 | 14 | 270 | 678 | 963 | 0 | 117 | 3561 | 1045 | 0 | 27 | 94 | 81 | 27 | 13 | 27 | 135 | 175 | 710 | 0 |
| SRR7412340 | 0 | 24 | 0 | 0 | 19 | 0 | 0 | 113 | 113 | 113 | 20919 | 17584 | 0 | 0 | 0 | 0 | 0 | 327 | 32 | 622 | 401 | 641 | 0 | 25 | 262 | 970 | 0 | 24 | 36 | 24 | 0 | 0 | 72 | 72 | 120 | 3675 | 0 |
| SRR7412341 | 0 | 0 | 37 | 0 | 17 | 0 | 0 | 55 | 110 | 166 | 18383 | 15954 | 0 | 0 | 0 | 55 | 0 | 337 | 16 | 1435 | 366 | 565 | 0 | 0 | 203 | 545 | 0 | 37 | 84 | 66 | 103 | 19 | 28 | 56 | 94 | 2650 | 0 |
| SRR7412342 | 0 | 0 | 0 | 0 | 11 | 0 | 0 | 488 | 0 | 139 | 276 | 14977 | 0 | 70 | 70 | 139 | 0 | 318 | 0 | 139 | 845 | 1051 | 0 | 26 | 2533 | 883 | 0 | 15 | 15 | 0 | 0 | 0 | 0 | 30 | 45 | 836 | 0 |
| SRR7412343 | 0 | 25 | 0 | 0 | 20 | 0 | 0 | 500 | 600 | 300 | 21906 | 18105 | 0 | 0 | 0 | 0 | 0 | 148 | 13 | 550 | 466 | 1111 | 0 | 22 | 2405 | 1005 | 0 | 33 | 58 | 25 | 0 | 0 | 17 | 75 | 100 | 2501 | 0 |
| SRR7412344 | 0 | 0 | 45 | 0 | 29 | 0 | 0 | 0 | 235 | 202 | 512 | 10146 | 0 | 34 | 0 | 67 | 0 | 293 | 10 | 504 | 504 | 699 | 0 | 25 | 1788 | 584 | 15 | 22 | 19 | 12 | 30 | 0 | 22 | 18 | 14 | 1545 | 0 |
| SRR7412345 | 0 | 78 | 20 | 0 | 44 | 0 | 0 | 468 | 156 | 234 | 32917 | 24493 | 0 | 0 | 0 | 156 | 0 | 417 | 21 | 78 | 666 | 1232 | 0 | 12 | 1256 | 766 | 0 | 0 | 78 | 78 | 39 | 20 | 20 | 21 | 13 | 468 | 0 |
| SRR7466734 | 0 | 43 | 14 | 0 | 0 | 0 | 0 | 609 | 171 | 5118 | 94 | 2226 | 0 | 4795 | 0 | 0 | 0 | 69 | 68 | 1503 | 3815 | 7804 | 16 | 7630 | 6894 | 3339 | 0 | 119 | 141 | 223 | 161 | 36 | 146 | 264 | 59 | 3767 | 27 |
| SRR7466735 | 0 | 60 | 17 | 0 | 0 | 0 | 0 | 547 | 98 | 5722 | 114 | 1797 | 0 | 5703 | 20 | 0 | 0 | 57 | 48 | 1601 | 3199 | 6975 | 16 | 7732 | 6427 | 3275 | 0 | 132 | 137 | 221 | 160 | 34 | 136 | 277 | 60 | 3476 | 27 |
| SRR7466736 | 0 | 26 | 42 | 0 | 0 | 0 | 0 | 290 | 375 | 8192 | 1792 | 2133 | 0 | 10718 | 17 | 0 | 0 | 69 | 44 | 1331 | 2777 | 5608 | 19 | 10093 | 13 | 3558 | 0 | 891 | 735 | 1074 | 440 | 100 | 668 | 1113 | 43 | 5683 | 34 |
| SRR7466737 | 0 | 36 | 31 | 0 | 0 | 0 | 0 | 379 | 224 | 7776 | 1776 | 1897 | 0 | 11069 | 0 | 0 | 0 | 44 | 46 | 1517 | 2276 | 5017 | 24 | 10307 | 9013 | 3489 | 0 | 953 | 719 | 989 | 528 | 129 | 636 | 1144 | 33 | 6862 | 47 |
| SRR7466738 | 0 | 30 | 12 | 0 | 0 | 0 | 0 | 295 | 263 | 8006 | 2215 | 2379 | 0 | 10664 | 0 | 0 | 0 | 60 | 57 | 1116 | 2740 | 5767 | 15 | 9967 | 17 | 3463 | 0 | 295 | 217 | 314 | 142 | 32 | 196 | 336 | 27 | 5873 | 46 |
| SRR7466739 | 0 | 21 | 24 | 0 | 0 | 13 | 0 | 437 | 269 | 7880 | 1966 | 2134 | 0 | 10904 | 0 | 17 | 0 | 58 | 42 | 1714 | 2331 | 5021 | 24 | 10217 | 11 | 3608 | 0 | 968 | 698 | 968 | 464 | 118 | 573 | 1181 | 31 | 5763 | 29 |
| SRR7466740 | 0 | 23 | 41 | 0 | 12 | 0 | 0 | 626 | 383 | 10717 | 1837 | 1938 | 0 | 9224 | 0 | 0 | 0 | 54 | 63 | 1917 | 3262 | 6496 | 26 | 8160 | 15 | 2275 | 0 | 212 | 318 | 893 | 561 | 71 | 483 | 1162 | 24 | 6479 | 32 |
| SRR7466741 | 0 | 21 | 31 | 0 | 0 | 0 | 0 | 554 | 349 | 10902 | 1786 | 2012 | 0 | 8911 | 0 | 21 | 0 | 52 | 42 | 1766 | 2668 | 5714 | 19 | 8087 | 8327 | 2296 | 0 | 215 | 275 | 833 | 598 | 73 | 500 | 1252 | 33 | 6960 | 37 |
| SRR7466742 | 0 | 27 | 12 | 0 | 0 | 0 | 0 | 410 | 181 | 8814 | 102 | 1723 | 0 | 7780 | 0 | 16 | 0 | 48 | 50 | 1789 | 3326 | 6509 | 39 | 8024 | 8952 | 2312 | 0 | 44 | 89 | 223 | 141 | 22 | 116 | 315 | 22 | 4989 | 47 |
| SRR7466743 | 0 | 24 | 27 | 0 | 0 | 0 | 0 | 504 | 343 | 11346 | 1773 | 2297 | 0 | 9250 | 0 | 20 | 0 | 36 | 48 | 2056 | 2801 | 5692 | 31 | 8198 | 8533 | 2301 | 0 | 184 | 318 | 957 | 615 | 80 | 495 | 1136 | 37 | 7355 | 27 |
| SRR7466744 | 0 | 24 | 32 | 0 | 0 | 0 | 0 | 200 | 109 | 8855 | 98 | 2255 | 0 | 7455 | 18 | 0 | 0 | 43 | 43 | 1618 | 2553 | 5424 | 24 | 4292 | 3412 | 2517 | 0 | 72 | 86 | 215 | 176 | 28 | 184 | 392 | 42 | 5237 | 38 |
| SRR7466745 | 0 | 42 | 47 | 0 | 0 | 0 | 0 | 418 | 286 | 10774 | 2023 | 2089 | 0 | 9366 | 0 | 0 | 0 | 50 | 49 | 1605 | 3120 | 6204 | 13 | 4214 | 3565 | 2407 | 0 | 361 | 416 | 1205 | 1033 | 191 | 777 | 2183 | 44 | 6442 | 17 |
| SRR7466746 | 0 | 48 | 90 | 0 | 0 | 0 | 0 | 347 | 154 | 5642 | 116 | 2561 | 0 | 5546 | 0 | 0 | 0 | 73 | 57 | 2311 | 3094 | 6197 | 18 | 6127 | 5239 | 2291 | 0 | 354 | 568 | 1520 | 1093 | 195 | 973 | 2172 | 42 | 6085 | 26 |
| SRR7466747 | 0 | 54 | 71 | 0 | 0 | 0 | 0 | 496 | 153 | 5935 | 101 | 2233 | 0 | 5592 | 19 | 19 | 0 | 76 | 65 | 2271 | 3716 | 10 | 14 | 6085 | 13 | 2333 | 0 | 336 | 631 | 1503 | 1125 | 137 | 1024 | 2030 | 30 | 5553 | 30 |
| SRR7466748 | 0 | 62 | 138 | 0 | 0 | 0 | 0 | 370 | 87 | 6515 | 2419 | 3029 | 0 | 6864 | 0 | 22 | 0 | 61 | 77 | 2179 | 3183 | 6509 | 23 | 6365 | 5374 | 2409 | 0 | 464 | 614 | 1502 | 1087 | 188 | 1087 | 2113 | 59 | 6472 | 26 |
| SRR7466749 | 0 | 54 | 14 | 0 | 0 | 0 | 0 | 472 | 193 | 7395 | 2508 | 3001 | 0 | 7245 | 0 | 21 | 0 | 68 | 72 | 2551 | 3697 | 7445 | 13 | 6063 | 5724 | 2297 | 0 | 50 | 83 | 197 | 147 | 29 | 136 | 298 | 57 | 6816 | 20 |
| SRR7466750 | 0 | 20 | 24 | 0 | 0 | 0 | 0 | 477 | 204 | 8040 | 71 | 1522 | 0 | 8290 | 0 | 23 | 0 | 49 | 40 | 1862 | 2599 | 6351 | 14 | 6705 | 5648 | 2277 | 0 | 157 | 234 | 724 | 449 | 59 | 478 | 1180 | 29 | 8835 | 25 |
| SRR7466751 | 0 | 27 | 30 | 0 | 0 | 0 | 0 | 579 | 132 | 9978 | 1685 | 2132 | 0 | 9162 | 0 | 0 | 0 | 32 | 60 | 2238 | 3240 | 7404 | 0 | 6521 | 5862 | 2269 | 0 | 56 | 67 | 199 | 150 | 21 | 124 | 300 | 12 | 10267 | 31 |
| SRR7466752 | 0 | 15 | 12 | 0 | 0 | 0 | 0 | 628 | 471 | 9133 | 2146 | 2826 | 0 | 10258 | 0 | 0 | 0 | 34 | 40 | 2905 | 2764 | 6385 | 15 | 6679 | 5949 | 2284 | 0 | 60 | 80 | 206 | 131 | 17 | 129 | 326 | 20 | 11305 | 34 |
| SRR7466753 | 0 | 11 | 29 | 0 | 0 | 0 | 0 | 432 | 305 | 9733 | 2059 | 1855 | 0 | 9683 | 25 | 25 | 0 | 41 | 60 | 2516 | 3216 | 7275 | 20 | 6515 | 5977 | 2285 | 0 | 63 | 66 | 216 | 144 | 20 | 126 | 328 | 34 | 11538 | 29 |
| SRR7466754 | 0 | 37 | 49 | 0 | 0 | 0 | 0 | 549 | 264 | 8984 | 1582 | 1582 | 0 | 12060 | 22 | 0 | 0 | 33 | 61 | 1933 | 3262 | 7218 | 18 | 7414 | 13 | 3451 | 0 | 293 | 242 | 347 | 191 | 34 | 217 | 422 | 12 | 4986 | 33 |
| SRR7466755 | 0 | 37 | 12 | 0 | 0 | 0 | 0 | 624 | 223 | 10790 | 1605 | 1917 | 0 | 11459 | 22 | 0 | 0 | 36 | 32 | 1271 | 2680 | 6186 | 15 | 7677 | 6281 | 3561 | 0 | 279 | 243 | 353 | 188 | 39 | 236 | 419 | 50 | 5283 | 40 |
| SRR7466756 | 0 | 33 | 71 | 0 | 0 | 0 | 0 | 365 | 116 | 7692 | 54 | 1479 | 0 | 10051 | 17 | 0 | 0 | 44 | 53 | 1096 | 3266 | 6966 | 19 | 7825 | 12 | 3566 | 0 | 1623 | 1323 | 1934 | 1017 | 252 | 1276 | 2450 | 92 | 3572 | 34 |
| SRR7466757 | 0 | 30 | 55 | 0 | 0 | 0 | 0 | 377 | 205 | 7873 | 78 | 1369 | 0 | 9345 | 0 | 17 | 0 | 47 | 50 | 1027 | 2822 | 6205 | 19 | 7758 | 13 | 3535 | 0 | 1747 | 1381 | 1915 | 1110 | 247 | 1323 | 2405 | 58 | 4245 | 19 |
| SRR7466758 | 0 | 13 | 19 | 0 | 0 | 0 | 0 | 344 | 57 | 8897 | 43 | 1579 | 0 | 10361 | 0 | 0 | 0 | 33 | 60 | 1263 | 3105 | 7412 | 10 | 6355 | 5812 | 3823 | 0 | 212 | 205 | 326 | 162 | 35 | 200 | 419 | 67 | 3272 | 19 |
| SRR7466759 | 0 | 54 | 16 | 0 | 0 | 0 | 0 | 630 | 148 | 11012 | 1483 | 2484 | 0 | 10753 | 0 | 0 | 0 | 34 | 42 | 1187 | 2771 | 6616 | 25 | 6354 | 12 | 3798 | 0 | 216 | 182 | 311 | 179 | 40 | 213 | 424 | 67 | 5302 | 11 |
| SRR7466760 | 0 | 19 | 103 | 0 | 0 | 0 | 0 | 581 | 83 | 8662 | 38 | 1882 | 0 | 9603 | 0 | 0 | 0 | 26 | 40 | 1245 | 3330 | 7431 | 28 | 6431 | 6024 | 3745 | 0 | 1613 | 1465 | 2493 | 1291 | 270 | 1664 | 3007 | 71 | 4040 | 16 |
| SRR7466761 | 0 | 105 | 138 | 0 | 0 | 0 | 0 | 544 | 115 | 9080 | 44 | 1690 | 0 | 9911 | 29 | 0 | 0 | 33 | 41 | 1232 | 2733 | 6706 | 19 | 6259 | 17 | 3949 | 0 | 1578 | 1532 | 2163 | 1216 | 302 | 1663 | 3215 | 72 | 3695 | 16 |
| SRR7466762 | 0 | 74 | 13 | 0 | 0 | 0 | 0 | 470 | 20 | 5169 | 101 | 2056 | 0 | 5169 | 0 | 20 | 0 | 77 | 66 | 1704 | 3691 | 7803 | 20 | 7688 | 6871 | 3320 | 0 | 119 | 141 | 223 | 147 | 28 | 127 | 247 | 69 | 3799 | 33 |
| SRR7466763 | 0 | 58 | 11 | 0 | 0 | 0 | 0 | 747 | 289 | 6895 | 1832 | 2580 | 0 | 6027 | 0 | 0 | 0 | 58 | 52 | 1519 | 3075 | 11 | 21 | 7673 | 15 | 3222 | 0 | 116 | 137 | 237 | 151 | 34 | 129 | 270 | 66 | 4677 | 31 |
| SRR7466764 | 0 | 10 | 10 | 0 | 0 | 0 | 0 | 512 | 235 | 10542 | 1985 | 2390 | 0 | 9944 | 0 | 21 | 0 | 37 | 64 | 1750 | 3146 | 6414 | 17 | 4346 | 3717 | 2557 | 0 | 58 | 74 | 222 | 163 | 25 | 171 | 408 | 48 | 6189 | 23 |
| SRR7466765 | 0 | 33 | 46 | 0 | 0 | 0 | 0 | 477 | 238 | 10723 | 2318 | 2188 | 0 | 10203 | 22 | 0 | 0 | 47 | 61 | 1885 | 2687 | 5658 | 0 | 4312 | 3495 | 2591 | 0 | 61 | 65 | 209 | 213 | 27 | 170 | 435 | 36 | 6152 | 30 |
| SRR7541164 | 0 | 28 | 13 | 0 | 0 | 0 | 0 | 435 | 79 | 435 | 5080 | 9294 | 0 | 80 | 218 | 1278 | 0 | 81 | 19 | 763 | 620 | 2581 | 0 | 80 | 3262 | 0 | 0 | 0 | 17 | 211 | 140 | 0 | 14 | 23 | 0 | 1090 | 15 |
| SRR7541165 | 0 | 0 | 0 | 0 | 0 | 0 | 0 | 219 | 61 | 838 | 2610 | 16326 | 0 | 2997 | 12 | 33 | 0 | 173 | 21 | 1087 | 225 | 613 | 0 | 88 | 1625 | 13 | 0 | 0 | 0 | 17 | 12 | 0 | 0 | 13 | 0 | 3518 | 0 |
| SRR7541166 | 0 | 17 | 0 | 0 | 10 | 0 | 0 | 1090 | 301 | 362 | 814 | 13311 | 0 | 342 | 51 | 146 | 0 | 173 | 0 | 2612 | 1008 | 890 | 23 | 578 | 6795 | 0 | 0 | 0 | 23 | 448 | 234 | 0 | 28 | 25 | 0 | 5245 | 45 |
| SRR7541167 | 0 | 32 | 0 | 0 | 0 | 0 | 0 | 620 | 104 | 396 | 1912 | 10172 | 0 | 297 | 0 | 32 | 0 | 146 | 0 | 1451 | 1367 | 1611 | 19 | 537 | 8801 | 0 | 0 | 0 | 0 | 38 | 21 | 0 | 0 | 0 | 0 | 3402 | 13 |
| SRR7541168 | 0 | 20 | 20 | 0 | 0 | 0 | 0 | 206 | 67 | 379 | 5923 | 15813 | 0 | 872 | 444 | 306 | 0 | 121 | 39 | 899 | 1080 | 1282 | 20 | 868 | 5083 | 0 | 14 | 81 | 192 | 678 | 511 | 59 | 158 | 250 | 12 | 2939 | 17 |
| SRR7541169 | 0 | 30 | 14 | 0 | 0 | 0 | 0 | 288 | 39 | 409 | 7184 | 29417 | 0 | 144 | 112 | 166 | 15 | 122 | 0 | 669 | 1283 | 608 | 38 | 541 | 3102 | 0 | 22 | 0 | 19 | 85 | 31 | 0 | 40 | 11 | 0 | 3662 | 14 |
| SRR7541170 | 0 | 15 | 0 | 0 | 15 | 0 | 0 | 470 | 160 | 525 | 5877 | 19685 | 0 | 374 | 17 | 46 | 0 | 248 | 0 | 1356 | 969 | 2093 | 101 | 1629 | 11005 | 0 | 13 | 0 | 19 | 38 | 17 | 0 | 0 | 17 | 0 | 4584 | 47 |
| SRR7541171 | 0 | 32 | 32 | 0 | 0 | 0 | 0 | 209 | 127 | 1058 | 11309 | 34447 | 0 | 127 | 46 | 109 | 0 | 217 | 0 | 590 | 1304 | 2453 | 637 | 4717 | 8553 | 0 | 0 | 20 | 64 | 51 | 44 | 20 | 61 | 60 | 0 | 957 | 0 |
| SRR7541172 | 0 | 13 | 13 | 0 | 0 | 0 | 0 | 702 | 169 | 1149 | 13344 | 31964 | 0 | 145 | 67 | 43 | 0 | 130 | 0 | 612 | 580 | 851 | 57 | 1938 | 5136 | 19 | 0 | 0 | 59 | 68 | 41 | 0 | 18 | 25 | 0 | 1396 | 0 |
| SRR7541173 | 0 | 22 | 44 | 0 | 72 | 0 | 0 | 258 | 61 | 346 | 2821 | 17428 | 0 | 442 | 51 | 45 | 0 | 690 | 13 | 1220 | 1462 | 2589 | 42 | 858 | 13660 | 0 | 0 | 45 | 121 | 579 | 201 | 20 | 29 | 41 | 0 | 2995 | 20 |
| SRR7541174 | 0 | 22 | 0 | 0 | 0 | 0 | 0 | 423 | 75 | 694 | 7171 | 28091 | 0 | 109 | 118 | 526 | 0 | 518 | 37 | 1412 | 86 | 3780 | 88 | 2496 | 18108 | 0 | 0 | 0 | 25 | 47 | 29 | 0 | 23 | 44 | 0 | 2478 | 75 |
| SRR7541175 | 0 | 42 | 16 | 0 | 0 | 0 | 0 | 976 | 192 | 2046 | 15026 | 24863 | 0 | 137 | 17 | 25 | 0 | 184 | 0 | 555 | 658 | 2564 | 274 | 5631 | 11529 | 0 | 0 | 0 | 14 | 51 | 20 | 0 | 25 | 31 | 0 | 856 | 0 |
| SRR7541176 | 0 | 0 | 0 | 0 | 0 | 0 | 0 | 354 | 73 | 150 | 1526 | 28472 | 0 | 136 | 29 | 131 | 0 | 170 | 13 | 917 | 1600 | 2192 | 375 | 3056 | 5191 | 0 | 0 | 14 | 13 | 67 | 44 | 20 | 92 | 109 | 0 | 2061 | 35 |
| SRR7541177 | 0 | 30 | 43 | 0 | 0 | 0 | 0 | 385 | 84 | 1302 | 16680 | 27119 | 0 | 56 | 61 | 105 | 0 | 80 | 11 | 720 | 369 | 1205 | 182 | 4798 | 7398 | 0 | 16 | 23 | 560 | 2050 | 822 | 24 | 95 | 126 | 0 | 1011 | 29 |
| SRR7541178 | 0 | 59 | 67 | 0 | 26 | 0 | 0 | 609 | 136 | 1127 | 6436 | 25061 | 0 | 280 | 0 | 10 | 0 | 298 | 0 | 0 | 431 | 2420 | 23 | 379 | 8140 | 0 | 0 | 19 | 15 | 54 | 19 | 0 | 29 | 46 | 0 | 0 | 0 |
| SRR7541179 | 0 | 34 | 0 | 0 | 22 | 0 | 0 | 283 | 61 | 438 | 2127 | 14473 | 0 | 39 | 11 | 35 | 0 | 148 | 0 | 1021 | 341 | 1393 | 52 | 916 | 9187 | 0 | 0 | 20 | 44 | 173 | 85 | 0 | 0 | 14 | 0 | 1586 | 0 |
| SRR7541180 | 0 | 37 | 59 | 0 | 0 | 0 | 0 | 619 | 255 | 1499 | 2888 | 2574 | 0 | 1782 | 0 | 0 | 14 | 149 | 11 | 421 | 357 | 719 | 147 | 2996 | 2637 | 0 | 0 | 33 | 41 | 148 | 49 | 42 | 15 | 24 | 0 | 715 | 14 |
| SRR7541181 | 10 | 14 | 19 | 0 | 0 | 0 | 0 | 499 | 125 | 328 | 21054 | 28277 | 0 | 223 | 20 | 72 | 0 | 164 | 0 | 722 | 121 | 851 | 196 | 1511 | 3027 | 0 | 0 | 142 | 437 | 1256 | 292 | 27 | 50 | 72 | 21 | 3953 | 0 |
| SRR7541182 | 0 | 46 | 20 | 0 | 0 | 0 | 0 | 759 | 230 | 810 | 5928 | 12730 | 0 | 272 | 0 | 0 | 0 | 119 | 0 | 503 | 1459 | 2050 | 139 | 2644 | 7765 | 0 | 0 | 34 | 479 | 1840 | 917 | 27 | 93 | 127 | 0 | 851 | 0 |
| SRR7541183 | 0 | 54 | 18 | 0 | 0 | 0 | 0 | 511 | 77 | 406 | 1809 | 26874 | 0 | 272 | 0 | 10 | 0 | 183 | 10 | 564 | 680 | 995 | 11 | 330 | 10384 | 0 | 0 | 19 | 69 | 478 | 278 | 0 | 13 | 26 | 0 | 813 | 39 |
| SRR7541184 | 0 | 36 | 28 | 0 | 21 | 0 | 0 | 350 | 144 | 725 | 2595 | 12755 | 0 | 587 | 0 | 26 | 0 | 189 | 0 | 278 | 1381 | 1403 | 50 | 1072 | 8278 | 0 | 0 | 17 | 118 | 95 | 50 | 13 | 29 | 33 | 0 | 452 | 0 |
| SRR7541185 | 0 | 156 | 132 | 0 | 0 | 0 | 0 | 305 | 237 | 421 | 7694 | 16313 | 0 | 1022 | 14 | 10 | 0 | 648 | 0 | 234 | 5305 | 1597 | 0 | 78 | 5687 | 0 | 0 | 32 | 42 | 101 | 37 | 0 | 22 | 57 | 0 | 244 | 11 |
| SRR7541186 | 0 | 20 | 16 | 0 | 0 | 0 | 0 | 331 | 58 | 380 | 14641 | 26958 | 0 | 0 | 0 | 0 | 0 | 144 | 0 | 306 | 557 | 1162 | 0 | 164 | 8990 | 0 | 0 | 0 | 12 | 108 | 60 | 0 | 22 | 28 | 0 | 676 | 37 |
| SRR7541187 | 0 | 0 | 0 | 0 | 13 | 0 | 0 | 247 | 62 | 836 | 2266 | 21729 | 0 | 39 | 29 | 0 | 12 | 153 | 0 | 314 | 300 | 2189 | 0 | 326 | 15066 | 0 | 24 | 0 | 13 | 36 | 17 | 0 | 0 | 0 | 0 | 623 | 17 |
| SRR7541188 | 0 | 14 | 0 | 0 | 17 | 0 | 0 | 550 | 183 | 1291 | 2631 | 21185 | 0 | 79 | 0 | 0 | 0 | 133 | 0 | 354 | 108 | 2730 | 1001 | 11580 | 11056 | 0 | 0 | 0 | 30 | 233 | 92 | 0 | 10 | 0 | 0 | 399 | 18 |
| SRR7541189 | 0 | 35 | 18 | 0 | 36 | 0 | 0 | 997 | 561 | 155 | 6365 | 18140 | 0 | 126 | 37 | 58 | 0 | 213 | 0 | 1886 | 864 | 1315 | 0 | 33 | 11888 | 0 | 0 | 125 | 779 | 1335 | 434 | 29 | 79 | 121 | 0 | 2477 | 23 |
| SRR7541190 | 0 | 23 | 25 | 0 | 15 | 0 | 0 | 344 | 93 | 747 | 17700 | 23366 | 0 | 120 | 0 | 16 | 13 | 87 | 0 | 536 | 1841 | 1240 | 0 | 0 | 4173 | 0 | 40 | 24 | 296 | 922 | 233 | 0 | 46 | 52 | 0 | 1461 | 83 |
| SRR7541191 | 0 | 52 | 17 | 0 | 0 | 0 | 0 | 380 | 259 | 742 | 1231 | 6555 | 0 | 1013 | 0 | 0 | 0 | 513 | 0 | 733 | 1876 | 2831 | 48 | 824 | 7814 | 13 | 0 | 0 | 56 | 24 | 121 | 0 | 12 | 21 | 0 | 1565 | 0 |
| SRR7541192 | 0 | 38 | 0 | 0 | 10 | 0 | 0 | 597 | 184 | 1473 | 3420 | 23440 | 0 | 157 | 10 | 0 | 0 | 179 | 0 | 460 | 1278 | 4053 | 1044 | 14239 | 20884 | 0 | 15 | 0 | 56 | 367 | 133 | 0 | 19 | 26 | 0 | 586 | 21 |
| SRR7541193 | 0 | 20 | 0 | 0 | 26 | 0 | 0 | 965 | 293 | 2424 | 7568 | 6666 | 0 | 261 | 59 | 24 | 0 | 96 | 0 | 1127 | 465 | 1666 | 80 | 795 | 7485 | 0 | 21 | 11 | 100 | 346 | 87 | 0 | 16 | 16 | 0 | 9663 | 0 |
| SRR7547853 | 0 | 404 | 24 | 528 | 2360 | 0 | 0 | 3634 | 15578 | 21415 | 399 | 558 | 0 | 572 | 366 | 1250 | 13 | 853 | 86 | 2956 | 1114 | 1645 | 156 | 7089 | 5250 | 0 | 0 | 0 | 23 | 12 | 21 | 16 | 0 | 0 | 23 | 1543 | 516 |
| SRR7547854 | 0 | 424 | 74 | 232 | 387 | 0 | 0 | 3274 | 9716 | 17329 | 508 | 3331 | 0 | 583 | 18233 | 24801 | 0 | 1100 | 50 | 2380 | 603 | 2411 | 302 | 10548 | 3747 | 0 | 0 | 0 | 34 | 18 | 24 | 22 | 0 | 0 | 12 | 1193 | 829 |
| SRR7547855 | 0 | 1011 | 265 | 101 | 371 | 0 | 13 | 3988 | 13269 | 17435 | 170 | 328 | 0 | 1125 | 395 | 1082 | 0 | 2415 | 17 | 1249 | 618 | 1377 | 116 | 3889 | 2250 | 0 | 0 | 0 | 33 | 16 | 31 | 10 | 0 | 0 | 23 | 822 | 31 |
| SRR7547856 | 0 | 1320 | 101 | 249 | 1299 | 0 | 0 | 2567 | 8884 | 18582 | 152 | 340 | 0 | 1916 | 2597 | 4041 | 10 | 2463 | 130 | 687 | 2287 | 2097 | 136 | 10514 | 4463 | 0 | 0 | 0 | 119 | 87 | 96 | 17 | 0 | 0 | 19 | 376 | 1182 |
| SRR7547857 | 0 | 1507 | 125 | 193 | 1698 | 11 | 0 | 330 | 1238 | 5522 | 66 | 117 | 0 | 2103 | 177 | 245 | 0 | 990 | 66 | 119 | 314 | 876 | 43 | 2194 | 2903 | 0 | 0 | 0 | 115 | 72 | 101 | 27 | 0 | 0 | 35 | 57 | 810 |
| SRR7547858 | 0 | 1642 | 129 | 219 | 1174 | 0 | 0 | 1743 | 7052 | 13602 | 471 | 787 | 0 | 882 | 544 | 2417 | 0 | 2135 | 43 | 1325 | 685 | 1887 | 229 | 8717 | 2906 | 0 | 0 | 0 | 48 | 45 | 82 | 0 | 0 | 0 | 31 | 563 | 414 |
| SRR7547859 | 0 | 139 | 15 | 94 | 1870 | 0 | 11 | 2997 | 15445 | 21968 | 26 | 23 | 0 | 163 | 27 | 65 | 12 | 2218 | 60 | 145 | 354 | 460 | 39 | 2842 | 2567 | 0 | 0 | 0 | 80 | 27 | 70 | 16 | 0 | 0 | 36 | 173 | 388 |
| SRR7547860 | 0 | 1620 | 150 | 32 | 787 | 0 | 13 | 2692 | 7898 | 14553 | 735 | 933 | 0 | 1648 | 853 | 1661 | 11 | 1402 | 63 | 1914 | 422 | 1818 | 207 | 5783 | 2786 | 0 | 0 | 0 | 80 | 59 | 67 | 18 | 0 | 0 | 30 | 1267 | 225 |
| SRR7547861 | 0 | 308 | 55 | 77 | 525 | 0 | 0 | 1527 | 4892 | 13802 | 454 | 1605 | 0 | 1902 | 8903 | 20283 | 12 | 1797 | 46 | 1640 | 278 | 2996 | 104 | 13694 | 5525 | 0 | 0 | 12 | 118 | 40 | 71 | 21 | 19 | 29 | 25 | 775 | 220 |
| SRR7547862 | 0 | 498 | 14 | 367 | 3462 | 0 | 0 | 2000 | 12554 | 29342 | 478 | 793 | 0 | 682 | 307 | 1010 | 17 | 1163 | 19 | 713 | 1330 | 1070 | 51 | 1131 | 2969 | 0 | 0 | 0 | 40 | 37 | 52 | 0 | 0 | 0 | 17 | 979 | 437 |
| SRR7547863 | 0 | 552 | 13 | 102 | 1107 | 0 | 0 | 1794 | 10966 | 13410 | 510 | 374 | 0 | 717 | 1054 | 1085 | 0 | 1610 | 49 | 603 | 1146 | 2402 | 241 | 10334 | 6020 | 12 | 0 | 14 | 40 | 13 | 20 | 36 | 10 | 18 | 30 | 973 | 1054 |
| SRR7547864 | 0 | 3323 | 141 | 499 | 477 | 27 | 34 | 1546 | 6358 | 13482 | 654 | 5508 | 0 | 793 | 512 | 893 | 40 | 2004 | 24 | 2071 | 1065 | 1465 | 251 | 4758 | 2055 | 0 | 19 | 12 | 79 | 37 | 44 | 0 | 0 | 0 | 0 | 718 | 350 |
| SRR7547865 | 0 | 239 | 24 | 858 | 1074 | 0 | 0 | 5393 | 23901 | 55108 | 139 | 695 | 0 | 549 | 38 | 158 | 20 | 1208 | 41 | 615 | 1345 | 1523 | 66 | 2706 | 6549 | 0 | 0 | 0 | 14 | 58 | 14 | 18 | 0 | 0 | 28 | 392 | 28 |
| SRR7547866 | 0 | 430 | 20 | 285 | 3084 | 0 | 0 | 3141 | 15895 | 28561 | 285 | 1685 | 0 | 705 | 827 | 1616 | 11 | 1682 | 24 | 1293 | 1182 | 2224 | 380 | 7653 | 2502 | 0 | 0 | 11 | 24 | 16 | 27 | 28 | 0 | 0 | 22 | 709 | 388 |
| SRR7547867 | 0 | 1868 | 84 | 144 | 1833 | 0 | 0 | 2190 | 8185 | 16450 | 268 | 617 | 0 | 1165 | 358 | 725 | 26 | 2420 | 36 | 594 | 358 | 3018 | 252 | 11266 | 3612 | 0 | 0 | 19 | 90 | 59 | 120 | 0 | 12 | 0 | 12 | 744 | 1026 |
| SRR7547868 | 0 | 1336 | 215 | 133 | 1361 | 0 | 0 | 4263 | 22244 | 12945 | 543 | 665 | 0 | 663 | 592 | 2073 | 12 | 1900 | 39 | 2591 | 1269 | 2266 | 367 | 8965 | 1628 | 0 | 0 | 0 | 38 | 36 | 57 | 20 | 0 | 0 | 40 | 1117 | 347 |
| SRR7547869 | 0 | 1302 | 101 | 267 | 1599 | 0 | 0 | 1686 | 7747 | 9809 | 277 | 565 | 0 | 865 | 4150 | 4896 | 0 | 1063 | 81 | 888 | 645 | 3132 | 476 | 25799 | 4794 | 0 | 0 | 0 | 74 | 60 | 75 | 17 | 0 | 0 | 29 | 681 | 353 |
| SRR7547870 | 0 | 229 | 26 | 162 | 1508 | 0 | 0 | 2878 | 10492 | 16551 | 276 | 382 | 0 | 845 | 779 | 518 | 14 | 1235 | 49 | 588 | 771 | 1311 | 73 | 4905 | 2988 | 0 | 0 | 0 | 21 | 12 | 20 | 16 | 0 | 0 | 20 | 383 | 909 |
| SRR7547871 | 0 | 1003 | 169 | 150 | 2021 | 0 | 0 | 1757 | 7595 | 21000 | 153 | 480 | 0 | 941 | 699 | 1201 | 0 | 1017 | 44 | 1232 | 1596 | 2127 | 365 | 11575 | 2183 | 0 | 0 | 0 | 72 | 86 | 129 | 11 | 0 | 0 | 45 | 903 | 293 |
| SRR7547872 | 0 | 232 | 14 | 110 | 2174 | 0 | 0 | 1918 | 6550 | 19304 | 280 | 221 | 0 | 865 | 129 | 132 | 12 | 1509 | 52 | 149 | 394 | 1097 | 117 | 6019 | 1905 | 0 | 0 | 0 | 59 | 54 | 67 | 0 | 0 | 0 | 19 | 152 | 292 |
| SRR7547873 | 0 | 306 | 19 | 157 | 1019 | 0 | 10 | 985 | 4096 | 7709 | 582 | 1077 | 0 | 589 | 542 | 774 | 12 | 1225 | 50 | 500 | 1362 | 3861 | 569 | 19319 | 5263 | 0 | 0 | 25 | 58 | 40 | 64 | 14 | 19 | 17 | 17 | 261 | 631 |
| SRR7547874 | 0 | 1268 | 144 | 152 | 948 | 0 | 0 | 2363 | 7975 | 17990 | 418 | 459 | 0 | 949 | 455 | 672 | 0 | 1870 | 22 | 936 | 743 | 2831 | 470 | 16342 | 2070 | 0 | 0 | 0 | 117 | 100 | 124 | 28 | 0 | 0 | 41 | 349 | 468 |
| SRR7547875 | 0 | 1065 | 139 | 218 | 1901 | 0 | 0 | 2255 | 9562 | 29616 | 332 | 459 | 0 | 1161 | 298 | 462 | 0 | 2109 | 14 | 504 | 552 | 630 | 57 | 1835 | 1960 | 0 | 0 | 0 | 109 | 70 | 103 | 24 | 0 | 0 | 22 | 256 | 219 |
| SRR7547876 | 0 | 609 | 13 | 319 | 2199 | 21 | 38 | 3766 | 13831 | 15777 | 1025 | 5961 | 0 | 960 | 338 | 658 | 18 | 2411 | 29 | 1063 | 931 | 1372 | 87 | 1586 | 2361 | 0 | 13 | 30 | 13 | 25 | 30 | 0 | 0 | 0 | 0 | 563 | 355 |
| SRR7547877 | 0 | 1907 | 84 | 145 | 1488 | 0 | 0 | 4215 | 16026 | 33006 | 203 | 1574 | 0 | 551 | 4801 | 3798 | 15 | 1322 | 15 | 3022 | 686 | 1483 | 145 | 5052 | 2427 | 0 | 0 | 0 | 140 | 91 | 124 | 22 | 10 | 0 | 32 | 1221 | 619 |
| SRR7547878 | 0 | 382 | 16 | 128 | 733 | 0 | 28 | 1732 | 5723 | 8559 | 409 | 1266 | 0 | 720 | 1723 | 2365 | 30 | 1256 | 0 | 602 | 443 | 941 | 32 | 3317 | 6077 | 0 | 0 | 23 | 35 | 10 | 12 | 18 | 0 | 0 | 15 | 541 | 1345 |
| SRR7547879 | 0 | 246 | 11 | 367 | 1273 | 0 | 24 | 1846 | 6704 | 12336 | 696 | 2000 | 0 | 1606 | 1442 | 2148 | 46 | 3805 | 35 | 301 | 1259 | 626 | 183 | 3943 | 2050 | 0 | 16 | 28 | 36 | 13 | 56 | 24 | 0 | 0 | 14 | 404 | 133 |
| SRR7547880 | 0 | 1796 | 60 | 86 | 572 | 28 | 70 | 3972 | 12603 | 11580 | 1408 | 1857 | 0 | 837 | 126 | 337 | 60 | 2036 | 109 | 731 | 678 | 1257 | 150 | 4427 | 1845 | 20 | 24 | 19 | 75 | 25 | 25 | 0 | 0 | 0 | 0 | 510 | 377 |
| SRR7547881 | 0 | 3040 | 149 | 123 | 978 | 18 | 32 | 3236 | 9721 | 9810 | 1832 | 6714 | 0 | 652 | 157 | 310 | 36 | 2934 | 22 | 582 | 1061 | 815 | 83 | 3650 | 1963 | 0 | 20 | 39 | 214 | 61 | 57 | 14 | 0 | 0 | 12 | 386 | 211 |
| SRR7547882 | 0 | 368 | 64 | 105 | 26 | 0 | 0 | 2990 | 9586 | 12683 | 223 | 2827 | 0 | 1303 | 696 | 2466 | 0 | 1397 | 27 | 2794 | 760 | 4233 | 81 | 1635 | 279 | 0 | 0 | 0 | 14 | 20 | 18 | 10 | 0 | 0 | 36 | 2360 | 14 |
| SRR7547883 | 0 | 599 | 61 | 100 | 76 | 0 | 0 | 2230 | 6341 | 12149 | 14 | 45 | 0 | 5815 | 50 | 297 | 0 | 2138 | 18 | 656 | 322 | 4365 | 29 | 1535 | 586 | 18 | 0 | 0 | 16 | 0 | 18 | 0 | 0 | 0 | 0 | 1010 | 47 |
| SRR7547884 | 0 | 320 | 34 | 184 | 35 | 0 | 0 | 1455 | 7477 | 32089 | 840 | 1581 | 0 | 1035 | 34 | 284 | 0 | 2036 | 24 | 1106 | 794 | 4639 | 88 | 2654 | 398 | 0 | 0 | 0 | 14 | 10 | 21 | 14 | 0 | 0 | 22 | 769 | 29 |
| SRR7547885 | 0 | 1851 | 52 | 280 | 1509 | 0 | 19 | 744 | 2563 | 6531 | 1051 | 1289 | 0 | 1234 | 2893 | 2085 | 37 | 1815 | 168 | 467 | 2086 | 1906 | 107 | 6705 | 4518 | 0 | 21 | 27 | 296 | 89 | 76 | 23 | 0 | 0 | 20 | 230 | 532 |
| SRR7547886 | 0 | 322 | 29 | 295 | 1118 | 0 | 25 | 1479 | 4803 | 9959 | 1412 | 6452 | 0 | 1871 | 2589 | 5570 | 34 | 7313 | 35 | 1219 | 889 | 2181 | 47 | 3330 | 7505 | 13 | 22 | 36 | 108 | 21 | 16 | 22 | 0 | 11 | 17 | 3293 | 184 |
| SRR7547887 | 0 | 340 | 50 | 232 | 1046 | 0 | 25 | 1466 | 4750 | 8265 | 2581 | 2548 | 0 | 812 | 1603 | 977 | 46 | 1619 | 78 | 817 | 856 | 1522 | 70 | 4002 | 4418 | 0 | 15 | 28 | 31 | 13 | 11 | 14 | 0 | 0 | 10 | 406 | 1584 |
| SRR7547888 | 0 | 1310 | 123 | 570 | 1187 | 0 | 28 | 1973 | 5216 | 9207 | 705 | 4811 | 0 | 1254 | 471 | 362 | 64 | 3137 | 34 | 442 | 742 | 939 | 0 | 472 | 4654 | 0 | 32 | 106 | 522 | 86 | 70 | 20 | 0 | 0 | 10 | 304 | 250 |
| SRR7547889 | 0 | 1287 | 60 | 231 | 891 | 0 | 20 | 2786 | 6971 | 16373 | 6469 | 5767 | 0 | 862 | 4787 | 8795 | 49 | 1663 | 45 | 1051 | 471 | 2402 | 85 | 7550 | 7086 | 0 | 15 | 16 | 124 | 38 | 47 | 13 | 0 | 0 | 0 | 1039 | 827 |
| SRR7547890 | 0 | 343 | 11 | 234 | 795 | 0 | 27 | 2043 | 6500 | 10113 | 2590 | 3479 | 0 | 1018 | 177 | 691 | 11 | 1389 | 94 | 1406 | 637 | 796 | 11 | 774 | 3236 | 0 | 26 | 33 | 41 | 14 | 12 | 19 | 0 | 0 | 13 | 1248 | 1859 |
| SRR7547891 | 0 | 2828 | 68 | 123 | 1578 | 16 | 35 | 3213 | 10102 | 15286 | 798 | 4837 | 0 | 1268 | 1898 | 4550 | 50 | 2400 | 42 | 1804 | 1427 | 970 | 45 | 1722 | 1741 | 0 | 24 | 11 | 45 | 20 | 23 | 0 | 0 | 0 | 0 | 775 | 220 |
| SRR7547892 | 0 | 800 | 19 | 149 | 900 | 21 | 32 | 1951 | 6699 | 10425 | 2166 | 3071 | 0 | 1405 | 4296 | 4403 | 14 | 2118 | 55 | 342 | 2244 | 1385 | 173 | 5691 | 4596 | 0 | 16 | 33 | 37 | 16 | 22 | 17 | 0 | 11 | 14 | 202 | 529 |
| SRR7547893 | 0 | 281 | 13 | 216 | 1964 | 0 | 24 | 2250 | 6955 | 9788 | 538 | 4944 | 0 | 870 | 2885 | 1301 | 37 | 748 | 20 | 1536 | 659 | 748 | 78 | 4490 | 1865 | 0 | 18 | 19 | 33 | 59 | 50 | 12 | 10 | 0 | 12 | 1488 | 447 |
| SRR7547894 | 0 | 2020 | 82 | 356 | 808 | 15 | 19 | 802 | 2778 | 5909 | 421 | 436 | 0 | 866 | 0 | 0 | 28 | 2165 | 70 | 177 | 1065 | 1742 | 25 | 1036 | 9070 | 0 | 16 | 14 | 61 | 28 | 28 | 0 | 0 | 0 | 0 | 116 | 691 |
| SRR7547895 | 0 | 445 | 10 | 244 | 1229 | 0 | 29 | 652 | 2076 | 3041 | 4628 | 6503 | 0 | 2291 | 645 | 470 | 41 | 1613 | 142 | 825 | 447 | 1108 | 19 | 1036 | 3778 | 0 | 23 | 18 | 23 | 11 | 51 | 17 | 0 | 0 | 0 | 424 | 1924 |
| SRR7547896 | 0 | 3129 | 87 | 187 | 1736 | 16 | 26 | 2604 | 8580 | 12726 | 903 | 10647 | 0 | 884 | 105 | 149 | 67 | 1175 | 67 | 951 | 828 | 1055 | 52 | 1391 | 2360 | 0 | 11 | 30 | 122 | 46 | 48 | 0 | 0 | 0 | 0 | 746 | 1911 |

### Supp Table S8.4.Columns_DI_to_ES

| | | | | | | | | | | | | | | | | | | | | | | | | | | | | | | | | | | | | | |
|---|---|---|---|---|---|---|---|---|---|---|---|---|---|---|---|---|---|---|---|---|---|---|---|---|---|---|---|---|---|---|---|---|---|---|---|---|---|
| SRR7547897 | 0 | 387 | 25 | 386 | 536 | 13 | 23 | 1842 | 4479 | 6033 | 1533 | 14427 | 0 | 365 | 1201 | 4485 | 20 | 969 | 41 | 2537 | 1287 | 1554 | 168 | 3482 | 4804 | 0 | 15 | 30 | 30 | 55 | 51 | 18 | 0 | 0 | 17 | 1389 | 829 |
| SRR7547898 | 0 | 5908 | 74 | 442 | 2262 | 18 | 19 | 2307 | 6920 | 12289 | 1550 | 15786 | 0 | 335 | 62 | 384 | 52 | 981 | 72 | 4241 | 1387 | 1629 | 37 | 1552 | 5585 | 0 | 10 | 43 | 178 | 93 | 110 | 28 | 0 | 0 | 20 | 1761 | 286 |
| SRR7547899 | 0 | 3629 | 75 | 332 | 772 | 23 | 19 | 574 | 1850 | 2564 | 1541 | 9765 | 0 | 324 | 1419 | 2390 | 33 | 1226 | 70 | 6530 | 480 | 1752 | 352 | 13377 | 4066 | 0 | 0 | 15 | 119 | 71 | 61 | 12 | 0 | 0 | 12 | 1773 | 1740 |
| SRR7547900 | 0 | 1761 | 57 | 287 | 2131 | 0 | 31 | 2431 | 5605 | 9522 | 1216 | 6004 | 0 | 1522 | 2801 | 2750 | 54 | 2063 | 24 | 1704 | 493 | 636 | 55 | 1585 | 1776 | 0 | 21 | 17 | 92 | 43 | 49 | 12 | 0 | 0 | 0 | 852 | 244 |
| SRR7547901 | 0 | 4731 | 181 | 173 | 1376 | 18 | 36 | 1733 | 7881 | 4816 | 822 | 1365 | 0 | 654 | 11 | 56 | 56 | 1598 | 43 | 853 | 535 | 734 | 39 | 1592 | 2888 | 0 | 11 | 36 | 161 | 70 | 68 | 12 | 0 | 0 | 22 | 435 | 461 |
| SRR7547902 | 0 | 4486 | 68 | 82 | 1145 | 12 | 23 | 1102 | 5785 | 5253 | 362 | 691 | 0 | 267 | 559 | 554 | 34 | 1506 | 96 | 1239 | 1148 | 936 | 152 | 5349 | 2813 | 0 | 15 | 45 | 169 | 64 | 74 | 29 | 0 | 0 | 16 | 4985 | 1046 |
| SRR7547903 | 0 | 798 | 12 | 767 | 1715 | 13 | 25 | 2723 | 9401 | 15944 | 1118 | 7836 | 0 | 341 | 2299 | 3060 | 36 | 983 | 53 | 2390 | 750 | 1252 | 105 | 3715 | 4643 | 0 | 16 | 26 | 20 | 50 | 47 | 12 | 0 | 0 | 11 | 925 | 1422 |
| SRR7547904 | 0 | 4257 | 67 | 151 | 3485 | 32 | 47 | 3045 | 11264 | 11941 | 1244 | 4712 | 0 | 905 | 221 | 513 | 65 | 1224 | 45 | 533 | 1248 | 1517 | 346 | 8331 | 1428 | 0 | 13 | 23 | 73 | 53 | 36 | 0 | 0 | 0 | 0 | 377 | 288 |
| SRR7547905 | 0 | 4109 | 85 | 99 | 1063 | 17 | 31 | 3678 | 9373 | 12299 | 4664 | 6160 | 0 | 995 | 0 | 15 | 49 | 1833 | 35 | 1087 | 728 | 996 | 164 | 5836 | 1824 | 0 | 18 | 23 | 224 | 45 | 37 | 0 | 0 | 0 | 0 | 530 | 262 |
| SRR7547906 | 0 | 1419 | 59 | 394 | 64 | 0 | 23 | 1617 | 3582 | 10369 | 1194 | 1795 | 0 | 2096 | 254 | 620 | 26 | 1892 | 39 | 2247 | 440 | 2456 | 82 | 1789 | 497 | 0 | 0 | 0 | 22 | 13 | 12 | 0 | 0 | 0 | 0 | 2848 | 39 |
| SRR7547907 | 0 | 2130 | 76 | 106 | 35 | 0 | 22 | 2077 | 6124 | 9976 | 2277 | 3943 | 0 | 2368 | 19 | 171 | 45 | 1990 | 21 | 1078 | 301 | 2261 | 33 | 842 | 246 | 0 | 0 | 0 | 34 | 24 | 27 | 0 | 0 | 0 | 18 | 2887 | 13 |
| SRR7547908 | 0 | 4673 | 179 | 215 | 78 | 0 | 13 | 1026 | 5799 | 11938 | 3137 | 10132 | 0 | 619 | 153 | 492 | 24 | 1710 | 15 | 2442 | 1210 | 3132 | 255 | 2503 | 310 | 0 | 11 | 15 | 125 | 50 | 66 | 20 | 0 | 0 | 23 | 1755 | 28 |
| SRR7547909 | 0 | 2045 | 245 | 406 | 1125 | 14 | 32 | 1914 | 6760 | 13620 | 646 | 788 | 0 | 407 | 313 | 397 | 35 | 1894 | 55 | 191 | 1181 | 1101 | 214 | 10136 | 2565 | 0 | 12 | 50 | 250 | 79 | 71 | 15 | 0 | 0 | 12 | 110 | 1692 |
| SRR7547910 | 0 | 1430 | 134 | 442 | 1021 | 0 | 18 | 1932 | 6036 | 13564 | 274 | 522 | 0 | 563 | 276 | 423 | 21 | 1093 | 61 | 534 | 4221 | 695 | 37 | 1340 | 3645 | 0 | 15 | 20 | 184 | 68 | 68 | 17 | 0 | 0 | 18 | 247 | 736 |
| SRR7547911 | 0 | 339 | 48 | 607 | 1135 | 11 | 37 | 4354 | 13804 | 18845 | 1265 | 2067 | 0 | 463 | 312 | 438 | 10 | 1225 | 40 | 1531 | 2983 | 2125 | 290 | 15408 | 4311 | 0 | 23 | 19 | 17 | 10 | 40 | 0 | 0 | 0 | 0 | 759 | 671 |
| SRR7547912 | 0 | 1794 | 142 | 235 | 844 | 18 | 21 | 2413 | 6507 | 10582 | 988 | 2976 | 0 | 332 | 485 | 1054 | 22 | 1846 | 49 | 1796 | 479 | 939 | 77 | 2033 | 2524 | 0 | 15 | 21 | 128 | 74 | 54 | 20 | 0 | 0 | 19 | 685 | 367 |
| SRR7547913 | 0 | 395 | 18 | 176 | 1169 | 12 | 31 | 3157 | 8551 | 11385 | 1625 | 9260 | 0 | 390 | 1269 | 1161 | 40 | 1616 | 71 | 2372 | 607 | 1092 | 80 | 3270 | 2319 | 0 | 20 | 14 | 24 | 10 | 41 | 11 | 0 | 0 | 11 | 1054 | 1315 |
| SRR7547914 | 0 | 1812 | 117 | 237 | 962 | 15 | 46 | 3383 | 9672 | 12445 | 1747 | 3314 | 0 | 400 | 88 | 91 | 50 | 1779 | 23 | 1490 | 736 | 1125 | 118 | 3335 | 2365 | 0 | 16 | 15 | 100 | 56 | 62 | 13 | 0 | 0 | 13 | 577 | 874 |
| SRR7547915 | 0 | 502 | 14 | 250 | 1831 | 11 | 35 | 1078 | 5207 | 6220 | 675 | 648 | 0 | 1200 | 95 | 151 | 32 | 1637 | 32 | 450 | 1206 | 1689 | 271 | 11872 | 3459 | 0 | 16 | 29 | 16 | 37 | 51 | 10 | 0 | 0 | 12 | 377 | 772 |
| SRR7547916 | 0 | 3062 | 51 | 273 | 2336 | 50 | 43 | 2801 | 8133 | 11213 | 3227 | 3361 | 0 | 365 | 212 | 339 | 47 | 2244 | 146 | 1093 | 1116 | 2032 | 166 | 4746 | 4224 | 0 | 15 | 10 | 46 | 22 | 18 | 0 | 0 | 0 | 0 | 574 | 531 |
| SRR7547917 | 0 | 3187 | 127 | 258 | 36 | 0 | 17 | 1280 | 5615 | 14647 | 1743 | 2766 | 0 | 1753 | 61 | 409 | 31 | 1439 | 20 | 1799 | 1002 | 2513 | 138 | 1752 | 214 | 0 | 25 | 14 | 69 | 39 | 58 | 10 | 0 | 0 | 27 | 3143 | 19 |
| SRR7547918 | 0 | 2630 | 80 | 519 | 41 | 0 | 34 | 3244 | 9726 | 16668 | 1151 | 3244 | 0 | 1804 | 14 | 154 | 74 | 1957 | 15 | 1395 | 668 | 2116 | 42 | 1231 | 257 | 0 | 10 | 0 | 21 | 0 | 10 | 0 | 0 | 0 | 0 | 5507 | 35 |
| SRR7547919 | 0 | 1778 | 57 | 234 | 42 | 0 | 36 | 2323 | 5966 | 8549 | 3477 | 5987 | 0 | 6174 | 83 | 405 | 65 | 1634 | 19 | 2188 | 304 | 2605 | 30 | 1162 | 323 | 12 | 11 | 0 | 21 | 0 | 14 | 0 | 0 | 0 | 0 | 2126 | 26 |
| SRR7547920 | 0 | 1178 | 46 | 230 | 68 | 14 | 15 | 3007 | 9901 | 17686 | 17395 | 48800 | 0 | 770 | 1500 | 2838 | 26 | 1588 | 20 | 7610 | 1051 | 2583 | 204 | 2599 | 348 | 0 | 18 | 11 | 24 | 27 | 18 | 11 | 0 | 0 | 10 | 5467 | 38 |
| SRR7585373 | 0 | 13 | 0 | 0 | 0 | 0 | 0 | 982 | 180 | 1000 | 74134 | 9581 | 13 | 190 | 260 | 1130 | 0 | 252 | 0 | 672 | 5398 | 2679 | 0 | 302 | 4528 | 0 | 52 | 0 | 36 | 55 | 55 | 0 | 18 | 52 | 42 | 2908 | 812 |
| SRR7586110 | 0 | 649 | 148 | 264 | 373 | 0 | 11 | 554 | 1684 | 5142 | 15239 | 27918 | 0 | 2652 | 148 | 375 | 33 | 270 | 26 | 2242 | 61 | 1316 | 0 | 0 | 1748 | 0 | 0 | 19 | 114 | 184 | 187 | 15 | 11 | 10 | 72 | 3075 | 0 |
| SRR7586111 | 0 | 237 | 66 | 264 | 411 | 0 | 11 | 434 | 1603 | 4376 | 2696 | 24333 | 0 | 2323 | 148 | 327 | 12 | 258 | 29 | 1946 | 70 | 1311 | 0 | 11 | 1819 | 0 | 0 | 18 | 36 | 59 | 57 | 19 | 17 | 25 | 31 | 2691 | 0 |
| SRR7586113 | 0 | 491 | 156 | 401 | 317 | 0 | 14 | 630 | 1469 | 3044 | 24816 | 40869 | 0 | 2166 | 36 | 175 | 46 | 263 | 73 | 2442 | 67 | 1452 | 0 | 13 | 2850 | 0 | 0 | 20 | 97 | 186 | 193 | 25 | 0 | 0 | 94 | 2478 | 0 |
| SRR7586114 | 0 | 488 | 169 | 380 | 319 | 0 | 14 | 590 | 1320 | 2631 | 5488 | 35739 | 0 | 1677 | 36 | 140 | 47 | 253 | 68 | 2062 | 56 | 1482 | 0 | 14 | 3146 | 0 | 14 | 22 | 91 | 173 | 187 | 27 | 25 | 32 | 115 | 2266 | 0 |
| SRR7586116 | 0 | 302 | 51 | 48 | 525 | 0 | 13 | 376 | 806 | 1508 | 9695 | 41477 | 0 | 1220 | 181 | 363 | 35 | 197 | 53 | 6209 | 32 | 1847 | 0 | 37 | 2888 | 0 | 0 | 21 | 113 | 96 | 153 | 31 | 31 | 11 | 63 | 7101 | 0 |
| SRR7586117 | 0 | 1158 | 202 | 64 | 547 | 0 | 12 | 365 | 788 | 1506 | 7566 | 41121 | 0 | 1140 | 145 | 385 | 33 | 193 | 63 | 5891 | 28 | 1927 | 0 | 30 | 3208 | 0 | 0 | 79 | 444 | 340 | 544 | 122 | 159 | 162 | 328 | 6911 | 0 |
| SRR7586118 | 0 | 312 | 55 | 58 | 536 | 0 | 12 | 446 | 1022 | 1775 | 25074 | 50439 | 0 | 1495 | 203 | 474 | 32 | 198 | 65 | 7013 | 33 | 1947 | 0 | 40 | 3238 | 0 | 0 | 22 | 113 | 91 | 145 | 33 | 45 | 45 | 84 | 8488 | 0 |
| SRR7586119 | 0 | 681 | 77 | 155 | 637 | 0 | 0 | 618 | 2146 | 4653 | 6691 | 30857 | 0 | 1853 | 41 | 105 | 11 | 228 | 88 | 3677 | 222 | 1850 | 0 | 15 | 3100 | 0 | 0 | 10 | 71 | 97 | 147 | 20 | 15 | 19 | 30 | 3731 | 28 |
| SRR7586120 | 0 | 1828 | 232 | 156 | 619 | 0 | 13 | 714 | 2204 | 4347 | 4016 | 31272 | 0 | 1623 | 44 | 103 | 28 | 227 | 95 | 3944 | 225 | 1945 | 0 | 16 | 3257 | 0 | 0 | 30 | 181 | 247 | 372 | 63 | 62 | 66 | 96 | 3493 | 24 |
| SRR7586122 | 0 | 767 | 242 | 55 | 228 | 0 | 12 | 423 | 1112 | 1254 | 7797 | 28791 | 0 | 1428 | 157 | 722 | 24 | 186 | 163 | 4899 | 134 | 1186 | 0 | 136 | 2630 | 0 | 0 | 29 | 146 | 200 | 297 | 49 | 16 | 23 | 230 | 6443 | 0 |
| SRR7586123 | 0 | 733 | 255 | 59 | 227 | 0 | 0 | 456 | 1282 | 1382 | 21843 | 34247 | 0 | 2023 | 180 | 946 | 27 | 200 | 153 | 5629 | 123 | 1251 | 0 | 121 | 2915 | 0 | 0 | 31 | 128 | 172 | 282 | 58 | 66 | 66 | 295 | 7612 | 0 |
| SRR7586125 | 0 | 300 | 122 | 117 | 459 | 0 | 17 | 720 | 2295 | 3533 | 5465 | 24712 | 0 | 1963 | 82 | 290 | 13 | 230 | 82 | 5393 | 152 | 1274 | 0 | 47 | 2590 | 0 | 0 | 11 | 60 | 96 | 81 | 11 | 0 | 0 | 21 | 6926 | 12 |
| SRR7586126 | 0 | 849 | 358 | 143 | 420 | 0 | 13 | 826 | 2254 | 3088 | 1794 | 24524 | 0 | 2331 | 103 | 241 | 40 | 211 | 71 | 5695 | 151 | 1299 | 0 | 48 | 2761 | 0 | 0 | 28 | 155 | 258 | 199 | 34 | 21 | 28 | 87 | 6985 | 0 |
| SRR7586128 | 0 | 374 | 86 | 72 | 215 | 0 | 18 | 573 | 1236 | 2369 | 7755 | 21699 | 0 | 2213 | 70 | 166 | 12 | 278 | 77 | 9685 | 51 | 1305 | 0 | 14 | 2704 | 0 | 0 | 24 | 64 | 89 | 123 | 29 | 16 | 25 | 66 | 14653 | 0 |
| SRR7586129 | 0 | 368 | 91 | 78 | 204 | 0 | 13 | 586 | 1322 | 2691 | 17198 | 24325 | 0 | 2439 | 75 | 194 | 11 | 290 | 78 | 11036 | 43 | 1400 | 0 | 16 | 3038 | 0 | 0 | 24 | 64 | 87 | 123 | 30 | 34 | 37 | 84 | 15894 | 0 |
| SRR7586131 | 0 | 164 | 60 | 97 | 181 | 0 | 19 | 529 | 1486 | 1299 | 4238 | 24408 | 0 | 1515 | 34 | 115 | 16 | 164 | 111 | 5566 | 85 | 872 | 0 | 98 | 2068 | 0 | 11 | 28 | 46 | 55 | 85 | 17 | 11 | 18 | 68 | 8639 | 0 |
| SRR7586132 | 0 | 519 | 220 | 108 | 181 | 0 | 17 | 616 | 1554 | 1427 | 23421 | 27444 | 0 | 1486 | 45 | 199 | 54 | 149 | 93 | 5881 | 85 | 1002 | 0 | 76 | 2292 | 0 | 11 | 31 | 138 | 142 | 266 | 63 | 55 | 63 | 276 | 9465 | 0 |
| SRR7586134 | 0 | 340 | 110 | 75 | 242 | 0 | 14 | 625 | 1717 | 3148 | 15635 | 19010 | 0 | 3537 | 88 | 273 | 15 | 192 | 82 | 6985 | 68 | 1337 | 0 | 17 | 2512 | 0 | 0 | 10 | 59 | 77 | 114 | 26 | 15 | 10 | 69 | 7874 | 0 |
| SRR7586135 | 0 | 329 | 119 | 84 | 215 | 0 | 13 | 661 | 1616 | 3281 | 15807 | 19132 | 0 | 3812 | 89 | 192 | 15 | 171 | 103 | 6459 | 55 | 1370 | 0 | 14 | 2739 | 0 | 0 | 22 | 61 | 71 | 105 | 24 | 30 | 37 | 88 | 7986 | 0 |
| SRR7586137 | 0 | 369 | 146 | 294 | 411 | 0 | 0 | 489 | 1752 | 3124 | 1733 | 9814 | 0 | 1955 | 73 | 130 | 28 | 215 | 80 | 2854 | 108 | 1196 | 0 | 22 | 2501 | 0 | 0 | 19 | 73 | 127 | 171 | 21 | 0 | 0 | 151 | 2568 | 0 |
| SRR7586138 | 0 | 102 | 40 | 366 | 423 | 0 | 0 | 567 | 1773 | 3267 | 1250 | 9225 | 0 | 1802 | 79 | 115 | 27 | 230 | 98 | 2807 | 105 | 1306 | 0 | 24 | 2775 | 0 | 0 | 20 | 18 | 31 | 44 | 21 | 19 | 34 | 46 | 2477 | 0 |
| SRR7586140 | 0 | 273 | 110 | 98 | 229 | 0 | 12 | 956 | 2509 | 5408 | 2346 | 5968 | 0 | 3227 | 61 | 134 | 10 | 222 | 51 | 5958 | 89 | 987 | 0 | 34 | 2483 | 0 | 0 | 18 | 60 | 60 | 111 | 22 | 10 | 10 | 56 | 6982 | 0 |
| SRR7586141 | 0 | 258 | 120 | 119 | 222 | 0 | 12 | 1026 | 2650 | 5823 | 6425 | 6229 | 0 | 3730 | 60 | 155 | 12 | 200 | 53 | 6327 | 74 | 1036 | 0 | 25 | 2718 | 0 | 0 | 19 | 59 | 58 | 108 | 25 | 28 | 39 | 74 | 7676 | 0 |
| SRR7586143 | 0 | 257 | 87 | 59 | 166 | 0 | 19 | 873 | 1581 | 2574 | 4865 | 30362 | 0 | 1908 | 79 | 158 | 22 | 203 | 142 | 5032 | 77 | 940 | 0 | 45 | 2757 | 0 | 0 | 10 | 72 | 88 | 108 | 23 | 0 | 18 | 58 | 5595 | 0 |
| SRR7586144 | 0 | 645 | 242 | 59 | 179 | 0 | 16 | 802 | 1526 | 2518 | 6080 | 30681 | 0 | 1740 | 76 | 234 | 65 | 181 | 140 | 4960 | 80 | 966 | 0 | 36 | 3038 | 0 | 0 | 21 | 174 | 211 | 274 | 69 | 51 | 64 | 197 | 5534 | 0 |
| SRR7586146 | 0 | 371 | 108 | 225 | 376 | 0 | 21 | 378 | 1017 | 2577 | 4529 | 42981 | 0 | 1592 | 11 | 69 | 15 | 291 | 182 | 4446 | 151 | 2308 | 0 | 22 | 3422 | 0 | 0 | 32 | 40 | 84 | 137 | 12 | 0 | 11 | 0 | 5851 | 0 |
| SRR7586147 | 0 | 1200 | 352 | 243 | 425 | 0 | 20 | 424 | 962 | 2763 | 25172 | 46582 | 0 | 1616 | 32 | 108 | 56 | 282 | 185 | 4903 | 164 | 2301 | 0 | 18 | 3756 | 0 | 12 | 26 | 129 | 263 | 439 | 41 | 40 | 36 | 10 | 5946 | 0 |
| SRR7586149 | 0 | 935 | 138 | 413 | 350 | 0 | 15 | 478 | 1157 | 4276 | 6404 | 33670 | 0 | 2198 | 43 | 96 | 81 | 324 | 78 | 2736 | 159 | 944 | 0 | 26 | 2915 | 0 | 12 | 49 | 188 | 382 | 551 | 43 | 0 | 15 | 66 | 2809 | 0 |
| SRR7586150 | 0 | 324 | 50 | 427 | 372 | 0 | 20 | 534 | 1243 | 4178 | 4302 | 32702 | 0 | 2255 | 46 | 113 | 29 | 293 | 79 | 2832 | 160 | 1058 | 0 | 28 | 3058 | 0 | 16 | 17 | 65 | 124 | 188 | 18 | 16 | 14 | 28 | 2702 | 0 |
| SRR7586152 | 0 | 998 | 203 | 181 | 336 | 0 | 21 | 615 | 1811 | 6214 | 2508 | 21149 | 0 | 2365 | 112 | 292 | 57 | 375 | 114 | 3157 | 44 | 1907 | 0 | 12 | 3562 | 0 | 0 | 21 | 115 | 279 | 654 | 54 | 12 | 15 | 37 | 3657 | 0 |
| SRR7586153 | 0 | 333 | 72 | 195 | 360 | 0 | 21 | 700 | 1954 | 6728 | 10881 | 23053 | 0 | 2760 | 184 | 392 | 22 | 339 | 123 | 3479 | 47 | 2002 | 0 | 17 | 3810 | 0 | 0 | 19 | 34 | 92 | 214 | 19 | 14 | 15 | 18 | 3780 | 0 |
| SRR7586155 | 0 | 389 | 43 | 250 | 272 | 0 | 31 | 363 | 923 | 3093 | 29330 | 49265 | 0 | 2982 | 71 | 116 | 30 | 357 | 95 | 3572 | 38 | 1516 | 0 | 0 | 2405 | 0 | 17 | 13 | 63 | 99 | 133 | 14 | 0 | 17 | 23 | 4576 | 0 |
| SRR7586156 | 0 | 375 | 40 | 290 | 266 | 0 | 22 | 304 | 974 | 2965 | 5625 | 45190 | 0 | 2856 | 19 | 105 | 27 | 371 | 121 | 3088 | 47 | 1528 | 0 | 15 | 2721 | 0 | 17 | 13 | 55 | 91 | 138 | 13 | 35 | 13 | 33 | 4314 | 0 |
| SRR7586158 | 0 | 1489 | 253 | 201 | 313 | 0 | 0 | 448 | 1479 | 6344 | 10712 | 21824 | 0 | 3406 | 89 | 312 | 36 | 371 | 98 | 7056 | 134 | 1498 | 0 | 15 | 3042 | 0 | 0 | 28 | 212 | 458 | 1159 | 103 | 25 | 29 | 138 | 7333 | 0 |
| SRR7586159 | 0 | 1426 | 254 | 208 | 328 | 0 | 0 | 523 | 1532 | 6181 | 10136 | 21351 | 0 | 3554 | 118 | 360 | 31 | 349 | 99 | 6410 | 148 | 1529 | 0 | 17 | 3403 | 0 | 0 | 25 | 220 | 468 | 1124 | 102 | 121 | 112 | 186 | 7468 | 0 |
| SRR7609258 | 0 | 32 | 0 | 339 | 630 | 12 | 0 | 2086 | 2410 | 5985 | 2146 | 5223 | 0 | 214 | 0 | 0 | 0 | 532 | 108 | 169 | 1301 | 476 | 16 | 255 | 2274 | 0 | 22 | 90 | 18 | 17 | 83 | 0 | 0 | 0 | 0 | 259 | 80 |
| SRR7609259 | 0 | 123 | 35 | 34 | 551 | 42 | 42 | 2907 | 3377 | 4416 | 1690 | 2444 | 23 | 1248 | 0 | 64 | 27 | 530 | 70 | 226 | 126 | 379 | 130 | 1501 | 697 | 0 | 99 | 61 | 11 | 11 | 24 | 60 | 50 | 21 | 0 | 318 | 91 |
| SRR7609260 | 0 | 202 | 50 | 36 | 339 | 34 | 37 | 944 | 1129 | 1199 | 1209 | 1383 | 27 | 1503 | 0 | 0 | 27 | 182 | 62 | 84 | 169 | 756 | 265 | 3942 | 1125 | 0 | 236 | 36 | 178 | 64 | 197 | 40 | 0 | 0 | 0 | 114 | 498 |
| SRR7609261 | 0 | 250 | 0 | 33 | 299 | 0 | 11 | 1001 | 1175 | 1074 | 1410 | 1524 | 0 | 1141 | 11 | 22 | 0 | 252 | 12 | 378 | 453 | 769 | 709 | 3095 | 681 | 0 | 308 | 25 | 55 | 26 | 44 | 0 | 0 | 0 | 0 | 1151 | 466 |
| SRR7609262 | 0 | 157 | 0 | 0 | 110 | 0 | 54 | 5531 | 5978 | 2595 | 580 | 2201 | 0 | 1421 | 0 | 0 | 168 | 143 | 15 | 76 | 245 | 412 | 14 | 242 | 1157 | 29 | 11 | 20 | 71 | 14 | 55 | 0 | 0 | 0 | 0 | 127 | 195 |
| SRR7609263 | 0 | 116 | 12 | 28 | 490 | 33 | 21 | 497 | 315 | 895 | 1126 | 6579 | 0 | 1152 | 41 | 547 | 16 | 354 | 71 | 33 | 343 | 450 | 129 | 1845 | 1013 | 0 | 147 | 109 | 191 | 40 | 65 | 0 | 0 | 0 | 0 | 21 | 1166 |
| SRR7609264 | 0 | 14 | 0 | 26 | 532 | 25 | 15 | 573 | 674 | 1218 | 5863 | 7912 | 0 | 1096 | 21 | 135 | 28 | 238 | 57 | 30 | 132 | 221 | 78 | 674 | 629 | 0 | 62 | 34 | 77 | 23 | 65 | 0 | 0 | 0 | 0 | 35 | 115 |
| SRR7609265 | 0 | 20 | 0 | 28 | 160 | 20 | 13 | 359 | 304 | 365 | 6035 | 21671 | 0 | 526 | 32 | 143 | 11 | 187 | 85 | 38 | 146 | 401 | 218 | 2539 | 539 | 0 | 132 | 73 | 115 | 21 | 46 | 0 | 0 | 0 | 0 | 0 | 366 |
| SRR7609266 | 0 | 90 | 0 | 86 | 373 | 303 | 0 | 706 | 718 | 1022 | 3440 | 3983 | 0 | 1130 | 0 | 39 | 47 | 264 | 64 | 101 | 281 | 380 | 290 | 1353 | 504 | 0 | 32 | 12 | 84 | 35 | 49 | 0 | 0 | 0 | 0 | 47 | 485 |
| SRR7609267 | 12 | 103 | 0 | 46 | 624 | 117 | 62 | 1920 | 2270 | 2292 | 1253 | 1342 | 30 | 1364 | 0 | 0 | 71 | 489 | 25 | 50 | 422 | 456 | 117 | 871 | 630 | 0 | 15 | 25 | 117 | 36 | 44 | 0 | 0 | 0 | 0 | 59 | 559 |
| SRR7609268 | 0 | 20 | 0 | 38 | 74 | 12 | 17 | 485 | 481 | 693 | 1489 | 3597 | 0 | 1426 | 0 | 14 | 27 | 187 | 36 | 49 | 163 | 446 | 213 | 981 | 449 | 0 | 83 | 29 | 79 | 26 | 64 | 0 | 0 | 0 | 0 | 44 | 260 |
| SRR7609269 | 0 | 17 | 0 | 47 | 392 | 0 | 0 | 1627 | 1879 | 1675 | 832 | 2495 | 0 | 820 | 0 | 18 | 0 | 445 | 120 | 31 | 1133 | 391 | 149 | 2555 | 987 | 0 | 20 | 41 | 20 | 41 | 75 | 0 | 0 | 0 | 0 | 29 | 313 |
| SRR7609270 | 11 | 80 | 0 | 38 | 414 | 16 | 52 | 1282 | 1097 | 602 | 3611 | 5019 | 0 | 1042 | 39 | 75 | 40 | 152 | 34 | 29 | 80 | 208 | 21 | 135 | 909 | 0 | 104 | 107 | 364 | 143 | 268 | 18 | 0 | 0 | 0 | 36 | 106 |
| SRR7609271 | 0 | 15 | 0 | 45 | 261 | 16 | 11 | 1434 | 1310 | 916 | 4857 | 11922 | 0 | 1349 | 0 | 33 | 20 | 356 | 76 | 35 | 954 | 417 | 356 | 1185 | 492 | 0 | 52 | 49 | 105 | 28 | 55 | 0 | 0 | 0 | 0 | 0 | 380 |
| SRR7609272 | 0 | 30 | 0 | 24 | 63 | 0 | 0 | 631 | 657 | 715 | 6804 | 8717 | 0 | 1198 | 21 | 11 | 11 | 152 | 30 | 83 | 136 | 534 | 229 | 2515 | 465 | 0 | 34 | 19 | 64 | 21 | 56 | 0 | 0 | 0 | 0 | 86 | 197 |
| SRR7609273 | 0 | 278 | 0 | 77 | 545 | 0 | 17 | 712 | 673 | 1627 | 2243 | 5595 | 0 | 767 | 59 | 69 | 68 | 240 | 82 | 129 | 401 | 437 | 138 | 1898 | 795 | 11 | 65 | 21 | 70 | 25 | 43 | 0 | 0 | 0 | 0 | 153 | 671 |
| SRR7609274 | 0 | 13 | 0 | 29 | 490 | 0 | 11 | 1997 | 2440 | 3569 | 1606 | 7152 | 0 | 878 | 0 | 57 | 16 | 584 | 25 | 30 | 325 | 208 | 55 | 460 | 433 | 0 | 162 | 55 | 98 | 25 | 41 | 0 | 0 | 0 | 0 | 20 | 354 |
| SRR7609275 | 0 | 245 | 13 | 21 | 187 | 14 | 19 | 781 | 764 | 1172 | 2990 | 5500 | 0 | 1452 | 0 | 0 | 0 | 241 | 110 | 64 | 344 | 203 | 31 | 226 | 508 | 0 | 103 | 59 | 356 | 84 | 141 | 23 | 19 | 0 | 0 | 37 | 345 |
| SRR7609276 | 0 | 254 | 0 | 19 | 47 | 0 | 14 | 658 | 595 | 635 | 1350 | 12107 | 0 | 539 | 0 | 24 | 14 | 107 | 10 | 151 | 82 | 307 | 55 | 406 | 399 | 0 | 47 | 12 | 25 | 0 | 12 | 0 | 0 | 0 | 0 | 103 | 0 |
| SRR7609277 | 10 | 11 | 0 | 11 | 259 | 16 | 28 | 1441 | 1589 | 2204 | 1579 | 2146 | 10 | 1939 | 0 | 0 | 26 | 267 | 177 | 64 | 459 | 417 | 146 | 2508 | 647 | 0 | 134 | 29 | 15 | 112 | 15 | 35 | 14 | 0 | 0 | 61 | 107 |
| SRR7609278 | 0 | 13 | 0 | 23 | 450 | 0 | 0 | 435 | 474 | 706 | 12814 | 12182 | 0 | 622 | 0 | 49 | 0 | 268 | 138 | 109 | 117 | 299 | 145 | 1819 | 469 | 0 | 60 | 22 | 84 | 20 | 43 | 0 | 0 | 0 | 0 | 138 | 197 |
| SRR7609279 | 0 | 141 | 0 | 52 | 202 | 0 | 11 | 1441 | 1633 | 1244 | 3806 | 8140 | 0 | 909 | 0 | 0 | 0 | 362 | 35 | 70 | 236 | 307 | 137 | 860 | 700 | 0 | 14 | 46 | 23 | 81 | 120 | 16 | 0 | 0 | 0 | 67 | 469 |
| SRR7609280 | 0 | 18 | 0 | 34 | 154 | 20 | 15 | 685 | 565 | 761 | 3257 | 5670 | 0 | 839 | 0 | 0 | 0 | 203 | 45 | 32 | 154 | 553 | 197 | 1489 | 605 | 0 | 74 | 10 | 76 | 31 | 37 | 0 | 0 | 0 | 0 | 32 | 514 |
| SRR7609281 | 0 | 176 | 0 | 16 | 619 | 22 | 27 | 1605 | 1891 | 2908 | 1596 | 5429 | 0 | 1764 | 0 | 0 | 0 | 215 | 95 | 140 | 169 | 205 | 118 | 961 | 425 | 0 | 115 | 37 | 139 | 22 | 0 | 0 | 0 | 0 | 0 | 312 | 344 |
| SRR7609282 | 0 | 183 | 16 | 45 | 744 | 14 | 29 | 1025 | 905 | 979 | 1811 | 2437 | 0 | 2048 | 0 | 44 | 0 | 135 | 67 | 75 | 277 | 198 | 262 | 1027 | 366 | 0 | 78 | 90 | 1209 | 203 | 369 | 57 | 18 | 0 | 0 | 42 | 358 |
| SRR7609283 | 0 | 110 | 17 | 21 | 123 | 51 | 33 | 1154 | 1239 | 1185 | 3421 | 4423 | 0 | 2531 | 0 | 52 | 0 | 471 | 33 | 273 | 184 | 408 | 179 | 824 | 596 | 0 | 112 | 25 | 221 | 62 | 172 | 29 | 12 | 0 | 0 | 125 | 294 |
| SRR7609284 | 0 | 27 | 0 | 31 | 239 | 0 | 0 | 263 | 331 | 733 | 2659 | 5991 | 0 | 1124 | 31 | 137 | 11 | 145 | 38 | 85 | 385 | 370 | 52 | 329 | 837 | 0 | 60 | 41 | 124 | 43 | 106 | 24 | 0 | 0 | 0 | 83 | 479 |
| SRR7609285 | 0 | 157 | 0 | 20 | 175 | 28 | 41 | 609 | 514 | 542 | 2104 | 5409 | 0 | 3061 | 0 | 47 | 0 | 459 | 43 | 122 | 115 | 388 | 140 | 833 | 753 | 0 | 130 | 35 | 270 | 89 | 160 | 28 | 0 | 0 | 0 | 82 | 616 |
| SRR7609286 | 0 | 100 | 13 | 38 | 101 | 15 | 29 | 2051 | 2243 | 2616 | 1035 | 4741 | 0 | 1710 | 0 | 90 | 13 | 554 | 28 | 290 | 170 | 464 | 201 | 2032 | 728 | 0 | 68 | 28 | 98 | 32 | 56 | 0 | 0 | 0 | 0 | 306 | 111 |
| SRR7609287 | 0 | 139 | 11 | 27 | 1047 | 34 | 23 | 5792 | 7821 | 8550 | 370 | 401 | 0 | 1309 | 0 | 0 | 20 | 461 | 30 | 59 | 224 | 172 | 136 | 1294 | 540 | 0 | 136 | 37 | 372 | 139 | 264 | 54 | 23 | 34 | 0 | 71 | 149 |
| SRR7609288 | 0 | 13 | 0 | 40 | 509 | 14 | 26 | 607 | 663 | 1934 | 770 | 2392 | 10 | 5100 | 13 | 26 | 0 | 588 | 257 | 66 | 3485 | 931 | 363 | 5938 | 1562 | 0 | 129 | 33 | 11 | 117 | 21 | 78 | 24 | 26 | 0 | 144 | 1213 |
| SRR7609289 | 10 | 35 | 0 | 28 | 665 | 32 | 11 | 2077 | 2340 | 1191 | 522 | 651 | 0 | 1495 | 0 | 0 | 16 | 337 | 45 | 147 | 542 | 409 | 129 | 1084 | 710 | 0 | 137 | 17 | 52 | 24 | 41 | 0 | 0 | 0 | 0 | 246 | 446 |
| SRR7609290 | 12 | 297 | 0 | 53 | 512 | 43 | 24 | 2008 | 2210 | 2422 | 696 | 687 | 0 | 834 | 0 | 0 | 24 | 247 | 35 | 25 | 343 | 555 | 231 | 1557 | 1093 | 0 | 122 | 92 | 284 | 156 | 272 | 12 | 21 | 18 | 0 | 39 | 1305 |
| SRR7609291 | 0 | 20 | 0 | 47 | 159 | 0 | 15 | 2868 | 3854 | 1753 | 81 | 261 | 0 | 296 | 0 | 0 | 63 | 431 | 96 | 301 | 194 | 100 | 155 | 2076 | 2325 | 30 | 1036 | 0 | 17 | 24 | 42 | 0 | 0 | 0 | 0 | 972 | 322 |
| SRR7609292 | 0 | 19 | 0 | 54 | 138 | 21 | 11 | 751 | 674 | 749 | 1735 | 2390 | 0 | 1200 | 0 | 0 | 58 | 285 | 39 | 43 | 234 | 464 | 318 | 1509 | 504 | 0 | 99 | 30 | 95 | 36 | 67 | 0 | 0 | 0 | 0 | 69 | 541 |
| SRR7609293 | 0 | 212 | 15 | 26 | 397 | 0 | 34 | 2137 | 2681 | 1891 | 3506 | 6736 | 0 | 1458 | 0 | 0 | 16 | 361 | 49 | 28 | 255 | 290 | 43 | 217 | 612 | 0 | 179 | 107 | 794 | 149 | 187 | 24 | 15 | 18 | 0 | 53 | 155 |
| SRR7609294 | 0 | 72 | 15 | 35 | 150 | 13 | 35 | 1535 | 1701 | 2172 | 739 | 976 | 14 | 1191 | 0 | 11 | 18 | 720 | 20 | 467 | 210 | 292 | 139 | 663 | 527 | 0 | 37 | 22 | 17 | 63 | 12 | 32 | 20 | 16 | 0 | 1487 | 550 |
| SRR7609295 | 14 | 66 | 0 | 25 | 544 | 0 | 46 | 371 | 538 | 1701 | 1329 | 5660 | 0 | 1576 | 0 | 21 | 18 | 208 | 120 | 18 | 154 | 235 | 97 | 1880 | 452 | 0 | 289 | 88 | 30 | 24 | 78 | 14 | 0 | 12 | 24 | 18 | 421 |
| SRR7609296 | 0 | 38 | 0 | 52 | 367 | 23 | 21 | 921 | 931 | 3218 | 803 | 10674 | 0 | 337 | 109 | 703 | 14 | 416 | 13 | 129 | 299 | 240 | 275 | 2722 | 426 | 0 | 112 | 11 | 14 | 0 | 0 | 0 | 0 | 0 | 0 | 366 | 162 |
| SRR7609297 | 10 | 56 | 0 | 24 | 145 | 22 | 14 | 1100 | 1051 | 3506 | 825 | 3230 | 15 | 1428 | 31 | 83 | 12 | 140 | 51 | 169 | 271 | 371 | 310 | 1577 | 410 | 0 | 181 | 29 | 84 | 14 | 20 | 0 | 22 | 14 | 10 | 190 | 155 |
| SRR7609298 | 0 | 68 | 0 | 38 | 112 | 12 | 12 | 206 | 214 | 580 | 12044 | 14472 | 0 | 855 | 909 | 6171 | 0 | 182 | 22 | 244 | 244 | 659 | 1331 | 3458 | 403 | 0 | 64 | 56 | 29 | 14 | 31 | 18 | 0 | 0 | 0 | 160 | 133 |
| SRR7609299 | 0 | 38 | 0 | 27 | 295 | 14 | 16 | 623 | 860 | 2045 | 2225 | 4673 | 0 | 1933 | 13 | 39 | 16 | 206 | 48 | 0 | 266 | 248 | 113 | 949 | 496 | 0 | 155 | 48 | 78 | 22 | 105 | 0 | 14 | 18 | 0 | 29 | 283 |
| SRR7609300 | 0 | 108 | 26 | 56 | 546 | 0 | 23 | 3591 | 5026 | 9728 | 2982 | 3758 | 0 | 652 | 0 | 37 | 32 | 263 | 25 | 93 | 230 | 81 | 55 | 258 | 308 | 0 | 94 | 35 | 53 | 20 | 88 | 20 | 29 | 23 | 0 | 603 | 254 |
| SRR7609301 | 0 | 47 | 0 | 13 | 379 | 0 | 27 | 523 | 695 | 1707 | 974 | 1393 | 12 | 1907 | 21 | 103 | 22 | 317 | 122 | 24 | 1355 | 198 | 77 | 563 | 397 | 0 | 123 | 29 | 12 | 56 | 113 | 39 | 37 | 32 | 17 | 24 | 154 |
| SRR7609302 | 0 | 37 | 40 | 60 | 92 | 12 | 12 | 332 | 373 | 1046 | 399 | 2364 | 0 | 1627 | 168 | 814 | 15 | 215 | 35 | 118 | 283 | 209 | 388 | 721 | 274 | 0 | 113 | 12 | 49 | 15 | 74 | 18 | 15 | 21 | 28 | 95 | 541 |
| SRR7644105 | 0 | 0 | 0 | 0 | 0 | 11 | 0 | 0 | 0 | 1314 | 0 | 0 | 0 | 6132 | 0 | 1314 | 0 | 14 | 0 | 0 | 0 | 11 | 18 | 43 | 139 | 75 | 300 | 0 | 0 | 0 | 0 | 0 | 0 | 0 | 0 | 1752 | 0 |
| SRR7644106 | 0 | 0 | 0 | 0 | 0 | 163 | 0 | 0 | 0 | 479 | 0 | 0 | 0 | 5750 | 0 | 479 | 0 | 0 | 0 | 0 | 0 | 11 | 11 | 15 | 116 | 60 | 553 | 0 | 0 | 0 | 0 | 0 | 0 | 0 | 0 | 958 | 0 |
| SRR7644107 | 0 | 30 | 0 | 0 | 0 | 335 | 0 | 0 | 0 | 441 | 0 | 0 | 0 | 8826 | 0 | 1324 | 0 | 0 | 0 | 0 | 0 | 18 | 22 | 32 | 151 | 97 | 548 | 0 | 30 | 0 | 0 | 0 | 0 | 0 | 0 | 1765 | 0 |
| SRR7644108 | 0 | 32 | 0 | 0 | 0 | 225 | 0 | 967 | 0 | 484 | 0 | 0 | 0 | 5319 | 0 | 484 | 0 | 0 | 0 | 0 | 0 | 19 | 30 | 15 | 152 | 49 | 10 | 0 | 32 | 0 | 0 | 0 | 0 | 0 | 0 | 2418 | 0 |
| SRR7644109 | 0 | 0 | 0 | 0 | 0 | 53 | 0 | 1467 | 1956 | 0 | 0 | 0 | 0 | 5868 | 0 | 978 | 0 | 0 | 0 | 0 | 0 | 14 | 11 | 107 | 210 | 96 | 294 | 0 | 0 | 0 | 0 | 0 | 0 | 0 | 0 | 0 | 0 |
| SRR7644110 | 0 | 29 | 0 | 0 | 0 | 202 | 0 | 0 | 0 | 524 | 524 | 0 | 0 | 1571 | 0 | 524 | 0 | 12 | 12 | 0 | 0 | 19 | 12 | 89 | 185 | 104 | 260 | 0 | 0 | 0 | 0 | 0 | 0 | 0 | 0 | 524 | 0 |
| SRR7644111 | 0 | 0 | 0 | 0 | 0 | 272 | 0 | 1464 | 1464 | 488 | 490 | 0 | 0 | 2928 | 0 | 0 | 0 | 11 | 19 | 0 | 0 | 30 | 26 | 119 | 263 | 122 | 10 | 0 | 0 | 0 | 0 | 0 | 0 | 0 | 0 | 0 | 0 |
| SRR7644112 | 0 | 29 | 0 | 0 | 0 | 114 | 0 | 1557 | 519 | 0 | 1558 | 0 | 0 | 5189 | 0 | 0 | 0 | 0 | 0 | 0 | 0 | 0 | 0 | 121 | 180 | 121 | 427 | 0 | 0 | 0 | 0 | 0 | 0 | 0 | 0 | 519 | 0 |
| SRR7644113 | 0 | 0 | 0 | 0 | 0 | 89 | 0 | 0 | 0 | 292 | 583 | 0 | 0 | 3792 | 0 | 0 | 0 | 0 | 17 | 583 | 0 | 17 | 15 | 86 | 306 | 116 | 13 | 0 | 0 | 0 | 0 | 0 | 0 | 0 | 0 | 3501 | 0 |
| SRR7644114 | 0 | 0 | 0 | 0 | 0 | 283 | 0 | 0 | 0 | 1280 | 321 | 0 | 0 | 3519 | 0 | 0 | 0 | 0 | 16 | 0 | 0 | 23 | 16 | 130 | 331 | 115 | 377 | 0 | 0 | 0 | 0 | 0 | 0 | 0 | 0 | 2239 | 0 |
| SRR7644115 | 0 | 0 | 0 | 0 | 0 | 149 | 0 | 0 | 0 | 1245 | 311 | 0 | 0 | 3111 | 0 | 0 | 0 | 15 | 22 | 933 | 0 | 30 | 25 | 89 | 325 | 194 | 20 | 0 | 0 | 0 | 0 | 0 | 0 | 0 | 0 | 1556 | 0 |
| SRR7644116 | 0 | 0 | 0 | 0 | 0 | 125 | 0 | 0 | 0 | 649 | 325 | 0 | 0 | 3895 | 0 | 0 | 0 | 21 | 16 | 649 | 0 | 21 | 18 | 94 | 294 | 163 | 312 | 0 | 0 | 0 | 0 | 0 | 0 | 0 | 0 | 3570 | 0 |
| SRR7644117 | 0 | 0 | 0 | 0 | 0 | 53 | 0 | 0 | 0 | 1835 | 0 | 0 | 0 | 3670 | 0 | 0 | 0 | 12 | 35 | 0 | 0 | 35 | 0 | 93 | 356 | 198 | 15 | 0 | 0 | 0 | 0 | 0 | 0 | 0 | 0 | 917 | 0 |
| SRR7644118 | 0 | 0 | 0 | 0 | 0 | 255 | 0 | 0 | 0 | 996 | 0 | 0 | 0 | 1494 | 0 | 0 | 0 | 12 | 37 | 1494 | 0 | 50 | 12 | 50 | 218 | 143 | 566 | 0 | 0 | 0 | 0 | 0 | 0 | 0 | 0 | 2988 | 0 |
| SRR7644119 | 0 | 0 | 0 | 0 | 0 | 186 | 0 | 0 | 0 | 1367 | 0 | 0 | 0 | 3646 | 0 | 456 | 0 | 0 | 36 | 1823 | 0 | 24 | 0 | 95 | 320 | 178 | 266 | 0 | 0 | 0 | 0 | 0 | 0 | 0 | 0 | 2735 | 0 |
| SRR7644120 | 0 | 0 | 0 | 0 | 0 | 196 | 0 | 0 | 0 | 3005 | 0 | 0 | 0 | 3005 | 0 | 501 | 0 | 19 | 56 | 0 | 0 | 13 | 0 | 94 | 364 | 176 | 309 | 0 | 0 | 0 | 0 | 0 | 0 | 0 | 0 | 2003 | 0 |
| SRR7644121 | 0 | 0 | 0 | 0 | 0 | 26 | 0 | 0 | 284 | 1701 | 0 | 0 | 0 | 1985 | 0 | 284 | 0 | 25 | 39 | 1701 | 0 | 18 | 32 | 118 | 357 | 121 | 472 | 0 | 0 | 0 | 0 | 0 | 0 | 0 | 0 | 1985 | 0 |
| SRR7644122 | 0 | 0 | 0 | 0 | 0 | 86 | 0 | 601 | 300 | 1201 | 0 | 0 | 0 | 1201 | 0 | 901 | 0 | 12 | 0 | 601 | 0 | 39 | 31 | 93 | 344 | 186 | 690 | 0 | 0 | 0 | 0 | 0 | 0 | 0 | 0 | 1201 | 0 |
| SRR7644123 | 0 | 0 | 0 | 0 | 0 | 214 | 0 | 277 | 0 | 277 | 0 | 0 | 0 | 2217 | 0 | 0 | 0 | 40 | 18 | 1663 | 0 | 55 | 15 | 121 | 318 | 172 | 322 | 0 | 0 | 0 | 0 | 0 | 0 | 0 | 0 | 3325 | 0 |

### Supp Table S8.4.Columns_DI_to_ES

| | | | | | | | | | | | | | | | | | | | | | | | | | | | | | | | | | | | | | |
|---|---|---|---|---|---|---|---|---|---|---|---|---|---|---|---|---|---|---|---|---|---|---|---|---|---|---|---|---|---|---|---|---|---|---|---|---|---|
| SRR7644124 | 0 | 0 | 0 | 0 | 0 | 57 | 0 | 0 | 592 | 1776 | 0 | 0 | 0 | 1776 | 0 | 0 | 0 | 27 | 23 | 592 | 0 | 19 | 31 | 109 | 372 | 159 | 537 | 0 | 0 | 0 | 0 | 0 | 0 | 0 | 0 | 2664 | 0 |
| SRR7644125 | 0 | 0 | 0 | 0 | 0 | 69 | 0 | 519 | 519 | 1297 | 0 | 0 | 0 | 1556 | 0 | 0 | 0 | 18 | 25 | 1297 | 0 | 12 | 34 | 59 | 197 | 237 | 458 | 0 | 0 | 0 | 0 | 0 | 0 | 0 | 0 | 4668 | 0 |
| SRR7644126 | 0 | 0 | 0 | 0 | 0 | 124 | 0 | 282 | 0 | 1126 | 0 | 0 | 0 | 845 | 282 | 282 | 0 | 17 | 37 | 1126 | 0 | 30 | 20 | 76 | 222 | 239 | 14 | 0 | 0 | 0 | 0 | 0 | 0 | 0 | 0 | 2534 | 0 |
| SRR7644128 | 0 | 46 | 0 | 0 | 0 | 115 | 0 | 262 | 262 | 523 | 0 | 0 | 0 | 2355 | 262 | 0 | 0 | 25 | 25 | 785 | 0 | 44 | 0 | 66 | 226 | 251 | 598 | 0 | 0 | 0 | 0 | 0 | 0 | 0 | 0 | 3141 | 0 |
| SRR7644129 | 0 | 0 | 0 | 0 | 0 | 74 | 0 | 0 | 283 | 566 | 0 | 0 | 0 | 1416 | 283 | 566 | 0 | 13 | 37 | 1699 | 0 | 13 | 24 | 54 | 209 | 188 | 297 | 0 | 0 | 0 | 0 | 0 | 0 | 0 | 0 | 3398 | 0 |
| SRR7663662 | 45 | 0 | 0 | 0 | 0 | 23 | 17 | 40530 | 1806 | 152 | 1186 | 968 | 0 | 76 | 0 | 0 | 30 | 11 | 0 | 0 | 639 | 360 | 0 | 66 | 1845 | 0 | 0 | 0 | 0 | 0 | 0 | 0 | 0 | 0 | 0 | 0 | 0 |
| SRR7663663 | 18 | 0 | 0 | 0 | 0 | 24 | 13 | 117173 | 3660 | 234 | 265 | 1153 | 0 | 62 | 0 | 0 | 0 | 0 | 0 | 0 | 757 | 131 | 0 | 17 | 1218 | 0 | 0 | 0 | 0 | 0 | 0 | 0 | 0 | 0 | 0 | 0 | 0 |
| SRR7664346 | 0 | 121 | 0 | 0 | 18 | 0 | 0 | 408 | 370 | 26516 | 5724 | 2489 | 0 | 1901 | 0 | 0 | 0 | 948 | 14 | 708 | 932 | 1545 | 41 | 21335 | 6302 | 23 | 0 | 30 | 47 | 91 | 50 | 12 | 11 | 16 | 0 | 14592 | 0 |
| SRR7664347 | 0 | 133 | 0 | 0 | 19 | 0 | 0 | 233 | 191 | 11286 | 22881 | 7504 | 0 | 1058 | 0 | 0 | 0 | 786 | 14 | 305 | 1040 | 1175 | 37 | 15976 | 4863 | 29 | 0 | 60 | 70 | 91 | 42 | 0 | 0 | 0 | 0 | 5334 | 0 |
| SRR7690697 | 0 | 333 | 40 | 0 | 0 | 0 | 0 | 604 | 164 | 2206 | 2560 | 1956 | 0 | 197 | 1153 | 2579 | 0 | 377 | 0 | 1018 | 1936 | 2675 | 82 | 1749 | 5361 | 71 | 0 | 0 | 0 | 18 | 11 | 0 | 0 | 20 | 28 | 2668 | 0 |
| SRR7690698 | 0 | 115 | 10 | 0 | 0 | 0 | 0 | 517 | 177 | 2345 | 3605 | 2142 | 0 | 247 | 1010 | 2450 | 0 | 184 | 0 | 1062 | 2076 | 3368 | 79 | 1604 | 5362 | 47 | 0 | 0 | 0 | 10 | 0 | 0 | 0 | 12 | 17 | 2252 | 0 |
| SRR7690699 | 0 | 204 | 54 | 0 | 0 | 0 | 0 | 926 | 565 | 1633 | 2618 | 1860 | 0 | 205 | 1147 | 1891 | 0 | 497 | 0 | 1208 | 1910 | 2157 | 69 | 938 | 2411 | 39 | 0 | 0 | 0 | 15 | 12 | 0 | 0 | 17 | 22 | 2272 | 0 |
| SRR7690700 | 0 | 2701 | 104 | 392 | 1339 | 0 | 23 | 3086 | 11666 | 15388 | 1380 | 6061 | 0 | 799 | 243 | 469 | 77 | 1337 | 25 | 2200 | 724 | 800 | 30 | 508 | 1927 | 16 | 21 | 15 | 76 | 65 | 72 | 13 | 0 | 0 | 0 | 902 | 211 |
| SRR7690701 | 0 | 2885 | 40 | 134 | 911 | 0 | 30 | 6674 | 21032 | 19117 | 4772 | 7296 | 0 | 603 | 47 | 100 | 74 | 1422 | 28 | 1952 | 790 | 780 | 47 | 610 | 1114 | 28 | 16 | 18 | 62 | 44 | 36 | 0 | 0 | 0 | 0 | 1480 | 111 |
| SRR7690702 | 0 | 2956 | 101 | 191 | 877 | 0 | 30 | 4884 | 16286 | 15666 | 1342 | 10814 | 0 | 340 | 304 | 589 | 29 | 1138 | 33 | 3135 | 809 | 834 | 45 | 673 | 2326 | 0 | 20 | 16 | 80 | 40 | 34 | 0 | 0 | 0 | 0 | 1862 | 85 |
| SRR7690703 | 0 | 1519 | 45 | 217 | 574 | 0 | 42 | 10048 | 26593 | 25841 | 108 | 406 | 0 | 523 | 188 | 305 | 58 | 1584 | 33 | 1601 | 1081 | 714 | 55 | 693 | 1864 | 16 | 16 | 0 | 26 | 21 | 20 | 0 | 0 | 0 | 0 | 1765 | 97 |
| SRR7690704 | 0 | 293 | 12 | 285 | 859 | 0 | 30 | 4152 | 12836 | 15367 | 1090 | 10220 | 0 | 491 | 61 | 123 | 49 | 1410 | 22 | 4809 | 826 | 900 | 41 | 781 | 2225 | 15 | 25 | 12 | 16 | 16 | 17 | 25 | 0 | 0 | 0 | 1849 | 114 |
| SRR7690705 | 0 | 1653 | 31 | 429 | 758 | 0 | 32 | 7378 | 15177 | 12977 | 1047 | 8716 | 0 | 672 | 18 | 61 | 44 | 1446 | 16 | 2231 | 480 | 1004 | 30 | 715 | 2876 | 17 | 24 | 13 | 46 | 29 | 36 | 0 | 0 | 0 | 0 | 978 | 87 |
| SRR7690706 | 0 | 2936 | 55 | 513 | 250 | 0 | 26 | 4562 | 12252 | 11736 | 295 | 3344 | 0 | 756 | 66 | 264 | 40 | 2670 | 42 | 3350 | 1115 | 1381 | 53 | 1132 | 2946 | 0 | 19 | 10 | 44 | 20 | 34 | 0 | 0 | 0 | 0 | 2876 | 80 |
| SRR7690707 | 0 | 79 | 48 | 0 | 0 | 0 | 0 | 766 | 194 | 1267 | 4284 | 2743 | 0 | 132 | 2095 | 16526 | 0 | 352 | 0 | 1423 | 2436 | 4387 | 53 | 1185 | 5666 | 24 | 0 | 0 | 0 | 0 | 0 | 0 | 0 | 0 | 23 | 3594 | 0 |
| SRR7690708 | 0 | 2228 | 87 | 435 | 605 | 0 | 28 | 4018 | 11950 | 13029 | 9052 | 11445 | 0 | 519 | 159 | 242 | 26 | 1917 | 29 | 6120 | 963 | 1186 | 58 | 1131 | 2289 | 0 | 26 | 0 | 64 | 29 | 49 | 13 | 11 | 0 | 0 | 3458 | 96 |
| SRR7690713 | 0 | 2017 | 138 | 523 | 610 | 0 | 24 | 3174 | 11088 | 14990 | 511 | 5003 | 0 | 723 | 359 | 1042 | 31 | 2155 | 35 | 3354 | 1131 | 1190 | 52 | 982 | 2848 | 0 | 25 | 14 | 78 | 50 | 77 | 18 | 0 | 0 | 0 | 2522 | 55 |
| SRR7690714 | 0 | 2069 | 135 | 608 | 280 | 11 | 18 | 5198 | 12048 | 9785 | 684 | 4756 | 0 | 1025 | 480 | 1227 | 37 | 2280 | 42 | 5116 | 726 | 1634 | 38 | 1095 | 3308 | 0 | 21 | 0 | 42 | 50 | 45 | 13 | 0 | 0 | 0 | 4211 | 61 |
| SRR7690715 | 0 | 27 | 13 | 0 | 0 | 0 | 0 | 824 | 142 | 2307 | 2854 | 1505 | 0 | 180 | 1273 | 3288 | 0 | 240 | 0 | 1828 | 2445 | 2970 | 70 | 1508 | 4248 | 26 | 0 | 0 | 0 | 0 | 0 | 0 | 0 | 0 | 15 | 4277 | 0 |
| SRR7690716 | 0 | 27 | 11 | 0 | 0 | 0 | 0 | 514 | 119 | 1657 | 633 | 2095 | 0 | 308 | 832 | 3104 | 0 | 283 | 0 | 1317 | 1846 | 2008 | 83 | 1263 | 4354 | 21 | 0 | 0 | 0 | 0 | 0 | 0 | 0 | 10 | 25 | 4179 | 0 |
| SRR7690717 | 0 | 3332 | 79 | 201 | 983 | 0 | 27 | 5086 | 17138 | 16128 | 1031 | 4336 | 0 | 521 | 29 | 122 | 26 | 1564 | 35 | 3942 | 1047 | 1605 | 16 | 409 | 1920 | 10 | 16 | 0 | 91 | 83 | 112 | 18 | 0 | 0 | 0 | 2256 | 131 |
| SRR7690718 | 0 | 901 | 59 | 277 | 539 | 0 | 51 | 3778 | 11297 | 13870 | 255 | 1432 | 0 | 956 | 60 | 286 | 47 | 1166 | 48 | 1287 | 1036 | 1808 | 12 | 215 | 1155 | 0 | 23 | 0 | 14 | 11 | 17 | 0 | 0 | 0 | 0 | 1670 | 29 |
| SRR7690719 | 0 | 810 | 20 | 658 | 1299 | 0 | 21 | 6147 | 22109 | 21601 | 3783 | 4953 | 0 | 620 | 83 | 75 | 38 | 2175 | 25 | 2564 | 728 | 1213 | 45 | 1089 | 2231 | 22 | 0 | 13 | 18 | 17 | 21 | 13 | 0 | 0 | 0 | 1386 | 91 |
| SRR7690720 | 0 | 361 | 16 | 505 | 712 | 0 | 12 | 6411 | 20262 | 16122 | 6709 | 8697 | 0 | 514 | 149 | 164 | 29 | 2004 | 48 | 3693 | 1244 | 1024 | 38 | 875 | 3219 | 0 | 19 | 16 | 22 | 16 | 18 | 20 | 12 | 12 | 0 | 2011 | 75 |
| SRR7690721 | 0 | 350 | 19 | 484 | 684 | 0 | 15 | 3913 | 11309 | 9899 | 4681 | 7427 | 0 | 643 | 125 | 337 | 39 | 1925 | 70 | 3114 | 851 | 797 | 54 | 1085 | 2606 | 0 | 22 | 21 | 18 | 10 | 10 | 0 | 0 | 0 | 0 | 1666 | 50 |
| SRR7690722 | 0 | 13 | 20 | 60 | 239 | 58 | 30 | 1369 | 4542 | 16047 | 3680 | 6081 | 17 | 1845 | 316 | 526 | 0 | 672 | 33 | 1394 | 400 | 509 | 113 | 585 | 431 | 0 | 127 | 19 | 11 | 19 | 25 | 12 | 144 | 109 | 0 | 2100 | 52 |
| SRR7690723 | 17 | 214 | 90 | 75 | 168 | 69 | 44 | 2868 | 8803 | 17706 | 8033 | 13939 | 25 | 2012 | 71 | 185 | 18 | 523 | 34 | 2811 | 417 | 638 | 88 | 378 | 371 | 0 | 75 | 15 | 126 | 350 | 459 | 132 | 40 | 20 | 0 | 3110 | 38 |
| SRR7690724 | 0 | 149 | 166 | 87 | 345 | 57 | 55 | 535 | 1515 | 18213 | 560 | 2865 | 15 | 1505 | 0 | 49 | 27 | 526 | 35 | 652 | 574 | 342 | 95 | 526 | 646 | 0 | 156 | 20 | 206 | 13 | 27 | 11 | 104 | 97 | 0 | 1048 | 135 |
| SRR7690726 | 10 | 654 | 200 | 81 | 333 | 38 | 18 | 3292 | 11041 | 38694 | 1701 | 3266 | 18 | 1517 | 49 | 176 | 0 | 874 | 46 | 608 | 656 | 680 | 95 | 583 | 528 | 0 | 81 | 28 | 176 | 322 | 500 | 210 | 95 | 69 | 0 | 833 | 54 |
| SRR7690727 | 0 | 1802 | 87 | 196 | 732 | 0 | 30 | 5847 | 19054 | 22653 | 2883 | 5323 | 0 | 752 | 126 | 211 | 76 | 1519 | 43 | 2455 | 1125 | 2044 | 24 | 329 | 1020 | 0 | 17 | 0 | 39 | 41 | 67 | 11 | 0 | 0 | 0 | 1664 | 40 |
| SRR7690728 | 0 | 2214 | 48 | 63 | 336 | 0 | 52 | 3321 | 9692 | 7593 | 1330 | 9771 | 0 | 516 | 141 | 494 | 62 | 1414 | 48 | 2762 | 1091 | 2256 | 11 | 145 | 290 | 0 | 22 | 21 | 50 | 32 | 38 | 0 | 0 | 0 | 0 | 1594 | 17 |
| SRR7690729 | 0 | 436 | 29 | 227 | 1072 | 0 | 35 | 5888 | 18276 | 20946 | 6125 | 10869 | 0 | 659 | 64 | 214 | 50 | 1613 | 32 | 3265 | 797 | 1237 | 28 | 579 | 1627 | 25 | 12 | 34 | 19 | 25 | 36 | 35 | 0 | 0 | 0 | 1283 | 154 |
| SRR7690730 | 0 | 1209 | 69 | 217 | 390 | 12 | 35 | 2094 | 6498 | 5053 | 1497 | 19518 | 0 | 1616 | 159 | 899 | 54 | 1378 | 28 | 1502 | 682 | 1372 | 22 | 500 | 2312 | 0 | 52 | 0 | 32 | 46 | 51 | 15 | 0 | 0 | 0 | 1036 | 35 |
| SRR7690731 | 0 | 2364 | 33 | 302 | 478 | 0 | 50 | 7455 | 22254 | 15235 | 455 | 3841 | 0 | 516 | 25 | 46 | 44 | 1587 | 36 | 1509 | 1584 | 1353 | 48 | 864 | 3487 | 0 | 42 | 0 | 47 | 26 | 49 | 0 | 0 | 0 | 0 | 1040 | 112 |
| SRR7690732 | 0 | 255 | 17 | 670 | 572 | 0 | 28 | 3894 | 9508 | 8070 | 3480 | 5614 | 0 | 1024 | 267 | 545 | 11 | 1672 | 45 | 1857 | 681 | 1095 | 45 | 969 | 3131 | 0 | 35 | 0 | 14 | 41 | 44 | 0 | 0 | 0 | 0 | 1661 | 43 |
| SRR7690733 | 0 | 342 | 12 | 463 | 986 | 13 | 32 | 2884 | 6937 | 5212 | 1023 | 12993 | 0 | 279 | 81 | 225 | 44 | 1511 | 45 | 2444 | 576 | 931 | 36 | 864 | 3278 | 0 | 33 | 19 | 19 | 13 | 16 | 13 | 0 | 0 | 0 | 1869 | 51 |
| SRR7690734 | 0 | 125 | 119 | 134 | 425 | 18 | 131 | 2193 | 4188 | 12988 | 1942 | 2352 | 12 | 647 | 40 | 132 | 41 | 611 | 47 | 3713 | 627 | 509 | 73 | 492 | 621 | 14 | 529 | 0 | 62 | 273 | 233 | 50 | 0 | 0 | 0 | 10399 | 120 |
| SRR7690736 | 0 | 600 | 474 | 37 | 206 | 18 | 21 | 4928 | 10761 | 25443 | 2810 | 3957 | 13 | 1736 | 537 | 883 | 0 | 570 | 45 | 4310 | 351 | 834 | 190 | 1209 | 612 | 15 | 30 | 33 | 241 | 628 | 708 | 265 | 74 | 52 | 15 | 7694 | 57 |
| SRR7690737 | 0 | 458 | 235 | 84 | 450 | 23 | 31 | 1571 | 3225 | 9607 | 7045 | 12809 | 11 | 1043 | 60 | 37 | 0 | 612 | 35 | 648 | 338 | 562 | 67 | 332 | 397 | 16 | 48 | 85 | 328 | 693 | 682 | 133 | 91 | 23 | 0 | 1214 | 45 |
| SRR7690739 | 0 | 540 | 528 | 69 | 255 | 15 | 0 | 1317 | 5557 | 17590 | 5792 | 9070 | 15 | 1399 | 70 | 76 | 0 | 589 | 44 | 696 | 403 | 662 | 111 | 582 | 471 | 15 | 45 | 63 | 238 | 573 | 733 | 208 | 187 | 130 | 15 | 722 | 52 |
| SRR7690740 | 0 | 608 | 512 | 68 | 265 | 0 | 0 | 1671 | 5985 | 17990 | 825 | 7661 | 10 | 1784 | 20 | 59 | 0 | 599 | 44 | 482 | 408 | 609 | 93 | 574 | 409 | 16 | 63 | 71 | 207 | 507 | 820 | 199 | 141 | 108 | 13 | 786 | 38 |
| SRR7690741 | 0 | 60 | 33 | 0 | 10 | 0 | 0 | 674 | 218 | 1082 | 3335 | 2937 | 0 | 152 | 1586 | 3217 | 0 | 309 | 0 | 1013 | 1924 | 3268 | 87 | 1518 | 5481 | 17 | 0 | 0 | 0 | 18 | 13 | 0 | 0 | 21 | 27 | 1648 | 0 |
| SRR7690742 | 0 | 35 | 34 | 0 | 0 | 0 | 0 | 730 | 104 | 708 | 1463 | 2918 | 0 | 190 | 742 | 1463 | 0 | 355 | 0 | 1189 | 1988 | 2813 | 97 | 1677 | 6106 | 13 | 0 | 0 | 0 | 10 | 10 | 0 | 0 | 12 | 21 | 2346 | 0 |
| SRR7690743 | 0 | 91 | 45 | 0 | 12 | 0 | 0 | 659 | 140 | 867 | 874 | 2908 | 0 | 123 | 1743 | 3308 | 0 | 296 | 0 | 1203 | 1993 | 2809 | 84 | 1328 | 5034 | 0 | 0 | 0 | 0 | 0 | 0 | 0 | 0 | 0 | 14 | 2011 | 0 |
| SRR7690744 | 0 | 440 | 36 | 87 | 425 | 0 | 13 | 3538 | 12859 | 8722 | 3846 | 7980 | 0 | 731 | 520 | 1066 | 23 | 1574 | 29 | 951 | 852 | 1039 | 26 | 526 | 3494 | 0 | 13 | 21 | 23 | 50 | 13 | 11 | 0 | 0 | 0 | 470 | 88 |
| SRR7690745 | 0 | 353 | 12 | 164 | 823 | 0 | 29 | 6461 | 20796 | 12069 | 3777 | 7468 | 0 | 658 | 62 | 196 | 25 | 1510 | 29 | 2351 | 851 | 1059 | 56 | 655 | 2713 | 0 | 13 | 17 | 17 | 67 | 53 | 0 | 0 | 0 | 0 | 1326 | 80 |
| SRR7690746 | 0 | 284 | 30 | 102 | 393 | 0 | 15 | 9143 | 27766 | 18022 | 464 | 3711 | 0 | 808 | 1225 | 1937 | 14 | 1336 | 38 | 2964 | 444 | 746 | 35 | 849 | 3068 | 0 | 0 | 11 | 31 | 14 | 18 | 0 | 0 | 0 | 0 | 1857 | 44 |
| SRR7690747 | 0 | 286 | 43 | 371 | 1004 | 0 | 23 | 7057 | 22904 | 15072 | 644 | 5198 | 0 | 861 | 35 | 71 | 31 | 1747 | 26 | 1110 | 766 | 1599 | 35 | 704 | 3278 | 0 | 16 | 15 | 11 | 11 | 12 | 0 | 0 | 0 | 0 | 562 | 26 |
| SRR7690748 | 0 | 2034 | 104 | 286 | 305 | 0 | 25 | 6359 | 19923 | 18329 | 752 | 4499 | 0 | 630 | 342 | 529 | 24 | 2204 | 26 | 5141 | 739 | 1472 | 31 | 1028 | 2223 | 0 | 24 | 0 | 56 | 22 | 60 | 10 | 0 | 0 | 0 | 3860 | 250 |
| SRR7690749 | 0 | 2599 | 104 | 158 | 35 | 0 | 21 | 4438 | 10246 | 8770 | 1754 | 1972 | 0 | 584 | 249 | 342 | 35 | 2149 | 61 | 2687 | 344 | 1898 | 17 | 1090 | 3076 | 11 | 24 | 0 | 38 | 22 | 23 | 10 | 0 | 0 | 0 | 3662 | 71 |
| SRR7690750 | 0 | 2137 | 104 | 142 | 662 | 0 | 29 | 7398 | 24952 | 27022 | 930 | 4442 | 0 | 761 | 118 | 280 | 55 | 1524 | 33 | 2276 | 1181 | 1403 | 44 | 435 | 987 | 0 | 0 | 12 | 49 | 37 | 40 | 0 | 0 | 0 | 0 | 1386 | 31 |
| SRR7690751 | 0 | 688 | 34 | 106 | 423 | 0 | 31 | 7657 | 25554 | 26218 | 2487 | 5976 | 0 | 708 | 379 | 1483 | 10 | 1318 | 36 | 2294 | 1023 | 1122 | 33 | 557 | 1448 | 0 | 14 | 14 | 14 | 26 | 11 | 0 | 0 | 0 | 0 | 1355 | 60 |
| SRR7690752 | 0 | 562 | 38 | 242 | 622 | 0 | 27 | 6880 | 20889 | 17617 | 650 | 7563 | 0 | 631 | 46 | 307 | 10 | 1433 | 28 | 2583 | 1159 | 1077 | 57 | 636 | 1758 | 0 | 18 | 14 | 15 | 43 | 11 | 15 | 0 | 0 | 0 | 1152 | 54 |
| SRR7690753 | 0 | 636 | 40 | 132 | 384 | 0 | 17 | 6216 | 24109 | 17732 | 3357 | 5305 | 0 | 498 | 526 | 687 | 25 | 2111 | 47 | 4883 | 1534 | 1092 | 64 | 1277 | 2730 | 0 | 19 | 13 | 25 | 12 | 15 | 14 | 11 | 0 | 0 | 3333 | 52 |
| SRR7690754 | 0 | 767 | 29 | 399 | 512 | 0 | 22 | 6482 | 25878 | 23668 | 6355 | 10621 | 0 | 862 | 217 | 268 | 13 | 2007 | 50 | 5633 | 1171 | 1586 | 53 | 1086 | 2522 | 0 | 17 | 0 | 24 | 13 | 21 | 20 | 14 | 0 | 0 | 2829 | 31 |
| SRR7690755 | 0 | 1218 | 74 | 452 | 643 | 0 | 0 | 3720 | 12873 | 9985 | 942 | 9082 | 0 | 555 | 405 | 427 | 41 | 2127 | 112 | 5682 | 1212 | 779 | 59 | 881 | 2615 | 0 | 36 | 17 | 152 | 71 | 78 | 25 | 17 | 16 | 0 | 3328 | 71 |
| SRR7690756 | 0 | 954 | 62 | 473 | 856 | 0 | 29 | 4527 | 15574 | 10975 | 669 | 10709 | 0 | 317 | 102 | 143 | 31 | 2219 | 46 | 2156 | 1570 | 1096 | 74 | 1418 | 3344 | 0 | 38 | 17 | 17 | 11 | 11 | 18 | 0 | 0 | 0 | 1093 | 90 |
| SRR7690757 | 0 | 2076 | 75 | 387 | 1116 | 0 | 26 | 4116 | 16020 | 13452 | 7039 | 8677 | 0 | 586 | 383 | 980 | 30 | 1269 | 51 | 2100 | 1055 | 837 | 42 | 863 | 2982 | 0 | 28 | 13 | 98 | 48 | 56 | 11 | 0 | 0 | 0 | 1627 | 55 |
| SRR7690758 | 0 | 300 | 304 | 32 | 135 | 0 | 0 | 662 | 2683 | 11663 | 781 | 1323 | 38 | 1874 | 0 | 35 | 17 | 529 | 47 | 252 | 368 | 395 | 169 | 717 | 487 | 10 | 12 | 35 | 15 | 11 | 21 | 251 | 154 | 94 | 14 | 715 | 33 |
| SRR7690759 | 0 | 20 | 37 | 39 | 96 | 33 | 53 | 3921 | 11065 | 23568 | 709 | 9130 | 27 | 1678 | 374 | 515 | 0 | 608 | 67 | 5823 | 530 | 876 | 233 | 1115 | 427 | 14 | 40 | 33 | 13 | 42 | 60 | 26 | 11 | 110 | 0 | 9670 | 24 |
| SRR7690762 | 11 | 20 | 12 | 59 | 253 | 40 | 11 | 2032 | 6846 | 19334 | 708 | 11244 | 0 | 1759 | 38 | 76 | 11 | 560 | 50 | 584 | 412 | 668 | 126 | 660 | 449 | 0 | 65 | 57 | 194 | 17 | 26 | 208 | 122 | 71 | 0 | 1266 | 32 |
| SRR7690764 | 0 | 32 | 14 | 0 | 0 | 0 | 0 | 550 | 147 | 1172 | 671 | 1774 | 0 | 283 | 1404 | 9226 | 0 | 453 | 0 | 1088 | 2747 | 3848 | 85 | 1458 | 6852 | 20 | 0 | 0 | 0 | 0 | 0 | 0 | 0 | 13 | 11 | 2357 | 0 |
| SRR7690765 | 0 | 109 | 54 | 0 | 13 | 12 | 0 | 505 | 136 | 1972 | 1233 | 1379 | 0 | 306 | 452 | 2171 | 0 | 325 | 0 | 1097 | 2383 | 2557 | 84 | 1550 | 5628 | 0 | 0 | 0 | 0 | 0 | 0 | 0 | 0 | 0 | 35 | 2117 | 0 |
| SRR7690766 | 0 | 494 | 15 | 282 | 948 | 0 | 27 | 4572 | 14835 | 18908 | 2661 | 4827 | 0 | 787 | 130 | 580 | 41 | 1694 | 35 | 2138 | 1469 | 1127 | 44 | 686 | 2753 | 0 | 15 | 12 | 13 | 10 | 14 | 11 | 0 | 0 | 0 | 1517 | 83 |
| SRR7690767 | 0 | 2221 | 63 | 181 | 711 | 0 | 26 | 5184 | 17652 | 21374 | 572 | 4322 | 0 | 665 | 59 | 219 | 57 | 1338 | 41 | 2227 | 1105 | 1318 | 31 | 492 | 1960 | 0 | 24 | 17 | 66 | 35 | 43 | 0 | 0 | 0 | 0 | 1393 | 67 |
| SRR7690768 | 0 | 322 | 13 | 463 | 591 | 0 | 29 | 5149 | 17438 | 13083 | 3663 | 7819 | 0 | 692 | 167 | 416 | 27 | 1931 | 34 | 2863 | 1147 | 1124 | 55 | 996 | 3718 | 0 | 24 | 13 | 22 | 11 | 11 | 16 | 0 | 0 | 0 | 1960 | 56 |
| SRR7690769 | 11 | 11 | 12 | 99 | 282 | 66 | 46 | 2231 | 6342 | 20324 | 371 | 9725 | 18 | 1473 | 627 | 1152 | 18 | 631 | 43 | 3382 | 589 | 642 | 143 | 614 | 523 | 0 | 103 | 39 | 112 | 18 | 26 | 132 | 50 | 50 | 0 | 4942 | 42 |
| SRR7690770 | 0 | 315 | 442 | 102 | 306 | 62 | 38 | 2017 | 6220 | 23785 | 2279 | 4204 | 20 | 1796 | 198 | 350 | 13 | 745 | 33 | 1331 | 729 | 628 | 123 | 688 | 698 | 0 | 71 | 36 | 193 | 490 | 783 | 226 | 158 | 91 | 0 | 2095 | 34 |
| SRR7690773 | 11 | 24 | 144 | 47 | 220 | 53 | 53 | 2929 | 6111 | 13667 | 3735 | 6569 | 19 | 1295 | 134 | 245 | 0 | 551 | 55 | 1011 | 640 | 390 | 133 | 507 | 594 | 0 | 72 | 76 | 19 | 13 | 25 | 235 | 140 | 65 | 0 | 1579 | 43 |
| SRR7690775 | 0 | 2132 | 96 | 544 | 1479 | 0 | 14 | 4289 | 21824 | 10727 | 616 | 6150 | 0 | 663 | 152 | 632 | 32 | 2180 | 58 | 1340 | 742 | 953 | 50 | 757 | 3857 | 0 | 15 | 24 | 94 | 55 | 43 | 0 | 0 | 0 | 0 | 1396 | 57 |
| SRR7690777 | 0 | 1636 | 127 | 375 | 982 | 0 | 20 | 3928 | 14554 | 16557 | 4038 | 6554 | 0 | 801 | 144 | 805 | 30 | 1358 | 29 | 1469 | 855 | 1186 | 63 | 978 | 4388 | 0 | 0 | 19 | 143 | 100 | 85 | 0 | 0 | 0 | 0 | 748 | 146 |
| SRR7690778 | 0 | 1218 | 61 | 647 | 166 | 0 | 17 | 4624 | 12869 | 9640 | 2411 | 2941 | 0 | 739 | 162 | 375 | 15 | 1936 | 41 | 3233 | 420 | 1516 | 55 | 1479 | 5122 | 0 | 14 | 0 | 14 | 14 | 15 | 0 | 0 | 0 | 0 | 4411 | 41 |
| SRR7690779 | 0 | 1099 | 40 | 158 | 176 | 0 | 17 | 4175 | 10452 | 7867 | 4621 | 5104 | 0 | 639 | 260 | 357 | 133 | 1434 | 273 | 12473 | 484 | 305 | 47 | 1040 | 1323 | 0 | 25 | 30 | 191 | 47 | 46 | 0 | 0 | 0 | 0 | 6849 | 21 |
| SRR7690780 | 0 | 749 | 22 | 413 | 777 | 0 | 16 | 4387 | 18304 | 14379 | 8451 | 12508 | 0 | 496 | 144 | 431 | 28 | 2647 | 54 | 3486 | 824 | 1267 | 47 | 885 | 2318 | 0 | 20 | 22 | 24 | 14 | 13 | 0 | 0 | 0 | 0 | 1588 | 37 |
| SRR7690781 | 0 | 417 | 37 | 367 | 508 | 0 | 20 | 6228 | 22048 | 23060 | 1373 | 2017 | 0 | 1394 | 364 | 475 | 27 | 2032 | 30 | 2845 | 966 | 1048 | 64 | 1545 | 2719 | 0 | 12 | 11 | 22 | 12 | 18 | 29 | 23 | 13 | 0 | 1479 | 75 |
| SRR7690783 | 0 | 447 | 277 | 41 | 185 | 25 | 15 | 2795 | 7722 | 20196 | 883 | 9916 | 11 | 1424 | 338 | 668 | 0 | 493 | 54 | 3092 | 417 | 602 | 159 | 751 | 518 | 0 | 44 | 36 | 142 | 341 | 589 | 282 | 100 | 80 | 0 | 3665 | 34 |
| SRR7690788 | 0 | 76 | 28 | 141 | 188 | 0 | 22 | 4873 | 15324 | 57753 | 389 | 3152 | 0 | 2071 | 195 | 575 | 0 | 939 | 56 | 1211 | 698 | 792 | 199 | 1438 | 810 | 0 | 19 | 17 | 12 | 36 | 39 | 12 | 63 | 48 | 0 | 1542 | 28 |
| SRR7690789 | 0 | 13 | 16 | 61 | 272 | 23 | 10 | 1022 | 3842 | 21292 | 259 | 495 | 13 | 1552 | 68 | 149 | 10 | 725 | 46 | 456 | 383 | 636 | 162 | 917 | 641 | 0 | 31 | 0 | 10 | 14 | 41 | 10 | 151 | 149 | 13 | 521 | 23 |
| SRR7690790 | 0 | 1582 | 237 | 530 | 1095 | 0 | 0 | 3819 | 21683 | 14587 | 1131 | 2291 | 0 | 1065 | 363 | 1783 | 10 | 2552 | 39 | 2167 | 812 | 1308 | 45 | 816 | 4459 | 0 | 0 | 0 | 56 | 47 | 65 | 0 | 0 | 0 | 0 | 2553 | 64 |
| SRR7690791 | 0 | 2475 | 153 | 183 | 663 | 0 | 19 | 4438 | 15779 | 17723 | 3466 | 6071 | 0 | 756 | 203 | 1162 | 13 | 2141 | 36 | 2039 | 1370 | 2001 | 22 | 211 | 1071 | 0 | 0 | 24 | 84 | 42 | 60 | 12 | 0 | 0 | 0 | 1090 | 30 |
| SRR7690792 | 12 | 1394 | 93 | 305 | 964 | 13 | 37 | 2234 | 7340 | 13375 | 694 | 2969 | 0 | 950 | 72 | 593 | 16 | 1155 | 25 | 1009 | 1025 | 821 | 40 | 613 | 4135 | 0 | 21 | 19 | 133 | 64 | 47 | 0 | 0 | 0 | 0 | 707 | 144 |
| SRR7690793 | 0 | 5032 | 100 | 111 | 119 | 0 | 25 | 5844 | 13277 | 10247 | 1121 | 6724 | 0 | 608 | 165 | 467 | 42 | 1742 | 58 | 5384 | 755 | 1552 | 27 | 1050 | 4045 | 0 | 20 | 0 | 34 | 31 | 34 | 18 | 0 | 0 | 0 | 3305 | 45 |
| SRR7690794 | 0 | 924 | 38 | 179 | 22 | 0 | 22 | 5128 | 9943 | 5327 | 6136 | 9539 | 0 | 673 | 16 | 88 | 10 | 1823 | 20 | 4259 | 314 | 1860 | 0 | 622 | 1764 | 0 | 11 | 0 | 16 | 11 | 11 | 0 | 0 | 0 | 0 | 3687 | 27 |
| SRR7690795 | 0 | 663 | 83 | 180 | 52 | 0 | 15 | 3187 | 7841 | 6494 | 2788 | 5021 | 0 | 786 | 437 | 1605 | 13 | 2005 | 22 | 4974 | 482 | 2406 | 13 | 595 | 1877 | 0 | 16 | 0 | 12 | 19 | 14 | 0 | 0 | 0 | 0 | 5009 | 23 |
| SRR7690796 | 0 | 2348 | 133 | 374 | 532 | 0 | 24 | 5297 | 20574 | 22186 | 641 | 4295 | 0 | 861 | 522 | 596 | 24 | 2402 | 27 | 3446 | 1283 | 1166 | 51 | 1037 | 3092 | 0 | 17 | 14 | 95 | 62 | 65 | 17 | 11 | 0 | 0 | 2077 | 44 |
| SRR7690797 | 0 | 11 | 29 | 15 | 89 | 30 | 11 | 3509 | 9578 | 19129 | 496 | 9011 | 11 | 1979 | 1689 | 2784 | 0 | 533 | 41 | 2889 | 252 | 658 | 145 | 589 | 456 | 0 | 52 | 70 | 15 | 13 | 29 | 299 | 258 | 196 | 15 | 4697 | 23 |
| SRR7690798 | 0 | 13 | 11 | 98 | 237 | 50 | 36 | 1997 | 4465 | 18225 | 4002 | 6424 | 33 | 2126 | 197 | 342 | 0 | 607 | 39 | 1990 | 678 | 735 | 132 | 698 | 482 | 0 | 173 | 33 | 159 | 16 | 31 | 287 | 137 | 106 | 0 | 2635 | 25 |
| SRR7690800 | 10 | 12 | 11 | 51 | 145 | 65 | 61 | 4668 | 12774 | 34162 | 580 | 11470 | 27 | 1865 | 43 | 108 | 0 | 731 | 35 | 3482 | 586 | 612 | 139 | 671 | 382 | 0 | 73 | 37 | 95 | 19 | 30 | 14 | 107 | 75 | 0 | 6306 | 20 |
| SRR7690801 | 0 | 732 | 233 | 110 | 430 | 25 | 13 | 2314 | 6631 | 20636 | 434 | 5539 | 13 | 1373 | 23 | 82 | 17 | 606 | 36 | 269 | 480 | 475 | 121 | 695 | 690 | 0 | 55 | 89 | 562 | 922 | 1121 | 190 | 110 | 85 | 0 | 654 | 49 |
| SRR7690802 | 15 | 23 | 76 | 41 | 175 | 24 | 50 | 4294 | 10260 | 14965 | 596 | 20300 | 22 | 865 | 44 | 44 | 11 | 587 | 44 | 2096 | 560 | 676 | 118 | 511 | 414 | 0 | 74 | 54 | 155 | 15 | 13 | 120 | 41 | 26 | 0 | 1964 | 17 |
| SRR7690803 | 17 | 620 | 314 | 50 | 192 | 41 | 34 | 4683 | 13389 | 34795 | 461 | 9747 | 17 | 1794 | 190 | 471 | 0 | 879 | 37 | 2546 | 688 | 710 | 115 | 535 | 485 | 0 | 73 | 49 | 171 | 282 | 321 | 160 | 88 | 64 | 0 | 3080 | 19 |
| SRR7819267 | 0 | 31 | 0 | 33 | 799 | 67 | 0 | 0 | 66 | 0 | 77 | 16673 | 10 | 0 | 0 | 0 | 77 | 225 | 80 | 66 | 21 | 68 | 47 | 54 | 495 | 1251 | 426 | 0 | 15 | 0 | 0 | 0 | 0 | 0 | 0 | 0 | 0 |
| SRR7819269 | 0 | 29 | 0 | 31 | 1404 | 29 | 57 | 118 | 0 | 118 | 29 | 2831 | 29 | 59 | 0 | 0 | 57 | 533 | 120 | 59 | 43 | 120 | 17 | 110 | 3458 | 1581 | 940 | 0 | 0 | 0 | 0 | 0 | 0 | 0 | 0 | 236 | 0 |
| SRR7819272 | 0 | 59 | 0 | 30 | 793 | 0 | 0 | 124 | 41 | 289 | 1695 | 3266 | 0 | 413 | 0 | 0 | 131 | 412 | 205 | 496 | 24 | 73 | 25 | 74 | 1449 | 2102 | 288 | 0 | 13 | 0 | 0 | 0 | 0 | 0 | 0 | 331 | 0 |
| SRR7851651 | 0 | 2608 | 0 | 11 | 238 | 39 | 128 | 1397 | 184 | 265 | 53081 | 24280 | 15 | 81 | 0 | 0 | 614 | 3353 | 50 | 11 | 1347 | 1484 | 0 | 2486 | 2284 | 0 | 52 | 0 | 15 | 0 | 0 | 0 | 0 | 0 | 0 | 38 | 0 |
| SRR7851652 | 0 | 479 | 0 | 0 | 208 | 18 | 52 | 1873 | 172 | 292 | 10734 | 23192 | 17 | 19 | 0 | 0 | 525 | 3231 | 47 | 0 | 1348 | 1564 | 0 | 2522 | 2571 | 0 | 27 | 0 | 0 | 0 | 0 | 0 | 0 | 0 | 0 | 23 | 0 |
| SRR7851653 | 0 | 395 | 0 | 16 | 184 | 15 | 46 | 534 | 83 | 143 | 11366 | 27261 | 10 | 23 | 0 | 0 | 824 | 3232 | 49 | 18 | 1685 | 1010 | 0 | 1830 | 1863 | 0 | 39 | 0 | 0 | 0 | 0 | 0 | 0 | 0 | 0 | 46 | 0 |
| SRR7851655 | 0 | 1526 | 0 | 0 | 198 | 50 | 164 | 1385 | 226 | 283 | 9497 | 24051 | 16 | 42 | 0 | 0 | 1128 | 3206 | 46 | 0 | 1277 | 1415 | 0 | 2494 | 2464 | 0 | 50 | 0 | 0 | 0 | 0 | 0 | 0 | 0 | 0 | 16 | 0 |
| SRR7851656 | 0 | 609 | 0 | 0 | 206 | 27 | 32 | 783 | 131 | 444 | 70414 | 34737 | 11 | 59 | 0 | 0 | 246 | 3345 | 49 | 13 | 1288 | 1310 | 0 | 2172 | 2102 | 0 | 13 | 11 | 12 | 0 | 0 | 0 | 0 | 0 | 0 | 20 | 0 |
| SRR7901057 | 0 | 187 | 24 | 0 | 16 | 0 | 0 | 129 | 28 | 110 | 11422 | 8461 | 0 | 2817 | 0 | 0 | 11 | 332 | 1641 | 1496 | 12 | 219 | 0 | 1012 | 26 | 0 | 30 | 0 | 22 | 19 | 125 | 75 | 63 | 183 | 27 | 2315 | 35 |
| SRR7901058 | 0 | 834 | 20 | 0 | 0 | 0 | 0 | 394 | 33 | 126 | 17902 | 14720 | 0 | 2453 | 0 | 0 | 20 | 426 | 1479 | 1295 | 0 | 283 | 0 | 819 | 25 | 0 | 375 | 0 | 35 | 55 | 275 | 177 | 143 | 368 | 17 | 1947 | 25 |
| SRR7901059 | 0 | 246 | 19 | 0 | 14 | 0 | 0 | 103 | 61 | 94 | 11472 | 11297 | 0 | 2442 | 0 | 15 | 12 | 362 | 1331 | 1336 | 15 | 214 | 0 | 1185 | 52 | 0 | 16 | 0 | 15 | 18 | 102 | 53 | 43 | 98 | 12 | 1833 | 34 |
| SRR7903701 | 0 | 904 | 741 | 12 | 566 | 0 | 0 | 892 | 2042 | 18766 | 41229 | 49513 | 0 | 1021 | 0 | 0 | 0 | 180 | 300 | 6064 | 2455 | 538 | 0 | 206 | 733 | 19 | 80 | 136 | 555 | 977 | 1892 | 479 | 533 | 719 | 244 | 3578 | 520 |
| SRR7903702 | 0 | 787 | 784 | 15 | 529 | 0 | 0 | 793 | 1851 | 27211 | 37118 | 51357 | 0 | 1619 | 0 | 0 | 0 | 148 | 264 | 6441 | 1936 | 522 | 0 | 200 | 786 | 20 | 21 | 152 | 576 | 1122 | 1726 | 414 | 391 | 513 | 264 | 3293 | 496 |
| SRR7903703 | 0 | 939 | 668 | 14 | 617 | 0 | 0 | 958 | 2143 | 21757 | 39428 | 47920 | 0 | 1237 | 0 | 0 | 0 | 172 | 311 | 5536 | 2448 | 539 | 0 | 217 | 781 | 21 | 21 | 139 | 538 | 1007 | 1649 | 371 | 347 | 461 | 215 | 3019 | 490 |
| SRR7903704 | 0 | 2217 | 130 | 0 | 106 | 0 | 0 | 63 | 122 | 479 | 5195 | 4391 | 0 | 1149 | 0 | 0 | 0 | 116 | 908 | 1972 | 189 | 311 | 0 | 59 | 50 | 31 | 15 | 16 | 50 | 145 | 238 | 107 | 81 | 156 | 179 | 1084 | 440 |
| SRR7903705 | 0 | 7083 | 134 | 0 | 73 | 0 | 0 | 88 | 137 | 642 | 9762 | 3122 | 0 | 1452 | 0 | 0 | 0 | 109 | 1202 | 3115 | 175 | 366 | 0 | 50 | 46 | 32 | 0 | 0 | 55 | 176 | 253 | 75 | 79 | 151 | 153 | 1553 | 309 |
| SRR7903706 | 0 | 8352 | 97 | 0 | 125 | 0 | 0 | 68 | 139 | 429 | 4666 | 2774 | 0 | 863 | 0 | 0 | 0 | 144 | 1561 | 1900 | 242 | 300 | 0 | 44 | 71 | 18 | 0 | 16 | 59 | 147 | 195 | 67 | 69 | 170 | 191 | 1007 | 399 |
| SRR8063607 | 0 | 3280 | 237 | 12 | 82 | 0 | 0 | 398 | 1398 | 3254 | 12377 | 24476 | 0 | 482 | 0 | 24 | 0 | 81 | 1670 | 253 | 897 | 1822 | 98 | 8513 | 4761 | 0 | 0 | 423 | 1137 | 6425 | 3692 | 590 | 423 | 204 | 0 | 1928 | 242 |
| SRR8063608 | 0 | 334 | 25 | 0 | 95 | 0 | 0 | 294 | 1176 | 3470 | 374 | 20144 | 0 | 294 | 0 | 29 | 0 | 79 | 1687 | 765 | 869 | 1891 | 77 | 8528 | 4896 | 0 | 0 | 45 | 132 | 689 | 412 | 83 | 45 | 26 | 0 | 7175 | 237 |
| SRR8063609 | 0 | 354 | 34 | 0 | 80 | 0 | 0 | 344 | 1757 | 3808 | 12709 | 23470 | 0 | 611 | 0 | 0 | 0 | 95 | 1704 | 408 | 882 | 1863 | 89 | 8968 | 5067 | 0 | 0 | 52 | 125 | 614 | 410 | 94 | 71 | 46 | 0 | 3362 | 217 |
| SRR8063610 | 0 | 132 | 23 | 0 | 51 | 0 | 0 | 111 | 382 | 1211 | 490 | 1598 | 40 | 134 | 0 | 19 | 0 | 2438 | 1348 | 25973 | 969 | 1564 | 131 | 8167 | 4234 | 0 | 29 | 55 | 33 | 69 | 88 | 20 | 27 | 13 | 0 | 318282 | 216 |
| SRR8063611 | 0 | 146 | 40 | 0 | 71 | 0 | 13 | 93 | 346 | 1482 | 925 | 1673 | 39 | 137 | 0 | 16 | 11 | 2547 | 1412 | 24920 | 1020 | 1621 | 112 | 7832 | 4037 | 0 | 24 | 12 | 28 | 66 | 79 | 16 | 18 | 11 | 0 | 317694 | 218 |
| SRR8063612 | 0 | 125 | 44 | 13 | 75 | 0 | 19 | 115 | 304 | 1602 | 1005 | 1494 | 44 | 134 | 0 | 26 | 17 | 2368 | 1401 | 25434 | 1069 | 1612 | 136 | 7819 | 4185 | 0 | 24 | 40 | 26 | 52 | 70 | 14 | 18 | 11 | 0 | 323782 | 274 |
| SRR8063613 | 0 | 84 | 0 | 0 | 0 | 0 | 17 | 192 | 256 | 703 | 639 | 1535 | 17 | 1215 | 0 | 64 | 50 | 1621 | 2502 | 35363 | 98 | 1765 | 11 | 1689 | 354 | 0 | 285 | 17 | 0 | 117 | 168 | 117 | 84 | 101 | 0 | 278169 | 119 |
| SRR8063614 | 0 | 12 | 12 | 0 | 0 | 0 | 34 | 88 | 141 | 651 | 106 | 1900 | 0 | 862 | 158 | 1091 | 0 | 1780 | 2361 | 14320 | 161 | 2395 | 32 | 5038 | 1755 | 0 | 15 | 12 | 0 | 40 | 22 | 12 | 0 | 0 | 0 | 130392 | 60 |
| SRR8063615 | 0 | 11 | 0 | 0 | 0 | 0 | 25 | 114 | 57 | 739 | 49 | 2331 | 0 | 796 | 0 | 0 | 25 | 1598 | 2292 | 20750 | 159 | 2539 | 39 | 4814 | 1744 | 0 | 18 | 25 | 0 | 49 | 57 | 0 | 0 | 0 | 0 | 159864 | 74 |
| SRR8074870 | 0 | 14 | 0 | 0 | 12 | 0 | 88 | 90 | 12 | 12 | 1625 | 1016 | 88 | 198 | 0 | 0 | 135 | 857 | 1252 | 71 | 324 | 9322 | 89 | 45067 | 9427 | 47 | 0 | 46 | 116 | 0 | 0 | 0 | 0 | 0 | 0 | 60 | 154 |
| SRR8074871 | 0 | 18 | 0 | 0 | 14 | 0 | 81 | 112 | 14 | 0 | 1678 | 1084 | 65 | 195 | 0 | 0 | 136 | 847 | 1234 | 74 | 318 | 9412 | 87 | 44447 | 9273 | 42 | 0 | 47 | 152 | 0 | 11 | 0 | 0 | 0 | 0 | 55 | 157 |

### Supp Table S8.4.Columns_DI_to_ES

| | | | | | | | | | | | | | | | | | | | | | | | | | | | | | | | | | | | | | |
|---|---|---|---|---|---|---|---|---|---|---|---|---|---|---|---|---|---|---|---|---|---|---|---|---|---|---|---|---|---|---|---|---|---|---|---|---|---|
| SRR8074873 | 0 | 0 | 0 | 0 | 13 | 0 | 44 | 306 | 75 | 25 | 468 | 191 | 54 | 285 | 0 | 0 | 65 | 1101 | 1228 | 181 | 250 | 12 | 67 | 60083 | 11797 | 30 | 0 | 11 | 30 | 0 | 29 | 0 | 0 | 0 | 0 | 97 | 255 |
| SRR8074874 | 0 | 0 | 0 | 0 | 13 | 0 | 15 | 329 | 93 | 76 | 520 | 196 | 38 | 984 | 0 | 0 | 30 | 900 | 1107 | 297 | 361 | 11 | 67 | 52780 | 11891 | 18 | 0 | 19 | 105 | 0 | 0 | 0 | 0 | 0 | 0 | 224 | 347 |
| SRR8074875 | 0 | 0 | 0 | 0 | 14 | 0 | 17 | 323 | 106 | 75 | 496 | 207 | 21 | 906 | 0 | 0 | 35 | 903 | 1140 | 245 | 373 | 9344 | 63 | 51592 | 11687 | 18 | 0 | 15 | 94 | 0 | 10 | 0 | 0 | 0 | 0 | 251 | 332 |
| SRR8094756 | 33 | 52 | 13 | 99 | 737 | 0 | 27 | 3415 | 7814 | 63574 | 26726 | 29866 | 30 | 3690 | 0 | 0 | 22 | 2594 | 67 | 1337 | 434 | 659 | 48 | 2216 | 326 | 0 | 50 | 37 | 72 | 14 | 59 | 28 | 21 | 21 | 0 | 1427 | 0 |
| SRR8094757 | 0 | 58 | 68 | 132 | 993 | 0 | 47 | 9533 | 2323 | 12646 | 556 | 831 | 0 | 2171 | 0 | 0 | 42 | 1411 | 84 | 1954 | 696 | 1195 | 32 | 2194 | 239 | 16 | 23 | 0 | 17 | 195 | 982 | 255 | 301 | 281 | 40 | 3752 | 0 |
| SRR8094758 | 68 | 41 | 16 | 22 | 298 | 44 | 24 | 5301 | 5489 | 31296 | 2433 | 7795 | 27 | 5176 | 0 | 0 | 41 | 864 | 50 | 549 | 260 | 306 | 13 | 1439 | 113 | 0 | 129 | 51 | 26 | 26 | 182 | 84 | 103 | 84 | 12 | 1807 | 0 |
| SRR8094759 | 0 | 28 | 32 | 60 | 159 | 15 | 14 | 404 | 2324 | 8583 | 120 | 291 | 0 | 401 | 294 | 390 | 0 | 1733 | 11 | 1901 | 5876 | 2007 | 398 | 3262 | 2192 | 0 | 16 | 0 | 16 | 19 | 59 | 29 | 24 | 16 | 0 | 3476 | 0 |
| SRR8094760 | 0 | 18 | 21 | 24 | 104 | 19 | 30 | 840 | 5543 | 9335 | 476 | 8724 | 11 | 240 | 10 | 50 | 22 | 892 | 0 | 11366 | 2309 | 800 | 360 | 1906 | 1384 | 0 | 36 | 0 | 69 | 30 | 82 | 23 | 31 | 20 | 0 | 11926 | 13 |
| SRR8094761 | 0 | 29 | 31 | 39 | 155 | 0 | 24 | 393 | 1667 | 2125 | 4472 | 21848 | 0 | 69 | 53 | 50 | 21 | 997 | 0 | 8877 | 5051 | 1258 | 272 | 2062 | 1305 | 0 | 16 | 36 | 22 | 39 | 118 | 26 | 39 | 32 | 0 | 40579 | 0 |
| SRR8094762 | 0 | 84 | 826 | 37 | 120 | 0 | 38 | 365 | 1922 | 12144 | 1228 | 10768 | 0 | 361 | 473 | 817 | 18 | 1030 | 14 | 7153 | 3499 | 1274 | 255 | 2051 | 1501 | 0 | 840 | 18 | 613 | 2342 | 5110 | 1061 | 1755 | 1464 | 50 | 38338 | 25 |
| SRR8094763 | 0 | 336 | 305 | 145 | 1563 | 0 | 0 | 705 | 1948 | 21102 | 1075 | 2684 | 0 | 841 | 971 | 846 | 14 | 297 | 15 | 822 | 1266 | 1123 | 0 | 34 | 1973 | 0 | 161 | 25 | 553 | 632 | 1252 | 180 | 262 | 261 | 44 | 2084 | 11 |
| SRR8094764 | 0 | 60 | 55 | 117 | 1409 | 0 | 15 | 2066 | 973 | 21148 | 1406 | 3193 | 28 | 182 | 173 | 164 | 35 | 688 | 58 | 106 | 2806 | 1904 | 0 | 53 | 2136 | 0 | 38 | 51 | 55 | 93 | 177 | 30 | 41 | 29 | 11 | 272 | 129 |
| SRR8094765 | 0 | 21 | 14 | 234 | 2960 | 0 | 35 | 2635 | 1627 | 16440 | 628 | 2782 | 16 | 269 | 433 | 610 | 19 | 466 | 40 | 339 | 2076 | 1171 | 0 | 40 | 2426 | 0 | 40 | 23 | 20 | 41 | 55 | 179 | 16 | 12 | 33 | 610 | 90 |
| SRR8094766 | 0 | 26 | 37 | 218 | 1906 | 0 | 25 | 1275 | 1345 | 26806 | 1709 | 3614 | 0 | 897 | 41 | 62 | 34 | 883 | 57 | 226 | 2407 | 2738 | 0 | 127 | 1905 | 0 | 219 | 23 | 45 | 98 | 155 | 15 | 23 | 21 | 0 | 579 | 0 |
| SRR8094767 | 23 | 13 | 13 | 182 | 2147 | 0 | 29 | 18702 | 32978 | 11757 | 817 | 8759 | 17 | 221 | 171 | 207 | 17 | 1313 | 52 | 128 | 2108 | 1621 | 0 | 31 | 1882 | 0 | 23 | 87 | 20 | 26 | 36 | 413 | 506 | 465 | 23 | 307 | 181 |
| SRR8094768 | 0 | 62 | 127 | 129 | 1186 | 0 | 0 | 1563 | 880 | 17312 | 1747 | 3989 | 11 | 304 | 353 | 353 | 0 | 919 | 51 | 95 | 3083 | 2496 | 0 | 77 | 3176 | 0 | 12 | 70 | 61 | 101 | 175 | 35 | 53 | 43 | 43 | 370 | 82 |
| SRR8094769 | 0 | 493 | 1327 | 276 | 1644 | 0 | 0 | 770 | 604 | 12383 | 1666 | 11469 | 0 | 382 | 337 | 435 | 0 | 742 | 66 | 56 | 2606 | 1516 | 0 | 85 | 3601 | 0 | 105 | 118 | 698 | 3021 | 4703 | 554 | 597 | 570 | 12 | 219 | 37 |
| SRR8094770 | 0 | 371 | 91 | 20 | 462 | 0 | 37 | 331 | 48 | 1265 | 2009 | 5488 | 41 | 93 | 0 | 0 | 34 | 219 | 92 | 152 | 1758 | 857 | 0 | 34 | 2465 | 0 | 1302 | 2725 | 1077 | 2212 | 4973 | 2143 | 2747 | 2500 | 14 | 336 | 1289 |
| SRR8137033 | 0 | 36 | 101 | 0 | 16 | 0 | 0 | 213 | 53 | 2233 | 279 | 55027 | 0 | 478 | 0 | 160 | 0 | 160 | 694 | 213 | 523 | 1021 | 56 | 5151 | 2322 | 0 | 20 | 25 | 25 | 35 | 180 | 51 | 24 | 11 | 61 | 585 | 563 |
| SRR8137034 | 0 | 39 | 53 | 0 | 0 | 0 | 0 | 247 | 288 | 5719 | 619 | 29621 | 0 | 1604 | 41 | 391 | 0 | 199 | 637 | 350 | 273 | 1572 | 16 | 1946 | 2429 | 0 | 0 | 0 | 106 | 68 | 198 | 83 | 79 | 88 | 155 | 535 | 966 |
| SRR8137035 | 0 | 17 | 12 | 0 | 0 | 0 | 0 | 87 | 216 | 4932 | 361 | 54424 | 0 | 1817 | 43 | 87 | 0 | 18 | 353 | 2163 | 26 | 139 | 18 | 1119 | 315 | 0 | 0 | 0 | 42 | 42 | 198 | 102 | 51 | 21 | 103 | 2509 | 250 |
| SRR8137036 | 10 | 262 | 80 | 0 | 0 | 10 | 0 | 0 | 214 | 2388 | 675 | 96696 | 0 | 642 | 0 | 36 | 0 | 39 | 548 | 428 | 130 | 415 | 24 | 2655 | 1082 | 0 | 0 | 121 | 161 | 171 | 1257 | 473 | 151 | 70 | 221 | 178 | 394 |
| SRR8137037 | 0 | 0 | 22 | 446 | 590 | 0 | 0 | 2891 | 13051 | 42789 | 5700 | 9252 | 0 | 4543 | 0 | 83 | 0 | 2386 | 336 | 1404 | 336 | 2441 | 267 | 7925 | 878 | 27 | 0 | 0 | 91 | 71 | 205 | 189 | 355 | 164 | 71 | 5287 | 69 |
| SRR8203922 | 0 | 0 | 0 | 0 | 0 | 0 | 0 | 2491 | 104 | 0 | 487 | 225 | 0 | 112 | 0 | 0 | 0 | 86 | 102 | 0 | 1953 | 1764 | 0 | 10425 | 5922 | 0 | 0 | 0 | 0 | 0 | 0 | 0 | 0 | 11 | 0 | 0 | 26 |
| SRR8203923 | 0 | 0 | 0 | 0 | 0 | 0 | 0 | 2530 | 110 | 0 | 513 | 364 | 0 | 85 | 0 | 0 | 0 | 94 | 124 | 0 | 2139 | 1607 | 0 | 10516 | 5707 | 0 | 0 | 0 | 0 | 0 | 0 | 0 | 0 | 11 | 0 | 0 | 23 |
| SRR8203924 | 0 | 0 | 0 | 0 | 0 | 0 | 0 | 2536 | 124 | 0 | 3887 | 232 | 0 | 141 | 0 | 0 | 0 | 79 | 115 | 0 | 2159 | 1608 | 16 | 10607 | 5902 | 0 | 0 | 0 | 0 | 0 | 0 | 0 | 0 | 17 | 0 | 33 | 30 |
| SRR8203925 | 0 | 0 | 0 | 0 | 0 | 0 | 0 | 2568 | 118 | 0 | 4224 | 296 | 0 | 51 | 0 | 0 | 0 | 114 | 151 | 0 | 2081 | 1610 | 23 | 10159 | 5324 | 0 | 0 | 0 | 0 | 0 | 0 | 0 | 0 | 11 | 0 | 0 | 44 |
| SRR8203926 | 0 | 10 | 0 | 0 | 0 | 0 | 0 | 1673 | 111 | 27 | 798 | 408 | 0 | 133 | 0 | 13 | 0 | 120 | 213 | 0 | 2471 | 1606 | 32 | 9941 | 7588 | 0 | 0 | 0 | 0 | 0 | 0 | 0 | 0 | 0 | 0 | 0 | 24 |
| SRR8203927 | 0 | 22 | 0 | 0 | 0 | 0 | 0 | 1620 | 164 | 42 | 3704 | 352 | 0 | 99 | 0 | 19 | 0 | 106 | 193 | 0 | 2522 | 1528 | 11 | 9819 | 7197 | 0 | 0 | 0 | 0 | 0 | 0 | 0 | 0 | 19 | 0 | 28 | 22 |
| SRR8203928 | 0 | 11 | 0 | 0 | 0 | 0 | 0 | 1702 | 178 | 32 | 3618 | 338 | 0 | 119 | 27 | 0 | 0 | 79 | 241 | 0 | 2619 | 1566 | 14 | 9914 | 7319 | 0 | 0 | 0 | 0 | 0 | 0 | 0 | 0 | 18 | 0 | 27 | 30 |
| SRR8203929 | 0 | 19 | 0 | 0 | 0 | 0 | 0 | 1822 | 192 | 37 | 3916 | 350 | 0 | 154 | 0 | 0 | 0 | 87 | 207 | 0 | 2563 | 1521 | 14 | 9647 | 7132 | 0 | 0 | 0 | 0 | 0 | 0 | 0 | 0 | 13 | 0 | 0 | 14 |
| SRR8203930 | 0 | 24 | 0 | 0 | 0 | 0 | 0 | 1528 | 90 | 19 | 3508 | 404 | 0 | 120 | 0 | 0 | 0 | 88 | 206 | 0 | 2534 | 964 | 15 | 15217 | 11055 | 0 | 0 | 0 | 0 | 0 | 0 | 0 | 0 | 11 | 0 | 0 | 0 |
| SRR8203931 | 0 | 27 | 0 | 0 | 0 | 0 | 0 | 1594 | 97 | 12 | 867 | 350 | 0 | 113 | 0 | 0 | 0 | 115 | 146 | 0 | 2506 | 946 | 16 | 14902 | 10492 | 0 | 0 | 0 | 0 | 0 | 0 | 0 | 0 | 17 | 0 | 19 | 20 |
| SRR8203932 | 0 | 27 | 0 | 0 | 0 | 0 | 0 | 1689 | 129 | 15 | 945 | 338 | 0 | 129 | 0 | 0 | 0 | 105 | 163 | 0 | 2470 | 948 | 17 | 14939 | 11112 | 0 | 0 | 0 | 0 | 0 | 0 | 0 | 0 | 0 | 0 | 0 | 16 |
| SRR8203933 | 0 | 36 | 0 | 0 | 0 | 0 | 0 | 1588 | 106 | 0 | 3744 | 335 | 0 | 122 | 0 | 0 | 0 | 76 | 170 | 0 | 2515 | 1035 | 16 | 14816 | 10200 | 0 | 0 | 0 | 0 | 0 | 0 | 0 | 0 | 0 | 0 | 32 | 16 |
| SRR8203934 | 0 | 19 | 0 | 0 | 0 | 0 | 0 | 2032 | 151 | 27 | 2923 | 285 | 0 | 116 | 0 | 0 | 0 | 127 | 242 | 0 | 1855 | 1581 | 14 | 14297 | 12787 | 0 | 0 | 0 | 0 | 0 | 0 | 0 | 0 | 0 | 0 | 0 | 23 |
| SRR8203935 | 0 | 20 | 0 | 0 | 0 | 0 | 0 | 2030 | 122 | 20 | 2950 | 295 | 0 | 131 | 0 | 0 | 0 | 147 | 223 | 0 | 1638 | 1463 | 15 | 14503 | 12031 | 0 | 0 | 0 | 0 | 0 | 0 | 0 | 0 | 0 | 0 | 0 | 25 |
| SRR8203936 | 0 | 19 | 0 | 0 | 0 | 0 | 0 | 2167 | 141 | 24 | 2643 | 334 | 0 | 115 | 0 | 0 | 0 | 154 | 243 | 0 | 1671 | 1462 | 13 | 14274 | 12887 | 0 | 0 | 0 | 0 | 0 | 0 | 0 | 0 | 0 | 0 | 0 | 22 |
| SRR8203937 | 0 | 33 | 0 | 0 | 0 | 0 | 0 | 1951 | 142 | 25 | 2632 | 325 | 0 | 104 | 0 | 0 | 0 | 160 | 249 | 0 | 1656 | 1515 | 11 | 14371 | 11612 | 0 | 0 | 0 | 0 | 0 | 0 | 0 | 0 | 0 | 0 | 0 | 18 |
| SRR8203938 | 0 | 28 | 0 | 0 | 0 | 0 | 0 | 2423 | 142 | 35 | 3283 | 381 | 0 | 160 | 0 | 0 | 0 | 132 | 261 | 0 | 2699 | 1816 | 21 | 15584 | 14727 | 0 | 0 | 0 | 0 | 0 | 0 | 0 | 0 | 0 | 0 | 0 | 21 |
| SRR8203939 | 0 | 31 | 0 | 0 | 0 | 0 | 0 | 2577 | 152 | 47 | 3429 | 390 | 0 | 146 | 0 | 0 | 0 | 144 | 263 | 0 | 2606 | 1752 | 21 | 15851 | 13986 | 0 | 0 | 0 | 0 | 0 | 0 | 0 | 0 | 0 | 0 | 0 | 22 |
| SRR8203940 | 0 | 32 | 0 | 0 | 0 | 0 | 0 | 2525 | 136 | 32 | 1946 | 353 | 0 | 144 | 0 | 0 | 0 | 156 | 242 | 0 | 2654 | 1892 | 23 | 15628 | 14775 | 0 | 0 | 0 | 0 | 0 | 0 | 0 | 0 | 0 | 0 | 0 | 16 |
| SRR8203941 | 0 | 39 | 0 | 0 | 0 | 0 | 0 | 2638 | 183 | 65 | 1828 | 343 | 0 | 165 | 0 | 0 | 0 | 138 | 264 | 0 | 2664 | 1719 | 33 | 15780 | 13834 | 0 | 0 | 0 | 0 | 0 | 0 | 0 | 0 | 0 | 0 | 10 | 29 |
| SRR8203942 | 0 | 10 | 0 | 0 | 0 | 0 | 0 | 1760 | 212 | 53 | 2361 | 188 | 0 | 124 | 0 | 0 | 0 | 112 | 362 | 0 | 1112 | 1255 | 20 | 25406 | 5570 | 0 | 10 | 0 | 0 | 0 | 0 | 0 | 0 | 0 | 0 | 12 | 0 |
| SRR8203943 | 0 | 0 | 0 | 0 | 0 | 0 | 0 | 1718 | 218 | 31 | 2004 | 193 | 0 | 162 | 19 | 0 | 0 | 85 | 411 | 0 | 1190 | 1334 | 11 | 25885 | 5442 | 0 | 0 | 0 | 0 | 0 | 0 | 0 | 0 | 0 | 0 | 25 | 27 |
| SRR8203944 | 10 | 21 | 0 | 0 | 0 | 0 | 0 | 1658 | 175 | 30 | 336 | 157 | 0 | 121 | 0 | 0 | 0 | 94 | 366 | 0 | 1167 | 1146 | 10 | 25299 | 5575 | 0 | 16 | 0 | 0 | 0 | 0 | 0 | 0 | 0 | 0 | 0 | 0 |
| SRR8203945 | 0 | 0 | 0 | 0 | 0 | 0 | 0 | 1685 | 154 | 68 | 2314 | 241 | 0 | 105 | 0 | 0 | 0 | 74 | 318 | 0 | 1225 | 1262 | 11 | 26329 | 5246 | 0 | 21 | 0 | 0 | 11 | 0 | 0 | 0 | 0 | 0 | 25 | 0 |
| SRR8203946 | 0 | 21 | 0 | 0 | 0 | 0 | 0 | 1222 | 104 | 60 | 1034 | 403 | 0 | 185 | 0 | 0 | 0 | 114 | 355 | 0 | 2005 | 1384 | 0 | 27072 | 8912 | 0 | 21 | 0 | 0 | 0 | 0 | 0 | 0 | 0 | 0 | 40 | 17 |
| SRR8203947 | 0 | 19 | 0 | 0 | 0 | 0 | 0 | 1292 | 117 | 60 | 3419 | 340 | 0 | 173 | 0 | 0 | 0 | 137 | 367 | 0 | 1908 | 1376 | 15 | 27515 | 8849 | 0 | 12 | 0 | 0 | 0 | 0 | 0 | 0 | 0 | 0 | 35 | 13 |
| SRR8203948 | 0 | 22 | 0 | 0 | 0 | 0 | 0 | 1387 | 117 | 62 | 1013 | 285 | 0 | 199 | 10 | 0 | 0 | 99 | 370 | 0 | 1790 | 1222 | 13 | 27194 | 9056 | 0 | 22 | 0 | 0 | 0 | 0 | 0 | 0 | 0 | 0 | 31 | 11 |
| SRR8203949 | 0 | 33 | 0 | 0 | 0 | 0 | 0 | 1259 | 99 | 60 | 1010 | 370 | 0 | 212 | 0 | 0 | 0 | 84 | 465 | 0 | 1978 | 1262 | 0 | 27709 | 8722 | 0 | 15 | 0 | 0 | 0 | 0 | 0 | 0 | 0 | 0 | 35 | 20 |
| SRR8248787 | 0 | 270 | 110 | 239 | 302 | 0 | 0 | 239 | 722 | 13754 | 18263 | 22212 | 0 | 23735 | 0 | 0 | 0 | 840 | 240 | 852 | 71 | 501 | 41 | 737 | 79 | 92 | 0 | 318 | 242 | 338 | 1068 | 844 | 1354 | 1223 | 772 | 3198 | 0 |
| SRR8248788 | 0 | 440 | 218 | 183 | 298 | 0 | 0 | 296 | 541 | 11860 | 20533 | 25740 | 0 | 22123 | 0 | 0 | 23 | 798 | 233 | 1761 | 55 | 445 | 35 | 595 | 65 | 102 | 49 | 724 | 589 | 703 | 2464 | 2260 | 3897 | 3269 | 356 | 4907 | 0 |
| SRR8248789 | 0 | 277 | 153 | 179 | 261 | 0 | 0 | 234 | 507 | 9550 | 20248 | 24520 | 0 | 24529 | 0 | 0 | 14 | 746 | 243 | 1705 | 29 | 392 | 30 | 571 | 60 | 104 | 28 | 478 | 430 | 542 | 2136 | 1865 | 2976 | 2647 | 665 | 4980 | 0 |
| SRR8248790 | 0 | 67 | 34 | 129 | 168 | 0 | 0 | 264 | 449 | 10497 | 19066 | 23149 | 0 | 35448 | 32 | 11 | 0 | 516 | 196 | 1362 | 29 | 318 | 19 | 381 | 36 | 79 | 0 | 92 | 62 | 73 | 404 | 519 | 610 | 580 | 769 | 4841 | 0 |
| SRR8248791 | 0 | 33 | 101 | 171 | 230 | 0 | 0 | 393 | 696 | 9346 | 8127 | 31473 | 0 | 17739 | 0 | 0 | 19 | 651 | 271 | 1322 | 31 | 279 | 34 | 589 | 63 | 87 | 41 | 58 | 47 | 59 | 205 | 147 | 175 | 136 | 122 | 3189 | 0 |
| SRR8248792 | 0 | 413 | 69 | 85 | 139 | 25 | 0 | 336 | 578 | 9596 | 20922 | 29547 | 0 | 23918 | 0 | 0 | 44 | 364 | 304 | 1613 | 21 | 131 | 25 | 649 | 31 | 132 | 76 | 519 | 303 | 533 | 3954 | 3425 | 3096 | 2543 | 217 | 6453 | 0 |
| SRR8272058 | 0 | 0 | 0 | 0 | 11 | 32 | 0 | 0 | 0 | 0 | 424 | 78 | 0 | 0 | 0 | 0 | 28 | 33 | 24 | 0 | 76 | 326 | 154 | 87545 | 1166 | 65 | 0 | 14 | 0 | 27 | 0 | 0 | 0 | 0 | 0 | 14 | 79 |
| SRR8280405 | 0 | 0 | 0 | 0 | 0 | 0 | 0 | 146 | 0 | 1533 | 0 | 0 | 0 | 1168 | 0 | 0 | 0 | 0 | 0 | 0 | 0 | 19 | 17 | 33 | 193 | 46 | 44 | 0 | 0 | 0 | 0 | 0 | 0 | 0 | 0 | 511 | 0 |
| SRR8280408 | 0 | 0 | 0 | 0 | 0 | 0 | 0 | 184 | 0 | 1652 | 0 | 184 | 0 | 826 | 92 | 184 | 0 | 0 | 40 | 0 | 0 | 35 | 0 | 44 | 436 | 88 | 0 | 0 | 0 | 0 | 0 | 0 | 0 | 0 | 0 | 1010 | 0 |
| SRR8280414 | 22 | 0 | 0 | 0 | 0 | 0 | 0 | 439 | 0 | 731 | 219 | 0 | 0 | 1828 | 0 | 73 | 22 | 0 | 11 | 0 | 21 | 15 | 96 | 140 | 659 | 28 | 45 | 0 | 0 | 0 | 0 | 0 | 0 | 0 | 0 | 804 | 0 |
| SRR8280415 | 0 | 0 | 0 | 0 | 0 | 611 | 0 | 0 | 0 | 2485 | 0 | 0 | 0 | 2209 | 0 | 0 | 0 | 0 | 0 | 0 | 12 | 55 | 24 | 57 | 336 | 45 | 0 | 0 | 0 | 0 | 0 | 0 | 0 | 0 | 0 | 736 | 0 |
| SRR8280416 | 194 | 0 | 0 | 0 | 0 | 194 | 0 | 153 | 0 | 3069 | 0 | 153 | 0 | 1995 | 0 | 0 | 0 | 0 | 11 | 0 | 16 | 41 | 41 | 30 | 291 | 19 | 0 | 0 | 0 | 0 | 0 | 0 | 0 | 0 | 0 | 153 | 0 |
| SRR8280417 | 0 | 51 | 0 | 0 | 0 | 171 | 0 | 1169 | 146 | 828 | 244 | 0 | 0 | 1169 | 0 | 0 | 0 | 0 | 0 | 0 | 35 | 50 | 152 | 300 | 905 | 48 | 0 | 0 | 0 | 0 | 0 | 0 | 0 | 0 | 0 | 487 | 0 |
| SRR8280423 | 0 | 0 | 0 | 0 | 0 | 0 | 0 | 0 | 0 | 726 | 0 | 0 | 0 | 2418 | 0 | 0 | 0 | 0 | 12 | 0 | 0 | 0 | 0 | 0 | 103 | 26 | 0 | 0 | 0 | 0 | 0 | 0 | 0 | 0 | 0 | 1451 | 0 |
| SRR8280425 | 68 | 0 | 0 | 0 | 0 | 0 | 0 | 0 | 0 | 673 | 0 | 0 | 0 | 2244 | 0 | 0 | 0 | 0 | 0 | 0 | 0 | 22 | 11 | 26 | 118 | 19 | 68 | 0 | 0 | 0 | 0 | 0 | 0 | 0 | 0 | 224 | 0 |
| SRR8280432 | 0 | 0 | 0 | 0 | 0 | 0 | 0 | 801 | 0 | 1247 | 0 | 0 | 0 | 1781 | 0 | 178 | 0 | 0 | 0 | 89 | 0 | 0 | 0 | 22 | 151 | 27 | 0 | 0 | 0 | 0 | 0 | 0 | 0 | 0 | 0 | 801 | 0 |
| SRR8280440 | 0 | 0 | 0 | 0 | 0 | 0 | 0 | 0 | 0 | 932 | 0 | 0 | 0 | 414 | 0 | 0 | 0 | 0 | 105 | 0 | 20 | 0 | 32 | 44 | 332 | 15 | 34 | 0 | 0 | 0 | 0 | 0 | 0 | 0 | 0 | 1277 | 0 |
| SRR8280442 | 0 | 0 | 0 | 0 | 0 | 0 | 0 | 297 | 54 | 1000 | 0 | 0 | 0 | 487 | 108 | 135 | 0 | 0 | 0 | 27 | 0 | 20 | 24 | 50 | 354 | 31 | 0 | 0 | 0 | 0 | 0 | 0 | 0 | 0 | 0 | 595 | 0 |
| SRR8280452 | 0 | 0 | 0 | 0 | 0 | 0 | 0 | 0 | 0 | 364 | 0 | 0 | 0 | 1092 | 0 | 0 | 0 | 0 | 0 | 0 | 0 | 0 | 0 | 0 | 70 | 0 | 0 | 0 | 0 | 0 | 0 | 0 | 0 | 0 | 0 | 0 | 0 |
| SRR8280460 | 14 | 0 | 0 | 0 | 0 | 0 | 0 | 221 | 0 | 1635 | 0 | 0 | 0 | 884 | 0 | 88 | 0 | 0 | 0 | 44 | 0 | 13 | 30 | 60 | 153 | 28 | 0 | 0 | 0 | 0 | 0 | 0 | 0 | 0 | 14 | 795 | 0 |
| SRR8280461 | 0 | 0 | 0 | 0 | 0 | 0 | 0 | 0 | 0 | 325 | 0 | 163 | 0 | 325 | 0 | 163 | 0 | 0 | 13 | 0 | 0 | 0 | 0 | 15 | 53 | 18 | 0 | 0 | 0 | 0 | 0 | 0 | 0 | 0 | 0 | 0 | 0 |
| SRR8280463 | 0 | 0 | 0 | 0 | 0 | 0 | 0 | 167 | 83 | 917 | 0 | 0 | 0 | 208 | 0 | 0 | 0 | 0 | 0 | 0 | 0 | 0 | 0 | 0 | 72 | 19 | 0 | 0 | 0 | 0 | 0 | 0 | 0 | 0 | 32 | 0 | 0 |
| SRR8280464 | 0 | 0 | 0 | 0 | 0 | 0 | 0 | 106 | 0 | 1005 | 0 | 53 | 0 | 476 | 0 | 53 | 0 | 0 | 11 | 0 | 16 | 72 | 25 | 56 | 433 | 228 | 0 | 0 | 0 | 0 | 0 | 0 | 0 | 0 | 0 | 0 | 0 |
| SRR8280467 | 0 | 0 | 0 | 0 | 0 | 0 | 0 | 73 | 0 | 915 | 24 | 0 | 0 | 232 | 0 | 122 | 0 | 0 | 0 | 159 | 19 | 22 | 11 | 33 | 259 | 65 | 62 | 0 | 0 | 0 | 0 | 0 | 0 | 0 | 12 | 2660 | 0 |
| SRR8280468 | 0 | 0 | 0 | 0 | 0 | 0 | 0 | 0 | 0 | 1753 | 0 | 0 | 0 | 701 | 0 | 0 | 0 | 0 | 18 | 0 | 0 | 13 | 0 | 26 | 70 | 61 | 0 | 0 | 0 | 0 | 0 | 0 | 0 | 0 | 0 | 351 | 0 |
| SRR8280472 | 0 | 0 | 0 | 0 | 0 | 0 | 0 | 331 | 0 | 331 | 0 | 0 | 0 | 662 | 0 | 0 | 0 | 0 | 0 | 0 | 0 | 0 | 0 | 0 | 203 | 23 | 0 | 0 | 0 | 0 | 0 | 0 | 0 | 0 | 0 | 1324 | 0 |
| SRR8280473 | 0 | 0 | 0 | 0 | 0 | 96 | 0 | 0 | 0 | 995 | 398 | 0 | 0 | 597 | 0 | 199 | 0 | 0 | 0 | 199 | 11 | 31 | 0 | 0 | 157 | 97 | 48 | 0 | 0 | 0 | 0 | 0 | 0 | 0 | 0 | 1592 | 0 |
| SRR8280476 | 0 | 0 | 0 | 0 | 0 | 0 | 0 | 706 | 177 | 1236 | 0 | 0 | 0 | 177 | 0 | 0 | 0 | 0 | 12 | 0 | 0 | 13 | 0 | 19 | 165 | 29 | 0 | 0 | 0 | 0 | 0 | 0 | 0 | 0 | 0 | 1236 | 0 |
| SRR8280479 | 0 | 0 | 0 | 0 | 0 | 0 | 0 | 248 | 496 | 1487 | 0 | 0 | 0 | 496 | 0 | 0 | 0 | 0 | 0 | 496 | 15 | 20 | 0 | 33 | 397 | 0 | 221 | 0 | 0 | 0 | 0 | 0 | 0 | 0 | 0 | 0 | 0 |
| SRR8280481 | 129 | 0 | 0 | 0 | 0 | 387 | 0 | 0 | 0 | 181 | 181 | 0 | 0 | 181 | 0 | 0 | 0 | 0 | 0 | 0 | 0 | 26 | 0 | 39 | 336 | 22 | 0 | 0 | 0 | 0 | 0 | 0 | 0 | 0 | 0 | 181 | 0 |
| SRR8280483 | 68 | 0 | 0 | 0 | 0 | 204 | 0 | 1094 | 1368 | 274 | 0 | 0 | 0 | 547 | 0 | 0 | 68 | 0 | 0 | 0 | 15 | 0 | 69 | 57 | 446 | 12 | 340 | 0 | 0 | 0 | 0 | 0 | 0 | 0 | 0 | 0 | 0 |
| SRR8280485 | 0 | 0 | 0 | 0 | 0 | 299 | 0 | 268 | 375 | 321 | 0 | 54 | 0 | 1232 | 0 | 0 | 0 | 0 | 0 | 0 | 0 | 15 | 11 | 16 | 255 | 0 | 326 | 0 | 0 | 0 | 0 | 0 | 0 | 0 | 0 | 0 | 0 |
| SRR8280487 | 0 | 0 | 0 | 0 | 0 | 0 | 0 | 299 | 0 | 1197 | 0 | 0 | 0 | 3290 | 0 | 0 | 0 | 0 | 12 | 0 | 0 | 0 | 0 | 12 | 209 | 64 | 0 | 0 | 0 | 0 | 0 | 0 | 0 | 0 | 0 | 0 | 0 |
| SRR8280490 | 0 | 0 | 0 | 0 | 0 | 0 | 0 | 0 | 0 | 0 | 0 | 0 | 0 | 571 | 0 | 0 | 0 | 0 | 0 | 0 | 13 | 27 | 0 | 40 | 426 | 240 | 0 | 0 | 0 | 0 | 0 | 0 | 0 | 0 | 0 | 1713 | 0 |
| SRR8280494 | 0 | 0 | 0 | 0 | 0 | 0 | 0 | 0 | 0 | 1512 | 0 | 0 | 0 | 0 | 0 | 0 | 0 | 0 | 0 | 0 | 0 | 0 | 0 | 19 | 110 | 19 | 0 | 0 | 0 | 0 | 0 | 0 | 0 | 0 | 0 | 0 | 0 |
| SRR8280497 | 22 | 0 | 0 | 0 | 0 | 0 | 0 | 405 | 45 | 1620 | 0 | 45 | 0 | 900 | 45 | 0 | 11 | 0 | 0 | 90 | 29 | 30 | 80 | 99 | 375 | 47 | 32 | 0 | 0 | 0 | 0 | 0 | 0 | 0 | 0 | 2024 | 0 |
| SRR8280498 | 0 | 0 | 0 | 0 | 0 | 85 | 0 | 999 | 286 | 1285 | 0 | 0 | 0 | 857 | 428 | 0 | 0 | 0 | 0 | 0 | 16 | 46 | 62 | 76 | 545 | 28 | 28 | 0 | 0 | 0 | 0 | 0 | 0 | 0 | 0 | 286 | 0 |
| SRR8280500 | 61 | 0 | 0 | 0 | 0 | 290 | 0 | 3481 | 591 | 1642 | 0 | 0 | 31 | 755 | 0 | 0 | 0 | 0 | 0 | 0 | 54 | 83 | 118 | 171 | 1046 | 42 | 168 | 0 | 0 | 0 | 0 | 0 | 0 | 0 | 0 | 66 | 0 |
| SRR8280501 | 0 | 0 | 0 | 0 | 0 | 0 | 0 | 331 | 16 | 1339 | 0 | 0 | 0 | 281 | 53 | 42 | 0 | 817 | 203 | 105 | 1072 | 536 | 78 | 170 | 3790 | 59 | 25 | 0 | 0 | 0 | 0 | 0 | 0 | 0 | 0 | 3068 | 85 |
| SRR8280502 | 104 | 0 | 0 | 0 | 0 | 0 | 0 | 0 | 0 | 2505 | 0 | 0 | 0 | 2147 | 0 | 0 | 0 | 0 | 0 | 0 | 23 | 23 | 16 | 62 | 494 | 26 | 0 | 0 | 0 | 0 | 0 | 0 | 0 | 0 | 0 | 0 | 0 |
| SRR8280503 | 0 | 0 | 0 | 0 | 17 | 0 | 0 | 352 | 0 | 1379 | 0 | 0 | 0 | 447 | 46 | 34 | 0 | 471 | 212 | 44 | 898 | 359 | 85 | 178 | 3238 | 68 | 0 | 0 | 0 | 0 | 0 | 0 | 0 | 0 | 20 | 1294 | 46 |
| SRR8280504 | 0 | 0 | 0 | 0 | 0 | 0 | 0 | 1141 | 88 | 2721 | 0 | 0 | 0 | 1141 | 0 | 88 | 0 | 0 | 0 | 0 | 17 | 57 | 77 | 130 | 720 | 64 | 54 | 0 | 0 | 0 | 0 | 0 | 0 | 0 | 0 | 176 | 0 |
| SRR8280505 | 0 | 0 | 0 | 0 | 0 | 0 | 0 | 31 | 0 | 1200 | 0 | 0 | 0 | 69 | 13 | 14 | 0 | 1237 | 202 | 67 | 2663 | 542 | 18 | 78 | 4397 | 29 | 19 | 0 | 0 | 0 | 0 | 0 | 0 | 0 | 20 | 1098 | 163 |
| SRR8280506 | 0 | 0 | 0 | 0 | 0 | 43 | 0 | 1025 | 205 | 2153 | 0 | 205 | 0 | 923 | 0 | 0 | 43 | 0 | 0 | 0 | 14 | 19 | 29 | 59 | 406 | 27 | 43 | 0 | 0 | 0 | 0 | 0 | 0 | 0 | 0 | 103 | 0 |
| SRR8280507 | 0 | 0 | 0 | 0 | 0 | 0 | 0 | 130 | 13 | 1265 | 0 | 0 | 0 | 318 | 74 | 67 | 0 | 995 | 117 | 68 | 1447 | 508 | 48 | 165 | 3718 | 55 | 0 | 0 | 0 | 0 | 0 | 0 | 0 | 0 | 0 | 2119 | 48 |
| SRR8280509 | 0 | 0 | 0 | 0 | 0 | 0 | 0 | 105 | 0 | 1463 | 0 | 0 | 0 | 226 | 18 | 23 | 0 | 1127 | 222 | 62 | 2793 | 1153 | 48 | 79 | 3682 | 26 | 19 | 0 | 0 | 0 | 0 | 0 | 0 | 0 | 0 | 1314 | 69 |
| SRR8280510 | 0 | 0 | 0 | 0 | 0 | 0 | 0 | 270 | 1080 | 1890 | 0 | 0 | 0 | 1350 | 0 | 0 | 0 | 0 | 14 | 0 | 0 | 31 | 76 | 111 | 639 | 10 | 0 | 0 | 0 | 0 | 0 | 0 | 0 | 0 | 0 | 0 | 0 |
| SRR8280511 | 0 | 0 | 0 | 0 | 0 | 0 | 0 | 50 | 0 | 594 | 0 | 0 | 0 | 193 | 17 | 27 | 0 | 778 | 142 | 44 | 3830 | 2730 | 73 | 132 | 3558 | 26 | 36 | 0 | 0 | 0 | 0 | 0 | 0 | 0 | 0 | 912 | 47 |
| SRR8280512 | 0 | 0 | 0 | 0 | 0 | 139 | 0 | 1352 | 0 | 2287 | 0 | 0 | 0 | 1144 | 0 | 0 | 0 | 0 | 0 | 104 | 12 | 47 | 61 | 89 | 374 | 25 | 46 | 0 | 0 | 0 | 0 | 0 | 0 | 0 | 0 | 520 | 0 |
| SRR8280513 | 0 | 0 | 0 | 0 | 0 | 0 | 0 | 38 | 0 | 938 | 0 | 0 | 0 | 177 | 79 | 189 | 0 | 567 | 228 | 340 | 825 | 231 | 86 | 188 | 2125 | 56 | 0 | 0 | 0 | 0 | 0 | 0 | 0 | 0 | 0 | 7445 | 26 |
| SRR8280515 | 123 | 0 | 0 | 0 | 0 | 245 | 0 | 658 | 94 | 3197 | 0 | 0 | 0 | 376 | 188 | 0 | 0 | 0 | 0 | 0 | 17 | 29 | 45 | 50 | 395 | 12 | 245 | 0 | 0 | 0 | 0 | 0 | 0 | 0 | 0 | 470 | 0 |
| SRR8280519 | 0 | 0 | 0 | 0 | 0 | 37 | 0 | 395 | 0 | 2921 | 0 | 0 | 0 | 0 | 0 | 0 | 183 | 0 | 24 | 0 | 30 | 40 | 57 | 48 | 956 | 23 | 37 | 0 | 0 | 0 | 0 | 0 | 0 | 0 | 0 | 711 | 0 |
| SRR8280521 | 0 | 0 | 0 | 0 | 0 | 0 | 0 | 282 | 0 | 864 | 0 | 19 | 0 | 244 | 19 | 94 | 0 | 0 | 23 | 75 | 14 | 84 | 30 | 157 | 912 | 98 | 0 | 0 | 0 | 0 | 0 | 0 | 0 | 0 | 0 | 657 | 0 |
| SRR8280525 | 0 | 0 | 0 | 0 | 0 | 0 | 0 | 667 | 0 | 3468 | 0 | 0 | 0 | 667 | 0 | 0 | 0 | 12 | 12 | 267 | 12 | 24 | 18 | 54 | 330 | 30 | 0 | 0 | 0 | 0 | 0 | 0 | 0 | 0 | 0 | 267 | 0 |
| SRR8280531 | 0 | 0 | 0 | 0 | 0 | 0 | 0 | 395 | 99 | 1974 | 0 | 0 | 0 | 888 | 0 | 0 | 0 | 0 | 0 | 0 | 0 | 0 | 0 | 35 | 204 | 33 | 0 | 0 | 0 | 0 | 0 | 0 | 0 | 0 | 0 | 1086 | 0 |
| SRR8280533 | 0 | 0 | 0 | 0 | 0 | 0 | 0 | 495 | 82 | 1484 | 0 | 0 | 0 | 165 | 0 | 0 | 0 | 0 | 0 | 82 | 0 | 18 | 19 | 40 | 260 | 33 | 0 | 0 | 0 | 0 | 0 | 0 | 0 | 0 | 0 | 907 | 0 |
| SRR8280536 | 0 | 0 | 0 | 0 | 0 | 0 | 47 | 623 | 0 | 1345 | 66 | 0 | 0 | 1739 | 33 | 33 | 0 | 0 | 0 | 131 | 0 | 0 | 14 | 22 | 106 | 23 | 0 | 0 | 0 | 0 | 0 | 0 | 0 | 0 | 0 | 1411 | 0 |
| SRR8280539 | 117 | 0 | 0 | 0 | 0 | 0 | 0 | 344 | 0 | 1618 | 0 | 0 | 0 | 384 | 0 | 61 | 0 | 0 | 17 | 20 | 12 | 26 | 15 | 41 | 282 | 84 | 0 | 0 | 0 | 0 | 0 | 0 | 0 | 0 | 0 | 930 | 0 |
| SRR8280541 | 0 | 0 | 0 | 0 | 0 | 0 | 0 | 345 | 0 | 1240 | 138 | 138 | 0 | 586 | 0 | 241 | 0 | 0 | 0 | 69 | 0 | 54 | 19 | 20 | 323 | 87 | 73 | 0 | 0 | 0 | 0 | 0 | 0 | 0 | 0 | 1103 | 0 |
| SRR8280543 | 256 | 0 | 0 | 0 | 0 | 0 | 0 | 0 | 0 | 0 | 0 | 0 | 0 | 6061 | 0 | 0 | 0 | 0 | 31 | 0 | 0 | 0 | 31 | 69 | 169 | 0 | 0 | 0 | 0 | 0 | 0 | 0 | 0 | 0 | 0 | 0 | 0 |
| SRR8280544 | 63 | 0 | 0 | 0 | 0 | 0 | 0 | 540 | 0 | 810 | 0 | 0 | 0 | 1350 | 0 | 0 | 0 | 0 | 0 | 0 | 0 | 10 | 0 | 26 | 96 | 38 | 63 | 0 | 0 | 0 | 0 | 0 | 0 | 0 | 0 | 1890 | 0 |
| SRR8280546 | 118 | 0 | 0 | 0 | 0 | 0 | 0 | 936 | 0 | 2807 | 626 | 0 | 0 | 2183 | 0 | 0 | 0 | 0 | 0 | 0 | 0 | 0 | 0 | 33 | 137 | 20 | 0 | 0 | 0 | 0 | 0 | 0 | 0 | 0 | 0 | 624 | 0 |
| SRR8280548 | 0 | 0 | 0 | 0 | 0 | 0 | 0 | 0 | 0 | 0 | 0 | 0 | 0 | 0 | 0 | 0 | 0 | 0 | 0 | 0 | 0 | 0 | 29 | 29 | 167 | 23 | 580 | 0 | 0 | 0 | 0 | 0 | 0 | 0 | 0 | 0 | 0 |
| SRR8280550 | 123 | 0 | 0 | 0 | 0 | 0 | 0 | 36 | 0 | 859 | 0 | 36 | 0 | 465 | 0 | 0 | 0 | 0 | 0 | 0 | 0 | 0 | 14 | 17 | 133 | 25 | 0 | 0 | 0 | 0 | 0 | 0 | 0 | 0 | 0 | 0 | 0 |
| SRR8280551 | 0 | 0 | 0 | 0 | 0 | 0 | 0 | 0 | 0 | 1485 | 0 | 0 | 0 | 165 | 0 | 0 | 0 | 0 | 0 | 0 | 0 | 0 | 14 | 11 | 80 | 0 | 0 | 0 | 0 | 0 | 0 | 0 | 0 | 0 | 0 | 413 | 0 |
| SRR8280552 | 0 | 0 | 0 | 0 | 0 | 0 | 0 | 0 | 0 | 1823 | 0 | 0 | 0 | 1139 | 0 | 0 | 0 | 0 | 17 | 228 | 0 | 17 | 0 | 0 | 77 | 28 | 0 | 0 | 0 | 0 | 0 | 0 | 0 | 0 | 0 | 684 | 0 |
| SRR8280554 | 0 | 0 | 0 | 0 | 0 | 0 | 0 | 633 | 0 | 886 | 0 | 0 | 0 | 506 | 0 | 127 | 0 | 0 | 0 | 0 | 0 | 37 | 0 | 21 | 292 | 209 | 0 | 0 | 0 | 0 | 0 | 0 | 0 | 0 | 0 | 506 | 0 |
| SRR8280558 | 219 | 0 | 0 | 0 | 0 | 0 | 0 | 1824 | 0 | 782 | 0 | 261 | 0 | 261 | 0 | 0 | 0 | 0 | 0 | 0 | 0 | 15 | 0 | 12 | 127 | 58 | 0 | 0 | 0 | 0 | 0 | 0 | 0 | 0 | 0 | 1303 | 0 |
| SRR8280560 | 37 | 0 | 0 | 0 | 0 | 0 | 110 | 0 | 0 | 1197 | 50 | 50 | 0 | 1197 | 0 | 150 | 0 | 0 | 0 | 0 | 0 | 0 | 10 | 17 | 63 | 16 | 0 | 0 | 0 | 0 | 0 | 0 | 0 | 0 | 0 | 0 | 0 |
| SRR8280561 | 0 | 0 | 0 | 0 | 0 | 0 | 0 | 293 | 0 | 1537 | 0 | 0 | 0 | 659 | 0 | 73 | 0 | 0 | 0 | 0 | 0 | 0 | 0 | 11 | 69 | 27 | 0 | 0 | 0 | 0 | 0 | 0 | 0 | 0 | 0 | 0 | 0 |
| SRR8280562 | 0 | 0 | 0 | 0 | 0 | 0 | 0 | 299 | 0 | 2844 | 0 | 0 | 0 | 1497 | 0 | 0 | 0 | 0 | 12 | 200 | 0 | 23 | 42 | 92 | 377 | 71 | 0 | 0 | 0 | 0 | 0 | 0 | 0 | 0 | 0 | 1098 | 0 |
| SRR8280564 | 0 | 0 | 0 | 0 | 0 | 0 | 0 | 307 | 103 | 2409 | 0 | 0 | 0 | 512 | 51 | 0 | 0 | 0 | 0 | 103 | 0 | 11 | 11 | 27 | 190 | 33 | 49 | 0 | 0 | 0 | 0 | 0 | 0 | 0 | 0 | 512 | 0 |
| SRR8280566 | 0 | 0 | 0 | 0 | 0 | 0 | 0 | 1088 | 263 | 1726 | 0 | 0 | 0 | 113 | 0 | 38 | 0 | 0 | 0 | 0 | 0 | 19 | 12 | 35 | 402 | 28 | 78 | 0 | 0 | 0 | 0 | 0 | 0 | 0 | 0 | 600 | 0 |

### Supp Table S8.4.Columns_DI_to_ES

| | | | | | | | | | | | | | | | | | | | | | | | | | | | | | | | | | | | | | |
|---|---|---|---|---|---|---|---|---|---|---|---|---|---|---|---|---|---|---|---|---|---|---|---|---|---|---|---|---|---|---|---|---|---|---|---|---|---|
| SRR8280568 | 0 | 0 | 0 | 0 | 0 | 43 | 0 | 606 | 303 | 1818 | 0 | 202 | 0 | 404 | 0 | 202 | 0 | 0 | 0 | 0 | 0 | 14 | 22 | 27 | 211 | 29 | 43 | 0 | 0 | 0 | 0 | 0 | 0 | 0 | 43 | 1616 | 0 |
| SRR8280572 | 157 | 0 | 0 | 0 | 0 | 0 | 0 | 305 | 0 | 1100 | 428 | 122 | 0 | 3604 | 0 | 0 | 0 | 0 | 0 | 0 | 10 | 11 | 77 | 88 | 284 | 16 | 57 | 0 | 0 | 0 | 0 | 0 | 0 | 0 | 0 | 550 | 0 |
| SRR8280573 | 38 | 0 | 0 | 0 | 0 | 154 | 0 | 1742 | 134 | 3260 | 0 | 89 | 0 | 1384 | 179 | 134 | 0 | 0 | 0 | 89 | 46 | 80 | 32 | 109 | 639 | 62 | 38 | 0 | 0 | 0 | 0 | 0 | 0 | 0 | 0 | 1161 | 0 |
| SRR8280574 | 0 | 0 | 0 | 0 | 0 | 76 | 0 | 196 | 196 | 785 | 0 | 0 | 0 | 3927 | 0 | 0 | 0 | 0 | 10 | 0 | 23 | 61 | 13 | 53 | 396 | 13 | 0 | 0 | 0 | 0 | 0 | 0 | 0 | 0 | 0 | 0 | 0 |
| SRR8280576 | 0 | 0 | 0 | 0 | 0 | 0 | 0 | 4148 | 0 | 1091 | 0 | 0 | 0 | 1746 | 0 | 0 | 0 | 0 | 0 | 0 | 13 | 0 | 111 | 86 | 230 | 15 | 349 | 0 | 0 | 0 | 0 | 0 | 0 | 0 | 0 | 0 | 0 |
| SRR8280580 | 81 | 0 | 0 | 0 | 0 | 40 | 0 | 550 | 0 | 1884 | 79 | 236 | 0 | 314 | 157 | 0 | 0 | 0 | 0 | 0 | 13 | 11 | 16 | 44 | 528 | 15 | 121 | 0 | 0 | 0 | 0 | 0 | 0 | 0 | 0 | 79 | 0 |
| SRR8280582 | 0 | 0 | 0 | 0 | 0 | 0 | 0 | 1973 | 172 | 2831 | 0 | 257 | 0 | 257 | 0 | 86 | 0 | 0 | 0 | 0 | 35 | 45 | 65 | 126 | 557 | 45 | 0 | 0 | 0 | 0 | 0 | 0 | 0 | 0 | 0 | 172 | 0 |
| SRR8280583 | 0 | 0 | 0 | 0 | 0 | 0 | 0 | 78 | 0 | 487 | 0 | 0 | 0 | 284 | 0 | 0 | 12 | 396 | 185 | 0 | 1496 | 323 | 51 | 153 | 2043 | 54 | 24 | 0 | 0 | 0 | 0 | 0 | 0 | 0 | 0 | 161 | 42 |
| SRR8280584 | 0 | 0 | 0 | 0 | 0 | 294 | 0 | 699 | 299 | 3094 | 0 | 100 | 0 | 399 | 0 | 100 | 0 | 0 | 0 | 100 | 0 | 24 | 56 | 85 | 386 | 40 | 49 | 0 | 0 | 0 | 0 | 0 | 0 | 0 | 0 | 399 | 0 |
| SRR8280589 | 0 | 0 | 0 | 0 | 0 | 0 | 0 | 496 | 0 | 496 | 0 | 0 | 0 | 695 | 0 | 0 | 0 | 0 | 0 | 0 | 0 | 0 | 22 | 20 | 109 | 27 | 0 | 0 | 0 | 0 | 0 | 0 | 0 | 0 | 0 | 298 | 0 |
| SRR8280590 | 0 | 0 | 0 | 0 | 0 | 0 | 0 | 0 | 0 | 1521 | 0 | 0 | 0 | 811 | 101 | 203 | 0 | 0 | 0 | 0 | 0 | 36 | 13 | 11 | 306 | 58 | 0 | 0 | 0 | 0 | 0 | 0 | 0 | 0 | 0 | 609 | 0 |
| SRR8280592 | 0 | 0 | 0 | 0 | 0 | 0 | 0 | 0 | 0 | 917 | 689 | 0 | 0 | 917 | 0 | 0 | 0 | 0 | 0 | 0 | 0 | 28 | 16 | 85 | 133 | 12 | 116 | 0 | 0 | 0 | 0 | 0 | 0 | 0 | 0 | 229 | 0 |
| SRR8280593 | 0 | 0 | 0 | 0 | 0 | 33 | 0 | 217 | 0 | 977 | 0 | 163 | 0 | 923 | 0 | 0 | 0 | 0 | 0 | 0 | 0 | 0 | 0 | 18 | 91 | 40 | 0 | 0 | 0 | 0 | 0 | 0 | 0 | 0 | 33 | 814 | 0 |
| SRR8311061 | 61 | 0 | 0 | 0 | 0 | 92 | 0 | 0 | 859 | 859 | 0 | 0 | 0 | 0 | 0 | 0 | 0 | 0 | 0 | 0 | 0 | 0 | 11 | 32 | 804 | 205 | 176 | 0 | 0 | 0 | 0 | 0 | 0 | 0 | 0 | 0 | 0 |
| SRR8311062 | 18 | 0 | 0 | 0 | 0 | 474 | 0 | 0 | 0 | 0 | 0 | 0 | 0 | 3189 | 0 | 0 | 0 | 41 | 0 | 0 | 0 | 14 | 78 | 253 | 613 | 192 | 0 | 0 | 0 | 0 | 0 | 0 | 0 | 0 | 0 | 0 | 0 |
| SRR8311063 | 0 | 0 | 0 | 0 | 37 | 151 | 0 | 0 | 0 | 0 | 0 | 0 | 0 | 0 | 0 | 0 | 0 | 88 | 30 | 752 | 28 | 32 | 204 | 715 | 833 | 306 | 0 | 0 | 0 | 0 | 0 | 0 | 0 | 0 | 0 | 0 | 0 |
| SRR8311064 | 0 | 0 | 0 | 0 | 0 | 0 | 0 | 0 | 0 | 0 | 0 | 0 | 0 | 0 | 0 | 0 | 0 | 72 | 14 | 0 | 28 | 27 | 53 | 140 | 697 | 548 | 0 | 0 | 0 | 0 | 0 | 0 | 0 | 0 | 0 | 187 | 0 |
| SRR8311070 | 0 | 0 | 298 | 0 | 18 | 0 | 0 | 0 | 0 | 0 | 0 | 0 | 0 | 0 | 0 | 3226 | 0 | 107 | 0 | 0 | 20 | 16 | 175 | 175 | 646 | 21 | 0 | 0 | 0 | 0 | 0 | 0 | 0 | 0 | 0 | 0 | 0 |
| SRR8311071 | 0 | 0 | 0 | 0 | 0 | 0 | 0 | 0 | 0 | 0 | 0 | 0 | 0 | 0 | 0 | 0 | 0 | 55 | 0 | 0 | 14 | 17 | 92 | 314 | 760 | 255 | 154 | 0 | 0 | 0 | 0 | 0 | 0 | 0 | 0 | 0 | 0 |
| SRR8311072 | 0 | 0 | 51 | 0 | 0 | 96 | 0 | 0 | 0 | 553 | 0 | 1056 | 0 | 25 | 0 | 0 | 0 | 36 | 27 | 0 | 0 | 22 | 33 | 92 | 594 | 284 | 68 | 0 | 0 | 0 | 0 | 0 | 0 | 0 | 0 | 0 | 0 |
| SRR8311073 | 0 | 0 | 0 | 0 | 19 | 0 | 0 | 983 | 0 | 0 | 0 | 0 | 0 | 3811 | 0 | 1065 | 0 | 38 | 33 | 0 | 21 | 79 | 91 | 423 | 676 | 198 | 216 | 0 | 0 | 0 | 0 | 0 | 0 | 0 | 0 | 1065 | 0 |
| SRR8311074 | 0 | 0 | 202 | 0 | 0 | 0 | 0 | 0 | 0 | 4062 | 2998 | 3772 | 0 | 0 | 0 | 0 | 0 | 33 | 11 | 0 | 0 | 70 | 310 | 641 | 1025 | 185 | 0 | 0 | 0 | 0 | 0 | 0 | 0 | 0 | 0 | 2128 | 0 |
| SRR8311076 | 0 | 0 | 45 | 0 | 0 | 348 | 0 | 0 | 0 | 0 | 0 | 0 | 0 | 2166 | 0 | 0 | 0 | 24 | 20 | 0 | 0 | 20 | 35 | 70 | 636 | 342 | 0 | 0 | 0 | 0 | 0 | 0 | 0 | 0 | 0 | 0 | 0 |
| SRR8311078 | 0 | 0 | 0 | 0 | 0 | 228 | 0 | 0 | 0 | 948 | 0 | 0 | 0 | 0 | 0 | 0 | 0 | 32 | 15 | 0 | 0 | 30 | 54 | 211 | 681 | 247 | 205 | 0 | 0 | 0 | 0 | 0 | 0 | 0 | 0 | 0 | 0 |
| SRR8311080 | 0 | 0 | 0 | 0 | 0 | 0 | 0 | 0 | 0 | 0 | 0 | 0 | 0 | 829 | 0 | 319 | 0 | 0 | 18 | 0 | 0 | 0 | 30 | 58 | 542 | 179 | 211 | 0 | 0 | 0 | 0 | 0 | 0 | 0 | 0 | 0 | 0 |
| SRR8311081 | 0 | 0 | 0 | 0 | 0 | 0 | 0 | 0 | 1788 | 0 | 0 | 0 | 0 | 3260 | 0 | 0 | 0 | 0 | 0 | 0 | 0 | 0 | 42 | 59 | 602 | 154 | 0 | 0 | 0 | 0 | 0 | 0 | 0 | 0 | 0 | 0 | 0 |
| SRR8311082 | 0 | 0 | 0 | 0 | 0 | 552 | 0 | 0 | 0 | 0 | 0 | 0 | 0 | 1218 | 0 | 0 | 0 | 20 | 0 | 0 | 11 | 20 | 82 | 309 | 351 | 252 | 127 | 0 | 0 | 0 | 0 | 0 | 0 | 0 | 0 | 0 | 0 |
| SRR8311085 | 0 | 116 | 0 | 0 | 12 | 650 | 0 | 0 | 0 | 1118 | 0 | 0 | 0 | 578 | 0 | 0 | 0 | 37 | 0 | 0 | 0 | 48 | 117 | 262 | 838 | 190 | 0 | 0 | 0 | 0 | 0 | 0 | 0 | 0 | 0 | 0 | 0 |
| SRR8311087 | 0 | 0 | 0 | 0 | 14 | 0 | 0 | 0 | 0 | 0 | 0 | 1960 | 0 | 3731 | 0 | 0 | 0 | 39 | 0 | 0 | 23 | 53 | 92 | 385 | 1333 | 283 | 0 | 0 | 0 | 0 | 0 | 0 | 0 | 0 | 0 | 0 | 0 |
| SRR8311089 | 0 | 0 | 0 | 0 | 0 | 531 | 0 | 0 | 0 | 0 | 97 | 0 | 0 | 0 | 2317 | 0 | 0 | 40 | 0 | 0 | 45 | 33 | 55 | 111 | 415 | 146 | 0 | 0 | 0 | 0 | 0 | 0 | 0 | 0 | 0 | 2414 | 0 |
| SRR8311092 | 0 | 111 | 0 | 0 | 37 | 0 | 0 | 0 | 0 | 0 | 0 | 0 | 0 | 48 | 0 | 0 | 0 | 84 | 0 | 0 | 37 | 19 | 177 | 602 | 1120 | 528 | 0 | 0 | 0 | 0 | 0 | 0 | 0 | 0 | 0 | 0 | 0 |
| SRR8311093 | 0 | 0 | 0 | 0 | 0 | 151 | 0 | 0 | 0 | 0 | 0 | 1297 | 0 | 2545 | 0 | 0 | 0 | 0 | 13 | 1148 | 0 | 23 | 31 | 146 | 644 | 186 | 174 | 0 | 0 | 0 | 0 | 0 | 0 | 0 | 0 | 1347 | 0 |
| SRR8311095 | 0 | 0 | 0 | 0 | 0 | 523 | 0 | 0 | 0 | 0 | 0 | 0 | 0 | 0 | 0 | 0 | 0 | 15 | 0 | 0 | 0 | 13 | 81 | 182 | 500 | 122 | 114 | 0 | 0 | 0 | 0 | 0 | 0 | 0 | 0 | 0 | 0 |
| SRR8311098 | 0 | 0 | 0 | 0 | 0 | 157 | 0 | 0 | 0 | 0 | 0 | 0 | 0 | 2498 | 0 | 0 | 0 | 103 | 15 | 0 | 20 | 46 | 336 | 725 | 1093 | 133 | 0 | 0 | 0 | 0 | 0 | 0 | 0 | 0 | 0 | 1318 | 0 |
| SRR8311105 | 0 | 0 | 0 | 0 | 21 | 1187 | 0 | 0 | 0 | 1348 | 0 | 0 | 0 | 0 | 0 | 0 | 0 | 15 | 15 | 0 | 0 | 11 | 97 | 181 | 723 | 278 | 0 | 0 | 0 | 0 | 0 | 0 | 0 | 0 | 0 | 0 | 0 |
| SRR8311106 | 0 | 0 | 0 | 0 | 28 | 0 | 0 | 0 | 0 | 0 | 0 | 0 | 0 | 2597 | 0 | 0 | 0 | 19 | 0 | 0 | 25 | 135 | 295 | 775 | 802 | 203 | 203 | 0 | 0 | 0 | 0 | 0 | 0 | 0 | 0 | 0 | 0 |
| SRR835324 | 0 | 1412 | 28 | 106 | 0 | 0 | 28 | 2573 | 9070 | 4521 | 2168 | 2658 | 0 | 384 | 12 | 0 | 29 | 34 | 0 | 3063 | 42 | 118 | 13 | 308 | 101 | 0 | 29 | 16 | 34 | 15 | 12 | 25 | 0 | 0 | 0 | 3525 | 0 |
| SRR835325 | 0 | 1195 | 11 | 59 | 0 | 0 | 25 | 7159 | 31077 | 12353 | 56 | 337 | 0 | 272 | 43 | 18 | 27 | 26 | 13 | 201 | 39 | 131 | 0 | 424 | 163 | 0 | 12 | 13 | 10 | 28 | 23 | 10 | 0 | 0 | 0 | 304 | 0 |
| SRR8354354 | 0 | 263 | 15 | 104 | 773 | 0 | 0 | 3121 | 7943 | 16170 | 0 | 7660 | 0 | 0 | 0 | 0 | 91 | 817 | 0 | 0 | 357 | 1375 | 82 | 5715 | 1419 | 0 | 30 | 0 | 0 | 0 | 0 | 0 | 0 | 0 | 0 | 0 | 1167 |
| SRR837775 | 0 | 22 | 15 | 0 | 0 | 0 | 11 | 314 | 21 | 352 | 4347 | 6939 | 0 | 0 | 0 | 0 | 0 | 133 | 197 | 354 | 1048 | 2725 | 20 | 4808 | 16 | 68 | 0 | 246 | 268 | 502 | 492 | 51 | 108 | 190 | 23 | 649 | 108 |
| SRR837776 | 0 | 18 | 0 | 0 | 0 | 0 | 14 | 223 | 0 | 372 | 1879 | 7103 | 0 | 0 | 0 | 0 | 0 | 269 | 93 | 283 | 1397 | 2684 | 44 | 3271 | 18 | 24 | 29 | 45 | 28 | 40 | 24 | 38 | 113 | 171 | 0 | 457 | 90 |
| SRR837777 | 0 | 19 | 0 | 0 | 28 | 0 | 0 | 36 | 16 | 158 | 13022 | 25738 | 0 | 0 | 0 | 25 | 0 | 540 | 28 | 96 | 1808 | 2390 | 17 | 14482 | 4078 | 214 | 0 | 14 | 11 | 35 | 18 | 0 | 0 | 0 | 0 | 254 | 14 |
| SRR837778 | 0 | 19 | 0 | 0 | 12 | 0 | 0 | 100 | 0 | 320 | 10242 | 22310 | 0 | 0 | 0 | 0 | 0 | 347 | 0 | 94 | 363 | 1475 | 11 | 16910 | 2880 | 34 | 0 | 14 | 0 | 15 | 12 | 0 | 0 | 0 | 0 | 70 | 0 |
| SRR837779 | 0 | 18 | 0 | 0 | 0 | 0 | 14 | 223 | 0 | 372 | 1879 | 7103 | 0 | 0 | 0 | 0 | 0 | 269 | 93 | 283 | 1397 | 2684 | 44 | 3271 | 18 | 24 | 29 | 45 | 28 | 40 | 24 | 38 | 113 | 171 | 0 | 457 | 90 |
| SRR837780 | 0 | 22 | 15 | 0 | 0 | 0 | 11 | 305 | 20 | 343 | 2285 | 6754 | 0 | 0 | 0 | 0 | 0 | 133 | 197 | 344 | 1048 | 2725 | 20 | 4808 | 16 | 68 | 0 | 246 | 268 | 502 | 492 | 51 | 108 | 190 | 23 | 631 | 108 |
| SRR8380550 | 0 | 264 | 0 | 456 | 1629 | 0 | 0 | 1119 | 2917 | 10370 | 11 | 29 | 0 | 243 | 0 | 0 | 0 | 1010 | 0 | 18 | 1300 | 82 | 265 | 1831 | 1210 | 0 | 17 | 0 | 0 | 0 | 0 | 0 | 0 | 0 | 0 | 18 | 0 |
| SRR8380552 | 0 | 219 | 11 | 0 | 82 | 0 | 46 | 483 | 840 | 2562 | 146 | 265 | 0 | 1100 | 0 | 14 | 0 | 190 | 0 | 2639 | 320 | 977 | 82 | 1108 | 125 | 0 | 436 | 0 | 58 | 63 | 105 | 30 | 12 | 0 | 0 | 7457 | 0 |
| SRR8380553 | 0 | 120 | 0 | 0 | 36 | 0 | 84 | 259 | 662 | 2848 | 121 | 97 | 0 | 2142 | 0 | 14 | 0 | 572 | 34 | 1392 | 653 | 1796 | 86 | 1495 | 146 | 0 | 1260 | 0 | 29 | 62 | 122 | 29 | 0 | 0 | 0 | 3655 | 313 |
| SRR8380554 | 0 | 66 | 0 | 12 | 0 | 0 | 24 | 3698 | 7307 | 16288 | 12 | 13 | 0 | 1237 | 0 | 0 | 0 | 246 | 14 | 116 | 727 | 1610 | 106 | 1101 | 79 | 0 | 257 | 0 | 0 | 0 | 0 | 0 | 0 | 0 | 0 | 403 | 0 |
| SRR8380555 | 0 | 2623 | 0 | 289 | 2360 | 0 | 0 | 1501 | 3419 | 11786 | 301 | 1110 | 0 | 366 | 0 | 0 | 0 | 548 | 25 | 280 | 14 | 548 | 35 | 391 | 416 | 0 | 52 | 0 | 0 | 14 | 0 | 0 | 0 | 0 | 0 | 698 | 0 |
| SRR8380722 | 0 | 660 | 133 | 252 | 155 | 73 | 13 | 487 | 1635 | 47221 | 85 | 100 | 0 | 3580 | 58 | 110 | 16 | 743 | 26 | 796 | 1804 | 1034 | 509 | 6817 | 1309 | 229 | 74 | 0 | 71 | 111 | 508 | 156 | 122 | 105 | 34 | 1854 | 33 |
| SRR8380723 | 0 | 3102 | 35 | 368 | 236 | 61 | 0 | 1027 | 3552 | 37148 | 342 | 201 | 10 | 3746 | 13 | 20 | 25 | 1156 | 59 | 631 | 1961 | 1823 | 539 | 8846 | 2279 | 148 | 20 | 46 | 96 | 117 | 299 | 15 | 20 | 10 | 0 | 826 | 0 |
| SRR8380724 | 0 | 1027 | 240 | 158 | 86 | 0 | 0 | 498 | 1733 | 39595 | 66 | 56 | 0 | 3656 | 173 | 342 | 0 | 937 | 44 | 346 | 1880 | 2692 | 551 | 11346 | 1927 | 84 | 16 | 0 | 42 | 77 | 259 | 48 | 54 | 67 | 45 | 677 | 0 |
| SRR8380725 | 0 | 1248 | 35 | 780 | 217 | 89 | 0 | 674 | 3712 | 30363 | 2181 | 3061 | 0 | 2975 | 0 | 35 | 27 | 729 | 33 | 401 | 1782 | 1221 | 469 | 4573 | 2243 | 221 | 54 | 19 | 154 | 88 | 150 | 35 | 21 | 44 | 0 | 328 | 13 |
| SRR8380726 | 0 | 29 | 54 | 329 | 114 | 63 | 15 | 621 | 1906 | 26521 | 869 | 1015 | 0 | 5151 | 11 | 13 | 28 | 655 | 43 | 1071 | 1691 | 892 | 276 | 3369 | 1212 | 151 | 86 | 18 | 136 | 160 | 18 | 105 | 95 | 63 | 0 | 1182 | 0 |
| SRR8380727 | 0 | 788 | 33 | 228 | 162 | 126 | 24 | 874 | 3591 | 53111 | 142 | 253 | 0 | 3838 | 0 | 0 | 31 | 1075 | 40 | 518 | 2042 | 1549 | 419 | 4182 | 959 | 193 | 102 | 11 | 58 | 79 | 158 | 22 | 26 | 20 | 0 | 950 | 11 |
| SRR8400006 | 77 | 491 | 50 | 0 | 235 | 22 | 67 | 482 | 1103 | 2943 | 475 | 131 | 0 | 274 | 0 | 0 | 0 | 5325 | 0 | 0 | 674 | 314 | 0 | 105 | 129 | 0 | 16 | 13 | 19 | 33 | 20 | 0 | 15 | 18 | 0 | 43 | 1658 |
| SRR8400007 | 127 | 2449 | 0 | 0 | 420 | 20 | 70 | 46 | 48 | 154 | 2324 | 867 | 0 | 0 | 0 | 0 | 0 | 8697 | 0 | 0 | 1173 | 310 | 20 | 269 | 140 | 0 | 19 | 17 | 76 | 0 | 0 | 0 | 0 | 0 | 0 | 12 | 945 |
| SRR8400008 | 48 | 10011 | 15 | 0 | 324 | 33 | 36 | 373 | 462 | 2237 | 1861 | 423 | 0 | 44 | 0 | 0 | 0 | 7813 | 0 | 0 | 624 | 255 | 11 | 161 | 105 | 0 | 0 | 76 | 779 | 48 | 16 | 10 | 13 | 0 | 0 | 106 | 740 |
| SRR8400009 | 61 | 1097 | 18 | 0 | 185 | 18 | 27 | 198 | 501 | 16097 | 663 | 360 | 0 | 23 | 0 | 0 | 0 | 5855 | 0 | 22 | 1513 | 734 | 38 | 436 | 281 | 0 | 11 | 0 | 107 | 13 | 0 | 0 | 0 | 0 | 0 | 192 | 1544 |
| SRR8400010 | 15 | 120 | 0 | 0 | 110 | 0 | 18 | 130 | 192 | 1087 | 4138 | 1771 | 0 | 0 | 0 | 0 | 0 | 7159 | 0 | 0 | 1289 | 654 | 29 | 317 | 77 | 0 | 15 | 10 | 251 | 39 | 0 | 0 | 0 | 0 | 0 | 254 | 1201 |
| SRR8417876 | 0 | 13 | 14 | 0 | 13 | 0 | 0 | 2274 | 1067 | 17920 | 3670 | 1670 | 0 | 3547 | 73 | 70 | 0 | 239 | 0 | 623 | 306 | 1944 | 16 | 34723 | 3125 | 258 | 0 | 287 | 52 | 138 | 188 | 56 | 164 | 409 | 126 | 2587 | 0 |
| SRR8417877 | 0 | 39 | 13 | 0 | 0 | 0 | 0 | 2680 | 876 | 12961 | 4797 | 1898 | 0 | 3467 | 114 | 46 | 0 | 339 | 10 | 553 | 381 | 2291 | 15 | 34318 | 3710 | 210 | 0 | 48 | 34 | 49 | 60 | 43 | 77 | 19 | 10 | 2136 | 0 |
| SRR8417878 | 0 | 29 | 33 | 0 | 10 | 0 | 0 | 2661 | 819 | 11870 | 3516 | 1484 | 0 | 4827 | 20 | 0 | 0 | 359 | 12 | 454 | 337 | 2056 | 14 | 38388 | 4303 | 201 | 0 | 85 | 16 | 41 | 66 | 30 | 43 | 97 | 25 | 1856 | 0 |
| SRR8435171 | 26 | 899 | 0 | 185 | 2116 | 25 | 217 | 928 | 2193 | 1834 | 18044 | 25440 | 0 | 33 | 0 | 0 | 361 | 824 | 47 | 77 | 1132 | 1145 | 60 | 3269 | 1782 | 0 | 578 | 42 | 169 | 178 | 161 | 0 | 0 | 0 | 0 | 149 | 113 |
| SRR8435172 | 53 | 87 | 0 | 149 | 1698 | 70 | 573 | 1156 | 2621 | 1792 | 2425 | 17286 | 12 | 19 | 0 | 0 | 798 | 742 | 139 | 39 | 1181 | 1172 | 91 | 3360 | 1017 | 0 | 1976 | 0 | 32 | 38 | 44 | 0 | 0 | 0 | 0 | 51 | 99 |
| SRR8481810 | 0 | 16 | 18 | 0 | 54 | 0 | 0 | 616 | 369 | 1724 | 16575 | 10773 | 0 | 139 | 292 | 154 | 0 | 949 | 29 | 769 | 1982 | 4079 | 0 | 227 | 74 | 0 | 0 | 15 | 14 | 18 | 14 | 15 | 61 | 12 | 37 | 1170 | 0 |
| SRR8481811 | 0 | 42 | 41 | 0 | 61 | 0 | 0 | 2246 | 2847 | 1960 | 1682 | 8790 | 0 | 130 | 174 | 155 | 0 | 586 | 20 | 1421 | 2040 | 3701 | 0 | 294 | 7161 | 0 | 10 | 32 | 39 | 14 | 59 | 0 | 21 | 56 | 0 | 16793 | 0 |
| SRR8481812 | 0 | 12 | 120 | 0 | 65 | 0 | 0 | 503 | 286 | 4020 | 1546 | 13397 | 0 | 148 | 331 | 114 | 0 | 657 | 18 | 1051 | 2254 | 3947 | 0 | 236 | 89 | 0 | 0 | 33 | 101 | 13 | 131 | 26 | 61 | 96 | 12 | 1096 | 0 |
| SRR8481813 | 0 | 0 | 0 | 0 | 86 | 0 | 0 | 4153 | 4910 | 1613 | 2040 | 1504 | 0 | 310 | 0 | 32 | 13 | 395 | 19 | 2558 | 1518 | 2421 | 0 | 437 | 13 | 0 | 10 | 18 | 14 | 28 | 16 | 0 | 12 | 29 | 0 | 38818 | 0 |
| SRR8481814 | 0 | 95 | 21 | 0 | 79 | 0 | 0 | 509 | 307 | 2296 | 18138 | 16928 | 0 | 259 | 279 | 317 | 0 | 399 | 14 | 769 | 1565 | 3159 | 0 | 228 | 60 | 0 | 0 | 25 | 62 | 15 | 105 | 19 | 41 | 78 | 12 | 1038 | 0 |
| SRR8481815 | 0 | 55 | 25 | 0 | 79 | 0 | 0 | 2068 | 2648 | 2205 | 13745 | 10865 | 0 | 169 | 169 | 95 | 10 | 662 | 22 | 1097 | 1521 | 3075 | 0 | 267 | 6715 | 0 | 0 | 43 | 57 | 86 | 55 | 0 | 37 | 61 | 16 | 14779 | 0 |
| SRR8481816 | 0 | 108 | 29 | 0 | 77 | 0 | 0 | 465 | 248 | 1613 | 2798 | 12461 | 0 | 96 | 1383 | 1087 | 0 | 605 | 31 | 848 | 1743 | 3052 | 0 | 305 | 7565 | 0 | 0 | 22 | 70 | 182 | 179 | 15 | 58 | 134 | 48 | 1483 | 0 |
| SRR8481817 | 0 | 0 | 0 | 0 | 84 | 0 | 0 | 4321 | 4900 | 1376 | 1992 | 1120 | 0 | 210 | 135 | 233 | 0 | 366 | 21 | 1526 | 1392 | 2616 | 0 | 683 | 6845 | 0 | 0 | 18 | 12 | 11 | 18 | 0 | 16 | 13 | 11 | 46183 | 0 |
| SRR8481818 | 0 | 32 | 37 | 0 | 62 | 0 | 0 | 1128 | 369 | 2118 | 1723 | 9418 | 0 | 130 | 1807 | 1438 | 0 | 497 | 19 | 1373 | 1801 | 4104 | 0 | 193 | 9458 | 0 | 0 | 21 | 54 | 73 | 76 | 21 | 52 | 91 | 62 | 3021 | 0 |
| SRR8481819 | 0 | 44 | 42 | 0 | 74 | 0 | 0 | 2101 | 1440 | 2345 | 12187 | 7569 | 0 | 118 | 1729 | 1403 | 0 | 741 | 19 | 933 | 1474 | 3685 | 0 | 264 | 9653 | 0 | 0 | 56 | 92 | 139 | 113 | 36 | 86 | 14 | 10 | 10729 | 0 |
| SRR8481820 | 0 | 27 | 49 | 0 | 100 | 0 | 0 | 1630 | 466 | 1862 | 828 | 7045 | 0 | 147 | 1336 | 943 | 13 | 810 | 22 | 257 | 1406 | 3152 | 0 | 287 | 7956 | 0 | 0 | 58 | 134 | 161 | 170 | 27 | 116 | 156 | 116 | 1005 | 0 |
| SRR8481821 | 0 | 0 | 0 | 0 | 88 | 0 | 0 | 7470 | 5105 | 1018 | 1230 | 917 | 0 | 98 | 104 | 150 | 0 | 426 | 11 | 1948 | 936 | 1950 | 0 | 457 | 6269 | 0 | 0 | 21 | 0 | 23 | 0 | 0 | 16 | 12 | 12 | 42542 | 0 |
| SRR8485055 | 0 | 120 | 31 | 0 | 194 | 0 | 0 | 629 | 13160 | 33342 | 281 | 76 | 0 | 509 | 5299 | 13298 | 0 | 4285 | 25 | 116 | 466 | 1198 | 111 | 1695 | 969 | 17 | 0 | 40 | 14 | 28 | 35 | 0 | 29 | 50 | 12 | 900 | 0 |
| SRR8485056 | 0 | 1244 | 290 | 0 | 105 | 0 | 0 | 1559 | 12813 | 23352 | 191 | 583 | 0 | 339 | 203 | 668 | 0 | 3377 | 41 | 182 | 298 | 1209 | 174 | 1862 | 1332 | 24 | 0 | 41 | 183 | 37 | 39 | 22 | 24 | 37 | 21 | 743 | 0 |
| SRR8485057 | 0 | 371 | 15 | 0 | 81 | 0 | 0 | 3295 | 54955 | 21886 | 323 | 479 | 0 | 908 | 615 | 1664 | 0 | 2127 | 35 | 260 | 303 | 1527 | 42 | 3627 | 1308 | 23 | 17 | 54 | 46 | 30 | 23 | 12 | 25 | 36 | 19 | 2390 | 0 |
| SRR8485058 | 0 | 1179 | 217 | 0 | 88 | 0 | 0 | 190 | 1809 | 4811 | 624 | 416 | 0 | 902 | 117 | 310 | 0 | 2850 | 29 | 111 | 231 | 1013 | 61 | 1582 | 664 | 24 | 52 | 34 | 136 | 17 | 10 | 0 | 13 | 12 | 0 | 804 | 0 |
| SRR8485059 | 0 | 313 | 67 | 0 | 85 | 0 | 0 | 1151 | 19142 | 12043 | 769 | 297 | 0 | 244 | 197 | 898 | 0 | 1247 | 34 | 32 | 345 | 921 | 42 | 1595 | 769 | 20 | 0 | 68 | 39 | 36 | 26 | 12 | 39 | 71 | 49 | 308 | 0 |
| SRR8485060 | 0 | 546 | 46 | 0 | 102 | 0 | 0 | 983 | 6323 | 28055 | 1932 | 3286 | 0 | 184 | 7219 | 8783 | 0 | 3478 | 31 | 873 | 823 | 856 | 65 | 1417 | 455 | 30 | 0 | 11 | 50 | 20 | 17 | 0 | 12 | 13 | 11 | 6391 | 0 |
| SRR8485061 | 0 | 1044 | 261 | 0 | 107 | 0 | 0 | 750 | 8829 | 5938 | 246 | 395 | 0 | 373 | 74 | 112 | 0 | 2665 | 37 | 140 | 471 | 1767 | 153 | 2401 | 1927 | 17 | 0 | 70 | 295 | 51 | 46 | 30 | 43 | 57 | 24 | 1254 | 0 |
| SRR8485062 | 0 | 174 | 18 | 0 | 64 | 0 | 0 | 1102 | 8615 | 20651 | 471 | 971 | 0 | 374 | 606 | 2246 | 0 | 3043 | 21 | 154 | 644 | 1388 | 163 | 2064 | 1471 | 11 | 0 | 29 | 12 | 24 | 23 | 13 | 24 | 36 | 15 | 1313 | 0 |
| SRR8485063 | 0 | 1855 | 86 | 0 | 184 | 0 | 0 | 497 | 3981 | 6110 | 1350 | 2110 | 0 | 382 | 7018 | 13424 | 0 | 3361 | 33 | 180 | 360 | 681 | 89 | 2340 | 1648 | 42 | 0 | 63 | 237 | 41 | 34 | 16 | 34 | 55 | 12 | 1512 | 0 |
| SRR8485064 | 0 | 1593 | 218 | 0 | 212 | 0 | 0 | 198 | 1330 | 5988 | 899 | 814 | 0 | 352 | 15007 | 26904 | 0 | 3759 | 17 | 221 | 417 | 1672 | 75 | 2439 | 1916 | 13 | 0 | 62 | 348 | 70 | 45 | 21 | 46 | 81 | 18 | 1833 | 0 |
| SRR8485065 | 0 | 202 | 48 | 0 | 115 | 0 | 0 | 553 | 5674 | 14795 | 17 | 19 | 0 | 469 | 1005 | 4204 | 0 | 2770 | 34 | 134 | 1046 | 1336 | 103 | 2657 | 1466 | 16 | 0 | 28 | 11 | 19 | 19 | 0 | 11 | 15 | 0 | 1363 | 0 |
| SRR8485066 | 0 | 2054 | 85 | 0 | 72 | 0 | 0 | 2681 | 42120 | 15468 | 874 | 815 | 0 | 171 | 1946 | 5436 | 0 | 3095 | 43 | 156 | 759 | 1596 | 149 | 1664 | 1764 | 15 | 11 | 38 | 112 | 25 | 21 | 16 | 25 | 28 | 13 | 1052 | 0 |
| SRR8485067 | 0 | 78 | 137 | 0 | 250 | 0 | 0 | 633 | 9637 | 4530 | 285 | 23 | 0 | 83 | 15330 | 24446 | 0 | 4208 | 48 | 79 | 959 | 1039 | 155 | 2927 | 1744 | 14 | 0 | 28 | 13 | 75 | 52 | 39 | 60 | 75 | 0 | 467 | 0 |
| SRR8487217 | 0 | 391 | 23 | 0 | 178 | 0 | 0 | 54 | 348 | 1559 | 5143 | 3519 | 0 | 54 | 512 | 2267 | 0 | 3377 | 31 | 2925 | 620 | 937 | 162 | 3406 | 1192 | 34 | 11 | 48 | 130 | 50 | 33 | 10 | 19 | 28 | 14 | 31418 | 0 |
| SRR8487218 | 0 | 105 | 12 | 0 | 143 | 0 | 0 | 292 | 3573 | 6053 | 142 | 235 | 0 | 227 | 658 | 734 | 0 | 2251 | 47 | 34 | 136 | 401 | 56 | 929 | 712 | 14 | 0 | 21 | 22 | 46 | 62 | 19 | 35 | 47 | 0 | 402 | 0 |
| SRR8487219 | 0 | 296 | 45 | 0 | 192 | 0 | 0 | 833 | 3856 | 8795 | 765 | 767 | 0 | 166 | 230 | 555 | 0 | 2592 | 24 | 339 | 167 | 551 | 65 | 2124 | 1031 | 17 | 25 | 38 | 20 | 12 | 110 | 32 | 46 | 93 | 56 | 6269 | 0 |
| SRR8487220 | 0 | 205 | 74 | 0 | 244 | 0 | 0 | 310 | 6070 | 3179 | 452 | 39 | 0 | 40 | 606 | 1238 | 0 | 2782 | 60 | 70 | 690 | 840 | 107 | 2888 | 2521 | 17 | 11 | 55 | 10 | 48 | 44 | 21 | 34 | 54 | 15 | 744 | 0 |
| SRR8487221 | 0 | 1337 | 33 | 0 | 215 | 0 | 0 | 384 | 1301 | 4437 | 1664 | 626 | 0 | 122 | 688 | 2949 | 0 | 1852 | 35 | 243 | 560 | 186 | 114 | 4326 | 1101 | 71 | 0 | 19 | 85 | 11 | 35 | 17 | 27 | 34 | 22 | 1870 | 0 |
| SRR8487222 | 0 | 3698 | 162 | 0 | 132 | 0 | 0 | 537 | 3798 | 3369 | 4271 | 2134 | 0 | 108 | 370 | 1245 | 0 | 1314 | 36 | 316 | 400 | 262 | 235 | 3415 | 2583 | 22 | 0 | 107 | 339 | 57 | 35 | 28 | 32 | 42 | 15 | 2209 | 0 |
| SRR8487223 | 0 | 488 | 15 | 0 | 358 | 0 | 0 | 311 | 920 | 1500 | 350 | 417 | 0 | 109 | 1406 | 1565 | 45 | 1016 | 102 | 508 | 75 | 82 | 106 | 3209 | 1994 | 40 | 16 | 14 | 73 | 21 | 15 | 24 | 27 | 42 | 45 | 8133 | 0 |
| SRR8487224 | 0 | 672 | 17 | 0 | 261 | 0 | 0 | 523 | 4133 | 7867 | 96 | 182 | 0 | 105 | 18 | 32 | 34 | 2185 | 19 | 456 | 428 | 652 | 236 | 4364 | 1850 | 12 | 143 | 0 | 31 | 10 | 11 | 0 | 0 | 0 | 0 | 6799 | 0 |
| SRR8586304 | 0 | 43 | 0 | 0 | 28 | 0 | 0 | 375 | 857 | 83475 | 20 | 15 | 0 | 2937 | 0 | 0 | 0 | 373 | 56 | 0 | 390 | 1236 | 123 | 2027 | 1342 | 0 | 0 | 0 | 0 | 0 | 0 | 0 | 0 | 0 | 0 | 0 | 17 |
| SRR871609 | 0 | 22 | 0 | 0 | 0 | 0 | 0 | 59 | 560 | 134 | 837 | 7796 | 0 | 0 | 0 | 0 | 0 | 9274 | 0 | 301 | 803 | 447 | 263 | 820 | 285 | 0 | 12 | 35 | 22 | 0 | 0 | 0 | 0 | 0 | 0 | 376 | 69 |
| SRR871671 | 0 | 0 | 0 | 0 | 0 | 0 | 0 | 0 | 0 | 0 | 271 | 34965 | 0 | 0 | 163 | 0 | 0 | 2083 | 0 | 1471 | 55 | 22 | 0 | 27 | 12 | 0 | 83 | 15 | 38 | 15 | 0 | 0 | 0 | 0 | 0 | 54 | 13 |
| SRR873381 | 0 | 0 | 43 | 0 | 11 | 0 | 0 | 747 | 601 | 10286 | 2913 | 5275 | 0 | 383 | 20 | 32 | 0 | 83 | 0 | 1358 | 404 | 113 | 182 | 1824 | 1619 | 0 | 0 | 0 | 18 | 40 | 170 | 15 | 29 | 50 | 0 | 3899 | 14 |
| SRR873401 | 0 | 0 | 38 | 0 | 0 | 0 | 0 | 792 | 296 | 7506 | 7169 | 15353 | 0 | 319 | 132 | 237 | 0 | 22 | 0 | 2103 | 339 | 135 | 85 | 401 | 224 | 0 | 18 | 0 | 35 | 130 | 214 | 33 | 130 | 294 | 0 | 8658 | 20 |
| SRR873410 | 0 | 0 | 28 | 0 | 0 | 0 | 0 | 651 | 214 | 5491 | 2317 | 10589 | 0 | 138 | 387 | 199 | 12 | 39 | 0 | 1508 | 862 | 295 | 85 | 572 | 1150 | 0 | 79 | 0 | 15 | 119 | 146 | 19 | 51 | 136 | 0 | 6804 | 34 |
| SRR8834226 | 0 | 0 | 0 | 178 | 0 | 0 | 0 | 0 | 0 | 12 | 0 | 0 | 0 | 0 | 0 | 0 | 0 | 6007 | 41 | 0 | 0 | 160 | 0 | 0 | 0 | 0 | 0 | 0 | 0 | 0 | 0 | 0 | 0 | 0 | 0 | 0 | 0 |
| SRR8834227 | 0 | 0 | 0 | 294 | 32 | 0 | 0 | 0 | 0 | 0 | 0 | 0 | 0 | 0 | 0 | 42 | 0 | 19484 | 200 | 0 | 32 | 53 | 0 | 0 | 0 | 0 | 0 | 0 | 0 | 0 | 0 | 0 | 0 | 0 | 0 | 0 | 0 |
| SRR8839811 | 0 | 0 | 0 | 29 | 571 | 27 | 0 | 217 | 217 | 4978 | 24 | 61 | 0 | 42710 | 0 | 18 | 24 | 117 | 196 | 37 | 144 | 401 | 116 | 2478 | 59 | 29 | 11 | 0 | 0 | 0 | 0 | 0 | 0 | 0 | 0 | 122 | 214 |
| SRR8839812 | 0 | 0 | 0 | 36 | 462 | 0 | 0 | 1487 | 7167 | 13054 | 0 | 0 | 0 | 4014 | 0 | 0 | 0 | 213 | 213 | 10 | 36 | 213 | 213 | 10308 | 178 | 71 | 11 | 0 | 0 | 0 | 11 | 0 | 0 | 0 | 0 | 64 | 36 |
| SRR8850891 | 0 | 1581 | 26 | 229 | 1256 | 0 | 0 | 4994 | 12717 | 28326 | 12764 | 11103 | 0 | 8745 | 35 | 116 | 0 | 780 | 142 | 1022 | 120 | 849 | 41 | 1563 | 221 | 0 | 15 | 70 | 131 | 128 | 196 | 90 | 111 | 126 | 131 | 1161 | 0 |
| SRR8850892 | 0 | 247 | 69 | 244 | 1559 | 0 | 0 | 5657 | 12652 | 31530 | 2208 | 12083 | 0 | 7089 | 29 | 135 | 0 | 804 | 115 | 857 | 153 | 867 | 61 | 2168 | 347 | 0 | 0 | 97 | 17 | 16 | 31 | 13 | 20 | 19 | 18 | 1150 | 0 |
| SRR8850893 | 0 | 4173 | 70 | 237 | 1823 | 0 | 0 | 2893 | 7592 | 31082 | 1333 | 9272 | 0 | 4588 | 19 | 76 | 0 | 877 | 95 | 2564 | 161 | 743 | 52 | 2096 | 381 | 0 | 0 | 62 | 241 | 393 | 720 | 281 | 419 | 575 | 242 | 2698 | 0 |
| SRR8850894 | 0 | 316 | 45 | 250 | 3014 | 0 | 0 | 2093 | 5931 | 21007 | 4075 | 18485 | 0 | 2483 | 21 | 99 | 12 | 813 | 88 | 2728 | 178 | 835 | 63 | 2293 | 407 | 0 | 15 | 88 | 24 | 27 | 54 | 19 | 30 | 37 | 24 | 2820 | 0 |
| SRR8850895 | 0 | 183 | 37 | 349 | 2367 | 0 | 0 | 5201 | 12192 | 17803 | 2633 | 17386 | 0 | 7666 | 23 | 106 | 0 | 829 | 112 | 341 | 155 | 621 | 48 | 1210 | 255 | 0 | 0 | 72 | 16 | 16 | 25 | 84 | 15 | 14 | 12 | 584 | 0 |
| SRR8850896 | 0 | 891 | 24 | 346 | 1974 | 0 | 0 | 5271 | 12124 | 14375 | 19105 | 12893 | 0 | 7551 | 57 | 128 | 0 | 844 | 134 | 541 | 228 | 590 | 48 | 1428 | 327 | 0 | 11 | 45 | 102 | 89 | 181 | 66 | 106 | 92 | 119 | 613 | 0 |
| SRR8850897 | 0 | 607 | 31 | 255 | 704 | 0 | 0 | 4382 | 9174 | 13113 | 2561 | 22537 | 0 | 3246 | 38 | 186 | 11 | 758 | 97 | 982 | 107 | 629 | 42 | 1688 | 459 | 0 | 30 | 34 | 105 | 111 | 359 | 220 | 329 | 352 | 238 | 1007 | 0 |
| SRR8850898 | 0 | 481 | 43 | 392 | 2487 | 0 | 0 | 3433 | 8942 | 19020 | 9484 | 11285 | 0 | 3049 | 107 | 261 | 0 | 1000 | 94 | 506 | 229 | 619 | 63 | 1938 | 507 | 0 | 12 | 10 | 23 | 30 | 51 | 15 | 25 | 33 | 21 | 680 | 26 |
| SRR8850899 | 0 | 496 | 43 | 296 | 1031 | 11 | 0 | 3782 | 7972 | 30283 | 1945 | 10487 | 0 | 10102 | 13 | 13 | 15 | 873 | 101 | 377 | 126 | 993 | 25 | 1347 | 190 | 12 | 13 | 96 | 17 | 24 | 40 | 14 | 15 | 14 | 73 | 587 | 0 |
| SRR8850900 | 0 | 5936 | 31 | 517 | 2600 | 0 | 0 | 3043 | 7134 | 35713 | 9749 | 13557 | 0 | 10810 | 0 | 24 | 0 | 1043 | 95 | 271 | 150 | 1184 | 46 | 1393 | 211 | 11 | 0 | 115 | 138 | 264 | 423 | 82 | 92 | 109 | 59 | 410 | 40 |
| SRR8850901 | 0 | 6077 | 42 | 331 | 1335 | 0 | 0 | 3022 | 7304 | 28295 | 10851 | 18470 | 0 | 3646 | 12 | 18 | 23 | 908 | 98 | 770 | 134 | 857 | 31 | 2497 | 470 | 15 | 24 | 98 | 155 | 273 | 497 | 145 | 239 | 231 | 132 | 1114 | 14 |
| SRR8850902 | 0 | 577 | 26 | 401 | 1370 | 0 | 0 | 1937 | 4518 | 17345 | 6495 | 11212 | 0 | 2541 | 0 | 21 | 23 | 962 | 92 | 783 | 143 | 884 | 38 | 2471 | 474 | 13 | 28 | 75 | 15 | 26 | 48 | 15 | 17 | 20 | 40 | 961 | 0 |
| SRR8858246 | 0 | 0 | 0 | 0 | 18 | 0 | 0 | 100 | 0 | 0 | 408 | 2295 | 0 | 0 | 0 | 0 | 21 | 0 | 0 | 0 | 439 | 603 | 0 | 317 | 2238 | 0 | 484 | 46 | 162 | 2077 | 33 | 0 | 31 | 0 | 0 | 0 | 42 |
| SRR8858247 | 0 | 0 | 0 | 0 | 20 | 0 | 0 | 743 | 0 | 0 | 626 | 1185 | 0 | 0 | 0 | 0 | 0 | 0 | 0 | 0 | 528 | 713 | 34 | 1791 | 3209 | 0 | 87 | 0 | 85 | 67 | 199 | 0 | 0 | 0 | 0 | 209 | 84 |
| SRR8858248 | 0 | 0 | 0 | 0 | 19 | 157 | 19 | 259 | 0 | 0 | 3551 | 1946 | 0 | 0 | 0 | 0 | 0 | 0 | 0 | 0 | 511 | 532 | 40 | 1177 | 2795 | 0 | 10 | 0 | 0 | 13 | 0 | 0 | 0 | 0 | 0 | 114 | 77 |

### Supp Table S8.4.Columns_DI_to_ES

| | | | | | | | | | | | | | | | | | | | | | | | | | | | | | | | | | | | | | |
|---|---|---|---|---|---|---|---|---|---|---|---|---|---|---|---|---|---|---|---|---|---|---|---|---|---|---|---|---|---|---|---|---|---|---|---|---|---|
| SRR8858254 | 0 | 0 | 0 | 0 | 0 | 0 | 0 | 1627 | 0 | 1264 | 2684 | 871 | 0 | 0 | 0 | 0 | 0 | 25 | 0 | 0 | 449 | 692 | 23 | 545 | 2437 | 0 | 15 | 0 | 0 | 0 | 81 | 0 | 0 | 0 | 0 | 0 | 39 |
| SRR8858258 | 0 | 0 | 0 | 0 | 0 | 90 | 46 | 0 | 0 | 0 | 202 | 0 | 0 | 0 | 0 | 0 | 0 | 0 | 0 | 0 | 371 | 395 | 53 | 1628 | 992 | 0 | 135 | 47 | 0 | 90 | 0 | 0 | 0 | 0 | 0 | 0 | 53 |
| SRR8907824 | 0 | 0 | 0 | 0 | 27 | 0 | 45 | 2687 | 722 | 43 | 0 | 0 | 40 | 241 | 14 | 0 | 0 | 137 | 80 | 115 | 0 | 164 | 31 | 3175 | 62 | 106 | 19 | 0 | 0 | 0 | 26 | 0 | 0 | 0 | 0 | 159 | 0 |
| SRR8932122 | 0 | 32 | 0 | 0 | 50 | 21 | 0 | 5053 | 453 | 3395 | 184 | 64 | 0 | 374 | 0 | 0 | 0 | 365 | 0 | 102 | 363 | 277 | 99 | 2365 | 3929 | 12 | 18 | 0 | 0 | 11 | 0 | 0 | 0 | 0 | 0 | 251 | 0 |
| SRR8932123 | 12 | 28 | 0 | 0 | 0 | 18 | 0 | 8073 | 527 | 1674 | 495 | 731 | 0 | 381 | 0 | 14 | 11 | 172 | 0 | 406 | 502 | 261 | 134 | 1127 | 1095 | 0 | 19 | 0 | 0 | 0 | 0 | 0 | 0 | 0 | 0 | 1023 | 0 |
| SRR8932124 | 17 | 29 | 0 | 0 | 20 | 25 | 0 | 1509 | 286 | 2043 | 2310 | 545 | 0 | 181 | 49 | 43 | 11 | 269 | 0 | 498 | 580 | 335 | 148 | 1403 | 2582 | 0 | 19 | 0 | 0 | 0 | 0 | 0 | 0 | 0 | 0 | 1241 | 26 |
| SRR8932125 | 13 | 16 | 0 | 0 | 13 | 35 | 0 | 1671 | 284 | 3734 | 1909 | 1179 | 0 | 187 | 428 | 762 | 0 | 135 | 0 | 2656 | 849 | 489 | 150 | 2448 | 3432 | 0 | 18 | 0 | 0 | 0 | 0 | 0 | 0 | 0 | 0 | 6105 | 0 |
| SRR8944248 | 0 | 0 | 0 | 0 | 0 | 0 | 0 | 6531 | 1451 | 3628 | 0 | 0 | 0 | 363 | 0 | 0 | 0 | 0 | 0 | 6894 | 0 | 25 | 0 | 16 | 148 | 48 | 0 | 0 | 0 | 0 | 0 | 0 | 0 | 0 | 0 | 5080 | 0 |
| SRR8944250 | 0 | 0 | 0 | 0 | 0 | 0 | 0 | 17863 | 1751 | 0 | 352 | 0 | 0 | 0 | 0 | 0 | 0 | 0 | 0 | 19965 | 0 | 61 | 0 | 19 | 214 | 119 | 0 | 0 | 0 | 0 | 0 | 0 | 0 | 0 | 0 | 32574 | 0 |
| SRR8944251 | 0 | 0 | 0 | 0 | 0 | 0 | 0 | 14661 | 2728 | 0 | 3409 | 0 | 0 | 2046 | 0 | 0 | 0 | 0 | 0 | 0 | 0 | 44 | 0 | 26 | 166 | 161 | 0 | 0 | 0 | 0 | 0 | 0 | 0 | 0 | 0 | 3750 | 0 |
| SRR8944252 | 0 | 0 | 0 | 0 | 0 | 0 | 0 | 4872 | 0 | 1154 | 0 | 1538 | 0 | 1154 | 0 | 0 | 0 | 15 | 0 | 11667 | 16 | 112 | 0 | 53 | 376 | 466 | 0 | 0 | 0 | 0 | 0 | 0 | 0 | 0 | 0 | 24487 | 0 |
| SRR8944257 | 0 | 0 | 0 | 0 | 0 | 0 | 0 | 20891 | 4730 | 0 | 0 | 0 | 0 | 0 | 0 | 0 | 0 | 0 | 0 | 11431 | 0 | 24 | 0 | 58 | 112 | 134 | 0 | 0 | 0 | 0 | 0 | 0 | 0 | 0 | 0 | 11037 | 0 |
| SRR8944259 | 0 | 0 | 0 | 0 | 0 | 0 | 0 | 9966 | 2677 | 0 | 892 | 1339 | 0 | 149 | 0 | 0 | 0 | 17 | 0 | 12792 | 0 | 85 | 0 | 69 | 429 | 390 | 0 | 0 | 0 | 0 | 0 | 0 | 0 | 0 | 0 | 17552 | 0 |
| SRR8956086 | 0 | 0 | 0 | 109 | 195 | 0 | 0 | 112 | 315 | 13266 | 23 | 13 | 0 | 47403 | 0 | 0 | 0 | 633 | 516 | 88 | 32 | 1487 | 249 | 11267 | 788 | 34 | 0 | 0 | 0 | 0 | 12 | 0 | 0 | 0 | 0 | 362 | 72 |
| SRR8956087 | 0 | 0 | 0 | 95 | 92 | 0 | 0 | 126 | 344 | 21526 | 13 | 0 | 0 | 55037 | 0 | 0 | 0 | 905 | 636 | 133 | 47 | 1346 | 142 | 10736 | 828 | 52 | 0 | 0 | 0 | 0 | 12 | 0 | 0 | 0 | 0 | 585 | 72 |
| SRR8956089 | 0 | 12 | 28 | 221 | 371 | 0 | 0 | 338 | 1104 | 35942 | 96 | 50 | 0 | 34625 | 0 | 0 | 0 | 824 | 547 | 65 | 99 | 1816 | 742 | 21156 | 1203 | 11 | 0 | 0 | 12 | 18 | 35 | 0 | 0 | 0 | 0 | 173 | 120 |
| SRR8956090 | 0 | 15 | 31 | 153 | 247 | 0 | 0 | 425 | 1352 | 52458 | 148 | 76 | 0 | 26451 | 0 | 0 | 0 | 503 | 479 | 128 | 113 | 1164 | 850 | 28651 | 825 | 13 | 0 | 0 | 14 | 17 | 68 | 15 | 0 | 0 | 0 | 339 | 119 |
| SRR8956091 | 0 | 14 | 31 | 175 | 287 | 0 | 0 | 318 | 976 | 32634 | 61 | 62 | 0 | 30380 | 0 | 0 | 12 | 600 | 554 | 86 | 96 | 1358 | 776 | 26502 | 884 | 19 | 0 | 0 | 10 | 17 | 39 | 11 | 0 | 0 | 0 | 228 | 117 |
| SRR8992368 | 0 | 0 | 0 | 151 | 86 | 12 | 102 | 878 | 3479 | 5406 | 0 | 0 | 0 | 681 | 0 | 0 | 0 | 170 | 0 | 0 | 15 | 17 | 247 | 2347 | 1940 | 0 | 70 | 0 | 0 | 0 | 0 | 0 | 0 | 0 | 0 | 0 | 0 |
| SRR8992369 | 0 | 0 | 0 | 547 | 120 | 0 | 48 | 714 | 2421 | 6354 | 0 | 0 | 0 | 3144 | 0 | 0 | 20 | 135 | 0 | 0 | 59 | 21 | 324 | 4653 | 4202 | 0 | 60 | 0 | 0 | 0 | 0 | 0 | 0 | 0 | 0 | 0 | 0 |
| SRR8992370 | 0 | 0 | 0 | 219 | 108 | 0 | 48 | 824 | 1775 | 5657 | 0 | 0 | 0 | 4366 | 0 | 0 | 22 | 125 | 0 | 0 | 60 | 28 | 262 | 3141 | 2228 | 0 | 243 | 0 | 0 | 0 | 0 | 0 | 0 | 0 | 0 | 0 | 0 |
| SRR8992371 | 0 | 0 | 0 | 436 | 124 | 0 | 20 | 1102 | 2740 | 6219 | 0 | 0 | 0 | 2941 | 0 | 0 | 37 | 154 | 0 | 0 | 96 | 52 | 293 | 4232 | 3620 | 0 | 178 | 0 | 0 | 0 | 0 | 0 | 0 | 0 | 0 | 0 | 0 |
| SRR8992378 | 0 | 0 | 0 | 247 | 107 | 0 | 38 | 1063 | 2574 | 7098 | 0 | 0 | 0 | 4190 | 0 | 0 | 16 | 128 | 13 | 0 | 58 | 57 | 265 | 3815 | 2571 | 0 | 11 | 0 | 0 | 0 | 0 | 0 | 0 | 0 | 0 | 0 | 0 |
| SRR8992379 | 0 | 0 | 0 | 153 | 124 | 33 | 94 | 691 | 2414 | 4705 | 0 | 0 | 0 | 771 | 0 | 0 | 11 | 159 | 0 | 0 | 11 | 0 | 250 | 1750 | 1673 | 0 | 88 | 0 | 0 | 0 | 0 | 0 | 0 | 0 | 0 | 0 | 0 |
| SRR8992380 | 0 | 0 | 0 | 348 | 126 | 0 | 27 | 1110 | 2927 | 7583 | 0 | 0 | 0 | 3130 | 0 | 0 | 27 | 119 | 0 | 13 | 102 | 40 | 280 | 5140 | 3639 | 0 | 54 | 0 | 0 | 0 | 0 | 0 | 0 | 0 | 0 | 19 | 0 |
| SRR8992381 | 0 | 0 | 0 | 213 | 71 | 18 | 67 | 602 | 2703 | 15961 | 0 | 0 | 0 | 406 | 0 | 0 | 0 | 192 | 0 | 0 | 20 | 14 | 228 | 2148 | 1754 | 0 | 31 | 0 | 0 | 0 | 0 | 0 | 0 | 0 | 0 | 0 | 0 |
| SRR8992382 | 0 | 0 | 0 | 148 | 75 | 0 | 79 | 725 | 2347 | 7441 | 0 | 0 | 0 | 901 | 0 | 0 | 0 | 86 | 0 | 0 | 0 | 0 | 256 | 1817 | 1722 | 11 | 36 | 0 | 0 | 0 | 0 | 0 | 0 | 0 | 0 | 0 | 0 |
| SRR8992383 | 0 | 0 | 0 | 198 | 68 | 12 | 77 | 481 | 2123 | 10893 | 0 | 0 | 0 | 825 | 0 | 0 | 0 | 158 | 0 | 0 | 18 | 19 | 204 | 2160 | 1751 | 0 | 35 | 0 | 0 | 0 | 0 | 0 | 0 | 0 | 0 | 0 | 0 |
| SRR8992384 | 0 | 0 | 0 | 226 | 104 | 0 | 44 | 1660 | 3851 | 9478 | 0 | 0 | 0 | 3350 | 0 | 0 | 12 | 156 | 0 | 0 | 58 | 43 | 372 | 4247 | 2966 | 18 | 35 | 0 | 0 | 0 | 0 | 0 | 0 | 0 | 0 | 0 | 0 |
| SRR8992385 | 0 | 0 | 0 | 194 | 84 | 0 | 51 | 1789 | 3755 | 9149 | 0 | 0 | 0 | 3186 | 0 | 0 | 29 | 95 | 0 | 0 | 33 | 17 | 334 | 4183 | 2775 | 0 | 86 | 0 | 0 | 0 | 0 | 0 | 0 | 0 | 0 | 0 | 0 |
| SRR8992387 | 0 | 0 | 0 | 265 | 98 | 0 | 39 | 1633 | 3326 | 8374 | 0 | 0 | 0 | 3531 | 0 | 0 | 24 | 127 | 0 | 0 | 51 | 33 | 266 | 3443 | 2847 | 0 | 92 | 0 | 0 | 0 | 0 | 0 | 0 | 0 | 0 | 0 | 0 |
| SRR8992388 | 0 | 0 | 0 | 264 | 147 | 0 | 25 | 1616 | 3488 | 9767 | 0 | 0 | 0 | 3216 | 0 | 0 | 43 | 73 | 19 | 0 | 42 | 22 | 418 | 5043 | 2785 | 0 | 84 | 0 | 0 | 0 | 0 | 0 | 0 | 0 | 0 | 13 | 0 |
| SRR8992389 | 0 | 0 | 0 | 355 | 180 | 0 | 30 | 1789 | 4422 | 8906 | 0 | 0 | 0 | 2242 | 0 | 0 | 36 | 126 | 0 | 0 | 65 | 39 | 330 | 4333 | 3466 | 0 | 11 | 0 | 0 | 0 | 0 | 0 | 0 | 0 | 0 | 0 | 0 |
| SRR8992390 | 0 | 0 | 0 | 302 | 177 | 0 | 30 | 1191 | 2614 | 7205 | 0 | 0 | 0 | 4277 | 0 | 0 | 25 | 121 | 0 | 16 | 46 | 51 | 314 | 3579 | 3182 | 0 | 56 | 0 | 0 | 0 | 0 | 0 | 0 | 0 | 0 | 20 | 0 |
| SRR8992391 | 0 | 0 | 0 | 280 | 151 | 0 | 48 | 1059 | 3090 | 6006 | 0 | 0 | 0 | 2283 | 0 | 0 | 25 | 116 | 14 | 0 | 31 | 35 | 189 | 1559 | 1682 | 19 | 72 | 0 | 0 | 0 | 0 | 0 | 0 | 0 | 0 | 0 | 0 |
| SRR9003330 | 0 | 49 | 33 | 21 | 169 | 0 | 22 | 675 | 1270 | 7546 | 982 | 2760 | 0 | 3146 | 626 | 1291 | 11 | 123 | 825 | 729 | 92 | 1854 | 0 | 15 | 4606 | 0 | 34 | 98 | 42 | 58 | 70 | 29 | 12 | 151 | 13 | 1990 | 240 |
| SRR9003332 | 0 | 12 | 0 | 112 | 178 | 0 | 15 | 0 | 0 | 0 | 2895 | 5009 | 0 | 0 | 0 | 0 | 19 | 565 | 645 | 0 | 415 | 6312 | 0 | 28 | 14882 | 0 | 17 | 64 | 203 | 69 | 0 | 0 | 0 | 0 | 0 | 0 | 852 |
| SRR9003333 | 0 | 48 | 625 | 34 | 259 | 0 | 24 | 537 | 979 | 6444 | 693 | 2977 | 0 | 2594 | 221 | 398 | 11 | 147 | 898 | 906 | 179 | 1229 | 0 | 19 | 6603 | 0 | 48 | 3138 | 1114 | 1355 | 1721 | 674 | 228 | 194 | 24 | 2248 | 306 |
| SRR9003334 | 0 | 16 | 0 | 70 | 125 | 0 | 28 | 0 | 0 | 0 | 3479 | 5756 | 25 | 0 | 0 | 0 | 56 | 436 | 446 | 0 | 280 | 4739 | 0 | 23 | 12319 | 0 | 34 | 3240 | 5212 | 311 | 0 | 0 | 0 | 0 | 0 | 0 | 718 |
| SRR9003336 | 0 | 55 | 544 | 24 | 163 | 0 | 28 | 682 | 1238 | 5673 | 1041 | 2142 | 0 | 2506 | 302 | 620 | 14 | 152 | 1067 | 1123 | 132 | 1608 | 0 | 21 | 6368 | 0 | 63 | 2589 | 1200 | 1630 | 1980 | 949 | 308 | 250 | 13 | 2643 | 261 |
| SRR9003338 | 0 | 28 | 0 | 67 | 118 | 0 | 26 | 0 | 0 | 0 | 8952 | 15105 | 26 | 0 | 0 | 0 | 46 | 427 | 791 | 0 | 220 | 6351 | 0 | 27 | 14063 | 0 | 55 | 4115 | 4912 | 529 | 0 | 0 | 0 | 0 | 0 | 0 | 867 |
| SRR9003339 | 0 | 58 | 31 | 25 | 192 | 0 | 31 | 748 | 1487 | 7717 | 638 | 1802 | 0 | 3401 | 417 | 1098 | 13 | 162 | 618 | 891 | 168 | 1525 | 0 | 17 | 5049 | 0 | 55 | 138 | 59 | 79 | 109 | 42 | 16 | 16 | 48 | 2494 | 221 |
| SRR9003341 | 0 | 66 | 0 | 73 | 126 | 0 | 0 | 0 | 0 | 0 | 10515 | 15932 | 12 | 12 | 0 | 0 | 35 | 484 | 722 | 12 | 229 | 29 | 0 | 33 | 17678 | 0 | 48 | 3050 | 5822 | 920 | 23 | 0 | 0 | 0 | 0 | 0 | 851 |
| SRR9003342 | 0 | 61 | 37 | 21 | 145 | 0 | 19 | 649 | 1351 | 6828 | 1450 | 3206 | 0 | 2961 | 412 | 516 | 11 | 145 | 770 | 1098 | 130 | 1630 | 0 | 17 | 5150 | 0 | 54 | 125 | 43 | 61 | 84 | 41 | 16 | 14 | 20 | 2476 | 239 |
| SRR9018258 | 47 | 3412 | 45 | 18 | 60 | 0 | 17 | 18123 | 13613 | 2262 | 3696 | 10780 | 0 | 187 | 0 | 29 | 29 | 55 | 1287 | 71 | 850 | 1244 | 65 | 6330 | 3552 | 0 | 18 | 234 | 523 | 3614 | 2007 | 479 | 319 | 218 | 20 | 134 | 217 |
| SRR9018259 | 0 | 4060 | 91 | 13 | 32 | 0 | 0 | 4440 | 4226 | 4079 | 16877 | 25207 | 0 | 869 | 76 | 133 | 0 | 16 | 775 | 256 | 953 | 1324 | 74 | 6893 | 4125 | 0 | 16 | 208 | 223 | 4782 | 2721 | 454 | 125 | 70 | 0 | 275 | 190 |
| SRR9018260 | 0 | 155 | 0 | 0 | 0 | 0 | 19 | 1168 | 278 | 246 | 1286 | 10147 | 42 | 556 | 0 | 48 | 77 | 0 | 1796 | 238 | 56 | 1186 | 11 | 983 | 445 | 0 | 23 | 60 | 106 | 367 | 982 | 263 | 164 | 148 | 396 | 580 | 96 |
| SRR9018261 | 27 | 28 | 21 | 0 | 0 | 0 | 31 | 10531 | 5535 | 859 | 4871 | 9353 | 11 | 213 | 137 | 430 | 35 | 0 | 988 | 75 | 329 | 1083 | 43 | 4087 | 1404 | 0 | 0 | 39 | 27 | 76 | 153 | 45 | 34 | 27 | 21 | 221 | 276 |
| SRR952248 | 0 | 45 | 67 | 0 | 13 | 0 | 0 | 686 | 377 | 6061 | 4110 | 5542 | 0 | 1227 | 0 | 0 | 0 | 395 | 15 | 3101 | 825 | 2025 | 0 | 3914 | 6564 | 383 | 11 | 16 | 117 | 116 | 15 | 48 | 16 | 32 | 27 | 22594 | 63 |
| SRR952249 | 0 | 12 | 45 | 0 | 16 | 0 | 0 | 832 | 381 | 5581 | 4015 | 3037 | 0 | 947 | 0 | 0 | 0 | 384 | 12 | 4584 | 1122 | 1773 | 10 | 5309 | 5459 | 257 | 0 | 100 | 104 | 89 | 141 | 30 | 115 | 231 | 314 | 34544 | 72 |
| SRR952288 | 0 | 14 | 42 | 0 | 11 | 0 | 0 | 888 | 341 | 5852 | 4070 | 3048 | 0 | 917 | 0 | 0 | 0 | 392 | 13 | 4751 | 1127 | 1783 | 10 | 5326 | 5543 | 254 | 11 | 13 | 12 | 76 | 18 | 33 | 15 | 31 | 41 | 34710 | 72 |
| SRR952289 | 0 | 24 | 51 | 0 | 20 | 0 | 0 | 723 | 352 | 5518 | 5955 | 6499 | 0 | 873 | 0 | 0 | 0 | 425 | 11 | 5237 | 1158 | 1809 | 14 | 5131 | 5659 | 262 | 24 | 18 | 11 | 14 | 24 | 84 | 22 | 52 | 54 | 35252 | 64 |
| SRR952290 | 0 | 25 | 53 | 0 | 16 | 0 | 0 | 688 | 314 | 5325 | 14013 | 6260 | 0 | 864 | 0 | 0 | 0 | 423 | 11 | 4981 | 1157 | 1806 | 14 | 5193 | 5701 | 256 | 29 | 18 | 11 | 14 | 25 | 89 | 24 | 52 | 52 | 33878 | 66 |
| SRR952309 | 0 | 116 | 62 | 0 | 10 | 0 | 0 | 278 | 99 | 1949 | 17190 | 16587 | 0 | 396 | 0 | 0 | 0 | 315 | 0 | 1375 | 1591 | 3284 | 0 | 4765 | 7705 | 717 | 0 | 31 | 14 | 22 | 14 | 56 | 121 | 13 | 34 | 6715 | 64 |
| SRR952310 | 0 | 187 | 89 | 0 | 11 | 0 | 0 | 571 | 236 | 3730 | 10907 | 12183 | 0 | 669 | 0 | 0 | 0 | 418 | 0 | 2502 | 1981 | 4174 | 0 | 4633 | 8020 | 627 | 0 | 252 | 113 | 209 | 144 | 39 | 138 | 175 | 57 | 13033 | 53 |
| SRR952311 | 0 | 117 | 69 | 0 | 12 | 0 | 0 | 380 | 190 | 4551 | 13291 | 10308 | 0 | 1240 | 0 | 0 | 0 | 487 | 0 | 2520 | 1857 | 3630 | 0 | 5741 | 8644 | 718 | 23 | 329 | 165 | 261 | 158 | 53 | 152 | 180 | 45 | 6742 | 83 |
| SRR952312 | 0 | 0 | 98 | 0 | 0 | 0 | 0 | 1493 | 163 | 1719 | 372 | 3068 | 0 | 950 | 0 | 0 | 0 | 198 | 0 | 4534 | 330 | 466 | 0 | 1252 | 1928 | 99 | 18 | 210 | 84 | 154 | 151 | 91 | 84 | 203 | 151 | 15240 | 60 |
| SRR952313 | 0 | 0 | 73 | 0 | 0 | 0 | 0 | 1336 | 361 | 1678 | 1564 | 2057 | 0 | 744 | 0 | 0 | 0 | 195 | 0 | 5371 | 352 | 397 | 11 | 1514 | 1837 | 109 | 0 | 204 | 61 | 78 | 147 | 81 | 187 | 333 | 309 | 20945 | 57 |
| SRR952314 | 0 | 17 | 202 | 0 | 0 | 0 | 0 | 886 | 149 | 1995 | 3836 | 4690 | 0 | 296 | 0 | 0 | 0 | 173 | 15 | 944 | 621 | 914 | 0 | 3974 | 3664 | 0 | 11 | 308 | 154 | 164 | 261 | 109 | 353 | 578 | 401 | 10362 | 29 |
| SRR952315 | 0 | 0 | 125 | 0 | 0 | 0 | 0 | 857 | 191 | 1893 | 4126 | 4665 | 0 | 284 | 0 | 0 | 0 | 180 | 0 | 946 | 518 | 1041 | 0 | 3075 | 4069 | 0 | 0 | 13 | 95 | 150 | 12 | 71 | 11 | 17 | 20 | 8813 | 34 |
| SRR952316 | 0 | 0 | 57 | 0 | 0 | 0 | 0 | 2263 | 52 | 6378 | 2380 | 2719 | 0 | 2843 | 0 | 0 | 0 | 4551 | 0 | 78 | 0 | 77 | 0 | 393 | 106 | 18006 | 0 | 0 | 0 | 0 | 0 | 0 | 0 | 0 | 0 | 251 | 249 |
| SRR952317 | 0 | 0 | 312 | 0 | 0 | 0 | 0 | 3383 | 1043 | 5653 | 1599 | 2111 | 0 | 2704 | 0 | 0 | 0 | 2767 | 128 | 141 | 322 | 344 | 0 | 665 | 790 | 15440 | 0 | 0 | 0 | 0 | 0 | 0 | 0 | 0 | 0 | 232 | 126 |
| SRR952318 | 0 | 0 | 631 | 0 | 0 | 0 | 0 | 1481 | 272 | 4277 | 1292 | 2903 | 0 | 893 | 0 | 0 | 0 | 2657 | 207 | 0 | 292 | 302 | 0 | 1032 | 843 | 511 | 0 | 0 | 0 | 0 | 0 | 0 | 0 | 0 | 0 | 14 | 0 |
| SRR952319 | 0 | 0 | 416 | 0 | 0 | 0 | 0 | 863 | 316 | 3888 | 1453 | 3343 | 0 | 947 | 0 | 0 | 0 | 2965 | 0 | 75 | 0 | 557 | 0 | 263 | 705 | 0 | 0 | 0 | 0 | 0 | 0 | 0 | 0 | 0 | 0 | 25 | 269 |
| SRR9588764 | 0 | 1861 | 24 | 733 | 2319 | 0 | 35 | 2565 | 11988 | 35696 | 84 | 73 | 0 | 1644 | 0 | 59 | 121 | 1228 | 105 | 440 | 1254 | 1627 | 150 | 10534 | 3492 | 129 | 0 | 0 | 19 | 22 | 0 | 0 | 0 | 0 | 0 | 527 | 0 |
| SRR9588765 | 0 | 1539 | 31 | 876 | 222 | 11 | 11 | 2050 | 7999 | 29921 | 131 | 449 | 0 | 2192 | 96 | 226 | 59 | 864 | 62 | 2092 | 2342 | 2407 | 65 | 8105 | 3807 | 53 | 0 | 0 | 35 | 32 | 14 | 0 | 0 | 0 | 0 | 2027 | 113 |
| SRR9588766 | 0 | 257 | 15 | 783 | 3099 | 0 | 12 | 2423 | 14163 | 29727 | 45 | 163 | 0 | 4267 | 0 | 16 | 30 | 1034 | 103 | 1195 | 2047 | 1135 | 156 | 8081 | 2329 | 0 | 10 | 0 | 16 | 0 | 0 | 0 | 0 | 0 | 0 | 2060 | 12 |
| SRR9588767 | 0 | 1800 | 67 | 445 | 135 | 0 | 16 | 5624 | 22271 | 45515 | 110 | 221 | 0 | 989 | 11 | 15 | 106 | 486 | 60 | 740 | 2004 | 1109 | 143 | 6948 | 1683 | 0 | 0 | 18 | 71 | 60 | 26 | 0 | 0 | 0 | 0 | 777 | 30 |
| SRR9618218 | 0 | 24 | 35 | 0 | 16 | 0 | 0 | 1397 | 530 | 724 | 17500 | 26592 | 0 | 0 | 39 | 91 | 0 | 481 | 15 | 479 | 1774 | 5504 | 0 | 507 | 13874 | 56 | 49 | 14 | 46 | 14 | 70 | 0 | 46 | 73 | 65 | 1125 | 0 |
| SRR9618219 | 0 | 224 | 32 | 0 | 51 | 0 | 0 | 1208 | 534 | 449 | 6947 | 11838 | 0 | 0 | 0 | 11 | 16 | 463 | 115 | 866 | 1450 | 5495 | 0 | 63 | 18597 | 91 | 419 | 37 | 288 | 430 | 242 | 64 | 64 | 62 | 0 | 390 | 0 |
| SRR9618220 | 0 | 20 | 40 | 0 | 29 | 0 | 0 | 1463 | 1024 | 799 | 780 | 11770 | 0 | 0 | 56 | 203 | 0 | 365 | 13 | 461 | 1390 | 5561 | 0 | 405 | 15522 | 47 | 43 | 13 | 70 | 214 | 114 | 33 | 100 | 187 | 77 | 1800 | 0 |
| SRR9618221 | 0 | 24 | 31 | 0 | 23 | 0 | 0 | 1448 | 557 | 612 | 763 | 15290 | 0 | 0 | 33 | 33 | 0 | 410 | 20 | 757 | 1547 | 5553 | 0 | 488 | 14590 | 61 | 48 | 0 | 22 | 14 | 45 | 0 | 48 | 57 | 26 | 1927 | 0 |
| SRR9618222 | 0 | 13 | 0 | 0 | 53 | 0 | 34 | 1187 | 400 | 320 | 1010 | 19787 | 17 | 53 | 0 | 0 | 0 | 229 | 53 | 680 | 381 | 2788 | 0 | 38 | 12858 | 88 | 98 | 17 | 283 | 304 | 351 | 69 | 124 | 351 | 56 | 1693 | 0 |
| SRR9627609 | 0 | 0 | 22 | 0 | 23 | 0 | 0 | 308 | 706 | 199 | 3765 | 5303 | 0 | 181 | 72 | 18 | 0 | 455 | 11 | 3457 | 1260 | 12 | 0 | 17 | 1782 | 33 | 0 | 11 | 12 | 10 | 21 | 0 | 0 | 53 | 18 | 3765 | 0 |
| SRR9627610 | 0 | 0 | 50 | 0 | 21 | 0 | 0 | 170 | 536 | 170 | 4136 | 5493 | 0 | 152 | 0 | 36 | 0 | 451 | 0 | 1867 | 1206 | 1857 | 0 | 19 | 1624 | 28 | 0 | 0 | 13 | 28 | 22 | 0 | 0 | 21 | 0 | 2394 | 0 |
| SRR9627611 | 0 | 0 | 70 | 0 | 16 | 0 | 0 | 249 | 598 | 532 | 182 | 4136 | 0 | 150 | 50 | 17 | 0 | 560 | 0 | 2791 | 855 | 2116 | 0 | 25 | 1857 | 44 | 0 | 0 | 56 | 31 | 17 | 0 | 0 | 26 | 0 | 2774 | 0 |
| SRR9627612 | 0 | 15 | 23 | 0 | 17 | 0 | 0 | 178 | 487 | 132 | 702 | 5624 | 0 | 147 | 54 | 39 | 0 | 518 | 15 | 3489 | 1115 | 1773 | 0 | 23 | 2189 | 20 | 0 | 0 | 14 | 35 | 15 | 0 | 0 | 28 | 11 | 30316 | 0 |
| SRR9627613 | 0 | 0 | 17 | 0 | 12 | 0 | 0 | 382 | 340 | 180 | 851 | 5865 | 0 | 153 | 14 | 49 | 0 | 521 | 0 | 3832 | 1037 | 1739 | 0 | 23 | 2109 | 27 | 0 | 0 | 16 | 31 | 0 | 0 | 0 | 30 | 15 | 29271 | 0 |
| SRR9627614 | 0 | 10 | 47 | 0 | 21 | 0 | 0 | 188 | 338 | 125 | 5729 | 7567 | 0 | 75 | 13 | 0 | 0 | 424 | 13 | 3527 | 1308 | 1588 | 0 | 24 | 2030 | 16 | 0 | 14 | 20 | 58 | 25 | 0 | 0 | 15 | 0 | 22652 | 0 |
| SRR9627651 | 0 | 46 | 47 | 0 | 58 | 0 | 0 | 102 | 526 | 879 | 12890 | 18783 | 0 | 110 | 0 | 0 | 0 | 1265 | 20 | 1012 | 253 | 1467 | 0 | 0 | 724 | 22 | 0 | 0 | 15 | 39 | 59 | 14 | 27 | 40 | 12 | 267 | 0 |
| SRR9627653 | 0 | 61 | 141 | 0 | 53 | 12 | 0 | 31 | 184 | 314 | 2251 | 24139 | 0 | 38 | 0 | 0 | 0 | 923 | 15 | 390 | 168 | 1072 | 0 | 0 | 478 | 17 | 0 | 11 | 18 | 40 | 41 | 11 | 12 | 13 | 0 | 99 | 0 |
| SRR9627654 | 0 | 661 | 255 | 0 | 54 | 0 | 0 | 49 | 367 | 407 | 2360 | 24065 | 0 | 90 | 0 | 0 | 0 | 1086 | 12 | 1222 | 184 | 1157 | 0 | 0 | 704 | 14 | 0 | 0 | 239 | 40 | 39 | 0 | 12 | 25 | 0 | 1581 | 0 |
| SRR9627655 | 0 | 708 | 120 | 0 | 47 | 0 | 0 | 36 | 189 | 108 | 26237 | 28871 | 0 | 36 | 0 | 0 | 0 | 972 | 14 | 530 | 151 | 1052 | 0 | 0 | 542 | 12 | 0 | 0 | 245 | 28 | 21 | 0 | 0 | 0 | 0 | 503 | 0 |
| SRR9627656 | 0 | 58 | 51 | 0 | 50 | 0 | 0 | 27 | 440 | 513 | 2051 | 16381 | 0 | 160 | 0 | 0 | 0 | 1035 | 14 | 1953 | 174 | 1133 | 0 | 0 | 745 | 14 | 0 | 0 | 26 | 35 | 37 | 0 | 20 | 35 | 13 | 2499 | 0 |
| SRR9627693 | 0 | 23 | 13 | 0 | 40 | 0 | 0 | 415 | 776 | 2284 | 206 | 3873 | 0 | 460 | 0 | 54 | 0 | 697 | 15 | 172 | 158 | 608 | 0 | 0 | 589 | 77 | 0 | 23 | 22 | 17 | 0 | 0 | 0 | 19 | 29 | 1815 | 0 |
| SRR9627694 | 0 | 37 | 13 | 0 | 44 | 0 | 0 | 487 | 1175 | 2695 | 218 | 4711 | 0 | 564 | 29 | 48 | 0 | 783 | 17 | 153 | 200 | 798 | 0 | 0 | 548 | 64 | 0 | 34 | 19 | 13 | 10 | 0 | 0 | 20 | 36 | 1596 | 0 |
| SRR9627695 | 0 | 28 | 13 | 0 | 40 | 0 | 0 | 542 | 1139 | 2398 | 3011 | 5067 | 0 | 518 | 16 | 40 | 0 | 760 | 13 | 127 | 178 | 749 | 0 | 0 | 551 | 65 | 0 | 28 | 19 | 12 | 13 | 0 | 0 | 28 | 37 | 1418 | 0 |
| SRR9627696 | 0 | 67 | 55 | 0 | 43 | 0 | 0 | 495 | 547 | 1252 | 211 | 3610 | 0 | 442 | 32 | 11 | 0 | 550 | 0 | 389 | 107 | 509 | 0 | 0 | 552 | 52 | 0 | 0 | 10 | 0 | 0 | 0 | 0 | 13 | 10 | 21216 | 0 |
| SRR9627697 | 0 | 70 | 60 | 0 | 39 | 0 | 0 | 417 | 417 | 871 | 1969 | 3054 | 0 | 576 | 0 | 12 | 0 | 594 | 11 | 245 | 101 | 511 | 0 | 0 | 708 | 51 | 0 | 11 | 75 | 11 | 0 | 0 | 0 | 16 | 64 | 20022 | 0 |
| SRR9627698 | 0 | 12 | 75 | 0 | 40 | 0 | 0 | 386 | 492 | 424 | 295 | 3599 | 0 | 652 | 0 | 30 | 0 | 473 | 0 | 356 | 117 | 440 | 0 | 0 | 669 | 49 | 0 | 23 | 16 | 10 | 0 | 0 | 0 | 29 | 10 | 28837 | 0 |
| SRR9648387 | 0 | 0 | 0 | 22 | 220 | 0 | 0 | 699 | 1696 | 2472 | 0 | 0 | 0 | 8936 | 0 | 0 | 0 | 77 | 220 | 19 | 0 | 33 | 11 | 363 | 0 | 0 | 12 | 0 | 0 | 0 | 0 | 0 | 0 | 0 | 0 | 69 | 11 |
| SRR9648388 | 0 | 0 | 0 | 0 | 0 | 0 | 0 | 12432 | 8022 | 91 | 0 | 0 | 0 | 107 | 0 | 0 | 0 | 0 | 0 | 31 | 0 | 0 | 0 | 167 | 0 | 0 | 0 | 0 | 0 | 0 | 0 | 0 | 0 | 0 | 0 | 118 | 0 |
| SRR9648389 | 0 | 0 | 0 | 62 | 124 | 0 | 0 | 8130 | 6155 | 152 | 0 | 0 | 0 | 30 | 0 | 0 | 0 | 62 | 124 | 0 | 0 | 0 | 62 | 1555 | 0 | 0 | 0 | 0 | 0 | 0 | 0 | 0 | 0 | 0 | 0 | 15 | 0 |
| SRR965653 | 0 | 0 | 0 | 0 | 19 | 0 | 13 | 209 | 38 | 1897 | 4117 | 2068 | 0 | 2296 | 209 | 247 | 0 | 0 | 12 | 13983 | 290 | 1468 | 0 | 290 | 2694 | 0 | 35 | 0 | 0 | 0 | 0 | 0 | 0 | 16 | 0 | 64147 | 0 |
| SRR965655 | 0 | 0 | 37 | 60 | 23 | 0 | 0 | 536 | 583 | 5213 | 431 | 942 | 0 | 364 | 800 | 533 | 0 | 16 | 0 | 1297 | 1251 | 2424 | 0 | 810 | 6772 | 0 | 0 | 19 | 13 | 25 | 15 | 66 | 14 | 40 | 107 | 4524 | 0 |
| SRR9833150 | 0 | 0 | 0 | 0 | 0 | 0 | 12 | 1044 | 235 | 4958 | 2557 | 14218 | 0 | 1213 | 0 | 0 | 15 | 106 | 38 | 1883 | 1747 | 11845 | 10 | 6010 | 13951 | 0 | 93 | 0 | 12 | 90 | 43 | 13 | 15 | 29 | 0 | 5423 | 15 |
| SRR9833151 | 0 | 0 | 0 | 0 | 0 | 0 | 0 | 796 | 117 | 5316 | 4486 | 18691 | 0 | 306 | 11 | 0 | 11 | 82 | 23 | 607 | 1447 | 9525 | 13 | 5924 | 10025 | 12 | 50 | 0 | 0 | 36 | 27 | 0 | 0 | 0 | 0 | 2688 | 15 |
| SRR9833152 | 0 | 12 | 0 | 0 | 0 | 0 | 12 | 1144 | 180 | 5423 | 2686 | 16711 | 0 | 819 | 0 | 0 | 13 | 90 | 31 | 1672 | 1828 | 12436 | 0 | 6198 | 14621 | 16 | 55 | 0 | 13 | 31 | 17 | 0 | 15 | 19 | 0 | 5232 | 16 |
| SRR9833153 | 0 | 0 | 0 | 0 | 0 | 0 | 20 | 983 | 160 | 3742 | 2435 | 19474 | 0 | 891 | 0 | 0 | 17 | 32 | 30 | 694 | 824 | 5875 | 0 | 3898 | 10747 | 12 | 91 | 0 | 30 | 154 | 152 | 38 | 37 | 62 | 0 | 2464 | 20 |
| SRR9833154 | 0 | 11 | 0 | 0 | 0 | 0 | 12 | 707 | 114 | 4401 | 29261 | 22907 | 0 | 228 | 0 | 0 | 13 | 98 | 43 | 528 | 1318 | 8553 | 0 | 4913 | 10026 | 15 | 25 | 0 | 17 | 31 | 27 | 0 | 0 | 15 | 0 | 1501 | 17 |
| SRR9833155 | 0 | 0 | 0 | 0 | 0 | 0 | 18 | 1768 | 319 | 6441 | 2670 | 16088 | 0 | 1945 | 0 | 0 | 15 | 98 | 43 | 1689 | 1268 | 7357 | 14 | 4146 | 11199 | 17 | 127 | 0 | 53 | 164 | 276 | 107 | 115 | 199 | 0 | 4957 | 21 |
| SRR9833156 | 0 | 0 | 0 | 0 | 0 | 0 | 0 | 2142 | 572 | 18749 | 4729 | 25679 | 0 | 1738 | 0 | 0 | 0 | 54 | 44 | 2900 | 1565 | 11061 | 11 | 4689 | 10747 | 11 | 36 | 0 | 16 | 42 | 85 | 28 | 78 | 189 | 0 | 7067 | 27 |
| SRR9833157 | 0 | 0 | 11 | 0 | 0 | 0 | 0 | 2039 | 402 | 20080 | 3960 | 12791 | 0 | 751 | 0 | 0 | 0 | 93 | 31 | 3134 | 1735 | 12719 | 0 | 7598 | 10385 | 16 | 12 | 0 | 0 | 20 | 20 | 0 | 18 | 22 | 0 | 9914 | 17 |
| SRR9833158 | 0 | 0 | 12 | 0 | 0 | 0 | 19 | 1534 | 287 | 8847 | 1945 | 9491 | 0 | 1298 | 107 | 144 | 13 | 86 | 24 | 4268 | 1949 | 15609 | 0 | 6925 | 16221 | 13 | 78 | 0 | 0 | 34 | 34 | 15 | 30 | 97 | 21 | 19881 | 18 |
| SRR9833159 | 0 | 0 | 0 | 0 | 0 | 0 | 39 | 1055 | 78 | 2509 | 2119 | 3850 | 0 | 374 | 283 | 803 | 33 | 84 | 20 | 3221 | 2806 | 19472 | 20 | 10040 | 26746 | 23 | 166 | 0 | 0 | 11 | 12 | 0 | 0 | 22 | 10 | 49596 | 0 |
| SRR9833160 | 0 | 0 | 29 | 0 | 0 | 0 | 31 | 945 | 65 | 3114 | 4742 | 4432 | 0 | 680 | 177 | 508 | 51 | 74 | 25 | 4391 | 2065 | 16807 | 21 | 13154 | 28576 | 0 | 213 | 0 | 0 | 24 | 18 | 0 | 14 | 30 | 11 | 43533 | 0 |
| SRR9833161 | 0 | 0 | 0 | 0 | 0 | 0 | 25 | 376 | 23 | 1543 | 560 | 1927 | 0 | 437 | 182 | 354 | 16 | 61 | 53 | 4748 | 636 | 5058 | 12 | 13331 | 16353 | 19 | 116 | 0 | 0 | 18 | 23 | 0 | 12 | 12 | 14 | 44833 | 0 |
| SRR9833162 | 0 | 0 | 21 | 0 | 0 | 0 | 37 | 1764 | 97 | 3432 | 9196 | 6867 | 0 | 433 | 62 | 200 | 67 | 92 | 18 | 3504 | 2073 | 16159 | 14 | 10801 | 21123 | 22 | 244 | 0 | 14 | 27 | 26 | 0 | 12 | 35 | 21 | 37007 | 0 |
| SRR9833163 | 0 | 0 | 12 | 0 | 0 | 0 | 55 | 930 | 82 | 2505 | 2026 | 5658 | 0 | 562 | 141 | 155 | 74 | 109 | 27 | 2965 | 2036 | 13408 | 18 | 9280 | 20979 | 22 | 326 | 0 | 0 | 22 | 20 | 0 | 10 | 35 | 14 | 54274 | 0 |
| SRR9833164 | 0 | 0 | 20 | 0 | 0 | 0 | 25 | 1212 | 57 | 1672 | 10055 | 8208 | 0 | 474 | 163 | 260 | 36 | 108 | 20 | 4184 | 2382 | 15186 | 17 | 11303 | 19779 | 0 | 187 | 0 | 12 | 35 | 21 | 0 | 15 | 36 | 15 | 59358 | 12 |
| SRR9856157 | 0 | 65 | 42 | 40 | 125 | 67 | 18 | 538 | 1850 | 4302 | 700 | 5450 | 0 | 147 | 196 | 196 | 16 | 901 | 0 | 12681 | 9897 | 3098 | 133 | 3907 | 1754 | 13 | 131 | 23 | 51 | 72 | 190 | 81 | 29 | 26 | 0 | 12475 | 0 |
| SRR9856158 | 0 | 1290 | 55 | 160 | 342 | 116 | 12 | 80 | 685 | 6902 | 1036 | 2252 | 0 | 299 | 42 | 49 | 0 | 1309 | 0 | 2513 | 7538 | 3475 | 290 | 4040 | 2762 | 0 | 147 | 0 | 119 | 236 | 496 | 92 | 46 | 18 | 0 | 4581 | 0 |
| SRR9856159 | 0 | 58 | 118 | 88 | 326 | 47 | 0 | 847 | 3041 | 7720 | 2097 | 4212 | 0 | 345 | 131 | 162 | 0 | 1563 | 0 | 3662 | 6509 | 2100 | 372 | 6016 | 2011 | 18 | 92 | 22 | 93 | 127 | 13 | 105 | 66 | 45 | 0 | 8857 | 0 |
| SRR9856160 | 0 | 226 | 19 | 20 | 11 | 149 | 0 | 206 | 411 | 4696 | 265 | 359 | 12 | 180 | 98 | 41 | 30 | 627 | 0 | 58943 | 28 | 1488 | 73 | 6689 | 966 | 0 | 454 | 0 | 26 | 33 | 10 | 137 | 37 | 26 | 0 | 53630 | 0 |
| SRR9856161 | 0 | 1700 | 82 | 136 | 280 | 75 | 0 | 267 | 1056 | 6253 | 1700 | 3751 | 0 | 447 | 26 | 39 | 18 | 1339 | 0 | 3917 | 14035 | 4626 | 289 | 4569 | 2257 | 0 | 124 | 15 | 42 | 145 | 296 | 39 | 21 | 18 | 0 | 4794 | 0 |
| SRR9856162 | 0 | 1582 | 61 | 109 | 187 | 23 | 0 | 68 | 448 | 2193 | 1955 | 4928 | 0 | 170 | 41 | 27 | 0 | 1388 | 0 | 2070 | 38 | 10 | 319 | 3888 | 3124 | 0 | 103 | 0 | 112 | 234 | 389 | 47 | 51 | 19 | 0 | 5831 | 0 |
| SRR9856163 | 0 | 1287 | 67 | 153 | 285 | 28 | 0 | 268 | 970 | 7971 | 288 | 2273 | 0 | 231 | 61 | 61 | 0 | 1209 | 0 | 3032 | 12661 | 4097 | 274 | 2568 | 1562 | 0 | 64 | 0 | 79 | 205 | 385 | 70 | 31 | 0 | 0 | 9043 | 0 |
| SRR9856164 | 0 | 26 | 51 | 159 | 655 | 89 | 0 | 345 | 1454 | 7296 | 1007 | 1989 | 0 | 458 | 48 | 40 | 22 | 1658 | 0 | 4869 | 20 | 3576 | 238 | 3968 | 2019 | 0 | 85 | 14 | 51 | 80 | 232 | 49 | 22 | 13 | 0 | 5140 | 0 |
| SRR9856165 | 0 | 26 | 61 | 77 | 138 | 47 | 0 | 236 | 863 | 3557 | 895 | 12989 | 0 | 214 | 44 | 81 | 31 | 846 | 0 | 7380 | 10207 | 2784 | 185 | 2979 | 1424 | 0 | 122 | 17 | 47 | 112 | 441 | 51 | 44 | 34 | 0 | 8605 | 0 |
| SRR9856166 | 0 | 21 | 65 | 59 | 142 | 87 | 0 | 209 | 702 | 2620 | 1877 | 4254 | 0 | 203 | 27 | 68 | 16 | 836 | 0 | 3903 | 17 | 2686 | 156 | 2156 | 1104 | 0 | 50 | 19 | 22 | 56 | 131 | 19 | 12 | 0 | 0 | 5942 | 0 |
| SRR9856167 | 0 | 26 | 33 | 55 | 138 | 49 | 15 | 210 | 751 | 3615 | 3228 | 4956 | 0 | 171 | 270 | 237 | 29 | 1211 | 0 | 10209 | 24 | 2215 | 224 | 5635 | 1858 | 0 | 92 | 15 | 51 | 92 | 14 | 40 | 26 | 18 | 0 | 11247 | 0 |
| SRR9856168 | 0 | 1032 | 50 | 70 | 334 | 43 | 0 | 272 | 867 | 7376 | 251 | 2772 | 0 | 361 | 132 | 77 | 12 | 1998 | 0 | 2933 | 25 | 3495 | 311 | 4194 | 1539 | 18 | 54 | 19 | 78 | 177 | 473 | 62 | 38 | 14 | 0 | 5837 | 0 |
| SRR9856169 | 0 | 40 | 43 | 75 | 521 | 61 | 0 | 247 | 969 | 3988 | 846 | 2796 | 0 | 360 | 49 | 61 | 23 | 1857 | 0 | 5740 | 36 | 3503 | 398 | 5246 | 2064 | 14 | 95 | 0 | 77 | 195 | 17 | 59 | 25 | 24 | 0 | 12561 | 0 |
| SRR9856170 | 0 | 698 | 35 | 242 | 313 | 132 | 0 | 111 | 485 | 2527 | 1804 | 3757 | 0 | 72 | 0 | 0 | 30 | 3903 | 0 | 3667 | 16815 | 4327 | 346 | 4357 | 2789 | 12 | 409 | 19 | 106 | 282 | 616 | 124 | 35 | 23 | 0 | 4416 | 0 |

### Supp Table S8.4.Columns_DI_to_ES

| | | | | | | | | | | | | | | | | | | | | | | | | | | | | | | | | | | | | | |
|---|---|---|---|---|---|---|---|---|---|---|---|---|---|---|---|---|---|---|---|---|---|---|---|---|---|---|---|---|---|---|---|---|---|---|---|---|---|
| SRR9856171 | 0 | 26 | 0 | 150 | 226 | 118 | 13 | 47 | 91 | 227 | 1420 | 4259 | 0 | 0 | 0 | 16 | 40 | 2606 | 0 | 4572 | 42 | 5301 | 535 | 5888 | 4729 | 17 | 16 | 0 | 25 | 66 | 12 | 29 | 10 | 0 | 0 | 3568 | 0 |
| SRR9856172 | 0 | 28 | 65 | 117 | 472 | 57 | 0 | 300 | 800 | 3148 | 5888 | 16857 | 0 | 123 | 108 | 23 | 22 | 1644 | 0 | 1978 | 19183 | 5538 | 487 | 4653 | 3618 | 12 | 74 | 12 | 125 | 232 | 503 | 74 | 31 | 34 | 0 | 3133 | 0 |
| SRR9856173 | 0 | 739 | 29 | 126 | 738 | 85 | 12 | 583 | 1920 | 20229 | 410 | 3428 | 0 | 370 | 66 | 63 | 33 | 1615 | 0 | 4087 | 9041 | 2814 | 499 | 7025 | 2494 | 16 | 114 | 0 | 36 | 77 | 195 | 27 | 16 | 12 | 0 | 9542 | 0 |
| SRR9856174 | 0 | 31 | 39 | 77 | 381 | 172 | 0 | 453 | 2159 | 10645 | 2195 | 4359 | 0 | 562 | 46 | 36 | 59 | 2470 | 0 | 11938 | 18727 | 2843 | 240 | 6006 | 2389 | 12 | 173 | 12 | 118 | 222 | 35 | 120 | 64 | 40 | 0 | 11263 | 0 |
| SRR9856175 | 0 | 842 | 49 | 94 | 422 | 58 | 0 | 245 | 1047 | 6728 | 544 | 858 | 0 | 435 | 133 | 112 | 21 | 1399 | 0 | 6279 | 12532 | 4219 | 291 | 5768 | 1871 | 17 | 139 | 15 | 51 | 142 | 556 | 66 | 30 | 28 | 0 | 15558 | 0 |
| SRR9856176 | 0 | 75 | 71 | 75 | 267 | 77 | 18 | 891 | 3027 | 6662 | 2498 | 6335 | 0 | 368 | 184 | 234 | 19 | 1022 | 0 | 6039 | 6798 | 2515 | 291 | 5637 | 2039 | 21 | 146 | 0 | 61 | 60 | 14 | 75 | 39 | 32 | 0 | 9238 | 0 |
| SRR9856177 | 0 | 598 | 29 | 100 | 1522 | 46 | 13 | 805 | 3872 | 8611 | 735 | 2156 | 0 | 359 | 16 | 20 | 22 | 1860 | 0 | 3196 | 11 | 1406 | 334 | 4517 | 1802 | 13 | 101 | 10 | 36 | 36 | 141 | 41 | 18 | 17 | 0 | 8662 | 0 |
| SRR9856178 | 0 | 22 | 38 | 28 | 0 | 71 | 0 | 564 | 909 | 7261 | 195 | 2707 | 0 | 115 | 1024 | 497 | 38 | 279 | 0 | 19880 | 4545 | 1267 | 85 | 7300 | 1710 | 0 | 125 | 0 | 20 | 56 | 20 | 56 | 41 | 25 | 0 | 17680 | 0 |
| SRR9856179 | 0 | 887 | 40 | 127 | 95 | 125 | 0 | 44 | 305 | 1262 | 256 | 6502 | 0 | 78 | 296 | 279 | 13 | 829 | 0 | 3055 | 15818 | 4948 | 251 | 2466 | 1953 | 0 | 113 | 0 | 32 | 56 | 148 | 25 | 27 | 17 | 0 | 4700 | 0 |
| SRR9856180 | 0 | 20 | 52 | 102 | 122 | 65 | 0 | 99 | 482 | 1793 | 3176 | 8712 | 0 | 106 | 362 | 532 | 0 | 764 | 0 | 6883 | 32 | 12 | 223 | 2497 | 1823 | 0 | 65 | 19 | 45 | 168 | 530 | 52 | 29 | 13 | 0 | 4976 | 0 |
| SRR9856181 | 0 | 2187 | 63 | 98 | 125 | 42 | 0 | 115 | 508 | 2228 | 1878 | 4230 | 0 | 235 | 175 | 102 | 21 | 775 | 0 | 6389 | 11355 | 4038 | 216 | 2879 | 1434 | 0 | 84 | 17 | 110 | 291 | 868 | 122 | 42 | 42 | 0 | 5732 | 0 |
| SRR9856182 | 0 | 31 | 61 | 83 | 75 | 52 | 0 | 49 | 183 | 790 | 1166 | 2058 | 0 | 60 | 330 | 200 | 0 | 518 | 0 | 6185 | 10643 | 4260 | 204 | 3404 | 1762 | 0 | 81 | 0 | 45 | 135 | 384 | 45 | 26 | 39 | 0 | 7436 | 0 |
| SRR9856183 | 0 | 45 | 12 | 108 | 146 | 93 | 0 | 114 | 386 | 2166 | 397 | 3003 | 0 | 155 | 34 | 49 | 29 | 1331 | 0 | 4169 | 14791 | 5827 | 270 | 5524 | 2626 | 0 | 21 | 0 | 17 | 32 | 100 | 0 | 10 | 0 | 0 | 5612 | 0 |
| SRR9856184 | 0 | 23 | 48 | 62 | 75 | 60 | 0 | 122 | 240 | 3074 | 298 | 3615 | 0 | 96 | 1284 | 1270 | 0 | 952 | 0 | 6047 | 10487 | 6394 | 202 | 4129 | 1690 | 11 | 88 | 14 | 41 | 111 | 292 | 51 | 49 | 26 | 0 | 7806 | 0 |
| SRR9856185 | 0 | 1357 | 65 | 14 | 14 | 58 | 15 | 203 | 1115 | 3750 | 85 | 5540 | 0 | 203 | 439 | 304 | 44 | 542 | 0 | 32733 | 10 | 1068 | 115 | 2499 | 892 | 0 | 232 | 15 | 116 | 116 | 595 | 167 | 58 | 51 | 0 | 19795 | 0 |
| SRR9856186 | 0 | 1073 | 27 | 45 | 49 | 61 | 0 | 100 | 561 | 5079 | 707 | 5809 | 0 | 221 | 159 | 95 | 19 | 1161 | 0 | 17793 | 18 | 4081 | 205 | 4892 | 2831 | 16 | 305 | 0 | 65 | 291 | 1201 | 235 | 157 | 99 | 0 | 10321 | 0 |
| SRR990360 | 0 | 27 | 0 | 0 | 25 | 0 | 0 | 0 | 0 | 0 | 3394 | 2388 | 0 | 0 | 0 | 0 | 59 | 5139 | 0 | 0 | 6096 | 3440 | 272 | 2643 | 2834 | 0 | 13 | 12 | 0 | 0 | 0 | 0 | 0 | 0 | 0 | 0 | 63 |
| SRR9924500 | 0 | 61 | 27 | 0 | 29 | 0 | 0 | 36198 | 3727 | 4532 | 613 | 2769 | 0 | 383 | 43 | 57 | 0 | 232 | 0 | 819 | 2162 | 4347 | 0 | 153 | 10236 | 0 | 0 | 155 | 35 | 74 | 66 | 14 | 54 | 93 | 28 | 2018 | 285 |
| SRR9924502 | 0 | 71 | 18 | 0 | 26 | 13 | 0 | 22129 | 2572 | 5060 | 701 | 3505 | 0 | 335 | 24 | 0 | 0 | 208 | 0 | 1001 | 2410 | 4136 | 0 | 185 | 9805 | 0 | 0 | 16 | 24 | 45 | 35 | 0 | 21 | 45 | 0 | 2400 | 263 |
| SRR9924511 | 0 | 68 | 36 | 0 | 45 | 0 | 0 | 17364 | 1999 | 5402 | 2024 | 9625 | 0 | 725 | 365 | 216 | 0 | 224 | 0 | 1023 | 1468 | 2623 | 0 | 111 | 8080 | 0 | 0 | 111 | 11 | 24 | 36 | 0 | 24 | 43 | 26 | 2560 | 150 |
| SRR9924512 | 0 | 47 | 22 | 0 | 41 | 11 | 0 | 22129 | 2143 | 4737 | 1353 | 6694 | 0 | 579 | 167 | 48 | 0 | 238 | 0 | 770 | 1314 | 2120 | 0 | 123 | 7959 | 0 | 0 | 76 | 12 | 28 | 31 | 0 | 10 | 46 | 0 | 2990 | 168 |
| SRR9924514 | 0 | 11 | 29 | 0 | 67 | 26 | 0 | 4568 | 668 | 6848 | 9160 | 11778 | 0 | 399 | 221 | 123 | 11 | 200 | 0 | 1251 | 1478 | 2593 | 0 | 118 | 8273 | 0 | 0 | 11 | 0 | 22 | 23 | 0 | 18 | 38 | 21 | 3243 | 155 |
| SRR9924515 | 0 | 47 | 21 | 0 | 49 | 0 | 0 | 10969 | 1217 | 3848 | 1665 | 9193 | 0 | 506 | 183 | 105 | 0 | 216 | 0 | 1030 | 1451 | 2362 | 0 | 119 | 7622 | 0 | 0 | 59 | 0 | 15 | 11 | 0 | 17 | 27 | 0 | 3320 | 135 |
| SRR9924516 | 0 | 11 | 12 | 0 | 56 | 18 | 0 | 3265 | 479 | 2929 | 2399 | 17265 | 0 | 267 | 19 | 0 | 11 | 166 | 0 | 858 | 1590 | 2585 | 0 | 118 | 8057 | 0 | 0 | 11 | 12 | 22 | 26 | 0 | 13 | 26 | 0 | 1922 | 156 |
| SRR9924517 | 0 | 13 | 25 | 0 | 62 | 16 | 0 | 3658 | 503 | 5097 | 10983 | 13413 | 0 | 595 | 92 | 38 | 0 | 182 | 0 | 1412 | 1471 | 2590 | 0 | 119 | 7973 | 0 | 0 | 11 | 15 | 35 | 34 | 0 | 14 | 44 | 13 | 3501 | 157 |
| SRR9924518 | 0 | 47 | 24 | 0 | 29 | 0 | 0 | 50017 | 4438 | 2478 | 2162 | 3716 | 0 | 239 | 49 | 30 | 0 | 215 | 0 | 390 | 2627 | 4051 | 0 | 134 | 8929 | 0 | 0 | 13 | 22 | 34 | 42 | 0 | 18 | 48 | 0 | 1099 | 277 |
| SRR9924519 | 0 | 46 | 14 | 0 | 27 | 12 | 0 | 49214 | 4537 | 3849 | 4127 | 3292 | 0 | 211 | 124 | 112 | 0 | 187 | 0 | 526 | 2501 | 4065 | 0 | 153 | 9509 | 0 | 0 | 50 | 0 | 26 | 18 | 0 | 18 | 29 | 20 | 1810 | 304 |
| SRR9924520 | 0 | 79 | 25 | 0 | 24 | 17 | 0 | 6010 | 1154 | 7544 | 883 | 5499 | 0 | 476 | 216 | 137 | 0 | 169 | 0 | 1379 | 2662 | 4717 | 0 | 162 | 9309 | 0 | 0 | 90 | 29 | 37 | 39 | 0 | 26 | 83 | 36 | 4296 | 309 |
| SRR9924521 | 0 | 33 | 22 | 0 | 17 | 11 | 0 | 49803 | 4445 | 3451 | 1943 | 3277 | 0 | 192 | 93 | 65 | 0 | 174 | 0 | 342 | 2417 | 4176 | 0 | 137 | 8405 | 0 | 0 | 13 | 20 | 34 | 47 | 0 | 23 | 56 | 14 | 1295 | 250 |
| SRR9924522 | 0 | 39 | 0 | 0 | 20 | 25 | 0 | 10019 | 1142 | 5345 | 6552 | 8550 | 0 | 434 | 24 | 12 | 0 | 148 | 0 | 1094 | 2508 | 4211 | 0 | 158 | 8873 | 0 | 0 | 81 | 28 | 31 | 22 | 0 | 14 | 42 | 0 | 2878 | 278 |
| SRR9924523 | 0 | 23 | 15 | 0 | 23 | 0 | 0 | 10931 | 1657 | 5714 | 749 | 5741 | 0 | 326 | 112 | 86 | 0 | 175 | 0 | 1262 | 2713 | 4213 | 0 | 172 | 9003 | 0 | 0 | 14 | 15 | 31 | 37 | 0 | 18 | 56 | 11 | 2790 | 255 |
| SRR9924524 | 0 | 28 | 25 | 0 | 23 | 0 | 0 | 25281 | 2869 | 7992 | 1592 | 4006 | 0 | 472 | 135 | 98 | 0 | 231 | 0 | 881 | 2089 | 3296 | 0 | 188 | 9232 | 0 | 0 | 41 | 22 | 24 | 22 | 0 | 12 | 25 | 0 | 2541 | 323 |
| SRR9924525 | 0 | 63 | 37 | 0 | 31 | 0 | 0 | 42981 | 3260 | 4172 | 3289 | 3042 | 0 | 335 | 157 | 87 | 0 | 217 | 0 | 688 | 2111 | 3377 | 0 | 154 | 8716 | 0 | 0 | 145 | 43 | 69 | 53 | 10 | 27 | 82 | 69 | 2004 | 252 |
| SRR9924526 | 0 | 42 | 35 | 0 | 29 | 0 | 0 | 34261 | 4029 | 8277 | 1916 | 3802 | 0 | 579 | 198 | 146 | 0 | 231 | 0 | 544 | 2059 | 3302 | 0 | 183 | 8864 | 0 | 0 | 11 | 22 | 31 | 22 | 0 | 13 | 25 | 21 | 1476 | 303 |
| SRR9924527 | 0 | 44 | 33 | 0 | 21 | 14 | 0 | 31983 | 3845 | 8797 | 4722 | 4964 | 0 | 444 | 145 | 68 | 0 | 196 | 0 | 944 | 2347 | 3941 | 0 | 170 | 8849 | 0 | 0 | 64 | 27 | 30 | 17 | 0 | 18 | 43 | 10 | 2229 | 267 |
