## Supplementary material for "Single-nucleotide Differences and Cell Type Decide the Subcellular Localization of miRNA Isoforms (isomiRs), tRNA-derived Fragments (tRFs) and rRNA-derived Fragments (rRFs)": Supp. Table S8 (5 of 5)

### Supp Table S8.5.Columns_ET_to_FD

**Supplemental Table S8:** The abundances

#### 11 5´ ETS rRF (Figure 6)

| | >hg38_21_+_8433172_8446622_RNA45SN1_45S_50ntFlanks@475.524.50\|CTTCGTGATCGATGTGGTGACGTCGTGCTCTCCCGGGCCGGGTCCGAGCC | >hg38_21_+_8433172_8446622_RNA45SN1_45S_50ntFlanks@475.517.43\|CTTCGTGATCGATGTGGTGACGTCGTGCTCTCCCGGGCCGGGT | >hg38_21_+_8433172_8446622_RNA45SN1_45S_50ntFlanks@475.516.42\|CTTCGTGATCGATGTGGTGACGTCGTGCTCTCCCGGGCCGGG | >hg38_21_+_8433172_8446622_RNA45SN1_45S_50ntFlanks@475.515.41\|CTTCGTGATCGATGTGGTGACGTCGTGCTCTCCCGGGCCGG | >hg38_21_+_8433172_8446622_RNA45SN1_45S_50ntFlanks@475.506.32\|CTTCGTGATCGATGTGGTGACGTCGTGCTCTC | >hg38_21_+_8433172_8446622_RNA45SN1_45S_50ntFlanks@475.505.31\|CTTCGTGATCGATGTGGTGACGTCGTGCTCT | >hg38_21_+_8433172_8446622_RNA45SN1_45S_50ntFlanks@475.504.30\|CTTCGTGATCGATGTGGTGACGTCGTGCTC | >hg38_21_+_8433172_8446622_RNA45SN1_45S_50ntFlanks@475.503.29\|CTTCGTGATCGATGTGGTGACGTCGTGCT | >hg38_21_+_8433172_8446622_RNA45SN1_45S_50ntFlanks@475.498.24\|CTTCGTGATCGATGTGGTGACGTC | >hg38_21_+_8433172_8446622_RNA45SN1_45S_50ntFlanks@475.497.23\|CTTCGTGATCGATGTGGTGACGT | >hg38_21_+_8433172_8446622_RNA45SN1_45S_50ntFlanks@475.496.22\|CTTCGTGATCGATGTGGTGACG |
|---|---|---|---|---|---|---|---|---|---|---|---|
| SRR10024150 | 0 | 31 | 138 | 102 | 634 | 976 | 82 | 465 | 128 | 0 | 118 |
| SRR10024151 | 0 | 28 | 246 | 133 | 1173 | 2545 | 199 | 1665 | 301 | 25 | 182 |
| SRR10024152 | 0 | 10 | 82 | 46 | 406 | 812 | 51 | 180 | 180 | 0 | 77 |
| SRR10024153 | 0 | 0 | 25 | 25 | 597 | 1733 | 80 | 1284 | 299 | 30 | 229 |
| SRR10024154 | 0 | 16 | 22 | 41 | 168 | 383 | 25 | 136 | 51 | 16 | 32 |
| SRR10024155 | 0 | 0 | 16 | 0 | 222 | 556 | 38 | 279 | 72 | 16 | 99 |
| SRR10024156 | 0 | 20 | 81 | 23 | 516 | 1463 | 112 | 783 | 183 | 13 | 91 |
| SRR10024158 | 0 | 0 | 0 | 0 | 133 | 338 | 34 | 307 | 104 | 0 | 87 |
| SRR10024159 | 0 | 0 | 0 | 0 | 380 | 1127 | 70 | 654 | 313 | 17 | 157 |
| SRR10024161 | 0 | 15 | 15 | 15 | 244 | 527 | 57 | 584 | 134 | 19 | 118 |
| SRR10024162 | 0 | 25 | 146 | 53 | 1006 | 2843 | 190 | 1647 | 536 | 62 | 330 |
| SRR10024163 | 0 | 0 | 210 | 98 | 1199 | 1735 | 131 | 760 | 434 | 23 | 131 |
| SRR10024164 | 0 | 0 | 0 | 35 | 702 | 1282 | 121 | 139 | 52 | 0 | 43 |
| SRR10024166 | 0 | 0 | 41 | 0 | 1400 | 2783 | 235 | 950 | 562 | 29 | 182 |
| SRR10024167 | 0 | 0 | 0 | 0 | 92 | 294 | 51 | 277 | 215 | 0 | 113 |
| SRR10024168 | 0 | 0 | 19 | 15 | 237 | 305 | 29 | 121 | 53 | 0 | 19 |
| SRR10024169 | 0 | 0 | 0 | 13 | 534 | 1540 | 109 | 1558 | 439 | 47 | 182 |
| SRR10024170 | 0 | 0 | 22 | 30 | 525 | 1340 | 121 | 273 | 191 | 22 | 95 |
| SRR10024171 | 0 | 0 | 72 | 47 | 294 | 447 | 89 | 291 | 160 | 0 | 86 |
| SRR10024173 | 0 | 0 | 0 | 23 | 545 | 653 | 90 | 108 | 45 | 23 | 45 |
| SRR10024176 | 0 | 0 | 0 | 0 | 67 | 147 | 24 | 218 | 70 | 0 | 39 |
| SRR10024178 | 0 | 0 | 33 | 17 | 214 | 412 | 64 | 228 | 150 | 14 | 53 |
| SRR10024179 | 0 | 0 | 0 | 0 | 89 | 215 | 17 | 169 | 93 | 0 | 28 |
| SRR10024180 | 0 | 0 | 0 | 0 | 272 | 655 | 70 | 409 | 199 | 0 | 110 |
| SRR10024181 | 0 | 35 | 154 | 75 | 367 | 1033 | 91 | 640 | 210 | 19 | 163 |
| SRR10024182 | 0 | 0 | 150 | 48 | 970 | 2269 | 101 | 830 | 227 | 19 | 193 |
| SRR10024183 | 0 | 0 | 89 | 46 | 348 | 931 | 86 | 205 | 214 | 21 | 92 |
| SRR10024185 | 0 | 0 | 57 | 48 | 602 | 1380 | 111 | 464 | 139 | 0 | 111 |
| SRR10024186 | 0 | 0 | 22 | 11 | 149 | 352 | 39 | 344 | 154 | 17 | 69 |
| SRR10024187 | 0 | 0 | 25 | 31 | 1304 | 1820 | 183 | 82 | 38 | 13 | 82 |
| SRR10024188 | 0 | 0 | 32 | 16 | 782 | 1915 | 165 | 210 | 347 | 20 | 56 |
| SRR10024189 | 0 | 0 | 17 | 0 | 813 | 1901 | 125 | 700 | 275 | 13 | 146 |
| SRR10024190 | 0 | 0 | 105 | 12 | 2673 | 4921 | 277 | 125 | 515 | 0 | 52 |
| SRR10024191 | 0 | 0 | 0 | 33 | 434 | 662 | 98 | 206 | 119 | 54 | 54 |
| SRR10024192 | 0 | 11 | 64 | 64 | 1399 | 3374 | 171 | 438 | 320 | 43 | 85 |
| SRR10024193 | 0 | 0 | 69 | 69 | 547 | 1218 | 120 | 1225 | 308 | 43 | 236 |
| SRR10024194 | 0 | 0 | 15 | 0 | 173 | 352 | 41 | 138 | 120 | 15 | 47 |
| SRR10024195 | 0 | 12 | 169 | 31 | 397 | 994 | 55 | 534 | 193 | 12 | 118 |
| SRR10024196 | 0 | 0 | 39 | 31 | 884 | 1432 | 180 | 509 | 376 | 16 | 86 |
| SRR10024197 | 0 | 0 | 35 | 30 | 195 | 311 | 52 | 216 | 52 | 0 | 48 |
| SRR10024198 | 0 | 0 | 50 | 13 | 165 | 326 | 48 | 202 | 143 | 13 | 48 |
| SRR10024199 | 0 | 13 | 52 | 26 | 626 | 988 | 65 | 155 | 168 | 19 | 19 |
| SRR10024201 | 0 | 0 | 41 | 23 | 429 | 1222 | 99 | 1037 | 347 | 27 | 257 |
| SRR10024202 | 0 | 0 | 40 | 16 | 1204 | 1674 | 191 | 151 | 263 | 16 | 24 |
| SRR10024203 | 0 | 23 | 97 | 28 | 506 | 1674 | 143 | 1357 | 244 | 23 | 322 |
| SRR10024204 | 0 | 0 | 0 | 0 | 306 | 1620 | 78 | 888 | 371 | 26 | 205 |
| SRR10024205 | 0 | 0 | 70 | 23 | 1649 | 4437 | 244 | 709 | 290 | 46 | 348 |
| SRR10024206 | 0 | 0 | 46 | 19 | 316 | 843 | 84 | 774 | 167 | 31 | 128 |
| SRR10024207 | 0 | 0 | 0 | 0 | 265 | 566 | 46 | 260 | 204 | 25 | 87 |
| SRR10024208 | 0 | 0 | 36 | 0 | 375 | 682 | 36 | 207 | 88 | 0 | 76 |
| SRR1003025 | 0 | 39 | 145 | 137 | 85 | 534 | 237 | 811 | 775 | 179 | 1522 |
| SRR1005875 | 0 | 0 | 0 | 0 | 0 | 0 | 0 | 0 | 0 | 0 | 0 |
| SRR10072377 | 0 | 0 | 0 | 0 | 16 | 42 | 0 | 34 | 65 | 15 | 11 |
| SRR10072378 | 0 | 0 | 0 | 0 | 0 | 38 | 15 | 31 | 62 | 0 | 19 |
| SRR10072379 | 0 | 0 | 0 | 13 | 22 | 63 | 13 | 46 | 121 | 14 | 24 |
| SRR10072380 | 0 | 0 | 0 | 12 | 28 | 67 | 13 | 43 | 110 | 0 | 28 |
| SRR10072381 | 0 | 0 | 0 | 17 | 0 | 61 | 16 | 38 | 66 | 0 | 0 |
| SRR10072382 | 0 | 0 | 13 | 24 | 13 | 52 | 10 | 24 | 81 | 0 | 0 |
| SRR10072383 | 0 | 0 | 0 | 21 | 45 | 124 | 33 | 99 | 178 | 40 | 33 |
| SRR10072384 | 0 | 0 | 0 | 21 | 72 | 148 | 28 | 97 | 171 | 25 | 25 |
| SRR10072385 | 0 | 0 | 14 | 22 | 30 | 119 | 30 | 55 | 109 | 19 | 18 |
| SRR10072386 | 0 | 0 | 0 | 20 | 26 | 90 | 29 | 67 | 108 | 18 | 23 |
| SRR10072387 | 0 | 0 | 0 | 0 | 12 | 44 | 12 | 21 | 94 | 0 | 0 |
| SRR10072388 | 0 | 0 | 0 | 15 | 0 | 39 | 15 | 27 | 81 | 0 | 0 |
| SRR10072389 | 0 | 0 | 0 | 61 | 293 | 789 | 229 | 455 | 312 | 140 | 93 |
| SRR10072390 | 0 | 0 | 0 | 63 | 255 | 821 | 175 | 372 | 284 | 131 | 87 |
| SRR10072391 | 0 | 0 | 0 | 17 | 35 | 96 | 36 | 59 | 183 | 23 | 23 |
| SRR10072392 | 0 | 0 | 0 | 11 | 34 | 77 | 30 | 68 | 184 | 28 | 23 |
| SRR10072393 | 0 | 0 | 0 | 21 | 30 | 50 | 16 | 44 | 121 | 11 | 25 |
| SRR10072394 | 0 | 0 | 12 | 21 | 44 | 72 | 28 | 51 | 153 | 16 | 32 |
| SRR10072395 | 0 | 0 | 0 | 48 | 429 | 524 | 109 | 225 | 129 | 41 | 54 |
| SRR10072396 | 0 | 0 | 0 | 27 | 342 | 456 | 141 | 235 | 121 | 47 | 60 |
| SRR10072397 | 0 | 0 | 20 | 12 | 79 | 204 | 51 | 74 | 78 | 34 | 27 |
| SRR10072398 | 0 | 0 | 0 | 15 | 60 | 224 | 32 | 103 | 101 | 30 | 25 |
| SRR10072399 | 0 | 0 | 0 | 14 | 13 | 39 | 0 | 32 | 142 | 12 | 23 |
| SRR10072400 | 0 | 0 | 10 | 14 | 0 | 36 | 12 | 36 | 144 | 0 | 14 |
| SRR10077412 | 14 | 0 | 14 | 19 | 29 | 319 | 53 | 608 | 980 | 541 | 2100 |
| SRR10077413 | 0 | 0 | 0 | 15 | 20 | 414 | 79 | 755 | 1051 | 533 | 2077 |
| SRR10077414 | 0 | 0 | 0 | 21 | 26 | 395 | 90 | 706 | 880 | 580 | 1898 |
| SRR10077415 | 0 | 0 | 25 | 15 | 0 | 303 | 60 | 776 | 975 | 473 | 1975 |
| SRR10077416 | 0 | 0 | 0 | 14 | 0 | 106 | 14 | 207 | 948 | 428 | 1752 |
| SRR10077417 | 0 | 0 | 0 | 0 | 0 | 88 | 0 | 205 | 895 | 548 | 1883 |
| SRR10077418 | 0 | 0 | 0 | 0 | 0 | 58 | 21 | 189 | 886 | 503 | 1866 |
| SRR10077419 | 0 | 0 | 0 | 0 | 0 | 49 | 25 | 168 | 998 | 548 | 1754 |
| SRR10077420 | 0 | 0 | 0 | 0 | 0 | 99 | 17 | 248 | 618 | 335 | 1376 |
| SRR10077421 | 0 | 0 | 0 | 0 | 0 | 54 | 21 | 228 | 624 | 360 | 1311 |
| SRR10077422 | 0 | 0 | 0 | 0 | 0 | 61 | 19 | 166 | 642 | 415 | 1342 |
| SRR10077423 | 0 | 0 | 0 | 0 | 0 | 78 | 24 | 175 | 667 | 317 | 1285 |

### Supp Table S8.5.Columns_ET_to_FD

| | | | | | | | | | | | |
|---|---|---|---|---|---|---|---|---|---|---|---|
| SRR10077424 | 0 | 0 | 0 | 0 | 18 | 191 | 51 | 350 | 927 | 790 | 2364 |
| SRR10077425 | 0 | 0 | 0 | 0 | 0 | 180 | 44 | 468 | 788 | 685 | 2214 |
| SRR10077426 | 0 | 0 | 0 | 0 | 0 | 161 | 59 | 448 | 919 | 825 | 2165 |
| SRR10077427 | 0 | 0 | 0 | 0 | 0 | 144 | 48 | 369 | 971 | 831 | 2141 |
| SRR10077428 | 0 | 0 | 14 | 0 | 71 | 595 | 177 | 928 | 857 | 644 | 1707 |
| SRR10077429 | 0 | 0 | 29 | 0 | 22 | 803 | 153 | 825 | 1044 | 540 | 1606 |
| SRR10077430 | 0 | 0 | 23 | 16 | 47 | 774 | 141 | 828 | 719 | 555 | 1688 |
| SRR10077431 | 15 | 15 | 0 | 0 | 87 | 597 | 87 | 764 | 837 | 713 | 1564 |
| SRR10077432 | 0 | 0 | 11 | 0 | 21 | 274 | 81 | 561 | 1126 | 870 | 2565 |
| SRR10077433 | 0 | 15 | 0 | 0 | 33 | 294 | 105 | 707 | 1120 | 866 | 2385 |
| SRR10077434 | 0 | 0 | 0 | 0 | 12 | 270 | 104 | 747 | 1052 | 790 | 2454 |
| SRR10077435 | 0 | 0 | 0 | 0 | 29 | 318 | 91 | 712 | 1008 | 902 | 2392 |
| SRR10116271 | 0 | 0 | 0 | 0 | 0 | 37 | 0 | 260 | 917 | 1167 | 292 |
| SRR10116272 | 0 | 0 | 0 | 0 | 0 | 40 | 0 | 83 | 123 | 218 | 90 |
| SRR10116273 | 0 | 0 | 0 | 0 | 0 | 72 | 10 | 386 | 3914 | 5718 | 1011 |
| SRR10116274 | 0 | 0 | 0 | 0 | 14 | 60 | 26 | 201 | 295 | 275 | 116 |
| SRR10116275 | 0 | 0 | 0 | 0 | 0 | 73 | 17 | 204 | 389 | 435 | 391 |
| SRR10119011 | 0 | 0 | 0 | 0 | 0 | 13 | 0 | 0 | 0 | 0 | 0 |
| SRR10119012 | 0 | 0 | 0 | 0 | 0 | 0 | 0 | 0 | 0 | 0 | 0 |
| SRR10119014 | 0 | 0 | 0 | 0 | 0 | 0 | 0 | 0 | 0 | 0 | 0 |
| SRR10119022 | 0 | 0 | 38 | 16 | 0 | 0 | 0 | 0 | 0 | 0 | 0 |
| SRR10119033 | 0 | 0 | 0 | 0 | 0 | 0 | 0 | 0 | 0 | 0 | 0 |
| SRR10119034 | 0 | 0 | 0 | 0 | 0 | 0 | 0 | 0 | 0 | 0 | 0 |
| SRR10119035 | 0 | 17 | 48 | 23 | 0 | 0 | 0 | 0 | 0 | 0 | 0 |
| SRR10119037 | 0 | 0 | 0 | 0 | 0 | 0 | 0 | 0 | 0 | 0 | 0 |
| SRR10119040 | 0 | 0 | 0 | 0 | 0 | 0 | 0 | 0 | 0 | 0 | 0 |
| SRR10119044 | 0 | 0 | 0 | 0 | 0 | 0 | 0 | 0 | 0 | 0 | 0 |
| SRR10119046 | 0 | 12 | 34 | 41 | 0 | 0 | 0 | 0 | 0 | 0 | 0 |
| SRR10119048 | 0 | 0 | 0 | 0 | 0 | 0 | 0 | 0 | 0 | 0 | 0 |
| SRR10119051 | 0 | 0 | 0 | 0 | 0 | 0 | 0 | 0 | 0 | 0 | 0 |
| SRR10119052 | 0 | 0 | 0 | 0 | 0 | 0 | 0 | 0 | 0 | 0 | 0 |
| SRR10119056 | 0 | 0 | 0 | 0 | 0 | 0 | 0 | 0 | 0 | 0 | 0 |
| SRR10119057 | 0 | 0 | 27 | 17 | 0 | 0 | 0 | 0 | 0 | 0 | 0 |
| SRR10119058 | 0 | 0 | 22 | 25 | 0 | 0 | 0 | 0 | 0 | 0 | 0 |
| SRR10119059 | 0 | 0 | 11 | 11 | 0 | 22 | 11 | 0 | 0 | 0 | 11 |
| SRR10119063 | 0 | 0 | 0 | 0 | 0 | 0 | 0 | 0 | 0 | 12 | 0 |
| SRR10119066 | 0 | 11 | 46 | 33 | 0 | 0 | 0 | 0 | 13 | 0 | 0 |
| SRR10119067 | 0 | 13 | 48 | 45 | 0 | 0 | 0 | 0 | 0 | 0 | 0 |
| SRR10119068 | 0 | 12 | 37 | 22 | 0 | 0 | 0 | 0 | 0 | 0 | 0 |
| SRR10119069 | 0 | 12 | 49 | 20 | 0 | 0 | 0 | 0 | 0 | 0 | 0 |
| SRR10119070 | 0 | 0 | 14 | 0 | 0 | 0 | 0 | 0 | 0 | 0 | 0 |
| SRR10119071 | 0 | 0 | 20 | 10 | 0 | 0 | 0 | 0 | 0 | 0 | 0 |
| SRR10119072 | 0 | 0 | 0 | 0 | 0 | 0 | 0 | 0 | 0 | 0 | 0 |
| SRR10119073 | 0 | 0 | 0 | 0 | 0 | 0 | 0 | 0 | 0 | 0 | 0 |
| SRR10119074 | 0 | 0 | 0 | 0 | 0 | 0 | 0 | 0 | 0 | 0 | 0 |
| SRR10119075 | 0 | 0 | 37 | 15 | 0 | 0 | 0 | 10 | 0 | 0 | 0 |
| SRR10119076 | 0 | 0 | 21 | 19 | 0 | 0 | 0 | 0 | 0 | 0 | 0 |
| SRR10119077 | 0 | 13 | 33 | 24 | 0 | 0 | 0 | 0 | 0 | 0 | 0 |
| SRR10119078 | 0 | 21 | 53 | 26 | 0 | 0 | 0 | 0 | 0 | 0 | 0 |
| SRR10119079 | 0 | 12 | 36 | 25 | 0 | 0 | 0 | 16 | 0 | 0 | 0 |
| SRR10119080 | 0 | 14 | 48 | 14 | 0 | 12 | 0 | 0 | 0 | 0 | 0 |
| SRR10119081 | 0 | 0 | 12 | 0 | 0 | 0 | 0 | 0 | 0 | 0 | 0 |
| SRR10119082 | 0 | 0 | 0 | 0 | 0 | 0 | 0 | 0 | 0 | 0 | 0 |
| SRR1014666 | 0 | 0 | 0 | 0 | 0 | 0 | 17 | 1100 | 1401 | 907 | 960 |
| SRR1014667 | 0 | 0 | 0 | 0 | 0 | 11 | 10 | 44 | 27 | 18 | 13 |
| SRR1014669 | 0 | 0 | 0 | 0 | 0 | 0 | 0 | 12 | 30 | 38 | 29 |
| SRR10197184 | 0 | 0 | 0 | 0 | 37 | 97 | 62 | 52 | 38 | 18 | 83 |
| SRR10197185 | 0 | 0 | 99 | 0 | 143 | 326 | 52 | 0 | 238 | 0 | 68 |
| SRR10197186 | 0 | 0 | 0 | 0 | 30 | 99 | 40 | 19 | 607 | 48 | 40 |
| SRR10197187 | 0 | 0 | 0 | 0 | 0 | 37 | 16 | 0 | 212 | 23 | 32 |
| SRR10197188 | 0 | 0 | 0 | 0 | 0 | 28 | 13 | 10 | 57 | 0 | 26 |
| SRR10197189 | 0 | 0 | 0 | 0 | 13 | 51 | 17 | 0 | 275 | 17 | 10 |
| SRR10197190 | 0 | 0 | 0 | 0 | 0 | 11 | 0 | 0 | 13 | 0 | 19 |
| SRR10197191 | 0 | 0 | 0 | 0 | 0 | 14 | 0 | 0 | 31 | 0 | 11 |
| SRR10197192 | 0 | 0 | 0 | 0 | 0 | 15 | 0 | 0 | 0 | 0 | 0 |
| SRR10225092 | 1577 | 526 | 1577 | 2278 | 88 | 1139 | 263 | 876 | 2541 | 1226 | 613 |
| SRR10225093 | 0 | 0 | 0 | 0 | 0 | 0 | 0 | 0 | 0 | 0 | 0 |
| SRR10225094 | 220 | 529 | 717 | 496 | 55 | 198 | 55 | 198 | 132 | 121 | 121 |
| SRR10225096 | 29 | 127 | 362 | 274 | 0 | 49 | 0 | 59 | 166 | 117 | 68 |
| SRR10282881 | 0 | 0 | 0 | 0 | 0 | 0 | 0 | 0 | 0 | 0 | 0 |
| SRR10282882 | 0 | 0 | 0 | 0 | 0 | 0 | 0 | 0 | 0 | 0 | 0 |
| SRR10282883 | 0 | 0 | 0 | 0 | 0 | 0 | 0 | 0 | 0 | 0 | 0 |
| SRR10282884 | 0 | 0 | 0 | 0 | 0 | 0 | 0 | 0 | 0 | 0 | 0 |
| SRR10282885 | 0 | 0 | 0 | 0 | 0 | 0 | 0 | 0 | 0 | 0 | 0 |
| SRR10282886 | 0 | 0 | 0 | 0 | 0 | 0 | 0 | 0 | 0 | 0 | 0 |
| SRR10282890 | 0 | 0 | 0 | 0 | 0 | 15 | 0 | 15 | 0 | 0 | 0 |
| SRR10282891 | 0 | 0 | 0 | 0 | 0 | 0 | 0 | 15 | 0 | 0 | 0 |
| SRR10282892 | 0 | 0 | 0 | 0 | 0 | 14 | 0 | 28 | 0 | 0 | 0 |
| SRR10282893 | 0 | 0 | 0 | 0 | 0 | 12 | 0 | 28 | 0 | 0 | 0 |
| SRR10282894 | 0 | 0 | 0 | 0 | 0 | 14 | 0 | 11 | 0 | 0 | 0 |
| SRR10282895 | 0 | 0 | 0 | 0 | 0 | 12 | 0 | 0 | 0 | 0 | 0 |
| SRR10323931 | 0 | 0 | 0 | 0 | 1941 | 4512 | 1891 | 418 | 1374 | 1807 | 1192 |
| SRR10323932 | 0 | 0 | 0 | 0 | 2192 | 4219 | 1605 | 442 | 745 | 1073 | 1101 |
| SRR10323933 | 0 | 11 | 40 | 48 | 2739 | 4294 | 2003 | 391 | 608 | 590 | 572 |
| SRR10323939 | 0 | 0 | 0 | 0 | 599 | 1683 | 841 | 383 | 774 | 867 | 855 |
| SRR10323940 | 0 | 0 | 0 | 0 | 1309 | 3837 | 1215 | 596 | 675 | 1353 | 759 |
| SRR10323941 | 0 | 0 | 0 | 0 | 714 | 2063 | 703 | 480 | 484 | 760 | 1306 |
| SRR10385125 | 0 | 0 | 0 | 0 | 513 | 883 | 393 | 795 | 2491 | 723 | 730 |
| SRR10385126 | 0 | 31 | 40 | 32 | 1032 | 1747 | 647 | 1565 | 7778 | 2096 | 1677 |
| SRR10416439 | 0 | 0 | 0 | 0 | 227 | 798 | 56 | 995 | 0 | 23 | 50 |
| SRR10416440 | 0 | 0 | 0 | 0 | 118 | 407 | 17 | 514 | 0 | 12 | 62 |
| SRR10416441 | 0 | 0 | 0 | 0 | 147 | 464 | 30 | 693 | 10 | 19 | 48 |
| SRR10416442 | 0 | 0 | 0 | 0 | 285 | 1121 | 311 | 1051 | 13 | 27 | 76 |
| SRR10416496 | 0 | 0 | 0 | 0 | 0 | 0 | 0 | 0 | 0 | 10 | 0 |
| SRR10416497 | 0 | 0 | 0 | 0 | 0 | 18 | 0 | 0 | 0 | 18 | 0 |
| SRR10416498 | 0 | 0 | 0 | 0 | 0 | 0 | 0 | 0 | 0 | 0 | 0 |
| SRR10416499 | 0 | 0 | 0 | 0 | 0 | 0 | 0 | 0 | 0 | 0 | 0 |
| SRR10416500 | 0 | 0 | 0 | 15 | 0 | 0 | 0 | 0 | 0 | 0 | 0 |
| SRR10416501 | 0 | 0 | 0 | 0 | 0 | 15 | 0 | 0 | 0 | 15 | 0 |
| SRR10416502 | 0 | 0 | 0 | 20 | 0 | 0 | 0 | 0 | 0 | 0 | 0 |
| SRR10416503 | 0 | 0 | 0 | 0 | 0 | 0 | 0 | 22 | 0 | 22 | 0 |
| SRR10416504 | 0 | 0 | 0 | 0 | 0 | 0 | 0 | 0 | 0 | 42 | 0 |
| SRR10416505 | 0 | 0 | 0 | 0 | 0 | 0 | 0 | 0 | 0 | 0 | 0 |
| SRR10416506 | 0 | 0 | 18 | 0 | 35 | 0 | 0 | 0 | 0 | 0 | 0 |
| SRR10416507 | 0 | 0 | 0 | 0 | 0 | 0 | 0 | 0 | 0 | 15 | 0 |
| SRR10416508 | 0 | 0 | 0 | 23 | 0 | 0 | 0 | 0 | 0 | 0 | 0 |
| SRR10416509 | 0 | 0 | 0 | 0 | 0 | 0 | 0 | 0 | 0 | 0 | 0 |
| SRR10416510 | 31 | 0 | 0 | 0 | 0 | 0 | 0 | 0 | 0 | 0 | 0 |
| SRR10416513 | 0 | 0 | 0 | 0 | 0 | 0 | 0 | 0 | 0 | 0 | 0 |
| SRR10416514 | 23 | 0 | 0 | 0 | 0 | 0 | 0 | 23 | 0 | 0 | 23 |
| SRR10416515 | 23 | 0 | 45 | 0 | 0 | 0 | 0 | 0 | 0 | 0 | 0 |
| SRR10416516 | 0 | 0 | 0 | 0 | 0 | 0 | 0 | 0 | 0 | 0 | 0 |
| SRR10416517 | 0 | 0 | 15 | 46 | 0 | 0 | 0 | 0 | 0 | 0 | 0 |
| SRR10416518 | 0 | 0 | 0 | 0 | 0 | 0 | 0 | 0 | 0 | 0 | 0 |
| SRR10416519 | 97 | 0 | 16 | 32 | 0 | 16 | 0 | 0 | 32 | 0 | 0 |
| SRR10416520 | 0 | 0 | 39 | 39 | 0 | 0 | 0 | 0 | 0 | 0 | 0 |
| SRR10416521 | 0 | 0 | 0 | 0 | 0 | 0 | 0 | 0 | 0 | 0 | 0 |
| SRR10416522 | 0 | 0 | 0 | 0 | 0 | 0 | 0 | 0 | 0 | 0 | 0 |
| SRR10416523 | 0 | 0 | 0 | 0 | 0 | 21 | 0 | 0 | 0 | 0 | 0 |

### Supp Table S8.5.Columns_ET_to_FD

| | | | | | | | | | | | |
|---|---|---|---|---|---|---|---|---|---|---|---|
| SRR10416524 | 0 | 0 | 0 | 30 | 0 | 0 | 0 | 0 | 0 | 0 | 0 |
| SRR10416525 | 0 | 0 | 0 | 0 | 0 | 0 | 0 | 0 | 0 | 0 | 0 |
| SRR10416526 | 0 | 39 | 0 | 39 | 0 | 0 | 0 | 0 | 0 | 0 | 0 |
| SRR10416527 | 0 | 0 | 0 | 0 | 0 | 0 | 0 | 0 | 0 | 0 | 0 |
| SRR10416528 | 0 | 0 | 0 | 81 | 0 | 0 | 0 | 0 | 0 | 0 | 0 |
| SRR10416529 | 0 | 0 | 0 | 0 | 0 | 0 | 0 | 0 | 55 | 0 | 0 |
| SRR10416530 | 0 | 0 | 0 | 60 | 0 | 0 | 0 | 0 | 0 | 0 | 0 |
| SRR10416531 | 0 | 0 | 0 | 0 | 0 | 0 | 0 | 0 | 0 | 0 | 0 |
| SRR10416532 | 0 | 0 | 69 | 0 | 0 | 0 | 0 | 0 | 0 | 0 | 0 |
| SRR10416533 | 0 | 0 | 0 | 44 | 0 | 0 | 0 | 0 | 0 | 0 | 0 |
| SRR10416534 | 0 | 0 | 0 | 0 | 0 | 0 | 0 | 0 | 0 | 0 | 0 |
| SRR10416535 | 0 | 0 | 0 | 0 | 0 | 0 | 0 | 0 | 0 | 0 | 0 |
| SRR10416536 | 44 | 0 | 44 | 0 | 0 | 0 | 0 | 0 | 0 | 0 | 0 |
| SRR10416537 | 0 | 0 | 0 | 0 | 0 | 0 | 0 | 0 | 0 | 0 | 0 |
| SRR10416538 | 0 | 0 | 0 | 0 | 0 | 0 | 0 | 0 | 0 | 0 | 34 |
| SRR10416539 | 0 | 0 | 0 | 0 | 0 | 0 | 0 | 0 | 0 | 0 | 0 |
| SRR10416540 | 0 | 0 | 0 | 0 | 0 | 0 | 0 | 0 | 0 | 0 | 0 |
| SRR10416541 | 0 | 0 | 0 | 0 | 0 | 0 | 0 | 228 | 0 | 0 | 0 |
| SRR10416542 | 0 | 0 | 0 | 0 | 0 | 0 | 0 | 0 | 0 | 0 | 0 |
| SRR10416543 | 0 | 0 | 0 | 0 | 0 | 0 | 0 | 0 | 0 | 0 | 0 |
| SRR10416544 | 0 | 0 | 0 | 0 | 0 | 0 | 0 | 0 | 78 | 0 | 0 |
| SRR10416545 | 0 | 0 | 19 | 38 | 0 | 0 | 0 | 0 | 0 | 0 | 0 |
| SRR10416546 | 0 | 0 | 66 | 0 | 0 | 0 | 0 | 0 | 0 | 0 | 0 |
| SRR10416547 | 0 | 0 | 0 | 0 | 0 | 0 | 0 | 0 | 48 | 0 | 0 |
| SRR10416549 | 0 | 0 | 0 | 45 | 0 | 0 | 0 | 0 | 0 | 0 | 0 |
| SRR10416550 | 0 | 0 | 0 | 21 | 0 | 0 | 0 | 21 | 0 | 0 | 0 |
| SRR10416551 | 0 | 0 | 0 | 27 | 0 | 0 | 0 | 0 | 0 | 0 | 0 |
| SRR10416553 | 0 | 0 | 0 | 130 | 0 | 0 | 0 | 0 | 0 | 0 | 0 |
| SRR10416554 | 0 | 0 | 71 | 0 | 0 | 0 | 0 | 0 | 0 | 0 | 0 |
| SRR10416555 | 0 | 0 | 0 | 0 | 0 | 0 | 0 | 0 | 0 | 0 | 0 |
| SRR10416557 | 0 | 0 | 0 | 115 | 0 | 0 | 0 | 0 | 0 | 0 | 0 |
| SRR10416558 | 0 | 0 | 0 | 0 | 0 | 0 | 0 | 0 | 0 | 0 | 0 |
| SRR10416559 | 0 | 0 | 39 | 0 | 0 | 0 | 0 | 0 | 0 | 0 | 0 |
| SRR10416560 | 0 | 0 | 23 | 0 | 0 | 0 | 0 | 0 | 0 | 0 | 0 |
| SRR10416562 | 0 | 0 | 0 | 0 | 0 | 0 | 0 | 0 | 0 | 0 | 0 |
| SRR10416563 | 30 | 0 | 30 | 15 | 0 | 0 | 0 | 0 | 0 | 0 | 0 |
| SRR10416564 | 0 | 0 | 0 | 0 | 0 | 0 | 0 | 0 | 0 | 0 | 0 |
| SRR10416565 | 0 | 0 | 0 | 0 | 0 | 0 | 0 | 0 | 0 | 0 | 0 |
| SRR10416566 | 0 | 0 | 0 | 72 | 0 | 0 | 0 | 0 | 0 | 0 | 0 |
| SRR10416567 | 0 | 0 | 0 | 0 | 0 | 0 | 0 | 0 | 0 | 0 | 0 |
| SRR10416568 | 0 | 0 | 0 | 0 | 0 | 0 | 0 | 0 | 0 | 0 | 0 |
| SRR10416569 | 0 | 0 | 0 | 0 | 0 | 0 | 0 | 0 | 0 | 0 | 0 |
| SRR10416570 | 0 | 0 | 53 | 0 | 0 | 53 | 0 | 0 | 0 | 0 | 0 |
| SRR10416571 | 0 | 0 | 61 | 61 | 0 | 0 | 0 | 0 | 0 | 0 | 0 |
| SRR10416572 | 0 | 44 | 0 | 131 | 0 | 0 | 0 | 0 | 0 | 0 | 0 |
| SRR10416573 | 0 | 0 | 0 | 204 | 0 | 0 | 0 | 0 | 0 | 0 | 0 |
| SRR10416574 | 0 | 0 | 0 | 0 | 0 | 0 | 0 | 0 | 0 | 0 | 0 |
| SRR10416575 | 42 | 0 | 0 | 0 | 0 | 0 | 0 | 0 | 0 | 0 | 0 |
| SRR10416576 | 141 | 0 | 28 | 0 | 0 | 0 | 0 | 0 | 0 | 0 | 0 |
| SRR10416577 | 0 | 0 | 0 | 0 | 0 | 0 | 0 | 0 | 0 | 0 | 0 |
| SRR10416578 | 0 | 0 | 0 | 0 | 0 | 0 | 0 | 0 | 0 | 0 | 0 |
| SRR10416579 | 0 | 0 | 0 | 0 | 0 | 0 | 0 | 0 | 0 | 0 | 0 |
| SRR10416580 | 0 | 0 | 0 | 0 | 0 | 0 | 0 | 0 | 0 | 0 | 0 |
| SRR10416581 | 0 | 0 | 0 | 20 | 0 | 0 | 0 | 0 | 0 | 0 | 0 |
| SRR10416582 | 0 | 0 | 0 | 0 | 0 | 0 | 0 | 0 | 0 | 0 | 0 |
| SRR10416583 | 0 | 0 | 0 | 0 | 0 | 0 | 0 | 0 | 0 | 0 | 0 |
| SRR10416584 | 0 | 0 | 0 | 0 | 0 | 0 | 0 | 0 | 0 | 0 | 0 |
| SRR10416585 | 0 | 0 | 0 | 0 | 0 | 0 | 0 | 0 | 0 | 0 | 0 |
| SRR10416586 | 0 | 0 | 0 | 0 | 0 | 0 | 0 | 0 | 0 | 0 | 0 |
| SRR10416587 | 0 | 0 | 0 | 0 | 0 | 0 | 0 | 0 | 0 | 0 | 0 |
| SRR10416588 | 78 | 0 | 0 | 0 | 0 | 0 | 0 | 0 | 0 | 0 | 0 |
| SRR10416589 | 0 | 0 | 0 | 0 | 0 | 0 | 0 | 0 | 0 | 0 | 0 |
| SRR10416590 | 0 | 0 | 0 | 0 | 0 | 0 | 0 | 0 | 0 | 0 | 0 |
| SRR10416591 | 0 | 0 | 0 | 0 | 0 | 0 | 0 | 0 | 0 | 0 | 0 |
| SRR10416592 | 0 | 0 | 0 | 0 | 0 | 0 | 0 | 0 | 0 | 0 | 0 |
| SRR10416593 | 47 | 0 | 47 | 140 | 0 | 0 | 0 | 0 | 47 | 0 | 0 |
| SRR10416594 | 0 | 0 | 0 | 0 | 0 | 0 | 0 | 0 | 0 | 0 | 0 |
| SRR10416595 | 0 | 0 | 0 | 42 | 0 | 0 | 0 | 0 | 0 | 0 | 0 |
| SRR10416596 | 19 | 0 | 19 | 39 | 0 | 0 | 0 | 0 | 0 | 0 | 0 |
| SRR10416597 | 0 | 0 | 0 | 27 | 0 | 0 | 0 | 0 | 0 | 0 | 0 |
| SRR10416598 | 28 | 0 | 0 | 0 | 0 | 0 | 0 | 0 | 0 | 0 | 0 |
| SRR10416599 | 0 | 0 | 0 | 0 | 0 | 0 | 0 | 0 | 0 | 0 | 0 |
| SRR10416600 | 0 | 0 | 0 | 20 | 0 | 0 | 0 | 0 | 0 | 0 | 0 |
| SRR10428553 | 0 | 0 | 0 | 0 | 0 | 0 | 0 | 0 | 13 | 0 | 0 |
| SRR10428554 | 0 | 0 | 17 | 14 | 0 | 25 | 0 | 53 | 57 | 45 | 59 |
| SRR10428555 | 0 | 0 | 0 | 0 | 0 | 15 | 0 | 13 | 0 | 0 | 19 |
| SRR10428556 | 0 | 0 | 0 | 12 | 0 | 29 | 0 | 54 | 0 | 14 | 48 |
| SRR10428557 | 0 | 0 | 0 | 0 | 0 | 10 | 0 | 0 | 0 | 0 | 0 |
| SRR10428558 | 0 | 0 | 0 | 0 | 0 | 0 | 0 | 0 | 0 | 0 | 21 |
| SRR10428559 | 0 | 0 | 0 | 0 | 0 | 31 | 0 | 37 | 12 | 16 | 38 |
| SRR10428560 | 0 | 0 | 0 | 0 | 0 | 29 | 0 | 46 | 29 | 24 | 66 |
| SRR10446207 | 0 | 0 | 0 | 11 | 97 | 43 | 11 | 136 | 452 | 26 | 432 |
| SRR10446208 | 0 | 0 | 0 | 0 | 48 | 42 | 26 | 83 | 208 | 19 | 182 |
| SRR10446209 | 0 | 0 | 0 | 39 | 128 | 167 | 54 | 275 | 206 | 25 | 447 |
| SRR10446210 | 0 | 0 | 0 | 26 | 131 | 138 | 52 | 282 | 820 | 171 | 1037 |
| SRR10446211 | 0 | 0 | 0 | 0 | 23 | 27 | 15 | 29 | 239 | 0 | 62 |
| SRR10446212 | 0 | 0 | 0 | 13 | 95 | 108 | 64 | 156 | 681 | 32 | 411 |
| SRR10531363 | 0 | 0 | 0 | 0 | 0 | 21 | 0 | 0 | 0 | 0 | 0 |
| SRR10531364 | 0 | 0 | 0 | 0 | 0 | 0 | 0 | 0 | 0 | 0 | 0 |
| SRR10531365 | 0 | 0 | 0 | 0 | 0 | 0 | 0 | 0 | 0 | 0 | 0 |
| SRR10531367 | 0 | 0 | 0 | 0 | 0 | 0 | 0 | 0 | 0 | 0 | 0 |
| SRR10531368 | 0 | 0 | 0 | 0 | 0 | 0 | 0 | 0 | 0 | 0 | 0 |
| SRR10531369 | 0 | 0 | 0 | 0 | 0 | 0 | 0 | 0 | 0 | 0 | 0 |
| SRR10531370 | 0 | 0 | 0 | 0 | 0 | 0 | 0 | 0 | 0 | 0 | 0 |
| SRR10544939 | 0 | 0 | 0 | 0 | 0 | 0 | 0 | 0 | 13 | 11 | 16 |
| SRR10544940 | 0 | 0 | 0 | 0 | 0 | 0 | 0 | 0 | 13 | 11 | 19 |
| SRR10544941 | 0 | 0 | 0 | 0 | 24 | 19 | 15 | 0 | 34 | 28 | 52 |
| SRR10544942 | 0 | 0 | 0 | 0 | 0 | 0 | 0 | 0 | 10 | 0 | 21 |
| SRR10544943 | 0 | 0 | 0 | 0 | 0 | 0 | 0 | 0 | 0 | 0 | 12 |
| SRR10544944 | 0 | 0 | 0 | 0 | 0 | 0 | 0 | 0 | 0 | 0 | 0 |
| SRR10544945 | 0 | 0 | 0 | 0 | 0 | 0 | 0 | 0 | 0 | 0 | 0 |
| SRR10544946 | 0 | 0 | 0 | 0 | 0 | 0 | 0 | 0 | 0 | 0 | 0 |
| SRR10544947 | 0 | 0 | 0 | 0 | 0 | 0 | 0 | 0 | 0 | 0 | 0 |
| SRR10544948 | 0 | 0 | 0 | 0 | 0 | 0 | 0 | 0 | 0 | 0 | 0 |
| SRR10544949 | 0 | 0 | 0 | 0 | 0 | 0 | 0 | 0 | 0 | 0 | 0 |
| SRR10544950 | 0 | 0 | 0 | 0 | 0 | 0 | 0 | 0 | 0 | 0 | 0 |
| SRR10756575 | 0 | 237 | 762 | 133 | 1229 | 3761 | 1892 | 2581 | 642 | 96 | 434 |
| SRR10756576 | 0 | 118 | 346 | 66 | 1461 | 4197 | 3430 | 3645 | 780 | 155 | 590 |
| SRR10756577 | 0 | 43 | 93 | 15 | 1009 | 3552 | 2207 | 3781 | 1004 | 227 | 544 |
| SRR10756578 | 0 | 157 | 363 | 126 | 889 | 2668 | 1784 | 3271 | 504 | 102 | 342 |
| SRR10756579 | 0 | 127 | 408 | 81 | 1678 | 4667 | 3240 | 4369 | 1099 | 220 | 448 |
| SRR10756581 | 0 | 237 | 657 | 171 | 5175 | 14069 | 8531 | 8270 | 894 | 119 | 398 |
| SRR10756582 | 0 | 1050 | 2669 | 384 | 4044 | 12421 | 7401 | 5634 | 673 | 229 | 569 |
| SRR10756583 | 0 | 201 | 369 | 151 | 1215 | 6149 | 3728 | 5939 | 1349 | 151 | 662 |
| SRR10875785 | 0 | 0 | 78 | 76 | 3582 | 9410 | 5201 | 1683 | 268 | 198 | 298 |
| SRR10875786 | 0 | 0 | 0 | 0 | 238 | 420 | 198 | 53 | 12 | 0 | 13 |
| SRR10953234 | 0 | 221 | 567 | 210 | 225 | 654 | 259 | 253 | 93 | 10 | 44 |
| SRR10953235 | 0 | 231 | 540 | 172 | 226 | 640 | 329 | 240 | 109 | 14 | 40 |
| SRR10953236 | 0 | 193 | 467 | 148 | 189 | 597 | 250 | 284 | 127 | 14 | 44 |

### Supp Table S8.5.Columns_ET_to_FD

| | | | | | | | | | | | |
|---|---|---|---|---|---|---|---|---|---|---|---|
| SRR10953237 | 0 | 188 | 425 | 136 | 258 | 757 | 406 | 307 | 121 | 14 | 42 |
| SRR10953238 | 0 | 224 | 512 | 157 | 200 | 593 | 260 | 259 | 135 | 13 | 52 |
| SRR10953239 | 0 | 218 | 464 | 118 | 246 | 752 | 414 | 305 | 193 | 19 | 49 |
| SRR1107492 | 0 | 0 | 21 | 12 | 73 | 635 | 60 | 993 | 125 | 104 | 425 |
| SRR1107493 | 0 | 0 | 0 | 0 | 14 | 158 | 0 | 191 | 89 | 34 | 170 |
| SRR1107494 | 0 | 34 | 133 | 87 | 28 | 198 | 24 | 395 | 65 | 42 | 165 |
| SRR1107495 | 0 | 52 | 150 | 96 | 32 | 185 | 34 | 512 | 84 | 41 | 179 |
| SRR1107496 | 0 | 57 | 127 | 148 | 25 | 160 | 29 | 396 | 80 | 50 | 209 |
| SRR1107497 | 0 | 72 | 144 | 186 | 47 | 186 | 21 | 483 | 59 | 47 | 305 |
| SRR1159512 | 0 | 115 | 257 | 168 | 133 | 1099 | 287 | 1850 | 7906 | 7890 | 13145 |
| SRR1207050 | 0 | 0 | 0 | 0 | 0 | 0 | 0 | 0 | 0 | 0 | 15 |
| SRR1207051 | 0 | 0 | 0 | 15 | 25 | 0 | 22 | 34 | 1027 | 329 | 228 |
| SRR1207053 | 0 | 0 | 0 | 0 | 14 | 0 | 0 | 20 | 793 | 205 | 213 |
| SRR1207054 | 0 | 0 | 0 | 0 | 0 | 0 | 0 | 0 | 0 | 0 | 0 |
| SRR1207055 | 0 | 0 | 0 | 0 | 0 | 0 | 0 | 0 | 109 | 27 | 11 |
| SRR1232072 | 0 | 0 | 0 | 0 | 0 | 0 | 0 | 0 | 0 | 0 | 0 |
| SRR1232073 | 0 | 0 | 22 | 11 | 0 | 17 | 0 | 54 | 11 | 0 | 0 |
| SRR1240812 | 0 | 208 | 383 | 752 | 259 | 1293 | 237 | 2653 | 11033 | 15571 | 6107 |
| SRR1240813 | 0 | 191 | 353 | 632 | 136 | 708 | 142 | 1256 | 4997 | 7281 | 3223 |
| SRR1240814 | 0 | 130 | 295 | 357 | 117 | 808 | 129 | 874 | 1062 | 715 | 543 |
| SRR1240815 | 0 | 105 | 261 | 328 | 106 | 834 | 156 | 966 | 1169 | 833 | 600 |
| SRR1240816 | 0 | 191 | 390 | 886 | 234 | 1531 | 517 | 2349 | 2301 | 3653 | 2627 |
| SRR1240817 | 0 | 233 | 380 | 943 | 240 | 1529 | 567 | 2437 | 2536 | 4011 | 2861 |
| SRR1240818 | 0 | 21 | 48 | 32 | 22 | 143 | 22 | 193 | 449 | 708 | 245 |
| SRR1241600 | 0 | 0 | 11 | 0 | 0 | 16 | 39 | 71 | 932 | 606 | 1007 |
| SRR1241601 | 0 | 27 | 66 | 36 | 42 | 13 | 39 | 17 | 279 | 503 | 2294 |
| SRR1241602 | 0 | 0 | 13 | 0 | 13 | 0 | 14 | 0 | 694 | 1028 | 1682 |
| SRR1241603 | 0 | 39 | 142 | 94 | 290 | 3314 | 798 | 8458 | 68194 | 49021 | 11262 |
| SRR1241604 | 0 | 65 | 211 | 160 | 199 | 1934 | 452 | 4986 | 35858 | 31833 | 8675 |
| SRR1241605 | 0 | 48 | 156 | 154 | 304 | 2292 | 675 | 5528 | 33064 | 25533 | 7455 |
| SRR1264353 | 0 | 0 | 0 | 0 | 26 | 61 | 31 | 33 | 43 | 23 | 16 |
| SRR1264354 | 0 | 0 | 0 | 0 | 40 | 80 | 39 | 92 | 60 | 18 | 13 |
| SRR1270153 | 0 | 0 | 0 | 0 | 0 | 0 | 0 | 0 | 12 | 0 | 0 |
| SRR1290784 | 0 | 49 | 76 | 40 | 252 | 5663 | 897 | 1638 | 1525 | 971 | 830 |
| SRR1290785 | 20 | 47 | 133 | 32 | 122 | 2750 | 378 | 742 | 928 | 667 | 483 |
| SRR1290786 | 13 | 37 | 66 | 24 | 113 | 3119 | 371 | 816 | 964 | 570 | 429 |
| SRR1290787 | 0 | 16 | 55 | 16 | 131 | 2951 | 330 | 611 | 540 | 257 | 207 |
| SRR1290788 | 0 | 18 | 42 | 24 | 106 | 2653 | 325 | 666 | 825 | 434 | 383 |
| SRR1290789 | 0 | 0 | 18 | 0 | 164 | 4477 | 547 | 1297 | 1025 | 588 | 610 |
| SRR1290790 | 0 | 20 | 80 | 26 | 95 | 2329 | 330 | 678 | 781 | 550 | 473 |
| SRR1290791 | 0 | 46 | 81 | 42 | 405 | 7443 | 1139 | 1699 | 1159 | 737 | 757 |
| SRR1290792 | 0 | 41 | 111 | 54 | 242 | 4582 | 664 | 1230 | 819 | 446 | 482 |
| SRR1290793 | 0 | 0 | 55 | 32 | 192 | 4432 | 606 | 1665 | 1645 | 887 | 486 |
| SRR1290794 | 0 | 10 | 19 | 11 | 131 | 1916 | 337 | 543 | 570 | 223 | 127 |
| SRR1290795 | 0 | 0 | 26 | 0 | 107 | 2206 | 316 | 710 | 571 | 262 | 115 |
| SRR1290796 | 14 | 0 | 25 | 11 | 49 | 1739 | 221 | 794 | 498 | 213 | 181 |
| SRR1290797 | 0 | 28 | 89 | 41 | 84 | 2361 | 290 | 1178 | 846 | 385 | 295 |
| SRR1290798 | 32 | 111 | 307 | 89 | 201 | 6672 | 870 | 3603 | 1748 | 1069 | 734 |
| SRR1290799 | 0 | 55 | 123 | 65 | 70 | 1082 | 116 | 409 | 476 | 180 | 81 |
| SRR1290800 | 14 | 76 | 172 | 42 | 43 | 585 | 64 | 225 | 314 | 98 | 68 |
| SRR1290801 | 0 | 44 | 123 | 45 | 50 | 749 | 52 | 237 | 230 | 60 | 34 |
| SRR1290802 | 18 | 45 | 103 | 55 | 133 | 3048 | 565 | 2073 | 1195 | 855 | 687 |
| SRR1290803 | 28 | 69 | 137 | 67 | 91 | 2061 | 332 | 1266 | 890 | 621 | 423 |
| SRR1290804 | 38 | 63 | 120 | 85 | 201 | 3476 | 690 | 1828 | 1049 | 804 | 644 |
| SRR1290805 | 0 | 56 | 180 | 69 | 167 | 4236 | 548 | 1729 | 802 | 576 | 518 |
| SRR1290806 | 0 | 12 | 77 | 31 | 202 | 4174 | 811 | 2616 | 942 | 789 | 591 |
| SRR1290807 | 0 | 39 | 94 | 44 | 241 | 4559 | 824 | 2497 | 1063 | 793 | 547 |
| SRR1290808 | 0 | 59 | 114 | 53 | 321 | 4142 | 758 | 2060 | 1168 | 803 | 725 |
| SRR1290809 | 0 | 92 | 252 | 114 | 248 | 4950 | 877 | 2602 | 1281 | 1015 | 717 |
| SRR1292461 | 0 | 0 | 0 | 0 | 15 | 0 | 48 | 523 | 6132 | 1703 | 3618 |
| SRR1304304 | 0 | 0 | 0 | 0 | 61 | 887 | 276 | 1269 | 46956 | 65615 | 65245 |
| SRR1304307 | 0 | 0 | 0 | 0 | 0 | 24 | 17 | 511 | 2698 | 1328 | 2840 |
| SRR1304309 | 0 | 0 | 0 | 0 | 13 | 176 | 169 | 1906 | 18137 | 6519 | 23363 |
| SRR1304313 | 0 | 93 | 239 | 349 | 102 | 814 | 139 | 1356 | 3481 | 1514 | 7645 |
| SRR1393733 | 0 | 0 | 0 | 0 | 512 | 5891 | 1873 | 20041 | 18585 | 32052 | 16229 |
| SRR1510077 | 0 | 0 | 17 | 21 | 0 | 11 | 0 | 0 | 0 | 0 | 32 |
| SRR1510078 | 0 | 0 | 0 | 0 | 0 | 0 | 0 | 13 | 0 | 0 | 0 |
| SRR1510079 | 0 | 14 | 68 | 64 | 0 | 0 | 0 | 14 | 0 | 0 | 27 |
| SRR1510080 | 0 | 0 | 12 | 14 | 0 | 12 | 0 | 14 | 0 | 0 | 0 |
| SRR1510081 | 0 | 0 | 19 | 21 | 0 | 0 | 0 | 17 | 0 | 0 | 11 |
| SRR1510082 | 0 | 12 | 29 | 17 | 0 | 23 | 0 | 12 | 0 | 0 | 0 |
| SRR1510083 | 0 | 13 | 22 | 23 | 0 | 12 | 0 | 10 | 16 | 0 | 0 |
| SRR1510084 | 0 | 0 | 14 | 25 | 0 | 37 | 0 | 45 | 37 | 20 | 20 |
| SRR1510085 | 0 | 0 | 0 | 0 | 0 | 0 | 0 | 16 | 0 | 0 | 0 |
| SRR1510086 | 0 | 13 | 38 | 10 | 0 | 10 | 0 | 20 | 13 | 23 | 10 |
| SRR1510087 | 0 | 40 | 88 | 53 | 0 | 25 | 0 | 21 | 14 | 0 | 0 |
| SRR1510088 | 0 | 0 | 15 | 10 | 0 | 27 | 0 | 67 | 41 | 27 | 18 |
| SRR1510089 | 0 | 0 | 36 | 23 | 0 | 11 | 0 | 25 | 13 | 0 | 11 |
| SRR1510090 | 0 | 10 | 36 | 52 | 0 | 44 | 0 | 42 | 23 | 0 | 16 |
| SRR1510091 | 0 | 12 | 19 | 12 | 0 | 19 | 0 | 12 | 0 | 0 | 0 |
| SRR1510092 | 0 | 0 | 26 | 10 | 0 | 0 | 0 | 0 | 0 | 0 | 0 |
| SRR1510093 | 0 | 0 | 18 | 14 | 0 | 20 | 0 | 10 | 0 | 0 | 0 |
| SRR1510094 | 0 | 14 | 10 | 17 | 0 | 27 | 0 | 14 | 27 | 0 | 14 |
| SRR1510095 | 0 | 11 | 52 | 71 | 0 | 26 | 0 | 56 | 37 | 19 | 37 |
| SRR1510096 | 0 | 0 | 30 | 30 | 0 | 18 | 0 | 21 | 12 | 0 | 0 |
| SRR1510097 | 0 | 14 | 41 | 45 | 0 | 50 | 0 | 18 | 0 | 14 | 14 |
| SRR1510098 | 0 | 30 | 55 | 78 | 0 | 45 | 0 | 43 | 20 | 0 | 13 |
| SRR1510099 | 0 | 0 | 20 | 16 | 0 | 32 | 0 | 11 | 14 | 0 | 0 |
| SRR1510100 | 0 | 13 | 21 | 14 | 0 | 14 | 0 | 10 | 32 | 0 | 11 |
| SRR1510101 | 0 | 14 | 63 | 33 | 16 | 24 | 0 | 24 | 11 | 0 | 0 |
| SRR1510102 | 0 | 14 | 27 | 15 | 0 | 22 | 0 | 35 | 18 | 0 | 13 |
| SRR1510103 | 0 | 19 | 46 | 45 | 0 | 41 | 0 | 77 | 49 | 17 | 21 |
| SRR1510104 | 0 | 27 | 26 | 23 | 0 | 0 | 0 | 0 | 14 | 0 | 0 |
| SRR1510105 | 0 | 0 | 0 | 24 | 0 | 12 | 0 | 32 | 0 | 12 | 20 |
| SRR1510106 | 0 | 0 | 12 | 19 | 0 | 21 | 0 | 26 | 19 | 0 | 0 |
| SRR1510107 | 0 | 0 | 36 | 14 | 0 | 22 | 0 | 12 | 22 | 12 | 0 |
| SRR1510108 | 0 | 0 | 29 | 17 | 0 | 11 | 0 | 14 | 0 | 0 | 0 |
| SRR1510109 | 0 | 0 | 33 | 34 | 0 | 26 | 0 | 25 | 19 | 10 | 14 |
| SRR1510110 | 0 | 17 | 18 | 24 | 0 | 15 | 0 | 0 | 13 | 0 | 0 |
| SRR1510111 | 0 | 30 | 30 | 43 | 0 | 38 | 0 | 64 | 21 | 13 | 30 |
| SRR1510112 | 0 | 24 | 59 | 40 | 11 | 35 | 0 | 48 | 16 | 0 | 13 |
| SRR1510113 | 0 | 15 | 48 | 46 | 0 | 41 | 0 | 72 | 31 | 11 | 33 |
| SRR1510114 | 0 | 14 | 0 | 12 | 0 | 27 | 0 | 20 | 18 | 0 | 18 |
| SRR1510115 | 0 | 0 | 0 | 11 | 0 | 29 | 0 | 19 | 0 | 0 | 0 |
| SRR1510116 | 0 | 0 | 30 | 37 | 0 | 23 | 0 | 14 | 0 | 0 | 0 |
| SRR1510117 | 0 | 24 | 78 | 57 | 0 | 13 | 0 | 0 | 0 | 0 | 0 |
| SRR1510118 | 0 | 14 | 61 | 43 | 0 | 38 | 0 | 0 | 0 | 0 | 11 |
| SRR1510119 | 0 | 19 | 63 | 24 | 0 | 30 | 0 | 31 | 24 | 0 | 19 |
| SRR1510120 | 0 | 11 | 37 | 23 | 0 | 37 | 0 | 66 | 34 | 0 | 26 |
| SRR1510121 | 0 | 32 | 83 | 60 | 0 | 51 | 14 | 58 | 46 | 39 | 81 |
| SRR1519906 | 0 | 0 | 0 | 0 | 0 | 0 | 0 | 0 | 0 | 0 | 0 |
| SRR1519919 | 0 | 0 | 0 | 0 | 0 | 0 | 0 | 0 | 21 | 0 | 0 |
| SRR1525198 | 0 | 78 | 180 | 106 | 353 | 4262 | 1384 | 17100 | 43402 | 66936 | 22051 |
| SRR1525199 | 0 | 98 | 156 | 123 | 226 | 3974 | 1325 | 18489 | 40818 | 76522 | 28379 |
| SRR1525200 | 0 | 116 | 184 | 82 | 307 | 6011 | 1940 | 17513 | 22267 | 32492 | 13388 |
| SRR1525201 | 0 | 155 | 200 | 145 | 291 | 3968 | 1453 | 14516 | 22342 | 36750 | 14987 |
| SRR1525202 | 0 | 17 | 18 | 15 | 29 | 583 | 190 | 1658 | 2606 | 4141 | 1759 |
| SRR1525203 | 0 | 107 | 129 | 121 | 324 | 4973 | 1610 | 16759 | 24105 | 43093 | 18415 |
| SRR1563015 | 0 | 0 | 0 | 0 | 123 | 1637 | 229 | 2694 | 326 | 239 | 267 |

### Supp Table S8.5.Columns_ET_to_FD

| | | | | | | | | | | | |
|---|---|---|---|---|---|---|---|---|---|---|---|
| SRR1563018 | 13 | 82 | 102 | 105 | 470 | 8648 | 1068 | 9112 | 3609 | 1367 | 4027 |
| SRR1563057 | 62 | 124 | 228 | 41 | 1068 | 18520 | 3401 | 21392 | 10390 | 3847 | 20272 |
| SRR1563058 | 0 | 0 | 22 | 0 | 0 | 21 | 0 | 32 | 34 | 16 | 24 |
| SRR1563059 | 0 | 98 | 378 | 233 | 371 | 4230 | 563 | 11487 | 2676 | 1193 | 6613 |
| SRR1563060 | 0 | 0 | 0 | 0 | 0 | 20 | 0 | 38 | 27 | 17 | 21 |
| SRR1563061 | 0 | 0 | 14 | 0 | 0 | 267 | 33 | 664 | 310 | 144 | 365 |
| SRR1563062 | 0 | 0 | 11 | 11 | 0 | 0 | 0 | 14 | 29 | 22 | 25 |
| SRR1563063 | 0 | 84 | 215 | 192 | 801 | 3812 | 1092 | 6241 | 3639 | 2907 | 5866 |
| SRR1563064 | 0 | 0 | 12 | 0 | 0 | 26 | 0 | 46 | 41 | 19 | 23 |
| SRR1575598 | 0 | 0 | 0 | 0 | 0 | 0 | 0 | 0 | 560 | 1144 | 1389 |
| SRR1575599 | 0 | 0 | 0 | 0 | 0 | 188 | 48 | 460 | 636 | 640 | 607 |
| SRR1575601 | 0 | 0 | 0 | 0 | 0 | 0 | 0 | 0 | 173 | 916 | 1631 |
| SRR1575602 | 0 | 0 | 0 | 0 | 70 | 705 | 103 | 1122 | 2243 | 2421 | 2340 |
| SRR1592244 | 0 | 118 | 314 | 96 | 0 | 24 | 0 | 29 | 0 | 0 | 0 |
| SRR1592245 | 0 | 16 | 41 | 0 | 0 | 0 | 0 | 0 | 0 | 0 | 0 |
| SRR1592246 | 0 | 19 | 37 | 15 | 0 | 0 | 0 | 0 | 0 | 0 | 0 |
| SRR1592247 | 0 | 14 | 54 | 12 | 0 | 0 | 0 | 0 | 0 | 0 | 0 |
| SRR1592248 | 0 | 18 | 68 | 25 | 0 | 0 | 0 | 0 | 0 | 0 | 0 |
| SRR1596226 | 0 | 0 | 0 | 0 | 0 | 18 | 0 | 18 | 23 | 66 | 21 |
| SRR1596227 | 0 | 0 | 0 | 22 | 0 | 29 | 0 | 22 | 26 | 77 | 15 |
| SRR1596228 | 0 | 0 | 0 | 0 | 0 | 0 | 0 | 17 | 40 | 27 | 0 |
| SRR1596229 | 0 | 0 | 0 | 0 | 0 | 0 | 0 | 0 | 0 | 28 | 21 |
| SRR1596230 | 0 | 0 | 0 | 20 | 24 | 14 | 10 | 47 | 64 | 64 | 44 |
| SRR1596231 | 0 | 0 | 0 | 33 | 0 | 41 | 24 | 65 | 73 | 81 | 77 |
| SRR1608128 | 0 | 0 | 0 | 0 | 0 | 10 | 12 | 0 | 0 | 123 | 42 |
| SRR1608138 | 0 | 0 | 0 | 0 | 0 | 0 | 10 | 0 | 21 | 175 | 18 |
| SRR1608143 | 0 | 0 | 0 | 0 | 0 | 0 | 0 | 0 | 0 | 36 | 0 |
| SRR1608150 | 0 | 0 | 0 | 0 | 0 | 0 | 0 | 0 | 10 | 132 | 15 |
| SRR1608158 | 0 | 0 | 0 | 0 | 0 | 0 | 0 | 0 | 0 | 90 | 0 |
| SRR1608160 | 0 | 0 | 0 | 0 | 0 | 0 | 0 | 0 | 0 | 55 | 0 |
| SRR1608163 | 0 | 0 | 0 | 0 | 0 | 0 | 0 | 0 | 0 | 109 | 33 |
| SRR1608165 | 0 | 0 | 0 | 0 | 0 | 0 | 0 | 0 | 22 | 272 | 51 |
| SRR1608166 | 0 | 0 | 0 | 0 | 0 | 0 | 0 | 0 | 0 | 158 | 13 |
| SRR1608167 | 0 | 0 | 0 | 0 | 0 | 0 | 0 | 0 | 0 | 130 | 13 |
| SRR1608168 | 0 | 0 | 0 | 0 | 0 | 0 | 0 | 0 | 0 | 136 | 18 |
| SRR1608169 | 0 | 0 | 0 | 0 | 0 | 0 | 0 | 0 | 0 | 53 | 0 |
| SRR1608170 | 0 | 0 | 0 | 0 | 0 | 0 | 0 | 0 | 0 | 34 | 0 |
| SRR1608171 | 0 | 0 | 0 | 0 | 0 | 0 | 0 | 0 | 0 | 148 | 14 |
| SRR1608172 | 0 | 0 | 0 | 0 | 0 | 0 | 0 | 0 | 0 | 41 | 0 |
| SRR1608174 | 0 | 0 | 0 | 0 | 0 | 0 | 12 | 18 | 14 | 147 | 100 |
| SRR1608175 | 0 | 0 | 0 | 0 | 0 | 0 | 0 | 0 | 15 | 243 | 31 |
| SRR1608177 | 0 | 0 | 0 | 0 | 0 | 0 | 0 | 0 | 0 | 65 | 0 |
| SRR1608178 | 0 | 0 | 0 | 0 | 0 | 12 | 35 | 19 | 23 | 206 | 43 |
| SRR1608179 | 0 | 0 | 0 | 0 | 0 | 0 | 19 | 0 | 27 | 317 | 80 |
| SRR1608180 | 0 | 0 | 0 | 0 | 0 | 14 | 14 | 0 | 0 | 235 | 39 |
| SRR1608181 | 0 | 0 | 0 | 0 | 0 | 0 | 0 | 0 | 0 | 120 | 18 |
| SRR1608182 | 0 | 0 | 0 | 0 | 0 | 0 | 0 | 0 | 0 | 107 | 18 |
| SRR1608184 | 0 | 0 | 0 | 0 | 0 | 0 | 14 | 0 | 0 | 278 | 35 |
| SRR1608185 | 0 | 0 | 0 | 0 | 0 | 0 | 0 | 0 | 0 | 92 | 11 |
| SRR1608186 | 0 | 0 | 0 | 0 | 0 | 0 | 14 | 0 | 10 | 156 | 20 |
| SRR1608189 | 0 | 0 | 0 | 0 | 0 | 0 | 0 | 0 | 0 | 45 | 16 |
| SRR1608190 | 0 | 0 | 0 | 0 | 0 | 0 | 12 | 0 | 0 | 266 | 17 |
| SRR1608191 | 0 | 0 | 0 | 0 | 0 | 0 | 12 | 0 | 15 | 229 | 15 |
| SRR1608192 | 0 | 0 | 0 | 0 | 0 | 0 | 0 | 0 | 0 | 105 | 0 |
| SRR1608193 | 0 | 0 | 0 | 0 | 0 | 0 | 0 | 0 | 16 | 154 | 34 |
| SRR1608194 | 0 | 0 | 0 | 0 | 0 | 0 | 26 | 0 | 15 | 245 | 19 |
| SRR1608196 | 0 | 0 | 0 | 0 | 0 | 0 | 0 | 0 | 0 | 92 | 0 |
| SRR1608201 | 0 | 0 | 0 | 0 | 0 | 0 | 23 | 0 | 18 | 230 | 54 |
| SRR1608208 | 0 | 0 | 0 | 0 | 0 | 0 | 0 | 0 | 14 | 109 | 25 |
| SRR1608213 | 0 | 0 | 0 | 0 | 0 | 0 | 14 | 19 | 48 | 354 | 137 |
| SRR1608214 | 0 | 0 | 0 | 0 | 0 | 0 | 0 | 0 | 0 | 110 | 12 |
| SRR1608222 | 0 | 0 | 0 | 0 | 0 | 0 | 0 | 0 | 0 | 0 | 0 |
| SRR1608232 | 0 | 0 | 0 | 0 | 0 | 0 | 31 | 10 | 0 | 80 | 28 |
| SRR1608237 | 0 | 0 | 0 | 0 | 0 | 0 | 0 | 0 | 0 | 61 | 12 |
| SRR1608254 | 0 | 0 | 0 | 0 | 0 | 0 | 0 | 0 | 0 | 35 | 0 |
| SRR1608255 | 0 | 0 | 0 | 0 | 0 | 0 | 0 | 0 | 0 | 32 | 0 |
| SRR1608256 | 0 | 0 | 0 | 0 | 0 | 0 | 0 | 0 | 0 | 0 | 0 |
| SRR1608257 | 0 | 0 | 0 | 0 | 0 | 0 | 0 | 0 | 0 | 36 | 0 |
| SRR1608260 | 0 | 0 | 0 | 0 | 0 | 0 | 0 | 0 | 0 | 42 | 0 |
| SRR1608263 | 0 | 0 | 0 | 0 | 0 | 0 | 0 | 0 | 0 | 34 | 0 |
| SRR1608264 | 0 | 0 | 0 | 0 | 0 | 11 | 11 | 11 | 0 | 38 | 11 |
| SRR1608266 | 0 | 0 | 0 | 0 | 0 | 0 | 14 | 0 | 0 | 35 | 0 |
| SRR1608267 | 0 | 0 | 0 | 0 | 0 | 12 | 0 | 0 | 0 | 30 | 20 |
| SRR1608269 | 0 | 0 | 0 | 0 | 0 | 0 | 22 | 0 | 0 | 106 | 13 |
| SRR1608270 | 0 | 0 | 0 | 0 | 0 | 33 | 37 | 0 | 0 | 71 | 33 |
| SRR1608271 | 0 | 0 | 0 | 0 | 0 | 0 | 0 | 0 | 0 | 34 | 0 |
| SRR1608274 | 0 | 0 | 0 | 0 | 0 | 13 | 0 | 0 | 0 | 109 | 20 |
| SRR1608276 | 14 | 0 | 0 | 0 | 0 | 28 | 19 | 0 | 0 | 94 | 14 |
| SRR1608277 | 16 | 0 | 0 | 0 | 0 | 14 | 16 | 0 | 0 | 78 | 0 |
| SRR1608278 | 0 | 0 | 0 | 0 | 0 | 0 | 0 | 0 | 0 | 27 | 0 |
| SRR1608281 | 0 | 0 | 0 | 0 | 0 | 0 | 15 | 0 | 0 | 88 | 18 |
| SRR1608282 | 0 | 0 | 0 | 0 | 0 | 0 | 11 | 0 | 0 | 45 | 0 |
| SRR1608283 | 0 | 0 | 0 | 0 | 0 | 0 | 12 | 0 | 0 | 35 | 0 |
| SRR1608286 | 0 | 0 | 0 | 0 | 0 | 0 | 0 | 0 | 0 | 25 | 11 |
| SRR1608287 | 0 | 0 | 0 | 0 | 0 | 0 | 0 | 0 | 0 | 30 | 15 |
| SRR1608289 | 0 | 0 | 0 | 0 | 0 | 0 | 17 | 0 | 0 | 25 | 30 |
| SRR1608290 | 0 | 0 | 0 | 0 | 0 | 0 | 22 | 0 | 0 | 32 | 16 |
| SRR1608302 | 0 | 0 | 0 | 0 | 0 | 0 | 0 | 0 | 0 | 52 | 0 |
| SRR1608307 | 0 | 0 | 0 | 0 | 0 | 0 | 0 | 0 | 0 | 38 | 21 |
| SRR1608308 | 0 | 0 | 0 | 0 | 0 | 0 | 11 | 0 | 0 | 131 | 0 |
| SRR1634379 | 0 | 0 | 0 | 0 | 0 | 11 | 0 | 13 | 18 | 12 | 0 |
| SRR1634381 | 0 | 0 | 0 | 0 | 15 | 23 | 15 | 23 | 31 | 31 | 0 |
| SRR1634384 | 0 | 0 | 0 | 0 | 0 | 0 | 0 | 0 | 0 | 0 | 0 |
| SRR1634388 | 0 | 0 | 0 | 0 | 0 | 0 | 0 | 0 | 0 | 0 | 0 |
| SRR1634390 | 0 | 0 | 0 | 0 | 0 | 33 | 11 | 42 | 30 | 47 | 14 |
| SRR1634392 | 0 | 14 | 0 | 0 | 10 | 18 | 0 | 18 | 26 | 17 | 0 |
| SRR1634394 | 0 | 0 | 0 | 0 | 0 | 13 | 0 | 0 | 13 | 12 | 0 |
| SRR1634395 | 0 | 0 | 0 | 0 | 21 | 38 | 13 | 78 | 59 | 82 | 19 |
| SRR1634398 | 0 | 0 | 0 | 0 | 15 | 28 | 12 | 56 | 44 | 73 | 21 |
| SRR1634399 | 0 | 0 | 0 | 0 | 11 | 33 | 10 | 41 | 34 | 45 | 14 |
| SRR1634400 | 0 | 0 | 0 | 0 | 11 | 30 | 18 | 45 | 35 | 48 | 11 |
| SRR1634406 | 0 | 0 | 0 | 0 | 13 | 24 | 0 | 23 | 30 | 16 | 0 |
| SRR1634409 | 0 | 0 | 0 | 0 | 21 | 36 | 14 | 65 | 80 | 120 | 25 |
| SRR1634411 | 0 | 0 | 0 | 0 | 15 | 39 | 16 | 56 | 40 | 50 | 20 |
| SRR1634412 | 0 | 0 | 0 | 12 | 21 | 34 | 16 | 61 | 35 | 40 | 29 |
| SRR1634414 | 0 | 0 | 0 | 0 | 11 | 29 | 13 | 44 | 51 | 78 | 23 |
| SRR1634415 | 0 | 0 | 0 | 0 | 16 | 27 | 14 | 41 | 43 | 45 | 17 |
| SRR1634417 | 0 | 0 | 0 | 0 | 15 | 27 | 15 | 58 | 49 | 75 | 25 |
| SRR1636959 | 30 | 0 | 61 | 43 | 0 | 121 | 22 | 146 | 22 | 43 | 45 |
| SRR1636960 | 16 | 73 | 141 | 49 | 0 | 179 | 35 | 290 | 0 | 57 | 52 |
| SRR1636961 | 0 | 42 | 72 | 28 | 11 | 141 | 11 | 285 | 11 | 30 | 39 |
| SRR1636962 | 47 | 81 | 152 | 115 | 47 | 149 | 40 | 301 | 127 | 230 | 99 |
| SRR1636963 | 32 | 69 | 205 | 244 | 59 | 159 | 27 | 271 | 112 | 255 | 147 |
| SRR1636964 | 51 | 81 | 257 | 257 | 48 | 147 | 21 | 215 | 90 | 233 | 182 |
| SRR1636965 | 13 | 0 | 26 | 18 | 0 | 35 | 0 | 78 | 0 | 23 | 40 |
| SRR1636966 | 0 | 0 | 23 | 31 | 0 | 0 | 0 | 12 | 0 | 0 | 15 |
| SRR1636967 | 0 | 0 | 78 | 83 | 0 | 14 | 14 | 46 | 0 | 0 | 14 |
| SRR1636968 | 84 | 100 | 230 | 116 | 131 | 699 | 81 | 1051 | 30 | 101 | 97 |
| SRR1636969 | 59 | 130 | 265 | 121 | 164 | 830 | 108 | 1320 | 27 | 114 | 104 |

### Supp Table S8.5.Columns_ET_to_FD

| | | | | | | | | | | | |
|---|---|---|---|---|---|---|---|---|---|---|---|
| SRR1636970 | 115 | 237 | 472 | 179 | 183 | 968 | 130 | 1360 | 55 | 114 | 158 |
| SRR1642941 | 0 | 0 | 0 | 0 | 0 | 0 | 0 | 0 | 39 | 0 | 0 |
| SRR1642942 | 0 | 0 | 12 | 0 | 13 | 29 | 13 | 16 | 166 | 0 | 46 |
| SRR1642943 | 0 | 0 | 0 | 0 | 0 | 0 | 0 | 0 | 20 | 0 | 0 |
| SRR1642944 | 0 | 0 | 11 | 0 | 10 | 24 | 10 | 12 | 96 | 0 | 21 |
| SRR1642945 | 0 | 0 | 0 | 0 | 0 | 0 | 0 | 0 | 12 | 0 | 0 |
| SRR1642946 | 0 | 0 | 0 | 0 | 0 | 0 | 0 | 0 | 31 | 0 | 0 |
| SRR1642947 | 0 | 0 | 0 | 0 | 0 | 0 | 0 | 0 | 21 | 0 | 0 |
| SRR1642948 | 0 | 0 | 0 | 0 | 0 | 15 | 0 | 13 | 71 | 0 | 15 |
| SRR1642949 | 0 | 0 | 0 | 0 | 0 | 0 | 0 | 0 | 24 | 0 | 0 |
| SRR1642950 | 0 | 0 | 0 | 0 | 0 | 0 | 0 | 0 | 37 | 0 | 11 |
| SRR1642951 | 0 | 0 | 0 | 0 | 0 | 10 | 0 | 0 | 28 | 0 | 10 |
| SRR1642952 | 0 | 0 | 0 | 0 | 0 | 10 | 0 | 0 | 41 | 0 | 17 |
| SRR1642953 | 0 | 0 | 0 | 0 | 0 | 14 | 0 | 0 | 33 | 0 | 0 |
| SRR1642954 | 0 | 0 | 0 | 0 | 13 | 36 | 13 | 18 | 100 | 0 | 42 |
| SRR1642955 | 0 | 0 | 0 | 0 | 0 | 0 | 0 | 0 | 11 | 0 | 0 |
| SRR1642956 | 0 | 0 | 0 | 0 | 21 | 55 | 21 | 23 | 96 | 0 | 51 |
| SRR1642957 | 0 | 0 | 0 | 0 | 0 | 34 | 19 | 24 | 75 | 0 | 29 |
| SRR1642958 | 0 | 0 | 0 | 0 | 0 | 15 | 11 | 14 | 45 | 0 | 17 |
| SRR1642959 | 0 | 0 | 0 | 0 | 0 | 11 | 0 | 0 | 38 | 0 | 0 |
| SRR1642960 | 0 | 0 | 0 | 0 | 0 | 14 | 0 | 0 | 49 | 0 | 13 |
| SRR1642961 | 0 | 0 | 0 | 0 | 0 | 13 | 0 | 0 | 25 | 0 | 0 |
| SRR1642962 | 0 | 0 | 0 | 0 | 0 | 17 | 0 | 0 | 52 | 0 | 25 |
| SRR1642963 | 0 | 0 | 0 | 0 | 0 | 12 | 0 | 10 | 37 | 0 | 0 |
| SRR1642964 | 0 | 0 | 0 | 0 | 16 | 41 | 11 | 15 | 97 | 0 | 20 |
| SRR1642965 | 0 | 0 | 0 | 0 | 0 | 0 | 0 | 0 | 63 | 0 | 22 |
| SRR1642966 | 0 | 0 | 0 | 0 | 0 | 14 | 0 | 0 | 93 | 0 | 37 |
| SRR1642967 | 0 | 0 | 0 | 0 | 0 | 0 | 0 | 0 | 35 | 0 | 18 |
| SRR1642968 | 0 | 0 | 0 | 0 | 0 | 0 | 0 | 0 | 73 | 0 | 23 |
| SRR1642969 | 0 | 0 | 0 | 0 | 0 | 0 | 0 | 0 | 26 | 0 | 0 |
| SRR1642970 | 0 | 0 | 0 | 0 | 0 | 24 | 13 | 18 | 124 | 0 | 33 |
| SRR1642971 | 0 | 0 | 0 | 0 | 0 | 11 | 0 | 0 | 45 | 0 | 23 |
| SRR1642972 | 0 | 0 | 0 | 0 | 0 | 21 | 11 | 14 | 100 | 0 | 30 |
| SRR1642973 | 0 | 0 | 0 | 0 | 0 | 10 | 0 | 0 | 56 | 0 | 25 |
| SRR1642974 | 0 | 0 | 0 | 0 | 0 | 20 | 0 | 13 | 120 | 0 | 38 |
| SRR1642975 | 0 | 0 | 0 | 0 | 0 | 12 | 0 | 14 | 89 | 0 | 25 |
| SRR1642976 | 0 | 0 | 0 | 0 | 0 | 12 | 0 | 12 | 61 | 0 | 24 |
| SRR1642977 | 0 | 0 | 0 | 0 | 16 | 32 | 13 | 12 | 72 | 0 | 11 |
| SRR1642978 | 0 | 0 | 0 | 0 | 0 | 24 | 15 | 18 | 120 | 0 | 28 |
| SRR1642979 | 0 | 0 | 0 | 0 | 11 | 32 | 19 | 17 | 50 | 0 | 22 |
| SRR1642980 | 0 | 0 | 0 | 0 | 18 | 64 | 30 | 39 | 250 | 11 | 43 |
| SRR1642981 | 0 | 0 | 0 | 0 | 0 | 0 | 0 | 0 | 73 | 0 | 11 |
| SRR1642982 | 0 | 0 | 0 | 0 | 14 | 42 | 13 | 24 | 125 | 0 | 33 |
| SRR1642983 | 0 | 0 | 0 | 0 | 0 | 12 | 0 | 0 | 39 | 0 | 0 |
| SRR1642984 | 0 | 0 | 0 | 0 | 12 | 33 | 0 | 14 | 68 | 0 | 22 |
| SRR1642985 | 0 | 0 | 0 | 0 | 0 | 15 | 12 | 11 | 82 | 0 | 22 |
| SRR1642986 | 0 | 0 | 0 | 0 | 0 | 19 | 0 | 12 | 78 | 0 | 15 |
| SRR1642987 | 0 | 0 | 0 | 0 | 0 | 13 | 0 | 0 | 48 | 0 | 0 |
| SRR1642988 | 0 | 0 | 0 | 0 | 17 | 34 | 18 | 18 | 113 | 0 | 24 |
| SRR1657194 | 0 | 18 | 45 | 65 | 69 | 1014 | 129 | 2105 | 6313 | 11373 | 6119 |
| SRR1657195 | 0 | 47 | 112 | 89 | 218 | 2767 | 338 | 7142 | 72510 | 118362 | 23400 |
| SRR1657196 | 0 | 42 | 86 | 56 | 139 | 1483 | 211 | 3952 | 9647 | 17190 | 6385 |
| SRR1657197 | 0 | 235 | 525 | 235 | 180 | 746 | 138 | 1382 | 4352 | 7724 | 3468 |
| SRR1657198 | 0 | 0 | 16 | 0 | 21 | 367 | 48 | 1074 | 4423 | 7692 | 3103 |
| SRR1686806 | 0 | 0 | 0 | 0 | 0 | 45 | 73 | 46 | 11 | 0 | 0 |
| SRR1707574 | 0 | 0 | 0 | 86 | 0 | 24 | 0 | 36 | 60 | 0 | 0 |
| SRR1707575 | 0 | 0 | 49 | 0 | 13 | 0 | 86 | 0 | 145 | 0 | 577 |
| SRR1708906 | 0 | 16 | 63 | 18 | 220 | 3564 | 693 | 2194 | 186 | 112 | 735 |
| SRR1708907 | 0 | 0 | 0 | 0 | 106 | 2280 | 334 | 1545 | 145 | 68 | 454 |
| SRR1708908 | 0 | 0 | 0 | 0 | 50 | 1119 | 202 | 840 | 151 | 67 | 547 |
| SRR1708909 | 0 | 0 | 72 | 65 | 37 | 755 | 90 | 745 | 167 | 45 | 235 |
| SRR1708910 | 0 | 0 | 0 | 0 | 57 | 859 | 235 | 548 | 291 | 59 | 548 |
| SRR1708911 | 0 | 0 | 0 | 0 | 45 | 717 | 175 | 655 | 350 | 147 | 847 |
| SRR1708912 | 0 | 0 | 0 | 0 | 64 | 1097 | 323 | 727 | 271 | 52 | 693 |
| SRR1708913 | 0 | 0 | 0 | 0 | 42 | 733 | 206 | 386 | 316 | 77 | 662 |
| SRR1708914 | 0 | 0 | 0 | 0 | 104 | 1023 | 243 | 1033 | 263 | 144 | 1143 |
| SRR1708915 | 0 | 0 | 30 | 19 | 85 | 784 | 165 | 503 | 257 | 159 | 570 |
| SRR1708916 | 0 | 0 | 0 | 0 | 46 | 992 | 293 | 848 | 266 | 111 | 656 |
| SRR1708917 | 0 | 0 | 0 | 0 | 71 | 931 | 231 | 608 | 266 | 165 | 768 |
| SRR1708918 | 0 | 0 | 0 | 0 | 26 | 725 | 163 | 418 | 337 | 88 | 1055 |
| SRR1708919 | 0 | 0 | 0 | 0 | 69 | 916 | 158 | 680 | 290 | 50 | 976 |
| SRR1708920 | 0 | 0 | 0 | 0 | 32 | 435 | 100 | 362 | 139 | 47 | 488 |
| SRR1708921 | 0 | 0 | 0 | 16 | 47 | 576 | 107 | 475 | 257 | 49 | 548 |
| SRR1708922 | 0 | 0 | 0 | 0 | 49 | 963 | 195 | 434 | 225 | 79 | 798 |
| SRR1708923 | 0 | 0 | 0 | 12 | 76 | 770 | 122 | 411 | 216 | 85 | 666 |
| SRR1708924 | 0 | 0 | 0 | 0 | 173 | 2465 | 514 | 1855 | 146 | 100 | 420 |
| SRR1708925 | 0 | 0 | 21 | 11 | 75 | 2297 | 423 | 2431 | 177 | 104 | 501 |
| SRR1708926 | 0 | 45 | 97 | 73 | 133 | 1807 | 376 | 837 | 112 | 56 | 477 |
| SRR1708927 | 0 | 0 | 20 | 16 | 116 | 1666 | 342 | 1974 | 204 | 102 | 816 |
| SRR1708928 | 0 | 0 | 0 | 0 | 112 | 3057 | 524 | 2236 | 239 | 167 | 832 |
| SRR1708929 | 0 | 0 | 0 | 0 | 82 | 1084 | 238 | 868 | 187 | 77 | 393 |
| SRR1708930 | 0 | 0 | 22 | 0 | 56 | 934 | 204 | 1271 | 222 | 17 | 672 |
| SRR1708931 | 0 | 0 | 0 | 0 | 67 | 1346 | 187 | 1082 | 146 | 51 | 513 |
| SRR1708932 | 0 | 0 | 0 | 0 | 48 | 738 | 133 | 1587 | 342 | 74 | 959 |
| SRR1708933 | 0 | 0 | 0 | 0 | 57 | 1212 | 278 | 2161 | 462 | 351 | 1324 |
| SRR1708934 | 0 | 0 | 19 | 24 | 102 | 2163 | 404 | 2870 | 408 | 93 | 860 |
| SRR1708935 | 0 | 0 | 0 | 0 | 31 | 698 | 129 | 758 | 245 | 137 | 957 |
| SRR1708936 | 0 | 0 | 49 | 43 | 43 | 707 | 122 | 718 | 171 | 56 | 572 |
| SRR1708937 | 0 | 0 | 0 | 0 | 0 | 133 | 36 | 396 | 169 | 70 | 647 |
| SRR1708938 | 0 | 0 | 0 | 0 | 19 | 412 | 84 | 718 | 310 | 74 | 663 |
| SRR1708939 | 0 | 0 | 0 | 0 | 39 | 654 | 120 | 891 | 452 | 107 | 654 |
| SRR1708940 | 0 | 0 | 0 | 0 | 33 | 904 | 193 | 1111 | 214 | 56 | 616 |
| SRR1708941 | 0 | 0 | 0 | 0 | 41 | 697 | 153 | 598 | 303 | 88 | 722 |
| SRR1708942 | 0 | 0 | 0 | 0 | 158 | 2641 | 598 | 2606 | 175 | 103 | 609 |
| SRR1708943 | 0 | 0 | 0 | 0 | 31 | 409 | 135 | 648 | 103 | 44 | 235 |
| SRR1708944 | 0 | 0 | 0 | 0 | 99 | 1325 | 204 | 744 | 185 | 91 | 392 |
| SRR1708945 | 0 | 13 | 53 | 32 | 101 | 1153 | 313 | 1280 | 194 | 82 | 474 |
| SRR1708946 | 0 | 0 | 0 | 0 | 66 | 1263 | 253 | 939 | 132 | 89 | 345 |
| SRR1708947 | 0 | 0 | 27 | 20 | 83 | 1052 | 167 | 573 | 141 | 86 | 283 |
| SRR1708948 | 0 | 0 | 0 | 0 | 30 | 800 | 146 | 1004 | 240 | 68 | 554 |
| SRR1708949 | 0 | 0 | 11 | 0 | 65 | 1325 | 230 | 1106 | 298 | 71 | 599 |
| SRR1708950 | 0 | 0 | 0 | 0 | 43 | 610 | 174 | 800 | 88 | 46 | 381 |
| SRR1708951 | 0 | 0 | 0 | 14 | 54 | 800 | 172 | 1261 | 197 | 156 | 698 |
| SRR1708952 | 0 | 0 | 0 | 0 | 37 | 1140 | 267 | 2016 | 211 | 98 | 689 |
| SRR1708953 | 0 | 0 | 0 | 0 | 39 | 951 | 147 | 1204 | 224 | 92 | 631 |
| SRR1708954 | 0 | 0 | 0 | 0 | 29 | 611 | 124 | 772 | 250 | 93 | 766 |
| SRR1708955 | 0 | 0 | 0 | 0 | 24 | 450 | 71 | 676 | 141 | 67 | 617 |
| SRR1708956 | 0 | 0 | 0 | 0 | 17 | 425 | 64 | 898 | 211 | 93 | 632 |
| SRR1708957 | 0 | 0 | 18 | 51 | 30 | 437 | 106 | 842 | 154 | 94 | 727 |
| SRR1708958 | 0 | 0 | 0 | 0 | 0 | 449 | 54 | 988 | 163 | 53 | 421 |
| SRR1708959 | 0 | 0 | 0 | 0 | 32 | 563 | 90 | 645 | 215 | 64 | 615 |
| SRR1708960 | 0 | 18 | 24 | 27 | 159 | 2073 | 536 | 1804 | 139 | 110 | 449 |
| SRR1708961 | 0 | 0 | 22 | 25 | 171 | 2558 | 454 | 2216 | 146 | 130 | 363 |
| SRR1708962 | 0 | 0 | 0 | 0 | 80 | 1390 | 290 | 867 | 108 | 88 | 448 |
| SRR1708963 | 0 | 0 | 0 | 0 | 99 | 1401 | 297 | 1713 | 165 | 101 | 374 |
| SRR1708964 | 0 | 0 | 15 | 0 | 192 | 1879 | 546 | 1461 | 237 | 256 | 934 |
| SRR1708965 | 0 | 0 | 0 | 0 | 45 | 849 | 189 | 904 | 126 | 114 | 488 |
| SRR1708966 | 0 | 0 | 11 | 15 | 65 | 860 | 163 | 994 | 149 | 31 | 360 |
| SRR1708967 | 0 | 0 | 0 | 0 | 23 | 459 | 106 | 820 | 88 | 46 | 422 |

### Supp Table S8.5.Columns_ET_to_FD

| | | | | | | | | | | | |
|---|---|---|---|---|---|---|---|---|---|---|---|
| SRR1708968 | 0 | 0 | 0 | 0 | 16 | 616 | 141 | 1853 | 228 | 109 | 734 |
| SRR1708969 | 0 | 0 | 0 | 0 | 49 | 1014 | 158 | 1709 | 294 | 207 | 769 |
| SRR1708970 | 0 | 0 | 0 | 0 | 35 | 740 | 189 | 1341 | 184 | 106 | 513 |
| SRR1708971 | 0 | 0 | 0 | 0 | 22 | 752 | 149 | 1335 | 569 | 210 | 1274 |
| SRR1708972 | 0 | 0 | 0 | 0 | 18 | 405 | 52 | 808 | 167 | 75 | 664 |
| SRR1708973 | 0 | 0 | 0 | 0 | 0 | 78 | 15 | 322 | 150 | 72 | 700 |
| SRR1708974 | 0 | 0 | 0 | 0 | 17 | 261 | 36 | 817 | 151 | 77 | 447 |
| SRR1708975 | 0 | 0 | 0 | 0 | 32 | 384 | 76 | 897 | 209 | 57 | 596 |
| SRR1708976 | 0 | 0 | 0 | 0 | 17 | 616 | 74 | 1167 | 139 | 52 | 438 |
| SRR1708977 | 0 | 0 | 15 | 0 | 31 | 464 | 88 | 498 | 125 | 76 | 555 |
| SRR1708978 | 0 | 0 | 0 | 0 | 103 | 2193 | 571 | 2260 | 175 | 75 | 437 |
| SRR1708979 | 0 | 0 | 0 | 0 | 123 | 1613 | 330 | 1529 | 127 | 78 | 291 |
| SRR1708980 | 0 | 0 | 0 | 0 | 65 | 777 | 217 | 653 | 129 | 81 | 616 |
| SRR1708981 | 0 | 0 | 0 | 0 | 63 | 1311 | 276 | 1064 | 133 | 36 | 356 |
| SRR1708982 | 0 | 0 | 0 | 0 | 95 | 1845 | 323 | 1068 | 222 | 144 | 921 |
| SRR1708983 | 0 | 0 | 0 | 11 | 79 | 879 | 188 | 426 | 111 | 67 | 206 |
| SRR1708984 | 0 | 0 | 0 | 0 | 85 | 1345 | 225 | 634 | 280 | 77 | 755 |
| SRR1708985 | 0 | 11 | 35 | 36 | 60 | 993 | 220 | 656 | 137 | 23 | 511 |
| SRR1708986 | 0 | 0 | 13 | 10 | 84 | 1402 | 276 | 1086 | 266 | 71 | 895 |
| SRR1708987 | 0 | 0 | 0 | 0 | 79 | 1174 | 195 | 1493 | 229 | 75 | 885 |
| SRR1708988 | 0 | 0 | 0 | 0 | 26 | 832 | 85 | 1576 | 219 | 69 | 632 |
| SRR1708989 | 0 | 0 | 0 | 0 | 51 | 1307 | 173 | 2221 | 538 | 220 | 1687 |
| SRR1708990 | 0 | 0 | 18 | 18 | 60 | 950 | 202 | 547 | 212 | 93 | 742 |
| SRR1708991 | 0 | 0 | 0 | 0 | 14 | 767 | 118 | 596 | 171 | 61 | 692 |
| SRR1708992 | 0 | 0 | 0 | 58 | 52 | 663 | 133 | 757 | 207 | 87 | 492 |
| SRR1708993 | 0 | 0 | 0 | 0 | 12 | 583 | 47 | 836 | 221 | 37 | 577 |
| SRR1708994 | 0 | 0 | 0 | 0 | 17 | 528 | 67 | 801 | 145 | 52 | 415 |
| SRR1708995 | 0 | 0 | 0 | 0 | 38 | 738 | 86 | 758 | 225 | 65 | 758 |
| SRR1708996 | 0 | 0 | 0 | 0 | 98 | 1668 | 409 | 1777 | 113 | 74 | 452 |
| SRR1708997 | 0 | 25 | 75 | 50 | 92 | 1110 | 271 | 884 | 57 | 53 | 262 |
| SRR1708998 | 0 | 0 | 13 | 18 | 103 | 1748 | 405 | 1211 | 194 | 77 | 819 |
| SRR1708999 | 0 | 0 | 0 | 0 | 33 | 945 | 234 | 1072 | 211 | 64 | 414 |
| SRR1709000 | 0 | 0 | 11 | 0 | 142 | 1577 | 331 | 997 | 315 | 50 | 690 |
| SRR1709001 | 0 | 0 | 0 | 0 | 73 | 1097 | 241 | 883 | 197 | 29 | 598 |
| SRR1709002 | 0 | 19 | 66 | 38 | 75 | 1324 | 174 | 1405 | 202 | 100 | 516 |
| SRR1709003 | 0 | 0 | 0 | 0 | 59 | 762 | 166 | 1427 | 246 | 95 | 679 |
| SRR1709004 | 0 | 0 | 0 | 0 | 192 | 2904 | 611 | 2250 | 348 | 130 | 524 |
| SRR1709005 | 0 | 0 | 17 | 24 | 137 | 1906 | 484 | 1683 | 108 | 55 | 405 |
| SRR1709006 | 0 | 0 | 24 | 65 | 99 | 1426 | 396 | 1055 | 177 | 147 | 763 |
| SRR1709007 | 0 | 13 | 35 | 27 | 110 | 2094 | 507 | 1709 | 99 | 67 | 556 |
| SRR1709008 | 0 | 0 | 0 | 0 | 79 | 1303 | 278 | 872 | 162 | 114 | 557 |
| SRR1709009 | 0 | 0 | 0 | 0 | 99 | 1535 | 327 | 1019 | 186 | 162 | 572 |
| SRR1709010 | 0 | 0 | 0 | 0 | 61 | 1238 | 186 | 1586 | 539 | 61 | 1248 |
| SRR1709011 | 0 | 0 | 13 | 15 | 96 | 1618 | 177 | 1886 | 250 | 36 | 899 |
| SRR1709012 | 0 | 0 | 34 | 15 | 76 | 1005 | 162 | 1653 | 182 | 71 | 717 |
| SRR1709013 | 0 | 0 | 0 | 0 | 36 | 791 | 97 | 1806 | 245 | 82 | 761 |
| SRR1709014 | 0 | 0 | 12 | 0 | 83 | 2428 | 396 | 4926 | 446 | 151 | 1586 |
| SRR1709015 | 0 | 0 | 0 | 0 | 73 | 1341 | 145 | 1448 | 343 | 161 | 1046 |
| SRR1709016 | 0 | 0 | 0 | 0 | 65 | 806 | 187 | 1244 | 204 | 75 | 1059 |
| SRR1709017 | 0 | 0 | 0 | 0 | 34 | 674 | 102 | 1227 | 414 | 81 | 1026 |
| SRR1709018 | 0 | 0 | 0 | 0 | 16 | 314 | 51 | 1234 | 220 | 73 | 663 |
| SRR1709019 | 0 | 0 | 0 | 0 | 42 | 656 | 78 | 1210 | 353 | 94 | 1131 |
| SRR1709020 | 0 | 0 | 0 | 16 | 36 | 822 | 99 | 1062 | 151 | 57 | 407 |
| SRR1709021 | 0 | 0 | 0 | 0 | 29 | 711 | 148 | 916 | 424 | 80 | 1142 |
| SRR1732015 | 0 | 0 | 12 | 12 | 17 | 48 | 15 | 62 | 447 | 299 | 211 |
| SRR1732016 | 0 | 0 | 0 | 0 | 0 | 0 | 0 | 0 | 13 | 0 | 16 |
| SRR1748132 | 0 | 0 | 0 | 0 | 0 | 59 | 16 | 247 | 33 | 42 | 59 |
| SRR1748133 | 0 | 0 | 0 | 0 | 0 | 22 | 0 | 40 | 26 | 27 | 34 |
| SRR1759212 | 0 | 0 | 20 | 16 | 0 | 16 | 0 | 25 | 30 | 0 | 0 |
| SRR1759213 | 0 | 0 | 0 | 0 | 0 | 0 | 0 | 0 | 0 | 0 | 0 |
| SRR1759214 | 0 | 0 | 17 | 0 | 0 | 21 | 0 | 26 | 15 | 0 | 0 |
| SRR1759215 | 0 | 0 | 11 | 0 | 0 | 0 | 0 | 24 | 18 | 0 | 0 |
| SRR1759216 | 0 | 0 | 0 | 18 | 0 | 17 | 0 | 28 | 15 | 0 | 0 |
| SRR1759217 | 0 | 0 | 0 | 13 | 0 | 11 | 0 | 18 | 0 | 0 | 0 |
| SRR1759218 | 0 | 0 | 15 | 11 | 0 | 31 | 0 | 31 | 14 | 0 | 0 |
| SRR1759219 | 0 | 0 | 21 | 19 | 0 | 19 | 0 | 25 | 13 | 0 | 0 |
| SRR1759220 | 0 | 0 | 0 | 19 | 0 | 19 | 0 | 35 | 31 | 0 | 0 |
| SRR1759221 | 0 | 0 | 42 | 62 | 0 | 50 | 0 | 90 | 48 | 11 | 21 |
| SRR1759222 | 0 | 0 | 0 | 0 | 0 | 37 | 0 | 80 | 37 | 17 | 51 |
| SRR1759223 | 0 | 11 | 42 | 56 | 0 | 46 | 0 | 71 | 26 | 0 | 13 |
| SRR1759224 | 0 | 0 | 53 | 33 | 0 | 70 | 0 | 141 | 50 | 18 | 0 |
| SRR1759225 | 0 | 0 | 0 | 0 | 0 | 43 | 0 | 100 | 21 | 10 | 19 |
| SRR1759226 | 0 | 0 | 18 | 0 | 0 | 33 | 0 | 72 | 29 | 0 | 20 |
| SRR1759227 | 0 | 0 | 43 | 31 | 0 | 43 | 0 | 86 | 48 | 11 | 25 |
| SRR1759228 | 0 | 0 | 0 | 21 | 0 | 60 | 0 | 88 | 45 | 35 | 25 |
| SRR1759229 | 0 | 11 | 72 | 42 | 0 | 29 | 10 | 55 | 21 | 14 | 14 |
| SRR1759230 | 0 | 0 | 0 | 0 | 0 | 31 | 0 | 77 | 36 | 12 | 21 |
| SRR1759231 | 0 | 0 | 41 | 24 | 0 | 43 | 0 | 100 | 18 | 0 | 20 |
| SRR1759232 | 0 | 0 | 50 | 41 | 0 | 113 | 0 | 213 | 30 | 33 | 61 |
| SRR1759233 | 0 | 0 | 0 | 0 | 0 | 57 | 0 | 106 | 51 | 20 | 47 |
| SRR1759234 | 0 | 19 | 40 | 25 | 0 | 37 | 0 | 60 | 18 | 0 | 12 |
| SRR1759235 | 0 | 0 | 34 | 14 | 0 | 19 | 0 | 44 | 39 | 0 | 0 |
| SRR1759236 | 0 | 0 | 83 | 73 | 0 | 42 | 0 | 72 | 40 | 20 | 28 |
| SRR1759237 | 0 | 32 | 93 | 107 | 0 | 52 | 0 | 73 | 47 | 21 | 0 |
| SRR1759238 | 0 | 30 | 100 | 113 | 0 | 72 | 0 | 94 | 64 | 30 | 40 |
| SRR1759239 | 0 | 0 | 0 | 0 | 11 | 34 | 21 | 67 | 44 | 40 | 23 |
| SRR1759240 | 0 | 23 | 62 | 48 | 0 | 52 | 0 | 66 | 33 | 26 | 25 |
| SRR1759241 | 0 | 14 | 43 | 50 | 0 | 76 | 0 | 110 | 59 | 19 | 21 |
| SRR1759242 | 0 | 35 | 87 | 99 | 15 | 101 | 16 | 104 | 95 | 41 | 16 |
| SRR1759243 | 0 | 0 | 45 | 30 | 0 | 30 | 13 | 117 | 60 | 15 | 12 |
| SRR1759244 | 0 | 0 | 12 | 0 | 0 | 12 | 0 | 20 | 20 | 0 | 0 |
| SRR1759245 | 0 | 0 | 17 | 18 | 0 | 15 | 0 | 25 | 25 | 0 | 0 |
| SRR1759246 | 0 | 15 | 58 | 39 | 0 | 23 | 0 | 55 | 31 | 16 | 28 |
| SRR1759247 | 0 | 19 | 28 | 33 | 0 | 54 | 0 | 96 | 33 | 13 | 41 |
| SRR1759248 | 0 | 19 | 66 | 89 | 0 | 32 | 0 | 74 | 42 | 24 | 32 |
| SRR1759249 | 0 | 11 | 31 | 42 | 0 | 30 | 0 | 32 | 23 | 13 | 0 |
| SRR1759250 | 0 | 0 | 0 | 0 | 0 | 18 | 0 | 29 | 25 | 0 | 0 |
| SRR1759251 | 0 | 16 | 27 | 24 | 0 | 11 | 0 | 26 | 20 | 13 | 0 |
| SRR1759252 | 0 | 0 | 51 | 69 | 0 | 39 | 0 | 90 | 25 | 0 | 16 |
| SRR1759253 | 0 | 13 | 58 | 30 | 0 | 30 | 0 | 51 | 32 | 17 | 22 |
| SRR1759254 | 0 | 0 | 29 | 38 | 0 | 15 | 0 | 24 | 16 | 0 | 0 |
| SRR1759255 | 0 | 0 | 20 | 0 | 0 | 0 | 0 | 22 | 0 | 0 | 0 |
| SRR1759256 | 0 | 0 | 38 | 38 | 0 | 88 | 12 | 127 | 85 | 22 | 54 |
| SRR1759257 | 0 | 0 | 17 | 0 | 0 | 42 | 0 | 37 | 53 | 101 | 0 |
| SRR1759258 | 0 | 123 | 449 | 475 | 28 | 228 | 40 | 414 | 92 | 98 | 266 |
| SRR1759259 | 0 | 55 | 238 | 288 | 25 | 202 | 22 | 276 | 108 | 65 | 195 |
| SRR1759260 | 0 | 0 | 0 | 0 | 0 | 14 | 0 | 61 | 26 | 0 | 0 |
| SRR1759261 | 0 | 22 | 59 | 36 | 0 | 49 | 0 | 116 | 66 | 30 | 28 |
| SRR1759262 | 0 | 0 | 15 | 15 | 0 | 13 | 0 | 42 | 21 | 0 | 0 |
| SRR1759263 | 0 | 0 | 0 | 0 | 0 | 58 | 0 | 85 | 46 | 0 | 19 |
| SRR1759264 | 0 | 0 | 35 | 25 | 0 | 37 | 0 | 87 | 23 | 0 | 25 |
| SRR1759265 | 0 | 0 | 12 | 0 | 0 | 24 | 0 | 39 | 24 | 78 | 20 |
| SRR1759266 | 0 | 0 | 18 | 18 | 0 | 13 | 0 | 0 | 22 | 33 | 0 |
| SRR1759267 | 0 | 0 | 0 | 0 | 0 | 52 | 11 | 110 | 47 | 23 | 23 |
| SRR1759268 | 0 | 0 | 24 | 22 | 0 | 51 | 0 | 90 | 34 | 19 | 22 |
| SRR1759269 | 0 | 0 | 0 | 18 | 0 | 16 | 0 | 43 | 18 | 13 | 0 |
| SRR1759270 | 0 | 0 | 25 | 23 | 0 | 32 | 0 | 37 | 25 | 15 | 0 |
| SRR1759271 | 0 | 0 | 10 | 10 | 0 | 40 | 18 | 79 | 26 | 14 | 0 |
| SRR1759272 | 0 | 0 | 23 | 23 | 0 | 89 | 11 | 114 | 25 | 0 | 25 |

### Supp Table S8.5.Columns_ET_to_FD

| | | | | | | | | | | | |
|---|---|---|---|---|---|---|---|---|---|---|---|
| SRR1759273 | 0 | 0 | 25 | 22 | 11 | 37 | 0 | 59 | 33 | 11 | 12 |
| SRR1759274 | 0 | 0 | 0 | 0 | 0 | 18 | 0 | 80 | 46 | 30 | 10 |
| SRR1759275 | 0 | 0 | 43 | 33 | 0 | 27 | 0 | 59 | 16 | 39 | 0 |
| SRR1777167 | 0 | 12 | 107 | 0 | 117 | 165 | 51 | 37 | 180 | 15 | 26 |
| SRR1820679 | 0 | 0 | 19 | 0 | 20 | 23 | 12 | 14 | 86 | 0 | 0 |
| SRR1820680 | 0 | 0 | 0 | 0 | 0 | 14 | 0 | 0 | 19 | 0 | 0 |
| SRR1820681 | 0 | 0 | 16 | 0 | 0 | 13 | 0 | 0 | 62 | 0 | 0 |
| SRR1820682 | 0 | 0 | 0 | 0 | 0 | 21 | 10 | 14 | 113 | 0 | 0 |
| SRR1820683 | 0 | 0 | 17 | 0 | 12 | 17 | 0 | 14 | 55 | 0 | 0 |
| SRR1820684 | 0 | 0 | 0 | 0 | 20 | 23 | 21 | 33 | 258 | 0 | 23 |
| SRR1820685 | 0 | 12 | 31 | 0 | 0 | 19 | 0 | 12 | 81 | 0 | 0 |
| SRR1820686 | 0 | 0 | 0 | 0 | 0 | 18 | 0 | 18 | 123 | 0 | 0 |
| SRR1820687 | 0 | 0 | 0 | 0 | 10 | 21 | 12 | 12 | 229 | 0 | 0 |
| SRR1820688 | 0 | 0 | 0 | 0 | 0 | 19 | 0 | 0 | 156 | 0 | 0 |
| SRR1820689 | 0 | 0 | 0 | 0 | 0 | 15 | 11 | 0 | 245 | 0 | 0 |
| SRR1820690 | 0 | 0 | 0 | 0 | 0 | 0 | 0 | 0 | 151 | 0 | 0 |
| SRR1853804 | 0 | 94 | 251 | 109 | 0 | 118 | 42 | 230 | 0 | 0 | 54 |
| SRR1853805 | 0 | 48 | 162 | 53 | 0 | 88 | 31 | 197 | 35 | 0 | 39 |
| SRR1853806 | 0 | 113 | 317 | 244 | 17 | 158 | 117 | 334 | 29 | 0 | 64 |
| SRR1853807 | 0 | 23 | 162 | 181 | 17 | 231 | 109 | 229 | 13 | 0 | 19 |
| SRR1853808 | 0 | 58 | 478 | 888 | 48 | 388 | 109 | 221 | 29 | 19 | 67 |
| SRR1853809 | 0 | 105 | 665 | 580 | 15 | 346 | 95 | 279 | 27 | 0 | 37 |
| SRR1917323 | 20 | 16 | 164 | 52 | 4001 | 5472 | 3139 | 367 | 138 | 911 | 402 |
| SRR1917324 | 12 | 20 | 144 | 24 | 4488 | 6227 | 3167 | 455 | 318 | 754 | 450 |
| SRR1917325 | 0 | 0 | 69 | 32 | 5438 | 8686 | 3758 | 546 | 107 | 717 | 247 |
| SRR1917326 | 0 | 0 | 0 | 0 | 0 | 0 | 0 | 0 | 14 | 11 | 0 |
| SRR1917327 | 0 | 0 | 0 | 0 | 0 | 0 | 0 | 0 | 0 | 0 | 0 |
| SRR1917328 | 0 | 0 | 0 | 0 | 0 | 0 | 0 | 0 | 0 | 0 | 0 |
| SRR1917329 | 34 | 10 | 81 | 57 | 2821 | 3684 | 1882 | 415 | 132 | 284 | 175 |
| SRR1917330 | 17 | 15 | 139 | 68 | 4082 | 5351 | 2353 | 422 | 120 | 307 | 163 |
| SRR1917331 | 32 | 42 | 359 | 167 | 1721 | 2671 | 976 | 275 | 92 | 200 | 118 |
| SRR1917332 | 18 | 0 | 0 | 0 | 0 | 0 | 0 | 0 | 0 | 0 | 0 |
| SRR1917333 | 0 | 0 | 0 | 0 | 0 | 0 | 0 | 0 | 0 | 0 | 0 |
| SRR1917334 | 26 | 0 | 0 | 0 | 0 | 0 | 0 | 0 | 13 | 0 | 0 |
| SRR1917335 | 44 | 0 | 87 | 49 | 3881 | 5031 | 2647 | 386 | 125 | 574 | 293 |
| SRR1917336 | 24 | 0 | 70 | 30 | 5540 | 8356 | 4170 | 810 | 187 | 1771 | 660 |
| SRR1917337 | 16 | 0 | 58 | 27 | 2356 | 3745 | 1741 | 566 | 136 | 571 | 237 |
| SRR1917338 | 0 | 0 | 0 | 0 | 0 | 0 | 0 | 0 | 0 | 0 | 0 |
| SRR1917339 | 37 | 0 | 0 | 0 | 15 | 22 | 0 | 0 | 0 | 0 | 0 |
| SRR1917340 | 13 | 0 | 0 | 0 | 14 | 21 | 0 | 0 | 0 | 0 | 0 |
| SRR1922516 | 0 | 0 | 0 | 0 | 0 | 25 | 0 | 31 | 35 | 53 | 41 |
| SRR1922517 | 0 | 0 | 0 | 0 | 0 | 17 | 0 | 23 | 25 | 39 | 31 |
| SRR1922518 | 0 | 0 | 0 | 0 | 0 | 25 | 0 | 33 | 32 | 51 | 36 |
| SRR1922519 | 0 | 0 | 0 | 0 | 0 | 25 | 0 | 35 | 40 | 69 | 52 |
| SRR1922520 | 0 | 0 | 0 | 0 | 0 | 27 | 0 | 29 | 27 | 43 | 30 |
| SRR1922521 | 0 | 0 | 0 | 0 | 0 | 18 | 0 | 22 | 22 | 36 | 28 |
| SRR1922522 | 0 | 0 | 0 | 0 | 0 | 35 | 0 | 41 | 40 | 69 | 48 |
| SRR1922523 | 0 | 0 | 0 | 0 | 0 | 26 | 0 | 29 | 33 | 51 | 43 |
| SRR1922524 | 0 | 0 | 0 | 0 | 0 | 37 | 0 | 50 | 41 | 69 | 51 |
| SRR1922525 | 0 | 0 | 0 | 0 | 0 | 29 | 0 | 40 | 38 | 64 | 45 |
| SRR1922526 | 0 | 0 | 0 | 0 | 0 | 47 | 0 | 66 | 88 | 138 | 92 |
| SRR1922527 | 0 | 0 | 0 | 0 | 0 | 32 | 0 | 50 | 56 | 92 | 57 |
| SRR1973407 | 212 | 74 | 201 | 106 | 21 | 106 | 21 | 201 | 106 | 127 | 180 |
| SRR1973408 | 106 | 47 | 130 | 35 | 47 | 35 | 12 | 154 | 142 | 83 | 367 |
| SRR1973409 | 187 | 49 | 226 | 118 | 59 | 148 | 39 | 256 | 148 | 148 | 384 |
| SRR1973410 | 109 | 20 | 99 | 139 | 40 | 109 | 30 | 238 | 119 | 139 | 377 |
| SRR1973411 | 186 | 31 | 93 | 155 | 0 | 93 | 0 | 186 | 31 | 186 | 340 |
| SRR1973412 | 135 | 45 | 90 | 225 | 30 | 150 | 0 | 285 | 255 | 360 | 585 |
| SRR1973413 | 191 | 11 | 34 | 112 | 22 | 90 | 0 | 382 | 112 | 180 | 393 |
| SRR1973414 | 176 | 0 | 44 | 293 | 15 | 103 | 44 | 396 | 366 | 410 | 879 |
| SRR1973415 | 193 | 16 | 0 | 129 | 32 | 129 | 16 | 548 | 290 | 403 | 935 |
| SRR1973952 | 555 | 232 | 297 | 1496 | 8757 | 29443 | 11117 | 1444 | 335 | 3044 | 1315 |
| SRR1973953 | 45 | 56 | 61 | 67 | 108 | 201 | 138 | 131 | 301 | 94 | 89 |
| SRR1973956 | 0 | 0 | 0 | 0 | 0 | 0 | 0 | 0 | 0 | 0 | 0 |
| SRR1973958 | 0 | 0 | 0 | 0 | 0 | 0 | 0 | 0 | 0 | 0 | 0 |
| SRR1973964 | 0 | 0 | 0 | 0 | 0 | 0 | 0 | 0 | 0 | 0 | 0 |
| SRR1973966 | 16 | 0 | 0 | 20 | 33 | 53 | 33 | 37 | 62 | 13 | 52 |
| SRR1973967 | 21 | 79 | 61 | 56 | 49 | 137 | 87 | 116 | 479 | 81 | 116 |
| SRR1979367 | 0 | 0 | 0 | 0 | 33 | 0 | 17 | 0 | 555 | 1093 | 1400 |
| SRR1988280 | 0 | 26 | 41 | 21 | 70 | 521 | 161 | 799 | 85 | 73 | 164 |
| SRR1988281 | 0 | 33 | 47 | 25 | 51 | 574 | 176 | 762 | 73 | 83 | 175 |
| SRR1988282 | 0 | 22 | 42 | 18 | 49 | 469 | 182 | 842 | 67 | 75 | 161 |
| SRR1988283 | 0 | 31 | 38 | 25 | 43 | 406 | 77 | 534 | 903 | 1282 | 1323 |
| SRR1988284 | 0 | 26 | 32 | 25 | 35 | 404 | 55 | 513 | 719 | 838 | 1051 |
| SRR1988285 | 0 | 24 | 36 | 21 | 38 | 340 | 53 | 440 | 722 | 861 | 981 |
| SRR1988286 | 0 | 23 | 45 | 27 | 42 | 351 | 55 | 470 | 759 | 1000 | 1160 |
| SRR1988287 | 0 | 15 | 51 | 24 | 64 | 285 | 48 | 356 | 1040 | 817 | 466 |
| SRR1988288 | 0 | 23 | 46 | 26 | 77 | 285 | 63 | 360 | 1005 | 811 | 474 |
| SRR1988289 | 0 | 20 | 35 | 20 | 60 | 258 | 36 | 272 | 601 | 534 | 306 |
| SRR1988290 | 0 | 18 | 27 | 17 | 60 | 206 | 34 | 221 | 562 | 512 | 286 |
| SRR1988291 | 0 | 22 | 53 | 19 | 45 | 574 | 148 | 1254 | 102 | 97 | 181 |
| SRR1988292 | 0 | 32 | 66 | 28 | 46 | 651 | 169 | 1530 | 95 | 88 | 196 |
| SRR1988293 | 0 | 28 | 59 | 35 | 44 | 596 | 191 | 1322 | 114 | 86 | 226 |
| SRR1988294 | 0 | 32 | 46 | 20 | 47 | 604 | 160 | 1227 | 94 | 76 | 180 |
| SRR1988295 | 0 | 28 | 54 | 15 | 65 | 400 | 32 | 305 | 170 | 124 | 139 |
| SRR1988296 | 0 | 17 | 53 | 22 | 74 | 347 | 26 | 262 | 151 | 129 | 135 |
| SRR1988297 | 0 | 32 | 58 | 24 | 66 | 371 | 35 | 295 | 157 | 145 | 147 |
| SRR1988298 | 0 | 21 | 28 | 12 | 43 | 222 | 16 | 209 | 132 | 115 | 118 |
| SRR1988299 | 0 | 79 | 117 | 83 | 52 | 713 | 222 | 1305 | 81 | 83 | 258 |
| SRR1988300 | 0 | 48 | 82 | 40 | 77 | 826 | 216 | 1773 | 88 | 92 | 314 |
| SRR1988301 | 0 | 81 | 170 | 94 | 88 | 762 | 312 | 1885 | 116 | 121 | 304 |
| SRR1988302 | 0 | 38 | 94 | 64 | 63 | 754 | 194 | 1666 | 80 | 96 | 283 |
| SRR1988303 | 0 | 42 | 104 | 51 | 58 | 603 | 161 | 1288 | 122 | 103 | 242 |
| SRR1988304 | 0 | 71 | 97 | 66 | 67 | 764 | 190 | 1616 | 119 | 107 | 298 |
| SRR1988305 | 0 | 60 | 100 | 54 | 41 | 648 | 168 | 1373 | 106 | 109 | 270 |
| SRR1988306 | 0 | 47 | 82 | 38 | 53 | 776 | 165 | 1474 | 80 | 69 | 247 |
| SRR1988307 | 0 | 69 | 114 | 51 | 90 | 1391 | 293 | 1910 | 259 | 214 | 340 |
| SRR1988308 | 0 | 48 | 115 | 83 | 67 | 762 | 153 | 982 | 67 | 54 | 141 |
| SRR1988309 | 0 | 65 | 140 | 72 | 94 | 1293 | 275 | 1923 | 194 | 159 | 277 |
| SRR1988310 | 0 | 55 | 128 | 50 | 99 | 998 | 282 | 2438 | 150 | 149 | 283 |
| SRR1988311 | 0 | 37 | 82 | 55 | 72 | 1054 | 231 | 2152 | 162 | 182 | 254 |
| SRR1988312 | 0 | 46 | 98 | 46 | 77 | 1065 | 251 | 2225 | 211 | 194 | 278 |
| SRR2052873 | 0 | 49 | 211 | 110 | 242 | 3247 | 368 | 7931 | 6953 | 8852 | 6743 |
| SRR2052874 | 0 | 45 | 211 | 102 | 179 | 2852 | 293 | 7597 | 6932 | 8417 | 6682 |
| SRR2052875 | 0 | 45 | 171 | 109 | 178 | 2465 | 360 | 7665 | 6703 | 7498 | 7090 |
| SRR2052877 | 0 | 19 | 105 | 77 | 140 | 1967 | 260 | 4630 | 3948 | 4386 | 3646 |
| SRR2052878 | 0 | 32 | 217 | 105 | 264 | 3454 | 410 | 7709 | 6342 | 6736 | 5430 |
| SRR2053027 | 0 | 0 | 79 | 68 | 137 | 1900 | 251 | 4324 | 2506 | 2857 | 2592 |
| SRR2053028 | 0 | 17 | 87 | 78 | 167 | 2220 | 277 | 5973 | 3125 | 3241 | 2771 |
| SRR2053029 | 0 | 29 | 156 | 106 | 185 | 2300 | 283 | 5055 | 2903 | 3010 | 2891 |
| SRR2061931 | 0 | 0 | 71 | 67 | 0 | 25 | 0 | 63 | 42 | 0 | 134 |
| SRR2061932 | 0 | 14 | 14 | 14 | 0 | 0 | 0 | 27 | 27 | 27 | 54 |
| SRR2061933 | 0 | 13 | 35 | 30 | 0 | 0 | 0 | 0 | 13 | 0 | 17 |
| SRR2061934 | 0 | 23 | 92 | 46 | 0 | 0 | 0 | 0 | 0 | 0 | 92 |
| SRR2061935 | 0 | 25 | 76 | 51 | 0 | 25 | 0 | 25 | 0 | 0 | 126 |
| SRR2061936 | 0 | 0 | 43 | 48 | 0 | 11 | 0 | 32 | 11 | 0 | 21 |
| SRR2061937 | 0 | 0 | 50 | 0 | 0 | 0 | 0 | 36 | 0 | 0 | 14 |
| SRR2061938 | 0 | 0 | 98 | 0 | 0 | 0 | 0 | 24 | 39 | 0 | 49 |
| SRR2061940 | 0 | 0 | 0 | 0 | 0 | 0 | 0 | 24 | 24 | 0 | 24 |

### Supp Table S8.5.Columns_ET_to_FD

| | | | | | | | | | | | |
|---|---|---|---|---|---|---|---|---|---|---|---|
| SRR2061941 | 0 | 73 | 352 | 219 | 24 | 364 | 49 | 522 | 498 | 121 | 559 |
| SRR2061942 | 0 | 74 | 248 | 159 | 22 | 314 | 44 | 503 | 432 | 203 | 392 |
| SRR2061943 | 0 | 69 | 216 | 95 | 31 | 283 | 46 | 440 | 620 | 337 | 682 |
| SRR2061944 | 0 | 64 | 148 | 89 | 15 | 173 | 0 | 183 | 213 | 49 | 247 |
| SRR2061945 | 0 | 80 | 173 | 111 | 10 | 277 | 24 | 304 | 270 | 159 | 380 |
| SRR2061946 | 0 | 32 | 77 | 55 | 0 | 82 | 0 | 156 | 318 | 155 | 502 |
| SRR2061947 | 0 | 47 | 102 | 93 | 51 | 275 | 33 | 312 | 377 | 168 | 433 |
| SRR2061948 | 0 | 59 | 110 | 75 | 12 | 161 | 20 | 271 | 275 | 126 | 298 |
| SRR2061949 | 0 | 45 | 92 | 53 | 17 | 56 | 0 | 165 | 268 | 195 | 430 |
| SRR2061950 | 0 | 131 | 348 | 208 | 54 | 678 | 154 | 926 | 411 | 425 | 872 |
| SRR2061951 | 0 | 215 | 426 | 287 | 45 | 685 | 163 | 1021 | 391 | 578 | 944 |
| SRR2061952 | 0 | 149 | 319 | 243 | 40 | 442 | 91 | 1042 | 1309 | 1977 | 2075 |
| SRR2061953 | 0 | 125 | 394 | 223 | 111 | 1136 | 162 | 1934 | 914 | 1387 | 1702 |
| SRR2061954 | 0 | 231 | 348 | 190 | 44 | 801 | 190 | 1043 | 554 | 740 | 869 |
| SRR2061955 | 0 | 116 | 222 | 247 | 53 | 434 | 49 | 962 | 1269 | 1805 | 1861 |
| SRR2061956 | 0 | 137 | 444 | 281 | 91 | 925 | 144 | 1640 | 752 | 1225 | 1353 |
| SRR2061957 | 0 | 172 | 423 | 296 | 75 | 780 | 150 | 1447 | 791 | 1173 | 1244 |
| SRR2061958 | 0 | 133 | 238 | 212 | 52 | 384 | 46 | 817 | 1292 | 1507 | 1624 |
| SRR2069629 | 0 | 0 | 0 | 0 | 0 | 17 | 10 | 17 | 52 | 35 | 16 |
| SRR2069630 | 0 | 0 | 0 | 0 | 0 | 15 | 0 | 0 | 17 | 149 | 239 |
| SRR2084358 | 0 | 0 | 392 | 392 | 392 | 4071 | 834 | 2747 | 2011 | 2306 | 294 |
| SRR2084359 | 0 | 35 | 247 | 211 | 317 | 4758 | 987 | 3665 | 2220 | 1656 | 141 |
| SRR2085173 | 0 | 0 | 0 | 0 | 0 | 0 | 40 | 105 | 1212 | 226 | 8520 |
| SRR2103379 | 0 | 26 | 63 | 37 | 74 | 684 | 142 | 1112 | 1449 | 1360 | 1254 |
| SRR2103380 | 0 | 49 | 99 | 54 | 63 | 646 | 117 | 987 | 1257 | 1279 | 1278 |
| SRR2103381 | 0 | 44 | 105 | 54 | 81 | 728 | 135 | 1114 | 1423 | 1450 | 1376 |
| SRR2103382 | 0 | 87 | 181 | 123 | 96 | 797 | 277 | 2475 | 632 | 843 | 666 |
| SRR2103383 | 0 | 64 | 145 | 95 | 100 | 752 | 273 | 2377 | 684 | 787 | 658 |
| SRR2103384 | 0 | 75 | 196 | 126 | 92 | 1168 | 359 | 4234 | 3629 | 4289 | 2822 |
| SRR2105940 | 0 | 0 | 0 | 0 | 0 | 0 | 0 | 55 | 76 | 36 | 33 |
| SRR2105943 | 0 | 0 | 0 | 0 | 0 | 0 | 0 | 40 | 0 | 20 | 0 |
| SRR2105944 | 0 | 0 | 0 | 0 | 0 | 0 | 0 | 45 | 29 | 10 | 23 |
| SRR2105949 | 0 | 0 | 0 | 0 | 0 | 0 | 0 | 0 | 28 | 0 | 12 |
| SRR2105951 | 0 | 0 | 0 | 0 | 0 | 0 | 0 | 38 | 0 | 0 | 13 |
| SRR2106873 | 0 | 0 | 25 | 18 | 88 | 1600 | 217 | 1522 | 618 | 673 | 630 |
| SRR2106874 | 0 | 0 | 0 | 0 | 33 | 1080 | 202 | 1564 | 470 | 340 | 290 |
| SRR2106875 | 0 | 15 | 21 | 48 | 70 | 1007 | 112 | 1252 | 363 | 365 | 230 |
| SRR2106876 | 0 | 35 | 97 | 93 | 12 | 186 | 50 | 337 | 500 | 411 | 217 |
| SRR2106877 | 0 | 104 | 195 | 213 | 98 | 1213 | 219 | 841 | 598 | 628 | 622 |
| SRR2106878 | 0 | 160 | 510 | 362 | 49 | 1259 | 141 | 1481 | 590 | 424 | 534 |
| SRR2106879 | 0 | 0 | 15 | 18 | 15 | 311 | 44 | 362 | 370 | 282 | 282 |
| SRR2106880 | 0 | 0 | 13 | 17 | 29 | 445 | 88 | 773 | 437 | 248 | 185 |
| SRR2106881 | 0 | 0 | 21 | 24 | 24 | 465 | 48 | 562 | 680 | 408 | 381 |
| SRR2106882 | 0 | 0 | 0 | 0 | 43 | 577 | 72 | 609 | 276 | 138 | 131 |
| SRR2106883 | 0 | 39 | 148 | 106 | 46 | 476 | 72 | 616 | 956 | 485 | 413 |
| SRR2106884 | 0 | 18 | 64 | 49 | 43 | 544 | 69 | 583 | 552 | 285 | 270 |
| SRR2124760 | 0 | 0 | 27 | 22 | 0 | 0 | 0 | 0 | 0 | 0 | 14 |
| SRR2124761 | 0 | 0 | 23 | 24 | 0 | 0 | 0 | 14 | 0 | 0 | 0 |
| SRR2124762 | 0 | 0 | 16 | 15 | 0 | 0 | 0 | 0 | 0 | 0 | 11 |
| SRR2124763 | 0 | 0 | 20 | 46 | 0 | 18 | 0 | 14 | 0 | 0 | 14 |
| SRR2124764 | 0 | 0 | 15 | 0 | 0 | 0 | 0 | 10 | 0 | 0 | 11 |
| SRR2124765 | 0 | 0 | 20 | 11 | 0 | 0 | 0 | 0 | 0 | 0 | 0 |
| SRR2124766 | 0 | 0 | 14 | 16 | 0 | 0 | 0 | 0 | 0 | 0 | 0 |
| SRR2124767 | 0 | 0 | 15 | 23 | 0 | 10 | 0 | 11 | 0 | 0 | 0 |
| SRR2124768 | 0 | 0 | 16 | 26 | 0 | 0 | 0 | 10 | 0 | 0 | 0 |
| SRR2124769 | 0 | 0 | 23 | 21 | 0 | 12 | 0 | 0 | 0 | 0 | 10 |
| SRR2124770 | 0 | 0 | 0 | 0 | 0 | 0 | 0 | 0 | 0 | 0 | 0 |
| SRR2124771 | 0 | 0 | 0 | 0 | 0 | 0 | 0 | 10 | 0 | 0 | 0 |
| SRR2124772 | 0 | 0 | 0 | 0 | 0 | 0 | 0 | 0 | 0 | 0 | 0 |
| SRR2124773 | 0 | 0 | 15 | 35 | 0 | 0 | 0 | 0 | 0 | 0 | 11 |
| SRR2124774 | 0 | 0 | 20 | 34 | 0 | 0 | 0 | 12 | 0 | 0 | 16 |
| SRR2124775 | 0 | 0 | 26 | 47 | 0 | 25 | 0 | 28 | 10 | 0 | 14 |
| SRR2124776 | 0 | 0 | 23 | 41 | 0 | 0 | 0 | 0 | 0 | 0 | 0 |
| SRR2124777 | 0 | 0 | 43 | 50 | 0 | 14 | 0 | 31 | 0 | 0 | 0 |
| SRR2124778 | 0 | 0 | 0 | 14 | 0 | 0 | 0 | 0 | 0 | 0 | 0 |
| SRR2124779 | 0 | 0 | 22 | 20 | 0 | 11 | 0 | 0 | 0 | 0 | 0 |
| SRR2126527 | 0 | 1073 | 1599 | 1281 | 245 | 1645 | 248 | 2745 | 570 | 474 | 312 |
| SRR2126528 | 0 | 1190 | 1834 | 1492 | 352 | 1899 | 328 | 3035 | 658 | 571 | 364 |
| SRR2126529 | 0 | 1114 | 1667 | 1325 | 321 | 2131 | 351 | 3218 | 629 | 534 | 364 |
| SRR2126530 | 0 | 1032 | 1660 | 1109 | 180 | 1096 | 137 | 1695 | 430 | 488 | 371 |
| SRR2126531 | 0 | 989 | 1545 | 1216 | 181 | 1101 | 169 | 1719 | 442 | 461 | 366 |
| SRR2126532 | 0 | 1199 | 2013 | 1540 | 328 | 1991 | 289 | 2832 | 690 | 673 | 454 |
| SRR2135530 | 0 | 0 | 11 | 0 | 326 | 682 | 169 | 295 | 1099 | 99 | 110 |
| SRR2135531 | 0 | 37 | 140 | 46 | 438 | 452 | 210 | 169 | 334 | 66 | 140 |
| SRR2135532 | 0 | 0 | 0 | 0 | 421 | 428 | 214 | 184 | 355 | 99 | 206 |
| SRR2229584 | 0 | 14 | 69 | 26 | 0 | 48 | 16 | 34 | 832 | 406 | 490 |
| SRR2229585 | 0 | 41 | 137 | 51 | 10 | 58 | 0 | 75 | 517 | 308 | 247 |
| SRR2229586 | 0 | 22 | 0 | 18 | 0 | 22 | 0 | 0 | 11 | 14 | 93 |
| SRR2229587 | 0 | 13 | 0 | 0 | 0 | 0 | 0 | 0 | 0 | 0 | 52 |
| SRR2229588 | 0 | 147 | 152 | 24 | 29 | 49 | 24 | 34 | 147 | 44 | 78 |
| SRR2229589 | 0 | 65 | 77 | 24 | 37 | 69 | 16 | 16 | 65 | 41 | 114 |
| SRR2287868 | 0 | 26 | 153 | 119 | 0 | 39 | 21 | 73 | 35 | 28 | 76 |
| SRR2289624 | 0 | 0 | 0 | 0 | 0 | 0 | 0 | 0 | 0 | 0 | 0 |
| SRR2289625 | 0 | 0 | 18 | 18 | 0 | 50 | 0 | 50 | 22 | 16 | 28 |
| SRR2289626 | 0 | 0 | 0 | 0 | 0 | 0 | 0 | 0 | 14 | 42 | 0 |
| SRR2294377 | 0 | 0 | 33 | 44 | 0 | 41 | 0 | 31 | 0 | 14 | 25 |
| SRR2294380 | 0 | 0 | 22 | 40 | 30 | 79 | 23 | 100 | 60 | 396 | 45 |
| SRR2294739 | 0 | 29 | 120 | 80 | 11 | 44 | 0 | 74 | 30 | 41 | 114 |
| SRR2294742 | 0 | 43 | 102 | 109 | 14 | 72 | 24 | 92 | 39 | 46 | 35 |
| SRR2294746 | 0 | 0 | 70 | 48 | 11 | 33 | 0 | 74 | 22 | 37 | 26 |
| SRR2294747 | 0 | 0 | 0 | 0 | 0 | 51 | 0 | 16 | 0 | 260 | 0 |
| SRR2294748 | 0 | 26 | 115 | 70 | 0 | 34 | 0 | 33 | 0 | 25 | 15 |
| SRR2294820 | 0 | 29 | 95 | 81 | 15 | 103 | 27 | 110 | 49 | 95 | 68 |
| SRR2296788 | 0 | 345 | 624 | 583 | 24 | 309 | 36 | 660 | 208 | 202 | 190 |
| SRR2296789 | 0 | 593 | 943 | 1172 | 76 | 730 | 183 | 1248 | 335 | 259 | 304 |
| SRR2296790 | 0 | 133 | 282 | 349 | 17 | 465 | 66 | 730 | 332 | 199 | 249 |
| SRR2296791 | 0 | 263 | 711 | 659 | 40 | 408 | 40 | 975 | 369 | 132 | 316 |
| SRR2296792 | 0 | 248 | 867 | 706 | 87 | 657 | 74 | 1053 | 74 | 74 | 149 |
| SRR2296793 | 0 | 124 | 227 | 405 | 27 | 247 | 48 | 515 | 110 | 144 | 117 |
| SRR2296794 | 0 | 179 | 497 | 457 | 46 | 219 | 40 | 629 | 172 | 139 | 126 |
| SRR2296795 | 0 | 118 | 371 | 434 | 36 | 353 | 36 | 651 | 109 | 154 | 81 |
| SRR2296797 | 0 | 99 | 334 | 386 | 24 | 268 | 52 | 485 | 99 | 71 | 113 |
| SRR2296798 | 0 | 177 | 594 | 481 | 25 | 316 | 51 | 645 | 240 | 354 | 240 |
| SRR2296799 | 0 | 203 | 567 | 416 | 31 | 380 | 89 | 755 | 234 | 172 | 213 |
| SRR2296800 | 0 | 323 | 914 | 1181 | 70 | 590 | 84 | 1124 | 211 | 84 | 267 |
| SRR2296801 | 0 | 128 | 313 | 360 | 0 | 133 | 0 | 337 | 90 | 62 | 57 |
| SRR2296802 | 0 | 24 | 435 | 435 | 24 | 217 | 48 | 507 | 290 | 72 | 97 |
| SRR2296805 | 0 | 208 | 654 | 612 | 45 | 399 | 52 | 709 | 283 | 258 | 292 |
| SRR2296806 | 0 | 290 | 830 | 698 | 13 | 738 | 79 | 1291 | 237 | 250 | 264 |
| SRR2296807 | 0 | 237 | 558 | 615 | 36 | 265 | 32 | 506 | 193 | 169 | 193 |
| SRR2296808 | 0 | 330 | 972 | 625 | 69 | 417 | 17 | 556 | 313 | 122 | 139 |
| SRR2296809 | 0 | 73 | 192 | 202 | 0 | 149 | 23 | 245 | 43 | 46 | 23 |
| SRR2296810 | 0 | 113 | 435 | 792 | 44 | 348 | 87 | 739 | 252 | 130 | 113 |
| SRR2353417 | 0 | 0 | 0 | 0 | 0 | 0 | 0 | 17 | 14 | 0 | 0 |
| SRR2353418 | 0 | 0 | 0 | 0 | 0 | 0 | 0 | 14 | 10 | 0 | 0 |
| SRR2353419 | 0 | 0 | 16 | 35 | 0 | 22 | 0 | 35 | 0 | 0 | 0 |
| SRR2353420 | 0 | 0 | 0 | 0 | 0 | 18 | 0 | 16 | 18 | 11 | 0 |
| SRR2353421 | 0 | 0 | 17 | 11 | 0 | 21 | 0 | 43 | 11 | 0 | 0 |
| SRR2353422 | 0 | 0 | 0 | 0 | 0 | 0 | 0 | 14 | 12 | 0 | 0 |

| | | | | | | | | | | | |
|---|---|---|---|---|---|---|---|---|---|---|---|
| SRR2353423 | 0 | 0 | 12 | 11 | 0 | 23 | 0 | 20 | 25 | 11 | 0 |
| SRR2353424 | 0 | 0 | 0 | 0 | 0 | 12 | 0 | 20 | 0 | 0 | 0 |
| SRR2353425 | 0 | 0 | 0 | 19 | 0 | 48 | 0 | 45 | 15 | 15 | 22 |
| SRR2353426 | 0 | 0 | 25 | 12 | 0 | 33 | 0 | 35 | 14 | 0 | 0 |
| SRR2353427 | 0 | 0 | 0 | 0 | 0 | 11 | 0 | 31 | 26 | 0 | 0 |
| SRR2353428 | 0 | 0 | 0 | 0 | 0 | 0 | 0 | 45 | 13 | 0 | 0 |
| SRR2353429 | 0 | 0 | 11 | 0 | 0 | 15 | 0 | 35 | 12 | 0 | 0 |
| SRR2353430 | 0 | 0 | 0 | 0 | 0 | 0 | 0 | 20 | 11 | 0 | 0 |
| SRR2353431 | 0 | 0 | 13 | 12 | 0 | 24 | 0 | 45 | 43 | 17 | 0 |
| SRR2353432 | 0 | 0 | 11 | 0 | 0 | 14 | 11 | 69 | 30 | 0 | 0 |
| SRR2353433 | 0 | 0 | 0 | 0 | 0 | 0 | 0 | 40 | 22 | 0 | 0 |
| SRR2353434 | 0 | 0 | 23 | 10 | 0 | 22 | 0 | 36 | 25 | 0 | 0 |
| SRR2353435 | 0 | 0 | 10 | 19 | 0 | 21 | 0 | 50 | 38 | 15 | 12 |
| SRR2353436 | 0 | 0 | 0 | 0 | 0 | 0 | 0 | 17 | 10 | 0 | 0 |
| SRR2353437 | 0 | 0 | 0 | 0 | 0 | 13 | 0 | 16 | 0 | 0 | 0 |
| SRR2353438 | 0 | 0 | 15 | 0 | 0 | 0 | 0 | 28 | 25 | 13 | 0 |
| SRR2353439 | 0 | 0 | 0 | 0 | 0 | 0 | 0 | 0 | 0 | 0 | 0 |
| SRR2353440 | 0 | 0 | 13 | 14 | 0 | 16 | 0 | 16 | 22 | 0 | 0 |
| SRR2353441 | 0 | 0 | 37 | 22 | 0 | 22 | 0 | 32 | 17 | 0 | 0 |
| SRR2353442 | 0 | 0 | 0 | 0 | 0 | 0 | 0 | 14 | 16 | 0 | 0 |
| SRR2353443 | 0 | 14 | 23 | 26 | 0 | 25 | 0 | 43 | 40 | 14 | 11 |
| SRR2353444 | 0 | 0 | 0 | 0 | 0 | 12 | 0 | 31 | 0 | 0 | 0 |
| SRR2353445 | 0 | 0 | 13 | 13 | 0 | 17 | 0 | 20 | 15 | 0 | 0 |
| SRR2422969 | 255 | 32 | 64 | 32 | 191 | 159 | 223 | 287 | 987 | 1273 | 860 |
| SRR2422970 | 366 | 33 | 0 | 166 | 133 | 133 | 67 | 333 | 1365 | 932 | 832 |
| SRR2422971 | 333 | 0 | 30 | 60 | 91 | 242 | 181 | 484 | 1573 | 1089 | 1059 |
| SRR2422972 | 341 | 62 | 0 | 62 | 62 | 434 | 279 | 310 | 1147 | 930 | 961 |
| SRR2454328 | 0 | 0 | 0 | 0 | 44 | 368 | 101 | 876 | 1811 | 2549 | 2648 |
| SRR2454329 | 0 | 0 | 0 | 0 | 35 | 338 | 95 | 850 | 1549 | 2226 | 2279 |
| SRR2454330 | 0 | 0 | 0 | 0 | 38 | 407 | 94 | 947 | 2947 | 3137 | 2731 |
| SRR2454331 | 0 | 0 | 0 | 0 | 50 | 369 | 87 | 900 | 2231 | 2437 | 2131 |
| SRR2454332 | 0 | 0 | 17 | 17 | 165 | 1205 | 225 | 1827 | 2390 | 2695 | 2470 |
| SRR2454333 | 0 | 0 | 0 | 12 | 150 | 1160 | 271 | 2253 | 2720 | 3396 | 3118 |
| SRR2454334 | 0 | 0 | 0 | 14 | 115 | 1186 | 216 | 2333 | 2776 | 4273 | 4885 |
| SRR2454335 | 0 | 0 | 0 | 11 | 64 | 613 | 164 | 1920 | 4061 | 6437 | 7823 |
| SRR2454336 | 0 | 0 | 0 | 0 | 59 | 595 | 154 | 1599 | 3488 | 5461 | 5587 |
| SRR2454337 | 0 | 0 | 0 | 0 | 21 | 544 | 187 | 1566 | 3503 | 5543 | 5498 |
| SRR2487412 | 0 | 0 | 0 | 11 | 0 | 64 | 11 | 243 | 48 | 21 | 164 |
| SRR2487413 | 0 | 0 | 0 | 0 | 0 | 79 | 12 | 200 | 36 | 24 | 115 |
| SRR2487414 | 0 | 0 | 0 | 0 | 0 | 62 | 21 | 202 | 31 | 41 | 155 |
| SRR2487415 | 0 | 0 | 14 | 42 | 14 | 347 | 28 | 1026 | 42 | 69 | 873 |
| SRR2487416 | 0 | 0 | 16 | 0 | 0 | 594 | 32 | 1269 | 96 | 96 | 916 |
| SRR2487417 | 0 | 0 | 14 | 27 | 0 | 438 | 55 | 1135 | 123 | 123 | 1026 |
| SRR2487418 | 20 | 0 | 0 | 20 | 0 | 121 | 0 | 502 | 20 | 20 | 161 |
| SRR2487419 | 0 | 0 | 0 | 23 | 0 | 92 | 0 | 575 | 115 | 23 | 138 |
| SRR2487420 | 20 | 0 | 0 | 0 | 20 | 99 | 0 | 418 | 80 | 0 | 279 |
| SRR2487422 | 0 | 0 | 0 | 0 | 49 | 196 | 0 | 588 | 98 | 245 | 662 |
| SRR2487423 | 0 | 0 | 0 | 0 | 28 | 55 | 0 | 469 | 110 | 83 | 828 |
| SRR2487424 | 0 | 0 | 0 | 24 | 0 | 215 | 24 | 741 | 120 | 215 | 741 |
| SRR2487426 | 0 | 0 | 0 | 0 | 0 | 236 | 0 | 579 | 319 | 95 | 177 |
| SRR2487427 | 0 | 0 | 13 | 0 | 0 | 296 | 13 | 350 | 336 | 134 | 551 |
| SRR2487428 | 0 | 0 | 0 | 0 | 0 | 222 | 12 | 491 | 257 | 82 | 409 |
| SRR2487430 | 0 | 0 | 0 | 0 | 0 | 263 | 16 | 675 | 115 | 33 | 922 |
| SRR2487431 | 0 | 0 | 0 | 0 | 38 | 244 | 19 | 825 | 38 | 38 | 881 |
| SRR2487432 | 0 | 0 | 0 | 16 | 0 | 292 | 0 | 812 | 81 | 162 | 942 |
| SRR2487434 | 0 | 0 | 0 | 0 | 0 | 41 | 0 | 223 | 41 | 47 | 108 |
| SRR2487435 | 0 | 0 | 0 | 0 | 0 | 62 | 16 | 311 | 62 | 47 | 109 |
| SRR2487436 | 0 | 0 | 0 | 0 | 0 | 60 | 0 | 219 | 40 | 33 | 146 |
| SRR2487438 | 0 | 0 | 0 | 11 | 11 | 100 | 0 | 391 | 67 | 22 | 201 |
| SRR2487439 | 0 | 13 | 0 | 0 | 0 | 140 | 13 | 178 | 76 | 51 | 191 |
| SRR2487441 | 0 | 13 | 13 | 26 | 0 | 157 | 0 | 222 | 13 | 52 | 261 |
| SRR2487443 | 0 | 0 | 0 | 67 | 0 | 161 | 0 | 403 | 13 | 54 | 242 |
| SRR2487445 | 0 | 0 | 34 | 34 | 0 | 168 | 0 | 303 | 17 | 84 | 135 |
| SRR2487447 | 0 | 33 | 17 | 33 | 0 | 133 | 17 | 283 | 0 | 50 | 183 |
| SRR2487449 | 0 | 22 | 78 | 112 | 11 | 358 | 11 | 381 | 112 | 90 | 370 |
| SRR2487451 | 0 | 11 | 69 | 57 | 11 | 332 | 11 | 447 | 103 | 92 | 344 |
| SRR2487453 | 0 | 0 | 43 | 72 | 0 | 158 | 14 | 459 | 72 | 72 | 431 |
| SRR2487455 | 0 | 14 | 128 | 113 | 14 | 255 | 0 | 383 | 128 | 85 | 383 |
| SRR2487457 | 0 | 0 | 0 | 27 | 0 | 122 | 14 | 203 | 41 | 27 | 95 |
| SRR2487459 | 0 | 0 | 0 | 14 | 0 | 157 | 0 | 185 | 28 | 43 | 71 |
| SRR2487461 | 0 | 0 | 18 | 36 | 0 | 90 | 0 | 162 | 36 | 18 | 108 |
| SRR2487463 | 0 | 0 | 0 | 0 | 0 | 141 | 18 | 141 | 35 | 18 | 141 |
| SRR2487465 | 0 | 0 | 66 | 83 | 33 | 249 | 0 | 581 | 66 | 66 | 282 |
| SRR2487467 | 0 | 0 | 17 | 69 | 34 | 172 | 51 | 961 | 103 | 137 | 206 |
| SRR2487469 | 0 | 0 | 21 | 43 | 43 | 301 | 0 | 301 | 21 | 64 | 258 |
| SRR2487471 | 0 | 0 | 21 | 21 | 21 | 256 | 21 | 449 | 43 | 85 | 363 |
| SRR2487473 | 0 | 17 | 17 | 51 | 17 | 475 | 17 | 865 | 153 | 220 | 509 |
| SRR2487475 | 0 | 17 | 0 | 104 | 17 | 383 | 17 | 818 | 244 | 104 | 679 |
| SRR2487477 | 0 | 0 | 109 | 88 | 22 | 350 | 22 | 1094 | 306 | 131 | 416 |
| SRR2487479 | 0 | 0 | 22 | 43 | 0 | 239 | 22 | 1043 | 65 | 65 | 456 |
| SRR2487481 | 0 | 14 | 28 | 112 | 28 | 266 | 0 | 644 | 140 | 84 | 434 |
| SRR2487483 | 0 | 29 | 72 | 57 | 0 | 244 | 29 | 675 | 129 | 72 | 445 |
| SRR2487485 | 0 | 0 | 54 | 90 | 0 | 288 | 0 | 523 | 72 | 36 | 433 |
| SRR2487487 | 0 | 0 | 72 | 36 | 0 | 251 | 18 | 412 | 72 | 36 | 376 |
| SRR2487489 | 0 | 0 | 16 | 0 | 0 | 62 | 0 | 91 | 47 | 22 | 56 |
| SRR2487491 | 0 | 0 | 13 | 0 | 0 | 71 | 0 | 58 | 29 | 23 | 32 |
| SRR2487493 | 0 | 0 | 0 | 0 | 0 | 43 | 0 | 82 | 20 | 31 | 55 |
| SRR2487495 | 0 | 0 | 0 | 0 | 0 | 52 | 0 | 68 | 28 | 44 | 44 |
| SRR2487497 | 0 | 0 | 34 | 34 | 0 | 86 | 17 | 207 | 69 | 69 | 146 |
| SRR2487499 | 0 | 0 | 44 | 18 | 0 | 106 | 0 | 142 | 35 | 71 | 257 |
| SRR2487501 | 0 | 22 | 33 | 44 | 0 | 122 | 22 | 144 | 111 | 66 | 210 |
| SRR2487503 | 0 | 0 | 55 | 11 | 0 | 154 | 11 | 154 | 33 | 33 | 253 |
| SRR2487504 | 0 | 0 | 0 | 0 | 60 | 120 | 0 | 360 | 0 | 60 | 300 |
| SRR2487505 | 0 | 41 | 0 | 41 | 0 | 124 | 0 | 289 | 83 | 41 | 248 |
| SRR2487507 | 0 | 0 | 83 | 0 | 0 | 83 | 41 | 248 | 124 | 0 | 165 |
| SRR2487508 | 0 | 0 | 0 | 63 | 0 | 253 | 63 | 443 | 63 | 63 | 126 |
| SRR2487509 | 0 | 0 | 0 | 0 | 0 | 132 | 0 | 330 | 66 | 66 | 264 |
| SRR2487510 | 0 | 0 | 135 | 0 | 0 | 135 | 0 | 473 | 135 | 0 | 0 |
| SRR2487512 | 0 | 0 | 32 | 64 | 0 | 96 | 0 | 288 | 64 | 64 | 192 |
| SRR2487514 | 0 | 0 | 0 | 33 | 33 | 166 | 33 | 299 | 166 | 0 | 33 |
| SRR2487516 | 0 | 0 | 0 | 151 | 0 | 151 | 0 | 452 | 101 | 0 | 352 |
| SRR2487517 | 0 | 0 | 65 | 33 | 65 | 392 | 0 | 457 | 0 | 33 | 261 |
| SRR2487519 | 0 | 0 | 32 | 32 | 0 | 227 | 0 | 259 | 0 | 65 | 194 |
| SRR2487520 | 0 | 0 | 0 | 103 | 51 | 51 | 0 | 411 | 0 | 51 | 154 |
| SRR2487521 | 0 | 0 | 0 | 0 | 56 | 223 | 0 | 724 | 0 | 56 | 390 |
| SRR2487522 | 0 | 0 | 0 | 0 | 0 | 281 | 0 | 450 | 281 | 56 | 169 |
| SRR2487524 | 0 | 0 | 0 | 76 | 0 | 127 | 0 | 508 | 51 | 25 | 508 |
| SRR2487526 | 0 | 26 | 104 | 52 | 26 | 260 | 0 | 313 | 52 | 26 | 104 |
| SRR2487529 | 0 | 0 | 27 | 44 | 0 | 231 | 53 | 559 | 53 | 35 | 89 |
| SRR2487531 | 0 | 0 | 55 | 37 | 27 | 293 | 0 | 732 | 64 | 64 | 220 |
| SRR2487533 | 0 | 0 | 23 | 34 | 23 | 249 | 11 | 590 | 136 | 34 | 102 |
| SRR2487535 | 0 | 0 | 22 | 22 | 11 | 336 | 56 | 639 | 90 | 34 | 235 |
| SRR2487537 | 0 | 0 | 30 | 30 | 15 | 258 | 30 | 334 | 0 | 46 | 273 |
| SRR2487539 | 0 | 0 | 32 | 0 | 0 | 221 | 47 | 299 | 32 | 79 | 536 |
| SRR2487541 | 0 | 59 | 39 | 0 | 20 | 176 | 20 | 352 | 20 | 20 | 391 |
| SRR2487543 | 0 | 0 | 39 | 0 | 39 | 136 | 58 | 446 | 19 | 39 | 310 |
| SRR2487545 | 0 | 0 | 31 | 122 | 122 | 397 | 31 | 764 | 31 | 31 | 183 |
| SRR2487547 | 0 | 0 | 125 | 93 | 31 | 280 | 31 | 748 | 0 | 62 | 125 |
| SRR2487549 | 0 | 0 | 118 | 0 | 78 | 392 | 0 | 824 | 118 | 0 | 235 |

### Supp Table S8.5.Columns_ET_to_FD

| | | | | | | | | | | | |
|---|---|---|---|---|---|---|---|---|---|---|---|
| SRR2487551 | 0 | 0 | 39 | 78 | 0 | 428 | 39 | 1052 | 156 | 0 | 273 |
| SRR2487553 | 0 | 61 | 81 | 283 | 20 | 465 | 0 | 445 | 0 | 0 | 40 |
| SRR2487555 | 0 | 21 | 228 | 145 | 41 | 559 | 62 | 269 | 0 | 0 | 62 |
| SRR2487560 | 0 | 0 | 0 | 0 | 0 | 138 | 0 | 277 | 138 | 138 | 484 |
| SRR2487561 | 0 | 0 | 170 | 38 | 0 | 189 | 19 | 151 | 57 | 0 | 0 |
| SRR2487563 | 0 | 19 | 39 | 233 | 0 | 155 | 19 | 117 | 39 | 0 | 97 |
| SRR2487564 | 0 | 0 | 0 | 0 | 0 | 223 | 0 | 744 | 74 | 0 | 819 |
| SRR2487565 | 0 | 0 | 77 | 230 | 77 | 460 | 0 | 384 | 77 | 153 | 767 |
| SRR2487566 | 0 | 0 | 0 | 0 | 79 | 79 | 79 | 629 | 236 | 79 | 786 |
| SRR2487568 | 0 | 0 | 103 | 147 | 0 | 250 | 29 | 220 | 0 | 15 | 44 |
| SRR2487570 | 15 | 0 | 61 | 76 | 0 | 137 | 15 | 213 | 15 | 0 | 61 |
| SRR2487572 | 0 | 0 | 65 | 0 | 0 | 260 | 32 | 941 | 32 | 65 | 812 |
| SRR2487573 | 0 | 30 | 179 | 268 | 30 | 447 | 0 | 298 | 0 | 0 | 0 |
| SRR2487575 | 0 | 30 | 240 | 210 | 0 | 360 | 60 | 570 | 30 | 0 | 0 |
| SRR2487576 | 0 | 34 | 67 | 34 | 34 | 436 | 34 | 804 | 34 | 0 | 570 |
| SRR2487577 | 0 | 36 | 36 | 36 | 36 | 399 | 36 | 761 | 72 | 145 | 797 |
| SRR2487578 | 0 | 0 | 0 | 36 | 0 | 292 | 0 | 766 | 73 | 109 | 547 |
| SRR2487580 | 23 | 23 | 181 | 136 | 23 | 136 | 23 | 295 | 0 | 0 | 0 |
| SRR2487582 | 23 | 46 | 185 | 116 | 70 | 93 | 23 | 301 | 23 | 0 | 46 |
| SRR2487584 | 0 | 0 | 0 | 0 | 0 | 33 | 0 | 47 | 58 | 33 | 54 |
| SRR2487585 | 0 | 0 | 0 | 32 | 0 | 32 | 0 | 32 | 0 | 0 | 0 |
| SRR2487587 | 0 | 0 | 23 | 37 | 0 | 28 | 0 | 37 | 14 | 0 | 0 |
| SRR2487588 | 0 | 0 | 19 | 0 | 0 | 43 | 0 | 66 | 66 | 54 | 66 |
| SRR2487589 | 0 | 0 | 0 | 0 | 0 | 37 | 0 | 57 | 61 | 37 | 81 |
| SRR2487590 | 0 | 0 | 0 | 0 | 0 | 12 | 0 | 70 | 74 | 79 | 91 |
| SRR2487592 | 0 | 0 | 21 | 32 | 0 | 25 | 0 | 32 | 0 | 0 | 0 |
| SRR2487594 | 0 | 15 | 11 | 26 | 0 | 29 | 0 | 29 | 11 | 0 | 11 |
| SRR2487597 | 0 | 11 | 85 | 42 | 32 | 254 | 21 | 454 | 32 | 42 | 285 |
| SRR2487598 | 0 | 0 | 11 | 56 | 0 | 215 | 0 | 373 | 102 | 113 | 215 |
| SRR2487599 | 0 | 0 | 34 | 23 | 11 | 205 | 11 | 421 | 34 | 125 | 285 |
| SRR2487600 | 0 | 16 | 173 | 126 | 0 | 205 | 32 | 268 | 0 | 0 | 32 |
| SRR2487602 | 0 | 16 | 243 | 195 | 32 | 195 | 0 | 211 | 16 | 0 | 0 |
| SRR2487604 | 41 | 0 | 163 | 204 | 20 | 183 | 0 | 163 | 20 | 0 | 0 |
| SRR2487606 | 0 | 21 | 186 | 249 | 0 | 186 | 0 | 331 | 0 | 0 | 41 |
| SRR2487607 | 0 | 10 | 60 | 40 | 10 | 201 | 50 | 503 | 91 | 40 | 292 |
| SRR2487609 | 0 | 0 | 12 | 35 | 23 | 207 | 12 | 507 | 127 | 127 | 311 |
| SRR2487610 | 0 | 0 | 12 | 12 | 0 | 93 | 0 | 439 | 139 | 139 | 139 |
| SRR2487611 | 0 | 0 | 12 | 0 | 12 | 188 | 23 | 528 | 176 | 82 | 235 |
| SRR2487612 | 34 | 34 | 34 | 201 | 0 | 134 | 0 | 302 | 34 | 34 | 67 |
| SRR2487614 | 0 | 0 | 35 | 70 | 0 | 210 | 0 | 385 | 0 | 35 | 0 |
| SRR2487616 | 0 | 0 | 0 | 136 | 0 | 91 | 45 | 318 | 45 | 0 | 0 |
| SRR2487618 | 0 | 46 | 229 | 137 | 0 | 46 | 0 | 274 | 46 | 0 | 0 |
| SRR2487619 | 0 | 0 | 21 | 11 | 0 | 159 | 42 | 541 | 159 | 95 | 201 |
| SRR2487621 | 0 | 0 | 0 | 0 | 0 | 77 | 0 | 307 | 26 | 154 | 435 |
| SRR2487622 | 0 | 0 | 27 | 0 | 0 | 191 | 0 | 327 | 218 | 109 | 654 |
| SRR2487623 | 0 | 0 | 0 | 28 | 0 | 249 | 28 | 415 | 138 | 55 | 277 |
| SRR2487624 | 0 | 15 | 77 | 107 | 0 | 77 | 0 | 107 | 0 | 0 | 15 |
| SRR2487626 | 0 | 0 | 47 | 79 | 0 | 126 | 0 | 110 | 0 | 0 | 79 |
| SRR2487628 | 0 | 0 | 119 | 60 | 0 | 79 | 40 | 198 | 0 | 0 | 40 |
| SRR2487630 | 0 | 20 | 60 | 60 | 0 | 60 | 0 | 199 | 0 | 20 | 0 |
| SRR2487631 | 0 | 0 | 0 | 48 | 0 | 121 | 0 | 338 | 169 | 72 | 411 |
| SRR2487633 | 0 | 37 | 203 | 221 | 37 | 221 | 18 | 369 | 37 | 37 | 0 |
| SRR2487635 | 0 | 0 | 154 | 193 | 77 | 367 | 19 | 425 | 19 | 19 | 0 |
| SRR2487637 | 0 | 71 | 119 | 95 | 24 | 167 | 0 | 238 | 0 | 0 | 24 |
| SRR2487639 | 0 | 24 | 72 | 48 | 48 | 338 | 0 | 290 | 24 | 0 | 72 |
| SRR2487641 | 0 | 23 | 252 | 229 | 0 | 206 | 46 | 435 | 46 | 0 | 69 |
| SRR2487643 | 0 | 47 | 285 | 142 | 71 | 285 | 24 | 427 | 47 | 47 | 214 |
| SRR2487645 | 0 | 0 | 242 | 242 | 0 | 454 | 0 | 303 | 30 | 0 | 91 |
| SRR2487647 | 0 | 31 | 276 | 307 | 31 | 369 | 92 | 461 | 0 | 0 | 31 |
| SRR2487648 | 0 | 0 | 21 | 0 | 21 | 314 | 0 | 670 | 440 | 105 | 419 |
| SRR2487649 | 24 | 24 | 215 | 72 | 0 | 119 | 48 | 334 | 0 | 0 | 0 |
| SRR2487651 | 0 | 48 | 143 | 214 | 48 | 143 | 0 | 309 | 48 | 24 | 24 |
| SRR2487652 | 0 | 0 | 0 | 42 | 42 | 337 | 0 | 695 | 211 | 84 | 337 |
| SRR2487653 | 0 | 0 | 0 | 23 | 23 | 229 | 0 | 549 | 229 | 114 | 343 |
| SRR2487654 | 0 | 0 | 23 | 0 | 0 | 275 | 23 | 665 | 207 | 138 | 482 |
| SRR2487656 | 0 | 0 | 19 | 93 | 19 | 185 | 0 | 222 | 19 | 37 | 19 |
| SRR2487658 | 0 | 19 | 57 | 76 | 0 | 134 | 0 | 497 | 38 | 0 | 19 |
| SRR2487661 | 0 | 0 | 85 | 57 | 0 | 254 | 28 | 424 | 28 | 0 | 57 |
| SRR2487663 | 0 | 29 | 87 | 0 | 0 | 233 | 0 | 204 | 0 | 0 | 29 |
| SRR2487665 | 0 | 0 | 0 | 148 | 37 | 334 | 0 | 185 | 37 | 0 | 111 |
| SRR2487667 | 0 | 0 | 75 | 75 | 37 | 262 | 0 | 262 | 75 | 0 | 75 |
| SRR2487668 | 0 | 0 | 0 | 0 | 36 | 291 | 109 | 182 | 146 | 109 | 182 |
| SRR2487669 | 0 | 33 | 33 | 165 | 66 | 232 | 0 | 66 | 33 | 0 | 33 |
| SRR2487671 | 0 | 0 | 300 | 133 | 133 | 167 | 0 | 167 | 0 | 67 | 33 |
| SRR2487672 | 0 | 0 | 38 | 0 | 38 | 263 | 0 | 338 | 188 | 0 | 338 |
| SRR2487673 | 0 | 0 | 0 | 0 | 40 | 316 | 79 | 79 | 118 | 0 | 316 |
| SRR2487674 | 0 | 0 | 80 | 40 | 0 | 120 | 0 | 399 | 80 | 40 | 240 |
| SRR2487676 | 26 | 78 | 104 | 78 | 26 | 235 | 0 | 313 | 52 | 0 | 52 |
| SRR2487678 | 27 | 0 | 81 | 188 | 27 | 296 | 27 | 161 | 0 | 0 | 27 |
| SRR2487688 | 0 | 0 | 91 | 68 | 0 | 272 | 68 | 725 | 159 | 113 | 589 |
| SRR2487689 | 0 | 59 | 177 | 118 | 0 | 294 | 0 | 530 | 59 | 0 | 118 |
| SRR2487691 | 0 | 119 | 179 | 299 | 0 | 418 | 0 | 478 | 60 | 119 | 179 |
| SRR2487692 | 0 | 0 | 23 | 23 | 0 | 369 | 23 | 806 | 46 | 92 | 852 |
| SRR2487693 | 26 | 26 | 26 | 26 | 26 | 282 | 26 | 642 | 51 | 77 | 642 |
| SRR2487694 | 0 | 0 | 51 | 76 | 0 | 459 | 25 | 663 | 102 | 102 | 1147 |
| SRR2487696 | 0 | 91 | 273 | 228 | 0 | 364 | 0 | 683 | 0 | 0 | 91 |
| SRR2487698 | 0 | 0 | 276 | 276 | 0 | 368 | 0 | 506 | 0 | 0 | 138 |
| SRR2487700 | 0 | 0 | 26 | 13 | 26 | 130 | 13 | 626 | 52 | 52 | 274 |
| SRR2487701 | 0 | 0 | 73 | 171 | 24 | 146 | 0 | 219 | 24 | 0 | 0 |
| SRR2487703 | 0 | 0 | 49 | 122 | 0 | 49 | 0 | 219 | 49 | 0 | 73 |
| SRR2487704 | 0 | 0 | 0 | 26 | 39 | 210 | 0 | 406 | 13 | 52 | 288 |
| SRR2487705 | 0 | 0 | 0 | 43 | 29 | 200 | 0 | 472 | 100 | 29 | 200 |
| SRR2487706 | 0 | 0 | 0 | 43 | 14 | 215 | 14 | 545 | 86 | 86 | 330 |
| SRR2487708 | 0 | 0 | 113 | 75 | 19 | 132 | 19 | 302 | 0 | 0 | 0 |
| SRR2487710 | 0 | 78 | 97 | 58 | 39 | 116 | 0 | 291 | 0 | 0 | 0 |
| SRR2529523 | 0 | 0 | 13 | 0 | 0 | 24 | 0 | 44 | 21 | 0 | 50 |
| SRR2529524 | 0 | 0 | 0 | 0 | 0 | 32 | 0 | 45 | 20 | 0 | 65 |
| SRR2529525 | 0 | 17 | 14 | 20 | 14 | 165 | 49 | 178 | 53 | 32 | 136 |
| SRR2529526 | 0 | 0 | 31 | 21 | 21 | 134 | 40 | 203 | 63 | 35 | 116 |
| SRR2529527 | 0 | 14 | 24 | 13 | 25 | 211 | 58 | 299 | 105 | 86 | 220 |
| SRR2529528 | 0 | 16 | 36 | 14 | 19 | 225 | 60 | 291 | 135 | 70 | 210 |
| SRR2529529 | 0 | 0 | 16 | 0 | 0 | 32 | 0 | 46 | 11 | 11 | 66 |
| SRR2529530 | 0 | 12 | 16 | 0 | 0 | 56 | 0 | 80 | 0 | 0 | 56 |
| SRR2529531 | 0 | 0 | 35 | 17 | 12 | 130 | 51 | 228 | 46 | 17 | 69 |
| SRR2529532 | 0 | 14 | 28 | 24 | 13 | 149 | 60 | 256 | 36 | 16 | 105 |
| SRR2529533 | 0 | 0 | 20 | 0 | 12 | 54 | 20 | 113 | 22 | 11 | 108 |
| SRR2529534 | 0 | 0 | 29 | 17 | 11 | 61 | 11 | 109 | 28 | 21 | 79 |
| SRR2529535 | 0 | 0 | 0 | 0 | 0 | 11 | 0 | 38 | 0 | 0 | 0 |
| SRR2529536 | 0 | 0 | 0 | 0 | 0 | 13 | 0 | 35 | 0 | 0 | 0 |
| SRR2529537 | 0 | 0 | 20 | 36 | 11 | 54 | 18 | 231 | 12 | 0 | 78 |
| SRR2529538 | 0 | 0 | 28 | 29 | 0 | 89 | 22 | 224 | 26 | 0 | 64 |
| SRR2529539 | 0 | 0 | 26 | 33 | 0 | 87 | 22 | 255 | 20 | 20 | 122 |
| SRR2529540 | 0 | 10 | 28 | 34 | 0 | 123 | 16 | 262 | 29 | 11 | 133 |
| SRR2529541 | 0 | 0 | 0 | 19 | 0 | 30 | 0 | 60 | 11 | 0 | 0 |
| SRR2529542 | 0 | 0 | 16 | 21 | 0 | 21 | 0 | 47 | 0 | 0 | 0 |
| SRR2529543 | 0 | 0 | 11 | 23 | 0 | 15 | 0 | 114 | 13 | 0 | 17 |
| SRR2529544 | 0 | 0 | 14 | 25 | 0 | 20 | 0 | 103 | 0 | 0 | 35 |
| SRR2529545 | 0 | 0 | 18 | 20 | 0 | 29 | 0 | 65 | 0 | 0 | 0 |
| SRR2529546 | 0 | 0 | 16 | 23 | 0 | 44 | 0 | 71 | 14 | 0 | 16 |

### Supp Table S8.5.Columns_ET_to_FD

| | | | | | | | | | | | |
|---|---|---|---|---|---|---|---|---|---|---|---|
| SRR2529547 | 0 | 0 | 12 | 0 | 0 | 95 | 20 | 107 | 43 | 20 | 174 |
| SRR2529548 | 0 | 0 | 15 | 12 | 0 | 89 | 15 | 92 | 45 | 30 | 196 |
| SRR2529549 | 0 | 15 | 42 | 31 | 18 | 118 | 35 | 215 | 74 | 28 | 175 |
| SRR2529550 | 0 | 11 | 25 | 18 | 0 | 176 | 34 | 212 | 106 | 41 | 178 |
| SRR2529551 | 0 | 14 | 48 | 17 | 19 | 220 | 44 | 260 | 116 | 70 | 227 |
| SRR2529552 | 0 | 0 | 23 | 17 | 27 | 238 | 38 | 240 | 118 | 73 | 242 |
| SRR2529553 | 0 | 0 | 16 | 16 | 10 | 49 | 13 | 57 | 13 | 13 | 104 |
| SRR2529554 | 0 | 0 | 13 | 0 | 0 | 37 | 0 | 61 | 13 | 11 | 93 |
| SRR2529555 | 0 | 11 | 49 | 21 | 16 | 148 | 36 | 246 | 41 | 24 | 122 |
| SRR2529556 | 0 | 0 | 44 | 38 | 0 | 142 | 33 | 211 | 35 | 22 | 102 |
| SRR2529557 | 0 | 0 | 23 | 39 | 0 | 107 | 12 | 174 | 28 | 14 | 146 |
| SRR2529558 | 0 | 14 | 29 | 41 | 12 | 136 | 17 | 191 | 41 | 12 | 155 |
| SRR2529559 | 0 | 0 | 0 | 0 | 0 | 25 | 0 | 59 | 21 | 0 | 32 |
| SRR2529560 | 0 | 0 | 0 | 0 | 0 | 30 | 0 | 65 | 15 | 0 | 33 |
| SRR2529561 | 0 | 0 | 14 | 23 | 0 | 100 | 17 | 93 | 42 | 16 | 116 |
| SRR2529562 | 0 | 0 | 10 | 20 | 0 | 94 | 16 | 110 | 39 | 17 | 80 |
| SRR2529563 | 0 | 0 | 12 | 11 | 0 | 145 | 36 | 97 | 66 | 54 | 157 |
| SRR2529564 | 0 | 0 | 20 | 14 | 0 | 122 | 30 | 97 | 77 | 31 | 139 |
| SRR2529565 | 0 | 0 | 0 | 0 | 0 | 19 | 0 | 31 | 0 | 0 | 31 |
| SRR2529566 | 0 | 0 | 13 | 0 | 0 | 19 | 0 | 35 | 13 | 0 | 13 |
| SRR2529567 | 0 | 15 | 13 | 21 | 0 | 55 | 0 | 93 | 19 | 0 | 49 |
| SRR2529568 | 0 | 0 | 26 | 15 | 0 | 28 | 0 | 90 | 24 | 0 | 66 |
| SRR2529569 | 0 | 0 | 0 | 0 | 0 | 0 | 0 | 0 | 16 | 0 | 28 |
| SRR2529570 | 0 | 0 | 0 | 0 | 0 | 0 | 0 | 19 | 16 | 0 | 23 |
| SRR2529571 | 0 | 0 | 12 | 12 | 0 | 37 | 0 | 85 | 16 | 0 | 75 |
| SRR2529572 | 0 | 0 | 14 | 20 | 0 | 56 | 0 | 71 | 16 | 0 | 65 |
| SRR2529573 | 0 | 14 | 27 | 27 | 13 | 199 | 47 | 228 | 49 | 36 | 143 |
| SRR2529574 | 0 | 19 | 26 | 22 | 0 | 161 | 33 | 278 | 63 | 26 | 178 |
| SRR2529575 | 0 | 21 | 27 | 12 | 28 | 200 | 59 | 271 | 110 | 59 | 182 |
| SRR2529576 | 0 | 15 | 20 | 15 | 35 | 225 | 38 | 295 | 102 | 49 | 190 |
| SRR2529577 | 0 | 0 | 0 | 0 | 0 | 66 | 11 | 55 | 15 | 0 | 63 |
| SRR2529578 | 0 | 0 | 13 | 0 | 0 | 40 | 13 | 91 | 15 | 0 | 53 |
| SRR2529579 | 0 | 15 | 30 | 31 | 15 | 142 | 18 | 325 | 50 | 13 | 81 |
| SRR2529580 | 0 | 11 | 33 | 20 | 12 | 111 | 40 | 330 | 46 | 15 | 73 |
| SRR2529581 | 0 | 0 | 23 | 0 | 0 | 76 | 18 | 161 | 28 | 0 | 102 |
| SRR2529582 | 0 | 0 | 15 | 19 | 0 | 71 | 19 | 159 | 19 | 14 | 125 |
| SRR2529583 | 0 | 0 | 0 | 0 | 0 | 14 | 0 | 74 | 16 | 0 | 19 |
| SRR2529584 | 0 | 0 | 0 | 0 | 0 | 15 | 0 | 51 | 16 | 0 | 27 |
| SRR2529585 | 0 | 0 | 24 | 17 | 0 | 76 | 12 | 287 | 35 | 17 | 169 |
| SRR2529586 | 0 | 0 | 23 | 22 | 0 | 83 | 14 | 297 | 36 | 20 | 171 |
| SRR2529587 | 0 | 0 | 30 | 13 | 0 | 108 | 14 | 341 | 65 | 25 | 364 |
| SRR2529588 | 0 | 0 | 24 | 21 | 0 | 111 | 18 | 324 | 81 | 20 | 347 |
| SRR2529589 | 0 | 0 | 0 | 11 | 0 | 27 | 0 | 80 | 0 | 0 | 18 |
| SRR2529590 | 0 | 0 | 0 | 0 | 0 | 23 | 0 | 73 | 0 | 0 | 27 |
| SRR2529591 | 0 | 16 | 27 | 21 | 0 | 53 | 11 | 128 | 13 | 0 | 56 |
| SRR2529592 | 0 | 0 | 22 | 30 | 0 | 49 | 0 | 145 | 16 | 0 | 27 |
| SRR2529593 | 0 | 0 | 0 | 15 | 0 | 21 | 0 | 77 | 15 | 0 | 47 |
| SRR2529594 | 0 | 0 | 15 | 18 | 0 | 33 | 0 | 67 | 18 | 0 | 49 |
| SRR2648184 | 0 | 0 | 0 | 0 | 0 | 78 | 15 | 216 | 172 | 166 | 323 |
| SRR2648185 | 0 | 0 | 0 | 0 | 0 | 51 | 14 | 137 | 121 | 125 | 254 |
| SRR2648186 | 0 | 0 | 0 | 0 | 0 | 51 | 12 | 118 | 124 | 113 | 182 |
| SRR2648187 | 0 | 0 | 0 | 19 | 25 | 484 | 63 | 1778 | 645 | 526 | 1184 |
| SRR2648188 | 0 | 0 | 0 | 20 | 31 | 579 | 99 | 1944 | 705 | 608 | 1280 |
| SRR2648189 | 0 | 0 | 0 | 14 | 35 | 415 | 57 | 1239 | 492 | 442 | 907 |
| SRR2728221 | 0 | 0 | 0 | 0 | 0 | 0 | 0 | 0 | 0 | 23 | 0 |
| SRR2731278 | 0 | 0 | 0 | 0 | 0 | 17 | 0 | 0 | 18 | 0 | 0 |
| SRR2731279 | 0 | 13 | 21 | 0 | 24 | 52 | 23 | 20 | 47 | 0 | 0 |
| SRR2731280 | 0 | 0 | 11 | 0 | 19 | 36 | 17 | 21 | 55 | 11 | 0 |
| SRR2731281 | 0 | 0 | 23 | 12 | 26 | 57 | 26 | 41 | 96 | 0 | 0 |
| SRR2731282 | 0 | 17 | 35 | 12 | 29 | 83 | 28 | 44 | 50 | 0 | 0 |
| SRR2731283 | 0 | 0 | 0 | 0 | 15 | 27 | 10 | 15 | 27 | 0 | 0 |
| SRR2731284 | 0 | 0 | 13 | 0 | 13 | 27 | 12 | 13 | 39 | 0 | 0 |
| SRR2731285 | 0 | 0 | 0 | 0 | 13 | 29 | 10 | 19 | 36 | 0 | 0 |
| SRR2731286 | 0 | 0 | 11 | 0 | 13 | 39 | 14 | 22 | 111 | 0 | 0 |
| SRR2731287 | 0 | 16 | 26 | 0 | 14 | 65 | 18 | 24 | 159 | 0 | 0 |
| SRR2731288 | 0 | 0 | 22 | 0 | 26 | 94 | 33 | 54 | 189 | 16 | 16 |
| SRR2731289 | 0 | 23 | 84 | 0 | 56 | 256 | 53 | 47 | 363 | 0 | 0 |
| SRR2731290 | 0 | 12 | 41 | 0 | 62 | 131 | 40 | 31 | 187 | 0 | 0 |
| SRR2731291 | 0 | 17 | 66 | 0 | 43 | 256 | 48 | 69 | 470 | 12 | 0 |
| SRR2731292 | 0 | 25 | 103 | 11 | 103 | 514 | 80 | 59 | 275 | 0 | 0 |
| SRR2731293 | 0 | 0 | 0 | 0 | 12 | 28 | 0 | 12 | 18 | 0 | 0 |
| SRR2731294 | 0 | 0 | 17 | 0 | 27 | 49 | 28 | 29 | 92 | 0 | 0 |
| SRR2731295 | 0 | 15 | 34 | 16 | 84 | 147 | 74 | 78 | 223 | 24 | 30 |
| SRR2731296 | 0 | 11 | 17 | 13 | 11 | 47 | 14 | 26 | 32 | 0 | 0 |
| SRR2731297 | 0 | 0 | 0 | 0 | 0 | 16 | 0 | 0 | 17 | 0 | 0 |
| SRR2731298 | 0 | 0 | 84 | 19 | 133 | 223 | 110 | 78 | 463 | 58 | 49 |
| SRR2731299 | 0 | 0 | 24 | 0 | 15 | 24 | 0 | 18 | 13 | 0 | 0 |
| SRR2731300 | 0 | 0 | 0 | 0 | 14 | 30 | 14 | 19 | 39 | 0 | 0 |
| SRR2731301 | 0 | 0 | 15 | 0 | 21 | 34 | 15 | 22 | 85 | 0 | 0 |
| SRR2731302 | 0 | 13 | 36 | 15 | 28 | 79 | 29 | 41 | 61 | 0 | 0 |
| SRR2731303 | 0 | 0 | 19 | 0 | 37 | 41 | 19 | 15 | 66 | 0 | 0 |
| SRR2731304 | 0 | 0 | 18 | 11 | 24 | 48 | 33 | 35 | 49 | 0 | 0 |
| SRR2731305 | 0 | 0 | 12 | 0 | 10 | 29 | 0 | 13 | 19 | 0 | 0 |
| SRR2731306 | 0 | 14 | 12 | 0 | 14 | 42 | 14 | 19 | 23 | 0 | 0 |
| SRR2731307 | 0 | 11 | 17 | 0 | 21 | 58 | 24 | 29 | 51 | 0 | 0 |
| SRR2731308 | 0 | 10 | 26 | 0 | 20 | 47 | 14 | 29 | 43 | 0 | 0 |
| SRR2731309 | 0 | 11 | 15 | 0 | 11 | 34 | 11 | 18 | 30 | 0 | 0 |
| SRR2731310 | 0 | 21 | 40 | 17 | 30 | 92 | 29 | 45 | 65 | 0 | 0 |
| SRR2731311 | 0 | 0 | 11 | 0 | 11 | 33 | 12 | 19 | 39 | 0 | 0 |
| SRR2731312 | 0 | 0 | 0 | 0 | 21 | 45 | 14 | 22 | 42 | 0 | 0 |
| SRR2731313 | 0 | 0 | 0 | 0 | 16 | 36 | 12 | 19 | 53 | 0 | 0 |
| SRR2731314 | 0 | 0 | 0 | 0 | 0 | 24 | 0 | 12 | 33 | 0 | 0 |
| SRR2731315 | 0 | 0 | 25 | 18 | 32 | 65 | 28 | 35 | 32 | 0 | 0 |
| SRR2731316 | 0 | 0 | 0 | 0 | 15 | 40 | 14 | 16 | 24 | 0 | 0 |
| SRR2731317 | 0 | 0 | 11 | 0 | 16 | 59 | 17 | 35 | 44 | 0 | 0 |
| SRR2731318 | 0 | 0 | 18 | 18 | 105 | 112 | 80 | 50 | 54 | 12 | 0 |
| SRR2731319 | 0 | 0 | 19 | 0 | 17 | 32 | 17 | 23 | 49 | 0 | 0 |
| SRR2731320 | 0 | 17 | 18 | 0 | 58 | 154 | 48 | 65 | 82 | 19 | 0 |
| SRR2731321 | 0 | 0 | 12 | 0 | 81 | 111 | 66 | 55 | 91 | 0 | 10 |
| SRR2731322 | 0 | 0 | 0 | 0 | 30 | 43 | 29 | 16 | 25 | 0 | 0 |
| SRR2731323 | 0 | 10 | 0 | 0 | 29 | 66 | 27 | 41 | 57 | 0 | 0 |
| SRR2731324 | 0 | 0 | 0 | 0 | 0 | 23 | 0 | 16 | 27 | 0 | 0 |
| SRR2731325 | 0 | 0 | 0 | 0 | 12 | 36 | 16 | 19 | 44 | 0 | 0 |
| SRR2731326 | 0 | 0 | 10 | 0 | 15 | 40 | 13 | 23 | 39 | 0 | 0 |
| SRR2731327 | 0 | 0 | 0 | 0 | 13 | 24 | 11 | 19 | 38 | 0 | 0 |
| SRR2731328 | 0 | 0 | 0 | 0 | 11 | 30 | 0 | 18 | 22 | 0 | 0 |
| SRR2731329 | 0 | 29 | 60 | 23 | 56 | 186 | 51 | 111 | 146 | 11 | 11 |
| SRR2731330 | 0 | 96 | 109 | 96 | 199 | 492 | 157 | 271 | 234 | 53 | 19 |
| SRR2731331 | 0 | 0 | 0 | 19 | 196 | 231 | 87 | 192 | 174 | 58 | 23 |
| SRR2731332 | 0 | 0 | 20 | 0 | 39 | 84 | 30 | 53 | 32 | 11 | 0 |
| SRR2731333 | 0 | 15 | 19 | 20 | 126 | 279 | 104 | 165 | 99 | 36 | 13 |
| SRR2731335 | 0 | 0 | 0 | 0 | 14 | 22 | 13 | 16 | 29 | 0 | 0 |
| SRR2731336 | 0 | 0 | 16 | 18 | 113 | 135 | 42 | 124 | 50 | 22 | 0 |
| SRR2731337 | 0 | 0 | 0 | 12 | 38 | 79 | 30 | 113 | 111 | 21 | 11 |
| SRR2731338 | 0 | 0 | 12 | 22 | 343 | 338 | 160 | 171 | 84 | 49 | 10 |
| SRR2731339 | 0 | 0 | 0 | 0 | 23 | 44 | 22 | 26 | 42 | 21 | 0 |
| SRR2731340 | 0 | 34 | 38 | 26 | 105 | 237 | 90 | 149 | 97 | 18 | 19 |
| SRR2731341 | 0 | 0 | 0 | 0 | 11 | 31 | 11 | 16 | 28 | 0 | 0 |
| SRR2731342 | 0 | 0 | 0 | 0 | 0 | 18 | 0 | 0 | 30 | 0 | 0 |

### Supp Table S8.5.Columns_ET_to_FD

| | | | | | | | | | | | |
|---|---|---|---|---|---|---|---|---|---|---|---|
| SRR2731343 | 0 | 0 | 0 | 0 | 0 | 13 | 0 | 0 | 25 | 0 | 0 |
| SRR2731344 | 0 | 0 | 15 | 14 | 23 | 79 | 31 | 60 | 111 | 13 | 12 |
| SRR2731345 | 0 | 0 | 0 | 0 | 19 | 61 | 19 | 17 | 84 | 15 | 12 |
| SRR2731346 | 0 | 0 | 17 | 0 | 16 | 44 | 22 | 29 | 75 | 0 | 0 |
| SRR2731347 | 0 | 0 | 0 | 0 | 17 | 55 | 18 | 28 | 54 | 0 | 0 |
| SRR2731348 | 0 | 0 | 12 | 0 | 12 | 32 | 14 | 22 | 57 | 0 | 0 |
| SRR2731349 | 0 | 0 | 0 | 0 | 35 | 38 | 16 | 25 | 78 | 0 | 20 |
| SRR2731350 | 0 | 0 | 16 | 0 | 15 | 59 | 17 | 29 | 27 | 0 | 0 |
| SRR2910706 | 0 | 47 | 117 | 123 | 26 | 196 | 40 | 531 | 61 | 24 | 168 |
| SRR2910737 | 0 | 32 | 89 | 73 | 34 | 213 | 28 | 509 | 88 | 52 | 209 |
| SRR2910738 | 0 | 43 | 143 | 125 | 28 | 212 | 21 | 428 | 87 | 54 | 235 |
| SRR2912507 | 0 | 0 | 0 | 0 | 72 | 285 | 338 | 750 | 3972 | 151 | 426 |
| SRR2912508 | 0 | 0 | 0 | 0 | 259 | 624 | 862 | 2038 | 7564 | 424 | 1018 |
| SRR2912510 | 0 | 0 | 0 | 0 | 0 | 0 | 0 | 18 | 179 | 59 | 19 |
| SRR2912511 | 0 | 0 | 0 | 0 | 0 | 11 | 0 | 29 | 193 | 47 | 23 |
| SRR2912513 | 0 | 0 | 0 | 0 | 0 | 0 | 0 | 11 | 40 | 0 | 0 |
| SRR2912514 | 0 | 0 | 0 | 0 | 66 | 206 | 330 | 577 | 1302 | 67 | 190 |
| SRR2912515 | 0 | 0 | 0 | 0 | 0 | 0 | 0 | 13 | 67 | 27 | 10 |
| SRR2913202 | 0 | 0 | 17 | 18 | 0 | 0 | 0 | 0 | 0 | 0 | 0 |
| SRR2913203 | 0 | 16 | 45 | 27 | 0 | 11 | 0 | 0 | 0 | 0 | 0 |
| SRR2913204 | 0 | 0 | 22 | 21 | 0 | 0 | 0 | 0 | 0 | 0 | 0 |
| SRR2913205 | 0 | 0 | 15 | 15 | 0 | 0 | 0 | 0 | 0 | 0 | 0 |
| SRR2913206 | 0 | 0 | 12 | 0 | 0 | 0 | 0 | 0 | 0 | 0 | 0 |
| SRR2913207 | 0 | 11 | 25 | 29 | 0 | 15 | 0 | 28 | 0 | 0 | 0 |
| SRR2913208 | 0 | 0 | 16 | 13 | 0 | 0 | 0 | 0 | 0 | 0 | 0 |
| SRR2913209 | 0 | 0 | 18 | 19 | 0 | 0 | 0 | 0 | 0 | 0 | 0 |
| SRR2913210 | 0 | 0 | 0 | 14 | 0 | 0 | 0 | 0 | 0 | 0 | 0 |
| SRR2913211 | 0 | 0 | 16 | 13 | 0 | 0 | 0 | 0 | 0 | 0 | 18 |
| SRR2913212 | 0 | 0 | 0 | 11 | 0 | 0 | 0 | 0 | 0 | 0 | 0 |
| SRR2913213 | 0 | 0 | 16 | 13 | 0 | 0 | 0 | 0 | 0 | 0 | 0 |
| SRR2913214 | 0 | 0 | 18 | 16 | 0 | 0 | 0 | 0 | 0 | 0 | 0 |
| SRR2913215 | 0 | 0 | 12 | 13 | 0 | 0 | 0 | 0 | 0 | 0 | 0 |
| SRR2913216 | 0 | 0 | 49 | 39 | 0 | 0 | 0 | 0 | 0 | 0 | 0 |
| SRR2913217 | 0 | 0 | 22 | 17 | 0 | 0 | 0 | 0 | 0 | 0 | 11 |
| SRR2913218 | 0 | 0 | 0 | 0 | 0 | 0 | 0 | 0 | 0 | 0 | 0 |
| SRR2913219 | 0 | 0 | 12 | 13 | 0 | 0 | 0 | 0 | 0 | 0 | 0 |
| SRR2913220 | 0 | 0 | 18 | 15 | 0 | 0 | 0 | 0 | 0 | 0 | 0 |
| SRR2913221 | 0 | 0 | 0 | 11 | 0 | 0 | 0 | 0 | 0 | 0 | 0 |
| SRR2913222 | 0 | 0 | 13 | 17 | 0 | 0 | 0 | 0 | 0 | 0 | 0 |
| SRR2913223 | 0 | 0 | 18 | 17 | 0 | 0 | 0 | 12 | 0 | 0 | 0 |
| SRR2913224 | 0 | 0 | 12 | 23 | 0 | 0 | 0 | 0 | 0 | 0 | 0 |
| SRR2913225 | 0 | 0 | 16 | 25 | 0 | 0 | 0 | 10 | 0 | 0 | 0 |
| SRR2915342 | 0 | 15 | 42 | 0 | 73 | 163 | 55 | 281 | 458 | 20 | 104 |
| SRR2915343 | 0 | 26 | 69 | 13 | 92 | 185 | 79 | 246 | 437 | 22 | 98 |
| SRR2915344 | 0 | 26 | 70 | 10 | 95 | 209 | 64 | 336 | 263 | 18 | 83 |
| SRR2915345 | 0 | 23 | 99 | 23 | 316 | 680 | 882 | 831 | 550 | 149 | 186 |
| SRR2915346 | 0 | 32 | 83 | 21 | 132 | 299 | 342 | 371 | 320 | 55 | 83 |
| SRR2915347 | 12 | 43 | 110 | 30 | 186 | 387 | 506 | 568 | 398 | 80 | 132 |
| SRR2915348 | 11 | 69 | 199 | 27 | 148 | 249 | 270 | 286 | 194 | 44 | 78 |
| SRR2915349 | 0 | 20 | 60 | 0 | 88 | 168 | 178 | 284 | 303 | 55 | 111 |
| SRR2915350 | 0 | 28 | 114 | 18 | 167 | 270 | 315 | 349 | 282 | 56 | 101 |
| SRR2915905 | 0 | 0 | 49 | 34 | 0 | 106 | 0 | 142 | 90 | 135 | 274 |
| SRR2915915 | 0 | 0 | 17 | 20 | 0 | 172 | 17 | 271 | 17 | 17 | 72 |
| SRR2915918 | 0 | 30 | 89 | 72 | 10 | 129 | 12 | 191 | 52 | 25 | 47 |
| SRR2915921 | 0 | 0 | 0 | 0 | 0 | 14 | 0 | 22 | 0 | 0 | 0 |
| SRR2915923 | 0 | 0 | 12 | 0 | 0 | 23 | 0 | 34 | 19 | 23 | 38 |
| SRR2915926 | 0 | 16 | 34 | 18 | 0 | 40 | 0 | 67 | 68 | 74 | 55 |
| SRR2915928 | 0 | 0 | 17 | 0 | 0 | 35 | 0 | 90 | 27 | 58 | 49 |
| SRR2960039 | 0 | 0 | 0 | 0 | 0 | 0 | 101 | 308 | 1499 | 168 | 459 |
| SRR2976280 | 0 | 0 | 0 | 0 | 0 | 0 | 0 | 0 | 0 | 0 | 0 |
| SRR2976305 | 0 | 0 | 0 | 0 | 0 | 0 | 0 | 0 | 22 | 0 | 17 |
| SRR3092114 | 0 | 0 | 0 | 0 | 190 | 875 | 925 | 611 | 60 | 351 | 85 |
| SRR3092115 | 0 | 0 | 0 | 0 | 144 | 666 | 668 | 420 | 55 | 185 | 49 |
| SRR3103222 | 26 | 0 | 0 | 10 | 10 | 76 | 31 | 193 | 99 | 146 | 480 |
| SRR3103223 | 43 | 14 | 43 | 28 | 0 | 71 | 0 | 142 | 14 | 71 | 114 |
| SRR3103224 | 13 | 0 | 0 | 13 | 0 | 77 | 13 | 90 | 58 | 32 | 180 |
| SRR3103225 | 406 | 61 | 90 | 61 | 0 | 57 | 0 | 128 | 28 | 24 | 109 |
| SRR3103226 | 21 | 0 | 42 | 0 | 0 | 42 | 0 | 106 | 42 | 85 | 127 |
| SRR3103227 | 141 | 10 | 20 | 71 | 0 | 151 | 10 | 192 | 30 | 10 | 182 |
| SRR3103228 | 40 | 0 | 17 | 17 | 0 | 44 | 17 | 54 | 40 | 10 | 87 |
| SRR3103229 | 154 | 25 | 88 | 56 | 11 | 105 | 11 | 151 | 46 | 46 | 175 |
| SRR3103230 | 67 | 0 | 42 | 42 | 17 | 67 | 0 | 109 | 59 | 34 | 93 |
| SRR3103231 | 191 | 18 | 50 | 46 | 0 | 59 | 0 | 77 | 14 | 36 | 146 |
| SRR3103232 | 36 | 0 | 54 | 18 | 0 | 54 | 0 | 125 | 0 | 36 | 161 |
| SRR3103233 | 83 | 0 | 0 | 12 | 0 | 59 | 12 | 112 | 0 | 12 | 166 |
| SRR3103234 | 104 | 33 | 48 | 56 | 0 | 45 | 19 | 152 | 0 | 15 | 122 |
| SRR3103235 | 148 | 0 | 32 | 48 | 0 | 55 | 13 | 97 | 26 | 32 | 129 |
| SRR3103236 | 71 | 10 | 24 | 40 | 0 | 40 | 0 | 114 | 27 | 0 | 185 |
| SRR3103237 | 119 | 19 | 52 | 67 | 0 | 109 | 29 | 133 | 19 | 29 | 204 |
| SRR3103238 | 15 | 12 | 29 | 50 | 29 | 90 | 26 | 140 | 123 | 341 | 823 |
| SRR3103239 | 1039 | 19 | 56 | 79 | 0 | 56 | 0 | 100 | 14 | 21 | 61 |
| SRR3103240 | 76 | 30 | 60 | 72 | 0 | 72 | 12 | 125 | 65 | 113 | 229 |
| SRR3103241 | 126 | 0 | 114 | 101 | 0 | 177 | 0 | 151 | 151 | 296 | 448 |
| SRR3103242 | 88 | 70 | 130 | 97 | 19 | 167 | 37 | 357 | 534 | 687 | 844 |
| SRR3103243 | 0 | 0 | 0 | 0 | 0 | 87 | 0 | 277 | 695 | 786 | 2314 |
| SRR3103244 | 148 | 24 | 71 | 53 | 0 | 214 | 42 | 344 | 89 | 95 | 196 |
| SRR3103245 | 54 | 15 | 27 | 36 | 12 | 185 | 15 | 324 | 307 | 158 | 610 |
| SRR3103246 | 36 | 22 | 91 | 86 | 12 | 138 | 11 | 226 | 90 | 160 | 273 |
| SRR3103247 | 122 | 20 | 78 | 78 | 15 | 161 | 44 | 377 | 465 | 333 | 710 |
| SRR3103248 | 78 | 11 | 23 | 13 | 0 | 55 | 0 | 105 | 78 | 61 | 78 |
| SRR3103249 | 24 | 14 | 65 | 27 | 0 | 53 | 0 | 81 | 198 | 294 | 312 |
| SRR3103250 | 52 | 0 | 27 | 33 | 11 | 107 | 0 | 224 | 235 | 238 | 580 |
| SRR3103251 | 32 | 0 | 50 | 72 | 0 | 158 | 25 | 369 | 523 | 660 | 1036 |
| SRR3103252 | 92 | 0 | 15 | 46 | 0 | 92 | 0 | 139 | 200 | 108 | 216 |
| SRR3103253 | 48 | 0 | 0 | 0 | 17 | 39 | 17 | 78 | 35 | 43 | 182 |
| SRR3103254 | 365 | 103 | 219 | 87 | 23 | 126 | 23 | 377 | 494 | 1003 | 2926 |
| SRR3103255 | 161 | 0 | 51 | 30 | 21 | 179 | 24 | 419 | 438 | 414 | 1414 |
| SRR3103256 | 150 | 19 | 30 | 30 | 14 | 145 | 44 | 235 | 96 | 156 | 437 |
| SRR3103257 | 53 | 0 | 34 | 19 | 0 | 72 | 12 | 116 | 155 | 131 | 401 |
| SRR3103258 | 921 | 181 | 519 | 488 | 86 | 1193 | 181 | 2226 | 2633 | 6586 | 15720 |
| SRR3103259 | 154 | 0 | 18 | 31 | 0 | 39 | 0 | 67 | 26 | 18 | 67 |
| SRR3103260 | 155 | 28 | 85 | 51 | 11 | 136 | 0 | 216 | 246 | 216 | 244 |
| SRR3103261 | 360 | 73 | 163 | 107 | 31 | 258 | 28 | 289 | 85 | 118 | 165 |
| SRR3103262 | 73 | 19 | 54 | 39 | 19 | 135 | 23 | 270 | 305 | 745 | 1058 |
| SRR3103263 | 0 | 0 | 0 | 0 | 23 | 185 | 19 | 629 | 398 | 366 | 1883 |
| SRR3103264 | 198 | 12 | 41 | 41 | 0 | 18 | 0 | 33 | 12 | 0 | 0 |
| SRR3103265 | 1098 | 23 | 100 | 46 | 0 | 103 | 0 | 138 | 86 | 72 | 249 |
| SRR3103266 | 115 | 29 | 182 | 153 | 57 | 239 | 29 | 191 | 220 | 574 | 459 |
| SRR3103267 | 42 | 12 | 33 | 33 | 24 | 209 | 33 | 391 | 421 | 631 | 1225 |
| SRR3103268 | 104 | 26 | 49 | 26 | 20 | 108 | 20 | 189 | 98 | 108 | 26 |
| SRR3103269 | 70 | 28 | 186 | 186 | 28 | 446 | 65 | 888 | 711 | 600 | 446 |
| SRR3103270 | 72 | 12 | 84 | 108 | 24 | 108 | 0 | 335 | 36 | 72 | 227 |
| SRR3103271 | 32 | 19 | 97 | 45 | 13 | 317 | 52 | 362 | 446 | 517 | 1215 |
| SRR3103272 | 76 | 0 | 38 | 19 | 0 | 38 | 0 | 57 | 95 | 114 | 362 |
| SRR3103273 | 69 | 26 | 52 | 69 | 0 | 95 | 17 | 190 | 104 | 61 | 286 |
| SRR3103274 | 49 | 52 | 92 | 60 | 19 | 190 | 27 | 418 | 361 | 565 | 1783 |
| SRR3103275 | 175 | 37 | 83 | 42 | 20 | 121 | 20 | 302 | 227 | 381 | 812 |
| SRR3103276 | 68 | 0 | 30 | 21 | 12 | 94 | 28 | 246 | 117 | 161 | 463 |
| SRR3103277 | 54 | 18 | 24 | 24 | 0 | 125 | 0 | 137 | 214 | 262 | 590 |

### Supp Table S8.5.Columns_ET_to_FD

| | | | | | | | | | | | |
|---|---|---|---|---|---|---|---|---|---|---|---|
| SRR3103278 | 483 | 276 | 742 | 793 | 198 | 828 | 155 | 2182 | 1776 | 6665 | 16564 |
| SRR3103279 | 92 | 20 | 54 | 44 | 10 | 49 | 0 | 106 | 18 | 29 | 143 |
| SRR3103280 | 243 | 97 | 335 | 411 | 28 | 441 | 81 | 1136 | 2337 | 4935 | 10268 |
| SRR3103281 | 921 | 185 | 630 | 738 | 56 | 577 | 67 | 1803 | 4180 | 8147 | 16136 |
| SRR3103386 | 0 | 0 | 97 | 65 | 0 | 97 | 32 | 389 | 0 | 0 | 65 |
| SRR3103387 | 0 | 63 | 32 | 32 | 32 | 126 | 32 | 284 | 0 | 32 | 63 |
| SRR3103388 | 0 | 0 | 0 | 0 | 44 | 88 | 0 | 131 | 0 | 0 | 131 |
| SRR3103389 | 0 | 32 | 63 | 63 | 0 | 0 | 0 | 222 | 0 | 32 | 32 |
| SRR3103390 | 0 | 0 | 56 | 28 | 0 | 111 | 28 | 195 | 0 | 0 | 84 |
| SRR3103391 | 0 | 0 | 83 | 83 | 0 | 166 | 42 | 125 | 42 | 21 | 104 |
| SRR3103392 | 0 | 97 | 271 | 348 | 77 | 252 | 97 | 465 | 58 | 39 | 97 |
| SRR3103393 | 0 | 0 | 0 | 172 | 86 | 86 | 0 | 86 | 0 | 0 | 86 |
| SRR3103394 | 0 | 17 | 50 | 75 | 0 | 159 | 42 | 394 | 17 | 0 | 75 |
| SRR3103395 | 0 | 21 | 42 | 169 | 42 | 63 | 21 | 42 | 0 | 0 | 148 |
| SRR3103396 | 0 | 24 | 48 | 71 | 48 | 95 | 48 | 119 | 0 | 0 | 48 |
| SRR3103397 | 0 | 116 | 47 | 116 | 23 | 70 | 0 | 70 | 0 | 0 | 23 |
| SRR3103400 | 0 | 41 | 166 | 41 | 41 | 166 | 41 | 1037 | 124 | 41 | 124 |
| SRR3103401 | 0 | 109 | 0 | 437 | 0 | 875 | 0 | 1093 | 109 | 219 | 219 |
| SRR3103402 | 0 | 0 | 487 | 81 | 0 | 649 | 0 | 1380 | 0 | 0 | 244 |
| SRR3103403 | 0 | 0 | 162 | 270 | 54 | 431 | 0 | 809 | 0 | 54 | 108 |
| SRR3103404 | 0 | 114 | 285 | 171 | 0 | 114 | 57 | 742 | 171 | 57 | 628 |
| SRR3103405 | 0 | 54 | 268 | 161 | 214 | 375 | 321 | 964 | 54 | 0 | 0 |
| SRR3103406 | 0 | 0 | 33 | 0 | 0 | 200 | 67 | 534 | 0 | 67 | 33 |
| SRR3103407 | 0 | 58 | 58 | 0 | 117 | 58 | 0 | 700 | 0 | 0 | 58 |
| SRR3103408 | 0 | 0 | 114 | 0 | 114 | 228 | 0 | 685 | 0 | 0 | 114 |
| SRR3103409 | 0 | 0 | 0 | 62 | 0 | 124 | 62 | 249 | 0 | 0 | 62 |
| SRR3138807 | 0 | 0 | 0 | 0 | 0 | 20 | 49 | 11 | 0 | 0 | 18 |
| SRR3138808 | 0 | 0 | 0 | 0 | 0 | 16 | 22 | 0 | 0 | 23 | 15 |
| SRR3138809 | 0 | 0 | 0 | 0 | 0 | 17 | 24 | 0 | 0 | 28 | 22 |
| SRR3138810 | 0 | 0 | 0 | 0 | 0 | 16 | 20 | 0 | 0 | 0 | 0 |
| SRR3199254 | 0 | 0 | 0 | 0 | 286 | 471 | 354 | 342 | 388 | 231 | 116 |
| SRR3199255 | 0 | 0 | 0 | 18 | 286 | 1038 | 737 | 804 | 82 | 44 | 60 |
| SRR3199256 | 0 | 0 | 0 | 0 | 232 | 323 | 328 | 261 | 862 | 328 | 456 |
| SRR3199257 | 0 | 0 | 0 | 10 | 471 | 742 | 1083 | 4553 | 522 | 130 | 110 |
| SRR3199258 | 0 | 0 | 0 | 0 | 215 | 67 | 115 | 84 | 151 | 20 | 0 |
| SRR3199259 | 0 | 0 | 0 | 0 | 89 | 85 | 64 | 79 | 28 | 14 | 0 |
| SRR3199260 | 0 | 0 | 0 | 0 | 157 | 240 | 73 | 156 | 18 | 0 | 0 |
| SRR3199261 | 0 | 0 | 0 | 0 | 64 | 34 | 43 | 46 | 40 | 0 | 15 |
| SRR3199262 | 0 | 0 | 0 | 0 | 230 | 129 | 86 | 148 | 20 | 23 | 16 |
| SRR3199263 | 0 | 0 | 0 | 0 | 178 | 934 | 178 | 3070 | 222 | 311 | 133 |
| SRR3235777 | 0 | 0 | 0 | 0 | 35 | 85 | 131 | 174 | 660 | 394 | 62 |
| SRR3235778 | 0 | 0 | 0 | 0 | 73 | 165 | 195 | 253 | 583 | 460 | 100 |
| SRR3235779 | 0 | 0 | 0 | 0 | 290 | 397 | 423 | 605 | 135 | 353 | 77 |
| SRR3235780 | 0 | 0 | 0 | 0 | 80 | 114 | 106 | 234 | 250 | 236 | 64 |
| SRR3235781 | 10 | 24 | 21 | 31 | 312 | 593 | 538 | 333 | 604 | 458 | 302 |
| SRR3279553 | 0 | 0 | 0 | 0 | 0 | 0 | 0 | 0 | 18 | 0 | 0 |
| SRR3279554 | 0 | 0 | 0 | 0 | 0 | 0 | 0 | 59 | 88 | 29 | 88 |
| SRR3279555 | 0 | 0 | 0 | 0 | 0 | 0 | 0 | 0 | 0 | 0 | 22 |
| SRR3279556 | 0 | 0 | 0 | 0 | 10 | 0 | 27 | 27 | 57 | 51 | 17 |
| SRR3279557 | 0 | 0 | 0 | 0 | 31 | 57 | 39 | 159 | 183 | 284 | 194 |
| SRR3279558 | 0 | 0 | 0 | 0 | 0 | 39 | 29 | 48 | 67 | 144 | 77 |
| SRR3279559 | 0 | 0 | 0 | 0 | 18 | 48 | 18 | 85 | 54 | 66 | 36 |
| SRR3279560 | 0 | 0 | 0 | 0 | 0 | 12 | 12 | 0 | 25 | 12 | 135 |
| SRR3279561 | 0 | 0 | 0 | 0 | 16 | 40 | 16 | 88 | 64 | 96 | 116 |
| SRR3279562 | 0 | 0 | 0 | 0 | 0 | 0 | 0 | 43 | 43 | 62 | 130 |
| SRR3279563 | 0 | 0 | 0 | 0 | 0 | 0 | 0 | 70 | 23 | 12 | 23 |
| SRR3279564 | 0 | 0 | 0 | 0 | 14 | 48 | 0 | 41 | 55 | 65 | 31 |
| SRR3279565 | 0 | 0 | 0 | 0 | 0 | 17 | 11 | 14 | 28 | 23 | 26 |
| SRR3279566 | 0 | 0 | 0 | 0 | 0 | 0 | 0 | 0 | 0 | 11 | 0 |
| SRR3279567 | 0 | 0 | 0 | 0 | 61 | 209 | 152 | 388 | 743 | 764 | 449 |
| SRR3279568 | 0 | 0 | 0 | 0 | 0 | 0 | 0 | 56 | 14 | 14 | 28 |
| SRR3279569 | 0 | 0 | 0 | 0 | 0 | 32 | 0 | 64 | 0 | 64 | 0 |
| SRR3279570 | 0 | 0 | 0 | 0 | 0 | 0 | 0 | 0 | 12 | 28 | 0 |
| SRR3279571 | 0 | 0 | 0 | 0 | 0 | 41 | 0 | 41 | 54 | 82 | 109 |
| SRR3279572 | 0 | 0 | 0 | 0 | 17 | 51 | 22 | 163 | 107 | 275 | 343 |
| SRR3279573 | 0 | 0 | 0 | 0 | 0 | 0 | 0 | 16 | 14 | 0 | 0 |
| SRR3279574 | 0 | 0 | 0 | 0 | 0 | 15 | 15 | 0 | 0 | 0 | 0 |
| SRR3279575 | 0 | 0 | 0 | 0 | 11 | 18 | 11 | 34 | 66 | 103 | 29 |
| SRR3279576 | 0 | 0 | 0 | 0 | 0 | 15 | 0 | 15 | 15 | 44 | 15 |
| SRR3279577 | 0 | 0 | 0 | 0 | 0 | 16 | 0 | 24 | 47 | 51 | 43 |
| SRR3279578 | 0 | 0 | 0 | 0 | 0 | 24 | 18 | 52 | 0 | 0 | 21 |
| SRR3279579 | 0 | 0 | 0 | 0 | 0 | 0 | 0 | 0 | 23 | 0 | 0 |
| SRR3279580 | 0 | 0 | 0 | 0 | 0 | 18 | 12 | 45 | 48 | 54 | 42 |
| SRR3279581 | 0 | 0 | 0 | 0 | 0 | 21 | 13 | 77 | 39 | 56 | 51 |
| SRR3279582 | 0 | 0 | 0 | 0 | 0 | 11 | 0 | 0 | 0 | 17 | 0 |
| SRR3279583 | 0 | 0 | 0 | 0 | 0 | 23 | 0 | 51 | 28 | 46 | 41 |
| SRR3279584 | 0 | 0 | 0 | 12 | 0 | 19 | 11 | 30 | 22 | 33 | 34 |
| SRR3279585 | 0 | 0 | 0 | 0 | 0 | 0 | 0 | 0 | 0 | 0 | 0 |
| SRR3279586 | 0 | 0 | 0 | 0 | 0 | 0 | 0 | 0 | 0 | 0 | 0 |
| SRR3317221 | 0 | 0 | 0 | 0 | 179 | 286 | 251 | 501 | 394 | 286 | 107 |
| SRR3477269 | 0 | 0 | 0 | 0 | 0 | 0 | 0 | 0 | 0 | 0 | 0 |
| SRR3477271 | 0 | 0 | 0 | 0 | 0 | 0 | 0 | 0 | 0 | 0 | 0 |
| SRR3490374 | 189 | 0 | 0 | 0 | 63 | 63 | 0 | 0 | 0 | 0 | 189 |
| SRR3490375 | 3189 | 290 | 290 | 0 | 0 | 1160 | 0 | 1160 | 290 | 0 | 2030 |
| SRR3490376 | 0 | 0 | 15 | 30 | 15 | 44 | 0 | 44 | 0 | 59 | 207 |
| SRR3490377 | 0 | 0 | 0 | 0 | 0 | 0 | 0 | 0 | 0 | 0 | 18 |
| SRR3490378 | 0 | 0 | 24 | 0 | 0 | 24 | 0 | 24 | 24 | 48 | 24 |
| SRR3490379 | 0 | 92 | 0 | 275 | 0 | 92 | 0 | 183 | 0 | 0 | 366 |
| SRR3490380 | 0 | 0 | 86 | 0 | 0 | 43 | 0 | 0 | 0 | 86 | 599 |
| SRR3490382 | 2620 | 403 | 403 | 202 | 0 | 0 | 202 | 0 | 0 | 0 | 1612 |
| SRR3490383 | 13 | 0 | 25 | 0 | 0 | 13 | 0 | 63 | 25 | 0 | 189 |
| SRR3490384 | 0 | 47 | 140 | 0 | 0 | 47 | 0 | 93 | 93 | 0 | 326 |
| SRR3490385 | 0 | 0 | 152 | 101 | 51 | 507 | 51 | 507 | 51 | 51 | 355 |
| SRR3490386 | 50 | 50 | 0 | 0 | 0 | 50 | 0 | 300 | 0 | 0 | 300 |
| SRR3490387 | 36 | 36 | 109 | 145 | 0 | 145 | 0 | 218 | 36 | 0 | 545 |
| SRR3490388 | 0 | 400 | 200 | 200 | 0 | 200 | 0 | 600 | 0 | 0 | 200 |
| SRR3490389 | 1762 | 77 | 153 | 306 | 77 | 230 | 153 | 230 | 153 | 0 | 690 |
| SRR3490390 | 1426 | 324 | 130 | 65 | 0 | 65 | 0 | 130 | 0 | 0 | 454 |
| SRR3490391 | 3868 | 0 | 0 | 0 | 0 | 242 | 0 | 0 | 0 | 0 | 725 |
| SRR3490392 | 17 | 0 | 17 | 34 | 17 | 69 | 17 | 120 | 0 | 51 | 240 |
| SRR3490393 | 0 | 23 | 47 | 70 | 0 | 141 | 0 | 94 | 47 | 47 | 141 |
| SRR3490394 | 33 | 33 | 66 | 266 | 0 | 299 | 0 | 531 | 100 | 33 | 565 |
| SRR3493372 | 0 | 0 | 0 | 0 | 0 | 0 | 0 | 0 | 0 | 0 | 0 |
| SRR3493373 | 0 | 0 | 0 | 0 | 0 | 0 | 0 | 0 | 0 | 0 | 0 |
| SRR3534933 | 11 | 55 | 144 | 54 | 89 | 1002 | 179 | 2394 | 1764 | 1582 | 1587 |
| SRR3534934 | 21 | 29 | 79 | 37 | 95 | 1004 | 136 | 2188 | 1861 | 1736 | 1831 |
| SRR3534935 | 0 | 42 | 111 | 43 | 62 | 652 | 125 | 1285 | 1307 | 1341 | 1399 |
| SRR3534936 | 23 | 38 | 117 | 42 | 110 | 931 | 188 | 1308 | 2045 | 2043 | 2184 |
| SRR3534937 | 0 | 65 | 129 | 72 | 46 | 408 | 58 | 629 | 461 | 162 | 135 |
| SRR3534938 | 0 | 35 | 79 | 39 | 89 | 785 | 132 | 898 | 515 | 209 | 183 |
| SRR3566024 | 0 | 17 | 0 | 0 | 51 | 244 | 67 | 202 | 93 | 59 | 17 |
| SRR3566026 | 0 | 0 | 0 | 0 | 152 | 284 | 114 | 379 | 208 | 47 | 28 |
| SRR3566150 | 0 | 0 | 0 | 10 | 21 | 131 | 42 | 125 | 47 | 0 | 0 |
| SRR3566151 | 0 | 0 | 0 | 0 | 22 | 164 | 33 | 88 | 93 | 11 | 44 |
| SRR3566152 | 0 | 0 | 0 | 0 | 16 | 16 | 23 | 70 | 62 | 0 | 31 |
| SRR3566153 | 0 | 0 | 0 | 0 | 0 | 60 | 0 | 80 | 133 | 0 | 60 |
| SRR3566154 | 0 | 0 | 0 | 0 | 0 | 0 | 0 | 0 | 0 | 0 | 0 |
| SRR3566155 | 0 | 0 | 0 | 0 | 0 | 0 | 0 | 0 | 0 | 0 | 0 |
| SRR3582392 | 0 | 35 | 0 | 0 | 248 | 548 | 265 | 672 | 248 | 106 | 35 |

### Supp Table S8.5.Columns_ET_to_FD

| | | | | | | | | | | | |
|---|---|---|---|---|---|---|---|---|---|---|---|
| SRR3582393 | 0 | 19 | 19 | 0 | 212 | 714 | 463 | 792 | 347 | 39 | 77 |
| SRR3623374 | 0 | 0 | 0 | 0 | 0 | 0 | 0 | 0 | 0 | 0 | 0 |
| SRR3624069 | 0 | 0 | 10 | 0 | 150 | 370 | 71 | 55 | 399 | 36 | 174 |
| SRR3624070 | 0 | 30 | 34 | 11 | 995 | 2336 | 334 | 150 | 739 | 122 | 439 |
| SRR3624071 | 0 | 0 | 16 | 0 | 627 | 1779 | 281 | 138 | 1010 | 106 | 415 |
| SRR3624072 | 0 | 53 | 64 | 15 | 1168 | 3492 | 664 | 218 | 630 | 100 | 256 |
| SRR3624073 | 0 | 0 | 10 | 0 | 408 | 1089 | 180 | 140 | 708 | 55 | 231 |
| SRR3624074 | 0 | 0 | 15 | 0 | 1072 | 3206 | 678 | 242 | 679 | 85 | 258 |
| SRR3675003 | 0 | 0 | 53 | 0 | 0 | 51 | 12 | 112 | 180 | 314 | 86 |
| SRR3675004 | 0 | 13 | 74 | 12 | 0 | 56 | 19 | 107 | 166 | 260 | 94 |
| SRR3675005 | 0 | 34 | 120 | 86 | 231 | 2112 | 349 | 4398 | 7720 | 10060 | 1835 |
| SRR3675006 | 0 | 15 | 81 | 14 | 0 | 66 | 27 | 124 | 229 | 291 | 115 |
| SRR3675007 | 0 | 17 | 94 | 15 | 12 | 74 | 31 | 139 | 168 | 277 | 102 |
| SRR3675008 | 0 | 13 | 96 | 18 | 0 | 35 | 12 | 65 | 106 | 196 | 73 |
| SRR3713941 | 0 | 0 | 15 | 0 | 55 | 117 | 110 | 66 | 601 | 45 | 64 |
| SRR3713942 | 0 | 0 | 0 | 0 | 0 | 0 | 0 | 0 | 0 | 0 | 0 |
| SRR3713943 | 0 | 0 | 0 | 0 | 25 | 37 | 52 | 45 | 73 | 0 | 0 |
| SRR3713945 | 0 | 0 | 19 | 0 | 30 | 44 | 37 | 17 | 91 | 65 | 89 |
| SRR3713946 | 0 | 0 | 0 | 14 | 29 | 50 | 51 | 21 | 163 | 236 | 149 |
| SRR3879751 | 0 | 0 | 0 | 0 | 0 | 0 | 0 | 0 | 20 | 11 | 0 |
| SRR3879752 | 0 | 0 | 0 | 0 | 0 | 0 | 0 | 0 | 0 | 0 | 0 |
| SRR3879754 | 0 | 0 | 0 | 0 | 0 | 0 | 0 | 0 | 0 | 0 | 0 |
| SRR3879755 | 0 | 0 | 0 | 0 | 0 | 0 | 0 | 0 | 12 | 0 | 0 |
| SRR3879756 | 0 | 0 | 0 | 0 | 0 | 0 | 0 | 0 | 12 | 0 | 0 |
| SRR3879757 | 0 | 0 | 0 | 0 | 0 | 0 | 0 | 0 | 0 | 0 | 0 |
| SRR3879758 | 0 | 0 | 0 | 0 | 0 | 0 | 0 | 0 | 0 | 0 | 0 |
| SRR3879759 | 0 | 0 | 0 | 0 | 0 | 0 | 0 | 0 | 0 | 0 | 0 |
| SRR3879760 | 0 | 0 | 0 | 0 | 0 | 0 | 0 | 0 | 0 | 10 | 0 |
| SRR3879761 | 0 | 0 | 0 | 0 | 0 | 0 | 0 | 0 | 0 | 0 | 0 |
| SRR3879766 | 0 | 0 | 0 | 0 | 153 | 438 | 172 | 306 | 885 | 406 | 220 |
| SRR3879767 | 0 | 0 | 0 | 0 | 34 | 297 | 16 | 0 | 204 | 40 | 16 |
| SRR3879768 | 0 | 0 | 0 | 0 | 80 | 237 | 63 | 109 | 758 | 256 | 93 |
| SRR3879769 | 0 | 0 | 0 | 0 | 73 | 225 | 41 | 27 | 932 | 300 | 180 |
| SRR3879770 | 0 | 0 | 0 | 0 | 195 | 1143 | 139 | 0 | 221 | 34 | 18 |
| SRR3923813 | 0 | 0 | 0 | 0 | 0 | 0 | 0 | 11 | 20 | 0 | 0 |
| SRR3931357 | 0 | 36 | 85 | 113 | 113 | 1602 | 265 | 1996 | 6622 | 8113 | 3755 |
| SRR3931358 | 0 | 38 | 102 | 176 | 146 | 1663 | 369 | 2034 | 7988 | 10789 | 6643 |
| SRR3931359 | 0 | 114 | 304 | 677 | 213 | 1622 | 207 | 1047 | 2197 | 3458 | 2467 |
| SRR3931360 | 0 | 136 | 378 | 843 | 184 | 1642 | 220 | 1167 | 2143 | 3297 | 2422 |
| SRR3931984 | 0 | 0 | 0 | 0 | 0 | 218 | 69 | 1055 | 6087 | 7247 | 7416 |
| SRR3931987 | 0 | 0 | 0 | 0 | 0 | 21 | 0 | 182 | 2300 | 2883 | 4044 |
| SRR3931988 | 0 | 0 | 0 | 0 | 17 | 270 | 71 | 1460 | 10389 | 10858 | 11660 |
| SRR3992213 | 0 | 0 | 62 | 0 | 233 | 725 | 88 | 109 | 268 | 15 | 18 |
| SRR3992215 | 0 | 0 | 0 | 0 | 29 | 124 | 38 | 68 | 209 | 0 | 32 |
| SRR3992474 | 0 | 0 | 0 | 0 | 0 | 0 | 0 | 0 | 0 | 0 | 0 |
| SRR3992475 | 0 | 0 | 0 | 0 | 0 | 0 | 0 | 0 | 0 | 0 | 0 |
| SRR3992476 | 109 | 82 | 154 | 95 | 29 | 126 | 38 | 187 | 320 | 440 | 448 |
| SRR3992477 | 382 | 210 | 389 | 232 | 47 | 306 | 103 | 470 | 1060 | 1383 | 1308 |
| SRR3996364 | 248 | 54 | 13 | 13 | 34 | 322 | 27 | 657 | 1039 | 355 | 577 |
| SRR3996365 | 1841 | 93 | 23 | 15 | 0 | 209 | 70 | 913 | 1648 | 418 | 650 |
| SRR3996366 | 2577 | 93 | 42 | 13 | 34 | 388 | 110 | 962 | 2640 | 856 | 886 |
| SRR3996367 | 414 | 76 | 55 | 28 | 35 | 449 | 124 | 1402 | 3211 | 766 | 1056 |
| SRR3996368 | 137 | 25 | 0 | 0 | 21 | 75 | 17 | 150 | 191 | 83 | 254 |
| SRR3996369 | 701 | 18 | 0 | 0 | 0 | 94 | 25 | 258 | 280 | 94 | 243 |
| SRR4000625 | 0 | 0 | 0 | 0 | 42 | 58 | 0 | 0 | 0 | 0 | 0 |
| SRR4000626 | 0 | 0 | 0 | 0 | 34 | 60 | 0 | 0 | 0 | 0 | 0 |
| SRR4000627 | 0 | 0 | 0 | 0 | 0 | 0 | 0 | 0 | 0 | 0 | 0 |
| SRR4000628 | 0 | 0 | 0 | 0 | 0 | 0 | 0 | 0 | 0 | 0 | 0 |
| SRR4000629 | 0 | 0 | 0 | 0 | 68 | 126 | 35 | 0 | 0 | 0 | 0 |
| SRR4000630 | 0 | 0 | 0 | 0 | 77 | 139 | 36 | 0 | 0 | 0 | 0 |
| SRR4000631 | 0 | 0 | 0 | 0 | 53 | 68 | 12 | 0 | 0 | 0 | 0 |
| SRR4000632 | 0 | 0 | 0 | 0 | 59 | 71 | 12 | 0 | 0 | 0 | 0 |
| SRR4000633 | 0 | 0 | 0 | 0 | 45 | 91 | 15 | 0 | 0 | 0 | 0 |
| SRR4000634 | 0 | 0 | 0 | 0 | 39 | 94 | 17 | 0 | 0 | 0 | 0 |
| SRR4000635 | 0 | 0 | 0 | 0 | 10 | 18 | 0 | 0 | 0 | 0 | 0 |
| SRR4000636 | 0 | 0 | 0 | 0 | 12 | 23 | 0 | 0 | 0 | 0 | 0 |
| SRR4000638 | 0 | 0 | 0 | 0 | 0 | 0 | 0 | 0 | 0 | 0 | 0 |
| SRR4000639 | 0 | 0 | 0 | 0 | 0 | 0 | 0 | 0 | 0 | 0 | 0 |
| SRR4000640 | 0 | 0 | 0 | 0 | 0 | 0 | 0 | 0 | 0 | 0 | 0 |
| SRR4000641 | 0 | 0 | 0 | 0 | 0 | 0 | 0 | 0 | 0 | 0 | 0 |
| SRR4000642 | 0 | 0 | 0 | 0 | 0 | 0 | 0 | 0 | 0 | 0 | 0 |
| SRR4000643 | 0 | 0 | 0 | 0 | 0 | 0 | 0 | 0 | 0 | 0 | 0 |
| SRR4000644 | 0 | 0 | 0 | 0 | 0 | 0 | 0 | 0 | 0 | 0 | 0 |
| SRR4000645 | 0 | 0 | 0 | 0 | 0 | 0 | 0 | 0 | 0 | 0 | 0 |
| SRR4000649 | 0 | 0 | 0 | 0 | 0 | 0 | 0 | 0 | 0 | 0 | 0 |
| SRR4000650 | 0 | 0 | 0 | 0 | 0 | 0 | 0 | 0 | 0 | 0 | 0 |
| SRR4011144 | 0 | 0 | 0 | 0 | 822 | 0 | 0 | 1287 | 0 | 0 | 0 |
| SRR4044175 | 0 | 0 | 16 | 0 | 0 | 53 | 0 | 113 | 159 | 230 | 113 |
| SRR4044176 | 0 | 0 | 11 | 0 | 0 | 39 | 0 | 87 | 240 | 392 | 316 |
| SRR4044177 | 0 | 0 | 28 | 12 | 12 | 78 | 0 | 122 | 94 | 185 | 135 |
| SRR4044178 | 0 | 0 | 12 | 0 | 0 | 75 | 15 | 158 | 248 | 343 | 190 |
| SRR4044179 | 0 | 13 | 33 | 13 | 0 | 51 | 0 | 107 | 261 | 418 | 310 |
| SRR4044180 | 0 | 0 | 0 | 0 | 0 | 47 | 0 | 77 | 93 | 180 | 136 |
| SRR4044206 | 0 | 0 | 0 | 0 | 0 | 0 | 0 | 0 | 0 | 0 | 0 |
| SRR4044210 | 0 | 0 | 0 | 0 | 0 | 0 | 0 | 0 | 0 | 0 | 0 |
| SRR4044211 | 0 | 13 | 17 | 0 | 0 | 31 | 0 | 0 | 52 | 0 | 0 |
| SRR4270461 | 0 | 129 | 199 | 71 | 208 | 447 | 253 | 322 | 1088 | 292 | 216 |
| SRR4270462 | 0 | 129 | 249 | 65 | 299 | 603 | 389 | 371 | 1669 | 522 | 437 |
| SRR4292754 | 0 | 13 | 20 | 0 | 0 | 66 | 24 | 148 | 1414 | 1938 | 1219 |
| SRR4292755 | 0 | 0 | 13 | 10 | 0 | 72 | 19 | 141 | 1485 | 1798 | 1364 |
| SRR4292756 | 0 | 0 | 12 | 11 | 11 | 57 | 19 | 129 | 1125 | 1242 | 1783 |
| SRR4292757 | 0 | 0 | 10 | 11 | 0 | 67 | 27 | 147 | 955 | 961 | 1191 |
| SRR4296311 | 0 | 280 | 692 | 197 | 2176 | 5543 | 1773 | 1730 | 3420 | 898 | 1935 |
| SRR4296312 | 360 | 214 | 462 | 192 | 2095 | 5666 | 1633 | 1667 | 3537 | 901 | 2005 |
| SRR4296313 | 0 | 206 | 368 | 263 | 1334 | 2999 | 1035 | 1661 | 2987 | 881 | 2146 |
| SRR4296314 | 236 | 193 | 364 | 187 | 964 | 2714 | 974 | 1638 | 2671 | 889 | 2088 |
| SRR4296315 | 0 | 178 | 657 | 278 | 2550 | 5684 | 1592 | 1659 | 1720 | 1002 | 2505 |
| SRR4296316 | 166 | 177 | 521 | 222 | 2438 | 5397 | 1651 | 1795 | 1574 | 909 | 2715 |
| SRR4296317 | 0 | 613 | 1248 | 315 | 3969 | 8148 | 2401 | 1971 | 2070 | 834 | 1871 |
| SRR4296318 | 436 | 594 | 990 | 261 | 3429 | 7413 | 2107 | 1631 | 1829 | 1022 | 1647 |
| SRR4296319 | 0 | 177 | 244 | 57 | 994 | 2596 | 602 | 550 | 889 | 292 | 554 |
| SRR4296320 | 41 | 119 | 199 | 81 | 882 | 2413 | 639 | 569 | 952 | 356 | 624 |
| SRR4296321 | 0 | 98 | 192 | 94 | 823 | 1795 | 483 | 477 | 679 | 287 | 559 |
| SRR4296322 | 68 | 58 | 146 | 73 | 792 | 2152 | 505 | 340 | 714 | 350 | 442 |
| SRR4296979 | 0 | 49 | 118 | 29 | 26 | 309 | 52 | 225 | 182 | 139 | 219 |
| SRR4296985 | 0 | 0 | 0 | 0 | 0 | 0 | 0 | 0 | 0 | 0 | 0 |
| SRR4296986 | 0 | 0 | 0 | 0 | 0 | 0 | 0 | 0 | 0 | 0 | 0 |
| SRR4296987 | 0 | 0 | 0 | 0 | 0 | 0 | 0 | 0 | 0 | 0 | 0 |
| SRR4296988 | 0 | 0 | 0 | 0 | 0 | 0 | 0 | 0 | 0 | 0 | 0 |
| SRR4328127 | 0 | 0 | 0 | 0 | 0 | 0 | 0 | 0 | 0 | 0 | 0 |
| SRR4328130 | 0 | 0 | 0 | 0 | 0 | 0 | 0 | 0 | 0 | 0 | 0 |
| SRR4328135 | 0 | 0 | 0 | 0 | 0 | 0 | 0 | 0 | 0 | 0 | 0 |
| SRR4328136 | 0 | 0 | 0 | 0 | 0 | 0 | 0 | 0 | 0 | 0 | 0 |
| SRR4328137 | 0 | 0 | 0 | 0 | 0 | 0 | 0 | 0 | 0 | 0 | 0 |
| SRR4409201 | 0 | 13 | 35 | 60 | 12 | 35 | 0 | 28 | 0 | 0 | 45 |
| SRR4409202 | 0 | 402 | 510 | 210 | 0 | 44 | 0 | 18 | 21 | 16 | 24 |
| SRR4409203 | 0 | 15 | 39 | 59 | 0 | 24 | 0 | 21 | 13 | 0 | 51 |
| SRR4409204 | 0 | 11 | 25 | 0 | 0 | 0 | 0 | 0 | 0 | 0 | 0 |
| SRR4417245 | 0 | 0 | 0 | 0 | 0 | 0 | 0 | 0 | 0 | 0 | 0 |

### Supp Table S8.5.Columns_ET_to_FD

| | | | | | | | | | | | |
|---|---|---|---|---|---|---|---|---|---|---|---|
| SRR4417246 | 0 | 0 | 0 | 0 | 0 | 0 | 0 | 0 | 15 | 0 | 0 |
| SRR4417247 | 0 | 0 | 0 | 0 | 0 | 0 | 0 | 0 | 0 | 0 | 0 |
| SRR4417248 | 0 | 0 | 0 | 0 | 0 | 0 | 0 | 0 | 0 | 0 | 0 |
| SRR4417249 | 0 | 0 | 0 | 0 | 0 | 0 | 0 | 0 | 0 | 0 | 0 |
| SRR4417250 | 0 | 0 | 0 | 0 | 0 | 0 | 0 | 0 | 0 | 0 | 0 |
| SRR4423247 | 0 | 262 | 637 | 559 | 114 | 880 | 272 | 3429 | 1023 | 941 | 1459 |
| SRR4423248 | 0 | 158 | 487 | 401 | 82 | 430 | 56 | 556 | 289 | 218 | 358 |
| SRR4423250 | 0 | 232 | 559 | 327 | 364 | 4896 | 2465 | 11865 | 796 | 661 | 701 |
| SRR4423251 | 0 | 416 | 908 | 386 | 619 | 7143 | 1932 | 5745 | 1315 | 1049 | 785 |
| SRR4473670 | 56 | 90 | 186 | 76 | 48 | 188 | 35 | 97 | 115 | 78 | 91 |
| SRR4734676 | 0 | 0 | 0 | 0 | 13 | 228 | 66 | 1242 | 91 | 43 | 294 |
| SRR4734677 | 0 | 0 | 0 | 17 | 17 | 520 | 134 | 3235 | 201 | 84 | 821 |
| SRR4734678 | 0 | 0 | 0 | 40 | 13 | 106 | 40 | 835 | 27 | 40 | 172 |
| SRR4734679 | 0 | 0 | 14 | 14 | 27 | 178 | 82 | 561 | 41 | 0 | 178 |
| SRR4734680 | 0 | 0 | 34 | 17 | 34 | 84 | 0 | 421 | 84 | 17 | 202 |
| SRR4734681 | 0 | 0 | 15 | 23 | 38 | 437 | 153 | 2155 | 153 | 54 | 798 |
| SRR4734682 | 0 | 0 | 0 | 0 | 0 | 154 | 48 | 646 | 32 | 0 | 226 |
| SRR4734683 | 0 | 0 | 0 | 11 | 13 | 122 | 38 | 468 | 36 | 0 | 96 |
| SRR4734684 | 0 | 0 | 15 | 0 | 49 | 434 | 157 | 1874 | 61 | 18 | 342 |
| SRR4734685 | 0 | 0 | 0 | 0 | 26 | 174 | 47 | 507 | 17 | 0 | 107 |
| SRR4734686 | 0 | 0 | 0 | 13 | 63 | 717 | 202 | 3478 | 139 | 63 | 780 |
| SRR4787084 | 0 | 0 | 0 | 0 | 0 | 0 | 0 | 0 | 0 | 0 | 0 |
| SRR4787085 | 0 | 0 | 0 | 36 | 0 | 0 | 0 | 36 | 0 | 71 | 0 |
| SRR4787086 | 0 | 0 | 0 | 0 | 129 | 0 | 0 | 0 | 64 | 0 | 0 |
| SRR4787087 | 0 | 0 | 0 | 0 | 0 | 134 | 45 | 0 | 90 | 0 | 0 |
| SRR4787088 | 0 | 0 | 0 | 0 | 0 | 0 | 0 | 0 | 0 | 0 | 66 |
| SRR4787089 | 0 | 0 | 0 | 0 | 0 | 0 | 0 | 0 | 0 | 89 | 0 |
| SRR4787090 | 18 | 0 | 18 | 0 | 13 | 27 | 13 | 18 | 148 | 18 | 18 |
| SRR4787091 | 0 | 0 | 0 | 0 | 157 | 0 | 0 | 0 | 0 | 0 | 0 |
| SRR4787092 | 14 | 19 | 38 | 0 | 19 | 34 | 14 | 0 | 207 | 0 | 24 |
| SRR4787093 | 23 | 0 | 37 | 0 | 23 | 18 | 14 | 23 | 179 | 14 | 32 |
| SRR4787095 | 0 | 13 | 0 | 17 | 26 | 99 | 39 | 112 | 619 | 21 | 125 |
| SRR4787096 | 0 | 0 | 0 | 10 | 82 | 108 | 82 | 97 | 272 | 0 | 97 |
| SRR4787097 | 0 | 0 | 0 | 0 | 26 | 13 | 0 | 0 | 0 | 0 | 0 |
| SRR4787098 | 0 | 0 | 0 | 0 | 0 | 16 | 10 | 0 | 13 | 0 | 0 |
| SRR4787099 | 0 | 0 | 0 | 0 | 13 | 0 | 0 | 0 | 0 | 0 | 22 |
| SRR4787100 | 0 | 0 | 0 | 0 | 46 | 55 | 18 | 25 | 15 | 0 | 0 |
| SRR4787101 | 0 | 0 | 0 | 0 | 40 | 91 | 0 | 24 | 16 | 0 | 16 |
| SRR4787102 | 0 | 0 | 0 | 0 | 39 | 51 | 17 | 22 | 15 | 0 | 0 |
| SRR4787103 | 10 | 0 | 0 | 0 | 91 | 131 | 61 | 106 | 40 | 0 | 25 |
| SRR4787104 | 0 | 0 | 0 | 0 | 85 | 106 | 42 | 56 | 67 | 0 | 46 |
| SRR4787105 | 0 | 0 | 0 | 0 | 0 | 75 | 0 | 83 | 0 | 0 | 17 |
| SRR4787106 | 0 | 0 | 25 | 0 | 13 | 102 | 0 | 121 | 13 | 0 | 0 |
| SRR4787107 | 0 | 0 | 12 | 0 | 0 | 61 | 0 | 70 | 0 | 0 | 0 |
| SRR4787108 | 0 | 0 | 0 | 0 | 0 | 53 | 0 | 44 | 0 | 0 | 0 |
| SRR4787109 | 0 | 0 | 18 | 18 | 0 | 75 | 0 | 97 | 0 | 0 | 14 |
| SRR4787110 | 0 | 0 | 12 | 0 | 0 | 51 | 0 | 116 | 30 | 0 | 0 |
| SRR4787118 | 0 | 0 | 0 | 0 | 0 | 0 | 0 | 0 | 0 | 0 | 0 |
| SRR4787119 | 0 | 0 | 0 | 5152 | 0 | 0 | 0 | 0 | 0 | 0 | 0 |
| SRR4787120 | 0 | 0 | 0 | 0 | 101 | 101 | 74 | 129 | 345 | 23 | 106 |
| SRR4787121 | 0 | 0 | 0 | 0 | 11 | 15 | 0 | 0 | 191 | 0 | 18 |
| SRR4787122 | 0 | 0 | 0 | 0 | 18 | 0 | 13 | 0 | 75 | 0 | 11 |
| SRR4787123 | 0 | 0 | 0 | 0 | 0 | 0 | 0 | 16 | 36 | 0 | 14 |
| SRR4787124 | 0 | 0 | 0 | 0 | 0 | 0 | 0 | 0 | 451 | 0 | 0 |
| SRR4787125 | 0 | 0 | 0 | 0 | 0 | 0 | 0 | 0 | 0 | 0 | 0 |
| SRR4787126 | 0 | 0 | 0 | 0 | 0 | 0 | 0 | 0 | 583 | 0 | 0 |
| SRR4787127 | 0 | 0 | 0 | 0 | 0 | 0 | 0 | 0 | 0 | 0 | 0 |
| SRR4787132 | 0 | 0 | 24 | 47 | 0 | 0 | 0 | 24 | 0 | 24 | 0 |
| SRR4787133 | 0 | 0 | 0 | 0 | 0 | 0 | 0 | 0 | 0 | 100 | 0 |
| SRR4787134 | 0 | 0 | 0 | 0 | 0 | 0 | 0 | 35 | 199 | 0 | 0 |
| SRR4787135 | 0 | 0 | 0 | 0 | 79 | 111 | 23 | 69 | 28 | 0 | 51 |
| SRR4787136 | 0 | 0 | 0 | 0 | 11 | 37 | 50 | 13 | 395 | 0 | 74 |
| SRR4787137 | 0 | 0 | 0 | 0 | 0 | 0 | 0 | 13 | 310 | 17 | 30 |
| SRR4787138 | 0 | 0 | 0 | 0 | 42 | 25 | 11 | 34 | 700 | 12 | 17 |
| SRR4787139 | 0 | 0 | 0 | 0 | 26 | 46 | 33 | 50 | 503 | 0 | 21 |
| SRR4787140 | 0 | 0 | 0 | 0 | 45 | 90 | 36 | 60 | 729 | 28 | 66 |
| SRR4787141 | 0 | 0 | 0 | 0 | 28 | 46 | 27 | 42 | 422 | 14 | 56 |
| SRR4787142 | 0 | 0 | 0 | 0 | 105 | 128 | 67 | 65 | 648 | 21 | 51 |
| SRR4787143 | 0 | 0 | 0 | 0 | 88 | 88 | 11 | 66 | 808 | 11 | 66 |
| SRR4836291 | 0 | 26 | 82 | 15 | 10 | 131 | 10 | 193 | 577 | 512 | 322 |
| SRR4836293 | 0 | 60 | 54 | 16 | 25 | 123 | 16 | 177 | 606 | 543 | 265 |
| SRR4836294 | 0 | 45 | 56 | 30 | 26 | 160 | 23 | 216 | 557 | 535 | 316 |
| SRR4836295 | 0 | 88 | 129 | 36 | 129 | 350 | 175 | 206 | 175 | 299 | 541 |
| SRR4836297 | 0 | 94 | 126 | 94 | 157 | 372 | 126 | 189 | 208 | 315 | 453 |
| SRR4836299 | 0 | 120 | 131 | 71 | 157 | 426 | 183 | 307 | 213 | 243 | 527 |
| SRR4836301 | 0 | 59 | 44 | 12 | 23 | 224 | 23 | 293 | 959 | 1782 | 1252 |
| SRR4836302 | 0 | 83 | 67 | 24 | 33 | 219 | 39 | 306 | 1108 | 1794 | 1182 |
| SRR4836303 | 0 | 50 | 67 | 26 | 44 | 217 | 54 | 314 | 1104 | 1941 | 1381 |
| SRR4836305 | 0 | 69 | 99 | 44 | 25 | 247 | 74 | 341 | 519 | 909 | 1615 |
| SRR4836307 | 0 | 115 | 90 | 72 | 18 | 368 | 36 | 325 | 452 | 934 | 1374 |
| SRR4836308 | 0 | 73 | 109 | 54 | 47 | 327 | 47 | 425 | 570 | 969 | 1582 |
| SRR4836310 | 0 | 97 | 206 | 86 | 51 | 405 | 47 | 810 | 1508 | 2111 | 1807 |
| SRR4836311 | 0 | 85 | 161 | 66 | 33 | 450 | 52 | 849 | 1659 | 2024 | 1797 |
| SRR4836312 | 0 | 114 | 261 | 97 | 43 | 366 | 63 | 861 | 1741 | 2389 | 2034 |
| SRR4836313 | 0 | 66 | 95 | 59 | 133 | 878 | 145 | 2046 | 1364 | 1910 | 2473 |
| SRR4836314 | 0 | 68 | 93 | 49 | 137 | 1018 | 153 | 2061 | 1307 | 2089 | 2495 |
| SRR4836315 | 0 | 67 | 105 | 61 | 169 | 937 | 126 | 2026 | 1309 | 1997 | 2487 |
| SRR4836316 | 0 | 23 | 52 | 12 | 52 | 452 | 45 | 677 | 900 | 1249 | 2627 |
| SRR4836317 | 0 | 29 | 49 | 25 | 47 | 445 | 36 | 741 | 1044 | 1408 | 2565 |
| SRR4836318 | 0 | 29 | 42 | 29 | 46 | 448 | 46 | 731 | 1002 | 1331 | 2670 |
| SRR4836320 | 0 | 95 | 162 | 91 | 83 | 667 | 87 | 1748 | 1628 | 1922 | 6624 |
| SRR4836321 | 0 | 76 | 236 | 84 | 104 | 516 | 72 | 1663 | 1659 | 1983 | 7083 |
| SRR4836323 | 0 | 68 | 166 | 74 | 68 | 549 | 74 | 1534 | 1612 | 1774 | 6684 |
| SRR4836325 | 0 | 46 | 92 | 43 | 248 | 1837 | 207 | 2246 | 3919 | 5718 | 7331 |
| SRR4836326 | 0 | 75 | 52 | 47 | 282 | 1713 | 217 | 2370 | 3899 | 5659 | 7579 |
| SRR4836327 | 0 | 35 | 93 | 55 | 251 | 1612 | 209 | 2218 | 3920 | 5915 | 7094 |
| SRR4836328 | 0 | 19 | 23 | 15 | 12 | 133 | 23 | 198 | 101 | 103 | 232 |
| SRR4836330 | 0 | 11 | 32 | 11 | 13 | 107 | 20 | 203 | 96 | 101 | 228 |
| SRR4836332 | 0 | 22 | 16 | 16 | 33 | 121 | 16 | 170 | 92 | 97 | 209 |
| SRR4836334 | 0 | 57 | 82 | 34 | 69 | 713 | 67 | 1748 | 7134 | 8408 | 16131 |
| SRR4836335 | 0 | 44 | 67 | 38 | 61 | 692 | 85 | 1712 | 7171 | 8733 | 16308 |
| SRR4836336 | 0 | 38 | 105 | 50 | 71 | 678 | 88 | 1751 | 6997 | 8443 | 15837 |
| SRR4836337 | 0 | 49 | 60 | 17 | 12 | 154 | 28 | 221 | 311 | 492 | 600 |
| SRR4836339 | 0 | 49 | 42 | 32 | 12 | 131 | 23 | 203 | 226 | 362 | 644 |
| SRR4836341 | 0 | 46 | 72 | 25 | 21 | 179 | 0 | 259 | 303 | 454 | 727 |
| SRR4836343 | 0 | 40 | 83 | 28 | 24 | 157 | 22 | 275 | 301 | 483 | 692 |
| SRR4836344 | 0 | 102 | 199 | 49 | 88 | 905 | 126 | 1649 | 506 | 486 | 754 |
| SRR4836345 | 0 | 92 | 203 | 81 | 92 | 773 | 97 | 1501 | 341 | 417 | 672 |
| SRR4836346 | 0 | 130 | 218 | 65 | 84 | 910 | 122 | 1827 | 439 | 397 | 715 |
| SRR4836347 | 0 | 128 | 190 | 43 | 66 | 816 | 104 | 1690 | 425 | 414 | 615 |
| SRR4836348 | 0 | 143 | 110 | 49 | 98 | 489 | 41 | 530 | 1756 | 1508 | 1190 |
| SRR4836349 | 0 | 107 | 111 | 30 | 60 | 506 | 34 | 480 | 1702 | 1471 | 1218 |
| SRR4836350 | 0 | 106 | 158 | 45 | 87 | 518 | 26 | 534 | 2044 | 1614 | 1366 |
| SRR4836351 | 0 | 126 | 132 | 13 | 122 | 582 | 60 | 516 | 1833 | 1469 | 1228 |
| SRR4836352 | 0 | 73 | 95 | 38 | 50 | 394 | 43 | 835 | 914 | 1224 | 3332 |
| SRR4836353 | 0 | 74 | 79 | 22 | 45 | 298 | 67 | 774 | 846 | 1113 | 3192 |
| SRR4836354 | 0 | 98 | 99 | 51 | 38 | 372 | 52 | 864 | 851 | 1147 | 3517 |
| SRR4836355 | 0 | 97 | 107 | 49 | 32 | 355 | 33 | 846 | 899 | 1166 | 3374 |
| SRR4836356 | 0 | 54 | 60 | 18 | 23 | 211 | 40 | 321 | 127 | 139 | 234 |

### Supp Table S8.5.Columns_ET_to_FD

| | | | | | | | | | | | |
|---|---|---|---|---|---|---|---|---|---|---|---|
| SRR4836357 | 0 | 40 | 60 | 21 | 24 | 215 | 39 | 303 | 115 | 141 | 245 |
| SRR4836359 | 0 | 44 | 81 | 28 | 29 | 216 | 43 | 320 | 128 | 153 | 239 |
| SRR4836361 | 0 | 45 | 64 | 18 | 27 | 230 | 42 | 297 | 115 | 134 | 245 |
| SRR4836362 | 0 | 48 | 71 | 27 | 28 | 205 | 21 | 397 | 210 | 318 | 687 |
| SRR4836364 | 0 | 52 | 79 | 22 | 29 | 177 | 27 | 334 | 229 | 279 | 579 |
| SRR4836366 | 0 | 57 | 76 | 33 | 32 | 198 | 24 | 433 | 197 | 345 | 662 |
| SRR4836368 | 0 | 52 | 71 | 36 | 32 | 171 | 23 | 326 | 209 | 319 | 597 |
| SRR4836369 | 0 | 66 | 91 | 22 | 103 | 510 | 25 | 412 | 1421 | 911 | 1421 |
| SRR4836370 | 0 | 51 | 64 | 26 | 102 | 562 | 36 | 446 | 1439 | 1106 | 1484 |
| SRR4836371 | 0 | 81 | 84 | 33 | 79 | 504 | 28 | 423 | 1421 | 1046 | 1453 |
| SRR4836373 | 0 | 87 | 75 | 30 | 103 | 548 | 38 | 479 | 1535 | 1132 | 1645 |
| SRR4836375 | 0 | 126 | 132 | 62 | 55 | 536 | 32 | 661 | 333 | 209 | 733 |
| SRR4836376 | 0 | 98 | 142 | 54 | 48 | 487 | 48 | 642 | 291 | 226 | 686 |
| SRR4836378 | 0 | 79 | 121 | 50 | 52 | 485 | 28 | 640 | 340 | 250 | 767 |
| SRR4836380 | 0 | 101 | 118 | 55 | 88 | 547 | 57 | 676 | 315 | 259 | 785 |
| SRR4836382 | 0 | 30 | 45 | 18 | 19 | 125 | 18 | 242 | 444 | 573 | 1145 |
| SRR4836383 | 0 | 35 | 35 | 21 | 12 | 115 | 0 | 239 | 456 | 621 | 1182 |
| SRR4836384 | 0 | 23 | 34 | 19 | 16 | 119 | 23 | 220 | 477 | 617 | 1098 |
| SRR4836386 | 0 | 23 | 47 | 13 | 13 | 140 | 17 | 263 | 461 | 682 | 1250 |
| SRR4836388 | 0 | 115 | 185 | 60 | 57 | 569 | 49 | 753 | 2441 | 1554 | 4929 |
| SRR4836390 | 0 | 153 | 176 | 89 | 69 | 626 | 26 | 846 | 2479 | 1516 | 5370 |
| SRR4836391 | 0 | 143 | 193 | 81 | 58 | 586 | 62 | 866 | 2367 | 1471 | 5022 |
| SRR4836392 | 0 | 143 | 222 | 63 | 74 | 637 | 61 | 905 | 2711 | 1603 | 5586 |
| SRR4836394 | 0 | 23 | 26 | 18 | 19 | 123 | 20 | 204 | 58 | 50 | 168 |
| SRR4836396 | 0 | 23 | 34 | 17 | 16 | 132 | 29 | 198 | 62 | 60 | 163 |
| SRR4836398 | 0 | 25 | 28 | 15 | 10 | 136 | 28 | 208 | 71 | 53 | 157 |
| SRR4836399 | 0 | 32 | 29 | 20 | 0 | 160 | 28 | 193 | 80 | 42 | 164 |
| SRR4836400 | 0 | 300 | 369 | 196 | 138 | 978 | 144 | 2080 | 6972 | 8873 | 12035 |
| SRR4836402 | 0 | 241 | 410 | 223 | 144 | 949 | 137 | 2246 | 7606 | 9439 | 12464 |
| SRR4836404 | 0 | 263 | 363 | 149 | 155 | 898 | 93 | 2159 | 7485 | 9016 | 11966 |
| SRR4836406 | 0 | 348 | 440 | 203 | 134 | 1020 | 148 | 2199 | 8170 | 9755 | 12659 |
| SRR4836407 | 0 | 68 | 91 | 41 | 82 | 579 | 41 | 343 | 694 | 871 | 893 |
| SRR4836408 | 0 | 63 | 94 | 44 | 96 | 551 | 30 | 405 | 814 | 1002 | 956 |
| SRR4836410 | 0 | 62 | 87 | 39 | 92 | 586 | 47 | 376 | 794 | 1041 | 937 |
| SRR4836412 | 0 | 34 | 98 | 23 | 17 | 153 | 44 | 297 | 149 | 410 | 440 |
| SRR4836414 | 0 | 41 | 99 | 33 | 23 | 146 | 41 | 311 | 196 | 501 | 530 |
| SRR4836415 | 0 | 43 | 106 | 27 | 12 | 133 | 22 | 335 | 216 | 383 | 470 |
| SRR4836416 | 0 | 90 | 141 | 43 | 47 | 292 | 37 | 497 | 1532 | 2605 | 3502 |
| SRR4836417 | 0 | 117 | 164 | 54 | 41 | 310 | 55 | 612 | 1778 | 3040 | 4113 |
| SRR4836419 | 0 | 105 | 145 | 47 | 41 | 291 | 42 | 593 | 1713 | 2993 | 3957 |
| SRR4836421 | 0 | 135 | 156 | 66 | 31 | 224 | 25 | 420 | 343 | 544 | 683 |
| SRR4836422 | 0 | 116 | 164 | 65 | 32 | 238 | 30 | 476 | 455 | 613 | 879 |
| SRR4836423 | 0 | 118 | 158 | 60 | 43 | 263 | 41 | 425 | 354 | 597 | 783 |
| SRR4836425 | 0 | 90 | 110 | 43 | 51 | 332 | 63 | 794 | 164 | 90 | 141 |
| SRR4836427 | 0 | 87 | 155 | 42 | 34 | 340 | 49 | 1034 | 143 | 151 | 245 |
| SRR4836428 | 0 | 84 | 131 | 38 | 38 | 381 | 56 | 1004 | 178 | 84 | 172 |
| SRR4836429 | 0 | 104 | 150 | 55 | 37 | 265 | 29 | 334 | 277 | 326 | 706 |
| SRR4836431 | 0 | 124 | 189 | 71 | 42 | 333 | 37 | 426 | 313 | 407 | 907 |
| SRR4836433 | 0 | 116 | 175 | 50 | 40 | 325 | 50 | 365 | 317 | 365 | 910 |
| SRR4836434 | 0 | 11 | 11 | 11 | 24 | 106 | 16 | 175 | 82 | 85 | 289 |
| SRR4836435 | 0 | 18 | 18 | 0 | 16 | 89 | 24 | 225 | 73 | 94 | 330 |
| SRR4836437 | 0 | 13 | 30 | 17 | 19 | 78 | 15 | 262 | 85 | 74 | 368 |
| SRR4836439 | 0 | 11 | 13 | 17 | 21 | 98 | 23 | 228 | 81 | 79 | 358 |
| SRR4836440 | 0 | 67 | 125 | 58 | 240 | 1353 | 144 | 1593 | 1487 | 1228 | 1699 |
| SRR4836441 | 0 | 66 | 94 | 28 | 272 | 1612 | 159 | 1556 | 1631 | 1125 | 1622 |
| SRR4836443 | 0 | 69 | 192 | 61 | 184 | 1426 | 207 | 1633 | 1472 | 1388 | 1756 |
| SRR4836445 | 0 | 84 | 84 | 53 | 306 | 1773 | 176 | 1735 | 1544 | 1368 | 1475 |
| SRR4836446 | 0 | 27 | 48 | 24 | 43 | 582 | 46 | 854 | 1166 | 636 | 4261 |
| SRR4836447 | 0 | 37 | 58 | 48 | 66 | 538 | 50 | 927 | 1260 | 768 | 4512 |
| SRR4836448 | 0 | 26 | 60 | 30 | 60 | 537 | 36 | 837 | 1178 | 685 | 4306 |
| SRR4836450 | 0 | 37 | 53 | 40 | 99 | 603 | 55 | 944 | 1298 | 746 | 4306 |
| SRR4836452 | 0 | 28 | 0 | 0 | 48 | 336 | 56 | 220 | 476 | 468 | 516 |
| SRR4836453 | 0 | 23 | 55 | 12 | 55 | 301 | 27 | 227 | 551 | 481 | 684 |
| SRR4836454 | 0 | 0 | 13 | 13 | 51 | 289 | 35 | 286 | 410 | 502 | 636 |
| SRR4836456 | 0 | 26 | 32 | 0 | 48 | 309 | 48 | 261 | 544 | 521 | 627 |
| SRR4836458 | 0 | 11 | 11 | 0 | 54 | 504 | 57 | 688 | 341 | 444 | 817 |
| SRR4836459 | 0 | 0 | 14 | 11 | 45 | 492 | 51 | 628 | 349 | 431 | 884 |
| SRR4836461 | 0 | 11 | 18 | 0 | 64 | 445 | 80 | 651 | 336 | 454 | 868 |
| SRR4836463 | 0 | 14 | 16 | 14 | 72 | 580 | 55 | 709 | 354 | 485 | 964 |
| SRR4836464 | 0 | 33 | 44 | 68 | 23 | 242 | 45 | 607 | 573 | 361 | 1626 |
| SRR4836466 | 0 | 28 | 50 | 55 | 13 | 251 | 27 | 622 | 625 | 407 | 1740 |
| SRR4836468 | 0 | 23 | 58 | 64 | 23 | 250 | 35 | 600 | 588 | 375 | 1715 |
| SRR4836469 | 0 | 30 | 40 | 71 | 34 | 265 | 28 | 597 | 630 | 398 | 1741 |
| SRR4836471 | 0 | 33 | 38 | 46 | 81 | 644 | 57 | 1032 | 739 | 717 | 1882 |
| SRR4836473 | 0 | 22 | 58 | 28 | 101 | 640 | 86 | 1134 | 825 | 762 | 2295 |
| SRR4836474 | 0 | 45 | 47 | 56 | 72 | 808 | 74 | 1231 | 883 | 891 | 2357 |
| SRR4836475 | 0 | 21 | 54 | 21 | 63 | 447 | 84 | 580 | 1286 | 914 | 1432 |
| SRR4836477 | 0 | 22 | 43 | 0 | 52 | 597 | 77 | 766 | 1505 | 1086 | 1742 |
| SRR4836479 | 0 | 0 | 59 | 13 | 66 | 713 | 138 | 710 | 1611 | 1117 | 1796 |
| SRR4836481 | 0 | 12 | 66 | 21 | 58 | 433 | 78 | 870 | 1583 | 2556 | 2696 |
| SRR4836482 | 0 | 37 | 52 | 24 | 61 | 428 | 131 | 1002 | 1800 | 2983 | 3093 |
| SRR4836483 | 0 | 47 | 47 | 13 | 76 | 517 | 101 | 1043 | 1762 | 3004 | 3190 |
| SRR4836484 | 0 | 80 | 114 | 69 | 251 | 1497 | 171 | 3862 | 4696 | 2674 | 8102 |
| SRR4836485 | 0 | 189 | 137 | 52 | 189 | 1940 | 223 | 4223 | 5527 | 3270 | 9209 |
| SRR4836487 | 0 | 139 | 182 | 35 | 199 | 2219 | 208 | 4266 | 5973 | 3485 | 9346 |
| SRR4836489 | 0 | 15 | 13 | 0 | 28 | 220 | 30 | 258 | 747 | 863 | 781 |
| SRR4836491 | 0 | 13 | 21 | 13 | 35 | 235 | 35 | 294 | 860 | 1001 | 873 |
| SRR4836492 | 0 | 0 | 21 | 0 | 32 | 291 | 21 | 323 | 958 | 1067 | 944 |
| SRR4836493 | 0 | 18 | 30 | 12 | 65 | 499 | 74 | 572 | 421 | 451 | 778 |
| SRR4836494 | 0 | 18 | 23 | 16 | 76 | 610 | 89 | 678 | 426 | 490 | 861 |
| SRR4836495 | 0 | 20 | 23 | 19 | 67 | 609 | 101 | 710 | 435 | 541 | 904 |
| SRR4836496 | 0 | 48 | 87 | 29 | 65 | 396 | 67 | 691 | 1008 | 1099 | 1096 |
| SRR4836498 | 0 | 50 | 68 | 29 | 66 | 402 | 98 | 724 | 944 | 1160 | 1274 |
| SRR4836500 | 0 | 59 | 80 | 29 | 54 | 477 | 83 | 723 | 978 | 1173 | 1167 |
| SRR4836502 | 0 | 48 | 98 | 25 | 78 | 508 | 85 | 831 | 1210 | 1411 | 1405 |
| SRR4836503 | 0 | 34 | 30 | 20 | 21 | 245 | 52 | 317 | 151 | 240 | 249 |
| SRR4836504 | 0 | 27 | 47 | 11 | 11 | 223 | 52 | 380 | 164 | 234 | 292 |
| SRR4836505 | 0 | 24 | 45 | 0 | 19 | 208 | 54 | 420 | 187 | 232 | 300 |
| SRR4836507 | 0 | 45 | 52 | 10 | 22 | 292 | 40 | 446 | 196 | 290 | 364 |
| SRR4836509 | 0 | 72 | 134 | 37 | 37 | 278 | 65 | 566 | 1933 | 5527 | 4428 |
| SRR4836511 | 0 | 60 | 118 | 29 | 37 | 244 | 76 | 556 | 2002 | 5895 | 4593 |
| SRR4836512 | 0 | 72 | 128 | 36 | 43 | 248 | 64 | 607 | 1935 | 5591 | 4559 |
| SRR4836513 | 0 | 67 | 142 | 40 | 56 | 282 | 79 | 729 | 2278 | 6944 | 5487 |
| SRR4836514 | 0 | 94 | 166 | 60 | 125 | 876 | 43 | 803 | 1386 | 780 | 6787 |
| SRR4836515 | 0 | 105 | 141 | 53 | 138 | 830 | 53 | 777 | 1446 | 779 | 7029 |
| SRR4836517 | 0 | 98 | 135 | 62 | 110 | 872 | 48 | 758 | 1437 | 783 | 6958 |
| SRR4836519 | 0 | 133 | 187 | 76 | 144 | 1089 | 53 | 977 | 1670 | 952 | 8468 |
| SRR4836521 | 0 | 45 | 80 | 40 | 25 | 194 | 22 | 482 | 265 | 430 | 542 |
| SRR4836522 | 0 | 60 | 51 | 43 | 11 | 289 | 26 | 493 | 232 | 420 | 601 |
| SRR4836523 | 0 | 62 | 102 | 30 | 19 | 261 | 16 | 398 | 261 | 412 | 573 |
| SRR4836524 | 0 | 74 | 102 | 46 | 26 | 293 | 59 | 511 | 280 | 503 | 750 |
| SRR4836525 | 0 | 155 | 184 | 77 | 128 | 883 | 47 | 947 | 3397 | 1835 | 15946 |
| SRR4836526 | 0 | 151 | 167 | 72 | 142 | 973 | 38 | 1034 | 3608 | 1960 | 16211 |
| SRR4836528 | 0 | 149 | 208 | 86 | 141 | 884 | 55 | 1080 | 3660 | 1875 | 16016 |
| SRR4836529 | 0 | 153 | 184 | 112 | 189 | 994 | 49 | 1156 | 4195 | 2204 | 19695 |
| SRR4836531 | 0 | 34 | 54 | 20 | 51 | 366 | 45 | 723 | 970 | 1108 | 1613 |
| SRR4836533 | 0 | 28 | 47 | 25 | 61 | 457 | 58 | 723 | 929 | 1358 | 1960 |
| SRR4836534 | 0 | 30 | 46 | 41 | 61 | 458 | 61 | 741 | 952 | 1324 | 1921 |
| SRR4836535 | 0 | 23 | 31 | 0 | 39 | 182 | 14 | 303 | 558 | 571 | 475 |

### Supp Table S8.5.Columns_ET_to_FD

| | | | | | | | | | | | |
|---|---|---|---|---|---|---|---|---|---|---|---|
| SRR4836536 | 0 | 20 | 26 | 0 | 26 | 218 | 26 | 353 | 588 | 633 | 498 |
| SRR4836537 | 0 | 20 | 14 | 12 | 33 | 205 | 15 | 331 | 610 | 550 | 477 |
| SRR4836539 | 0 | 0 | 11 | 0 | 0 | 39 | 11 | 68 | 33 | 67 | 122 |
| SRR4836541 | 0 | 0 | 0 | 0 | 11 | 58 | 18 | 92 | 27 | 68 | 164 |
| SRR4836543 | 0 | 0 | 0 | 0 | 0 | 40 | 0 | 72 | 34 | 64 | 123 |
| SRR4836544 | 0 | 41 | 50 | 12 | 118 | 883 | 168 | 1492 | 1242 | 1304 | 1981 |
| SRR4836545 | 0 | 37 | 35 | 20 | 84 | 1012 | 208 | 1661 | 1422 | 1621 | 2270 |
| SRR4836546 | 0 | 42 | 44 | 25 | 65 | 1012 | 177 | 1651 | 1359 | 1569 | 2153 |
| SRR4836548 | 0 | 0 | 40 | 49 | 85 | 687 | 147 | 1771 | 500 | 326 | 1731 |
| SRR4836550 | 0 | 13 | 44 | 49 | 49 | 848 | 132 | 2301 | 618 | 362 | 1877 |
| SRR4836552 | 0 | 38 | 38 | 33 | 85 | 582 | 112 | 1961 | 486 | 281 | 1898 |
| SRR4836553 | 0 | 30 | 30 | 35 | 56 | 502 | 95 | 1380 | 1380 | 2016 | 3076 |
| SRR4836554 | 0 | 25 | 42 | 47 | 68 | 520 | 110 | 1662 | 1764 | 2313 | 3337 |
| SRR4836555 | 0 | 34 | 61 | 34 | 63 | 561 | 58 | 1558 | 1776 | 2161 | 3434 |
| SRR4836556 | 0 | 0 | 0 | 0 | 92 | 932 | 257 | 2704 | 903 | 1417 | 2238 |
| SRR4836560 | 0 | 0 | 0 | 0 | 89 | 860 | 225 | 2522 | 879 | 1464 | 2425 |
| SRR4836561 | 0 | 0 | 23 | 0 | 102 | 916 | 204 | 2792 | 901 | 1558 | 2443 |
| SRR4836562 | 0 | 12 | 24 | 0 | 56 | 685 | 56 | 1418 | 1851 | 709 | 7783 |
| SRR4836563 | 0 | 12 | 24 | 0 | 36 | 550 | 51 | 1428 | 1756 | 748 | 8070 |
| SRR4836565 | 0 | 0 | 23 | 0 | 61 | 630 | 45 | 1479 | 2020 | 659 | 7921 |
| SRR4836567 | 0 | 0 | 20 | 0 | 59 | 686 | 49 | 1421 | 2126 | 741 | 8279 |
| SRR4836569 | 0 | 0 | 18 | 0 | 211 | 3825 | 429 | 3112 | 517 | 575 | 681 |
| SRR4836570 | 0 | 0 | 0 | 0 | 297 | 3711 | 419 | 3382 | 616 | 588 | 699 |
| SRR4836571 | 0 | 12 | 0 | 0 | 254 | 3934 | 349 | 3186 | 563 | 664 | 716 |
| SRR4836572 | 0 | 0 | 0 | 0 | 287 | 3951 | 500 | 3184 | 597 | 650 | 597 |
| SRR4896694 | 0 | 12 | 24 | 34 | 101 | 237 | 46 | 301 | 151 | 304 | 259 |
| SRR4896695 | 0 | 0 | 0 | 36 | 24 | 313 | 36 | 526 | 518 | 1313 | 2611 |
| SRR4896698 | 0 | 0 | 0 | 21 | 64 | 182 | 43 | 236 | 1110 | 1244 | 1136 |
| SRR4896701 | 0 | 35 | 105 | 158 | 80 | 353 | 103 | 706 | 3229 | 8760 | 19666 |
| SRR4896708 | 0 | 0 | 0 | 18 | 180 | 586 | 240 | 757 | 2674 | 4339 | 2875 |
| SRR5022482 | 0 | 24 | 44 | 208 | 262 | 437 | 146 | 496 | 621 | 406 | 446 |
| SRR5022483 | 0 | 0 | 0 | 0 | 0 | 13 | 0 | 0 | 15 | 0 | 0 |
| SRR5022484 | 0 | 0 | 38 | 26 | 70 | 525 | 96 | 964 | 4619 | 3044 | 2185 |
| SRR5022485 | 0 | 0 | 0 | 0 | 0 | 41 | 0 | 23 | 35 | 0 | 11 |
| SRR5029397 | 0 | 0 | 0 | 0 | 33 | 236 | 40 | 153 | 291 | 110 | 40 |
| SRR5029398 | 0 | 0 | 0 | 0 | 0 | 37 | 14 | 32 | 118 | 12 | 0 |
| SRR5029399 | 0 | 0 | 0 | 0 | 166 | 451 | 94 | 155 | 174 | 37 | 37 |
| SRR5029400 | 0 | 0 | 0 | 0 | 42 | 324 | 52 | 270 | 621 | 144 | 92 |
| SRR5029401 | 0 | 0 | 0 | 0 | 35 | 139 | 22 | 73 | 227 | 64 | 36 |
| SRR5029402 | 0 | 0 | 0 | 0 | 39 | 198 | 32 | 95 | 489 | 115 | 48 |
| SRR5029403 | 0 | 0 | 0 | 0 | 0 | 50 | 13 | 86 | 93 | 13 | 28 |
| SRR5029404 | 0 | 0 | 0 | 0 | 0 | 46 | 0 | 46 | 323 | 76 | 32 |
| SRR5029405 | 0 | 0 | 0 | 0 | 27 | 171 | 22 | 121 | 478 | 45 | 14 |
| SRR5029406 | 0 | 0 | 0 | 0 | 190 | 894 | 150 | 341 | 968 | 117 | 105 |
| SRR5029407 | 0 | 0 | 0 | 0 | 10 | 33 | 11 | 36 | 216 | 57 | 57 |
| SRR5029408 | 0 | 0 | 0 | 0 | 0 | 14 | 0 | 28 | 89 | 14 | 38 |
| SRR5029409 | 0 | 0 | 0 | 0 | 36 | 143 | 67 | 98 | 207 | 42 | 50 |
| SRR5029410 | 0 | 0 | 0 | 0 | 164 | 569 | 95 | 225 | 235 | 0 | 42 |
| SRR5029411 | 0 | 0 | 0 | 0 | 12 | 144 | 15 | 105 | 266 | 81 | 48 |
| SRR5029412 | 0 | 0 | 0 | 0 | 59 | 178 | 52 | 181 | 113 | 34 | 38 |
| SRR5029414 | 0 | 0 | 0 | 0 | 0 | 58 | 0 | 46 | 240 | 42 | 24 |
| SRR5029415 | 0 | 0 | 0 | 0 | 0 | 48 | 0 | 48 | 211 | 48 | 48 |
| SRR5029416 | 0 | 0 | 0 | 0 | 112 | 373 | 81 | 149 | 94 | 18 | 16 |
| SRR5029417 | 0 | 0 | 0 | 0 | 10 | 50 | 18 | 55 | 277 | 119 | 61 |
| SRR5029418 | 0 | 0 | 0 | 0 | 70 | 306 | 54 | 99 | 267 | 59 | 30 |
| SRR5029419 | 0 | 0 | 0 | 0 | 0 | 40 | 0 | 21 | 46 | 11 | 0 |
| SRR5029420 | 0 | 0 | 0 | 0 | 62 | 173 | 113 | 111 | 194 | 19 | 43 |
| SRR5029422 | 0 | 0 | 0 | 0 | 113 | 360 | 113 | 240 | 307 | 193 | 127 |
| SRR5029424 | 0 | 0 | 0 | 0 | 0 | 41 | 0 | 23 | 0 | 0 | 0 |
| SRR5029425 | 0 | 0 | 0 | 0 | 12 | 52 | 14 | 46 | 166 | 72 | 0 |
| SRR5029426 | 0 | 0 | 0 | 0 | 26 | 88 | 24 | 58 | 109 | 19 | 46 |
| SRR5029427 | 0 | 0 | 0 | 0 | 121 | 449 | 100 | 256 | 406 | 108 | 79 |
| SRR5029428 | 0 | 0 | 0 | 0 | 355 | 839 | 133 | 107 | 0 | 0 | 0 |
| SRR5029429 | 0 | 0 | 0 | 0 | 41 | 160 | 72 | 57 | 145 | 52 | 16 |
| SRR5029430 | 0 | 0 | 0 | 0 | 69 | 312 | 69 | 217 | 269 | 116 | 53 |
| SRR5029431 | 0 | 0 | 0 | 0 | 0 | 32 | 0 | 31 | 152 | 22 | 35 |
| SRR5029432 | 0 | 0 | 0 | 0 | 82 | 310 | 82 | 180 | 212 | 51 | 44 |
| SRR5029433 | 0 | 0 | 0 | 0 | 0 | 33 | 12 | 23 | 237 | 75 | 29 |
| SRR5029434 | 0 | 0 | 0 | 0 | 64 | 457 | 48 | 173 | 858 | 90 | 75 |
| SRR5029435 | 0 | 0 | 0 | 0 | 50 | 247 | 53 | 87 | 209 | 62 | 31 |
| SRR5029436 | 0 | 0 | 0 | 0 | 163 | 661 | 92 | 435 | 624 | 89 | 80 |
| SRR5029437 | 0 | 0 | 0 | 0 | 24 | 181 | 91 | 66 | 24 | 0 | 12 |
| SRR5029438 | 0 | 0 | 0 | 0 | 55 | 193 | 62 | 83 | 228 | 38 | 48 |
| SRR5029439 | 0 | 0 | 0 | 0 | 27 | 171 | 24 | 123 | 284 | 45 | 92 |
| SRR5029440 | 0 | 0 | 0 | 0 | 42 | 294 | 57 | 78 | 29 | 0 | 18 |
| SRR5029441 | 0 | 0 | 0 | 0 | 41 | 217 | 80 | 126 | 390 | 145 | 58 |
| SRR5029442 | 0 | 0 | 0 | 0 | 53 | 166 | 48 | 132 | 326 | 97 | 128 |
| SRR5029444 | 0 | 0 | 0 | 0 | 0 | 62 | 31 | 108 | 263 | 62 | 124 |
| SRR5029446 | 0 | 0 | 0 | 0 | 11 | 22 | 11 | 18 | 0 | 0 | 0 |
| SRR5029447 | 0 | 0 | 0 | 0 | 0 | 80 | 33 | 88 | 253 | 62 | 36 |
| SRR5029448 | 0 | 0 | 0 | 0 | 24 | 145 | 26 | 174 | 172 | 55 | 55 |
| SRR5029452 | 0 | 0 | 0 | 0 | 23 | 147 | 23 | 34 | 49 | 0 | 11 |
| SRR5029455 | 0 | 0 | 0 | 0 | 22 | 69 | 38 | 94 | 63 | 34 | 0 |
| SRR5029456 | 0 | 0 | 0 | 0 | 22 | 139 | 32 | 123 | 857 | 89 | 171 |
| SRR5029457 | 0 | 0 | 0 | 0 | 16 | 62 | 15 | 47 | 189 | 21 | 16 |
| SRR5029459 | 0 | 0 | 0 | 0 | 160 | 589 | 110 | 200 | 516 | 134 | 118 |
| SRR5029460 | 0 | 0 | 0 | 0 | 10 | 56 | 22 | 59 | 226 | 30 | 44 |
| SRR5029462 | 0 | 0 | 0 | 0 | 81 | 348 | 78 | 163 | 252 | 34 | 27 |
| SRR5029463 | 0 | 0 | 0 | 0 | 0 | 94 | 29 | 53 | 211 | 18 | 29 |
| SRR5029465 | 0 | 0 | 0 | 0 | 15 | 24 | 0 | 0 | 0 | 0 | 0 |
| SRR5029466 | 0 | 0 | 0 | 0 | 93 | 362 | 29 | 267 | 485 | 60 | 72 |
| SRR5029467 | 0 | 0 | 0 | 0 | 0 | 33 | 11 | 27 | 278 | 53 | 55 |
| SRR5029469 | 0 | 0 | 0 | 0 | 0 | 59 | 0 | 66 | 480 | 61 | 48 |
| SRR5029471 | 0 | 0 | 0 | 0 | 16 | 69 | 16 | 56 | 351 | 27 | 38 |
| SRR5029472 | 0 | 0 | 0 | 0 | 124 | 509 | 72 | 207 | 661 | 120 | 101 |
| SRR5029473 | 0 | 0 | 0 | 0 | 194 | 663 | 142 | 204 | 541 | 166 | 90 |
| SRR5029474 | 0 | 0 | 0 | 0 | 123 | 335 | 100 | 158 | 383 | 117 | 44 |
| SRR5029476 | 0 | 0 | 0 | 0 | 118 | 458 | 101 | 214 | 456 | 96 | 54 |
| SRR5029477 | 0 | 0 | 0 | 0 | 0 | 90 | 25 | 108 | 941 | 136 | 210 |
| SRR5029478 | 0 | 0 | 0 | 0 | 40 | 176 | 34 | 150 | 691 | 78 | 71 |
| SRR5029480 | 0 | 0 | 0 | 0 | 29 | 172 | 33 | 110 | 84 | 29 | 18 |
| SRR5029481 | 0 | 0 | 0 | 0 | 54 | 172 | 36 | 107 | 360 | 77 | 36 |
| SRR5029482 | 0 | 0 | 0 | 0 | 0 | 55 | 13 | 50 | 331 | 69 | 57 |
| SRR5029483 | 0 | 0 | 0 | 0 | 22 | 92 | 42 | 82 | 671 | 58 | 91 |
| SRR5029484 | 0 | 0 | 0 | 0 | 205 | 374 | 83 | 163 | 224 | 86 | 60 |
| SRR5029486 | 0 | 0 | 0 | 0 | 33 | 154 | 36 | 103 | 352 | 83 | 66 |
| SRR5029487 | 0 | 0 | 0 | 0 | 14 | 61 | 19 | 61 | 340 | 89 | 99 |
| SRR5029488 | 0 | 0 | 0 | 0 | 15 | 177 | 0 | 191 | 294 | 44 | 59 |
| SRR5029489 | 0 | 0 | 0 | 0 | 117 | 423 | 109 | 202 | 500 | 84 | 66 |
| SRR5029490 | 0 | 0 | 0 | 0 | 36 | 167 | 36 | 82 | 277 | 39 | 40 |
| SRR5029491 | 0 | 0 | 0 | 0 | 61 | 214 | 65 | 205 | 690 | 137 | 144 |
| SRR5029492 | 0 | 0 | 0 | 0 | 18 | 108 | 25 | 83 | 317 | 53 | 20 |
| SRR5029493 | 0 | 0 | 0 | 0 | 28 | 103 | 69 | 120 | 291 | 84 | 35 |
| SRR5029494 | 0 | 0 | 0 | 0 | 51 | 284 | 41 | 225 | 519 | 99 | 63 |
| SRR5029495 | 0 | 0 | 0 | 0 | 88 | 446 | 96 | 352 | 1137 | 88 | 106 |
| SRR5029496 | 0 | 0 | 0 | 0 | 26 | 110 | 16 | 106 | 485 | 83 | 33 |
| SRR5029497 | 0 | 0 | 0 | 0 | 39 | 196 | 26 | 218 | 534 | 122 | 109 |
| SRR5029498 | 0 | 0 | 0 | 0 | 52 | 308 | 74 | 284 | 1645 | 248 | 293 |
| SRR5029499 | 0 | 0 | 0 | 0 | 25 | 98 | 29 | 104 | 598 | 92 | 70 |

### Supp Table S8.5.Columns_ET_to_FD

| | | | | | | | | | | | |
|---|---|---|---|---|---|---|---|---|---|---|---|
| SRR5029500 | 0 | 0 | 0 | 0 | 11 | 106 | 21 | 138 | 540 | 116 | 85 |
| SRR5029501 | 0 | 0 | 0 | 0 | 17 | 49 | 13 | 44 | 357 | 75 | 98 |
| SRR5029502 | 0 | 0 | 0 | 0 | 89 | 334 | 22 | 267 | 456 | 211 | 67 |
| SRR5029503 | 0 | 0 | 0 | 0 | 21 | 120 | 32 | 148 | 473 | 81 | 74 |
| SRR5029504 | 0 | 0 | 0 | 0 | 24 | 137 | 35 | 131 | 919 | 137 | 97 |
| SRR5029505 | 0 | 0 | 0 | 0 | 20 | 106 | 41 | 135 | 510 | 129 | 42 |
| SRR5029506 | 0 | 0 | 0 | 0 | 184 | 455 | 137 | 214 | 372 | 97 | 57 |
| SRR5029507 | 0 | 0 | 0 | 0 | 43 | 344 | 71 | 166 | 354 | 57 | 83 |
| SRR5029508 | 0 | 0 | 0 | 0 | 29 | 320 | 53 | 227 | 1903 | 127 | 124 |
| SRR5029509 | 0 | 0 | 0 | 0 | 32 | 153 | 17 | 90 | 411 | 111 | 95 |
| SRR5029510 | 0 | 0 | 0 | 0 | 51 | 232 | 73 | 116 | 369 | 93 | 57 |
| SRR5029511 | 0 | 0 | 0 | 0 | 60 | 307 | 70 | 161 | 189 | 68 | 33 |
| SRR5029512 | 0 | 0 | 0 | 0 | 88 | 253 | 84 | 149 | 464 | 59 | 70 |
| SRR5029513 | 0 | 0 | 0 | 0 | 12 | 138 | 28 | 109 | 561 | 147 | 93 |
| SRR5029514 | 0 | 0 | 0 | 0 | 14 | 55 | 12 | 77 | 390 | 82 | 57 |
| SRR5029515 | 0 | 0 | 0 | 0 | 12 | 35 | 17 | 50 | 601 | 111 | 114 |
| SRR5029516 | 0 | 0 | 0 | 0 | 77 | 229 | 68 | 130 | 285 | 77 | 25 |
| SRR5029870 | 0 | 14 | 83 | 0 | 208 | 3884 | 569 | 8087 | 291 | 83 | 264 |
| SRR5029871 | 0 | 19 | 19 | 57 | 151 | 4044 | 661 | 8447 | 548 | 189 | 416 |
| SRR5029872 | 0 | 46 | 77 | 107 | 123 | 3828 | 567 | 8085 | 352 | 77 | 199 |
| SRR5029873 | 0 | 0 | 20 | 39 | 157 | 4013 | 590 | 8558 | 393 | 118 | 571 |
| SRR5029874 | 0 | 0 | 85 | 57 | 100 | 3441 | 597 | 7665 | 156 | 43 | 356 |
| SRR5029875 | 19 | 0 | 39 | 19 | 135 | 4043 | 481 | 8413 | 539 | 39 | 327 |
| SRR5029876 | 0 | 30 | 135 | 75 | 240 | 4106 | 719 | 7538 | 405 | 45 | 255 |
| SRR5029877 | 0 | 20 | 20 | 20 | 159 | 4246 | 655 | 8155 | 476 | 119 | 536 |
| SRR5029878 | 0 | 30 | 161 | 81 | 444 | 4894 | 616 | 8678 | 192 | 141 | 353 |
| SRR5029879 | 0 | 0 | 101 | 151 | 202 | 4342 | 555 | 8532 | 404 | 101 | 505 |
| SRR5029880 | 0 | 54 | 108 | 86 | 292 | 4449 | 842 | 7980 | 151 | 173 | 216 |
| SRR5029882 | 0 | 10 | 62 | 103 | 381 | 4669 | 638 | 7764 | 175 | 123 | 278 |
| SRR5029883 | 0 | 0 | 153 | 153 | 102 | 4937 | 662 | 7075 | 458 | 356 | 254 |
| SRR5029884 | 0 | 31 | 73 | 126 | 366 | 4968 | 847 | 8272 | 188 | 199 | 366 |
| SRR5029885 | 0 | 0 | 105 | 52 | 262 | 4349 | 838 | 7859 | 105 | 314 | 419 |
| SRR5029886 | 0 | 52 | 135 | 125 | 552 | 12905 | 1979 | 18268 | 167 | 73 | 177 |
| SRR5029887 | 0 | 18 | 74 | 0 | 525 | 10623 | 1795 | 17876 | 331 | 166 | 313 |
| SRR5029888 | 0 | 78 | 111 | 56 | 511 | 12468 | 1821 | 18374 | 200 | 89 | 211 |
| SRR5029889 | 0 | 19 | 48 | 0 | 420 | 10766 | 1844 | 17329 | 325 | 182 | 287 |
| SRR5029890 | 0 | 64 | 117 | 53 | 584 | 13250 | 2187 | 17529 | 202 | 127 | 297 |
| SRR5029891 | 0 | 0 | 19 | 28 | 378 | 10815 | 1625 | 17511 | 387 | 151 | 274 |
| SRR5029892 | 0 | 22 | 131 | 76 | 555 | 12400 | 2110 | 17947 | 152 | 98 | 261 |
| SRR5029893 | 0 | 38 | 48 | 38 | 548 | 10567 | 1844 | 17723 | 346 | 134 | 346 |
| SRR5029894 | 0 | 25 | 92 | 12 | 227 | 2654 | 270 | 3938 | 111 | 12 | 123 |
| SRR5029895 | 0 | 0 | 15 | 15 | 260 | 2701 | 367 | 5027 | 214 | 54 | 222 |
| SRR5029896 | 0 | 0 | 46 | 20 | 251 | 2547 | 357 | 4089 | 106 | 33 | 119 |
| SRR5029897 | 0 | 16 | 24 | 16 | 176 | 2606 | 264 | 4684 | 152 | 80 | 272 |
| SRR5029898 | 0 | 31 | 57 | 19 | 208 | 2749 | 264 | 3926 | 113 | 63 | 101 |
| SRR5029899 | 0 | 0 | 39 | 31 | 218 | 2630 | 164 | 4729 | 203 | 70 | 257 |
| SRR5029900 | 0 | 13 | 64 | 19 | 238 | 2484 | 290 | 3880 | 71 | 19 | 122 |
| SRR5029901 | 0 | 0 | 24 | 16 | 160 | 2505 | 319 | 4826 | 128 | 48 | 271 |
| SRR5029902 | 0 | 0 | 48 | 48 | 164 | 3263 | 198 | 4817 | 198 | 48 | 327 |
| SRR5029904 | 0 | 37 | 60 | 37 | 201 | 2949 | 320 | 4751 | 216 | 67 | 320 |
| SRR5029905 | 0 | 0 | 0 | 0 | 335 | 2788 | 335 | 4740 | 112 | 56 | 669 |
| SRR5029906 | 0 | 28 | 85 | 42 | 282 | 3423 | 352 | 4747 | 183 | 106 | 261 |
| SRR5029908 | 0 | 29 | 51 | 58 | 210 | 3046 | 420 | 4587 | 137 | 130 | 232 |
| SRR5029910 | 0 | 41 | 102 | 102 | 61 | 2847 | 574 | 11861 | 389 | 61 | 492 |
| SRR5029911 | 0 | 0 | 42 | 42 | 148 | 2755 | 466 | 14667 | 360 | 233 | 339 |
| SRR5029912 | 0 | 45 | 45 | 134 | 156 | 3029 | 468 | 13141 | 312 | 223 | 267 |
| SRR5029913 | 0 | 22 | 0 | 0 | 111 | 2972 | 577 | 13972 | 421 | 266 | 421 |
| SRR5029914 | 0 | 21 | 21 | 85 | 107 | 3182 | 406 | 12427 | 192 | 85 | 171 |
| SRR5029915 | 0 | 0 | 0 | 0 | 109 | 3352 | 675 | 14564 | 414 | 109 | 283 |
| SRR5029916 | 0 | 43 | 64 | 64 | 64 | 3201 | 580 | 12569 | 279 | 86 | 344 |
| SRR5029917 | 0 | 44 | 44 | 22 | 131 | 3192 | 612 | 14102 | 481 | 153 | 372 |
| SRR5029918 | 0 | 17 | 133 | 17 | 216 | 4487 | 349 | 7295 | 249 | 116 | 216 |
| SRR5029919 | 0 | 24 | 32 | 0 | 315 | 4116 | 444 | 8288 | 266 | 178 | 315 |
| SRR5029920 | 0 | 124 | 142 | 53 | 478 | 4924 | 319 | 7173 | 213 | 142 | 266 |
| SRR5029921 | 0 | 0 | 42 | 0 | 285 | 3979 | 453 | 8026 | 327 | 227 | 353 |
| SRR5029922 | 0 | 51 | 102 | 17 | 237 | 4645 | 542 | 8222 | 322 | 136 | 271 |
| SRR5029923 | 0 | 16 | 49 | 0 | 458 | 4349 | 392 | 7855 | 401 | 139 | 311 |
| SRR5029924 | 0 | 17 | 87 | 17 | 399 | 5071 | 556 | 7224 | 191 | 52 | 208 |
| SRR5029925 | 0 | 0 | 25 | 0 | 276 | 4373 | 435 | 8395 | 259 | 159 | 318 |
| SRR5029926 | 0 | 0 | 0 | 24 | 63 | 1237 | 111 | 2388 | 167 | 71 | 167 |
| SRR5029927 | 0 | 0 | 28 | 0 | 56 | 1471 | 183 | 3287 | 239 | 106 | 178 |
| SRR5029928 | 0 | 0 | 34 | 17 | 60 | 1300 | 120 | 2246 | 138 | 0 | 86 |
| SRR5029929 | 0 | 0 | 0 | 0 | 75 | 1331 | 197 | 3009 | 266 | 81 | 203 |
| SRR5029930 | 0 | 24 | 49 | 0 | 65 | 1201 | 203 | 2426 | 81 | 41 | 130 |
| SRR5029931 | 0 | 0 | 0 | 0 | 101 | 1275 | 326 | 3168 | 286 | 107 | 247 |
| SRR5029932 | 0 | 0 | 42 | 33 | 109 | 1136 | 159 | 2346 | 200 | 42 | 150 |
| SRR5029933 | 0 | 0 | 0 | 0 | 98 | 1371 | 226 | 3009 | 318 | 75 | 284 |
| SRR5029934 | 0 | 0 | 53 | 40 | 212 | 4464 | 543 | 7696 | 212 | 66 | 119 |
| SRR5029935 | 0 | 15 | 30 | 15 | 180 | 3082 | 389 | 7212 | 165 | 60 | 150 |
| SRR5029936 | 0 | 0 | 129 | 58 | 244 | 4212 | 460 | 7273 | 316 | 58 | 129 |
| SRR5029937 | 0 | 16 | 16 | 31 | 125 | 3367 | 485 | 6733 | 313 | 63 | 188 |
| SRR5029938 | 0 | 14 | 27 | 0 | 314 | 4602 | 382 | 7838 | 150 | 109 | 178 |
| SRR5029939 | 0 | 15 | 15 | 15 | 107 | 3621 | 520 | 7396 | 168 | 61 | 214 |
| SRR5029940 | 0 | 56 | 42 | 0 | 211 | 4537 | 534 | 7992 | 309 | 56 | 126 |
| SRR5029941 | 0 | 16 | 0 | 0 | 156 | 3582 | 438 | 6585 | 360 | 16 | 156 |
| SRR5029942 | 0 | 28 | 98 | 70 | 14 | 799 | 98 | 1864 | 280 | 140 | 98 |
| SRR5029943 | 0 | 0 | 28 | 0 | 44 | 694 | 116 | 2004 | 314 | 275 | 297 |
| SRR5029944 | 15 | 0 | 77 | 31 | 46 | 944 | 93 | 2090 | 186 | 139 | 62 |
| SRR5029945 | 0 | 0 | 0 | 34 | 46 | 667 | 109 | 1920 | 282 | 270 | 328 |
| SRR5029946 | 0 | 29 | 102 | 102 | 58 | 902 | 73 | 2153 | 233 | 160 | 73 |
| SRR5029947 | 0 | 0 | 17 | 11 | 50 | 757 | 67 | 1866 | 407 | 200 | 312 |
| SRR5029948 | 0 | 30 | 61 | 45 | 45 | 698 | 45 | 1592 | 303 | 152 | 152 |
| SRR5029949 | 0 | 0 | 17 | 17 | 58 | 656 | 63 | 2176 | 449 | 201 | 242 |
| SRR5029950 | 0 | 28 | 107 | 121 | 89 | 2657 | 312 | 6531 | 415 | 61 | 210 |
| SRR5029951 | 0 | 0 | 85 | 0 | 0 | 2393 | 555 | 6708 | 427 | 0 | 214 |
| SRR5029952 | 0 | 20 | 200 | 90 | 70 | 2965 | 351 | 6165 | 361 | 70 | 265 |
| SRR5029953 | 0 | 0 | 44 | 0 | 44 | 3216 | 308 | 6079 | 308 | 132 | 220 |
| SRR5029954 | 0 | 53 | 91 | 91 | 81 | 3074 | 263 | 5985 | 345 | 81 | 220 |
| SRR5029955 | 0 | 0 | 131 | 87 | 87 | 2354 | 654 | 6191 | 567 | 131 | 262 |
| SRR5029956 | 0 | 20 | 146 | 156 | 68 | 2758 | 322 | 6044 | 405 | 78 | 244 |
| SRR5029957 | 0 | 0 | 88 | 88 | 44 | 2826 | 221 | 5388 | 530 | 44 | 132 |
| SRR5029958 | 0 | 44 | 103 | 44 | 471 | 8070 | 913 | 11590 | 236 | 74 | 147 |
| SRR5029959 | 0 | 0 | 41 | 16 | 319 | 5850 | 743 | 11022 | 376 | 188 | 212 |
| SRR5029960 | 0 | 63 | 158 | 111 | 521 | 7388 | 789 | 11256 | 537 | 63 | 47 |
| SRR5029961 | 0 | 0 | 0 | 0 | 314 | 6304 | 763 | 11363 | 305 | 119 | 161 |
| SRR5029962 | 0 | 60 | 210 | 60 | 540 | 8236 | 870 | 11476 | 345 | 45 | 75 |
| SRR5029963 | 0 | 0 | 33 | 17 | 224 | 6102 | 739 | 11672 | 357 | 116 | 174 |
| SRR5029964 | 0 | 46 | 185 | 46 | 432 | 7903 | 772 | 12687 | 170 | 123 | 93 |
| SRR5029965 | 0 | 25 | 17 | 25 | 305 | 6043 | 797 | 11526 | 364 | 136 | 102 |
| SRR5029966 | 0 | 53 | 276 | 202 | 223 | 4986 | 976 | 13781 | 329 | 42 | 117 |
| SRR5029967 | 0 | 17 | 33 | 44 | 160 | 4699 | 893 | 14775 | 458 | 171 | 320 |
| SRR5029968 | 0 | 35 | 257 | 163 | 222 | 5624 | 910 | 14654 | 245 | 93 | 105 |
| SRR5029969 | 0 | 0 | 41 | 17 | 186 | 4688 | 946 | 15138 | 383 | 116 | 209 |
| SRR5029970 | 0 | 44 | 309 | 176 | 287 | 5379 | 1069 | 14978 | 242 | 88 | 132 |
| SRR5029971 | 0 | 0 | 68 | 23 | 141 | 4615 | 904 | 15200 | 384 | 119 | 249 |
| SRR5029972 | 0 | 57 | 318 | 181 | 250 | 5875 | 1009 | 14346 | 193 | 91 | 136 |
| SRR5029973 | 0 | 0 | 64 | 41 | 210 | 4640 | 875 | 15332 | 438 | 158 | 268 |
| SRR5029974 | 0 | 16 | 66 | 41 | 280 | 3406 | 543 | 5998 | 107 | 49 | 66 |
| SRR5029975 | 0 | 0 | 0 | 0 | 125 | 2586 | 446 | 5576 | 137 | 77 | 113 |

### Supp Table S8.5.Columns_ET_to_FD

| | | | | | | | | | | | |
|---|---|---|---|---|---|---|---|---|---|---|---|
| SRR5029976 | 0 | 18 | 54 | 36 | 180 | 3928 | 658 | 6280 | 108 | 18 | 45 |
| SRR5029977 | 0 | 0 | 12 | 0 | 136 | 2874 | 421 | 5865 | 211 | 74 | 87 |
| SRR5029978 | 0 | 17 | 60 | 52 | 233 | 3462 | 611 | 5899 | 112 | 112 | 103 |
| SRR5029979 | 0 | 12 | 0 | 0 | 165 | 2876 | 447 | 5734 | 196 | 80 | 110 |
| SRR5029980 | 0 | 44 | 78 | 26 | 226 | 4102 | 496 | 6785 | 113 | 78 | 96 |
| SRR5029981 | 0 | 0 | 19 | 0 | 149 | 3036 | 584 | 5656 | 137 | 99 | 124 |
| SRR5029982 | 0 | 0 | 55 | 55 | 124 | 3453 | 511 | 5636 | 138 | 124 | 249 |
| SRR5029983 | 0 | 0 | 0 | 0 | 94 | 3835 | 374 | 5332 | 187 | 187 | 281 |
| SRR5029984 | 0 | 15 | 30 | 45 | 120 | 2755 | 434 | 5406 | 240 | 120 | 225 |
| SRR5029986 | 0 | 0 | 28 | 71 | 198 | 2760 | 396 | 5860 | 283 | 85 | 241 |
| SRR5029988 | 0 | 0 | 44 | 29 | 131 | 3126 | 424 | 5681 | 278 | 146 | 161 |
| SRR5029990 | 0 | 44 | 208 | 102 | 91 | 1368 | 121 | 2092 | 282 | 102 | 150 |
| SRR5029991 | 0 | 0 | 85 | 50 | 70 | 1008 | 55 | 2116 | 339 | 130 | 245 |
| SRR5029992 | 0 | 32 | 212 | 124 | 60 | 1438 | 116 | 2127 | 264 | 148 | 132 |
| SRR5029993 | 0 | 0 | 58 | 21 | 63 | 1024 | 79 | 2043 | 326 | 194 | 226 |
| SRR5029994 | 0 | 60 | 177 | 151 | 83 | 1318 | 94 | 2189 | 260 | 132 | 143 |
| SRR5029995 | 0 | 15 | 66 | 41 | 81 | 1194 | 112 | 2042 | 335 | 168 | 223 |
| SRR5029996 | 0 | 43 | 253 | 171 | 86 | 1262 | 105 | 2251 | 257 | 132 | 101 |
| SRR5029997 | 0 | 21 | 58 | 31 | 37 | 1125 | 84 | 1936 | 324 | 157 | 225 |
| SRR5029998 | 0 | 18 | 168 | 44 | 319 | 2021 | 186 | 2854 | 186 | 44 | 80 |
| SRR5029999 | 0 | 0 | 37 | 16 | 235 | 2067 | 235 | 3065 | 283 | 96 | 139 |
| SRR5030000 | 0 | 0 | 170 | 38 | 330 | 2263 | 198 | 2810 | 170 | 132 | 160 |
| SRR5030001 | 0 | 0 | 33 | 0 | 223 | 1683 | 268 | 3255 | 240 | 117 | 212 |
| SRR5030002 | 0 | 18 | 99 | 72 | 271 | 2212 | 235 | 3070 | 235 | 27 | 117 |
| SRR5030003 | 0 | 22 | 44 | 33 | 234 | 1758 | 250 | 3048 | 234 | 114 | 169 |
| SRR5030004 | 0 | 27 | 128 | 82 | 192 | 2454 | 357 | 3086 | 247 | 119 | 64 |
| SRR5030005 | 0 | 0 | 67 | 17 | 194 | 1937 | 333 | 3181 | 272 | 122 | 155 |
| SRR5030006 | 0 | 33 | 25 | 42 | 234 | 8066 | 1237 | 16750 | 493 | 167 | 318 |
| SRR5030008 | 0 | 0 | 99 | 45 | 198 | 7459 | 1198 | 16585 | 423 | 234 | 369 |
| SRR5030009 | 0 | 0 | 0 | 0 | 236 | 6363 | 1100 | 16184 | 786 | 314 | 628 |
| SRR5030010 | 0 | 0 | 77 | 69 | 249 | 7562 | 1298 | 16524 | 421 | 284 | 361 |
| SRR5030011 | 0 | 0 | 0 | 232 | 155 | 6897 | 852 | 15885 | 930 | 77 | 542 |
| SRR5030012 | 0 | 0 | 88 | 26 | 263 | 7549 | 1403 | 16475 | 386 | 289 | 473 |
| SRR5030014 | 14 | 28 | 110 | 152 | 303 | 4767 | 730 | 5125 | 207 | 83 | 55 |
| SRR5030015 | 0 | 0 | 27 | 27 | 150 | 4071 | 533 | 4673 | 492 | 178 | 150 |
| SRR5030016 | 0 | 0 | 137 | 106 | 456 | 4893 | 547 | 5045 | 137 | 106 | 91 |
| SRR5030017 | 0 | 29 | 43 | 0 | 272 | 4295 | 487 | 4939 | 215 | 186 | 129 |
| SRR5030018 | 0 | 0 | 201 | 86 | 272 | 4684 | 731 | 4885 | 258 | 57 | 57 |
| SRR5030019 | 0 | 14 | 14 | 14 | 209 | 3912 | 473 | 5276 | 404 | 153 | 167 |
| SRR5030020 | 0 | 15 | 206 | 103 | 295 | 4863 | 634 | 5217 | 192 | 88 | 147 |
| SRR5030021 | 0 | 0 | 29 | 57 | 216 | 4066 | 589 | 5187 | 244 | 187 | 216 |
| SRR5030022 | 0 | 0 | 36 | 36 | 97 | 799 | 109 | 1501 | 194 | 109 | 194 |
| SRR5030023 | 0 | 0 | 22 | 22 | 179 | 627 | 134 | 1499 | 134 | 90 | 336 |
| SRR5030024 | 0 | 13 | 65 | 26 | 39 | 751 | 104 | 1360 | 130 | 65 | 207 |
| SRR5030025 | 0 | 0 | 23 | 0 | 23 | 930 | 47 | 1628 | 140 | 70 | 302 |
| SRR5030026 | 0 | 12 | 87 | 50 | 99 | 758 | 87 | 1343 | 199 | 124 | 224 |
| SRR5030027 | 0 | 23 | 23 | 0 | 90 | 810 | 158 | 1620 | 203 | 68 | 158 |
| SRR5030028 | 0 | 25 | 38 | 25 | 101 | 896 | 114 | 1312 | 202 | 114 | 240 |
| SRR5030030 | 0 | 11 | 124 | 90 | 112 | 4848 | 416 | 5883 | 202 | 67 | 360 |
| SRR5030031 | 0 | 0 | 12 | 23 | 127 | 3613 | 475 | 5836 | 232 | 185 | 405 |
| SRR5030032 | 0 | 0 | 121 | 12 | 230 | 4796 | 521 | 5704 | 157 | 85 | 327 |
| SRR5030033 | 0 | 0 | 24 | 0 | 60 | 3820 | 301 | 6121 | 289 | 205 | 422 |
| SRR5030034 | 0 | 58 | 58 | 35 | 196 | 4872 | 587 | 6657 | 207 | 58 | 265 |
| SRR5030035 | 0 | 0 | 24 | 0 | 94 | 3671 | 295 | 5583 | 319 | 153 | 401 |
| SRR5030036 | 0 | 12 | 82 | 59 | 176 | 5033 | 447 | 5280 | 306 | 47 | 318 |
| SRR5030038 | 0 | 0 | 76 | 51 | 203 | 4825 | 775 | 14323 | 394 | 140 | 368 |
| SRR5030039 | 0 | 0 | 141 | 0 | 141 | 4092 | 611 | 14581 | 635 | 235 | 588 |
| SRR5030040 | 0 | 0 | 148 | 13 | 188 | 4351 | 698 | 13551 | 322 | 175 | 336 |
| SRR5030041 | 0 | 0 | 97 | 24 | 170 | 4041 | 803 | 15727 | 463 | 122 | 463 |
| SRR5030042 | 0 | 26 | 116 | 52 | 259 | 4797 | 595 | 13913 | 388 | 194 | 375 |
| SRR5030043 | 0 | 0 | 24 | 71 | 238 | 4575 | 739 | 14749 | 548 | 191 | 381 |
| SRR5030044 | 0 | 0 | 92 | 105 | 237 | 4722 | 671 | 14770 | 263 | 132 | 447 |
| SRR5030045 | 0 | 0 | 0 | 0 | 195 | 4585 | 561 | 14510 | 536 | 293 | 415 |
| SRR5030046 | 0 | 45 | 141 | 119 | 111 | 2823 | 208 | 5705 | 141 | 74 | 119 |
| SRR5030048 | 0 | 0 | 105 | 80 | 137 | 2623 | 193 | 5519 | 185 | 72 | 24 |
| SRR5030050 | 0 | 15 | 69 | 122 | 107 | 2774 | 251 | 5548 | 114 | 30 | 91 |
| SRR5030052 | 0 | 24 | 165 | 87 | 181 | 2929 | 276 | 5543 | 134 | 71 | 31 |
| SRR5030053 | 0 | 0 | 0 | 46 | 46 | 2922 | 137 | 5341 | 137 | 137 | 91 |
| SRR5030054 | 0 | 29 | 117 | 87 | 539 | 6510 | 714 | 10514 | 772 | 262 | 583 |
| SRR5030055 | 0 | 48 | 48 | 48 | 433 | 4999 | 721 | 9326 | 481 | 192 | 865 |
| SRR5030056 | 0 | 0 | 157 | 31 | 314 | 6711 | 674 | 10568 | 737 | 188 | 564 |
| SRR5030057 | 0 | 0 | 51 | 0 | 204 | 6275 | 969 | 10408 | 918 | 153 | 867 |
| SRR5030058 | 0 | 90 | 135 | 60 | 420 | 6961 | 840 | 9857 | 420 | 195 | 630 |
| SRR5030059 | 0 | 0 | 49 | 148 | 295 | 6200 | 837 | 11465 | 837 | 394 | 935 |
| SRR5030060 | 0 | 46 | 31 | 77 | 430 | 6468 | 845 | 10047 | 645 | 215 | 538 |
| SRR5030061 | 0 | 0 | 50 | 0 | 302 | 6341 | 1006 | 10719 | 856 | 252 | 956 |
| SRR5030062 | 0 | 13 | 78 | 26 | 65 | 688 | 104 | 2999 | 909 | 208 | 325 |
| SRR5030064 | 0 | 14 | 14 | 42 | 70 | 683 | 56 | 2859 | 823 | 209 | 432 |
| SRR5030066 | 0 | 27 | 40 | 13 | 94 | 936 | 67 | 2994 | 749 | 254 | 361 |
| SRR5030068 | 0 | 0 | 81 | 27 | 108 | 753 | 135 | 3188 | 942 | 229 | 309 |
| SRR5030069 | 0 | 0 | 0 | 0 | 250 | 563 | 125 | 2000 | 563 | 250 | 563 |
| SRR5030070 | 0 | 0 | 50 | 149 | 129 | 2188 | 259 | 1999 | 139 | 60 | 199 |
| SRR5030071 | 0 | 0 | 0 | 21 | 185 | 2178 | 288 | 2404 | 247 | 82 | 123 |
| SRR5030072 | 0 | 11 | 54 | 43 | 193 | 2116 | 290 | 2008 | 129 | 86 | 161 |
| SRR5030073 | 0 | 0 | 0 | 21 | 170 | 1870 | 340 | 2422 | 191 | 85 | 340 |
| SRR5030074 | 0 | 10 | 61 | 31 | 204 | 2033 | 358 | 2023 | 123 | 92 | 143 |
| SRR5030076 | 0 | 31 | 83 | 21 | 198 | 2282 | 417 | 2282 | 104 | 63 | 271 |
| SRR5030077 | 0 | 0 | 21 | 0 | 149 | 2376 | 170 | 2185 | 127 | 106 | 233 |
| SRR5030078 | 0 | 42 | 116 | 32 | 180 | 1491 | 63 | 2327 | 264 | 201 | 106 |
| SRR5030079 | 0 | 17 | 0 | 17 | 17 | 1271 | 172 | 2146 | 326 | 137 | 137 |
| SRR5030080 | 0 | 0 | 80 | 114 | 126 | 1431 | 114 | 2472 | 275 | 149 | 126 |
| SRR5030081 | 0 | 18 | 18 | 18 | 89 | 1466 | 125 | 2414 | 340 | 250 | 161 |
| SRR5030082 | 0 | 22 | 66 | 98 | 55 | 1572 | 131 | 2271 | 186 | 131 | 109 |
| SRR5030083 | 0 | 35 | 53 | 35 | 70 | 1002 | 70 | 2303 | 387 | 229 | 123 |
| SRR5030084 | 0 | 0 | 90 | 34 | 168 | 1817 | 168 | 2232 | 404 | 157 | 123 |
| SRR5030085 | 0 | 0 | 36 | 0 | 53 | 1371 | 125 | 2314 | 356 | 196 | 160 |
| SRR5030086 | 0 | 0 | 20 | 15 | 59 | 931 | 124 | 1837 | 59 | 25 | 114 |
| SRR5030087 | 0 | 0 | 17 | 0 | 119 | 1048 | 153 | 2232 | 85 | 0 | 204 |
| SRR5030088 | 0 | 0 | 21 | 16 | 63 | 853 | 94 | 1815 | 115 | 58 | 84 |
| SRR5030090 | 0 | 0 | 25 | 20 | 55 | 917 | 85 | 1810 | 70 | 15 | 85 |
| SRR5030091 | 0 | 0 | 0 | 26 | 103 | 964 | 112 | 2204 | 103 | 17 | 138 |
| SRR5030092 | 0 | 0 | 0 | 41 | 81 | 829 | 137 | 1811 | 97 | 56 | 127 |
| SRR5030093 | 0 | 0 | 0 | 0 | 88 | 866 | 124 | 2554 | 150 | 62 | 115 |
| SRR5030094 | 0 | 0 | 20 | 0 | 12 | 296 | 45 | 1246 | 69 | 15 | 11 |
| SRR5030095 | 0 | 0 | 0 | 0 | 21 | 329 | 41 | 1254 | 144 | 0 | 82 |
| SRR5030096 | 0 | 0 | 18 | 0 | 18 | 352 | 48 | 1193 | 60 | 15 | 15 |
| SRR5030097 | 0 | 0 | 64 | 0 | 43 | 447 | 64 | 1597 | 106 | 21 | 0 |
| SRR5030098 | 0 | 0 | 16 | 0 | 27 | 331 | 63 | 1254 | 58 | 22 | 19 |
| SRR5030100 | 0 | 0 | 21 | 0 | 19 | 336 | 49 | 1318 | 31 | 0 | 11 |
| SRR5030101 | 0 | 0 | 0 | 0 | 85 | 426 | 64 | 1577 | 21 | 0 | 64 |
| SRR5030102 | 15 | 46 | 131 | 62 | 193 | 4477 | 624 | 9338 | 200 | 69 | 108 |
| SRR5030103 | 0 | 0 | 28 | 0 | 235 | 3716 | 488 | 9844 | 206 | 160 | 178 |
| SRR5030104 | 0 | 0 | 193 | 34 | 159 | 4479 | 646 | 9436 | 226 | 75 | 176 |
| SRR5030105 | 0 | 0 | 59 | 29 | 156 | 3822 | 585 | 9613 | 254 | 146 | 234 |
| SRR5030106 | 0 | 32 | 119 | 40 | 206 | 4061 | 603 | 9303 | 143 | 79 | 190 |
| SRR5030107 | 0 | 19 | 76 | 29 | 133 | 3716 | 523 | 10084 | 314 | 152 | 228 |
| SRR5030108 | 0 | 16 | 156 | 66 | 180 | 4483 | 647 | 9580 | 246 | 139 | 139 |
| SRR5030109 | 0 | 20 | 98 | 20 | 137 | 4057 | 441 | 9300 | 265 | 59 | 216 |
| SRR5030110 | 0 | 30 | 90 | 20 | 169 | 2417 | 318 | 6425 | 298 | 90 | 298 |

### Supp Table S8.5.Columns_ET_to_FD

| | | | | | | | | | | | |
|---|---|---|---|---|---|---|---|---|---|---|---|
| SRR5030111 | 0 | 0 | 22 | 0 | 106 | 2072 | 329 | 6228 | 557 | 134 | 474 |
| SRR5030112 | 0 | 21 | 128 | 64 | 223 | 2149 | 287 | 6650 | 404 | 138 | 234 |
| SRR5030113 | 0 | 12 | 0 | 0 | 121 | 1857 | 317 | 6363 | 519 | 104 | 375 |
| SRR5030114 | 0 | 10 | 71 | 61 | 143 | 2739 | 316 | 6680 | 377 | 81 | 153 |
| SRR5030115 | 0 | 0 | 28 | 0 | 85 | 2104 | 339 | 6415 | 662 | 85 | 328 |
| SRR5030116 | 10 | 41 | 73 | 83 | 114 | 2375 | 322 | 6680 | 415 | 124 | 176 |
| SRR5030117 | 0 | 0 | 40 | 0 | 132 | 2249 | 287 | 6340 | 551 | 138 | 373 |
| SRR5030118 | 0 | 0 | 17 | 14 | 83 | 1090 | 111 | 2957 | 73 | 69 | 52 |
| SRR5030119 | 0 | 0 | 13 | 16 | 73 | 1151 | 193 | 3479 | 121 | 70 | 127 |
| SRR5030120 | 0 | 0 | 26 | 15 | 45 | 1187 | 182 | 3056 | 100 | 26 | 74 |
| SRR5030121 | 0 | 0 | 0 | 0 | 73 | 1049 | 185 | 3524 | 89 | 40 | 103 |
| SRR5030122 | 0 | 18 | 43 | 14 | 86 | 1209 | 189 | 3149 | 93 | 50 | 82 |
| SRR5030123 | 0 | 0 | 13 | 0 | 80 | 1062 | 199 | 3632 | 97 | 48 | 68 |
| SRR5030124 | 0 | 0 | 25 | 36 | 73 | 1044 | 185 | 3121 | 87 | 18 | 83 |
| SRR5030125 | 0 | 0 | 13 | 16 | 66 | 1092 | 197 | 3503 | 95 | 49 | 98 |
| SRR5030126 | 0 | 0 | 0 | 15 | 88 | 1616 | 191 | 3916 | 220 | 96 | 73 |
| SRR5030127 | 0 | 0 | 14 | 0 | 117 | 1944 | 310 | 5431 | 234 | 179 | 159 |
| SRR5030128 | 0 | 0 | 31 | 0 | 70 | 1675 | 235 | 3946 | 164 | 63 | 94 |
| SRR5030130 | 0 | 15 | 45 | 15 | 90 | 1671 | 187 | 3544 | 157 | 120 | 75 |
| SRR5030131 | 0 | 0 | 0 | 0 | 91 | 1936 | 274 | 5176 | 323 | 154 | 281 |
| SRR5030132 | 0 | 0 | 31 | 15 | 46 | 1658 | 123 | 3593 | 92 | 69 | 131 |
| SRR5030133 | 0 | 0 | 0 | 0 | 122 | 2142 | 323 | 4996 | 237 | 122 | 173 |
| SRR5030134 | 0 | 0 | 12 | 0 | 38 | 832 | 71 | 1865 | 30 | 0 | 18 |
| SRR5030135 | 0 | 0 | 0 | 0 | 41 | 840 | 79 | 2012 | 72 | 0 | 38 |
| SRR5030136 | 0 | 0 | 15 | 0 | 56 | 747 | 73 | 1896 | 26 | 0 | 23 |
| SRR5030137 | 0 | 0 | 0 | 0 | 57 | 817 | 71 | 2126 | 67 | 18 | 25 |
| SRR5030138 | 0 | 0 | 12 | 0 | 49 | 827 | 94 | 1895 | 29 | 0 | 15 |
| SRR5030139 | 0 | 0 | 0 | 0 | 87 | 748 | 55 | 2037 | 45 | 28 | 24 |
| SRR5030140 | 0 | 0 | 16 | 0 | 47 | 873 | 77 | 1856 | 20 | 0 | 17 |
| SRR5030141 | 0 | 0 | 0 | 0 | 53 | 786 | 92 | 2175 | 64 | 11 | 21 |
| SRR5030142 | 0 | 21 | 45 | 10 | 90 | 2017 | 372 | 4904 | 125 | 17 | 21 |
| SRR5030143 | 0 | 0 | 60 | 60 | 150 | 1830 | 210 | 5639 | 90 | 30 | 30 |
| SRR5030144 | 0 | 11 | 46 | 23 | 69 | 1907 | 325 | 4885 | 80 | 31 | 19 |
| SRR5030145 | 0 | 32 | 0 | 0 | 64 | 1882 | 319 | 4434 | 32 | 0 | 64 |
| SRR5030146 | 0 | 0 | 47 | 43 | 57 | 1903 | 344 | 4821 | 129 | 18 | 29 |
| SRR5030147 | 0 | 0 | 31 | 0 | 124 | 1804 | 311 | 4448 | 124 | 31 | 62 |
| SRR5030148 | 0 | 22 | 74 | 30 | 108 | 1936 | 408 | 5201 | 111 | 41 | 37 |
| SRR5030149 | 0 | 0 | 63 | 31 | 126 | 1446 | 566 | 5185 | 189 | 31 | 63 |
| SRR5030150 | 0 | 0 | 14 | 0 | 48 | 1026 | 146 | 2268 | 89 | 23 | 30 |
| SRR5030151 | 0 | 0 | 20 | 0 | 20 | 1143 | 197 | 2128 | 79 | 0 | 59 |
| SRR5030152 | 0 | 0 | 22 | 0 | 67 | 1021 | 141 | 2233 | 67 | 22 | 30 |
| SRR5030153 | 0 | 0 | 41 | 20 | 41 | 963 | 164 | 2847 | 20 | 0 | 102 |
| SRR5030154 | 0 | 0 | 24 | 14 | 40 | 1035 | 129 | 2267 | 66 | 28 | 52 |
| SRR5030155 | 0 | 0 | 20 | 0 | 81 | 1197 | 101 | 2476 | 61 | 20 | 41 |
| SRR5030156 | 0 | 0 | 17 | 0 | 75 | 1061 | 142 | 2303 | 70 | 41 | 43 |
| SRR5030157 | 0 | 21 | 0 | 21 | 41 | 1512 | 145 | 2279 | 83 | 0 | 41 |
| SRR5030158 | 0 | 16 | 49 | 21 | 165 | 3024 | 346 | 5463 | 144 | 37 | 95 |
| SRR5030159 | 0 | 0 | 0 | 23 | 140 | 2372 | 210 | 5841 | 129 | 58 | 181 |
| SRR5030160 | 0 | 0 | 49 | 18 | 208 | 2604 | 305 | 5596 | 137 | 35 | 137 |
| SRR5030161 | 0 | 0 | 24 | 24 | 104 | 2247 | 269 | 5964 | 190 | 61 | 153 |
| SRR5030162 | 0 | 29 | 84 | 29 | 185 | 2829 | 324 | 5843 | 135 | 46 | 130 |
| SRR5030163 | 0 | 0 | 30 | 18 | 154 | 2553 | 326 | 5521 | 178 | 41 | 142 |
| SRR5030164 | 0 | 34 | 60 | 30 | 176 | 2568 | 301 | 5837 | 142 | 22 | 116 |
| SRR5030165 | 0 | 0 | 0 | 12 | 146 | 2179 | 323 | 5602 | 165 | 73 | 177 |
| SRR5030166 | 0 | 26 | 91 | 65 | 456 | 9438 | 963 | 12862 | 143 | 13 | 117 |
| SRR5030167 | 0 | 0 | 29 | 0 | 391 | 8915 | 916 | 11883 | 210 | 67 | 153 |
| SRR5030168 | 0 | 71 | 127 | 14 | 566 | 9704 | 891 | 11670 | 141 | 85 | 99 |
| SRR5030169 | 0 | 10 | 30 | 30 | 381 | 8714 | 782 | 11823 | 160 | 120 | 191 |
| SRR5030170 | 0 | 27 | 108 | 27 | 552 | 9885 | 1010 | 11528 | 148 | 148 | 108 |
| SRR5030171 | 0 | 0 | 19 | 0 | 429 | 8992 | 740 | 11729 | 195 | 107 | 244 |
| SRR5030172 | 0 | 14 | 69 | 83 | 513 | 9589 | 958 | 11393 | 180 | 83 | 56 |
| SRR5030173 | 0 | 0 | 0 | 30 | 296 | 8269 | 918 | 11515 | 227 | 128 | 247 |
| SRR5030174 | 0 | 0 | 0 | 0 | 47 | 499 | 48 | 1207 | 42 | 11 | 17 |
| SRR5030175 | 0 | 0 | 0 | 0 | 62 | 494 | 66 | 1250 | 74 | 28 | 19 |
| SRR5030176 | 0 | 0 | 0 | 0 | 37 | 498 | 58 | 1086 | 47 | 0 | 21 |
| SRR5030177 | 0 | 0 | 0 | 0 | 37 | 469 | 52 | 1313 | 59 | 15 | 0 |
| SRR5030178 | 0 | 0 | 0 | 0 | 46 | 514 | 68 | 1162 | 35 | 11 | 15 |
| SRR5030179 | 0 | 0 | 0 | 0 | 29 | 518 | 72 | 1283 | 55 | 22 | 36 |
| SRR5030180 | 0 | 0 | 0 | 0 | 39 | 521 | 63 | 1148 | 33 | 16 | 13 |
| SRR5030181 | 0 | 0 | 0 | 0 | 34 | 553 | 86 | 1255 | 61 | 12 | 17 |
| SRR5030182 | 0 | 27 | 96 | 82 | 384 | 4818 | 645 | 6946 | 233 | 110 | 261 |
| SRR5030183 | 0 | 0 | 12 | 46 | 161 | 4573 | 587 | 7245 | 380 | 127 | 633 |
| SRR5030184 | 0 | 30 | 134 | 134 | 298 | 4873 | 730 | 6319 | 268 | 119 | 417 |
| SRR5030185 | 0 | 0 | 48 | 12 | 275 | 5055 | 526 | 7134 | 370 | 84 | 693 |
| SRR5030186 | 0 | 57 | 85 | 99 | 255 | 5285 | 652 | 7184 | 326 | 85 | 397 |
| SRR5030187 | 0 | 12 | 47 | 23 | 186 | 4732 | 466 | 7214 | 420 | 175 | 618 |
| SRR5030188 | 0 | 29 | 87 | 58 | 305 | 4815 | 626 | 6983 | 407 | 116 | 305 |
| SRR5030189 | 0 | 0 | 48 | 12 | 179 | 4740 | 669 | 6924 | 466 | 215 | 573 |
| SRR5030190 | 0 | 0 | 16 | 0 | 56 | 489 | 56 | 590 | 26 | 16 | 14 |
| SRR5030191 | 0 | 0 | 0 | 0 | 89 | 558 | 18 | 823 | 27 | 0 | 18 |
| SRR5030192 | 0 | 0 | 15 | 0 | 40 | 496 | 52 | 615 | 25 | 12 | 20 |
| SRR5030193 | 0 | 0 | 0 | 0 | 55 | 605 | 64 | 734 | 18 | 0 | 0 |
| SRR5030194 | 0 | 0 | 0 | 0 | 60 | 484 | 36 | 682 | 17 | 17 | 19 |
| SRR5030196 | 0 | 0 | 0 | 0 | 56 | 537 | 41 | 637 | 27 | 12 | 15 |
| SRR5030197 | 0 | 0 | 0 | 0 | 101 | 541 | 28 | 752 | 28 | 0 | 0 |
| SRR5030198 | 0 | 0 | 18 | 0 | 123 | 2554 | 296 | 3078 | 146 | 59 | 96 |
| SRR5030199 | 0 | 0 | 22 | 0 | 153 | 2188 | 240 | 3365 | 160 | 124 | 65 |
| SRR5030200 | 0 | 15 | 0 | 0 | 201 | 2432 | 318 | 2965 | 108 | 64 | 88 |
| SRR5030201 | 0 | 0 | 15 | 0 | 144 | 2216 | 357 | 3461 | 182 | 68 | 76 |
| SRR5030202 | 0 | 0 | 23 | 0 | 163 | 2512 | 280 | 2880 | 140 | 93 | 135 |
| SRR5030203 | 0 | 0 | 22 | 0 | 156 | 2353 | 319 | 3177 | 126 | 89 | 156 |
| SRR5030204 | 0 | 0 | 24 | 19 | 195 | 2623 | 323 | 3017 | 123 | 85 | 138 |
| SRR5030205 | 0 | 0 | 0 | 0 | 152 | 2546 | 349 | 3478 | 99 | 61 | 182 |
| SRR5030206 | 0 | 0 | 19 | 0 | 81 | 1135 | 299 | 7119 | 262 | 131 | 156 |
| SRR5030207 | 0 | 0 | 0 | 0 | 79 | 950 | 277 | 7283 | 237 | 119 | 158 |
| SRR5030208 | 0 | 20 | 40 | 13 | 86 | 1229 | 213 | 7082 | 286 | 140 | 153 |
| SRR5030209 | 0 | 0 | 41 | 41 | 122 | 1102 | 286 | 7101 | 367 | 122 | 245 |
| SRR5030210 | 0 | 13 | 19 | 0 | 51 | 1159 | 243 | 7192 | 295 | 109 | 109 |
| SRR5030211 | 0 | 0 | 0 | 0 | 41 | 1511 | 204 | 6493 | 286 | 163 | 123 |
| SRR5030212 | 0 | 0 | 45 | 19 | 52 | 1194 | 310 | 6546 | 252 | 161 | 110 |
| SRR5030213 | 0 | 0 | 0 | 42 | 42 | 1081 | 332 | 7813 | 332 | 125 | 208 |
| SRR5030214 | 0 | 14 | 11 | 14 | 25 | 701 | 93 | 1740 | 72 | 29 | 93 |
| SRR5030216 | 0 | 15 | 23 | 27 | 31 | 669 | 77 | 1801 | 65 | 31 | 85 |
| SRR5030217 | 0 | 0 | 0 | 0 | 0 | 811 | 74 | 2139 | 37 | 0 | 74 |
| SRR5030218 | 0 | 0 | 26 | 26 | 30 | 653 | 140 | 1719 | 33 | 0 | 81 |
| SRR5030220 | 0 | 0 | 30 | 19 | 34 | 669 | 142 | 1768 | 71 | 11 | 64 |
| SRR5030221 | 0 | 0 | 0 | 0 | 74 | 670 | 186 | 1786 | 37 | 37 | 112 |
| SRR5030222 | 0 | 0 | 33 | 0 | 147 | 2883 | 508 | 4636 | 344 | 33 | 180 |
| SRR5030223 | 0 | 0 | 0 | 0 | 119 | 2484 | 509 | 5868 | 301 | 89 | 136 |
| SRR5030224 | 0 | 35 | 70 | 88 | 105 | 2853 | 455 | 5145 | 228 | 35 | 88 |
| SRR5030225 | 0 | 0 | 0 | 0 | 180 | 2407 | 465 | 5914 | 254 | 75 | 127 |
| SRR5030226 | 0 | 33 | 133 | 33 | 217 | 2534 | 483 | 5317 | 267 | 67 | 117 |
| SRR5030227 | 0 | 0 | 22 | 0 | 155 | 2499 | 436 | 5759 | 302 | 56 | 147 |
| SRR5030229 | 0 | 0 | 13 | 0 | 167 | 2558 | 399 | 5615 | 259 | 61 | 127 |
| SRR5030230 | 0 | 34 | 102 | 68 | 305 | 5826 | 610 | 12534 | 474 | 136 | 102 |
| SRR5030231 | 0 | 0 | 25 | 34 | 254 | 4597 | 663 | 11771 | 452 | 152 | 266 |
| SRR5030232 | 0 | 36 | 181 | 145 | 363 | 5619 | 653 | 12399 | 363 | 109 | 218 |
| SRR5030233 | 0 | 0 | 0 | 22 | 296 | 4629 | 562 | 11912 | 445 | 109 | 231 |
| SRR5030234 | 0 | 35 | 138 | 138 | 380 | 5937 | 587 | 11355 | 242 | 69 | 35 |

### Supp Table S8.5.Columns_ET_to_FD

| | | | | | | | | | | | |
|---|---|---|---|---|---|---|---|---|---|---|---|
| SRR5030235 | 0 | 0 | 26 | 30 | 284 | 4769 | 624 | 11404 | 533 | 142 | 206 |
| SRR5030236 | 0 | 0 | 105 | 141 | 176 | 4253 | 668 | 11670 | 316 | 70 | 176 |
| SRR5030237 | 0 | 0 | 35 | 22 | 249 | 4890 | 655 | 11928 | 393 | 166 | 231 |
| SRR5030238 | 0 | 29 | 86 | 43 | 237 | 3787 | 818 | 14933 | 287 | 86 | 122 |
| SRR5030239 | 0 | 0 | 37 | 25 | 211 | 3647 | 905 | 16881 | 372 | 112 | 149 |
| SRR5030240 | 0 | 15 | 85 | 54 | 223 | 3774 | 647 | 15420 | 262 | 77 | 131 |
| SRR5030241 | 0 | 0 | 39 | 64 | 219 | 4159 | 695 | 17034 | 348 | 103 | 167 |
| SRR5030242 | 0 | 0 | 81 | 22 | 198 | 3567 | 769 | 15784 | 330 | 95 | 146 |
| SRR5030243 | 0 | 25 | 38 | 25 | 253 | 4032 | 809 | 17343 | 228 | 101 | 164 |
| SRR5030244 | 0 | 23 | 90 | 45 | 308 | 3638 | 825 | 15031 | 255 | 98 | 188 |
| SRR5030245 | 0 | 13 | 39 | 26 | 335 | 3903 | 889 | 17505 | 399 | 103 | 193 |
| SRR5030246 | 0 | 32 | 75 | 43 | 64 | 1776 | 246 | 3949 | 128 | 64 | 118 |
| SRR5030247 | 0 | 0 | 0 | 0 | 97 | 1540 | 166 | 4254 | 214 | 76 | 228 |
| SRR5030248 | 0 | 23 | 125 | 57 | 114 | 1788 | 364 | 3564 | 80 | 114 | 171 |
| SRR5030249 | 0 | 0 | 29 | 0 | 72 | 1632 | 266 | 4630 | 216 | 93 | 237 |
| SRR5030250 | 0 | 0 | 216 | 32 | 151 | 1795 | 151 | 4239 | 195 | 54 | 76 |
| SRR5030251 | 0 | 0 | 35 | 14 | 56 | 1388 | 273 | 4627 | 273 | 112 | 203 |
| SRR5030252 | 0 | 11 | 134 | 22 | 111 | 1769 | 211 | 4517 | 189 | 122 | 111 |
| SRR5030253 | 0 | 0 | 36 | 14 | 86 | 1741 | 251 | 4343 | 222 | 93 | 208 |
| SRR5030254 | 0 | 0 | 0 | 0 | 0 | 52 | 0 | 94 | 186 | 94 | 214 |
| SRR5030255 | 0 | 0 | 0 | 0 | 0 | 38 | 0 | 108 | 154 | 77 | 189 |
| SRR5030256 | 0 | 0 | 0 | 0 | 0 | 54 | 0 | 68 | 135 | 103 | 215 |
| SRR5030257 | 0 | 0 | 0 | 0 | 0 | 54 | 0 | 62 | 155 | 85 | 197 |
| SRR5030258 | 0 | 0 | 0 | 0 | 14 | 127 | 31 | 164 | 199 | 127 | 257 |
| SRR5030259 | 0 | 0 | 0 | 0 | 0 | 119 | 0 | 105 | 177 | 126 | 216 |
| SRR5030260 | 0 | 0 | 0 | 0 | 0 | 101 | 10 | 188 | 129 | 101 | 286 |
| SRR5030261 | 0 | 0 | 0 | 0 | 0 | 94 | 0 | 101 | 195 | 97 | 167 |
| SRR5030262 | 0 | 0 | 0 | 0 | 87 | 575 | 76 | 890 | 448 | 310 | 1028 |
| SRR5030263 | 0 | 0 | 0 | 0 | 53 | 666 | 63 | 857 | 375 | 254 | 1005 |
| SRR5030265 | 0 | 10 | 10 | 0 | 62 | 671 | 36 | 996 | 330 | 387 | 1053 |
| SRR5030266 | 0 | 0 | 0 | 0 | 0 | 182 | 30 | 470 | 1449 | 979 | 774 |
| SRR5030267 | 0 | 0 | 0 | 0 | 24 | 102 | 31 | 386 | 1448 | 811 | 945 |
| SRR5030268 | 0 | 0 | 0 | 0 | 31 | 231 | 31 | 285 | 1242 | 956 | 787 |
| SRR5030270 | 0 | 0 | 0 | 0 | 33 | 342 | 53 | 887 | 953 | 473 | 605 |
| SRR5030271 | 0 | 0 | 0 | 0 | 21 | 368 | 49 | 805 | 916 | 569 | 666 |
| SRR5030272 | 0 | 0 | 0 | 0 | 27 | 381 | 40 | 936 | 990 | 535 | 622 |
| SRR5030273 | 0 | 0 | 20 | 0 | 13 | 341 | 67 | 948 | 855 | 534 | 554 |
| SRR5030274 | 0 | 0 | 0 | 0 | 20 | 258 | 0 | 185 | 563 | 318 | 172 |
| SRR5030275 | 0 | 0 | 0 | 0 | 28 | 326 | 35 | 194 | 666 | 333 | 215 |
| SRR5030276 | 0 | 0 | 0 | 0 | 34 | 291 | 14 | 189 | 629 | 223 | 230 |
| SRR5030277 | 0 | 0 | 0 | 0 | 27 | 257 | 14 | 176 | 467 | 264 | 223 |
| SRR5030278 | 0 | 0 | 0 | 0 | 32 | 224 | 29 | 302 | 356 | 183 | 173 |
| SRR5030279 | 0 | 0 | 0 | 0 | 21 | 190 | 18 | 310 | 366 | 210 | 233 |
| SRR5030280 | 0 | 0 | 12 | 0 | 37 | 179 | 17 | 323 | 370 | 204 | 154 |
| SRR5030281 | 0 | 0 | 0 | 0 | 17 | 241 | 27 | 328 | 363 | 214 | 164 |
| SRR5030282 | 0 | 0 | 0 | 0 | 36 | 623 | 99 | 2602 | 68 | 26 | 109 |
| SRR5030283 | 0 | 0 | 0 | 0 | 27 | 619 | 60 | 2535 | 65 | 22 | 76 |
| SRR5030284 | 0 | 0 | 0 | 0 | 21 | 582 | 101 | 2571 | 85 | 48 | 79 |
| SRR5030285 | 0 | 0 | 0 | 0 | 16 | 580 | 116 | 2310 | 69 | 32 | 127 |
| SRR5030287 | 0 | 0 | 0 | 0 | 0 | 491 | 123 | 2893 | 175 | 96 | 193 |
| SRR5030289 | 0 | 0 | 0 | 0 | 0 | 752 | 85 | 2735 | 180 | 162 | 231 |
| SRR5030290 | 0 | 0 | 10 | 0 | 89 | 1271 | 303 | 4221 | 241 | 115 | 298 |
| SRR5030291 | 0 | 0 | 16 | 0 | 87 | 1324 | 358 | 4462 | 244 | 136 | 353 |
| SRR5030292 | 0 | 0 | 0 | 0 | 59 | 1342 | 319 | 4297 | 197 | 176 | 346 |
| SRR5030293 | 0 | 0 | 11 | 0 | 90 | 1331 | 308 | 4428 | 260 | 133 | 323 |
| SRR5030294 | 0 | 0 | 0 | 0 | 113 | 1665 | 272 | 2627 | 232 | 73 | 411 |
| SRR5030295 | 0 | 0 | 0 | 0 | 70 | 1502 | 362 | 2859 | 202 | 153 | 577 |
| SRR5030296 | 0 | 13 | 0 | 13 | 88 | 1523 | 377 | 2520 | 249 | 148 | 539 |
| SRR5030297 | 0 | 0 | 14 | 14 | 101 | 1317 | 270 | 2837 | 182 | 115 | 385 |
| SRR5034616 | 0 | 0 | 0 | 0 | 0 | 0 | 0 | 0 | 0 | 0 | 0 |
| SRR5034617 | 0 | 0 | 52 | 17 | 0 | 48 | 0 | 69 | 0 | 0 | 0 |
| SRR5034618 | 0 | 22 | 31 | 15 | 0 | 19 | 0 | 47 | 0 | 0 | 0 |
| SRR5034619 | 0 | 0 | 22 | 0 | 0 | 15 | 0 | 0 | 0 | 0 | 0 |
| SRR5034620 | 0 | 37 | 0 | 12 | 0 | 99 | 0 | 43 | 0 | 0 | 0 |
| SRR5034621 | 0 | 70 | 120 | 32 | 0 | 82 | 0 | 73 | 0 | 0 | 0 |
| SRR5034622 | 0 | 12 | 39 | 15 | 0 | 17 | 0 | 34 | 0 | 0 | 0 |
| SRR5034623 | 0 | 0 | 0 | 0 | 0 | 0 | 0 | 0 | 0 | 0 | 0 |
| SRR5034625 | 0 | 0 | 13 | 0 | 0 | 0 | 0 | 0 | 0 | 0 | 0 |
| SRR5034626 | 0 | 31 | 44 | 0 | 0 | 11 | 0 | 28 | 0 | 0 | 0 |
| SRR5034627 | 0 | 0 | 0 | 0 | 0 | 0 | 0 | 0 | 0 | 0 | 0 |
| SRR5034628 | 0 | 0 | 0 | 0 | 0 | 48 | 0 | 48 | 0 | 0 | 32 |
| SRR5034629 | 0 | 11 | 16 | 22 | 0 | 22 | 0 | 0 | 0 | 0 | 11 |
| SRR5034630 | 0 | 0 | 24 | 0 | 0 | 0 | 0 | 24 | 0 | 0 | 0 |
| SRR5034632 | 0 | 0 | 0 | 0 | 0 | 0 | 0 | 0 | 0 | 0 | 0 |
| SRR5034634 | 0 | 0 | 0 | 0 | 0 | 0 | 0 | 0 | 0 | 0 | 0 |
| SRR5034635 | 0 | 0 | 0 | 0 | 0 | 0 | 0 | 0 | 0 | 0 | 0 |
| SRR5034636 | 0 | 0 | 0 | 0 | 0 | 0 | 0 | 0 | 0 | 0 | 0 |
| SRR5034637 | 0 | 0 | 0 | 0 | 0 | 0 | 0 | 0 | 0 | 0 | 0 |
| SRR5034639 | 0 | 0 | 0 | 0 | 0 | 0 | 0 | 0 | 0 | 0 | 0 |
| SRR5034641 | 0 | 0 | 0 | 0 | 0 | 0 | 0 | 0 | 0 | 0 | 0 |
| SRR5034642 | 0 | 0 | 0 | 0 | 0 | 0 | 0 | 22 | 0 | 0 | 0 |
| SRR5034643 | 0 | 0 | 0 | 0 | 0 | 29 | 0 | 26 | 0 | 0 | 0 |
| SRR5034644 | 0 | 12 | 0 | 0 | 0 | 0 | 0 | 23 | 0 | 0 | 0 |
| SRR5034645 | 0 | 0 | 0 | 0 | 0 | 0 | 0 | 0 | 0 | 0 | 0 |
| SRR5034646 | 0 | 0 | 0 | 0 | 0 | 0 | 0 | 0 | 12 | 0 | 0 |
| SRR5034647 | 0 | 0 | 0 | 0 | 0 | 0 | 0 | 0 | 0 | 0 | 0 |
| SRR5034648 | 0 | 0 | 15 | 0 | 0 | 0 | 0 | 0 | 0 | 0 | 0 |
| SRR5034649 | 0 | 0 | 0 | 0 | 0 | 0 | 0 | 0 | 0 | 0 | 0 |
| SRR5034650 | 0 | 129 | 144 | 70 | 0 | 20 | 0 | 224 | 35 | 15 | 0 |
| SRR5034651 | 0 | 14 | 56 | 0 | 0 | 0 | 0 | 0 | 0 | 0 | 0 |
| SRR5034652 | 0 | 13 | 22 | 22 | 0 | 13 | 0 | 0 | 0 | 0 | 0 |
| SRR5034653 | 0 | 0 | 21 | 0 | 0 | 0 | 0 | 0 | 0 | 0 | 0 |
| SRR5034654 | 0 | 0 | 20 | 0 | 0 | 11 | 0 | 24 | 13 | 0 | 0 |
| SRR5034655 | 0 | 0 | 0 | 0 | 0 | 0 | 0 | 0 | 0 | 0 | 0 |
| SRR5034656 | 0 | 0 | 0 | 0 | 0 | 0 | 0 | 0 | 0 | 0 | 0 |
| SRR5034657 | 0 | 0 | 0 | 0 | 0 | 32 | 0 | 26 | 0 | 0 | 0 |
| SRR5034658 | 0 | 0 | 22 | 0 | 0 | 0 | 0 | 0 | 0 | 0 | 0 |
| SRR5034659 | 0 | 0 | 0 | 0 | 0 | 0 | 0 | 0 | 0 | 0 | 0 |
| SRR5034660 | 0 | 0 | 0 | 0 | 0 | 0 | 0 | 0 | 0 | 0 | 0 |
| SRR5034661 | 0 | 0 | 0 | 0 | 0 | 11 | 0 | 11 | 0 | 0 | 0 |
| SRR5034662 | 0 | 0 | 0 | 0 | 0 | 0 | 0 | 0 | 0 | 0 | 0 |
| SRR5034663 | 0 | 0 | 0 | 0 | 0 | 14 | 0 | 19 | 0 | 0 | 0 |
| SRR5054304 | 0 | 74 | 180 | 29 | 12280 | 39195 | 20460 | 9587 | 10970 | 18663 | 11028 |
| SRR5054305 | 0 | 674 | 1218 | 125 | 20111 | 102562 | 19627 | 18534 | 12727 | 29010 | 10114 |
| SRR5087508 | 0 | 0 | 12 | 22 | 14 | 69 | 11 | 164 | 70 | 77 | 92 |
| SRR5087510 | 0 | 0 | 15 | 25 | 17 | 80 | 16 | 171 | 81 | 88 | 121 |
| SRR5087512 | 0 | 0 | 11 | 22 | 13 | 72 | 11 | 177 | 86 | 82 | 119 |
| SRR5087516 | 0 | 0 | 0 | 11 | 11 | 52 | 0 | 121 | 53 | 66 | 40 |
| SRR5087517 | 0 | 0 | 0 | 0 | 0 | 50 | 0 | 100 | 32 | 56 | 48 |
| SRR5087519 | 0 | 0 | 0 | 12 | 11 | 41 | 13 | 123 | 53 | 65 | 53 |
| SRR5087522 | 0 | 32 | 75 | 117 | 13 | 244 | 27 | 519 | 220 | 178 | 498 |
| SRR5087527 | 0 | 36 | 42 | 74 | 30 | 219 | 43 | 495 | 207 | 148 | 464 |
| SRR5087554 | 0 | 21 | 80 | 114 | 24 | 226 | 21 | 467 | 206 | 155 | 512 |
| SRR5090283 | 0 | 0 | 13 | 0 | 17 | 30 | 21 | 12 | 178 | 0 | 11 |
| SRR5092303 | 0 | 0 | 0 | 0 | 0 | 39 | 0 | 0 | 20 | 20 | 0 |
| SRR5092304 | 0 | 0 | 0 | 0 | 0 | 93 | 0 | 23 | 23 | 93 | 0 |
| SRR5092305 | 0 | 0 | 0 | 0 | 0 | 22 | 0 | 22 | 0 | 0 | 0 |
| SRR5092836 | 0 | 38 | 0 | 38 | 0 | 0 | 0 | 190 | 114 | 38 | 76 |
| SRR5100734 | 0 | 0 | 0 | 0 | 0 | 12 | 0 | 0 | 0 | 0 | 0 |

### Supp Table S8.5.Columns_ET_to_FD

| | | | | | | | | | | | |
|---|---|---|---|---|---|---|---|---|---|---|---|
| SRR5100735 | 0 | 155 | 217 | 10 | 304 | 544 | 224 | 72 | 343 | 148 | 126 |
| SRR5104206 | 0 | 0 | 23 | 32 | 0 | 82 | 12 | 130 | 71 | 22 | 92 |
| SRR5104207 | 0 | 0 | 13 | 18 | 0 | 80 | 20 | 121 | 50 | 18 | 169 |
| SRR5104208 | 0 | 0 | 18 | 19 | 13 | 79 | 21 | 88 | 48 | 12 | 128 |
| SRR5104209 | 0 | 0 | 13 | 18 | 0 | 62 | 0 | 74 | 39 | 18 | 115 |
| SRR5104210 | 0 | 0 | 13 | 0 | 0 | 64 | 21 | 118 | 11 | 15 | 63 |
| SRR5104211 | 0 | 0 | 0 | 12 | 0 | 18 | 0 | 65 | 15 | 0 | 29 |
| SRR5104212 | 0 | 0 | 11 | 0 | 0 | 39 | 12 | 55 | 13 | 18 | 71 |
| SRR5104213 | 0 | 0 | 12 | 0 | 0 | 49 | 21 | 82 | 0 | 0 | 79 |
| SRR5104214 | 0 | 0 | 16 | 0 | 0 | 0 | 0 | 31 | 0 | 0 | 25 |
| SRR5104215 | 0 | 0 | 0 | 15 | 0 | 18 | 0 | 58 | 15 | 0 | 46 |
| SRR5104216 | 0 | 0 | 0 | 15 | 0 | 39 | 11 | 90 | 18 | 0 | 79 |
| SRR5104217 | 0 | 0 | 0 | 0 | 0 | 25 | 12 | 64 | 10 | 25 | 39 |
| SRR5104218 | 0 | 0 | 0 | 0 | 0 | 21 | 13 | 43 | 0 | 0 | 21 |
| SRR5104219 | 0 | 0 | 0 | 10 | 0 | 23 | 16 | 60 | 10 | 13 | 57 |
| SRR5104220 | 0 | 0 | 0 | 0 | 0 | 36 | 18 | 94 | 25 | 21 | 80 |
| SRR5104221 | 0 | 0 | 12 | 0 | 0 | 16 | 11 | 60 | 12 | 0 | 16 |
| SRR5104222 | 0 | 0 | 0 | 10 | 0 | 28 | 0 | 46 | 0 | 10 | 28 |
| SRR5104223 | 0 | 0 | 17 | 0 | 0 | 13 | 0 | 35 | 10 | 0 | 19 |
| SRR5104224 | 0 | 0 | 0 | 0 | 0 | 0 | 0 | 20 | 0 | 0 | 14 |
| SRR5104225 | 0 | 0 | 23 | 0 | 0 | 35 | 0 | 68 | 0 | 12 | 56 |
| SRR5116489 | 0 | 0 | 0 | 0 | 73 | 660 | 73 | 1247 | 734 | 1577 | 844 |
| SRR5116490 | 92 | 79 | 119 | 26 | 26 | 777 | 53 | 1093 | 1040 | 1225 | 1185 |
| SRR5116491 | 21 | 0 | 21 | 85 | 149 | 831 | 43 | 1151 | 298 | 1087 | 661 |
| SRR5116492 | 55 | 32 | 60 | 55 | 156 | 1011 | 151 | 1529 | 2549 | 3048 | 3245 |
| SRR5116493 | 16 | 16 | 0 | 0 | 63 | 803 | 47 | 1385 | 1370 | 1684 | 1322 |
| SRR5116494 | 0 | 22 | 34 | 0 | 34 | 1010 | 101 | 2042 | 3423 | 2940 | 3636 |
| SRR5116495 | 19 | 0 | 14 | 24 | 62 | 835 | 81 | 1150 | 1408 | 1632 | 1026 |
| SRR5116496 | 0 | 0 | 0 | 0 | 49 | 697 | 59 | 1620 | 3005 | 3889 | 2494 |
| SRR5121477 | 22 | 25 | 53 | 21 | 0 | 112 | 10 | 204 | 369 | 148 | 239 |
| SRR5121478 | 51 | 51 | 88 | 31 | 86 | 770 | 65 | 370 | 421 | 149 | 137 |
| SRR5121479 | 0 | 156 | 286 | 225 | 439 | 2397 | 621 | 4283 | 15664 | 26768 | 44955 |
| SRR5121480 | 34 | 118 | 158 | 83 | 197 | 1120 | 154 | 483 | 2082 | 1048 | 3345 |
| SRR5121481 | 0 | 122 | 263 | 175 | 63 | 554 | 88 | 1063 | 1207 | 368 | 2180 |
| SRR5121482 | 13 | 196 | 592 | 248 | 285 | 1417 | 256 | 995 | 807 | 582 | 861 |
| SRR5121483 | 0 | 0 | 0 | 0 | 48 | 380 | 14 | 74 | 34 | 0 | 21 |
| SRR5121484 | 207 | 279 | 711 | 539 | 314 | 2437 | 253 | 1944 | 4044 | 3123 | 4177 |
| SRR5121485 | 154 | 604 | 1540 | 1314 | 340 | 7204 | 1331 | 10676 | 1115 | 788 | 8656 |
| SRR5121486 | 0 | 0 | 0 | 0 | 0 | 54 | 0 | 142 | 29 | 0 | 48 |
| SRR5121487 | 257 | 370 | 1382 | 1327 | 166 | 2660 | 326 | 2428 | 764 | 560 | 5208 |
| SRR5127200 | 0 | 54 | 136 | 61 | 16 | 390 | 51 | 923 | 50 | 15 | 106 |
| SRR5127201 | 0 | 16 | 42 | 16 | 115 | 917 | 236 | 1326 | 246 | 206 | 365 |
| SRR5127202 | 0 | 39 | 177 | 86 | 137 | 2008 | 366 | 2231 | 10661 | 9623 | 5436 |
| SRR5127203 | 0 | 82 | 234 | 129 | 262 | 3129 | 610 | 3031 | 2597 | 1901 | 1476 |
| SRR5127204 | 0 | 25 | 54 | 54 | 68 | 787 | 129 | 674 | 706 | 491 | 810 |
| SRR5127205 | 0 | 18 | 64 | 72 | 77 | 1247 | 174 | 444 | 432 | 267 | 947 |
| SRR5127206 | 0 | 51 | 116 | 57 | 77 | 1014 | 216 | 972 | 524 | 388 | 1002 |
| SRR5127207 | 0 | 0 | 0 | 0 | 0 | 35 | 0 | 22 | 0 | 0 | 92 |
| SRR5127208 | 0 | 21 | 48 | 28 | 236 | 3295 | 467 | 1358 | 965 | 790 | 1343 |
| SRR5127209 | 0 | 36 | 107 | 83 | 85 | 861 | 130 | 773 | 270 | 369 | 433 |
| SRR5127210 | 0 | 106 | 349 | 219 | 256 | 3303 | 456 | 2433 | 1914 | 1589 | 1473 |
| SRR5127211 | 0 | 15 | 31 | 11 | 0 | 44 | 0 | 36 | 31 | 22 | 60 |
| SRR5127212 | 0 | 28 | 61 | 30 | 57 | 711 | 103 | 1137 | 354 | 331 | 475 |
| SRR5127213 | 0 | 92 | 303 | 131 | 216 | 2485 | 341 | 3104 | 3434 | 2821 | 2527 |
| SRR5127214 | 0 | 32 | 87 | 52 | 68 | 1290 | 278 | 1125 | 457 | 310 | 456 |
| SRR5127215 | 0 | 51 | 135 | 56 | 71 | 1303 | 291 | 1455 | 414 | 283 | 922 |
| SRR5127216 | 0 | 17 | 26 | 18 | 102 | 1059 | 153 | 810 | 511 | 466 | 546 |
| SRR5127217 | 0 | 34 | 106 | 58 | 52 | 764 | 108 | 575 | 653 | 411 | 708 |
| SRR5127218 | 0 | 23 | 64 | 36 | 98 | 1186 | 217 | 1060 | 732 | 588 | 944 |
| SRR5127219 | 0 | 16 | 61 | 25 | 0 | 50 | 0 | 102 | 0 | 12 | 11 |
| SRR5127220 | 0 | 33 | 116 | 47 | 156 | 2357 | 486 | 2265 | 5308 | 5527 | 5678 |
| SRR5127221 | 0 | 56 | 174 | 71 | 194 | 2704 | 518 | 2246 | 840 | 728 | 1129 |
| SRR5127222 | 0 | 40 | 70 | 48 | 93 | 1794 | 294 | 1300 | 1091 | 1202 | 1543 |
| SRR5127223 | 0 | 64 | 165 | 72 | 159 | 2018 | 318 | 1943 | 3230 | 2778 | 3380 |
| SRR5127224 | 0 | 28 | 55 | 37 | 114 | 1289 | 208 | 1807 | 3828 | 3894 | 2114 |
| SRR5127225 | 0 | 124 | 383 | 166 | 297 | 3648 | 543 | 4732 | 2654 | 1847 | 2217 |
| SRR5127226 | 0 | 51 | 118 | 52 | 141 | 1644 | 323 | 1322 | 519 | 421 | 977 |
| SRR5127227 | 0 | 42 | 124 | 77 | 98 | 1343 | 247 | 1060 | 999 | 1035 | 1814 |
| SRR5127228 | 0 | 237 | 599 | 201 | 104 | 937 | 287 | 1116 | 397 | 317 | 383 |
| SRR5127229 | 0 | 41 | 162 | 68 | 138 | 2027 | 295 | 1221 | 1105 | 757 | 1492 |
| SRR5127230 | 0 | 41 | 135 | 86 | 110 | 1365 | 213 | 1354 | 869 | 498 | 1136 |
| SRR5127231 | 0 | 12 | 38 | 17 | 0 | 64 | 12 | 96 | 111 | 103 | 85 |
| SRR5127232 | 0 | 0 | 0 | 0 | 0 | 14 | 0 | 25 | 0 | 0 | 17 |
| SRR5127233 | 0 | 19 | 56 | 37 | 53 | 689 | 148 | 563 | 135 | 95 | 382 |
| SRR5127234 | 0 | 13 | 52 | 56 | 36 | 417 | 52 | 402 | 123 | 179 | 179 |
| SRR5127235 | 0 | 84 | 306 | 136 | 190 | 2559 | 504 | 2392 | 1951 | 1561 | 1641 |
| SRR5127236 | 0 | 0 | 0 | 0 | 11 | 127 | 16 | 59 | 45 | 46 | 74 |
| SRR5136285 | 0 | 0 | 0 | 0 | 0 | 0 | 0 | 0 | 0 | 0 | 0 |
| SRR5136286 | 0 | 0 | 0 | 0 | 0 | 0 | 0 | 0 | 0 | 0 | 0 |
| SRR5136287 | 0 | 0 | 0 | 0 | 0 | 0 | 0 | 0 | 0 | 0 | 0 |
| SRR5136288 | 0 | 0 | 0 | 0 | 0 | 0 | 0 | 0 | 0 | 0 | 0 |
| SRR5136289 | 0 | 0 | 0 | 0 | 0 | 0 | 0 | 0 | 0 | 0 | 0 |
| SRR5136290 | 0 | 0 | 0 | 0 | 0 | 0 | 0 | 0 | 0 | 0 | 0 |
| SRR5136291 | 0 | 0 | 0 | 0 | 0 | 0 | 0 | 0 | 0 | 0 | 0 |
| SRR5136292 | 0 | 0 | 0 | 0 | 0 | 0 | 0 | 0 | 0 | 0 | 0 |
| SRR5136293 | 0 | 0 | 0 | 0 | 0 | 0 | 0 | 0 | 0 | 0 | 0 |
| SRR5136294 | 0 | 0 | 0 | 0 | 0 | 0 | 0 | 0 | 0 | 0 | 0 |
| SRR5136295 | 0 | 0 | 0 | 0 | 0 | 0 | 0 | 0 | 0 | 0 | 0 |
| SRR5136296 | 0 | 0 | 0 | 0 | 0 | 0 | 0 | 0 | 0 | 0 | 0 |
| SRR5136297 | 0 | 0 | 0 | 0 | 0 | 0 | 0 | 0 | 0 | 0 | 0 |
| SRR5136298 | 0 | 0 | 0 | 0 | 0 | 0 | 0 | 0 | 0 | 0 | 0 |
| SRR5136299 | 0 | 0 | 0 | 0 | 0 | 0 | 0 | 0 | 0 | 0 | 0 |
| SRR5136300 | 0 | 0 | 0 | 0 | 0 | 0 | 0 | 0 | 0 | 0 | 0 |
| SRR5139121 | 0 | 63 | 172 | 104 | 130 | 1377 | 255 | 2174 | 1201 | 986 | 1474 |
| SRR5145238 | 0 | 63 | 103 | 49 | 214 | 596 | 295 | 489 | 1306 | 334 | 205 |
| SRR5145239 | 0 | 75 | 321 | 95 | 1050 | 2126 | 1511 | 922 | 427 | 328 | 191 |
| SRR5145240 | 0 | 0 | 0 | 0 | 82 | 134 | 62 | 102 | 136 | 42 | 35 |
| SRR5145241 | 0 | 13 | 37 | 36 | 773 | 1077 | 404 | 383 | 440 | 74 | 79 |
| SRR5145242 | 0 | 0 | 0 | 0 | 26 | 45 | 25 | 30 | 150 | 59 | 38 |
| SRR5145243 | 0 | 0 | 0 | 0 | 70 | 155 | 120 | 129 | 1148 | 155 | 308 |
| SRR5145244 | 0 | 0 | 0 | 0 | 18 | 44 | 17 | 53 | 154 | 59 | 34 |
| SRR5145245 | 0 | 58 | 147 | 66 | 225 | 553 | 365 | 339 | 363 | 58 | 70 |
| SRR5163971 | 0 | 65 | 67 | 71 | 85 | 331 | 71 | 115 | 28 | 12 | 24 |
| SRR5163972 | 0 | 43 | 55 | 82 | 118 | 314 | 106 | 122 | 24 | 20 | 31 |
| SRR5163973 | 0 | 68 | 79 | 38 | 251 | 815 | 180 | 142 | 59 | 64 | 28 |
| SRR5163974 | 0 | 58 | 77 | 39 | 281 | 896 | 194 | 106 | 44 | 63 | 29 |
| SRR5163975 | 0 | 48 | 117 | 44 | 547 | 1481 | 456 | 241 | 367 | 666 | 294 |
| SRR5163976 | 0 | 39 | 133 | 65 | 667 | 1467 | 398 | 194 | 389 | 657 | 236 |
| SRR5163977 | 0 | 72 | 85 | 68 | 285 | 1083 | 248 | 220 | 120 | 153 | 158 |
| SRR5163978 | 0 | 73 | 127 | 71 | 251 | 1036 | 220 | 223 | 152 | 157 | 195 |
| SRR5163979 | 0 | 26 | 25 | 33 | 53 | 273 | 65 | 43 | 25 | 0 | 28 |
| SRR5163980 | 0 | 27 | 16 | 22 | 70 | 216 | 35 | 51 | 32 | 27 | 27 |
| SRR5163981 | 0 | 78 | 90 | 83 | 138 | 361 | 101 | 92 | 19 | 0 | 13 |
| SRR5163982 | 0 | 42 | 121 | 74 | 180 | 691 | 143 | 87 | 61 | 61 | 68 |
| SRR5163983 | 0 | 29 | 85 | 80 | 137 | 478 | 76 | 66 | 29 | 10 | 14 |
| SRR5163984 | 0 | 29 | 88 | 75 | 130 | 453 | 67 | 92 | 13 | 0 | 17 |
| SRR5163985 | 0 | 18 | 29 | 18 | 117 | 427 | 123 | 82 | 47 | 0 | 23 |
| SRR5163986 | 0 | 28 | 37 | 19 | 140 | 438 | 88 | 140 | 47 | 19 | 28 |
| SRR5163987 | 0 | 72 | 98 | 123 | 196 | 838 | 154 | 130 | 26 | 0 | 40 |

### Supp Table S8.5.Columns_ET_to_FD

| | | | | | | | | | | | |
|---|---|---|---|---|---|---|---|---|---|---|---|
| SRR5168444 | 10 | 14 | 15 | 0 | 44 | 134 | 44 | 89 | 61 | 18 | 41 |
| SRR5168445 | 0 | 18 | 29 | 0 | 11 | 13 | 0 | 0 | 65 | 0 | 0 |
| SRR5168447 | 0 | 0 | 0 | 0 | 11 | 39 | 0 | 47 | 157 | 22 | 22 |
| SRR5168448 | 0 | 0 | 12 | 0 | 58 | 247 | 44 | 203 | 76 | 247 | 26 |
| SRR5168451 | 0 | 0 | 0 | 0 | 0 | 19 | 0 | 15 | 28 | 0 | 0 |
| SRR5168454 | 0 | 0 | 27 | 0 | 16 | 26 | 20 | 22 | 348 | 11 | 27 |
| SRR5168458 | 0 | 0 | 20 | 0 | 10 | 17 | 0 | 16 | 196 | 0 | 14 |
| SRR5168460 | 0 | 0 | 11 | 0 | 0 | 30 | 14 | 24 | 92 | 19 | 26 |
| SRR5168465 | 0 | 0 | 13 | 0 | 22 | 71 | 22 | 74 | 2101 | 52 | 65 |
| SRR5168467 | 0 | 0 | 0 | 0 | 0 | 14 | 0 | 0 | 28 | 18 | 0 |
| SRR5168471 | 0 | 0 | 29 | 0 | 12 | 34 | 28 | 30 | 392 | 17 | 29 |
| SRR5168475 | 0 | 0 | 0 | 0 | 0 | 0 | 0 | 0 | 82 | 0 | 0 |
| SRR5171728 | 0 | 101 | 99 | 97 | 73 | 126 | 90 | 84 | 162 | 147 | 376 |
| SRR5171729 | 0 | 125 | 197 | 168 | 96 | 170 | 119 | 92 | 295 | 237 | 535 |
| SRR5171730 | 0 | 119 | 183 | 97 | 49 | 97 | 70 | 59 | 334 | 162 | 237 |
| SRR5171731 | 0 | 70 | 130 | 155 | 65 | 155 | 88 | 73 | 165 | 140 | 245 |
| SRR5171732 | 0 | 73 | 113 | 126 | 100 | 119 | 20 | 80 | 153 | 133 | 126 |
| SRR5171733 | 0 | 138 | 170 | 238 | 145 | 191 | 128 | 109 | 113 | 108 | 206 |
| SRR5195030 | 0 | 0 | 0 | 0 | 0 | 87 | 18 | 103 | 145 | 111 | 191 |
| SRR5195032 | 0 | 0 | 0 | 0 | 0 | 29 | 0 | 45 | 72 | 36 | 119 |
| SRR5195033 | 0 | 0 | 11 | 0 | 0 | 26 | 0 | 23 | 78 | 62 | 103 |
| SRR5195034 | 0 | 0 | 33 | 16 | 0 | 46 | 0 | 51 | 50 | 33 | 59 |
| SRR5224010 | 0 | 26 | 52 | 10 | 80 | 278 | 50 | 154 | 463 | 30 | 34 |
| SRR5224011 | 0 | 26 | 45 | 0 | 63 | 178 | 44 | 99 | 406 | 33 | 29 |
| SRR5224012 | 12 | 106 | 170 | 29 | 186 | 518 | 121 | 181 | 335 | 86 | 52 |
| SRR5224013 | 0 | 17 | 28 | 0 | 72 | 284 | 50 | 221 | 1220 | 58 | 73 |
| SRR5224014 | 0 | 31 | 49 | 0 | 94 | 359 | 61 | 168 | 1176 | 75 | 68 |
| SRR5224015 | 0 | 11 | 28 | 0 | 66 | 172 | 37 | 136 | 1241 | 41 | 66 |
| SRR5224016 | 0 | 21 | 45 | 12 | 91 | 257 | 53 | 224 | 1910 | 81 | 150 |
| SRR5224017 | 0 | 24 | 51 | 0 | 163 | 773 | 106 | 274 | 2184 | 226 | 133 |
| SRR5224018 | 0 | 19 | 58 | 13 | 152 | 381 | 76 | 282 | 1905 | 100 | 171 |
| SRR5224019 | 0 | 19 | 34 | 0 | 65 | 231 | 32 | 211 | 1433 | 78 | 74 |
| SRR5224020 | 0 | 19 | 43 | 0 | 96 | 240 | 46 | 197 | 1370 | 53 | 99 |
| SRR5224021 | 0 | 22 | 44 | 0 | 93 | 306 | 54 | 232 | 1521 | 100 | 86 |
| SRR5224022 | 13 | 26 | 60 | 10 | 140 | 339 | 77 | 264 | 1804 | 81 | 128 |
| SRR5224023 | 0 | 21 | 36 | 0 | 74 | 367 | 36 | 320 | 1755 | 119 | 118 |
| SRR5224024 | 0 | 35 | 78 | 12 | 115 | 277 | 88 | 182 | 1537 | 67 | 147 |
| SRR5230550 | 0 | 0 | 0 | 0 | 0 | 0 | 0 | 0 | 0 | 0 | 0 |
| SRR5230551 | 0 | 0 | 0 | 0 | 0 | 0 | 0 | 0 | 0 | 0 | 0 |
| SRR5230552 | 0 | 0 | 0 | 0 | 0 | 0 | 0 | 0 | 16 | 0 | 0 |
| SRR5230553 | 0 | 0 | 0 | 0 | 0 | 0 | 0 | 0 | 0 | 0 | 0 |
| SRR5230554 | 0 | 0 | 0 | 0 | 0 | 0 | 0 | 0 | 0 | 0 | 0 |
| SRR5230555 | 0 | 0 | 0 | 0 | 0 | 0 | 0 | 0 | 0 | 0 | 0 |
| SRR5230556 | 0 | 0 | 0 | 0 | 0 | 0 | 0 | 0 | 0 | 0 | 0 |
| SRR5230557 | 0 | 0 | 0 | 0 | 0 | 0 | 0 | 0 | 0 | 0 | 0 |
| SRR5230558 | 0 | 0 | 0 | 0 | 0 | 0 | 0 | 0 | 20 | 0 | 0 |
| SRR5230559 | 0 | 0 | 0 | 0 | 0 | 0 | 0 | 0 | 0 | 0 | 0 |
| SRR5230560 | 0 | 0 | 0 | 0 | 0 | 0 | 0 | 0 | 0 | 0 | 0 |
| SRR5230561 | 0 | 0 | 0 | 0 | 0 | 0 | 0 | 0 | 0 | 0 | 0 |
| SRR5230562 | 0 | 0 | 0 | 0 | 0 | 0 | 0 | 0 | 0 | 0 | 0 |
| SRR5230563 | 0 | 0 | 0 | 0 | 0 | 0 | 0 | 0 | 0 | 0 | 0 |
| SRR5230564 | 0 | 0 | 0 | 0 | 0 | 0 | 0 | 0 | 0 | 0 | 0 |
| SRR5230565 | 0 | 0 | 0 | 0 | 0 | 0 | 0 | 0 | 0 | 0 | 0 |
| SRR5230566 | 0 | 0 | 0 | 0 | 0 | 0 | 0 | 0 | 0 | 0 | 0 |
| SRR5230567 | 0 | 0 | 0 | 0 | 0 | 0 | 0 | 0 | 13 | 0 | 0 |
| SRR5230568 | 0 | 0 | 0 | 0 | 0 | 0 | 0 | 0 | 0 | 0 | 0 |
| SRR5230569 | 0 | 0 | 0 | 0 | 0 | 0 | 0 | 0 | 0 | 0 | 0 |
| SRR5230570 | 0 | 0 | 0 | 0 | 0 | 0 | 0 | 0 | 0 | 0 | 0 |
| SRR5230571 | 0 | 0 | 0 | 0 | 0 | 0 | 0 | 0 | 10 | 0 | 0 |
| SRR5230572 | 0 | 0 | 0 | 0 | 0 | 0 | 0 | 0 | 0 | 0 | 0 |
| SRR5230573 | 0 | 0 | 0 | 0 | 0 | 0 | 0 | 0 | 0 | 0 | 0 |
| SRR5230574 | 0 | 0 | 0 | 0 | 0 | 0 | 0 | 0 | 0 | 0 | 0 |
| SRR5230575 | 0 | 0 | 0 | 0 | 0 | 0 | 0 | 0 | 0 | 0 | 0 |
| SRR5230576 | 0 | 0 | 0 | 0 | 0 | 0 | 0 | 0 | 0 | 0 | 0 |
| SRR5230577 | 0 | 0 | 0 | 0 | 0 | 0 | 0 | 0 | 0 | 0 | 0 |
| SRR5230578 | 0 | 0 | 0 | 0 | 0 | 0 | 0 | 0 | 0 | 0 | 0 |
| SRR5230579 | 0 | 0 | 0 | 0 | 11 | 0 | 0 | 0 | 0 | 0 | 0 |
| SRR5230580 | 0 | 0 | 0 | 0 | 0 | 0 | 0 | 0 | 0 | 0 | 0 |
| SRR5230581 | 0 | 0 | 0 | 0 | 0 | 0 | 0 | 0 | 0 | 0 | 0 |
| SRR5230582 | 0 | 16 | 0 | 0 | 0 | 32 | 0 | 0 | 0 | 0 | 0 |
| SRR5230583 | 0 | 16 | 0 | 0 | 0 | 47 | 0 | 0 | 0 | 0 | 0 |
| SRR5230584 | 0 | 0 | 0 | 0 | 0 | 0 | 0 | 0 | 0 | 0 | 0 |
| SRR5230585 | 0 | 0 | 0 | 0 | 0 | 0 | 0 | 0 | 0 | 0 | 0 |
| SRR5230586 | 0 | 0 | 0 | 0 | 0 | 0 | 0 | 0 | 0 | 0 | 0 |
| SRR5230587 | 0 | 0 | 0 | 0 | 0 | 0 | 0 | 0 | 0 | 0 | 0 |
| SRR5230588 | 0 | 0 | 0 | 0 | 0 | 0 | 0 | 0 | 0 | 0 | 0 |
| SRR5230589 | 0 | 0 | 0 | 0 | 0 | 15 | 0 | 0 | 0 | 0 | 0 |
| SRR5230590 | 0 | 0 | 0 | 0 | 0 | 0 | 0 | 0 | 0 | 0 | 0 |
| SRR5230591 | 0 | 0 | 0 | 0 | 0 | 0 | 0 | 0 | 0 | 0 | 0 |
| SRR5230592 | 0 | 0 | 0 | 0 | 0 | 0 | 0 | 0 | 0 | 0 | 0 |
| SRR5230593 | 0 | 0 | 0 | 0 | 0 | 0 | 0 | 0 | 0 | 0 | 0 |
| SRR5230594 | 0 | 0 | 0 | 0 | 0 | 0 | 0 | 0 | 21 | 0 | 0 |
| SRR5230595 | 0 | 0 | 0 | 0 | 0 | 17 | 0 | 0 | 0 | 0 | 0 |
| SRR5230596 | 0 | 0 | 0 | 0 | 0 | 0 | 0 | 0 | 0 | 0 | 0 |
| SRR5230597 | 0 | 0 | 0 | 0 | 0 | 0 | 0 | 0 | 0 | 0 | 0 |
| SRR5230600 | 0 | 0 | 0 | 0 | 0 | 0 | 0 | 0 | 0 | 0 | 0 |
| SRR5230601 | 0 | 0 | 0 | 0 | 0 | 0 | 0 | 0 | 0 | 0 | 0 |
| SRR5230602 | 0 | 0 | 0 | 0 | 0 | 14 | 0 | 0 | 0 | 0 | 0 |
| SRR5230603 | 0 | 0 | 0 | 0 | 0 | 0 | 0 | 0 | 0 | 0 | 0 |
| SRR5230604 | 0 | 0 | 0 | 0 | 0 | 0 | 0 | 0 | 0 | 0 | 0 |
| SRR5230605 | 0 | 0 | 0 | 0 | 0 | 0 | 0 | 0 | 0 | 0 | 0 |
| SRR5230606 | 0 | 0 | 0 | 0 | 0 | 0 | 0 | 0 | 0 | 0 | 0 |
| SRR5230607 | 0 | 0 | 0 | 0 | 0 | 0 | 0 | 0 | 0 | 0 | 0 |
| SRR5230608 | 0 | 0 | 0 | 0 | 0 | 0 | 0 | 0 | 0 | 0 | 0 |
| SRR5230609 | 0 | 0 | 0 | 0 | 0 | 0 | 0 | 0 | 12 | 0 | 0 |
| SRR5230610 | 0 | 0 | 0 | 0 | 0 | 0 | 0 | 0 | 0 | 0 | 0 |
| SRR5230611 | 0 | 0 | 0 | 0 | 0 | 0 | 0 | 0 | 0 | 0 | 0 |
| SRR5230613 | 0 | 0 | 0 | 0 | 0 | 0 | 0 | 0 | 0 | 0 | 0 |
| SRR5230614 | 0 | 0 | 0 | 0 | 0 | 0 | 0 | 0 | 0 | 0 | 0 |
| SRR5230615 | 0 | 0 | 0 | 0 | 0 | 0 | 0 | 0 | 0 | 0 | 0 |
| SRR5230617 | 0 | 0 | 0 | 0 | 0 | 0 | 0 | 0 | 0 | 0 | 0 |
| SRR5230618 | 0 | 0 | 0 | 0 | 0 | 0 | 0 | 0 | 0 | 0 | 0 |
| SRR5230619 | 0 | 0 | 0 | 0 | 0 | 0 | 0 | 0 | 17 | 0 | 0 |
| SRR5230620 | 0 | 0 | 0 | 0 | 0 | 0 | 0 | 0 | 0 | 0 | 0 |
| SRR5230621 | 0 | 0 | 0 | 0 | 0 | 0 | 0 | 0 | 0 | 0 | 0 |
| SRR5230622 | 0 | 0 | 0 | 0 | 0 | 0 | 0 | 0 | 0 | 0 | 0 |
| SRR5230623 | 0 | 0 | 0 | 0 | 0 | 0 | 0 | 0 | 0 | 0 | 0 |
| SRR5230624 | 0 | 0 | 0 | 0 | 0 | 0 | 0 | 0 | 0 | 0 | 0 |
| SRR5230625 | 0 | 0 | 0 | 0 | 0 | 0 | 0 | 0 | 0 | 0 | 0 |
| SRR5230626 | 0 | 0 | 0 | 0 | 0 | 0 | 0 | 0 | 18 | 0 | 0 |
| SRR5230627 | 0 | 0 | 0 | 0 | 0 | 0 | 0 | 0 | 0 | 0 | 0 |
| SRR5230628 | 0 | 0 | 0 | 0 | 0 | 0 | 0 | 0 | 0 | 0 | 0 |
| SRR5230629 | 0 | 0 | 0 | 0 | 0 | 0 | 0 | 0 | 0 | 0 | 0 |
| SRR5230630 | 0 | 0 | 0 | 0 | 0 | 0 | 0 | 0 | 0 | 0 | 0 |
| SRR5230631 | 0 | 0 | 0 | 0 | 0 | 0 | 0 | 0 | 0 | 0 | 0 |
| SRR5230632 | 0 | 0 | 0 | 0 | 0 | 0 | 0 | 0 | 0 | 0 | 0 |
| SRR5230633 | 0 | 0 | 0 | 0 | 0 | 0 | 0 | 0 | 0 | 0 | 0 |
| SRR5230634 | 0 | 0 | 0 | 0 | 0 | 0 | 0 | 0 | 0 | 0 | 0 |
| SRR5230635 | 0 | 0 | 0 | 0 | 0 | 0 | 0 | 0 | 0 | 0 | 0 |

### Supp Table S8.5.Columns_ET_to_FD

| | | | | | | | | | | | |
|---|---|---|---|---|---|---|---|---|---|---|---|
| SRR5230636 | 0 | 0 | 0 | 0 | 0 | 0 | 0 | 0 | 0 | 0 | 0 |
| SRR5230637 | 0 | 0 | 0 | 0 | 0 | 0 | 0 | 0 | 0 | 0 | 0 |
| SRR5230638 | 0 | 0 | 0 | 0 | 0 | 0 | 0 | 0 | 16 | 0 | 0 |
| SRR5230639 | 0 | 0 | 0 | 0 | 0 | 0 | 0 | 0 | 0 | 0 | 0 |
| SRR5230640 | 0 | 0 | 0 | 0 | 0 | 0 | 0 | 0 | 0 | 0 | 0 |
| SRR5230641 | 0 | 0 | 0 | 0 | 0 | 0 | 0 | 0 | 0 | 0 | 0 |
| SRR5230642 | 0 | 0 | 0 | 0 | 0 | 0 | 0 | 0 | 0 | 0 | 0 |
| SRR5230643 | 0 | 0 | 0 | 0 | 0 | 0 | 0 | 0 | 0 | 0 | 0 |
| SRR5230644 | 0 | 0 | 0 | 0 | 0 | 0 | 0 | 0 | 0 | 0 | 0 |
| SRR5230645 | 0 | 0 | 0 | 0 | 0 | 0 | 0 | 0 | 0 | 0 | 0 |
| SRR5230646 | 0 | 0 | 0 | 0 | 0 | 0 | 0 | 0 | 0 | 0 | 0 |
| SRR5230647 | 0 | 0 | 0 | 0 | 0 | 0 | 0 | 0 | 0 | 0 | 0 |
| SRR5230648 | 0 | 0 | 10 | 0 | 0 | 15 | 0 | 0 | 0 | 0 | 0 |
| SRR5230649 | 0 | 0 | 0 | 0 | 0 | 0 | 0 | 0 | 0 | 13 | 0 |
| SRR5230650 | 0 | 0 | 0 | 0 | 0 | 0 | 0 | 0 | 0 | 0 | 0 |
| SRR5230651 | 0 | 0 | 0 | 0 | 0 | 0 | 0 | 0 | 0 | 0 | 0 |
| SRR5230652 | 0 | 0 | 0 | 0 | 0 | 0 | 0 | 0 | 0 | 0 | 0 |
| SRR5230653 | 0 | 0 | 0 | 0 | 0 | 0 | 0 | 0 | 0 | 0 | 0 |
| SRR5230654 | 0 | 0 | 0 | 0 | 0 | 0 | 0 | 0 | 0 | 0 | 0 |
| SRR5230655 | 0 | 0 | 0 | 0 | 0 | 0 | 0 | 0 | 0 | 0 | 0 |
| SRR5230656 | 0 | 0 | 0 | 0 | 0 | 19 | 0 | 0 | 0 | 0 | 0 |
| SRR5230657 | 0 | 0 | 0 | 0 | 0 | 0 | 0 | 0 | 0 | 0 | 0 |
| SRR5230658 | 0 | 0 | 0 | 0 | 0 | 0 | 0 | 0 | 0 | 0 | 0 |
| SRR5230659 | 0 | 0 | 0 | 0 | 0 | 0 | 0 | 0 | 11 | 0 | 0 |
| SRR5230660 | 0 | 0 | 0 | 0 | 0 | 14 | 0 | 0 | 0 | 0 | 0 |
| SRR5230661 | 0 | 0 | 0 | 0 | 0 | 0 | 0 | 0 | 0 | 0 | 0 |
| SRR5230662 | 0 | 0 | 0 | 0 | 0 | 0 | 0 | 0 | 0 | 0 | 0 |
| SRR5230663 | 0 | 0 | 0 | 0 | 0 | 0 | 0 | 0 | 0 | 0 | 0 |
| SRR5230664 | 0 | 0 | 0 | 0 | 0 | 0 | 0 | 0 | 0 | 0 | 0 |
| SRR5230665 | 0 | 0 | 0 | 0 | 0 | 0 | 0 | 0 | 0 | 0 | 0 |
| SRR5230666 | 0 | 0 | 0 | 17 | 0 | 0 | 0 | 0 | 0 | 0 | 0 |
| SRR5230667 | 0 | 0 | 0 | 0 | 0 | 0 | 0 | 0 | 0 | 0 | 0 |
| SRR5230668 | 0 | 0 | 0 | 0 | 0 | 0 | 0 | 0 | 0 | 0 | 0 |
| SRR5230669 | 0 | 12 | 0 | 0 | 0 | 16 | 14 | 0 | 0 | 0 | 0 |
| SRR5230670 | 0 | 0 | 0 | 0 | 0 | 0 | 0 | 0 | 0 | 0 | 0 |
| SRR5230671 | 0 | 0 | 0 | 0 | 0 | 0 | 0 | 0 | 0 | 0 | 0 |
| SRR5230673 | 0 | 0 | 0 | 0 | 0 | 0 | 0 | 0 | 0 | 0 | 0 |
| SRR5230674 | 0 | 0 | 0 | 0 | 0 | 0 | 0 | 0 | 0 | 0 | 0 |
| SRR5230675 | 0 | 0 | 0 | 0 | 0 | 0 | 0 | 0 | 0 | 0 | 0 |
| SRR5230676 | 0 | 0 | 0 | 0 | 0 | 0 | 0 | 0 | 0 | 0 | 0 |
| SRR5230677 | 0 | 0 | 0 | 0 | 0 | 0 | 0 | 0 | 0 | 0 | 0 |
| SRR5230678 | 0 | 19 | 0 | 0 | 0 | 0 | 0 | 0 | 0 | 0 | 0 |
| SRR5230679 | 0 | 0 | 0 | 0 | 0 | 0 | 0 | 0 | 0 | 0 | 0 |
| SRR5230680 | 0 | 0 | 0 | 0 | 0 | 0 | 0 | 0 | 0 | 0 | 0 |
| SRR5230681 | 0 | 0 | 0 | 0 | 0 | 0 | 0 | 0 | 0 | 0 | 0 |
| SRR5230682 | 0 | 0 | 0 | 0 | 0 | 0 | 0 | 0 | 0 | 0 | 0 |
| SRR5230683 | 0 | 0 | 0 | 0 | 0 | 0 | 0 | 0 | 0 | 0 | 0 |
| SRR5230684 | 0 | 0 | 0 | 0 | 0 | 0 | 0 | 0 | 0 | 0 | 0 |
| SRR5230685 | 0 | 0 | 0 | 0 | 0 | 0 | 0 | 0 | 0 | 0 | 0 |
| SRR5230686 | 0 | 0 | 0 | 0 | 14 | 20 | 0 | 0 | 0 | 0 | 0 |
| SRR5230687 | 0 | 0 | 0 | 0 | 0 | 0 | 0 | 0 | 0 | 0 | 0 |
| SRR5230688 | 0 | 0 | 0 | 0 | 0 | 0 | 0 | 0 | 0 | 0 | 0 |
| SRR5230689 | 0 | 0 | 0 | 0 | 0 | 0 | 0 | 0 | 0 | 0 | 0 |
| SRR5230690 | 0 | 0 | 0 | 0 | 0 | 0 | 0 | 0 | 0 | 0 | 0 |
| SRR5230691 | 0 | 0 | 0 | 0 | 0 | 0 | 0 | 0 | 0 | 0 | 0 |
| SRR5230692 | 0 | 0 | 0 | 0 | 0 | 0 | 0 | 0 | 0 | 0 | 0 |
| SRR5230693 | 0 | 0 | 0 | 0 | 0 | 0 | 0 | 0 | 0 | 0 | 0 |
| SRR5230694 | 0 | 0 | 0 | 0 | 0 | 0 | 0 | 0 | 0 | 0 | 0 |
| SRR5230695 | 0 | 0 | 15 | 0 | 0 | 0 | 0 | 0 | 0 | 0 | 0 |
| SRR5230696 | 0 | 0 | 0 | 0 | 0 | 0 | 0 | 0 | 0 | 0 | 0 |
| SRR5230697 | 0 | 0 | 0 | 0 | 0 | 0 | 0 | 0 | 0 | 0 | 0 |
| SRR5230698 | 0 | 14 | 0 | 0 | 0 | 0 | 0 | 0 | 0 | 0 | 0 |
| SRR5230699 | 0 | 0 | 0 | 0 | 0 | 0 | 0 | 0 | 0 | 0 | 0 |
| SRR5230700 | 0 | 0 | 0 | 0 | 0 | 0 | 0 | 0 | 0 | 0 | 0 |
| SRR5230701 | 0 | 0 | 0 | 0 | 0 | 0 | 0 | 0 | 0 | 0 | 0 |
| SRR5230702 | 0 | 0 | 0 | 0 | 0 | 0 | 0 | 0 | 0 | 0 | 0 |
| SRR5230703 | 0 | 0 | 0 | 0 | 0 | 0 | 0 | 0 | 0 | 0 | 0 |
| SRR5230704 | 0 | 0 | 0 | 0 | 0 | 0 | 0 | 0 | 0 | 0 | 0 |
| SRR5230705 | 0 | 0 | 0 | 0 | 0 | 0 | 0 | 0 | 0 | 0 | 0 |
| SRR5230706 | 0 | 0 | 0 | 0 | 0 | 0 | 0 | 0 | 0 | 0 | 0 |
| SRR5230707 | 0 | 0 | 0 | 0 | 0 | 0 | 0 | 0 | 0 | 0 | 0 |
| SRR5230708 | 0 | 0 | 0 | 0 | 0 | 0 | 0 | 0 | 0 | 0 | 0 |
| SRR5230709 | 0 | 0 | 0 | 0 | 0 | 0 | 0 | 0 | 0 | 0 | 0 |
| SRR5230710 | 0 | 0 | 0 | 0 | 0 | 0 | 0 | 0 | 0 | 0 | 0 |
| SRR5230711 | 0 | 0 | 0 | 0 | 0 | 0 | 0 | 0 | 0 | 0 | 0 |
| SRR5230712 | 0 | 0 | 0 | 0 | 0 | 0 | 0 | 0 | 0 | 0 | 0 |
| SRR5230713 | 0 | 0 | 0 | 0 | 0 | 0 | 0 | 0 | 0 | 0 | 0 |
| SRR5230714 | 0 | 0 | 0 | 0 | 0 | 0 | 0 | 0 | 0 | 0 | 0 |
| SRR5230716 | 0 | 0 | 0 | 0 | 0 | 0 | 0 | 0 | 0 | 0 | 0 |
| SRR5230717 | 0 | 0 | 0 | 0 | 0 | 0 | 0 | 0 | 0 | 0 | 0 |
| SRR5230718 | 0 | 0 | 0 | 0 | 0 | 0 | 0 | 0 | 0 | 0 | 0 |
| SRR5230719 | 0 | 0 | 0 | 0 | 0 | 0 | 0 | 0 | 0 | 0 | 0 |
| SRR5230720 | 0 | 0 | 0 | 0 | 0 | 0 | 0 | 0 | 0 | 0 | 0 |
| SRR5230721 | 0 | 0 | 0 | 0 | 0 | 0 | 0 | 0 | 0 | 0 | 0 |
| SRR5230722 | 0 | 0 | 0 | 0 | 0 | 0 | 0 | 0 | 0 | 0 | 0 |
| SRR5230723 | 0 | 0 | 0 | 0 | 0 | 0 | 0 | 0 | 0 | 0 | 0 |
| SRR5230724 | 0 | 0 | 0 | 0 | 0 | 0 | 0 | 0 | 0 | 0 | 0 |
| SRR5230725 | 0 | 0 | 0 | 0 | 0 | 0 | 0 | 0 | 0 | 0 | 0 |
| SRR5230726 | 0 | 0 | 0 | 0 | 0 | 0 | 0 | 0 | 0 | 0 | 0 |
| SRR5230727 | 0 | 0 | 0 | 0 | 0 | 0 | 0 | 0 | 0 | 0 | 0 |
| SRR5230728 | 0 | 0 | 0 | 0 | 0 | 0 | 0 | 0 | 0 | 0 | 0 |
| SRR5230729 | 0 | 0 | 0 | 0 | 0 | 0 | 0 | 0 | 0 | 0 | 0 |
| SRR5230732 | 0 | 0 | 0 | 0 | 0 | 0 | 0 | 0 | 0 | 0 | 0 |
| SRR5230733 | 0 | 0 | 0 | 0 | 0 | 0 | 0 | 0 | 0 | 0 | 0 |
| SRR5230734 | 0 | 0 | 0 | 0 | 0 | 0 | 0 | 0 | 0 | 0 | 0 |
| SRR5230735 | 0 | 0 | 0 | 0 | 0 | 0 | 0 | 0 | 0 | 0 | 0 |
| SRR5230736 | 0 | 0 | 0 | 0 | 0 | 0 | 0 | 0 | 0 | 0 | 0 |
| SRR5230737 | 0 | 0 | 0 | 0 | 0 | 0 | 0 | 0 | 20 | 0 | 0 |
| SRR5230738 | 0 | 0 | 0 | 0 | 0 | 0 | 0 | 0 | 0 | 0 | 0 |
| SRR5230739 | 0 | 0 | 0 | 0 | 0 | 0 | 0 | 0 | 0 | 0 | 0 |
| SRR5230740 | 0 | 0 | 0 | 0 | 0 | 0 | 0 | 0 | 0 | 0 | 0 |
| SRR5230741 | 0 | 0 | 0 | 0 | 0 | 0 | 0 | 0 | 0 | 0 | 0 |
| SRR5230742 | 0 | 0 | 0 | 0 | 0 | 0 | 0 | 0 | 0 | 0 | 0 |
| SRR5230743 | 0 | 0 | 0 | 0 | 0 | 0 | 0 | 0 | 0 | 0 | 0 |
| SRR5230744 | 0 | 0 | 0 | 0 | 0 | 0 | 0 | 0 | 0 | 0 | 0 |
| SRR5230745 | 0 | 0 | 0 | 0 | 13 | 0 | 0 | 0 | 0 | 0 | 0 |
| SRR5230746 | 0 | 0 | 0 | 0 | 0 | 0 | 0 | 0 | 0 | 0 | 0 |
| SRR5230747 | 0 | 0 | 0 | 0 | 0 | 0 | 0 | 0 | 0 | 0 | 0 |
| SRR5230748 | 0 | 0 | 0 | 0 | 0 | 11 | 0 | 0 | 0 | 0 | 0 |
| SRR5230749 | 0 | 0 | 0 | 0 | 0 | 0 | 0 | 19 | 0 | 0 | 0 |
| SRR5230750 | 0 | 0 | 0 | 0 | 0 | 0 | 0 | 0 | 0 | 0 | 0 |
| SRR5230751 | 0 | 0 | 0 | 0 | 0 | 0 | 0 | 0 | 0 | 0 | 0 |
| SRR5230752 | 0 | 0 | 0 | 0 | 0 | 0 | 0 | 0 | 0 | 0 | 0 |
| SRR5230753 | 0 | 0 | 0 | 0 | 0 | 0 | 0 | 0 | 0 | 0 | 0 |
| SRR5231309 | 0 | 0 | 28 | 11 | 0 | 17 | 0 | 31 | 15 | 16 | 12 |
| SRR5231310 | 0 | 0 | 11 | 0 | 0 | 17 | 0 | 26 | 21 | 34 | 27 |
| SRR5231311 | 0 | 0 | 10 | 0 | 0 | 0 | 0 | 10 | 0 | 10 | 0 |
| SRR5231312 | 0 | 0 | 0 | 0 | 0 | 0 | 0 | 15 | 12 | 15 | 12 |
| SRR5231313 | 0 | 0 | 13 | 0 | 0 | 16 | 0 | 39 | 19 | 18 | 11 |

### Supp Table S8.5.Columns_ET_to_FD

| | | | | | | | | | | | |
|---|---|---|---|---|---|---|---|---|---|---|---|
| SRR5231314 | 0 | 0 | 0 | 0 | 0 | 0 | 0 | 14 | 0 | 11 | 0 |
| SRR5239182 | 0 | 0 | 0 | 0 | 0 | 82 | 16 | 184 | 264 | 193 | 94 |
| SRR5239183 | 0 | 79 | 141 | 44 | 0 | 51 | 0 | 60 | 107 | 49 | 48 |
| SRR5239184 | 0 | 33 | 74 | 25 | 10 | 60 | 0 | 89 | 142 | 118 | 78 |
| SRR5239185 | 0 | 0 | 0 | 0 | 0 | 61 | 14 | 98 | 67 | 57 | 40 |
| SRR5239186 | 0 | 0 | 0 | 0 | 0 | 90 | 19 | 132 | 193 | 162 | 91 |
| SRR5239187 | 0 | 25 | 71 | 28 | 17 | 81 | 15 | 98 | 155 | 121 | 104 |
| SRR5239188 | 0 | 0 | 18 | 11 | 0 | 115 | 17 | 184 | 124 | 101 | 67 |
| SRR5275577 | 0 | 0 | 18 | 0 | 31 | 468 | 59 | 262 | 382 | 460 | 220 |
| SRR5275578 | 0 | 0 | 14 | 0 | 23 | 393 | 52 | 224 | 452 | 599 | 295 |
| SRR5275579 | 0 | 0 | 14 | 0 | 26 | 436 | 59 | 260 | 369 | 443 | 215 |
| SRR5275580 | 0 | 11 | 44 | 16 | 88 | 1082 | 166 | 1077 | 1515 | 1929 | 792 |
| SRR5275581 | 0 | 0 | 16 | 0 | 63 | 831 | 133 | 913 | 1462 | 1876 | 754 |
| SRR5275582 | 0 | 32 | 104 | 45 | 107 | 1243 | 172 | 936 | 875 | 1060 | 415 |
| SRR5284195 | 0 | 0 | 10 | 0 | 273 | 398 | 279 | 77 | 325 | 26 | 42 |
| SRR5284196 | 0 | 17 | 37 | 21 | 132 | 156 | 124 | 79 | 279 | 12 | 18 |
| SRR5284197 | 0 | 20 | 33 | 24 | 271 | 644 | 271 | 86 | 208 | 24 | 16 |
| SRR5284198 | 0 | 10 | 24 | 13 | 153 | 158 | 119 | 53 | 229 | 0 | 14 |
| SRR5284199 | 0 | 0 | 25 | 10 | 51 | 67 | 55 | 50 | 223 | 10 | 13 |
| SRR5284200 | 0 | 0 | 26 | 20 | 119 | 118 | 68 | 40 | 232 | 0 | 0 |
| SRR5284201 | 0 | 0 | 15 | 12 | 88 | 128 | 49 | 55 | 160 | 17 | 0 |
| SRR5284202 | 0 | 0 | 0 | 0 | 53 | 138 | 55 | 45 | 180 | 13 | 0 |
| SRR5284203 | 0 | 23 | 35 | 25 | 55 | 163 | 50 | 69 | 150 | 13 | 0 |
| SRR5284204 | 0 | 0 | 18 | 17 | 135 | 120 | 79 | 52 | 353 | 16 | 17 |
| SRR5284205 | 0 | 15 | 24 | 12 | 334 | 588 | 211 | 126 | 348 | 38 | 22 |
| SRR5284206 | 0 | 13 | 21 | 0 | 288 | 564 | 202 | 83 | 274 | 35 | 12 |
| SRR5317818 | 0 | 0 | 0 | 0 | 0 | 32 | 0 | 36 | 21 | 31 | 68 |
| SRR5317819 | 0 | 0 | 29 | 57 | 12 | 48 | 0 | 77 | 29 | 33 | 46 |
| SRR5317820 | 0 | 0 | 0 | 0 | 0 | 28 | 0 | 41 | 26 | 24 | 66 |
| SRR5317821 | 0 | 0 | 0 | 0 | 0 | 0 | 0 | 0 | 0 | 0 | 18 |
| SRR5317822 | 0 | 0 | 0 | 0 | 0 | 27 | 0 | 36 | 25 | 27 | 166 |
| SRR5317823 | 0 | 0 | 0 | 0 | 0 | 18 | 0 | 20 | 0 | 13 | 16 |
| SRR5317824 | 0 | 0 | 0 | 0 | 0 | 16 | 0 | 36 | 26 | 27 | 20 |
| SRR5317825 | 0 | 0 | 0 | 11 | 0 | 15 | 0 | 25 | 0 | 11 | 0 |
| SRR5317826 | 0 | 0 | 0 | 0 | 0 | 35 | 0 | 40 | 24 | 43 | 21 |
| SRR5317827 | 0 | 0 | 0 | 17 | 0 | 13 | 0 | 14 | 11 | 20 | 16 |
| SRR5387779 | 0 | 0 | 0 | 14 | 0 | 0 | 0 | 0 | 0 | 14 | 0 |
| SRR5387780 | 0 | 0 | 23 | 0 | 0 | 15 | 0 | 0 | 0 | 0 | 15 |
| SRR5387781 | 0 | 0 | 18 | 0 | 0 | 0 | 0 | 0 | 0 | 0 | 0 |
| SRR5387782 | 0 | 0 | 22 | 0 | 0 | 0 | 0 | 0 | 0 | 0 | 22 |
| SRR5387783 | 0 | 0 | 25 | 0 | 0 | 0 | 0 | 0 | 0 | 0 | 0 |
| SRR5387784 | 0 | 0 | 0 | 0 | 0 | 0 | 0 | 0 | 0 | 0 | 0 |
| SRR5387785 | 0 | 0 | 0 | 0 | 0 | 0 | 0 | 13 | 0 | 0 | 0 |
| SRR5387786 | 0 | 0 | 0 | 0 | 0 | 0 | 0 | 0 | 0 | 0 | 0 |
| SRR5387787 | 0 | 0 | 0 | 0 | 0 | 0 | 0 | 0 | 0 | 0 | 0 |
| SRR5387788 | 0 | 0 | 21 | 14 | 0 | 0 | 0 | 0 | 0 | 0 | 0 |
| SRR5387789 | 0 | 0 | 0 | 0 | 0 | 0 | 0 | 0 | 0 | 0 | 0 |
| SRR5387790 | 0 | 0 | 14 | 0 | 0 | 14 | 0 | 0 | 0 | 0 | 0 |
| SRR5387791 | 0 | 0 | 18 | 0 | 0 | 0 | 0 | 18 | 0 | 0 | 0 |
| SRR5387792 | 0 | 0 | 10 | 31 | 0 | 10 | 0 | 0 | 0 | 0 | 0 |
| SRR5387793 | 0 | 14 | 43 | 29 | 0 | 0 | 0 | 14 | 0 | 0 | 0 |
| SRR5387794 | 0 | 13 | 13 | 0 | 0 | 0 | 0 | 0 | 13 | 0 | 13 |
| SRR5387795 | 0 | 11 | 0 | 22 | 0 | 0 | 0 | 0 | 0 | 0 | 0 |
| SRR5387796 | 0 | 0 | 0 | 0 | 0 | 0 | 0 | 0 | 0 | 0 | 0 |
| SRR5387797 | 0 | 0 | 0 | 0 | 0 | 0 | 0 | 16 | 0 | 0 | 0 |
| SRR5387798 | 0 | 0 | 22 | 44 | 0 | 0 | 0 | 0 | 0 | 0 | 0 |
| SRR5387799 | 0 | 0 | 0 | 0 | 0 | 0 | 0 | 0 | 0 | 0 | 0 |
| SRR5387800 | 0 | 0 | 33 | 0 | 0 | 0 | 0 | 0 | 0 | 0 | 0 |
| SRR5387801 | 0 | 0 | 0 | 19 | 0 | 0 | 0 | 0 | 0 | 0 | 0 |
| SRR5387802 | 0 | 0 | 0 | 13 | 0 | 0 | 0 | 13 | 0 | 0 | 13 |
| SRR5387803 | 0 | 0 | 0 | 20 | 0 | 0 | 0 | 0 | 0 | 0 | 0 |
| SRR5387804 | 0 | 0 | 16 | 16 | 0 | 0 | 0 | 0 | 0 | 0 | 0 |
| SRR5387805 | 0 | 0 | 0 | 0 | 0 | 0 | 0 | 0 | 0 | 0 | 0 |
| SRR5387806 | 0 | 0 | 0 | 32 | 0 | 0 | 0 | 0 | 0 | 0 | 0 |
| SRR5387807 | 0 | 0 | 24 | 59 | 0 | 0 | 0 | 0 | 0 | 0 | 12 |
| SRR5387808 | 0 | 0 | 30 | 0 | 0 | 0 | 0 | 0 | 20 | 0 | 0 |
| SRR5387809 | 0 | 0 | 35 | 18 | 0 | 18 | 0 | 18 | 0 | 0 | 0 |
| SRR5387810 | 0 | 0 | 0 | 0 | 0 | 0 | 0 | 0 | 0 | 0 | 0 |
| SRR5387811 | 0 | 0 | 0 | 31 | 0 | 0 | 0 | 0 | 0 | 0 | 0 |
| SRR5387812 | 0 | 0 | 0 | 0 | 0 | 0 | 0 | 0 | 0 | 0 | 0 |
| SRR5387813 | 0 | 0 | 0 | 30 | 0 | 0 | 0 | 0 | 0 | 0 | 0 |
| SRR5387814 | 0 | 0 | 18 | 0 | 0 | 35 | 0 | 0 | 0 | 0 | 0 |
| SRR5387815 | 0 | 0 | 28 | 0 | 0 | 0 | 0 | 0 | 0 | 0 | 14 |
| SRR5387816 | 0 | 0 | 56 | 19 | 0 | 0 | 0 | 0 | 0 | 0 | 0 |
| SRR5387817 | 0 | 0 | 13 | 0 | 0 | 0 | 0 | 13 | 0 | 0 | 0 |
| SRR5387818 | 0 | 0 | 0 | 0 | 0 | 0 | 0 | 0 | 0 | 0 | 0 |
| SRR5387819 | 0 | 0 | 0 | 0 | 0 | 0 | 0 | 34 | 0 | 0 | 17 |
| SRR5387820 | 0 | 0 | 29 | 19 | 0 | 0 | 0 | 0 | 0 | 0 | 0 |
| SRR5387821 | 0 | 0 | 0 | 0 | 0 | 0 | 0 | 0 | 0 | 0 | 0 |
| SRR5387822 | 0 | 0 | 13 | 0 | 0 | 13 | 0 | 0 | 0 | 0 | 0 |
| SRR5387823 | 0 | 0 | 16 | 0 | 0 | 0 | 0 | 0 | 0 | 0 | 0 |
| SRR5387824 | 0 | 0 | 0 | 0 | 0 | 0 | 0 | 0 | 0 | 0 | 0 |
| SRR5387825 | 0 | 0 | 0 | 0 | 0 | 0 | 0 | 0 | 0 | 0 | 0 |
| SRR5387826 | 0 | 0 | 0 | 0 | 0 | 0 | 0 | 0 | 12 | 0 | 0 |
| SRR5387827 | 0 | 0 | 32 | 16 | 0 | 16 | 0 | 16 | 32 | 0 | 16 |
| SRR5387828 | 0 | 0 | 0 | 0 | 0 | 0 | 0 | 0 | 0 | 0 | 0 |
| SRR5387829 | 0 | 10 | 30 | 30 | 0 | 0 | 0 | 0 | 0 | 0 | 0 |
| SRR5387830 | 0 | 0 | 0 | 32 | 0 | 0 | 0 | 0 | 0 | 0 | 0 |
| SRR5387831 | 0 | 0 | 0 | 0 | 0 | 0 | 0 | 0 | 0 | 0 | 0 |
| SRR5387832 | 0 | 0 | 10 | 31 | 0 | 0 | 0 | 10 | 0 | 0 | 0 |
| SRR5387833 | 0 | 0 | 23 | 0 | 0 | 12 | 0 | 0 | 12 | 0 | 12 |
| SRR5387834 | 0 | 0 | 0 | 0 | 0 | 0 | 0 | 0 | 0 | 0 | 0 |
| SRR5387835 | 0 | 0 | 0 | 0 | 0 | 0 | 0 | 0 | 0 | 0 | 0 |
| SRR5387836 | 0 | 0 | 22 | 0 | 0 | 0 | 0 | 15 | 0 | 0 | 0 |
| SRR5387837 | 0 | 0 | 15 | 0 | 0 | 0 | 0 | 0 | 0 | 0 | 15 |
| SRR5387838 | 0 | 0 | 38 | 19 | 0 | 0 | 0 | 0 | 0 | 0 | 0 |
| SRR5387839 | 0 | 0 | 11 | 32 | 0 | 0 | 0 | 0 | 0 | 0 | 32 |
| SRR5387840 | 0 | 0 | 29 | 29 | 14 | 0 | 0 | 14 | 0 | 0 | 14 |
| SRR5387841 | 0 | 0 | 16 | 16 | 0 | 0 | 0 | 0 | 0 | 0 | 0 |
| SRR5387842 | 0 | 0 | 0 | 0 | 0 | 0 | 0 | 0 | 0 | 0 | 0 |
| SRR5387843 | 0 | 0 | 20 | 13 | 0 | 13 | 0 | 0 | 0 | 0 | 0 |
| SRR5387844 | 0 | 0 | 18 | 0 | 0 | 0 | 0 | 0 | 0 | 0 | 0 |
| SRR5387845 | 0 | 0 | 20 | 0 | 0 | 0 | 0 | 0 | 0 | 0 | 0 |
| SRR5387846 | 0 | 0 | 0 | 25 | 0 | 0 | 0 | 0 | 0 | 0 | 0 |
| SRR5387847 | 0 | 0 | 0 | 0 | 0 | 0 | 0 | 0 | 0 | 0 | 0 |
| SRR5387848 | 0 | 0 | 14 | 34 | 0 | 0 | 0 | 0 | 0 | 0 | 0 |
| SRR5387849 | 0 | 0 | 23 | 0 | 0 | 0 | 0 | 0 | 0 | 0 | 0 |
| SRR5387850 | 0 | 0 | 27 | 0 | 0 | 0 | 0 | 0 | 0 | 0 | 0 |
| SRR5387851 | 0 | 0 | 0 | 0 | 0 | 0 | 0 | 14 | 0 | 14 | 14 |
| SRR5387852 | 0 | 0 | 0 | 18 | 0 | 18 | 0 | 0 | 0 | 0 | 0 |
| SRR5387853 | 0 | 0 | 10 | 10 | 0 | 0 | 0 | 0 | 0 | 0 | 0 |
| SRR5387854 | 0 | 0 | 0 | 0 | 0 | 0 | 0 | 0 | 0 | 0 | 14 |
| SRR5387855 | 0 | 18 | 18 | 28 | 0 | 0 | 0 | 0 | 0 | 0 | 0 |
| SRR5387856 | 0 | 0 | 0 | 14 | 0 | 0 | 0 | 0 | 0 | 0 | 0 |
| SRR5387857 | 0 | 0 | 11 | 11 | 0 | 0 | 0 | 0 | 0 | 0 | 0 |
| SRR5387858 | 0 | 0 | 0 | 25 | 0 | 0 | 0 | 0 | 0 | 0 | 0 |
| SRR5387859 | 0 | 0 | 13 | 0 | 0 | 0 | 0 | 0 | 0 | 0 | 13 |
| SRR5387860 | 0 | 0 | 11 | 11 | 0 | 0 | 0 | 11 | 0 | 0 | 0 |
| SRR5387861 | 0 | 0 | 20 | 20 | 0 | 20 | 0 | 20 | 0 | 0 | 0 |

| | | | | | | | | | | | |
|---|---|---|---|---|---|---|---|---|---|---|---|
| SRR5387862 | 0 | 0 | 0 | 17 | 0 | 0 | 0 | 0 | 0 | 0 | 0 |
| SRR5387863 | 0 | 0 | 19 | 0 | 0 | 0 | 0 | 19 | 0 | 0 | 0 |
| SRR5387864 | 0 | 0 | 15 | 15 | 0 | 0 | 0 | 0 | 0 | 0 | 0 |
| SRR5387865 | 0 | 0 | 13 | 0 | 0 | 0 | 0 | 0 | 20 | 0 | 0 |
| SRR5387866 | 0 | 0 | 0 | 0 | 0 | 0 | 0 | 0 | 0 | 0 | 0 |
| SRR5387867 | 0 | 0 | 0 | 0 | 0 | 16 | 0 | 0 | 24 | 0 | 0 |
| SRR5387868 | 0 | 0 | 21 | 10 | 0 | 0 | 0 | 0 | 0 | 0 | 0 |
| SRR5387869 | 0 | 0 | 32 | 0 | 0 | 0 | 0 | 0 | 0 | 0 | 0 |
| SRR5387870 | 0 | 0 | 15 | 15 | 0 | 0 | 0 | 0 | 0 | 0 | 0 |
| SRR5387871 | 0 | 0 | 0 | 12 | 0 | 0 | 0 | 0 | 12 | 0 | 0 |
| SRR5387872 | 0 | 0 | 0 | 0 | 0 | 0 | 0 | 20 | 0 | 0 | 0 |
| SRR5387873 | 0 | 0 | 0 | 36 | 0 | 0 | 0 | 18 | 18 | 0 | 18 |
| SRR5387874 | 0 | 0 | 0 | 0 | 0 | 0 | 0 | 0 | 0 | 0 | 25 |
| SRR5387875 | 0 | 0 | 0 | 0 | 0 | 0 | 0 | 0 | 0 | 0 | 0 |
| SRR5387876 | 0 | 0 | 15 | 0 | 0 | 0 | 0 | 0 | 0 | 0 | 0 |
| SRR5387877 | 0 | 0 | 23 | 23 | 0 | 0 | 0 | 0 | 11 | 0 | 0 |
| SRR5387878 | 0 | 0 | 0 | 0 | 0 | 0 | 0 | 0 | 0 | 0 | 0 |
| SRR5387879 | 0 | 0 | 31 | 0 | 0 | 0 | 0 | 0 | 0 | 0 | 0 |
| SRR5387880 | 0 | 0 | 0 | 0 | 0 | 0 | 0 | 0 | 0 | 0 | 0 |
| SRR5387881 | 0 | 0 | 15 | 0 | 0 | 0 | 0 | 0 | 0 | 15 | 0 |
| SRR5387882 | 0 | 0 | 0 | 0 | 0 | 0 | 0 | 0 | 0 | 0 | 0 |
| SRR5387883 | 0 | 0 | 0 | 0 | 0 | 0 | 0 | 15 | 0 | 0 | 15 |
| SRR5387884 | 0 | 0 | 26 | 0 | 0 | 0 | 0 | 0 | 0 | 0 | 0 |
| SRR5387885 | 0 | 0 | 0 | 0 | 0 | 10 | 0 | 10 | 0 | 0 | 21 |
| SRR5387886 | 0 | 0 | 0 | 18 | 0 | 0 | 0 | 0 | 0 | 0 | 0 |
| SRR5387887 | 0 | 0 | 0 | 0 | 0 | 11 | 0 | 11 | 0 | 0 | 0 |
| SRR5387888 | 0 | 0 | 15 | 0 | 0 | 0 | 0 | 0 | 0 | 0 | 0 |
| SRR5387889 | 0 | 0 | 0 | 0 | 0 | 0 | 0 | 0 | 0 | 0 | 0 |
| SRR5387890 | 0 | 0 | 28 | 0 | 0 | 0 | 0 | 0 | 0 | 0 | 0 |
| SRR5387891 | 0 | 0 | 0 | 0 | 0 | 0 | 0 | 19 | 0 | 0 | 0 |
| SRR5387892 | 0 | 0 | 14 | 0 | 0 | 0 | 0 | 0 | 0 | 0 | 0 |
| SRR5387893 | 0 | 0 | 12 | 0 | 0 | 12 | 0 | 0 | 0 | 0 | 12 |
| SRR5387894 | 0 | 19 | 0 | 0 | 0 | 0 | 0 | 0 | 0 | 0 | 0 |
| SRR5387895 | 0 | 0 | 18 | 18 | 0 | 0 | 0 | 0 | 0 | 0 | 0 |
| SRR5387896 | 0 | 13 | 13 | 33 | 0 | 0 | 0 | 0 | 0 | 0 | 0 |
| SRR5387897 | 0 | 11 | 11 | 32 | 0 | 11 | 0 | 11 | 0 | 0 | 21 |
| SRR5387898 | 0 | 0 | 17 | 0 | 0 | 0 | 0 | 0 | 0 | 0 | 0 |
| SRR5387899 | 0 | 0 | 0 | 28 | 0 | 0 | 0 | 0 | 0 | 0 | 18 |
| SRR5387900 | 0 | 0 | 0 | 17 | 0 | 0 | 0 | 0 | 0 | 0 | 0 |
| SRR5387901 | 0 | 0 | 0 | 0 | 0 | 0 | 0 | 0 | 0 | 0 | 0 |
| SRR5387902 | 0 | 0 | 0 | 0 | 0 | 0 | 0 | 0 | 0 | 0 | 0 |
| SRR5387903 | 0 | 0 | 15 | 0 | 0 | 0 | 0 | 0 | 0 | 0 | 0 |
| SRR5387904 | 0 | 0 | 18 | 18 | 0 | 0 | 0 | 0 | 0 | 0 | 0 |
| SRR5387905 | 0 | 12 | 24 | 0 | 0 | 12 | 0 | 0 | 0 | 0 | 0 |
| SRR5387906 | 0 | 0 | 23 | 0 | 0 | 0 | 0 | 0 | 0 | 0 | 0 |
| SRR5387907 | 0 | 0 | 0 | 0 | 0 | 0 | 0 | 0 | 0 | 0 | 0 |
| SRR5387908 | 0 | 0 | 0 | 0 | 0 | 0 | 0 | 0 | 15 | 0 | 0 |
| SRR5387909 | 0 | 0 | 18 | 18 | 0 | 18 | 0 | 0 | 0 | 0 | 0 |
| SRR5387910 | 0 | 17 | 0 | 0 | 0 | 0 | 0 | 0 | 0 | 0 | 0 |
| SRR5387911 | 0 | 0 | 0 | 17 | 0 | 0 | 0 | 0 | 0 | 0 | 42 |
| SRR5387912 | 0 | 0 | 23 | 15 | 0 | 0 | 0 | 15 | 0 | 0 | 0 |
| SRR5387913 | 0 | 12 | 23 | 23 | 0 | 12 | 0 | 12 | 0 | 0 | 35 |
| SRR5387914 | 0 | 0 | 39 | 59 | 0 | 0 | 0 | 0 | 0 | 0 | 0 |
| SRR5387915 | 0 | 0 | 0 | 23 | 0 | 0 | 0 | 0 | 0 | 0 | 0 |
| SRR5387916 | 0 | 0 | 62 | 27 | 0 | 0 | 0 | 0 | 0 | 0 | 0 |
| SRR5387917 | 0 | 0 | 32 | 0 | 0 | 0 | 0 | 0 | 0 | 0 | 16 |
| SRR5387918 | 0 | 0 | 25 | 0 | 0 | 0 | 0 | 0 | 0 | 0 | 0 |
| SRR5387919 | 0 | 0 | 0 | 20 | 0 | 0 | 0 | 20 | 0 | 0 | 0 |
| SRR5387920 | 0 | 12 | 12 | 12 | 0 | 0 | 0 | 0 | 0 | 0 | 0 |
| SRR5387921 | 0 | 0 | 0 | 0 | 0 | 0 | 0 | 0 | 0 | 0 | 0 |
| SRR5387922 | 0 | 0 | 40 | 20 | 0 | 0 | 0 | 0 | 0 | 0 | 0 |
| SRR5387923 | 0 | 0 | 34 | 0 | 11 | 0 | 0 | 11 | 0 | 0 | 0 |
| SRR5387924 | 0 | 0 | 0 | 0 | 0 | 0 | 0 | 0 | 0 | 0 | 0 |
| SRR5387925 | 0 | 0 | 0 | 0 | 0 | 0 | 0 | 0 | 0 | 0 | 0 |
| SRR5387926 | 0 | 0 | 0 | 0 | 0 | 0 | 0 | 0 | 0 | 0 | 0 |
| SRR5387927 | 0 | 0 | 28 | 28 | 0 | 0 | 0 | 38 | 0 | 0 | 0 |
| SRR5387928 | 0 | 16 | 16 | 0 | 0 | 0 | 0 | 0 | 16 | 0 | 0 |
| SRR5387929 | 0 | 0 | 15 | 38 | 0 | 0 | 0 | 0 | 0 | 0 | 31 |
| SRR5387930 | 0 | 12 | 25 | 37 | 0 | 0 | 0 | 0 | 0 | 12 | 0 |
| SRR5387931 | 0 | 0 | 0 | 0 | 0 | 0 | 0 | 0 | 0 | 17 | 0 |
| SRR5387932 | 0 | 16 | 0 | 0 | 0 | 0 | 0 | 0 | 0 | 0 | 0 |
| SRR5387933 | 0 | 0 | 0 | 20 | 0 | 0 | 0 | 0 | 0 | 0 | 0 |
| SRR5387934 | 0 | 0 | 0 | 0 | 0 | 0 | 0 | 0 | 0 | 0 | 0 |
| SRR5387935 | 0 | 21 | 0 | 0 | 0 | 21 | 0 | 10 | 0 | 0 | 21 |
| SRR5387936 | 0 | 0 | 17 | 17 | 0 | 0 | 0 | 0 | 0 | 0 | 0 |
| SRR5387937 | 0 | 0 | 0 | 14 | 0 | 0 | 0 | 0 | 0 | 0 | 0 |
| SRR5387938 | 0 | 0 | 0 | 29 | 0 | 0 | 0 | 14 | 0 | 0 | 0 |
| SRR5387939 | 0 | 0 | 0 | 0 | 0 | 0 | 0 | 0 | 0 | 0 | 0 |
| SRR5387940 | 0 | 0 | 0 | 0 | 0 | 0 | 0 | 0 | 0 | 0 | 0 |
| SRR5387941 | 0 | 0 | 29 | 37 | 0 | 0 | 0 | 0 | 0 | 0 | 0 |
| SRR5387942 | 0 | 0 | 0 | 0 | 0 | 0 | 0 | 0 | 0 | 0 | 0 |
| SRR5387943 | 0 | 0 | 0 | 0 | 0 | 0 | 0 | 0 | 0 | 0 | 0 |
| SRR5387944 | 0 | 16 | 16 | 24 | 0 | 0 | 0 | 0 | 16 | 0 | 24 |
| SRR5387945 | 0 | 0 | 0 | 0 | 0 | 0 | 0 | 0 | 0 | 0 | 0 |
| SRR5387946 | 0 | 0 | 0 | 0 | 0 | 0 | 0 | 0 | 0 | 0 | 0 |
| SRR5387947 | 0 | 0 | 22 | 11 | 0 | 11 | 0 | 0 | 0 | 0 | 11 |
| SRR5387948 | 0 | 0 | 0 | 0 | 0 | 0 | 0 | 0 | 0 | 0 | 0 |
| SRR5387949 | 0 | 0 | 13 | 0 | 0 | 0 | 0 | 0 | 0 | 13 | 13 |
| SRR5387950 | 0 | 23 | 23 | 0 | 0 | 0 | 0 | 0 | 0 | 0 | 15 |
| SRR5387951 | 0 | 0 | 0 | 22 | 0 | 0 | 0 | 0 | 0 | 0 | 11 |
| SRR5387952 | 0 | 0 | 41 | 0 | 0 | 0 | 0 | 0 | 0 | 0 | 10 |
| SRR5387953 | 0 | 0 | 0 | 0 | 0 | 0 | 0 | 19 | 0 | 0 | 0 |
| SRR5387954 | 0 | 0 | 12 | 0 | 0 | 0 | 0 | 0 | 0 | 0 | 0 |
| SRR5387955 | 0 | 0 | 0 | 22 | 0 | 22 | 0 | 22 | 0 | 0 | 11 |
| SRR5387956 | 0 | 0 | 15 | 15 | 0 | 0 | 0 | 0 | 0 | 0 | 0 |
| SRR5387957 | 0 | 0 | 0 | 0 | 0 | 0 | 0 | 0 | 0 | 0 | 0 |
| SRR5387958 | 0 | 0 | 0 | 10 | 0 | 0 | 0 | 0 | 0 | 0 | 0 |
| SRR5387959 | 0 | 0 | 19 | 0 | 0 | 0 | 0 | 0 | 0 | 0 | 0 |
| SRR5387960 | 0 | 0 | 0 | 15 | 0 | 0 | 0 | 0 | 0 | 0 | 0 |
| SRR5387961 | 0 | 0 | 0 | 27 | 0 | 0 | 0 | 0 | 0 | 0 | 0 |
| SRR5387962 | 0 | 0 | 12 | 0 | 12 | 12 | 0 | 0 | 0 | 0 | 0 |
| SRR5387963 | 0 | 0 | 0 | 25 | 0 | 0 | 0 | 0 | 0 | 0 | 0 |
| SRR5387964 | 0 | 0 | 0 | 0 | 0 | 0 | 0 | 0 | 0 | 0 | 0 |
| SRR5387965 | 0 | 0 | 0 | 0 | 0 | 0 | 0 | 0 | 0 | 0 | 0 |
| SRR5387966 | 0 | 0 | 16 | 0 | 0 | 0 | 0 | 31 | 0 | 0 | 0 |
| SRR5387967 | 0 | 0 | 0 | 14 | 0 | 22 | 0 | 0 | 0 | 0 | 0 |
| SRR5387968 | 0 | 0 | 0 | 0 | 0 | 0 | 0 | 0 | 0 | 0 | 0 |
| SRR5387969 | 0 | 24 | 0 | 24 | 0 | 0 | 0 | 0 | 0 | 0 | 16 |
| SRR5387970 | 0 | 0 | 34 | 34 | 0 | 0 | 0 | 0 | 0 | 0 | 0 |
| SRR5387971 | 0 | 0 | 31 | 21 | 0 | 21 | 0 | 21 | 0 | 0 | 0 |
| SRR5387972 | 0 | 0 | 15 | 0 | 0 | 15 | 0 | 0 | 0 | 0 | 0 |
| SRR5387973 | 0 | 10 | 51 | 21 | 0 | 10 | 0 | 0 | 0 | 0 | 0 |
| SRR5387974 | 0 | 0 | 20 | 0 | 0 | 0 | 0 | 0 | 0 | 0 | 0 |
| SRR5387975 | 0 | 0 | 0 | 18 | 0 | 0 | 0 | 0 | 0 | 0 | 0 |
| SRR5387976 | 0 | 0 | 0 | 17 | 0 | 0 | 0 | 0 | 0 | 0 | 0 |
| SRR5387977 | 0 | 0 | 0 | 0 | 0 | 0 | 0 | 0 | 0 | 0 | 0 |
| SRR5387978 | 0 | 0 | 19 | 76 | 0 | 28 | 0 | 0 | 0 | 0 | 0 |
| SRR5387979 | 0 | 0 | 12 | 12 | 0 | 0 | 0 | 0 | 0 | 0 | 0 |
| SRR5387980 | 0 | 0 | 0 | 45 | 0 | 18 | 0 | 0 | 0 | 0 | 0 |

| | | | | | | | | | | | |
|---|---|---|---|---|---|---|---|---|---|---|---|
| SRR5387981 | 0 | 0 | 0 | 0 | 0 | 0 | 0 | 14 | 14 | 0 | 0 |
| SRR5387982 | 0 | 11 | 0 | 11 | 0 | 11 | 0 | 0 | 0 | 0 | 0 |
| SRR5387983 | 0 | 16 | 0 | 16 | 16 | 0 | 0 | 0 | 0 | 0 | 16 |
| SRR5387984 | 0 | 0 | 16 | 16 | 0 | 0 | 0 | 0 | 0 | 0 | 0 |
| SRR5387985 | 0 | 0 | 0 | 19 | 0 | 0 | 0 | 0 | 0 | 0 | 0 |
| SRR5387986 | 0 | 0 | 0 | 15 | 0 | 0 | 0 | 0 | 0 | 0 | 0 |
| SRR5387987 | 0 | 0 | 30 | 20 | 0 | 0 | 0 | 0 | 0 | 0 | 20 |
| SRR5387988 | 0 | 0 | 17 | 26 | 0 | 0 | 0 | 0 | 0 | 0 | 0 |
| SRR5387989 | 0 | 0 | 0 | 17 | 0 | 11 | 0 | 0 | 0 | 0 | 0 |
| SRR5387990 | 0 | 0 | 0 | 0 | 0 | 0 | 0 | 0 | 0 | 0 | 0 |
| SRR5387991 | 0 | 0 | 0 | 15 | 0 | 0 | 0 | 0 | 0 | 0 | 0 |
| SRR5387992 | 0 | 0 | 35 | 17 | 0 | 0 | 0 | 0 | 0 | 0 | 0 |
| SRR5387993 | 0 | 0 | 0 | 43 | 0 | 0 | 0 | 0 | 0 | 0 | 0 |
| SRR5387994 | 0 | 0 | 15 | 20 | 0 | 0 | 0 | 0 | 0 | 0 | 0 |
| SRR5387995 | 0 | 0 | 43 | 17 | 0 | 0 | 0 | 17 | 0 | 0 | 35 |
| SRR5387996 | 0 | 0 | 21 | 0 | 0 | 0 | 0 | 0 | 0 | 0 | 0 |
| SRR5387997 | 0 | 0 | 0 | 0 | 0 | 0 | 0 | 0 | 0 | 0 | 0 |
| SRR5387998 | 0 | 11 | 0 | 0 | 0 | 0 | 0 | 0 | 11 | 0 | 0 |
| SRR5387999 | 0 | 0 | 0 | 0 | 0 | 0 | 0 | 0 | 0 | 0 | 23 |
| SRR5388000 | 0 | 0 | 0 | 0 | 0 | 0 | 0 | 0 | 0 | 0 | 0 |
| SRR5388001 | 0 | 0 | 22 | 0 | 0 | 0 | 0 | 0 | 0 | 0 | 0 |
| SRR5388002 | 0 | 0 | 0 | 0 | 0 | 0 | 0 | 0 | 0 | 0 | 0 |
| SRR5388003 | 0 | 13 | 13 | 53 | 0 | 0 | 0 | 0 | 0 | 0 | 0 |
| SRR5388004 | 0 | 0 | 0 | 0 | 0 | 0 | 0 | 0 | 0 | 0 | 0 |
| SRR5388005 | 0 | 0 | 0 | 0 | 0 | 0 | 0 | 0 | 0 | 0 | 42 |
| SRR5388006 | 0 | 0 | 16 | 24 | 0 | 0 | 0 | 0 | 0 | 0 | 0 |
| SRR5388007 | 0 | 0 | 0 | 0 | 0 | 0 | 0 | 0 | 0 | 0 | 0 |
| SRR5388008 | 0 | 0 | 0 | 0 | 0 | 0 | 0 | 0 | 0 | 0 | 0 |
| SRR5388009 | 0 | 0 | 10 | 0 | 0 | 0 | 0 | 10 | 0 | 0 | 0 |
| SRR5388010 | 0 | 0 | 23 | 0 | 0 | 0 | 0 | 0 | 0 | 0 | 0 |
| SRR5388011 | 0 | 0 | 11 | 0 | 0 | 11 | 0 | 11 | 0 | 0 | 0 |
| SRR5388012 | 0 | 0 | 10 | 10 | 0 | 10 | 0 | 10 | 10 | 0 | 0 |
| SRR5388013 | 0 | 0 | 0 | 14 | 0 | 0 | 0 | 0 | 0 | 0 | 0 |
| SRR5388014 | 0 | 0 | 12 | 47 | 0 | 0 | 0 | 0 | 0 | 0 | 12 |
| SRR5388015 | 0 | 0 | 0 | 0 | 0 | 0 | 0 | 0 | 26 | 0 | 0 |
| SRR5388016 | 0 | 0 | 18 | 0 | 0 | 12 | 0 | 0 | 0 | 0 | 18 |
| SRR5388017 | 0 | 0 | 0 | 0 | 0 | 0 | 0 | 0 | 0 | 0 | 0 |
| SRR5388018 | 0 | 0 | 0 | 0 | 0 | 0 | 0 | 0 | 0 | 0 | 0 |
| SRR5388019 | 0 | 11 | 0 | 11 | 0 | 0 | 0 | 22 | 0 | 0 | 11 |
| SRR5388020 | 0 | 0 | 28 | 28 | 0 | 0 | 0 | 0 | 0 | 0 | 0 |
| SRR5388021 | 0 | 0 | 0 | 0 | 0 | 0 | 0 | 0 | 17 | 17 | 0 |
| SRR5388022 | 0 | 0 | 13 | 19 | 0 | 0 | 0 | 0 | 0 | 0 | 0 |
| SRR5388023 | 0 | 0 | 31 | 0 | 0 | 0 | 0 | 0 | 10 | 0 | 0 |
| SRR5388024 | 0 | 0 | 25 | 16 | 0 | 0 | 0 | 0 | 0 | 0 | 0 |
| SRR5388025 | 0 | 0 | 0 | 16 | 0 | 0 | 0 | 16 | 16 | 0 | 0 |
| SRR5388026 | 0 | 0 | 0 | 46 | 0 | 0 | 0 | 15 | 0 | 0 | 15 |
| SRR5388027 | 0 | 0 | 12 | 0 | 0 | 12 | 0 | 0 | 0 | 0 | 12 |
| SRR5388028 | 0 | 0 | 0 | 0 | 0 | 0 | 0 | 0 | 0 | 0 | 0 |
| SRR5388029 | 0 | 0 | 0 | 0 | 0 | 0 | 0 | 0 | 0 | 0 | 0 |
| SRR5388030 | 0 | 0 | 0 | 0 | 0 | 0 | 0 | 17 | 0 | 0 | 0 |
| SRR5388031 | 0 | 0 | 0 | 0 | 0 | 0 | 0 | 0 | 0 | 0 | 12 |
| SRR5388032 | 0 | 0 | 0 | 0 | 0 | 0 | 0 | 0 | 0 | 0 | 0 |
| SRR5388033 | 0 | 0 | 39 | 39 | 0 | 0 | 0 | 0 | 0 | 0 | 29 |
| SRR5388034 | 0 | 0 | 0 | 0 | 0 | 0 | 0 | 0 | 0 | 0 | 0 |
| SRR5388035 | 0 | 24 | 35 | 35 | 0 | 0 | 0 | 0 | 0 | 0 | 0 |
| SRR5388036 | 0 | 0 | 29 | 29 | 0 | 0 | 0 | 0 | 0 | 0 | 0 |
| SRR5388037 | 0 | 0 | 0 | 0 | 0 | 0 | 0 | 0 | 0 | 0 | 0 |
| SRR5388038 | 0 | 0 | 33 | 33 | 0 | 0 | 0 | 11 | 0 | 0 | 11 |
| SRR5388039 | 0 | 14 | 14 | 0 | 0 | 0 | 0 | 0 | 0 | 0 | 0 |
| SRR5388040 | 0 | 0 | 37 | 22 | 0 | 15 | 0 | 0 | 0 | 0 | 0 |
| SRR5388041 | 0 | 0 | 0 | 0 | 0 | 0 | 0 | 0 | 0 | 0 | 0 |
| SRR5388042 | 0 | 0 | 34 | 17 | 17 | 0 | 0 | 0 | 0 | 0 | 0 |
| SRR5388043 | 0 | 0 | 0 | 0 | 0 | 0 | 0 | 0 | 0 | 0 | 0 |
| SRR5388044 | 0 | 0 | 10 | 20 | 0 | 0 | 0 | 0 | 0 | 0 | 10 |
| SRR5388045 | 0 | 0 | 16 | 16 | 0 | 0 | 0 | 0 | 0 | 0 | 16 |
| SRR5388046 | 0 | 0 | 0 | 17 | 0 | 0 | 0 | 0 | 0 | 0 | 0 |
| SRR5388047 | 0 | 0 | 0 | 0 | 0 | 0 | 0 | 0 | 0 | 0 | 0 |
| SRR5388048 | 0 | 0 | 0 | 25 | 0 | 16 | 0 | 0 | 0 | 0 | 0 |
| SRR5388049 | 0 | 0 | 0 | 30 | 0 | 0 | 0 | 0 | 0 | 0 | 0 |
| SRR5388050 | 0 | 0 | 0 | 19 | 0 | 12 | 0 | 12 | 0 | 0 | 0 |
| SRR5388051 | 0 | 0 | 14 | 0 | 0 | 0 | 0 | 0 | 0 | 0 | 0 |
| SRR5388052 | 0 | 0 | 23 | 17 | 0 | 0 | 0 | 29 | 0 | 11 | 0 |
| SRR5388053 | 0 | 0 | 37 | 12 | 0 | 12 | 0 | 0 | 0 | 0 | 19 |
| SRR5388054 | 0 | 0 | 0 | 0 | 0 | 0 | 0 | 0 | 0 | 0 | 0 |
| SRR5388055 | 0 | 0 | 19 | 38 | 0 | 0 | 0 | 0 | 0 | 0 | 0 |
| SRR5388056 | 0 | 0 | 0 | 31 | 0 | 0 | 0 | 0 | 0 | 0 | 0 |
| SRR5388057 | 0 | 0 | 18 | 0 | 0 | 0 | 0 | 0 | 0 | 0 | 0 |
| SRR5388058 | 0 | 0 | 0 | 0 | 0 | 12 | 0 | 0 | 0 | 0 | 0 |
| SRR5388059 | 0 | 0 | 36 | 0 | 0 | 0 | 0 | 0 | 14 | 0 | 0 |
| SRR5388060 | 0 | 0 | 0 | 13 | 0 | 0 | 0 | 0 | 0 | 0 | 0 |
| SRR5388061 | 0 | 0 | 23 | 0 | 0 | 0 | 0 | 0 | 0 | 0 | 0 |
| SRR5388062 | 0 | 0 | 0 | 0 | 0 | 0 | 0 | 0 | 0 | 0 | 0 |
| SRR5388063 | 0 | 21 | 0 | 32 | 0 | 0 | 0 | 0 | 0 | 0 | 0 |
| SRR5388064 | 0 | 0 | 0 | 0 | 0 | 0 | 0 | 18 | 0 | 0 | 0 |
| SRR5388065 | 0 | 0 | 28 | 0 | 0 | 0 | 0 | 14 | 0 | 0 | 0 |
| SRR5388066 | 0 | 0 | 18 | 0 | 0 | 0 | 0 | 0 | 0 | 0 | 0 |
| SRR5388067 | 0 | 15 | 0 | 15 | 0 | 0 | 0 | 15 | 15 | 0 | 0 |
| SRR5388068 | 0 | 0 | 17 | 17 | 0 | 0 | 0 | 0 | 0 | 17 | 0 |
| SRR5388069 | 0 | 0 | 24 | 24 | 0 | 0 | 0 | 0 | 0 | 0 | 0 |
| SRR5388070 | 0 | 0 | 40 | 40 | 0 | 0 | 0 | 0 | 0 | 0 | 16 |
| SRR5388071 | 0 | 0 | 12 | 0 | 0 | 0 | 0 | 12 | 0 | 0 | 0 |
| SRR5388072 | 0 | 0 | 0 | 20 | 0 | 0 | 0 | 0 | 0 | 0 | 0 |
| SRR5388073 | 0 | 0 | 27 | 18 | 0 | 0 | 0 | 0 | 0 | 0 | 0 |
| SRR5388074 | 0 | 0 | 0 | 0 | 0 | 0 | 0 | 0 | 0 | 0 | 0 |
| SRR5388075 | 0 | 0 | 0 | 0 | 0 | 0 | 0 | 0 | 0 | 0 | 0 |
| SRR5388076 | 0 | 0 | 34 | 34 | 0 | 25 | 0 | 0 | 0 | 0 | 0 |
| SRR5388077 | 0 | 0 | 21 | 10 | 0 | 0 | 0 | 10 | 0 | 10 | 0 |
| SRR5388078 | 0 | 0 | 0 | 0 | 0 | 0 | 0 | 0 | 0 | 0 | 0 |
| SRR5388079 | 0 | 0 | 0 | 19 | 0 | 0 | 0 | 0 | 0 | 0 | 0 |
| SRR5388080 | 0 | 0 | 0 | 17 | 0 | 0 | 0 | 0 | 0 | 0 | 0 |
| SRR5388081 | 0 | 0 | 17 | 0 | 0 | 0 | 0 | 0 | 0 | 0 | 0 |
| SRR5388082 | 0 | 0 | 0 | 20 | 0 | 13 | 0 | 0 | 0 | 0 | 0 |
| SRR5388083 | 0 | 0 | 0 | 0 | 0 | 0 | 0 | 0 | 0 | 0 | 0 |
| SRR5388084 | 0 | 0 | 28 | 0 | 0 | 0 | 0 | 0 | 0 | 0 | 0 |
| SRR5388085 | 0 | 10 | 0 | 0 | 0 | 0 | 0 | 0 | 0 | 0 | 0 |
| SRR5388086 | 0 | 0 | 0 | 17 | 0 | 0 | 0 | 17 | 0 | 0 | 0 |
| SRR5388087 | 0 | 15 | 23 | 0 | 0 | 0 | 0 | 15 | 0 | 0 | 0 |
| SRR5388088 | 0 | 0 | 20 | 13 | 0 | 0 | 0 | 13 | 0 | 0 | 20 |
| SRR5388089 | 0 | 0 | 0 | 0 | 0 | 0 | 0 | 0 | 0 | 0 | 0 |
| SRR5388090 | 0 | 0 | 12 | 0 | 0 | 0 | 0 | 0 | 0 | 0 | 0 |
| SRR5388091 | 0 | 0 | 0 | 0 | 0 | 0 | 0 | 0 | 0 | 0 | 0 |
| SRR5388092 | 0 | 0 | 0 | 0 | 0 | 0 | 0 | 0 | 13 | 0 | 0 |
| SRR5388093 | 0 | 0 | 0 | 11 | 0 | 0 | 0 | 0 | 0 | 0 | 0 |
| SRR5388094 | 0 | 0 | 0 | 0 | 0 | 0 | 0 | 0 | 0 | 0 | 0 |
| SRR5388095 | 0 | 12 | 12 | 37 | 0 | 12 | 0 | 12 | 0 | 0 | 0 |
| SRR5388096 | 0 | 0 | 22 | 0 | 0 | 0 | 0 | 0 | 0 | 0 | 0 |
| SRR5388097 | 0 | 0 | 0 | 0 | 0 | 0 | 0 | 0 | 0 | 0 | 19 |
| SRR5388098 | 0 | 0 | 23 | 0 | 0 | 0 | 0 | 0 | 0 | 0 | 0 |
| SRR5388099 | 0 | 0 | 0 | 0 | 0 | 0 | 0 | 0 | 0 | 0 | 0 |

| | | | | | | | | | | | |
|---|---|---|---|---|---|---|---|---|---|---|---|
| SRR5388100 | 0 | 0 | 11 | 0 | 0 | 0 | 0 | 0 | 22 | 0 | 0 |
| SRR5388101 | 0 | 0 | 0 | 0 | 0 | 0 | 0 | 0 | 0 | 0 | 0 |
| SRR5388102 | 0 | 0 | 0 | 0 | 0 | 0 | 0 | 0 | 0 | 0 | 0 |
| SRR5388103 | 0 | 11 | 11 | 0 | 0 | 0 | 0 | 0 | 0 | 0 | 11 |
| SRR5388104 | 0 | 0 | 22 | 0 | 0 | 0 | 0 | 0 | 15 | 0 | 0 |
| SRR5388105 | 0 | 0 | 0 | 25 | 0 | 0 | 0 | 0 | 0 | 0 | 0 |
| SRR5388106 | 0 | 0 | 0 | 0 | 0 | 0 | 0 | 0 | 0 | 0 | 0 |
| SRR5388107 | 0 | 12 | 0 | 12 | 0 | 0 | 0 | 0 | 0 | 0 | 12 |
| SRR5388108 | 0 | 0 | 14 | 0 | 0 | 0 | 0 | 0 | 0 | 0 | 0 |
| SRR5388109 | 0 | 0 | 0 | 11 | 0 | 23 | 0 | 0 | 0 | 0 | 0 |
| SRR5388110 | 0 | 0 | 0 | 14 | 0 | 0 | 0 | 0 | 0 | 0 | 0 |
| SRR5388111 | 0 | 0 | 0 | 12 | 0 | 12 | 0 | 0 | 0 | 0 | 0 |
| SRR5388112 | 0 | 0 | 0 | 11 | 0 | 16 | 0 | 0 | 0 | 0 | 0 |
| SRR5388113 | 0 | 0 | 0 | 23 | 0 | 0 | 0 | 0 | 0 | 0 | 0 |
| SRR5388114 | 0 | 0 | 12 | 12 | 0 | 0 | 0 | 0 | 0 | 0 | 0 |
| SRR5388115 | 0 | 16 | 16 | 33 | 0 | 0 | 0 | 16 | 0 | 0 | 0 |
| SRR5388116 | 0 | 0 | 19 | 38 | 0 | 0 | 0 | 0 | 0 | 0 | 0 |
| SRR5388117 | 0 | 0 | 0 | 0 | 0 | 0 | 0 | 0 | 0 | 0 | 0 |
| SRR5388118 | 0 | 0 | 12 | 12 | 0 | 0 | 0 | 0 | 0 | 0 | 0 |
| SRR5388119 | 0 | 0 | 18 | 0 | 0 | 0 | 0 | 18 | 0 | 0 | 0 |
| SRR5388120 | 0 | 0 | 0 | 0 | 0 | 0 | 0 | 0 | 0 | 0 | 0 |
| SRR5388121 | 0 | 0 | 0 | 0 | 0 | 15 | 0 | 0 | 0 | 0 | 0 |
| SRR5388122 | 0 | 0 | 17 | 0 | 0 | 0 | 0 | 0 | 0 | 0 | 0 |
| SRR5388123 | 0 | 0 | 0 | 12 | 0 | 0 | 0 | 12 | 0 | 0 | 12 |
| SRR5388124 | 0 | 0 | 0 | 0 | 0 | 0 | 0 | 0 | 0 | 0 | 0 |
| SRR5388125 | 0 | 0 | 0 | 0 | 0 | 0 | 0 | 0 | 0 | 0 | 0 |
| SRR5388126 | 0 | 0 | 25 | 37 | 0 | 0 | 0 | 25 | 0 | 0 | 0 |
| SRR5388127 | 0 | 11 | 22 | 32 | 0 | 22 | 0 | 0 | 22 | 11 | 11 |
| SRR5388128 | 0 | 0 | 23 | 0 | 0 | 16 | 0 | 0 | 0 | 0 | 23 |
| SRR5388129 | 0 | 0 | 0 | 10 | 0 | 0 | 0 | 0 | 0 | 0 | 0 |
| SRR5388130 | 0 | 0 | 0 | 54 | 0 | 0 | 0 | 0 | 0 | 0 | 0 |
| SRR5388131 | 0 | 14 | 0 | 29 | 0 | 14 | 0 | 0 | 14 | 0 | 0 |
| SRR5388132 | 0 | 0 | 13 | 0 | 0 | 0 | 0 | 0 | 0 | 0 | 0 |
| SRR5388133 | 0 | 0 | 0 | 0 | 0 | 0 | 0 | 0 | 0 | 0 | 0 |
| SRR5388134 | 0 | 0 | 0 | 13 | 0 | 0 | 0 | 0 | 0 | 0 | 0 |
| SRR5388135 | 0 | 0 | 0 | 23 | 0 | 0 | 0 | 0 | 0 | 0 | 16 |
| SRR5388136 | 0 | 0 | 0 | 0 | 0 | 0 | 0 | 0 | 0 | 0 | 0 |
| SRR5388137 | 0 | 0 | 12 | 0 | 0 | 0 | 0 | 0 | 0 | 0 | 0 |
| SRR5388138 | 0 | 0 | 0 | 0 | 0 | 0 | 0 | 0 | 0 | 0 | 0 |
| SRR5388139 | 0 | 0 | 37 | 55 | 0 | 0 | 0 | 0 | 0 | 0 | 18 |
| SRR5388140 | 0 | 0 | 26 | 0 | 0 | 0 | 0 | 0 | 0 | 0 | 13 |
| SRR5388141 | 0 | 0 | 11 | 32 | 0 | 21 | 0 | 0 | 11 | 0 | 21 |
| SRR5388142 | 0 | 0 | 0 | 0 | 0 | 0 | 0 | 0 | 0 | 0 | 0 |
| SRR5388143 | 0 | 10 | 10 | 20 | 0 | 10 | 0 | 0 | 0 | 0 | 20 |
| SRR5388144 | 0 | 0 | 0 | 12 | 0 | 36 | 0 | 0 | 0 | 12 | 12 |
| SRR5388145 | 0 | 0 | 0 | 33 | 0 | 11 | 0 | 0 | 11 | 0 | 0 |
| SRR5388146 | 0 | 0 | 11 | 11 | 0 | 0 | 0 | 0 | 0 | 0 | 0 |
| SRR5388147 | 0 | 17 | 33 | 17 | 0 | 17 | 0 | 0 | 17 | 0 | 0 |
| SRR5388148 | 0 | 0 | 0 | 15 | 0 | 0 | 0 | 0 | 0 | 0 | 0 |
| SRR5388149 | 0 | 0 | 26 | 0 | 0 | 0 | 0 | 0 | 0 | 0 | 0 |
| SRR5388150 | 0 | 0 | 16 | 0 | 0 | 0 | 0 | 0 | 0 | 0 | 0 |
| SRR5388151 | 0 | 0 | 0 | 0 | 0 | 0 | 0 | 0 | 0 | 0 | 0 |
| SRR5388152 | 0 | 0 | 35 | 18 | 0 | 0 | 0 | 0 | 18 | 0 | 18 |
| SRR5388153 | 0 | 0 | 0 | 33 | 0 | 0 | 0 | 0 | 0 | 0 | 0 |
| SRR5388154 | 0 | 0 | 0 | 38 | 0 | 0 | 0 | 0 | 0 | 0 | 0 |
| SRR5388155 | 0 | 0 | 0 | 0 | 0 | 0 | 0 | 0 | 0 | 0 | 14 |
| SRR5388156 | 0 | 0 | 0 | 0 | 0 | 0 | 0 | 11 | 0 | 0 | 0 |
| SRR5388157 | 0 | 0 | 23 | 0 | 0 | 0 | 0 | 0 | 0 | 12 | 12 |
| SRR5388158 | 0 | 0 | 24 | 0 | 0 | 0 | 0 | 0 | 0 | 0 | 0 |
| SRR5388159 | 0 | 10 | 10 | 31 | 0 | 10 | 0 | 0 | 0 | 0 | 0 |
| SRR5388160 | 0 | 0 | 0 | 0 | 0 | 0 | 0 | 0 | 16 | 0 | 0 |
| SRR5388161 | 0 | 0 | 0 | 0 | 0 | 0 | 0 | 0 | 0 | 0 | 0 |
| SRR5388162 | 0 | 0 | 0 | 28 | 0 | 0 | 0 | 0 | 0 | 0 | 0 |
| SRR5388163 | 0 | 0 | 26 | 19 | 0 | 13 | 0 | 0 | 13 | 0 | 0 |
| SRR5388164 | 0 | 0 | 12 | 25 | 0 | 0 | 12 | 0 | 0 | 0 | 0 |
| SRR5388165 | 0 | 0 | 11 | 27 | 0 | 0 | 0 | 0 | 0 | 0 | 0 |
| SRR5388166 | 0 | 0 | 11 | 22 | 0 | 0 | 0 | 0 | 11 | 0 | 11 |
| SRR5388167 | 0 | 0 | 0 | 0 | 0 | 0 | 0 | 0 | 0 | 0 | 0 |
| SRR5388168 | 0 | 0 | 0 | 0 | 0 | 0 | 0 | 0 | 0 | 0 | 0 |
| SRR5388169 | 0 | 0 | 0 | 0 | 0 | 0 | 0 | 0 | 0 | 0 | 0 |
| SRR5388170 | 0 | 0 | 0 | 0 | 0 | 0 | 0 | 0 | 0 | 0 | 0 |
| SRR5388171 | 0 | 0 | 0 | 0 | 0 | 0 | 0 | 0 | 0 | 0 | 0 |
| SRR5388172 | 0 | 0 | 0 | 0 | 0 | 0 | 0 | 0 | 0 | 0 | 0 |
| SRR5388173 | 0 | 0 | 20 | 0 | 0 | 0 | 0 | 0 | 0 | 0 | 0 |
| SRR5388174 | 0 | 0 | 12 | 24 | 0 | 12 | 0 | 0 | 0 | 0 | 12 |
| SRR5388175 | 0 | 0 | 0 | 0 | 0 | 0 | 0 | 0 | 0 | 0 | 0 |
| SRR5388176 | 0 | 0 | 0 | 10 | 0 | 0 | 0 | 0 | 0 | 0 | 0 |
| SRR5388177 | 0 | 0 | 0 | 10 | 0 | 0 | 0 | 0 | 10 | 10 | 0 |
| SRR5388178 | 0 | 0 | 11 | 11 | 0 | 0 | 0 | 0 | 0 | 0 | 0 |
| SRR5388179 | 0 | 0 | 0 | 0 | 0 | 0 | 0 | 0 | 0 | 0 | 0 |
| SRR5388180 | 0 | 0 | 0 | 0 | 0 | 0 | 0 | 0 | 0 | 0 | 0 |
| SRR5388181 | 0 | 0 | 10 | 10 | 0 | 0 | 0 | 0 | 0 | 0 | 0 |
| SRR5388182 | 0 | 0 | 21 | 0 | 0 | 0 | 0 | 0 | 0 | 0 | 0 |
| SRR5388183 | 0 | 0 | 0 | 14 | 0 | 0 | 0 | 0 | 0 | 0 | 14 |
| SRR5388184 | 0 | 0 | 44 | 11 | 11 | 0 | 0 | 0 | 0 | 0 | 0 |
| SRR5388185 | 0 | 0 | 0 | 0 | 0 | 0 | 0 | 0 | 0 | 0 | 0 |
| SRR5388186 | 0 | 0 | 17 | 17 | 0 | 0 | 0 | 0 | 0 | 0 | 0 |
| SRR5388187 | 0 | 0 | 12 | 36 | 12 | 0 | 0 | 0 | 0 | 0 | 0 |
| SRR5388188 | 0 | 0 | 14 | 28 | 0 | 0 | 0 | 0 | 0 | 0 | 0 |
| SRR5388189 | 0 | 0 | 0 | 18 | 0 | 0 | 0 | 0 | 0 | 0 | 0 |
| SRR5388190 | 0 | 0 | 0 | 14 | 0 | 0 | 0 | 0 | 0 | 0 | 14 |
| SRR5388191 | 0 | 0 | 0 | 35 | 0 | 0 | 0 | 0 | 0 | 0 | 18 |
| SRR5388192 | 0 | 0 | 0 | 0 | 0 | 0 | 0 | 0 | 0 | 0 | 0 |
| SRR5388193 | 0 | 0 | 0 | 25 | 0 | 0 | 0 | 0 | 0 | 0 | 0 |
| SRR5388194 | 0 | 0 | 0 | 37 | 0 | 19 | 0 | 0 | 0 | 0 | 0 |
| SRR5388195 | 0 | 0 | 0 | 20 | 0 | 0 | 0 | 0 | 0 | 0 | 0 |
| SRR5388196 | 0 | 0 | 0 | 0 | 0 | 0 | 0 | 0 | 0 | 0 | 0 |
| SRR5388197 | 0 | 0 | 14 | 21 | 0 | 0 | 0 | 0 | 0 | 0 | 14 |
| SRR5388198 | 0 | 0 | 0 | 37 | 0 | 0 | 0 | 12 | 0 | 0 | 0 |
| SRR5388199 | 0 | 0 | 0 | 0 | 0 | 0 | 0 | 0 | 0 | 0 | 0 |
| SRR5388200 | 0 | 0 | 0 | 0 | 0 | 0 | 0 | 0 | 0 | 0 | 0 |
| SRR5388201 | 0 | 0 | 0 | 0 | 0 | 0 | 0 | 0 | 0 | 0 | 16 |
| SRR5388202 | 0 | 0 | 22 | 0 | 0 | 0 | 0 | 0 | 0 | 0 | 0 |
| SRR5388203 | 0 | 0 | 25 | 17 | 0 | 0 | 0 | 0 | 0 | 0 | 0 |
| SRR5388204 | 0 | 12 | 12 | 24 | 0 | 12 | 0 | 12 | 0 | 0 | 0 |
| SRR5388205 | 0 | 0 | 0 | 0 | 0 | 0 | 0 | 0 | 0 | 0 | 0 |
| SRR5388206 | 0 | 0 | 24 | 24 | 0 | 0 | 0 | 0 | 0 | 0 | 0 |
| SRR5388207 | 0 | 0 | 0 | 16 | 0 | 0 | 0 | 0 | 0 | 0 | 0 |
| SRR5388208 | 0 | 0 | 19 | 0 | 0 | 0 | 0 | 0 | 0 | 0 | 0 |
| SRR5388209 | 0 | 14 | 0 | 28 | 0 | 0 | 0 | 0 | 0 | 0 | 0 |
| SRR5388210 | 0 | 0 | 0 | 22 | 0 | 0 | 0 | 0 | 0 | 0 | 0 |
| SRR5388211 | 0 | 0 | 0 | 0 | 0 | 0 | 0 | 0 | 0 | 0 | 0 |
| SRR5388212 | 0 | 0 | 0 | 0 | 0 | 0 | 0 | 0 | 0 | 0 | 0 |
| SRR5388213 | 0 | 0 | 0 | 0 | 0 | 0 | 0 | 0 | 0 | 0 | 0 |
| SRR5388214 | 0 | 0 | 11 | 21 | 0 | 0 | 0 | 11 | 0 | 0 | 0 |
| SRR5388215 | 0 | 21 | 0 | 0 | 0 | 10 | 0 | 21 | 0 | 0 | 0 |
| SRR5388216 | 0 | 0 | 0 | 0 | 0 | 0 | 0 | 0 | 12 | 0 | 0 |
| SRR5388217 | 0 | 18 | 0 | 18 | 0 | 0 | 0 | 18 | 0 | 18 | 18 |
| SRR5388218 | 0 | 0 | 0 | 0 | 0 | 0 | 0 | 0 | 0 | 0 | 0 |

### Supp Table S8.5.Columns_ET_to_FD

| | | | | | | | | | | | |
|---|---|---|---|---|---|---|---|---|---|---|---|
| SRR5388219 | 0 | 0 | 0 | 18 | 0 | 0 | 0 | 0 | 0 | 0 | 0 |
| SRR5388220 | 0 | 0 | 0 | 0 | 0 | 0 | 0 | 0 | 0 | 0 | 0 |
| SRR5388221 | 0 | 0 | 0 | 0 | 0 | 0 | 0 | 0 | 10 | 0 | 0 |
| SRR5388222 | 0 | 10 | 10 | 0 | 0 | 0 | 0 | 0 | 0 | 0 | 0 |
| SRR5388223 | 0 | 0 | 0 | 22 | 0 | 0 | 0 | 0 | 11 | 11 | 0 |
| SRR5388224 | 0 | 10 | 10 | 0 | 0 | 0 | 0 | 10 | 0 | 10 | 0 |
| SRR5388225 | 0 | 0 | 18 | 18 | 0 | 0 | 0 | 0 | 0 | 0 | 0 |
| SRR5388226 | 0 | 0 | 0 | 28 | 0 | 0 | 0 | 0 | 0 | 0 | 0 |
| SRR5388227 | 0 | 0 | 0 | 0 | 0 | 0 | 0 | 0 | 0 | 0 | 0 |
| SRR5388228 | 0 | 0 | 0 | 0 | 0 | 0 | 0 | 11 | 0 | 0 | 0 |
| SRR5388229 | 0 | 11 | 11 | 22 | 0 | 11 | 0 | 0 | 11 | 0 | 0 |
| SRR5388230 | 0 | 0 | 0 | 0 | 0 | 0 | 0 | 0 | 0 | 0 | 0 |
| SRR5388231 | 0 | 0 | 0 | 0 | 0 | 0 | 0 | 16 | 0 | 0 | 25 |
| SRR5388232 | 0 | 0 | 27 | 0 | 0 | 0 | 0 | 0 | 0 | 0 | 0 |
| SRR5388233 | 0 | 0 | 16 | 0 | 0 | 0 | 0 | 0 | 0 | 0 | 0 |
| SRR5388234 | 0 | 0 | 24 | 0 | 0 | 0 | 0 | 0 | 0 | 0 | 24 |
| SRR5388235 | 0 | 0 | 0 | 0 | 0 | 0 | 0 | 14 | 0 | 0 | 0 |
| SRR5388236 | 0 | 0 | 0 | 12 | 0 | 0 | 0 | 0 | 0 | 0 | 0 |
| SRR5388237 | 0 | 0 | 0 | 0 | 0 | 0 | 0 | 0 | 0 | 0 | 0 |
| SRR5388238 | 0 | 0 | 17 | 0 | 0 | 0 | 0 | 0 | 0 | 0 | 0 |
| SRR5388239 | 0 | 0 | 0 | 19 | 0 | 0 | 0 | 0 | 0 | 0 | 0 |
| SRR5388240 | 0 | 0 | 18 | 26 | 0 | 0 | 0 | 0 | 0 | 0 | 0 |
| SRR5388241 | 0 | 0 | 0 | 0 | 0 | 0 | 0 | 0 | 11 | 0 | 0 |
| SRR5388242 | 0 | 0 | 0 | 0 | 0 | 0 | 0 | 0 | 0 | 0 | 0 |
| SRR5388243 | 0 | 0 | 0 | 0 | 0 | 0 | 0 | 0 | 0 | 0 | 0 |
| SRR5388244 | 0 | 0 | 61 | 41 | 0 | 0 | 0 | 0 | 10 | 0 | 10 |
| SRR5388245 | 0 | 0 | 0 | 0 | 0 | 0 | 0 | 0 | 0 | 0 | 0 |
| SRR5388246 | 0 | 0 | 12 | 0 | 0 | 0 | 0 | 0 | 0 | 0 | 0 |
| SRR5388247 | 0 | 0 | 22 | 0 | 0 | 0 | 0 | 0 | 0 | 0 | 0 |
| SRR5388248 | 0 | 0 | 0 | 0 | 0 | 0 | 0 | 0 | 0 | 0 | 0 |
| SRR5388249 | 0 | 19 | 0 | 0 | 0 | 0 | 0 | 0 | 0 | 0 | 0 |
| SRR5388250 | 0 | 0 | 0 | 0 | 0 | 0 | 0 | 0 | 0 | 0 | 0 |
| SRR5388251 | 0 | 0 | 0 | 18 | 0 | 0 | 0 | 0 | 0 | 0 | 0 |
| SRR5388252 | 0 | 0 | 0 | 0 | 0 | 0 | 0 | 0 | 0 | 0 | 0 |
| SRR5388253 | 0 | 0 | 30 | 30 | 0 | 0 | 0 | 0 | 0 | 0 | 40 |
| SRR5388254 | 0 | 0 | 0 | 0 | 0 | 0 | 0 | 0 | 0 | 0 | 0 |
| SRR5388255 | 0 | 0 | 0 | 0 | 0 | 10 | 0 | 0 | 0 | 0 | 10 |
| SRR5388256 | 0 | 0 | 0 | 21 | 0 | 0 | 0 | 0 | 0 | 0 | 0 |
| SRR5388257 | 0 | 0 | 0 | 22 | 0 | 0 | 0 | 11 | 11 | 0 | 11 |
| SRR5388258 | 0 | 0 | 0 | 12 | 0 | 0 | 0 | 12 | 23 | 12 | 35 |
| SRR5388259 | 0 | 0 | 0 | 41 | 0 | 0 | 0 | 0 | 0 | 0 | 0 |
| SRR5388260 | 0 | 0 | 0 | 28 | 0 | 0 | 0 | 0 | 0 | 0 | 0 |
| SRR5388261 | 0 | 0 | 0 | 0 | 0 | 0 | 0 | 0 | 0 | 0 | 0 |
| SRR5388262 | 0 | 0 | 0 | 0 | 0 | 0 | 0 | 0 | 13 | 0 | 0 |
| SRR5388263 | 0 | 20 | 29 | 0 | 0 | 0 | 0 | 0 | 0 | 0 | 0 |
| SRR5388264 | 0 | 0 | 11 | 0 | 0 | 0 | 0 | 11 | 0 | 0 | 0 |
| SRR5388265 | 0 | 0 | 19 | 0 | 0 | 0 | 0 | 0 | 0 | 0 | 0 |
| SRR5388266 | 0 | 0 | 15 | 0 | 0 | 0 | 0 | 15 | 0 | 0 | 0 |
| SRR5388267 | 0 | 0 | 0 | 0 | 0 | 0 | 0 | 0 | 0 | 0 | 0 |
| SRR5388268 | 0 | 0 | 13 | 0 | 0 | 0 | 0 | 0 | 0 | 0 | 0 |
| SRR5388269 | 0 | 0 | 0 | 18 | 0 | 0 | 0 | 0 | 0 | 0 | 0 |
| SRR5388270 | 0 | 0 | 28 | 0 | 0 | 28 | 0 | 0 | 0 | 0 | 19 |
| SRR5388271 | 0 | 0 | 0 | 0 | 0 | 0 | 0 | 0 | 0 | 0 | 0 |
| SRR5388272 | 0 | 0 | 29 | 0 | 0 | 19 | 0 | 0 | 0 | 0 | 19 |
| SRR5388273 | 0 | 0 | 0 | 0 | 0 | 24 | 0 | 0 | 0 | 0 | 0 |
| SRR5388274 | 0 | 0 | 0 | 12 | 0 | 0 | 0 | 0 | 0 | 0 | 0 |
| SRR5388275 | 0 | 0 | 0 | 0 | 0 | 0 | 0 | 0 | 0 | 0 | 0 |
| SRR5388276 | 0 | 0 | 0 | 0 | 0 | 0 | 0 | 0 | 0 | 0 | 0 |
| SRR5388277 | 0 | 0 | 0 | 10 | 0 | 0 | 0 | 0 | 0 | 0 | 0 |
| SRR5388278 | 0 | 0 | 0 | 25 | 0 | 0 | 0 | 0 | 0 | 0 | 0 |
| SRR5388279 | 0 | 0 | 0 | 0 | 0 | 0 | 0 | 18 | 0 | 0 | 0 |
| SRR5388280 | 0 | 0 | 0 | 0 | 0 | 0 | 0 | 0 | 0 | 0 | 0 |
| SRR5388281 | 0 | 0 | 24 | 24 | 0 | 0 | 0 | 48 | 0 | 0 | 24 |
| SRR5388282 | 0 | 0 | 28 | 28 | 0 | 0 | 0 | 14 | 0 | 0 | 14 |
| SRR5388283 | 0 | 0 | 13 | 0 | 0 | 0 | 0 | 0 | 0 | 0 | 0 |
| SRR5388284 | 0 | 14 | 0 | 0 | 0 | 0 | 0 | 0 | 0 | 0 | 0 |
| SRR5388285 | 0 | 14 | 0 | 0 | 0 | 0 | 0 | 14 | 0 | 0 | 0 |
| SRR5388286 | 0 | 0 | 15 | 0 | 0 | 15 | 0 | 0 | 0 | 0 | 0 |
| SRR5388287 | 0 | 0 | 11 | 17 | 0 | 0 | 0 | 0 | 0 | 0 | 0 |
| SRR5388288 | 0 | 0 | 0 | 0 | 0 | 0 | 0 | 0 | 0 | 0 | 0 |
| SRR5388289 | 0 | 0 | 12 | 47 | 0 | 0 | 0 | 0 | 0 | 0 | 0 |
| SRR5388290 | 0 | 0 | 0 | 38 | 0 | 0 | 0 | 0 | 0 | 0 | 0 |
| SRR5388291 | 0 | 0 | 12 | 0 | 0 | 0 | 0 | 0 | 12 | 0 | 0 |
| SRR5388292 | 0 | 0 | 30 | 0 | 0 | 0 | 0 | 0 | 0 | 0 | 0 |
| SRR5388293 | 0 | 0 | 0 | 0 | 0 | 0 | 0 | 16 | 0 | 0 | 0 |
| SRR5388294 | 0 | 0 | 20 | 13 | 0 | 0 | 0 | 0 | 0 | 0 | 0 |
| SRR5476966 | 0 | 0 | 340 | 681 | 0 | 340 | 0 | 340 | 681 | 1702 | 681 |
| SRR5476967 | 0 | 0 | 0 | 0 | 0 | 0 | 0 | 0 | 301 | 1803 | 301 |
| SRR5476969 | 0 | 0 | 0 | 0 | 0 | 0 | 0 | 0 | 933 | 4664 | 5597 |
| SRR5517463 | 0 | 0 | 0 | 0 | 51 | 183 | 214 | 233 | 3552 | 490 | 541 |
| SRR5517464 | 0 | 0 | 0 | 0 | 236 | 963 | 827 | 1162 | 1393 | 977 | 436 |
| SRR5517465 | 0 | 0 | 15 | 0 | 149 | 482 | 467 | 628 | 1337 | 379 | 244 |
| SRR5517466 | 0 | 0 | 0 | 0 | 31 | 141 | 132 | 256 | 1598 | 264 | 251 |
| SRR5517467 | 0 | 0 | 0 | 0 | 109 | 413 | 359 | 577 | 2087 | 563 | 607 |
| SRR5517468 | 0 | 0 | 0 | 0 | 248 | 996 | 1025 | 1483 | 1336 | 804 | 579 |
| SRR5517469 | 0 | 0 | 0 | 0 | 60 | 284 | 313 | 443 | 1395 | 305 | 438 |
| SRR5517470 | 0 | 0 | 0 | 0 | 142 | 595 | 473 | 915 | 1131 | 413 | 266 |
| SRR5575011 | 55 | 15 | 593 | 60 | 0 | 20 | 30 | 17 | 82 | 70 | 760 |
| SRR5575012 | 31 | 16 | 618 | 38 | 0 | 16 | 33 | 29 | 84 | 64 | 760 |
| SRR5575013 | 52 | 45 | 623 | 73 | 10 | 17 | 24 | 31 | 194 | 128 | 813 |
| SRR5575014 | 83 | 43 | 608 | 70 | 11 | 26 | 24 | 43 | 61 | 50 | 509 |
| SRR5575015 | 86 | 56 | 627 | 60 | 32 | 56 | 80 | 86 | 91 | 58 | 739 |
| SRR5575016 | 42 | 27 | 681 | 48 | 15 | 24 | 27 | 30 | 96 | 118 | 705 |
| SRR5575017 | 59 | 55 | 399 | 59 | 29 | 18 | 59 | 29 | 88 | 66 | 553 |
| SRR5575018 | 104 | 45 | 512 | 120 | 24 | 41 | 88 | 104 | 75 | 61 | 502 |
| SRR5575019 | 63 | 24 | 439 | 73 | 10 | 16 | 47 | 24 | 146 | 104 | 797 |
| SRR5644067 | 0 | 30 | 89 | 74 | 37 | 130 | 30 | 231 | 93 | 168 | 246 |
| SRR5644068 | 0 | 15 | 30 | 61 | 61 | 243 | 0 | 396 | 259 | 502 | 396 |
| SRR5644069 | 0 | 47 | 79 | 55 | 0 | 47 | 0 | 94 | 47 | 47 | 47 |
| SRR5644070 | 0 | 0 | 10 | 18 | 13 | 48 | 10 | 61 | 10 | 15 | 23 |
| SRR5644071 | 0 | 0 | 22 | 67 | 0 | 76 | 0 | 138 | 218 | 330 | 254 |
| SRR5644072 | 0 | 0 | 13 | 13 | 0 | 28 | 0 | 61 | 18 | 10 | 23 |
| SRR5644073 | 0 | 35 | 41 | 57 | 0 | 32 | 0 | 54 | 0 | 0 | 13 |
| SRR5644074 | 0 | 0 | 0 | 23 | 0 | 51 | 0 | 112 | 39 | 14 | 28 |
| SRR5644075 | 0 | 42 | 72 | 48 | 0 | 36 | 0 | 60 | 12 | 0 | 18 |
| SRR5644076 | 0 | 0 | 0 | 0 | 0 | 54 | 0 | 132 | 93 | 36 | 95 |
| SRR5644077 | 0 | 0 | 0 | 0 | 13 | 73 | 0 | 95 | 110 | 20 | 28 |
| SRR5644078 | 0 | 0 | 0 | 0 | 0 | 16 | 0 | 38 | 31 | 0 | 30 |
| SRR5644085 | 0 | 18 | 76 | 112 | 18 | 108 | 11 | 155 | 313 | 435 | 302 |
| SRR5644086 | 0 | 13 | 29 | 28 | 0 | 38 | 0 | 33 | 0 | 13 | 15 |
| SRR5644087 | 0 | 22 | 104 | 82 | 49 | 121 | 16 | 143 | 16 | 16 | 66 |
| SRR5644088 | 0 | 31 | 78 | 51 | 27 | 71 | 10 | 82 | 20 | 27 | 27 |
| SRR5657537 | 0 | 0 | 0 | 24 | 0 | 88 | 10 | 120 | 30 | 44 | 83 |
| SRR5657538 | 0 | 0 | 0 | 0 | 0 | 22 | 0 | 34 | 25 | 30 | 32 |
| SRR5657539 | 0 | 0 | 0 | 0 | 0 | 21 | 0 | 42 | 21 | 33 | 30 |
| SRR5657540 | 0 | 0 | 0 | 0 | 0 | 38 | 0 | 68 | 30 | 47 | 102 |
| SRR5657541 | 0 | 0 | 0 | 17 | 14 | 56 | 19 | 98 | 22 | 22 | 68 |
| SRR5657542 | 0 | 0 | 0 | 0 | 0 | 38 | 10 | 70 | 33 | 44 | 65 |
| SRR5657543 | 0 | 0 | 0 | 0 | 0 | 20 | 0 | 38 | 30 | 41 | 43 |

### Supp Table S8.5.Columns_ET_to_FD

| | | | | | | | | | | | |
|---|---|---|---|---|---|---|---|---|---|---|---|
| SRR5657544 | 0 | 0 | 0 | 11 | 11 | 24 | 0 | 24 | 24 | 50 | 83 |
| SRR5657545 | 0 | 0 | 0 | 0 | 0 | 13 | 0 | 24 | 63 | 57 | 36 |
| SRR5657546 | 0 | 0 | 0 | 0 | 12 | 27 | 0 | 46 | 89 | 129 | 66 |
| SRR5658693 | 0 | 26 | 63 | 91 | 0 | 81 | 16 | 141 | 35 | 29 | 165 |
| SRR5658694 | 0 | 36 | 132 | 196 | 14 | 223 | 21 | 310 | 40 | 21 | 159 |
| SRR5658695 | 0 | 25 | 76 | 82 | 17 | 243 | 28 | 360 | 31 | 17 | 134 |
| SRR5658696 | 0 | 26 | 58 | 113 | 19 | 263 | 41 | 375 | 32 | 29 | 239 |
| SRR5658697 | 0 | 21 | 82 | 100 | 30 | 436 | 34 | 656 | 57 | 40 | 292 |
| SRR5687455 | 0 | 0 | 0 | 0 | 86 | 1120 | 83 | 802 | 766 | 588 | 1388 |
| SRR5687456 | 0 | 0 | 0 | 0 | 86 | 1413 | 113 | 1635 | 798 | 557 | 1245 |
| SRR5687457 | 0 | 0 | 0 | 0 | 93 | 1582 | 108 | 2032 | 954 | 726 | 1412 |
| SRR5687458 | 0 | 0 | 0 | 0 | 87 | 1427 | 146 | 1658 | 937 | 668 | 1630 |
| SRR5687459 | 0 | 0 | 0 | 0 | 47 | 1163 | 117 | 1043 | 966 | 656 | 1658 |
| SRR5687461 | 0 | 0 | 0 | 0 | 26 | 551 | 59 | 835 | 384 | 373 | 369 |
| SRR5687462 | 0 | 0 | 0 | 0 | 40 | 650 | 59 | 904 | 416 | 334 | 363 |
| SRR5687463 | 0 | 0 | 0 | 0 | 37 | 861 | 89 | 1216 | 444 | 351 | 437 |
| SRR5687464 | 0 | 0 | 0 | 0 | 20 | 351 | 35 | 516 | 445 | 384 | 492 |
| SRR5687465 | 0 | 0 | 0 | 0 | 61 | 1011 | 89 | 1360 | 275 | 296 | 327 |
| SRR5687466 | 0 | 0 | 0 | 0 | 40 | 883 | 76 | 1407 | 345 | 337 | 468 |
| SRR5687467 | 0 | 0 | 0 | 0 | 58 | 995 | 85 | 1510 | 358 | 342 | 538 |
| SRR5687468 | 0 | 0 | 0 | 0 | 21 | 741 | 72 | 1605 | 277 | 352 | 452 |
| SRR5687469 | 0 | 0 | 0 | 0 | 41 | 1006 | 106 | 1769 | 363 | 382 | 584 |
| SRR5753916 | 252 | 901 | 1212 | 379 | 2454 | 9833 | 6788 | 15495 | 21097 | 14365 | 14346 |
| SRR5755813 | 0 | 10 | 81 | 71 | 0 | 267 | 35 | 1173 | 60 | 30 | 297 |
| SRR5755814 | 0 | 26 | 78 | 0 | 26 | 391 | 104 | 990 | 78 | 104 | 261 |
| SRR5755815 | 0 | 0 | 70 | 105 | 0 | 628 | 175 | 2687 | 140 | 0 | 523 |
| SRR5755818 | 0 | 12 | 39 | 23 | 0 | 104 | 19 | 464 | 31 | 19 | 81 |
| SRR5755819 | 0 | 22 | 67 | 67 | 0 | 202 | 11 | 986 | 235 | 67 | 146 |
| SRR5755820 | 0 | 0 | 88 | 0 | 29 | 353 | 29 | 1678 | 265 | 0 | 236 |
| SRR5755821 | 0 | 40 | 169 | 129 | 50 | 279 | 100 | 1424 | 30 | 40 | 129 |
| SRR5755824 | 0 | 28 | 99 | 42 | 0 | 113 | 0 | 480 | 28 | 0 | 99 |
| SRR5755828 | 0 | 27 | 119 | 86 | 40 | 254 | 52 | 1354 | 59 | 15 | 90 |
| SRR5755830 | 0 | 43 | 184 | 33 | 11 | 434 | 11 | 4199 | 553 | 174 | 347 |
| SRR5755831 | 0 | 52 | 90 | 62 | 0 | 114 | 0 | 509 | 117 | 34 | 76 |
| SRR5755833 | 0 | 59 | 135 | 88 | 18 | 441 | 29 | 1111 | 112 | 0 | 241 |
| SRR5755835 | 0 | 31 | 238 | 150 | 38 | 582 | 69 | 1726 | 38 | 44 | 156 |
| SRR5755837 | 0 | 66 | 103 | 33 | 23 | 281 | 23 | 1292 | 52 | 0 | 187 |
| SRR5755838 | 0 | 38 | 54 | 33 | 17 | 197 | 13 | 761 | 33 | 13 | 84 |
| SRR5755842 | 0 | 60 | 255 | 135 | 15 | 405 | 75 | 1471 | 45 | 45 | 240 |
| SRR5755844 | 0 | 33 | 150 | 50 | 50 | 300 | 0 | 1466 | 83 | 33 | 200 |
| SRR5755847 | 0 | 0 | 187 | 75 | 0 | 149 | 37 | 1307 | 224 | 75 | 299 |
| SRR5755849 | 0 | 31 | 280 | 171 | 62 | 497 | 47 | 1444 | 47 | 47 | 47 |
| SRR5755851 | 0 | 71 | 142 | 24 | 0 | 213 | 24 | 1067 | 24 | 0 | 95 |
| SRR5755854 | 0 | 0 | 103 | 34 | 0 | 342 | 17 | 1316 | 103 | 17 | 205 |
| SRR5755855 | 0 | 20 | 203 | 0 | 41 | 344 | 41 | 1580 | 20 | 20 | 162 |
| SRR5755856 | 0 | 95 | 285 | 272 | 54 | 489 | 109 | 1969 | 27 | 41 | 136 |
| SRR5755858 | 0 | 42 | 52 | 52 | 0 | 268 | 23 | 1720 | 94 | 0 | 169 |
| SRR5755861 | 0 | 44 | 137 | 77 | 16 | 198 | 27 | 670 | 82 | 16 | 159 |
| SRR5755862 | 0 | 0 | 84 | 76 | 0 | 143 | 17 | 749 | 59 | 25 | 151 |
| SRR5755863 | 0 | 25 | 202 | 218 | 67 | 470 | 59 | 1848 | 34 | 34 | 151 |
| SRR5755865 | 0 | 45 | 109 | 39 | 0 | 193 | 13 | 1050 | 116 | 13 | 180 |
| SRR5755866 | 0 | 29 | 236 | 141 | 0 | 271 | 24 | 677 | 53 | 18 | 94 |
| SRR5755868 | 0 | 26 | 144 | 26 | 0 | 157 | 0 | 262 | 92 | 65 | 157 |
| SRR5755870 | 0 | 20 | 242 | 268 | 20 | 471 | 20 | 1001 | 46 | 20 | 183 |
| SRR5755875 | 0 | 104 | 231 | 45 | 15 | 254 | 0 | 813 | 127 | 90 | 216 |
| SRR5755876 | 0 | 29 | 221 | 110 | 22 | 133 | 15 | 618 | 59 | 15 | 118 |
| SRR5755877 | 0 | 57 | 257 | 224 | 51 | 630 | 84 | 1485 | 51 | 39 | 194 |
| SRR5755879 | 0 | 22 | 67 | 41 | 0 | 157 | 22 | 797 | 84 | 36 | 183 |
| SRR5755880 | 0 | 21 | 170 | 59 | 0 | 191 | 0 | 725 | 47 | 38 | 85 |
| SRR5755882 | 0 | 38 | 306 | 64 | 13 | 229 | 26 | 739 | 26 | 51 | 127 |
| SRR5755884 | 0 | 65 | 273 | 243 | 33 | 642 | 75 | 1843 | 65 | 45 | 93 |
| SRR5755887 | 0 | 32 | 81 | 0 | 0 | 113 | 16 | 694 | 81 | 16 | 81 |
| SRR5755889 | 0 | 21 | 103 | 27 | 27 | 486 | 68 | 1135 | 103 | 34 | 212 |
| SRR5755890 | 0 | 46 | 62 | 31 | 31 | 364 | 46 | 1570 | 132 | 70 | 232 |
| SRR5755891 | 0 | 31 | 301 | 160 | 68 | 891 | 80 | 1236 | 49 | 37 | 221 |
| SRR5755894 | 0 | 18 | 46 | 46 | 46 | 392 | 37 | 457 | 23 | 14 | 111 |
| SRR5755896 | 0 | 64 | 176 | 106 | 23 | 410 | 41 | 2105 | 223 | 88 | 311 |
| SRR5755897 | 0 | 40 | 144 | 72 | 16 | 359 | 24 | 1445 | 216 | 64 | 112 |
| SRR5755898 | 0 | 43 | 201 | 244 | 72 | 747 | 129 | 2844 | 43 | 57 | 101 |
| SRR5755901 | 0 | 44 | 159 | 62 | 0 | 406 | 57 | 1298 | 49 | 31 | 93 |
| SRR5755903 | 0 | 23 | 59 | 35 | 16 | 286 | 26 | 852 | 115 | 28 | 218 |
| SRR5755905 | 0 | 53 | 185 | 170 | 79 | 519 | 83 | 1526 | 42 | 64 | 140 |
| SRR5755908 | 0 | 16 | 88 | 95 | 0 | 127 | 16 | 422 | 64 | 24 | 80 |
| SRR5755910 | 0 | 40 | 79 | 41 | 33 | 194 | 16 | 660 | 84 | 35 | 149 |
| SRR5755911 | 0 | 55 | 117 | 78 | 0 | 243 | 47 | 759 | 94 | 23 | 164 |
| SRR5755912 | 0 | 41 | 263 | 138 | 41 | 491 | 86 | 1535 | 83 | 55 | 183 |
| SRR5755914 | 0 | 21 | 88 | 33 | 0 | 151 | 0 | 883 | 67 | 13 | 96 |
| SRR5755915 | 0 | 68 | 182 | 74 | 28 | 222 | 17 | 586 | 63 | 28 | 125 |
| SRR5755917 | 0 | 79 | 251 | 265 | 79 | 782 | 136 | 2072 | 93 | 22 | 595 |
| SRR5755918 | 0 | 20 | 156 | 88 | 20 | 482 | 34 | 1282 | 122 | 34 | 231 |
| SRR5755919 | 0 | 58 | 204 | 126 | 49 | 865 | 78 | 2302 | 97 | 68 | 223 |
| SRR5755922 | 0 | 40 | 137 | 109 | 17 | 337 | 29 | 1063 | 23 | 34 | 97 |
| SRR5755924 | 0 | 39 | 157 | 83 | 0 | 228 | 0 | 1103 | 98 | 39 | 228 |
| SRR5755925 | 0 | 11 | 97 | 91 | 34 | 324 | 0 | 1263 | 80 | 51 | 171 |
| SRR5755926 | 0 | 42 | 315 | 257 | 53 | 615 | 74 | 1408 | 32 | 21 | 142 |
| SRR5755928 | 0 | 21 | 81 | 38 | 0 | 210 | 11 | 817 | 43 | 21 | 97 |
| SRR5755929 | 0 | 0 | 161 | 12 | 25 | 223 | 12 | 929 | 12 | 12 | 62 |
| SRR5755931 | 0 | 75 | 198 | 44 | 22 | 287 | 18 | 869 | 84 | 31 | 181 |
| SRR5755933 | 0 | 50 | 252 | 191 | 0 | 423 | 20 | 1350 | 50 | 71 | 91 |
| SRR5755936 | 0 | 64 | 198 | 70 | 23 | 294 | 26 | 616 | 28 | 26 | 36 |
| SRR5755938 | 0 | 26 | 218 | 152 | 0 | 561 | 40 | 1663 | 119 | 66 | 238 |
| SRR5755939 | 0 | 100 | 232 | 111 | 27 | 520 | 22 | 1679 | 71 | 33 | 149 |
| SRR5755940 | 0 | 125 | 384 | 266 | 118 | 1129 | 180 | 2665 | 141 | 47 | 329 |
| SRR5755943 | 0 | 76 | 192 | 119 | 13 | 527 | 46 | 1611 | 43 | 23 | 129 |
| SRR5755946 | 0 | 61 | 251 | 69 | 0 | 304 | 38 | 1720 | 99 | 15 | 266 |
| SRR5755947 | 0 | 49 | 544 | 453 | 25 | 1021 | 124 | 3254 | 82 | 33 | 272 |
| SRR5755950 | 0 | 31 | 186 | 72 | 52 | 238 | 21 | 1147 | 93 | 31 | 93 |
| SRR5755952 | 0 | 122 | 169 | 75 | 19 | 424 | 19 | 1450 | 104 | 38 | 480 |
| SRR5755953 | 0 | 93 | 166 | 124 | 26 | 285 | 26 | 1134 | 47 | 36 | 181 |
| SRR5755954 | 0 | 83 | 266 | 183 | 83 | 515 | 166 | 1662 | 66 | 50 | 183 |
| SRR5755956 | 0 | 22 | 182 | 85 | 0 | 186 | 30 | 985 | 45 | 22 | 197 |
| SRR5755957 | 0 | 85 | 231 | 109 | 0 | 122 | 49 | 815 | 85 | 12 | 24 |
| SRR5755959 | 0 | 55 | 137 | 82 | 22 | 724 | 82 | 2133 | 164 | 66 | 422 |
| SRR5755960 | 0 | 35 | 150 | 62 | 0 | 308 | 18 | 2186 | 79 | 35 | 194 |
| SRR5755961 | 0 | 59 | 364 | 238 | 42 | 807 | 142 | 2957 | 33 | 67 | 222 |
| SRR5755963 | 0 | 67 | 208 | 74 | 0 | 402 | 20 | 2494 | 80 | 27 | 261 |
| SRR5755964 | 0 | 98 | 321 | 187 | 0 | 366 | 80 | 1890 | 53 | 18 | 178 |
| SRR5755966 | 0 | 80 | 216 | 80 | 15 | 351 | 25 | 1215 | 65 | 25 | 196 |
| SRR5755967 | 0 | 71 | 206 | 87 | 24 | 333 | 48 | 1134 | 79 | 71 | 111 |
| SRR5755968 | 0 | 40 | 115 | 86 | 35 | 403 | 75 | 1612 | 46 | 58 | 138 |
| SRR5755971 | 0 | 21 | 57 | 45 | 18 | 169 | 12 | 722 | 54 | 21 | 94 |
| SRR5755973 | 0 | 53 | 99 | 114 | 23 | 258 | 38 | 1220 | 91 | 114 | 318 |
| SRR5755974 | 0 | 25 | 62 | 66 | 0 | 115 | 0 | 611 | 62 | 21 | 201 |
| SRR5755975 | 0 | 37 | 214 | 317 | 78 | 697 | 95 | 1686 | 91 | 70 | 284 |
| SRR5755978 | 0 | 50 | 50 | 50 | 0 | 226 | 17 | 888 | 42 | 42 | 151 |
| SRR5755980 | 0 | 27 | 121 | 52 | 12 | 225 | 46 | 1071 | 112 | 70 | 349 |
| SRR5755981 | 0 | 40 | 134 | 65 | 15 | 203 | 0 | 1335 | 149 | 69 | 382 |
| SRR5755982 | 0 | 33 | 223 | 211 | 39 | 451 | 74 | 1677 | 68 | 74 | 202 |
| SRR5755984 | 0 | 15 | 83 | 77 | 0 | 169 | 28 | 1238 | 117 | 31 | 323 |

### Supp Table S8.5.Columns_ET_to_FD

| | | | | | | | | | | | |
|---|---|---|---|---|---|---|---|---|---|---|---|
| SRR5755985 | 0 | 40 | 178 | 109 | 20 | 226 | 20 | 1023 | 85 | 12 | 133 |
| SRR5755987 | 0 | 22 | 106 | 57 | 27 | 199 | 27 | 929 | 137 | 66 | 371 |
| SRR5755988 | 0 | 29 | 151 | 58 | 15 | 93 | 29 | 585 | 97 | 54 | 151 |
| SRR5755989 | 0 | 120 | 443 | 508 | 74 | 942 | 148 | 2624 | 102 | 120 | 499 |
| SRR5755991 | 0 | 17 | 110 | 44 | 0 | 154 | 0 | 1028 | 91 | 42 | 293 |
| SRR5755992 | 0 | 21 | 107 | 73 | 0 | 141 | 0 | 527 | 47 | 30 | 103 |
| SRR5755994 | 0 | 117 | 352 | 285 | 45 | 586 | 106 | 2754 | 84 | 50 | 492 |
| SRR5755995 | 0 | 28 | 177 | 88 | 0 | 326 | 23 | 1811 | 182 | 42 | 289 |
| SRR5755996 | 0 | 53 | 284 | 287 | 64 | 977 | 146 | 4187 | 109 | 58 | 308 |
| SRR5755997 | 0 | 0 | 0 | 0 | 0 | 0 | 0 | 28 | 19 | 0 | 28 |
| SRR5755998 | 0 | 33 | 137 | 45 | 0 | 304 | 19 | 2269 | 152 | 37 | 245 |
| SRR5755999 | 0 | 56 | 106 | 128 | 11 | 378 | 44 | 1889 | 128 | 22 | 178 |
| SRR5756000 | 0 | 0 | 0 | 0 | 0 | 0 | 0 | 0 | 0 | 0 | 0 |
| SRR5756001 | 0 | 65 | 178 | 121 | 16 | 242 | 0 | 1292 | 97 | 77 | 266 |
| SRR5756002 | 0 | 65 | 186 | 81 | 0 | 283 | 40 | 1943 | 162 | 40 | 364 |
| SRR5756003 | 0 | 65 | 276 | 402 | 96 | 870 | 115 | 3127 | 123 | 111 | 295 |
| SRR5756005 | 0 | 15 | 139 | 45 | 0 | 203 | 15 | 1469 | 119 | 30 | 178 |
| SRR5756006 | 0 | 60 | 179 | 192 | 33 | 498 | 73 | 2051 | 173 | 93 | 252 |
| SRR5756007 | 0 | 0 | 0 | 0 | 0 | 0 | 0 | 0 | 0 | 0 | 0 |
| SRR5756008 | 0 | 67 | 152 | 110 | 0 | 213 | 24 | 694 | 104 | 12 | 189 |
| SRR5756009 | 0 | 27 | 352 | 227 | 32 | 536 | 70 | 1294 | 65 | 87 | 217 |
| SRR5756011 | 0 | 21 | 101 | 52 | 0 | 185 | 21 | 1031 | 101 | 35 | 228 |
| SRR5756012 | 0 | 53 | 251 | 90 | 0 | 155 | 12 | 496 | 34 | 15 | 81 |
| SRR5756014 | 0 | 54 | 198 | 58 | 0 | 184 | 31 | 790 | 112 | 54 | 216 |
| SRR5756016 | 0 | 37 | 295 | 267 | 75 | 464 | 98 | 1445 | 34 | 18 | 121 |
| SRR5756017 | 0 | 0 | 0 | 0 | 0 | 41 | 20 | 0 | 0 | 20 | 0 |
| SRR5756018 | 0 | 0 | 69 | 26 | 0 | 135 | 0 | 902 | 65 | 26 | 191 |
| SRR5756019 | 0 | 26 | 259 | 181 | 13 | 194 | 0 | 544 | 104 | 39 | 156 |
| SRR5756021 | 0 | 11 | 157 | 45 | 11 | 210 | 34 | 814 | 112 | 60 | 274 |
| SRR5756022 | 0 | 25 | 123 | 55 | 0 | 178 | 12 | 522 | 86 | 25 | 86 |
| SRR5756023 | 0 | 38 | 121 | 151 | 15 | 454 | 45 | 1438 | 106 | 30 | 182 |
| SRR5756025 | 0 | 0 | 116 | 0 | 0 | 161 | 27 | 884 | 80 | 54 | 143 |
| SRR5756026 | 0 | 38 | 82 | 69 | 19 | 151 | 38 | 509 | 69 | 38 | 31 |
| SRR5756027 | 0 | 17 | 57 | 24 | 0 | 101 | 24 | 415 | 101 | 91 | 294 |
| SRR5756028 | 0 | 31 | 31 | 0 | 0 | 116 | 0 | 318 | 70 | 85 | 256 |
| SRR5756029 | 0 | 22 | 106 | 53 | 28 | 285 | 53 | 601 | 56 | 50 | 221 |
| SRR5756031 | 0 | 14 | 43 | 0 | 0 | 121 | 11 | 515 | 82 | 34 | 234 |
| SRR5756032 | 0 | 13 | 99 | 46 | 0 | 132 | 0 | 357 | 73 | 46 | 112 |
| SRR5756033 | 0 | 53 | 127 | 53 | 13 | 347 | 27 | 1417 | 100 | 33 | 394 |
| SRR5756034 | 0 | 0 | 83 | 61 | 0 | 162 | 24 | 1177 | 91 | 77 | 243 |
| SRR5756035 | 0 | 24 | 120 | 160 | 20 | 385 | 44 | 1528 | 40 | 72 | 176 |
| SRR5756037 | 0 | 22 | 75 | 32 | 12 | 159 | 17 | 1387 | 104 | 30 | 249 |
| SRR5756038 | 0 | 27 | 48 | 48 | 0 | 42 | 11 | 626 | 69 | 42 | 85 |
| SRR5756040 | 0 | 0 | 73 | 26 | 0 | 141 | 13 | 906 | 137 | 51 | 355 |
| SRR5756041 | 0 | 11 | 80 | 23 | 0 | 97 | 11 | 517 | 68 | 28 | 136 |
| SRR5756042 | 0 | 0 | 172 | 55 | 23 | 414 | 16 | 1616 | 70 | 31 | 226 |
| SRR5756044 | 0 | 0 | 49 | 18 | 0 | 153 | 0 | 1014 | 55 | 12 | 208 |
| SRR5756045 | 0 | 0 | 49 | 63 | 0 | 161 | 21 | 455 | 49 | 35 | 105 |
| SRR5756047 | 0 | 46 | 229 | 147 | 23 | 399 | 50 | 1316 | 105 | 32 | 472 |
| SRR5756048 | 0 | 28 | 208 | 60 | 0 | 333 | 19 | 1036 | 176 | 32 | 278 |
| SRR5756049 | 0 | 35 | 268 | 146 | 42 | 582 | 70 | 1929 | 77 | 52 | 122 |
| SRR5756051 | 0 | 39 | 264 | 66 | 0 | 268 | 19 | 1797 | 210 | 39 | 345 |
| SRR5756052 | 0 | 53 | 99 | 64 | 0 | 187 | 0 | 613 | 64 | 18 | 53 |
| SRR5756054 | 0 | 32 | 77 | 58 | 26 | 336 | 39 | 1659 | 103 | 26 | 413 |
| SRR5756055 | 0 | 0 | 56 | 0 | 0 | 212 | 28 | 1384 | 56 | 28 | 169 |
| SRR5756056 | 0 | 44 | 144 | 105 | 33 | 503 | 44 | 1504 | 94 | 61 | 122 |
| SRR5756058 | 0 | 21 | 82 | 43 | 0 | 189 | 18 | 1138 | 116 | 27 | 222 |
| SRR5756059 | 0 | 0 | 65 | 28 | 19 | 261 | 19 | 1138 | 56 | 0 | 112 |
| SRR5756061 | 0 | 49 | 160 | 102 | 0 | 253 | 27 | 1046 | 84 | 49 | 292 |
| SRR5756062 | 0 | 14 | 123 | 58 | 0 | 96 | 14 | 492 | 55 | 20 | 85 |
| SRR5756063 | 0 | 19 | 174 | 167 | 13 | 367 | 52 | 1482 | 58 | 45 | 110 |
| SRR5756065 | 0 | 26 | 86 | 34 | 0 | 170 | 14 | 1250 | 112 | 49 | 259 |
| SRR5756066 | 0 | 29 | 103 | 57 | 0 | 211 | 20 | 927 | 64 | 41 | 100 |
| SRR5756068 | 0 | 13 | 138 | 48 | 0 | 193 | 13 | 718 | 132 | 42 | 232 |
| SRR5756069 | 0 | 16 | 212 | 98 | 12 | 183 | 20 | 876 | 134 | 49 | 281 |
| SRR5756070 | 0 | 65 | 351 | 351 | 65 | 865 | 171 | 3225 | 90 | 49 | 376 |
| SRR5756072 | 0 | 15 | 93 | 21 | 0 | 93 | 0 | 617 | 81 | 45 | 186 |
| SRR5756073 | 0 | 29 | 182 | 73 | 15 | 285 | 0 | 781 | 88 | 58 | 153 |
| SRR5756075 | 0 | 38 | 163 | 75 | 0 | 207 | 13 | 745 | 119 | 38 | 507 |
| SRR5756076 | 0 | 17 | 106 | 42 | 0 | 91 | 0 | 403 | 119 | 25 | 147 |
| SRR5756077 | 0 | 41 | 400 | 276 | 69 | 717 | 97 | 1917 | 83 | 41 | 469 |
| SRR5756079 | 0 | 0 | 90 | 13 | 0 | 202 | 13 | 880 | 99 | 45 | 229 |
| SRR5756080 | 0 | 0 | 71 | 41 | 0 | 127 | 11 | 352 | 64 | 14 | 136 |
| SRR5756082 | 0 | 10 | 51 | 41 | 0 | 97 | 15 | 439 | 77 | 36 | 240 |
| SRR5756083 | 0 | 0 | 61 | 25 | 0 | 106 | 0 | 439 | 47 | 33 | 189 |
| SRR5756084 | 0 | 85 | 340 | 239 | 39 | 471 | 39 | 1374 | 77 | 139 | 371 |
| SRR5756086 | 0 | 21 | 82 | 24 | 15 | 122 | 0 | 550 | 116 | 49 | 290 |
| SRR5756087 | 0 | 12 | 46 | 17 | 0 | 121 | 29 | 345 | 81 | 12 | 75 |
| SRR5756089 | 0 | 29 | 216 | 70 | 18 | 222 | 35 | 714 | 105 | 64 | 392 |
| SRR5756090 | 0 | 30 | 156 | 62 | 0 | 152 | 15 | 478 | 126 | 54 | 233 |
| SRR5756091 | 0 | 34 | 174 | 123 | 17 | 292 | 30 | 1161 | 72 | 97 | 199 |
| SRR5756093 | 0 | 24 | 111 | 59 | 0 | 122 | 11 | 629 | 103 | 49 | 157 |
| SRR5756094 | 0 | 14 | 113 | 70 | 0 | 137 | 0 | 492 | 60 | 38 | 113 |
| SRR5756096 | 0 | 43 | 82 | 53 | 0 | 146 | 18 | 758 | 137 | 50 | 252 |
| SRR5756097 | 0 | 36 | 217 | 72 | 0 | 127 | 18 | 742 | 72 | 54 | 199 |
| SRR5756098 | 0 | 34 | 214 | 227 | 36 | 371 | 71 | 1588 | 69 | 76 | 312 |
| SRR5756100 | 0 | 22 | 98 | 28 | 0 | 174 | 17 | 1088 | 108 | 43 | 322 |
| SRR5756101 | 0 | 42 | 113 | 48 | 12 | 125 | 0 | 549 | 78 | 42 | 84 |
| SRR5756103 | 0 | 59 | 164 | 104 | 59 | 401 | 89 | 1457 | 119 | 59 | 587 |
| SRR5756104 | 0 | 34 | 118 | 91 | 0 | 444 | 46 | 1784 | 259 | 108 | 499 |
| SRR5756105 | 0 | 97 | 350 | 593 | 53 | 1093 | 120 | 4209 | 237 | 160 | 803 |
| SRR5756106 | 0 | 0 | 18 | 18 | 0 | 18 | 0 | 407 | 0 | 0 | 36 |
| SRR5756107 | 0 | 54 | 149 | 76 | 0 | 331 | 0 | 1629 | 225 | 76 | 571 |
| SRR5756108 | 0 | 31 | 331 | 212 | 26 | 635 | 36 | 2464 | 201 | 88 | 439 |
| SRR5756110 | 0 | 0 | 0 | 35 | 0 | 0 | 0 | 0 | 35 | 0 | 0 |
| SRR5756114 | 0 | 0 | 0 | 47 | 0 | 0 | 0 | 24 | 0 | 0 | 24 |
| SRR5756115 | 0 | 0 | 0 | 0 | 0 | 0 | 0 | 0 | 0 | 0 | 0 |
| SRR5756116 | 0 | 0 | 0 | 0 | 0 | 0 | 0 | 0 | 68 | 0 | 0 |
| SRR5756118 | 0 | 0 | 0 | 0 | 0 | 0 | 0 | 0 | 0 | 0 | 0 |
| SRR5756120 | 0 | 0 | 0 | 0 | 0 | 0 | 0 | 0 | 0 | 0 | 54 |
| SRR5756121 | 0 | 0 | 44 | 0 | 0 | 0 | 0 | 0 | 0 | 0 | 0 |
| SRR5756122 | 0 | 0 | 0 | 0 | 0 | 0 | 0 | 27 | 0 | 27 | 80 |
| SRR5756123 | 0 | 0 | 0 | 57 | 0 | 0 | 0 | 0 | 57 | 0 | 86 |
| SRR5756125 | 0 | 0 | 0 | 0 | 0 | 0 | 0 | 16 | 32 | 0 | 32 |
| SRR5756128 | 0 | 0 | 0 | 46 | 0 | 0 | 0 | 0 | 0 | 0 | 0 |
| SRR5756130 | 0 | 0 | 0 | 0 | 0 | 0 | 0 | 0 | 0 | 0 | 0 |
| SRR5756131 | 0 | 0 | 0 | 0 | 26 | 0 | 0 | 26 | 26 | 0 | 0 |
| SRR5756132 | 0 | 0 | 0 | 0 | 0 | 0 | 0 | 0 | 0 | 0 | 0 |
| SRR5756133 | 0 | 0 | 36 | 0 | 0 | 36 | 0 | 73 | 109 | 0 | 0 |
| SRR5756135 | 0 | 0 | 0 | 37 | 0 | 37 | 0 | 37 | 0 | 0 | 37 |
| SRR5756136 | 0 | 0 | 0 | 0 | 0 | 0 | 0 | 0 | 36 | 0 | 0 |
| SRR5756137 | 0 | 18 | 36 | 36 | 0 | 36 | 0 | 18 | 0 | 0 | 125 |
| SRR5756138 | 0 | 0 | 29 | 0 | 0 | 0 | 0 | 0 | 29 | 0 | 57 |
| SRR5756139 | 0 | 0 | 69 | 35 | 0 | 35 | 0 | 0 | 0 | 0 | 0 |
| SRR5756140 | 0 | 0 | 29 | 29 | 0 | 29 | 0 | 0 | 29 | 0 | 0 |
| SRR5756141 | 0 | 0 | 32 | 16 | 0 | 0 | 0 | 0 | 16 | 0 | 32 |
| SRR5756142 | 0 | 0 | 36 | 55 | 0 | 0 | 0 | 36 | 0 | 0 | 36 |
| SRR5756143 | 0 | 0 | 0 | 0 | 0 | 0 | 0 | 0 | 18 | 0 | 0 |
| SRR5756144 | 0 | 0 | 0 | 61 | 0 | 0 | 0 | 0 | 0 | 0 | 20 |

### Supp Table S8.5.Columns_ET_to_FD

| | | | | | | | | | | | |
|---|---|---|---|---|---|---|---|---|---|---|---|
| SRR5756145 | 0 | 0 | 0 | 76 | 0 | 0 | 0 | 0 | 0 | 0 | 38 |
| SRR5756146 | 0 | 0 | 0 | 40 | 0 | 0 | 0 | 0 | 0 | 0 | 40 |
| SRR5756147 | 0 | 0 | 0 | 78 | 0 | 0 | 0 | 0 | 39 | 0 | 0 |
| SRR5756148 | 0 | 0 | 27 | 0 | 0 | 0 | 0 | 0 | 0 | 0 | 0 |
| SRR5756149 | 0 | 0 | 0 | 38 | 0 | 0 | 0 | 0 | 0 | 0 | 38 |
| SRR5756150 | 0 | 0 | 19 | 0 | 0 | 0 | 0 | 39 | 0 | 0 | 19 |
| SRR5756151 | 0 | 0 | 77 | 0 | 0 | 0 | 0 | 0 | 0 | 0 | 115 |
| SRR5756152 | 0 | 0 | 0 | 17 | 0 | 0 | 0 | 17 | 0 | 0 | 0 |
| SRR5756153 | 0 | 0 | 0 | 0 | 0 | 0 | 0 | 0 | 0 | 0 | 24 |
| SRR5756154 | 0 | 0 | 45 | 90 | 45 | 45 | 0 | 0 | 0 | 0 | 90 |
| SRR5756155 | 0 | 0 | 0 | 0 | 0 | 0 | 0 | 14 | 0 | 0 | 0 |
| SRR5756156 | 0 | 0 | 21 | 41 | 0 | 21 | 0 | 21 | 0 | 0 | 41 |
| SRR5756157 | 0 | 0 | 0 | 19 | 0 | 0 | 0 | 0 | 38 | 0 | 0 |
| SRR5756158 | 0 | 0 | 0 | 45 | 0 | 0 | 0 | 23 | 23 | 0 | 23 |
| SRR5756159 | 0 | 21 | 42 | 0 | 0 | 21 | 0 | 0 | 0 | 0 | 21 |
| SRR5756160 | 0 | 0 | 0 | 0 | 0 | 0 | 0 | 53 | 0 | 0 | 53 |
| SRR5756162 | 0 | 0 | 0 | 0 | 0 | 0 | 0 | 58 | 0 | 0 | 58 |
| SRR5756164 | 0 | 0 | 0 | 0 | 0 | 22 | 0 | 0 | 0 | 0 | 22 |
| SRR5756166 | 0 | 0 | 0 | 0 | 0 | 25 | 0 | 0 | 50 | 0 | 0 |
| SRR5756167 | 0 | 0 | 0 | 0 | 0 | 30 | 30 | 0 | 0 | 0 | 59 |
| SRR5756168 | 0 | 0 | 0 | 0 | 0 | 0 | 0 | 0 | 0 | 0 | 0 |
| SRR5756177 | 0 | 0 | 0 | 0 | 110 | 0 | 0 | 0 | 110 | 0 | 0 |
| SRR5756178 | 0 | 0 | 0 | 89 | 0 | 0 | 0 | 0 | 0 | 0 | 0 |
| SRR5756179 | 0 | 0 | 0 | 0 | 0 | 0 | 0 | 0 | 0 | 0 | 83 |
| SRR5756180 | 0 | 0 | 0 | 0 | 0 | 28 | 0 | 0 | 0 | 0 | 0 |
| SRR5756182 | 0 | 0 | 0 | 0 | 0 | 0 | 0 | 0 | 0 | 0 | 0 |
| SRR5756183 | 0 | 0 | 0 | 34 | 11 | 23 | 0 | 0 | 23 | 0 | 45 |
| SRR5756184 | 0 | 0 | 0 | 30 | 0 | 0 | 0 | 30 | 30 | 0 | 90 |
| SRR5756185 | 0 | 0 | 17 | 70 | 0 | 0 | 0 | 0 | 0 | 0 | 17 |
| SRR5756186 | 0 | 0 | 0 | 82 | 0 | 0 | 0 | 49 | 16 | 0 | 16 |
| SRR5804909 | 0 | 0 | 0 | 0 | 0 | 0 | 0 | 0 | 0 | 0 | 0 |
| SRR5804912 | 0 | 0 | 0 | 0 | 0 | 0 | 0 | 0 | 0 | 0 | 0 |
| SRR5804914 | 0 | 0 | 0 | 0 | 0 | 0 | 0 | 0 | 0 | 0 | 0 |
| SRR5817623 | 0 | 61 | 73 | 52 | 437 | 559 | 566 | 249 | 11138 | 455 | 1229 |
| SRR5817624 | 0 | 39 | 28 | 48 | 265 | 340 | 350 | 157 | 4457 | 195 | 728 |
| SRR5817625 | 0 | 84 | 75 | 38 | 398 | 514 | 570 | 248 | 11174 | 492 | 1156 |
| SRR5817626 | 0 | 74 | 62 | 58 | 324 | 448 | 392 | 197 | 7266 | 291 | 922 |
| SRR5821803 | 0 | 0 | 19 | 0 | 11 | 75 | 12 | 276 | 1278 | 1344 | 660 |
| SRR5821804 | 0 | 0 | 0 | 0 | 0 | 65 | 19 | 120 | 206 | 245 | 442 |
| SRR5821805 | 0 | 12 | 33 | 22 | 116 | 806 | 157 | 1434 | 5140 | 7929 | 7718 |
| SRR5832003 | 6031 | 37 | 74 | 37 | 0 | 37 | 0 | 74 | 0 | 37 | 0 |
| SRR5832004 | 3029 | 31 | 125 | 94 | 0 | 125 | 0 | 62 | 94 | 0 | 0 |
| SRR5832005 | 1063 | 70 | 28 | 0 | 0 | 0 | 0 | 70 | 98 | 56 | 42 |
| SRR5832006 | 2409 | 0 | 86 | 172 | 0 | 0 | 0 | 86 | 0 | 0 | 86 |
| SRR5832007 | 1218 | 0 | 84 | 0 | 42 | 126 | 42 | 252 | 42 | 0 | 0 |
| SRR5832008 | 724 | 12 | 0 | 25 | 12 | 12 | 0 | 123 | 25 | 98 | 98 |
| SRR5832009 | 3285 | 36 | 71 | 0 | 0 | 0 | 0 | 179 | 71 | 71 | 71 |
| SRR5832010 | 2178 | 0 | 0 | 28 | 0 | 28 | 0 | 28 | 28 | 0 | 28 |
| SRR5832011 | 930 | 0 | 0 | 58 | 0 | 116 | 0 | 58 | 174 | 0 | 116 |
| SRR5832012 | 3381 | 0 | 0 | 0 | 0 | 130 | 0 | 65 | 65 | 0 | 0 |
| SRR5832013 | 4896 | 42 | 42 | 167 | 0 | 0 | 0 | 251 | 84 | 126 | 0 |
| SRR5832014 | 2798 | 122 | 122 | 61 | 0 | 61 | 0 | 122 | 122 | 122 | 0 |
| SRR5832015 | 1578 | 0 | 0 | 0 | 0 | 73 | 37 | 110 | 0 | 0 | 0 |
| SRR5832016 | 743 | 0 | 57 | 29 | 29 | 29 | 0 | 143 | 57 | 29 | 57 |
| SRR5832017 | 251 | 0 | 101 | 0 | 0 | 50 | 0 | 0 | 50 | 302 | 101 |
| SRR5832018 | 948 | 0 | 75 | 25 | 0 | 0 | 25 | 25 | 25 | 0 | 50 |
| SRR5832019 | 1056 | 0 | 182 | 146 | 36 | 36 | 0 | 36 | 0 | 0 | 146 |
| SRR5832020 | 1126 | 31 | 31 | 63 | 0 | 31 | 0 | 94 | 94 | 0 | 63 |
| SRR5832021 | 167 | 0 | 0 | 56 | 0 | 0 | 0 | 56 | 0 | 0 | 0 |
| SRR5832022 | 260 | 26 | 26 | 0 | 0 | 0 | 0 | 52 | 52 | 130 | 0 |
| SRR5832023 | 413 | 0 | 44 | 22 | 0 | 22 | 0 | 65 | 87 | 0 | 0 |
| SRR5832024 | 3517 | 0 | 56 | 0 | 0 | 0 | 0 | 223 | 0 | 223 | 0 |
| SRR5832025 | 623 | 0 | 11 | 11 | 0 | 45 | 0 | 57 | 11 | 57 | 34 |
| SRR5832026 | 1240 | 0 | 16 | 16 | 0 | 48 | 0 | 161 | 64 | 16 | 0 |
| SRR5832027 | 424 | 0 | 33 | 33 | 0 | 98 | 0 | 33 | 65 | 33 | 98 |
| SRR5832028 | 191 | 14 | 14 | 0 | 14 | 55 | 0 | 82 | 0 | 27 | 41 |
| SRR5832029 | 295 | 49 | 0 | 0 | 0 | 0 | 0 | 0 | 0 | 49 | 0 |
| SRR5868085 | 0 | 50 | 255 | 107 | 6516 | 22088 | 10503 | 1994 | 656 | 407 | 822 |
| SRR5901584 | 0 | 0 | 60 | 20 | 27 | 231 | 22 | 415 | 342 | 412 | 139 |
| SRR5901585 | 0 | 0 | 50 | 14 | 22 | 205 | 18 | 351 | 266 | 360 | 131 |
| SRR5901586 | 0 | 0 | 57 | 20 | 30 | 363 | 29 | 501 | 612 | 597 | 299 |
| SRR5901587 | 0 | 0 | 20 | 0 | 32 | 345 | 28 | 580 | 938 | 907 | 274 |
| SRR5901588 | 0 | 0 | 22 | 0 | 36 | 358 | 35 | 569 | 695 | 707 | 255 |
| SRR5901589 | 0 | 0 | 14 | 0 | 28 | 394 | 35 | 633 | 987 | 1025 | 303 |
| SRR5901590 | 0 | 0 | 0 | 0 | 59 | 977 | 131 | 2192 | 778 | 1263 | 419 |
| SRR5901591 | 0 | 0 | 19 | 0 | 64 | 810 | 94 | 1954 | 649 | 1084 | 364 |
| SRR5901592 | 0 | 0 | 0 | 0 | 27 | 606 | 70 | 1268 | 486 | 908 | 298 |
| SRR5901593 | 0 | 0 | 0 | 0 | 50 | 1029 | 96 | 1576 | 767 | 1057 | 450 |
| SRR5901594 | 0 | 0 | 0 | 0 | 34 | 870 | 57 | 1324 | 1373 | 1722 | 521 |
| SRR5901595 | 0 | 0 | 0 | 0 | 29 | 1135 | 52 | 1750 | 778 | 1222 | 455 |
| SRR5922753 | 0 | 0 | 0 | 16 | 159 | 320 | 308 | 398 | 742 | 94 | 487 |
| SRR5922754 | 0 | 0 | 0 | 0 | 21 | 106 | 86 | 129 | 982 | 73 | 386 |
| SRR5922756 | 0 | 0 | 0 | 0 | 0 | 0 | 0 | 14 | 138 | 13 | 45 |
| SRR5925650 | 0 | 0 | 17 | 12 | 0 | 24 | 0 | 29 | 52 | 40 | 67 |
| SRR5925651 | 0 | 0 | 0 | 0 | 0 | 23 | 0 | 50 | 126 | 96 | 123 |
| SRR5925652 | 0 | 15 | 34 | 28 | 27 | 175 | 46 | 218 | 1024 | 896 | 860 |
| SRR5925665 | 0 | 386 | 337 | 74 | 134 | 1075 | 183 | 936 | 708 | 337 | 520 |
| SRR5925666 | 0 | 419 | 299 | 70 | 116 | 1192 | 221 | 1009 | 683 | 237 | 439 |
| SRR5925667 | 0 | 717 | 624 | 130 | 292 | 1919 | 449 | 1899 | 1126 | 539 | 956 |
| SRR5925668 | 0 | 507 | 449 | 105 | 223 | 1430 | 331 | 1393 | 970 | 502 | 733 |
| SRR5925669 | 0 | 604 | 477 | 118 | 216 | 1835 | 363 | 1844 | 950 | 436 | 840 |
| SRR5925670 | 0 | 526 | 653 | 114 | 202 | 1389 | 364 | 1617 | 1004 | 495 | 793 |
| SRR5925671 | 0 | 745 | 604 | 172 | 126 | 1311 | 271 | 1609 | 1295 | 688 | 1139 |
| SRR5925672 | 0 | 627 | 595 | 189 | 217 | 1411 | 361 | 1833 | 1238 | 873 | 1091 |
| SRR5925673 | 0 | 490 | 363 | 121 | 118 | 858 | 198 | 784 | 380 | 180 | 525 |
| SRR5925674 | 0 | 590 | 450 | 52 | 118 | 657 | 177 | 627 | 435 | 133 | 516 |
| SRR5925675 | 0 | 193 | 213 | 67 | 23 | 181 | 53 | 295 | 173 | 53 | 149 |
| SRR5925676 | 0 | 397 | 222 | 101 | 77 | 427 | 89 | 581 | 347 | 119 | 350 |
| SRR5925677 | 0 | 437 | 397 | 120 | 94 | 571 | 76 | 517 | 780 | 366 | 669 |
| SRR5925678 | 0 | 552 | 414 | 128 | 118 | 700 | 155 | 811 | 771 | 263 | 532 |
| SRR5925679 | 0 | 536 | 364 | 169 | 100 | 513 | 73 | 636 | 812 | 318 | 762 |
| SRR5925680 | 0 | 436 | 317 | 126 | 126 | 498 | 100 | 743 | 752 | 314 | 781 |
| SRR5925681 | 0 | 377 | 352 | 96 | 117 | 549 | 117 | 813 | 674 | 272 | 855 |
| SRR5925682 | 0 | 396 | 350 | 92 | 138 | 685 | 138 | 911 | 969 | 323 | 808 |
| SRR5925683 | 0 | 666 | 447 | 130 | 143 | 720 | 130 | 1230 | 1833 | 854 | 1364 |
| SRR5925684 | 0 | 630 | 403 | 103 | 264 | 698 | 134 | 1028 | 1421 | 687 | 1297 |
| SRR5925685 | 0 | 366 | 309 | 167 | 82 | 504 | 57 | 618 | 1672 | 863 | 1019 |
| SRR5925686 | 0 | 386 | 333 | 151 | 79 | 416 | 83 | 730 | 1665 | 772 | 938 |
| SRR5925687 | 0 | 467 | 406 | 144 | 79 | 734 | 105 | 935 | 1944 | 856 | 887 |
| SRR5925688 | 0 | 439 | 382 | 126 | 98 | 509 | 61 | 806 | 1969 | 899 | 1074 |
| SRR5925689 | 0 | 479 | 364 | 148 | 97 | 454 | 65 | 666 | 2135 | 972 | 936 |
| SRR5925690 | 0 | 455 | 367 | 113 | 55 | 516 | 70 | 748 | 1677 | 748 | 861 |
| SRR5925691 | 0 | 501 | 378 | 123 | 112 | 631 | 112 | 813 | 2701 | 1328 | 1142 |
| SRR5925692 | 0 | 501 | 435 | 128 | 144 | 527 | 75 | 749 | 2572 | 1447 | 1087 |
| SRR5925693 | 0 | 378 | 294 | 181 | 97 | 627 | 81 | 849 | 1263 | 827 | 895 |
| SRR5925694 | 0 | 415 | 283 | 132 | 99 | 569 | 93 | 788 | 1285 | 656 | 857 |
| SRR5925695 | 0 | 491 | 368 | 132 | 90 | 543 | 80 | 699 | 1614 | 699 | 916 |
| SRR5925696 | 0 | 554 | 394 | 146 | 103 | 930 | 178 | 756 | 1395 | 822 | 911 |
| SRR5925697 | 0 | 587 | 398 | 230 | 88 | 749 | 101 | 1057 | 2589 | 1161 | 1426 |

### Supp Table S8.5.Columns_ET_to_FD

| | | | | | | | | | | | |
|---|---|---|---|---|---|---|---|---|---|---|---|
| SRR5925698 | 0 | 617 | 385 | 183 | 84 | 947 | 143 | 982 | 2408 | 1317 | 1406 |
| SRR5925699 | 0 | 495 | 370 | 174 | 124 | 544 | 68 | 710 | 1977 | 1022 | 1109 |
| SRR5925700 | 0 | 543 | 459 | 173 | 109 | 711 | 84 | 838 | 2003 | 1010 | 981 |
| SRR5925701 | 0 | 464 | 278 | 116 | 54 | 569 | 72 | 698 | 2188 | 856 | 1046 |
| SRR5925702 | 0 | 444 | 308 | 97 | 117 | 619 | 86 | 752 | 1881 | 822 | 1110 |
| SRR5925703 | 0 | 555 | 314 | 146 | 92 | 682 | 115 | 816 | 1789 | 766 | 977 |
| SRR5925704 | 0 | 534 | 472 | 210 | 122 | 822 | 137 | 777 | 1744 | 880 | 1118 |
| SRR5925705 | 0 | 69 | 78 | 20 | 93 | 629 | 160 | 426 | 500 | 170 | 287 |
| SRR5925706 | 0 | 32 | 0 | 0 | 57 | 1411 | 144 | 996 | 522 | 157 | 262 |
| SRR5925707 | 0 | 50 | 74 | 13 | 128 | 948 | 231 | 732 | 841 | 249 | 449 |
| SRR5925708 | 0 | 29 | 16 | 0 | 41 | 1048 | 144 | 908 | 459 | 148 | 214 |
| SRR5925709 | 0 | 303 | 330 | 80 | 149 | 829 | 223 | 1648 | 1042 | 441 | 1254 |
| SRR5925710 | 0 | 0 | 0 | 463 | 0 | 0 | 463 | 926 | 463 | 0 | 926 |
| SRR5925711 | 0 | 39 | 28 | 0 | 35 | 683 | 75 | 731 | 176 | 50 | 291 |
| SRR5925712 | 0 | 42 | 189 | 25 | 60 | 232 | 97 | 239 | 119 | 51 | 356 |
| SRR5925713 | 0 | 74 | 59 | 15 | 114 | 718 | 184 | 515 | 142 | 41 | 171 |
| SRR5925714 | 0 | 68 | 65 | 21 | 108 | 664 | 159 | 480 | 152 | 43 | 128 |
| SRR5925715 | 0 | 31 | 11 | 0 | 60 | 1163 | 127 | 864 | 234 | 79 | 139 |
| SRR5925716 | 0 | 16 | 16 | 0 | 27 | 673 | 101 | 529 | 145 | 41 | 89 |
| SRR5925717 | 0 | 0 | 0 | 0 | 37 | 769 | 77 | 511 | 39 | 0 | 19 |
| SRR5925718 | 0 | 0 | 0 | 0 | 31 | 821 | 92 | 554 | 32 | 0 | 28 |
| SRR5925719 | 0 | 20 | 12 | 0 | 63 | 743 | 55 | 834 | 2693 | 688 | 908 |
| SRR5925720 | 0 | 24 | 10 | 0 | 56 | 807 | 56 | 825 | 2781 | 731 | 983 |
| SRR5925721 | 0 | 19 | 0 | 0 | 35 | 328 | 27 | 389 | 987 | 276 | 318 |
| SRR5925722 | 0 | 17 | 12 | 0 | 38 | 290 | 28 | 389 | 1076 | 285 | 356 |
| SRR5925723 | 0 | 10 | 0 | 0 | 45 | 392 | 36 | 209 | 602 | 180 | 283 |
| SRR5925724 | 0 | 16 | 0 | 0 | 26 | 346 | 38 | 188 | 667 | 211 | 302 |
| SRR5925725 | 0 | 10 | 0 | 0 | 29 | 315 | 28 | 160 | 298 | 61 | 194 |
| SRR5925726 | 0 | 0 | 0 | 0 | 33 | 242 | 27 | 145 | 341 | 106 | 211 |
| SRR5925727 | 0 | 0 | 0 | 0 | 0 | 0 | 0 | 0 | 0 | 1828 | 0 |
| SRR5925728 | 0 | 28 | 18 | 0 | 69 | 480 | 39 | 272 | 451 | 98 | 149 |
| SRR5925729 | 0 | 22 | 0 | 0 | 35 | 469 | 22 | 585 | 2124 | 800 | 897 |
| SRR5925730 | 0 | 12 | 0 | 0 | 22 | 410 | 30 | 465 | 2276 | 699 | 884 |
| SRR5925731 | 0 | 47 | 22 | 13 | 84 | 932 | 54 | 815 | 768 | 205 | 322 |
| SRR5925732 | 0 | 41 | 0 | 0 | 66 | 1119 | 84 | 803 | 1031 | 291 | 377 |
| SRR5930976 | 0 | 49 | 105 | 80 | 102 | 993 | 154 | 1319 | 4369 | 2948 | 2671 |
| SRR5930977 | 0 | 20 | 35 | 25 | 0 | 11 | 0 | 16 | 16 | 15 | 27 |
| SRR5930978 | 0 | 0 | 0 | 0 | 0 | 62 | 0 | 116 | 156 | 124 | 195 |
| SRR5936591 | 0 | 33 | 79 | 36 | 30 | 140 | 54 | 281 | 2236 | 1591 | 1329 |
| SRR5936592 | 0 | 0 | 10 | 0 | 0 | 23 | 0 | 31 | 187 | 271 | 296 |
| SRR5951878 | 10 | 0 | 14 | 0 | 1727 | 2206 | 2483 | 538 | 831 | 6264 | 2076 |
| SRR5951879 | 0 | 0 | 0 | 0 | 143 | 225 | 185 | 246 | 464 | 562 | 398 |
| SRR5951880 | 0 | 0 | 0 | 0 | 0 | 0 | 0 | 0 | 0 | 0 | 0 |
| SRR5951881 | 21 | 0 | 25 | 0 | 1502 | 1672 | 1550 | 313 | 373 | 1647 | 716 |
| SRR5951882 | 16 | 0 | 0 | 0 | 311 | 515 | 231 | 194 | 352 | 540 | 485 |
| SRR5951883 | 0 | 0 | 0 | 0 | 0 | 32 | 0 | 0 | 0 | 0 | 0 |
| SRR5951884 | 41 | 22 | 57 | 18 | 1767 | 4263 | 3452 | 1660 | 215 | 1250 | 240 |
| SRR5951885 | 55 | 20 | 52 | 13 | 1570 | 3896 | 3481 | 1584 | 215 | 910 | 201 |
| SRR5951886 | 40 | 25 | 101 | 0 | 1714 | 3667 | 2615 | 1482 | 428 | 1546 | 256 |
| SRR5951887 | 58 | 26 | 87 | 23 | 3731 | 8447 | 8413 | 4769 | 837 | 3199 | 919 |
| SRR5951888 | 69 | 36 | 117 | 27 | 4053 | 10533 | 9614 | 5667 | 1160 | 4991 | 1141 |
| SRR5951889 | 93 | 38 | 112 | 33 | 4426 | 9962 | 10293 | 4851 | 937 | 4800 | 1110 |
| SRR5951890 | 146 | 37 | 103 | 24 | 4829 | 9593 | 10248 | 3808 | 503 | 3736 | 789 |
| SRR5951891 | 106 | 29 | 142 | 20 | 3997 | 8797 | 8839 | 3793 | 437 | 2350 | 547 |
| SRR5951892 | 120 | 17 | 107 | 24 | 6418 | 13817 | 14461 | 3980 | 339 | 3325 | 614 |
| SRR5951893 | 30 | 23 | 103 | 0 | 1222 | 2634 | 1579 | 831 | 325 | 1592 | 321 |
| SRR5951894 | 111 | 43 | 208 | 0 | 1522 | 2896 | 1618 | 1195 | 612 | 2313 | 389 |
| SRR5951895 | 51 | 19 | 92 | 20 | 1316 | 2762 | 1730 | 694 | 273 | 1821 | 295 |
| SRR5951896 | 33 | 0 | 36 | 11 | 304 | 424 | 463 | 313 | 91 | 123 | 130 |
| SRR5951897 | 35 | 0 | 21 | 40 | 300 | 341 | 268 | 153 | 33 | 77 | 268 |
| SRR5951898 | 0 | 0 | 0 | 0 | 0 | 0 | 0 | 0 | 0 | 0 | 0 |
| SRR5951899 | 64 | 22 | 53 | 59 | 436 | 615 | 665 | 245 | 90 | 109 | 296 |
| SRR5951900 | 30 | 17 | 25 | 25 | 335 | 451 | 423 | 244 | 39 | 119 | 453 |
| SRR5951901 | 0 | 0 | 0 | 0 | 0 | 0 | 0 | 0 | 0 | 0 | 0 |
| SRR5951902 | 32 | 0 | 45 | 28 | 221 | 354 | 449 | 294 | 98 | 63 | 136 |
| SRR5951903 | 10 | 0 | 0 | 13 | 169 | 251 | 214 | 250 | 38 | 77 | 184 |
| SRR5951904 | 0 | 0 | 0 | 0 | 0 | 11 | 0 | 0 | 0 | 0 | 0 |
| SRR5951905 | 29 | 16 | 78 | 23 | 124 | 254 | 228 | 170 | 239 | 83 | 192 |
| SRR5951906 | 18 | 10 | 19 | 10 | 165 | 333 | 241 | 305 | 77 | 30 | 109 |
| SRR5951907 | 0 | 0 | 0 | 0 | 0 | 0 | 0 | 0 | 0 | 0 | 0 |
| SRR5951908 | 115 | 150 | 335 | 84 | 1757 | 4602 | 3889 | 2436 | 69 | 151 | 192 |
| SRR5951909 | 66 | 47 | 238 | 36 | 1128 | 3024 | 2473 | 1937 | 49 | 73 | 82 |
| SRR5951910 | 65 | 69 | 270 | 46 | 1510 | 4453 | 3160 | 2297 | 31 | 78 | 78 |
| SRR5951911 | 33 | 19 | 110 | 10 | 519 | 1282 | 702 | 521 | 33 | 93 | 45 |
| SRR5951912 | 29 | 25 | 73 | 14 | 300 | 900 | 440 | 437 | 24 | 71 | 29 |
| SRR5951913 | 37 | 48 | 112 | 33 | 507 | 1456 | 717 | 489 | 35 | 95 | 60 |
| SRR6020460 | 0 | 0 | 0 | 0 | 0 | 0 | 0 | 0 | 12 | 14 | 0 |
| SRR6020461 | 0 | 0 | 0 | 0 | 0 | 14 | 0 | 21 | 0 | 0 | 0 |
| SRR6020462 | 0 | 0 | 0 | 0 | 0 | 0 | 0 | 0 | 12 | 14 | 0 |
| SRR6020463 | 0 | 0 | 0 | 0 | 0 | 11 | 0 | 24 | 10 | 27 | 20 |
| SRR6020464 | 0 | 0 | 0 | 0 | 0 | 12 | 0 | 26 | 35 | 37 | 29 |
| SRR6020465 | 0 | 0 | 0 | 0 | 0 | 0 | 0 | 12 | 0 | 18 | 0 |
| SRR6020466 | 0 | 0 | 0 | 0 | 0 | 0 | 0 | 11 | 0 | 14 | 13 |
| SRR6020467 | 0 | 0 | 0 | 0 | 0 | 0 | 0 | 0 | 11 | 21 | 0 |
| SRR6020468 | 0 | 0 | 0 | 0 | 0 | 0 | 0 | 0 | 19 | 40 | 19 |
| SRR6020469 | 0 | 0 | 0 | 0 | 0 | 0 | 0 | 11 | 0 | 13 | 13 |
| SRR6020470 | 0 | 0 | 0 | 0 | 0 | 0 | 0 | 0 | 19 | 37 | 15 |
| SRR6020471 | 0 | 0 | 0 | 0 | 0 | 0 | 0 | 0 | 0 | 0 | 0 |
| SRR6020472 | 0 | 0 | 0 | 0 | 0 | 0 | 0 | 0 | 0 | 27 | 27 |
| SRR6020473 | 0 | 0 | 0 | 0 | 0 | 0 | 0 | 24 | 51 | 37 | 33 |
| SRR6020474 | 0 | 0 | 0 | 0 | 0 | 0 | 0 | 10 | 14 | 28 | 0 |
| SRR6020475 | 0 | 0 | 0 | 0 | 0 | 0 | 0 | 10 | 14 | 20 | 13 |
| SRR6020476 | 0 | 0 | 0 | 0 | 0 | 0 | 0 | 10 | 0 | 0 | 0 |
| SRR6020477 | 0 | 0 | 0 | 0 | 0 | 0 | 0 | 12 | 24 | 15 | 0 |
| SRR6020478 | 0 | 0 | 0 | 0 | 0 | 0 | 0 | 0 | 12 | 21 | 11 |
| SRR6020479 | 0 | 0 | 0 | 0 | 0 | 0 | 0 | 18 | 24 | 21 | 25 |
| SRR6020480 | 0 | 0 | 0 | 0 | 0 | 0 | 0 | 13 | 0 | 13 | 0 |
| SRR6020481 | 0 | 0 | 0 | 0 | 0 | 0 | 0 | 16 | 0 | 18 | 0 |
| SRR6020482 | 0 | 0 | 0 | 0 | 0 | 0 | 0 | 11 | 11 | 0 | 13 |
| SRR6020483 | 0 | 0 | 0 | 0 | 0 | 0 | 0 | 0 | 19 | 30 | 22 |
| SRR6020484 | 0 | 0 | 0 | 0 | 0 | 0 | 0 | 0 | 16 | 31 | 13 |
| SRR6020485 | 0 | 0 | 0 | 0 | 0 | 0 | 0 | 15 | 12 | 23 | 13 |
| SRR6020486 | 0 | 0 | 0 | 0 | 0 | 0 | 0 | 0 | 0 | 26 | 0 |
| SRR6020487 | 0 | 0 | 0 | 0 | 0 | 0 | 0 | 11 | 28 | 38 | 35 |
| SRR6020488 | 0 | 0 | 0 | 0 | 0 | 0 | 0 | 0 | 0 | 12 | 0 |
| SRR6020489 | 0 | 0 | 0 | 0 | 0 | 0 | 0 | 0 | 0 | 0 | 0 |
| SRR6020490 | 0 | 0 | 0 | 0 | 0 | 0 | 0 | 12 | 0 | 15 | 12 |
| SRR6020491 | 0 | 0 | 0 | 0 | 0 | 0 | 0 | 0 | 0 | 13 | 0 |
| SRR6020492 | 0 | 0 | 0 | 10 | 0 | 14 | 0 | 15 | 0 | 0 | 16 |
| SRR6020493 | 0 | 0 | 0 | 0 | 0 | 0 | 0 | 0 | 17 | 20 | 15 |
| SRR6020494 | 0 | 0 | 0 | 0 | 0 | 0 | 0 | 0 | 0 | 12 | 0 |
| SRR6020495 | 0 | 0 | 0 | 0 | 0 | 0 | 0 | 0 | 13 | 18 | 15 |
| SRR6020496 | 0 | 0 | 0 | 0 | 0 | 0 | 0 | 0 | 17 | 17 | 23 |
| SRR6020497 | 0 | 0 | 0 | 0 | 0 | 0 | 0 | 0 | 14 | 21 | 0 |
| SRR6020498 | 0 | 0 | 21 | 25 | 0 | 34 | 0 | 40 | 19 | 17 | 11 |
| SRR6020499 | 0 | 0 | 0 | 0 | 0 | 0 | 0 | 0 | 14 | 16 | 0 |
| SRR6020500 | 0 | 0 | 0 | 0 | 0 | 0 | 0 | 17 | 0 | 10 | 0 |
| SRR6020501 | 0 | 0 | 0 | 0 | 0 | 0 | 0 | 18 | 25 | 29 | 26 |
| SRR6020502 | 0 | 0 | 0 | 0 | 0 | 0 | 0 | 0 | 0 | 0 | 0 |

### Supp Table S8.5.Columns_ET_to_FD

| | | | | | | | | | | | |
|---|---|---|---|---|---|---|---|---|---|---|---|
| SRR6020503 | 0 | 0 | 0 | 0 | 0 | 0 | 0 | 10 | 18 | 24 | 20 |
| SRR6020504 | 0 | 0 | 0 | 0 | 0 | 0 | 0 | 23 | 51 | 57 | 41 |
| SRR6020505 | 0 | 0 | 0 | 0 | 0 | 0 | 0 | 15 | 19 | 40 | 17 |
| SRR6020506 | 0 | 0 | 0 | 0 | 0 | 0 | 0 | 0 | 0 | 15 | 0 |
| SRR6020507 | 0 | 0 | 0 | 0 | 0 | 0 | 0 | 0 | 11 | 11 | 11 |
| SRR6020508 | 0 | 0 | 0 | 0 | 0 | 0 | 0 | 13 | 12 | 20 | 0 |
| SRR6020509 | 0 | 0 | 0 | 0 | 0 | 0 | 0 | 11 | 11 | 20 | 15 |
| SRR6020510 | 0 | 0 | 0 | 0 | 0 | 0 | 0 | 0 | 16 | 16 | 22 |
| SRR6020511 | 0 | 0 | 0 | 0 | 0 | 0 | 0 | 0 | 0 | 19 | 13 |
| SRR6020512 | 0 | 0 | 0 | 0 | 0 | 0 | 0 | 0 | 12 | 26 | 22 |
| SRR6020513 | 0 | 0 | 0 | 0 | 0 | 0 | 0 | 0 | 26 | 22 | 19 |
| SRR6020514 | 0 | 0 | 0 | 0 | 0 | 0 | 0 | 0 | 0 | 0 | 0 |
| SRR6020515 | 0 | 0 | 0 | 0 | 0 | 0 | 0 | 13 | 19 | 27 | 24 |
| SRR6020516 | 0 | 0 | 0 | 0 | 0 | 0 | 0 | 13 | 0 | 24 | 0 |
| SRR6020517 | 0 | 0 | 0 | 0 | 0 | 0 | 0 | 11 | 0 | 10 | 0 |
| SRR6020518 | 0 | 0 | 0 | 0 | 0 | 0 | 0 | 0 | 21 | 18 | 0 |
| SRR6020519 | 0 | 0 | 0 | 0 | 0 | 0 | 0 | 18 | 0 | 24 | 18 |
| SRR6020520 | 0 | 0 | 0 | 0 | 0 | 0 | 0 | 0 | 23 | 34 | 17 |
| SRR6020521 | 0 | 0 | 0 | 0 | 0 | 0 | 0 | 0 | 0 | 23 | 12 |
| SRR6020522 | 0 | 0 | 0 | 0 | 0 | 0 | 0 | 0 | 21 | 19 | 11 |
| SRR6020523 | 0 | 0 | 0 | 0 | 0 | 0 | 0 | 13 | 12 | 19 | 27 |
| SRR6020524 | 0 | 0 | 0 | 0 | 0 | 0 | 0 | 0 | 15 | 31 | 15 |
| SRR6020525 | 0 | 0 | 0 | 0 | 0 | 0 | 0 | 0 | 0 | 14 | 19 |
| SRR6020526 | 0 | 0 | 0 | 0 | 0 | 0 | 0 | 0 | 25 | 23 | 22 |
| SRR6020527 | 0 | 0 | 0 | 0 | 0 | 0 | 0 | 14 | 0 | 0 | 0 |
| SRR6020528 | 0 | 0 | 15 | 0 | 0 | 0 | 0 | 12 | 10 | 10 | 0 |
| SRR6020529 | 0 | 0 | 0 | 0 | 0 | 0 | 0 | 0 | 0 | 10 | 12 |
| SRR6020530 | 0 | 0 | 0 | 0 | 0 | 0 | 0 | 0 | 0 | 12 | 10 |
| SRR6020531 | 0 | 0 | 0 | 0 | 0 | 0 | 0 | 0 | 0 | 11 | 0 |
| SRR6020532 | 0 | 0 | 0 | 0 | 0 | 0 | 0 | 14 | 17 | 21 | 12 |
| SRR6025030 | 324 | 18 | 26 | 0 | 48 | 138 | 31 | 44 | 72 | 78 | 12 |
| SRR6025031 | 0 | 0 | 0 | 0 | 0 | 0 | 13 | 0 | 0 | 0 | 0 |
| SRR6025033 | 0 | 0 | 0 | 0 | 0 | 0 | 0 | 11 | 0 | 0 | 11 |
| SRR6025035 | 27 | 0 | 21 | 12 | 0 | 30 | 0 | 44 | 94 | 74 | 23 |
| SRR6025036 | 27 | 0 | 20 | 15 | 0 | 18 | 0 | 40 | 49 | 16 | 14 |
| SRR6025037 | 448 | 0 | 18 | 13 | 0 | 28 | 0 | 40 | 54 | 38 | 11 |
| SRR6025038 | 0 | 0 | 0 | 0 | 0 | 0 | 0 | 0 | 0 | 0 | 0 |
| SRR6025041 | 0 | 0 | 0 | 0 | 13 | 59 | 13 | 15 | 406 | 0 | 12 |
| SRR6025042 | 0 | 0 | 0 | 10 | 0 | 10 | 0 | 0 | 0 | 0 | 0 |
| SRR6025043 | 0 | 0 | 18 | 0 | 0 | 35 | 0 | 71 | 111 | 80 | 35 |
| SRR6025044 | 0 | 0 | 0 | 0 | 0 | 44 | 0 | 64 | 94 | 13 | 17 |
| SRR6025047 | 0 | 0 | 0 | 0 | 34 | 83 | 18 | 41 | 386 | 23 | 17 |
| SRR6025048 | 0 | 12 | 41 | 0 | 29 | 123 | 15 | 11 | 262 | 0 | 0 |
| SRR6025050 | 350 | 22 | 58 | 15 | 54 | 219 | 36 | 24 | 179 | 71 | 36 |
| SRR6025051 | 10 | 0 | 0 | 12 | 0 | 31 | 0 | 95 | 73 | 19 | 27 |
| SRR6025053 | 246 | 0 | 17 | 10 | 0 | 15 | 0 | 28 | 0 | 0 | 0 |
| SRR6025055 | 207 | 0 | 55 | 11 | 83 | 138 | 28 | 61 | 41 | 11 | 0 |
| SRR6025056 | 88 | 0 | 15 | 18 | 62 | 106 | 32 | 65 | 18 | 12 | 0 |
| SRR6025058 | 103 | 0 | 24 | 19 | 0 | 24 | 0 | 31 | 20 | 0 | 0 |
| SRR6025061 | 0 | 10 | 28 | 42 | 0 | 23 | 0 | 40 | 18 | 13 | 0 |
| SRR6025062 | 62 | 10 | 40 | 23 | 0 | 38 | 0 | 61 | 28 | 12 | 0 |
| SRR6025065 | 12 | 11 | 50 | 19 | 0 | 34 | 0 | 72 | 64 | 102 | 44 |
| SRR6025066 | 32 | 49 | 66 | 113 | 26 | 35 | 16 | 46 | 10 | 0 | 29 |
| SRR6025067 | 415 | 16 | 74 | 36 | 0 | 51 | 0 | 78 | 25 | 61 | 20 |
| SRR6025068 | 167 | 0 | 27 | 30 | 0 | 37 | 0 | 63 | 14 | 25 | 30 |
| SRR6025069 | 12 | 16 | 35 | 41 | 11 | 29 | 0 | 73 | 24 | 32 | 51 |
| SRR6025070 | 0 | 0 | 19 | 0 | 25 | 132 | 12 | 61 | 287 | 42 | 18 |
| SRR6025071 | 0 | 16 | 51 | 0 | 44 | 133 | 13 | 16 | 188 | 11 | 12 |
| SRR6025072 | 138 | 0 | 0 | 10 | 0 | 23 | 0 | 33 | 0 | 0 | 15 |
| SRR6025073 | 107 | 0 | 0 | 0 | 0 | 21 | 0 | 37 | 0 | 11 | 0 |
| SRR6025075 | 81 | 0 | 13 | 11 | 0 | 40 | 0 | 48 | 12 | 14 | 39 |
| SRR6025079 | 0 | 32 | 92 | 11 | 61 | 194 | 23 | 28 | 58 | 0 | 0 |
| SRR6025080 | 283 | 13 | 33 | 25 | 60 | 129 | 46 | 41 | 261 | 21 | 21 |
| SRR6025082 | 0 | 11 | 33 | 0 | 36 | 70 | 24 | 24 | 125 | 10 | 0 |
| SRR6025083 | 381 | 21 | 67 | 17 | 65 | 187 | 43 | 72 | 373 | 89 | 29 |
| SRR6025084 | 24 | 0 | 33 | 0 | 62 | 159 | 42 | 51 | 387 | 24 | 13 |
| SRR6025085 | 0 | 11 | 36 | 0 | 33 | 66 | 22 | 25 | 201 | 25 | 15 |
| SRR6025086 | 46 | 45 | 75 | 146 | 32 | 64 | 31 | 95 | 20 | 16 | 69 |
| SRR6025087 | 20 | 49 | 87 | 58 | 30 | 39 | 18 | 48 | 26 | 20 | 52 |
| SRR6025088 | 0 | 0 | 13 | 0 | 21 | 49 | 14 | 16 | 107 | 0 | 0 |
| SRR6025091 | 22 | 0 | 17 | 0 | 19 | 90 | 14 | 41 | 238 | 0 | 13 |
| SRR6025092 | 240 | 0 | 21 | 0 | 0 | 25 | 0 | 25 | 13 | 26 | 12 |
| SRR6025093 | 0 | 15 | 25 | 0 | 23 | 59 | 16 | 49 | 195 | 21 | 17 |
| SRR6025094 | 53 | 0 | 0 | 10 | 0 | 26 | 0 | 39 | 18 | 41 | 15 |
| SRR6025095 | 0 | 0 | 30 | 0 | 65 | 226 | 26 | 67 | 186 | 24 | 24 |
| SRR6025096 | 16 | 0 | 30 | 0 | 35 | 187 | 16 | 76 | 370 | 20 | 18 |
| SRR6025097 | 0 | 21 | 33 | 0 | 37 | 99 | 23 | 54 | 216 | 10 | 12 |
| SRR6025099 | 0 | 0 | 0 | 0 | 0 | 0 | 10 | 0 | 0 | 0 | 0 |
| SRR6025101 | 0 | 32 | 92 | 0 | 62 | 189 | 25 | 23 | 77 | 0 | 0 |
| SRR6025103 | 0 | 0 | 29 | 0 | 38 | 124 | 15 | 14 | 331 | 0 | 0 |
| SRR6025104 | 0 | 0 | 0 | 0 | 14 | 38 | 0 | 0 | 235 | 0 | 0 |
| SRR6025105 | 0 | 13 | 17 | 0 | 0 | 30 | 0 | 28 | 20 | 28 | 17 |
| SRR6025106 | 0 | 13 | 52 | 0 | 35 | 122 | 16 | 20 | 247 | 0 | 0 |
| SRR6025108 | 0 | 0 | 0 | 0 | 0 | 0 | 0 | 0 | 0 | 0 | 0 |
| SRR6025111 | 197 | 29 | 25 | 17 | 76 | 139 | 25 | 46 | 67 | 13 | 0 |
| SRR6025113 | 395 | 0 | 0 | 0 | 26 | 99 | 18 | 29 | 29 | 11 | 0 |
| SRR6025114 | 471 | 0 | 32 | 11 | 43 | 72 | 43 | 29 | 32 | 14 | 0 |
| SRR6025116 | 478 | 13 | 162 | 0 | 62 | 241 | 21 | 0 | 313 | 38 | 11 |
| SRR6025119 | 0 | 19 | 50 | 0 | 33 | 129 | 14 | 17 | 168 | 11 | 0 |
| SRR6025120 | 98 | 0 | 0 | 11 | 0 | 41 | 0 | 36 | 11 | 0 | 11 |
| SRR6025121 | 0 | 0 | 12 | 12 | 0 | 19 | 0 | 47 | 22 | 15 | 35 |
| SRR6025123 | 91 | 14 | 25 | 0 | 24 | 78 | 11 | 26 | 227 | 13 | 0 |
| SRR6025125 | 0 | 10 | 41 | 0 | 29 | 99 | 17 | 14 | 211 | 0 | 0 |
| SRR6025126 | 0 | 0 | 31 | 0 | 32 | 97 | 13 | 0 | 262 | 0 | 0 |
| SRR6025128 | 214 | 24 | 40 | 0 | 22 | 103 | 0 | 53 | 214 | 30 | 25 |
| SRR6040044 | 45 | 31 | 63 | 28 | 3787 | 8303 | 4895 | 3330 | 3327 | 6705 | 2862 |
| SRR6040045 | 80 | 20 | 57 | 20 | 5501 | 10543 | 6945 | 3958 | 4178 | 7416 | 3781 |
| SRR6076852 | 0 | 0 | 0 | 0 | 0 | 39 | 31 | 0 | 55 | 0 | 0 |
| SRR6076853 | 0 | 0 | 0 | 0 | 0 | 55 | 0 | 21 | 0 | 14 | 0 |
| SRR6076854 | 0 | 11 | 0 | 0 | 64 | 96 | 64 | 43 | 64 | 32 | 0 |
| SRR6076855 | 0 | 0 | 0 | 0 | 0 | 22 | 22 | 0 | 30 | 0 | 0 |
| SRR6076856 | 0 | 0 | 0 | 0 | 0 | 22 | 0 | 17 | 34 | 11 | 17 |
| SRR6076858 | 0 | 0 | 0 | 0 | 0 | 0 | 0 | 17 | 0 | 26 | 0 |
| SRR6076861 | 0 | 0 | 0 | 0 | 16 | 43 | 38 | 0 | 32 | 0 | 11 |
| SRR6153965 | 0 | 0 | 0 | 0 | 0 | 0 | 0 | 0 | 0 | 0 | 0 |
| SRR6153968 | 0 | 0 | 0 | 0 | 0 | 0 | 0 | 0 | 0 | 0 | 0 |
| SRR6153976 | 0 | 0 | 0 | 0 | 0 | 0 | 0 | 0 | 0 | 0 | 0 |
| SRR6153984 | 0 | 0 | 0 | 0 | 0 | 0 | 0 | 0 | 0 | 0 | 0 |
| SRR6153998 | 0 | 0 | 0 | 0 | 0 | 0 | 0 | 0 | 0 | 0 | 0 |
| SRR6154002 | 0 | 0 | 0 | 0 | 0 | 0 | 0 | 0 | 0 | 0 | 0 |
| SRR6154003 | 0 | 0 | 0 | 0 | 0 | 0 | 0 | 0 | 0 | 0 | 0 |
| SRR6154004 | 0 | 0 | 0 | 0 | 0 | 0 | 0 | 0 | 0 | 0 | 0 |
| SRR6154005 | 0 | 0 | 0 | 0 | 0 | 0 | 0 | 0 | 0 | 0 | 0 |
| SRR6154006 | 0 | 0 | 12 | 0 | 0 | 0 | 0 | 0 | 0 | 0 | 0 |
| SRR6203722 | 0 | 0 | 25 | 12 | 0 | 12 | 0 | 12 | 12 | 25 | 12 |
| SRR6203757 | 0 | 23 | 46 | 23 | 0 | 0 | 23 | 46 | 46 | 115 | 46 |
| SRR6203760 | 0 | 0 | 0 | 0 | 0 | 0 | 0 | 0 | 0 | 0 | 0 |
| SRR6203761 | 0 | 176 | 0 | 0 | 0 | 0 | 0 | 0 | 0 | 0 | 0 |
| SRR6203772 | 0 | 19 | 0 | 0 | 0 | 0 | 0 | 28 | 0 | 57 | 0 |

### Supp Table S8.5.Columns_ET_to_FD

| | | | | | | | | | | | |
|---|---|---|---|---|---|---|---|---|---|---|---|
| SRR6203807 | 0 | 0 | 0 | 0 | 0 | 0 | 0 | 0 | 0 | 0 | 0 |
| SRR6224558 | 0 | 0 | 0 | 0 | 0 | 0 | 0 | 0 | 0 | 12 | 0 |
| SRR6226650 | 0 | 0 | 0 | 0 | 163 | 628 | 564 | 236 | 88 | 97 | 325 |
| SRR6226651 | 0 | 0 | 0 | 0 | 884 | 2864 | 3420 | 1003 | 298 | 154 | 321 |
| SRR6226652 | 0 | 0 | 0 | 0 | 204 | 539 | 518 | 251 | 77 | 28 | 49 |
| SRR6226653 | 0 | 0 | 0 | 0 | 55 | 163 | 117 | 89 | 90 | 16 | 39 |
| SRR6226654 | 0 | 0 | 0 | 0 | 317 | 780 | 554 | 288 | 86 | 62 | 135 |
| SRR6226655 | 0 | 0 | 0 | 0 | 667 | 874 | 507 | 188 | 89 | 69 | 151 |
| SRR6226656 | 0 | 0 | 0 | 0 | 354 | 756 | 572 | 104 | 20 | 51 | 40 |
| SRR6226657 | 0 | 0 | 0 | 0 | 501 | 931 | 547 | 89 | 14 | 16 | 32 |
| SRR6230411 | 0 | 124 | 238 | 56 | 5509 | 21858 | 7876 | 6633 | 14577 | 21872 | 14016 |
| SRR6230412 | 0 | 101 | 400 | 348 | 13434 | 41713 | 10722 | 6515 | 4356 | 13123 | 5660 |
| SRR6230413 | 0 | 41 | 133 | 76 | 8413 | 21909 | 8671 | 6865 | 5437 | 14674 | 6767 |
| SRR6246305 | 0 | 44 | 122 | 268 | 43 | 189 | 32 | 171 | 88 | 236 | 94 |
| SRR6246306 | 0 | 30 | 111 | 226 | 53 | 180 | 41 | 182 | 87 | 234 | 101 |
| SRR6246307 | 0 | 29 | 119 | 237 | 46 | 182 | 46 | 181 | 101 | 225 | 118 |
| SRR6246308 | 0 | 33 | 99 | 246 | 53 | 177 | 41 | 189 | 96 | 235 | 103 |
| SRR6246309 | 0 | 0 | 0 | 11 | 0 | 32 | 0 | 35 | 0 | 12 | 10 |
| SRR6246310 | 0 | 0 | 0 | 12 | 10 | 28 | 0 | 42 | 0 | 14 | 0 |
| SRR6246311 | 0 | 0 | 0 | 12 | 0 | 19 | 0 | 35 | 13 | 17 | 0 |
| SRR6246312 | 0 | 0 | 0 | 10 | 0 | 21 | 0 | 36 | 15 | 0 | 0 |
| SRR6246313 | 0 | 0 | 0 | 0 | 0 | 0 | 0 | 12 | 0 | 0 | 0 |
| SRR6246314 | 0 | 0 | 0 | 0 | 0 | 0 | 0 | 0 | 0 | 0 | 0 |
| SRR6246315 | 0 | 0 | 0 | 0 | 0 | 0 | 0 | 0 | 0 | 0 | 0 |
| SRR6246316 | 0 | 0 | 0 | 0 | 0 | 0 | 0 | 10 | 0 | 0 | 0 |
| SRR6246317 | 0 | 0 | 0 | 0 | 0 | 0 | 0 | 0 | 0 | 0 | 0 |
| SRR6246318 | 0 | 0 | 0 | 0 | 0 | 0 | 0 | 0 | 0 | 0 | 0 |
| SRR6246319 | 0 | 0 | 0 | 0 | 0 | 0 | 0 | 0 | 0 | 0 | 0 |
| SRR6246320 | 0 | 0 | 0 | 0 | 0 | 0 | 0 | 0 | 0 | 0 | 0 |
| SRR6246321 | 12 | 25 | 85 | 101 | 0 | 36 | 0 | 107 | 12 | 37 | 22 |
| SRR6246322 | 0 | 23 | 75 | 61 | 0 | 44 | 0 | 100 | 22 | 49 | 18 |
| SRR6246323 | 0 | 25 | 58 | 97 | 0 | 41 | 0 | 107 | 15 | 30 | 13 |
| SRR6246324 | 13 | 29 | 54 | 74 | 0 | 50 | 0 | 97 | 14 | 37 | 22 |
| SRR6246325 | 0 | 0 | 0 | 11 | 0 | 11 | 0 | 0 | 0 | 0 | 0 |
| SRR6246326 | 0 | 0 | 0 | 0 | 0 | 0 | 0 | 16 | 0 | 0 | 0 |
| SRR6246327 | 0 | 0 | 0 | 0 | 0 | 12 | 0 | 0 | 0 | 0 | 0 |
| SRR6246328 | 0 | 0 | 0 | 0 | 0 | 19 | 0 | 0 | 0 | 0 | 0 |
| SRR6246329 | 0 | 0 | 0 | 0 | 0 | 0 | 0 | 13 | 0 | 0 | 0 |
| SRR6246330 | 0 | 0 | 0 | 0 | 0 | 0 | 0 | 0 | 0 | 0 | 0 |
| SRR6246331 | 0 | 0 | 0 | 0 | 0 | 0 | 0 | 13 | 0 | 0 | 0 |
| SRR6246332 | 0 | 0 | 0 | 0 | 0 | 0 | 0 | 0 | 0 | 0 | 0 |
| SRR6246333 | 0 | 0 | 0 | 0 | 0 | 0 | 0 | 0 | 0 | 0 | 0 |
| SRR6246334 | 0 | 0 | 0 | 0 | 0 | 0 | 0 | 0 | 0 | 0 | 0 |
| SRR6246335 | 0 | 0 | 0 | 0 | 0 | 0 | 0 | 0 | 0 | 0 | 0 |
| SRR6246336 | 0 | 0 | 0 | 0 | 0 | 0 | 0 | 0 | 0 | 0 | 0 |
| SRR6246337 | 0 | 0 | 18 | 12 | 0 | 12 | 0 | 26 | 0 | 0 | 0 |
| SRR6246338 | 12 | 0 | 18 | 13 | 0 | 0 | 0 | 28 | 0 | 0 | 0 |
| SRR6246339 | 0 | 0 | 26 | 0 | 0 | 0 | 0 | 21 | 0 | 0 | 0 |
| SRR6246340 | 0 | 0 | 19 | 0 | 0 | 12 | 0 | 26 | 0 | 0 | 0 |
| SRR6246341 | 0 | 0 | 0 | 0 | 0 | 0 | 0 | 0 | 0 | 0 | 0 |
| SRR6246342 | 0 | 0 | 0 | 0 | 0 | 0 | 0 | 0 | 0 | 0 | 0 |
| SRR6246343 | 0 | 0 | 0 | 0 | 0 | 0 | 0 | 0 | 0 | 0 | 0 |
| SRR6246344 | 0 | 0 | 0 | 0 | 0 | 0 | 0 | 0 | 0 | 0 | 0 |
| SRR6246345 | 0 | 0 | 0 | 0 | 0 | 0 | 0 | 0 | 0 | 0 | 0 |
| SRR6246346 | 0 | 0 | 0 | 0 | 0 | 0 | 0 | 0 | 0 | 0 | 0 |
| SRR6246347 | 0 | 0 | 0 | 0 | 0 | 0 | 0 | 0 | 0 | 0 | 0 |
| SRR6246348 | 0 | 0 | 0 | 0 | 0 | 0 | 0 | 0 | 0 | 0 | 0 |
| SRR6246349 | 0 | 0 | 0 | 0 | 0 | 0 | 0 | 0 | 0 | 0 | 0 |
| SRR6246350 | 0 | 0 | 0 | 0 | 0 | 0 | 0 | 0 | 0 | 0 | 0 |
| SRR6246351 | 0 | 0 | 0 | 0 | 0 | 0 | 0 | 0 | 0 | 0 | 0 |
| SRR6246352 | 0 | 0 | 0 | 0 | 0 | 0 | 0 | 0 | 0 | 0 | 0 |
| SRR6246353 | 0 | 0 | 0 | 0 | 17 | 28 | 15 | 24 | 40 | 45 | 11 |
| SRR6246354 | 0 | 0 | 0 | 0 | 17 | 25 | 0 | 26 | 30 | 31 | 0 |
| SRR6246355 | 0 | 0 | 0 | 0 | 24 | 24 | 13 | 22 | 39 | 40 | 11 |
| SRR6246356 | 0 | 0 | 0 | 0 | 13 | 25 | 11 | 18 | 35 | 42 | 0 |
| SRR6246358 | 0 | 0 | 11 | 0 | 0 | 0 | 0 | 0 | 0 | 0 | 0 |
| SRR6246359 | 0 | 0 | 0 | 0 | 0 | 0 | 0 | 0 | 11 | 0 | 0 |
| SRR6246360 | 0 | 0 | 0 | 0 | 0 | 0 | 0 | 0 | 11 | 0 | 0 |
| SRR6246361 | 0 | 0 | 12 | 0 | 0 | 0 | 12 | 0 | 0 | 0 | 0 |
| SRR6246362 | 0 | 12 | 0 | 0 | 0 | 0 | 0 | 0 | 12 | 0 | 12 |
| SRR6246363 | 0 | 0 | 0 | 0 | 0 | 12 | 0 | 0 | 0 | 0 | 12 |
| SRR6246364 | 0 | 0 | 0 | 0 | 0 | 0 | 12 | 0 | 12 | 0 | 0 |
| SRR6246365 | 0 | 0 | 0 | 0 | 0 | 29 | 0 | 0 | 0 | 0 | 0 |
| SRR6246366 | 0 | 0 | 0 | 0 | 0 | 0 | 0 | 0 | 0 | 0 | 0 |
| SRR6246367 | 0 | 0 | 0 | 0 | 0 | 0 | 0 | 0 | 29 | 0 | 0 |
| SRR6246368 | 0 | 0 | 0 | 0 | 0 | 30 | 0 | 0 | 0 | 0 | 0 |
| SRR6246369 | 0 | 0 | 0 | 0 | 13 | 28 | 0 | 27 | 32 | 138 | 11 |
| SRR6246370 | 0 | 0 | 0 | 0 | 13 | 27 | 0 | 17 | 35 | 137 | 13 |
| SRR6246371 | 0 | 0 | 0 | 0 | 0 | 32 | 0 | 24 | 30 | 147 | 10 |
| SRR6246372 | 0 | 0 | 0 | 0 | 11 | 27 | 0 | 25 | 31 | 138 | 0 |
| SRR6246373 | 0 | 0 | 0 | 0 | 0 | 0 | 0 | 0 | 0 | 0 | 0 |
| SRR6246374 | 0 | 0 | 0 | 0 | 0 | 0 | 0 | 0 | 0 | 0 | 0 |
| SRR6246375 | 0 | 0 | 0 | 0 | 0 | 0 | 0 | 0 | 0 | 0 | 0 |
| SRR6246376 | 0 | 0 | 0 | 0 | 0 | 0 | 0 | 0 | 0 | 0 | 0 |
| SRR6246379 | 0 | 0 | 0 | 0 | 0 | 0 | 0 | 0 | 0 | 0 | 0 |
| SRR6246385 | 0 | 0 | 15 | 13 | 11 | 34 | 22 | 37 | 49 | 40 | 24 |
| SRR6246386 | 0 | 0 | 15 | 15 | 13 | 38 | 17 | 41 | 61 | 35 | 25 |
| SRR6246387 | 0 | 0 | 15 | 14 | 11 | 32 | 16 | 38 | 58 | 37 | 21 |
| SRR6246388 | 0 | 0 | 12 | 13 | 16 | 29 | 16 | 41 | 62 | 37 | 23 |
| SRR6246389 | 0 | 0 | 0 | 0 | 0 | 0 | 0 | 0 | 0 | 0 | 0 |
| SRR6246390 | 0 | 0 | 0 | 0 | 0 | 0 | 0 | 0 | 0 | 0 | 0 |
| SRR6246391 | 0 | 0 | 0 | 0 | 0 | 0 | 0 | 0 | 0 | 0 | 0 |
| SRR6246392 | 0 | 0 | 0 | 0 | 0 | 0 | 0 | 0 | 0 | 0 | 0 |
| SRR6246393 | 0 | 0 | 0 | 0 | 0 | 0 | 0 | 0 | 0 | 0 | 14 |
| SRR6246394 | 0 | 0 | 0 | 0 | 0 | 0 | 0 | 0 | 0 | 0 | 0 |
| SRR6246395 | 0 | 0 | 0 | 0 | 0 | 0 | 0 | 0 | 0 | 0 | 0 |
| SRR6246396 | 0 | 0 | 15 | 0 | 0 | 0 | 0 | 0 | 0 | 0 | 0 |
| SRR6246397 | 0 | 0 | 0 | 0 | 0 | 0 | 0 | 0 | 0 | 0 | 0 |
| SRR6246398 | 0 | 0 | 0 | 0 | 0 | 0 | 0 | 0 | 0 | 0 | 0 |
| SRR6246399 | 0 | 0 | 0 | 0 | 0 | 0 | 0 | 0 | 0 | 0 | 0 |
| SRR6246400 | 0 | 0 | 0 | 0 | 0 | 0 | 0 | 0 | 0 | 0 | 0 |
| SRR6265480 | 0 | 17 | 41 | 27 | 29 | 181 | 20 | 216 | 110 | 81 | 187 |
| SRR6265481 | 0 | 56 | 87 | 48 | 0 | 53 | 0 | 76 | 39 | 19 | 117 |
| SRR6265482 | 0 | 0 | 18 | 12 | 18 | 177 | 24 | 212 | 302 | 280 | 494 |
| SRR6265483 | 0 | 12 | 25 | 20 | 27 | 112 | 30 | 155 | 150 | 85 | 125 |
| SRR6265484 | 0 | 18 | 49 | 20 | 11 | 87 | 13 | 83 | 71 | 44 | 117 |
| SRR6265485 | 0 | 42 | 135 | 45 | 24 | 88 | 13 | 193 | 45 | 49 | 112 |
| SRR6265486 | 0 | 0 | 16 | 0 | 29 | 333 | 47 | 676 | 1076 | 765 | 905 |
| SRR6265487 | 0 | 18 | 35 | 21 | 0 | 94 | 0 | 174 | 74 | 29 | 356 |
| SRR6265488 | 0 | 28 | 62 | 54 | 15 | 188 | 25 | 280 | 131 | 67 | 216 |
| SRR6265489 | 0 | 0 | 0 | 0 | 0 | 0 | 0 | 14 | 0 | 0 | 17 |
| SRR6265490 | 28 | 124 | 297 | 100 | 28 | 210 | 35 | 239 | 129 | 113 | 217 |
| SRR6265491 | 0 | 12 | 15 | 14 | 12 | 99 | 0 | 115 | 26 | 23 | 124 |
| SRR6306014 | 0 | 0 | 17 | 0 | 114 | 1711 | 257 | 2500 | 10065 | 5501 | 3591 |
| SRR6306015 | 0 | 0 | 0 | 15 | 137 | 1988 | 198 | 2701 | 13250 | 7153 | 3455 |
| SRR6313948 | 0 | 17 | 157 | 40 | 3262 | 5637 | 2967 | 1086 | 1106 | 1152 | 1127 |
| SRR6313949 | 0 | 0 | 0 | 0 | 1310 | 2456 | 1221 | 752 | 355 | 280 | 349 |
| SRR6317622 | 0 | 42 | 268 | 127 | 14 | 408 | 49 | 577 | 838 | 648 | 1000 |
| SRR6317623 | 0 | 43 | 137 | 99 | 24 | 393 | 66 | 591 | 1286 | 1291 | 1258 |

### Supp Table S8.5.Columns_ET_to_FD

| | | | | | | | | | | | |
|---|---|---|---|---|---|---|---|---|---|---|---|
| SRR6317624 | 0 | 34 | 109 | 109 | 50 | 474 | 88 | 508 | 772 | 579 | 969 |
| SRR6317625 | 0 | 57 | 121 | 107 | 50 | 597 | 135 | 540 | 391 | 348 | 874 |
| SRR6317626 | 0 | 41 | 78 | 35 | 0 | 97 | 16 | 168 | 181 | 73 | 216 |
| SRR6317627 | 0 | 18 | 36 | 18 | 0 | 31 | 0 | 50 | 41 | 38 | 36 |
| SRR6317628 | 0 | 40 | 100 | 43 | 0 | 246 | 33 | 219 | 309 | 173 | 545 |
| SRR6317629 | 0 | 21 | 52 | 63 | 0 | 79 | 0 | 146 | 213 | 152 | 319 |
| SRR6317630 | 0 | 123 | 212 | 164 | 38 | 434 | 31 | 724 | 953 | 946 | 980 |
| SRR6317631 | 0 | 90 | 156 | 146 | 16 | 339 | 72 | 342 | 451 | 367 | 647 |
| SRR6317632 | 0 | 62 | 173 | 123 | 25 | 440 | 78 | 617 | 822 | 691 | 839 |
| SRR6317633 | 0 | 19 | 112 | 84 | 0 | 284 | 56 | 326 | 391 | 298 | 470 |
| SRR6317634 | 0 | 25 | 57 | 54 | 0 | 94 | 15 | 178 | 185 | 116 | 232 |
| SRR6317635 | 0 | 22 | 42 | 36 | 0 | 41 | 0 | 55 | 33 | 20 | 36 |
| SRR6317636 | 0 | 57 | 144 | 105 | 11 | 305 | 45 | 274 | 285 | 283 | 585 |
| SRR6317637 | 0 | 17 | 17 | 28 | 0 | 65 | 0 | 83 | 163 | 131 | 195 |
| SRR6317638 | 0 | 58 | 111 | 82 | 24 | 269 | 43 | 548 | 846 | 755 | 1029 |
| SRR6317639 | 0 | 53 | 143 | 128 | 34 | 414 | 45 | 561 | 1106 | 1068 | 967 |
| SRR6317640 | 0 | 50 | 109 | 59 | 24 | 423 | 56 | 535 | 591 | 544 | 617 |
| SRR6317641 | 0 | 18 | 83 | 28 | 37 | 275 | 83 | 316 | 316 | 261 | 555 |
| SRR6317642 | 0 | 22 | 83 | 52 | 12 | 89 | 12 | 148 | 213 | 114 | 240 |
| SRR6317643 | 0 | 53 | 66 | 46 | 12 | 68 | 17 | 134 | 124 | 83 | 169 |
| SRR6317644 | 0 | 98 | 158 | 84 | 24 | 402 | 33 | 283 | 266 | 215 | 522 |
| SRR6317645 | 0 | 43 | 70 | 68 | 13 | 70 | 0 | 140 | 163 | 113 | 256 |
| SRR6324194 | 0 | 0 | 0 | 0 | 12 | 55 | 24 | 155 | 679 | 627 | 1367 |
| SRR6324195 | 0 | 19 | 61 | 35 | 54 | 142 | 57 | 149 | 370 | 460 | 1117 |
| SRR6363052 | 104 | 452 | 835 | 438 | 1086 | 2865 | 1758 | 3290 | 6071 | 8119 | 6915 |
| SRR6363053 | 78 | 245 | 605 | 290 | 875 | 2563 | 1368 | 3364 | 5397 | 7293 | 6664 |
| SRR6363054 | 21 | 129 | 302 | 166 | 588 | 1870 | 1053 | 2710 | 6413 | 8716 | 8588 |
| SRR6370846 | 0 | 0 | 0 | 0 | 0 | 0 | 0 | 0 | 0 | 0 | 11 |
| SRR6370847 | 0 | 0 | 0 | 0 | 17 | 67 | 17 | 25 | 0 | 0 | 17 |
| SRR6370848 | 0 | 0 | 0 | 0 | 0 | 30 | 0 | 12 | 0 | 0 | 0 |
| SRR6370849 | 10 | 10 | 0 | 0 | 27 | 94 | 27 | 57 | 13 | 0 | 17 |
| SRR6370852 | 0 | 0 | 0 | 18 | 109 | 173 | 146 | 55 | 18 | 18 | 0 |
| SRR6370853 | 0 | 0 | 0 | 0 | 0 | 0 | 0 | 13 | 0 | 0 | 39 |
| SRR6370856 | 0 | 16 | 38 | 27 | 38 | 112 | 46 | 30 | 27 | 0 | 19 |
| SRR6372849 | 0 | 16 | 20 | 13 | 142 | 360 | 81 | 147 | 1704 | 253 | 142 |
| SRR6372850 | 0 | 0 | 0 | 0 | 72 | 149 | 108 | 128 | 457 | 98 | 60 |
| SRR6372851 | 0 | 0 | 0 | 0 | 111 | 158 | 127 | 69 | 1832 | 131 | 106 |
| SRR6372852 | 0 | 42 | 90 | 21 | 189 | 695 | 370 | 1100 | 720 | 190 | 129 |
| SRR6372853 | 0 | 26 | 66 | 21 | 73 | 183 | 176 | 541 | 558 | 53 | 91 |
| SRR6372854 | 0 | 0 | 0 | 0 | 144 | 733 | 336 | 1526 | 4957 | 713 | 352 |
| SRR6372855 | 0 | 0 | 0 | 0 | 0 | 0 | 0 | 11 | 120 | 0 | 0 |
| SRR6372856 | 0 | 0 | 0 | 0 | 19 | 34 | 16 | 39 | 258 | 10 | 35 |
| SRR6372857 | 0 | 0 | 0 | 0 | 32 | 83 | 37 | 56 | 376 | 26 | 50 |
| SRR6389795 | 0 | 28 | 90 | 71 | 521 | 4435 | 403 | 5115 | 524 | 316 | 183 |
| SRR6389796 | 0 | 40 | 88 | 63 | 557 | 4537 | 544 | 7783 | 2343 | 1139 | 482 |
| SRR6389797 | 0 | 138 | 282 | 194 | 966 | 7372 | 609 | 11514 | 5632 | 3561 | 1213 |
| SRR6389798 | 0 | 18 | 38 | 16 | 236 | 3287 | 434 | 7722 | 421 | 292 | 225 |
| SRR6389799 | 0 | 52 | 114 | 85 | 644 | 5819 | 627 | 8099 | 804 | 377 | 280 |
| SRR6389800 | 0 | 17 | 31 | 31 | 587 | 8573 | 950 | 14844 | 1564 | 670 | 430 |
| SRR6389801 | 0 | 0 | 0 | 0 | 399 | 4323 | 858 | 9573 | 926 | 391 | 735 |
| SRR6389802 | 0 | 0 | 0 | 0 | 2123 | 20410 | 3858 | 33229 | 20653 | 9823 | 4888 |
| SRR6389803 | 0 | 10 | 14 | 20 | 1796 | 21827 | 2906 | 34288 | 1992 | 877 | 738 |
| SRR6389804 | 0 | 0 | 0 | 14 | 1844 | 23510 | 3269 | 25640 | 7559 | 3724 | 2320 |
| SRR6389805 | 0 | 0 | 15 | 14 | 671 | 5863 | 692 | 9199 | 1958 | 1224 | 667 |
| SRR6389806 | 0 | 73 | 114 | 63 | 608 | 11054 | 1634 | 14828 | 1013 | 643 | 457 |
| SRR6389807 | 0 | 0 | 32 | 29 | 473 | 4127 | 429 | 7916 | 2508 | 2095 | 1547 |
| SRR6389808 | 0 | 41 | 211 | 164 | 1325 | 12168 | 1882 | 21643 | 5317 | 2353 | 694 |
| SRR6389809 | 0 | 0 | 19 | 31 | 789 | 5955 | 600 | 11497 | 1175 | 848 | 943 |
| SRR6389810 | 0 | 15 | 54 | 37 | 485 | 4176 | 546 | 7955 | 2164 | 1458 | 1469 |
| SRR6389811 | 0 | 38 | 105 | 69 | 349 | 3084 | 278 | 3448 | 1455 | 1050 | 473 |
| SRR6389812 | 0 | 0 | 0 | 0 | 271 | 3128 | 332 | 4339 | 2262 | 1502 | 798 |
| SRR6389813 | 0 | 13 | 26 | 26 | 1571 | 14901 | 1843 | 24321 | 8073 | 6618 | 3431 |
| SRR6389814 | 0 | 0 | 12 | 0 | 204 | 2205 | 228 | 2393 | 455 | 267 | 295 |
| SRR6389815 | 0 | 0 | 0 | 0 | 470 | 3778 | 307 | 4422 | 287 | 153 | 119 |
| SRR6389816 | 0 | 0 | 34 | 26 | 1016 | 13548 | 1431 | 23450 | 8722 | 8663 | 4522 |
| SRR6389817 | 0 | 0 | 19 | 0 | 1326 | 21995 | 1894 | 22013 | 2358 | 774 | 1324 |
| SRR6389818 | 0 | 0 | 12 | 0 | 791 | 10425 | 1851 | 11892 | 1957 | 1161 | 1181 |
| SRR6389819 | 0 | 0 | 26 | 13 | 771 | 11983 | 1723 | 11106 | 4435 | 1986 | 2260 |
| SRR6389820 | 0 | 0 | 0 | 0 | 794 | 8180 | 998 | 17298 | 1250 | 801 | 493 |
| SRR6409751 | 0 | 0 | 0 | 0 | 0 | 0 | 0 | 0 | 0 | 0 | 0 |
| SRR6409760 | 0 | 0 | 0 | 0 | 0 | 0 | 0 | 0 | 0 | 0 | 0 |
| SRR6409798 | 0 | 0 | 0 | 362 | 0 | 0 | 0 | 0 | 895 | 0 | 0 |
| SRR6409800 | 0 | 0 | 0 | 0 | 0 | 0 | 0 | 0 | 0 | 0 | 0 |
| SRR6409827 | 0 | 0 | 0 | 0 | 0 | 0 | 0 | 0 | 0 | 0 | 0 |
| SRR6409832 | 0 | 0 | 0 | 0 | 0 | 0 | 0 | 0 | 0 | 0 | 0 |
| SRR6409833 | 0 | 0 | 159 | 0 | 0 | 0 | 0 | 0 | 0 | 0 | 0 |
| SRR6409834 | 0 | 0 | 0 | 0 | 0 | 0 | 0 | 0 | 0 | 0 | 0 |
| SRR6409845 | 0 | 0 | 0 | 0 | 0 | 0 | 0 | 0 | 0 | 0 | 0 |
| SRR6409851 | 0 | 0 | 29 | 27 | 0 | 526 | 0 | 0 | 0 | 0 | 0 |
| SRR6409853 | 0 | 0 | 0 | 29 | 0 | 0 | 0 | 0 | 0 | 0 | 0 |
| SRR6409859 | 0 | 0 | 0 | 187 | 0 | 0 | 0 | 0 | 0 | 0 | 0 |
| SRR6409860 | 0 | 115 | 0 | 0 | 0 | 0 | 0 | 0 | 0 | 0 | 0 |
| SRR6409863 | 0 | 0 | 0 | 0 | 0 | 0 | 0 | 0 | 0 | 0 | 0 |
| SRR6409866 | 0 | 0 | 184 | 0 | 0 | 0 | 0 | 0 | 0 | 0 | 0 |
| SRR6409867 | 0 | 0 | 103 | 0 | 0 | 0 | 0 | 0 | 0 | 0 | 0 |
| SRR6409868 | 0 | 0 | 0 | 0 | 0 | 0 | 0 | 0 | 0 | 0 | 0 |
| SRR6450851 | 0 | 112 | 0 | 112 | 223 | 1338 | 223 | 334 | 3010 | 2230 | 2118 |
| SRR6450852 | 71 | 29 | 88 | 53 | 370 | 958 | 523 | 388 | 1617 | 1235 | 1523 |
| SRR6450853 | 17 | 17 | 99 | 33 | 363 | 793 | 380 | 446 | 1751 | 1272 | 1718 |
| SRR6450854 | 12 | 23 | 46 | 46 | 267 | 814 | 546 | 244 | 2034 | 1569 | 1813 |
| SRR6450855 | 0 | 0 | 66 | 66 | 464 | 596 | 199 | 398 | 2849 | 1855 | 2253 |
| SRR6450856 | 36 | 24 | 204 | 60 | 240 | 852 | 468 | 588 | 2088 | 1560 | 1644 |
| SRR6450857 | 0 | 91 | 182 | 91 | 273 | 729 | 91 | 364 | 2278 | 1458 | 2187 |
| SRR6450858 | 58 | 116 | 58 | 70 | 395 | 1301 | 476 | 441 | 2079 | 1405 | 1765 |
| SRR6450859 | 0 | 111 | 166 | 111 | 553 | 664 | 609 | 332 | 2047 | 1771 | 1936 |
| SRR6450860 | 139 | 46 | 232 | 185 | 371 | 696 | 464 | 556 | 1623 | 1530 | 1762 |
| SRR6450861 | 0 | 0 | 31 | 31 | 343 | 685 | 374 | 405 | 2492 | 1526 | 2212 |
| SRR6450862 | 0 | 0 | 39 | 22 | 320 | 842 | 409 | 507 | 2077 | 1654 | 2060 |
| SRR6450863 | 24 | 31 | 94 | 24 | 384 | 971 | 533 | 243 | 1520 | 1167 | 2005 |
| SRR6450867 | 37 | 341 | 931 | 1244 | 322 | 332 | 396 | 673 | 9582 | 3068 | 10051 |
| SRR6450868 | 0 | 0 | 0 | 0 | 20 | 23 | 38 | 77 | 5067 | 696 | 22706 |
| SRR6450869 | 0 | 0 | 0 | 0 | 20 | 0 | 40 | 130 | 5390 | 559 | 22717 |
| SRR6453341 | 0 | 0 | 0 | 0 | 19 | 40 | 0 | 0 | 77 | 0 | 0 |
| SRR6453342 | 0 | 0 | 0 | 0 | 16 | 29 | 0 | 0 | 38 | 0 | 0 |
| SRR6453343 | 0 | 0 | 0 | 0 | 32 | 70 | 14 | 0 | 109 | 0 | 0 |
| SRR6453344 | 0 | 0 | 22 | 0 | 34 | 62 | 13 | 0 | 92 | 0 | 0 |
| SRR6453345 | 0 | 11 | 29 | 0 | 43 | 89 | 12 | 15 | 101 | 0 | 0 |
| SRR6453346 | 0 | 0 | 21 | 0 | 31 | 43 | 0 | 0 | 93 | 0 | 0 |
| SRR6453347 | 0 | 0 | 0 | 0 | 11 | 21 | 0 | 0 | 38 | 0 | 0 |
| SRR6453348 | 0 | 10 | 34 | 0 | 52 | 103 | 16 | 11 | 91 | 0 | 0 |
| SRR6453349 | 0 | 0 | 25 | 0 | 51 | 95 | 18 | 14 | 95 | 0 | 0 |
| SRR6453350 | 0 | 0 | 31 | 0 | 46 | 102 | 22 | 19 | 127 | 0 | 10 |
| SRR6453351 | 0 | 0 | 12 | 0 | 19 | 36 | 0 | 0 | 69 | 0 | 0 |
| SRR6453352 | 0 | 0 | 11 | 0 | 32 | 62 | 16 | 12 | 100 | 0 | 0 |
| SRR6453353 | 0 | 0 | 0 | 0 | 17 | 24 | 0 | 0 | 52 | 0 | 0 |
| SRR6453354 | 0 | 0 | 12 | 0 | 23 | 25 | 12 | 0 | 52 | 0 | 0 |
| SRR6453355 | 0 | 0 | 0 | 0 | 17 | 33 | 10 | 11 | 57 | 0 | 0 |
| SRR6453356 | 0 | 0 | 14 | 0 | 32 | 43 | 16 | 13 | 77 | 0 | 0 |
| SRR6453357 | 0 | 0 | 0 | 0 | 30 | 65 | 14 | 11 | 80 | 0 | 0 |

Supp Table S8.5.Columns_ET_to_FD

| | | | | | | | | | | | |
|---|---|---|---|---|---|---|---|---|---|---|---|
| SRR6453358 | 0 | 0 | 10 | 0 | 27 | 46 | 0 | 11 | 35 | 0 | 0 |
| SRR6453359 | 0 | 0 | 0 | 0 | 0 | 13 | 0 | 0 | 33 | 0 | 0 |
| SRR6453360 | 0 | 0 | 0 | 0 | 20 | 55 | 16 | 0 | 132 | 0 | 0 |
| SRR6453361 | 0 | 0 | 15 | 0 | 15 | 21 | 0 | 0 | 52 | 0 | 0 |
| SRR6453362 | 0 | 0 | 11 | 0 | 11 | 27 | 0 | 0 | 71 | 0 | 0 |
| SRR6453363 | 0 | 0 | 0 | 0 | 12 | 28 | 0 | 0 | 74 | 0 | 0 |
| SRR6453364 | 0 | 0 | 10 | 0 | 33 | 64 | 14 | 0 | 123 | 0 | 0 |
| SRR6453365 | 0 | 0 | 0 | 0 | 23 | 43 | 16 | 13 | 73 | 0 | 0 |
| SRR6453366 | 0 | 0 | 0 | 0 | 14 | 24 | 0 | 0 | 54 | 0 | 0 |
| SRR6453367 | 0 | 0 | 0 | 0 | 0 | 12 | 0 | 0 | 35 | 0 | 0 |
| SRR6453368 | 0 | 0 | 0 | 0 | 0 | 17 | 0 | 10 | 40 | 0 | 0 |
| SRR6453369 | 0 | 0 | 13 | 0 | 25 | 45 | 0 | 0 | 56 | 0 | 0 |
| SRR6453370 | 0 | 0 | 0 | 0 | 21 | 48 | 0 | 0 | 59 | 0 | 0 |
| SRR6453371 | 0 | 0 | 0 | 0 | 12 | 37 | 0 | 0 | 64 | 0 | 0 |
| SRR6453372 | 0 | 0 | 0 | 0 | 10 | 24 | 0 | 0 | 46 | 0 | 0 |
| SRR6453373 | 0 | 0 | 17 | 0 | 36 | 69 | 17 | 11 | 96 | 0 | 0 |
| SRR6453374 | 0 | 0 | 0 | 0 | 15 | 33 | 0 | 0 | 74 | 0 | 0 |
| SRR6453375 | 0 | 0 | 0 | 0 | 13 | 30 | 0 | 0 | 62 | 0 | 0 |
| SRR6453376 | 0 | 0 | 14 | 0 | 0 | 25 | 0 | 0 | 117 | 0 | 0 |
| SRR6453377 | 0 | 0 | 11 | 0 | 18 | 49 | 10 | 11 | 74 | 0 | 0 |
| SRR6453378 | 0 | 0 | 0 | 0 | 15 | 49 | 10 | 0 | 82 | 0 | 0 |
| SRR6453379 | 0 | 0 | 31 | 0 | 45 | 78 | 13 | 0 | 93 | 0 | 0 |
| SRR6453380 | 0 | 0 | 0 | 0 | 0 | 20 | 0 | 0 | 26 | 0 | 0 |
| SRR6453381 | 0 | 0 | 0 | 0 | 15 | 38 | 11 | 0 | 51 | 0 | 0 |
| SRR6453382 | 0 | 0 | 0 | 0 | 19 | 37 | 0 | 0 | 38 | 0 | 0 |
| SRR6453383 | 0 | 0 | 12 | 0 | 24 | 54 | 14 | 0 | 50 | 0 | 0 |
| SRR6453384 | 0 | 0 | 0 | 0 | 13 | 40 | 0 | 0 | 65 | 0 | 0 |
| SRR6453385 | 0 | 0 | 0 | 0 | 11 | 24 | 0 | 0 | 46 | 0 | 0 |
| SRR6453386 | 0 | 0 | 25 | 0 | 0 | 25 | 0 | 0 | 54 | 0 | 0 |
| SRR6453387 | 0 | 13 | 39 | 0 | 13 | 31 | 0 | 0 | 56 | 0 | 0 |
| SRR6453388 | 0 | 0 | 35 | 0 | 10 | 41 | 17 | 0 | 62 | 0 | 0 |
| SRR6453389 | 0 | 0 | 27 | 0 | 0 | 38 | 0 | 0 | 100 | 0 | 0 |
| SRR6453390 | 0 | 17 | 89 | 27 | 20 | 60 | 23 | 12 | 106 | 0 | 0 |
| SRR6453391 | 0 | 14 | 51 | 0 | 16 | 56 | 13 | 0 | 129 | 0 | 0 |
| SRR6453392 | 0 | 0 | 27 | 0 | 0 | 33 | 11 | 0 | 38 | 0 | 0 |
| SRR6453393 | 0 | 12 | 61 | 17 | 15 | 37 | 12 | 0 | 110 | 0 | 0 |
| SRR6453394 | 0 | 0 | 13 | 0 | 0 | 21 | 0 | 0 | 53 | 0 | 0 |
| SRR6453395 | 0 | 10 | 32 | 0 | 0 | 32 | 0 | 0 | 44 | 0 | 0 |
| SRR6453396 | 0 | 0 | 27 | 0 | 0 | 26 | 0 | 0 | 64 | 0 | 0 |
| SRR6453397 | 0 | 0 | 30 | 0 | 0 | 15 | 0 | 0 | 51 | 0 | 0 |
| SRR6453398 | 0 | 0 | 15 | 0 | 11 | 25 | 0 | 0 | 39 | 0 | 0 |
| SRR6453399 | 0 | 0 | 21 | 0 | 12 | 29 | 13 | 0 | 91 | 0 | 0 |
| SRR6453400 | 0 | 0 | 13 | 0 | 0 | 22 | 0 | 0 | 75 | 0 | 0 |
| SRR6453401 | 0 | 10 | 43 | 0 | 20 | 49 | 15 | 0 | 58 | 0 | 0 |
| SRR6453402 | 0 | 51 | 107 | 17 | 21 | 58 | 15 | 0 | 58 | 0 | 0 |
| SRR6453403 | 0 | 17 | 41 | 0 | 0 | 35 | 0 | 0 | 92 | 0 | 0 |
| SRR6453404 | 0 | 22 | 54 | 0 | 10 | 32 | 0 | 0 | 75 | 0 | 0 |
| SRR6453405 | 0 | 12 | 77 | 10 | 22 | 63 | 24 | 0 | 125 | 0 | 0 |
| SRR6453406 | 0 | 14 | 38 | 0 | 20 | 49 | 15 | 0 | 87 | 0 | 0 |
| SRR6453407 | 0 | 0 | 21 | 0 | 0 | 17 | 0 | 0 | 38 | 0 | 0 |
| SRR6453408 | 0 | 0 | 56 | 0 | 16 | 45 | 11 | 0 | 59 | 0 | 0 |
| SRR6453409 | 0 | 61 | 87 | 0 | 13 | 47 | 11 | 0 | 40 | 0 | 0 |
| SRR6453410 | 0 | 12 | 29 | 0 | 13 | 31 | 0 | 0 | 54 | 0 | 0 |
| SRR6453411 | 0 | 13 | 49 | 0 | 19 | 58 | 15 | 0 | 95 | 0 | 0 |
| SRR6453412 | 0 | 15 | 77 | 0 | 0 | 28 | 0 | 0 | 58 | 0 | 0 |
| SRR6453413 | 0 | 0 | 24 | 0 | 13 | 32 | 0 | 0 | 67 | 0 | 0 |
| SRR6453414 | 0 | 12 | 22 | 0 | 0 | 0 | 0 | 0 | 73 | 0 | 0 |
| SRR6453415 | 0 | 11 | 28 | 0 | 0 | 0 | 0 | 0 | 49 | 0 | 0 |
| SRR6453416 | 0 | 27 | 42 | 24 | 0 | 22 | 0 | 0 | 67 | 0 | 0 |
| SRR6453417 | 0 | 68 | 93 | 15 | 0 | 12 | 0 | 10 | 50 | 0 | 0 |
| SRR6453418 | 0 | 29 | 105 | 14 | 0 | 30 | 0 | 0 | 23 | 0 | 0 |
| SRR6453419 | 0 | 19 | 102 | 16 | 14 | 50 | 17 | 0 | 68 | 0 | 0 |
| SRR6453420 | 0 | 16 | 45 | 0 | 0 | 25 | 0 | 0 | 70 | 0 | 0 |
| SRR6453421 | 0 | 34 | 125 | 11 | 35 | 112 | 24 | 16 | 28 | 0 | 0 |
| SRR6453422 | 0 | 22 | 118 | 0 | 24 | 69 | 10 | 0 | 129 | 0 | 0 |
| SRR6453423 | 0 | 0 | 26 | 0 | 0 | 18 | 0 | 0 | 34 | 0 | 0 |
| SRR6453424 | 0 | 46 | 65 | 12 | 0 | 0 | 0 | 11 | 46 | 0 | 0 |
| SRR6453425 | 0 | 10 | 19 | 0 | 0 | 0 | 0 | 0 | 29 | 0 | 0 |
| SRR6453426 | 0 | 42 | 66 | 13 | 0 | 20 | 0 | 0 | 40 | 0 | 0 |
| SRR6453427 | 0 | 0 | 17 | 0 | 0 | 0 | 0 | 0 | 58 | 0 | 0 |
| SRR6453428 | 0 | 0 | 29 | 11 | 0 | 11 | 0 | 0 | 28 | 0 | 0 |
| SRR6454160 | 0 | 0 | 106 | 186 | 292 | 558 | 160 | 1223 | 186 | 53 | 585 |
| SRR6454161 | 0 | 0 | 0 | 0 | 15 | 107 | 15 | 314 | 36 | 36 | 137 |
| SRR6454162 | 0 | 0 | 0 | 92 | 0 | 31 | 0 | 0 | 0 | 0 | 0 |
| SRR6454163 | 0 | 0 | 10 | 10 | 114 | 1662 | 93 | 4243 | 196 | 372 | 681 |
| SRR6454164 | 0 | 0 | 0 | 0 | 0 | 31 | 0 | 61 | 0 | 0 | 61 |
| SRR6454165 | 0 | 15 | 15 | 92 | 84 | 1235 | 84 | 3260 | 330 | 338 | 1143 |
| SRR6454166 | 0 | 0 | 0 | 32 | 0 | 0 | 0 | 63 | 0 | 0 | 32 |
| SRR6454167 | 0 | 0 | 24 | 12 | 131 | 1399 | 120 | 4028 | 311 | 454 | 1136 |
| SRR6454169 | 0 | 0 | 21 | 82 | 92 | 1005 | 123 | 3610 | 328 | 308 | 892 |
| SRR6454170 | 0 | 0 | 0 | 20 | 0 | 20 | 0 | 0 | 20 | 0 | 0 |
| SRR6454171 | 0 | 18 | 44 | 80 | 18 | 550 | 44 | 1047 | 302 | 284 | 648 |
| SRR6454172 | 0 | 0 | 0 | 33 | 0 | 0 | 0 | 0 | 0 | 0 | 0 |
| SRR6454173 | 0 | 0 | 16 | 47 | 55 | 343 | 23 | 1146 | 133 | 117 | 553 |
| SRR6454174 | 0 | 32 | 16 | 0 | 0 | 16 | 16 | 0 | 48 | 0 | 16 |
| SRR6454175 | 0 | 0 | 32 | 64 | 44 | 413 | 57 | 699 | 25 | 114 | 305 |
| SRR6454176 | 0 | 13 | 0 | 13 | 0 | 13 | 0 | 13 | 39 | 0 | 0 |
| SRR6454177 | 0 | 26 | 13 | 143 | 182 | 1120 | 78 | 3244 | 430 | 274 | 782 |
| SRR6454178 | 0 | 0 | 0 | 0 | 0 | 0 | 0 | 17 | 0 | 0 | 35 |
| SRR6454179 | 0 | 0 | 13 | 126 | 38 | 440 | 0 | 1508 | 226 | 264 | 805 |
| SRR6454180 | 0 | 0 | 0 | 24 | 0 | 0 | 0 | 0 | 0 | 0 | 0 |
| SRR6454181 | 0 | 0 | 20 | 50 | 10 | 727 | 71 | 1383 | 172 | 151 | 384 |
| SRR6454182 | 0 | 0 | 0 | 27 | 0 | 0 | 0 | 0 | 0 | 0 | 0 |
| SRR6454183 | 0 | 0 | 0 | 35 | 21 | 331 | 35 | 733 | 134 | 134 | 367 |
| SRR6454184 | 0 | 0 | 0 | 16 | 0 | 0 | 0 | 16 | 0 | 0 | 0 |
| SRR6454185 | 0 | 26 | 65 | 92 | 59 | 350 | 23 | 612 | 88 | 62 | 357 |
| SRR6454186 | 0 | 16 | 47 | 31 | 0 | 0 | 0 | 31 | 16 | 0 | 0 |
| SRR6454187 | 0 | 18 | 0 | 62 | 26 | 423 | 62 | 886 | 123 | 194 | 547 |
| SRR6454188 | 0 | 0 | 0 | 0 | 0 | 0 | 0 | 21 | 21 | 0 | 0 |
| SRR6454189 | 0 | 0 | 0 | 22 | 0 | 581 | 45 | 3172 | 357 | 849 | 1162 |
| SRR6454190 | 0 | 0 | 0 | 0 | 0 | 0 | 0 | 0 | 0 | 0 | 0 |
| SRR6454191 | 0 | 15 | 23 | 53 | 15 | 198 | 15 | 959 | 137 | 221 | 404 |
| SRR6454193 | 0 | 14 | 14 | 98 | 14 | 810 | 56 | 2584 | 182 | 363 | 1006 |
| SRR6454194 | 0 | 25 | 0 | 74 | 0 | 0 | 0 | 0 | 49 | 0 | 0 |
| SRR6454195 | 0 | 18 | 18 | 53 | 53 | 942 | 53 | 2949 | 373 | 444 | 1777 |
| SRR6454196 | 0 | 0 | 0 | 0 | 0 | 0 | 0 | 0 | 0 | 0 | 0 |
| SRR6454197 | 0 | 28 | 49 | 247 | 148 | 643 | 92 | 861 | 92 | 169 | 487 |
| SRR6454198 | 0 | 0 | 93 | 40 | 0 | 0 | 0 | 13 | 0 | 13 | 13 |
| SRR6454199 | 0 | 12 | 74 | 283 | 99 | 443 | 43 | 622 | 68 | 105 | 468 |
| SRR6454201 | 0 | 29 | 109 | 299 | 37 | 526 | 15 | 1255 | 102 | 58 | 212 |
| SRR6454203 | 0 | 0 | 51 | 85 | 51 | 1236 | 17 | 2947 | 203 | 68 | 593 |
| SRR6454205 | 0 | 12 | 12 | 74 | 86 | 824 | 49 | 2877 | 344 | 332 | 1094 |
| SRR6454206 | 0 | 0 | 28 | 0 | 0 | 28 | 0 | 0 | 0 | 28 | 28 |
| SRR6454207 | 0 | 0 | 23 | 137 | 91 | 1093 | 91 | 2983 | 387 | 490 | 2300 |
| SRR6454208 | 0 | 0 | 0 | 58 | 0 | 0 | 0 | 0 | 29 | 0 | 0 |
| SRR6454209 | 0 | 0 | 0 | 42 | 34 | 681 | 67 | 2513 | 311 | 311 | 849 |
| SRR6454211 | 0 | 0 | 11 | 27 | 49 | 483 | 22 | 1063 | 255 | 282 | 656 |
| SRR6454213 | 0 | 0 | 0 | 68 | 23 | 734 | 68 | 2235 | 542 | 553 | 1185 |
| SRR6454214 | 0 | 0 | 0 | 54 | 0 | 0 | 0 | 0 | 54 | 0 | 0 |

### Supp Table S8.5.Columns_ET_to_FD

| | | | | | | | | | | | |
|---|---|---|---|---|---|---|---|---|---|---|---|
| SRR6454215 | 0 | 0 | 30 | 42 | 42 | 601 | 42 | 1351 | 190 | 238 | 595 |
| SRR6454216 | 0 | 0 | 0 | 0 | 49 | 0 | 0 | 98 | 0 | 0 | 0 |
| SRR6454217 | 0 | 17 | 100 | 324 | 100 | 1205 | 108 | 2401 | 141 | 108 | 565 |
| SRR6454218 | 0 | 0 | 0 | 0 | 0 | 27 | 0 | 0 | 0 | 0 | 54 |
| SRR6454219 | 0 | 17 | 17 | 122 | 70 | 504 | 61 | 991 | 70 | 148 | 313 |
| SRR6454220 | 0 | 0 | 0 | 22 | 0 | 0 | 0 | 22 | 22 | 0 | 22 |
| SRR6454221 | 0 | 23 | 34 | 147 | 158 | 1072 | 135 | 1906 | 147 | 102 | 384 |
| SRR6454222 | 0 | 0 | 50 | 0 | 0 | 0 | 0 | 50 | 0 | 25 | 25 |
| SRR6454223 | 0 | 13 | 52 | 287 | 98 | 732 | 111 | 1176 | 131 | 235 | 536 |
| SRR6454224 | 0 | 0 | 0 | 0 | 0 | 0 | 0 | 0 | 0 | 0 | 0 |
| SRR6454225 | 0 | 14 | 28 | 154 | 140 | 698 | 126 | 1061 | 70 | 154 | 335 |
| SRR6454226 | 0 | 0 | 0 | 24 | 0 | 0 | 0 | 0 | 24 | 0 | 0 |
| SRR6454227 | 0 | 41 | 65 | 247 | 114 | 426 | 57 | 904 | 77 | 134 | 373 |
| SRR6454228 | 0 | 0 | 0 | 35 | 0 | 0 | 0 | 0 | 17 | 0 | 17 |
| SRR6454229 | 0 | 0 | 17 | 233 | 67 | 1066 | 100 | 2016 | 83 | 267 | 583 |
| SRR6454230 | 0 | 0 | 0 | 0 | 0 | 0 | 0 | 19 | 38 | 0 | 38 |
| SRR6454231 | 0 | 26 | 26 | 155 | 52 | 699 | 117 | 1373 | 168 | 389 | 842 |
| SRR6454232 | 0 | 0 | 0 | 0 | 16 | 0 | 0 | 32 | 0 | 16 | 16 |
| SRR6454233 | 0 | 0 | 0 | 63 | 31 | 424 | 39 | 903 | 55 | 102 | 220 |
| SRR6454234 | 0 | 0 | 19 | 0 | 38 | 254 | 0 | 451 | 47 | 103 | 357 |
| SRR6454235 | 0 | 0 | 0 | 0 | 0 | 16 | 0 | 33 | 0 | 0 | 33 |
| SRR6454236 | 0 | 0 | 43 | 86 | 62 | 338 | 38 | 609 | 52 | 105 | 228 |
| SRR6454237 | 0 | 0 | 0 | 0 | 0 | 19 | 0 | 19 | 0 | 0 | 0 |
| SRR6454238 | 0 | 0 | 0 | 24 | 48 | 329 | 16 | 746 | 0 | 104 | 144 |
| SRR6454239 | 0 | 0 | 0 | 0 | 0 | 10 | 10 | 10 | 21 | 0 | 31 |
| SRR6454240 | 0 | 11 | 70 | 124 | 43 | 484 | 59 | 1177 | 48 | 97 | 102 |
| SRR6454241 | 0 | 0 | 0 | 0 | 0 | 0 | 0 | 0 | 12 | 0 | 12 |
| SRR6454242 | 0 | 10 | 20 | 55 | 20 | 346 | 50 | 801 | 90 | 155 | 260 |
| SRR6454244 | 0 | 14 | 28 | 131 | 83 | 723 | 117 | 1425 | 207 | 392 | 551 |
| SRR6454245 | 0 | 0 | 0 | 0 | 0 | 0 | 27 | 27 | 55 | 0 | 0 |
| SRR6454246 | 0 | 0 | 0 | 58 | 33 | 681 | 17 | 1519 | 100 | 191 | 249 |
| SRR6454247 | 0 | 0 | 21 | 48 | 75 | 301 | 27 | 541 | 75 | 151 | 329 |
| SRR6454249 | 0 | 0 | 26 | 92 | 39 | 564 | 39 | 971 | 26 | 223 | 276 |
| SRR6454251 | 0 | 0 | 33 | 42 | 17 | 543 | 33 | 994 | 25 | 100 | 84 |
| SRR6454252 | 0 | 0 | 0 | 0 | 0 | 0 | 0 | 0 | 0 | 0 | 0 |
| SRR6454253 | 0 | 14 | 14 | 105 | 56 | 482 | 70 | 1118 | 77 | 119 | 217 |
| SRR6454254 | 0 | 14 | 70 | 56 | 14 | 14 | 0 | 0 | 0 | 14 | 42 |
| SRR6454255 | 0 | 27 | 0 | 47 | 34 | 429 | 34 | 798 | 20 | 121 | 188 |
| SRR6454256 | 0 | 0 | 0 | 32 | 0 | 0 | 0 | 0 | 0 | 0 | 0 |
| SRR6454257 | 0 | 14 | 50 | 86 | 65 | 516 | 57 | 746 | 65 | 115 | 437 |
| SRR6454258 | 0 | 0 | 16 | 0 | 49 | 591 | 33 | 870 | 82 | 115 | 410 |
| SRR6454259 | 0 | 0 | 0 | 0 | 78 | 322 | 49 | 752 | 244 | 293 | 517 |
| SRR6454260 | 0 | 0 | 0 | 0 | 86 | 546 | 48 | 1045 | 58 | 125 | 182 |
| SRR6454261 | 0 | 0 | 0 | 26 | 38 | 300 | 26 | 536 | 57 | 147 | 287 |
| SRR6454262 | 0 | 0 | 13 | 0 | 25 | 177 | 25 | 443 | 38 | 266 | 291 |
| SRR6454263 | 0 | 13 | 25 | 101 | 32 | 291 | 70 | 594 | 95 | 114 | 291 |
| SRR6454264 | 0 | 0 | 0 | 36 | 18 | 303 | 89 | 571 | 71 | 107 | 392 |
| SRR6454265 | 0 | 0 | 15 | 30 | 30 | 345 | 45 | 706 | 60 | 360 | 330 |
| SRR6454266 | 0 | 0 | 0 | 0 | 13 | 233 | 40 | 654 | 40 | 161 | 228 |
| SRR6454267 | 0 | 0 | 18 | 32 | 55 | 407 | 18 | 770 | 60 | 100 | 247 |
| SRR6454268 | 0 | 0 | 0 | 27 | 54 | 401 | 54 | 696 | 80 | 214 | 589 |
| SRR6454269 | 0 | 0 | 0 | 53 | 26 | 132 | 26 | 688 | 26 | 264 | 159 |
| SRR6454270 | 0 | 0 | 0 | 28 | 28 | 293 | 14 | 376 | 28 | 209 | 195 |
| SRR6454271 | 0 | 60 | 213 | 1267 | 162 | 891 | 162 | 1505 | 60 | 150 | 764 |
| SRR6454272 | 0 | 48 | 111 | 865 | 103 | 547 | 127 | 1626 | 71 | 111 | 444 |
| SRR6454273 | 0 | 71 | 399 | 248 | 89 | 868 | 124 | 1692 | 53 | 27 | 213 |
| SRR6454274 | 0 | 124 | 304 | 219 | 38 | 361 | 57 | 1550 | 48 | 76 | 143 |
| SRR6454275 | 0 | 0 | 64 | 117 | 37 | 469 | 37 | 1119 | 80 | 170 | 218 |
| SRR6454276 | 0 | 15 | 50 | 166 | 80 | 744 | 45 | 1946 | 131 | 231 | 488 |
| SRR6454277 | 0 | 15 | 25 | 84 | 40 | 361 | 49 | 904 | 163 | 222 | 361 |
| SRR6454278 | 0 | 10 | 25 | 231 | 50 | 1140 | 55 | 3185 | 171 | 367 | 357 |
| SRR6454279 | 0 | 0 | 14 | 19 | 24 | 256 | 53 | 801 | 174 | 362 | 656 |
| SRR6454280 | 0 | 0 | 15 | 69 | 11 | 157 | 19 | 413 | 31 | 65 | 160 |
| SRR6454281 | 0 | 0 | 12 | 102 | 51 | 330 | 55 | 1213 | 39 | 247 | 228 |
| SRR6454282 | 0 | 0 | 0 | 51 | 37 | 370 | 14 | 1310 | 66 | 178 | 370 |
| SRR6454283 | 0 | 0 | 0 | 131 | 47 | 513 | 61 | 1889 | 65 | 210 | 313 |
| SRR6454284 | 0 | 16 | 43 | 162 | 70 | 686 | 32 | 2280 | 103 | 238 | 470 |
| SRR6454285 | 0 | 0 | 0 | 56 | 0 | 56 | 15 | 116 | 35 | 25 | 116 |
| SRR6454286 | 0 | 12 | 87 | 255 | 99 | 699 | 93 | 1568 | 66 | 99 | 330 |
| SRR6454287 | 0 | 54 | 0 | 243 | 27 | 675 | 54 | 1674 | 108 | 27 | 270 |
| SRR6454288 | 0 | 41 | 206 | 124 | 0 | 578 | 41 | 1857 | 41 | 0 | 83 |
| SRR6454289 | 0 | 101 | 185 | 84 | 67 | 370 | 50 | 1716 | 135 | 67 | 151 |
| SRR6454290 | 0 | 0 | 45 | 56 | 112 | 625 | 45 | 1809 | 123 | 123 | 268 |
| SRR6454291 | 0 | 10 | 91 | 81 | 40 | 514 | 91 | 1775 | 101 | 81 | 414 |
| SRR6454292 | 0 | 69 | 118 | 39 | 29 | 412 | 88 | 1550 | 49 | 78 | 206 |
| SRR6454293 | 0 | 70 | 35 | 70 | 35 | 105 | 70 | 1856 | 175 | 280 | 700 |
| SRR6454294 | 0 | 24 | 24 | 0 | 24 | 415 | 146 | 3563 | 317 | 146 | 732 |
| SRR6454295 | 0 | 26 | 53 | 66 | 13 | 722 | 105 | 1865 | 158 | 118 | 381 |
| SRR6454296 | 0 | 53 | 53 | 36 | 18 | 374 | 178 | 1692 | 107 | 36 | 249 |
| SRR6454298 | 0 | 0 | 0 | 0 | 85 | 714 | 101 | 2103 | 272 | 559 | 2017 |
| SRR6454299 | 0 | 0 | 0 | 0 | 0 | 221 | 14 | 1214 | 64 | 29 | 64 |
| SRR6454300 | 0 | 0 | 0 | 0 | 0 | 227 | 63 | 1729 | 75 | 48 | 102 |
| SRR6454301 | 0 | 17 | 21 | 141 | 51 | 525 | 55 | 2103 | 98 | 346 | 469 |
| SRR6454302 | 0 | 43 | 89 | 190 | 19 | 474 | 31 | 1143 | 190 | 218 | 599 |
| SRR6454303 | 0 | 0 | 22 | 49 | 27 | 374 | 27 | 916 | 42 | 135 | 177 |
| SRR6454304 | 0 | 0 | 38 | 59 | 21 | 156 | 0 | 389 | 124 | 158 | 381 |
| SRR6454305 | 0 | 0 | 11 | 16 | 27 | 188 | 11 | 441 | 75 | 108 | 161 |
| SRR6454306 | 0 | 0 | 12 | 44 | 14 | 112 | 15 | 288 | 34 | 41 | 109 |
| SRR6454307 | 0 | 0 | 0 | 31 | 17 | 283 | 31 | 770 | 94 | 234 | 324 |
| SRR6454308 | 0 | 0 | 22 | 39 | 0 | 248 | 44 | 740 | 63 | 102 | 425 |
| SRR6454309 | 0 | 25 | 73 | 228 | 76 | 469 | 41 | 1138 | 82 | 143 | 276 |
| SRR6454310 | 0 | 21 | 101 | 201 | 27 | 199 | 31 | 441 | 51 | 61 | 205 |
| SRR6454311 | 0 | 11 | 65 | 119 | 11 | 627 | 54 | 2574 | 87 | 22 | 389 |
| SRR6454312 | 0 | 81 | 180 | 90 | 33 | 390 | 70 | 1268 | 26 | 35 | 136 |
| SRR6454313 | 0 | 0 | 35 | 35 | 23 | 221 | 23 | 1149 | 46 | 93 | 383 |
| SRR6454314 | 0 | 85 | 62 | 39 | 31 | 216 | 69 | 710 | 15 | 0 | 131 |
| SRR6454315 | 0 | 19 | 13 | 13 | 19 | 493 | 102 | 2881 | 115 | 32 | 410 |
| SRR6454316 | 0 | 0 | 60 | 0 | 0 | 444 | 85 | 2678 | 94 | 17 | 367 |
| SRR6454317 | 0 | 13 | 33 | 20 | 13 | 429 | 86 | 2134 | 112 | 112 | 515 |
| SRR6454318 | 0 | 0 | 11 | 43 | 21 | 450 | 32 | 1608 | 193 | 75 | 375 |
| SRR6454319 | 0 | 0 | 74 | 22 | 30 | 864 | 216 | 4349 | 164 | 82 | 425 |
| SRR6454320 | 0 | 0 | 11 | 46 | 57 | 1005 | 320 | 5239 | 240 | 80 | 685 |
| SRR6454321 | 0 | 30 | 166 | 76 | 0 | 151 | 60 | 1420 | 30 | 15 | 181 |
| SRR6454322 | 0 | 53 | 106 | 66 | 0 | 253 | 53 | 705 | 53 | 80 | 160 |
| SRR6454323 | 0 | 125 | 325 | 135 | 42 | 371 | 65 | 1623 | 70 | 0 | 181 |
| SRR6454324 | 0 | 30 | 163 | 151 | 66 | 1189 | 217 | 3538 | 97 | 91 | 616 |
| SRR6454325 | 0 | 29 | 59 | 358 | 72 | 433 | 49 | 1152 | 104 | 299 | 495 |
| SRR6454326 | 0 | 23 | 92 | 234 | 38 | 276 | 39 | 704 | 91 | 73 | 250 |
| SRR6454331 | 0 | 32 | 182 | 182 | 21 | 374 | 32 | 1463 | 118 | 64 | 513 |
| SRR6454332 | 0 | 102 | 181 | 181 | 0 | 373 | 113 | 2021 | 0 | 34 | 158 |
| SRR6454335 | 0 | 0 | 68 | 0 | 102 | 819 | 137 | 2833 | 102 | 137 | 205 |
| SRR6454337 | 0 | 0 | 0 | 0 | 73 | 619 | 96 | 2425 | 191 | 223 | 855 |
| SRR6454338 | 0 | 0 | 0 | 0 | 82 | 783 | 170 | 3874 | 149 | 283 | 706 |
| SRR6454339 | 0 | 0 | 0 | 0 | 55 | 472 | 34 | 1842 | 89 | 172 | 258 |
| SRR6454340 | 0 | 0 | 0 | 0 | 17 | 213 | 34 | 331 | 449 | 241 | 1004 |
| SRR6454341 | 0 | 0 | 0 | 0 | 44 | 347 | 41 | 761 | 70 | 122 | 451 |
| SRR6454342 | 0 | 0 | 0 | 0 | 12 | 141 | 42 | 283 | 87 | 61 | 438 |
| SRR6454343 | 0 | 0 | 0 | 0 | 66 | 419 | 53 | 741 | 88 | 103 | 626 |
| SRR6454345 | 0 | 0 | 0 | 0 | 0 | 281 | 0 | 1347 | 41 | 33 | 74 |

### Supp Table S8.5.Columns_ET_to_FD

| | | | | | | | | | | | |
|---|---|---|---|---|---|---|---|---|---|---|---|
| SRR6454347 | 0 | 0 | 0 | 0 | 34 | 968 | 139 | 4572 | 116 | 75 | 229 |
| SRR6454348 | 0 | 0 | 0 | 0 | 29 | 714 | 154 | 4355 | 100 | 74 | 147 |
| SRR6454349 | 0 | 0 | 0 | 0 | 14 | 585 | 83 | 3163 | 58 | 25 | 127 |
| SRR6454350 | 0 | 0 | 0 | 0 | 28 | 199 | 47 | 1641 | 47 | 66 | 76 |
| SRR6454351 | 0 | 0 | 0 | 0 | 29 | 171 | 20 | 439 | 24 | 49 | 98 |
| SRR6454352 | 0 | 0 | 0 | 0 | 35 | 757 | 38 | 1334 | 41 | 114 | 59 |
| SRR6454354 | 0 | 0 | 0 | 0 | 12 | 418 | 19 | 758 | 12 | 43 | 35 |
| SRR6496456 | 0 | 11 | 16 | 14 | 94 | 190 | 232 | 176 | 148 | 50 | 21 |
| SRR6496457 | 0 | 0 | 0 | 0 | 30 | 67 | 108 | 112 | 117 | 29 | 11 |
| SRR6496458 | 0 | 0 | 0 | 0 | 60 | 117 | 151 | 126 | 226 | 65 | 25 |
| SRR6496459 | 0 | 0 | 0 | 0 | 33 | 62 | 107 | 88 | 135 | 38 | 13 |
| SRR6496460 | 0 | 0 | 0 | 0 | 74 | 124 | 182 | 109 | 290 | 74 | 30 |
| SRR6496461 | 0 | 38 | 158 | 58 | 0 | 38 | 0 | 61 | 25 | 28 | 17 |
| SRR6496462 | 0 | 60 | 187 | 63 | 0 | 40 | 0 | 55 | 17 | 22 | 15 |
| SRR6496463 | 0 | 56 | 136 | 87 | 0 | 30 | 0 | 57 | 21 | 29 | 23 |
| SRR6496464 | 0 | 41 | 134 | 63 | 0 | 35 | 0 | 65 | 19 | 32 | 19 |
| SRR6496465 | 0 | 66 | 128 | 86 | 0 | 48 | 17 | 69 | 31 | 31 | 24 |
| SRR6498854 | 26 | 24 | 49 | 69 | 18 | 207 | 37 | 353 | 195 | 185 | 623 |
| SRR6498855 | 34 | 26 | 90 | 69 | 21 | 430 | 55 | 557 | 387 | 312 | 918 |
| SRR6498856 | 36 | 31 | 59 | 64 | 17 | 235 | 22 | 439 | 263 | 267 | 883 |
| SRR6498857 | 29 | 37 | 80 | 74 | 21 | 385 | 39 | 419 | 383 | 331 | 905 |
| SRR6498858 | 69 | 73 | 158 | 107 | 28 | 325 | 40 | 650 | 255 | 351 | 539 |
| SRR6498860 | 17 | 25 | 73 | 71 | 33 | 396 | 54 | 379 | 254 | 244 | 577 |
| SRR6498861 | 0 | 13 | 46 | 29 | 29 | 684 | 26 | 916 | 664 | 684 | 710 |
| SRR6498862 | 49 | 32 | 99 | 42 | 27 | 288 | 30 | 379 | 271 | 261 | 545 |
| SRR6498863 | 0 | 44 | 60 | 48 | 32 | 559 | 46 | 556 | 428 | 536 | 582 |
| SRR6498864 | 17 | 77 | 109 | 91 | 49 | 460 | 79 | 625 | 323 | 410 | 545 |
| SRR6498865 | 26 | 106 | 62 | 70 | 0 | 458 | 106 | 660 | 519 | 343 | 370 |
| SRR6498866 | 33 | 29 | 70 | 81 | 16 | 188 | 26 | 302 | 142 | 97 | 358 |
| SRR6498867 | 70 | 47 | 99 | 55 | 0 | 95 | 15 | 156 | 125 | 74 | 161 |
| SRR6498868 | 50 | 31 | 87 | 119 | 17 | 139 | 24 | 258 | 104 | 50 | 221 |
| SRR6498869 | 42 | 37 | 127 | 78 | 22 | 240 | 23 | 398 | 237 | 240 | 443 |
| SRR6498870 | 53 | 30 | 92 | 62 | 18 | 175 | 19 | 486 | 162 | 196 | 198 |
| SRR6498871 | 0 | 19 | 55 | 61 | 19 | 203 | 40 | 298 | 195 | 169 | 370 |
| SRR6705804 | 0 | 0 | 0 | 0 | 0 | 0 | 0 | 25 | 0 | 0 | 0 |
| SRR6705805 | 0 | 0 | 0 | 0 | 0 | 0 | 0 | 0 | 0 | 0 | 0 |
| SRR6705806 | 0 | 0 | 0 | 0 | 0 | 0 | 0 | 0 | 0 | 0 | 0 |
| SRR6705808 | 0 | 0 | 0 | 0 | 59 | 86 | 43 | 123 | 11 | 0 | 0 |
| SRR6705809 | 0 | 0 | 0 | 0 | 40 | 103 | 70 | 308 | 20 | 0 | 70 |
| SRR6705810 | 0 | 0 | 0 | 0 | 81 | 102 | 29 | 322 | 22 | 22 | 132 |
| SRR6705811 | 0 | 0 | 0 | 0 | 0 | 0 | 0 | 0 | 0 | 0 | 0 |
| SRR6705812 | 0 | 0 | 0 | 0 | 0 | 0 | 0 | 0 | 0 | 0 | 11 |
| SRR6705813 | 0 | 0 | 0 | 0 | 0 | 0 | 0 | 0 | 0 | 0 | 0 |
| SRR6705814 | 0 | 0 | 0 | 0 | 20 | 0 | 0 | 0 | 0 | 0 | 33 |
| SRR6705815 | 0 | 0 | 13 | 0 | 52 | 87 | 48 | 117 | 61 | 0 | 65 |
| SRR6705816 | 0 | 0 | 0 | 0 | 35 | 39 | 14 | 44 | 12 | 0 | 30 |
| SRR6705817 | 0 | 0 | 0 | 0 | 43 | 58 | 38 | 108 | 66 | 0 | 151 |
| SRR6705818 | 0 | 22 | 17 | 15 | 20 | 30 | 0 | 82 | 22 | 0 | 37 |
| SRR6705819 | 0 | 92 | 90 | 59 | 67 | 59 | 39 | 167 | 74 | 0 | 213 |
| SRR6705820 | 0 | 0 | 0 | 0 | 0 | 10 | 0 | 13 | 0 | 0 | 0 |
| SRR6705821 | 0 | 0 | 0 | 17 | 60 | 66 | 35 | 63 | 17 | 0 | 58 |
| SRR6705822 | 0 | 0 | 0 | 0 | 68 | 118 | 69 | 204 | 19 | 20 | 51 |
| SRR6705823 | 0 | 0 | 0 | 0 | 53 | 58 | 51 | 87 | 16 | 0 | 47 |
| SRR6705824 | 0 | 0 | 0 | 0 | 0 | 14 | 0 | 59 | 0 | 0 | 0 |
| SRR6705825 | 0 | 0 | 0 | 0 | 49 | 185 | 58 | 185 | 26 | 0 | 69 |
| SRR6705826 | 0 | 0 | 0 | 0 | 21 | 17 | 0 | 49 | 0 | 0 | 24 |
| SRR6705827 | 0 | 0 | 0 | 0 | 109 | 59 | 101 | 185 | 0 | 0 | 67 |
| SRR6705828 | 0 | 0 | 0 | 0 | 0 | 0 | 17 | 19 | 0 | 0 | 64 |
| SRR6705829 | 0 | 0 | 0 | 0 | 79 | 73 | 99 | 101 | 22 | 30 | 30 |
| SRR6705830 | 0 | 0 | 0 | 0 | 0 | 0 | 0 | 0 | 0 | 0 | 0 |
| SRR6705831 | 0 | 0 | 0 | 0 | 0 | 15 | 14 | 12 | 0 | 0 | 0 |
| SRR6705832 | 0 | 0 | 0 | 0 | 0 | 0 | 0 | 0 | 0 | 0 | 0 |
| SRR6705833 | 0 | 0 | 0 | 0 | 0 | 13 | 0 | 19 | 0 | 0 | 0 |
| SRR6705834 | 0 | 0 | 0 | 0 | 87 | 44 | 59 | 124 | 60 | 0 | 110 |
| SRR6705835 | 0 | 0 | 0 | 0 | 66 | 46 | 34 | 134 | 26 | 0 | 71 |
| SRR6705837 | 0 | 0 | 0 | 0 | 0 | 0 | 0 | 15 | 0 | 0 | 0 |
| SRR6705838 | 0 | 0 | 0 | 0 | 50 | 21 | 21 | 30 | 12 | 0 | 38 |
| SRR6705839 | 0 | 0 | 0 | 0 | 25 | 19 | 16 | 29 | 10 | 0 | 0 |
| SRR6705840 | 0 | 0 | 0 | 0 | 32 | 14 | 0 | 59 | 19 | 0 | 41 |
| SRR6705841 | 0 | 0 | 0 | 0 | 27 | 36 | 29 | 43 | 16 | 0 | 29 |
| SRR6705842 | 0 | 0 | 0 | 0 | 44 | 37 | 22 | 63 | 20 | 0 | 34 |
| SRR6705843 | 0 | 0 | 0 | 0 | 32 | 29 | 0 | 60 | 0 | 0 | 13 |
| SRR6705844 | 0 | 0 | 0 | 0 | 0 | 0 | 0 | 0 | 0 | 0 | 12 |
| SRR6705845 | 0 | 19 | 22 | 22 | 58 | 33 | 20 | 86 | 19 | 10 | 45 |
| SRR6705846 | 0 | 12 | 12 | 17 | 17 | 18 | 18 | 33 | 14 | 0 | 20 |
| SRR6705847 | 0 | 0 | 0 | 0 | 0 | 18 | 14 | 35 | 0 | 0 | 22 |
| SRR6705848 | 0 | 0 | 0 | 0 | 0 | 0 | 0 | 11 | 0 | 0 | 0 |
| SRR6705849 | 0 | 0 | 0 | 0 | 0 | 11 | 0 | 0 | 0 | 0 | 0 |
| SRR6705850 | 0 | 15 | 18 | 0 | 0 | 11 | 0 | 15 | 0 | 0 | 18 |
| SRR6705851 | 0 | 0 | 0 | 0 | 0 | 0 | 0 | 21 | 0 | 0 | 27 |
| SRR6705852 | 0 | 0 | 0 | 11 | 0 | 0 | 0 | 19 | 0 | 0 | 19 |
| SRR6705853 | 0 | 0 | 0 | 0 | 26 | 19 | 21 | 24 | 0 | 0 | 24 |
| SRR6705854 | 0 | 0 | 0 | 0 | 16 | 0 | 13 | 40 | 0 | 0 | 0 |
| SRR6705855 | 0 | 0 | 0 | 0 | 12 | 10 | 0 | 16 | 0 | 0 | 0 |
| SRR6705856 | 0 | 0 | 0 | 0 | 210 | 155 | 152 | 243 | 43 | 0 | 109 |
| SRR6705857 | 0 | 13 | 0 | 29 | 42 | 64 | 48 | 71 | 0 | 0 | 48 |
| SRR6705859 | 0 | 0 | 0 | 15 | 25 | 0 | 15 | 54 | 0 | 0 | 35 |
| SRR6705860 | 0 | 0 | 0 | 0 | 17 | 21 | 10 | 55 | 17 | 0 | 10 |
| SRR6705861 | 0 | 0 | 0 | 0 | 24 | 19 | 0 | 11 | 0 | 0 | 0 |
| SRR6705862 | 0 | 0 | 0 | 0 | 24 | 14 | 0 | 20 | 16 | 0 | 0 |
| SRR6705863 | 0 | 0 | 0 | 0 | 27 | 30 | 27 | 41 | 30 | 0 | 22 |
| SRR6705866 | 0 | 0 | 0 | 0 | 0 | 12 | 12 | 0 | 12 | 0 | 37 |
| SRR6705867 | 0 | 0 | 0 | 0 | 23 | 16 | 0 | 47 | 23 | 0 | 23 |
| SRR6705868 | 0 | 0 | 0 | 0 | 0 | 0 | 0 | 0 | 11 | 0 | 0 |
| SRR6705869 | 0 | 0 | 0 | 0 | 38 | 31 | 19 | 38 | 0 | 0 | 25 |
| SRR6705870 | 0 | 0 | 0 | 0 | 0 | 0 | 0 | 31 | 0 | 0 | 13 |
| SRR6705871 | 0 | 16 | 0 | 16 | 24 | 28 | 12 | 35 | 28 | 0 | 45 |
| SRR6705874 | 0 | 0 | 0 | 0 | 0 | 0 | 0 | 0 | 0 | 0 | 0 |
| SRR6705875 | 0 | 0 | 0 | 0 | 0 | 0 | 0 | 0 | 0 | 0 | 0 |
| SRR6705876 | 0 | 0 | 0 | 0 | 0 | 0 | 0 | 0 | 0 | 0 | 0 |
| SRR6705877 | 0 | 0 | 0 | 0 | 0 | 0 | 0 | 0 | 0 | 0 | 0 |
| SRR6705878 | 0 | 0 | 0 | 0 | 0 | 17 | 0 | 14 | 0 | 0 | 0 |
| SRR6705879 | 0 | 0 | 0 | 0 | 0 | 0 | 0 | 11 | 0 | 0 | 12 |
| SRR6705880 | 0 | 0 | 0 | 0 | 0 | 0 | 0 | 0 | 0 | 0 | 0 |
| SRR6705881 | 0 | 0 | 0 | 0 | 0 | 0 | 0 | 0 | 0 | 0 | 12 |
| SRR6705882 | 0 | 0 | 0 | 0 | 0 | 0 | 0 | 0 | 0 | 0 | 0 |
| SRR6705883 | 0 | 0 | 0 | 0 | 0 | 0 | 0 | 0 | 0 | 0 | 0 |
| SRR6705884 | 0 | 0 | 13 | 0 | 0 | 0 | 0 | 19 | 12 | 0 | 14 |
| SRR6705885 | 0 | 0 | 0 | 0 | 0 | 0 | 0 | 13 | 13 | 0 | 27 |
| SRR6705886 | 0 | 0 | 0 | 0 | 18 | 0 | 12 | 12 | 0 | 0 | 0 |
| SRR6705887 | 0 | 0 | 0 | 0 | 15 | 17 | 10 | 30 | 17 | 0 | 23 |
| SRR6705888 | 0 | 0 | 0 | 0 | 21 | 23 | 20 | 31 | 14 | 0 | 20 |
| SRR6705889 | 0 | 0 | 0 | 0 | 0 | 0 | 0 | 17 | 0 | 0 | 0 |
| SRR6705890 | 0 | 0 | 29 | 36 | 69 | 51 | 14 | 97 | 29 | 11 | 51 |
| SRR6705891 | 0 | 10 | 15 | 0 | 27 | 20 | 12 | 36 | 0 | 0 | 10 |
| SRR6705892 | 0 | 0 | 0 | 0 | 13 | 47 | 0 | 22 | 22 | 0 | 38 |
| SRR6705893 | 0 | 0 | 0 | 0 | 10 | 0 | 10 | 29 | 0 | 0 | 14 |
| SRR6705894 | 0 | 0 | 0 | 0 | 16 | 49 | 31 | 45 | 0 | 0 | 14 |
| SRR6705895 | 0 | 0 | 0 | 0 | 104 | 195 | 202 | 248 | 0 | 0 | 26 |

### Supp Table S8.5.Columns_ET_to_FD

| | | | | | | | | | | | |
|---|---|---|---|---|---|---|---|---|---|---|---|
| SRR6705897 | 0 | 0 | 13 | 0 | 172 | 395 | 293 | 312 | 32 | 0 | 76 |
| SRR6705898 | 0 | 0 | 0 | 0 | 0 | 0 | 0 | 0 | 0 | 0 | 0 |
| SRR6705899 | 0 | 0 | 0 | 0 | 0 | 0 | 0 | 11 | 0 | 0 | 0 |
| SRR6705900 | 0 | 0 | 0 | 0 | 0 | 0 | 0 | 0 | 0 | 0 | 0 |
| SRR6705901 | 0 | 0 | 0 | 0 | 28 | 51 | 17 | 12 | 0 | 0 | 0 |
| SRR6705902 | 0 | 0 | 0 | 0 | 39 | 14 | 14 | 21 | 14 | 0 | 0 |
| SRR6705903 | 0 | 0 | 0 | 0 | 47 | 22 | 11 | 26 | 26 | 0 | 25 |
| SRR6726498 | 0 | 30 | 58 | 19 | 18 | 71 | 54 | 53 | 70 | 77 | 361 |
| SRR6726499 | 0 | 50 | 399 | 179 | 0 | 209 | 0 | 837 | 329 | 339 | 2631 |
| SRR6757230 | 0 | 237 | 442 | 106 | 92 | 750 | 151 | 1021 | 505 | 366 | 487 |
| SRR6757231 | 0 | 214 | 400 | 98 | 87 | 687 | 140 | 893 | 487 | 344 | 425 |
| SRR6757232 | 0 | 82 | 211 | 55 | 43 | 396 | 54 | 557 | 182 | 103 | 133 |
| SRR6757233 | 0 | 69 | 213 | 54 | 42 | 367 | 55 | 582 | 157 | 85 | 172 |
| SRR6757234 | 0 | 183 | 330 | 81 | 47 | 325 | 67 | 584 | 167 | 131 | 236 |
| SRR6757235 | 0 | 170 | 324 | 77 | 30 | 276 | 57 | 574 | 162 | 120 | 217 |
| SRR6757236 | 0 | 43 | 205 | 53 | 125 | 991 | 168 | 1313 | 330 | 340 | 433 |
| SRR6757237 | 0 | 46 | 176 | 36 | 86 | 929 | 168 | 1267 | 326 | 265 | 357 |
| SRR6757238 | 0 | 309 | 670 | 155 | 81 | 873 | 150 | 951 | 124 | 58 | 57 |
| SRR6757239 | 0 | 322 | 638 | 150 | 101 | 745 | 146 | 884 | 100 | 57 | 53 |
| SRR6757240 | 0 | 448 | 947 | 258 | 136 | 896 | 223 | 1087 | 96 | 53 | 59 |
| SRR6757241 | 0 | 446 | 901 | 234 | 133 | 879 | 186 | 1063 | 95 | 44 | 44 |
| SRR6757242 | 0 | 278 | 636 | 185 | 82 | 561 | 92 | 517 | 172 | 124 | 163 |
| SRR6757243 | 0 | 277 | 612 | 190 | 80 | 556 | 91 | 511 | 164 | 111 | 140 |
| SRR6757244 | 0 | 303 | 624 | 166 | 90 | 811 | 142 | 849 | 333 | 202 | 212 |
| SRR6757245 | 0 | 340 | 608 | 152 | 77 | 787 | 173 | 824 | 315 | 177 | 199 |
| SRR6757246 | 0 | 172 | 290 | 80 | 51 | 415 | 94 | 408 | 318 | 204 | 332 |
| SRR6757247 | 0 | 151 | 283 | 74 | 65 | 389 | 80 | 342 | 308 | 191 | 341 |
| SRR6757248 | 0 | 202 | 408 | 95 | 56 | 340 | 61 | 503 | 115 | 62 | 80 |
| SRR6757249 | 0 | 157 | 372 | 95 | 53 | 302 | 67 | 465 | 107 | 71 | 88 |
| SRR6757250 | 0 | 201 | 474 | 137 | 65 | 418 | 74 | 530 | 310 | 175 | 283 |
| SRR6757251 | 0 | 194 | 475 | 103 | 56 | 453 | 72 | 529 | 298 | 186 | 284 |
| SRR6757252 | 0 | 258 | 509 | 130 | 70 | 573 | 111 | 665 | 288 | 190 | 205 |
| SRR6757253 | 0 | 229 | 547 | 127 | 79 | 541 | 108 | 606 | 279 | 166 | 204 |
| SRR6757254 | 0 | 245 | 513 | 159 | 112 | 1025 | 134 | 1531 | 152 | 99 | 215 |
| SRR6757255 | 0 | 248 | 441 | 149 | 111 | 985 | 131 | 1409 | 155 | 90 | 198 |
| SRR6757256 | 0 | 353 | 691 | 181 | 123 | 791 | 84 | 795 | 105 | 55 | 159 |
| SRR6757257 | 0 | 309 | 642 | 166 | 98 | 727 | 84 | 754 | 110 | 56 | 132 |
| SRR6757258 | 0 | 222 | 376 | 120 | 56 | 430 | 70 | 695 | 66 | 27 | 101 |
| SRR6757259 | 0 | 200 | 364 | 104 | 49 | 434 | 61 | 673 | 67 | 32 | 104 |
| SRR6757260 | 0 | 468 | 904 | 190 | 145 | 1440 | 174 | 1728 | 397 | 186 | 488 |
| SRR6757261 | 0 | 403 | 850 | 209 | 160 | 1281 | 176 | 1589 | 403 | 182 | 435 |
| SRR6757262 | 0 | 570 | 1003 | 245 | 286 | 2543 | 344 | 2722 | 424 | 184 | 289 |
| SRR6757263 | 0 | 531 | 905 | 226 | 239 | 2389 | 353 | 2593 | 373 | 196 | 281 |
| SRR6757264 | 0 | 604 | 1087 | 322 | 208 | 1721 | 219 | 1655 | 154 | 70 | 286 |
| SRR6757265 | 0 | 534 | 1050 | 279 | 212 | 1672 | 183 | 1578 | 179 | 71 | 265 |
| SRR6757266 | 0 | 421 | 768 | 183 | 115 | 1148 | 141 | 1052 | 151 | 69 | 109 |
| SRR6757267 | 0 | 408 | 788 | 190 | 113 | 1102 | 117 | 858 | 131 | 61 | 77 |
| SRR6757268 | 0 | 289 | 502 | 140 | 75 | 747 | 96 | 798 | 328 | 165 | 283 |
| SRR6757269 | 0 | 281 | 468 | 114 | 95 | 779 | 91 | 765 | 303 | 178 | 294 |
| SRR6757270 | 0 | 222 | 418 | 101 | 60 | 572 | 74 | 506 | 277 | 164 | 344 |
| SRR6757271 | 0 | 196 | 417 | 101 | 66 | 524 | 74 | 501 | 247 | 174 | 330 |
| SRR6757272 | 0 | 283 | 521 | 160 | 62 | 519 | 64 | 527 | 197 | 126 | 229 |
| SRR6757273 | 0 | 258 | 497 | 128 | 76 | 478 | 74 | 520 | 197 | 106 | 215 |
| SRR6757274 | 0 | 319 | 771 | 208 | 110 | 940 | 95 | 807 | 194 | 119 | 243 |
| SRR6757275 | 0 | 338 | 761 | 198 | 113 | 894 | 81 | 789 | 170 | 101 | 254 |
| SRR6757276 | 0 | 299 | 636 | 152 | 72 | 517 | 75 | 621 | 381 | 165 | 440 |
| SRR6757277 | 0 | 345 | 551 | 145 | 76 | 537 | 60 | 577 | 295 | 168 | 460 |
| SRR6757373 | 0 | 0 | 0 | 0 | 0 | 12 | 0 | 26 | 31 | 26 | 187 |
| SRR6757374 | 0 | 0 | 0 | 0 | 0 | 12 | 0 | 50 | 121 | 55 | 451 |
| SRR6757375 | 0 | 0 | 0 | 0 | 0 | 0 | 0 | 0 | 18 | 17 | 139 |
| SRR6757376 | 0 | 0 | 0 | 0 | 0 | 0 | 0 | 38 | 134 | 71 | 444 |
| SRR6757377 | 0 | 0 | 0 | 0 | 0 | 0 | 0 | 0 | 30 | 13 | 146 |
| SRR6757378 | 0 | 0 | 0 | 0 | 0 | 36 | 0 | 166 | 188 | 83 | 342 |
| SRR6757379 | 0 | 0 | 0 | 0 | 0 | 21 | 0 | 89 | 24 | 11 | 64 |
| SRR6757380 | 0 | 0 | 0 | 0 | 0 | 31 | 0 | 126 | 67 | 46 | 165 |
| SRR6757381 | 0 | 0 | 0 | 0 | 0 | 70 | 12 | 127 | 25 | 14 | 84 |
| SRR6757382 | 0 | 0 | 0 | 0 | 0 | 33 | 0 | 121 | 118 | 57 | 228 |
| SRR6757383 | 0 | 0 | 0 | 0 | 0 | 0 | 0 | 35 | 41 | 24 | 150 |
| SRR6757384 | 0 | 0 | 0 | 0 | 0 | 18 | 0 | 35 | 140 | 75 | 813 |
| SRR6757385 | 0 | 0 | 0 | 0 | 0 | 11 | 0 | 24 | 39 | 30 | 138 |
| SRR6757386 | 0 | 0 | 0 | 0 | 0 | 12 | 0 | 36 | 76 | 69 | 243 |
| SRR6757387 | 0 | 0 | 0 | 0 | 0 | 0 | 0 | 27 | 89 | 67 | 567 |
| SRR6757388 | 0 | 0 | 0 | 0 | 0 | 27 | 0 | 106 | 487 | 568 | 654 |
| SRR6757389 | 0 | 0 | 0 | 0 | 0 | 53 | 0 | 85 | 21 | 16 | 115 |
| SRR6757390 | 0 | 0 | 10 | 23 | 19 | 104 | 17 | 140 | 64 | 44 | 238 |
| SRR6757391 | 0 | 0 | 0 | 0 | 0 | 0 | 0 | 48 | 45 | 24 | 223 |
| SRR6757392 | 0 | 0 | 0 | 0 | 0 | 18 | 0 | 93 | 180 | 112 | 770 |
| SRR6757393 | 0 | 0 | 0 | 0 | 0 | 0 | 0 | 0 | 22 | 27 | 196 |
| SRR6757394 | 0 | 0 | 0 | 0 | 0 | 0 | 0 | 15 | 94 | 101 | 689 |
| SRR6757395 | 0 | 0 | 0 | 0 | 0 | 65 | 17 | 130 | 23 | 13 | 76 |
| SRR6757396 | 0 | 0 | 0 | 0 | 0 | 36 | 0 | 95 | 66 | 32 | 175 |
| SRR6757397 | 0 | 0 | 0 | 0 | 0 | 14 | 0 | 40 | 12 | 0 | 53 |
| SRR6757398 | 0 | 0 | 0 | 0 | 0 | 34 | 0 | 113 | 156 | 73 | 440 |
| SRR6757399 | 0 | 0 | 0 | 0 | 0 | 0 | 0 | 37 | 19 | 17 | 96 |
| SRR6757400 | 0 | 0 | 0 | 0 | 0 | 57 | 13 | 208 | 105 | 87 | 264 |
| SRR6757401 | 0 | 0 | 0 | 0 | 0 | 25 | 0 | 71 | 40 | 24 | 174 |
| SRR6757402 | 0 | 0 | 0 | 0 | 0 | 19 | 0 | 65 | 58 | 32 | 273 |
| SRR6757403 | 0 | 0 | 0 | 0 | 0 | 0 | 0 | 33 | 16 | 17 | 126 |
| SRR6757404 | 0 | 0 | 0 | 0 | 0 | 50 | 12 | 218 | 137 | 79 | 500 |
| SRR6757405 | 0 | 0 | 0 | 0 | 0 | 19 | 0 | 63 | 17 | 10 | 79 |
| SRR6757406 | 0 | 0 | 0 | 0 | 0 | 12 | 0 | 43 | 23 | 10 | 103 |
| SRR6757407 | 0 | 0 | 0 | 0 | 0 | 0 | 0 | 45 | 13 | 12 | 90 |
| SRR6757408 | 0 | 0 | 0 | 0 | 0 | 0 | 0 | 14 | 23 | 17 | 106 |
| SRR6757409 | 0 | 0 | 0 | 0 | 0 | 0 | 0 | 18 | 32 | 35 | 155 |
| SRR6757410 | 0 | 0 | 0 | 0 | 0 | 17 | 0 | 100 | 72 | 63 | 426 |
| SRR6757411 | 0 | 0 | 0 | 0 | 0 | 24 | 0 | 70 | 27 | 14 | 120 |
| SRR6757412 | 0 | 0 | 13 | 10 | 0 | 131 | 16 | 390 | 133 | 86 | 479 |
| SRR6757413 | 0 | 0 | 0 | 0 | 17 | 78 | 17 | 229 | 58 | 55 | 153 |
| SRR6757414 | 0 | 0 | 0 | 0 | 0 | 0 | 0 | 40 | 104 | 66 | 478 |
| SRR6757415 | 0 | 0 | 0 | 0 | 0 | 0 | 0 | 24 | 16 | 11 | 77 |
| SRR6757416 | 0 | 0 | 0 | 0 | 0 | 0 | 0 | 16 | 35 | 28 | 245 |
| SRR6757417 | 0 | 0 | 0 | 0 | 0 | 35 | 0 | 74 | 24 | 12 | 123 |
| SRR6757418 | 0 | 0 | 0 | 0 | 0 | 124 | 18 | 339 | 91 | 54 | 232 |
| SRR6757419 | 0 | 0 | 0 | 0 | 0 | 68 | 0 | 145 | 36 | 22 | 147 |
| SRR6757420 | 0 | 0 | 0 | 0 | 0 | 82 | 12 | 140 | 69 | 33 | 319 |
| SRR6757421 | 0 | 0 | 0 | 0 | 0 | 0 | 0 | 28 | 31 | 0 | 130 |
| SRR6757422 | 0 | 0 | 0 | 0 | 0 | 0 | 0 | 25 | 45 | 24 | 175 |
| SRR6757423 | 0 | 0 | 0 | 0 | 0 | 11 | 0 | 23 | 13 | 0 | 40 |
| SRR6757424 | 0 | 12 | 28 | 26 | 0 | 67 | 16 | 204 | 78 | 43 | 187 |
| SRR6757425 | 0 | 0 | 0 | 0 | 0 | 18 | 0 | 42 | 28 | 28 | 145 |
| SRR6757426 | 0 | 0 | 0 | 0 | 0 | 29 | 0 | 129 | 115 | 46 | 142 |
| SRR6757427 | 0 | 0 | 17 | 23 | 0 | 31 | 11 | 87 | 40 | 29 | 135 |
| SRR6757428 | 0 | 0 | 0 | 0 | 0 | 84 | 0 | 237 | 191 | 114 | 281 |
| SRR6757429 | 0 | 0 | 0 | 0 | 0 | 0 | 0 | 26 | 30 | 11 | 66 |
| SRR6757430 | 0 | 0 | 0 | 0 | 0 | 0 | 0 | 29 | 19 | 10 | 50 |
| SRR6757431 | 0 | 0 | 0 | 0 | 0 | 0 | 0 | 26 | 22 | 31 | 173 |
| SRR6757432 | 0 | 0 | 0 | 0 | 0 | 83 | 16 | 350 | 153 | 56 | 202 |
| SRR6757433 | 0 | 0 | 0 | 0 | 0 | 0 | 0 | 17 | 12 | 0 | 87 |
| SRR6757434 | 0 | 0 | 0 | 0 | 0 | 18 | 0 | 85 | 101 | 142 | 206 |

### Supp Table S8.5.Columns_ET_to_FD

| | | | | | | | | | | | |
|---|---|---|---|---|---|---|---|---|---|---|---|
| SRR6757435 | 0 | 0 | 0 | 0 | 0 | 38 | 0 | 121 | 48 | 21 | 94 |
| SRR6757436 | 0 | 0 | 0 | 0 | 0 | 48 | 0 | 194 | 132 | 77 | 244 |
| SRR6757437 | 0 | 0 | 0 | 0 | 0 | 39 | 0 | 119 | 33 | 15 | 67 |
| SRR6757438 | 0 | 0 | 0 | 0 | 0 | 124 | 22 | 347 | 131 | 71 | 173 |
| SRR6757439 | 0 | 0 | 0 | 0 | 0 | 113 | 18 | 245 | 41 | 20 | 60 |
| SRR6757440 | 0 | 0 | 0 | 0 | 0 | 137 | 13 | 292 | 86 | 38 | 97 |
| SRR6757441 | 0 | 0 | 0 | 0 | 12 | 127 | 16 | 257 | 55 | 13 | 80 |
| SRR6757442 | 0 | 0 | 0 | 0 | 0 | 33 | 0 | 99 | 108 | 48 | 256 |
| SRR6757443 | 0 | 0 | 0 | 0 | 0 | 19 | 0 | 74 | 26 | 0 | 64 |
| SRR6757444 | 0 | 0 | 0 | 0 | 0 | 90 | 0 | 283 | 150 | 60 | 230 |
| SRR6757445 | 0 | 0 | 0 | 0 | 0 | 38 | 0 | 130 | 42 | 16 | 172 |
| SRR6757446 | 0 | 0 | 0 | 0 | 0 | 0 | 0 | 55 | 80 | 43 | 255 |
| SRR6757447 | 0 | 0 | 0 | 0 | 0 | 25 | 0 | 79 | 32 | 14 | 103 |
| SRR6757448 | 0 | 0 | 0 | 0 | 0 | 76 | 11 | 326 | 344 | 362 | 475 |
| SRR6757449 | 0 | 0 | 0 | 0 | 0 | 0 | 0 | 12 | 26 | 18 | 157 |
| SRR6757450 | 0 | 0 | 0 | 0 | 0 | 11 | 0 | 69 | 192 | 163 | 1472 |
| SRR6757451 | 0 | 0 | 0 | 22 | 11 | 64 | 0 | 185 | 40 | 26 | 192 |
| SRR6757452 | 0 | 0 | 0 | 0 | 24 | 251 | 47 | 819 | 750 | 999 | 860 |
| SRR6757453 | 0 | 0 | 0 | 0 | 0 | 35 | 11 | 104 | 13 | 18 | 90 |
| SRR6757454 | 0 | 0 | 0 | 0 | 0 | 51 | 13 | 132 | 60 | 31 | 258 |
| SRR6757455 | 0 | 0 | 0 | 0 | 0 | 0 | 0 | 29 | 25 | 14 | 138 |
| SRR6757456 | 0 | 0 | 0 | 0 | 0 | 31 | 0 | 98 | 214 | 172 | 413 |
| SRR6757457 | 0 | 0 | 0 | 0 | 0 | 0 | 0 | 19 | 25 | 17 | 102 |
| SRR6757458 | 0 | 0 | 0 | 0 | 0 | 0 | 0 | 0 | 29 | 30 | 158 |
| SRR6757459 | 0 | 0 | 0 | 0 | 0 | 18 | 0 | 42 | 23 | 11 | 74 |
| SRR6757460 | 0 | 0 | 0 | 0 | 0 | 0 | 0 | 34 | 104 | 70 | 290 |
| SRR6757461 | 0 | 0 | 0 | 0 | 0 | 0 | 0 | 15 | 15 | 34 | 86 |
| SRR6757462 | 0 | 0 | 0 | 0 | 0 | 0 | 0 | 24 | 59 | 49 | 169 |
| SRR6757463 | 0 | 0 | 0 | 0 | 0 | 14 | 0 | 72 | 29 | 35 | 160 |
| SRR6757464 | 0 | 0 | 0 | 0 | 0 | 56 | 0 | 208 | 275 | 334 | 426 |
| SRR6757465 | 0 | 0 | 0 | 0 | 0 | 21 | 0 | 65 | 36 | 28 | 120 |
| SRR6757466 | 0 | 0 | 0 | 0 | 0 | 86 | 20 | 322 | 147 | 157 | 289 |
| SRR6757467 | 0 | 0 | 0 | 0 | 0 | 0 | 0 | 21 | 25 | 33 | 339 |
| SRR6757468 | 0 | 0 | 0 | 0 | 0 | 0 | 0 | 26 | 50 | 40 | 484 |
| SRR6761159 | 0 | 0 | 0 | 0 | 0 | 0 | 0 | 0 | 0 | 0 | 0 |
| SRR6761162 | 0 | 0 | 0 | 0 | 0 | 0 | 0 | 0 | 0 | 0 | 0 |
| SRR6761163 | 0 | 0 | 0 | 0 | 0 | 0 | 0 | 0 | 0 | 0 | 0 |
| SRR6761165 | 0 | 0 | 0 | 0 | 0 | 0 | 0 | 0 | 0 | 0 | 0 |
| SRR6761167 | 0 | 0 | 0 | 0 | 0 | 0 | 0 | 0 | 0 | 0 | 0 |
| SRR6761175 | 0 | 0 | 0 | 0 | 0 | 0 | 0 | 0 | 0 | 0 | 0 |
| SRR6761179 | 0 | 0 | 0 | 0 | 0 | 0 | 0 | 0 | 0 | 0 | 0 |
| SRR6761181 | 0 | 0 | 0 | 0 | 0 | 0 | 0 | 0 | 0 | 0 | 0 |
| SRR6761182 | 0 | 0 | 0 | 0 | 0 | 0 | 0 | 0 | 0 | 0 | 0 |
| SRR6761183 | 0 | 0 | 0 | 0 | 0 | 0 | 0 | 0 | 0 | 0 | 0 |
| SRR6761184 | 0 | 0 | 0 | 0 | 0 | 0 | 0 | 0 | 0 | 0 | 0 |
| SRR6761188 | 0 | 0 | 0 | 0 | 0 | 0 | 0 | 0 | 0 | 0 | 0 |
| SRR6761192 | 0 | 0 | 0 | 0 | 0 | 0 | 0 | 0 | 0 | 0 | 0 |
| SRR6761198 | 0 | 0 | 0 | 0 | 0 | 0 | 0 | 0 | 0 | 0 | 0 |
| SRR6767691 | 0 | 0 | 0 | 0 | 0 | 0 | 0 | 0 | 0 | 0 | 0 |
| SRR6767701 | 0 | 0 | 0 | 0 | 0 | 0 | 0 | 0 | 0 | 0 | 0 |
| SRR6767712 | 0 | 0 | 0 | 0 | 0 | 0 | 0 | 0 | 0 | 0 | 0 |
| SRR6767713 | 0 | 0 | 0 | 0 | 0 | 0 | 0 | 0 | 0 | 0 | 0 |
| SRR6821085 | 0 | 17 | 33 | 47 | 21 | 89 | 22 | 64 | 101 | 54 | 87 |
| SRR6821086 | 0 | 0 | 22 | 46 | 43 | 50 | 37 | 48 | 66 | 110 | 242 |
| SRR6821087 | 0 | 0 | 0 | 0 | 13 | 34 | 17 | 29 | 43 | 95 | 89 |
| SRR6821088 | 0 | 0 | 23 | 55 | 13 | 123 | 16 | 182 | 71 | 16 | 62 |
| SRR6821089 | 0 | 0 | 10 | 0 | 0 | 26 | 0 | 19 | 25 | 0 | 0 |
| SRR6821090 | 0 | 20 | 45 | 41 | 0 | 63 | 0 | 76 | 22 | 29 | 22 |
| SRR6823984 | 0 | 222 | 475 | 272 | 3481 | 9929 | 2565 | 1878 | 5159 | 5233 | 4043 |
| SRR6823985 | 0 | 56 | 160 | 53 | 3304 | 8398 | 2113 | 1530 | 6130 | 6000 | 5664 |
| SRR6823986 | 0 | 332 | 590 | 312 | 5541 | 12793 | 3538 | 6699 | 18039 | 23065 | 27005 |
| SRR6823987 | 0 | 375 | 550 | 129 | 2864 | 7799 | 1971 | 4253 | 8753 | 12974 | 14315 |
| SRR6823990 | 0 | 251 | 389 | 217 | 2957 | 5716 | 1190 | 2062 | 3770 | 2078 | 1974 |
| SRR6823991 | 0 | 83 | 189 | 109 | 1482 | 3457 | 728 | 1275 | 1176 | 1316 | 1780 |
| SRR6829958 | 0 | 126 | 411 | 103 | 12151 | 33097 | 25261 | 4729 | 822 | 211 | 569 |
| SRR6829959 | 0 | 40 | 139 | 151 | 2099 | 5395 | 2233 | 824 | 123 | 20 | 46 |
| SRR6829960 | 0 | 146 | 569 | 190 | 18193 | 56513 | 48034 | 8753 | 1930 | 516 | 1291 |
| SRR6829961 | 0 | 45 | 134 | 109 | 3241 | 8708 | 4387 | 1644 | 310 | 54 | 92 |
| SRR6829962 | 0 | 64 | 273 | 78 | 11127 | 35110 | 30249 | 4907 | 2185 | 321 | 867 |
| SRR6829963 | 0 | 15 | 52 | 48 | 4330 | 11709 | 4616 | 773 | 299 | 38 | 155 |
| SRR6831609 | 0 | 0 | 0 | 0 | 0 | 22 | 0 | 0 | 22 | 22 | 22 |
| SRR6831611 | 0 | 0 | 0 | 0 | 0 | 0 | 0 | 0 | 0 | 0 | 0 |
| SRR6831613 | 0 | 0 | 0 | 0 | 0 | 0 | 0 | 0 | 29 | 0 | 0 |
| SRR6837863 | 0 | 0 | 0 | 0 | 0 | 90 | 33 | 438 | 1415 | 915 | 1086 |
| SRR6837867 | 0 | 0 | 0 | 0 | 0 | 29 | 12 | 128 | 800 | 565 | 654 |
| SRR6837868 | 0 | 0 | 0 | 0 | 0 | 75 | 28 | 244 | 910 | 664 | 750 |
| SRR6880326 | 0 | 0 | 0 | 0 | 0 | 0 | 0 | 0 | 0 | 0 | 0 |
| SRR6880328 | 0 | 0 | 0 | 0 | 0 | 0 | 0 | 0 | 0 | 0 | 0 |
| SRR6880329 | 0 | 0 | 0 | 0 | 0 | 0 | 0 | 0 | 0 | 0 | 0 |
| SRR6880330 | 0 | 0 | 0 | 0 | 0 | 0 | 0 | 0 | 0 | 0 | 0 |
| SRR6880331 | 0 | 0 | 0 | 0 | 0 | 0 | 0 | 0 | 0 | 0 | 0 |
| SRR6880333 | 0 | 0 | 0 | 0 | 0 | 20 | 0 | 14 | 0 | 0 | 0 |
| SRR6880334 | 0 | 0 | 0 | 0 | 0 | 20 | 0 | 19 | 0 | 0 | 0 |
| SRR6880335 | 0 | 0 | 0 | 0 | 0 | 23 | 0 | 20 | 0 | 0 | 0 |
| SRR6880336 | 0 | 0 | 0 | 0 | 0 | 0 | 0 | 0 | 0 | 0 | 0 |
| SRR6880337 | 0 | 0 | 0 | 0 | 0 | 0 | 0 | 0 | 0 | 0 | 0 |
| SRR6880338 | 0 | 0 | 0 | 0 | 0 | 0 | 0 | 0 | 0 | 0 | 0 |
| SRR6880339 | 0 | 0 | 0 | 0 | 0 | 0 | 0 | 0 | 0 | 0 | 0 |
| SRR6880341 | 0 | 0 | 0 | 0 | 0 | 0 | 0 | 0 | 0 | 0 | 0 |
| SRR6880342 | 0 | 0 | 0 | 0 | 0 | 0 | 0 | 0 | 0 | 0 | 0 |
| SRR6880343 | 0 | 0 | 0 | 0 | 0 | 0 | 0 | 0 | 0 | 0 | 0 |
| SRR6880345 | 0 | 0 | 0 | 0 | 0 | 0 | 0 | 0 | 0 | 0 | 0 |
| SRR6880351 | 0 | 0 | 0 | 0 | 0 | 0 | 0 | 0 | 0 | 0 | 0 |
| SRR6880356 | 0 | 0 | 0 | 0 | 0 | 0 | 0 | 0 | 0 | 0 | 0 |
| SRR6880357 | 0 | 0 | 0 | 0 | 0 | 0 | 0 | 0 | 0 | 0 | 0 |
| SRR6898464 | 0 | 27 | 30 | 57 | 0 | 46 | 12 | 91 | 62 | 22 | 61 |
| SRR6914752 | 0 | 63 | 84 | 168 | 10834 | 25447 | 9784 | 3464 | 1491 | 3422 | 1575 |
| SRR6914753 | 0 | 48 | 32 | 87 | 17988 | 14756 | 17610 | 6785 | 8957 | 2073 | 1362 |
| SRR6914754 | 0 | 100 | 160 | 186 | 3419 | 5701 | 3080 | 1284 | 725 | 652 | 512 |
| SRR6914755 | 0 | 17 | 11 | 64 | 3835 | 4730 | 5355 | 1554 | 1667 | 539 | 581 |
| SRR6928219 | 0 | 131 | 315 | 290 | 3057 | 3560 | 3364 | 1036 | 5763 | 3918 | 1097 |
| SRR6928220 | 0 | 248 | 715 | 496 | 4846 | 4806 | 4478 | 1612 | 12063 | 10887 | 4440 |
| SRR6928221 | 0 | 63 | 89 | 131 | 1019 | 3928 | 2852 | 1035 | 855 | 538 | 189 |
| SRR6928222 | 0 | 37 | 124 | 105 | 836 | 1224 | 907 | 357 | 1254 | 501 | 420 |
| SRR6928223 | 0 | 167 | 519 | 428 | 7104 | 11512 | 11341 | 2445 | 11793 | 10958 | 1721 |
| SRR6928224 | 0 | 200 | 520 | 406 | 2948 | 3020 | 3090 | 1194 | 11525 | 13099 | 6037 |
| SRR6928225 | 0 | 67 | 141 | 227 | 1770 | 5776 | 5071 | 1600 | 971 | 508 | 155 |
| SRR6928226 | 0 | 48 | 122 | 104 | 627 | 1133 | 841 | 365 | 977 | 487 | 413 |
| SRR6930936 | 0 | 0 | 66 | 0 | 223 | 423 | 377 | 75 | 152 | 137 | 55 |
| SRR6930937 | 0 | 0 | 68 | 0 | 116 | 545 | 268 | 191 | 196 | 184 | 56 |
| SRR6930939 | 0 | 0 | 0 | 0 | 21 | 114 | 34 | 98 | 155 | 126 | 66 |
| SRR6930941 | 0 | 0 | 43 | 0 | 227 | 975 | 487 | 268 | 248 | 165 | 54 |
| SRR6932941 | 0 | 0 | 0 | 0 | 0 | 220 | 149 | 1200 | 24142 | 18893 | 8228 |
| SRR6932942 | 0 | 0 | 0 | 0 | 0 | 159 | 278 | 1044 | 8637 | 8120 | 4343 |
| SRR6932943 | 0 | 0 | 0 | 0 | 90 | 785 | 269 | 1316 | 9938 | 10005 | 2946 |
| SRR6932946 | 0 | 0 | 0 | 0 | 175 | 867 | 433 | 1037 | 12790 | 10577 | 5251 |
| SRR6932949 | 0 | 0 | 0 | 0 | 0 | 0 | 0 | 0 | 5776 | 5998 | 2989 |
| SRR6932950 | 0 | 0 | 0 | 0 | 0 | 0 | 15 | 0 | 5162 | 6082 | 3175 |
| SRR6932951 | 0 | 0 | 0 | 0 | 0 | 0 | 10 | 0 | 4128 | 6538 | 3001 |

### Supp Table S8.5.Columns_ET_to_FD

| | | | | | | | | | | | |
|---|---|---|---|---|---|---|---|---|---|---|---|
| SRR6932952 | 0 | 16 | 0 | 0 | 2749 | 6230 | 2223 | 7178 | 4468 | 4121 | 4495 |
| SRR6932953 | 0 | 15 | 26 | 26 | 2571 | 4162 | 965 | 6836 | 11480 | 8858 | 5604 |
| SRR6932954 | 0 | 0 | 0 | 11 | 3389 | 2409 | 341 | 7784 | 21464 | 22541 | 10020 |
| SRR6932955 | 0 | 0 | 0 | 0 | 383 | 585 | 192 | 974 | 10485 | 10561 | 3164 |
| SRR7062776 | 0 | 0 | 0 | 12 | 32 | 74 | 31 | 94 | 143 | 70 | 110 |
| SRR7062777 | 0 | 0 | 0 | 14 | 31 | 49 | 23 | 70 | 131 | 58 | 108 |
| SRR7062778 | 0 | 0 | 0 | 0 | 31 | 53 | 15 | 66 | 72 | 50 | 78 |
| SRR7062779 | 0 | 0 | 0 | 16 | 37 | 45 | 23 | 75 | 76 | 41 | 70 |
| SRR7062780 | 0 | 0 | 0 | 11 | 27 | 48 | 28 | 75 | 127 | 71 | 96 |
| SRR7062781 | 0 | 0 | 0 | 13 | 34 | 46 | 28 | 80 | 129 | 83 | 130 |
| SRR7081766 | 0 | 0 | 0 | 0 | 0 | 0 | 0 | 0 | 30 | 0 | 0 |
| SRR7081767 | 0 | 0 | 0 | 0 | 0 | 0 | 0 | 0 | 0 | 0 | 0 |
| SRR7081768 | 0 | 0 | 0 | 0 | 0 | 0 | 0 | 0 | 0 | 0 | 20 |
| SRR7081769 | 0 | 0 | 0 | 0 | 0 | 0 | 0 | 0 | 0 | 0 | 0 |
| SRR7081770 | 0 | 0 | 0 | 0 | 0 | 0 | 0 | 98 | 0 | 0 | 0 |
| SRR7081771 | 0 | 0 | 0 | 0 | 0 | 0 | 0 | 21 | 0 | 0 | 0 |
| SRR7081772 | 0 | 0 | 0 | 0 | 0 | 0 | 0 | 0 | 0 | 0 | 0 |
| SRR7081773 | 0 | 0 | 0 | 0 | 0 | 0 | 0 | 13 | 0 | 0 | 0 |
| SRR7081774 | 0 | 0 | 0 | 0 | 0 | 0 | 0 | 0 | 0 | 0 | 0 |
| SRR7081775 | 0 | 0 | 0 | 0 | 0 | 0 | 0 | 0 | 0 | 0 | 0 |
| SRR7081776 | 0 | 0 | 0 | 0 | 0 | 0 | 0 | 0 | 0 | 0 | 0 |
| SRR7081777 | 0 | 0 | 0 | 0 | 0 | 0 | 0 | 0 | 0 | 0 | 0 |
| SRR7107416 | 0 | 0 | 263 | 229 | 162 | 0 | 2776 | 0 | 3868 | 0 | 4677 |
| SRR7107417 | 0 | 0 | 148 | 111 | 191 | 0 | 3624 | 0 | 5029 | 0 | 6315 |
| SRR7107418 | 0 | 0 | 0 | 29 | 40 | 757 | 105 | 968 | 731 | 381 | 0 |
| SRR7107419 | 0 | 0 | 0 | 36 | 39 | 832 | 109 | 1122 | 693 | 362 | 0 |
| SRR7107420 | 0 | 0 | 127 | 199 | 95 | 0 | 2693 | 0 | 4134 | 0 | 7229 |
| SRR7107421 | 0 | 0 | 143 | 204 | 82 | 0 | 2377 | 0 | 3692 | 0 | 6777 |
| SRR7126083 | 0 | 0 | 15 | 0 | 15 | 44 | 29 | 0 | 0 | 15 | 0 |
| SRR7126084 | 0 | 629 | 1317 | 265 | 4728 | 14021 | 10790 | 5086 | 3386 | 2820 | 1247 |
| SRR7126085 | 0 | 760 | 1830 | 390 | 6703 | 17851 | 15262 | 5893 | 2915 | 2071 | 1642 |
| SRR7126086 | 0 | 597 | 1924 | 408 | 5512 | 15271 | 13090 | 5320 | 2942 | 1762 | 1687 |
| SRR7126087 | 0 | 464 | 783 | 223 | 2190 | 6147 | 5647 | 4839 | 1387 | 1001 | 810 |
| SRR7126088 | 0 | 517 | 1702 | 302 | 3755 | 12885 | 10583 | 4373 | 1656 | 1040 | 1055 |
| SRR7126089 | 0 | 656 | 1541 | 269 | 3972 | 11460 | 9442 | 3270 | 1258 | 676 | 819 |
| SRR7126090 | 0 | 1051 | 2331 | 256 | 6548 | 17814 | 12794 | 4471 | 1686 | 1099 | 881 |
| SRR7126091 | 0 | 643 | 1469 | 277 | 6295 | 18084 | 14398 | 6303 | 2805 | 2129 | 1405 |
| SRR7126092 | 0 | 526 | 1187 | 211 | 3310 | 10074 | 8526 | 4233 | 1430 | 958 | 948 |
| SRR7126093 | 0 | 634 | 2085 | 365 | 5531 | 16995 | 15637 | 6778 | 4273 | 2553 | 1905 |
| SRR7153253 | 0 | 48 | 84 | 84 | 0 | 34 | 0 | 83 | 28 | 65 | 96 |
| SRR7153254 | 0 | 23 | 45 | 77 | 12 | 68 | 0 | 138 | 16 | 23 | 48 |
| SRR7153255 | 0 | 28 | 45 | 120 | 0 | 63 | 0 | 117 | 15 | 17 | 42 |
| SRR7153256 | 0 | 72 | 199 | 327 | 12 | 118 | 0 | 238 | 50 | 65 | 101 |
| SRR7153257 | 0 | 22 | 64 | 69 | 0 | 22 | 0 | 54 | 11 | 16 | 34 |
| SRR7153258 | 0 | 86 | 192 | 217 | 0 | 53 | 0 | 101 | 0 | 14 | 23 |
| SRR7153259 | 0 | 39 | 81 | 94 | 0 | 44 | 0 | 74 | 31 | 29 | 69 |
| SRR7153260 | 0 | 48 | 135 | 155 | 0 | 52 | 0 | 100 | 21 | 22 | 51 |
| SRR7153261 | 0 | 73 | 189 | 158 | 0 | 38 | 0 | 81 | 19 | 29 | 33 |
| SRR7153262 | 0 | 21 | 29 | 34 | 0 | 24 | 0 | 51 | 30 | 46 | 61 |
| SRR7153263 | 0 | 38 | 110 | 159 | 0 | 49 | 0 | 130 | 23 | 36 | 55 |
| SRR7153264 | 0 | 91 | 188 | 226 | 10 | 114 | 0 | 217 | 51 | 105 | 144 |
| SRR7153265 | 0 | 48 | 104 | 145 | 0 | 58 | 0 | 122 | 43 | 50 | 83 |
| SRR7153266 | 0 | 70 | 219 | 244 | 0 | 44 | 0 | 69 | 0 | 12 | 21 |
| SRR7153267 | 0 | 28 | 68 | 99 | 0 | 42 | 0 | 81 | 19 | 21 | 32 |
| SRR7153268 | 0 | 93 | 249 | 296 | 0 | 42 | 0 | 54 | 0 | 0 | 0 |
| SRR7153269 | 0 | 13 | 46 | 74 | 0 | 51 | 0 | 83 | 35 | 30 | 71 |
| SRR7153270 | 0 | 39 | 101 | 121 | 0 | 47 | 0 | 54 | 12 | 0 | 23 |
| SRR7153271 | 0 | 26 | 81 | 98 | 0 | 32 | 0 | 40 | 0 | 0 | 18 |
| SRR7153272 | 0 | 21 | 61 | 59 | 0 | 23 | 0 | 60 | 21 | 14 | 34 |
| SRR7153273 | 0 | 17 | 69 | 70 | 0 | 71 | 0 | 192 | 35 | 32 | 24 |
| SRR7153274 | 0 | 21 | 48 | 72 | 0 | 34 | 0 | 86 | 35 | 22 | 48 |
| SRR7153275 | 0 | 88 | 215 | 252 | 10 | 86 | 12 | 198 | 33 | 34 | 54 |
| SRR7153276 | 0 | 18 | 55 | 92 | 0 | 29 | 0 | 62 | 15 | 10 | 34 |
| SRR7153277 | 0 | 17 | 19 | 50 | 0 | 18 | 0 | 44 | 10 | 11 | 19 |
| SRR7153278 | 0 | 22 | 44 | 57 | 0 | 23 | 0 | 38 | 12 | 16 | 15 |
| SRR7153279 | 0 | 12 | 36 | 45 | 0 | 52 | 0 | 115 | 42 | 39 | 44 |
| SRR7153280 | 0 | 17 | 68 | 82 | 0 | 35 | 0 | 58 | 11 | 12 | 15 |
| SRR7153281 | 0 | 56 | 113 | 113 | 0 | 42 | 0 | 106 | 26 | 25 | 32 |
| SRR7153282 | 0 | 22 | 60 | 60 | 0 | 27 | 0 | 44 | 30 | 36 | 52 |
| SRR7153283 | 0 | 38 | 65 | 113 | 0 | 50 | 0 | 118 | 24 | 29 | 58 |
| SRR7153284 | 0 | 26 | 61 | 36 | 0 | 16 | 0 | 23 | 0 | 0 | 16 |
| SRR7153285 | 0 | 10 | 27 | 53 | 0 | 42 | 0 | 75 | 16 | 18 | 45 |
| SRR7153286 | 0 | 16 | 32 | 68 | 0 | 30 | 0 | 75 | 35 | 53 | 75 |
| SRR7153287 | 0 | 63 | 175 | 193 | 0 | 15 | 0 | 25 | 0 | 0 | 0 |
| SRR7153288 | 0 | 26 | 76 | 119 | 0 | 48 | 0 | 119 | 25 | 27 | 82 |
| SRR7153289 | 0 | 21 | 57 | 59 | 0 | 30 | 0 | 51 | 19 | 34 | 43 |
| SRR7153290 | 0 | 31 | 54 | 82 | 0 | 50 | 0 | 94 | 15 | 37 | 50 |
| SRR7153291 | 0 | 0 | 29 | 28 | 0 | 67 | 0 | 102 | 54 | 57 | 92 |
| SRR7153292 | 0 | 18 | 43 | 48 | 0 | 58 | 0 | 102 | 58 | 63 | 56 |
| SRR7153293 | 0 | 23 | 84 | 68 | 0 | 55 | 0 | 82 | 30 | 27 | 27 |
| SRR7153294 | 0 | 24 | 51 | 57 | 0 | 11 | 0 | 22 | 0 | 11 | 28 |
| SRR7153295 | 0 | 53 | 103 | 101 | 0 | 29 | 0 | 35 | 17 | 27 | 19 |
| SRR7153296 | 0 | 13 | 33 | 37 | 0 | 19 | 0 | 27 | 0 | 0 | 17 |
| SRR7153297 | 0 | 0 | 0 | 11 | 0 | 0 | 0 | 12 | 0 | 0 | 0 |
| SRR7153298 | 0 | 28 | 64 | 89 | 0 | 21 | 0 | 59 | 14 | 14 | 41 |
| SRR7153299 | 0 | 0 | 71 | 85 | 10 | 68 | 0 | 135 | 47 | 51 | 80 |
| SRR7153300 | 0 | 25 | 77 | 91 | 0 | 54 | 0 | 82 | 25 | 39 | 67 |
| SRR7153301 | 0 | 12 | 20 | 32 | 0 | 14 | 0 | 30 | 0 | 0 | 0 |
| SRR7153302 | 0 | 12 | 50 | 34 | 0 | 25 | 0 | 40 | 23 | 28 | 30 |
| SRR7153303 | 0 | 0 | 19 | 30 | 0 | 76 | 0 | 169 | 62 | 53 | 54 |
| SRR7153304 | 0 | 39 | 67 | 74 | 0 | 33 | 0 | 46 | 0 | 0 | 25 |
| SRR7153305 | 0 | 36 | 80 | 83 | 0 | 29 | 0 | 60 | 18 | 15 | 28 |
| SRR7153306 | 0 | 32 | 59 | 47 | 0 | 23 | 0 | 48 | 23 | 27 | 26 |
| SRR7153307 | 0 | 41 | 79 | 129 | 0 | 40 | 0 | 93 | 27 | 28 | 65 |
| SRR7153308 | 0 | 22 | 47 | 57 | 0 | 29 | 0 | 71 | 33 | 31 | 49 |
| SRR7153309 | 0 | 28 | 56 | 73 | 0 | 25 | 0 | 47 | 19 | 18 | 50 |
| SRR7153310 | 0 | 0 | 0 | 0 | 0 | 55 | 0 | 121 | 54 | 56 | 92 |
| SRR7153311 | 0 | 22 | 49 | 66 | 0 | 30 | 0 | 66 | 24 | 26 | 57 |
| SRR7153312 | 0 | 18 | 70 | 74 | 0 | 83 | 0 | 187 | 81 | 80 | 77 |
| SRR7153313 | 0 | 16 | 66 | 64 | 0 | 55 | 11 | 126 | 45 | 54 | 80 |
| SRR7153314 | 0 | 20 | 60 | 62 | 0 | 35 | 0 | 56 | 25 | 28 | 55 |
| SRR7153315 | 0 | 28 | 63 | 79 | 0 | 31 | 0 | 50 | 14 | 20 | 26 |
| SRR7153316 | 0 | 31 | 90 | 91 | 0 | 23 | 0 | 38 | 0 | 0 | 10 |
| SRR7153317 | 0 | 0 | 0 | 16 | 0 | 95 | 0 | 190 | 95 | 64 | 91 |
| SRR7153318 | 0 | 17 | 41 | 28 | 0 | 20 | 0 | 28 | 0 | 0 | 0 |
| SRR7153319 | 0 | 0 | 18 | 15 | 0 | 53 | 0 | 95 | 37 | 39 | 50 |
| SRR7153320 | 0 | 22 | 56 | 75 | 0 | 82 | 0 | 158 | 67 | 53 | 79 |
| SRR7153321 | 0 | 28 | 96 | 90 | 0 | 99 | 0 | 127 | 46 | 40 | 56 |
| SRR7153322 | 0 | 0 | 0 | 12 | 15 | 180 | 11 | 299 | 122 | 142 | 138 |
| SRR7153323 | 0 | 10 | 41 | 28 | 0 | 46 | 0 | 94 | 72 | 63 | 78 |
| SRR7153324 | 0 | 21 | 47 | 68 | 0 | 42 | 0 | 92 | 28 | 33 | 54 |
| SRR7153325 | 0 | 0 | 10 | 12 | 0 | 20 | 0 | 49 | 33 | 13 | 10 |
| SRR7153326 | 0 | 0 | 31 | 40 | 0 | 58 | 0 | 135 | 31 | 23 | 43 |
| SRR7153327 | 0 | 0 | 34 | 45 | 0 | 100 | 0 | 206 | 103 | 163 | 115 |
| SRR7153328 | 0 | 13 | 31 | 29 | 0 | 34 | 0 | 89 | 51 | 58 | 55 |
| SRR7153329 | 0 | 0 | 26 | 38 | 0 | 71 | 0 | 114 | 44 | 30 | 56 |
| SRR7153330 | 0 | 10 | 35 | 29 | 0 | 31 | 0 | 84 | 32 | 26 | 37 |
| SRR7153331 | 0 | 0 | 36 | 38 | 0 | 49 | 0 | 100 | 65 | 60 | 71 |
| SRR7162919 | 0 | 0 | 0 | 0 | 29 | 72 | 43 | 237 | 295 | 323 | 129 |

### Supp Table S8.5.Columns_ET_to_FD

| | | | | | | | | | | | |
|---|---|---|---|---|---|---|---|---|---|---|---|
| SRR7162920 | 0 | 0 | 0 | 0 | 25 | 50 | 41 | 115 | 174 | 145 | 176 |
| SRR7162921 | 0 | 0 | 0 | 0 | 11 | 51 | 114 | 205 | 495 | 233 | 171 |
| SRR7178585 | 0 | 25 | 95 | 71 | 14 | 67 | 0 | 74 | 49 | 32 | 71 |
| SRR7178586 | 0 | 36 | 119 | 119 | 13 | 106 | 0 | 82 | 26 | 26 | 96 |
| SRR7178587 | 0 | 0 | 16 | 32 | 32 | 16 | 16 | 32 | 0 | 32 | 48 |
| SRR7178588 | 0 | 35 | 35 | 106 | 18 | 0 | 0 | 18 | 53 | 18 | 35 |
| SRR7178589 | 0 | 0 | 18 | 23 | 11 | 45 | 0 | 93 | 124 | 68 | 187 |
| SRR7178590 | 0 | 0 | 25 | 23 | 0 | 59 | 0 | 114 | 133 | 57 | 226 |
| SRR7178591 | 0 | 0 | 0 | 0 | 0 | 37 | 0 | 56 | 75 | 66 | 112 |
| SRR7178592 | 0 | 11 | 0 | 11 | 0 | 44 | 0 | 44 | 33 | 0 | 98 |
| SRR7178593 | 0 | 44 | 93 | 186 | 0 | 80 | 13 | 177 | 106 | 88 | 199 |
| SRR7178594 | 0 | 51 | 146 | 137 | 0 | 116 | 0 | 137 | 69 | 34 | 240 |
| SRR7178595 | 0 | 17 | 86 | 156 | 0 | 35 | 0 | 104 | 35 | 35 | 35 |
| SRR7178596 | 0 | 48 | 24 | 48 | 0 | 24 | 24 | 121 | 24 | 0 | 48 |
| SRR7178597 | 0 | 16 | 52 | 36 | 0 | 41 | 0 | 109 | 107 | 57 | 284 |
| SRR7178598 | 0 | 0 | 35 | 28 | 0 | 44 | 0 | 102 | 67 | 39 | 155 |
| SRR7178599 | 0 | 0 | 22 | 22 | 0 | 46 | 0 | 101 | 53 | 43 | 137 |
| SRR7178600 | 0 | 0 | 0 | 0 | 0 | 20 | 0 | 33 | 83 | 18 | 37 |
| SRR7178601 | 0 | 0 | 0 | 0 | 0 | 0 | 0 | 36 | 77 | 24 | 38 |
| SRR7178602 | 0 | 0 | 0 | 0 | 0 | 16 | 0 | 53 | 75 | 16 | 21 |
| SRR7178603 | 0 | 0 | 0 | 0 | 0 | 0 | 0 | 44 | 51 | 19 | 44 |
| SRR7178604 | 0 | 16 | 51 | 29 | 0 | 32 | 0 | 29 | 22 | 16 | 0 |
| SRR7178605 | 0 | 34 | 15 | 37 | 15 | 46 | 0 | 28 | 12 | 0 | 40 |
| SRR7178606 | 0 | 42 | 28 | 14 | 28 | 28 | 0 | 28 | 0 | 14 | 0 |
| SRR7178607 | 0 | 19 | 0 | 19 | 0 | 39 | 0 | 58 | 0 | 0 | 0 |
| SRR7178608 | 0 | 19 | 54 | 47 | 0 | 47 | 0 | 75 | 65 | 35 | 208 |
| SRR7178609 | 0 | 22 | 57 | 63 | 0 | 39 | 0 | 105 | 81 | 54 | 176 |
| SRR7178610 | 0 | 19 | 28 | 76 | 0 | 0 | 0 | 0 | 28 | 28 | 142 |
| SRR7178611 | 0 | 11 | 0 | 43 | 0 | 33 | 0 | 43 | 87 | 11 | 87 |
| SRR7178612 | 0 | 44 | 56 | 65 | 0 | 63 | 0 | 118 | 100 | 65 | 255 |
| SRR7178613 | 0 | 18 | 65 | 97 | 13 | 54 | 0 | 119 | 72 | 65 | 337 |
| SRR7178614 | 0 | 19 | 86 | 67 | 0 | 48 | 0 | 96 | 29 | 38 | 105 |
| SRR7178615 | 0 | 66 | 66 | 40 | 0 | 40 | 0 | 27 | 27 | 53 | 106 |
| SRR7178616 | 0 | 15 | 34 | 43 | 0 | 27 | 0 | 59 | 41 | 38 | 106 |
| SRR7178617 | 0 | 17 | 37 | 46 | 0 | 32 | 0 | 58 | 41 | 22 | 104 |
| SRR7178618 | 0 | 0 | 18 | 41 | 0 | 29 | 0 | 53 | 24 | 18 | 71 |
| SRR7178619 | 0 | 0 | 27 | 34 | 0 | 14 | 0 | 48 | 27 | 0 | 82 |
| SRR7178620 | 0 | 19 | 24 | 21 | 0 | 38 | 0 | 45 | 50 | 21 | 57 |
| SRR7178621 | 0 | 0 | 30 | 16 | 0 | 0 | 0 | 50 | 28 | 23 | 48 |
| SRR7178622 | 0 | 0 | 0 | 11 | 0 | 33 | 0 | 11 | 44 | 22 | 44 |
| SRR7178623 | 0 | 0 | 30 | 0 | 0 | 15 | 0 | 76 | 30 | 15 | 61 |
| SRR7178624 | 0 | 40 | 26 | 37 | 0 | 57 | 0 | 83 | 89 | 75 | 187 |
| SRR7178625 | 0 | 34 | 77 | 40 | 0 | 58 | 0 | 103 | 87 | 74 | 201 |
| SRR7178626 | 0 | 0 | 25 | 25 | 13 | 63 | 0 | 114 | 89 | 114 | 89 |
| SRR7178627 | 0 | 0 | 56 | 28 | 0 | 83 | 0 | 125 | 14 | 42 | 125 |
| SRR7178628 | 0 | 0 | 23 | 21 | 0 | 21 | 0 | 29 | 26 | 23 | 83 |
| SRR7178629 | 0 | 0 | 24 | 17 | 0 | 22 | 0 | 15 | 15 | 17 | 87 |
| SRR7178630 | 0 | 11 | 23 | 16 | 0 | 20 | 0 | 27 | 39 | 20 | 66 |
| SRR7178631 | 0 | 14 | 18 | 21 | 0 | 16 | 0 | 39 | 35 | 28 | 28 |
| SRR7178632 | 0 | 0 | 19 | 19 | 0 | 13 | 0 | 41 | 41 | 17 | 62 |
| SRR7178633 | 0 | 0 | 0 | 10 | 0 | 10 | 0 | 30 | 10 | 20 | 50 |
| SRR7178634 | 0 | 0 | 11 | 23 | 0 | 34 | 0 | 23 | 0 | 23 | 57 |
| SRR7178635 | 0 | 14 | 36 | 43 | 0 | 41 | 0 | 88 | 74 | 55 | 221 |
| SRR7178636 | 0 | 14 | 35 | 38 | 0 | 43 | 0 | 105 | 64 | 67 | 251 |
| SRR7178637 | 0 | 0 | 27 | 40 | 0 | 40 | 0 | 27 | 60 | 40 | 181 |
| SRR7178638 | 0 | 18 | 18 | 0 | 0 | 18 | 0 | 83 | 55 | 37 | 222 |
| SRR7178639 | 0 | 21 | 93 | 145 | 10 | 49 | 13 | 114 | 75 | 75 | 256 |
| SRR7178640 | 0 | 16 | 74 | 119 | 0 | 68 | 0 | 123 | 74 | 55 | 177 |
| SRR7178641 | 0 | 27 | 47 | 104 | 10 | 67 | 0 | 104 | 80 | 94 | 258 |
| SRR7178642 | 0 | 15 | 39 | 52 | 0 | 17 | 0 | 72 | 33 | 48 | 153 |
| SRR7178643 | 0 | 14 | 31 | 40 | 0 | 14 | 0 | 61 | 38 | 38 | 145 |
| SRR7178644 | 0 | 12 | 39 | 32 | 0 | 22 | 0 | 39 | 61 | 44 | 132 |
| SRR7178645 | 0 | 23 | 63 | 56 | 0 | 26 | 0 | 46 | 66 | 36 | 159 |
| SRR7178646 | 0 | 34 | 28 | 65 | 0 | 40 | 0 | 37 | 65 | 53 | 161 |
| SRR7178647 | 0 | 0 | 40 | 67 | 0 | 27 | 0 | 27 | 40 | 0 | 161 |
| SRR7178648 | 0 | 15 | 76 | 15 | 0 | 15 | 0 | 15 | 30 | 0 | 91 |
| SRR7178649 | 0 | 58 | 101 | 130 | 0 | 76 | 0 | 112 | 76 | 40 | 90 |
| SRR7178650 | 0 | 29 | 110 | 48 | 0 | 62 | 14 | 57 | 62 | 29 | 77 |
| SRR7178651 | 0 | 35 | 96 | 70 | 20 | 30 | 0 | 86 | 55 | 35 | 106 |
| SRR7178652 | 0 | 0 | 20 | 22 | 0 | 27 | 0 | 42 | 52 | 37 | 57 |
| SRR7178653 | 0 | 17 | 33 | 19 | 0 | 31 | 0 | 48 | 43 | 41 | 55 |
| SRR7178654 | 0 | 0 | 10 | 10 | 0 | 21 | 0 | 21 | 52 | 31 | 73 |
| SRR7178655 | 0 | 0 | 0 | 58 | 0 | 15 | 0 | 15 | 44 | 15 | 87 |
| SRR7178656 | 0 | 18 | 41 | 76 | 0 | 51 | 0 | 132 | 84 | 124 | 221 |
| SRR7178657 | 0 | 22 | 78 | 63 | 0 | 58 | 0 | 117 | 71 | 83 | 227 |
| SRR7178658 | 0 | 11 | 55 | 77 | 0 | 33 | 11 | 77 | 88 | 66 | 131 |
| SRR7178659 | 0 | 15 | 30 | 45 | 0 | 60 | 0 | 30 | 45 | 45 | 90 |
| SRR7178660 | 0 | 55 | 107 | 127 | 0 | 79 | 13 | 203 | 79 | 63 | 153 |
| SRR7178661 | 0 | 45 | 99 | 111 | 0 | 68 | 0 | 203 | 52 | 40 | 104 |
| SRR7178662 | 0 | 39 | 127 | 110 | 17 | 59 | 0 | 159 | 54 | 32 | 135 |
| SRR7178663 | 0 | 20 | 45 | 41 | 0 | 41 | 0 | 65 | 77 | 0 | 73 |
| SRR7178664 | 0 | 15 | 15 | 57 | 0 | 38 | 0 | 76 | 42 | 11 | 121 |
| SRR7178665 | 0 | 0 | 37 | 0 | 0 | 19 | 0 | 75 | 75 | 19 | 0 |
| SRR7178666 | 0 | 44 | 44 | 22 | 0 | 44 | 0 | 0 | 22 | 0 | 0 |
| SRR7178667 | 0 | 12 | 19 | 33 | 0 | 24 | 0 | 69 | 29 | 33 | 102 |
| SRR7178668 | 0 | 11 | 13 | 42 | 0 | 24 | 0 | 49 | 47 | 29 | 75 |
| SRR7178669 | 0 | 0 | 11 | 42 | 21 | 53 | 11 | 63 | 11 | 53 | 42 |
| SRR7178670 | 0 | 0 | 24 | 49 | 0 | 24 | 0 | 49 | 37 | 37 | 37 |
| SRR7178671 | 0 | 35 | 60 | 76 | 11 | 22 | 0 | 65 | 65 | 60 | 169 |
| SRR7178672 | 0 | 42 | 74 | 92 | 0 | 47 | 0 | 79 | 68 | 71 | 184 |
| SRR7178673 | 0 | 11 | 22 | 33 | 0 | 11 | 0 | 76 | 22 | 22 | 87 |
| SRR7178674 | 0 | 15 | 44 | 44 | 0 | 15 | 0 | 44 | 29 | 0 | 73 |
| SRR7178675 | 0 | 32 | 69 | 96 | 0 | 46 | 0 | 106 | 106 | 78 | 248 |
| SRR7178676 | 0 | 47 | 98 | 98 | 0 | 30 | 0 | 90 | 102 | 77 | 264 |
| SRR7178677 | 0 | 39 | 39 | 0 | 0 | 39 | 0 | 78 | 78 | 78 | 214 |
| SRR7178678 | 0 | 0 | 111 | 133 | 0 | 22 | 0 | 44 | 67 | 111 | 155 |
| SRR7178679 | 0 | 91 | 246 | 255 | 22 | 69 | 13 | 268 | 129 | 79 | 306 |
| SRR7178680 | 0 | 89 | 217 | 224 | 15 | 108 | 15 | 271 | 89 | 66 | 364 |
| SRR7178681 | 0 | 80 | 196 | 228 | 0 | 100 | 20 | 244 | 60 | 116 | 285 |
| SRR7178682 | 0 | 20 | 83 | 83 | 0 | 44 | 11 | 103 | 44 | 38 | 162 |
| SRR7178683 | 0 | 32 | 83 | 76 | 0 | 39 | 0 | 133 | 48 | 36 | 237 |
| SRR7178684 | 0 | 22 | 51 | 109 | 0 | 15 | 0 | 80 | 29 | 36 | 138 |
| SRR7178685 | 0 | 25 | 34 | 51 | 0 | 25 | 0 | 76 | 42 | 34 | 93 |
| SRR7178686 | 0 | 16 | 51 | 55 | 12 | 53 | 16 | 112 | 25 | 0 | 59 |
| SRR7178687 | 0 | 17 | 40 | 55 | 0 | 53 | 0 | 119 | 19 | 0 | 66 |
| SRR7178688 | 0 | 25 | 25 | 42 | 0 | 51 | 0 | 76 | 17 | 0 | 25 |
| SRR7178689 | 0 | 0 | 12 | 23 | 0 | 12 | 0 | 59 | 12 | 0 | 47 |
| SRR7178690 | 0 | 11 | 44 | 46 | 0 | 26 | 0 | 72 | 44 | 29 | 77 |
| SRR7178691 | 0 | 12 | 52 | 41 | 12 | 49 | 0 | 99 | 73 | 41 | 238 |
| SRR7178692 | 0 | 27 | 33 | 51 | 15 | 27 | 0 | 87 | 72 | 33 | 235 |
| SRR7178693 | 0 | 0 | 41 | 50 | 0 | 56 | 0 | 79 | 61 | 45 | 238 |
| SRR7178694 | 0 | 24 | 59 | 55 | 0 | 41 | 12 | 67 | 71 | 59 | 219 |
| SRR7178695 | 0 | 0 | 26 | 26 | 0 | 26 | 0 | 68 | 68 | 26 | 136 |
| SRR7178696 | 0 | 0 | 24 | 24 | 0 | 35 | 0 | 35 | 47 | 35 | 71 |
| SRR7178697 | 0 | 11 | 16 | 26 | 0 | 30 | 0 | 98 | 63 | 58 | 156 |
| SRR7178698 | 0 | 16 | 51 | 35 | 0 | 21 | 0 | 60 | 39 | 31 | 72 |
| SRR7178699 | 0 | 17 | 55 | 28 | 0 | 25 | 0 | 68 | 30 | 15 | 104 |
| SRR7178700 | 0 | 22 | 43 | 65 | 0 | 36 | 0 | 132 | 53 | 38 | 84 |
| SRR7178701 | 0 | 16 | 62 | 85 | 0 | 25 | 0 | 124 | 48 | 37 | 83 |

### Supp Table S8.5.Columns_ET_to_FD

| | | | | | | | | | | | |
|---|---|---|---|---|---|---|---|---|---|---|---|
| SRR7178702 | 0 | 0 | 74 | 42 | 0 | 42 | 0 | 74 | 32 | 42 | 84 |
| SRR7178703 | 0 | 29 | 15 | 29 | 15 | 29 | 0 | 44 | 15 | 0 | 103 |
| SRR7178704 | 0 | 20 | 50 | 34 | 0 | 42 | 0 | 50 | 36 | 0 | 67 |
| SRR7178705 | 0 | 25 | 50 | 50 | 0 | 27 | 0 | 32 | 22 | 15 | 52 |
| SRR7178706 | 0 | 18 | 52 | 36 | 0 | 26 | 10 | 49 | 23 | 10 | 75 |
| SRR7178707 | 0 | 13 | 19 | 57 | 0 | 13 | 0 | 89 | 57 | 57 | 134 |
| SRR7178708 | 0 | 12 | 61 | 18 | 0 | 68 | 0 | 86 | 74 | 31 | 172 |
| SRR7178709 | 0 | 0 | 25 | 0 | 0 | 0 | 0 | 25 | 25 | 50 | 101 |
| SRR7178710 | 0 | 0 | 34 | 34 | 0 | 68 | 0 | 34 | 34 | 0 | 68 |
| SRR7178711 | 0 | 65 | 191 | 273 | 32 | 122 | 0 | 288 | 61 | 76 | 342 |
| SRR7178712 | 0 | 35 | 173 | 301 | 18 | 128 | 0 | 257 | 84 | 49 | 381 |
| SRR7178713 | 0 | 50 | 146 | 283 | 14 | 105 | 23 | 251 | 50 | 78 | 329 |
| SRR7178714 | 0 | 11 | 34 | 41 | 0 | 71 | 15 | 86 | 75 | 56 | 209 |
| SRR7178715 | 0 | 0 | 63 | 70 | 18 | 35 | 0 | 116 | 85 | 88 | 240 |
| SRR7178716 | 0 | 0 | 16 | 62 | 0 | 31 | 0 | 16 | 47 | 93 | 264 |
| SRR7178717 | 0 | 0 | 0 | 70 | 0 | 35 | 0 | 88 | 140 | 35 | 158 |
| SRR7178718 | 0 | 37 | 98 | 105 | 0 | 37 | 0 | 105 | 56 | 56 | 260 |
| SRR7178719 | 0 | 42 | 100 | 97 | 0 | 42 | 0 | 106 | 37 | 44 | 204 |
| SRR7178720 | 0 | 34 | 94 | 118 | 0 | 43 | 0 | 123 | 53 | 36 | 202 |
| SRR7178721 | 0 | 104 | 208 | 256 | 0 | 84 | 0 | 260 | 128 | 124 | 364 |
| SRR7178722 | 0 | 74 | 214 | 311 | 23 | 93 | 12 | 241 | 155 | 155 | 342 |
| SRR7178723 | 0 | 47 | 140 | 187 | 0 | 78 | 0 | 249 | 93 | 140 | 156 |
| SRR7178724 | 0 | 129 | 129 | 215 | 22 | 43 | 22 | 194 | 86 | 65 | 129 |
| SRR7213733 | 0 | 0 | 0 | 0 | 0 | 0 | 0 | 0 | 0 | 0 | 0 |
| SRR7213734 | 0 | 0 | 0 | 0 | 0 | 15 | 0 | 0 | 22 | 0 | 0 |
| SRR7213735 | 0 | 0 | 0 | 0 | 0 | 0 | 0 | 0 | 40 | 0 | 0 |
| SRR7226512 | 0 | 0 | 0 | 0 | 89 | 140 | 82 | 132 | 605 | 27 | 81 |
| SRR7226513 | 0 | 17 | 0 | 14 | 21 | 29 | 25 | 33 | 121 | 0 | 14 |
| SRR7226514 | 0 | 41 | 19 | 24 | 555 | 953 | 615 | 870 | 2465 | 132 | 1025 |
| SRR7226515 | 0 | 0 | 0 | 0 | 68 | 147 | 90 | 182 | 2261 | 59 | 267 |
| SRR7226516 | 0 | 0 | 0 | 0 | 211 | 276 | 148 | 231 | 1259 | 67 | 433 |
| SRR7226517 | 0 | 0 | 0 | 0 | 168 | 368 | 231 | 606 | 384 | 30 | 127 |
| SRR7226518 | 0 | 11 | 0 | 0 | 127 | 257 | 135 | 480 | 200 | 18 | 141 |
| SRR7226519 | 0 | 0 | 0 | 0 | 71 | 155 | 69 | 221 | 223 | 14 | 127 |
| SRR7226520 | 0 | 14 | 0 | 16 | 112 | 171 | 132 | 225 | 403 | 20 | 58 |
| SRR7226521 | 0 | 13 | 0 | 16 | 161 | 295 | 254 | 373 | 1077 | 61 | 165 |
| SRR7226522 | 0 | 0 | 0 | 12 | 186 | 289 | 192 | 220 | 613 | 28 | 102 |
| SRR7226523 | 0 | 13 | 0 | 16 | 522 | 657 | 458 | 375 | 1695 | 101 | 392 |
| SRR7226524 | 0 | 0 | 0 | 0 | 38 | 53 | 29 | 48 | 147 | 0 | 46 |
| SRR7226525 | 0 | 73 | 30 | 46 | 643 | 1074 | 719 | 1205 | 1933 | 172 | 888 |
| SRR7226526 | 0 | 121 | 47 | 58 | 859 | 1074 | 867 | 1476 | 1978 | 174 | 796 |
| SRR7233553 | 0 | 0 | 0 | 0 | 77 | 229 | 50 | 38 | 457 | 12 | 15 |
| SRR7233554 | 0 | 0 | 0 | 0 | 55 | 125 | 42 | 21 | 261 | 0 | 15 |
| SRR7233556 | 0 | 0 | 0 | 0 | 44 | 84 | 27 | 0 | 167 | 0 | 0 |
| SRR7233557 | 0 | 0 | 0 | 0 | 43 | 104 | 25 | 20 | 199 | 0 | 18 |
| SRR7233558 | 0 | 0 | 0 | 0 | 54 | 113 | 30 | 16 | 156 | 0 | 14 |
| SRR7233559 | 0 | 0 | 0 | 0 | 58 | 145 | 36 | 25 | 323 | 15 | 17 |
| SRR7233560 | 0 | 0 | 0 | 0 | 30 | 80 | 20 | 17 | 233 | 11 | 0 |
| SRR7233561 | 0 | 0 | 0 | 0 | 127 | 291 | 73 | 49 | 541 | 15 | 26 |
| SRR7233562 | 0 | 0 | 0 | 0 | 67 | 166 | 44 | 34 | 280 | 11 | 14 |
| SRR7233563 | 0 | 0 | 0 | 0 | 121 | 258 | 62 | 28 | 305 | 10 | 13 |
| SRR7233564 | 0 | 0 | 0 | 0 | 133 | 350 | 65 | 40 | 417 | 11 | 19 |
| SRR7233565 | 0 | 0 | 0 | 0 | 173 | 352 | 90 | 38 | 281 | 19 | 22 |
| SRR7233566 | 0 | 0 | 0 | 0 | 192 | 390 | 92 | 49 | 328 | 13 | 31 |
| SRR7233568 | 0 | 0 | 0 | 0 | 55 | 110 | 26 | 18 | 144 | 0 | 0 |
| SRR7233570 | 0 | 0 | 0 | 0 | 46 | 103 | 29 | 12 | 157 | 0 | 0 |
| SRR7233571 | 0 | 0 | 0 | 0 | 89 | 176 | 45 | 20 | 354 | 0 | 17 |
| SRR7233572 | 0 | 0 | 0 | 0 | 55 | 125 | 38 | 20 | 216 | 0 | 16 |
| SRR7262826 | 0 | 0 | 0 | 0 | 98 | 218 | 74 | 77 | 139 | 32 | 49 |
| SRR7262827 | 0 | 0 | 0 | 0 | 116 | 256 | 93 | 79 | 243 | 40 | 55 |
| SRR7262828 | 0 | 0 | 13 | 16 | 193 | 297 | 140 | 102 | 208 | 34 | 39 |
| SRR7262829 | 0 | 0 | 32 | 35 | 164 | 206 | 96 | 46 | 115 | 19 | 38 |
| SRR7262830 | 0 | 0 | 0 | 0 | 120 | 278 | 112 | 109 | 281 | 44 | 41 |
| SRR7288164 | 0 | 0 | 0 | 0 | 0 | 0 | 0 | 0 | 0 | 0 | 0 |
| SRR7288166 | 0 | 0 | 0 | 0 | 0 | 0 | 0 | 0 | 0 | 0 | 0 |
| SRR7288167 | 0 | 0 | 0 | 0 | 0 | 0 | 0 | 0 | 0 | 0 | 0 |
| SRR7288236 | 0 | 0 | 0 | 0 | 0 | 0 | 0 | 0 | 0 | 0 | 0 |
| SRR7288237 | 0 | 0 | 0 | 0 | 0 | 0 | 0 | 0 | 0 | 0 | 0 |
| SRR7288238 | 0 | 0 | 0 | 0 | 0 | 0 | 0 | 0 | 0 | 0 | 0 |
| SRR7288239 | 0 | 0 | 0 | 0 | 0 | 0 | 0 | 0 | 0 | 0 | 0 |
| SRR7288244 | 0 | 0 | 0 | 0 | 0 | 0 | 0 | 0 | 0 | 0 | 0 |
| SRR7288264 | 0 | 0 | 0 | 0 | 0 | 0 | 0 | 0 | 0 | 0 | 0 |
| SRR7288265 | 0 | 0 | 0 | 0 | 0 | 0 | 0 | 0 | 0 | 0 | 0 |
| SRR7288266 | 0 | 0 | 0 | 0 | 0 | 0 | 0 | 0 | 0 | 0 | 0 |
| SRR7288267 | 0 | 0 | 0 | 0 | 0 | 0 | 0 | 0 | 0 | 0 | 0 |
| SRR7288277 | 0 | 0 | 0 | 0 | 0 | 0 | 0 | 0 | 0 | 0 | 0 |
| SRR7288300 | 0 | 0 | 0 | 0 | 0 | 0 | 0 | 0 | 0 | 0 | 0 |
| SRR7288301 | 0 | 0 | 0 | 0 | 0 | 0 | 0 | 0 | 0 | 0 | 0 |
| SRR7288302 | 0 | 0 | 0 | 0 | 0 | 0 | 0 | 0 | 0 | 0 | 0 |
| SRR7288303 | 0 | 0 | 0 | 0 | 0 | 0 | 0 | 0 | 0 | 0 | 0 |
| SRR7288308 | 0 | 0 | 0 | 0 | 0 | 0 | 0 | 0 | 0 | 0 | 0 |
| SRR7288309 | 0 | 0 | 0 | 0 | 0 | 0 | 0 | 0 | 0 | 0 | 0 |
| SRR7288310 | 0 | 0 | 0 | 0 | 0 | 0 | 0 | 0 | 0 | 0 | 0 |
| SRR7288311 | 0 | 0 | 0 | 0 | 0 | 0 | 0 | 0 | 0 | 0 | 0 |
| SRR7288320 | 0 | 0 | 0 | 0 | 0 | 0 | 0 | 0 | 0 | 0 | 0 |
| SRR7288321 | 0 | 0 | 0 | 0 | 0 | 0 | 0 | 0 | 0 | 0 | 0 |
| SRR7288322 | 0 | 0 | 0 | 0 | 0 | 0 | 0 | 0 | 0 | 0 | 0 |
| SRR7288323 | 0 | 0 | 0 | 0 | 0 | 0 | 0 | 0 | 0 | 0 | 0 |
| SRR7288376 | 0 | 0 | 0 | 0 | 0 | 0 | 0 | 0 | 0 | 0 | 0 |
| SRR7288377 | 0 | 0 | 0 | 0 | 0 | 0 | 0 | 0 | 0 | 0 | 0 |
| SRR7288378 | 0 | 0 | 0 | 0 | 0 | 0 | 0 | 0 | 0 | 0 | 0 |
| SRR7288379 | 0 | 0 | 0 | 0 | 0 | 0 | 0 | 0 | 0 | 0 | 0 |
| SRR7288380 | 0 | 0 | 0 | 0 | 0 | 0 | 0 | 0 | 0 | 0 | 0 |
| SRR7288381 | 0 | 0 | 0 | 0 | 0 | 0 | 0 | 0 | 0 | 0 | 0 |
| SRR7288382 | 0 | 0 | 0 | 0 | 0 | 0 | 0 | 0 | 0 | 0 | 0 |
| SRR7288383 | 0 | 0 | 0 | 0 | 0 | 0 | 0 | 0 | 0 | 0 | 0 |
| SRR7288384 | 0 | 0 | 0 | 0 | 0 | 0 | 0 | 0 | 0 | 0 | 0 |
| SRR7288385 | 0 | 0 | 0 | 0 | 0 | 0 | 0 | 0 | 0 | 0 | 0 |
| SRR7288386 | 0 | 0 | 0 | 0 | 0 | 0 | 0 | 0 | 0 | 0 | 0 |
| SRR7288387 | 0 | 0 | 0 | 0 | 0 | 0 | 0 | 0 | 0 | 0 | 0 |
| SRR7288400 | 0 | 0 | 0 | 0 | 0 | 0 | 0 | 0 | 0 | 0 | 0 |
| SRR7288402 | 0 | 0 | 0 | 0 | 0 | 0 | 0 | 0 | 0 | 0 | 0 |
| SRR7288472 | 0 | 0 | 0 | 0 | 0 | 0 | 0 | 0 | 0 | 0 | 0 |
| SRR7288474 | 0 | 0 | 0 | 0 | 0 | 0 | 0 | 0 | 0 | 0 | 0 |
| SRR7288475 | 0 | 0 | 0 | 0 | 0 | 0 | 0 | 0 | 0 | 0 | 0 |
| SRR7288512 | 0 | 0 | 0 | 0 | 0 | 0 | 0 | 0 | 0 | 0 | 0 |
| SRR7288513 | 0 | 0 | 0 | 0 | 0 | 0 | 0 | 0 | 0 | 0 | 0 |
| SRR7288514 | 0 | 0 | 0 | 0 | 0 | 0 | 0 | 0 | 0 | 0 | 0 |
| SRR7288515 | 0 | 0 | 0 | 0 | 0 | 0 | 0 | 0 | 0 | 0 | 0 |
| SRR7293222 | 0 | 0 | 16 | 16 | 37 | 673 | 94 | 1747 | 1484 | 1519 | 1364 |
| SRR7293223 | 0 | 0 | 0 | 17 | 25 | 598 | 66 | 1252 | 811 | 911 | 767 |
| SRR7293224 | 0 | 13 | 21 | 21 | 94 | 1223 | 109 | 1748 | 472 | 476 | 460 |
| SRR7293225 | 0 | 0 | 21 | 85 | 51 | 677 | 63 | 1803 | 320 | 302 | 507 |
| SRR7293226 | 0 | 0 | 12 | 14 | 34 | 506 | 74 | 1315 | 1775 | 1857 | 1802 |
| SRR7293227 | 0 | 0 | 0 | 13 | 19 | 278 | 28 | 719 | 888 | 915 | 995 |
| SRR7293228 | 0 | 0 | 0 | 0 | 69 | 999 | 136 | 2835 | 1924 | 2305 | 1926 |
| SRR7293229 | 0 | 0 | 0 | 0 | 67 | 910 | 93 | 2548 | 302 | 333 | 848 |
| SRR7293230 | 0 | 0 | 0 | 17 | 12 | 108 | 25 | 192 | 703 | 379 | 492 |
| SRR7293231 | 0 | 0 | 0 | 0 | 0 | 106 | 16 | 170 | 391 | 237 | 344 |

### Supp Table S8.5.Columns_ET_to_FD

| | | | | | | | | | | | |
|---|---|---|---|---|---|---|---|---|---|---|---|
| SRR7293232 | 0 | 0 | 0 | 0 | 78 | 971 | 66 | 628 | 359 | 222 | 271 |
| SRR7293233 | 0 | 0 | 0 | 24 | 0 | 74 | 0 | 161 | 79 | 66 | 240 |
| SRR7343626 | 0 | 0 | 0 | 21 | 0 | 0 | 0 | 0 | 0 | 0 | 0 |
| SRR7343627 | 0 | 0 | 0 | 0 | 0 | 0 | 0 | 50 | 0 | 0 | 0 |
| SRR7343629 | 0 | 0 | 0 | 0 | 0 | 0 | 0 | 0 | 0 | 0 | 0 |
| SRR7343630 | 0 | 0 | 0 | 0 | 0 | 0 | 0 | 0 | 0 | 0 | 0 |
| SRR7343631 | 0 | 0 | 0 | 0 | 0 | 0 | 0 | 0 | 0 | 0 | 0 |
| SRR7343632 | 0 | 0 | 0 | 0 | 0 | 0 | 0 | 0 | 0 | 0 | 0 |
| SRR7343633 | 0 | 0 | 0 | 0 | 0 | 0 | 0 | 0 | 0 | 0 | 0 |
| SRR7343635 | 0 | 0 | 0 | 0 | 0 | 39 | 0 | 0 | 0 | 0 | 0 |
| SRR7343636 | 0 | 0 | 0 | 0 | 0 | 24 | 0 | 0 | 0 | 0 | 0 |
| SRR7343637 | 0 | 0 | 0 | 0 | 0 | 0 | 0 | 0 | 0 | 0 | 0 |
| SRR7343638 | 0 | 0 | 0 | 0 | 0 | 0 | 0 | 0 | 0 | 0 | 0 |
| SRR7343639 | 0 | 0 | 0 | 0 | 0 | 44 | 0 | 44 | 44 | 0 | 0 |
| SRR7343640 | 0 | 0 | 0 | 19 | 0 | 0 | 0 | 0 | 0 | 0 | 19 |
| SRR7412243 | 0 | 21 | 50 | 43 | 14 | 21 | 21 | 71 | 36 | 21 | 270 |
| SRR7412244 | 0 | 65 | 147 | 286 | 18 | 74 | 18 | 65 | 74 | 37 | 341 |
| SRR7412245 | 0 | 39 | 140 | 171 | 0 | 101 | 31 | 31 | 78 | 16 | 156 |
| SRR7412246 | 0 | 51 | 77 | 147 | 45 | 147 | 71 | 45 | 32 | 13 | 199 |
| SRR7412247 | 0 | 32 | 64 | 131 | 12 | 91 | 20 | 119 | 48 | 36 | 266 |
| SRR7412248 | 0 | 108 | 140 | 248 | 19 | 65 | 19 | 168 | 94 | 42 | 402 |
| SRR7412249 | 0 | 42 | 150 | 243 | 14 | 89 | 28 | 136 | 80 | 33 | 388 |
| SRR7412250 | 0 | 14 | 48 | 178 | 0 | 69 | 34 | 75 | 69 | 69 | 322 |
| SRR7412251 | 0 | 17 | 149 | 149 | 17 | 116 | 0 | 248 | 17 | 50 | 479 |
| SRR7412252 | 0 | 20 | 82 | 160 | 0 | 29 | 0 | 29 | 45 | 49 | 303 |
| SRR7412253 | 0 | 81 | 137 | 223 | 12 | 149 | 50 | 366 | 81 | 87 | 745 |
| SRR7412254 | 0 | 79 | 211 | 158 | 0 | 79 | 79 | 158 | 79 | 66 | 435 |
| SRR7412255 | 0 | 41 | 186 | 243 | 10 | 72 | 36 | 83 | 98 | 36 | 434 |
| SRR7412256 | 0 | 21 | 73 | 239 | 31 | 62 | 31 | 135 | 52 | 10 | 239 |
| SRR7412257 | 0 | 95 | 155 | 182 | 14 | 95 | 27 | 142 | 47 | 41 | 345 |
| SRR7412258 | 0 | 36 | 144 | 72 | 0 | 108 | 0 | 0 | 36 | 108 | 216 |
| SRR7412259 | 0 | 32 | 128 | 186 | 38 | 51 | 19 | 289 | 71 | 32 | 456 |
| SRR7412260 | 0 | 72 | 90 | 18 | 0 | 90 | 72 | 72 | 0 | 18 | 288 |
| SRR7412261 | 0 | 35 | 125 | 236 | 21 | 62 | 49 | 97 | 90 | 28 | 430 |
| SRR7412262 | 0 | 43 | 34 | 111 | 0 | 60 | 17 | 128 | 68 | 34 | 383 |
| SRR7412263 | 0 | 95 | 284 | 213 | 47 | 95 | 95 | 402 | 189 | 118 | 1135 |
| SRR7412264 | 0 | 53 | 158 | 368 | 39 | 197 | 39 | 289 | 105 | 66 | 723 |
| SRR7412265 | 0 | 93 | 295 | 286 | 17 | 152 | 25 | 244 | 118 | 51 | 682 |
| SRR7412266 | 0 | 74 | 170 | 273 | 0 | 140 | 30 | 502 | 598 | 133 | 1277 |
| SRR7412267 | 0 | 60 | 140 | 160 | 13 | 80 | 0 | 133 | 187 | 47 | 620 |
| SRR7412268 | 0 | 56 | 144 | 194 | 19 | 107 | 13 | 150 | 69 | 31 | 382 |
| SRR7412269 | 0 | 60 | 218 | 236 | 0 | 133 | 24 | 248 | 109 | 67 | 466 |
| SRR7412270 | 0 | 89 | 283 | 297 | 0 | 134 | 15 | 268 | 59 | 15 | 491 |
| SRR7412271 | 0 | 57 | 126 | 91 | 23 | 80 | 46 | 137 | 46 | 69 | 343 |
| SRR7412272 | 0 | 48 | 96 | 145 | 19 | 39 | 19 | 116 | 29 | 29 | 251 |
| SRR7412273 | 0 | 57 | 77 | 130 | 49 | 97 | 36 | 357 | 61 | 32 | 162 |
| SRR7412274 | 0 | 59 | 166 | 186 | 0 | 146 | 49 | 244 | 98 | 49 | 459 |
| SRR7412275 | 0 | 129 | 233 | 181 | 0 | 97 | 32 | 194 | 97 | 45 | 252 |
| SRR7412276 | 0 | 27 | 63 | 89 | 18 | 54 | 27 | 152 | 89 | 0 | 339 |
| SRR7412277 | 0 | 56 | 153 | 121 | 32 | 72 | 16 | 266 | 24 | 16 | 137 |
| SRR7412278 | 0 | 42 | 99 | 113 | 56 | 0 | 71 | 42 | 28 | 14 | 127 |
| SRR7412279 | 0 | 57 | 123 | 90 | 0 | 57 | 25 | 25 | 57 | 0 | 262 |
| SRR7412280 | 0 | 55 | 154 | 227 | 0 | 73 | 55 | 109 | 109 | 55 | 754 |
| SRR7412281 | 0 | 79 | 98 | 236 | 20 | 79 | 49 | 148 | 89 | 59 | 246 |
| SRR7412282 | 0 | 68 | 91 | 137 | 11 | 80 | 11 | 103 | 80 | 114 | 662 |
| SRR7412283 | 0 | 61 | 61 | 306 | 0 | 114 | 53 | 114 | 0 | 0 | 280 |
| SRR7412284 | 0 | 81 | 45 | 125 | 45 | 63 | 36 | 98 | 0 | 0 | 125 |
| SRR7412285 | 0 | 152 | 199 | 450 | 47 | 82 | 47 | 99 | 53 | 35 | 321 |
| SRR7412286 | 0 | 56 | 137 | 201 | 0 | 40 | 24 | 32 | 16 | 16 | 217 |
| SRR7412287 | 0 | 34 | 155 | 74 | 13 | 101 | 13 | 94 | 20 | 20 | 161 |
| SRR7412288 | 0 | 109 | 253 | 545 | 30 | 134 | 84 | 322 | 99 | 64 | 684 |
| SRR7412289 | 0 | 82 | 113 | 205 | 10 | 41 | 41 | 72 | 92 | 10 | 235 |
| SRR7412290 | 0 | 28 | 56 | 42 | 14 | 42 | 42 | 56 | 0 | 0 | 42 |
| SRR7412291 | 0 | 40 | 88 | 104 | 16 | 32 | 16 | 48 | 32 | 24 | 120 |
| SRR7412292 | 0 | 0 | 27 | 165 | 37 | 46 | 27 | 55 | 0 | 18 | 220 |
| SRR7412293 | 0 | 46 | 92 | 212 | 0 | 83 | 37 | 184 | 64 | 64 | 663 |
| SRR7412294 | 0 | 135 | 114 | 285 | 21 | 36 | 29 | 78 | 0 | 64 | 200 |
| SRR7412295 | 0 | 219 | 271 | 691 | 18 | 149 | 122 | 306 | 122 | 44 | 779 |
| SRR7412296 | 0 | 24 | 61 | 73 | 12 | 97 | 49 | 402 | 61 | 12 | 402 |
| SRR7412297 | 0 | 0 | 55 | 65 | 22 | 71 | 16 | 224 | 38 | 16 | 207 |
| SRR7412298 | 0 | 108 | 125 | 274 | 25 | 50 | 42 | 75 | 25 | 25 | 357 |
| SRR7412299 | 0 | 0 | 36 | 125 | 0 | 36 | 18 | 160 | 53 | 18 | 267 |
| SRR7412300 | 0 | 56 | 134 | 189 | 45 | 45 | 67 | 279 | 45 | 45 | 591 |
| SRR7412301 | 0 | 102 | 125 | 238 | 23 | 79 | 45 | 136 | 34 | 34 | 272 |
| SRR7412302 | 0 | 47 | 205 | 295 | 13 | 73 | 56 | 98 | 94 | 81 | 608 |
| SRR7412303 | 0 | 37 | 116 | 128 | 18 | 134 | 67 | 306 | 147 | 18 | 1008 |
| SRR7412304 | 0 | 0 | 134 | 134 | 0 | 100 | 33 | 159 | 75 | 33 | 493 |
| SRR7412305 | 0 | 57 | 195 | 321 | 34 | 69 | 57 | 126 | 46 | 57 | 459 |
| SRR7412306 | 0 | 10 | 92 | 173 | 0 | 31 | 61 | 143 | 143 | 71 | 621 |
| SRR7412307 | 0 | 49 | 49 | 115 | 49 | 33 | 66 | 66 | 33 | 16 | 115 |
| SRR7412308 | 0 | 29 | 124 | 181 | 19 | 29 | 19 | 86 | 38 | 29 | 382 |
| SRR7412309 | 0 | 50 | 83 | 225 | 25 | 117 | 50 | 83 | 25 | 0 | 217 |
| SRR7412310 | 0 | 38 | 107 | 222 | 19 | 61 | 27 | 84 | 38 | 15 | 291 |
| SRR7412311 | 0 | 16 | 99 | 181 | 41 | 74 | 33 | 49 | 58 | 16 | 460 |
| SRR7412312 | 0 | 34 | 119 | 204 | 25 | 34 | 17 | 68 | 93 | 25 | 407 |
| SRR7412313 | 0 | 38 | 124 | 76 | 0 | 86 | 0 | 67 | 105 | 48 | 525 |
| SRR7412314 | 0 | 52 | 74 | 74 | 22 | 45 | 22 | 37 | 0 | 15 | 208 |
| SRR7412315 | 0 | 59 | 129 | 94 | 24 | 59 | 59 | 188 | 71 | 35 | 306 |
| SRR7412316 | 0 | 109 | 337 | 492 | 0 | 164 | 46 | 382 | 228 | 127 | 1512 |
| SRR7412317 | 0 | 29 | 92 | 121 | 12 | 17 | 12 | 23 | 29 | 29 | 219 |
| SRR7412318 | 0 | 64 | 51 | 115 | 0 | 13 | 38 | 51 | 38 | 0 | 165 |
| SRR7412319 | 0 | 28 | 114 | 220 | 43 | 128 | 28 | 149 | 50 | 14 | 426 |
| SRR7412320 | 0 | 60 | 186 | 714 | 30 | 181 | 151 | 377 | 206 | 106 | 1096 |
| SRR7412321 | 0 | 57 | 95 | 123 | 19 | 161 | 28 | 209 | 104 | 57 | 521 |
| SRR7412322 | 0 | 74 | 143 | 212 | 11 | 109 | 29 | 149 | 132 | 52 | 722 |
| SRR7412323 | 0 | 139 | 287 | 615 | 16 | 205 | 49 | 197 | 271 | 107 | 1435 |
| SRR7412324 | 0 | 36 | 164 | 400 | 36 | 100 | 73 | 109 | 73 | 27 | 801 |
| SRR7412325 | 0 | 11 | 163 | 314 | 0 | 108 | 43 | 54 | 33 | 65 | 423 |
| SRR7412326 | 0 | 36 | 109 | 189 | 0 | 60 | 68 | 117 | 68 | 40 | 439 |
| SRR7412327 | 0 | 46 | 255 | 380 | 13 | 144 | 33 | 111 | 59 | 20 | 590 |
| SRR7412328 | 0 | 46 | 167 | 179 | 23 | 237 | 81 | 473 | 110 | 23 | 312 |
| SRR7412329 | 0 | 94 | 103 | 122 | 19 | 65 | 19 | 168 | 103 | 75 | 243 |
| SRR7412330 | 0 | 168 | 455 | 239 | 0 | 112 | 24 | 359 | 144 | 48 | 112 |
| SRR7412331 | 0 | 96 | 167 | 287 | 32 | 135 | 24 | 430 | 96 | 56 | 757 |
| SRR7412332 | 0 | 66 | 284 | 256 | 28 | 66 | 76 | 199 | 95 | 47 | 644 |
| SRR7412333 | 0 | 44 | 44 | 117 | 0 | 97 | 39 | 97 | 49 | 0 | 336 |
| SRR7412334 | 0 | 56 | 91 | 77 | 0 | 126 | 0 | 356 | 91 | 14 | 258 |
| SRR7412335 | 0 | 105 | 120 | 195 | 0 | 75 | 15 | 195 | 30 | 0 | 361 |
| SRR7412336 | 0 | 66 | 206 | 169 | 19 | 159 | 38 | 300 | 94 | 56 | 572 |
| SRR7412337 | 0 | 20 | 79 | 395 | 40 | 198 | 79 | 395 | 49 | 0 | 444 |
| SRR7412338 | 0 | 90 | 466 | 481 | 75 | 226 | 60 | 406 | 135 | 90 | 1460 |
| SRR7412339 | 0 | 27 | 162 | 202 | 13 | 94 | 0 | 148 | 94 | 0 | 229 |
| SRR7412340 | 0 | 60 | 60 | 180 | 48 | 96 | 72 | 96 | 144 | 36 | 241 |
| SRR7412341 | 0 | 47 | 56 | 141 | 37 | 253 | 75 | 112 | 215 | 0 | 150 |
| SRR7412342 | 0 | 30 | 23 | 60 | 0 | 83 | 0 | 105 | 30 | 15 | 256 |
| SRR7412343 | 0 | 42 | 125 | 142 | 17 | 92 | 33 | 209 | 100 | 25 | 392 |
| SRR7412344 | 0 | 0 | 165 | 90 | 15 | 82 | 22 | 195 | 157 | 60 | 636 |
| SRR7412345 | 0 | 59 | 117 | 215 | 39 | 117 | 20 | 215 | 20 | 0 | 293 |
| SRR7466734 | 0 | 47 | 196 | 133 | 184 | 3562 | 422 | 9779 | 17486 | 12157 | 17216 |

### Supp Table S8.5.Columns_ET_to_FD

| | | | | | | | | | | | |
|---|---|---|---|---|---|---|---|---|---|---|---|
| SRR7466735 | 0 | 52 | 192 | 112 | 192 | 3624 | 344 | 9877 | 17746 | 12030 | 16622 |
| SRR7466736 | 0 | 19 | 68 | 47 | 81 | 1289 | 119 | 3681 | 8107 | 7434 | 10426 |
| SRR7466737 | 0 | 15 | 93 | 78 | 82 | 1242 | 97 | 3826 | 8037 | 7518 | 10407 |
| SRR7466738 | 0 | 20 | 70 | 49 | 79 | 1251 | 112 | 3849 | 7971 | 7450 | 10654 |
| SRR7466739 | 0 | 24 | 60 | 60 | 85 | 1181 | 125 | 3904 | 8030 | 7385 | 10752 |
| SRR7466740 | 0 | 11 | 78 | 50 | 112 | 1950 | 173 | 5087 | 4722 | 2931 | 4865 |
| SRR7466741 | 0 | 21 | 85 | 51 | 117 | 2129 | 161 | 5229 | 4671 | 2927 | 4803 |
| SRR7466742 | 0 | 31 | 78 | 56 | 92 | 2035 | 162 | 5082 | 4892 | 2904 | 4853 |
| SRR7466743 | 0 | 35 | 83 | 46 | 86 | 2034 | 157 | 5258 | 4858 | 2872 | 4822 |
| SRR7466744 | 0 | 42 | 158 | 100 | 160 | 3562 | 313 | 7597 | 5825 | 3028 | 5478 |
| SRR7466745 | 0 | 58 | 136 | 94 | 212 | 3694 | 272 | 7373 | 5851 | 2800 | 5498 |
| SRR7466746 | 0 | 57 | 153 | 90 | 204 | 4280 | 366 | 10245 | 10011 | 5572 | 8677 |
| SRR7466747 | 0 | 57 | 152 | 104 | 247 | 3938 | 408 | 9997 | 9550 | 5394 | 8690 |
| SRR7466748 | 0 | 62 | 159 | 97 | 250 | 4170 | 370 | 9829 | 9876 | 5639 | 8772 |
| SRR7466749 | 0 | 49 | 192 | 100 | 198 | 4240 | 352 | 10341 | 9794 | 5694 | 8766 |
| SRR7466750 | 0 | 25 | 89 | 27 | 81 | 1722 | 151 | 4472 | 3817 | 2197 | 3599 |
| SRR7466751 | 0 | 10 | 87 | 52 | 81 | 1723 | 143 | 4463 | 3809 | 2337 | 3650 |
| SRR7466752 | 0 | 35 | 58 | 38 | 93 | 1755 | 165 | 4537 | 3793 | 2268 | 3608 |
| SRR7466753 | 0 | 23 | 77 | 39 | 72 | 1727 | 156 | 4551 | 3712 | 2229 | 3886 |
| SRR7466754 | 0 | 37 | 133 | 124 | 167 | 2842 | 263 | 8445 | 15146 | 11784 | 19243 |
| SRR7466755 | 0 | 37 | 144 | 134 | 187 | 2902 | 256 | 8600 | 15244 | 11683 | 19295 |
| SRR7466756 | 0 | 27 | 125 | 71 | 193 | 2806 | 225 | 8718 | 15470 | 11812 | 18857 |
| SRR7466757 | 0 | 49 | 131 | 119 | 198 | 2902 | 259 | 8600 | 15454 | 11707 | 18795 |
| SRR7466758 | 0 | 27 | 235 | 141 | 195 | 3789 | 410 | 11959 | 20854 | 13175 | 22365 |
| SRR7466759 | 0 | 74 | 169 | 175 | 263 | 3955 | 317 | 11473 | 20037 | 13686 | 21522 |
| SRR7466760 | 0 | 39 | 212 | 77 | 263 | 3990 | 296 | 11738 | 20090 | 14089 | 22493 |
| SRR7466761 | 0 | 39 | 184 | 112 | 217 | 3589 | 335 | 11859 | 21285 | 13713 | 22850 |
| SRR7466762 | 12 | 49 | 188 | 106 | 213 | 3744 | 339 | 9308 | 16711 | 11977 | 17017 |
| SRR7466763 | 0 | 66 | 227 | 83 | 248 | 3702 | 331 | 9642 | 17190 | 11886 | 16851 |
| SRR7466764 | 0 | 43 | 140 | 120 | 196 | 3617 | 356 | 7575 | 5884 | 2862 | 5719 |
| SRR7466765 | 0 | 49 | 142 | 95 | 183 | 3513 | 335 | 7592 | 5757 | 2790 | 5878 |
| SRR7541164 | 0 | 0 | 0 | 0 | 20 | 391 | 54 | 1376 | 44 | 21 | 133 |
| SRR7541165 | 0 | 0 | 0 | 0 | 0 | 20 | 0 | 61 | 13 | 0 | 19 |
| SRR7541166 | 0 | 0 | 0 | 0 | 17 | 700 | 44 | 2211 | 40 | 0 | 80 |
| SRR7541167 | 0 | 0 | 0 | 0 | 0 | 48 | 0 | 204 | 20 | 0 | 61 |
| SRR7541168 | 0 | 0 | 28 | 15 | 225 | 1744 | 232 | 3973 | 1182 | 1284 | 1920 |
| SRR7541169 | 0 | 0 | 0 | 0 | 38 | 720 | 47 | 2782 | 436 | 139 | 383 |
| SRR7541170 | 0 | 0 | 0 | 0 | 11 | 195 | 18 | 897 | 59 | 20 | 110 |
| SRR7541171 | 0 | 0 | 10 | 12 | 147 | 2458 | 309 | 5036 | 317 | 123 | 1001 |
| SRR7541172 | 0 | 0 | 0 | 0 | 23 | 918 | 71 | 2810 | 405 | 89 | 303 |
| SRR7541173 | 0 | 0 | 0 | 0 | 43 | 603 | 125 | 2890 | 670 | 211 | 1155 |
| SRR7541174 | 0 | 0 | 0 | 0 | 33 | 651 | 74 | 2126 | 90 | 22 | 109 |
| SRR7541175 | 0 | 0 | 15 | 0 | 33 | 643 | 105 | 2764 | 554 | 219 | 374 |
| SRR7541176 | 0 | 0 | 0 | 0 | 78 | 1756 | 381 | 5004 | 114 | 44 | 520 |
| SRR7541177 | 0 | 10 | 33 | 15 | 114 | 2612 | 336 | 10172 | 1157 | 408 | 669 |
| SRR7541178 | 0 | 0 | 22 | 10 | 24 | 555 | 85 | 2862 | 533 | 180 | 500 |
| SRR7541179 | 0 | 0 | 0 | 0 | 12 | 231 | 41 | 1021 | 198 | 67 | 498 |
| SRR7541180 | 0 | 0 | 0 | 0 | 119 | 1966 | 363 | 9559 | 1714 | 538 | 1533 |
| SRR7541181 | 0 | 0 | 31 | 35 | 72 | 804 | 225 | 5073 | 2445 | 616 | 3830 |
| SRR7541182 | 0 | 0 | 17 | 19 | 97 | 2352 | 359 | 8363 | 654 | 326 | 804 |
| SRR7541183 | 0 | 0 | 0 | 0 | 46 | 1062 | 110 | 3619 | 312 | 94 | 805 |
| SRR7541184 | 0 | 0 | 0 | 0 | 89 | 1768 | 176 | 6482 | 254 | 94 | 798 |
| SRR7541185 | 0 | 0 | 0 | 0 | 52 | 2467 | 289 | 9726 | 927 | 181 | 1112 |
| SRR7541186 | 0 | 0 | 0 | 0 | 31 | 1055 | 98 | 3798 | 152 | 48 | 324 |
| SRR7541187 | 0 | 0 | 0 | 0 | 0 | 232 | 19 | 905 | 114 | 30 | 128 |
| SRR7541188 | 0 | 0 | 0 | 0 | 15 | 395 | 61 | 1997 | 102 | 36 | 169 |
| SRR7541189 | 0 | 0 | 18 | 20 | 54 | 983 | 145 | 5968 | 1990 | 686 | 3711 |
| SRR7541190 | 0 | 0 | 23 | 19 | 44 | 1099 | 214 | 6323 | 754 | 194 | 1714 |
| SRR7541191 | 0 | 0 | 0 | 0 | 14 | 310 | 59 | 2000 | 166 | 59 | 359 |
| SRR7541192 | 0 | 0 | 13 | 12 | 41 | 542 | 127 | 2538 | 194 | 66 | 376 |
| SRR7541193 | 0 | 0 | 0 | 0 | 0 | 302 | 43 | 2456 | 862 | 157 | 686 |
| SRR7547853 | 0 | 22 | 47 | 35 | 42 | 117 | 48 | 62 | 151 | 12 | 35 |
| SRR7547854 | 0 | 35 | 102 | 33 | 42 | 75 | 40 | 84 | 493 | 12 | 27 |
| SRR7547855 | 0 | 21 | 43 | 28 | 16 | 24 | 18 | 19 | 171 | 0 | 12 |
| SRR7547856 | 0 | 22 | 48 | 19 | 32 | 100 | 37 | 75 | 160 | 18 | 25 |
| SRR7547857 | 0 | 17 | 31 | 15 | 28 | 64 | 17 | 48 | 146 | 13 | 13 |
| SRR7547858 | 0 | 25 | 48 | 31 | 34 | 68 | 19 | 46 | 184 | 0 | 14 |
| SRR7547859 | 0 | 31 | 50 | 16 | 24 | 57 | 32 | 24 | 196 | 22 | 20 |
| SRR7547860 | 0 | 16 | 33 | 10 | 16 | 35 | 15 | 35 | 177 | 12 | 16 |
| SRR7547861 | 0 | 56 | 123 | 34 | 117 | 187 | 100 | 117 | 332 | 60 | 141 |
| SRR7547862 | 0 | 15 | 35 | 31 | 42 | 89 | 35 | 44 | 108 | 0 | 31 |
| SRR7547863 | 0 | 17 | 49 | 15 | 75 | 121 | 81 | 55 | 188 | 25 | 54 |
| SRR7547864 | 0 | 0 | 24 | 17 | 23 | 55 | 27 | 47 | 387 | 17 | 73 |
| SRR7547865 | 0 | 21 | 34 | 19 | 34 | 93 | 36 | 71 | 96 | 13 | 18 |
| SRR7547866 | 0 | 27 | 62 | 59 | 163 | 364 | 108 | 95 | 73 | 51 | 61 |
| SRR7547867 | 0 | 15 | 33 | 10 | 57 | 143 | 65 | 52 | 87 | 19 | 30 |
| SRR7547868 | 0 | 44 | 91 | 46 | 43 | 89 | 30 | 44 | 131 | 0 | 24 |
| SRR7547869 | 0 | 36 | 46 | 20 | 42 | 89 | 28 | 61 | 183 | 20 | 17 |
| SRR7547870 | 0 | 24 | 39 | 19 | 21 | 65 | 16 | 42 | 116 | 0 | 14 |
| SRR7547871 | 0 | 54 | 84 | 33 | 58 | 136 | 31 | 91 | 336 | 22 | 21 |
| SRR7547872 | 0 | 23 | 32 | 17 | 18 | 50 | 22 | 41 | 134 | 0 | 11 |
| SRR7547873 | 0 | 56 | 105 | 40 | 133 | 301 | 127 | 118 | 133 | 52 | 66 |
| SRR7547874 | 0 | 41 | 70 | 51 | 57 | 164 | 48 | 137 | 172 | 12 | 13 |
| SRR7547875 | 0 | 31 | 41 | 14 | 56 | 123 | 44 | 77 | 118 | 0 | 19 |
| SRR7547876 | 0 | 0 | 14 | 0 | 16 | 32 | 17 | 22 | 125 | 20 | 72 |
| SRR7547877 | 0 | 38 | 94 | 59 | 71 | 209 | 70 | 117 | 160 | 20 | 38 |
| SRR7547878 | 0 | 10 | 30 | 12 | 26 | 45 | 22 | 36 | 445 | 30 | 72 |
| SRR7547879 | 0 | 0 | 24 | 0 | 33 | 61 | 35 | 44 | 335 | 44 | 89 |
| SRR7547880 | 0 | 0 | 11 | 0 | 0 | 14 | 15 | 14 | 219 | 12 | 48 |
| SRR7547881 | 0 | 0 | 23 | 0 | 21 | 38 | 20 | 39 | 291 | 34 | 96 |
| SRR7547882 | 0 | 64 | 155 | 32 | 57 | 76 | 38 | 31 | 166 | 0 | 0 |
| SRR7547883 | 0 | 0 | 14 | 0 | 19 | 37 | 12 | 20 | 62 | 0 | 0 |
| SRR7547884 | 0 | 72 | 162 | 59 | 60 | 139 | 48 | 48 | 333 | 18 | 0 |
| SRR7547885 | 0 | 18 | 57 | 27 | 43 | 87 | 44 | 86 | 450 | 37 | 89 |
| SRR7547886 | 0 | 22 | 52 | 24 | 31 | 84 | 36 | 110 | 629 | 45 | 101 |
| SRR7547887 | 0 | 12 | 27 | 0 | 24 | 40 | 27 | 40 | 475 | 26 | 60 |
| SRR7547888 | 0 | 10 | 25 | 0 | 29 | 59 | 29 | 63 | 508 | 140 | 258 |
| SRR7547889 | 0 | 0 | 22 | 0 | 22 | 45 | 26 | 45 | 245 | 18 | 36 |
| SRR7547890 | 0 | 13 | 47 | 0 | 39 | 66 | 30 | 70 | 432 | 39 | 75 |
| SRR7547891 | 0 | 0 | 15 | 0 | 14 | 27 | 15 | 25 | 257 | 16 | 44 |
| SRR7547892 | 0 | 0 | 16 | 0 | 31 | 54 | 25 | 32 | 255 | 21 | 61 |
| SRR7547893 | 0 | 12 | 36 | 0 | 19 | 47 | 25 | 50 | 363 | 24 | 50 |
| SRR7547894 | 0 | 0 | 0 | 0 | 10 | 20 | 0 | 11 | 74 | 0 | 23 |
| SRR7547895 | 0 | 13 | 30 | 0 | 28 | 46 | 28 | 60 | 434 | 36 | 80 |
| SRR7547896 | 0 | 0 | 23 | 10 | 47 | 67 | 48 | 47 | 178 | 26 | 94 |
| SRR7547897 | 0 | 0 | 38 | 15 | 27 | 52 | 32 | 49 | 250 | 22 | 83 |
| SRR7547898 | 0 | 11 | 41 | 15 | 80 | 122 | 68 | 70 | 220 | 51 | 143 |
| SRR7547899 | 0 | 0 | 25 | 0 | 17 | 50 | 26 | 60 | 377 | 24 | 48 |
| SRR7547900 | 0 | 0 | 18 | 0 | 13 | 30 | 14 | 27 | 358 | 13 | 36 |
| SRR7547901 | 0 | 11 | 42 | 13 | 60 | 93 | 73 | 71 | 284 | 39 | 127 |
| SRR7547902 | 0 | 0 | 25 | 0 | 31 | 44 | 28 | 34 | 277 | 13 | 51 |
| SRR7547903 | 0 | 0 | 28 | 11 | 22 | 49 | 28 | 35 | 147 | 18 | 57 |
| SRR7547904 | 0 | 0 | 15 | 0 | 0 | 26 | 15 | 16 | 121 | 13 | 32 |
| SRR7547905 | 0 | 0 | 24 | 0 | 15 | 25 | 29 | 35 | 335 | 12 | 57 |
| SRR7547906 | 0 | 0 | 0 | 0 | 16 | 43 | 18 | 32 | 286 | 0 | 17 |
| SRR7547907 | 0 | 15 | 48 | 0 | 34 | 56 | 31 | 41 | 342 | 10 | 26 |
| SRR7547908 | 0 | 16 | 84 | 41 | 113 | 145 | 107 | 81 | 609 | 36 | 108 |
| SRR7547909 | 0 | 17 | 43 | 0 | 40 | 75 | 54 | 66 | 741 | 87 | 230 |
| SRR7547910 | 0 | 0 | 41 | 22 | 36 | 68 | 37 | 62 | 683 | 33 | 94 |

### Supp Table S8.5.Columns_ET_to_FD

| | | | | | | | | | | | |
|---|---|---|---|---|---|---|---|---|---|---|---|
| SRR7547911 | 0 | 0 | 15 | 0 | 20 | 37 | 21 | 31 | 266 | 19 | 50 |
| SRR7547912 | 0 | 0 | 35 | 0 | 30 | 46 | 29 | 44 | 616 | 22 | 66 |
| SRR7547913 | 0 | 0 | 38 | 0 | 25 | 44 | 28 | 46 | 343 | 19 | 54 |
| SRR7547914 | 0 | 0 | 18 | 0 | 18 | 36 | 16 | 28 | 258 | 14 | 45 |
| SRR7547915 | 0 | 0 | 24 | 0 | 29 | 54 | 35 | 48 | 279 | 29 | 108 |
| SRR7547916 | 0 | 0 | 0 | 0 | 0 | 16 | 19 | 19 | 157 | 12 | 42 |
| SRR7547917 | 0 | 24 | 75 | 22 | 107 | 122 | 57 | 50 | 474 | 31 | 57 |
| SRR7547918 | 0 | 0 | 23 | 0 | 17 | 28 | 18 | 18 | 279 | 13 | 24 |
| SRR7547919 | 0 | 0 | 21 | 0 | 15 | 32 | 13 | 21 | 262 | 12 | 22 |
| SRR7547920 | 0 | 11 | 71 | 29 | 87 | 146 | 84 | 65 | 507 | 26 | 61 |
| SRR7585373 | 0 | 21 | 67 | 66 | 14 | 145 | 15 | 203 | 23 | 27 | 32 |
| SRR7586110 | 11 | 108 | 171 | 14 | 103 | 298 | 131 | 299 | 750 | 30 | 98 |
| SRR7586111 | 23 | 128 | 223 | 22 | 96 | 305 | 136 | 299 | 701 | 37 | 107 |
| SRR7586113 | 12 | 129 | 214 | 21 | 133 | 435 | 181 | 370 | 1560 | 54 | 138 |
| SRR7586114 | 27 | 165 | 263 | 34 | 127 | 445 | 185 | 356 | 1484 | 50 | 132 |
| SRR7586116 | 52 | 325 | 980 | 98 | 556 | 1050 | 541 | 587 | 2016 | 193 | 688 |
| SRR7586117 | 120 | 416 | 1245 | 163 | 545 | 1057 | 558 | 577 | 1956 | 179 | 652 |
| SRR7586118 | 127 | 431 | 1277 | 161 | 571 | 1071 | 568 | 585 | 1971 | 192 | 659 |
| SRR7586119 | 13 | 103 | 345 | 59 | 360 | 869 | 560 | 508 | 1280 | 49 | 206 |
| SRR7586120 | 30 | 132 | 422 | 86 | 368 | 885 | 550 | 497 | 1224 | 51 | 212 |
| SRR7586122 | 44 | 254 | 601 | 37 | 265 | 626 | 317 | 390 | 1820 | 55 | 242 |
| SRR7586123 | 95 | 352 | 755 | 61 | 288 | 617 | 306 | 352 | 1794 | 48 | 219 |
| SRR7586125 | 0 | 86 | 160 | 12 | 169 | 505 | 222 | 564 | 2496 | 72 | 262 |
| SRR7586126 | 0 | 93 | 178 | 19 | 171 | 465 | 230 | 531 | 2386 | 63 | 274 |
| SRR7586128 | 18 | 211 | 547 | 41 | 362 | 700 | 453 | 472 | 2332 | 66 | 213 |
| SRR7586129 | 38 | 279 | 668 | 57 | 360 | 672 | 454 | 464 | 2373 | 58 | 225 |
| SRR7586131 | 44 | 277 | 621 | 33 | 379 | 750 | 402 | 384 | 1792 | 67 | 238 |
| SRR7586132 | 115 | 402 | 771 | 50 | 362 | 772 | 406 | 368 | 1790 | 68 | 208 |
| SRR7586134 | 13 | 215 | 517 | 39 | 342 | 638 | 381 | 352 | 1681 | 51 | 210 |
| SRR7586135 | 21 | 284 | 639 | 56 | 354 | 647 | 392 | 349 | 1650 | 50 | 201 |
| SRR7586137 | 27 | 210 | 271 | 18 | 138 | 442 | 131 | 264 | 1137 | 48 | 137 |
| SRR7586138 | 52 | 281 | 331 | 32 | 144 | 444 | 134 | 277 | 1120 | 53 | 129 |
| SRR7586140 | 0 | 122 | 312 | 23 | 241 | 487 | 243 | 245 | 1717 | 44 | 186 |
| SRR7586141 | 16 | 167 | 414 | 38 | 254 | 494 | 242 | 226 | 1664 | 39 | 190 |
| SRR7586143 | 10 | 167 | 395 | 25 | 323 | 674 | 370 | 476 | 2627 | 76 | 217 |
| SRR7586144 | 24 | 224 | 487 | 46 | 335 | 674 | 380 | 470 | 2494 | 69 | 185 |
| SRR7586146 | 0 | 16 | 44 | 0 | 243 | 672 | 304 | 419 | 1039 | 66 | 168 |
| SRR7586147 | 0 | 23 | 48 | 0 | 234 | 665 | 281 | 422 | 1032 | 65 | 161 |
| SRR7586149 | 0 | 68 | 138 | 15 | 204 | 919 | 226 | 590 | 2234 | 87 | 322 |
| SRR7586150 | 0 | 104 | 163 | 19 | 214 | 945 | 258 | 594 | 2166 | 85 | 319 |
| SRR7586152 | 0 | 60 | 110 | 0 | 351 | 1312 | 426 | 557 | 1929 | 78 | 158 |
| SRR7586153 | 0 | 87 | 139 | 15 | 339 | 1309 | 455 | 534 | 1827 | 80 | 161 |
| SRR7586155 | 0 | 33 | 77 | 11 | 303 | 929 | 397 | 648 | 3093 | 146 | 316 |
| SRR7586156 | 0 | 52 | 94 | 18 | 307 | 922 | 438 | 662 | 3055 | 139 | 314 |
| SRR7586158 | 0 | 216 | 448 | 35 | 710 | 2429 | 860 | 983 | 1905 | 128 | 216 |
| SRR7586159 | 0 | 273 | 575 | 56 | 740 | 2454 | 853 | 974 | 1808 | 122 | 202 |
| SRR7609258 | 0 | 0 | 0 | 0 | 71 | 246 | 206 | 563 | 2278 | 191 | 399 |
| SRR7609259 | 0 | 0 | 0 | 0 | 444 | 1050 | 646 | 383 | 2466 | 90 | 185 |
| SRR7609260 | 0 | 0 | 0 | 0 | 147 | 316 | 160 | 151 | 2106 | 76 | 276 |
| SRR7609261 | 0 | 0 | 0 | 0 | 31 | 111 | 66 | 42 | 765 | 42 | 103 |
| SRR7609262 | 0 | 0 | 0 | 0 | 46 | 117 | 81 | 59 | 957 | 95 | 149 |
| SRR7609263 | 0 | 0 | 0 | 0 | 40 | 91 | 30 | 44 | 3772 | 128 | 228 |
| SRR7609264 | 0 | 0 | 0 | 0 | 25 | 65 | 29 | 39 | 1009 | 35 | 135 |
| SRR7609265 | 0 | 0 | 0 | 0 | 14 | 72 | 42 | 24 | 2229 | 90 | 219 |
| SRR7609266 | 0 | 0 | 0 | 0 | 34 | 118 | 63 | 58 | 774 | 25 | 79 |
| SRR7609267 | 0 | 0 | 0 | 0 | 33 | 71 | 53 | 61 | 460 | 39 | 77 |
| SRR7609268 | 0 | 0 | 0 | 0 | 34 | 128 | 89 | 52 | 285 | 24 | 60 |
| SRR7609269 | 0 | 0 | 0 | 0 | 33 | 81 | 51 | 45 | 1001 | 57 | 138 |
| SRR7609270 | 0 | 0 | 0 | 0 | 62 | 220 | 186 | 127 | 2189 | 68 | 297 |
| SRR7609271 | 0 | 0 | 0 | 0 | 33 | 92 | 68 | 42 | 3038 | 48 | 201 |
| SRR7609272 | 0 | 0 | 0 | 0 | 39 | 108 | 72 | 23 | 828 | 22 | 64 |
| SRR7609273 | 0 | 0 | 0 | 0 | 16 | 48 | 25 | 25 | 663 | 39 | 110 |
| SRR7609274 | 0 | 0 | 0 | 0 | 44 | 87 | 54 | 36 | 1468 | 60 | 201 |
| SRR7609275 | 0 | 0 | 13 | 0 | 103 | 305 | 242 | 132 | 2614 | 125 | 289 |
| SRR7609276 | 0 | 0 | 0 | 0 | 0 | 0 | 0 | 0 | 580 | 0 | 47 |
| SRR7609277 | 0 | 0 | 0 | 0 | 170 | 422 | 279 | 150 | 1039 | 73 | 112 |
| SRR7609278 | 0 | 0 | 0 | 0 | 27 | 72 | 20 | 25 | 1500 | 68 | 197 |
| SRR7609279 | 0 | 0 | 0 | 0 | 72 | 209 | 114 | 108 | 872 | 67 | 223 |
| SRR7609280 | 0 | 0 | 0 | 0 | 24 | 72 | 78 | 35 | 398 | 20 | 53 |
| SRR7609281 | 0 | 0 | 0 | 0 | 15 | 12 | 12 | 54 | 2458 | 76 | 405 |
| SRR7609282 | 0 | 0 | 0 | 0 | 142 | 468 | 287 | 291 | 3331 | 113 | 271 |
| SRR7609283 | 0 | 0 | 0 | 0 | 129 | 324 | 163 | 137 | 2235 | 42 | 316 |
| SRR7609284 | 0 | 0 | 12 | 0 | 101 | 213 | 113 | 98 | 2138 | 84 | 128 |
| SRR7609285 | 0 | 0 | 0 | 0 | 89 | 230 | 117 | 94 | 1798 | 67 | 242 |
| SRR7609286 | 0 | 0 | 0 | 0 | 76 | 199 | 80 | 75 | 2849 | 95 | 281 |
| SRR7609287 | 0 | 0 | 11 | 0 | 216 | 523 | 134 | 153 | 2197 | 168 | 284 |
| SRR7609288 | 0 | 0 | 0 | 0 | 332 | 715 | 269 | 152 | 2908 | 181 | 125 |
| SRR7609289 | 0 | 0 | 0 | 0 | 28 | 58 | 45 | 42 | 361 | 39 | 38 |
| SRR7609290 | 0 | 0 | 0 | 0 | 55 | 303 | 135 | 159 | 1028 | 95 | 223 |
| SRR7609291 | 0 | 0 | 0 | 0 | 21 | 77 | 62 | 35 | 121 | 56 | 12 |
| SRR7609292 | 0 | 0 | 0 | 0 | 38 | 160 | 125 | 69 | 426 | 15 | 71 |
| SRR7609293 | 0 | 0 | 22 | 0 | 226 | 498 | 209 | 316 | 3676 | 221 | 1175 |
| SRR7609294 | 0 | 0 | 0 | 0 | 185 | 373 | 157 | 125 | 630 | 36 | 200 |
| SRR7609295 | 38 | 12 | 30 | 0 | 128 | 235 | 80 | 12 | 622 | 82 | 311 |
| SRR7609296 | 0 | 0 | 0 | 0 | 21 | 23 | 16 | 0 | 546 | 25 | 133 |
| SRR7609297 | 14 | 22 | 12 | 14 | 48 | 107 | 43 | 24 | 1046 | 31 | 179 |
| SRR7609298 | 12 | 0 | 18 | 0 | 31 | 70 | 19 | 19 | 908 | 43 | 126 |
| SRR7609299 | 14 | 0 | 24 | 0 | 58 | 129 | 103 | 40 | 1056 | 52 | 274 |
| SRR7609300 | 29 | 18 | 61 | 44 | 117 | 199 | 117 | 67 | 880 | 15 | 208 |
| SRR7609301 | 49 | 39 | 98 | 22 | 159 | 358 | 113 | 83 | 613 | 76 | 199 |
| SRR7609302 | 12 | 31 | 34 | 12 | 61 | 110 | 43 | 34 | 221 | 15 | 80 |
| SRR7644105 | 0 | 0 | 0 | 0 | 0 | 0 | 0 | 0 | 0 | 0 | 0 |
| SRR7644106 | 0 | 0 | 0 | 0 | 0 | 0 | 0 | 0 | 0 | 0 | 0 |
| SRR7644107 | 0 | 0 | 0 | 0 | 0 | 0 | 0 | 0 | 0 | 0 | 0 |
| SRR7644108 | 0 | 0 | 0 | 0 | 0 | 0 | 0 | 0 | 0 | 0 | 0 |
| SRR7644109 | 0 | 0 | 0 | 0 | 0 | 0 | 0 | 0 | 0 | 0 | 0 |
| SRR7644110 | 0 | 0 | 0 | 0 | 0 | 0 | 0 | 0 | 0 | 0 | 0 |
| SRR7644111 | 0 | 0 | 0 | 0 | 0 | 0 | 0 | 0 | 0 | 0 | 0 |
| SRR7644112 | 0 | 0 | 0 | 0 | 0 | 0 | 0 | 0 | 0 | 0 | 0 |
| SRR7644113 | 0 | 0 | 0 | 0 | 0 | 0 | 30 | 0 | 30 | 0 | 0 |
| SRR7644114 | 0 | 0 | 0 | 0 | 0 | 0 | 31 | 0 | 94 | 0 | 0 |
| SRR7644115 | 0 | 0 | 0 | 0 | 0 | 0 | 0 | 0 | 149 | 0 | 0 |
| SRR7644116 | 0 | 0 | 0 | 0 | 0 | 0 | 0 | 0 | 125 | 0 | 0 |
| SRR7644117 | 0 | 0 | 0 | 0 | 0 | 0 | 0 | 0 | 0 | 0 | 0 |
| SRR7644118 | 0 | 0 | 0 | 0 | 0 | 0 | 0 | 0 | 0 | 0 | 0 |
| SRR7644119 | 0 | 0 | 0 | 0 | 0 | 0 | 0 | 0 | 0 | 0 | 0 |
| SRR7644120 | 0 | 0 | 0 | 0 | 0 | 0 | 0 | 0 | 0 | 0 | 0 |
| SRR7644121 | 0 | 0 | 0 | 0 | 0 | 0 | 0 | 0 | 0 | 0 | 0 |
| SRR7644122 | 0 | 0 | 0 | 0 | 0 | 0 | 0 | 0 | 0 | 0 | 0 |
| SRR7644123 | 0 | 0 | 0 | 0 | 0 | 0 | 0 | 0 | 0 | 0 | 0 |
| SRR7644124 | 0 | 0 | 0 | 0 | 0 | 0 | 0 | 0 | 0 | 0 | 0 |
| SRR7644125 | 0 | 0 | 0 | 0 | 0 | 0 | 0 | 0 | 0 | 0 | 0 |
| SRR7644126 | 0 | 0 | 0 | 0 | 0 | 0 | 0 | 0 | 0 | 0 | 0 |
| SRR7644128 | 0 | 0 | 0 | 0 | 0 | 0 | 0 | 0 | 0 | 0 | 0 |
| SRR7644129 | 0 | 0 | 0 | 0 | 0 | 0 | 0 | 0 | 0 | 0 | 0 |
| SRR7663662 | 0 | 0 | 0 | 0 | 0 | 0 | 0 | 0 | 0 | 0 | 0 |
| SRR7663663 | 0 | 0 | 0 | 0 | 0 | 0 | 0 | 0 | 0 | 0 | 0 |
| SRR7664346 | 0 | 0 | 0 | 0 | 16 | 136 | 32 | 311 | 1206 | 1489 | 2017 |
| SRR7664347 | 0 | 0 | 0 | 0 | 14 | 102 | 22 | 284 | 1175 | 1378 | 3240 |

### Supp Table S8.5.Columns_ET_to_FD

| | | | | | | | | | | | |
|---|---|---|---|---|---|---|---|---|---|---|---|
| SRR7690697 | 0 | 11 | 40 | 31 | 0 | 24 | 0 | 55 | 79 | 22 | 47 |
| SRR7690698 | 0 | 10 | 29 | 24 | 0 | 11 | 0 | 21 | 30 | 0 | 13 |
| SRR7690699 | 0 | 12 | 35 | 38 | 0 | 26 | 0 | 48 | 107 | 13 | 58 |
| SRR7690700 | 0 | 0 | 0 | 0 | 34 | 70 | 42 | 78 | 348 | 28 | 42 |
| SRR7690701 | 0 | 0 | 0 | 0 | 24 | 52 | 34 | 51 | 187 | 18 | 47 |
| SRR7690702 | 0 | 0 | 0 | 0 | 23 | 42 | 30 | 39 | 261 | 11 | 31 |
| SRR7690703 | 0 | 0 | 0 | 0 | 13 | 19 | 14 | 17 | 79 | 0 | 11 |
| SRR7690704 | 0 | 0 | 0 | 0 | 40 | 110 | 39 | 107 | 365 | 19 | 39 |
| SRR7690705 | 0 | 0 | 0 | 0 | 15 | 26 | 17 | 22 | 204 | 11 | 18 |
| SRR7690706 | 0 | 0 | 0 | 0 | 31 | 53 | 20 | 21 | 419 | 22 | 28 |
| SRR7690707 | 0 | 18 | 40 | 46 | 0 | 11 | 0 | 18 | 24 | 0 | 18 |
| SRR7690708 | 0 | 0 | 0 | 0 | 33 | 70 | 30 | 46 | 670 | 31 | 39 |
| SRR7690713 | 0 | 0 | 0 | 0 | 66 | 106 | 38 | 75 | 365 | 27 | 43 |
| SRR7690714 | 0 | 0 | 0 | 0 | 35 | 59 | 24 | 41 | 294 | 16 | 28 |
| SRR7690715 | 0 | 0 | 31 | 15 | 0 | 0 | 0 | 12 | 35 | 0 | 19 |
| SRR7690716 | 0 | 16 | 58 | 34 | 0 | 11 | 0 | 26 | 22 | 11 | 19 |
| SRR7690717 | 0 | 0 | 0 | 0 | 25 | 67 | 34 | 44 | 167 | 11 | 17 |
| SRR7690718 | 0 | 0 | 0 | 0 | 0 | 16 | 0 | 0 | 40 | 0 | 0 |
| SRR7690719 | 0 | 0 | 0 | 0 | 43 | 86 | 45 | 58 | 237 | 0 | 24 |
| SRR7690720 | 0 | 0 | 0 | 0 | 62 | 102 | 47 | 89 | 599 | 29 | 48 |
| SRR7690721 | 0 | 0 | 0 | 0 | 32 | 53 | 44 | 57 | 602 | 46 | 84 |
| SRR7690722 | 0 | 0 | 28 | 41 | 523 | 1253 | 421 | 579 | 775 | 64 | 358 |
| SRR7690723 | 0 | 0 | 0 | 0 | 243 | 683 | 166 | 494 | 618 | 49 | 206 |
| SRR7690724 | 0 | 0 | 12 | 20 | 576 | 1474 | 484 | 571 | 876 | 141 | 370 |
| SRR7690726 | 0 | 0 | 12 | 22 | 474 | 1068 | 342 | 569 | 915 | 87 | 318 |
| SRR7690727 | 0 | 0 | 0 | 0 | 38 | 73 | 29 | 54 | 145 | 0 | 17 |
| SRR7690728 | 0 | 0 | 0 | 0 | 32 | 51 | 41 | 37 | 209 | 0 | 21 |
| SRR7690729 | 0 | 0 | 0 | 0 | 99 | 255 | 106 | 186 | 482 | 47 | 90 |
| SRR7690730 | 0 | 0 | 0 | 0 | 38 | 89 | 23 | 76 | 287 | 14 | 18 |
| SRR7690731 | 0 | 0 | 0 | 0 | 27 | 69 | 32 | 35 | 257 | 14 | 37 |
| SRR7690732 | 0 | 0 | 0 | 0 | 18 | 42 | 19 | 39 | 254 | 0 | 19 |
| SRR7690733 | 0 | 0 | 0 | 0 | 40 | 87 | 36 | 101 | 645 | 38 | 61 |
| SRR7690734 | 0 | 0 | 0 | 0 | 175 | 541 | 142 | 483 | 324 | 38 | 139 |
| SRR7690736 | 0 | 0 | 28 | 20 | 657 | 1542 | 564 | 1042 | 1621 | 56 | 361 |
| SRR7690737 | 0 | 0 | 14 | 0 | 515 | 1565 | 586 | 1273 | 3573 | 246 | 1149 |
| SRR7690739 | 0 | 0 | 63 | 36 | 723 | 1827 | 609 | 892 | 2225 | 115 | 787 |
| SRR7690740 | 0 | 0 | 61 | 43 | 754 | 1934 | 580 | 973 | 2345 | 144 | 711 |
| SRR7690741 | 0 | 16 | 54 | 43 | 0 | 31 | 0 | 64 | 65 | 21 | 50 |
| SRR7690742 | 0 | 0 | 40 | 39 | 0 | 25 | 0 | 45 | 55 | 0 | 30 |
| SRR7690743 | 0 | 0 | 28 | 38 | 0 | 27 | 0 | 43 | 45 | 0 | 35 |
| SRR7690744 | 0 | 0 | 0 | 0 | 59 | 98 | 50 | 56 | 818 | 16 | 65 |
| SRR7690745 | 0 | 0 | 0 | 0 | 39 | 74 | 47 | 114 | 750 | 25 | 90 |
| SRR7690746 | 0 | 0 | 0 | 0 | 27 | 47 | 27 | 33 | 684 | 11 | 28 |
| SRR7690747 | 0 | 0 | 0 | 0 | 20 | 55 | 26 | 43 | 703 | 28 | 34 |
| SRR7690748 | 0 | 0 | 0 | 0 | 68 | 113 | 49 | 40 | 572 | 18 | 39 |
| SRR7690749 | 0 | 0 | 0 | 0 | 0 | 23 | 14 | 23 | 166 | 0 | 16 |
| SRR7690750 | 0 | 0 | 0 | 0 | 37 | 56 | 34 | 50 | 168 | 0 | 26 |
| SRR7690751 | 0 | 0 | 0 | 0 | 17 | 37 | 27 | 28 | 196 | 11 | 35 |
| SRR7690752 | 0 | 0 | 0 | 0 | 26 | 56 | 30 | 50 | 311 | 15 | 52 |
| SRR7690753 | 0 | 0 | 0 | 0 | 45 | 79 | 45 | 67 | 580 | 27 | 61 |
| SRR7690754 | 0 | 0 | 0 | 0 | 68 | 141 | 63 | 92 | 541 | 23 | 54 |
| SRR7690755 | 0 | 0 | 0 | 14 | 57 | 97 | 55 | 68 | 686 | 35 | 72 |
| SRR7690756 | 0 | 0 | 0 | 0 | 36 | 88 | 47 | 76 | 645 | 56 | 88 |
| SRR7690757 | 0 | 0 | 0 | 12 | 36 | 64 | 28 | 63 | 497 | 36 | 69 |
| SRR7690758 | 0 | 0 | 10 | 0 | 946 | 1759 | 680 | 736 | 1179 | 115 | 464 |
| SRR7690759 | 0 | 0 | 43 | 29 | 977 | 2161 | 633 | 1081 | 936 | 99 | 508 |
| SRR7690762 | 0 | 14 | 65 | 37 | 635 | 1691 | 518 | 974 | 2146 | 179 | 734 |
| SRR7690764 | 0 | 30 | 52 | 52 | 0 | 17 | 0 | 21 | 31 | 0 | 23 |
| SRR7690765 | 0 | 18 | 53 | 34 | 0 | 0 | 0 | 15 | 35 | 14 | 35 |
| SRR7690766 | 0 | 0 | 0 | 0 | 23 | 47 | 24 | 39 | 132 | 12 | 24 |
| SRR7690767 | 0 | 0 | 0 | 0 | 20 | 42 | 32 | 29 | 142 | 18 | 28 |
| SRR7690768 | 0 | 0 | 0 | 0 | 39 | 88 | 43 | 87 | 399 | 29 | 50 |
| SRR7690769 | 0 | 0 | 32 | 18 | 397 | 1279 | 283 | 641 | 732 | 89 | 221 |
| SRR7690770 | 0 | 0 | 16 | 20 | 648 | 1706 | 488 | 745 | 590 | 82 | 375 |
| SRR7690773 | 0 | 0 | 27 | 11 | 782 | 1621 | 797 | 824 | 2782 | 129 | 888 |
| SRR7690775 | 0 | 0 | 0 | 0 | 19 | 59 | 31 | 68 | 1104 | 46 | 92 |
| SRR7690777 | 0 | 0 | 0 | 0 | 45 | 119 | 34 | 129 | 1041 | 31 | 68 |
| SRR7690778 | 0 | 0 | 0 | 0 | 0 | 16 | 19 | 14 | 122 | 0 | 13 |
| SRR7690779 | 0 | 0 | 15 | 13 | 36 | 45 | 49 | 44 | 2326 | 78 | 204 |
| SRR7690780 | 0 | 0 | 0 | 0 | 34 | 65 | 38 | 58 | 572 | 29 | 47 |
| SRR7690781 | 0 | 0 | 0 | 0 | 70 | 145 | 62 | 97 | 504 | 29 | 59 |
| SRR7690783 | 0 | 0 | 74 | 38 | 680 | 1385 | 487 | 678 | 1290 | 80 | 464 |
| SRR7690788 | 0 | 0 | 39 | 35 | 382 | 1117 | 288 | 768 | 793 | 66 | 282 |
| SRR7690789 | 0 | 10 | 101 | 78 | 1024 | 2087 | 813 | 561 | 855 | 44 | 228 |
| SRR7690790 | 0 | 0 | 0 | 0 | 66 | 102 | 46 | 53 | 608 | 13 | 38 |
| SRR7690791 | 0 | 0 | 0 | 0 | 36 | 80 | 57 | 56 | 673 | 21 | 59 |
| SRR7690792 | 0 | 0 | 0 | 0 | 26 | 73 | 29 | 90 | 751 | 25 | 58 |
| SRR7690793 | 0 | 0 | 0 | 0 | 19 | 32 | 22 | 18 | 102 | 11 | 14 |
| SRR7690794 | 0 | 0 | 0 | 0 | 0 | 24 | 16 | 32 | 168 | 11 | 16 |
| SRR7690795 | 0 | 0 | 0 | 0 | 0 | 0 | 0 | 14 | 82 | 0 | 0 |
| SRR7690796 | 0 | 0 | 0 | 0 | 43 | 103 | 33 | 83 | 374 | 24 | 61 |
| SRR7690797 | 0 | 22 | 103 | 85 | 1147 | 1959 | 801 | 679 | 1708 | 59 | 295 |
| SRR7690798 | 0 | 17 | 39 | 31 | 769 | 2017 | 513 | 822 | 939 | 72 | 340 |
| SRR7690800 | 0 | 0 | 34 | 0 | 458 | 1113 | 247 | 594 | 795 | 89 | 383 |
| SRR7690801 | 0 | 0 | 17 | 13 | 478 | 1844 | 643 | 1091 | 2491 | 207 | 651 |
| SRR7690802 | 0 | 0 | 0 | 0 | 274 | 564 | 235 | 438 | 634 | 74 | 438 |
| SRR7690803 | 0 | 0 | 0 | 23 | 385 | 667 | 269 | 415 | 724 | 43 | 327 |
| SRR7819267 | 0 | 0 | 0 | 0 | 0 | 0 | 0 | 0 | 210 | 92 | 31 |
| SRR7819269 | 0 | 0 | 0 | 0 | 0 | 0 | 0 | 0 | 171 | 114 | 0 |
| SRR7819272 | 0 | 0 | 0 | 0 | 0 | 0 | 20 | 13 | 301 | 79 | 0 |
| SRR7851651 | 0 | 0 | 0 | 0 | 0 | 0 | 0 | 0 | 108 | 22 | 67 |
| SRR7851652 | 0 | 0 | 0 | 0 | 0 | 0 | 0 | 0 | 83 | 16 | 36 |
| SRR7851653 | 0 | 0 | 0 | 0 | 0 | 0 | 0 | 0 | 34 | 0 | 32 |
| SRR7851655 | 0 | 0 | 0 | 0 | 0 | 0 | 0 | 0 | 107 | 29 | 52 |
| SRR7851656 | 0 | 0 | 0 | 0 | 0 | 0 | 0 | 0 | 126 | 30 | 99 |
| SRR7901057 | 0 | 97 | 143 | 0 | 61 | 119 | 38 | 43 | 85 | 0 | 0 |
| SRR7901058 | 0 | 47 | 62 | 0 | 71 | 107 | 48 | 45 | 77 | 0 | 0 |
| SRR7901059 | 0 | 81 | 122 | 0 | 55 | 99 | 38 | 50 | 48 | 0 | 0 |
| SRR7903701 | 0 | 279 | 485 | 166 | 2138 | 4325 | 1923 | 1609 | 9388 | 1191 | 986 |
| SRR7903702 | 0 | 317 | 556 | 140 | 1716 | 3924 | 1490 | 1841 | 9480 | 1273 | 952 |
| SRR7903703 | 0 | 279 | 448 | 153 | 1708 | 3760 | 1684 | 1780 | 9168 | 1262 | 863 |
| SRR7903704 | 0 | 360 | 254 | 14 | 212 | 413 | 125 | 258 | 757 | 35 | 38 |
| SRR7903705 | 0 | 286 | 185 | 21 | 231 | 392 | 139 | 288 | 539 | 31 | 43 |
| SRR7903706 | 0 | 381 | 227 | 16 | 198 | 428 | 120 | 281 | 475 | 33 | 35 |
| SRR8063607 | 0 | 11 | 26 | 22 | 2431 | 9654 | 1764 | 10861 | 3794 | 8437 | 3262 |
| SRR8063608 | 0 | 0 | 29 | 19 | 2326 | 9489 | 1898 | 10900 | 3465 | 8097 | 3338 |
| SRR8063609 | 0 | 0 | 81 | 106 | 2569 | 8544 | 2085 | 9308 | 3296 | 7194 | 3274 |
| SRR8063610 | 0 | 0 | 0 | 0 | 493 | 1105 | 258 | 560 | 390 | 832 | 249 |
| SRR8063611 | 0 | 0 | 0 | 0 | 463 | 982 | 243 | 531 | 385 | 835 | 223 |
| SRR8063612 | 0 | 0 | 0 | 0 | 417 | 852 | 238 | 463 | 341 | 758 | 231 |
| SRR8063613 | 0 | 0 | 0 | 0 | 184 | 251 | 101 | 168 | 285 | 134 | 17 |
| SRR8063614 | 0 | 26 | 30 | 14 | 26 | 82 | 22 | 137 | 70 | 24 | 24 |
| SRR8063615 | 0 | 0 | 0 | 0 | 57 | 139 | 25 | 237 | 123 | 49 | 49 |
| SRR8074870 | 0 | 0 | 0 | 0 | 12 | 13 | 0 | 0 | 503 | 258 | 76 |
| SRR8074871 | 0 | 0 | 0 | 0 | 0 | 18 | 0 | 12 | 495 | 181 | 69 |
| SRR8074873 | 0 | 0 | 0 | 0 | 22 | 48 | 0 | 0 | 800 | 236 | 76 |
| SRR8074874 | 0 | 0 | 0 | 0 | 0 | 30 | 0 | 14 | 608 | 132 | 33 |
| SRR8074875 | 0 | 0 | 0 | 0 | 13 | 27 | 0 | 0 | 578 | 135 | 24 |
| SRR8094756 | 0 | 14 | 58 | 39 | 2710 | 4201 | 4670 | 716 | 276 | 110 | 347 |

### Supp Table S8.5.Columns_ET_to_FD

| | | | | | | | | | | | |
|---|---|---|---|---|---|---|---|---|---|---|---|
| SRR8094757 | 0 | 89 | 335 | 81 | 1938 | 3977 | 1807 | 408 | 60 | 52 | 79 |
| SRR8094758 | 0 | 25 | 159 | 35 | 3372 | 5312 | 2782 | 397 | 62 | 46 | 176 |
| SRR8094759 | 0 | 0 | 0 | 0 | 190 | 284 | 156 | 41 | 0 | 14 | 15 |
| SRR8094760 | 0 | 0 | 0 | 13 | 819 | 1234 | 633 | 240 | 23 | 24 | 84 |
| SRR8094761 | 0 | 0 | 98 | 83 | 4291 | 7617 | 3970 | 1509 | 146 | 146 | 294 |
| SRR8094762 | 0 | 64 | 453 | 587 | 6150 | 12077 | 7446 | 2506 | 30 | 43 | 250 |
| SRR8094763 | 0 | 44 | 221 | 286 | 922 | 2453 | 730 | 687 | 89 | 71 | 302 |
| SRR8094764 | 0 | 20 | 140 | 353 | 3015 | 6755 | 2529 | 2124 | 140 | 167 | 677 |
| SRR8094765 | 0 | 45 | 190 | 273 | 818 | 1730 | 626 | 665 | 70 | 90 | 305 |
| SRR8094766 | 0 | 38 | 160 | 189 | 2011 | 7397 | 2135 | 2427 | 99 | 137 | 469 |
| SRR8094767 | 0 | 23 | 87 | 192 | 1518 | 3949 | 1501 | 1745 | 326 | 413 | 913 |
| SRR8094768 | 0 | 22 | 395 | 926 | 4356 | 9944 | 3877 | 3215 | 229 | 155 | 857 |
| SRR8094769 | 0 | 32 | 294 | 400 | 2804 | 9922 | 3110 | 3143 | 217 | 323 | 1603 |
| SRR8094770 | 0 | 22 | 245 | 93 | 9517 | 11489 | 7521 | 2062 | 33627 | 41011 | 29256 |
| SRR8137033 | 0 | 263 | 365 | 182 | 1864 | 3404 | 871 | 567 | 1013 | 993 | 973 |
| SRR8137034 | 0 | 307 | 466 | 85 | 374 | 627 | 266 | 167 | 554 | 237 | 190 |
| SRR8137035 | 0 | 822 | 938 | 75 | 1059 | 2114 | 428 | 79 | 656 | 116 | 71 |
| SRR8137036 | 0 | 412 | 433 | 171 | 1016 | 2766 | 513 | 292 | 1318 | 251 | 221 |
| SRR8137037 | 0 | 720 | 1689 | 203 | 1349 | 2089 | 524 | 504 | 995 | 530 | 557 |
| SRR8203922 | 0 | 0 | 0 | 0 | 0 | 0 | 0 | 0 | 0 | 16 | 0 |
| SRR8203923 | 0 | 0 | 0 | 0 | 0 | 0 | 11 | 11 | 0 | 20 | 0 |
| SRR8203924 | 0 | 0 | 0 | 0 | 0 | 0 | 0 | 0 | 0 | 22 | 0 |
| SRR8203925 | 0 | 0 | 0 | 0 | 0 | 0 | 0 | 0 | 0 | 0 | 0 |
| SRR8203926 | 0 | 0 | 0 | 0 | 0 | 13 | 0 | 21 | 0 | 23 | 0 |
| SRR8203927 | 0 | 0 | 14 | 0 | 11 | 16 | 0 | 11 | 14 | 33 | 0 |
| SRR8203928 | 0 | 0 | 0 | 0 | 0 | 26 | 0 | 16 | 0 | 32 | 0 |
| SRR8203929 | 0 | 0 | 11 | 0 | 0 | 0 | 0 | 19 | 0 | 19 | 0 |
| SRR8203930 | 0 | 0 | 0 | 0 | 0 | 0 | 0 | 0 | 0 | 36 | 0 |
| SRR8203931 | 0 | 0 | 0 | 0 | 0 | 0 | 0 | 10 | 0 | 22 | 0 |
| SRR8203932 | 0 | 0 | 0 | 0 | 0 | 0 | 0 | 13 | 0 | 30 | 0 |
| SRR8203933 | 0 | 0 | 0 | 0 | 0 | 0 | 0 | 10 | 12 | 27 | 0 |
| SRR8203934 | 0 | 0 | 0 | 0 | 0 | 0 | 0 | 0 | 0 | 0 | 0 |
| SRR8203935 | 0 | 0 | 0 | 0 | 0 | 0 | 0 | 11 | 0 | 0 | 0 |
| SRR8203936 | 0 | 0 | 0 | 0 | 0 | 0 | 0 | 12 | 0 | 13 | 0 |
| SRR8203937 | 0 | 0 | 0 | 0 | 0 | 0 | 0 | 0 | 0 | 0 | 0 |
| SRR8203938 | 0 | 0 | 0 | 0 | 0 | 0 | 0 | 11 | 0 | 20 | 0 |
| SRR8203939 | 0 | 0 | 0 | 0 | 0 | 0 | 0 | 13 | 0 | 23 | 0 |
| SRR8203940 | 0 | 0 | 0 | 0 | 0 | 0 | 0 | 0 | 0 | 16 | 0 |
| SRR8203941 | 0 | 0 | 0 | 0 | 0 | 0 | 0 | 0 | 0 | 30 | 0 |
| SRR8203942 | 0 | 0 | 0 | 0 | 0 | 0 | 0 | 0 | 0 | 0 | 0 |
| SRR8203943 | 0 | 0 | 0 | 0 | 0 | 0 | 0 | 0 | 11 | 0 | 11 |
| SRR8203944 | 0 | 0 | 0 | 0 | 0 | 0 | 0 | 0 | 16 | 0 | 0 |
| SRR8203945 | 0 | 0 | 11 | 11 | 0 | 0 | 0 | 0 | 0 | 16 | 0 |
| SRR8203946 | 0 | 0 | 0 | 0 | 0 | 0 | 0 | 0 | 12 | 12 | 0 |
| SRR8203947 | 0 | 0 | 0 | 0 | 0 | 0 | 0 | 0 | 10 | 10 | 0 |
| SRR8203948 | 0 | 0 | 0 | 0 | 0 | 0 | 0 | 0 | 12 | 29 | 0 |
| SRR8203949 | 0 | 0 | 0 | 0 | 0 | 0 | 0 | 0 | 21 | 14 | 0 |
| SRR8248787 | 631 | 4049 | 4814 | 491 | 1988 | 3486 | 1537 | 1098 | 4381 | 1530 | 1241 |
| SRR8248788 | 64 | 1859 | 3536 | 393 | 5019 | 8342 | 3691 | 2177 | 8922 | 3335 | 3128 |
| SRR8248789 | 114 | 3353 | 4880 | 406 | 4259 | 7063 | 3000 | 1625 | 7263 | 2530 | 2489 |
| SRR8248790 | 2393 | 3934 | 3998 | 199 | 1152 | 1526 | 674 | 292 | 1400 | 321 | 338 |
| SRR8248791 | 37 | 549 | 1027 | 86 | 4668 | 8311 | 4062 | 2136 | 14756 | 4056 | 4222 |
| SRR8248792 | 34 | 1134 | 1783 | 150 | 8718 | 13916 | 5483 | 1869 | 7913 | 1772 | 1628 |
| SRR8272058 | 0 | 0 | 0 | 0 | 0 | 0 | 0 | 28 | 717 | 358 | 305 |
| SRR8280405 | 0 | 0 | 0 | 0 | 0 | 0 | 0 | 0 | 0 | 0 | 0 |
| SRR8280408 | 0 | 0 | 0 | 0 | 0 | 0 | 0 | 0 | 0 | 0 | 0 |
| SRR8280414 | 0 | 0 | 0 | 0 | 0 | 0 | 0 | 0 | 0 | 0 | 0 |
| SRR8280415 | 0 | 0 | 0 | 0 | 0 | 0 | 0 | 0 | 0 | 0 | 0 |
| SRR8280416 | 0 | 49 | 49 | 0 | 0 | 0 | 0 | 0 | 0 | 0 | 0 |
| SRR8280417 | 0 | 0 | 0 | 0 | 0 | 0 | 0 | 0 | 0 | 0 | 0 |
| SRR8280423 | 0 | 0 | 0 | 0 | 0 | 0 | 0 | 0 | 0 | 0 | 0 |
| SRR8280425 | 0 | 0 | 0 | 0 | 0 | 0 | 0 | 0 | 0 | 0 | 0 |
| SRR8280432 | 0 | 37 | 0 | 0 | 0 | 0 | 0 | 0 | 0 | 0 | 0 |
| SRR8280440 | 0 | 0 | 0 | 0 | 0 | 0 | 0 | 0 | 0 | 0 | 0 |
| SRR8280442 | 0 | 18 | 0 | 0 | 0 | 0 | 0 | 0 | 0 | 0 | 0 |
| SRR8280452 | 0 | 0 | 0 | 90 | 0 | 0 | 0 | 0 | 0 | 0 | 0 |
| SRR8280460 | 0 | 14 | 0 | 43 | 0 | 0 | 0 | 0 | 0 | 0 | 0 |
| SRR8280461 | 0 | 0 | 0 | 0 | 0 | 0 | 0 | 0 | 0 | 0 | 0 |
| SRR8280463 | 0 | 0 | 0 | 0 | 0 | 0 | 0 | 0 | 0 | 0 | 0 |
| SRR8280464 | 0 | 0 | 0 | 0 | 0 | 0 | 0 | 0 | 0 | 0 | 0 |
| SRR8280467 | 0 | 25 | 0 | 0 | 0 | 0 | 0 | 0 | 0 | 0 | 0 |
| SRR8280468 | 0 | 0 | 0 | 0 | 0 | 0 | 0 | 0 | 0 | 0 | 0 |
| SRR8280472 | 0 | 0 | 0 | 0 | 0 | 0 | 0 | 0 | 0 | 0 | 0 |
| SRR8280473 | 0 | 0 | 0 | 0 | 0 | 0 | 0 | 0 | 0 | 0 | 0 |
| SRR8280476 | 0 | 0 | 0 | 0 | 0 | 0 | 0 | 0 | 0 | 0 | 0 |
| SRR8280479 | 0 | 0 | 0 | 0 | 0 | 0 | 0 | 0 | 0 | 0 | 0 |
| SRR8280481 | 0 | 0 | 0 | 0 | 0 | 0 | 0 | 0 | 0 | 0 | 0 |
| SRR8280483 | 0 | 0 | 0 | 0 | 0 | 0 | 0 | 0 | 0 | 0 | 0 |
| SRR8280485 | 0 | 0 | 0 | 0 | 0 | 0 | 0 | 0 | 0 | 0 | 0 |
| SRR8280487 | 0 | 0 | 0 | 0 | 0 | 0 | 0 | 0 | 0 | 0 | 0 |
| SRR8280490 | 0 | 0 | 0 | 0 | 0 | 0 | 0 | 0 | 0 | 0 | 0 |
| SRR8280494 | 0 | 0 | 0 | 0 | 0 | 0 | 0 | 0 | 0 | 0 | 0 |
| SRR8280497 | 0 | 0 | 11 | 0 | 0 | 0 | 0 | 0 | 0 | 0 | 0 |
| SRR8280498 | 0 | 0 | 0 | 0 | 0 | 0 | 0 | 0 | 0 | 0 | 0 |
| SRR8280500 | 0 | 0 | 0 | 0 | 0 | 0 | 0 | 0 | 0 | 0 | 0 |
| SRR8280501 | 0 | 0 | 0 | 0 | 0 | 0 | 0 | 0 | 0 | 0 | 0 |
| SRR8280502 | 0 | 0 | 0 | 0 | 0 | 0 | 0 | 0 | 0 | 0 | 0 |
| SRR8280503 | 0 | 0 | 0 | 0 | 0 | 0 | 0 | 0 | 0 | 0 | 0 |
| SRR8280504 | 0 | 0 | 0 | 0 | 0 | 0 | 0 | 0 | 0 | 0 | 0 |
| SRR8280505 | 0 | 0 | 0 | 0 | 0 | 0 | 0 | 0 | 0 | 0 | 0 |
| SRR8280506 | 0 | 0 | 0 | 0 | 0 | 0 | 0 | 0 | 0 | 0 | 0 |
| SRR8280507 | 0 | 0 | 0 | 0 | 0 | 0 | 0 | 0 | 0 | 0 | 0 |
| SRR8280509 | 0 | 0 | 0 | 0 | 0 | 0 | 0 | 0 | 0 | 0 | 0 |
| SRR8280510 | 0 | 0 | 0 | 0 | 0 | 0 | 0 | 0 | 0 | 0 | 0 |
| SRR8280511 | 0 | 0 | 0 | 0 | 0 | 0 | 0 | 0 | 0 | 0 | 0 |
| SRR8280512 | 0 | 0 | 0 | 0 | 0 | 0 | 0 | 0 | 0 | 0 | 0 |
| SRR8280513 | 0 | 0 | 0 | 0 | 0 | 0 | 0 | 0 | 0 | 0 | 0 |
| SRR8280515 | 0 | 0 | 0 | 0 | 0 | 0 | 0 | 0 | 0 | 0 | 0 |
| SRR8280519 | 0 | 0 | 0 | 0 | 0 | 0 | 0 | 0 | 0 | 0 | 0 |
| SRR8280521 | 0 | 0 | 0 | 0 | 0 | 0 | 0 | 0 | 0 | 0 | 0 |
| SRR8280525 | 0 | 0 | 0 | 0 | 0 | 0 | 0 | 0 | 0 | 0 | 0 |
| SRR8280531 | 0 | 0 | 0 | 0 | 0 | 0 | 0 | 0 | 0 | 0 | 0 |
| SRR8280533 | 0 | 0 | 0 | 0 | 0 | 0 | 0 | 0 | 0 | 0 | 0 |
| SRR8280536 | 0 | 0 | 0 | 0 | 0 | 0 | 0 | 0 | 0 | 0 | 0 |
| SRR8280539 | 0 | 0 | 0 | 0 | 0 | 0 | 0 | 0 | 0 | 0 | 0 |
| SRR8280541 | 0 | 0 | 0 | 0 | 0 | 0 | 0 | 0 | 0 | 0 | 0 |
| SRR8280543 | 0 | 0 | 0 | 0 | 0 | 0 | 0 | 0 | 0 | 0 | 0 |
| SRR8280544 | 0 | 0 | 0 | 0 | 0 | 0 | 0 | 0 | 63 | 0 | 0 |
| SRR8280546 | 0 | 0 | 0 | 0 | 0 | 0 | 0 | 0 | 0 | 0 | 0 |
| SRR8280548 | 0 | 0 | 0 | 0 | 0 | 0 | 0 | 0 | 0 | 0 | 0 |
| SRR8280550 | 0 | 0 | 0 | 0 | 0 | 0 | 0 | 0 | 0 | 0 | 0 |
| SRR8280551 | 0 | 0 | 0 | 0 | 0 | 0 | 0 | 0 | 0 | 0 | 0 |
| SRR8280552 | 0 | 0 | 0 | 0 | 0 | 0 | 0 | 0 | 0 | 0 | 0 |
| SRR8280554 | 0 | 0 | 0 | 0 | 0 | 0 | 0 | 0 | 0 | 0 | 0 |
| SRR8280558 | 0 | 0 | 0 | 0 | 0 | 0 | 0 | 0 | 0 | 0 | 0 |
| SRR8280560 | 0 | 0 | 0 | 0 | 0 | 0 | 0 | 0 | 0 | 0 | 0 |
| SRR8280561 | 0 | 0 | 0 | 0 | 0 | 0 | 0 | 0 | 0 | 0 | 0 |
| SRR8280562 | 0 | 0 | 0 | 0 | 0 | 0 | 0 | 0 | 0 | 0 | 0 |
| SRR8280564 | 0 | 0 | 0 | 0 | 0 | 0 | 0 | 0 | 0 | 0 | 0 |

### Supp Table S8.5.Columns_ET_to_FD

| | | | | | | | | | | | |
|---|---|---|---|---|---|---|---|---|---|---|---|
| SRR8280566 | 0 | 52 | 0 | 0 | 0 | 0 | 0 | 0 | 0 | 0 | 0 |
| SRR8280568 | 0 | 0 | 0 | 0 | 0 | 0 | 0 | 0 | 0 | 0 | 0 |
| SRR8280572 | 0 | 0 | 0 | 0 | 0 | 43 | 0 | 0 | 0 | 0 | 0 |
| SRR8280573 | 0 | 0 | 0 | 0 | 0 | 0 | 0 | 0 | 0 | 0 | 0 |
| SRR8280574 | 0 | 0 | 0 | 0 | 0 | 0 | 0 | 0 | 0 | 0 | 0 |
| SRR8280576 | 0 | 0 | 0 | 0 | 0 | 0 | 0 | 0 | 0 | 0 | 0 |
| SRR8280580 | 0 | 0 | 0 | 0 | 0 | 0 | 0 | 0 | 0 | 0 | 0 |
| SRR8280582 | 0 | 0 | 0 | 0 | 0 | 0 | 0 | 0 | 0 | 0 | 0 |
| SRR8280583 | 0 | 0 | 0 | 0 | 0 | 0 | 0 | 0 | 0 | 0 | 0 |
| SRR8280584 | 0 | 0 | 0 | 0 | 0 | 0 | 0 | 0 | 0 | 0 | 0 |
| SRR8280589 | 0 | 0 | 0 | 0 | 0 | 0 | 0 | 0 | 0 | 0 | 0 |
| SRR8280590 | 0 | 0 | 0 | 0 | 0 | 0 | 0 | 0 | 0 | 0 | 0 |
| SRR8280592 | 0 | 0 | 0 | 0 | 0 | 0 | 0 | 0 | 0 | 0 | 0 |
| SRR8280593 | 0 | 0 | 0 | 0 | 0 | 0 | 0 | 0 | 0 | 0 | 0 |
| SRR8311061 | 0 | 0 | 0 | 0 | 0 | 0 | 0 | 0 | 0 | 0 | 0 |
| SRR8311062 | 0 | 0 | 0 | 0 | 0 | 0 | 0 | 0 | 0 | 0 | 0 |
| SRR8311063 | 0 | 0 | 0 | 0 | 0 | 0 | 0 | 0 | 0 | 0 | 0 |
| SRR8311064 | 0 | 0 | 0 | 0 | 0 | 0 | 0 | 0 | 0 | 0 | 0 |
| SRR8311070 | 0 | 0 | 0 | 0 | 0 | 0 | 0 | 0 | 0 | 0 | 0 |
| SRR8311071 | 0 | 0 | 0 | 0 | 0 | 0 | 0 | 0 | 0 | 0 | 0 |
| SRR8311072 | 0 | 0 | 0 | 0 | 0 | 0 | 0 | 0 | 0 | 0 | 0 |
| SRR8311073 | 0 | 0 | 0 | 0 | 0 | 0 | 0 | 0 | 0 | 0 | 0 |
| SRR8311074 | 0 | 0 | 0 | 0 | 0 | 0 | 0 | 0 | 0 | 0 | 0 |
| SRR8311076 | 0 | 0 | 0 | 0 | 0 | 0 | 0 | 0 | 0 | 0 | 0 |
| SRR8311078 | 0 | 0 | 0 | 0 | 0 | 0 | 0 | 0 | 0 | 0 | 0 |
| SRR8311080 | 0 | 0 | 0 | 0 | 0 | 0 | 0 | 0 | 0 | 0 | 0 |
| SRR8311081 | 0 | 0 | 0 | 0 | 0 | 0 | 0 | 0 | 0 | 0 | 0 |
| SRR8311082 | 0 | 0 | 0 | 0 | 0 | 0 | 0 | 0 | 0 | 76 | 0 |
| SRR8311085 | 0 | 0 | 0 | 0 | 0 | 0 | 0 | 0 | 0 | 0 | 0 |
| SRR8311087 | 0 | 0 | 0 | 0 | 0 | 0 | 0 | 0 | 0 | 0 | 0 |
| SRR8311089 | 0 | 0 | 0 | 0 | 0 | 0 | 0 | 0 | 0 | 0 | 0 |
| SRR8311092 | 0 | 0 | 0 | 0 | 0 | 0 | 0 | 0 | 0 | 0 | 0 |
| SRR8311093 | 0 | 0 | 0 | 0 | 0 | 0 | 0 | 0 | 0 | 0 | 0 |
| SRR8311095 | 0 | 0 | 0 | 0 | 0 | 0 | 0 | 0 | 0 | 0 | 0 |
| SRR8311098 | 0 | 0 | 0 | 0 | 0 | 0 | 0 | 0 | 0 | 0 | 0 |
| SRR8311105 | 0 | 0 | 0 | 0 | 0 | 0 | 0 | 0 | 0 | 0 | 0 |
| SRR8311106 | 0 | 0 | 0 | 0 | 0 | 0 | 0 | 0 | 0 | 0 | 0 |
| SRR835324 | 0 | 0 | 0 | 0 | 18 | 69 | 20 | 50 | 32 | 13 | 40 |
| SRR835325 | 0 | 0 | 0 | 0 | 40 | 123 | 34 | 91 | 74 | 14 | 46 |
| SRR8354354 | 0 | 0 | 0 | 0 | 0 | 0 | 0 | 0 | 0 | 0 | 0 |
| SRR837775 | 0 | 26 | 41 | 23 | 112 | 1304 | 310 | 2036 | 2984 | 2831 | 3478 |
| SRR837776 | 0 | 0 | 18 | 15 | 84 | 969 | 161 | 2576 | 4418 | 4470 | 9436 |
| SRR837777 | 35 | 0 | 0 | 0 | 0 | 19 | 0 | 46 | 445 | 478 | 285 |
| SRR837778 | 0 | 0 | 0 | 0 | 0 | 22 | 0 | 30 | 144 | 79 | 144 |
| SRR837779 | 0 | 0 | 18 | 15 | 84 | 969 | 161 | 2576 | 4418 | 4470 | 9436 |
| SRR837780 | 0 | 26 | 41 | 23 | 112 | 1304 | 310 | 2036 | 2984 | 2831 | 3478 |
| SRR8380550 | 0 | 0 | 0 | 0 | 0 | 0 | 0 | 0 | 0 | 0 | 0 |
| SRR8380552 | 0 | 0 | 0 | 0 | 125 | 293 | 80 | 143 | 135 | 18 | 35 |
| SRR8380553 | 0 | 0 | 0 | 0 | 94 | 277 | 80 | 175 | 120 | 15 | 42 |
| SRR8380554 | 0 | 0 | 0 | 0 | 0 | 0 | 0 | 0 | 0 | 0 | 0 |
| SRR8380555 | 0 | 0 | 0 | 0 | 0 | 15 | 0 | 20 | 50 | 0 | 10 |
| SRR8380722 | 0 | 30 | 98 | 62 | 529 | 1035 | 312 | 124 | 123 | 86 | 45 |
| SRR8380723 | 0 | 0 | 0 | 0 | 193 | 502 | 167 | 132 | 132 | 243 | 61 |
| SRR8380724 | 0 | 29 | 96 | 109 | 285 | 458 | 173 | 115 | 67 | 22 | 22 |
| SRR8380725 | 0 | 0 | 0 | 0 | 147 | 226 | 96 | 173 | 236 | 161 | 104 |
| SRR8380726 | 0 | 0 | 0 | 0 | 231 | 441 | 199 | 114 | 292 | 91 | 67 |
| SRR8380727 | 0 | 0 | 0 | 0 | 112 | 267 | 99 | 92 | 125 | 142 | 54 |
| SRR8400006 | 0 | 0 | 0 | 0 | 0 | 14 | 13 | 44 | 108 | 184 | 730 |
| SRR8400007 | 0 | 0 | 0 | 0 | 0 | 0 | 0 | 0 | 257 | 329 | 2143 |
| SRR8400008 | 0 | 0 | 0 | 0 | 0 | 22 | 20 | 102 | 1145 | 1626 | 5178 |
| SRR8400009 | 0 | 0 | 0 | 0 | 0 | 0 | 0 | 18 | 39 | 32 | 548 |
| SRR8400010 | 0 | 0 | 0 | 0 | 0 | 23 | 26 | 55 | 120 | 228 | 2413 |
| SRR8417876 | 33 | 140 | 368 | 138 | 101 | 563 | 81 | 882 | 7035 | 6780 | 4300 |
| SRR8417877 | 25 | 94 | 250 | 111 | 47 | 298 | 50 | 373 | 2145 | 1905 | 1812 |
| SRR8417878 | 0 | 32 | 98 | 47 | 41 | 197 | 31 | 272 | 1140 | 1149 | 1740 |
| SRR8435171 | 0 | 0 | 0 | 0 | 30 | 143 | 122 | 135 | 120 | 346 | 195 |
| SRR8435172 | 0 | 0 | 0 | 0 | 15 | 57 | 40 | 43 | 17 | 52 | 44 |
| SRR8481810 | 0 | 21 | 49 | 24 | 28 | 388 | 73 | 461 | 752 | 422 | 608 |
| SRR8481811 | 0 | 0 | 16 | 16 | 0 | 126 | 17 | 207 | 239 | 205 | 239 |
| SRR8481812 | 0 | 12 | 28 | 38 | 14 | 239 | 33 | 363 | 579 | 455 | 579 |
| SRR8481813 | 0 | 0 | 0 | 0 | 0 | 21 | 0 | 47 | 53 | 87 | 54 |
| SRR8481814 | 0 | 0 | 31 | 0 | 21 | 288 | 35 | 390 | 598 | 359 | 458 |
| SRR8481815 | 0 | 0 | 27 | 20 | 16 | 158 | 27 | 268 | 336 | 299 | 364 |
| SRR8481816 | 0 | 35 | 99 | 63 | 28 | 497 | 61 | 568 | 480 | 392 | 403 |
| SRR8481817 | 0 | 0 | 16 | 0 | 0 | 40 | 0 | 64 | 72 | 105 | 76 |
| SRR8481818 | 0 | 40 | 76 | 73 | 11 | 211 | 28 | 322 | 308 | 242 | 526 |
| SRR8481819 | 0 | 47 | 119 | 157 | 30 | 376 | 74 | 480 | 611 | 451 | 1061 |
| SRR8481820 | 0 | 107 | 205 | 183 | 49 | 415 | 54 | 639 | 464 | 469 | 1161 |
| SRR8481821 | 0 | 14 | 20 | 11 | 0 | 26 | 0 | 51 | 45 | 81 | 90 |
| SRR8485055 | 0 | 12 | 0 | 17 | 35 | 89 | 45 | 91 | 516 | 25 | 106 |
| SRR8485056 | 0 | 12 | 0 | 18 | 91 | 118 | 92 | 79 | 278 | 27 | 122 |
| SRR8485057 | 0 | 16 | 0 | 21 | 34 | 59 | 36 | 92 | 1014 | 28 | 138 |
| SRR8485058 | 0 | 0 | 0 | 0 | 24 | 28 | 22 | 47 | 804 | 19 | 70 |
| SRR8485059 | 0 | 38 | 14 | 59 | 60 | 108 | 84 | 170 | 1259 | 43 | 305 |
| SRR8485060 | 0 | 0 | 0 | 17 | 25 | 35 | 38 | 64 | 733 | 19 | 76 |
| SRR8485061 | 0 | 0 | 0 | 19 | 55 | 97 | 42 | 108 | 250 | 26 | 183 |
| SRR8485062 | 0 | 11 | 0 | 23 | 58 | 75 | 37 | 84 | 282 | 12 | 114 |
| SRR8485063 | 0 | 14 | 0 | 19 | 61 | 97 | 50 | 117 | 698 | 30 | 238 |
| SRR8485064 | 0 | 13 | 0 | 23 | 68 | 111 | 66 | 168 | 688 | 41 | 194 |
| SRR8485065 | 41 | 0 | 0 | 19 | 23 | 37 | 23 | 50 | 186 | 14 | 58 |
| SRR8485066 | 31 | 0 | 0 | 26 | 33 | 48 | 29 | 60 | 225 | 18 | 121 |
| SRR8485067 | 28 | 0 | 0 | 28 | 86 | 143 | 46 | 203 | 208 | 26 | 162 |
| SRR8487217 | 0 | 49 | 17 | 63 | 299 | 632 | 274 | 1038 | 463 | 42 | 288 |
| SRR8487218 | 0 | 15 | 0 | 0 | 128 | 200 | 133 | 156 | 350 | 16 | 59 |
| SRR8487219 | 0 | 56 | 16 | 52 | 164 | 324 | 244 | 412 | 293 | 29 | 184 |
| SRR8487220 | 0 | 0 | 0 | 31 | 95 | 108 | 67 | 136 | 651 | 52 | 262 |
| SRR8487221 | 0 | 10 | 0 | 62 | 56 | 112 | 67 | 170 | 1335 | 58 | 299 |
| SRR8487222 | 0 | 0 | 0 | 52 | 81 | 111 | 80 | 160 | 755 | 53 | 486 |
| SRR8487223 | 0 | 34 | 0 | 55 | 130 | 249 | 172 | 437 | 499 | 62 | 257 |
| SRR8487224 | 0 | 0 | 0 | 0 | 20 | 35 | 16 | 27 | 66 | 0 | 33 |
| SRR8586304 | 0 | 0 | 0 | 0 | 0 | 0 | 0 | 0 | 0 | 0 | 0 |
| SRR871609 | 0 | 0 | 0 | 0 | 0 | 0 | 0 | 0 | 40 | 15 | 531 |
| SRR871671 | 0 | 0 | 0 | 0 | 0 | 0 | 38 | 61 | 30 | 0 | 38 |
| SRR873381 | 0 | 0 | 0 | 0 | 48 | 521 | 62 | 198 | 52 | 48 | 148 |
| SRR873401 | 0 | 0 | 0 | 0 | 81 | 1029 | 157 | 932 | 90 | 52 | 205 |
| SRR873410 | 0 | 0 | 0 | 0 | 19 | 428 | 65 | 803 | 49 | 30 | 64 |
| SRR8834226 | 0 | 0 | 0 | 0 | 0 | 0 | 0 | 0 | 0 | 0 | 0 |
| SRR8834227 | 0 | 0 | 0 | 0 | 0 | 0 | 0 | 0 | 0 | 0 | 0 |
| SRR8839811 | 0 | 0 | 0 | 0 | 0 | 0 | 0 | 0 | 0 | 0 | 0 |
| SRR8839812 | 0 | 0 | 0 | 0 | 0 | 18 | 0 | 0 | 0 | 0 | 0 |
| SRR8850891 | 0 | 381 | 830 | 100 | 465 | 602 | 412 | 250 | 1435 | 315 | 415 |
| SRR8850892 | 0 | 475 | 1031 | 181 | 690 | 1033 | 639 | 396 | 2603 | 587 | 518 |
| SRR8850893 | 0 | 590 | 1277 | 260 | 2512 | 2703 | 2341 | 899 | 2379 | 629 | 495 |
| SRR8850894 | 0 | 691 | 1307 | 200 | 1766 | 2057 | 1807 | 769 | 2032 | 511 | 462 |
| SRR8850895 | 0 | 291 | 494 | 81 | 426 | 658 | 380 | 282 | 1470 | 434 | 470 |
| SRR8850896 | 0 | 394 | 447 | 66 | 306 | 483 | 251 | 176 | 962 | 211 | 250 |
| SRR8850897 | 0 | 674 | 1445 | 149 | 1409 | 1308 | 1184 | 286 | 756 | 220 | 278 |
| SRR8850898 | 0 | 375 | 566 | 163 | 839 | 1114 | 1028 | 427 | 1168 | 331 | 256 |
| SRR8850899 | 0 | 182 | 363 | 125 | 626 | 1201 | 543 | 538 | 2579 | 613 | 445 |

### Supp Table S8.5.Columns_ET_to_FD

| | | | | | | | | | | | |
|---|---|---|---|---|---|---|---|---|---|---|---|
| SRR8850900 | 0 | 195 | 245 | 77 | 404 | 1214 | 328 | 656 | 2282 | 785 | 333 |
| SRR8850901 | 0 | 313 | 542 | 126 | 1083 | 1683 | 1029 | 687 | 1863 | 533 | 304 |
| SRR8850902 | 0 | 97 | 195 | 61 | 879 | 1562 | 901 | 705 | 1889 | 582 | 260 |
| SRR8858246 | 0 | 0 | 0 | 0 | 0 | 16 | 0 | 3879 | 1022 | 534 | 920 |
| SRR8858247 | 0 | 0 | 0 | 0 | 0 | 33 | 92 | 5618 | 242 | 67 | 375 |
| SRR8858248 | 0 | 0 | 0 | 0 | 0 | 34 | 0 | 1282 | 740 | 266 | 600 |
| SRR8858254 | 0 | 0 | 0 | 0 | 0 | 0 | 0 | 0 | 185 | 0 | 290 |
| SRR8858258 | 0 | 0 | 0 | 0 | 0 | 0 | 0 | 82 | 145 | 65 | 226 |
| SRR8907824 | 0 | 0 | 0 | 0 | 18 | 78 | 15 | 0 | 17 | 0 | 18 |
| SRR8932122 | 0 | 0 | 0 | 13 | 0 | 20 | 0 | 42 | 42 | 63 | 57 |
| SRR8932123 | 0 | 0 | 0 | 0 | 0 | 0 | 0 | 10 | 13 | 20 | 20 |
| SRR8932124 | 0 | 0 | 0 | 0 | 0 | 12 | 0 | 20 | 34 | 48 | 49 |
| SRR8932125 | 0 | 0 | 0 | 0 | 0 | 10 | 0 | 20 | 25 | 25 | 24 |
| SRR8944248 | 0 | 0 | 0 | 0 | 0 | 0 | 0 | 0 | 0 | 0 | 0 |
| SRR8944250 | 0 | 0 | 0 | 0 | 0 | 0 | 0 | 0 | 0 | 0 | 0 |
| SRR8944251 | 0 | 0 | 0 | 0 | 0 | 0 | 0 | 0 | 0 | 0 | 0 |
| SRR8944252 | 0 | 0 | 0 | 0 | 0 | 0 | 0 | 0 | 0 | 0 | 0 |
| SRR8944257 | 0 | 0 | 0 | 0 | 0 | 0 | 0 | 0 | 0 | 0 | 0 |
| SRR8944259 | 0 | 0 | 0 | 0 | 0 | 0 | 0 | 0 | 0 | 0 | 0 |
| SRR8956086 | 0 | 0 | 0 | 0 | 14 | 22 | 0 | 16 | 16 | 0 | 0 |
| SRR8956087 | 0 | 0 | 0 | 0 | 14 | 22 | 0 | 18 | 0 | 0 | 0 |
| SRR8956089 | 0 | 0 | 0 | 0 | 25 | 57 | 20 | 44 | 46 | 11 | 0 |
| SRR8956090 | 0 | 0 | 0 | 0 | 57 | 121 | 47 | 34 | 49 | 16 | 0 |
| SRR8956091 | 0 | 0 | 0 | 0 | 43 | 93 | 27 | 30 | 45 | 14 | 0 |
| SRR8992368 | 0 | 0 | 0 | 0 | 0 | 0 | 0 | 0 | 0 | 0 | 0 |
| SRR8992369 | 0 | 0 | 0 | 0 | 0 | 0 | 0 | 0 | 0 | 0 | 0 |
| SRR8992370 | 0 | 0 | 0 | 0 | 0 | 0 | 0 | 0 | 0 | 0 | 0 |
| SRR8992371 | 0 | 0 | 0 | 0 | 0 | 0 | 0 | 0 | 0 | 0 | 0 |
| SRR8992378 | 0 | 0 | 0 | 0 | 0 | 0 | 0 | 0 | 0 | 0 | 0 |
| SRR8992379 | 0 | 0 | 0 | 0 | 0 | 0 | 0 | 0 | 0 | 0 | 0 |
| SRR8992380 | 0 | 0 | 0 | 0 | 0 | 0 | 0 | 0 | 0 | 0 | 0 |
| SRR8992381 | 0 | 0 | 0 | 0 | 0 | 0 | 0 | 0 | 0 | 0 | 0 |
| SRR8992382 | 0 | 0 | 0 | 0 | 0 | 0 | 0 | 0 | 0 | 0 | 0 |
| SRR8992383 | 0 | 0 | 0 | 0 | 0 | 0 | 0 | 0 | 0 | 0 | 0 |
| SRR8992384 | 0 | 0 | 0 | 0 | 0 | 0 | 0 | 0 | 0 | 0 | 0 |
| SRR8992385 | 0 | 0 | 0 | 0 | 0 | 0 | 0 | 0 | 0 | 0 | 0 |
| SRR8992387 | 0 | 0 | 0 | 0 | 0 | 0 | 0 | 0 | 0 | 0 | 0 |
| SRR8992388 | 0 | 0 | 0 | 0 | 0 | 0 | 0 | 0 | 0 | 0 | 0 |
| SRR8992389 | 0 | 0 | 0 | 0 | 0 | 0 | 0 | 0 | 0 | 0 | 0 |
| SRR8992390 | 0 | 0 | 0 | 0 | 0 | 0 | 0 | 0 | 0 | 0 | 0 |
| SRR8992391 | 0 | 0 | 0 | 0 | 0 | 0 | 0 | 0 | 0 | 0 | 0 |
| SRR9003330 | 0 | 190 | 501 | 298 | 1531 | 2503 | 1416 | 1841 | 3969 | 2027 | 2357 |
| SRR9003332 | 0 | 0 | 0 | 0 | 0 | 0 | 0 | 392 | 3366 | 1984 | 1690 |
| SRR9003333 | 0 | 28 | 74 | 90 | 2161 | 3039 | 1931 | 1889 | 5574 | 2822 | 2750 |
| SRR9003334 | 0 | 0 | 0 | 0 | 0 | 0 | 0 | 0 | 17605 | 8405 | 6970 |
| SRR9003336 | 0 | 13 | 38 | 70 | 2615 | 3684 | 2504 | 2685 | 5998 | 3074 | 2811 |
| SRR9003338 | 0 | 0 | 0 | 0 | 0 | 0 | 0 | 13 | 21574 | 12337 | 8863 |
| SRR9003339 | 18 | 35 | 143 | 195 | 2570 | 3795 | 2509 | 2564 | 5449 | 2882 | 2772 |
| SRR9003341 | 0 | 0 | 0 | 0 | 46 | 46 | 46 | 46 | 18606 | 10122 | 7861 |
| SRR9003342 | 0 | 17 | 35 | 45 | 2341 | 3338 | 2032 | 2133 | 5089 | 2588 | 3219 |
| SRR9018258 | 0 | 62 | 164 | 143 | 1875 | 5002 | 1425 | 6635 | 1104 | 1953 | 2039 |
| SRR9018259 | 0 | 25 | 54 | 63 | 1666 | 6846 | 730 | 8075 | 823 | 1063 | 2411 |
| SRR9018260 | 360 | 1771 | 958 | 113 | 1977 | 4103 | 2345 | 1147 | 764 | 283 | 122 |
| SRR9018261 | 720 | 320 | 691 | 131 | 1463 | 2842 | 1226 | 867 | 581 | 455 | 390 |
| SRR952248 | 0 | 504 | 752 | 281 | 92 | 811 | 156 | 835 | 2244 | 2830 | 3932 |
| SRR952249 | 0 | 346 | 501 | 215 | 71 | 554 | 125 | 619 | 2300 | 2939 | 3394 |
| SRR952288 | 0 | 316 | 498 | 241 | 70 | 553 | 138 | 616 | 2335 | 2947 | 3390 |
| SRR952289 | 0 | 94 | 199 | 94 | 234 | 1249 | 241 | 1168 | 3029 | 3760 | 5056 |
| SRR952290 | 0 | 95 | 199 | 88 | 230 | 1285 | 231 | 1159 | 3011 | 3758 | 5089 |
| SRR952309 | 0 | 55 | 106 | 61 | 125 | 1069 | 153 | 1785 | 5251 | 7272 | 8426 |
| SRR952310 | 0 | 81 | 163 | 82 | 105 | 774 | 109 | 1129 | 3232 | 4436 | 4713 |
| SRR952311 | 0 | 65 | 134 | 93 | 103 | 825 | 93 | 1465 | 4051 | 5058 | 5421 |
| SRR952312 | 0 | 186 | 375 | 224 | 56 | 434 | 186 | 686 | 3347 | 4453 | 5745 |
| SRR952313 | 0 | 491 | 979 | 400 | 113 | 445 | 151 | 444 | 2538 | 3226 | 4750 |
| SRR952314 | 0 | 437 | 849 | 422 | 191 | 918 | 343 | 918 | 3856 | 5540 | 7288 |
| SRR952315 | 0 | 286 | 842 | 336 | 119 | 742 | 209 | 635 | 3273 | 5026 | 7410 |
| SRR952316 | 0 | 0 | 0 | 0 | 0 | 32 | 66 | 0 | 0 | 0 | 57 |
| SRR952317 | 0 | 23 | 65 | 13 | 0 | 0 | 0 | 12 | 0 | 47 | 29 |
| SRR952318 | 0 | 20 | 27 | 12 | 0 | 0 | 0 | 0 | 0 | 14 | 76 |
| SRR952319 | 0 | 0 | 0 | 0 | 0 | 0 | 0 | 0 | 41 | 60 | 0 |
| SRR9588764 | 0 | 0 | 0 | 0 | 0 | 0 | 0 | 0 | 23 | 13 | 0 |
| SRR9588765 | 0 | 0 | 0 | 0 | 0 | 0 | 0 | 15 | 71 | 11 | 0 |
| SRR9588766 | 0 | 0 | 0 | 0 | 0 | 0 | 0 | 0 | 67 | 0 | 0 |
| SRR9588767 | 0 | 18 | 0 | 0 | 12 | 20 | 0 | 43 | 250 | 32 | 33 |
| SRR9618218 | 0 | 43 | 136 | 165 | 30 | 244 | 24 | 626 | 87 | 41 | 295 |
| SRR9618219 | 0 | 0 | 34 | 101 | 176 | 958 | 396 | 2141 | 718 | 185 | 1468 |
| SRR9618220 | 0 | 80 | 221 | 337 | 37 | 408 | 114 | 1176 | 114 | 40 | 304 |
| SRR9618221 | 0 | 0 | 67 | 72 | 19 | 199 | 45 | 608 | 91 | 41 | 299 |
| SRR9618222 | 0 | 17 | 107 | 265 | 150 | 813 | 235 | 1267 | 180 | 120 | 788 |
| SRR9627609 | 0 | 0 | 21 | 15 | 0 | 28 | 14 | 87 | 53 | 13 | 28 |
| SRR9627610 | 0 | 0 | 0 | 0 | 0 | 23 | 11 | 50 | 46 | 13 | 33 |
| SRR9627611 | 0 | 0 | 0 | 0 | 0 | 16 | 0 | 39 | 48 | 24 | 18 |
| SRR9627612 | 0 | 0 | 11 | 17 | 0 | 17 | 0 | 49 | 42 | 15 | 29 |
| SRR9627613 | 0 | 0 | 13 | 14 | 0 | 11 | 0 | 32 | 49 | 18 | 40 |
| SRR9627614 | 0 | 0 | 0 | 0 | 0 | 19 | 0 | 65 | 79 | 27 | 61 |
| SRR9627651 | 0 | 0 | 18 | 0 | 21 | 68 | 60 | 137 | 81 | 29 | 39 |
| SRR9627653 | 0 | 0 | 0 | 0 | 16 | 49 | 44 | 136 | 117 | 26 | 40 |
| SRR9627654 | 0 | 0 | 0 | 0 | 23 | 41 | 43 | 95 | 91 | 19 | 64 |
| SRR9627655 | 0 | 0 | 0 | 0 | 0 | 43 | 28 | 76 | 100 | 26 | 65 |
| SRR9627656 | 0 | 0 | 22 | 0 | 13 | 59 | 49 | 143 | 102 | 17 | 40 |
| SRR9627693 | 0 | 14 | 0 | 0 | 16 | 27 | 22 | 57 | 1694 | 13 | 44 |
| SRR9627694 | 0 | 19 | 0 | 0 | 12 | 41 | 19 | 62 | 1402 | 14 | 43 |
| SRR9627695 | 11 | 16 | 0 | 0 | 24 | 29 | 26 | 48 | 1381 | 15 | 36 |
| SRR9627696 | 15 | 27 | 0 | 15 | 0 | 12 | 0 | 34 | 727 | 0 | 26 |
| SRR9627697 | 17 | 26 | 0 | 16 | 0 | 15 | 11 | 29 | 1131 | 0 | 23 |
| SRR9627698 | 19 | 23 | 0 | 22 | 0 | 0 | 0 | 41 | 2125 | 0 | 41 |
| SRR9648387 | 0 | 0 | 0 | 0 | 0 | 11 | 0 | 0 | 0 | 0 | 0 |
| SRR9648388 | 0 | 0 | 0 | 0 | 0 | 0 | 0 | 0 | 0 | 0 | 0 |
| SRR9648389 | 0 | 0 | 0 | 0 | 0 | 0 | 0 | 0 | 0 | 0 | 0 |
| SRR965653 | 0 | 0 | 0 | 0 | 0 | 18 | 0 | 27 | 60 | 16 | 63 |
| SRR965655 | 0 | 86 | 264 | 367 | 108 | 834 | 149 | 1322 | 3517 | 1642 | 7606 |
| SRR9833150 | 0 | 0 | 0 | 0 | 14 | 150 | 34 | 604 | 38 | 40 | 52 |
| SRR9833151 | 0 | 0 | 0 | 0 | 0 | 102 | 18 | 228 | 34 | 26 | 77 |
| SRR9833152 | 0 | 0 | 0 | 0 | 16 | 117 | 27 | 398 | 23 | 24 | 44 |
| SRR9833153 | 0 | 0 | 0 | 0 | 44 | 365 | 44 | 842 | 84 | 86 | 67 |
| SRR9833154 | 0 | 0 | 0 | 0 | 20 | 177 | 32 | 444 | 66 | 53 | 109 |
| SRR9833155 | 0 | 0 | 0 | 0 | 132 | 729 | 71 | 785 | 65 | 67 | 78 |
| SRR9833156 | 0 | 0 | 12 | 0 | 40 | 254 | 28 | 289 | 29 | 29 | 68 |
| SRR9833157 | 0 | 0 | 20 | 13 | 19 | 156 | 24 | 256 | 22 | 19 | 59 |
| SRR9833158 | 0 | 15 | 49 | 19 | 17 | 144 | 16 | 263 | 21 | 23 | 44 |
| SRR9833159 | 0 | 0 | 17 | 13 | 0 | 39 | 0 | 47 | 0 | 0 | 17 |
| SRR9833160 | 0 | 13 | 16 | 11 | 0 | 55 | 0 | 127 | 0 | 0 | 23 |
| SRR9833161 | 0 | 16 | 16 | 11 | 12 | 59 | 0 | 98 | 10 | 0 | 15 |
| SRR9833162 | 0 | 11 | 36 | 23 | 0 | 90 | 0 | 172 | 16 | 12 | 27 |
| SRR9833163 | 0 | 0 | 15 | 0 | 0 | 61 | 0 | 80 | 0 | 11 | 20 |
| SRR9833164 | 0 | 0 | 35 | 29 | 0 | 65 | 0 | 135 | 0 | 0 | 23 |
| SRR9856157 | 0 | 0 | 0 | 18 | 153 | 167 | 107 | 48 | 30 | 22 | 18 |
| SRR9856158 | 0 | 0 | 0 | 0 | 218 | 551 | 162 | 211 | 49 | 89 | 40 |
| SRR9856159 | 0 | 0 | 0 | 24 | 283 | 327 | 147 | 95 | 38 | 31 | 60 |

| | | | | | | | | | | | |
|---|---|---|---|---|---|---|---|---|---|---|---|
| SRR9856160 | 0 | 0 | 0 | 0 | 191 | 619 | 86 | 19 | 102 | 0 | 0 |
| SRR9856161 | 0 | 0 | 0 | 0 | 145 | 305 | 97 | 121 | 30 | 39 | 21 |
| SRR9856162 | 0 | 0 | 0 | 0 | 150 | 398 | 98 | 159 | 70 | 61 | 28 |
| SRR9856163 | 0 | 0 | 0 | 21 | 101 | 287 | 70 | 131 | 28 | 37 | 21 |
| SRR9856164 | 0 | 0 | 0 | 0 | 92 | 210 | 38 | 69 | 24 | 20 | 11 |
| SRR9856165 | 0 | 0 | 0 | 0 | 173 | 360 | 112 | 78 | 34 | 37 | 17 |
| SRR9856166 | 0 | 0 | 0 | 0 | 59 | 100 | 41 | 78 | 22 | 34 | 0 |
| SRR9856167 | 0 | 0 | 14 | 0 | 93 | 301 | 68 | 56 | 19 | 29 | 0 |
| SRR9856168 | 0 | 0 | 0 | 14 | 196 | 478 | 99 | 106 | 45 | 57 | 26 |
| SRR9856169 | 0 | 0 | 0 | 15 | 118 | 324 | 56 | 125 | 31 | 34 | 0 |
| SRR9856170 | 0 | 0 | 0 | 0 | 322 | 841 | 195 | 261 | 54 | 162 | 57 |
| SRR9856171 | 0 | 0 | 0 | 0 | 63 | 138 | 49 | 49 | 16 | 37 | 12 |
| SRR9856172 | 0 | 0 | 0 | 14 | 184 | 441 | 96 | 194 | 74 | 74 | 31 |
| SRR9856173 | 0 | 0 | 0 | 0 | 61 | 160 | 41 | 44 | 28 | 27 | 10 |
| SRR9856174 | 0 | 0 | 12 | 17 | 449 | 1867 | 409 | 194 | 22 | 91 | 32 |
| SRR9856175 | 0 | 0 | 10 | 0 | 194 | 500 | 122 | 84 | 27 | 35 | 12 |
| SRR9856176 | 0 | 0 | 0 | 18 | 170 | 199 | 94 | 41 | 27 | 24 | 18 |
| SRR9856177 | 0 | 0 | 0 | 0 | 85 | 103 | 63 | 28 | 23 | 15 | 0 |
| SRR9856178 | 0 | 0 | 12 | 0 | 108 | 727 | 215 | 59 | 81 | 16 | 12 |
| SRR9856179 | 0 | 0 | 0 | 11 | 89 | 167 | 46 | 51 | 21 | 24 | 16 |
| SRR9856180 | 0 | 0 | 0 | 19 | 123 | 517 | 207 | 181 | 42 | 45 | 0 |
| SRR9856181 | 0 | 0 | 0 | 21 | 265 | 788 | 265 | 160 | 13 | 34 | 13 |
| SRR9856182 | 0 | 0 | 0 | 13 | 119 | 277 | 87 | 52 | 35 | 29 | 0 |
| SRR9856183 | 0 | 0 | 0 | 0 | 40 | 118 | 34 | 27 | 18 | 16 | 0 |
| SRR9856184 | 0 | 0 | 15 | 41 | 148 | 376 | 93 | 79 | 23 | 29 | 14 |
| SRR9856185 | 0 | 0 | 0 | 0 | 210 | 769 | 254 | 44 | 22 | 58 | 15 |
| SRR9856186 | 0 | 0 | 27 | 0 | 482 | 1758 | 889 | 196 | 15 | 34 | 29 |
| SRR990360 | 0 | 0 | 0 | 0 | 0 | 0 | 0 | 0 | 0 | 0 | 12 |
| SRR9924500 | 0 | 39 | 108 | 61 | 21 | 212 | 24 | 350 | 1048 | 1278 | 1460 |
| SRR9924502 | 0 | 0 | 25 | 15 | 11 | 136 | 0 | 196 | 679 | 889 | 884 |
| SRR9924511 | 0 | 23 | 79 | 39 | 14 | 159 | 26 | 257 | 700 | 973 | 1425 |
| SRR9924512 | 0 | 18 | 63 | 33 | 11 | 125 | 18 | 267 | 892 | 1434 | 1403 |
| SRR9924514 | 0 | 17 | 43 | 32 | 11 | 125 | 22 | 167 | 461 | 596 | 839 |
| SRR9924515 | 0 | 14 | 30 | 22 | 0 | 96 | 10 | 158 | 380 | 627 | 826 |
| SRR9924516 | 0 | 0 | 0 | 0 | 0 | 122 | 16 | 242 | 678 | 990 | 1360 |
| SRR9924517 | 0 | 13 | 34 | 22 | 12 | 138 | 22 | 245 | 611 | 860 | 1155 |
| SRR9924518 | 0 | 17 | 37 | 30 | 0 | 115 | 0 | 184 | 554 | 804 | 745 |
| SRR9924519 | 0 | 15 | 37 | 18 | 0 | 70 | 0 | 117 | 301 | 407 | 375 |
| SRR9924520 | 0 | 29 | 90 | 34 | 14 | 143 | 14 | 184 | 670 | 901 | 768 |
| SRR9924521 | 0 | 18 | 44 | 24 | 11 | 93 | 12 | 143 | 458 | 552 | 576 |
| SRR9924522 | 0 | 0 | 15 | 0 | 0 | 104 | 0 | 150 | 574 | 778 | 748 |
| SRR9924523 | 0 | 13 | 35 | 15 | 12 | 86 | 0 | 141 | 567 | 709 | 699 |
| SRR9924524 | 0 | 0 | 34 | 16 | 0 | 48 | 0 | 103 | 250 | 316 | 383 |
| SRR9924525 | 0 | 57 | 161 | 98 | 16 | 141 | 25 | 242 | 879 | 1140 | 1278 |
| SRR9924526 | 0 | 19 | 47 | 28 | 0 | 63 | 14 | 112 | 257 | 357 | 435 |
| SRR9924527 | 0 | 0 | 30 | 17 | 0 | 74 | 0 | 118 | 317 | 400 | 486 |
