## Supplementary material for "Single-nucleotide Differences and Cell Type Decide the Subcellular Localization of miRNA Isoforms (isomiRs), tRNA-derived Fragments (tRFs) and rRNA-derived Fragments (rRFs)": Supp. Table S9 (1 of 4)

**Supplemental Table S9:** The average reconstructed abundances of each of the short RNA in the cell fractions of BT-20, MDA-MB-231, and MDA-MB468 cells.

| Column | Name | Description |
| --- | --- | --- |
| A | Full Name | Full name of short RNA. Includes all instances of RNA when mapped to isomiR, tRF, and rRF space |
| B | Short Name | Shortened name of short RNA |
| C | Start | Start position of short RNA in precursor molecule |
| D | End | End position of short RNA in precursor molecule |
| E | Length | Length of short RNA |
| F | Additional Nucleotides | Additional nucleotides (isomiRs only) |
| G | group | Short RNA group (isomiRs, nuclear tRF, mitochondrial tRF, nuclear rRF 45S, nuclear rRF 5S rRF, mitochondrial rRF 12S & 16S |
| H | Outlier | Short RNA is (YES) or isn't (NO) an outlier in one or more replicates |
| I | AVE NUC | Average abundance of reconstructed short RNA in nuclear fraction |
| J | AVE CYTO | Average abundance of reconstructed short RNA in cytosolic fraction |
| K | AVE MITO | Average abundance of reconstructed short RNA in mitochondrial fraction |
| L | AVE MP | Average abundance of reconstructed short RNA in mitoplast fraction |
