## Supplementary material for "Single-nucleotide Differences and Cell Type Decide the Subcellular Localization of miRNA Isoforms (isomiRs), tRNA-derived Fragments (tRFs) and rRNA-derived Fragments (rRFs)": Supp. Table S9 (2 of 4)

# BT-20

<table>
<tr><th>Full Name</th><th>Short Name</th><th>Start</th><th>End</th><th>Length</th><th>Additional Nucleotides</th><th>group</th><th>Outlier</th><th>AVE NUC</th><th>AVE CYTO</th><th>AVE MITO</th><th>AVE MP</th></tr>
<tr><td>>MI0000061|hsa-let-7a-2&WithFlank&11|-|122146516|122146599@11.28.18;MI0000064|hsa-let-7c&WithFlank&21|+|16539822|16539917@17.34.18;MI0000062|hsa-let-7a-3&WithFlank&22|+|46112743|46112828@10.27.18;MI0000060|hsa-let-7a-1&WithFlank&9|+|94175951|94176042@12.29.18&[MIMAT0000062&hsa-let-7a-5p&offsets|0|-4;m-6&11|-|122146568|122146589&offsets|0|-4];[MIMAT0000064&hsa-let-7c-5p&offsets|0|-4;m-37&21|+|16539838|16539859&offsets|0|-4];[MIMAT0000062_1&hsa-let-7a-5p&offsets|0|-4;m-5&22|+|46112752|46112773&offsets|0|-4];[MIMAT0000062_2&hsa-let-7a-5p&offsets|0|-4;m-4&9|+|94175962|94175983&offsets|0|-4]|TGAGGTAGTAGGTTGTAT</td><td>>MI0000061|hsa-let-7a-2&WithFlank&11|-|122146516|122146599</td><td>11</td><td>28</td><td>18</td><td></td><td>isomiR</td><td>NO</td><td>125.86</td><td>280.03</td><td>77.36</td><td>39.57</td></tr>
<tr><td>>MI0000061|hsa-let-7a-2&WithFlank&11|-|122146516|122146599@11.29.19;MI0000062|hsa-let-7a-3&WithFlank&22|+|46112743|46112828@10.28.19;MI0000060|hsa-let-7a-1&WithFlank&9|+|94175951|94176042@12.30.19&[MIMAT0000062&hsa-let-7a-5p&offsets|0|-3;m-6&11|-|122146568|122146589&offsets|0|-3];[MIMAT0000062_1&hsa-let-7a-5p&offsets|0|-3;m-5&22|+|46112752|46112773&offsets|0|-3];[MIMAT0000062_2&hsa-let-7a-5p&offsets|0|-3;m-4&9|+|94175962|94175983&offsets|0|-3]|TGAGGTAGTAGGTTGTATA</td><td>>MI0000061|hsa-let-7a-2&WithFlank&11|-|122146516|122146599</td><td>11</td><td>29</td><td>19</td><td></td><td>isomiR</td><td>NO</td><td>27.68</td><td>272.23</td><td>0.00</td><td>0.00</td></tr>
<tr><td>>MI0000061|hsa-let-7a-2&WithFlank&11|-|122146516|122146599@11.30.20(+1C);MI0000062|hsa-let-7a-3&WithFlank&22|+|46112743|46112828@10.29.20(+1C);MI0000060|hsa-let-7a-1&WithFlank&9|+|94175951|94176042@12.31.20(+1C)&[MIMAT0000062&hsa-let-7a-5p&offsets|0|-2(+1C);m-6&11|-|122146568|122146589&offsets|0|-2(+1C)];[MIMAT0000062_1&hsa-let-7a-5p&offsets|0|-2(+1C);m-5&22|+|46112752|46112773&offsets|0|-2(+1C)];[MIMAT0000062_2&hsa-let-7a-5p&offsets|0|-2(+1C);m-4&9|+|94175962|94175983&offsets|0|-2(+1C)]|TGAGGTAGTAGGTTGTATAGC</td><td>>MI0000061|hsa-let-7a-2&WithFlank&11|-|122146516|122146599</td><td>11</td><td>30</td><td>20</td><td>(+1C)</td><td>isomiR</td><td>NO</td><td>39.78</td><td>92.26</td><td>0.00</td><td>0.00</td></tr>
<tr><td>>MI0000061|hsa-let-7a-2&WithFlank&11|-|122146516|122146599@11.31.21(+1A);MI0000062|hsa-let-7a-3&WithFlank&22|+|46112743|46112828@10.30.21(+1A);MI0000060|hsa-let-7a-1&WithFlank&9|+|94175951|94176042@12.32.21(+1A)&[MIMAT0000062&hsa-let-7a-5p&offsets|0|-1(+1A);m-6&11|-|122146568|122146589&offsets|0|-1(+1A)];[MIMAT0000062_1&hsa-let-7a-5p&offsets|0|-1(+1A);m-5&22|+|46112752|46112773&offsets|0|-1(+1A)];[MIMAT0000062_2&hsa-let-7a-5p&offsets|0|-1(+1A);m-4&9|+|94175962|94175983&offsets|0|-1(+1A)]|TGAGGTAGTAGGTTGTATAGTA</td><td>>MI0000061|hsa-let-7a-2&WithFlank&11|-|122146516|122146599</td><td>11</td><td>31</td><td>21</td><td>(+1A)</td><td>isomiR</td><td>NO</td><td>94.93</td><td>295.62</td><td>10.96</td><td>28.83</td></tr>
<tr><td>>MI0000061|hsa-let-7a-2&WithFlank&11|-|122146516|122146599@11.31.21(+1C);MI0000062|hsa-let-7a-3&WithFlank&22|+|46112743|46112828@10.30.21(+1C);MI0000060|hsa-let-7a-1&WithFlank&9|+|94175951|94176042@12.32.21(+1C)&[MIMAT0000062&hsa-let-7a-5p&offsets|0|-1(+1C);m-6&11|-|122146568|122146589&offsets|0|-1(+1C)];[MIMAT0000062_1&hsa-let-7a-5p&offsets|0|-1(+1C);m-5&22|+|46112752|46112773&offsets|0|-1(+1C)];[MIMAT0000062_2&hsa-let-7a-5p&offsets|0|-1(+1C);m-4&9|+|94175962|94175983&offsets|0|-1(+1C)]|TGAGGTAGTAGGTTGTATAGTC</td><td>>MI0000061|hsa-let-7a-2&WithFlank&11|-|122146516|122146599</td><td>11</td><td>31</td><td>21</td><td>(+1C)</td><td>isomiR</td><td>NO</td><td>414.90</td><td>889.65</td><td>142.87</td><td>192.55</td></tr>
<tr><td>>MI0000061|hsa-let-7a-2&WithFlank&11|-|122146516|122146599@11.31.21(+1G);MI0000062|hsa-let-7a-3&WithFlank&22|+|46112743|46112828@10.30.21(+1G);MI0000060|hsa-let-7a-1&WithFlank&9|+|94175951|94176042@12.32.21(+1G)&[MIMAT0000062&hsa-let-7a-5p&offsets|0|-1(+1G);m-6&11|-|122146568|122146589&offsets|0|-1(+1G)];[MIMAT0000062_1&hsa-let-7a-5p&offsets|0|-1(+1G);m-5&22|+|46112752|46112773&offsets|0|-1(+1G)];[MIMAT0000062_2&hsa-let-7a-5p&offsets|0|-1(+1G);m-4&9|+|94175962|94175983&offsets|0|-1(+1G)]|TGAGGTAGTAGGTTGTATAGTG</td><td>>MI0000061|hsa-let-7a-2&WithFlank&11|-|122146516|122146599</td><td>11</td><td>31</td><td>21</td><td>(+1G)</td><td>isomiR</td><td>NO</td><td>41.56</td><td>122.11</td><td>0.00</td><td>0.00</td></tr>
<tr><td>>MI0000061|hsa-let-7a-2&WithFlank&11|-|122146516|122146599@11.31.21;MI0000062|hsa-let-7a-3&WithFlank&22|+|46112743|46112828@10.30.21;MI0000060|hsa-let-7a-1&WithFlank&9|+|94175951|94176042@12.32.21&[MIMAT0000062&hsa-let-7a-5p&offsets|0|-1;m-6&11|-|122146568|122146589&offsets|0|-1];[MIMAT0000062_1&hsa-let-7a-5p&offsets|0|-1;m-5&22|+|46112752|46112773&offsets|0|-1];[MIMAT0000062_2&hsa-let-7a-5p&offsets|0|-1;m-4&9|+|94175962|94175983&offsets|0|-1]|TGAGGTAGTAGGTTGTATAGT</td><td>>MI0000061|hsa-let-7a-2&WithFlank&11|-|122146516|122146599</td><td>11</td><td>31</td><td>21</td><td></td><td>isomiR</td><td>NO</td><td>292.99</td><td>642.38</td><td>138.11</td><td>125.07</td></tr>
<tr><td>>MI0000061|hsa-let-7a-2&WithFlank&11|-|122146516|122146599@11.32.22(+1A);MI0000062|hsa-let-7a-3&WithFlank&22|+|46112743|46112828@10.31.22(+1A);MI0000060|hsa-let-7a-1&WithFlank&9|+|94175951|94176042@12.33.22(+1A)&[MIMAT0000062&hsa-let-7a-5p&offsets|0|0(+1A);m-6&11|-|122146568|122146589&offsets|0|0(+1A)];[MIMAT0000062_1&hsa-let-7a-5p&offsets|0|0(+1A);m-5&22|+|46112752|46112773&offsets|0|0(+1A)];[MIMAT0000062_2&hsa-let-7a-5p&offsets|0|0(+1A);m-4&9|+|94175962|94175983&offsets|0|0(+1A)]|TGAGGTAGTAGGTTGTATAGTTA</td><td>>MI0000061|hsa-let-7a-2&WithFlank&11|-|122146516|122146599</td><td>11</td><td>32</td><td>22</td><td>(+1A)</td><td>isomiR</td><td>NO</td><td>71.47</td><td>142.32</td><td>88.87</td><td>0.00</td></tr>
<tr><td>>MI0000061|hsa-let-7a-2&WithFlank&11|-|122146516|122146599@11.32.22;MI0000062|hsa-let-7a-3&WithFlank&22|+|46112743|46112828@10.31.22;MI0000060|hsa-let-7a-1&WithFlank&9|+|94175951|94176042@12.33.22&[MIMAT0000062&hsa-let-7a-5p&offsets|0|0;m-6&11|-|122146568|122146589&offsets|0|0];[MIMAT0000062_1&hsa-let-7a-5p&offsets|0|0;m-5&22|+|46112752|46112773&offsets|0|0];[MIMAT0000062_2&hsa-let-7a-5p&offsets|0|0;m-4&9|+|94175962|94175983&offsets|0|0]|TGAGGTAGTAGGTTGTATAGTT</td><td>>MI0000061|hsa-let-7a-2&WithFlank&11|-|122146516|122146599</td><td>11</td><td>32</td><td>22</td><td></td><td>isomiR</td><td>YES</td><td>1960.49</td><td>5670.71</td><td>769.17</td><td>919.65</td></tr>
<tr><td>>MI0000061|hsa-let-7a-2&WithFlank&11|-|122146516|122146599@11.33.23;MI0000062|hsa-let-7a-3&WithFlank&22|+|46112743|46112828@10.32.23;MI0000060|hsa-let-7a-1&WithFlank&9|+|94175951|94176042@12.34.23&[MIMAT0000062&hsa-let-7a-5p&offsets|0|+1;m-6&11|-|122146568|122146589&offsets|0|+1];[MIMAT0000062_1&hsa-let-7a-5p&offsets|0|+1;m-5&22|+|46112752|46112773&offsets|0|+1];[MIMAT0000062_2&hsa-let-7a-5p&offsets|0|+1;m-4&9|+|94175962|94175983&offsets|0|+1]|TGAGGTAGTAGGTTGTATAGTTT</td><td>>MI0000061|hsa-let-7a-2&WithFlank&11|-|122146516|122146599</td><td>11</td><td>33</td><td>23</td><td></td><td>isomiR</td><td>NO</td><td>96.67</td><td>163.08</td><td>96.78</td><td>28.28</td></tr>
</table>

# BT-20

| | | | | | | | | | | | |
|---|---|---|---|---|---|---|---|---|---|---|---|
| >MI0000062\|hsa-let-7a-3&WithFlank&22\|+\|46112743\|46112828@58.78.21(+1U);MI0000060\|hsa-let-7a-1&WithFlank&9\|+\|94175951\|94176042@63.83.21(+1U)&[MIMAT0004481&hsa-let-7a-3p&offsets\|0\|0(+1U);m-180&22\|+\|46112800\|46112821&offsets\|0\|-1(+1U)];[MIMAT0004481_1&hsa-let-7a-3p&offsets\|0\|0(+1U);m-181&9\|+\|94176013\|94176034&offsets\|0\|-1(+1U)]\|CTATACAATCTACTGTCTTTCT | >MI0000062\|hsa-let-7a-3&WithFlank&22\|+\|46112743\|46112828 | 58 | 78 | 21 | (+1U) | isomiR | NO | 28.65 | 30.63 | 12.05 | 0.00 |
| >MI0000063\|hsa-let-7b&WithFlank&22\|+\|46113680\|46113774@12.29.18&[MIMAT0000063&hsa-let-7b-5p&offsets\|0\|-4;m-14&22\|+\|46113691\|46113712&offsets\|0\|-4]\|TGAGGTAGTAGGTTGTGT | >MI0000063\|hsa-let-7b&WithFlank&22\|+\|46113680\|46113774 | 12 | 29 | 18 | | isomiR | NO | 0.00 | 31.29 | 67.01 | 0.00 |
| >MI0000063\|hsa-let-7b&WithFlank&22\|+\|46113680\|46113774@12.30.19&[MIMAT0000063&hsa-let-7b-5p&offsets\|0\|-3;m-14&22\|+\|46113691\|46113712&offsets\|0\|-3]\|TGAGGTAGTAGGTTGTGTG | >MI0000063\|hsa-let-7b&WithFlank&22\|+\|46113680\|46113774 | 12 | 30 | 19 | | isomiR | NO | 0.00 | 0.00 | 10.66 | 0.00 |
| >MI0000063\|hsa-let-7b&WithFlank&22\|+\|46113680\|46113774@12.30.19(+1A)&[MIMAT0000063&hsa-let-7b-5p&offsets\|0\|-3(+1A);m-14&22\|+\|46113691\|46113712&offsets\|0\|-3(+1A)]\|TGAGGTAGTAGGTTGTGTGA | >MI0000063\|hsa-let-7b&WithFlank&22\|+\|46113680\|46113774 | 12 | 30 | 19 | (+1A) | isomiR | NO | 0.00 | 0.00 | 11.50 | 0.00 |
| >MI0000063\|hsa-let-7b&WithFlank&22\|+\|46113680\|46113774@12.31.20&[MIMAT0000063&hsa-let-7b-5p&offsets\|0\|-2;m-14&22\|+\|46113691\|46113712&offsets\|0\|-2]\|TGAGGTAGTAGGTTGTGTGG | >MI0000063\|hsa-let-7b&WithFlank&22\|+\|46113680\|46113774 | 12 | 31 | 20 | | isomiR | NO | 32.44 | 107.49 | 166.84 | 0.00 |
| >MI0000063\|hsa-let-7b&WithFlank&22\|+\|46113680\|46113774@12.31.20(+1A)&[MIMAT0000063&hsa-let-7b-5p&offsets\|0\|-2(+1A);m-14&22\|+\|46113691\|46113712&offsets\|0\|-2(+1A)]\|TGAGGTAGTAGGTTGTGTGGA | >MI0000063\|hsa-let-7b&WithFlank&22\|+\|46113680\|46113774 | 12 | 31 | 20 | (+1A) | isomiR | NO | 0.00 | 32.80 | 11.24 | 0.00 |
| >MI0000063\|hsa-let-7b&WithFlank&22\|+\|46113680\|46113774@12.31.20(+1C)&[MIMAT0000063&hsa-let-7b-5p&offsets\|0\|-2(+1C);m-14&22\|+\|46113691\|46113712&offsets\|0\|-2(+1C)]\|TGAGGTAGTAGGTTGTGTGGC | >MI0000063\|hsa-let-7b&WithFlank&22\|+\|46113680\|46113774 | 12 | 31 | 20 | (+1C) | isomiR | NO | 33.62 | 105.34 | 101.92 | 0.00 |
| >MI0000063\|hsa-let-7b&WithFlank&22\|+\|46113680\|46113774@12.32.21&[MIMAT0000063&hsa-let-7b-5p&offsets\|0\|-1;m-14&22\|+\|46113691\|46113712&offsets\|0\|-1]\|TGAGGTAGTAGGTTGTGTGGT | >MI0000063\|hsa-let-7b&WithFlank&22\|+\|46113680\|46113774 | 12 | 32 | 21 | | isomiR | NO | 236.79 | 689.71 | 552.90 | 87.25 |
| >MI0000063\|hsa-let-7b&WithFlank&22\|+\|46113680\|46113774@12.32.21(+1A)&[MIMAT0000063&hsa-let-7b-5p&offsets\|0\|-1(+1A);m-14&22\|+\|46113691\|46113712&offsets\|0\|-1(+1A)]\|TGAGGTAGTAGGTTGTGTGGTA | >MI0000063\|hsa-let-7b&WithFlank&22\|+\|46113680\|46113774 | 12 | 32 | 21 | (+1A) | isomiR | NO | 104.35 | 392.58 | 149.91 | 24.96 |
| >MI0000063\|hsa-let-7b&WithFlank&22\|+\|46113680\|46113774@12.32.21(+1C)&[MIMAT0000063&hsa-let-7b-5p&offsets\|0\|-1(+1C);m-14&22\|+\|46113691\|46113712&offsets\|0\|-1(+1C)]\|TGAGGTAGTAGGTTGTGTGGTC | >MI0000063\|hsa-let-7b&WithFlank&22\|+\|46113680\|46113774 | 12 | 32 | 21 | (+1C) | isomiR | NO | 278.20 | 764.52 | 410.23 | 104.36 |
| >MI0000063\|hsa-let-7b&WithFlank&22\|+\|46113680\|46113774@12.32.21(+1G)&[MIMAT0000063&hsa-let-7b-5p&offsets\|0\|-1(+1G);m-14&22\|+\|46113691\|46113712&offsets\|0\|-1(+1G)]\|TGAGGTAGTAGGTTGTGTGGTG | >MI0000063\|hsa-let-7b&WithFlank&22\|+\|46113680\|46113774 | 12 | 32 | 21 | (+1G) | isomiR | NO | 38.21 | 134.49 | 15.58 | 0.00 |
| >MI0000063\|hsa-let-7b&WithFlank&22\|+\|46113680\|46113774@12.33.22&[MIMAT0000063&hsa-let-7b-5p&offsets\|0\|0;m-14&22\|+\|46113691\|46113712&offsets\|0\|0]\|TGAGGTAGTAGGTTGTGTGGTT | >MI0000063\|hsa-let-7b&WithFlank&22\|+\|46113680\|46113774 | 12 | 33 | 22 | | isomiR | YES | 1305.19 | 4799.22 | 2230.35 | 512.48 |
| >MI0000063\|hsa-let-7b&WithFlank&22\|+\|46113680\|46113774@12.33.22(+1A)&[MIMAT0000063&hsa-let-7b-5p&offsets\|0\|0(+1A);m-14&22\|+\|46113691\|46113712&offsets\|0\|0(+1A)]\|TGAGGTAGTAGGTTGTGTGGTTA | >MI0000063\|hsa-let-7b&WithFlank&22\|+\|46113680\|46113774 | 12 | 33 | 22 | (+1A) | isomiR | NO | 112.04 | 299.83 | 326.78 | 31.95 |
| >MI0000063\|hsa-let-7b&WithFlank&22\|+\|46113680\|46113774@12.33.22(+1C)&[MIMAT0000063&hsa-let-7b-5p&offsets\|0\|0(+1C);m-14&22\|+\|46113691\|46113712&offsets\|0\|0(+1C)]\|TGAGGTAGTAGGTTGTGTGGTTC | >MI0000063\|hsa-let-7b&WithFlank&22\|+\|46113680\|46113774 | 12 | 33 | 22 | (+1C) | isomiR | NO | 31.71 | 108.02 | 124.42 | 0.00 |
| >MI0000063\|hsa-let-7b&WithFlank&22\|+\|46113680\|46113774@12.33.22(+1G)&[MIMAT0000063&hsa-let-7b-5p&offsets\|0\|0(+1G);m-14&22\|+\|46113691\|46113712&offsets\|0\|0(+1G)]\|TGAGGTAGTAGGTTGTGTGGTTG | >MI0000063\|hsa-let-7b&WithFlank&22\|+\|46113680\|46113774 | 12 | 33 | 22 | (+1G) | isomiR | NO | 0.00 | 33.97 | 59.86 | 0.00 |
| >MI0000063\|hsa-let-7b&WithFlank&22\|+\|46113680\|46113774@12.33.22(+2A)&[MIMAT0000063&hsa-let-7b-5p&offsets\|0\|0(+2A);m-14&22\|+\|46113691\|46113712&offsets\|0\|0(+2A)]\|TGAGGTAGTAGGTTGTGTGGTTAA | >MI0000063\|hsa-let-7b&WithFlank&22\|+\|46113680\|46113774 | 12 | 33 | 22 | (+2A) | isomiR | NO | 0.00 | 23.32 | 12.40 | 0.00 |
| >MI0000063\|hsa-let-7b&WithFlank&22\|+\|46113680\|46113774@12.34.23&[MIMAT0000063&hsa-let-7b-5p&offsets\|0\|+1;m-14&22\|+\|46113691\|46113712&offsets\|0\|+1]\|TGAGGTAGTAGGTTGTGTGGTTT | >MI0000063\|hsa-let-7b&WithFlank&22\|+\|46113680\|46113774 | 12 | 34 | 23 | | isomiR | NO | 245.42 | 728.04 | 695.46 | 106.79 |
| >MI0000063\|hsa-let-7b&WithFlank&22\|+\|46113680\|46113774@12.34.23(+1U)&[MIMAT0000063&hsa-let-7b-5p&offsets\|0\|+1(+1U);m-14&22\|+\|46113691\|46113712&offsets\|0\|+1(+1U)]\|TGAGGTAGTAGGTTGTGTGGTTTT | >MI0000063\|hsa-let-7b&WithFlank&22\|+\|46113680\|46113774 | 12 | 34 | 23 | (+1U) | isomiR | NO | 10.19 | 92.58 | 61.30 | 0.00 |
| >MI0000063\|hsa-let-7b&WithFlank&22\|+\|46113680\|46113774@66.86.21(+1U)&[MIMAT0004482&hsa-let-7b-3p&offsets\|0\|-1(+1U);m-171&22\|+\|46113745\|46113766&offsets\|0\|-1(+1U)]\|CTATACAACCTACTGCCTTCCT | >MI0000063\|hsa-let-7b&WithFlank&22\|+\|46113680\|46113774 | 66 | 86 | 21 | (+1U) | isomiR | NO | 0.00 | 9.96 | 12.78 | 0.00 |
| >MI0000064\|hsa-let-7c&WithFlank&21\|+\|16539822\|16539917@17.38.22&[MIMAT0000064&hsa-let-7c-5p&offsets\|0\|0;m-37&21\|+\|16539838\|16539859&offsets\|0\|0]\|TGAGGTAGTAGGTTGTATGGTT | >MI0000064\|hsa-let-7c&WithFlank&21\|+\|16539822\|16539917 | 17 | 38 | 22 | | isomiR | NO | 0.00 | 38.98 | 0.00 | 0.00 |
| >MI0000065\|hsa-let-7d&WithFlank&9\|+\|94178828\|94178926@14.31.18&[MIMAT0000065&hsa-let-7d-5p&offsets\|0\|-4;m-90&9\|+\|94178841\|94178862&offsets\|0\|-4]\|AGAGGTAGTAGGTTGCAT | >MI0000065\|hsa-let-7d&WithFlank&9\|+\|94178828\|94178926 | 14 | 31 | 18 | | isomiR | NO | 0.00 | 32.56 | 0.00 | 0.00 |

# BT-20

<table>
<tr><th></th><th></th><th></th><th></th><th></th><th></th><th></th><th></th><th></th><th></th><th></th><th></th></tr>
<tr><td>>MI0000065|hsa-let-7d&WithFlank&9|+|94178828|94178926@14.34.21&[MIMAT0000065&hsa-let-7d-5p&offsets|0|-1;m-90&9|+|94178841|94178862&offsets|0|-1]|AGAGGTAGTAGGTTGCATAGT</td><td>>MI0000065|hsa-let-7d&WithFlank&9|+|94178828|94178926</td><td>14</td><td>34</td><td>21</td><td></td><td>isomiR</td><td>NO</td><td>95.04</td><td>150.71</td><td>13.95</td><td>26.79</td></tr>
<tr><td>>MI0000065|hsa-let-7d&WithFlank&9|+|94178828|94178926@14.34.21(+1A)&[MIMAT0000065&hsa-let-7d-5p&offsets|0|-1(+1A);m-90&9|+|94178841|94178862&offsets|0|-1(+1A)]|AGAGGTAGTAGGTTGCATAGTA</td><td>>MI0000065|hsa-let-7d&WithFlank&9|+|94178828|94178926</td><td>14</td><td>34</td><td>21</td><td>(+1A)</td><td>isomiR</td><td>NO</td><td>0.00</td><td>33.09</td><td>0.00</td><td>0.00</td></tr>
<tr><td>>MI0000065|hsa-let-7d&WithFlank&9|+|94178828|94178926@14.34.21(+1C)&[MIMAT0000065&hsa-let-7d-5p&offsets|0|-1(+1C);m-90&9|+|94178841|94178862&offsets|0|-1(+1C)]|AGAGGTAGTAGGTTGCATAGTC</td><td>>MI0000065|hsa-let-7d&WithFlank&9|+|94178828|94178926</td><td>14</td><td>34</td><td>21</td><td>(+1C)</td><td>isomiR</td><td>NO</td><td>89.22</td><td>145.84</td><td>13.46</td><td>23.67</td></tr>
<tr><td>>MI0000065|hsa-let-7d&WithFlank&9|+|94178828|94178926@14.35.22&[MIMAT0000065&hsa-let-7d-5p&offsets|0|0;m-90&9|+|94178841|94178862&offsets|0|0]|AGAGGTAGTAGGTTGCATAGTT</td><td>>MI0000065|hsa-let-7d&WithFlank&9|+|94178828|94178926</td><td>14</td><td>35</td><td>22</td><td></td><td>isomiR</td><td>NO</td><td>439.31</td><td>929.50</td><td>211.01</td><td>175.66</td></tr>
<tr><td>>MI0000065|hsa-let-7d&WithFlank&9|+|94178828|94178926@14.36.23&[MIMAT0000065&hsa-let-7d-5p&offsets|0|+1;m-90&9|+|94178841|94178862&offsets|0|+1]|AGAGGTAGTAGGTTGCATAGTTT</td><td>>MI0000065|hsa-let-7d&WithFlank&9|+|94178828|94178926</td><td>14</td><td>36</td><td>23</td><td></td><td>isomiR</td><td>NO</td><td>9.55</td><td>28.71</td><td>12.38</td><td>0.00</td></tr>
<tr><td>>MI0000065|hsa-let-7d&WithFlank&9|+|94178828|94178926@68.88.21&[MIMAT0004484&hsa-let-7d-3p&offsets|0|-1;m-98&9|+|94178895|94178916&offsets|0|-1]|CTATACGACCTGCTGCCTTTC</td><td>>MI0000065|hsa-let-7d&WithFlank&9|+|94178828|94178926</td><td>68</td><td>88</td><td>21</td><td></td><td>isomiR</td><td>NO</td><td>9.54</td><td>32.54</td><td>72.39</td><td>0.00</td></tr>
<tr><td>>MI0000065|hsa-let-7d&WithFlank&9|+|94178828|94178926@68.88.21(+1A)&[MIMAT0004484&hsa-let-7d-3p&offsets|0|-1(+1A);m-98&9|+|94178895|94178916&offsets|0|-1(+1A)]|CTATACGACCTGCTGCCTTTCA</td><td>>MI0000065|hsa-let-7d&WithFlank&9|+|94178828|94178926</td><td>68</td><td>88</td><td>21</td><td>(+1A)</td><td>isomiR</td><td>NO</td><td>0.00</td><td>27.37</td><td>0.00</td><td>0.00</td></tr>
<tr><td>>MI0000065|hsa-let-7d&WithFlank&9|+|94178828|94178926@68.88.21(+1C)&[MIMAT0004484&hsa-let-7d-3p&offsets|0|-1(+1C);m-98&9|+|94178895|94178916&offsets|0|-1(+1C)]|CTATACGACCTGCTGCCTTTCC</td><td>>MI0000065|hsa-let-7d&WithFlank&9|+|94178828|94178926</td><td>68</td><td>88</td><td>21</td><td>(+1C)</td><td>isomiR</td><td>NO</td><td>23.93</td><td>38.89</td><td>15.42</td><td>0.00</td></tr>
<tr><td>>MI0000065|hsa-let-7d&WithFlank&9|+|94178828|94178926@68.89.22&[MIMAT0004484&hsa-let-7d-3p&offsets|0|0;m-98&9|+|94178895|94178916&offsets|0|0]|CTATACGACCTGCTGCCTTTCT</td><td>>MI0000065|hsa-let-7d&WithFlank&9|+|94178828|94178926</td><td>68</td><td>89</td><td>22</td><td></td><td>isomiR</td><td>NO</td><td>154.56</td><td>323.39</td><td>226.61</td><td>89.51</td></tr>
<tr><td>>MI0000066|hsa-let-7e&WithFlank&19|+|51692780|51692870@14.31.18&[MIMAT0000066&hsa-let-7e-5p&offsets|0|-4;m-50&19|+|51692793|51692814&offsets|0|-4]|TGAGGTAGGAGGTTGTAT</td><td>>MI0000066|hsa-let-7e&WithFlank&19|+|51692780|51692870</td><td>14</td><td>31</td><td>18</td><td></td><td>isomiR</td><td>NO</td><td>0.00</td><td>90.96</td><td>0.00</td><td>0.00</td></tr>
<tr><td>>MI0000066|hsa-let-7e&WithFlank&19|+|51692780|51692870@14.32.19&[MIMAT0000066&hsa-let-7e-5p&offsets|0|-3;m-50&19|+|51692793|51692814&offsets|0|-3]|TGAGGTAGGAGGTTGTATA</td><td>>MI0000066|hsa-let-7e&WithFlank&19|+|51692780|51692870</td><td>14</td><td>32</td><td>19</td><td></td><td>isomiR</td><td>NO</td><td>0.00</td><td>67.19</td><td>0.00</td><td>0.00</td></tr>
<tr><td>>MI0000066|hsa-let-7e&WithFlank&19|+|51692780|51692870@14.33.20(+1C)&[MIMAT0000066&hsa-let-7e-5p&offsets|0|-2(+1C);m-50&19|+|51692793|51692814&offsets|0|-2(+1C)]|TGAGGTAGGAGGTTGTATAGC</td><td>>MI0000066|hsa-let-7e&WithFlank&19|+|51692780|51692870</td><td>14</td><td>33</td><td>20</td><td>(+1C)</td><td>isomiR</td><td>NO</td><td>0.00</td><td>36.32</td><td>0.00</td><td>0.00</td></tr>
<tr><td>>MI0000066|hsa-let-7e&WithFlank&19|+|51692780|51692870@14.34.21&[MIMAT0000066&hsa-let-7e-5p&offsets|0|-1;m-50&19|+|51692793|51692814&offsets|0|-1]|TGAGGTAGGAGGTTGTATAGT</td><td>>MI0000066|hsa-let-7e&WithFlank&19|+|51692780|51692870</td><td>14</td><td>34</td><td>21</td><td></td><td>isomiR</td><td>NO</td><td>76.38</td><td>357.98</td><td>14.86</td><td>26.98</td></tr>
<tr><td>>MI0000066|hsa-let-7e&WithFlank&19|+|51692780|51692870@14.34.21(+1A)&[MIMAT0000066&hsa-let-7e-5p&offsets|0|-1(+1A);m-50&19|+|51692793|51692814&offsets|0|-1(+1A)]|TGAGGTAGGAGGTTGTATAGTA</td><td>>MI0000066|hsa-let-7e&WithFlank&19|+|51692780|51692870</td><td>14</td><td>34</td><td>21</td><td>(+1A)</td><td>isomiR</td><td>NO</td><td>0.00</td><td>98.57</td><td>0.00</td><td>0.00</td></tr>
<tr><td>>MI0000066|hsa-let-7e&WithFlank&19|+|51692780|51692870@14.34.21(+1C)&[MIMAT0000066&hsa-let-7e-5p&offsets|0|-1(+1C);m-50&19|+|51692793|51692814&offsets|0|-1(+1C)]|TGAGGTAGGAGGTTGTATAGTC</td><td>>MI0000066|hsa-let-7e&WithFlank&19|+|51692780|51692870</td><td>14</td><td>34</td><td>21</td><td>(+1C)</td><td>isomiR</td><td>NO</td><td>72.01</td><td>286.97</td><td>10.11</td><td>29.53</td></tr>
<tr><td>>MI0000066|hsa-let-7e&WithFlank&19|+|51692780|51692870@14.34.21(+1G)&[MIMAT0000066&hsa-let-7e-5p&offsets|0|-1(+1G);m-50&19|+|51692793|51692814&offsets|0|-1(+1G)]|TGAGGTAGGAGGTTGTATAGTG</td><td>>MI0000066|hsa-let-7e&WithFlank&19|+|51692780|51692870</td><td>14</td><td>34</td><td>21</td><td>(+1G)</td><td>isomiR</td><td>NO</td><td>0.00</td><td>27.12</td><td>0.00</td><td>0.00</td></tr>
<tr><td>>MI0000066|hsa-let-7e&WithFlank&19|+|51692780|51692870@14.35.22&[MIMAT0000066&hsa-let-7e-5p&offsets|0|0;m-50&19|+|51692793|51692814&offsets|0|0]|TGAGGTAGGAGGTTGTATAGTT</td><td>>MI0000066|hsa-let-7e&WithFlank&19|+|51692780|51692870</td><td>14</td><td>35</td><td>22</td><td></td><td>isomiR</td><td>NO</td><td>348.02</td><td>1814.90</td><td>159.27</td><td>226.77</td></tr>
<tr><td>>MI0000066|hsa-let-7e&WithFlank&19|+|51692780|51692870@14.35.22(+1A)&[MIMAT0000066&hsa-let-7e-5p&offsets|0|0(+1A);m-50&19|+|51692793|51692814&offsets|0|0(+1A)]|TGAGGTAGGAGGTTGTATAGTTA</td><td>>MI0000066|hsa-let-7e&WithFlank&19|+|51692780|51692870</td><td>14</td><td>35</td><td>22</td><td>(+1A)</td><td>isomiR</td><td>NO</td><td>0.00</td><td>85.85</td><td>0.00</td><td>0.00</td></tr>
<tr><td>>MI0000066|hsa-let-7e&WithFlank&19|+|51692780|51692870@14.35.22(+1U)&[MIMAT0000066&hsa-let-7e-5p&offsets|0|0(+1U);m-50&19|+|51692793|51692814&offsets|0|0(+1U)]|TGAGGTAGGAGGTTGTATAGTTT</td><td>>MI0000066|hsa-let-7e&WithFlank&19|+|51692780|51692870</td><td>14</td><td>35</td><td>22</td><td>(+1U)</td><td>isomiR</td><td>NO</td><td>24.02</td><td>125.57</td><td>10.82</td><td>0.00</td></tr>
<tr><td>>MI0000066|hsa-let-7e&WithFlank&19|+|51692780|51692870@14.36.23&[MIMAT0000066&hsa-let-7e-5p&offsets|0|+1;m-50&19|+|51692793|51692814&offsets|0|+1]|TGAGGTAGGAGGTTGTATAGTTG</td><td>>MI0000066|hsa-let-7e&WithFlank&19|+|51692780|51692870</td><td>14</td><td>36</td><td>23</td><td></td><td>isomiR</td><td>NO</td><td>0.00</td><td>99.77</td><td>0.00</td><td>0.00</td></tr>
<tr><td>>MI0000067|hsa-let-7f-1&WithFlank&9|+|94176341|94176439@13.30.18;MI0000068|hsa-let-7f-2&WithFlank&X|-|53557186|53557280@14.31.18&[MIMAT0000067&hsa-let-7f-5p&offsets|0|-4;m-20&9|+|94176353|94176374&offsets|0|-4];[MIMAT0000067_1&hsa-let-7f-5p&offsets|0|-4;m-19&X|-|53557246|53557267&offsets|0|-4]|TGAGGTAGTAGATTGTAT</td><td>>MI0000067|hsa-let-7f-1&WithFlank&9|+|94176341|94176439</td><td>13</td><td>30</td><td>18</td><td></td><td>isomiR</td><td>NO</td><td>205.25</td><td>346.96</td><td>94.55</td><td>84.21</td></tr>
</table>

# BT-20

<table>
<tr><th></th><th></th><th></th><th></th><th></th><th></th><th></th><th></th><th></th><th></th><th></th><th></th></tr>
<tr><td>>MI0000067|hsa-let-7f-1&WithFlank&9|+|94176341|94176439@13.31.19;MI0000068|hsa-let-7f-2&WithFlank&X|-|53557186|53557280@14.32.19&[MIMAT0000067&hsa-let-7f-5p&offsets|0|-3;m-20&9|+|94176353|94176374&offsets|0|-3];[MIMAT0000067_1&hsa-let-7f-5p&offsets|0|-3;m-19&X|-|53557246|53557267&offsets|0|-3]|TGAGGTAGTAGATTGTATA</td><td>>MI0000067|hsa-let-7f-1&WithFlank&9|+|94176341|94176439</td><td>13</td><td>31</td><td>19</td><td></td><td>isomiR</td><td>NO</td><td>42.53</td><td>246.76</td><td>0.00</td><td>0.00</td></tr>
<tr><td>>MI0000067|hsa-let-7f-1&WithFlank&9|+|94176341|94176439@13.32.20(+1C);MI0000068|hsa-let-7f-2&WithFlank&X|-|53557186|53557280@14.33.20(+1C)&[MIMAT0000067&hsa-let-7f-5p&offsets|0|-2(+1C);m-20&9|+|94176353|94176374&offsets|0|-2(+1C)];[MIMAT0000067_1&hsa-let-7f-5p&offsets|0|-2(+1C);m-19&X|-|53557246|53557267&offsets|0|-2(+1C)]|TGAGGTAGTAGATTGTATAGC</td><td>>MI0000067|hsa-let-7f-1&WithFlank&9|+|94176341|94176439</td><td>13</td><td>32</td><td>20</td><td>(+1C)</td><td>isomiR</td><td>NO</td><td>72.48</td><td>97.32</td><td>0.00</td><td>0.00</td></tr>
<tr><td>>MI0000067|hsa-let-7f-1&WithFlank&9|+|94176341|94176439@13.33.21(+1A);MI0000068|hsa-let-7f-2&WithFlank&X|-|53557186|53557280@14.34.21(+1A)&[MIMAT0000067&hsa-let-7f-5p&offsets|0|-1(+1A);m-20&9|+|94176353|94176374&offsets|0|-1(+1A)];[MIMAT0000067_1&hsa-let-7f-5p&offsets|0|-1(+1A);m-19&X|-|53557246|53557267&offsets|0|-1(+1A)]|TGAGGTAGTAGATTGTATAGTA</td><td>>MI0000067|hsa-let-7f-1&WithFlank&9|+|94176341|94176439</td><td>13</td><td>33</td><td>21</td><td>(+1A)</td><td>isomiR</td><td>NO</td><td>140.88</td><td>346.07</td><td>9.51</td><td>34.21</td></tr>
<tr><td>>MI0000067|hsa-let-7f-1&WithFlank&9|+|94176341|94176439@13.33.21(+1C);MI0000068|hsa-let-7f-2&WithFlank&X|-|53557186|53557280@14.34.21(+1C)&[MIMAT0000067&hsa-let-7f-5p&offsets|0|-1(+1C);m-20&9|+|94176353|94176374&offsets|0|-1(+1C)];[MIMAT0000067_1&hsa-let-7f-5p&offsets|0|-1(+1C);m-19&X|-|53557246|53557267&offsets|0|-1(+1C)]|TGAGGTAGTAGATTGTATAGTC</td><td>>MI0000067|hsa-let-7f-1&WithFlank&9|+|94176341|94176439</td><td>13</td><td>33</td><td>21</td><td>(+1C)</td><td>isomiR</td><td>NO</td><td>624.40</td><td>1061.98</td><td>129.33</td><td>244.34</td></tr>
<tr><td>>MI0000067|hsa-let-7f-1&WithFlank&9|+|94176341|94176439@13.33.21(+1G);MI0000068|hsa-let-7f-2&WithFlank&X|-|53557186|53557280@14.34.21(+1G)&[MIMAT0000067&hsa-let-7f-5p&offsets|0|-1(+1G);m-20&9|+|94176353|94176374&offsets|0|-1(+1G)];[MIMAT0000067_1&hsa-let-7f-5p&offsets|0|-1(+1G);m-19&X|-|53557246|53557267&offsets|0|-1(+1G)]|TGAGGTAGTAGATTGTATAGTG</td><td>>MI0000067|hsa-let-7f-1&WithFlank&9|+|94176341|94176439</td><td>13</td><td>33</td><td>21</td><td>(+1G)</td><td>isomiR</td><td>NO</td><td>88.89</td><td>141.35</td><td>0.00</td><td>0.00</td></tr>
<tr><td>>MI0000067|hsa-let-7f-1&WithFlank&9|+|94176341|94176439@13.33.21;MI0000068|hsa-let-7f-2&WithFlank&X|-|53557186|53557280@14.34.21&[MIMAT0000067&hsa-let-7f-5p&offsets|0|-1;m-20&9|+|94176353|94176374&offsets|0|-1];[MIMAT0000067_1&hsa-let-7f-5p&offsets|0|-1;m-19&X|-|53557246|53557267&offsets|0|-1]|TGAGGTAGTAGATTGTATAGT</td><td>>MI0000067|hsa-let-7f-1&WithFlank&9|+|94176341|94176439</td><td>13</td><td>33</td><td>21</td><td></td><td>isomiR</td><td>NO</td><td>369.52</td><td>673.72</td><td>100.83</td><td>141.09</td></tr>
<tr><td>>MI0000067|hsa-let-7f-1&WithFlank&9|+|94176341|94176439@13.34.22(+1A);MI0000068|hsa-let-7f-2&WithFlank&X|-|53557186|53557280@14.35.22(+1A)&[MIMAT0000067&hsa-let-7f-5p&offsets|0|0(+1A);m-20&9|+|94176353|94176374&offsets|0|0(+1A)];[MIMAT0000067_1&hsa-let-7f-5p&offsets|0|0(+1A);m-19&X|-|53557246|53557267&offsets|0|0(+1A)]|TGAGGTAGTAGATTGTATAGTTA</td><td>>MI0000067|hsa-let-7f-1&WithFlank&9|+|94176341|94176439</td><td>13</td><td>34</td><td>22</td><td>(+1A)</td><td>isomiR</td><td>NO</td><td>108.34</td><td>173.81</td><td>139.81</td><td>23.67</td></tr>
<tr><td>>MI0000067|hsa-let-7f-1&WithFlank&9|+|94176341|94176439@13.34.22;MI0000068|hsa-let-7f-2&WithFlank&X|-|53557186|53557280@14.35.22&[MIMAT0000067&hsa-let-7f-5p&offsets|0|0;m-20&9|+|94176353|94176374&offsets|0|0];[MIMAT0000067_1&hsa-let-7f-5p&offsets|0|0;m-19&X|-|53557246|53557267&offsets|0|0]|TGAGGTAGTAGATTGTATAGTT</td><td>>MI0000067|hsa-let-7f-1&WithFlank&9|+|94176341|94176439</td><td>13</td><td>34</td><td>22</td><td></td><td>isomiR</td><td>YES</td><td>2982.46</td><td>6788.06</td><td>701.94</td><td>1202.10</td></tr>
<tr><td>>MI0000067|hsa-let-7f-1&WithFlank&9|+|94176341|94176439@13.35.23&[MIMAT0000067&hsa-let-7f-5p&offsets|0|+1;m-20&9|+|94176353|94176374&offsets|0|+1]|TGAGGTAGTAGATTGTATAGTTG</td><td>>MI0000067|hsa-let-7f-1&WithFlank&9|+|94176341|94176439</td><td>13</td><td>35</td><td>23</td><td></td><td>isomiR</td><td>NO</td><td>0.00</td><td>25.46</td><td>0.00</td><td>0.00</td></tr>
<tr><td>>MI0000068|hsa-let-7f-2&WithFlank&X|-|53557186|53557280@14.36.23&[MIMAT0000067_1&hsa-let-7f-5p&offsets|0|+1;m-19&X|-|53557246|53557267&offsets|0|+1]|TGAGGTAGTAGATTGTATAGTTT</td><td>>MI0000068|hsa-let-7f-2&WithFlank&X|-|53557186|53557280</td><td>14</td><td>36</td><td>23</td><td></td><td>isomiR</td><td>NO</td><td>102.23</td><td>172.93</td><td>147.73</td><td>31.50</td></tr>
<tr><td>>MI0000070|hsa-mir-16-1&WithFlank&13|-|50048967|50049067@20.39.20(+1U);MI0000115|hsa-mir-16-2&WithFlank&3|+|160404739|160404831@16.35.20(+1U)&[MIMAT0000069&hsa-miR-16-5p&offsets|0|-2(+1U);m-62&13|-|50049027|50049048&offsets|0|-2(+1U)];[MIMAT0000069_1&hsa-miR-16-5p&offsets|0|-2(+1U);m-61&3|+|160404754|160404775&offsets|0|-2(+1U)]|TAGCAGCACGTAAATATTGGT</td><td>>MI0000070|hsa-mir-16-1&WithFlank&13|-|50048967|50049067</td><td>20</td><td>39</td><td>20</td><td>(+1U)</td><td>isomiR</td><td>NO</td><td>0.00</td><td>36.82</td><td>10.24</td><td>0.00</td></tr>
<tr><td>>MI0000070|hsa-mir-16-1&WithFlank&13|-|50048967|50049067@20.39.20;MI0000115|hsa-mir-16-2&WithFlank&3|+|160404739|160404831@16.35.20&[MIMAT0000069&hsa-miR-16-5p&offsets|0|-2;m-62&13|-|50049027|50049048&offsets|0|-2];[MIMAT0000069_1&hsa-miR-16-5p&offsets|0|-2;m-61&3|+|160404754|160404775&offsets|0|-2]|TAGCAGCACGTAAATATTGG</td><td>>MI0000070|hsa-mir-16-1&WithFlank&13|-|50048967|50049067</td><td>20</td><td>39</td><td>20</td><td></td><td>isomiR</td><td>NO</td><td>0.00</td><td>24.28</td><td>11.66</td><td>0.00</td></tr>
<tr><td>>MI0000070|hsa-mir-16-1&WithFlank&13|-|50048967|50049067@20.40.21(+1A);MI0000115|hsa-mir-16-2&WithFlank&3|+|160404739|160404831@16.36.21(+1A)&[MIMAT0000069&hsa-miR-16-5p&offsets|0|-1(+1A);m-62&13|-|50049027|50049048&offsets|0|-1(+1A)];[MIMAT0000069_1&hsa-miR-16-5p&offsets|0|-1(+1A);m-61&3|+|160404754|160404775&offsets|0|-1(+1A)]|TAGCAGCACGTAAATATTGGCA</td><td>>MI0000070|hsa-mir-16-1&WithFlank&13|-|50048967|50049067</td><td>20</td><td>40</td><td>21</td><td>(+1A)</td><td>isomiR</td><td>NO</td><td>73.99</td><td>118.90</td><td>80.60</td><td>0.00</td></tr>
<tr><td>>MI0000070|hsa-mir-16-1&WithFlank&13|-|50048967|50049067@20.40.21;MI0000115|hsa-mir-16-2&WithFlank&3|+|160404739|160404831@16.36.21&[MIMAT0000069&hsa-miR-16-5p&offsets|0|-1;m-62&13|-|50049027|50049048&offsets|0|-1];[MIMAT0000069_1&hsa-miR-16-5p&offsets|0|-1;m-61&3|+|160404754|160404775&offsets|0|-1]|TAGCAGCACGTAAATATTGGC</td><td>>MI0000070|hsa-mir-16-1&WithFlank&13|-|50048967|50049067</td><td>20</td><td>40</td><td>21</td><td></td><td>isomiR</td><td>NO</td><td>71.66</td><td>141.57</td><td>80.02</td><td>0.00</td></tr>
<tr><td>>MI0000070|hsa-mir-16-1&WithFlank&13|-|50048967|50049067@20.41.22;MI0000115|hsa-mir-16-2&WithFlank&3|+|160404739|160404831@16.37.22&[MIMAT0000069&hsa-miR-16-5p&offsets|0|0;m-62&13|-|50049027|50049048&offsets|0|0];[MIMAT0000069_1&hsa-miR-16-5p&offsets|0|0;m-61&3|+|160404754|160404775&offsets|0|0]|TAGCAGCACGTAAATATTGGCG</td><td>>MI0000070|hsa-mir-16-1&WithFlank&13|-|50048967|50049067</td><td>20</td><td>41</td><td>22</td><td></td><td>isomiR</td><td>NO</td><td>232.30</td><td>350.84</td><td>335.60</td><td>0.00</td></tr>
<tr><td>>MI0000070|hsa-mir-16-1&WithFlank&13|-|50048967|50049067@20.42.23;MI0000115|hsa-mir-16-2&WithFlank&3|+|160404739|160404831@16.38.23&[MIMAT0000069&hsa-miR-16-5p&offsets|0|+1;m-62&13|-|50049027|50049048&offsets|0|+1];[MIMAT0000069_1&hsa-miR-16-5p&offsets|0|+1;m-61&3|+|160404754|160404775&offsets|0|+1]|TAGCAGCACGTAAATATTGGCGT</td><td>>MI0000070|hsa-mir-16-1&WithFlank&13|-|50048967|50049067</td><td>20</td><td>42</td><td>23</td><td></td><td>isomiR</td><td>NO</td><td>123.30</td><td>174.44</td><td>125.68</td><td>0.00</td></tr>
<tr><td>>MI0000071|hsa-mir-17&WithFlank&13|+|91350599|91350694@20.41.22(+1A)&[MIMAT0000070&hsa-miR-17-5p&offsets|0|-1(+1A);m-88&13|+|91350618|91350640&offsets|0|-1(+1A)]|CAAAGTGCTTACAGTGCAGGTAA</td><td>>MI0000071|hsa-mir-17&WithFlank&13|+|91350599|91350694</td><td>20</td><td>41</td><td>22</td><td>(+1A)</td><td>isomiR</td><td>NO</td><td>0.00</td><td>34.84</td><td>0.00</td><td>0.00</td></tr>
</table>

# BT-20

| | | | | | | | | | | | |
|---|---|---|---|---|---|---|---|---|---|---|---|
| >MI0000071\|hsa-mir-17&WithFlank&13\|+\|91350599\|91350694@20.42.23&[MIMAT0000070&hsa-miR-17-5p&offsets\|0\|0;m-88&13\|+\|91350618\|91350640&offsets\|0\|0]\|CAAAGTGCTTACAGTGCAGGTAG | >MI0000071\|hsa-mir-17&WithFlank&13\|+\|91350599\|91350694 | 20 | 42 | 23 | | isomiR | NO | 161.92 | 276.17 | 111.08 | 29.26 |
| >MI0000071\|hsa-mir-17&WithFlank&13\|+\|91350599\|91350694@20.42.23(+1A)&[MIMAT0000070&hsa-miR-17-5p&offsets\|0\|0(+1A);m-88&13\|+\|91350618\|91350640&offsets\|0\|0(+1A)]\|CAAAGTGCTTACAGTGCAGGTAGA | >MI0000071\|hsa-mir-17&WithFlank&13\|+\|91350599\|91350694 | 20 | 42 | 23 | (+1A) | isomiR | NO | 39.73 | 196.97 | 0.00 | 9.92 |
| >MI0000071\|hsa-mir-17&WithFlank&13\|+\|91350599\|91350694@20.43.24&[MIMAT0000070&hsa-miR-17-5p&offsets\|0\|+1;m-88&13\|+\|91350618\|91350640&offsets\|0\|+1]\|CAAAGTGCTTACAGTGCAGGTAGT | >MI0000071\|hsa-mir-17&WithFlank&13\|+\|91350599\|91350694 | 20 | 43 | 24 | | isomiR | NO | 9.59 | 36.52 | 0.00 | 0.00 |
| >MI0000074\|hsa-mir-19b-1&WithFlank&13\|+\|91351186\|91351284@60.81.22;MI0000075\|hsa-mir-19b-2&WithFlank&X\|-\|134169665\|134169772@68.89.22&[MIMAT0000074&hsa-miR-19b-3p&offsets\|0\|-1;m-122&13\|+\|91351245\|91351267&offsets\|0\|-1];[MIMAT0000074_1&hsa-miR-19b-3p&offsets\|0\|-1;m-121&X\|-\|134169683\|134169705&offsets\|0\|-1]\|TGTGCAAATCCATGCAAAACTG | >MI0000074\|hsa-mir-19b-1&WithFlank&13\|+\|91351186\|91351284 | 60 | 81 | 22 | | isomiR | NO | 0.00 | 0.00 | 10.12 | 0.00 |
| >MI0000074\|hsa-mir-19b-1&WithFlank&13\|+\|91351186\|91351284@60.82.23;MI0000075\|hsa-mir-19b-2&WithFlank&X\|-\|134169665\|134169772@68.90.23&[MIMAT0000074&hsa-miR-19b-3p&offsets\|0\|0;m-122&13\|+\|91351245\|91351267&offsets\|0\|0];[MIMAT0000074_1&hsa-miR-19b-3p&offsets\|0\|0;m-121&X\|-\|134169683\|134169705&offsets\|0\|0]\|TGTGCAAATCCATGCAAAACTGA | >MI0000074\|hsa-mir-19b-1&WithFlank&13\|+\|91351186\|91351284 | 60 | 82 | 23 | | isomiR | NO | 11.16 | 0.00 | 0.00 | 0.00 |
| >MI0000076\|hsa-mir-20a&WithFlank&13\|+\|91351059\|91351141@14.35.22(+1A)&[MIMAT0000075&hsa-miR-20a-5p&offsets\|0\|-1(+1A);m-96&13\|+\|91351072\|91351094&offsets\|0\|-1(+1A)]\|TAAAGTGCTTATAGTGCAGGTAA | >MI0000076\|hsa-mir-20a&WithFlank&13\|+\|91351059\|91351141 | 14 | 35 | 22 | (+1A) | isomiR | NO | 9.48 | 22.99 | 0.00 | 0.00 |
| >MI0000076\|hsa-mir-20a&WithFlank&13\|+\|91351059\|91351141@14.36.23&[MIMAT0000075&hsa-miR-20a-5p&offsets\|0\|0;m-96&13\|+\|91351072\|91351094&offsets\|0\|0]\|TAAAGTGCTTATAGTGCAGGTAG | >MI0000076\|hsa-mir-20a&WithFlank&13\|+\|91351059\|91351141 | 14 | 36 | 23 | | isomiR | NO | 175.34 | 187.99 | 95.90 | 26.93 |
| >MI0000076\|hsa-mir-20a&WithFlank&13\|+\|91351059\|91351141@14.36.23(+1A)&[MIMAT0000075&hsa-miR-20a-5p&offsets\|0\|0(+1A);m-96&13\|+\|91351072\|91351094&offsets\|0\|0(+1A)]\|TAAAGTGCTTATAGTGCAGGTAGA | >MI0000076\|hsa-mir-20a&WithFlank&13\|+\|91351059\|91351141 | 14 | 36 | 23 | (+1A) | isomiR | NO | 29.45 | 72.29 | 0.00 | 0.00 |
| >MI0000077\|hsa-mir-21&WithFlank&17\|+\|59841260\|59841343@13.35.23&[MIMAT0000076&hsa-miR-21-5p&offsets\|-1\|0;m-1&17\|+\|59841273\|59841294&offsets\|-1\|0]\|GTAGCTTATCAGACTGATGTTGA | >MI0000077\|hsa-mir-21&WithFlank&17\|+\|59841260\|59841343 | 13 | 35 | 23 | | isomiR | NO | 24.29 | 43.15 | 96.58 | 0.00 |
| >MI0000077\|hsa-mir-21&WithFlank&17\|+\|59841260\|59841343@14.31.18&[MIMAT0000076&hsa-miR-21-5p&offsets\|0\|-4;m-1&17\|+\|59841273\|59841294&offsets\|0\|-4]\|TAGCTTATCAGACTGATG | >MI0000077\|hsa-mir-21&WithFlank&17\|+\|59841260\|59841343 | 14 | 31 | 18 | | isomiR | NO | 69.71 | 93.71 | 14.79 | 0.00 |
| >MI0000077\|hsa-mir-21&WithFlank&17\|+\|59841260\|59841343@14.32.19&[MIMAT0000076&hsa-miR-21-5p&offsets\|0\|-3;m-1&17\|+\|59841273\|59841294&offsets\|0\|-3]\|TAGCTTATCAGACTGATGT | >MI0000077\|hsa-mir-21&WithFlank&17\|+\|59841260\|59841343 | 14 | 32 | 19 | | isomiR | NO | 65.90 | 112.03 | 114.87 | 0.00 |
| >MI0000077\|hsa-mir-21&WithFlank&17\|+\|59841260\|59841343@14.33.20&[MIMAT0000076&hsa-miR-21-5p&offsets\|0\|-2;m-1&17\|+\|59841273\|59841294&offsets\|0\|-2]\|TAGCTTATCAGACTGATGTT | >MI0000077\|hsa-mir-21&WithFlank&17\|+\|59841260\|59841343 | 14 | 33 | 20 | | isomiR | NO | 77.24 | 109.77 | 79.15 | 0.00 |
| >MI0000077\|hsa-mir-21&WithFlank&17\|+\|59841260\|59841343@14.33.20(+1A)&[MIMAT0000076&hsa-miR-21-5p&offsets\|0\|-2(+1A);m-1&17\|+\|59841273\|59841294&offsets\|0\|-2(+1A)]\|TAGCTTATCAGACTGATGTTA | >MI0000077\|hsa-mir-21&WithFlank&17\|+\|59841260\|59841343 | 14 | 33 | 20 | (+1A) | isomiR | NO | 75.80 | 197.50 | 102.25 | 0.00 |
| >MI0000077\|hsa-mir-21&WithFlank&17\|+\|59841260\|59841343@14.34.21&[MIMAT0000076&hsa-miR-21-5p&offsets\|0\|-1;m-1&17\|+\|59841273\|59841294&offsets\|0\|-1]\|TAGCTTATCAGACTGATGTTG | >MI0000077\|hsa-mir-21&WithFlank&17\|+\|59841260\|59841343 | 14 | 34 | 21 | | isomiR | NO | 419.65 | 984.33 | 646.67 | 117.86 |
| >MI0000077\|hsa-mir-21&WithFlank&17\|+\|59841260\|59841343@14.34.21(+1G)&[MIMAT0000076&hsa-miR-21-5p&offsets\|0\|-1(+1G);m-1&17\|+\|59841273\|59841294&offsets\|0\|-1(+1G)]\|TAGCTTATCAGACTGATGTTGG | >MI0000077\|hsa-mir-21&WithFlank&17\|+\|59841260\|59841343 | 14 | 34 | 21 | (+1G) | isomiR | YES | 145.69 | 238.81 | 159.21 | 0.00 |
| >MI0000077\|hsa-mir-21&WithFlank&17\|+\|59841260\|59841343@14.34.21(+1U)&[MIMAT0000076&hsa-miR-21-5p&offsets\|0\|-1(+1U);m-1&17\|+\|59841273\|59841294&offsets\|0\|-1(+1U)]\|TAGCTTATCAGACTGATGTTGT | >MI0000077\|hsa-mir-21&WithFlank&17\|+\|59841260\|59841343 | 14 | 34 | 21 | (+1U) | isomiR | NO | 106.44 | 189.15 | 136.97 | 0.00 |
| >MI0000077\|hsa-mir-21&WithFlank&17\|+\|59841260\|59841343@14.34.21(+2C)&[MIMAT0000076&hsa-miR-21-5p&offsets\|0\|-1(+2C);m-1&17\|+\|59841273\|59841294&offsets\|0\|-1(+2C)]\|TAGCTTATCAGACTGATGTTGCC | >MI0000077\|hsa-mir-21&WithFlank&17\|+\|59841260\|59841343 | 14 | 34 | 21 | (+2C) | isomiR | NO | 0.00 | 28.74 | 0.00 | 0.00 |
| >MI0000077\|hsa-mir-21&WithFlank&17\|+\|59841260\|59841343@14.35.22&[MIMAT0000076&hsa-miR-21-5p&offsets\|0\|0;m-1&17\|+\|59841273\|59841294&offsets\|0\|0]\|TAGCTTATCAGACTGATGTTGA | >MI0000077\|hsa-mir-21&WithFlank&17\|+\|59841260\|59841343 | 14 | 35 | 22 | | isomiR | YES | 2559.37 | 6595.63 | 4818.81 | 517.81 |
| >MI0000077\|hsa-mir-21&WithFlank&17\|+\|59841260\|59841343@14.35.22(+1A)&[MIMAT0000076&hsa-miR-21-5p&offsets\|0\|0(+1A);m-1&17\|+\|59841273\|59841294&offsets\|0\|0(+1A)]\|TAGCTTATCAGACTGATGTTGAA | >MI0000077\|hsa-mir-21&WithFlank&17\|+\|59841260\|59841343 | 14 | 35 | 22 | (+1A) | isomiR | YES | 472.13 | 2051.59 | 151.68 | 161.24 |
| >MI0000077\|hsa-mir-21&WithFlank&17\|+\|59841260\|59841343@14.35.22(+1G)&[MIMAT0000076&hsa-miR-21-5p&offsets\|0\|0(+1G);m-1&17\|+\|59841273\|59841294&offsets\|0\|0(+1G)]\|TAGCTTATCAGACTGATGTTGAG | >MI0000077\|hsa-mir-21&WithFlank&17\|+\|59841260\|59841343 | 14 | 35 | 22 | (+1G) | isomiR | NO | 65.65 | 191.63 | 0.00 | 0.00 |
| >MI0000077\|hsa-mir-21&WithFlank&17\|+\|59841260\|59841343@14.35.22(+1U)&[MIMAT0000076&hsa-miR-21-5p&offsets\|0\|0(+1U);m-1&17\|+\|59841273\|59841294&offsets\|0\|0(+1U)]\|TAGCTTATCAGACTGATGTTGAT | >MI0000077\|hsa-mir-21&WithFlank&17\|+\|59841260\|59841343 | 14 | 35 | 22 | (+1U) | isomiR | YES | 1769.85 | 8123.59 | 573.27 | 626.07 |

# BT-20

<table>
<tr><td>>MI0000077|hsa-mir-21&WithFlank&17|+|59841260|59841343@14.35.22(+2A)&[MIMAT0000076&hsa-miR-21-5p&offsets|0|0(+2A);m-1&17|+|59841273|59841294&offsets|0|0(+2A)]|TAGCTTATCAGACTGATGTTGAAA</td><td>>MI0000077|hsa-mir-21&WithFlank&17|+|59841260|59841343</td><td>14</td><td>35</td><td>22</td><td>(+2A)</td><td>isomiR</td><td>NO</td><td>32.20</td><td>99.77</td><td>0.00</td><td>0.00</td></tr>
<tr><td>>MI0000077|hsa-mir-21&WithFlank&17|+|59841260|59841343@14.35.22(+2U)&[MIMAT0000076&hsa-miR-21-5p&offsets|0|0(+2U);m-1&17|+|59841273|59841294&offsets|0|0(+2U)]|TAGCTTATCAGACTGATGTTGATT</td><td>>MI0000077|hsa-mir-21&WithFlank&17|+|59841260|59841343</td><td>14</td><td>35</td><td>22</td><td>(+2U)</td><td>isomiR</td><td>NO</td><td>10.02</td><td>28.69</td><td>0.00</td><td>0.00</td></tr>
<tr><td>>MI0000077|hsa-mir-21&WithFlank&17|+|59841260|59841343@14.36.23&[MIMAT0000076&hsa-miR-21-5p&offsets|0|+1;m-1&17|+|59841273|59841294&offsets|0|+1]|TAGCTTATCAGACTGATGTTGAC</td><td>>MI0000077|hsa-mir-21&WithFlank&17|+|59841260|59841343</td><td>14</td><td>36</td><td>23</td><td></td><td>isomiR</td><td>YES</td><td>8809.01</td><td>45838.51</td><td>3205.02</td><td>3500.89</td></tr>
<tr><td>>MI0000077|hsa-mir-21&WithFlank&17|+|59841260|59841343@14.36.23(+1A)&[MIMAT0000076&hsa-miR-21-5p&offsets|0|+1(+1A);m-1&17|+|59841273|59841294&offsets|0|+1(+1A)]|TAGCTTATCAGACTGATGTTGACA</td><td>>MI0000077|hsa-mir-21&WithFlank&17|+|59841260|59841343</td><td>14</td><td>36</td><td>23</td><td>(+1A)</td><td>isomiR</td><td>YES</td><td>1893.67</td><td>8999.02</td><td>686.08</td><td>1018.40</td></tr>
<tr><td>>MI0000077|hsa-mir-21&WithFlank&17|+|59841260|59841343@14.36.23(+1C)&[MIMAT0000076&hsa-miR-21-5p&offsets|0|+1(+1C);m-1&17|+|59841273|59841294&offsets|0|+1(+1C)]|TAGCTTATCAGACTGATGTTGACC</td><td>>MI0000077|hsa-mir-21&WithFlank&17|+|59841260|59841343</td><td>14</td><td>36</td><td>23</td><td>(+1C)</td><td>isomiR</td><td>YES</td><td>750.52</td><td>2143.43</td><td>231.95</td><td>381.56</td></tr>
<tr><td>>MI0000077|hsa-mir-21&WithFlank&17|+|59841260|59841343@14.36.23(+1G)&[MIMAT0000076&hsa-miR-21-5p&offsets|0|+1(+1G);m-1&17|+|59841273|59841294&offsets|0|+1(+1G)]|TAGCTTATCAGACTGATGTTGACG</td><td>>MI0000077|hsa-mir-21&WithFlank&17|+|59841260|59841343</td><td>14</td><td>36</td><td>23</td><td>(+1G)</td><td>isomiR</td><td>YES</td><td>360.70</td><td>978.27</td><td>77.16</td><td>103.55</td></tr>
<tr><td>>MI0000077|hsa-mir-21&WithFlank&17|+|59841260|59841343@14.37.24&[MIMAT0000076&hsa-miR-21-5p&offsets|0|+2;m-1&17|+|59841273|59841294&offsets|0|+2]|TAGCTTATCAGACTGATGTTGACT</td><td>>MI0000077|hsa-mir-21&WithFlank&17|+|59841260|59841343</td><td>14</td><td>37</td><td>24</td><td></td><td>isomiR</td><td>YES</td><td>2671.81</td><td>7967.78</td><td>836.98</td><td>1394.33</td></tr>
<tr><td>>MI0000077|hsa-mir-21&WithFlank&17|+|59841260|59841343@14.39.26(+1U)&[MIMAT0000076&hsa-miR-21-5p&offsets|0|+4(+1U);m-1&17|+|59841273|59841294&offsets|0|+4(+1U)]|TAGCTTATCAGACTGATGTTGACTGTT</td><td>>MI0000077|hsa-mir-21&WithFlank&17|+|59841260|59841343</td><td>14</td><td>39</td><td>26</td><td>(+1U)</td><td>isomiR</td><td>NO</td><td>10.20</td><td>0.00</td><td>0.00</td><td>0.00</td></tr>
<tr><td>>MI0000077|hsa-mir-21&WithFlank&17|+|59841260|59841343@15.36.22&[MIMAT0000076&hsa-miR-21-5p&offsets|+1|+1;m-1&17|+|59841273|59841294&offsets|+1|+1]|AGCTTATCAGACTGATGTTGAC</td><td>>MI0000077|hsa-mir-21&WithFlank&17|+|59841260|59841343</td><td>15</td><td>36</td><td>22</td><td></td><td>isomiR</td><td>NO</td><td>65.07</td><td>79.49</td><td>0.00</td><td>30.34</td></tr>
<tr><td>>MI0000077|hsa-mir-21&WithFlank&17|+|59841260|59841343@15.36.22(+1A)&[MIMAT0000076&hsa-miR-21-5p&offsets|+1|+1(+1A);m-1&17|+|59841273|59841294&offsets|+1|+1(+1A)]|AGCTTATCAGACTGATGTTGACA</td><td>>MI0000077|hsa-mir-21&WithFlank&17|+|59841260|59841343</td><td>15</td><td>36</td><td>22</td><td>(+1A)</td><td>isomiR</td><td>NO</td><td>0.00</td><td>10.14</td><td>0.00</td><td>0.00</td></tr>
<tr><td>>MI0000077|hsa-mir-21&WithFlank&17|+|59841260|59841343@16.36.21&[MIMAT0000076&hsa-miR-21-5p&offsets|+2|+1;m-1&17|+|59841273|59841294&offsets|+2|+1]|GCTTATCAGACTGATGTTGAC</td><td>>MI0000077|hsa-mir-21&WithFlank&17|+|59841260|59841343</td><td>16</td><td>36</td><td>21</td><td></td><td>isomiR</td><td>NO</td><td>0.00</td><td>30.88</td><td>0.00</td><td>0.00</td></tr>
<tr><td>>MI0000077|hsa-mir-21&WithFlank&17|+|59841260|59841343@52.72.21&[MIMAT0004494&hsa-miR-21-3p&offsets|0|0;m-52&17|+|59841311|59841332&offsets|0|-1]|CAACACCAGTCGATGGGCTGT</td><td>>MI0000077|hsa-mir-21&WithFlank&17|+|59841260|59841343</td><td>52</td><td>72</td><td>21</td><td></td><td>isomiR</td><td>NO</td><td>0.00</td><td>9.97</td><td>83.07</td><td>0.00</td></tr>
<tr><td>>MI0000077|hsa-mir-21&WithFlank&17|+|59841260|59841343@52.73.22&[MIMAT0004494&hsa-miR-21-3p&offsets|0|+1;m-52&17|+|59841311|59841332&offsets|0|0]|CAACACCAGTCGATGGGCTGTC</td><td>>MI0000077|hsa-mir-21&WithFlank&17|+|59841260|59841343</td><td>52</td><td>73</td><td>22</td><td></td><td>isomiR</td><td>NO</td><td>0.00</td><td>0.00</td><td>67.21</td><td>0.00</td></tr>
<tr><td>>MI0000078|hsa-mir-22&WithFlank&17|-|1713897|1713993@21.41.21&[MIMAT0004495&hsa-miR-22-5p&offsets|0|-1;m-217&17|-|1713953|1713973&offsets|0|0]|AGTTCTTCAGTGGCAAGCTTT</td><td>>MI0000078|hsa-mir-22&WithFlank&17|-|1713897|1713993</td><td>21</td><td>41</td><td>21</td><td></td><td>isomiR</td><td>NO</td><td>0.00</td><td>29.75</td><td>66.60</td><td>0.00</td></tr>
<tr><td>>MI0000078|hsa-mir-22&WithFlank&17|-|1713897|1713993@59.78.20&[MIMAT0000077&hsa-miR-22-3p&offsets|0|-2;m-2&17|-|1713914|1713935&offsets|0|-2]|AAGCTGCCAGTTGAAGAACT</td><td>>MI0000078|hsa-mir-22&WithFlank&17|-|1713897|1713993</td><td>59</td><td>78</td><td>20</td><td></td><td>isomiR</td><td>NO</td><td>11.16</td><td>75.26</td><td>13.17</td><td>0.00</td></tr>
<tr><td>>MI0000078|hsa-mir-22&WithFlank&17|-|1713897|1713993@59.79.21(+1A)&[MIMAT0000077&hsa-miR-22-3p&offsets|0|-1(+1A);m-2&17|-|1713914|1713935&offsets|0|-1(+1A)]|AAGCTGCCAGTTGAAGAACTGA</td><td>>MI0000078|hsa-mir-22&WithFlank&17|-|1713897|1713993</td><td>59</td><td>79</td><td>21</td><td>(+1A)</td><td>isomiR</td><td>NO</td><td>0.00</td><td>31.55</td><td>38.77</td><td>0.00</td></tr>
<tr><td>>MI0000078|hsa-mir-22&WithFlank&17|-|1713897|1713993@59.79.21(+1C)&[MIMAT0000077&hsa-miR-22-3p&offsets|0|-1(+1C);m-2&17|-|1713914|1713935&offsets|0|-1(+1C)]|AAGCTGCCAGTTGAAGAACTGC</td><td>>MI0000078|hsa-mir-22&WithFlank&17|-|1713897|1713993</td><td>59</td><td>79</td><td>21</td><td>(+1C)</td><td>isomiR</td><td>NO</td><td>78.81</td><td>127.35</td><td>185.23</td><td>10.50</td></tr>
<tr><td>>MI0000078|hsa-mir-22&WithFlank&17|-|1713897|1713993@59.80.22&[MIMAT0000077&hsa-miR-22-3p&offsets|0|0;m-2&17|-|1713914|1713935&offsets|0|0]|AAGCTGCCAGTTGAAGAACTGT</td><td>>MI0000078|hsa-mir-22&WithFlank&17|-|1713897|1713993</td><td>59</td><td>80</td><td>22</td><td></td><td>isomiR</td><td>NO</td><td>366.86</td><td>812.31</td><td>979.38</td><td>166.35</td></tr>
<tr><td>>MI0000078|hsa-mir-22&WithFlank&17|-|1713897|1713993@59.80.22(+1A)&[MIMAT0000077&hsa-miR-22-3p&offsets|0|0(+1A);m-2&17|-|1713914|1713935&offsets|0|0(+1A)]|AAGCTGCCAGTTGAAGAACTGTA</td><td>>MI0000078|hsa-mir-22&WithFlank&17|-|1713897|1713993</td><td>59</td><td>80</td><td>22</td><td>(+1A)</td><td>isomiR</td><td>NO</td><td>0.00</td><td>26.32</td><td>11.28</td><td>0.00</td></tr>
<tr><td>>MI0000079|hsa-mir-23a&WithFlank&19|-|13836581|13836665@54.71.18&[MIMAT0000078&hsa-miR-23a-3p&offsets|+3|0;m-31&19|-|13836595|13836615&offsets|+3|0]|ACATTGCCAGGGATTTCC</td><td>>MI0000079|hsa-mir-23a&WithFlank&19|-|13836581|13836665</td><td>54</td><td>71</td><td>18</td><td></td><td>isomiR</td><td>NO</td><td>164.04</td><td>671.01</td><td>234.75</td><td>33.98</td></tr>
<tr><td>>MI0000079|hsa-mir-23a&WithFlank&19|-|13836581|13836665@54.71.18(+1G)&[MIMAT0000078&hsa-miR-23a-3p&offsets|+3|0(+1G);m-31&19|-|13836595|13836615&offsets|+3|0(+1G)]|ACATTGCCAGGGATTTCCG</td><td>>MI0000079|hsa-mir-23a&WithFlank&19|-|13836581|13836665</td><td>54</td><td>71</td><td>18</td><td>(+1G)</td><td>isomiR</td><td>NO</td><td>0.00</td><td>24.62</td><td>0.00</td><td>0.00</td></tr>
<tr><td>>MI0000079|hsa-mir-23a&WithFlank&19|-|13836581|13836665@54.72.19&[MIMAT0000078&hsa-miR-23a-3p&offsets|+3|+1;m-31&19|-|13836595|13836615&offsets|+3|+1]|ACATTGCCAGGGATTTCCA</td><td>>MI0000079|hsa-mir-23a&WithFlank&19|-|13836581|13836665</td><td>54</td><td>72</td><td>19</td><td></td><td>isomiR</td><td>NO</td><td>223.94</td><td>758.19</td><td>98.61</td><td>86.79</td></tr>
<tr><td>>MI0000079|hsa-mir-23a&WithFlank&19|-|13836581|13836665@54.72.19(+1U)&[MIMAT0000078&hsa-miR-23a-3p&offsets|+3|+1(+1U);m-31&19|-|13836595|13836615&offsets|+3|+1(+1U)]|ACATTGCCAGGGATTTCCAT</td><td>>MI0000079|hsa-mir-23a&WithFlank&19|-|13836581|13836665</td><td>54</td><td>72</td><td>19</td><td>(+1U)</td><td>isomiR</td><td>NO</td><td>10.35</td><td>42.39</td><td>0.00</td><td>0.00</td></tr>
</table>

# BT-20

<table>
<tr><td>>MI0000079|hsa-mir-23a&WithFlank&19|-|13836581|13836665@54.73.20&[MIMAT0000078&hsa-miR-23a-3p&offsets|+3|+2;m-31&19|-|13836595|13836615&offsets|+3|+2]|ACATTGCCAGGGATTTCCAA</td><td>>MI0000079|hsa-mir-23a&WithFlank&19|-|13836581|13836665</td><td>54</td><td>73</td><td>20</td><td></td><td>isomiR</td><td>NO</td><td>79.54</td><td>214.82</td><td>0.00</td><td>0.00</td></tr>
<tr><td>>MI0000079|hsa-mir-23a&WithFlank&19|-|13836581|13836665@54.73.20(+1U)&[MIMAT0000078&hsa-miR-23a-3p&offsets|+3|+2(+1U);m-31&19|-|13836595|13836615&offsets|+3|+2(+1U)]|ACATTGCCAGGGATTTCCAAT</td><td>>MI0000079|hsa-mir-23a&WithFlank&19|-|13836581|13836665</td><td>54</td><td>73</td><td>20</td><td>(+1U)</td><td>isomiR</td><td>NO</td><td>32.53</td><td>101.80</td><td>0.00</td><td>0.00</td></tr>
<tr><td>>MI0000081|hsa-mir-24-2&WithFlank&19|-|13836281|13836365@56.74.19(+1U);MI0000080|hsa-mir-24-1&WithFlank&9|+|95086015|95086094@50.68.19(+1U)&[MIMAT0000080&hsa-miR-24-3p&offsets|0|-3(+1U);m-36&19|-|13836288|13836310&offsets|0|-4(+1U)];[MIMAT0000080_1&hsa-miR-24-3p&offsets|0|-3(+1U);m-35&9|+|95086064|95086086&offsets|0|-4(+1U)]|TGGCTCAGTTCAGCAGGAAT</td><td>>MI0000081|hsa-mir-24-2&WithFlank&19|-|13836281|13836365</td><td>56</td><td>74</td><td>19</td><td>(+1U)</td><td>isomiR</td><td>NO</td><td>37.37</td><td>81.83</td><td>139.79</td><td>23.81</td></tr>
<tr><td>>MI0000081|hsa-mir-24-2&WithFlank&19|-|13836281|13836365@56.75.20;MI0000080|hsa-mir-24-1&WithFlank&9|+|95086015|95086094@50.69.20&[MIMAT0000080&hsa-miR-24-3p&offsets|0|-2;m-36&19|-|13836288|13836310&offsets|0|-3];[MIMAT0000080_1&hsa-miR-24-3p&offsets|0|-2;m-35&9|+|95086064|95086086&offsets|0|-3]|TGGCTCAGTTCAGCAGGAAC</td><td>>MI0000081|hsa-mir-24-2&WithFlank&19|-|13836281|13836365</td><td>56</td><td>75</td><td>20</td><td></td><td>isomiR</td><td>NO</td><td>377.78</td><td>611.84</td><td>1076.88</td><td>271.53</td></tr>
<tr><td>>MI0000081|hsa-mir-24-2&WithFlank&19|-|13836281|13836365@56.76.21(+1A);MI0000080|hsa-mir-24-1&WithFlank&9|+|95086015|95086094@50.70.21(+1A)&[MIMAT0000080&hsa-miR-24-3p&offsets|0|-1(+1A);m-36&19|-|13836288|13836310&offsets|0|-2(+1A)];[MIMAT0000080_1&hsa-miR-24-3p&offsets|0|-1(+1A);m-35&9|+|95086064|95086086&offsets|0|-2(+1A)]|TGGCTCAGTTCAGCAGGAACAA</td><td>>MI0000081|hsa-mir-24-2&WithFlank&19|-|13836281|13836365</td><td>56</td><td>76</td><td>21</td><td>(+1A)</td><td>isomiR</td><td>NO</td><td>42.35</td><td>92.66</td><td>244.36</td><td>25.36</td></tr>
<tr><td>>MI0000081|hsa-mir-24-2&WithFlank&19|-|13836281|13836365@56.76.21(+1U);MI0000080|hsa-mir-24-1&WithFlank&9|+|95086015|95086094@50.70.21(+1U)&[MIMAT0000080&hsa-miR-24-3p&offsets|0|-1(+1U);m-36&19|-|13836288|13836310&offsets|0|-2(+1U)];[MIMAT0000080_1&hsa-miR-24-3p&offsets|0|-1(+1U);m-35&9|+|95086064|95086086&offsets|0|-2(+1U)]|TGGCTCAGTTCAGCAGGAACAT</td><td>>MI0000081|hsa-mir-24-2&WithFlank&19|-|13836281|13836365</td><td>56</td><td>76</td><td>21</td><td>(+1U)</td><td>isomiR</td><td>NO</td><td>0.00</td><td>0.00</td><td>14.78</td><td>0.00</td></tr>
<tr><td>>MI0000081|hsa-mir-24-2&WithFlank&19|-|13836281|13836365@56.76.21;MI0000080|hsa-mir-24-1&WithFlank&9|+|95086015|95086094@50.70.21&[MIMAT0000080&hsa-miR-24-3p&offsets|0|-1;m-36&19|-|13836288|13836310&offsets|0|-2];[MIMAT0000080_1&hsa-miR-24-3p&offsets|0|-1;m-35&9|+|95086064|95086086&offsets|0|-2]|TGGCTCAGTTCAGCAGGAACA</td><td>>MI0000081|hsa-mir-24-2&WithFlank&19|-|13836281|13836365</td><td>56</td><td>76</td><td>21</td><td></td><td>isomiR</td><td>NO</td><td>110.79</td><td>150.39</td><td>374.73</td><td>112.94</td></tr>
<tr><td>>MI0000081|hsa-mir-24-2&WithFlank&19|-|13836281|13836365@56.77.22(+1A);MI0000080|hsa-mir-24-1&WithFlank&9|+|95086015|95086094@50.71.22(+1A)&[MIMAT0000080&hsa-miR-24-3p&offsets|0|0(+1A);m-36&19|-|13836288|13836310&offsets|0|-1(+1A)];[MIMAT0000080_1&hsa-miR-24-3p&offsets|0|0(+1A);m-35&9|+|95086064|95086086&offsets|0|-1(+1A)]|TGGCTCAGTTCAGCAGGAACAGA</td><td>>MI0000081|hsa-mir-24-2&WithFlank&19|-|13836281|13836365</td><td>56</td><td>77</td><td>22</td><td>(+1A)</td><td>isomiR</td><td>NO</td><td>31.74</td><td>25.42</td><td>0.00</td><td>0.00</td></tr>
<tr><td>>MI0000081|hsa-mir-24-2&WithFlank&19|-|13836281|13836365@56.77.22(+1C);MI0000080|hsa-mir-24-1&WithFlank&9|+|95086015|95086094@50.71.22(+1C)&[MIMAT0000080&hsa-miR-24-3p&offsets|0|0(+1C);m-36&19|-|13836288|13836310&offsets|0|-1(+1C)];[MIMAT0000080_1&hsa-miR-24-3p&offsets|0|0(+1C);m-35&9|+|95086064|95086086&offsets|0|-1(+1C)]|TGGCTCAGTTCAGCAGGAACAGC</td><td>>MI0000081|hsa-mir-24-2&WithFlank&19|-|13836281|13836365</td><td>56</td><td>77</td><td>22</td><td>(+1C)</td><td>isomiR</td><td>NO</td><td>35.17</td><td>24.02</td><td>0.00</td><td>0.00</td></tr>
<tr><td>>MI0000081|hsa-mir-24-2&WithFlank&19|-|13836281|13836365@56.77.22(+1U);MI0000080|hsa-mir-24-1&WithFlank&9|+|95086015|95086094@50.71.22(+1U)&[MIMAT0000080&hsa-miR-24-3p&offsets|0|0(+1U);m-36&19|-|13836288|13836310&offsets|0|-1(+1U)];[MIMAT0000080_1&hsa-miR-24-3p&offsets|0|0(+1U);m-35&9|+|95086064|95086086&offsets|0|-1(+1U)]|TGGCTCAGTTCAGCAGGAACAGT</td><td>>MI0000081|hsa-mir-24-2&WithFlank&19|-|13836281|13836365</td><td>56</td><td>77</td><td>22</td><td>(+1U)</td><td>isomiR</td><td>NO</td><td>260.12</td><td>239.89</td><td>95.69</td><td>99.46</td></tr>
<tr><td>>MI0000081|hsa-mir-24-2&WithFlank&19|-|13836281|13836365@56.77.22;MI0000080|hsa-mir-24-1&WithFlank&9|+|95086015|95086094@50.71.22&[MIMAT0000080&hsa-miR-24-3p&offsets|0|0;m-36&19|-|13836288|13836310&offsets|0|-1];[MIMAT0000080_1&hsa-miR-24-3p&offsets|0|0;m-35&9|+|95086064|95086086&offsets|0|-1]|TGGCTCAGTTCAGCAGGAACAG</td><td>>MI0000081|hsa-mir-24-2&WithFlank&19|-|13836281|13836365</td><td>56</td><td>77</td><td>22</td><td></td><td>isomiR</td><td>NO</td><td>339.97</td><td>476.09</td><td>1568.07</td><td>228.55</td></tr>
<tr><td>>MI0000082|hsa-mir-25&WithFlank&7|-|100093554|100093649@58.77.20&[MIMAT0000081&hsa-miR-25-3p&offsets|0|-2;m-23&7|-|100093571|100093592&offsets|0|-2]|CATTGCACTTGTCTCGGTCT</td><td>>MI0000082|hsa-mir-25&WithFlank&7|-|100093554|100093649</td><td>58</td><td>77</td><td>20</td><td></td><td>isomiR</td><td>NO</td><td>0.00</td><td>30.24</td><td>0.00</td><td>0.00</td></tr>
<tr><td>>MI0000082|hsa-mir-25&WithFlank&7|-|100093554|100093649@58.77.20(+1A)&[MIMAT0000081&hsa-miR-25-3p&offsets|0|-2(+1A);m-23&7|-|100093571|100093592&offsets|0|-2(+1A)]|CATTGCACTTGTCTCGGTCTA</td><td>>MI0000082|hsa-mir-25&WithFlank&7|-|100093554|100093649</td><td>58</td><td>77</td><td>20</td><td>(+1A)</td><td>isomiR</td><td>NO</td><td>11.04</td><td>85.31</td><td>58.74</td><td>0.00</td></tr>
<tr><td>>MI0000082|hsa-mir-25&WithFlank&7|-|100093554|100093649@58.78.21&[MIMAT0000081&hsa-miR-25-3p&offsets|0|-1;m-23&7|-|100093571|100093592&offsets|0|-1]|CATTGCACTTGTCTCGGTCTG</td><td>>MI0000082|hsa-mir-25&WithFlank&7|-|100093554|100093649</td><td>58</td><td>78</td><td>21</td><td></td><td>isomiR</td><td>NO</td><td>170.50</td><td>406.75</td><td>356.21</td><td>136.87</td></tr>
<tr><td>>MI0000082|hsa-mir-25&WithFlank&7|-|100093554|100093649@58.78.21(+1C)&[MIMAT0000081&hsa-miR-25-3p&offsets|0|-1(+1C);m-23&7|-|100093571|100093592&offsets|0|-1(+1C)]|CATTGCACTTGTCTCGGTCTGC</td><td>>MI0000082|hsa-mir-25&WithFlank&7|-|100093554|100093649</td><td>58</td><td>78</td><td>21</td><td>(+1C)</td><td>isomiR</td><td>NO</td><td>0.00</td><td>32.59</td><td>9.85</td><td>0.00</td></tr>
<tr><td>>MI0000082|hsa-mir-25&WithFlank&7|-|100093554|100093649@58.78.21(+1G)&[MIMAT0000081&hsa-miR-25-3p&offsets|0|-1(+1G);m-23&7|-|100093571|100093592&offsets|0|-1(+1G)]|CATTGCACTTGTCTCGGTCTGG</td><td>>MI0000082|hsa-mir-25&WithFlank&7|-|100093554|100093649</td><td>58</td><td>78</td><td>21</td><td>(+1G)</td><td>isomiR</td><td>NO</td><td>34.47</td><td>134.08</td><td>16.31</td><td>0.00</td></tr>
<tr><td>>MI0000082|hsa-mir-25&WithFlank&7|-|100093554|100093649@58.78.21(+1U)&[MIMAT0000081&hsa-miR-25-3p&offsets|0|-1(+1U);m-23&7|-|100093571|100093592&offsets|0|-1(+1U)]|CATTGCACTTGTCTCGGTCTGT</td><td>>MI0000082|hsa-mir-25&WithFlank&7|-|100093554|100093649</td><td>58</td><td>78</td><td>21</td><td>(+1U)</td><td>isomiR</td><td>NO</td><td>102.35</td><td>319.22</td><td>156.28</td><td>66.66</td></tr>
<tr><td>>MI0000082|hsa-mir-25&WithFlank&7|-|100093554|100093649@58.79.22&[MIMAT0000081&hsa-miR-25-3p&offsets|0|0;m-23&7|-|100093571|100093592&offsets|0|0]|CATTGCACTTGTCTCGGTCTGA</td><td>>MI0000082|hsa-mir-25&WithFlank&7|-|100093554|100093649</td><td>58</td><td>79</td><td>22</td><td></td><td>isomiR</td><td>YES</td><td>862.02</td><td>3627.56</td><td>1515.47</td><td>643.68</td></tr>
</table>

# BT-20

| | | | | | | | | | | | |
|---|---|---|---|---|---|---|---|---|---|---|---|
| >MI0000082\|hsa-mir-25&WithFlank&7\|-\|100093554\|100093649@58.79.22(+1A)&[MIMAT0000081&hsa-miR-25-3p&offsets\|0\|0(+1A);m-23&7\|-\|100093571\|100093592&offsets\|0\|0(+1A)]\|CATTGCACTTGTCTCGGTCTGAA | >MI0000082\|hsa-mir-25&WithFlank&7\|-\|100093554\|100093649 | 58 | 79 | 22 | (+1A) | isomiR | NO | 32.07 | 193.92 | 12.27 | 10.70 |
| >MI0000082\|hsa-mir-25&WithFlank&7\|-\|100093554\|100093649@58.79.22(+1U)&[MIMAT0000081&hsa-miR-25-3p&offsets\|0\|0(+1U);m-23&7\|-\|100093571\|100093592&offsets\|0\|0(+1U)]\|CATTGCACTTGTCTCGGTCTGAT | >MI0000082\|hsa-mir-25&WithFlank&7\|-\|100093554\|100093649 | 58 | 79 | 22 | (+1U) | isomiR | NO | 10.11 | 116.25 | 0.00 | 0.00 |
| >MI0000082\|hsa-mir-25&WithFlank&7\|-\|100093554\|100093649@58.79.22(+2A)&[MIMAT0000081&hsa-miR-25-3p&offsets\|0\|0(+2A);m-23&7\|-\|100093571\|100093592&offsets\|0\|0(+2A)]\|CATTGCACTTGTCTCGGTCTGAAA | >MI0000082\|hsa-mir-25&WithFlank&7\|-\|100093554\|100093649 | 58 | 79 | 22 | (+2A) | isomiR | NO | 0.00 | 69.65 | 0.00 | 0.00 |
| >MI0000084\|hsa-mir-26b&WithFlank&2\|+\|218402640\|218402728@18.38.21&[MIMAT0000083&hsa-miR-26b-5p&offsets\|0\|0;m-53&2\|+\|218402657\|218402678&offsets\|0\|-1]\|TTCAAGTAATTCAGGATAGGT | >MI0000084\|hsa-mir-26b&WithFlank&2\|+\|218402640\|218402728 | 18 | 38 | 21 | | isomiR | NO | 41.25 | 88.73 | 10.00 | 0.00 |
| >MI0000084\|hsa-mir-26b&WithFlank&2\|+\|218402640\|218402728@18.38.21(+1A)&[MIMAT0000083&hsa-miR-26b-5p&offsets\|0\|0(+1A);m-53&2\|+\|218402657\|218402678&offsets\|0\|-1(+1A)]\|TTCAAGTAATTCAGGATAGGTA | >MI0000084\|hsa-mir-26b&WithFlank&2\|+\|218402640\|218402728 | 18 | 38 | 21 | (+1A) | isomiR | NO | 0.00 | 24.69 | 0.00 | 0.00 |
| >MI0000084\|hsa-mir-26b&WithFlank&2\|+\|218402640\|218402728@18.38.21(+1C)&[MIMAT0000083&hsa-miR-26b-5p&offsets\|0\|0(+1C);m-53&2\|+\|218402657\|218402678&offsets\|0\|-1(+1C)]\|TTCAAGTAATTCAGGATAGGTC | >MI0000084\|hsa-mir-26b&WithFlank&2\|+\|218402640\|218402728 | 18 | 38 | 21 | (+1C) | isomiR | NO | 43.45 | 81.16 | 0.00 | 0.00 |
| >MI0000084\|hsa-mir-26b&WithFlank&2\|+\|218402640\|218402728@18.39.22&[MIMAT0000083&hsa-miR-26b-5p&offsets\|0\|+1;m-53&2\|+\|218402657\|218402678&offsets\|0\|0]\|TTCAAGTAATTCAGGATAGGTT | >MI0000084\|hsa-mir-26b&WithFlank&2\|+\|218402640\|218402728 | 18 | 39 | 22 | | isomiR | NO | 315.27 | 513.48 | 57.62 | 65.70 |
| >MI0000085\|hsa-mir-27a&WithFlank&19\|-\|13836434\|13836523@16.36.21&[MIMAT0004501&hsa-miR-27a-5p&offsets\|0\|-1;m-270&19\|-\|13836487\|13836508&offsets\|0\|-1]\|AGGGCTTAGCTGCTTGTGAGC | >MI0000085\|hsa-mir-27a&WithFlank&19\|-\|13836434\|13836523 | 16 | 36 | 21 | | isomiR | NO | 0.00 | 27.95 | 0.00 | 0.00 |
| >MI0000085\|hsa-mir-27a&WithFlank&19\|-\|13836434\|13836523@16.37.22&[MIMAT0004501&hsa-miR-27a-5p&offsets\|0\|0;m-270&19\|-\|13836487\|13836508&offsets\|0\|0]\|AGGGCTTAGCTGCTTGTGAGCA | >MI0000085\|hsa-mir-27a&WithFlank&19\|-\|13836434\|13836523 | 16 | 37 | 22 | | isomiR | NO | 0.00 | 93.63 | 0.00 | 0.00 |
| >MI0000085\|hsa-mir-27a&WithFlank&19\|-\|13836434\|13836523@57.74.18;MI0000440\|hsa-mir-27b&WithFlank&9\|+\|95085439\|95085547@67.84.18&[MIMAT0000084&hsa-miR-27a-3p&offsets\|0\|-3;m-43&19\|-\|13836447\|13836467&offsets\|0\|-3];[MIMAT0000419&hsa-miR-27b-3p&offsets\|0\|-3;m-39&9\|+\|95085505\|95085525&offsets\|0\|-3]\|TTCACAGTGGCTAAGTTC | >MI0000085\|hsa-mir-27a&WithFlank&19\|-\|13836434\|13836523 | 57 | 74 | 18 | | isomiR | NO | 0.00 | 10.47 | 11.76 | 0.00 |
| >MI0000085\|hsa-mir-27a&WithFlank&19\|-\|13836434\|13836523@57.75.19&[MIMAT0000084&hsa-miR-27a-3p&offsets\|0\|-2;m-43&19\|-\|13836447\|13836467&offsets\|0\|-2]\|TTCACAGTGGCTAAGTTCC | >MI0000085\|hsa-mir-27a&WithFlank&19\|-\|13836434\|13836523 | 57 | 75 | 19 | | isomiR | NO | 0.00 | 25.13 | 12.91 | 0.00 |
| >MI0000085\|hsa-mir-27a&WithFlank&19\|-\|13836434\|13836523@57.75.19(+1A)&[MIMAT0000084&hsa-miR-27a-3p&offsets\|0\|-2(+1A);m-43&19\|-\|13836447\|13836467&offsets\|0\|-2(+1A)]\|TTCACAGTGGCTAAGTTCCA | >MI0000085\|hsa-mir-27a&WithFlank&19\|-\|13836434\|13836523 | 57 | 75 | 19 | (+1A) | isomiR | NO | 28.72 | 76.71 | 59.18 | 0.00 |
| >MI0000085\|hsa-mir-27a&WithFlank&19\|-\|13836434\|13836523@57.76.20&[MIMAT0000084&hsa-miR-27a-3p&offsets\|0\|-1;m-43&19\|-\|13836447\|13836467&offsets\|0\|-1]\|TTCACAGTGGCTAAGTTCCG | >MI0000085\|hsa-mir-27a&WithFlank&19\|-\|13836434\|13836523 | 57 | 76 | 20 | | isomiR | NO | 154.76 | 270.71 | 263.38 | 79.62 |
| >MI0000085\|hsa-mir-27a&WithFlank&19\|-\|13836434\|13836523@57.76.20(+1A)&[MIMAT0000084&hsa-miR-27a-3p&offsets\|0\|-1(+1A);m-43&19\|-\|13836447\|13836467&offsets\|0\|-1(+1A)]\|TTCACAGTGGCTAAGTTCCGA | >MI0000085\|hsa-mir-27a&WithFlank&19\|-\|13836434\|13836523 | 57 | 76 | 20 | (+1A) | isomiR | NO | 39.89 | 96.67 | 112.83 | 9.50 |
| >MI0000085\|hsa-mir-27a&WithFlank&19\|-\|13836434\|13836523@57.76.20(+1U)&[MIMAT0000084&hsa-miR-27a-3p&offsets\|0\|-1(+1U);m-43&19\|-\|13836447\|13836467&offsets\|0\|-1(+1U)]\|TTCACAGTGGCTAAGTTCCGT | >MI0000085\|hsa-mir-27a&WithFlank&19\|-\|13836434\|13836523 | 57 | 76 | 20 | (+1U) | isomiR | NO | 96.15 | 107.38 | 113.31 | 27.06 |
| >MI0000085\|hsa-mir-27a&WithFlank&19\|-\|13836434\|13836523@57.76.20(+2U)&[MIMAT0000084&hsa-miR-27a-3p&offsets\|0\|-1(+2U);m-43&19\|-\|13836447\|13836467&offsets\|0\|-1(+2U)]\|TTCACAGTGGCTAAGTTCCGTT | >MI0000085\|hsa-mir-27a&WithFlank&19\|-\|13836434\|13836523 | 57 | 76 | 20 | (+2U) | isomiR | NO | 26.09 | 32.32 | 12.27 | 0.00 |
| >MI0000085\|hsa-mir-27a&WithFlank&19\|-\|13836434\|13836523@57.77.21&[MIMAT0000084&hsa-miR-27a-3p&offsets\|0\|0;m-43&19\|-\|13836447\|13836467&offsets\|0\|0]\|TTCACAGTGGCTAAGTTCCGC | >MI0000085\|hsa-mir-27a&WithFlank&19\|-\|13836434\|13836523 | 57 | 77 | 21 | | isomiR | NO | 36.89 | 37.17 | 100.21 | 0.00 |
| >MI0000086\|hsa-mir-28&WithFlank&3\|+\|188688775\|188688872@20.40.21&[MIMAT0000085&hsa-miR-28-5p&offsets\|0\|-1;m-133&3\|+\|188688794\|188688815&offsets\|0\|-1]\|AAGGAGCTCACAGTCTATTGA | >MI0000086\|hsa-mir-28&WithFlank&3\|+\|188688775\|188688872 | 20 | 40 | 21 | | isomiR | NO | 0.00 | 23.30 | 13.03 | 0.00 |
| >MI0000086\|hsa-mir-28&WithFlank&3\|+\|188688775\|188688872@20.41.22&[MIMAT0000085&hsa-miR-28-5p&offsets\|0\|0;m-133&3\|+\|188688794\|188688815&offsets\|0\|0]\|AAGGAGCTCACAGTCTATTGAG | >MI0000086\|hsa-mir-28&WithFlank&3\|+\|188688775\|188688872 | 20 | 41 | 22 | | isomiR | NO | 30.78 | 34.76 | 55.47 | 0.00 |
| >MI0000086\|hsa-mir-28&WithFlank&3\|+\|188688775\|188688872@60.79.20&[MIMAT0004502&hsa-miR-28-3p&offsets\|0\|-2;m-28&3\|+\|188688834\|188688855&offsets\|0\|-2]\|CACTAGATTGTGAGCTCCTG | >MI0000086\|hsa-mir-28&WithFlank&3\|+\|188688775\|188688872 | 60 | 79 | 20 | | isomiR | NO | 0.00 | 10.17 | 0.00 | 0.00 |
| >MI0000086\|hsa-mir-28&WithFlank&3\|+\|188688775\|188688872@60.79.20(+1A)&[MIMAT0004502&hsa-miR-28-3p&offsets\|0\|-2(+1A);m-28&3\|+\|188688834\|188688855&offsets\|0\|-2(+1A)]\|CACTAGATTGTGAGCTCCTGA | >MI0000086\|hsa-mir-28&WithFlank&3\|+\|188688775\|188688872 | 60 | 79 | 20 | (+1A) | isomiR | NO | 0.00 | 42.94 | 0.00 | 0.00 |
| >MI0000086\|hsa-mir-28&WithFlank&3\|+\|188688775\|188688872@60.80.21&[MIMAT0004502&hsa-miR-28-3p&offsets\|0\|-1;m-28&3\|+\|188688834\|188688855&offsets\|0\|-1]\|CACTAGATTGTGAGCTCCTGG | >MI0000086\|hsa-mir-28&WithFlank&3\|+\|188688775\|188688872 | 60 | 80 | 21 | | isomiR | NO | 30.69 | 281.24 | 126.28 | 0.00 |
| >MI0000086\|hsa-mir-28&WithFlank&3\|+\|188688775\|188688872@60.80.21(+1G)&[MIMAT0004502&hsa-miR-28-3p&offsets\|0\|-1(+1G);m-28&3\|+\|188688834\|188688855&offsets\|0\|-1(+1G)]\|CACTAGATTGTGAGCTCCTGGG | >MI0000086\|hsa-mir-28&WithFlank&3\|+\|188688775\|188688872 | 60 | 80 | 21 | (+1G) | isomiR | NO | 0.00 | 45.06 | 0.00 | 0.00 |
| >MI0000086\|hsa-mir-28&WithFlank&3\|+\|188688775\|188688872@60.80.21(+1U)&[MIMAT0004502&hsa-miR-28-3p&offsets\|0\|-1(+1U);m-28&3\|+\|188688834\|188688855&offsets\|0\|-1(+1U)]\|CACTAGATTGTGAGCTCCTGGT | >MI0000086\|hsa-mir-28&WithFlank&3\|+\|188688775\|188688872 | 60 | 80 | 21 | (+1U) | isomiR | NO | 0.00 | 86.45 | 0.00 | 0.00 |

# BT-20

<table>
<tr><td></td><td></td><td></td><td></td><td></td><td></td><td></td><td></td><td></td><td></td><td></td><td></td></tr>
<tr><td>>MI0000086|hsa-mir-28&WithFlank&3|+|188688775|188688872@60.81.22&[MIMAT0004502&hsa-miR-28-3p&offsets|0|0;m-28&3|+|188688834|188688855&offsets|0|0]|CACTAGATTGTGAGCTCCTGGA</td><td>>MI0000086|hsa-mir-28&WithFlank&3|+|188688775|188688872</td><td>60</td><td>81</td><td>22</td><td></td><td>isomiR</td><td>NO</td><td>180.64</td><td>1825.41</td><td>404.97</td><td>131.71</td></tr>
<tr><td>>MI0000086|hsa-mir-28&WithFlank&3|+|188688775|188688872@60.81.22(+1A)&[MIMAT0004502&hsa-miR-28-3p&offsets|0|0(+1A);m-28&3|+|188688834|188688855&offsets|0|0(+1A)]|CACTAGATTGTGAGCTCCTGGAA</td><td>>MI0000086|hsa-mir-28&WithFlank&3|+|188688775|188688872</td><td>60</td><td>81</td><td>22</td><td>(+1A)</td><td>isomiR</td><td>NO</td><td>33.15</td><td>345.90</td><td>79.57</td><td>23.63</td></tr>
<tr><td>>MI0000086|hsa-mir-28&WithFlank&3|+|188688775|188688872@60.81.22(+1U)&[MIMAT0004502&hsa-miR-28-3p&offsets|0|0(+1U);m-28&3|+|188688834|188688855&offsets|0|0(+1U)]|CACTAGATTGTGAGCTCCTGGAT</td><td>>MI0000086|hsa-mir-28&WithFlank&3|+|188688775|188688872</td><td>60</td><td>81</td><td>22</td><td>(+1U)</td><td>isomiR</td><td>NO</td><td>0.00</td><td>81.76</td><td>0.00</td><td>0.00</td></tr>
<tr><td>>MI0000086|hsa-mir-28&WithFlank&3|+|188688775|188688872@60.81.22(+2A)&[MIMAT0004502&hsa-miR-28-3p&offsets|0|0(+2A);m-28&3|+|188688834|188688855&offsets|0|0(+2A)]|CACTAGATTGTGAGCTCCTGGAAA</td><td>>MI0000086|hsa-mir-28&WithFlank&3|+|188688775|188688872</td><td>60</td><td>81</td><td>22</td><td>(+2A)</td><td>isomiR</td><td>NO</td><td>0.00</td><td>30.86</td><td>0.00</td><td>0.00</td></tr>
<tr><td>>MI0000086|hsa-mir-28&WithFlank&3|+|188688775|188688872@60.82.23&[MIMAT0004502&hsa-miR-28-3p&offsets|0|+1;m-28&3|+|188688834|188688855&offsets|0|+1]|CACTAGATTGTGAGCTCCTGGAG</td><td>>MI0000086|hsa-mir-28&WithFlank&3|+|188688775|188688872</td><td>60</td><td>82</td><td>23</td><td></td><td>isomiR</td><td>NO</td><td>0.00</td><td>138.37</td><td>37.31</td><td>0.00</td></tr>
<tr><td>>MI0000086|hsa-mir-28&WithFlank&3|+|188688775|188688872@60.82.23(+1U)&[MIMAT0004502&hsa-miR-28-3p&offsets|0|+1(+1U);m-28&3|+|188688834|188688855&offsets|0|+1(+1U)]|CACTAGATTGTGAGCTCCTGGAGT</td><td>>MI0000086|hsa-mir-28&WithFlank&3|+|188688775|188688872</td><td>60</td><td>82</td><td>23</td><td>(+1U)</td><td>isomiR</td><td>NO</td><td>0.00</td><td>23.46</td><td>11.79</td><td>0.00</td></tr>
<tr><td>>MI0000086|hsa-mir-28&WithFlank&3|+|188688775|188688872@61.82.22&[MIMAT0004502&hsa-miR-28-3p&offsets|+1|+1;m-28&3|+|188688834|188688855&offsets|+1|+1]|ACTAGATTGTGAGCTCCTGGAG</td><td>>MI0000086|hsa-mir-28&WithFlank&3|+|188688775|188688872</td><td>61</td><td>82</td><td>22</td><td></td><td>isomiR</td><td>NO</td><td>0.00</td><td>104.76</td><td>38.59</td><td>0.00</td></tr>
<tr><td>>MI0000086|hsa-mir-28&WithFlank&3|+|188688775|188688872@61.82.22(+1U)&[MIMAT0004502&hsa-miR-28-3p&offsets|+1|+1(+1U);m-28&3|+|188688834|188688855&offsets|+1|+1(+1U)]|ACTAGATTGTGAGCTCCTGGAGT</td><td>>MI0000086|hsa-mir-28&WithFlank&3|+|188688775|188688872</td><td>61</td><td>82</td><td>22</td><td>(+1U)</td><td>isomiR</td><td>NO</td><td>0.00</td><td>38.69</td><td>39.82</td><td>0.00</td></tr>
<tr><td>>MI0000086|hsa-mir-28&WithFlank&3|+|188688775|188688872@62.82.21&[MIMAT0004502&hsa-miR-28-3p&offsets|+2|+1;m-28&3|+|188688834|188688855&offsets|+2|+1]|CTAGATTGTGAGCTCCTGGAG</td><td>>MI0000086|hsa-mir-28&WithFlank&3|+|188688775|188688872</td><td>62</td><td>82</td><td>21</td><td></td><td>isomiR</td><td>NO</td><td>0.00</td><td>29.99</td><td>12.12</td><td>0.00</td></tr>
<tr><td>>MI0000086|hsa-mir-28&WithFlank&3|+|188688775|188688872@62.82.21(+1U)&[MIMAT0004502&hsa-miR-28-3p&offsets|+2|+1(+1U);m-28&3|+|188688834|188688855&offsets|+2|+1(+1U)]|CTAGATTGTGAGCTCCTGGAGT</td><td>>MI0000086|hsa-mir-28&WithFlank&3|+|188688775|188688872</td><td>62</td><td>82</td><td>21</td><td>(+1U)</td><td>isomiR</td><td>NO</td><td>0.00</td><td>34.91</td><td>72.39</td><td>0.00</td></tr>
<tr><td>>am_MI0000086|hsa-mir-28&WithFlank&3|+|188688775|188688872@20.39.20&[MIMAT0000085&hsa-miR-28-5p&offsets|0|-2;m-133&3|+|188688794|188688815&offsets|0|-2]|AAGGAGCTCACAGTCTATTG</td><td>>MI0000086|hsa-mir-28&WithFlank&3|+|188688775|188688872.am</td><td>20</td><td>39</td><td>20</td><td></td><td>isomiR</td><td>NO</td><td>0.00</td><td>0.00</td><td>38.54</td><td>0.00</td></tr>
<tr><td>>am_MI0000086|hsa-mir-28&WithFlank&3|+|188688775|188688872@60.78.19&[MIMAT0004502&hsa-miR-28-3p&offsets|0|-3;m-28&3|+|188688834|188688855&offsets|0|-3]|CACTAGATTGTGAGCTCCT</td><td>>MI0000086|hsa-mir-28&WithFlank&3|+|188688775|188688872.am</td><td>60</td><td>78</td><td>19</td><td></td><td>isomiR</td><td>NO</td><td>0.00</td><td>27.48</td><td>11.05</td><td>0.00</td></tr>
<tr><td>>MI0000087|hsa-mir-29a&WithFlank&7|-|130876741|130876816@47.67.21(+1A)&[MIMAT0000086&hsa-miR-29a-3p&offsets|-1|-2(+1A);m-26&7|-|130876749|130876769&offsets|-1|-1(+1A)]|CTAGCACCATCTGAAATCGGTA</td><td>>MI0000087|hsa-mir-29a&WithFlank&7|-|130876741|130876816</td><td>47</td><td>67</td><td>21</td><td>(+1A)</td><td>isomiR</td><td>NO</td><td>0.00</td><td>24.42</td><td>10.75</td><td>0.00</td></tr>
<tr><td>>MI0000087|hsa-mir-29a&WithFlank&7|-|130876741|130876816@47.67.21(+1C)&[MIMAT0000086&hsa-miR-29a-3p&offsets|-1|-2(+1C);m-26&7|-|130876749|130876769&offsets|-1|-1(+1C)]|CTAGCACCATCTGAAATCGGTC</td><td>>MI0000087|hsa-mir-29a&WithFlank&7|-|130876741|130876816</td><td>47</td><td>67</td><td>21</td><td>(+1C)</td><td>isomiR</td><td>NO</td><td>72.21</td><td>83.82</td><td>108.61</td><td>28.68</td></tr>
<tr><td>>MI0000087|hsa-mir-29a&WithFlank&7|-|130876741|130876816@47.68.22&[MIMAT0000086&hsa-miR-29a-3p&offsets|-1|-1;m-26&7|-|130876749|130876769&offsets|-1|0]|CTAGCACCATCTGAAATCGGTT</td><td>>MI0000087|hsa-mir-29a&WithFlank&7|-|130876741|130876816</td><td>47</td><td>68</td><td>22</td><td></td><td>isomiR</td><td>NO</td><td>356.90</td><td>564.01</td><td>618.39</td><td>222.63</td></tr>
<tr><td>>MI0000087|hsa-mir-29a&WithFlank&7|-|130876741|130876816@47.68.22(+1U)&[MIMAT0000086&hsa-miR-29a-3p&offsets|-1|-1(+1U);m-26&7|-|130876749|130876769&offsets|-1|0(+1U)]|CTAGCACCATCTGAAATCGGTTT</td><td>>MI0000087|hsa-mir-29a&WithFlank&7|-|130876741|130876816</td><td>47</td><td>68</td><td>22</td><td>(+1U)</td><td>isomiR</td><td>NO</td><td>24.41</td><td>25.31</td><td>114.96</td><td>0.00</td></tr>
<tr><td>>MI0000087|hsa-mir-29a&WithFlank&7|-|130876741|130876816@47.69.23&[MIMAT0000086&hsa-miR-29a-3p&offsets|-1|0;m-26&7|-|130876749|130876769&offsets|-1|+1]|CTAGCACCATCTGAAATCGGTTA</td><td>>MI0000087|hsa-mir-29a&WithFlank&7|-|130876741|130876816</td><td>47</td><td>69</td><td>23</td><td></td><td>isomiR</td><td>NO</td><td>30.41</td><td>32.51</td><td>102.37</td><td>0.00</td></tr>
<tr><td>>MI0000087|hsa-mir-29a&WithFlank&7|-|130876741|130876816@48.67.20(+1C)&[MIMAT0000086&hsa-miR-29a-3p&offsets|0|-2(+1C);m-26&7|-|130876749|130876769&offsets|0|-1(+1C)]|TAGCACCATCTGAAATCGGTC</td><td>>MI0000087|hsa-mir-29a&WithFlank&7|-|130876741|130876816</td><td>48</td><td>67</td><td>20</td><td>(+1C)</td><td>isomiR</td><td>NO</td><td>32.49</td><td>40.38</td><td>79.41</td><td>0.00</td></tr>
<tr><td>>MI0000087|hsa-mir-29a&WithFlank&7|-|130876741|130876816@48.68.21&[MIMAT0000086&hsa-miR-29a-3p&offsets|0|-1;m-26&7|-|130876749|130876769&offsets|0|0]|TAGCACCATCTGAAATCGGTT</td><td>>MI0000087|hsa-mir-29a&WithFlank&7|-|130876741|130876816</td><td>48</td><td>68</td><td>21</td><td></td><td>isomiR</td><td>NO</td><td>238.32</td><td>376.03</td><td>413.26</td><td>68.94</td></tr>
<tr><td>>MI0000087|hsa-mir-29a&WithFlank&7|-|130876741|130876816@48.68.21(+1U)&[MIMAT0000086&hsa-miR-29a-3p&offsets|0|-1(+1U);m-26&7|-|130876749|130876769&offsets|0|0(+1U)]|TAGCACCATCTGAAATCGGTTT</td><td>>MI0000087|hsa-mir-29a&WithFlank&7|-|130876741|130876816</td><td>48</td><td>68</td><td>21</td><td>(+1U)</td><td>isomiR</td><td>NO</td><td>0.00</td><td>0.00</td><td>14.04</td><td>0.00</td></tr>
<tr><td>>MI0000087|hsa-mir-29a&WithFlank&7|-|130876741|130876816@48.69.22&[MIMAT0000086&hsa-miR-29a-3p&offsets|0|0;m-26&7|-|130876749|130876769&offsets|0|+1]|TAGCACCATCTGAAATCGGTTA</td><td>>MI0000087|hsa-mir-29a&WithFlank&7|-|130876741|130876816</td><td>48</td><td>69</td><td>22</td><td></td><td>isomiR</td><td>NO</td><td>215.60</td><td>233.57</td><td>541.81</td><td>78.43</td></tr>
<tr><td>>MI0000088|hsa-mir-30a&WithFlank&6|-|71403545|71403627@12.31.20&[MIMAT0000087&hsa-miR-30a-5p&offsets|0|-2;m-15&6|-|71403595|71403616&offsets|0|-2]|TGTAAACATCCTCGACTGGA</td><td>>MI0000088|hsa-mir-30a&WithFlank&6|-|71403545|71403627</td><td>12</td><td>31</td><td>20</td><td></td><td>isomiR</td><td>NO</td><td>31.02</td><td>36.39</td><td>179.21</td><td>0.00</td></tr>
<tr><td>>MI0000088|hsa-mir-30a&WithFlank&6|-|71403545|71403627@12.32.21&[MIMAT0000087&hsa-miR-30a-5p&offsets|0|-1;m-15&6|-|71403595|71403616&offsets|0|-1]|TGTAAACATCCTCGACTGGAA</td><td>>MI0000088|hsa-mir-30a&WithFlank&6|-|71403545|71403627</td><td>12</td><td>32</td><td>21</td><td></td><td>isomiR</td><td>NO</td><td>0.00</td><td>0.00</td><td>13.02</td><td>0.00</td></tr>
</table>

# BT-20

| | | | | | | | | | | | |
|---|---|---|---|---|---|---|---|---|---|---|---|
| >MI0000088\|hsa-mir-30a&WithFlank&6\|-\|71403545\|71403627@12.33.22&[MIMAT0000087&hsa-miR-30a-5p&offsets\|0\|0;m-15&6\|-\|71403595\|71403616&offsets\|0\|0]\|TGTAAACATCCTCGACTGGAAG | >MI0000088\|hsa-mir-30a&WithFlank&6\|-\|71403545\|71403627 | 12 | 33 | 22 | | isomiR | NO | 111.20 | 140.16 | 199.19 | 10.98 |
| >MI0000088\|hsa-mir-30a&WithFlank&6\|-\|71403545\|71403627@12.33.22(+1A)&[MIMAT0000087&hsa-miR-30a-5p&offsets\|0\|0(+1A);m-15&6\|-\|71403595\|71403616&offsets\|0\|0(+1A)]\|TGTAAACATCCTCGACTGGAAGA | >MI0000088\|hsa-mir-30a&WithFlank&6\|-\|71403545\|71403627 | 12 | 33 | 22 | (+1A) | isomiR | NO | 35.64 | 77.29 | 95.66 | 0.00 |
| >MI0000088\|hsa-mir-30a&WithFlank&6\|-\|71403545\|71403627@12.33.22(+1U)&[MIMAT0000087&hsa-miR-30a-5p&offsets\|0\|0(+1U);m-15&6\|-\|71403595\|71403616&offsets\|0\|0(+1U)]\|TGTAAACATCCTCGACTGGAAGT | >MI0000088\|hsa-mir-30a&WithFlank&6\|-\|71403545\|71403627 | 12 | 33 | 22 | (+1U) | isomiR | YES | 177.49 | 218.68 | 301.36 | 63.48 |
| >MI0000088\|hsa-mir-30a&WithFlank&6\|-\|71403545\|71403627@12.34.23&[MIMAT0000087&hsa-miR-30a-5p&offsets\|0\|+1;m-15&6\|-\|71403595\|71403616&offsets\|0\|+1]\|TGTAAACATCCTCGACTGGAAGC | >MI0000088\|hsa-mir-30a&WithFlank&6\|-\|71403545\|71403627 | 12 | 34 | 23 | | isomiR | YES | 467.32 | 642.89 | 924.67 | 189.17 |
| >MI0000088\|hsa-mir-30a&WithFlank&6\|-\|71403545\|71403627@12.34.23(+1A)&[MIMAT0000087&hsa-miR-30a-5p&offsets\|0\|+1(+1A);m-15&6\|-\|71403595\|71403616&offsets\|0\|+1(+1A)]\|TGTAAACATCCTCGACTGGAAGCA | >MI0000088\|hsa-mir-30a&WithFlank&6\|-\|71403545\|71403627 | 12 | 34 | 23 | (+1A) | isomiR | YES | 115.26 | 195.30 | 274.16 | 35.76 |
| >MI0000088\|hsa-mir-30a&WithFlank&6\|-\|71403545\|71403627@12.34.23(+1C)&[MIMAT0000087&hsa-miR-30a-5p&offsets\|0\|+1(+1C);m-15&6\|-\|71403595\|71403616&offsets\|0\|+1(+1C)]\|TGTAAACATCCTCGACTGGAAGCC | >MI0000088\|hsa-mir-30a&WithFlank&6\|-\|71403545\|71403627 | 12 | 34 | 23 | (+1C) | isomiR | YES | 176.87 | 218.51 | 400.80 | 79.18 |
| >MI0000088\|hsa-mir-30a&WithFlank&6\|-\|71403545\|71403627@12.34.23(+1G)&[MIMAT0000087&hsa-miR-30a-5p&offsets\|0\|+1(+1G);m-15&6\|-\|71403595\|71403616&offsets\|0\|+1(+1G)]\|TGTAAACATCCTCGACTGGAAGCG | >MI0000088\|hsa-mir-30a&WithFlank&6\|-\|71403545\|71403627 | 12 | 34 | 23 | (+1G) | isomiR | NO | 37.67 | 42.86 | 85.62 | 0.00 |
| >MI0000088\|hsa-mir-30a&WithFlank&6\|-\|71403545\|71403627@12.35.24&[MIMAT0000087&hsa-miR-30a-5p&offsets\|0\|+2;m-15&6\|-\|71403595\|71403616&offsets\|0\|+2]\|TGTAAACATCCTCGACTGGAAGCT | >MI0000088\|hsa-mir-30a&WithFlank&6\|-\|71403545\|71403627 | 12 | 35 | 24 | | isomiR | YES | 706.43 | 1202.03 | 1957.79 | 347.78 |
| >MI0000088\|hsa-mir-30a&WithFlank&6\|-\|71403545\|71403627@12.35.24(+1U)&[MIMAT0000087&hsa-miR-30a-5p&offsets\|0\|+2(+1U);m-15&6\|-\|71403595\|71403616&offsets\|0\|+2(+1U)]\|TGTAAACATCCTCGACTGGAAGCTT | >MI0000088\|hsa-mir-30a&WithFlank&6\|-\|71403545\|71403627 | 12 | 35 | 24 | (+1U) | isomiR | NO | 10.70 | 26.65 | 116.93 | 0.00 |
| >MI0000088\|hsa-mir-30a&WithFlank&6\|-\|71403545\|71403627@13.35.23&[MIMAT0000087&hsa-miR-30a-5p&offsets\|+1\|+2;m-15&6\|-\|71403595\|71403616&offsets\|+1\|+2]\|GTAAACATCCTCGACTGGAAGCT | >MI0000088\|hsa-mir-30a&WithFlank&6\|-\|71403545\|71403627 | 13 | 35 | 23 | | isomiR | NO | 23.80 | 24.90 | 121.72 | 0.00 |
| >MI0000088\|hsa-mir-30a&WithFlank&6\|-\|71403545\|71403627@53.71.19&[MIMAT0000088&hsa-miR-30a-3p&offsets\|0\|-3;m-38&6\|-\|71403554\|71403575&offsets\|0\|-3]\|CTTTCAGTCGGATGTTTGC | >MI0000088\|hsa-mir-30a&WithFlank&6\|-\|71403545\|71403627 | 53 | 71 | 19 | | isomiR | NO | 23.51 | 84.59 | 12.65 | 0.00 |
| >MI0000088\|hsa-mir-30a&WithFlank&6\|-\|71403545\|71403627@53.73.21&[MIMAT0000088&hsa-miR-30a-3p&offsets\|0\|-1;m-38&6\|-\|71403554\|71403575&offsets\|0\|-1]\|CTTTCAGTCGGATGTTTGCAG | >MI0000088\|hsa-mir-30a&WithFlank&6\|-\|71403545\|71403627 | 53 | 73 | 21 | | isomiR | NO | 0.00 | 41.19 | 11.27 | 0.00 |
| >MI0000088\|hsa-mir-30a&WithFlank&6\|-\|71403545\|71403627@53.73.21(+1A)&[MIMAT0000088&hsa-miR-30a-3p&offsets\|0\|-1(+1A);m-38&6\|-\|71403554\|71403575&offsets\|0\|-1(+1A)]\|CTTTCAGTCGGATGTTTGCAGA | >MI0000088\|hsa-mir-30a&WithFlank&6\|-\|71403545\|71403627 | 53 | 73 | 21 | (+1A) | isomiR | NO | 0.00 | 44.52 | 0.00 | 0.00 |
| >MI0000088\|hsa-mir-30a&WithFlank&6\|-\|71403545\|71403627@53.73.21(+1U)&[MIMAT0000088&hsa-miR-30a-3p&offsets\|0\|-1(+1U);m-38&6\|-\|71403554\|71403575&offsets\|0\|-1(+1U)]\|CTTTCAGTCGGATGTTTGCAGT | >MI0000088\|hsa-mir-30a&WithFlank&6\|-\|71403545\|71403627 | 53 | 73 | 21 | (+1U) | isomiR | NO | 150.83 | 415.00 | 58.45 | 63.85 |
| >MI0000088\|hsa-mir-30a&WithFlank&6\|-\|71403545\|71403627@53.73.21(+2U)&[MIMAT0000088&hsa-miR-30a-3p&offsets\|0\|-1(+2U);m-38&6\|-\|71403554\|71403575&offsets\|0\|-1(+2U)]\|CTTTCAGTCGGATGTTTGCAGTT | >MI0000088\|hsa-mir-30a&WithFlank&6\|-\|71403545\|71403627 | 53 | 73 | 21 | (+2U) | isomiR | NO | 0.00 | 10.30 | 0.00 | 0.00 |
| >MI0000088\|hsa-mir-30a&WithFlank&6\|-\|71403545\|71403627@53.74.22&[MIMAT0000088&hsa-miR-30a-3p&offsets\|0\|0;m-38&6\|-\|71403554\|71403575&offsets\|0\|0]\|CTTTCAGTCGGATGTTTGCAGC | >MI0000088\|hsa-mir-30a&WithFlank&6\|-\|71403545\|71403627 | 53 | 74 | 22 | | isomiR | NO | 163.59 | 576.75 | 72.07 | 72.99 |
| >MI0000088\|hsa-mir-30a&WithFlank&6\|-\|71403545\|71403627@53.74.22(+1A)&[MIMAT0000088&hsa-miR-30a-3p&offsets\|0\|0(+1A);m-38&6\|-\|71403554\|71403575&offsets\|0\|0(+1A)]\|CTTTCAGTCGGATGTTTGCAGCA | >MI0000088\|hsa-mir-30a&WithFlank&6\|-\|71403545\|71403627 | 53 | 74 | 22 | (+1A) | isomiR | NO | 0.00 | 24.55 | 0.00 | 0.00 |
| >MI0000088\|hsa-mir-30a&WithFlank&6\|-\|71403545\|71403627@54.71.18&[MIMAT0000088&hsa-miR-30a-3p&offsets\|+1\|-3;m-38&6\|-\|71403554\|71403575&offsets\|+1\|-3]\|TTTCAGTCGGATGTTTGC | >MI0000088\|hsa-mir-30a&WithFlank&6\|-\|71403545\|71403627 | 54 | 71 | 18 | | isomiR | NO | 0.00 | 27.78 | 0.00 | 0.00 |
| >MI0000088\|hsa-mir-30a&WithFlank&6\|-\|71403545\|71403627@54.73.20(+1U)&[MIMAT0000088&hsa-miR-30a-3p&offsets\|+1\|-1(+1U);m-38&6\|-\|71403554\|71403575&offsets\|+1\|-1(+1U)]\|TTTCAGTCGGATGTTTGCAGT | >MI0000088\|hsa-mir-30a&WithFlank&6\|-\|71403545\|71403627 | 54 | 73 | 20 | (+1U) | isomiR | NO | 73.17 | 187.61 | 13.57 | 23.60 |
| >MI0000088\|hsa-mir-30a&WithFlank&6\|-\|71403545\|71403627@54.73.20(+2U)&[MIMAT0000088&hsa-miR-30a-3p&offsets\|+1\|-1(+2U);m-38&6\|-\|71403554\|71403575&offsets\|+1\|-1(+2U)]\|TTTCAGTCGGATGTTTGCAGTT | >MI0000088\|hsa-mir-30a&WithFlank&6\|-\|71403545\|71403627 | 54 | 73 | 20 | (+2U) | isomiR | NO | 0.00 | 25.22 | 0.00 | 0.00 |
| >MI0000088\|hsa-mir-30a&WithFlank&6\|-\|71403545\|71403627@54.74.21&[MIMAT0000088&hsa-miR-30a-3p&offsets\|+1\|0;m-38&6\|-\|71403554\|71403575&offsets\|+1\|0]\|TTTCAGTCGGATGTTTGCAGC | >MI0000088\|hsa-mir-30a&WithFlank&6\|-\|71403545\|71403627 | 54 | 74 | 21 | | isomiR | NO | 72.26 | 222.85 | 15.26 | 25.16 |
| >MI0000088\|hsa-mir-30a&WithFlank&6\|-\|71403545\|71403627@54.75.22&[MIMAT0000088&hsa-miR-30a-3p&offsets\|+1\|+1;m-38&6\|-\|71403554\|71403575&offsets\|+1\|+1]\|TTTCAGTCGGATGTTTGCAGCT | >MI0000088\|hsa-mir-30a&WithFlank&6\|-\|71403545\|71403627 | 54 | 75 | 22 | | isomiR | NO | 0.00 | 31.26 | 0.00 | 0.00 |
| >MI0000089\|hsa-mir-31&WithFlank&9\|-\|21512109\|21512191@14.34.21&[MIMAT0000089&hsa-miR-31-5p&offsets\|0\|0;m-360&9\|-\|21512157\|21512178&offsets\|0\|-1]\|AGGCAAGATGCTGGCATAGCT | >MI0000089\|hsa-mir-31&WithFlank&9\|-\|21512109\|21512191 | 14 | 34 | 21 | | isomiR | NO | 24.01 | 0.00 | 0.00 | 0.00 |
| >MI0000089\|hsa-mir-31&WithFlank&9\|-\|21512109\|21512191@14.35.22&[MIMAT0000089&hsa-miR-31-5p&offsets\|0\|+1;m-360&9\|-\|21512157\|21512178&offsets\|0\|0]\|AGGCAAGATGCTGGCATAGCTG | >MI0000089\|hsa-mir-31&WithFlank&9\|-\|21512109\|21512191 | 14 | 35 | 22 | | isomiR | NO | 41.95 | 43.84 | 10.21 | 0.00 |
| >MI0000089\|hsa-mir-31&WithFlank&9\|-\|21512109\|21512191@14.35.22(+1A)&[MIMAT0000089&hsa-miR-31-5p&offsets\|0\|+1(+1A);m-360&9\|-\|21512157\|21512178&offsets\|0\|0(+1A)]\|AGGCAAGATGCTGGCATAGCTGA | >MI0000089\|hsa-mir-31&WithFlank&9\|-\|21512109\|21512191 | 14 | 35 | 22 | (+1A) | isomiR | NO | 0.00 | 10.25 | 0.00 | 0.00 |
| >MI0000089\|hsa-mir-31&WithFlank&9\|-\|21512109\|21512191@14.36.23&[MIMAT0000089&hsa-miR-31-5p&offsets\|0\|+2;m-360&9\|-\|21512157\|21512178&offsets\|0\|+1]\|AGGCAAGATGCTGGCATAGCTGT | >MI0000089\|hsa-mir-31&WithFlank&9\|-\|21512109\|21512191 | 14 | 36 | 23 | | isomiR | NO | 75.64 | 118.54 | 73.79 | 0.00 |

# BT-20

<table>
<tr><td>>MI0000093|hsa-mir-92a-1&WithFlank&13|+|91351308|91351397@54.74.21(+1A);MI0000094|hsa-mir-92a-2&WithFlank&X|-|134169532|134169618@54.74.21(+1A)&[MIMAT0000092&hsa-miR-92a-3p&offsets|0|-1(+1A);m-9&13|+|91351361|91351382&offsets|0|-1(+1A)];[MIMAT0000092_1&hsa-miR-92a-3p&offsets|0|-1(+1A);m-11&X|-|134169544|134169565&offsets|0|-1(+1A)]|TATTGCACTTGTCCCGGCCTGA</td><td>>MI0000093|hsa-mir-92a-1&WithFlank&13|+|91351308|91351397</td><td>54</td><td>74</td><td>21</td><td>(+1A)</td><td>isomiR</td><td>NO</td><td>38.22</td><td>217.23</td><td>15.42</td><td>10.94</td></tr>
<tr><td>>MI0000093|hsa-mir-92a-1&WithFlank&13|+|91351308|91351397@54.74.21(+1C);MI0000094|hsa-mir-92a-2&WithFlank&X|-|134169532|134169618@54.74.21(+1C)&[MIMAT0000092&hsa-miR-92a-3p&offsets|0|-1(+1C);m-9&13|+|91351361|91351382&offsets|0|-1(+1C)];[MIMAT0000092_1&hsa-miR-92a-3p&offsets|0|-1(+1C);m-11&X|-|134169544|134169565&offsets|0|-1(+1C)]|TATTGCACTTGTCCCGGCCTGC</td><td>>MI0000093|hsa-mir-92a-1&WithFlank&13|+|91351308|91351397</td><td>54</td><td>74</td><td>21</td><td>(+1C)</td><td>isomiR</td><td>NO</td><td>175.40</td><td>501.35</td><td>143.44</td><td>110.77</td></tr>
<tr><td>>MI0000093|hsa-mir-92a-1&WithFlank&13|+|91351308|91351397@54.74.21(+1G);MI0000094|hsa-mir-92a-2&WithFlank&X|-|134169532|134169618@54.74.21(+1G)&[MIMAT0000092&hsa-miR-92a-3p&offsets|0|-1(+1G);m-9&13|+|91351361|91351382&offsets|0|-1(+1G)];[MIMAT0000092_1&hsa-miR-92a-3p&offsets|0|-1(+1G);m-11&X|-|134169544|134169565&offsets|0|-1(+1G)]|TATTGCACTTGTCCCGGCCTGG</td><td>>MI0000093|hsa-mir-92a-1&WithFlank&13|+|91351308|91351397</td><td>54</td><td>74</td><td>21</td><td>(+1G)</td><td>isomiR</td><td>NO</td><td>29.39</td><td>106.01</td><td>0.00</td><td>0.00</td></tr>
<tr><td>>MI0000093|hsa-mir-92a-1&WithFlank&13|+|91351308|91351397@54.74.21;MI0000094|hsa-mir-92a-2&WithFlank&X|-|134169532|134169618@54.74.21&[MIMAT0000092&hsa-miR-92a-3p&offsets|0|-1;m-9&13|+|91351361|91351382&offsets|0|-1];[MIMAT0000092_1&hsa-miR-92a-3p&offsets|0|-1;m-11&X|-|134169544|134169565&offsets|0|-1]|TATTGCACTTGTCCCGGCCTG</td><td>>MI0000093|hsa-mir-92a-1&WithFlank&13|+|91351308|91351397</td><td>54</td><td>74</td><td>21</td><td></td><td>isomiR</td><td>NO</td><td>0.00</td><td>36.35</td><td>10.04</td><td>0.00</td></tr>
<tr><td>>MI0000093|hsa-mir-92a-1&WithFlank&13|+|91351308|91351397@54.75.22(+1A);MI0000094|hsa-mir-92a-2&WithFlank&X|-|134169532|134169618@54.75.22(+1A)&[MIMAT0000092&hsa-miR-92a-3p&offsets|0|0(+1A);m-9&13|+|91351361|91351382&offsets|0|0(+1A)];[MIMAT0000092_1&hsa-miR-92a-3p&offsets|0|0(+1A);m-11&X|-|134169544|134169565&offsets|0|0(+1A)]|TATTGCACTTGTCCCGGCCTGTA</td><td>>MI0000093|hsa-mir-92a-1&WithFlank&13|+|91351308|91351397</td><td>54</td><td>75</td><td>22</td><td>(+1A)</td><td>isomiR</td><td>NO</td><td>168.70</td><td>747.05</td><td>105.18</td><td>217.96</td></tr>
<tr><td>>MI0000093|hsa-mir-92a-1&WithFlank&13|+|91351308|91351397@54.75.22(+1C);MI0000094|hsa-mir-92a-2&WithFlank&X|-|134169532|134169618@54.75.22(+1C)&[MIMAT0000092&hsa-miR-92a-3p&offsets|0|0(+1C);m-9&13|+|91351361|91351382&offsets|0|0(+1C)];[MIMAT0000092_1&hsa-miR-92a-3p&offsets|0|0(+1C);m-11&X|-|134169544|134169565&offsets|0|0(+1C)]|TATTGCACTTGTCCCGGCCTGTC</td><td>>MI0000093|hsa-mir-92a-1&WithFlank&13|+|91351308|91351397</td><td>54</td><td>75</td><td>22</td><td>(+1C)</td><td>isomiR</td><td>NO</td><td>0.00</td><td>26.79</td><td>0.00</td><td>0.00</td></tr>
<tr><td>>MI0000093|hsa-mir-92a-1&WithFlank&13|+|91351308|91351397@54.75.22(+2A);MI0000094|hsa-mir-92a-2&WithFlank&X|-|134169532|134169618@54.75.22(+2A)&[MIMAT0000092&hsa-miR-92a-3p&offsets|0|0(+2A);m-9&13|+|91351361|91351382&offsets|0|0(+2A)];[MIMAT0000092_1&hsa-miR-92a-3p&offsets|0|0(+2A);m-11&X|-|134169544|134169565&offsets|0|0(+2A)]|TATTGCACTTGTCCCGGCCTGTAA</td><td>>MI0000093|hsa-mir-92a-1&WithFlank&13|+|91351308|91351397</td><td>54</td><td>75</td><td>22</td><td>(+2A)</td><td>isomiR</td><td>NO</td><td>0.00</td><td>37.06</td><td>0.00</td><td>0.00</td></tr>
<tr><td>>MI0000093|hsa-mir-92a-1&WithFlank&13|+|91351308|91351397@54.75.22;MI0000094|hsa-mir-92a-2&WithFlank&X|-|134169532|134169618@54.75.22&[MIMAT0000092&hsa-miR-92a-3p&offsets|0|0;m-9&13|+|91351361|91351382&offsets|0|0];[MIMAT0000092_1&hsa-miR-92a-3p&offsets|0|0;m-11&X|-|134169544|134169565&offsets|0|0]|TATTGCACTTGTCCCGGCCTGT</td><td>>MI0000093|hsa-mir-92a-1&WithFlank&13|+|91351308|91351397</td><td>54</td><td>75</td><td>22</td><td></td><td>isomiR</td><td>YES</td><td>779.11</td><td>3058.34</td><td>745.50</td><td>519.69</td></tr>
<tr><td>>MI0000093|hsa-mir-92a-1&WithFlank&13|+|91351308|91351397@54.76.23&[MIMAT0000092&hsa-miR-92a-3p&offsets|0|+1;m-9&13|+|91351361|91351382&offsets|0|+1]|TATTGCACTTGTCCCGGCCTGTT</td><td>>MI0000093|hsa-mir-92a-1&WithFlank&13|+|91351308|91351397</td><td>54</td><td>76</td><td>23</td><td></td><td>isomiR</td><td>NO</td><td>66.09</td><td>252.40</td><td>11.97</td><td>41.30</td></tr>
<tr><td>>MI0000093|hsa-mir-92a-1&WithFlank&13|+|91351308|91351397@55.75.21;MI0000094|hsa-mir-92a-2&WithFlank&X|-|134169532|134169618@55.75.21&[MIMAT0000092&hsa-miR-92a-3p&offsets|+1|0;m-9&13|+|91351361|91351382&offsets|+1|0];[MIMAT0000092_1&hsa-miR-92a-3p&offsets|+1|0;m-11&X|-|134169544|134169565&offsets|+1|0]|ATTGCACTTGTCCCGGCCTGT</td><td>>MI0000093|hsa-mir-92a-1&WithFlank&13|+|91351308|91351397</td><td>55</td><td>75</td><td>21</td><td></td><td>isomiR</td><td>NO</td><td>0.00</td><td>44.10</td><td>12.96</td><td>0.00</td></tr>
<tr><td>>MI0000095|hsa-mir-93&WithFlank&7|-|100093762|100093853@17.37.21&[MIMAT0000093&hsa-miR-93-5p&offsets|0|-2;m-29&7|-|100093815|100093837&offsets|0|-2]|CAAAGTGCTGTTCGTGCAGGT</td><td>>MI0000095|hsa-mir-93&WithFlank&7|-|100093762|100093853</td><td>17</td><td>37</td><td>21</td><td></td><td>isomiR</td><td>NO</td><td>31.65</td><td>37.52</td><td>90.32</td><td>0.00</td></tr>
<tr><td>>MI0000095|hsa-mir-93&WithFlank&7|-|100093762|100093853@17.38.22&[MIMAT0000093&hsa-miR-93-5p&offsets|0|-1;m-29&7|-|100093815|100093837&offsets|0|-1]|CAAAGTGCTGTTCGTGCAGGTA</td><td>>MI0000095|hsa-mir-93&WithFlank&7|-|100093762|100093853</td><td>17</td><td>38</td><td>22</td><td></td><td>isomiR</td><td>NO</td><td>9.71</td><td>32.87</td><td>0.00</td><td>0.00</td></tr>
<tr><td>>MI0000095|hsa-mir-93&WithFlank&7|-|100093762|100093853@17.38.22(+1A)&[MIMAT0000093&hsa-miR-93-5p&offsets|0|-1(+1A);m-29&7|-|100093815|100093837&offsets|0|-1(+1A)]|CAAAGTGCTGTTCGTGCAGGTAA</td><td>>MI0000095|hsa-mir-93&WithFlank&7|-|100093762|100093853</td><td>17</td><td>38</td><td>22</td><td>(+1A)</td><td>isomiR</td><td>NO</td><td>36.13</td><td>123.30</td><td>89.32</td><td>0.00</td></tr>
<tr><td>>MI0000095|hsa-mir-93&WithFlank&7|-|100093762|100093853@17.39.23&[MIMAT0000093&hsa-miR-93-5p&offsets|0|0;m-29&7|-|100093815|100093837&offsets|0|0]|CAAAGTGCTGTTCGTGCAGGTAG</td><td>>MI0000095|hsa-mir-93&WithFlank&7|-|100093762|100093853</td><td>17</td><td>39</td><td>23</td><td></td><td>isomiR</td><td>NO</td><td>306.49</td><td>675.48</td><td>614.23</td><td>101.16</td></tr>
<tr><td>>MI0000095|hsa-mir-93&WithFlank&7|-|100093762|100093853@17.39.23(+1A)&[MIMAT0000093&hsa-miR-93-5p&offsets|0|0(+1A);m-29&7|-|100093815|100093837&offsets|0|0(+1A)]|CAAAGTGCTGTTCGTGCAGGTAGA</td><td>>MI0000095|hsa-mir-93&WithFlank&7|-|100093762|100093853</td><td>17</td><td>39</td><td>23</td><td>(+1A)</td><td>isomiR</td><td>NO</td><td>73.25</td><td>232.41</td><td>12.43</td><td>26.91</td></tr>
<tr><td>>MI0000095|hsa-mir-93&WithFlank&7|-|100093762|100093853@17.40.24&[MIMAT0000093&hsa-miR-93-5p&offsets|0|+1;m-29&7|-|100093815|100093837&offsets|0|+1]|CAAAGTGCTGTTCGTGCAGGTAGT</td><td>>MI0000095|hsa-mir-93&WithFlank&7|-|100093762|100093853</td><td>17</td><td>40</td><td>24</td><td></td><td>isomiR</td><td>NO</td><td>42.87</td><td>129.61</td><td>0.00</td><td>0.00</td></tr>
<tr><td>>MI0000098|hsa-mir-96&WithFlank&7|-|129774686|129774775@15.36.22(+1A)&[MIMAT0000095&hsa-miR-96-5p&offsets|0|-1(+1A);m-248&7|-|129774739|129774761&offsets|0|-1(+1A)]|TTTGGCACTAGCACATTTTTGCA</td><td>>MI0000098|hsa-mir-96&WithFlank&7|-|129774686|129774775</td><td>15</td><td>36</td><td>22</td><td>(+1A)</td><td>isomiR</td><td>NO</td><td>26.87</td><td>26.88</td><td>0.00</td><td>0.00</td></tr>
<tr><td>>MI0000098|hsa-mir-96&WithFlank&7|-|129774686|129774775@15.36.22(+1C)&[MIMAT0000095&hsa-miR-96-5p&offsets|0|-1(+1C);m-248&7|-|129774739|129774761&offsets|0|-1(+1C)]|TTTGGCACTAGCACATTTTTGCC</td><td>>MI0000098|hsa-mir-96&WithFlank&7|-|129774686|129774775</td><td>15</td><td>36</td><td>22</td><td>(+1C)</td><td>isomiR</td><td>NO</td><td>39.17</td><td>27.83</td><td>0.00</td><td>0.00</td></tr>
<tr><td>>MI0000098|hsa-mir-96&WithFlank&7|-|129774686|129774775@15.37.23&[MIMAT0000095&hsa-miR-96-5p&offsets|0|0;m-248&7|-|129774739|129774761&offsets|0|0]|TTTGGCACTAGCACATTTTTGCT</td><td>>MI0000098|hsa-mir-96&WithFlank&7|-|129774686|129774775</td><td>15</td><td>37</td><td>23</td><td></td><td>isomiR</td><td>NO</td><td>247.09</td><td>233.39</td><td>67.59</td><td>107.87</td></tr>
</table>

# BT-20

<table>
<tr><td>>MI0000100|hsa-mir-98&WithFlank&X|-|53556217|53556347@28.48.21(+1C)&[MIMAT0000096&hsa-miR-98-5p&offsets|0|-1(+1C);m-143&X|-|53556299|53556320&offsets|0|-1(+1C)]|TGAGGTAGTAAGTTGTATTGTC</td><td>>MI0000100|hsa-mir-98&WithFlank&X|-|53556217|53556347</td><td>28</td><td>48</td><td>21</td><td>(+1C)</td><td>isomiR</td><td>NO</td><td>9.90</td><td>33.89</td><td>0.00</td><td>0.00</td></tr>
<tr><td>>MI0000100|hsa-mir-98&WithFlank&X|-|53556217|53556347@28.49.22&[MIMAT0000096&hsa-miR-98-5p&offsets|0|0;m-143&X|-|53556299|53556320&offsets|0|0]|TGAGGTAGTAAGTTGTATTGTT</td><td>>MI0000100|hsa-mir-98&WithFlank&X|-|53556217|53556347</td><td>28</td><td>49</td><td>22</td><td></td><td>isomiR</td><td>NO</td><td>137.70</td><td>308.69</td><td>73.00</td><td>38.01</td></tr>
<tr><td>>MI0000102|hsa-mir-100&WithFlank&11|-|122152223|122152314@19.38.20&[MIMAT0000098&hsa-miR-100-5p&offsets|0|-2;m-27&11|-|122152276|122152296&offsets|0|-1]|AACCCGTAGATCCGAACTTG</td><td>>MI0000102|hsa-mir-100&WithFlank&11|-|122152223|122152314</td><td>19</td><td>38</td><td>20</td><td></td><td>isomiR</td><td>NO</td><td>0.00</td><td>37.49</td><td>11.45</td><td>0.00</td></tr>
<tr><td>>MI0000102|hsa-mir-100&WithFlank&11|-|122152223|122152314@19.38.20(+1C)&[MIMAT0000098&hsa-miR-100-5p&offsets|0|-2(+1C);m-27&11|-|122152276|122152296&offsets|0|-1(+1C)]|AACCCGTAGATCCGAACTTGC</td><td>>MI0000102|hsa-mir-100&WithFlank&11|-|122152223|122152314</td><td>19</td><td>38</td><td>20</td><td>(+1C)</td><td>isomiR</td><td>NO</td><td>0.00</td><td>33.08</td><td>0.00</td><td>0.00</td></tr>
<tr><td>>MI0000102|hsa-mir-100&WithFlank&11|-|122152223|122152314@19.39.21&[MIMAT0000098&hsa-miR-100-5p&offsets|0|-1;m-27&11|-|122152276|122152296&offsets|0|0]|AACCCGTAGATCCGAACTTGT</td><td>>MI0000102|hsa-mir-100&WithFlank&11|-|122152223|122152314</td><td>19</td><td>39</td><td>21</td><td></td><td>isomiR</td><td>NO</td><td>36.19</td><td>289.39</td><td>106.15</td><td>0.00</td></tr>
<tr><td>>MI0000102|hsa-mir-100&WithFlank&11|-|122152223|122152314@19.39.21(+1A)&[MIMAT0000098&hsa-miR-100-5p&offsets|0|-1(+1A);m-27&11|-|122152276|122152296&offsets|0|0(+1A)]|AACCCGTAGATCCGAACTTGTA</td><td>>MI0000102|hsa-mir-100&WithFlank&11|-|122152223|122152314</td><td>19</td><td>39</td><td>21</td><td>(+1A)</td><td>isomiR</td><td>NO</td><td>26.50</td><td>241.82</td><td>11.09</td><td>0.00</td></tr>
<tr><td>>MI0000102|hsa-mir-100&WithFlank&11|-|122152223|122152314@19.40.22&[MIMAT0000098&hsa-miR-100-5p&offsets|0|0;m-27&11|-|122152276|122152296&offsets|0|+1]|AACCCGTAGATCCGAACTTGTG</td><td>>MI0000102|hsa-mir-100&WithFlank&11|-|122152223|122152314</td><td>19</td><td>40</td><td>22</td><td></td><td>isomiR</td><td>NO</td><td>193.01</td><td>1136.59</td><td>215.00</td><td>108.97</td></tr>
<tr><td>>MI0000102|hsa-mir-100&WithFlank&11|-|122152223|122152314@19.40.22(+1A)&[MIMAT0000098&hsa-miR-100-5p&offsets|0|0(+1A);m-27&11|-|122152276|122152296&offsets|0|+1(+1A)]|AACCCGTAGATCCGAACTTGTGA</td><td>>MI0000102|hsa-mir-100&WithFlank&11|-|122152223|122152314</td><td>19</td><td>40</td><td>22</td><td>(+1A)</td><td>isomiR</td><td>NO</td><td>0.00</td><td>72.81</td><td>0.00</td><td>0.00</td></tr>
<tr><td>>MI0000102|hsa-mir-100&WithFlank&11|-|122152223|122152314@19.40.22(+1U)&[MIMAT0000098&hsa-miR-100-5p&offsets|0|0(+1U);m-27&11|-|122152276|122152296&offsets|0|+1(+1U)]|AACCCGTAGATCCGAACTTGTGT</td><td>>MI0000102|hsa-mir-100&WithFlank&11|-|122152223|122152314</td><td>19</td><td>40</td><td>22</td><td>(+1U)</td><td>isomiR</td><td>NO</td><td>0.00</td><td>97.24</td><td>0.00</td><td>0.00</td></tr>
<tr><td>>MI0000103|hsa-mir-101-1&WithFlank&1|-|65058428|65058514@52.72.21;MI0000739|hsa-mir-101-2&WithFlank&9|+|4850291|4850381@54.74.21&[MIMAT0000099&hsa-miR-101-3p&offsets|-1|-1;m-17&1|-|65058443|65058462&offsets|-1|0];[MIMAT0000099_1&hsa-miR-101-3p&offsets|-1|-1;m-16&9|+|4850345|4850364&offsets|-1|0]|GTACAGTACTGTGATAACTGA</td><td>>MI0000103|hsa-mir-101-1&WithFlank&1|-|65058428|65058514</td><td>52</td><td>72</td><td>21</td><td></td><td>isomiR</td><td>NO</td><td>28.67</td><td>0.00</td><td>0.00</td><td>0.00</td></tr>
<tr><td>>MI0000103|hsa-mir-101-1&WithFlank&1|-|65058428|65058514@52.73.22;MI0000739|hsa-mir-101-2&WithFlank&9|+|4850291|4850381@54.75.22&[MIMAT0000099&hsa-miR-101-3p&offsets|-1|0;m-17&1|-|65058443|65058462&offsets|-1|+1];[MIMAT0000099_1&hsa-miR-101-3p&offsets|-1|0;m-16&9|+|4850345|4850364&offsets|-1|+1]|GTACAGTACTGTGATAACTGAA</td><td>>MI0000103|hsa-mir-101-1&WithFlank&1|-|65058428|65058514</td><td>52</td><td>73</td><td>22</td><td></td><td>isomiR</td><td>NO</td><td>70.50</td><td>27.13</td><td>15.23</td><td>0.00</td></tr>
<tr><td>>MI0000103|hsa-mir-101-1&WithFlank&1|-|65058428|65058514@53.72.20;MI0000739|hsa-mir-101-2&WithFlank&9|+|4850291|4850381@55.74.20&[MIMAT0000099&hsa-miR-101-3p&offsets|0|-1;m-17&1|-|65058443|65058462&offsets|0|0];[MIMAT0000099_1&hsa-miR-101-3p&offsets|0|-1;m-16&9|+|4850345|4850364&offsets|0|0]|TACAGTACTGTGATAACTGA</td><td>>MI0000103|hsa-mir-101-1&WithFlank&1|-|65058428|65058514</td><td>53</td><td>72</td><td>20</td><td></td><td>isomiR</td><td>NO</td><td>34.58</td><td>32.37</td><td>68.43</td><td>0.00</td></tr>
<tr><td>>MI0000103|hsa-mir-101-1&WithFlank&1|-|65058428|65058514@53.74.22;MI0000739|hsa-mir-101-2&WithFlank&9|+|4850291|4850381@55.76.22&[MIMAT0000099&hsa-miR-101-3p&offsets|0|+1;m-17&1|-|65058443|65058462&offsets|0|+2];[MIMAT0000099_1&hsa-miR-101-3p&offsets|0|+1;m-16&9|+|4850345|4850364&offsets|0|+2]|TACAGTACTGTGATAACTGAAG</td><td>>MI0000103|hsa-mir-101-1&WithFlank&1|-|65058428|65058514</td><td>53</td><td>74</td><td>22</td><td></td><td>isomiR</td><td>NO</td><td>103.38</td><td>70.52</td><td>39.67</td><td>0.00</td></tr>
<tr><td>>MI0000107|hsa-mir-29b-2&WithFlank&1|-|207802437|207802529@58.80.23;MI0000105|hsa-mir-29b-1&WithFlank&7|-|130877453|130877545@57.79.23&[MIMAT0000100&hsa-miR-29b-3p&offsets|0|0;m-81&1|-|207802450|207802472&offsets|0|0];[MIMAT0000100_1&hsa-miR-29b-3p&offsets|0|0;m-82&7|-|130877467|130877489&offsets|0|0]|TAGCACCATTTGAAATCAGTGTT</td><td>>MI0000107|hsa-mir-29b-2&WithFlank&1|-|207802437|207802529</td><td>58</td><td>80</td><td>23</td><td></td><td>isomiR</td><td>NO</td><td>40.41</td><td>0.00</td><td>0.00</td><td>0.00</td></tr>
<tr><td>>MI0000108|hsa-mir-103a-2&WithFlank&20|+|3917488|3917577@54.75.22(+1G);MI0000109|hsa-mir-103a-1&WithFlank&5|-|168560890|168560979@54.75.22(+1G)&[MIMAT0000101&hsa-miR-103a-3p&offsets|0|-1(+1G);m-12&20|+|3917541|3917563&offsets|0|-1(+1G)];[MIMAT0000101_1&hsa-miR-103a-3p&offsets|0|-1(+1G);m-13&5|-|168560904|168560926&offsets|0|-1(+1G);m-44220&5|-|168560908|168560926&offsets|0|+3(+1G)]|AGCAGCATTGTACAGGGCTATGG</td><td>>MI0000108|hsa-mir-103a-2&WithFlank&20|+|3917488|3917577</td><td>54</td><td>75</td><td>22</td><td>(+1G)</td><td>isomiR</td><td>NO</td><td>28.06</td><td>38.33</td><td>0.00</td><td>0.00</td></tr>
<tr><td>>MI0000108|hsa-mir-103a-2&WithFlank&20|+|3917488|3917577@54.75.22(+1U);MI0000109|hsa-mir-103a-1&WithFlank&5|-|168560890|168560979@54.75.22(+1U)&[MIMAT0000101&hsa-miR-103a-3p&offsets|0|-1(+1U);m-12&20|+|3917541|3917563&offsets|0|-1(+1U)];[MIMAT0000101_1&hsa-miR-103a-3p&offsets|0|-1(+1U);m-13&5|-|168560904|168560926&offsets|0|-1(+1U);m-44220&5|-|168560908|168560926&offsets|0|+3(+1U)]|AGCAGCATTGTACAGGGCTATGT</td><td>>MI0000108|hsa-mir-103a-2&WithFlank&20|+|3917488|3917577</td><td>54</td><td>75</td><td>22</td><td>(+1U)</td><td>isomiR</td><td>NO</td><td>40.01</td><td>85.07</td><td>0.00</td><td>0.00</td></tr>
<tr><td>>MI0000108|hsa-mir-103a-2&WithFlank&20|+|3917488|3917577@54.75.22;MI0000109|hsa-mir-103a-1&WithFlank&5|-|168560890|168560979@54.75.22&[MIMAT0000101&hsa-miR-103a-3p&offsets|0|-1;m-12&20|+|3917541|3917563&offsets|0|-1];[MIMAT0000101_1&hsa-miR-103a-3p&offsets|0|-1;m-13&5|-|168560904|168560926&offsets|0|-1;m-44220&5|-|168560908|168560926&offsets|0|+3]|AGCAGCATTGTACAGGGCTATG</td><td>>MI0000108|hsa-mir-103a-2&WithFlank&20|+|3917488|3917577</td><td>54</td><td>75</td><td>22</td><td></td><td>isomiR</td><td>NO</td><td>209.88</td><td>326.72</td><td>108.31</td><td>117.76</td></tr>
<tr><td>>MI0000108|hsa-mir-103a-2&WithFlank&20|+|3917488|3917577@54.76.23(+1U);MI0000109|hsa-mir-103a-1&WithFlank&5|-|168560890|168560979@54.76.23(+1U)&[MIMAT0000101&hsa-miR-103a-3p&offsets|0|0(+1U);m-12&20|+|3917541|3917563&offsets|0|0(+1U)];[MIMAT0000101_1&hsa-miR-103a-3p&offsets|0|0(+1U);m-13&5|-|168560904|168560926&offsets|0|0(+1U);m-44220&5|-|168560908|168560926&offsets|0|+4(+1U)]|AGCAGCATTGTACAGGGCTATGAT</td><td>>MI0000108|hsa-mir-103a-2&WithFlank&20|+|3917488|3917577</td><td>54</td><td>76</td><td>23</td><td>(+1U)</td><td>isomiR</td><td>NO</td><td>92.02</td><td>166.02</td><td>0.00</td><td>34.28</td></tr>
<tr><td>>MI0000108|hsa-mir-103a-2&WithFlank&20|+|3917488|3917577@54.76.23(+2U);MI0000109|hsa-mir-103a-1&WithFlank&5|-|168560890|168560979@54.76.23(+2U)&[MIMAT0000101&hsa-miR-103a-3p&offsets|0|0(+2U);m-12&20|+|3917541|3917563&offsets|0|0(+2U)];[MIMAT0000101_1&hsa-miR-103a-3p&offsets|0|0(+2U);m-13&5|-|168560904|168560926&offsets|0|0(+2U);m-44220&5|-|168560908|168560926&offsets|0|+4(+2U)]|AGCAGCATTGTACAGGGCTATGATT</td><td>>MI0000108|hsa-mir-103a-2&WithFlank&20|+|3917488|3917577</td><td>54</td><td>76</td><td>23</td><td>(+2U)</td><td>isomiR</td><td>NO</td><td>0.00</td><td>26.71</td><td>0.00</td><td>0.00</td></tr>
</table>

# BT-20

<table>
<tr><th></th><th></th><th></th><th></th><th></th><th></th><th></th><th></th><th></th><th></th><th></th><th></th></tr>
<tr><td>>MI0000108|hsa-mir-103a-2&WithFlank&20|+|3917488|3917577@54.76.23;MI0000109|hsa-mir-103a-1&WithFlank&5|-|168560890|168560979@54.76.23&[MIMAT0000101&hsa-miR-103a-3p&offsets|0|0;m-12&20|+|3917541|3917563&offsets|0|0];[MIMAT0000101_1&hsa-miR-103a-3p&offsets|0|0;m-13&5|-|168560904|168560926&offsets|0|0;m-44220&5|-|168560908|168560926&offsets|0|+4]|AGCAGCATTGTACAGGGCTATGA</td><td>>MI0000108|hsa-mir-103a-2&WithFlank&20|+|3917488|3917577</td><td>54</td><td>76</td><td>23</td><td></td><td>isomiR</td><td>YES</td><td>735.54</td><td>1585.57</td><td>285.73</td><td>351.04</td></tr>
<tr><td>>MI0000108|hsa-mir-103a-2&WithFlank&20|+|3917488|3917577@54.77.24;MI0000109|hsa-mir-103a-1&WithFlank&5|-|168560890|168560979@54.77.24&[MIMAT0000101&hsa-miR-103a-3p&offsets|0|+1;m-12&20|+|3917541|3917563&offsets|0|+1];[MIMAT0000101_1&hsa-miR-103a-3p&offsets|0|+1;m-13&5|-|168560904|168560926&offsets|0|+1;m-44220&5|-|168560908|168560926&offsets|0|+5]|AGCAGCATTGTACAGGGCTATGAA</td><td>>MI0000108|hsa-mir-103a-2&WithFlank&20|+|3917488|3917577</td><td>54</td><td>77</td><td>24</td><td></td><td>isomiR</td><td>NO</td><td>38.33</td><td>80.38</td><td>0.00</td><td>0.00</td></tr>
<tr><td>>MI0000111|hsa-mir-105-1&WithFlank&X|-|152392213|152392305@19.40.22;MI0000112|hsa-mir-105-2&WithFlank&X|-|152394406|152394498@19.40.22&[MIMAT0000102&hsa-miR-105-5p&offsets|0|-1;m-735&X|-|152392266|152392287&offsets|0|0];[MIMAT0000102_1&hsa-miR-105-5p&offsets|0|-1;m-735&X|-|152394459|152394480&offsets|0|0]|TCAAATGCTCAGACTCCTGTGG</td><td>>MI0000111|hsa-mir-105-1&WithFlank&X|-|152392213|152392305</td><td>19</td><td>40</td><td>22</td><td></td><td>isomiR</td><td>NO</td><td>0.00</td><td>32.59</td><td>12.23</td><td>0.00</td></tr>
<tr><td>>MI0000114|hsa-mir-107&WithFlank&10|-|89592741|89592833@56.76.21(+1A);MI0000108|hsa-mir-103a-2&WithFlank&20|+|3917488|3917577@54.74.21(+1A);MI0000109|hsa-mir-103a-1&WithFlank&5|-|168560890|168560979@54.74.21(+1A)&[MIMAT0000104&hsa-miR-107&offsets|0|-2(+1A);m-78&10|-|89592758|89592778&offsets|0|0(+1A)];[MIMAT0000101&hsa-miR-103a-3p&offsets|0|-2(+1A);m-12&20|+|3917541|3917563&offsets|0|-2(+1A)];[MIMAT0000101_1&hsa-miR-103a-3p&offsets|0|-2(+1A);m-13&5|-|168560904|168560926&offsets|0|-2(+1A);m-44220&5|-|168560908|168560926&offsets|0|+2(+1A)]|AGCAGCATTGTACAGGGCTATA</td><td>>MI0000114|hsa-mir-107&WithFlank&10|-|89592741|89592833</td><td>56</td><td>76</td><td>21</td><td>(+1A)</td><td>isomiR</td><td>NO</td><td>23.88</td><td>42.20</td><td>0.00</td><td>0.00</td></tr>
<tr><td>>MI0000114|hsa-mir-107&WithFlank&10|-|89592741|89592833@56.76.21;MI0000108|hsa-mir-103a-2&WithFlank&20|+|3917488|3917577@54.74.21;MI0000109|hsa-mir-103a-1&WithFlank&5|-|168560890|168560979@54.74.21&[MIMAT0000104&hsa-miR-107&offsets|0|-2;m-78&10|-|89592758|89592778&offsets|0|0];[MIMAT0000101&hsa-miR-103a-3p&offsets|0|-2;m-12&20|+|3917541|3917563&offsets|0|-2];[MIMAT0000101_1&hsa-miR-103a-3p&offsets|0|-2;m-13&5|-|168560904|168560926&offsets|0|-2;m-44220&5|-|168560908|168560926&offsets|0|+2]|AGCAGCATTGTACAGGGCTAT</td><td>>MI0000114|hsa-mir-107&WithFlank&10|-|89592741|89592833</td><td>56</td><td>76</td><td>21</td><td></td><td>isomiR</td><td>NO</td><td>26.83</td><td>30.07</td><td>55.03</td><td>0.00</td></tr>
<tr><td>>MI0000114|hsa-mir-107&WithFlank&10|-|89592741|89592833@56.77.22&[MIMAT0000104&hsa-miR-107&offsets|0|-1;m-78&10|-|89592758|89592778&offsets|0|+1]|AGCAGCATTGTACAGGGCTATC</td><td>>MI0000114|hsa-mir-107&WithFlank&10|-|89592741|89592833</td><td>56</td><td>77</td><td>22</td><td></td><td>isomiR</td><td>NO</td><td>0.00</td><td>26.17</td><td>0.00</td><td>0.00</td></tr>
<tr><td>>MI0000114|hsa-mir-107&WithFlank&10|-|89592741|89592833@56.78.23&[MIMAT0000104&hsa-miR-107&offsets|0|0;m-78&10|-|89592758|89592778&offsets|0|+2]|AGCAGCATTGTACAGGGCTATCA</td><td>>MI0000114|hsa-mir-107&WithFlank&10|-|89592741|89592833</td><td>56</td><td>78</td><td>23</td><td></td><td>isomiR</td><td>NO</td><td>0.00</td><td>28.82</td><td>0.00</td><td>0.00</td></tr>
<tr><td>>MI0000115|hsa-mir-16-2&WithFlank&3|+|160404739|160404831@58.79.22&[MIMAT0004518&hsa-miR-16-2-3p&offsets|-1|-1;m-252&3|+|160404796|160404816&offsets|0|+1]|ACCAATATTACTGTGCTGCTTT</td><td>>MI0000115|hsa-mir-16-2&WithFlank&3|+|160404739|160404831</td><td>58</td><td>79</td><td>22</td><td></td><td>isomiR</td><td>NO</td><td>25.99</td><td>86.75</td><td>13.19</td><td>0.00</td></tr>
<tr><td>>MI0000234|hsa-mir-192&WithFlank&11|-|64891131|64891252@30.49.20(+1U)&[MIMAT0000222&hsa-miR-192-5p&offsets|0|-1(+1U);m-59&11|-|64891203|64891223&offsets|0|-1(+1U)]|CTGACCTATGAATTGACAGCT</td><td>>MI0000234|hsa-mir-192&WithFlank&11|-|64891131|64891252</td><td>30</td><td>49</td><td>20</td><td>(+1U)</td><td>isomiR</td><td>NO</td><td>0.00</td><td>10.16</td><td>11.61</td><td>0.00</td></tr>
<tr><td>>MI0000234|hsa-mir-192&WithFlank&11|-|64891131|64891252@30.50.21&[MIMAT0000222&hsa-miR-192-5p&offsets|0|0;m-59&11|-|64891203|64891223&offsets|0|0]|CTGACCTATGAATTGACAGCC</td><td>>MI0000234|hsa-mir-192&WithFlank&11|-|64891131|64891252</td><td>30</td><td>50</td><td>21</td><td></td><td>isomiR</td><td>NO</td><td>27.33</td><td>38.44</td><td>61.90</td><td>0.00</td></tr>
<tr><td>>MI0000239|hsa-mir-197&WithFlank&1|+|109598887|109598973@54.72.19&[MIMAT0000227&hsa-miR-197-3p&offsets|0|-3;m-99&1|+|109598940|109598961&offsets|0|-3]|TTCACCACCTTCTCCACCC</td><td>>MI0000239|hsa-mir-197&WithFlank&1|+|109598887|109598973</td><td>54</td><td>72</td><td>19</td><td></td><td>isomiR</td><td>NO</td><td>0.00</td><td>35.29</td><td>0.00</td><td>0.00</td></tr>
<tr><td>>MI0000239|hsa-mir-197&WithFlank&1|+|109598887|109598973@54.74.21&[MIMAT0000227&hsa-miR-197-3p&offsets|0|-1;m-99&1|+|109598940|109598961&offsets|0|-1]|TTCACCACCTTCTCCACCCAG</td><td>>MI0000239|hsa-mir-197&WithFlank&1|+|109598887|109598973</td><td>54</td><td>74</td><td>21</td><td></td><td>isomiR</td><td>NO</td><td>0.00</td><td>45.39</td><td>0.00</td><td>0.00</td></tr>
<tr><td>>MI0000239|hsa-mir-197&WithFlank&1|+|109598887|109598973@54.74.21(+1A)&[MIMAT0000227&hsa-miR-197-3p&offsets|0|-1(+1A);m-99&1|+|109598940|109598961&offsets|0|-1(+1A)]|TTCACCACCTTCTCCACCCAGA</td><td>>MI0000239|hsa-mir-197&WithFlank&1|+|109598887|109598973</td><td>54</td><td>74</td><td>21</td><td>(+1A)</td><td>isomiR</td><td>NO</td><td>26.88</td><td>138.28</td><td>0.00</td><td>0.00</td></tr>
<tr><td>>MI0000239|hsa-mir-197&WithFlank&1|+|109598887|109598973@54.74.21(+1U)&[MIMAT0000227&hsa-miR-197-3p&offsets|0|-1(+1U);m-99&1|+|109598940|109598961&offsets|0|-1(+1U)]|TTCACCACCTTCTCCACCCAGT</td><td>>MI0000239|hsa-mir-197&WithFlank&1|+|109598887|109598973</td><td>54</td><td>74</td><td>21</td><td>(+1U)</td><td>isomiR</td><td>NO</td><td>130.12</td><td>385.13</td><td>0.00</td><td>90.90</td></tr>
<tr><td>>MI0000239|hsa-mir-197&WithFlank&1|+|109598887|109598973@54.75.22&[MIMAT0000227&hsa-miR-197-3p&offsets|0|0;m-99&1|+|109598940|109598961&offsets|0|0]|TTCACCACCTTCTCCACCCAGC</td><td>>MI0000239|hsa-mir-197&WithFlank&1|+|109598887|109598973</td><td>54</td><td>75</td><td>22</td><td></td><td>isomiR</td><td>NO</td><td>359.28</td><td>1185.64</td><td>82.92</td><td>293.00</td></tr>
<tr><td>>MI0000239|hsa-mir-197&WithFlank&1|+|109598887|109598973@54.75.22(+1U)&[MIMAT0000227&hsa-miR-197-3p&offsets|0|0(+1U);m-99&1|+|109598940|109598961&offsets|0|0(+1U)]|TTCACCACCTTCTCCACCCAGCT</td><td>>MI0000239|hsa-mir-197&WithFlank&1|+|109598887|109598973</td><td>54</td><td>75</td><td>22</td><td>(+1U)</td><td>isomiR</td><td>NO</td><td>0.00</td><td>23.63</td><td>0.00</td><td>0.00</td></tr>
<tr><td>>MI0000239|hsa-mir-197&WithFlank&1|+|109598887|109598973@54.76.23&[MIMAT0000227&hsa-miR-197-3p&offsets|0|+1;m-99&1|+|109598940|109598961&offsets|0|+1]|TTCACCACCTTCTCCACCCAGCA</td><td>>MI0000239|hsa-mir-197&WithFlank&1|+|109598887|109598973</td><td>54</td><td>76</td><td>23</td><td></td><td>isomiR</td><td>NO</td><td>0.00</td><td>28.51</td><td>0.00</td><td>0.00</td></tr>
<tr><td>>am_MI0000239|hsa-mir-197&WithFlank&1|+|109598887|109598973@54.71.18(+1U)&[MIMAT0000227&hsa-miR-197-3p&offsets|0|-4(+1U);m-99&1|+|109598940|109598961&offsets|0|-4(+1U)]|TTCACCACCTTCTCCACCT</td><td>>MI0000239|hsa-mir-197&WithFlank&1|+|109598887|109598973.am</td><td>54</td><td>71</td><td>18</td><td>(+1U)</td><td>isomiR</td><td>NO</td><td>0.00</td><td>24.56</td><td>0.00</td><td>0.00</td></tr>
</table>

# BT-20

<table>
<tr><th></th><th></th><th></th><th></th><th></th><th></th><th></th><th></th><th></th><th></th><th></th><th></th></tr>
<tr><td>>MI0000253|hsa-mir-148a&WithFlank&7|-|25949913|25949992@12.33.22&[MIMAT0004549&hsa-miR-148a-5p&offsets|0|0;m-189&7|-|25949960|25949981&offsets|0|0]|AAAGTTCTGAGACACTCCGACT</td><td>>MI0000253|hsa-mir-148a&WithFlank&7|-|25949913|25949992</td><td>12</td><td>33</td><td>22</td><td></td><td>isomiR</td><td>NO</td><td>26.15</td><td>71.03</td><td>12.56</td><td>0.00</td></tr>
<tr><td>>MI0000253|hsa-mir-148a&WithFlank&7|-|25949913|25949992@50.70.21(+1A)&[MIMAT0000243&hsa-miR-148a-3p&offsets|0|-1(+1A);m-7&7|-|25949922|25949943&offsets|0|-1(+1A)]|TCAGTGCACTACAGAACTTTGA</td><td>>MI0000253|hsa-mir-148a&WithFlank&7|-|25949913|25949992</td><td>50</td><td>70</td><td>21</td><td>(+1A)</td><td>isomiR</td><td>NO</td><td>172.44</td><td>483.21</td><td>429.22</td><td>69.61</td></tr>
<tr><td>>MI0000253|hsa-mir-148a&WithFlank&7|-|25949913|25949992@50.70.21(+1C)&[MIMAT0000243&hsa-miR-148a-3p&offsets|0|-1(+1C);m-7&7|-|25949922|25949943&offsets|0|-1(+1C)]|TCAGTGCACTACAGAACTTTGC</td><td>>MI0000253|hsa-mir-148a&WithFlank&7|-|25949913|25949992</td><td>50</td><td>70</td><td>21</td><td>(+1C)</td><td>isomiR</td><td>YES</td><td>726.40</td><td>1391.65</td><td>1728.62</td><td>299.98</td></tr>
<tr><td>>MI0000253|hsa-mir-148a&WithFlank&7|-|25949913|25949992@50.70.21(+1G)&[MIMAT0000243&hsa-miR-148a-3p&offsets|0|-1(+1G);m-7&7|-|25949922|25949943&offsets|0|-1(+1G)]|TCAGTGCACTACAGAACTTTGG</td><td>>MI0000253|hsa-mir-148a&WithFlank&7|-|25949913|25949992</td><td>50</td><td>70</td><td>21</td><td>(+1G)</td><td>isomiR</td><td>YES</td><td>182.06</td><td>314.58</td><td>245.29</td><td>10.78</td></tr>
<tr><td>>MI0000253|hsa-mir-148a&WithFlank&7|-|25949913|25949992@50.71.22&[MIMAT0000243&hsa-miR-148a-3p&offsets|0|0;m-7&7|-|25949922|25949943&offsets|0|0]|TCAGTGCACTACAGAACTTTGT</td><td>>MI0000253|hsa-mir-148a&WithFlank&7|-|25949913|25949992</td><td>50</td><td>71</td><td>22</td><td></td><td>isomiR</td><td>YES</td><td>3513.72</td><td>9182.41</td><td>9675.09</td><td>1515.78</td></tr>
<tr><td>>MI0000253|hsa-mir-148a&WithFlank&7|-|25949913|25949992@50.71.22(+1U)&[MIMAT0000243&hsa-miR-148a-3p&offsets|0|0(+1U);m-7&7|-|25949922|25949943&offsets|0|0(+1U)]|TCAGTGCACTACAGAACTTTGTT</td><td>>MI0000253|hsa-mir-148a&WithFlank&7|-|25949913|25949992</td><td>50</td><td>71</td><td>22</td><td>(+1U)</td><td>isomiR</td><td>NO</td><td>30.51</td><td>68.80</td><td>0.00</td><td>0.00</td></tr>
<tr><td>>MI0000253|hsa-mir-148a&WithFlank&7|-|25949913|25949992@50.72.23&[MIMAT0000243&hsa-miR-148a-3p&offsets|0|+1;m-7&7|-|25949922|25949943&offsets|0|+1]|TCAGTGCACTACAGAACTTTGTC</td><td>>MI0000253|hsa-mir-148a&WithFlank&7|-|25949913|25949992</td><td>50</td><td>72</td><td>23</td><td></td><td>isomiR</td><td>NO</td><td>112.16</td><td>200.10</td><td>66.37</td><td>33.06</td></tr>
<tr><td>>MI0000254|hsa-mir-30c-2&WithFlank&6|-|71376954|71377037@53.74.22&[MIMAT0004550&hsa-miR-30c-2-3p&offsets|0|0;m-206&6|-|71376964|71376985&offsets|0|0]|CTGGGAGAAGGCTGTTTACTCT</td><td>>MI0000254|hsa-mir-30c-2&WithFlank&6|-|71376954|71377037</td><td>53</td><td>74</td><td>22</td><td></td><td>isomiR</td><td>NO</td><td>27.37</td><td>165.01</td><td>0.00</td><td>0.00</td></tr>
<tr><td>>MI0000255|hsa-mir-30d&WithFlank&8|-|134804870|134804951@12.31.20&[MIMAT0000245&hsa-miR-30d-5p&offsets|0|-2;m-21&8|-|134804919|134804940&offsets|0|-2]|TGTAAACATCCCCGACTGGA</td><td>>MI0000255|hsa-mir-30d&WithFlank&8|-|134804870|134804951</td><td>12</td><td>31</td><td>20</td><td></td><td>isomiR</td><td>NO</td><td>80.53</td><td>175.58</td><td>186.43</td><td>34.16</td></tr>
<tr><td>>MI0000255|hsa-mir-30d&WithFlank&8|-|134804870|134804951@12.32.21&[MIMAT0000245&hsa-miR-30d-5p&offsets|0|-1;m-21&8|-|134804919|134804940&offsets|0|-1]|TGTAAACATCCCCGACTGGAA</td><td>>MI0000255|hsa-mir-30d&WithFlank&8|-|134804870|134804951</td><td>12</td><td>32</td><td>21</td><td></td><td>isomiR</td><td>NO</td><td>38.51</td><td>111.26</td><td>10.52</td><td>23.33</td></tr>
<tr><td>>MI0000255|hsa-mir-30d&WithFlank&8|-|134804870|134804951@12.32.21(+1A)&[MIMAT0000245&hsa-miR-30d-5p&offsets|0|-1(+1A);m-21&8|-|134804919|134804940&offsets|0|-1(+1A)]|TGTAAACATCCCCGACTGGAAA</td><td>>MI0000255|hsa-mir-30d&WithFlank&8|-|134804870|134804951</td><td>12</td><td>32</td><td>21</td><td>(+1A)</td><td>isomiR</td><td>NO</td><td>25.97</td><td>86.93</td><td>0.00</td><td>0.00</td></tr>
<tr><td>>MI0000255|hsa-mir-30d&WithFlank&8|-|134804870|134804951@12.33.22&[MIMAT0000245&hsa-miR-30d-5p&offsets|0|0;m-21&8|-|134804919|134804940&offsets|0|0]|TGTAAACATCCCCGACTGGAAG</td><td>>MI0000255|hsa-mir-30d&WithFlank&8|-|134804870|134804951</td><td>12</td><td>33</td><td>22</td><td></td><td>isomiR</td><td>NO</td><td>217.29</td><td>421.37</td><td>119.13</td><td>122.90</td></tr>
<tr><td>>MI0000255|hsa-mir-30d&WithFlank&8|-|134804870|134804951@12.33.22(+1A)&[MIMAT0000245&hsa-miR-30d-5p&offsets|0|0(+1A);m-21&8|-|134804919|134804940&offsets|0|0(+1A)]|TGTAAACATCCCCGACTGGAAGA</td><td>>MI0000255|hsa-mir-30d&WithFlank&8|-|134804870|134804951</td><td>12</td><td>33</td><td>22</td><td>(+1A)</td><td>isomiR</td><td>NO</td><td>100.79</td><td>285.30</td><td>71.73</td><td>84.55</td></tr>
<tr><td>>MI0000255|hsa-mir-30d&WithFlank&8|-|134804870|134804951@12.33.22(+1U)&[MIMAT0000245&hsa-miR-30d-5p&offsets|0|0(+1U);m-21&8|-|134804919|134804940&offsets|0|0(+1U)]|TGTAAACATCCCCGACTGGAAGT</td><td>>MI0000255|hsa-mir-30d&WithFlank&8|-|134804870|134804951</td><td>12</td><td>33</td><td>22</td><td>(+1U)</td><td>isomiR</td><td>NO</td><td>358.85</td><td>905.79</td><td>254.44</td><td>304.46</td></tr>
<tr><td>>MI0000255|hsa-mir-30d&WithFlank&8|-|134804870|134804951@12.34.23&[MIMAT0000245&hsa-miR-30d-5p&offsets|0|+1;m-21&8|-|134804919|134804940&offsets|0|+1]|TGTAAACATCCCCGACTGGAAGC</td><td>>MI0000255|hsa-mir-30d&WithFlank&8|-|134804870|134804951</td><td>12</td><td>34</td><td>23</td><td></td><td>isomiR</td><td>YES</td><td>947.66</td><td>2604.96</td><td>758.95</td><td>892.97</td></tr>
<tr><td>>MI0000255|hsa-mir-30d&WithFlank&8|-|134804870|134804951@12.34.23(+1A)&[MIMAT0000245&hsa-miR-30d-5p&offsets|0|+1(+1A);m-21&8|-|134804919|134804940&offsets|0|+1(+1A)]|TGTAAACATCCCCGACTGGAAGCA</td><td>>MI0000255|hsa-mir-30d&WithFlank&8|-|134804870|134804951</td><td>12</td><td>34</td><td>23</td><td>(+1A)</td><td>isomiR</td><td>NO</td><td>233.77</td><td>671.52</td><td>146.58</td><td>144.00</td></tr>
<tr><td>>MI0000255|hsa-mir-30d&WithFlank&8|-|134804870|134804951@12.34.23(+1C)&[MIMAT0000245&hsa-miR-30d-5p&offsets|0|+1(+1C);m-21&8|-|134804919|134804940&offsets|0|+1(+1C)]|TGTAAACATCCCCGACTGGAAGCC</td><td>>MI0000255|hsa-mir-30d&WithFlank&8|-|134804870|134804951</td><td>12</td><td>34</td><td>23</td><td>(+1C)</td><td>isomiR</td><td>NO</td><td>364.42</td><td>764.87</td><td>230.65</td><td>228.02</td></tr>
<tr><td>>MI0000255|hsa-mir-30d&WithFlank&8|-|134804870|134804951@12.34.23(+1G)&[MIMAT0000245&hsa-miR-30d-5p&offsets|0|+1(+1G);m-21&8|-|134804919|134804940&offsets|0|+1(+1G)]|TGTAAACATCCCCGACTGGAAGCG</td><td>>MI0000255|hsa-mir-30d&WithFlank&8|-|134804870|134804951</td><td>12</td><td>34</td><td>23</td><td>(+1G)</td><td>isomiR</td><td>NO</td><td>112.23</td><td>210.90</td><td>14.17</td><td>23.56</td></tr>
<tr><td>>MI0000255|hsa-mir-30d&WithFlank&8|-|134804870|134804951@12.35.24&[MIMAT0000245&hsa-miR-30d-5p&offsets|0|+2;m-21&8|-|134804919|134804940&offsets|0|+2]|TGTAAACATCCCCGACTGGAAGCT</td><td>>MI0000255|hsa-mir-30d&WithFlank&8|-|134804870|134804951</td><td>12</td><td>35</td><td>24</td><td></td><td>isomiR</td><td>YES</td><td>1487.42</td><td>4221.06</td><td>1110.64</td><td>967.33</td></tr>
<tr><td>>MI0000255|hsa-mir-30d&WithFlank&8|-|134804870|134804951@12.35.24(+1A)&[MIMAT0000245&hsa-miR-30d-5p&offsets|0|+2(+1A);m-21&8|-|134804919|134804940&offsets|0|+2(+1A)]|TGTAAACATCCCCGACTGGAAGCTA</td><td>>MI0000255|hsa-mir-30d&WithFlank&8|-|134804870|134804951</td><td>12</td><td>35</td><td>24</td><td>(+1A)</td><td>isomiR</td><td>NO</td><td>0.00</td><td>23.86</td><td>0.00</td><td>0.00</td></tr>
<tr><td>>MI0000255|hsa-mir-30d&WithFlank&8|-|134804870|134804951@12.35.24(+1U)&[MIMAT0000245&hsa-miR-30d-5p&offsets|0|+2(+1U);m-21&8|-|134804919|134804940&offsets|0|+2(+1U)]|TGTAAACATCCCCGACTGGAAGCTT</td><td>>MI0000255|hsa-mir-30d&WithFlank&8|-|134804870|134804951</td><td>12</td><td>35</td><td>24</td><td>(+1U)</td><td>isomiR</td><td>NO</td><td>75.58</td><td>139.25</td><td>60.47</td><td>32.03</td></tr>
<tr><td>>MI0000261|hsa-mir-139&WithFlank&11|-|72615057|72615136@13.35.23&[MIMAT0000250&hsa-miR-139-5p&offsets|0|0;m-186&11|-|72615102|72615124&offsets|0|0]|TCTACAGTGCACGTGTCTCCAGT</td><td>>MI0000261|hsa-mir-139&WithFlank&11|-|72615057|72615136</td><td>13</td><td>35</td><td>23</td><td></td><td>isomiR</td><td>NO</td><td>0.00</td><td>76.22</td><td>10.70</td><td>0.00</td></tr>
<tr><td>>MI0000263|hsa-mir-7-1&WithFlank&9|-|83969742|83969863@30.54.25&[MIMAT0000252_2&hsa-miR-7-5p&offsets|0|+1;m-401&9|-|83969811|83969834&offsets|0|+1]|TGGAAGACTAGTGATTTTGTTGTTT</td><td>>MI0000263|hsa-mir-7-1&WithFlank&9|-|83969742|83969863</td><td>30</td><td>54</td><td>25</td><td></td><td>isomiR</td><td>NO</td><td>30.73</td><td>67.14</td><td>0.00</td><td>9.82</td></tr>
</table>

# BT-20

<table>
<tr><th></th><th></th><th></th><th></th><th></th><th></th><th></th><th></th><th></th><th></th><th></th><th></th></tr>
<tr><td>>MI0000264|hsa-mir-7-2&WithFlank&15|+|88611819|88611940@38.58.21;MI0000265|hsa-mir-7-3&WithFlank&19|+|4770664|4770785@37.57.21;MI0000263|hsa-mir-7-1&WithFlank&9|-|83969742|83969863@30.50.21&[MIMAT0000252&hsa-miR-7-5p&offsets|0|-3;m-400&15|+|88611856|88611879&offsets|0|-3];[MIMAT0000252_1&hsa-miR-7-5p&offsets|0|-3;m-402&19|+|4770700|4770723&offsets|0|-3];[MIMAT0000252_2&hsa-miR-7-5p&offsets|0|-3;m-401&9|-|83969811|83969834&offsets|0|-3]|TGGAAGACTAGTGATTTTGTT</td><td>>MI0000264|hsa-mir-7-2&WithFlank&15|+|88611819|88611940</td><td>38</td><td>58</td><td>21</td><td></td><td>isomiR</td><td>NO</td><td>39.89</td><td>71.44</td><td>12.54</td><td>0.00</td></tr>
<tr><td>>MI0000264|hsa-mir-7-2&WithFlank&15|+|88611819|88611940@38.59.22(+1A);MI0000265|hsa-mir-7-3&WithFlank&19|+|4770664|4770785@37.58.22(+1A);MI0000263|hsa-mir-7-1&WithFlank&9|-|83969742|83969863@30.51.22(+1A)&[MIMAT0000252&hsa-miR-7-5p&offsets|0|-2(+1A);m-400&15|+|88611856|88611879&offsets|0|-2(+1A)];[MIMAT0000252_1&hsa-miR-7-5p&offsets|0|-2(+1A);m-402&19|+|4770700|4770723&offsets|0|-2(+1A)];[MIMAT0000252_2&hsa-miR-7-5p&offsets|0|-2(+1A);m-401&9|-|83969811|83969834&offsets|0|-2(+1A)]|TGGAAGACTAGTGATTTTGTTGA</td><td>>MI0000264|hsa-mir-7-2&WithFlank&15|+|88611819|88611940</td><td>38</td><td>59</td><td>22</td><td>(+1A)</td><td>isomiR</td><td>NO</td><td>0.00</td><td>25.98</td><td>0.00</td><td>0.00</td></tr>
<tr><td>>MI0000264|hsa-mir-7-2&WithFlank&15|+|88611819|88611940@38.59.22(+1C);MI0000265|hsa-mir-7-3&WithFlank&19|+|4770664|4770785@37.58.22(+1C);MI0000263|hsa-mir-7-1&WithFlank&9|-|83969742|83969863@30.51.22(+1C)&[MIMAT0000252&hsa-miR-7-5p&offsets|0|-2(+1C);m-400&15|+|88611856|88611879&offsets|0|-2(+1C)];[MIMAT0000252_1&hsa-miR-7-5p&offsets|0|-2(+1C);m-402&19|+|4770700|4770723&offsets|0|-2(+1C)];[MIMAT0000252_2&hsa-miR-7-5p&offsets|0|-2(+1C);m-401&9|-|83969811|83969834&offsets|0|-2(+1C)]|TGGAAGACTAGTGATTTTGTTGC</td><td>>MI0000264|hsa-mir-7-2&WithFlank&15|+|88611819|88611940</td><td>38</td><td>59</td><td>22</td><td>(+1C)</td><td>isomiR</td><td>NO</td><td>69.83</td><td>91.63</td><td>38.58</td><td>0.00</td></tr>
<tr><td>>MI0000264|hsa-mir-7-2&WithFlank&15|+|88611819|88611940@38.59.22;MI0000265|hsa-mir-7-3&WithFlank&19|+|4770664|4770785@37.58.22;MI0000263|hsa-mir-7-1&WithFlank&9|-|83969742|83969863@30.51.22&[MIMAT0000252&hsa-miR-7-5p&offsets|0|-2;m-400&15|+|88611856|88611879&offsets|0|-2];[MIMAT0000252_1&hsa-miR-7-5p&offsets|0|-2;m-402&19|+|4770700|4770723&offsets|0|-2];[MIMAT0000252_2&hsa-miR-7-5p&offsets|0|-2;m-401&9|-|83969811|83969834&offsets|0|-2]|TGGAAGACTAGTGATTTTGTTG</td><td>>MI0000264|hsa-mir-7-2&WithFlank&15|+|88611819|88611940</td><td>38</td><td>59</td><td>22</td><td></td><td>isomiR</td><td>NO</td><td>39.17</td><td>42.21</td><td>54.24</td><td>0.00</td></tr>
<tr><td>>MI0000264|hsa-mir-7-2&WithFlank&15|+|88611819|88611940@38.60.23(+1A);MI0000265|hsa-mir-7-3&WithFlank&19|+|4770664|4770785@37.59.23(+1A);MI0000263|hsa-mir-7-1&WithFlank&9|-|83969742|83969863@30.52.23(+1A)&[MIMAT0000252&hsa-miR-7-5p&offsets|0|-1(+1A);m-400&15|+|88611856|88611879&offsets|0|-1(+1A)];[MIMAT0000252_1&hsa-miR-7-5p&offsets|0|-1(+1A);m-402&19|+|4770700|4770723&offsets|0|-1(+1A)];[MIMAT0000252_2&hsa-miR-7-5p&offsets|0|-1(+1A);m-401&9|-|83969811|83969834&offsets|0|-1(+1A)]|TGGAAGACTAGTGATTTTGTTGTA</td><td>>MI0000264|hsa-mir-7-2&WithFlank&15|+|88611819|88611940</td><td>38</td><td>60</td><td>23</td><td>(+1A)</td><td>isomiR</td><td>NO</td><td>94.18</td><td>299.74</td><td>12.06</td><td>67.23</td></tr>
<tr><td>>MI0000264|hsa-mir-7-2&WithFlank&15|+|88611819|88611940@38.60.23(+1C);MI0000265|hsa-mir-7-3&WithFlank&19|+|4770664|4770785@37.59.23(+1C);MI0000263|hsa-mir-7-1&WithFlank&9|-|83969742|83969863@30.52.23(+1C)&[MIMAT0000252&hsa-miR-7-5p&offsets|0|-1(+1C);m-400&15|+|88611856|88611879&offsets|0|-1(+1C)];[MIMAT0000252_1&hsa-miR-7-5p&offsets|0|-1(+1C);m-402&19|+|4770700|4770723&offsets|0|-1(+1C)];[MIMAT0000252_2&hsa-miR-7-5p&offsets|0|-1(+1C);m-401&9|-|83969811|83969834&offsets|0|-1(+1C)]|TGGAAGACTAGTGATTTTGTTGTC</td><td>>MI0000264|hsa-mir-7-2&WithFlank&15|+|88611819|88611940</td><td>38</td><td>60</td><td>23</td><td>(+1C)</td><td>isomiR</td><td>NO</td><td>340.41</td><td>761.33</td><td>135.40</td><td>258.63</td></tr>
<tr><td>>MI0000264|hsa-mir-7-2&WithFlank&15|+|88611819|88611940@38.60.23(+1G);MI0000265|hsa-mir-7-3&WithFlank&19|+|4770664|4770785@37.59.23(+1G);MI0000263|hsa-mir-7-1&WithFlank&9|-|83969742|83969863@30.52.23(+1G)&[MIMAT0000252&hsa-miR-7-5p&offsets|0|-1(+1G);m-400&15|+|88611856|88611879&offsets|0|-1(+1G)];[MIMAT0000252_1&hsa-miR-7-5p&offsets|0|-1(+1G);m-402&19|+|4770700|4770723&offsets|0|-1(+1G)];[MIMAT0000252_2&hsa-miR-7-5p&offsets|0|-1(+1G);m-401&9|-|83969811|83969834&offsets|0|-1(+1G)]|TGGAAGACTAGTGATTTTGTTGTG</td><td>>MI0000264|hsa-mir-7-2&WithFlank&15|+|88611819|88611940</td><td>38</td><td>60</td><td>23</td><td>(+1G)</td><td>isomiR</td><td>NO</td><td>42.95</td><td>130.93</td><td>0.00</td><td>0.00</td></tr>
<tr><td>>MI0000264|hsa-mir-7-2&WithFlank&15|+|88611819|88611940@38.60.23;MI0000265|hsa-mir-7-3&WithFlank&19|+|4770664|4770785@37.59.23;MI0000263|hsa-mir-7-1&WithFlank&9|-|83969742|83969863@30.52.23&[MIMAT0000252&hsa-miR-7-5p&offsets|0|-1;m-400&15|+|88611856|88611879&offsets|0|-1];[MIMAT0000252_1&hsa-miR-7-5p&offsets|0|-1;m-402&19|+|4770700|4770723&offsets|0|-1];[MIMAT0000252_2&hsa-miR-7-5p&offsets|0|-1;m-401&9|-|83969811|83969834&offsets|0|-1]|TGGAAGACTAGTGATTTTGTTGT</td><td>>MI0000264|hsa-mir-7-2&WithFlank&15|+|88611819|88611940</td><td>38</td><td>60</td><td>23</td><td></td><td>isomiR</td><td>YES</td><td>321.35</td><td>601.51</td><td>265.95</td><td>116.58</td></tr>
<tr><td>>MI0000264|hsa-mir-7-2&WithFlank&15|+|88611819|88611940@38.61.24(+1A);MI0000265|hsa-mir-7-3&WithFlank&19|+|4770664|4770785@37.60.24(+1A);MI0000263|hsa-mir-7-1&WithFlank&9|-|83969742|83969863@30.53.24(+1A)&[MIMAT0000252&hsa-miR-7-5p&offsets|0|0(+1A);m-400&15|+|88611856|88611879&offsets|0|0(+1A)];[MIMAT0000252_1&hsa-miR-7-5p&offsets|0|0(+1A);m-402&19|+|4770700|4770723&offsets|0|0(+1A)];[MIMAT0000252_2&hsa-miR-7-5p&offsets|0|0(+1A);m-401&9|-|83969811|83969834&offsets|0|0(+1A)]|TGGAAGACTAGTGATTTTGTTGTTA</td><td>>MI0000264|hsa-mir-7-2&WithFlank&15|+|88611819|88611940</td><td>38</td><td>61</td><td>24</td><td>(+1A)</td><td>isomiR</td><td>NO</td><td>10.85</td><td>69.91</td><td>0.00</td><td>0.00</td></tr>
<tr><td>>MI0000264|hsa-mir-7-2&WithFlank&15|+|88611819|88611940@38.61.24;MI0000265|hsa-mir-7-3&WithFlank&19|+|4770664|4770785@37.60.24;MI0000263|hsa-mir-7-1&WithFlank&9|-|83969742|83969863@30.53.24&[MIMAT0000252&hsa-miR-7-5p&offsets|0|0;m-400&15|+|88611856|88611879&offsets|0|0];[MIMAT0000252_1&hsa-miR-7-5p&offsets|0|0;m-402&19|+|4770700|4770723&offsets|0|0];[MIMAT0000252_2&hsa-miR-7-5p&offsets|0|0;m-401&9|-|83969811|83969834&offsets|0|0]|TGGAAGACTAGTGATTTTGTTGTT</td><td>>MI0000264|hsa-mir-7-2&WithFlank&15|+|88611819|88611940</td><td>38</td><td>61</td><td>24</td><td></td><td>isomiR</td><td>YES</td><td>1621.26</td><td>4775.37</td><td>722.72</td><td>1271.82</td></tr>
<tr><td>>MI0000266|hsa-mir-10a&WithFlank&17|-|48579832|48579953@28.48.21&[MIMAT0000253&hsa-miR-10a-5p&offsets|0|-2;m-18&17|-|48579905|48579926&offsets|0|-1]|TACCCTGTAGATCCGAATTTG</td><td>>MI0000266|hsa-mir-10a&WithFlank&17|-|48579832|48579953</td><td>28</td><td>48</td><td>21</td><td></td><td>isomiR</td><td>NO</td><td>0.00</td><td>76.77</td><td>11.80</td><td>0.00</td></tr>
</table>

# BT-20

| | | | | | | | | | | | |
|---|---|---|---|---|---|---|---|---|---|---|---|
| >MI0000266\|hsa-mir-10a&WithFlank&17\|-\|48579832\|48579953@28.48.21(+1A)&[MIMAT0000253&hsa-miR-10a-5p&offsets\|0\|-2(+1A);m-18&17\|-\|48579905\|48579926&offsets\|0\|-1(+1A)]\|TACCCTGTAGATCCGAATTTGA | >MI0000266\|hsa-mir-10a&WithFlank&17\|-\|48579832\|48579953 | 28 | 48 | 21 | (+1A) | isomiR | NO | 0.00 | 122.15 | 11.16 | 0.00 |
| >MI0000266\|hsa-mir-10a&WithFlank&17\|-\|48579832\|48579953@28.48.21(+1C)&[MIMAT0000253&hsa-miR-10a-5p&offsets\|0\|-2(+1C);m-18&17\|-\|48579905\|48579926&offsets\|0\|-1(+1C)]\|TACCCTGTAGATCCGAATTTGC | >MI0000266\|hsa-mir-10a&WithFlank&17\|-\|48579832\|48579953 | 28 | 48 | 21 | (+1C) | isomiR | NO | 80.55 | 337.88 | 121.24 | 11.04 |
| >MI0000266\|hsa-mir-10a&WithFlank&17\|-\|48579832\|48579953@28.48.21(+1G)&[MIMAT0000253&hsa-miR-10a-5p&offsets\|0\|-2(+1G);m-18&17\|-\|48579905\|48579926&offsets\|0\|-1(+1G)]\|TACCCTGTAGATCCGAATTTGG | >MI0000266\|hsa-mir-10a&WithFlank&17\|-\|48579832\|48579953 | 28 | 48 | 21 | (+1G) | isomiR | NO | 0.00 | 72.03 | 0.00 | 0.00 |
| >MI0000266\|hsa-mir-10a&WithFlank&17\|-\|48579832\|48579953@28.49.22&[MIMAT0000253&hsa-miR-10a-5p&offsets\|0\|-1;m-18&17\|-\|48579905\|48579926&offsets\|0\|0]\|TACCCTGTAGATCCGAATTTGT | >MI0000266\|hsa-mir-10a&WithFlank&17\|-\|48579832\|48579953 | 28 | 49 | 22 | | isomiR | NO | 366.32 | 2142.10 | 656.55 | 169.54 |
| >MI0000266\|hsa-mir-10a&WithFlank&17\|-\|48579832\|48579953@28.49.22(+1A)&[MIMAT0000253&hsa-miR-10a-5p&offsets\|0\|-1(+1A);m-18&17\|-\|48579905\|48579926&offsets\|0\|0(+1A)]\|TACCCTGTAGATCCGAATTTGTA | >MI0000266\|hsa-mir-10a&WithFlank&17\|-\|48579832\|48579953 | 28 | 49 | 22 | (+1A) | isomiR | NO | 36.68 | 269.03 | 54.51 | 0.00 |
| >MI0000266\|hsa-mir-10a&WithFlank&17\|-\|48579832\|48579953@28.49.22(+1U)&[MIMAT0000253&hsa-miR-10a-5p&offsets\|0\|-1(+1U);m-18&17\|-\|48579905\|48579926&offsets\|0\|0(+1U)]\|TACCCTGTAGATCCGAATTTGTT | >MI0000266\|hsa-mir-10a&WithFlank&17\|-\|48579832\|48579953 | 28 | 49 | 22 | (+1U) | isomiR | NO | 0.00 | 23.46 | 0.00 | 0.00 |
| >MI0000266\|hsa-mir-10a&WithFlank&17\|-\|48579832\|48579953@28.50.23&[MIMAT0000253&hsa-miR-10a-5p&offsets\|0\|0;m-18&17\|-\|48579905\|48579926&offsets\|0\|+1]\|TACCCTGTAGATCCGAATTTGTG | >MI0000266\|hsa-mir-10a&WithFlank&17\|-\|48579832\|48579953 | 28 | 50 | 23 | | isomiR | NO | 268.94 | 1246.14 | 332.13 | 165.60 |
| >MI0000266\|hsa-mir-10a&WithFlank&17\|-\|48579832\|48579953@28.50.23(+1A)&[MIMAT0000253&hsa-miR-10a-5p&offsets\|0\|0(+1A);m-18&17\|-\|48579905\|48579926&offsets\|0\|+1(+1A)]\|TACCCTGTAGATCCGAATTTGTGA | >MI0000266\|hsa-mir-10a&WithFlank&17\|-\|48579832\|48579953 | 28 | 50 | 23 | (+1A) | isomiR | NO | 27.68 | 125.65 | 11.67 | 23.69 |
| >MI0000266\|hsa-mir-10a&WithFlank&17\|-\|48579832\|48579953@28.51.24&[MIMAT0000253&hsa-miR-10a-5p&offsets\|0\|+1;m-18&17\|-\|48579905\|48579926&offsets\|0\|+2]\|TACCCTGTAGATCCGAATTTGTGT | >MI0000266\|hsa-mir-10a&WithFlank&17\|-\|48579832\|48579953 | 28 | 51 | 24 | | isomiR | NO | 25.80 | 152.35 | 0.00 | 0.00 |
| >MI0000266\|hsa-mir-10a&WithFlank&17\|-\|48579832\|48579953@29.48.20(+1C)&[MIMAT0000253&hsa-miR-10a-5p&offsets\|+1\|-2(+1C);m-18&17\|-\|48579905\|48579926&offsets\|+1\|-1(+1C)]\|ACCCTGTAGATCCGAATTTGC | >MI0000266\|hsa-mir-10a&WithFlank&17\|-\|48579832\|48579953 | 29 | 48 | 20 | (+1C) | isomiR | NO | 0.00 | 36.73 | 0.00 | 0.00 |
| >MI0000266\|hsa-mir-10a&WithFlank&17\|-\|48579832\|48579953@29.49.21&[MIMAT0000253&hsa-miR-10a-5p&offsets\|+1\|-1;m-18&17\|-\|48579905\|48579926&offsets\|+1\|0]\|ACCCTGTAGATCCGAATTTGT | >MI0000266\|hsa-mir-10a&WithFlank&17\|-\|48579832\|48579953 | 29 | 49 | 21 | | isomiR | NO | 72.36 | 332.17 | 116.80 | 26.55 |
| >MI0000266\|hsa-mir-10a&WithFlank&17\|-\|48579832\|48579953@29.49.21(+1A)&[MIMAT0000253&hsa-miR-10a-5p&offsets\|+1\|-1(+1A);m-18&17\|-\|48579905\|48579926&offsets\|+1\|0(+1A)]\|ACCCTGTAGATCCGAATTTGTA | >MI0000266\|hsa-mir-10a&WithFlank&17\|-\|48579832\|48579953 | 29 | 49 | 21 | (+1A) | isomiR | NO | 28.32 | 194.77 | 10.01 | 0.00 |
| >MI0000266\|hsa-mir-10a&WithFlank&17\|-\|48579832\|48579953@29.50.22&[MIMAT0000253&hsa-miR-10a-5p&offsets\|+1\|0;m-18&17\|-\|48579905\|48579926&offsets\|+1\|+1]\|ACCCTGTAGATCCGAATTTGTG | >MI0000266\|hsa-mir-10a&WithFlank&17\|-\|48579832\|48579953 | 29 | 50 | 22 | | isomiR | NO | 204.12 | 894.03 | 175.24 | 135.94 |
| >MI0000266\|hsa-mir-10a&WithFlank&17\|-\|48579832\|48579953@29.50.22(+1A)&[MIMAT0000253&hsa-miR-10a-5p&offsets\|+1\|0(+1A);m-18&17\|-\|48579905\|48579926&offsets\|+1\|+1(+1A)]\|ACCCTGTAGATCCGAATTTGTGA | >MI0000266\|hsa-mir-10a&WithFlank&17\|-\|48579832\|48579953 | 29 | 50 | 22 | (+1A) | isomiR | NO | 28.47 | 124.81 | 10.44 | 26.60 |
| >MI0000266\|hsa-mir-10a&WithFlank&17\|-\|48579832\|48579953@29.51.23&[MIMAT0000253&hsa-miR-10a-5p&offsets\|+1\|+1;m-18&17\|-\|48579905\|48579926&offsets\|+1\|+2]\|ACCCTGTAGATCCGAATTTGTGT | >MI0000266\|hsa-mir-10a&WithFlank&17\|-\|48579832\|48579953 | 29 | 51 | 23 | | isomiR | NO | 28.07 | 152.33 | 0.00 | 0.00 |
| >MI0000269\|hsa-mir-181a-2&WithFlank&9\|+\|124692436\|124692557@83.103.21(+1A)&[MIMAT0004558&hsa-miR-181a-2-3p&offsets\|0\|-1(+1A);m-71&9\|+\|124692518\|124692539&offsets\|0\|-1(+1A)]\|ACCACTGACCGTTGACTGTACA | >MI0000269\|hsa-mir-181a-2&WithFlank&9\|+\|124692436\|124692557 | 83 | 103 | 21 | (+1A) | isomiR | NO | 0.00 | 24.52 | 0.00 | 0.00 |
| >MI0000269\|hsa-mir-181a-2&WithFlank&9\|+\|124692436\|124692557@83.103.21(+1U)&[MIMAT0004558&hsa-miR-181a-2-3p&offsets\|0\|-1(+1U);m-71&9\|+\|124692518\|124692539&offsets\|0\|-1(+1U)]\|ACCACTGACCGTTGACTGTACT | >MI0000269\|hsa-mir-181a-2&WithFlank&9\|+\|124692436\|124692557 | 83 | 103 | 21 | (+1U) | isomiR | NO | 0.00 | 70.16 | 0.00 | 0.00 |
| >MI0000269\|hsa-mir-181a-2&WithFlank&9\|+\|124692436\|124692557@83.104.22&[MIMAT0004558&hsa-miR-181a-2-3p&offsets\|0\|0;m-71&9\|+\|124692518\|124692539&offsets\|0\|0]\|ACCACTGACCGTTGACTGTACC | >MI0000269\|hsa-mir-181a-2&WithFlank&9\|+\|124692436\|124692557 | 83 | 104 | 22 | | isomiR | NO | 0.00 | 155.17 | 0.00 | 0.00 |
| >MI0000270\|hsa-mir-181b-1&WithFlank&1\|-\|198858867\|198858988@42.63.22(+1A);MI0000683\|hsa-mir-181b-2&WithFlank&9\|+\|124693704\|124693804@22.43.22(+1A)&[MIMAT0000257&hsa-miR-181b-5p&offsets\|0\|-1(+1A);m-73&1\|-\|198858925\|198858947&offsets\|0\|-1(+1A)];[MIMAT0000257_1&hsa-miR-181b-5p&offsets\|0\|-1(+1A);m-72&9\|+\|124693725\|124693747&offsets\|0\|-1(+1A)]\|AACATTCATTGCTGTCGGTGGGA | >MI0000270\|hsa-mir-181b-1&WithFlank&1\|-\|198858867\|198858988 | 42 | 63 | 22 | (+1A) | isomiR | NO | 0.00 | 10.17 | 0.00 | 0.00 |
| >MI0000270\|hsa-mir-181b-1&WithFlank&1\|-\|198858867\|198858988@42.64.23;MI0000683\|hsa-mir-181b-2&WithFlank&9\|+\|124693704\|124693804@22.44.23&[MIMAT0000257&hsa-miR-181b-5p&offsets\|0\|0;m-73&1\|-\|198858925\|198858947&offsets\|0\|0];[MIMAT0000257_1&hsa-miR-181b-5p&offsets\|0\|0;m-72&9\|+\|124693725\|124693747&offsets\|0\|0]\|AACATTCATTGCTGTCGGTGGGT | >MI0000270\|hsa-mir-181b-1&WithFlank&1\|-\|198858867\|198858988 | 42 | 64 | 23 | | isomiR | NO | 34.21 | 114.73 | 38.02 | 23.29 |
| >MI0000270\|hsa-mir-181b-1&WithFlank&1\|-\|198858867\|198858988@42.65.24;MI0000683\|hsa-mir-181b-2&WithFlank&9\|+\|124693704\|124693804@22.45.24&[MIMAT0000257&hsa-miR-181b-5p&offsets\|0\|+1;m-73&1\|-\|198858925\|198858947&offsets\|0\|+1];[MIMAT0000257_1&hsa-miR-181b-5p&offsets\|0\|+1;m-72&9\|+\|124693725\|124693747&offsets\|0\|+1]\|AACATTCATTGCTGTCGGTGGGTT | >MI0000270\|hsa-mir-181b-1&WithFlank&1\|-\|198858867\|198858988 | 42 | 65 | 24 | | isomiR | NO | 31.25 | 39.19 | 38.29 | 0.00 |
| >MI0000272\|hsa-mir-182&WithFlank&7\|-\|129770377\|129770498@29.49.21&[MIMAT0000259&hsa-miR-182-5p&offsets\|0\|-3;m-22&7\|-\|129770449\|129770470&offsets\|0\|-1]\|TTTGGCAATGGTAGAACTCAC | >MI0000272\|hsa-mir-182&WithFlank&7\|-\|129770377\|129770498 | 29 | 49 | 21 | | isomiR | NO | 10.60 | 89.91 | 63.25 | 0.00 |
| >MI0000272\|hsa-mir-182&WithFlank&7\|-\|129770377\|129770498@29.50.22&[MIMAT0000259&hsa-miR-182-5p&offsets\|0\|-2;m-22&7\|-\|129770449\|129770470&offsets\|0\|0]\|TTTGGCAATGGTAGAACTCACA | >MI0000272\|hsa-mir-182&WithFlank&7\|-\|129770377\|129770498 | 29 | 50 | 22 | | isomiR | NO | 144.68 | 452.16 | 95.09 | 38.06 |

# BT-20

<table>
<tr><td>>MI0000272|hsa-mir-182&WithFlank&7|-|129770377|129770498@29.50.22(+1U)&[MIMAT0000259&hsa-miR-182-5p&offsets|0|-2(+1U);m-22&7|-|129770449|129770470&offsets|0|0(+1U)]|TTTGGCAATGGTAGAACTCACAT</td><td>>MI0000272|hsa-mir-182&WithFlank&7|-|129770377|129770498</td><td>29</td><td>50</td><td>22</td><td>(+1U)</td><td>isomiR</td><td>NO</td><td>26.39</td><td>72.63</td><td>0.00</td><td>0.00</td></tr>
<tr><td>>MI0000272|hsa-mir-182&WithFlank&7|-|129770377|129770498@29.51.23&[MIMAT0000259&hsa-miR-182-5p&offsets|0|-1;m-22&7|-|129770449|129770470&offsets|0|+1]|TTTGGCAATGGTAGAACTCACAC</td><td>>MI0000272|hsa-mir-182&WithFlank&7|-|129770377|129770498</td><td>29</td><td>51</td><td>23</td><td></td><td>isomiR</td><td>NO</td><td>148.21</td><td>311.38</td><td>67.46</td><td>35.58</td></tr>
<tr><td>>MI0000272|hsa-mir-182&WithFlank&7|-|129770377|129770498@29.51.23(+1A)&[MIMAT0000259&hsa-miR-182-5p&offsets|0|-1(+1A);m-22&7|-|129770449|129770470&offsets|0|+1(+1A)]|TTTGGCAATGGTAGAACTCACACA</td><td>>MI0000272|hsa-mir-182&WithFlank&7|-|129770377|129770498</td><td>29</td><td>51</td><td>23</td><td>(+1A)</td><td>isomiR</td><td>NO</td><td>148.75</td><td>691.71</td><td>38.38</td><td>39.03</td></tr>
<tr><td>>MI0000272|hsa-mir-182&WithFlank&7|-|129770377|129770498@29.51.23(+1C)&[MIMAT0000259&hsa-miR-182-5p&offsets|0|-1(+1C);m-22&7|-|129770449|129770470&offsets|0|+1(+1C)]|TTTGGCAATGGTAGAACTCACACC</td><td>>MI0000272|hsa-mir-182&WithFlank&7|-|129770377|129770498</td><td>29</td><td>51</td><td>23</td><td>(+1C)</td><td>isomiR</td><td>NO</td><td>289.82</td><td>1014.10</td><td>99.65</td><td>118.53</td></tr>
<tr><td>>MI0000272|hsa-mir-182&WithFlank&7|-|129770377|129770498@29.51.23(+1G)&[MIMAT0000259&hsa-miR-182-5p&offsets|0|-1(+1G);m-22&7|-|129770449|129770470&offsets|0|+1(+1G)]|TTTGGCAATGGTAGAACTCACACG</td><td>>MI0000272|hsa-mir-182&WithFlank&7|-|129770377|129770498</td><td>29</td><td>51</td><td>23</td><td>(+1G)</td><td>isomiR</td><td>NO</td><td>82.38</td><td>260.00</td><td>0.00</td><td>0.00</td></tr>
<tr><td>>MI0000272|hsa-mir-182&WithFlank&7|-|129770377|129770498@29.52.24&[MIMAT0000259&hsa-miR-182-5p&offsets|0|0;m-22&7|-|129770449|129770470&offsets|0|+2]|TTTGGCAATGGTAGAACTCACACT</td><td>>MI0000272|hsa-mir-182&WithFlank&7|-|129770377|129770498</td><td>29</td><td>52</td><td>24</td><td></td><td>isomiR</td><td>YES</td><td>1322.41</td><td>6120.53</td><td>516.19</td><td>550.47</td></tr>
<tr><td>>MI0000272|hsa-mir-182&WithFlank&7|-|129770377|129770498@29.52.24(+1A)&[MIMAT0000259&hsa-miR-182-5p&offsets|0|0(+1A);m-22&7|-|129770449|129770470&offsets|0|+2(+1A)]|TTTGGCAATGGTAGAACTCACACTA</td><td>>MI0000272|hsa-mir-182&WithFlank&7|-|129770377|129770498</td><td>29</td><td>52</td><td>24</td><td>(+1A)</td><td>isomiR</td><td>NO</td><td>69.69</td><td>220.73</td><td>0.00</td><td>24.96</td></tr>
<tr><td>>MI0000272|hsa-mir-182&WithFlank&7|-|129770377|129770498@29.52.24(+1U)&[MIMAT0000259&hsa-miR-182-5p&offsets|0|0(+1U);m-22&7|-|129770449|129770470&offsets|0|+2(+1U)]|TTTGGCAATGGTAGAACTCACACTT</td><td>>MI0000272|hsa-mir-182&WithFlank&7|-|129770377|129770498</td><td>29</td><td>52</td><td>24</td><td>(+1U)</td><td>isomiR</td><td>NO</td><td>0.00</td><td>29.26</td><td>0.00</td><td>0.00</td></tr>
<tr><td>>MI0000272|hsa-mir-182&WithFlank&7|-|129770377|129770498@29.53.25&[MIMAT0000259&hsa-miR-182-5p&offsets|0|+1;m-22&7|-|129770449|129770470&offsets|0|+3]|TTTGGCAATGGTAGAACTCACACTG</td><td>>MI0000272|hsa-mir-182&WithFlank&7|-|129770377|129770498</td><td>29</td><td>53</td><td>25</td><td></td><td>isomiR</td><td>NO</td><td>344.16</td><td>1000.47</td><td>122.14</td><td>201.43</td></tr>
<tr><td>>MI0000272|hsa-mir-182&WithFlank&7|-|129770377|129770498@29.53.25(+1A)&[MIMAT0000259&hsa-miR-182-5p&offsets|0|+1(+1A);m-22&7|-|129770449|129770470&offsets|0|+3(+1A)]|TTTGGCAATGGTAGAACTCACACTGA</td><td>>MI0000272|hsa-mir-182&WithFlank&7|-|129770377|129770498</td><td>29</td><td>53</td><td>25</td><td>(+1A)</td><td>isomiR</td><td>NO</td><td>37.13</td><td>122.78</td><td>0.00</td><td>0.00</td></tr>
<tr><td>>MI0000272|hsa-mir-182&WithFlank&7|-|129770377|129770498@29.53.25(+1C)&[MIMAT0000259&hsa-miR-182-5p&offsets|0|+1(+1C);m-22&7|-|129770449|129770470&offsets|0|+3(+1C)]|TTTGGCAATGGTAGAACTCACACTGC</td><td>>MI0000272|hsa-mir-182&WithFlank&7|-|129770377|129770498</td><td>29</td><td>53</td><td>25</td><td>(+1C)</td><td>isomiR</td><td>NO</td><td>0.00</td><td>29.14</td><td>0.00</td><td>0.00</td></tr>
<tr><td>>MI0000272|hsa-mir-182&WithFlank&7|-|129770377|129770498@29.53.25(+1U)&[MIMAT0000259&hsa-miR-182-5p&offsets|0|+1(+1U);m-22&7|-|129770449|129770470&offsets|0|+3(+1U)]|TTTGGCAATGGTAGAACTCACACTGT</td><td>>MI0000272|hsa-mir-182&WithFlank&7|-|129770377|129770498</td><td>29</td><td>53</td><td>25</td><td>(+1U)</td><td>isomiR</td><td>NO</td><td>72.26</td><td>242.28</td><td>0.00</td><td>29.20</td></tr>
<tr><td>>MI0000272|hsa-mir-182&WithFlank&7|-|129770377|129770498@29.54.26&[MIMAT0000259&hsa-miR-182-5p&offsets|0|+2;m-22&7|-|129770449|129770470&offsets|0|+4]|TTTGGCAATGGTAGAACTCACACTGG</td><td>>MI0000272|hsa-mir-182&WithFlank&7|-|129770377|129770498</td><td>29</td><td>54</td><td>26</td><td></td><td>isomiR</td><td>NO</td><td>206.45</td><td>436.29</td><td>90.81</td><td>121.70</td></tr>
<tr><td>>MI0000272|hsa-mir-182&WithFlank&7|-|129770377|129770498@29.54.26(+1U)&[MIMAT0000259&hsa-miR-182-5p&offsets|0|+2(+1U);m-22&7|-|129770449|129770470&offsets|0|+4(+1U)]|TTTGGCAATGGTAGAACTCACACTGGT</td><td>>MI0000272|hsa-mir-182&WithFlank&7|-|129770377|129770498</td><td>29</td><td>54</td><td>26</td><td>(+1U)</td><td>isomiR</td><td>NO</td><td>23.79</td><td>79.61</td><td>0.00</td><td>0.00</td></tr>
<tr><td>>MI0000273|hsa-mir-183&WithFlank&7|-|129774899|129775020@33.53.21&[MIMAT0000261&hsa-miR-183-5p&offsets|0|-1;m-41&7|-|129774966|129774987&offsets|-1|-2]|TATGGCACTGGTAGAATTCAC</td><td>>MI0000273|hsa-mir-183&WithFlank&7|-|129774899|129775020</td><td>33</td><td>53</td><td>21</td><td></td><td>isomiR</td><td>NO</td><td>0.00</td><td>26.49</td><td>11.47</td><td>0.00</td></tr>
<tr><td>>MI0000273|hsa-mir-183&WithFlank&7|-|129774899|129775020@33.53.21(+1A)&[MIMAT0000261&hsa-miR-183-5p&offsets|0|-1(+1A);m-41&7|-|129774966|129774987&offsets|-1|-2(+1A)]|TATGGCACTGGTAGAATTCACA</td><td>>MI0000273|hsa-mir-183&WithFlank&7|-|129774899|129775020</td><td>33</td><td>53</td><td>21</td><td>(+1A)</td><td>isomiR</td><td>NO</td><td>24.10</td><td>112.64</td><td>0.00</td><td>0.00</td></tr>
<tr><td>>MI0000273|hsa-mir-183&WithFlank&7|-|129774899|129775020@33.53.21(+1C)&[MIMAT0000261&hsa-miR-183-5p&offsets|0|-1(+1C);m-41&7|-|129774966|129774987&offsets|-1|-2(+1C)]|TATGGCACTGGTAGAATTCACC</td><td>>MI0000273|hsa-mir-183&WithFlank&7|-|129774899|129775020</td><td>33</td><td>53</td><td>21</td><td>(+1C)</td><td>isomiR</td><td>NO</td><td>84.37</td><td>192.19</td><td>14.53</td><td>37.22</td></tr>
<tr><td>>MI0000273|hsa-mir-183&WithFlank&7|-|129774899|129775020@33.53.21(+1G)&[MIMAT0000261&hsa-miR-183-5p&offsets|0|-1(+1G);m-41&7|-|129774966|129774987&offsets|-1|-2(+1G)]|TATGGCACTGGTAGAATTCACG</td><td>>MI0000273|hsa-mir-183&WithFlank&7|-|129774899|129775020</td><td>33</td><td>53</td><td>21</td><td>(+1G)</td><td>isomiR</td><td>NO</td><td>0.00</td><td>31.79</td><td>0.00</td><td>0.00</td></tr>
<tr><td>>MI0000273|hsa-mir-183&WithFlank&7|-|129774899|129775020@33.54.22&[MIMAT0000261&hsa-miR-183-5p&offsets|0|0;m-41&7|-|129774966|129774987&offsets|-1|-1]|TATGGCACTGGTAGAATTCACT</td><td>>MI0000273|hsa-mir-183&WithFlank&7|-|129774899|129775020</td><td>33</td><td>54</td><td>22</td><td></td><td>isomiR</td><td>NO</td><td>396.18</td><td>1213.28</td><td>222.11</td><td>267.64</td></tr>
<tr><td>>MI0000273|hsa-mir-183&WithFlank&7|-|129774899|129775020@33.54.22(+1A)&[MIMAT0000261&hsa-miR-183-5p&offsets|0|0(+1A);m-41&7|-|129774966|129774987&offsets|-1|-1(+1A)]|TATGGCACTGGTAGAATTCACTA</td><td>>MI0000273|hsa-mir-183&WithFlank&7|-|129774899|129775020</td><td>33</td><td>54</td><td>22</td><td>(+1A)</td><td>isomiR</td><td>NO</td><td>10.73</td><td>80.98</td><td>10.48</td><td>0.00</td></tr>
<tr><td>>MI0000273|hsa-mir-183&WithFlank&7|-|129774899|129775020@33.55.23&[MIMAT0000261&hsa-miR-183-5p&offsets|0|+1;m-41&7|-|129774966|129774987&offsets|-1|0]|TATGGCACTGGTAGAATTCACTG</td><td>>MI0000273|hsa-mir-183&WithFlank&7|-|129774899|129775020</td><td>33</td><td>55</td><td>23</td><td></td><td>isomiR</td><td>NO</td><td>161.81</td><td>370.89</td><td>184.48</td><td>100.06</td></tr>
<tr><td>>MI0000273|hsa-mir-183&WithFlank&7|-|129774899|129775020@33.56.24&[MIMAT0000261&hsa-miR-183-5p&offsets|0|+2;m-41&7|-|129774966|129774987&offsets|-1|+1]|TATGGCACTGGTAGAATTCACTGT</td><td>>MI0000273|hsa-mir-183&WithFlank&7|-|129774899|129775020</td><td>33</td><td>56</td><td>24</td><td></td><td>isomiR</td><td>NO</td><td>9.60</td><td>133.09</td><td>0.00</td><td>0.00</td></tr>
<tr><td>>MI0000273|hsa-mir-183&WithFlank&7|-|129774899|129775020@34.53.20(+1A)&[MIMAT0000261&hsa-miR-183-5p&offsets|+1|-1(+1A);m-41&7|-|129774966|129774987&offsets|0|-2(+1A)]|ATGGCACTGGTAGAATTCACA</td><td>>MI0000273|hsa-mir-183&WithFlank&7|-|129774899|129775020</td><td>33</td><td>53</td><td>20</td><td>(+1A)</td><td>isomiR</td><td>NO</td><td>0.00</td><td>26.20</td><td>0.00</td><td>0.00</td></tr>
</table>

# BT-20

<table>
<tr><td>>MI0000273|hsa-mir-183&WithFlank&7|-|129774899|129775020@34.53.20(+1C)&[MIMAT0000261&hsa-miR-183-5p&offsets|+1|-1(+1C);m-41&7|-|129774966|129774987&offsets|0|-2(+1C)]|ATGGCACTGGTAGAATTCACC</td><td>>MI0000273|hsa-mir-183&WithFlank&7|-|129774899|129775020</td><td>34</td><td>53</td><td>20</td><td>(+1C)</td><td>isomiR</td><td>NO</td><td>10.62</td><td>45.16</td><td>0.00</td><td>9.40</td></tr>
<tr><td>>MI0000273|hsa-mir-183&WithFlank&7|-|129774899|129775020@34.54.21&[MIMAT0000261&hsa-miR-183-5p&offsets|+1|0;m-41&7|-|129774966|129774987&offsets|0|-1]|ATGGCACTGGTAGAATTCACT</td><td>>MI0000273|hsa-mir-183&WithFlank&7|-|129774899|129775020</td><td>34</td><td>54</td><td>21</td><td></td><td>isomiR</td><td>NO</td><td>153.33</td><td>396.19</td><td>107.89</td><td>146.49</td></tr>
<tr><td>>MI0000273|hsa-mir-183&WithFlank&7|-|129774899|129775020@34.54.21(+1A)&[MIMAT0000261&hsa-miR-183-5p&offsets|+1|0(+1A);m-41&7|-|129774966|129774987&offsets|0|-1(+1A)]|ATGGCACTGGTAGAATTCACTA</td><td>>MI0000273|hsa-mir-183&WithFlank&7|-|129774899|129775020</td><td>34</td><td>54</td><td>21</td><td>(+1A)</td><td>isomiR</td><td>NO</td><td>9.67</td><td>73.83</td><td>10.85</td><td>0.00</td></tr>
<tr><td>>MI0000273|hsa-mir-183&WithFlank&7|-|129774899|129775020@34.55.22&[MIMAT0000261&hsa-miR-183-5p&offsets|+1|+1;m-41&7|-|129774966|129774987&offsets|0|0]|ATGGCACTGGTAGAATTCACTG</td><td>>MI0000273|hsa-mir-183&WithFlank&7|-|129774899|129775020</td><td>34</td><td>55</td><td>22</td><td></td><td>isomiR</td><td>NO</td><td>150.98</td><td>344.28</td><td>179.81</td><td>127.33</td></tr>
<tr><td>>MI0000273|hsa-mir-183&WithFlank&7|-|129774899|129775020@34.55.22(+1C)&[MIMAT0000261&hsa-miR-183-5p&offsets|+1|+1(+1C);m-41&7|-|129774966|129774987&offsets|0|0(+1C)]|ATGGCACTGGTAGAATTCACTGC</td><td>>MI0000273|hsa-mir-183&WithFlank&7|-|129774899|129775020</td><td>34</td><td>55</td><td>22</td><td>(+1C)</td><td>isomiR</td><td>NO</td><td>0.00</td><td>10.07</td><td>0.00</td><td>0.00</td></tr>
<tr><td>>MI0000273|hsa-mir-183&WithFlank&7|-|129774899|129775020@34.56.23&[MIMAT0000261&hsa-miR-183-5p&offsets|+1|+2;m-41&7|-|129774966|129774987&offsets|0|+1]|ATGGCACTGGTAGAATTCACTGT</td><td>>MI0000273|hsa-mir-183&WithFlank&7|-|129774899|129775020</td><td>34</td><td>56</td><td>23</td><td></td><td>isomiR</td><td>NO</td><td>30.07</td><td>202.13</td><td>13.52</td><td>9.82</td></tr>
<tr><td>>MI0000279|hsa-mir-196a-2&WithFlank&12|+|53991732|53991853@31.51.21(+1A);MI0000238|hsa-mir-196a-1&WithFlank&17|-|48632484|48632565@13.33.21(+1A)&[MIMAT0000226&hsa-miR-196a-5p&offsets|0|-1(+1A);m-261&12|+|53991762|53991783&offsets|0|-1(+1A)];[MIMAT0000226_1&hsa-miR-196a-5p&offsets|0|-1(+1A);m-268&17|-|48632532|48632553&offsets|0|-1(+1A)]|TAGGTAGTTTCATGTTGTTGGA</td><td>>MI0000279|hsa-mir-196a-2&WithFlank&12|+|53991732|53991853</td><td>31</td><td>51</td><td>21</td><td>(+1A)</td><td>isomiR</td><td>NO</td><td>27.59</td><td>96.99</td><td>12.65</td><td>0.00</td></tr>
<tr><td>>MI0000279|hsa-mir-196a-2&WithFlank&12|+|53991732|53991853@31.51.21;MI0000238|hsa-mir-196a-1&WithFlank&17|-|48632484|48632565@13.33.21&[MIMAT0000226&hsa-miR-196a-5p&offsets|0|-1;m-261&12|+|53991762|53991783&offsets|0|-1];[MIMAT0000226_1&hsa-miR-196a-5p&offsets|0|-1;m-268&17|-|48632532|48632553&offsets|0|-1]|TAGGTAGTTTCATGTTGTTGG</td><td>>MI0000279|hsa-mir-196a-2&WithFlank&12|+|53991732|53991853</td><td>31</td><td>51</td><td>21</td><td></td><td>isomiR</td><td>NO</td><td>38.29</td><td>125.11</td><td>15.03</td><td>0.00</td></tr>
<tr><td>>MI0000279|hsa-mir-196a-2&WithFlank&12|+|53991732|53991853@31.52.22(+1U);MI0000238|hsa-mir-196a-1&WithFlank&17|-|48632484|48632565@13.34.22(+1U)&[MIMAT0000226&hsa-miR-196a-5p&offsets|0|0(+1U);m-261&12|+|53991762|53991783&offsets|0|0(+1U)];[MIMAT0000226_1&hsa-miR-196a-5p&offsets|0|0(+1U);m-268&17|-|48632532|48632553&offsets|0|0(+1U)]|TAGGTAGTTTCATGTTGTTGGGT</td><td>>MI0000279|hsa-mir-196a-2&WithFlank&12|+|53991732|53991853</td><td>31</td><td>52</td><td>22</td><td>(+1U)</td><td>isomiR</td><td>NO</td><td>10.87</td><td>34.02</td><td>14.07</td><td>0.00</td></tr>
<tr><td>>MI0000279|hsa-mir-196a-2&WithFlank&12|+|53991732|53991853@31.52.22;MI0000238|hsa-mir-196a-1&WithFlank&17|-|48632484|48632565@13.34.22&[MIMAT0000226&hsa-miR-196a-5p&offsets|0|0;m-261&12|+|53991762|53991783&offsets|0|0];[MIMAT0000226_1&hsa-miR-196a-5p&offsets|0|0;m-268&17|-|48632532|48632553&offsets|0|0]|TAGGTAGTTTCATGTTGTTGGG</td><td>>MI0000279|hsa-mir-196a-2&WithFlank&12|+|53991732|53991853</td><td>31</td><td>52</td><td>22</td><td></td><td>isomiR</td><td>NO</td><td>175.08</td><td>428.90</td><td>219.18</td><td>29.54</td></tr>
<tr><td>>MI0000279|hsa-mir-196a-2&WithFlank&12|+|53991732|53991853@31.53.23&[MIMAT0000226&hsa-miR-196a-5p&offsets|0|+1;m-261&12|+|53991762|53991783&offsets|0|+1]|TAGGTAGTTTCATGTTGTTGGGA</td><td>>MI0000279|hsa-mir-196a-2&WithFlank&12|+|53991732|53991853</td><td>31</td><td>53</td><td>23</td><td></td><td>isomiR</td><td>NO</td><td>80.80</td><td>197.01</td><td>78.95</td><td>0.00</td></tr>
<tr><td>>MI0000283|hsa-mir-203a&WithFlank&14|+|104117399|104117520@71.90.20&[MIMAT0000264&hsa-miR-203a-3p&offsets|0|-2;m-79&14|+|104117469|104117490&offsets|0|-2]|GTGAAATGTTTAGGACCACT</td><td>>MI0000283|hsa-mir-203a&WithFlank&14|+|104117399|104117520</td><td>71</td><td>90</td><td>20</td><td></td><td>isomiR</td><td>NO</td><td>10.51</td><td>44.75</td><td>12.15</td><td>0.00</td></tr>
<tr><td>>MI0000283|hsa-mir-203a&WithFlank&14|+|104117399|104117520@71.91.21(+1A)&[MIMAT0000264&hsa-miR-203a-3p&offsets|0|-1(+1A);m-79&14|+|104117469|104117490&offsets|0|-1(+1A)]|GTGAAATGTTTAGGACCACTAA</td><td>>MI0000283|hsa-mir-203a&WithFlank&14|+|104117399|104117520</td><td>71</td><td>91</td><td>21</td><td>(+1A)</td><td>isomiR</td><td>NO</td><td>0.00</td><td>25.62</td><td>0.00</td><td>0.00</td></tr>
<tr><td>>MI0000283|hsa-mir-203a&WithFlank&14|+|104117399|104117520@71.92.22&[MIMAT0000264&hsa-miR-203a-3p&offsets|0|0;m-79&14|+|104117469|104117490&offsets|0|0]|GTGAAATGTTTAGGACCACTAG</td><td>>MI0000283|hsa-mir-203a&WithFlank&14|+|104117399|104117520</td><td>71</td><td>92</td><td>22</td><td></td><td>isomiR</td><td>NO</td><td>72.22</td><td>205.07</td><td>52.72</td><td>0.00</td></tr>
<tr><td>>MI0000283|hsa-mir-203a&WithFlank&14|+|104117399|104117520@71.93.23&[MIMAT0000264&hsa-miR-203a-3p&offsets|0|+1;m-79&14|+|104117469|104117490&offsets|0|+1]|GTGAAATGTTTAGGACCACTAGA</td><td>>MI0000283|hsa-mir-203a&WithFlank&14|+|104117399|104117520</td><td>71</td><td>93</td><td>23</td><td></td><td>isomiR</td><td>NO</td><td>0.00</td><td>33.40</td><td>0.00</td><td>0.00</td></tr>
<tr><td>>MI0000283|hsa-mir-203a&WithFlank&14|+|104117399|104117520@72.90.19&[MIMAT0000264&hsa-miR-203a-3p&offsets|+1|-2;m-79&14|+|104117469|104117490&offsets|+1|-2]|TGAAATGTTTAGGACCACT</td><td>>MI0000283|hsa-mir-203a&WithFlank&14|+|104117399|104117520</td><td>72</td><td>90</td><td>19</td><td></td><td>isomiR</td><td>NO</td><td>10.06</td><td>78.06</td><td>11.31</td><td>0.00</td></tr>
<tr><td>>MI0000283|hsa-mir-203a&WithFlank&14|+|104117399|104117520@72.91.20(+1A)&[MIMAT0000264&hsa-miR-203a-3p&offsets|+1|-1(+1A);m-79&14|+|104117469|104117490&offsets|+1|-1(+1A)]|TGAAATGTTTAGGACCACTAA</td><td>>MI0000283|hsa-mir-203a&WithFlank&14|+|104117399|104117520</td><td>72</td><td>91</td><td>20</td><td>(+1A)</td><td>isomiR</td><td>NO</td><td>0.00</td><td>28.93</td><td>0.00</td><td>0.00</td></tr>
<tr><td>>MI0000283|hsa-mir-203a&WithFlank&14|+|104117399|104117520@72.92.21&[MIMAT0000264&hsa-miR-203a-3p&offsets|+1|0;m-79&14|+|104117469|104117490&offsets|+1|0]|TGAAATGTTTAGGACCACTAG</td><td>>MI0000283|hsa-mir-203a&WithFlank&14|+|104117399|104117520</td><td>72</td><td>92</td><td>21</td><td></td><td>isomiR</td><td>NO</td><td>80.26</td><td>231.56</td><td>55.67</td><td>0.00</td></tr>
<tr><td>>MI0000283|hsa-mir-203a&WithFlank&14|+|104117399|104117520@72.92.21(+1U)&[MIMAT0000264&hsa-miR-203a-3p&offsets|+1|0(+1U);m-79&14|+|104117469|104117490&offsets|+1|0(+1U)]|TGAAATGTTTAGGACCACTAGT</td><td>>MI0000283|hsa-mir-203a&WithFlank&14|+|104117399|104117520</td><td>72</td><td>92</td><td>21</td><td>(+1U)</td><td>isomiR</td><td>NO</td><td>33.84</td><td>106.20</td><td>0.00</td><td>0.00</td></tr>
</table>

# BT-20

<table>
<tr><td>>MI0000283|hsa-mir-203a&WithFlank&14|+|104117399|104117520@72.93.22&[MIMAT0000264&hsa-miR-203a-3p&offsets|+1|+1;m-79&14|+|104117469|104117490&offsets|+1|+1]|TGAAATGTTTAGGACCACTAGA</td><td>>MI0000283|hsa-mir-203a&WithFlank&14|+|104117399|104117520</td><td>72</td><td>93</td><td>22</td><td></td><td>isomiR</td><td>NO</td><td>27.53</td><td>78.07</td><td>0.00</td><td>0.00</td></tr>
<tr><td>>MI0000285|hsa-mir-205&WithFlank&1|+|209432127|209432248@40.59.20&[MIMAT0000266&hsa-miR-205-5p&offsets|0|-2;m-351&1|+|209432166|209432187&offsets|0|-2]|TCCTTCATTCCACCGGAGTC</td><td>>MI0000285|hsa-mir-205&WithFlank&1|+|209432127|209432248</td><td>40</td><td>59</td><td>20</td><td></td><td>isomiR</td><td>NO</td><td>0.00</td><td>31.48</td><td>0.00</td><td>0.00</td></tr>
<tr><td>>MI0000285|hsa-mir-205&WithFlank&1|+|209432127|209432248@40.59.20(+1A)&[MIMAT0000266&hsa-miR-205-5p&offsets|0|-2(+1A);m-351&1|+|209432166|209432187&offsets|0|-2(+1A)]|TCCTTCATTCCACCGGAGTCA</td><td>>MI0000285|hsa-mir-205&WithFlank&1|+|209432127|209432248</td><td>40</td><td>59</td><td>20</td><td>(+1A)</td><td>isomiR</td><td>NO</td><td>25.70</td><td>95.95</td><td>10.18</td><td>0.00</td></tr>
<tr><td>>MI0000285|hsa-mir-205&WithFlank&1|+|209432127|209432248@40.59.20(+1C)&[MIMAT0000266&hsa-miR-205-5p&offsets|0|-2(+1C);m-351&1|+|209432166|209432187&offsets|0|-2(+1C)]|TCCTTCATTCCACCGGAGTCC</td><td>>MI0000285|hsa-mir-205&WithFlank&1|+|209432127|209432248</td><td>40</td><td>59</td><td>20</td><td>(+1C)</td><td>isomiR</td><td>NO</td><td>36.54</td><td>95.69</td><td>13.31</td><td>0.00</td></tr>
<tr><td>>MI0000285|hsa-mir-205&WithFlank&1|+|209432127|209432248@40.60.21&[MIMAT0000266&hsa-miR-205-5p&offsets|0|-1;m-351&1|+|209432166|209432187&offsets|0|-1]|TCCTTCATTCCACCGGAGTCT</td><td>>MI0000285|hsa-mir-205&WithFlank&1|+|209432127|209432248</td><td>40</td><td>60</td><td>21</td><td></td><td>isomiR</td><td>NO</td><td>226.89</td><td>551.55</td><td>189.55</td><td>86.82</td></tr>
<tr><td>>MI0000285|hsa-mir-205&WithFlank&1|+|209432127|209432248@40.60.21(+1A)&[MIMAT0000266&hsa-miR-205-5p&offsets|0|-1(+1A);m-351&1|+|209432166|209432187&offsets|0|-1(+1A)]|TCCTTCATTCCACCGGAGTCTA</td><td>>MI0000285|hsa-mir-205&WithFlank&1|+|209432127|209432248</td><td>40</td><td>60</td><td>21</td><td>(+1A)</td><td>isomiR</td><td>NO</td><td>153.45</td><td>660.30</td><td>204.64</td><td>72.55</td></tr>
<tr><td>>MI0000285|hsa-mir-205&WithFlank&1|+|209432127|209432248@40.60.21(+1U)&[MIMAT0000266&hsa-miR-205-5p&offsets|0|-1(+1U);m-351&1|+|209432166|209432187&offsets|0|-1(+1U)]|TCCTTCATTCCACCGGAGTCTT</td><td>>MI0000285|hsa-mir-205&WithFlank&1|+|209432127|209432248</td><td>40</td><td>60</td><td>21</td><td>(+1U)</td><td>isomiR</td><td>NO</td><td>29.50</td><td>124.60</td><td>55.66</td><td>0.00</td></tr>
<tr><td>>MI0000285|hsa-mir-205&WithFlank&1|+|209432127|209432248@40.61.22&[MIMAT0000266&hsa-miR-205-5p&offsets|0|0;m-351&1|+|209432166|209432187&offsets|0|0]|TCCTTCATTCCACCGGAGTCTG</td><td>>MI0000285|hsa-mir-205&WithFlank&1|+|209432127|209432248</td><td>40</td><td>61</td><td>22</td><td></td><td>isomiR</td><td>YES</td><td>713.21</td><td>2830.18</td><td>1110.71</td><td>377.37</td></tr>
<tr><td>>MI0000285|hsa-mir-205&WithFlank&1|+|209432127|209432248@40.61.22(+1A)&[MIMAT0000266&hsa-miR-205-5p&offsets|0|0(+1A);m-351&1|+|209432166|209432187&offsets|0|0(+1A)]|TCCTTCATTCCACCGGAGTCTGA</td><td>>MI0000285|hsa-mir-205&WithFlank&1|+|209432127|209432248</td><td>40</td><td>61</td><td>22</td><td>(+1A)</td><td>isomiR</td><td>YES</td><td>812.33</td><td>4257.84</td><td>211.03</td><td>889.86</td></tr>
<tr><td>>MI0000285|hsa-mir-205&WithFlank&1|+|209432127|209432248@40.61.22(+1C)&[MIMAT0000266&hsa-miR-205-5p&offsets|0|0(+1C);m-351&1|+|209432166|209432187&offsets|0|0(+1C)]|TCCTTCATTCCACCGGAGTCTGC</td><td>>MI0000285|hsa-mir-205&WithFlank&1|+|209432127|209432248</td><td>40</td><td>61</td><td>22</td><td>(+1C)</td><td>isomiR</td><td>NO</td><td>216.90</td><td>745.33</td><td>39.04</td><td>131.90</td></tr>
<tr><td>>MI0000285|hsa-mir-205&WithFlank&1|+|209432127|209432248@40.61.22(+1G)&[MIMAT0000266&hsa-miR-205-5p&offsets|0|0(+1G);m-351&1|+|209432166|209432187&offsets|0|0(+1G)]|TCCTTCATTCCACCGGAGTCTGG</td><td>>MI0000285|hsa-mir-205&WithFlank&1|+|209432127|209432248</td><td>40</td><td>61</td><td>22</td><td>(+1G)</td><td>isomiR</td><td>NO</td><td>88.64</td><td>296.60</td><td>0.00</td><td>24.11</td></tr>
<tr><td>>MI0000285|hsa-mir-205&WithFlank&1|+|209432127|209432248@40.61.22(+2A)&[MIMAT0000266&hsa-miR-205-5p&offsets|0|0(+2A);m-351&1|+|209432166|209432187&offsets|0|0(+2A)]|TCCTTCATTCCACCGGAGTCTGAA</td><td>>MI0000285|hsa-mir-205&WithFlank&1|+|209432127|209432248</td><td>40</td><td>61</td><td>22</td><td>(+2A)</td><td>isomiR</td><td>NO</td><td>34.13</td><td>247.18</td><td>0.00</td><td>30.42</td></tr>
<tr><td>>MI0000285|hsa-mir-205&WithFlank&1|+|209432127|209432248@40.61.22(+3A)&[MIMAT0000266&hsa-miR-205-5p&offsets|0|0(+3A);m-351&1|+|209432166|209432187&offsets|0|0(+3A)]|TCCTTCATTCCACCGGAGTCTGAAA</td><td>>MI0000285|hsa-mir-205&WithFlank&1|+|209432127|209432248</td><td>40</td><td>61</td><td>22</td><td>(+3A)</td><td>isomiR</td><td>NO</td><td>0.00</td><td>31.71</td><td>0.00</td><td>0.00</td></tr>
<tr><td>>MI0000285|hsa-mir-205&WithFlank&1|+|209432127|209432248@40.62.23&[MIMAT0000266&hsa-miR-205-5p&offsets|0|+1;m-351&1|+|209432166|209432187&offsets|0|+1]|TCCTTCATTCCACCGGAGTCTGT</td><td>>MI0000285|hsa-mir-205&WithFlank&1|+|209432127|209432248</td><td>40</td><td>62</td><td>23</td><td></td><td>isomiR</td><td>YES</td><td>961.55</td><td>4625.48</td><td>260.77</td><td>616.34</td></tr>
<tr><td>>MI0000285|hsa-mir-205&WithFlank&1|+|209432127|209432248@40.62.23(+1A)&[MIMAT0000266&hsa-miR-205-5p&offsets|0|+1(+1A);m-351&1|+|209432166|209432187&offsets|0|+1(+1A)]|TCCTTCATTCCACCGGAGTCTGTA</td><td>>MI0000285|hsa-mir-205&WithFlank&1|+|209432127|209432248</td><td>40</td><td>62</td><td>23</td><td>(+1A)</td><td>isomiR</td><td>NO</td><td>40.00</td><td>285.87</td><td>0.00</td><td>34.30</td></tr>
<tr><td>>MI0000285|hsa-mir-205&WithFlank&1|+|209432127|209432248@40.62.23(+1U)&[MIMAT0000266&hsa-miR-205-5p&offsets|0|+1(+1U);m-351&1|+|209432166|209432187&offsets|0|+1(+1U)]|TCCTTCATTCCACCGGAGTCTGTT</td><td>>MI0000285|hsa-mir-205&WithFlank&1|+|209432127|209432248</td><td>40</td><td>62</td><td>23</td><td>(+1U)</td><td>isomiR</td><td>NO</td><td>33.48</td><td>151.33</td><td>0.00</td><td>10.12</td></tr>
<tr><td>>MI0000285|hsa-mir-205&WithFlank&1|+|209432127|209432248@40.63.24&[MIMAT0000266&hsa-miR-205-5p&offsets|0|+2;m-351&1|+|209432166|209432187&offsets|0|+2]|TCCTTCATTCCACCGGAGTCTGTC</td><td>>MI0000285|hsa-mir-205&WithFlank&1|+|209432127|209432248</td><td>40</td><td>63</td><td>24</td><td></td><td>isomiR</td><td>NO</td><td>0.00</td><td>10.39</td><td>0.00</td><td>0.00</td></tr>
<tr><td>>MI0000286|hsa-mir-210&WithFlank&11|-|568083|568204@72.92.21(+1U)&[MIMAT0000267&hsa-miR-210-3p&offsets|0|-1(+1U);m-102&11|-|568112|568133&offsets|0|-1(+1U)]|CTGTGCGTGTGACAGCGGCTGT</td><td>>MI0000286|hsa-mir-210&WithFlank&11|-|568083|568204</td><td>72</td><td>92</td><td>21</td><td>(+1U)</td><td>isomiR</td><td>NO</td><td>0.00</td><td>0.00</td><td>11.91</td><td>0.00</td></tr>
<tr><td>>MI0000286|hsa-mir-210&WithFlank&11|-|568083|568204@72.93.22&[MIMAT0000267&hsa-miR-210-3p&offsets|0|0;m-102&11|-|568112|568133&offsets|0|0]|CTGTGCGTGTGACAGCGGCTGA</td><td>>MI0000286|hsa-mir-210&WithFlank&11|-|568083|568204</td><td>72</td><td>93</td><td>22</td><td></td><td>isomiR</td><td>NO</td><td>28.06</td><td>40.39</td><td>109.19</td><td>0.00</td></tr>
<tr><td>>MI0000286|hsa-mir-210&WithFlank&11|-|568083|568204@72.93.22(+1A)&[MIMAT0000267&hsa-miR-210-3p&offsets|0|0(+1A);m-102&11|-|568112|568133&offsets|0|0(+1A)]|CTGTGCGTGTGACAGCGGCTGAA</td><td>>MI0000286|hsa-mir-210&WithFlank&11|-|568083|568204</td><td>72</td><td>93</td><td>22</td><td>(+1A)</td><td>isomiR</td><td>NO</td><td>27.32</td><td>35.96</td><td>83.13</td><td>0.00</td></tr>
<tr><td>>MI0000286|hsa-mir-210&WithFlank&11|-|568083|568204@72.93.22(+2A)&[MIMAT0000267&hsa-miR-210-3p&offsets|0|0(+2A);m-102&11|-|568112|568133&offsets|0|0(+2A)]|CTGTGCGTGTGACAGCGGCTGAAA</td><td>>MI0000286|hsa-mir-210&WithFlank&11|-|568083|568204</td><td>72</td><td>93</td><td>22</td><td>(+2A)</td><td>isomiR</td><td>NO</td><td>106.26</td><td>151.19</td><td>190.20</td><td>116.29</td></tr>
</table>

# BT-20

<table>
<tr><th></th><th></th><th></th><th></th><th></th><th></th><th></th><th></th><th></th><th></th><th></th><th></th></tr>
<tr><td>>MI0000286|hsa-mir-210&WithFlank&11|-|568083|568204@72.94.23&[MIMAT0000267&hsa-miR-210-3p&offsets|0|+1;m-102&11|-|568112|568133&offsets|0|+1]|CTGTGCGTGTGACAGCGGCTGAT</td><td>>MI0000286|hsa-mir-210&WithFlank&11|-|568083|568204</td><td>72</td><td>94</td><td>23</td><td></td><td>isomiR</td><td>NO</td><td>0.00</td><td>0.00</td><td>12.05</td><td>0.00</td></tr>
<tr><td>>MI0000289|hsa-mir-181a-1&WithFlank&1|-|198859038|198859159@30.52.23;MI0000269|hsa-mir-181a-2&WithFlank&9|+|124692436|124692557@45.67.23&[MIMAT0000256&hsa-miR-181a-5p&offsets|0|0;m-32&1|-|198859108|198859130&offsets|0|0];[MIMAT0000256_1&hsa-miR-181a-5p&offsets|0|0;m-32&9|+|124692480|124692502&offsets|0|0]|AACATTCAACGCTGTCGGTGAGT</td><td>>MI0000289|hsa-mir-181a-1&WithFlank&1|-|198859038|198859159</td><td>30</td><td>52</td><td>23</td><td></td><td>isomiR</td><td>NO</td><td>65.01</td><td>114.16</td><td>86.20</td><td>26.00</td></tr>
<tr><td>>MI0000298|hsa-mir-221&WithFlank&X|-|45746151|45746272@31.54.24&[MIMAT0004568&hsa-miR-221-5p&offsets|0|+2;m-405&X|-|45746220|45746242&offsets|0|+1]|ACCTGGCATACAATGTAGATTTCT</td><td>>MI0000298|hsa-mir-221&WithFlank&X|-|45746151|45746272</td><td>31</td><td>54</td><td>24</td><td></td><td>isomiR</td><td>NO</td><td>0.00</td><td>30.77</td><td>0.00</td><td>0.00</td></tr>
<tr><td>>MI0000298|hsa-mir-221&WithFlank&X|-|45746151|45746272@31.56.26&[MIMAT0004568&hsa-miR-221-5p&offsets|0|+4;m-405&X|-|45746220|45746242&offsets|0|+3]|ACCTGGCATACAATGTAGATTTCTGT</td><td>>MI0000298|hsa-mir-221&WithFlank&X|-|45746151|45746272</td><td>31</td><td>56</td><td>26</td><td></td><td>isomiR</td><td>NO</td><td>0.00</td><td>71.32</td><td>0.00</td><td>0.00</td></tr>
<tr><td>>MI0000298|hsa-mir-221&WithFlank&X|-|45746151|45746272@71.90.20&[MIMAT0000278&hsa-miR-221-3p&offsets|0|-3;m-91&X|-|45746181|45746202&offsets|0|-2]|AGCTACATTGTCTGCTGGGT</td><td>>MI0000298|hsa-mir-221&WithFlank&X|-|45746151|45746272</td><td>71</td><td>90</td><td>20</td><td></td><td>isomiR</td><td>NO</td><td>0.00</td><td>24.96</td><td>13.12</td><td>0.00</td></tr>
<tr><td>>MI0000298|hsa-mir-221&WithFlank&X|-|45746151|45746272@71.91.21&[MIMAT0000278&hsa-miR-221-3p&offsets|0|-2;m-91&X|-|45746181|45746202&offsets|0|-1]|AGCTACATTGTCTGCTGGGTT</td><td>>MI0000298|hsa-mir-221&WithFlank&X|-|45746151|45746272</td><td>71</td><td>91</td><td>21</td><td></td><td>isomiR</td><td>NO</td><td>23.95</td><td>41.59</td><td>89.62</td><td>0.00</td></tr>
<tr><td>>MI0000298|hsa-mir-221&WithFlank&X|-|45746151|45746272@71.91.21(+1A)&[MIMAT0000278&hsa-miR-221-3p&offsets|0|-2(+1A);m-91&X|-|45746181|45746202&offsets|0|-1(+1A)]|AGCTACATTGTCTGCTGGGTTA</td><td>>MI0000298|hsa-mir-221&WithFlank&X|-|45746151|45746272</td><td>71</td><td>91</td><td>21</td><td>(+1A)</td><td>isomiR</td><td>NO</td><td>0.00</td><td>10.51</td><td>9.76</td><td>0.00</td></tr>
<tr><td>>MI0000298|hsa-mir-221&WithFlank&X|-|45746151|45746272@71.91.21(+1C)&[MIMAT0000278&hsa-miR-221-3p&offsets|0|-2(+1C);m-91&X|-|45746181|45746202&offsets|0|-1(+1C)]|AGCTACATTGTCTGCTGGGTTC</td><td>>MI0000298|hsa-mir-221&WithFlank&X|-|45746151|45746272</td><td>71</td><td>91</td><td>21</td><td>(+1C)</td><td>isomiR</td><td>NO</td><td>23.98</td><td>80.48</td><td>93.52</td><td>0.00</td></tr>
<tr><td>>MI0000298|hsa-mir-221&WithFlank&X|-|45746151|45746272@71.92.22&[MIMAT0000278&hsa-miR-221-3p&offsets|0|-1;m-91&X|-|45746181|45746202&offsets|0|0]|AGCTACATTGTCTGCTGGGTTT</td><td>>MI0000298|hsa-mir-221&WithFlank&X|-|45746151|45746272</td><td>71</td><td>92</td><td>22</td><td></td><td>isomiR</td><td>NO</td><td>181.46</td><td>495.15</td><td>469.34</td><td>135.61</td></tr>
<tr><td>>MI0000298|hsa-mir-221&WithFlank&X|-|45746151|45746272@71.92.22(+1A)&[MIMAT0000278&hsa-miR-221-3p&offsets|0|-1(+1A);m-91&X|-|45746181|45746202&offsets|0|0(+1A)]|AGCTACATTGTCTGCTGGGTTTA</td><td>>MI0000298|hsa-mir-221&WithFlank&X|-|45746151|45746272</td><td>71</td><td>92</td><td>22</td><td>(+1A)</td><td>isomiR</td><td>NO</td><td>0.00</td><td>68.72</td><td>11.28</td><td>0.00</td></tr>
<tr><td>>MI0000298|hsa-mir-221&WithFlank&X|-|45746151|45746272@71.92.22(+1U)&[MIMAT0000278&hsa-miR-221-3p&offsets|0|-1(+1U);m-91&X|-|45746181|45746202&offsets|0|0(+1U)]|AGCTACATTGTCTGCTGGGTTTT</td><td>>MI0000298|hsa-mir-221&WithFlank&X|-|45746151|45746272</td><td>71</td><td>92</td><td>22</td><td>(+1U)</td><td>isomiR</td><td>NO</td><td>30.61</td><td>142.23</td><td>111.43</td><td>11.05</td></tr>
<tr><td>>MI0000298|hsa-mir-221&WithFlank&X|-|45746151|45746272@71.93.23&[MIMAT0000278&hsa-miR-221-3p&offsets|0|0;m-91&X|-|45746181|45746202&offsets|0|+1]|AGCTACATTGTCTGCTGGGTTTC</td><td>>MI0000298|hsa-mir-221&WithFlank&X|-|45746151|45746272</td><td>71</td><td>93</td><td>23</td><td></td><td>isomiR</td><td>NO</td><td>255.86</td><td>1161.74</td><td>628.32</td><td>222.78</td></tr>
<tr><td>>MI0000298|hsa-mir-221&WithFlank&X|-|45746151|45746272@71.93.23(+1C)&[MIMAT0000278&hsa-miR-221-3p&offsets|0|0(+1C);m-91&X|-|45746181|45746202&offsets|0|+1(+1C)]|AGCTACATTGTCTGCTGGGTTTCC</td><td>>MI0000298|hsa-mir-221&WithFlank&X|-|45746151|45746272</td><td>71</td><td>93</td><td>23</td><td>(+1C)</td><td>isomiR</td><td>NO</td><td>0.00</td><td>76.94</td><td>0.00</td><td>0.00</td></tr>
<tr><td>>MI0000298|hsa-mir-221&WithFlank&X|-|45746151|45746272@71.93.23(+1G)&[MIMAT0000278&hsa-miR-221-3p&offsets|0|0(+1G);m-91&X|-|45746181|45746202&offsets|0|+1(+1G)]|AGCTACATTGTCTGCTGGGTTTCG</td><td>>MI0000298|hsa-mir-221&WithFlank&X|-|45746151|45746272</td><td>71</td><td>93</td><td>23</td><td>(+1G)</td><td>isomiR</td><td>NO</td><td>0.00</td><td>23.50</td><td>0.00</td><td>0.00</td></tr>
<tr><td>>MI0000298|hsa-mir-221&WithFlank&X|-|45746151|45746272@71.93.23(+1U)&[MIMAT0000278&hsa-miR-221-3p&offsets|0|0(+1U);m-91&X|-|45746181|45746202&offsets|0|+1(+1U)]|AGCTACATTGTCTGCTGGGTTTCT</td><td>>MI0000298|hsa-mir-221&WithFlank&X|-|45746151|45746272</td><td>71</td><td>93</td><td>23</td><td>(+1U)</td><td>isomiR</td><td>NO</td><td>84.85</td><td>448.85</td><td>73.15</td><td>87.32</td></tr>
<tr><td>>MI0000298|hsa-mir-221&WithFlank&X|-|45746151|45746272@71.94.24&[MIMAT0000278&hsa-miR-221-3p&offsets|0|+1;m-91&X|-|45746181|45746202&offsets|0|+2]|AGCTACATTGTCTGCTGGGTTTCA</td><td>>MI0000298|hsa-mir-221&WithFlank&X|-|45746151|45746272</td><td>71</td><td>94</td><td>24</td><td></td><td>isomiR</td><td>NO</td><td>0.00</td><td>118.92</td><td>0.00</td><td>0.00</td></tr>
<tr><td>>MI0000299|hsa-mir-222&WithFlank&X|-|45747009|45747130@75.94.20(+1C)&[MIMAT0000279&hsa-miR-222-3p&offsets|0|-1(+1C);m-140&X|-|45747033|45747056&offsets|0|-4(+1C)]|AGCTACATCTGGCTACTGGGC</td><td>>MI0000299|hsa-mir-222&WithFlank&X|-|45747009|45747130</td><td>75</td><td>94</td><td>20</td><td>(+1C)</td><td>isomiR</td><td>NO</td><td>0.00</td><td>0.00</td><td>13.23</td><td>0.00</td></tr>
<tr><td>>MI0000299|hsa-mir-222&WithFlank&X|-|45747009|45747130@75.95.21&[MIMAT0000279&hsa-miR-222-3p&offsets|0|0;m-140&X|-|45747033|45747056&offsets|0|-3]|AGCTACATCTGGCTACTGGGT</td><td>>MI0000299|hsa-mir-222&WithFlank&X|-|45747009|45747130</td><td>75</td><td>95</td><td>21</td><td></td><td>isomiR</td><td>NO</td><td>39.95</td><td>85.48</td><td>197.28</td><td>10.94</td></tr>
<tr><td>>MI0000299|hsa-mir-222&WithFlank&X|-|45747009|45747130@75.96.22&[MIMAT0000279&hsa-miR-222-3p&offsets|0|+1;m-140&X|-|45747033|45747056&offsets|0|-2]|AGCTACATCTGGCTACTGGGTC</td><td>>MI0000299|hsa-mir-222&WithFlank&X|-|45747009|45747130</td><td>75</td><td>96</td><td>22</td><td></td><td>isomiR</td><td>NO</td><td>24.53</td><td>73.47</td><td>63.10</td><td>9.81</td></tr>
<tr><td>>MI0000299|hsa-mir-222&WithFlank&X|-|45747009|45747130@75.96.22(+1A)&[MIMAT0000279&hsa-miR-222-3p&offsets|0|+1(+1A);m-140&X|-|45747033|45747056&offsets|0|-2(+1A)]|AGCTACATCTGGCTACTGGGTCA</td><td>>MI0000299|hsa-mir-222&WithFlank&X|-|45747009|45747130</td><td>75</td><td>96</td><td>22</td><td>(+1A)</td><td>isomiR</td><td>NO</td><td>0.00</td><td>27.19</td><td>0.00</td><td>0.00</td></tr>
<tr><td>>MI0000299|hsa-mir-222&WithFlank&X|-|45747009|45747130@75.96.22(+1C)&[MIMAT0000279&hsa-miR-222-3p&offsets|0|+1(+1C);m-140&X|-|45747033|45747056&offsets|0|-2(+1C)]|AGCTACATCTGGCTACTGGGTCC</td><td>>MI0000299|hsa-mir-222&WithFlank&X|-|45747009|45747130</td><td>75</td><td>96</td><td>22</td><td>(+1C)</td><td>isomiR</td><td>NO</td><td>0.00</td><td>25.70</td><td>0.00</td><td>0.00</td></tr>
<tr><td>>MI0000299|hsa-mir-222&WithFlank&X|-|45747009|45747130@75.97.23&[MIMAT0000279&hsa-miR-222-3p&offsets|0|+2;m-140&X|-|45747033|45747056&offsets|0|-1]|AGCTACATCTGGCTACTGGGTCT</td><td>>MI0000299|hsa-mir-222&WithFlank&X|-|45747009|45747130</td><td>75</td><td>97</td><td>23</td><td></td><td>isomiR</td><td>NO</td><td>95.36</td><td>203.97</td><td>116.10</td><td>105.01</td></tr>
<tr><td>>MI0000299|hsa-mir-222&WithFlank&X|-|45747009|45747130@75.97.23(+1A)&[MIMAT0000279&hsa-miR-222-3p&offsets|0|+2(+1A);m-140&X|-|45747033|45747056&offsets|0|-1(+1A)]|AGCTACATCTGGCTACTGGGTCTA</td><td>>MI0000299|hsa-mir-222&WithFlank&X|-|45747009|45747130</td><td>75</td><td>97</td><td>23</td><td>(+1A)</td><td>isomiR</td><td>NO</td><td>0.00</td><td>44.08</td><td>0.00</td><td>0.00</td></tr>
<tr><td>>MI0000299|hsa-mir-222&WithFlank&X|-|45747009|45747130@75.97.23(+1U)&[MIMAT0000279&hsa-miR-222-3p&offsets|0|+2(+1U);m-140&X|-|45747033|45747056&offsets|0|-1(+1U)]|AGCTACATCTGGCTACTGGGTCTT</td><td>>MI0000299|hsa-mir-222&WithFlank&X|-|45747009|45747130</td><td>75</td><td>97</td><td>23</td><td>(+1U)</td><td>isomiR</td><td>NO</td><td>28.72</td><td>126.86</td><td>39.89</td><td>31.50</td></tr>
<tr><td>>MI0000299|hsa-mir-222&WithFlank&X|-|45747009|45747130@75.98.24&[MIMAT0000279&hsa-miR-222-3p&offsets|0|+3;m-140&X|-|45747033|45747056&offsets|0|0]|AGCTACATCTGGCTACTGGGTCTC</td><td>>MI0000299|hsa-mir-222&WithFlank&X|-|45747009|45747130</td><td>75</td><td>98</td><td>24</td><td></td><td>isomiR</td><td>NO</td><td>141.74</td><td>530.52</td><td>199.58</td><td>178.25</td></tr>
</table>

# BT-20

| | | | | | | | | | | | |
|---|---|---|---|---|---|---|---|---|---|---|---|
| >MI0000299\|hsa-mir-222&WithFlank&X\|-\|45747009\|45747130@75.98.24(+1A)&[MIMAT0000279&hsa-miR-222-3p&offsets\|0\|+3(+1A);m-140&X\|-\|45747033\|45747056&offsets\|0\|0(+1A)]\|AGCTACATCTGGCTACTGGGTCTCA | >MI0000299\|hsa-mir-222&WithFlank&X\|-\|45747009\|45747130 | 75 | 98 | 24 | (+1A) | isomiR | NO | 24.42 | 193.98 | 9.43 | 29.23 |
| >MI0000299\|hsa-mir-222&WithFlank&X\|-\|45747009\|45747130@75.98.24(+1C)&[MIMAT0000279&hsa-miR-222-3p&offsets\|0\|+3(+1C);m-140&X\|-\|45747033\|45747056&offsets\|0\|0(+1C)]\|AGCTACATCTGGCTACTGGGTCTCC | >MI0000299\|hsa-mir-222&WithFlank&X\|-\|45747009\|45747130 | 75 | 98 | 24 | (+1C) | isomiR | NO | 36.88 | 241.37 | 15.80 | 79.91 |
| >MI0000299\|hsa-mir-222&WithFlank&X\|-\|45747009\|45747130@75.98.24(+1G)&[MIMAT0000279&hsa-miR-222-3p&offsets\|0\|+3(+1G);m-140&X\|-\|45747033\|45747056&offsets\|0\|0(+1G)]\|AGCTACATCTGGCTACTGGGTCTCG | >MI0000299\|hsa-mir-222&WithFlank&X\|-\|45747009\|45747130 | 75 | 98 | 24 | (+1G) | isomiR | NO | 0.00 | 39.68 | 0.00 | 0.00 |
| >MI0000299\|hsa-mir-222&WithFlank&X\|-\|45747009\|45747130@75.99.25&[MIMAT0000279&hsa-miR-222-3p&offsets\|0\|+4;m-140&X\|-\|45747033\|45747056&offsets\|0\|+1]\|AGCTACATCTGGCTACTGGGTCTCT | >MI0000299\|hsa-mir-222&WithFlank&X\|-\|45747009\|45747130 | 75 | 99 | 25 | | isomiR | NO | 181.58 | 1077.30 | 188.93 | 235.39 |
| >MI0000299\|hsa-mir-222&WithFlank&X\|-\|45747009\|45747130@75.99.25(+1U)&[MIMAT0000279&hsa-miR-222-3p&offsets\|0\|+4(+1U);m-140&X\|-\|45747033\|45747056&offsets\|0\|+1(+1U)]\|AGCTACATCTGGCTACTGGGTCTCTT | >MI0000299\|hsa-mir-222&WithFlank&X\|-\|45747009\|45747130 | 75 | 99 | 25 | (+1U) | isomiR | NO | 0.00 | 130.19 | 0.00 | 11.01 |
| >MI0000299\|hsa-mir-222&WithFlank&X\|-\|45747009\|45747130@75.99.25(+2U)&[MIMAT0000279&hsa-miR-222-3p&offsets\|0\|+4(+2U);m-140&X\|-\|45747033\|45747056&offsets\|0\|+1(+2U)]\|AGCTACATCTGGCTACTGGGTCTCTTT | >MI0000299\|hsa-mir-222&WithFlank&X\|-\|45747009\|45747130 | 75 | 99 | 25 | (+2U) | isomiR | NO | 0.00 | 33.08 | 0.00 | 0.00 |
| >MI0000301\|hsa-mir-224&WithFlank&X\|-\|151958572\|151958664@14.34.21&[MIMAT0000281&hsa-miR-224-5p&offsets\|+1\|-3;m-238&X\|-\|151958630\|151958651&offsets\|0\|-1]\|CAAGTCACTAGTGGTTCCGTT | >MI0000301\|hsa-mir-224&WithFlank&X\|-\|151958572\|151958664 | 14 | 34 | 21 | | isomiR | NO | 43.25 | 123.30 | 57.43 | 25.32 |
| >MI0000301\|hsa-mir-224&WithFlank&X\|-\|151958572\|151958664@14.34.21(+1A)&[MIMAT0000281&hsa-miR-224-5p&offsets\|+1\|-3(+1A);m-238&X\|-\|151958630\|151958651&offsets\|0\|-1(+1A)]\|CAAGTCACTAGTGGTTCCGTTA | >MI0000301\|hsa-mir-224&WithFlank&X\|-\|151958572\|151958664 | 14 | 34 | 21 | (+1A) | isomiR | NO | 0.00 | 33.63 | 0.00 | 0.00 |
| >MI0000301\|hsa-mir-224&WithFlank&X\|-\|151958572\|151958664@14.34.21(+1C)&[MIMAT0000281&hsa-miR-224-5p&offsets\|+1\|-3(+1C);m-238&X\|-\|151958630\|151958651&offsets\|0\|-1(+1C)]\|CAAGTCACTAGTGGTTCCGTTC | >MI0000301\|hsa-mir-224&WithFlank&X\|-\|151958572\|151958664 | 14 | 34 | 21 | (+1C) | isomiR | NO | 36.66 | 130.41 | 59.18 | 0.00 |
| >MI0000301\|hsa-mir-224&WithFlank&X\|-\|151958572\|151958664@14.35.22&[MIMAT0000281&hsa-miR-224-5p&offsets\|+1\|-2;m-238&X\|-\|151958630\|151958651&offsets\|0\|0]\|CAAGTCACTAGTGGTTCCGTTT | >MI0000301\|hsa-mir-224&WithFlank&X\|-\|151958572\|151958664 | 14 | 35 | 22 | | isomiR | NO | 304.21 | 992.28 | 380.29 | 164.09 |
| >MI0000301\|hsa-mir-224&WithFlank&X\|-\|151958572\|151958664@14.36.23&[MIMAT0000281&hsa-miR-224-5p&offsets\|+1\|-1;m-238&X\|-\|151958630\|151958651&offsets\|0\|+1]\|CAAGTCACTAGTGGTTCCGTTTA | >MI0000301\|hsa-mir-224&WithFlank&X\|-\|151958572\|151958664 | 14 | 36 | 23 | | isomiR | NO | 65.70 | 196.25 | 0.00 | 27.21 |
| >MI0000301\|hsa-mir-224&WithFlank&X\|-\|151958572\|151958664@14.36.23(+1A)&[MIMAT0000281&hsa-miR-224-5p&offsets\|+1\|-1(+1A);m-238&X\|-\|151958630\|151958651&offsets\|0\|+1(+1A)]\|CAAGTCACTAGTGGTTCCGTTTAA | >MI0000301\|hsa-mir-224&WithFlank&X\|-\|151958572\|151958664 | 14 | 36 | 23 | (+1A) | isomiR | NO | 0.00 | 29.60 | 0.00 | 0.00 |
| >MI0000301\|hsa-mir-224&WithFlank&X\|-\|151958572\|151958664@14.37.24&[MIMAT0000281&hsa-miR-224-5p&offsets\|+1\|0;m-238&X\|-\|151958630\|151958651&offsets\|0\|+2]\|CAAGTCACTAGTGGTTCCGTTTAG | >MI0000301\|hsa-mir-224&WithFlank&X\|-\|151958572\|151958664 | 14 | 37 | 24 | | isomiR | NO | 118.30 | 308.95 | 59.93 | 115.78 |
| >MI0000301\|hsa-mir-224&WithFlank&X\|-\|151958572\|151958664@14.38.25&[MIMAT0000281&hsa-miR-224-5p&offsets\|+1\|+1;m-238&X\|-\|151958630\|151958651&offsets\|0\|+3]\|CAAGTCACTAGTGGTTCCGTTTAGT | >MI0000301\|hsa-mir-224&WithFlank&X\|-\|151958572\|151958664 | 14 | 38 | 25 | | isomiR | NO | 0.00 | 69.35 | 0.00 | 0.00 |
| >MI0000342\|hsa-mir-200b&WithFlank&1\|+\|1167098\|1167204@63.84.22&[MIMAT0000318&hsa-miR-200b-3p&offsets\|0\|0;m-93&1\|+\|1167160\|1167182&offsets\|0\|-1]\|TAATACTGCCTGGTAATGATGA | >MI0000342\|hsa-mir-200b&WithFlank&1\|+\|1167098\|1167204 | 63 | 84 | 22 | | isomiR | NO | 0.00 | 24.24 | 0.00 | 0.00 |
| >MI0000342\|hsa-mir-200b&WithFlank&1\|+\|1167098\|1167204@63.85.23&[MIMAT0000318&hsa-miR-200b-3p&offsets\|0\|+1;m-93&1\|+\|1167160\|1167182&offsets\|0\|0]\|TAATACTGCCTGGTAATGATGAC | >MI0000342\|hsa-mir-200b&WithFlank&1\|+\|1167098\|1167204 | 63 | 85 | 23 | | isomiR | NO | 0.00 | 112.61 | 0.00 | 0.00 |
| >MI0000342\|hsa-mir-200b&WithFlank&1\|+\|1167098\|1167204@63.85.23(+1U)&[MIMAT0000318&hsa-miR-200b-3p&offsets\|0\|+1(+1U);m-93&1\|+\|1167160\|1167182&offsets\|0\|0(+1U)]\|TAATACTGCCTGGTAATGATGACT | >MI0000342\|hsa-mir-200b&WithFlank&1\|+\|1167098\|1167204 | 63 | 85 | 23 | (+1U) | isomiR | NO | 0.00 | 28.18 | 0.00 | 0.00 |
| >MI0000433\|hsa-let-7g&WithFlank&3\|-\|52268272\|52268367@11.28.18&[MIMAT0000414&hsa-let-7g-5p&offsets\|0\|-4;m-58&3\|-\|52268336\|52268357&offsets\|0\|-4]\|TGAGGTAGTAGTTTGTAC | >MI0000433\|hsa-let-7g&WithFlank&3\|-\|52268272\|52268367 | 11 | 28 | 18 | | isomiR | NO | 139.70 | 167.70 | 73.99 | 70.44 |
| >MI0000433\|hsa-let-7g&WithFlank&3\|-\|52268272\|52268367@11.29.19&[MIMAT0000414&hsa-let-7g-5p&offsets\|0\|-3;m-58&3\|-\|52268336\|52268357&offsets\|0\|-3]\|TGAGGTAGTAGTTTGTACA | >MI0000433\|hsa-let-7g&WithFlank&3\|-\|52268272\|52268367 | 11 | 29 | 19 | | isomiR | NO | 0.00 | 30.79 | 0.00 | 0.00 |
| >MI0000433\|hsa-let-7g&WithFlank&3\|-\|52268272\|52268367@11.30.20(+1C)&[MIMAT0000414&hsa-let-7g-5p&offsets\|0\|-2(+1C);m-58&3\|-\|52268336\|52268357&offsets\|0\|-2(+1C)]\|TGAGGTAGTAGTTTGTACAGC | >MI0000433\|hsa-let-7g&WithFlank&3\|-\|52268272\|52268367 | 11 | 30 | 20 | (+1C) | isomiR | NO | 28.13 | 29.13 | 0.00 | 0.00 |
| >MI0000433\|hsa-let-7g&WithFlank&3\|-\|52268272\|52268367@11.31.21&[MIMAT0000414&hsa-let-7g-5p&offsets\|0\|-1;m-58&3\|-\|52268336\|52268357&offsets\|0\|-1]\|TGAGGTAGTAGTTTGTACAGT | >MI0000433\|hsa-let-7g&WithFlank&3\|-\|52268272\|52268367 | 11 | 31 | 21 | | isomiR | NO | 209.63 | 293.73 | 117.72 | 85.50 |
| >MI0000433\|hsa-let-7g&WithFlank&3\|-\|52268272\|52268367@11.31.21(+1A)&[MIMAT0000414&hsa-let-7g-5p&offsets\|0\|-1(+1A);m-58&3\|-\|52268336\|52268357&offsets\|0\|-1(+1A)]\|TGAGGTAGTAGTTTGTACAGTA | >MI0000433\|hsa-let-7g&WithFlank&3\|-\|52268272\|52268367 | 11 | 31 | 21 | (+1A) | isomiR | NO | 132.01 | 219.69 | 12.67 | 69.84 |
| >MI0000433\|hsa-let-7g&WithFlank&3\|-\|52268272\|52268367@11.31.21(+1C)&[MIMAT0000414&hsa-let-7g-5p&offsets\|0\|-1(+1C);m-58&3\|-\|52268336\|52268357&offsets\|0\|-1(+1C)]\|TGAGGTAGTAGTTTGTACAGTC | >MI0000433\|hsa-let-7g&WithFlank&3\|-\|52268272\|52268367 | 11 | 31 | 21 | (+1C) | isomiR | NO | 463.21 | 563.39 | 131.85 | 253.72 |
| >MI0000433\|hsa-let-7g&WithFlank&3\|-\|52268272\|52268367@11.31.21(+1G)&[MIMAT0000414&hsa-let-7g-5p&offsets\|0\|-1(+1G);m-58&3\|-\|52268336\|52268357&offsets\|0\|-1(+1G)]\|TGAGGTAGTAGTTTGTACAGTG | >MI0000433\|hsa-let-7g&WithFlank&3\|-\|52268272\|52268367 | 11 | 31 | 21 | (+1G) | isomiR | NO | 77.20 | 83.83 | 0.00 | 0.00 |
| >MI0000433\|hsa-let-7g&WithFlank&3\|-\|52268272\|52268367@11.32.22&[MIMAT0000414&hsa-let-7g-5p&offsets\|0\|0;m-58&3\|-\|52268336\|52268357&offsets\|0\|0]\|TGAGGTAGTAGTTTGTACAGTT | >MI0000433\|hsa-let-7g&WithFlank&3\|-\|52268272\|52268367 | 11 | 32 | 22 | | isomiR | YES | 2176.51 | 3469.40 | 713.30 | 1214.51 |
| >MI0000433\|hsa-let-7g&WithFlank&3\|-\|52268272\|52268367@11.32.22(+1A)&[MIMAT0000414&hsa-let-7g-5p&offsets\|0\|0(+1A);m-58&3\|-\|52268336\|52268357&offsets\|0\|0(+1A)]\|TGAGGTAGTAGTTTGTACAGTTA | >MI0000433\|hsa-let-7g&WithFlank&3\|-\|52268272\|52268367 | 11 | 32 | 22 | (+1A) | isomiR | NO | 77.06 | 110.88 | 126.61 | 25.56 |

# BT-20

<table>
<tr><td>>MI0000433|hsa-let-7g&WithFlank&3|-|52268272|52268367@11.33.23&[MIMAT0000414&hsa-let-7g-5p&offsets|0|+1;m-58&3|-|52268336|52268357&offsets|0|+1]|TGAGGTAGTAGTTTGTACAGTTT</td><td>>MI0000433|hsa-let-7g&WithFlank&3|-|52268272|52268367</td><td>11</td><td>33</td><td>23</td><td></td><td>isomiR</td><td>NO</td><td>65.97</td><td>74.16</td><td>73.35</td><td>0.00</td></tr>
<tr><td>>MI0000434|hsa-let-7i&WithFlank&12|+|62603680|62603775@12.31.20(+1C)&[MIMAT0000415&hsa-let-7i-5p&offsets|0|-2(+1C);m-49&12|+|62603691|62603712&offsets|0|-2(+1C)]|TGAGGTAGTAGTTTGTGCTGC</td><td>>MI0000434|hsa-let-7i&WithFlank&12|+|62603680|62603775</td><td>12</td><td>31</td><td>20</td><td>(+1C)</td><td>isomiR</td><td>NO</td><td>10.19</td><td>28.36</td><td>11.95</td><td>0.00</td></tr>
<tr><td>>MI0000434|hsa-let-7i&WithFlank&12|+|62603680|62603775@12.32.21&[MIMAT0000415&hsa-let-7i-5p&offsets|0|-1;m-49&12|+|62603691|62603712&offsets|0|-1]|TGAGGTAGTAGTTTGTGCTGT</td><td>>MI0000434|hsa-let-7i&WithFlank&12|+|62603680|62603775</td><td>12</td><td>32</td><td>21</td><td></td><td>isomiR</td><td>NO</td><td>146.08</td><td>271.04</td><td>191.58</td><td>67.55</td></tr>
<tr><td>>MI0000434|hsa-let-7i&WithFlank&12|+|62603680|62603775@12.32.21(+1A)&[MIMAT0000415&hsa-let-7i-5p&offsets|0|-1(+1A);m-49&12|+|62603691|62603712&offsets|0|-1(+1A)]|TGAGGTAGTAGTTTGTGCTGTA</td><td>>MI0000434|hsa-let-7i&WithFlank&12|+|62603680|62603775</td><td>12</td><td>32</td><td>21</td><td>(+1A)</td><td>isomiR</td><td>NO</td><td>132.22</td><td>311.02</td><td>144.69</td><td>75.06</td></tr>
<tr><td>>MI0000434|hsa-let-7i&WithFlank&12|+|62603680|62603775@12.32.21(+1C)&[MIMAT0000415&hsa-let-7i-5p&offsets|0|-1(+1C);m-49&12|+|62603691|62603712&offsets|0|-1(+1C)]|TGAGGTAGTAGTTTGTGCTGTC</td><td>>MI0000434|hsa-let-7i&WithFlank&12|+|62603680|62603775</td><td>12</td><td>32</td><td>21</td><td>(+1C)</td><td>isomiR</td><td>NO</td><td>396.04</td><td>672.96</td><td>461.89</td><td>238.26</td></tr>
<tr><td>>MI0000434|hsa-let-7i&WithFlank&12|+|62603680|62603775@12.32.21(+1G)&[MIMAT0000415&hsa-let-7i-5p&offsets|0|-1(+1G);m-49&12|+|62603691|62603712&offsets|0|-1(+1G)]|TGAGGTAGTAGTTTGTGCTGTG</td><td>>MI0000434|hsa-let-7i&WithFlank&12|+|62603680|62603775</td><td>12</td><td>32</td><td>21</td><td>(+1G)</td><td>isomiR</td><td>NO</td><td>77.23</td><td>123.36</td><td>55.16</td><td>0.00</td></tr>
<tr><td>>MI0000434|hsa-let-7i&WithFlank&12|+|62603680|62603775@12.33.22&[MIMAT0000415&hsa-let-7i-5p&offsets|0|0;m-49&12|+|62603691|62603712&offsets|0|0]|TGAGGTAGTAGTTTGTGCTGTT</td><td>>MI0000434|hsa-let-7i&WithFlank&12|+|62603680|62603775</td><td>12</td><td>33</td><td>22</td><td></td><td>isomiR</td><td>YES</td><td>1942.32</td><td>4475.08</td><td>2632.00</td><td>1211.32</td></tr>
<tr><td>>MI0000434|hsa-let-7i&WithFlank&12|+|62603680|62603775@12.33.22(+1A)&[MIMAT0000415&hsa-let-7i-5p&offsets|0|0(+1A);m-49&12|+|62603691|62603712&offsets|0|0(+1A)]|TGAGGTAGTAGTTTGTGCTGTTA</td><td>>MI0000434|hsa-let-7i&WithFlank&12|+|62603680|62603775</td><td>12</td><td>33</td><td>22</td><td>(+1A)</td><td>isomiR</td><td>NO</td><td>164.10</td><td>267.27</td><td>457.86</td><td>98.32</td></tr>
<tr><td>>MI0000434|hsa-let-7i&WithFlank&12|+|62603680|62603775@12.33.22(+1C)&[MIMAT0000415&hsa-let-7i-5p&offsets|0|0(+1C);m-49&12|+|62603691|62603712&offsets|0|0(+1C)]|TGAGGTAGTAGTTTGTGCTGTTC</td><td>>MI0000434|hsa-let-7i&WithFlank&12|+|62603680|62603775</td><td>12</td><td>33</td><td>22</td><td>(+1C)</td><td>isomiR</td><td>NO</td><td>0.00</td><td>9.40</td><td>39.08</td><td>0.00</td></tr>
<tr><td>>MI0000434|hsa-let-7i&WithFlank&12|+|62603680|62603775@12.33.22(+1U)&[MIMAT0000415&hsa-let-7i-5p&offsets|0|0(+1U);m-49&12|+|62603691|62603712&offsets|0|0(+1U)]|TGAGGTAGTAGTTTGTGCTGTTT</td><td>>MI0000434|hsa-let-7i&WithFlank&12|+|62603680|62603775</td><td>12</td><td>33</td><td>22</td><td>(+1U)</td><td>isomiR</td><td>NO</td><td>135.00</td><td>171.83</td><td>281.90</td><td>72.84</td></tr>
<tr><td>>MI0000434|hsa-let-7i&WithFlank&12|+|62603680|62603775@12.34.23&[MIMAT0000415&hsa-let-7i-5p&offsets|0|+1;m-49&12|+|62603691|62603712&offsets|0|+1]|TGAGGTAGTAGTTTGTGCTGTTG</td><td>>MI0000434|hsa-let-7i&WithFlank&12|+|62603680|62603775</td><td>12</td><td>34</td><td>23</td><td></td><td>isomiR</td><td>NO</td><td>0.00</td><td>0.00</td><td>14.16</td><td>0.00</td></tr>
<tr><td>>MI0000438|hsa-mir-15b&WithFlank&3|+|160404582|160404691@26.46.21&[MIMAT0000417&hsa-miR-15b-5p&offsets|0|-1;m-110&3|+|160404607|160404628&offsets|0|-1]|TAGCAGCACATCATGGTTTAC</td><td>>MI0000438|hsa-mir-15b&WithFlank&3|+|160404582|160404691</td><td>26</td><td>46</td><td>21</td><td></td><td>isomiR</td><td>NO</td><td>28.11</td><td>0.00</td><td>0.00</td><td>0.00</td></tr>
<tr><td>>MI0000438|hsa-mir-15b&WithFlank&3|+|160404582|160404691@26.47.22&[MIMAT0000417&hsa-miR-15b-5p&offsets|0|0;m-110&3|+|160404607|160404628&offsets|0|0]|TAGCAGCACATCATGGTTTACA</td><td>>MI0000438|hsa-mir-15b&WithFlank&3|+|160404582|160404691</td><td>26</td><td>47</td><td>22</td><td></td><td>isomiR</td><td>NO</td><td>145.16</td><td>103.21</td><td>0.00</td><td>0.00</td></tr>
<tr><td>>MI0000438|hsa-mir-15b&WithFlank&3|+|160404582|160404691@64.84.21&[MIMAT0004586&hsa-miR-15b-3p&offsets|0|-1;m-204&3|+|160404645|160404665&offsets|0|0]|CGAATCATTATTTGCTGCTCT</td><td>>MI0000438|hsa-mir-15b&WithFlank&3|+|160404582|160404691</td><td>64</td><td>84</td><td>21</td><td></td><td>isomiR</td><td>NO</td><td>0.00</td><td>10.24</td><td>0.00</td><td>0.00</td></tr>
<tr><td>>MI0000439|hsa-mir-23b&WithFlank&9|+|95085202|95085310@67.84.18&[MIMAT0000418&hsa-miR-23b-3p&offsets|+3|-2;m-42&9|+|95085265|95085285&offsets|+3|0]|ACATTGCCAGGGATTACC</td><td>>MI0000439|hsa-mir-23b&WithFlank&9|+|95085202|95085310</td><td>67</td><td>84</td><td>18</td><td></td><td>isomiR</td><td>NO</td><td>41.49</td><td>188.20</td><td>9.48</td><td>0.00</td></tr>
<tr><td>>MI0000439|hsa-mir-23b&WithFlank&9|+|95085202|95085310@67.85.19&[MIMAT0000418&hsa-miR-23b-3p&offsets|+3|-1;m-42&9|+|95085265|95085285&offsets|+3|+1]|ACATTGCCAGGGATTACCA</td><td>>MI0000439|hsa-mir-23b&WithFlank&9|+|95085202|95085310</td><td>67</td><td>85</td><td>19</td><td></td><td>isomiR</td><td>NO</td><td>27.49</td><td>97.87</td><td>0.00</td><td>0.00</td></tr>
<tr><td>>MI0000439|hsa-mir-23b&WithFlank&9|+|95085202|95085310@67.85.19(+1U)&[MIMAT0000418&hsa-miR-23b-3p&offsets|+3|-1(+1U);m-42&9|+|95085265|95085285&offsets|+3|+1(+1U)]|ACATTGCCAGGGATTACCAT</td><td>>MI0000439|hsa-mir-23b&WithFlank&9|+|95085202|95085310</td><td>67</td><td>85</td><td>19</td><td>(+1U)</td><td>isomiR</td><td>NO</td><td>25.02</td><td>139.65</td><td>0.00</td><td>0.00</td></tr>
<tr><td>>MI0000439|hsa-mir-23b&WithFlank&9|+|95085202|95085310@67.85.19(+2U)&[MIMAT0000418&hsa-miR-23b-3p&offsets|+3|-1(+2U);m-42&9|+|95085265|95085285&offsets|+3|+1(+2U)]|ACATTGCCAGGGATTACCATT</td><td>>MI0000439|hsa-mir-23b&WithFlank&9|+|95085202|95085310</td><td>67</td><td>85</td><td>19</td><td>(+2U)</td><td>isomiR</td><td>NO</td><td>0.00</td><td>39.71</td><td>0.00</td><td>0.00</td></tr>
<tr><td>>MI0000439|hsa-mir-23b&WithFlank&9|+|95085202|95085310@67.86.20&[MIMAT0000418&hsa-miR-23b-3p&offsets|+3|0;m-42&9|+|95085265|95085285&offsets|+3|+2]|ACATTGCCAGGGATTACCAC</td><td>>MI0000439|hsa-mir-23b&WithFlank&9|+|95085202|95085310</td><td>67</td><td>86</td><td>20</td><td></td><td>isomiR</td><td>NO</td><td>98.22</td><td>448.20</td><td>0.00</td><td>30.35</td></tr>
<tr><td>>MI0000439|hsa-mir-23b&WithFlank&9|+|95085202|95085310@67.86.20(+1A)&[MIMAT0000418&hsa-miR-23b-3p&offsets|+3|0(+1A);m-42&9|+|95085265|95085285&offsets|+3|+2(+1A)]|ACATTGCCAGGGATTACCACA</td><td>>MI0000439|hsa-mir-23b&WithFlank&9|+|95085202|95085310</td><td>67</td><td>86</td><td>20</td><td>(+1A)</td><td>isomiR</td><td>NO</td><td>10.36</td><td>91.31</td><td>0.00</td><td>0.00</td></tr>
<tr><td>>MI0000439|hsa-mir-23b&WithFlank&9|+|95085202|95085310@67.86.20(+1C)&[MIMAT0000418&hsa-miR-23b-3p&offsets|+3|0(+1C);m-42&9|+|95085265|95085285&offsets|+3|+2(+1C)]|ACATTGCCAGGGATTACCACC</td><td>>MI0000439|hsa-mir-23b&WithFlank&9|+|95085202|95085310</td><td>67</td><td>86</td><td>20</td><td>(+1C)</td><td>isomiR</td><td>NO</td><td>31.35</td><td>105.60</td><td>0.00</td><td>10.73</td></tr>
<tr><td>>MI0000439|hsa-mir-23b&WithFlank&9|+|95085202|95085310@67.86.20(+1U)&[MIMAT0000418&hsa-miR-23b-3p&offsets|+3|0(+1U);m-42&9|+|95085265|95085285&offsets|+3|+2(+1U)]|ACATTGCCAGGGATTACCACT</td><td>>MI0000439|hsa-mir-23b&WithFlank&9|+|95085202|95085310</td><td>67</td><td>86</td><td>20</td><td>(+1U)</td><td>isomiR</td><td>NO</td><td>197.25</td><td>583.07</td><td>110.78</td><td>151.59</td></tr>
</table>

# BT-20

| | | | | | | | | | | | |
|---|---|---|---|---|---|---|---|---|---|---|---|
| >MI0000440\|hsa-mir-27b&WithFlank&9\|+\|95085439\|95085547@25.46.22&[MIMAT0004588&hsa-miR-27b-5p&offsets\|0\|0;m-240&9\|+\|95085463\|95085484&offsets\|0\|0]\|AGAGCTTAGCTGATTGGTGAAC | >MI0000440\|hsa-mir-27b&WithFlank&9\|+\|95085439\|95085547 | 25 | 46 | 22 | | isomiR | NO | 0.00 | 10.36 | 0.00 | 0.00 |
| >MI0000440\|hsa-mir-27b&WithFlank&9\|+\|95085439\|95085547@66.87.22&[MIMAT0000419&hsa-miR-27b-3p&offsets\|-1\|0;m-39&9\|+\|95085505\|95085525&offsets\|-1\|0]\|GTTCACAGTGGCTAAGTTCTGC | >MI0000440\|hsa-mir-27b&WithFlank&9\|+\|95085439\|95085547 | 66 | 87 | 22 | | isomiR | NO | 9.63 | 75.95 | 0.00 | 0.00 |
| >MI0000440\|hsa-mir-27b&WithFlank&9\|+\|95085439\|95085547@67.85.19&[MIMAT0000419&hsa-miR-27b-3p&offsets\|0\|-2;m-39&9\|+\|95085505\|95085525&offsets\|0\|-2]\|TTCACAGTGGCTAAGTTCT | >MI0000440\|hsa-mir-27b&WithFlank&9\|+\|95085439\|95085547 | 67 | 85 | 19 | | isomiR | NO | 73.09 | 143.85 | 112.03 | 23.42 |
| >MI0000440\|hsa-mir-27b&WithFlank&9\|+\|95085439\|95085547@67.85.19(+1A)&[MIMAT0000419&hsa-miR-27b-3p&offsets\|0\|-2(+1A);m-39&9\|+\|95085505\|95085525&offsets\|0\|-2(+1A)]\|TTCACAGTGGCTAAGTTCTA | >MI0000440\|hsa-mir-27b&WithFlank&9\|+\|95085439\|95085547 | 67 | 85 | 19 | (+1A) | isomiR | NO | 28.92 | 143.26 | 14.90 | 0.00 |
| >MI0000440\|hsa-mir-27b&WithFlank&9\|+\|95085439\|95085547@67.86.20&[MIMAT0000419&hsa-miR-27b-3p&offsets\|0\|-1;m-39&9\|+\|95085505\|95085525&offsets\|0\|-1]\|TTCACAGTGGCTAAGTTCTG | >MI0000440\|hsa-mir-27b&WithFlank&9\|+\|95085439\|95085547 | 67 | 86 | 20 | | isomiR | NO | 252.93 | 758.99 | 286.20 | 99.00 |
| >MI0000440\|hsa-mir-27b&WithFlank&9\|+\|95085439\|95085547@67.86.20(+1A)&[MIMAT0000419&hsa-miR-27b-3p&offsets\|0\|-1(+1A);m-39&9\|+\|95085505\|95085525&offsets\|0\|-1(+1A)]\|TTCACAGTGGCTAAGTTCTGA | >MI0000440\|hsa-mir-27b&WithFlank&9\|+\|95085439\|95085547 | 67 | 86 | 20 | (+1A) | isomiR | NO | 124.63 | 379.78 | 78.82 | 35.58 |
| >MI0000440\|hsa-mir-27b&WithFlank&9\|+\|95085439\|95085547@67.86.20(+1U)&[MIMAT0000419&hsa-miR-27b-3p&offsets\|0\|-1(+1U);m-39&9\|+\|95085505\|95085525&offsets\|0\|-1(+1U)]\|TTCACAGTGGCTAAGTTCTGT | >MI0000440\|hsa-mir-27b&WithFlank&9\|+\|95085439\|95085547 | 67 | 86 | 20 | (+1U) | isomiR | YES | 371.34 | 931.69 | 207.82 | 162.09 |
| >MI0000440\|hsa-mir-27b&WithFlank&9\|+\|95085439\|95085547@67.87.21&[MIMAT0000419&hsa-miR-27b-3p&offsets\|0\|0;m-39&9\|+\|95085505\|95085525&offsets\|0\|0]\|TTCACAGTGGCTAAGTTCTGC | >MI0000440\|hsa-mir-27b&WithFlank&9\|+\|95085439\|95085547 | 67 | 87 | 21 | | isomiR | YES | 1162.33 | 3248.29 | 774.50 | 589.71 |
| >MI0000440\|hsa-mir-27b&WithFlank&9\|+\|95085439\|95085547@67.87.21(+1U)&[MIMAT0000419&hsa-miR-27b-3p&offsets\|0\|0(+1U);m-39&9\|+\|95085505\|95085525&offsets\|0\|0(+1U)]\|TTCACAGTGGCTAAGTTCTGCT | >MI0000440\|hsa-mir-27b&WithFlank&9\|+\|95085439\|95085547 | 67 | 87 | 21 | (+1U) | isomiR | NO | 33.45 | 36.68 | 0.00 | 0.00 |
| >MI0000440\|hsa-mir-27b&WithFlank&9\|+\|95085439\|95085547@67.88.22&[MIMAT0000419&hsa-miR-27b-3p&offsets\|0\|+1;m-39&9\|+\|95085505\|95085525&offsets\|0\|+1]\|TTCACAGTGGCTAAGTTCTGCA | >MI0000440\|hsa-mir-27b&WithFlank&9\|+\|95085439\|95085547 | 67 | 88 | 22 | | isomiR | NO | 182.85 | 578.66 | 14.68 | 139.41 |
| >MI0000440\|hsa-mir-27b&WithFlank&9\|+\|95085439\|95085547@67.88.22(+1A)&[MIMAT0000419&hsa-miR-27b-3p&offsets\|0\|+1(+1A);m-39&9\|+\|95085505\|95085525&offsets\|0\|+1(+1A)]\|TTCACAGTGGCTAAGTTCTGCAA | >MI0000440\|hsa-mir-27b&WithFlank&9\|+\|95085439\|95085547 | 67 | 88 | 22 | (+1A) | isomiR | NO | 0.00 | 39.37 | 0.00 | 0.00 |
| >MI0000440\|hsa-mir-27b&WithFlank&9\|+\|95085439\|95085547@67.88.22(+1U)&[MIMAT0000419&hsa-miR-27b-3p&offsets\|0\|+1(+1U);m-39&9\|+\|95085505\|95085525&offsets\|0\|+1(+1U)]\|TTCACAGTGGCTAAGTTCTGCAT | >MI0000440\|hsa-mir-27b&WithFlank&9\|+\|95085439\|95085547 | 67 | 88 | 22 | (+1U) | isomiR | NO | 0.00 | 27.16 | 0.00 | 0.00 |
| >MI0000440\|hsa-mir-27b&WithFlank&9\|+\|95085439\|95085547@70.87.18&[MIMAT0000419&hsa-miR-27b-3p&offsets\|+3\|0;m-39&9\|+\|95085505\|95085525&offsets\|+3\|0]\|ACAGTGGCTAAGTTCTGC | >MI0000440\|hsa-mir-27b&WithFlank&9\|+\|95085439\|95085547 | 70 | 87 | 18 | | isomiR | NO | 30.52 | 80.17 | 0.00 | 0.00 |
| >MI0000441\|hsa-mir-30b&WithFlank&8\|-\|134800514\|134800613@23.43.21(+1A)&[MIMAT0000420&hsa-miR-30b-5p&offsets\|0\|-1(+1A);m-76&8\|-\|134800570\|134800591&offsets\|0\|-1(+1A)]\|TGTAAACATCCTACACTCAGCA | >MI0000441\|hsa-mir-30b&WithFlank&8\|-\|134800514\|134800613 | 23 | 43 | 21 | (+1A) | isomiR | NO | 26.73 | 23.84 | 0.00 | 0.00 |
| >MI0000441\|hsa-mir-30b&WithFlank&8\|-\|134800514\|134800613@23.43.21(+1C)&[MIMAT0000420&hsa-miR-30b-5p&offsets\|0\|-1(+1C);m-76&8\|-\|134800570\|134800591&offsets\|0\|-1(+1C)]\|TGTAAACATCCTACACTCAGCC | >MI0000441\|hsa-mir-30b&WithFlank&8\|-\|134800514\|134800613 | 23 | 43 | 21 | (+1C) | isomiR | NO | 37.78 | 26.18 | 0.00 | 0.00 |
| >MI0000441\|hsa-mir-30b&WithFlank&8\|-\|134800514\|134800613@23.44.22&[MIMAT0000420&hsa-miR-30b-5p&offsets\|0\|0;m-76&8\|-\|134800570\|134800591&offsets\|0\|0]\|TGTAAACATCCTACACTCAGCT | >MI0000441\|hsa-mir-30b&WithFlank&8\|-\|134800514\|134800613 | 23 | 44 | 22 | | isomiR | YES | 239.17 | 213.62 | 10.58 | 10.95 |
| >MI0000446\|hsa-mir-125b-1&WithFlank&11\|-\|122099751\|122099850@21.41.21;MI0000470\|hsa-mir-125b-2&WithFlank&21\|+\|16590231\|16590331@23.43.21&[MIMAT0000423&hsa-miR-125b-5p&offsets\|0\|-1;m-67&11\|-\|122099809\|122099830&offsets\|0\|-1];[MIMAT0000423_1&hsa-miR-125b-5p&offsets\|0\|-1;m-68&21\|+\|16590253\|16590274&offsets\|0\|-1]\|TCCCTGAGACCCTAACTTGTG | >MI0000446\|hsa-mir-125b-1&WithFlank&11\|-\|122099751\|122099850 | 21 | 41 | 21 | | isomiR | YES | 0.00 | 30.26 | 0.00 | 0.00 |
| >MI0000446\|hsa-mir-125b-1&WithFlank&11\|-\|122099751\|122099850@21.42.22;MI0000470\|hsa-mir-125b-2&WithFlank&21\|+\|16590231\|16590331@23.44.22&[MIMAT0000423&hsa-miR-125b-5p&offsets\|0\|0;m-67&11\|-\|122099809\|122099830&offsets\|0\|0];[MIMAT0000423_1&hsa-miR-125b-5p&offsets\|0\|0;m-68&21\|+\|16590253\|16590274&offsets\|0\|0]\|TCCCTGAGACCCTAACTTGTGA | >MI0000446\|hsa-mir-125b-1&WithFlank&11\|-\|122099751\|122099850 | 21 | 42 | 22 | | isomiR | YES | 0.00 | 33.45 | 0.00 | 0.00 |
| >MI0000447\|hsa-mir-128-1&WithFlank&2\|+\|135665391\|135665484@56.75.20(+1A);MI0000727\|hsa-mir-128-2&WithFlank&3\|+\|35744470\|35744565@58.77.20(+1A)&[MIMAT0000424&hsa-miR-128-3p&offsets\|0\|-1(+1A);m-108&2\|+\|135665446\|135665466&offsets\|0\|-1(+1A)];[MIMAT0000424_1&hsa-miR-128-3p&offsets\|0\|-1(+1A);m-117&3\|+\|35744527\|35744547&offsets\|0\|-1(+1A)]\|TCACAGTGAACCGGTCTCTTA | >MI0000447\|hsa-mir-128-1&WithFlank&2\|+\|135665391\|135665484 | 56 | 75 | 20 | (+1A) | isomiR | NO | 0.00 | 24.24 | 0.00 | 0.00 |
| >MI0000447\|hsa-mir-128-1&WithFlank&2\|+\|135665391\|135665484@56.75.20(+1C);MI0000727\|hsa-mir-128-2&WithFlank&3\|+\|35744470\|35744565@58.77.20(+1C)&[MIMAT0000424&hsa-miR-128-3p&offsets\|0\|-1(+1C);m-108&2\|+\|135665446\|135665466&offsets\|0\|-1(+1C)];[MIMAT0000424_1&hsa-miR-128-3p&offsets\|0\|-1(+1C);m-117&3\|+\|35744527\|35744547&offsets\|0\|-1(+1C)]\|TCACAGTGAACCGGTCTCTTC | >MI0000447\|hsa-mir-128-1&WithFlank&2\|+\|135665391\|135665484 | 56 | 75 | 20 | (+1C) | isomiR | NO | 34.38 | 94.68 | 57.57 | 27.11 |

# BT-20

| | | | | | | | | | | | |
|---|---|---|---|---|---|---|---|---|---|---|---|
| >MI0000447\|hsa-mir-128-1&WithFlank&2\|+\|135665391\|135665484@56.75.20;MI0000727\|hsa-mir-128-2&WithFlank&3\|+\|35744470\|35744565@58.77.20&[MIMAT0000424&hsa-miR-128-3p&offsets\|0\|-1;m-108&2\|+\|135665446\|135665466&offsets\|0\|-1];[MIMAT0000424_1&hsa-miR-128-3p&offsets\|0\|-1;m-117&3\|+\|35744527\|35744547&offsets\|0\|-1]\|TCACAGTGAACCGGTCTCTT | >MI0000447\|hsa-mir-128-1&WithFlank&2\|+\|135665391\|135665484 | 56 | 75 | 20 | | isomiR | NO | 27.79 | 85.06 | 15.60 | 10.46 |
| >MI0000447\|hsa-mir-128-1&WithFlank&2\|+\|135665391\|135665484@56.76.21(+1A);MI0000727\|hsa-mir-128-2&WithFlank&3\|+\|35744470\|35744565@58.78.21(+1A)&[MIMAT0000424&hsa-miR-128-3p&offsets\|0\|0(+1A);m-108&2\|+\|135665446\|135665466&offsets\|0\|0(+1A)];[MIMAT0000424_1&hsa-miR-128-3p&offsets\|0\|0(+1A);m-117&3\|+\|35744527\|35744547&offsets\|0\|0(+1A)]\|TCACAGTGAACCGGTCTCTTTA | >MI0000447\|hsa-mir-128-1&WithFlank&2\|+\|135665391\|135665484 | 56 | 76 | 21 | (+1A) | isomiR | NO | 39.02 | 130.94 | 15.65 | 33.68 |
| >MI0000447\|hsa-mir-128-1&WithFlank&2\|+\|135665391\|135665484@56.76.21(+2A);MI0000727\|hsa-mir-128-2&WithFlank&3\|+\|35744470\|35744565@58.78.21(+2A)&[MIMAT0000424&hsa-miR-128-3p&offsets\|0\|0(+2A);m-108&2\|+\|135665446\|135665466&offsets\|0\|0(+2A)];[MIMAT0000424_1&hsa-miR-128-3p&offsets\|0\|0(+2A);m-117&3\|+\|35744527\|35744547&offsets\|0\|0(+2A)]\|TCACAGTGAACCGGTCTCTTTAA | >MI0000447\|hsa-mir-128-1&WithFlank&2\|+\|135665391\|135665484 | 56 | 76 | 21 | (+2A) | isomiR | NO | 11.05 | 72.63 | 10.92 | 0.00 |
| >MI0000447\|hsa-mir-128-1&WithFlank&2\|+\|135665391\|135665484@56.76.21;MI0000727\|hsa-mir-128-2&WithFlank&3\|+\|35744470\|35744565@58.78.21&[MIMAT0000424&hsa-miR-128-3p&offsets\|0\|0;m-108&2\|+\|135665446\|135665466&offsets\|0\|0];[MIMAT0000424_1&hsa-miR-128-3p&offsets\|0\|0;m-117&3\|+\|35744527\|35744547&offsets\|0\|0]\|TCACAGTGAACCGGTCTCTTT | >MI0000447\|hsa-mir-128-1&WithFlank&2\|+\|135665391\|135665484 | 56 | 76 | 21 | | isomiR | NO | 278.99 | 682.64 | 338.93 | 229.90 |
| >MI0000447\|hsa-mir-128-1&WithFlank&2\|+\|135665391\|135665484@56.77.22&[MIMAT0000424&hsa-miR-128-3p&offsets\|0\|+1;m-108&2\|+\|135665446\|135665466&offsets\|0\|+1]\|TCACAGTGAACCGGTCTCTTTT | >MI0000447\|hsa-mir-128-1&WithFlank&2\|+\|135665391\|135665484 | 56 | 77 | 22 | | isomiR | NO | 90.17 | 225.91 | 81.73 | 78.38 |
| >MI0000449\|hsa-mir-132&WithFlank&17\|-\|2049902\|2050014@65.86.22&[MIMAT0000426&hsa-miR-132-3p&offsets\|0\|0;m-138&17\|-\|2049929\|2049950&offsets\|0\|0]\|TAACAGTCTACAGCCATGGTCG | >MI0000449\|hsa-mir-132&WithFlank&17\|-\|2049902\|2050014 | 65 | 86 | 22 | | isomiR | NO | 0.00 | 66.70 | 14.04 | 0.00 |
| >MI0000449\|hsa-mir-132&WithFlank&17\|-\|2049902\|2050014@65.86.22(+1U)&[MIMAT0000426&hsa-miR-132-3p&offsets\|0\|0(+1U);m-138&17\|-\|2049929\|2049950&offsets\|0\|0(+1U)]\|TAACAGTCTACAGCCATGGTCGT | >MI0000449\|hsa-mir-132&WithFlank&17\|-\|2049902\|2050014 | 65 | 86 | 22 | (+1U) | isomiR | NO | 0.00 | 37.43 | 0.00 | 0.00 |
| >MI0000456\|hsa-mir-140&WithFlank&16\|+\|69933075\|69933186@29.50.22&[MIMAT0000431&hsa-miR-140-5p&offsets\|0\|0;m-173&16\|+\|69933103\|69933124&offsets\|0\|0]\|CAGTGGTTTTACCCTATGGTAG | >MI0000456\|hsa-mir-140&WithFlank&16\|+\|69933075\|69933186 | 29 | 50 | 22 | | isomiR | NO | 0.00 | 0.00 | 16.19 | 0.00 |
| >MI0000456\|hsa-mir-140&WithFlank&16\|+\|69933075\|69933186@68.88.21&[MIMAT0004597&hsa-miR-140-3p&offsets\|0\|0;m-45&16\|+\|69933143\|69933164&offsets\|-1\|-2]\|TACCACAGGGTAGAACCACGG | >MI0000456\|hsa-mir-140&WithFlank&16\|+\|69933075\|69933186 | 68 | 88 | 21 | | isomiR | NO | 0.00 | 26.21 | 0.00 | 0.00 |
| >MI0000456\|hsa-mir-140&WithFlank&16\|+\|69933075\|69933186@68.89.22&[MIMAT0004597&hsa-miR-140-3p&offsets\|0\|+1;m-45&16\|+\|69933143\|69933164&offsets\|-1\|-1]\|TACCACAGGGTAGAACCACGGA | >MI0000456\|hsa-mir-140&WithFlank&16\|+\|69933075\|69933186 | 68 | 89 | 22 | | isomiR | NO | 0.00 | 153.30 | 10.62 | 0.00 |
| >MI0000456\|hsa-mir-140&WithFlank&16\|+\|69933075\|69933186@68.90.23&[MIMAT0004597&hsa-miR-140-3p&offsets\|0\|+2;m-45&16\|+\|69933143\|69933164&offsets\|-1\|0]\|TACCACAGGGTAGAACCACGGAC | >MI0000456\|hsa-mir-140&WithFlank&16\|+\|69933075\|69933186 | 68 | 90 | 23 | | isomiR | NO | 0.00 | 108.13 | 0.00 | 0.00 |
| >MI0000456\|hsa-mir-140&WithFlank&16\|+\|69933075\|69933186@68.90.23(+1U)&[MIMAT0004597&hsa-miR-140-3p&offsets\|0\|+2(+1U);m-45&16\|+\|69933143\|69933164&offsets\|-1\|0(+1U)]\|TACCACAGGGTAGAACCACGGACT | >MI0000456\|hsa-mir-140&WithFlank&16\|+\|69933075\|69933186 | 68 | 90 | 23 | (+1U) | isomiR | NO | 0.00 | 23.80 | 0.00 | 0.00 |
| >MI0000456\|hsa-mir-140&WithFlank&16\|+\|69933075\|69933186@68.91.24&[MIMAT0004597&hsa-miR-140-3p&offsets\|0\|+3;m-45&16\|+\|69933143\|69933164&offsets\|-1\|+1]\|TACCACAGGGTAGAACCACGGACA | >MI0000456\|hsa-mir-140&WithFlank&16\|+\|69933075\|69933186 | 68 | 91 | 24 | | isomiR | NO | 0.00 | 43.62 | 0.00 | 0.00 |
| >MI0000456\|hsa-mir-140&WithFlank&16\|+\|69933075\|69933186@69.88.20&[MIMAT0004597&hsa-miR-140-3p&offsets\|+1\|0;m-45&16\|+\|69933143\|69933164&offsets\|0\|-2]\|ACCACAGGGTAGAACCACGG | >MI0000456\|hsa-mir-140&WithFlank&16\|+\|69933075\|69933186 | 69 | 88 | 20 | | isomiR | NO | 0.00 | 0.00 | 10.62 | 0.00 |
| >MI0000456\|hsa-mir-140&WithFlank&16\|+\|69933075\|69933186@69.89.21&[MIMAT0004597&hsa-miR-140-3p&offsets\|+1\|+1;m-45&16\|+\|69933143\|69933164&offsets\|0\|-1]\|ACCACAGGGTAGAACCACGGA | >MI0000456\|hsa-mir-140&WithFlank&16\|+\|69933075\|69933186 | 69 | 89 | 21 | | isomiR | NO | 26.22 | 129.47 | 59.62 | 0.00 |
| >MI0000456\|hsa-mir-140&WithFlank&16\|+\|69933075\|69933186@69.89.21(+1U)&[MIMAT0004597&hsa-miR-140-3p&offsets\|+1\|+1(+1U);m-45&16\|+\|69933143\|69933164&offsets\|0\|-1(+1U)]\|ACCACAGGGTAGAACCACGGAT | >MI0000456\|hsa-mir-140&WithFlank&16\|+\|69933075\|69933186 | 69 | 89 | 21 | (+1U) | isomiR | NO | 0.00 | 27.27 | 0.00 | 0.00 |
| >MI0000456\|hsa-mir-140&WithFlank&16\|+\|69933075\|69933186@69.90.22&[MIMAT0004597&hsa-miR-140-3p&offsets\|+1\|+2;m-45&16\|+\|69933143\|69933164&offsets\|0\|0]\|ACCACAGGGTAGAACCACGGAC | >MI0000456\|hsa-mir-140&WithFlank&16\|+\|69933075\|69933186 | 69 | 90 | 22 | | isomiR | NO | 36.11 | 193.41 | 70.12 | 0.00 |
| >MI0000456\|hsa-mir-140&WithFlank&16\|+\|69933075\|69933186@69.90.22(+1U)&[MIMAT0004597&hsa-miR-140-3p&offsets\|+1\|+2(+1U);m-45&16\|+\|69933143\|69933164&offsets\|0\|0(+1U)]\|ACCACAGGGTAGAACCACGGACT | >MI0000456\|hsa-mir-140&WithFlank&16\|+\|69933075\|69933186 | 69 | 90 | 22 | (+1U) | isomiR | NO | 0.00 | 81.12 | 53.75 | 0.00 |
| >MI0000456\|hsa-mir-140&WithFlank&16\|+\|69933075\|69933186@69.91.23&[MIMAT0004597&hsa-miR-140-3p&offsets\|+1\|+3;m-45&16\|+\|69933143\|69933164&offsets\|0\|+1]\|ACCACAGGGTAGAACCACGGACA | >MI0000456\|hsa-mir-140&WithFlank&16\|+\|69933075\|69933186 | 69 | 91 | 23 | | isomiR | NO | 10.59 | 125.90 | 54.22 | 0.00 |

# BT-20

<table>
<tr><td>>MI0000457|hsa-mir-141&WithFlank&12|+|6964091|6964197@65.85.21&[MIMAT0000432&hsa-miR-141-3p&offsets|0|-1;m-134&12|+|6964155|6964175&offsets|0|0]|TAACACTGTCTGGTAAAGATG</td><td>>MI0000457|hsa-mir-141&WithFlank&12|+|6964091|6964197</td><td>65</td><td>85</td><td>21</td><td></td><td>isomiR</td><td>NO</td><td>10.77</td><td>24.29</td><td>69.10</td><td>0.00</td></tr>
<tr><td>>MI0000462|hsa-mir-152&WithFlank&17|-|48037155|48037253@60.80.21&[MIMAT0000438&hsa-miR-152-3p&offsets|0|0;m-97&17|-|48037174|48037194&offsets|0|0]|TCAGTGCATGACAGAACTTGG</td><td>>MI0000462|hsa-mir-152&WithFlank&17|-|48037155|48037253</td><td>60</td><td>80</td><td>21</td><td></td><td>isomiR</td><td>NO</td><td>38.71</td><td>69.96</td><td>53.23</td><td>0.00</td></tr>
<tr><td>>MI0000462|hsa-mir-152&WithFlank&17|-|48037155|48037253@60.80.21(+1U)&[MIMAT0000438&hsa-miR-152-3p&offsets|0|0(+1U);m-97&17|-|48037174|48037194&offsets|0|0(+1U)]|TCAGTGCATGACAGAACTTGGT</td><td>>MI0000462|hsa-mir-152&WithFlank&17|-|48037155|48037253</td><td>60</td><td>80</td><td>21</td><td>(+1U)</td><td>isomiR</td><td>NO</td><td>0.00</td><td>10.24</td><td>0.00</td><td>0.00</td></tr>
<tr><td>>MI0000462|hsa-mir-152&WithFlank&17|-|48037155|48037253@60.81.22&[MIMAT0000438&hsa-miR-152-3p&offsets|0|+1;m-97&17|-|48037174|48037194&offsets|0|+1]|TCAGTGCATGACAGAACTTGGG</td><td>>MI0000462|hsa-mir-152&WithFlank&17|-|48037155|48037253</td><td>60</td><td>81</td><td>22</td><td></td><td>isomiR</td><td>NO</td><td>28.53</td><td>75.47</td><td>82.80</td><td>0.00</td></tr>
<tr><td>>MI0000462|hsa-mir-152&WithFlank&17|-|48037155|48037253@60.81.22(+1A)&[MIMAT0000438&hsa-miR-152-3p&offsets|0|+1(+1A);m-97&17|-|48037174|48037194&offsets|0|+1(+1A)]|TCAGTGCATGACAGAACTTGGGA</td><td>>MI0000462|hsa-mir-152&WithFlank&17|-|48037155|48037253</td><td>60</td><td>81</td><td>22</td><td>(+1A)</td><td>isomiR</td><td>NO</td><td>0.00</td><td>32.83</td><td>0.00</td><td>0.00</td></tr>
<tr><td>>MI0000462|hsa-mir-152&WithFlank&17|-|48037155|48037253@60.81.22(+1U)&[MIMAT0000438&hsa-miR-152-3p&offsets|0|+1(+1U);m-97&17|-|48037174|48037194&offsets|0|+1(+1U)]|TCAGTGCATGACAGAACTTGGGT</td><td>>MI0000462|hsa-mir-152&WithFlank&17|-|48037155|48037253</td><td>60</td><td>81</td><td>22</td><td>(+1U)</td><td>isomiR</td><td>NO</td><td>0.00</td><td>41.98</td><td>9.75</td><td>0.00</td></tr>
<tr><td>>MI0000465|hsa-mir-191&WithFlank&3|-|49020612|49020715@22.42.21&[MIMAT0000440&hsa-miR-191-5p&offsets|0|-2;m-48&3|-|49020672|49020694&offsets|0|-2]|CAACGGAATCCCAAAAGCAGC</td><td>>MI0000465|hsa-mir-191&WithFlank&3|-|49020612|49020715</td><td>22</td><td>42</td><td>21</td><td></td><td>isomiR</td><td>NO</td><td>30.72</td><td>79.51</td><td>11.12</td><td>0.00</td></tr>
<tr><td>>MI0000465|hsa-mir-191&WithFlank&3|-|49020612|49020715@22.42.21(+1A)&[MIMAT0000440&hsa-miR-191-5p&offsets|0|-2(+1A);m-48&3|-|49020672|49020694&offsets|0|-2(+1A)]|CAACGGAATCCCAAAAGCAGCA</td><td>>MI0000465|hsa-mir-191&WithFlank&3|-|49020612|49020715</td><td>22</td><td>42</td><td>21</td><td>(+1A)</td><td>isomiR</td><td>NO</td><td>33.68</td><td>106.02</td><td>0.00</td><td>0.00</td></tr>
<tr><td>>MI0000465|hsa-mir-191&WithFlank&3|-|49020612|49020715@22.42.21(+1C)&[MIMAT0000440&hsa-miR-191-5p&offsets|0|-2(+1C);m-48&3|-|49020672|49020694&offsets|0|-2(+1C)]|CAACGGAATCCCAAAAGCAGCC</td><td>>MI0000465|hsa-mir-191&WithFlank&3|-|49020612|49020715</td><td>22</td><td>42</td><td>21</td><td>(+1C)</td><td>isomiR</td><td>NO</td><td>80.32</td><td>125.47</td><td>14.52</td><td>25.15</td></tr>
<tr><td>>MI0000465|hsa-mir-191&WithFlank&3|-|49020612|49020715@22.42.21(+1G)&[MIMAT0000440&hsa-miR-191-5p&offsets|0|-2(+1G);m-48&3|-|49020672|49020694&offsets|0|-2(+1G)]|CAACGGAATCCCAAAAGCAGCG</td><td>>MI0000465|hsa-mir-191&WithFlank&3|-|49020612|49020715</td><td>22</td><td>42</td><td>21</td><td>(+1G)</td><td>isomiR</td><td>NO</td><td>0.00</td><td>24.76</td><td>0.00</td><td>0.00</td></tr>
<tr><td>>MI0000465|hsa-mir-191&WithFlank&3|-|49020612|49020715@22.43.22&[MIMAT0000440&hsa-miR-191-5p&offsets|0|-1;m-48&3|-|49020672|49020694&offsets|0|-1]|CAACGGAATCCCAAAAGCAGCT</td><td>>MI0000465|hsa-mir-191&WithFlank&3|-|49020612|49020715</td><td>22</td><td>43</td><td>22</td><td></td><td>isomiR</td><td>NO</td><td>348.58</td><td>728.59</td><td>211.49</td><td>174.76</td></tr>
<tr><td>>MI0000465|hsa-mir-191&WithFlank&3|-|49020612|49020715@22.43.22(+1A)&[MIMAT0000440&hsa-miR-191-5p&offsets|0|-1(+1A);m-48&3|-|49020672|49020694&offsets|0|-1(+1A)]|CAACGGAATCCCAAAAGCAGCTA</td><td>>MI0000465|hsa-mir-191&WithFlank&3|-|49020612|49020715</td><td>22</td><td>43</td><td>22</td><td>(+1A)</td><td>isomiR</td><td>NO</td><td>79.98</td><td>225.19</td><td>96.23</td><td>33.42</td></tr>
<tr><td>>MI0000465|hsa-mir-191&WithFlank&3|-|49020612|49020715@22.43.22(+1U)&[MIMAT0000440&hsa-miR-191-5p&offsets|0|-1(+1U);m-48&3|-|49020672|49020694&offsets|0|-1(+1U)]|CAACGGAATCCCAAAAGCAGCTT</td><td>>MI0000465|hsa-mir-191&WithFlank&3|-|49020612|49020715</td><td>22</td><td>43</td><td>22</td><td>(+1U)</td><td>isomiR</td><td>NO</td><td>0.00</td><td>33.51</td><td>0.00</td><td>0.00</td></tr>
<tr><td>>MI0000465|hsa-mir-191&WithFlank&3|-|49020612|49020715@22.44.23&[MIMAT0000440&hsa-miR-191-5p&offsets|0|0;m-48&3|-|49020672|49020694&offsets|0|0]|CAACGGAATCCCAAAAGCAGCTG</td><td>>MI0000465|hsa-mir-191&WithFlank&3|-|49020612|49020715</td><td>22</td><td>44</td><td>23</td><td></td><td>isomiR</td><td>NO</td><td>427.82</td><td>1107.45</td><td>631.59</td><td>307.68</td></tr>
<tr><td>>MI0000465|hsa-mir-191&WithFlank&3|-|49020612|49020715@22.44.23(+1A)&[MIMAT0000440&hsa-miR-191-5p&offsets|0|0(+1A);m-48&3|-|49020672|49020694&offsets|0|0(+1A)]|CAACGGAATCCCAAAAGCAGCTGA</td><td>>MI0000465|hsa-mir-191&WithFlank&3|-|49020612|49020715</td><td>22</td><td>44</td><td>23</td><td>(+1A)</td><td>isomiR</td><td>NO</td><td>0.00</td><td>30.27</td><td>0.00</td><td>0.00</td></tr>
<tr><td>>MI0000465|hsa-mir-191&WithFlank&3|-|49020612|49020715@22.45.24&[MIMAT0000440&hsa-miR-191-5p&offsets|0|+1;m-48&3|-|49020672|49020694&offsets|0|+1]|CAACGGAATCCCAAAAGCAGCTGT</td><td>>MI0000465|hsa-mir-191&WithFlank&3|-|49020612|49020715</td><td>22</td><td>45</td><td>24</td><td></td><td>isomiR</td><td>NO</td><td>115.78</td><td>183.86</td><td>100.68</td><td>26.09</td></tr>
<tr><td>>MI0000465|hsa-mir-191&WithFlank&3|-|49020612|49020715@23.44.22&[MIMAT0000440&hsa-miR-191-5p&offsets|+1|0;m-48&3|-|49020672|49020694&offsets|+1|0]|AACGGAATCCCAAAAGCAGCTG</td><td>>MI0000465|hsa-mir-191&WithFlank&3|-|49020612|49020715</td><td>23</td><td>44</td><td>22</td><td></td><td>isomiR</td><td>NO</td><td>0.00</td><td>25.60</td><td>10.36</td><td>0.00</td></tr>
<tr><td>>MI0000466|hsa-mir-9-1&WithFlank&1|-|156420335|156420435@22.43.22;MI0000468|hsa-mir-9-3&WithFlank&15|+|89368011|89368112@22.43.22;MI0000467|hsa-mir-9-2&WithFlank&5|-|88666847|88666945@22.43.22&[MIMAT0000441&hsa-miR-9-5p&offsets|0|-1;m-104&1|-|156420392|156420414&offsets|0|-1];[MIMAT0000441_1&hsa-miR-9-5p&offsets|0|-1;m-105&15|+|89368032|89368054&offsets|0|-1];[MIMAT0000441_2&hsa-miR-9-5p&offsets|0|-1;m-104&5|-|88666902|88666924&offsets|0|-1]|TCTTTGGTTATCTAGCTGTATG</td><td>>MI0000466|hsa-mir-9-1&WithFlank&1|-|156420335|156420435</td><td>22</td><td>43</td><td>22</td><td></td><td>isomiR</td><td>NO</td><td>41.13</td><td>114.03</td><td>0.00</td><td>26.92</td></tr>
<tr><td>>MI0000466|hsa-mir-9-1&WithFlank&1|-|156420335|156420435@22.44.23;MI0000468|hsa-mir-9-3&WithFlank&15|+|89368011|89368112@22.44.23;MI0000467|hsa-mir-9-2&WithFlank&5|-|88666847|88666945@22.44.23&[MIMAT0000441&hsa-miR-9-5p&offsets|0|0;m-104&1|-|156420392|156420414&offsets|0|0];[MIMAT0000441_1&hsa-miR-9-5p&offsets|0|0;m-105&15|+|89368032|89368054&offsets|0|0];[MIMAT0000441_2&hsa-miR-9-5p&offsets|0|0;m-104&5|-|88666902|88666924&offsets|0|0]|TCTTTGGTTATCTAGCTGTATGA</td><td>>MI0000466|hsa-mir-9-1&WithFlank&1|-|156420335|156420435</td><td>22</td><td>44</td><td>23</td><td></td><td>isomiR</td><td>NO</td><td>83.40</td><td>100.82</td><td>10.79</td><td>36.11</td></tr>
<tr><td>>MI0000469|hsa-mir-125a&WithFlank&19|+|51693248|51693345@21.41.21&[MIMAT0000443&hsa-miR-125a-5p&offsets|0|-3;m-63&19|+|51693268|51693289&offsets|0|-1]|TCCCTGAGACCCTTTAACCTG</td><td>>MI0000469|hsa-mir-125a&WithFlank&19|+|51693248|51693345</td><td>21</td><td>41</td><td>21</td><td></td><td>isomiR</td><td>NO</td><td>0.00</td><td>29.96</td><td>0.00</td><td>0.00</td></tr>
<tr><td>>MI0000469|hsa-mir-125a&WithFlank&19|+|51693248|51693345@21.41.21(+1C)&[MIMAT0000443&hsa-miR-125a-5p&offsets|0|-3(+1C);m-63&19|+|51693268|51693289&offsets|0|-1(+1C)]|TCCCTGAGACCCTTTAACCTGC</td><td>>MI0000469|hsa-mir-125a&WithFlank&19|+|51693248|51693345</td><td>21</td><td>41</td><td>21</td><td>(+1C)</td><td>isomiR</td><td>NO</td><td>26.52</td><td>81.45</td><td>0.00</td><td>24.48</td></tr>
<tr><td>>MI0000469|hsa-mir-125a&WithFlank&19|+|51693248|51693345@21.42.22&[MIMAT0000443&hsa-miR-125a-5p&offsets|0|-2;m-63&19|+|51693268|51693289&offsets|0|0]|TCCCTGAGACCCTTTAACCTGT</td><td>>MI0000469|hsa-mir-125a&WithFlank&19|+|51693248|51693345</td><td>21</td><td>42</td><td>22</td><td></td><td>isomiR</td><td>NO</td><td>184.96</td><td>537.41</td><td>13.82</td><td>190.78</td></tr>
<tr><td>>MI0000469|hsa-mir-125a&WithFlank&19|+|51693248|51693345@21.42.22(+1A)&[MIMAT0000443&hsa-miR-125a-5p&offsets|0|-2(+1A);m-63&19|+|51693268|51693289&offsets|0|0(+1A)]|TCCCTGAGACCCTTTAACCTGTA</td><td>>MI0000469|hsa-mir-125a&WithFlank&19|+|51693248|51693345</td><td>21</td><td>42</td><td>22</td><td>(+1A)</td><td>isomiR</td><td>NO</td><td>107.30</td><td>373.92</td><td>10.71</td><td>130.10</td></tr>
</table>

# BT-20

| | | | | | | | | | | | |
|---|---|---|---|---|---|---|---|---|---|---|---|
| >MI0000469\|hsa-mir-125a&WithFlank&19\|+\|51693248\|51693345@21.42.22(+1U)&[MIMAT0000443&hsa-miR-125a-5p&offsets\|0\|-2(+1U);m-63&19\|+\|51693268\|51693289&offsets\|0\|0(+1U)]\|TCCCTGAGACCCTTTAACCTGTT | >MI0000469\|hsa-mir-125a&WithFlank&19\|+\|51693248\|51693345 | 21 | 42 | 22 | (+1U) | isomiR | NO | 0.00 | 28.40 | 0.00 | 0.00 |
| >MI0000469\|hsa-mir-125a&WithFlank&19\|+\|51693248\|51693345@21.43.23&[MIMAT0000443&hsa-miR-125a-5p&offsets\|0\|-1;m-63&19\|+\|51693268\|51693289&offsets\|0\|+1]\|TCCCTGAGACCCTTTAACCTGTG | >MI0000469\|hsa-mir-125a&WithFlank&19\|+\|51693248\|51693345 | 21 | 43 | 23 | | isomiR | YES | 606.35 | 1947.17 | 218.75 | 820.96 |
| >MI0000469\|hsa-mir-125a&WithFlank&19\|+\|51693248\|51693345@21.43.23(+1U)&[MIMAT0000443&hsa-miR-125a-5p&offsets\|0\|-1(+1U);m-63&19\|+\|51693268\|51693289&offsets\|0\|+1(+1U)]\|TCCCTGAGACCCTTTAACCTGTGT | >MI0000469\|hsa-mir-125a&WithFlank&19\|+\|51693248\|51693345 | 21 | 43 | 23 | (+1U) | isomiR | NO | 30.36 | 105.07 | 0.00 | 35.82 |
| >MI0000469\|hsa-mir-125a&WithFlank&19\|+\|51693248\|51693345@21.44.24&[MIMAT0000443&hsa-miR-125a-5p&offsets\|0\|0;m-63&19\|+\|51693268\|51693289&offsets\|0\|+2]\|TCCCTGAGACCCTTTAACCTGTGA | >MI0000469\|hsa-mir-125a&WithFlank&19\|+\|51693248\|51693345 | 21 | 44 | 24 | | isomiR | NO | 185.48 | 457.47 | 11.87 | 166.99 |
| >MI0000469\|hsa-mir-125a&WithFlank&19\|+\|51693248\|51693345@59.79.21&[MIMAT0004602&hsa-miR-125a-3p&offsets\|0\|-1;m-333&19\|+\|51693306\|51693326&offsets\|0\|0]\|ACAGGTGAGGTTCTTGGGAGC | >MI0000469\|hsa-mir-125a&WithFlank&19\|+\|51693248\|51693345 | 59 | 79 | 21 | | isomiR | NO | 0.00 | 76.04 | 39.10 | 0.00 |
| >MI0000471\|hsa-mir-126&WithFlank&9\|+\|136670596\|136670692@58.77.20&[MIMAT0000445&hsa-miR-126-3p&offsets\|0\|-2;m-40&9\|+\|136670653\|136670674&offsets\|0\|-2]\|TCGTACCGTGAGTAATAATG | >MI0000471\|hsa-mir-126&WithFlank&9\|+\|136670596\|136670692 | 58 | 77 | 20 | | isomiR | NO | 0.00 | 9.45 | 10.52 | 0.00 |
| >MI0000471\|hsa-mir-126&WithFlank&9\|+\|136670596\|136670692@58.77.20(+1U)&[MIMAT0000445&hsa-miR-126-3p&offsets\|0\|-2(+1U);m-40&9\|+\|136670653\|136670674&offsets\|0\|-2(+1U)]\|TCGTACCGTGAGTAATAATGT | >MI0000471\|hsa-mir-126&WithFlank&9\|+\|136670596\|136670692 | 58 | 77 | 20 | (+1U) | isomiR | NO | 10.37 | 30.35 | 13.27 | 0.00 |
| >MI0000471\|hsa-mir-126&WithFlank&9\|+\|136670596\|136670692@58.78.21&[MIMAT0000445&hsa-miR-126-3p&offsets\|0\|-1;m-40&9\|+\|136670653\|136670674&offsets\|0\|-1]\|TCGTACCGTGAGTAATAATGC | >MI0000471\|hsa-mir-126&WithFlank&9\|+\|136670596\|136670692 | 58 | 78 | 21 | | isomiR | NO | 86.91 | 125.46 | 123.70 | 41.03 |
| >MI0000471\|hsa-mir-126&WithFlank&9\|+\|136670596\|136670692@58.78.21(+1A)&[MIMAT0000445&hsa-miR-126-3p&offsets\|0\|-1(+1A);m-40&9\|+\|136670653\|136670674&offsets\|0\|-1(+1A)]\|TCGTACCGTGAGTAATAATGCA | >MI0000471\|hsa-mir-126&WithFlank&9\|+\|136670596\|136670692 | 58 | 78 | 21 | (+1A) | isomiR | NO | 32.19 | 70.92 | 16.01 | 0.00 |
| >MI0000471\|hsa-mir-126&WithFlank&9\|+\|136670596\|136670692@58.79.22&[MIMAT0000445&hsa-miR-126-3p&offsets\|0\|0;m-40&9\|+\|136670653\|136670674&offsets\|0\|0]\|TCGTACCGTGAGTAATAATGCG | >MI0000471\|hsa-mir-126&WithFlank&9\|+\|136670596\|136670692 | 58 | 79 | 22 | | isomiR | NO | 160.79 | 223.22 | 205.73 | 105.48 |
| >MI0000471\|hsa-mir-126&WithFlank&9\|+\|136670596\|136670692@59.78.20&[MIMAT0000445&hsa-miR-126-3p&offsets\|+1\|-1;m-40&9\|+\|136670653\|136670674&offsets\|+1\|-1]\|CGTACCGTGAGTAATAATGC | >MI0000471\|hsa-mir-126&WithFlank&9\|+\|136670596\|136670692 | 59 | 78 | 20 | | isomiR | NO | 0.00 | 32.67 | 15.73 | 0.00 |
| >MI0000471\|hsa-mir-126&WithFlank&9\|+\|136670596\|136670692@59.79.21&[MIMAT0000445&hsa-miR-126-3p&offsets\|+1\|0;m-40&9\|+\|136670653\|136670674&offsets\|+1\|0]\|CGTACCGTGAGTAATAATGCG | >MI0000471\|hsa-mir-126&WithFlank&9\|+\|136670596\|136670692 | 59 | 79 | 21 | | isomiR | NO | 0.00 | 0.00 | 11.78 | 0.00 |
| >MI0000478\|hsa-mir-149&WithFlank&2\|+\|240455995\|240456095@21.41.21&[MIMAT0000450&hsa-miR-149-5p&offsets\|0\|-2;m-195&2\|+\|240456015\|240456037&offsets\|0\|-2]\|TCTGGCTCCGTGTCTTCACTC | >MI0000478\|hsa-mir-149&WithFlank&2\|+\|240455995\|240456095 | 21 | 41 | 21 | | isomiR | NO | 0.00 | 32.07 | 0.00 | 0.00 |
| >MI0000478\|hsa-mir-149&WithFlank&2\|+\|240455995\|240456095@21.41.21(+1A)&[MIMAT0000450&hsa-miR-149-5p&offsets\|0\|-2(+1A);m-195&2\|+\|240456015\|240456037&offsets\|0\|-2(+1A)]\|TCTGGCTCCGTGTCTTCACTCA | >MI0000478\|hsa-mir-149&WithFlank&2\|+\|240455995\|240456095 | 21 | 41 | 21 | (+1A) | isomiR | NO | 0.00 | 29.89 | 0.00 | 0.00 |
| >MI0000478\|hsa-mir-149&WithFlank&2\|+\|240455995\|240456095@21.41.21(+1U)&[MIMAT0000450&hsa-miR-149-5p&offsets\|0\|-2(+1U);m-195&2\|+\|240456015\|240456037&offsets\|0\|-2(+1U)]\|TCTGGCTCCGTGTCTTCACTCT | >MI0000478\|hsa-mir-149&WithFlank&2\|+\|240455995\|240456095 | 21 | 41 | 21 | (+1U) | isomiR | NO | 10.56 | 89.94 | 11.64 | 0.00 |
| >MI0000478\|hsa-mir-149&WithFlank&2\|+\|240455995\|240456095@21.42.22&[MIMAT0000450&hsa-miR-149-5p&offsets\|0\|-1;m-195&2\|+\|240456015\|240456037&offsets\|0\|-1]\|TCTGGCTCCGTGTCTTCACTCC | >MI0000478\|hsa-mir-149&WithFlank&2\|+\|240455995\|240456095 | 21 | 42 | 22 | | isomiR | NO | 43.49 | 173.26 | 74.53 | 34.66 |
| >MI0000478\|hsa-mir-149&WithFlank&2\|+\|240455995\|240456095@21.42.22(+1A)&[MIMAT0000450&hsa-miR-149-5p&offsets\|0\|-1(+1A);m-195&2\|+\|240456015\|240456037&offsets\|0\|-1(+1A)]\|TCTGGCTCCGTGTCTTCACTCCA | >MI0000478\|hsa-mir-149&WithFlank&2\|+\|240455995\|240456095 | 21 | 42 | 22 | (+1A) | isomiR | NO | 0.00 | 44.79 | 0.00 | 0.00 |
| >MI0000478\|hsa-mir-149&WithFlank&2\|+\|240455995\|240456095@21.42.22(+1U)&[MIMAT0000450&hsa-miR-149-5p&offsets\|0\|-1(+1U);m-195&2\|+\|240456015\|240456037&offsets\|0\|-1(+1U)]\|TCTGGCTCCGTGTCTTCACTCCT | >MI0000478\|hsa-mir-149&WithFlank&2\|+\|240455995\|240456095 | 21 | 42 | 22 | (+1U) | isomiR | NO | 39.63 | 129.60 | 13.43 | 30.37 |
| >MI0000478\|hsa-mir-149&WithFlank&2\|+\|240455995\|240456095@21.43.23&[MIMAT0000450&hsa-miR-149-5p&offsets\|0\|0;m-195&2\|+\|240456015\|240456037&offsets\|0\|0]\|TCTGGCTCCGTGTCTTCACTCCC | >MI0000478\|hsa-mir-149&WithFlank&2\|+\|240455995\|240456095 | 21 | 43 | 23 | | isomiR | NO | 92.09 | 220.55 | 73.32 | 81.75 |
| >MI0000481\|hsa-mir-184&WithFlank&15\|+\|79209782\|79209877@59.80.22&[MIMAT0000454&hsa-miR-184&offsets\|0\|0;m-460&15\|+\|79209840\|79209861&offsets\|0\|0]\|TGGACGGAGAACTGATAAGGGT | >MI0000481\|hsa-mir-184&WithFlank&15\|+\|79209782\|79209877 | 59 | 80 | 22 | | isomiR | NO | 0.00 | 9.76 | 53.90 | 0.00 |
| >MI0000482\|hsa-mir-185&WithFlank&22\|+\|20033133\|20033226@21.42.22&[MIMAT0000455&hsa-miR-185-5p&offsets\|0\|0;m-145&22\|+\|20033153\|20033174&offsets\|0\|0]\|TGGAGAGAAAGGCAGTTCCTGA | >MI0000482\|hsa-mir-185&WithFlank&22\|+\|20033133\|20033226 | 21 | 42 | 22 | | isomiR | NO | 80.61 | 164.19 | 84.89 | 41.40 |

# BT-20

<table>
<tr><th></th><th></th><th></th><th></th><th></th><th></th><th></th><th></th><th></th><th></th><th></th><th></th></tr>
<tr><td>>MI0000482|hsa-mir-185&WithFlank&22|+|20033133|20033226@21.42.22(+1A)&[MIMAT0000455&hsa-miR-185-5p&offsets|0|0(+1A);m-145&22|+|20033153|20033174&offsets|0|0(+1A)]|TGGAGAGAAAGGCAGTTCCTGAA</td><td>>MI0000482|hsa-mir-185&WithFlank&22|+|20033133|20033226</td><td>21</td><td>42</td><td>22</td><td>(+1A)</td><td>isomiR</td><td>NO</td><td>0.00</td><td>27.15</td><td>0.00</td><td>0.00</td></tr>
<tr><td>>MI0000482|hsa-mir-185&WithFlank&22|+|20033133|20033226@21.43.23&[MIMAT0000455&hsa-miR-185-5p&offsets|0|+1;m-145&22|+|20033153|20033174&offsets|0|+1]|TGGAGAGAAAGGCAGTTCCTGAT</td><td>>MI0000482|hsa-mir-185&WithFlank&22|+|20033133|20033226</td><td>21</td><td>43</td><td>23</td><td></td><td>isomiR</td><td>NO</td><td>0.00</td><td>30.76</td><td>0.00</td><td>0.00</td></tr>
<tr><td>>MI0000483|hsa-mir-186&WithFlank&1|-|71067625|71067722@21.41.21&[MIMAT0000456&hsa-miR-186-5p&offsets|0|-1;m-84&1|-|71067681|71067702&offsets|0|-1]|CAAAGAATTCTCCTTTTGGGC</td><td>>MI0000483|hsa-mir-186&WithFlank&1|-|71067625|71067722</td><td>21</td><td>41</td><td>21</td><td></td><td>isomiR</td><td>NO</td><td>0.00</td><td>27.22</td><td>0.00</td><td>0.00</td></tr>
<tr><td>>MI0000483|hsa-mir-186&WithFlank&1|-|71067625|71067722@21.42.22&[MIMAT0000456&hsa-miR-186-5p&offsets|0|0;m-84&1|-|71067681|71067702&offsets|0|0]|CAAAGAATTCTCCTTTTGGGCT</td><td>>MI0000483|hsa-mir-186&WithFlank&1|-|71067625|71067722</td><td>21</td><td>42</td><td>22</td><td></td><td>isomiR</td><td>NO</td><td>99.93</td><td>122.75</td><td>40.09</td><td>30.20</td></tr>
<tr><td>>MI0000483|hsa-mir-186&WithFlank&1|-|71067625|71067722@21.43.23&[MIMAT0000456&hsa-miR-186-5p&offsets|0|+1;m-84&1|-|71067681|71067702&offsets|0|+1]|CAAAGAATTCTCCTTTTGGGCTT</td><td>>MI0000483|hsa-mir-186&WithFlank&1|-|71067625|71067722</td><td>21</td><td>43</td><td>23</td><td></td><td>isomiR</td><td>NO</td><td>43.47</td><td>32.24</td><td>12.30</td><td>0.00</td></tr>
<tr><td>>MI0000487|hsa-mir-193a&WithFlank&17|+|31559990|31560089@27.47.21&[MIMAT0004614&hsa-miR-193a-5p&offsets|0|-1;m-123&17|+|31560016|31560037&offsets|0|-1]|TGGGTCTTTGCGGGCGAGATG</td><td>>MI0000487|hsa-mir-193a&WithFlank&17|+|31559990|31560089</td><td>27</td><td>47</td><td>21</td><td></td><td>isomiR</td><td>NO</td><td>0.00</td><td>28.98</td><td>69.85</td><td>0.00</td></tr>
<tr><td>>MI0000487|hsa-mir-193a&WithFlank&17|+|31559990|31560089@27.48.22&[MIMAT0004614&hsa-miR-193a-5p&offsets|0|0;m-123&17|+|31560016|31560037&offsets|0|0]|TGGGTCTTTGCGGGCGAGATGA</td><td>>MI0000487|hsa-mir-193a&WithFlank&17|+|31559990|31560089</td><td>27</td><td>48</td><td>22</td><td></td><td>isomiR</td><td>NO</td><td>0.00</td><td>31.59</td><td>71.66</td><td>0.00</td></tr>
<tr><td>>MI0000488|hsa-mir-194-1&WithFlank&1|-|220118151|220118247@21.42.22;MI0000732|hsa-mir-194-2&WithFlank&11|-|64891349|64891445@21.42.22&[MIMAT0000460&hsa-miR-194-5p&offsets|0|0;m-131&1|-|220118206|220118227&offsets|0|0];[MIMAT0000460_1&hsa-miR-194-5p&offsets|0|0;m-126&11|-|64891404|64891425&offsets|0|0]|TGTAACAGCAACTCCATGTGGA</td><td>>MI0000488|hsa-mir-194-1&WithFlank&1|-|220118151|220118247</td><td>21</td><td>42</td><td>22</td><td></td><td>isomiR</td><td>NO</td><td>0.00</td><td>29.19</td><td>0.00</td><td>0.00</td></tr>
<tr><td>>MI0000542|hsa-mir-320a&WithFlank&8|-|22244960|22245043@47.69.23&[MIMAT0000510&hsa-miR-320a-3p&offsets|-1|0;m-60&8|-|22244975|22244996&offsets|-1|0]|GAAAAGCTGGGTTGAGAGGGCGA</td><td>>MI0000542|hsa-mir-320a&WithFlank&8|-|22244960|22245043</td><td>47</td><td>69</td><td>23</td><td></td><td>isomiR</td><td>NO</td><td>0.00</td><td>23.89</td><td>38.73</td><td>0.00</td></tr>
<tr><td>>MI0000542|hsa-mir-320a&WithFlank&8|-|22244960|22245043@48.68.21&[MIMAT0000510&hsa-miR-320a-3p&offsets|0|-1;m-60&8|-|22244975|22244996&offsets|0|-1]|AAAAGCTGGGTTGAGAGGGCG</td><td>>MI0000542|hsa-mir-320a&WithFlank&8|-|22244960|22245043</td><td>48</td><td>68</td><td>21</td><td></td><td>isomiR</td><td>NO</td><td>144.75</td><td>1196.46</td><td>1151.76</td><td>78.54</td></tr>
<tr><td>>MI0000542|hsa-mir-320a&WithFlank&8|-|22244960|22245043@48.68.21(+1C)&[MIMAT0000510&hsa-miR-320a-3p&offsets|0|-1(+1C);m-60&8|-|22244975|22244996&offsets|0|-1(+1C)]|AAAAGCTGGGTTGAGAGGGCGC</td><td>>MI0000542|hsa-mir-320a&WithFlank&8|-|22244960|22245043</td><td>48</td><td>68</td><td>21</td><td>(+1C)</td><td>isomiR</td><td>NO</td><td>9.42</td><td>313.91</td><td>319.00</td><td>0.00</td></tr>
<tr><td>>MI0000542|hsa-mir-320a&WithFlank&8|-|22244960|22245043@48.68.21(+1G)&[MIMAT0000510&hsa-miR-320a-3p&offsets|0|-1(+1G);m-60&8|-|22244975|22244996&offsets|0|-1(+1G)]|AAAAGCTGGGTTGAGAGGGCGG</td><td>>MI0000542|hsa-mir-320a&WithFlank&8|-|22244960|22245043</td><td>48</td><td>68</td><td>21</td><td>(+1G)</td><td>isomiR</td><td>NO</td><td>0.00</td><td>177.41</td><td>136.65</td><td>0.00</td></tr>
<tr><td>>MI0000542|hsa-mir-320a&WithFlank&8|-|22244960|22245043@48.68.21(+1U)&[MIMAT0000510&hsa-miR-320a-3p&offsets|0|-1(+1U);m-60&8|-|22244975|22244996&offsets|0|-1(+1U)]|AAAAGCTGGGTTGAGAGGGCGT</td><td>>MI0000542|hsa-mir-320a&WithFlank&8|-|22244960|22245043</td><td>48</td><td>68</td><td>21</td><td>(+1U)</td><td>isomiR</td><td>NO</td><td>134.89</td><td>1979.83</td><td>1704.82</td><td>28.26</td></tr>
<tr><td>>MI0000542|hsa-mir-320a&WithFlank&8|-|22244960|22245043@48.68.21(+2U)&[MIMAT0000510&hsa-miR-320a-3p&offsets|0|-1(+2U);m-60&8|-|22244975|22244996&offsets|0|-1(+2U)]|AAAAGCTGGGTTGAGAGGGCGTT</td><td>>MI0000542|hsa-mir-320a&WithFlank&8|-|22244960|22245043</td><td>48</td><td>68</td><td>21</td><td>(+2U)</td><td>isomiR</td><td>NO</td><td>94.22</td><td>899.59</td><td>357.57</td><td>31.23</td></tr>
<tr><td>>MI0000542|hsa-mir-320a&WithFlank&8|-|22244960|22245043@48.68.21(+3U)&[MIMAT0000510&hsa-miR-320a-3p&offsets|0|-1(+3U);m-60&8|-|22244975|22244996&offsets|0|-1(+3U)]|AAAAGCTGGGTTGAGAGGGCGTTT</td><td>>MI0000542|hsa-mir-320a&WithFlank&8|-|22244960|22245043</td><td>48</td><td>68</td><td>21</td><td>(+3U)</td><td>isomiR</td><td>NO</td><td>9.42</td><td>394.83</td><td>95.19</td><td>0.00</td></tr>
<tr><td>>MI0000542|hsa-mir-320a&WithFlank&8|-|22244960|22245043@48.68.21(+4U)&[MIMAT0000510&hsa-miR-320a-3p&offsets|0|-1(+4U);m-60&8|-|22244975|22244996&offsets|0|-1(+4U)]|AAAAGCTGGGTTGAGAGGGCGTTTT</td><td>>MI0000542|hsa-mir-320a&WithFlank&8|-|22244960|22245043</td><td>48</td><td>68</td><td>21</td><td>(+4U)</td><td>isomiR</td><td>NO</td><td>0.00</td><td>40.06</td><td>0.00</td><td>0.00</td></tr>
<tr><td>>MI0000542|hsa-mir-320a&WithFlank&8|-|22244960|22245043@48.69.22&[MIMAT0000510&hsa-miR-320a-3p&offsets|0|0;m-60&8|-|22244975|22244996&offsets|0|0]|AAAAGCTGGGTTGAGAGGGCGA</td><td>>MI0000542|hsa-mir-320a&WithFlank&8|-|22244960|22245043</td><td>48</td><td>69</td><td>22</td><td></td><td>isomiR</td><td>YES</td><td>367.82</td><td>3628.70</td><td>3549.77</td><td>133.75</td></tr>
<tr><td>>MI0000542|hsa-mir-320a&WithFlank&8|-|22244960|22245043@48.69.22(+1C)&[MIMAT0000510&hsa-miR-320a-3p&offsets|0|0(+1C);m-60&8|-|22244975|22244996&offsets|0|0(+1C)]|AAAAGCTGGGTTGAGAGGGCGAC</td><td>>MI0000542|hsa-mir-320a&WithFlank&8|-|22244960|22245043</td><td>48</td><td>69</td><td>22</td><td>(+1C)</td><td>isomiR</td><td>NO</td><td>32.44</td><td>331.06</td><td>113.85</td><td>0.00</td></tr>
<tr><td>>MI0000542|hsa-mir-320a&WithFlank&8|-|22244960|22245043@48.69.22(+1G)&[MIMAT0000510&hsa-miR-320a-3p&offsets|0|0(+1G);m-60&8|-|22244975|22244996&offsets|0|0(+1G)]|AAAAGCTGGGTTGAGAGGGCGAG</td><td>>MI0000542|hsa-mir-320a&WithFlank&8|-|22244960|22245043</td><td>48</td><td>69</td><td>22</td><td>(+1G)</td><td>isomiR</td><td>NO</td><td>0.00</td><td>177.55</td><td>66.26</td><td>0.00</td></tr>
<tr><td>>MI0000542|hsa-mir-320a&WithFlank&8|-|22244960|22245043@48.69.22(+1U)&[MIMAT0000510&hsa-miR-320a-3p&offsets|0|0(+1U);m-60&8|-|22244975|22244996&offsets|0|0(+1U)]|AAAAGCTGGGTTGAGAGGGCGAT</td><td>>MI0000542|hsa-mir-320a&WithFlank&8|-|22244960|22245043</td><td>48</td><td>69</td><td>22</td><td>(+1U)</td><td>isomiR</td><td>NO</td><td>234.56</td><td>2215.87</td><td>592.25</td><td>89.85</td></tr>
<tr><td>>MI0000542|hsa-mir-320a&WithFlank&8|-|22244960|22245043@48.69.22(+2U)&[MIMAT0000510&hsa-miR-320a-3p&offsets|0|0(+2U);m-60&8|-|22244975|22244996&offsets|0|0(+2U)]|AAAAGCTGGGTTGAGAGGGCGATT</td><td>>MI0000542|hsa-mir-320a&WithFlank&8|-|22244960|22245043</td><td>48</td><td>69</td><td>22</td><td>(+2U)</td><td>isomiR</td><td>NO</td><td>86.44</td><td>1247.63</td><td>137.55</td><td>0.00</td></tr>
<tr><td>>MI0000542|hsa-mir-320a&WithFlank&8|-|22244960|22245043@48.69.22(+3U)&[MIMAT0000510&hsa-miR-320a-3p&offsets|0|0(+3U);m-60&8|-|22244975|22244996&offsets|0|0(+3U)]|AAAAGCTGGGTTGAGAGGGCGATTT</td><td>>MI0000542|hsa-mir-320a&WithFlank&8|-|22244960|22245043</td><td>48</td><td>69</td><td>22</td><td>(+3U)</td><td>isomiR</td><td>NO</td><td>0.00</td><td>69.76</td><td>0.00</td><td>0.00</td></tr>
<tr><td>>MI0000542|hsa-mir-320a&WithFlank&8|-|22244960|22245043@48.70.23&[MIMAT0000510&hsa-miR-320a-3p&offsets|0|+1;m-60&8|-|22244975|22244996&offsets|0|+1]|AAAAGCTGGGTTGAGAGGGCGAA</td><td>>MI0000542|hsa-mir-320a&WithFlank&8|-|22244960|22245043</td><td>48</td><td>70</td><td>23</td><td></td><td>isomiR</td><td>NO</td><td>229.88</td><td>1881.42</td><td>751.04</td><td>106.12</td></tr>
<tr><td>>MI0000542|hsa-mir-320a&WithFlank&8|-|22244960|22245043@48.70.23(+1C)&[MIMAT0000510&hsa-miR-320a-3p&offsets|0|+1(+1C);m-60&8|-|22244975|22244996&offsets|0|+1(+1C)]|AAAAGCTGGGTTGAGAGGGCGAAC</td><td>>MI0000542|hsa-mir-320a&WithFlank&8|-|22244960|22245043</td><td>48</td><td>70</td><td>23</td><td>(+1C)</td><td>isomiR</td><td>NO</td><td>10.42</td><td>272.52</td><td>111.19</td><td>0.00</td></tr>
</table>

# BT-20

<table>
<tr><td></td><td></td><td></td><td></td><td></td><td></td><td></td><td></td><td></td><td></td><td></td><td></td></tr>
<tr><td>>MI0000542|hsa-mir-320a&WithFlank&8|-|22244960|22245043@48.70.23(+1G)&[MIMAT0000510&hsa-miR-320a-3p&offsets|0|+1(+1G);m-60&8|-|22244975|22244996&offsets|0|+1(+1G)]|AAAAGCTGGGTTGAGAGGGCGAAG</td><td>>MI0000542|hsa-mir-320a&WithFlank&8|-|22244960|22245043</td><td>48</td><td>70</td><td>23</td><td>(+1G)</td><td>isomiR</td><td>NO</td><td>0.00</td><td>175.62</td><td>94.58</td><td>0.00</td></tr>
<tr><td>>MI0000542|hsa-mir-320a&WithFlank&8|-|22244960|22245043@48.70.23(+1U)&[MIMAT0000510&hsa-miR-320a-3p&offsets|0|+1(+1U);m-60&8|-|22244975|22244996&offsets|0|+1(+1U)]|AAAAGCTGGGTTGAGAGGGCGAAT</td><td>>MI0000542|hsa-mir-320a&WithFlank&8|-|22244960|22245043</td><td>48</td><td>70</td><td>23</td><td>(+1U)</td><td>isomiR</td><td>NO</td><td>149.77</td><td>1648.65</td><td>558.53</td><td>87.07</td></tr>
<tr><td>>MI0000542|hsa-mir-320a&WithFlank&8|-|22244960|22245043@48.70.23(+2U)&[MIMAT0000510&hsa-miR-320a-3p&offsets|0|+1(+2U);m-60&8|-|22244975|22244996&offsets|0|+1(+2U)]|AAAAGCTGGGTTGAGAGGGCGAATT</td><td>>MI0000542|hsa-mir-320a&WithFlank&8|-|22244960|22245043</td><td>48</td><td>70</td><td>23</td><td>(+2U)</td><td>isomiR</td><td>NO</td><td>0.00</td><td>139.77</td><td>38.10</td><td>0.00</td></tr>
<tr><td>>MI0000542|hsa-mir-320a&WithFlank&8|-|22244960|22245043@48.71.24&[MIMAT0000510&hsa-miR-320a-3p&offsets|0|+2;m-60&8|-|22244975|22244996&offsets|0|+2]|AAAAGCTGGGTTGAGAGGGCGAAA</td><td>>MI0000542|hsa-mir-320a&WithFlank&8|-|22244960|22245043</td><td>48</td><td>71</td><td>24</td><td></td><td>isomiR</td><td>NO</td><td>83.52</td><td>830.48</td><td>218.98</td><td>23.46</td></tr>
<tr><td>>MI0000542|hsa-mir-320a&WithFlank&8|-|22244960|22245043@48.71.24(+1U)&[MIMAT0000510&hsa-miR-320a-3p&offsets|0|+2(+1U);m-60&8|-|22244975|22244996&offsets|0|+2(+1U)]|AAAAGCTGGGTTGAGAGGGCGAAAT</td><td>>MI0000542|hsa-mir-320a&WithFlank&8|-|22244960|22245043</td><td>48</td><td>71</td><td>24</td><td>(+1U)</td><td>isomiR</td><td>NO</td><td>0.00</td><td>157.89</td><td>100.78</td><td>0.00</td></tr>
<tr><td>>MI0000542|hsa-mir-320a&WithFlank&8|-|22244960|22245043@48.72.25&[MIMAT0000510&hsa-miR-320a-3p&offsets|0|+3;m-60&8|-|22244975|22244996&offsets|0|+3]|AAAAGCTGGGTTGAGAGGGCGAAAA</td><td>>MI0000542|hsa-mir-320a&WithFlank&8|-|22244960|22245043</td><td>48</td><td>72</td><td>25</td><td></td><td>isomiR</td><td>NO</td><td>0.00</td><td>70.21</td><td>11.65</td><td>0.00</td></tr>
<tr><td>>MI0000542|hsa-mir-320a&WithFlank&8|-|22244960|22245043@49.68.20&[MIMAT0000510&hsa-miR-320a-3p&offsets|+1|-1;m-60&8|-|22244975|22244996&offsets|+1|-1]|AAAGCTGGGTTGAGAGGGCG</td><td>>MI0000542|hsa-mir-320a&WithFlank&8|-|22244960|22245043</td><td>49</td><td>68</td><td>20</td><td></td><td>isomiR</td><td>NO</td><td>0.00</td><td>75.61</td><td>93.12</td><td>0.00</td></tr>
<tr><td>>MI0000542|hsa-mir-320a&WithFlank&8|-|22244960|22245043@49.68.20(+1U)&[MIMAT0000510&hsa-miR-320a-3p&offsets|+1|-1(+1U);m-60&8|-|22244975|22244996&offsets|+1|-1(+1U)]|AAAGCTGGGTTGAGAGGGCGT</td><td>>MI0000542|hsa-mir-320a&WithFlank&8|-|22244960|22245043</td><td>49</td><td>68</td><td>20</td><td>(+1U)</td><td>isomiR</td><td>NO</td><td>0.00</td><td>75.36</td><td>128.03</td><td>0.00</td></tr>
<tr><td>>MI0000542|hsa-mir-320a&WithFlank&8|-|22244960|22245043@49.68.20(+2U)&[MIMAT0000510&hsa-miR-320a-3p&offsets|+1|-1(+2U);m-60&8|-|22244975|22244996&offsets|+1|-1(+2U)]|AAAGCTGGGTTGAGAGGGCGTT</td><td>>MI0000542|hsa-mir-320a&WithFlank&8|-|22244960|22245043</td><td>49</td><td>68</td><td>20</td><td>(+2U)</td><td>isomiR</td><td>NO</td><td>0.00</td><td>34.32</td><td>39.18</td><td>0.00</td></tr>
<tr><td>>MI0000542|hsa-mir-320a&WithFlank&8|-|22244960|22245043@49.69.21&[MIMAT0000510&hsa-miR-320a-3p&offsets|+1|0;m-60&8|-|22244975|22244996&offsets|+1|0]|AAAGCTGGGTTGAGAGGGCGA</td><td>>MI0000542|hsa-mir-320a&WithFlank&8|-|22244960|22245043</td><td>49</td><td>69</td><td>21</td><td></td><td>isomiR</td><td>NO</td><td>10.83</td><td>127.55</td><td>257.98</td><td>0.00</td></tr>
<tr><td>>MI0000542|hsa-mir-320a&WithFlank&8|-|22244960|22245043@49.69.21(+1U)&[MIMAT0000510&hsa-miR-320a-3p&offsets|+1|0(+1U);m-60&8|-|22244975|22244996&offsets|+1|0(+1U)]|AAAGCTGGGTTGAGAGGGCGAT</td><td>>MI0000542|hsa-mir-320a&WithFlank&8|-|22244960|22245043</td><td>49</td><td>69</td><td>21</td><td>(+1U)</td><td>isomiR</td><td>NO</td><td>0.00</td><td>96.71</td><td>70.92</td><td>0.00</td></tr>
<tr><td>>MI0000542|hsa-mir-320a&WithFlank&8|-|22244960|22245043@49.69.21(+2U)&[MIMAT0000510&hsa-miR-320a-3p&offsets|+1|0(+2U);m-60&8|-|22244975|22244996&offsets|+1|0(+2U)]|AAAGCTGGGTTGAGAGGGCGATT</td><td>>MI0000542|hsa-mir-320a&WithFlank&8|-|22244960|22245043</td><td>49</td><td>69</td><td>21</td><td>(+2U)</td><td>isomiR</td><td>NO</td><td>0.00</td><td>32.60</td><td>0.00</td><td>0.00</td></tr>
<tr><td>>MI0000542|hsa-mir-320a&WithFlank&8|-|22244960|22245043@49.70.22&[MIMAT0000510&hsa-miR-320a-3p&offsets|+1|+1;m-60&8|-|22244975|22244996&offsets|+1|+1]|AAAGCTGGGTTGAGAGGGCGAA</td><td>>MI0000542|hsa-mir-320a&WithFlank&8|-|22244960|22245043</td><td>49</td><td>70</td><td>22</td><td></td><td>isomiR</td><td>NO</td><td>0.00</td><td>95.42</td><td>99.57</td><td>0.00</td></tr>
<tr><td>>MI0000542|hsa-mir-320a&WithFlank&8|-|22244960|22245043@49.70.22(+1U)&[MIMAT0000510&hsa-miR-320a-3p&offsets|+1|+1(+1U);m-60&8|-|22244975|22244996&offsets|+1|+1(+1U)]|AAAGCTGGGTTGAGAGGGCGAAT</td><td>>MI0000542|hsa-mir-320a&WithFlank&8|-|22244960|22245043</td><td>49</td><td>70</td><td>22</td><td>(+1U)</td><td>isomiR</td><td>NO</td><td>0.00</td><td>106.49</td><td>101.71</td><td>0.00</td></tr>
<tr><td>>MI0000542|hsa-mir-320a&WithFlank&8|-|22244960|22245043@49.71.23&[MIMAT0000510&hsa-miR-320a-3p&offsets|+1|+2;m-60&8|-|22244975|22244996&offsets|+1|+2]|AAAGCTGGGTTGAGAGGGCGAAA</td><td>>MI0000542|hsa-mir-320a&WithFlank&8|-|22244960|22245043</td><td>49</td><td>71</td><td>23</td><td></td><td>isomiR</td><td>NO</td><td>0.00</td><td>40.07</td><td>13.97</td><td>0.00</td></tr>
<tr><td>>MI0000542|hsa-mir-320a&WithFlank&8|-|22244960|22245043@50.69.20&[MIMAT0000510&hsa-miR-320a-3p&offsets|+2|0;m-60&8|-|22244975|22244996&offsets|+2|0]|AAGCTGGGTTGAGAGGGCGA</td><td>>MI0000542|hsa-mir-320a&WithFlank&8|-|22244960|22245043</td><td>50</td><td>69</td><td>20</td><td></td><td>isomiR</td><td>NO</td><td>0.00</td><td>0.00</td><td>15.53</td><td>0.00</td></tr>
<tr><td>>MI0000542|hsa-mir-320a&WithFlank&8|-|22244960|22245043@50.70.21&[MIMAT0000510&hsa-miR-320a-3p&offsets|+2|+1;m-60&8|-|22244975|22244996&offsets|+2|+1]|AAGCTGGGTTGAGAGGGCGAA</td><td>>MI0000542|hsa-mir-320a&WithFlank&8|-|22244960|22245043</td><td>50</td><td>70</td><td>21</td><td></td><td>isomiR</td><td>NO</td><td>0.00</td><td>0.00</td><td>15.51</td><td>0.00</td></tr>
<tr><td>>MI0000542|hsa-mir-320a&WithFlank&8|-|22244960|22245043@50.70.21(+1U)&[MIMAT0000510&hsa-miR-320a-3p&offsets|+2|+1(+1U);m-60&8|-|22244975|22244996&offsets|+2|+1(+1U)]|AAGCTGGGTTGAGAGGGCGAAT</td><td>>MI0000542|hsa-mir-320a&WithFlank&8|-|22244960|22245043</td><td>50</td><td>70</td><td>21</td><td>(+1U)</td><td>isomiR</td><td>NO</td><td>0.00</td><td>23.16</td><td>55.33</td><td>0.00</td></tr>
<tr><td>>MI0000542|hsa-mir-320a&WithFlank&8|-|22244960|22245043@50.71.22&[MIMAT0000510&hsa-miR-320a-3p&offsets|+2|+2;m-60&8|-|22244975|22244996&offsets|+2|+2]|AAGCTGGGTTGAGAGGGCGAAA</td><td>>MI0000542|hsa-mir-320a&WithFlank&8|-|22244960|22245043</td><td>50</td><td>71</td><td>22</td><td></td><td>isomiR</td><td>NO</td><td>0.00</td><td>9.52</td><td>11.81</td><td>0.00</td></tr>
<tr><td>>MI0000650|hsa-mir-200c&WithFlank&12|+|6963693|6963772@49.70.22&[MIMAT0000617&hsa-miR-200c-3p&offsets|-1|-2;m-54&12|+|6963742|6963764&offsets|-1|-2]|CTAATACTGCCGGGTAATGATG</td><td>>MI0000650|hsa-mir-200c&WithFlank&12|+|6963693|6963772</td><td>49</td><td>70</td><td>22</td><td></td><td>isomiR</td><td>NO</td><td>0.00</td><td>34.19</td><td>0.00</td><td>0.00</td></tr>
<tr><td>>MI0000650|hsa-mir-200c&WithFlank&12|+|6963693|6963772@49.71.23&[MIMAT0000617&hsa-miR-200c-3p&offsets|-1|-1;m-54&12|+|6963742|6963764&offsets|-1|-1]|CTAATACTGCCGGGTAATGATGG</td><td>>MI0000650|hsa-mir-200c&WithFlank&12|+|6963693|6963772</td><td>49</td><td>71</td><td>23</td><td></td><td>isomiR</td><td>NO</td><td>10.80</td><td>42.96</td><td>0.00</td><td>0.00</td></tr>
<tr><td>>MI0000650|hsa-mir-200c&WithFlank&12|+|6963693|6963772@49.72.24&[MIMAT0000617&hsa-miR-200c-3p&offsets|-1|0;m-54&12|+|6963742|6963764&offsets|-1|0]|CTAATACTGCCGGGTAATGATGGA</td><td>>MI0000650|hsa-mir-200c&WithFlank&12|+|6963693|6963772</td><td>49</td><td>72</td><td>24</td><td></td><td>isomiR</td><td>NO</td><td>9.66</td><td>39.43</td><td>0.00</td><td>0.00</td></tr>
<tr><td>>MI0000650|hsa-mir-200c&WithFlank&12|+|6963693|6963772@50.69.20&[MIMAT0000617&hsa-miR-200c-3p&offsets|0|-3;m-54&12|+|6963742|6963764&offsets|0|-3]|TAATACTGCCGGGTAATGAT</td><td>>MI0000650|hsa-mir-200c&WithFlank&12|+|6963693|6963772</td><td>50</td><td>69</td><td>20</td><td></td><td>isomiR</td><td>NO</td><td>90.12</td><td>150.46</td><td>113.30</td><td>69.67</td></tr>
</table>

# BT-20

<table>
<tr><td>>MI0000650|hsa-mir-200c&WithFlank&12|+|6963693|6963772@50.69.20(+1A)&[MIMAT0000617&hsa-miR-200c-3p&offsets|0|-3(+1A);m-54&12|+|6963742|6963764&offsets|0|-3(+1A)]|TAATACTGCCGGGTAATGATA</td><td>>MI0000650|hsa-mir-200c&WithFlank&12|+|6963693|6963772</td><td>50</td><td>69</td><td>20</td><td>(+1A)</td><td>isomiR</td><td>NO</td><td>0.00</td><td>70.48</td><td>14.71</td><td>0.00</td></tr>
<tr><td>>MI0000650|hsa-mir-200c&WithFlank&12|+|6963693|6963772@50.70.21&[MIMAT0000617&hsa-miR-200c-3p&offsets|0|-2;m-54&12|+|6963742|6963764&offsets|0|-2]|TAATACTGCCGGGTAATGATG</td><td>>MI0000650|hsa-mir-200c&WithFlank&12|+|6963693|6963772</td><td>50</td><td>70</td><td>21</td><td></td><td>isomiR</td><td>NO</td><td>139.39</td><td>360.44</td><td>286.32</td><td>104.01</td></tr>
<tr><td>>MI0000650|hsa-mir-200c&WithFlank&12|+|6963693|6963772@50.70.21(+1A)&[MIMAT0000617&hsa-miR-200c-3p&offsets|0|-2(+1A);m-54&12|+|6963742|6963764&offsets|0|-2(+1A)]|TAATACTGCCGGGTAATGATGA</td><td>>MI0000650|hsa-mir-200c&WithFlank&12|+|6963693|6963772</td><td>50</td><td>70</td><td>21</td><td>(+1A)</td><td>isomiR</td><td>NO</td><td>135.67</td><td>355.00</td><td>169.40</td><td>36.10</td></tr>
<tr><td>>MI0000650|hsa-mir-200c&WithFlank&12|+|6963693|6963772@50.70.21(+1U)&[MIMAT0000617&hsa-miR-200c-3p&offsets|0|-2(+1U);m-54&12|+|6963742|6963764&offsets|0|-2(+1U)]|TAATACTGCCGGGTAATGATGT</td><td>>MI0000650|hsa-mir-200c&WithFlank&12|+|6963693|6963772</td><td>50</td><td>70</td><td>21</td><td>(+1U)</td><td>isomiR</td><td>NO</td><td>24.55</td><td>45.32</td><td>14.67</td><td>0.00</td></tr>
<tr><td>>MI0000650|hsa-mir-200c&WithFlank&12|+|6963693|6963772@50.71.22&[MIMAT0000617&hsa-miR-200c-3p&offsets|0|-1;m-54&12|+|6963742|6963764&offsets|0|-1]|TAATACTGCCGGGTAATGATGG</td><td>>MI0000650|hsa-mir-200c&WithFlank&12|+|6963693|6963772</td><td>50</td><td>71</td><td>22</td><td></td><td>isomiR</td><td>NO</td><td>607.19</td><td>1615.63</td><td>988.06</td><td>270.62</td></tr>
<tr><td>>MI0000650|hsa-mir-200c&WithFlank&12|+|6963693|6963772@50.71.22(+1C)&[MIMAT0000617&hsa-miR-200c-3p&offsets|0|-1(+1C);m-54&12|+|6963742|6963764&offsets|0|-1(+1C)]|TAATACTGCCGGGTAATGATGGC</td><td>>MI0000650|hsa-mir-200c&WithFlank&12|+|6963693|6963772</td><td>50</td><td>71</td><td>22</td><td>(+1C)</td><td>isomiR</td><td>NO</td><td>10.16</td><td>23.83</td><td>0.00</td><td>0.00</td></tr>
<tr><td>>MI0000650|hsa-mir-200c&WithFlank&12|+|6963693|6963772@50.71.22(+1G)&[MIMAT0000617&hsa-miR-200c-3p&offsets|0|-1(+1G);m-54&12|+|6963742|6963764&offsets|0|-1(+1G)]|TAATACTGCCGGGTAATGATGGG</td><td>>MI0000650|hsa-mir-200c&WithFlank&12|+|6963693|6963772</td><td>50</td><td>71</td><td>22</td><td>(+1G)</td><td>isomiR</td><td>NO</td><td>193.07</td><td>307.41</td><td>108.47</td><td>67.79</td></tr>
<tr><td>>MI0000650|hsa-mir-200c&WithFlank&12|+|6963693|6963772@50.71.22(+1U)&[MIMAT0000617&hsa-miR-200c-3p&offsets|0|-1(+1U);m-54&12|+|6963742|6963764&offsets|0|-1(+1U)]|TAATACTGCCGGGTAATGATGGT</td><td>>MI0000650|hsa-mir-200c&WithFlank&12|+|6963693|6963772</td><td>50</td><td>71</td><td>22</td><td>(+1U)</td><td>isomiR</td><td>NO</td><td>160.36</td><td>264.31</td><td>137.95</td><td>74.47</td></tr>
<tr><td>>MI0000650|hsa-mir-200c&WithFlank&12|+|6963693|6963772@50.72.23&[MIMAT0000617&hsa-miR-200c-3p&offsets|0|0;m-54&12|+|6963742|6963764&offsets|0|0]|TAATACTGCCGGGTAATGATGGA</td><td>>MI0000650|hsa-mir-200c&WithFlank&12|+|6963693|6963772</td><td>50</td><td>72</td><td>23</td><td></td><td>isomiR</td><td>YES</td><td>3212.69</td><td>8482.85</td><td>3419.72</td><td>2194.32</td></tr>
<tr><td>>MI0000650|hsa-mir-200c&WithFlank&12|+|6963693|6963772@50.72.23(+1A)&[MIMAT0000617&hsa-miR-200c-3p&offsets|0|0(+1A);m-54&12|+|6963742|6963764&offsets|0|0(+1A)]|TAATACTGCCGGGTAATGATGGAA</td><td>>MI0000650|hsa-mir-200c&WithFlank&12|+|6963693|6963772</td><td>50</td><td>72</td><td>23</td><td>(+1A)</td><td>isomiR</td><td>NO</td><td>452.41</td><td>1316.23</td><td>249.31</td><td>363.66</td></tr>
<tr><td>>MI0000650|hsa-mir-200c&WithFlank&12|+|6963693|6963772@50.72.23(+1C)&[MIMAT0000617&hsa-miR-200c-3p&offsets|0|0(+1C);m-54&12|+|6963742|6963764&offsets|0|0(+1C)]|TAATACTGCCGGGTAATGATGGAC</td><td>>MI0000650|hsa-mir-200c&WithFlank&12|+|6963693|6963772</td><td>50</td><td>72</td><td>23</td><td>(+1C)</td><td>isomiR</td><td>NO</td><td>68.69</td><td>160.47</td><td>10.65</td><td>35.74</td></tr>
<tr><td>>MI0000650|hsa-mir-200c&WithFlank&12|+|6963693|6963772@50.72.23(+1U)&[MIMAT0000617&hsa-miR-200c-3p&offsets|0|0(+1U);m-54&12|+|6963742|6963764&offsets|0|0(+1U)]|TAATACTGCCGGGTAATGATGGAT</td><td>>MI0000650|hsa-mir-200c&WithFlank&12|+|6963693|6963772</td><td>50</td><td>72</td><td>23</td><td>(+1U)</td><td>isomiR</td><td>NO</td><td>302.43</td><td>889.01</td><td>153.48</td><td>241.95</td></tr>
<tr><td>>MI0000650|hsa-mir-200c&WithFlank&12|+|6963693|6963772@50.72.23(+2A)&[MIMAT0000617&hsa-miR-200c-3p&offsets|0|0(+2A);m-54&12|+|6963742|6963764&offsets|0|0(+2A)]|TAATACTGCCGGGTAATGATGGAAA</td><td>>MI0000650|hsa-mir-200c&WithFlank&12|+|6963693|6963772</td><td>50</td><td>72</td><td>23</td><td>(+2A)</td><td>isomiR</td><td>NO</td><td>32.21</td><td>125.18</td><td>0.00</td><td>29.24</td></tr>
<tr><td>>MI0000650|hsa-mir-200c&WithFlank&12|+|6963693|6963772@50.72.23(+2U)&[MIMAT0000617&hsa-miR-200c-3p&offsets|0|0(+2U);m-54&12|+|6963742|6963764&offsets|0|0(+2U)]|TAATACTGCCGGGTAATGATGGATT</td><td>>MI0000650|hsa-mir-200c&WithFlank&12|+|6963693|6963772</td><td>50</td><td>72</td><td>23</td><td>(+2U)</td><td>isomiR</td><td>NO</td><td>0.00</td><td>34.58</td><td>0.00</td><td>0.00</td></tr>
<tr><td>>MI0000650|hsa-mir-200c&WithFlank&12|+|6963693|6963772@50.73.24&[MIMAT0000617&hsa-miR-200c-3p&offsets|0|+1;m-54&12|+|6963742|6963764&offsets|0|+1]|TAATACTGCCGGGTAATGATGGAG</td><td>>MI0000650|hsa-mir-200c&WithFlank&12|+|6963693|6963772</td><td>50</td><td>73</td><td>24</td><td></td><td>isomiR</td><td>NO</td><td>10.83</td><td>70.44</td><td>0.00</td><td>0.00</td></tr>
<tr><td>>MI0000734|hsa-mir-106b&WithFlank&7|-|100093987|100094080@18.38.21&[MIMAT0000680&hsa-miR-106b-5p&offsets|0|0;m-103&7|-|100094043|100094063&offsets|0|0]|TAAAGTGCTGACAGTGCAGAT</td><td>>MI0000734|hsa-mir-106b&WithFlank&7|-|100093987|100094080</td><td>18</td><td>38</td><td>21</td><td></td><td>isomiR</td><td>NO</td><td>74.23</td><td>0.00</td><td>0.00</td><td>0.00</td></tr>
<tr><td>>MI0000734|hsa-mir-106b&WithFlank&7|-|100093987|100094080@18.39.22&[MIMAT0000680&hsa-miR-106b-5p&offsets|0|+1;m-103&7|-|100094043|100094063&offsets|0|+1]|TAAAGTGCTGACAGTGCAGATA</td><td>>MI0000734|hsa-mir-106b&WithFlank&7|-|100093987|100094080</td><td>18</td><td>39</td><td>22</td><td></td><td>isomiR</td><td>NO</td><td>107.96</td><td>31.49</td><td>0.00</td><td>0.00</td></tr>
<tr><td>>MI0000734|hsa-mir-106b&WithFlank&7|-|100093987|100094080@56.77.22&[MIMAT0004672&hsa-miR-106b-3p&offsets|-2|-2;m-87&7|-|100094003|100094023&offsets|-2|-1]|TACCGCACTGTGGGTACTTGCT</td><td>>MI0000734|hsa-mir-106b&WithFlank&7|-|100093987|100094080</td><td>56</td><td>77</td><td>22</td><td></td><td>isomiR</td><td>NO</td><td>30.18</td><td>131.31</td><td>14.86</td><td>0.00</td></tr>
<tr><td>>MI0000734|hsa-mir-106b&WithFlank&7|-|100093987|100094080@58.76.19&[MIMAT0004672&hsa-miR-106b-3p&offsets|0|-3;m-87&7|-|100094003|100094023&offsets|0|-2]|CCGCACTGTGGGTACTTGC</td><td>>MI0000734|hsa-mir-106b&WithFlank&7|-|100093987|100094080</td><td>58</td><td>76</td><td>19</td><td></td><td>isomiR</td><td>NO</td><td>0.00</td><td>26.42</td><td>11.46</td><td>0.00</td></tr>
<tr><td>>MI0000734|hsa-mir-106b&WithFlank&7|-|100093987|100094080@58.77.20&[MIMAT0004672&hsa-miR-106b-3p&offsets|0|-2;m-87&7|-|100094003|100094023&offsets|0|-1]|CCGCACTGTGGGTACTTGCT</td><td>>MI0000734|hsa-mir-106b&WithFlank&7|-|100093987|100094080</td><td>58</td><td>77</td><td>20</td><td></td><td>isomiR</td><td>NO</td><td>32.41</td><td>156.90</td><td>78.57</td><td>10.41</td></tr>
<tr><td>>MI0000734|hsa-mir-106b&WithFlank&7|-|100093987|100094080@58.77.20(+1A)&[MIMAT0004672&hsa-miR-106b-3p&offsets|0|-2(+1A);m-87&7|-|100094003|100094023&offsets|0|-1(+1A)]|CCGCACTGTGGGTACTTGCTA</td><td>>MI0000734|hsa-mir-106b&WithFlank&7|-|100093987|100094080</td><td>58</td><td>77</td><td>20</td><td>(+1A)</td><td>isomiR</td><td>NO</td><td>0.00</td><td>75.08</td><td>0.00</td><td>0.00</td></tr>
<tr><td>>MI0000734|hsa-mir-106b&WithFlank&7|-|100093987|100094080@58.78.21&[MIMAT0004672&hsa-miR-106b-3p&offsets|0|-1;m-87&7|-|100094003|100094023&offsets|0|0]|CCGCACTGTGGGTACTTGCTG</td><td>>MI0000734|hsa-mir-106b&WithFlank&7|-|100093987|100094080</td><td>58</td><td>78</td><td>21</td><td></td><td>isomiR</td><td>NO</td><td>85.41</td><td>389.70</td><td>169.86</td><td>39.59</td></tr>
</table>

# BT-20

| | | | | | | | | | | | |
|---|---|---|---|---|---|---|---|---|---|---|---|
| >MI0000734\|hsa-mir-106b&WithFlank&7\|-\|100093987\|100094080@58.78.21(+1A)&[MIMAT0004672&hsa-miR-106b-3p&offsets\|0\|-1(+1A);m-87&7\|-\|100094003\|100094023&offsets\|0\|0(+1A)]\|CCGCACTGTGGGTACTTGCTGA | >MI0000734\|hsa-mir-106b&WithFlank&7\|-\|100093987\|100094080 | 58 | 78 | 21 | (+1A) | isomiR | NO | 0.00 | 81.41 | 12.19 | 0.00 |
| >MI0000734\|hsa-mir-106b&WithFlank&7\|-\|100093987\|100094080@58.78.21(+1U)&[MIMAT0004672&hsa-miR-106b-3p&offsets\|0\|-1(+1U);m-87&7\|-\|100094003\|100094023&offsets\|0\|0(+1U)]\|CCGCACTGTGGGTACTTGCTGT | >MI0000734\|hsa-mir-106b&WithFlank&7\|-\|100093987\|100094080 | 58 | 78 | 21 | (+1U) | isomiR | NO | 66.63 | 240.93 | 122.55 | 25.51 |
| >MI0000734\|hsa-mir-106b&WithFlank&7\|-\|100093987\|100094080@58.79.22&[MIMAT0004672&hsa-miR-106b-3p&offsets\|0\|0;m-87&7\|-\|100094003\|100094023&offsets\|0\|+1]\|CCGCACTGTGGGTACTTGCTGC | >MI0000734\|hsa-mir-106b&WithFlank&7\|-\|100093987\|100094080 | 58 | 79 | 22 | | isomiR | NO | 165.37 | 667.14 | 349.02 | 105.83 |
| >MI0000734\|hsa-mir-106b&WithFlank&7\|-\|100093987\|100094080@58.79.22(+1A)&[MIMAT0004672&hsa-miR-106b-3p&offsets\|0\|0(+1A);m-87&7\|-\|100094003\|100094023&offsets\|0\|+1(+1A)]\|CCGCACTGTGGGTACTTGCTGCA | >MI0000734\|hsa-mir-106b&WithFlank&7\|-\|100093987\|100094080 | 58 | 79 | 22 | (+1A) | isomiR | NO | 0.00 | 25.47 | 0.00 | 0.00 |
| >MI0000736\|hsa-mir-30c-1&WithFlank&1\|+\|40757278\|40757378@23.42.20;MI0000254\|hsa-mir-30c-2&WithFlank&6\|-\|71376954\|71377037@13.32.20&[MIMAT0000244&hsa-miR-30c-5p&offsets\|0\|-3;m-57&1\|+\|40757300\|40757323&offsets\|0\|-4];[MIMAT0000244_1&hsa-miR-30c-5p&offsets\|0\|-3;m-56&6\|-\|71377002\|71377025&offsets\|0\|-4]\|TGTAAACATCCTACACTCTC | >MI0000736\|hsa-mir-30c-1&WithFlank&1\|+\|40757278\|40757378 | 23 | 42 | 20 | | isomiR | NO | 23.60 | 30.77 | 0.00 | 0.00 |
| >MI0000736\|hsa-mir-30c-1&WithFlank&1\|+\|40757278\|40757378@23.44.22(+1A);MI0000254\|hsa-mir-30c-2&WithFlank&6\|-\|71376954\|71377037@13.34.22(+1A)&[MIMAT0000244&hsa-miR-30c-5p&offsets\|0\|-1(+1A);m-57&1\|+\|40757300\|40757323&offsets\|0\|-2(+1A)];[MIMAT0000244_1&hsa-miR-30c-5p&offsets\|0\|-1(+1A);m-56&6\|-\|71377002\|71377025&offsets\|0\|-2(+1A)]\|TGTAAACATCCTACACTCTCAGA | >MI0000736\|hsa-mir-30c-1&WithFlank&1\|+\|40757278\|40757378 | 23 | 44 | 22 | (+1A) | isomiR | NO | 10.41 | 72.05 | 0.00 | 0.00 |
| >MI0000736\|hsa-mir-30c-1&WithFlank&1\|+\|40757278\|40757378@23.44.22(+1U);MI0000254\|hsa-mir-30c-2&WithFlank&6\|-\|71376954\|71377037@13.34.22(+1U)&[MIMAT0000244&hsa-miR-30c-5p&offsets\|0\|-1(+1U);m-57&1\|+\|40757300\|40757323&offsets\|0\|-2(+1U)];[MIMAT0000244_1&hsa-miR-30c-5p&offsets\|0\|-1(+1U);m-56&6\|-\|71377002\|71377025&offsets\|0\|-2(+1U)]\|TGTAAACATCCTACACTCTCAGT | >MI0000736\|hsa-mir-30c-1&WithFlank&1\|+\|40757278\|40757378 | 23 | 44 | 22 | (+1U) | isomiR | NO | 113.33 | 216.42 | 11.17 | 29.99 |
| >MI0000736\|hsa-mir-30c-1&WithFlank&1\|+\|40757278\|40757378@23.45.23(+1A);MI0000254\|hsa-mir-30c-2&WithFlank&6\|-\|71376954\|71377037@13.35.23(+1A)&[MIMAT0000244&hsa-miR-30c-5p&offsets\|0\|0(+1A);m-57&1\|+\|40757300\|40757323&offsets\|0\|-1(+1A)];[MIMAT0000244_1&hsa-miR-30c-5p&offsets\|0\|0(+1A);m-56&6\|-\|71377002\|71377025&offsets\|0\|-1(+1A)]\|TGTAAACATCCTACACTCTCAGCA | >MI0000736\|hsa-mir-30c-1&WithFlank&1\|+\|40757278\|40757378 | 23 | 45 | 23 | (+1A) | isomiR | NO | 73.93 | 119.04 | 0.00 | 0.00 |
| >MI0000736\|hsa-mir-30c-1&WithFlank&1\|+\|40757278\|40757378@23.45.23(+1C);MI0000254\|hsa-mir-30c-2&WithFlank&6\|-\|71376954\|71377037@13.35.23(+1C)&[MIMAT0000244&hsa-miR-30c-5p&offsets\|0\|0(+1C);m-57&1\|+\|40757300\|40757323&offsets\|0\|-1(+1C)];[MIMAT0000244_1&hsa-miR-30c-5p&offsets\|0\|0(+1C);m-56&6\|-\|71377002\|71377025&offsets\|0\|-1(+1C)]\|TGTAAACATCCTACACTCTCAGCC | >MI0000736\|hsa-mir-30c-1&WithFlank&1\|+\|40757278\|40757378 | 23 | 45 | 23 | (+1C) | isomiR | NO | 97.52 | 115.29 | 0.00 | 24.13 |
| >MI0000736\|hsa-mir-30c-1&WithFlank&1\|+\|40757278\|40757378@23.45.23(+1G);MI0000254\|hsa-mir-30c-2&WithFlank&6\|-\|71376954\|71377037@13.35.23(+1G)&[MIMAT0000244&hsa-miR-30c-5p&offsets\|0\|0(+1G);m-57&1\|+\|40757300\|40757323&offsets\|0\|-1(+1G)];[MIMAT0000244_1&hsa-miR-30c-5p&offsets\|0\|0(+1G);m-56&6\|-\|71377002\|71377025&offsets\|0\|-1(+1G)]\|TGTAAACATCCTACACTCTCAGCG | >MI0000736\|hsa-mir-30c-1&WithFlank&1\|+\|40757278\|40757378 | 23 | 45 | 23 | (+1G) | isomiR | NO | 24.02 | 23.86 | 0.00 | 0.00 |
| >MI0000736\|hsa-mir-30c-1&WithFlank&1\|+\|40757278\|40757378@23.45.23;MI0000254\|hsa-mir-30c-2&WithFlank&6\|-\|71376954\|71377037@13.35.23&[MIMAT0000244&hsa-miR-30c-5p&offsets\|0\|0;m-57&1\|+\|40757300\|40757323&offsets\|0\|-1];[MIMAT0000244_1&hsa-miR-30c-5p&offsets\|0\|0;m-56&6\|-\|71377002\|71377025&offsets\|0\|-1]\|TGTAAACATCCTACACTCTCAGC | >MI0000736\|hsa-mir-30c-1&WithFlank&1\|+\|40757278\|40757378 | 23 | 45 | 23 | | isomiR | NO | 311.10 | 651.16 | 103.33 | 142.64 |
| >MI0000736\|hsa-mir-30c-1&WithFlank&1\|+\|40757278\|40757378@23.46.24;MI0000254\|hsa-mir-30c-2&WithFlank&6\|-\|71376954\|71377037@13.36.24&[MIMAT0000244&hsa-miR-30c-5p&offsets\|0\|+1;m-57&1\|+\|40757300\|40757323&offsets\|0\|0];[MIMAT0000244_1&hsa-miR-30c-5p&offsets\|0\|+1;m-56&6\|-\|71377002\|71377025&offsets\|0\|0]\|TGTAAACATCCTACACTCTCAGCT | >MI0000736\|hsa-mir-30c-1&WithFlank&1\|+\|40757278\|40757378 | 23 | 46 | 24 | | isomiR | NO | 421.41 | 655.52 | 137.22 | 157.37 |
| >MI0000746\|hsa-mir-99b&WithFlank&19\|+\|51692606\|51692687@13.31.19&[MIMAT0000689&hsa-miR-99b-5p&offsets\|0\|-3;m-10&19\|+\|51692618\|51692639&offsets\|0\|-3]\|CACCCGTAGAACCGACCTT | >MI0000746\|hsa-mir-99b&WithFlank&19\|+\|51692606\|51692687 | 13 | 31 | 19 | | isomiR | NO | 0.00 | 23.27 | 0.00 | 0.00 |
| >MI0000746\|hsa-mir-99b&WithFlank&19\|+\|51692606\|51692687@13.31.19(+1A)&[MIMAT0000689&hsa-miR-99b-5p&offsets\|0\|-3(+1A);m-10&19\|+\|51692618\|51692639&offsets\|0\|-3(+1A)]\|CACCCGTAGAACCGACCTTA | >MI0000746\|hsa-mir-99b&WithFlank&19\|+\|51692606\|51692687 | 13 | 31 | 19 | (+1A) | isomiR | NO | 0.00 | 38.17 | 0.00 | 0.00 |
| >MI0000746\|hsa-mir-99b&WithFlank&19\|+\|51692606\|51692687@13.32.20&[MIMAT0000689&hsa-miR-99b-5p&offsets\|0\|-2;m-10&19\|+\|51692618\|51692639&offsets\|0\|-2]\|CACCCGTAGAACCGACCTTG | >MI0000746\|hsa-mir-99b&WithFlank&19\|+\|51692606\|51692687 | 13 | 32 | 20 | | isomiR | NO | 84.69 | 318.50 | 174.70 | 37.88 |
| >MI0000746\|hsa-mir-99b&WithFlank&19\|+\|51692606\|51692687@13.32.20(+1A)&[MIMAT0000689&hsa-miR-99b-5p&offsets\|0\|-2(+1A);m-10&19\|+\|51692618\|51692639&offsets\|0\|-2(+1A)]\|CACCCGTAGAACCGACCTTGA | >MI0000746\|hsa-mir-99b&WithFlank&19\|+\|51692606\|51692687 | 13 | 32 | 20 | (+1A) | isomiR | NO | 10.69 | 217.88 | 15.62 | 0.00 |
| >MI0000746\|hsa-mir-99b&WithFlank&19\|+\|51692606\|51692687@13.32.20(+1G)&[MIMAT0000689&hsa-miR-99b-5p&offsets\|0\|-2(+1G);m-10&19\|+\|51692618\|51692639&offsets\|0\|-2(+1G)]\|CACCCGTAGAACCGACCTTGG | >MI0000746\|hsa-mir-99b&WithFlank&19\|+\|51692606\|51692687 | 13 | 32 | 20 | (+1G) | isomiR | NO | 0.00 | 71.72 | 0.00 | 0.00 |
| >MI0000746\|hsa-mir-99b&WithFlank&19\|+\|51692606\|51692687@13.32.20(+1U)&[MIMAT0000689&hsa-miR-99b-5p&offsets\|0\|-2(+1U);m-10&19\|+\|51692618\|51692639&offsets\|0\|-2(+1U)]\|CACCCGTAGAACCGACCTTGT | >MI0000746\|hsa-mir-99b&WithFlank&19\|+\|51692606\|51692687 | 13 | 32 | 20 | (+1U) | isomiR | NO | 314.98 | 2098.86 | 464.10 | 141.73 |

# BT-20

<table>
<tr><td></td><td></td><td></td><td></td><td></td><td></td><td></td><td></td><td></td><td></td><td></td><td></td></tr>
<tr><td>>MI0000746|hsa-mir-99b&WithFlank&19|+|51692606|51692687@13.32.20(+2U)&[MIMAT0000689&hsa-miR-99b-5p&offsets|0|-2(+2U);m-10&19|+|51692618|51692639&offsets|0|-2(+2U)]|CACCCGTAGAACCGACCTTGTT</td><td>>MI0000746|hsa-mir-99b&WithFlank&19|+|51692606|51692687</td><td>13</td><td>32</td><td>20</td><td>(+2U)</td><td>isomiR</td><td>NO</td><td>28.13</td><td>182.16</td><td>13.83</td><td>0.00</td></tr>
<tr><td>>MI0000746|hsa-mir-99b&WithFlank&19|+|51692606|51692687@13.33.21&[MIMAT0000689&hsa-miR-99b-5p&offsets|0|-1;m-10&19|+|51692618|51692639&offsets|0|-1]|CACCCGTAGAACCGACCTTGC</td><td>>MI0000746|hsa-mir-99b&WithFlank&19|+|51692606|51692687</td><td>13</td><td>33</td><td>21</td><td></td><td>isomiR</td><td>NO</td><td>249.75</td><td>1335.03</td><td>406.50</td><td>157.17</td></tr>
<tr><td>>MI0000746|hsa-mir-99b&WithFlank&19|+|51692606|51692687@13.33.21(+1A)&[MIMAT0000689&hsa-miR-99b-5p&offsets|0|-1(+1A);m-10&19|+|51692618|51692639&offsets|0|-1(+1A)]|CACCCGTAGAACCGACCTTGCA</td><td>>MI0000746|hsa-mir-99b&WithFlank&19|+|51692606|51692687</td><td>13</td><td>33</td><td>21</td><td>(+1A)</td><td>isomiR</td><td>NO</td><td>168.74</td><td>1358.90</td><td>125.06</td><td>99.43</td></tr>
<tr><td>>MI0000746|hsa-mir-99b&WithFlank&19|+|51692606|51692687@13.33.21(+1C)&[MIMAT0000689&hsa-miR-99b-5p&offsets|0|-1(+1C);m-10&19|+|51692618|51692639&offsets|0|-1(+1C)]|CACCCGTAGAACCGACCTTGCC</td><td>>MI0000746|hsa-mir-99b&WithFlank&19|+|51692606|51692687</td><td>13</td><td>33</td><td>21</td><td>(+1C)</td><td>isomiR</td><td>NO</td><td>0.00</td><td>43.58</td><td>0.00</td><td>0.00</td></tr>
<tr><td>>MI0000746|hsa-mir-99b&WithFlank&19|+|51692606|51692687@13.33.21(+1U)&[MIMAT0000689&hsa-miR-99b-5p&offsets|0|-1(+1U);m-10&19|+|51692618|51692639&offsets|0|-1(+1U)]|CACCCGTAGAACCGACCTTGCT</td><td>>MI0000746|hsa-mir-99b&WithFlank&19|+|51692606|51692687</td><td>13</td><td>33</td><td>21</td><td>(+1U)</td><td>isomiR</td><td>NO</td><td>10.29</td><td>166.26</td><td>0.00</td><td>0.00</td></tr>
<tr><td>>MI0000746|hsa-mir-99b&WithFlank&19|+|51692606|51692687@13.34.22&[MIMAT0000689&hsa-miR-99b-5p&offsets|0|0;m-10&19|+|51692618|51692639&offsets|0|0]|CACCCGTAGAACCGACCTTGCG</td><td>>MI0000746|hsa-mir-99b&WithFlank&19|+|51692606|51692687</td><td>13</td><td>34</td><td>22</td><td></td><td>isomiR</td><td>YES</td><td>589.76</td><td>4553.71</td><td>557.43</td><td>398.46</td></tr>
<tr><td>>MI0000746|hsa-mir-99b&WithFlank&19|+|51692606|51692687@13.34.22(+1A)&[MIMAT0000689&hsa-miR-99b-5p&offsets|0|0(+1A);m-10&19|+|51692618|51692639&offsets|0|0(+1A)]|CACCCGTAGAACCGACCTTGCGA</td><td>>MI0000746|hsa-mir-99b&WithFlank&19|+|51692606|51692687</td><td>13</td><td>34</td><td>22</td><td>(+1A)</td><td>isomiR</td><td>NO</td><td>42.39</td><td>297.82</td><td>13.67</td><td>27.11</td></tr>
<tr><td>>MI0000746|hsa-mir-99b&WithFlank&19|+|51692606|51692687@13.34.22(+1C)&[MIMAT0000689&hsa-miR-99b-5p&offsets|0|0(+1C);m-10&19|+|51692618|51692639&offsets|0|0(+1C)]|CACCCGTAGAACCGACCTTGCGC</td><td>>MI0000746|hsa-mir-99b&WithFlank&19|+|51692606|51692687</td><td>13</td><td>34</td><td>22</td><td>(+1C)</td><td>isomiR</td><td>NO</td><td>0.00</td><td>38.37</td><td>0.00</td><td>0.00</td></tr>
<tr><td>>MI0000746|hsa-mir-99b&WithFlank&19|+|51692606|51692687@13.34.22(+1U)&[MIMAT0000689&hsa-miR-99b-5p&offsets|0|0(+1U);m-10&19|+|51692618|51692639&offsets|0|0(+1U)]|CACCCGTAGAACCGACCTTGCGT</td><td>>MI0000746|hsa-mir-99b&WithFlank&19|+|51692606|51692687</td><td>13</td><td>34</td><td>22</td><td>(+1U)</td><td>isomiR</td><td>NO</td><td>87.62</td><td>345.43</td><td>57.37</td><td>35.68</td></tr>
<tr><td>>MI0000746|hsa-mir-99b&WithFlank&19|+|51692606|51692687@14.34.21&[MIMAT0000689&hsa-miR-99b-5p&offsets|+1|0;m-10&19|+|51692618|51692639&offsets|+1|0]|ACCCGTAGAACCGACCTTGCG</td><td>>MI0000746|hsa-mir-99b&WithFlank&19|+|51692606|51692687</td><td>14</td><td>34</td><td>21</td><td></td><td>isomiR</td><td>NO</td><td>0.00</td><td>35.42</td><td>0.00</td><td>0.00</td></tr>
<tr><td>>MI0000746|hsa-mir-99b&WithFlank&19|+|51692606|51692687@51.70.20&[MIMAT0004678&hsa-miR-99b-3p&offsets|0|-2;m-151&19|+|51692656|51692678&offsets|0|-3]|CAAGCTCGTGTCTGTGGGTC</td><td>>MI0000746|hsa-mir-99b&WithFlank&19|+|51692606|51692687</td><td>51</td><td>70</td><td>20</td><td></td><td>isomiR</td><td>NO</td><td>0.00</td><td>44.80</td><td>13.11</td><td>0.00</td></tr>
<tr><td>>MI0000746|hsa-mir-99b&WithFlank&19|+|51692606|51692687@51.70.20(+1U)&[MIMAT0004678&hsa-miR-99b-3p&offsets|0|-2(+1U);m-151&19|+|51692656|51692678&offsets|0|-3(+1U)]|CAAGCTCGTGTCTGTGGGTCT</td><td>>MI0000746|hsa-mir-99b&WithFlank&19|+|51692606|51692687</td><td>51</td><td>70</td><td>20</td><td>(+1U)</td><td>isomiR</td><td>NO</td><td>0.00</td><td>33.79</td><td>12.23</td><td>0.00</td></tr>
<tr><td>>MI0000746|hsa-mir-99b&WithFlank&19|+|51692606|51692687@51.71.21&[MIMAT0004678&hsa-miR-99b-3p&offsets|0|-1;m-151&19|+|51692656|51692678&offsets|0|-2]|CAAGCTCGTGTCTGTGGGTCC</td><td>>MI0000746|hsa-mir-99b&WithFlank&19|+|51692606|51692687</td><td>51</td><td>71</td><td>21</td><td></td><td>isomiR</td><td>NO</td><td>0.00</td><td>72.03</td><td>54.28</td><td>0.00</td></tr>
<tr><td>>MI0000746|hsa-mir-99b&WithFlank&19|+|51692606|51692687@51.71.21(+1A)&[MIMAT0004678&hsa-miR-99b-3p&offsets|0|-1(+1A);m-151&19|+|51692656|51692678&offsets|0|-2(+1A)]|CAAGCTCGTGTCTGTGGGTCCA</td><td>>MI0000746|hsa-mir-99b&WithFlank&19|+|51692606|51692687</td><td>51</td><td>71</td><td>21</td><td>(+1A)</td><td>isomiR</td><td>NO</td><td>0.00</td><td>70.09</td><td>10.80</td><td>0.00</td></tr>
<tr><td>>MI0000746|hsa-mir-99b&WithFlank&19|+|51692606|51692687@51.72.22&[MIMAT0004678&hsa-miR-99b-3p&offsets|0|0;m-151&19|+|51692656|51692678&offsets|0|-1]|CAAGCTCGTGTCTGTGGGTCCG</td><td>>MI0000746|hsa-mir-99b&WithFlank&19|+|51692606|51692687</td><td>51</td><td>72</td><td>22</td><td></td><td>isomiR</td><td>NO</td><td>66.27</td><td>205.17</td><td>129.35</td><td>41.48</td></tr>
<tr><td>>MI0000746|hsa-mir-99b&WithFlank&19|+|51692606|51692687@51.72.22(+1A)&[MIMAT0004678&hsa-miR-99b-3p&offsets|0|0(+1A);m-151&19|+|51692656|51692678&offsets|0|-1(+1A)]|CAAGCTCGTGTCTGTGGGTCCGA</td><td>>MI0000746|hsa-mir-99b&WithFlank&19|+|51692606|51692687</td><td>51</td><td>72</td><td>22</td><td>(+1A)</td><td>isomiR</td><td>NO</td><td>9.61</td><td>151.81</td><td>62.55</td><td>0.00</td></tr>
<tr><td>>MI0000746|hsa-mir-99b&WithFlank&19|+|51692606|51692687@51.72.22(+1C)&[MIMAT0004678&hsa-miR-99b-3p&offsets|0|0(+1C);m-151&19|+|51692656|51692678&offsets|0|-1(+1C)]|CAAGCTCGTGTCTGTGGGTCCGC</td><td>>MI0000746|hsa-mir-99b&WithFlank&19|+|51692606|51692687</td><td>51</td><td>72</td><td>22</td><td>(+1C)</td><td>isomiR</td><td>NO</td><td>0.00</td><td>24.13</td><td>0.00</td><td>0.00</td></tr>
<tr><td>>MI0000746|hsa-mir-99b&WithFlank&19|+|51692606|51692687@51.73.23&[MIMAT0004678&hsa-miR-99b-3p&offsets|0|+1;m-151&19|+|51692656|51692678&offsets|0|0]|CAAGCTCGTGTCTGTGGGTCCGT</td><td>>MI0000746|hsa-mir-99b&WithFlank&19|+|51692606|51692687</td><td>51</td><td>73</td><td>23</td><td></td><td>isomiR</td><td>NO</td><td>29.49</td><td>204.33</td><td>55.41</td><td>10.01</td></tr>
<tr><td>>MI0000746|hsa-mir-99b&WithFlank&19|+|51692606|51692687@51.73.23(+1U)&[MIMAT0004678&hsa-miR-99b-3p&offsets|0|+1(+1U);m-151&19|+|51692656|51692678&offsets|0|0(+1U)]|CAAGCTCGTGTCTGTGGGTCCGTT</td><td>>MI0000746|hsa-mir-99b&WithFlank&19|+|51692606|51692687</td><td>51</td><td>73</td><td>23</td><td>(+1U)</td><td>isomiR</td><td>NO</td><td>0.00</td><td>31.40</td><td>0.00</td><td>0.00</td></tr>
<tr><td>>MI0000748|hsa-mir-130b&WithFlank&22|+|21653298|21653391@19.40.22&[MIMAT0004680&hsa-miR-130b-5p&offsets|0|+1;m-234&22|+|21653316|21653337&offsets|0|0]|ACTCTTTCCCTGTTGCACTACT</td><td>>MI0000748|hsa-mir-130b&WithFlank&22|+|21653298|21653391</td><td>19</td><td>40</td><td>22</td><td></td><td>isomiR</td><td>NO</td><td>0.00</td><td>110.48</td><td>0.00</td><td>0.00</td></tr>
<tr><td>>am_MI0000748|hsa-mir-130b&WithFlank&22|+|21653298|21653391@19.39.21&[MIMAT0004680&hsa-miR-130b-5p&offsets|0|0;m-234&22|+|21653316|21653337&offsets|0|-1]|ACTCTTTCCCTGTTGCACTAC</td><td>>MI0000748|hsa-mir-130b&WithFlank&22|+|21653298|21653391.am</td><td>19</td><td>39</td><td>21</td><td></td><td>isomiR</td><td>NO</td><td>0.00</td><td>28.76</td><td>0.00</td><td>0.00</td></tr>
</table>

# BT-20

<table>
<tr><th></th><th></th><th></th><th></th><th></th><th></th><th></th><th></th><th></th><th></th><th></th><th></th></tr>
<tr><td>>MI0000749|hsa-mir-30e&WithFlank&1|+|40754349|40754452@23.45.23(+1A)&[MIMAT0000692&hsa-miR-30e-5p&offsets|0|+1(+1A);m-24&1|+|40754371|40754394&offsets|0|-1(+1A)]|TGTAAACATCCTTGACTGGAAGCA</td><td>>MI0000749|hsa-mir-30e&WithFlank&1|+|40754349|40754452</td><td>23</td><td>45</td><td>23</td><td>(+1A)</td><td>isomiR</td><td>NO</td><td>0.00</td><td>0.00</td><td>15.86</td><td>0.00</td></tr>
<tr><td>>MI0000749|hsa-mir-30e&WithFlank&1|+|40754349|40754452@23.45.23(+1C)&[MIMAT0000692&hsa-miR-30e-5p&offsets|0|+1(+1C);m-24&1|+|40754371|40754394&offsets|0|-1(+1C)]|TGTAAACATCCTTGACTGGAAGCC</td><td>>MI0000749|hsa-mir-30e&WithFlank&1|+|40754349|40754452</td><td>23</td><td>45</td><td>23</td><td>(+1C)</td><td>isomiR</td><td>NO</td><td>0.00</td><td>0.00</td><td>73.69</td><td>0.00</td></tr>
<tr><td>>MI0000749|hsa-mir-30e&WithFlank&1|+|40754349|40754452@23.46.24&[MIMAT0000692&hsa-miR-30e-5p&offsets|0|+2;m-24&1|+|40754371|40754394&offsets|0|0]|TGTAAACATCCTTGACTGGAAGCT</td><td>>MI0000749|hsa-mir-30e&WithFlank&1|+|40754349|40754452</td><td>23</td><td>46</td><td>24</td><td></td><td>isomiR</td><td>NO</td><td>105.94</td><td>115.84</td><td>335.39</td><td>33.69</td></tr>
<tr><td>>MI0000749|hsa-mir-30e&WithFlank&1|+|40754349|40754452@24.46.23&[MIMAT0000692&hsa-miR-30e-5p&offsets|+1|+2;m-24&1|+|40754371|40754394&offsets|+1|0]|GTAAACATCCTTGACTGGAAGCT</td><td>>MI0000749|hsa-mir-30e&WithFlank&1|+|40754349|40754452</td><td>24</td><td>46</td><td>23</td><td></td><td>isomiR</td><td>NO</td><td>0.00</td><td>0.00</td><td>56.15</td><td>0.00</td></tr>
<tr><td>>MI0000749|hsa-mir-30e&WithFlank&1|+|40754349|40754452@65.83.19&[MIMAT0000693&hsa-miR-30e-3p&offsets|0|-3;m-25&1|+|40754413|40754434&offsets|0|-3]|CTTTCAGTCGGATGTTTAC</td><td>>MI0000749|hsa-mir-30e&WithFlank&1|+|40754349|40754452</td><td>65</td><td>83</td><td>19</td><td></td><td>isomiR</td><td>NO</td><td>0.00</td><td>38.03</td><td>0.00</td><td>0.00</td></tr>
<tr><td>>MI0000749|hsa-mir-30e&WithFlank&1|+|40754349|40754452@65.85.21&[MIMAT0000693&hsa-miR-30e-3p&offsets|0|-1;m-25&1|+|40754413|40754434&offsets|0|-1]|CTTTCAGTCGGATGTTTACAG</td><td>>MI0000749|hsa-mir-30e&WithFlank&1|+|40754349|40754452</td><td>65</td><td>85</td><td>21</td><td></td><td>isomiR</td><td>NO</td><td>0.00</td><td>23.79</td><td>0.00</td><td>0.00</td></tr>
<tr><td>>MI0000749|hsa-mir-30e&WithFlank&1|+|40754349|40754452@65.85.21(+1U)&[MIMAT0000693&hsa-miR-30e-3p&offsets|0|-1(+1U);m-25&1|+|40754413|40754434&offsets|0|-1(+1U)]|CTTTCAGTCGGATGTTTACAGT</td><td>>MI0000749|hsa-mir-30e&WithFlank&1|+|40754349|40754452</td><td>65</td><td>85</td><td>21</td><td>(+1U)</td><td>isomiR</td><td>NO</td><td>109.26</td><td>164.18</td><td>10.26</td><td>23.74</td></tr>
<tr><td>>MI0000749|hsa-mir-30e&WithFlank&1|+|40754349|40754452@65.85.21(+2U)&[MIMAT0000693&hsa-miR-30e-3p&offsets|0|-1(+2U);m-25&1|+|40754413|40754434&offsets|0|-1(+2U)]|CTTTCAGTCGGATGTTTACAGTT</td><td>>MI0000749|hsa-mir-30e&WithFlank&1|+|40754349|40754452</td><td>65</td><td>85</td><td>21</td><td>(+2U)</td><td>isomiR</td><td>NO</td><td>0.00</td><td>36.37</td><td>0.00</td><td>0.00</td></tr>
<tr><td>>MI0000749|hsa-mir-30e&WithFlank&1|+|40754349|40754452@65.86.22&[MIMAT0000693&hsa-miR-30e-3p&offsets|0|0;m-25&1|+|40754413|40754434&offsets|0|0]|CTTTCAGTCGGATGTTTACAGC</td><td>>MI0000749|hsa-mir-30e&WithFlank&1|+|40754349|40754452</td><td>65</td><td>86</td><td>22</td><td></td><td>isomiR</td><td>NO</td><td>43.31</td><td>154.49</td><td>0.00</td><td>0.00</td></tr>
<tr><td>>MI0000750|hsa-mir-26a-2&WithFlank&12|-|57824603|57824698@20.39.20(+1U);MI0000083|hsa-mir-26a-1&WithFlank&3|+|37969398|37969486@16.35.20(+1U)&[MIMAT0000082&hsa-miR-26a-5p&offsets|0|-2(+1U);m-33&12|-|57824658|57824679&offsets|0|-2(+1U)];[MIMAT0000082_1&hsa-miR-26a-5p&offsets|0|-2(+1U);m-34&3|+|37969413|37969434&offsets|0|-2(+1U)]|TTCAAGTAATCCAGGATAGGT</td><td>>MI0000750|hsa-mir-26a-2&WithFlank&12|-|57824603|57824698</td><td>20</td><td>39</td><td>20</td><td>(+1U)</td><td>isomiR</td><td>NO</td><td>0.00</td><td>25.78</td><td>0.00</td><td>0.00</td></tr>
<tr><td>>MI0000750|hsa-mir-26a-2&WithFlank&12|-|57824603|57824698@20.39.20;MI0000083|hsa-mir-26a-1&WithFlank&3|+|37969398|37969486@16.35.20&[MIMAT0000082&hsa-miR-26a-5p&offsets|0|-2;m-33&12|-|57824658|57824679&offsets|0|-2];[MIMAT0000082_1&hsa-miR-26a-5p&offsets|0|-2;m-34&3|+|37969413|37969434&offsets|0|-2]|TTCAAGTAATCCAGGATAGG</td><td>>MI0000750|hsa-mir-26a-2&WithFlank&12|-|57824603|57824698</td><td>20</td><td>39</td><td>20</td><td></td><td>isomiR</td><td>NO</td><td>0.00</td><td>70.18</td><td>0.00</td><td>0.00</td></tr>
<tr><td>>MI0000750|hsa-mir-26a-2&WithFlank&12|-|57824603|57824698@20.40.21(+1A);MI0000083|hsa-mir-26a-1&WithFlank&3|+|37969398|37969486@16.36.21(+1A)&[MIMAT0000082&hsa-miR-26a-5p&offsets|0|-1(+1A);m-33&12|-|57824658|57824679&offsets|0|-1(+1A)];[MIMAT0000082_1&hsa-miR-26a-5p&offsets|0|-1(+1A);m-34&3|+|37969413|37969434&offsets|0|-1(+1A)]|TTCAAGTAATCCAGGATAGGCA</td><td>>MI0000750|hsa-mir-26a-2&WithFlank&12|-|57824603|57824698</td><td>20</td><td>40</td><td>21</td><td>(+1A)</td><td>isomiR</td><td>NO</td><td>180.20</td><td>564.52</td><td>89.24</td><td>128.96</td></tr>
<tr><td>>MI0000750|hsa-mir-26a-2&WithFlank&12|-|57824603|57824698@20.40.21(+1C);MI0000083|hsa-mir-26a-1&WithFlank&3|+|37969398|37969486@16.36.21(+1C)&[MIMAT0000082&hsa-miR-26a-5p&offsets|0|-1(+1C);m-33&12|-|57824658|57824679&offsets|0|-1(+1C)];[MIMAT0000082_1&hsa-miR-26a-5p&offsets|0|-1(+1C);m-34&3|+|37969413|37969434&offsets|0|-1(+1C)]|TTCAAGTAATCCAGGATAGGCC</td><td>>MI0000750|hsa-mir-26a-2&WithFlank&12|-|57824603|57824698</td><td>20</td><td>40</td><td>21</td><td>(+1C)</td><td>isomiR</td><td>NO</td><td>268.09</td><td>620.04</td><td>140.35</td><td>199.50</td></tr>
<tr><td>>MI0000750|hsa-mir-26a-2&WithFlank&12|-|57824603|57824698@20.40.21(+1G);MI0000083|hsa-mir-26a-1&WithFlank&3|+|37969398|37969486@16.36.21(+1G)&[MIMAT0000082&hsa-miR-26a-5p&offsets|0|-1(+1G);m-33&12|-|57824658|57824679&offsets|0|-1(+1G)];[MIMAT0000082_1&hsa-miR-26a-5p&offsets|0|-1(+1G);m-34&3|+|37969413|37969434&offsets|0|-1(+1G)]|TTCAAGTAATCCAGGATAGGCG</td><td>>MI0000750|hsa-mir-26a-2&WithFlank&12|-|57824603|57824698</td><td>20</td><td>40</td><td>21</td><td>(+1G)</td><td>isomiR</td><td>NO</td><td>88.30</td><td>181.50</td><td>0.00</td><td>10.54</td></tr>
<tr><td>>MI0000750|hsa-mir-26a-2&WithFlank&12|-|57824603|57824698@20.40.21;MI0000083|hsa-mir-26a-1&WithFlank&3|+|37969398|37969486@16.36.21&[MIMAT0000082&hsa-miR-26a-5p&offsets|0|-1;m-33&12|-|57824658|57824679&offsets|0|-1];[MIMAT0000082_1&hsa-miR-26a-5p&offsets|0|-1;m-34&3|+|37969413|37969434&offsets|0|-1]|TTCAAGTAATCCAGGATAGGC</td><td>>MI0000750|hsa-mir-26a-2&WithFlank&12|-|57824603|57824698</td><td>20</td><td>40</td><td>21</td><td></td><td>isomiR</td><td>NO</td><td>39.96</td><td>112.57</td><td>0.00</td><td>0.00</td></tr>
<tr><td>>MI0000750|hsa-mir-26a-2&WithFlank&12|-|57824603|57824698@20.41.22(+1A);MI0000083|hsa-mir-26a-1&WithFlank&3|+|37969398|37969486@16.37.22(+1A)&[MIMAT0000082&hsa-miR-26a-5p&offsets|0|0(+1A);m-33&12|-|57824658|57824679&offsets|0|0(+1A)];[MIMAT0000082_1&hsa-miR-26a-5p&offsets|0|0(+1A);m-34&3|+|37969413|37969434&offsets|0|0(+1A)]|TTCAAGTAATCCAGGATAGGCTA</td><td>>MI0000750|hsa-mir-26a-2&WithFlank&12|-|57824603|57824698</td><td>20</td><td>41</td><td>22</td><td>(+1A)</td><td>isomiR</td><td>NO</td><td>32.24</td><td>102.86</td><td>12.30</td><td>0.00</td></tr>
<tr><td>>MI0000750|hsa-mir-26a-2&WithFlank&12|-|57824603|57824698@20.41.22(+1C);MI0000083|hsa-mir-26a-1&WithFlank&3|+|37969398|37969486@16.37.22(+1C)&[MIMAT0000082&hsa-miR-26a-5p&offsets|0|0(+1C);m-33&12|-|57824658|57824679&offsets|0|0(+1C)];[MIMAT0000082_1&hsa-miR-26a-5p&offsets|0|0(+1C);m-34&3|+|37969413|37969434&offsets|0|0(+1C)]|TTCAAGTAATCCAGGATAGGCTC</td><td>>MI0000750|hsa-mir-26a-2&WithFlank&12|-|57824603|57824698</td><td>20</td><td>41</td><td>22</td><td>(+1C)</td><td>isomiR</td><td>NO</td><td>0.00</td><td>29.74</td><td>0.00</td><td>0.00</td></tr>
</table>

# BT-20

<table>
<tr><td>>MI0000750|hsa-mir-26a-2&WithFlank&12|-|57824603|57824698@20.41.22(+1U);MI0000083|hsa-mir-26a-1&WithFlank&3|+|37969398|37969486@16.37.22(+1U)&[MIMAT0000082&hsa-miR-26a-5p&offsets|0|0(+1U);m-33&12|-|57824658|57824679&offsets|0|0(+1U)];[MIMAT0000082_1&hsa-miR-26a-5p&offsets|0|0(+1U);m-34&3|+|37969413|37969434&offsets|0|0(+1U)]|TTCAAGTAATCCAGGATAGGCTT</td><td>>MI0000750|hsa-mir-26a-2&WithFlank&12|-|57824603|57824698</td><td>20</td><td>41</td><td>22</td><td>(+1U)</td><td>isomiR</td><td>NO</td><td>116.90</td><td>254.87</td><td>84.64</td><td>27.70</td></tr>
<tr><td>>MI0000750|hsa-mir-26a-2&WithFlank&12|-|57824603|57824698@20.41.22(+2U);MI0000083|hsa-mir-26a-1&WithFlank&3|+|37969398|37969486@16.37.22(+2U)&[MIMAT0000082&hsa-miR-26a-5p&offsets|0|0(+2U);m-33&12|-|57824658|57824679&offsets|0|0(+2U)];[MIMAT0000082_1&hsa-miR-26a-5p&offsets|0|0(+2U);m-34&3|+|37969413|37969434&offsets|0|0(+2U)]|TTCAAGTAATCCAGGATAGGCTTT</td><td>>MI0000750|hsa-mir-26a-2&WithFlank&12|-|57824603|57824698</td><td>20</td><td>41</td><td>22</td><td>(+2U)</td><td>isomiR</td><td>NO</td><td>38.77</td><td>121.61</td><td>14.03</td><td>0.00</td></tr>
<tr><td>>MI0000750|hsa-mir-26a-2&WithFlank&12|-|57824603|57824698@20.41.22;MI0000083|hsa-mir-26a-1&WithFlank&3|+|37969398|37969486@16.37.22&[MIMAT0000082&hsa-miR-26a-5p&offsets|0|0;m-33&12|-|57824658|57824679&offsets|0|0];[MIMAT0000082_1&hsa-miR-26a-5p&offsets|0|0;m-34&3|+|37969413|37969434&offsets|0|0]|TTCAAGTAATCCAGGATAGGCT</td><td>>MI0000750|hsa-mir-26a-2&WithFlank&12|-|57824603|57824698</td><td>20</td><td>41</td><td>22</td><td></td><td>isomiR</td><td>YES</td><td>1178.04</td><td>3690.55</td><td>722.82</td><td>918.31</td></tr>
<tr><td>>MI0000750|hsa-mir-26a-2&WithFlank&12|-|57824603|57824698@23.41.19;MI0000083|hsa-mir-26a-1&WithFlank&3|+|37969398|37969486@19.37.19&[MIMAT0000082&hsa-miR-26a-5p&offsets|+3|0;m-33&12|-|57824658|57824679&offsets|+3|0];[MIMAT0000082_1&hsa-miR-26a-5p&offsets|+3|0;m-34&3|+|37969413|37969434&offsets|+3|0]|AAGTAATCCAGGATAGGCT</td><td>>MI0000750|hsa-mir-26a-2&WithFlank&12|-|57824603|57824698</td><td>23</td><td>41</td><td>19</td><td></td><td>isomiR</td><td>NO</td><td>28.88</td><td>82.02</td><td>0.00</td><td>10.74</td></tr>
<tr><td>>MI0000760|hsa-mir-361&WithFlank&X|-|85903630|85903713@51.73.23&[MIMAT0004682&hsa-miR-361-3p&offsets|0|0;m-124&X|-|85903642|85903663&offsets|0|+1]|TCCCCCAGGTGTGATTCTGATTT</td><td>>MI0000760|hsa-mir-361&WithFlank&X|-|85903630|85903713</td><td>51</td><td>73</td><td>23</td><td></td><td>isomiR</td><td>NO</td><td>0.00</td><td>75.52</td><td>0.00</td><td>0.00</td></tr>
<tr><td>>MI0000760|hsa-mir-361&WithFlank&X|-|85903630|85903713@51.74.24&[MIMAT0004682&hsa-miR-361-3p&offsets|0|+1;m-124&X|-|85903642|85903663&offsets|0|+2]|TCCCCCAGGTGTGATTCTGATTTG</td><td>>MI0000760|hsa-mir-361&WithFlank&X|-|85903630|85903713</td><td>51</td><td>74</td><td>24</td><td></td><td>isomiR</td><td>NO</td><td>0.00</td><td>37.33</td><td>0.00</td><td>0.00</td></tr>
<tr><td>>MI0000760|hsa-mir-361&WithFlank&X|-|85903630|85903713@52.75.24&[MIMAT0004682&hsa-miR-361-3p&offsets|+1|+2;m-124&X|-|85903642|85903663&offsets|+1|+3]|CCCCCAGGTGTGATTCTGATTTGC</td><td>>MI0000760|hsa-mir-361&WithFlank&X|-|85903630|85903713</td><td>52</td><td>75</td><td>24</td><td></td><td>isomiR</td><td>NO</td><td>0.00</td><td>34.38</td><td>0.00</td><td>0.00</td></tr>
<tr><td>>MI0000762|hsa-mir-362&WithFlank&X|+|50008958|50009034@11.34.24&[MIMAT0000705&hsa-miR-362-5p&offsets|0|0;m-267&X|+|50008968|50008991&offsets|0|0]|AATCCTTGGAACCTAGGTGTGAGT</td><td>>MI0000762|hsa-mir-362&WithFlank&X|+|50008958|50009034</td><td>11</td><td>34</td><td>24</td><td></td><td>isomiR</td><td>NO</td><td>0.00</td><td>24.30</td><td>0.00</td><td>0.00</td></tr>
<tr><td>>MI0000764|hsa-mir-363&WithFlank&X|-|134169372|134169458@56.77.22&[MIMAT0000707&hsa-miR-363-3p&offsets|0|0;m-178&X|-|134169381|134169403&offsets|0|-1]|AATTGCACGGTATCCATCTGTA</td><td>>MI0000764|hsa-mir-363&WithFlank&X|-|134169372|134169458</td><td>56</td><td>77</td><td>22</td><td></td><td>isomiR</td><td>NO</td><td>42.79</td><td>74.72</td><td>115.39</td><td>37.43</td></tr>
<tr><td>>MI0000767|hsa-mir-365a&WithFlank&16|+|14309279|14309377@22.43.22&[MIMAT0009199&hsa-miR-365a-5p&offsets|0|-1;m-595&16|+|14309300|14309322&offsets|0|-1]|AGGGACTTTTGGGGGCAGATGT</td><td>>MI0000767|hsa-mir-365a&WithFlank&16|+|14309279|14309377</td><td>22</td><td>43</td><td>22</td><td></td><td>isomiR</td><td>NO</td><td>0.00</td><td>0.00</td><td>13.52</td><td>0.00</td></tr>
<tr><td>>MI0000767|hsa-mir-365a&WithFlank&16|+|14309279|14309377@62.81.20(+2U);MI0000769|hsa-mir-365b&WithFlank&17|+|31575405|31575527@74.93.20(+2U)&[MIMAT0000710&hsa-miR-365a-3p&offsets|0|-2(+2U);m-176&16|+|14309340|14309361&offsets|0|-2(+2U)];[MIMAT0022834&hsa-miR-365b-3p&offsets|0|-2(+2U);m-175&17|+|31575478|31575499&offsets|0|-2(+2U)]|TAATGCCCCTAAAAATCCTTTT</td><td>>MI0000767|hsa-mir-365a&WithFlank&16|+|14309279|14309377</td><td>62</td><td>81</td><td>20</td><td>(+2U)</td><td>isomiR</td><td>NO</td><td>0.00</td><td>34.25</td><td>0.00</td><td>0.00</td></tr>
<tr><td>>MI0000767|hsa-mir-365a&WithFlank&16|+|14309279|14309377@62.82.21;MI0000769|hsa-mir-365b&WithFlank&17|+|31575405|31575527@74.94.21&[MIMAT0000710&hsa-miR-365a-3p&offsets|0|-1;m-176&16|+|14309340|14309361&offsets|0|-1];[MIMAT0022834&hsa-miR-365b-3p&offsets|0|-1;m-175&17|+|31575478|31575499&offsets|0|-1]|TAATGCCCCTAAAAATCCTTA</td><td>>MI0000767|hsa-mir-365a&WithFlank&16|+|14309279|14309377</td><td>62</td><td>82</td><td>21</td><td></td><td>isomiR</td><td>NO</td><td>9.85</td><td>36.92</td><td>0.00</td><td>0.00</td></tr>
<tr><td>>MI0000767|hsa-mir-365a&WithFlank&16|+|14309279|14309377@62.83.22;MI0000769|hsa-mir-365b&WithFlank&17|+|31575405|31575527@74.95.22&[MIMAT0000710&hsa-miR-365a-3p&offsets|0|0;m-176&16|+|14309340|14309361&offsets|0|0];[MIMAT0022834&hsa-miR-365b-3p&offsets|0|0;m-175&17|+|31575478|31575499&offsets|0|0]|TAATGCCCCTAAAAATCCTTAT</td><td>>MI0000767|hsa-mir-365a&WithFlank&16|+|14309279|14309377</td><td>62</td><td>83</td><td>22</td><td></td><td>isomiR</td><td>NO</td><td>99.08</td><td>181.02</td><td>9.95</td><td>67.01</td></tr>
<tr><td>>MI0000767|hsa-mir-365a&WithFlank&16|+|14309279|14309377@62.84.23;MI0000769|hsa-mir-365b&WithFlank&17|+|31575405|31575527@74.96.23&[MIMAT0000710&hsa-miR-365a-3p&offsets|0|+1;m-176&16|+|14309340|14309361&offsets|0|+1];[MIMAT0022834&hsa-miR-365b-3p&offsets|0|+1;m-175&17|+|31575478|31575499&offsets|0|+1]|TAATGCCCCTAAAAATCCTTATT</td><td>>MI0000767|hsa-mir-365a&WithFlank&16|+|14309279|14309377</td><td>62</td><td>84</td><td>23</td><td></td><td>isomiR</td><td>NO</td><td>25.52</td><td>74.53</td><td>0.00</td><td>0.00</td></tr>
<tr><td>>am_MI0000786|hsa-mir-378a&WithFlank&5|+|149732819|149732896@49.69.21&[MIMAT0000732&hsa-miR-378a-3p&offsets|0|-1;m-44&5|+|149732867|149732888&offsets|0|-1]|ACTGGACTTGGAGTCAGAAGG</td><td>>MI0000786|hsa-mir-378a&WithFlank&5|+|149732819|149732896.an</td><td>49</td><td>69</td><td>21</td><td></td><td>isomiR</td><td>NO</td><td>103.83</td><td>181.00</td><td>230.46</td><td>29.91</td></tr>
<tr><td>>am_MI0000786|hsa-mir-378a&WithFlank&5|+|149732819|149732896@49.69.21(+1A)&[MIMAT0000732&hsa-miR-378a-3p&offsets|0|-1(+1A);m-44&5|+|149732867|149732888&offsets|0|-1(+1A)]|ACTGGACTTGGAGTCAGAAGGA</td><td>>MI0000786|hsa-mir-378a&WithFlank&5|+|149732819|149732896.an</td><td>49</td><td>69</td><td>21</td><td>(+1A)</td><td>isomiR</td><td>NO</td><td>35.72</td><td>198.76</td><td>165.12</td><td>0.00</td></tr>
<tr><td>>am_MI0000786|hsa-mir-378a&WithFlank&5|+|149732819|149732896@49.69.21(+1U)&[MIMAT0000732&hsa-miR-378a-3p&offsets|0|-1(+1U);m-44&5|+|149732867|149732888&offsets|0|-1(+1U)]|ACTGGACTTGGAGTCAGAAGGT</td><td>>MI0000786|hsa-mir-378a&WithFlank&5|+|149732819|149732896.an</td><td>49</td><td>69</td><td>21</td><td>(+1U)</td><td>isomiR</td><td>NO</td><td>133.09</td><td>479.90</td><td>478.16</td><td>69.48</td></tr>
<tr><td>>am_MI0000786|hsa-mir-378a&WithFlank&5|+|149732819|149732896@49.70.22&[MIMAT0000732&hsa-miR-378a-3p&offsets|0|0;m-44&5|+|149732867|149732888&offsets|0|0]|ACTGGACTTGGAGTCAGAAGGC</td><td>>MI0000786|hsa-mir-378a&WithFlank&5|+|149732819|149732896.an</td><td>49</td><td>70</td><td>22</td><td></td><td>isomiR</td><td>NO</td><td>332.56</td><td>1357.81</td><td>1405.24</td><td>198.47</td></tr>
<tr><td>>am_MI0000786|hsa-mir-378a&WithFlank&5|+|149732819|149732896@49.70.22(+1A)&[MIMAT0000732&hsa-miR-378a-3p&offsets|0|0(+1A);m-44&5|+|149732867|149732888&offsets|0|0(+1A)]|ACTGGACTTGGAGTCAGAAGGCA</td><td>>MI0000786|hsa-mir-378a&WithFlank&5|+|149732819|149732896.an</td><td>49</td><td>70</td><td>22</td><td>(+1A)</td><td>isomiR</td><td>NO</td><td>39.60</td><td>264.70</td><td>271.63</td><td>9.94</td></tr>
</table>

# BT-20

<table>
<tr><td>>am_MI0000786|hsa-mir-378a&WithFlank&5|+|149732819|149732896@49.70.22(+1G)&[MIMAT0000732&hsa-miR-378a-3p&offsets|0|0(+1G);m-44&5|+|149732867|149732888&offsets|0|0(+1G)]|ACTGGACTTGGAGTCAGAAGGCG</td><td>>MI0000786|hsa-mir-378a&WithFlank&5|+|149732819|149732896.an</td><td>49</td><td>70</td><td>22</td><td>(+1G)</td><td>isomiR</td><td>NO</td><td>0.00</td><td>33.26</td><td>102.28</td><td>0.00</td></tr>
<tr><td>>am_MI0000786|hsa-mir-378a&WithFlank&5|+|149732819|149732896@49.70.22(+1U)&[MIMAT0000732&hsa-miR-378a-3p&offsets|0|0(+1U);m-44&5|+|149732867|149732888&offsets|0|0(+1U)]|ACTGGACTTGGAGTCAGAAGGCT</td><td>>MI0000786|hsa-mir-378a&WithFlank&5|+|149732819|149732896.an</td><td>49</td><td>70</td><td>22</td><td>(+1U)</td><td>isomiR</td><td>NO</td><td>27.27</td><td>164.10</td><td>335.03</td><td>10.38</td></tr>
<tr><td>>am_MI0000786|hsa-mir-378a&WithFlank&5|+|149732819|149732896@49.71.23&[MIMAT0000732&hsa-miR-378a-3p&offsets|0|+1;m-44&5|+|149732867|149732888&offsets|0|+1]|ACTGGACTTGGAGTCAGAAGGCC</td><td>>MI0000786|hsa-mir-378a&WithFlank&5|+|149732819|149732896.an</td><td>49</td><td>71</td><td>23</td><td></td><td>isomiR</td><td>NO</td><td>0.00</td><td>9.84</td><td>72.34</td><td>0.00</td></tr>
<tr><td>>am_MI0000786|hsa-mir-378a&WithFlank&5|+|149732819|149732896@50.69.20&[MIMAT0000732&hsa-miR-378a-3p&offsets|+1|-1;m-44&5|+|149732867|149732888&offsets|+1|-1]|CTGGACTTGGAGTCAGAAGG</td><td>>MI0000786|hsa-mir-378a&WithFlank&5|+|149732819|149732896.an</td><td>50</td><td>69</td><td>20</td><td></td><td>isomiR</td><td>NO</td><td>0.00</td><td>0.00</td><td>10.49</td><td>0.00</td></tr>
<tr><td>>am_MI0000786|hsa-mir-378a&WithFlank&5|+|149732819|149732896@50.69.20(+1U)&[MIMAT0000732&hsa-miR-378a-3p&offsets|+1|-1(+1U);m-44&5|+|149732867|149732888&offsets|+1|-1(+1U)]|CTGGACTTGGAGTCAGAAGGT</td><td>>MI0000786|hsa-mir-378a&WithFlank&5|+|149732819|149732896.an</td><td>50</td><td>69</td><td>20</td><td>(+1U)</td><td>isomiR</td><td>NO</td><td>0.00</td><td>74.16</td><td>81.67</td><td>0.00</td></tr>
<tr><td>>am_MI0000786|hsa-mir-378a&WithFlank&5|+|149732819|149732896@50.70.21&[MIMAT0000732&hsa-miR-378a-3p&offsets|+1|0;m-44&5|+|149732867|149732888&offsets|+1|0]|CTGGACTTGGAGTCAGAAGGC</td><td>>MI0000786|hsa-mir-378a&WithFlank&5|+|149732819|149732896.an</td><td>50</td><td>70</td><td>21</td><td></td><td>isomiR</td><td>NO</td><td>35.98</td><td>211.11</td><td>242.93</td><td>10.54</td></tr>
<tr><td>>am_MI0000786|hsa-mir-378a&WithFlank&5|+|149732819|149732896@50.70.21(+1A)&[MIMAT0000732&hsa-miR-378a-3p&offsets|+1|0(+1A);m-44&5|+|149732867|149732888&offsets|+1|0(+1A)]|CTGGACTTGGAGTCAGAAGGCA</td><td>>MI0000786|hsa-mir-378a&WithFlank&5|+|149732819|149732896.an</td><td>50</td><td>70</td><td>21</td><td>(+1A)</td><td>isomiR</td><td>NO</td><td>0.00</td><td>31.26</td><td>62.08</td><td>0.00</td></tr>
<tr><td>>am_MI0000786|hsa-mir-378a&WithFlank&5|+|149732819|149732896@50.70.21(+1U)&[MIMAT0000732&hsa-miR-378a-3p&offsets|+1|0(+1U);m-44&5|+|149732867|149732888&offsets|+1|0(+1U)]|CTGGACTTGGAGTCAGAAGGCT</td><td>>MI0000786|hsa-mir-378a&WithFlank&5|+|149732819|149732896.an</td><td>50</td><td>70</td><td>21</td><td>(+1U)</td><td>isomiR</td><td>NO</td><td>0.00</td><td>78.92</td><td>178.54</td><td>0.00</td></tr>
<tr><td>>MI0000802|hsa-mir-340&WithFlank&5|-|180015297|180015403@22.43.22&[MIMAT0004692&hsa-miR-340-5p&offsets|0|0;m-220&5|-|180015361|180015382&offsets|0|0]|TTATAAAGCAATGAGACTGATT</td><td>>MI0000802|hsa-mir-340&WithFlank&5|-|180015297|180015403</td><td>22</td><td>43</td><td>22</td><td></td><td>isomiR</td><td>NO</td><td>160.88</td><td>91.34</td><td>37.36</td><td>0.00</td></tr>
<tr><td>>MI0000803|hsa-mir-330&WithFlank&19|-|45638988|45639093@63.85.23&[MIMAT0000751&hsa-miR-330-3p&offsets|0|0;m-347&19|-|45639009|45639030&offsets|-1|0]|GCAAAGCACACGGCCTGCAGAGA</td><td>>MI0000803|hsa-mir-330&WithFlank&19|-|45638988|45639093</td><td>63</td><td>85</td><td>23</td><td></td><td>isomiR</td><td>NO</td><td>0.00</td><td>24.20</td><td>0.00</td><td>0.00</td></tr>
<tr><td>>MI0000803|hsa-mir-330&WithFlank&19|-|45638988|45639093@63.86.24(+1U)&[MIMAT0000751&hsa-miR-330-3p&offsets|0|+1(+1U);m-347&19|-|45639009|45639030&offsets|-1|+1(+1U)]|GCAAAGCACACGGCCTGCAGAGAGT</td><td>>MI0000803|hsa-mir-330&WithFlank&19|-|45638988|45639093</td><td>63</td><td>86</td><td>24</td><td>(+1U)</td><td>isomiR</td><td>NO</td><td>0.00</td><td>35.95</td><td>0.00</td><td>0.00</td></tr>
<tr><td>>MI0000803|hsa-mir-330&WithFlank&19|-|45638988|45639093@64.86.23(+1U)&[MIMAT0000751&hsa-miR-330-3p&offsets|+1|+1(+1U);m-347&19|-|45639009|45639030&offsets|0|+1(+1U)]|CAAAGCACACGGCCTGCAGAGAGT</td><td>>MI0000803|hsa-mir-330&WithFlank&19|-|45638988|45639093</td><td>64</td><td>86</td><td>23</td><td>(+1U)</td><td>isomiR</td><td>NO</td><td>0.00</td><td>10.18</td><td>0.00</td><td>0.00</td></tr>
<tr><td>>MI0000804|hsa-mir-328&WithFlank&16|-|67202315|67202401@54.75.22&[MIMAT0000752&hsa-miR-328-3p&offsets|0|0;m-160&16|-|67202327|67202348&offsets|0|0]|CTGGCCCTCTCTGCCCTTCCGT</td><td>>MI0000804|hsa-mir-328&WithFlank&16|-|67202315|67202401</td><td>54</td><td>75</td><td>22</td><td></td><td>isomiR</td><td>NO</td><td>25.50</td><td>143.89</td><td>0.00</td><td>24.90</td></tr>
<tr><td>>MI0000805|hsa-mir-342&WithFlank&14|+|100109649|100109759@67.90.24&[MIMAT0000753&hsa-miR-342-3p&offsets|0|+1;m-128&14|+|100109715|100109737&offsets|0|+1]|TCTCACACAGAAATCGCACCCGTC</td><td>>MI0000805|hsa-mir-342&WithFlank&14|+|100109649|100109759</td><td>67</td><td>90</td><td>24</td><td></td><td>isomiR</td><td>NO</td><td>0.00</td><td>86.02</td><td>0.00</td><td>0.00</td></tr>
<tr><td>>MI0000809|hsa-mir-151a&WithFlank&8|-|140732558|140732659@51.72.22&[MIMAT0000757&hsa-miR-151a-3p&offsets|-2|-1;m-30&8|-|140732587|140732607&offsets|-2|-1]|TACTAGACTGAAGCTCCTTGAG</td><td>>MI0000809|hsa-mir-151a&WithFlank&8|-|140732558|140732659</td><td>51</td><td>72</td><td>22</td><td></td><td>isomiR</td><td>NO</td><td>0.00</td><td>90.18</td><td>67.75</td><td>0.00</td></tr>
<tr><td>>MI0000809|hsa-mir-151a&WithFlank&8|-|140732558|140732659@51.72.22(+1A)&[MIMAT0000757&hsa-miR-151a-3p&offsets|-2|-1(+1A);m-30&8|-|140732587|140732607&offsets|-2|-1(+1A)]|TACTAGACTGAAGCTCCTTGAGA</td><td>>MI0000809|hsa-mir-151a&WithFlank&8|-|140732558|140732659</td><td>51</td><td>72</td><td>22</td><td>(+1A)</td><td>isomiR</td><td>NO</td><td>0.00</td><td>124.45</td><td>12.26</td><td>0.00</td></tr>
<tr><td>>MI0000809|hsa-mir-151a&WithFlank&8|-|140732558|140732659@51.72.22(+1U)&[MIMAT0000757&hsa-miR-151a-3p&offsets|-2|-1(+1U);m-30&8|-|140732587|140732607&offsets|-2|-1(+1U)]|TACTAGACTGAAGCTCCTTGAGT</td><td>>MI0000809|hsa-mir-151a&WithFlank&8|-|140732558|140732659</td><td>51</td><td>72</td><td>22</td><td>(+1U)</td><td>isomiR</td><td>NO</td><td>0.00</td><td>35.19</td><td>0.00</td><td>0.00</td></tr>
<tr><td>>MI0000809|hsa-mir-151a&WithFlank&8|-|140732558|140732659@51.73.23&[MIMAT0000757&hsa-miR-151a-3p&offsets|-2|0;m-30&8|-|140732587|140732607&offsets|-2|0]|TACTAGACTGAAGCTCCTTGAGG</td><td>>MI0000809|hsa-mir-151a&WithFlank&8|-|140732558|140732659</td><td>51</td><td>73</td><td>23</td><td></td><td>isomiR</td><td>NO</td><td>104.57</td><td>551.98</td><td>194.81</td><td>114.06</td></tr>
<tr><td>>MI0000809|hsa-mir-151a&WithFlank&8|-|140732558|140732659@51.73.23(+1U)&[MIMAT0000757&hsa-miR-151a-3p&offsets|-2|0(+1U);m-30&8|-|140732587|140732607&offsets|-2|0(+1U)]|TACTAGACTGAAGCTCCTTGAGGT</td><td>>MI0000809|hsa-mir-151a&WithFlank&8|-|140732558|140732659</td><td>51</td><td>73</td><td>23</td><td>(+1U)</td><td>isomiR</td><td>NO</td><td>0.00</td><td>27.72</td><td>0.00</td><td>0.00</td></tr>
<tr><td>>MI0000809|hsa-mir-151a&WithFlank&8|-|140732558|140732659@53.74.22&[MIMAT0000757&hsa-miR-151a-3p&offsets|0|+1;m-30&8|-|140732587|140732607&offsets|0|+1]|CTAGACTGAAGCTCCTTGAGGA</td><td>>MI0000809|hsa-mir-151a&WithFlank&8|-|140732558|140732659</td><td>53</td><td>74</td><td>22</td><td></td><td>isomiR</td><td>NO</td><td>392.47</td><td>1614.76</td><td>753.51</td><td>414.55</td></tr>
<tr><td>>MI0000809|hsa-mir-151a&WithFlank&8|-|140732558|140732659@53.74.22(+1A)&[MIMAT0000757&hsa-miR-151a-3p&offsets|0|+1(+1A);m-30&8|-|140732587|140732607&offsets|0|+1(+1A)]|CTAGACTGAAGCTCCTTGAGGAA</td><td>>MI0000809|hsa-mir-151a&WithFlank&8|-|140732558|140732659</td><td>53</td><td>74</td><td>22</td><td>(+1A)</td><td>isomiR</td><td>NO</td><td>167.46</td><td>581.34</td><td>360.68</td><td>205.35</td></tr>
<tr><td>>MI0000809|hsa-mir-151a&WithFlank&8|-|140732558|140732659@53.74.22(+1U)&[MIMAT0000757&hsa-miR-151a-3p&offsets|0|+1(+1U);m-30&8|-|140732587|140732607&offsets|0|+1(+1U)]|CTAGACTGAAGCTCCTTGAGGAT</td><td>>MI0000809|hsa-mir-151a&WithFlank&8|-|140732558|140732659</td><td>53</td><td>74</td><td>22</td><td>(+1U)</td><td>isomiR</td><td>NO</td><td>40.46</td><td>146.77</td><td>92.91</td><td>65.25</td></tr>
</table>

# BT-20

<table>
<tr><th></th><th></th><th></th><th></th><th></th><th></th><th></th><th></th><th></th><th></th><th></th><th></th></tr>
<tr><td>>MI0000809|hsa-mir-151a&WithFlank&8|-|140732558|140732659@53.74.22(+2A)&[MIMAT0000757&hsa-miR-151a-3p&offsets|0|+1(+2A);m-30&8|-|140732587|140732607&offsets|0|+1(+2A)]|CTAGACTGAAGCTCCTTGAGGAAA</td><td>>MI0000809|hsa-mir-151a&WithFlank&8|-|140732558|140732659</td><td>53</td><td>74</td><td>22</td><td>(+2A)</td><td>isomiR</td><td>NO</td><td>26.98</td><td>126.64</td><td>75.51</td><td>30.35</td></tr>
<tr><td>>MI0000809|hsa-mir-151a&WithFlank&8|-|140732558|140732659@53.74.22(+2U)&[MIMAT0000757&hsa-miR-151a-3p&offsets|0|+1(+2U);m-30&8|-|140732587|140732607&offsets|0|+1(+2U)]|CTAGACTGAAGCTCCTTGAGGATT</td><td>>MI0000809|hsa-mir-151a&WithFlank&8|-|140732558|140732659</td><td>53</td><td>74</td><td>22</td><td>(+2U)</td><td>isomiR</td><td>NO</td><td>0.00</td><td>36.94</td><td>16.24</td><td>0.00</td></tr>
<tr><td>>am_MI0000809|hsa-mir-151a&WithFlank&8|-|140732558|140732659@17.37.21&[MIMAT0004697&hsa-miR-151a-5p&offsets|0|0;m-75&8|-|140732623|140732643&offsets|0|0]|TCGAGGAGCTCACAGTCTAGT</td><td>>MI0000809|hsa-mir-151a&WithFlank&8|-|140732558|140732659.am</td><td>17</td><td>37</td><td>21</td><td></td><td>isomiR</td><td>NO</td><td>145.17</td><td>246.22</td><td>14.62</td><td>0.00</td></tr>
<tr><td>>am_MI0000809|hsa-mir-151a&WithFlank&8|-|140732558|140732659@17.38.22&[MIMAT0004697&hsa-miR-151a-5p&offsets|0|+1;m-75&8|-|140732623|140732643&offsets|0|+1]|TCGAGGAGCTCACAGTCTAGTA</td><td>>MI0000809|hsa-mir-151a&WithFlank&8|-|140732558|140732659.am</td><td>17</td><td>38</td><td>22</td><td></td><td>isomiR</td><td>NO</td><td>65.24</td><td>92.16</td><td>0.00</td><td>0.00</td></tr>
<tr><td>>am_MI0000809|hsa-mir-151a&WithFlank&8|-|140732558|140732659@52.72.21(+1A)&[MIMAT0000757&hsa-miR-151a-3p&offsets|-1|-1(+1A);m-30&8|-|140732587|140732607&offsets|-1|-1(+1A)]|ACTAGACTGAAGCTCCTTGAGA</td><td>>MI0000809|hsa-mir-151a&WithFlank&8|-|140732558|140732659.am</td><td>52</td><td>72</td><td>21</td><td>(+1A)</td><td>isomiR</td><td>NO</td><td>0.00</td><td>31.94</td><td>0.00</td><td>0.00</td></tr>
<tr><td>>am_MI0000809|hsa-mir-151a&WithFlank&8|-|140732558|140732659@52.73.22&[MIMAT0000757&hsa-miR-151a-3p&offsets|-1|0;m-30&8|-|140732587|140732607&offsets|-1|0]|ACTAGACTGAAGCTCCTTGAGG</td><td>>MI0000809|hsa-mir-151a&WithFlank&8|-|140732558|140732659.am</td><td>52</td><td>73</td><td>22</td><td></td><td>isomiR</td><td>NO</td><td>41.97</td><td>222.20</td><td>91.40</td><td>36.19</td></tr>
<tr><td>>am_MI0000809|hsa-mir-151a&WithFlank&8|-|140732558|140732659@53.70.18&[MIMAT0000757&hsa-miR-151a-3p&offsets|0|-3;m-30&8|-|140732587|140732607&offsets|0|-3]|CTAGACTGAAGCTCCTTG</td><td>>MI0000809|hsa-mir-151a&WithFlank&8|-|140732558|140732659.am</td><td>53</td><td>70</td><td>18</td><td></td><td>isomiR</td><td>NO</td><td>66.30</td><td>149.32</td><td>193.73</td><td>40.76</td></tr>
<tr><td>>am_MI0000809|hsa-mir-151a&WithFlank&8|-|140732558|140732659@53.70.18(+3G)&[MIMAT0000757&hsa-miR-151a-3p&offsets|0|-3(+3G);m-30&8|-|140732587|140732607&offsets|0|-3(+3G)]|CTAGACTGAAGCTCCTTGGGG</td><td>>MI0000809|hsa-mir-151a&WithFlank&8|-|140732558|140732659.am</td><td>53</td><td>70</td><td>18</td><td>(+3G)</td><td>isomiR</td><td>NO</td><td>0.00</td><td>36.16</td><td>15.24</td><td>0.00</td></tr>
<tr><td>>am_MI0000809|hsa-mir-151a&WithFlank&8|-|140732558|140732659@53.71.19&[MIMAT0000757&hsa-miR-151a-3p&offsets|0|-2;m-30&8|-|140732587|140732607&offsets|0|-2]|CTAGACTGAAGCTCCTTGA</td><td>>MI0000809|hsa-mir-151a&WithFlank&8|-|140732558|140732659.am</td><td>53</td><td>71</td><td>19</td><td></td><td>isomiR</td><td>NO</td><td>38.63</td><td>103.88</td><td>125.63</td><td>33.47</td></tr>
<tr><td>>am_MI0000809|hsa-mir-151a&WithFlank&8|-|140732558|140732659@53.71.19(+1A)&[MIMAT0000757&hsa-miR-151a-3p&offsets|0|-2(+1A);m-30&8|-|140732587|140732607&offsets|0|-2(+1A)]|CTAGACTGAAGCTCCTTGAA</td><td>>MI0000809|hsa-mir-151a&WithFlank&8|-|140732558|140732659.am</td><td>53</td><td>71</td><td>19</td><td>(+1A)</td><td>isomiR</td><td>NO</td><td>0.00</td><td>38.37</td><td>39.14</td><td>0.00</td></tr>
<tr><td>>am_MI0000809|hsa-mir-151a&WithFlank&8|-|140732558|140732659@53.72.20&[MIMAT0000757&hsa-miR-151a-3p&offsets|0|-1;m-30&8|-|140732587|140732607&offsets|0|-1]|CTAGACTGAAGCTCCTTGAG</td><td>>MI0000809|hsa-mir-151a&WithFlank&8|-|140732558|140732659.am</td><td>53</td><td>72</td><td>20</td><td></td><td>isomiR</td><td>NO</td><td>130.78</td><td>315.29</td><td>356.09</td><td>84.97</td></tr>
<tr><td>>am_MI0000809|hsa-mir-151a&WithFlank&8|-|140732558|140732659@53.72.20(+1A)&[MIMAT0000757&hsa-miR-151a-3p&offsets|0|-1(+1A);m-30&8|-|140732587|140732607&offsets|0|-1(+1A)]|CTAGACTGAAGCTCCTTGAGA</td><td>>MI0000809|hsa-mir-151a&WithFlank&8|-|140732558|140732659.am</td><td>53</td><td>72</td><td>20</td><td>(+1A)</td><td>isomiR</td><td>NO</td><td>325.85</td><td>1549.56</td><td>703.86</td><td>261.95</td></tr>
<tr><td>>am_MI0000809|hsa-mir-151a&WithFlank&8|-|140732558|140732659@53.72.20(+1C)&[MIMAT0000757&hsa-miR-151a-3p&offsets|0|-1(+1C);m-30&8|-|140732587|140732607&offsets|0|-1(+1C)]|CTAGACTGAAGCTCCTTGAGC</td><td>>MI0000809|hsa-mir-151a&WithFlank&8|-|140732558|140732659.am</td><td>53</td><td>72</td><td>20</td><td>(+1C)</td><td>isomiR</td><td>NO</td><td>29.73</td><td>100.87</td><td>73.95</td><td>0.00</td></tr>
<tr><td>>am_MI0000809|hsa-mir-151a&WithFlank&8|-|140732558|140732659@53.72.20(+1U)&[MIMAT0000757&hsa-miR-151a-3p&offsets|0|-1(+1U);m-30&8|-|140732587|140732607&offsets|0|-1(+1U)]|CTAGACTGAAGCTCCTTGAGT</td><td>>MI0000809|hsa-mir-151a&WithFlank&8|-|140732558|140732659.am</td><td>53</td><td>72</td><td>20</td><td>(+1U)</td><td>isomiR</td><td>NO</td><td>163.26</td><td>449.11</td><td>298.08</td><td>124.65</td></tr>
<tr><td>>am_MI0000809|hsa-mir-151a&WithFlank&8|-|140732558|140732659@53.73.21&[MIMAT0000757&hsa-miR-151a-3p&offsets|0|0;m-30&8|-|140732587|140732607&offsets|0|0]|CTAGACTGAAGCTCCTTGAGG</td><td>>MI0000809|hsa-mir-151a&WithFlank&8|-|140732558|140732659.am</td><td>53</td><td>73</td><td>21</td><td></td><td>isomiR</td><td>YES</td><td>1525.01</td><td>7396.83</td><td>4172.97</td><td>1332.48</td></tr>
<tr><td>>am_MI0000809|hsa-mir-151a&WithFlank&8|-|140732558|140732659@53.73.21(+1C)&[MIMAT0000757&hsa-miR-151a-3p&offsets|0|0(+1C);m-30&8|-|140732587|140732607&offsets|0|0(+1C)]|CTAGACTGAAGCTCCTTGAGGC</td><td>>MI0000809|hsa-mir-151a&WithFlank&8|-|140732558|140732659.am</td><td>53</td><td>73</td><td>21</td><td>(+1C)</td><td>isomiR</td><td>NO</td><td>40.82</td><td>179.18</td><td>91.54</td><td>38.18</td></tr>
<tr><td>>am_MI0000809|hsa-mir-151a&WithFlank&8|-|140732558|140732659@53.73.21(+1G)&[MIMAT0000757&hsa-miR-151a-3p&offsets|0|0(+1G);m-30&8|-|140732587|140732607&offsets|0|0(+1G)]|CTAGACTGAAGCTCCTTGAGGG</td><td>>MI0000809|hsa-mir-151a&WithFlank&8|-|140732558|140732659.am</td><td>53</td><td>73</td><td>21</td><td>(+1G)</td><td>isomiR</td><td>NO</td><td>9.98</td><td>78.56</td><td>10.30</td><td>0.00</td></tr>
<tr><td>>am_MI0000809|hsa-mir-151a&WithFlank&8|-|140732558|140732659@53.73.21(+1U)&[MIMAT0000757&hsa-miR-151a-3p&offsets|0|0(+1U);m-30&8|-|140732587|140732607&offsets|0|0(+1U)]|CTAGACTGAAGCTCCTTGAGGT</td><td>>MI0000809|hsa-mir-151a&WithFlank&8|-|140732558|140732659.am</td><td>53</td><td>73</td><td>21</td><td>(+1U)</td><td>isomiR</td><td>NO</td><td>262.94</td><td>982.58</td><td>447.05</td><td>250.30</td></tr>
<tr><td>>am_MI0000809|hsa-mir-151a&WithFlank&8|-|140732558|140732659@53.73.21(+2U)&[MIMAT0000757&hsa-miR-151a-3p&offsets|0|0(+2U);m-30&8|-|140732587|140732607&offsets|0|0(+2U)]|CTAGACTGAAGCTCCTTGAGGTT</td><td>>MI0000809|hsa-mir-151a&WithFlank&8|-|140732558|140732659.am</td><td>53</td><td>73</td><td>21</td><td>(+2U)</td><td>isomiR</td><td>NO</td><td>26.86</td><td>101.49</td><td>39.57</td><td>9.65</td></tr>
<tr><td>>am_MI0000809|hsa-mir-151a&WithFlank&8|-|140732558|140732659@53.73.21(+3U)&[MIMAT0000757&hsa-miR-151a-3p&offsets|0|0(+3U);m-30&8|-|140732587|140732607&offsets|0|0(+3U)]|CTAGACTGAAGCTCCTTGAGGTTT</td><td>>MI0000809|hsa-mir-151a&WithFlank&8|-|140732558|140732659.am</td><td>53</td><td>73</td><td>21</td><td>(+3U)</td><td>isomiR</td><td>NO</td><td>0.00</td><td>9.95</td><td>37.27</td><td>0.00</td></tr>
<tr><td>>MI0000810|hsa-mir-135b&WithFlank&1|-|205448296|205448404@22.44.23&[MIMAT0000758&hsa-miR-135b-5p&offsets|0|0;m-321&1|-|205448361|205448383&offsets|0|0]|TATGGCTTTTCATTCCTATGTGA</td><td>>MI0000810|hsa-mir-135b&WithFlank&1|-|205448296|205448404</td><td>22</td><td>44</td><td>23</td><td></td><td>isomiR</td><td>NO</td><td>65.55</td><td>0.00</td><td>60.40</td><td>0.00</td></tr>
<tr><td>>MI0000811|hsa-mir-148b&WithFlank&12|+|54337210|54337320@69.89.21&[MIMAT0000759&hsa-miR-148b-3p&offsets|0|-1;m-77&12|+|54337278|54337299&offsets|0|-1]|TCAGTGCATCACAGAACTTTG</td><td>>MI0000811|hsa-mir-148b&WithFlank&12|+|54337210|54337320</td><td>69</td><td>89</td><td>21</td><td></td><td>isomiR</td><td>NO</td><td>0.00</td><td>0.00</td><td>12.97</td><td>0.00</td></tr>
<tr><td>>MI0000811|hsa-mir-148b&WithFlank&12|+|54337210|54337320@69.89.21(+1C)&[MIMAT0000759&hsa-miR-148b-3p&offsets|0|-1(+1C);m-77&12|+|54337278|54337299&offsets|0|-1(+1C)]|TCAGTGCATCACAGAACTTTGC</td><td>>MI0000811|hsa-mir-148b&WithFlank&12|+|54337210|54337320</td><td>69</td><td>89</td><td>21</td><td>(+1C)</td><td>isomiR</td><td>NO</td><td>27.53</td><td>75.19</td><td>55.52</td><td>0.00</td></tr>
</table>

# BT-20

| | | | | | | | | | | | |
|---|---|---|---|---|---|---|---|---|---|---|---|
| >MI0000811\|hsa-mir-148b&WithFlank&12\|+\|54337210\|54337320@69.90.22&[MIMAT0000759&hsa-miR-148b-3p&offsets\|0\|0;m-77&12\|+\|54337278\|54337299&offsets\|0\|0]\|TCAGTGCATCACAGAACTTTGT | >MI0000811\|hsa-mir-148b&WithFlank&12\|+\|54337210\|54337320 | 69 | 90 | 22 | | isomiR | NO | 201.64 | 496.05 | 292.55 | 34.40 |
| >MI0000811\|hsa-mir-148b&WithFlank&12\|+\|54337210\|54337320@69.90.22(+1A)&[MIMAT0000759&hsa-miR-148b-3p&offsets\|0\|0(+1A);m-77&12\|+\|54337278\|54337299&offsets\|0\|0(+1A)]\|TCAGTGCATCACAGAACTTTGTA | >MI0000811\|hsa-mir-148b&WithFlank&12\|+\|54337210\|54337320 | 69 | 90 | 22 | (+1A) | isomiR | NO | 0.00 | 10.07 | 0.00 | 0.00 |
| >MI0000811\|hsa-mir-148b&WithFlank&12\|+\|54337210\|54337320@69.90.22(+1U)&[MIMAT0000759&hsa-miR-148b-3p&offsets\|0\|0(+1U);m-77&12\|+\|54337278\|54337299&offsets\|0\|0(+1U)]\|TCAGTGCATCACAGAACTTTGTT | >MI0000811\|hsa-mir-148b&WithFlank&12\|+\|54337210\|54337320 | 69 | 90 | 22 | (+1U) | isomiR | NO | 136.34 | 360.74 | 11.00 | 35.19 |
| >MI0000811\|hsa-mir-148b&WithFlank&12\|+\|54337210\|54337320@69.91.23&[MIMAT0000759&hsa-miR-148b-3p&offsets\|0\|+1;m-77&12\|+\|54337278\|54337299&offsets\|0\|+1]\|TCAGTGCATCACAGAACTTTGTC | >MI0000811\|hsa-mir-148b&WithFlank&12\|+\|54337210\|54337320 | 69 | 91 | 23 | | isomiR | NO | 0.00 | 38.90 | 0.00 | 0.00 |
| >MI0000812\|hsa-mir-331&WithFlank&12\|+\|95308414\|95308519@67.87.21(+1U)&[MIMAT0000760&hsa-miR-331-3p&offsets\|0\|0(+1U);m-192&12\|+\|95308480\|95308501&offsets\|0\|-1(+1U)]\|GCCCCTGGGCCTATCCTAGAAT | >MI0000812\|hsa-mir-331&WithFlank&12\|+\|95308414\|95308519 | 67 | 87 | 21 | (+1U) | isomiR | NO | 35.32 | 34.02 | 0.00 | 0.00 |
| >MI0000815\|hsa-mir-339&WithFlank&7\|-\|1022927\|1023032@21.43.23&[MIMAT0000764&hsa-miR-339-5p&offsets\|0\|0;m-163&7\|-\|1022990\|1023012&offsets\|0\|0]\|TCCCTGTCCTCCAGGAGCTCACG | >MI0000815\|hsa-mir-339&WithFlank&7\|-\|1022927\|1023032 | 21 | 43 | 23 | | isomiR | NO | 10.75 | 134.37 | 0.00 | 0.00 |
| >MI0000815\|hsa-mir-339&WithFlank&7\|-\|1022927\|1023032@56.76.21&[MIMAT0004702&hsa-miR-339-3p&offsets\|0\|-2;m-235&7\|-\|1022957\|1022977&offsets\|0\|0]\|TGAGCGCCTCGACGACAGAGC | >MI0000815\|hsa-mir-339&WithFlank&7\|-\|1022927\|1023032 | 56 | 76 | 21 | | isomiR | NO | 0.00 | 24.73 | 84.73 | 0.00 |
| >MI0000815\|hsa-mir-339&WithFlank&7\|-\|1022927\|1023032@56.78.23&[MIMAT0004702&hsa-miR-339-3p&offsets\|0\|0;m-235&7\|-\|1022957\|1022977&offsets\|0\|+2]\|TGAGCGCCTCGACGACAGAGCCG | >MI0000815\|hsa-mir-339&WithFlank&7\|-\|1022927\|1023032 | 56 | 78 | 23 | | isomiR | NO | 0.00 | 0.00 | 12.37 | 0.00 |
| >am_MI0000815\|hsa-mir-339&WithFlank&7\|-\|1022927\|1023032@21.42.22(+1A)&[MIMAT0000764&hsa-miR-339-5p&offsets\|0\|-1(+1A);m-163&7\|-\|1022990\|1023012&offsets\|0\|-1(+1A)]\|TCCCTGTCCTCCAGGAGCTCACA | >MI0000815\|hsa-mir-339&WithFlank&7\|-\|1022927\|1023032.am | 21 | 42 | 22 | (+1A) | isomiR | NO | 0.00 | 32.46 | 0.00 | 0.00 |
| >MI0000816\|hsa-mir-335&WithFlank&7\|+\|130496105\|130496210@58.78.21(+1A)&[MIMAT0004703&hsa-miR-335-3p&offsets\|0\|-1(+1A);m-159&7\|+\|130496162\|130496183&offsets\|0\|-1(+1A)]\|TTTTTCATTATTGCTCCTGACA | >MI0000816\|hsa-mir-335&WithFlank&7\|+\|130496105\|130496210 | 58 | 78 | 21 | (+1A) | isomiR | NO | 0.00 | 28.02 | 0.00 | 0.00 |
| >MI0000816\|hsa-mir-335&WithFlank&7\|+\|130496105\|130496210@58.78.21(+1U)&[MIMAT0004703&hsa-miR-335-3p&offsets\|0\|-1(+1U);m-159&7\|+\|130496162\|130496183&offsets\|0\|-1(+1U)]\|TTTTTCATTATTGCTCCTGACT | >MI0000816\|hsa-mir-335&WithFlank&7\|+\|130496105\|130496210 | 58 | 78 | 21 | (+1U) | isomiR | NO | 0.00 | 73.09 | 0.00 | 0.00 |
| >MI0000816\|hsa-mir-335&WithFlank&7\|+\|130496105\|130496210@58.79.22&[MIMAT0004703&hsa-miR-335-3p&offsets\|0\|0;m-159&7\|+\|130496162\|130496183&offsets\|0\|0]\|TTTTTCATTATTGCTCCTGACC | >MI0000816\|hsa-mir-335&WithFlank&7\|+\|130496105\|130496210 | 58 | 79 | 22 | | isomiR | NO | 11.16 | 147.31 | 0.00 | 0.00 |
| >MI0001445\|hsa-mir-423&WithFlank&17\|+\|30117073\|30117178@23.42.20(+1A)&[MIMAT0004748&hsa-miR-423-5p&offsets\|0\|-3(+1A);m-141&17\|+\|30117095\|30117117&offsets\|0\|-3(+1A)]\|TGAGGGGCAGAGAGCGAGACA | >MI0001445\|hsa-mir-423&WithFlank&17\|+\|30117073\|30117178 | 23 | 42 | 20 | (+1A) | isomiR | NO | 0.00 | 24.82 | 90.71 | 0.00 |
| >MI0001445\|hsa-mir-423&WithFlank&17\|+\|30117073\|30117178@23.42.20(+1C)&[MIMAT0004748&hsa-miR-423-5p&offsets\|0\|-3(+1C);m-141&17\|+\|30117095\|30117117&offsets\|0\|-3(+1C)]\|TGAGGGGCAGAGAGCGAGACC | >MI0001445\|hsa-mir-423&WithFlank&17\|+\|30117073\|30117178 | 23 | 42 | 20 | (+1C) | isomiR | NO | 0.00 | 25.57 | 109.34 | 0.00 |
| >MI0001445\|hsa-mir-423&WithFlank&17\|+\|30117073\|30117178@23.43.21&[MIMAT0004748&hsa-miR-423-5p&offsets\|0\|-2;m-141&17\|+\|30117095\|30117117&offsets\|0\|-2]\|TGAGGGGCAGAGAGCGAGACT | >MI0001445\|hsa-mir-423&WithFlank&17\|+\|30117073\|30117178 | 23 | 43 | 21 | | isomiR | NO | 10.36 | 219.57 | 554.85 | 0.00 |
| >MI0001445\|hsa-mir-423&WithFlank&17\|+\|30117073\|30117178@23.44.22&[MIMAT0004748&hsa-miR-423-5p&offsets\|0\|-1;m-141&17\|+\|30117095\|30117117&offsets\|0\|-1]\|TGAGGGGCAGAGAGCGAGACTT | >MI0001445\|hsa-mir-423&WithFlank&17\|+\|30117073\|30117178 | 23 | 44 | 22 | | isomiR | NO | 0.00 | 153.04 | 377.54 | 0.00 |
| >MI0001445\|hsa-mir-423&WithFlank&17\|+\|30117073\|30117178@23.44.22(+1A)&[MIMAT0004748&hsa-miR-423-5p&offsets\|0\|-1(+1A);m-141&17\|+\|30117095\|30117117&offsets\|0\|-1(+1A)]\|TGAGGGGCAGAGAGCGAGACTTA | >MI0001445\|hsa-mir-423&WithFlank&17\|+\|30117073\|30117178 | 23 | 44 | 22 | (+1A) | isomiR | NO | 0.00 | 10.25 | 107.25 | 0.00 |
| >MI0001445\|hsa-mir-423&WithFlank&17\|+\|30117073\|30117178@23.44.22(+1C)&[MIMAT0004748&hsa-miR-423-5p&offsets\|0\|-1(+1C);m-141&17\|+\|30117095\|30117117&offsets\|0\|-1(+1C)]\|TGAGGGGCAGAGAGCGAGACTTC | >MI0001445\|hsa-mir-423&WithFlank&17\|+\|30117073\|30117178 | 23 | 44 | 22 | (+1C) | isomiR | NO | 0.00 | 112.52 | 475.84 | 0.00 |
| >MI0001445\|hsa-mir-423&WithFlank&17\|+\|30117073\|30117178@23.44.22(+1G)&[MIMAT0004748&hsa-miR-423-5p&offsets\|0\|-1(+1G);m-141&17\|+\|30117095\|30117117&offsets\|0\|-1(+1G)]\|TGAGGGGCAGAGAGCGAGACTTG | >MI0001445\|hsa-mir-423&WithFlank&17\|+\|30117073\|30117178 | 23 | 44 | 22 | (+1G) | isomiR | NO | 0.00 | 0.00 | 11.13 | 0.00 |
| >MI0001445\|hsa-mir-423&WithFlank&17\|+\|30117073\|30117178@23.45.23&[MIMAT0004748&hsa-miR-423-5p&offsets\|0\|0;m-141&17\|+\|30117095\|30117117&offsets\|0\|0]\|TGAGGGGCAGAGAGCGAGACTTT | >MI0001445\|hsa-mir-423&WithFlank&17\|+\|30117073\|30117178 | 23 | 45 | 23 | | isomiR | YES | 134.67 | 859.95 | 3070.80 | 39.06 |
| >MI0001445\|hsa-mir-423&WithFlank&17\|+\|30117073\|30117178@23.46.24&[MIMAT0004748&hsa-miR-423-5p&offsets\|0\|+1;m-141&17\|+\|30117095\|30117117&offsets\|0\|+1]\|TGAGGGGCAGAGAGCGAGACTTTT | >MI0001445\|hsa-mir-423&WithFlank&17\|+\|30117073\|30117178 | 23 | 46 | 24 | | isomiR | NO | 26.02 | 224.90 | 380.57 | 0.00 |

# BT-20

<table>
<tr><td>>MI0001445|hsa-mir-423&WithFlank&17|+|30117073|30117178@58.78.21&[MIMAT0001340&hsa-miR-423-3p&offsets|-1|-3;m-100&17|+|30117131|30117153&offsets|-1|-3]|AAGCTCGGTCTGAGGCCCCTC</td><td>>MI0001445|hsa-mir-423&WithFlank&17|+|30117073|30117178</td><td>58</td><td>78</td><td>21</td><td></td><td>isomiR</td><td>NO</td><td>27.27</td><td>90.56</td><td>69.52</td><td>73.45</td></tr>
<tr><td>>MI0001445|hsa-mir-423&WithFlank&17|+|30117073|30117178@58.79.22&[MIMAT0001340&hsa-miR-423-3p&offsets|-1|-2;m-100&17|+|30117131|30117153&offsets|-1|-2]|AAGCTCGGTCTGAGGCCCCTCA</td><td>>MI0001445|hsa-mir-423&WithFlank&17|+|30117073|30117178</td><td>58</td><td>79</td><td>22</td><td></td><td>isomiR</td><td>NO</td><td>0.00</td><td>36.81</td><td>0.00</td><td>0.00</td></tr>
<tr><td>>MI0001445|hsa-mir-423&WithFlank&17|+|30117073|30117178@58.80.23(+1C)&[MIMAT0001340&hsa-miR-423-3p&offsets|-1|-1(+1C);m-100&17|+|30117131|30117153&offsets|-1|-1(+1C)]|AAGCTCGGTCTGAGGCCCCTCAGC</td><td>>MI0001445|hsa-mir-423&WithFlank&17|+|30117073|30117178</td><td>58</td><td>80</td><td>23</td><td>(+1C)</td><td>isomiR</td><td>NO</td><td>0.00</td><td>31.97</td><td>0.00</td><td>24.59</td></tr>
<tr><td>>MI0001445|hsa-mir-423&WithFlank&17|+|30117073|30117178@58.81.24&[MIMAT0001340&hsa-miR-423-3p&offsets|-1|0;m-100&17|+|30117131|30117153&offsets|-1|0]|AAGCTCGGTCTGAGGCCCCTCAGT</td><td>>MI0001445|hsa-mir-423&WithFlank&17|+|30117073|30117178</td><td>58</td><td>81</td><td>24</td><td></td><td>isomiR</td><td>NO</td><td>118.09</td><td>311.15</td><td>145.94</td><td>186.56</td></tr>
<tr><td>>MI0001445|hsa-mir-423&WithFlank&17|+|30117073|30117178@59.77.19(+1U)&[MIMAT0001340&hsa-miR-423-3p&offsets|0|-4(+1U);m-100&17|+|30117131|30117153&offsets|0|-4(+1U)]|AGCTCGGTCTGAGGCCCCTT</td><td>>MI0001445|hsa-mir-423&WithFlank&17|+|30117073|30117178</td><td>59</td><td>77</td><td>19</td><td>(+1U)</td><td>isomiR</td><td>NO</td><td>0.00</td><td>39.70</td><td>14.03</td><td>30.82</td></tr>
<tr><td>>MI0001445|hsa-mir-423&WithFlank&17|+|30117073|30117178@59.78.20&[MIMAT0001340&hsa-miR-423-3p&offsets|0|-3;m-100&17|+|30117131|30117153&offsets|0|-3]|AGCTCGGTCTGAGGCCCCTC</td><td>>MI0001445|hsa-mir-423&WithFlank&17|+|30117073|30117178</td><td>59</td><td>78</td><td>20</td><td></td><td>isomiR</td><td>NO</td><td>126.50</td><td>299.21</td><td>192.63</td><td>238.55</td></tr>
<tr><td>>MI0001445|hsa-mir-423&WithFlank&17|+|30117073|30117178@59.79.21&[MIMAT0001340&hsa-miR-423-3p&offsets|0|-2;m-100&17|+|30117131|30117153&offsets|0|-2]|AGCTCGGTCTGAGGCCCCTCA</td><td>>MI0001445|hsa-mir-423&WithFlank&17|+|30117073|30117178</td><td>59</td><td>79</td><td>21</td><td></td><td>isomiR</td><td>NO</td><td>0.00</td><td>105.28</td><td>52.77</td><td>31.17</td></tr>
<tr><td>>MI0001445|hsa-mir-423&WithFlank&17|+|30117073|30117178@59.80.22&[MIMAT0001340&hsa-miR-423-3p&offsets|0|-1;m-100&17|+|30117131|30117153&offsets|0|-1]|AGCTCGGTCTGAGGCCCCTCAG</td><td>>MI0001445|hsa-mir-423&WithFlank&17|+|30117073|30117178</td><td>59</td><td>80</td><td>22</td><td></td><td>isomiR</td><td>NO</td><td>29.17</td><td>126.08</td><td>80.37</td><td>74.28</td></tr>
<tr><td>>MI0001445|hsa-mir-423&WithFlank&17|+|30117073|30117178@59.80.22(+1A)&[MIMAT0001340&hsa-miR-423-3p&offsets|0|-1(+1A);m-100&17|+|30117131|30117153&offsets|0|-1(+1A)]|AGCTCGGTCTGAGGCCCCTCAGA</td><td>>MI0001445|hsa-mir-423&WithFlank&17|+|30117073|30117178</td><td>59</td><td>80</td><td>22</td><td>(+1A)</td><td>isomiR</td><td>NO</td><td>26.53</td><td>120.62</td><td>65.45</td><td>38.17</td></tr>
<tr><td>>MI0001445|hsa-mir-423&WithFlank&17|+|30117073|30117178@59.80.22(+1C)&[MIMAT0001340&hsa-miR-423-3p&offsets|0|-1(+1C);m-100&17|+|30117131|30117153&offsets|0|-1(+1C)]|AGCTCGGTCTGAGGCCCCTCAGC</td><td>>MI0001445|hsa-mir-423&WithFlank&17|+|30117073|30117178</td><td>59</td><td>80</td><td>22</td><td>(+1C)</td><td>isomiR</td><td>NO</td><td>111.62</td><td>261.50</td><td>113.82</td><td>153.73</td></tr>
<tr><td>>MI0001445|hsa-mir-423&WithFlank&17|+|30117073|30117178@59.80.22(+1G)&[MIMAT0001340&hsa-miR-423-3p&offsets|0|-1(+1G);m-100&17|+|30117131|30117153&offsets|0|-1(+1G)]|AGCTCGGTCTGAGGCCCCTCAGG</td><td>>MI0001445|hsa-mir-423&WithFlank&17|+|30117073|30117178</td><td>59</td><td>80</td><td>22</td><td>(+1G)</td><td>isomiR</td><td>NO</td><td>0.00</td><td>38.75</td><td>0.00</td><td>0.00</td></tr>
<tr><td>>MI0001445|hsa-mir-423&WithFlank&17|+|30117073|30117178@59.81.23&[MIMAT0001340&hsa-miR-423-3p&offsets|0|0;m-100&17|+|30117131|30117153&offsets|0|0]|AGCTCGGTCTGAGGCCCCTCAGT</td><td>>MI0001445|hsa-mir-423&WithFlank&17|+|30117073|30117178</td><td>59</td><td>81</td><td>23</td><td></td><td>isomiR</td><td>NO</td><td>540.67</td><td>1734.66</td><td>644.09</td><td>774.88</td></tr>
<tr><td>>MI0001445|hsa-mir-423&WithFlank&17|+|30117073|30117178@59.81.23(+1A)&[MIMAT0001340&hsa-miR-423-3p&offsets|0|0(+1A);m-100&17|+|30117131|30117153&offsets|0|0(+1A)]|AGCTCGGTCTGAGGCCCCTCAGTA</td><td>>MI0001445|hsa-mir-423&WithFlank&17|+|30117073|30117178</td><td>59</td><td>81</td><td>23</td><td>(+1A)</td><td>isomiR</td><td>NO</td><td>32.21</td><td>114.12</td><td>0.00</td><td>99.23</td></tr>
<tr><td>>MI0001445|hsa-mir-423&WithFlank&17|+|30117073|30117178@59.81.23(+1U)&[MIMAT0001340&hsa-miR-423-3p&offsets|0|0(+1U);m-100&17|+|30117131|30117153&offsets|0|0(+1U)]|AGCTCGGTCTGAGGCCCCTCAGTT</td><td>>MI0001445|hsa-mir-423&WithFlank&17|+|30117073|30117178</td><td>59</td><td>81</td><td>23</td><td>(+1U)</td><td>isomiR</td><td>NO</td><td>39.90</td><td>143.94</td><td>11.17</td><td>114.11</td></tr>
<tr><td>>MI0001446|hsa-mir-424&WithFlank&X|-|134546608|134546717@54.74.21&[MIMAT0004749&hsa-miR-424-3p&offsets|0|0;m-305&X|-|134546644|134546664&offsets|0|0]|CAAAACGTGAGGCGCTGCTAT</td><td>>MI0001446|hsa-mir-424&WithFlank&X|-|134546608|134546717</td><td>54</td><td>74</td><td>21</td><td></td><td>isomiR</td><td>NO</td><td>0.00</td><td>100.42</td><td>13.08</td><td>0.00</td></tr>
<tr><td>>MI0001446|hsa-mir-424&WithFlank&X|-|134546608|134546717@54.75.22&[MIMAT0004749&hsa-miR-424-3p&offsets|0|+1;m-305&X|-|134546644|134546664&offsets|0|+1]|CAAAACGTGAGGCGCTGCTATA</td><td>>MI0001446|hsa-mir-424&WithFlank&X|-|134546608|134546717</td><td>54</td><td>75</td><td>22</td><td></td><td>isomiR</td><td>NO</td><td>0.00</td><td>33.23</td><td>0.00</td><td>0.00</td></tr>
<tr><td>>MI0001448|hsa-mir-425&WithFlank&3|-|49020142|49020240@20.41.22&[MIMAT0003393&hsa-miR-425-5p&offsets|0|-1;m-95&3|-|49020200|49020221&offsets|0|0]|AATGACACGATCACTCCCGTTG</td><td>>MI0001448|hsa-mir-425&WithFlank&3|-|49020142|49020240</td><td>20</td><td>41</td><td>22</td><td></td><td>isomiR</td><td>NO</td><td>0.00</td><td>33.66</td><td>0.00</td><td>0.00</td></tr>
<tr><td>>MI0001448|hsa-mir-425&WithFlank&3|-|49020142|49020240@20.42.23&[MIMAT0003393&hsa-miR-425-5p&offsets|0|0;m-95&3|-|49020200|49020221&offsets|0|+1]|AATGACACGATCACTCCCGTTGA</td><td>>MI0001448|hsa-mir-425&WithFlank&3|-|49020142|49020240</td><td>20</td><td>42</td><td>23</td><td></td><td>isomiR</td><td>NO</td><td>0.00</td><td>35.12</td><td>0.00</td><td>0.00</td></tr>
<tr><td>>MI0001448|hsa-mir-425&WithFlank&3|-|49020142|49020240@20.43.24&[MIMAT0003393&hsa-miR-425-5p&offsets|0|+1;m-95&3|-|49020200|49020221&offsets|0|+2]|AATGACACGATCACTCCCGTTGAG</td><td>>MI0001448|hsa-mir-425&WithFlank&3|-|49020142|49020240</td><td>20</td><td>43</td><td>24</td><td></td><td>isomiR</td><td>NO</td><td>36.74</td><td>145.25</td><td>0.00</td><td>0.00</td></tr>
<tr><td>>MI0001448|hsa-mir-425&WithFlank&3|-|49020142|49020240@20.43.24(+1C)&[MIMAT0003393&hsa-miR-425-5p&offsets|0|+1(+1C);m-95&3|-|49020200|49020221&offsets|0|+2(+1C)]|AATGACACGATCACTCCCGTTGAGC</td><td>>MI0001448|hsa-mir-425&WithFlank&3|-|49020142|49020240</td><td>20</td><td>43</td><td>24</td><td>(+1C)</td><td>isomiR</td><td>NO</td><td>0.00</td><td>72.16</td><td>0.00</td><td>0.00</td></tr>
<tr><td>>MI0001448|hsa-mir-425&WithFlank&3|-|49020142|49020240@20.44.25&[MIMAT0003393&hsa-miR-425-5p&offsets|0|+2;m-95&3|-|49020200|49020221&offsets|0|+3]|AATGACACGATCACTCCCGTTGAGT</td><td>>MI0001448|hsa-mir-425&WithFlank&3|-|49020142|49020240</td><td>20</td><td>44</td><td>25</td><td></td><td>isomiR</td><td>NO</td><td>125.67</td><td>479.03</td><td>12.34</td><td>28.98</td></tr>
<tr><td>>MI0001448|hsa-mir-425&WithFlank&3|-|49020142|49020240@21.44.24&[MIMAT0003393&hsa-miR-425-5p&offsets|+1|+2;m-95&3|-|49020200|49020221&offsets|+1|+3]|ATGACACGATCACTCCCGTTGAGT</td><td>>MI0001448|hsa-mir-425&WithFlank&3|-|49020142|49020240</td><td>21</td><td>44</td><td>24</td><td></td><td>isomiR</td><td>NO</td><td>0.00</td><td>42.42</td><td>0.00</td><td>0.00</td></tr>
</table>

# BT-20

<table>
<tr><td>>MI0001448|hsa-mir-425&WithFlank&3|-|49020142|49020240@60.81.22&[MIMAT0001343&hsa-miR-425-3p&offsets|-1|-1;m-258&3|-|49020160|49020181&offsets|0|0]|CATCGGGAATGTCGTGTCCGCC</td><td>>MI0001448|hsa-mir-425&WithFlank&3|-|49020142|49020240</td><td>60</td><td>81</td><td>22</td><td></td><td>isomiR</td><td>NO</td><td>0.00</td><td>0.00</td><td>12.23</td><td>0.00</td></tr>
<tr><td>>MI0001733|hsa-mir-452&WithFlank&X|-|151959622|151959718@20.41.22&[MIMAT0001635&hsa-miR-452-5p&offsets|0|0;m-188&X|-|151959677|151959699&offsets|0|-1]|AACTGTTTGCAGAGGAAACTGA</td><td>>MI0001733|hsa-mir-452&WithFlank&X|-|151959622|151959718</td><td>20</td><td>41</td><td>22</td><td></td><td>isomiR</td><td>NO</td><td>10.62</td><td>0.00</td><td>0.00</td><td>0.00</td></tr>
<tr><td>>MI0001733|hsa-mir-452&WithFlank&X|-|151959622|151959718@20.42.23&[MIMAT0001635&hsa-miR-452-5p&offsets|0|+1;m-188&X|-|151959677|151959699&offsets|0|0]|AACTGTTTGCAGAGGAAACTGAG</td><td>>MI0001733|hsa-mir-452&WithFlank&X|-|151959622|151959718</td><td>20</td><td>42</td><td>23</td><td></td><td>isomiR</td><td>NO</td><td>69.77</td><td>110.99</td><td>15.17</td><td>23.72</td></tr>
<tr><td>>MI0001733|hsa-mir-452&WithFlank&X|-|151959622|151959718@20.43.24&[MIMAT0001635&hsa-miR-452-5p&offsets|0|+2;m-188&X|-|151959677|151959699&offsets|0|+1]|AACTGTTTGCAGAGGAAACTGAGA</td><td>>MI0001733|hsa-mir-452&WithFlank&X|-|151959622|151959718</td><td>20</td><td>43</td><td>24</td><td></td><td>isomiR</td><td>NO</td><td>64.77</td><td>103.46</td><td>0.00</td><td>9.99</td></tr>
<tr><td>>MI0001733|hsa-mir-452&WithFlank&X|-|151959622|151959718@23.43.21&[MIMAT0001635&hsa-miR-452-5p&offsets|+3|+2;m-188&X|-|151959677|151959699&offsets|+3|+1]|TGTTTGCAGAGGAAACTGAGA</td><td>>MI0001733|hsa-mir-452&WithFlank&X|-|151959622|151959718</td><td>23</td><td>43</td><td>21</td><td></td><td>isomiR</td><td>NO</td><td>28.57</td><td>73.97</td><td>12.72</td><td>0.00</td></tr>
<tr><td>>MI0001733|hsa-mir-452&WithFlank&X|-|151959622|151959718@23.44.22&[MIMAT0001635&hsa-miR-452-5p&offsets|+3|+3;m-188&X|-|151959677|151959699&offsets|+3|+2]|TGTTTGCAGAGGAAACTGAGAC</td><td>>MI0001733|hsa-mir-452&WithFlank&X|-|151959622|151959718</td><td>23</td><td>44</td><td>22</td><td></td><td>isomiR</td><td>NO</td><td>0.00</td><td>9.68</td><td>91.18</td><td>0.00</td></tr>
<tr><td>>MI0002468|hsa-mir-484&WithFlank&16|+|15643288|15643378@14.35.22&[MIMAT0002174&hsa-miR-484&offsets|0|0;m-127&16|+|15643301|15643321&offsets|0|+1]|TCAGGCTCAGTCCCCTCCCGAT</td><td>>MI0002468|hsa-mir-484&WithFlank&16|+|15643288|15643378</td><td>14</td><td>35</td><td>22</td><td></td><td>isomiR</td><td>NO</td><td>25.39</td><td>115.31</td><td>0.00</td><td>32.35</td></tr>
<tr><td>>MI0002468|hsa-mir-484&WithFlank&16|+|15643288|15643378@14.35.22(+1U)&[MIMAT0002174&hsa-miR-484&offsets|0|0(+1U);m-127&16|+|15643301|15643321&offsets|0|+1(+1U)]|TCAGGCTCAGTCCCCTCCCGATT</td><td>>MI0002468|hsa-mir-484&WithFlank&16|+|15643288|15643378</td><td>14</td><td>35</td><td>22</td><td>(+1U)</td><td>isomiR</td><td>NO</td><td>0.00</td><td>36.10</td><td>0.00</td><td>0.00</td></tr>
<tr><td>>MI0003129|hsa-mir-146b&WithFlank&10|+|102436506|102436590@15.38.24&[MIMAT0002809&hsa-miR-146b-5p&offsets|0|+1;m-70&10|+|102436520|102436543&offsets|0|0]|TGAGAACTGAATTCCATAGGCTGT</td><td>>MI0003129|hsa-mir-146b&WithFlank&10|+|102436506|102436590</td><td>15</td><td>38</td><td>24</td><td></td><td>isomiR</td><td>NO</td><td>0.00</td><td>10.48</td><td>0.00</td><td>0.00</td></tr>
<tr><td>>MI0003137|hsa-mir-193b&WithFlank&16|+|14303961|14304055@57.77.21&[MIMAT0002819&hsa-miR-193b-3p&offsets|0|-1;m-166&16|+|14304017|14304038&offsets|0|-1]|AACTGGCCCTCAAAGTCCCGC</td><td>>MI0003137|hsa-mir-193b&WithFlank&16|+|14303961|14304055</td><td>57</td><td>77</td><td>21</td><td></td><td>isomiR</td><td>NO</td><td>0.00</td><td>0.00</td><td>84.67</td><td>0.00</td></tr>
<tr><td>>MI0003137|hsa-mir-193b&WithFlank&16|+|14303961|14304055@57.77.21(+1A)&[MIMAT0002819&hsa-miR-193b-3p&offsets|0|-1(+1A);m-166&16|+|14304017|14304038&offsets|0|-1(+1A)]|AACTGGCCCTCAAAGTCCCGCA</td><td>>MI0003137|hsa-mir-193b&WithFlank&16|+|14303961|14304055</td><td>57</td><td>77</td><td>21</td><td>(+1A)</td><td>isomiR</td><td>NO</td><td>11.01</td><td>24.60</td><td>14.94</td><td>0.00</td></tr>
<tr><td>>MI0003137|hsa-mir-193b&WithFlank&16|+|14303961|14304055@57.77.21(+1C)&[MIMAT0002819&hsa-miR-193b-3p&offsets|0|-1(+1C);m-166&16|+|14304017|14304038&offsets|0|-1(+1C)]|AACTGGCCCTCAAAGTCCCGCC</td><td>>MI0003137|hsa-mir-193b&WithFlank&16|+|14303961|14304055</td><td>57</td><td>77</td><td>21</td><td>(+1C)</td><td>isomiR</td><td>NO</td><td>37.51</td><td>29.54</td><td>72.05</td><td>0.00</td></tr>
<tr><td>>MI0003137|hsa-mir-193b&WithFlank&16|+|14303961|14304055@57.78.22&[MIMAT0002819&hsa-miR-193b-3p&offsets|0|0;m-166&16|+|14304017|14304038&offsets|0|0]|AACTGGCCCTCAAAGTCCCGCT</td><td>>MI0003137|hsa-mir-193b&WithFlank&16|+|14303961|14304055</td><td>57</td><td>78</td><td>22</td><td></td><td>isomiR</td><td>NO</td><td>224.81</td><td>241.25</td><td>352.13</td><td>70.44</td></tr>
<tr><td>>MI0003137|hsa-mir-193b&WithFlank&16|+|14303961|14304055@57.79.23&[MIMAT0002819&hsa-miR-193b-3p&offsets|0|+1;m-166&16|+|14304017|14304038&offsets|0|+1]|AACTGGCCCTCAAAGTCCCGCTT</td><td>>MI0003137|hsa-mir-193b&WithFlank&16|+|14303961|14304055</td><td>57</td><td>79</td><td>23</td><td></td><td>isomiR</td><td>NO</td><td>0.00</td><td>0.00</td><td>90.76</td><td>0.00</td></tr>
<tr><td>>MI0003188|hsa-mir-503&WithFlank&X|-|134546322|134546404@12.31.20&[MIMAT0002874&hsa-miR-503-5p&offsets|0|-3;m-297&X|-|134546371|134546393&offsets|0|-3]|TAGCAGCGGGAACAGTTCTG</td><td>>MI0003188|hsa-mir-503&WithFlank&X|-|134546322|134546404</td><td>12</td><td>31</td><td>20</td><td></td><td>isomiR</td><td>NO</td><td>0.00</td><td>30.66</td><td>65.79</td><td>0.00</td></tr>
<tr><td>>MI0003205|hsa-mir-532&WithFlank&X|+|50003142|50003244@26.46.21(+1C)&[MIMAT0002888&hsa-miR-532-5p&offsets|0|-1(+1C);m-65&X|+|50003167|50003188&offsets|0|-1(+1C)]|CATGCCTTGAGTGTAGGACCGC</td><td>>MI0003205|hsa-mir-532&WithFlank&X|+|50003142|50003244</td><td>26</td><td>46</td><td>21</td><td>(+1C)</td><td>isomiR</td><td>NO</td><td>0.00</td><td>38.12</td><td>56.06</td><td>0.00</td></tr>
<tr><td>>MI0003205|hsa-mir-532&WithFlank&X|+|50003142|50003244@26.47.22&[MIMAT0002888&hsa-miR-532-5p&offsets|0|0;m-65&X|+|50003167|50003188&offsets|0|0]|CATGCCTTGAGTGTAGGACCGT</td><td>>MI0003205|hsa-mir-532&WithFlank&X|+|50003142|50003244</td><td>26</td><td>47</td><td>22</td><td></td><td>isomiR</td><td>NO</td><td>101.97</td><td>348.01</td><td>308.44</td><td>39.80</td></tr>
<tr><td>>MI0003205|hsa-mir-532&WithFlank&X|+|50003142|50003244@27.47.21&[MIMAT0002888&hsa-miR-532-5p&offsets|+1|0;m-65&X|+|50003167|50003188&offsets|+1|0]|ATGCCTTGAGTGTAGGACCGT</td><td>>MI0003205|hsa-mir-532&WithFlank&X|+|50003142|50003244</td><td>27</td><td>47</td><td>21</td><td></td><td>isomiR</td><td>NO</td><td>0.00</td><td>0.00</td><td>10.49</td><td>0.00</td></tr>
<tr><td>>MI0003560|hsa-mir-92b&WithFlank&1|+|155195171|155195278@67.86.20&[MIMAT0003218&hsa-miR-92b-3p&offsets|0|-2;m-115&1|+|155195237|155195258&offsets|0|-2]|TATTGCACTCGTCCCGGCCT</td><td>>MI0003560|hsa-mir-92b&WithFlank&1|+|155195171|155195278</td><td>67</td><td>86</td><td>20</td><td></td><td>isomiR</td><td>NO</td><td>0.00</td><td>44.67</td><td>9.53</td><td>0.00</td></tr>
<tr><td>>MI0003560|hsa-mir-92b&WithFlank&1|+|155195171|155195278@67.86.20(+1U)&[MIMAT0003218&hsa-miR-92b-3p&offsets|0|-2(+1U);m-115&1|+|155195237|155195258&offsets|0|-2(+1U)]|TATTGCACTCGTCCCGGCCTT</td><td>>MI0003560|hsa-mir-92b&WithFlank&1|+|155195171|155195278</td><td>67</td><td>86</td><td>20</td><td>(+1U)</td><td>isomiR</td><td>NO</td><td>0.00</td><td>32.12</td><td>0.00</td><td>0.00</td></tr>
<tr><td>>MI0003560|hsa-mir-92b&WithFlank&1|+|155195171|155195278@67.87.21&[MIMAT0003218&hsa-miR-92b-3p&offsets|0|-1;m-115&1|+|155195237|155195258&offsets|0|-1]|TATTGCACTCGTCCCGGCCTC</td><td>>MI0003560|hsa-mir-92b&WithFlank&1|+|155195171|155195278</td><td>67</td><td>87</td><td>21</td><td></td><td>isomiR</td><td>NO</td><td>30.16</td><td>127.25</td><td>71.54</td><td>0.00</td></tr>
<tr><td>>MI0003560|hsa-mir-92b&WithFlank&1|+|155195171|155195278@67.87.21(+1A)&[MIMAT0003218&hsa-miR-92b-3p&offsets|0|-1(+1A);m-115&1|+|155195237|155195258&offsets|0|-1(+1A)]|TATTGCACTCGTCCCGGCCTCA</td><td>>MI0003560|hsa-mir-92b&WithFlank&1|+|155195171|155195278</td><td>67</td><td>87</td><td>21</td><td>(+1A)</td><td>isomiR</td><td>NO</td><td>31.04</td><td>203.95</td><td>54.16</td><td>10.20</td></tr>
</table>

# BT-20

<table>
<tr><th></th><th></th><th></th><th></th><th></th><th></th><th></th><th></th><th></th><th></th><th></th><th></th></tr>
<tr><td>>MI0003560|hsa-mir-92b&WithFlank&1|+|155195171|155195278@67.87.21(+1U)&[MIMAT0003218&hsa-miR-92b-3p&offsets|0|-1(+1U);m-115&1|+|155195237|155195258&offsets|0|-1(+1U)]|TATTGCACTCGTCCCGGCCTCT</td><td>>MI0003560|hsa-mir-92b&WithFlank&1|+|155195171|155195278</td><td>67</td><td>87</td><td>21</td><td>(+1U)</td><td>isomiR</td><td>NO</td><td>94.98</td><td>387.95</td><td>138.25</td><td>65.86</td></tr>
<tr><td>>MI0003560|hsa-mir-92b&WithFlank&1|+|155195171|155195278@67.88.22&[MIMAT0003218&hsa-miR-92b-3p&offsets|0|0;m-115&1|+|155195237|155195258&offsets|0|0]|TATTGCACTCGTCCCGGCCTCC</td><td>>MI0003560|hsa-mir-92b&WithFlank&1|+|155195171|155195278</td><td>67</td><td>88</td><td>22</td><td></td><td>isomiR</td><td>NO</td><td>149.84</td><td>597.08</td><td>244.19</td><td>109.09</td></tr>
<tr><td>>MI0003560|hsa-mir-92b&WithFlank&1|+|155195171|155195278@67.88.22(+1A)&[MIMAT0003218&hsa-miR-92b-3p&offsets|0|0(+1A);m-115&1|+|155195237|155195258&offsets|0|0(+1A)]|TATTGCACTCGTCCCGGCCTCCA</td><td>>MI0003560|hsa-mir-92b&WithFlank&1|+|155195171|155195278</td><td>67</td><td>88</td><td>22</td><td>(+1A)</td><td>isomiR</td><td>NO</td><td>0.00</td><td>42.10</td><td>0.00</td><td>0.00</td></tr>
<tr><td>>MI0003560|hsa-mir-92b&WithFlank&1|+|155195171|155195278@67.88.22(+1U)&[MIMAT0003218&hsa-miR-92b-3p&offsets|0|0(+1U);m-115&1|+|155195237|155195258&offsets|0|0(+1U)]|TATTGCACTCGTCCCGGCCTCCT</td><td>>MI0003560|hsa-mir-92b&WithFlank&1|+|155195171|155195278</td><td>67</td><td>88</td><td>22</td><td>(+1U)</td><td>isomiR</td><td>NO</td><td>0.00</td><td>82.22</td><td>0.00</td><td>0.00</td></tr>
<tr><td>>MI0003581|hsa-mir-574&WithFlank&4|+|38868026|38868133@67.88.22&[MIMAT0003239&hsa-miR-574-3p&offsets|0|0;m-149&4|+|38868092|38868113&offsets|0|0]|CACGCTCATGCACACACCCACA</td><td>>MI0003581|hsa-mir-574&WithFlank&4|+|38868026|38868133</td><td>67</td><td>88</td><td>22</td><td></td><td>isomiR</td><td>NO</td><td>9.89</td><td>202.17</td><td>0.00</td><td>0.00</td></tr>
<tr><td>>am_MI0003581|hsa-mir-574&WithFlank&4|+|38868026|38868133@31.53.23&[MIMAT0004795&hsa-miR-574-5p&offsets|0|0;m-213&4|+|38868056|38868078&offsets|0|0]|TGAGTGTGTGTGTGTGAGTGTGT</td><td>>MI0003581|hsa-mir-574&WithFlank&4|+|38868026|38868133.am</td><td>31</td><td>53</td><td>23</td><td></td><td>isomiR</td><td>NO</td><td>0.00</td><td>68.31</td><td>72.92</td><td>0.00</td></tr>
<tr><td>>am_MI0003583|hsa-mir-576&WithFlank&4|+|109488692|109488801@61.83.23&[MIMAT0004796&hsa-miR-576-3p&offsets|0|+1;m-524&4|+|109488752|109488772&offsets|0|+2]|AAGATGTGGAAAAATTGGAATCC</td><td>>MI0003583|hsa-mir-576&WithFlank&4|+|109488692|109488801.am</td><td>61</td><td>83</td><td>23</td><td></td><td>isomiR</td><td>NO</td><td>0.00</td><td>0.00</td><td>70.26</td><td>0.00</td></tr>
<tr><td>>MI0003591|hsa-mir-584&WithFlank&5|-|149062307|149062415@22.40.19&[MIMAT0003249&hsa-miR-584-5p&offsets|0|-3;m-167&5|-|149062374|149062394&offsets|0|-2]|TTATGGTTTGCCTGGGACT</td><td>>MI0003591|hsa-mir-584&WithFlank&5|-|149062307|149062415</td><td>22</td><td>40</td><td>19</td><td></td><td>isomiR</td><td>NO</td><td>0.00</td><td>31.78</td><td>100.49</td><td>0.00</td></tr>
<tr><td>>MI0003591|hsa-mir-584&WithFlank&5|-|149062307|149062415@22.41.20&[MIMAT0003249&hsa-miR-584-5p&offsets|0|-2;m-167&5|-|149062374|149062394&offsets|0|-1]|TTATGGTTTGCCTGGGACTG</td><td>>MI0003591|hsa-mir-584&WithFlank&5|-|149062307|149062415</td><td>22</td><td>41</td><td>20</td><td></td><td>isomiR</td><td>NO</td><td>0.00</td><td>0.00</td><td>91.04</td><td>0.00</td></tr>
<tr><td>>MI0003591|hsa-mir-584&WithFlank&5|-|149062307|149062415@22.41.20(+1U)&[MIMAT0003249&hsa-miR-584-5p&offsets|0|-2(+1U);m-167&5|-|149062374|149062394&offsets|0|-1(+1U)]|TTATGGTTTGCCTGGGACTGT</td><td>>MI0003591|hsa-mir-584&WithFlank&5|-|149062307|149062415</td><td>22</td><td>41</td><td>20</td><td>(+1U)</td><td>isomiR</td><td>NO</td><td>10.62</td><td>38.34</td><td>99.56</td><td>9.84</td></tr>
<tr><td>>MI0003591|hsa-mir-584&WithFlank&5|-|149062307|149062415@22.42.21&[MIMAT0003249&hsa-miR-584-5p&offsets|0|-1;m-167&5|-|149062374|149062394&offsets|0|0]|TTATGGTTTGCCTGGGACTGA</td><td>>MI0003591|hsa-mir-584&WithFlank&5|-|149062307|149062415</td><td>22</td><td>42</td><td>21</td><td></td><td>isomiR</td><td>NO</td><td>87.64</td><td>119.06</td><td>344.77</td><td>84.85</td></tr>
<tr><td>>MI0003599|hsa-mir-589&WithFlank&7|-|5495813|5495923@30.50.21&[MIMAT0004799&hsa-miR-589-5p&offsets|0|-1;m-162&7|-|5495874|5495894&offsets|0|0]|TGAGAACCACGTCTGCTCTGA</td><td>>MI0003599|hsa-mir-589&WithFlank&7|-|5495813|5495923</td><td>30</td><td>50</td><td>21</td><td></td><td>isomiR</td><td>NO</td><td>0.00</td><td>9.95</td><td>12.12</td><td>0.00</td></tr>
<tr><td>>MI0003599|hsa-mir-589&WithFlank&7|-|5495813|5495923@30.51.22(+1U)&[MIMAT0004799&hsa-miR-589-5p&offsets|0|0(+1U);m-162&7|-|5495874|5495894&offsets|0|+1(+1U)]|TGAGAACCACGTCTGCTCTGAGT</td><td>>MI0003599|hsa-mir-589&WithFlank&7|-|5495813|5495923</td><td>30</td><td>51</td><td>22</td><td>(+1U)</td><td>isomiR</td><td>NO</td><td>0.00</td><td>25.25</td><td>0.00</td><td>0.00</td></tr>
<tr><td>>MI0003599|hsa-mir-589&WithFlank&7|-|5495813|5495923@30.52.23&[MIMAT0004799&hsa-miR-589-5p&offsets|0|+1;m-162&7|-|5495874|5495894&offsets|0|+2]|TGAGAACCACGTCTGCTCTGAGC</td><td>>MI0003599|hsa-mir-589&WithFlank&7|-|5495813|5495923</td><td>30</td><td>52</td><td>23</td><td></td><td>isomiR</td><td>NO</td><td>29.58</td><td>135.57</td><td>13.59</td><td>9.67</td></tr>
<tr><td>>MI0003628|hsa-mir-615&WithFlank&12|+|54033944|54034051@67.86.20&[MIMAT0003283&hsa-miR-615-3p&offsets|0|-2;m-495&12|+|54034010|54034030&offsets|0|-1]|TCCGAGCCTGGGTCTCCCTC</td><td>>MI0003628|hsa-mir-615&WithFlank&12|+|54033944|54034051</td><td>67</td><td>86</td><td>20</td><td></td><td>isomiR</td><td>NO</td><td>24.86</td><td>41.30</td><td>0.00</td><td>36.50</td></tr>
<tr><td>>MI0003628|hsa-mir-615&WithFlank&12|+|54033944|54034051@67.86.20(+1A)&[MIMAT0003283&hsa-miR-615-3p&offsets|0|-2(+1A);m-495&12|+|54034010|54034030&offsets|0|-1(+1A)]|TCCGAGCCTGGGTCTCCCTCA</td><td>>MI0003628|hsa-mir-615&WithFlank&12|+|54033944|54034051</td><td>67</td><td>86</td><td>20</td><td>(+1A)</td><td>isomiR</td><td>NO</td><td>0.00</td><td>24.22</td><td>0.00</td><td>0.00</td></tr>
<tr><td>>MI0003628|hsa-mir-615&WithFlank&12|+|54033944|54034051@67.86.20(+1C)&[MIMAT0003283&hsa-miR-615-3p&offsets|0|-2(+1C);m-495&12|+|54034010|54034030&offsets|0|-1(+1C)]|TCCGAGCCTGGGTCTCCCTCC</td><td>>MI0003628|hsa-mir-615&WithFlank&12|+|54033944|54034051</td><td>67</td><td>86</td><td>20</td><td>(+1C)</td><td>isomiR</td><td>NO</td><td>0.00</td><td>10.02</td><td>0.00</td><td>24.86</td></tr>
<tr><td>>MI0003628|hsa-mir-615&WithFlank&12|+|54033944|54034051@67.87.21&[MIMAT0003283&hsa-miR-615-3p&offsets|0|-1;m-495&12|+|54034010|54034030&offsets|0|0]|TCCGAGCCTGGGTCTCCCTCT</td><td>>MI0003628|hsa-mir-615&WithFlank&12|+|54033944|54034051</td><td>67</td><td>87</td><td>21</td><td></td><td>isomiR</td><td>NO</td><td>92.98</td><td>187.02</td><td>11.33</td><td>178.14</td></tr>
<tr><td>>MI0003628|hsa-mir-615&WithFlank&12|+|54033944|54034051@67.88.22&[MIMAT0003283&hsa-miR-615-3p&offsets|0|0;m-495&12|+|54034010|54034030&offsets|0|+1]|TCCGAGCCTGGGTCTCCCTCTT</td><td>>MI0003628|hsa-mir-615&WithFlank&12|+|54033944|54034051</td><td>67</td><td>88</td><td>22</td><td></td><td>isomiR</td><td>NO</td><td>64.68</td><td>160.89</td><td>11.99</td><td>108.72</td></tr>
<tr><td>>MI0003643|hsa-mir-629&WithFlank&15|-|70079366|70079474@28.48.21&[MIMAT0004810&hsa-miR-629-5p&offsets|0|0;m-150&15|-|70079427|70079447&offsets|0|0]|TGGGTTTACGTTGGGAGAACT</td><td>>MI0003643|hsa-mir-629&WithFlank&15|-|70079366|70079474</td><td>28</td><td>48</td><td>21</td><td></td><td>isomiR</td><td>NO</td><td>24.12</td><td>72.81</td><td>297.24</td><td>0.00</td></tr>
<tr><td>>MI0003643|hsa-mir-629&WithFlank&15|-|70079366|70079474@28.48.21(+1A)&[MIMAT0004810&hsa-miR-629-5p&offsets|0|0(+1A);m-150&15|-|70079427|70079447&offsets|0|0(+1A)]|TGGGTTTACGTTGGGAGAACTA</td><td>>MI0003643|hsa-mir-629&WithFlank&15|-|70079366|70079474</td><td>28</td><td>48</td><td>21</td><td>(+1A)</td><td>isomiR</td><td>NO</td><td>0.00</td><td>0.00</td><td>10.70</td><td>0.00</td></tr>
<tr><td>>MI0003643|hsa-mir-629&WithFlank&15|-|70079366|70079474@28.48.21(+1C)&[MIMAT0004810&hsa-miR-629-5p&offsets|0|0(+1C);m-150&15|-|70079427|70079447&offsets|0|0(+1C)]|TGGGTTTACGTTGGGAGAACTC</td><td>>MI0003643|hsa-mir-629&WithFlank&15|-|70079366|70079474</td><td>28</td><td>48</td><td>21</td><td>(+1C)</td><td>isomiR</td><td>NO</td><td>0.00</td><td>28.17</td><td>102.84</td><td>0.00</td></tr>
<tr><td>>MI0003643|hsa-mir-629&WithFlank&15|-|70079366|70079474@28.49.22&[MIMAT0004810&hsa-miR-629-5p&offsets|0|+1;m-150&15|-|70079427|70079447&offsets|0|+1]|TGGGTTTACGTTGGGAGAACTT</td><td>>MI0003643|hsa-mir-629&WithFlank&15|-|70079366|70079474</td><td>28</td><td>49</td><td>22</td><td></td><td>isomiR</td><td>NO</td><td>86.51</td><td>264.13</td><td>546.82</td><td>65.11</td></tr>
</table>

# BT-20

| | | | | | | | | | | | |
|---|---|---|---|---|---|---|---|---|---|---|---|
| >MI0003667\|hsa-mir-652&WithFlank&X\|+\|110055323\|110055432@67.86.20&[MIMAT0003322&hsa-miR-652-3p&offsets\|0\|-1;m-148&X\|+\|110055389\|110055410&offsets\|0\|-2]\|AATGGCGCCACTAGGGTTGT | >MI0003667\|hsa-mir-652&WithFlank&X\|+\|110055323\|110055432 | 67 | 86 | 20 | | isomiR | NO | 0.00 | 0.00 | 13.62 | 0.00 |
| >MI0003667\|hsa-mir-652&WithFlank&X\|+\|110055323\|110055432@67.87.21&[MIMAT0003322&hsa-miR-652-3p&offsets\|0\|0;m-148&X\|+\|110055389\|110055410&offsets\|0\|-1]\|AATGGCGCCACTAGGGTTGTG | >MI0003667\|hsa-mir-652&WithFlank&X\|+\|110055323\|110055432 | 67 | 87 | 21 | | isomiR | NO | 0.00 | 26.72 | 88.81 | 0.00 |
| >MI0003667\|hsa-mir-652&WithFlank&X\|+\|110055323\|110055432@67.87.21(+1U)&[MIMAT0003322&hsa-miR-652-3p&offsets\|0\|0(+1U);m-148&X\|+\|110055389\|110055410&offsets\|0\|-1(+1U)]\|AATGGCGCCACTAGGGTTGTGT | >MI0003667\|hsa-mir-652&WithFlank&X\|+\|110055323\|110055432 | 67 | 87 | 21 | (+1U) | isomiR | NO | 23.62 | 111.43 | 80.73 | 0.00 |
| >MI0003763\|hsa-mir-767&WithFlank&X\|-\|152393415\|152393535@33.55.23&[MIMAT0003882&hsa-miR-767-5p&offsets\|0\|0;m-788&X\|-\|152393481\|152393503&offsets\|0\|0]\|TGCACCATGGTTGTCTGAGCATG | >MI0003763\|hsa-mir-767&WithFlank&X\|-\|152393415\|152393535 | 33 | 55 | 23 | | isomiR | NO | 0.00 | 25.59 | 0.00 | 0.00 |
| >MI0003763\|hsa-mir-767&WithFlank&X\|-\|152393415\|152393535@33.56.24&[MIMAT0003882&hsa-miR-767-5p&offsets\|0\|+1;m-788&X\|-\|152393481\|152393503&offsets\|0\|+1]\|TGCACCATGGTTGTCTGAGCATGC | >MI0003763\|hsa-mir-767&WithFlank&X\|-\|152393415\|152393535 | 33 | 56 | 24 | | isomiR | NO | 0.00 | 45.06 | 0.00 | 0.00 |
| >am_MI0003772\|hsa-mir-151b&WithFlank&14\|-\|100109413\|100109520@66.85.20(+1C);MI0000809\|hsa-mir-151a&WithFlank&8\|-\|140732558\|140732659@17.36.20(+1C)&[MIMAT0010214&hsa-miR-151b&offsets\|0\|+2(+1C);m-179&14\|-\|100109436\|100109455&offsets\|0\|0(+1C)];[MIMAT0004697&hsa-miR-151a-5p&offsets\|0\|-1(+1C);m-75&8\|-\|140732623\|140732643&offsets\|0\|-1(+1C)]\|TCGAGGAGCTCACAGTCTAGC | >MI0003772\|hsa-mir-151b&WithFlank&14\|-\|100109413\|100109520.a | 66 | 85 | 20 | (+1C) | isomiR | NO | 0.00 | 24.83 | 0.00 | 0.00 |
| >MI0003776\|hsa-mir-320b-1&WithFlank&1\|+\|116671740\|116671823@48.66.19(+1A);MI0003839\|hsa-mir-320b-2&WithFlank&1\|-\|224257034\|224257116@47.65.19(+1A);MI0003778\|hsa-mir-320c-1&WithFlank&18\|+\|21683512\|21683595@48.66.19(+1A);MI0008191\|hsa-mir-320c-2&WithFlank&18\|+\|24321669\|24321752@48.66.19(+1A);MI0000542\|hsa-mir-320a&WithFlank&8\|-\|22244960\|22245043@48.66.19(+1A)&[MIMAT0005792&hsa-miR-320b&offsets\|0\|-3(+1A);m-183&1\|+\|116671787\|116671806&offsets\|0\|-1(+1A)];[MIMAT0005792_1&hsa-miR-320b&offsets\|0\|-3(+1A);m-183&1\|-\|224257051\|224257070&offsets\|0\|-1(+1A)];[MIMAT0005793&hsa-miR-320c&offsets\|0\|-1(+1A);m-222&18\|+\|21683559\|21683578&offsets\|0\|-1(+1A)];[MIMAT0005793_1&hsa-miR-320c&offsets\|0\|-1(+1A);m-219&18\|+\|24321716\|24321735&offsets\|0\|-1(+1A)];[MIMAT0000510&hsa-miR-320a-3p&offsets\|0\|-3(+1A);m-60&8\|-\|22244975\|22244996&offsets\|0\|-3(+1A)]\|AAAAGCTGGGTTGAGAGGGA | >MI0003776\|hsa-mir-320b-1&WithFlank&1\|+\|116671740\|116671823 | 48 | 66 | 19 | (+1A) | isomiR | NO | 0.00 | 83.74 | 125.24 | 0.00 |
| >MI0003776\|hsa-mir-320b-1&WithFlank&1\|+\|116671740\|116671823@48.66.19;MI0003839\|hsa-mir-320b-2&WithFlank&1\|-\|224257034\|224257116@47.65.19;MI0003778\|hsa-mir-320c-1&WithFlank&18\|+\|21683512\|21683595@48.66.19;MI0008191\|hsa-mir-320c-2&WithFlank&18\|+\|24321669\|24321752@48.66.19;MI0000542\|hsa-mir-320a&WithFlank&8\|-\|22244960\|22245043@48.66.19&[MIMAT0005792&hsa-miR-320b&offsets\|0\|-3;m-183&1\|+\|116671787\|116671806&offsets\|0\|-1];[MIMAT0005792_1&hsa-miR-320b&offsets\|0\|-3;m-183&1\|-\|224257051\|224257070&offsets\|0\|-1];[MIMAT0005793&hsa-miR-320c&offsets\|0\|-1;m-222&18\|+\|21683559\|21683578&offsets\|0\|-1];[MIMAT0005793_1&hsa-miR-320c&offsets\|0\|-1;m-219&18\|+\|24321716\|24321735&offsets\|0\|-1];[MIMAT0000510&hsa-miR-320a-3p&offsets\|0\|-3;m-60&8\|-\|22244975\|22244996&offsets\|0\|-3]\|AAAAGCTGGGTTGAGAGGG | >MI0003776\|hsa-mir-320b-1&WithFlank&1\|+\|116671740\|116671823 | 48 | 66 | 19 | | isomiR | NO | 0.00 | 40.33 | 150.18 | 0.00 |
| >MI0003776\|hsa-mir-320b-1&WithFlank&1\|+\|116671740\|116671823@48.67.20(+1U);MI0003839\|hsa-mir-320b-2&WithFlank&1\|-\|224257034\|224257116@47.66.20(+1U);MI0000542\|hsa-mir-320a&WithFlank&8\|-\|22244960\|22245043@48.67.20(+1U)&[MIMAT0005792&hsa-miR-320b&offsets\|0\|-2(+1U);m-183&1\|+\|116671787\|116671806&offsets\|0\|0(+1U)];[MIMAT0005792_1&hsa-miR-320b&offsets\|0\|-2(+1U);m-183&1\|-\|224257051\|224257070&offsets\|0\|0(+1U)];[MIMAT0000510&hsa-miR-320a-3p&offsets\|0\|-2(+1U);m-60&8\|-\|22244975\|22244996&offsets\|0\|-2(+1U)]\|AAAAGCTGGGTTGAGAGGGCT | >MI0003776\|hsa-mir-320b-1&WithFlank&1\|+\|116671740\|116671823 | 48 | 67 | 20 | (+1U) | isomiR | NO | 0.00 | 112.99 | 126.66 | 0.00 |
| >MI0003776\|hsa-mir-320b-1&WithFlank&1\|+\|116671740\|116671823@48.67.20;MI0003839\|hsa-mir-320b-2&WithFlank&1\|-\|224257034\|224257116@47.66.20;MI0000542\|hsa-mir-320a&WithFlank&8\|-\|22244960\|22245043@48.67.20&[MIMAT0005792&hsa-miR-320b&offsets\|0\|-2;m-183&1\|+\|116671787\|116671806&offsets\|0\|0];[MIMAT0005792_1&hsa-miR-320b&offsets\|0\|-2;m-183&1\|-\|224257051\|224257070&offsets\|0\|0];[MIMAT0000510&hsa-miR-320a-3p&offsets\|0\|-2;m-60&8\|-\|22244975\|22244996&offsets\|0\|-2]\|AAAAGCTGGGTTGAGAGGGC | >MI0003776\|hsa-mir-320b-1&WithFlank&1\|+\|116671740\|116671823 | 48 | 67 | 20 | | isomiR | NO | 42.91 | 382.41 | 683.11 | 9.60 |
| >MI0003776\|hsa-mir-320b-1&WithFlank&1\|+\|116671740\|116671823@48.68.21;MI0003839\|hsa-mir-320b-2&WithFlank&1\|-\|224257034\|224257116@47.67.21&[MIMAT0005792&hsa-miR-320b&offsets\|0\|-1;m-183&1\|+\|116671787\|116671806&offsets\|0\|+1];[MIMAT0005792_1&hsa-miR-320b&offsets\|0\|-1;m-183&1\|-\|224257051\|224257070&offsets\|0\|+1]\|AAAAGCTGGGTTGAGAGGGCA | >MI0003776\|hsa-mir-320b-1&WithFlank&1\|+\|116671740\|116671823 | 48 | 68 | 21 | | isomiR | NO | 30.69 | 391.55 | 294.75 | 0.00 |

# BT-20

| | | | | | | | | | | | |
|---|---|---|---|---|---|---|---|---|---|---|---|
| >am_MI0003776\|hsa-mir-320b-1&WithFlank&1\|+\|116671740\|116671823@48.65.18;MI0003839\|hsa-mir-320b-2&WithFlank&1\|-\|224257034\|224257116@47.64.18;MI0008190\|hsa-mir-320d-1&WithFlank&13\|-\|40727810\|40727893@48.65.18;MI0003778\|hsa-mir-320c-1&WithFlank&18\|+\|21683512\|21683595@48.65.18;MI0008191\|hsa-mir-320c-2&WithFlank&18\|+\|24321669\|24321752@48.65.18;MI0000542\|hsa-mir-320a&WithFlank&8\|-\|22244960\|22245043@48.65.18;MI0008192\|hsa-mir-320d-2&WithFlank&X\|-\|140926154\|140926237@48.65.18&[MIMAT0005792&hsa-miR-320b&offsets\|0\|-4;m-183&1\|+\|116671787\|116671806&offsets\|0\|-2];[MIMAT0005792_1&hsa-miR-320b&offsets\|0\|-4;m-183&1\|-\|224257051\|224257070&offsets\|0\|-2];[MIMAT0006764&hsa-miR-320d&offsets\|0\|-1;m-246&13\|-\|40727829\|40727846&offsets\|0\|0];[MIMAT0005793&hsa-miR-320c&offsets\|0\|-2;m-222&18\|+\|21683559\|21683578&offsets\|0\|-2];[MIMAT0005793_1&hsa-miR-320c&offsets\|0\|-2;m-219&18\|+\|24321716\|24321735&offsets\|0\|-2];[MIMAT0000510&hsa-miR-320a-3p&offsets\|0\|-4;m-60&8\|-\|22244975\|22244996&offsets\|0\|-4];[MIMAT0006764_1&hsa-miR-320d&offsets\|0\|-1;m-253&X\|-\|140926173\|140926190&offsets\|0\|0]\|AAAAGCTGGGTTGAGAGG | >MI0003776\|hsa-mir-320b-1&WithFlank&1\|+\|116671740\|116671823.a | 48 | 65 | 18 | | isomiR | NO | 0.00 | 90.82 | 159.46 | 0.00 |
| >MI0003778\|hsa-mir-320c-1&WithFlank&18\|+\|21683512\|21683595@48.67.20;MI0008191\|hsa-mir-320c-2&WithFlank&18\|+\|24321669\|24321752@48.67.20&[MIMAT0005793&hsa-miR-320c&offsets\|0\|0;m-222&18\|+\|21683559\|21683578&offsets\|0\|0];[MIMAT0005793_1&hsa-miR-320c&offsets\|0\|0;m-219&18\|+\|24321716\|24321735&offsets\|0\|0]\|AAAAGCTGGGTTGAGAGGGT | >MI0003778\|hsa-mir-320c-1&WithFlank&18\|+\|21683512\|21683595 | 48 | 67 | 20 | | isomiR | NO | 9.55 | 157.60 | 251.30 | 0.00 |
| >MI0003780\|hsa-mir-1296&WithFlank&10\|-\|63372951\|63373054@22.43.22&[MIMAT0005794&hsa-miR-1296-5p&offsets\|0\|0;m-304&10\|-\|63373012\|63373033&offsets\|0\|0]\|TTAGGGCCCTGGCTCCATCTCC | >MI0003780\|hsa-mir-1296&WithFlank&10\|-\|63372951\|63373054 | 22 | 43 | 22 | | isomiR | NO | 0.00 | 26.75 | 10.52 | 0.00 |
| >MI0003820\|hsa-mir-454&WithFlank&17\|-\|59137752\|59137878@70.92.23&[MIMAT0003885&hsa-miR-454-3p&offsets\|0\|0;m-272&17\|-\|59137787\|59137809&offsets\|0\|0]\|TAGTGCAATATTGCTTATAGGGT | >MI0003820\|hsa-mir-454&WithFlank&17\|-\|59137752\|59137878 | 70 | 92 | 23 | | isomiR | NO | 0.00 | 10.08 | 0.00 | 0.00 |
| >MI0003820\|hsa-mir-454&WithFlank&17\|-\|59137752\|59137878@70.93.24&[MIMAT0003885&hsa-miR-454-3p&offsets\|0\|+1;m-272&17\|-\|59137787\|59137809&offsets\|0\|+1]\|TAGTGCAATATTGCTTATAGGGTT | >MI0003820\|hsa-mir-454&WithFlank&17\|-\|59137752\|59137878 | 70 | 93 | 24 | | isomiR | NO | 24.21 | 10.48 | 0.00 | 0.00 |
| >MI0003834\|hsa-mir-769&WithFlank&19\|+\|46018926\|46019055@36.57.22&[MIMAT0003886&hsa-miR-769-5p&offsets\|0\|0;m-216&19\|+\|46018961\|46018982&offsets\|0\|0]\|TGAGACCTCTGGGTTCTGAGCT | >MI0003834\|hsa-mir-769&WithFlank&19\|+\|46018926\|46019055 | 36 | 57 | 22 | | isomiR | NO | 37.86 | 112.23 | 81.51 | 0.00 |
| >MI0005531\|hsa-mir-450b&WithFlank&X\|-\|134540179\|134540268@17.37.21&[MIMAT0004909&hsa-miR-450b-5p&offsets\|0\|-1;m-266&X\|-\|134540232\|134540252&offsets\|0\|0]\|TTTTGCAATATGTTCCTGAAT | >MI0005531\|hsa-mir-450b&WithFlank&X\|-\|134540179\|134540268 | 17 | 37 | 21 | | isomiR | NO | 0.00 | 32.61 | 0.00 | 0.00 |
| >MI0005559\|hsa-mir-744&WithFlank&17\|+\|12081893\|12082002@17.34.18&[MIMAT0004945&hsa-miR-744-5p&offsets\|0\|-4;m-172&17\|+\|12081909\|12081929&offsets\|0\|-3]\|TGCGGGGCTAGGGCTAAC | >MI0005559\|hsa-mir-744&WithFlank&17\|+\|12081893\|12082002 | 17 | 34 | 18 | | isomiR | NO | 0.00 | 0.00 | 13.23 | 0.00 |
| >MI0005559\|hsa-mir-744&WithFlank&17\|+\|12081893\|12082002@17.36.20(+1U)&[MIMAT0004945&hsa-miR-744-5p&offsets\|0\|-2(+1U);m-172&17\|+\|12081909\|12081929&offsets\|0\|-1(+1U)]\|TGCGGGGCTAGGGCTAACAGT | >MI0005559\|hsa-mir-744&WithFlank&17\|+\|12081893\|12082002 | 17 | 36 | 20 | (+1U) | isomiR | NO | 0.00 | 73.28 | 67.56 | 10.89 |
| >MI0005559\|hsa-mir-744&WithFlank&17\|+\|12081893\|12082002@17.37.21&[MIMAT0004945&hsa-miR-744-5p&offsets\|0\|-1;m-172&17\|+\|12081909\|12081929&offsets\|0\|0]\|TGCGGGGCTAGGGCTAACAGC | >MI0005559\|hsa-mir-744&WithFlank&17\|+\|12081893\|12082002 | 17 | 37 | 21 | | isomiR | NO | 77.34 | 279.88 | 256.36 | 131.47 |
| >MI0005559\|hsa-mir-744&WithFlank&17\|+\|12081893\|12082002@17.38.22&[MIMAT0004945&hsa-miR-744-5p&offsets\|0\|0;m-172&17\|+\|12081909\|12081929&offsets\|0\|+1]\|TGCGGGGCTAGGGCTAACAGCA | >MI0005559\|hsa-mir-744&WithFlank&17\|+\|12081893\|12082002 | 17 | 38 | 22 | | isomiR | NO | 24.68 | 86.91 | 56.47 | 32.49 |
| >MI0005561\|hsa-mir-877&WithFlank&6\|+\|30584326\|30584423@7.30.24&[MIMAT0004949&hsa-miR-877-5p&offsets\|0\|+4;m-382&6\|+\|30584332\|30584354&offsets\|0\|+1]\|GTAGAGGAGATGGCGCAGGGGACA | >MI0005561\|hsa-mir-877&WithFlank&6\|+\|30584326\|30584423 | 7 | 30 | 24 | | isomiR | NO | 0.00 | 10.16 | 0.00 | 0.00 |
| >MI0005561\|hsa-mir-877&WithFlank&6\|+\|30584326\|30584423@72.92.21(+1U)&[MIMAT0004950&hsa-miR-877-3p&offsets\|0\|0(+1U);m-607&6\|+\|30584397\|30584418&offsets\|0\|-1(+1U)]\|TCCTCTTCTCCCTCCTCCCAGT | >MI0005561\|hsa-mir-877&WithFlank&6\|+\|30584326\|30584423 | 72 | 92 | 21 | (+1U) | isomiR | NO | 0.00 | 32.66 | 0.00 | 0.00 |
| >MI0005566\|hsa-mir-374b&WithFlank&X\|-\|74218541\|74218624@17.37.21&[MIMAT0004955&hsa-miR-374b-5p&offsets\|0\|-1;m-153&X\|-\|74218587\|74218608&offsets\|0\|-1]\|ATATAATACAACCTGCTAAGT | >MI0005566\|hsa-mir-374b&WithFlank&X\|-\|74218541\|74218624 | 17 | 37 | 21 | | isomiR | NO | 25.89 | 42.81 | 0.00 | 0.00 |
| >MI0005566\|hsa-mir-374b&WithFlank&X\|-\|74218541\|74218624@17.38.22&[MIMAT0004955&hsa-miR-374b-5p&offsets\|0\|0;m-153&X\|-\|74218587\|74218608&offsets\|0\|0]\|ATATAATACAACCTGCTAAGTG | >MI0005566\|hsa-mir-374b&WithFlank&X\|-\|74218541\|74218624 | 17 | 38 | 22 | | isomiR | NO | 28.95 | 34.54 | 0.00 | 0.00 |
| >MI0005756\|hsa-mir-934&WithFlank&X\|+\|136550872\|136550966@21.40.20(+1A)&[MIMAT0004977&hsa-miR-934&offsets\|0\|-2(+1A);m-943&X\|+\|136550892\|136550913&offsets\|0\|-2(+1A)]\|TGTCTACTACTGGAGACACTA | >MI0005756\|hsa-mir-934&WithFlank&X\|+\|136550872\|136550966 | 21 | 40 | 20 | (+1A) | isomiR | NO | 0.00 | 0.00 | 12.43 | 0.00 |
| >MI0005756\|hsa-mir-934&WithFlank&X\|+\|136550872\|136550966@21.41.21&[MIMAT0004977&hsa-miR-934&offsets\|0\|-1;m-943&X\|+\|136550892\|136550913&offsets\|0\|-1]\|TGTCTACTACTGGAGACACTG | >MI0005756\|hsa-mir-934&WithFlank&X\|+\|136550872\|136550966 | 21 | 41 | 21 | | isomiR | NO | 93.90 | 87.81 | 211.54 | 71.33 |
| >MI0005756\|hsa-mir-934&WithFlank&X\|+\|136550872\|136550966@21.42.22&[MIMAT0004977&hsa-miR-934&offsets\|0\|0;m-943&X\|+\|136550892\|136550913&offsets\|0\|0]\|TGTCTACTACTGGAGACACTGG | >MI0005756\|hsa-mir-934&WithFlank&X\|+\|136550872\|136550966 | 21 | 42 | 22 | | isomiR | NO | 115.33 | 103.37 | 62.18 | 39.43 |
| >MI0005756\|hsa-mir-934&WithFlank&X\|+\|136550872\|136550966@21.42.22(+1A)&[MIMAT0004977&hsa-miR-934&offsets\|0\|0(+1A);m-943&X\|+\|136550892\|136550913&offsets\|0\|0(+1A)]\|TGTCTACTACTGGAGACACTGGA | >MI0005756\|hsa-mir-934&WithFlank&X\|+\|136550872\|136550966 | 21 | 42 | 22 | (+1A) | isomiR | NO | 94.94 | 136.31 | 12.02 | 28.88 |

# BT-20

<table>
<tr><td>>MI0005756|hsa-mir-934&WithFlank&X|+|136550872|136550966@21.42.22(+2A)&[MIMAT0004977&hsa-miR-934&offsets|0|0(+2A);m-943&X|+|136550892|136550913&offsets|0|0(+2A)]|TGTCTACTACTGGAGACACTGGAA</td><td>>MI0005756|hsa-mir-934&WithFlank&X|+|136550872|136550966</td><td>21</td><td>42</td><td>22</td><td>(+2A)</td><td>isomiR</td><td>NO</td><td>25.88</td><td>32.14</td><td>0.00</td><td>0.00</td></tr>
<tr><td>>MI0005756|hsa-mir-934&WithFlank&X|+|136550872|136550966@21.43.23&[MIMAT0004977&hsa-miR-934&offsets|0|+1;m-943&X|+|136550892|136550913&offsets|0|+1]|TGTCTACTACTGGAGACACTGGT</td><td>>MI0005756|hsa-mir-934&WithFlank&X|+|136550872|136550966</td><td>21</td><td>43</td><td>23</td><td></td><td>isomiR</td><td>NO</td><td>69.31</td><td>132.52</td><td>65.04</td><td>38.75</td></tr>
<tr><td>>am_MI0005763|hsa-mir-941-1&WithFlank&20|+|63919443|63919526@53.74.22(+1A);MI0005764|hsa-mir-941-2&WithFlank&20|+|63919499|63919582@53.74.22(+1A);MI0005765|hsa-mir-941-3&WithFlank&20|+|63919555|63919638@53.74.22(+1A);MI0005766|hsa-mir-941-4&WithFlank&20|+|63919750|63919833@53.74.22(+1A);MI0031520|hsa-mir-941-5&WithFlank&20|+|63919862|63919945@53.74.22(+1A)&[MIMAT0004984&hsa-miR-941&offsets|0|-1(+1A);m-263&20|+|63919495|63919517&offsets|0|-1(+1A)];[MIMAT0004984_1&hsa-miR-941&offsets|0|-1(+1A);m-263&20|+|63919551|63919573&offsets|0|-1(+1A)];[MIMAT0004984_2&hsa-miR-941&offsets|0|-1(+1A);m-263&20|+|63919607|63919629&offsets|0|-1(+1A)];[MIMAT0004984_3&hsa-miR-941&offsets|0|-1(+1A);m-263&20|+|63919802|63919824&offsets|0|-1(+1A)];[MIMAT0004984_4&hsa-miR-941&offsets|0|-1(+1A);m-263&20|+|63919914|63919936&offsets|0|-1(+1A)]|CACCCGGCTGTGTGCACATGTGA</td><td>>MI0005763|hsa-mir-941-1&WithFlank&20|+|63919443|63919526.am</td><td>53</td><td>74</td><td>22</td><td>(+1A)</td><td>isomiR</td><td>NO</td><td>0.00</td><td>159.04</td><td>13.89</td><td>10.21</td></tr>
<tr><td>>am_MI0005763|hsa-mir-941-1&WithFlank&20|+|63919443|63919526@53.74.22(+1U);MI0005764|hsa-mir-941-2&WithFlank&20|+|63919499|63919582@53.74.22(+1U);MI0005765|hsa-mir-941-3&WithFlank&20|+|63919555|63919638@53.74.22(+1U);MI0005766|hsa-mir-941-4&WithFlank&20|+|63919750|63919833@53.74.22(+1U);MI0031520|hsa-mir-941-5&WithFlank&20|+|63919862|63919945@53.74.22(+1U)&[MIMAT0004984&hsa-miR-941&offsets|0|-1(+1U);m-263&20|+|63919495|63919517&offsets|0|-1(+1U)];[MIMAT0004984_1&hsa-miR-941&offsets|0|-1(+1U);m-263&20|+|63919551|63919573&offsets|0|-1(+1U)];[MIMAT0004984_2&hsa-miR-941&offsets|0|-1(+1U);m-263&20|+|63919607|63919629&offsets|0|-1(+1U)];[MIMAT0004984_3&hsa-miR-941&offsets|0|-1(+1U);m-263&20|+|63919802|63919824&offsets|0|-1(+1U)];[MIMAT0004984_4&hsa-miR-941&offsets|0|-1(+1U);m-263&20|+|63919914|63919936&offsets|0|-1(+1U)]|CACCCGGCTGTGTGCACATGTGT</td><td>>MI0005763|hsa-mir-941-1&WithFlank&20|+|63919443|63919526.am</td><td>53</td><td>74</td><td>22</td><td>(+1U)</td><td>isomiR</td><td>NO</td><td>92.18</td><td>450.48</td><td>130.21</td><td>109.20</td></tr>
<tr><td>>am_MI0005763|hsa-mir-941-1&WithFlank&20|+|63919443|63919526@53.74.22;MI0005764|hsa-mir-941-2&WithFlank&20|+|63919499|63919582@53.74.22;MI0005765|hsa-mir-941-3&WithFlank&20|+|63919555|63919638@53.74.22;MI0005766|hsa-mir-941-4&WithFlank&20|+|63919750|63919833@53.74.22;MI0031520|hsa-mir-941-5&WithFlank&20|+|63919862|63919945@53.74.22&[MIMAT0004984&hsa-miR-941&offsets|0|-1;m-263&20|+|63919495|63919517&offsets|0|-1];[MIMAT0004984_1&hsa-miR-941&offsets|0|-1;m-263&20|+|63919551|63919573&offsets|0|-1];[MIMAT0004984_2&hsa-miR-941&offsets|0|-1;m-263&20|+|63919607|63919629&offsets|0|-1];[MIMAT0004984_3&hsa-miR-941&offsets|0|-1;m-263&20|+|63919802|63919824&offsets|0|-1];[MIMAT0004984_4&hsa-miR-941&offsets|0|-1;m-263&20|+|63919914|63919936&offsets|0|-1]|CACCCGGCTGTGTGCACATGTG</td><td>>MI0005763|hsa-mir-941-1&WithFlank&20|+|63919443|63919526.am</td><td>53</td><td>74</td><td>22</td><td></td><td>isomiR</td><td>NO</td><td>0.00</td><td>110.50</td><td>14.64</td><td>0.00</td></tr>
<tr><td>>am_MI0005763|hsa-mir-941-1&WithFlank&20|+|63919443|63919526@53.75.23(+1U);MI0005764|hsa-mir-941-2&WithFlank&20|+|63919499|63919582@53.75.23(+1U);MI0005765|hsa-mir-941-3&WithFlank&20|+|63919555|63919638@53.75.23(+1U);MI0005766|hsa-mir-941-4&WithFlank&20|+|63919750|63919833@53.75.23(+1U);MI0031520|hsa-mir-941-5&WithFlank&20|+|63919862|63919945@53.75.23(+1U)&[MIMAT0004984&hsa-miR-941&offsets|0|0(+1U);m-263&20|+|63919495|63919517&offsets|0|0(+1U)];[MIMAT0004984_1&hsa-miR-941&offsets|0|0(+1U);m-263&20|+|63919551|63919573&offsets|0|0(+1U)];[MIMAT0004984_2&hsa-miR-941&offsets|0|0(+1U);m-263&20|+|63919607|63919629&offsets|0|0(+1U)];[MIMAT0004984_3&hsa-miR-941&offsets|0|0(+1U);m-263&20|+|63919802|63919824&offsets|0|0(+1U)];[MIMAT0004984_4&hsa-miR-941&offsets|0|0(+1U);m-263&20|+|63919914|63919936&offsets|0|0(+1U)]|CACCCGGCTGTGTGCACATGTGCT</td><td>>MI0005763|hsa-mir-941-1&WithFlank&20|+|63919443|63919526.am</td><td>53</td><td>75</td><td>23</td><td>(+1U)</td><td>isomiR</td><td>NO</td><td>0.00</td><td>31.42</td><td>0.00</td><td>0.00</td></tr>
<tr><td>>am_MI0005763|hsa-mir-941-1&WithFlank&20|+|63919443|63919526@53.75.23;MI0005764|hsa-mir-941-2&WithFlank&20|+|63919499|63919582@53.75.23;MI0005765|hsa-mir-941-3&WithFlank&20|+|63919555|63919638@53.75.23;MI0005766|hsa-mir-941-4&WithFlank&20|+|63919750|63919833@53.75.23;MI0031520|hsa-mir-941-5&WithFlank&20|+|63919862|63919945@53.75.23&[MIMAT0004984&hsa-miR-941&offsets|0|0;m-263&20|+|63919495|63919517&offsets|0|0];[MIMAT0004984_1&hsa-miR-941&offsets|0|0;m-263&20|+|63919551|63919573&offsets|0|0];[MIMAT0004984_2&hsa-miR-941&offsets|0|0;m-263&20|+|63919607|63919629&offsets|0|0];[MIMAT0004984_3&hsa-miR-941&offsets|0|0;m-263&20|+|63919802|63919824&offsets|0|0];[MIMAT0004984_4&hsa-miR-941&offsets|0|0;m-263&20|+|63919914|63919936&offsets|0|0]|CACCCGGCTGTGTGCACATGTGC</td><td>>MI0005763|hsa-mir-941-1&WithFlank&20|+|63919443|63919526.am</td><td>53</td><td>75</td><td>23</td><td></td><td>isomiR</td><td>NO</td><td>235.45</td><td>1223.23</td><td>381.84</td><td>330.09</td></tr>
<tr><td>>MI0006273|hsa-mir-1180&WithFlank&17|-|19344500|19344580@47.66.20&[MIMAT0005825&hsa-miR-1180-3p&offsets|0|-2;m-214&17|-|19344513|19344534&offsets|0|-2]|TTTCCGGCTCGCGTGGGTGT</td><td>>MI0006273|hsa-mir-1180&WithFlank&17|-|19344500|19344580</td><td>47</td><td>66</td><td>20</td><td></td><td>isomiR</td><td>NO</td><td>0.00</td><td>85.43</td><td>16.11</td><td>0.00</td></tr>
</table>

# BT-20

| | | | | | | | | | | | |
|---|---|---|---|---|---|---|---|---|---|---|---|
| >MI0006273\|hsa-mir-1180&WithFlank&17\|-\|19344500\|19344580@47.67.21(+1C)&[MIMAT0005825&hsa-miR-1180-3p&offsets\|0\|-1(+1C);m-214&17\|-\|19344513\|19344534&offsets\|0\|-1(+1C)]\|TTTCCGGCTCGCGTGGGTGTGC | >MI0006273\|hsa-mir-1180&WithFlank&17\|-\|19344500\|19344580 | 47 | 67 | 21 | (+1C) | isomiR | NO | 0.00 | 33.71 | 0.00 | 0.00 |
| >MI0006273\|hsa-mir-1180&WithFlank&17\|-\|19344500\|19344580@47.68.22&[MIMAT0005825&hsa-miR-1180-3p&offsets\|0\|0;m-214&17\|-\|19344513\|19344534&offsets\|0\|0]\|TTTCCGGCTCGCGTGGGTGTGT | >MI0006273\|hsa-mir-1180&WithFlank&17\|-\|19344500\|19344580 | 47 | 68 | 22 | | isomiR | NO | 65.19 | 289.74 | 141.12 | 32.22 |
| >MI0006444\|hsa-mir-1307&WithFlank&10\|-\|103394247\|103394407@86.107.22&[MIMAT0005951&hsa-miR-1307-3p&offsets\|0\|0;m-66&10\|-\|103394300\|103394322&offsets\|0\|-1]\|ACTCGGCGTGGCGTCGGTCGTG | >MI0006444\|hsa-mir-1307&WithFlank&10\|-\|103394247\|103394407 | 86 | 107 | 22 | | isomiR | NO | 0.00 | 45.37 | 10.59 | 0.00 |
| >MI0006444\|hsa-mir-1307&WithFlank&10\|-\|103394247\|103394407@86.107.22(+1A)&[MIMAT0005951&hsa-miR-1307-3p&offsets\|0\|0(+1A);m-66&10\|-\|103394300\|103394322&offsets\|0\|-1(+1A)]\|ACTCGGCGTGGCGTCGGTCGTGA | >MI0006444\|hsa-mir-1307&WithFlank&10\|-\|103394247\|103394407 | 86 | 107 | 22 | (+1A) | isomiR | NO | 0.00 | 114.95 | 0.00 | 0.00 |
| >MI0006444\|hsa-mir-1307&WithFlank&10\|-\|103394247\|103394407@86.107.22(+1U)&[MIMAT0005951&hsa-miR-1307-3p&offsets\|0\|0(+1U);m-66&10\|-\|103394300\|103394322&offsets\|0\|-1(+1U)]\|ACTCGGCGTGGCGTCGGTCGTGT | >MI0006444\|hsa-mir-1307&WithFlank&10\|-\|103394247\|103394407 | 86 | 107 | 22 | (+1U) | isomiR | NO | 0.00 | 104.18 | 10.19 | 0.00 |
| >MI0006444\|hsa-mir-1307&WithFlank&10\|-\|103394247\|103394407@86.108.23&[MIMAT0005951&hsa-miR-1307-3p&offsets\|0\|+1;m-66&10\|-\|103394300\|103394322&offsets\|0\|0]\|ACTCGGCGTGGCGTCGGTCGTGG | >MI0006444\|hsa-mir-1307&WithFlank&10\|-\|103394247\|103394407 | 86 | 108 | 23 | | isomiR | NO | 72.72 | 392.14 | 112.42 | 95.89 |
| >MI0006444\|hsa-mir-1307&WithFlank&10\|-\|103394247\|103394407@86.108.23(+1A)&[MIMAT0005951&hsa-miR-1307-3p&offsets\|0\|+1(+1A);m-66&10\|-\|103394300\|103394322&offsets\|0\|0(+1A)]\|ACTCGGCGTGGCGTCGGTCGTGGA | >MI0006444\|hsa-mir-1307&WithFlank&10\|-\|103394247\|103394407 | 86 | 108 | 23 | (+1A) | isomiR | NO | 0.00 | 30.64 | 0.00 | 0.00 |
| >MI0006444\|hsa-mir-1307&WithFlank&10\|-\|103394247\|103394407@86.109.24&[MIMAT0005951&hsa-miR-1307-3p&offsets\|0\|+2;m-66&10\|-\|103394300\|103394322&offsets\|0\|+1]\|ACTCGGCGTGGCGTCGGTCGTGGT | >MI0006444\|hsa-mir-1307&WithFlank&10\|-\|103394247\|103394407 | 86 | 109 | 24 | | isomiR | NO | 0.00 | 38.57 | 0.00 | 0.00 |
| >MI0006444\|hsa-mir-1307&WithFlank&10\|-\|103394247\|103394407@87.107.21(+1A)&[MIMAT0005951&hsa-miR-1307-3p&offsets\|+1\|0(+1A);m-66&10\|-\|103394300\|103394322&offsets\|+1\|-1(+1A)]\|CTCGGCGTGGCGTCGGTCGTGA | >MI0006444\|hsa-mir-1307&WithFlank&10\|-\|103394247\|103394407 | 87 | 107 | 21 | (+1A) | isomiR | NO | 0.00 | 24.23 | 0.00 | 0.00 |
| >MI0006444\|hsa-mir-1307&WithFlank&10\|-\|103394247\|103394407@87.107.21(+1U)&[MIMAT0005951&hsa-miR-1307-3p&offsets\|+1\|0(+1U);m-66&10\|-\|103394300\|103394322&offsets\|+1\|-1(+1U)]\|CTCGGCGTGGCGTCGGTCGTGT | >MI0006444\|hsa-mir-1307&WithFlank&10\|-\|103394247\|103394407 | 87 | 107 | 21 | (+1U) | isomiR | NO | 0.00 | 22.88 | 0.00 | 0.00 |
| >MI0006444\|hsa-mir-1307&WithFlank&10\|-\|103394247\|103394407@87.108.22&[MIMAT0005951&hsa-miR-1307-3p&offsets\|+1\|+1;m-66&10\|-\|103394300\|103394322&offsets\|+1\|0]\|CTCGGCGTGGCGTCGGTCGTGG | >MI0006444\|hsa-mir-1307&WithFlank&10\|-\|103394247\|103394407 | 87 | 108 | 22 | | isomiR | NO | 27.07 | 127.36 | 85.49 | 33.11 |
| >MI0006444\|hsa-mir-1307&WithFlank&10\|-\|103394247\|103394407@87.109.23&[MIMAT0005951&hsa-miR-1307-3p&offsets\|+1\|+2;m-66&10\|-\|103394300\|103394322&offsets\|+1\|+1]\|CTCGGCGTGGCGTCGGTCGTGGT | >MI0006444\|hsa-mir-1307&WithFlank&10\|-\|103394247\|103394407 | 87 | 109 | 23 | | isomiR | NO | 0.00 | 23.12 | 0.00 | 0.00 |
| >am_MI0008190\|hsa-mir-320d-1&WithFlank&13\|-\|40727810\|40727893@48.66.19;MI0008192\|hsa-mir-320d-2&WithFlank&X\|-\|140926154\|140926237@48.66.19&[MIMAT0006764&hsa-miR-320d&offsets\|0\|0;m-246&13\|-\|40727829\|40727846&offsets\|0\|+1];[MIMAT0006764_1&hsa-miR-320d&offsets\|0\|0;m-253&X\|-\|140926173\|140926190&offsets\|0\|+1]\|AAAAGCTGGGTTGAGAGGA | >MI0008190\|hsa-mir-320d-1&WithFlank&13\|-\|40727810\|40727893.an | 48 | 66 | 19 | | isomiR | NO | 0.00 | 0.00 | 14.70 | 0.00 |
| >am_MI0015825\|hsa-mir-378c&WithFlank&10\|-\|130962582\|130962674@17.34.18;MI0016749\|hsa-mir-378d-1&WithFlank&4\|-\|5923269\|5923334@37.54.18;MI0000786\|hsa-mir-378a&WithFlank&5\|+\|149732819\|149732896@49.66.18;MI0003840\|hsa-mir-378d-2&WithFlank&8\|-\|93916016\|93916125@28.45.18&[MIMAT0016847&hsa-miR-378c&offsets\|0\|-7;m-130&10\|-\|130962639\|130962658&offsets\|0\|-2];[MIMAT0018926&hsa-miR-378d&offsets\|0\|-2;m-228&4\|-\|5923281\|5923298&offsets\|0\|0];[MIMAT0000732&hsa-miR-378a-3p&offsets\|0\|-4;m-44&5\|+\|149732867\|149732888&offsets\|0\|-4];[MIMAT0018926_1&hsa-miR-378d&offsets\|0\|-2;m-227&8\|-\|93916081\|93916098&offsets\|0\|0]\|ACTGGACTTGGAGTCAGA | >MI0015825\|hsa-mir-378c&WithFlank&10\|-\|130962582\|130962674.ar | 17 | 34 | 18 | | isomiR | NO | 0.00 | 23.85 | 59.58 | 0.00 |
| >am_MI0015825\|hsa-mir-378c&WithFlank&10\|-\|130962582\|130962674@17.35.19;MI0016749\|hsa-mir-378d-1&WithFlank&4\|-\|5923269\|5923334@37.55.19;MI0000786\|hsa-mir-378a&WithFlank&5\|+\|149732819\|149732896@49.67.19;MI0003840\|hsa-mir-378d-2&WithFlank&8\|-\|93916016\|93916125@28.46.19&[MIMAT0016847&hsa-miR-378c&offsets\|0\|-6;m-130&10\|-\|130962639\|130962658&offsets\|0\|-1];[MIMAT0018926&hsa-miR-378d&offsets\|0\|-1;m-228&4\|-\|5923281\|5923298&offsets\|0\|+1];[MIMAT0000732&hsa-miR-378a-3p&offsets\|0\|-3;m-44&5\|+\|149732867\|149732888&offsets\|0\|-3];[MIMAT0018926_1&hsa-miR-378d&offsets\|0\|-1;m-227&8\|-\|93916081\|93916098&offsets\|0\|+1]\|ACTGGACTTGGAGTCAGAA | >MI0015825\|hsa-mir-378c&WithFlank&10\|-\|130962582\|130962674.ar | 17 | 35 | 19 | | isomiR | NO | 0.00 | 0.00 | 66.50 | 0.00 |
| >am_MI0015825\|hsa-mir-378c&WithFlank&10\|-\|130962582\|130962674@17.37.21&[MIMAT0016847&hsa-miR-378c&offsets\|0\|-4;m-130&10\|-\|130962639\|130962658&offsets\|0\|+1]\|ACTGGACTTGGAGTCAGAAGA | >MI0015825\|hsa-mir-378c&WithFlank&10\|-\|130962582\|130962674.ar | 17 | 37 | 21 | | isomiR | NO | 0.00 | 25.90 | 13.32 | 0.00 |
| >MI0015866\|hsa-mir-4326&WithFlank&20\|+\|63286802\|63286872@17.39.23&[MIMAT0016888&hsa-miR-4326&offsets\|0\|+3;m-339&20\|+\|63286818\|63286841&offsets\|0\|-1]\|TGTTCCTCTGTCTCCCAGACTCT | >MI0015866\|hsa-mir-4326&WithFlank&20\|+\|63286802\|63286872 | 17 | 39 | 23 | | isomiR | NO | 0.00 | 28.27 | 0.00 | 0.00 |
| >MI0015866\|hsa-mir-4326&WithFlank&20\|+\|63286802\|63286872@17.40.24&[MIMAT0016888&hsa-miR-4326&offsets\|0\|+4;m-339&20\|+\|63286818\|63286841&offsets\|0\|0]\|TGTTCCTCTGTCTCCCAGACTCTG | >MI0015866\|hsa-mir-4326&WithFlank&20\|+\|63286802\|63286872 | 17 | 40 | 24 | | isomiR | NO | 0.00 | 70.89 | 15.46 | 0.00 |

# BT-20

| | | | | | | | | | | | |
|---|---|---|---|---|---|---|---|---|---|---|---|
| >MI0016005\|hsa-mir-3615&WithFlank&17\|+\|74748607\|74748705@57.78.22&[MIMAT0017994&hsa-miR-3615&offsets\|0\|+1;m-271&17\|+\|74748663\|74748684&offsets\|0\|0]\|TCTCTCGGCTCCTCGCGGCTCG | >MI0016005\|hsa-mir-3615&WithFlank&17\|+\|74748607\|74748705 | 57 | 78 | 22 | | isomiR | NO | 0.00 | 94.00 | 0.00 | 0.00 |
| >MI0016746\|hsa-mir-548o-2&WithFlank&20\|+\|38516557\|38516638@51.72.22;MI0006402\|hsa-mir-548o&WithFlank&7\|-\|102405736\|102405861@93.114.22&[MIMAT0005919&hsa-miR-548o-3p&offsets\|0\|0;m-431&20\|+\|38516607\|38516628&offsets\|0\|0];[MIMAT0005919_1&hsa-miR-548o-3p&offsets\|0\|0;m-430&7\|-\|102405748\|102405769&offsets\|0\|0]\|CCAAAACTGCAGTTACTTTTGC | >MI0016746\|hsa-mir-548o-2&WithFlank&20\|+\|38516557\|38516638 | 51 | 72 | 22 | | isomiR | NO | 0.00 | 123.03 | 14.91 | 0.00 |
| >am_MI0016746\|hsa-mir-548o-2&WithFlank&20\|+\|38516557\|38516638@51.71.21(+1U);MI0006402\|hsa-mir-548o&WithFlank&7\|-\|102405736\|102405861@93.113.21(+1U)&[MIMAT0005919&hsa-miR-548o-3p&offsets\|0\|-1(+1U);m-431&20\|+\|38516607\|38516628&offsets\|0\|-1(+1U)];[MIMAT0005919_1&hsa-miR-548o-3p&offsets\|0\|-1(+1U);m-430&7\|-\|102405748\|102405769&offsets\|0\|-1(+1U)]\|CCAAAACTGCAGTTACTTTTGT | >MI0016746\|hsa-mir-548o-2&WithFlank&20\|+\|38516557\|38516638.a | 51 | 71 | 21 | (+1U) | isomiR | NO | 0.00 | 28.82 | 0.00 | 0.00 |
| >MI0016887\|hsa-mir-4521&WithFlank&17\|+\|8186939\|8187010@10.34.25&[MIMAT0019058&hsa-miR-4521&offsets\|0\|+3;m-765&17\|+\|8186948\|8186969&offsets\|0\|+3]\|GCTAAGGAAGTCCTGTGCTCAGTTT | >MI0016887\|hsa-mir-4521&WithFlank&17\|+\|8186939\|8187010 | 10 | 34 | 25 | | isomiR | NO | 0.00 | 154.71 | 0.00 | 0.00 |
| >MI0016887\|hsa-mir-4521&WithFlank&17\|+\|8186939\|8187010@10.35.26&[MIMAT0019058&hsa-miR-4521&offsets\|0\|+4;m-765&17\|+\|8186948\|8186969&offsets\|0\|+4]\|GCTAAGGAAGTCCTGTGCTCAGTTTT | >MI0016887\|hsa-mir-4521&WithFlank&17\|+\|8186939\|8187010 | 10 | 35 | 26 | | isomiR | NO | 0.00 | 90.88 | 0.00 | 0.00 |
| >MI0016887\|hsa-mir-4521&WithFlank&17\|+\|8186939\|8187010@13.34.22&[MIMAT0019058&hsa-miR-4521&offsets\|+3\|+3;m-765&17\|+\|8186948\|8186969&offsets\|+3\|+3]\|AAGGAAGTCCTGTGCTCAGTTT | >MI0016887\|hsa-mir-4521&WithFlank&17\|+\|8186939\|8187010 | 13 | 34 | 22 | | isomiR | NO | 0.00 | 29.26 | 0.00 | 0.00 |
| >MI0016887\|hsa-mir-4521&WithFlank&17\|+\|8186939\|8187010@13.35.23&[MIMAT0019058&hsa-miR-4521&offsets\|+3\|+4;m-765&17\|+\|8186948\|8186969&offsets\|+3\|+4]\|AAGGAAGTCCTGTGCTCAGTTTT | >MI0016887\|hsa-mir-4521&WithFlank&17\|+\|8186939\|8187010 | 13 | 35 | 23 | | isomiR | NO | 0.00 | 25.73 | 0.00 | 0.00 |
| >MI0016887\|hsa-mir-4521&WithFlank&17\|+\|8186939\|8187010@14.34.21&[MIMAT0019058&hsa-miR-4521&offsets\|+4\|+3;m-765&17\|+\|8186948\|8186969&offsets\|+4\|+3]\|AGGAAGTCCTGTGCTCAGTTT | >MI0016887\|hsa-mir-4521&WithFlank&17\|+\|8186939\|8187010 | 14 | 34 | 21 | | isomiR | NO | 0.00 | 30.17 | 0.00 | 0.00 |
| >MI0025242\|hsa-mir-7706&WithFlank&15\|+\|85380590\|85380668@50.71.22&[MIMAT0030021&hsa-miR-7706&offsets\|0\|-2;m-390&15\|+\|85380639\|85380662&offsets\|0\|-2]\|TGAAGCGCCTGTGCTCTGCCGA | >MI0025242\|hsa-mir-7706&WithFlank&15\|+\|85380590\|85380668 | 50 | 71 | 22 | | isomiR | NO | 0.00 | 25.34 | 0.00 | 0.00 |
| >MI0025242\|hsa-mir-7706&WithFlank&15\|+\|85380590\|85380668@50.72.23&[MIMAT0030021&hsa-miR-7706&offsets\|0\|-1;m-390&15\|+\|85380639\|85380662&offsets\|0\|-1]\|TGAAGCGCCTGTGCTCTGCCGAG | >MI0025242\|hsa-mir-7706&WithFlank&15\|+\|85380590\|85380668 | 50 | 72 | 23 | | isomiR | NO | 0.00 | 24.45 | 0.00 | 0.00 |
| >am_MI0039740\|hsa-mir-12136&WithFlank&1\|-\|632609\|632691@7.31.25&[MIMAT0049032&hsa-miR-12136&offsets\|0\|+7]\|GAAAAAGTCATGGAGGCCATGGGGT | >MI0039740\|hsa-mir-12136&WithFlank&1\|-\|632609\|632691.am | 7 | 31 | 25 | | isomiR | NO | 9.80 | 0.00 | 74.43 | 0.00 |
| >am_MI0039740\|hsa-mir-12136&WithFlank&1\|-\|632609\|632691@7.32.26&[MIMAT0049032&hsa-miR-12136&offsets\|0\|+8]\|GAAAAAGTCATGGAGGCCATGGGGTT | >MI0039740\|hsa-mir-12136&WithFlank&1\|-\|632609\|632691.am | 7 | 32 | 26 | | isomiR | NO | 66.25 | 0.00 | 119.93 | 0.00 |
| >hg38_MT_+_1621_3279_RNR2_16S_50ntFlanks@51.90.40\|GCTAAACCTAGCCCCAAACCCACTCCACCTTACTACCAGA | >hg38_MT_+_1621_3279_RNR2_16S_50ntFlanks | 51 | 90 | 40 | | rRF 12S and 16S | NO | 0.00 | 0.00 | 112.62 | 24.74 |
| >hg38_MT_+_1621_3279_RNR2_16S_50ntFlanks@51.91.41\|GCTAAACCTAGCCCCAAACCCACTCCACCTTACTACCAGAC | >hg38_MT_+_1621_3279_RNR2_16S_50ntFlanks | 51 | 91 | 41 | | rRF 12S and 16S | NO | 0.00 | 0.00 | 13.07 | 23.38 |
| >hg38_MT_+_1621_3279_RNR2_16S_50ntFlanks@51.92.42\|GCTAAACCTAGCCCCAAACCCACTCCACCTTACTACCAGACA | >hg38_MT_+_1621_3279_RNR2_16S_50ntFlanks | 51 | 92 | 42 | | rRF 12S and 16S | NO | 0.00 | 0.00 | 59.73 | 37.44 |
| >hg38_MT_+_1621_3279_RNR2_16S_50ntFlanks@51.94.44\|GCTAAACCTAGCCCCAAACCCACTCCACCTTACTACCAGACAAC | >hg38_MT_+_1621_3279_RNR2_16S_50ntFlanks | 51 | 94 | 44 | | rRF 12S and 16S | NO | 0.00 | 0.00 | 15.60 | 116.39 |
| >hg38_MT_+_1621_3279_RNR2_16S_50ntFlanks@51.95.45\|GCTAAACCTAGCCCCAAACCCACTCCACCTTACTACCAGACAACC | >hg38_MT_+_1621_3279_RNR2_16S_50ntFlanks | 51 | 95 | 45 | | rRF 12S and 16S | NO | 0.00 | 0.00 | 73.02 | 196.65 |
| >hg38_MT_+_1621_3279_RNR2_16S_50ntFlanks@622.652.31\|TATAACTGAACTCCTCACACCCAATTGGACC | >hg38_MT_+_1621_3279_RNR2_16S_50ntFlanks | 622 | 652 | 31 | | rRF 12S and 16S | NO | 0.00 | 0.00 | 13.39 | 0.00 |
| >hg38_MT_+_1621_3279_RNR2_16S_50ntFlanks@1184.1215.32\|AATTTCGGTTGGGGCGACCTCGGAGCAGAACC | >hg38_MT_+_1621_3279_RNR2_16S_50ntFlanks | 1184 | 1215 | 32 | | rRF 12S and 16S | NO | 0.00 | 0.00 | 13.03 | 0.00 |
| >hg38_MT_+_1621_3279_RNR2_16S_50ntFlanks@1184.1216.33\|AATTTCGGTTGGGGCGACCTCGGAGCAGAACCC | >hg38_MT_+_1621_3279_RNR2_16S_50ntFlanks | 1184 | 1216 | 33 | | rRF 12S and 16S | NO | 0.00 | 0.00 | 11.29 | 0.00 |
| >hg38_MT_+_1621_3279_RNR2_16S_50ntFlanks@1185.1215.31\|ATTTCGGTTGGGGCGACCTCGGAGCAGAACC | >hg38_MT_+_1621_3279_RNR2_16S_50ntFlanks | 1185 | 1215 | 31 | | rRF 12S and 16S | NO | 0.00 | 0.00 | 10.18 | 0.00 |
| >hg38_MT_+_1621_3279_RNR2_16S_50ntFlanks@1348.1385.38\|AACAATAGGGTTTACGACCTCGATGTTGGATCAGGACA | >hg38_MT_+_1621_3279_RNR2_16S_50ntFlanks | 1348 | 1385 | 38 | | rRF 12S and 16S | NO | 0.00 | 0.00 | 11.31 | 0.00 |
| >hg38_MT_+_1621_3279_RNR2_16S_50ntFlanks@1367.1396.30\|TCGATGTTGGATCAGGACATCCCGATGGTG | >hg38_MT_+_1621_3279_RNR2_16S_50ntFlanks | 1367 | 1396 | 30 | | rRF 12S and 16S | NO | 0.00 | 0.00 | 15.00 | 0.00 |
| >hg38_MT_+_1621_3279_RNR2_16S_50ntFlanks@1370.1396.27\|ATGTTGGATCAGGACATCCCGATGGTG | >hg38_MT_+_1621_3279_RNR2_16S_50ntFlanks | 1370 | 1396 | 27 | | rRF 12S and 16S | NO | 0.00 | 0.00 | 81.25 | 0.00 |
| >hg38_MT_+_1621_3279_RNR2_16S_50ntFlanks@1438.1477.40\|TACGTGATCTGAGTTCAGACCGGAGTAATCCAGGTCGGTT | >hg38_MT_+_1621_3279_RNR2_16S_50ntFlanks | 1438 | 1477 | 40 | | rRF 12S and 16S | NO | 0.00 | 0.00 | 11.47 | 9.73 |
| >multi-hg38_MT_+_1621_3279_RNR2_16S_50ntFlanks@51.70.20\|GCTAAACCTAGCCCCAAACC | >hg38_MT_+_1621_3279_RNR2_16S_50ntFlanks.multi | 51 | 70 | 20 | | rRF 12S and 16S | NO | 0.00 | 0.00 | 16.24 | 0.00 |

# BT-20

| | | | | | | | | | | |
|---|---|---|---|---|---|---|---|---|---|---|
| >multi-hg38_MT_+_1621_3279_RNR2_16S_50ntFlanks@51.71.21\|GCTAAACCTAGCCCCAAACCC | >hg38_MT_+_1621_3279_RNR2_16S_50ntFlanks.multi | 51 | 71 | 21 | rRF 12S and 16S | NO | 23.79 | 0.00 | 160.94 | 86.70 |
| >multi-hg38_MT_+_1621_3279_RNR2_16S_50ntFlanks@51.72.22\|GCTAAACCTAGCCCCAAACCCA | >hg38_MT_+_1621_3279_RNR2_16S_50ntFlanks.multi | 51 | 72 | 22 | rRF 12S and 16S | NO | 10.88 | 0.00 | 15.27 | 121.60 |
| >multi-hg38_MT_+_1621_3279_RNR2_16S_50ntFlanks@51.73.23\|GCTAAACCTAGCCCCAAACCCAC | >hg38_MT_+_1621_3279_RNR2_16S_50ntFlanks.multi | 51 | 73 | 23 | rRF 12S and 16S | NO | 0.00 | 0.00 | 37.39 | 67.46 |
| >multi-hg38_MT_+_1621_3279_RNR2_16S_50ntFlanks@51.74.24\|GCTAAACCTAGCCCCAAACCCACT | >hg38_MT_+_1621_3279_RNR2_16S_50ntFlanks.multi | 51 | 74 | 24 | rRF 12S and 16S | NO | 42.56 | 0.00 | 218.55 | 164.85 |
| >multi-hg38_MT_+_1621_3279_RNR2_16S_50ntFlanks@51.76.26\|GCTAAACCTAGCCCCAAACCCACTCC | >hg38_MT_+_1621_3279_RNR2_16S_50ntFlanks.multi | 51 | 76 | 26 | rRF 12S and 16S | NO | 0.00 | 0.00 | 37.62 | 25.49 |
| >multi-hg38_MT_+_1621_3279_RNR2_16S_50ntFlanks@51.79.29\|GCTAAACCTAGCCCCAAACCCACTCCACC | >hg38_MT_+_1621_3279_RNR2_16S_50ntFlanks.multi | 51 | 79 | 29 | rRF 12S and 16S | NO | 23.53 | 0.00 | 299.27 | 163.21 |
| >multi-hg38_MT_+_1621_3279_RNR2_16S_50ntFlanks@51.80.30\|GCTAAACCTAGCCCCAAACCCACTCCACCT | >hg38_MT_+_1621_3279_RNR2_16S_50ntFlanks.multi | 51 | 80 | 30 | rRF 12S and 16S | NO | 0.00 | 0.00 | 70.92 | 39.46 |
| >multi-hg38_MT_+_1621_3279_RNR2_16S_50ntFlanks@51.81.31\|GCTAAACCTAGCCCCAAACCCACTCCACCTT | >hg38_MT_+_1621_3279_RNR2_16S_50ntFlanks.multi | 51 | 81 | 31 | rRF 12S and 16S | NO | 25.67 | 0.00 | 124.48 | 85.07 |
| >multi-hg38_MT_+_1621_3279_RNR2_16S_50ntFlanks@51.82.32\|GCTAAACCTAGCCCCAAACCCACTCCACCTTA | >hg38_MT_+_1621_3279_RNR2_16S_50ntFlanks.multi | 51 | 82 | 32 | rRF 12S and 16S | NO | 0.00 | 0.00 | 317.62 | 34.66 |
| >multi-hg38_MT_+_1621_3279_RNR2_16S_50ntFlanks@51.84.34\|GCTAAACCTAGCCCCAAACCCACTCCACCTTACT | >hg38_MT_+_1621_3279_RNR2_16S_50ntFlanks.multi | 51 | 84 | 34 | rRF 12S and 16S | NO | 27.24 | 0.00 | 431.33 | 214.78 |
| >multi-hg38_MT_+_1621_3279_RNR2_16S_50ntFlanks@51.85.35\|GCTAAACCTAGCCCCAAACCCACTCCACCTTACTA | >hg38_MT_+_1621_3279_RNR2_16S_50ntFlanks.multi | 51 | 85 | 35 | rRF 12S and 16S | NO | 0.00 | 0.00 | 61.48 | 35.96 |
| >multi-hg38_MT_+_1621_3279_RNR2_16S_50ntFlanks@51.87.37\|GCTAAACCTAGCCCCAAACCCACTCCACCTTACTACC | >hg38_MT_+_1621_3279_RNR2_16S_50ntFlanks.multi | 51 | 87 | 37 | rRF 12S and 16S | NO | 0.00 | 0.00 | 461.25 | 93.89 |
| >multi-hg38_MT_+_1621_3279_RNR2_16S_50ntFlanks@51.88.38\|GCTAAACCTAGCCCCAAACCCACTCCACCTTACTACCA | >hg38_MT_+_1621_3279_RNR2_16S_50ntFlanks.multi | 51 | 88 | 38 | rRF 12S and 16S | NO | 0.00 | 0.00 | 146.23 | 80.62 |
| >multi-hg38_MT_+_1621_3279_RNR2_16S_50ntFlanks@292.311.20\|AGACGAGCTACCTAAGAACA | >hg38_MT_+_1621_3279_RNR2_16S_50ntFlanks.multi | 292 | 311 | 20 | rRF 12S and 16S | NO | 0.00 | 0.00 | 54.67 | 0.00 |
| >multi-hg38_MT_+_1621_3279_RNR2_16S_50ntFlanks@459.492.34\|CTTGTAAATTTAACTGTTAGTCCAAAGAGGAACA | >hg38_MT_+_1621_3279_RNR2_16S_50ntFlanks.multi | 459 | 492 | 34 | rRF 12S and 16S | NO | 0.00 | 0.00 | 10.07 | 0.00 |
| >multi-hg38_MT_+_1621_3279_RNR2_16S_50ntFlanks@1095.1120.26\|AGACGAGAAGACCCTATGGAGCTTTA | >hg38_MT_+_1621_3279_RNR2_16S_50ntFlanks.multi | 1095 | 1120 | 26 | rRF 12S and 16S | NO | 0.00 | 0.00 | 15.37 | 0.00 |
| >multi-hg38_MT_+_1621_3279_RNR2_16S_50ntFlanks@1428.1458.31\|ATTAAAGTCCTACGTGATCTGAGTTCAGACC | >hg38_MT_+_1621_3279_RNR2_16S_50ntFlanks.multi | 1428 | 1458 | 31 | rRF 12S and 16S | NO | 0.00 | 0.00 | 11.60 | 0.00 |
| >hg38_MT_+_598_1651_RNR1_12S_50ntFlanks@916.946.31\|AAGGACATTTAACTAAAACCCCTACGCATTT | >hg38_MT_+_598_1651_RNR1_12S_50ntFlanks | 916 | 946 | 31 | rRF 12S and 16S | NO | 0.00 | 0.00 | 131.10 | 0.00 |
| >multi-hg38_MT_+_598_1651_RNR1_12S_50ntFlanks@51.88.38\|AATAGGTTTGGTCCTAGCCTTTCTATTAGCTCTTAGTA | >hg38_MT_+_598_1651_RNR1_12S_50ntFlanks.multi | 51 | 88 | 38 | rRF 12S and 16S | NO | 0.00 | 0.00 | 280.02 | 0.00 |
| >multi-hg38_MT_+_598_1651_RNR1_12S_50ntFlanks@52.70.19\|ATAGGTTTGGTCCTAGCCT | >hg38_MT_+_598_1651_RNR1_12S_50ntFlanks.multi | 52 | 70 | 19 | rRF 12S and 16S | NO | 0.00 | 0.00 | 59.22 | 0.00 |
| >multi-hg38_MT_+_598_1651_RNR1_12S_50ntFlanks@52.72.21\|ATAGGTTTGGTCCTAGCCTTT | >hg38_MT_+_598_1651_RNR1_12S_50ntFlanks.multi | 52 | 72 | 21 | rRF 12S and 16S | NO | 0.00 | 0.00 | 52.71 | 0.00 |
| >multi-hg38_MT_+_598_1651_RNR1_12S_50ntFlanks@52.74.23\|ATAGGTTTGGTCCTAGCCTTTCT | >hg38_MT_+_598_1651_RNR1_12S_50ntFlanks.multi | 52 | 74 | 23 | rRF 12S and 16S | NO | 0.00 | 0.00 | 162.36 | 0.00 |
| >multi-hg38_MT_+_598_1651_RNR1_12S_50ntFlanks@52.77.26\|ATAGGTTTGGTCCTAGCCTTTCTATT | >hg38_MT_+_598_1651_RNR1_12S_50ntFlanks.multi | 52 | 77 | 26 | rRF 12S and 16S | NO | 0.00 | 0.00 | 806.73 | 112.71 |
| >multi-hg38_MT_+_598_1651_RNR1_12S_50ntFlanks@52.81.30\|ATAGGTTTGGTCCTAGCCTTTCTATTAGCT | >hg38_MT_+_598_1651_RNR1_12S_50ntFlanks.multi | 52 | 81 | 30 | rRF 12S and 16S | NO | 0.00 | 0.00 | 94.98 | 0.00 |
| >multi-hg38_MT_+_598_1651_RNR1_12S_50ntFlanks@52.83.32\|ATAGGTTTGGTCCTAGCCTTTCTATTAGCTCT | >hg38_MT_+_598_1651_RNR1_12S_50ntFlanks.multi | 52 | 83 | 32 | rRF 12S and 16S | NO | 0.00 | 0.00 | 165.04 | 23.71 |
| >multi-hg38_MT_+_598_1651_RNR1_12S_50ntFlanks@52.84.33\|ATAGGTTTGGTCCTAGCCTTTCTATTAGCTCTT | >hg38_MT_+_598_1651_RNR1_12S_50ntFlanks.multi | 52 | 84 | 33 | rRF 12S and 16S | NO | 0.00 | 0.00 | 445.50 | 28.20 |
| >multi-hg38_MT_+_598_1651_RNR1_12S_50ntFlanks@52.85.34\|ATAGGTTTGGTCCTAGCCTTTCTATTAGCTCTTA | >hg38_MT_+_598_1651_RNR1_12S_50ntFlanks.multi | 52 | 85 | 34 | rRF 12S and 16S | NO | 0.00 | 0.00 | 373.26 | 0.00 |
| >multi-hg38_MT_+_598_1651_RNR1_12S_50ntFlanks@52.87.36\|ATAGGTTTGGTCCTAGCCTTTCTATTAGCTCTTAGT | >hg38_MT_+_598_1651_RNR1_12S_50ntFlanks.multi | 52 | 87 | 36 | rRF 12S and 16S | NO | 9.59 | 0.00 | 354.21 | 72.18 |

# BT-20

| | | | | | | | | | | |
|---|---|---|---|---|---|---|---|---|---|---|
| >multi-hg38_MT_+_598_1651_RNR1_12S_50ntFlanks@52.88.37\|ATAGGTTTGGTCCTAGCCTTTCTATTAGCTCTTAGTA | >hg38_MT_+_598_1651_RNR1_12S_50ntFlanks.multi | 52 | 88 | 37 | rRF 12S and 16S | NO | 34.75 | 0.00 | 2287.95 | 247.37 |
| >multi-hg38_MT_+_598_1651_RNR1_12S_50ntFlanks@52.90.39\|ATAGGTTTGGTCCTAGCCTTTCTATTAGCTCTTAGTAAG | >hg38_MT_+_598_1651_RNR1_12S_50ntFlanks.multi | 52 | 90 | 39 | rRF 12S and 16S | NO | 0.00 | 0.00 | 15.96 | 0.00 |
| >multi-hg38_MT_+_598_1651_RNR1_12S_50ntFlanks@52.93.42\|ATAGGTTTGGTCCTAGCCTTTCTATTAGCTCTTAGTAAGATT | >hg38_MT_+_598_1651_RNR1_12S_50ntFlanks.multi | 52 | 93 | 42 | rRF 12S and 16S | NO | 0.00 | 0.00 | 76.96 | 0.00 |
| >multi-hg38_MT_+_598_1651_RNR1_12S_50ntFlanks@368.404.37\|CAATAAAGCTAAAACTCACCTGAGTTGTAAAAAACTC | >hg38_MT_+_598_1651_RNR1_12S_50ntFlanks.multi | 368 | 404 | 37 | rRF 12S and 16S | NO | 0.00 | 0.00 | 12.61 | 28.53 |
| >multi-hg38_MT_+_598_1651_RNR1_12S_50ntFlanks@369.404.36\|AATAAAGCTAAAACTCACCTGAGTTGTAAAAAACTC | >hg38_MT_+_598_1651_RNR1_12S_50ntFlanks.multi | 369 | 404 | 36 | rRF 12S and 16S | NO | 0.00 | 0.00 | 39.83 | 35.82 |
| >multi-hg38_MT_+_598_1651_RNR1_12S_50ntFlanks@370.404.35\|ATAAAGCTAAAACTCACCTGAGTTGTAAAAAACTC | >hg38_MT_+_598_1651_RNR1_12S_50ntFlanks.multi | 370 | 404 | 35 | rRF 12S and 16S | NO | 0.00 | 0.00 | 12.32 | 27.35 |
| >multi-hg38_MT_+_598_1651_RNR1_12S_50ntFlanks@373.404.32\|AAGCTAAAACTCACCTGAGTTGTAAAAAACTC | >hg38_MT_+_598_1651_RNR1_12S_50ntFlanks.multi | 373 | 404 | 32 | rRF 12S and 16S | NO | 0.00 | 0.00 | 10.05 | 0.00 |
| >multi-hg38_MT_+_598_1651_RNR1_12S_50ntFlanks@425.455.31\|ACGAAAGTGGCTTTAACATATCTGAACACAC | >hg38_MT_+_598_1651_RNR1_12S_50ntFlanks.multi | 425 | 455 | 31 | rRF 12S and 16S | NO | 0.00 | 0.00 | 11.24 | 0.00 |
| >multi-hg38_MT_+_598_1651_RNR1_12S_50ntFlanks@428.452.25\|AAAGTGGCTTTAACATATCTGAACA | >hg38_MT_+_598_1651_RNR1_12S_50ntFlanks.multi | 428 | 452 | 25 | rRF 12S and 16S | NO | 0.00 | 0.00 | 15.30 | 0.00 |
| >multi-hg38_MT_+_598_1651_RNR1_12S_50ntFlanks@428.455.28\|AAAGTGGCTTTAACATATCTGAACACAC | >hg38_MT_+_598_1651_RNR1_12S_50ntFlanks.multi | 428 | 455 | 28 | rRF 12S and 16S | NO | 0.00 | 0.00 | 37.32 | 0.00 |
| >multi-hg38_MT_+_598_1651_RNR1_12S_50ntFlanks@428.468.41\|AAAGTGGCTTTAACATATCTGAACACACAATAGCTAAGACC | >hg38_MT_+_598_1651_RNR1_12S_50ntFlanks.multi | 428 | 468 | 41 | rRF 12S and 16S | NO | 0.00 | 0.00 | 55.80 | 25.80 |
| >multi-hg38_MT_+_598_1651_RNR1_12S_50ntFlanks@428.469.42\|AAAGTGGCTTTAACATATCTGAACACACAATAGCTAAGACCC | >hg38_MT_+_598_1651_RNR1_12S_50ntFlanks.multi | 428 | 469 | 42 | rRF 12S and 16S | NO | 0.00 | 0.00 | 14.24 | 10.54 |
| >multi-hg38_MT_+_598_1651_RNR1_12S_50ntFlanks@905.929.25\|AGTATACTTCAAAGGACATTTAACT | >hg38_MT_+_598_1651_RNR1_12S_50ntFlanks.multi | 905 | 929 | 25 | rRF 12S and 16S | NO | 0.00 | 0.00 | 15.23 | 0.00 |
| >hg38_21_+_8433172_8446622_RNA45SN1_45S_50ntFlanks@474.504.31\|TCTTCGTGATCGATGTGGTGACGTCGTGCTC | >hg38_21_+_8433172_8446622_RNA45SN1_45S_50ntFlanks | 474 | 504 | 31 | rRF 45S | NO | 24.28 | 0.00 | 0.00 | 0.00 |
| >hg38_21_+_8433172_8446622_RNA45SN1_45S_50ntFlanks@474.505.32\|TCTTCGTGATCGATGTGGTGACGTCGTGCTCT | >hg38_21_+_8433172_8446622_RNA45SN1_45S_50ntFlanks | 474 | 505 | 32 | rRF 45S | NO | 37.13 | 0.00 | 0.00 | 0.00 |
| >hg38_21_+_8433172_8446622_RNA45SN1_45S_50ntFlanks@474.506.33\|TCTTCGTGATCGATGTGGTGACGTCGTGCTCTC | >hg38_21_+_8433172_8446622_RNA45SN1_45S_50ntFlanks | 474 | 506 | 33 | rRF 45S | NO | 10.22 | 0.00 | 0.00 | 0.00 |
| >hg38_21_+_8433172_8446622_RNA45SN1_45S_50ntFlanks@475.492.18\|CTTCGTGATCGATGTGGT | >hg38_21_+_8433172_8446622_RNA45SN1_45S_50ntFlanks | 475 | 492 | 18 | rRF 45S | NO | 141.39 | 0.00 | 0.00 | 0.00 |
| >hg38_21_+_8433172_8446622_RNA45SN1_45S_50ntFlanks@475.493.19\|CTTCGTGATCGATGTGGTG | >hg38_21_+_8433172_8446622_RNA45SN1_45S_50ntFlanks | 475 | 493 | 19 | rRF 45S | NO | 10.99 | 0.00 | 0.00 | 0.00 |
| >hg38_21_+_8433172_8446622_RNA45SN1_45S_50ntFlanks@475.495.21\|CTTCGTGATCGATGTGGTGAC | >hg38_21_+_8433172_8446622_RNA45SN1_45S_50ntFlanks | 475 | 495 | 21 | rRF 45S | NO | 84.65 | 0.00 | 0.00 | 0.00 |
| >hg38_21_+_8433172_8446622_RNA45SN1_45S_50ntFlanks@475.496.22\|CTTCGTGATCGATGTGGTGACG | >hg38_21_+_8433172_8446622_RNA45SN1_45S_50ntFlanks | 475 | 496 | 22 | rRF 45S | NO | 200.40 | 9.82 | 60.47 | 40.25 |
| >hg38_21_+_8433172_8446622_RNA45SN1_45S_50ntFlanks@475.497.23\|CTTCGTGATCGATGTGGTGACGT | >hg38_21_+_8433172_8446622_RNA45SN1_45S_50ntFlanks | 475 | 497 | 23 | rRF 45S | NO | 309.26 | 26.81 | 92.19 | 95.86 |
| >hg38_21_+_8433172_8446622_RNA45SN1_45S_50ntFlanks@475.498.24\|CTTCGTGATCGATGTGGTGACGTC | >hg38_21_+_8433172_8446622_RNA45SN1_45S_50ntFlanks | 475 | 498 | 24 | rRF 45S | NO | 677.33 | 95.14 | 288.84 | 250.22 |
| >hg38_21_+_8433172_8446622_RNA45SN1_45S_50ntFlanks@475.499.25\|CTTCGTGATCGATGTGGTGACGTCG | >hg38_21_+_8433172_8446622_RNA45SN1_45S_50ntFlanks | 475 | 499 | 25 | rRF 45S | NO | 24.83 | 0.00 | 0.00 | 0.00 |
| >hg38_21_+_8433172_8446622_RNA45SN1_45S_50ntFlanks@475.501.27\|CTTCGTGATCGATGTGGTGACGTCGTG | >hg38_21_+_8433172_8446622_RNA45SN1_45S_50ntFlanks | 475 | 501 | 27 | rRF 45S | NO | 93.42 | 0.00 | 0.00 | 24.24 |
| >hg38_21_+_8433172_8446622_RNA45SN1_45S_50ntFlanks@475.502.28\|CTTCGTGATCGATGTGGTGACGTCGTGC | >hg38_21_+_8433172_8446622_RNA45SN1_45S_50ntFlanks | 475 | 502 | 28 | rRF 45S | NO | 90.22 | 0.00 | 0.00 | 0.00 |
| >hg38_21_+_8433172_8446622_RNA45SN1_45S_50ntFlanks@475.503.29\|CTTCGTGATCGATGTGGTGACGTCGTGCT | >hg38_21_+_8433172_8446622_RNA45SN1_45S_50ntFlanks | 475 | 503 | 29 | rRF 45S | NO | 293.41 | 71.98 | 55.44 | 86.88 |
| >hg38_21_+_8433172_8446622_RNA45SN1_45S_50ntFlanks@475.504.30\|CTTCGTGATCGATGTGGTGACGTCGTGCTC | >hg38_21_+_8433172_8446622_RNA45SN1_45S_50ntFlanks | 475 | 504 | 30 | rRF 45S | NO | 548.30 | 290.12 | 122.19 | 188.25 |
| >hg38_21_+_8433172_8446622_RNA45SN1_45S_50ntFlanks@475.505.31\|CTTCGTGATCGATGTGGTGACGTCGTGCTCT | >hg38_21_+_8433172_8446622_RNA45SN1_45S_50ntFlanks | 475 | 505 | 31 | rRF 45S | NO | 952.52 | 461.97 | 153.64 | 226.31 |
| >hg38_21_+_8433172_8446622_RNA45SN1_45S_50ntFlanks@475.506.32\|CTTCGTGATCGATGTGGTGACGTCGTGCTCTC | >hg38_21_+_8433172_8446622_RNA45SN1_45S_50ntFlanks | 475 | 506 | 32 | rRF 45S | NO | 548.27 | 344.71 | 53.53 | 174.93 |
| >hg38_21_+_8433172_8446622_RNA45SN1_45S_50ntFlanks@475.507.33\|CTTCGTGATCGATGTGGTGACGTCGTGCTCTCC | >hg38_21_+_8433172_8446622_RNA45SN1_45S_50ntFlanks | 475 | 507 | 33 | rRF 45S | NO | 220.50 | 100.25 | 0.00 | 88.81 |
| >hg38_21_+_8433172_8446622_RNA45SN1_45S_50ntFlanks@475.508.34\|CTTCGTGATCGATGTGGTGACGTCGTGCTCTCCC | >hg38_21_+_8433172_8446622_RNA45SN1_45S_50ntFlanks | 475 | 508 | 34 | rRF 45S | NO | 91.46 | 24.80 | 0.00 | 0.00 |
| >hg38_21_+_8433172_8446622_RNA45SN1_45S_50ntFlanks@475.509.35\|CTTCGTGATCGATGTGGTGACGTCGTGCTCTCCCG | >hg38_21_+_8433172_8446622_RNA45SN1_45S_50ntFlanks | 475 | 509 | 35 | rRF 45S | NO | 215.34 | 87.59 | 0.00 | 87.00 |

# BT-20

| | | | | | | | | | | |
|---|---|---|---|---|---|---|---|---|---|---|
| >hg38_21_+_8433172_8446622_RNA45SN1_45S_50ntFlanks@475.510.36\|CTTCGTGATCGATG<br>TGGTGACGTCGTGCTCTCCCGG | >hg38_21_+_8433172_8446622_RNA45SN1_45S_50ntFlanks | 475 | 510 | 36 | rRF 45S | NO | 252.25 | 139.54 | 0.00 | 119.00 |
| >hg38_21_+_8433172_8446622_RNA45SN1_45S_50ntFlanks@475.511.37\|CTTCGTGATCGATG<br>TGGTGACGTCGTGCTCTCCCGGG | >hg38_21_+_8433172_8446622_RNA45SN1_45S_50ntFlanks | 475 | 511 | 37 | rRF 45S | NO | 175.09 | 85.33 | 0.00 | 74.73 |
| >hg38_21_+_8433172_8446622_RNA45SN1_45S_50ntFlanks@475.512.38\|CTTCGTGATCGATG<br>TGGTGACGTCGTGCTCTCCCGGGC | >hg38_21_+_8433172_8446622_RNA45SN1_45S_50ntFlanks | 475 | 512 | 38 | rRF 45S | NO | 42.28 | 0.00 | 0.00 | 0.00 |
| >hg38_21_+_8433172_8446622_RNA45SN1_45S_50ntFlanks@475.513.39\|CTTCGTGATCGATG<br>TGGTGACGTCGTGCTCTCCCGGGCC | >hg38_21_+_8433172_8446622_RNA45SN1_45S_50ntFlanks | 475 | 513 | 39 | rRF 45S | NO | 65.34 | 0.00 | 0.00 | 0.00 |
| >hg38_21_+_8433172_8446622_RNA45SN1_45S_50ntFlanks@475.514.40\|CTTCGTGATCGATG<br>TGGTGACGTCGTGCTCTCCCGGGCCG | >hg38_21_+_8433172_8446622_RNA45SN1_45S_50ntFlanks | 475 | 514 | 40 | rRF 45S | NO | 158.86 | 36.42 | 0.00 | 0.00 |
| >hg38_21_+_8433172_8446622_RNA45SN1_45S_50ntFlanks@475.515.41\|CTTCGTGATCGATG<br>TGGTGACGTCGTGCTCTCCCGGGCCGG | >hg38_21_+_8433172_8446622_RNA45SN1_45S_50ntFlanks | 475 | 515 | 41 | rRF 45S | NO | 105.75 | 0.00 | 0.00 | 0.00 |
| >hg38_21_+_8433172_8446622_RNA45SN1_45S_50ntFlanks@475.516.42\|CTTCGTGATCGATG<br>TGGTGACGTCGTGCTCTCCCGGGCCGGG | >hg38_21_+_8433172_8446622_RNA45SN1_45S_50ntFlanks | 475 | 516 | 42 | rRF 45S | NO | 306.80 | 23.09 | 0.00 | 134.18 |
| >hg38_21_+_8433172_8446622_RNA45SN1_45S_50ntFlanks@475.517.43\|CTTCGTGATCGATG<br>TGGTGACGTCGTGCTCTCCCGGGCCGGGT | >hg38_21_+_8433172_8446622_RNA45SN1_45S_50ntFlanks | 475 | 517 | 43 | rRF 45S | NO | 222.92 | 0.00 | 0.00 | 163.15 |
| >hg38_21_+_8433172_8446622_RNA45SN1_45S_50ntFlanks@475.518.44\|CTTCGTGATCGATG<br>TGGTGACGTCGTGCTCTCCCGGGCCGGGTC | >hg38_21_+_8433172_8446622_RNA45SN1_45S_50ntFlanks | 475 | 518 | 44 | rRF 45S | NO | 133.17 | 0.00 | 0.00 | 97.66 |
| >hg38_21_+_8433172_8446622_RNA45SN1_45S_50ntFlanks@475.519.45\|CTTCGTGATCGATG<br>TGGTGACGTCGTGCTCTCCCGGGCCGGGTCC | >hg38_21_+_8433172_8446622_RNA45SN1_45S_50ntFlanks | 475 | 519 | 45 | rRF 45S | NO | 10.11 | 0.00 | 0.00 | 0.00 |
| >hg38_21_+_8433172_8446622_RNA45SN1_45S_50ntFlanks@475.520.46\|CTTCGTGATCGATG<br>TGGTGACGTCGTGCTCTCCCGGGCCGGGTCCG | >hg38_21_+_8433172_8446622_RNA45SN1_45S_50ntFlanks | 475 | 520 | 46 | rRF 45S | NO | 40.46 | 0.00 | 0.00 | 0.00 |
| >hg38_21_+_8433172_8446622_RNA45SN1_45S_50ntFlanks@475.521.47\|CTTCGTGATCGATG<br>TGGTGACGTCGTGCTCTCCCGGGCCGGGTCCGA | >hg38_21_+_8433172_8446622_RNA45SN1_45S_50ntFlanks | 475 | 521 | 47 | rRF 45S | NO | 92.20 | 0.00 | 0.00 | 27.06 |
| >hg38_21_+_8433172_8446622_RNA45SN1_45S_50ntFlanks@475.522.48\|CTTCGTGATCGATG<br>TGGTGACGTCGTGCTCTCCCGGGCCGGGTCCGAG | >hg38_21_+_8433172_8446622_RNA45SN1_45S_50ntFlanks | 475 | 522 | 48 | rRF 45S | NO | 32.47 | 0.00 | 0.00 | 0.00 |
| >hg38_21_+_8433172_8446622_RNA45SN1_45S_50ntFlanks@475.523.49\|CTTCGTGATCGATG<br>TGGTGACGTCGTGCTCTCCCGGGCCGGGTCCGAGC | >hg38_21_+_8433172_8446622_RNA45SN1_45S_50ntFlanks | 475 | 523 | 49 | rRF 45S | NO | 28.60 | 0.00 | 0.00 | 0.00 |
| >hg38_21_+_8433172_8446622_RNA45SN1_45S_50ntFlanks@476.495.20\|TTCGTGATCGATGT<br>GGTGAC | >hg38_21_+_8433172_8446622_RNA45SN1_45S_50ntFlanks | 476 | 495 | 20 | rRF 45S | NO | 39.77 | 0.00 | 0.00 | 0.00 |
| >hg38_21_+_8433172_8446622_RNA45SN1_45S_50ntFlanks@476.496.21\|TTCGTGATCGATGT<br>GGTGACG | >hg38_21_+_8433172_8446622_RNA45SN1_45S_50ntFlanks | 476 | 496 | 21 | rRF 45S | NO | 26.37 | 0.00 | 0.00 | 0.00 |
| >hg38_21_+_8433172_8446622_RNA45SN1_45S_50ntFlanks@476.497.22\|TTCGTGATCGATGT<br>GGTGACGT | >hg38_21_+_8433172_8446622_RNA45SN1_45S_50ntFlanks | 476 | 497 | 22 | rRF 45S | NO | 29.32 | 0.00 | 0.00 | 0.00 |
| >hg38_21_+_8433172_8446622_RNA45SN1_45S_50ntFlanks@476.498.23\|TTCGTGATCGATGT<br>GGTGACGTC | >hg38_21_+_8433172_8446622_RNA45SN1_45S_50ntFlanks | 476 | 498 | 23 | rRF 45S | NO | 84.04 | 0.00 | 0.00 | 0.00 |
| >hg38_21_+_8433172_8446622_RNA45SN1_45S_50ntFlanks@476.499.24\|TTCGTGATCGATGT<br>GGTGACGTCG | >hg38_21_+_8433172_8446622_RNA45SN1_45S_50ntFlanks | 476 | 499 | 24 | rRF 45S | NO | 138.31 | 30.77 | 0.00 | 39.57 |
| >hg38_21_+_8433172_8446622_RNA45SN1_45S_50ntFlanks@476.501.26\|TTCGTGATCGATGT<br>GGTGACGTCGTG | >hg38_21_+_8433172_8446622_RNA45SN1_45S_50ntFlanks | 476 | 501 | 26 | rRF 45S | NO | 119.69 | 0.00 | 0.00 | 9.57 |
| >hg38_21_+_8433172_8446622_RNA45SN1_45S_50ntFlanks@476.502.27\|TTCGTGATCGATGT<br>GGTGACGTCGTGC | >hg38_21_+_8433172_8446622_RNA45SN1_45S_50ntFlanks | 476 | 502 | 27 | rRF 45S | NO | 72.51 | 0.00 | 0.00 | 0.00 |
| >hg38_21_+_8433172_8446622_RNA45SN1_45S_50ntFlanks@476.503.28\|TTCGTGATCGATGT<br>GGTGACGTCGTGCT | >hg38_21_+_8433172_8446622_RNA45SN1_45S_50ntFlanks | 476 | 503 | 28 | rRF 45S | NO | 158.53 | 0.00 | 0.00 | 10.76 |
| >hg38_21_+_8433172_8446622_RNA45SN1_45S_50ntFlanks@476.504.29\|TTCGTGATCGATGT<br>GGTGACGTCGTGCTC | >hg38_21_+_8433172_8446622_RNA45SN1_45S_50ntFlanks | 476 | 504 | 29 | rRF 45S | NO | 206.10 | 38.22 | 0.00 | 62.48 |
| >hg38_21_+_8433172_8446622_RNA45SN1_45S_50ntFlanks@476.505.30\|TTCGTGATCGATGT<br>GGTGACGTCGTGCTCT | >hg38_21_+_8433172_8446622_RNA45SN1_45S_50ntFlanks | 476 | 505 | 30 | rRF 45S | NO | 357.77 | 96.06 | 10.02 | 87.91 |
| >hg38_21_+_8433172_8446622_RNA45SN1_45S_50ntFlanks@476.506.31\|TTCGTGATCGATGT<br>GGTGACGTCGTGCTCTC | >hg38_21_+_8433172_8446622_RNA45SN1_45S_50ntFlanks | 476 | 506 | 31 | rRF 45S | NO | 221.92 | 79.62 | 0.00 | 40.78 |
| >hg38_21_+_8433172_8446622_RNA45SN1_45S_50ntFlanks@476.507.32\|TTCGTGATCGATGT<br>GGTGACGTCGTGCTCTCC | >hg38_21_+_8433172_8446622_RNA45SN1_45S_50ntFlanks | 476 | 507 | 32 | rRF 45S | NO | 116.78 | 25.26 | 0.00 | 0.00 |
| >hg38_21_+_8433172_8446622_RNA45SN1_45S_50ntFlanks@476.508.33\|TTCGTGATCGATGT<br>GGTGACGTCGTGCTCTCCC | >hg38_21_+_8433172_8446622_RNA45SN1_45S_50ntFlanks | 476 | 508 | 33 | rRF 45S | NO | 30.47 | 0.00 | 0.00 | 0.00 |
| >hg38_21_+_8433172_8446622_RNA45SN1_45S_50ntFlanks@476.509.34\|TTCGTGATCGATGT<br>GGTGACGTCGTGCTCTCCCG | >hg38_21_+_8433172_8446622_RNA45SN1_45S_50ntFlanks | 476 | 509 | 34 | rRF 45S | NO | 114.75 | 10.38 | 0.00 | 0.00 |
| >hg38_21_+_8433172_8446622_RNA45SN1_45S_50ntFlanks@476.510.35\|TTCGTGATCGATGT<br>GGTGACGTCGTGCTCTCCCGG | >hg38_21_+_8433172_8446622_RNA45SN1_45S_50ntFlanks | 476 | 510 | 35 | rRF 45S | NO | 99.28 | 27.74 | 0.00 | 0.00 |
| >hg38_21_+_8433172_8446622_RNA45SN1_45S_50ntFlanks@476.511.36\|TTCGTGATCGATGT<br>GGTGACGTCGTGCTCTCCCGGG | >hg38_21_+_8433172_8446622_RNA45SN1_45S_50ntFlanks | 476 | 511 | 36 | rRF 45S | NO | 85.68 | 10.19 | 0.00 | 0.00 |
| >hg38_21_+_8433172_8446622_RNA45SN1_45S_50ntFlanks@476.512.37\|TTCGTGATCGATGT<br>GGTGACGTCGTGCTCTCCCGGGC | >hg38_21_+_8433172_8446622_RNA45SN1_45S_50ntFlanks | 476 | 512 | 37 | rRF 45S | NO | 34.29 | 0.00 | 0.00 | 0.00 |
| >hg38_21_+_8433172_8446622_RNA45SN1_45S_50ntFlanks@476.513.38\|TTCGTGATCGATGT<br>GGTGACGTCGTGCTCTCCCGGGCC | >hg38_21_+_8433172_8446622_RNA45SN1_45S_50ntFlanks | 476 | 513 | 38 | rRF 45S | NO | 26.66 | 0.00 | 0.00 | 0.00 |
| >hg38_21_+_8433172_8446622_RNA45SN1_45S_50ntFlanks@476.514.39\|TTCGTGATCGATGT<br>GGTGACGTCGTGCTCTCCCGGGCCG | >hg38_21_+_8433172_8446622_RNA45SN1_45S_50ntFlanks | 476 | 514 | 39 | rRF 45S | NO | 10.68 | 0.00 | 0.00 | 0.00 |
| >hg38_21_+_8433172_8446622_RNA45SN1_45S_50ntFlanks@476.516.41\|TTCGTGATCGATGT<br>GGTGACGTCGTGCTCTCCCGGGCCGGG | >hg38_21_+_8433172_8446622_RNA45SN1_45S_50ntFlanks | 476 | 516 | 41 | rRF 45S | NO | 184.19 | 0.00 | 0.00 | 0.00 |
| >hg38_21_+_8433172_8446622_RNA45SN1_45S_50ntFlanks@476.517.42\|TTCGTGATCGATGT<br>GGTGACGTCGTGCTCTCCCGGGCCGGGT | >hg38_21_+_8433172_8446622_RNA45SN1_45S_50ntFlanks | 476 | 517 | 42 | rRF 45S | NO | 136.29 | 0.00 | 0.00 | 0.00 |
| >hg38_21_+_8433172_8446622_RNA45SN1_45S_50ntFlanks@476.518.43\|TTCGTGATCGATGT<br>GGTGACGTCGTGCTCTCCCGGGCCGGGTC | >hg38_21_+_8433172_8446622_RNA45SN1_45S_50ntFlanks | 476 | 518 | 43 | rRF 45S | NO | 82.71 | 0.00 | 0.00 | 10.49 |
| >hg38_21_+_8433172_8446622_RNA45SN1_45S_50ntFlanks@476.520.45\|TTCGTGATCGATGT<br>GGTGACGTCGTGCTCTCCCGGGCCGGGTCCG | >hg38_21_+_8433172_8446622_RNA45SN1_45S_50ntFlanks | 476 | 520 | 45 | rRF 45S | NO | 33.39 | 0.00 | 0.00 | 0.00 |
| >hg38_21_+_8433172_8446622_RNA45SN1_45S_50ntFlanks@476.521.46\|TTCGTGATCGATGT<br>GGTGACGTCGTGCTCTCCCGGGCCGGGTCCGA | >hg38_21_+_8433172_8446622_RNA45SN1_45S_50ntFlanks | 476 | 521 | 46 | rRF 45S | NO | 34.95 | 0.00 | 0.00 | 0.00 |

# BT-20

| | | | | | | | | | | |
|---|---|---|---|---|---|---|---|---|---|---|
| >hg38_21_+_8433172_8446622_RNA45SN1_45S_50ntFlanks@476.522.47\|TTCGTGATCGATGTGGTGACGTCGTGCTCTCCCGGGCCGGGTCCGAG | >hg38_21_+_8433172_8446622_RNA45SN1_45S_50ntFlanks | 476 | 522 | 47 | rRF 45S | YES | 10.42 | 0.00 | 0.00 | 0.00 |
| >hg38_21_+_8433172_8446622_RNA45SN1_45S_50ntFlanks@476.523.48\|TTCGTGATCGATGTGGTGACGTCGTGCTCTCCCGGGCCGGGTCCGAGC | >hg38_21_+_8433172_8446622_RNA45SN1_45S_50ntFlanks | 476 | 523 | 48 | rRF 45S | NO | 31.26 | 0.00 | 0.00 | 0.00 |
| >hg38_21_+_8433172_8446622_RNA45SN1_45S_50ntFlanks@483.505.23\|TCGATGTGGTGACGTCGTGCTCT | >hg38_21_+_8433172_8446622_RNA45SN1_45S_50ntFlanks | 483 | 505 | 23 | rRF 45S | NO | 82.68 | 0.00 | 0.00 | 0.00 |
| >hg38_21_+_8433172_8446622_RNA45SN1_45S_50ntFlanks@483.506.24\|TCGATGTGGTGACGTCGTGCTCTC | >hg38_21_+_8433172_8446622_RNA45SN1_45S_50ntFlanks | 483 | 506 | 24 | rRF 45S | NO | 24.86 | 0.00 | 0.00 | 0.00 |
| >hg38_21_+_8433172_8446622_RNA45SN1_45S_50ntFlanks@483.532.50\|TCGATGTGGTGACGTCGTGCTCTCCCGGGCCGGGTCCGAGCCGCGACGGG | >hg38_21_+_8433172_8446622_RNA45SN1_45S_50ntFlanks | 483 | 532 | 50 | rRF 45S | YES | 28.62 | 0.00 | 0.00 | 0.00 |
| >hg38_21_+_8433172_8446622_RNA45SN1_45S_50ntFlanks@485.504.20\|GATGTGGTGACGTCGTGCTC | >hg38_21_+_8433172_8446622_RNA45SN1_45S_50ntFlanks | 485 | 504 | 20 | rRF 45S | NO | 67.19 | 0.00 | 16.05 | 0.00 |
| >hg38_21_+_8433172_8446622_RNA45SN1_45S_50ntFlanks@485.505.21\|GATGTGGTGACGTCGTGCTCT | >hg38_21_+_8433172_8446622_RNA45SN1_45S_50ntFlanks | 485 | 505 | 21 | rRF 45S | NO | 634.92 | 92.41 | 325.55 | 92.49 |
| >hg38_21_+_8433172_8446622_RNA45SN1_45S_50ntFlanks@485.506.22\|GATGTGGTGACGTCGTGCTCTC | >hg38_21_+_8433172_8446622_RNA45SN1_45S_50ntFlanks | 485 | 506 | 22 | rRF 45S | NO | 361.90 | 110.80 | 64.31 | 79.78 |
| >hg38_21_+_8433172_8446622_RNA45SN1_45S_50ntFlanks@485.507.23\|GATGTGGTGACGTCGTGCTCTCC | >hg38_21_+_8433172_8446622_RNA45SN1_45S_50ntFlanks | 485 | 507 | 23 | rRF 45S | NO | 239.67 | 33.88 | 0.00 | 33.28 |
| >hg38_21_+_8433172_8446622_RNA45SN1_45S_50ntFlanks@485.508.24\|GATGTGGTGACGTCGTGCTCTCCC | >hg38_21_+_8433172_8446622_RNA45SN1_45S_50ntFlanks | 485 | 508 | 24 | rRF 45S | NO | 90.15 | 0.00 | 0.00 | 0.00 |
| >hg38_21_+_8433172_8446622_RNA45SN1_45S_50ntFlanks@485.509.25\|GATGTGGTGACGTCGTGCTCTCCCG | >hg38_21_+_8433172_8446622_RNA45SN1_45S_50ntFlanks | 485 | 509 | 25 | rRF 45S | NO | 95.00 | 0.00 | 0.00 | 0.00 |
| >hg38_21_+_8433172_8446622_RNA45SN1_45S_50ntFlanks@485.510.26\|GATGTGGTGACGTCGTGCTCTCCCGG | >hg38_21_+_8433172_8446622_RNA45SN1_45S_50ntFlanks | 485 | 510 | 26 | rRF 45S | NO | 98.52 | 0.00 | 0.00 | 0.00 |
| >hg38_21_+_8433172_8446622_RNA45SN1_45S_50ntFlanks@485.511.27\|GATGTGGTGACGTCGTGCTCTCCCGGG | >hg38_21_+_8433172_8446622_RNA45SN1_45S_50ntFlanks | 485 | 511 | 27 | rRF 45S | NO | 78.64 | 0.00 | 0.00 | 0.00 |
| >hg38_21_+_8433172_8446622_RNA45SN1_45S_50ntFlanks@485.513.29\|GATGTGGTGACGTCGTGCTCTCCCGGGCC | >hg38_21_+_8433172_8446622_RNA45SN1_45S_50ntFlanks | 485 | 513 | 29 | rRF 45S | NO | 23.59 | 0.00 | 0.00 | 0.00 |
| >hg38_21_+_8433172_8446622_RNA45SN1_45S_50ntFlanks@485.514.30\|GATGTGGTGACGTCGTGCTCTCCCGGGCCG | >hg38_21_+_8433172_8446622_RNA45SN1_45S_50ntFlanks | 485 | 514 | 30 | rRF 45S | NO | 106.58 | 0.00 | 0.00 | 0.00 |
| >hg38_21_+_8433172_8446622_RNA45SN1_45S_50ntFlanks@485.515.31\|GATGTGGTGACGTCGTGCTCTCCCGGGCCGG | >hg38_21_+_8433172_8446622_RNA45SN1_45S_50ntFlanks | 485 | 515 | 31 | rRF 45S | NO | 125.38 | 0.00 | 0.00 | 0.00 |
| >hg38_21_+_8433172_8446622_RNA45SN1_45S_50ntFlanks@485.516.32\|GATGTGGTGACGTCGTGCTCTCCCGGGCCGGG | >hg38_21_+_8433172_8446622_RNA45SN1_45S_50ntFlanks | 485 | 516 | 32 | rRF 45S | NO | 130.38 | 0.00 | 0.00 | 0.00 |
| >hg38_21_+_8433172_8446622_RNA45SN1_45S_50ntFlanks@485.517.33\|GATGTGGTGACGTCGTGCTCTCCCGGGCCGGGT | >hg38_21_+_8433172_8446622_RNA45SN1_45S_50ntFlanks | 485 | 517 | 33 | rRF 45S | NO | 163.30 | 0.00 | 9.51 | 29.74 |
| >hg38_21_+_8433172_8446622_RNA45SN1_45S_50ntFlanks@485.518.34\|GATGTGGTGACGTCGTGCTCTCCCGGGCCGGGTC | >hg38_21_+_8433172_8446622_RNA45SN1_45S_50ntFlanks | 485 | 518 | 34 | rRF 45S | NO | 87.77 | 0.00 | 0.00 | 0.00 |
| >hg38_21_+_8433172_8446622_RNA45SN1_45S_50ntFlanks@485.519.35\|GATGTGGTGACGTCGTGCTCTCCCGGGCCGGGTCC | >hg38_21_+_8433172_8446622_RNA45SN1_45S_50ntFlanks | 485 | 519 | 35 | rRF 45S | NO | 38.68 | 0.00 | 0.00 | 0.00 |
| >hg38_21_+_8433172_8446622_RNA45SN1_45S_50ntFlanks@485.520.36\|GATGTGGTGACGTCGTGCTCTCCCGGGCCGGGTCCG | >hg38_21_+_8433172_8446622_RNA45SN1_45S_50ntFlanks | 485 | 520 | 36 | rRF 45S | NO | 121.94 | 0.00 | 0.00 | 0.00 |
| >hg38_21_+_8433172_8446622_RNA45SN1_45S_50ntFlanks@485.521.37\|GATGTGGTGACGTCGTGCTCTCCCGGGCCGGGTCCGA | >hg38_21_+_8433172_8446622_RNA45SN1_45S_50ntFlanks | 485 | 521 | 37 | rRF 45S | NO | 170.73 | 0.00 | 0.00 | 24.54 |
| >hg38_21_+_8433172_8446622_RNA45SN1_45S_50ntFlanks@485.522.38\|GATGTGGTGACGTCGTGCTCTCCCGGGCCGGGTCCGAG | >hg38_21_+_8433172_8446622_RNA45SN1_45S_50ntFlanks | 485 | 522 | 38 | rRF 45S | NO | 171.82 | 0.00 | 0.00 | 0.00 |
| >hg38_21_+_8433172_8446622_RNA45SN1_45S_50ntFlanks@485.524.40\|GATGTGGTGACGTCGTGCTCTCCCGGGCCGGGTCCGAGCC | >hg38_21_+_8433172_8446622_RNA45SN1_45S_50ntFlanks | 485 | 524 | 40 | rRF 45S | NO | 89.56 | 0.00 | 0.00 | 0.00 |
| >hg38_21_+_8433172_8446622_RNA45SN1_45S_50ntFlanks@485.525.41\|GATGTGGTGACGTCGTGCTCTCCCGGGCCGGGTCCGAGCCG | >hg38_21_+_8433172_8446622_RNA45SN1_45S_50ntFlanks | 485 | 525 | 41 | rRF 45S | NO | 126.03 | 0.00 | 0.00 | 0.00 |
| >hg38_21_+_8433172_8446622_RNA45SN1_45S_50ntFlanks@485.526.42\|GATGTGGTGACGTCGTGCTCTCCCGGGCCGGGTCCGAGCCGC | >hg38_21_+_8433172_8446622_RNA45SN1_45S_50ntFlanks | 485 | 526 | 42 | rRF 45S | NO | 69.57 | 0.00 | 0.00 | 0.00 |
| >hg38_21_+_8433172_8446622_RNA45SN1_45S_50ntFlanks@485.527.43\|GATGTGGTGACGTCGTGCTCTCCCGGGCCGGGTCCGAGCCGCG | >hg38_21_+_8433172_8446622_RNA45SN1_45S_50ntFlanks | 485 | 527 | 43 | rRF 45S | NO | 106.27 | 0.00 | 0.00 | 0.00 |
| >hg38_21_+_8433172_8446622_RNA45SN1_45S_50ntFlanks@485.528.44\|GATGTGGTGACGTCGTGCTCTCCCGGGCCGGGTCCGAGCCGCGA | >hg38_21_+_8433172_8446622_RNA45SN1_45S_50ntFlanks | 485 | 528 | 44 | rRF 45S | YES | 37.84 | 0.00 | 0.00 | 0.00 |
| >hg38_21_+_8433172_8446622_RNA45SN1_45S_50ntFlanks@485.529.45\|GATGTGGTGACGTCGTGCTCTCCCGGGCCGGGTCCGAGCCGCGAC | >hg38_21_+_8433172_8446622_RNA45SN1_45S_50ntFlanks | 485 | 529 | 45 | rRF 45S | NO | 65.88 | 0.00 | 0.00 | 0.00 |
| >hg38_21_+_8433172_8446622_RNA45SN1_45S_50ntFlanks@485.530.46\|GATGTGGTGACGTCGTGCTCTCCCGGGCCGGGTCCGAGCCGCGACG | >hg38_21_+_8433172_8446622_RNA45SN1_45S_50ntFlanks | 485 | 530 | 46 | rRF 45S | NO | 32.40 | 0.00 | 0.00 | 0.00 |
| >hg38_21_+_8433172_8446622_RNA45SN1_45S_50ntFlanks@485.531.47\|GATGTGGTGACGTCGTGCTCTCCCGGGCCGGGTCCGAGCCGCGACGG | >hg38_21_+_8433172_8446622_RNA45SN1_45S_50ntFlanks | 485 | 531 | 47 | rRF 45S | YES | 29.26 | 0.00 | 0.00 | 0.00 |
| >hg38_21_+_8433172_8446622_RNA45SN1_45S_50ntFlanks@485.532.48\|GATGTGGTGACGTCGTGCTCTCCCGGGCCGGGTCCGAGCCGCGACGGG | >hg38_21_+_8433172_8446622_RNA45SN1_45S_50ntFlanks | 485 | 532 | 48 | rRF 45S | NO | 133.75 | 0.00 | 0.00 | 0.00 |
| >hg38_21_+_8433172_8446622_RNA45SN1_45S_50ntFlanks@3704.3725.22\|CTACCTGGTTGATCCTGCCAGT | >hg38_21_+_8433172_8446622_RNA45SN1_45S_50ntFlanks | 3704 | 3725 | 22 | rRF 45S | NO | 0.00 | 24.54 | 0.00 | 0.00 |
| >hg38_21_+_8433172_8446622_RNA45SN1_45S_50ntFlanks@3705.3722.18\|TACCTGGTTGATCCTGCC | >hg38_21_+_8433172_8446622_RNA45SN1_45S_50ntFlanks | 3705 | 3722 | 18 | rRF 45S | NO | 248.53 | 1630.73 | 120.67 | 178.69 |
| >hg38_21_+_8433172_8446622_RNA45SN1_45S_50ntFlanks@3705.3723.19\|TACCTGGTTGATCCTGCCA | >hg38_21_+_8433172_8446622_RNA45SN1_45S_50ntFlanks | 3705 | 3723 | 19 | rRF 45S | NO | 10.92 | 105.55 | 0.00 | 0.00 |
| >hg38_21_+_8433172_8446622_RNA45SN1_45S_50ntFlanks@3705.3725.21\|TACCTGGTTGATCCTGCCAGT | >hg38_21_+_8433172_8446622_RNA45SN1_45S_50ntFlanks | 3705 | 3725 | 21 | rRF 45S | NO | 184.31 | 1283.90 | 14.91 | 86.39 |
| >hg38_21_+_8433172_8446622_RNA45SN1_45S_50ntFlanks@3705.3726.22\|TACCTGGTTGATCCTGCCAGTA | >hg38_21_+_8433172_8446622_RNA45SN1_45S_50ntFlanks | 3705 | 3726 | 22 | rRF 45S | NO | 0.00 | 67.36 | 0.00 | 0.00 |
| >hg38_21_+_8433172_8446622_RNA45SN1_45S_50ntFlanks@3705.3728.24\|TACCTGGTTGATCCTGCCAGTAGC | >hg38_21_+_8433172_8446622_RNA45SN1_45S_50ntFlanks | 3705 | 3728 | 24 | rRF 45S | NO | 25.17 | 516.78 | 0.00 | 0.00 |

# BT-20

| | | | | | | | | | | |
|---|---|---|---|---|---|---|---|---|---|---|
| >hg38_21_+_8433172_8446622_RNA45SN1_45S_50ntFlanks@3705.3729.25\|TACCTGGTTGAT CCTGCCAGTAGCA | >hg38_21_+_8433172_8446622_RNA45SN1_45S_50ntFlanks | 3705 | 3729 | 25 | rRF 45S | NO | 0.00 | 10.03 | 0.00 | 0.00 |
| >hg38_21_+_8433172_8446622_RNA45SN1_45S_50ntFlanks@3705.3735.31\|TACCTGGTTGAT CCTGCCAGTAGCATATGCT | >hg38_21_+_8433172_8446622_RNA45SN1_45S_50ntFlanks | 3705 | 3735 | 31 | rRF 45S | NO | 0.00 | 146.95 | 0.00 | 0.00 |
| >hg38_21_+_8433172_8446622_RNA45SN1_45S_50ntFlanks@3705.3736.32\|TACCTGGTTGAT CCTGCCAGTAGCATATGCTT | >hg38_21_+_8433172_8446622_RNA45SN1_45S_50ntFlanks | 3705 | 3736 | 32 | rRF 45S | NO | 34.00 | 166.46 | 0.00 | 35.40 |
| >hg38_21_+_8433172_8446622_RNA45SN1_45S_50ntFlanks@3705.3737.33\|TACCTGGTTGAT CCTGCCAGTAGCATATGCTTG | >hg38_21_+_8433172_8446622_RNA45SN1_45S_50ntFlanks | 3705 | 3737 | 33 | rRF 45S | NO | 0.00 | 29.23 | 0.00 | 0.00 |
| >hg38_21_+_8433172_8446622_RNA45SN1_45S_50ntFlanks@3705.3739.35\|TACCTGGTTGAT CCTGCCAGTAGCATATGCTTGTC | >hg38_21_+_8433172_8446622_RNA45SN1_45S_50ntFlanks | 3705 | 3739 | 35 | rRF 45S | NO | 84.70 | 234.09 | 0.00 | 10.03 |
| >hg38_21_+_8433172_8446622_RNA45SN1_45S_50ntFlanks@3705.3740.36\|TACCTGGTTGAT CCTGCCAGTAGCATATGCTTGTCT | >hg38_21_+_8433172_8446622_RNA45SN1_45S_50ntFlanks | 3705 | 3740 | 36 | rRF 45S | NO | 0.00 | 43.60 | 0.00 | 0.00 |
| >hg38_21_+_8433172_8446622_RNA45SN1_45S_50ntFlanks@3705.3741.37\|TACCTGGTTGAT CCTGCCAGTAGCATATGCTTGTCTC | >hg38_21_+_8433172_8446622_RNA45SN1_45S_50ntFlanks | 3705 | 3741 | 37 | rRF 45S | NO | 90.54 | 367.58 | 0.00 | 29.37 |
| >hg38_21_+_8433172_8446622_RNA45SN1_45S_50ntFlanks@3705.3742.38\|TACCTGGTTGAT CCTGCCAGTAGCATATGCTTGTCTCA | >hg38_21_+_8433172_8446622_RNA45SN1_45S_50ntFlanks | 3705 | 3742 | 38 | rRF 45S | NO | 89.81 | 412.13 | 0.00 | 26.69 |
| >hg38_21_+_8433172_8446622_RNA45SN1_45S_50ntFlanks@3705.3743.39\|TACCTGGTTGAT CCTGCCAGTAGCATATGCTTGTCTCAA | >hg38_21_+_8433172_8446622_RNA45SN1_45S_50ntFlanks | 3705 | 3743 | 39 | rRF 45S | NO | 77.87 | 294.40 | 9.52 | 27.12 |
| >hg38_21_+_8433172_8446622_RNA45SN1_45S_50ntFlanks@3705.3744.40\|TACCTGGTTGAT CCTGCCAGTAGCATATGCTTGTCTCAAA | >hg38_21_+_8433172_8446622_RNA45SN1_45S_50ntFlanks | 3705 | 3744 | 40 | rRF 45S | NO | 73.89 | 102.62 | 0.00 | 0.00 |
| >hg38_21_+_8433172_8446622_RNA45SN1_45S_50ntFlanks@3705.3745.41\|TACCTGGTTGAT CCTGCCAGTAGCATATGCTTGTCTCAAAG | >hg38_21_+_8433172_8446622_RNA45SN1_45S_50ntFlanks | 3705 | 3745 | 41 | rRF 45S | NO | 29.61 | 91.38 | 0.00 | 0.00 |
| >hg38_21_+_8433172_8446622_RNA45SN1_45S_50ntFlanks@3705.3746.42\|TACCTGGTTGAT CCTGCCAGTAGCATATGCTTGTCTCAAAGA | >hg38_21_+_8433172_8446622_RNA45SN1_45S_50ntFlanks | 3705 | 3746 | 42 | rRF 45S | NO | 124.67 | 178.02 | 0.00 | 31.53 |
| >hg38_21_+_8433172_8446622_RNA45SN1_45S_50ntFlanks@3705.3747.43\|TACCTGGTTGAT CCTGCCAGTAGCATATGCTTGTCTCAAAGAT | >hg38_21_+_8433172_8446622_RNA45SN1_45S_50ntFlanks | 3705 | 3747 | 43 | rRF 45S | NO | 25.30 | 33.85 | 0.00 | 0.00 |
| >hg38_21_+_8433172_8446622_RNA45SN1_45S_50ntFlanks@3705.3748.44\|TACCTGGTTGAT CCTGCCAGTAGCATATGCTTGTCTCAAAGATT | >hg38_21_+_8433172_8446622_RNA45SN1_45S_50ntFlanks | 3705 | 3748 | 44 | rRF 45S | NO | 141.09 | 171.18 | 0.00 | 9.50 |
| >hg38_21_+_8433172_8446622_RNA45SN1_45S_50ntFlanks@3839.3860.22\|TCTCCTACTTGG ATAACTGTGG | >hg38_21_+_8433172_8446622_RNA45SN1_45S_50ntFlanks | 3839 | 3860 | 22 | rRF 45S | NO | 0.00 | 29.72 | 0.00 | 0.00 |
| >hg38_21_+_8433172_8446622_RNA45SN1_45S_50ntFlanks@3839.3861.23\|TCTCCTACTTGG ATAACTGTGGT | >hg38_21_+_8433172_8446622_RNA45SN1_45S_50ntFlanks | 3839 | 3861 | 23 | rRF 45S | NO | 0.00 | 24.03 | 0.00 | 0.00 |
| >hg38_21_+_8433172_8446622_RNA45SN1_45S_50ntFlanks@3839.3862.24\|TCTCCTACTTGG ATAACTGTGGTA | >hg38_21_+_8433172_8446622_RNA45SN1_45S_50ntFlanks | 3839 | 3862 | 24 | rRF 45S | NO | 0.00 | 37.27 | 0.00 | 0.00 |
| >hg38_21_+_8433172_8446622_RNA45SN1_45S_50ntFlanks@3839.3868.30\|TCTCCTACTTGG ATAACTGTGGTAATTCTA | >hg38_21_+_8433172_8446622_RNA45SN1_45S_50ntFlanks | 3839 | 3868 | 30 | rRF 45S | NO | 0.00 | 26.38 | 0.00 | 0.00 |
| >hg38_21_+_8433172_8446622_RNA45SN1_45S_50ntFlanks@3840.3859.20\|CTCCTACTTGGA TAACTGTG | >hg38_21_+_8433172_8446622_RNA45SN1_45S_50ntFlanks | 3840 | 3859 | 20 | rRF 45S | NO | 0.00 | 24.44 | 0.00 | 0.00 |
| >hg38_21_+_8433172_8446622_RNA45SN1_45S_50ntFlanks@3840.3860.21\|CTCCTACTTGGA TAACTGTGG | >hg38_21_+_8433172_8446622_RNA45SN1_45S_50ntFlanks | 3840 | 3860 | 21 | rRF 45S | NO | 0.00 | 96.52 | 0.00 | 0.00 |
| >hg38_21_+_8433172_8446622_RNA45SN1_45S_50ntFlanks@3840.3861.22\|CTCCTACTTGGA TAACTGTGGT | >hg38_21_+_8433172_8446622_RNA45SN1_45S_50ntFlanks | 3840 | 3861 | 22 | rRF 45S | NO | 0.00 | 87.64 | 0.00 | 0.00 |
| >hg38_21_+_8433172_8446622_RNA45SN1_45S_50ntFlanks@3840.3862.23\|CTCCTACTTGGA TAACTGTGGTA | >hg38_21_+_8433172_8446622_RNA45SN1_45S_50ntFlanks | 3840 | 3862 | 23 | rRF 45S | NO | 0.00 | 161.90 | 0.00 | 0.00 |
| >hg38_21_+_8433172_8446622_RNA45SN1_45S_50ntFlanks@3840.3864.25\|CTCCTACTTGGA TAACTGTGGTAAT | >hg38_21_+_8433172_8446622_RNA45SN1_45S_50ntFlanks | 3840 | 3864 | 25 | rRF 45S | NO | 0.00 | 10.39 | 0.00 | 0.00 |
| >hg38_21_+_8433172_8446622_RNA45SN1_45S_50ntFlanks@3840.3865.26\|CTCCTACTTGGA TAACTGTGGTAATT | >hg38_21_+_8433172_8446622_RNA45SN1_45S_50ntFlanks | 3840 | 3865 | 26 | rRF 45S | NO | 0.00 | 43.03 | 0.00 | 0.00 |
| >hg38_21_+_8433172_8446622_RNA45SN1_45S_50ntFlanks@3840.3867.28\|CTCCTACTTGGA TAACTGTGGTAATTCT | >hg38_21_+_8433172_8446622_RNA45SN1_45S_50ntFlanks | 3840 | 3867 | 28 | rRF 45S | NO | 0.00 | 26.63 | 0.00 | 0.00 |
| >hg38_21_+_8433172_8446622_RNA45SN1_45S_50ntFlanks@3840.3868.29\|CTCCTACTTGGA TAACTGTGGTAATTCTA | >hg38_21_+_8433172_8446622_RNA45SN1_45S_50ntFlanks | 3840 | 3868 | 29 | rRF 45S | NO | 24.52 | 105.73 | 0.00 | 0.00 |
| >hg38_21_+_8433172_8446622_RNA45SN1_45S_50ntFlanks@3840.3871.32\|CTCCTACTTGGA TAACTGTGGTAATTCTAGAG | >hg38_21_+_8433172_8446622_RNA45SN1_45S_50ntFlanks | 3840 | 3871 | 32 | rRF 45S | NO | 0.00 | 25.53 | 0.00 | 0.00 |
| >hg38_21_+_8433172_8446622_RNA45SN1_45S_50ntFlanks@3840.3872.33\|CTCCTACTTGGA TAACTGTGGTAATTCTAGAGC | >hg38_21_+_8433172_8446622_RNA45SN1_45S_50ntFlanks | 3840 | 3872 | 33 | rRF 45S | NO | 0.00 | 27.40 | 0.00 | 0.00 |
| >hg38_21_+_8433172_8446622_RNA45SN1_45S_50ntFlanks@3840.3873.34\|CTCCTACTTGGA TAACTGTGGTAATTCTAGAGCT | >hg38_21_+_8433172_8446622_RNA45SN1_45S_50ntFlanks | 3840 | 3873 | 34 | rRF 45S | NO | 0.00 | 39.46 | 0.00 | 0.00 |
| >hg38_21_+_8433172_8446622_RNA45SN1_45S_50ntFlanks@3840.3879.40\|CTCCTACTTGGA TAACTGTGGTAATTCTAGAGCTAATACA | >hg38_21_+_8433172_8446622_RNA45SN1_45S_50ntFlanks | 3840 | 3879 | 40 | rRF 45S | NO | 0.00 | 42.99 | 0.00 | 0.00 |
| >hg38_21_+_8433172_8446622_RNA45SN1_45S_50ntFlanks@3841.3868.28\|TCCTACTTGGAT AACTGTGGTAATTCTA | >hg38_21_+_8433172_8446622_RNA45SN1_45S_50ntFlanks | 3841 | 3868 | 28 | rRF 45S | NO | 0.00 | 36.17 | 0.00 | 0.00 |
| >hg38_21_+_8433172_8446622_RNA45SN1_45S_50ntFlanks@3841.3879.39\|TCCTACTTGGAT AACTGTGGTAATTCTAGAGCTAATACA | >hg38_21_+_8433172_8446622_RNA45SN1_45S_50ntFlanks | 3841 | 3879 | 39 | rRF 45S | NO | 0.00 | 26.53 | 0.00 | 0.00 |
| >hg38_21_+_8433172_8446622_RNA45SN1_45S_50ntFlanks@3842.3868.27\|CCTACTTGGATA ACTGTGGTAATTCTA | >hg38_21_+_8433172_8446622_RNA45SN1_45S_50ntFlanks | 3842 | 3868 | 27 | rRF 45S | NO | 0.00 | 25.71 | 0.00 | 0.00 |
| >hg38_21_+_8433172_8446622_RNA45SN1_45S_50ntFlanks@3844.3868.25\|TACTTGGATAAC TGTGGTAATTCTA | >hg38_21_+_8433172_8446622_RNA45SN1_45S_50ntFlanks | 3844 | 3868 | 25 | rRF 45S | NO | 0.00 | 23.02 | 0.00 | 0.00 |
| >hg38_21_+_8433172_8446622_RNA45SN1_45S_50ntFlanks@3845.3868.24\|ACTTGGATAACT GTGGTAATTCTA | >hg38_21_+_8433172_8446622_RNA45SN1_45S_50ntFlanks | 3845 | 3868 | 24 | rRF 45S | NO | 0.00 | 28.88 | 0.00 | 0.00 |
| >hg38_21_+_8433172_8446622_RNA45SN1_45S_50ntFlanks@3848.3866.19\|TGGATAACTGTG GTAATTC | >hg38_21_+_8433172_8446622_RNA45SN1_45S_50ntFlanks | 3848 | 3866 | 19 | rRF 45S | NO | 0.00 | 10.54 | 0.00 | 0.00 |
| >hg38_21_+_8433172_8446622_RNA45SN1_45S_50ntFlanks@3853.3884.32\|AACTGTGGTAAT TCTAGAGCTAATACATGCCG | >hg38_21_+_8433172_8446622_RNA45SN1_45S_50ntFlanks | 3853 | 3884 | 32 | rRF 45S | NO | 0.00 | 10.38 | 0.00 | 0.00 |
| >hg38_21_+_8433172_8446622_RNA45SN1_45S_50ntFlanks@3986.4023.38\|GGCTTTGGTGAC TCTAGATAACCTCGGGCCGATCGCAC | >hg38_21_+_8433172_8446622_RNA45SN1_45S_50ntFlanks | 3986 | 4023 | 38 | rRF 45S | NO | 0.00 | 97.64 | 0.00 | 0.00 |

# BT-20

| | | | | | | | | | | |
|---|---|---|---|---|---|---|---|---|---|---|
| >hg38_21_+_8433172_8446622_RNA45SN1_45S_50ntFlanks@3987.4023.37\|GCTTTGGTGACTCTAGATAACCTCGGGCCGATCGCAC | >hg38_21_+_8433172_8446622_RNA45SN1_45S_50ntFlanks | 3987 | 4023 | 37 | rRF 45S | NO | 0.00 | 81.06 | 14.56 | 0.00 |
| >hg38_21_+_8433172_8446622_RNA45SN1_45S_50ntFlanks@3988.4023.36\|CTTTGGTGACTCTAGATAACCTCGGGCCGATCGCAC | >hg38_21_+_8433172_8446622_RNA45SN1_45S_50ntFlanks | 3988 | 4023 | 36 | rRF 45S | NO | 24.74 | 159.67 | 64.18 | 9.96 |
| >hg38_21_+_8433172_8446622_RNA45SN1_45S_50ntFlanks@3989.4023.35\|TTTGGTGACTCTAGATAACCTCGGGCCGATCGCAC | >hg38_21_+_8433172_8446622_RNA45SN1_45S_50ntFlanks | 3989 | 4023 | 35 | rRF 45S | NO | 0.00 | 24.17 | 0.00 | 0.00 |
| >hg38_21_+_8433172_8446622_RNA45SN1_45S_50ntFlanks@3990.4023.34\|TTGGTGACTCTAGATAACCTCGGGCCGATCGCAC | >hg38_21_+_8433172_8446622_RNA45SN1_45S_50ntFlanks | 3990 | 4023 | 34 | rRF 45S | NO | 0.00 | 74.35 | 0.00 | 0.00 |
| >hg38_21_+_8433172_8446622_RNA45SN1_45S_50ntFlanks@3991.4023.33\|TGGTGACTCTAGATAACCTCGGGCCGATCGCAC | >hg38_21_+_8433172_8446622_RNA45SN1_45S_50ntFlanks | 3991 | 4023 | 33 | rRF 45S | NO | 0.00 | 28.86 | 0.00 | 0.00 |
| >hg38_21_+_8433172_8446622_RNA45SN1_45S_50ntFlanks@3992.4023.32\|GGTGACTCTAGATAACCTCGGGCCGATCGCAC | >hg38_21_+_8433172_8446622_RNA45SN1_45S_50ntFlanks | 3992 | 4023 | 32 | rRF 45S | NO | 0.00 | 30.77 | 0.00 | 0.00 |
| >hg38_21_+_8433172_8446622_RNA45SN1_45S_50ntFlanks@3993.4023.31\|GTGACTCTAGATAACCTCGGGCCGATCGCAC | >hg38_21_+_8433172_8446622_RNA45SN1_45S_50ntFlanks | 3993 | 4023 | 31 | rRF 45S | NO | 0.00 | 68.70 | 0.00 | 0.00 |
| >hg38_21_+_8433172_8446622_RNA45SN1_45S_50ntFlanks@4005.4023.19\|AACCTCGGGCCGATCGCAC | >hg38_21_+_8433172_8446622_RNA45SN1_45S_50ntFlanks | 4005 | 4023 | 19 | rRF 45S | NO | 0.00 | 29.29 | 0.00 | 0.00 |
| >hg38_21_+_8433172_8446622_RNA45SN1_45S_50ntFlanks@4006.4023.18\|ACCTCGGGCCGATCGCAC | >hg38_21_+_8433172_8446622_RNA45SN1_45S_50ntFlanks | 4006 | 4023 | 18 | rRF 45S | NO | 0.00 | 23.58 | 0.00 | 0.00 |
| >hg38_21_+_8433172_8446622_RNA45SN1_45S_50ntFlanks@4046.4066.21\|CATTCGAACGTCTGCCCTATC | >hg38_21_+_8433172_8446622_RNA45SN1_45S_50ntFlanks | 4046 | 4066 | 21 | rRF 45S | NO | 0.00 | 10.07 | 0.00 | 0.00 |
| >hg38_21_+_8433172_8446622_RNA45SN1_45S_50ntFlanks@4498.4520.23\|AAGCGTTTACTTTGAAAAAATTA | >hg38_21_+_8433172_8446622_RNA45SN1_45S_50ntFlanks | 4498 | 4520 | 23 | rRF 45S | NO | 0.00 | 40.41 | 0.00 | 0.00 |
| >hg38_21_+_8433172_8446622_RNA45SN1_45S_50ntFlanks@4565.4589.25\|AATAATGGAATAGGACCGCGGTTCT | >hg38_21_+_8433172_8446622_RNA45SN1_45S_50ntFlanks | 4565 | 4589 | 25 | rRF 45S | NO | 0.00 | 0.00 | 12.36 | 0.00 |
| >hg38_21_+_8433172_8446622_RNA45SN1_45S_50ntFlanks@4727.4758.32\|AATCAAGAACGAAAGTCGGAGGTTCGAAGACG | >hg38_21_+_8433172_8446622_RNA45SN1_45S_50ntFlanks | 4727 | 4758 | 32 | rRF 45S | NO | 0.00 | 33.55 | 39.61 | 0.00 |
| >hg38_21_+_8433172_8446622_RNA45SN1_45S_50ntFlanks@4727.4761.35\|AATCAAGAACGAAAGTCGGAGGTTCGAAGACGATC | >hg38_21_+_8433172_8446622_RNA45SN1_45S_50ntFlanks | 4727 | 4761 | 35 | rRF 45S | NO | 0.00 | 69.93 | 11.42 | 0.00 |
| >hg38_21_+_8433172_8446622_RNA45SN1_45S_50ntFlanks@4727.4765.39\|AATCAAGAACGAAAGTCGGAGGTTCGAAGACGATCAGAT | >hg38_21_+_8433172_8446622_RNA45SN1_45S_50ntFlanks | 4727 | 4765 | 39 | rRF 45S | NO | 10.49 | 125.09 | 74.96 | 0.00 |
| >hg38_21_+_8433172_8446622_RNA45SN1_45S_50ntFlanks@4727.4767.41\|AATCAAGAACGAAAGTCGGAGGTTCGAAGACGATCAGATAC | >hg38_21_+_8433172_8446622_RNA45SN1_45S_50ntFlanks | 4727 | 4767 | 41 | rRF 45S | NO | 0.00 | 99.94 | 0.00 | 0.00 |
| >hg38_21_+_8433172_8446622_RNA45SN1_45S_50ntFlanks@4730.4758.29\|CAAGAACGAAAGTCGGAGGTTCGAAGACG | >hg38_21_+_8433172_8446622_RNA45SN1_45S_50ntFlanks | 4730 | 4758 | 29 | rRF 45S | NO | 0.00 | 36.07 | 68.66 | 0.00 |
| >hg38_21_+_8433172_8446622_RNA45SN1_45S_50ntFlanks@4730.4761.32\|CAAGAACGAAAGTCGGAGGTTCGAAGACGATC | >hg38_21_+_8433172_8446622_RNA45SN1_45S_50ntFlanks | 4730 | 4761 | 32 | rRF 45S | NO | 0.00 | 32.15 | 0.00 | 0.00 |
| >hg38_21_+_8433172_8446622_RNA45SN1_45S_50ntFlanks@4730.4765.36\|CAAGAACGAAAGTCGGAGGTTCGAAGACGATCAGAT | >hg38_21_+_8433172_8446622_RNA45SN1_45S_50ntFlanks | 4730 | 4765 | 36 | rRF 45S | NO | 0.00 | 73.54 | 10.56 | 0.00 |
| >hg38_21_+_8433172_8446622_RNA45SN1_45S_50ntFlanks@4730.4767.38\|CAAGAACGAAAGTCGGAGGTTCGAAGACGATCAGATAC | >hg38_21_+_8433172_8446622_RNA45SN1_45S_50ntFlanks | 4730 | 4767 | 38 | rRF 45S | NO | 0.00 | 109.65 | 0.00 | 0.00 |
| >hg38_21_+_8433172_8446622_RNA45SN1_45S_50ntFlanks@4731.4751.21\|AAGAACGAAAGTCGGAGGTTC | >hg38_21_+_8433172_8446622_RNA45SN1_45S_50ntFlanks | 4731 | 4751 | 21 | rRF 45S | NO | 0.00 | 25.12 | 10.58 | 0.00 |
| >hg38_21_+_8433172_8446622_RNA45SN1_45S_50ntFlanks@4731.4757.27\|AAGAACGAAAGTCGGAGGTTCGAAGAC | >hg38_21_+_8433172_8446622_RNA45SN1_45S_50ntFlanks | 4731 | 4757 | 27 | rRF 45S | NO | 0.00 | 33.29 | 0.00 | 0.00 |
| >hg38_21_+_8433172_8446622_RNA45SN1_45S_50ntFlanks@4731.4758.28\|AAGAACGAAAGTCGGAGGTTCGAAGACG | >hg38_21_+_8433172_8446622_RNA45SN1_45S_50ntFlanks | 4731 | 4758 | 28 | rRF 45S | NO | 0.00 | 181.08 | 145.09 | 0.00 |
| >hg38_21_+_8433172_8446622_RNA45SN1_45S_50ntFlanks@4731.4761.31\|AAGAACGAAAGTCGGAGGTTCGAAGACGATC | >hg38_21_+_8433172_8446622_RNA45SN1_45S_50ntFlanks | 4731 | 4761 | 31 | rRF 45S | NO | 24.33 | 179.73 | 96.52 | 0.00 |
| >hg38_21_+_8433172_8446622_RNA45SN1_45S_50ntFlanks@4731.4762.32\|AAGAACGAAAGTCGGAGGTTCGAAGACGATCA | >hg38_21_+_8433172_8446622_RNA45SN1_45S_50ntFlanks | 4731 | 4762 | 32 | rRF 45S | NO | 0.00 | 39.68 | 0.00 | 0.00 |
| >hg38_21_+_8433172_8446622_RNA45SN1_45S_50ntFlanks@4731.4765.35\|AAGAACGAAAGTCGGAGGTTCGAAGACGATCAGAT | >hg38_21_+_8433172_8446622_RNA45SN1_45S_50ntFlanks | 4731 | 4765 | 35 | rRF 45S | NO | 24.73 | 210.40 | 91.69 | 0.00 |
| >hg38_21_+_8433172_8446622_RNA45SN1_45S_50ntFlanks@4731.4766.36\|AAGAACGAAAGTCGGAGGTTCGAAGACGATCAGATA | >hg38_21_+_8433172_8446622_RNA45SN1_45S_50ntFlanks | 4731 | 4766 | 36 | rRF 45S | NO | 0.00 | 158.54 | 0.00 | 0.00 |
| >hg38_21_+_8433172_8446622_RNA45SN1_45S_50ntFlanks@4731.4767.37\|AAGAACGAAAGTCGGAGGTTCGAAGACGATCAGATAC | >hg38_21_+_8433172_8446622_RNA45SN1_45S_50ntFlanks | 4731 | 4767 | 37 | rRF 45S | NO | 0.00 | 274.19 | 11.38 | 0.00 |
| >hg38_21_+_8433172_8446622_RNA45SN1_45S_50ntFlanks@4731.4768.38\|AAGAACGAAAGTCGGAGGTTCGAAGACGATCAGATACC | >hg38_21_+_8433172_8446622_RNA45SN1_45S_50ntFlanks | 4731 | 4768 | 38 | rRF 45S | NO | 0.00 | 158.47 | 0.00 | 0.00 |
| >hg38_21_+_8433172_8446622_RNA45SN1_45S_50ntFlanks@4732.4758.27\|AGAACGAAAGTCGGAGGTTCGAAGACG | >hg38_21_+_8433172_8446622_RNA45SN1_45S_50ntFlanks | 4732 | 4758 | 27 | rRF 45S | NO | 0.00 | 78.58 | 63.11 | 0.00 |
| >hg38_21_+_8433172_8446622_RNA45SN1_45S_50ntFlanks@4732.4761.30\|AGAACGAAAGTCGGAGGTTCGAAGACGATC | >hg38_21_+_8433172_8446622_RNA45SN1_45S_50ntFlanks | 4732 | 4761 | 30 | rRF 45S | NO | 0.00 | 98.37 | 82.78 | 0.00 |
| >hg38_21_+_8433172_8446622_RNA45SN1_45S_50ntFlanks@4732.4765.34\|AGAACGAAAGTCGGAGGTTCGAAGACGATCAGAT | >hg38_21_+_8433172_8446622_RNA45SN1_45S_50ntFlanks | 4732 | 4765 | 34 | rRF 45S | NO | 0.00 | 99.99 | 13.36 | 0.00 |
| >hg38_21_+_8433172_8446622_RNA45SN1_45S_50ntFlanks@4732.4766.35\|AGAACGAAAGTCGGAGGTTCGAAGACGATCAGATA | >hg38_21_+_8433172_8446622_RNA45SN1_45S_50ntFlanks | 4732 | 4766 | 35 | rRF 45S | NO | 0.00 | 99.42 | 0.00 | 0.00 |
| >hg38_21_+_8433172_8446622_RNA45SN1_45S_50ntFlanks@4732.4767.36\|AGAACGAAAGTCGGAGGTTCGAAGACGATCAGATAC | >hg38_21_+_8433172_8446622_RNA45SN1_45S_50ntFlanks | 4732 | 4767 | 36 | rRF 45S | NO | 0.00 | 163.30 | 0.00 | 0.00 |
| >hg38_21_+_8433172_8446622_RNA45SN1_45S_50ntFlanks@4732.4768.37\|AGAACGAAAGTCGGAGGTTCGAAGACGATCAGATACC | >hg38_21_+_8433172_8446622_RNA45SN1_45S_50ntFlanks | 4732 | 4768 | 37 | rRF 45S | NO | 0.00 | 89.72 | 0.00 | 0.00 |
| >hg38_21_+_8433172_8446622_RNA45SN1_45S_50ntFlanks@4733.4758.26\|GAACGAAAGTCGGAGGTTCGAAGACG | >hg38_21_+_8433172_8446622_RNA45SN1_45S_50ntFlanks | 4733 | 4758 | 26 | rRF 45S | NO | 0.00 | 32.50 | 109.13 | 0.00 |
| >hg38_21_+_8433172_8446622_RNA45SN1_45S_50ntFlanks@4733.4761.29\|GAACGAAAGTCGGAGGTTCGAAGACGATC | >hg38_21_+_8433172_8446622_RNA45SN1_45S_50ntFlanks | 4733 | 4761 | 29 | rRF 45S | NO | 0.00 | 39.53 | 81.45 | 0.00 |
| >hg38_21_+_8433172_8446622_RNA45SN1_45S_50ntFlanks@4733.4765.33\|GAACGAAAGTCGGAGGTTCGAAGACGATCAGAT | >hg38_21_+_8433172_8446622_RNA45SN1_45S_50ntFlanks | 4733 | 4765 | 33 | rRF 45S | NO | 0.00 | 43.47 | 56.96 | 0.00 |

# BT-20

| | | | | | | | | | | |
|---|---|---|---|---|---|---|---|---|---|---|
| >hg38_21_+_8433172_8446622_RNA45SN1_45S_50ntFlanks@4733.4767.35\|GAACGAAAGTC GGAGGTTCGAAGACGATCAGATAC | >hg38_21_+_8433172_8446622_RNA45SN1_45S_50ntFlanks | 4733 | 4767 | 35 | rRF 45S | NO | 0.00 | 98.72 | 0.00 | 0.00 |
| >hg38_21_+_8433172_8446622_RNA45SN1_45S_50ntFlanks@4733.4768.36\|GAACGAAAGTC GGAGGTTCGAAGACGATCAGATACC | >hg38_21_+_8433172_8446622_RNA45SN1_45S_50ntFlanks | 4733 | 4768 | 36 | rRF 45S | NO | 0.00 | 26.08 | 0.00 | 0.00 |
| >hg38_21_+_8433172_8446622_RNA45SN1_45S_50ntFlanks@4734.4758.25\|AACGAAAGTCG GAGGTTCGAAGACG | >hg38_21_+_8433172_8446622_RNA45SN1_45S_50ntFlanks | 4734 | 4758 | 25 | rRF 45S | NO | 0.00 | 34.65 | 243.11 | 0.00 |
| >hg38_21_+_8433172_8446622_RNA45SN1_45S_50ntFlanks@4734.4761.28\|AACGAAAGTCG GAGGTTCGAAGACGATC | >hg38_21_+_8433172_8446622_RNA45SN1_45S_50ntFlanks | 4734 | 4761 | 28 | rRF 45S | NO | 0.00 | 34.22 | 87.91 | 0.00 |
| >hg38_21_+_8433172_8446622_RNA45SN1_45S_50ntFlanks@4734.4765.32\|AACGAAAGTCG GAGGTTCGAAGACGATCAGAT | >hg38_21_+_8433172_8446622_RNA45SN1_45S_50ntFlanks | 4734 | 4765 | 32 | rRF 45S | NO | 0.00 | 30.33 | 67.35 | 0.00 |
| >hg38_21_+_8433172_8446622_RNA45SN1_45S_50ntFlanks@4734.4767.34\|AACGAAAGTCG GAGGTTCGAAGACGATCAGATAC | >hg38_21_+_8433172_8446622_RNA45SN1_45S_50ntFlanks | 4734 | 4767 | 34 | rRF 45S | NO | 0.00 | 68.08 | 0.00 | 0.00 |
| >hg38_21_+_8433172_8446622_RNA45SN1_45S_50ntFlanks@4735.4758.24\|ACGAAAGTCGG AGGTTCGAAGACG | >hg38_21_+_8433172_8446622_RNA45SN1_45S_50ntFlanks | 4735 | 4758 | 24 | rRF 45S | NO | 0.00 | 29.57 | 12.89 | 0.00 |
| >hg38_21_+_8433172_8446622_RNA45SN1_45S_50ntFlanks@4735.4761.27\|ACGAAAGTCGG AGGTTCGAAGACGATC | >hg38_21_+_8433172_8446622_RNA45SN1_45S_50ntFlanks | 4735 | 4761 | 27 | rRF 45S | NO | 0.00 | 34.68 | 0.00 | 0.00 |
| >hg38_21_+_8433172_8446622_RNA45SN1_45S_50ntFlanks@4735.4765.31\|ACGAAAGTCGG AGGTTCGAAGACGATCAGAT | >hg38_21_+_8433172_8446622_RNA45SN1_45S_50ntFlanks | 4735 | 4765 | 31 | rRF 45S | NO | 0.00 | 10.45 | 0.00 | 0.00 |
| >hg38_21_+_8433172_8446622_RNA45SN1_45S_50ntFlanks@4738.4758.21\|AAAGTCGGAGGT TCGAAGACG | >hg38_21_+_8433172_8446622_RNA45SN1_45S_50ntFlanks | 4738 | 4758 | 21 | rRF 45S | NO | 0.00 | 74.98 | 162.24 | 0.00 |
| >hg38_21_+_8433172_8446622_RNA45SN1_45S_50ntFlanks@4738.4761.24\|AAAGTCGGAGGT TCGAAGACGATC | >hg38_21_+_8433172_8446622_RNA45SN1_45S_50ntFlanks | 4738 | 4761 | 24 | rRF 45S | NO | 0.00 | 66.63 | 69.07 | 0.00 |
| >hg38_21_+_8433172_8446622_RNA45SN1_45S_50ntFlanks@4738.4765.28\|AAAGTCGGAGGT TCGAAGACGATCAGAT | >hg38_21_+_8433172_8446622_RNA45SN1_45S_50ntFlanks | 4738 | 4765 | 28 | rRF 45S | NO | 0.00 | 28.33 | 14.78 | 0.00 |
| >hg38_21_+_8433172_8446622_RNA45SN1_45S_50ntFlanks@4738.4766.29\|AAAGTCGGAGGT TCGAAGACGATCAGATA | >hg38_21_+_8433172_8446622_RNA45SN1_45S_50ntFlanks | 4738 | 4766 | 29 | rRF 45S | NO | 0.00 | 39.01 | 0.00 | 0.00 |
| >hg38_21_+_8433172_8446622_RNA45SN1_45S_50ntFlanks@4738.4767.30\|AAAGTCGGAGGT TCGAAGACGATCAGATAC | >hg38_21_+_8433172_8446622_RNA45SN1_45S_50ntFlanks | 4738 | 4767 | 30 | rRF 45S | NO | 0.00 | 101.39 | 0.00 | 0.00 |
| >hg38_21_+_8433172_8446622_RNA45SN1_45S_50ntFlanks@4738.4768.31\|AAAGTCGGAGGT TCGAAGACGATCAGATACC | >hg38_21_+_8433172_8446622_RNA45SN1_45S_50ntFlanks | 4738 | 4768 | 31 | rRF 45S | NO | 0.00 | 106.75 | 10.03 | 0.00 |
| >hg38_21_+_8433172_8446622_RNA45SN1_45S_50ntFlanks@4975.5001.27\|CCCGGACACGG ACAGGATTGACAGATT | >hg38_21_+_8433172_8446622_RNA45SN1_45S_50ntFlanks | 4975 | 5001 | 27 | rRF 45S | NO | 10.09 | 0.00 | 0.00 | 0.00 |
| >hg38_21_+_8433172_8446622_RNA45SN1_45S_50ntFlanks@4976.4993.18\|CCGGACACGGA CAGGATT | >hg38_21_+_8433172_8446622_RNA45SN1_45S_50ntFlanks | 4976 | 4993 | 18 | rRF 45S | NO | 0.00 | 28.20 | 0.00 | 0.00 |
| >hg38_21_+_8433172_8446622_RNA45SN1_45S_50ntFlanks@4976.5001.26\|CCGGACACGGA CAGGATTGACAGATT | >hg38_21_+_8433172_8446622_RNA45SN1_45S_50ntFlanks | 4976 | 5001 | 26 | rRF 45S | NO | 0.00 | 26.86 | 0.00 | 0.00 |
| >hg38_21_+_8433172_8446622_RNA45SN1_45S_50ntFlanks@5055.5081.27\|GGAGCGATTTGT CTGGTTAATTCCGAT | >hg38_21_+_8433172_8446622_RNA45SN1_45S_50ntFlanks | 5055 | 5081 | 27 | rRF 45S | NO | 0.00 | 0.00 | 82.80 | 0.00 |
| >hg38_21_+_8433172_8446622_RNA45SN1_45S_50ntFlanks@5057.5081.25\|AGCGATTTGTCT GGTTAATTCCGAT | >hg38_21_+_8433172_8446622_RNA45SN1_45S_50ntFlanks | 5057 | 5081 | 25 | rRF 45S | NO | 0.00 | 0.00 | 37.58 | 0.00 |
| >hg38_21_+_8433172_8446622_RNA45SN1_45S_50ntFlanks@5082.5104.23\|AACGAACGAGA CTCTGGCATGCT | >hg38_21_+_8433172_8446622_RNA45SN1_45S_50ntFlanks | 5082 | 5104 | 23 | rRF 45S | NO | 0.00 | 10.22 | 16.33 | 0.00 |
| >hg38_21_+_8433172_8446622_RNA45SN1_45S_50ntFlanks@5083.5100.18\|ACGAACGAGAC TCTGGCA | >hg38_21_+_8433172_8446622_RNA45SN1_45S_50ntFlanks | 5083 | 5100 | 18 | rRF 45S | NO | 0.00 | 24.83 | 0.00 | 0.00 |
| >hg38_21_+_8433172_8446622_RNA45SN1_45S_50ntFlanks@5086.5104.19\|AACGAGACTCTG GCATGCT | >hg38_21_+_8433172_8446622_RNA45SN1_45S_50ntFlanks | 5086 | 5104 | 19 | rRF 45S | NO | 0.00 | 0.00 | 10.45 | 0.00 |
| >hg38_21_+_8433172_8446622_RNA45SN1_45S_50ntFlanks@5125.5145.21\|AGCGGTCGGCG TCCCCCAACT | >hg38_21_+_8433172_8446622_RNA45SN1_45S_50ntFlanks | 5125 | 5145 | 21 | rRF 45S | NO | 0.00 | 33.98 | 0.00 | 0.00 |
| >hg38_21_+_8433172_8446622_RNA45SN1_45S_50ntFlanks@5125.5158.34\|AGCGGTCGGCG TCCCCCAACTTCTTAGAGGGACA | >hg38_21_+_8433172_8446622_RNA45SN1_45S_50ntFlanks | 5125 | 5158 | 34 | rRF 45S | NO | 0.00 | 123.06 | 0.00 | 0.00 |
| >hg38_21_+_8433172_8446622_RNA45SN1_45S_50ntFlanks@5237.5255.19\|ACTGACTGGCTC AGCGTGT | >hg38_21_+_8433172_8446622_RNA45SN1_45S_50ntFlanks | 5237 | 5255 | 19 | rRF 45S | NO | 0.00 | 0.00 | 56.63 | 0.00 |
| >hg38_21_+_8433172_8446622_RNA45SN1_45S_50ntFlanks@5444.5466.23\|CTCGGATCGGC CCCGCCGGGGTC | >hg38_21_+_8433172_8446622_RNA45SN1_45S_50ntFlanks | 5444 | 5466 | 23 | rRF 45S | NO | 0.00 | 36.73 | 0.00 | 0.00 |
| >hg38_21_+_8433172_8446622_RNA45SN1_45S_50ntFlanks@5471.5501.31\|CACGGCCCTGG CGGAGCGCTGAGAAGACGGT | >hg38_21_+_8433172_8446622_RNA45SN1_45S_50ntFlanks | 5471 | 5501 | 31 | rRF 45S | NO | 0.00 | 39.35 | 0.00 | 0.00 |
| >hg38_21_+_8433172_8446622_RNA45SN1_45S_50ntFlanks@5472.5489.18\|ACGGCCCTGGC GGAGCGC | >hg38_21_+_8433172_8446622_RNA45SN1_45S_50ntFlanks | 5472 | 5489 | 18 | rRF 45S | NO | 0.00 | 141.69 | 0.00 | 0.00 |
| >hg38_21_+_8433172_8446622_RNA45SN1_45S_50ntFlanks@5472.5490.19\|ACGGCCCTGGC GGAGCGCT | >hg38_21_+_8433172_8446622_RNA45SN1_45S_50ntFlanks | 5472 | 5490 | 19 | rRF 45S | NO | 0.00 | 114.49 | 0.00 | 0.00 |
| >hg38_21_+_8433172_8446622_RNA45SN1_45S_50ntFlanks@5472.5491.20\|ACGGCCCTGGC GGAGCGCTG | >hg38_21_+_8433172_8446622_RNA45SN1_45S_50ntFlanks | 5472 | 5491 | 20 | rRF 45S | NO | 0.00 | 72.79 | 0.00 | 0.00 |
| >hg38_21_+_8433172_8446622_RNA45SN1_45S_50ntFlanks@5472.5494.23\|ACGGCCCTGGC GGAGCGCTGAGA | >hg38_21_+_8433172_8446622_RNA45SN1_45S_50ntFlanks | 5472 | 5494 | 23 | rRF 45S | NO | 0.00 | 185.68 | 9.48 | 0.00 |
| >hg38_21_+_8433172_8446622_RNA45SN1_45S_50ntFlanks@5472.5498.27\|ACGGCCCTGGC GGAGCGCTGAGAAGAC | >hg38_21_+_8433172_8446622_RNA45SN1_45S_50ntFlanks | 5472 | 5498 | 27 | rRF 45S | NO | 0.00 | 91.59 | 0.00 | 0.00 |
| >hg38_21_+_8433172_8446622_RNA45SN1_45S_50ntFlanks@5472.5499.28\|ACGGCCCTGGC GGAGCGCTGAGAAGACG | >hg38_21_+_8433172_8446622_RNA45SN1_45S_50ntFlanks | 5472 | 5499 | 28 | rRF 45S | NO | 0.00 | 245.71 | 62.97 | 10.46 |
| >hg38_21_+_8433172_8446622_RNA45SN1_45S_50ntFlanks@5472.5500.29\|ACGGCCCTGGC GGAGCGCTGAGAAGACGG | >hg38_21_+_8433172_8446622_RNA45SN1_45S_50ntFlanks | 5472 | 5500 | 29 | rRF 45S | NO | 0.00 | 74.74 | 9.84 | 0.00 |
| >hg38_21_+_8433172_8446622_RNA45SN1_45S_50ntFlanks@5472.5501.30\|ACGGCCCTGGC GGAGCGCTGAGAAGACGGT | >hg38_21_+_8433172_8446622_RNA45SN1_45S_50ntFlanks | 5472 | 5501 | 30 | rRF 45S | NO | 10.25 | 496.68 | 16.20 | 27.70 |
| >hg38_21_+_8433172_8446622_RNA45SN1_45S_50ntFlanks@5472.5502.31\|ACGGCCCTGGC GGAGCGCTGAGAAGACGGTC | >hg38_21_+_8433172_8446622_RNA45SN1_45S_50ntFlanks | 5472 | 5502 | 31 | rRF 45S | NO | 0.00 | 169.34 | 12.49 | 0.00 |
| >hg38_21_+_8433172_8446622_RNA45SN1_45S_50ntFlanks@5472.5506.35\|ACGGCCCTGGC GGAGCGCTGAGAAGACGGTCGAAC | >hg38_21_+_8433172_8446622_RNA45SN1_45S_50ntFlanks | 5472 | 5506 | 35 | rRF 45S | NO | 0.00 | 106.63 | 0.00 | 0.00 |

# BT-20

| | | | | | | | | | | |
|---|---|---|---|---|---|---|---|---|---|---|
| >hg38_21_+_8433172_8446622_RNA45SN1_45S_50ntFlanks@5472.5507.36\|ACGGCCCTGGCGGAGCGCTGAGAAGACGGTCGAACT | >hg38_21_+_8433172_8446622_RNA45SN1_45S_50ntFlanks | 5472 | 5507 | 36 | rRF 45S | NO | 0.00 | 143.83 | 0.00 | 0.00 |
| >hg38_21_+_8433172_8446622_RNA45SN1_45S_50ntFlanks@5472.5509.38\|ACGGCCCTGGCGGAGCGCTGAGAAGACGGTCGAACTTG | >hg38_21_+_8433172_8446622_RNA45SN1_45S_50ntFlanks | 5472 | 5509 | 38 | rRF 45S | NO | 23.99 | 220.89 | 10.12 | 23.89 |
| >hg38_21_+_8433172_8446622_RNA45SN1_45S_50ntFlanks@5472.5510.39\|ACGGCCCTGGCGGAGCGCTGAGAAGACGGTCGAACTTGA | >hg38_21_+_8433172_8446622_RNA45SN1_45S_50ntFlanks | 5472 | 5510 | 39 | rRF 45S | NO | 0.00 | 33.45 | 0.00 | 0.00 |
| >hg38_21_+_8433172_8446622_RNA45SN1_45S_50ntFlanks@5472.5512.41\|ACGGCCCTGGCGGAGCGCTGAGAAGACGGTCGAACTTGACT | >hg38_21_+_8433172_8446622_RNA45SN1_45S_50ntFlanks | 5472 | 5512 | 41 | rRF 45S | NO | 28.90 | 220.86 | 13.63 | 25.38 |
| >hg38_21_+_8433172_8446622_RNA45SN1_45S_50ntFlanks@5472.5516.45\|ACGGCCCTGGCGGAGCGCTGAGAAGACGGTCGAACTTGACTATCT | >hg38_21_+_8433172_8446622_RNA45SN1_45S_50ntFlanks | 5472 | 5516 | 45 | rRF 45S | NO | 10.40 | 268.87 | 79.69 | 200.39 |
| >hg38_21_+_8433172_8446622_RNA45SN1_45S_50ntFlanks@5473.5490.18\|CGGCCCTGGCGGAGCGCT | >hg38_21_+_8433172_8446622_RNA45SN1_45S_50ntFlanks | 5473 | 5490 | 18 | rRF 45S | NO | 0.00 | 28.55 | 0.00 | 0.00 |
| >hg38_21_+_8433172_8446622_RNA45SN1_45S_50ntFlanks@5473.5494.22\|CGGCCCTGGCGGAGCGCTGAGA | >hg38_21_+_8433172_8446622_RNA45SN1_45S_50ntFlanks | 5473 | 5494 | 22 | rRF 45S | NO | 0.00 | 76.67 | 0.00 | 0.00 |
| >hg38_21_+_8433172_8446622_RNA45SN1_45S_50ntFlanks@5473.5499.27\|CGGCCCTGGCGGAGCGCTGAGAAGACG | >hg38_21_+_8433172_8446622_RNA45SN1_45S_50ntFlanks | 5473 | 5499 | 27 | rRF 45S | NO | 0.00 | 161.45 | 13.70 | 0.00 |
| >hg38_21_+_8433172_8446622_RNA45SN1_45S_50ntFlanks@5473.5501.29\|CGGCCCTGGCGGAGCGCTGAGAAGACGGT | >hg38_21_+_8433172_8446622_RNA45SN1_45S_50ntFlanks | 5473 | 5501 | 29 | rRF 45S | NO | 0.00 | 189.11 | 0.00 | 0.00 |
| >hg38_21_+_8433172_8446622_RNA45SN1_45S_50ntFlanks@5473.5502.30\|CGGCCCTGGCGGAGCGCTGAGAAGACGGTC | >hg38_21_+_8433172_8446622_RNA45SN1_45S_50ntFlanks | 5473 | 5502 | 30 | rRF 45S | NO | 0.00 | 82.34 | 0.00 | 0.00 |
| >hg38_21_+_8433172_8446622_RNA45SN1_45S_50ntFlanks@5473.5505.33\|CGGCCCTGGCGGAGCGCTGAGAAGACGGTCGAA | >hg38_21_+_8433172_8446622_RNA45SN1_45S_50ntFlanks | 5473 | 5505 | 33 | rRF 45S | NO | 0.00 | 79.87 | 0.00 | 0.00 |
| >hg38_21_+_8433172_8446622_RNA45SN1_45S_50ntFlanks@5473.5507.35\|CGGCCCTGGCGGAGCGCTGAGAAGACGGTCGAACT | >hg38_21_+_8433172_8446622_RNA45SN1_45S_50ntFlanks | 5473 | 5507 | 35 | rRF 45S | NO | 0.00 | 39.23 | 0.00 | 0.00 |
| >hg38_21_+_8433172_8446622_RNA45SN1_45S_50ntFlanks@5473.5509.37\|CGGCCCTGGCGGAGCGCTGAGAAGACGGTCGAACTTG | >hg38_21_+_8433172_8446622_RNA45SN1_45S_50ntFlanks | 5473 | 5509 | 37 | rRF 45S | NO | 0.00 | 82.70 | 0.00 | 0.00 |
| >hg38_21_+_8433172_8446622_RNA45SN1_45S_50ntFlanks@5473.5512.40\|CGGCCCTGGCGGAGCGCTGAGAAGACGGTCGAACTTGACT | >hg38_21_+_8433172_8446622_RNA45SN1_45S_50ntFlanks | 5473 | 5512 | 40 | rRF 45S | NO | 0.00 | 95.93 | 0.00 | 0.00 |
| >hg38_21_+_8433172_8446622_RNA45SN1_45S_50ntFlanks@5473.5516.44\|CGGCCCTGGCGGAGCGCTGAGAAGACGGTCGAACTTGACTATCT | >hg38_21_+_8433172_8446622_RNA45SN1_45S_50ntFlanks | 5473 | 5516 | 44 | rRF 45S | NO | 0.00 | 162.39 | 69.00 | 143.94 |
| >hg38_21_+_8433172_8446622_RNA45SN1_45S_50ntFlanks@5475.5516.42\|GCCCTGGCGGAGCGCTGAGAAGACGGTCGAACTTGACTATCT | >hg38_21_+_8433172_8446622_RNA45SN1_45S_50ntFlanks | 5475 | 5516 | 42 | rRF 45S | NO | 0.00 | 22.73 | 0.00 | 64.58 |
| >hg38_21_+_8433172_8446622_RNA45SN1_45S_50ntFlanks@5476.5516.41\|CCCTGGCGGAGCGCTGAGAAGACGGTCGAACTTGACTATCT | >hg38_21_+_8433172_8446622_RNA45SN1_45S_50ntFlanks | 5476 | 5516 | 41 | rRF 45S | NO | 0.00 | 83.07 | 0.00 | 63.02 |
| >hg38_21_+_8433172_8446622_RNA45SN1_45S_50ntFlanks@5477.5516.40\|CCTGGCGGAGCGCTGAGAAGACGGTCGAACTTGACTATCT | >hg38_21_+_8433172_8446622_RNA45SN1_45S_50ntFlanks | 5477 | 5516 | 40 | rRF 45S | NO | 0.00 | 95.01 | 11.58 | 30.95 |
| >hg38_21_+_8433172_8446622_RNA45SN1_45S_50ntFlanks@5485.5507.23\|AGCGCTGAGAAGACGGTCGAACT | >hg38_21_+_8433172_8446622_RNA45SN1_45S_50ntFlanks | 5485 | 5507 | 23 | rRF 45S | NO | 0.00 | 24.50 | 0.00 | 0.00 |
| >hg38_21_+_8433172_8446622_RNA45SN1_45S_50ntFlanks@5485.5512.28\|AGCGCTGAGAAGACGGTCGAACTTGACT | >hg38_21_+_8433172_8446622_RNA45SN1_45S_50ntFlanks | 5485 | 5512 | 28 | rRF 45S | NO | 0.00 | 37.58 | 0.00 | 0.00 |
| >hg38_21_+_8433172_8446622_RNA45SN1_45S_50ntFlanks@5485.5516.32\|AGCGCTGAGAAGACGGTCGAACTTGACTATCT | >hg38_21_+_8433172_8446622_RNA45SN1_45S_50ntFlanks | 5485 | 5516 | 32 | rRF 45S | NO | 65.60 | 367.03 | 153.22 | 153.97 |
| >hg38_21_+_8433172_8446622_RNA45SN1_45S_50ntFlanks@5486.5512.27\|GCGCTGAGAAGACGGTCGAACTTGACT | >hg38_21_+_8433172_8446622_RNA45SN1_45S_50ntFlanks | 5486 | 5512 | 27 | rRF 45S | NO | 0.00 | 37.83 | 0.00 | 0.00 |
| >hg38_21_+_8433172_8446622_RNA45SN1_45S_50ntFlanks@5486.5516.31\|GCGCTGAGAAGACGGTCGAACTTGACTATCT | >hg38_21_+_8433172_8446622_RNA45SN1_45S_50ntFlanks | 5486 | 5516 | 31 | rRF 45S | NO | 0.00 | 142.65 | 0.00 | 9.66 |
| >hg38_21_+_8433172_8446622_RNA45SN1_45S_50ntFlanks@5487.5516.30\|CGCTGAGAAGACGGTCGAACTTGACTATCT | >hg38_21_+_8433172_8446622_RNA45SN1_45S_50ntFlanks | 5487 | 5516 | 30 | rRF 45S | NO | 0.00 | 410.33 | 65.22 | 71.12 |
| >hg38_21_+_8433172_8446622_RNA45SN1_45S_50ntFlanks@5488.5516.29\|GCTGAGAAGACGGTCGAACTTGACTATCT | >hg38_21_+_8433172_8446622_RNA45SN1_45S_50ntFlanks | 5488 | 5516 | 29 | rRF 45S | NO | 0.00 | 342.57 | 15.03 | 26.00 |
| >hg38_21_+_8433172_8446622_RNA45SN1_45S_50ntFlanks@5489.5512.24\|CTGAGAAGACGGTCGAACTTGACT | >hg38_21_+_8433172_8446622_RNA45SN1_45S_50ntFlanks | 5489 | 5512 | 24 | rRF 45S | NO | 0.00 | 42.12 | 0.00 | 0.00 |
| >hg38_21_+_8433172_8446622_RNA45SN1_45S_50ntFlanks@5489.5516.28\|CTGAGAAGACGGTCGAACTTGACTATCT | >hg38_21_+_8433172_8446622_RNA45SN1_45S_50ntFlanks | 5489 | 5516 | 28 | rRF 45S | NO | 34.98 | 941.73 | 205.06 | 123.33 |
| >hg38_21_+_8433172_8446622_RNA45SN1_45S_50ntFlanks@5490.5512.23\|TGAGAAGACGGTCGAACTTGACT | >hg38_21_+_8433172_8446622_RNA45SN1_45S_50ntFlanks | 5490 | 5512 | 23 | rRF 45S | NO | 0.00 | 72.54 | 0.00 | 0.00 |
| >hg38_21_+_8433172_8446622_RNA45SN1_45S_50ntFlanks@5490.5516.27\|TGAGAAGACGGTCGAACTTGACTATCT | >hg38_21_+_8433172_8446622_RNA45SN1_45S_50ntFlanks | 5490 | 5516 | 27 | rRF 45S | NO | 26.82 | 596.15 | 112.31 | 106.60 |
| >hg38_21_+_8433172_8446622_RNA45SN1_45S_50ntFlanks@5491.5512.22\|GAGAAGACGGTCGAACTTGACT | >hg38_21_+_8433172_8446622_RNA45SN1_45S_50ntFlanks | 5491 | 5512 | 22 | rRF 45S | NO | 0.00 | 107.33 | 0.00 | 0.00 |
| >hg38_21_+_8433172_8446622_RNA45SN1_45S_50ntFlanks@5491.5515.25\|GAGAAGACGGTCGAACTTGACTATC | >hg38_21_+_8433172_8446622_RNA45SN1_45S_50ntFlanks | 5491 | 5515 | 25 | rRF 45S | NO | 0.00 | 23.99 | 0.00 | 0.00 |
| >hg38_21_+_8433172_8446622_RNA45SN1_45S_50ntFlanks@5491.5516.26\|GAGAAGACGGTCGAACTTGACTATCT | >hg38_21_+_8433172_8446622_RNA45SN1_45S_50ntFlanks | 5491 | 5516 | 26 | rRF 45S | YES | 122.59 | 1664.40 | 278.05 | 278.59 |
| >hg38_21_+_8433172_8446622_RNA45SN1_45S_50ntFlanks@5491.5525.35\|GAGAAGACGGTCGAACTTGACTATCTAGAGGAAGT | >hg38_21_+_8433172_8446622_RNA45SN1_45S_50ntFlanks | 5491 | 5525 | 35 | rRF 45S | NO | 0.00 | 37.17 | 11.32 | 0.00 |
| >hg38_21_+_8433172_8446622_RNA45SN1_45S_50ntFlanks@5492.5512.21\|AGAAGACGGTCGAACTTGACT | >hg38_21_+_8433172_8446622_RNA45SN1_45S_50ntFlanks | 5492 | 5512 | 21 | rRF 45S | NO | 0.00 | 69.11 | 0.00 | 0.00 |
| >hg38_21_+_8433172_8446622_RNA45SN1_45S_50ntFlanks@5492.5516.25\|AGAAGACGGTCGAACTTGACTATCT | >hg38_21_+_8433172_8446622_RNA45SN1_45S_50ntFlanks | 5492 | 5516 | 25 | rRF 45S | YES | 174.81 | 1521.32 | 241.17 | 300.09 |
| >hg38_21_+_8433172_8446622_RNA45SN1_45S_50ntFlanks@5492.5525.34\|AGAAGACGGTCGAACTTGACTATCTAGAGGAAGT | >hg38_21_+_8433172_8446622_RNA45SN1_45S_50ntFlanks | 5492 | 5525 | 34 | rRF 45S | NO | 0.00 | 35.75 | 10.42 | 0.00 |
| >hg38_21_+_8433172_8446622_RNA45SN1_45S_50ntFlanks@5492.5532.41\|AGAAGACGGTCGAACTTGACTATCTAGAGGAAGTAAAAGTC | >hg38_21_+_8433172_8446622_RNA45SN1_45S_50ntFlanks | 5492 | 5532 | 41 | rRF 45S | NO | 0.00 | 25.36 | 0.00 | 0.00 |
| >hg38_21_+_8433172_8446622_RNA45SN1_45S_50ntFlanks@5493.5516.24\|GAAGACGGTCGAACTTGACTATCT | >hg38_21_+_8433172_8446622_RNA45SN1_45S_50ntFlanks | 5493 | 5516 | 24 | rRF 45S | NO | 0.00 | 173.76 | 38.60 | 35.15 |

# BT-20

| | | | | | | | | | | |
|---|---|---|---|---|---|---|---|---|---|---|
| >hg38_21_+_8433172_8446622_RNA45SN1_45S_50ntFlanks@5494.5516.23\|AAGACGGTCGAACTTGACTATCT | >hg38_21_+_8433172_8446622_RNA45SN1_45S_50ntFlanks | 5494 | 5516 | 23 | rRF 45S | NO | 9.70 | 298.65 | 82.63 | 84.74 |
| >hg38_21_+_8433172_8446622_RNA45SN1_45S_50ntFlanks@5495.5516.22\|AGACGGTCGAACTTGACTATCT | >hg38_21_+_8433172_8446622_RNA45SN1_45S_50ntFlanks | 5495 | 5516 | 22 | rRF 45S | NO | 0.00 | 125.59 | 14.72 | 28.80 |
| >hg38_21_+_8433172_8446622_RNA45SN1_45S_50ntFlanks@5496.5516.21\|GACGGTCGAACTTGACTATCT | >hg38_21_+_8433172_8446622_RNA45SN1_45S_50ntFlanks | 5496 | 5516 | 21 | rRF 45S | NO | 0.00 | 34.20 | 9.56 | 0.00 |
| >hg38_21_+_8433172_8446622_RNA45SN1_45S_50ntFlanks@6644.6665.22\|CCTCGTACGACTCTTAGCGGTG | >hg38_21_+_8433172_8446622_RNA45SN1_45S_50ntFlanks | 6644 | 6665 | 22 | rRF 45S | NO | 28.42 | 0.00 | 61.73 | 40.50 |
| >hg38_21_+_8433172_8446622_RNA45SN1_45S_50ntFlanks@6645.6662.18\|CTCGTACGACTCTTAGCG | >hg38_21_+_8433172_8446622_RNA45SN1_45S_50ntFlanks | 6645 | 6662 | 18 | rRF 45S | NO | 151.80 | 291.37 | 521.31 | 250.12 |
| >hg38_21_+_8433172_8446622_RNA45SN1_45S_50ntFlanks@6645.6663.19\|CTCGTACGACTCTTAGCGG | >hg38_21_+_8433172_8446622_RNA45SN1_45S_50ntFlanks | 6645 | 6663 | 19 | rRF 45S | NO | 102.49 | 159.97 | 280.92 | 190.71 |
| >hg38_21_+_8433172_8446622_RNA45SN1_45S_50ntFlanks@6645.6664.20\|CTCGTACGACTCTTAGCGGT | >hg38_21_+_8433172_8446622_RNA45SN1_45S_50ntFlanks | 6645 | 6664 | 20 | rRF 45S | NO | 127.86 | 241.83 | 420.62 | 269.35 |
| >hg38_21_+_8433172_8446622_RNA45SN1_45S_50ntFlanks@6645.6665.21\|CTCGTACGACTCTTAGCGGTG | >hg38_21_+_8433172_8446622_RNA45SN1_45S_50ntFlanks | 6645 | 6665 | 21 | rRF 45S | NO | 315.80 | 704.93 | 1190.32 | 591.67 |
| >hg38_21_+_8433172_8446622_RNA45SN1_45S_50ntFlanks@6645.6666.22\|CTCGTACGACTCTTAGCGGTGG | >hg38_21_+_8433172_8446622_RNA45SN1_45S_50ntFlanks | 6645 | 6666 | 22 | rRF 45S | NO | 113.85 | 208.26 | 257.58 | 304.81 |
| >hg38_21_+_8433172_8446622_RNA45SN1_45S_50ntFlanks@6645.6667.23\|CTCGTACGACTCTTAGCGGTGGA | >hg38_21_+_8433172_8446622_RNA45SN1_45S_50ntFlanks | 6645 | 6667 | 23 | rRF 45S | NO | 93.77 | 240.63 | 163.94 | 291.50 |
| >hg38_21_+_8433172_8446622_RNA45SN1_45S_50ntFlanks@6645.6668.24\|CTCGTACGACTCTTAGCGGTGGAT | >hg38_21_+_8433172_8446622_RNA45SN1_45S_50ntFlanks | 6645 | 6668 | 24 | rRF 45S | NO | 86.66 | 153.09 | 109.08 | 241.25 |
| >hg38_21_+_8433172_8446622_RNA45SN1_45S_50ntFlanks@6645.6669.25\|CTCGTACGACTCTTAGCGGTGGATC | >hg38_21_+_8433172_8446622_RNA45SN1_45S_50ntFlanks | 6645 | 6669 | 25 | rRF 45S | NO | 209.68 | 351.29 | 183.03 | 681.87 |
| >hg38_21_+_8433172_8446622_RNA45SN1_45S_50ntFlanks@6645.6672.28\|CTCGTACGACTCTTAGCGGTGGATCACT | >hg38_21_+_8433172_8446622_RNA45SN1_45S_50ntFlanks | 6645 | 6672 | 28 | rRF 45S | NO | 74.02 | 95.85 | 12.39 | 174.73 |
| >hg38_21_+_8433172_8446622_RNA45SN1_45S_50ntFlanks@6645.6673.29\|CTCGTACGACTCTTAGCGGTGGATCACTC | >hg38_21_+_8433172_8446622_RNA45SN1_45S_50ntFlanks | 6645 | 6673 | 29 | rRF 45S | NO | 0.00 | 0.00 | 13.27 | 27.83 |
| >hg38_21_+_8433172_8446622_RNA45SN1_45S_50ntFlanks@6645.6674.30\|CTCGTACGACTCTTAGCGGTGGATCACTCG | >hg38_21_+_8433172_8446622_RNA45SN1_45S_50ntFlanks | 6645 | 6674 | 30 | rRF 45S | NO | 26.70 | 23.22 | 69.22 | 150.16 |
| >hg38_21_+_8433172_8446622_RNA45SN1_45S_50ntFlanks@6645.6675.31\|CTCGTACGACTCTTAGCGGTGGATCACTCGG | >hg38_21_+_8433172_8446622_RNA45SN1_45S_50ntFlanks | 6645 | 6675 | 31 | rRF 45S | NO | 0.00 | 0.00 | 12.99 | 39.82 |
| >hg38_21_+_8433172_8446622_RNA45SN1_45S_50ntFlanks@6645.6676.32\|CTCGTACGACTCTTAGCGGTGGATCACTCGGC | >hg38_21_+_8433172_8446622_RNA45SN1_45S_50ntFlanks | 6645 | 6676 | 32 | rRF 45S | NO | 26.69 | 36.23 | 79.48 | 112.64 |
| >hg38_21_+_8433172_8446622_RNA45SN1_45S_50ntFlanks@6645.6677.33\|CTCGTACGACTCTTAGCGGTGGATCACTCGGCT | >hg38_21_+_8433172_8446622_RNA45SN1_45S_50ntFlanks | 6645 | 6677 | 33 | rRF 45S | NO | 33.47 | 26.27 | 73.33 | 97.48 |
| >hg38_21_+_8433172_8446622_RNA45SN1_45S_50ntFlanks@6645.6678.34\|CTCGTACGACTCTTAGCGGTGGATCACTCGGCTC | >hg38_21_+_8433172_8446622_RNA45SN1_45S_50ntFlanks | 6645 | 6678 | 34 | rRF 45S | NO | 31.02 | 0.00 | 67.47 | 90.22 |
| >hg38_21_+_8433172_8446622_RNA45SN1_45S_50ntFlanks@6645.6679.35\|CTCGTACGACTCTTAGCGGTGGATCACTCGGCTCG | >hg38_21_+_8433172_8446622_RNA45SN1_45S_50ntFlanks | 6645 | 6679 | 35 | rRF 45S | NO | 9.77 | 25.11 | 13.14 | 111.61 |
| >hg38_21_+_8433172_8446622_RNA45SN1_45S_50ntFlanks@6645.6680.36\|CTCGTACGACTCTTAGCGGTGGATCACTCGGCTCGT | >hg38_21_+_8433172_8446622_RNA45SN1_45S_50ntFlanks | 6645 | 6680 | 36 | rRF 45S | NO | 30.51 | 32.36 | 72.32 | 97.90 |
| >hg38_21_+_8433172_8446622_RNA45SN1_45S_50ntFlanks@6645.6681.37\|CTCGTACGACTCTTAGCGGTGGATCACTCGGCTCGTG | >hg38_21_+_8433172_8446622_RNA45SN1_45S_50ntFlanks | 6645 | 6681 | 37 | rRF 45S | NO | 10.70 | 0.00 | 60.16 | 35.68 |
| >hg38_21_+_8433172_8446622_RNA45SN1_45S_50ntFlanks@6645.6685.41\|CTCGTACGACTCTTAGCGGTGGATCACTCGGCTCGTGCGTC | >hg38_21_+_8433172_8446622_RNA45SN1_45S_50ntFlanks | 6645 | 6685 | 41 | rRF 45S | NO | 27.55 | 10.39 | 78.22 | 0.00 |
| >hg38_21_+_8433172_8446622_RNA45SN1_45S_50ntFlanks@6646.6663.18\|TCGTACGACTCTTAGCGG | >hg38_21_+_8433172_8446622_RNA45SN1_45S_50ntFlanks | 6646 | 6663 | 18 | rRF 45S | NO | 546.60 | 862.34 | 1011.98 | 1094.28 |
| >hg38_21_+_8433172_8446622_RNA45SN1_45S_50ntFlanks@6646.6664.19\|TCGTACGACTCTTAGCGGT | >hg38_21_+_8433172_8446622_RNA45SN1_45S_50ntFlanks | 6646 | 6664 | 19 | rRF 45S | YES | 957.96 | 1654.05 | 1746.27 | 2490.16 |
| >hg38_21_+_8433172_8446622_RNA45SN1_45S_50ntFlanks@6646.6665.20\|TCGTACGACTCTTAGCGGTG | >hg38_21_+_8433172_8446622_RNA45SN1_45S_50ntFlanks | 6646 | 6665 | 20 | rRF 45S | YES | 1163.29 | 2712.07 | 3081.85 | 2314.28 |
| >hg38_21_+_8433172_8446622_RNA45SN1_45S_50ntFlanks@6646.6666.21\|TCGTACGACTCTTAGCGGTGG | >hg38_21_+_8433172_8446622_RNA45SN1_45S_50ntFlanks | 6646 | 6666 | 21 | rRF 45S | NO | 426.83 | 811.23 | 851.76 | 1120.21 |
| >hg38_21_+_8433172_8446622_RNA45SN1_45S_50ntFlanks@6646.6667.22\|TCGTACGACTCTTAGCGGTGGA | >hg38_21_+_8433172_8446622_RNA45SN1_45S_50ntFlanks | 6646 | 6667 | 22 | rRF 45S | NO | 349.99 | 904.43 | 547.67 | 1010.96 |
| >hg38_21_+_8433172_8446622_RNA45SN1_45S_50ntFlanks@6646.6668.23\|TCGTACGACTCTTAGCGGTGGAT | >hg38_21_+_8433172_8446622_RNA45SN1_45S_50ntFlanks | 6646 | 6668 | 23 | rRF 45S | NO | 352.87 | 668.48 | 355.76 | 949.74 |
| >hg38_21_+_8433172_8446622_RNA45SN1_45S_50ntFlanks@6646.6669.24\|TCGTACGACTCTTAGCGGTGGATC | >hg38_21_+_8433172_8446622_RNA45SN1_45S_50ntFlanks | 6646 | 6669 | 24 | rRF 45S | NO | 901.33 | 1725.00 | 773.24 | 2633.96 |
| >hg38_21_+_8433172_8446622_RNA45SN1_45S_50ntFlanks@6646.6670.25\|TCGTACGACTCTTAGCGGTGGATCA | >hg38_21_+_8433172_8446622_RNA45SN1_45S_50ntFlanks | 6646 | 6670 | 25 | rRF 45S | NO | 77.51 | 101.34 | 9.70 | 240.84 |
| >hg38_21_+_8433172_8446622_RNA45SN1_45S_50ntFlanks@6646.6671.26\|TCGTACGACTCTTAGCGGTGGATCAC | >hg38_21_+_8433172_8446622_RNA45SN1_45S_50ntFlanks | 6646 | 6671 | 26 | rRF 45S | NO | 10.71 | 39.44 | 0.00 | 88.41 |
| >hg38_21_+_8433172_8446622_RNA45SN1_45S_50ntFlanks@6646.6672.27\|TCGTACGACTCTTAGCGGTGGATCACT | >hg38_21_+_8433172_8446622_RNA45SN1_45S_50ntFlanks | 6646 | 6672 | 27 | rRF 45S | NO | 343.70 | 562.83 | 146.06 | 796.24 |
| >hg38_21_+_8433172_8446622_RNA45SN1_45S_50ntFlanks@6646.6673.28\|TCGTACGACTCTTAGCGGTGGATCACTC | >hg38_21_+_8433172_8446622_RNA45SN1_45S_50ntFlanks | 6646 | 6673 | 28 | rRF 45S | NO | 92.52 | 96.48 | 117.30 | 189.79 |
| >hg38_21_+_8433172_8446622_RNA45SN1_45S_50ntFlanks@6646.6674.29\|TCGTACGACTCTTAGCGGTGGATCACTCG | >hg38_21_+_8433172_8446622_RNA45SN1_45S_50ntFlanks | 6646 | 6674 | 29 | rRF 45S | NO | 94.18 | 94.26 | 145.49 | 324.51 |
| >hg38_21_+_8433172_8446622_RNA45SN1_45S_50ntFlanks@6646.6675.30\|TCGTACGACTCTTAGCGGTGGATCACTCGG | >hg38_21_+_8433172_8446622_RNA45SN1_45S_50ntFlanks | 6646 | 6675 | 30 | rRF 45S | NO | 100.55 | 83.35 | 140.82 | 262.60 |
| >hg38_21_+_8433172_8446622_RNA45SN1_45S_50ntFlanks@6646.6676.31\|TCGTACGACTCTTAGCGGTGGATCACTCGGC | >hg38_21_+_8433172_8446622_RNA45SN1_45S_50ntFlanks | 6646 | 6676 | 31 | rRF 45S | NO | 160.94 | 186.62 | 244.17 | 410.93 |
| >hg38_21_+_8433172_8446622_RNA45SN1_45S_50ntFlanks@6646.6677.32\|TCGTACGACTCTTAGCGGTGGATCACTCGGCT | >hg38_21_+_8433172_8446622_RNA45SN1_45S_50ntFlanks | 6646 | 6677 | 32 | rRF 45S | NO | 207.87 | 196.35 | 271.90 | 429.27 |

# BT-20

| | | | | | | | | | | |
|---|---|---|---|---|---|---|---|---|---|---|
| >hg38_21_+_8433172_8446622_RNA45SN1_45S_50ntFlanks@6646.6678.33\|TCGTACGACTCTTAGCGGTGGATCACTCGGCTC | >hg38_21_+_8433172_8446622_RNA45SN1_45S_50ntFlanks | 6646 | 6678 | 33 | rRF 45S | NO | 165.96 | 122.69 | 233.93 | 326.51 |
| >hg38_21_+_8433172_8446622_RNA45SN1_45S_50ntFlanks@6646.6679.34\|TCGTACGACTCTTAGCGGTGGATCACTCGGCTCG | >hg38_21_+_8433172_8446622_RNA45SN1_45S_50ntFlanks | 6646 | 6679 | 34 | rRF 45S | NO | 114.83 | 108.45 | 147.16 | 379.33 |
| >hg38_21_+_8433172_8446622_RNA45SN1_45S_50ntFlanks@6646.6680.35\|TCGTACGACTCTTAGCGGTGGATCACTCGGCTCGT | >hg38_21_+_8433172_8446622_RNA45SN1_45S_50ntFlanks | 6646 | 6680 | 35 | rRF 45S | NO | 191.21 | 265.69 | 236.21 | 399.89 |
| >hg38_21_+_8433172_8446622_RNA45SN1_45S_50ntFlanks@6646.6681.36\|TCGTACGACTCTTAGCGGTGGATCACTCGGCTCGTG | >hg38_21_+_8433172_8446622_RNA45SN1_45S_50ntFlanks | 6646 | 6681 | 36 | rRF 45S | NO | 229.15 | 105.12 | 259.43 | 418.40 |
| >hg38_21_+_8433172_8446622_RNA45SN1_45S_50ntFlanks@6646.6682.37\|TCGTACGACTCTTAGCGGTGGATCACTCGGCTCGTGC | >hg38_21_+_8433172_8446622_RNA45SN1_45S_50ntFlanks | 6646 | 6682 | 37 | rRF 45S | NO | 9.91 | 0.00 | 38.77 | 24.15 |
| >hg38_21_+_8433172_8446622_RNA45SN1_45S_50ntFlanks@6646.6683.38\|TCGTACGACTCTTAGCGGTGGATCACTCGGCTCGTGCG | >hg38_21_+_8433172_8446622_RNA45SN1_45S_50ntFlanks | 6646 | 6683 | 38 | rRF 45S | NO | 76.00 | 39.57 | 98.89 | 26.31 |
| >hg38_21_+_8433172_8446622_RNA45SN1_45S_50ntFlanks@6646.6685.40\|TCGTACGACTCTTAGCGGTGGATCACTCGGCTCGTGCGTC | >hg38_21_+_8433172_8446622_RNA45SN1_45S_50ntFlanks | 6646 | 6685 | 40 | rRF 45S | NO | 636.87 | 420.86 | 831.56 | 498.33 |
| >hg38_21_+_8433172_8446622_RNA45SN1_45S_50ntFlanks@6646.6688.43\|TCGTACGACTCTTAGCGGTGGATCACTCGGCTCGTGCGTCGAT | >hg38_21_+_8433172_8446622_RNA45SN1_45S_50ntFlanks | 6646 | 6688 | 43 | rRF 45S | NO | 10.68 | 0.00 | 10.35 | 36.87 |
| >hg38_21_+_8433172_8446622_RNA45SN1_45S_50ntFlanks@6646.6689.44\|TCGTACGACTCTTAGCGGTGGATCACTCGGCTCGTGCGTCGATG | >hg38_21_+_8433172_8446622_RNA45SN1_45S_50ntFlanks | 6646 | 6689 | 44 | rRF 45S | NO | 226.76 | 90.77 | 265.27 | 517.13 |
| >hg38_21_+_8433172_8446622_RNA45SN1_45S_50ntFlanks@6646.6690.45\|TCGTACGACTCTTAGCGGTGGATCACTCGGCTCGTGCGTCGATGA | >hg38_21_+_8433172_8446622_RNA45SN1_45S_50ntFlanks | 6646 | 6690 | 45 | rRF 45S | NO | 10.52 | 0.00 | 58.57 | 35.60 |
| >hg38_21_+_8433172_8446622_RNA45SN1_45S_50ntFlanks@6646.6695.50\|TCGTACGACTCTTAGCGGTGGATCACTCGGCTCGTGCGTCGATGAAGAAC | >hg38_21_+_8433172_8446622_RNA45SN1_45S_50ntFlanks | 6646 | 6695 | 50 | rRF 45S | NO | 33.89 | 0.00 | 65.48 | 283.17 |
| >hg38_21_+_8433172_8446622_RNA45SN1_45S_50ntFlanks@6647.6664.18\|CGTACGACTCTTAGCGGT | >hg38_21_+_8433172_8446622_RNA45SN1_45S_50ntFlanks | 6647 | 6664 | 18 | rRF 45S | NO | 189.20 | 196.90 | 436.72 | 451.20 |
| >hg38_21_+_8433172_8446622_RNA45SN1_45S_50ntFlanks@6647.6665.19\|CGTACGACTCTTAGCGGTG | >hg38_21_+_8433172_8446622_RNA45SN1_45S_50ntFlanks | 6647 | 6665 | 19 | rRF 45S | NO | 279.87 | 368.83 | 968.11 | 548.34 |
| >hg38_21_+_8433172_8446622_RNA45SN1_45S_50ntFlanks@6647.6666.20\|CGTACGACTCTTAGCGGTGG | >hg38_21_+_8433172_8446622_RNA45SN1_45S_50ntFlanks | 6647 | 6666 | 20 | rRF 45S | NO | 86.95 | 102.05 | 236.94 | 205.01 |
| >hg38_21_+_8433172_8446622_RNA45SN1_45S_50ntFlanks@6647.6667.21\|CGTACGACTCTTAGCGGTGGA | >hg38_21_+_8433172_8446622_RNA45SN1_45S_50ntFlanks | 6647 | 6667 | 21 | rRF 45S | NO | 83.93 | 144.60 | 194.30 | 213.58 |
| >hg38_21_+_8433172_8446622_RNA45SN1_45S_50ntFlanks@6647.6668.22\|CGTACGACTCTTAGCGGTGGAT | >hg38_21_+_8433172_8446622_RNA45SN1_45S_50ntFlanks | 6647 | 6668 | 22 | rRF 45S | NO | 97.53 | 118.70 | 164.82 | 233.44 |
| >hg38_21_+_8433172_8446622_RNA45SN1_45S_50ntFlanks@6647.6669.23\|CGTACGACTCTTAGCGGTGGATC | >hg38_21_+_8433172_8446622_RNA45SN1_45S_50ntFlanks | 6647 | 6669 | 23 | rRF 45S | NO | 207.81 | 251.87 | 205.95 | 556.26 |
| >hg38_21_+_8433172_8446622_RNA45SN1_45S_50ntFlanks@6647.6672.26\|CGTACGACTCTTAGCGGTGGATCACT | >hg38_21_+_8433172_8446622_RNA45SN1_45S_50ntFlanks | 6647 | 6672 | 26 | rRF 45S | NO | 78.68 | 77.42 | 11.75 | 160.90 |
| >hg38_21_+_8433172_8446622_RNA45SN1_45S_50ntFlanks@6647.6673.27\|CGTACGACTCTTAGCGGTGGATCACTC | >hg38_21_+_8433172_8446622_RNA45SN1_45S_50ntFlanks | 6647 | 6673 | 27 | rRF 45S | NO | 0.00 | 0.00 | 10.41 | 26.56 |
| >hg38_21_+_8433172_8446622_RNA45SN1_45S_50ntFlanks@6647.6674.28\|CGTACGACTCTTAGCGGTGGATCACTCG | >hg38_21_+_8433172_8446622_RNA45SN1_45S_50ntFlanks | 6647 | 6674 | 28 | rRF 45S | NO | 0.00 | 0.00 | 11.51 | 34.18 |
| >hg38_21_+_8433172_8446622_RNA45SN1_45S_50ntFlanks@6647.6676.30\|CGTACGACTCTTAGCGGTGGATCACTCGGC | >hg38_21_+_8433172_8446622_RNA45SN1_45S_50ntFlanks | 6647 | 6676 | 30 | rRF 45S | NO | 31.63 | 28.31 | 73.80 | 121.10 |
| >hg38_21_+_8433172_8446622_RNA45SN1_45S_50ntFlanks@6647.6677.31\|CGTACGACTCTTAGCGGTGGATCACTCGGCT | >hg38_21_+_8433172_8446622_RNA45SN1_45S_50ntFlanks | 6647 | 6677 | 31 | rRF 45S | NO | 34.28 | 0.00 | 70.10 | 94.84 |
| >hg38_21_+_8433172_8446622_RNA45SN1_45S_50ntFlanks@6647.6678.32\|CGTACGACTCTTAGCGGTGGATCACTCGGCTC | >hg38_21_+_8433172_8446622_RNA45SN1_45S_50ntFlanks | 6647 | 6678 | 32 | rRF 45S | NO | 27.09 | 0.00 | 62.53 | 82.64 |
| >hg38_21_+_8433172_8446622_RNA45SN1_45S_50ntFlanks@6647.6680.34\|CGTACGACTCTTAGCGGTGGATCACTCGGCTCGT | >hg38_21_+_8433172_8446622_RNA45SN1_45S_50ntFlanks | 6647 | 6680 | 34 | rRF 45S | NO | 28.82 | 27.36 | 58.83 | 76.24 |
| >hg38_21_+_8433172_8446622_RNA45SN1_45S_50ntFlanks@6647.6681.35\|CGTACGACTCTTAGCGGTGGATCACTCGGCTCGTG | >hg38_21_+_8433172_8446622_RNA45SN1_45S_50ntFlanks | 6647 | 6681 | 35 | rRF 45S | NO | 109.04 | 0.00 | 70.22 | 91.87 |
| >hg38_21_+_8433172_8446622_RNA45SN1_45S_50ntFlanks@6647.6689.43\|CGTACGACTCTTAGCGGTGGATCACTCGGCTCGTGCGTCGATG | >hg38_21_+_8433172_8446622_RNA45SN1_45S_50ntFlanks | 6647 | 6689 | 43 | rRF 45S | NO | 26.51 | 0.00 | 37.84 | 68.93 |
| >hg38_21_+_8433172_8446622_RNA45SN1_45S_50ntFlanks@6648.6665.18\|GTACGACTCTTAGCGGTG | >hg38_21_+_8433172_8446622_RNA45SN1_45S_50ntFlanks | 6648 | 6665 | 18 | rRF 45S | NO | 39.96 | 27.27 | 136.07 | 126.99 |
| >hg38_21_+_8433172_8446622_RNA45SN1_45S_50ntFlanks@6648.6666.19\|GTACGACTCTTAGCGGTGG | >hg38_21_+_8433172_8446622_RNA45SN1_45S_50ntFlanks | 6648 | 6666 | 19 | rRF 45S | NO | 0.00 | 0.00 | 10.85 | 33.98 |
| >hg38_21_+_8433172_8446622_RNA45SN1_45S_50ntFlanks@6648.6667.20\|GTACGACTCTTAGCGGTGGA | >hg38_21_+_8433172_8446622_RNA45SN1_45S_50ntFlanks | 6648 | 6667 | 20 | rRF 45S | NO | 0.00 | 0.00 | 15.27 | 39.49 |
| >hg38_21_+_8433172_8446622_RNA45SN1_45S_50ntFlanks@6648.6668.21\|GTACGACTCTTAGCGGTGGAT | >hg38_21_+_8433172_8446622_RNA45SN1_45S_50ntFlanks | 6648 | 6668 | 21 | rRF 45S | NO | 71.36 | 39.06 | 78.50 | 199.54 |
| >hg38_21_+_8433172_8446622_RNA45SN1_45S_50ntFlanks@6648.6669.22\|GTACGACTCTTAGCGGTGGATC | >hg38_21_+_8433172_8446622_RNA45SN1_45S_50ntFlanks | 6648 | 6669 | 22 | rRF 45S | NO | 82.28 | 72.71 | 78.88 | 211.45 |
| >hg38_21_+_8433172_8446622_RNA45SN1_45S_50ntFlanks@6648.6672.25\|GTACGACTCTTAGCGGTGGATCACT | >hg38_21_+_8433172_8446622_RNA45SN1_45S_50ntFlanks | 6648 | 6672 | 25 | rRF 45S | NO | 27.54 | 0.00 | 0.00 | 96.37 |
| >hg38_21_+_8433172_8446622_RNA45SN1_45S_50ntFlanks@6648.6676.29\|GTACGACTCTTAGCGGTGGATCACTCGGC | >hg38_21_+_8433172_8446622_RNA45SN1_45S_50ntFlanks | 6648 | 6676 | 29 | rRF 45S | NO | 37.92 | 31.35 | 72.77 | 190.99 |
| >hg38_21_+_8433172_8446622_RNA45SN1_45S_50ntFlanks@6648.6677.30\|GTACGACTCTTAGCGGTGGATCACTCGGCT | >hg38_21_+_8433172_8446622_RNA45SN1_45S_50ntFlanks | 6648 | 6677 | 30 | rRF 45S | NO | 29.05 | 0.00 | 15.91 | 119.76 |
| >hg38_21_+_8433172_8446622_RNA45SN1_45S_50ntFlanks@6648.6678.31\|GTACGACTCTTAGCGGTGGATCACTCGGCTC | >hg38_21_+_8433172_8446622_RNA45SN1_45S_50ntFlanks | 6648 | 6678 | 31 | rRF 45S | NO | 34.01 | 0.00 | 38.79 | 161.72 |
| >hg38_21_+_8433172_8446622_RNA45SN1_45S_50ntFlanks@6648.6679.32\|GTACGACTCTTAGCGGTGGATCACTCGGCTCG | >hg38_21_+_8433172_8446622_RNA45SN1_45S_50ntFlanks | 6648 | 6679 | 32 | rRF 45S | NO | 10.74 | 0.00 | 11.42 | 124.70 |
| >hg38_21_+_8433172_8446622_RNA45SN1_45S_50ntFlanks@6648.6680.33\|GTACGACTCTTAGCGGTGGATCACTCGGCTCGT | >hg38_21_+_8433172_8446622_RNA45SN1_45S_50ntFlanks | 6648 | 6680 | 33 | rRF 45S | NO | 31.70 | 30.42 | 15.87 | 102.45 |
| >hg38_21_+_8433172_8446622_RNA45SN1_45S_50ntFlanks@6648.6681.34\|GTACGACTCTTAGCGGTGGATCACTCGGCTCGTG | >hg38_21_+_8433172_8446622_RNA45SN1_45S_50ntFlanks | 6648 | 6681 | 34 | rRF 45S | NO | 37.85 | 0.00 | 13.36 | 84.99 |

# BT-20

| | | | | | | | | | | |
|---|---|---|---|---|---|---|---|---|---|---|
| >hg38_21_+_8433172_8446622_RNA45SN1_45S_50ntFlanks@6648.6689.42\|GTACGACTCTTAGCGGTGGATCACTCGGCTCGTGCGTCGATG | >hg38_21_+_8433172_8446622_RNA45SN1_45S_50ntFlanks | 6648 | 6689 | 42 | rRF 45S | NO | 68.74 | 0.00 | 63.79 | 174.24 |
| >hg38_21_+_8433172_8446622_RNA45SN1_45S_50ntFlanks@6649.6668.20\|TACGACTCTTAGCGGTGGAT | >hg38_21_+_8433172_8446622_RNA45SN1_45S_50ntFlanks | 6649 | 6668 | 20 | rRF 45S | NO | 29.74 | 9.44 | 10.38 | 141.77 |
| >hg38_21_+_8433172_8446622_RNA45SN1_45S_50ntFlanks@6649.6669.21\|TACGACTCTTAGCGGTGGATC | >hg38_21_+_8433172_8446622_RNA45SN1_45S_50ntFlanks | 6649 | 6669 | 21 | rRF 45S | NO | 38.29 | 25.70 | 39.70 | 173.77 |
| >hg38_21_+_8433172_8446622_RNA45SN1_45S_50ntFlanks@6649.6677.29\|TACGACTCTTAGCGGTGGATCACTCGGCT | >hg38_21_+_8433172_8446622_RNA45SN1_45S_50ntFlanks | 6649 | 6677 | 29 | rRF 45S | NO | 10.42 | 0.00 | 9.64 | 78.48 |
| >hg38_21_+_8433172_8446622_RNA45SN1_45S_50ntFlanks@6649.6683.35\|TACGACTCTTAGCGGTGGATCACTCGGCTCGTGCG | >hg38_21_+_8433172_8446622_RNA45SN1_45S_50ntFlanks | 6649 | 6683 | 35 | rRF 45S | NO | 33.60 | 0.00 | 0.00 | 0.00 |
| >hg38_21_+_8433172_8446622_RNA45SN1_45S_50ntFlanks@6649.6689.41\|TACGACTCTTAGCGGTGGATCACTCGGCTCGTGCGTCGATG | >hg38_21_+_8433172_8446622_RNA45SN1_45S_50ntFlanks | 6649 | 6689 | 41 | rRF 45S | NO | 23.48 | 0.00 | 0.00 | 66.30 |
| >hg38_21_+_8433172_8446622_RNA45SN1_45S_50ntFlanks@6650.6667.18\|ACGACTCTTAGCGGTGGA | >hg38_21_+_8433172_8446622_RNA45SN1_45S_50ntFlanks | 6650 | 6667 | 18 | rRF 45S | NO | 0.00 | 0.00 | 10.68 | 35.76 |
| >hg38_21_+_8433172_8446622_RNA45SN1_45S_50ntFlanks@6650.6668.19\|ACGACTCTTAGCGGTGGAT | >hg38_21_+_8433172_8446622_RNA45SN1_45S_50ntFlanks | 6650 | 6668 | 19 | rRF 45S | NO | 101.37 | 45.46 | 82.54 | 228.52 |
| >hg38_21_+_8433172_8446622_RNA45SN1_45S_50ntFlanks@6650.6669.20\|ACGACTCTTAGCGGTGGATC | >hg38_21_+_8433172_8446622_RNA45SN1_45S_50ntFlanks | 6650 | 6669 | 20 | rRF 45S | NO | 122.00 | 68.47 | 122.20 | 278.53 |
| >hg38_21_+_8433172_8446622_RNA45SN1_45S_50ntFlanks@6650.6672.23\|ACGACTCTTAGCGGTGGATCACT | >hg38_21_+_8433172_8446622_RNA45SN1_45S_50ntFlanks | 6650 | 6672 | 23 | rRF 45S | NO | 69.13 | 0.00 | 0.00 | 144.40 |
| >hg38_21_+_8433172_8446622_RNA45SN1_45S_50ntFlanks@6650.6674.25\|ACGACTCTTAGCGGTGGATCACTCG | >hg38_21_+_8433172_8446622_RNA45SN1_45S_50ntFlanks | 6650 | 6674 | 25 | rRF 45S | NO | 27.21 | 0.00 | 16.13 | 143.21 |
| >hg38_21_+_8433172_8446622_RNA45SN1_45S_50ntFlanks@6650.6675.26\|ACGACTCTTAGCGGTGGATCACTCGG | >hg38_21_+_8433172_8446622_RNA45SN1_45S_50ntFlanks | 6650 | 6675 | 26 | rRF 45S | NO | 26.39 | 0.00 | 59.44 | 74.51 |
| >hg38_21_+_8433172_8446622_RNA45SN1_45S_50ntFlanks@6650.6676.27\|ACGACTCTTAGCGGTGGATCACTCGGC | >hg38_21_+_8433172_8446622_RNA45SN1_45S_50ntFlanks | 6650 | 6676 | 27 | rRF 45S | NO | 38.83 | 67.14 | 90.01 | 167.31 |
| >hg38_21_+_8433172_8446622_RNA45SN1_45S_50ntFlanks@6650.6677.28\|ACGACTCTTAGCGGTGGATCACTCGGCT | >hg38_21_+_8433172_8446622_RNA45SN1_45S_50ntFlanks | 6650 | 6677 | 28 | rRF 45S | NO | 86.95 | 0.00 | 73.51 | 157.77 |
| >hg38_21_+_8433172_8446622_RNA45SN1_45S_50ntFlanks@6650.6678.29\|ACGACTCTTAGCGGTGGATCACTCGGCTC | >hg38_21_+_8433172_8446622_RNA45SN1_45S_50ntFlanks | 6650 | 6678 | 29 | rRF 45S | NO | 75.32 | 10.31 | 81.86 | 172.07 |
| >hg38_21_+_8433172_8446622_RNA45SN1_45S_50ntFlanks@6650.6679.30\|ACGACTCTTAGCGGTGGATCACTCGGCTCG | >hg38_21_+_8433172_8446622_RNA45SN1_45S_50ntFlanks | 6650 | 6679 | 30 | rRF 45S | NO | 37.54 | 35.59 | 67.07 | 156.11 |
| >hg38_21_+_8433172_8446622_RNA45SN1_45S_50ntFlanks@6650.6680.31\|ACGACTCTTAGCGGTGGATCACTCGGCTCGT | >hg38_21_+_8433172_8446622_RNA45SN1_45S_50ntFlanks | 6650 | 6680 | 31 | rRF 45S | NO | 0.00 | 0.00 | 10.19 | 0.00 |
| >hg38_21_+_8433172_8446622_RNA45SN1_45S_50ntFlanks@6650.6681.32\|ACGACTCTTAGCGGTGGATCACTCGGCTCGTG | >hg38_21_+_8433172_8446622_RNA45SN1_45S_50ntFlanks | 6650 | 6681 | 32 | rRF 45S | NO | 89.28 | 0.00 | 69.76 | 113.66 |
| >hg38_21_+_8433172_8446622_RNA45SN1_45S_50ntFlanks@6650.6683.34\|ACGACTCTTAGCGGTGGATCACTCGGCTCGTGCG | >hg38_21_+_8433172_8446622_RNA45SN1_45S_50ntFlanks | 6650 | 6683 | 34 | rRF 45S | NO | 92.34 | 0.00 | 0.00 | 9.47 |
| >hg38_21_+_8433172_8446622_RNA45SN1_45S_50ntFlanks@6650.6689.40\|ACGACTCTTAGCGGTGGATCACTCGGCTCGTGCGTCGATG | >hg38_21_+_8433172_8446622_RNA45SN1_45S_50ntFlanks | 6650 | 6689 | 40 | rRF 45S | NO | 81.67 | 0.00 | 66.62 | 107.04 |
| >hg38_21_+_8433172_8446622_RNA45SN1_45S_50ntFlanks@6650.6692.43\|ACGACTCTTAGCGGTGGATCACTCGGCTCGTGCGTCGATGAAG | >hg38_21_+_8433172_8446622_RNA45SN1_45S_50ntFlanks | 6650 | 6692 | 43 | rRF 45S | NO | 24.96 | 0.00 | 0.00 | 30.07 |
| >hg38_21_+_8433172_8446622_RNA45SN1_45S_50ntFlanks@6650.6695.46\|ACGACTCTTAGCGGTGGATCACTCGGCTCGTGCGTCGATGAAGAAC | >hg38_21_+_8433172_8446622_RNA45SN1_45S_50ntFlanks | 6650 | 6695 | 46 | rRF 45S | NO | 30.16 | 0.00 | 9.75 | 114.23 |
| >hg38_21_+_8433172_8446622_RNA45SN1_45S_50ntFlanks@6650.6696.47\|ACGACTCTTAGCGGTGGATCACTCGGCTCGTGCGTCGATGAAGAACG | >hg38_21_+_8433172_8446622_RNA45SN1_45S_50ntFlanks | 6650 | 6696 | 47 | rRF 45S | NO | 24.15 | 0.00 | 0.00 | 34.46 |
| >hg38_21_+_8433172_8446622_RNA45SN1_45S_50ntFlanks@6650.6698.49\|ACGACTCTTAGCGGTGGATCACTCGGCTCGTGCGTCGATGAAGAACGCA | >hg38_21_+_8433172_8446622_RNA45SN1_45S_50ntFlanks | 6650 | 6698 | 49 | rRF 45S | NO | 9.41 | 0.00 | 11.52 | 102.12 |
| >hg38_21_+_8433172_8446622_RNA45SN1_45S_50ntFlanks@6651.6668.18\|CGACTCTTAGCGGTGGAT | >hg38_21_+_8433172_8446622_RNA45SN1_45S_50ntFlanks | 6651 | 6668 | 18 | rRF 45S | NO | 546.54 | 670.59 | 346.68 | 1486.06 |
| >hg38_21_+_8433172_8446622_RNA45SN1_45S_50ntFlanks@6651.6669.19\|CGACTCTTAGCGGTGGATC | >hg38_21_+_8433172_8446622_RNA45SN1_45S_50ntFlanks | 6651 | 6669 | 19 | rRF 45S | NO | 556.06 | 570.59 | 539.16 | 1350.89 |
| >hg38_21_+_8433172_8446622_RNA45SN1_45S_50ntFlanks@6651.6670.20\|CGACTCTTAGCGGTGGATCA | >hg38_21_+_8433172_8446622_RNA45SN1_45S_50ntFlanks | 6651 | 6670 | 20 | rRF 45S | NO | 28.01 | 0.00 | 0.00 | 78.82 |
| >hg38_21_+_8433172_8446622_RNA45SN1_45S_50ntFlanks@6651.6672.22\|CGACTCTTAGCGGTGGATCACT | >hg38_21_+_8433172_8446622_RNA45SN1_45S_50ntFlanks | 6651 | 6672 | 22 | rRF 45S | NO | 340.21 | 259.24 | 113.37 | 719.50 |
| >hg38_21_+_8433172_8446622_RNA45SN1_45S_50ntFlanks@6651.6673.23\|CGACTCTTAGCGGTGGATCACTC | >hg38_21_+_8433172_8446622_RNA45SN1_45S_50ntFlanks | 6651 | 6673 | 23 | rRF 45S | NO | 149.60 | 107.87 | 127.98 | 366.20 |
| >hg38_21_+_8433172_8446622_RNA45SN1_45S_50ntFlanks@6651.6674.24\|CGACTCTTAGCGGTGGATCACTCG | >hg38_21_+_8433172_8446622_RNA45SN1_45S_50ntFlanks | 6651 | 6674 | 24 | rRF 45S | NO | 117.65 | 93.47 | 136.12 | 429.99 |
| >hg38_21_+_8433172_8446622_RNA45SN1_45S_50ntFlanks@6651.6675.25\|CGACTCTTAGCGGTGGATCACTCGG | >hg38_21_+_8433172_8446622_RNA45SN1_45S_50ntFlanks | 6651 | 6675 | 25 | rRF 45S | NO | 138.55 | 118.71 | 135.47 | 307.62 |
| >hg38_21_+_8433172_8446622_RNA45SN1_45S_50ntFlanks@6651.6676.26\|CGACTCTTAGCGGTGGATCACTCGGC | >hg38_21_+_8433172_8446622_RNA45SN1_45S_50ntFlanks | 6651 | 6676 | 26 | rRF 45S | NO | 373.32 | 819.60 | 428.66 | 1265.21 |
| >hg38_21_+_8433172_8446622_RNA45SN1_45S_50ntFlanks@6651.6677.27\|CGACTCTTAGCGGTGGATCACTCGGCT | >hg38_21_+_8433172_8446622_RNA45SN1_45S_50ntFlanks | 6651 | 6677 | 27 | rRF 45S | NO | 428.01 | 253.51 | 366.20 | 954.18 |
| >hg38_21_+_8433172_8446622_RNA45SN1_45S_50ntFlanks@6651.6678.28\|CGACTCTTAGCGGTGGATCACTCGGCTC | >hg38_21_+_8433172_8446622_RNA45SN1_45S_50ntFlanks | 6651 | 6678 | 28 | rRF 45S | NO | 436.78 | 304.39 | 652.15 | 1188.97 |
| >hg38_21_+_8433172_8446622_RNA45SN1_45S_50ntFlanks@6651.6679.29\|CGACTCTTAGCGGTGGATCACTCGGCTCG | >hg38_21_+_8433172_8446622_RNA45SN1_45S_50ntFlanks | 6651 | 6679 | 29 | rRF 45S | NO | 145.71 | 226.31 | 154.74 | 450.82 |
| >hg38_21_+_8433172_8446622_RNA45SN1_45S_50ntFlanks@6651.6680.30\|CGACTCTTAGCGGTGGATCACTCGGCTCGT | >hg38_21_+_8433172_8446622_RNA45SN1_45S_50ntFlanks | 6651 | 6680 | 30 | rRF 45S | NO | 170.82 | 121.48 | 177.95 | 328.53 |
| >hg38_21_+_8433172_8446622_RNA45SN1_45S_50ntFlanks@6651.6681.31\|CGACTCTTAGCGGTGGATCACTCGGCTCGTG | >hg38_21_+_8433172_8446622_RNA45SN1_45S_50ntFlanks | 6651 | 6681 | 31 | rRF 45S | NO | 544.78 | 194.54 | 389.78 | 936.34 |
| >hg38_21_+_8433172_8446622_RNA45SN1_45S_50ntFlanks@6651.6682.32\|CGACTCTTAGCGGTGGATCACTCGGCTCGTGC | >hg38_21_+_8433172_8446622_RNA45SN1_45S_50ntFlanks | 6651 | 6682 | 32 | rRF 45S | NO | 156.84 | 90.73 | 128.20 | 355.59 |

# BT-20

| | | | | | | | | | | |
|---|---|---|---|---|---|---|---|---|---|---|
| >hg38_21_+_8433172_8446622_RNA45SN1_45S_50ntFlanks@6651.6683.33\|CGACTCTTAGCGGTGGATCACTCGGCTCGTGCG | >hg38_21_+_8433172_8446622_RNA45SN1_45S_50ntFlanks | 6651 | 6683 | 33 | rRF 45S | NO | 260.46 | 28.00 | 40.06 | 66.15 |
| >hg38_21_+_8433172_8446622_RNA45SN1_45S_50ntFlanks@6651.6684.34\|CGACTCTTAGCGGTGGATCACTCGGCTCGTGCGT | >hg38_21_+_8433172_8446622_RNA45SN1_45S_50ntFlanks | 6651 | 6684 | 34 | rRF 45S | NO | 271.96 | 154.63 | 84.67 | 235.81 |
| >hg38_21_+_8433172_8446622_RNA45SN1_45S_50ntFlanks@6651.6685.35\|CGACTCTTAGCGGTGGATCACTCGGCTCGTGCGTC | >hg38_21_+_8433172_8446622_RNA45SN1_45S_50ntFlanks | 6651 | 6685 | 35 | rRF 45S | NO | 159.23 | 81.30 | 503.08 | 151.62 |
| >hg38_21_+_8433172_8446622_RNA45SN1_45S_50ntFlanks@6651.6686.36\|CGACTCTTAGCGGTGGATCACTCGGCTCGTGCGTCG | >hg38_21_+_8433172_8446622_RNA45SN1_45S_50ntFlanks | 6651 | 6686 | 36 | rRF 45S | NO | 34.45 | 0.00 | 0.00 | 0.00 |
| >hg38_21_+_8433172_8446622_RNA45SN1_45S_50ntFlanks@6651.6687.37\|CGACTCTTAGCGGTGGATCACTCGGCTCGTGCGTCGA | >hg38_21_+_8433172_8446622_RNA45SN1_45S_50ntFlanks | 6651 | 6687 | 37 | rRF 45S | NO | 104.49 | 25.46 | 0.00 | 66.21 |
| >hg38_21_+_8433172_8446622_RNA45SN1_45S_50ntFlanks@6651.6688.38\|CGACTCTTAGCGGTGGATCACTCGGCTCGTGCGTCGAT | >hg38_21_+_8433172_8446622_RNA45SN1_45S_50ntFlanks | 6651 | 6688 | 38 | rRF 45S | NO | 262.80 | 103.14 | 120.31 | 321.36 |
| >hg38_21_+_8433172_8446622_RNA45SN1_45S_50ntFlanks@6651.6689.39\|CGACTCTTAGCGGTGGATCACTCGGCTCGTGCGTCGATG | >hg38_21_+_8433172_8446622_RNA45SN1_45S_50ntFlanks | 6651 | 6689 | 39 | rRF 45S | NO | 513.55 | 251.30 | 389.33 | 799.50 |
| >hg38_21_+_8433172_8446622_RNA45SN1_45S_50ntFlanks@6651.6690.40\|CGACTCTTAGCGGTGGATCACTCGGCTCGTGCGTCGATGA | >hg38_21_+_8433172_8446622_RNA45SN1_45S_50ntFlanks | 6651 | 6690 | 40 | rRF 45S | NO | 134.77 | 45.61 | 107.38 | 186.87 |
| >hg38_21_+_8433172_8446622_RNA45SN1_45S_50ntFlanks@6651.6691.41\|CGACTCTTAGCGGTGGATCACTCGGCTCGTGCGTCGATGAA | >hg38_21_+_8433172_8446622_RNA45SN1_45S_50ntFlanks | 6651 | 6691 | 41 | rRF 45S | NO | 33.62 | 0.00 | 0.00 | 28.47 |
| >hg38_21_+_8433172_8446622_RNA45SN1_45S_50ntFlanks@6651.6692.42\|CGACTCTTAGCGGTGGATCACTCGGCTCGTGCGTCGATGAAG | >hg38_21_+_8433172_8446622_RNA45SN1_45S_50ntFlanks | 6651 | 6692 | 42 | rRF 45S | NO | 188.23 | 137.62 | 98.86 | 286.99 |
| >hg38_21_+_8433172_8446622_RNA45SN1_45S_50ntFlanks@6651.6693.43\|CGACTCTTAGCGGTGGATCACTCGGCTCGTGCGTCGATGAAGA | >hg38_21_+_8433172_8446622_RNA45SN1_45S_50ntFlanks | 6651 | 6693 | 43 | rRF 45S | NO | 212.23 | 106.00 | 73.33 | 449.11 |
| >hg38_21_+_8433172_8446622_RNA45SN1_45S_50ntFlanks@6651.6694.44\|CGACTCTTAGCGGTGGATCACTCGGCTCGTGCGTCGATGAAGAA | >hg38_21_+_8433172_8446622_RNA45SN1_45S_50ntFlanks | 6651 | 6694 | 44 | rRF 45S | NO | 121.52 | 30.44 | 10.65 | 165.18 |
| >hg38_21_+_8433172_8446622_RNA45SN1_45S_50ntFlanks@6651.6695.45\|CGACTCTTAGCGGTGGATCACTCGGCTCGTGCGTCGATGAAGAAC | >hg38_21_+_8433172_8446622_RNA45SN1_45S_50ntFlanks | 6651 | 6695 | 45 | rRF 45S | NO | 246.34 | 123.10 | 149.47 | 577.28 |
| >hg38_21_+_8433172_8446622_RNA45SN1_45S_50ntFlanks@6651.6696.46\|CGACTCTTAGCGGTGGATCACTCGGCTCGTGCGTCGATGAAGAACG | >hg38_21_+_8433172_8446622_RNA45SN1_45S_50ntFlanks | 6651 | 6696 | 46 | rRF 45S | NO | 271.03 | 44.44 | 110.68 | 476.23 |
| >hg38_21_+_8433172_8446622_RNA45SN1_45S_50ntFlanks@6651.6697.47\|CGACTCTTAGCGGTGGATCACTCGGCTCGTGCGTCGATGAAGAACGC | >hg38_21_+_8433172_8446622_RNA45SN1_45S_50ntFlanks | 6651 | 6697 | 47 | rRF 45S | NO | 95.13 | 26.07 | 86.37 | 303.38 |
| >hg38_21_+_8433172_8446622_RNA45SN1_45S_50ntFlanks@6651.6698.48\|CGACTCTTAGCGGTGGATCACTCGGCTCGTGCGTCGATGAAGAACGCA | >hg38_21_+_8433172_8446622_RNA45SN1_45S_50ntFlanks | 6651 | 6698 | 48 | rRF 45S | NO | 105.19 | 10.32 | 125.67 | 386.62 |
| >hg38_21_+_8433172_8446622_RNA45SN1_45S_50ntFlanks@6651.6699.49\|CGACTCTTAGCGGTGGATCACTCGGCTCGTGCGTCGATGAAGAACGCAG | >hg38_21_+_8433172_8446622_RNA45SN1_45S_50ntFlanks | 6651 | 6699 | 49 | rRF 45S | NO | 143.83 | 0.00 | 55.90 | 227.88 |
| >hg38_21_+_8433172_8446622_RNA45SN1_45S_50ntFlanks@6651.6700.50\|CGACTCTTAGCGGTGGATCACTCGGCTCGTGCGTCGATGAAGAACGCAGC | >hg38_21_+_8433172_8446622_RNA45SN1_45S_50ntFlanks | 6651 | 6700 | 50 | rRF 45S | NO | 79.67 | 0.00 | 176.75 | 416.15 |
| >hg38_21_+_8433172_8446622_RNA45SN1_45S_50ntFlanks@6652.6669.18\|GACTCTTAGCGGTGGATC | >hg38_21_+_8433172_8446622_RNA45SN1_45S_50ntFlanks | 6652 | 6669 | 18 | rRF 45S | NO | 10.72 | 26.47 | 16.33 | 33.44 |
| >hg38_21_+_8433172_8446622_RNA45SN1_45S_50ntFlanks@6652.6676.25\|GACTCTTAGCGGTGGATCACTCGGC | >hg38_21_+_8433172_8446622_RNA45SN1_45S_50ntFlanks | 6652 | 6676 | 25 | rRF 45S | NO | 40.61 | 136.84 | 222.05 | 175.74 |
| >hg38_21_+_8433172_8446622_RNA45SN1_45S_50ntFlanks@6652.6677.26\|GACTCTTAGCGGTGGATCACTCGGCT | >hg38_21_+_8433172_8446622_RNA45SN1_45S_50ntFlanks | 6652 | 6677 | 26 | rRF 45S | NO | 42.33 | 33.46 | 266.97 | 118.64 |
| >hg38_21_+_8433172_8446622_RNA45SN1_45S_50ntFlanks@6652.6678.27\|GACTCTTAGCGGTGGATCACTCGGCTC | >hg38_21_+_8433172_8446622_RNA45SN1_45S_50ntFlanks | 6652 | 6678 | 27 | rRF 45S | NO | 99.71 | 80.29 | 249.68 | 286.15 |
| >hg38_21_+_8433172_8446622_RNA45SN1_45S_50ntFlanks@6652.6679.28\|GACTCTTAGCGGTGGATCACTCGGCTCG | >hg38_21_+_8433172_8446622_RNA45SN1_45S_50ntFlanks | 6652 | 6679 | 28 | rRF 45S | NO | 27.85 | 69.38 | 71.97 | 150.36 |
| >hg38_21_+_8433172_8446622_RNA45SN1_45S_50ntFlanks@6652.6680.29\|GACTCTTAGCGGTGGATCACTCGGCTCGT | >hg38_21_+_8433172_8446622_RNA45SN1_45S_50ntFlanks | 6652 | 6680 | 29 | rRF 45S | NO | 0.00 | 0.00 | 76.72 | 25.77 |
| >hg38_21_+_8433172_8446622_RNA45SN1_45S_50ntFlanks@6652.6681.30\|GACTCTTAGCGGTGGATCACTCGGCTCGTG | >hg38_21_+_8433172_8446622_RNA45SN1_45S_50ntFlanks | 6652 | 6681 | 30 | rRF 45S | NO | 111.23 | 24.11 | 195.22 | 115.14 |
| >hg38_21_+_8433172_8446622_RNA45SN1_45S_50ntFlanks@6652.6682.31\|GACTCTTAGCGGTGGATCACTCGGCTCGTGC | >hg38_21_+_8433172_8446622_RNA45SN1_45S_50ntFlanks | 6652 | 6682 | 31 | rRF 45S | NO | 33.06 | 0.00 | 90.11 | 94.65 |
| >hg38_21_+_8433172_8446622_RNA45SN1_45S_50ntFlanks@6652.6683.32\|GACTCTTAGCGGTGGATCACTCGGCTCGTGCG | >hg38_21_+_8433172_8446622_RNA45SN1_45S_50ntFlanks | 6652 | 6683 | 32 | rRF 45S | NO | 154.34 | 0.00 | 11.20 | 72.44 |
| >hg38_21_+_8433172_8446622_RNA45SN1_45S_50ntFlanks@6652.6684.33\|GACTCTTAGCGGTGGATCACTCGGCTCGTGCGT | >hg38_21_+_8433172_8446622_RNA45SN1_45S_50ntFlanks | 6652 | 6684 | 33 | rRF 45S | NO | 72.89 | 29.74 | 52.35 | 67.55 |
| >hg38_21_+_8433172_8446622_RNA45SN1_45S_50ntFlanks@6652.6686.35\|GACTCTTAGCGGTGGATCACTCGGCTCGTGCGTCG | >hg38_21_+_8433172_8446622_RNA45SN1_45S_50ntFlanks | 6652 | 6686 | 35 | rRF 45S | NO | 70.81 | 0.00 | 16.08 | 66.90 |
| >hg38_21_+_8433172_8446622_RNA45SN1_45S_50ntFlanks@6652.6687.36\|GACTCTTAGCGGTGGATCACTCGGCTCGTGCGTCGA | >hg38_21_+_8433172_8446622_RNA45SN1_45S_50ntFlanks | 6652 | 6687 | 36 | rRF 45S | NO | 115.36 | 0.00 | 13.63 | 39.02 |
| >hg38_21_+_8433172_8446622_RNA45SN1_45S_50ntFlanks@6652.6688.37\|GACTCTTAGCGGTGGATCACTCGGCTCGTGCGTCGAT | >hg38_21_+_8433172_8446622_RNA45SN1_45S_50ntFlanks | 6652 | 6688 | 37 | rRF 45S | NO | 198.68 | 37.70 | 138.82 | 259.32 |
| >hg38_21_+_8433172_8446622_RNA45SN1_45S_50ntFlanks@6652.6689.38\|GACTCTTAGCGGTGGATCACTCGGCTCGTGCGTCGATG | >hg38_21_+_8433172_8446622_RNA45SN1_45S_50ntFlanks | 6652 | 6689 | 38 | rRF 45S | NO | 205.99 | 89.82 | 429.34 | 387.16 |
| >hg38_21_+_8433172_8446622_RNA45SN1_45S_50ntFlanks@6652.6690.39\|GACTCTTAGCGGTGGATCACTCGGCTCGTGCGTCGATGA | >hg38_21_+_8433172_8446622_RNA45SN1_45S_50ntFlanks | 6652 | 6690 | 39 | rRF 45S | NO | 37.63 | 0.00 | 110.38 | 72.37 |
| >hg38_21_+_8433172_8446622_RNA45SN1_45S_50ntFlanks@6652.6692.41\|GACTCTTAGCGGTGGATCACTCGGCTCGTGCGTCGATGAAG | >hg38_21_+_8433172_8446622_RNA45SN1_45S_50ntFlanks | 6652 | 6692 | 41 | rRF 45S | NO | 42.46 | 31.59 | 84.72 | 130.73 |
| >hg38_21_+_8433172_8446622_RNA45SN1_45S_50ntFlanks@6652.6693.42\|GACTCTTAGCGGTGGATCACTCGGCTCGTGCGTCGATGAAGA | >hg38_21_+_8433172_8446622_RNA45SN1_45S_50ntFlanks | 6652 | 6693 | 42 | rRF 45S | NO | 42.31 | 0.00 | 38.82 | 141.52 |
| >hg38_21_+_8433172_8446622_RNA45SN1_45S_50ntFlanks@6652.6694.43\|GACTCTTAGCGGTGGATCACTCGGCTCGTGCGTCGATGAAGAA | >hg38_21_+_8433172_8446622_RNA45SN1_45S_50ntFlanks | 6652 | 6694 | 43 | rRF 45S | NO | 26.09 | 0.00 | 0.00 | 70.97 |
| >hg38_21_+_8433172_8446622_RNA45SN1_45S_50ntFlanks@6652.6695.44\|GACTCTTAGCGGTGGATCACTCGGCTCGTGCGTCGATGAAGAAC | >hg38_21_+_8433172_8446622_RNA45SN1_45S_50ntFlanks | 6652 | 6695 | 44 | rRF 45S | NO | 84.97 | 28.21 | 161.23 | 228.29 |
| >hg38_21_+_8433172_8446622_RNA45SN1_45S_50ntFlanks@6652.6696.45\|GACTCTTAGCGGTGGATCACTCGGCTCGTGCGTCGATGAAGAACG | >hg38_21_+_8433172_8446622_RNA45SN1_45S_50ntFlanks | 6652 | 6696 | 45 | rRF 45S | NO | 74.04 | 0.00 | 79.25 | 118.63 |

# BT-20

| | | | | | | | | | | | |
|---|---|---|---|---|---|---|---|---|---|---|---|
| >hg38_21_+_8433172_8446622_RNA45SN1_45S_50ntFlanks@6652.6697.46\|GACTCTTAGCGG TGGATCACTCGGCTCGTGCGTCGATGAAGAACGC | >hg38_21_+_8433172_8446622_RNA45SN1_45S_50ntFlanks | 6652 | 6697 | 46 | | rRF 45S | NO | 10.56 | 0.00 | 97.94 | 105.75 |
| >hg38_21_+_8433172_8446622_RNA45SN1_45S_50ntFlanks@6652.6698.47\|GACTCTTAGCGG TGGATCACTCGGCTCGTGCGTCGATGAAGAACGCA | >hg38_21_+_8433172_8446622_RNA45SN1_45S_50ntFlanks | 6652 | 6698 | 47 | | rRF 45S | NO | 68.21 | 0.00 | 241.67 | 201.44 |
| >hg38_21_+_8433172_8446622_RNA45SN1_45S_50ntFlanks@6652.6699.48\|GACTCTTAGCGG TGGATCACTCGGCTCGTGCGTCGATGAAGAACGCAG | >hg38_21_+_8433172_8446622_RNA45SN1_45S_50ntFlanks | 6652 | 6699 | 48 | | rRF 45S | NO | 33.11 | 0.00 | 16.00 | 40.97 |
| >hg38_21_+_8433172_8446622_RNA45SN1_45S_50ntFlanks@6652.6700.49\|GACTCTTAGCGG TGGATCACTCGGCTCGTGCGTCGATGAAGAACGCAGC | >hg38_21_+_8433172_8446622_RNA45SN1_45S_50ntFlanks | 6652 | 6700 | 49 | | rRF 45S | NO | 9.77 | 0.00 | 212.86 | 119.58 |
| >hg38_21_+_8433172_8446622_RNA45SN1_45S_50ntFlanks@6652.6701.50\|GACTCTTAGCGG TGGATCACTCGGCTCGTGCGTCGATGAAGAACGCAGCT | >hg38_21_+_8433172_8446622_RNA45SN1_45S_50ntFlanks | 6652 | 6701 | 50 | | rRF 45S | NO | 0.00 | 0.00 | 70.94 | 111.63 |
| >hg38_21_+_8433172_8446622_RNA45SN1_45S_50ntFlanks@6653.6676.24\|ACTCTTAGCGGT GGATCACTCGGC | >hg38_21_+_8433172_8446622_RNA45SN1_45S_50ntFlanks | 6653 | 6676 | 24 | | rRF 45S | NO | 0.00 | 23.74 | 0.00 | 9.54 |
| >hg38_21_+_8433172_8446622_RNA45SN1_45S_50ntFlanks@6653.6681.29\|ACTCTTAGCGGT GGATCACTCGGCTCGTG | >hg38_21_+_8433172_8446622_RNA45SN1_45S_50ntFlanks | 6653 | 6681 | 29 | | rRF 45S | NO | 37.31 | 0.00 | 9.61 | 0.00 |
| >hg38_21_+_8433172_8446622_RNA45SN1_45S_50ntFlanks@6653.6683.31\|ACTCTTAGCGGT GGATCACTCGGCTCGTGCG | >hg38_21_+_8433172_8446622_RNA45SN1_45S_50ntFlanks | 6653 | 6683 | 31 | | rRF 45S | NO | 41.73 | 0.00 | 0.00 | 0.00 |
| >hg38_21_+_8433172_8446622_RNA45SN1_45S_50ntFlanks@6653.6688.36\|ACTCTTAGCGGT GGATCACTCGGCTCGTGCGTCGAT | >hg38_21_+_8433172_8446622_RNA45SN1_45S_50ntFlanks | 6653 | 6688 | 36 | | rRF 45S | NO | 35.16 | 0.00 | 0.00 | 119.27 |
| >hg38_21_+_8433172_8446622_RNA45SN1_45S_50ntFlanks@6653.6689.37\|ACTCTTAGCGGT GGATCACTCGGCTCGTGCGTCGATG | >hg38_21_+_8433172_8446622_RNA45SN1_45S_50ntFlanks | 6653 | 6689 | 37 | | rRF 45S | NO | 25.17 | 0.00 | 54.49 | 71.03 |
| >hg38_21_+_8433172_8446622_RNA45SN1_45S_50ntFlanks@6655.6698.44\|TCTTAGCGGTGG ATCACTCGGCTCGTGCGTCGATGAAGAACGCA | >hg38_21_+_8433172_8446622_RNA45SN1_45S_50ntFlanks | 6655 | 6698 | 44 | | rRF 45S | NO | 0.00 | 0.00 | 13.18 | 0.00 |
| >hg38_21_+_8433172_8446622_RNA45SN1_45S_50ntFlanks@6660.6688.29\|GCGGTGGATCA CTCGGCTCGTGCGTCGAT | >hg38_21_+_8433172_8446622_RNA45SN1_45S_50ntFlanks | 6660 | 6688 | 29 | | rRF 45S | NO | 0.00 | 0.00 | 10.99 | 0.00 |
| >hg38_21_+_8433172_8446622_RNA45SN1_45S_50ntFlanks@6660.6689.30\|GCGGTGGATCA CTCGGCTCGTGCGTCGATG | >hg38_21_+_8433172_8446622_RNA45SN1_45S_50ntFlanks | 6660 | 6689 | 30 | | rRF 45S | NO | 0.00 | 0.00 | 57.63 | 0.00 |
| >hg38_21_+_8433172_8446622_RNA45SN1_45S_50ntFlanks@6661.6678.18\|CGGTGGATCACT CGGCTC | >hg38_21_+_8433172_8446622_RNA45SN1_45S_50ntFlanks | 6661 | 6678 | 18 | | rRF 45S | NO | 37.10 | 40.40 | 148.01 | 83.82 |
| >hg38_21_+_8433172_8446622_RNA45SN1_45S_50ntFlanks@6661.6679.19\|CGGTGGATCACT CGGCTCG | >hg38_21_+_8433172_8446622_RNA45SN1_45S_50ntFlanks | 6661 | 6679 | 19 | | rRF 45S | NO | 0.00 | 29.56 | 13.09 | 0.00 |
| >hg38_21_+_8433172_8446622_RNA45SN1_45S_50ntFlanks@6661.6680.20\|CGGTGGATCACT CGGCTCGT | >hg38_21_+_8433172_8446622_RNA45SN1_45S_50ntFlanks | 6661 | 6680 | 20 | | rRF 45S | NO | 0.00 | 0.00 | 66.18 | 24.12 |
| >hg38_21_+_8433172_8446622_RNA45SN1_45S_50ntFlanks@6661.6681.21\|CGGTGGATCACT CGGCTCGTG | >hg38_21_+_8433172_8446622_RNA45SN1_45S_50ntFlanks | 6661 | 6681 | 21 | | rRF 45S | NO | 33.32 | 25.54 | 110.95 | 40.09 |
| >hg38_21_+_8433172_8446622_RNA45SN1_45S_50ntFlanks@6661.6682.22\|CGGTGGATCACT CGGCTCGTGC | >hg38_21_+_8433172_8446622_RNA45SN1_45S_50ntFlanks | 6661 | 6682 | 22 | | rRF 45S | NO | 0.00 | 0.00 | 14.49 | 0.00 |
| >hg38_21_+_8433172_8446622_RNA45SN1_45S_50ntFlanks@6661.6683.23\|CGGTGGATCACT CGGCTCGTGCG | >hg38_21_+_8433172_8446622_RNA45SN1_45S_50ntFlanks | 6661 | 6683 | 23 | | rRF 45S | NO | 117.10 | 0.00 | 10.63 | 30.92 |
| >hg38_21_+_8433172_8446622_RNA45SN1_45S_50ntFlanks@6661.6684.24\|CGGTGGATCACT CGGCTCGTGCGT | >hg38_21_+_8433172_8446622_RNA45SN1_45S_50ntFlanks | 6661 | 6684 | 24 | | rRF 45S | NO | 25.11 | 0.00 | 15.92 | 10.37 |
| >hg38_21_+_8433172_8446622_RNA45SN1_45S_50ntFlanks@6661.6685.25\|CGGTGGATCACT CGGCTCGTGCGTC | >hg38_21_+_8433172_8446622_RNA45SN1_45S_50ntFlanks | 6661 | 6685 | 25 | | rRF 45S | NO | 70.92 | 26.06 | 543.36 | 176.22 |
| >hg38_21_+_8433172_8446622_RNA45SN1_45S_50ntFlanks@6661.6686.26\|CGGTGGATCACT CGGCTCGTGCGTCG | >hg38_21_+_8433172_8446622_RNA45SN1_45S_50ntFlanks | 6661 | 6686 | 26 | | rRF 45S | NO | 43.42 | 0.00 | 57.89 | 87.85 |
| >hg38_21_+_8433172_8446622_RNA45SN1_45S_50ntFlanks@6661.6687.27\|CGGTGGATCACT CGGCTCGTGCGTCGA | >hg38_21_+_8433172_8446622_RNA45SN1_45S_50ntFlanks | 6661 | 6687 | 27 | | rRF 45S | NO | 90.58 | 0.00 | 63.53 | 78.49 |
| >hg38_21_+_8433172_8446622_RNA45SN1_45S_50ntFlanks@6661.6688.28\|CGGTGGATCACT CGGCTCGTGCGTCGAT | >hg38_21_+_8433172_8446622_RNA45SN1_45S_50ntFlanks | 6661 | 6688 | 28 | | rRF 45S | NO | 149.81 | 32.54 | 227.89 | 310.94 |
| >hg38_21_+_8433172_8446622_RNA45SN1_45S_50ntFlanks@6661.6689.29\|CGGTGGATCACT CGGCTCGTGCGTCGATG | >hg38_21_+_8433172_8446622_RNA45SN1_45S_50ntFlanks | 6661 | 6689 | 29 | | rRF 45S | NO | 164.86 | 91.27 | 582.03 | 406.89 |
| >hg38_21_+_8433172_8446622_RNA45SN1_45S_50ntFlanks@6661.6690.30\|CGGTGGATCACT CGGCTCGTGCGTCGATGA | >hg38_21_+_8433172_8446622_RNA45SN1_45S_50ntFlanks | 6661 | 6690 | 30 | | rRF 45S | NO | 0.00 | 0.00 | 72.93 | 0.00 |
| >hg38_21_+_8433172_8446622_RNA45SN1_45S_50ntFlanks@6661.6692.32\|CGGTGGATCACT CGGCTCGTGCGTCGATGAAG | >hg38_21_+_8433172_8446622_RNA45SN1_45S_50ntFlanks | 6661 | 6692 | 32 | | rRF 45S | NO | 0.00 | 0.00 | 39.52 | 23.28 |
| >hg38_21_+_8433172_8446622_RNA45SN1_45S_50ntFlanks@6661.6693.33\|CGGTGGATCACT CGGCTCGTGCGTCGATGAAGA | >hg38_21_+_8433172_8446622_RNA45SN1_45S_50ntFlanks | 6661 | 6693 | 33 | | rRF 45S | NO | 24.33 | 0.00 | 82.09 | 32.31 |
| >hg38_21_+_8433172_8446622_RNA45SN1_45S_50ntFlanks@6661.6695.35\|CGGTGGATCACT CGGCTCGTGCGTCGATGAAGAAC | >hg38_21_+_8433172_8446622_RNA45SN1_45S_50ntFlanks | 6661 | 6695 | 35 | | rRF 45S | NO | 86.06 | 39.21 | 231.93 | 133.66 |
| >hg38_21_+_8433172_8446622_RNA45SN1_45S_50ntFlanks@6661.6696.36\|CGGTGGATCACT CGGCTCGTGCGTCGATGAAGAACG | >hg38_21_+_8433172_8446622_RNA45SN1_45S_50ntFlanks | 6661 | 6696 | 36 | | rRF 45S | NO | 36.33 | 0.00 | 110.55 | 29.23 |
| >hg38_21_+_8433172_8446622_RNA45SN1_45S_50ntFlanks@6661.6697.37\|CGGTGGATCACT CGGCTCGTGCGTCGATGAAGAACGC | >hg38_21_+_8433172_8446622_RNA45SN1_45S_50ntFlanks | 6661 | 6697 | 37 | | rRF 45S | NO | 33.23 | 26.85 | 153.75 | 62.74 |
| >hg38_21_+_8433172_8446622_RNA45SN1_45S_50ntFlanks@6661.6698.38\|CGGTGGATCACT CGGCTCGTGCGTCGATGAAGAACGCA | >hg38_21_+_8433172_8446622_RNA45SN1_45S_50ntFlanks | 6661 | 6698 | 38 | | rRF 45S | NO | 10.73 | 0.00 | 132.49 | 24.62 |
| >hg38_21_+_8433172_8446622_RNA45SN1_45S_50ntFlanks@6661.6699.39\|CGGTGGATCACT CGGCTCGTGCGTCGATGAAGAACGCAG | >hg38_21_+_8433172_8446622_RNA45SN1_45S_50ntFlanks | 6661 | 6699 | 39 | | rRF 45S | NO | 82.55 | 10.38 | 221.89 | 32.08 |
| >hg38_21_+_8433172_8446622_RNA45SN1_45S_50ntFlanks@6661.6700.40\|CGGTGGATCACT CGGCTCGTGCGTCGATGAAGAACGCAGC | >hg38_21_+_8433172_8446622_RNA45SN1_45S_50ntFlanks | 6661 | 6700 | 40 | | rRF 45S | NO | 79.21 | 39.76 | 401.74 | 95.90 |
| >hg38_21_+_8433172_8446622_RNA45SN1_45S_50ntFlanks@6661.6701.41\|CGGTGGATCACT CGGCTCGTGCGTCGATGAAGAACGCAGCT | >hg38_21_+_8433172_8446622_RNA45SN1_45S_50ntFlanks | 6661 | 6701 | 41 | | rRF 45S | NO | 80.45 | 43.51 | 121.36 | 169.86 |
| >hg38_21_+_8433172_8446622_RNA45SN1_45S_50ntFlanks@6661.6706.46\|CGGTGGATCACT CGGCTCGTGCGTCGATGAAGAACGCAGCTAGCTG | >hg38_21_+_8433172_8446622_RNA45SN1_45S_50ntFlanks | 6661 | 6706 | 46 | | rRF 45S | NO | 83.93 | 36.37 | 11.92 | 374.72 |
| >hg38_21_+_8433172_8446622_RNA45SN1_45S_50ntFlanks@6661.6707.47\|CGGTGGATCACT CGGCTCGTGCGTCGATGAAGAACGCAGCTAGCTGC | >hg38_21_+_8433172_8446622_RNA45SN1_45S_50ntFlanks | 6661 | 6707 | 47 | | rRF 45S | NO | 39.27 | 0.00 | 15.35 | 411.17 |
| >hg38_21_+_8433172_8446622_RNA45SN1_45S_50ntFlanks@6698.6720.23\|AGCTAGCTGCG AGAATTAATGTG | >hg38_21_+_8433172_8446622_RNA45SN1_45S_50ntFlanks | 6698 | 6720 | 23 | | rRF 45S | NO | 0.00 | 29.83 | 92.73 | 0.00 |

# BT-20

| | | | | | | | | | | |
|---|---|---|---|---|---|---|---|---|---|---|
| >hg38_21_+_8433172_8446622_RNA45SN1_45S_50ntFlanks@6699.6720.22\|GCTAGCTGCGAGAATTAATGTG | >hg38_21_+_8433172_8446622_RNA45SN1_45S_50ntFlanks | 6699 | 6720 | 22 | rRF 45S | NO | 0.00 | 32.39 | 107.52 | 0.00 |
| >hg38_21_+_8433172_8446622_RNA45SN1_45S_50ntFlanks@6725.6747.23\|GCAGGACACATTGATCATCGACA | >hg38_21_+_8433172_8446622_RNA45SN1_45S_50ntFlanks | 6725 | 6747 | 23 | rRF 45S | NO | 0.00 | 23.86 | 0.00 | 0.00 |
| >hg38_21_+_8433172_8446622_RNA45SN1_45S_50ntFlanks@6732.6773.42\|ACATTGATCATCGACACTTCGAACGCACTTGCGGCCCCGGGT | >hg38_21_+_8433172_8446622_RNA45SN1_45S_50ntFlanks | 6732 | 6773 | 42 | rRF 45S | NO | 24.41 | 25.85 | 0.00 | 111.47 |
| >hg38_21_+_8433172_8446622_RNA45SN1_45S_50ntFlanks@6734.6773.40\|ATTGATCATCGACACTTCGAACGCACTTGCGGCCCCGGGT | >hg38_21_+_8433172_8446622_RNA45SN1_45S_50ntFlanks | 6734 | 6773 | 40 | rRF 45S | NO | 33.44 | 32.50 | 0.00 | 137.29 |
| >hg38_21_+_8433172_8446622_RNA45SN1_45S_50ntFlanks@6734.6774.41\|ATTGATCATCGACACTTCGAACGCACTTGCGGCCCCGGGTT | >hg38_21_+_8433172_8446622_RNA45SN1_45S_50ntFlanks | 6734 | 6774 | 41 | rRF 45S | NO | 25.11 | 9.57 | 0.00 | 101.47 |
| >hg38_21_+_8433172_8446622_RNA45SN1_45S_50ntFlanks@6773.6806.34\|TTCCTCCCGGGGCTACGCCTGTCTGAGCGTCGCT | >hg38_21_+_8433172_8446622_RNA45SN1_45S_50ntFlanks | 6773 | 6806 | 34 | rRF 45S | NO | 28.75 | 35.59 | 0.00 | 94.88 |
| >hg38_21_+_8433172_8446622_RNA45SN1_45S_50ntFlanks@6774.6806.33\|TCCTCCCGGGGCTACGCCTGTCTGAGCGTCGCT | >hg38_21_+_8433172_8446622_RNA45SN1_45S_50ntFlanks | 6774 | 6806 | 33 | rRF 45S | NO | 128.23 | 156.31 | 14.10 | 232.93 |
| >hg38_21_+_8433172_8446622_RNA45SN1_45S_50ntFlanks@6775.6806.32\|CCTCCCGGGGCTACGCCTGTCTGAGCGTCGCT | >hg38_21_+_8433172_8446622_RNA45SN1_45S_50ntFlanks | 6775 | 6806 | 32 | rRF 45S | NO | 10.74 | 26.49 | 0.00 | 37.76 |
| >hg38_21_+_8433172_8446622_RNA45SN1_45S_50ntFlanks@6776.6806.31\|CTCCCGGGGCTACGCCTGTCTGAGCGTCGCT | >hg38_21_+_8433172_8446622_RNA45SN1_45S_50ntFlanks | 6776 | 6806 | 31 | rRF 45S | NO | 66.37 | 91.04 | 14.49 | 194.80 |
| >hg38_21_+_8433172_8446622_RNA45SN1_45S_50ntFlanks@6778.6802.25\|CCCGGGGCTACGCCTGTCTGAGCGT | >hg38_21_+_8433172_8446622_RNA45SN1_45S_50ntFlanks | 6778 | 6802 | 25 | rRF 45S | NO | 27.75 | 34.18 | 0.00 | 93.20 |
| >hg38_21_+_8433172_8446622_RNA45SN1_45S_50ntFlanks@6778.6806.29\|CCCGGGGCTACGCCTGTCTGAGCGTCGCT | >hg38_21_+_8433172_8446622_RNA45SN1_45S_50ntFlanks | 6778 | 6806 | 29 | rRF 45S | NO | 36.72 | 71.37 | 14.24 | 176.74 |
| >hg38_21_+_8433172_8446622_RNA45SN1_45S_50ntFlanks@6779.6806.28\|CCGGGGCTACGCCTGTCTGAGCGTCGCT | >hg38_21_+_8433172_8446622_RNA45SN1_45S_50ntFlanks | 6779 | 6806 | 28 | rRF 45S | NO | 0.00 | 26.96 | 38.13 | 76.41 |
| >hg38_21_+_8433172_8446622_RNA45SN1_45S_50ntFlanks@6780.6802.23\|CGGGGCTACGCCTGTCTGAGCGT | >hg38_21_+_8433172_8446622_RNA45SN1_45S_50ntFlanks | 6780 | 6802 | 23 | rRF 45S | NO | 0.00 | 0.00 | 39.59 | 0.00 |
| >hg38_21_+_8433172_8446622_RNA45SN1_45S_50ntFlanks@6780.6806.27\|CGGGGCTACGCCTGTCTGAGCGTCGCT | >hg38_21_+_8433172_8446622_RNA45SN1_45S_50ntFlanks | 6780 | 6806 | 27 | rRF 45S | NO | 0.00 | 0.00 | 12.64 | 28.81 |
| >hg38_21_+_8433172_8446622_RNA45SN1_45S_50ntFlanks@6785.6806.22\|CTACGCCTGTCTGAGCGTCGCT | >hg38_21_+_8433172_8446622_RNA45SN1_45S_50ntFlanks | 6785 | 6806 | 22 | rRF 45S | NO | 67.42 | 107.82 | 87.50 | 133.67 |
| >hg38_21_+_8433172_8446622_RNA45SN1_45S_50ntFlanks@6786.6806.21\|TACGCCTGTCTGAGCGTCGCT | >hg38_21_+_8433172_8446622_RNA45SN1_45S_50ntFlanks | 6786 | 6806 | 21 | rRF 45S | NO | 72.10 | 106.16 | 79.44 | 140.42 |
| >hg38_21_+_8433172_8446622_RNA45SN1_45S_50ntFlanks@6787.6806.20\|ACGCCTGTCTGAGCGTCGCT | >hg38_21_+_8433172_8446622_RNA45SN1_45S_50ntFlanks | 6787 | 6806 | 20 | rRF 45S | NO | 42.41 | 104.04 | 86.47 | 110.59 |
| >hg38_21_+_8433172_8446622_RNA45SN1_45S_50ntFlanks@6788.6806.19\|CGCCTGTCTGAGCGTCGCT | >hg38_21_+_8433172_8446622_RNA45SN1_45S_50ntFlanks | 6788 | 6806 | 19 | rRF 45S | NO | 91.88 | 244.91 | 128.65 | 267.52 |
| >hg38_21_+_8433172_8446622_RNA45SN1_45S_50ntFlanks@6788.6807.20\|CGCCTGTCTGAGCGTCGCTT | >hg38_21_+_8433172_8446622_RNA45SN1_45S_50ntFlanks | 6788 | 6807 | 20 | rRF 45S | NO | 24.28 | 66.85 | 13.70 | 104.63 |
| >hg38_21_+_8433172_8446622_RNA45SN1_45S_50ntFlanks@6789.6806.18\|GCCTGTCTGAGCGTCGCT | >hg38_21_+_8433172_8446622_RNA45SN1_45S_50ntFlanks | 6789 | 6806 | 18 | rRF 45S | NO | 37.18 | 96.43 | 15.92 | 167.18 |
| >hg38_21_+_8433172_8446622_RNA45SN1_45S_50ntFlanks@6790.6807.18\|CCTGTCTGAGCGTCGCTT | >hg38_21_+_8433172_8446622_RNA45SN1_45S_50ntFlanks | 6790 | 6807 | 18 | rRF 45S | NO | 39.35 | 76.30 | 14.85 | 183.59 |
| >hg38_21_+_8433172_8446622_RNA45SN1_45S_50ntFlanks@7968.7988.21\|TCCGAGACGCGACCTCAGATC | >hg38_21_+_8433172_8446622_RNA45SN1_45S_50ntFlanks | 7968 | 7988 | 21 | rRF 45S | NO | 92.57 | 0.00 | 0.00 | 0.00 |
| >hg38_21_+_8433172_8446622_RNA45SN1_45S_50ntFlanks@7968.7993.26\|TCCGAGACGCGACCTCAGATCAGACG | >hg38_21_+_8433172_8446622_RNA45SN1_45S_50ntFlanks | 7968 | 7993 | 26 | rRF 45S | NO | 11.14 | 0.00 | 0.00 | 0.00 |
| >hg38_21_+_8433172_8446622_RNA45SN1_45S_50ntFlanks@7968.7994.27\|TCCGAGACGCGACCTCAGATCAGACGT | >hg38_21_+_8433172_8446622_RNA45SN1_45S_50ntFlanks | 7968 | 7994 | 27 | rRF 45S | NO | 71.31 | 0.00 | 0.00 | 0.00 |
| >hg38_21_+_8433172_8446622_RNA45SN1_45S_50ntFlanks@7968.8010.43\|TCCGAGACGCGACCTCAGATCAGACGTGGCGACCCGCTGAATT | >hg38_21_+_8433172_8446622_RNA45SN1_45S_50ntFlanks | 7968 | 8010 | 43 | rRF 45S | NO | 77.18 | 0.00 | 0.00 | 0.00 |
| >hg38_21_+_8433172_8446622_RNA45SN1_45S_50ntFlanks@7971.7988.18\|GAGACGCGACCTCAGATC | >hg38_21_+_8433172_8446622_RNA45SN1_45S_50ntFlanks | 7971 | 7988 | 18 | rRF 45S | NO | 81.98 | 0.00 | 73.43 | 0.00 |
| >hg38_21_+_8433172_8446622_RNA45SN1_45S_50ntFlanks@7973.7991.19\|GACGCGACCTCAGATCAGA | >hg38_21_+_8433172_8446622_RNA45SN1_45S_50ntFlanks | 7973 | 7991 | 19 | rRF 45S | NO | 80.46 | 0.00 | 16.33 | 101.21 |
| >hg38_21_+_8433172_8446622_RNA45SN1_45S_50ntFlanks@7973.7992.20\|GACGCGACCTCAGATCAGAC | >hg38_21_+_8433172_8446622_RNA45SN1_45S_50ntFlanks | 7973 | 7992 | 20 | rRF 45S | NO | 115.93 | 0.00 | 62.55 | 137.26 |
| >hg38_21_+_8433172_8446622_RNA45SN1_45S_50ntFlanks@7973.7993.21\|GACGCGACCTCAGATCAGACG | >hg38_21_+_8433172_8446622_RNA45SN1_45S_50ntFlanks | 7973 | 7993 | 21 | rRF 45S | NO | 114.60 | 0.00 | 112.41 | 117.75 |
| >hg38_21_+_8433172_8446622_RNA45SN1_45S_50ntFlanks@7973.7994.22\|GACGCGACCTCAGATCAGACGT | >hg38_21_+_8433172_8446622_RNA45SN1_45S_50ntFlanks | 7973 | 7994 | 22 | rRF 45S | NO | 93.76 | 0.00 | 63.57 | 26.77 |
| >hg38_21_+_8433172_8446622_RNA45SN1_45S_50ntFlanks@7973.7995.23\|GACGCGACCTCAGATCAGACGTG | >hg38_21_+_8433172_8446622_RNA45SN1_45S_50ntFlanks | 7973 | 7995 | 23 | rRF 45S | NO | 41.61 | 0.00 | 9.61 | 28.51 |
| >hg38_21_+_8433172_8446622_RNA45SN1_45S_50ntFlanks@7973.7997.25\|GACGCGACCTCAGATCAGACGTGGC | >hg38_21_+_8433172_8446622_RNA45SN1_45S_50ntFlanks | 7973 | 7997 | 25 | rRF 45S | NO | 73.54 | 0.00 | 0.00 | 26.00 |
| >hg38_21_+_8433172_8446622_RNA45SN1_45S_50ntFlanks@7973.8005.33\|GACGCGACCTCAGATCAGACGTGGCGACCCGCT | >hg38_21_+_8433172_8446622_RNA45SN1_45S_50ntFlanks | 7973 | 8005 | 33 | rRF 45S | NO | 347.25 | 0.00 | 76.11 | 188.63 |
| >hg38_21_+_8433172_8446622_RNA45SN1_45S_50ntFlanks@7973.8006.34\|GACGCGACCTCAGATCAGACGTGGCGACCCGCTG | >hg38_21_+_8433172_8446622_RNA45SN1_45S_50ntFlanks | 7973 | 8006 | 34 | rRF 45S | NO | 111.24 | 0.00 | 0.00 | 40.72 |
| >hg38_21_+_8433172_8446622_RNA45SN1_45S_50ntFlanks@7973.8007.35\|GACGCGACCTCAGATCAGACGTGGCGACCCGCTGA | >hg38_21_+_8433172_8446622_RNA45SN1_45S_50ntFlanks | 7973 | 8007 | 35 | rRF 45S | NO | 67.67 | 0.00 | 0.00 | 25.08 |
| >hg38_21_+_8433172_8446622_RNA45SN1_45S_50ntFlanks@7973.8008.36\|GACGCGACCTCAGATCAGACGTGGCGACCCGCTGAA | >hg38_21_+_8433172_8446622_RNA45SN1_45S_50ntFlanks | 7973 | 8008 | 36 | rRF 45S | NO | 112.82 | 31.30 | 14.69 | 173.08 |
| >hg38_21_+_8433172_8446622_RNA45SN1_45S_50ntFlanks@7973.8009.37\|GACGCGACCTCAGATCAGACGTGGCGACCCGCTGAAT | >hg38_21_+_8433172_8446622_RNA45SN1_45S_50ntFlanks | 7973 | 8009 | 37 | rRF 45S | NO | 292.09 | 106.13 | 113.64 | 261.39 |
| >hg38_21_+_8433172_8446622_RNA45SN1_45S_50ntFlanks@7973.8010.38\|GACGCGACCTCAGATCAGACGTGGCGACCCGCTGAATT | >hg38_21_+_8433172_8446622_RNA45SN1_45S_50ntFlanks | 7973 | 8010 | 38 | rRF 45S | NO | 449.69 | 71.53 | 12.00 | 245.32 |

# BT-20

| | | | | | | | | | | |
|---|---|---|---|---|---|---|---|---|---|---|
| >hg38_21_+_8433172_8446622_RNA45SN1_45S_50ntFlanks@7973.8011.39\|GACGCGACCTCAGATCAGACGTGGCGACCCGCTGAATTT | >hg38_21_+_8433172_8446622_RNA45SN1_45S_50ntFlanks | 7973 | 8011 | 39 | rRF 45S | NO | 383.75 | 23.86 | 0.00 | 216.84 |
| >hg38_21_+_8433172_8446622_RNA45SN1_45S_50ntFlanks@7973.8012.40\|GACGCGACCTCAGATCAGACGTGGCGACCCGCTGAATTTA | >hg38_21_+_8433172_8446622_RNA45SN1_45S_50ntFlanks | 7973 | 8012 | 40 | rRF 45S | NO | 24.02 | 0.00 | 0.00 | 0.00 |
| >hg38_21_+_8433172_8446622_RNA45SN1_45S_50ntFlanks@7974.7991.18\|ACGCGACCTCAGATCAGA | >hg38_21_+_8433172_8446622_RNA45SN1_45S_50ntFlanks | 7974 | 7991 | 18 | rRF 45S | NO | 177.86 | 105.90 | 168.18 | 469.50 |
| >hg38_21_+_8433172_8446622_RNA45SN1_45S_50ntFlanks@7974.7992.19\|ACGCGACCTCAGATCAGAC | >hg38_21_+_8433172_8446622_RNA45SN1_45S_50ntFlanks | 7974 | 7992 | 19 | rRF 45S | NO | 198.67 | 31.84 | 193.31 | 515.90 |
| >hg38_21_+_8433172_8446622_RNA45SN1_45S_50ntFlanks@7974.7993.20\|ACGCGACCTCAGATCAGACG | >hg38_21_+_8433172_8446622_RNA45SN1_45S_50ntFlanks | 7974 | 7993 | 20 | rRF 45S | NO | 275.90 | 152.17 | 517.55 | 466.07 |
| >hg38_21_+_8433172_8446622_RNA45SN1_45S_50ntFlanks@7974.7994.21\|ACGCGACCTCAGATCAGACGT | >hg38_21_+_8433172_8446622_RNA45SN1_45S_50ntFlanks | 7974 | 7994 | 21 | rRF 45S | NO | 238.69 | 35.13 | 208.32 | 179.36 |
| >hg38_21_+_8433172_8446622_RNA45SN1_45S_50ntFlanks@7974.7995.22\|ACGCGACCTCAGATCAGACGTG | >hg38_21_+_8433172_8446622_RNA45SN1_45S_50ntFlanks | 7974 | 7995 | 22 | rRF 45S | NO | 160.07 | 126.65 | 155.88 | 225.18 |
| >hg38_21_+_8433172_8446622_RNA45SN1_45S_50ntFlanks@7974.7996.23\|ACGCGACCTCAGATCAGACGTGG | >hg38_21_+_8433172_8446622_RNA45SN1_45S_50ntFlanks | 7974 | 7996 | 23 | rRF 45S | NO | 9.92 | 23.38 | 0.00 | 25.63 |
| >hg38_21_+_8433172_8446622_RNA45SN1_45S_50ntFlanks@7974.7997.24\|ACGCGACCTCAGATCAGACGTGGC | >hg38_21_+_8433172_8446622_RNA45SN1_45S_50ntFlanks | 7974 | 7997 | 24 | rRF 45S | NO | 380.77 | 82.45 | 208.95 | 469.89 |
| >hg38_21_+_8433172_8446622_RNA45SN1_45S_50ntFlanks@7974.8001.28\|ACGCGACCTCAGATCAGACGTGGCGACC | >hg38_21_+_8433172_8446622_RNA45SN1_45S_50ntFlanks | 7974 | 8001 | 28 | rRF 45S | NO | 24.82 | 0.00 | 0.00 | 0.00 |
| >hg38_21_+_8433172_8446622_RNA45SN1_45S_50ntFlanks@7974.8002.29\|ACGCGACCTCAGATCAGACGTGGCGACCC | >hg38_21_+_8433172_8446622_RNA45SN1_45S_50ntFlanks | 7974 | 8002 | 29 | rRF 45S | NO | 90.07 | 0.00 | 121.70 | 29.84 |
| >hg38_21_+_8433172_8446622_RNA45SN1_45S_50ntFlanks@7974.8004.31\|ACGCGACCTCAGATCAGACGTGGCGACCCGC | >hg38_21_+_8433172_8446622_RNA45SN1_45S_50ntFlanks | 7974 | 8004 | 31 | rRF 45S | NO | 203.43 | 90.27 | 123.12 | 202.56 |
| >hg38_21_+_8433172_8446622_RNA45SN1_45S_50ntFlanks@7974.8005.32\|ACGCGACCTCAGATCAGACGTGGCGACCCGCT | >hg38_21_+_8433172_8446622_RNA45SN1_45S_50ntFlanks | 7974 | 8005 | 32 | rRF 45S | NO | 1074.97 | 234.73 | 407.57 | 1199.74 |
| >hg38_21_+_8433172_8446622_RNA45SN1_45S_50ntFlanks@7974.8006.33\|ACGCGACCTCAGATCAGACGTGGCGACCCGCTG | >hg38_21_+_8433172_8446622_RNA45SN1_45S_50ntFlanks | 7974 | 8006 | 33 | rRF 45S | NO | 403.19 | 118.57 | 138.94 | 545.23 |
| >hg38_21_+_8433172_8446622_RNA45SN1_45S_50ntFlanks@7974.8007.34\|ACGCGACCTCAGATCAGACGTGGCGACCCGCTGA | >hg38_21_+_8433172_8446622_RNA45SN1_45S_50ntFlanks | 7974 | 8007 | 34 | rRF 45S | NO | 219.78 | 73.89 | 76.28 | 277.54 |
| >hg38_21_+_8433172_8446622_RNA45SN1_45S_50ntFlanks@7974.8008.35\|ACGCGACCTCAGATCAGACGTGGCGACCCGCTGAA | >hg38_21_+_8433172_8446622_RNA45SN1_45S_50ntFlanks | 7974 | 8008 | 35 | rRF 45S | NO | 411.23 | 520.04 | 211.05 | 981.39 |
| >hg38_21_+_8433172_8446622_RNA45SN1_45S_50ntFlanks@7974.8009.36\|ACGCGACCTCAGATCAGACGTGGCGACCCGCTGAAT | >hg38_21_+_8433172_8446622_RNA45SN1_45S_50ntFlanks | 7974 | 8009 | 36 | rRF 45S | NO | 1082.95 | 1021.97 | 586.13 | 1511.72 |
| >hg38_21_+_8433172_8446622_RNA45SN1_45S_50ntFlanks@7974.8010.37\|ACGCGACCTCAGATCAGACGTGGCGACCCGCTGAATT | >hg38_21_+_8433172_8446622_RNA45SN1_45S_50ntFlanks | 7974 | 8010 | 37 | rRF 45S | NO | 1671.76 | 686.71 | 228.47 | 1483.37 |
| >hg38_21_+_8433172_8446622_RNA45SN1_45S_50ntFlanks@7974.8011.38\|ACGCGACCTCAGATCAGACGTGGCGACCCGCTGAATTT | >hg38_21_+_8433172_8446622_RNA45SN1_45S_50ntFlanks | 7974 | 8011 | 38 | rRF 45S | NO | 1516.37 | 343.98 | 139.34 | 1364.94 |
| >hg38_21_+_8433172_8446622_RNA45SN1_45S_50ntFlanks@7974.8012.39\|ACGCGACCTCAGATCAGACGTGGCGACCCGCTGAATTTA | >hg38_21_+_8433172_8446622_RNA45SN1_45S_50ntFlanks | 7974 | 8012 | 39 | rRF 45S | NO | 116.27 | 27.13 | 0.00 | 172.00 |
| >hg38_21_+_8433172_8446622_RNA45SN1_45S_50ntFlanks@7974.8013.40\|ACGCGACCTCAGATCAGACGTGGCGACCCGCTGAATTTAA | >hg38_21_+_8433172_8446622_RNA45SN1_45S_50ntFlanks | 7974 | 8013 | 40 | rRF 45S | NO | 30.58 | 0.00 | 0.00 | 110.94 |
| >hg38_21_+_8433172_8446622_RNA45SN1_45S_50ntFlanks@7974.8014.41\|ACGCGACCTCAGATCAGACGTGGCGACCCGCTGAATTTAAG | >hg38_21_+_8433172_8446622_RNA45SN1_45S_50ntFlanks | 7974 | 8014 | 41 | rRF 45S | NO | 30.80 | 0.00 | 0.00 | 120.11 |
| >hg38_21_+_8433172_8446622_RNA45SN1_45S_50ntFlanks@7974.8015.42\|ACGCGACCTCAGATCAGACGTGGCGACCCGCTGAATTTAAGC | >hg38_21_+_8433172_8446622_RNA45SN1_45S_50ntFlanks | 7974 | 8015 | 42 | rRF 45S | NO | 129.54 | 0.00 | 10.55 | 278.85 |
| >hg38_21_+_8433172_8446622_RNA45SN1_45S_50ntFlanks@7974.8016.43\|ACGCGACCTCAGATCAGACGTGGCGACCCGCTGAATTTAAGCA | >hg38_21_+_8433172_8446622_RNA45SN1_45S_50ntFlanks | 7974 | 8016 | 43 | rRF 45S | NO | 42.20 | 0.00 | 0.00 | 270.27 |
| >hg38_21_+_8433172_8446622_RNA45SN1_45S_50ntFlanks@7975.7992.18\|CGCGACCTCAGATCAGAC | >hg38_21_+_8433172_8446622_RNA45SN1_45S_50ntFlanks | 7975 | 7992 | 18 | rRF 45S | NO | 851.47 | 367.68 | 931.73 | 2782.23 |
| >hg38_21_+_8433172_8446622_RNA45SN1_45S_50ntFlanks@7975.7993.19\|CGCGACCTCAGATCAGACG | >hg38_21_+_8433172_8446622_RNA45SN1_45S_50ntFlanks | 7975 | 7993 | 19 | rRF 45S | NO | 888.58 | 923.01 | 1672.86 | 1937.72 |
| >hg38_21_+_8433172_8446622_RNA45SN1_45S_50ntFlanks@7975.7994.20\|CGCGACCTCAGATCAGACGT | >hg38_21_+_8433172_8446622_RNA45SN1_45S_50ntFlanks | 7975 | 7994 | 20 | rRF 45S | YES | 4151.99 | 1530.91 | 4903.55 | 7463.01 |
| >hg38_21_+_8433172_8446622_RNA45SN1_45S_50ntFlanks@7975.7995.21\|CGCGACCTCAGATCAGACGTG | >hg38_21_+_8433172_8446622_RNA45SN1_45S_50ntFlanks | 7975 | 7995 | 21 | rRF 45S | NO | 535.99 | 615.60 | 524.16 | 703.17 |
| >hg38_21_+_8433172_8446622_RNA45SN1_45S_50ntFlanks@7975.7996.22\|CGCGACCTCAGATCAGACGTGG | >hg38_21_+_8433172_8446622_RNA45SN1_45S_50ntFlanks | 7975 | 7996 | 22 | rRF 45S | NO | 236.53 | 266.73 | 259.66 | 488.98 |
| >hg38_21_+_8433172_8446622_RNA45SN1_45S_50ntFlanks@7975.7997.23\|CGCGACCTCAGATCAGACGTGGC | >hg38_21_+_8433172_8446622_RNA45SN1_45S_50ntFlanks | 7975 | 7997 | 23 | rRF 45S | NO | 2208.41 | 636.65 | 1061.74 | 3346.83 |
| >hg38_21_+_8433172_8446622_RNA45SN1_45S_50ntFlanks@7975.7998.24\|CGCGACCTCAGATCAGACGTGGCG | >hg38_21_+_8433172_8446622_RNA45SN1_45S_50ntFlanks | 7975 | 7998 | 24 | rRF 45S | NO | 86.83 | 23.59 | 13.39 | 27.12 |
| >hg38_21_+_8433172_8446622_RNA45SN1_45S_50ntFlanks@7975.8000.26\|CGCGACCTCAGATCAGACGTGGCGAC | >hg38_21_+_8433172_8446622_RNA45SN1_45S_50ntFlanks | 7975 | 8000 | 26 | rRF 45S | NO | 102.54 | 10.06 | 15.08 | 82.43 |
| >hg38_21_+_8433172_8446622_RNA45SN1_45S_50ntFlanks@7975.8001.27\|CGCGACCTCAGATCAGACGTGGCGACC | >hg38_21_+_8433172_8446622_RNA45SN1_45S_50ntFlanks | 7975 | 8001 | 27 | rRF 45S | NO | 148.76 | 39.96 | 82.58 | 80.11 |
| >hg38_21_+_8433172_8446622_RNA45SN1_45S_50ntFlanks@7975.8002.28\|CGCGACCTCAGATCAGACGTGGCGACCC | >hg38_21_+_8433172_8446622_RNA45SN1_45S_50ntFlanks | 7975 | 8002 | 28 | rRF 45S | NO | 366.80 | 101.95 | 583.63 | 249.43 |
| >hg38_21_+_8433172_8446622_RNA45SN1_45S_50ntFlanks@7975.8003.29\|CGCGACCTCAGATCAGACGTGGCGACCCG | >hg38_21_+_8433172_8446622_RNA45SN1_45S_50ntFlanks | 7975 | 8003 | 29 | rRF 45S | NO | 188.29 | 124.84 | 157.54 | 161.48 |
| >hg38_21_+_8433172_8446622_RNA45SN1_45S_50ntFlanks@7975.8004.30\|CGCGACCTCAGATCAGACGTGGCGACCCGC | >hg38_21_+_8433172_8446622_RNA45SN1_45S_50ntFlanks | 7975 | 8004 | 30 | rRF 45S | NO | 906.18 | 591.85 | 559.10 | 1160.29 |
| >hg38_21_+_8433172_8446622_RNA45SN1_45S_50ntFlanks@7975.8005.31\|CGCGACCTCAGATCAGACGTGGCGACCCGCT | >hg38_21_+_8433172_8446622_RNA45SN1_45S_50ntFlanks | 7975 | 8005 | 31 | rRF 45S | YES | 5299.78 | 1737.10 | 2089.72 | 8048.39 |
| >hg38_21_+_8433172_8446622_RNA45SN1_45S_50ntFlanks@7975.8006.32\|CGCGACCTCAGATCAGACGTGGCGACCCGCTG | >hg38_21_+_8433172_8446622_RNA45SN1_45S_50ntFlanks | 7975 | 8006 | 32 | rRF 45S | NO | 2063.97 | 741.13 | 680.12 | 3354.25 |

# BT-20

| | | | | | | | | | | |
|---|---|---|---|---|---|---|---|---|---|---|
| >hg38_21_+_8433172_8446622_RNA45SN1_45S_50ntFlanks@7975.8007.33\|CGCGACCTCAG ATCAGACGTGGCGACCCGCTGA | >hg38_21_+_8433172_8446622_RNA45SN1_45S_50ntFlanks | 7975 | 8007 | 33 | rRF 45S | NO | 1097.48 | 500.78 | 378.77 | 1801.69 |
| >hg38_21_+_8433172_8446622_RNA45SN1_45S_50ntFlanks@7975.8008.34\|CGCGACCTCAG ATCAGACGTGGCGACCCGCTGAA | >hg38_21_+_8433172_8446622_RNA45SN1_45S_50ntFlanks | 7975 | 8008 | 34 | rRF 45S | YES | 1887.43 | 2923.90 | 1146.03 | 5431.45 |
| >hg38_21_+_8433172_8446622_RNA45SN1_45S_50ntFlanks@7975.8009.35\|CGCGACCTCAG ATCAGACGTGGCGACCCGCTGAAT | >hg38_21_+_8433172_8446622_RNA45SN1_45S_50ntFlanks | 7975 | 8009 | 35 | rRF 45S | YES | 5440.68 | 6202.68 | 2759.08 | 9680.45 |
| >hg38_21_+_8433172_8446622_RNA45SN1_45S_50ntFlanks@7975.8010.36\|CGCGACCTCAG ATCAGACGTGGCGACCCGCTGAATT | >hg38_21_+_8433172_8446622_RNA45SN1_45S_50ntFlanks | 7975 | 8010 | 36 | rRF 45S | YES | 8391.78 | 5010.80 | 1490.24 | 9242.23 |
| >hg38_21_+_8433172_8446622_RNA45SN1_45S_50ntFlanks@7975.8011.37\|CGCGACCTCAG ATCAGACGTGGCGACCCGCTGAATTT | >hg38_21_+_8433172_8446622_RNA45SN1_45S_50ntFlanks | 7975 | 8011 | 37 | rRF 45S | YES | 8379.17 | 2950.98 | 951.62 | 9093.73 |
| >hg38_21_+_8433172_8446622_RNA45SN1_45S_50ntFlanks@7975.8012.38\|CGCGACCTCAG ATCAGACGTGGCGACCCGCTGAATTTA | >hg38_21_+_8433172_8446622_RNA45SN1_45S_50ntFlanks | 7975 | 8012 | 38 | rRF 45S | NO | 589.69 | 348.93 | 137.62 | 1122.38 |
| >hg38_21_+_8433172_8446622_RNA45SN1_45S_50ntFlanks@7975.8013.39\|CGCGACCTCAG ATCAGACGTGGCGACCCGCTGAATTTAA | >hg38_21_+_8433172_8446622_RNA45SN1_45S_50ntFlanks | 7975 | 8013 | 39 | rRF 45S | NO | 223.20 | 181.89 | 80.44 | 665.90 |
| >hg38_21_+_8433172_8446622_RNA45SN1_45S_50ntFlanks@7975.8014.40\|CGCGACCTCAG ATCAGACGTGGCGACCCGCTGAATTTAAG | >hg38_21_+_8433172_8446622_RNA45SN1_45S_50ntFlanks | 7975 | 8014 | 40 | rRF 45S | NO | 287.97 | 145.53 | 68.75 | 981.64 |
| >hg38_21_+_8433172_8446622_RNA45SN1_45S_50ntFlanks@7975.8015.41\|CGCGACCTCAG ATCAGACGTGGCGACCCGCTGAATTTAAGC | >hg38_21_+_8433172_8446622_RNA45SN1_45S_50ntFlanks | 7975 | 8015 | 41 | rRF 45S | NO | 682.62 | 220.44 | 200.93 | 1595.49 |
| >hg38_21_+_8433172_8446622_RNA45SN1_45S_50ntFlanks@7975.8016.42\|CGCGACCTCAG ATCAGACGTGGCGACCCGCTGAATTTAAGCA | >hg38_21_+_8433172_8446622_RNA45SN1_45S_50ntFlanks | 7975 | 8016 | 42 | rRF 45S | NO | 368.15 | 226.81 | 150.71 | 1960.63 |
| >hg38_21_+_8433172_8446622_RNA45SN1_45S_50ntFlanks@7975.8017.43\|CGCGACCTCAG ATCAGACGTGGCGACCCGCTGAATTTAAGCAT | >hg38_21_+_8433172_8446622_RNA45SN1_45S_50ntFlanks | 7975 | 8017 | 43 | rRF 45S | NO | 173.64 | 27.36 | 12.63 | 569.38 |
| >hg38_21_+_8433172_8446622_RNA45SN1_45S_50ntFlanks@7975.8018.44\|CGCGACCTCAG ATCAGACGTGGCGACCCGCTGAATTTAAGCATA | >hg38_21_+_8433172_8446622_RNA45SN1_45S_50ntFlanks | 7975 | 8018 | 44 | rRF 45S | NO | 71.43 | 0.00 | 54.34 | 690.20 |
| >hg38_21_+_8433172_8446622_RNA45SN1_45S_50ntFlanks@7976.8009.34\|GCGACCTCAGA TCAGACGTGGCGACCCGCTGAAT | >hg38_21_+_8433172_8446622_RNA45SN1_45S_50ntFlanks | 7976 | 8009 | 34 | rRF 45S | NO | 39.57 | 25.22 | 100.22 | 84.95 |
| >hg38_21_+_8433172_8446622_RNA45SN1_45S_50ntFlanks@7976.8010.35\|GCGACCTCAGA TCAGACGTGGCGACCCGCTGAATT | >hg38_21_+_8433172_8446622_RNA45SN1_45S_50ntFlanks | 7976 | 8010 | 35 | rRF 45S | NO | 86.67 | 0.00 | 12.65 | 95.12 |
| >hg38_21_+_8433172_8446622_RNA45SN1_45S_50ntFlanks@7976.8011.36\|GCGACCTCAGA TCAGACGTGGCGACCCGCTGAATTT | >hg38_21_+_8433172_8446622_RNA45SN1_45S_50ntFlanks | 7976 | 8011 | 36 | rRF 45S | NO | 88.94 | 0.00 | 12.70 | 93.26 |
| >hg38_21_+_8433172_8446622_RNA45SN1_45S_50ntFlanks@7977.8009.33\|CGACCTCAGAT CAGACGTGGCGACCCGCTGAAT | >hg38_21_+_8433172_8446622_RNA45SN1_45S_50ntFlanks | 7977 | 8009 | 33 | rRF 45S | NO | 0.00 | 0.00 | 14.39 | 0.00 |
| >hg38_21_+_8433172_8446622_RNA45SN1_45S_50ntFlanks@7977.8010.34\|CGACCTCAGAT CAGACGTGGCGACCCGCTGAATT | >hg38_21_+_8433172_8446622_RNA45SN1_45S_50ntFlanks | 7977 | 8010 | 34 | rRF 45S | NO | 24.16 | 0.00 | 0.00 | 10.07 |
| >hg38_21_+_8433172_8446622_RNA45SN1_45S_50ntFlanks@7977.8011.35\|CGACCTCAGAT CAGACGTGGCGACCCGCTGAATTT | >hg38_21_+_8433172_8446622_RNA45SN1_45S_50ntFlanks | 7977 | 8011 | 35 | rRF 45S | NO | 32.19 | 10.22 | 12.72 | 33.83 |
| >hg38_21_+_8433172_8446622_RNA45SN1_45S_50ntFlanks@7978.8009.32\|GACCTCAGATCA GACGTGGCGACCCGCTGAAT | >hg38_21_+_8433172_8446622_RNA45SN1_45S_50ntFlanks | 7978 | 8009 | 32 | rRF 45S | NO | 0.00 | 0.00 | 10.05 | 0.00 |
| >hg38_21_+_8433172_8446622_RNA45SN1_45S_50ntFlanks@7978.8011.34\|GACCTCAGATCA GACGTGGCGACCCGCTGAATTT | >hg38_21_+_8433172_8446622_RNA45SN1_45S_50ntFlanks | 7978 | 8011 | 34 | rRF 45S | NO | 10.97 | 0.00 | 0.00 | 28.73 |
| >hg38_21_+_8433172_8446622_RNA45SN1_45S_50ntFlanks@7979.8010.32\|ACCTCAGATCAG ACGTGGCGACCCGCTGAATT | >hg38_21_+_8433172_8446622_RNA45SN1_45S_50ntFlanks | 7979 | 8010 | 32 | rRF 45S | NO | 10.22 | 0.00 | 0.00 | 10.43 |
| >hg38_21_+_8433172_8446622_RNA45SN1_45S_50ntFlanks@7979.8011.33\|ACCTCAGATCAG ACGTGGCGACCCGCTGAATTT | >hg38_21_+_8433172_8446622_RNA45SN1_45S_50ntFlanks | 7979 | 8011 | 33 | rRF 45S | NO | 25.02 | 0.00 | 0.00 | 25.45 |
| >hg38_21_+_8433172_8446622_RNA45SN1_45S_50ntFlanks@7988.8009.22\|CAGACGTGGCG ACCCGCTGAAT | >hg38_21_+_8433172_8446622_RNA45SN1_45S_50ntFlanks | 7988 | 8009 | 22 | rRF 45S | NO | 0.00 | 9.99 | 10.53 | 0.00 |
| >hg38_21_+_8433172_8446622_RNA45SN1_45S_50ntFlanks@7988.8011.24\|CAGACGTGGCG ACCCGCTGAATTT | >hg38_21_+_8433172_8446622_RNA45SN1_45S_50ntFlanks | 7988 | 8011 | 24 | rRF 45S | NO | 40.98 | 0.00 | 0.00 | 0.00 |
| >hg38_21_+_8433172_8446622_RNA45SN1_45S_50ntFlanks@7989.8007.19\|AGACGTGGCGA CCCGCTGA | >hg38_21_+_8433172_8446622_RNA45SN1_45S_50ntFlanks | 7989 | 8007 | 19 | rRF 45S | NO | 0.00 | 0.00 | 64.17 | 0.00 |
| >hg38_21_+_8433172_8446622_RNA45SN1_45S_50ntFlanks@7989.8008.20\|AGACGTGGCGA CCCGCTGAA | >hg38_21_+_8433172_8446622_RNA45SN1_45S_50ntFlanks | 7989 | 8008 | 20 | rRF 45S | NO | 25.32 | 24.41 | 81.68 | 0.00 |
| >hg38_21_+_8433172_8446622_RNA45SN1_45S_50ntFlanks@7989.8009.21\|AGACGTGGCGA CCCGCTGAAT | >hg38_21_+_8433172_8446622_RNA45SN1_45S_50ntFlanks | 7989 | 8009 | 21 | rRF 45S | NO | 90.56 | 45.15 | 150.43 | 23.39 |
| >hg38_21_+_8433172_8446622_RNA45SN1_45S_50ntFlanks@7989.8010.22\|AGACGTGGCGA CCCGCTGAATT | >hg38_21_+_8433172_8446622_RNA45SN1_45S_50ntFlanks | 7989 | 8010 | 22 | rRF 45S | NO | 81.04 | 36.87 | 55.36 | 25.21 |
| >hg38_21_+_8433172_8446622_RNA45SN1_45S_50ntFlanks@7989.8011.23\|AGACGTGGCGA CCCGCTGAATTT | >hg38_21_+_8433172_8446622_RNA45SN1_45S_50ntFlanks | 7989 | 8011 | 23 | rRF 45S | NO | 156.09 | 86.05 | 141.64 | 63.54 |
| >hg38_21_+_8433172_8446622_RNA45SN1_45S_50ntFlanks@7989.8012.24\|AGACGTGGCGA CCCGCTGAATTTA | >hg38_21_+_8433172_8446622_RNA45SN1_45S_50ntFlanks | 7989 | 8012 | 24 | rRF 45S | NO | 0.00 | 0.00 | 15.08 | 0.00 |
| >hg38_21_+_8433172_8446622_RNA45SN1_45S_50ntFlanks@7989.8015.27\|AGACGTGGCGA CCCGCTGAATTTAAGC | >hg38_21_+_8433172_8446622_RNA45SN1_45S_50ntFlanks | 7989 | 8015 | 27 | rRF 45S | NO | 27.57 | 0.00 | 0.00 | 0.00 |
| >hg38_21_+_8433172_8446622_RNA45SN1_45S_50ntFlanks@7990.8008.19\|GACGTGGCGAC CCGCTGAA | >hg38_21_+_8433172_8446622_RNA45SN1_45S_50ntFlanks | 7990 | 8008 | 19 | rRF 45S | NO | 0.00 | 0.00 | 13.27 | 0.00 |
| >hg38_21_+_8433172_8446622_RNA45SN1_45S_50ntFlanks@7990.8009.20\|GACGTGGCGAC CCGCTGAAT | >hg38_21_+_8433172_8446622_RNA45SN1_45S_50ntFlanks | 7990 | 8009 | 20 | rRF 45S | NO | 10.44 | 0.00 | 87.34 | 0.00 |
| >hg38_21_+_8433172_8446622_RNA45SN1_45S_50ntFlanks@7990.8010.21\|GACGTGGCGAC CCGCTGAATT | >hg38_21_+_8433172_8446622_RNA45SN1_45S_50ntFlanks | 7990 | 8010 | 21 | rRF 45S | NO | 10.38 | 0.00 | 11.02 | 0.00 |
| >hg38_21_+_8433172_8446622_RNA45SN1_45S_50ntFlanks@7990.8011.22\|GACGTGGCGAC CCGCTGAATTT | >hg38_21_+_8433172_8446622_RNA45SN1_45S_50ntFlanks | 7990 | 8011 | 22 | rRF 45S | NO | 41.10 | 23.56 | 84.92 | 26.09 |
| >hg38_21_+_8433172_8446622_RNA45SN1_45S_50ntFlanks@7991.8009.19\|ACGTGGCGACC CGCTGAAT | >hg38_21_+_8433172_8446622_RNA45SN1_45S_50ntFlanks | 7991 | 8009 | 19 | rRF 45S | NO | 28.16 | 0.00 | 37.92 | 0.00 |
| >hg38_21_+_8433172_8446622_RNA45SN1_45S_50ntFlanks@7991.8010.20\|ACGTGGCGACC CGCTGAATT | >hg38_21_+_8433172_8446622_RNA45SN1_45S_50ntFlanks | 7991 | 8010 | 20 | rRF 45S | NO | 24.57 | 0.00 | 0.00 | 0.00 |
| >hg38_21_+_8433172_8446622_RNA45SN1_45S_50ntFlanks@7991.8011.21\|ACGTGGCGACC CGCTGAATTT | >hg38_21_+_8433172_8446622_RNA45SN1_45S_50ntFlanks | 7991 | 8011 | 21 | rRF 45S | NO | 74.26 | 9.65 | 62.15 | 31.43 |

# BT-20

| | | | | | | | | | | |
|---|---|---|---|---|---|---|---|---|---|---|
| >hg38_21_+_8433172_8446622_RNA45SN1_45S_50ntFlanks@7992.8009.18\|CGTGGCGACCCGCTGAAT | >hg38_21_+_8433172_8446622_RNA45SN1_45S_50ntFlanks | 7992 | 8009 | 18 | rRF 45S | NO | 10.62 | 0.00 | 0.00 | 0.00 |
| >hg38_21_+_8433172_8446622_RNA45SN1_45S_50ntFlanks@7992.8010.19\|CGTGGCGACCCGCTGAATT | >hg38_21_+_8433172_8446622_RNA45SN1_45S_50ntFlanks | 7992 | 8010 | 19 | rRF 45S | NO | 23.58 | 0.00 | 0.00 | 0.00 |
| >hg38_21_+_8433172_8446622_RNA45SN1_45S_50ntFlanks@7992.8011.20\|CGTGGCGACCCGCTGAATTT | >hg38_21_+_8433172_8446622_RNA45SN1_45S_50ntFlanks | 7992 | 8011 | 20 | rRF 45S | NO | 38.99 | 0.00 | 0.00 | 0.00 |
| >hg38_21_+_8433172_8446622_RNA45SN1_45S_50ntFlanks@8104.8141.38\|CCGCGGCGGGGCGCGGGACATGTGGCGTACGGAAGACC | >hg38_21_+_8433172_8446622_RNA45SN1_45S_50ntFlanks | 8104 | 8141 | 38 | rRF 45S | NO | 0.00 | 0.00 | 13.84 | 0.00 |
| >hg38_21_+_8433172_8446622_RNA45SN1_45S_50ntFlanks@8115.8141.27\|CGCGGGACATGTGGCGTACGGAAGACC | >hg38_21_+_8433172_8446622_RNA45SN1_45S_50ntFlanks | 8115 | 8141 | 27 | rRF 45S | NO | 0.00 | 0.00 | 14.51 | 0.00 |
| >hg38_21_+_8433172_8446622_RNA45SN1_45S_50ntFlanks@8117.8134.18\|CGGGACATGTGGCGTACG | >hg38_21_+_8433172_8446622_RNA45SN1_45S_50ntFlanks | 8117 | 8134 | 18 | rRF 45S | NO | 0.00 | 0.00 | 13.39 | 0.00 |
| >hg38_21_+_8433172_8446622_RNA45SN1_45S_50ntFlanks@8174.8202.29\|TCCTTCTGATCGAGGCCCAGCCCGTGGAC | >hg38_21_+_8433172_8446622_RNA45SN1_45S_50ntFlanks | 8174 | 8202 | 29 | rRF 45S | NO | 0.00 | 24.58 | 0.00 | 0.00 |
| >hg38_21_+_8433172_8446622_RNA45SN1_45S_50ntFlanks@8175.8202.28\|CCTTCTGATCGAGGCCCAGCCCGTGGAC | >hg38_21_+_8433172_8446622_RNA45SN1_45S_50ntFlanks | 8175 | 8202 | 28 | rRF 45S | NO | 0.00 | 24.14 | 0.00 | 0.00 |
| >hg38_21_+_8433172_8446622_RNA45SN1_45S_50ntFlanks@8175.8216.42\|CCTTCTGATCGAGGCCCAGCCCGTGGACGGTGTGAGGCCGGT | >hg38_21_+_8433172_8446622_RNA45SN1_45S_50ntFlanks | 8175 | 8216 | 42 | rRF 45S | NO | 0.00 | 0.00 | 72.41 | 9.55 |
| >hg38_21_+_8433172_8446622_RNA45SN1_45S_50ntFlanks@8176.8195.20\|CTTCTGATCGAGGCCCAGCC | >hg38_21_+_8433172_8446622_RNA45SN1_45S_50ntFlanks | 8176 | 8195 | 20 | rRF 45S | NO | 0.00 | 169.30 | 0.00 | 0.00 |
| >hg38_21_+_8433172_8446622_RNA45SN1_45S_50ntFlanks@8176.8202.27\|CTTCTGATCGAGGCCCAGCCCGTGGAC | >hg38_21_+_8433172_8446622_RNA45SN1_45S_50ntFlanks | 8176 | 8202 | 27 | rRF 45S | NO | 0.00 | 120.75 | 0.00 | 10.79 |
| >hg38_21_+_8433172_8446622_RNA45SN1_45S_50ntFlanks@8177.8195.19\|TTCTGATCGAGGCCCAGCC | >hg38_21_+_8433172_8446622_RNA45SN1_45S_50ntFlanks | 8177 | 8195 | 19 | rRF 45S | NO | 0.00 | 33.72 | 0.00 | 0.00 |
| >hg38_21_+_8433172_8446622_RNA45SN1_45S_50ntFlanks@8195.8213.19\|CCGTGGACGGTGTGAGGCC | >hg38_21_+_8433172_8446622_RNA45SN1_45S_50ntFlanks | 8195 | 8213 | 19 | rRF 45S | NO | 0.00 | 97.65 | 85.40 | 0.00 |
| >hg38_21_+_8433172_8446622_RNA45SN1_45S_50ntFlanks@8196.8213.18\|CGTGGACGGTGTGAGGCC | >hg38_21_+_8433172_8446622_RNA45SN1_45S_50ntFlanks | 8196 | 8213 | 18 | rRF 45S | NO | 0.00 | 138.97 | 134.63 | 0.00 |
| >hg38_21_+_8433172_8446622_RNA45SN1_45S_50ntFlanks@8342.8365.24\|CGTAAGGGAAAGTTGAAAAGAACT | >hg38_21_+_8433172_8446622_RNA45SN1_45S_50ntFlanks | 8342 | 8365 | 24 | rRF 45S | NO | 0.00 | 90.65 | 15.00 | 0.00 |
| >hg38_21_+_8433172_8446622_RNA45SN1_45S_50ntFlanks@8342.8378.37\|CGTAAGGGAAAGTTGAAAAGAACTTTGAAGAGAGAGT | >hg38_21_+_8433172_8446622_RNA45SN1_45S_50ntFlanks | 8342 | 8378 | 37 | rRF 45S | NO | 0.00 | 35.56 | 12.91 | 0.00 |
| >hg38_21_+_8433172_8446622_RNA45SN1_45S_50ntFlanks@8342.8379.38\|CGTAAGGGAAAGTTGAAAAGAACTTTGAAGAGAGAGTT | >hg38_21_+_8433172_8446622_RNA45SN1_45S_50ntFlanks | 8342 | 8379 | 38 | rRF 45S | NO | 0.00 | 42.32 | 0.00 | 0.00 |
| >hg38_21_+_8433172_8446622_RNA45SN1_45S_50ntFlanks@9546.9571.26\|TGGAGGTCCGTAGCGGTCCTGACGTG | >hg38_21_+_8433172_8446622_RNA45SN1_45S_50ntFlanks | 9546 | 9571 | 26 | rRF 45S | NO | 0.00 | 24.79 | 0.00 | 0.00 |
| >hg38_21_+_8433172_8446622_RNA45SN1_45S_50ntFlanks@9549.9570.22\|AGGTCCGTAGCGGTCCTGACGT | >hg38_21_+_8433172_8446622_RNA45SN1_45S_50ntFlanks | 9549 | 9570 | 22 | rRF 45S | NO | 0.00 | 0.00 | 11.75 | 0.00 |
| >hg38_21_+_8433172_8446622_RNA45SN1_45S_50ntFlanks@9554.9571.18\|CGTAGCGGTCCTGACGTG | >hg38_21_+_8433172_8446622_RNA45SN1_45S_50ntFlanks | 9554 | 9571 | 18 | rRF 45S | NO | 0.00 | 139.33 | 14.37 | 10.12 |
| >hg38_21_+_8433172_8446622_RNA45SN1_45S_50ntFlanks@9554.9572.19\|CGTAGCGGTCCTGACGTGC | >hg38_21_+_8433172_8446622_RNA45SN1_45S_50ntFlanks | 9554 | 9572 | 19 | rRF 45S | NO | 0.00 | 38.80 | 10.56 | 0.00 |
| >hg38_21_+_8433172_8446622_RNA45SN1_45S_50ntFlanks@9554.9573.20\|CGTAGCGGTCCTGACGTGCA | >hg38_21_+_8433172_8446622_RNA45SN1_45S_50ntFlanks | 9554 | 9573 | 20 | rRF 45S | NO | 0.00 | 108.20 | 0.00 | 0.00 |
| >hg38_21_+_8433172_8446622_RNA45SN1_45S_50ntFlanks@9554.9576.23\|CGTAGCGGTCCTGACGTGCAAAT | >hg38_21_+_8433172_8446622_RNA45SN1_45S_50ntFlanks | 9554 | 9576 | 23 | rRF 45S | NO | 0.00 | 76.38 | 0.00 | 0.00 |
| >hg38_21_+_8433172_8446622_RNA45SN1_45S_50ntFlanks@9554.9577.24\|CGTAGCGGTCCTGACGTGCAAATC | >hg38_21_+_8433172_8446622_RNA45SN1_45S_50ntFlanks | 9554 | 9577 | 24 | rRF 45S | NO | 0.00 | 44.62 | 0.00 | 0.00 |
| >hg38_21_+_8433172_8446622_RNA45SN1_45S_50ntFlanks@9573.9603.31\|AAATCGGTCGTCCGACCTGGGTATAGGGGCG | >hg38_21_+_8433172_8446622_RNA45SN1_45S_50ntFlanks | 9573 | 9603 | 31 | rRF 45S | NO | 0.00 | 0.00 | 13.53 | 0.00 |
| >hg38_21_+_8433172_8446622_RNA45SN1_45S_50ntFlanks@9574.9601.28\|AATCGGTCGTCCGACCTGGGTATAGGGG | >hg38_21_+_8433172_8446622_RNA45SN1_45S_50ntFlanks | 9574 | 9601 | 28 | rRF 45S | NO | 0.00 | 0.00 | 16.17 | 0.00 |
| >hg38_21_+_8433172_8446622_RNA45SN1_45S_50ntFlanks@9574.9602.29\|AATCGGTCGTCCGACCTGGGTATAGGGGC | >hg38_21_+_8433172_8446622_RNA45SN1_45S_50ntFlanks | 9574 | 9602 | 29 | rRF 45S | NO | 0.00 | 0.00 | 55.57 | 0.00 |
| >hg38_21_+_8433172_8446622_RNA45SN1_45S_50ntFlanks@9574.9603.30\|AATCGGTCGTCCGACCTGGGTATAGGGGCG | >hg38_21_+_8433172_8446622_RNA45SN1_45S_50ntFlanks | 9574 | 9603 | 30 | rRF 45S | NO | 0.00 | 0.00 | 111.98 | 25.43 |
| >hg38_21_+_8433172_8446622_RNA45SN1_45S_50ntFlanks@9574.9605.32\|AATCGGTCGTCCGACCTGGGTATAGGGGCGAA | >hg38_21_+_8433172_8446622_RNA45SN1_45S_50ntFlanks | 9574 | 9605 | 32 | rRF 45S | NO | 0.00 | 0.00 | 73.86 | 0.00 |
| >hg38_21_+_8433172_8446622_RNA45SN1_45S_50ntFlanks@9578.9603.26\|GGTCGTCCGACCTGGGTATAGGGGCG | >hg38_21_+_8433172_8446622_RNA45SN1_45S_50ntFlanks | 9578 | 9603 | 26 | rRF 45S | NO | 0.00 | 0.00 | 56.80 | 0.00 |
| >hg38_21_+_8433172_8446622_RNA45SN1_45S_50ntFlanks@9937.9955.19\|CAGCAGGACGGTGGCCATG | >hg38_21_+_8433172_8446622_RNA45SN1_45S_50ntFlanks | 9937 | 9955 | 19 | rRF 45S | NO | 0.00 | 0.00 | 11.74 | 0.00 |
| >hg38_21_+_8433172_8446622_RNA45SN1_45S_50ntFlanks@10240.10259.20\|CTACGCCGCGACGAGTAGGA | >hg38_21_+_8433172_8446622_RNA45SN1_45S_50ntFlanks | 10240 | 10259 | 20 | rRF 45S | NO | 9.88 | 0.00 | 295.49 | 28.56 |
| >hg38_21_+_8433172_8446622_RNA45SN1_45S_50ntFlanks@10319.10340.22\|GCAGATCTTGGTGGTAGTAGCA | >hg38_21_+_8433172_8446622_RNA45SN1_45S_50ntFlanks | 10319 | 10340 | 22 | rRF 45S | NO | 0.00 | 0.00 | 10.86 | 0.00 |
| >hg38_21_+_8433172_8446622_RNA45SN1_45S_50ntFlanks@10348.10369.22\|AAACGAGAACTTTGAAGGCCGA | >hg38_21_+_8433172_8446622_RNA45SN1_45S_50ntFlanks | 10348 | 10369 | 22 | rRF 45S | NO | 0.00 | 0.00 | 52.57 | 0.00 |
| >hg38_21_+_8433172_8446622_RNA45SN1_45S_50ntFlanks@10348.10371.24\|AAACGAGAACTTTGAAGGCCGAAG | >hg38_21_+_8433172_8446622_RNA45SN1_45S_50ntFlanks | 10348 | 10371 | 24 | rRF 45S | NO | 0.00 | 0.00 | 61.96 | 0.00 |
| >hg38_21_+_8433172_8446622_RNA45SN1_45S_50ntFlanks@10348.10372.25\|AAACGAGAACTTTGAAGGCCGAAGT | >hg38_21_+_8433172_8446622_RNA45SN1_45S_50ntFlanks | 10348 | 10372 | 25 | rRF 45S | NO | 0.00 | 0.00 | 39.44 | 0.00 |
| >hg38_21_+_8433172_8446622_RNA45SN1_45S_50ntFlanks@10348.10382.35\|AAACGAGAACTTTGAAGGCCGAAGTGGAGAAGGGT | >hg38_21_+_8433172_8446622_RNA45SN1_45S_50ntFlanks | 10348 | 10382 | 35 | rRF 45S | NO | 0.00 | 34.46 | 12.66 | 0.00 |
| >hg38_21_+_8433172_8446622_RNA45SN1_45S_50ntFlanks@10349.10369.21\|AACGAGAACTTTGAAGGCCGA | >hg38_21_+_8433172_8446622_RNA45SN1_45S_50ntFlanks | 10349 | 10369 | 21 | rRF 45S | NO | 0.00 | 0.00 | 66.24 | 10.73 |

# BT-20

<table>
<tr><th></th><th></th><th></th><th></th><th></th><th></th><th></th><th></th><th></th><th></th><th></th></tr>
<tr><td>>hg38_21_+_8433172_8446622_RNA45SN1_45S_50ntFlanks@10349.10371.23|AACGAGAACTTTGAAGGCCGAAG</td><td>>hg38_21_+_8433172_8446622_RNA45SN1_45S_50ntFlanks</td><td>10349</td><td>10371</td><td>23</td><td>rRF 45S</td><td>NO</td><td>0.00</td><td>0.00</td><td>60.95</td><td>0.00</td></tr>
<tr><td>>hg38_21_+_8433172_8446622_RNA45SN1_45S_50ntFlanks@10349.10372.24|AACGAGAACTTTGAAGGCCGAAGT</td><td>>hg38_21_+_8433172_8446622_RNA45SN1_45S_50ntFlanks</td><td>10349</td><td>10372</td><td>24</td><td>rRF 45S</td><td>NO</td><td>0.00</td><td>0.00</td><td>59.60</td><td>10.44</td></tr>
<tr><td>>hg38_21_+_8433172_8446622_RNA45SN1_45S_50ntFlanks@10349.10382.34|AACGAGAACTTTGAAGGCCGAAGTGGAGAAGGGT</td><td>>hg38_21_+_8433172_8446622_RNA45SN1_45S_50ntFlanks</td><td>10349</td><td>10382</td><td>34</td><td>rRF 45S</td><td>NO</td><td>0.00</td><td>68.56</td><td>14.97</td><td>0.00</td></tr>
<tr><td>>hg38_21_+_8433172_8446622_RNA45SN1_45S_50ntFlanks@10349.10383.35|AACGAGAACTTTGAAGGCCGAAGTGGAGAAGGGTT</td><td>>hg38_21_+_8433172_8446622_RNA45SN1_45S_50ntFlanks</td><td>10349</td><td>10383</td><td>35</td><td>rRF 45S</td><td>NO</td><td>0.00</td><td>26.02</td><td>16.09</td><td>0.00</td></tr>
<tr><td>>hg38_21_+_8433172_8446622_RNA45SN1_45S_50ntFlanks@10350.10372.23|ACGAGAACTTTGAAGGCCGAAGT</td><td>>hg38_21_+_8433172_8446622_RNA45SN1_45S_50ntFlanks</td><td>10350</td><td>10372</td><td>23</td><td>rRF 45S</td><td>NO</td><td>0.00</td><td>0.00</td><td>10.40</td><td>0.00</td></tr>
<tr><td>>hg38_21_+_8433172_8446622_RNA45SN1_45S_50ntFlanks@10381.10424.44|GTTCCATGTGAACAGCAGTTGAACATGGGTCAGTCGGTCCTGAG</td><td>>hg38_21_+_8433172_8446622_RNA45SN1_45S_50ntFlanks</td><td>10381</td><td>10424</td><td>44</td><td>rRF 45S</td><td>NO</td><td>0.00</td><td>0.00</td><td>72.62</td><td>184.18</td></tr>
<tr><td>>hg38_21_+_8433172_8446622_RNA45SN1_45S_50ntFlanks@10382.10424.43|TTCCATGTGAACAGCAGTTGAACATGGGTCAGTCGGTCCTGAG</td><td>>hg38_21_+_8433172_8446622_RNA45SN1_45S_50ntFlanks</td><td>10382</td><td>10424</td><td>43</td><td>rRF 45S</td><td>NO</td><td>0.00</td><td>9.76</td><td>107.32</td><td>221.54</td></tr>
<tr><td>>hg38_21_+_8433172_8446622_RNA45SN1_45S_50ntFlanks@10383.10424.42|TCCATGTGAACAGCAGTTGAACATGGGTCAGTCGGTCCTGAG</td><td>>hg38_21_+_8433172_8446622_RNA45SN1_45S_50ntFlanks</td><td>10383</td><td>10424</td><td>42</td><td>rRF 45S</td><td>NO</td><td>30.44</td><td>35.55</td><td>77.75</td><td>223.52</td></tr>
<tr><td>>hg38_21_+_8433172_8446622_RNA45SN1_45S_50ntFlanks@10385.10424.40|CATGTGAACAGCAGTTGAACATGGGTCAGTCGGTCCTGAG</td><td>>hg38_21_+_8433172_8446622_RNA45SN1_45S_50ntFlanks</td><td>10385</td><td>10424</td><td>40</td><td>rRF 45S</td><td>NO</td><td>0.00</td><td>0.00</td><td>14.94</td><td>79.21</td></tr>
<tr><td>>hg38_21_+_8433172_8446622_RNA45SN1_45S_50ntFlanks@10386.10424.39|ATGTGAACAGCAGTTGAACATGGGTCAGTCGGTCCTGAG</td><td>>hg38_21_+_8433172_8446622_RNA45SN1_45S_50ntFlanks</td><td>10386</td><td>10424</td><td>39</td><td>rRF 45S</td><td>NO</td><td>42.57</td><td>86.02</td><td>320.12</td><td>396.36</td></tr>
<tr><td>>hg38_21_+_8433172_8446622_RNA45SN1_45S_50ntFlanks@10387.10424.38|TGTGAACAGCAGTTGAACATGGGTCAGTCGGTCCTGAG</td><td>>hg38_21_+_8433172_8446622_RNA45SN1_45S_50ntFlanks</td><td>10387</td><td>10424</td><td>38</td><td>rRF 45S</td><td>NO</td><td>41.00</td><td>106.27</td><td>298.96</td><td>400.02</td></tr>
<tr><td>>hg38_21_+_8433172_8446622_RNA45SN1_45S_50ntFlanks@10388.10424.37|GTGAACAGCAGTTGAACATGGGTCAGTCGGTCCTGAG</td><td>>hg38_21_+_8433172_8446622_RNA45SN1_45S_50ntFlanks</td><td>10388</td><td>10424</td><td>37</td><td>rRF 45S</td><td>NO</td><td>76.88</td><td>140.42</td><td>474.13</td><td>507.88</td></tr>
<tr><td>>hg38_21_+_8433172_8446622_RNA45SN1_45S_50ntFlanks@10389.10424.36|TGAACAGCAGTTGAACATGGGTCAGTCGGTCCTGAG</td><td>>hg38_21_+_8433172_8446622_RNA45SN1_45S_50ntFlanks</td><td>10389</td><td>10424</td><td>36</td><td>rRF 45S</td><td>NO</td><td>43.32</td><td>140.82</td><td>527.85</td><td>471.15</td></tr>
<tr><td>>hg38_21_+_8433172_8446622_RNA45SN1_45S_50ntFlanks@10390.10424.35|GAACAGCAGTTGAACATGGGTCAGTCGGTCCTGAG</td><td>>hg38_21_+_8433172_8446622_RNA45SN1_45S_50ntFlanks</td><td>10390</td><td>10424</td><td>35</td><td>rRF 45S</td><td>NO</td><td>0.00</td><td>32.75</td><td>178.16</td><td>196.99</td></tr>
<tr><td>>hg38_21_+_8433172_8446622_RNA45SN1_45S_50ntFlanks@10391.10424.34|AACAGCAGTTGAACATGGGTCAGTCGGTCCTGAG</td><td>>hg38_21_+_8433172_8446622_RNA45SN1_45S_50ntFlanks</td><td>10391</td><td>10424</td><td>34</td><td>rRF 45S</td><td>NO</td><td>109.99</td><td>145.68</td><td>468.31</td><td>581.50</td></tr>
<tr><td>>hg38_21_+_8433172_8446622_RNA45SN1_45S_50ntFlanks@10391.10429.39|AACAGCAGTTGAACATGGGTCAGTCGGTCCTGAGAGATG</td><td>>hg38_21_+_8433172_8446622_RNA45SN1_45S_50ntFlanks</td><td>10391</td><td>10429</td><td>39</td><td>rRF 45S</td><td>NO</td><td>0.00</td><td>25.03</td><td>0.00</td><td>0.00</td></tr>
<tr><td>>hg38_21_+_8433172_8446622_RNA45SN1_45S_50ntFlanks@10392.10424.33|ACAGCAGTTGAACATGGGTCAGTCGGTCCTGAG</td><td>>hg38_21_+_8433172_8446622_RNA45SN1_45S_50ntFlanks</td><td>10392</td><td>10424</td><td>33</td><td>rRF 45S</td><td>NO</td><td>26.19</td><td>41.16</td><td>324.04</td><td>320.87</td></tr>
<tr><td>>hg38_21_+_8433172_8446622_RNA45SN1_45S_50ntFlanks@10393.10424.32|CAGCAGTTGAACATGGGTCAGTCGGTCCTGAG</td><td>>hg38_21_+_8433172_8446622_RNA45SN1_45S_50ntFlanks</td><td>10393</td><td>10424</td><td>32</td><td>rRF 45S</td><td>NO</td><td>33.75</td><td>42.88</td><td>365.93</td><td>329.56</td></tr>
<tr><td>>hg38_21_+_8433172_8446622_RNA45SN1_45S_50ntFlanks@10394.10424.31|AGCAGTTGAACATGGGTCAGTCGGTCCTGAG</td><td>>hg38_21_+_8433172_8446622_RNA45SN1_45S_50ntFlanks</td><td>10394</td><td>10424</td><td>31</td><td>rRF 45S</td><td>NO</td><td>29.62</td><td>33.81</td><td>436.06</td><td>279.48</td></tr>
<tr><td>>hg38_21_+_8433172_8446622_RNA45SN1_45S_50ntFlanks@10395.10424.30|GCAGTTGAACATGGGTCAGTCGGTCCTGAG</td><td>>hg38_21_+_8433172_8446622_RNA45SN1_45S_50ntFlanks</td><td>10395</td><td>10424</td><td>30</td><td>rRF 45S</td><td>NO</td><td>33.08</td><td>39.51</td><td>403.33</td><td>339.56</td></tr>
<tr><td>>hg38_21_+_8433172_8446622_RNA45SN1_45S_50ntFlanks@10396.10424.29|CAGTTGAACATGGGTCAGTCGGTCCTGAG</td><td>>hg38_21_+_8433172_8446622_RNA45SN1_45S_50ntFlanks</td><td>10396</td><td>10424</td><td>29</td><td>rRF 45S</td><td>NO</td><td>105.41</td><td>195.41</td><td>591.70</td><td>550.69</td></tr>
<tr><td>>hg38_21_+_8433172_8446622_RNA45SN1_45S_50ntFlanks@10397.10424.28|AGTTGAACATGGGTCAGTCGGTCCTGAG</td><td>>hg38_21_+_8433172_8446622_RNA45SN1_45S_50ntFlanks</td><td>10397</td><td>10424</td><td>28</td><td>rRF 45S</td><td>NO</td><td>0.00</td><td>0.00</td><td>126.25</td><td>123.51</td></tr>
<tr><td>>hg38_21_+_8433172_8446622_RNA45SN1_45S_50ntFlanks@10398.10424.27|GTTGAACATGGGTCAGTCGGTCCTGAG</td><td>>hg38_21_+_8433172_8446622_RNA45SN1_45S_50ntFlanks</td><td>10398</td><td>10424</td><td>27</td><td>rRF 45S</td><td>NO</td><td>69.65</td><td>110.03</td><td>443.34</td><td>394.87</td></tr>
<tr><td>>hg38_21_+_8433172_8446622_RNA45SN1_45S_50ntFlanks@10399.10424.26|TTGAACATGGGTCAGTCGGTCCTGAG</td><td>>hg38_21_+_8433172_8446622_RNA45SN1_45S_50ntFlanks</td><td>10399</td><td>10424</td><td>26</td><td>rRF 45S</td><td>NO</td><td>0.00</td><td>0.00</td><td>100.77</td><td>106.92</td></tr>
<tr><td>>hg38_21_+_8433172_8446622_RNA45SN1_45S_50ntFlanks@10400.10424.25|TGAACATGGGTCAGTCGGTCCTGAG</td><td>>hg38_21_+_8433172_8446622_RNA45SN1_45S_50ntFlanks</td><td>10400</td><td>10424</td><td>25</td><td>rRF 45S</td><td>NO</td><td>40.26</td><td>95.53</td><td>380.15</td><td>363.65</td></tr>
<tr><td>>hg38_21_+_8433172_8446622_RNA45SN1_45S_50ntFlanks@10401.10424.24|GAACATGGGTCAGTCGGTCCTGAG</td><td>>hg38_21_+_8433172_8446622_RNA45SN1_45S_50ntFlanks</td><td>10401</td><td>10424</td><td>24</td><td>rRF 45S</td><td>NO</td><td>78.15</td><td>107.80</td><td>541.94</td><td>418.60</td></tr>
<tr><td>>hg38_21_+_8433172_8446622_RNA45SN1_45S_50ntFlanks@10402.10424.23|AACATGGGTCAGTCGGTCCTGAG</td><td>>hg38_21_+_8433172_8446622_RNA45SN1_45S_50ntFlanks</td><td>10402</td><td>10424</td><td>23</td><td>rRF 45S</td><td>NO</td><td>72.04</td><td>104.94</td><td>472.12</td><td>390.76</td></tr>
<tr><td>>hg38_21_+_8433172_8446622_RNA45SN1_45S_50ntFlanks@10403.10424.22|ACATGGGTCAGTCGGTCCTGAG</td><td>>hg38_21_+_8433172_8446622_RNA45SN1_45S_50ntFlanks</td><td>10403</td><td>10424</td><td>22</td><td>rRF 45S</td><td>NO</td><td>36.00</td><td>75.17</td><td>508.45</td><td>289.39</td></tr>
<tr><td>>hg38_21_+_8433172_8446622_RNA45SN1_45S_50ntFlanks@10404.10424.21|CATGGGTCAGTCGGTCCTGAG</td><td>>hg38_21_+_8433172_8446622_RNA45SN1_45S_50ntFlanks</td><td>10404</td><td>10424</td><td>21</td><td>rRF 45S</td><td>NO</td><td>33.89</td><td>84.30</td><td>504.79</td><td>276.97</td></tr>
<tr><td>>hg38_21_+_8433172_8446622_RNA45SN1_45S_50ntFlanks@10405.10424.20|ATGGGTCAGTCGGTCCTGAG</td><td>>hg38_21_+_8433172_8446622_RNA45SN1_45S_50ntFlanks</td><td>10405</td><td>10424</td><td>20</td><td>rRF 45S</td><td>NO</td><td>70.27</td><td>88.64</td><td>883.23</td><td>288.63</td></tr>
<tr><td>>hg38_21_+_8433172_8446622_RNA45SN1_45S_50ntFlanks@10406.10424.19|TGGGTCAGTCGGTCCTGAG</td><td>>hg38_21_+_8433172_8446622_RNA45SN1_45S_50ntFlanks</td><td>10406</td><td>10424</td><td>19</td><td>rRF 45S</td><td>NO</td><td>0.00</td><td>0.00</td><td>173.69</td><td>108.87</td></tr>
<tr><td>>hg38_21_+_8433172_8446622_RNA45SN1_45S_50ntFlanks@10677.10707.31|GCGGTTCCGGCGGCGTCCGGTGAGCTCTCGC</td><td>>hg38_21_+_8433172_8446622_RNA45SN1_45S_50ntFlanks</td><td>10677</td><td>10707</td><td>31</td><td>rRF 45S</td><td>NO</td><td>0.00</td><td>0.00</td><td>11.48</td><td>0.00</td></tr>
<tr><td>>hg38_21_+_8433172_8446622_RNA45SN1_45S_50ntFlanks@10677.10708.32|GCGGTTCCGGCGGCGTCCGGTGAGCTCTCGCT</td><td>>hg38_21_+_8433172_8446622_RNA45SN1_45S_50ntFlanks</td><td>10677</td><td>10708</td><td>32</td><td>rRF 45S</td><td>NO</td><td>0.00</td><td>0.00</td><td>10.01</td><td>0.00</td></tr>
<tr><td>>hg38_21_+_8433172_8446622_RNA45SN1_45S_50ntFlanks@10678.10708.31|CGGTTCCGGCGGCGTCCGGTGAGCTCTCGCT</td><td>>hg38_21_+_8433172_8446622_RNA45SN1_45S_50ntFlanks</td><td>10678</td><td>10708</td><td>31</td><td>rRF 45S</td><td>NO</td><td>0.00</td><td>0.00</td><td>10.56</td><td>0.00</td></tr>
<tr><td>>hg38_21_+_8433172_8446622_RNA45SN1_45S_50ntFlanks@10678.10710.33|CGGTTCCGGCGGCGTCCGGTGAGCTCTCGCTGG</td><td>>hg38_21_+_8433172_8446622_RNA45SN1_45S_50ntFlanks</td><td>10678</td><td>10710</td><td>33</td><td>rRF 45S</td><td>NO</td><td>0.00</td><td>91.43</td><td>60.00</td><td>33.60</td></tr>
<tr><td>>hg38_21_+_8433172_8446622_RNA45SN1_45S_50ntFlanks@10678.10711.34|CGGTTCCGGCGGCGTCCGGTGAGCTCTCGCTGGC</td><td>>hg38_21_+_8433172_8446622_RNA45SN1_45S_50ntFlanks</td><td>10678</td><td>10711</td><td>34</td><td>rRF 45S</td><td>NO</td><td>0.00</td><td>23.95</td><td>0.00</td><td>0.00</td></tr>
<tr><td>>hg38_21_+_8433172_8446622_RNA45SN1_45S_50ntFlanks@10686.10707.22|GCGGCGTCCGGTGAGCTCTCGC</td><td>>hg38_21_+_8433172_8446622_RNA45SN1_45S_50ntFlanks</td><td>10686</td><td>10707</td><td>22</td><td>rRF 45S</td><td>NO</td><td>0.00</td><td>0.00</td><td>10.72</td><td>0.00</td></tr>
<tr><td>>hg38_21_+_8433172_8446622_RNA45SN1_45S_50ntFlanks@10841.10867.27|CGTAACTTCGGGATAAGGATTGGCTCT</td><td>>hg38_21_+_8433172_8446622_RNA45SN1_45S_50ntFlanks</td><td>10841</td><td>10867</td><td>27</td><td>rRF 45S</td><td>NO</td><td>25.07</td><td>167.84</td><td>0.00</td><td>0.00</td></tr>
</table>

# BT-20

| | | | | | | | | | | |
|---|---|---|---|---|---|---|---|---|---|---|
| >hg38_21_+_8433172_8446622_RNA45SN1_45S_50ntFlanks@10841.10868.28\|CGTAACTTCGGGATAAGGATTGGCTCTA | >hg38_21_+_8433172_8446622_RNA45SN1_45S_50ntFlanks | 10841 | 10868 | 28 | rRF 45S | NO | 0.00 | 36.88 | 0.00 | 0.00 |
| >hg38_21_+_8433172_8446622_RNA45SN1_45S_50ntFlanks@11566.11584.19\|TAGCAGCCGACTTAGAACT | >hg38_21_+_8433172_8446622_RNA45SN1_45S_50ntFlanks | 11566 | 11584 | 19 | rRF 45S | NO | 0.00 | 42.66 | 0.00 | 0.00 |
| >hg38_21_+_8433172_8446622_RNA45SN1_45S_50ntFlanks@11566.11611.46\|TAGCAGCCGACTTAGAACTGGTGCGGACCAGGGGAATCCGACTGTT | >hg38_21_+_8433172_8446622_RNA45SN1_45S_50ntFlanks | 11566 | 11611 | 46 | rRF 45S | NO | 0.00 | 0.00 | 10.08 | 127.61 |
| >hg38_21_+_8433172_8446622_RNA45SN1_45S_50ntFlanks@11567.11584.18\|AGCAGCCGACTTAGAACT | >hg38_21_+_8433172_8446622_RNA45SN1_45S_50ntFlanks | 11567 | 11584 | 18 | rRF 45S | NO | 40.01 | 140.92 | 37.82 | 72.53 |
| >hg38_21_+_8433172_8446622_RNA45SN1_45S_50ntFlanks@11567.11611.45\|AGCAGCCGACTTAGAACTGGTGCGGACCAGGGGAATCCGACTGTT | >hg38_21_+_8433172_8446622_RNA45SN1_45S_50ntFlanks | 11567 | 11611 | 45 | rRF 45S | NO | 0.00 | 0.00 | 10.19 | 132.58 |
| >hg38_21_+_8433172_8446622_RNA45SN1_45S_50ntFlanks@11568.11587.20\|GCAGCCGACTTAGAACTGGT | >hg38_21_+_8433172_8446622_RNA45SN1_45S_50ntFlanks | 11568 | 11587 | 20 | rRF 45S | NO | 0.00 | 0.00 | 10.08 | 0.00 |
| >hg38_21_+_8433172_8446622_RNA45SN1_45S_50ntFlanks@11568.11611.44\|GCAGCCGACTTAGAACTGGTGCGGACCAGGGGAATCCGACTGTT | >hg38_21_+_8433172_8446622_RNA45SN1_45S_50ntFlanks | 11568 | 11611 | 44 | rRF 45S | NO | 83.33 | 74.42 | 15.90 | 334.04 |
| >hg38_21_+_8433172_8446622_RNA45SN1_45S_50ntFlanks@11578.11613.36\|TAGAACTGGTGCGGACCAGGGGAATCCGACTGTTTA | >hg38_21_+_8433172_8446622_RNA45SN1_45S_50ntFlanks | 11578 | 11613 | 36 | rRF 45S | NO | 43.20 | 0.00 | 0.00 | 83.77 |
| >hg38_21_+_8433172_8446622_RNA45SN1_45S_50ntFlanks@11580.11613.34\|GAACTGGTGCGGACCAGGGGAATCCGACTGTTTA | >hg38_21_+_8433172_8446622_RNA45SN1_45S_50ntFlanks | 11580 | 11613 | 34 | rRF 45S | NO | 42.67 | 74.43 | 11.57 | 146.39 |
| >hg38_21_+_8433172_8446622_RNA45SN1_45S_50ntFlanks@11582.11602.21\|ACTGGTGCGGACCAGGGGAAT | >hg38_21_+_8433172_8446622_RNA45SN1_45S_50ntFlanks | 11582 | 11602 | 21 | rRF 45S | NO | 0.00 | 24.45 | 0.00 | 0.00 |
| >hg38_21_+_8433172_8446622_RNA45SN1_45S_50ntFlanks@11582.11611.30\|ACTGGTGCGGACCAGGGGAATCCGACTGTT | >hg38_21_+_8433172_8446622_RNA45SN1_45S_50ntFlanks | 11582 | 11611 | 30 | rRF 45S | NO | 0.00 | 0.00 | 10.00 | 0.00 |
| >hg38_21_+_8433172_8446622_RNA45SN1_45S_50ntFlanks@11582.11616.35\|ACTGGTGCGGACCAGGGGAATCCGACTGTTTAATT | >hg38_21_+_8433172_8446622_RNA45SN1_45S_50ntFlanks | 11582 | 11616 | 35 | rRF 45S | NO | 0.00 | 0.00 | 11.44 | 0.00 |
| >hg38_21_+_8433172_8446622_RNA45SN1_45S_50ntFlanks@11583.11611.29\|CTGGTGCGGACCAGGGGAATCCGACTGTT | >hg38_21_+_8433172_8446622_RNA45SN1_45S_50ntFlanks | 11583 | 11611 | 29 | rRF 45S | NO | 40.14 | 9.44 | 9.85 | 32.22 |
| >hg38_21_+_8433172_8446622_RNA45SN1_45S_50ntFlanks@11583.11612.30\|CTGGTGCGGACCAGGGGAATCCGACTGTTT | >hg38_21_+_8433172_8446622_RNA45SN1_45S_50ntFlanks | 11583 | 11612 | 30 | rRF 45S | NO | 27.72 | 0.00 | 0.00 | 0.00 |
| >hg38_21_+_8433172_8446622_RNA45SN1_45S_50ntFlanks@11584.11611.28\|TGGTGCGGACCAGGGGAATCCGACTGTT | >hg38_21_+_8433172_8446622_RNA45SN1_45S_50ntFlanks | 11584 | 11611 | 28 | rRF 45S | NO | 38.86 | 0.00 | 0.00 | 33.65 |
| >hg38_21_+_8433172_8446622_RNA45SN1_45S_50ntFlanks@11584.11612.29\|TGGTGCGGACCAGGGGAATCCGACTGTTT | >hg38_21_+_8433172_8446622_RNA45SN1_45S_50ntFlanks | 11584 | 11612 | 29 | rRF 45S | NO | 27.04 | 0.00 | 0.00 | 0.00 |
| >hg38_21_+_8433172_8446622_RNA45SN1_45S_50ntFlanks@11584.11616.33\|TGGTGCGGACCAGGGGAATCCGACTGTTTAATT | >hg38_21_+_8433172_8446622_RNA45SN1_45S_50ntFlanks | 11584 | 11616 | 33 | rRF 45S | NO | 23.77 | 0.00 | 0.00 | 0.00 |
| >hg38_21_+_8433172_8446622_RNA45SN1_45S_50ntFlanks@11604.11623.20\|CGACTGTTTAATTAAAACAA | >hg38_21_+_8433172_8446622_RNA45SN1_45S_50ntFlanks | 11604 | 11623 | 20 | rRF 45S | NO | 0.00 | 70.33 | 53.79 | 160.85 |
| >hg38_21_+_8433172_8446622_RNA45SN1_45S_50ntFlanks@11666.11685.20\|TCTGCCCAGTGCTCTGAATG | >hg38_21_+_8433172_8446622_RNA45SN1_45S_50ntFlanks | 11666 | 11685 | 20 | rRF 45S | NO | 0.00 | 0.00 | 11.54 | 0.00 |
| >hg38_21_+_8433172_8446622_RNA45SN1_45S_50ntFlanks@11666.11687.22\|TCTGCCCAGTGCTCTGAATGTC | >hg38_21_+_8433172_8446622_RNA45SN1_45S_50ntFlanks | 11666 | 11687 | 22 | rRF 45S | NO | 0.00 | 32.63 | 10.64 | 0.00 |
| >hg38_21_+_8433172_8446622_RNA45SN1_45S_50ntFlanks@11666.11702.37\|TCTGCCCAGTGCTCTGAATGTCAAAGTGAAGAAATTC | >hg38_21_+_8433172_8446622_RNA45SN1_45S_50ntFlanks | 11666 | 11702 | 37 | rRF 45S | NO | 0.00 | 28.64 | 0.00 | 0.00 |
| >hg38_21_+_8433172_8446622_RNA45SN1_45S_50ntFlanks@11667.11687.21\|CTGCCCAGTGCTCTGAATGTC | >hg38_21_+_8433172_8446622_RNA45SN1_45S_50ntFlanks | 11667 | 11687 | 21 | rRF 45S | NO | 0.00 | 28.59 | 0.00 | 0.00 |
| >hg38_21_+_8433172_8446622_RNA45SN1_45S_50ntFlanks@11667.11691.25\|CTGCCCAGTGCTCTGAATGTCAAAG | >hg38_21_+_8433172_8446622_RNA45SN1_45S_50ntFlanks | 11667 | 11691 | 25 | rRF 45S | NO | 0.00 | 25.25 | 0.00 | 9.77 |
| >hg38_21_+_8433172_8446622_RNA45SN1_45S_50ntFlanks@11667.11702.36\|CTGCCCAGTGCTCTGAATGTCAAAGTGAAGAAATTC | >hg38_21_+_8433172_8446622_RNA45SN1_45S_50ntFlanks | 11667 | 11702 | 36 | rRF 45S | NO | 0.00 | 22.97 | 0.00 | 0.00 |
| >hg38_21_+_8433172_8446622_RNA45SN1_45S_50ntFlanks@11668.11687.20\|TGCCCAGTGCTCTGAATGTC | >hg38_21_+_8433172_8446622_RNA45SN1_45S_50ntFlanks | 11668 | 11687 | 20 | rRF 45S | NO | 0.00 | 38.30 | 0.00 | 0.00 |
| >hg38_21_+_8433172_8446622_RNA45SN1_45S_50ntFlanks@11668.11702.35\|TGCCCAGTGCTCTGAATGTCAAAGTGAAGAAATTC | >hg38_21_+_8433172_8446622_RNA45SN1_45S_50ntFlanks | 11668 | 11702 | 35 | rRF 45S | NO | 0.00 | 87.59 | 0.00 | 0.00 |
| >hg38_21_+_8433172_8446622_RNA45SN1_45S_50ntFlanks@11668.11703.36\|TGCCCAGTGCTCTGAATGTCAAAGTGAAGAAATTCA | >hg38_21_+_8433172_8446622_RNA45SN1_45S_50ntFlanks | 11668 | 11703 | 36 | rRF 45S | NO | 0.00 | 26.24 | 0.00 | 0.00 |
| >hg38_21_+_8433172_8446622_RNA45SN1_45S_50ntFlanks@11668.11706.39\|TGCCCAGTGCTCTGAATGTCAAAGTGAAGAAATTCAATG | >hg38_21_+_8433172_8446622_RNA45SN1_45S_50ntFlanks | 11668 | 11706 | 39 | rRF 45S | NO | 0.00 | 29.79 | 0.00 | 0.00 |
| >hg38_21_+_8433172_8446622_RNA45SN1_45S_50ntFlanks@11669.11687.19\|GCCCAGTGCTCTGAATGTC | >hg38_21_+_8433172_8446622_RNA45SN1_45S_50ntFlanks | 11669 | 11687 | 19 | rRF 45S | NO | 0.00 | 38.53 | 0.00 | 0.00 |
| >hg38_21_+_8433172_8446622_RNA45SN1_45S_50ntFlanks@11669.11702.34\|GCCCAGTGCTCTGAATGTCAAAGTGAAGAAATTC | >hg38_21_+_8433172_8446622_RNA45SN1_45S_50ntFlanks | 11669 | 11702 | 34 | rRF 45S | NO | 0.00 | 69.67 | 0.00 | 0.00 |
| >hg38_21_+_8433172_8446622_RNA45SN1_45S_50ntFlanks@11669.11706.38\|GCCCAGTGCTCTGAATGTCAAAGTGAAGAAATTCAATG | >hg38_21_+_8433172_8446622_RNA45SN1_45S_50ntFlanks | 11669 | 11706 | 38 | rRF 45S | NO | 0.00 | 34.71 | 0.00 | 0.00 |
| >hg38_21_+_8433172_8446622_RNA45SN1_45S_50ntFlanks@11675.11702.28\|TGCTCTGAATGTCAAAGTGAAGAAATTC | >hg38_21_+_8433172_8446622_RNA45SN1_45S_50ntFlanks | 11675 | 11702 | 28 | rRF 45S | NO | 0.00 | 26.46 | 0.00 | 0.00 |
| >hg38_21_+_8433172_8446622_RNA45SN1_45S_50ntFlanks@11675.11706.32\|TGCTCTGAATGTCAAAGTGAAGAAATTCAATG | >hg38_21_+_8433172_8446622_RNA45SN1_45S_50ntFlanks | 11675 | 11706 | 32 | rRF 45S | NO | 0.00 | 28.42 | 0.00 | 0.00 |
| >hg38_21_+_8433172_8446622_RNA45SN1_45S_50ntFlanks@11770.11794.25\|AATTAGTGACGCGCATGAATGGATG | >hg38_21_+_8433172_8446622_RNA45SN1_45S_50ntFlanks | 11770 | 11794 | 25 | rRF 45S | NO | 0.00 | 24.80 | 0.00 | 0.00 |
| >hg38_21_+_8433172_8446622_RNA45SN1_45S_50ntFlanks@12004.12026.23\|CGGGCCGCCGGTGAAATACCACT | >hg38_21_+_8433172_8446622_RNA45SN1_45S_50ntFlanks | 12004 | 12026 | 23 | rRF 45S | NO | 0.00 | 0.00 | 11.00 | 0.00 |
| >hg38_21_+_8433172_8446622_RNA45SN1_45S_50ntFlanks@12004.12033.30\|CGGGCCGCCGGTGAAATACCACTACTCTGA | >hg38_21_+_8433172_8446622_RNA45SN1_45S_50ntFlanks | 12004 | 12033 | 30 | rRF 45S | NO | 0.00 | 32.55 | 0.00 | 79.66 |
| >hg38_21_+_8433172_8446622_RNA45SN1_45S_50ntFlanks@12004.12035.32\|CGGGCCGCCGGTGAAATACCACTACTCTGATC | >hg38_21_+_8433172_8446622_RNA45SN1_45S_50ntFlanks | 12004 | 12035 | 32 | rRF 45S | NO | 11.02 | 27.38 | 70.04 | 103.58 |
| >hg38_21_+_8433172_8446622_RNA45SN1_45S_50ntFlanks@12396.12414.19\|GCGACGTCGCTTTTTGATC | >hg38_21_+_8433172_8446622_RNA45SN1_45S_50ntFlanks | 12396 | 12414 | 19 | rRF 45S | NO | 0.00 | 10.45 | 0.00 | 0.00 |

# BT-20

| | | | | | | | | | | |
|---|---|---|---|---|---|---|---|---|---|---|
| >hg38_21_+_8433172_8446622_RNA45SN1_45S_50ntFlanks@12397.12418.22\|CGACGTCGCTTTTGATCCTTC | >hg38_21_+_8433172_8446622_RNA45SN1_45S_50ntFlanks | 12397 | 12418 | 22 | rRF 45S | NO | 0.00 | 10.40 | 0.00 | 0.00 |
| >hg38_21_+_8433172_8446622_RNA45SN1_45S_50ntFlanks@12407.12433.27\|TTTTGATCCTTCGATGTCGGCTCTTCC | >hg38_21_+_8433172_8446622_RNA45SN1_45S_50ntFlanks | 12407 | 12433 | 27 | rRF 45S | NO | 0.00 | 10.13 | 0.00 | 0.00 |
| >hg38_21_+_8433172_8446622_RNA45SN1_45S_50ntFlanks@12409.12434.26\|TTGATCCTTCGATGTCGGCTCTTCCT | >hg38_21_+_8433172_8446622_RNA45SN1_45S_50ntFlanks | 12409 | 12434 | 26 | rRF 45S | NO | 11.17 | 0.00 | 0.00 | 0.00 |
| >hg38_21_+_8433172_8446622_RNA45SN1_45S_50ntFlanks@12415.12433.19\|CTTCGATGTCGGCTCTTCC | >hg38_21_+_8433172_8446622_RNA45SN1_45S_50ntFlanks | 12415 | 12433 | 19 | rRF 45S | NO | 0.00 | 70.32 | 0.00 | 0.00 |
| >hg38_21_+_8433172_8446622_RNA45SN1_45S_50ntFlanks@12415.12434.20\|CTTCGATGTCGGCTCTTCCT | >hg38_21_+_8433172_8446622_RNA45SN1_45S_50ntFlanks | 12415 | 12434 | 20 | rRF 45S | NO | 112.49 | 97.10 | 0.00 | 27.36 |
| >hg38_21_+_8433172_8446622_RNA45SN1_45S_50ntFlanks@12415.12437.23\|CTTCGATGTCGGCTCTTCCTATC | >hg38_21_+_8433172_8446622_RNA45SN1_45S_50ntFlanks | 12415 | 12437 | 23 | rRF 45S | NO | 0.00 | 35.88 | 0.00 | 25.16 |
| >hg38_21_+_8433172_8446622_RNA45SN1_45S_50ntFlanks@12415.12438.24\|CTTCGATGTCGGCTCTTCCTATCA | >hg38_21_+_8433172_8446622_RNA45SN1_45S_50ntFlanks | 12415 | 12438 | 24 | rRF 45S | NO | 11.21 | 10.28 | 0.00 | 0.00 |
| >hg38_21_+_8433172_8446622_RNA45SN1_45S_50ntFlanks@12415.12439.25\|CTTCGATGTCGGCTCTTCCTATCAT | >hg38_21_+_8433172_8446622_RNA45SN1_45S_50ntFlanks | 12415 | 12439 | 25 | rRF 45S | NO | 0.00 | 24.26 | 0.00 | 0.00 |
| >hg38_21_+_8433172_8446622_RNA45SN1_45S_50ntFlanks@12415.12440.26\|CTTCGATGTCGGCTCTTCCTATCATT | >hg38_21_+_8433172_8446622_RNA45SN1_45S_50ntFlanks | 12415 | 12440 | 26 | rRF 45S | NO | 0.00 | 10.13 | 0.00 | 0.00 |
| >hg38_21_+_8433172_8446622_RNA45SN1_45S_50ntFlanks@12415.12442.28\|CTTCGATGTCGGCTCTTCCTATCATTGT | >hg38_21_+_8433172_8446622_RNA45SN1_45S_50ntFlanks | 12415 | 12442 | 28 | rRF 45S | NO | 0.00 | 23.17 | 0.00 | 0.00 |
| >hg38_21_+_8433172_8446622_RNA45SN1_45S_50ntFlanks@12415.12443.29\|CTTCGATGTCGGCTCTTCCTATCATTGTG | >hg38_21_+_8433172_8446622_RNA45SN1_45S_50ntFlanks | 12415 | 12443 | 29 | rRF 45S | NO | 0.00 | 24.86 | 0.00 | 0.00 |
| >hg38_21_+_8433172_8446622_RNA45SN1_45S_50ntFlanks@12415.12444.30\|CTTCGATGTCGGCTCTTCCTATCATTGTGA | >hg38_21_+_8433172_8446622_RNA45SN1_45S_50ntFlanks | 12415 | 12444 | 30 | rRF 45S | NO | 0.00 | 27.88 | 0.00 | 0.00 |
| >hg38_21_+_8433172_8446622_RNA45SN1_45S_50ntFlanks@12415.12448.34\|CTTCGATGTCGGCTCTTCCTATCATTGTGAAGCA | >hg38_21_+_8433172_8446622_RNA45SN1_45S_50ntFlanks | 12415 | 12448 | 34 | rRF 45S | NO | 0.00 | 43.41 | 0.00 | 41.24 |
| >hg38_21_+_8433172_8446622_RNA45SN1_45S_50ntFlanks@12416.12433.18\|TTCGATGTCGGCTCTTCC | >hg38_21_+_8433172_8446622_RNA45SN1_45S_50ntFlanks | 12416 | 12433 | 18 | rRF 45S | NO | 0.00 | 121.31 | 0.00 | 0.00 |
| >hg38_21_+_8433172_8446622_RNA45SN1_45S_50ntFlanks@12416.12434.19\|TTCGATGTCGGCTCTTCCT | >hg38_21_+_8433172_8446622_RNA45SN1_45S_50ntFlanks | 12416 | 12434 | 19 | rRF 45S | NO | 0.00 | 109.88 | 0.00 | 0.00 |
| >hg38_21_+_8433172_8446622_RNA45SN1_45S_50ntFlanks@12416.12437.22\|TTCGATGTCGGCTCTTCCTATC | >hg38_21_+_8433172_8446622_RNA45SN1_45S_50ntFlanks | 12416 | 12437 | 22 | rRF 45S | NO | 0.00 | 98.05 | 0.00 | 0.00 |
| >hg38_21_+_8433172_8446622_RNA45SN1_45S_50ntFlanks@12416.12438.23\|TTCGATGTCGGCTCTTCCTATCA | >hg38_21_+_8433172_8446622_RNA45SN1_45S_50ntFlanks | 12416 | 12438 | 23 | rRF 45S | NO | 0.00 | 67.83 | 0.00 | 10.21 |
| >hg38_21_+_8433172_8446622_RNA45SN1_45S_50ntFlanks@12416.12439.24\|TTCGATGTCGGCTCTTCCTATCAT | >hg38_21_+_8433172_8446622_RNA45SN1_45S_50ntFlanks | 12416 | 12439 | 24 | rRF 45S | NO | 0.00 | 39.27 | 0.00 | 0.00 |
| >hg38_21_+_8433172_8446622_RNA45SN1_45S_50ntFlanks@12416.12440.25\|TTCGATGTCGGCTCTTCCTATCATT | >hg38_21_+_8433172_8446622_RNA45SN1_45S_50ntFlanks | 12416 | 12440 | 25 | rRF 45S | NO | 0.00 | 69.49 | 0.00 | 0.00 |
| >hg38_21_+_8433172_8446622_RNA45SN1_45S_50ntFlanks@12416.12441.26\|TTCGATGTCGGCTCTTCCTATCATTG | >hg38_21_+_8433172_8446622_RNA45SN1_45S_50ntFlanks | 12416 | 12441 | 26 | rRF 45S | NO | 0.00 | 26.33 | 0.00 | 0.00 |
| >hg38_21_+_8433172_8446622_RNA45SN1_45S_50ntFlanks@12416.12442.27\|TTCGATGTCGGCTCTTCCTATCATTGT | >hg38_21_+_8433172_8446622_RNA45SN1_45S_50ntFlanks | 12416 | 12442 | 27 | rRF 45S | NO | 0.00 | 40.68 | 0.00 | 0.00 |
| >hg38_21_+_8433172_8446622_RNA45SN1_45S_50ntFlanks@12416.12443.28\|TTCGATGTCGGCTCTTCCTATCATTGTG | >hg38_21_+_8433172_8446622_RNA45SN1_45S_50ntFlanks | 12416 | 12443 | 28 | rRF 45S | NO | 0.00 | 31.74 | 0.00 | 0.00 |
| >hg38_21_+_8433172_8446622_RNA45SN1_45S_50ntFlanks@12416.12444.29\|TTCGATGTCGGCTCTTCCTATCATTGTGA | >hg38_21_+_8433172_8446622_RNA45SN1_45S_50ntFlanks | 12416 | 12444 | 29 | rRF 45S | NO | 0.00 | 68.25 | 0.00 | 0.00 |
| >hg38_21_+_8433172_8446622_RNA45SN1_45S_50ntFlanks@12416.12446.31\|TTCGATGTCGGCTCTTCCTATCATTGTGAAG | >hg38_21_+_8433172_8446622_RNA45SN1_45S_50ntFlanks | 12416 | 12446 | 31 | rRF 45S | NO | 0.00 | 10.08 | 0.00 | 0.00 |
| >hg38_21_+_8433172_8446622_RNA45SN1_45S_50ntFlanks@12416.12447.32\|TTCGATGTCGGCTCTTCCTATCATTGTGAAGC | >hg38_21_+_8433172_8446622_RNA45SN1_45S_50ntFlanks | 12416 | 12447 | 32 | rRF 45S | NO | 0.00 | 25.57 | 0.00 | 0.00 |
| >hg38_21_+_8433172_8446622_RNA45SN1_45S_50ntFlanks@12416.12448.33\|TTCGATGTCGGCTCTTCCTATCATTGTGAAGCA | >hg38_21_+_8433172_8446622_RNA45SN1_45S_50ntFlanks | 12416 | 12448 | 33 | rRF 45S | NO | 0.00 | 103.81 | 0.00 | 0.00 |
| >hg38_21_+_8433172_8446622_RNA45SN1_45S_50ntFlanks@12417.12434.18\|TCGATGTCGGCTCTTCCT | >hg38_21_+_8433172_8446622_RNA45SN1_45S_50ntFlanks | 12417 | 12434 | 18 | rRF 45S | NO | 11.04 | 40.81 | 0.00 | 0.00 |
| >hg38_21_+_8433172_8446622_RNA45SN1_45S_50ntFlanks@12417.12437.21\|TCGATGTCGGCTCTTCCTATC | >hg38_21_+_8433172_8446622_RNA45SN1_45S_50ntFlanks | 12417 | 12437 | 21 | rRF 45S | NO | 0.00 | 29.22 | 0.00 | 0.00 |
| >hg38_21_+_8433172_8446622_RNA45SN1_45S_50ntFlanks@12417.12438.22\|TCGATGTCGGCTCTTCCTATCA | >hg38_21_+_8433172_8446622_RNA45SN1_45S_50ntFlanks | 12417 | 12438 | 22 | rRF 45S | NO | 0.00 | 24.80 | 0.00 | 0.00 |
| >hg38_21_+_8433172_8446622_RNA45SN1_45S_50ntFlanks@12417.12448.32\|TCGATGTCGGCTCTTCCTATCATTGTGAAGCA | >hg38_21_+_8433172_8446622_RNA45SN1_45S_50ntFlanks | 12417 | 12448 | 32 | rRF 45S | NO | 0.00 | 31.22 | 0.00 | 0.00 |
| >hg38_21_+_8433172_8446622_RNA45SN1_45S_50ntFlanks@12418.12437.20\|CGATGTCGGCTCTTCCTATC | >hg38_21_+_8433172_8446622_RNA45SN1_45S_50ntFlanks | 12418 | 12437 | 20 | rRF 45S | NO | 0.00 | 24.92 | 0.00 | 0.00 |
| >hg38_21_+_8433172_8446622_RNA45SN1_45S_50ntFlanks@12418.12439.22\|CGATGTCGGCTCTTCCTATCAT | >hg38_21_+_8433172_8446622_RNA45SN1_45S_50ntFlanks | 12418 | 12439 | 22 | rRF 45S | NO | 0.00 | 10.18 | 0.00 | 0.00 |
| >hg38_21_+_8433172_8446622_RNA45SN1_45S_50ntFlanks@12418.12448.31\|CGATGTCGGCTCTTCCTATCATTGTGAAGCA | >hg38_21_+_8433172_8446622_RNA45SN1_45S_50ntFlanks | 12418 | 12448 | 31 | rRF 45S | NO | 0.00 | 25.06 | 0.00 | 0.00 |
| >hg38_21_+_8433172_8446622_RNA45SN1_45S_50ntFlanks@12419.12437.19\|GATGTCGGCTCTTCCTATC | >hg38_21_+_8433172_8446622_RNA45SN1_45S_50ntFlanks | 12419 | 12437 | 19 | rRF 45S | NO | 24.34 | 152.51 | 0.00 | 27.12 |
| >hg38_21_+_8433172_8446622_RNA45SN1_45S_50ntFlanks@12419.12438.20\|GATGTCGGCTCTTCCTATCA | >hg38_21_+_8433172_8446622_RNA45SN1_45S_50ntFlanks | 12419 | 12438 | 20 | rRF 45S | NO | 0.00 | 71.17 | 0.00 | 28.06 |
| >hg38_21_+_8433172_8446622_RNA45SN1_45S_50ntFlanks@12419.12439.21\|GATGTCGGCTCTTCCTATCAT | >hg38_21_+_8433172_8446622_RNA45SN1_45S_50ntFlanks | 12419 | 12439 | 21 | rRF 45S | NO | 0.00 | 76.23 | 0.00 | 0.00 |
| >hg38_21_+_8433172_8446622_RNA45SN1_45S_50ntFlanks@12419.12440.22\|GATGTCGGCTCTTCCTATCATT | >hg38_21_+_8433172_8446622_RNA45SN1_45S_50ntFlanks | 12419 | 12440 | 22 | rRF 45S | NO | 0.00 | 31.66 | 0.00 | 0.00 |
| >hg38_21_+_8433172_8446622_RNA45SN1_45S_50ntFlanks@12419.12441.23\|GATGTCGGCTCTTCCTATCATTG | >hg38_21_+_8433172_8446622_RNA45SN1_45S_50ntFlanks | 12419 | 12441 | 23 | rRF 45S | NO | 0.00 | 27.71 | 0.00 | 0.00 |

# BT-20

| | | | | | | | | | | |
|---|---|---|---|---|---|---|---|---|---|---|
| >hg38_21_+_8433172_8446622_RNA45SN1_45S_50ntFlanks@12419.12442.24\|GATGTCGGCTCTTCCTATCATTGT | >hg38_21_+_8433172_8446622_RNA45SN1_45S_50ntFlanks | 12419 | 12442 | 24 | rRF 45S | NO | 0.00 | 69.78 | 0.00 | 0.00 |
| >hg38_21_+_8433172_8446622_RNA45SN1_45S_50ntFlanks@12419.12443.25\|GATGTCGGCTCTTCCTATCATTGTG | >hg38_21_+_8433172_8446622_RNA45SN1_45S_50ntFlanks | 12419 | 12443 | 25 | rRF 45S | NO | 0.00 | 30.49 | 0.00 | 0.00 |
| >hg38_21_+_8433172_8446622_RNA45SN1_45S_50ntFlanks@12419.12444.26\|GATGTCGGCTCTTCCTATCATTGTGA | >hg38_21_+_8433172_8446622_RNA45SN1_45S_50ntFlanks | 12419 | 12444 | 26 | rRF 45S | NO | 0.00 | 33.59 | 0.00 | 0.00 |
| >hg38_21_+_8433172_8446622_RNA45SN1_45S_50ntFlanks@12419.12447.29\|GATGTCGGCTCTTCCTATCATTGTGAAGC | >hg38_21_+_8433172_8446622_RNA45SN1_45S_50ntFlanks | 12419 | 12447 | 29 | rRF 45S | NO | 0.00 | 27.00 | 0.00 | 0.00 |
| >hg38_21_+_8433172_8446622_RNA45SN1_45S_50ntFlanks@12419.12448.30\|GATGTCGGCTCTTCCTATCATTGTGAAGCA | >hg38_21_+_8433172_8446622_RNA45SN1_45S_50ntFlanks | 12419 | 12448 | 30 | rRF 45S | NO | 0.00 | 105.84 | 0.00 | 0.00 |
| >hg38_21_+_8433172_8446622_RNA45SN1_45S_50ntFlanks@12419.12449.31\|GATGTCGGCTCTTCCTATCATTGTGAAGCAG | >hg38_21_+_8433172_8446622_RNA45SN1_45S_50ntFlanks | 12419 | 12449 | 31 | rRF 45S | NO | 0.00 | 27.40 | 0.00 | 0.00 |
| >hg38_21_+_8433172_8446622_RNA45SN1_45S_50ntFlanks@12419.12457.39\|GATGTCGGCTCTTCCTATCATTGTGAAGCAGAATTCACC | >hg38_21_+_8433172_8446622_RNA45SN1_45S_50ntFlanks | 12419 | 12457 | 39 | rRF 45S | NO | 0.00 | 23.14 | 0.00 | 0.00 |
| >hg38_21_+_8433172_8446622_RNA45SN1_45S_50ntFlanks@12420.12437.18\|ATGTCGGCTCTTCCTATC | >hg38_21_+_8433172_8446622_RNA45SN1_45S_50ntFlanks | 12420 | 12437 | 18 | rRF 45S | NO | 0.00 | 38.31 | 0.00 | 0.00 |
| >hg38_21_+_8433172_8446622_RNA45SN1_45S_50ntFlanks@12420.12439.20\|ATGTCGGCTCTTCCTATCAT | >hg38_21_+_8433172_8446622_RNA45SN1_45S_50ntFlanks | 12420 | 12439 | 20 | rRF 45S | NO | 0.00 | 26.72 | 0.00 | 0.00 |
| >hg38_21_+_8433172_8446622_RNA45SN1_45S_50ntFlanks@12420.12448.29\|ATGTCGGCTCTTCCTATCATTGTGAAGCA | >hg38_21_+_8433172_8446622_RNA45SN1_45S_50ntFlanks | 12420 | 12448 | 29 | rRF 45S | NO | 0.00 | 28.64 | 0.00 | 0.00 |
| >hg38_21_+_8433172_8446622_RNA45SN1_45S_50ntFlanks@12449.12480.32\|GAATTCACCAAGCGTTGGATTGTTCACCCACT | >hg38_21_+_8433172_8446622_RNA45SN1_45S_50ntFlanks | 12449 | 12480 | 32 | rRF 45S | NO | 0.00 | 40.65 | 0.00 | 0.00 |
| >hg38_21_+_8433172_8446622_RNA45SN1_45S_50ntFlanks@12450.12480.31\|AATTCACCAAGCGTTGGATTGTTCACCCACT | >hg38_21_+_8433172_8446622_RNA45SN1_45S_50ntFlanks | 12450 | 12480 | 31 | rRF 45S | NO | 0.00 | 36.97 | 0.00 | 0.00 |
| >hg38_21_+_8433172_8446622_RNA45SN1_45S_50ntFlanks@12458.12476.19\|AAGCGTTGGATTGTTCACC | >hg38_21_+_8433172_8446622_RNA45SN1_45S_50ntFlanks | 12458 | 12476 | 19 | rRF 45S | NO | 0.00 | 32.35 | 10.24 | 0.00 |
| >hg38_21_+_8433172_8446622_RNA45SN1_45S_50ntFlanks@12463.12496.34\|TTGGATTGTTCACCCACTAATAGGGAACGTGAGC | >hg38_21_+_8433172_8446622_RNA45SN1_45S_50ntFlanks | 12463 | 12496 | 34 | rRF 45S | NO | 0.00 | 23.76 | 0.00 | 0.00 |
| >hg38_21_+_8433172_8446622_RNA45SN1_45S_50ntFlanks@12474.12496.23\|ACCCACTAATAGGGAACGTGAGC | >hg38_21_+_8433172_8446622_RNA45SN1_45S_50ntFlanks | 12474 | 12496 | 23 | rRF 45S | NO | 0.00 | 23.55 | 0.00 | 0.00 |
| >hg38_21_+_8433172_8446622_RNA45SN1_45S_50ntFlanks@12477.12496.20\|CACTAATAGGGAACGTGAGC | >hg38_21_+_8433172_8446622_RNA45SN1_45S_50ntFlanks | 12477 | 12496 | 20 | rRF 45S | NO | 0.00 | 117.31 | 0.00 | 0.00 |
| >hg38_21_+_8433172_8446622_RNA45SN1_45S_50ntFlanks@12477.12497.21\|CACTAATAGGGAACGTGAGCT | >hg38_21_+_8433172_8446622_RNA45SN1_45S_50ntFlanks | 12477 | 12497 | 21 | rRF 45S | NO | 0.00 | 114.99 | 0.00 | 0.00 |
| >hg38_21_+_8433172_8446622_RNA45SN1_45S_50ntFlanks@12478.12496.19\|ACTAATAGGGAACGTGAGC | >hg38_21_+_8433172_8446622_RNA45SN1_45S_50ntFlanks | 12478 | 12496 | 19 | rRF 45S | NO | 0.00 | 94.45 | 9.58 | 0.00 |
| >hg38_21_+_8433172_8446622_RNA45SN1_45S_50ntFlanks@12478.12497.20\|ACTAATAGGGAACGTGAGCT | >hg38_21_+_8433172_8446622_RNA45SN1_45S_50ntFlanks | 12478 | 12497 | 20 | rRF 45S | NO | 0.00 | 122.13 | 14.05 | 0.00 |
| >hg38_21_+_8433172_8446622_RNA45SN1_45S_50ntFlanks@12479.12496.18\|CTAATAGGGAACGTGAGC | >hg38_21_+_8433172_8446622_RNA45SN1_45S_50ntFlanks | 12479 | 12496 | 18 | rRF 45S | NO | 0.00 | 95.69 | 0.00 | 0.00 |
| >hg38_21_+_8433172_8446622_RNA45SN1_45S_50ntFlanks@12479.12497.19\|CTAATAGGGAACGTGAGCT | >hg38_21_+_8433172_8446622_RNA45SN1_45S_50ntFlanks | 12479 | 12497 | 19 | rRF 45S | NO | 0.00 | 144.49 | 11.05 | 0.00 |
| >hg38_21_+_8433172_8446622_RNA45SN1_45S_50ntFlanks@12480.12497.18\|TAATAGGGAACGTGAGCT | >hg38_21_+_8433172_8446622_RNA45SN1_45S_50ntFlanks | 12480 | 12497 | 18 | rRF 45S | NO | 0.00 | 89.75 | 0.00 | 0.00 |
| >hg38_21_+_8433172_8446622_RNA45SN1_45S_50ntFlanks@12941.12960.20\|ATTCGTAGACGACCTGCTTC | >hg38_21_+_8433172_8446622_RNA45SN1_45S_50ntFlanks | 12941 | 12960 | 20 | rRF 45S | NO | 0.00 | 31.99 | 0.00 | 0.00 |
| >hg38_21_+_8433172_8446622_RNA45SN1_45S_50ntFlanks@12959.12993.35\|TCTGGGTCGGGGTTTCGTACGTAGCAGAGCAGCTC | >hg38_21_+_8433172_8446622_RNA45SN1_45S_50ntFlanks | 12959 | 12993 | 35 | rRF 45S | NO | 0.00 | 23.46 | 0.00 | 0.00 |
| >hg38_21_+_8433172_8446622_RNA45SN1_45S_50ntFlanks@12993.13039.47\|CCCTCGCTGCGATCTATTGAAAGTCAGCCCTCGACACAAGGGTTTGT | >hg38_21_+_8433172_8446622_RNA45SN1_45S_50ntFlanks | 12993 | 13039 | 47 | rRF 45S | NO | 0.00 | 31.05 | 0.00 | 242.68 |
| >hg38_21_+_8433172_8446622_RNA45SN1_45S_50ntFlanks@12994.13039.46\|CCTCGCTGCGATCTATTGAAAGTCAGCCCTCGACACAAGGGTTTGT | >hg38_21_+_8433172_8446622_RNA45SN1_45S_50ntFlanks | 12994 | 13039 | 46 | rRF 45S | NO | 0.00 | 136.49 | 10.43 | 411.17 |
| >hg38_21_+_8433172_8446622_RNA45SN1_45S_50ntFlanks@12995.13013.19\|CTCGCTGCGATCTATTGAA | >hg38_21_+_8433172_8446622_RNA45SN1_45S_50ntFlanks | 12995 | 13013 | 19 | rRF 45S | NO | 0.00 | 28.86 | 0.00 | 0.00 |
| >hg38_21_+_8433172_8446622_RNA45SN1_45S_50ntFlanks@12995.13016.22\|CTCGCTGCGATCTATTGAAAGT | >hg38_21_+_8433172_8446622_RNA45SN1_45S_50ntFlanks | 12995 | 13016 | 22 | rRF 45S | NO | 0.00 | 45.17 | 0.00 | 0.00 |
| >hg38_21_+_8433172_8446622_RNA45SN1_45S_50ntFlanks@12995.13017.23\|CTCGCTGCGATCTATTGAAAGTC | >hg38_21_+_8433172_8446622_RNA45SN1_45S_50ntFlanks | 12995 | 13017 | 23 | rRF 45S | NO | 0.00 | 32.06 | 0.00 | 0.00 |
| >hg38_21_+_8433172_8446622_RNA45SN1_45S_50ntFlanks@12995.13036.42\|CTCGCTGCGATCTATTGAAAGTCAGCCCTCGACACAAGGGTT | >hg38_21_+_8433172_8446622_RNA45SN1_45S_50ntFlanks | 12995 | 13036 | 42 | rRF 45S | NO | 25.90 | 188.21 | 0.00 | 100.91 |
| >hg38_21_+_8433172_8446622_RNA45SN1_45S_50ntFlanks@12995.13038.44\|CTCGCTGCGATCTATTGAAAGTCAGCCCTCGACACAAGGGTTTG | >hg38_21_+_8433172_8446622_RNA45SN1_45S_50ntFlanks | 12995 | 13038 | 44 | rRF 45S | NO | 0.00 | 28.60 | 0.00 | 30.04 |
| >hg38_21_+_8433172_8446622_RNA45SN1_45S_50ntFlanks@12995.13039.45\|CTCGCTGCGATCTATTGAAAGTCAGCCCTCGACACAAGGGTTTGT | >hg38_21_+_8433172_8446622_RNA45SN1_45S_50ntFlanks | 12995 | 13039 | 45 | rRF 45S | YES | 400.62 | 3334.88 | 321.76 | 2853.34 |
| >hg38_21_+_8433172_8446622_RNA45SN1_45S_50ntFlanks@12996.13013.18\|TCGCTGCGATCTATTGAA | >hg38_21_+_8433172_8446622_RNA45SN1_45S_50ntFlanks | 12996 | 13013 | 18 | rRF 45S | NO | 0.00 | 163.97 | 0.00 | 0.00 |
| >hg38_21_+_8433172_8446622_RNA45SN1_45S_50ntFlanks@12996.13014.19\|TCGCTGCGATCTATTGAAA | >hg38_21_+_8433172_8446622_RNA45SN1_45S_50ntFlanks | 12996 | 13014 | 19 | rRF 45S | NO | 0.00 | 89.81 | 0.00 | 0.00 |
| >hg38_21_+_8433172_8446622_RNA45SN1_45S_50ntFlanks@12996.13017.22\|TCGCTGCGATCTATTGAAAGTC | >hg38_21_+_8433172_8446622_RNA45SN1_45S_50ntFlanks | 12996 | 13017 | 22 | rRF 45S | NO | 0.00 | 84.48 | 0.00 | 0.00 |
| >hg38_21_+_8433172_8446622_RNA45SN1_45S_50ntFlanks@12996.13036.41\|TCGCTGCGATCTATTGAAAGTCAGCCCTCGACACAAGGGTT | >hg38_21_+_8433172_8446622_RNA45SN1_45S_50ntFlanks | 12996 | 13036 | 41 | rRF 45S | NO | 38.66 | 163.00 | 0.00 | 101.39 |
| >hg38_21_+_8433172_8446622_RNA45SN1_45S_50ntFlanks@12996.13038.43\|TCGCTGCGATCTATTGAAAGTCAGCCCTCGACACAAGGGTTTG | >hg38_21_+_8433172_8446622_RNA45SN1_45S_50ntFlanks | 12996 | 13038 | 43 | rRF 45S | NO | 0.00 | 24.83 | 0.00 | 0.00 |
| >hg38_21_+_8433172_8446622_RNA45SN1_45S_50ntFlanks@12996.13039.44\|TCGCTGCGATCTATTGAAAGTCAGCCCTCGACACAAGGGTTTGT | >hg38_21_+_8433172_8446622_RNA45SN1_45S_50ntFlanks | 12996 | 13039 | 44 | rRF 45S | YES | 499.93 | 2022.16 | 310.13 | 2119.68 |

# BT-20

| | | | | | | | | | | |
|---|---|---|---|---|---|---|---|---|---|---|
| >hg38_21_+_8433172_8446622_RNA45SN1_45S_50ntFlanks@12997.13014.18\|CGCTGCGATCTATTGAAA | >hg38_21_+_8433172_8446622_RNA45SN1_45S_50ntFlanks | 12997 | 13014 | 18 | rRF 45S | NO | 0.00 | 164.50 | 0.00 | 0.00 |
| >hg38_21_+_8433172_8446622_RNA45SN1_45S_50ntFlanks@12997.13015.19\|CGCTGCGATCTATTGAAAG | >hg38_21_+_8433172_8446622_RNA45SN1_45S_50ntFlanks | 12997 | 13015 | 19 | rRF 45S | NO | 0.00 | 24.43 | 0.00 | 0.00 |
| >hg38_21_+_8433172_8446622_RNA45SN1_45S_50ntFlanks@12997.13017.21\|CGCTGCGATCTATTGAAAGTC | >hg38_21_+_8433172_8446622_RNA45SN1_45S_50ntFlanks | 12997 | 13017 | 21 | rRF 45S | NO | 0.00 | 100.46 | 0.00 | 0.00 |
| >hg38_21_+_8433172_8446622_RNA45SN1_45S_50ntFlanks@12997.13036.40\|CGCTGCGATCTATTGAAAGTCAGCCCTCGACACAAGGGTT | >hg38_21_+_8433172_8446622_RNA45SN1_45S_50ntFlanks | 12997 | 13036 | 40 | rRF 45S | NO | 37.61 | 154.28 | 0.00 | 86.49 |
| >hg38_21_+_8433172_8446622_RNA45SN1_45S_50ntFlanks@12997.13038.42\|CGCTGCGATCTATTGAAAGTCAGCCCTCGACACAAGGGTTTG | >hg38_21_+_8433172_8446622_RNA45SN1_45S_50ntFlanks | 12997 | 13038 | 42 | rRF 45S | NO | 0.00 | 22.96 | 0.00 | 10.20 |
| >hg38_21_+_8433172_8446622_RNA45SN1_45S_50ntFlanks@12997.13039.43\|CGCTGCGATCTATTGAAAGTCAGCCCTCGACACAAGGGTTTGT | >hg38_21_+_8433172_8446622_RNA45SN1_45S_50ntFlanks | 12997 | 13039 | 43 | rRF 45S | YES | 547.82 | 2084.39 | 322.17 | 1892.09 |
| >hg38_21_+_8433172_8446622_RNA45SN1_45S_50ntFlanks@12998.13039.42\|GCTGCGATCTATTGAAAGTCAGCCCTCGACACAAGGGTTTGT | >hg38_21_+_8433172_8446622_RNA45SN1_45S_50ntFlanks | 12998 | 13039 | 42 | rRF 45S | NO | 119.49 | 152.67 | 82.91 | 225.35 |
| >hg38_21_+_8433172_8446622_RNA45SN1_45S_50ntFlanks@12999.13039.41\|CTGCGATCTATTGAAAGTCAGCCCTCGACACAAGGGTTTGT | >hg38_21_+_8433172_8446622_RNA45SN1_45S_50ntFlanks | 12999 | 13039 | 41 | rRF 45S | NO | 69.20 | 150.09 | 67.53 | 177.74 |
| >hg38_21_+_8433172_8446622_RNA45SN1_45S_50ntFlanks@13006.13039.34\|CTATTGAAAGTCAGCCCTCGACACAAGGGTTTGT | >hg38_21_+_8433172_8446622_RNA45SN1_45S_50ntFlanks | 13006 | 13039 | 34 | rRF 45S | NO | 32.57 | 41.82 | 52.67 | 82.38 |
| >hg38_21_+_8433172_8446622_RNA45SN1_45S_50ntFlanks@13007.13039.33\|TATTGAAAGTCAGCCCTCGACACAAGGGTTTGT | >hg38_21_+_8433172_8446622_RNA45SN1_45S_50ntFlanks | 13007 | 13039 | 33 | rRF 45S | NO | 40.20 | 113.20 | 62.79 | 159.69 |
| >hg38_21_+_8433172_8446622_RNA45SN1_45S_50ntFlanks@13008.13039.32\|ATTGAAAGTCAGCCCTCGACACAAGGGTTTGT | >hg38_21_+_8433172_8446622_RNA45SN1_45S_50ntFlanks | 13008 | 13039 | 32 | rRF 45S | NO | 156.07 | 211.77 | 135.43 | 277.85 |
| >hg38_21_+_8433172_8446622_RNA45SN1_45S_50ntFlanks@13022.13039.18\|CTCGACACAAGGGTTTGT | >hg38_21_+_8433172_8446622_RNA45SN1_45S_50ntFlanks | 13022 | 13039 | 18 | rRF 45S | NO | 0.00 | 36.90 | 0.00 | 30.93 |
| >hg38_1_-_228634819_228635039_RNA5S12_5S_50ntFlanks@51.74.24\|GTCTACGGCCATACCACCCTGAAC | >hg38_1_-_228634819_228635039_RNA5S12_5S_50ntFlanks | 51 | 74 | 24 | rRF 5S | NO | 0.00 | 79.13 | 93.33 | 0.00 |
| >hg38_1_-_228634819_228635039_RNA5S12_5S_50ntFlanks@51.75.25\|GTCTACGGCCATACCACCCTGAACG | >hg38_1_-_228634819_228635039_RNA5S12_5S_50ntFlanks | 51 | 75 | 25 | rRF 5S | NO | 0.00 | 94.59 | 82.67 | 0.00 |
| >hg38_1_-_228634819_228635039_RNA5S12_5S_50ntFlanks@51.76.26\|GTCTACGGCCATACCACCCTGAACGC | >hg38_1_-_228634819_228635039_RNA5S12_5S_50ntFlanks | 51 | 76 | 26 | rRF 5S | NO | 0.00 | 68.40 | 166.94 | 0.00 |
| >hg38_1_-_228634819_228635039_RNA5S12_5S_50ntFlanks@51.78.28\|GTCTACGGCCATACCACCCTGAACGCGC | >hg38_1_-_228634819_228635039_RNA5S12_5S_50ntFlanks | 51 | 78 | 28 | rRF 5S | NO | 0.00 | 42.23 | 0.00 | 0.00 |
| >hg38_1_-_228634819_228635039_RNA5S12_5S_50ntFlanks@51.79.29\|GTCTACGGCCATACCACCCTGAACGCGCC | >hg38_1_-_228634819_228635039_RNA5S12_5S_50ntFlanks | 51 | 79 | 29 | rRF 5S | NO | 27.13 | 108.52 | 109.90 | 0.00 |
| >hg38_1_-_228634819_228635039_RNA5S12_5S_50ntFlanks@51.80.30\|GTCTACGGCCATACCACCCTGAACGCGCCC | >hg38_1_-_228634819_228635039_RNA5S12_5S_50ntFlanks | 51 | 80 | 30 | rRF 5S | NO | 0.00 | 111.47 | 164.88 | 0.00 |
| >hg38_1_-_228634819_228635039_RNA5S12_5S_50ntFlanks@51.81.31\|GTCTACGGCCATACCACCCTGAACGCGCCCG | >hg38_1_-_228634819_228635039_RNA5S12_5S_50ntFlanks | 51 | 81 | 31 | rRF 5S | NO | 11.24 | 101.90 | 70.18 | 0.00 |
| >hg38_1_-_228634819_228635039_RNA5S12_5S_50ntFlanks@51.82.32\|GTCTACGGCCATACCACCCTGAACGCGCCCGA | >hg38_1_-_228634819_228635039_RNA5S12_5S_50ntFlanks | 51 | 82 | 32 | rRF 5S | NO | 23.85 | 153.11 | 14.82 | 0.00 |
| >hg38_1_-_228634819_228635039_RNA5S12_5S_50ntFlanks@51.83.33\|GTCTACGGCCATACCACCCTGAACGCGCCCGAT | >hg38_1_-_228634819_228635039_RNA5S12_5S_50ntFlanks | 51 | 83 | 33 | rRF 5S | NO | 35.25 | 193.22 | 121.20 | 0.00 |
| >hg38_1_-_228634819_228635039_RNA5S12_5S_50ntFlanks@51.84.34\|GTCTACGGCCATACCACCCTGAACGCGCCCGATC | >hg38_1_-_228634819_228635039_RNA5S12_5S_50ntFlanks | 51 | 84 | 34 | rRF 5S | NO | 189.02 | 2012.95 | 656.42 | 98.37 |
| >hg38_1_-_228634819_228635039_RNA5S12_5S_50ntFlanks@51.85.35\|GTCTACGGCCATACCACCCTGAACGCGCCCGATCT | >hg38_1_-_228634819_228635039_RNA5S12_5S_50ntFlanks | 51 | 85 | 35 | rRF 5S | NO | 28.72 | 150.51 | 58.25 | 0.00 |
| >hg38_1_-_228634819_228635039_RNA5S12_5S_50ntFlanks@51.86.36\|GTCTACGGCCATACCACCCTGAACGCGCCCGATCTC | >hg38_1_-_228634819_228635039_RNA5S12_5S_50ntFlanks | 51 | 86 | 36 | rRF 5S | NO | 30.43 | 198.23 | 127.73 | 0.00 |
| >hg38_1_-_228634819_228635039_RNA5S12_5S_50ntFlanks@51.87.37\|GTCTACGGCCATACCACCCTGAACGCGCCCGATCTCG | >hg38_1_-_228634819_228635039_RNA5S12_5S_50ntFlanks | 51 | 87 | 37 | rRF 5S | NO | 74.09 | 206.57 | 11.74 | 0.00 |
| >hg38_1_-_228634819_228635039_RNA5S12_5S_50ntFlanks@51.88.38\|GTCTACGGCCATACCACCCTGAACGCGCCCGATCTCGT | >hg38_1_-_228634819_228635039_RNA5S12_5S_50ntFlanks | 51 | 88 | 38 | rRF 5S | NO | 126.87 | 386.72 | 88.18 | 62.47 |
| >hg38_1_-_228634819_228635039_RNA5S12_5S_50ntFlanks@51.89.39\|GTCTACGGCCATACCACCCTGAACGCGCCCGATCTCGTC | >hg38_1_-_228634819_228635039_RNA5S12_5S_50ntFlanks | 51 | 89 | 39 | rRF 5S | NO | 601.25 | 2305.89 | 387.86 | 423.59 |
| >hg38_1_-_228634819_228635039_RNA5S12_5S_50ntFlanks@51.90.40\|GTCTACGGCCATACCACCCTGAACGCGCCCGATCTCGTCT | >hg38_1_-_228634819_228635039_RNA5S12_5S_50ntFlanks | 51 | 90 | 40 | rRF 5S | NO | 38.09 | 94.42 | 0.00 | 0.00 |
| >hg38_1_-_228634819_228635039_RNA5S12_5S_50ntFlanks@51.91.41\|GTCTACGGCCATACCACCCTGAACGCGCCCGATCTCGTCTG | >hg38_1_-_228634819_228635039_RNA5S12_5S_50ntFlanks | 51 | 91 | 41 | rRF 5S | NO | 33.76 | 183.23 | 326.55 | 0.00 |
| >hg38_1_-_228634819_228635039_RNA5S12_5S_50ntFlanks@51.92.42\|GTCTACGGCCATACCACCCTGAACGCGCCCGATCTCGTCTGA | >hg38_1_-_228634819_228635039_RNA5S12_5S_50ntFlanks | 51 | 92 | 42 | rRF 5S | NO | 26.62 | 41.04 | 0.00 | 26.51 |

# BT-20

| | | | | | | | | | | |
|---|---|---|---|---|---|---|---|---|---|---|
| >hg38_1_-_228634819_228635039_RNA5S12_5S_50ntFlanks@51.94.44\|GTCTACGGCCATACCACCCTGAACGCGCCCGATCTCGTCTGATC | >hg38_1_-_228634819_228635039_RNA5S12_5S_50ntFlanks | 51 | 94 | 44 | rRF 5S | NO | 0.00 | 82.71 | 13.18 | 0.00 |
| >hg38_1_-_228634819_228635039_RNA5S12_5S_50ntFlanks@125.170.46\|GTACTTGGATGGGAGACCGCCTGGGAATACCGGGTGCTGTAGGCTT | >hg38_1_-_228634819_228635039_RNA5S12_5S_50ntFlanks | 125 | 170 | 46 | rRF 5S | NO | 0.00 | 0.00 | 14.99 | 9.43 |
| >hg38_1_-_228634819_228635039_RNA5S12_5S_50ntFlanks@126.169.44\|TACTTGGATGGGAGACCGCCTGGGAATACCGGGTGCTGTAGGCT | >hg38_1_-_228634819_228635039_RNA5S12_5S_50ntFlanks | 126 | 169 | 44 | rRF 5S | NO | 0.00 | 10.43 | 0.00 | 0.00 |
| >hg38_1_-_228634819_228635039_RNA5S12_5S_50ntFlanks@126.170.45\|TACTTGGATGGGAGACCGCCTGGGAATACCGGGTGCTGTAGGCTT | >hg38_1_-_228634819_228635039_RNA5S12_5S_50ntFlanks | 126 | 170 | 45 | rRF 5S | NO | 0.00 | 34.03 | 65.26 | 10.80 |
| >hg38_1_-_228634819_228635039_RNA5S12_5S_50ntFlanks@127.169.43\|ACTTGGATGGGAGACCGCCTGGGAATACCGGGTGCTGTAGGCT | >hg38_1_-_228634819_228635039_RNA5S12_5S_50ntFlanks | 127 | 169 | 43 | rRF 5S | NO | 0.00 | 28.65 | 13.21 | 0.00 |
| >hg38_1_-_228634819_228635039_RNA5S12_5S_50ntFlanks@127.170.44\|ACTTGGATGGGAGACCGCCTGGGAATACCGGGTGCTGTAGGCTT | >hg38_1_-_228634819_228635039_RNA5S12_5S_50ntFlanks | 127 | 170 | 44 | rRF 5S | NO | 0.00 | 42.03 | 85.24 | 32.25 |
| >hg38_1_-_228634819_228635039_RNA5S12_5S_50ntFlanks@128.169.42\|CTTGGATGGGAGACCGCCTGGGAATACCGGGTGCTGTAGGCT | >hg38_1_-_228634819_228635039_RNA5S12_5S_50ntFlanks | 128 | 169 | 42 | rRF 5S | NO | 0.00 | 41.20 | 15.73 | 0.00 |
| >hg38_1_-_228634819_228635039_RNA5S12_5S_50ntFlanks@128.170.43\|CTTGGATGGGAGACCGCCTGGGAATACCGGGTGCTGTAGGCTT | >hg38_1_-_228634819_228635039_RNA5S12_5S_50ntFlanks | 128 | 170 | 43 | rRF 5S | NO | 0.00 | 91.01 | 122.72 | 30.69 |
| >hg38_1_-_228634819_228635039_RNA5S12_5S_50ntFlanks@128.171.44\|CTTGGATGGGAGACCGCCTGGGAATACCGGGTGCTGTAGGCTTT | >hg38_1_-_228634819_228635039_RNA5S12_5S_50ntFlanks | 128 | 171 | 44 | rRF 5S | NO | 0.00 | 26.79 | 12.57 | 0.00 |
| >hg38_1_-_228634819_228635039_RNA5S12_5S_50ntFlanks@129.169.41\|TTGGATGGGAGACCGCCTGGGAATACCGGGTGCTGTAGGCT | >hg38_1_-_228634819_228635039_RNA5S12_5S_50ntFlanks | 129 | 169 | 41 | rRF 5S | NO | 0.00 | 83.58 | 75.30 | 0.00 |
| >hg38_1_-_228634819_228635039_RNA5S12_5S_50ntFlanks@129.170.42\|TTGGATGGGAGACCGCCTGGGAATACCGGGTGCTGTAGGCTT | >hg38_1_-_228634819_228635039_RNA5S12_5S_50ntFlanks | 129 | 170 | 42 | rRF 5S | NO | 0.00 | 154.44 | 307.56 | 66.78 |
| >hg38_1_-_228634819_228635039_RNA5S12_5S_50ntFlanks@129.171.43\|TTGGATGGGAGACCGCCTGGGAATACCGGGTGCTGTAGGCTTT | >hg38_1_-_228634819_228635039_RNA5S12_5S_50ntFlanks | 129 | 171 | 43 | rRF 5S | NO | 0.00 | 37.30 | 68.02 | 10.78 |
| >hg38_1_-_228634819_228635039_RNA5S12_5S_50ntFlanks@130.169.40\|TGGATGGGAGACCGCCTGGGAATACCGGGTGCTGTAGGCT | >hg38_1_-_228634819_228635039_RNA5S12_5S_50ntFlanks | 130 | 169 | 40 | rRF 5S | NO | 0.00 | 127.31 | 114.52 | 0.00 |
| >hg38_1_-_228634819_228635039_RNA5S12_5S_50ntFlanks@130.170.41\|TGGATGGGAGACCGCCTGGGAATACCGGGTGCTGTAGGCTT | >hg38_1_-_228634819_228635039_RNA5S12_5S_50ntFlanks | 130 | 170 | 41 | rRF 5S | NO | 10.23 | 256.89 | 487.45 | 80.98 |
| >hg38_1_-_228634819_228635039_RNA5S12_5S_50ntFlanks@130.171.42\|TGGATGGGAGACCGCCTGGGAATACCGGGTGCTGTAGGCTTT | >hg38_1_-_228634819_228635039_RNA5S12_5S_50ntFlanks | 130 | 171 | 42 | rRF 5S | NO | 0.00 | 94.95 | 104.23 | 27.58 |
| >hg38_1_-_228634819_228635039_RNA5S12_5S_50ntFlanks@131.169.39\|GGATGGGAGACCGCCTGGGAATACCGGGTGCTGTAGGCT | >hg38_1_-_228634819_228635039_RNA5S12_5S_50ntFlanks | 131 | 169 | 39 | rRF 5S | NO | 0.00 | 70.96 | 100.03 | 0.00 |
| >hg38_1_-_228634819_228635039_RNA5S12_5S_50ntFlanks@131.170.40\|GGATGGGAGACCGCCTGGGAATACCGGGTGCTGTAGGCTT | >hg38_1_-_228634819_228635039_RNA5S12_5S_50ntFlanks | 131 | 170 | 40 | rRF 5S | NO | 0.00 | 135.63 | 384.33 | 34.51 |
| >hg38_1_-_228634819_228635039_RNA5S12_5S_50ntFlanks@131.171.41\|GGATGGGAGACCGCCTGGGAATACCGGGTGCTGTAGGCTTT | >hg38_1_-_228634819_228635039_RNA5S12_5S_50ntFlanks | 131 | 171 | 41 | rRF 5S | NO | 0.00 | 68.67 | 98.00 | 0.00 |
| >hg38_1_-_228634819_228635039_RNA5S12_5S_50ntFlanks@132.169.38\|GATGGGAGACCGCCTGGGAATACCGGGTGCTGTAGGCT | >hg38_1_-_228634819_228635039_RNA5S12_5S_50ntFlanks | 132 | 169 | 38 | rRF 5S | NO | 38.62 | 616.73 | 711.23 | 90.21 |
| >hg38_1_-_228634819_228635039_RNA5S12_5S_50ntFlanks@132.170.39\|GATGGGAGACCGCCTGGGAATACCGGGTGCTGTAGGCTT | >hg38_1_-_228634819_228635039_RNA5S12_5S_50ntFlanks | 132 | 170 | 39 | rRF 5S | YES | 322.89 | 2071.24 | 5199.34 | 583.23 |
| >hg38_1_-_228634819_228635039_RNA5S12_5S_50ntFlanks@132.171.40\|GATGGGAGACCGCCTGGGAATACCGGGTGCTGTAGGCTTT | >hg38_1_-_228634819_228635039_RNA5S12_5S_50ntFlanks | 132 | 171 | 40 | rRF 5S | NO | 107.07 | 520.90 | 964.26 | 185.15 |
| >hg38_1_-_228634819_228635039_RNA5S12_5S_50ntFlanks@133.169.37\|ATGGGAGACCGCCTGGGAATACCGGGTGCTGTAGGCT | >hg38_1_-_228634819_228635039_RNA5S12_5S_50ntFlanks | 133 | 169 | 37 | rRF 5S | NO | 30.67 | 420.07 | 305.73 | 41.25 |
| >hg38_1_-_228634819_228635039_RNA5S12_5S_50ntFlanks@133.170.38\|ATGGGAGACCGCCTGGGAATACCGGGTGCTGTAGGCTT | >hg38_1_-_228634819_228635039_RNA5S12_5S_50ntFlanks | 133 | 170 | 38 | rRF 5S | NO | 194.97 | 891.34 | 2023.33 | 342.60 |
| >hg38_1_-_228634819_228635039_RNA5S12_5S_50ntFlanks@133.171.39\|ATGGGAGACCGCCTGGGAATACCGGGTGCTGTAGGCTTT | >hg38_1_-_228634819_228635039_RNA5S12_5S_50ntFlanks | 133 | 171 | 39 | rRF 5S | NO | 74.24 | 309.83 | 366.67 | 113.16 |
| >hg38_1_-_228634819_228635039_RNA5S12_5S_50ntFlanks@134.168.35\|TGGGAGACCGCCTGGGAATACCGGGTGCTGTAGGC | >hg38_1_-_228634819_228635039_RNA5S12_5S_50ntFlanks | 134 | 168 | 35 | rRF 5S | NO | 0.00 | 26.63 | 0.00 | 0.00 |
| >hg38_1_-_228634819_228635039_RNA5S12_5S_50ntFlanks@134.169.36\|TGGGAGACCGCCTGGGAATACCGGGTGCTGTAGGCT | >hg38_1_-_228634819_228635039_RNA5S12_5S_50ntFlanks | 134 | 169 | 36 | rRF 5S | NO | 117.15 | 871.94 | 242.07 | 134.16 |

# BT-20

| | | | | | | | | | | |
|---|---|---|---|---|---|---|---|---|---|---|
| >hg38_1_-_228634819_228635039_RNA5S12_5S_50ntFlanks@134.170.37\|TGGGAGACCGCCTGGGAATACCGGGTGCTGTAGGCTT | >hg38_1_-_228634819_228635039_RNA5S12_5S_50ntFlanks | 134 | 170 | 37 | rRF 5S | YES | 550.85 | 2338.47 | 2282.45 | 791.42 |
| >hg38_1_-_228634819_228635039_RNA5S12_5S_50ntFlanks@134.171.38\|TGGGAGACCGCCTGGGAATACCGGGTGCTGTAGGCTTT | >hg38_1_-_228634819_228635039_RNA5S12_5S_50ntFlanks | 134 | 171 | 38 | rRF 5S | NO | 170.06 | 724.03 | 405.11 | 219.40 |
| >hg38_1_-_228634819_228635039_RNA5S12_5S_50ntFlanks@135.169.35\|GGGAGACCGCCTGGGAATACCGGGTGCTGTAGGCT | >hg38_1_-_228634819_228635039_RNA5S12_5S_50ntFlanks | 135 | 169 | 35 | rRF 5S | NO | 30.79 | 253.84 | 107.88 | 29.41 |
| >hg38_1_-_228634819_228635039_RNA5S12_5S_50ntFlanks@135.170.36\|GGGAGACCGCCTGGGAATACCGGGTGCTGTAGGCTT | >hg38_1_-_228634819_228635039_RNA5S12_5S_50ntFlanks | 135 | 170 | 36 | rRF 5S | NO | 343.22 | 1209.71 | 1757.72 | 423.16 |
| >hg38_1_-_228634819_228635039_RNA5S12_5S_50ntFlanks@135.171.37\|GGGAGACCGCCTGGGAATACCGGGTGCTGTAGGCTTT | >hg38_1_-_228634819_228635039_RNA5S12_5S_50ntFlanks | 135 | 171 | 37 | rRF 5S | NO | 201.84 | 650.20 | 365.08 | 150.13 |
| >hg38_1_-_228634819_228635039_RNA5S12_5S_50ntFlanks@136.169.34\|GGAGACCGCCTGGGAATACCGGGTGCTGTAGGCT | >hg38_1_-_228634819_228635039_RNA5S12_5S_50ntFlanks | 136 | 169 | 34 | rRF 5S | NO | 0.00 | 0.00 | 95.09 | 0.00 |
| >hg38_1_-_228634819_228635039_RNA5S12_5S_50ntFlanks@136.170.35\|GGAGACCGCCTGGGAATACCGGGTGCTGTAGGCTT | >hg38_1_-_228634819_228635039_RNA5S12_5S_50ntFlanks | 136 | 170 | 35 | rRF 5S | NO | 43.51 | 121.38 | 547.29 | 81.43 |
| >hg38_1_-_228634819_228635039_RNA5S12_5S_50ntFlanks@136.171.36\|GGAGACCGCCTGGGAATACCGGGTGCTGTAGGCTTT | >hg38_1_-_228634819_228635039_RNA5S12_5S_50ntFlanks | 136 | 171 | 36 | rRF 5S | NO | 38.02 | 110.38 | 370.51 | 89.35 |
| >hg38_1_-_228634819_228635039_RNA5S12_5S_50ntFlanks@137.169.33\|GAGACCGCCTGGGAATACCGGGTGCTGTAGGCT | >hg38_1_-_228634819_228635039_RNA5S12_5S_50ntFlanks | 137 | 169 | 33 | rRF 5S | NO | 0.00 | 26.57 | 72.52 | 0.00 |
| >hg38_1_-_228634819_228635039_RNA5S12_5S_50ntFlanks@137.170.34\|GAGACCGCCTGGGAATACCGGGTGCTGTAGGCTT | >hg38_1_-_228634819_228635039_RNA5S12_5S_50ntFlanks | 137 | 170 | 34 | rRF 5S | NO | 0.00 | 41.78 | 84.33 | 0.00 |
| >hg38_1_-_228634819_228635039_RNA5S12_5S_50ntFlanks@137.171.35\|GAGACCGCCTGGGAATACCGGGTGCTGTAGGCTTT | >hg38_1_-_228634819_228635039_RNA5S12_5S_50ntFlanks | 137 | 171 | 35 | rRF 5S | NO | 0.00 | 99.14 | 134.67 | 0.00 |
| >hg38_1_-_228634819_228635039_RNA5S12_5S_50ntFlanks@138.169.32\|AGACCGCCTGGGAATACCGGGTGCTGTAGGCT | >hg38_1_-_228634819_228635039_RNA5S12_5S_50ntFlanks | 138 | 169 | 32 | rRF 5S | NO | 0.00 | 42.88 | 102.13 | 0.00 |
| >hg38_1_-_228634819_228635039_RNA5S12_5S_50ntFlanks@138.170.33\|AGACCGCCTGGGAATACCGGGTGCTGTAGGCTT | >hg38_1_-_228634819_228635039_RNA5S12_5S_50ntFlanks | 138 | 170 | 33 | rRF 5S | NO | 0.00 | 98.03 | 168.35 | 0.00 |
| >hg38_1_-_228634819_228635039_RNA5S12_5S_50ntFlanks@138.171.34\|AGACCGCCTGGGAATACCGGGTGCTGTAGGCTTT | >hg38_1_-_228634819_228635039_RNA5S12_5S_50ntFlanks | 138 | 171 | 34 | rRF 5S | NO | 0.00 | 83.86 | 94.75 | 0.00 |
| >hg38_1_-_228634819_228635039_RNA5S12_5S_50ntFlanks@139.169.31\|GACCGCCTGGGAATACCGGGTGCTGTAGGCT | >hg38_1_-_228634819_228635039_RNA5S12_5S_50ntFlanks | 139 | 169 | 31 | rRF 5S | NO | 0.00 | 113.15 | 159.98 | 0.00 |
| >hg38_1_-_228634819_228635039_RNA5S12_5S_50ntFlanks@139.170.32\|GACCGCCTGGGAATACCGGGTGCTGTAGGCTT | >hg38_1_-_228634819_228635039_RNA5S12_5S_50ntFlanks | 139 | 170 | 32 | rRF 5S | NO | 0.00 | 112.81 | 189.38 | 0.00 |
| >hg38_1_-_228634819_228635039_RNA5S12_5S_50ntFlanks@139.171.33\|GACCGCCTGGGAATACCGGGTGCTGTAGGCTTT | >hg38_1_-_228634819_228635039_RNA5S12_5S_50ntFlanks | 139 | 171 | 33 | rRF 5S | NO | 0.00 | 101.70 | 93.64 | 0.00 |
| >hg38_1_-_228634819_228635039_RNA5S12_5S_50ntFlanks@140.169.30\|ACCGCCTGGGAATACCGGGTGCTGTAGGCT | >hg38_1_-_228634819_228635039_RNA5S12_5S_50ntFlanks | 140 | 169 | 30 | rRF 5S | NO | 35.28 | 421.81 | 280.86 | 76.47 |
| >hg38_1_-_228634819_228635039_RNA5S12_5S_50ntFlanks@140.170.31\|ACCGCCTGGGAATACCGGGTGCTGTAGGCTT | >hg38_1_-_228634819_228635039_RNA5S12_5S_50ntFlanks | 140 | 170 | 31 | rRF 5S | NO | 127.99 | 815.95 | 641.54 | 174.28 |
| >hg38_1_-_228634819_228635039_RNA5S12_5S_50ntFlanks@140.171.32\|ACCGCCTGGGAATACCGGGTGCTGTAGGCTTT | >hg38_1_-_228634819_228635039_RNA5S12_5S_50ntFlanks | 140 | 171 | 32 | rRF 5S | NO | 26.35 | 326.58 | 162.65 | 30.73 |
| >hg38_1_-_228634819_228635039_RNA5S12_5S_50ntFlanks@141.169.29\|CCGCCTGGGAATACCGGGTGCTGTAGGCT | >hg38_1_-_228634819_228635039_RNA5S12_5S_50ntFlanks | 141 | 169 | 29 | rRF 5S | NO | 0.00 | 146.73 | 111.00 | 0.00 |
| >hg38_1_-_228634819_228635039_RNA5S12_5S_50ntFlanks@141.170.30\|CCGCCTGGGAATACCGGGTGCTGTAGGCTT | >hg38_1_-_228634819_228635039_RNA5S12_5S_50ntFlanks | 141 | 170 | 30 | rRF 5S | NO | 26.93 | 223.29 | 165.95 | 9.75 |
| >hg38_1_-_228634819_228635039_RNA5S12_5S_50ntFlanks@141.171.31\|CCGCCTGGGAATACCGGGTGCTGTAGGCTTT | >hg38_1_-_228634819_228635039_RNA5S12_5S_50ntFlanks | 141 | 171 | 31 | rRF 5S | NO | 0.00 | 220.75 | 97.73 | 0.00 |
| >hg38_1_-_228634819_228635039_RNA5S12_5S_50ntFlanks@142.169.28\|CGCCTGGGAATACCGGGTGCTGTAGGCT | >hg38_1_-_228634819_228635039_RNA5S12_5S_50ntFlanks | 142 | 169 | 28 | rRF 5S | NO | 0.00 | 35.79 | 0.00 | 0.00 |
| >hg38_1_-_228634819_228635039_RNA5S12_5S_50ntFlanks@142.170.29\|CGCCTGGGAATACCGGGTGCTGTAGGCTT | >hg38_1_-_228634819_228635039_RNA5S12_5S_50ntFlanks | 142 | 170 | 29 | rRF 5S | NO | 0.00 | 99.99 | 80.65 | 0.00 |
| >hg38_1_-_228634819_228635039_RNA5S12_5S_50ntFlanks@142.171.30\|CGCCTGGGAATACCGGGTGCTGTAGGCTTT | >hg38_1_-_228634819_228635039_RNA5S12_5S_50ntFlanks | 142 | 171 | 30 | rRF 5S | NO | 0.00 | 92.02 | 64.74 | 0.00 |

# BT-20

| | | | | | | | | | | |
|---|---|---|---|---|---|---|---|---|---|---|
| >hg38_1_-_228634819_228635039_RNA5S12_5S_50ntFlanks@144.169.26\|CCTGGGAATACCGGGTGCTGTAGGCT | >hg38_1_-_228634819_228635039_RNA5S12_5S_50ntFlanks | 144 | 169 | 26 | rRF 5S | NO | 0.00 | 36.68 | 0.00 | 0.00 |
| >hg38_1_-_228634819_228635039_RNA5S12_5S_50ntFlanks@144.170.27\|CCTGGGAATACCGGGTGCTGTAGGCTT | >hg38_1_-_228634819_228635039_RNA5S12_5S_50ntFlanks | 144 | 170 | 27 | rRF 5S | NO | 78.70 | 169.19 | 127.53 | 9.88 |
| >hg38_1_-_228634819_228635039_RNA5S12_5S_50ntFlanks@144.171.28\|CCTGGGAATACCGGGTGCTGTAGGCTTT | >hg38_1_-_228634819_228635039_RNA5S12_5S_50ntFlanks | 144 | 171 | 28 | rRF 5S | NO | 40.69 | 161.28 | 79.03 | 10.21 |
| >hg38_1_-_228634819_228635039_RNA5S12_5S_50ntFlanks@145.169.25\|CTGGGAATACCGGGTGCTGTAGGCT | >hg38_1_-_228634819_228635039_RNA5S12_5S_50ntFlanks | 145 | 169 | 25 | rRF 5S | NO | 0.00 | 87.78 | 13.33 | 0.00 |
| >hg38_1_-_228634819_228635039_RNA5S12_5S_50ntFlanks@145.170.26\|CTGGGAATACCGGGTGCTGTAGGCTT | >hg38_1_-_228634819_228635039_RNA5S12_5S_50ntFlanks | 145 | 170 | 26 | rRF 5S | NO | 23.94 | 176.19 | 132.63 | 9.70 |
| >hg38_1_-_228634819_228635039_RNA5S12_5S_50ntFlanks@145.171.27\|CTGGGAATACCGGGTGCTGTAGGCTTT | >hg38_1_-_228634819_228635039_RNA5S12_5S_50ntFlanks | 145 | 171 | 27 | rRF 5S | NO | 0.00 | 135.73 | 61.71 | 0.00 |
| >hg38_1_-_228634819_228635039_RNA5S12_5S_50ntFlanks@146.169.24\|TGGGAATACCGGGTGCTGTAGGCT | >hg38_1_-_228634819_228635039_RNA5S12_5S_50ntFlanks | 146 | 169 | 24 | rRF 5S | NO | 0.00 | 160.97 | 120.48 | 0.00 |
| >hg38_1_-_228634819_228635039_RNA5S12_5S_50ntFlanks@146.170.25\|TGGGAATACCGGGTGCTGTAGGCTT | >hg38_1_-_228634819_228635039_RNA5S12_5S_50ntFlanks | 146 | 170 | 25 | rRF 5S | NO | 37.32 | 423.24 | 381.38 | 71.41 |
| >hg38_1_-_228634819_228635039_RNA5S12_5S_50ntFlanks@146.171.26\|TGGGAATACCGGGTGCTGTAGGCTTT | >hg38_1_-_228634819_228635039_RNA5S12_5S_50ntFlanks | 146 | 171 | 26 | rRF 5S | NO | 9.48 | 236.40 | 145.36 | 24.54 |
| >hg38_1_-_228634819_228635039_RNA5S12_5S_50ntFlanks@147.169.23\|GGGAATACCGGGTGCTGTAGGCT | >hg38_1_-_228634819_228635039_RNA5S12_5S_50ntFlanks | 147 | 169 | 23 | rRF 5S | NO | 0.00 | 157.27 | 390.31 | 0.00 |
| >hg38_1_-_228634819_228635039_RNA5S12_5S_50ntFlanks@147.170.24\|GGGAATACCGGGTGCTGTAGGCTT | >hg38_1_-_228634819_228635039_RNA5S12_5S_50ntFlanks | 147 | 170 | 24 | rRF 5S | NO | 76.29 | 413.10 | 1139.16 | 85.32 |
| >hg38_1_-_228634819_228635039_RNA5S12_5S_50ntFlanks@147.171.25\|GGGAATACCGGGTGCTGTAGGCTTT | >hg38_1_-_228634819_228635039_RNA5S12_5S_50ntFlanks | 147 | 171 | 25 | rRF 5S | NO | 32.54 | 234.13 | 332.90 | 31.76 |
| >hg38_1_-_228634819_228635039_RNA5S12_5S_50ntFlanks@148.169.22\|GGAATACCGGGTGCTGTAGGCT | >hg38_1_-_228634819_228635039_RNA5S12_5S_50ntFlanks | 148 | 169 | 22 | rRF 5S | NO | 24.76 | 178.89 | 471.43 | 9.55 |
| >hg38_1_-_228634819_228635039_RNA5S12_5S_50ntFlanks@148.170.23\|GGAATACCGGGTGCTGTAGGCTT | >hg38_1_-_228634819_228635039_RNA5S12_5S_50ntFlanks | 148 | 170 | 23 | rRF 5S | NO | 80.24 | 323.80 | 943.31 | 71.93 |
| >hg38_1_-_228634819_228635039_RNA5S12_5S_50ntFlanks@148.171.24\|GGAATACCGGGTGCTGTAGGCTTT | >hg38_1_-_228634819_228635039_RNA5S12_5S_50ntFlanks | 148 | 171 | 24 | rRF 5S | NO | 42.21 | 291.83 | 449.02 | 32.95 |
| >hg38_1_-_228634819_228635039_RNA5S12_5S_50ntFlanks@149.169.21\|GAATACCGGGTGCTGTAGGCT | >hg38_1_-_228634819_228635039_RNA5S12_5S_50ntFlanks | 149 | 169 | 21 | rRF 5S | NO | 0.00 | 123.44 | 173.64 | 0.00 |
| >hg38_1_-_228634819_228635039_RNA5S12_5S_50ntFlanks@149.170.22\|GAATACCGGGTGCTGTAGGCTT | >hg38_1_-_228634819_228635039_RNA5S12_5S_50ntFlanks | 149 | 170 | 22 | rRF 5S | NO | 26.12 | 292.05 | 480.97 | 30.98 |
| >hg38_1_-_228634819_228635039_RNA5S12_5S_50ntFlanks@149.171.23\|GAATACCGGGTGCTGTAGGCTTT | >hg38_1_-_228634819_228635039_RNA5S12_5S_50ntFlanks | 149 | 171 | 23 | rRF 5S | NO | 9.75 | 228.67 | 182.03 | 26.16 |
| >hg38_1_-_228634819_228635039_RNA5S12_5S_50ntFlanks@150.169.20\|AATACCGGGTGCTGTAGGCT | >hg38_1_-_228634819_228635039_RNA5S12_5S_50ntFlanks | 150 | 169 | 20 | rRF 5S | NO | 0.00 | 122.42 | 198.91 | 0.00 |
| >hg38_1_-_228634819_228635039_RNA5S12_5S_50ntFlanks@150.170.21\|AATACCGGGTGCTGTAGGCTT | >hg38_1_-_228634819_228635039_RNA5S12_5S_50ntFlanks | 150 | 170 | 21 | rRF 5S | NO | 25.36 | 268.48 | 568.90 | 10.83 |
| >hg38_1_-_228634819_228635039_RNA5S12_5S_50ntFlanks@150.171.22\|AATACCGGGTGCTGTAGGCTTT | >hg38_1_-_228634819_228635039_RNA5S12_5S_50ntFlanks | 150 | 171 | 22 | rRF 5S | NO | 10.26 | 211.16 | 157.00 | 24.98 |
| >hg38_1_-_228634819_228635039_RNA5S12_5S_50ntFlanks@151.169.19\|ATACCGGGTGCTGTAGGCT | >hg38_1_-_228634819_228635039_RNA5S12_5S_50ntFlanks | 151 | 169 | 19 | rRF 5S | NO | 0.00 | 102.35 | 209.16 | 0.00 |
| >hg38_1_-_228634819_228635039_RNA5S12_5S_50ntFlanks@151.170.20\|ATACCGGGTGCTGTAGGCTT | >hg38_1_-_228634819_228635039_RNA5S12_5S_50ntFlanks | 151 | 170 | 20 | rRF 5S | NO | 10.77 | 221.62 | 607.50 | 24.06 |
| >hg38_1_-_228634819_228635039_RNA5S12_5S_50ntFlanks@151.171.21\|ATACCGGGTGCTGTAGGCTTT | >hg38_1_-_228634819_228635039_RNA5S12_5S_50ntFlanks | 151 | 171 | 21 | rRF 5S | NO | 0.00 | 157.82 | 124.34 | 9.80 |
| >hg38_1_-_228634819_228635039_RNA5S12_5S_50ntFlanks@152.169.18\|TACCGGGTGCTGTAGGCT | >hg38_1_-_228634819_228635039_RNA5S12_5S_50ntFlanks | 152 | 169 | 18 | rRF 5S | NO | 0.00 | 98.39 | 114.54 | 0.00 |
| >hg38_1_-_228634819_228635039_RNA5S12_5S_50ntFlanks@152.170.19\|TACCGGGTGCTGTAGGCTT | >hg38_1_-_228634819_228635039_RNA5S12_5S_50ntFlanks | 152 | 170 | 19 | rRF 5S | NO | 10.11 | 228.13 | 328.37 | 25.72 |

# BT-20

<table>
<tr><th></th><th></th><th></th><th></th><th></th><th></th><th></th><th></th><th></th><th></th><th></th></tr>
<tr><td>>hg38_1_-_228634819_228635039_RNA5S12_5S_50ntFlanks@152.171.20|TACCGGGTGCTGTAGGCTTT</td><td>>hg38_1_-_228634819_228635039_RNA5S12_5S_50ntFlanks</td><td>152</td><td>171</td><td>20</td><td>rRF 5S</td><td>NO</td><td>0.00</td><td>182.91</td><td>94.89</td><td>10.46</td></tr>
<tr><td>>hg38_1_-_228634819_228635039_RNA5S12_5S_50ntFlanks@153.170.18|ACCGGGTGCTGTAGGCTT</td><td>>hg38_1_-_228634819_228635039_RNA5S12_5S_50ntFlanks</td><td>153</td><td>170</td><td>18</td><td>rRF 5S</td><td>YES</td><td>242.10</td><td>1186.90</td><td>2206.15</td><td>273.78</td></tr>
<tr><td>>hg38_1_-_228634819_228635039_RNA5S12_5S_50ntFlanks@153.171.19|ACCGGGTGCTGTAGGCTTT</td><td>>hg38_1_-_228634819_228635039_RNA5S12_5S_50ntFlanks</td><td>153</td><td>171</td><td>19</td><td>rRF 5S</td><td>YES</td><td>165.06</td><td>791.25</td><td>833.88</td><td>202.54</td></tr>
<tr><td>>hg38_1_-_228634819_228635039_RNA5S12_5S_50ntFlanks@154.171.18|CCGGGTGCTGTAGGCTTT</td><td>>hg38_1_-_228634819_228635039_RNA5S12_5S_50ntFlanks</td><td>154</td><td>171</td><td>18</td><td>rRF 5S</td><td>NO</td><td>31.66</td><td>297.60</td><td>274.49</td><td>26.55</td></tr>
<tr><td>>multi-trnaMT_GlnTTG_MT_-_4329_4400@1.26.26,trnalookalike2_GlnTTG_1_-_564879_564950@1.26.26|TAGGATGGGGTGTGATAGGTGGCACG</td><td>>GlnTTG.trnaMT_MT_-_4329_4400.multi</td><td>1</td><td>26</td><td>26</td><td>tRF lookalike</td><td>NO</td><td>0.00</td><td>0.00</td><td>71.22</td><td>0.00</td></tr>
<tr><td>>multi-trnaMT_GlnTTG_MT_-_4329_4400@1.29.29,trnalookalike2_GlnTTG_1_-_564879_564950@1.29.29|TAGGATGGGGTGTGATAGGTGGCACGGAG</td><td>>GlnTTG.trnaMT_MT_-_4329_4400.multi</td><td>1</td><td>29</td><td>29</td><td>tRF lookalike</td><td>NO</td><td>0.00</td><td>0.00</td><td>39.07</td><td>0.00</td></tr>
<tr><td>>multi-trnaMT_GlnTTG_MT_-_4329_4400@1.30.30,trnalookalike2_GlnTTG_1_-_564879_564950@1.30.30|TAGGATGGGGTGTGATAGGTGGCACGGAGA</td><td>>GlnTTG.trnaMT_MT_-_4329_4400.multi</td><td>1</td><td>30</td><td>30</td><td>tRF lookalike</td><td>NO</td><td>0.00</td><td>0.00</td><td>105.09</td><td>0.00</td></tr>
<tr><td>>multi-trnaMT_GlnTTG_MT_-_4329_4400@1.32.32,trnalookalike2_GlnTTG_1_-_564879_564950@1.32.32|TAGGATGGGGTGTGATAGGTGGCACGGAGAAT</td><td>>GlnTTG.trnaMT_MT_-_4329_4400.multi</td><td>1</td><td>32</td><td>32</td><td>tRF lookalike</td><td>NO</td><td>0.00</td><td>0.00</td><td>12.50</td><td>0.00</td></tr>
<tr><td>>multi-trnaMT_GlnTTG_MT_-_4329_4400@1.33.33,trnalookalike2_GlnTTG_1_-_564879_564950@1.33.33|TAGGATGGGGTGTGATAGGTGGCACGGAGAATT</td><td>>GlnTTG.trnaMT_MT_-_4329_4400.multi</td><td>1</td><td>33</td><td>33</td><td>tRF lookalike</td><td>NO</td><td>0.00</td><td>26.61</td><td>39.01</td><td>0.00</td></tr>
<tr><td>>multi-trnaMT_GlnTTG_MT_-_4329_4400@1.34.34,trnalookalike2_GlnTTG_1_-_564879_564950@1.34.34|TAGGATGGGGTGTGATAGGTGGCACGGAGAATTT</td><td>>GlnTTG.trnaMT_MT_-_4329_4400.multi</td><td>1</td><td>34</td><td>34</td><td>tRF lookalike</td><td>NO</td><td>0.00</td><td>0.00</td><td>67.22</td><td>0.00</td></tr>
<tr><td>>multi-trnaMT_GlnTTG_MT_-_4329_4400@1.36.36,trnalookalike2_GlnTTG_1_-_564879_564950@1.36.36|TAGGATGGGGTGTGATAGGTGGCACGGAGAATTTTG</td><td>>GlnTTG.trnaMT_MT_-_4329_4400.multi</td><td>1</td><td>36</td><td>36</td><td>tRF lookalike</td><td>NO</td><td>0.00</td><td>0.00</td><td>14.24</td><td>0.00</td></tr>
<tr><td>>trnaMT_GluTTC_MT_-_14674_14742@31.72.42,trnalookalike8_GluTTC_5_-_93905172_93905240@31.72.42|TTCATATCATTGGTCGTGGTTGTAGTCCGTGCGAGAATACCA</td><td>>GluTTC.trnaMT_MT_-_14674_14742</td><td>31</td><td>72</td><td>42</td><td>tRF lookalike</td><td>NO</td><td>0.00</td><td>0.00</td><td>11.38</td><td>9.84</td></tr>
<tr><td>>trnaMT_GluTTC_MT_-_14674_14742@32.72.41,trnalookalike8_GluTTC_5_-_93905172_93905240@32.72.41|TCATATCATTGGTCGTGGTTGTAGTCCGTGCGAGAATACCA</td><td>>GluTTC.trnaMT_MT_-_14674_14742</td><td>32</td><td>72</td><td>41</td><td>tRF lookalike</td><td>NO</td><td>0.00</td><td>0.00</td><td>59.77</td><td>25.64</td></tr>
<tr><td>>trnaMT_GluTTC_MT_-_14674_14742@33.72.40,trnalookalike8_GluTTC_5_-_93905172_93905240@33.72.40|CATATCATTGGTCGTGGTTGTAGTCCGTGCGAGAATACCA</td><td>>GluTTC.trnaMT_MT_-_14674_14742</td><td>33</td><td>72</td><td>40</td><td>tRF lookalike</td><td>NO</td><td>0.00</td><td>0.00</td><td>57.37</td><td>25.77</td></tr>
<tr><td>>trnaMT_GluTTC_MT_-_14674_14742@34.71.38,trnalookalike8_GluTTC_5_-_93905172_93905240@34.71.38|ATATCATTGGTCGTGGTTGTAGTCCGTGCGAGAATACC</td><td>>GluTTC.trnaMT_MT_-_14674_14742</td><td>34</td><td>71</td><td>38</td><td>tRF lookalike</td><td>NO</td><td>0.00</td><td>0.00</td><td>11.65</td><td>0.00</td></tr>
<tr><td>>trnaMT_GluTTC_MT_-_14674_14742@34.72.39,trnalookalike8_GluTTC_5_-_93905172_93905240@34.72.39|ATATCATTGGTCGTGGTTGTAGTCCGTGCGAGAATACCA</td><td>>GluTTC.trnaMT_MT_-_14674_14742</td><td>34</td><td>72</td><td>39</td><td>tRF lookalike</td><td>NO</td><td>0.00</td><td>23.79</td><td>220.24</td><td>134.40</td></tr>
<tr><td>>trnaMT_GluTTC_MT_-_14674_14742@35.72.38,trnalookalike8_GluTTC_5_-_93905172_93905240@35.72.38|TATCATTGGTCGTGGTTGTAGTCCGTGCGAGAATACCA</td><td>>GluTTC.trnaMT_MT_-_14674_14742</td><td>35</td><td>72</td><td>38</td><td>tRF lookalike</td><td>NO</td><td>0.00</td><td>34.52</td><td>201.90</td><td>114.30</td></tr>
<tr><td>>trnaMT_GluTTC_MT_-_14674_14742@37.72.36,trnalookalike8_GluTTC_5_-_93905172_93905240@37.72.36|TCATTGGTCGTGGTTGTAGTCCGTGCGAGAATACCA</td><td>>GluTTC.trnaMT_MT_-_14674_14742</td><td>37</td><td>72</td><td>36</td><td>tRF lookalike</td><td>NO</td><td>0.00</td><td>80.08</td><td>135.16</td><td>74.76</td></tr>
<tr><td>>trnaMT_GluTTC_MT_-_14674_14742@38.71.34,trnalookalike8_GluTTC_5_-_93905172_93905240@38.71.34|CATTGGTCGTGGTTGTAGTCCGTGCGAGAATACC</td><td>>GluTTC.trnaMT_MT_-_14674_14742</td><td>38</td><td>71</td><td>34</td><td>tRF lookalike</td><td>NO</td><td>0.00</td><td>0.00</td><td>39.65</td><td>23.47</td></tr>
<tr><td>>trnaMT_GluTTC_MT_-_14674_14742@38.72.35,trnalookalike8_GluTTC_5_-_93905172_93905240@38.72.35|CATTGGTCGTGGTTGTAGTCCGTGCGAGAATACCA</td><td>>GluTTC.trnaMT_MT_-_14674_14742</td><td>38</td><td>72</td><td>35</td><td>tRF lookalike</td><td>NO</td><td>32.24</td><td>40.59</td><td>558.70</td><td>270.65</td></tr>
<tr><td>>trnaMT_GluTTC_MT_-_14674_14742@39.70.32,trnalookalike8_GluTTC_5_-_93905172_93905240@39.70.32|ATTGGTCGTGGTTGTAGTCCGTGCGAGAATAC</td><td>>GluTTC.trnaMT_MT_-_14674_14742</td><td>39</td><td>70</td><td>32</td><td>tRF lookalike</td><td>NO</td><td>0.00</td><td>0.00</td><td>14.67</td><td>0.00</td></tr>
<tr><td>>trnaMT_GluTTC_MT_-_14674_14742@39.71.33,trnalookalike8_GluTTC_5_-_93905172_93905240@39.71.33|ATTGGTCGTGGTTGTAGTCCGTGCGAGAATACC</td><td>>GluTTC.trnaMT_MT_-_14674_14742</td><td>39</td><td>71</td><td>33</td><td>tRF lookalike</td><td>NO</td><td>0.00</td><td>0.00</td><td>168.26</td><td>74.09</td></tr>
<tr><td>>trnaMT_GluTTC_MT_-_14674_14742@39.72.34,trnalookalike8_GluTTC_5_-_93905172_93905240@39.72.34|ATTGGTCGTGGTTGTAGTCCGTGCGAGAATACCA</td><td>>GluTTC.trnaMT_MT_-_14674_14742</td><td>39</td><td>72</td><td>34</td><td>tRF lookalike</td><td>NO</td><td>147.13</td><td>196.05</td><td>1581.87</td><td>557.27</td></tr>
<tr><td>>trnaMT_GluTTC_MT_-_14674_14742@40.70.31,trnalookalike8_GluTTC_5_-_93905172_93905240@40.70.31|TTGGTCGTGGTTGTAGTCCGTGCGAGAATAC</td><td>>GluTTC.trnaMT_MT_-_14674_14742</td><td>40</td><td>70</td><td>31</td><td>tRF lookalike</td><td>NO</td><td>0.00</td><td>0.00</td><td>14.51</td><td>0.00</td></tr>
<tr><td>>trnaMT_GluTTC_MT_-_14674_14742@40.71.32,trnalookalike8_GluTTC_5_-_93905172_93905240@40.71.32|TTGGTCGTGGTTGTAGTCCGTGCGAGAATACC</td><td>>GluTTC.trnaMT_MT_-_14674_14742</td><td>40</td><td>71</td><td>32</td><td>tRF lookalike</td><td>NO</td><td>0.00</td><td>0.00</td><td>123.14</td><td>38.70</td></tr>
<tr><td>>trnaMT_GluTTC_MT_-_14674_14742@40.72.33,trnalookalike8_GluTTC_5_-_93905172_93905240@40.72.33|TTGGTCGTGGTTGTAGTCCGTGCGAGAATACCA</td><td>>GluTTC.trnaMT_MT_-_14674_14742</td><td>40</td><td>72</td><td>33</td><td>tRF lookalike</td><td>NO</td><td>113.90</td><td>115.01</td><td>1478.88</td><td>556.92</td></tr>
<tr><td>>trnaMT_GluTTC_MT_-_14674_14742@41.71.31,trnalookalike8_GluTTC_5_-_93905172_93905240@41.71.31|TGGTCGTGGTTGTAGTCCGTGCGAGAATACC</td><td>>GluTTC.trnaMT_MT_-_14674_14742</td><td>41</td><td>71</td><td>31</td><td>tRF lookalike</td><td>NO</td><td>0.00</td><td>0.00</td><td>14.55</td><td>0.00</td></tr>
<tr><td>>trnaMT_GluTTC_MT_-_14674_14742@41.72.32,trnalookalike8_GluTTC_5_-_93905172_93905240@41.72.32|TGGTCGTGGTTGTAGTCCGTGCGAGAATACCA</td><td>>GluTTC.trnaMT_MT_-_14674_14742</td><td>41</td><td>72</td><td>32</td><td>tRF lookalike</td><td>NO</td><td>28.61</td><td>32.04</td><td>561.64</td><td>219.57</td></tr>
<tr><td>>trnaMT_GluTTC_MT_-_14674_14742@44.72.29,trnalookalike8_GluTTC_5_-_93905172_93905240@44.72.29|TCGTGGTTGTAGTCCGTGCGAGAATACCA</td><td>>GluTTC.trnaMT_MT_-_14674_14742</td><td>44</td><td>72</td><td>29</td><td>tRF lookalike</td><td>NO</td><td>0.00</td><td>0.00</td><td>72.43</td><td>32.10</td></tr>
<tr><td>>trnaMT_GluTTC_MT_-_14674_14742@45.72.28,trnalookalike8_GluTTC_5_-_93905172_93905240@45.72.28|CGTGGTTGTAGTCCGTGCGAGAATACCA</td><td>>GluTTC.trnaMT_MT_-_14674_14742</td><td>45</td><td>72</td><td>28</td><td>tRF lookalike</td><td>NO</td><td>0.00</td><td>0.00</td><td>57.78</td><td>10.19</td></tr>
<tr><td>>trnaMT_GluTTC_MT_-_14674_14742@46.72.27,trnalookalike8_GluTTC_5_-_93905172_93905240@46.72.27|GTGGTTGTAGTCCGTGCGAGAATACCA</td><td>>GluTTC.trnaMT_MT_-_14674_14742</td><td>46</td><td>72</td><td>27</td><td>tRF lookalike</td><td>NO</td><td>0.00</td><td>0.00</td><td>73.32</td><td>25.53</td></tr>
<tr><td>>multi-trnaMT_GluTTC_MT_-_14674_14742@21.58.38,trnalookalike8_GluTTC_5_-_93905172_93905240@21.58.38|ACGATGGTTTTTCATATCATTGGTCGTGGTTGTAGTCC</td><td>>GluTTC.trnaMT_MT_-_14674_14742.multi</td><td>21</td><td>58</td><td>38</td><td>tRF lookalike</td><td>NO</td><td>0.00</td><td>0.00</td><td>53.70</td><td>9.77</td></tr>
<tr><td>>multi-trnaMT_GluTTC_MT_-_14674_14742@24.58.35,trnalookalike8_GluTTC_5_-_93905172_93905240@24.58.35|ATGGTTTTTCATATCATTGGTCGTGGTTGTAGTCC</td><td>>GluTTC.trnaMT_MT_-_14674_14742.multi</td><td>24</td><td>58</td><td>35</td><td>tRF lookalike</td><td>NO</td><td>0.00</td><td>0.00</td><td>54.44</td><td>0.00</td></tr>
<tr><td>>multi-trnaMT_GluTTC_MT_-_14674_14742@26.58.33,trnalookalike8_GluTTC_5_-_93905172_93905240@26.58.33|GGTTTTTCATATCATTGGTCGTGGTTGTAGTCC</td><td>>GluTTC.trnaMT_MT_-_14674_14742.multi</td><td>26</td><td>58</td><td>33</td><td>tRF lookalike</td><td>NO</td><td>0.00</td><td>0.00</td><td>13.91</td><td>0.00</td></tr>
<tr><td>>multi-trnaMT_GluTTC_MT_-_14674_14742@26.61.36,trnalookalike8_GluTTC_5_-_93905172_93905240@26.61.36|GGTTTTTCATATCATTGGTCGTGGTTGTAGTCCGTG</td><td>>GluTTC.trnaMT_MT_-_14674_14742.multi</td><td>26</td><td>61</td><td>36</td><td>tRF lookalike</td><td>NO</td><td>0.00</td><td>0.00</td><td>11.73</td><td>0.00</td></tr>
<tr><td>>multi-trnaMT_GluTTC_MT_-_14674_14742@38.58.21,trnalookalike8_GluTTC_5_-_93905172_93905240@38.58.21|CATTGGTCGTGGTTGTAGTCC</td><td>>GluTTC.trnaMT_MT_-_14674_14742.multi</td><td>38</td><td>58</td><td>21</td><td>tRF lookalike</td><td>NO</td><td>0.00</td><td>0.00</td><td>11.57</td><td>0.00</td></tr>
<tr><td>>multi-trnaMT_GluTTC_MT_-_14674_14742@38.61.24,trnalookalike8_GluTTC_5_-_93905172_93905240@38.61.24|CATTGGTCGTGGTTGTAGTCCGTG</td><td>>GluTTC.trnaMT_MT_-_14674_14742.multi</td><td>38</td><td>61</td><td>24</td><td>tRF lookalike</td><td>NO</td><td>0.00</td><td>0.00</td><td>14.02</td><td>0.00</td></tr>
</table>

# BT-20

| | | | | | | | | | | |
|---|---|---|---|---|---|---|---|---|---|---|
| >multi-trnaMT_GluTTC_MT_-_14674_14742@38.66.29,trnalookalike8_GluTTC_5_-_93905172_93905240@38.66.29\|CATTGGTCGTGGTTGTAGTCCGTGCGAGA | >GluTTC.trnaMT_MT_-_14674_14742.multi | 38 | 66 | 29 | tRF lookalike | NO | 0.00 | 0.00 | 12.76 | 10.03 |
| >multi-trnaMT_GluTTC_MT_-_14674_14742@39.57.19,trnalookalike8_GluTTC_5_-_93905172_93905240@39.57.19\|ATTGGTCGTGGTTGTAGTC | >GluTTC.trnaMT_MT_-_14674_14742.multi | 39 | 57 | 19 | tRF lookalike | NO | 0.00 | 0.00 | 67.76 | 0.00 |
| >multi-trnaMT_GluTTC_MT_-_14674_14742@39.58.20,trnalookalike8_GluTTC_5_-_93905172_93905240@39.58.20\|ATTGGTCGTGGTTGTAGTCC | >GluTTC.trnaMT_MT_-_14674_14742.multi | 39 | 58 | 20 | tRF lookalike | NO | 0.00 | 10.48 | 129.57 | 31.19 |
| >multi-trnaMT_GluTTC_MT_-_14674_14742@39.61.23,trnalookalike8_GluTTC_5_-_93905172_93905240@39.61.23\|ATTGGTCGTGGTTGTAGTCCGTG | >GluTTC.trnaMT_MT_-_14674_14742.multi | 39 | 61 | 23 | tRF lookalike | NO | 0.00 | 0.00 | 86.32 | 10.98 |
| >multi-trnaMT_GluTTC_MT_-_14674_14742@39.63.25,trnalookalike8_GluTTC_5_-_93905172_93905240@39.63.25\|ATTGGTCGTGGTTGTAGTCCGTGCG | >GluTTC.trnaMT_MT_-_14674_14742.multi | 39 | 63 | 25 | tRF lookalike | NO | 0.00 | 0.00 | 88.79 | 26.35 |
| >multi-trnaMT_GluTTC_MT_-_14674_14742@39.65.27,trnalookalike8_GluTTC_5_-_93905172_93905240@39.65.27\|ATTGGTCGTGGTTGTAGTCCGTGCGAG | >GluTTC.trnaMT_MT_-_14674_14742.multi | 39 | 65 | 27 | tRF lookalike | NO | 0.00 | 0.00 | 15.24 | 9.87 |
| >multi-trnaMT_GluTTC_MT_-_14674_14742@39.66.28,trnalookalike8_GluTTC_5_-_93905172_93905240@39.66.28\|ATTGGTCGTGGTTGTAGTCCGTGCGAGA | >GluTTC.trnaMT_MT_-_14674_14742.multi | 39 | 66 | 28 | tRF lookalike | NO | 0.00 | 0.00 | 94.33 | 29.58 |
| >multi-trnaMT_GluTTC_MT_-_14674_14742@39.68.30,trnalookalike8_GluTTC_5_-_93905172_93905240@39.68.30\|ATTGGTCGTGGTTGTAGTCCGTGCGAGAAT | >GluTTC.trnaMT_MT_-_14674_14742.multi | 39 | 68 | 30 | tRF lookalike | NO | 0.00 | 0.00 | 15.64 | 0.00 |
| >multi-trnaMT_GluTTC_MT_-_14674_14742@39.69.31,trnalookalike8_GluTTC_5_-_93905172_93905240@39.69.31\|ATTGGTCGTGGTTGTAGTCCGTGCGAGAATA | >GluTTC.trnaMT_MT_-_14674_14742.multi | 39 | 69 | 31 | tRF lookalike | NO | 0.00 | 0.00 | 10.98 | 0.00 |
| >multi-trnaMT_GluTTC_MT_-_14674_14742@40.57.18,trnalookalike8_GluTTC_5_-_93905172_93905240@40.57.18\|TTGGTCGTGGTTGTAGTC | >GluTTC.trnaMT_MT_-_14674_14742.multi | 40 | 57 | 18 | tRF lookalike | NO | 0.00 | 0.00 | 14.98 | 0.00 |
| >multi-trnaMT_GluTTC_MT_-_14674_14742@40.58.19,trnalookalike8_GluTTC_5_-_93905172_93905240@40.58.19\|TTGGTCGTGGTTGTAGTCC | >GluTTC.trnaMT_MT_-_14674_14742.multi | 40 | 58 | 19 | tRF lookalike | NO | 0.00 | 30.54 | 92.27 | 28.34 |
| >multi-trnaMT_GluTTC_MT_-_14674_14742@40.61.22,trnalookalike8_GluTTC_5_-_93905172_93905240@40.61.22\|TTGGTCGTGGTTGTAGTCCGTG | >GluTTC.trnaMT_MT_-_14674_14742.multi | 40 | 61 | 22 | tRF lookalike | NO | 0.00 | 0.00 | 60.84 | 0.00 |
| >multi-trnaMT_GluTTC_MT_-_14674_14742@40.63.24,trnalookalike8_GluTTC_5_-_93905172_93905240@40.63.24\|TTGGTCGTGGTTGTAGTCCGTGCG | >GluTTC.trnaMT_MT_-_14674_14742.multi | 40 | 63 | 24 | tRF lookalike | NO | 0.00 | 0.00 | 65.14 | 9.82 |
| >multi-trnaMT_GluTTC_MT_-_14674_14742@40.65.26,trnalookalike8_GluTTC_5_-_93905172_93905240@40.65.26\|TTGGTCGTGGTTGTAGTCCGTGCGAG | >GluTTC.trnaMT_MT_-_14674_14742.multi | 40 | 65 | 26 | tRF lookalike | NO | 0.00 | 0.00 | 10.53 | 0.00 |
| >multi-trnaMT_GluTTC_MT_-_14674_14742@40.66.27,trnalookalike8_GluTTC_5_-_93905172_93905240@40.66.27\|TTGGTCGTGGTTGTAGTCCGTGCGAGA | >GluTTC.trnaMT_MT_-_14674_14742.multi | 40 | 66 | 27 | tRF lookalike | NO | 0.00 | 0.00 | 77.09 | 30.34 |
| >multi-am_trnaMT_GluTTC_MT_-_14674_14742@21.54.34,trnalookalike8_GluTTC_5_-_93905172_93905240@21.54.34\|ACGATGGTTTTTCATATCATTGGTCGTGGTTGTA | >GluTTC.trnaMT_MT_-_14674_14742.multi.am | 21 | 54 | 34 | tRF lookalike | NO | 0.00 | 0.00 | 13.98 | 0.00 |
| >multi-am_trnaMT_GluTTC_MT_-_14674_14742@41.58.18,trnalookalike8_GluTTC_5_-_93905172_93905240@41.58.18\|TGGTCGTGGTTGTAGTCC | >GluTTC.trnaMT_MT_-_14674_14742.multi.am | 41 | 58 | 18 | tRF lookalike | NO | 0.00 | 0.00 | 13.62 | 0.00 |
| >multi-trnaMT_LysTTT_MT_+_8295_8364@1.23.23,trnalookalike7_LysTTT_1_+_568844_568913@1.23.23\|CACTGTAAAGCTAACTTAGCATT | >LysTTT.trnaMT_MT_+_8295_8364.multi | 1 | 23 | 23 | tRF lookalike | NO | 0.00 | 0.00 | 15.12 | 0.00 |
| >multi-trnaMT_LysTTT_MT_+_8295_8364@1.26.26,trnalookalike7_LysTTT_1_+_568844_568913@1.26.26\|CACTGTAAAGCTAACTTAGCATTAAC | >LysTTT.trnaMT_MT_+_8295_8364.multi | 1 | 26 | 26 | tRF lookalike | NO | 0.00 | 0.00 | 13.54 | 0.00 |
| >multi-trnaMT_LysTTT_MT_+_8295_8364@1.36.36,trnalookalike7_LysTTT_1_+_568844_568913@1.36.36\|CACTGTAAAGCTAACTTAGCATTAACCTTTTAAGTT | >LysTTT.trnaMT_MT_+_8295_8364.multi | 1 | 36 | 36 | tRF lookalike | NO | 0.00 | 0.00 | 13.31 | 0.00 |
| >multi-trnaMT_LysTTT_MT_+_8295_8364@1.38.38,trnalookalike7_LysTTT_1_+_568844_568913@1.38.38\|CACTGTAAAGCTAACTTAGCATTAACCTTTTAAGTTAA | >LysTTT.trnaMT_MT_+_8295_8364.multi | 1 | 38 | 38 | tRF lookalike | NO | 0.00 | 0.00 | 39.66 | 0.00 |
| >multi-am_trnaMT_LysTTT_MT_+_8295_8364@28.52.25,trnalookalike7_LysTTT_1_+_568844_568913@28.52.25\|TTTTAAGTTAAAGATTAAGAGAACC | >LysTTT.trnaMT_MT_+_8295_8364.multi.am | 28 | 52 | 25 | tRF lookalike | NO | 0.00 | 24.12 | 0.00 | 0.00 |
| >multi-am_trnaMT_LysTTT_MT_+_8295_8364@29.52.24,trnalookalike7_LysTTT_1_+_568844_568913@29.52.24\|TTTAAGTTAAAGATTAAGAGAACC | >LysTTT.trnaMT_MT_+_8295_8364.multi.am | 29 | 52 | 24 | tRF lookalike | NO | 0.00 | 26.36 | 0.00 | 0.00 |
| >multi-am_trnaMT_LysTTT_MT_+_8295_8364@30.52.23,trnalookalike7_LysTTT_1_+_568844_568913@30.52.23\|TTAAGTTAAAGATTAAGAGAACC | >LysTTT.trnaMT_MT_+_8295_8364.multi.am | 30 | 52 | 23 | tRF lookalike | NO | 0.00 | 31.31 | 0.00 | 0.00 |
| >multi-am_trnaMT_LysTTT_MT_+_8295_8364@34.52.19,trnalookalike7_LysTTT_1_+_568844_568913@34.52.19\|GTTAAAGATTAAGAGAACC | >LysTTT.trnaMT_MT_+_8295_8364.multi.am | 34 | 52 | 19 | tRF lookalike | NO | 0.00 | 37.11 | 0.00 | 0.00 |
| >multi-trnaMT_SerTGA_MT_-_7446_7514@1.25.25,trnalookalike6_SerTGA_1_-_567997_568065@1.25.25\|GAAAAAGTCATGGAGGCCATGGGGT | >SerTGA.trnaMT_MT_-_7446_7514.multi | 1 | 25 | 25 | tRF lookalike | NO | 0.00 | 0.00 | 82.06 | 0.00 |
| >multi-trnaMT_SerTGA_MT_-_7446_7514@1.26.26,trnalookalike6_SerTGA_1_-_567997_568065@1.26.26\|GAAAAAGTCATGGAGGCCATGGGGTT | >SerTGA.trnaMT_MT_-_7446_7514.multi | 1 | 26 | 26 | tRF lookalike | NO | 39.92 | 0.00 | 132.12 | 0.00 |
| >trnaMT_HisGTG_MT_+_12138_12206@26.63.38\|AGATTGTGAATCTGACAACAGAGGCTTACGACCCCTTA | >HisGTG.trnaMT_MT_+_12138_12206 | 26 | 63 | 38 | tRF MT | NO | 0.00 | 0.00 | 11.22 | 0.00 |
| >trnaMT_HisGTG_MT_+_12138_12206@32.63.32\|TGAATCTGACAACAGAGGCTTACGACCCCTTA | >HisGTG.trnaMT_MT_+_12138_12206 | 32 | 63 | 32 | tRF MT | NO | 0.00 | 0.00 | 11.25 | 0.00 |
| >trnaMT_HisGTG_MT_+_12138_12206@32.70.39\|TGAATCTGACAACAGAGGCTTACGACCCCTTATTTACCC | >HisGTG.trnaMT_MT_+_12138_12206 | 32 | 70 | 39 | tRF MT | NO | 0.00 | 0.00 | 13.45 | 0.00 |
| >trnaMT_HisGTG_MT_+_12138_12206@32.71.40\|TGAATCTGACAACAGAGGCTTACGACCCCTTATTTACCCC | >HisGTG.trnaMT_MT_+_12138_12206 | 32 | 71 | 40 | tRF MT | NO | 0.00 | 0.00 | 107.32 | 36.62 |
| >trnaMT_HisGTG_MT_+_12138_12206@32.72.41\|TGAATCTGACAACAGAGGCTTACGACCCCTTATTTACCCCA | >HisGTG.trnaMT_MT_+_12138_12206 | 32 | 72 | 41 | tRF MT | NO | 0.00 | 0.00 | 60.25 | 23.36 |
| >multi-am_trnaMT_HisGTG_MT_+_12138_12206@18.51.34\|AAAACATCAGATTGTGAATCTGACAACAGAGGCT | >HisGTG.trnaMT_MT_+_12138_12206.multi.am | 18 | 51 | 34 | tRF MT | NO | 0.00 | 0.00 | 75.20 | 9.52 |

# BT-20

| | | | | | | | | | | |
|---|---|---|---|---|---|---|---|---|---|---|
| >multi-am_trnaMT_HisGTG_MT_+_12138_12206@19.51.33\|AAACATCAGATTGTGAATCTGACAACAGAGGCT | >HisGTG.trnaMT_MT_+_12138_12206.multi.am | 19 | 51 | 33 | tRF MT | NO | 0.00 | 0.00 | 79.96 | 10.35 |
| >multi-am_trnaMT_HisGTG_MT_+_12138_12206@20.51.32\|AACATCAGATTGTGAATCTGACAACAGAGGCT | >HisGTG.trnaMT_MT_+_12138_12206.multi.am | 20 | 51 | 32 | tRF MT | NO | 0.00 | 0.00 | 67.37 | 9.62 |
| >trnaMT_LeuTAA_MT_+_3230_3304@1.25.25\|GTTAAGATGGCAGAGCCCGGTAATC | >LeuTAA.trnaMT_MT_+_3230_3304 | 1 | 25 | 25 | tRF MT | NO | 0.00 | 0.00 | 10.67 | 0.00 |
| >trnaMT_LeuTAA_MT_+_3230_3304@1.30.30\|GTTAAGATGGCAGAGCCCGGTAATCGCATA | >LeuTAA.trnaMT_MT_+_3230_3304 | 1 | 30 | 30 | tRF MT | NO | 0.00 | 0.00 | 115.42 | 0.00 |
| >trnaMT_LeuTAA_MT_+_3230_3304@18.42.25\|CGGTAATCGCATAAAACTTAAAACT | >LeuTAA.trnaMT_MT_+_3230_3304 | 18 | 42 | 25 | tRF MT | NO | 0.00 | 25.99 | 0.00 | 0.00 |
| >trnaMT_LeuTAA_MT_+_3230_3304@18.44.27\|CGGTAATCGCATAAAACTTAAAACTTT | >LeuTAA.trnaMT_MT_+_3230_3304 | 18 | 44 | 27 | tRF MT | NO | 0.00 | 30.75 | 0.00 | 0.00 |
| >trnaMT_LeuTAA_MT_+_3230_3304@19.44.26\|GGTAATCGCATAAAACTTAAAACTTT | >LeuTAA.trnaMT_MT_+_3230_3304 | 19 | 44 | 26 | tRF MT | NO | 0.00 | 23.57 | 0.00 | 0.00 |
| >trnaMT_MetCAT_MT_+_4402_4469@37.71.35\|CCGAAAATGTTGGTTATACCCTTCCCGTACTACCA | >MetCAT.trnaMT_MT_+_4402_4469 | 37 | 71 | 35 | tRF MT | NO | 0.00 | 0.00 | 39.64 | 9.75 |
| >trnaMT_MetCAT_MT_+_4402_4469@38.71.34\|CGAAAATGTTGGTTATACCCTTCCCGTACTACCA | >MetCAT.trnaMT_MT_+_4402_4469 | 38 | 71 | 34 | tRF MT | NO | 0.00 | 0.00 | 12.31 | 0.00 |
| >trnaMT_MetCAT_MT_+_4402_4469@40.71.32\|AAAATGTTGGTTATACCCTTCCCGTACTACCA | >MetCAT.trnaMT_MT_+_4402_4469 | 40 | 71 | 32 | tRF MT | NO | 0.00 | 0.00 | 10.28 | 0.00 |
| >trnaMT_MetCAT_MT_+_4402_4469@42.71.30\|AATGTTGGTTATACCCTTCCCGTACTACCA | >MetCAT.trnaMT_MT_+_4402_4469 | 42 | 71 | 30 | tRF MT | NO | 0.00 | 0.00 | 10.46 | 0.00 |
| >trnaMT_MetCAT_MT_+_4402_4469@52.71.20\|ATACCCTTCCCGTACTACCA | >MetCAT.trnaMT_MT_+_4402_4469 | 52 | 71 | 20 | tRF MT | NO | 0.00 | 27.31 | 0.00 | 0.00 |
| >trnaMT_MetCAT_MT_+_4402_4469@53.71.19\|TACCCTTCCCGTACTACCA | >MetCAT.trnaMT_MT_+_4402_4469 | 53 | 71 | 19 | tRF MT | NO | 0.00 | 24.66 | 0.00 | 0.00 |
| >trnaMT_MetCAT_MT_+_4402_4469@54.71.18\|ACCCTTCCCGTACTACCA | >MetCAT.trnaMT_MT_+_4402_4469 | 54 | 71 | 18 | tRF MT | NO | 0.00 | 32.88 | 0.00 | 0.00 |
| >multi-am_trnaMT_MetCAT_MT_+_4402_4469@1.21.21\|AGTAAGGTCAGCTAAATAAGC | >MetCAT.trnaMT_MT_+_4402_4469.multi.am | 1 | 21 | 21 | tRF MT | NO | 0.00 | 0.00 | 60.21 | 0.00 |
| >multi-am_trnaMT_MetCAT_MT_+_4402_4469@1.22.22\|AGTAAGGTCAGCTAAATAAGCT | >MetCAT.trnaMT_MT_+_4402_4469.multi.am | 1 | 22 | 22 | tRF MT | NO | 0.00 | 0.00 | 11.10 | 0.00 |
| >multi-am_trnaMT_MetCAT_MT_+_4402_4469@1.47.47\|AGTAAGGTCAGCTAAATAAGCTATCGGGCCCATACCCCGAAAATGTT | >MetCAT.trnaMT_MT_+_4402_4469.multi.am | 1 | 47 | 47 | tRF MT | NO | 0.00 | 0.00 | 15.60 | 0.00 |
| >trnaMT_PheGAA_MT_+_577_647@1.21.21\|GTTTATGTAGCTTACCTCCTC | >PheGAA.trnaMT_MT_+_577_647 | 1 | 21 | 21 | tRF MT | NO | 0.00 | 0.00 | 76.20 | 0.00 |
| >trnaMT_PheGAA_MT_+_577_647@1.30.30\|GTTTATGTAGCTTACCTCCTCAAAGCAATA | >PheGAA.trnaMT_MT_+_577_647 | 1 | 30 | 30 | tRF MT | NO | 0.00 | 0.00 | 11.58 | 0.00 |
| >trnaMT_PheGAA_MT_+_577_647@35.53.19\|GAAAATGTTTAGACGGGCT | >PheGAA.trnaMT_MT_+_577_647 | 35 | 53 | 19 | tRF MT | NO | 0.00 | 0.00 | 60.37 | 0.00 |
| >trnaMT_PheGAA_MT_+_577_647@35.54.20\|GAAAATGTTTAGACGGGCTC | >PheGAA.trnaMT_MT_+_577_647 | 35 | 54 | 20 | tRF MT | NO | 0.00 | 0.00 | 71.79 | 30.48 |
| >trnaMT_PheGAA_MT_+_577_647@35.55.21\|GAAAATGTTTAGACGGGCTCA | >PheGAA.trnaMT_MT_+_577_647 | 35 | 55 | 21 | tRF MT | NO | 0.00 | 0.00 | 152.61 | 77.89 |
| >trnaMT_PheGAA_MT_+_577_647@35.56.22\|GAAAATGTTTAGACGGGCTCAC | >PheGAA.trnaMT_MT_+_577_647 | 35 | 56 | 22 | tRF MT | NO | 0.00 | 0.00 | 90.74 | 39.47 |
| >trnaMT_PheGAA_MT_+_577_647@35.57.23\|GAAAATGTTTAGACGGGCTCACA | >PheGAA.trnaMT_MT_+_577_647 | 35 | 57 | 23 | tRF MT | NO | 42.99 | 27.00 | 149.73 | 111.51 |
| >trnaMT_PheGAA_MT_+_577_647@35.59.25\|GAAAATGTTTAGACGGGCTCACATC | >PheGAA.trnaMT_MT_+_577_647 | 35 | 59 | 25 | tRF MT | NO | 0.00 | 0.00 | 54.26 | 28.23 |
| >trnaMT_PheGAA_MT_+_577_647@35.60.26\|GAAAATGTTTAGACGGGCTCACATCA | >PheGAA.trnaMT_MT_+_577_647 | 35 | 60 | 26 | tRF MT | NO | 0.00 | 0.00 | 38.24 | 30.13 |
| >trnaMT_PheGAA_MT_+_577_647@35.62.28\|GAAAATGTTTAGACGGGCTCACATCACC | >PheGAA.trnaMT_MT_+_577_647 | 35 | 62 | 28 | tRF MT | NO | 0.00 | 0.00 | 58.37 | 25.02 |
| >trnaMT_PheGAA_MT_+_577_647@36.54.19\|AAAATGTTTAGACGGGCTC | >PheGAA.trnaMT_MT_+_577_647 | 36 | 54 | 19 | tRF MT | NO | 0.00 | 0.00 | 16.22 | 0.00 |
| >trnaMT_PheGAA_MT_+_577_647@36.55.20\|AAAATGTTTAGACGGGCTCA | >PheGAA.trnaMT_MT_+_577_647 | 36 | 55 | 20 | tRF MT | NO | 0.00 | 0.00 | 73.94 | 25.03 |
| >trnaMT_PheGAA_MT_+_577_647@36.56.21\|AAAATGTTTAGACGGGCTCAC | >PheGAA.trnaMT_MT_+_577_647 | 36 | 56 | 21 | tRF MT | NO | 0.00 | 0.00 | 54.12 | 27.04 |
| >trnaMT_PheGAA_MT_+_577_647@36.57.22\|AAAATGTTTAGACGGGCTCACA | >PheGAA.trnaMT_MT_+_577_647 | 36 | 57 | 22 | tRF MT | NO | 0.00 | 0.00 | 81.40 | 38.18 |
| >trnaMT_PheGAA_MT_+_577_647@36.62.27\|AAAATGTTTAGACGGGCTCACATCACC | >PheGAA.trnaMT_MT_+_577_647 | 36 | 62 | 27 | tRF MT | NO | 0.00 | 0.00 | 13.76 | 10.27 |
| >trnaMT_PheGAA_MT_+_577_647@38.56.19\|AATGTTTAGACGGGCTCAC | >PheGAA.trnaMT_MT_+_577_647 | 38 | 56 | 19 | tRF MT | NO | 0.00 | 0.00 | 15.48 | 31.40 |
| >trnaMT_PheGAA_MT_+_577_647@39.56.18\|ATGTTTAGACGGGCTCAC | >PheGAA.trnaMT_MT_+_577_647 | 39 | 56 | 18 | tRF MT | NO | 31.55 | 93.39 | 297.94 | 292.49 |
| >trnaMT_PheGAA_MT_+_577_647@39.57.19\|ATGTTTAGACGGGCTCACA | >PheGAA.trnaMT_MT_+_577_647 | 39 | 57 | 19 | tRF MT | NO | 0.00 | 24.83 | 63.10 | 65.34 |
| >trnaMT_PheGAA_MT_+_577_647@39.59.21\|ATGTTTAGACGGGCTCACATC | >PheGAA.trnaMT_MT_+_577_647 | 39 | 59 | 21 | tRF MT | NO | 0.00 | 0.00 | 12.55 | 10.41 |
| >trnaMT_PheGAA_MT_+_577_647@39.62.24\|ATGTTTAGACGGGCTCACATCACC | >PheGAA.trnaMT_MT_+_577_647 | 39 | 62 | 24 | tRF MT | NO | 0.00 | 0.00 | 13.53 | 9.96 |
| >trnaMT_PheGAA_MT_+_577_647@42.74.33\|TTTAGACGGGCTCACATCACCCCATAAACACCA | >PheGAA.trnaMT_MT_+_577_647 | 42 | 74 | 33 | tRF MT | NO | 0.00 | 0.00 | 59.09 | 0.00 |
| >trnaMT_PheGAA_MT_+_577_647@43.74.32\|TTAGACGGGCTCACATCACCCCATAAACACCA | >PheGAA.trnaMT_MT_+_577_647 | 43 | 74 | 32 | tRF MT | NO | 0.00 | 0.00 | 11.13 | 0.00 |
| >trnaMT_PheGAA_MT_+_577_647@44.74.31\|TAGACGGGCTCACATCACCCCATAAACACCA | >PheGAA.trnaMT_MT_+_577_647 | 44 | 74 | 31 | tRF MT | NO | 24.47 | 0.00 | 10.15 | 0.00 |
| >trnaMT_PheGAA_MT_+_577_647@56.74.19\|CATCACCCCATAAACACCA | >PheGAA.trnaMT_MT_+_577_647 | 56 | 74 | 19 | tRF MT | NO | 0.00 | 31.63 | 0.00 | 28.62 |
| >multi-am_trnaMT_PheGAA_MT_+_577_647@40.57.18\|TGTTTAGACGGGCTCACA | >PheGAA.trnaMT_MT_+_577_647.multi.am | 40 | 57 | 18 | tRF MT | NO | 0.00 | 26.87 | 53.94 | 40.76 |
| >multi-am_trnaMT_PheGAA_MT_+_577_647@41.58.18\|GTTTAGACGGGCTCACAT | >PheGAA.trnaMT_MT_+_577_647.multi.am | 41 | 58 | 18 | tRF MT | NO | 0.00 | 0.00 | 63.74 | 81.96 |
| >multi-am_trnaMT_PheGAA_MT_+_577_647@41.59.19\|GTTTAGACGGGCTCACATC | >PheGAA.trnaMT_MT_+_577_647.multi.am | 41 | 59 | 19 | tRF MT | NO | 0.00 | 31.17 | 53.11 | 0.00 |
| >multi-am_trnaMT_PheGAA_MT_+_577_647@41.62.22\|GTTTAGACGGGCTCACATCACC | >PheGAA.trnaMT_MT_+_577_647.multi.am | 41 | 62 | 22 | tRF MT | NO | 0.00 | 45.61 | 60.34 | 10.32 |
| >multi-am_trnaMT_PheGAA_MT_+_577_647@42.62.21\|TTTAGACGGGCTCACATCACC | >PheGAA.trnaMT_MT_+_577_647.multi.am | 42 | 62 | 21 | tRF MT | NO | 0.00 | 26.80 | 9.68 | 0.00 |
| >multi-am_trnaMT_PheGAA_MT_+_577_647@44.62.19\|TAGACGGGCTCACATCACC | >PheGAA.trnaMT_MT_+_577_647.multi.am | 44 | 62 | 19 | tRF MT | NO | 0.00 | 71.54 | 52.90 | 9.91 |
| >multi-am_trnaMT_PheGAA_MT_+_577_647@45.62.18\|AGACGGGCTCACATCACC | >PheGAA.trnaMT_MT_+_577_647.multi.am | 45 | 62 | 18 | tRF MT | NO | 0.00 | 35.16 | 10.11 | 0.00 |
| >trnaMT_ProTGG_MT_-_15956_16023@40.58.19\|AATGGTGGAGTTAAAGACT | >ProTGG.trnaMT_MT_-_15956_16023 | 40 | 58 | 19 | tRF MT | NO | 0.00 | 0.00 | 54.88 | 10.55 |
| >multi-am_trnaMT_ProTGG_MT_-_15956_16023@52.70.19\|AAAGACTTTTTCTCTGACC | >ProTGG.trnaMT_MT_-_15956_16023.multi.am | 52 | 70 | 19 | tRF MT | NO | 0.00 | 0.00 | 54.46 | 10.75 |
| >multi-am_trnaMT_ProTGG_MT_-_15956_16023@52.71.20\|AAAGACTTTTTCTCTGACCA | >ProTGG.trnaMT_MT_-_15956_16023.multi.am | 52 | 71 | 20 | tRF MT | NO | 25.57 | 104.51 | 70.26 | 85.01 |
| >multi-am_trnaMT_ProTGG_MT_-_15956_16023@53.70.18\|AAGACTTTTTCTCTGACC | >ProTGG.trnaMT_MT_-_15956_16023.multi.am | 53 | 70 | 18 | tRF MT | NO | 0.00 | 26.63 | 13.28 | 10.64 |
| >multi-am_trnaMT_ProTGG_MT_-_15956_16023@53.71.19\|AAGACTTTTTCTCTGACCA | >ProTGG.trnaMT_MT_-_15956_16023.multi.am | 53 | 71 | 19 | tRF MT | NO | 24.12 | 141.36 | 59.47 | 95.85 |
| >multi-am_trnaMT_ProTGG_MT_-_15956_16023@54.71.18\|AGACTTTTTCTCTGACCA | >ProTGG.trnaMT_MT_-_15956_16023.multi.am | 54 | 71 | 18 | tRF MT | NO | 26.60 | 184.84 | 39.27 | 71.61 |
| >trnaMT_SerGCT_MT_+_12207_12265@1.18.18\|GAGAAAGCTCACAAGAAC | >SerGCT.trnaMT_MT_+_12207_12265 | 1 | 18 | 18 | tRF MT | YES | 271.76 | 123.56 | 661.29 | 143.32 |
| >trnaMT_SerGCT_MT_+_12207_12265@1.19.19\|GAGAAAGCTCACAAGAACT | >SerGCT.trnaMT_MT_+_12207_12265 | 1 | 19 | 19 | tRF MT | NO | 218.12 | 81.36 | 607.57 | 111.88 |
| >trnaMT_SerGCT_MT_+_12207_12265@1.20.20\|GAGAAAGCTCACAAGAACTG | >SerGCT.trnaMT_MT_+_12207_12265 | 1 | 20 | 20 | tRF MT | NO | 0.00 | 0.00 | 146.02 | 10.38 |
| >trnaMT_SerGCT_MT_+_12207_12265@1.21.21\|GAGAAAGCTCACAAGAACTGC | >SerGCT.trnaMT_MT_+_12207_12265 | 1 | 21 | 21 | tRF MT | NO | 0.00 | 0.00 | 163.57 | 37.62 |
| >trnaMT_SerGCT_MT_+_12207_12265@1.22.22\|GAGAAAGCTCACAAGAACTGCT | >SerGCT.trnaMT_MT_+_12207_12265 | 1 | 22 | 22 | tRF MT | NO | 109.14 | 42.55 | 580.75 | 182.24 |
| >trnaMT_SerGCT_MT_+_12207_12265@1.23.23\|GAGAAAGCTCACAAGAACTGCTA | >SerGCT.trnaMT_MT_+_12207_12265 | 1 | 23 | 23 | tRF MT | NO | 0.00 | 0.00 | 80.93 | 0.00 |
| >trnaMT_SerGCT_MT_+_12207_12265@1.24.24\|GAGAAAGCTCACAAGAACTGCTAA | >SerGCT.trnaMT_MT_+_12207_12265 | 1 | 24 | 24 | tRF MT | NO | 0.00 | 0.00 | 53.95 | 0.00 |
| >trnaMT_SerGCT_MT_+_12207_12265@1.25.25\|GAGAAAGCTCACAAGAACTGCTAAC | >SerGCT.trnaMT_MT_+_12207_12265 | 1 | 25 | 25 | tRF MT | NO | 117.86 | 163.05 | 745.13 | 474.09 |
| >trnaMT_SerGCT_MT_+_12207_12265@1.26.26\|GAGAAAGCTCACAAGAACTGCTAACT | >SerGCT.trnaMT_MT_+_12207_12265 | 1 | 26 | 26 | tRF MT | NO | 80.66 | 45.06 | 753.11 | 222.66 |
| >trnaMT_SerGCT_MT_+_12207_12265@13.62.50\|AAGAACTGCTAACTCATGCCCCCATGTCTAACAACATGGCTTTCTCACCA | >SerGCT.trnaMT_MT_+_12207_12265 | 13 | 62 | 50 | tRF MT | NO | 0.00 | 0.00 | 12.75 | 0.00 |
| >trnaMT_SerGCT_MT_+_12207_12265@14.41.28\|AGAACTGCTAACTCATGCCCCCATGTCT | >SerGCT.trnaMT_MT_+_12207_12265 | 14 | 41 | 28 | tRF MT | NO | 0.00 | 0.00 | 10.43 | 0.00 |

# BT-20

| | | | | | | | | | | |
|---|---|---|---|---|---|---|---|---|---|---|
| >trnaMT_SerGCT_MT_+_12207_12265@14.62.49\|AGAACTGCTAACTCATGCCCCCATGTCTAACAACATGGCTTTCTCACCA | >SerGCT.trnaMT_MT_+_12207_12265 | 14 | 62 | 49 | tRF MT | NO | 0.00 | 0.00 | 11.23 | 25.24 |
| >multi-am_trnaMT_SerGCT_MT_+_12207_12265@14.31.18\|AGAACTGCTAACTCATGC | >SerGCT.trnaMT_MT_+_12207_12265.multi.am | 14 | 31 | 18 | tRF MT | NO | 0.00 | 29.02 | 0.00 | 0.00 |
| >multi-am_trnaMT_SerGCT_MT_+_12207_12265@14.32.19\|AGAACTGCTAACTCATGCC | >SerGCT.trnaMT_MT_+_12207_12265.multi.am | 14 | 32 | 19 | tRF MT | NO | 0.00 | 33.87 | 0.00 | 0.00 |
| >trnaMT_ThrTGT_MT_+_15888_15953@31.69.39\|TTGTAAACCGGAGATGAAAACCTTTTTCCAAGGACACCA | >ThrTGT.trnaMT_MT_+_15888_15953 | 31 | 69 | 39 | tRF MT | NO | 0.00 | 0.00 | 15.77 | 0.00 |
| >trnaMT_ThrTGT_MT_+_15888_15953@37.69.33\|ACCGGAGATGAAAACCTTTTTCCAAGGACACCA | >ThrTGT.trnaMT_MT_+_15888_15953 | 37 | 69 | 33 | tRF MT | NO | 0.00 | 0.00 | 15.94 | 0.00 |
| >trnaMT_ValTAC_MT_+_1602_1670@28.72.45\|AACTTACACTTAGGAGATTTCAACTTAACTTGACCGCTCTGACCA | >ValTAC.trnaMT_MT_+_1602_1670 | 28 | 72 | 45 | tRF MT | NO | 0.00 | 0.00 | 15.74 | 30.57 |
| >trnaMT_ValTAC_MT_+_1602_1670@29.72.44\|ACTTACACTTAGGAGATTTCAACTTAACTTGACCGCTCTGACCA | >ValTAC.trnaMT_MT_+_1602_1670 | 29 | 72 | 44 | tRF MT | NO | 0.00 | 0.00 | 11.74 | 10.34 |
| >trnaMT_ValTAC_MT_+_1602_1670@39.72.34\|AGGAGATTTCAACTTAACTTGACCGCTCTGACCA | >ValTAC.trnaMT_MT_+_1602_1670 | 39 | 72 | 34 | tRF MT | NO | 0.00 | 0.00 | 10.35 | 0.00 |
| >trnaMT_ValTAC_MT_+_1602_1670@41.72.32\|GAGATTTCAACTTAACTTGACCGCTCTGACCA | >ValTAC.trnaMT_MT_+_1602_1670 | 41 | 72 | 32 | tRF MT | NO | 0.00 | 0.00 | 11.23 | 0.00 |
| >trnaMT_ValTAC_MT_+_1602_1670@42.71.30\|AGATTTCAACTTAACTTGACCGCTCTGACC | >ValTAC.trnaMT_MT_+_1602_1670 | 42 | 71 | 30 | tRF MT | NO | 0.00 | 0.00 | 95.23 | 109.22 |
| >trnaMT_ValTAC_MT_+_1602_1670@42.72.31\|AGATTTCAACTTAACTTGACCGCTCTGACCA | >ValTAC.trnaMT_MT_+_1602_1670 | 42 | 72 | 31 | tRF MT | NO | 36.77 | 0.00 | 138.20 | 286.29 |
| >trnaMT_ValTAC_MT_+_1602_1670@43.72.30\|GATTTCAACTTAACTTGACCGCTCTGACCA | >ValTAC.trnaMT_MT_+_1602_1670 | 43 | 72 | 30 | tRF MT | NO | 0.00 | 0.00 | 12.03 | 82.60 |
| >trnaMT_ValTAC_MT_+_1602_1670@44.72.29\|ATTTCAACTTAACTTGACCGCTCTGACCA | >ValTAC.trnaMT_MT_+_1602_1670 | 44 | 72 | 29 | tRF MT | NO | 0.00 | 0.00 | 15.91 | 126.18 |
| >trnaMT_ValTAC_MT_+_1602_1670@48.70.23\|CAACTTAACTTGACCGCTCTGAC | >ValTAC.trnaMT_MT_+_1602_1670 | 48 | 70 | 23 | tRF MT | NO | 0.00 | 33.45 | 0.00 | 0.00 |
| >trnaMT_ValTAC_MT_+_1602_1670@48.71.24\|CAACTTAACTTGACCGCTCTGACC | >ValTAC.trnaMT_MT_+_1602_1670 | 48 | 71 | 24 | tRF MT | NO | 0.00 | 73.86 | 11.13 | 24.17 |
| >trnaMT_ValTAC_MT_+_1602_1670@48.72.25\|CAACTTAACTTGACCGCTCTGACCA | >ValTAC.trnaMT_MT_+_1602_1670 | 48 | 72 | 25 | tRF MT | NO | 35.29 | 160.82 | 39.60 | 105.05 |
| >trnaMT_ValTAC_MT_+_1602_1670@49.71.23\|AACTTAACTTGACCGCTCTGACC | >ValTAC.trnaMT_MT_+_1602_1670 | 49 | 71 | 23 | tRF MT | NO | 0.00 | 23.63 | 10.66 | 26.86 |
| >trnaMT_ValTAC_MT_+_1602_1670@49.72.24\|AACTTAACTTGACCGCTCTGACCA | >ValTAC.trnaMT_MT_+_1602_1670 | 49 | 72 | 24 | tRF MT | NO | 24.13 | 85.37 | 76.09 | 118.81 |
| >trnaMT_ValTAC_MT_+_1602_1670@50.70.21\|ACTTAACTTGACCGCTCTGAC | >ValTAC.trnaMT_MT_+_1602_1670 | 50 | 70 | 21 | tRF MT | NO | 0.00 | 10.04 | 0.00 | 0.00 |
| >trnaMT_ValTAC_MT_+_1602_1670@50.71.22\|ACTTAACTTGACCGCTCTGACC | >ValTAC.trnaMT_MT_+_1602_1670 | 50 | 71 | 22 | tRF MT | NO | 0.00 | 37.27 | 13.40 | 31.04 |
| >trnaMT_ValTAC_MT_+_1602_1670@50.72.23\|ACTTAACTTGACCGCTCTGACCA | >ValTAC.trnaMT_MT_+_1602_1670 | 50 | 72 | 23 | tRF MT | NO | 33.11 | 160.80 | 97.56 | 164.25 |
| >trnaMT_ValTAC_MT_+_1602_1670@51.72.22\|CTTAACTTGACCGCTCTGACCA | >ValTAC.trnaMT_MT_+_1602_1670 | 51 | 72 | 22 | tRF MT | NO | 0.00 | 10.53 | 0.00 | 31.65 |
| >trnaMT_ValTAC_MT_+_1602_1670@52.72.21\|TTAACTTGACCGCTCTGACCA | >ValTAC.trnaMT_MT_+_1602_1670 | 52 | 72 | 21 | tRF MT | NO | 0.00 | 25.15 | 11.88 | 71.88 |
| >trnaMT_ValTAC_MT_+_1602_1670@53.72.20\|TAACTTGACCGCTCTGACCA | >ValTAC.trnaMT_MT_+_1602_1670 | 53 | 72 | 20 | tRF MT | NO | 30.14 | 95.65 | 58.85 | 125.78 |
| >trnaMT_ValTAC_MT_+_1602_1670@54.71.18\|AACTTGACCGCTCTGACC | >ValTAC.trnaMT_MT_+_1602_1670 | 54 | 71 | 18 | tRF MT | NO | 0.00 | 30.01 | 9.60 | 24.34 |
| >trnaMT_ValTAC_MT_+_1602_1670@54.72.19\|AACTTGACCGCTCTGACCA | >ValTAC.trnaMT_MT_+_1602_1670 | 54 | 72 | 19 | tRF MT | NO | 90.67 | 122.00 | 75.72 | 140.87 |
| >trnaMT_ValTAC_MT_+_1602_1670@55.72.18\|ACTTGACCGCTCTGACCA | >ValTAC.trnaMT_MT_+_1602_1670 | 55 | 72 | 18 | tRF MT | NO | 81.34 | 154.33 | 81.60 | 179.60 |
| >multi-am_trnaMT_ValTAC_MT_+_1602_1670@29.52.24\|ACTTACACTTAGGAGATTTCAACT | >ValTAC.trnaMT_MT_+_1602_1670.multi.am | 29 | 52 | 24 | tRF MT | NO | 0.00 | 10.12 | 0.00 | 0.00 |
| >multi-am_trnaMT_ValTAC_MT_+_1602_1670@46.67.22\|TTCAACTTAACTTGACCGCTCT | >ValTAC.trnaMT_MT_+_1602_1670.multi.am | 46 | 67 | 22 | tRF MT | NO | 0.00 | 25.75 | 0.00 | 0.00 |
| >multi-am_trnaMT_ValTAC_MT_+_1602_1670@48.66.19\|CAACTTAACTTGACCGCTC | >ValTAC.trnaMT_MT_+_1602_1670.multi.am | 48 | 66 | 19 | tRF MT | NO | 0.00 | 40.49 | 0.00 | 0.00 |
| >multi-am_trnaMT_ValTAC_MT_+_1602_1670@48.67.20\|CAACTTAACTTGACCGCTCT | >ValTAC.trnaMT_MT_+_1602_1670.multi.am | 48 | 67 | 20 | tRF MT | NO | 94.28 | 724.85 | 0.00 | 38.89 |
| >multi-am_trnaMT_ValTAC_MT_+_1602_1670@48.68.21\|CAACTTAACTTGACCGCTCTG | >ValTAC.trnaMT_MT_+_1602_1670.multi.am | 48 | 68 | 21 | tRF MT | NO | 0.00 | 40.25 | 0.00 | 0.00 |
| >multi-am_trnaMT_ValTAC_MT_+_1602_1670@49.67.19\|AACTTAACTTGACCGCTCT | >ValTAC.trnaMT_MT_+_1602_1670.multi.am | 49 | 67 | 19 | tRF MT | NO | 30.31 | 303.13 | 0.00 | 29.64 |
| >multi-am_trnaMT_ValTAC_MT_+_1602_1670@50.67.18\|ACTTAACTTGACCGCTCT | >ValTAC.trnaMT_MT_+_1602_1670.multi.am | 50 | 67 | 18 | tRF MT | NO | 28.82 | 330.46 | 0.00 | 32.59 |
| >multi-am_trnaMT_ValTAC_MT_+_1602_1670@50.68.19\|ACTTAACTTGACCGCTCTG | >ValTAC.trnaMT_MT_+_1602_1670.multi.am | 50 | 68 | 19 | tRF MT | NO | 0.00 | 25.34 | 0.00 | 0.00 |
| >trna101_AlaAGC_6_-_28831462_28831533@58.75.18,trna102_AlaAGC_6_-_28806221_28806292@58.75.18,trna110_AlaTGC_6_-_28757547_28757618@58.75.18,trna113_AlaTGC_6_-_28726141_28726212@58.75.18,trna65_AlaAGC_6_+_28574933_28575004@58.75.18\|TCCCCGGCACCTCCACCA | >AlaAGC.trna101_6_-_28831462_28831533 | 58 | 75 | 18 | tRF nuclear | NO | 25.14 | 1032.83 | 0.00 | 0.00 |
| >trna108_AlaAGC_6_-_28763741_28763812@58.75.18\|TCCCCAGTACCTCCACCA | >AlaAGC.trna108_6_-_28763741_28763812 | 58 | 75 | 18 | tRF nuclear | NO | 0.00 | 234.00 | 0.00 | 0.00 |
| >trna10_AlaCGC_6_+_26553731_26553802@54.75.22,trna119_AlaCGC_6_-_28641613_28641684@54.75.22,trna13_AlaTGC_12_-_125406301_125406372@54.75.22,trna66_AlaTGC_6_+_28611222_28611293@54.75.22,trna8_AlaTGC_5_+_180633868_180633939@54.75.22\|TCGATCCCCGGCATCTCCACCA | >AlaCGC.trna10_6_+_26553731_26553802 | 54 | 75 | 22 | tRF nuclear | NO | 0.00 | 32.91 | 0.00 | 0.00 |
| >trna10_AlaCGC_6_+_26553731_26553802@58.75.18,trna119_AlaCGC_6_-_28641613_28641684@58.75.18,trna13_AlaCGC_2_+_157257281_157257352@58.75.18,trna13_AlaTGC_12_-_125406301_125406372@58.75.18,trna66_AlaTGC_6_+_28611222_28611293@58.75.18,trna8_AlaTGC_12_+_125424512_125424583@58.75.18,trna8_AlaTGC_5_+_180633868_180633939@58.75.18\|TCCCCGGCATCTCCACCA | >AlaCGC.trna10_6_+_26553731_26553802 | 58 | 75 | 18 | tRF nuclear | NO | 0.00 | 1199.45 | 0.00 | 0.00 |
| >trna70_AlaCGC_6_+_28697092_28697163@58.75.18\|CCCCCGGCTCCTCCACCA | >AlaCGC.trna70_6_+_28697092_28697163 | 58 | 75 | 18 | tRF nuclear | NO | 0.00 | 35.96 | 0.00 | 0.00 |
| >trna12_ArgCCT_16_+_3243918_3243990@35.54.20,trna18_ArgCCT_17_+_73030001_73030073@35.54.20,trna21_ArgCCT_17_-_73030526_73030598@35.54.20,trna2_ArgCCT_16_+_3202901_3202973@35.54.20,trna3_ArgCCT_7_+_139025446_139025518@35.54.20\|CTAAGCCAGGGATTGTGGGT | >ArgCCT.trna12_16_+_3243918_3243990 | 35 | 54 | 20 | tRF nuclear | NO | 0.00 | 85.81 | 0.00 | 0.00 |
| >trna10_AspGTC_12_-_125424193_125424264@31.53.23,trna12_AspGTC_12_-_125411891_125411962@31.53.23,trna38_AspGTC_17_-_8125556_8125627@31.53.23,trna45_AspGTC_6_+_27447453_27447524@31.53.23,trna48_AspGTC_6_+_27471523_27471594@31.53.23,trna4_AspGTC_12_+_96429799_96429870@31.53.23,trna5_AspGTC_12_+_98897281_98897352@31.53.23,trna69_AspGTC_1_-_161440205_161440276@31.53.23,trna6_AspGTC_9_-_77517990_77518061@31.53.23,trna72_AspGTC_1_-_161432824_161432895@31.53.23,trna75_AspGTC_1_-_161425414_161425485@31.53.23,trna78_AspGTC_1_-_161418033_161418104@31.53.23,trna81_AspGTC_1_-_161410615_161410686@31.53.23\|CCTGTCACGCGGGAGACCGGGT | >AspGTC.trna10_12_-_125424193_125424264 | 31 | 53 | 23 | tRF nuclear | NO | 0.00 | 93.75 | 0.00 | 0.00 |

# BT-20

| | | | | | | | | | | |
|---|---|---|---|---|---|---|---|---|---|---|
| >trna10_AspGTC_12_-_125424193_125424264@35.75.41,trna12_AspGTC_12_-_125411891_125411962@35.75.41,trna144_AspGTC_6_-_27551236_27551307@35.75.41,trna38_AspGTC_17_-_8125556_8125627@35.75.41,trna45_AspGTC_6_+_27447453_27447524@35.75.41,trna48_AspGTC_6_+_27471523_27471594@35.75.41,trna4_AspGTC_12_+_96429799_96429870@35.75.41,trna69_AspGTC_1_-_161440205_161440276@35.75.41,trna72_AspGTC_1_-_161432824_161432895@35.75.41,trna75_AspGTC_1_-_161425414_161425485@35.75.41,trna78_AspGTC_1_-_161418033_161418104@35.75.41,trna81_AspGTC_1_-_161410615_161410686@35.75.41\|TCACGCGGGAGACCGGGGTTCGATTCCCCGACGGGGAGCCA | >AspGTC.trna10_12_-_125424193_125424264 | 35 | 75 | 41 | tRF nuclear | NO | 0.00 | 39.16 | 0.00 | 0.00 |
| >trna10_AspGTC_12_-_125424193_125424264@36.75.40,trna12_AspGTC_12_-_125411891_125411962@36.75.40,trna144_AspGTC_6_-_27551236_27551307@36.75.40,trna38_AspGTC_17_-_8125556_8125627@36.75.40,trna45_AspGTC_6_+_27447453_27447524@36.75.40,trna48_AspGTC_6_+_27471523_27471594@36.75.40,trna4_AspGTC_12_+_96429799_96429870@36.75.40,trna69_AspGTC_1_-_161440205_161440276@36.75.40,trna72_AspGTC_1_-_161432824_161432895@36.75.40,trna75_AspGTC_1_-_161425414_161425485@36.75.40,trna78_AspGTC_1_-_161418033_161418104@36.75.40,trna81_AspGTC_1_-_161410615_161410686@36.75.40\|CACGCGGGAGACCGGGGTTCGATTCCCCGACGGGGAGCCA | >AspGTC.trna10_12_-_125424193_125424264 | 36 | 75 | 40 | tRF nuclear | NO | 0.00 | 144.63 | 0.00 | 0.00 |
| >trna10_AspGTC_12_-_125424193_125424264@38.69.32,trna12_AspGTC_12_-_125411891_125411962@38.69.32,trna144_AspGTC_6_-_27551236_27551307@38.69.32,trna38_AspGTC_17_-_8125556_8125627@38.69.32,trna45_AspGTC_6_+_27447453_27447524@38.69.32,trna48_AspGTC_6_+_27471523_27471594@38.69.32,trna4_AspGTC_12_+_96429799_96429870@38.69.32,trna69_AspGTC_1_-_161440205_161440276@38.69.32,trna72_AspGTC_1_-_161432824_161432895@38.69.32,trna75_AspGTC_1_-_161425414_161425485@38.69.32,trna78_AspGTC_1_-_161418033_161418104@38.69.32,trna81_AspGTC_1_-_161410615_161410686@38.69.32\|CGCGGGAGACCGGGGTTCGATTCCCCGACGGG | >AspGTC.trna10_12_-_125424193_125424264 | 38 | 69 | 32 | tRF nuclear | NO | 0.00 | 0.00 | 74.00 | 0.00 |
| >trna10_AspGTC_12_-_125424193_125424264@39.74.36,trna12_AspGTC_12_-_125411891_125411962@39.74.36,trna144_AspGTC_6_-_27551236_27551307@39.74.36,trna38_AspGTC_17_-_8125556_8125627@39.74.36,trna45_AspGTC_6_+_27447453_27447524@39.74.36,trna48_AspGTC_6_+_27471523_27471594@39.74.36,trna4_AspGTC_12_+_96429799_96429870@39.74.36,trna69_AspGTC_1_-_161440205_161440276@39.74.36,trna72_AspGTC_1_-_161432824_161432895@39.74.36,trna75_AspGTC_1_-_161425414_161425485@39.74.36,trna78_AspGTC_1_-_161418033_161418104@39.74.36,trna81_AspGTC_1_-_161410615_161410686@39.74.36\|GCGGGAGACCGGGGTTCGATTCCCCGACGGGGAGCC | >AspGTC.trna10_12_-_125424193_125424264 | 39 | 74 | 36 | tRF nuclear | NO | 0.00 | 30.15 | 0.00 | 0.00 |
| >trna10_AspGTC_12_-_125424193_125424264@39.75.37,trna12_AspGTC_12_-_125411891_125411962@39.75.37,trna144_AspGTC_6_-_27551236_27551307@39.75.37,trna38_AspGTC_17_-_8125556_8125627@39.75.37,trna45_AspGTC_6_+_27447453_27447524@39.75.37,trna48_AspGTC_6_+_27471523_27471594@39.75.37,trna4_AspGTC_12_+_96429799_96429870@39.75.37,trna69_AspGTC_1_-_161440205_161440276@39.75.37,trna72_AspGTC_1_-_161432824_161432895@39.75.37,trna75_AspGTC_1_-_161425414_161425485@39.75.37,trna78_AspGTC_1_-_161418033_161418104@39.75.37,trna81_AspGTC_1_-_161410615_161410686@39.75.37\|GCGGGAGACCGGGGTTCGATTCCCCGACGGGGAGCCA | >AspGTC.trna10_12_-_125424193_125424264 | 39 | 75 | 37 | tRF nuclear | YES | 0.00 | 110.27 | 0.00 | 0.00 |
| >trna146_GlnCTG_6_-_27515531_27515602@17.53.37,trna1_GlnCTG_6_+_18836402_18836473@17.53.37,trna3_GlnCTG_17_+_8023070_8023141@17.53.37,trna49_GlnCTG_6_+_27487308_27487379@17.53.37,trna7_GlnCTG_15_-_66161400_66161471@17.53.37,trna99_GlnCTG_6_-_28909378_28909449@17.53.37\|GGTTAGCACTCTGGACTCTGAATCCAGCGATCCGAGT | >GlnCTG.trna146_6_-_27515531_27515602 | 17 | 53 | 37 | tRF nuclear | NO | 0.00 | 10.14 | 0.00 | 0.00 |
| >trna146_GlnCTG_6_-_27515531_27515602@17.54.38,trna1_GlnCTG_6_+_18836402_18836473@17.54.38,trna3_GlnCTG_17_+_8023070_8023141@17.54.38,trna49_GlnCTG_6_+_27487308_27487379@17.54.38,trna7_GlnCTG_15_-_66161400_66161471@17.54.38,trna99_GlnCTG_6_-_28909378_28909449@17.54.38\|GGTTAGCACTCTGGACTCTGAATCCAGCGATCCGAGTT | >GlnCTG.trna146_6_-_27515531_27515602 | 17 | 54 | 38 | tRF nuclear | NO | 0.00 | 41.08 | 0.00 | 0.00 |
| >trna10_GluCTC_3_-_103879524_103879595@16.33.18,trna116_GluCTC_1_-_145399233_145399304@16.33.18,trna25_GluCTC_2_-_71273488_71273560@16.33.18,trna59_GluCTC_1_+_249168447_249168518@16.33.18,trna71_GluCTC_1_-_161439189_161439260@16.33.18,trna74_GluCTC_1_-_161431809_161431880@16.33.18,trna77_GluCTC_1_-_161424398_161424469@16.33.18,trna77_GluCTC_6_+_28949976_28950047@16.33.18,trna80_GluCTC_1_-_161417018_161417089@16.33.18,trna87_GluCTC_6_-_126101393_126101464@16.33.18\|TGGTTAGGATTCGGCGCT | >GluCTC.trna10_3_-_103879524_103879595 | 16 | 33 | 18 | tRF nuclear | NO | 0.00 | 82.25 | 0.00 | 0.00 |
| >trna10_GluCTC_3_-_103879524_103879595@16.39.24,trna116_GluCTC_1_-_145399233_145399304@16.39.24,trna59_GluCTC_1_+_249168447_249168518@16.39.24,trna71_GluCTC_1_-_161439189_161439260@16.39.24,trna74_GluCTC_1_-_161431809_161431880@16.39.24,trna77_GluCTC_1_-_161424398_161424469@16.39.24,trna77_GluCTC_6_+_28949976_28950047@16.39.24,trna80_GluCTC_1_-_161417018_161417089@16.39.24,trna87_GluCTC_6_-_126101393_126101464@16.39.24\|TGGTTAGGATTCGGCGCTCTCACC | >GluCTC.trna10_3_-_103879524_103879595 | 16 | 39 | 24 | tRF nuclear | NO | 0.00 | 38.86 | 0.00 | 0.00 |

# BT-20

| | | | | | | | | | | |
|---|---|---|---|---|---|---|---|---|---|---|
| >trna10_GluCTC_3_-_103879524_103879595@16.42.27,trna116_GluCTC_1_-_145399233_145399304@16.42.27,trna59_GluCTC_1_+_249168447_249168518@16.42.27,trna71_GluCTC_1_-_161439189_161439260@16.42.27,trna74_GluCTC_1_-_161431809_161431880@16.42.27,trna77_GluCTC_1_-_161424398_161424469@16.42.27,trna77_GluCTC_6_+_28949976_28950047@16.42.27,trna80_GluCTC_1_-_161417018_161417089@16.42.27,trna87_GluCTC_6_-_126101393_126101464@16.42.27\|TGGTTAGGATTCGGCGCTCTCACCGCC | >GluCTC.trna10_3_-_103879524_103879595 | 16 | 42 | 27 | tRF nuclear | NO | 0.00 | 89.81 | 0.00 | 0.00 |
| >trna116_GluCTC_1_-_145399233_145399304@1.31.31,trna59_GluCTC_1_+_249168447_249168518@1.31.31,trna71_GluCTC_1_-_161439189_161439260@1.31.31,trna74_GluCTC_1_-_161431809_161431880@1.31.31,trna77_GluCTC_1_-_161424398_161424469@1.31.31,trna77_GluCTC_6_+_28949976_28950047@1.31.31,trna80_GluCTC_1_-_161417018_161417089@1.31.31,trna87_GluCTC_6_-_126101393_126101464@1.31.31\|TCCCTGGTGGTCTAGTGGTTAGGATTCGGCG | >GluCTC.trna116_1_-_145399233_145399304 | 1 | 31 | 31 | tRF nuclear | NO | 0.00 | 481.63 | 102.08 | 73.46 |
| >trna116_GluCTC_1_-_145399233_145399304@1.32.32,trna59_GluCTC_1_+_249168447_249168518@1.32.32,trna71_GluCTC_1_-_161439189_161439260@1.32.32,trna74_GluCTC_1_-_161431809_161431880@1.32.32,trna77_GluCTC_1_-_161424398_161424469@1.32.32,trna77_GluCTC_6_+_28949976_28950047@1.32.32,trna80_GluCTC_1_-_161417018_161417089@1.32.32,trna87_GluCTC_6_-_126101393_126101464@1.32.32\|TCCCTGGTGGTCTAGTGGTTAGGATTCGGCGC | >GluCTC.trna116_1_-_145399233_145399304 | 1 | 32 | 32 | tRF nuclear | NO | 26.32 | 499.11 | 65.70 | 113.43 |
| >trna116_GluCTC_1_-_145399233_145399304@1.33.33,trna59_GluCTC_1_+_249168447_249168518@1.33.33,trna71_GluCTC_1_-_161439189_161439260@1.33.33,trna74_GluCTC_1_-_161431809_161431880@1.33.33,trna77_GluCTC_1_-_161424398_161424469@1.33.33,trna77_GluCTC_6_+_28949976_28950047@1.33.33,trna80_GluCTC_1_-_161417018_161417089@1.33.33,trna87_GluCTC_6_-_126101393_126101464@1.33.33\|TCCCTGGTGGTCTAGTGGTTAGGATTCGGCGCT | >GluCTC.trna116_1_-_145399233_145399304 | 1 | 33 | 33 | tRF nuclear | YES | 281.77 | 5755.30 | 342.64 | 666.03 |
| >trna116_GluCTC_1_-_145399233_145399304@1.34.34,trna59_GluCTC_1_+_249168447_249168518@1.34.34,trna71_GluCTC_1_-_161439189_161439260@1.34.34,trna74_GluCTC_1_-_161431809_161431880@1.34.34,trna77_GluCTC_1_-_161424398_161424469@1.34.34,trna77_GluCTC_6_+_28949976_28950047@1.34.34,trna80_GluCTC_1_-_161417018_161417089@1.34.34,trna87_GluCTC_6_-_126101393_126101464@1.34.34\|TCCCTGGTGGTCTAGTGGTTAGGATTCGGCGCTC | >GluCTC.trna116_1_-_145399233_145399304 | 1 | 34 | 34 | tRF nuclear | NO | 99.65 | 956.97 | 135.50 | 274.57 |
| >trna116_GluCTC_1_-_145399233_145399304@1.35.35,trna59_GluCTC_1_+_249168447_249168518@1.35.35,trna71_GluCTC_1_-_161439189_161439260@1.35.35,trna74_GluCTC_1_-_161431809_161431880@1.35.35,trna77_GluCTC_1_-_161424398_161424469@1.35.35,trna77_GluCTC_6_+_28949976_28950047@1.35.35,trna80_GluCTC_1_-_161417018_161417089@1.35.35,trna87_GluCTC_6_-_126101393_126101464@1.35.35\|TCCCTGGTGGTCTAGTGGTTAGGATTCGGCGCTCT | >GluCTC.trna116_1_-_145399233_145399304 | 1 | 35 | 35 | tRF nuclear | NO | 94.04 | 1568.09 | 96.18 | 182.63 |
| >trna116_GluCTC_1_-_145399233_145399304@1.36.36,trna59_GluCTC_1_+_249168447_249168518@1.36.36,trna71_GluCTC_1_-_161439189_161439260@1.36.36,trna74_GluCTC_1_-_161431809_161431880@1.36.36,trna77_GluCTC_1_-_161424398_161424469@1.36.36,trna77_GluCTC_6_+_28949976_28950047@1.36.36,trna80_GluCTC_1_-_161417018_161417089@1.36.36,trna87_GluCTC_6_-_126101393_126101464@1.36.36\|TCCCTGGTGGTCTAGTGGTTAGGATTCGGCGCTCTC | >GluCTC.trna116_1_-_145399233_145399304 | 1 | 36 | 36 | tRF nuclear | NO | 31.01 | 488.60 | 63.81 | 94.53 |
| >trna116_GluCTC_1_-_145399233_145399304@1.37.37,trna59_GluCTC_1_+_249168447_249168518@1.37.37,trna71_GluCTC_1_-_161439189_161439260@1.37.37,trna74_GluCTC_1_-_161431809_161431880@1.37.37,trna77_GluCTC_1_-_161424398_161424469@1.37.37,trna77_GluCTC_6_+_28949976_28950047@1.37.37,trna80_GluCTC_1_-_161417018_161417089@1.37.37,trna87_GluCTC_6_-_126101393_126101464@1.37.37\|TCCCTGGTGGTCTAGTGGTTAGGATTCGGCGCTCTCA | >GluCTC.trna116_1_-_145399233_145399304 | 1 | 37 | 37 | tRF nuclear | NO | 0.00 | 23.44 | 0.00 | 0.00 |
| >trna116_GluCTC_1_-_145399233_145399304@1.39.39,trna59_GluCTC_1_+_249168447_249168518@1.39.39,trna71_GluCTC_1_-_161439189_161439260@1.39.39,trna74_GluCTC_1_-_161431809_161431880@1.39.39,trna77_GluCTC_1_-_161424398_161424469@1.39.39,trna77_GluCTC_6_+_28949976_28950047@1.39.39,trna80_GluCTC_1_-_161417018_161417089@1.39.39,trna87_GluCTC_6_-_126101393_126101464@1.39.39\|TCCCTGGTGGTCTAGTGGTTAGGATTCGGCGCTCTCACC | >GluCTC.trna116_1_-_145399233_145399304 | 1 | 39 | 39 | tRF nuclear | NO | 10.96 | 655.03 | 54.10 | 183.84 |
| >trna116_GluCTC_1_-_145399233_145399304@1.42.42,trna59_GluCTC_1_+_249168447_249168518@1.42.42,trna71_GluCTC_1_-_161439189_161439260@1.42.42,trna74_GluCTC_1_-_161431809_161431880@1.42.42,trna77_GluCTC_1_-_161424398_161424469@1.42.42,trna77_GluCTC_6_+_28949976_28950047@1.42.42,trna80_GluCTC_1_-_161417018_161417089@1.42.42,trna87_GluCTC_6_-_126101393_126101464@1.42.42\|TCCCTGGTGGTCTAGTGGTTAGGATTCGGCGCTCTCACCGCC | >GluCTC.trna116_1_-_145399233_145399304 | 1 | 42 | 42 | tRF nuclear | NO | 0.00 | 100.83 | 0.00 | 0.00 |

# BT-20

| | | | | | | | | | | |
|---|---|---|---|---|---|---|---|---|---|---|
| >trna116_GluCTC_1_-_145399233_145399304@2.33.32,trna59_GluCTC_1_+_249168447_249168518@2.33.32,trna71_GluCTC_1_-_161439189_161439260@2.33.32,trna74_GluCTC_1_-_161431809_161431880@2.33.32,trna77_GluCTC_1_-_161424398_161424469@2.33.32,trna77_GluCTC_6_+_28949976_28950047@2.33.32,trna80_GluCTC_1_-_161417018_161417089@2.33.32,trna87_GluCTC_6_-_126101393_126101464@2.33.32\|CCCTGGTGGTCTAGTGGTTAGGATTCGGCGCT | >GluCTC.trna116_1_-_145399233_145399304 | 2 | 33 | 32 | tRF nuclear | NO | 0.00 | 32.52 | 0.00 | 0.00 |
| >trna116_GluCTC_1_-_145399233_145399304@36.75.40,trna134_GluTTC_1_-_16861774_16861845@36.75.40,trna71_GluCTC_1_-_161439189_161439260@36.75.40,trna74_GluCTC_1_-_161431809_161431880@36.75.40,trna77_GluCTC_1_-_161424398_161424469@36.75.40,trna77_GluCTC_6_+_28949976_28950047@36.75.40,trna80_GluCTC_1_-_161417018_161417089@36.75.40,trna84_GluTTC_1_-_161391883_161391954@36.75.40,trna87_GluCTC_6_-_126101393_126101464@36.75.40\|CACCGCCGCGGCCCGGGTTCGATTCCCGGTCAGGGAACCA | >GluCTC.trna116_1_-_145399233_145399304 | 36 | 75 | 40 | tRF nuclear | NO | 0.00 | 107.10 | 0.00 | 0.00 |
| >trna116_GluCTC_1_-_145399233_145399304@37.74.38,trna134_GluTTC_1_-_16861774_16861845@37.74.38,trna71_GluCTC_1_-_161439189_161439260@37.74.38,trna74_GluCTC_1_-_161431809_161431880@37.74.38,trna77_GluCTC_1_-_161424398_161424469@37.74.38,trna77_GluCTC_6_+_28949976_28950047@37.74.38,trna80_GluCTC_1_-_161417018_161417089@37.74.38,trna84_GluTTC_1_-_161391883_161391954@37.74.38,trna87_GluCTC_6_-_126101393_126101464@37.74.38\|ACCGCCGCGGCCCGGGTTCGATTCCCGGTCAGGGAACC | >GluCTC.trna116_1_-_145399233_145399304 | 37 | 74 | 38 | tRF nuclear | NO | 0.00 | 30.72 | 0.00 | 0.00 |
| >trna116_GluCTC_1_-_145399233_145399304@37.75.39,trna134_GluTTC_1_-_16861774_16861845@37.75.39,trna71_GluCTC_1_-_161439189_161439260@37.75.39,trna74_GluCTC_1_-_161431809_161431880@37.75.39,trna77_GluCTC_1_-_161424398_161424469@37.75.39,trna77_GluCTC_6_+_28949976_28950047@37.75.39,trna80_GluCTC_1_-_161417018_161417089@37.75.39,trna84_GluTTC_1_-_161391883_161391954@37.75.39,trna87_GluCTC_6_-_126101393_126101464@37.75.39\|ACCGCCGCGGCCCGGGTTCGATTCCCGGTCAGGGAACCA | >GluCTC.trna116_1_-_145399233_145399304 | 37 | 75 | 39 | tRF nuclear | NO | 0.00 | 93.71 | 0.00 | 0.00 |
| >am_trna116_GluCTC_1_-_145399233_145399304@1.26.26,trna59_GluCTC_1_+_249168447_249168518@1.26.26,trna71_GluCTC_1_-_161439189_161439260@1.26.26,trna74_GluCTC_1_-_161431809_161431880@1.26.26,trna77_GluCTC_1_-_161424398_161424469@1.26.26,trna77_GluCTC_6_+_28949976_28950047@1.26.26,trna80_GluCTC_1_-_161417018_161417089@1.26.26,trna87_GluCTC_6_-_126101393_126101464@1.26.26\|TCCCTGGTGGTCTAGTGGTTAGGATT | >GluCTC.trna116_1_-_145399233_145399304.am | 1 | 26 | 26 | tRF nuclear | NO | 0.00 | 384.45 | 70.47 | 0.00 |
| >am_trna116_GluCTC_1_-_145399233_145399304@1.27.27,trna59_GluCTC_1_+_249168447_249168518@1.27.27,trna71_GluCTC_1_-_161439189_161439260@1.27.27,trna74_GluCTC_1_-_161431809_161431880@1.27.27,trna77_GluCTC_1_-_161424398_161424469@1.27.27,trna77_GluCTC_6_+_28949976_28950047@1.27.27,trna80_GluCTC_1_-_161417018_161417089@1.27.27,trna87_GluCTC_6_-_126101393_126101464@1.27.27\|TCCCTGGTGGTCTAGTGGTTAGGATTC | >GluCTC.trna116_1_-_145399233_145399304.am | 1 | 27 | 27 | tRF nuclear | NO | 0.00 | 269.31 | 74.80 | 0.00 |
| >am_trna116_GluCTC_1_-_145399233_145399304@1.28.28,trna59_GluCTC_1_+_249168447_249168518@1.28.28,trna71_GluCTC_1_-_161439189_161439260@1.28.28,trna74_GluCTC_1_-_161431809_161431880@1.28.28,trna77_GluCTC_1_-_161424398_161424469@1.28.28,trna77_GluCTC_6_+_28949976_28950047@1.28.28,trna80_GluCTC_1_-_161417018_161417089@1.28.28,trna87_GluCTC_6_-_126101393_126101464@1.28.28\|TCCCTGGTGGTCTAGTGGTTAGGATTCG | >GluCTC.trna116_1_-_145399233_145399304.am | 1 | 28 | 28 | tRF nuclear | NO | 0.00 | 137.72 | 84.38 | 10.97 |
| >am_trna116_GluCTC_1_-_145399233_145399304@1.29.29,trna59_GluCTC_1_+_249168447_249168518@1.29.29,trna71_GluCTC_1_-_161439189_161439260@1.29.29,trna74_GluCTC_1_-_161431809_161431880@1.29.29,trna77_GluCTC_1_-_161424398_161424469@1.29.29,trna77_GluCTC_6_+_28949976_28950047@1.29.29,trna80_GluCTC_1_-_161417018_161417089@1.29.29,trna87_GluCTC_6_-_126101393_126101464@1.29.29\|TCCCTGGTGGTCTAGTGGTTAGGATTCGG | >GluCTC.trna116_1_-_145399233_145399304.am | 1 | 29 | 29 | tRF nuclear | NO | 0.00 | 196.95 | 71.97 | 23.66 |
| >am_trna116_GluCTC_1_-_145399233_145399304@1.30.30,trna59_GluCTC_1_+_249168447_249168518@1.30.30,trna71_GluCTC_1_-_161439189_161439260@1.30.30,trna74_GluCTC_1_-_161431809_161431880@1.30.30,trna77_GluCTC_1_-_161424398_161424469@1.30.30,trna77_GluCTC_6_+_28949976_28950047@1.30.30,trna80_GluCTC_1_-_161417018_161417089@1.30.30,trna87_GluCTC_6_-_126101393_126101464@1.30.30\|TCCCTGGTGGTCTAGTGGTTAGGATTCGGC | >GluCTC.trna116_1_-_145399233_145399304.am | 1 | 30 | 30 | tRF nuclear | NO | 10.58 | 1026.32 | 182.21 | 84.77 |
| >am_trna116_GluCTC_1_-_145399233_145399304@13.30.18,trna25_GluCTC_2_-_71273488_71273560@13.30.18,trna59_GluCTC_1_+_249168447_249168518@13.30.18,trna71_GluCTC_1_-_161439189_161439260@13.30.18,trna74_GluCTC_1_-_161431809_161431880@13.30.18,trna77_GluCTC_1_-_161424398_161424469@13.30.18,trna77_GluCTC_6_+_28949976_28950047@13.30.18,trna80_GluCTC_1_-_161417018_161417089@13.30.18,trna87_GluCTC_6_-_126101393_126101464@13.30.18\|TAGTGGTTAGGATTCGGC | >GluCTC.trna116_1_-_145399233_145399304.am | 13 | 30 | 18 | tRF nuclear | NO | 0.00 | 118.80 | 0.00 | 0.00 |

# BT-20

| | | | | | | | | | | |
|---|---|---|---|---|---|---|---|---|---|---|
| >trna11_GluTTC_15_-_26327381_26327452@1.29.29,trna3_GluTTC_13_-_45492062_45492133@1.29.29\|TCCCACATGGTCTAGCGGTTAGGATTCCT | >GluTTC.trna11_15_-_26327381_26327452 | 1 | 29 | 29 | tRF nuclear | NO | 0.00 | 183.76 | 0.00 | 0.00 |
| >trna11_GluTTC_15_-_26327381_26327452@1.32.32,trna3_GluTTC_13_-_45492062_45492133@1.32.32\|TCCCACATGGTCTAGCGGTTAGGATTCCTGGT | >GluTTC.trna11_15_-_26327381_26327452 | 1 | 32 | 32 | tRF nuclear | NO | 0.00 | 23.78 | 0.00 | 0.00 |
| >trna11_GluTTC_15_-_26327381_26327452@1.33.33,trna3_GluTTC_13_-_45492062_45492133@1.33.33\|TCCCACATGGTCTAGCGGTTAGGATTCCTGGTT | >GluTTC.trna11_15_-_26327381_26327452 | 1 | 33 | 33 | tRF nuclear | NO | 11.23 | 528.58 | 56.15 | 31.74 |
| >trna11_GluTTC_15_-_26327381_26327452@1.34.34,trna3_GluTTC_13_-_45492062_45492133@1.34.34\|TCCCACATGGTCTAGCGGTTAGGATTCCTGGTTT | >GluTTC.trna11_15_-_26327381_26327452 | 1 | 34 | 34 | tRF nuclear | NO | 0.00 | 30.61 | 0.00 | 0.00 |
| >trna11_GluTTC_15_-_26327381_26327452@1.35.35,trna3_GluTTC_13_-_45492062_45492133@1.35.35\|TCCCACATGGTCTAGCGGTTAGGATTCCTGGTTTT | >GluTTC.trna11_15_-_26327381_26327452 | 1 | 35 | 35 | tRF nuclear | NO | 0.00 | 10.07 | 0.00 | 0.00 |
| >trna11_GluTTC_15_-_26327381_26327452@1.36.36,trna3_GluTTC_13_-_45492062_45492133@1.36.36\|TCCCACATGGTCTAGCGGTTAGGATTCCTGGTTTTC | >GluTTC.trna11_15_-_26327381_26327452 | 1 | 36 | 36 | tRF nuclear | NO | 0.00 | 24.83 | 0.00 | 0.00 |
| >trna134_GluTTC_1_-_16861774_16861845@1.33.33,trna5_GluTTC_1_+_17199078_17199149@1.33.33,trna84_GluTTC_1_-_161391883_161391954@1.33.33,trna94_GluTTC_1_-_149664355_149664427@1.33.33\|TCCCTGGTGGTCTAGTGGCTAGGATTCGGCGCT | >GluTTC.trna134_1_-_16861774_16861845 | 1 | 33 | 33 | tRF nuclear | NO | 0.00 | 556.37 | 0.00 | 28.96 |
| >trna134_GluTTC_1_-_16861774_16861845@1.34.34,trna5_GluTTC_1_+_17199078_17199149@1.34.34,trna84_GluTTC_1_-_161391883_161391954@1.34.34,trna94_GluTTC_1_-_149664355_149664427@1.34.34\|TCCCTGGTGGTCTAGTGGCTAGGATTCGGCGCTT | >GluTTC.trna134_1_-_16861774_16861845 | 1 | 34 | 34 | tRF nuclear | NO | 0.00 | 39.22 | 0.00 | 0.00 |
| >am_trna134_GluTTC_1_-_16861774_16861845@1.30.30,trna5_GluTTC_1_+_17199078_17199149@1.30.30,trna84_GluTTC_1_-_161391883_161391954@1.30.30,trna94_GluTTC_1_-_149664355_149664427@1.30.30\|TCCCTGGTGGTCTAGTGGCTAGGATTCGGC | >GluTTC.trna134_1_-_16861774_16861845.am | 1 | 30 | 30 | tRF nuclear | NO | 0.00 | 95.70 | 0.00 | 0.00 |
| >trna20_GluTTC_2_-_131094701_131094772@1.33.33,trna5_GluTTC_13_-_41634874_41634945@1.33.33\|TCCCATATGGTCTAGCGGTTAGGATTCCTGGTT | >GluTTC.trna20_2_-_131094701_131094772 | 1 | 33 | 33 | tRF nuclear | NO | 0.00 | 80.86 | 0.00 | 0.00 |
| >trna128_GlyGCC_6_-_27870686_27870756@16.52.37,trna19_GlyGCC_16_+_70823410_70823480@16.52.37,trna19_GlyGCC_2_-_157257659_157257729@16.52.37,trna24_GlyGCC_16_-_70812942_70813012@16.52.37,trna25_GlyGCC_16_-_70812114_70812184@16.52.37,trna2_GlyGCC_21_-_18827107_18827177@16.52.37,trna35_GlyGCC_1_+_161413094_161413164@16.52.37,trna37_GlyGCC_1_+_161420467_161420537@16.52.37,trna39_GlyGCC_1_+_161427898_161427968@16.52.37,trna41_GlyGCC_1_+_161435258_161435328@16.52.37,trna5_GlyGCC_17_+_8029064_8029134@16.52.37,trna68_GlyGCC_1_-_161493637_161493707@16.52.37\|TGGTAGAATTCTCGCCTGCCACGCGGGAGGCCCGGGT | >GlyGCC.trna128_6_-_27870686_27870756 | 16 | 52 | 37 | tRF nuclear | NO | 0.00 | 40.12 | 0.00 | 0.00 |
| >trna128_GlyGCC_6_-_27870686_27870756@19.38.20,trna19_GlyGCC_16_+_70823410_70823480@19.38.20,trna19_GlyGCC_2_-_157257659_157257729@19.38.20,trna24_GlyGCC_16_-_70812942_70813012@19.38.20,trna25_GlyGCC_16_-_70812114_70812184@19.38.20,trna2_GlyGCC_21_-_18827107_18827177@19.38.20,trna35_GlyGCC_1_+_161413094_161413164@19.38.20,trna37_GlyGCC_1_+_161420467_161420537@19.38.20,trna39_GlyGCC_1_+_161427898_161427968@19.38.20,trna41_GlyGCC_1_+_161435258_161435328@19.38.20,trna5_GlyGCC_17_+_8029064_8029134@19.38.20,trna68_GlyGCC_1_-_161493637_161493707@19.38.20\|TAGAATTCTCGCCTGCCACG | >GlyGCC.trna128_6_-_27870686_27870756 | 19 | 38 | 20 | tRF nuclear | NO | 0.00 | 199.62 | 0.00 | 0.00 |
| >am_trna128_GlyGCC_6_-_27870686_27870756@1.25.25,trna133_GlyCCC_1_-_16872434_16872504@1.25.25,trna18_GlyGCC_16_+_70822597_70822667@1.25.25,trna19_GlyGCC_16_+_70823410_70823480@1.25.25,trna19_GlyGCC_2_-_157257659_157257729@1.25.25,trna24_GlyGCC_16_-_70812942_70813012@1.25.25,trna25_GlyGCC_16_-_70812114_70812184@1.25.25,trna4_GlyCCC_1_+_17188416_17188486@1.25.25,trna5_GlyGCC_17_+_8029064_8029134@1.25.25,trna68_GlyGCC_1_-_161493637_161493707@1.25.25\|GCATTGGTGGTTCAGTGGTAGAATT | >GlyGCC.trna128_6_-_27870686_27870756.am | 1 | 25 | 25 | tRF nuclear | NO | 0.00 | 82.98 | 13.10 | 0.00 |
| >am_trna128_GlyGCC_6_-_27870686_27870756@1.27.27,trna133_GlyCCC_1_-_16872434_16872504@1.27.27,trna18_GlyGCC_16_+_70822597_70822667@1.27.27,trna19_GlyGCC_16_+_70823410_70823480@1.27.27,trna19_GlyGCC_2_-_157257659_157257729@1.27.27,trna24_GlyGCC_16_-_70812942_70813012@1.27.27,trna25_GlyGCC_16_-_70812114_70812184@1.27.27,trna4_GlyCCC_1_+_17188416_17188486@1.27.27,trna5_GlyGCC_17_+_8029064_8029134@1.27.27,trna68_GlyGCC_1_-_161493637_161493707@1.27.27\|GCATTGGTGGTTCAGTGGTAGAATTCT | >GlyGCC.trna128_6_-_27870686_27870756.am | 1 | 27 | 27 | tRF nuclear | NO | 0.00 | 44.24 | 12.46 | 0.00 |
| >am_trna128_GlyGCC_6_-_27870686_27870756@1.28.28,trna133_GlyCCC_1_-_16872434_16872504@1.28.28,trna18_GlyGCC_16_+_70822597_70822667@1.28.28,trna19_GlyGCC_16_+_70823410_70823480@1.28.28,trna19_GlyGCC_2_-_157257659_157257729@1.28.28,trna24_GlyGCC_16_-_70812942_70813012@1.28.28,trna25_GlyGCC_16_-_70812114_70812184@1.28.28,trna4_GlyCCC_1_+_17188416_17188486@1.28.28,trna5_GlyGCC_17_+_8029064_8029134@1.28.28,trna68_GlyGCC_1_-_161493637_161493707@1.28.28\|GCATTGGTGGTTCAGTGGTAGAATTCTC | >GlyGCC.trna128_6_-_27870686_27870756.am | 1 | 28 | 28 | tRF nuclear | NO | 0.00 | 169.85 | 174.40 | 0.00 |

# BT-20

| | | | | | | | | | | |
|---|---|---|---|---|---|---|---|---|---|---|
| >am_trna128_GlyGCC_6_-_27870686_27870756@1.29.29,trna133_GlyCCC_1_-_16872434_16872504@1.29.29,trna18_GlyGCC_16_+_70822597_70822667@1.29.29,trna19_GlyGCC_16_+_70823410_70823480@1.29.29,trna19_GlyGCC_2_-_157257659_157257729@1.29.29,trna24_GlyGCC_16_-_70812942_70813012@1.29.29,trna25_GlyGCC_16_-_70812114_70812184@1.29.29,trna4_GlyCCC_1_+_17188416_17188486@1.29.29,trna5_GlyGCC_17_+_8029064_8029134@1.29.29,trna68_GlyGCC_1_-_161493637_161493707@1.29.29\|GCATTGGTGGTTCAGTGGTAGAATTCTCG | >GlyGCC.trna128_6_-_27870686_27870756.am | 1 | 29 | 29 | tRF nuclear | NO | 0.00 | 32.30 | 11.31 | 0.00 |
| >am_trna128_GlyGCC_6_-_27870686_27870756@1.30.30,trna133_GlyCCC_1_-_16872434_16872504@1.30.30,trna18_GlyGCC_16_+_70822597_70822667@1.30.30,trna19_GlyGCC_16_+_70823410_70823480@1.30.30,trna19_GlyGCC_2_-_157257659_157257729@1.30.30,trna24_GlyGCC_16_-_70812942_70813012@1.30.30,trna25_GlyGCC_16_-_70812114_70812184@1.30.30,trna4_GlyCCC_1_+_17188416_17188486@1.30.30,trna5_GlyGCC_17_+_8029064_8029134@1.30.30,trna68_GlyGCC_1_-_161493637_161493707@1.30.30\|GCATTGGTGGTTCAGTGGTAGAATTCTCGC | >GlyGCC.trna128_6_-_27870686_27870756.am | 1 | 30 | 30 | tRF nuclear | NO | 0.00 | 128.13 | 176.92 | 0.00 |
| >am_trna128_GlyGCC_6_-_27870686_27870756@1.31.31,trna133_GlyCCC_1_-_16872434_16872504@1.31.31,trna18_GlyGCC_16_+_70822597_70822667@1.31.31,trna19_GlyGCC_16_+_70823410_70823480@1.31.31,trna19_GlyGCC_2_-_157257659_157257729@1.31.31,trna24_GlyGCC_16_-_70812942_70813012@1.31.31,trna25_GlyGCC_16_-_70812114_70812184@1.31.31,trna4_GlyCCC_1_+_17188416_17188486@1.31.31,trna5_GlyGCC_17_+_8029064_8029134@1.31.31,trna68_GlyGCC_1_-_161493637_161493707@1.31.31\|GCATTGGTGGTTCAGTGGTAGAATTCTCGCC | >GlyGCC.trna128_6_-_27870686_27870756.am | 1 | 31 | 31 | tRF nuclear | NO | 77.92 | 1536.88 | 867.14 | 229.24 |
| >am_trna128_GlyGCC_6_-_27870686_27870756@1.32.32,trna133_GlyCCC_1_-_16872434_16872504@1.32.32,trna18_GlyGCC_16_+_70822597_70822667@1.32.32,trna19_GlyGCC_16_+_70823410_70823480@1.32.32,trna19_GlyGCC_2_-_157257659_157257729@1.32.32,trna24_GlyGCC_16_-_70812942_70813012@1.32.32,trna25_GlyGCC_16_-_70812114_70812184@1.32.32,trna4_GlyCCC_1_+_17188416_17188486@1.32.32,trna5_GlyGCC_17_+_8029064_8029134@1.32.32,trna68_GlyGCC_1_-_161493637_161493707@1.32.32\|GCATTGGTGGTTCAGTGGTAGAATTCTCGCCT | >GlyGCC.trna128_6_-_27870686_27870756.am | 1 | 32 | 32 | tRF nuclear | NO | 36.03 | 501.58 | 300.17 | 129.58 |
| >am_trna128_GlyGCC_6_-_27870686_27870756@1.33.33,trna18_GlyGCC_16_+_70822597_70822667@1.33.33,trna19_GlyGCC_16_+_70823410_70823480@1.33.33,trna19_GlyGCC_2_-_157257659_157257729@1.33.33,trna24_GlyGCC_16_-_70812942_70813012@1.33.33,trna25_GlyGCC_16_-_70812114_70812184@1.33.33,trna5_GlyGCC_17_+_8029064_8029134@1.33.33,trna68_GlyGCC_1_-_161493637_161493707@1.33.33\|GCATTGGTGGTTCAGTGGTAGAATTCTCGCCTG | >GlyGCC.trna128_6_-_27870686_27870756.am | 1 | 33 | 33 | tRF nuclear | NO | 0.00 | 208.68 | 97.43 | 27.43 |
| >am_trna128_GlyGCC_6_-_27870686_27870756@1.35.35,trna18_GlyGCC_16_+_70822597_70822667@1.35.35,trna19_GlyGCC_16_+_70823410_70823480@1.35.35,trna19_GlyGCC_2_-_157257659_157257729@1.35.35,trna24_GlyGCC_16_-_70812942_70813012@1.35.35,trna25_GlyGCC_16_-_70812114_70812184@1.35.35,trna5_GlyGCC_17_+_8029064_8029134@1.35.35,trna68_GlyGCC_1_-_161493637_161493707@1.35.35\|GCATTGGTGGTTCAGTGGTAGAATTCTCGCCTGCC | >GlyGCC.trna128_6_-_27870686_27870756.am | 1 | 35 | 35 | tRF nuclear | NO | 0.00 | 126.01 | 92.21 | 0.00 |
| >am_trna128_GlyGCC_6_-_27870686_27870756@1.36.36,trna18_GlyGCC_16_+_70822597_70822667@1.36.36,trna19_GlyGCC_16_+_70823410_70823480@1.36.36,trna19_GlyGCC_2_-_157257659_157257729@1.36.36,trna24_GlyGCC_16_-_70812942_70813012@1.36.36,trna25_GlyGCC_16_-_70812114_70812184@1.36.36,trna5_GlyGCC_17_+_8029064_8029134@1.36.36,trna68_GlyGCC_1_-_161493637_161493707@1.36.36\|GCATTGGTGGTTCAGTGGTAGAATTCTCGCCTGCCA | >GlyGCC.trna128_6_-_27870686_27870756.am | 1 | 36 | 36 | tRF nuclear | NO | 0.00 | 0.00 | 11.10 | 0.00 |
| >am_trna128_GlyGCC_6_-_27870686_27870756@19.37.19,trna19_GlyGCC_16_+_70823410_70823480@19.37.19,trna19_GlyGCC_2_-_157257659_157257729@19.37.19,trna24_GlyGCC_16_-_70812942_70813012@19.37.19,trna25_GlyGCC_16_-_70812114_70812184@19.37.19,trna2_GlyGCC_21_-_18827107_18827177@19.37.19,trna35_GlyGCC_1_+_161413094_161413164@19.37.19,trna37_GlyGCC_1_+_161420467_161420537@19.37.19,trna39_GlyGCC_1_+_161427898_161427968@19.37.19,trna41_GlyGCC_1_+_161435258_161435328@19.37.19,trna5_GlyGCC_17_+_8029064_8029134@19.37.19,trna68_GlyGCC_1_-_161493637_161493707@19.37.19\|TAGAATTCTCGCCTGCCAC | >GlyGCC.trna128_6_-_27870686_27870756.am | 19 | 37 | 19 | tRF nuclear | NO | 0.00 | 68.13 | 0.00 | 0.00 |
| >trna2_GlyGCC_21_-_18827107_18827177@1.28.28,trna35_GlyGCC_1_+_161413094_161413164@1.28.28,trna37_GlyGCC_1_+_161420467_161420537@1.28.28,trna39_GlyGCC_1_+_161427898_161427968@1.28.28,trna41_GlyGCC_1_+_161435258_161435328@1.28.28\|GCATGGGTGGTTCAGTGGTAGAATTCTC | >GlyGCC.trna2_21_-_18827107_18827177 | 1 | 28 | 28 | tRF nuclear | NO | 0.00 | 399.92 | 353.05 | 10.30 |
| >trna2_GlyGCC_21_-_18827107_18827177@1.29.29,trna35_GlyGCC_1_+_161413094_161413164@1.29.29,trna37_GlyGCC_1_+_161420467_161420537@1.29.29,trna39_GlyGCC_1_+_161427898_161427968@1.29.29,trna41_GlyGCC_1_+_161435258_161435328@1.29.29\|GCATGGGTGGTTCAGTGGTAGAATTCTCG | >GlyGCC.trna2_21_-_18827107_18827177 | 1 | 29 | 29 | tRF nuclear | NO | 0.00 | 147.27 | 77.76 | 0.00 |

# BT-20

<table>
<tr><th></th><th></th><th></th><th></th><th></th><th></th><th></th><th></th><th></th><th></th><th></th></tr>
<tr><td>>trna2_GlyGCC_21_-_18827107_18827177@1.30.30,trna35_GlyGCC_1_+_161413094_161413164@1.30.30,trna37_GlyGCC_1_+_161420467_161420537@1.30.30,trna39_GlyGCC_1_+_161427898_161427968@1.30.30,trna41_GlyGCC_1_+_161435258_161435328@1.30.30|GCATGGGTGGTTCAGTGGTAGAATTCTCGC</td><td>>GlyGCC.trna2_21_-_18827107_18827177</td><td>1</td><td>30</td><td>30</td><td>tRF nuclear</td><td>NO</td><td>0.00</td><td>245.66</td><td>262.81</td><td>0.00</td></tr>
<tr><td>>trna2_GlyGCC_21_-_18827107_18827177@1.31.31,trna35_GlyGCC_1_+_161413094_161413164@1.31.31,trna37_GlyGCC_1_+_161420467_161420537@1.31.31,trna39_GlyGCC_1_+_161427898_161427968@1.31.31,trna41_GlyGCC_1_+_161435258_161435328@1.31.31|GCATGGGTGGTTCAGTGGTAGAATTCTCGCC</td><td>>GlyGCC.trna2_21_-_18827107_18827177</td><td>1</td><td>31</td><td>31</td><td>tRF nuclear</td><td>YES</td><td>148.76</td><td>3211.17</td><td>1661.10</td><td>389.20</td></tr>
<tr><td>>trna2_GlyGCC_21_-_18827107_18827177@1.32.32,trna35_GlyGCC_1_+_161413094_161413164@1.32.32,trna37_GlyGCC_1_+_161420467_161420537@1.32.32,trna39_GlyGCC_1_+_161427898_161427968@1.32.32,trna41_GlyGCC_1_+_161435258_161435328@1.32.32|GCATGGGTGGTTCAGTGGTAGAATTCTCGCCT</td><td>>GlyGCC.trna2_21_-_18827107_18827177</td><td>1</td><td>32</td><td>32</td><td>tRF nuclear</td><td>NO</td><td>67.35</td><td>696.69</td><td>401.66</td><td>111.97</td></tr>
<tr><td>>trna2_GlyGCC_21_-_18827107_18827177@1.33.33,trna35_GlyGCC_1_+_161413094_161413164@1.33.33,trna37_GlyGCC_1_+_161420467_161420537@1.33.33,trna39_GlyGCC_1_+_161427898_161427968@1.33.33,trna41_GlyGCC_1_+_161435258_161435328@1.33.33|GCATGGGTGGTTCAGTGGTAGAATTCTCGCCTG</td><td>>GlyGCC.trna2_21_-_18827107_18827177</td><td>1</td><td>33</td><td>33</td><td>tRF nuclear</td><td>NO</td><td>0.00</td><td>147.34</td><td>70.50</td><td>0.00</td></tr>
<tr><td>>am_trna2_GlyGCC_21_-_18827107_18827177@1.25.25,trna35_GlyGCC_1_+_161413094_161413164@1.25.25,trna37_GlyGCC_1_+_161420467_161420537@1.25.25,trna39_GlyGCC_1_+_161427898_161427968@1.25.25,trna41_GlyGCC_1_+_161435258_161435328@1.25.25|GCATGGGTGGTTCAGTGGTAGAATT</td><td>>GlyGCC.trna2_21_-_18827107_18827177.am</td><td>1</td><td>25</td><td>25</td><td>tRF nuclear</td><td>NO</td><td>0.00</td><td>143.99</td><td>209.91</td><td>0.00</td></tr>
<tr><td>>am_trna2_GlyGCC_21_-_18827107_18827177@1.26.26,trna35_GlyGCC_1_+_161413094_161413164@1.26.26,trna37_GlyGCC_1_+_161420467_161420537@1.26.26,trna39_GlyGCC_1_+_161427898_161427968@1.26.26,trna41_GlyGCC_1_+_161435258_161435328@1.26.26|GCATGGGTGGTTCAGTGGTAGAATTC</td><td>>GlyGCC.trna2_21_-_18827107_18827177.am</td><td>1</td><td>26</td><td>26</td><td>tRF nuclear</td><td>NO</td><td>0.00</td><td>38.34</td><td>156.88</td><td>0.00</td></tr>
<tr><td>>am_trna2_GlyGCC_21_-_18827107_18827177@1.27.27,trna35_GlyGCC_1_+_161413094_161413164@1.27.27,trna37_GlyGCC_1_+_161420467_161420537@1.27.27,trna39_GlyGCC_1_+_161427898_161427968@1.27.27,trna41_GlyGCC_1_+_161435258_161435328@1.27.27|GCATGGGTGGTTCAGTGGTAGAATTCT</td><td>>GlyGCC.trna2_21_-_18827107_18827177.am</td><td>1</td><td>27</td><td>27</td><td>tRF nuclear</td><td>NO</td><td>0.00</td><td>188.29</td><td>158.51</td><td>30.81</td></tr>
<tr><td>>am_trna2_GlyGCC_21_-_18827107_18827177@57.74.18,trna35_GlyGCC_1_+_161413094_161413164@57.74.18,trna37_GlyGCC_1_+_161420467_161420537@57.74.18,trna39_GlyGCC_1_+_161427898_161427968@57.74.18,trna41_GlyGCC_1_+_161435258_161435328@57.74.18|TTCCCGGCCCATGCACCA</td><td>>GlyGCC.trna2_21_-_18827107_18827177.am</td><td>57</td><td>74</td><td>18</td><td>tRF nuclear</td><td>NO</td><td>0.00</td><td>131.18</td><td>0.00</td><td>0.00</td></tr>
<tr><td>>am_trna117_GlyTCC_1_-_145397864_145397935@1.32.32,trna45_GlyTCC_1_+_161500903_161500974@1.32.32,trna70_GlyTCC_1_-_161439547_161439618@1.32.32,trna73_GlyTCC_1_-_161432166_161432237@1.32.32,trna76_GlyTCC_1_-_161424756_161424827@1.32.32,trna79_GlyTCC_1_-_161417375_161417446@1.32.32|GCGTTGGTGGTATAGTGGTGAGCATAGCTGCC</td><td>>GlyTCC.trna117_1_-_145397864_145397935.am</td><td>1</td><td>32</td><td>32</td><td>tRF nuclear</td><td>NO</td><td>0.00</td><td>23.34</td><td>0.00</td><td>0.00</td></tr>
<tr><td>>trna111_HisGTG_1_-_147774845_147774916@1.30.30,trna118_HisGTG_1_-_145396881_145396952@1.30.30,trna16_HisGTG_1_+_146544773_146544844@1.30.30,trna1_HisGTG_15_+_45493349_45493420@1.30.30,trna21_HisGTG_1_+_147753471_147753542@1.30.30,trna33_HisGTG_6_+_27125906_27125977@1.30.30,trna7_HisGTG_9_-_14433938_14434009@1.30.30,trna8_HisGTG_15_-_45492611_45492682@1.30.30,trna9_HisGTG_15_-_45490804_45490875@1.30.30|GCCGTGATCGTATAGTGGTTAGTACTCTGC</td><td>>HisGTG.trna111_1_-_147774845_147774916</td><td>1</td><td>30</td><td>30</td><td>tRF nuclear</td><td>NO</td><td>0.00</td><td>84.91</td><td>0.00</td><td>0.00</td></tr>
<tr><td>>trna111_HisGTG_1_-_147774845_147774916@1.31.31,trna118_HisGTG_1_-_145396881_145396952@1.31.31,trna16_HisGTG_1_+_146544773_146544844@1.31.31,trna1_HisGTG_15_+_45493349_45493420@1.31.31,trna21_HisGTG_1_+_147753471_147753542@1.31.31,trna33_HisGTG_6_+_27125906_27125977@1.31.31,trna7_HisGTG_9_-_14433938_14434009@1.31.31,trna8_HisGTG_15_-_45492611_45492682@1.31.31,trna9_HisGTG_15_-_45490804_45490875@1.31.31|GCCGTGATCGTATAGTGGTTAGTACTCTGCG</td><td>>HisGTG.trna111_1_-_147774845_147774916</td><td>1</td><td>31</td><td>31</td><td>tRF nuclear</td><td>NO</td><td>0.00</td><td>193.57</td><td>10.43</td><td>0.00</td></tr>
<tr><td>>trna111_HisGTG_1_-_147774845_147774916@1.32.32,trna118_HisGTG_1_-_145396881_145396952@1.32.32,trna16_HisGTG_1_+_146544773_146544844@1.32.32,trna1_HisGTG_15_+_45493349_45493420@1.32.32,trna21_HisGTG_1_+_147753471_147753542@1.32.32,trna33_HisGTG_6_+_27125906_27125977@1.32.32,trna7_HisGTG_9_-_14433938_14434009@1.32.32,trna8_HisGTG_15_-_45492611_45492682@1.32.32,trna9_HisGTG_15_-_45490804_45490875@1.32.32|GCCGTGATCGTATAGTGGTTAGTACTCTGCGT</td><td>>HisGTG.trna111_1_-_147774845_147774916</td><td>1</td><td>32</td><td>32</td><td>tRF nuclear</td><td>NO</td><td>0.00</td><td>37.31</td><td>0.00</td><td>0.00</td></tr>
<tr><td>>trna111_HisGTG_1_-_147774845_147774916@1.33.33,trna118_HisGTG_1_-_145396881_145396952@1.33.33,trna16_HisGTG_1_+_146544773_146544844@1.33.33,trna1_HisGTG_15_+_45493349_45493420@1.33.33,trna21_HisGTG_1_+_147753471_147753542@1.33.33,trna33_HisGTG_6_+_27125906_27125977@1.33.33,trna7_HisGTG_9_-_14433938_14434009@1.33.33,trna8_HisGTG_15_-_45492611_45492682@1.33.33,trna9_HisGTG_15_-_45490804_45490875@1.33.33|GCCGTGATCGTATAGTGGTTAGTACTCTGCGTT</td><td>>HisGTG.trna111_1_-_147774845_147774916</td><td>1</td><td>33</td><td>33</td><td>tRF nuclear</td><td>NO</td><td>99.95</td><td>753.92</td><td>89.46</td><td>24.99</td></tr>
</table>

# BT-20

| | | | | | | | | | | |
|---|---|---|---|---|---|---|---|---|---|---|
| >trna111_HisGTG_1_-_147774845_147774916@1.34.34,trna118_HisGTG_1_-_145396881_145396952@1.34.34,trna16_HisGTG_1_+_146544773_146544844@1.34.34,trna1_HisGTG_15_+_45493349_45493420@1.34.34,trna21_HisGTG_1_+_147753471_147753542@1.34.34,trna33_HisGTG_6_+_27125906_27125977@1.34.34,trna7_HisGTG_9_-_14433938_14434009@1.34.34,trna8_HisGTG_15_-_45492611_45492682@1.34.34,trna9_HisGTG_15_-_45490804_45490875@1.34.34\|GCCGTGATCGTATAGTGGTTAGTACTCTGCGTTG | >HisGTG.trna111_1_-_147774845_147774916 | 1 | 34 | 34 | tRF nuclear | NO | 9.58 | 278.04 | 9.96 | 39.90 |
| >trna111_HisGTG_1_-_147774845_147774916@1.35.35,trna118_HisGTG_1_-_145396881_145396952@1.35.35,trna16_HisGTG_1_+_146544773_146544844@1.35.35,trna1_HisGTG_15_+_45493349_45493420@1.35.35,trna21_HisGTG_1_+_147753471_147753542@1.35.35,trna33_HisGTG_6_+_27125906_27125977@1.35.35,trna7_HisGTG_9_-_14433938_14434009@1.35.35,trna8_HisGTG_15_-_45492611_45492682@1.35.35,trna9_HisGTG_15_-_45490804_45490875@1.35.35\|GCCGTGATCGTATAGTGGTTAGTACTCTGCGTTGT | >HisGTG.trna111_1_-_147774845_147774916 | 1 | 35 | 35 | tRF nuclear | NO | 0.00 | 99.78 | 0.00 | 0.00 |
| >trna111_HisGTG_1_-_147774845_147774916@1.36.36,trna118_HisGTG_1_-_145396881_145396952@1.36.36,trna16_HisGTG_1_+_146544773_146544844@1.36.36,trna1_HisGTG_15_+_45493349_45493420@1.36.36,trna21_HisGTG_1_+_147753471_147753542@1.36.36,trna33_HisGTG_6_+_27125906_27125977@1.36.36,trna7_HisGTG_9_-_14433938_14434009@1.36.36,trna8_HisGTG_15_-_45492611_45492682@1.36.36,trna9_HisGTG_15_-_45490804_45490875@1.36.36\|GCCGTGATCGTATAGTGGTTAGTACTCTGCGTTGTG | >HisGTG.trna111_1_-_147774845_147774916 | 1 | 36 | 36 | tRF nuclear | NO | 0.00 | 32.90 | 0.00 | 0.00 |
| >trna111_HisGTG_1_-_147774845_147774916@2.30.29,trna118_HisGTG_1_-_145396881_145396952@2.30.29,trna16_HisGTG_1_+_146544773_146544844@2.30.29,trna1_HisGTG_15_+_45493349_45493420@2.30.29,trna21_HisGTG_1_+_147753471_147753542@2.30.29,trna33_HisGTG_6_+_27125906_27125977@2.30.29,trna7_HisGTG_9_-_14433938_14434009@2.30.29,trna8_HisGTG_15_-_45492611_45492682@2.30.29,trna9_HisGTG_15_-_45490804_45490875@2.30.29\|CCGTGATCGTATAGTGGTTAGTACTCTGC | >HisGTG.trna111_1_-_147774845_147774916 | 2 | 30 | 29 | tRF nuclear | NO | 0.00 | 175.60 | 9.67 | 0.00 |
| >trna111_HisGTG_1_-_147774845_147774916@2.31.30,trna118_HisGTG_1_-_145396881_145396952@2.31.30,trna16_HisGTG_1_+_146544773_146544844@2.31.30,trna1_HisGTG_15_+_45493349_45493420@2.31.30,trna21_HisGTG_1_+_147753471_147753542@2.31.30,trna33_HisGTG_6_+_27125906_27125977@2.31.30,trna7_HisGTG_9_-_14433938_14434009@2.31.30,trna8_HisGTG_15_-_45492611_45492682@2.31.30,trna9_HisGTG_15_-_45490804_45490875@2.31.30\|CCGTGATCGTATAGTGGTTAGTACTCTGCG | >HisGTG.trna111_1_-_147774845_147774916 | 2 | 31 | 30 | tRF nuclear | NO | 0.00 | 77.06 | 0.00 | 0.00 |
| >trna111_HisGTG_1_-_147774845_147774916@3.30.28,trna118_HisGTG_1_-_145396881_145396952@3.30.28,trna16_HisGTG_1_+_146544773_146544844@3.30.28,trna1_HisGTG_15_+_45493349_45493420@3.30.28,trna21_HisGTG_1_+_147753471_147753542@3.30.28,trna33_HisGTG_6_+_27125906_27125977@3.30.28,trna7_HisGTG_9_-_14433938_14434009@3.30.28,trna8_HisGTG_15_-_45492611_45492682@3.30.28,trna9_HisGTG_15_-_45490804_45490875@3.30.28\|CGTGATCGTATAGTGGTTAGTACTCTGC | >HisGTG.trna111_1_-_147774845_147774916 | 3 | 30 | 28 | tRF nuclear | NO | 0.00 | 41.53 | 0.00 | 0.00 |
| >trna111_HisGTG_1_-_147774845_147774916@16.33.18,trna118_HisGTG_1_-_145396881_145396952@16.33.18,trna16_HisGTG_1_+_146544773_146544844@16.33.18,trna1_HisGTG_15_+_45493349_45493420@16.33.18,trna21_HisGTG_1_+_147753471_147753542@16.33.18,trna33_HisGTG_6_+_27125906_27125977@16.33.18,trna7_HisGTG_9_-_14433938_14434009@16.33.18,trna8_HisGTG_15_-_45492611_45492682@16.33.18,trna9_HisGTG_15_-_45490804_45490875@16.33.18\|TGGTTAGTACTCTGCGTT | >HisGTG.trna111_1_-_147774845_147774916 | 16 | 33 | 18 | tRF nuclear | NO | 0.00 | 34.73 | 0.00 | 0.00 |
| >trna111_HisGTG_1_-_147774845_147774916@-1A.33.34,trna118_HisGTG_1_-_145396881_145396952@-1A.33.34,trna16_HisGTG_1_+_146544773_146544844@-1A.33.34,trna1_HisGTG_15_+_45493349_45493420@-1A.33.34,trna21_HisGTG_1_+_147753471_147753542@-1A.33.34,trna33_HisGTG_6_+_27125906_27125977@-1A.33.34,trna7_HisGTG_9_-_14433938_14434009@-1A.33.34,trna8_HisGTG_15_-_45492611_45492682@-1A.33.34,trna9_HisGTG_15_-_45490804_45490875@-1A.33.34\|AGCCGTGATCGTATAGTGGTTAGTACTCTGCGTT | >HisGTG.trna111_1_-_147774845_147774916 | -1A | 33 | 34 | tRF nuclear | NO | 0.00 | 23.32 | 0.00 | 0.00 |
| >trna111_HisGTG_1_-_147774845_147774916@-1G.30.31,trna118_HisGTG_1_-_145396881_145396952@-1G.30.31,trna16_HisGTG_1_+_146544773_146544844@-1G.30.31,trna1_HisGTG_15_+_45493349_45493420@-1G.30.31,trna21_HisGTG_1_+_147753471_147753542@-1G.30.31,trna33_HisGTG_6_+_27125906_27125977@-1G.30.31,trna7_HisGTG_9_-_14433938_14434009@-1G.30.31,trna8_HisGTG_15_-_45492611_45492682@-1G.30.31,trna9_HisGTG_15_-_45490804_45490875@-1G.30.31\|GGCCGTGATCGTATAGTGGTTAGTACTCTGC | >HisGTG.trna111_1_-_147774845_147774916 | -1G | 30 | 31 | tRF nuclear | NO | 0.00 | 32.94 | 0.00 | 0.00 |
| >trna111_HisGTG_1_-_147774845_147774916@-1G.31.32,trna118_HisGTG_1_-_145396881_145396952@-1G.31.32,trna16_HisGTG_1_+_146544773_146544844@-1G.31.32,trna1_HisGTG_15_+_45493349_45493420@-1G.31.32,trna21_HisGTG_1_+_147753471_147753542@-1G.31.32,trna33_HisGTG_6_+_27125906_27125977@-1G.31.32,trna7_HisGTG_9_-_14433938_14434009@-1G.31.32,trna8_HisGTG_15_-_45492611_45492682@-1G.31.32,trna9_HisGTG_15_-_45490804_45490875@-1G.31.32\|GGCCGTGATCGTATAGTGGTTAGTACTCTGCG | >HisGTG.trna111_1_-_147774845_147774916 | -1G | 31 | 32 | tRF nuclear | NO | 0.00 | 70.94 | 63.85 | 0.00 |

# BT-20

| | | | | | | | | | | |
|---|---|---|---|---|---|---|---|---|---|---|
| >trna111_HisGTG_1_-_147774845_147774916@-1G.33.34,trna118_HisGTG_1_-_145396881_145396952@-1G.33.34,trna16_HisGTG_1_+_146544773_146544844@-1G.33.34,trna1_HisGTG_15_+_45493349_45493420@-1G.33.34,trna21_HisGTG_1_+_147753471_147753542@-1G.33.34,trna33_HisGTG_6_+_27125906_27125977@-1G.33.34,trna7_HisGTG_9_-_14433938_14434009@-1G.33.34,trna8_HisGTG_15_-_45492611_45492682@-1G.33.34,trna9_HisGTG_15_-_45490804_45490875@-1G.33.34\|GGCCGTGATCGTATAGTGGTTAGTACTCTGCGTT | >HisGTG.trna111_1_-_147774845_147774916 | -1G | 33 | 34 | tRF nuclear | NO | 31.50 | 253.99 | 120.75 | 0.00 |
| >trna111_HisGTG_1_-_147774845_147774916@-1G.34.35,trna118_HisGTG_1_-_145396881_145396952@-1G.34.35,trna16_HisGTG_1_+_146544773_146544844@-1G.34.35,trna1_HisGTG_15_+_45493349_45493420@-1G.34.35,trna21_HisGTG_1_+_147753471_147753542@-1G.34.35,trna33_HisGTG_6_+_27125906_27125977@-1G.34.35,trna7_HisGTG_9_-_14433938_14434009@-1G.34.35,trna8_HisGTG_15_-_45492611_45492682@-1G.34.35,trna9_HisGTG_15_-_45490804_45490875@-1G.34.35\|GGCCGTGATCGTATAGTGGTTAGTACTCTGCGTTG | >HisGTG.trna111_1_-_147774845_147774916 | -1G | 34 | 35 | tRF nuclear | NO | 0.00 | 74.31 | 0.00 | 0.00 |
| >trna111_HisGTG_1_-_147774845_147774916@-1G.35.36,trna118_HisGTG_1_-_145396881_145396952@-1G.35.36,trna16_HisGTG_1_+_146544773_146544844@-1G.35.36,trna1_HisGTG_15_+_45493349_45493420@-1G.35.36,trna21_HisGTG_1_+_147753471_147753542@-1G.35.36,trna33_HisGTG_6_+_27125906_27125977@-1G.35.36,trna7_HisGTG_9_-_14433938_14434009@-1G.35.36,trna8_HisGTG_15_-_45492611_45492682@-1G.35.36,trna9_HisGTG_15_-_45490804_45490875@-1G.35.36\|GGCCGTGATCGTATAGTGGTTAGTACTCTGCGTTGT | >HisGTG.trna111_1_-_147774845_147774916 | -1G | 35 | 36 | tRF nuclear | NO | 0.00 | 24.68 | 0.00 | 0.00 |
| >trna111_HisGTG_1_-_147774845_147774916@-1T.26.27,trna118_HisGTG_1_-_145396881_145396952@-1T.26.27,trna16_HisGTG_1_+_146544773_146544844@-1T.26.27,trna1_HisGTG_15_+_45493349_45493420@-1T.26.27,trna21_HisGTG_1_+_147753471_147753542@-1T.26.27,trna33_HisGTG_6_+_27125906_27125977@-1T.26.27,trna7_HisGTG_9_-_14433938_14434009@-1T.26.27,trna8_HisGTG_15_-_45492611_45492682@-1T.26.27,trna9_HisGTG_15_-_45490804_45490875@-1T.26.27\|TGCCGTGATCGTATAGTGGTTAGTACT | >HisGTG.trna111_1_-_147774845_147774916 | -1T | 26 | 27 | tRF nuclear | NO | 0.00 | 44.00 | 0.00 | 0.00 |
| >trna111_HisGTG_1_-_147774845_147774916@-1T.31.32,trna118_HisGTG_1_-_145396881_145396952@-1T.31.32,trna16_HisGTG_1_+_146544773_146544844@-1T.31.32,trna1_HisGTG_15_+_45493349_45493420@-1T.31.32,trna21_HisGTG_1_+_147753471_147753542@-1T.31.32,trna33_HisGTG_6_+_27125906_27125977@-1T.31.32,trna7_HisGTG_9_-_14433938_14434009@-1T.31.32,trna8_HisGTG_15_-_45492611_45492682@-1T.31.32,trna9_HisGTG_15_-_45490804_45490875@-1T.31.32\|TGCCGTGATCGTATAGTGGTTAGTACTCTGCG | >HisGTG.trna111_1_-_147774845_147774916 | -1T | 31 | 32 | tRF nuclear | NO | 0.00 | 72.90 | 0.00 | 0.00 |
| >trna111_HisGTG_1_-_147774845_147774916@-1T.33.34,trna118_HisGTG_1_-_145396881_145396952@-1T.33.34,trna16_HisGTG_1_+_146544773_146544844@-1T.33.34,trna1_HisGTG_15_+_45493349_45493420@-1T.33.34,trna21_HisGTG_1_+_147753471_147753542@-1T.33.34,trna33_HisGTG_6_+_27125906_27125977@-1T.33.34,trna7_HisGTG_9_-_14433938_14434009@-1T.33.34,trna8_HisGTG_15_-_45492611_45492682@-1T.33.34,trna9_HisGTG_15_-_45490804_45490875@-1T.33.34\|TGCCGTGATCGTATAGTGGTTAGTACTCTGCGTT | >HisGTG.trna111_1_-_147774845_147774916 | -1T | 33 | 34 | tRF nuclear | NO | 0.00 | 154.11 | 0.00 | 0.00 |
| >trna111_HisGTG_1_-_147774845_147774916@-1T.34.35,trna118_HisGTG_1_-_145396881_145396952@-1T.34.35,trna16_HisGTG_1_+_146544773_146544844@-1T.34.35,trna1_HisGTG_15_+_45493349_45493420@-1T.34.35,trna21_HisGTG_1_+_147753471_147753542@-1T.34.35,trna33_HisGTG_6_+_27125906_27125977@-1T.34.35,trna7_HisGTG_9_-_14433938_14434009@-1T.34.35,trna8_HisGTG_15_-_45492611_45492682@-1T.34.35,trna9_HisGTG_15_-_45490804_45490875@-1T.34.35\|TGCCGTGATCGTATAGTGGTTAGTACTCTGCGTTG | >HisGTG.trna111_1_-_147774845_147774916 | -1T | 34 | 35 | tRF nuclear | NO | 0.00 | 66.93 | 0.00 | 0.00 |
| >am_trna111_HisGTG_1_-_147774845_147774916@1.26.26,trna118_HisGTG_1_-_145396881_145396952@1.26.26,trna16_HisGTG_1_+_146544773_146544844@1.26.26,trna1_HisGTG_15_+_45493349_45493420@1.26.26,trna21_HisGTG_1_+_147753471_147753542@1.26.26,trna33_HisGTG_6_+_27125906_27125977@1.26.26,trna7_HisGTG_9_-_14433938_14434009@1.26.26,trna8_HisGTG_15_-_45492611_45492682@1.26.26,trna9_HisGTG_15_-_45490804_45490875@1.26.26\|GCCGTGATCGTATAGTGGTTAGTACT | >HisGTG.trna111_1_-_147774845_147774916.am | 1 | 26 | 26 | tRF nuclear | NO | 0.00 | 41.59 | 0.00 | 0.00 |
| >am_trna111_HisGTG_1_-_147774845_147774916@1.28.28,trna118_HisGTG_1_-_145396881_145396952@1.28.28,trna16_HisGTG_1_+_146544773_146544844@1.28.28,trna1_HisGTG_15_+_45493349_45493420@1.28.28,trna21_HisGTG_1_+_147753471_147753542@1.28.28,trna33_HisGTG_6_+_27125906_27125977@1.28.28,trna7_HisGTG_9_-_14433938_14434009@1.28.28,trna8_HisGTG_15_-_45492611_45492682@1.28.28,trna9_HisGTG_15_-_45490804_45490875@1.28.28\|GCCGTGATCGTATAGTGGTTAGTACTCT | >HisGTG.trna111_1_-_147774845_147774916.am | 1 | 28 | 28 | tRF nuclear | NO | 0.00 | 23.23 | 0.00 | 0.00 |

# BT-20

| | | | | | | | | | | |
|---|---|---|---|---|---|---|---|---|---|---|
| >am_trna111_HisGTG_1_-_147774845_147774916@-1G.26.27,trna118_HisGTG_1_-_145396881_145396952@-1G.26.27,trna16_HisGTG_1_+_146544773_146544844@-1G.26.27,trna1_HisGTG_15_+_45493349_45493420@-1G.26.27,trna21_HisGTG_1_+_147753471_147753542@-1G.26.27,trna33_HisGTG_6_+_27125906_27125977@-1G.26.27,trna7_HisGTG_9_-_14433938_14434009@-1G.26.27,trna8_HisGTG_15_-_45492611_45492682@-1G.26.27,trna9_HisGTG_15_-_45490804_45490875@-1G.26.27\|GGCCGTGATCGTATAGTGGTTAGTACT | >HisGTG.trna111_1_-_147774845_147774916.am | -1G | 26 | 27 | tRF nuclear | NO | 0.00 | 23.88 | 13.79 | 0.00 |
| >am_trna111_HisGTG_1_-_147774845_147774916@-1G.28.29,trna118_HisGTG_1_-_145396881_145396952@-1G.28.29,trna16_HisGTG_1_+_146544773_146544844@-1G.28.29,trna1_HisGTG_15_+_45493349_45493420@-1G.28.29,trna21_HisGTG_1_+_147753471_147753542@-1G.28.29,trna33_HisGTG_6_+_27125906_27125977@-1G.28.29,trna7_HisGTG_9_-_14433938_14434009@-1G.28.29,trna8_HisGTG_15_-_45492611_45492682@-1G.28.29,trna9_HisGTG_15_-_45490804_45490875@-1G.28.29\|GGCCGTGATCGTATAGTGGTTAGTACTCT | >HisGTG.trna111_1_-_147774845_147774916.am | -1G | 28 | 29 | tRF nuclear | NO | 0.00 | 0.00 | 15.03 | 0.00 |
| >trna10_LysCTT_16_+_3241501_3241573@1.33.33,trna13_LysCTT_14_-_58706613_58706685@1.33.33,trna2_LysCTT_15_+_79152904_79152976@1.33.33\|GCCCGGCTAGCTCAGTCGGTAGAGCATGGGACT | >LysCTT.trna10_16_+_3241501_3241573 | 1 | 33 | 33 | tRF nuclear | NO | 0.00 | 68.37 | 0.00 | 0.00 |
| >trna10_LysCTT_16_+_3241501_3241573@1.34.34,trna13_LysCTT_14_-_58706613_58706685@1.34.34,trna2_LysCTT_15_+_79152904_79152976@1.34.34\|GCCCGGCTAGCTCAGTCGGTAGAGCATGGGACTC | >LysCTT.trna10_16_+_3241501_3241573 | 1 | 34 | 34 | tRF nuclear | NO | 0.00 | 122.11 | 79.71 | 0.00 |
| >trna10_LysCTT_16_+_3241501_3241573@1.35.35,trna13_LysCTT_14_-_58706613_58706685@1.35.35,trna2_LysCTT_15_+_79152904_79152976@1.35.35\|GCCCGGCTAGCTCAGTCGGTAGAGCATGGGACTCT | >LysCTT.trna10_16_+_3241501_3241573 | 1 | 35 | 35 | tRF nuclear | YES | 91.88 | 7784.52 | 157.10 | 31.36 |
| >trna10_LysCTT_16_+_3241501_3241573@1.36.36,trna13_LysCTT_14_-_58706613_58706685@1.36.36,trna2_LysCTT_15_+_79152904_79152976@1.36.36\|GCCCGGCTAGCTCAGTCGGTAGAGCATGGGACTCTT | >LysCTT.trna10_16_+_3241501_3241573 | 1 | 36 | 36 | tRF nuclear | NO | 0.00 | 469.47 | 140.97 | 0.00 |
| >trna10_LysCTT_16_+_3241501_3241573@1.40.40,trna13_LysCTT_14_-_58706613_58706685@1.40.40,trna2_LysCTT_15_+_79152904_79152976@1.40.40\|GCCCGGCTAGCTCAGTCGGTAGAGCATGGGACTCTTAATC | >LysCTT.trna10_16_+_3241501_3241573 | 1 | 40 | 40 | tRF nuclear | NO | 0.00 | 34.80 | 0.00 | 0.00 |
| >trna119_LysCTT_1_-_145395522_145395594@1.32.32,trna11_LysCTT_5_-_180648979_180649051@1.32.32,trna13_LysCTT_6_+_26556774_26556846@1.32.32,trna32_LysCTT_16_-_3207406_3207478@1.32.32,trna7_LysCTT_16_+_3225692_3225764@1.32.32,trna9_LysCTT_5_+_180634755_180634827@1.32.32\|GCCCGGCTAGCTCAGTCGGTAGAGCATGAGAC | >LysCTT.trna119_1_-_145395522_145395594 | 1 | 32 | 32 | tRF nuclear | NO | 0.00 | 72.09 | 0.00 | 0.00 |
| >trna119_LysCTT_1_-_145395522_145395594@1.33.33,trna11_LysCTT_5_-_180648979_180649051@1.33.33,trna13_LysCTT_6_+_26556774_26556846@1.33.33,trna7_LysCTT_16_+_3225692_3225764@1.33.33,trna9_LysCTT_5_+_180634755_180634827@1.33.33\|GCCCGGCTAGCTCAGTCGGTAGAGCATGAGACT | >LysCTT.trna119_1_-_145395522_145395594 | 1 | 33 | 33 | tRF nuclear | NO | 0.00 | 323.74 | 14.41 | 9.46 |
| >trna119_LysCTT_1_-_145395522_145395594@1.34.34,trna11_LysCTT_5_-_180648979_180649051@1.34.34,trna13_LysCTT_6_+_26556774_26556846@1.34.34,trna7_LysCTT_16_+_3225692_3225764@1.34.34,trna9_LysCTT_5_+_180634755_180634827@1.34.34\|GCCCGGCTAGCTCAGTCGGTAGAGCATGAGACTC | >LysCTT.trna119_1_-_145395522_145395594 | 1 | 34 | 34 | tRF nuclear | NO | 0.00 | 220.08 | 0.00 | 0.00 |
| >trna119_LysCTT_1_-_145395522_145395594@1.35.35,trna11_LysCTT_5_-_180648979_180649051@1.35.35,trna13_LysCTT_6_+_26556774_26556846@1.35.35,trna7_LysCTT_16_+_3225692_3225764@1.35.35,trna9_LysCTT_5_+_180634755_180634827@1.35.35\|GCCCGGCTAGCTCAGTCGGTAGAGCATGAGACTCT | >LysCTT.trna119_1_-_145395522_145395594 | 1 | 35 | 35 | tRF nuclear | NO | 24.42 | 1580.63 | 15.17 | 9.53 |
| >trna119_LysCTT_1_-_145395522_145395594@1.36.36,trna11_LysCTT_5_-_180648979_180649051@1.36.36,trna13_LysCTT_6_+_26556774_26556846@1.36.36,trna7_LysCTT_16_+_3225692_3225764@1.36.36,trna9_LysCTT_5_+_180634755_180634827@1.36.36\|GCCCGGCTAGCTCAGTCGGTAGAGCATGAGACTCTT | >LysCTT.trna119_1_-_145395522_145395594 | 1 | 36 | 36 | tRF nuclear | NO | 0.00 | 260.90 | 0.00 | 0.00 |
| >am_trna119_LysCTT_1_-_145395522_145395594@14.32.19,trna11_LysCTT_5_-_180648979_180649051@14.32.19,trna128_LysCTT_1_-_55423542_55423614@14.32.19,trna13_LysCTT_6_+_26556774_26556846@14.32.19,trna32_LysCTT_16_-_3207406_3207478@14.32.19,trna6_LysCTT_19_-_52425396_52425468@14.32.19,trna7_LysCTT_16_+_3225692_3225764@14.32.19,trna9_LysCTT_5_+_180634755_180634827@14.32.19\|AGTCGGTAGAGCATGAGAC | >LysCTT.trna119_1_-_145395522_145395594.am | 14 | 32 | 19 | tRF nuclear | NO | 0.00 | 34.98 | 0.00 | 0.00 |
| >trna32_LysCTT_16_-_3207406_3207478@1.33.33\|GCCCGGCTAGCTCAGTCGGTAGAGCATGAGACC | >LysCTT.trna32_16_-_3207406_3207478 | 1 | 33 | 33 | tRF nuclear | NO | 0.00 | 38.65 | 0.00 | 0.00 |
| >trna32_LysCTT_16_-_3207406_3207478@1.35.35\|GCCCGGCTAGCTCAGTCGGTAGAGCATGAGACCCT | >LysCTT.trna32_16_-_3207406_3207478 | 1 | 35 | 35 | tRF nuclear | NO | 0.00 | 10.32 | 0.00 | 0.00 |
| >trna14_LysTTT_11_-_59327808_59327880@1.33.33,trna2_LysTTT_17_+_8022473_8022545@1.33.33,trna54_LysTTT_1_+_204475655_204475727@1.33.33,trna5_LysTTT_11_+_59323902_59323974@1.33.33,trna62_LysTTT_1_-_204476158_204476230@1.33.33,trna76_LysTTT_6_+_28918806_28918878@1.33.33\|GCCCGGATAGCTCAGTCGGTAGAGCATCAGACT | >LysTTT.trna14_11_-_59327808_59327880 | 1 | 33 | 33 | tRF nuclear | NO | 9.77 | 218.87 | 87.83 | 0.00 |
| >trna14_LysTTT_11_-_59327808_59327880@1.36.36,trna2_LysTTT_17_+_8022473_8022545@1.36.36,trna54_LysTTT_1_+_204475655_204475727@1.36.36,trna5_LysTTT_11_+_59323902_59323974@1.36.36,trna62_LysTTT_1_-_204476158_204476230@1.36.36,trna76_LysTTT_6_+_28918806_28918878@1.36.36\|GCCCGGATAGCTCAGTCGGTAGAGCATCAGACTTTT | >LysTTT.trna14_11_-_59327808_59327880 | 1 | 36 | 36 | tRF nuclear | NO | 0.00 | 30.54 | 0.00 | 0.00 |

# BT-20

| | | | | | | | | | | |
|---|---|---|---|---|---|---|---|---|---|---|
| >trna143_LysTTT_6_-_27559593_27559665@14.31.18,trna149_LysTTT_6_-_27302769_27302841@14.31.18,trna14_LysTTT_11_-_59327808_59327880@14.31.18,trna2_LysTTT_17_+_8022473_8022545@14.31.18,trna54_LysTTT_1_+_204475655_204475727@14.31.18,trna5_LysTTT_11_+_59323902_59323974@14.31.18,trna62_LysTTT_1_-_204476158_204476230@14.31.18,trna76_LysTTT_6_+_28918806_28918878@14.31.18\|AGTCGGTAGAGCATCAGA | >LysTTT.trna143_6_-_27559593_27559665 | 14 | 31 | 18 | tRF nuclear | NO | 0.00 | 42.08 | 0.00 | 0.00 |
| >am_trna162_MetCAT_6_-_26758550_26758622@34.54.21,trna164_MetCAT_6_-_26735574_26735646@34.54.21,trna20_MetCAT_16_+_71460396_71460468@34.54.21,trna21_MetCAT_6_+_26701712_26701784@34.54.21,trna27_MetCAT_6_+_26766444_26766516@34.54.21,trna75_MetCAT_6_+_28912352_28912424@34.54.21,trna97_MetCAT_6_-_28921042_28921114@34.54.21\|CATAATCTGAAGGTCCTGAGT | >MetCAT.trna162_6_-_26758550_26758622.am | 34 | 54 | 21 | tRF nuclear | NO | 0.00 | 41.01 | 0.00 | 0.00 |
| >am_trna11_ProAGG_16_+_3241989_3242060@1.31.31,trna12_ProAGG_6_+_26555498_26555569@1.31.31,trna14_ProTGG_5_-_180615854_180615925@1.31.31,trna22_ProAGG_14_-_21081560_21081631@1.31.31,trna23_ProAGG_14_-_21077495_21077566@1.31.31,trna28_ProTGG_16_-_3234133_3234204@1.31.31,trna29_ProAGG_16_-_3232635_3232706@1.31.31,trna2_ProAGG_7_+_128423504_128423575@1.31.31,trna30_ProCGG_6_+_27059521_27059592@1.31.31,trna37_ProCGG_17_-_8126151_8126222@1.31.31,trna3_ProTGG_16_+_3208923_3208994@1.31.31,trna52_ProCGG_1_+_167683962_167684033@1.31.31,trna65_ProAGG_1_-_167684725_167684796@1.31.31,trna6_ProCGG_16_+_3222049_3222120@1.31.31,trna6_ProTGG_14_+_21152175_21152246@1.31.31,trna8_ProTGG_16_+_3238094_3238165@1.31.31,trna9_ProAGG_11_+_75946557_75946628@1.31.31,trna9_ProAGG_16_+_3239634_3239705@1.31.31\|GGCTCGTTGGTCTAGGGGTATGATTCTCGCT | >ProAGG.trna11_16_+_3241989_3242060.am | 1 | 31 | 31 | tRF nuclear | YES | 0.00 | 28.42 | 0.00 | 0.00 |
| >trna16_TyrGTA_14_-_21131351_21131444@59.76.18,trna17_TyrGTA_14_-_21128117_21128210@59.76.18,trna19_TyrGTA_14_-_21121258_21121351@59.76.18,trna2_TyrGTA_2_+_27273650_27273738@59.76.18,trna4_TyrGTA_8_+_67025602_67025694@59.76.18,trna5_TyrGTA_14_+_21151432_21151520@59.76.18,trna5_TyrGTA_8_+_67026223_67026311@59.76.18\|TTCCGGCTCGAAGGACCA | >TyrGTA.trna16_14_-_21131351_21131444 | 59 | 76 | 18 | tRF nuclear | NO | 10.12 | 251.08 | 66.10 | 10.03 |
| >am_trna16_TyrGTA_14_-_21131351_21131444@1.25.25,trna17_TyrGTA_14_-_21128117_21128210@1.25.25,trna17_TyrGTA_6_+_26595102_26595190@1.25.25,trna18_TyrGTA_14_-_21125623_21125716@1.25.25,trna19_TyrGTA_14_-_21121258_21121351@1.25.25,trna4_TyrGTA_8_+_67025602_67025694@1.25.25,trna5_TyrGTA_14_+_21151432_21151520@1.25.25,trna5_TyrGTA_8_+_67026223_67026311@1.25.25\|CCTTCGATAGCTCAGCTGGTAGAGC | >TyrGTA.trna16_14_-_21131351_21131444.am | 1 | 25 | 25 | tRF nuclear | NO | 0.00 | 27.85 | 0.00 | 0.00 |
| >trna139_ValAAC_6_-_27618707_27618779@36.56.21\|CACGCGAAAGGTCCCTGGATC | >ValAAC.trna139_6_-_27618707_27618779 | 36 | 56 | 21 | tRF nuclear | NO | 0.00 | 29.67 | 10.31 | 0.00 |
| >trna15_ValAAC_5_-_180615416_180615488@1.23.23\|GTTTCCGTAGTGTAGTGGTCATC | >ValAAC.trna15_5_-_180615416_180615488 | 1 | 23 | 23 | tRF nuclear | NO | 0.00 | 10.32 | 56.96 | 0.00 |
| >trna15_ValAAC_5_-_180615416_180615488@1.32.32\|GTTTCCGTAGTGTAGTGGTCATCACGTTCGCC | >ValAAC.trna15_5_-_180615416_180615488 | 1 | 32 | 32 | tRF nuclear | NO | 0.00 | 26.90 | 0.00 | 0.00 |
| >am_trna10_ValCAC_5_-_180649395_180649467@1.23.23,trna12_ValAAC_5_-_180645270_180645342@1.23.23,trna132_ValAAC_6_-_27721179_27721251@1.23.23,trna136_ValAAC_6_-_27648885_27648957@1.23.23,trna139_ValAAC_6_-_27618707_27618779@1.23.23,trna18_ValCAC_5_-_180529253_180529325@1.23.23,trna2_ValAAC_3_+_169490018_169490090@1.23.23,trna2_ValCAC_5_+_180524070_180524142@1.23.23,trna37_ValAAC_6_+_27203288_27203360@1.23.23,trna4_ValAAC_5_+_180591154_180591226@1.23.23,trna5_ValAAC_5_+_180596610_180596682@1.23.23,trna6_ValCAC_5_+_180600650_180600722@1.23.23,trna85_ValCAC_1_-_161369490_161369562@1.23.23,trna90_ValCAC_1_-_149684088_149684161@1.23.23,trna98_ValCAC_1_-_149298555_149298627@1.23.23,trna9_ValCAC_6_+_26538282_26538354@1.23.23\|GTTTCCGTAGTGTAGTGGTTATC | >ValCAC.trna10_5_-_180649395_180649467.am | 1 | 23 | 23 | tRF nuclear | NO | 0.00 | 150.44 | 219.39 | 31.25 |
| >am_trna10_ValCAC_5_-_180649395_180649467@1.32.32,trna12_ValAAC_5_-_180645270_180645342@1.32.32,trna132_ValAAC_6_-_27721179_27721251@1.32.32,trna136_ValAAC_6_-_27648885_27648957@1.32.32,trna139_ValAAC_6_-_27618707_27618779@1.32.32,trna18_ValCAC_5_-_180529253_180529325@1.32.32,trna2_ValAAC_3_+_169490018_169490090@1.32.32,trna2_ValCAC_5_+_180524070_180524142@1.32.32,trna4_ValAAC_5_+_180591154_180591226@1.32.32,trna5_ValAAC_5_+_180596610_180596682@1.32.32,trna6_ValCAC_5_+_180600650_180600722@1.32.32,trna85_ValCAC_1_-_161369490_161369562@1.32.32,trna90_ValCAC_1_-_149684088_149684161@1.32.32,trna98_ValCAC_1_-_149298555_149298627@1.32.32,trna9_ValCAC_6_+_26538282_26538354@1.32.32\|GTTTCCGTAGTGTAGTGGTTATCACGTTCGCC | >ValCAC.trna10_5_-_180649395_180649467.am | 1 | 32 | 32 | tRF nuclear | NO | 0.00 | 257.13 | 37.80 | 38.10 |

# BT-20

| | | | | | | | | | | |
|---|---|---|---|---|---|---|---|---|---|---|
| >am_trna10_ValCAC_5_-_180649395_180649467@1.33.33,trna12_ValAAC_5_-_180645270_180645342@1.33.33,trna132_ValAAC_6_-_27721179_27721251@1.33.33,trna136_ValAAC_6_-_27648885_27648957@1.33.33,trna139_ValAAC_6_-_27618707_27618779@1.33.33,trna18_ValCAC_5_-_180529253_180529325@1.33.33,trna2_ValAAC_3_+_169490018_169490090@1.33.33,trna2_ValCAC_5_+_180524070_180524142@1.33.33,trna4_ValAAC_5_+_180591154_180591226@1.33.33,trna5_ValAAC_5_+_180596610_180596682@1.33.33,trna6_ValCAC_5_+_180600650_180600722@1.33.33,trna85_ValCAC_1_-_161369490_161369562@1.33.33,trna90_ValCAC_1_-_149684088_149684161@1.33.33,trna98_ValCAC_1_-_149298555_149298627@1.33.33,trna9_ValCAC_6_+_26538282_26538354@1.33.33\|GTTTCCGTAGTGTAGTGGTTATCACGTTCGCCT | >ValCAC.trna10_5_-_180649395_180649467.am | 1 | 33 | 33 | | tRF nuclear | NO | 0.00 | 103.53 | 11.83 | 24.02 |
| >am_trna10_ValCAC_5_-_180649395_180649467@1.34.34,trna18_ValCAC_5_-_180529253_180529325@1.34.34,trna2_ValCAC_5_+_180524070_180524142@1.34.34,trna6_ValCAC_5_+_180600650_180600722@1.34.34,trna85_ValCAC_1_-_161369490_161369562@1.34.34,trna90_ValCAC_1_-_149684088_149684161@1.34.34,trna98_ValCAC_1_-_149298555_149298627@1.34.34,trna9_ValCAC_6_+_26538282_26538354@1.34.34\|GTTTCCGTAGTGTAGTGGTTATCACGTTCGCCTC | >ValCAC.trna10_5_-_180649395_180649467.am | 1 | 34 | 34 | | tRF nuclear | NO | 0.00 | 26.85 | 11.60 | 0.00 |
| >trna152_ValCAC_6_-_27248049_27248121@1.23.23\|GCTTCTGTAGTGTAGTGGTTATC | >ValCAC.trna152_6_-_27248049_27248121 | 1 | 23 | 23 | | tRF nuclear | NO | 0.00 | 0.00 | 14.44 | 0.00 |
| >trna152_ValCAC_6_-_27248049_27248121@1.32.32\|GCTTCTGTAGTGTAGTGGTTATCACGTTCGCC | >ValCAC.trna152_6_-_27248049_27248121 | 1 | 32 | 32 | | tRF nuclear | NO | 0.00 | 75.19 | 13.66 | 0.00 |
| >trna152_ValCAC_6_-_27248049_27248121@1.33.33\|GCTTCTGTAGTGTAGTGGTTATCACGTTCGCCT | >ValCAC.trna152_6_-_27248049_27248121 | 1 | 33 | 33 | | tRF nuclear | NO | 0.00 | 34.09 | 0.00 | 0.00 |
| >trna152_ValCAC_6_-_27248049_27248121@1.34.34\|GCTTCTGTAGTGTAGTGGTTATCACGTTCGCCTC | >ValCAC.trna152_6_-_27248049_27248121 | 1 | 34 | 34 | | tRF nuclear | NO | 0.00 | 74.55 | 15.96 | 0.00 |
