## Supplementary material for "Single-nucleotide Differences and Cell Type Decide the Subcellular Localization of miRNA Isoforms (isomiRs), tRNA-derived Fragments (tRFs) and rRNA-derived Fragments (rRFs)": Supp. Table S9 (3 of 4)

### MDA-MB-231

<table>
<tr><th>Full Name</th><th>Short Name</th><th>Start</th><th>End</th><th>Length</th><th>Additional Nucleotides</th><th>group</th><th>Outlier</th><th>AVE NUC</th><th>AVE CYTO</th><th>AVE MITO</th><th>AVE MP</th></tr>
<tr><td>>MI0000061|hsa-let-7a-2&WithFlank&11|-|122146516|122146599@11.28.18;MI0000064|hsa-let-7c&WithFlank&21|+|16539822|16539917@17.34.18;MI0000062|hsa-let-7a-3&WithFlank&22|+|46112743|46112828@10.27.18;MI0000060|hsa-let-7a-1&WithFlank&9|+|94175951|94176042@12.29.18&[MIMAT0000062&hsa-let-7a-5p&offsets|0|-4;m-6&11|-|122146568|122146589&offsets|0|-4];[MIMAT0000064&hsa-let-7c-5p&offsets|0|-4;m-37&21|+|16539838|16539859&offsets|0|-4];[MIMAT0000062_1&hsa-let-7a-5p&offsets|0|-4;m-5&22|+|46112752|46112773&offsets|0|-4];[MIMAT0000062_2&hsa-let-7a-5p&offsets|0|-4;m-4&9|+|94175962|94175983&offsets|0|-4]|TGAGGTAGTAGGTTGTAT</td><td>>MI0000061|hsa-let-7a-2&WithFlank&11|-|122146516|122146599</td><td>11</td><td>28</td><td>18</td><td></td><td>isomiR</td><td>NO</td><td>150.29</td><td>144.94</td><td>177.52</td><td>19.61</td></tr>
<tr><td>>MI0000061|hsa-let-7a-2&WithFlank&11|-|122146516|122146599@11.30.20(+1C);MI0000062|hsa-let-7a-3&WithFlank&22|+|46112743|46112828@10.29.20(+1C);MI0000060|hsa-let-7a-1&WithFlank&9|+|94175951|94176042@12.31.20(+1C)&[MIMAT0000062&hsa-let-7a-5p&offsets|0|-2(+1C);m-6&11|-|122146568|122146589&offsets|0|-2(+1C)];[MIMAT0000062_1&hsa-let-7a-5p&offsets|0|-2(+1C);m-5&22|+|46112752|46112773&offsets|0|-2(+1C)];[MIMAT0000062_2&hsa-let-7a-5p&offsets|0|-2(+1C);m-4&9|+|94175962|94175983&offsets|0|-2(+1C)]|TGAGGTAGTAGGTTGTATAGC</td><td>>MI0000061|hsa-let-7a-2&WithFlank&11|-|122146516|122146599</td><td>11</td><td>30</td><td>20</td><td>(+1C)</td><td>isomiR</td><td>NO</td><td>76.74</td><td>32.36</td><td>103.15</td><td>14.48</td></tr>
<tr><td>>MI0000061|hsa-let-7a-2&WithFlank&11|-|122146516|122146599@11.30.20;MI0000062|hsa-let-7a-3&WithFlank&22|+|46112743|46112828@10.29.20;MI0000060|hsa-let-7a-1&WithFlank&9|+|94175951|94176042@12.31.20&[MIMAT0000062&hsa-let-7a-5p&offsets|0|-2;m-6&11|-|122146568|122146589&offsets|0|-2];[MIMAT0000062_1&hsa-let-7a-5p&offsets|0|-2;m-5&22|+|46112752|46112773&offsets|0|-2];[MIMAT0000062_2&hsa-let-7a-5p&offsets|0|-2;m-4&9|+|94175962|94175983&offsets|0|-2]|TGAGGTAGTAGGTTGTATAG</td><td>>MI0000061|hsa-let-7a-2&WithFlank&11|-|122146516|122146599</td><td>11</td><td>30</td><td>20</td><td></td><td>isomiR</td><td>NO</td><td>0.00</td><td>0.00</td><td>12.06</td><td>0.00</td></tr>
<tr><td>>MI0000061|hsa-let-7a-2&WithFlank&11|-|122146516|122146599@11.31.21(+1A);MI0000062|hsa-let-7a-3&WithFlank&22|+|46112743|46112828@10.30.21(+1A);MI0000060|hsa-let-7a-1&WithFlank&9|+|94175951|94176042@12.32.21(+1A)&[MIMAT0000062&hsa-let-7a-5p&offsets|0|-1(+1A);m-6&11|-|122146568|122146589&offsets|0|-1(+1A)];[MIMAT0000062_1&hsa-let-7a-5p&offsets|0|-1(+1A);m-5&22|+|46112752|46112773&offsets|0|-1(+1A)];[MIMAT0000062_2&hsa-let-7a-5p&offsets|0|-1(+1A);m-4&9|+|94175962|94175983&offsets|0|-1(+1A)]|TGAGGTAGTAGGTTGTATAGTA</td><td>>MI0000061|hsa-let-7a-2&WithFlank&11|-|122146516|122146599</td><td>11</td><td>31</td><td>21</td><td>(+1A)</td><td>isomiR</td><td>NO</td><td>100.57</td><td>98.71</td><td>175.92</td><td>20.51</td></tr>
<tr><td>>MI0000061|hsa-let-7a-2&WithFlank&11|-|122146516|122146599@11.31.21(+1C);MI0000062|hsa-let-7a-3&WithFlank&22|+|46112743|46112828@10.30.21(+1C);MI0000060|hsa-let-7a-1&WithFlank&9|+|94175951|94176042@12.32.21(+1C)&[MIMAT0000062&hsa-let-7a-5p&offsets|0|-1(+1C);m-6&11|-|122146568|122146589&offsets|0|-1(+1C)];[MIMAT0000062_1&hsa-let-7a-5p&offsets|0|-1(+1C);m-5&22|+|46112752|46112773&offsets|0|-1(+1C)];[MIMAT0000062_2&hsa-let-7a-5p&offsets|0|-1(+1C);m-4&9|+|94175962|94175983&offsets|0|-1(+1C)]|TGAGGTAGTAGGTTGTATAGTC</td><td>>MI0000061|hsa-let-7a-2&WithFlank&11|-|122146516|122146599</td><td>11</td><td>31</td><td>21</td><td>(+1C)</td><td>isomiR</td><td>NO</td><td>472.84</td><td>367.49</td><td>664.30</td><td>160.56</td></tr>
<tr><td>>MI0000061|hsa-let-7a-2&WithFlank&11|-|122146516|122146599@11.31.21(+1G);MI0000062|hsa-let-7a-3&WithFlank&22|+|46112743|46112828@10.30.21(+1G);MI0000060|hsa-let-7a-1&WithFlank&9|+|94175951|94176042@12.32.21(+1G)&[MIMAT0000062&hsa-let-7a-5p&offsets|0|-1(+1G);m-6&11|-|122146568|122146589&offsets|0|-1(+1G)];[MIMAT0000062_1&hsa-let-7a-5p&offsets|0|-1(+1G);m-5&22|+|46112752|46112773&offsets|0|-1(+1G)];[MIMAT0000062_2&hsa-let-7a-5p&offsets|0|-1(+1G);m-4&9|+|94175962|94175983&offsets|0|-1(+1G)]|TGAGGTAGTAGGTTGTATAGTG</td><td>>MI0000061|hsa-let-7a-2&WithFlank&11|-|122146516|122146599</td><td>11</td><td>31</td><td>21</td><td>(+1G)</td><td>isomiR</td><td>NO</td><td>70.13</td><td>25.86</td><td>55.81</td><td>0.00</td></tr>
<tr><td>>MI0000061|hsa-let-7a-2&WithFlank&11|-|122146516|122146599@11.31.21;MI0000062|hsa-let-7a-3&WithFlank&22|+|46112743|46112828@10.30.21;MI0000060|hsa-let-7a-1&WithFlank&9|+|94175951|94176042@12.32.21&[MIMAT0000062&hsa-let-7a-5p&offsets|0|-1;m-6&11|-|122146568|122146589&offsets|0|-1];[MIMAT0000062_1&hsa-let-7a-5p&offsets|0|-1;m-5&22|+|46112752|46112773&offsets|0|-1];[MIMAT0000062_2&hsa-let-7a-5p&offsets|0|-1;m-4&9|+|94175962|94175983&offsets|0|-1]|TGAGGTAGTAGGTTGTATAGT</td><td>>MI0000061|hsa-let-7a-2&WithFlank&11|-|122146516|122146599</td><td>11</td><td>31</td><td>21</td><td></td><td>isomiR</td><td>NO</td><td>478.11</td><td>357.67</td><td>515.62</td><td>135.29</td></tr>
<tr><td>>MI0000061|hsa-let-7a-2&WithFlank&11|-|122146516|122146599@11.32.22(+1A);MI0000062|hsa-let-7a-3&WithFlank&22|+|46112743|46112828@10.31.22(+1A);MI0000060|hsa-let-7a-1&WithFlank&9|+|94175951|94176042@12.33.22(+1A)&[MIMAT0000062&hsa-let-7a-5p&offsets|0|0(+1A);m-6&11|-|122146568|122146589&offsets|0|0(+1A)];[MIMAT0000062_1&hsa-let-7a-5p&offsets|0|0(+1A);m-5&22|+|46112752|46112773&offsets|0|0(+1A)];[MIMAT0000062_2&hsa-let-7a-5p&offsets|0|0(+1A);m-4&9|+|94175962|94175983&offsets|0|0(+1A)]|TGAGGTAGTAGGTTGTATAGTTA</td><td>>MI0000061|hsa-let-7a-2&WithFlank&11|-|122146516|122146599</td><td>11</td><td>32</td><td>22</td><td>(+1A)</td><td>isomiR</td><td>NO</td><td>76.76</td><td>36.19</td><td>171.86</td><td>11.68</td></tr>
<tr><td>>MI0000061|hsa-let-7a-2&WithFlank&11|-|122146516|122146599@11.32.22(+1C);MI0000062|hsa-let-7a-3&WithFlank&22|+|46112743|46112828@10.31.22(+1C);MI0000060|hsa-let-7a-1&WithFlank&9|+|94175951|94176042@12.33.22(+1C)&[MIMAT0000062&hsa-let-7a-5p&offsets|0|0(+1C);m-6&11|-|122146568|122146589&offsets|0|0(+1C)];[MIMAT0000062_1&hsa-let-7a-5p&offsets|0|0(+1C);m-5&22|+|46112752|46112773&offsets|0|0(+1C)];[MIMAT0000062_2&hsa-let-7a-5p&offsets|0|0(+1C);m-4&9|+|94175962|94175983&offsets|0|0(+1C)]|TGAGGTAGTAGGTTGTATAGTTC</td><td>>MI0000061|hsa-let-7a-2&WithFlank&11|-|122146516|122146599</td><td>11</td><td>32</td><td>22</td><td>(+1C)</td><td>isomiR</td><td>NO</td><td>0.00</td><td>0.00</td><td>100.11</td><td>0.00</td></tr>
</table>

### MDA-MB-231

| | | | | | | | | | | | |
|---|---|---|---|---|---|---|---|---|---|---|---|
| >MI0000061\|hsa-let-7a-2&WithFlank&11\|-\|122146516\|122146599@11.32.22(+1G);MI0000062\|hsa-let-7a-3&WithFlank&22\|+\|46112743\|46112828@10.31.22(+1G);MI0000060\|hsa-let-7a-1&WithFlank&9\|+\|94175951\|94176042@12.33.22(+1G)&[MIMAT0000062&hsa-let-7a-5p&offsets\|0\|0(+1G);m-6&11\|-\|122146568\|122146589&offsets\|0\|0(+1G)];[MIMAT0000062_1&hsa-let-7a-5p&offsets\|0\|0(+1G);m-5&22\|+\|46112752\|46112773&offsets\|0\|0(+1G)];[MIMAT0000062_2&hsa-let-7a-5p&offsets\|0\|0(+1G);m-4&9\|+\|94175962\|94175983&offsets\|0\|0(+1G)]\|TGAGGTAGTAGGTTGTATAGTTG | >MI0000061\|hsa-let-7a-2&WithFlank&11\|-\|122146516\|122146599 | 11 | 32 | 22 | (+1G) | isomiR | NO | 0.00 | 0.00 | 10.73 | 0.00 |
| >MI0000061\|hsa-let-7a-2&WithFlank&11\|-\|122146516\|122146599@11.32.22;MI0000062\|hsa-let-7a-3&WithFlank&22\|+\|46112743\|46112828@10.31.22;MI0000060\|hsa-let-7a-1&WithFlank&9\|+\|94175951\|94176042@12.33.22&[MIMAT0000062&hsa-let-7a-5p&offsets\|0\|0;m-6&11\|-\|122146568\|122146589&offsets\|0\|0];[MIMAT0000062_1&hsa-let-7a-5p&offsets\|0\|0;m-5&22\|+\|46112752\|46112773&offsets\|0\|0];[MIMAT0000062_2&hsa-let-7a-5p&offsets\|0\|0;m-4&9\|+\|94175962\|94175983&offsets\|0\|0]\|TGAGGTAGTAGGTTGTATAGTT | >MI0000061\|hsa-let-7a-2&WithFlank&11\|-\|122146516\|122146599 | 11 | 32 | 22 | | isomiR | YES | 2776.22 | 2523.15 | 2963.96 | 598.82 |
| >MI0000061\|hsa-let-7a-2&WithFlank&11\|-\|122146516\|122146599@11.33.23;MI0000062\|hsa-let-7a-3&WithFlank&22\|+\|46112743\|46112828@10.32.23;MI0000060\|hsa-let-7a-1&WithFlank&9\|+\|94175951\|94176042@12.34.23&[MIMAT0000062&hsa-let-7a-5p&offsets\|0\|+1;m-6&11\|-\|122146568\|122146589&offsets\|0\|+1];[MIMAT0000062_1&hsa-let-7a-5p&offsets\|0\|+1;m-5&22\|+\|46112752\|46112773&offsets\|0\|+1];[MIMAT0000062_2&hsa-let-7a-5p&offsets\|0\|+1;m-4&9\|+\|94175962\|94175983&offsets\|0\|+1]\|TGAGGTAGTAGGTTGTATAGTTT | >MI0000061\|hsa-let-7a-2&WithFlank&11\|-\|122146516\|122146599 | 11 | 33 | 23 | | isomiR | NO | 166.97 | 140.09 | 471.86 | 57.44 |
| >MI0000062\|hsa-let-7a-3&WithFlank&22\|+\|46112743\|46112828@58.78.21(+1U);MI0000060\|hsa-let-7a-1&WithFlank&9\|+\|94175951\|94176042@63.83.21(+1U)&[MIMAT0004481&hsa-let-7a-3p&offsets\|0\|0(+1U);m-180&22\|+\|46112800\|46112821&offsets\|0\|-1(+1U)];[MIMAT0004481_1&hsa-let-7a-3p&offsets\|0\|0(+1U);m-181&9\|+\|94176013\|94176034&offsets\|0\|-1(+1U)]\|CTATACAATCTACTGTCTTTCT | >MI0000062\|hsa-let-7a-3&WithFlank&22\|+\|46112743\|46112828 | 58 | 78 | 21 | (+1U) | isomiR | NO | 100.16 | 24.78 | 99.09 | 54.81 |
| >MI0000062\|hsa-let-7a-3&WithFlank&22\|+\|46112743\|46112828@58.79.22;MI0000060\|hsa-let-7a-1&WithFlank&9\|+\|94175951\|94176042@63.84.22&[MIMAT0004481&hsa-let-7a-3p&offsets\|0\|+1;m-180&22\|+\|46112800\|46112821&offsets\|0\|0];[MIMAT0004481_1&hsa-let-7a-3p&offsets\|0\|+1;m-181&9\|+\|94176013\|94176034&offsets\|0\|0]\|CTATACAATCTACTGTCTTTCC | >MI0000062\|hsa-let-7a-3&WithFlank&22\|+\|46112743\|46112828 | 58 | 79 | 22 | | isomiR | NO | 0.00 | 0.00 | 12.97 | 0.00 |
| >MI0000063\|hsa-let-7b&WithFlank&22\|+\|46113680\|46113774@12.31.20&[MIMAT0000063&hsa-let-7b-5p&offsets\|0\|-2;m-14&22\|+\|46113691\|46113712&offsets\|0\|-2]\|TGAGGTAGTAGGTTGTGTGG | >MI0000063\|hsa-let-7b&WithFlank&22\|+\|46113680\|46113774 | 12 | 31 | 20 | | isomiR | NO | 0.00 | 0.00 | 35.55 | 0.00 |
| >MI0000063\|hsa-let-7b&WithFlank&22\|+\|46113680\|46113774@12.31.20(+1C)&[MIMAT0000063&hsa-let-7b-5p&offsets\|0\|-2(+1C);m-14&22\|+\|46113691\|46113712&offsets\|0\|-2(+1C)]\|TGAGGTAGTAGGTTGTGTGGC | >MI0000063\|hsa-let-7b&WithFlank&22\|+\|46113680\|46113774 | 12 | 31 | 20 | (+1C) | isomiR | NO | 0.00 | 0.00 | 87.51 | 0.00 |
| >MI0000063\|hsa-let-7b&WithFlank&22\|+\|46113680\|46113774@12.32.21&[MIMAT0000063&hsa-let-7b-5p&offsets\|0\|-1;m-14&22\|+\|46113691\|46113712&offsets\|0\|-1]\|TGAGGTAGTAGGTTGTGTGGT | >MI0000063\|hsa-let-7b&WithFlank&22\|+\|46113680\|46113774 | 12 | 32 | 21 | | isomiR | NO | 85.48 | 75.18 | 418.93 | 0.00 |
| >MI0000063\|hsa-let-7b&WithFlank&22\|+\|46113680\|46113774@12.32.21(+1A)&[MIMAT0000063&hsa-let-7b-5p&offsets\|0\|-1(+1A);m-14&22\|+\|46113691\|46113712&offsets\|0\|-1(+1A)]\|TGAGGTAGTAGGTTGTGTGGTA | >MI0000063\|hsa-let-7b&WithFlank&22\|+\|46113680\|46113774 | 12 | 32 | 21 | (+1A) | isomiR | NO | 0.00 | 0.00 | 150.27 | 0.00 |
| >MI0000063\|hsa-let-7b&WithFlank&22\|+\|46113680\|46113774@12.32.21(+1C)&[MIMAT0000063&hsa-let-7b-5p&offsets\|0\|-1(+1C);m-14&22\|+\|46113691\|46113712&offsets\|0\|-1(+1C)]\|TGAGGTAGTAGGTTGTGTGGTC | >MI0000063\|hsa-let-7b&WithFlank&22\|+\|46113680\|46113774 | 12 | 32 | 21 | (+1C) | isomiR | NO | 70.67 | 67.47 | 389.03 | 10.89 |
| >MI0000063\|hsa-let-7b&WithFlank&22\|+\|46113680\|46113774@12.32.21(+1G)&[MIMAT0000063&hsa-let-7b-5p&offsets\|0\|-1(+1G);m-14&22\|+\|46113691\|46113712&offsets\|0\|-1(+1G)]\|TGAGGTAGTAGGTTGTGTGGTG | >MI0000063\|hsa-let-7b&WithFlank&22\|+\|46113680\|46113774 | 12 | 32 | 21 | (+1G) | isomiR | NO | 0.00 | 0.00 | 16.32 | 0.00 |
| >MI0000063\|hsa-let-7b&WithFlank&22\|+\|46113680\|46113774@12.33.22&[MIMAT0000063&hsa-let-7b-5p&offsets\|0\|0;m-14&22\|+\|46113691\|46113712&offsets\|0\|0]\|TGAGGTAGTAGGTTGTGTGGTT | >MI0000063\|hsa-let-7b&WithFlank&22\|+\|46113680\|46113774 | 12 | 33 | 22 | | isomiR | NO | 408.07 | 459.83 | 1754.73 | 95.56 |
| >MI0000063\|hsa-let-7b&WithFlank&22\|+\|46113680\|46113774@12.33.22(+1A)&[MIMAT0000063&hsa-let-7b-5p&offsets\|0\|0(+1A);m-14&22\|+\|46113691\|46113712&offsets\|0\|0(+1A)]\|TGAGGTAGTAGGTTGTGTGGTTA | >MI0000063\|hsa-let-7b&WithFlank&22\|+\|46113680\|46113774 | 12 | 33 | 22 | (+1A) | isomiR | NO | 10.43 | 0.00 | 212.26 | 0.00 |
| >MI0000063\|hsa-let-7b&WithFlank&22\|+\|46113680\|46113774@12.33.22(+1C)&[MIMAT0000063&hsa-let-7b-5p&offsets\|0\|0(+1C);m-14&22\|+\|46113691\|46113712&offsets\|0\|0(+1C)]\|TGAGGTAGTAGGTTGTGTGGTTC | >MI0000063\|hsa-let-7b&WithFlank&22\|+\|46113680\|46113774 | 12 | 33 | 22 | (+1C) | isomiR | NO | 0.00 | 0.00 | 118.39 | 0.00 |
| >MI0000063\|hsa-let-7b&WithFlank&22\|+\|46113680\|46113774@12.34.23&[MIMAT0000063&hsa-let-7b-5p&offsets\|0\|+1;m-14&22\|+\|46113691\|46113712&offsets\|0\|+1]\|TGAGGTAGTAGGTTGTGTGGTTT | >MI0000063\|hsa-let-7b&WithFlank&22\|+\|46113680\|46113774 | 12 | 34 | 23 | | isomiR | NO | 106.56 | 117.83 | 556.90 | 14.75 |
| >MI0000063\|hsa-let-7b&WithFlank&22\|+\|46113680\|46113774@12.34.23(+1U)&[MIMAT0000063&hsa-let-7b-5p&offsets\|0\|+1(+1U);m-14&22\|+\|46113691\|46113712&offsets\|0\|+1(+1U)]\|TGAGGTAGTAGGTTGTGTGGTTTT | >MI0000063\|hsa-let-7b&WithFlank&22\|+\|46113680\|46113774 | 12 | 34 | 23 | (+1U) | isomiR | NO | 0.00 | 0.00 | 84.22 | 0.00 |
| >MI0000064\|hsa-let-7c&WithFlank&21\|+\|16539822\|16539917@17.38.22&[MIMAT0000064&hsa-let-7c-5p&offsets\|0\|0;m-37&21\|+\|16539838\|16539859&offsets\|0\|0]\|TGAGGTAGTAGGTTGTATGGTT | >MI0000064\|hsa-let-7c&WithFlank&21\|+\|16539822\|16539917 | 17 | 38 | 22 | | isomiR | NO | 0.00 | 0.00 | 59.27 | 0.00 |
| >MI0000065\|hsa-let-7d&WithFlank&9\|+\|94178828\|94178926@14.31.18&[MIMAT0000065&hsa-let-7d-5p&offsets\|0\|-4;m-90&9\|+\|94178841\|94178862&offsets\|0\|-4]\|AGAGGTAGTAGGTTGCAT | >MI0000065\|hsa-let-7d&WithFlank&9\|+\|94178828\|94178926 | 14 | 31 | 18 | | isomiR | NO | 0.00 | 0.00 | 17.44 | 0.00 |

### MDA-MB-231

| | | | | | | | | | | | |
|---|---|---|---|---|---|---|---|---|---|---|---|
| >MI0000065\|hsa-let-7d&WithFlank&9\|+\|94178828\|94178926@14.34.21&[MIMAT0000065&hsa-let-7d-5p&offsets\|0\|-1;m-90&9\|+\|94178841\|94178862&offsets\|0\|-1]\|AGAGGTAGTAGGTTGCATAGT | >MI0000065\|hsa-let-7d&WithFlank&9\|+\|94178828\|94178926 | 14 | 34 | 21 | | isomiR | NO | 75.76 | 36.65 | 101.88 | 13.58 |
| >MI0000065\|hsa-let-7d&WithFlank&9\|+\|94178828\|94178926@14.34.21(+1C)&[MIMAT0000065&hsa-let-7d-5p&offsets\|0\|-1(+1C);m-90&9\|+\|94178841\|94178862&offsets\|0\|-1(+1C)]\|AGAGGTAGTAGGTTGCATAGTC | >MI0000065\|hsa-let-7d&WithFlank&9\|+\|94178828\|94178926 | 14 | 34 | 21 | (+1C) | isomiR | NO | 52.91 | 13.15 | 94.95 | 0.00 |
| >MI0000065\|hsa-let-7d&WithFlank&9\|+\|94178828\|94178926@14.35.22&[MIMAT0000065&hsa-let-7d-5p&offsets\|0\|0;m-90&9\|+\|94178841\|94178862&offsets\|0\|0]\|AGAGGTAGTAGGTTGCATAGTT | >MI0000065\|hsa-let-7d&WithFlank&9\|+\|94178828\|94178926 | 14 | 35 | 22 | | isomiR | NO | 306.37 | 225.73 | 440.79 | 87.71 |
| >MI0000065\|hsa-let-7d&WithFlank&9\|+\|94178828\|94178926@14.36.23&[MIMAT0000065&hsa-let-7d-5p&offsets\|0\|+1;m-90&9\|+\|94178841\|94178862&offsets\|0\|+1]\|AGAGGTAGTAGGTTGCATAGTTT | >MI0000065\|hsa-let-7d&WithFlank&9\|+\|94178828\|94178926 | 14 | 36 | 23 | | isomiR | NO | 0.00 | 0.00 | 99.48 | 0.00 |
| >MI0000065\|hsa-let-7d&WithFlank&9\|+\|94178828\|94178926@68.88.21&[MIMAT0004484&hsa-let-7d-3p&offsets\|0\|-1;m-98&9\|+\|94178895\|94178916&offsets\|0\|-1]\|CTATACGACCTGCTGCCTTTC | >MI0000065\|hsa-let-7d&WithFlank&9\|+\|94178828\|94178926 | 68 | 88 | 21 | | isomiR | NO | 45.22 | 0.00 | 70.02 | 0.00 |
| >MI0000065\|hsa-let-7d&WithFlank&9\|+\|94178828\|94178926@68.88.21(+1A)&[MIMAT0004484&hsa-let-7d-3p&offsets\|0\|-1(+1A);m-98&9\|+\|94178895\|94178916&offsets\|0\|-1(+1A)]\|CTATACGACCTGCTGCCTTTCA | >MI0000065\|hsa-let-7d&WithFlank&9\|+\|94178828\|94178926 | 68 | 88 | 21 | (+1A) | isomiR | NO | 0.00 | 0.00 | 13.63 | 0.00 |
| >MI0000065\|hsa-let-7d&WithFlank&9\|+\|94178828\|94178926@68.88.21(+1C)&[MIMAT0004484&hsa-let-7d-3p&offsets\|0\|-1(+1C);m-98&9\|+\|94178895\|94178916&offsets\|0\|-1(+1C)]\|CTATACGACCTGCTGCCTTTCC | >MI0000065\|hsa-let-7d&WithFlank&9\|+\|94178828\|94178926 | 68 | 88 | 21 | (+1C) | isomiR | NO | 0.00 | 0.00 | 64.01 | 0.00 |
| >MI0000065\|hsa-let-7d&WithFlank&9\|+\|94178828\|94178926@68.89.22&[MIMAT0004484&hsa-let-7d-3p&offsets\|0\|0;m-98&9\|+\|94178895\|94178916&offsets\|0\|0]\|CTATACGACCTGCTGCCTTTCT | >MI0000065\|hsa-let-7d&WithFlank&9\|+\|94178828\|94178926 | 68 | 89 | 22 | | isomiR | NO | 139.01 | 109.17 | 250.56 | 64.47 |
| >MI0000066\|hsa-let-7e&WithFlank&19\|+\|51692780\|51692870@14.34.21&[MIMAT0000066&hsa-let-7e-5p&offsets\|0\|-1;m-50&19\|+\|51692793\|51692814&offsets\|0\|-1]\|TGAGGTAGGAGGTTGTATAGT | >MI0000066\|hsa-let-7e&WithFlank&19\|+\|51692780\|51692870 | 14 | 34 | 21 | | isomiR | NO | 64.26 | 109.22 | 56.39 | 11.60 |
| >MI0000066\|hsa-let-7e&WithFlank&19\|+\|51692780\|51692870@14.34.21(+1C)&[MIMAT0000066&hsa-let-7e-5p&offsets\|0\|-1(+1C);m-50&19\|+\|51692793\|51692814&offsets\|0\|-1(+1C)]\|TGAGGTAGGAGGTTGTATAGTC | >MI0000066\|hsa-let-7e&WithFlank&19\|+\|51692780\|51692870 | 14 | 34 | 21 | (+1C) | isomiR | NO | 24.80 | 63.35 | 16.71 | 11.96 |
| >MI0000066\|hsa-let-7e&WithFlank&19\|+\|51692780\|51692870@14.35.22&[MIMAT0000066&hsa-let-7e-5p&offsets\|0\|0;m-50&19\|+\|51692793\|51692814&offsets\|0\|0]\|TGAGGTAGGAGGTTGTATAGTT | >MI0000066\|hsa-let-7e&WithFlank&19\|+\|51692780\|51692870 | 14 | 35 | 22 | | isomiR | NO | 251.00 | 433.78 | 190.80 | 107.07 |
| >MI0000066\|hsa-let-7e&WithFlank&19\|+\|51692780\|51692870@14.35.22(+1U)&[MIMAT0000066&hsa-let-7e-5p&offsets\|0\|0(+1U);m-50&19\|+\|51692793\|51692814&offsets\|0\|0(+1U)]\|TGAGGTAGGAGGTTGTATAGTTT | >MI0000066\|hsa-let-7e&WithFlank&19\|+\|51692780\|51692870 | 14 | 35 | 22 | (+1U) | isomiR | NO | 0.00 | 36.20 | 63.02 | 0.00 |
| >MI0000066\|hsa-let-7e&WithFlank&19\|+\|51692780\|51692870@14.36.23&[MIMAT0000066&hsa-let-7e-5p&offsets\|0\|+1;m-50&19\|+\|51692793\|51692814&offsets\|0\|+1]\|TGAGGTAGGAGGTTGTATAGTTG | >MI0000066\|hsa-let-7e&WithFlank&19\|+\|51692780\|51692870 | 14 | 36 | 23 | | isomiR | NO | 0.00 | 0.00 | 10.75 | 0.00 |
| >MI0000067\|hsa-let-7f-1&WithFlank&9\|+\|94176341\|94176439@13.30.18;MI0000068\|hsa-let-7f-2&WithFlank&X\|-\|53557186\|53557280@14.31.18&[MIMAT0000067&hsa-let-7f-5p&offsets\|0\|-4;m-20&9\|+\|94176353\|94176374&offsets\|0\|-4];[MIMAT0000067_1&hsa-let-7f-5p&offsets\|0\|-4;m-19&X\|-\|53557246\|53557267&offsets\|0\|-4]\|TGAGGTAGTAGATTGTAT | >MI0000067\|hsa-let-7f-1&WithFlank&9\|+\|94176341\|94176439 | 13 | 30 | 18 | | isomiR | NO | 219.10 | 99.29 | 288.29 | 16.45 |
| >MI0000067\|hsa-let-7f-1&WithFlank&9\|+\|94176341\|94176439@13.32.20(+1C);MI0000068\|hsa-let-7f-2&WithFlank&X\|-\|53557186\|53557280@14.33.20(+1C)&[MIMAT0000067&hsa-let-7f-5p&offsets\|0\|-2(+1C);m-20&9\|+\|94176353\|94176374&offsets\|0\|-2(+1C)];[MIMAT0000067_1&hsa-let-7f-5p&offsets\|0\|-2(+1C);m-19&X\|-\|53557246\|53557267&offsets\|0\|-2(+1C)]\|TGAGGTAGTAGATTGTATAGC | >MI0000067\|hsa-let-7f-1&WithFlank&9\|+\|94176341\|94176439 | 13 | 32 | 20 | (+1C) | isomiR | NO | 86.98 | 0.00 | 95.74 | 0.00 |
| >MI0000067\|hsa-let-7f-1&WithFlank&9\|+\|94176341\|94176439@13.32.20;MI0000068\|hsa-let-7f-2&WithFlank&X\|-\|53557186\|53557280@14.33.20&[MIMAT0000067&hsa-let-7f-5p&offsets\|0\|-2;m-20&9\|+\|94176353\|94176374&offsets\|0\|-2];[MIMAT0000067_1&hsa-let-7f-5p&offsets\|0\|-2;m-19&X\|-\|53557246\|53557267&offsets\|0\|-2]\|TGAGGTAGTAGATTGTATAG | >MI0000067\|hsa-let-7f-1&WithFlank&9\|+\|94176341\|94176439 | 13 | 32 | 20 | | isomiR | NO | 0.00 | 0.00 | 12.97 | 0.00 |
| >MI0000067\|hsa-let-7f-1&WithFlank&9\|+\|94176341\|94176439@13.33.21(+1A);MI0000068\|hsa-let-7f-2&WithFlank&X\|-\|53557186\|53557280@14.34.21(+1A)&[MIMAT0000067&hsa-let-7f-5p&offsets\|0\|-1(+1A);m-20&9\|+\|94176353\|94176374&offsets\|0\|-1(+1A)];[MIMAT0000067_1&hsa-let-7f-5p&offsets\|0\|-1(+1A);m-19&X\|-\|53557246\|53557267&offsets\|0\|-1(+1A)]\|TGAGGTAGTAGATTGTATAGTA | >MI0000067\|hsa-let-7f-1&WithFlank&9\|+\|94176341\|94176439 | 13 | 33 | 21 | (+1A) | isomiR | NO | 121.78 | 72.43 | 191.58 | 13.87 |
| >MI0000067\|hsa-let-7f-1&WithFlank&9\|+\|94176341\|94176439@13.33.21(+1C);MI0000068\|hsa-let-7f-2&WithFlank&X\|-\|53557186\|53557280@14.34.21(+1C)&[MIMAT0000067&hsa-let-7f-5p&offsets\|0\|-1(+1C);m-20&9\|+\|94176353\|94176374&offsets\|0\|-1(+1C)];[MIMAT0000067_1&hsa-let-7f-5p&offsets\|0\|-1(+1C);m-19&X\|-\|53557246\|53557267&offsets\|0\|-1(+1C)]\|TGAGGTAGTAGATTGTATAGTC | >MI0000067\|hsa-let-7f-1&WithFlank&9\|+\|94176341\|94176439 | 13 | 33 | 21 | (+1C) | isomiR | NO | 592.73 | 284.20 | 752.50 | 112.54 |
| >MI0000067\|hsa-let-7f-1&WithFlank&9\|+\|94176341\|94176439@13.33.21(+1G);MI0000068\|hsa-let-7f-2&WithFlank&X\|-\|53557186\|53557280@14.34.21(+1G)&[MIMAT0000067&hsa-let-7f-5p&offsets\|0\|-1(+1G);m-20&9\|+\|94176353\|94176374&offsets\|0\|-1(+1G)];[MIMAT0000067_1&hsa-let-7f-5p&offsets\|0\|-1(+1G);m-19&X\|-\|53557246\|53557267&offsets\|0\|-1(+1G)]\|TGAGGTAGTAGATTGTATAGTG | >MI0000067\|hsa-let-7f-1&WithFlank&9\|+\|94176341\|94176439 | 13 | 33 | 21 | (+1G) | isomiR | NO | 85.78 | 0.00 | 63.40 | 0.00 |
| >MI0000067\|hsa-let-7f-1&WithFlank&9\|+\|94176341\|94176439@13.33.21;MI0000068\|hsa-let-7f-2&WithFlank&X\|-\|53557186\|53557280@14.34.21&[MIMAT0000067&hsa-let-7f-5p&offsets\|0\|-1;m-20&9\|+\|94176353\|94176374&offsets\|0\|-1];[MIMAT0000067_1&hsa-let-7f-5p&offsets\|0\|-1;m-19&X\|-\|53557246\|53557267&offsets\|0\|-1]\|TGAGGTAGTAGATTGTATAGT | >MI0000067\|hsa-let-7f-1&WithFlank&9\|+\|94176341\|94176439 | 13 | 33 | 21 | | isomiR | NO | 544.87 | 242.63 | 487.36 | 91.34 |

### MDA-MB-231

| | | | | | | | | | | | |
|---|---|---|---|---|---|---|---|---|---|---|---|
| >MI0000067\|hsa-let-7f-1&WithFlank&9\|+\|94176341\|94176439@13.34.22(+1A);MI0000068\|hsa-let-7f-2&WithFlank&X\|-\|53557186\|53557280@14.35.22(+1A)&[MIMAT0000067&hsa-let-7f-5p&offsets\|0\|0(+1A);m-20&9\|+\|94176353\|94176374&offsets\|0\|0(+1A)];[MIMAT0000067_1&hsa-let-7f-5p&offsets\|0\|0(+1A);m-19&X\|-\|53557246\|53557267&offsets\|0\|0(+1A)]\|TGAGGTAGTAGATTGTATAGTTA | >MI0000067\|hsa-let-7f-1&WithFlank&9\|+\|94176341\|94176439 | 13 | 34 | 22 | (+1A) | isomiR | NO | 86.25 | 27.29 | 205.25 | 0.00 |
| >MI0000067\|hsa-let-7f-1&WithFlank&9\|+\|94176341\|94176439@13.34.22(+1C);MI0000068\|hsa-let-7f-2&WithFlank&X\|-\|53557186\|53557280@14.35.22(+1C)&[MIMAT0000067&hsa-let-7f-5p&offsets\|0\|0(+1C);m-20&9\|+\|94176353\|94176374&offsets\|0\|0(+1C)];[MIMAT0000067_1&hsa-let-7f-5p&offsets\|0\|0(+1C);m-19&X\|-\|53557246\|53557267&offsets\|0\|0(+1C)]\|TGAGGTAGTAGATTGTATAGTTC | >MI0000067\|hsa-let-7f-1&WithFlank&9\|+\|94176341\|94176439 | 13 | 34 | 22 | (+1C) | isomiR | NO | 0.00 | 0.00 | 94.63 | 0.00 |
| >MI0000067\|hsa-let-7f-1&WithFlank&9\|+\|94176341\|94176439@13.34.22;MI0000068\|hsa-let-7f-2&WithFlank&X\|-\|53557186\|53557280@14.35.22&[MIMAT0000067&hsa-let-7f-5p&offsets\|0\|0;m-20&9\|+\|94176353\|94176374&offsets\|0\|0];[MIMAT0000067_1&hsa-let-7f-5p&offsets\|0\|0;m-19&X\|-\|53557246\|53557267&offsets\|0\|0]\|TGAGGTAGTAGATTGTATAGTT | >MI0000067\|hsa-let-7f-1&WithFlank&9\|+\|94176341\|94176439 | 13 | 34 | 22 | | isomiR | YES | 3516.58 | 1938.23 | 3374.54 | 425.87 |
| >MI0000067\|hsa-let-7f-1&WithFlank&9\|+\|94176341\|94176439@13.35.23&[MIMAT0000067&hsa-let-7f-5p&offsets\|0\|+1;m-20&9\|+\|94176353\|94176374&offsets\|0\|+1]\|TGAGGTAGTAGATTGTATAGTTG | >MI0000067\|hsa-let-7f-1&WithFlank&9\|+\|94176341\|94176439 | 13 | 35 | 23 | | isomiR | NO | 0.00 | 0.00 | 12.48 | 0.00 |
| >MI0000067\|hsa-let-7f-1&WithFlank&9\|+\|94176341\|94176439@69.89.21(+1U)&[MIMAT0004486&hsa-let-7f-1-3p&offsets\|0\|-1(+1U);m-275&9\|+\|94176409\|94176430&offsets\|0\|-1(+1U)]\|CTATACAATCTATTGCCTTCCT | >MI0000067\|hsa-let-7f-1&WithFlank&9\|+\|94176341\|94176439 | 69 | 89 | 21 | (+1U) | isomiR | NO | 0.00 | 0.00 | 65.32 | 0.00 |
| >MI0000068\|hsa-let-7f-2&WithFlank&X\|-\|53557186\|53557280@14.36.23&[MIMAT0000067_1&hsa-let-7f-5p&offsets\|0\|+1;m-19&X\|-\|53557246\|53557267&offsets\|0\|+1]\|TGAGGTAGTAGATTGTATAGTTT | >MI0000068\|hsa-let-7f-2&WithFlank&X\|-\|53557186\|53557280 | 14 | 36 | 23 | | isomiR | NO | 163.99 | 117.47 | 458.31 | 13.23 |
| >MI0000068\|hsa-let-7f-2&WithFlank&X\|-\|53557186\|53557280@64.84.21(+1U)&[MIMAT0004487&hsa-let-7f-2-3p&offsets\|0\|-1(+1U);m-453&X\|-\|53557196\|53557217&offsets\|0\|-1(+1U)]\|CTATACAGTCTACTGTCTTTCT | >MI0000068\|hsa-let-7f-2&WithFlank&X\|-\|53557186\|53557280 | 64 | 84 | 21 | (+1U) | isomiR | NO | 0.00 | 0.00 | 13.49 | 0.00 |
| >MI0000070\|hsa-mir-16-1&WithFlank&13\|-\|50048967\|50049067@20.39.20(+1U);MI0000115\|hsa-mir-16-2&WithFlank&3\|+\|160404739\|160404831@16.35.20(+1U)&[MIMAT0000069&hsa-miR-16-5p&offsets\|0\|-2(+1U);m-62&13\|-\|50049027\|50049048&offsets\|0\|-2(+1U)];[MIMAT0000069_1&hsa-miR-16-5p&offsets\|0\|-2(+1U);m-61&3\|+\|160404754\|160404775&offsets\|0\|-2(+1U)]\|TAGCAGCACGTAAATATTGGT | >MI0000070\|hsa-mir-16-1&WithFlank&13\|-\|50048967\|50049067 | 20 | 39 | 20 | (+1U) | isomiR | NO | 0.00 | 0.00 | 111.87 | 0.00 |
| >MI0000070\|hsa-mir-16-1&WithFlank&13\|-\|50048967\|50049067@20.39.20;MI0000115\|hsa-mir-16-2&WithFlank&3\|+\|160404739\|160404831@16.35.20&[MIMAT0000069&hsa-miR-16-5p&offsets\|0\|-2;m-62&13\|-\|50049027\|50049048&offsets\|0\|-2];[MIMAT0000069_1&hsa-miR-16-5p&offsets\|0\|-2;m-61&3\|+\|160404754\|160404775&offsets\|0\|-2]\|TAGCAGCACGTAAATATTGG | >MI0000070\|hsa-mir-16-1&WithFlank&13\|-\|50048967\|50049067 | 20 | 39 | 20 | | isomiR | NO | 0.00 | 0.00 | 68.09 | 0.00 |
| >MI0000070\|hsa-mir-16-1&WithFlank&13\|-\|50048967\|50049067@20.40.21(+1A);MI0000115\|hsa-mir-16-2&WithFlank&3\|+\|160404739\|160404831@16.36.21(+1A)&[MIMAT0000069&hsa-miR-16-5p&offsets\|0\|-1(+1A);m-62&13\|-\|50049027\|50049048&offsets\|0\|-1(+1A)];[MIMAT0000069_1&hsa-miR-16-5p&offsets\|0\|-1(+1A);m-61&3\|+\|160404754\|160404775&offsets\|0\|-1(+1A)]\|TAGCAGCACGTAAATATTGGCA | >MI0000070\|hsa-mir-16-1&WithFlank&13\|-\|50048967\|50049067 | 20 | 40 | 21 | (+1A) | isomiR | NO | 0.00 | 0.00 | 242.97 | 0.00 |
| >MI0000070\|hsa-mir-16-1&WithFlank&13\|-\|50048967\|50049067@20.40.21;MI0000115\|hsa-mir-16-2&WithFlank&3\|+\|160404739\|160404831@16.36.21&[MIMAT0000069&hsa-miR-16-5p&offsets\|0\|-1;m-62&13\|-\|50049027\|50049048&offsets\|0\|-1];[MIMAT0000069_1&hsa-miR-16-5p&offsets\|0\|-1;m-61&3\|+\|160404754\|160404775&offsets\|0\|-1]\|TAGCAGCACGTAAATATTGGC | >MI0000070\|hsa-mir-16-1&WithFlank&13\|-\|50048967\|50049067 | 20 | 40 | 21 | | isomiR | NO | 65.86 | 0.00 | 256.62 | 10.70 |
| >MI0000070\|hsa-mir-16-1&WithFlank&13\|-\|50048967\|50049067@20.41.22(+1C);MI0000115\|hsa-mir-16-2&WithFlank&3\|+\|160404739\|160404831@16.37.22(+1C)&[MIMAT0000069&hsa-miR-16-5p&offsets\|0\|0(+1C);m-62&13\|-\|50049027\|50049048&offsets\|0\|0(+1C)];[MIMAT0000069_1&hsa-miR-16-5p&offsets\|0\|0(+1C);m-61&3\|+\|160404754\|160404775&offsets\|0\|0(+1C)]\|TAGCAGCACGTAAATATTGGCGC | >MI0000070\|hsa-mir-16-1&WithFlank&13\|-\|50048967\|50049067 | 20 | 41 | 22 | (+1C) | isomiR | NO | 0.00 | 0.00 | 83.48 | 0.00 |
| >MI0000070\|hsa-mir-16-1&WithFlank&13\|-\|50048967\|50049067@20.41.22;MI0000115\|hsa-mir-16-2&WithFlank&3\|+\|160404739\|160404831@16.37.22&[MIMAT0000069&hsa-miR-16-5p&offsets\|0\|0;m-62&13\|-\|50049027\|50049048&offsets\|0\|0];[MIMAT0000069_1&hsa-miR-16-5p&offsets\|0\|0;m-61&3\|+\|160404754\|160404775&offsets\|0\|0]\|TAGCAGCACGTAAATATTGGCG | >MI0000070\|hsa-mir-16-1&WithFlank&13\|-\|50048967\|50049067 | 20 | 41 | 22 | | isomiR | NO | 132.23 | 33.48 | 770.37 | 13.85 |
| >MI0000070\|hsa-mir-16-1&WithFlank&13\|-\|50048967\|50049067@20.42.23;MI0000115\|hsa-mir-16-2&WithFlank&3\|+\|160404739\|160404831@16.38.23&[MIMAT0000069&hsa-miR-16-5p&offsets\|0\|+1;m-62&13\|-\|50049027\|50049048&offsets\|0\|+1];[MIMAT0000069_1&hsa-miR-16-5p&offsets\|0\|+1;m-61&3\|+\|160404754\|160404775&offsets\|0\|+1]\|TAGCAGCACGTAAATATTGGCGT | >MI0000070\|hsa-mir-16-1&WithFlank&13\|-\|50048967\|50049067 | 20 | 42 | 23 | | isomiR | NO | 91.86 | 0.00 | 393.01 | 14.38 |
| >MI0000070\|hsa-mir-16-1&WithFlank&13\|-\|50048967\|50049067@20.43.24&[MIMAT0000069&hsa-miR-16-5p&offsets\|0\|+2;m-62&13\|-\|50049027\|50049048&offsets\|0\|+2]\|TAGCAGCACGTAAATATTGGCGTT | >MI0000070\|hsa-mir-16-1&WithFlank&13\|-\|50048967\|50049067 | 20 | 43 | 24 | | isomiR | NO | 0.00 | 0.00 | 13.58 | 0.00 |
| >MI0000071\|hsa-mir-17&WithFlank&13\|+\|91350599\|91350694@19.42.24&[MIMAT0000070&hsa-miR-17-5p&offsets\|-1\|0;m-88&13\|+\|91350618\|91350640&offsets\|-1\|0]\|TCAAAGTGCTTACAGTGCAGGTAG | >MI0000071\|hsa-mir-17&WithFlank&13\|+\|91350599\|91350694 | 19 | 42 | 24 | | isomiR | NO | 0.00 | 0.00 | 11.12 | 0.00 |
| >MI0000071\|hsa-mir-17&WithFlank&13\|+\|91350599\|91350694@20.39.20(+1C)&[MIMAT0000070&hsa-miR-17-5p&offsets\|0\|-3(+1C);m-88&13\|+\|91350618\|91350640&offsets\|0\|-3(+1C)]\|CAAAGTGCTTACAGTGCAGGC | >MI0000071\|hsa-mir-17&WithFlank&13\|+\|91350599\|91350694 | 20 | 39 | 20 | (+1C) | isomiR | NO | 0.00 | 0.00 | 36.01 | 0.00 |
| >MI0000071\|hsa-mir-17&WithFlank&13\|+\|91350599\|91350694@20.40.21&[MIMAT0000070&hsa-miR-17-5p&offsets\|0\|-2;m-88&13\|+\|91350618\|91350640&offsets\|0\|-2]\|CAAAGTGCTTACAGTGCAGGT | >MI0000071\|hsa-mir-17&WithFlank&13\|+\|91350599\|91350694 | 20 | 40 | 21 | | isomiR | NO | 64.70 | 0.00 | 234.03 | 11.92 |
| >MI0000071\|hsa-mir-17&WithFlank&13\|+\|91350599\|91350694@20.41.22&[MIMAT0000070&hsa-miR-17-5p&offsets\|0\|-1;m-88&13\|+\|91350618\|91350640&offsets\|0\|-1]\|CAAAGTGCTTACAGTGCAGGTA | >MI0000071\|hsa-mir-17&WithFlank&13\|+\|91350599\|91350694 | 20 | 41 | 22 | | isomiR | NO | 25.94 | 0.00 | 125.32 | 0.00 |

### MDA-MB-231

<table>
<tr><td>>MI0000071|hsa-mir-17&WithFlank&13|+|91350599|91350694@20.41.22(+1A)&[MIMAT0000070&hsa-miR-17-5p&offsets|0|-1(+1A);m-88&13|+|91350618|91350640&offsets|0|-1(+1A)]|CAAAGTGCTTACAGTGCAGGTAA</td><td>>MI0000071|hsa-mir-17&WithFlank&13|+|91350599|91350694</td><td>20</td><td>41</td><td>22</td><td>(+1A)</td><td>isomiR</td><td>NO</td><td>46.80</td><td>0.00</td><td>231.34</td><td>12.30</td></tr>
<tr><td>>MI0000071|hsa-mir-17&WithFlank&13|+|91350599|91350694@20.42.23&[MIMAT0000070&hsa-miR-17-5p&offsets|0|0;m-88&13|+|91350618|91350640&offsets|0|0]|CAAAGTGCTTACAGTGCAGGTAG</td><td>>MI0000071|hsa-mir-17&WithFlank&13|+|91350599|91350694</td><td>20</td><td>42</td><td>23</td><td></td><td>isomiR</td><td>NO</td><td>414.26</td><td>94.99</td><td>1166.78</td><td>132.31</td></tr>
<tr><td>>MI0000071|hsa-mir-17&WithFlank&13|+|91350599|91350694@20.42.23(+1A)&[MIMAT0000070&hsa-miR-17-5p&offsets|0|0(+1A);m-88&13|+|91350618|91350640&offsets|0|0(+1A)]|CAAAGTGCTTACAGTGCAGGTAGA</td><td>>MI0000071|hsa-mir-17&WithFlank&13|+|91350599|91350694</td><td>20</td><td>42</td><td>23</td><td>(+1A)</td><td>isomiR</td><td>NO</td><td>160.25</td><td>76.09</td><td>180.38</td><td>88.13</td></tr>
<tr><td>>MI0000071|hsa-mir-17&WithFlank&13|+|91350599|91350694@20.42.23(+1C)&[MIMAT0000070&hsa-miR-17-5p&offsets|0|0(+1C);m-88&13|+|91350618|91350640&offsets|0|0(+1C)]|CAAAGTGCTTACAGTGCAGGTAGC</td><td>>MI0000071|hsa-mir-17&WithFlank&13|+|91350599|91350694</td><td>20</td><td>42</td><td>23</td><td>(+1C)</td><td>isomiR</td><td>NO</td><td>0.00</td><td>0.00</td><td>10.54</td><td>0.00</td></tr>
<tr><td>>MI0000071|hsa-mir-17&WithFlank&13|+|91350599|91350694@20.43.24&[MIMAT0000070&hsa-miR-17-5p&offsets|0|+1;m-88&13|+|91350618|91350640&offsets|0|+1]|CAAAGTGCTTACAGTGCAGGTAGT</td><td>>MI0000071|hsa-mir-17&WithFlank&13|+|91350599|91350694</td><td>20</td><td>43</td><td>24</td><td></td><td>isomiR</td><td>NO</td><td>130.34</td><td>13.42</td><td>119.52</td><td>67.88</td></tr>
<tr><td>>MI0000072|hsa-mir-18a&WithFlank&13|+|91350745|91350827@12.34.23&[MIMAT0000072&hsa-miR-18a-5p&offsets|0|0;m-262&13|+|91350756|91350778&offsets|0|0]|TAAGGTGCATCTAGTGCAGATAG</td><td>>MI0000072|hsa-mir-18a&WithFlank&13|+|91350745|91350827</td><td>12</td><td>34</td><td>23</td><td></td><td>isomiR</td><td>NO</td><td>10.50</td><td>0.00</td><td>109.33</td><td>0.00</td></tr>
<tr><td>>MI0000073|hsa-mir-19a&WithFlank&13|+|91350885|91350978@54.76.23(+1U)&[MIMAT0000073&hsa-miR-19a-3p&offsets|-1|-1(+1U);m-201&13|+|91350939|91350961&offsets|-1|-1(+1U)]|TTGTGCAAATCTATGCAAAACTGT</td><td>>MI0000073|hsa-mir-19a&WithFlank&13|+|91350885|91350978</td><td>54</td><td>76</td><td>23</td><td>(+1U)</td><td>isomiR</td><td>NO</td><td>0.00</td><td>0.00</td><td>35.54</td><td>0.00</td></tr>
<tr><td>>MI0000073|hsa-mir-19a&WithFlank&13|+|91350885|91350978@55.76.22(+1U)&[MIMAT0000073&hsa-miR-19a-3p&offsets|0|-1(+1U);m-201&13|+|91350939|91350961&offsets|0|-1(+1U)]|TGTGCAAATCTATGCAAAACTGT</td><td>>MI0000073|hsa-mir-19a&WithFlank&13|+|91350885|91350978</td><td>55</td><td>76</td><td>22</td><td>(+1U)</td><td>isomiR</td><td>NO</td><td>0.00</td><td>0.00</td><td>112.78</td><td>0.00</td></tr>
<tr><td>>MI0000073|hsa-mir-19a&WithFlank&13|+|91350885|91350978@55.78.24&[MIMAT0000073&hsa-miR-19a-3p&offsets|0|+1;m-201&13|+|91350939|91350961&offsets|0|+1]|TGTGCAAATCTATGCAAAACTGAT</td><td>>MI0000073|hsa-mir-19a&WithFlank&13|+|91350885|91350978</td><td>55</td><td>78</td><td>24</td><td></td><td>isomiR</td><td>NO</td><td>0.00</td><td>0.00</td><td>88.23</td><td>0.00</td></tr>
<tr><td>>MI0000074|hsa-mir-19b-1&WithFlank&13|+|91351186|91351284@60.81.22(+1U);MI0000075|hsa-mir-19b-2&WithFlank&X|-|134169665|134169772@68.89.22(+1U)&[MIMAT0000074&hsa-miR-19b-3p&offsets|0|-1(+1U);m-122&13|+|91351245|91351267&offsets|0|-1(+1U)];[MIMAT0000074_1&hsa-miR-19b-3p&offsets|0|-1(+1U);m-121&X|-|134169683|134169705&offsets|0|-1(+1U)]|TGTGCAAATCCATGCAAAACTGT</td><td>>MI0000074|hsa-mir-19b-1&WithFlank&13|+|91351186|91351284</td><td>60</td><td>81</td><td>22</td><td>(+1U)</td><td>isomiR</td><td>NO</td><td>0.00</td><td>0.00</td><td>59.09</td><td>0.00</td></tr>
<tr><td>>MI0000074|hsa-mir-19b-1&WithFlank&13|+|91351186|91351284@60.81.22;MI0000075|hsa-mir-19b-2&WithFlank&X|-|134169665|134169772@68.89.22&[MIMAT0000074&hsa-miR-19b-3p&offsets|0|-1;m-122&13|+|91351245|91351267&offsets|0|-1];[MIMAT0000074_1&hsa-miR-19b-3p&offsets|0|-1;m-121&X|-|134169683|134169705&offsets|0|-1]|TGTGCAAATCCATGCAAAACTG</td><td>>MI0000074|hsa-mir-19b-1&WithFlank&13|+|91351186|91351284</td><td>60</td><td>81</td><td>22</td><td></td><td>isomiR</td><td>NO</td><td>0.00</td><td>0.00</td><td>112.86</td><td>0.00</td></tr>
<tr><td>>MI0000074|hsa-mir-19b-1&WithFlank&13|+|91351186|91351284@60.82.23;MI0000075|hsa-mir-19b-2&WithFlank&X|-|134169665|134169772@68.90.23&[MIMAT0000074&hsa-miR-19b-3p&offsets|0|0;m-122&13|+|91351245|91351267&offsets|0|0];[MIMAT0000074_1&hsa-miR-19b-3p&offsets|0|0;m-121&X|-|134169683|134169705&offsets|0|0]|TGTGCAAATCCATGCAAAACTGA</td><td>>MI0000074|hsa-mir-19b-1&WithFlank&13|+|91351186|91351284</td><td>60</td><td>82</td><td>23</td><td></td><td>isomiR</td><td>NO</td><td>49.15</td><td>0.00</td><td>158.40</td><td>0.00</td></tr>
<tr><td>>MI0000076|hsa-mir-20a&WithFlank&13|+|91351059|91351141@14.33.20(+1C)&[MIMAT0000075&hsa-miR-20a-5p&offsets|0|-3(+1C);m-96&13|+|91351072|91351094&offsets|0|-3(+1C)]|TAAAGTGCTTATAGTGCAGGC</td><td>>MI0000076|hsa-mir-20a&WithFlank&13|+|91351059|91351141</td><td>14</td><td>33</td><td>20</td><td>(+1C)</td><td>isomiR</td><td>NO</td><td>0.00</td><td>0.00</td><td>55.02</td><td>0.00</td></tr>
<tr><td>>MI0000076|hsa-mir-20a&WithFlank&13|+|91351059|91351141@14.34.21&[MIMAT0000075&hsa-miR-20a-5p&offsets|0|-2;m-96&13|+|91351072|91351094&offsets|0|-2]|TAAAGTGCTTATAGTGCAGGT</td><td>>MI0000076|hsa-mir-20a&WithFlank&13|+|91351059|91351141</td><td>14</td><td>34</td><td>21</td><td></td><td>isomiR</td><td>NO</td><td>74.36</td><td>0.00</td><td>259.40</td><td>16.70</td></tr>
<tr><td>>MI0000076|hsa-mir-20a&WithFlank&13|+|91351059|91351141@14.35.22&[MIMAT0000075&hsa-miR-20a-5p&offsets|0|-1;m-96&13|+|91351072|91351094&offsets|0|-1]|TAAAGTGCTTATAGTGCAGGTA</td><td>>MI0000076|hsa-mir-20a&WithFlank&13|+|91351059|91351141</td><td>14</td><td>35</td><td>22</td><td></td><td>isomiR</td><td>NO</td><td>0.00</td><td>0.00</td><td>76.10</td><td>0.00</td></tr>
<tr><td>>MI0000076|hsa-mir-20a&WithFlank&13|+|91351059|91351141@14.35.22(+1A)&[MIMAT0000075&hsa-miR-20a-5p&offsets|0|-1(+1A);m-96&13|+|91351072|91351094&offsets|0|-1(+1A)]|TAAAGTGCTTATAGTGCAGGTAA</td><td>>MI0000076|hsa-mir-20a&WithFlank&13|+|91351059|91351141</td><td>14</td><td>35</td><td>22</td><td>(+1A)</td><td>isomiR</td><td>NO</td><td>82.51</td><td>0.00</td><td>334.66</td><td>20.68</td></tr>
<tr><td>>MI0000076|hsa-mir-20a&WithFlank&13|+|91351059|91351141@14.35.22(+1U)&[MIMAT0000075&hsa-miR-20a-5p&offsets|0|-1(+1U);m-96&13|+|91351072|91351094&offsets|0|-1(+1U)]|TAAAGTGCTTATAGTGCAGGTAT</td><td>>MI0000076|hsa-mir-20a&WithFlank&13|+|91351059|91351141</td><td>14</td><td>35</td><td>22</td><td>(+1U)</td><td>isomiR</td><td>NO</td><td>0.00</td><td>0.00</td><td>32.33</td><td>0.00</td></tr>
<tr><td>>MI0000076|hsa-mir-20a&WithFlank&13|+|91351059|91351141@14.36.23&[MIMAT0000075&hsa-miR-20a-5p&offsets|0|0;m-96&13|+|91351072|91351094&offsets|0|0]|TAAAGTGCTTATAGTGCAGGTAG</td><td>>MI0000076|hsa-mir-20a&WithFlank&13|+|91351059|91351141</td><td>14</td><td>36</td><td>23</td><td></td><td>isomiR</td><td>YES</td><td>737.21</td><td>83.89</td><td>1708.03</td><td>223.63</td></tr>
<tr><td>>MI0000076|hsa-mir-20a&WithFlank&13|+|91351059|91351141@14.36.23(+1A)&[MIMAT0000075&hsa-miR-20a-5p&offsets|0|0(+1A);m-96&13|+|91351072|91351094&offsets|0|0(+1A)]|TAAAGTGCTTATAGTGCAGGTAGA</td><td>>MI0000076|hsa-mir-20a&WithFlank&13|+|91351059|91351141</td><td>14</td><td>36</td><td>23</td><td>(+1A)</td><td>isomiR</td><td>NO</td><td>148.01</td><td>0.00</td><td>129.69</td><td>97.14</td></tr>
</table>

### MDA-MB-231

| | | | | | | | | | | | |
|---|---|---|---|---|---|---|---|---|---|---|---|
| >MI0000076\|hsa-mir-20a&WithFlank&13\|+\|91351059\|91351141@14.37.24&[MIMAT0000075&hsa-miR-20a-5p&offsets\|0\|+1;m-96&13\|+\|91351072\|91351094&offsets\|0\|+1]\|TAAAGTGCTTATAGTGCAGGTAGT | >MI0000076\|hsa-mir-20a&WithFlank&13\|+\|91351059\|91351141 | 14 | 37 | 24 | | isomiR | NO | 84.21 | 0.00 | 70.33 | 59.39 |
| >MI0000077\|hsa-mir-21&WithFlank&17\|+\|59841260\|59841343@13.34.22&[MIMAT0000076&hsa-miR-21-5p&offsets\|-1\|-1;m-1&17\|+\|59841273\|59841294&offsets\|-1\|-1]\|GTAGCTTATCAGACTGATGTTG | >MI0000077\|hsa-mir-21&WithFlank&17\|+\|59841260\|59841343 | 13 | 34 | 22 | | isomiR | NO | 0.00 | 0.00 | 33.40 | 0.00 |
| >MI0000077\|hsa-mir-21&WithFlank&17\|+\|59841260\|59841343@13.35.23&[MIMAT0000076&hsa-miR-21-5p&offsets\|-1\|0;m-1&17\|+\|59841273\|59841294&offsets\|-1\|0]\|GTAGCTTATCAGACTGATGTTGA | >MI0000077\|hsa-mir-21&WithFlank&17\|+\|59841260\|59841343 | 13 | 35 | 23 | | isomiR | NO | 24.54 | 0.00 | 339.44 | 0.00 |
| >MI0000077\|hsa-mir-21&WithFlank&17\|+\|59841260\|59841343@13.36.24&[MIMAT0000076&hsa-miR-21-5p&offsets\|-1\|+1;m-1&17\|+\|59841273\|59841294&offsets\|-1\|+1]\|GTAGCTTATCAGACTGATGTTGAC | >MI0000077\|hsa-mir-21&WithFlank&17\|+\|59841260\|59841343 | 13 | 36 | 24 | | isomiR | NO | 0.00 | 0.00 | 15.66 | 0.00 |
| >MI0000077\|hsa-mir-21&WithFlank&17\|+\|59841260\|59841343@14.31.18&[MIMAT0000076&hsa-miR-21-5p&offsets\|0\|-4;m-1&17\|+\|59841273\|59841294&offsets\|0\|-4]\|TAGCTTATCAGACTGATG | >MI0000077\|hsa-mir-21&WithFlank&17\|+\|59841260\|59841343 | 14 | 31 | 18 | | isomiR | NO | 48.81 | 0.00 | 106.33 | 0.00 |
| >MI0000077\|hsa-mir-21&WithFlank&17\|+\|59841260\|59841343@14.32.19&[MIMAT0000076&hsa-miR-21-5p&offsets\|0\|-3;m-1&17\|+\|59841273\|59841294&offsets\|0\|-3]\|TAGCTTATCAGACTGATGT | >MI0000077\|hsa-mir-21&WithFlank&17\|+\|59841260\|59841343 | 14 | 32 | 19 | | isomiR | NO | 46.13 | 0.00 | 351.32 | 0.00 |
| >MI0000077\|hsa-mir-21&WithFlank&17\|+\|59841260\|59841343@14.33.20&[MIMAT0000076&hsa-miR-21-5p&offsets\|0\|-2;m-1&17\|+\|59841273\|59841294&offsets\|0\|-2]\|TAGCTTATCAGACTGATGTT | >MI0000077\|hsa-mir-21&WithFlank&17\|+\|59841260\|59841343 | 14 | 33 | 20 | | isomiR | NO | 52.08 | 0.00 | 267.41 | 0.00 |
| >MI0000077\|hsa-mir-21&WithFlank&17\|+\|59841260\|59841343@14.33.20(+1A)&[MIMAT0000076&hsa-miR-21-5p&offsets\|0\|-2(+1A);m-1&17\|+\|59841273\|59841294&offsets\|0\|-2(+1A)]\|TAGCTTATCAGACTGATGTTA | >MI0000077\|hsa-mir-21&WithFlank&17\|+\|59841260\|59841343 | 14 | 33 | 20 | (+1A) | isomiR | NO | 53.43 | 0.00 | 421.53 | 11.26 |
| >MI0000077\|hsa-mir-21&WithFlank&17\|+\|59841260\|59841343@14.33.20(+1U)&[MIMAT0000076&hsa-miR-21-5p&offsets\|0\|-2(+1U);m-1&17\|+\|59841273\|59841294&offsets\|0\|-2(+1U)]\|TAGCTTATCAGACTGATGTTT | >MI0000077\|hsa-mir-21&WithFlank&17\|+\|59841260\|59841343 | 14 | 33 | 20 | (+1U) | isomiR | NO | 0.00 | 0.00 | 83.08 | 0.00 |
| >MI0000077\|hsa-mir-21&WithFlank&17\|+\|59841260\|59841343@14.34.21&[MIMAT0000076&hsa-miR-21-5p&offsets\|0\|-1;m-1&17\|+\|59841273\|59841294&offsets\|0\|-1]\|TAGCTTATCAGACTGATGTTG | >MI0000077\|hsa-mir-21&WithFlank&17\|+\|59841260\|59841343 | 14 | 34 | 21 | | isomiR | NO | 417.37 | 112.55 | 1967.04 | 111.97 |
| >MI0000077\|hsa-mir-21&WithFlank&17\|+\|59841260\|59841343@14.34.21(+1C)&[MIMAT0000076&hsa-miR-21-5p&offsets\|0\|-1(+1C);m-1&17\|+\|59841273\|59841294&offsets\|0\|-1(+1C)]\|TAGCTTATCAGACTGATGTTGC | >MI0000077\|hsa-mir-21&WithFlank&17\|+\|59841260\|59841343 | 14 | 34 | 21 | (+1C) | isomiR | NO | 0.00 | 0.00 | 125.52 | 0.00 |
| >MI0000077\|hsa-mir-21&WithFlank&17\|+\|59841260\|59841343@14.34.21(+1G)&[MIMAT0000076&hsa-miR-21-5p&offsets\|0\|-1(+1G);m-1&17\|+\|59841273\|59841294&offsets\|0\|-1(+1G)]\|TAGCTTATCAGACTGATGTTGG | >MI0000077\|hsa-mir-21&WithFlank&17\|+\|59841260\|59841343 | 14 | 34 | 21 | (+1G) | isomiR | NO | 86.19 | 0.00 | 745.75 | 11.04 |
| >MI0000077\|hsa-mir-21&WithFlank&17\|+\|59841260\|59841343@14.34.21(+1U)&[MIMAT0000076&hsa-miR-21-5p&offsets\|0\|-1(+1U);m-1&17\|+\|59841273\|59841294&offsets\|0\|-1(+1U)]\|TAGCTTATCAGACTGATGTTGT | >MI0000077\|hsa-mir-21&WithFlank&17\|+\|59841260\|59841343 | 14 | 34 | 21 | (+1U) | isomiR | NO | 65.37 | 0.00 | 764.66 | 15.84 |
| >MI0000077\|hsa-mir-21&WithFlank&17\|+\|59841260\|59841343@14.34.21(+2C)&[MIMAT0000076&hsa-miR-21-5p&offsets\|0\|-1(+2C);m-1&17\|+\|59841273\|59841294&offsets\|0\|-1(+2C)]\|TAGCTTATCAGACTGATGTTGCC | >MI0000077\|hsa-mir-21&WithFlank&17\|+\|59841260\|59841343 | 14 | 34 | 21 | (+2C) | isomiR | NO | 127.18 | 29.48 | 125.87 | 53.64 |
| >MI0000077\|hsa-mir-21&WithFlank&17\|+\|59841260\|59841343@14.35.22&[MIMAT0000076&hsa-miR-21-5p&offsets\|0\|0;m-1&17\|+\|59841273\|59841294&offsets\|0\|0]\|TAGCTTATCAGACTGATGTTGA | >MI0000077\|hsa-mir-21&WithFlank&17\|+\|59841260\|59841343 | 14 | 35 | 22 | | isomiR | YES | 2060.89 | 590.08 | 22600.23 | 467.84 |
| >MI0000077\|hsa-mir-21&WithFlank&17\|+\|59841260\|59841343@14.35.22(+1A)&[MIMAT0000076&hsa-miR-21-5p&offsets\|0\|0(+1A);m-1&17\|+\|59841273\|59841294&offsets\|0\|0(+1A)]\|TAGCTTATCAGACTGATGTTGAA | >MI0000077\|hsa-mir-21&WithFlank&17\|+\|59841260\|59841343 | 14 | 35 | 22 | (+1A) | isomiR | YES | 521.43 | 256.44 | 1117.61 | 209.90 |
| >MI0000077\|hsa-mir-21&WithFlank&17\|+\|59841260\|59841343@14.35.22(+1G)&[MIMAT0000076&hsa-miR-21-5p&offsets\|0\|0(+1G);m-1&17\|+\|59841273\|59841294&offsets\|0\|0(+1G)]\|TAGCTTATCAGACTGATGTTGAG | >MI0000077\|hsa-mir-21&WithFlank&17\|+\|59841260\|59841343 | 14 | 35 | 22 | (+1G) | isomiR | NO | 106.29 | 24.15 | 146.81 | 19.76 |
| >MI0000077\|hsa-mir-21&WithFlank&17\|+\|59841260\|59841343@14.35.22(+1U)&[MIMAT0000076&hsa-miR-21-5p&offsets\|0\|0(+1U);m-1&17\|+\|59841273\|59841294&offsets\|0\|0(+1U)]\|TAGCTTATCAGACTGATGTTGAT | >MI0000077\|hsa-mir-21&WithFlank&17\|+\|59841260\|59841343 | 14 | 35 | 22 | (+1U) | isomiR | YES | 2024.77 | 1102.42 | 4605.27 | 650.54 |
| >MI0000077\|hsa-mir-21&WithFlank&17\|+\|59841260\|59841343@14.35.22(+2A)&[MIMAT0000076&hsa-miR-21-5p&offsets\|0\|0(+2A);m-1&17\|+\|59841273\|59841294&offsets\|0\|0(+2A)]\|TAGCTTATCAGACTGATGTTGAAA | >MI0000077\|hsa-mir-21&WithFlank&17\|+\|59841260\|59841343 | 14 | 35 | 22 | (+2A) | isomiR | NO | 49.01 | 0.00 | 10.64 | 11.64 |
| >MI0000077\|hsa-mir-21&WithFlank&17\|+\|59841260\|59841343@14.35.22(+2U)&[MIMAT0000076&hsa-miR-21-5p&offsets\|0\|0(+2U);m-1&17\|+\|59841273\|59841294&offsets\|0\|0(+2U)]\|TAGCTTATCAGACTGATGTTGATT | >MI0000077\|hsa-mir-21&WithFlank&17\|+\|59841260\|59841343 | 14 | 35 | 22 | (+2U) | isomiR | NO | 48.35 | 0.00 | 0.00 | 0.00 |
| >MI0000077\|hsa-mir-21&WithFlank&17\|+\|59841260\|59841343@14.36.23&[MIMAT0000076&hsa-miR-21-5p&offsets\|0\|+1;m-1&17\|+\|59841273\|59841294&offsets\|0\|+1]\|TAGCTTATCAGACTGATGTTGAC | >MI0000077\|hsa-mir-21&WithFlank&17\|+\|59841260\|59841343 | 14 | 36 | 23 | | isomiR | YES | 14756.82 | 8116.49 | 18818.50 | 2724.63 |

### MDA-MB-231

<table>
<tr><td>>MI0000077|hsa-mir-21&WithFlank&17|+|59841260|59841343@14.36.23(+1A)&[MIMAT0000076&hsa-miR-21-5p&offsets|0|+1(+1A);m-1&17|+|59841273|59841294&offsets|0|+1(+1A)]|TAGCTTATCAGACTGATGTTGACA</td><td>>MI0000077|hsa-mir-21&WithFlank&17|+|59841260|59841343</td><td>14</td><td>36</td><td>23</td><td>(+1A)</td><td>isomiR</td><td>YES</td><td>2507.57</td><td>1606.04</td><td>1429.46</td><td>693.64</td></tr>
<tr><td>>MI0000077|hsa-mir-21&WithFlank&17|+|59841260|59841343@14.36.23(+1C)&[MIMAT0000076&hsa-miR-21-5p&offsets|0|+1(+1C);m-1&17|+|59841273|59841294&offsets|0|+1(+1C)]|TAGCTTATCAGACTGATGTTGACC</td><td>>MI0000077|hsa-mir-21&WithFlank&17|+|59841260|59841343</td><td>14</td><td>36</td><td>23</td><td>(+1C)</td><td>isomiR</td><td>YES</td><td>1368.10</td><td>631.30</td><td>699.13</td><td>400.70</td></tr>
<tr><td>>MI0000077|hsa-mir-21&WithFlank&17|+|59841260|59841343@14.36.23(+1G)&[MIMAT0000076&hsa-miR-21-5p&offsets|0|+1(+1G);m-1&17|+|59841273|59841294&offsets|0|+1(+1G)]|TAGCTTATCAGACTGATGTTGACG</td><td>>MI0000077|hsa-mir-21&WithFlank&17|+|59841260|59841343</td><td>14</td><td>36</td><td>23</td><td>(+1G)</td><td>isomiR</td><td>YES</td><td>429.33</td><td>137.63</td><td>195.49</td><td>108.34</td></tr>
<tr><td>>MI0000077|hsa-mir-21&WithFlank&17|+|59841260|59841343@14.37.24&[MIMAT0000076&hsa-miR-21-5p&offsets|0|+2;m-1&17|+|59841273|59841294&offsets|0|+2]|TAGCTTATCAGACTGATGTTGACT</td><td>>MI0000077|hsa-mir-21&WithFlank&17|+|59841260|59841343</td><td>14</td><td>37</td><td>24</td><td></td><td>isomiR</td><td>YES</td><td>5422.79</td><td>2195.97</td><td>2623.26</td><td>1122.68</td></tr>
<tr><td>>MI0000077|hsa-mir-21&WithFlank&17|+|59841260|59841343@15.35.21(+1U)&[MIMAT0000076&hsa-miR-21-5p&offsets|+1|0(+1U);m-1&17|+|59841273|59841294&offsets|+1|0(+1U)]|AGCTTATCAGACTGATGTTGAT</td><td>>MI0000077|hsa-mir-21&WithFlank&17|+|59841260|59841343</td><td>15</td><td>35</td><td>21</td><td>(+1U)</td><td>isomiR</td><td>NO</td><td>0.00</td><td>0.00</td><td>12.01</td><td>0.00</td></tr>
<tr><td>>MI0000077|hsa-mir-21&WithFlank&17|+|59841260|59841343@15.36.22&[MIMAT0000076&hsa-miR-21-5p&offsets|+1|+1;m-1&17|+|59841273|59841294&offsets|+1|+1]|AGCTTATCAGACTGATGTTGAC</td><td>>MI0000077|hsa-mir-21&WithFlank&17|+|59841260|59841343</td><td>15</td><td>36</td><td>22</td><td></td><td>isomiR</td><td>NO</td><td>69.82</td><td>0.00</td><td>128.01</td><td>17.71</td></tr>
<tr><td>>MI0000077|hsa-mir-21&WithFlank&17|+|59841260|59841343@15.37.23&[MIMAT0000076&hsa-miR-21-5p&offsets|+1|+2;m-1&17|+|59841273|59841294&offsets|+1|+2]|AGCTTATCAGACTGATGTTGACT</td><td>>MI0000077|hsa-mir-21&WithFlank&17|+|59841260|59841343</td><td>15</td><td>37</td><td>23</td><td></td><td>isomiR</td><td>NO</td><td>0.00</td><td>0.00</td><td>12.28</td><td>0.00</td></tr>
<tr><td>>MI0000077|hsa-mir-21&WithFlank&17|+|59841260|59841343@52.72.21&[MIMAT0004494&hsa-miR-21-3p&offsets|0|0;m-52&17|+|59841311|59841332&offsets|0|-1]|CAACACCAGTCGATGGGCTGT</td><td>>MI0000077|hsa-mir-21&WithFlank&17|+|59841260|59841343</td><td>52</td><td>72</td><td>21</td><td></td><td>isomiR</td><td>NO</td><td>0.00</td><td>0.00</td><td>195.14</td><td>0.00</td></tr>
<tr><td>>MI0000077|hsa-mir-21&WithFlank&17|+|59841260|59841343@52.73.22&[MIMAT0004494&hsa-miR-21-3p&offsets|0|+1;m-52&17|+|59841311|59841332&offsets|0|0]|CAACACCAGTCGATGGGCTGTC</td><td>>MI0000077|hsa-mir-21&WithFlank&17|+|59841260|59841343</td><td>52</td><td>73</td><td>22</td><td></td><td>isomiR</td><td>NO</td><td>0.00</td><td>0.00</td><td>218.76</td><td>0.00</td></tr>
<tr><td>>MI0000077|hsa-mir-21&WithFlank&17|+|59841260|59841343@52.74.23&[MIMAT0004494&hsa-miR-21-3p&offsets|0|+2;m-52&17|+|59841311|59841332&offsets|0|+1]|CAACACCAGTCGATGGGCTGTCT</td><td>>MI0000077|hsa-mir-21&WithFlank&17|+|59841260|59841343</td><td>52</td><td>74</td><td>23</td><td></td><td>isomiR</td><td>NO</td><td>0.00</td><td>0.00</td><td>108.89</td><td>0.00</td></tr>
<tr><td>>MI0000078|hsa-mir-22&WithFlank&17|-|1713897|1713993@21.41.21&[MIMAT0004495&hsa-miR-22-5p&offsets|0|-1;m-217&17|-|1713953|1713973&offsets|0|0]|AGTTCTTCAGTGGCAAGCTTT</td><td>>MI0000078|hsa-mir-22&WithFlank&17|-|1713897|1713993</td><td>21</td><td>41</td><td>21</td><td></td><td>isomiR</td><td>NO</td><td>50.19</td><td>0.00</td><td>345.12</td><td>0.00</td></tr>
<tr><td>>MI0000078|hsa-mir-22&WithFlank&17|-|1713897|1713993@21.42.22&[MIMAT0004495&hsa-miR-22-5p&offsets|0|0;m-217&17|-|1713953|1713973&offsets|0|+1]|AGTTCTTCAGTGGCAAGCTTTA</td><td>>MI0000078|hsa-mir-22&WithFlank&17|-|1713897|1713993</td><td>21</td><td>42</td><td>22</td><td></td><td>isomiR</td><td>NO</td><td>0.00</td><td>0.00</td><td>123.88</td><td>0.00</td></tr>
<tr><td>>MI0000078|hsa-mir-22&WithFlank&17|-|1713897|1713993@59.78.20&[MIMAT0000077&hsa-miR-22-3p&offsets|0|-2;m-2&17|-|1713914|1713935&offsets|0|-2]|AAGCTGCCAGTTGAAGAACT</td><td>>MI0000078|hsa-mir-22&WithFlank&17|-|1713897|1713993</td><td>59</td><td>78</td><td>20</td><td></td><td>isomiR</td><td>NO</td><td>24.62</td><td>0.00</td><td>60.12</td><td>10.73</td></tr>
<tr><td>>MI0000078|hsa-mir-22&WithFlank&17|-|1713897|1713993@59.79.21(+1A)&[MIMAT0000077&hsa-miR-22-3p&offsets|0|-1(+1A);m-2&17|-|1713914|1713935&offsets|0|-1(+1A)]|AAGCTGCCAGTTGAAGAACTGA</td><td>>MI0000078|hsa-mir-22&WithFlank&17|-|1713897|1713993</td><td>59</td><td>79</td><td>21</td><td>(+1A)</td><td>isomiR</td><td>NO</td><td>0.00</td><td>0.00</td><td>141.44</td><td>0.00</td></tr>
<tr><td>>MI0000078|hsa-mir-22&WithFlank&17|-|1713897|1713993@59.79.21(+1C)&[MIMAT0000077&hsa-miR-22-3p&offsets|0|-1(+1C);m-2&17|-|1713914|1713935&offsets|0|-1(+1C)]|AAGCTGCCAGTTGAAGAACTGC</td><td>>MI0000078|hsa-mir-22&WithFlank&17|-|1713897|1713993</td><td>59</td><td>79</td><td>21</td><td>(+1C)</td><td>isomiR</td><td>NO</td><td>74.72</td><td>0.00</td><td>500.06</td><td>16.07</td></tr>
<tr><td>>MI0000078|hsa-mir-22&WithFlank&17|-|1713897|1713993@59.79.21(+1G)&[MIMAT0000077&hsa-miR-22-3p&offsets|0|-1(+1G);m-2&17|-|1713914|1713935&offsets|0|-1(+1G)]|AAGCTGCCAGTTGAAGAACTGG</td><td>>MI0000078|hsa-mir-22&WithFlank&17|-|1713897|1713993</td><td>59</td><td>79</td><td>21</td><td>(+1G)</td><td>isomiR</td><td>NO</td><td>0.00</td><td>0.00</td><td>63.58</td><td>0.00</td></tr>
<tr><td>>MI0000078|hsa-mir-22&WithFlank&17|-|1713897|1713993@59.80.22&[MIMAT0000077&hsa-miR-22-3p&offsets|0|0;m-2&17|-|1713914|1713935&offsets|0|0]|AAGCTGCCAGTTGAAGAACTGT</td><td>>MI0000078|hsa-mir-22&WithFlank&17|-|1713897|1713993</td><td>59</td><td>80</td><td>22</td><td></td><td>isomiR</td><td>NO</td><td>415.61</td><td>134.19</td><td>2234.91</td><td>143.82</td></tr>
<tr><td>>MI0000078|hsa-mir-22&WithFlank&17|-|1713897|1713993@59.80.22(+1A)&[MIMAT0000077&hsa-miR-22-3p&offsets|0|0(+1A);m-2&17|-|1713914|1713935&offsets|0|0(+1A)]|AAGCTGCCAGTTGAAGAACTGTA</td><td>>MI0000078|hsa-mir-22&WithFlank&17|-|1713897|1713993</td><td>59</td><td>80</td><td>22</td><td>(+1A)</td><td>isomiR</td><td>NO</td><td>0.00</td><td>0.00</td><td>87.77</td><td>0.00</td></tr>
<tr><td>>MI0000078|hsa-mir-22&WithFlank&17|-|1713897|1713993@59.81.23&[MIMAT0000077&hsa-miR-22-3p&offsets|0|+1;m-2&17|-|1713914|1713935&offsets|0|+1]|AAGCTGCCAGTTGAAGAACTGTT</td><td>>MI0000078|hsa-mir-22&WithFlank&17|-|1713897|1713993</td><td>59</td><td>81</td><td>23</td><td></td><td>isomiR</td><td>NO</td><td>0.00</td><td>0.00</td><td>117.34</td><td>0.00</td></tr>
<tr><td>>MI0000079|hsa-mir-23a&WithFlank&19|-|13836581|13836665@54.71.18&[MIMAT0000078&hsa-miR-23a-3p&offsets|+3|0;m-31&19|-|13836595|13836615&offsets|+3|0]|ACATTGCCAGGGATTTCC</td><td>>MI0000079|hsa-mir-23a&WithFlank&19|-|13836581|13836665</td><td>54</td><td>71</td><td>18</td><td></td><td>isomiR</td><td>NO</td><td>50.02</td><td>135.55</td><td>89.41</td><td>0.00</td></tr>
<tr><td>>MI0000079|hsa-mir-23a&WithFlank&19|-|13836581|13836665@54.72.19&[MIMAT0000078&hsa-miR-23a-3p&offsets|+3|+1;m-31&19|-|13836595|13836615&offsets|+3|+1]|ACATTGCCAGGGATTTCCA</td><td>>MI0000079|hsa-mir-23a&WithFlank&19|-|13836581|13836665</td><td>54</td><td>72</td><td>19</td><td></td><td>isomiR</td><td>NO</td><td>56.29</td><td>124.83</td><td>60.03</td><td>0.00</td></tr>
<tr><td>>MI0000079|hsa-mir-23a&WithFlank&19|-|13836581|13836665@54.72.19(+1U)&[MIMAT0000078&hsa-miR-23a-3p&offsets|+3|+1(+1U);m-31&19|-|13836595|13836615&offsets|+3|+1(+1U)]|ACATTGCCAGGGATTTCCAT</td><td>>MI0000079|hsa-mir-23a&WithFlank&19|-|13836581|13836665</td><td>54</td><td>72</td><td>19</td><td>(+1U)</td><td>isomiR</td><td>NO</td><td>0.00</td><td>24.57</td><td>0.00</td><td>0.00</td></tr>
<tr><td>>MI0000079|hsa-mir-23a&WithFlank&19|-|13836581|13836665@54.72.19(+2U)&[MIMAT0000078&hsa-miR-23a-3p&offsets|+3|+1(+2U);m-31&19|-|13836595|13836615&offsets|+3|+1(+2U)]|ACATTGCCAGGGATTTCCATT</td><td>>MI0000079|hsa-mir-23a&WithFlank&19|-|13836581|13836665</td><td>54</td><td>72</td><td>19</td><td>(+2U)</td><td>isomiR</td><td>NO</td><td>0.00</td><td>12.88</td><td>0.00</td><td>0.00</td></tr>
</table>

### MDA-MB-231

| | | | | | | | | | | | |
|---|---|---|---|---|---|---|---|---|---|---|---|
| >MI0000079\|hsa-mir-23a&WithFlank&19\|-\|13836581\|13836665@54.73.20&[MIMAT0000078&hsa-miR-23a-3p&offsets\|+3\|+2;m-31&19\|-\|13836595\|13836615&offsets\|+3\|+2]\|ACATTGCCAGGGATTTCCAA | >MI0000079\|hsa-mir-23a&WithFlank&19\|-\|13836581\|13836665 | 54 | 73 | 20 | | isomiR | NO | 45.08 | 70.73 | 0.00 | 0.00 |
| >MI0000079\|hsa-mir-23a&WithFlank&19\|-\|13836581\|13836665@54.73.20(+1U)&[MIMAT0000078&hsa-miR-23a-3p&offsets\|+3\|+2(+1U);m-31&19\|-\|13836595\|13836615&offsets\|+3\|+2(+1U)]\|ACATTGCCAGGGATTTCCAAT | >MI0000079\|hsa-mir-23a&WithFlank&19\|-\|13836581\|13836665 | 54 | 73 | 20 | (+1U) | isomiR | NO | 25.11 | 89.46 | 0.00 | 10.48 |
| >MI0000081\|hsa-mir-24-2&WithFlank&19\|-\|13836281\|13836365@56.74.19(+1A);MI0000080\|hsa-mir-24-1&WithFlank&9\|+\|95086015\|95086094@50.68.19(+1A)&[MIMAT0000080&hsa-miR-24-3p&offsets\|0\|-3(+1A);m-36&19\|-\|13836288\|13836310&offsets\|0\|-4(+1A)];[MIMAT0000080_1&hsa-miR-24-3p&offsets\|0\|-3(+1A);m-35&9\|+\|95086064\|95086086&offsets\|0\|-4(+1A)]\|TGGCTCAGTTCAGCAGGAAA | >MI0000081\|hsa-mir-24-2&WithFlank&19\|-\|13836281\|13836365 | 56 | 74 | 19 | (+1A) | isomiR | NO | 0.00 | 0.00 | 62.06 | 0.00 |
| >MI0000081\|hsa-mir-24-2&WithFlank&19\|-\|13836281\|13836365@56.74.19(+1U);MI0000080\|hsa-mir-24-1&WithFlank&9\|+\|95086015\|95086094@50.68.19(+1U)&[MIMAT0000080&hsa-miR-24-3p&offsets\|0\|-3(+1U);m-36&19\|-\|13836288\|13836310&offsets\|0\|-4(+1U)];[MIMAT0000080_1&hsa-miR-24-3p&offsets\|0\|-3(+1U);m-35&9\|+\|95086064\|95086086&offsets\|0\|-4(+1U)]\|TGGCTCAGTTCAGCAGGAAT | >MI0000081\|hsa-mir-24-2&WithFlank&19\|-\|13836281\|13836365 | 56 | 74 | 19 | (+1U) | isomiR | NO | 24.70 | 0.00 | 342.65 | 11.73 |
| >MI0000081\|hsa-mir-24-2&WithFlank&19\|-\|13836281\|13836365@56.75.20;MI0000080\|hsa-mir-24-1&WithFlank&9\|+\|95086015\|95086094@50.69.20&[MIMAT0000080&hsa-miR-24-3p&offsets\|0\|-2;m-36&19\|-\|13836288\|13836310&offsets\|0\|-3];[MIMAT0000080_1&hsa-miR-24-3p&offsets\|0\|-2;m-35&9\|+\|95086064\|95086086&offsets\|0\|-3]\|TGGCTCAGTTCAGCAGGAAC | >MI0000081\|hsa-mir-24-2&WithFlank&19\|-\|13836281\|13836365 | 56 | 75 | 20 | | isomiR | NO | 438.87 | 201.41 | 1944.44 | 153.62 |
| >MI0000081\|hsa-mir-24-2&WithFlank&19\|-\|13836281\|13836365@56.76.21(+1A);MI0000080\|hsa-mir-24-1&WithFlank&9\|+\|95086015\|95086094@50.70.21(+1A)&[MIMAT0000080&hsa-miR-24-3p&offsets\|0\|-1(+1A);m-36&19\|-\|13836288\|13836310&offsets\|0\|-2(+1A)];[MIMAT0000080_1&hsa-miR-24-3p&offsets\|0\|-1(+1A);m-35&9\|+\|95086064\|95086086&offsets\|0\|-2(+1A)]\|TGGCTCAGTTCAGCAGGAACAA | >MI0000081\|hsa-mir-24-2&WithFlank&19\|-\|13836281\|13836365 | 56 | 76 | 21 | (+1A) | isomiR | NO | 64.29 | 0.00 | 543.91 | 13.46 |
| >MI0000081\|hsa-mir-24-2&WithFlank&19\|-\|13836281\|13836365@56.76.21(+1U);MI0000080\|hsa-mir-24-1&WithFlank&9\|+\|95086015\|95086094@50.70.21(+1U)&[MIMAT0000080&hsa-miR-24-3p&offsets\|0\|-1(+1U);m-36&19\|-\|13836288\|13836310&offsets\|0\|-2(+1U)];[MIMAT0000080_1&hsa-miR-24-3p&offsets\|0\|-1(+1U);m-35&9\|+\|95086064\|95086086&offsets\|0\|-2(+1U)]\|TGGCTCAGTTCAGCAGGAACAT | >MI0000081\|hsa-mir-24-2&WithFlank&19\|-\|13836281\|13836365 | 56 | 76 | 21 | (+1U) | isomiR | NO | 0.00 | 0.00 | 100.97 | 0.00 |
| >MI0000081\|hsa-mir-24-2&WithFlank&19\|-\|13836281\|13836365@56.76.21;MI0000080\|hsa-mir-24-1&WithFlank&9\|+\|95086015\|95086094@50.70.21&[MIMAT0000080&hsa-miR-24-3p&offsets\|0\|-1;m-36&19\|-\|13836288\|13836310&offsets\|0\|-2];[MIMAT0000080_1&hsa-miR-24-3p&offsets\|0\|-1;m-35&9\|+\|95086064\|95086086&offsets\|0\|-2]\|TGGCTCAGTTCAGCAGGAACA | >MI0000081\|hsa-mir-24-2&WithFlank&19\|-\|13836281\|13836365 | 56 | 76 | 21 | | isomiR | NO | 139.37 | 34.25 | 532.72 | 62.58 |
| >MI0000081\|hsa-mir-24-2&WithFlank&19\|-\|13836281\|13836365@56.77.22(+1A);MI0000080\|hsa-mir-24-1&WithFlank&9\|+\|95086015\|95086094@50.71.22(+1A)&[MIMAT0000080&hsa-miR-24-3p&offsets\|0\|0(+1A);m-36&19\|-\|13836288\|13836310&offsets\|0\|-1(+1A)];[MIMAT0000080_1&hsa-miR-24-3p&offsets\|0\|0(+1A);m-35&9\|+\|95086064\|95086086&offsets\|0\|-1(+1A)]\|TGGCTCAGTTCAGCAGGAACAGA | >MI0000081\|hsa-mir-24-2&WithFlank&19\|-\|13836281\|13836365 | 56 | 77 | 22 | (+1A) | isomiR | NO | 0.00 | 0.00 | 89.80 | 0.00 |
| >MI0000081\|hsa-mir-24-2&WithFlank&19\|-\|13836281\|13836365@56.77.22(+1C);MI0000080\|hsa-mir-24-1&WithFlank&9\|+\|95086015\|95086094@50.71.22(+1C)&[MIMAT0000080&hsa-miR-24-3p&offsets\|0\|0(+1C);m-36&19\|-\|13836288\|13836310&offsets\|0\|-1(+1C)];[MIMAT0000080_1&hsa-miR-24-3p&offsets\|0\|0(+1C);m-35&9\|+\|95086064\|95086086&offsets\|0\|-1(+1C)]\|TGGCTCAGTTCAGCAGGAACAGC | >MI0000081\|hsa-mir-24-2&WithFlank&19\|-\|13836281\|13836365 | 56 | 77 | 22 | (+1C) | isomiR | NO | 51.81 | 0.00 | 185.06 | 13.48 |
| >MI0000081\|hsa-mir-24-2&WithFlank&19\|-\|13836281\|13836365@56.77.22(+1U);MI0000080\|hsa-mir-24-1&WithFlank&9\|+\|95086015\|95086094@50.71.22(+1U)&[MIMAT0000080&hsa-miR-24-3p&offsets\|0\|0(+1U);m-36&19\|-\|13836288\|13836310&offsets\|0\|-1(+1U)];[MIMAT0000080_1&hsa-miR-24-3p&offsets\|0\|0(+1U);m-35&9\|+\|95086064\|95086086&offsets\|0\|-1(+1U)]\|TGGCTCAGTTCAGCAGGAACAGT | >MI0000081\|hsa-mir-24-2&WithFlank&19\|-\|13836281\|13836365 | 56 | 77 | 22 | (+1U) | isomiR | NO | 316.02 | 76.44 | 881.19 | 124.52 |
| >MI0000081\|hsa-mir-24-2&WithFlank&19\|-\|13836281\|13836365@56.77.22;MI0000080\|hsa-mir-24-1&WithFlank&9\|+\|95086015\|95086094@50.71.22&[MIMAT0000080&hsa-miR-24-3p&offsets\|0\|0;m-36&19\|-\|13836288\|13836310&offsets\|0\|-1];[MIMAT0000080_1&hsa-miR-24-3p&offsets\|0\|0;m-35&9\|+\|95086064\|95086086&offsets\|0\|-1]\|TGGCTCAGTTCAGCAGGAACAG | >MI0000081\|hsa-mir-24-2&WithFlank&19\|-\|13836281\|13836365 | 56 | 77 | 22 | | isomiR | YES | 506.39 | 182.93 | 2562.31 | 133.52 |
| >MI0000081\|hsa-mir-24-2&WithFlank&19\|-\|13836281\|13836365@56.78.23;MI0000080\|hsa-mir-24-1&WithFlank&9\|+\|95086015\|95086094@50.72.23&[MIMAT0000080&hsa-miR-24-3p&offsets\|0\|+1;m-36&19\|-\|13836288\|13836310&offsets\|0\|0];[MIMAT0000080_1&hsa-miR-24-3p&offsets\|0\|+1;m-35&9\|+\|95086064\|95086086&offsets\|0\|0]\|TGGCTCAGTTCAGCAGGAACAGG | >MI0000081\|hsa-mir-24-2&WithFlank&19\|-\|13836281\|13836365 | 56 | 78 | 23 | | isomiR | NO | 0.00 | 0.00 | 11.14 | 0.00 |
| >MI0000082\|hsa-mir-25&WithFlank&7\|-\|100093554\|100093649@58.77.20&[MIMAT0000081&hsa-miR-25-3p&offsets\|0\|-2;m-23&7\|-\|100093571\|100093592&offsets\|0\|-2]\|CATTGCACTTGTCTCGGTCT | >MI0000082\|hsa-mir-25&WithFlank&7\|-\|100093554\|100093649 | 58 | 77 | 20 | | isomiR | NO | 0.00 | 0.00 | 16.92 | 0.00 |
| >MI0000082\|hsa-mir-25&WithFlank&7\|-\|100093554\|100093649@58.77.20(+1A)&[MIMAT0000081&hsa-miR-25-3p&offsets\|0\|-2(+1A);m-23&7\|-\|100093571\|100093592&offsets\|0\|-2(+1A)]\|CATTGCACTTGTCTCGGTCTA | >MI0000082\|hsa-mir-25&WithFlank&7\|-\|100093554\|100093649 | 58 | 77 | 20 | (+1A) | isomiR | NO | 10.60 | 0.00 | 120.65 | 0.00 |
| >MI0000082\|hsa-mir-25&WithFlank&7\|-\|100093554\|100093649@58.78.21&[MIMAT0000081&hsa-miR-25-3p&offsets\|0\|-1;m-23&7\|-\|100093571\|100093592&offsets\|0\|-1]\|CATTGCACTTGTCTCGGTCTG | >MI0000082\|hsa-mir-25&WithFlank&7\|-\|100093554\|100093649 | 58 | 78 | 21 | | isomiR | NO | 258.70 | 210.85 | 526.17 | 81.72 |
| >MI0000082\|hsa-mir-25&WithFlank&7\|-\|100093554\|100093649@58.78.21(+1C)&[MIMAT0000081&hsa-miR-25-3p&offsets\|0\|-1(+1C);m-23&7\|-\|100093571\|100093592&offsets\|0\|-1(+1C)]\|CATTGCACTTGTCTCGGTCTGC | >MI0000082\|hsa-mir-25&WithFlank&7\|-\|100093554\|100093649 | 58 | 78 | 21 | (+1C) | isomiR | NO | 0.00 | 0.00 | 16.01 | 0.00 |

### MDA-MB-231

<table>
<tr><td>>MI0000082|hsa-mir-25&WithFlank&7|-|100093554|100093649@58.78.21(+1G)&[MIMAT0000081&hsa-miR-25-3p&offsets|0|-1(+1G);m-23&7|-|100093571|100093592&offsets|0|-1(+1G)]|CATTGCACTTGTCTCGGTCTGG</td><td>>MI0000082|hsa-mir-25&WithFlank&7|-|100093554|100093649</td><td>58</td><td>78</td><td>21</td><td>(+1G)</td><td>isomiR</td><td>NO</td><td>64.31</td><td>0.00</td><td>31.41</td><td>0.00</td></tr>
<tr><td>>MI0000082|hsa-mir-25&WithFlank&7|-|100093554|100093649@58.78.21(+1U)&[MIMAT0000081&hsa-miR-25-3p&offsets|0|-1(+1U);m-23&7|-|100093571|100093592&offsets|0|-1(+1U)]|CATTGCACTTGTCTCGGTCTGT</td><td>>MI0000082|hsa-mir-25&WithFlank&7|-|100093554|100093649</td><td>58</td><td>78</td><td>21</td><td>(+1U)</td><td>isomiR</td><td>NO</td><td>99.98</td><td>93.15</td><td>194.30</td><td>18.65</td></tr>
<tr><td>>MI0000082|hsa-mir-25&WithFlank&7|-|100093554|100093649@58.79.22&[MIMAT0000081&hsa-miR-25-3p&offsets|0|0;m-23&7|-|100093571|100093592&offsets|0|0]|CATTGCACTTGTCTCGGTCTGA</td><td>>MI0000082|hsa-mir-25&WithFlank&7|-|100093554|100093649</td><td>58</td><td>79</td><td>22</td><td></td><td>isomiR</td><td>YES</td><td>1456.25</td><td>1075.33</td><td>1377.95</td><td>242.27</td></tr>
<tr><td>>MI0000082|hsa-mir-25&WithFlank&7|-|100093554|100093649@58.79.22(+1A)&[MIMAT0000081&hsa-miR-25-3p&offsets|0|0(+1A);m-23&7|-|100093571|100093592&offsets|0|0(+1A)]|CATTGCACTTGTCTCGGTCTGAA</td><td>>MI0000082|hsa-mir-25&WithFlank&7|-|100093554|100093649</td><td>58</td><td>79</td><td>22</td><td>(+1A)</td><td>isomiR</td><td>NO</td><td>77.15</td><td>30.19</td><td>14.27</td><td>0.00</td></tr>
<tr><td>>MI0000082|hsa-mir-25&WithFlank&7|-|100093554|100093649@58.79.22(+1U)&[MIMAT0000081&hsa-miR-25-3p&offsets|0|0(+1U);m-23&7|-|100093571|100093592&offsets|0|0(+1U)]|CATTGCACTTGTCTCGGTCTGAT</td><td>>MI0000082|hsa-mir-25&WithFlank&7|-|100093554|100093649</td><td>58</td><td>79</td><td>22</td><td>(+1U)</td><td>isomiR</td><td>NO</td><td>60.92</td><td>70.79</td><td>35.51</td><td>12.04</td></tr>
<tr><td>>MI0000084|hsa-mir-26b&WithFlank&2|+|218402640|218402728@18.38.21&[MIMAT0000083&hsa-miR-26b-5p&offsets|0|0;m-53&2|+|218402657|218402678&offsets|0|-1]|TTCAAGTAATTCAGGATAGGT</td><td>>MI0000084|hsa-mir-26b&WithFlank&2|+|218402640|218402728</td><td>18</td><td>38</td><td>21</td><td></td><td>isomiR</td><td>NO</td><td>50.14</td><td>0.00</td><td>171.41</td><td>0.00</td></tr>
<tr><td>>MI0000084|hsa-mir-26b&WithFlank&2|+|218402640|218402728@18.38.21(+1A)&[MIMAT0000083&hsa-miR-26b-5p&offsets|0|0(+1A);m-53&2|+|218402657|218402678&offsets|0|-1(+1A)]|TTCAAGTAATTCAGGATAGGTA</td><td>>MI0000084|hsa-mir-26b&WithFlank&2|+|218402640|218402728</td><td>18</td><td>38</td><td>21</td><td>(+1A)</td><td>isomiR</td><td>NO</td><td>0.00</td><td>0.00</td><td>12.15</td><td>0.00</td></tr>
<tr><td>>MI0000084|hsa-mir-26b&WithFlank&2|+|218402640|218402728@18.38.21(+1C)&[MIMAT0000083&hsa-miR-26b-5p&offsets|0|0(+1C);m-53&2|+|218402657|218402678&offsets|0|-1(+1C)]|TTCAAGTAATTCAGGATAGGTC</td><td>>MI0000084|hsa-mir-26b&WithFlank&2|+|218402640|218402728</td><td>18</td><td>38</td><td>21</td><td>(+1C)</td><td>isomiR</td><td>NO</td><td>0.00</td><td>0.00</td><td>100.41</td><td>10.56</td></tr>
<tr><td>>MI0000084|hsa-mir-26b&WithFlank&2|+|218402640|218402728@18.39.22&[MIMAT0000083&hsa-miR-26b-5p&offsets|0|+1;m-53&2|+|218402657|218402678&offsets|0|0]|TTCAAGTAATTCAGGATAGGTT</td><td>>MI0000084|hsa-mir-26b&WithFlank&2|+|218402640|218402728</td><td>18</td><td>39</td><td>22</td><td></td><td>isomiR</td><td>NO</td><td>231.58</td><td>101.67</td><td>455.99</td><td>96.86</td></tr>
<tr><td>>MI0000084|hsa-mir-26b&WithFlank&2|+|218402640|218402728@18.39.22(+1U)&[MIMAT0000083&hsa-miR-26b-5p&offsets|0|+1(+1U);m-53&2|+|218402657|218402678&offsets|0|0(+1U)]|TTCAAGTAATTCAGGATAGGTTT</td><td>>MI0000084|hsa-mir-26b&WithFlank&2|+|218402640|218402728</td><td>18</td><td>39</td><td>22</td><td>(+1U)</td><td>isomiR</td><td>NO</td><td>0.00</td><td>0.00</td><td>135.42</td><td>0.00</td></tr>
<tr><td>>MI0000085|hsa-mir-27a&WithFlank&19|-|13836434|13836523@16.36.21&[MIMAT0004501&hsa-miR-27a-5p&offsets|0|-1;m-270&19|-|13836487|13836508&offsets|0|-1]|AGGGCTTAGCTGCTTGTGAGC</td><td>>MI0000085|hsa-mir-27a&WithFlank&19|-|13836434|13836523</td><td>16</td><td>36</td><td>21</td><td></td><td>isomiR</td><td>NO</td><td>0.00</td><td>70.66</td><td>15.18</td><td>0.00</td></tr>
<tr><td>>MI0000085|hsa-mir-27a&WithFlank&19|-|13836434|13836523@16.37.22&[MIMAT0004501&hsa-miR-27a-5p&offsets|0|0;m-270&19|-|13836487|13836508&offsets|0|0]|AGGGCTTAGCTGCTTGTGAGCA</td><td>>MI0000085|hsa-mir-27a&WithFlank&19|-|13836434|13836523</td><td>16</td><td>37</td><td>22</td><td></td><td>isomiR</td><td>NO</td><td>0.00</td><td>91.44</td><td>0.00</td><td>0.00</td></tr>
<tr><td>>MI0000085|hsa-mir-27a&WithFlank&19|-|13836434|13836523@57.74.18(+1A);MI0000440|hsa-mir-27b&WithFlank&9|+|95085439|95085547@67.84.18(+1A)&[MIMAT0000084&hsa-miR-27a-3p&offsets|0|-3(+1A);m-43&19|-|13836447|13836467&offsets|0|-3(+1A)];[MIMAT0000419&hsa-miR-27b-3p&offsets|0|-3(+1A);m-39&9|+|95085505|95085525&offsets|0|-3(+1A)]|TTCACAGTGGCTAAGTTCA</td><td>>MI0000085|hsa-mir-27a&WithFlank&19|-|13836434|13836523</td><td>57</td><td>74</td><td>18</td><td>(+1A)</td><td>isomiR</td><td>NO</td><td>0.00</td><td>0.00</td><td>15.37</td><td>0.00</td></tr>
<tr><td>>MI0000085|hsa-mir-27a&WithFlank&19|-|13836434|13836523@57.74.18;MI0000440|hsa-mir-27b&WithFlank&9|+|95085439|95085547@67.84.18&[MIMAT0000084&hsa-miR-27a-3p&offsets|0|-3;m-43&19|-|13836447|13836467&offsets|0|-3];[MIMAT0000419&hsa-miR-27b-3p&offsets|0|-3;m-39&9|+|95085505|95085525&offsets|0|-3]|TTCACAGTGGCTAAGTTC</td><td>>MI0000085|hsa-mir-27a&WithFlank&19|-|13836434|13836523</td><td>57</td><td>74</td><td>18</td><td></td><td>isomiR</td><td>NO</td><td>45.40</td><td>0.00</td><td>183.77</td><td>0.00</td></tr>
<tr><td>>MI0000085|hsa-mir-27a&WithFlank&19|-|13836434|13836523@57.75.19&[MIMAT0000084&hsa-miR-27a-3p&offsets|0|-2;m-43&19|-|13836447|13836467&offsets|0|-2]|TTCACAGTGGCTAAGTTCC</td><td>>MI0000085|hsa-mir-27a&WithFlank&19|-|13836434|13836523</td><td>57</td><td>75</td><td>19</td><td></td><td>isomiR</td><td>NO</td><td>49.97</td><td>0.00</td><td>128.93</td><td>0.00</td></tr>
<tr><td>>MI0000085|hsa-mir-27a&WithFlank&19|-|13836434|13836523@57.75.19(+1A)&[MIMAT0000084&hsa-miR-27a-3p&offsets|0|-2(+1A);m-43&19|-|13836447|13836467&offsets|0|-2(+1A)]|TTCACAGTGGCTAAGTTCCA</td><td>>MI0000085|hsa-mir-27a&WithFlank&19|-|13836434|13836523</td><td>57</td><td>75</td><td>19</td><td>(+1A)</td><td>isomiR</td><td>NO</td><td>106.31</td><td>13.51</td><td>306.89</td><td>67.39</td></tr>
<tr><td>>MI0000085|hsa-mir-27a&WithFlank&19|-|13836434|13836523@57.75.19(+1C)&[MIMAT0000084&hsa-miR-27a-3p&offsets|0|-2(+1C);m-43&19|-|13836447|13836467&offsets|0|-2(+1C)]|TTCACAGTGGCTAAGTTCCC</td><td>>MI0000085|hsa-mir-27a&WithFlank&19|-|13836434|13836523</td><td>57</td><td>75</td><td>19</td><td>(+1C)</td><td>isomiR</td><td>NO</td><td>0.00</td><td>0.00</td><td>12.14</td><td>0.00</td></tr>
<tr><td>>MI0000085|hsa-mir-27a&WithFlank&19|-|13836434|13836523@57.75.19(+1U)&[MIMAT0000084&hsa-miR-27a-3p&offsets|0|-2(+1U);m-43&19|-|13836447|13836467&offsets|0|-2(+1U)]|TTCACAGTGGCTAAGTTCCT</td><td>>MI0000085|hsa-mir-27a&WithFlank&19|-|13836434|13836523</td><td>57</td><td>75</td><td>19</td><td>(+1U)</td><td>isomiR</td><td>NO</td><td>0.00</td><td>0.00</td><td>60.92</td><td>0.00</td></tr>
<tr><td>>MI0000085|hsa-mir-27a&WithFlank&19|-|13836434|13836523@57.75.19(+3U)&[MIMAT0000084&hsa-miR-27a-3p&offsets|0|-2(+3U);m-43&19|-|13836447|13836467&offsets|0|-2(+3U)]|TTCACAGTGGCTAAGTTCCTTT</td><td>>MI0000085|hsa-mir-27a&WithFlank&19|-|13836434|13836523</td><td>57</td><td>75</td><td>19</td><td>(+3U)</td><td>isomiR</td><td>NO</td><td>0.00</td><td>0.00</td><td>12.14</td><td>0.00</td></tr>
<tr><td>>MI0000085|hsa-mir-27a&WithFlank&19|-|13836434|13836523@57.76.20&[MIMAT0000084&hsa-miR-27a-3p&offsets|0|-1;m-43&19|-|13836447|13836467&offsets|0|-1]|TTCACAGTGGCTAAGTTCCG</td><td>>MI0000085|hsa-mir-27a&WithFlank&19|-|13836434|13836523</td><td>57</td><td>76</td><td>20</td><td></td><td>isomiR</td><td>NO</td><td>538.97</td><td>164.51</td><td>1053.03</td><td>198.53</td></tr>
<tr><td>>MI0000085|hsa-mir-27a&WithFlank&19|-|13836434|13836523@57.76.20(+1A)&[MIMAT0000084&hsa-miR-27a-3p&offsets|0|-1(+1A);m-43&19|-|13836447|13836467&offsets|0|-1(+1A)]|TTCACAGTGGCTAAGTTCCGA</td><td>>MI0000085|hsa-mir-27a&WithFlank&19|-|13836434|13836523</td><td>57</td><td>76</td><td>20</td><td>(+1A)</td><td>isomiR</td><td>NO</td><td>150.02</td><td>29.93</td><td>433.67</td><td>65.09</td></tr>
<tr><td>>MI0000085|hsa-mir-27a&WithFlank&19|-|13836434|13836523@57.76.20(+1G)&[MIMAT0000084&hsa-miR-27a-3p&offsets|0|-1(+1G);m-43&19|-|13836447|13836467&offsets|0|-1(+1G)]|TTCACAGTGGCTAAGTTCCGG</td><td>>MI0000085|hsa-mir-27a&WithFlank&19|-|13836434|13836523</td><td>57</td><td>76</td><td>20</td><td>(+1G)</td><td>isomiR</td><td>YES</td><td>0.00</td><td>0.00</td><td>13.11</td><td>0.00</td></tr>
<tr><td>>MI0000085|hsa-mir-27a&WithFlank&19|-|13836434|13836523@57.76.20(+1U)&[MIMAT0000084&hsa-miR-27a-3p&offsets|0|-1(+1U);m-43&19|-|13836447|13836467&offsets|0|-1(+1U)]|TTCACAGTGGCTAAGTTCCGT</td><td>>MI0000085|hsa-mir-27a&WithFlank&19|-|13836434|13836523</td><td>57</td><td>76</td><td>20</td><td>(+1U)</td><td>isomiR</td><td>NO</td><td>281.51</td><td>67.22</td><td>956.45</td><td>113.57</td></tr>
</table>

### MDA-MB-231

| | | | | | | | | | | | |
|---|---|---|---|---|---|---|---|---|---|---|---|
| >MI0000085\|hsa-mir-27a&WithFlank&19\|-\|13836434\|13836523@57.76.20(+2U)&[MIMAT0000084&hsa-miR-27a-3p&offsets\|0\|-1(+2U);m-43&19\|-\|13836447\|13836467&offsets\|0\|-1(+2U)]\|TTCACAGTGGCTAAGTTCCGTT | >MI0000085\|hsa-mir-27a&WithFlank&19\|-\|13836434\|13836523 | 57 | 76 | 20 | (+2U) | isomiR | NO | 112.91 | 13.41 | 416.44 | 59.62 |
| >MI0000085\|hsa-mir-27a&WithFlank&19\|-\|13836434\|13836523@57.76.20(+3U)&[MIMAT0000084&hsa-miR-27a-3p&offsets\|0\|-1(+3U);m-43&19\|-\|13836447\|13836467&offsets\|0\|-1(+3U)]\|TTCACAGTGGCTAAGTTCCGTTT | >MI0000085\|hsa-mir-27a&WithFlank&19\|-\|13836434\|13836523 | 57 | 76 | 20 | (+3U) | isomiR | NO | 61.13 | 0.00 | 244.79 | 17.66 |
| >MI0000085\|hsa-mir-27a&WithFlank&19\|-\|13836434\|13836523@57.76.20(+4U)&[MIMAT0000084&hsa-miR-27a-3p&offsets\|0\|-1(+4U);m-43&19\|-\|13836447\|13836467&offsets\|0\|-1(+4U)]\|TTCACAGTGGCTAAGTTCCGTTTT | >MI0000085\|hsa-mir-27a&WithFlank&19\|-\|13836434\|13836523 | 57 | 76 | 20 | (+4U) | isomiR | NO | 0.00 | 0.00 | 11.32 | 0.00 |
| >MI0000085\|hsa-mir-27a&WithFlank&19\|-\|13836434\|13836523@57.77.21&[MIMAT0000084&hsa-miR-27a-3p&offsets\|0\|0;m-43&19\|-\|13836447\|13836467&offsets\|0\|0]\|TTCACAGTGGCTAAGTTCCGC | >MI0000085\|hsa-mir-27a&WithFlank&19\|-\|13836434\|13836523 | 57 | 77 | 21 | | isomiR | NO | 95.87 | 0.00 | 1141.38 | 21.40 |
| >MI0000085\|hsa-mir-27a&WithFlank&19\|-\|13836434\|13836523@57.77.21(+1A)&[MIMAT0000084&hsa-miR-27a-3p&offsets\|0\|0(+1A);m-43&19\|-\|13836447\|13836467&offsets\|0\|0(+1A)]\|TTCACAGTGGCTAAGTTCCGCA | >MI0000085\|hsa-mir-27a&WithFlank&19\|-\|13836434\|13836523 | 57 | 77 | 21 | (+1A) | isomiR | NO | 0.00 | 0.00 | 16.29 | 0.00 |
| >MI0000085\|hsa-mir-27a&WithFlank&19\|-\|13836434\|13836523@57.77.21(+1U)&[MIMAT0000084&hsa-miR-27a-3p&offsets\|0\|0(+1U);m-43&19\|-\|13836447\|13836467&offsets\|0\|0(+1U)]\|TTCACAGTGGCTAAGTTCCGCT | >MI0000085\|hsa-mir-27a&WithFlank&19\|-\|13836434\|13836523 | 57 | 77 | 21 | (+1U) | isomiR | NO | 0.00 | 0.00 | 101.43 | 0.00 |
| >MI0000086\|hsa-mir-28&WithFlank&3\|+\|188688775\|188688872@20.40.21&[MIMAT0000085&hsa-miR-28-5p&offsets\|0\|-1;m-133&3\|+\|188688794\|188688815&offsets\|0\|-1]\|AAGGAGCTCACAGTCTATTGA | >MI0000086\|hsa-mir-28&WithFlank&3\|+\|188688775\|188688872 | 20 | 40 | 21 | | isomiR | NO | 0.00 | 0.00 | 31.75 | 0.00 |
| >MI0000086\|hsa-mir-28&WithFlank&3\|+\|188688775\|188688872@20.41.22&[MIMAT0000085&hsa-miR-28-5p&offsets\|0\|0;m-133&3\|+\|188688794\|188688815&offsets\|0\|0]\|AAGGAGCTCACAGTCTATTGAG | >MI0000086\|hsa-mir-28&WithFlank&3\|+\|188688775\|188688872 | 20 | 41 | 22 | | isomiR | NO | 0.00 | 0.00 | 124.63 | 0.00 |
| >MI0000086\|hsa-mir-28&WithFlank&3\|+\|188688775\|188688872@60.79.20(+1A)&[MIMAT0004502&hsa-miR-28-3p&offsets\|0\|-2(+1A);m-28&3\|+\|188688834\|188688855&offsets\|0\|-2(+1A)]\|CACTAGATTGTGAGCTCCTGA | >MI0000086\|hsa-mir-28&WithFlank&3\|+\|188688775\|188688872 | 60 | 79 | 20 | (+1A) | isomiR | NO | 0.00 | 0.00 | 13.40 | 0.00 |
| >MI0000086\|hsa-mir-28&WithFlank&3\|+\|188688775\|188688872@60.80.21&[MIMAT0004502&hsa-miR-28-3p&offsets\|0\|-1;m-28&3\|+\|188688834\|188688855&offsets\|0\|-1]\|CACTAGATTGTGAGCTCCTGG | >MI0000086\|hsa-mir-28&WithFlank&3\|+\|188688775\|188688872 | 60 | 80 | 21 | | isomiR | NO | 55.44 | 74.60 | 153.19 | 0.00 |
| >MI0000086\|hsa-mir-28&WithFlank&3\|+\|188688775\|188688872@60.80.21(+1U)&[MIMAT0004502&hsa-miR-28-3p&offsets\|0\|-1(+1U);m-28&3\|+\|188688834\|188688855&offsets\|0\|-1(+1U)]\|CACTAGATTGTGAGCTCCTGGT | >MI0000086\|hsa-mir-28&WithFlank&3\|+\|188688775\|188688872 | 60 | 80 | 21 | (+1U) | isomiR | NO | 0.00 | 0.00 | 12.78 | 0.00 |
| >MI0000086\|hsa-mir-28&WithFlank&3\|+\|188688775\|188688872@60.81.22&[MIMAT0004502&hsa-miR-28-3p&offsets\|0\|0;m-28&3\|+\|188688834\|188688855&offsets\|0\|0]\|CACTAGATTGTGAGCTCCTGGA | >MI0000086\|hsa-mir-28&WithFlank&3\|+\|188688775\|188688872 | 60 | 81 | 22 | | isomiR | NO | 172.60 | 306.08 | 350.92 | 55.96 |
| >MI0000086\|hsa-mir-28&WithFlank&3\|+\|188688775\|188688872@60.81.22(+1A)&[MIMAT0004502&hsa-miR-28-3p&offsets\|0\|0(+1A);m-28&3\|+\|188688834\|188688855&offsets\|0\|0(+1A)]\|CACTAGATTGTGAGCTCCTGGAA | >MI0000086\|hsa-mir-28&WithFlank&3\|+\|188688775\|188688872 | 60 | 81 | 22 | (+1A) | isomiR | NO | 0.00 | 32.63 | 61.75 | 0.00 |
| >MI0000086\|hsa-mir-28&WithFlank&3\|+\|188688775\|188688872@60.81.22(+1U)&[MIMAT0004502&hsa-miR-28-3p&offsets\|0\|0(+1U);m-28&3\|+\|188688834\|188688855&offsets\|0\|0(+1U)]\|CACTAGATTGTGAGCTCCTGGAT | >MI0000086\|hsa-mir-28&WithFlank&3\|+\|188688775\|188688872 | 60 | 81 | 22 | (+1U) | isomiR | NO | 0.00 | 0.00 | 34.02 | 0.00 |
| >MI0000086\|hsa-mir-28&WithFlank&3\|+\|188688775\|188688872@60.82.23&[MIMAT0004502&hsa-miR-28-3p&offsets\|0\|+1;m-28&3\|+\|188688834\|188688855&offsets\|0\|+1]\|CACTAGATTGTGAGCTCCTGGAG | >MI0000086\|hsa-mir-28&WithFlank&3\|+\|188688775\|188688872 | 60 | 82 | 23 | | isomiR | NO | 0.00 | 0.00 | 13.77 | 0.00 |
| >MI0000086\|hsa-mir-28&WithFlank&3\|+\|188688775\|188688872@60.82.23(+1U)&[MIMAT0004502&hsa-miR-28-3p&offsets\|0\|+1(+1U);m-28&3\|+\|188688834\|188688855&offsets\|0\|+1(+1U)]\|CACTAGATTGTGAGCTCCTGGAGT | >MI0000086\|hsa-mir-28&WithFlank&3\|+\|188688775\|188688872 | 60 | 82 | 23 | (+1U) | isomiR | NO | 0.00 | 0.00 | 65.03 | 0.00 |
| >MI0000086\|hsa-mir-28&WithFlank&3\|+\|188688775\|188688872@61.82.22&[MIMAT0004502&hsa-miR-28-3p&offsets\|+1\|+1;m-28&3\|+\|188688834\|188688855&offsets\|+1\|+1]\|ACTAGATTGTGAGCTCCTGGAG | >MI0000086\|hsa-mir-28&WithFlank&3\|+\|188688775\|188688872 | 61 | 82 | 22 | | isomiR | NO | 0.00 | 0.00 | 13.05 | 0.00 |
| >MI0000086\|hsa-mir-28&WithFlank&3\|+\|188688775\|188688872@61.82.22(+1U)&[MIMAT0004502&hsa-miR-28-3p&offsets\|+1\|+1(+1U);m-28&3\|+\|188688834\|188688855&offsets\|+1\|+1(+1U)]\|ACTAGATTGTGAGCTCCTGGAGT | >MI0000086\|hsa-mir-28&WithFlank&3\|+\|188688775\|188688872 | 61 | 82 | 22 | (+1U) | isomiR | NO | 0.00 | 0.00 | 58.91 | 0.00 |
| >MI0000086\|hsa-mir-28&WithFlank&3\|+\|188688775\|188688872@62.82.21(+1U)&[MIMAT0004502&hsa-miR-28-3p&offsets\|+2\|+1(+1U);m-28&3\|+\|188688834\|188688855&offsets\|+2\|+1(+1U)]\|CTAGATTGTGAGCTCCTGGAGT | >MI0000086\|hsa-mir-28&WithFlank&3\|+\|188688775\|188688872 | 62 | 82 | 21 | (+1U) | isomiR | NO | 0.00 | 0.00 | 35.88 | 0.00 |
| >am_MI0000086\|hsa-mir-28&WithFlank&3\|+\|188688775\|188688872@20.39.20&[MIMAT0000085&hsa-miR-28-5p&offsets\|0\|-2;m-133&3\|+\|188688794\|188688815&offsets\|0\|-2]\|AAGGAGCTCACAGTCTATTG | >MI0000086\|hsa-mir-28&WithFlank&3\|+\|188688775\|188688872.am | 20 | 39 | 20 | | isomiR | NO | 0.00 | 0.00 | 81.64 | 0.00 |
| >MI0000087\|hsa-mir-29a&WithFlank&7\|-\|130876741\|130876816@47.67.21&[MIMAT0000086&hsa-miR-29a-3p&offsets\|-1\|-2;m-26&7\|-\|130876749\|130876769&offsets\|-1\|-1]\|CTAGCACCATCTGAAATCGGT | >MI0000087\|hsa-mir-29a&WithFlank&7\|-\|130876741\|130876816 | 47 | 67 | 21 | | isomiR | NO | 0.00 | 0.00 | 56.78 | 0.00 |
| >MI0000087\|hsa-mir-29a&WithFlank&7\|-\|130876741\|130876816@47.67.21(+1A)&[MIMAT0000086&hsa-miR-29a-3p&offsets\|-1\|-2(+1A);m-26&7\|-\|130876749\|130876769&offsets\|-1\|-1(+1A)]\|CTAGCACCATCTGAAATCGGTA | >MI0000087\|hsa-mir-29a&WithFlank&7\|-\|130876741\|130876816 | 47 | 67 | 21 | (+1A) | isomiR | NO | 0.00 | 0.00 | 113.62 | 0.00 |
| >MI0000087\|hsa-mir-29a&WithFlank&7\|-\|130876741\|130876816@47.67.21(+1C)&[MIMAT0000086&hsa-miR-29a-3p&offsets\|-1\|-2(+1C);m-26&7\|-\|130876749\|130876769&offsets\|-1\|-1(+1C)]\|CTAGCACCATCTGAAATCGGTC | >MI0000087\|hsa-mir-29a&WithFlank&7\|-\|130876741\|130876816 | 47 | 67 | 21 | (+1C) | isomiR | NO | 92.31 | 0.00 | 366.77 | 18.41 |

### MDA-MB-231

| | | | | | | | | | | | |
|---|---|---|---|---|---|---|---|---|---|---|---|
| >MI0000087\|hsa-mir-29a&WithFlank&7\|-\|130876741\|130876816@47.67.21(+1G)&[MIMAT0000086&hsa-miR-29a-3p&offsets\|-1\|-2(+1G);m-26&7\|-\|130876749\|130876769&offsets\|-1\|-1(+1G)]\|CTAGCACCATCTGAAATCGGTG | >MI0000087\|hsa-mir-29a&WithFlank&7\|-\|130876741\|130876816 | 47 | 67 | 21 | (+1G) | isomiR | NO | 0.00 | 0.00 | 14.86 | 0.00 |
| >MI0000087\|hsa-mir-29a&WithFlank&7\|-\|130876741\|130876816@47.68.22&[MIMAT0000086&hsa-miR-29a-3p&offsets\|-1\|-1;m-26&7\|-\|130876749\|130876769&offsets\|-1\|0]\|CTAGCACCATCTGAAATCGGTT | >MI0000087\|hsa-mir-29a&WithFlank&7\|-\|130876741\|130876816 | 47 | 68 | 22 | | isomiR | NO | 577.66 | 172.42 | 1718.01 | 168.70 |
| >MI0000087\|hsa-mir-29a&WithFlank&7\|-\|130876741\|130876816@47.68.22(+1C)&[MIMAT0000086&hsa-miR-29a-3p&offsets\|-1\|-1(+1C);m-26&7\|-\|130876749\|130876769&offsets\|-1\|0(+1C)]\|CTAGCACCATCTGAAATCGGTTC | >MI0000087\|hsa-mir-29a&WithFlank&7\|-\|130876741\|130876816 | 47 | 68 | 22 | (+1C) | isomiR | NO | 0.00 | 0.00 | 82.81 | 0.00 |
| >MI0000087\|hsa-mir-29a&WithFlank&7\|-\|130876741\|130876816@47.68.22(+1U)&[MIMAT0000086&hsa-miR-29a-3p&offsets\|-1\|-1(+1U);m-26&7\|-\|130876749\|130876769&offsets\|-1\|0(+1U)]\|CTAGCACCATCTGAAATCGGTTT | >MI0000087\|hsa-mir-29a&WithFlank&7\|-\|130876741\|130876816 | 47 | 68 | 22 | (+1U) | isomiR | NO | 84.79 | 0.00 | 398.55 | 15.02 |
| >MI0000087\|hsa-mir-29a&WithFlank&7\|-\|130876741\|130876816@47.69.23&[MIMAT0000086&hsa-miR-29a-3p&offsets\|-1\|0;m-26&7\|-\|130876749\|130876769&offsets\|-1\|+1]\|CTAGCACCATCTGAAATCGGTTA | >MI0000087\|hsa-mir-29a&WithFlank&7\|-\|130876741\|130876816 | 47 | 69 | 23 | | isomiR | NO | 78.40 | 0.00 | 546.46 | 17.08 |
| >MI0000087\|hsa-mir-29a&WithFlank&7\|-\|130876741\|130876816@47.70.24&[MIMAT0000086&hsa-miR-29a-3p&offsets\|-1\|+1;m-26&7\|-\|130876749\|130876769&offsets\|-1\|+2]\|CTAGCACCATCTGAAATCGGTTAT | >MI0000087\|hsa-mir-29a&WithFlank&7\|-\|130876741\|130876816 | 47 | 70 | 24 | | isomiR | NO | 0.00 | 0.00 | 87.78 | 0.00 |
| >MI0000087\|hsa-mir-29a&WithFlank&7\|-\|130876741\|130876816@48.66.19&[MIMAT0000086&hsa-miR-29a-3p&offsets\|0\|-3;m-26&7\|-\|130876749\|130876769&offsets\|0\|-2]\|TAGCACCATCTGAAATCGG | >MI0000087\|hsa-mir-29a&WithFlank&7\|-\|130876741\|130876816 | 48 | 66 | 19 | | isomiR | NO | 0.00 | 0.00 | 12.91 | 0.00 |
| >MI0000087\|hsa-mir-29a&WithFlank&7\|-\|130876741\|130876816@48.67.20&[MIMAT0000086&hsa-miR-29a-3p&offsets\|0\|-2;m-26&7\|-\|130876749\|130876769&offsets\|0\|-1]\|TAGCACCATCTGAAATCGGT | >MI0000087\|hsa-mir-29a&WithFlank&7\|-\|130876741\|130876816 | 48 | 67 | 20 | | isomiR | NO | 0.00 | 0.00 | 185.14 | 0.00 |
| >MI0000087\|hsa-mir-29a&WithFlank&7\|-\|130876741\|130876816@48.67.20(+1A)&[MIMAT0000086&hsa-miR-29a-3p&offsets\|0\|-2(+1A);m-26&7\|-\|130876749\|130876769&offsets\|0\|-1(+1A)]\|TAGCACCATCTGAAATCGGTA | >MI0000087\|hsa-mir-29a&WithFlank&7\|-\|130876741\|130876816 | 48 | 67 | 20 | (+1A) | isomiR | NO | 0.00 | 0.00 | 164.69 | 0.00 |
| >MI0000087\|hsa-mir-29a&WithFlank&7\|-\|130876741\|130876816@48.67.20(+1C)&[MIMAT0000086&hsa-miR-29a-3p&offsets\|0\|-2(+1C);m-26&7\|-\|130876749\|130876769&offsets\|0\|-1(+1C)]\|TAGCACCATCTGAAATCGGTC | >MI0000087\|hsa-mir-29a&WithFlank&7\|-\|130876741\|130876816 | 48 | 67 | 20 | (+1C) | isomiR | NO | 52.05 | 0.00 | 544.29 | 12.95 |
| >MI0000087\|hsa-mir-29a&WithFlank&7\|-\|130876741\|130876816@48.67.20(+1G)&[MIMAT0000086&hsa-miR-29a-3p&offsets\|0\|-2(+1G);m-26&7\|-\|130876749\|130876769&offsets\|0\|-1(+1G)]\|TAGCACCATCTGAAATCGGTG | >MI0000087\|hsa-mir-29a&WithFlank&7\|-\|130876741\|130876816 | 48 | 67 | 20 | (+1G) | isomiR | NO | 0.00 | 0.00 | 54.79 | 0.00 |
| >MI0000087\|hsa-mir-29a&WithFlank&7\|-\|130876741\|130876816@48.68.21&[MIMAT0000086&hsa-miR-29a-3p&offsets\|0\|-1;m-26&7\|-\|130876749\|130876769&offsets\|0\|0]\|TAGCACCATCTGAAATCGGTT | >MI0000087\|hsa-mir-29a&WithFlank&7\|-\|130876741\|130876816 | 48 | 68 | 21 | | isomiR | NO | 318.31 | 131.89 | 2515.92 | 121.85 |
| >MI0000087\|hsa-mir-29a&WithFlank&7\|-\|130876741\|130876816@48.68.21(+1G)&[MIMAT0000086&hsa-miR-29a-3p&offsets\|0\|-1(+1G);m-26&7\|-\|130876749\|130876769&offsets\|0\|0(+1G)]\|TAGCACCATCTGAAATCGGTTG | >MI0000087\|hsa-mir-29a&WithFlank&7\|-\|130876741\|130876816 | 48 | 68 | 21 | (+1G) | isomiR | NO | 0.00 | 0.00 | 17.74 | 0.00 |
| >MI0000087\|hsa-mir-29a&WithFlank&7\|-\|130876741\|130876816@48.68.21(+1U)&[MIMAT0000086&hsa-miR-29a-3p&offsets\|0\|-1(+1U);m-26&7\|-\|130876749\|130876769&offsets\|0\|0(+1U)]\|TAGCACCATCTGAAATCGGTTT | >MI0000087\|hsa-mir-29a&WithFlank&7\|-\|130876741\|130876816 | 48 | 68 | 21 | (+1U) | isomiR | NO | 0.00 | 0.00 | 266.07 | 0.00 |
| >MI0000087\|hsa-mir-29a&WithFlank&7\|-\|130876741\|130876816@48.69.22&[MIMAT0000086&hsa-miR-29a-3p&offsets\|0\|0;m-26&7\|-\|130876749\|130876769&offsets\|0\|+1]\|TAGCACCATCTGAAATCGGTTA | >MI0000087\|hsa-mir-29a&WithFlank&7\|-\|130876741\|130876816 | 48 | 69 | 22 | | isomiR | YES | 293.72 | 74.69 | 2925.97 | 133.69 |
| >MI0000087\|hsa-mir-29a&WithFlank&7\|-\|130876741\|130876816@48.69.22(+1A)&[MIMAT0000086&hsa-miR-29a-3p&offsets\|0\|0(+1A);m-26&7\|-\|130876749\|130876769&offsets\|0\|+1(+1A)]\|TAGCACCATCTGAAATCGGTTAA | >MI0000087\|hsa-mir-29a&WithFlank&7\|-\|130876741\|130876816 | 48 | 69 | 22 | (+1A) | isomiR | NO | 0.00 | 0.00 | 75.96 | 0.00 |
| >MI0000087\|hsa-mir-29a&WithFlank&7\|-\|130876741\|130876816@48.70.23&[MIMAT0000086&hsa-miR-29a-3p&offsets\|0\|+1;m-26&7\|-\|130876749\|130876769&offsets\|0\|+2]\|TAGCACCATCTGAAATCGGTTAT | >MI0000087\|hsa-mir-29a&WithFlank&7\|-\|130876741\|130876816 | 48 | 70 | 23 | | isomiR | NO | 0.00 | 0.00 | 120.22 | 0.00 |
| >MI0000088\|hsa-mir-30a&WithFlank&6\|-\|71403545\|71403627@12.30.19&[MIMAT0000087&hsa-miR-30a-5p&offsets\|0\|-3;m-15&6\|-\|71403595\|71403616&offsets\|0\|-3]\|TGTAAACATCCTCGACTGG | >MI0000088\|hsa-mir-30a&WithFlank&6\|-\|71403545\|71403627 | 12 | 30 | 19 | | isomiR | NO | 0.00 | 0.00 | 16.02 | 0.00 |
| >MI0000088\|hsa-mir-30a&WithFlank&6\|-\|71403545\|71403627@12.30.19(+1G)&[MIMAT0000087&hsa-miR-30a-5p&offsets\|0\|-3(+1G);m-15&6\|-\|71403595\|71403616&offsets\|0\|-3(+1G)]\|TGTAAACATCCTCGACTGGG | >MI0000088\|hsa-mir-30a&WithFlank&6\|-\|71403545\|71403627 | 12 | 30 | 19 | (+1G) | isomiR | NO | 0.00 | 0.00 | 14.16 | 0.00 |
| >MI0000088\|hsa-mir-30a&WithFlank&6\|-\|71403545\|71403627@12.31.20&[MIMAT0000087&hsa-miR-30a-5p&offsets\|0\|-2;m-15&6\|-\|71403595\|71403616&offsets\|0\|-2]\|TGTAAACATCCTCGACTGGA | >MI0000088\|hsa-mir-30a&WithFlank&6\|-\|71403545\|71403627 | 12 | 31 | 20 | | isomiR | NO | 130.02 | 0.00 | 1156.15 | 18.93 |
| >MI0000088\|hsa-mir-30a&WithFlank&6\|-\|71403545\|71403627@12.32.21&[MIMAT0000087&hsa-miR-30a-5p&offsets\|0\|-1;m-15&6\|-\|71403595\|71403616&offsets\|0\|-1]\|TGTAAACATCCTCGACTGGAA | >MI0000088\|hsa-mir-30a&WithFlank&6\|-\|71403545\|71403627 | 12 | 32 | 21 | | isomiR | NO | 58.75 | 0.00 | 249.75 | 11.59 |
| >MI0000088\|hsa-mir-30a&WithFlank&6\|-\|71403545\|71403627@12.32.21(+1A)&[MIMAT0000087&hsa-miR-30a-5p&offsets\|0\|-1(+1A);m-15&6\|-\|71403595\|71403616&offsets\|0\|-1(+1A)]\|TGTAAACATCCTCGACTGGAAA | >MI0000088\|hsa-mir-30a&WithFlank&6\|-\|71403545\|71403627 | 12 | 32 | 21 | (+1A) | isomiR | NO | 0.00 | 0.00 | 145.18 | 0.00 |
| >MI0000088\|hsa-mir-30a&WithFlank&6\|-\|71403545\|71403627@12.33.22&[MIMAT0000087&hsa-miR-30a-5p&offsets\|0\|0;m-15&6\|-\|71403595\|71403616&offsets\|0\|0]\|TGTAAACATCCTCGACTGGAAG | >MI0000088\|hsa-mir-30a&WithFlank&6\|-\|71403545\|71403627 | 12 | 33 | 22 | | isomiR | NO | 180.71 | 29.21 | 1004.86 | 58.73 |
| >MI0000088\|hsa-mir-30a&WithFlank&6\|-\|71403545\|71403627@12.33.22(+1A)&[MIMAT0000087&hsa-miR-30a-5p&offsets\|0\|0(+1A);m-15&6\|-\|71403595\|71403616&offsets\|0\|0(+1A)]\|TGTAAACATCCTCGACTGGAAGA | >MI0000088\|hsa-mir-30a&WithFlank&6\|-\|71403545\|71403627 | 12 | 33 | 22 | (+1A) | isomiR | NO | 71.57 | 0.00 | 433.88 | 14.52 |
| >MI0000088\|hsa-mir-30a&WithFlank&6\|-\|71403545\|71403627@12.33.22(+1G)&[MIMAT0000087&hsa-miR-30a-5p&offsets\|0\|0(+1G);m-15&6\|-\|71403595\|71403616&offsets\|0\|0(+1G)]\|TGTAAACATCCTCGACTGGAAGG | >MI0000088\|hsa-mir-30a&WithFlank&6\|-\|71403545\|71403627 | 12 | 33 | 22 | (+1G) | isomiR | NO | 0.00 | 0.00 | 11.15 | 0.00 |
| >MI0000088\|hsa-mir-30a&WithFlank&6\|-\|71403545\|71403627@12.33.22(+1U)&[MIMAT0000087&hsa-miR-30a-5p&offsets\|0\|0(+1U);m-15&6\|-\|71403595\|71403616&offsets\|0\|0(+1U)]\|TGTAAACATCCTCGACTGGAAGT | >MI0000088\|hsa-mir-30a&WithFlank&6\|-\|71403545\|71403627 | 12 | 33 | 22 | (+1U) | isomiR | NO | 225.26 | 60.19 | 1453.13 | 91.48 |

### MDA-MB-231

<table>
<tr><td>>MI0000088|hsa-mir-30a&WithFlank&6|-|71403545|71403627@12.33.22(+2U)&[MIMAT0000087&hsa-miR-30a-5p&offsets|0|0(+2U);m-15&6|-|71403595|71403616&offsets|0|0(+2U)]|TGTAAACATCCTCGACTGGAAGTT</td><td>>MI0000088|hsa-mir-30a&WithFlank&6|-|71403545|71403627</td><td>12</td><td>33</td><td>22</td><td>(+2U)</td><td>isomiR</td><td>NO</td><td>0.00</td><td>0.00</td><td>61.94</td><td>0.00</td></tr>
<tr><td>>MI0000088|hsa-mir-30a&WithFlank&6|-|71403545|71403627@12.34.23&[MIMAT0000087&hsa-miR-30a-5p&offsets|0|+1;m-15&6|-|71403595|71403616&offsets|0|+1]|TGTAAACATCCTCGACTGGAAGC</td><td>>MI0000088|hsa-mir-30a&WithFlank&6|-|71403545|71403627</td><td>12</td><td>34</td><td>23</td><td></td><td>isomiR</td><td>YES</td><td>871.62</td><td>241.72</td><td>3384.28</td><td>227.24</td></tr>
<tr><td>>MI0000088|hsa-mir-30a&WithFlank&6|-|71403545|71403627@12.34.23(+1A)&[MIMAT0000087&hsa-miR-30a-5p&offsets|0|+1(+1A);m-15&6|-|71403595|71403616&offsets|0|+1(+1A)]|TGTAAACATCCTCGACTGGAAGCA</td><td>>MI0000088|hsa-mir-30a&WithFlank&6|-|71403545|71403627</td><td>12</td><td>34</td><td>23</td><td>(+1A)</td><td>isomiR</td><td>NO</td><td>289.59</td><td>113.41</td><td>2533.05</td><td>120.92</td></tr>
<tr><td>>MI0000088|hsa-mir-30a&WithFlank&6|-|71403545|71403627@12.34.23(+1C)&[MIMAT0000087&hsa-miR-30a-5p&offsets|0|+1(+1C);m-15&6|-|71403595|71403616&offsets|0|+1(+1C)]|TGTAAACATCCTCGACTGGAAGCC</td><td>>MI0000088|hsa-mir-30a&WithFlank&6|-|71403545|71403627</td><td>12</td><td>34</td><td>23</td><td>(+1C)</td><td>isomiR</td><td>YES</td><td>477.90</td><td>165.85</td><td>3420.99</td><td>168.47</td></tr>
<tr><td>>MI0000088|hsa-mir-30a&WithFlank&6|-|71403545|71403627@12.34.23(+1G)&[MIMAT0000087&hsa-miR-30a-5p&offsets|0|+1(+1G);m-15&6|-|71403595|71403616&offsets|0|+1(+1G)]|TGTAAACATCCTCGACTGGAAGCG</td><td>>MI0000088|hsa-mir-30a&WithFlank&6|-|71403545|71403627</td><td>12</td><td>34</td><td>23</td><td>(+1G)</td><td>isomiR</td><td>YES</td><td>151.14</td><td>13.68</td><td>635.58</td><td>17.24</td></tr>
<tr><td>>MI0000088|hsa-mir-30a&WithFlank&6|-|71403545|71403627@12.35.24&[MIMAT0000087&hsa-miR-30a-5p&offsets|0|+2;m-15&6|-|71403595|71403616&offsets|0|+2]|TGTAAACATCCTCGACTGGAAGCT</td><td>>MI0000088|hsa-mir-30a&WithFlank&6|-|71403545|71403627</td><td>12</td><td>35</td><td>24</td><td></td><td>isomiR</td><td>YES</td><td>2329.57</td><td>958.75</td><td>14221.75</td><td>559.47</td></tr>
<tr><td>>MI0000088|hsa-mir-30a&WithFlank&6|-|71403545|71403627@12.35.24(+1A)&[MIMAT0000087&hsa-miR-30a-5p&offsets|0|+2(+1A);m-15&6|-|71403595|71403616&offsets|0|+2(+1A)]|TGTAAACATCCTCGACTGGAAGCTA</td><td>>MI0000088|hsa-mir-30a&WithFlank&6|-|71403545|71403627</td><td>12</td><td>35</td><td>24</td><td>(+1A)</td><td>isomiR</td><td>NO</td><td>0.00</td><td>0.00</td><td>232.11</td><td>0.00</td></tr>
<tr><td>>MI0000088|hsa-mir-30a&WithFlank&6|-|71403545|71403627@12.35.24(+1C)&[MIMAT0000087&hsa-miR-30a-5p&offsets|0|+2(+1C);m-15&6|-|71403595|71403616&offsets|0|+2(+1C)]|TGTAAACATCCTCGACTGGAAGCTC</td><td>>MI0000088|hsa-mir-30a&WithFlank&6|-|71403545|71403627</td><td>12</td><td>35</td><td>24</td><td>(+1C)</td><td>isomiR</td><td>NO</td><td>10.36</td><td>0.00</td><td>419.93</td><td>0.00</td></tr>
<tr><td>>MI0000088|hsa-mir-30a&WithFlank&6|-|71403545|71403627@12.35.24(+1U)&[MIMAT0000087&hsa-miR-30a-5p&offsets|0|+2(+1U);m-15&6|-|71403595|71403616&offsets|0|+2(+1U)]|TGTAAACATCCTCGACTGGAAGCTT</td><td>>MI0000088|hsa-mir-30a&WithFlank&6|-|71403545|71403627</td><td>12</td><td>35</td><td>24</td><td>(+1U)</td><td>isomiR</td><td>NO</td><td>230.90</td><td>74.50</td><td>1931.74</td><td>88.38</td></tr>
<tr><td>>MI0000088|hsa-mir-30a&WithFlank&6|-|71403545|71403627@12.35.24(+2U)&[MIMAT0000087&hsa-miR-30a-5p&offsets|0|+2(+2U);m-15&6|-|71403595|71403616&offsets|0|+2(+2U)]|TGTAAACATCCTCGACTGGAAGCTTT</td><td>>MI0000088|hsa-mir-30a&WithFlank&6|-|71403545|71403627</td><td>12</td><td>35</td><td>24</td><td>(+2U)</td><td>isomiR</td><td>NO</td><td>105.34</td><td>0.00</td><td>362.83</td><td>22.13</td></tr>
<tr><td>>MI0000088|hsa-mir-30a&WithFlank&6|-|71403545|71403627@12.35.24(+3U)&[MIMAT0000087&hsa-miR-30a-5p&offsets|0|+2(+3U);m-15&6|-|71403595|71403616&offsets|0|+2(+3U)]|TGTAAACATCCTCGACTGGAAGCTTTT</td><td>>MI0000088|hsa-mir-30a&WithFlank&6|-|71403545|71403627</td><td>12</td><td>35</td><td>24</td><td>(+3U)</td><td>isomiR</td><td>NO</td><td>0.00</td><td>0.00</td><td>18.01</td><td>0.00</td></tr>
<tr><td>>MI0000088|hsa-mir-30a&WithFlank&6|-|71403545|71403627@12.36.25&[MIMAT0000087&hsa-miR-30a-5p&offsets|0|+3;m-15&6|-|71403595|71403616&offsets|0|+3]|TGTAAACATCCTCGACTGGAAGCTG</td><td>>MI0000088|hsa-mir-30a&WithFlank&6|-|71403545|71403627</td><td>12</td><td>36</td><td>25</td><td></td><td>isomiR</td><td>NO</td><td>0.00</td><td>0.00</td><td>65.09</td><td>0.00</td></tr>
<tr><td>>MI0000088|hsa-mir-30a&WithFlank&6|-|71403545|71403627@13.34.22&[MIMAT0000087&hsa-miR-30a-5p&offsets|+1|+1;m-15&6|-|71403595|71403616&offsets|+1|+1]|GTAAACATCCTCGACTGGAAGC</td><td>>MI0000088|hsa-mir-30a&WithFlank&6|-|71403545|71403627</td><td>13</td><td>34</td><td>22</td><td></td><td>isomiR</td><td>NO</td><td>0.00</td><td>0.00</td><td>34.21</td><td>0.00</td></tr>
<tr><td>>MI0000088|hsa-mir-30a&WithFlank&6|-|71403545|71403627@13.34.22(+1A)&[MIMAT0000087&hsa-miR-30a-5p&offsets|+1|+1(+1A);m-15&6|-|71403595|71403616&offsets|+1|+1(+1A)]|GTAAACATCCTCGACTGGAAGCA</td><td>>MI0000088|hsa-mir-30a&WithFlank&6|-|71403545|71403627</td><td>13</td><td>34</td><td>22</td><td>(+1A)</td><td>isomiR</td><td>NO</td><td>0.00</td><td>0.00</td><td>73.97</td><td>0.00</td></tr>
<tr><td>>MI0000088|hsa-mir-30a&WithFlank&6|-|71403545|71403627@13.34.22(+1C)&[MIMAT0000087&hsa-miR-30a-5p&offsets|+1|+1(+1C);m-15&6|-|71403595|71403616&offsets|+1|+1(+1C)]|GTAAACATCCTCGACTGGAAGCC</td><td>>MI0000088|hsa-mir-30a&WithFlank&6|-|71403545|71403627</td><td>13</td><td>34</td><td>22</td><td>(+1C)</td><td>isomiR</td><td>NO</td><td>0.00</td><td>0.00</td><td>105.36</td><td>0.00</td></tr>
<tr><td>>MI0000088|hsa-mir-30a&WithFlank&6|-|71403545|71403627@13.35.23&[MIMAT0000087&hsa-miR-30a-5p&offsets|+1|+2;m-15&6|-|71403595|71403616&offsets|+1|+2]|GTAAACATCCTCGACTGGAAGCT</td><td>>MI0000088|hsa-mir-30a&WithFlank&6|-|71403545|71403627</td><td>13</td><td>35</td><td>23</td><td></td><td>isomiR</td><td>NO</td><td>88.23</td><td>0.00</td><td>455.29</td><td>13.40</td></tr>
<tr><td>>MI0000088|hsa-mir-30a&WithFlank&6|-|71403545|71403627@13.35.23(+1U)&[MIMAT0000087&hsa-miR-30a-5p&offsets|+1|+2(+1U);m-15&6|-|71403595|71403616&offsets|+1|+2(+1U)]|GTAAACATCCTCGACTGGAAGCTT</td><td>>MI0000088|hsa-mir-30a&WithFlank&6|-|71403545|71403627</td><td>13</td><td>35</td><td>23</td><td>(+1U)</td><td>isomiR</td><td>NO</td><td>0.00</td><td>0.00</td><td>144.89</td><td>0.00</td></tr>
<tr><td>>MI0000088|hsa-mir-30a&WithFlank&6|-|71403545|71403627@53.70.18(+1U)&[MIMAT0000088&hsa-miR-30a-3p&offsets|0|-4(+1U);m-38&6|-|71403554|71403575&offsets|0|-4(+1U)]|CTTTCAGTCGGATGTTTGT</td><td>>MI0000088|hsa-mir-30a&WithFlank&6|-|71403545|71403627</td><td>53</td><td>70</td><td>18</td><td>(+1U)</td><td>isomiR</td><td>NO</td><td>0.00</td><td>24.35</td><td>85.51</td><td>0.00</td></tr>
<tr><td>>MI0000088|hsa-mir-30a&WithFlank&6|-|71403545|71403627@53.71.19&[MIMAT0000088&hsa-miR-30a-3p&offsets|0|-3;m-38&6|-|71403554|71403575&offsets|0|-3]|CTTTCAGTCGGATGTTTGC</td><td>>MI0000088|hsa-mir-30a&WithFlank&6|-|71403545|71403627</td><td>53</td><td>71</td><td>19</td><td></td><td>isomiR</td><td>NO</td><td>148.08</td><td>149.64</td><td>233.80</td><td>14.55</td></tr>
<tr><td>>MI0000088|hsa-mir-30a&WithFlank&6|-|71403545|71403627@53.72.20&[MIMAT0000088&hsa-miR-30a-3p&offsets|0|-2;m-38&6|-|71403554|71403575&offsets|0|-2]|CTTTCAGTCGGATGTTTGCA</td><td>>MI0000088|hsa-mir-30a&WithFlank&6|-|71403545|71403627</td><td>53</td><td>72</td><td>20</td><td></td><td>isomiR</td><td>NO</td><td>0.00</td><td>0.00</td><td>33.23</td><td>0.00</td></tr>
<tr><td>>MI0000088|hsa-mir-30a&WithFlank&6|-|71403545|71403627@53.72.20(+1A)&[MIMAT0000088&hsa-miR-30a-3p&offsets|0|-2(+1A);m-38&6|-|71403554|71403575&offsets|0|-2(+1A)]|CTTTCAGTCGGATGTTTGCAA</td><td>>MI0000088|hsa-mir-30a&WithFlank&6|-|71403545|71403627</td><td>53</td><td>72</td><td>20</td><td>(+1A)</td><td>isomiR</td><td>NO</td><td>0.00</td><td>0.00</td><td>16.04</td><td>0.00</td></tr>
<tr><td>>MI0000088|hsa-mir-30a&WithFlank&6|-|71403545|71403627@53.73.21&[MIMAT0000088&hsa-miR-30a-3p&offsets|0|-1;m-38&6|-|71403554|71403575&offsets|0|-1]|CTTTCAGTCGGATGTTTGCAG</td><td>>MI0000088|hsa-mir-30a&WithFlank&6|-|71403545|71403627</td><td>53</td><td>73</td><td>21</td><td></td><td>isomiR</td><td>NO</td><td>90.00</td><td>75.70</td><td>212.22</td><td>0.00</td></tr>
<tr><td>>MI0000088|hsa-mir-30a&WithFlank&6|-|71403545|71403627@53.73.21(+1A)&[MIMAT0000088&hsa-miR-30a-3p&offsets|0|-1(+1A);m-38&6|-|71403554|71403575&offsets|0|-1(+1A)]|CTTTCAGTCGGATGTTTGCAGA</td><td>>MI0000088|hsa-mir-30a&WithFlank&6|-|71403545|71403627</td><td>53</td><td>73</td><td>21</td><td>(+1A)</td><td>isomiR</td><td>NO</td><td>26.59</td><td>33.89</td><td>77.15</td><td>0.00</td></tr>
<tr><td>>MI0000088|hsa-mir-30a&WithFlank&6|-|71403545|71403627@53.73.21(+1U)&[MIMAT0000088&hsa-miR-30a-3p&offsets|0|-1(+1U);m-38&6|-|71403554|71403575&offsets|0|-1(+1U)]|CTTTCAGTCGGATGTTTGCAGT</td><td>>MI0000088|hsa-mir-30a&WithFlank&6|-|71403545|71403627</td><td>53</td><td>73</td><td>21</td><td>(+1U)</td><td>isomiR</td><td>NO</td><td>401.70</td><td>370.32</td><td>523.04</td><td>112.68</td></tr>
<tr><td>>MI0000088|hsa-mir-30a&WithFlank&6|-|71403545|71403627@53.73.21(+2U)&[MIMAT0000088&hsa-miR-30a-3p&offsets|0|-1(+2U);m-38&6|-|71403554|71403575&offsets|0|-1(+2U)]|CTTTCAGTCGGATGTTTGCAGTT</td><td>>MI0000088|hsa-mir-30a&WithFlank&6|-|71403545|71403627</td><td>53</td><td>73</td><td>21</td><td>(+2U)</td><td>isomiR</td><td>NO</td><td>52.76</td><td>34.83</td><td>72.62</td><td>0.00</td></tr>
</table>

### MDA-MB-231

<table>
<tr><th></th><th></th><th></th><th></th><th></th><th></th><th></th><th></th><th></th><th></th><th></th><th></th></tr>
<tr><td>>MI0000088|hsa-mir-30a&WithFlank&6|-|71403545|71403627@53.74.22&[MIMAT0000088&hsa-miR-30a-3p&offsets|0|0;m-38&6|-|71403554|71403575&offsets|0|0]|CTTTCAGTCGGATGTTTGCAGC</td><td>>MI0000088|hsa-mir-30a&WithFlank&6|-|71403545|71403627</td><td>53</td><td>74</td><td>22</td><td></td><td>isomiR</td><td>NO</td><td>543.25</td><td>688.07</td><td>616.82</td><td>126.65</td></tr>
<tr><td>>MI0000088|hsa-mir-30a&WithFlank&6|-|71403545|71403627@53.74.22(+1A)&[MIMAT0000088&hsa-miR-30a-3p&offsets|0|0(+1A);m-38&6|-|71403554|71403575&offsets|0|0(+1A)]|CTTTCAGTCGGATGTTTGCAGCA</td><td>>MI0000088|hsa-mir-30a&WithFlank&6|-|71403545|71403627</td><td>53</td><td>74</td><td>22</td><td>(+1A)</td><td>isomiR</td><td>NO</td><td>0.00</td><td>13.19</td><td>0.00</td><td>0.00</td></tr>
<tr><td>>MI0000088|hsa-mir-30a&WithFlank&6|-|71403545|71403627@53.75.23&[MIMAT0000088&hsa-miR-30a-3p&offsets|0|+1;m-38&6|-|71403554|71403575&offsets|0|+1]|CTTTCAGTCGGATGTTTGCAGCT</td><td>>MI0000088|hsa-mir-30a&WithFlank&6|-|71403545|71403627</td><td>53</td><td>75</td><td>23</td><td></td><td>isomiR</td><td>NO</td><td>0.00</td><td>0.00</td><td>12.70</td><td>0.00</td></tr>
<tr><td>>MI0000088|hsa-mir-30a&WithFlank&6|-|71403545|71403627@54.71.18&[MIMAT0000088&hsa-miR-30a-3p&offsets|+1|-3;m-38&6|-|71403554|71403575&offsets|+1|-3]|TTTCAGTCGGATGTTTGC</td><td>>MI0000088|hsa-mir-30a&WithFlank&6|-|71403545|71403627</td><td>54</td><td>71</td><td>18</td><td></td><td>isomiR</td><td>NO</td><td>71.71</td><td>57.04</td><td>108.70</td><td>0.00</td></tr>
<tr><td>>MI0000088|hsa-mir-30a&WithFlank&6|-|71403545|71403627@54.73.20&[MIMAT0000088&hsa-miR-30a-3p&offsets|+1|-1;m-38&6|-|71403554|71403575&offsets|+1|-1]|TTTCAGTCGGATGTTTGCAG</td><td>>MI0000088|hsa-mir-30a&WithFlank&6|-|71403545|71403627</td><td>54</td><td>73</td><td>20</td><td></td><td>isomiR</td><td>NO</td><td>0.00</td><td>0.00</td><td>70.61</td><td>0.00</td></tr>
<tr><td>>MI0000088|hsa-mir-30a&WithFlank&6|-|71403545|71403627@54.73.20(+1A)&[MIMAT0000088&hsa-miR-30a-3p&offsets|+1|-1(+1A);m-38&6|-|71403554|71403575&offsets|+1|-1(+1A)]|TTTCAGTCGGATGTTTGCAGA</td><td>>MI0000088|hsa-mir-30a&WithFlank&6|-|71403545|71403627</td><td>54</td><td>73</td><td>20</td><td>(+1A)</td><td>isomiR</td><td>NO</td><td>0.00</td><td>0.00</td><td>14.34</td><td>0.00</td></tr>
<tr><td>>MI0000088|hsa-mir-30a&WithFlank&6|-|71403545|71403627@54.73.20(+1U)&[MIMAT0000088&hsa-miR-30a-3p&offsets|+1|-1(+1U);m-38&6|-|71403554|71403575&offsets|+1|-1(+1U)]|TTTCAGTCGGATGTTTGCAGT</td><td>>MI0000088|hsa-mir-30a&WithFlank&6|-|71403545|71403627</td><td>54</td><td>73</td><td>20</td><td>(+1U)</td><td>isomiR</td><td>NO</td><td>210.44</td><td>153.08</td><td>274.09</td><td>69.83</td></tr>
<tr><td>>MI0000088|hsa-mir-30a&WithFlank&6|-|71403545|71403627@54.73.20(+2U)&[MIMAT0000088&hsa-miR-30a-3p&offsets|+1|-1(+2U);m-38&6|-|71403554|71403575&offsets|+1|-1(+2U)]|TTTCAGTCGGATGTTTGCAGTT</td><td>>MI0000088|hsa-mir-30a&WithFlank&6|-|71403545|71403627</td><td>54</td><td>73</td><td>20</td><td>(+2U)</td><td>isomiR</td><td>NO</td><td>25.14</td><td>0.00</td><td>63.75</td><td>0.00</td></tr>
<tr><td>>MI0000088|hsa-mir-30a&WithFlank&6|-|71403545|71403627@54.74.21&[MIMAT0000088&hsa-miR-30a-3p&offsets|+1|0;m-38&6|-|71403554|71403575&offsets|+1|0]|TTTCAGTCGGATGTTTGCAGC</td><td>>MI0000088|hsa-mir-30a&WithFlank&6|-|71403545|71403627</td><td>54</td><td>74</td><td>21</td><td></td><td>isomiR</td><td>NO</td><td>219.30</td><td>215.17</td><td>286.63</td><td>63.89</td></tr>
<tr><td>>MI0000088|hsa-mir-30a&WithFlank&6|-|71403545|71403627@54.75.22&[MIMAT0000088&hsa-miR-30a-3p&offsets|+1|+1;m-38&6|-|71403554|71403575&offsets|+1|+1]|TTTCAGTCGGATGTTTGCAGCT</td><td>>MI0000088|hsa-mir-30a&WithFlank&6|-|71403545|71403627</td><td>54</td><td>75</td><td>22</td><td></td><td>isomiR</td><td>NO</td><td>59.88</td><td>33.91</td><td>84.32</td><td>10.67</td></tr>
<tr><td>>MI0000090|hsa-mir-32&WithFlank&9|-|109046223|109046304@12.32.21&[MIMAT0000090&hsa-miR-32-5p&offsets|0|-1;m-208&9|-|109046273|109046293&offsets|0|0]|TATTGCACATTACTAAGTTGC</td><td>>MI0000090|hsa-mir-32&WithFlank&9|-|109046223|109046304</td><td>12</td><td>32</td><td>21</td><td></td><td>isomiR</td><td>NO</td><td>0.00</td><td>0.00</td><td>15.14</td><td>0.00</td></tr>
<tr><td>>MI0000090|hsa-mir-32&WithFlank&9|-|109046223|109046304@12.33.22&[MIMAT0000090&hsa-miR-32-5p&offsets|0|0;m-208&9|-|109046273|109046293&offsets|0|+1]|TATTGCACATTACTAAGTTGCA</td><td>>MI0000090|hsa-mir-32&WithFlank&9|-|109046223|109046304</td><td>12</td><td>33</td><td>22</td><td></td><td>isomiR</td><td>NO</td><td>0.00</td><td>0.00</td><td>105.81</td><td>0.00</td></tr>
<tr><td>>MI0000093|hsa-mir-92a-1&WithFlank&13|+|91351308|91351397@54.73.20(+1A);MI0000094|hsa-mir-92a-2&WithFlank&X|-|134169532|134169618@54.73.20(+1A)&[MIMAT0000092&hsa-miR-92a-3p&offsets|0|-2(+1A);m-9&13|+|91351361|91351382&offsets|0|-2(+1A)];[MIMAT0000092_1&hsa-miR-92a-3p&offsets|0|-2(+1A);m-11&X|-|134169544|134169565&offsets|0|-2(+1A)]|TATTGCACTTGTCCCGGCCTA</td><td>>MI0000093|hsa-mir-92a-1&WithFlank&13|+|91351308|91351397</td><td>54</td><td>73</td><td>20</td><td>(+1A)</td><td>isomiR</td><td>NO</td><td>0.00</td><td>0.00</td><td>11.09</td><td>0.00</td></tr>
<tr><td>>MI0000093|hsa-mir-92a-1&WithFlank&13|+|91351308|91351397@54.73.20;MI0000094|hsa-mir-92a-2&WithFlank&X|-|134169532|134169618@54.73.20&[MIMAT0000092&hsa-miR-92a-3p&offsets|0|-2;m-9&13|+|91351361|91351382&offsets|0|-2];[MIMAT0000092_1&hsa-miR-92a-3p&offsets|0|-2;m-11&X|-|134169544|134169565&offsets|0|-2]|TATTGCACTTGTCCCGGCCT</td><td>>MI0000093|hsa-mir-92a-1&WithFlank&13|+|91351308|91351397</td><td>54</td><td>73</td><td>20</td><td></td><td>isomiR</td><td>NO</td><td>0.00</td><td>0.00</td><td>13.69</td><td>0.00</td></tr>
<tr><td>>MI0000093|hsa-mir-92a-1&WithFlank&13|+|91351308|91351397@54.74.21(+1A);MI0000094|hsa-mir-92a-2&WithFlank&X|-|134169532|134169618@54.74.21(+1A)&[MIMAT0000092&hsa-miR-92a-3p&offsets|0|-1(+1A);m-9&13|+|91351361|91351382&offsets|0|-1(+1A)];[MIMAT0000092_1&hsa-miR-92a-3p&offsets|0|-1(+1A);m-11&X|-|134169544|134169565&offsets|0|-1(+1A)]|TATTGCACTTGTCCCGGCCTGA</td><td>>MI0000093|hsa-mir-92a-1&WithFlank&13|+|91351308|91351397</td><td>54</td><td>74</td><td>21</td><td>(+1A)</td><td>isomiR</td><td>NO</td><td>107.63</td><td>124.12</td><td>175.20</td><td>59.06</td></tr>
<tr><td>>MI0000093|hsa-mir-92a-1&WithFlank&13|+|91351308|91351397@54.74.21(+1C);MI0000094|hsa-mir-92a-2&WithFlank&X|-|134169532|134169618@54.74.21(+1C)&[MIMAT0000092&hsa-miR-92a-3p&offsets|0|-1(+1C);m-9&13|+|91351361|91351382&offsets|0|-1(+1C)];[MIMAT0000092_1&hsa-miR-92a-3p&offsets|0|-1(+1C);m-11&X|-|134169544|134169565&offsets|0|-1(+1C)]|TATTGCACTTGTCCCGGCCTGC</td><td>>MI0000093|hsa-mir-92a-1&WithFlank&13|+|91351308|91351397</td><td>54</td><td>74</td><td>21</td><td>(+1C)</td><td>isomiR</td><td>NO</td><td>322.62</td><td>357.42</td><td>526.75</td><td>153.14</td></tr>
<tr><td>>MI0000093|hsa-mir-92a-1&WithFlank&13|+|91351308|91351397@54.74.21(+1G);MI0000094|hsa-mir-92a-2&WithFlank&X|-|134169532|134169618@54.74.21(+1G)&[MIMAT0000092&hsa-miR-92a-3p&offsets|0|-1(+1G);m-9&13|+|91351361|91351382&offsets|0|-1(+1G)];[MIMAT0000092_1&hsa-miR-92a-3p&offsets|0|-1(+1G);m-11&X|-|134169544|134169565&offsets|0|-1(+1G)]|TATTGCACTTGTCCCGGCCTGG</td><td>>MI0000093|hsa-mir-92a-1&WithFlank&13|+|91351308|91351397</td><td>54</td><td>74</td><td>21</td><td>(+1G)</td><td>isomiR</td><td>NO</td><td>78.88</td><td>62.76</td><td>71.39</td><td>12.65</td></tr>
<tr><td>>MI0000093|hsa-mir-92a-1&WithFlank&13|+|91351308|91351397@54.74.21;MI0000094|hsa-mir-92a-2&WithFlank&X|-|134169532|134169618@54.74.21&[MIMAT0000092&hsa-miR-92a-3p&offsets|0|-1;m-9&13|+|91351361|91351382&offsets|0|-1];[MIMAT0000092_1&hsa-miR-92a-3p&offsets|0|-1;m-11&X|-|134169544|134169565&offsets|0|-1]|TATTGCACTTGTCCCGGCCTG</td><td>>MI0000093|hsa-mir-92a-1&WithFlank&13|+|91351308|91351397</td><td>54</td><td>74</td><td>21</td><td></td><td>isomiR</td><td>NO</td><td>68.39</td><td>63.67</td><td>98.33</td><td>11.80</td></tr>
<tr><td>>MI0000093|hsa-mir-92a-1&WithFlank&13|+|91351308|91351397@54.75.22(+1A);MI0000094|hsa-mir-92a-2&WithFlank&X|-|134169532|134169618@54.75.22(+1A)&[MIMAT0000092&hsa-miR-92a-3p&offsets|0|0(+1A);m-9&13|+|91351361|91351382&offsets|0|0(+1A)];[MIMAT0000092_1&hsa-miR-92a-3p&offsets|0|0(+1A);m-11&X|-|134169544|134169565&offsets|0|0(+1A)]|TATTGCACTTGTCCCGGCCTGTA</td><td>>MI0000093|hsa-mir-92a-1&WithFlank&13|+|91351308|91351397</td><td>54</td><td>75</td><td>22</td><td>(+1A)</td><td>isomiR</td><td>NO</td><td>300.39</td><td>453.76</td><td>274.53</td><td>138.79</td></tr>
</table>

### MDA-MB-231

<table>
<tr><td>>MI0000093|hsa-mir-92a-1&WithFlank&13|+|91351308|91351397@54.75.22(+1C);MI0000094|hsa-mir-92a-2&WithFlank&X|-|134169532|134169618@54.75.22(+1C)&[MIMAT0000092&hsa-miR-92a-3p&offsets|0|0(+1C);m-9&13|+|91351361|91351382&offsets|0|0(+1C)];[MIMAT0000092_1&hsa-miR-92a-3p&offsets|0|0(+1C);m-11&X|-|134169544|134169565&offsets|0|0(+1C)]|TATTGCACTTGTCCCGGCCTGTC</td><td>>MI0000093|hsa-mir-92a-1&WithFlank&13|+|91351308|91351397</td><td>54</td><td>75</td><td>22</td><td>(+1C)</td><td>isomiR</td><td>NO</td><td>24.32</td><td>34.55</td><td>35.18</td><td>14.48</td></tr>
<tr><td>>MI0000093|hsa-mir-92a-1&WithFlank&13|+|91351308|91351397@54.75.22(+2A);MI0000094|hsa-mir-92a-2&WithFlank&X|-|134169532|134169618@54.75.22(+2A)&[MIMAT0000092&hsa-miR-92a-3p&offsets|0|0(+2A);m-9&13|+|91351361|91351382&offsets|0|0(+2A)];[MIMAT0000092_1&hsa-miR-92a-3p&offsets|0|0(+2A);m-11&X|-|134169544|134169565&offsets|0|0(+2A)]|TATTGCACTTGTCCCGGCCTGTAA</td><td>>MI0000093|hsa-mir-92a-1&WithFlank&13|+|91351308|91351397</td><td>54</td><td>75</td><td>22</td><td>(+2A)</td><td>isomiR</td><td>NO</td><td>49.96</td><td>26.50</td><td>0.00</td><td>12.19</td></tr>
<tr><td>>MI0000093|hsa-mir-92a-1&WithFlank&13|+|91351308|91351397@54.75.22;MI0000094|hsa-mir-92a-2&WithFlank&X|-|134169532|134169618@54.75.22&[MIMAT0000092&hsa-miR-92a-3p&offsets|0|0;m-9&13|+|91351361|91351382&offsets|0|0];[MIMAT0000092_1&hsa-miR-92a-3p&offsets|0|0;m-11&X|-|134169544|134169565&offsets|0|0]|TATTGCACTTGTCCCGGCCTGT</td><td>>MI0000093|hsa-mir-92a-1&WithFlank&13|+|91351308|91351397</td><td>54</td><td>75</td><td>22</td><td></td><td>isomiR</td><td>YES</td><td>1764.31</td><td>2334.50</td><td>2328.32</td><td>543.78</td></tr>
<tr><td>>MI0000093|hsa-mir-92a-1&WithFlank&13|+|91351308|91351397@54.76.23&[MIMAT0000092&hsa-miR-92a-3p&offsets|0|+1;m-9&13|+|91351361|91351382&offsets|0|+1]|TATTGCACTTGTCCCGGCCTGTT</td><td>>MI0000093|hsa-mir-92a-1&WithFlank&13|+|91351308|91351397</td><td>54</td><td>76</td><td>23</td><td></td><td>isomiR</td><td>NO</td><td>235.43</td><td>341.49</td><td>224.85</td><td>125.47</td></tr>
<tr><td>>MI0000093|hsa-mir-92a-1&WithFlank&13|+|91351308|91351397@54.76.23(+1U)&[MIMAT0000092&hsa-miR-92a-3p&offsets|0|+1(+1U);m-9&13|+|91351361|91351382&offsets|0|+1(+1U)]|TATTGCACTTGTCCCGGCCTGTTT</td><td>>MI0000093|hsa-mir-92a-1&WithFlank&13|+|91351308|91351397</td><td>54</td><td>76</td><td>23</td><td>(+1U)</td><td>isomiR</td><td>NO</td><td>0.00</td><td>26.25</td><td>11.41</td><td>10.56</td></tr>
<tr><td>>MI0000093|hsa-mir-92a-1&WithFlank&13|+|91351308|91351397@55.75.21;MI0000094|hsa-mir-92a-2&WithFlank&X|-|134169532|134169618@55.75.21&[MIMAT0000092&hsa-miR-92a-3p&offsets|+1|0;m-9&13|+|91351361|91351382&offsets|+1|0];[MIMAT0000092_1&hsa-miR-92a-3p&offsets|+1|0;m-11&X|-|134169544|134169565&offsets|+1|0]|ATTGCACTTGTCCCGGCCTGT</td><td>>MI0000093|hsa-mir-92a-1&WithFlank&13|+|91351308|91351397</td><td>55</td><td>75</td><td>21</td><td></td><td>isomiR</td><td>NO</td><td>62.97</td><td>58.36</td><td>84.59</td><td>14.98</td></tr>
<tr><td>>MI0000095|hsa-mir-93&WithFlank&7|-|100093762|100093853@17.36.20(+1C)&[MIMAT0000093&hsa-miR-93-5p&offsets|0|-3(+1C);m-29&7|-|100093815|100093837&offsets|0|-3(+1C)]|CAAAGTGCTGTTCGTGCAGGC</td><td>>MI0000095|hsa-mir-93&WithFlank&7|-|100093762|100093853</td><td>17</td><td>36</td><td>20</td><td>(+1C)</td><td>isomiR</td><td>NO</td><td>0.00</td><td>0.00</td><td>104.19</td><td>0.00</td></tr>
<tr><td>>MI0000095|hsa-mir-93&WithFlank&7|-|100093762|100093853@17.37.21&[MIMAT0000093&hsa-miR-93-5p&offsets|0|-2;m-29&7|-|100093815|100093837&offsets|0|-2]|CAAAGTGCTGTTCGTGCAGGT</td><td>>MI0000095|hsa-mir-93&WithFlank&7|-|100093762|100093853</td><td>17</td><td>37</td><td>21</td><td></td><td>isomiR</td><td>NO</td><td>65.72</td><td>0.00</td><td>465.17</td><td>13.07</td></tr>
<tr><td>>MI0000095|hsa-mir-93&WithFlank&7|-|100093762|100093853@17.38.22&[MIMAT0000093&hsa-miR-93-5p&offsets|0|-1;m-29&7|-|100093815|100093837&offsets|0|-1]|CAAAGTGCTGTTCGTGCAGGTA</td><td>>MI0000095|hsa-mir-93&WithFlank&7|-|100093762|100093853</td><td>17</td><td>38</td><td>22</td><td></td><td>isomiR</td><td>NO</td><td>0.00</td><td>0.00</td><td>171.61</td><td>0.00</td></tr>
<tr><td>>MI0000095|hsa-mir-93&WithFlank&7|-|100093762|100093853@17.38.22(+1A)&[MIMAT0000093&hsa-miR-93-5p&offsets|0|-1(+1A);m-29&7|-|100093815|100093837&offsets|0|-1(+1A)]|CAAAGTGCTGTTCGTGCAGGTAA</td><td>>MI0000095|hsa-mir-93&WithFlank&7|-|100093762|100093853</td><td>17</td><td>38</td><td>22</td><td>(+1A)</td><td>isomiR</td><td>NO</td><td>46.25</td><td>0.00</td><td>398.43</td><td>0.00</td></tr>
<tr><td>>MI0000095|hsa-mir-93&WithFlank&7|-|100093762|100093853@17.38.22(+1U)&[MIMAT0000093&hsa-miR-93-5p&offsets|0|-1(+1U);m-29&7|-|100093815|100093837&offsets|0|-1(+1U)]|CAAAGTGCTGTTCGTGCAGGTAT</td><td>>MI0000095|hsa-mir-93&WithFlank&7|-|100093762|100093853</td><td>17</td><td>38</td><td>22</td><td>(+1U)</td><td>isomiR</td><td>NO</td><td>0.00</td><td>0.00</td><td>84.08</td><td>0.00</td></tr>
<tr><td>>MI0000095|hsa-mir-93&WithFlank&7|-|100093762|100093853@17.39.23&[MIMAT0000093&hsa-miR-93-5p&offsets|0|0;m-29&7|-|100093815|100093837&offsets|0|0]|CAAAGTGCTGTTCGTGCAGGTAG</td><td>>MI0000095|hsa-mir-93&WithFlank&7|-|100093762|100093853</td><td>17</td><td>39</td><td>23</td><td></td><td>isomiR</td><td>NO</td><td>394.81</td><td>162.65</td><td>2021.11</td><td>106.48</td></tr>
<tr><td>>MI0000095|hsa-mir-93&WithFlank&7|-|100093762|100093853@17.39.23(+1A)&[MIMAT0000093&hsa-miR-93-5p&offsets|0|0(+1A);m-29&7|-|100093815|100093837&offsets|0|0(+1A)]|CAAAGTGCTGTTCGTGCAGGTAGA</td><td>>MI0000095|hsa-mir-93&WithFlank&7|-|100093762|100093853</td><td>17</td><td>39</td><td>23</td><td>(+1A)</td><td>isomiR</td><td>NO</td><td>83.64</td><td>35.90</td><td>90.94</td><td>17.51</td></tr>
<tr><td>>MI0000095|hsa-mir-93&WithFlank&7|-|100093762|100093853@17.40.24&[MIMAT0000093&hsa-miR-93-5p&offsets|0|+1;m-29&7|-|100093815|100093837&offsets|0|+1]|CAAAGTGCTGTTCGTGCAGGTAGT</td><td>>MI0000095|hsa-mir-93&WithFlank&7|-|100093762|100093853</td><td>17</td><td>40</td><td>24</td><td></td><td>isomiR</td><td>NO</td><td>145.93</td><td>62.56</td><td>122.19</td><td>69.31</td></tr>
<tr><td>>MI0000098|hsa-mir-96&WithFlank&7|-|129774686|129774775@15.37.23&[MIMAT0000095&hsa-miR-96-5p&offsets|0|0;m-248&7|-|129774739|129774761&offsets|0|0]|TTTGGCACTAGCACATTTTTGCT</td><td>>MI0000098|hsa-mir-96&WithFlank&7|-|129774686|129774775</td><td>15</td><td>37</td><td>23</td><td></td><td>isomiR</td><td>NO</td><td>72.40</td><td>0.00</td><td>99.09</td><td>14.42</td></tr>
<tr><td>>MI0000100|hsa-mir-98&WithFlank&X|-|53556217|53556347@28.48.21&[MIMAT0000096&hsa-miR-98-5p&offsets|0|-1;m-143&X|-|53556299|53556320&offsets|0|-1]|TGAGGTAGTAAGTTGTATTGT</td><td>>MI0000100|hsa-mir-98&WithFlank&X|-|53556217|53556347</td><td>28</td><td>48</td><td>21</td><td></td><td>isomiR</td><td>NO</td><td>0.00</td><td>0.00</td><td>65.07</td><td>0.00</td></tr>
<tr><td>>MI0000100|hsa-mir-98&WithFlank&X|-|53556217|53556347@28.48.21(+1A)&[MIMAT0000096&hsa-miR-98-5p&offsets|0|-1(+1A);m-143&X|-|53556299|53556320&offsets|0|-1(+1A)]|TGAGGTAGTAAGTTGTATTGTA</td><td>>MI0000100|hsa-mir-98&WithFlank&X|-|53556217|53556347</td><td>28</td><td>48</td><td>21</td><td>(+1A)</td><td>isomiR</td><td>NO</td><td>0.00</td><td>0.00</td><td>10.48</td><td>0.00</td></tr>
<tr><td>>MI0000100|hsa-mir-98&WithFlank&X|-|53556217|53556347@28.48.21(+1C)&[MIMAT0000096&hsa-miR-98-5p&offsets|0|-1(+1C);m-143&X|-|53556299|53556320&offsets|0|-1(+1C)]|TGAGGTAGTAAGTTGTATTGTC</td><td>>MI0000100|hsa-mir-98&WithFlank&X|-|53556217|53556347</td><td>28</td><td>48</td><td>21</td><td>(+1C)</td><td>isomiR</td><td>NO</td><td>0.00</td><td>0.00</td><td>91.61</td><td>0.00</td></tr>
<tr><td>>MI0000100|hsa-mir-98&WithFlank&X|-|53556217|53556347@28.49.22&[MIMAT0000096&hsa-miR-98-5p&offsets|0|0;m-143&X|-|53556299|53556320&offsets|0|0]|TGAGGTAGTAAGTTGTATTGTT</td><td>>MI0000100|hsa-mir-98&WithFlank&X|-|53556217|53556347</td><td>28</td><td>49</td><td>22</td><td></td><td>isomiR</td><td>NO</td><td>204.66</td><td>84.62</td><td>417.67</td><td>59.91</td></tr>
<tr><td>>MI0000101|hsa-mir-99a&WithFlank&21|+|16539083|16539175@19.39.21&[MIMAT0000097&hsa-miR-99a-5p&offsets|0|-1;m-64&21|+|16539101|16539121&offsets|0|0]|AACCCGTAGATCCGATCTTGT</td><td>>MI0000101|hsa-mir-99a&WithFlank&21|+|16539083|16539175</td><td>19</td><td>39</td><td>21</td><td></td><td>isomiR</td><td>NO</td><td>107.53</td><td>32.80</td><td>116.66</td><td>17.11</td></tr>
<tr><td>>MI0000101|hsa-mir-99a&WithFlank&21|+|16539083|16539175@19.40.22&[MIMAT0000097&hsa-miR-99a-5p&offsets|0|0;m-64&21|+|16539101|16539121&offsets|0|+1]|AACCCGTAGATCCGATCTTGTG</td><td>>MI0000101|hsa-mir-99a&WithFlank&21|+|16539083|16539175</td><td>19</td><td>40</td><td>22</td><td></td><td>isomiR</td><td>NO</td><td>97.41</td><td>36.89</td><td>104.60</td><td>16.12</td></tr>
<tr><td>>MI0000101|hsa-mir-99a&WithFlank&21|+|16539083|16539175@19.40.22(+1A)&[MIMAT0000097&hsa-miR-99a-5p&offsets|0|0(+1A);m-64&21|+|16539101|16539121&offsets|0|+1(+1A)]|AACCCGTAGATCCGATCTTGTGA</td><td>>MI0000101|hsa-mir-99a&WithFlank&21|+|16539083|16539175</td><td>19</td><td>40</td><td>22</td><td>(+1A)</td><td>isomiR</td><td>NO</td><td>24.55</td><td>0.00</td><td>12.47</td><td>0.00</td></tr>
</table>

### MDA-MB-231

<table>
<tr><th></th><th></th><th></th><th></th><th></th><th></th><th></th><th></th><th></th><th></th><th></th><th></th></tr>
<tr><td>>MI0000102|hsa-mir-100&WithFlank&11|-|122152223|122152314@19.36.18&[MIMAT0000098&hsa-miR-100-5p&offsets|0|-4;m-27&11|-|122152276|122152296&offsets|0|-3]|AACCCGTAGATCCGAACT</td><td>>MI0000102|hsa-mir-100&WithFlank&11|-|122152223|122152314</td><td>19</td><td>36</td><td>18</td><td></td><td>isomiR</td><td>NO</td><td>0.00</td><td>0.00</td><td>73.01</td><td>0.00</td></tr>
<tr><td>>MI0000102|hsa-mir-100&WithFlank&11|-|122152223|122152314@19.37.19&[MIMAT0000098&hsa-miR-100-5p&offsets|0|-3;m-27&11|-|122152276|122152296&offsets|0|-2]|AACCCGTAGATCCGAACTT</td><td>>MI0000102|hsa-mir-100&WithFlank&11|-|122152223|122152314</td><td>19</td><td>37</td><td>19</td><td></td><td>isomiR</td><td>NO</td><td>0.00</td><td>0.00</td><td>87.34</td><td>0.00</td></tr>
<tr><td>>MI0000102|hsa-mir-100&WithFlank&11|-|122152223|122152314@19.37.19(+1A)&[MIMAT0000098&hsa-miR-100-5p&offsets|0|-3(+1A);m-27&11|-|122152276|122152296&offsets|0|-2(+1A)]|AACCCGTAGATCCGAACTTA</td><td>>MI0000102|hsa-mir-100&WithFlank&11|-|122152223|122152314</td><td>19</td><td>37</td><td>19</td><td>(+1A)</td><td>isomiR</td><td>NO</td><td>101.58</td><td>88.70</td><td>370.31</td><td>19.14</td></tr>
<tr><td>>MI0000102|hsa-mir-100&WithFlank&11|-|122152223|122152314@19.37.19(+1U)&[MIMAT0000098&hsa-miR-100-5p&offsets|0|-3(+1U);m-27&11|-|122152276|122152296&offsets|0|-2(+1U)]|AACCCGTAGATCCGAACTTT</td><td>>MI0000102|hsa-mir-100&WithFlank&11|-|122152223|122152314</td><td>19</td><td>37</td><td>19</td><td>(+1U)</td><td>isomiR</td><td>NO</td><td>0.00</td><td>0.00</td><td>17.39</td><td>0.00</td></tr>
<tr><td>>MI0000102|hsa-mir-100&WithFlank&11|-|122152223|122152314@19.38.20&[MIMAT0000098&hsa-miR-100-5p&offsets|0|-2;m-27&11|-|122152276|122152296&offsets|0|-1]|AACCCGTAGATCCGAACTTG</td><td>>MI0000102|hsa-mir-100&WithFlank&11|-|122152223|122152314</td><td>19</td><td>38</td><td>20</td><td></td><td>isomiR</td><td>YES</td><td>836.32</td><td>649.44</td><td>1889.78</td><td>203.54</td></tr>
<tr><td>>MI0000102|hsa-mir-100&WithFlank&11|-|122152223|122152314@19.38.20(+1A)&[MIMAT0000098&hsa-miR-100-5p&offsets|0|-2(+1A);m-27&11|-|122152276|122152296&offsets|0|-1(+1A)]|AACCCGTAGATCCGAACTTGA</td><td>>MI0000102|hsa-mir-100&WithFlank&11|-|122152223|122152314</td><td>19</td><td>38</td><td>20</td><td>(+1A)</td><td>isomiR</td><td>NO</td><td>183.81</td><td>220.00</td><td>651.94</td><td>67.35</td></tr>
<tr><td>>MI0000102|hsa-mir-100&WithFlank&11|-|122152223|122152314@19.38.20(+1C)&[MIMAT0000098&hsa-miR-100-5p&offsets|0|-2(+1C);m-27&11|-|122152276|122152296&offsets|0|-1(+1C)]|AACCCGTAGATCCGAACTTGC</td><td>>MI0000102|hsa-mir-100&WithFlank&11|-|122152223|122152314</td><td>19</td><td>38</td><td>20</td><td>(+1C)</td><td>isomiR</td><td>YES</td><td>769.66</td><td>736.12</td><td>2034.04</td><td>197.34</td></tr>
<tr><td>>MI0000102|hsa-mir-100&WithFlank&11|-|122152223|122152314@19.38.20(+1G)&[MIMAT0000098&hsa-miR-100-5p&offsets|0|-2(+1G);m-27&11|-|122152276|122152296&offsets|0|-1(+1G)]|AACCCGTAGATCCGAACTTGG</td><td>>MI0000102|hsa-mir-100&WithFlank&11|-|122152223|122152314</td><td>19</td><td>38</td><td>20</td><td>(+1G)</td><td>isomiR</td><td>NO</td><td>175.30</td><td>123.13</td><td>269.63</td><td>15.01</td></tr>
<tr><td>>MI0000102|hsa-mir-100&WithFlank&11|-|122152223|122152314@19.38.20(+2G)&[MIMAT0000098&hsa-miR-100-5p&offsets|0|-2(+2G);m-27&11|-|122152276|122152296&offsets|0|-1(+2G)]|AACCCGTAGATCCGAACTTGGG</td><td>>MI0000102|hsa-mir-100&WithFlank&11|-|122152223|122152314</td><td>19</td><td>38</td><td>20</td><td>(+2G)</td><td>isomiR</td><td>NO</td><td>79.10</td><td>31.39</td><td>71.65</td><td>12.32</td></tr>
<tr><td>>MI0000102|hsa-mir-100&WithFlank&11|-|122152223|122152314@19.39.21&[MIMAT0000098&hsa-miR-100-5p&offsets|0|-1;m-27&11|-|122152276|122152296&offsets|0|0]|AACCCGTAGATCCGAACTTGT</td><td>>MI0000102|hsa-mir-100&WithFlank&11|-|122152223|122152314</td><td>19</td><td>39</td><td>21</td><td></td><td>isomiR</td><td>YES</td><td>4191.74</td><td>4870.96</td><td>9028.50</td><td>710.99</td></tr>
<tr><td>>MI0000102|hsa-mir-100&WithFlank&11|-|122152223|122152314@19.39.21(+1A)&[MIMAT0000098&hsa-miR-100-5p&offsets|0|-1(+1A);m-27&11|-|122152276|122152296&offsets|0|0(+1A)]|AACCCGTAGATCCGAACTTGTA</td><td>>MI0000102|hsa-mir-100&WithFlank&11|-|122152223|122152314</td><td>19</td><td>39</td><td>21</td><td>(+1A)</td><td>isomiR</td><td>YES</td><td>1755.56</td><td>2388.07</td><td>4530.63</td><td>524.89</td></tr>
<tr><td>>MI0000102|hsa-mir-100&WithFlank&11|-|122152223|122152314@19.39.21(+1C)&[MIMAT0000098&hsa-miR-100-5p&offsets|0|-1(+1C);m-27&11|-|122152276|122152296&offsets|0|0(+1C)]|AACCCGTAGATCCGAACTTGTC</td><td>>MI0000102|hsa-mir-100&WithFlank&11|-|122152223|122152314</td><td>19</td><td>39</td><td>21</td><td>(+1C)</td><td>isomiR</td><td>NO</td><td>117.28</td><td>92.14</td><td>240.08</td><td>21.87</td></tr>
<tr><td>>MI0000102|hsa-mir-100&WithFlank&11|-|122152223|122152314@19.39.21(+1U)&[MIMAT0000098&hsa-miR-100-5p&offsets|0|-1(+1U);m-27&11|-|122152276|122152296&offsets|0|0(+1U)]|AACCCGTAGATCCGAACTTGTT</td><td>>MI0000102|hsa-mir-100&WithFlank&11|-|122152223|122152314</td><td>19</td><td>39</td><td>21</td><td>(+1U)</td><td>isomiR</td><td>NO</td><td>279.83</td><td>252.36</td><td>628.02</td><td>100.68</td></tr>
<tr><td>>MI0000102|hsa-mir-100&WithFlank&11|-|122152223|122152314@19.40.22&[MIMAT0000098&hsa-miR-100-5p&offsets|0|0;m-27&11|-|122152276|122152296&offsets|0|+1]|AACCCGTAGATCCGAACTTGTG</td><td>>MI0000102|hsa-mir-100&WithFlank&11|-|122152223|122152314</td><td>19</td><td>40</td><td>22</td><td></td><td>isomiR</td><td>YES</td><td>12784.46</td><td>16001.85</td><td>21217.00</td><td>2077.28</td></tr>
<tr><td>>MI0000102|hsa-mir-100&WithFlank&11|-|122152223|122152314@19.40.22(+1A)&[MIMAT0000098&hsa-miR-100-5p&offsets|0|0(+1A);m-27&11|-|122152276|122152296&offsets|0|+1(+1A)]|AACCCGTAGATCCGAACTTGTGA</td><td>>MI0000102|hsa-mir-100&WithFlank&11|-|122152223|122152314</td><td>19</td><td>40</td><td>22</td><td>(+1A)</td><td>isomiR</td><td>YES</td><td>1562.86</td><td>1046.32</td><td>1995.56</td><td>335.83</td></tr>
<tr><td>>MI0000102|hsa-mir-100&WithFlank&11|-|122152223|122152314@19.40.22(+1C)&[MIMAT0000098&hsa-miR-100-5p&offsets|0|0(+1C);m-27&11|-|122152276|122152296&offsets|0|+1(+1C)]|AACCCGTAGATCCGAACTTGTGC</td><td>>MI0000102|hsa-mir-100&WithFlank&11|-|122152223|122152314</td><td>19</td><td>40</td><td>22</td><td>(+1C)</td><td>isomiR</td><td>NO</td><td>315.07</td><td>293.80</td><td>587.09</td><td>109.91</td></tr>
<tr><td>>MI0000102|hsa-mir-100&WithFlank&11|-|122152223|122152314@19.40.22(+1U)&[MIMAT0000098&hsa-miR-100-5p&offsets|0|0(+1U);m-27&11|-|122152276|122152296&offsets|0|+1(+1U)]|AACCCGTAGATCCGAACTTGTGT</td><td>>MI0000102|hsa-mir-100&WithFlank&11|-|122152223|122152314</td><td>19</td><td>40</td><td>22</td><td>(+1U)</td><td>isomiR</td><td>YES</td><td>1725.41</td><td>1837.43</td><td>2681.70</td><td>391.73</td></tr>
<tr><td>>MI0000102|hsa-mir-100&WithFlank&11|-|122152223|122152314@19.40.22(+2A)&[MIMAT0000098&hsa-miR-100-5p&offsets|0|0(+2A);m-27&11|-|122152276|122152296&offsets|0|+1(+2A)]|AACCCGTAGATCCGAACTTGTGAA</td><td>>MI0000102|hsa-mir-100&WithFlank&11|-|122152223|122152314</td><td>19</td><td>40</td><td>22</td><td>(+2A)</td><td>isomiR</td><td>NO</td><td>135.49</td><td>75.75</td><td>121.17</td><td>21.95</td></tr>
<tr><td>>MI0000102|hsa-mir-100&WithFlank&11|-|122152223|122152314@19.40.22(+2U)&[MIMAT0000098&hsa-miR-100-5p&offsets|0|0(+2U);m-27&11|-|122152276|122152296&offsets|0|+1(+2U)]|AACCCGTAGATCCGAACTTGTGTT</td><td>>MI0000102|hsa-mir-100&WithFlank&11|-|122152223|122152314</td><td>19</td><td>40</td><td>22</td><td>(+2U)</td><td>isomiR</td><td>NO</td><td>58.63</td><td>13.42</td><td>14.98</td><td>14.08</td></tr>
<tr><td>>MI0000102|hsa-mir-100&WithFlank&11|-|122152223|122152314@19.41.23&[MIMAT0000098&hsa-miR-100-5p&offsets|0|+1;m-27&11|-|122152276|122152296&offsets|0|+2]|AACCCGTAGATCCGAACTTGTGG</td><td>>MI0000102|hsa-mir-100&WithFlank&11|-|122152223|122152314</td><td>19</td><td>41</td><td>23</td><td></td><td>isomiR</td><td>NO</td><td>178.76</td><td>94.47</td><td>241.25</td><td>35.48</td></tr>
<tr><td>>MI0000102|hsa-mir-100&WithFlank&11|-|122152223|122152314@20.39.20&[MIMAT0000098&hsa-miR-100-5p&offsets|+1|-1;m-27&11|-|122152276|122152296&offsets|+1|0]|ACCCGTAGATCCGAACTTGT</td><td>>MI0000102|hsa-mir-100&WithFlank&11|-|122152223|122152314</td><td>20</td><td>39</td><td>20</td><td></td><td>isomiR</td><td>NO</td><td>0.00</td><td>0.00</td><td>17.65</td><td>0.00</td></tr>
<tr><td>>MI0000102|hsa-mir-100&WithFlank&11|-|122152223|122152314@20.39.20(+1A)&[MIMAT0000098&hsa-miR-100-5p&offsets|+1|-1(+1A);m-27&11|-|122152276|122152296&offsets|+1|0(+1A)]|ACCCGTAGATCCGAACTTGTA</td><td>>MI0000102|hsa-mir-100&WithFlank&11|-|122152223|122152314</td><td>20</td><td>39</td><td>20</td><td>(+1A)</td><td>isomiR</td><td>NO</td><td>0.00</td><td>0.00</td><td>11.95</td><td>0.00</td></tr>
<tr><td>>MI0000102|hsa-mir-100&WithFlank&11|-|122152223|122152314@20.40.21&[MIMAT0000098&hsa-miR-100-5p&offsets|+1|0;m-27&11|-|122152276|122152296&offsets|+1|+1]|ACCCGTAGATCCGAACTTGTG</td><td>>MI0000102|hsa-mir-100&WithFlank&11|-|122152223|122152314</td><td>20</td><td>40</td><td>21</td><td></td><td>isomiR</td><td>NO</td><td>92.65</td><td>98.31</td><td>143.59</td><td>13.35</td></tr>
<tr><td>>MI0000102|hsa-mir-100&WithFlank&11|-|122152223|122152314@22.40.19&[MIMAT0000098&hsa-miR-100-5p&offsets|+3|0;m-27&11|-|122152276|122152296&offsets|+3|+1]|CCGTAGATCCGAACTTGTG</td><td>>MI0000102|hsa-mir-100&WithFlank&11|-|122152223|122152314</td><td>22</td><td>40</td><td>19</td><td></td><td>isomiR</td><td>NO</td><td>0.00</td><td>0.00</td><td>14.66</td><td>0.00</td></tr>
</table>

### MDA-MB-231

<table>
<tr><td>>MI0000103|hsa-mir-101-1&WithFlank&1|-|65058428|65058514@52.72.21(+1C);MI0000739|hsa-mir-101-2&WithFlank&9|+|4850291|4850381@54.74.21(+1C)&[MIMAT0000099&hsa-miR-101-3p&offsets|-1|-1(+1C);m-17&1|-|65058443|65058462&offsets|-1|0(+1C)];[MIMAT0000099_1&hsa-miR-101-3p&offsets|-1|-1(+1C);m-16&9|+|4850345|4850364&offsets|-1|0(+1C)]|GTACAGTACTGTGATAACTGAC</td><td>>MI0000103|hsa-mir-101-1&WithFlank&1|-|65058428|65058514</td><td>52</td><td>72</td><td>21</td><td>(+1C)</td><td>isomiR</td><td>NO</td><td>0.00</td><td>0.00</td><td>13.66</td><td>0.00</td></tr>
<tr><td>>MI0000103|hsa-mir-101-1&WithFlank&1|-|65058428|65058514@52.72.21;MI0000739|hsa-mir-101-2&WithFlank&9|+|4850291|4850381@54.74.21&[MIMAT0000099&hsa-miR-101-3p&offsets|-1|-1;m-17&1|-|65058443|65058462&offsets|-1|0];[MIMAT0000099_1&hsa-miR-101-3p&offsets|-1|-1;m-16&9|+|4850345|4850364&offsets|-1|0]|GTACAGTACTGTGATAACTGA</td><td>>MI0000103|hsa-mir-101-1&WithFlank&1|-|65058428|65058514</td><td>52</td><td>72</td><td>21</td><td></td><td>isomiR</td><td>NO</td><td>24.49</td><td>0.00</td><td>96.67</td><td>0.00</td></tr>
<tr><td>>MI0000103|hsa-mir-101-1&WithFlank&1|-|65058428|65058514@52.73.22(+1A);MI0000739|hsa-mir-101-2&WithFlank&9|+|4850291|4850381@54.75.22(+1A)&[MIMAT0000099&hsa-miR-101-3p&offsets|-1|0(+1A);m-17&1|-|65058443|65058462&offsets|-1|+1(+1A)];[MIMAT0000099_1&hsa-miR-101-3p&offsets|-1|0(+1A);m-16&9|+|4850345|4850364&offsets|-1|+1(+1A)]|GTACAGTACTGTGATAACTGAAA</td><td>>MI0000103|hsa-mir-101-1&WithFlank&1|-|65058428|65058514</td><td>52</td><td>73</td><td>22</td><td>(+1A)</td><td>isomiR</td><td>NO</td><td>0.00</td><td>0.00</td><td>14.02</td><td>0.00</td></tr>
<tr><td>>MI0000103|hsa-mir-101-1&WithFlank&1|-|65058428|65058514@52.73.22;MI0000739|hsa-mir-101-2&WithFlank&9|+|4850291|4850381@54.75.22&[MIMAT0000099&hsa-miR-101-3p&offsets|-1|0;m-17&1|-|65058443|65058462&offsets|-1|+1];[MIMAT0000099_1&hsa-miR-101-3p&offsets|-1|0;m-16&9|+|4850345|4850364&offsets|-1|+1]|GTACAGTACTGTGATAACTGAA</td><td>>MI0000103|hsa-mir-101-1&WithFlank&1|-|65058428|65058514</td><td>52</td><td>73</td><td>22</td><td></td><td>isomiR</td><td>NO</td><td>89.47</td><td>0.00</td><td>246.51</td><td>12.94</td></tr>
<tr><td>>MI0000103|hsa-mir-101-1&WithFlank&1|-|65058428|65058514@53.72.20;MI0000739|hsa-mir-101-2&WithFlank&9|+|4850291|4850381@55.74.20&[MIMAT0000099&hsa-miR-101-3p&offsets|0|-1;m-17&1|-|65058443|65058462&offsets|0|0];[MIMAT0000099_1&hsa-miR-101-3p&offsets|0|-1;m-16&9|+|4850345|4850364&offsets|0|0]|TACAGTACTGTGATAACTGA</td><td>>MI0000103|hsa-mir-101-1&WithFlank&1|-|65058428|65058514</td><td>53</td><td>72</td><td>20</td><td></td><td>isomiR</td><td>NO</td><td>81.32</td><td>0.00</td><td>236.65</td><td>0.00</td></tr>
<tr><td>>MI0000103|hsa-mir-101-1&WithFlank&1|-|65058428|65058514@53.73.21(+1A);MI0000739|hsa-mir-101-2&WithFlank&9|+|4850291|4850381@55.75.21(+1A)&[MIMAT0000099&hsa-miR-101-3p&offsets|0|0(+1A);m-17&1|-|65058443|65058462&offsets|0|+1(+1A)];[MIMAT0000099_1&hsa-miR-101-3p&offsets|0|0(+1A);m-16&9|+|4850345|4850364&offsets|0|+1(+1A)]|TACAGTACTGTGATAACTGAAA</td><td>>MI0000103|hsa-mir-101-1&WithFlank&1|-|65058428|65058514</td><td>53</td><td>73</td><td>21</td><td>(+1A)</td><td>isomiR</td><td>NO</td><td>0.00</td><td>0.00</td><td>32.28</td><td>0.00</td></tr>
<tr><td>>MI0000103|hsa-mir-101-1&WithFlank&1|-|65058428|65058514@53.73.21(+1U);MI0000739|hsa-mir-101-2&WithFlank&9|+|4850291|4850381@55.75.21(+1U)&[MIMAT0000099&hsa-miR-101-3p&offsets|0|0(+1U);m-17&1|-|65058443|65058462&offsets|0|+1(+1U)];[MIMAT0000099_1&hsa-miR-101-3p&offsets|0|0(+1U);m-16&9|+|4850345|4850364&offsets|0|+1(+1U)]|TACAGTACTGTGATAACTGAAT</td><td>>MI0000103|hsa-mir-101-1&WithFlank&1|-|65058428|65058514</td><td>53</td><td>73</td><td>21</td><td>(+1U)</td><td>isomiR</td><td>NO</td><td>54.25</td><td>0.00</td><td>68.72</td><td>0.00</td></tr>
<tr><td>>MI0000103|hsa-mir-101-1&WithFlank&1|-|65058428|65058514@53.73.21;MI0000739|hsa-mir-101-2&WithFlank&9|+|4850291|4850381@55.75.21&[MIMAT0000099&hsa-miR-101-3p&offsets|0|0;m-17&1|-|65058443|65058462&offsets|0|+1];[MIMAT0000099_1&hsa-miR-101-3p&offsets|0|0;m-16&9|+|4850345|4850364&offsets|0|+1]|TACAGTACTGTGATAACTGAA</td><td>>MI0000103|hsa-mir-101-1&WithFlank&1|-|65058428|65058514</td><td>53</td><td>73</td><td>21</td><td></td><td>isomiR</td><td>NO</td><td>0.00</td><td>0.00</td><td>104.30</td><td>0.00</td></tr>
<tr><td>>MI0000103|hsa-mir-101-1&WithFlank&1|-|65058428|65058514@53.74.22(+1U);MI0000739|hsa-mir-101-2&WithFlank&9|+|4850291|4850381@55.76.22(+1U)&[MIMAT0000099&hsa-miR-101-3p&offsets|0|+1(+1U);m-17&1|-|65058443|65058462&offsets|0|+2(+1U)];[MIMAT0000099_1&hsa-miR-101-3p&offsets|0|+1(+1U);m-16&9|+|4850345|4850364&offsets|0|+2(+1U)]|TACAGTACTGTGATAACTGAAGT</td><td>>MI0000103|hsa-mir-101-1&WithFlank&1|-|65058428|65058514</td><td>53</td><td>74</td><td>22</td><td>(+1U)</td><td>isomiR</td><td>NO</td><td>24.53</td><td>0.00</td><td>75.29</td><td>0.00</td></tr>
<tr><td>>MI0000103|hsa-mir-101-1&WithFlank&1|-|65058428|65058514@53.74.22;MI0000739|hsa-mir-101-2&WithFlank&9|+|4850291|4850381@55.76.22&[MIMAT0000099&hsa-miR-101-3p&offsets|0|+1;m-17&1|-|65058443|65058462&offsets|0|+2];[MIMAT0000099_1&hsa-miR-101-3p&offsets|0|+1;m-16&9|+|4850345|4850364&offsets|0|+2]|TACAGTACTGTGATAACTGAAG</td><td>>MI0000103|hsa-mir-101-1&WithFlank&1|-|65058428|65058514</td><td>53</td><td>74</td><td>22</td><td></td><td>isomiR</td><td>NO</td><td>111.60</td><td>0.00</td><td>199.14</td><td>10.93</td></tr>
<tr><td>>MI0000107|hsa-mir-29b-2&WithFlank&1|-|207802437|207802529@58.79.22(+1C);MI0000105|hsa-mir-29b-1&WithFlank&7|-|130877453|130877545@57.78.22(+1C)&[MIMAT0000100&hsa-miR-29b-3p&offsets|0|-1(+1C);m-81&1|-|207802450|207802472&offsets|0|-1(+1C)];[MIMAT0000100_1&hsa-miR-29b-3p&offsets|0|-1(+1C);m-82&7|-|130877467|130877489&offsets|0|-1(+1C)]|TAGCACCATTTGAAATCAGTGTC</td><td>>MI0000107|hsa-mir-29b-2&WithFlank&1|-|207802437|207802529</td><td>58</td><td>79</td><td>22</td><td>(+1C)</td><td>isomiR</td><td>NO</td><td>0.00</td><td>0.00</td><td>13.47</td><td>0.00</td></tr>
<tr><td>>MI0000107|hsa-mir-29b-2&WithFlank&1|-|207802437|207802529@58.80.23;MI0000105|hsa-mir-29b-1&WithFlank&7|-|130877453|130877545@57.79.23&[MIMAT0000100&hsa-miR-29b-3p&offsets|0|0;m-81&1|-|207802450|207802472&offsets|0|0];[MIMAT0000100_1&hsa-miR-29b-3p&offsets|0|0;m-82&7|-|130877467|130877489&offsets|0|0]|TAGCACCATTTGAAATCAGTGTT</td><td>>MI0000107|hsa-mir-29b-2&WithFlank&1|-|207802437|207802529</td><td>58</td><td>80</td><td>23</td><td></td><td>isomiR</td><td>NO</td><td>60.67</td><td>0.00</td><td>170.53</td><td>0.00</td></tr>
<tr><td>>MI0000108|hsa-mir-103a-2&WithFlank&20|+|3917488|3917577@54.75.22(+1C);MI0000109|hsa-mir-103a-1&WithFlank&5|-|168560890|168560979@54.75.22(+1C)&[MIMAT0000101&hsa-miR-103a-3p&offsets|0|-1(+1C);m-12&20|+|3917541|3917563&offsets|0|-1(+1C)];[MIMAT0000101_1&hsa-miR-103a-3p&offsets|0|-1(+1C);m-13&5|-|168560904|168560926&offsets|0|-1(+1C);m-44220&5|-|168560908|168560926&offsets|0|+3(+1C)]|AGCAGCATTGTACAGGGCTATGC</td><td>>MI0000108|hsa-mir-103a-2&WithFlank&20|+|3917488|3917577</td><td>54</td><td>75</td><td>22</td><td>(+1C)</td><td>isomiR</td><td>NO</td><td>0.00</td><td>0.00</td><td>34.30</td><td>0.00</td></tr>
<tr><td>>MI0000108|hsa-mir-103a-2&WithFlank&20|+|3917488|3917577@54.75.22(+1G);MI0000109|hsa-mir-103a-1&WithFlank&5|-|168560890|168560979@54.75.22(+1G)&[MIMAT0000101&hsa-miR-103a-3p&offsets|0|-1(+1G);m-12&20|+|3917541|3917563&offsets|0|-1(+1G)];[MIMAT0000101_1&hsa-miR-103a-3p&offsets|0|-1(+1G);m-13&5|-|168560904|168560926&offsets|0|-1(+1G);m-44220&5|-|168560908|168560926&offsets|0|+3(+1G)]|AGCAGCATTGTACAGGGCTATGG</td><td>>MI0000108|hsa-mir-103a-2&WithFlank&20|+|3917488|3917577</td><td>54</td><td>75</td><td>22</td><td>(+1G)</td><td>isomiR</td><td>NO</td><td>0.00</td><td>0.00</td><td>73.96</td><td>0.00</td></tr>
</table>

### MDA-MB-231

<table>
<tr><th></th><th></th><th></th><th></th><th></th><th></th><th></th><th></th><th></th><th></th><th></th><th></th><th></th></tr>
<tr><td>>MI0000108|hsa-mir-103a-2&WithFlank&20|+|3917488|3917577@54.75.22(+1U);MI0000109|hsa-mir-103a-1&WithFlank&5|-|168560890|168560979@54.75.22(+1U)&[MIMAT0000101&hsa-miR-103a-3p&offsets|0|-1(+1U);m-12&20|+|3917541|3917563&offsets|0|-1(+1U)];[MIMAT0000101_1&hsa-miR-103a-3p&offsets|0|-1(+1U);m-13&5|-|168560904|168560926&offsets|0|-1(+1U);m-44220&5|-|168560908|168560926&offsets|0|+3(+1U)]|AGCAGCATTGTACAGGGCTATGT</td><td>>MI0000108|hsa-mir-103a-2&WithFlank&20|+|3917488|3917577</td><td>54</td><td>75</td><td>22</td><td>(+1U)</td><td>isomiR</td><td>NO</td><td>72.97</td><td>0.00</td><td>255.83</td><td>17.16</td></tr>
<tr><td>>MI0000108|hsa-mir-103a-2&WithFlank&20|+|3917488|3917577@54.75.22;MI0000109|hsa-mir-103a-1&WithFlank&5|-|168560890|168560979@54.75.22&[MIMAT0000101&hsa-miR-103a-3p&offsets|0|-1;m-12&20|+|3917541|3917563&offsets|0|-1];[MIMAT0000101_1&hsa-miR-103a-3p&offsets|0|-1;m-13&5|-|168560904|168560926&offsets|0|-1;m-44220&5|-|168560908|168560926&offsets|0|+3]|AGCAGCATTGTACAGGGCTATG</td><td>>MI0000108|hsa-mir-103a-2&WithFlank&20|+|3917488|3917577</td><td>54</td><td>75</td><td>22</td><td></td><td>isomiR</td><td>NO</td><td>271.41</td><td>101.01</td><td>729.02</td><td>112.34</td></tr>
<tr><td>>MI0000108|hsa-mir-103a-2&WithFlank&20|+|3917488|3917577@54.76.23(+1U);MI0000109|hsa-mir-103a-1&WithFlank&5|-|168560890|168560979@54.76.23(+1U)&[MIMAT0000101&hsa-miR-103a-3p&offsets|0|0(+1U);m-12&20|+|3917541|3917563&offsets|0|0(+1U)];[MIMAT0000101_1&hsa-miR-103a-3p&offsets|0|0(+1U);m-13&5|-|168560904|168560926&offsets|0|0(+1U);m-44220&5|-|168560908|168560926&offsets|0|+4(+1U)]|AGCAGCATTGTACAGGGCTATGAT</td><td>>MI0000108|hsa-mir-103a-2&WithFlank&20|+|3917488|3917577</td><td>54</td><td>76</td><td>23</td><td>(+1U)</td><td>isomiR</td><td>NO</td><td>150.69</td><td>85.71</td><td>87.06</td><td>110.89</td></tr>
<tr><td>>MI0000108|hsa-mir-103a-2&WithFlank&20|+|3917488|3917577@54.76.23(+2U);MI0000109|hsa-mir-103a-1&WithFlank&5|-|168560890|168560979@54.76.23(+2U)&[MIMAT0000101&hsa-miR-103a-3p&offsets|0|0(+2U);m-12&20|+|3917541|3917563&offsets|0|0(+2U)];[MIMAT0000101_1&hsa-miR-103a-3p&offsets|0|0(+2U);m-13&5|-|168560904|168560926&offsets|0|0(+2U);m-44220&5|-|168560908|168560926&offsets|0|+4(+2U)]|AGCAGCATTGTACAGGGCTATGATT</td><td>>MI0000108|hsa-mir-103a-2&WithFlank&20|+|3917488|3917577</td><td>54</td><td>76</td><td>23</td><td>(+2U)</td><td>isomiR</td><td>NO</td><td>52.98</td><td>33.57</td><td>31.56</td><td>14.88</td></tr>
<tr><td>>MI0000108|hsa-mir-103a-2&WithFlank&20|+|3917488|3917577@54.76.23;MI0000109|hsa-mir-103a-1&WithFlank&5|-|168560890|168560979@54.76.23&[MIMAT0000101&hsa-miR-103a-3p&offsets|0|0;m-12&20|+|3917541|3917563&offsets|0|0];[MIMAT0000101_1&hsa-miR-103a-3p&offsets|0|0;m-13&5|-|168560904|168560926&offsets|0|0;m-44220&5|-|168560908|168560926&offsets|0|+4]|AGCAGCATTGTACAGGGCTATGA</td><td>>MI0000108|hsa-mir-103a-2&WithFlank&20|+|3917488|3917577</td><td>54</td><td>76</td><td>23</td><td></td><td>isomiR</td><td>YES</td><td>846.22</td><td>390.88</td><td>2171.77</td><td>297.69</td></tr>
<tr><td>>MI0000108|hsa-mir-103a-2&WithFlank&20|+|3917488|3917577@54.77.24;MI0000109|hsa-mir-103a-1&WithFlank&5|-|168560890|168560979@54.77.24&[MIMAT0000101&hsa-miR-103a-3p&offsets|0|+1;m-12&20|+|3917541|3917563&offsets|0|+1];[MIMAT0000101_1&hsa-miR-103a-3p&offsets|0|+1;m-13&5|-|168560904|168560926&offsets|0|+1;m-44220&5|-|168560908|168560926&offsets|0|+5]|AGCAGCATTGTACAGGGCTATGAA</td><td>>MI0000108|hsa-mir-103a-2&WithFlank&20|+|3917488|3917577</td><td>54</td><td>77</td><td>24</td><td></td><td>isomiR</td><td>NO</td><td>58.05</td><td>0.00</td><td>13.63</td><td>11.89</td></tr>
<tr><td>>MI0000108|hsa-mir-103a-2&WithFlank&20|+|3917488|3917577@56.76.21;MI0000109|hsa-mir-103a-1&WithFlank&5|-|168560890|168560979@56.76.21&[MIMAT0000101&hsa-miR-103a-3p&offsets|+2|0;m-12&20|+|3917541|3917563&offsets|+2|0];[MIMAT0000101_1&hsa-miR-103a-3p&offsets|+2|0;m-13&5|-|168560904|168560926&offsets|+2|0;m-44220&5|-|168560908|168560926&offsets|+2|+4]|CAGCATTGTACAGGGCTATGA</td><td>>MI0000108|hsa-mir-103a-2&WithFlank&20|+|3917488|3917577</td><td>56</td><td>76</td><td>21</td><td></td><td>isomiR</td><td>NO</td><td>0.00</td><td>0.00</td><td>17.64</td><td>0.00</td></tr>
<tr><td>>MI0000114|hsa-mir-107&WithFlank&10|-|89592741|89592833@56.76.21(+1A);MI0000108|hsa-mir-103a-2&WithFlank&20|+|3917488|3917577@54.74.21(+1A);MI0000109|hsa-mir-103a-1&WithFlank&5|-|168560890|168560979@54.74.21(+1A)&[MIMAT0000104&hsa-miR-107&offsets|0|-2(+1A);m-78&10|-|89592758|89592778&offsets|0|0(+1A)];[MIMAT0000101&hsa-miR-103a-3p&offsets|0|-2(+1A);m-12&20|+|3917541|3917563&offsets|0|-2(+1A)];[MIMAT0000101_1&hsa-miR-103a-3p&offsets|0|-2(+1A);m-13&5|-|168560904|168560926&offsets|0|-2(+1A);m-44220&5|-|168560908|168560926&offsets|0|+2(+1A)]|AGCAGCATTGTACAGGGCTATA</td><td>>MI0000114|hsa-mir-107&WithFlank&10|-|89592741|89592833</td><td>56</td><td>76</td><td>21</td><td>(+1A)</td><td>isomiR</td><td>NO</td><td>0.00</td><td>0.00</td><td>151.47</td><td>0.00</td></tr>
<tr><td>>MI0000114|hsa-mir-107&WithFlank&10|-|89592741|89592833@56.76.21(+1U);MI0000108|hsa-mir-103a-2&WithFlank&20|+|3917488|3917577@54.74.21(+1U);MI0000109|hsa-mir-103a-1&WithFlank&5|-|168560890|168560979@54.74.21(+1U)&[MIMAT0000104&hsa-miR-107&offsets|0|-2(+1U);m-78&10|-|89592758|89592778&offsets|0|0(+1U)];[MIMAT0000101&hsa-miR-103a-3p&offsets|0|-2(+1U);m-12&20|+|3917541|3917563&offsets|0|-2(+1U)];[MIMAT0000101_1&hsa-miR-103a-3p&offsets|0|-2(+1U);m-13&5|-|168560904|168560926&offsets|0|-2(+1U);m-44220&5|-|168560908|168560926&offsets|0|+2(+1U)]|AGCAGCATTGTACAGGGCTATT</td><td>>MI0000114|hsa-mir-107&WithFlank&10|-|89592741|89592833</td><td>56</td><td>76</td><td>21</td><td>(+1U)</td><td>isomiR</td><td>NO</td><td>0.00</td><td>0.00</td><td>75.22</td><td>0.00</td></tr>
<tr><td>>MI0000114|hsa-mir-107&WithFlank&10|-|89592741|89592833@56.76.21;MI0000108|hsa-mir-103a-2&WithFlank&20|+|3917488|3917577@54.74.21;MI0000109|hsa-mir-103a-1&WithFlank&5|-|168560890|168560979@54.74.21&[MIMAT0000104&hsa-miR-107&offsets|0|-2;m-78&10|-|89592758|89592778&offsets|0|0];[MIMAT0000101&hsa-miR-103a-3p&offsets|0|-2;m-12&20|+|3917541|3917563&offsets|0|-2];[MIMAT0000101_1&hsa-miR-103a-3p&offsets|0|-2;m-13&5|-|168560904|168560926&offsets|0|-2;m-44220&5|-|168560908|168560926&offsets|0|+2]|AGCAGCATTGTACAGGGCTAT</td><td>>MI0000114|hsa-mir-107&WithFlank&10|-|89592741|89592833</td><td>56</td><td>76</td><td>21</td><td></td><td>isomiR</td><td>NO</td><td>50.64</td><td>0.00</td><td>340.53</td><td>11.13</td></tr>
<tr><td>>MI0000114|hsa-mir-107&WithFlank&10|-|89592741|89592833@56.77.22&[MIMAT0000104&hsa-miR-107&offsets|0|-1;m-78&10|-|89592758|89592778&offsets|0|+1]|AGCAGCATTGTACAGGGCTATC</td><td>>MI0000114|hsa-mir-107&WithFlank&10|-|89592741|89592833</td><td>56</td><td>77</td><td>22</td><td></td><td>isomiR</td><td>NO</td><td>24.97</td><td>0.00</td><td>173.33</td><td>0.00</td></tr>
<tr><td>>MI0000115|hsa-mir-16-2&WithFlank&3|+|160404739|160404831@58.79.22&[MIMAT0004518&hsa-miR-16-2-3p&offsets|-1|-1;m-252&3|+|160404796|160404816&offsets|0|+1]|ACCAATATTACTGTGCTGCTTT</td><td>>MI0000115|hsa-mir-16-2&WithFlank&3|+|160404739|160404831</td><td>58</td><td>79</td><td>22</td><td></td><td>isomiR</td><td>NO</td><td>57.62</td><td>13.45</td><td>106.68</td><td>15.89</td></tr>
<tr><td>>MI0000234|hsa-mir-192&WithFlank&11|-|64891131|64891252@30.49.20&[MIMAT0000222&hsa-miR-192-5p&offsets|0|-1;m-59&11|-|64891203|64891223&offsets|0|-1]|CTGACCTATGAATTGACAGC</td><td>>MI0000234|hsa-mir-192&WithFlank&11|-|64891131|64891252</td><td>30</td><td>49</td><td>20</td><td></td><td>isomiR</td><td>NO</td><td>0.00</td><td>0.00</td><td>15.00</td><td>0.00</td></tr>
<tr><td>>MI0000234|hsa-mir-192&WithFlank&11|-|64891131|64891252@30.49.20(+1A)&[MIMAT0000222&hsa-miR-192-5p&offsets|0|-1(+1A);m-59&11|-|64891203|64891223&offsets|0|-1(+1A)]|CTGACCTATGAATTGACAGCA</td><td>>MI0000234|hsa-mir-192&WithFlank&11|-|64891131|64891252</td><td>30</td><td>49</td><td>20</td><td>(+1A)</td><td>isomiR</td><td>NO</td><td>0.00</td><td>0.00</td><td>33.79</td><td>0.00</td></tr>
</table>

### MDA-MB-231

<table>
<tr><td>>MI0000234|hsa-mir-192&WithFlank&11|-|64891131|64891252@30.49.20(+1U)&[MIMAT0000222&hsa-miR-192-5p&offsets|0|-1(+1U);m-59&11|-|64891203|64891223&offsets|0|-1(+1U)]|CTGACCTATGAATTGACAGCT</td><td>>MI0000234|hsa-mir-192&WithFlank&11|-|64891131|64891252</td><td>30</td><td>49</td><td>20</td><td>(+1U)</td><td>isomiR</td><td>NO</td><td>0.00</td><td>0.00</td><td>119.98</td><td>0.00</td></tr>
<tr><td>>MI0000234|hsa-mir-192&WithFlank&11|-|64891131|64891252@30.50.21&[MIMAT0000222&hsa-miR-192-5p&offsets|0|0;m-59&11|-|64891203|64891223&offsets|0|0]|CTGACCTATGAATTGACAGCC</td><td>>MI0000234|hsa-mir-192&WithFlank&11|-|64891131|64891252</td><td>30</td><td>50</td><td>21</td><td></td><td>isomiR</td><td>NO</td><td>61.31</td><td>0.00</td><td>170.83</td><td>10.57</td></tr>
<tr><td>>MI0000234|hsa-mir-192&WithFlank&11|-|64891131|64891252@30.51.22&[MIMAT0000222&hsa-miR-192-5p&offsets|0|+1;m-59&11|-|64891203|64891223&offsets|0|+1]|CTGACCTATGAATTGACAGCCA</td><td>>MI0000234|hsa-mir-192&WithFlank&11|-|64891131|64891252</td><td>30</td><td>51</td><td>22</td><td></td><td>isomiR</td><td>NO</td><td>0.00</td><td>0.00</td><td>13.28</td><td>0.00</td></tr>
<tr><td>>MI0000234|hsa-mir-192&WithFlank&11|-|64891131|64891252@30.51.22(+1C)&[MIMAT0000222&hsa-miR-192-5p&offsets|0|+1(+1C);m-59&11|-|64891203|64891223&offsets|0|+1(+1C)]|CTGACCTATGAATTGACAGCCAC</td><td>>MI0000234|hsa-mir-192&WithFlank&11|-|64891131|64891252</td><td>30</td><td>51</td><td>22</td><td>(+1C)</td><td>isomiR</td><td>NO</td><td>0.00</td><td>0.00</td><td>10.63</td><td>0.00</td></tr>
<tr><td>>MI0000234|hsa-mir-192&WithFlank&11|-|64891131|64891252@30.51.22(+1U)&[MIMAT0000222&hsa-miR-192-5p&offsets|0|+1(+1U);m-59&11|-|64891203|64891223&offsets|0|+1(+1U)]|CTGACCTATGAATTGACAGCCAT</td><td>>MI0000234|hsa-mir-192&WithFlank&11|-|64891131|64891252</td><td>30</td><td>51</td><td>22</td><td>(+1U)</td><td>isomiR</td><td>NO</td><td>63.65</td><td>0.00</td><td>128.12</td><td>12.07</td></tr>
<tr><td>>MI0000234|hsa-mir-192&WithFlank&11|-|64891131|64891252@31.51.21(+1U)&[MIMAT0000222&hsa-miR-192-5p&offsets|+1|+1(+1U);m-59&11|-|64891203|64891223&offsets|+1|+1(+1U)]|TGACCTATGAATTGACAGCCAT</td><td>>MI0000234|hsa-mir-192&WithFlank&11|-|64891131|64891252</td><td>31</td><td>51</td><td>21</td><td>(+1U)</td><td>isomiR</td><td>NO</td><td>0.00</td><td>0.00</td><td>65.55</td><td>0.00</td></tr>
<tr><td>>MI0000234|hsa-mir-192&WithFlank&11|-|64891131|64891252@31.52.22&[MIMAT0000222&hsa-miR-192-5p&offsets|+1|+2;m-59&11|-|64891203|64891223&offsets|+1|+2]|TGACCTATGAATTGACAGCCAG</td><td>>MI0000234|hsa-mir-192&WithFlank&11|-|64891131|64891252</td><td>31</td><td>52</td><td>22</td><td></td><td>isomiR</td><td>NO</td><td>0.00</td><td>0.00</td><td>32.24</td><td>0.00</td></tr>
<tr><td>>MI0000234|hsa-mir-192&WithFlank&11|-|64891131|64891252@31.53.23&[MIMAT0000222&hsa-miR-192-5p&offsets|+1|+3;m-59&11|-|64891203|64891223&offsets|+1|+3]|TGACCTATGAATTGACAGCCAGT</td><td>>MI0000234|hsa-mir-192&WithFlank&11|-|64891131|64891252</td><td>31</td><td>53</td><td>23</td><td></td><td>isomiR</td><td>NO</td><td>0.00</td><td>0.00</td><td>120.59</td><td>0.00</td></tr>
<tr><td>>MI0000239|hsa-mir-197&WithFlank&1|+|109598887|109598973@54.74.21(+1U)&[MIMAT0000227&hsa-miR-197-3p&offsets|0|-1(+1U);m-99&1|+|109598940|109598961&offsets|0|-1(+1U)]|TTCACCACCTTCTCCACCCAGT</td><td>>MI0000239|hsa-mir-197&WithFlank&1|+|109598887|109598973</td><td>54</td><td>74</td><td>21</td><td>(+1U)</td><td>isomiR</td><td>NO</td><td>0.00</td><td>13.63</td><td>0.00</td><td>0.00</td></tr>
<tr><td>>MI0000239|hsa-mir-197&WithFlank&1|+|109598887|109598973@54.75.22&[MIMAT0000227&hsa-miR-197-3p&offsets|0|0;m-99&1|+|109598940|109598961&offsets|0|0]|TTCACCACCTTCTCCACCCAGC</td><td>>MI0000239|hsa-mir-197&WithFlank&1|+|109598887|109598973</td><td>54</td><td>75</td><td>22</td><td></td><td>isomiR</td><td>NO</td><td>67.90</td><td>133.63</td><td>0.00</td><td>56.63</td></tr>
<tr><td>>MI0000253|hsa-mir-148a&WithFlank&7|-|25949913|25949992@50.70.21(+1A)&[MIMAT0000243&hsa-miR-148a-3p&offsets|0|-1(+1A);m-7&7|-|25949922|25949943&offsets|0|-1(+1A)]|TCAGTGCACTACAGAACTTTGA</td><td>>MI0000253|hsa-mir-148a&WithFlank&7|-|25949913|25949992</td><td>50</td><td>70</td><td>21</td><td>(+1A)</td><td>isomiR</td><td>NO</td><td>0.00</td><td>0.00</td><td>93.48</td><td>0.00</td></tr>
<tr><td>>MI0000253|hsa-mir-148a&WithFlank&7|-|25949913|25949992@50.70.21(+1C)&[MIMAT0000243&hsa-miR-148a-3p&offsets|0|-1(+1C);m-7&7|-|25949922|25949943&offsets|0|-1(+1C)]|TCAGTGCACTACAGAACTTTGC</td><td>>MI0000253|hsa-mir-148a&WithFlank&7|-|25949913|25949992</td><td>50</td><td>70</td><td>21</td><td>(+1C)</td><td>isomiR</td><td>NO</td><td>72.62</td><td>24.87</td><td>314.21</td><td>10.91</td></tr>
<tr><td>>MI0000253|hsa-mir-148a&WithFlank&7|-|25949913|25949992@50.70.21(+1G)&[MIMAT0000243&hsa-miR-148a-3p&offsets|0|-1(+1G);m-7&7|-|25949922|25949943&offsets|0|-1(+1G)]|TCAGTGCACTACAGAACTTTGG</td><td>>MI0000253|hsa-mir-148a&WithFlank&7|-|25949913|25949992</td><td>50</td><td>70</td><td>21</td><td>(+1G)</td><td>isomiR</td><td>NO</td><td>0.00</td><td>0.00</td><td>16.23</td><td>0.00</td></tr>
<tr><td>>MI0000253|hsa-mir-148a&WithFlank&7|-|25949913|25949992@50.71.22&[MIMAT0000243&hsa-miR-148a-3p&offsets|0|0;m-7&7|-|25949922|25949943&offsets|0|0]|TCAGTGCACTACAGAACTTTGT</td><td>>MI0000253|hsa-mir-148a&WithFlank&7|-|25949913|25949992</td><td>50</td><td>71</td><td>22</td><td></td><td>isomiR</td><td>YES</td><td>430.10</td><td>269.40</td><td>1487.27</td><td>102.02</td></tr>
<tr><td>>MI0000253|hsa-mir-148a&WithFlank&7|-|25949913|25949992@50.72.23&[MIMAT0000243&hsa-miR-148a-3p&offsets|0|+1;m-7&7|-|25949922|25949943&offsets|0|+1]|TCAGTGCACTACAGAACTTTGTC</td><td>>MI0000253|hsa-mir-148a&WithFlank&7|-|25949913|25949992</td><td>50</td><td>72</td><td>23</td><td></td><td>isomiR</td><td>NO</td><td>0.00</td><td>0.00</td><td>10.62</td><td>0.00</td></tr>
<tr><td>>MI0000254|hsa-mir-30c-2&WithFlank&6|-|71376954|71377037@53.74.22&[MIMAT0004550&hsa-miR-30c-2-3p&offsets|0|0;m-206&6|-|71376964|71376985&offsets|0|0]|CTGGGAGAAGGCTGTTTACTCT</td><td>>MI0000254|hsa-mir-30c-2&WithFlank&6|-|71376954|71377037</td><td>53</td><td>74</td><td>22</td><td></td><td>isomiR</td><td>NO</td><td>68.14</td><td>122.01</td><td>34.55</td><td>12.92</td></tr>
<tr><td>>MI0000255|hsa-mir-30d&WithFlank&8|-|134804870|134804951@12.31.20&[MIMAT0000245&hsa-miR-30d-5p&offsets|0|-2;m-21&8|-|134804919|134804940&offsets|0|-2]|TGTAAACATCCCCGACTGGA</td><td>>MI0000255|hsa-mir-30d&WithFlank&8|-|134804870|134804951</td><td>12</td><td>31</td><td>20</td><td></td><td>isomiR</td><td>NO</td><td>80.20</td><td>0.00</td><td>122.17</td><td>13.58</td></tr>
<tr><td>>MI0000255|hsa-mir-30d&WithFlank&8|-|134804870|134804951@12.33.22&[MIMAT0000245&hsa-miR-30d-5p&offsets|0|0;m-21&8|-|134804919|134804940&offsets|0|0]|TGTAAACATCCCCGACTGGAAG</td><td>>MI0000255|hsa-mir-30d&WithFlank&8|-|134804870|134804951</td><td>12</td><td>33</td><td>22</td><td></td><td>isomiR</td><td>NO</td><td>117.61</td><td>24.92</td><td>111.10</td><td>22.33</td></tr>
<tr><td>>MI0000255|hsa-mir-30d&WithFlank&8|-|134804870|134804951@12.33.22(+1A)&[MIMAT0000245&hsa-miR-30d-5p&offsets|0|0(+1A);m-21&8|-|134804919|134804940&offsets|0|0(+1A)]|TGTAAACATCCCCGACTGGAAGA</td><td>>MI0000255|hsa-mir-30d&WithFlank&8|-|134804870|134804951</td><td>12</td><td>33</td><td>22</td><td>(+1A)</td><td>isomiR</td><td>NO</td><td>51.10</td><td>0.00</td><td>16.51</td><td>13.67</td></tr>
<tr><td>>MI0000255|hsa-mir-30d&WithFlank&8|-|134804870|134804951@12.33.22(+1U)&[MIMAT0000245&hsa-miR-30d-5p&offsets|0|0(+1U);m-21&8|-|134804919|134804940&offsets|0|0(+1U)]|TGTAAACATCCCCGACTGGAAGT</td><td>>MI0000255|hsa-mir-30d&WithFlank&8|-|134804870|134804951</td><td>12</td><td>33</td><td>22</td><td>(+1U)</td><td>isomiR</td><td>NO</td><td>176.44</td><td>83.61</td><td>142.79</td><td>90.36</td></tr>
<tr><td>>MI0000255|hsa-mir-30d&WithFlank&8|-|134804870|134804951@12.34.23&[MIMAT0000245&hsa-miR-30d-5p&offsets|0|+1;m-21&8|-|134804919|134804940&offsets|0|+1]|TGTAAACATCCCCGACTGGAAGC</td><td>>MI0000255|hsa-mir-30d&WithFlank&8|-|134804870|134804951</td><td>12</td><td>34</td><td>23</td><td></td><td>isomiR</td><td>NO</td><td>683.65</td><td>333.71</td><td>322.81</td><td>228.60</td></tr>
<tr><td>>MI0000255|hsa-mir-30d&WithFlank&8|-|134804870|134804951@12.34.23(+1A)&[MIMAT0000245&hsa-miR-30d-5p&offsets|0|+1(+1A);m-21&8|-|134804919|134804940&offsets|0|+1(+1A)]|TGTAAACATCCCCGACTGGAAGCA</td><td>>MI0000255|hsa-mir-30d&WithFlank&8|-|134804870|134804951</td><td>12</td><td>34</td><td>23</td><td>(+1A)</td><td>isomiR</td><td>NO</td><td>163.19</td><td>95.81</td><td>209.45</td><td>87.39</td></tr>
<tr><td>>MI0000255|hsa-mir-30d&WithFlank&8|-|134804870|134804951@12.34.23(+1C)&[MIMAT0000245&hsa-miR-30d-5p&offsets|0|+1(+1C);m-21&8|-|134804919|134804940&offsets|0|+1(+1C)]|TGTAAACATCCCCGACTGGAAGCC</td><td>>MI0000255|hsa-mir-30d&WithFlank&8|-|134804870|134804951</td><td>12</td><td>34</td><td>23</td><td>(+1C)</td><td>isomiR</td><td>NO</td><td>283.80</td><td>147.76</td><td>298.16</td><td>128.49</td></tr>
<tr><td>>MI0000255|hsa-mir-30d&WithFlank&8|-|134804870|134804951@12.34.23(+1G)&[MIMAT0000245&hsa-miR-30d-5p&offsets|0|+1(+1G);m-21&8|-|134804919|134804940&offsets|0|+1(+1G)]|TGTAAACATCCCCGACTGGAAGCG</td><td>>MI0000255|hsa-mir-30d&WithFlank&8|-|134804870|134804951</td><td>12</td><td>34</td><td>23</td><td>(+1G)</td><td>isomiR</td><td>NO</td><td>83.10</td><td>0.00</td><td>34.64</td><td>12.50</td></tr>
</table>

### MDA-MB-231

| | | | | | | | | | | | |
|---|---|---|---|---|---|---|---|---|---|---|---|
| >MI0000255\|hsa-mir-30d&WithFlank&8\|-\|134804870\|134804951@12.35.24&[MIMAT0000245&hsa-miR-30d-5p&offsets\|0\|+2;m-21&8\|-\|134804919\|134804940&offsets\|0\|+2]\|TGTAAACATCCCCGACTGGAAGCT | >MI0000255\|hsa-mir-30d&WithFlank&8\|-\|134804870\|134804951 | 12 | 35 | 24 | | isomiR | YES | 1346.88 | 828.89 | 1194.29 | 418.29 |
| >MI0000255\|hsa-mir-30d&WithFlank&8\|-\|134804870\|134804951@12.35.24(+1C)&[MIMAT0000245&hsa-miR-30d-5p&offsets\|0\|+2(+1C);m-21&8\|-\|134804919\|134804940&offsets\|0\|+2(+1C)]\|TGTAAACATCCCCGACTGGAAGCTC | >MI0000255\|hsa-mir-30d&WithFlank&8\|-\|134804870\|134804951 | 12 | 35 | 24 | (+1C) | isomiR | NO | 0.00 | 0.00 | 14.65 | 0.00 |
| >MI0000255\|hsa-mir-30d&WithFlank&8\|-\|134804870\|134804951@12.35.24(+1U)&[MIMAT0000245&hsa-miR-30d-5p&offsets\|0\|+2(+1U);m-21&8\|-\|134804919\|134804940&offsets\|0\|+2(+1U)]\|TGTAAACATCCCCGACTGGAAGCTT | >MI0000255\|hsa-mir-30d&WithFlank&8\|-\|134804870\|134804951 | 12 | 35 | 24 | (+1U) | isomiR | NO | 141.55 | 72.77 | 172.95 | 69.44 |
| >MI0000255\|hsa-mir-30d&WithFlank&8\|-\|134804870\|134804951@12.35.24(+2U)&[MIMAT0000245&hsa-miR-30d-5p&offsets\|0\|+2(+2U);m-21&8\|-\|134804919\|134804940&offsets\|0\|+2(+2U)]\|TGTAAACATCCCCGACTGGAAGCTTT | >MI0000255\|hsa-mir-30d&WithFlank&8\|-\|134804870\|134804951 | 12 | 35 | 24 | (+2U) | isomiR | NO | 26.72 | 0.00 | 0.00 | 12.62 |
| >MI0000261\|hsa-mir-139&WithFlank&11\|-\|72615057\|72615136@13.34.22(+1C)&[MIMAT0000250&hsa-miR-139-5p&offsets\|0\|-1(+1C);m-186&11\|-\|72615102\|72615124&offsets\|0\|-1(+1C)]\|TCTACAGTGCACGTGTCTCCAGC | >MI0000261\|hsa-mir-139&WithFlank&11\|-\|72615057\|72615136 | 13 | 34 | 22 | (+1C) | isomiR | NO | 0.00 | 0.00 | 13.02 | 0.00 |
| >MI0000261\|hsa-mir-139&WithFlank&11\|-\|72615057\|72615136@13.35.23&[MIMAT0000250&hsa-miR-139-5p&offsets\|0\|0;m-186&11\|-\|72615102\|72615124&offsets\|0\|0]\|TCTACAGTGCACGTGTCTCCAGT | >MI0000261\|hsa-mir-139&WithFlank&11\|-\|72615057\|72615136 | 13 | 35 | 23 | | isomiR | NO | 91.43 | 98.71 | 164.27 | 20.90 |
| >MI0000263\|hsa-mir-7-1&WithFlank&9\|-\|83969742\|83969863@30.54.25&[MIMAT0000252_2&hsa-miR-7-5p&offsets\|0\|+1;m-401&9\|-\|83969811\|83969834&offsets\|0\|+1]\|TGGAAGACTAGTGATTTTGTTGTTT | >MI0000263\|hsa-mir-7-1&WithFlank&9\|-\|83969742\|83969863 | 30 | 54 | 25 | | isomiR | NO | 98.01 | 0.00 | 268.72 | 19.94 |
| >MI0000264\|hsa-mir-7-2&WithFlank&15\|+\|88611819\|88611940@38.57.20(+1C);MI0000265\|hsa-mir-7-3&WithFlank&19\|+\|4770664\|4770785@37.56.20(+1C);MI0000263\|hsa-mir-7-1&WithFlank&9\|-\|83969742\|83969863@30.49.20(+1C)&[MIMAT0000252&hsa-miR-7-5p&offsets\|0\|-4(+1C);m-400&15\|+\|88611856\|88611879&offsets\|0\|-4(+1C)];[MIMAT0000252_1&hsa-miR-7-5p&offsets\|0\|-4(+1C);m-402&19\|+\|4770700\|4770723&offsets\|0\|-4(+1C)];[MIMAT0000252_2&hsa-miR-7-5p&offsets\|0\|-4(+1C);m-401&9\|-\|83969811\|83969834&offsets\|0\|-4(+1C)]\|TGGAAGACTAGTGATTTTGTC | >MI0000264\|hsa-mir-7-2&WithFlank&15\|+\|88611819\|88611940 | 38 | 57 | 20 | (+1C) | isomiR | NO | 0.00 | 0.00 | 55.09 | 0.00 |
| >MI0000264\|hsa-mir-7-2&WithFlank&15\|+\|88611819\|88611940@38.57.20;MI0000265\|hsa-mir-7-3&WithFlank&19\|+\|4770664\|4770785@37.56.20;MI0000263\|hsa-mir-7-1&WithFlank&9\|-\|83969742\|83969863@30.49.20&[MIMAT0000252&hsa-miR-7-5p&offsets\|0\|-4;m-400&15\|+\|88611856\|88611879&offsets\|0\|-4];[MIMAT0000252_1&hsa-miR-7-5p&offsets\|0\|-4;m-402&19\|+\|4770700\|4770723&offsets\|0\|-4];[MIMAT0000252_2&hsa-miR-7-5p&offsets\|0\|-4;m-401&9\|-\|83969811\|83969834&offsets\|0\|-4]\|TGGAAGACTAGTGATTTTGT | >MI0000264\|hsa-mir-7-2&WithFlank&15\|+\|88611819\|88611940 | 38 | 57 | 20 | | isomiR | NO | 0.00 | 0.00 | 16.79 | 0.00 |
| >MI0000264\|hsa-mir-7-2&WithFlank&15\|+\|88611819\|88611940@38.58.21(+1A);MI0000265\|hsa-mir-7-3&WithFlank&19\|+\|4770664\|4770785@37.57.21(+1A);MI0000263\|hsa-mir-7-1&WithFlank&9\|-\|83969742\|83969863@30.50.21(+1A)&[MIMAT0000252&hsa-miR-7-5p&offsets\|0\|-3(+1A);m-400&15\|+\|88611856\|88611879&offsets\|0\|-3(+1A)];[MIMAT0000252_1&hsa-miR-7-5p&offsets\|0\|-3(+1A);m-402&19\|+\|4770700\|4770723&offsets\|0\|-3(+1A)];[MIMAT0000252_2&hsa-miR-7-5p&offsets\|0\|-3(+1A);m-401&9\|-\|83969811\|83969834&offsets\|0\|-3(+1A)]\|TGGAAGACTAGTGATTTTGTTA | >MI0000264\|hsa-mir-7-2&WithFlank&15\|+\|88611819\|88611940 | 38 | 58 | 21 | (+1A) | isomiR | NO | 0.00 | 0.00 | 35.31 | 0.00 |
| >MI0000264\|hsa-mir-7-2&WithFlank&15\|+\|88611819\|88611940@38.58.21;MI0000265\|hsa-mir-7-3&WithFlank&19\|+\|4770664\|4770785@37.57.21;MI0000263\|hsa-mir-7-1&WithFlank&9\|-\|83969742\|83969863@30.50.21&[MIMAT0000252&hsa-miR-7-5p&offsets\|0\|-3;m-400&15\|+\|88611856\|88611879&offsets\|0\|-3];[MIMAT0000252_1&hsa-miR-7-5p&offsets\|0\|-3;m-402&19\|+\|4770700\|4770723&offsets\|0\|-3];[MIMAT0000252_2&hsa-miR-7-5p&offsets\|0\|-3;m-401&9\|-\|83969811\|83969834&offsets\|0\|-3]\|TGGAAGACTAGTGATTTTGTT | >MI0000264\|hsa-mir-7-2&WithFlank&15\|+\|88611819\|88611940 | 38 | 58 | 21 | | isomiR | NO | 89.02 | 0.00 | 261.17 | 15.01 |
| >MI0000264\|hsa-mir-7-2&WithFlank&15\|+\|88611819\|88611940@38.59.22(+1A);MI0000265\|hsa-mir-7-3&WithFlank&19\|+\|4770664\|4770785@37.58.22(+1A);MI0000263\|hsa-mir-7-1&WithFlank&9\|-\|83969742\|83969863@30.51.22(+1A)&[MIMAT0000252&hsa-miR-7-5p&offsets\|0\|-2(+1A);m-400&15\|+\|88611856\|88611879&offsets\|0\|-2(+1A)];[MIMAT0000252_1&hsa-miR-7-5p&offsets\|0\|-2(+1A);m-402&19\|+\|4770700\|4770723&offsets\|0\|-2(+1A)];[MIMAT0000252_2&hsa-miR-7-5p&offsets\|0\|-2(+1A);m-401&9\|-\|83969811\|83969834&offsets\|0\|-2(+1A)]\|TGGAAGACTAGTGATTTTGTTGA | >MI0000264\|hsa-mir-7-2&WithFlank&15\|+\|88611819\|88611940 | 38 | 59 | 22 | (+1A) | isomiR | NO | 0.00 | 0.00 | 67.84 | 0.00 |
| >MI0000264\|hsa-mir-7-2&WithFlank&15\|+\|88611819\|88611940@38.59.22(+1C);MI0000265\|hsa-mir-7-3&WithFlank&19\|+\|4770664\|4770785@37.58.22(+1C);MI0000263\|hsa-mir-7-1&WithFlank&9\|-\|83969742\|83969863@30.51.22(+1C)&[MIMAT0000252&hsa-miR-7-5p&offsets\|0\|-2(+1C);m-400&15\|+\|88611856\|88611879&offsets\|0\|-2(+1C)];[MIMAT0000252_1&hsa-miR-7-5p&offsets\|0\|-2(+1C);m-402&19\|+\|4770700\|4770723&offsets\|0\|-2(+1C)];[MIMAT0000252_2&hsa-miR-7-5p&offsets\|0\|-2(+1C);m-401&9\|-\|83969811\|83969834&offsets\|0\|-2(+1C)]\|TGGAAGACTAGTGATTTTGTTGC | >MI0000264\|hsa-mir-7-2&WithFlank&15\|+\|88611819\|88611940 | 38 | 59 | 22 | (+1C) | isomiR | NO | 50.14 | 0.00 | 196.99 | 11.02 |
| >MI0000264\|hsa-mir-7-2&WithFlank&15\|+\|88611819\|88611940@38.59.22;MI0000265\|hsa-mir-7-3&WithFlank&19\|+\|4770664\|4770785@37.58.22;MI0000263\|hsa-mir-7-1&WithFlank&9\|-\|83969742\|83969863@30.51.22&[MIMAT0000252&hsa-miR-7-5p&offsets\|0\|-2;m-400&15\|+\|88611856\|88611879&offsets\|0\|-2];[MIMAT0000252_1&hsa-miR-7-5p&offsets\|0\|-2;m-402&19\|+\|4770700\|4770723&offsets\|0\|-2];[MIMAT0000252_2&hsa-miR-7-5p&offsets\|0\|-2;m-401&9\|-\|83969811\|83969834&offsets\|0\|-2]\|TGGAAGACTAGTGATTTTGTTG | >MI0000264\|hsa-mir-7-2&WithFlank&15\|+\|88611819\|88611940 | 38 | 59 | 22 | | isomiR | NO | 74.00 | 0.00 | 253.03 | 12.84 |

### MDA-MB-231

<table>
<tr><th></th><th></th><th></th><th></th><th></th><th></th><th></th><th></th><th></th><th></th><th></th><th></th></tr>
<tr><td>>MI0000264|hsa-mir-7-2&WithFlank&15|+|88611819|88611940@38.60.23(+1A);MI0000265|hsa-mir-7-3&WithFlank&19|+|4770664|4770785@37.59.23(+1A);MI0000263|hsa-mir-7-1&WithFlank&9|-|83969742|83969863@30.52.23(+1A)&[MIMAT0000252&hsa-miR-7-5p&offsets|0|-1(+1A);m-400&15|+|88611856|88611879&offsets|0|-1(+1A)];[MIMAT0000252_1&hsa-miR-7-5p&offsets|0|-1(+1A);m-402&19|+|4770700|4770723&offsets|0|-1(+1A)];[MIMAT0000252_2&hsa-miR-7-5p&offsets|0|-1(+1A);m-401&9|-|83969811|83969834&offsets|0|-1(+1A)]|TGGAAGACTAGTGATTTTGTTGTA</td><td>>MI0000264|hsa-mir-7-2&WithFlank&15|+|88611819|88611940</td><td>38</td><td>60</td><td>23</td><td>(+1A)</td><td>isomiR</td><td>NO</td><td>64.12</td><td>0.00</td><td>184.19</td><td>16.02</td></tr>
<tr><td>>MI0000264|hsa-mir-7-2&WithFlank&15|+|88611819|88611940@38.60.23(+1C);MI0000265|hsa-mir-7-3&WithFlank&19|+|4770664|4770785@37.59.23(+1C);MI0000263|hsa-mir-7-1&WithFlank&9|-|83969742|83969863@30.52.23(+1C)&[MIMAT0000252&hsa-miR-7-5p&offsets|0|-1(+1C);m-400&15|+|88611856|88611879&offsets|0|-1(+1C)];[MIMAT0000252_1&hsa-miR-7-5p&offsets|0|-1(+1C);m-402&19|+|4770700|4770723&offsets|0|-1(+1C)];[MIMAT0000252_2&hsa-miR-7-5p&offsets|0|-1(+1C);m-401&9|-|83969811|83969834&offsets|0|-1(+1C)]|TGGAAGACTAGTGATTTTGTTGTC</td><td>>MI0000264|hsa-mir-7-2&WithFlank&15|+|88611819|88611940</td><td>38</td><td>60</td><td>23</td><td>(+1C)</td><td>isomiR</td><td>NO</td><td>247.15</td><td>95.39</td><td>579.82</td><td>106.58</td></tr>
<tr><td>>MI0000264|hsa-mir-7-2&WithFlank&15|+|88611819|88611940@38.60.23(+1G);MI0000265|hsa-mir-7-3&WithFlank&19|+|4770664|4770785@37.59.23(+1G);MI0000263|hsa-mir-7-1&WithFlank&9|-|83969742|83969863@30.52.23(+1G)&[MIMAT0000252&hsa-miR-7-5p&offsets|0|-1(+1G);m-400&15|+|88611856|88611879&offsets|0|-1(+1G)];[MIMAT0000252_1&hsa-miR-7-5p&offsets|0|-1(+1G);m-402&19|+|4770700|4770723&offsets|0|-1(+1G)];[MIMAT0000252_2&hsa-miR-7-5p&offsets|0|-1(+1G);m-401&9|-|83969811|83969834&offsets|0|-1(+1G)]|TGGAAGACTAGTGATTTTGTTGTG</td><td>>MI0000264|hsa-mir-7-2&WithFlank&15|+|88611819|88611940</td><td>38</td><td>60</td><td>23</td><td>(+1G)</td><td>isomiR</td><td>NO</td><td>26.56</td><td>0.00</td><td>64.11</td><td>0.00</td></tr>
<tr><td>>MI0000264|hsa-mir-7-2&WithFlank&15|+|88611819|88611940@38.60.23;MI0000265|hsa-mir-7-3&WithFlank&19|+|4770664|4770785@37.59.23;MI0000263|hsa-mir-7-1&WithFlank&9|-|83969742|83969863@30.52.23&[MIMAT0000252&hsa-miR-7-5p&offsets|0|-1;m-400&15|+|88611856|88611879&offsets|0|-1];[MIMAT0000252_1&hsa-miR-7-5p&offsets|0|-1;m-402&19|+|4770700|4770723&offsets|0|-1];[MIMAT0000252_2&hsa-miR-7-5p&offsets|0|-1;m-401&9|-|83969811|83969834&offsets|0|-1]|TGGAAGACTAGTGATTTTGTTGT</td><td>>MI0000264|hsa-mir-7-2&WithFlank&15|+|88611819|88611940</td><td>38</td><td>60</td><td>23</td><td></td><td>isomiR</td><td>NO</td><td>291.34</td><td>82.38</td><td>905.78</td><td>99.59</td></tr>
<tr><td>>MI0000264|hsa-mir-7-2&WithFlank&15|+|88611819|88611940@38.61.24(+1A);MI0000265|hsa-mir-7-3&WithFlank&19|+|4770664|4770785@37.60.24(+1A);MI0000263|hsa-mir-7-1&WithFlank&9|-|83969742|83969863@30.53.24(+1A)&[MIMAT0000252&hsa-miR-7-5p&offsets|0|0(+1A);m-400&15|+|88611856|88611879&offsets|0|0(+1A)];[MIMAT0000252_1&hsa-miR-7-5p&offsets|0|0(+1A);m-402&19|+|4770700|4770723&offsets|0|0(+1A)];[MIMAT0000252_2&hsa-miR-7-5p&offsets|0|0(+1A);m-401&9|-|83969811|83969834&offsets|0|0(+1A)]|TGGAAGACTAGTGATTTTGTTGTTA</td><td>>MI0000264|hsa-mir-7-2&WithFlank&15|+|88611819|88611940</td><td>38</td><td>61</td><td>24</td><td>(+1A)</td><td>isomiR</td><td>NO</td><td>0.00</td><td>0.00</td><td>81.60</td><td>0.00</td></tr>
<tr><td>>MI0000264|hsa-mir-7-2&WithFlank&15|+|88611819|88611940@38.61.24;MI0000265|hsa-mir-7-3&WithFlank&19|+|4770664|4770785@37.60.24;MI0000263|hsa-mir-7-1&WithFlank&9|-|83969742|83969863@30.53.24&[MIMAT0000252&hsa-miR-7-5p&offsets|0|0;m-400&15|+|88611856|88611879&offsets|0|0];[MIMAT0000252_1&hsa-miR-7-5p&offsets|0|0;m-402&19|+|4770700|4770723&offsets|0|0];[MIMAT0000252_2&hsa-miR-7-5p&offsets|0|0;m-401&9|-|83969811|83969834&offsets|0|0]|TGGAAGACTAGTGATTTTGTTGTT</td><td>>MI0000264|hsa-mir-7-2&WithFlank&15|+|88611819|88611940</td><td>38</td><td>61</td><td>24</td><td></td><td>isomiR</td><td>YES</td><td>1434.28</td><td>640.13</td><td>2607.92</td><td>393.02</td></tr>
<tr><td>>MI0000265|hsa-mir-7-3&WithFlank&19|+|4770664|4770785@37.61.25&[MIMAT0000252_1&hsa-miR-7-5p&offsets|0|+1;m-402&19|+|4770700|4770723&offsets|0|+1]|TGGAAGACTAGTGATTTTGTTGTTC</td><td>>MI0000265|hsa-mir-7-3&WithFlank&19|+|4770664|4770785</td><td>37</td><td>61</td><td>25</td><td></td><td>isomiR</td><td>NO</td><td>0.00</td><td>0.00</td><td>55.21</td><td>0.00</td></tr>
<tr><td>>MI0000266|hsa-mir-10a&WithFlank&17|-|48579832|48579953@28.48.21&[MIMAT0000253&hsa-miR-10a-5p&offsets|0|-2;m-18&17|-|48579905|48579926&offsets|0|-1]|TACCCTGTAGATCCGAATTTG</td><td>>MI0000266|hsa-mir-10a&WithFlank&17|-|48579832|48579953</td><td>28</td><td>48</td><td>21</td><td></td><td>isomiR</td><td>NO</td><td>0.00</td><td>0.00</td><td>16.11</td><td>0.00</td></tr>
<tr><td>>MI0000266|hsa-mir-10a&WithFlank&17|-|48579832|48579953@28.48.21(+1A)&[MIMAT0000253&hsa-miR-10a-5p&offsets|0|-2(+1A);m-18&17|-|48579905|48579926&offsets|0|-1(+1A)]|TACCCTGTAGATCCGAATTTGA</td><td>>MI0000266|hsa-mir-10a&WithFlank&17|-|48579832|48579953</td><td>28</td><td>48</td><td>21</td><td>(+1A)</td><td>isomiR</td><td>NO</td><td>0.00</td><td>0.00</td><td>16.37</td><td>0.00</td></tr>
<tr><td>>MI0000266|hsa-mir-10a&WithFlank&17|-|48579832|48579953@28.48.21(+1C)&[MIMAT0000253&hsa-miR-10a-5p&offsets|0|-2(+1C);m-18&17|-|48579905|48579926&offsets|0|-1(+1C)]|TACCCTGTAGATCCGAATTTGC</td><td>>MI0000266|hsa-mir-10a&WithFlank&17|-|48579832|48579953</td><td>28</td><td>48</td><td>21</td><td>(+1C)</td><td>isomiR</td><td>NO</td><td>63.45</td><td>57.03</td><td>134.17</td><td>13.60</td></tr>
<tr><td>>MI0000266|hsa-mir-10a&WithFlank&17|-|48579832|48579953@28.49.22&[MIMAT0000253&hsa-miR-10a-5p&offsets|0|-1;m-18&17|-|48579905|48579926&offsets|0|0]|TACCCTGTAGATCCGAATTTGT</td><td>>MI0000266|hsa-mir-10a&WithFlank&17|-|48579832|48579953</td><td>28</td><td>49</td><td>22</td><td></td><td>isomiR</td><td>NO</td><td>357.96</td><td>376.13</td><td>595.92</td><td>121.27</td></tr>
<tr><td>>MI0000266|hsa-mir-10a&WithFlank&17|-|48579832|48579953@28.49.22(+1A)&[MIMAT0000253&hsa-miR-10a-5p&offsets|0|-1(+1A);m-18&17|-|48579905|48579926&offsets|0|0(+1A)]|TACCCTGTAGATCCGAATTTGTA</td><td>>MI0000266|hsa-mir-10a&WithFlank&17|-|48579832|48579953</td><td>28</td><td>49</td><td>22</td><td>(+1A)</td><td>isomiR</td><td>NO</td><td>0.00</td><td>0.00</td><td>71.68</td><td>0.00</td></tr>
<tr><td>>MI0000266|hsa-mir-10a&WithFlank&17|-|48579832|48579953@28.50.23&[MIMAT0000253&hsa-miR-10a-5p&offsets|0|0;m-18&17|-|48579905|48579926&offsets|0|+1]|TACCCTGTAGATCCGAATTTGTG</td><td>>MI0000266|hsa-mir-10a&WithFlank&17|-|48579832|48579953</td><td>28</td><td>50</td><td>23</td><td></td><td>isomiR</td><td>NO</td><td>205.62</td><td>203.62</td><td>322.72</td><td>78.97</td></tr>
<tr><td>>MI0000266|hsa-mir-10a&WithFlank&17|-|48579832|48579953@28.50.23(+1A)&[MIMAT0000253&hsa-miR-10a-5p&offsets|0|0(+1A);m-18&17|-|48579905|48579926&offsets|0|+1(+1A)]|TACCCTGTAGATCCGAATTTGTGA</td><td>>MI0000266|hsa-mir-10a&WithFlank&17|-|48579832|48579953</td><td>28</td><td>50</td><td>23</td><td>(+1A)</td><td>isomiR</td><td>NO</td><td>26.45</td><td>0.00</td><td>55.44</td><td>0.00</td></tr>
<tr><td>>MI0000266|hsa-mir-10a&WithFlank&17|-|48579832|48579953@28.51.24&[MIMAT0000253&hsa-miR-10a-5p&offsets|0|+1;m-18&17|-|48579905|48579926&offsets|0|+2]|TACCCTGTAGATCCGAATTTGTGT</td><td>>MI0000266|hsa-mir-10a&WithFlank&17|-|48579832|48579953</td><td>28</td><td>51</td><td>24</td><td></td><td>isomiR</td><td>NO</td><td>46.40</td><td>31.93</td><td>55.14</td><td>11.94</td></tr>
<tr><td>>MI0000266|hsa-mir-10a&WithFlank&17|-|48579832|48579953@29.48.20(+1C)&[MIMAT0000253&hsa-miR-10a-5p&offsets|+1|-2(+1C);m-18&17|-|48579905|48579926&offsets|+1|-1(+1C)]|ACCCTGTAGATCCGAATTTGC</td><td>>MI0000266|hsa-mir-10a&WithFlank&17|-|48579832|48579953</td><td>29</td><td>48</td><td>20</td><td>(+1C)</td><td>isomiR</td><td>NO</td><td>0.00</td><td>0.00</td><td>11.03</td><td>0.00</td></tr>
</table>

### MDA-MB-231

<table>
<tr><th></th><th></th><th></th><th></th><th></th><th></th><th></th><th></th><th></th><th></th><th></th><th></th></tr>
<tr><td>>MI0000266|hsa-mir-10a&WithFlank&17|-|48579832|48579953@29.49.21&[MIMAT0000253&hsa-miR-10a-5p&offsets|+1|-1;m-18&17|-|48579905|48579926&offsets|+1|0]|ACCCTGTAGATCCGAATTTGT</td><td>>MI0000266|hsa-mir-10a&WithFlank&17|-|48579832|48579953</td><td>29</td><td>49</td><td>21</td><td></td><td>isomiR</td><td>NO</td><td>84.42</td><td>79.01</td><td>123.79</td><td>16.04</td></tr>
<tr><td>>MI0000266|hsa-mir-10a&WithFlank&17|-|48579832|48579953@29.49.21(+1A)&[MIMAT0000253&hsa-miR-10a-5p&offsets|+1|-1(+1A);m-18&17|-|48579905|48579926&offsets|+1|0(+1A)]|ACCCTGTAGATCCGAATTTGTA</td><td>>MI0000266|hsa-mir-10a&WithFlank&17|-|48579832|48579953</td><td>29</td><td>49</td><td>21</td><td>(+1A)</td><td>isomiR</td><td>NO</td><td>0.00</td><td>0.00</td><td>34.24</td><td>0.00</td></tr>
<tr><td>>MI0000266|hsa-mir-10a&WithFlank&17|-|48579832|48579953@29.50.22&[MIMAT0000253&hsa-miR-10a-5p&offsets|+1|0;m-18&17|-|48579905|48579926&offsets|+1|+1]|ACCCTGTAGATCCGAATTTGTG</td><td>>MI0000266|hsa-mir-10a&WithFlank&17|-|48579832|48579953</td><td>29</td><td>50</td><td>22</td><td></td><td>isomiR</td><td>NO</td><td>185.78</td><td>179.74</td><td>232.38</td><td>69.21</td></tr>
<tr><td>>MI0000266|hsa-mir-10a&WithFlank&17|-|48579832|48579953@29.50.22(+1A)&[MIMAT0000253&hsa-miR-10a-5p&offsets|+1|0(+1A);m-18&17|-|48579905|48579926&offsets|+1|+1(+1A)]|ACCCTGTAGATCCGAATTTGTGA</td><td>>MI0000266|hsa-mir-10a&WithFlank&17|-|48579832|48579953</td><td>29</td><td>50</td><td>22</td><td>(+1A)</td><td>isomiR</td><td>NO</td><td>24.57</td><td>0.00</td><td>17.57</td><td>0.00</td></tr>
<tr><td>>MI0000266|hsa-mir-10a&WithFlank&17|-|48579832|48579953@29.51.23&[MIMAT0000253&hsa-miR-10a-5p&offsets|+1|+1;m-18&17|-|48579905|48579926&offsets|+1|+2]|ACCCTGTAGATCCGAATTTGTGT</td><td>>MI0000266|hsa-mir-10a&WithFlank&17|-|48579832|48579953</td><td>29</td><td>51</td><td>23</td><td></td><td>isomiR</td><td>NO</td><td>51.06</td><td>30.11</td><td>33.86</td><td>12.11</td></tr>
<tr><td>>MI0000266|hsa-mir-10a&WithFlank&17|-|48579832|48579953@69.89.21&[MIMAT0004555&hsa-miR-10a-3p&offsets|0|-1;m-468&17|-|48579865|48579885&offsets|0|0]|CAAATTCGTATCTAGGGGAAT</td><td>>MI0000266|hsa-mir-10a&WithFlank&17|-|48579832|48579953</td><td>69</td><td>89</td><td>21</td><td></td><td>isomiR</td><td>NO</td><td>0.00</td><td>0.00</td><td>14.49</td><td>0.00</td></tr>
<tr><td>>MI0000269|hsa-mir-181a-2&WithFlank&9|+|124692436|124692557@83.103.21&[MIMAT0004558&hsa-miR-181a-2-3p&offsets|0|-1;m-71&9|+|124692518|124692539&offsets|0|-1]|ACCACTGACCGTTGACTGTAC</td><td>>MI0000269|hsa-mir-181a-2&WithFlank&9|+|124692436|124692557</td><td>83</td><td>103</td><td>21</td><td></td><td>isomiR</td><td>NO</td><td>0.00</td><td>0.00</td><td>10.57</td><td>0.00</td></tr>
<tr><td>>MI0000269|hsa-mir-181a-2&WithFlank&9|+|124692436|124692557@83.103.21(+1U)&[MIMAT0004558&hsa-miR-181a-2-3p&offsets|0|-1(+1U);m-71&9|+|124692518|124692539&offsets|0|-1(+1U)]|ACCACTGACCGTTGACTGTACT</td><td>>MI0000269|hsa-mir-181a-2&WithFlank&9|+|124692436|124692557</td><td>83</td><td>103</td><td>21</td><td>(+1U)</td><td>isomiR</td><td>NO</td><td>0.00</td><td>36.48</td><td>13.74</td><td>0.00</td></tr>
<tr><td>>MI0000269|hsa-mir-181a-2&WithFlank&9|+|124692436|124692557@83.104.22&[MIMAT0004558&hsa-miR-181a-2-3p&offsets|0|0;m-71&9|+|124692518|124692539&offsets|0|0]|ACCACTGACCGTTGACTGTACC</td><td>>MI0000269|hsa-mir-181a-2&WithFlank&9|+|124692436|124692557</td><td>83</td><td>104</td><td>22</td><td></td><td>isomiR</td><td>NO</td><td>75.66</td><td>138.65</td><td>62.24</td><td>16.08</td></tr>
<tr><td>>MI0000270|hsa-mir-181b-1&WithFlank&1|-|198858867|198858988@42.60.19;MI0000683|hsa-mir-181b-2&WithFlank&9|+|124693704|124693804@22.40.19&[MIMAT0000257&hsa-miR-181b-5p&offsets|0|-4;m-73&1|-|198858925|198858947&offsets|0|-4];[MIMAT0000257_1&hsa-miR-181b-5p&offsets|0|-4;m-72&9|+|124693725|124693747&offsets|0|-4]|AACATTCATTGCTGTCGGT</td><td>>MI0000270|hsa-mir-181b-1&WithFlank&1|-|198858867|198858988</td><td>42</td><td>60</td><td>19</td><td></td><td>isomiR</td><td>NO</td><td>0.00</td><td>0.00</td><td>10.52</td><td>0.00</td></tr>
<tr><td>>MI0000270|hsa-mir-181b-1&WithFlank&1|-|198858867|198858988@42.61.20;MI0000683|hsa-mir-181b-2&WithFlank&9|+|124693704|124693804@22.41.20&[MIMAT0000257&hsa-miR-181b-5p&offsets|0|-3;m-73&1|-|198858925|198858947&offsets|0|-3];[MIMAT0000257_1&hsa-miR-181b-5p&offsets|0|-3;m-72&9|+|124693725|124693747&offsets|0|-3]|AACATTCATTGCTGTCGGTG</td><td>>MI0000270|hsa-mir-181b-1&WithFlank&1|-|198858867|198858988</td><td>42</td><td>61</td><td>20</td><td></td><td>isomiR</td><td>NO</td><td>0.00</td><td>0.00</td><td>35.05</td><td>0.00</td></tr>
<tr><td>>MI0000270|hsa-mir-181b-1&WithFlank&1|-|198858867|198858988@42.62.21;MI0000683|hsa-mir-181b-2&WithFlank&9|+|124693704|124693804@22.42.21&[MIMAT0000257&hsa-miR-181b-5p&offsets|0|-2;m-73&1|-|198858925|198858947&offsets|0|-2];[MIMAT0000257_1&hsa-miR-181b-5p&offsets|0|-2;m-72&9|+|124693725|124693747&offsets|0|-2]|AACATTCATTGCTGTCGGTGG</td><td>>MI0000270|hsa-mir-181b-1&WithFlank&1|-|198858867|198858988</td><td>42</td><td>62</td><td>21</td><td></td><td>isomiR</td><td>NO</td><td>24.28</td><td>0.00</td><td>17.30</td><td>0.00</td></tr>
<tr><td>>MI0000270|hsa-mir-181b-1&WithFlank&1|-|198858867|198858988@42.63.22(+1A);MI0000683|hsa-mir-181b-2&WithFlank&9|+|124693704|124693804@22.43.22(+1A)&[MIMAT0000257&hsa-miR-181b-5p&offsets|0|-1(+1A);m-73&1|-|198858925|198858947&offsets|0|-1(+1A)];[MIMAT0000257_1&hsa-miR-181b-5p&offsets|0|-1(+1A);m-72&9|+|124693725|124693747&offsets|0|-1(+1A)]|AACATTCATTGCTGTCGGTGGGA</td><td>>MI0000270|hsa-mir-181b-1&WithFlank&1|-|198858867|198858988</td><td>42</td><td>63</td><td>22</td><td>(+1A)</td><td>isomiR</td><td>NO</td><td>0.00</td><td>0.00</td><td>12.05</td><td>0.00</td></tr>
<tr><td>>MI0000270|hsa-mir-181b-1&WithFlank&1|-|198858867|198858988@42.63.22(+1C);MI0000683|hsa-mir-181b-2&WithFlank&9|+|124693704|124693804@22.43.22(+1C)&[MIMAT0000257&hsa-miR-181b-5p&offsets|0|-1(+1C);m-73&1|-|198858925|198858947&offsets|0|-1(+1C)];[MIMAT0000257_1&hsa-miR-181b-5p&offsets|0|-1(+1C);m-72&9|+|124693725|124693747&offsets|0|-1(+1C)]|AACATTCATTGCTGTCGGTGGGC</td><td>>MI0000270|hsa-mir-181b-1&WithFlank&1|-|198858867|198858988</td><td>42</td><td>63</td><td>22</td><td>(+1C)</td><td>isomiR</td><td>NO</td><td>0.00</td><td>0.00</td><td>34.71</td><td>0.00</td></tr>
<tr><td>>MI0000270|hsa-mir-181b-1&WithFlank&1|-|198858867|198858988@42.63.22;MI0000683|hsa-mir-181b-2&WithFlank&9|+|124693704|124693804@22.43.22&[MIMAT0000257&hsa-miR-181b-5p&offsets|0|-1;m-73&1|-|198858925|198858947&offsets|0|-1];[MIMAT0000257_1&hsa-miR-181b-5p&offsets|0|-1;m-72&9|+|124693725|124693747&offsets|0|-1]|AACATTCATTGCTGTCGGTGGG</td><td>>MI0000270|hsa-mir-181b-1&WithFlank&1|-|198858867|198858988</td><td>42</td><td>63</td><td>22</td><td></td><td>isomiR</td><td>NO</td><td>0.00</td><td>0.00</td><td>10.86</td><td>0.00</td></tr>
<tr><td>>MI0000270|hsa-mir-181b-1&WithFlank&1|-|198858867|198858988@42.64.23(+1A);MI0000683|hsa-mir-181b-2&WithFlank&9|+|124693704|124693804@22.44.23(+1A)&[MIMAT0000257&hsa-miR-181b-5p&offsets|0|0(+1A);m-73&1|-|198858925|198858947&offsets|0|0(+1A)];[MIMAT0000257_1&hsa-miR-181b-5p&offsets|0|0(+1A);m-72&9|+|124693725|124693747&offsets|0|0(+1A)]|AACATTCATTGCTGTCGGTGGGTA</td><td>>MI0000270|hsa-mir-181b-1&WithFlank&1|-|198858867|198858988</td><td>42</td><td>64</td><td>23</td><td>(+1A)</td><td>isomiR</td><td>NO</td><td>0.00</td><td>0.00</td><td>15.63</td><td>0.00</td></tr>
<tr><td>>MI0000270|hsa-mir-181b-1&WithFlank&1|-|198858867|198858988@42.64.23(+1C);MI0000683|hsa-mir-181b-2&WithFlank&9|+|124693704|124693804@22.44.23(+1C)&[MIMAT0000257&hsa-miR-181b-5p&offsets|0|0(+1C);m-73&1|-|198858925|198858947&offsets|0|0(+1C)];[MIMAT0000257_1&hsa-miR-181b-5p&offsets|0|0(+1C);m-72&9|+|124693725|124693747&offsets|0|0(+1C)]|AACATTCATTGCTGTCGGTGGGTC</td><td>>MI0000270|hsa-mir-181b-1&WithFlank&1|-|198858867|198858988</td><td>42</td><td>64</td><td>23</td><td>(+1C)</td><td>isomiR</td><td>NO</td><td>0.00</td><td>0.00</td><td>88.15</td><td>0.00</td></tr>
<tr><td>>MI0000270|hsa-mir-181b-1&WithFlank&1|-|198858867|198858988@42.64.23;MI0000683|hsa-mir-181b-2&WithFlank&9|+|124693704|124693804@22.44.23&[MIMAT0000257&hsa-miR-181b-5p&offsets|0|0;m-73&1|-|198858925|198858947&offsets|0|0];[MIMAT0000257_1&hsa-miR-181b-5p&offsets|0|0;m-72&9|+|124693725|124693747&offsets|0|0]|AACATTCATTGCTGTCGGTGGGT</td><td>>MI0000270|hsa-mir-181b-1&WithFlank&1|-|198858867|198858988</td><td>42</td><td>64</td><td>23</td><td></td><td>isomiR</td><td>NO</td><td>153.41</td><td>82.12</td><td>225.93</td><td>56.18</td></tr>
</table>

### MDA-MB-231

<table>
<tr><th></th><th></th><th></th><th></th><th></th><th></th><th></th><th></th><th></th><th></th><th></th><th></th></tr>
<tr><td>>MI0000270|hsa-mir-181b-1&WithFlank&1|-|198858867|198858988@42.65.24(+1C);MI0000683|hsa-mir-181b-2&WithFlank&9|+|124693704|124693804@22.45.24(+1C)&[MIMAT0000257&hsa-miR-181b-5p&offsets|0|+1(+1C);m-73&1|-|198858925|198858947&offsets|0|+1(+1C)];[MIMAT0000257_1&hsa-miR-181b-5p&offsets|0|+1(+1C);m-72&9|+|124693725|124693747&offsets|0|+1(+1C)]|AACATTCATTGCTGTCGGTGGGTTC</td><td>>MI0000270|hsa-mir-181b-1&WithFlank&1|-|198858867|198858988</td><td>42</td><td>65</td><td>24</td><td>(+1C)</td><td>isomiR</td><td>NO</td><td>0.00</td><td>0.00</td><td>12.40</td><td>0.00</td></tr>
<tr><td>>MI0000270|hsa-mir-181b-1&WithFlank&1|-|198858867|198858988@42.65.24;MI0000683|hsa-mir-181b-2&WithFlank&9|+|124693704|124693804@22.45.24&[MIMAT0000257&hsa-miR-181b-5p&offsets|0|+1;m-73&1|-|198858925|198858947&offsets|0|+1];[MIMAT0000257_1&hsa-miR-181b-5p&offsets|0|+1;m-72&9|+|124693725|124693747&offsets|0|+1]|AACATTCATTGCTGTCGGTGGGTT</td><td>>MI0000270|hsa-mir-181b-1&WithFlank&1|-|198858867|198858988</td><td>42</td><td>65</td><td>24</td><td></td><td>isomiR</td><td>NO</td><td>104.91</td><td>26.25</td><td>400.96</td><td>56.45</td></tr>
<tr><td>>MI0000272|hsa-mir-182&WithFlank&7|-|129770377|129770498@29.50.22&[MIMAT0000259&hsa-miR-182-5p&offsets|0|-2;m-22&7|-|129770449|129770470&offsets|0|0]|TTTGGCAATGGTAGAACTCACA</td><td>>MI0000272|hsa-mir-182&WithFlank&7|-|129770377|129770498</td><td>29</td><td>50</td><td>22</td><td></td><td>isomiR</td><td>NO</td><td>0.00</td><td>0.00</td><td>72.03</td><td>0.00</td></tr>
<tr><td>>MI0000272|hsa-mir-182&WithFlank&7|-|129770377|129770498@29.51.23&[MIMAT0000259&hsa-miR-182-5p&offsets|0|-1;m-22&7|-|129770449|129770470&offsets|0|+1]|TTTGGCAATGGTAGAACTCACAC</td><td>>MI0000272|hsa-mir-182&WithFlank&7|-|129770377|129770498</td><td>29</td><td>51</td><td>23</td><td></td><td>isomiR</td><td>NO</td><td>0.00</td><td>0.00</td><td>14.99</td><td>0.00</td></tr>
<tr><td>>MI0000272|hsa-mir-182&WithFlank&7|-|129770377|129770498@29.51.23(+1C)&[MIMAT0000259&hsa-miR-182-5p&offsets|0|-1(+1C);m-22&7|-|129770449|129770470&offsets|0|+1(+1C)]|TTTGGCAATGGTAGAACTCACACC</td><td>>MI0000272|hsa-mir-182&WithFlank&7|-|129770377|129770498</td><td>29</td><td>51</td><td>23</td><td>(+1C)</td><td>isomiR</td><td>NO</td><td>46.76</td><td>24.61</td><td>15.02</td><td>0.00</td></tr>
<tr><td>>MI0000272|hsa-mir-182&WithFlank&7|-|129770377|129770498@29.52.24&[MIMAT0000259&hsa-miR-182-5p&offsets|0|0;m-22&7|-|129770449|129770470&offsets|0|+2]|TTTGGCAATGGTAGAACTCACACT</td><td>>MI0000272|hsa-mir-182&WithFlank&7|-|129770377|129770498</td><td>29</td><td>52</td><td>24</td><td></td><td>isomiR</td><td>NO</td><td>264.10</td><td>235.03</td><td>168.98</td><td>88.63</td></tr>
<tr><td>>MI0000272|hsa-mir-182&WithFlank&7|-|129770377|129770498@29.53.25&[MIMAT0000259&hsa-miR-182-5p&offsets|0|+1;m-22&7|-|129770449|129770470&offsets|0|+3]|TTTGGCAATGGTAGAACTCACACTG</td><td>>MI0000272|hsa-mir-182&WithFlank&7|-|129770377|129770498</td><td>29</td><td>53</td><td>25</td><td></td><td>isomiR</td><td>NO</td><td>95.72</td><td>89.24</td><td>65.66</td><td>20.09</td></tr>
<tr><td>>MI0000272|hsa-mir-182&WithFlank&7|-|129770377|129770498@29.53.25(+1U)&[MIMAT0000259&hsa-miR-182-5p&offsets|0|+1(+1U);m-22&7|-|129770449|129770470&offsets|0|+3(+1U)]|TTTGGCAATGGTAGAACTCACACTGT</td><td>>MI0000272|hsa-mir-182&WithFlank&7|-|129770377|129770498</td><td>29</td><td>53</td><td>25</td><td>(+1U)</td><td>isomiR</td><td>NO</td><td>0.00</td><td>12.95</td><td>12.75</td><td>10.54</td></tr>
<tr><td>>MI0000272|hsa-mir-182&WithFlank&7|-|129770377|129770498@29.54.26&[MIMAT0000259&hsa-miR-182-5p&offsets|0|+2;m-22&7|-|129770449|129770470&offsets|0|+4]|TTTGGCAATGGTAGAACTCACACTGG</td><td>>MI0000272|hsa-mir-182&WithFlank&7|-|129770377|129770498</td><td>29</td><td>54</td><td>26</td><td></td><td>isomiR</td><td>NO</td><td>55.13</td><td>27.44</td><td>16.71</td><td>14.86</td></tr>
<tr><td>>MI0000273|hsa-mir-183&WithFlank&7|-|129774899|129775020@33.53.21(+1C)&[MIMAT0000261&hsa-miR-183-5p&offsets|0|-1(+1C);m-41&7|-|129774966|129774987&offsets|-1|-2(+1C)]|TATGGCACTGGTAGAATTCACC</td><td>>MI0000273|hsa-mir-183&WithFlank&7|-|129774899|129775020</td><td>33</td><td>53</td><td>21</td><td>(+1C)</td><td>isomiR</td><td>NO</td><td>0.00</td><td>0.00</td><td>13.75</td><td>0.00</td></tr>
<tr><td>>MI0000273|hsa-mir-183&WithFlank&7|-|129774899|129775020@33.54.22&[MIMAT0000261&hsa-miR-183-5p&offsets|0|0;m-41&7|-|129774966|129774987&offsets|-1|-1]|TATGGCACTGGTAGAATTCACT</td><td>>MI0000273|hsa-mir-183&WithFlank&7|-|129774899|129775020</td><td>33</td><td>54</td><td>22</td><td></td><td>isomiR</td><td>NO</td><td>184.03</td><td>114.18</td><td>157.95</td><td>60.53</td></tr>
<tr><td>>MI0000273|hsa-mir-183&WithFlank&7|-|129774899|129775020@33.55.23&[MIMAT0000261&hsa-miR-183-5p&offsets|0|+1;m-41&7|-|129774966|129774987&offsets|-1|0]|TATGGCACTGGTAGAATTCACTG</td><td>>MI0000273|hsa-mir-183&WithFlank&7|-|129774899|129775020</td><td>33</td><td>55</td><td>23</td><td></td><td>isomiR</td><td>NO</td><td>54.21</td><td>0.00</td><td>107.88</td><td>0.00</td></tr>
<tr><td>>MI0000273|hsa-mir-183&WithFlank&7|-|129774899|129775020@34.54.21&[MIMAT0000261&hsa-miR-183-5p&offsets|+1|0;m-41&7|-|129774966|129774987&offsets|0|-1]|ATGGCACTGGTAGAATTCACT</td><td>>MI0000273|hsa-mir-183&WithFlank&7|-|129774899|129775020</td><td>34</td><td>54</td><td>21</td><td></td><td>isomiR</td><td>NO</td><td>24.75</td><td>0.00</td><td>34.62</td><td>0.00</td></tr>
<tr><td>>MI0000273|hsa-mir-183&WithFlank&7|-|129774899|129775020@34.55.22&[MIMAT0000261&hsa-miR-183-5p&offsets|+1|+1;m-41&7|-|129774966|129774987&offsets|0|0]|ATGGCACTGGTAGAATTCACTG</td><td>>MI0000273|hsa-mir-183&WithFlank&7|-|129774899|129775020</td><td>34</td><td>55</td><td>22</td><td></td><td>isomiR</td><td>NO</td><td>0.00</td><td>0.00</td><td>81.83</td><td>0.00</td></tr>
<tr><td>>MI0000286|hsa-mir-210&WithFlank&11|-|568083|568204@72.93.22&[MIMAT0000267&hsa-miR-210-3p&offsets|0|0;m-102&11|-|568112|568133&offsets|0|0]|CTGTGCGTGTGACAGCGGCTGA</td><td>>MI0000286|hsa-mir-210&WithFlank&11|-|568083|568204</td><td>72</td><td>93</td><td>22</td><td></td><td>isomiR</td><td>NO</td><td>0.00</td><td>0.00</td><td>142.89</td><td>0.00</td></tr>
<tr><td>>MI0000286|hsa-mir-210&WithFlank&11|-|568083|568204@72.93.22(+1A)&[MIMAT0000267&hsa-miR-210-3p&offsets|0|0(+1A);m-102&11|-|568112|568133&offsets|0|0(+1A)]|CTGTGCGTGTGACAGCGGCTGAA</td><td>>MI0000286|hsa-mir-210&WithFlank&11|-|568083|568204</td><td>72</td><td>93</td><td>22</td><td>(+1A)</td><td>isomiR</td><td>NO</td><td>0.00</td><td>0.00</td><td>73.80</td><td>0.00</td></tr>
<tr><td>>MI0000286|hsa-mir-210&WithFlank&11|-|568083|568204@72.93.22(+2A)&[MIMAT0000267&hsa-miR-210-3p&offsets|0|0(+2A);m-102&11|-|568112|568133&offsets|0|0(+2A)]|CTGTGCGTGTGACAGCGGCTGAAA</td><td>>MI0000286|hsa-mir-210&WithFlank&11|-|568083|568204</td><td>72</td><td>93</td><td>22</td><td>(+2A)</td><td>isomiR</td><td>NO</td><td>0.00</td><td>0.00</td><td>178.73</td><td>10.99</td></tr>
<tr><td>>MI0000286|hsa-mir-210&WithFlank&11|-|568083|568204@72.94.23&[MIMAT0000267&hsa-miR-210-3p&offsets|0|+1;m-102&11|-|568112|568133&offsets|0|+1]|CTGTGCGTGTGACAGCGGCTGAT</td><td>>MI0000286|hsa-mir-210&WithFlank&11|-|568083|568204</td><td>72</td><td>94</td><td>23</td><td></td><td>isomiR</td><td>NO</td><td>0.00</td><td>0.00</td><td>81.94</td><td>0.00</td></tr>
<tr><td>>MI0000289|hsa-mir-181a-1&WithFlank&1|-|198859038|198859159@30.49.20;MI0000269|hsa-mir-181a-2&WithFlank&9|+|124692436|124692557@45.64.20&[MIMAT0000256&hsa-miR-181a-5p&offsets|0|-3;m-32&1|-|198859108|198859130&offsets|0|-3];[MIMAT0000256_1&hsa-miR-181a-5p&offsets|0|-3;m-32&9|+|124692480|124692502&offsets|0|-3]|AACATTCAACGCTGTCGGTG</td><td>>MI0000289|hsa-mir-181a-1&WithFlank&1|-|198859038|198859159</td><td>30</td><td>49</td><td>20</td><td></td><td>isomiR</td><td>NO</td><td>25.68</td><td>0.00</td><td>108.70</td><td>0.00</td></tr>
<tr><td>>MI0000289|hsa-mir-181a-1&WithFlank&1|-|198859038|198859159@30.50.21;MI0000269|hsa-mir-181a-2&WithFlank&9|+|124692436|124692557@45.65.21&[MIMAT0000256&hsa-miR-181a-5p&offsets|0|-2;m-32&1|-|198859108|198859130&offsets|0|-2];[MIMAT0000256_1&hsa-miR-181a-5p&offsets|0|-2;m-32&9|+|124692480|124692502&offsets|0|-2]|AACATTCAACGCTGTCGGTGA</td><td>>MI0000289|hsa-mir-181a-1&WithFlank&1|-|198859038|198859159</td><td>30</td><td>50</td><td>21</td><td></td><td>isomiR</td><td>NO</td><td>25.88</td><td>0.00</td><td>73.45</td><td>0.00</td></tr>
<tr><td>>MI0000289|hsa-mir-181a-1&WithFlank&1|-|198859038|198859159@30.51.22(+1A);MI0000269|hsa-mir-181a-2&WithFlank&9|+|124692436|124692557@45.66.22(+1A)&[MIMAT0000256&hsa-miR-181a-5p&offsets|0|-1(+1A);m-32&1|-|198859108|198859130&offsets|0|-1(+1A)];[MIMAT0000256_1&hsa-miR-181a-5p&offsets|0|-1(+1A);m-32&9|+|124692480|124692502&offsets|0|-1(+1A)]|AACATTCAACGCTGTCGGTGAGA</td><td>>MI0000289|hsa-mir-181a-1&WithFlank&1|-|198859038|198859159</td><td>30</td><td>51</td><td>22</td><td>(+1A)</td><td>isomiR</td><td>NO</td><td>0.00</td><td>0.00</td><td>78.16</td><td>0.00</td></tr>
</table>

### MDA-MB-231

| | | | | | | | | | | | |
|---|---|---|---|---|---|---|---|---|---|---|---|
| >MI0000289\|hsa-mir-181a-1&WithFlank&1\|-\|198859038\|198859159@30.51.22(+1C);MI0000269\|hsa-mir-181a-2&WithFlank&9\|+\|124692436\|124692557@45.66.22(+1C)&[MIMAT0000256&hsa-miR-181a-5p&offsets\|0\|-1(+1C);m-32&1\|-\|198859108\|198859130&offsets\|0\|-1(+1C)];[MIMAT0000256_1&hsa-miR-181a-5p&offsets\|0\|-1(+1C);m-32&9\|+\|124692480\|124692502&offsets\|0\|-1(+1C)]\|AACATTCAACGCTGTCGGTGAGC | >MI0000289\|hsa-mir-181a-1&WithFlank&1\|-\|198859038\|198859159 | 30 | 51 | 22 | (+1C) | isomiR | NO | 0.00 | 0.00 | 232.07 | 0.00 |
| >MI0000289\|hsa-mir-181a-1&WithFlank&1\|-\|198859038\|198859159@30.51.22(+1G);MI0000269\|hsa-mir-181a-2&WithFlank&9\|+\|124692436\|124692557@45.66.22(+1G)&[MIMAT0000256&hsa-miR-181a-5p&offsets\|0\|-1(+1G);m-32&1\|-\|198859108\|198859130&offsets\|0\|-1(+1G)];[MIMAT0000256_1&hsa-miR-181a-5p&offsets\|0\|-1(+1G);m-32&9\|+\|124692480\|124692502&offsets\|0\|-1(+1G)]\|AACATTCAACGCTGTCGGTGAGG | >MI0000289\|hsa-mir-181a-1&WithFlank&1\|-\|198859038\|198859159 | 30 | 51 | 22 | (+1G) | isomiR | NO | 0.00 | 0.00 | 12.16 | 0.00 |
| >MI0000289\|hsa-mir-181a-1&WithFlank&1\|-\|198859038\|198859159@30.51.22;MI0000269\|hsa-mir-181a-2&WithFlank&9\|+\|124692436\|124692557@45.66.22&[MIMAT0000256&hsa-miR-181a-5p&offsets\|0\|-1;m-32&1\|-\|198859108\|198859130&offsets\|0\|-1];[MIMAT0000256_1&hsa-miR-181a-5p&offsets\|0\|-1;m-32&9\|+\|124692480\|124692502&offsets\|0\|-1]\|AACATTCAACGCTGTCGGTGAG | >MI0000289\|hsa-mir-181a-1&WithFlank&1\|-\|198859038\|198859159 | 30 | 51 | 22 | | isomiR | NO | 10.33 | 0.00 | 84.39 | 0.00 |
| >MI0000289\|hsa-mir-181a-1&WithFlank&1\|-\|198859038\|198859159@30.52.23(+1C);MI0000269\|hsa-mir-181a-2&WithFlank&9\|+\|124692436\|124692557@45.67.23(+1C)&[MIMAT0000256&hsa-miR-181a-5p&offsets\|0\|0(+1C);m-32&1\|-\|198859108\|198859130&offsets\|0\|0(+1C)];[MIMAT0000256_1&hsa-miR-181a-5p&offsets\|0\|0(+1C);m-32&9\|+\|124692480\|124692502&offsets\|0\|0(+1C)]\|AACATTCAACGCTGTCGGTGAGTC | >MI0000289\|hsa-mir-181a-1&WithFlank&1\|-\|198859038\|198859159 | 30 | 52 | 23 | (+1C) | isomiR | NO | 0.00 | 0.00 | 149.36 | 0.00 |
| >MI0000289\|hsa-mir-181a-1&WithFlank&1\|-\|198859038\|198859159@30.52.23;MI0000269\|hsa-mir-181a-2&WithFlank&9\|+\|124692436\|124692557@45.67.23&[MIMAT0000256&hsa-miR-181a-5p&offsets\|0\|0;m-32&1\|-\|198859108\|198859130&offsets\|0\|0];[MIMAT0000256_1&hsa-miR-181a-5p&offsets\|0\|0;m-32&9\|+\|124692480\|124692502&offsets\|0\|0]\|AACATTCAACGCTGTCGGTGAGT | >MI0000289\|hsa-mir-181a-1&WithFlank&1\|-\|198859038\|198859159 | 30 | 52 | 23 | | isomiR | NO | 216.26 | 83.11 | 1074.56 | 89.96 |
| >MI0000289\|hsa-mir-181a-1&WithFlank&1\|-\|198859038\|198859159@30.53.24(+1C);MI0000269\|hsa-mir-181a-2&WithFlank&9\|+\|124692436\|124692557@45.68.24(+1C)&[MIMAT0000256&hsa-miR-181a-5p&offsets\|0\|+1(+1C);m-32&1\|-\|198859108\|198859130&offsets\|0\|+1(+1C)];[MIMAT0000256_1&hsa-miR-181a-5p&offsets\|0\|+1(+1C);m-32&9\|+\|124692480\|124692502&offsets\|0\|+1(+1C)]\|AACATTCAACGCTGTCGGTGAGTTC | >MI0000289\|hsa-mir-181a-1&WithFlank&1\|-\|198859038\|198859159 | 30 | 53 | 24 | (+1C) | isomiR | NO | 0.00 | 0.00 | 118.80 | 0.00 |
| >MI0000289\|hsa-mir-181a-1&WithFlank&1\|-\|198859038\|198859159@30.53.24;MI0000269\|hsa-mir-181a-2&WithFlank&9\|+\|124692436\|124692557@45.68.24&[MIMAT0000256&hsa-miR-181a-5p&offsets\|0\|+1;m-32&1\|-\|198859108\|198859130&offsets\|0\|+1];[MIMAT0000256_1&hsa-miR-181a-5p&offsets\|0\|+1;m-32&9\|+\|124692480\|124692502&offsets\|0\|+1]\|AACATTCAACGCTGTCGGTGAGTT | >MI0000289\|hsa-mir-181a-1&WithFlank&1\|-\|198859038\|198859159 | 30 | 53 | 24 | | isomiR | NO | 59.25 | 0.00 | 667.98 | 17.37 |
| >MI0000289\|hsa-mir-181a-1&WithFlank&1\|-\|198859038\|198859159@30.54.25(+1U);MI0000269\|hsa-mir-181a-2&WithFlank&9\|+\|124692436\|124692557@45.69.25(+1U)&[MIMAT0000256&hsa-miR-181a-5p&offsets\|0\|+2(+1U);m-32&1\|-\|198859108\|198859130&offsets\|0\|+2(+1U)];[MIMAT0000256_1&hsa-miR-181a-5p&offsets\|0\|+2(+1U);m-32&9\|+\|124692480\|124692502&offsets\|0\|+2(+1U)]\|AACATTCAACGCTGTCGGTGAGTTTT | >MI0000289\|hsa-mir-181a-1&WithFlank&1\|-\|198859038\|198859159 | 30 | 54 | 25 | (+1U) | isomiR | NO | 0.00 | 0.00 | 131.83 | 10.69 |
| >MI0000289\|hsa-mir-181a-1&WithFlank&1\|-\|198859038\|198859159@30.54.25;MI0000269\|hsa-mir-181a-2&WithFlank&9\|+\|124692436\|124692557@45.69.25&[MIMAT0000256&hsa-miR-181a-5p&offsets\|0\|+2;m-32&1\|-\|198859108\|198859130&offsets\|0\|+2];[MIMAT0000256_1&hsa-miR-181a-5p&offsets\|0\|+2;m-32&9\|+\|124692480\|124692502&offsets\|0\|+2]\|AACATTCAACGCTGTCGGTGAGTTT | >MI0000289\|hsa-mir-181a-1&WithFlank&1\|-\|198859038\|198859159 | 30 | 54 | 25 | | isomiR | NO | 70.54 | 0.00 | 540.03 | 22.08 |
| >MI0000289\|hsa-mir-181a-1&WithFlank&1\|-\|198859038\|198859159@70.90.21(+1U)&[MIMAT0000270&hsa-miR-181a-3p&offsets\|0\|-1(+1U);m-125&1\|-\|198859069\|198859090&offsets\|0\|-1(+1U)]\|ACCATCGACCGTTGATTGTACT | >MI0000289\|hsa-mir-181a-1&WithFlank&1\|-\|198859038\|198859159 | 70 | 90 | 21 | (+1U) | isomiR | NO | 0.00 | 30.96 | 33.53 | 0.00 |
| >MI0000289\|hsa-mir-181a-1&WithFlank&1\|-\|198859038\|198859159@70.91.22&[MIMAT0000270&hsa-miR-181a-3p&offsets\|0\|0;m-125&1\|-\|198859069\|198859090&offsets\|0\|0]\|ACCATCGACCGTTGATTGTACC | >MI0000289\|hsa-mir-181a-1&WithFlank&1\|-\|198859038\|198859159 | 70 | 91 | 22 | | isomiR | NO | 52.10 | 123.11 | 76.43 | 14.12 |
| >MI0000294\|hsa-mir-218-1&WithFlank&4\|+\|20528269\|20528390@31.51.21;MI0000295\|hsa-mir-218-2&WithFlank&5\|-\|168768140\|168768261@31.51.21&[MIMAT0000275&hsa-miR-218-5p&offsets\|0\|0;m-185&4\|+\|20528299\|20528319&offsets\|0\|0];[MIMAT0000275_1&hsa-miR-218-5p&offsets\|0\|0;m-187&5\|-\|168768211\|168768231&offsets\|0\|0]\|TTGTGCTTGATCTAACCATGT | >MI0000294\|hsa-mir-218-1&WithFlank&4\|+\|20528269\|20528390 | 31 | 51 | 21 | | isomiR | NO | 0.00 | 0.00 | 92.37 | 0.00 |
| >MI0000294\|hsa-mir-218-1&WithFlank&4\|+\|20528269\|20528390@31.52.22(+1U);MI0000295\|hsa-mir-218-2&WithFlank&5\|-\|168768140\|168768261@31.52.22(+1U)&[MIMAT0000275&hsa-miR-218-5p&offsets\|0\|+1(+1U);m-185&4\|+\|20528299\|20528319&offsets\|0\|+1(+1U)];[MIMAT0000275_1&hsa-miR-218-5p&offsets\|0\|+1(+1U);m-187&5\|-\|168768211\|168768231&offsets\|0\|+1(+1U)]\|TTGTGCTTGATCTAACCATGTGT | >MI0000294\|hsa-mir-218-1&WithFlank&4\|+\|20528269\|20528390 | 31 | 52 | 22 | (+1U) | isomiR | NO | 0.00 | 0.00 | 14.21 | 0.00 |
| >MI0000294\|hsa-mir-218-1&WithFlank&4\|+\|20528269\|20528390@31.52.22;MI0000295\|hsa-mir-218-2&WithFlank&5\|-\|168768140\|168768261@31.52.22&[MIMAT0000275&hsa-miR-218-5p&offsets\|0\|+1;m-185&4\|+\|20528299\|20528319&offsets\|0\|+1];[MIMAT0000275_1&hsa-miR-218-5p&offsets\|0\|+1;m-187&5\|-\|168768211\|168768231&offsets\|0\|+1]\|TTGTGCTTGATCTAACCATGTG | >MI0000294\|hsa-mir-218-1&WithFlank&4\|+\|20528269\|20528390 | 31 | 52 | 22 | | isomiR | NO | 0.00 | 0.00 | 13.77 | 0.00 |
| >MI0000298\|hsa-mir-221&WithFlank&X\|-\|45746151\|45746272@31.53.23&[MIMAT0004568&hsa-miR-221-5p&offsets\|0\|+1;m-405&X\|-\|45746220\|45746242&offsets\|0\|0]\|ACCTGGCATACAATGTAGATTTC | >MI0000298\|hsa-mir-221&WithFlank&X\|-\|45746151\|45746272 | 31 | 53 | 23 | | isomiR | NO | 25.50 | 0.00 | 16.71 | 11.42 |

### MDA-MB-231

<table>
<tr><th></th><th></th><th></th><th></th><th></th><th></th><th></th><th></th><th></th><th></th><th></th><th></th></tr>
<tr><td>>MI0000298|hsa-mir-221&WithFlank&X|-|45746151|45746272@31.54.24&[MIMAT0004568&hsa-miR-221-5p&offsets|0|+2;m-405&X|-|45746220|45746242&offsets|0|+1]|ACCTGGCATACAATGTAGATTTCT</td><td>>MI0000298|hsa-mir-221&WithFlank&X|-|45746151|45746272</td><td>31</td><td>54</td><td>24</td><td></td><td>isomiR</td><td>NO</td><td>91.68</td><td>105.19</td><td>17.18</td><td>64.25</td></tr>
<tr><td>>MI0000298|hsa-mir-221&WithFlank&X|-|45746151|45746272@31.56.26&[MIMAT0004568&hsa-miR-221-5p&offsets|0|+4;m-405&X|-|45746220|45746242&offsets|0|+3]|ACCTGGCATACAATGTAGATTTCTGT</td><td>>MI0000298|hsa-mir-221&WithFlank&X|-|45746151|45746272</td><td>31</td><td>56</td><td>26</td><td></td><td>isomiR</td><td>NO</td><td>100.16</td><td>182.57</td><td>0.00</td><td>64.11</td></tr>
<tr><td>>MI0000298|hsa-mir-221&WithFlank&X|-|45746151|45746272@71.90.20&[MIMAT0000278&hsa-miR-221-3p&offsets|0|-3;m-91&X|-|45746181|45746202&offsets|0|-2]|AGCTACATTGTCTGCTGGGT</td><td>>MI0000298|hsa-mir-221&WithFlank&X|-|45746151|45746272</td><td>71</td><td>90</td><td>20</td><td></td><td>isomiR</td><td>NO</td><td>24.56</td><td>0.00</td><td>148.11</td><td>10.50</td></tr>
<tr><td>>MI0000298|hsa-mir-221&WithFlank&X|-|45746151|45746272@71.90.20(+1C)&[MIMAT0000278&hsa-miR-221-3p&offsets|0|-3(+1C);m-91&X|-|45746181|45746202&offsets|0|-2(+1C)]|AGCTACATTGTCTGCTGGGTC</td><td>>MI0000298|hsa-mir-221&WithFlank&X|-|45746151|45746272</td><td>71</td><td>90</td><td>20</td><td>(+1C)</td><td>isomiR</td><td>NO</td><td>0.00</td><td>0.00</td><td>134.93</td><td>0.00</td></tr>
<tr><td>>MI0000298|hsa-mir-221&WithFlank&X|-|45746151|45746272@71.91.21&[MIMAT0000278&hsa-miR-221-3p&offsets|0|-2;m-91&X|-|45746181|45746202&offsets|0|-1]|AGCTACATTGTCTGCTGGGTT</td><td>>MI0000298|hsa-mir-221&WithFlank&X|-|45746151|45746272</td><td>71</td><td>91</td><td>21</td><td></td><td>isomiR</td><td>NO</td><td>109.34</td><td>36.99</td><td>625.40</td><td>21.42</td></tr>
<tr><td>>MI0000298|hsa-mir-221&WithFlank&X|-|45746151|45746272@71.91.21(+1A)&[MIMAT0000278&hsa-miR-221-3p&offsets|0|-2(+1A);m-91&X|-|45746181|45746202&offsets|0|-1(+1A)]|AGCTACATTGTCTGCTGGGTTA</td><td>>MI0000298|hsa-mir-221&WithFlank&X|-|45746151|45746272</td><td>71</td><td>91</td><td>21</td><td>(+1A)</td><td>isomiR</td><td>NO</td><td>23.60</td><td>0.00</td><td>226.57</td><td>12.29</td></tr>
<tr><td>>MI0000298|hsa-mir-221&WithFlank&X|-|45746151|45746272@71.91.21(+1C)&[MIMAT0000278&hsa-miR-221-3p&offsets|0|-2(+1C);m-91&X|-|45746181|45746202&offsets|0|-1(+1C)]|AGCTACATTGTCTGCTGGGTTC</td><td>>MI0000298|hsa-mir-221&WithFlank&X|-|45746151|45746272</td><td>71</td><td>91</td><td>21</td><td>(+1C)</td><td>isomiR</td><td>NO</td><td>125.02</td><td>62.58</td><td>691.97</td><td>77.23</td></tr>
<tr><td>>MI0000298|hsa-mir-221&WithFlank&X|-|45746151|45746272@71.91.21(+1G)&[MIMAT0000278&hsa-miR-221-3p&offsets|0|-2(+1G);m-91&X|-|45746181|45746202&offsets|0|-1(+1G)]|AGCTACATTGTCTGCTGGGTTG</td><td>>MI0000298|hsa-mir-221&WithFlank&X|-|45746151|45746272</td><td>71</td><td>91</td><td>21</td><td>(+1G)</td><td>isomiR</td><td>NO</td><td>0.00</td><td>0.00</td><td>59.53</td><td>0.00</td></tr>
<tr><td>>MI0000298|hsa-mir-221&WithFlank&X|-|45746151|45746272@71.91.21(+2C)&[MIMAT0000278&hsa-miR-221-3p&offsets|0|-2(+2C);m-91&X|-|45746181|45746202&offsets|0|-1(+2C)]|AGCTACATTGTCTGCTGGGTTCC</td><td>>MI0000298|hsa-mir-221&WithFlank&X|-|45746151|45746272</td><td>71</td><td>91</td><td>21</td><td>(+2C)</td><td>isomiR</td><td>NO</td><td>0.00</td><td>0.00</td><td>10.85</td><td>0.00</td></tr>
<tr><td>>MI0000298|hsa-mir-221&WithFlank&X|-|45746151|45746272@71.92.22&[MIMAT0000278&hsa-miR-221-3p&offsets|0|-1;m-91&X|-|45746181|45746202&offsets|0|0]|AGCTACATTGTCTGCTGGGTTT</td><td>>MI0000298|hsa-mir-221&WithFlank&X|-|45746151|45746272</td><td>71</td><td>92</td><td>22</td><td></td><td>isomiR</td><td>YES</td><td>727.61</td><td>381.67</td><td>3049.95</td><td>290.29</td></tr>
<tr><td>>MI0000298|hsa-mir-221&WithFlank&X|-|45746151|45746272@71.92.22(+1A)&[MIMAT0000278&hsa-miR-221-3p&offsets|0|-1(+1A);m-91&X|-|45746181|45746202&offsets|0|0(+1A)]|AGCTACATTGTCTGCTGGGTTTA</td><td>>MI0000298|hsa-mir-221&WithFlank&X|-|45746151|45746272</td><td>71</td><td>92</td><td>22</td><td>(+1A)</td><td>isomiR</td><td>NO</td><td>50.28</td><td>30.44</td><td>255.14</td><td>12.91</td></tr>
<tr><td>>MI0000298|hsa-mir-221&WithFlank&X|-|45746151|45746272@71.92.22(+1U)&[MIMAT0000278&hsa-miR-221-3p&offsets|0|-1(+1U);m-91&X|-|45746181|45746202&offsets|0|0(+1U)]|AGCTACATTGTCTGCTGGGTTTT</td><td>>MI0000298|hsa-mir-221&WithFlank&X|-|45746151|45746272</td><td>71</td><td>92</td><td>22</td><td>(+1U)</td><td>isomiR</td><td>NO</td><td>194.88</td><td>122.02</td><td>943.23</td><td>99.65</td></tr>
<tr><td>>MI0000298|hsa-mir-221&WithFlank&X|-|45746151|45746272@71.92.22(+2U)&[MIMAT0000278&hsa-miR-221-3p&offsets|0|-1(+2U);m-91&X|-|45746181|45746202&offsets|0|0(+2U)]|AGCTACATTGTCTGCTGGGTTTTT</td><td>>MI0000298|hsa-mir-221&WithFlank&X|-|45746151|45746272</td><td>71</td><td>92</td><td>22</td><td>(+2U)</td><td>isomiR</td><td>NO</td><td>0.00</td><td>0.00</td><td>96.65</td><td>10.68</td></tr>
<tr><td>>MI0000298|hsa-mir-221&WithFlank&X|-|45746151|45746272@71.93.23&[MIMAT0000278&hsa-miR-221-3p&offsets|0|0;m-91&X|-|45746181|45746202&offsets|0|+1]|AGCTACATTGTCTGCTGGGTTTC</td><td>>MI0000298|hsa-mir-221&WithFlank&X|-|45746151|45746272</td><td>71</td><td>93</td><td>23</td><td></td><td>isomiR</td><td>YES</td><td>1076.25</td><td>1032.41</td><td>3579.76</td><td>309.29</td></tr>
<tr><td>>MI0000298|hsa-mir-221&WithFlank&X|-|45746151|45746272@71.93.23(+1C)&[MIMAT0000278&hsa-miR-221-3p&offsets|0|0(+1C);m-91&X|-|45746181|45746202&offsets|0|+1(+1C)]|AGCTACATTGTCTGCTGGGTTTCC</td><td>>MI0000298|hsa-mir-221&WithFlank&X|-|45746151|45746272</td><td>71</td><td>93</td><td>23</td><td>(+1C)</td><td>isomiR</td><td>NO</td><td>76.57</td><td>98.20</td><td>140.34</td><td>18.88</td></tr>
<tr><td>>MI0000298|hsa-mir-221&WithFlank&X|-|45746151|45746272@71.93.23(+1G)&[MIMAT0000278&hsa-miR-221-3p&offsets|0|0(+1G);m-91&X|-|45746181|45746202&offsets|0|+1(+1G)]|AGCTACATTGTCTGCTGGGTTTCG</td><td>>MI0000298|hsa-mir-221&WithFlank&X|-|45746151|45746272</td><td>71</td><td>93</td><td>23</td><td>(+1G)</td><td>isomiR</td><td>NO</td><td>0.00</td><td>13.35</td><td>12.81</td><td>0.00</td></tr>
<tr><td>>MI0000298|hsa-mir-221&WithFlank&X|-|45746151|45746272@71.93.23(+1U)&[MIMAT0000278&hsa-miR-221-3p&offsets|0|0(+1U);m-91&X|-|45746181|45746202&offsets|0|+1(+1U)]|AGCTACATTGTCTGCTGGGTTTCT</td><td>>MI0000298|hsa-mir-221&WithFlank&X|-|45746151|45746272</td><td>71</td><td>93</td><td>23</td><td>(+1U)</td><td>isomiR</td><td>NO</td><td>413.22</td><td>586.37</td><td>575.19</td><td>155.29</td></tr>
<tr><td>>MI0000298|hsa-mir-221&WithFlank&X|-|45746151|45746272@71.94.24&[MIMAT0000278&hsa-miR-221-3p&offsets|0|+1;m-91&X|-|45746181|45746202&offsets|0|+2]|AGCTACATTGTCTGCTGGGTTTCA</td><td>>MI0000298|hsa-mir-221&WithFlank&X|-|45746151|45746272</td><td>71</td><td>94</td><td>24</td><td></td><td>isomiR</td><td>NO</td><td>110.98</td><td>119.89</td><td>216.95</td><td>62.54</td></tr>
<tr><td>>MI0000299|hsa-mir-222&WithFlank&X|-|45747009|45747130@75.94.20&[MIMAT0000279&hsa-miR-222-3p&offsets|0|-1;m-140&X|-|45747033|45747056&offsets|0|-4]|AGCTACATCTGGCTACTGGG</td><td>>MI0000299|hsa-mir-222&WithFlank&X|-|45747009|45747130</td><td>75</td><td>94</td><td>20</td><td></td><td>isomiR</td><td>NO</td><td>0.00</td><td>0.00</td><td>79.87</td><td>0.00</td></tr>
<tr><td>>MI0000299|hsa-mir-222&WithFlank&X|-|45747009|45747130@75.94.20(+1C)&[MIMAT0000279&hsa-miR-222-3p&offsets|0|-1(+1C);m-140&X|-|45747033|45747056&offsets|0|-4(+1C)]|AGCTACATCTGGCTACTGGGC</td><td>>MI0000299|hsa-mir-222&WithFlank&X|-|45747009|45747130</td><td>75</td><td>94</td><td>20</td><td>(+1C)</td><td>isomiR</td><td>NO</td><td>0.00</td><td>0.00</td><td>170.28</td><td>0.00</td></tr>
<tr><td>>MI0000299|hsa-mir-222&WithFlank&X|-|45747009|45747130@75.95.21&[MIMAT0000279&hsa-miR-222-3p&offsets|0|0;m-140&X|-|45747033|45747056&offsets|0|-3]|AGCTACATCTGGCTACTGGGT</td><td>>MI0000299|hsa-mir-222&WithFlank&X|-|45747009|45747130</td><td>75</td><td>95</td><td>21</td><td></td><td>isomiR</td><td>NO</td><td>129.19</td><td>63.58</td><td>771.77</td><td>60.52</td></tr>
<tr><td>>MI0000299|hsa-mir-222&WithFlank&X|-|45747009|45747130@75.95.21(+1A)&[MIMAT0000279&hsa-miR-222-3p&offsets|0|0(+1A);m-140&X|-|45747033|45747056&offsets|0|-3(+1A)]|AGCTACATCTGGCTACTGGGTA</td><td>>MI0000299|hsa-mir-222&WithFlank&X|-|45747009|45747130</td><td>75</td><td>95</td><td>21</td><td>(+1A)</td><td>isomiR</td><td>NO</td><td>0.00</td><td>0.00</td><td>13.65</td><td>0.00</td></tr>
<tr><td>>MI0000299|hsa-mir-222&WithFlank&X|-|45747009|45747130@75.95.21(+1U)&[MIMAT0000279&hsa-miR-222-3p&offsets|0|0(+1U);m-140&X|-|45747033|45747056&offsets|0|-3(+1U)]|AGCTACATCTGGCTACTGGGTT</td><td>>MI0000299|hsa-mir-222&WithFlank&X|-|45747009|45747130</td><td>75</td><td>95</td><td>21</td><td>(+1U)</td><td>isomiR</td><td>NO</td><td>0.00</td><td>0.00</td><td>66.76</td><td>0.00</td></tr>
<tr><td>>MI0000299|hsa-mir-222&WithFlank&X|-|45747009|45747130@75.96.22&[MIMAT0000279&hsa-miR-222-3p&offsets|0|+1;m-140&X|-|45747033|45747056&offsets|0|-2]|AGCTACATCTGGCTACTGGGTC</td><td>>MI0000299|hsa-mir-222&WithFlank&X|-|45747009|45747130</td><td>75</td><td>96</td><td>22</td><td></td><td>isomiR</td><td>NO</td><td>110.57</td><td>95.78</td><td>191.19</td><td>22.45</td></tr>
<tr><td>>MI0000299|hsa-mir-222&WithFlank&X|-|45747009|45747130@75.96.22(+1A)&[MIMAT0000279&hsa-miR-222-3p&offsets|0|+1(+1A);m-140&X|-|45747033|45747056&offsets|0|-2(+1A)]|AGCTACATCTGGCTACTGGGTCA</td><td>>MI0000299|hsa-mir-222&WithFlank&X|-|45747009|45747130</td><td>75</td><td>96</td><td>22</td><td>(+1A)</td><td>isomiR</td><td>NO</td><td>25.36</td><td>0.00</td><td>126.18</td><td>14.03</td></tr>
<tr><td>>MI0000299|hsa-mir-222&WithFlank&X|-|45747009|45747130@75.96.22(+1C)&[MIMAT0000279&hsa-miR-222-3p&offsets|0|+1(+1C);m-140&X|-|45747033|45747056&offsets|0|-2(+1C)]|AGCTACATCTGGCTACTGGGTCC</td><td>>MI0000299|hsa-mir-222&WithFlank&X|-|45747009|45747130</td><td>75</td><td>96</td><td>22</td><td>(+1C)</td><td>isomiR</td><td>NO</td><td>54.52</td><td>24.65</td><td>143.27</td><td>17.38</td></tr>
</table>

### MDA-MB-231

<table>
<tr><td>>MI0000299|hsa-mir-222&WithFlank&X|-|45747009|45747130@75.97.23&[MIMAT0000279&hsa-miR-222-3p&offsets|0|+2;m-140&X|-|45747033|45747056&offsets|0|-1]|AGCTACATCTGGCTACTGGGTCT</td><td>>MI0000299|hsa-mir-222&WithFlank&X|-|45747009|45747130</td><td>75</td><td>97</td><td>23</td><td></td><td>isomiR</td><td>NO</td><td>276.11</td><td>209.82</td><td>596.66</td><td>137.50</td></tr>
<tr><td>>MI0000299|hsa-mir-222&WithFlank&X|-|45747009|45747130@75.97.23(+1A)&[MIMAT0000279&hsa-miR-222-3p&offsets|0|+2(+1A);m-140&X|-|45747033|45747056&offsets|0|-1(+1A)]|AGCTACATCTGGCTACTGGGTCTA</td><td>>MI0000299|hsa-mir-222&WithFlank&X|-|45747009|45747130</td><td>75</td><td>97</td><td>23</td><td>(+1A)</td><td>isomiR</td><td>NO</td><td>50.18</td><td>99.50</td><td>77.62</td><td>17.25</td></tr>
<tr><td>>MI0000299|hsa-mir-222&WithFlank&X|-|45747009|45747130@75.97.23(+1U)&[MIMAT0000279&hsa-miR-222-3p&offsets|0|+2(+1U);m-140&X|-|45747033|45747056&offsets|0|-1(+1U)]|AGCTACATCTGGCTACTGGGTCTT</td><td>>MI0000299|hsa-mir-222&WithFlank&X|-|45747009|45747130</td><td>75</td><td>97</td><td>23</td><td>(+1U)</td><td>isomiR</td><td>NO</td><td>134.10</td><td>199.45</td><td>218.94</td><td>96.24</td></tr>
<tr><td>>MI0000299|hsa-mir-222&WithFlank&X|-|45747009|45747130@75.97.23(+2U)&[MIMAT0000279&hsa-miR-222-3p&offsets|0|+2(+2U);m-140&X|-|45747033|45747056&offsets|0|-1(+2U)]|AGCTACATCTGGCTACTGGGTCTTT</td><td>>MI0000299|hsa-mir-222&WithFlank&X|-|45747009|45747130</td><td>75</td><td>97</td><td>23</td><td>(+2U)</td><td>isomiR</td><td>NO</td><td>68.22</td><td>35.66</td><td>17.17</td><td>20.41</td></tr>
<tr><td>>MI0000299|hsa-mir-222&WithFlank&X|-|45747009|45747130@75.98.24&[MIMAT0000279&hsa-miR-222-3p&offsets|0|+3;m-140&X|-|45747033|45747056&offsets|0|0]|AGCTACATCTGGCTACTGGGTCTC</td><td>>MI0000299|hsa-mir-222&WithFlank&X|-|45747009|45747130</td><td>75</td><td>98</td><td>24</td><td></td><td>isomiR</td><td>YES</td><td>581.37</td><td>1142.99</td><td>534.47</td><td>256.69</td></tr>
<tr><td>>MI0000299|hsa-mir-222&WithFlank&X|-|45747009|45747130@75.98.24(+1A)&[MIMAT0000279&hsa-miR-222-3p&offsets|0|+3(+1A);m-140&X|-|45747033|45747056&offsets|0|0(+1A)]|AGCTACATCTGGCTACTGGGTCTCA</td><td>>MI0000299|hsa-mir-222&WithFlank&X|-|45747009|45747130</td><td>75</td><td>98</td><td>24</td><td>(+1A)</td><td>isomiR</td><td>NO</td><td>145.19</td><td>283.73</td><td>153.41</td><td>104.89</td></tr>
<tr><td>>MI0000299|hsa-mir-222&WithFlank&X|-|45747009|45747130@75.98.24(+1C)&[MIMAT0000279&hsa-miR-222-3p&offsets|0|+3(+1C);m-140&X|-|45747033|45747056&offsets|0|0(+1C)]|AGCTACATCTGGCTACTGGGTCTCC</td><td>>MI0000299|hsa-mir-222&WithFlank&X|-|45747009|45747130</td><td>75</td><td>98</td><td>24</td><td>(+1C)</td><td>isomiR</td><td>NO</td><td>261.61</td><td>482.15</td><td>224.90</td><td>174.08</td></tr>
<tr><td>>MI0000299|hsa-mir-222&WithFlank&X|-|45747009|45747130@75.98.24(+1G)&[MIMAT0000279&hsa-miR-222-3p&offsets|0|+3(+1G);m-140&X|-|45747033|45747056&offsets|0|0(+1G)]|AGCTACATCTGGCTACTGGGTCTCG</td><td>>MI0000299|hsa-mir-222&WithFlank&X|-|45747009|45747130</td><td>75</td><td>98</td><td>24</td><td>(+1G)</td><td>isomiR</td><td>NO</td><td>53.19</td><td>76.24</td><td>15.10</td><td>12.41</td></tr>
<tr><td>>MI0000299|hsa-mir-222&WithFlank&X|-|45747009|45747130@75.99.25&[MIMAT0000279&hsa-miR-222-3p&offsets|0|+4;m-140&X|-|45747033|45747056&offsets|0|+1]|AGCTACATCTGGCTACTGGGTCTCT</td><td>>MI0000299|hsa-mir-222&WithFlank&X|-|45747009|45747130</td><td>75</td><td>99</td><td>25</td><td></td><td>isomiR</td><td>YES</td><td>854.51</td><td>1997.15</td><td>864.84</td><td>393.42</td></tr>
<tr><td>>MI0000299|hsa-mir-222&WithFlank&X|-|45747009|45747130@75.99.25(+1A)&[MIMAT0000279&hsa-miR-222-3p&offsets|0|+4(+1A);m-140&X|-|45747033|45747056&offsets|0|+1(+1A)]|AGCTACATCTGGCTACTGGGTCTCTA</td><td>>MI0000299|hsa-mir-222&WithFlank&X|-|45747009|45747130</td><td>75</td><td>99</td><td>25</td><td>(+1A)</td><td>isomiR</td><td>NO</td><td>0.00</td><td>24.06</td><td>0.00</td><td>0.00</td></tr>
<tr><td>>MI0000299|hsa-mir-222&WithFlank&X|-|45747009|45747130@75.99.25(+1C)&[MIMAT0000279&hsa-miR-222-3p&offsets|0|+4(+1C);m-140&X|-|45747033|45747056&offsets|0|+1(+1C)]|AGCTACATCTGGCTACTGGGTCTCTC</td><td>>MI0000299|hsa-mir-222&WithFlank&X|-|45747009|45747130</td><td>75</td><td>99</td><td>25</td><td>(+1C)</td><td>isomiR</td><td>NO</td><td>0.00</td><td>30.72</td><td>0.00</td><td>12.70</td></tr>
<tr><td>>MI0000299|hsa-mir-222&WithFlank&X|-|45747009|45747130@75.99.25(+1U)&[MIMAT0000279&hsa-miR-222-3p&offsets|0|+4(+1U);m-140&X|-|45747033|45747056&offsets|0|+1(+1U)]|AGCTACATCTGGCTACTGGGTCTCTT</td><td>>MI0000299|hsa-mir-222&WithFlank&X|-|45747009|45747130</td><td>75</td><td>99</td><td>25</td><td>(+1U)</td><td>isomiR</td><td>NO</td><td>150.99</td><td>297.12</td><td>70.31</td><td>116.30</td></tr>
<tr><td>>MI0000299|hsa-mir-222&WithFlank&X|-|45747009|45747130@75.99.25(+2U)&[MIMAT0000279&hsa-miR-222-3p&offsets|0|+4(+2U);m-140&X|-|45747033|45747056&offsets|0|+1(+2U)]|AGCTACATCTGGCTACTGGGTCTCTTT</td><td>>MI0000299|hsa-mir-222&WithFlank&X|-|45747009|45747130</td><td>75</td><td>99</td><td>25</td><td>(+2U)</td><td>isomiR</td><td>NO</td><td>102.27</td><td>196.16</td><td>16.24</td><td>81.10</td></tr>
<tr><td>>MI0000299|hsa-mir-222&WithFlank&X|-|45747009|45747130@75.99.25(+3U)&[MIMAT0000279&hsa-miR-222-3p&offsets|0|+4(+3U);m-140&X|-|45747033|45747056&offsets|0|+1(+3U)]|AGCTACATCTGGCTACTGGGTCTCTTTT</td><td>>MI0000299|hsa-mir-222&WithFlank&X|-|45747009|45747130</td><td>75</td><td>99</td><td>25</td><td>(+3U)</td><td>isomiR</td><td>NO</td><td>77.34</td><td>152.94</td><td>0.00</td><td>85.53</td></tr>
<tr><td>>MI0000299|hsa-mir-222&WithFlank&X|-|45747009|45747130@75.99.25(+4U)&[MIMAT0000279&hsa-miR-222-3p&offsets|0|+4(+4U);m-140&X|-|45747033|45747056&offsets|0|+1(+4U)]|AGCTACATCTGGCTACTGGGTCTCTTTTT</td><td>>MI0000299|hsa-mir-222&WithFlank&X|-|45747009|45747130</td><td>75</td><td>99</td><td>25</td><td>(+4U)</td><td>isomiR</td><td>NO</td><td>0.00</td><td>30.53</td><td>0.00</td><td>12.50</td></tr>
<tr><td>>MI0000301|hsa-mir-224&WithFlank&X|-|151958572|151958664@14.35.22&[MIMAT0000281&hsa-miR-224-5p&offsets|+1|-2;m-238&X|-|151958630|151958651&offsets|0|0]|CAAGTCACTAGTGGTTCCGTTT</td><td>>MI0000301|hsa-mir-224&WithFlank&X|-|151958572|151958664</td><td>14</td><td>35</td><td>22</td><td></td><td>isomiR</td><td>NO</td><td>96.76</td><td>80.48</td><td>83.92</td><td>18.37</td></tr>
<tr><td>>MI0000301|hsa-mir-224&WithFlank&X|-|151958572|151958664@14.37.24&[MIMAT0000281&hsa-miR-224-5p&offsets|+1|0;m-238&X|-|151958630|151958651&offsets|0|+2]|CAAGTCACTAGTGGTTCCGTTTAG</td><td>>MI0000301|hsa-mir-224&WithFlank&X|-|151958572|151958664</td><td>14</td><td>37</td><td>24</td><td></td><td>isomiR</td><td>NO</td><td>97.51</td><td>68.45</td><td>15.54</td><td>21.97</td></tr>
<tr><td>>MI0000301|hsa-mir-224&WithFlank&X|-|151958572|151958664@14.38.25&[MIMAT0000281&hsa-miR-224-5p&offsets|+1|+1;m-238&X|-|151958630|151958651&offsets|0|+3]|CAAGTCACTAGTGGTTCCGTTTAGT</td><td>>MI0000301|hsa-mir-224&WithFlank&X|-|151958572|151958664</td><td>14</td><td>38</td><td>25</td><td></td><td>isomiR</td><td>NO</td><td>24.73</td><td>0.00</td><td>0.00</td><td>0.00</td></tr>
<tr><td>>MI0000433|hsa-let-7g&WithFlank&3|-|52268272|52268367@11.28.18&[MIMAT0000414&hsa-let-7g-5p&offsets|0|-4;m-58&3|-|52268336|52268357&offsets|0|-4]|TGAGGTAGTAGTTTGTAC</td><td>>MI0000433|hsa-let-7g&WithFlank&3|-|52268272|52268367</td><td>11</td><td>28</td><td>18</td><td></td><td>isomiR</td><td>NO</td><td>193.60</td><td>63.04</td><td>190.76</td><td>12.85</td></tr>
<tr><td>>MI0000433|hsa-let-7g&WithFlank&3|-|52268272|52268367@11.30.20(+1C)&[MIMAT0000414&hsa-let-7g-5p&offsets|0|-2(+1C);m-58&3|-|52268336|52268357&offsets|0|-2(+1C)]|TGAGGTAGTAGTTTGTACAGC</td><td>>MI0000433|hsa-let-7g&WithFlank&3|-|52268272|52268367</td><td>11</td><td>30</td><td>20</td><td>(+1C)</td><td>isomiR</td><td>NO</td><td>46.27</td><td>0.00</td><td>17.19</td><td>0.00</td></tr>
<tr><td>>MI0000433|hsa-let-7g&WithFlank&3|-|52268272|52268367@11.31.21&[MIMAT0000414&hsa-let-7g-5p&offsets|0|-1;m-58&3|-|52268336|52268357&offsets|0|-1]|TGAGGTAGTAGTTTGTACAGT</td><td>>MI0000433|hsa-let-7g&WithFlank&3|-|52268272|52268367</td><td>11</td><td>31</td><td>21</td><td></td><td>isomiR</td><td>NO</td><td>287.20</td><td>115.37</td><td>202.32</td><td>15.08</td></tr>
<tr><td>>MI0000433|hsa-let-7g&WithFlank&3|-|52268272|52268367@11.31.21(+1A)&[MIMAT0000414&hsa-let-7g-5p&offsets|0|-1(+1A);m-58&3|-|52268336|52268357&offsets|0|-1(+1A)]|TGAGGTAGTAGTTTGTACAGTA</td><td>>MI0000433|hsa-let-7g&WithFlank&3|-|52268272|52268367</td><td>11</td><td>31</td><td>21</td><td>(+1A)</td><td>isomiR</td><td>NO</td><td>135.87</td><td>35.89</td><td>148.46</td><td>12.33</td></tr>
<tr><td>>MI0000433|hsa-let-7g&WithFlank&3|-|52268272|52268367@11.31.21(+1C)&[MIMAT0000414&hsa-let-7g-5p&offsets|0|-1(+1C);m-58&3|-|52268336|52268357&offsets|0|-1(+1C)]|TGAGGTAGTAGTTTGTACAGTC</td><td>>MI0000433|hsa-let-7g&WithFlank&3|-|52268272|52268367</td><td>11</td><td>31</td><td>21</td><td>(+1C)</td><td>isomiR</td><td>NO</td><td>510.95</td><td>162.57</td><td>455.68</td><td>82.22</td></tr>
<tr><td>>MI0000433|hsa-let-7g&WithFlank&3|-|52268272|52268367@11.31.21(+1G)&[MIMAT0000414&hsa-let-7g-5p&offsets|0|-1(+1G);m-58&3|-|52268336|52268357&offsets|0|-1(+1G)]|TGAGGTAGTAGTTTGTACAGTG</td><td>>MI0000433|hsa-let-7g&WithFlank&3|-|52268272|52268367</td><td>11</td><td>31</td><td>21</td><td>(+1G)</td><td>isomiR</td><td>NO</td><td>80.62</td><td>0.00</td><td>17.56</td><td>0.00</td></tr>
<tr><td>>MI0000433|hsa-let-7g&WithFlank&3|-|52268272|52268367@11.32.22&[MIMAT0000414&hsa-let-7g-5p&offsets|0|0;m-58&3|-|52268336|52268357&offsets|0|0]|TGAGGTAGTAGTTTGTACAGTT</td><td>>MI0000433|hsa-let-7g&WithFlank&3|-|52268272|52268367</td><td>11</td><td>32</td><td>22</td><td></td><td>isomiR</td><td>YES</td><td>2878.68</td><td>1077.96</td><td>2006.63</td><td>302.65</td></tr>
<tr><td>>MI0000433|hsa-let-7g&WithFlank&3|-|52268272|52268367@11.32.22(+1A)&[MIMAT0000414&hsa-let-7g-5p&offsets|0|0(+1A);m-58&3|-|52268336|52268357&offsets|0|0(+1A)]|TGAGGTAGTAGTTTGTACAGTTA</td><td>>MI0000433|hsa-let-7g&WithFlank&3|-|52268272|52268367</td><td>11</td><td>32</td><td>22</td><td>(+1A)</td><td>isomiR</td><td>NO</td><td>79.70</td><td>0.00</td><td>154.83</td><td>0.00</td></tr>
</table>

### MDA-MB-231

| | | | | | | | | | | | |
|---|---|---|---|---|---|---|---|---|---|---|---|
| >MI0000433\|hsa-let-7g&WithFlank&3\|-\|52268272\|52268367@11.32.22(+1C)&[MIMAT0000414&hsa-let-7g-5p&offsets\|0\|0(+1C);m-58&3\|-\|52268336\|52268357&offsets\|0\|0(+1C)]\|TGAGGTAGTAGTTTGTACAGTTC | >MI0000433\|hsa-let-7g&WithFlank&3\|-\|52268272\|52268367 | 11 | 32 | 22 | (+1C) | isomiR | NO | 0.00 | 0.00 | 31.61 | 0.00 |
| >MI0000433\|hsa-let-7g&WithFlank&3\|-\|52268272\|52268367@11.33.23&[MIMAT0000414&hsa-let-7g-5p&offsets\|0\|+1;m-58&3\|-\|52268336\|52268357&offsets\|0\|+1]\|TGAGGTAGTAGTTTGTACAGTTT | >MI0000433\|hsa-let-7g&WithFlank&3\|-\|52268272\|52268367 | 11 | 33 | 23 | | isomiR | NO | 129.58 | 32.94 | 197.37 | 0.00 |
| >MI0000434\|hsa-let-7i&WithFlank&12\|+\|62603680\|62603775@12.30.19&[MIMAT0000415&hsa-let-7i-5p&offsets\|0\|-3;m-49&12\|+\|62603691\|62603712&offsets\|0\|-3]\|TGAGGTAGTAGTTTGTGCT | >MI0000434\|hsa-let-7i&WithFlank&12\|+\|62603680\|62603775 | 12 | 30 | 19 | | isomiR | NO | 0.00 | 0.00 | 14.04 | 0.00 |
| >MI0000434\|hsa-let-7i&WithFlank&12\|+\|62603680\|62603775@12.31.20&[MIMAT0000415&hsa-let-7i-5p&offsets\|0\|-2;m-49&12\|+\|62603691\|62603712&offsets\|0\|-2]\|TGAGGTAGTAGTTTGTGCTG | >MI0000434\|hsa-let-7i&WithFlank&12\|+\|62603680\|62603775 | 12 | 31 | 20 | | isomiR | NO | 24.32 | 0.00 | 59.35 | 0.00 |
| >MI0000434\|hsa-let-7i&WithFlank&12\|+\|62603680\|62603775@12.31.20(+1A)&[MIMAT0000415&hsa-let-7i-5p&offsets\|0\|-2(+1A);m-49&12\|+\|62603691\|62603712&offsets\|0\|-2(+1A)]\|TGAGGTAGTAGTTTGTGCTGA | >MI0000434\|hsa-let-7i&WithFlank&12\|+\|62603680\|62603775 | 12 | 31 | 20 | (+1A) | isomiR | NO | 0.00 | 0.00 | 14.55 | 0.00 |
| >MI0000434\|hsa-let-7i&WithFlank&12\|+\|62603680\|62603775@12.31.20(+1C)&[MIMAT0000415&hsa-let-7i-5p&offsets\|0\|-2(+1C);m-49&12\|+\|62603691\|62603712&offsets\|0\|-2(+1C)]\|TGAGGTAGTAGTTTGTGCTGC | >MI0000434\|hsa-let-7i&WithFlank&12\|+\|62603680\|62603775 | 12 | 31 | 20 | (+1C) | isomiR | NO | 57.55 | 0.00 | 118.26 | 0.00 |
| >MI0000434\|hsa-let-7i&WithFlank&12\|+\|62603680\|62603775@12.32.21&[MIMAT0000415&hsa-let-7i-5p&offsets\|0\|-1;m-49&12\|+\|62603691\|62603712&offsets\|0\|-1]\|TGAGGTAGTAGTTTGTGCTGT | >MI0000434\|hsa-let-7i&WithFlank&12\|+\|62603680\|62603775 | 12 | 32 | 21 | | isomiR | NO | 331.96 | 188.16 | 545.16 | 15.57 |
| >MI0000434\|hsa-let-7i&WithFlank&12\|+\|62603680\|62603775@12.32.21(+1A)&[MIMAT0000415&hsa-let-7i-5p&offsets\|0\|-1(+1A);m-49&12\|+\|62603691\|62603712&offsets\|0\|-1(+1A)]\|TGAGGTAGTAGTTTGTGCTGTA | >MI0000434\|hsa-let-7i&WithFlank&12\|+\|62603680\|62603775 | 12 | 32 | 21 | (+1A) | isomiR | NO | 228.07 | 136.84 | 587.10 | 13.65 |
| >MI0000434\|hsa-let-7i&WithFlank&12\|+\|62603680\|62603775@12.32.21(+1C)&[MIMAT0000415&hsa-let-7i-5p&offsets\|0\|-1(+1C);m-49&12\|+\|62603691\|62603712&offsets\|0\|-1(+1C)]\|TGAGGTAGTAGTTTGTGCTGTC | >MI0000434\|hsa-let-7i&WithFlank&12\|+\|62603680\|62603775 | 12 | 32 | 21 | (+1C) | isomiR | YES | 756.90 | 362.04 | 1582.07 | 80.55 |
| >MI0000434\|hsa-let-7i&WithFlank&12\|+\|62603680\|62603775@12.32.21(+1G)&[MIMAT0000415&hsa-let-7i-5p&offsets\|0\|-1(+1G);m-49&12\|+\|62603691\|62603712&offsets\|0\|-1(+1G)]\|TGAGGTAGTAGTTTGTGCTGTG | >MI0000434\|hsa-let-7i&WithFlank&12\|+\|62603680\|62603775 | 12 | 32 | 21 | (+1G) | isomiR | NO | 151.97 | 34.02 | 187.05 | 0.00 |
| >MI0000434\|hsa-let-7i&WithFlank&12\|+\|62603680\|62603775@12.33.22&[MIMAT0000415&hsa-let-7i-5p&offsets\|0\|0;m-49&12\|+\|62603691\|62603712&offsets\|0\|0]\|TGAGGTAGTAGTTTGTGCTGTT | >MI0000434\|hsa-let-7i&WithFlank&12\|+\|62603680\|62603775 | 12 | 33 | 22 | | isomiR | YES | 4520.40 | 2561.60 | 7400.34 | 307.48 |
| >MI0000434\|hsa-let-7i&WithFlank&12\|+\|62603680\|62603775@12.33.22(+1A)&[MIMAT0000415&hsa-let-7i-5p&offsets\|0\|0(+1A);m-49&12\|+\|62603691\|62603712&offsets\|0\|0(+1A)]\|TGAGGTAGTAGTTTGTGCTGTTA | >MI0000434\|hsa-let-7i&WithFlank&12\|+\|62603680\|62603775 | 12 | 33 | 22 | (+1A) | isomiR | NO | 255.68 | 92.38 | 750.12 | 14.99 |
| >MI0000434\|hsa-let-7i&WithFlank&12\|+\|62603680\|62603775@12.33.22(+1C)&[MIMAT0000415&hsa-let-7i-5p&offsets\|0\|0(+1C);m-49&12\|+\|62603691\|62603712&offsets\|0\|0(+1C)]\|TGAGGTAGTAGTTTGTGCTGTTC | >MI0000434\|hsa-let-7i&WithFlank&12\|+\|62603680\|62603775 | 12 | 33 | 22 | (+1C) | isomiR | NO | 54.64 | 0.00 | 199.63 | 0.00 |
| >MI0000434\|hsa-let-7i&WithFlank&12\|+\|62603680\|62603775@12.33.22(+1U)&[MIMAT0000415&hsa-let-7i-5p&offsets\|0\|0(+1U);m-49&12\|+\|62603691\|62603712&offsets\|0\|0(+1U)]\|TGAGGTAGTAGTTTGTGCTGTTT | >MI0000434\|hsa-let-7i&WithFlank&12\|+\|62603680\|62603775 | 12 | 33 | 22 | (+1U) | isomiR | NO | 326.47 | 161.31 | 963.24 | 16.76 |
| >MI0000434\|hsa-let-7i&WithFlank&12\|+\|62603680\|62603775@12.33.22(+2U)&[MIMAT0000415&hsa-let-7i-5p&offsets\|0\|0(+2U);m-49&12\|+\|62603691\|62603712&offsets\|0\|0(+2U)]\|TGAGGTAGTAGTTTGTGCTGTTTT | >MI0000434\|hsa-let-7i&WithFlank&12\|+\|62603680\|62603775 | 12 | 33 | 22 | (+2U) | isomiR | NO | 0.00 | 0.00 | 32.93 | 0.00 |
| >MI0000434\|hsa-let-7i&WithFlank&12\|+\|62603680\|62603775@12.34.23&[MIMAT0000415&hsa-let-7i-5p&offsets\|0\|+1;m-49&12\|+\|62603691\|62603712&offsets\|0\|+1]\|TGAGGTAGTAGTTTGTGCTGTTG | >MI0000434\|hsa-let-7i&WithFlank&12\|+\|62603680\|62603775 | 12 | 34 | 23 | | isomiR | NO | 0.00 | 0.00 | 56.22 | 0.00 |
| >MI0000438\|hsa-mir-15b&WithFlank&3\|+\|160404582\|160404691@26.47.22&[MIMAT0000417&hsa-miR-15b-5p&offsets\|0\|0;m-110&3\|+\|160404607\|160404628&offsets\|0\|0]\|TAGCAGCACATCATGGTTTACA | >MI0000438\|hsa-mir-15b&WithFlank&3\|+\|160404582\|160404691 | 26 | 47 | 22 | | isomiR | NO | 135.66 | 0.00 | 12.07 | 17.52 |
| >MI0000438\|hsa-mir-15b&WithFlank&3\|+\|160404582\|160404691@64.84.21&[MIMAT0004586&hsa-miR-15b-3p&offsets\|0\|-1;m-204&3\|+\|160404645\|160404665&offsets\|0\|0]\|CGAATCATTATTTGCTGCTCT | >MI0000438\|hsa-mir-15b&WithFlank&3\|+\|160404582\|160404691 | 64 | 84 | 21 | | isomiR | NO | 0.00 | 0.00 | 67.85 | 0.00 |
| >MI0000438\|hsa-mir-15b&WithFlank&3\|+\|160404582\|160404691@64.85.22&[MIMAT0004586&hsa-miR-15b-3p&offsets\|0\|0;m-204&3\|+\|160404645\|160404665&offsets\|0\|+1]\|CGAATCATTATTTGCTGCTCTA | >MI0000438\|hsa-mir-15b&WithFlank&3\|+\|160404582\|160404691 | 64 | 85 | 22 | | isomiR | NO | 0.00 | 0.00 | 10.37 | 0.00 |
| >MI0000439\|hsa-mir-23b&WithFlank&9\|+\|95085202\|95085310@67.84.18&[MIMAT0000418&hsa-miR-23b-3p&offsets\|+3\|-2;m-42&9\|+\|95085265\|95085285&offsets\|+3\|0]\|ACATTGCCAGGGATTACC | >MI0000439\|hsa-mir-23b&WithFlank&9\|+\|95085202\|95085310 | 67 | 84 | 18 | | isomiR | NO | 0.00 | 25.22 | 0.00 | 0.00 |
| >MI0000439\|hsa-mir-23b&WithFlank&9\|+\|95085202\|95085310@67.86.20&[MIMAT0000418&hsa-miR-23b-3p&offsets\|+3\|0;m-42&9\|+\|95085265\|95085285&offsets\|+3\|+2]\|ACATTGCCAGGGATTACCAC | >MI0000439\|hsa-mir-23b&WithFlank&9\|+\|95085202\|95085310 | 67 | 86 | 20 | | isomiR | NO | 0.00 | 66.01 | 0.00 | 0.00 |
| >MI0000439\|hsa-mir-23b&WithFlank&9\|+\|95085202\|95085310@67.86.20(+1U)&[MIMAT0000418&hsa-miR-23b-3p&offsets\|+3\|0(+1U);m-42&9\|+\|95085265\|95085285&offsets\|+3\|+2(+1U)]\|ACATTGCCAGGGATTACCACT | >MI0000439\|hsa-mir-23b&WithFlank&9\|+\|95085202\|95085310 | 67 | 86 | 20 | (+1U) | isomiR | NO | 52.18 | 92.73 | 16.33 | 12.76 |
| >MI0000440\|hsa-mir-27b&WithFlank&9\|+\|95085439\|95085547@67.85.19&[MIMAT0000419&hsa-miR-27b-3p&offsets\|0\|-2;m-39&9\|+\|95085505\|95085525&offsets\|0\|-2]\|TTCACAGTGGCTAAGTTCT | >MI0000440\|hsa-mir-27b&WithFlank&9\|+\|95085439\|95085547 | 67 | 85 | 19 | | isomiR | NO | 109.80 | 36.55 | 179.49 | 17.14 |

### MDA-MB-231

<table>
<tr><td></td><td></td><td></td><td></td><td></td><td></td><td></td><td></td><td></td><td></td><td></td><td></td><td></td></tr>
<tr><td>>MI0000440|hsa-mir-27b&WithFlank&9|+|95085439|95085547@67.85.19(+1A)&[MIMAT0000419&hsa-miR-27b-3p&offsets|0|-2(+1A);m-39&9|+|95085505|95085525&offsets|0|-2(+1A)]|TTCACAGTGGCTAAGTTCTA</td><td>>MI0000440|hsa-mir-27b&WithFlank&9|+|95085439|95085547</td><td>67</td><td>85</td><td>19</td><td>(+1A)</td><td>isomiR</td><td>NO</td><td>0.00</td><td>0.00</td><td>94.77</td><td>0.00</td></tr>
<tr><td>>MI0000440|hsa-mir-27b&WithFlank&9|+|95085439|95085547@67.86.20&[MIMAT0000419&hsa-miR-27b-3p&offsets|0|-1;m-39&9|+|95085505|95085525&offsets|0|-1]|TTCACAGTGGCTAAGTTCTG</td><td>>MI0000440|hsa-mir-27b&WithFlank&9|+|95085439|95085547</td><td>67</td><td>86</td><td>20</td><td></td><td>isomiR</td><td>NO</td><td>254.25</td><td>159.22</td><td>469.49</td><td>90.17</td></tr>
<tr><td>>MI0000440|hsa-mir-27b&WithFlank&9|+|95085439|95085547@67.86.20(+1A)&[MIMAT0000419&hsa-miR-27b-3p&offsets|0|-1(+1A);m-39&9|+|95085505|95085525&offsets|0|-1(+1A)]|TTCACAGTGGCTAAGTTCTGA</td><td>>MI0000440|hsa-mir-27b&WithFlank&9|+|95085439|95085547</td><td>67</td><td>86</td><td>20</td><td>(+1A)</td><td>isomiR</td><td>NO</td><td>90.98</td><td>35.23</td><td>217.37</td><td>63.23</td></tr>
<tr><td>>MI0000440|hsa-mir-27b&WithFlank&9|+|95085439|95085547@67.86.20(+1U)&[MIMAT0000419&hsa-miR-27b-3p&offsets|0|-1(+1U);m-39&9|+|95085505|95085525&offsets|0|-1(+1U)]|TTCACAGTGGCTAAGTTCTGT</td><td>>MI0000440|hsa-mir-27b&WithFlank&9|+|95085439|95085547</td><td>67</td><td>86</td><td>20</td><td>(+1U)</td><td>isomiR</td><td>NO</td><td>256.18</td><td>125.77</td><td>584.71</td><td>154.53</td></tr>
<tr><td>>MI0000440|hsa-mir-27b&WithFlank&9|+|95085439|95085547@67.87.21&[MIMAT0000419&hsa-miR-27b-3p&offsets|0|0;m-39&9|+|95085505|95085525&offsets|0|0]|TTCACAGTGGCTAAGTTCTGC</td><td>>MI0000440|hsa-mir-27b&WithFlank&9|+|95085439|95085547</td><td>67</td><td>87</td><td>21</td><td></td><td>isomiR</td><td>YES</td><td>1141.44</td><td>572.89</td><td>1716.64</td><td>425.80</td></tr>
<tr><td>>MI0000440|hsa-mir-27b&WithFlank&9|+|95085439|95085547@67.87.21(+1C)&[MIMAT0000419&hsa-miR-27b-3p&offsets|0|0(+1C);m-39&9|+|95085505|95085525&offsets|0|0(+1C)]|TTCACAGTGGCTAAGTTCTGCC</td><td>>MI0000440|hsa-mir-27b&WithFlank&9|+|95085439|95085547</td><td>67</td><td>87</td><td>21</td><td>(+1C)</td><td>isomiR</td><td>NO</td><td>0.00</td><td>0.00</td><td>15.68</td><td>0.00</td></tr>
<tr><td>>MI0000440|hsa-mir-27b&WithFlank&9|+|95085439|95085547@67.87.21(+1U)&[MIMAT0000419&hsa-miR-27b-3p&offsets|0|0(+1U);m-39&9|+|95085505|95085525&offsets|0|0(+1U)]|TTCACAGTGGCTAAGTTCTGCT</td><td>>MI0000440|hsa-mir-27b&WithFlank&9|+|95085439|95085547</td><td>67</td><td>87</td><td>21</td><td>(+1U)</td><td>isomiR</td><td>NO</td><td>64.71</td><td>0.00</td><td>175.19</td><td>19.35</td></tr>
<tr><td>>MI0000440|hsa-mir-27b&WithFlank&9|+|95085439|95085547@67.87.21(+2U)&[MIMAT0000419&hsa-miR-27b-3p&offsets|0|0(+2U);m-39&9|+|95085505|95085525&offsets|0|0(+2U)]|TTCACAGTGGCTAAGTTCTGCTT</td><td>>MI0000440|hsa-mir-27b&WithFlank&9|+|95085439|95085547</td><td>67</td><td>87</td><td>21</td><td>(+2U)</td><td>isomiR</td><td>NO</td><td>0.00</td><td>0.00</td><td>71.11</td><td>11.45</td></tr>
<tr><td>>MI0000440|hsa-mir-27b&WithFlank&9|+|95085439|95085547@67.88.22&[MIMAT0000419&hsa-miR-27b-3p&offsets|0|+1;m-39&9|+|95085505|95085525&offsets|0|+1]|TTCACAGTGGCTAAGTTCTGCA</td><td>>MI0000440|hsa-mir-27b&WithFlank&9|+|95085439|95085547</td><td>67</td><td>88</td><td>22</td><td></td><td>isomiR</td><td>NO</td><td>282.83</td><td>93.24</td><td>107.34</td><td>96.07</td></tr>
<tr><td>>MI0000441|hsa-mir-30b&WithFlank&8|-|134800514|134800613@23.43.21&[MIMAT0000420&hsa-miR-30b-5p&offsets|0|-1;m-76&8|-|134800570|134800591&offsets|0|-1]|TGTAAACATCCTACACTCAGC</td><td>>MI0000441|hsa-mir-30b&WithFlank&8|-|134800514|134800613</td><td>23</td><td>43</td><td>21</td><td></td><td>isomiR</td><td>NO</td><td>0.00</td><td>0.00</td><td>31.66</td><td>0.00</td></tr>
<tr><td>>MI0000441|hsa-mir-30b&WithFlank&8|-|134800514|134800613@23.43.21(+1A)&[MIMAT0000420&hsa-miR-30b-5p&offsets|0|-1(+1A);m-76&8|-|134800570|134800591&offsets|0|-1(+1A)]|TGTAAACATCCTACACTCAGCA</td><td>>MI0000441|hsa-mir-30b&WithFlank&8|-|134800514|134800613</td><td>23</td><td>43</td><td>21</td><td>(+1A)</td><td>isomiR</td><td>NO</td><td>0.00</td><td>0.00</td><td>12.25</td><td>0.00</td></tr>
<tr><td>>MI0000441|hsa-mir-30b&WithFlank&8|-|134800514|134800613@23.43.21(+1C)&[MIMAT0000420&hsa-miR-30b-5p&offsets|0|-1(+1C);m-76&8|-|134800570|134800591&offsets|0|-1(+1C)]|TGTAAACATCCTACACTCAGCC</td><td>>MI0000441|hsa-mir-30b&WithFlank&8|-|134800514|134800613</td><td>23</td><td>43</td><td>21</td><td>(+1C)</td><td>isomiR</td><td>NO</td><td>0.00</td><td>0.00</td><td>15.79</td><td>12.00</td></tr>
<tr><td>>MI0000441|hsa-mir-30b&WithFlank&8|-|134800514|134800613@23.44.22&[MIMAT0000420&hsa-miR-30b-5p&offsets|0|0;m-76&8|-|134800570|134800591&offsets|0|0]|TGTAAACATCCTACACTCAGCT</td><td>>MI0000441|hsa-mir-30b&WithFlank&8|-|134800514|134800613</td><td>23</td><td>44</td><td>22</td><td></td><td>isomiR</td><td>NO</td><td>187.81</td><td>29.83</td><td>180.47</td><td>96.98</td></tr>
<tr><td>>MI0000446|hsa-mir-125b-1&WithFlank&11|-|122099751|122099850@21.39.19;MI0000470|hsa-mir-125b-2&WithFlank&21|+|16590231|16590331@23.41.19&[MIMAT0000423&hsa-miR-125b-5p&offsets|0|-3;m-67&11|-|122099809|122099830&offsets|0|-3];[MIMAT0000423_1&hsa-miR-125b-5p&offsets|0|-3;m-68&21|+|16590253|16590274&offsets|0|-3]|TCCCTGAGACCCTAACTTG</td><td>>MI0000446|hsa-mir-125b-1&WithFlank&11|-|122099751|12209985</td><td>21</td><td>39</td><td>19</td><td></td><td>isomiR</td><td>NO</td><td>72.89</td><td>67.97</td><td>16.65</td><td>17.48</td></tr>
<tr><td>>MI0000446|hsa-mir-125b-1&WithFlank&11|-|122099751|122099850@21.40.20(+1A);MI0000470|hsa-mir-125b-2&WithFlank&21|+|16590231|16590331@23.42.20(+1A)&[MIMAT0000423&hsa-miR-125b-5p&offsets|0|-2(+1A);m-67&11|-|122099809|122099830&offsets|0|-2(+1A)];[MIMAT0000423_1&hsa-miR-125b-5p&offsets|0|-2(+1A);m-68&21|+|16590253|16590274&offsets|0|-2(+1A)]|TCCCTGAGACCCTAACTTGTA</td><td>>MI0000446|hsa-mir-125b-1&WithFlank&11|-|122099751|12209985</td><td>21</td><td>40</td><td>20</td><td>(+1A)</td><td>isomiR</td><td>NO</td><td>51.07</td><td>70.17</td><td>102.68</td><td>14.70</td></tr>
<tr><td>>MI0000446|hsa-mir-125b-1&WithFlank&11|-|122099751|122099850@21.40.20;MI0000470|hsa-mir-125b-2&WithFlank&21|+|16590231|16590331@23.42.20&[MIMAT0000423&hsa-miR-125b-5p&offsets|0|-2;m-67&11|-|122099809|122099830&offsets|0|-2];[MIMAT0000423_1&hsa-miR-125b-5p&offsets|0|-2;m-68&21|+|16590253|16590274&offsets|0|-2]|TCCCTGAGACCCTAACTTGT</td><td>>MI0000446|hsa-mir-125b-1&WithFlank&11|-|122099751|12209985</td><td>21</td><td>40</td><td>20</td><td></td><td>isomiR</td><td>NO</td><td>120.79</td><td>125.26</td><td>64.19</td><td>61.43</td></tr>
<tr><td>>MI0000446|hsa-mir-125b-1&WithFlank&11|-|122099751|122099850@21.41.21(+1G);MI0000470|hsa-mir-125b-2&WithFlank&21|+|16590231|16590331@23.43.21(+1G)&[MIMAT0000423&hsa-miR-125b-5p&offsets|0|-1(+1G);m-67&11|-|122099809|122099830&offsets|0|-1(+1G)];[MIMAT0000423_1&hsa-miR-125b-5p&offsets|0|-1(+1G);m-68&21|+|16590253|16590274&offsets|0|-1(+1G)]|TCCCTGAGACCCTAACTTGTGG</td><td>>MI0000446|hsa-mir-125b-1&WithFlank&11|-|122099751|12209985</td><td>21</td><td>41</td><td>21</td><td>(+1G)</td><td>isomiR</td><td>NO</td><td>0.00</td><td>0.00</td><td>16.35</td><td>0.00</td></tr>
<tr><td>>MI0000446|hsa-mir-125b-1&WithFlank&11|-|122099751|122099850@21.41.21(+1U);MI0000470|hsa-mir-125b-2&WithFlank&21|+|16590231|16590331@23.43.21(+1U)&[MIMAT0000423&hsa-miR-125b-5p&offsets|0|-1(+1U);m-67&11|-|122099809|122099830&offsets|0|-1(+1U)];[MIMAT0000423_1&hsa-miR-125b-5p&offsets|0|-1(+1U);m-68&21|+|16590253|16590274&offsets|0|-1(+1U)]|TCCCTGAGACCCTAACTTGTGT</td><td>>MI0000446|hsa-mir-125b-1&WithFlank&11|-|122099751|12209985</td><td>21</td><td>41</td><td>21</td><td>(+1U)</td><td>isomiR</td><td>NO</td><td>61.94</td><td>31.40</td><td>63.70</td><td>16.50</td></tr>
</table>

### MDA-MB-231

<table>
<tr><td>>MI0000446|hsa-mir-125b-1&WithFlank&11|-|122099751|122099850@21.41.21;MI0000470|hsa-mir-125b-2&WithFlank&21|+|16590231|16590331@23.43.21&[MIMAT0000423&hsa-miR-125b-5p&offsets|0|-1;m-67&11|-|122099809|122099830&offsets|0|-1];[MIMAT0000423_1&hsa-miR-125b-5p&offsets|0|-1;m-68&21|+|16590253|16590274&offsets|0|-1]|TCCCTGAGACCCTAACTTGTG</td><td>>MI0000446|hsa-mir-125b-1&WithFlank&11|-|122099751|12209985</td><td>21</td><td>41</td><td>21</td><td></td><td>isomiR</td><td>NO</td><td>381.48</td><td>466.34</td><td>477.03</td><td>137.61</td></tr>
<tr><td>>MI0000446|hsa-mir-125b-1&WithFlank&11|-|122099751|122099850@21.42.22(+1A);MI0000470|hsa-mir-125b-2&WithFlank&21|+|16590231|16590331@23.44.22(+1A)&[MIMAT0000423&hsa-miR-125b-5p&offsets|0|0(+1A);m-67&11|-|122099809|122099830&offsets|0|0(+1A)];[MIMAT0000423_1&hsa-miR-125b-5p&offsets|0|0(+1A);m-68&21|+|16590253|16590274&offsets|0|0(+1A)]|TCCCTGAGACCCTAACTTGTGAA</td><td>>MI0000446|hsa-mir-125b-1&WithFlank&11|-|122099751|12209985</td><td>21</td><td>42</td><td>22</td><td>(+1A)</td><td>isomiR</td><td>NO</td><td>211.36</td><td>184.10</td><td>113.46</td><td>108.33</td></tr>
<tr><td>>MI0000446|hsa-mir-125b-1&WithFlank&11|-|122099751|122099850@21.42.22(+2A);MI0000470|hsa-mir-125b-2&WithFlank&21|+|16590231|16590331@23.44.22(+2A)&[MIMAT0000423&hsa-miR-125b-5p&offsets|0|0(+2A);m-67&11|-|122099809|122099830&offsets|0|0(+2A)];[MIMAT0000423_1&hsa-miR-125b-5p&offsets|0|0(+2A);m-68&21|+|16590253|16590274&offsets|0|0(+2A)]|TCCCTGAGACCCTAACTTGTGAAA</td><td>>MI0000446|hsa-mir-125b-1&WithFlank&11|-|122099751|12209985</td><td>21</td><td>42</td><td>22</td><td>(+2A)</td><td>isomiR</td><td>NO</td><td>51.21</td><td>32.37</td><td>0.00</td><td>17.88</td></tr>
<tr><td>>MI0000446|hsa-mir-125b-1&WithFlank&11|-|122099751|122099850@21.42.22;MI0000470|hsa-mir-125b-2&WithFlank&21|+|16590231|16590331@23.44.22&[MIMAT0000423&hsa-miR-125b-5p&offsets|0|0;m-67&11|-|122099809|122099830&offsets|0|0];[MIMAT0000423_1&hsa-miR-125b-5p&offsets|0|0;m-68&21|+|16590253|16590274&offsets|0|0]|TCCCTGAGACCCTAACTTGTGA</td><td>>MI0000446|hsa-mir-125b-1&WithFlank&11|-|122099751|12209985</td><td>21</td><td>42</td><td>22</td><td></td><td>isomiR</td><td>NO</td><td>713.71</td><td>476.29</td><td>1247.74</td><td>223.81</td></tr>
<tr><td>>MI0000446|hsa-mir-125b-1&WithFlank&11|-|122099751|122099850@21.43.23&[MIMAT0000423&hsa-miR-125b-5p&offsets|0|+1;m-67&11|-|122099809|122099830&offsets|0|+1]|TCCCTGAGACCCTAACTTGTGAT</td><td>>MI0000446|hsa-mir-125b-1&WithFlank&11|-|122099751|12209985</td><td>21</td><td>43</td><td>23</td><td></td><td>isomiR</td><td>NO</td><td>99.70</td><td>91.60</td><td>16.84</td><td>67.77</td></tr>
<tr><td>>MI0000446|hsa-mir-125b-1&WithFlank&11|-|122099751|122099850@61.80.20(+1U)&[MIMAT0004592&hsa-miR-125b-1-3p&offsets|0|-2(+1U);m-485&11|-|122099769|122099790&offsets|0|-2(+1U)]|ACGGGTTAGGCTCTTGGGAGT</td><td>>MI0000446|hsa-mir-125b-1&WithFlank&11|-|122099751|12209985</td><td>61</td><td>80</td><td>20</td><td>(+1U)</td><td>isomiR</td><td>NO</td><td>0.00</td><td>73.94</td><td>116.78</td><td>0.00</td></tr>
<tr><td>>MI0000446|hsa-mir-125b-1&WithFlank&11|-|122099751|122099850@61.81.21&[MIMAT0004592&hsa-miR-125b-1-3p&offsets|0|-1;m-485&11|-|122099769|122099790&offsets|0|-1]|ACGGGTTAGGCTCTTGGGAGC</td><td>>MI0000446|hsa-mir-125b-1&WithFlank&11|-|122099751|12209985</td><td>61</td><td>81</td><td>21</td><td></td><td>isomiR</td><td>NO</td><td>0.00</td><td>78.79</td><td>33.48</td><td>0.00</td></tr>
<tr><td>>MI0000446|hsa-mir-125b-1&WithFlank&11|-|122099751|122099850@61.81.21(+1C)&[MIMAT0004592&hsa-miR-125b-1-3p&offsets|0|-1(+1C);m-485&11|-|122099769|122099790&offsets|0|-1(+1C)]|ACGGGTTAGGCTCTTGGGAGCC</td><td>>MI0000446|hsa-mir-125b-1&WithFlank&11|-|122099751|12209985</td><td>61</td><td>81</td><td>21</td><td>(+1C)</td><td>isomiR</td><td>NO</td><td>0.00</td><td>0.00</td><td>13.41</td><td>0.00</td></tr>
<tr><td>>MI0000446|hsa-mir-125b-1&WithFlank&11|-|122099751|122099850@61.82.22&[MIMAT0004592&hsa-miR-125b-1-3p&offsets|0|0;m-485&11|-|122099769|122099790&offsets|0|0]|ACGGGTTAGGCTCTTGGGAGCT</td><td>>MI0000446|hsa-mir-125b-1&WithFlank&11|-|122099751|12209985</td><td>61</td><td>82</td><td>22</td><td></td><td>isomiR</td><td>NO</td><td>23.73</td><td>80.65</td><td>128.30</td><td>0.00</td></tr>
<tr><td>>MI0000446|hsa-mir-125b-1&WithFlank&11|-|122099751|122099850@61.82.22(+1U)&[MIMAT0004592&hsa-miR-125b-1-3p&offsets|0|0(+1U);m-485&11|-|122099769|122099790&offsets|0|0(+1U)]|ACGGGTTAGGCTCTTGGGAGCTT</td><td>>MI0000446|hsa-mir-125b-1&WithFlank&11|-|122099751|12209985</td><td>61</td><td>82</td><td>22</td><td>(+1U)</td><td>isomiR</td><td>NO</td><td>0.00</td><td>0.00</td><td>17.64</td><td>0.00</td></tr>
<tr><td>>MI0000447|hsa-mir-128-1&WithFlank&2|+|135665391|135665484@56.75.20(+1C);MI0000727|hsa-mir-128-2&WithFlank&3|+|35744470|35744565@58.77.20(+1C)&[MIMAT0000424&hsa-miR-128-3p&offsets|0|-1(+1C);m-108&2|+|135665446|135665466&offsets|0|-1(+1C)];[MIMAT0000424_1&hsa-miR-128-3p&offsets|0|-1(+1C);m-117&3|+|35744527|35744547&offsets|0|-1(+1C)]|TCACAGTGAACCGGTCTCTTC</td><td>>MI0000447|hsa-mir-128-1&WithFlank&2|+|135665391|135665484</td><td>56</td><td>75</td><td>20</td><td>(+1C)</td><td>isomiR</td><td>NO</td><td>26.22</td><td>0.00</td><td>12.42</td><td>10.90</td></tr>
<tr><td>>MI0000447|hsa-mir-128-1&WithFlank&2|+|135665391|135665484@56.75.20;MI0000727|hsa-mir-128-2&WithFlank&3|+|35744470|35744565@58.77.20&[MIMAT0000424&hsa-miR-128-3p&offsets|0|-1;m-108&2|+|135665446|135665466&offsets|0|-1];[MIMAT0000424_1&hsa-miR-128-3p&offsets|0|-1;m-117&3|+|35744527|35744547&offsets|0|-1]|TCACAGTGAACCGGTCTCTT</td><td>>MI0000447|hsa-mir-128-1&WithFlank&2|+|135665391|135665484</td><td>56</td><td>75</td><td>20</td><td></td><td>isomiR</td><td>NO</td><td>71.99</td><td>0.00</td><td>13.49</td><td>13.48</td></tr>
<tr><td>>MI0000447|hsa-mir-128-1&WithFlank&2|+|135665391|135665484@56.76.21(+1A);MI0000727|hsa-mir-128-2&WithFlank&3|+|35744470|35744565@58.78.21(+1A)&[MIMAT0000424&hsa-miR-128-3p&offsets|0|0(+1A);m-108&2|+|135665446|135665466&offsets|0|0(+1A)];[MIMAT0000424_1&hsa-miR-128-3p&offsets|0|0(+1A);m-117&3|+|35744527|35744547&offsets|0|0(+1A)]|TCACAGTGAACCGGTCTCTTTA</td><td>>MI0000447|hsa-mir-128-1&WithFlank&2|+|135665391|135665484</td><td>56</td><td>76</td><td>21</td><td>(+1A)</td><td>isomiR</td><td>NO</td><td>48.92</td><td>0.00</td><td>0.00</td><td>10.41</td></tr>
<tr><td>>MI0000447|hsa-mir-128-1&WithFlank&2|+|135665391|135665484@56.76.21;MI0000727|hsa-mir-128-2&WithFlank&3|+|35744470|35744565@58.78.21&[MIMAT0000424&hsa-miR-128-3p&offsets|0|0;m-108&2|+|135665446|135665466&offsets|0|0];[MIMAT0000424_1&hsa-miR-128-3p&offsets|0|0;m-117&3|+|35744527|35744547&offsets|0|0]|TCACAGTGAACCGGTCTCTTT</td><td>>MI0000447|hsa-mir-128-1&WithFlank&2|+|135665391|135665484</td><td>56</td><td>76</td><td>21</td><td></td><td>isomiR</td><td>NO</td><td>294.18</td><td>139.43</td><td>152.06</td><td>107.51</td></tr>
<tr><td>>MI0000447|hsa-mir-128-1&WithFlank&2|+|135665391|135665484@56.77.22&[MIMAT0000424&hsa-miR-128-3p&offsets|0|+1;m-108&2|+|135665446|135665466&offsets|0|+1]|TCACAGTGAACCGGTCTCTTTT</td><td>>MI0000447|hsa-mir-128-1&WithFlank&2|+|135665391|135665484</td><td>56</td><td>77</td><td>22</td><td></td><td>isomiR</td><td>NO</td><td>134.07</td><td>69.60</td><td>56.41</td><td>64.16</td></tr>
<tr><td>>MI0000448|hsa-mir-130a&WithFlank&11|+|57641192|57641292@61.83.23&[MIMAT0000425&hsa-miR-130a-3p&offsets|0|+1;m-139&11|+|57641252|57641273&offsets|0|+1]|CAGTGCAATGTTAAAAGGGCATT</td><td>>MI0000448|hsa-mir-130a&WithFlank&11|+|57641192|57641292</td><td>61</td><td>83</td><td>23</td><td></td><td>isomiR</td><td>NO</td><td>0.00</td><td>0.00</td><td>11.83</td><td>0.00</td></tr>
</table>

### MDA-MB-231

| | | | | | | | | | | | |
|---|---|---|---|---|---|---|---|---|---|---|---|
| >MI0000449\|hsa-mir-132&WithFlank&17\|-\|2049902\|2050014@65.86.22&[MIMAT0000426&hsa-miR-132-3p&offsets\|0\|0;m-138&17\|-\|2049929\|2049950&offsets\|0\|0]\|TAACAGTCTACAGCCATGGTCG | >MI0000449\|hsa-mir-132&WithFlank&17\|-\|2049902\|2050014 | 65 | 86 | 22 | | isomiR | NO | 0.00 | 0.00 | 13.21 | 0.00 |
| >MI0000449\|hsa-mir-132&WithFlank&17\|-\|2049902\|2050014@65.86.22(+1U)&[MIMAT0000426&hsa-miR-132-3p&offsets\|0\|0(+1U);m-138&17\|-\|2049929\|2049950&offsets\|0\|0(+1U)]\|TAACAGTCTACAGCCATGGTCGT | >MI0000449\|hsa-mir-132&WithFlank&17\|-\|2049902\|2050014 | 65 | 86 | 22 | (+1U) | isomiR | NO | 0.00 | 0.00 | 14.73 | 0.00 |
| >MI0000456\|hsa-mir-140&WithFlank&16\|+\|69933075\|69933186@29.50.22&[MIMAT0000431&hsa-miR-140-5p&offsets\|0\|0;m-173&16\|+\|69933103\|69933124&offsets\|0\|0]\|CAGTGGTTTTACCCTATGGTAG | >MI0000456\|hsa-mir-140&WithFlank&16\|+\|69933075\|69933186 | 29 | 50 | 22 | | isomiR | NO | 0.00 | 0.00 | 154.80 | 0.00 |
| >MI0000456\|hsa-mir-140&WithFlank&16\|+\|69933075\|69933186@68.88.21&[MIMAT0004597&hsa-miR-140-3p&offsets\|0\|0;m-45&16\|+\|69933143\|69933164&offsets\|-1\|-2]\|TACCACAGGGTAGAACCACGG | >MI0000456\|hsa-mir-140&WithFlank&16\|+\|69933075\|69933186 | 68 | 88 | 21 | | isomiR | NO | 0.00 | 0.00 | 55.96 | 0.00 |
| >MI0000456\|hsa-mir-140&WithFlank&16\|+\|69933075\|69933186@68.89.22&[MIMAT0004597&hsa-miR-140-3p&offsets\|0\|+1;m-45&16\|+\|69933143\|69933164&offsets\|-1\|-1]\|TACCACAGGGTAGAACCACGGA | >MI0000456\|hsa-mir-140&WithFlank&16\|+\|69933075\|69933186 | 68 | 89 | 22 | | isomiR | NO | 0.00 | 27.36 | 121.31 | 0.00 |
| >MI0000456\|hsa-mir-140&WithFlank&16\|+\|69933075\|69933186@68.90.23&[MIMAT0004597&hsa-miR-140-3p&offsets\|0\|+2;m-45&16\|+\|69933143\|69933164&offsets\|-1\|0]\|TACCACAGGGTAGAACCACGGAC | >MI0000456\|hsa-mir-140&WithFlank&16\|+\|69933075\|69933186 | 68 | 90 | 23 | | isomiR | NO | 0.00 | 24.59 | 62.27 | 0.00 |
| >MI0000456\|hsa-mir-140&WithFlank&16\|+\|69933075\|69933186@68.90.23(+1U)&[MIMAT0004597&hsa-miR-140-3p&offsets\|0\|+2(+1U);m-45&16\|+\|69933143\|69933164&offsets\|-1\|0(+1U)]\|TACCACAGGGTAGAACCACGGACT | >MI0000456\|hsa-mir-140&WithFlank&16\|+\|69933075\|69933186 | 68 | 90 | 23 | (+1U) | isomiR | NO | 0.00 | 0.00 | 59.41 | 0.00 |
| >MI0000456\|hsa-mir-140&WithFlank&16\|+\|69933075\|69933186@68.91.24&[MIMAT0004597&hsa-miR-140-3p&offsets\|0\|+3;m-45&16\|+\|69933143\|69933164&offsets\|-1\|+1]\|TACCACAGGGTAGAACCACGGACA | >MI0000456\|hsa-mir-140&WithFlank&16\|+\|69933075\|69933186 | 68 | 91 | 24 | | isomiR | NO | 0.00 | 0.00 | 15.07 | 0.00 |
| >MI0000456\|hsa-mir-140&WithFlank&16\|+\|69933075\|69933186@69.88.20&[MIMAT0004597&hsa-miR-140-3p&offsets\|+1\|0;m-45&16\|+\|69933143\|69933164&offsets\|0\|-2]\|ACCACAGGGTAGAACCACGG | >MI0000456\|hsa-mir-140&WithFlank&16\|+\|69933075\|69933186 | 69 | 88 | 20 | | isomiR | NO | 0.00 | 0.00 | 113.61 | 0.00 |
| >MI0000456\|hsa-mir-140&WithFlank&16\|+\|69933075\|69933186@69.89.21&[MIMAT0004597&hsa-miR-140-3p&offsets\|+1\|+1;m-45&16\|+\|69933143\|69933164&offsets\|0\|-1]\|ACCACAGGGTAGAACCACGGA | >MI0000456\|hsa-mir-140&WithFlank&16\|+\|69933075\|69933186 | 69 | 89 | 21 | | isomiR | NO | 45.06 | 32.83 | 220.24 | 0.00 |
| >MI0000456\|hsa-mir-140&WithFlank&16\|+\|69933075\|69933186@69.89.21(+1A)&[MIMAT0004597&hsa-miR-140-3p&offsets\|+1\|+1(+1A);m-45&16\|+\|69933143\|69933164&offsets\|0\|-1(+1A)]\|ACCACAGGGTAGAACCACGGAA | >MI0000456\|hsa-mir-140&WithFlank&16\|+\|69933075\|69933186 | 69 | 89 | 21 | (+1A) | isomiR | NO | 0.00 | 0.00 | 11.30 | 0.00 |
| >MI0000456\|hsa-mir-140&WithFlank&16\|+\|69933075\|69933186@69.89.21(+1U)&[MIMAT0004597&hsa-miR-140-3p&offsets\|+1\|+1(+1U);m-45&16\|+\|69933143\|69933164&offsets\|0\|-1(+1U)]\|ACCACAGGGTAGAACCACGGAT | >MI0000456\|hsa-mir-140&WithFlank&16\|+\|69933075\|69933186 | 69 | 89 | 21 | (+1U) | isomiR | NO | 0.00 | 0.00 | 65.79 | 0.00 |
| >MI0000456\|hsa-mir-140&WithFlank&16\|+\|69933075\|69933186@69.90.22&[MIMAT0004597&hsa-miR-140-3p&offsets\|+1\|+2;m-45&16\|+\|69933143\|69933164&offsets\|0\|0]\|ACCACAGGGTAGAACCACGGAC | >MI0000456\|hsa-mir-140&WithFlank&16\|+\|69933075\|69933186 | 69 | 90 | 22 | | isomiR | NO | 52.57 | 57.71 | 207.28 | 0.00 |
| >MI0000456\|hsa-mir-140&WithFlank&16\|+\|69933075\|69933186@69.90.22(+1C)&[MIMAT0004597&hsa-miR-140-3p&offsets\|+1\|+2(+1C);m-45&16\|+\|69933143\|69933164&offsets\|0\|0(+1C)]\|ACCACAGGGTAGAACCACGGACC | >MI0000456\|hsa-mir-140&WithFlank&16\|+\|69933075\|69933186 | 69 | 90 | 22 | (+1C) | isomiR | NO | 0.00 | 0.00 | 16.23 | 0.00 |
| >MI0000456\|hsa-mir-140&WithFlank&16\|+\|69933075\|69933186@69.90.22(+1U)&[MIMAT0004597&hsa-miR-140-3p&offsets\|+1\|+2(+1U);m-45&16\|+\|69933143\|69933164&offsets\|0\|0(+1U)]\|ACCACAGGGTAGAACCACGGACT | >MI0000456\|hsa-mir-140&WithFlank&16\|+\|69933075\|69933186 | 69 | 90 | 22 | (+1U) | isomiR | NO | 10.63 | 37.20 | 187.47 | 0.00 |
| >MI0000456\|hsa-mir-140&WithFlank&16\|+\|69933075\|69933186@69.91.23&[MIMAT0004597&hsa-miR-140-3p&offsets\|+1\|+3;m-45&16\|+\|69933143\|69933164&offsets\|0\|+1]\|ACCACAGGGTAGAACCACGGACA | >MI0000456\|hsa-mir-140&WithFlank&16\|+\|69933075\|69933186 | 69 | 91 | 23 | | isomiR | NO | 0.00 | 0.00 | 86.07 | 0.00 |
| >MI0000462\|hsa-mir-152&WithFlank&17\|-\|48037155\|48037253@60.79.20(+1A)&[MIMAT0000438&hsa-miR-152-3p&offsets\|0\|-1(+1A);m-97&17\|-\|48037174\|48037194&offsets\|0\|-1(+1A)]\|TCAGTGCATGACAGAACTTGA | >MI0000462\|hsa-mir-152&WithFlank&17\|-\|48037155\|48037253 | 60 | 79 | 20 | (+1A) | isomiR | NO | 0.00 | 0.00 | 15.32 | 0.00 |
| >MI0000462\|hsa-mir-152&WithFlank&17\|-\|48037155\|48037253@60.80.21&[MIMAT0000438&hsa-miR-152-3p&offsets\|0\|0;m-97&17\|-\|48037174\|48037194&offsets\|0\|0]\|TCAGTGCATGACAGAACTTGG | >MI0000462\|hsa-mir-152&WithFlank&17\|-\|48037155\|48037253 | 60 | 80 | 21 | | isomiR | NO | 53.60 | 0.00 | 172.58 | 0.00 |
| >MI0000462\|hsa-mir-152&WithFlank&17\|-\|48037155\|48037253@60.80.21(+1U)&[MIMAT0000438&hsa-miR-152-3p&offsets\|0\|0(+1U);m-97&17\|-\|48037174\|48037194&offsets\|0\|0(+1U)]\|TCAGTGCATGACAGAACTTGGT | >MI0000462\|hsa-mir-152&WithFlank&17\|-\|48037155\|48037253 | 60 | 80 | 21 | (+1U) | isomiR | NO | 0.00 | 0.00 | 100.23 | 0.00 |
| >MI0000462\|hsa-mir-152&WithFlank&17\|-\|48037155\|48037253@60.81.22&[MIMAT0000438&hsa-miR-152-3p&offsets\|0\|+1;m-97&17\|-\|48037174\|48037194&offsets\|0\|+1]\|TCAGTGCATGACAGAACTTGGG | >MI0000462\|hsa-mir-152&WithFlank&17\|-\|48037155\|48037253 | 60 | 81 | 22 | | isomiR | NO | 0.00 | 0.00 | 122.38 | 0.00 |
| >MI0000462\|hsa-mir-152&WithFlank&17\|-\|48037155\|48037253@60.81.22(+1U)&[MIMAT0000438&hsa-miR-152-3p&offsets\|0\|+1(+1U);m-97&17\|-\|48037174\|48037194&offsets\|0\|+1(+1U)]\|TCAGTGCATGACAGAACTTGGGT | >MI0000462\|hsa-mir-152&WithFlank&17\|-\|48037155\|48037253 | 60 | 81 | 22 | (+1U) | isomiR | NO | 0.00 | 0.00 | 78.05 | 0.00 |
| >MI0000465\|hsa-mir-191&WithFlank&3\|-\|49020612\|49020715@22.42.21&[MIMAT0000440&hsa-miR-191-5p&offsets\|0\|-2;m-48&3\|-\|49020672\|49020694&offsets\|0\|-2]\|CAACGGAATCCCAAAAGCAGC | >MI0000465\|hsa-mir-191&WithFlank&3\|-\|49020612\|49020715 | 22 | 42 | 21 | | isomiR | NO | 48.74 | 0.00 | 72.77 | 0.00 |

### MDA-MB-231

| | | | | | | | | | | | |
|---|---|---|---|---|---|---|---|---|---|---|---|
| >MI0000465\|hsa-mir-191&WithFlank&3\|-\|49020612\|49020715@22.42.21(+1A)&[MIMAT0000440&hsa-miR-191-5p&offsets\|0\|-2(+1A);m-48&3\|-\|49020672\|49020694&offsets\|0\|-2(+1A)]\|CAACGGAATCCCAAAAGCAGCA | >MI0000465\|hsa-mir-191&WithFlank&3\|-\|49020612\|49020715 | 22 | 42 | 21 | (+1A) | isomiR | NO | 0.00 | 0.00 | 89.17 | 0.00 |
| >MI0000465\|hsa-mir-191&WithFlank&3\|-\|49020612\|49020715@22.42.21(+1C)&[MIMAT0000440&hsa-miR-191-5p&offsets\|0\|-2(+1C);m-48&3\|-\|49020672\|49020694&offsets\|0\|-2(+1C)]\|CAACGGAATCCCAAAAGCAGCC | >MI0000465\|hsa-mir-191&WithFlank&3\|-\|49020612\|49020715 | 22 | 42 | 21 | (+1C) | isomiR | NO | 58.30 | 0.00 | 128.24 | 12.04 |
| >MI0000465\|hsa-mir-191&WithFlank&3\|-\|49020612\|49020715@22.43.22&[MIMAT0000440&hsa-miR-191-5p&offsets\|0\|-1;m-48&3\|-\|49020672\|49020694&offsets\|0\|-1]\|CAACGGAATCCCAAAAGCAGCT | >MI0000465\|hsa-mir-191&WithFlank&3\|-\|49020612\|49020715 | 22 | 43 | 22 | | isomiR | NO | 316.70 | 162.02 | 548.91 | 100.00 |
| >MI0000465\|hsa-mir-191&WithFlank&3\|-\|49020612\|49020715@22.43.22(+1A)&[MIMAT0000440&hsa-miR-191-5p&offsets\|0\|-1(+1A);m-48&3\|-\|49020672\|49020694&offsets\|0\|-1(+1A)]\|CAACGGAATCCCAAAAGCAGCTA | >MI0000465\|hsa-mir-191&WithFlank&3\|-\|49020612\|49020715 | 22 | 43 | 22 | (+1A) | isomiR | NO | 52.30 | 30.78 | 209.07 | 12.36 |
| >MI0000465\|hsa-mir-191&WithFlank&3\|-\|49020612\|49020715@22.43.22(+1U)&[MIMAT0000440&hsa-miR-191-5p&offsets\|0\|-1(+1U);m-48&3\|-\|49020672\|49020694&offsets\|0\|-1(+1U)]\|CAACGGAATCCCAAAAGCAGCTT | >MI0000465\|hsa-mir-191&WithFlank&3\|-\|49020612\|49020715 | 22 | 43 | 22 | (+1U) | isomiR | NO | 0.00 | 0.00 | 60.70 | 0.00 |
| >MI0000465\|hsa-mir-191&WithFlank&3\|-\|49020612\|49020715@22.44.23&[MIMAT0000440&hsa-miR-191-5p&offsets\|0\|0;m-48&3\|-\|49020672\|49020694&offsets\|0\|0]\|CAACGGAATCCCAAAAGCAGCTG | >MI0000465\|hsa-mir-191&WithFlank&3\|-\|49020612\|49020715 | 22 | 44 | 23 | | isomiR | NO | 399.94 | 297.28 | 992.17 | 121.70 |
| >MI0000465\|hsa-mir-191&WithFlank&3\|-\|49020612\|49020715@22.44.23(+1A)&[MIMAT0000440&hsa-miR-191-5p&offsets\|0\|0(+1A);m-48&3\|-\|49020672\|49020694&offsets\|0\|0(+1A)]\|CAACGGAATCCCAAAAGCAGCTGA | >MI0000465\|hsa-mir-191&WithFlank&3\|-\|49020612\|49020715 | 22 | 44 | 23 | (+1A) | isomiR | NO | 0.00 | 0.00 | 74.40 | 0.00 |
| >MI0000465\|hsa-mir-191&WithFlank&3\|-\|49020612\|49020715@22.44.23(+1C)&[MIMAT0000440&hsa-miR-191-5p&offsets\|0\|0(+1C);m-48&3\|-\|49020672\|49020694&offsets\|0\|0(+1C)]\|CAACGGAATCCCAAAAGCAGCTGC | >MI0000465\|hsa-mir-191&WithFlank&3\|-\|49020612\|49020715 | 22 | 44 | 23 | (+1C) | isomiR | NO | 0.00 | 0.00 | 139.20 | 0.00 |
| >MI0000465\|hsa-mir-191&WithFlank&3\|-\|49020612\|49020715@22.45.24&[MIMAT0000440&hsa-miR-191-5p&offsets\|0\|+1;m-48&3\|-\|49020672\|49020694&offsets\|0\|+1]\|CAACGGAATCCCAAAAGCAGCTGT | >MI0000465\|hsa-mir-191&WithFlank&3\|-\|49020612\|49020715 | 22 | 45 | 24 | | isomiR | NO | 200.81 | 97.46 | 620.84 | 78.18 |
| >MI0000465\|hsa-mir-191&WithFlank&3\|-\|49020612\|49020715@23.44.22&[MIMAT0000440&hsa-miR-191-5p&offsets\|+1\|0;m-48&3\|-\|49020672\|49020694&offsets\|+1\|0]\|AACGGAATCCCAAAAGCAGCTG | >MI0000465\|hsa-mir-191&WithFlank&3\|-\|49020612\|49020715 | 23 | 44 | 22 | | isomiR | NO | 0.00 | 0.00 | 61.44 | 0.00 |
| >MI0000466\|hsa-mir-9-1&WithFlank&1\|-\|156420335\|156420435@22.44.23;MI0000468\|hsa-mir-9-3&WithFlank&15\|+\|89368011\|89368112@22.44.23;MI0000467\|hsa-mir-9-2&WithFlank&5\|-\|88666847\|88666945@22.44.23&[MIMAT0000441&hsa-miR-9-5p&offsets\|0\|0;m-104&1\|-\|156420392\|156420414&offsets\|0\|0];[MIMAT0000441_1&hsa-miR-9-5p&offsets\|0\|0;m-105&15\|+\|89368032\|89368054&offsets\|0\|0];[MIMAT0000441_2&hsa-miR-9-5p&offsets\|0\|0;m-104&5\|-\|88666902\|88666924&offsets\|0\|0]\|TCTTTGGTTATCTAGCTGTATGA | >MI0000466\|hsa-mir-9-1&WithFlank&1\|-\|156420335\|156420435 | 22 | 44 | 23 | | isomiR | NO | 0.00 | 0.00 | 16.50 | 0.00 |
| >MI0000469\|hsa-mir-125a&WithFlank&19\|+\|51693248\|51693345@21.42.22&[MIMAT0000443&hsa-miR-125a-5p&offsets\|0\|-2;m-63&19\|+\|51693268\|51693289&offsets\|0\|0]\|TCCCTGAGACCCTTTAACCTGT | >MI0000469\|hsa-mir-125a&WithFlank&19\|+\|51693248\|51693345 | 21 | 42 | 22 | | isomiR | NO | 104.98 | 107.74 | 0.00 | 62.29 |
| >MI0000469\|hsa-mir-125a&WithFlank&19\|+\|51693248\|51693345@21.42.22(+1A)&[MIMAT0000443&hsa-miR-125a-5p&offsets\|0\|-2(+1A);m-63&19\|+\|51693268\|51693289&offsets\|0\|0(+1A)]\|TCCCTGAGACCCTTTAACCTGTA | >MI0000469\|hsa-mir-125a&WithFlank&19\|+\|51693248\|51693345 | 21 | 42 | 22 | (+1A) | isomiR | NO | 49.40 | 33.54 | 0.00 | 19.85 |
| >MI0000469\|hsa-mir-125a&WithFlank&19\|+\|51693248\|51693345@21.43.23&[MIMAT0000443&hsa-miR-125a-5p&offsets\|0\|-1;m-63&19\|+\|51693268\|51693289&offsets\|0\|+1]\|TCCCTGAGACCCTTTAACCTGTG | >MI0000469\|hsa-mir-125a&WithFlank&19\|+\|51693248\|51693345 | 21 | 43 | 23 | | isomiR | NO | 385.12 | 337.81 | 120.65 | 198.39 |
| >MI0000469\|hsa-mir-125a&WithFlank&19\|+\|51693248\|51693345@21.43.23(+1U)&[MIMAT0000443&hsa-miR-125a-5p&offsets\|0\|-1(+1U);m-63&19\|+\|51693268\|51693289&offsets\|0\|+1(+1U)]\|TCCCTGAGACCCTTTAACCTGTGT | >MI0000469\|hsa-mir-125a&WithFlank&19\|+\|51693248\|51693345 | 21 | 43 | 23 | (+1U) | isomiR | NO | 55.64 | 29.15 | 0.00 | 16.42 |
| >MI0000469\|hsa-mir-125a&WithFlank&19\|+\|51693248\|51693345@21.44.24&[MIMAT0000443&hsa-miR-125a-5p&offsets\|0\|0;m-63&19\|+\|51693268\|51693289&offsets\|0\|+2]\|TCCCTGAGACCCTTTAACCTGTGA | >MI0000469\|hsa-mir-125a&WithFlank&19\|+\|51693248\|51693345 | 21 | 44 | 24 | | isomiR | NO | 137.95 | 105.04 | 16.17 | 85.45 |
| >MI0000471\|hsa-mir-126&WithFlank&9\|+\|136670596\|136670692@21.41.21&[MIMAT0000444&hsa-miR-126-5p&offsets\|0\|0;m-114&9\|+\|136670616\|136670636&offsets\|0\|0]\|CATTATTACTTTTGGTACGCG | >MI0000471\|hsa-mir-126&WithFlank&9\|+\|136670596\|136670692 | 21 | 41 | 21 | | isomiR | NO | 0.00 | 0.00 | 31.24 | 0.00 |
| >MI0000471\|hsa-mir-126&WithFlank&9\|+\|136670596\|136670692@57.79.23&[MIMAT0000445&hsa-miR-126-3p&offsets\|-1\|0;m-40&9\|+\|136670653\|136670674&offsets\|-1\|0]\|CTCGTACCGTGAGTAATAATGCG | >MI0000471\|hsa-mir-126&WithFlank&9\|+\|136670596\|136670692 | 57 | 79 | 23 | | isomiR | NO | 25.20 | 0.00 | 35.35 | 0.00 |
| >MI0000471\|hsa-mir-126&WithFlank&9\|+\|136670596\|136670692@58.77.20&[MIMAT0000445&hsa-miR-126-3p&offsets\|0\|-2;m-40&9\|+\|136670653\|136670674&offsets\|0\|-2]\|TCGTACCGTGAGTAATAATG | >MI0000471\|hsa-mir-126&WithFlank&9\|+\|136670596\|136670692 | 58 | 77 | 20 | | isomiR | NO | 0.00 | 0.00 | 17.89 | 0.00 |
| >MI0000471\|hsa-mir-126&WithFlank&9\|+\|136670596\|136670692@58.77.20(+1A)&[MIMAT0000445&hsa-miR-126-3p&offsets\|0\|-2(+1A);m-40&9\|+\|136670653\|136670674&offsets\|0\|-2(+1A)]\|TCGTACCGTGAGTAATAATGA | >MI0000471\|hsa-mir-126&WithFlank&9\|+\|136670596\|136670692 | 58 | 77 | 20 | (+1A) | isomiR | NO | 0.00 | 0.00 | 35.70 | 0.00 |
| >MI0000471\|hsa-mir-126&WithFlank&9\|+\|136670596\|136670692@58.77.20(+1U)&[MIMAT0000445&hsa-miR-126-3p&offsets\|0\|-2(+1U);m-40&9\|+\|136670653\|136670674&offsets\|0\|-2(+1U)]\|TCGTACCGTGAGTAATAATGT | >MI0000471\|hsa-mir-126&WithFlank&9\|+\|136670596\|136670692 | 58 | 77 | 20 | (+1U) | isomiR | NO | 26.25 | 0.00 | 177.88 | 0.00 |
| >MI0000471\|hsa-mir-126&WithFlank&9\|+\|136670596\|136670692@58.78.21&[MIMAT0000445&hsa-miR-126-3p&offsets\|0\|-1;m-40&9\|+\|136670653\|136670674&offsets\|0\|-1]\|TCGTACCGTGAGTAATAATGC | >MI0000471\|hsa-mir-126&WithFlank&9\|+\|136670596\|136670692 | 58 | 78 | 21 | | isomiR | NO | 170.80 | 62.23 | 430.54 | 21.27 |

### MDA-MB-231

<table>
<tr><td>>MI0000471|hsa-mir-126&WithFlank&9|+|136670596|136670692@58.78.21(+1A)&[MIMAT0000445&hsa-miR-126-3p&offsets|0|-1(+1A);m-40&9|+|136670653|136670674&offsets|0|-1(+1A)]|TCGTACCGTGAGTAATAATGCA</td><td>>MI0000471|hsa-mir-126&WithFlank&9|+|136670596|136670692</td><td>58</td><td>78</td><td>21</td><td>(+1A)</td><td>isomiR</td><td>NO</td><td>10.40</td><td>0.00</td><td>180.55</td><td>0.00</td></tr>
<tr><td>>MI0000471|hsa-mir-126&WithFlank&9|+|136670596|136670692@58.79.22&[MIMAT0000445&hsa-miR-126-3p&offsets|0|0;m-40&9|+|136670653|136670674&offsets|0|0]|TCGTACCGTGAGTAATAATGCG</td><td>>MI0000471|hsa-mir-126&WithFlank&9|+|136670596|136670692</td><td>58</td><td>79</td><td>22</td><td></td><td>isomiR</td><td>NO</td><td>181.69</td><td>33.05</td><td>568.26</td><td>58.17</td></tr>
<tr><td>>MI0000471|hsa-mir-126&WithFlank&9|+|136670596|136670692@59.77.19(+1U)&[MIMAT0000445&hsa-miR-126-3p&offsets|+1|-2(+1U);m-40&9|+|136670653|136670674&offsets|+1|-2(+1U)]|CGTACCGTGAGTAATAATGT</td><td>>MI0000471|hsa-mir-126&WithFlank&9|+|136670596|136670692</td><td>59</td><td>77</td><td>19</td><td>(+1U)</td><td>isomiR</td><td>NO</td><td>0.00</td><td>0.00</td><td>18.01</td><td>0.00</td></tr>
<tr><td>>MI0000471|hsa-mir-126&WithFlank&9|+|136670596|136670692@59.78.20&[MIMAT0000445&hsa-miR-126-3p&offsets|+1|-1;m-40&9|+|136670653|136670674&offsets|+1|-1]|CGTACCGTGAGTAATAATGC</td><td>>MI0000471|hsa-mir-126&WithFlank&9|+|136670596|136670692</td><td>59</td><td>78</td><td>20</td><td></td><td>isomiR</td><td>NO</td><td>10.61</td><td>0.00</td><td>123.16</td><td>0.00</td></tr>
<tr><td>>MI0000471|hsa-mir-126&WithFlank&9|+|136670596|136670692@59.79.21&[MIMAT0000445&hsa-miR-126-3p&offsets|+1|0;m-40&9|+|136670653|136670674&offsets|+1|0]|CGTACCGTGAGTAATAATGCG</td><td>>MI0000471|hsa-mir-126&WithFlank&9|+|136670596|136670692</td><td>59</td><td>79</td><td>21</td><td></td><td>isomiR</td><td>NO</td><td>0.00</td><td>0.00</td><td>69.68</td><td>0.00</td></tr>
<tr><td>>MI0000476|hsa-mir-138-1&WithFlank&3|+|44114206|44114316@29.52.24&[MIMAT0000430_1&hsa-miR-138-5p&offsets|0|+1;m-497&3|+|44114234|44114256&offsets|0|+1]|AGCTGGTGTTGTGAATCAGGCCGT</td><td>>MI0000476|hsa-mir-138-1&WithFlank&3|+|44114206|44114316</td><td>29</td><td>52</td><td>24</td><td></td><td>isomiR</td><td>NO</td><td>0.00</td><td>0.00</td><td>166.73</td><td>0.00</td></tr>
<tr><td>>MI0000477|hsa-mir-146a&WithFlank&5|+|160485346|160485456@26.47.22&[MIMAT0000449&hsa-miR-146a-5p&offsets|-1|-1;m-129&5|+|160485372|160485393&offsets|-1|-1]|TTGAGAACTGAATTCCATGGGT</td><td>>MI0000477|hsa-mir-146a&WithFlank&5|+|160485346|160485456</td><td>26</td><td>47</td><td>22</td><td></td><td>isomiR</td><td>NO</td><td>0.00</td><td>0.00</td><td>17.85</td><td>0.00</td></tr>
<tr><td>>MI0000477|hsa-mir-146a&WithFlank&5|+|160485346|160485456@27.46.20(+1C)&[MIMAT0000449&hsa-miR-146a-5p&offsets|0|-2(+1C);m-129&5|+|160485372|160485393&offsets|0|-2(+1C)]|TGAGAACTGAATTCCATGGGC</td><td>>MI0000477|hsa-mir-146a&WithFlank&5|+|160485346|160485456</td><td>27</td><td>46</td><td>20</td><td>(+1C)</td><td>isomiR</td><td>NO</td><td>0.00</td><td>0.00</td><td>167.85</td><td>0.00</td></tr>
<tr><td>>MI0000477|hsa-mir-146a&WithFlank&5|+|160485346|160485456@27.47.21&[MIMAT0000449&hsa-miR-146a-5p&offsets|0|-1;m-129&5|+|160485372|160485393&offsets|0|-1]|TGAGAACTGAATTCCATGGGT</td><td>>MI0000477|hsa-mir-146a&WithFlank&5|+|160485346|160485456</td><td>27</td><td>47</td><td>21</td><td></td><td>isomiR</td><td>NO</td><td>121.86</td><td>65.38</td><td>808.43</td><td>14.82</td></tr>
<tr><td>>MI0000477|hsa-mir-146a&WithFlank&5|+|160485346|160485456@27.47.21(+1A)&[MIMAT0000449&hsa-miR-146a-5p&offsets|0|-1(+1A);m-129&5|+|160485372|160485393&offsets|0|-1(+1A)]|TGAGAACTGAATTCCATGGGTA</td><td>>MI0000477|hsa-mir-146a&WithFlank&5|+|160485346|160485456</td><td>27</td><td>47</td><td>21</td><td>(+1A)</td><td>isomiR</td><td>NO</td><td>24.15</td><td>0.00</td><td>462.12</td><td>0.00</td></tr>
<tr><td>>MI0000477|hsa-mir-146a&WithFlank&5|+|160485346|160485456@27.47.21(+1C)&[MIMAT0000449&hsa-miR-146a-5p&offsets|0|-1(+1C);m-129&5|+|160485372|160485393&offsets|0|-1(+1C)]|TGAGAACTGAATTCCATGGGTC</td><td>>MI0000477|hsa-mir-146a&WithFlank&5|+|160485346|160485456</td><td>27</td><td>47</td><td>21</td><td>(+1C)</td><td>isomiR</td><td>NO</td><td>124.04</td><td>32.90</td><td>1278.86</td><td>16.57</td></tr>
<tr><td>>MI0000477|hsa-mir-146a&WithFlank&5|+|160485346|160485456@27.47.21(+1G)&[MIMAT0000449&hsa-miR-146a-5p&offsets|0|-1(+1G);m-129&5|+|160485372|160485393&offsets|0|-1(+1G)]|TGAGAACTGAATTCCATGGGTG</td><td>>MI0000477|hsa-mir-146a&WithFlank&5|+|160485346|160485456</td><td>27</td><td>47</td><td>21</td><td>(+1G)</td><td>isomiR</td><td>NO</td><td>0.00</td><td>0.00</td><td>138.93</td><td>0.00</td></tr>
<tr><td>>MI0000477|hsa-mir-146a&WithFlank&5|+|160485346|160485456@27.48.22&[MIMAT0000449&hsa-miR-146a-5p&offsets|0|0;m-129&5|+|160485372|160485393&offsets|0|0]|TGAGAACTGAATTCCATGGGTT</td><td>>MI0000477|hsa-mir-146a&WithFlank&5|+|160485346|160485456</td><td>27</td><td>48</td><td>22</td><td></td><td>isomiR</td><td>YES</td><td>729.12</td><td>305.44</td><td>5854.48</td><td>148.07</td></tr>
<tr><td>>MI0000477|hsa-mir-146a&WithFlank&5|+|160485346|160485456@27.48.22(+1A)&[MIMAT0000449&hsa-miR-146a-5p&offsets|0|0(+1A);m-129&5|+|160485372|160485393&offsets|0|0(+1A)]|TGAGAACTGAATTCCATGGGTTA</td><td>>MI0000477|hsa-mir-146a&WithFlank&5|+|160485346|160485456</td><td>27</td><td>48</td><td>22</td><td>(+1A)</td><td>isomiR</td><td>NO</td><td>10.65</td><td>0.00</td><td>231.37</td><td>0.00</td></tr>
<tr><td>>MI0000477|hsa-mir-146a&WithFlank&5|+|160485346|160485456@27.48.22(+1U)&[MIMAT0000449&hsa-miR-146a-5p&offsets|0|0(+1U);m-129&5|+|160485372|160485393&offsets|0|0(+1U)]|TGAGAACTGAATTCCATGGGTTT</td><td>>MI0000477|hsa-mir-146a&WithFlank&5|+|160485346|160485456</td><td>27</td><td>48</td><td>22</td><td>(+1U)</td><td>isomiR</td><td>NO</td><td>66.22</td><td>0.00</td><td>751.64</td><td>0.00</td></tr>
<tr><td>>MI0000477|hsa-mir-146a&WithFlank&5|+|160485346|160485456@27.49.23&[MIMAT0000449&hsa-miR-146a-5p&offsets|0|+1;m-129&5|+|160485372|160485393&offsets|0|+1]|TGAGAACTGAATTCCATGGGTTG</td><td>>MI0000477|hsa-mir-146a&WithFlank&5|+|160485346|160485456</td><td>27</td><td>49</td><td>23</td><td></td><td>isomiR</td><td>NO</td><td>94.38</td><td>24.48</td><td>521.15</td><td>14.03</td></tr>
<tr><td>>MI0000477|hsa-mir-146a&WithFlank&5|+|160485346|160485456@27.50.24&[MIMAT0000449&hsa-miR-146a-5p&offsets|0|+2;m-129&5|+|160485372|160485393&offsets|0|+2]|TGAGAACTGAATTCCATGGGTTGT</td><td>>MI0000477|hsa-mir-146a&WithFlank&5|+|160485346|160485456</td><td>27</td><td>50</td><td>24</td><td></td><td>isomiR</td><td>NO</td><td>0.00</td><td>0.00</td><td>173.30</td><td>0.00</td></tr>
<tr><td>>MI0000482|hsa-mir-185&WithFlank&22|+|20033133|20033226@21.42.22&[MIMAT0000455&hsa-miR-185-5p&offsets|0|0;m-145&22|+|20033153|20033174&offsets|0|0]|TGGAGAGAAAGGCAGTTCCTGA</td><td>>MI0000482|hsa-mir-185&WithFlank&22|+|20033133|20033226</td><td>21</td><td>42</td><td>22</td><td></td><td>isomiR</td><td>NO</td><td>50.77</td><td>0.00</td><td>126.48</td><td>0.00</td></tr>
<tr><td>>MI0000483|hsa-mir-186&WithFlank&1|-|71067625|71067722@21.40.20&[MIMAT0000456&hsa-miR-186-5p&offsets|0|-2;m-84&1|-|71067681|71067702&offsets|0|-2]|CAAAGAATTCTCCTTTTGGG</td><td>>MI0000483|hsa-mir-186&WithFlank&1|-|71067625|71067722</td><td>21</td><td>40</td><td>20</td><td></td><td>isomiR</td><td>NO</td><td>0.00</td><td>0.00</td><td>32.09</td><td>0.00</td></tr>
<tr><td>>MI0000483|hsa-mir-186&WithFlank&1|-|71067625|71067722@21.41.21&[MIMAT0000456&hsa-miR-186-5p&offsets|0|-1;m-84&1|-|71067681|71067702&offsets|0|-1]|CAAAGAATTCTCCTTTTGGGC</td><td>>MI0000483|hsa-mir-186&WithFlank&1|-|71067625|71067722</td><td>21</td><td>41</td><td>21</td><td></td><td>isomiR</td><td>NO</td><td>0.00</td><td>0.00</td><td>17.84</td><td>0.00</td></tr>
<tr><td>>MI0000483|hsa-mir-186&WithFlank&1|-|71067625|71067722@21.41.21(+1A)&[MIMAT0000456&hsa-miR-186-5p&offsets|0|-1(+1A);m-84&1|-|71067681|71067702&offsets|0|-1(+1A)]|CAAAGAATTCTCCTTTTGGGCA</td><td>>MI0000483|hsa-mir-186&WithFlank&1|-|71067625|71067722</td><td>21</td><td>41</td><td>21</td><td>(+1A)</td><td>isomiR</td><td>NO</td><td>0.00</td><td>0.00</td><td>13.90</td><td>0.00</td></tr>
<tr><td>>MI0000483|hsa-mir-186&WithFlank&1|-|71067625|71067722@21.41.21(+1C)&[MIMAT0000456&hsa-miR-186-5p&offsets|0|-1(+1C);m-84&1|-|71067681|71067702&offsets|0|-1(+1C)]|CAAAGAATTCTCCTTTTGGGCC</td><td>>MI0000483|hsa-mir-186&WithFlank&1|-|71067625|71067722</td><td>21</td><td>41</td><td>21</td><td>(+1C)</td><td>isomiR</td><td>NO</td><td>0.00</td><td>0.00</td><td>15.77</td><td>0.00</td></tr>
</table>

### MDA-MB-231

<table>
<tr><th></th><th></th><th></th><th></th><th></th><th></th><th></th><th></th><th></th><th></th><th></th><th></th></tr>
<tr><td>>MI0000483|hsa-mir-186&WithFlank&1|-|71067625|71067722@21.42.22&[MIMAT0000456&hsa-miR-186-5p&offsets|0|0;m-84&1|-|71067681|71067702&offsets|0|0]|CAAAGAATTCTCCTTTTGGGCT</td><td>>MI0000483|hsa-mir-186&WithFlank&1|-|71067625|71067722</td><td>21</td><td>42</td><td>22</td><td></td><td>isomiR</td><td>NO</td><td>69.73</td><td>0.00</td><td>180.18</td><td>16.43</td></tr>
<tr><td>>MI0000483|hsa-mir-186&WithFlank&1|-|71067625|71067722@21.43.23&[MIMAT0000456&hsa-miR-186-5p&offsets|0|+1;m-84&1|-|71067681|71067702&offsets|0|+1]|CAAAGAATTCTCCTTTTGGGCTT</td><td>>MI0000483|hsa-mir-186&WithFlank&1|-|71067625|71067722</td><td>21</td><td>43</td><td>23</td><td></td><td>isomiR</td><td>NO</td><td>0.00</td><td>0.00</td><td>212.64</td><td>0.00</td></tr>
<tr><td>>MI0000483|hsa-mir-186&WithFlank&1|-|71067625|71067722@21.44.24&[MIMAT0000456&hsa-miR-186-5p&offsets|0|+2;m-84&1|-|71067681|71067702&offsets|0|+2]|CAAAGAATTCTCCTTTTGGGCTTT</td><td>>MI0000483|hsa-mir-186&WithFlank&1|-|71067625|71067722</td><td>21</td><td>44</td><td>24</td><td></td><td>isomiR</td><td>NO</td><td>0.00</td><td>0.00</td><td>247.07</td><td>0.00</td></tr>
<tr><td>>MI0000488|hsa-mir-194-1&WithFlank&1|-|220118151|220118247@21.42.22;MI0000732|hsa-mir-194-2&WithFlank&11|-|64891349|64891445@21.42.22&[MIMAT0000460&hsa-miR-194-5p&offsets|0|0;m-131&1|-|220118206|220118227&offsets|0|0];[MIMAT0000460_1&hsa-miR-194-5p&offsets|0|0;m-126&11|-|64891404|64891425&offsets|0|0]|TGTAACAGCAACTCCATGTGGA</td><td>>MI0000488|hsa-mir-194-1&WithFlank&1|-|220118151|220118247</td><td>21</td><td>42</td><td>22</td><td></td><td>isomiR</td><td>NO</td><td>0.00</td><td>0.00</td><td>63.33</td><td>0.00</td></tr>
<tr><td>>MI0000542|hsa-mir-320a&WithFlank&8|-|22244960|22245043@48.68.21&[MIMAT0000510&hsa-miR-320a-3p&offsets|0|-1;m-60&8|-|22244975|22244996&offsets|0|-1]|AAAAGCTGGGTTGAGAGGGCG</td><td>>MI0000542|hsa-mir-320a&WithFlank&8|-|22244960|22245043</td><td>48</td><td>68</td><td>21</td><td></td><td>isomiR</td><td>NO</td><td>54.36</td><td>72.93</td><td>176.95</td><td>0.00</td></tr>
<tr><td>>MI0000542|hsa-mir-320a&WithFlank&8|-|22244960|22245043@48.68.21(+1C)&[MIMAT0000510&hsa-miR-320a-3p&offsets|0|-1(+1C);m-60&8|-|22244975|22244996&offsets|0|-1(+1C)]|AAAAGCTGGGTTGAGAGGGCGC</td><td>>MI0000542|hsa-mir-320a&WithFlank&8|-|22244960|22245043</td><td>48</td><td>68</td><td>21</td><td>(+1C)</td><td>isomiR</td><td>NO</td><td>0.00</td><td>0.00</td><td>91.65</td><td>0.00</td></tr>
<tr><td>>MI0000542|hsa-mir-320a&WithFlank&8|-|22244960|22245043@48.68.21(+1G)&[MIMAT0000510&hsa-miR-320a-3p&offsets|0|-1(+1G);m-60&8|-|22244975|22244996&offsets|0|-1(+1G)]|AAAAGCTGGGTTGAGAGGGCGG</td><td>>MI0000542|hsa-mir-320a&WithFlank&8|-|22244960|22245043</td><td>48</td><td>68</td><td>21</td><td>(+1G)</td><td>isomiR</td><td>NO</td><td>0.00</td><td>0.00</td><td>11.44</td><td>0.00</td></tr>
<tr><td>>MI0000542|hsa-mir-320a&WithFlank&8|-|22244960|22245043@48.68.21(+1U)&[MIMAT0000510&hsa-miR-320a-3p&offsets|0|-1(+1U);m-60&8|-|22244975|22244996&offsets|0|-1(+1U)]|AAAAGCTGGGTTGAGAGGGCGT</td><td>>MI0000542|hsa-mir-320a&WithFlank&8|-|22244960|22245043</td><td>48</td><td>68</td><td>21</td><td>(+1U)</td><td>isomiR</td><td>NO</td><td>73.42</td><td>111.51</td><td>410.33</td><td>0.00</td></tr>
<tr><td>>MI0000542|hsa-mir-320a&WithFlank&8|-|22244960|22245043@48.68.21(+2U)&[MIMAT0000510&hsa-miR-320a-3p&offsets|0|-1(+2U);m-60&8|-|22244975|22244996&offsets|0|-1(+2U)]|AAAAGCTGGGTTGAGAGGGCGTT</td><td>>MI0000542|hsa-mir-320a&WithFlank&8|-|22244960|22245043</td><td>48</td><td>68</td><td>21</td><td>(+2U)</td><td>isomiR</td><td>NO</td><td>0.00</td><td>30.11</td><td>96.00</td><td>0.00</td></tr>
<tr><td>>MI0000542|hsa-mir-320a&WithFlank&8|-|22244960|22245043@48.68.21(+3U)&[MIMAT0000510&hsa-miR-320a-3p&offsets|0|-1(+3U);m-60&8|-|22244975|22244996&offsets|0|-1(+3U)]|AAAAGCTGGGTTGAGAGGGCGTTT</td><td>>MI0000542|hsa-mir-320a&WithFlank&8|-|22244960|22245043</td><td>48</td><td>68</td><td>21</td><td>(+3U)</td><td>isomiR</td><td>NO</td><td>0.00</td><td>0.00</td><td>15.51</td><td>0.00</td></tr>
<tr><td>>MI0000542|hsa-mir-320a&WithFlank&8|-|22244960|22245043@48.69.22&[MIMAT0000510&hsa-miR-320a-3p&offsets|0|0;m-60&8|-|22244975|22244996&offsets|0|0]|AAAAGCTGGGTTGAGAGGGCGA</td><td>>MI0000542|hsa-mir-320a&WithFlank&8|-|22244960|22245043</td><td>48</td><td>69</td><td>22</td><td></td><td>isomiR</td><td>NO</td><td>145.43</td><td>135.42</td><td>673.17</td><td>0.00</td></tr>
<tr><td>>MI0000542|hsa-mir-320a&WithFlank&8|-|22244960|22245043@48.69.22(+1C)&[MIMAT0000510&hsa-miR-320a-3p&offsets|0|0(+1C);m-60&8|-|22244975|22244996&offsets|0|0(+1C)]|AAAAGCTGGGTTGAGAGGGCGAC</td><td>>MI0000542|hsa-mir-320a&WithFlank&8|-|22244960|22245043</td><td>48</td><td>69</td><td>22</td><td>(+1C)</td><td>isomiR</td><td>NO</td><td>0.00</td><td>0.00</td><td>17.92</td><td>0.00</td></tr>
<tr><td>>MI0000542|hsa-mir-320a&WithFlank&8|-|22244960|22245043@48.69.22(+1U)&[MIMAT0000510&hsa-miR-320a-3p&offsets|0|0(+1U);m-60&8|-|22244975|22244996&offsets|0|0(+1U)]|AAAAGCTGGGTTGAGAGGGCGAT</td><td>>MI0000542|hsa-mir-320a&WithFlank&8|-|22244960|22245043</td><td>48</td><td>69</td><td>22</td><td>(+1U)</td><td>isomiR</td><td>NO</td><td>132.45</td><td>122.66</td><td>205.19</td><td>0.00</td></tr>
<tr><td>>MI0000542|hsa-mir-320a&WithFlank&8|-|22244960|22245043@48.69.22(+2U)&[MIMAT0000510&hsa-miR-320a-3p&offsets|0|0(+2U);m-60&8|-|22244975|22244996&offsets|0|0(+2U)]|AAAAGCTGGGTTGAGAGGGCGATT</td><td>>MI0000542|hsa-mir-320a&WithFlank&8|-|22244960|22245043</td><td>48</td><td>69</td><td>22</td><td>(+2U)</td><td>isomiR</td><td>NO</td><td>55.86</td><td>84.43</td><td>90.46</td><td>0.00</td></tr>
<tr><td>>MI0000542|hsa-mir-320a&WithFlank&8|-|22244960|22245043@48.70.23&[MIMAT0000510&hsa-miR-320a-3p&offsets|0|+1;m-60&8|-|22244975|22244996&offsets|0|+1]|AAAAGCTGGGTTGAGAGGGCGAA</td><td>>MI0000542|hsa-mir-320a&WithFlank&8|-|22244960|22245043</td><td>48</td><td>70</td><td>23</td><td></td><td>isomiR</td><td>NO</td><td>122.50</td><td>107.47</td><td>198.31</td><td>0.00</td></tr>
<tr><td>>MI0000542|hsa-mir-320a&WithFlank&8|-|22244960|22245043@48.70.23(+1C)&[MIMAT0000510&hsa-miR-320a-3p&offsets|0|+1(+1C);m-60&8|-|22244975|22244996&offsets|0|+1(+1C)]|AAAAGCTGGGTTGAGAGGGCGAAC</td><td>>MI0000542|hsa-mir-320a&WithFlank&8|-|22244960|22245043</td><td>48</td><td>70</td><td>23</td><td>(+1C)</td><td>isomiR</td><td>NO</td><td>0.00</td><td>0.00</td><td>32.17</td><td>0.00</td></tr>
<tr><td>>MI0000542|hsa-mir-320a&WithFlank&8|-|22244960|22245043@48.70.23(+1U)&[MIMAT0000510&hsa-miR-320a-3p&offsets|0|+1(+1U);m-60&8|-|22244975|22244996&offsets|0|+1(+1U)]|AAAAGCTGGGTTGAGAGGGCGAAT</td><td>>MI0000542|hsa-mir-320a&WithFlank&8|-|22244960|22245043</td><td>48</td><td>70</td><td>23</td><td>(+1U)</td><td>isomiR</td><td>NO</td><td>83.54</td><td>167.13</td><td>225.37</td><td>0.00</td></tr>
<tr><td>>MI0000542|hsa-mir-320a&WithFlank&8|-|22244960|22245043@48.70.23(+2U)&[MIMAT0000510&hsa-miR-320a-3p&offsets|0|+1(+2U);m-60&8|-|22244975|22244996&offsets|0|+1(+2U)]|AAAAGCTGGGTTGAGAGGGCGAATT</td><td>>MI0000542|hsa-mir-320a&WithFlank&8|-|22244960|22245043</td><td>48</td><td>70</td><td>23</td><td>(+2U)</td><td>isomiR</td><td>NO</td><td>0.00</td><td>0.00</td><td>11.45</td><td>0.00</td></tr>
<tr><td>>MI0000542|hsa-mir-320a&WithFlank&8|-|22244960|22245043@48.71.24&[MIMAT0000510&hsa-miR-320a-3p&offsets|0|+2;m-60&8|-|22244975|22244996&offsets|0|+2]|AAAAGCTGGGTTGAGAGGGCGAAA</td><td>>MI0000542|hsa-mir-320a&WithFlank&8|-|22244960|22245043</td><td>48</td><td>71</td><td>24</td><td></td><td>isomiR</td><td>NO</td><td>0.00</td><td>26.75</td><td>56.99</td><td>0.00</td></tr>
<tr><td>>MI0000542|hsa-mir-320a&WithFlank&8|-|22244960|22245043@48.71.24(+1U)&[MIMAT0000510&hsa-miR-320a-3p&offsets|0|+2(+1U);m-60&8|-|22244975|22244996&offsets|0|+2(+1U)]|AAAAGCTGGGTTGAGAGGGCGAAAT</td><td>>MI0000542|hsa-mir-320a&WithFlank&8|-|22244960|22245043</td><td>48</td><td>71</td><td>24</td><td>(+1U)</td><td>isomiR</td><td>NO</td><td>0.00</td><td>0.00</td><td>14.45</td><td>0.00</td></tr>
<tr><td>>MI0000542|hsa-mir-320a&WithFlank&8|-|22244960|22245043@49.69.21&[MIMAT0000510&hsa-miR-320a-3p&offsets|+1|0;m-60&8|-|22244975|22244996&offsets|+1|0]|AAAGCTGGGTTGAGAGGGCGA</td><td>>MI0000542|hsa-mir-320a&WithFlank&8|-|22244960|22245043</td><td>49</td><td>69</td><td>21</td><td></td><td>isomiR</td><td>NO</td><td>0.00</td><td>0.00</td><td>54.66</td><td>0.00</td></tr>
<tr><td>>MI0000542|hsa-mir-320a&WithFlank&8|-|22244960|22245043@49.70.22(+1U)&[MIMAT0000510&hsa-miR-320a-3p&offsets|+1|+1(+1U);m-60&8|-|22244975|22244996&offsets|+1|+1(+1U)]|AAAGCTGGGTTGAGAGGGCGAAT</td><td>>MI0000542|hsa-mir-320a&WithFlank&8|-|22244960|22245043</td><td>49</td><td>70</td><td>22</td><td>(+1U)</td><td>isomiR</td><td>NO</td><td>0.00</td><td>0.00</td><td>12.09</td><td>0.00</td></tr>
<tr><td>>MI0000683|hsa-mir-181b-2&WithFlank&9|+|124693704|124693804@22.46.25&[MIMAT0000257_1&hsa-miR-181b-5p&offsets|0|+2;m-72&9|+|124693725|124693747&offsets|0|+2]|AACATTCATTGCTGTCGGTGGGTTT</td><td>>MI0000683|hsa-mir-181b-2&WithFlank&9|+|124693704|124693804</td><td>22</td><td>46</td><td>25</td><td></td><td>isomiR</td><td>NO</td><td>26.24</td><td>0.00</td><td>141.28</td><td>18.04</td></tr>
</table>

### MDA-MB-231

<table>
<tr><td>>MI0000683|hsa-mir-181b-2&WithFlank&9|+|124693704|124693804@22.46.25(+1U)&[MIMAT0000257_1&hsa-miR-181b-5p&offsets|0|+2(+1U);m-72&9|+|124693725|124693747&offsets|0|+2(+1U)]|AACATTCATTGCTGTCGGTGGGTTTT</td><td>>MI0000683|hsa-mir-181b-2&WithFlank&9|+|124693704|124693804</td><td>22</td><td>46</td><td>25</td><td>(+1U)</td><td>isomiR</td><td>NO</td><td>0.00</td><td>0.00</td><td>74.71</td><td>0.00</td></tr>
<tr><td>>MI0000734|hsa-mir-106b&WithFlank&7|-|100093987|100094080@18.37.20&[MIMAT0000680&hsa-miR-106b-5p&offsets|0|-1;m-103&7|-|100094043|100094063&offsets|0|-1]|TAAAGTGCTGACAGTGCAGA</td><td>>MI0000734|hsa-mir-106b&WithFlank&7|-|100093987|100094080</td><td>18</td><td>37</td><td>20</td><td></td><td>isomiR</td><td>NO</td><td>0.00</td><td>0.00</td><td>14.71</td><td>0.00</td></tr>
<tr><td>>MI0000734|hsa-mir-106b&WithFlank&7|-|100093987|100094080@18.38.21&[MIMAT0000680&hsa-miR-106b-5p&offsets|0|0;m-103&7|-|100094043|100094063&offsets|0|0]|TAAAGTGCTGACAGTGCAGAT</td><td>>MI0000734|hsa-mir-106b&WithFlank&7|-|100093987|100094080</td><td>18</td><td>38</td><td>21</td><td></td><td>isomiR</td><td>NO</td><td>73.30</td><td>0.00</td><td>114.26</td><td>11.14</td></tr>
<tr><td>>MI0000734|hsa-mir-106b&WithFlank&7|-|100093987|100094080@18.39.22&[MIMAT0000680&hsa-miR-106b-5p&offsets|0|+1;m-103&7|-|100094043|100094063&offsets|0|+1]|TAAAGTGCTGACAGTGCAGATA</td><td>>MI0000734|hsa-mir-106b&WithFlank&7|-|100093987|100094080</td><td>18</td><td>39</td><td>22</td><td></td><td>isomiR</td><td>NO</td><td>89.22</td><td>0.00</td><td>226.30</td><td>14.28</td></tr>
<tr><td>>MI0000734|hsa-mir-106b&WithFlank&7|-|100093987|100094080@56.77.22&[MIMAT0004672&hsa-miR-106b-3p&offsets|-2|-2;m-87&7|-|100094003|100094023&offsets|-2|-1]|TACCGCACTGTGGGTACTTGCT</td><td>>MI0000734|hsa-mir-106b&WithFlank&7|-|100093987|100094080</td><td>56</td><td>77</td><td>22</td><td></td><td>isomiR</td><td>NO</td><td>51.30</td><td>31.81</td><td>67.37</td><td>10.78</td></tr>
<tr><td>>MI0000734|hsa-mir-106b&WithFlank&7|-|100093987|100094080@58.77.20&[MIMAT0004672&hsa-miR-106b-3p&offsets|0|-2;m-87&7|-|100094003|100094023&offsets|0|-1]|CCGCACTGTGGGTACTTGCT</td><td>>MI0000734|hsa-mir-106b&WithFlank&7|-|100093987|100094080</td><td>58</td><td>77</td><td>20</td><td></td><td>isomiR</td><td>NO</td><td>58.08</td><td>71.14</td><td>67.78</td><td>0.00</td></tr>
<tr><td>>MI0000734|hsa-mir-106b&WithFlank&7|-|100093987|100094080@58.78.21&[MIMAT0004672&hsa-miR-106b-3p&offsets|0|-1;m-87&7|-|100094003|100094023&offsets|0|0]|CCGCACTGTGGGTACTTGCTG</td><td>>MI0000734|hsa-mir-106b&WithFlank&7|-|100093987|100094080</td><td>58</td><td>78</td><td>21</td><td></td><td>isomiR</td><td>NO</td><td>90.35</td><td>156.59</td><td>130.24</td><td>10.92</td></tr>
<tr><td>>MI0000734|hsa-mir-106b&WithFlank&7|-|100093987|100094080@58.78.21(+1U)&[MIMAT0004672&hsa-miR-106b-3p&offsets|0|-1(+1U);m-87&7|-|100094003|100094023&offsets|0|0(+1U)]|CCGCACTGTGGGTACTTGCTGT</td><td>>MI0000734|hsa-mir-106b&WithFlank&7|-|100093987|100094080</td><td>58</td><td>78</td><td>21</td><td>(+1U)</td><td>isomiR</td><td>NO</td><td>82.35</td><td>101.51</td><td>93.01</td><td>10.70</td></tr>
<tr><td>>MI0000734|hsa-mir-106b&WithFlank&7|-|100093987|100094080@58.79.22&[MIMAT0004672&hsa-miR-106b-3p&offsets|0|0;m-87&7|-|100094003|100094023&offsets|0|+1]|CCGCACTGTGGGTACTTGCTGC</td><td>>MI0000734|hsa-mir-106b&WithFlank&7|-|100093987|100094080</td><td>58</td><td>79</td><td>22</td><td></td><td>isomiR</td><td>NO</td><td>263.85</td><td>328.98</td><td>187.03</td><td>57.01</td></tr>
<tr><td>>MI0000736|hsa-mir-30c-1&WithFlank&1|+|40757278|40757378@23.42.20;MI0000254|hsa-mir-30c-2&WithFlank&6|-|71376954|71377037@13.32.20&[MIMAT0000244&hsa-miR-30c-5p&offsets|0|-3;m-57&1|+|40757300|40757323&offsets|0|-4];[MIMAT0000244_1&hsa-miR-30c-5p&offsets|0|-3;m-56&6|-|71377002|71377025&offsets|0|-4]|TGTAAACATCCTACACTCTC</td><td>>MI0000736|hsa-mir-30c-1&WithFlank&1|+|40757278|40757378</td><td>23</td><td>42</td><td>20</td><td></td><td>isomiR</td><td>NO</td><td>116.07</td><td>0.00</td><td>87.01</td><td>63.27</td></tr>
<tr><td>>MI0000736|hsa-mir-30c-1&WithFlank&1|+|40757278|40757378@23.43.21;MI0000254|hsa-mir-30c-2&WithFlank&6|-|71376954|71377037@13.33.21&[MIMAT0000244&hsa-miR-30c-5p&offsets|0|-2;m-57&1|+|40757300|40757323&offsets|0|-3];[MIMAT0000244_1&hsa-miR-30c-5p&offsets|0|-2;m-56&6|-|71377002|71377025&offsets|0|-3]|TGTAAACATCCTACACTCTCA</td><td>>MI0000736|hsa-mir-30c-1&WithFlank&1|+|40757278|40757378</td><td>23</td><td>43</td><td>21</td><td></td><td>isomiR</td><td>NO</td><td>0.00</td><td>0.00</td><td>17.73</td><td>0.00</td></tr>
<tr><td>>MI0000736|hsa-mir-30c-1&WithFlank&1|+|40757278|40757378@23.44.22(+1A);MI0000254|hsa-mir-30c-2&WithFlank&6|-|71376954|71377037@13.34.22(+1A)&[MIMAT0000244&hsa-miR-30c-5p&offsets|0|-1(+1A);m-57&1|+|40757300|40757323&offsets|0|-2(+1A)];[MIMAT0000244_1&hsa-miR-30c-5p&offsets|0|-1(+1A);m-56&6|-|71377002|71377025&offsets|0|-2(+1A)]|TGTAAACATCCTACACTCTCAGA</td><td>>MI0000736|hsa-mir-30c-1&WithFlank&1|+|40757278|40757378</td><td>23</td><td>44</td><td>22</td><td>(+1A)</td><td>isomiR</td><td>NO</td><td>91.27</td><td>12.87</td><td>31.11</td><td>67.31</td></tr>
<tr><td>>MI0000736|hsa-mir-30c-1&WithFlank&1|+|40757278|40757378@23.44.22(+1U);MI0000254|hsa-mir-30c-2&WithFlank&6|-|71376954|71377037@13.34.22(+1U)&[MIMAT0000244&hsa-miR-30c-5p&offsets|0|-1(+1U);m-57&1|+|40757300|40757323&offsets|0|-2(+1U)];[MIMAT0000244_1&hsa-miR-30c-5p&offsets|0|-1(+1U);m-56&6|-|71377002|71377025&offsets|0|-2(+1U)]|TGTAAACATCCTACACTCTCAGT</td><td>>MI0000736|hsa-mir-30c-1&WithFlank&1|+|40757278|40757378</td><td>23</td><td>44</td><td>22</td><td>(+1U)</td><td>isomiR</td><td>NO</td><td>311.81</td><td>108.80</td><td>157.21</td><td>181.04</td></tr>
<tr><td>>MI0000736|hsa-mir-30c-1&WithFlank&1|+|40757278|40757378@23.44.22;MI0000254|hsa-mir-30c-2&WithFlank&6|-|71376954|71377037@13.34.22&[MIMAT0000244&hsa-miR-30c-5p&offsets|0|-1;m-57&1|+|40757300|40757323&offsets|0|-2];[MIMAT0000244_1&hsa-miR-30c-5p&offsets|0|-1;m-56&6|-|71377002|71377025&offsets|0|-2]|TGTAAACATCCTACACTCTCAG</td><td>>MI0000736|hsa-mir-30c-1&WithFlank&1|+|40757278|40757378</td><td>23</td><td>44</td><td>22</td><td></td><td>isomiR</td><td>NO</td><td>0.00</td><td>0.00</td><td>54.87</td><td>0.00</td></tr>
<tr><td>>MI0000736|hsa-mir-30c-1&WithFlank&1|+|40757278|40757378@23.45.23(+1A);MI0000254|hsa-mir-30c-2&WithFlank&6|-|71376954|71377037@13.35.23(+1A)&[MIMAT0000244&hsa-miR-30c-5p&offsets|0|0(+1A);m-57&1|+|40757300|40757323&offsets|0|-1(+1A)];[MIMAT0000244_1&hsa-miR-30c-5p&offsets|0|0(+1A);m-56&6|-|71377002|71377025&offsets|0|-1(+1A)]|TGTAAACATCCTACACTCTCAGCA</td><td>>MI0000736|hsa-mir-30c-1&WithFlank&1|+|40757278|40757378</td><td>23</td><td>45</td><td>23</td><td>(+1A)</td><td>isomiR</td><td>NO</td><td>131.85</td><td>36.70</td><td>101.61</td><td>96.31</td></tr>
<tr><td>>MI0000736|hsa-mir-30c-1&WithFlank&1|+|40757278|40757378@23.45.23(+1C);MI0000254|hsa-mir-30c-2&WithFlank&6|-|71376954|71377037@13.35.23(+1C)&[MIMAT0000244&hsa-miR-30c-5p&offsets|0|0(+1C);m-57&1|+|40757300|40757323&offsets|0|-1(+1C)];[MIMAT0000244_1&hsa-miR-30c-5p&offsets|0|0(+1C);m-56&6|-|71377002|71377025&offsets|0|-1(+1C)]|TGTAAACATCCTACACTCTCAGCC</td><td>>MI0000736|hsa-mir-30c-1&WithFlank&1|+|40757278|40757378</td><td>23</td><td>45</td><td>23</td><td>(+1C)</td><td>isomiR</td><td>NO</td><td>189.21</td><td>62.83</td><td>126.84</td><td>122.31</td></tr>
<tr><td>>MI0000736|hsa-mir-30c-1&WithFlank&1|+|40757278|40757378@23.45.23(+1G);MI0000254|hsa-mir-30c-2&WithFlank&6|-|71376954|71377037@13.35.23(+1G)&[MIMAT0000244&hsa-miR-30c-5p&offsets|0|0(+1G);m-57&1|+|40757300|40757323&offsets|0|-1(+1G)];[MIMAT0000244_1&hsa-miR-30c-5p&offsets|0|0(+1G);m-56&6|-|71377002|71377025&offsets|0|-1(+1G)]|TGTAAACATCCTACACTCTCAGCG</td><td>>MI0000736|hsa-mir-30c-1&WithFlank&1|+|40757278|40757378</td><td>23</td><td>45</td><td>23</td><td>(+1G)</td><td>isomiR</td><td>NO</td><td>60.75</td><td>0.00</td><td>0.00</td><td>12.72</td></tr>
<tr><td>>MI0000736|hsa-mir-30c-1&WithFlank&1|+|40757278|40757378@23.45.23;MI0000254|hsa-mir-30c-2&WithFlank&6|-|71376954|71377037@13.35.23&[MIMAT0000244&hsa-miR-30c-5p&offsets|0|0;m-57&1|+|40757300|40757323&offsets|0|-1];[MIMAT0000244_1&hsa-miR-30c-5p&offsets|0|0;m-56&6|-|71377002|71377025&offsets|0|-1]|TGTAAACATCCTACACTCTCAGC</td><td>>MI0000736|hsa-mir-30c-1&WithFlank&1|+|40757278|40757378</td><td>23</td><td>45</td><td>23</td><td></td><td>isomiR</td><td>YES</td><td>1188.72</td><td>433.37</td><td>361.29</td><td>451.08</td></tr>
</table>

### MDA-MB-231

<table>
<tr><td>>MI0000736|hsa-mir-30c-1&WithFlank&1|+|40757278|40757378@23.46.24(+1U);MI0000254|hsa-mir-30c-2&WithFlank&6|-|71376954|71377037@13.36.24(+1U)&[MIMAT0000244&hsa-miR-30c-5p&offsets|0|+1(+1U);m-57&1|+|40757300|40757323&offsets|0|0(+1U)];[MIMAT0000244_1&hsa-miR-30c-5p&offsets|0|+1(+1U);m-56&6|-|71377002|71377025&offsets|0|0(+1U)]|TGTAAACATCCTACACTCTCAGCTT</td><td>>MI0000736|hsa-mir-30c-1&WithFlank&1|+|40757278|40757378</td><td>23</td><td>46</td><td>24</td><td>(+1U)</td><td>isomiR</td><td>NO</td><td>81.12</td><td>0.00</td><td>35.32</td><td>20.97</td></tr>
<tr><td>>MI0000736|hsa-mir-30c-1&WithFlank&1|+|40757278|40757378@23.46.24;MI0000254|hsa-mir-30c-2&WithFlank&6|-|71376954|71377037@13.36.24&[MIMAT0000244&hsa-miR-30c-5p&offsets|0|+1;m-57&1|+|40757300|40757323&offsets|0|0];[MIMAT0000244_1&hsa-miR-30c-5p&offsets|0|+1;m-56&6|-|71377002|71377025&offsets|0|0]|TGTAAACATCCTACACTCTCAGCT</td><td>>MI0000736|hsa-mir-30c-1&WithFlank&1|+|40757278|40757378</td><td>23</td><td>46</td><td>24</td><td></td><td>isomiR</td><td>NO</td><td>951.19</td><td>390.70</td><td>555.40</td><td>414.93</td></tr>
<tr><td>>MI0000743|hsa-mir-34c&WithFlank&11|+|111513433|111513521@19.40.22(+1U)&[MIMAT0000686&hsa-miR-34c-5p&offsets|0|-1(+1U);m-336&11|+|111513451|111513473&offsets|0|-1(+1U)]|AGGCAGTGTAGTTAGCTGATTGT</td><td>>MI0000743|hsa-mir-34c&WithFlank&11|+|111513433|111513521</td><td>19</td><td>40</td><td>22</td><td>(+1U)</td><td>isomiR</td><td>NO</td><td>0.00</td><td>0.00</td><td>64.92</td><td>0.00</td></tr>
<tr><td>>MI0000743|hsa-mir-34c&WithFlank&11|+|111513433|111513521@19.41.23&[MIMAT0000686&hsa-miR-34c-5p&offsets|0|0;m-336&11|+|111513451|111513473&offsets|0|0]|AGGCAGTGTAGTTAGCTGATTGC</td><td>>MI0000743|hsa-mir-34c&WithFlank&11|+|111513433|111513521</td><td>19</td><td>41</td><td>23</td><td></td><td>isomiR</td><td>NO</td><td>0.00</td><td>0.00</td><td>155.72</td><td>0.00</td></tr>
<tr><td>>MI0000746|hsa-mir-99b&WithFlank&19|+|51692606|51692687@13.32.20&[MIMAT0000689&hsa-miR-99b-5p&offsets|0|-2;m-10&19|+|51692618|51692639&offsets|0|-2]|CACCCGTAGAACCGACCTTG</td><td>>MI0000746|hsa-mir-99b&WithFlank&19|+|51692606|51692687</td><td>13</td><td>32</td><td>20</td><td></td><td>isomiR</td><td>NO</td><td>0.00</td><td>36.23</td><td>17.13</td><td>0.00</td></tr>
<tr><td>>MI0000746|hsa-mir-99b&WithFlank&19|+|51692606|51692687@13.32.20(+1A)&[MIMAT0000689&hsa-miR-99b-5p&offsets|0|-2(+1A);m-10&19|+|51692618|51692639&offsets|0|-2(+1A)]|CACCCGTAGAACCGACCTTGA</td><td>>MI0000746|hsa-mir-99b&WithFlank&19|+|51692606|51692687</td><td>13</td><td>32</td><td>20</td><td>(+1A)</td><td>isomiR</td><td>NO</td><td>0.00</td><td>24.17</td><td>13.81</td><td>0.00</td></tr>
<tr><td>>MI0000746|hsa-mir-99b&WithFlank&19|+|51692606|51692687@13.32.20(+1U)&[MIMAT0000689&hsa-miR-99b-5p&offsets|0|-2(+1U);m-10&19|+|51692618|51692639&offsets|0|-2(+1U)]|CACCCGTAGAACCGACCTTGT</td><td>>MI0000746|hsa-mir-99b&WithFlank&19|+|51692606|51692687</td><td>13</td><td>32</td><td>20</td><td>(+1U)</td><td>isomiR</td><td>NO</td><td>83.21</td><td>100.53</td><td>100.51</td><td>16.79</td></tr>
<tr><td>>MI0000746|hsa-mir-99b&WithFlank&19|+|51692606|51692687@13.33.21&[MIMAT0000689&hsa-miR-99b-5p&offsets|0|-1;m-10&19|+|51692618|51692639&offsets|0|-1]|CACCCGTAGAACCGACCTTGC</td><td>>MI0000746|hsa-mir-99b&WithFlank&19|+|51692606|51692687</td><td>13</td><td>33</td><td>21</td><td></td><td>isomiR</td><td>NO</td><td>381.47</td><td>441.72</td><td>279.47</td><td>112.15</td></tr>
<tr><td>>MI0000746|hsa-mir-99b&WithFlank&19|+|51692606|51692687@13.33.21(+1A)&[MIMAT0000689&hsa-miR-99b-5p&offsets|0|-1(+1A);m-10&19|+|51692618|51692639&offsets|0|-1(+1A)]|CACCCGTAGAACCGACCTTGCA</td><td>>MI0000746|hsa-mir-99b&WithFlank&19|+|51692606|51692687</td><td>13</td><td>33</td><td>21</td><td>(+1A)</td><td>isomiR</td><td>NO</td><td>147.76</td><td>267.94</td><td>189.17</td><td>71.42</td></tr>
<tr><td>>MI0000746|hsa-mir-99b&WithFlank&19|+|51692606|51692687@13.33.21(+1U)&[MIMAT0000689&hsa-miR-99b-5p&offsets|0|-1(+1U);m-10&19|+|51692618|51692639&offsets|0|-1(+1U)]|CACCCGTAGAACCGACCTTGCT</td><td>>MI0000746|hsa-mir-99b&WithFlank&19|+|51692606|51692687</td><td>13</td><td>33</td><td>21</td><td>(+1U)</td><td>isomiR</td><td>NO</td><td>0.00</td><td>24.88</td><td>12.94</td><td>0.00</td></tr>
<tr><td>>MI0000746|hsa-mir-99b&WithFlank&19|+|51692606|51692687@13.34.22&[MIMAT0000689&hsa-miR-99b-5p&offsets|0|0;m-10&19|+|51692618|51692639&offsets|0|0]|CACCCGTAGAACCGACCTTGCG</td><td>>MI0000746|hsa-mir-99b&WithFlank&19|+|51692606|51692687</td><td>13</td><td>34</td><td>22</td><td></td><td>isomiR</td><td>YES</td><td>679.99</td><td>1246.51</td><td>618.03</td><td>204.15</td></tr>
<tr><td>>MI0000746|hsa-mir-99b&WithFlank&19|+|51692606|51692687@13.34.22(+1A)&[MIMAT0000689&hsa-miR-99b-5p&offsets|0|0(+1A);m-10&19|+|51692618|51692639&offsets|0|0(+1A)]|CACCCGTAGAACCGACCTTGCGA</td><td>>MI0000746|hsa-mir-99b&WithFlank&19|+|51692606|51692687</td><td>13</td><td>34</td><td>22</td><td>(+1A)</td><td>isomiR</td><td>NO</td><td>25.44</td><td>61.35</td><td>14.45</td><td>0.00</td></tr>
<tr><td>>MI0000746|hsa-mir-99b&WithFlank&19|+|51692606|51692687@13.34.22(+1U)&[MIMAT0000689&hsa-miR-99b-5p&offsets|0|0(+1U);m-10&19|+|51692618|51692639&offsets|0|0(+1U)]|CACCCGTAGAACCGACCTTGCGT</td><td>>MI0000746|hsa-mir-99b&WithFlank&19|+|51692606|51692687</td><td>13</td><td>34</td><td>22</td><td>(+1U)</td><td>isomiR</td><td>NO</td><td>102.24</td><td>151.06</td><td>93.62</td><td>19.66</td></tr>
<tr><td>>MI0000746|hsa-mir-99b&WithFlank&19|+|51692606|51692687@51.72.22&[MIMAT0004678&hsa-miR-99b-3p&offsets|0|0;m-151&19|+|51692656|51692678&offsets|0|-1]|CAAGCTCGTGTCTGTGGGTCCG</td><td>>MI0000746|hsa-mir-99b&WithFlank&19|+|51692606|51692687</td><td>51</td><td>72</td><td>22</td><td></td><td>isomiR</td><td>NO</td><td>25.92</td><td>28.91</td><td>10.59</td><td>0.00</td></tr>
<tr><td>>MI0000746|hsa-mir-99b&WithFlank&19|+|51692606|51692687@51.73.23&[MIMAT0004678&hsa-miR-99b-3p&offsets|0|+1;m-151&19|+|51692656|51692678&offsets|0|0]|CAAGCTCGTGTCTGTGGGTCCGT</td><td>>MI0000746|hsa-mir-99b&WithFlank&19|+|51692606|51692687</td><td>51</td><td>73</td><td>23</td><td></td><td>isomiR</td><td>NO</td><td>0.00</td><td>12.85</td><td>13.03</td><td>0.00</td></tr>
<tr><td>>MI0000748|hsa-mir-130b&WithFlank&22|+|21653298|21653391@19.40.22&[MIMAT0004680&hsa-miR-130b-5p&offsets|0|+1;m-234&22|+|21653316|21653337&offsets|0|0]|ACTCTTTCCCTGTTGCACTACT</td><td>>MI0000748|hsa-mir-130b&WithFlank&22|+|21653298|21653391</td><td>19</td><td>40</td><td>22</td><td></td><td>isomiR</td><td>NO</td><td>63.29</td><td>101.00</td><td>0.00</td><td>20.26</td></tr>
<tr><td>>am_MI0000748|hsa-mir-130b&WithFlank&22|+|21653298|21653391@19.39.21&[MIMAT0004680&hsa-miR-130b-5p&offsets|0|0;m-234&22|+|21653316|21653337&offsets|0|-1]|ACTCTTTCCCTGTTGCACTAC</td><td>>MI0000748|hsa-mir-130b&WithFlank&22|+|21653298|21653391.an</td><td>19</td><td>39</td><td>21</td><td></td><td>isomiR</td><td>NO</td><td>24.63</td><td>61.83</td><td>0.00</td><td>0.00</td></tr>
<tr><td>>MI0000749|hsa-mir-30e&WithFlank&1|+|40754349|40754452@23.45.23&[MIMAT0000692&hsa-miR-30e-5p&offsets|0|+1;m-24&1|+|40754371|40754394&offsets|0|-1]|TGTAAACATCCTTGACTGGAAGC</td><td>>MI0000749|hsa-mir-30e&WithFlank&1|+|40754349|40754452</td><td>23</td><td>45</td><td>23</td><td></td><td>isomiR</td><td>NO</td><td>0.00</td><td>0.00</td><td>87.02</td><td>0.00</td></tr>
<tr><td>>MI0000749|hsa-mir-30e&WithFlank&1|+|40754349|40754452@23.45.23(+1A)&[MIMAT0000692&hsa-miR-30e-5p&offsets|0|+1(+1A);m-24&1|+|40754371|40754394&offsets|0|-1(+1A)]|TGTAAACATCCTTGACTGGAAGCA</td><td>>MI0000749|hsa-mir-30e&WithFlank&1|+|40754349|40754452</td><td>23</td><td>45</td><td>23</td><td>(+1A)</td><td>isomiR</td><td>NO</td><td>0.00</td><td>0.00</td><td>259.55</td><td>0.00</td></tr>
<tr><td>>MI0000749|hsa-mir-30e&WithFlank&1|+|40754349|40754452@23.45.23(+1C)&[MIMAT0000692&hsa-miR-30e-5p&offsets|0|+1(+1C);m-24&1|+|40754371|40754394&offsets|0|-1(+1C)]|TGTAAACATCCTTGACTGGAAGCC</td><td>>MI0000749|hsa-mir-30e&WithFlank&1|+|40754349|40754452</td><td>23</td><td>45</td><td>23</td><td>(+1C)</td><td>isomiR</td><td>NO</td><td>0.00</td><td>0.00</td><td>367.64</td><td>0.00</td></tr>
</table>

### MDA-MB-231

<table>
<tr><th></th><th></th><th></th><th></th><th></th><th></th><th></th><th></th><th></th><th></th><th></th><th></th></tr>
<tr><td>>MI0000749|hsa-mir-30e&WithFlank&1|+|40754349|40754452@23.45.23(+1G)&[MIMAT0000692&hsa-miR-30e-5p&offsets|0|+1(+1G);m-24&1|+|40754371|40754394&offsets|0|-1(+1G)]|TGTAAACATCCTTGACTGGAAGCG</td><td>>MI0000749|hsa-mir-30e&WithFlank&1|+|40754349|40754452</td><td>23</td><td>45</td><td>23</td><td>(+1G)</td><td>isomiR</td><td>NO</td><td>0.00</td><td>0.00</td><td>61.29</td><td>0.00</td></tr>
<tr><td>>MI0000749|hsa-mir-30e&WithFlank&1|+|40754349|40754452@23.46.24&[MIMAT0000692&hsa-miR-30e-5p&offsets|0|+2;m-24&1|+|40754371|40754394&offsets|0|0]|TGTAAACATCCTTGACTGGAAGCT</td><td>>MI0000749|hsa-mir-30e&WithFlank&1|+|40754349|40754452</td><td>23</td><td>46</td><td>24</td><td></td><td>isomiR</td><td>NO</td><td>195.15</td><td>25.81</td><td>1535.92</td><td>73.68</td></tr>
<tr><td>>MI0000749|hsa-mir-30e&WithFlank&1|+|40754349|40754452@23.46.24(+1U)&[MIMAT0000692&hsa-miR-30e-5p&offsets|0|+2(+1U);m-24&1|+|40754371|40754394&offsets|0|0(+1U)]|TGTAAACATCCTTGACTGGAAGCTT</td><td>>MI0000749|hsa-mir-30e&WithFlank&1|+|40754349|40754452</td><td>23</td><td>46</td><td>24</td><td>(+1U)</td><td>isomiR</td><td>NO</td><td>0.00</td><td>0.00</td><td>198.54</td><td>0.00</td></tr>
<tr><td>>MI0000749|hsa-mir-30e&WithFlank&1|+|40754349|40754452@23.46.24(+2U)&[MIMAT0000692&hsa-miR-30e-5p&offsets|0|+2(+2U);m-24&1|+|40754371|40754394&offsets|0|0(+2U)]|TGTAAACATCCTTGACTGGAAGCTTT</td><td>>MI0000749|hsa-mir-30e&WithFlank&1|+|40754349|40754452</td><td>23</td><td>46</td><td>24</td><td>(+2U)</td><td>isomiR</td><td>NO</td><td>0.00</td><td>0.00</td><td>54.67</td><td>0.00</td></tr>
<tr><td>>MI0000749|hsa-mir-30e&WithFlank&1|+|40754349|40754452@24.46.23&[MIMAT0000692&hsa-miR-30e-5p&offsets|+1|+2;m-24&1|+|40754371|40754394&offsets|+1|0]|GTAAACATCCTTGACTGGAAGCT</td><td>>MI0000749|hsa-mir-30e&WithFlank&1|+|40754349|40754452</td><td>24</td><td>46</td><td>23</td><td></td><td>isomiR</td><td>NO</td><td>0.00</td><td>0.00</td><td>125.99</td><td>0.00</td></tr>
<tr><td>>MI0000749|hsa-mir-30e&WithFlank&1|+|40754349|40754452@65.83.19&[MIMAT0000693&hsa-miR-30e-3p&offsets|0|-3;m-25&1|+|40754413|40754434&offsets|0|-3]|CTTTCAGTCGGATGTTTAC</td><td>>MI0000749|hsa-mir-30e&WithFlank&1|+|40754349|40754452</td><td>65</td><td>83</td><td>19</td><td></td><td>isomiR</td><td>NO</td><td>69.28</td><td>31.28</td><td>64.40</td><td>0.00</td></tr>
<tr><td>>MI0000749|hsa-mir-30e&WithFlank&1|+|40754349|40754452@65.85.21&[MIMAT0000693&hsa-miR-30e-3p&offsets|0|-1;m-25&1|+|40754413|40754434&offsets|0|-1]|CTTTCAGTCGGATGTTTACAG</td><td>>MI0000749|hsa-mir-30e&WithFlank&1|+|40754349|40754452</td><td>65</td><td>85</td><td>21</td><td></td><td>isomiR</td><td>NO</td><td>0.00</td><td>0.00</td><td>12.92</td><td>0.00</td></tr>
<tr><td>>MI0000749|hsa-mir-30e&WithFlank&1|+|40754349|40754452@65.85.21(+1U)&[MIMAT0000693&hsa-miR-30e-3p&offsets|0|-1(+1U);m-25&1|+|40754413|40754434&offsets|0|-1(+1U)]|CTTTCAGTCGGATGTTTACAGT</td><td>>MI0000749|hsa-mir-30e&WithFlank&1|+|40754349|40754452</td><td>65</td><td>85</td><td>21</td><td>(+1U)</td><td>isomiR</td><td>NO</td><td>179.98</td><td>76.70</td><td>157.41</td><td>22.37</td></tr>
<tr><td>>MI0000749|hsa-mir-30e&WithFlank&1|+|40754349|40754452@65.85.21(+2U)&[MIMAT0000693&hsa-miR-30e-3p&offsets|0|-1(+2U);m-25&1|+|40754413|40754434&offsets|0|-1(+2U)]|CTTTCAGTCGGATGTTTACAGTT</td><td>>MI0000749|hsa-mir-30e&WithFlank&1|+|40754349|40754452</td><td>65</td><td>85</td><td>21</td><td>(+2U)</td><td>isomiR</td><td>NO</td><td>61.91</td><td>27.44</td><td>34.15</td><td>0.00</td></tr>
<tr><td>>MI0000749|hsa-mir-30e&WithFlank&1|+|40754349|40754452@65.86.22&[MIMAT0000693&hsa-miR-30e-3p&offsets|0|0;m-25&1|+|40754413|40754434&offsets|0|0]|CTTTCAGTCGGATGTTTACAGC</td><td>>MI0000749|hsa-mir-30e&WithFlank&1|+|40754349|40754452</td><td>65</td><td>86</td><td>22</td><td></td><td>isomiR</td><td>NO</td><td>139.72</td><td>99.51</td><td>121.95</td><td>18.34</td></tr>
<tr><td>>MI0000750|hsa-mir-26a-2&WithFlank&12|-|57824603|57824698@20.39.20(+1U);MI0000083|hsa-mir-26a-1&WithFlank&3|+|37969398|37969486@16.35.20(+1U)&[MIMAT0000082&hsa-miR-26a-5p&offsets|0|-2(+1U);m-33&12|-|57824658|57824679&offsets|0|-2(+1U)];[MIMAT0000082_1&hsa-miR-26a-5p&offsets|0|-2(+1U);m-34&3|+|37969413|37969434&offsets|0|-2(+1U)]|TTCAAGTAATCCAGGATAGGT</td><td>>MI0000750|hsa-mir-26a-2&WithFlank&12|-|57824603|57824698</td><td>20</td><td>39</td><td>20</td><td>(+1U)</td><td>isomiR</td><td>NO</td><td>0.00</td><td>0.00</td><td>62.06</td><td>0.00</td></tr>
<tr><td>>MI0000750|hsa-mir-26a-2&WithFlank&12|-|57824603|57824698@20.39.20;MI0000083|hsa-mir-26a-1&WithFlank&3|+|37969398|37969486@16.35.20&[MIMAT0000082&hsa-miR-26a-5p&offsets|0|-2;m-33&12|-|57824658|57824679&offsets|0|-2];[MIMAT0000082_1&hsa-miR-26a-5p&offsets|0|-2;m-34&3|+|37969413|37969434&offsets|0|-2]|TTCAAGTAATCCAGGATAGG</td><td>>MI0000750|hsa-mir-26a-2&WithFlank&12|-|57824603|57824698</td><td>20</td><td>39</td><td>20</td><td></td><td>isomiR</td><td>NO</td><td>0.00</td><td>0.00</td><td>18.00</td><td>0.00</td></tr>
<tr><td>>MI0000750|hsa-mir-26a-2&WithFlank&12|-|57824603|57824698@20.40.21(+1A);MI0000083|hsa-mir-26a-1&WithFlank&3|+|37969398|37969486@16.36.21(+1A)&[MIMAT0000082&hsa-miR-26a-5p&offsets|0|-1(+1A);m-33&12|-|57824658|57824679&offsets|0|-1(+1A)];[MIMAT0000082_1&hsa-miR-26a-5p&offsets|0|-1(+1A);m-34&3|+|37969413|37969434&offsets|0|-1(+1A)]|TTCAAGTAATCCAGGATAGGCA</td><td>>MI0000750|hsa-mir-26a-2&WithFlank&12|-|57824603|57824698</td><td>20</td><td>40</td><td>21</td><td>(+1A)</td><td>isomiR</td><td>NO</td><td>238.40</td><td>181.30</td><td>614.42</td><td>148.96</td></tr>
<tr><td>>MI0000750|hsa-mir-26a-2&WithFlank&12|-|57824603|57824698@20.40.21(+1C);MI0000083|hsa-mir-26a-1&WithFlank&3|+|37969398|37969486@16.36.21(+1C)&[MIMAT0000082&hsa-miR-26a-5p&offsets|0|-1(+1C);m-33&12|-|57824658|57824679&offsets|0|-1(+1C)];[MIMAT0000082_1&hsa-miR-26a-5p&offsets|0|-1(+1C);m-34&3|+|37969413|37969434&offsets|0|-1(+1C)]|TTCAAGTAATCCAGGATAGGCC</td><td>>MI0000750|hsa-mir-26a-2&WithFlank&12|-|57824603|57824698</td><td>20</td><td>40</td><td>21</td><td>(+1C)</td><td>isomiR</td><td>NO</td><td>386.48</td><td>251.68</td><td>831.37</td><td>209.03</td></tr>
<tr><td>>MI0000750|hsa-mir-26a-2&WithFlank&12|-|57824603|57824698@20.40.21(+1G);MI0000083|hsa-mir-26a-1&WithFlank&3|+|37969398|37969486@16.36.21(+1G)&[MIMAT0000082&hsa-miR-26a-5p&offsets|0|-1(+1G);m-33&12|-|57824658|57824679&offsets|0|-1(+1G)];[MIMAT0000082_1&hsa-miR-26a-5p&offsets|0|-1(+1G);m-34&3|+|37969413|37969434&offsets|0|-1(+1G)]|TTCAAGTAATCCAGGATAGGCG</td><td>>MI0000750|hsa-mir-26a-2&WithFlank&12|-|57824603|57824698</td><td>20</td><td>40</td><td>21</td><td>(+1G)</td><td>isomiR</td><td>NO</td><td>122.72</td><td>58.54</td><td>150.46</td><td>21.13</td></tr>
<tr><td>>MI0000750|hsa-mir-26a-2&WithFlank&12|-|57824603|57824698@20.40.21;MI0000083|hsa-mir-26a-1&WithFlank&3|+|37969398|37969486@16.36.21&[MIMAT0000082&hsa-miR-26a-5p&offsets|0|-1;m-33&12|-|57824658|57824679&offsets|0|-1];[MIMAT0000082_1&hsa-miR-26a-5p&offsets|0|-1;m-34&3|+|37969413|37969434&offsets|0|-1]|TTCAAGTAATCCAGGATAGGC</td><td>>MI0000750|hsa-mir-26a-2&WithFlank&12|-|57824603|57824698</td><td>20</td><td>40</td><td>21</td><td></td><td>isomiR</td><td>NO</td><td>90.73</td><td>30.27</td><td>154.58</td><td>15.02</td></tr>
<tr><td>>MI0000750|hsa-mir-26a-2&WithFlank&12|-|57824603|57824698@20.41.22(+1A);MI0000083|hsa-mir-26a-1&WithFlank&3|+|37969398|37969486@16.37.22(+1A)&[MIMAT0000082&hsa-miR-26a-5p&offsets|0|0(+1A);m-33&12|-|57824658|57824679&offsets|0|0(+1A)];[MIMAT0000082_1&hsa-miR-26a-5p&offsets|0|0(+1A);m-34&3|+|37969413|37969434&offsets|0|0(+1A)]|TTCAAGTAATCCAGGATAGGCTA</td><td>>MI0000750|hsa-mir-26a-2&WithFlank&12|-|57824603|57824698</td><td>20</td><td>41</td><td>22</td><td>(+1A)</td><td>isomiR</td><td>NO</td><td>48.93</td><td>0.00</td><td>133.06</td><td>13.76</td></tr>
</table>

### MDA-MB-231

<table>
<tr><td>>MI0000750|hsa-mir-26a-2&WithFlank&12|-|57824603|57824698@20.41.22(+1C);MI0000083|hsa-mir-26a-1&WithFlank&3|+|37969398|37969486@16.37.22(+1C)&[MIMAT0000082&hsa-miR-26a-5p&offsets|0|0(+1C);m-33&12|-|57824658|57824679&offsets|0|0(+1C)];[MIMAT0000082_1&hsa-miR-26a-5p&offsets|0|0(+1C);m-34&3|+|37969413|37969434&offsets|0|0(+1C)]|TTCAAGTAATCCAGGATAGGCTC</td><td>>MI0000750|hsa-mir-26a-2&WithFlank&12|-|57824603|57824698</td><td>20</td><td>41</td><td>22</td><td>(+1C)</td><td>isomiR</td><td>NO</td><td>0.00</td><td>0.00</td><td>97.08</td><td>0.00</td></tr>
<tr><td>>MI0000750|hsa-mir-26a-2&WithFlank&12|-|57824603|57824698@20.41.22(+1U);MI0000083|hsa-mir-26a-1&WithFlank&3|+|37969398|37969486@16.37.22(+1U)&[MIMAT0000082&hsa-miR-26a-5p&offsets|0|0(+1U);m-33&12|-|57824658|57824679&offsets|0|0(+1U)];[MIMAT0000082_1&hsa-miR-26a-5p&offsets|0|0(+1U);m-34&3|+|37969413|37969434&offsets|0|0(+1U)]|TTCAAGTAATCCAGGATAGGCTT</td><td>>MI0000750|hsa-mir-26a-2&WithFlank&12|-|57824603|57824698</td><td>20</td><td>41</td><td>22</td><td>(+1U)</td><td>isomiR</td><td>NO</td><td>184.28</td><td>73.39</td><td>436.22</td><td>85.10</td></tr>
<tr><td>>MI0000750|hsa-mir-26a-2&WithFlank&12|-|57824603|57824698@20.41.22(+2A);MI0000083|hsa-mir-26a-1&WithFlank&3|+|37969398|37969486@16.37.22(+2A)&[MIMAT0000082&hsa-miR-26a-5p&offsets|0|0(+2A);m-33&12|-|57824658|57824679&offsets|0|0(+2A)];[MIMAT0000082_1&hsa-miR-26a-5p&offsets|0|0(+2A);m-34&3|+|37969413|37969434&offsets|0|0(+2A)]|TTCAAGTAATCCAGGATAGGCTAA</td><td>>MI0000750|hsa-mir-26a-2&WithFlank&12|-|57824603|57824698</td><td>20</td><td>41</td><td>22</td><td>(+2A)</td><td>isomiR</td><td>NO</td><td>0.00</td><td>0.00</td><td>73.85</td><td>0.00</td></tr>
<tr><td>>MI0000750|hsa-mir-26a-2&WithFlank&12|-|57824603|57824698@20.41.22(+2U);MI0000083|hsa-mir-26a-1&WithFlank&3|+|37969398|37969486@16.37.22(+2U)&[MIMAT0000082&hsa-miR-26a-5p&offsets|0|0(+2U);m-33&12|-|57824658|57824679&offsets|0|0(+2U)];[MIMAT0000082_1&hsa-miR-26a-5p&offsets|0|0(+2U);m-34&3|+|37969413|37969434&offsets|0|0(+2U)]|TTCAAGTAATCCAGGATAGGCTTT</td><td>>MI0000750|hsa-mir-26a-2&WithFlank&12|-|57824603|57824698</td><td>20</td><td>41</td><td>22</td><td>(+2U)</td><td>isomiR</td><td>NO</td><td>149.86</td><td>71.62</td><td>293.30</td><td>89.40</td></tr>
<tr><td>>MI0000750|hsa-mir-26a-2&WithFlank&12|-|57824603|57824698@20.41.22(+3U);MI0000083|hsa-mir-26a-1&WithFlank&3|+|37969398|37969486@16.37.22(+3U)&[MIMAT0000082&hsa-miR-26a-5p&offsets|0|0(+3U);m-33&12|-|57824658|57824679&offsets|0|0(+3U)];[MIMAT0000082_1&hsa-miR-26a-5p&offsets|0|0(+3U);m-34&3|+|37969413|37969434&offsets|0|0(+3U)]|TTCAAGTAATCCAGGATAGGCTTTT</td><td>>MI0000750|hsa-mir-26a-2&WithFlank&12|-|57824603|57824698</td><td>20</td><td>41</td><td>22</td><td>(+3U)</td><td>isomiR</td><td>NO</td><td>0.00</td><td>0.00</td><td>14.51</td><td>0.00</td></tr>
<tr><td>>MI0000750|hsa-mir-26a-2&WithFlank&12|-|57824603|57824698@20.41.22;MI0000083|hsa-mir-26a-1&WithFlank&3|+|37969398|37969486@16.37.22&[MIMAT0000082&hsa-miR-26a-5p&offsets|0|0;m-33&12|-|57824658|57824679&offsets|0|0];[MIMAT0000082_1&hsa-miR-26a-5p&offsets|0|0;m-34&3|+|37969413|37969434&offsets|0|0]|TTCAAGTAATCCAGGATAGGCT</td><td>>MI0000750|hsa-mir-26a-2&WithFlank&12|-|57824603|57824698</td><td>20</td><td>41</td><td>22</td><td></td><td>isomiR</td><td>YES</td><td>2055.45</td><td>1583.67</td><td>3657.35</td><td>724.51</td></tr>
<tr><td>>MI0000750|hsa-mir-26a-2&WithFlank&12|-|57824603|57824698@20.42.23;MI0000083|hsa-mir-26a-1&WithFlank&3|+|37969398|37969486@16.38.23&[MIMAT0000082&hsa-miR-26a-5p&offsets|0|+1;m-33&12|-|57824658|57824679&offsets|0|+1];[MIMAT0000082_1&hsa-miR-26a-5p&offsets|0|+1;m-34&3|+|37969413|37969434&offsets|0|+1]|TTCAAGTAATCCAGGATAGGCTG</td><td>>MI0000750|hsa-mir-26a-2&WithFlank&12|-|57824603|57824698</td><td>20</td><td>42</td><td>23</td><td></td><td>isomiR</td><td>NO</td><td>0.00</td><td>0.00</td><td>64.21</td><td>0.00</td></tr>
<tr><td>>MI0000750|hsa-mir-26a-2&WithFlank&12|-|57824603|57824698@20.43.24;MI0000083|hsa-mir-26a-1&WithFlank&3|+|37969398|37969486@16.39.24&[MIMAT0000082&hsa-miR-26a-5p&offsets|0|+2;m-33&12|-|57824658|57824679&offsets|0|+2];[MIMAT0000082_1&hsa-miR-26a-5p&offsets|0|+2;m-34&3|+|37969413|37969434&offsets|0|+2]|TTCAAGTAATCCAGGATAGGCTGT</td><td>>MI0000750|hsa-mir-26a-2&WithFlank&12|-|57824603|57824698</td><td>20</td><td>43</td><td>24</td><td></td><td>isomiR</td><td>NO</td><td>0.00</td><td>0.00</td><td>13.73</td><td>0.00</td></tr>
<tr><td>>MI0000750|hsa-mir-26a-2&WithFlank&12|-|57824603|57824698@23.41.19;MI0000083|hsa-mir-26a-1&WithFlank&3|+|37969398|37969486@19.37.19&[MIMAT0000082&hsa-miR-26a-5p&offsets|+3|0;m-33&12|-|57824658|57824679&offsets|+3|0];[MIMAT0000082_1&hsa-miR-26a-5p&offsets|+3|0;m-34&3|+|37969413|37969434&offsets|+3|0]|AAGTAATCCAGGATAGGCT</td><td>>MI0000750|hsa-mir-26a-2&WithFlank&12|-|57824603|57824698</td><td>23</td><td>41</td><td>19</td><td></td><td>isomiR</td><td>NO</td><td>0.00</td><td>0.00</td><td>14.58</td><td>0.00</td></tr>
<tr><td>>MI0000760|hsa-mir-361&WithFlank&X|-|85903630|85903713@12.33.22&[MIMAT0000703&hsa-miR-361-5p&offsets|0|0;m-89&X|-|85903681|85903702&offsets|0|0]|TTATCAGAATCTCCAGGGGTAC</td><td>>MI0000760|hsa-mir-361&WithFlank&X|-|85903630|85903713</td><td>12</td><td>33</td><td>22</td><td></td><td>isomiR</td><td>NO</td><td>0.00</td><td>0.00</td><td>15.74</td><td>0.00</td></tr>
<tr><td>>MI0000760|hsa-mir-361&WithFlank&X|-|85903630|85903713@52.75.24&[MIMAT0004682&hsa-miR-361-3p&offsets|+1|+2;m-124&X|-|85903642|85903663&offsets|+1|+3]|CCCCCAGGTGTGATTCTGATTTGC</td><td>>MI0000760|hsa-mir-361&WithFlank&X|-|85903630|85903713</td><td>52</td><td>75</td><td>24</td><td></td><td>isomiR</td><td>NO</td><td>0.00</td><td>27.04</td><td>0.00</td><td>0.00</td></tr>
<tr><td>>MI0000782|hsa-mir-374a&WithFlank&X|-|74287280|74287363@18.38.21&[MIMAT0000727&hsa-miR-374a-5p&offsets|0|-1;m-184&X|-|74287325|74287346&offsets|0|-1]|TTATAATACAACCTGATAAGT</td><td>>MI0000782|hsa-mir-374a&WithFlank&X|-|74287280|74287363</td><td>18</td><td>38</td><td>21</td><td></td><td>isomiR</td><td>NO</td><td>0.00</td><td>0.00</td><td>59.43</td><td>0.00</td></tr>
<tr><td>>MI0000782|hsa-mir-374a&WithFlank&X|-|74287280|74287363@18.39.22&[MIMAT0000727&hsa-miR-374a-5p&offsets|0|0;m-184&X|-|74287325|74287346&offsets|0|0]|TTATAATACAACCTGATAAGTG</td><td>>MI0000782|hsa-mir-374a&WithFlank&X|-|74287280|74287363</td><td>18</td><td>39</td><td>22</td><td></td><td>isomiR</td><td>NO</td><td>0.00</td><td>0.00</td><td>62.47</td><td>0.00</td></tr>
<tr><td>>MI0000782|hsa-mir-374a&WithFlank&X|-|74287280|74287363@48.69.22&[MIMAT0004688&hsa-miR-374a-3p&offsets|0|0;m-74&X|-|74287295|74287316&offsets|0|0]|CTTATCAGATTGTATTGTAATT</td><td>>MI0000782|hsa-mir-374a&WithFlank&X|-|74287280|74287363</td><td>48</td><td>69</td><td>22</td><td></td><td>isomiR</td><td>NO</td><td>0.00</td><td>0.00</td><td>12.08</td><td>0.00</td></tr>
<tr><td>>am_MI0000786|hsa-mir-378a&WithFlank&5|+|149732819|149732896@49.69.21&[MIMAT0000732&hsa-miR-378a-3p&offsets|0|-1;m-44&5|+|149732867|149732888&offsets|0|-1]|ACTGGACTTGGAGTCAGAAGG</td><td>>MI0000786|hsa-mir-378a&WithFlank&5|+|149732819|149732896.a</td><td>49</td><td>69</td><td>21</td><td></td><td>isomiR</td><td>NO</td><td>120.44</td><td>35.81</td><td>689.47</td><td>14.99</td></tr>
<tr><td>>am_MI0000786|hsa-mir-378a&WithFlank&5|+|149732819|149732896@49.69.21(+1A)&[MIMAT0000732&hsa-miR-378a-3p&offsets|0|-1(+1A);m-44&5|+|149732867|149732888&offsets|0|-1(+1A)]|ACTGGACTTGGAGTCAGAAGGA</td><td>>MI0000786|hsa-mir-378a&WithFlank&5|+|149732819|149732896.a</td><td>49</td><td>69</td><td>21</td><td>(+1A)</td><td>isomiR</td><td>NO</td><td>26.82</td><td>0.00</td><td>330.60</td><td>0.00</td></tr>
<tr><td>>am_MI0000786|hsa-mir-378a&WithFlank&5|+|149732819|149732896@49.69.21(+1U)&[MIMAT0000732&hsa-miR-378a-3p&offsets|0|-1(+1U);m-44&5|+|149732867|149732888&offsets|0|-1(+1U)]|ACTGGACTTGGAGTCAGAAGGT</td><td>>MI0000786|hsa-mir-378a&WithFlank&5|+|149732819|149732896.a</td><td>49</td><td>69</td><td>21</td><td>(+1U)</td><td>isomiR</td><td>NO</td><td>114.06</td><td>86.35</td><td>981.10</td><td>17.22</td></tr>
<tr><td>>am_MI0000786|hsa-mir-378a&WithFlank&5|+|149732819|149732896@49.70.22&[MIMAT0000732&hsa-miR-378a-3p&offsets|0|0;m-44&5|+|149732867|149732888&offsets|0|0]|ACTGGACTTGGAGTCAGAAGGC</td><td>>MI0000786|hsa-mir-378a&WithFlank&5|+|149732819|149732896.a</td><td>49</td><td>70</td><td>22</td><td></td><td>isomiR</td><td>NO</td><td>396.45</td><td>319.71</td><td>2260.42</td><td>97.79</td></tr>
</table>

### MDA-MB-231

<table>
<tr><th></th><th></th><th></th><th></th><th></th><th></th><th></th><th></th><th></th><th></th><th></th><th></th><th></th></tr>
<tr><td>>am_MI0000786|hsa-mir-378a&WithFlank&5|+|149732819|149732896@49.70.22(+1A)&[MIMAT0000732&hsa-miR-378a-3p&offsets|0|0(+1A);m-44&5|+|149732867|149732888&offsets|0|0(+1A)]|ACTGGACTTGGAGTCAGAAGGCA</td><td>>MI0000786|hsa-mir-378a&WithFlank&5|+|149732819|149732896.a</td><td>49</td><td>70</td><td>22</td><td>(+1A)</td><td>isomiR</td><td>NO</td><td>45.44</td><td>33.63</td><td>290.84</td><td>0.00</td></tr>
<tr><td>>am_MI0000786|hsa-mir-378a&WithFlank&5|+|149732819|149732896@49.70.22(+1G)&[MIMAT0000732&hsa-miR-378a-3p&offsets|0|0(+1G);m-44&5|+|149732867|149732888&offsets|0|0(+1G)]|ACTGGACTTGGAGTCAGAAGGCG</td><td>>MI0000786|hsa-mir-378a&WithFlank&5|+|149732819|149732896.a</td><td>49</td><td>70</td><td>22</td><td>(+1G)</td><td>isomiR</td><td>NO</td><td>0.00</td><td>0.00</td><td>65.93</td><td>0.00</td></tr>
<tr><td>>am_MI0000786|hsa-mir-378a&WithFlank&5|+|149732819|149732896@49.70.22(+1U)&[MIMAT0000732&hsa-miR-378a-3p&offsets|0|0(+1U);m-44&5|+|149732867|149732888&offsets|0|0(+1U)]|ACTGGACTTGGAGTCAGAAGGCT</td><td>>MI0000786|hsa-mir-378a&WithFlank&5|+|149732819|149732896.a</td><td>49</td><td>70</td><td>22</td><td>(+1U)</td><td>isomiR</td><td>NO</td><td>75.76</td><td>66.14</td><td>486.11</td><td>0.00</td></tr>
<tr><td>>am_MI0000786|hsa-mir-378a&WithFlank&5|+|149732819|149732896@49.71.23&[MIMAT0000732&hsa-miR-378a-3p&offsets|0|+1;m-44&5|+|149732867|149732888&offsets|0|+1]|ACTGGACTTGGAGTCAGAAGGCC</td><td>>MI0000786|hsa-mir-378a&WithFlank&5|+|149732819|149732896.a</td><td>49</td><td>71</td><td>23</td><td></td><td>isomiR</td><td>NO</td><td>0.00</td><td>0.00</td><td>118.45</td><td>0.00</td></tr>
<tr><td>>am_MI0000786|hsa-mir-378a&WithFlank&5|+|149732819|149732896@50.69.20&[MIMAT0000732&hsa-miR-378a-3p&offsets|+1|-1;m-44&5|+|149732867|149732888&offsets|+1|-1]|CTGGACTTGGAGTCAGAAGG</td><td>>MI0000786|hsa-mir-378a&WithFlank&5|+|149732819|149732896.a</td><td>50</td><td>69</td><td>20</td><td></td><td>isomiR</td><td>NO</td><td>0.00</td><td>0.00</td><td>69.27</td><td>0.00</td></tr>
<tr><td>>am_MI0000786|hsa-mir-378a&WithFlank&5|+|149732819|149732896@50.69.20(+1U)&[MIMAT0000732&hsa-miR-378a-3p&offsets|+1|-1(+1U);m-44&5|+|149732867|149732888&offsets|+1|-1(+1U)]|CTGGACTTGGAGTCAGAAGGT</td><td>>MI0000786|hsa-mir-378a&WithFlank&5|+|149732819|149732896.a</td><td>50</td><td>69</td><td>20</td><td>(+1U)</td><td>isomiR</td><td>NO</td><td>0.00</td><td>0.00</td><td>126.65</td><td>0.00</td></tr>
<tr><td>>am_MI0000786|hsa-mir-378a&WithFlank&5|+|149732819|149732896@50.70.21&[MIMAT0000732&hsa-miR-378a-3p&offsets|+1|0;m-44&5|+|149732867|149732888&offsets|+1|0]|CTGGACTTGGAGTCAGAAGGC</td><td>>MI0000786|hsa-mir-378a&WithFlank&5|+|149732819|149732896.a</td><td>50</td><td>70</td><td>21</td><td></td><td>isomiR</td><td>NO</td><td>45.33</td><td>64.25</td><td>295.42</td><td>0.00</td></tr>
<tr><td>>am_MI0000786|hsa-mir-378a&WithFlank&5|+|149732819|149732896@50.70.21(+1U)&[MIMAT0000732&hsa-miR-378a-3p&offsets|+1|0(+1U);m-44&5|+|149732867|149732888&offsets|+1|0(+1U)]|CTGGACTTGGAGTCAGAAGGCT</td><td>>MI0000786|hsa-mir-378a&WithFlank&5|+|149732819|149732896.a</td><td>50</td><td>70</td><td>21</td><td>(+1U)</td><td>isomiR</td><td>NO</td><td>0.00</td><td>24.21</td><td>197.31</td><td>0.00</td></tr>
<tr><td>>MI0000802|hsa-mir-340&WithFlank&5|-|180015297|180015403@22.42.21&[MIMAT0004692&hsa-miR-340-5p&offsets|0|-1;m-220&5|-|180015361|180015382&offsets|0|-1]|TTATAAAGCAATGAGACTGAT</td><td>>MI0000802|hsa-mir-340&WithFlank&5|-|180015297|180015403</td><td>22</td><td>42</td><td>21</td><td></td><td>isomiR</td><td>NO</td><td>0.00</td><td>0.00</td><td>10.46</td><td>0.00</td></tr>
<tr><td>>MI0000802|hsa-mir-340&WithFlank&5|-|180015297|180015403@22.42.21(+1C)&[MIMAT0004692&hsa-miR-340-5p&offsets|0|-1(+1C);m-220&5|-|180015361|180015382&offsets|0|-1(+1C)]|TTATAAAGCAATGAGACTGATC</td><td>>MI0000802|hsa-mir-340&WithFlank&5|-|180015297|180015403</td><td>22</td><td>42</td><td>21</td><td>(+1C)</td><td>isomiR</td><td>NO</td><td>0.00</td><td>0.00</td><td>16.74</td><td>0.00</td></tr>
<tr><td>>MI0000802|hsa-mir-340&WithFlank&5|-|180015297|180015403@22.43.22&[MIMAT0004692&hsa-miR-340-5p&offsets|0|0;m-220&5|-|180015361|180015382&offsets|0|0]|TTATAAAGCAATGAGACTGATT</td><td>>MI0000802|hsa-mir-340&WithFlank&5|-|180015297|180015403</td><td>22</td><td>43</td><td>22</td><td></td><td>isomiR</td><td>NO</td><td>107.14</td><td>0.00</td><td>262.15</td><td>11.93</td></tr>
<tr><td>>MI0000803|hsa-mir-330&WithFlank&19|-|45638988|45639093@63.86.24(+1U)&[MIMAT0000751&hsa-miR-330-3p&offsets|0|+1(+1U);m-347&19|-|45639009|45639030&offsets|-1|+1(+1U)]|GCAAAGCACACGGCCTGCAGAGAGT</td><td>>MI0000803|hsa-mir-330&WithFlank&19|-|45638988|45639093</td><td>63</td><td>86</td><td>24</td><td>(+1U)</td><td>isomiR</td><td>NO</td><td>0.00</td><td>60.22</td><td>17.75</td><td>0.00</td></tr>
<tr><td>>MI0000803|hsa-mir-330&WithFlank&19|-|45638988|45639093@64.86.23(+1U)&[MIMAT0000751&hsa-miR-330-3p&offsets|+1|+1(+1U);m-347&19|-|45639009|45639030&offsets|0|+1(+1U)]|CAAAGCACACGGCCTGCAGAGAGT</td><td>>MI0000803|hsa-mir-330&WithFlank&19|-|45638988|45639093</td><td>64</td><td>86</td><td>23</td><td>(+1U)</td><td>isomiR</td><td>NO</td><td>0.00</td><td>0.00</td><td>13.05</td><td>0.00</td></tr>
<tr><td>>MI0000803|hsa-mir-330&WithFlank&19|-|45638988|45639093@64.87.24&[MIMAT0000751&hsa-miR-330-3p&offsets|+1|+2;m-347&19|-|45639009|45639030&offsets|0|+2]|CAAAGCACACGGCCTGCAGAGAGG</td><td>>MI0000803|hsa-mir-330&WithFlank&19|-|45638988|45639093</td><td>64</td><td>87</td><td>24</td><td></td><td>isomiR</td><td>NO</td><td>0.00</td><td>0.00</td><td>16.55</td><td>0.00</td></tr>
<tr><td>>MI0000809|hsa-mir-151a&WithFlank&8|-|140732558|140732659@51.70.20&[MIMAT0000757&hsa-miR-151a-3p&offsets|-2|-3;m-30&8|-|140732587|140732607&offsets|-2|-3]|TACTAGACTGAAGCTCCTTG</td><td>>MI0000809|hsa-mir-151a&WithFlank&8|-|140732558|140732659</td><td>51</td><td>70</td><td>20</td><td></td><td>isomiR</td><td>NO</td><td>0.00</td><td>0.00</td><td>10.91</td><td>0.00</td></tr>
<tr><td>>MI0000809|hsa-mir-151a&WithFlank&8|-|140732558|140732659@51.72.22&[MIMAT0000757&hsa-miR-151a-3p&offsets|-2|-1;m-30&8|-|140732587|140732607&offsets|-2|-1]|TACTAGACTGAAGCTCCTTGAG</td><td>>MI0000809|hsa-mir-151a&WithFlank&8|-|140732558|140732659</td><td>51</td><td>72</td><td>22</td><td></td><td>isomiR</td><td>NO</td><td>24.96</td><td>0.00</td><td>82.56</td><td>0.00</td></tr>
<tr><td>>MI0000809|hsa-mir-151a&WithFlank&8|-|140732558|140732659@51.72.22(+1A)&[MIMAT0000757&hsa-miR-151a-3p&offsets|-2|-1(+1A);m-30&8|-|140732587|140732607&offsets|-2|-1(+1A)]|TACTAGACTGAAGCTCCTTGAGA</td><td>>MI0000809|hsa-mir-151a&WithFlank&8|-|140732558|140732659</td><td>51</td><td>72</td><td>22</td><td>(+1A)</td><td>isomiR</td><td>NO</td><td>0.00</td><td>0.00</td><td>60.66</td><td>0.00</td></tr>
<tr><td>>MI0000809|hsa-mir-151a&WithFlank&8|-|140732558|140732659@51.72.22(+1U)&[MIMAT0000757&hsa-miR-151a-3p&offsets|-2|-1(+1U);m-30&8|-|140732587|140732607&offsets|-2|-1(+1U)]|TACTAGACTGAAGCTCCTTGAGT</td><td>>MI0000809|hsa-mir-151a&WithFlank&8|-|140732558|140732659</td><td>51</td><td>72</td><td>22</td><td>(+1U)</td><td>isomiR</td><td>NO</td><td>24.65</td><td>0.00</td><td>70.99</td><td>10.38</td></tr>
<tr><td>>MI0000809|hsa-mir-151a&WithFlank&8|-|140732558|140732659@51.73.23&[MIMAT0000757&hsa-miR-151a-3p&offsets|-2|0;m-30&8|-|140732587|140732607&offsets|-2|0]|TACTAGACTGAAGCTCCTTGAGG</td><td>>MI0000809|hsa-mir-151a&WithFlank&8|-|140732558|140732659</td><td>51</td><td>73</td><td>23</td><td></td><td>isomiR</td><td>NO</td><td>173.10</td><td>185.66</td><td>269.23</td><td>75.34</td></tr>
<tr><td>>MI0000809|hsa-mir-151a&WithFlank&8|-|140732558|140732659@51.73.23(+1U)&[MIMAT0000757&hsa-miR-151a-3p&offsets|-2|0(+1U);m-30&8|-|140732587|140732607&offsets|-2|0(+1U)]|TACTAGACTGAAGCTCCTTGAGGT</td><td>>MI0000809|hsa-mir-151a&WithFlank&8|-|140732558|140732659</td><td>51</td><td>73</td><td>23</td><td>(+1U)</td><td>isomiR</td><td>NO</td><td>0.00</td><td>0.00</td><td>11.21</td><td>0.00</td></tr>
<tr><td>>MI0000809|hsa-mir-151a&WithFlank&8|-|140732558|140732659@53.74.22&[MIMAT0000757&hsa-miR-151a-3p&offsets|0|+1;m-30&8|-|140732587|140732607&offsets|0|+1]|CTAGACTGAAGCTCCTTGAGGA</td><td>>MI0000809|hsa-mir-151a&WithFlank&8|-|140732558|140732659</td><td>53</td><td>74</td><td>22</td><td></td><td>isomiR</td><td>NO</td><td>382.65</td><td>348.70</td><td>701.93</td><td>131.28</td></tr>
<tr><td>>MI0000809|hsa-mir-151a&WithFlank&8|-|140732558|140732659@53.74.22(+1A)&[MIMAT0000757&hsa-miR-151a-3p&offsets|0|+1(+1A);m-30&8|-|140732587|140732607&offsets|0|+1(+1A)]|CTAGACTGAAGCTCCTTGAGGAA</td><td>>MI0000809|hsa-mir-151a&WithFlank&8|-|140732558|140732659</td><td>53</td><td>74</td><td>22</td><td>(+1A)</td><td>isomiR</td><td>NO</td><td>122.59</td><td>102.49</td><td>243.79</td><td>21.96</td></tr>
</table>

### MDA-MB-231

| | | | | | | | | | | | |
|---|---|---|---|---|---|---|---|---|---|---|---|
| >MI0000809\|hsa-mir-151a&WithFlank&8\|-\|140732558\|140732659@53.74.22(+1U)&[MIMAT0000757&hsa-miR-151a-3p&offsets\|0\|+1(+1U);m-30&8\|-\|140732587\|140732607&offsets\|0\|+1(+1U)]\|CTAGACTGAAGCTCCTTGAGGAT | >MI0000809\|hsa-mir-151a&WithFlank&8\|-\|140732558\|140732659 | 53 | 74 | 22 | (+1U) | isomiR | NO | 87.16 | 34.62 | 166.65 | 17.31 |
| >MI0000809\|hsa-mir-151a&WithFlank&8\|-\|140732558\|140732659@53.74.22(+2A)&[MIMAT0000757&hsa-miR-151a-3p&offsets\|0\|+1(+2A);m-30&8\|-\|140732587\|140732607&offsets\|0\|+1(+2A)]\|CTAGACTGAAGCTCCTTGAGGAAA | >MI0000809\|hsa-mir-151a&WithFlank&8\|-\|140732558\|140732659 | 53 | 74 | 22 | (+2A) | isomiR | NO | 0.00 | 0.00 | 33.38 | 0.00 |
| >MI0000809\|hsa-mir-151a&WithFlank&8\|-\|140732558\|140732659@53.74.22(+2U)&[MIMAT0000757&hsa-miR-151a-3p&offsets\|0\|+1(+2U);m-30&8\|-\|140732587\|140732607&offsets\|0\|+1(+2U)]\|CTAGACTGAAGCTCCTTGAGGATT | >MI0000809\|hsa-mir-151a&WithFlank&8\|-\|140732558\|140732659 | 53 | 74 | 22 | (+2U) | isomiR | NO | 48.75 | 0.00 | 108.73 | 0.00 |
| >MI0000809\|hsa-mir-151a&WithFlank&8\|-\|140732558\|140732659@53.75.23&[MIMAT0000757&hsa-miR-151a-3p&offsets\|0\|+2;m-30&8\|-\|140732587\|140732607&offsets\|0\|+2]\|CTAGACTGAAGCTCCTTGAGGAC | >MI0000809\|hsa-mir-151a&WithFlank&8\|-\|140732558\|140732659 | 53 | 75 | 23 | | isomiR | NO | 0.00 | 0.00 | 14.66 | 0.00 |
| >am_MI0000809\|hsa-mir-151a&WithFlank&8\|-\|140732558\|140732659@17.37.21&[MIMAT0004697&hsa-miR-151a-5p&offsets\|0\|0;m-75&8\|-\|140732623\|140732643&offsets\|0\|0]\|TCGAGGAGCTCACAGTCTAGT | >MI0000809\|hsa-mir-151a&WithFlank&8\|-\|140732558\|140732659.a | 17 | 37 | 21 | | isomiR | NO | 70.94 | 0.00 | 72.40 | 12.19 |
| >am_MI0000809\|hsa-mir-151a&WithFlank&8\|-\|140732558\|140732659@52.73.22&[MIMAT0000757&hsa-miR-151a-3p&offsets\|-1\|0;m-30&8\|-\|140732587\|140732607&offsets\|-1\|0]\|ACTAGACTGAAGCTCCTTGAGG | >MI0000809\|hsa-mir-151a&WithFlank&8\|-\|140732558\|140732659.a | 52 | 73 | 22 | | isomiR | NO | 74.25 | 59.16 | 100.37 | 12.21 |
| >am_MI0000809\|hsa-mir-151a&WithFlank&8\|-\|140732558\|140732659@53.70.18&[MIMAT0000757&hsa-miR-151a-3p&offsets\|0\|-3;m-30&8\|-\|140732587\|140732607&offsets\|0\|-3]\|CTAGACTGAAGCTCCTTG | >MI0000809\|hsa-mir-151a&WithFlank&8\|-\|140732558\|140732659.a | 53 | 70 | 18 | | isomiR | NO | 46.81 | 26.21 | 144.65 | 0.00 |
| >am_MI0000809\|hsa-mir-151a&WithFlank&8\|-\|140732558\|140732659@53.70.18(+3G)&[MIMAT0000757&hsa-miR-151a-3p&offsets\|0\|-3(+3G);m-30&8\|-\|140732587\|140732607&offsets\|0\|-3(+3G)]\|CTAGACTGAAGCTCCTTGGGG | >MI0000809\|hsa-mir-151a&WithFlank&8\|-\|140732558\|140732659.a | 53 | 70 | 18 | (+3G) | isomiR | NO | 0.00 | 0.00 | 10.59 | 0.00 |
| >am_MI0000809\|hsa-mir-151a&WithFlank&8\|-\|140732558\|140732659@53.71.19&[MIMAT0000757&hsa-miR-151a-3p&offsets\|0\|-2;m-30&8\|-\|140732587\|140732607&offsets\|0\|-2]\|CTAGACTGAAGCTCCTTGA | >MI0000809\|hsa-mir-151a&WithFlank&8\|-\|140732558\|140732659.a | 53 | 71 | 19 | | isomiR | NO | 0.00 | 0.00 | 67.00 | 0.00 |
| >am_MI0000809\|hsa-mir-151a&WithFlank&8\|-\|140732558\|140732659@53.71.19(+1A)&[MIMAT0000757&hsa-miR-151a-3p&offsets\|0\|-2(+1A);m-30&8\|-\|140732587\|140732607&offsets\|0\|-2(+1A)]\|CTAGACTGAAGCTCCTTGAA | >MI0000809\|hsa-mir-151a&WithFlank&8\|-\|140732558\|140732659.a | 53 | 71 | 19 | (+1A) | isomiR | NO | 0.00 | 0.00 | 15.91 | 0.00 |
| >am_MI0000809\|hsa-mir-151a&WithFlank&8\|-\|140732558\|140732659@53.72.20&[MIMAT0000757&hsa-miR-151a-3p&offsets\|0\|-1;m-30&8\|-\|140732587\|140732607&offsets\|0\|-1]\|CTAGACTGAAGCTCCTTGAG | >MI0000809\|hsa-mir-151a&WithFlank&8\|-\|140732558\|140732659.a | 53 | 72 | 20 | | isomiR | NO | 97.97 | 58.82 | 213.12 | 14.48 |
| >am_MI0000809\|hsa-mir-151a&WithFlank&8\|-\|140732558\|140732659@53.72.20(+1A)&[MIMAT0000757&hsa-miR-151a-3p&offsets\|0\|-1(+1A);m-30&8\|-\|140732587\|140732607&offsets\|0\|-1(+1A)]\|CTAGACTGAAGCTCCTTGAGA | >MI0000809\|hsa-mir-151a&WithFlank&8\|-\|140732558\|140732659.a | 53 | 72 | 20 | (+1A) | isomiR | NO | 189.80 | 242.03 | 529.09 | 81.98 |
| >am_MI0000809\|hsa-mir-151a&WithFlank&8\|-\|140732558\|140732659@53.72.20(+1C)&[MIMAT0000757&hsa-miR-151a-3p&offsets\|0\|-1(+1C);m-30&8\|-\|140732587\|140732607&offsets\|0\|-1(+1C)]\|CTAGACTGAAGCTCCTTGAGC | >MI0000809\|hsa-mir-151a&WithFlank&8\|-\|140732558\|140732659.a | 53 | 72 | 20 | (+1C) | isomiR | NO | 0.00 | 0.00 | 82.74 | 0.00 |
| >am_MI0000809\|hsa-mir-151a&WithFlank&8\|-\|140732558\|140732659@53.72.20(+1U)&[MIMAT0000757&hsa-miR-151a-3p&offsets\|0\|-1(+1U);m-30&8\|-\|140732587\|140732607&offsets\|0\|-1(+1U)]\|CTAGACTGAAGCTCCTTGAGT | >MI0000809\|hsa-mir-151a&WithFlank&8\|-\|140732558\|140732659.a | 53 | 72 | 20 | (+1U) | isomiR | NO | 164.10 | 106.02 | 310.51 | 64.41 |
| >am_MI0000809\|hsa-mir-151a&WithFlank&8\|-\|140732558\|140732659@53.72.20(+2U)&[MIMAT0000757&hsa-miR-151a-3p&offsets\|0\|-1(+2U);m-30&8\|-\|140732587\|140732607&offsets\|0\|-1(+2U)]\|CTAGACTGAAGCTCCTTGAGTT | >MI0000809\|hsa-mir-151a&WithFlank&8\|-\|140732558\|140732659.a | 53 | 72 | 20 | (+2U) | isomiR | NO | 0.00 | 0.00 | 11.91 | 0.00 |
| >am_MI0000809\|hsa-mir-151a&WithFlank&8\|-\|140732558\|140732659@53.73.21&[MIMAT0000757&hsa-miR-151a-3p&offsets\|0\|0;m-30&8\|-\|140732587\|140732607&offsets\|0\|0]\|CTAGACTGAAGCTCCTTGAGG | >MI0000809\|hsa-mir-151a&WithFlank&8\|-\|140732558\|140732659.a | 53 | 73 | 21 | | isomiR | YES | 1247.94 | 1632.65 | 2481.86 | 305.34 |
| >am_MI0000809\|hsa-mir-151a&WithFlank&8\|-\|140732558\|140732659@53.73.21(+1C)&[MIMAT0000757&hsa-miR-151a-3p&offsets\|0\|0(+1C);m-30&8\|-\|140732587\|140732607&offsets\|0\|0(+1C)]\|CTAGACTGAAGCTCCTTGAGGC | >MI0000809\|hsa-mir-151a&WithFlank&8\|-\|140732558\|140732659.a | 53 | 73 | 21 | (+1C) | isomiR | NO | 64.16 | 33.97 | 130.49 | 13.50 |
| >am_MI0000809\|hsa-mir-151a&WithFlank&8\|-\|140732558\|140732659@53.73.21(+1G)&[MIMAT0000757&hsa-miR-151a-3p&offsets\|0\|0(+1G);m-30&8\|-\|140732587\|140732607&offsets\|0\|0(+1G)]\|CTAGACTGAAGCTCCTTGAGGG | >MI0000809\|hsa-mir-151a&WithFlank&8\|-\|140732558\|140732659.a | 53 | 73 | 21 | (+1G) | isomiR | NO | 0.00 | 0.00 | 12.45 | 0.00 |
| >am_MI0000809\|hsa-mir-151a&WithFlank&8\|-\|140732558\|140732659@53.73.21(+1U)&[MIMAT0000757&hsa-miR-151a-3p&offsets\|0\|0(+1U);m-30&8\|-\|140732587\|140732607&offsets\|0\|0(+1U)]\|CTAGACTGAAGCTCCTTGAGGT | >MI0000809\|hsa-mir-151a&WithFlank&8\|-\|140732558\|140732659.a | 53 | 73 | 21 | (+1U) | isomiR | NO | 325.87 | 297.79 | 531.86 | 112.42 |
| >am_MI0000809\|hsa-mir-151a&WithFlank&8\|-\|140732558\|140732659@53.73.21(+2U)&[MIMAT0000757&hsa-miR-151a-3p&offsets\|0\|0(+2U);m-30&8\|-\|140732587\|140732607&offsets\|0\|0(+2U)]\|CTAGACTGAAGCTCCTTGAGGTT | >MI0000809\|hsa-mir-151a&WithFlank&8\|-\|140732558\|140732659.a | 53 | 73 | 21 | (+2U) | isomiR | NO | 64.29 | 24.98 | 105.23 | 12.70 |
| >am_MI0000809\|hsa-mir-151a&WithFlank&8\|-\|140732558\|140732659@53.73.21(+3U)&[MIMAT0000757&hsa-miR-151a-3p&offsets\|0\|0(+3U);m-30&8\|-\|140732587\|140732607&offsets\|0\|0(+3U)]\|CTAGACTGAAGCTCCTTGAGGTTT | >MI0000809\|hsa-mir-151a&WithFlank&8\|-\|140732558\|140732659.a | 53 | 73 | 21 | (+3U) | isomiR | NO | 0.00 | 0.00 | 133.11 | 0.00 |
| >MI0000810\|hsa-mir-135b&WithFlank&1\|-\|205448296\|205448404@22.44.23&[MIMAT0000758&hsa-miR-135b-5p&offsets\|0\|0;m-321&1\|-\|205448361\|205448383&offsets\|0\|0]\|TATGGCTTTTCATTCCTATGTGA | >MI0000810\|hsa-mir-135b&WithFlank&1\|-\|205448296\|205448404 | 22 | 44 | 23 | | isomiR | NO | 57.61 | 0.00 | 189.90 | 0.00 |
| >MI0000811\|hsa-mir-148b&WithFlank&12\|+\|54337210\|54337320@69.89.21&[MIMAT0000759&hsa-miR-148b-3p&offsets\|0\|-1;m-77&12\|+\|54337278\|54337299&offsets\|0\|-1]\|TCAGTGCATCACAGAACTTTG | >MI0000811\|hsa-mir-148b&WithFlank&12\|+\|54337210\|54337320 | 69 | 89 | 21 | | isomiR | NO | 0.00 | 0.00 | 132.39 | 0.00 |

### MDA-MB-231

<table>
<tr><td>>MI0000811|hsa-mir-148b&WithFlank&12|+|54337210|54337320@69.89.21(+1A)&[MIMAT0000759&hsa-miR-148b-3p&offsets|0|-1(+1A);m-77&12|+|54337278|54337299&offsets|0|-1(+1A)]|TCAGTGCATCACAGAACTTTGA</td><td>>MI0000811|hsa-mir-148b&WithFlank&12|+|54337210|54337320</td><td>69</td><td>89</td><td>21</td><td>(+1A)</td><td>isomiR</td><td>NO</td><td>0.00</td><td>0.00</td><td>87.62</td><td>0.00</td></tr>
<tr><td>>MI0000811|hsa-mir-148b&WithFlank&12|+|54337210|54337320@69.89.21(+1C)&[MIMAT0000759&hsa-miR-148b-3p&offsets|0|-1(+1C);m-77&12|+|54337278|54337299&offsets|0|-1(+1C)]|TCAGTGCATCACAGAACTTTGC</td><td>>MI0000811|hsa-mir-148b&WithFlank&12|+|54337210|54337320</td><td>69</td><td>89</td><td>21</td><td>(+1C)</td><td>isomiR</td><td>NO</td><td>0.00</td><td>0.00</td><td>297.80</td><td>0.00</td></tr>
<tr><td>>MI0000811|hsa-mir-148b&WithFlank&12|+|54337210|54337320@69.89.21(+1G)&[MIMAT0000759&hsa-miR-148b-3p&offsets|0|-1(+1G);m-77&12|+|54337278|54337299&offsets|0|-1(+1G)]|TCAGTGCATCACAGAACTTTGG</td><td>>MI0000811|hsa-mir-148b&WithFlank&12|+|54337210|54337320</td><td>69</td><td>89</td><td>21</td><td>(+1G)</td><td>isomiR</td><td>NO</td><td>0.00</td><td>0.00</td><td>15.28</td><td>0.00</td></tr>
<tr><td>>MI0000811|hsa-mir-148b&WithFlank&12|+|54337210|54337320@69.90.22&[MIMAT0000759&hsa-miR-148b-3p&offsets|0|0;m-77&12|+|54337278|54337299&offsets|0|0]|TCAGTGCATCACAGAACTTTGT</td><td>>MI0000811|hsa-mir-148b&WithFlank&12|+|54337210|54337320</td><td>69</td><td>90</td><td>22</td><td></td><td>isomiR</td><td>NO</td><td>189.17</td><td>192.77</td><td>1395.54</td><td>62.64</td></tr>
<tr><td>>MI0000811|hsa-mir-148b&WithFlank&12|+|54337210|54337320@69.90.22(+1A)&[MIMAT0000759&hsa-miR-148b-3p&offsets|0|0(+1A);m-77&12|+|54337278|54337299&offsets|0|0(+1A)]|TCAGTGCATCACAGAACTTTGTA</td><td>>MI0000811|hsa-mir-148b&WithFlank&12|+|54337210|54337320</td><td>69</td><td>90</td><td>22</td><td>(+1A)</td><td>isomiR</td><td>NO</td><td>0.00</td><td>0.00</td><td>11.92</td><td>0.00</td></tr>
<tr><td>>MI0000811|hsa-mir-148b&WithFlank&12|+|54337210|54337320@69.90.22(+1U)&[MIMAT0000759&hsa-miR-148b-3p&offsets|0|0(+1U);m-77&12|+|54337278|54337299&offsets|0|0(+1U)]|TCAGTGCATCACAGAACTTTGTT</td><td>>MI0000811|hsa-mir-148b&WithFlank&12|+|54337210|54337320</td><td>69</td><td>90</td><td>22</td><td>(+1U)</td><td>isomiR</td><td>NO</td><td>158.32</td><td>174.69</td><td>268.78</td><td>76.05</td></tr>
<tr><td>>MI0000811|hsa-mir-148b&WithFlank&12|+|54337210|54337320@69.90.22(+2U)&[MIMAT0000759&hsa-miR-148b-3p&offsets|0|0(+2U);m-77&12|+|54337278|54337299&offsets|0|0(+2U)]|TCAGTGCATCACAGAACTTTGTTT</td><td>>MI0000811|hsa-mir-148b&WithFlank&12|+|54337210|54337320</td><td>69</td><td>90</td><td>22</td><td>(+2U)</td><td>isomiR</td><td>NO</td><td>0.00</td><td>0.00</td><td>68.56</td><td>0.00</td></tr>
<tr><td>>MI0000811|hsa-mir-148b&WithFlank&12|+|54337210|54337320@69.90.22(+3U)&[MIMAT0000759&hsa-miR-148b-3p&offsets|0|0(+3U);m-77&12|+|54337278|54337299&offsets|0|0(+3U)]|TCAGTGCATCACAGAACTTTGTTTT</td><td>>MI0000811|hsa-mir-148b&WithFlank&12|+|54337210|54337320</td><td>69</td><td>90</td><td>22</td><td>(+3U)</td><td>isomiR</td><td>NO</td><td>0.00</td><td>0.00</td><td>13.98</td><td>0.00</td></tr>
<tr><td>>MI0000811|hsa-mir-148b&WithFlank&12|+|54337210|54337320@69.91.23&[MIMAT0000759&hsa-miR-148b-3p&offsets|0|+1;m-77&12|+|54337278|54337299&offsets|0|+1]|TCAGTGCATCACAGAACTTTGTC</td><td>>MI0000811|hsa-mir-148b&WithFlank&12|+|54337210|54337320</td><td>69</td><td>91</td><td>23</td><td></td><td>isomiR</td><td>NO</td><td>0.00</td><td>0.00</td><td>59.59</td><td>0.00</td></tr>
<tr><td>>MI0000812|hsa-mir-331&WithFlank&12|+|95308414|95308519@67.87.21(+1U)&[MIMAT0000760&hsa-miR-331-3p&offsets|0|0(+1U);m-192&12|+|95308480|95308501&offsets|0|-1(+1U)]|GCCCCTGGGCCTATCCTAGAAT</td><td>>MI0000812|hsa-mir-331&WithFlank&12|+|95308414|95308519</td><td>67</td><td>87</td><td>21</td><td>(+1U)</td><td>isomiR</td><td>NO</td><td>50.75</td><td>0.00</td><td>0.00</td><td>17.22</td></tr>
<tr><td>>MI0000815|hsa-mir-339&WithFlank&7|-|1022927|1023032@21.43.23&[MIMAT0000764&hsa-miR-339-5p&offsets|0|0;m-163&7|-|1022990|1023012&offsets|0|0]|TCCCTGTCCTCCAGGAGCTCACG</td><td>>MI0000815|hsa-mir-339&WithFlank&7|-|1022927|1023032</td><td>21</td><td>43</td><td>23</td><td></td><td>isomiR</td><td>NO</td><td>0.00</td><td>13.15</td><td>0.00</td><td>12.23</td></tr>
<tr><td>>MI0000815|hsa-mir-339&WithFlank&7|-|1022927|1023032@56.76.21&[MIMAT0004702&hsa-miR-339-3p&offsets|0|-2;m-235&7|-|1022957|1022977&offsets|0|0]|TGAGCGCCTCGACGACAGAGC</td><td>>MI0000815|hsa-mir-339&WithFlank&7|-|1022927|1023032</td><td>56</td><td>76</td><td>21</td><td></td><td>isomiR</td><td>NO</td><td>0.00</td><td>0.00</td><td>34.88</td><td>0.00</td></tr>
<tr><td>>MI0000816|hsa-mir-335&WithFlank&7|+|130496105|130496210@22.43.22&[MIMAT0000765&hsa-miR-335-5p&offsets|0|-1;m-249&7|+|130496126|130496147&offsets|0|0]|TCAAGAGCAATAACGAAAAATG</td><td>>MI0000816|hsa-mir-335&WithFlank&7|+|130496105|130496210</td><td>22</td><td>43</td><td>22</td><td></td><td>isomiR</td><td>NO</td><td>0.00</td><td>0.00</td><td>35.20</td><td>0.00</td></tr>
<tr><td>>MI0000816|hsa-mir-335&WithFlank&7|+|130496105|130496210@58.78.21&[MIMAT0004703&hsa-miR-335-3p&offsets|0|-1;m-159&7|+|130496162|130496183&offsets|0|-1]|TTTTTCATTATTGCTCCTGAC</td><td>>MI0000816|hsa-mir-335&WithFlank&7|+|130496105|130496210</td><td>58</td><td>78</td><td>21</td><td></td><td>isomiR</td><td>NO</td><td>0.00</td><td>0.00</td><td>10.36</td><td>0.00</td></tr>
<tr><td>>MI0000816|hsa-mir-335&WithFlank&7|+|130496105|130496210@58.78.21(+1U)&[MIMAT0004703&hsa-miR-335-3p&offsets|0|-1(+1U);m-159&7|+|130496162|130496183&offsets|0|-1(+1U)]|TTTTTCATTATTGCTCCTGACT</td><td>>MI0000816|hsa-mir-335&WithFlank&7|+|130496105|130496210</td><td>58</td><td>78</td><td>21</td><td>(+1U)</td><td>isomiR</td><td>NO</td><td>24.73</td><td>66.91</td><td>17.20</td><td>16.48</td></tr>
<tr><td>>MI0000816|hsa-mir-335&WithFlank&7|+|130496105|130496210@58.79.22&[MIMAT0004703&hsa-miR-335-3p&offsets|0|0;m-159&7|+|130496162|130496183&offsets|0|0]|TTTTTCATTATTGCTCCTGACC</td><td>>MI0000816|hsa-mir-335&WithFlank&7|+|130496105|130496210</td><td>58</td><td>79</td><td>22</td><td></td><td>isomiR</td><td>NO</td><td>112.79</td><td>188.39</td><td>71.55</td><td>71.41</td></tr>
<tr><td>>MI0000825|hsa-mir-345&WithFlank&14|+|100307853|100307962@24.44.21&[MIMAT0000772&hsa-miR-345-5p&offsets|0|-1;m-193&14|+|100307876|100307896&offsets|0|0]|GCTGACTCCTAGTCCAGGGCT</td><td>>MI0000825|hsa-mir-345&WithFlank&14|+|100307853|100307962</td><td>24</td><td>44</td><td>21</td><td></td><td>isomiR</td><td>NO</td><td>0.00</td><td>0.00</td><td>17.88</td><td>0.00</td></tr>
<tr><td>>MI0000825|hsa-mir-345&WithFlank&14|+|100307853|100307962@24.45.22&[MIMAT0000772&hsa-miR-345-5p&offsets|0|0;m-193&14|+|100307876|100307896&offsets|0|+1]|GCTGACTCCTAGTCCAGGGCTC</td><td>>MI0000825|hsa-mir-345&WithFlank&14|+|100307853|100307962</td><td>24</td><td>45</td><td>22</td><td></td><td>isomiR</td><td>NO</td><td>0.00</td><td>0.00</td><td>83.90</td><td>0.00</td></tr>
<tr><td>>MI0000825|hsa-mir-345&WithFlank&14|+|100307853|100307962@24.45.22(+1U)&[MIMAT0000772&hsa-miR-345-5p&offsets|0|0(+1U);m-193&14|+|100307876|100307896&offsets|0|+1(+1U)]|GCTGACTCCTAGTCCAGGGCTCT</td><td>>MI0000825|hsa-mir-345&WithFlank&14|+|100307853|100307962</td><td>24</td><td>45</td><td>22</td><td>(+1U)</td><td>isomiR</td><td>NO</td><td>0.00</td><td>0.00</td><td>13.59</td><td>0.00</td></tr>
<tr><td>>MI0001445|hsa-mir-423&WithFlank&17|+|30117073|30117178@23.43.21&[MIMAT0004748&hsa-miR-423-5p&offsets|0|-2;m-141&17|+|30117095|30117117&offsets|0|-2]|TGAGGGGCAGAGAGCGAGACT</td><td>>MI0001445|hsa-mir-423&WithFlank&17|+|30117073|30117178</td><td>23</td><td>43</td><td>21</td><td></td><td>isomiR</td><td>NO</td><td>24.44</td><td>111.68</td><td>319.09</td><td>0.00</td></tr>
<tr><td>>MI0001445|hsa-mir-423&WithFlank&17|+|30117073|30117178@23.44.22&[MIMAT0004748&hsa-miR-423-5p&offsets|0|-1;m-141&17|+|30117095|30117117&offsets|0|-1]|TGAGGGGCAGAGAGCGAGACTT</td><td>>MI0001445|hsa-mir-423&WithFlank&17|+|30117073|30117178</td><td>23</td><td>44</td><td>22</td><td></td><td>isomiR</td><td>NO</td><td>0.00</td><td>68.88</td><td>214.78</td><td>0.00</td></tr>
</table>

### MDA-MB-231

<table>
<tr><th></th><th></th><th></th><th></th><th></th><th></th><th></th><th></th><th></th><th></th><th></th><th></th></tr>
<tr><td>>MI0001445|hsa-mir-423&WithFlank&17|+|30117073|30117178@23.44.22(+1C)&[MIMAT0004748&hsa-miR-423-5p&offsets|0|-1(+1C);m-141&17|+|30117095|30117117&offsets|0|-1(+1C)]|TGAGGGGCAGAGAGCGAGACTTC</td><td>>MI0001445|hsa-mir-423&WithFlank&17|+|30117073|30117178</td><td>23</td><td>44</td><td>22</td><td>(+1C)</td><td>isomiR</td><td>NO</td><td>0.00</td><td>26.25</td><td>247.71</td><td>0.00</td></tr>
<tr><td>>MI0001445|hsa-mir-423&WithFlank&17|+|30117073|30117178@23.45.23&[MIMAT0004748&hsa-miR-423-5p&offsets|0|0;m-141&17|+|30117095|30117117&offsets|0|0]|TGAGGGGCAGAGAGCGAGACTTT</td><td>>MI0001445|hsa-mir-423&WithFlank&17|+|30117073|30117178</td><td>23</td><td>45</td><td>23</td><td></td><td>isomiR</td><td>NO</td><td>122.62</td><td>313.76</td><td>1262.84</td><td>0.00</td></tr>
<tr><td>>MI0001445|hsa-mir-423&WithFlank&17|+|30117073|30117178@23.46.24&[MIMAT0004748&hsa-miR-423-5p&offsets|0|+1;m-141&17|+|30117095|30117117&offsets|0|+1]|TGAGGGGCAGAGAGCGAGACTTTT</td><td>>MI0001445|hsa-mir-423&WithFlank&17|+|30117073|30117178</td><td>23</td><td>46</td><td>24</td><td></td><td>isomiR</td><td>NO</td><td>10.59</td><td>88.87</td><td>272.47</td><td>0.00</td></tr>
<tr><td>>MI0001445|hsa-mir-423&WithFlank&17|+|30117073|30117178@58.78.21&[MIMAT0001340&hsa-miR-423-3p&offsets|-1|-3;m-100&17|+|30117131|30117153&offsets|-1|-3]|AAGCTCGGTCTGAGGCCCCTC</td><td>>MI0001445|hsa-mir-423&WithFlank&17|+|30117073|30117178</td><td>58</td><td>78</td><td>21</td><td></td><td>isomiR</td><td>NO</td><td>26.27</td><td>25.25</td><td>10.95</td><td>11.55</td></tr>
<tr><td>>MI0001445|hsa-mir-423&WithFlank&17|+|30117073|30117178@58.81.24&[MIMAT0001340&hsa-miR-423-3p&offsets|-1|0;m-100&17|+|30117131|30117153&offsets|-1|0]|AAGCTCGGTCTGAGGCCCCTCAGT</td><td>>MI0001445|hsa-mir-423&WithFlank&17|+|30117073|30117178</td><td>58</td><td>81</td><td>24</td><td></td><td>isomiR</td><td>NO</td><td>92.24</td><td>94.12</td><td>86.69</td><td>85.73</td></tr>
<tr><td>>MI0001445|hsa-mir-423&WithFlank&17|+|30117073|30117178@59.77.19(+1U)&[MIMAT0001340&hsa-miR-423-3p&offsets|0|-4(+1U);m-100&17|+|30117131|30117153&offsets|0|-4(+1U)]|AGCTCGGTCTGAGGCCCCTT</td><td>>MI0001445|hsa-mir-423&WithFlank&17|+|30117073|30117178</td><td>59</td><td>77</td><td>19</td><td>(+1U)</td><td>isomiR</td><td>NO</td><td>0.00</td><td>0.00</td><td>11.30</td><td>0.00</td></tr>
<tr><td>>MI0001445|hsa-mir-423&WithFlank&17|+|30117073|30117178@59.78.20&[MIMAT0001340&hsa-miR-423-3p&offsets|0|-3;m-100&17|+|30117131|30117153&offsets|0|-3]|AGCTCGGTCTGAGGCCCCTC</td><td>>MI0001445|hsa-mir-423&WithFlank&17|+|30117073|30117178</td><td>59</td><td>78</td><td>20</td><td></td><td>isomiR</td><td>NO</td><td>156.75</td><td>140.32</td><td>99.34</td><td>75.34</td></tr>
<tr><td>>MI0001445|hsa-mir-423&WithFlank&17|+|30117073|30117178@59.79.21&[MIMAT0001340&hsa-miR-423-3p&offsets|0|-2;m-100&17|+|30117131|30117153&offsets|0|-2]|AGCTCGGTCTGAGGCCCCTCA</td><td>>MI0001445|hsa-mir-423&WithFlank&17|+|30117073|30117178</td><td>59</td><td>79</td><td>21</td><td></td><td>isomiR</td><td>NO</td><td>0.00</td><td>13.16</td><td>10.47</td><td>10.44</td></tr>
<tr><td>>MI0001445|hsa-mir-423&WithFlank&17|+|30117073|30117178@59.80.22&[MIMAT0001340&hsa-miR-423-3p&offsets|0|-1;m-100&17|+|30117131|30117153&offsets|0|-1]|AGCTCGGTCTGAGGCCCCTCAG</td><td>>MI0001445|hsa-mir-423&WithFlank&17|+|30117073|30117178</td><td>59</td><td>80</td><td>22</td><td></td><td>isomiR</td><td>NO</td><td>62.27</td><td>60.62</td><td>58.40</td><td>17.57</td></tr>
<tr><td>>MI0001445|hsa-mir-423&WithFlank&17|+|30117073|30117178@59.80.22(+1A)&[MIMAT0001340&hsa-miR-423-3p&offsets|0|-1(+1A);m-100&17|+|30117131|30117153&offsets|0|-1(+1A)]|AGCTCGGTCTGAGGCCCCTCAGA</td><td>>MI0001445|hsa-mir-423&WithFlank&17|+|30117073|30117178</td><td>59</td><td>80</td><td>22</td><td>(+1A)</td><td>isomiR</td><td>NO</td><td>0.00</td><td>0.00</td><td>16.46</td><td>11.75</td></tr>
<tr><td>>MI0001445|hsa-mir-423&WithFlank&17|+|30117073|30117178@59.80.22(+1C)&[MIMAT0001340&hsa-miR-423-3p&offsets|0|-1(+1C);m-100&17|+|30117131|30117153&offsets|0|-1(+1C)]|AGCTCGGTCTGAGGCCCCTCAGC</td><td>>MI0001445|hsa-mir-423&WithFlank&17|+|30117073|30117178</td><td>59</td><td>80</td><td>22</td><td>(+1C)</td><td>isomiR</td><td>NO</td><td>77.59</td><td>72.12</td><td>105.20</td><td>71.91</td></tr>
<tr><td>>MI0001445|hsa-mir-423&WithFlank&17|+|30117073|30117178@59.81.23&[MIMAT0001340&hsa-miR-423-3p&offsets|0|0;m-100&17|+|30117131|30117153&offsets|0|0]|AGCTCGGTCTGAGGCCCCTCAGT</td><td>>MI0001445|hsa-mir-423&WithFlank&17|+|30117073|30117178</td><td>59</td><td>81</td><td>23</td><td></td><td>isomiR</td><td>NO</td><td>444.57</td><td>501.20</td><td>493.73</td><td>272.42</td></tr>
<tr><td>>MI0001445|hsa-mir-423&WithFlank&17|+|30117073|30117178@59.81.23(+1A)&[MIMAT0001340&hsa-miR-423-3p&offsets|0|0(+1A);m-100&17|+|30117131|30117153&offsets|0|0(+1A)]|AGCTCGGTCTGAGGCCCCTCAGTA</td><td>>MI0001445|hsa-mir-423&WithFlank&17|+|30117073|30117178</td><td>59</td><td>81</td><td>23</td><td>(+1A)</td><td>isomiR</td><td>NO</td><td>24.66</td><td>0.00</td><td>0.00</td><td>15.51</td></tr>
<tr><td>>MI0001445|hsa-mir-423&WithFlank&17|+|30117073|30117178@59.81.23(+1U)&[MIMAT0001340&hsa-miR-423-3p&offsets|0|0(+1U);m-100&17|+|30117131|30117153&offsets|0|0(+1U)]|AGCTCGGTCTGAGGCCCCTCAGTT</td><td>>MI0001445|hsa-mir-423&WithFlank&17|+|30117073|30117178</td><td>59</td><td>81</td><td>23</td><td>(+1U)</td><td>isomiR</td><td>NO</td><td>73.80</td><td>60.77</td><td>12.41</td><td>64.88</td></tr>
<tr><td>>MI0001448|hsa-mir-425&WithFlank&3|-|49020142|49020240@20.41.22&[MIMAT0003393&hsa-miR-425-5p&offsets|0|-1;m-95&3|-|49020200|49020221&offsets|0|0]|AATGACACGATCACTCCCGTTG</td><td>>MI0001448|hsa-mir-425&WithFlank&3|-|49020142|49020240</td><td>20</td><td>41</td><td>22</td><td></td><td>isomiR</td><td>NO</td><td>0.00</td><td>0.00</td><td>12.42</td><td>0.00</td></tr>
<tr><td>>MI0001448|hsa-mir-425&WithFlank&3|-|49020142|49020240@20.43.24&[MIMAT0003393&hsa-miR-425-5p&offsets|0|+1;m-95&3|-|49020200|49020221&offsets|0|+2]|AATGACACGATCACTCCCGTTGAG</td><td>>MI0001448|hsa-mir-425&WithFlank&3|-|49020142|49020240</td><td>20</td><td>43</td><td>24</td><td></td><td>isomiR</td><td>NO</td><td>95.14</td><td>66.01</td><td>66.19</td><td>17.88</td></tr>
<tr><td>>MI0001448|hsa-mir-425&WithFlank&3|-|49020142|49020240@20.43.24(+1C)&[MIMAT0003393&hsa-miR-425-5p&offsets|0|+1(+1C);m-95&3|-|49020200|49020221&offsets|0|+2(+1C)]|AATGACACGATCACTCCCGTTGAGC</td><td>>MI0001448|hsa-mir-425&WithFlank&3|-|49020142|49020240</td><td>20</td><td>43</td><td>24</td><td>(+1C)</td><td>isomiR</td><td>NO</td><td>24.12</td><td>0.00</td><td>11.28</td><td>11.23</td></tr>
<tr><td>>MI0001448|hsa-mir-425&WithFlank&3|-|49020142|49020240@20.44.25&[MIMAT0003393&hsa-miR-425-5p&offsets|0|+2;m-95&3|-|49020200|49020221&offsets|0|+3]|AATGACACGATCACTCCCGTTGAGT</td><td>>MI0001448|hsa-mir-425&WithFlank&3|-|49020142|49020240</td><td>20</td><td>44</td><td>25</td><td></td><td>isomiR</td><td>NO</td><td>241.90</td><td>158.47</td><td>139.85</td><td>105.23</td></tr>
<tr><td>>MI0001448|hsa-mir-425&WithFlank&3|-|49020142|49020240@60.81.22&[MIMAT0001343&hsa-miR-425-3p&offsets|-1|-1;m-258&3|-|49020160|49020181&offsets|0|0]|CATCGGGAATGTCGTGTCCGCC</td><td>>MI0001448|hsa-mir-425&WithFlank&3|-|49020142|49020240</td><td>60</td><td>81</td><td>22</td><td></td><td>isomiR</td><td>NO</td><td>0.00</td><td>0.00</td><td>33.93</td><td>0.00</td></tr>
<tr><td>>MI0001733|hsa-mir-452&WithFlank&X|-|151959622|151959718@20.43.24&[MIMAT0001635&hsa-miR-452-5p&offsets|0|+2;m-188&X|-|151959677|151959699&offsets|0|+1]|AACTGTTTGCAGAGGAAACTGAGA</td><td>>MI0001733|hsa-mir-452&WithFlank&X|-|151959622|151959718</td><td>20</td><td>43</td><td>24</td><td></td><td>isomiR</td><td>NO</td><td>0.00</td><td>0.00</td><td>10.36</td><td>0.00</td></tr>
<tr><td>>MI0001733|hsa-mir-452&WithFlank&X|-|151959622|151959718@23.43.21&[MIMAT0001635&hsa-miR-452-5p&offsets|+3|+2;m-188&X|-|151959677|151959699&offsets|+3|+1]|TGTTTGCAGAGGAAACTGAGA</td><td>>MI0001733|hsa-mir-452&WithFlank&X|-|151959622|151959718</td><td>23</td><td>43</td><td>21</td><td></td><td>isomiR</td><td>NO</td><td>0.00</td><td>0.00</td><td>10.62</td><td>0.00</td></tr>
</table>

### MDA-MB-231

<table>
<tr><td>>MI0002468|hsa-mir-484&WithFlank&16|+|15643288|15643378@14.35.22&[MIMAT0002174&hsa-miR-484&offsets|0|0;m-127&16|+|15643301|15643321&offsets|0|+1]|TCAGGCTCAGTCCCCTCCCGAT</td><td>>MI0002468|hsa-mir-484&WithFlank&16|+|15643288|15643378</td><td>14</td><td>35</td><td>22</td><td></td><td>isomiR</td><td>NO</td><td>52.58</td><td>32.61</td><td>0.00</td><td>63.22</td></tr>
<tr><td>>MI0002470|hsa-mir-486-1&WithFlank&8|-|41660435|41660514@10.31.22(+1U);MI0023622|hsa-mir-486-2&WithFlank&8|+|41660438|41660513@7.28.22(+1U)&[MIMAT0002177&hsa-miR-486-5p&offsets|0|0(+1U);m-107&8|-|41660484|41660505&offsets|0|0(+1U)];[MIMAT0002177_1&hsa-miR-486-5p&offsets|0|0(+1U);TJU_CMC_MD2.ID03138.5p-miR&offsets|0|-2(+1U);m-106&8|+|41660444|41660465&offsets|0|0(+1U)]|TCCTGTACTGAGCTGCCCCGAGT</td><td>>MI0002470|hsa-mir-486-1&WithFlank&8|-|41660435|41660514</td><td>10</td><td>31</td><td>22</td><td>(+1U)</td><td>isomiR</td><td>NO</td><td>0.00</td><td>25.81</td><td>35.03</td><td>0.00</td></tr>
<tr><td>>MI0002470|hsa-mir-486-1&WithFlank&8|-|41660435|41660514@10.31.22;MI0023622|hsa-mir-486-2&WithFlank&8|+|41660438|41660513@7.28.22&[MIMAT0002177&hsa-miR-486-5p&offsets|0|0;m-107&8|-|41660484|41660505&offsets|0|0];[MIMAT0002177_1&hsa-miR-486-5p&offsets|0|0;TJU_CMC_MD2.ID03138.5p-miR&offsets|0|-2;m-106&8|+|41660444|41660465&offsets|0|0]|TCCTGTACTGAGCTGCCCCGAG</td><td>>MI0002470|hsa-mir-486-1&WithFlank&8|-|41660435|41660514</td><td>10</td><td>31</td><td>22</td><td></td><td>isomiR</td><td>NO</td><td>0.00</td><td>0.00</td><td>12.86</td><td>0.00</td></tr>
<tr><td>>MI0003129|hsa-mir-146b&WithFlank&10|+|102436506|102436590@15.38.24&[MIMAT0002809&hsa-miR-146b-5p&offsets|0|+1;m-70&10|+|102436520|102436543&offsets|0|0]|TGAGAACTGAATTCCATAGGCTGT</td><td>>MI0003129|hsa-mir-146b&WithFlank&10|+|102436506|102436590</td><td>15</td><td>38</td><td>24</td><td></td><td>isomiR</td><td>NO</td><td>0.00</td><td>0.00</td><td>99.27</td><td>0.00</td></tr>
<tr><td>>MI0003137|hsa-mir-193b&WithFlank&16|+|14303961|14304055@57.78.22&[MIMAT0002819&hsa-miR-193b-3p&offsets|0|0;m-166&16|+|14304017|14304038&offsets|0|0]|AACTGGCCCTCAAAGTCCCGCT</td><td>>MI0003137|hsa-mir-193b&WithFlank&16|+|14303961|14304055</td><td>57</td><td>78</td><td>22</td><td></td><td>isomiR</td><td>NO</td><td>0.00</td><td>0.00</td><td>81.62</td><td>0.00</td></tr>
<tr><td>>MI0003185|hsa-mir-501&WithFlank&X|+|50009716|50009811@57.78.22&[MIMAT0004774&hsa-miR-501-3p&offsets|0|0;m-168&X|+|50009772|50009793&offsets|0|0]|AATGCACCCGGGCAAGGATTCT</td><td>>MI0003185|hsa-mir-501&WithFlank&X|+|50009716|50009811</td><td>57</td><td>78</td><td>22</td><td></td><td>isomiR</td><td>NO</td><td>0.00</td><td>0.00</td><td>54.68</td><td>0.00</td></tr>
<tr><td>>MI0003188|hsa-mir-503&WithFlank&X|-|134546322|134546404@12.31.20&[MIMAT0002874&hsa-miR-503-5p&offsets|0|-3;m-297&X|-|134546371|134546393&offsets|0|-3]|TAGCAGCGGGAACAGTTCTG</td><td>>MI0003188|hsa-mir-503&WithFlank&X|-|134546322|134546404</td><td>12</td><td>31</td><td>20</td><td></td><td>isomiR</td><td>NO</td><td>0.00</td><td>0.00</td><td>58.03</td><td>0.00</td></tr>
<tr><td>>MI0003188|hsa-mir-503&WithFlank&X|-|134546322|134546404@12.31.20(+1A)&[MIMAT0002874&hsa-miR-503-5p&offsets|0|-3(+1A);m-297&X|-|134546371|134546393&offsets|0|-3(+1A)]|TAGCAGCGGGAACAGTTCTGA</td><td>>MI0003188|hsa-mir-503&WithFlank&X|-|134546322|134546404</td><td>12</td><td>31</td><td>20</td><td>(+1A)</td><td>isomiR</td><td>NO</td><td>0.00</td><td>0.00</td><td>12.85</td><td>0.00</td></tr>
<tr><td>>MI0003205|hsa-mir-532&WithFlank&X|+|50003142|50003244@26.46.21(+1C)&[MIMAT0002888&hsa-miR-532-5p&offsets|0|-1(+1C);m-65&X|+|50003167|50003188&offsets|0|-1(+1C)]|CATGCCTTGAGTGTAGGACCGC</td><td>>MI0003205|hsa-mir-532&WithFlank&X|+|50003142|50003244</td><td>26</td><td>46</td><td>21</td><td>(+1C)</td><td>isomiR</td><td>NO</td><td>0.00</td><td>0.00</td><td>73.82</td><td>0.00</td></tr>
<tr><td>>MI0003205|hsa-mir-532&WithFlank&X|+|50003142|50003244@26.47.22&[MIMAT0002888&hsa-miR-532-5p&offsets|0|0;m-65&X|+|50003167|50003188&offsets|0|0]|CATGCCTTGAGTGTAGGACCGT</td><td>>MI0003205|hsa-mir-532&WithFlank&X|+|50003142|50003244</td><td>26</td><td>47</td><td>22</td><td></td><td>isomiR</td><td>NO</td><td>142.47</td><td>81.29</td><td>341.04</td><td>17.11</td></tr>
<tr><td>>MI0003205|hsa-mir-532&WithFlank&X|+|50003142|50003244@26.48.23&[MIMAT0002888&hsa-miR-532-5p&offsets|0|+1;m-65&X|+|50003167|50003188&offsets|0|+1]|CATGCCTTGAGTGTAGGACCGTT</td><td>>MI0003205|hsa-mir-532&WithFlank&X|+|50003142|50003244</td><td>26</td><td>48</td><td>23</td><td></td><td>isomiR</td><td>NO</td><td>0.00</td><td>0.00</td><td>11.84</td><td>0.00</td></tr>
<tr><td>>MI0003205|hsa-mir-532&WithFlank&X|+|50003142|50003244@27.47.21&[MIMAT0002888&hsa-miR-532-5p&offsets|+1|0;m-65&X|+|50003167|50003188&offsets|+1|0]|ATGCCTTGAGTGTAGGACCGT</td><td>>MI0003205|hsa-mir-532&WithFlank&X|+|50003142|50003244</td><td>27</td><td>47</td><td>21</td><td></td><td>isomiR</td><td>NO</td><td>0.00</td><td>0.00</td><td>11.98</td><td>0.00</td></tr>
<tr><td>>MI0003513|hsa-mir-455&WithFlank&9|+|114209428|114209535@22.43.22&[MIMAT0003150&hsa-miR-455-5p&offsets|0|0;m-232&9|+|114209449|114209470&offsets|0|0]|TATGTGCCTTTGGACTACATCG</td><td>>MI0003513|hsa-mir-455&WithFlank&9|+|114209428|114209535</td><td>22</td><td>43</td><td>22</td><td></td><td>isomiR</td><td>NO</td><td>0.00</td><td>0.00</td><td>64.09</td><td>0.00</td></tr>
<tr><td>>MI0003513|hsa-mir-455&WithFlank&9|+|114209428|114209535@22.43.22(+1C)&[MIMAT0003150&hsa-miR-455-5p&offsets|0|0(+1C);m-232&9|+|114209449|114209470&offsets|0|0(+1C)]|TATGTGCCTTTGGACTACATCGC</td><td>>MI0003513|hsa-mir-455&WithFlank&9|+|114209428|114209535</td><td>22</td><td>43</td><td>22</td><td>(+1C)</td><td>isomiR</td><td>NO</td><td>0.00</td><td>0.00</td><td>13.63</td><td>0.00</td></tr>
<tr><td>>MI0003513|hsa-mir-455&WithFlank&9|+|114209428|114209535@22.44.23&[MIMAT0003150&hsa-miR-455-5p&offsets|0|+1;m-232&9|+|114209449|114209470&offsets|0|+1]|TATGTGCCTTTGGACTACATCGT</td><td>>MI0003513|hsa-mir-455&WithFlank&9|+|114209428|114209535</td><td>22</td><td>44</td><td>23</td><td></td><td>isomiR</td><td>NO</td><td>69.06</td><td>0.00</td><td>167.52</td><td>15.11</td></tr>
<tr><td>>MI0003560|hsa-mir-92b&WithFlank&1|+|155195171|155195278@67.86.20&[MIMAT0003218&hsa-miR-92b-3p&offsets|0|-2;m-115&1|+|155195237|155195258&offsets|0|-2]|TATTGCACTCGTCCCGGCCT</td><td>>MI0003560|hsa-mir-92b&WithFlank&1|+|155195171|155195278</td><td>67</td><td>86</td><td>20</td><td></td><td>isomiR</td><td>NO</td><td>0.00</td><td>33.65</td><td>17.41</td><td>0.00</td></tr>
<tr><td>>MI0003560|hsa-mir-92b&WithFlank&1|+|155195171|155195278@67.86.20(+1U)&[MIMAT0003218&hsa-miR-92b-3p&offsets|0|-2(+1U);m-115&1|+|155195237|155195258&offsets|0|-2(+1U)]|TATTGCACTCGTCCCGGCCTT</td><td>>MI0003560|hsa-mir-92b&WithFlank&1|+|155195171|155195278</td><td>67</td><td>86</td><td>20</td><td>(+1U)</td><td>isomiR</td><td>NO</td><td>0.00</td><td>0.00</td><td>17.07</td><td>0.00</td></tr>
<tr><td>>MI0003560|hsa-mir-92b&WithFlank&1|+|155195171|155195278@67.87.21&[MIMAT0003218&hsa-miR-92b-3p&offsets|0|-1;m-115&1|+|155195237|155195258&offsets|0|-1]|TATTGCACTCGTCCCGGCCTC</td><td>>MI0003560|hsa-mir-92b&WithFlank&1|+|155195171|155195278</td><td>67</td><td>87</td><td>21</td><td></td><td>isomiR</td><td>NO</td><td>26.90</td><td>86.00</td><td>78.63</td><td>0.00</td></tr>
<tr><td>>MI0003560|hsa-mir-92b&WithFlank&1|+|155195171|155195278@67.87.21(+1A)&[MIMAT0003218&hsa-miR-92b-3p&offsets|0|-1(+1A);m-115&1|+|155195237|155195258&offsets|0|-1(+1A)]|TATTGCACTCGTCCCGGCCTCA</td><td>>MI0003560|hsa-mir-92b&WithFlank&1|+|155195171|155195278</td><td>67</td><td>87</td><td>21</td><td>(+1A)</td><td>isomiR</td><td>NO</td><td>0.00</td><td>63.12</td><td>64.82</td><td>0.00</td></tr>
<tr><td>>MI0003560|hsa-mir-92b&WithFlank&1|+|155195171|155195278@67.87.21(+1U)&[MIMAT0003218&hsa-miR-92b-3p&offsets|0|-1(+1U);m-115&1|+|155195237|155195258&offsets|0|-1(+1U)]|TATTGCACTCGTCCCGGCCTCT</td><td>>MI0003560|hsa-mir-92b&WithFlank&1|+|155195171|155195278</td><td>67</td><td>87</td><td>21</td><td>(+1U)</td><td>isomiR</td><td>NO</td><td>72.79</td><td>140.28</td><td>158.11</td><td>18.03</td></tr>
</table>

### MDA-MB-231

| | | | | | | | | | | | |
|---|---|---|---|---|---|---|---|---|---|---|---|
| >MI0003560\|hsa-mir-92b&WithFlank&1\|+\|155195171\|155195278@67.88.22&[MIMAT0003218&hsa-miR-92b-3p&offsets\|0\|0;m-115&1\|+\|155195237\|155195258&offsets\|0\|0]\|TATTGCACTCGTCCCGGCCTCC | >MI0003560\|hsa-mir-92b&WithFlank&1\|+\|155195171\|155195278 | 67 | 88 | 22 | | isomiR | NO | 125.63 | 232.00 | 204.39 | 62.07 |
| >MI0003560\|hsa-mir-92b&WithFlank&1\|+\|155195171\|155195278@67.88.22(+1U)&[MIMAT0003218&hsa-miR-92b-3p&offsets\|0\|0(+1U);m-115&1\|+\|155195237\|155195258&offsets\|0\|0(+1U)]\|TATTGCACTCGTCCCGGCCTCCT | >MI0003560\|hsa-mir-92b&WithFlank&1\|+\|155195171\|155195278 | 67 | 88 | 22 | (+1U) | isomiR | NO | 0.00 | 30.13 | 11.28 | 0.00 |
| >MI0003581\|hsa-mir-574&WithFlank&4\|+\|38868026\|38868133@67.87.21&[MIMAT0003239&hsa-miR-574-3p&offsets\|0\|-1;m-149&4\|+\|38868092\|38868113&offsets\|0\|-1]\|CACGCTCATGCACACACCCAC | >MI0003581\|hsa-mir-574&WithFlank&4\|+\|38868026\|38868133 | 67 | 87 | 21 | | isomiR | NO | 0.00 | 0.00 | 34.49 | 0.00 |
| >MI0003581\|hsa-mir-574&WithFlank&4\|+\|38868026\|38868133@67.88.22&[MIMAT0003239&hsa-miR-574-3p&offsets\|0\|0;m-149&4\|+\|38868092\|38868113&offsets\|0\|0]\|CACGCTCATGCACACACCCACA | >MI0003581\|hsa-mir-574&WithFlank&4\|+\|38868026\|38868133 | 67 | 88 | 22 | | isomiR | NO | 26.05 | 183.84 | 92.20 | 13.52 |
| >am_MI0003581\|hsa-mir-574&WithFlank&4\|+\|38868026\|38868133@31.51.21&[MIMAT0004795&hsa-miR-574-5p&offsets\|0\|-2;m-213&4\|+\|38868056\|38868078&offsets\|0\|-2]\|TGAGTGTGTGTGTGTGAGTGT | >MI0003581\|hsa-mir-574&WithFlank&4\|+\|38868026\|38868133.am | 31 | 51 | 21 | | isomiR | NO | 0.00 | 0.00 | 75.45 | 0.00 |
| >am_MI0003581\|hsa-mir-574&WithFlank&4\|+\|38868026\|38868133@31.52.22&[MIMAT0004795&hsa-miR-574-5p&offsets\|0\|-1;m-213&4\|+\|38868056\|38868078&offsets\|0\|-1]\|TGAGTGTGTGTGTGTGAGTGTG | >MI0003581\|hsa-mir-574&WithFlank&4\|+\|38868026\|38868133.am | 31 | 52 | 22 | | isomiR | NO | 0.00 | 0.00 | 34.00 | 0.00 |
| >am_MI0003581\|hsa-mir-574&WithFlank&4\|+\|38868026\|38868133@31.53.23&[MIMAT0004795&hsa-miR-574-5p&offsets\|0\|0;m-213&4\|+\|38868056\|38868078&offsets\|0\|0]\|TGAGTGTGTGTGTGTGAGTGTGT | >MI0003581\|hsa-mir-574&WithFlank&4\|+\|38868026\|38868133.am | 31 | 53 | 23 | | isomiR | NO | 10.65 | 28.41 | 213.96 | 0.00 |
| >am_MI0003581\|hsa-mir-574&WithFlank&4\|+\|38868026\|38868133@31.54.24&[MIMAT0004795&hsa-miR-574-5p&offsets\|0\|+1;m-213&4\|+\|38868056\|38868078&offsets\|0\|+1]\|TGAGTGTGTGTGTGTGAGTGTGTG | >MI0003581\|hsa-mir-574&WithFlank&4\|+\|38868026\|38868133.am | 31 | 54 | 24 | | isomiR | NO | 0.00 | 0.00 | 83.07 | 0.00 |
| >am_MI0003581\|hsa-mir-574&WithFlank&4\|+\|38868026\|38868133@31.55.25&[MIMAT0004795&hsa-miR-574-5p&offsets\|0\|+2;m-213&4\|+\|38868056\|38868078&offsets\|0\|+2]\|TGAGTGTGTGTGTGTGAGTGTGTGT | >MI0003581\|hsa-mir-574&WithFlank&4\|+\|38868026\|38868133.am | 31 | 55 | 25 | | isomiR | NO | 0.00 | 0.00 | 108.31 | 11.24 |
| >am_MI0003583\|hsa-mir-576&WithFlank&4\|+\|109488692\|109488801@61.83.23&[MIMAT0004796&hsa-miR-576-3p&offsets\|0\|+1;m-524&4\|+\|109488752\|109488772&offsets\|0\|+2]\|AAGATGTGGAAAAATTGGAATCC | >MI0003583\|hsa-mir-576&WithFlank&4\|+\|109488692\|109488801.an | 61 | 83 | 23 | | isomiR | NO | 0.00 | 0.00 | 98.76 | 0.00 |
| >MI0003591\|hsa-mir-584&WithFlank&5\|-\|149062307\|149062415@22.40.19&[MIMAT0003249&hsa-miR-584-5p&offsets\|0\|-3;m-167&5\|-\|149062374\|149062394&offsets\|0\|-2]\|TTATGGTTTGCCTGGGACT | >MI0003591\|hsa-mir-584&WithFlank&5\|-\|149062307\|149062415 | 22 | 40 | 19 | | isomiR | NO | 25.03 | 0.00 | 100.95 | 0.00 |
| >MI0003591\|hsa-mir-584&WithFlank&5\|-\|149062307\|149062415@22.41.20&[MIMAT0003249&hsa-miR-584-5p&offsets\|0\|-2;m-167&5\|-\|149062374\|149062394&offsets\|0\|-1]\|TTATGGTTTGCCTGGGACTG | >MI0003591\|hsa-mir-584&WithFlank&5\|-\|149062307\|149062415 | 22 | 41 | 20 | | isomiR | NO | 0.00 | 0.00 | 83.31 | 0.00 |
| >MI0003591\|hsa-mir-584&WithFlank&5\|-\|149062307\|149062415@22.41.20(+1U)&[MIMAT0003249&hsa-miR-584-5p&offsets\|0\|-2(+1U);m-167&5\|-\|149062374\|149062394&offsets\|0\|-1(+1U)]\|TTATGGTTTGCCTGGGACTGT | >MI0003591\|hsa-mir-584&WithFlank&5\|-\|149062307\|149062415 | 22 | 41 | 20 | (+1U) | isomiR | NO | 24.09 | 0.00 | 116.85 | 0.00 |
| >MI0003591\|hsa-mir-584&WithFlank&5\|-\|149062307\|149062415@22.42.21&[MIMAT0003249&hsa-miR-584-5p&offsets\|0\|-1;m-167&5\|-\|149062374\|149062394&offsets\|0\|0]\|TTATGGTTTGCCTGGGACTGA | >MI0003591\|hsa-mir-584&WithFlank&5\|-\|149062307\|149062415 | 22 | 42 | 21 | | isomiR | NO | 96.48 | 26.79 | 304.74 | 19.54 |
| >MI0003591\|hsa-mir-584&WithFlank&5\|-\|149062307\|149062415@22.42.21(+1A)&[MIMAT0003249&hsa-miR-584-5p&offsets\|0\|-1(+1A);m-167&5\|-\|149062374\|149062394&offsets\|0\|0(+1A)]\|TTATGGTTTGCCTGGGACTGAA | >MI0003591\|hsa-mir-584&WithFlank&5\|-\|149062307\|149062415 | 22 | 42 | 21 | (+1A) | isomiR | NO | 0.00 | 0.00 | 11.16 | 0.00 |
| >MI0003591\|hsa-mir-584&WithFlank&5\|-\|149062307\|149062415@22.42.21(+1U)&[MIMAT0003249&hsa-miR-584-5p&offsets\|0\|-1(+1U);m-167&5\|-\|149062374\|149062394&offsets\|0\|0(+1U)]\|TTATGGTTTGCCTGGGACTGAT | >MI0003591\|hsa-mir-584&WithFlank&5\|-\|149062307\|149062415 | 22 | 42 | 21 | (+1U) | isomiR | NO | 0.00 | 0.00 | 12.42 | 0.00 |
| >MI0003591\|hsa-mir-584&WithFlank&5\|-\|149062307\|149062415@22.42.21(+2U)&[MIMAT0003249&hsa-miR-584-5p&offsets\|0\|-1(+2U);m-167&5\|-\|149062374\|149062394&offsets\|0\|0(+2U)]\|TTATGGTTTGCCTGGGACTGATT | >MI0003591\|hsa-mir-584&WithFlank&5\|-\|149062307\|149062415 | 22 | 42 | 21 | (+2U) | isomiR | NO | 0.00 | 0.00 | 11.19 | 0.00 |
| >MI0003591\|hsa-mir-584&WithFlank&5\|-\|149062307\|149062415@22.43.22&[MIMAT0003249&hsa-miR-584-5p&offsets\|0\|0;m-167&5\|-\|149062374\|149062394&offsets\|0\|+1]\|TTATGGTTTGCCTGGGACTGAG | >MI0003591\|hsa-mir-584&WithFlank&5\|-\|149062307\|149062415 | 22 | 43 | 22 | | isomiR | NO | 0.00 | 0.00 | 16.23 | 0.00 |
| >MI0003591\|hsa-mir-584&WithFlank&5\|-\|149062307\|149062415@22.43.22(+1U)&[MIMAT0003249&hsa-miR-584-5p&offsets\|0\|0(+1U);m-167&5\|-\|149062374\|149062394&offsets\|0\|+1(+1U)]\|TTATGGTTTGCCTGGGACTGAGT | >MI0003591\|hsa-mir-584&WithFlank&5\|-\|149062307\|149062415 | 22 | 43 | 22 | (+1U) | isomiR | NO | 0.00 | 0.00 | 14.11 | 0.00 |
| >MI0003591\|hsa-mir-584&WithFlank&5\|-\|149062307\|149062415@22.44.23&[MIMAT0003249&hsa-miR-584-5p&offsets\|0\|+1;m-167&5\|-\|149062374\|149062394&offsets\|0\|+2]\|TTATGGTTTGCCTGGGACTGAGG | >MI0003591\|hsa-mir-584&WithFlank&5\|-\|149062307\|149062415 | 22 | 44 | 23 | | isomiR | NO | 0.00 | 0.00 | 13.94 | 0.00 |
| >MI0003599\|hsa-mir-589&WithFlank&7\|-\|5495813\|5495923@30.50.21&[MIMAT0004799&hsa-miR-589-5p&offsets\|0\|-1;m-162&7\|-\|5495874\|5495894&offsets\|0\|0]\|TGAGAACCACGTCTGCTCTGA | >MI0003599\|hsa-mir-589&WithFlank&7\|-\|5495813\|5495923 | 30 | 50 | 21 | | isomiR | NO | 10.45 | 0.00 | 70.04 | 0.00 |
| >MI0003599\|hsa-mir-589&WithFlank&7\|-\|5495813\|5495923@30.52.23&[MIMAT0004799&hsa-miR-589-5p&offsets\|0\|+1;m-162&7\|-\|5495874\|5495894&offsets\|0\|+2]\|TGAGAACCACGTCTGCTCTGAGC | >MI0003599\|hsa-mir-589&WithFlank&7\|-\|5495813\|5495923 | 30 | 52 | 23 | | isomiR | NO | 25.40 | 58.31 | 11.94 | 0.00 |
| >MI0003602\|hsa-mir-590&WithFlank&7\|+\|74191192\|74191300@62.82.21&[MIMAT0004801&hsa-miR-590-3p&offsets\|0\|0;m-332&7\|+\|74191253\|74191273&offsets\|0\|0]\|TAATTTTATGTATAAGCTAGT | >MI0003602\|hsa-mir-590&WithFlank&7\|+\|74191192\|74191300 | 62 | 82 | 21 | | isomiR | NO | 0.00 | 0.00 | 11.26 | 0.00 |

### MDA-MB-231

<table>
<tr><th></th><th></th><th></th><th></th><th></th><th></th><th></th><th></th><th></th><th></th><th></th><th></th></tr>
<tr><td>>am_MI0003639|hsa-mir-625&WithFlank&14|+|65471096|65471192@58.78.21(+1U)&[MIMAT0004808&hsa-miR-625-3p&offsets|0|-1(+1U);m-137&14|+|65471153|65471174&offsets|0|-1(+1U)]|GACTATAGAACTTTCCCCCTCT</td><td>>MI0003639|hsa-mir-625&WithFlank&14|+|65471096|65471192.am</td><td>58</td><td>78</td><td>21</td><td>(+1U)</td><td>isomiR</td><td>NO</td><td>0.00</td><td>0.00</td><td>13.79</td><td>0.00</td></tr>
<tr><td>>am_MI0003639|hsa-mir-625&WithFlank&14|+|65471096|65471192@58.79.22&[MIMAT0004808&hsa-miR-625-3p&offsets|0|0;m-137&14|+|65471153|65471174&offsets|0|0]|GACTATAGAACTTTCCCCCTCA</td><td>>MI0003639|hsa-mir-625&WithFlank&14|+|65471096|65471192.am</td><td>58</td><td>79</td><td>22</td><td></td><td>isomiR</td><td>NO</td><td>0.00</td><td>0.00</td><td>15.53</td><td>0.00</td></tr>
<tr><td>>am_MI0003639|hsa-mir-625&WithFlank&14|+|65471096|65471192@58.80.23&[MIMAT0004808&hsa-miR-625-3p&offsets|0|+1;m-137&14|+|65471153|65471174&offsets|0|+1]|GACTATAGAACTTTCCCCCTCAT</td><td>>MI0003639|hsa-mir-625&WithFlank&14|+|65471096|65471192.am</td><td>58</td><td>80</td><td>23</td><td></td><td>isomiR</td><td>NO</td><td>10.36</td><td>37.53</td><td>57.37</td><td>0.00</td></tr>
<tr><td>>MI0003643|hsa-mir-629&WithFlank&15|-|70079366|70079474@28.48.21&[MIMAT0004810&hsa-miR-629-5p&offsets|0|0;m-150&15|-|70079427|70079447&offsets|0|0]|TGGGTTTACGTTGGGAGAACT</td><td>>MI0003643|hsa-mir-629&WithFlank&15|-|70079366|70079474</td><td>28</td><td>48</td><td>21</td><td></td><td>isomiR</td><td>NO</td><td>0.00</td><td>0.00</td><td>171.10</td><td>0.00</td></tr>
<tr><td>>MI0003643|hsa-mir-629&WithFlank&15|-|70079366|70079474@28.49.22&[MIMAT0004810&hsa-miR-629-5p&offsets|0|+1;m-150&15|-|70079427|70079447&offsets|0|+1]|TGGGTTTACGTTGGGAGAACTT</td><td>>MI0003643|hsa-mir-629&WithFlank&15|-|70079366|70079474</td><td>28</td><td>49</td><td>22</td><td></td><td>isomiR</td><td>NO</td><td>0.00</td><td>31.83</td><td>368.27</td><td>0.00</td></tr>
<tr><td>>MI0003667|hsa-mir-652&WithFlank&X|+|110055323|110055432@67.87.21(+1U)&[MIMAT0003322&hsa-miR-652-3p&offsets|0|0(+1U);m-148&X|+|110055389|110055410&offsets|0|-1(+1U)]|AATGGCGCCACTAGGGTTGTGT</td><td>>MI0003667|hsa-mir-652&WithFlank&X|+|110055323|110055432</td><td>67</td><td>87</td><td>21</td><td>(+1U)</td><td>isomiR</td><td>NO</td><td>0.00</td><td>0.00</td><td>64.66</td><td>0.00</td></tr>
<tr><td>>MI0003684|hsa-mir-660&WithFlank&X|+|50013235|50013343@22.44.23&[MIMAT0003338&hsa-miR-660-5p&offsets|0|+1;m-157&X|+|50013256|50013278&offsets|0|0]|TACCCATTGCATATCGGAGTTGT</td><td>>MI0003684|hsa-mir-660&WithFlank&X|+|50013235|50013343</td><td>22</td><td>44</td><td>23</td><td></td><td>isomiR</td><td>NO</td><td>0.00</td><td>0.00</td><td>90.93</td><td>0.00</td></tr>
<tr><td>>MI0003760|hsa-mir-671&WithFlank&7|+|151238415|151238544@35.58.24&[MIMAT0003880&hsa-miR-671-5p&offsets|0|+1;m-316&7|+|151238449|151238472&offsets|0|0]|AGGAAGCCCTGGAGGGGCTGGAGG</td><td>>MI0003760|hsa-mir-671&WithFlank&7|+|151238415|151238544</td><td>35</td><td>58</td><td>24</td><td></td><td>isomiR</td><td>NO</td><td>0.00</td><td>0.00</td><td>13.54</td><td>0.00</td></tr>
<tr><td>>MI0003776|hsa-mir-320b-1&WithFlank&1|+|116671740|116671823@48.67.20;MI0003839|hsa-mir-320b-2&WithFlank&1|-|224257034|224257116@47.66.20;MI0000542|hsa-mir-320a&WithFlank&8|-|22244960|22245043@48.67.20&[MIMAT0005792&hsa-miR-320b&offsets|0|-2;m-183&1|+|116671787|116671806&offsets|0|0];[MIMAT0005792_1&hsa-miR-320b&offsets|0|-2;m-183&1|-|224257051|224257070&offsets|0|0];[MIMAT0000510&hsa-miR-320a-3p&offsets|0|-2;m-60&8|-|22244975|22244996&offsets|0|-2]|AAAAGCTGGGTTGAGAGGGC</td><td>>MI0003776|hsa-mir-320b-1&WithFlank&1|+|116671740|116671823</td><td>48</td><td>67</td><td>20</td><td></td><td>isomiR</td><td>NO</td><td>0.00</td><td>0.00</td><td>96.70</td><td>0.00</td></tr>
<tr><td>>MI0003776|hsa-mir-320b-1&WithFlank&1|+|116671740|116671823@48.68.21;MI0003839|hsa-mir-320b-2&WithFlank&1|-|224257034|224257116@47.67.21&[MIMAT0005792&hsa-miR-320b&offsets|0|-1;m-183&1|+|116671787|116671806&offsets|0|+1];[MIMAT0005792_1&hsa-miR-320b&offsets|0|-1;m-183&1|-|224257051|224257070&offsets|0|+1]|AAAAGCTGGGTTGAGAGGGCA</td><td>>MI0003776|hsa-mir-320b-1&WithFlank&1|+|116671740|116671823</td><td>48</td><td>68</td><td>21</td><td></td><td>isomiR</td><td>NO</td><td>0.00</td><td>0.00</td><td>60.53</td><td>0.00</td></tr>
<tr><td>>MI0003780|hsa-mir-1296&WithFlank&10|-|63372951|63373054@22.43.22&[MIMAT0005794&hsa-miR-1296-5p&offsets|0|0;m-304&10|-|63373012|63373033&offsets|0|0]|TTAGGGCCCTGGCTCCATCTCC</td><td>>MI0003780|hsa-mir-1296&WithFlank&10|-|63372951|63373054</td><td>22</td><td>43</td><td>22</td><td></td><td>isomiR</td><td>NO</td><td>0.00</td><td>0.00</td><td>17.06</td><td>0.00</td></tr>
<tr><td>>MI0003820|hsa-mir-454&WithFlank&17|-|59137752|59137878@70.92.23&[MIMAT0003885&hsa-miR-454-3p&offsets|0|0;m-272&17|-|59137787|59137809&offsets|0|0]|TAGTGCAATATTGCTTATAGGGT</td><td>>MI0003820|hsa-mir-454&WithFlank&17|-|59137752|59137878</td><td>70</td><td>92</td><td>23</td><td></td><td>isomiR</td><td>NO</td><td>0.00</td><td>0.00</td><td>78.47</td><td>0.00</td></tr>
<tr><td>>MI0003820|hsa-mir-454&WithFlank&17|-|59137752|59137878@70.93.24&[MIMAT0003885&hsa-miR-454-3p&offsets|0|+1;m-272&17|-|59137787|59137809&offsets|0|+1]|TAGTGCAATATTGCTTATAGGGTT</td><td>>MI0003820|hsa-mir-454&WithFlank&17|-|59137752|59137878</td><td>70</td><td>93</td><td>24</td><td></td><td>isomiR</td><td>NO</td><td>25.26</td><td>0.00</td><td>108.93</td><td>10.82</td></tr>
<tr><td>>MI0003820|hsa-mir-454&WithFlank&17|-|59137752|59137878@70.94.25&[MIMAT0003885&hsa-miR-454-3p&offsets|0|+2;m-272&17|-|59137787|59137809&offsets|0|+2]|TAGTGCAATATTGCTTATAGGGTTT</td><td>>MI0003820|hsa-mir-454&WithFlank&17|-|59137752|59137878</td><td>70</td><td>94</td><td>25</td><td></td><td>isomiR</td><td>NO</td><td>47.49</td><td>0.00</td><td>110.25</td><td>16.38</td></tr>
<tr><td>>MI0003834|hsa-mir-769&WithFlank&19|+|46018926|46019055@36.56.21&[MIMAT0003886&hsa-miR-769-5p&offsets|0|-1;m-216&19|+|46018961|46018982&offsets|0|-1]|TGAGACCTCTGGGTTCTGAGC</td><td>>MI0003834|hsa-mir-769&WithFlank&19|+|46018926|46019055</td><td>36</td><td>56</td><td>21</td><td></td><td>isomiR</td><td>NO</td><td>0.00</td><td>0.00</td><td>74.30</td><td>0.00</td></tr>
<tr><td>>MI0003834|hsa-mir-769&WithFlank&19|+|46018926|46019055@36.56.21(+1A)&[MIMAT0003886&hsa-miR-769-5p&offsets|0|-1(+1A);m-216&19|+|46018961|46018982&offsets|0|-1(+1A)]|TGAGACCTCTGGGTTCTGAGCA</td><td>>MI0003834|hsa-mir-769&WithFlank&19|+|46018926|46019055</td><td>36</td><td>56</td><td>21</td><td>(+1A)</td><td>isomiR</td><td>NO</td><td>0.00</td><td>0.00</td><td>12.38</td><td>0.00</td></tr>
<tr><td>>MI0003834|hsa-mir-769&WithFlank&19|+|46018926|46019055@36.56.21(+1C)&[MIMAT0003886&hsa-miR-769-5p&offsets|0|-1(+1C);m-216&19|+|46018961|46018982&offsets|0|-1(+1C)]|TGAGACCTCTGGGTTCTGAGCC</td><td>>MI0003834|hsa-mir-769&WithFlank&19|+|46018926|46019055</td><td>36</td><td>56</td><td>21</td><td>(+1C)</td><td>isomiR</td><td>NO</td><td>0.00</td><td>0.00</td><td>17.89</td><td>0.00</td></tr>
<tr><td>>MI0003834|hsa-mir-769&WithFlank&19|+|46018926|46019055@36.57.22&[MIMAT0003886&hsa-miR-769-5p&offsets|0|0;m-216&19|+|46018961|46018982&offsets|0|0]|TGAGACCTCTGGGTTCTGAGCT</td><td>>MI0003834|hsa-mir-769&WithFlank&19|+|46018926|46019055</td><td>36</td><td>57</td><td>22</td><td></td><td>isomiR</td><td>NO</td><td>50.59</td><td>13.67</td><td>198.00</td><td>11.18</td></tr>
<tr><td>>MI0003834|hsa-mir-769&WithFlank&19|+|46018926|46019055@36.57.22(+1U)&[MIMAT0003886&hsa-miR-769-5p&offsets|0|0(+1U);m-216&19|+|46018961|46018982&offsets|0|0(+1U)]|TGAGACCTCTGGGTTCTGAGCTT</td><td>>MI0003834|hsa-mir-769&WithFlank&19|+|46018926|46019055</td><td>36</td><td>57</td><td>22</td><td>(+1U)</td><td>isomiR</td><td>NO</td><td>0.00</td><td>0.00</td><td>62.60</td><td>0.00</td></tr>
<tr><td>>MI0005559|hsa-mir-744&WithFlank&17|+|12081893|12082002@17.37.21&[MIMAT0004945&hsa-miR-744-5p&offsets|0|-1;m-172&17|+|12081909|12081929&offsets|0|0]|TGCGGGGCTAGGGCTAACAGC</td><td>>MI0005559|hsa-mir-744&WithFlank&17|+|12081893|12082002</td><td>17</td><td>37</td><td>21</td><td></td><td>isomiR</td><td>NO</td><td>48.13</td><td>92.56</td><td>82.37</td><td>0.00</td></tr>
<tr><td>>MI0005559|hsa-mir-744&WithFlank&17|+|12081893|12082002@17.38.22&[MIMAT0004945&hsa-miR-744-5p&offsets|0|0;m-172&17|+|12081909|12081929&offsets|0|+1]|TGCGGGGCTAGGGCTAACAGCA</td><td>>MI0005559|hsa-mir-744&WithFlank&17|+|12081893|12082002</td><td>17</td><td>38</td><td>22</td><td></td><td>isomiR</td><td>NO</td><td>0.00</td><td>0.00</td><td>54.87</td><td>0.00</td></tr>
</table>

### MDA-MB-231

<table>
<tr><td></td><td></td><td></td><td></td><td></td><td></td><td></td><td></td><td></td><td></td><td></td><td></td></tr>
<tr><td>>MI0005566|hsa-mir-374b&WithFlank&X|-|74218541|74218624@17.37.21&[MIMAT0004955&hsa-miR-374b-5p&offsets|0|-1;m-153&X|-|74218587|74218608&offsets|0|-1]|ATATAATACAACCTGCTAAGT</td><td>>MI0005566|hsa-mir-374b&WithFlank&X|-|74218541|74218624</td><td>17</td><td>37</td><td>21</td><td></td><td>isomiR</td><td>NO</td><td>63.39</td><td>0.00</td><td>54.26</td><td>18.82</td></tr>
<tr><td>>MI0005566|hsa-mir-374b&WithFlank&X|-|74218541|74218624@17.38.22&[MIMAT0004955&hsa-miR-374b-5p&offsets|0|0;m-153&X|-|74218587|74218608&offsets|0|0]|ATATAATACAACCTGCTAAGTG</td><td>>MI0005566|hsa-mir-374b&WithFlank&X|-|74218541|74218624</td><td>17</td><td>38</td><td>22</td><td></td><td>isomiR</td><td>NO</td><td>71.48</td><td>0.00</td><td>109.23</td><td>56.39</td></tr>
<tr><td>>MI0005566|hsa-mir-374b&WithFlank&X|-|74218541|74218624@17.39.23&[MIMAT0004955&hsa-miR-374b-5p&offsets|0|+1;m-153&X|-|74218587|74218608&offsets|0|+1]|ATATAATACAACCTGCTAAGTGT</td><td>>MI0005566|hsa-mir-374b&WithFlank&X|-|74218541|74218624</td><td>17</td><td>39</td><td>23</td><td></td><td>isomiR</td><td>NO</td><td>0.00</td><td>0.00</td><td>63.69</td><td>0.00</td></tr>
<tr><td>>am_MI0005763|hsa-mir-941-1&WithFlank&20|+|63919443|63919526@53.74.22(+1A);MI0005764|hsa-mir-941-2&WithFlank&20|+|63919499|63919582@53.74.22(+1A);MI0005765|hsa-mir-941-3&WithFlank&20|+|63919555|63919638@53.74.22(+1A);MI0005766|hsa-mir-941-4&WithFlank&20|+|63919750|63919833@53.74.22(+1A);MI0031520|hsa-mir-941-5&WithFlank&20|+|63919862|63919945@53.74.22(+1A)&[MIMAT0004984&hsa-miR-941&offsets|0|-1(+1A);m-263&20|+|63919495|63919517&offsets|0|-1(+1A)];[MIMAT0004984_1&hsa-miR-941&offsets|0|-1(+1A);m-263&20|+|63919551|63919573&offsets|0|-1(+1A)];[MIMAT0004984_2&hsa-miR-941&offsets|0|-1(+1A);m-263&20|+|63919607|63919629&offsets|0|-1(+1A)];[MIMAT0004984_3&hsa-miR-941&offsets|0|-1(+1A);m-263&20|+|63919802|63919824&offsets|0|-1(+1A)];[MIMAT0004984_4&hsa-miR-941&offsets|0|-1(+1A);m-263&20|+|63919914|63919936&offsets|0|-1(+1A)]|CACCCGGCTGTGTGCACATGTGA</td><td>>MI0005763|hsa-mir-941-1&WithFlank&20|+|63919443|63919526.a</td><td>53</td><td>74</td><td>22</td><td>(+1A)</td><td>isomiR</td><td>NO</td><td>0.00</td><td>62.32</td><td>16.66</td><td>0.00</td></tr>
<tr><td>>am_MI0005763|hsa-mir-941-1&WithFlank&20|+|63919443|63919526@53.74.22(+1U);MI0005764|hsa-mir-941-2&WithFlank&20|+|63919499|63919582@53.74.22(+1U);MI0005765|hsa-mir-941-3&WithFlank&20|+|63919555|63919638@53.74.22(+1U);MI0005766|hsa-mir-941-4&WithFlank&20|+|63919750|63919833@53.74.22(+1U);MI0031520|hsa-mir-941-5&WithFlank&20|+|63919862|63919945@53.74.22(+1U)&[MIMAT0004984&hsa-miR-941&offsets|0|-1(+1U);m-263&20|+|63919495|63919517&offsets|0|-1(+1U)];[MIMAT0004984_1&hsa-miR-941&offsets|0|-1(+1U);m-263&20|+|63919551|63919573&offsets|0|-1(+1U)];[MIMAT0004984_2&hsa-miR-941&offsets|0|-1(+1U);m-263&20|+|63919607|63919629&offsets|0|-1(+1U)];[MIMAT0004984_3&hsa-miR-941&offsets|0|-1(+1U);m-263&20|+|63919802|63919824&offsets|0|-1(+1U)];[MIMAT0004984_4&hsa-miR-941&offsets|0|-1(+1U);m-263&20|+|63919914|63919936&offsets|0|-1(+1U)]|CACCCGGCTGTGTGCACATGTGT</td><td>>MI0005763|hsa-mir-941-1&WithFlank&20|+|63919443|63919526.a</td><td>53</td><td>74</td><td>22</td><td>(+1U)</td><td>isomiR</td><td>NO</td><td>113.57</td><td>195.15</td><td>137.80</td><td>56.84</td></tr>
<tr><td>>am_MI0005763|hsa-mir-941-1&WithFlank&20|+|63919443|63919526@53.74.22;MI0005764|hsa-mir-941-2&WithFlank&20|+|63919499|63919582@53.74.22;MI0005765|hsa-mir-941-3&WithFlank&20|+|63919555|63919638@53.74.22;MI0005766|hsa-mir-941-4&WithFlank&20|+|63919750|63919833@53.74.22;MI0031520|hsa-mir-941-5&WithFlank&20|+|63919862|63919945@53.74.22&[MIMAT0004984&hsa-miR-941&offsets|0|-1;m-263&20|+|63919495|63919517&offsets|0|-1];[MIMAT0004984_1&hsa-miR-941&offsets|0|-1;m-263&20|+|63919551|63919573&offsets|0|-1];[MIMAT0004984_2&hsa-miR-941&offsets|0|-1;m-263&20|+|63919607|63919629&offsets|0|-1];[MIMAT0004984_3&hsa-miR-941&offsets|0|-1;m-263&20|+|63919802|63919824&offsets|0|-1];[MIMAT0004984_4&hsa-miR-941&offsets|0|-1;m-263&20|+|63919914|63919936&offsets|0|-1]|CACCCGGCTGTGTGCACATGTG</td><td>>MI0005763|hsa-mir-941-1&WithFlank&20|+|63919443|63919526.a</td><td>53</td><td>74</td><td>22</td><td></td><td>isomiR</td><td>NO</td><td>0.00</td><td>24.91</td><td>81.76</td><td>0.00</td></tr>
<tr><td>>am_MI0005763|hsa-mir-941-1&WithFlank&20|+|63919443|63919526@53.75.23;MI0005764|hsa-mir-941-2&WithFlank&20|+|63919499|63919582@53.75.23;MI0005765|hsa-mir-941-3&WithFlank&20|+|63919555|63919638@53.75.23;MI0005766|hsa-mir-941-4&WithFlank&20|+|63919750|63919833@53.75.23;MI0031520|hsa-mir-941-5&WithFlank&20|+|63919862|63919945@53.75.23&[MIMAT0004984&hsa-miR-941&offsets|0|0;m-263&20|+|63919495|63919517&offsets|0|0];[MIMAT0004984_1&hsa-miR-941&offsets|0|0;m-263&20|+|63919551|63919573&offsets|0|0];[MIMAT0004984_2&hsa-miR-941&offsets|0|0;m-263&20|+|63919607|63919629&offsets|0|0];[MIMAT0004984_3&hsa-miR-941&offsets|0|0;m-263&20|+|63919802|63919824&offsets|0|0];[MIMAT0004984_4&hsa-miR-941&offsets|0|0;m-263&20|+|63919914|63919936&offsets|0|0]|CACCCGGCTGTGTGCACATGTGC</td><td>>MI0005763|hsa-mir-941-1&WithFlank&20|+|63919443|63919526.a</td><td>53</td><td>75</td><td>23</td><td></td><td>isomiR</td><td>NO</td><td>391.51</td><td>674.41</td><td>312.82</td><td>128.32</td></tr>
<tr><td>>MI0006273|hsa-mir-1180&WithFlank&17|-|19344500|19344580@47.68.22&[MIMAT0005825&hsa-miR-1180-3p&offsets|0|0;m-214&17|-|19344513|19344534&offsets|0|0]|TTTCCGGCTCGCGTGGGTGTGT</td><td>>MI0006273|hsa-mir-1180&WithFlank&17|-|19344500|19344580</td><td>47</td><td>68</td><td>22</td><td></td><td>isomiR</td><td>NO</td><td>0.00</td><td>24.90</td><td>61.29</td><td>0.00</td></tr>
<tr><td>>MI0006349|hsa-mir-1287&WithFlank&10|-|98395212|98395313@22.43.22&[MIMAT0005878&hsa-miR-1287-5p&offsets|0|0;m-244&10|-|98395272|98395292&offsets|0|+1]|TGCTGGATCAGTGGTTCGAGTC</td><td>>MI0006349|hsa-mir-1287&WithFlank&10|-|98395212|98395313</td><td>22</td><td>43</td><td>22</td><td></td><td>isomiR</td><td>NO</td><td>0.00</td><td>0.00</td><td>60.00</td><td>0.00</td></tr>
<tr><td>>MI0006444|hsa-mir-1307&WithFlank&10|-|103394247|103394407@86.106.21&[MIMAT0005951&hsa-miR-1307-3p&offsets|0|-1;m-66&10|-|103394300|103394322&offsets|0|-2]|ACTCGGCGTGGCGTCGGTCGT</td><td>>MI0006444|hsa-mir-1307&WithFlank&10|-|103394247|103394407</td><td>86</td><td>106</td><td>21</td><td></td><td>isomiR</td><td>NO</td><td>0.00</td><td>0.00</td><td>10.56</td><td>0.00</td></tr>
<tr><td>>MI0006444|hsa-mir-1307&WithFlank&10|-|103394247|103394407@86.107.22&[MIMAT0005951&hsa-miR-1307-3p&offsets|0|0;m-66&10|-|103394300|103394322&offsets|0|-1]|ACTCGGCGTGGCGTCGGTCGTG</td><td>>MI0006444|hsa-mir-1307&WithFlank&10|-|103394247|103394407</td><td>86</td><td>107</td><td>22</td><td></td><td>isomiR</td><td>NO</td><td>50.94</td><td>32.29</td><td>17.26</td><td>12.22</td></tr>
<tr><td>>MI0006444|hsa-mir-1307&WithFlank&10|-|103394247|103394407@86.107.22(+1A)&[MIMAT0005951&hsa-miR-1307-3p&offsets|0|0(+1A);m-66&10|-|103394300|103394322&offsets|0|-1(+1A)]|ACTCGGCGTGGCGTCGGTCGTGA</td><td>>MI0006444|hsa-mir-1307&WithFlank&10|-|103394247|103394407</td><td>86</td><td>107</td><td>22</td><td>(+1A)</td><td>isomiR</td><td>NO</td><td>0.00</td><td>29.62</td><td>10.73</td><td>0.00</td></tr>
</table>

### MDA-MB-231

<table>
<tr><th></th><th></th><th></th><th></th><th></th><th></th><th></th><th></th><th></th><th></th><th></th><th></th></tr>
<tr><td>>MI0006444|hsa-mir-1307&WithFlank&10|-|103394247|103394407@86.107.22(+1U)&[MIMAT0005951&hsa-miR-1307-3p&offsets|0|0(+1U);m-66&10|-|103394300|103394322&offsets|0|-1(+1U)]|ACTCGGCGTGGCGTCGGTCGTGT</td><td>>MI0006444|hsa-mir-1307&WithFlank&10|-|103394247|103394407</td><td>86</td><td>107</td><td>22</td><td>(+1U)</td><td>isomiR</td><td>NO</td><td>51.21</td><td>37.47</td><td>35.44</td><td>14.37</td></tr>
<tr><td>>MI0006444|hsa-mir-1307&WithFlank&10|-|103394247|103394407@86.108.23&[MIMAT0005951&hsa-miR-1307-3p&offsets|0|+1;m-66&10|-|103394300|103394322&offsets|0|0]|ACTCGGCGTGGCGTCGGTCGTGG</td><td>>MI0006444|hsa-mir-1307&WithFlank&10|-|103394247|103394407</td><td>86</td><td>108</td><td>23</td><td></td><td>isomiR</td><td>NO</td><td>139.93</td><td>189.76</td><td>99.82</td><td>62.32</td></tr>
<tr><td>>MI0006444|hsa-mir-1307&WithFlank&10|-|103394247|103394407@87.107.21&[MIMAT0005951&hsa-miR-1307-3p&offsets|+1|0;m-66&10|-|103394300|103394322&offsets|+1|-1]|CTCGGCGTGGCGTCGGTCGTG</td><td>>MI0006444|hsa-mir-1307&WithFlank&10|-|103394247|103394407</td><td>87</td><td>107</td><td>21</td><td></td><td>isomiR</td><td>NO</td><td>0.00</td><td>0.00</td><td>12.21</td><td>0.00</td></tr>
<tr><td>>MI0006444|hsa-mir-1307&WithFlank&10|-|103394247|103394407@87.107.21(+1U)&[MIMAT0005951&hsa-miR-1307-3p&offsets|+1|0(+1U);m-66&10|-|103394300|103394322&offsets|+1|-1(+1U)]|CTCGGCGTGGCGTCGGTCGTGT</td><td>>MI0006444|hsa-mir-1307&WithFlank&10|-|103394247|103394407</td><td>87</td><td>107</td><td>21</td><td>(+1U)</td><td>isomiR</td><td>NO</td><td>0.00</td><td>0.00</td><td>10.57</td><td>0.00</td></tr>
<tr><td>>MI0006444|hsa-mir-1307&WithFlank&10|-|103394247|103394407@87.108.22&[MIMAT0005951&hsa-miR-1307-3p&offsets|+1|+1;m-66&10|-|103394300|103394322&offsets|+1|0]|CTCGGCGTGGCGTCGGTCGTGG</td><td>>MI0006444|hsa-mir-1307&WithFlank&10|-|103394247|103394407</td><td>87</td><td>108</td><td>22</td><td></td><td>isomiR</td><td>NO</td><td>90.86</td><td>91.30</td><td>69.76</td><td>11.57</td></tr>
<tr><td>>am_MI0015825|hsa-mir-378c&WithFlank&10|-|130962582|130962674@17.34.18;MI0016749|hsa-mir-378d-1&WithFlank&4|-|5923269|5923334@37.54.18;MI0000786|hsa-mir-378a&WithFlank&5|+|149732819|149732896@49.66.18;MI0003840|hsa-mir-378d-2&WithFlank&8|-|93916016|93916125@28.45.18&[MIMAT0016847&hsa-miR-378c&offsets|0|-7;m-130&10|-|130962639|130962658&offsets|0|-2];[MIMAT0018926&hsa-miR-378d&offsets|0|-2;m-228&4|-|5923281|5923298&offsets|0|0];[MIMAT0000732&hsa-miR-378a-3p&offsets|0|-4;m-44&5|+|149732867|149732888&offsets|0|-4];[MIMAT0018926_1&hsa-miR-378d&offsets|0|-2;m-227&8|-|93916081|93916098&offsets|0|0]|ACTGGACTTGGAGTCAGA</td><td>>MI0015825|hsa-mir-378c&WithFlank&10|-|130962582|130962674.</td><td>17</td><td>34</td><td>18</td><td></td><td>isomiR</td><td>NO</td><td>0.00</td><td>0.00</td><td>137.09</td><td>0.00</td></tr>
<tr><td>>am_MI0015825|hsa-mir-378c&WithFlank&10|-|130962582|130962674@17.35.19;MI0016749|hsa-mir-378d-1&WithFlank&4|-|5923269|5923334@37.55.19;MI0000786|hsa-mir-378a&WithFlank&5|+|149732819|149732896@49.67.19;MI0003840|hsa-mir-378d-2&WithFlank&8|-|93916016|93916125@28.46.19&[MIMAT0016847&hsa-miR-378c&offsets|0|-6;m-130&10|-|130962639|130962658&offsets|0|-1];[MIMAT0018926&hsa-miR-378d&offsets|0|-1;m-228&4|-|5923281|5923298&offsets|0|+1];[MIMAT0000732&hsa-miR-378a-3p&offsets|0|-3;m-44&5|+|149732867|149732888&offsets|0|-3];[MIMAT0018926_1&hsa-miR-378d&offsets|0|-1;m-227&8|-|93916081|93916098&offsets|0|+1]|ACTGGACTTGGAGTCAGAA</td><td>>MI0015825|hsa-mir-378c&WithFlank&10|-|130962582|130962674.</td><td>17</td><td>35</td><td>19</td><td></td><td>isomiR</td><td>NO</td><td>0.00</td><td>0.00</td><td>278.34</td><td>0.00</td></tr>
<tr><td>>am_MI0015825|hsa-mir-378c&WithFlank&10|-|130962582|130962674@17.36.20;MI0000786|hsa-mir-378a&WithFlank&5|+|149732819|149732896@49.68.20&[MIMAT0016847&hsa-miR-378c&offsets|0|-5;m-130&10|-|130962639|130962658&offsets|0|0];[MIMAT0000732&hsa-miR-378a-3p&offsets|0|-2;m-44&5|+|149732867|149732888&offsets|0|-2]|ACTGGACTTGGAGTCAGAAG</td><td>>MI0015825|hsa-mir-378c&WithFlank&10|-|130962582|130962674.</td><td>17</td><td>36</td><td>20</td><td></td><td>isomiR</td><td>NO</td><td>0.00</td><td>0.00</td><td>109.87</td><td>0.00</td></tr>
<tr><td>>am_MI0015825|hsa-mir-378c&WithFlank&10|-|130962582|130962674@17.37.21&[MIMAT0016847&hsa-miR-378c&offsets|0|-4;m-130&10|-|130962639|130962658&offsets|0|+1]|ACTGGACTTGGAGTCAGAAGA</td><td>>MI0015825|hsa-mir-378c&WithFlank&10|-|130962582|130962674.</td><td>17</td><td>37</td><td>21</td><td></td><td>isomiR</td><td>NO</td><td>0.00</td><td>0.00</td><td>151.06</td><td>0.00</td></tr>
<tr><td>>MI0015866|hsa-mir-4326&WithFlank&20|+|63286802|63286872@17.40.24&[MIMAT0016888&hsa-miR-4326&offsets|0|+4;m-339&20|+|63286818|63286841&offsets|0|0]|TGTTCCTCTGTCTCCCAGACTCTG</td><td>>MI0015866|hsa-mir-4326&WithFlank&20|+|63286802|63286872</td><td>17</td><td>40</td><td>24</td><td></td><td>isomiR</td><td>NO</td><td>0.00</td><td>0.00</td><td>62.25</td><td>0.00</td></tr>
<tr><td>>MI0016003|hsa-mir-3613&WithFlank&13|-|49996409|49996507@22.43.22&[MIMAT0017990&hsa-miR-3613-5p&offsets|0|0;m-259&13|-|49996465|49996486&offsets|0|0]|TGTTGTACTTTTTTTTTTGTTC</td><td>>MI0016003|hsa-mir-3613&WithFlank&13|-|49996409|49996507</td><td>22</td><td>43</td><td>22</td><td></td><td>isomiR</td><td>NO</td><td>0.00</td><td>0.00</td><td>104.12</td><td>0.00</td></tr>
<tr><td>>MI0016005|hsa-mir-3615&WithFlank&17|+|74748607|74748705@57.78.22&[MIMAT0017994&hsa-miR-3615&offsets|0|+1;m-271&17|+|74748663|74748684&offsets|0|0]|TCTCTCGGCTCCTCGCGGCTCG</td><td>>MI0016005|hsa-mir-3615&WithFlank&17|+|74748607|74748705</td><td>57</td><td>78</td><td>22</td><td></td><td>isomiR</td><td>NO</td><td>48.16</td><td>85.70</td><td>0.00</td><td>11.52</td></tr>
<tr><td>>MI0016746|hsa-mir-548o-2&WithFlank&20|+|38516557|38516638@51.72.22;MI0006402|hsa-mir-548o&WithFlank&7|-|102405736|102405861@93.114.22&[MIMAT0005919&hsa-miR-548o-3p&offsets|0|0;m-431&20|+|38516607|38516628&offsets|0|0];[MIMAT0005919_1&hsa-miR-548o-3p&offsets|0|0;m-430&7|-|102405748|102405769&offsets|0|0]|CCAAAACTGCAGTTACTTTTGC</td><td>>MI0016746|hsa-mir-548o-2&WithFlank&20|+|38516557|38516638</td><td>51</td><td>72</td><td>22</td><td></td><td>isomiR</td><td>NO</td><td>0.00</td><td>0.00</td><td>80.43</td><td>0.00</td></tr>
<tr><td>>MI0016887|hsa-mir-4521&WithFlank&17|+|8186939|8187010@10.34.25&[MIMAT0019058&hsa-miR-4521&offsets|0|+3;m-765&17|+|8186948|8186969&offsets|0|+3]|GCTAAGGAAGTCCTGTGCTCAGTTT</td><td>>MI0016887|hsa-mir-4521&WithFlank&17|+|8186939|8187010</td><td>10</td><td>34</td><td>25</td><td></td><td>isomiR</td><td>NO</td><td>0.00</td><td>25.47</td><td>0.00</td><td>0.00</td></tr>
<tr><td>>MI0016887|hsa-mir-4521&WithFlank&17|+|8186939|8187010@10.35.26&[MIMAT0019058&hsa-miR-4521&offsets|0|+4;m-765&17|+|8186948|8186969&offsets|0|+4]|GCTAAGGAAGTCCTGTGCTCAGTTTT</td><td>>MI0016887|hsa-mir-4521&WithFlank&17|+|8186939|8187010</td><td>10</td><td>35</td><td>26</td><td></td><td>isomiR</td><td>NO</td><td>0.00</td><td>26.97</td><td>0.00</td><td>0.00</td></tr>
<tr><td>>MI0025750|hsa-mir-7974&WithFlank&19|-|11495538|11495628@62.85.24&[MIMAT0031177&hsa-miR-7974&offsets|0|0;m-867&19|-|11495544|11495567&offsets|0|0]|AGGCTGTGATGCTCTCCTGAGCCC</td><td>>MI0025750|hsa-mir-7974&WithFlank&19|-|11495538|11495628</td><td>62</td><td>85</td><td>24</td><td></td><td>isomiR</td><td>NO</td><td>10.47</td><td>0.00</td><td>0.00</td><td>0.00</td></tr>
<tr><td>>MI0033423|hsa-mir-10399&WithFlank&7|-|139044094|139044163@43.63.21(+1U)&[MIMAT0041630&hsa-miR-10399-3p&offsets|-1|-1(+1U);TJU_CMC_MD2.ID03061.3p-miR&offsets|+3|+2(+1U);m-2657&7|-|139044103|139044124&offsets|+3|+2(+1U)]|TCTCTCGGACAAGCTGTAGGTT</td><td>>MI0033423|hsa-mir-10399&WithFlank&7|-|139044094|139044163</td><td>43</td><td>63</td><td>21</td><td>(+1U)</td><td>isomiR</td><td>NO</td><td>0.00</td><td>0.00</td><td>32.17</td><td>0.00</td></tr>
</table>

### MDA-MB-231

<table>
<tr><td>>MI0033423|hsa-mir-10399&WithFlank&7|-|139044094|139044163@45.63.19(+2U)&[MIMAT0041630&hsa-miR-10399-3p&offsets|+1|-1(+2U);TJU_CMC_MD2.ID03061.3p-miR&offsets|+5|+2(+2U);m-2657&7|-|139044103|139044124&offsets|+5|+2(+2U)]|TCTCGGACAAGCTGTAGGTTT</td><td>>MI0033423|hsa-mir-10399&WithFlank&7|-|139044094|139044163</td><td>45</td><td>63</td><td>19</td><td>(+2U)</td><td>isomiR</td><td>NO</td><td>0.00</td><td>0.00</td><td>34.22</td><td>0.00</td></tr>
<tr><td>>hg38_MT_+_1621_3279_RNR2_16S_50ntFlanks@51.89.39|GCTAAACCTAGCCCCAAACCCACTCCACCTTACTACCAG</td><td>>hg38_MT_+_1621_3279_RNR2_16S_50ntFlanks</td><td>51</td><td>89</td><td>39</td><td></td><td>rRF 12S and 16S</td><td>NO</td><td>0.00</td><td>0.00</td><td>32.48</td><td>14.14</td></tr>
<tr><td>>hg38_MT_+_1621_3279_RNR2_16S_50ntFlanks@51.90.40|GCTAAACCTAGCCCCAAACCCACTCCACCTTACTACCAGA</td><td>>hg38_MT_+_1621_3279_RNR2_16S_50ntFlanks</td><td>51</td><td>90</td><td>40</td><td></td><td>rRF 12S and 16S</td><td>NO</td><td>0.00</td><td>0.00</td><td>175.49</td><td>101.49</td></tr>
<tr><td>>hg38_MT_+_1621_3279_RNR2_16S_50ntFlanks@51.91.41|GCTAAACCTAGCCCCAAACCCACTCCACCTTACTACCAGAC</td><td>>hg38_MT_+_1621_3279_RNR2_16S_50ntFlanks</td><td>51</td><td>91</td><td>41</td><td></td><td>rRF 12S and 16S</td><td>NO</td><td>0.00</td><td>0.00</td><td>66.14</td><td>56.15</td></tr>
<tr><td>>hg38_MT_+_1621_3279_RNR2_16S_50ntFlanks@51.92.42|GCTAAACCTAGCCCCAAACCCACTCCACCTTACTACCAGACA</td><td>>hg38_MT_+_1621_3279_RNR2_16S_50ntFlanks</td><td>51</td><td>92</td><td>42</td><td></td><td>rRF 12S and 16S</td><td>NO</td><td>0.00</td><td>0.00</td><td>122.81</td><td>123.28</td></tr>
<tr><td>>hg38_MT_+_1621_3279_RNR2_16S_50ntFlanks@51.93.43|GCTAAACCTAGCCCCAAACCCACTCCACCTTACTACCAGACAA</td><td>>hg38_MT_+_1621_3279_RNR2_16S_50ntFlanks</td><td>51</td><td>93</td><td>43</td><td></td><td>rRF 12S and 16S</td><td>NO</td><td>0.00</td><td>0.00</td><td>14.70</td><td>35.46</td></tr>
<tr><td>>hg38_MT_+_1621_3279_RNR2_16S_50ntFlanks@51.94.44|GCTAAACCTAGCCCCAAACCCACTCCACCTTACTACCAGACAAC</td><td>>hg38_MT_+_1621_3279_RNR2_16S_50ntFlanks</td><td>51</td><td>94</td><td>44</td><td></td><td>rRF 12S and 16S</td><td>NO</td><td>0.00</td><td>0.00</td><td>79.26</td><td>135.31</td></tr>
<tr><td>>hg38_MT_+_1621_3279_RNR2_16S_50ntFlanks@51.95.45|GCTAAACCTAGCCCCAAACCCACTCCACCTTACTACCAGACAACC</td><td>>hg38_MT_+_1621_3279_RNR2_16S_50ntFlanks</td><td>51</td><td>95</td><td>45</td><td></td><td>rRF 12S and 16S</td><td>NO</td><td>10.37</td><td>0.00</td><td>84.57</td><td>212.23</td></tr>
<tr><td>>hg38_MT_+_1621_3279_RNR2_16S_50ntFlanks@617.642.26|AAACATATAACTGAACTCCTCACACC</td><td>>hg38_MT_+_1621_3279_RNR2_16S_50ntFlanks</td><td>617</td><td>642</td><td>26</td><td></td><td>rRF 12S and 16S</td><td>NO</td><td>0.00</td><td>0.00</td><td>11.21</td><td>0.00</td></tr>
<tr><td>>hg38_MT_+_1621_3279_RNR2_16S_50ntFlanks@644.676.33|AATTGGACCAATCTATCACCCTATAGAAGAACT</td><td>>hg38_MT_+_1621_3279_RNR2_16S_50ntFlanks</td><td>644</td><td>676</td><td>33</td><td></td><td>rRF 12S and 16S</td><td>NO</td><td>0.00</td><td>0.00</td><td>12.74</td><td>0.00</td></tr>
<tr><td>>hg38_MT_+_1621_3279_RNR2_16S_50ntFlanks@729.759.31|GATTAAAACACTGAACTGACAATTAACAGCC</td><td>>hg38_MT_+_1621_3279_RNR2_16S_50ntFlanks</td><td>729</td><td>759</td><td>31</td><td></td><td>rRF 12S and 16S</td><td>NO</td><td>0.00</td><td>0.00</td><td>12.76</td><td>0.00</td></tr>
<tr><td>>hg38_MT_+_1621_3279_RNR2_16S_50ntFlanks@732.759.28|TAAAACACTGAACTGACAATTAACAGCC</td><td>>hg38_MT_+_1621_3279_RNR2_16S_50ntFlanks</td><td>732</td><td>759</td><td>28</td><td></td><td>rRF 12S and 16S</td><td>NO</td><td>0.00</td><td>0.00</td><td>12.08</td><td>0.00</td></tr>
<tr><td>>hg38_MT_+_1621_3279_RNR2_16S_50ntFlanks@929.958.30|CAGTGACACATGTTTAACGGCCGCGGTACC</td><td>>hg38_MT_+_1621_3279_RNR2_16S_50ntFlanks</td><td>929</td><td>958</td><td>30</td><td></td><td>rRF 12S and 16S</td><td>NO</td><td>0.00</td><td>0.00</td><td>15.48</td><td>0.00</td></tr>
<tr><td>>hg38_MT_+_1621_3279_RNR2_16S_50ntFlanks@930.958.29|AGTGACACATGTTTAACGGCCGCGGTACC</td><td>>hg38_MT_+_1621_3279_RNR2_16S_50ntFlanks</td><td>930</td><td>958</td><td>29</td><td></td><td>rRF 12S and 16S</td><td>NO</td><td>0.00</td><td>0.00</td><td>55.41</td><td>0.00</td></tr>
<tr><td>>hg38_MT_+_1621_3279_RNR2_16S_50ntFlanks@930.959.30|AGTGACACATGTTTAACGGCCGCGGTACCC</td><td>>hg38_MT_+_1621_3279_RNR2_16S_50ntFlanks</td><td>930</td><td>959</td><td>30</td><td></td><td>rRF 12S and 16S</td><td>NO</td><td>0.00</td><td>0.00</td><td>12.82</td><td>0.00</td></tr>
<tr><td>>hg38_MT_+_1621_3279_RNR2_16S_50ntFlanks@999.1018.20|AGGGACCTGTATGAATGGCT</td><td>>hg38_MT_+_1621_3279_RNR2_16S_50ntFlanks</td><td>999</td><td>1018</td><td>20</td><td></td><td>rRF 12S and 16S</td><td>NO</td><td>0.00</td><td>0.00</td><td>14.94</td><td>0.00</td></tr>
<tr><td>>hg38_MT_+_1621_3279_RNR2_16S_50ntFlanks@1000.1018.19|GGGACCTGTATGAATGGCT</td><td>>hg38_MT_+_1621_3279_RNR2_16S_50ntFlanks</td><td>1000</td><td>1018</td><td>19</td><td></td><td>rRF 12S and 16S</td><td>NO</td><td>0.00</td><td>0.00</td><td>11.38</td><td>0.00</td></tr>
<tr><td>>hg38_MT_+_1621_3279_RNR2_16S_50ntFlanks@1092.1120.29|GCAAGACGAGAAGACCCTATGGAGCTTTA</td><td>>hg38_MT_+_1621_3279_RNR2_16S_50ntFlanks</td><td>1092</td><td>1120</td><td>29</td><td></td><td>rRF 12S and 16S</td><td>NO</td><td>0.00</td><td>0.00</td><td>14.93</td><td>0.00</td></tr>
<tr><td>>hg38_MT_+_1621_3279_RNR2_16S_50ntFlanks@1184.1215.32|AATTTCGGTTGGGGCGACCTCGGAGCAGAACC</td><td>>hg38_MT_+_1621_3279_RNR2_16S_50ntFlanks</td><td>1184</td><td>1215</td><td>32</td><td></td><td>rRF 12S and 16S</td><td>NO</td><td>0.00</td><td>0.00</td><td>12.70</td><td>0.00</td></tr>
<tr><td>>hg38_MT_+_1621_3279_RNR2_16S_50ntFlanks@1216.1248.33|CAACCTCCGAGCAGTACATGCTAAGACTTCACC</td><td>>hg38_MT_+_1621_3279_RNR2_16S_50ntFlanks</td><td>1216</td><td>1248</td><td>33</td><td></td><td>rRF 12S and 16S</td><td>NO</td><td>0.00</td><td>0.00</td><td>35.36</td><td>18.15</td></tr>
<tr><td>>hg38_MT_+_1621_3279_RNR2_16S_50ntFlanks@1216.1262.47|CAACCTCCGAGCAGTACATGCTAAGACTTCACCAGTCAAAGCGAACT</td><td>>hg38_MT_+_1621_3279_RNR2_16S_50ntFlanks</td><td>1216</td><td>1262</td><td>47</td><td></td><td>rRF 12S and 16S</td><td>NO</td><td>0.00</td><td>24.16</td><td>0.00</td><td>121.29</td></tr>
<tr><td>>hg38_MT_+_1621_3279_RNR2_16S_50ntFlanks@1249.1271.23|AGTCAAAGCGAACTACTATACTC</td><td>>hg38_MT_+_1621_3279_RNR2_16S_50ntFlanks</td><td>1249</td><td>1271</td><td>23</td><td></td><td>rRF 12S and 16S</td><td>NO</td><td>0.00</td><td>0.00</td><td>31.02</td><td>0.00</td></tr>
<tr><td>>hg38_MT_+_1621_3279_RNR2_16S_50ntFlanks@1249.1275.27|AGTCAAAGCGAACTACTATACTCAATT</td><td>>hg38_MT_+_1621_3279_RNR2_16S_50ntFlanks</td><td>1249</td><td>1275</td><td>27</td><td></td><td>rRF 12S and 16S</td><td>NO</td><td>0.00</td><td>0.00</td><td>15.42</td><td>0.00</td></tr>
<tr><td>>hg38_MT_+_1621_3279_RNR2_16S_50ntFlanks@1249.1279.31|AGTCAAAGCGAACTACTATACTCAATTGATC</td><td>>hg38_MT_+_1621_3279_RNR2_16S_50ntFlanks</td><td>1249</td><td>1279</td><td>31</td><td></td><td>rRF 12S and 16S</td><td>NO</td><td>0.00</td><td>0.00</td><td>14.92</td><td>0.00</td></tr>
<tr><td>>hg38_MT_+_1621_3279_RNR2_16S_50ntFlanks@1250.1279.30|GTCAAAGCGAACTACTATACTCAATTGATC</td><td>>hg38_MT_+_1621_3279_RNR2_16S_50ntFlanks</td><td>1250</td><td>1279</td><td>30</td><td></td><td>rRF 12S and 16S</td><td>NO</td><td>0.00</td><td>0.00</td><td>10.46</td><td>0.00</td></tr>
<tr><td>>hg38_MT_+_1621_3279_RNR2_16S_50ntFlanks@1263.1292.30|ACTATACTCAATTGATCCAATAACTTGACC</td><td>>hg38_MT_+_1621_3279_RNR2_16S_50ntFlanks</td><td>1263</td><td>1292</td><td>30</td><td></td><td>rRF 12S and 16S</td><td>NO</td><td>0.00</td><td>0.00</td><td>12.54</td><td>0.00</td></tr>
<tr><td>>hg38_MT_+_1621_3279_RNR2_16S_50ntFlanks@1348.1385.38|AACAATAGGGTTTACGACCTCGATGTTGGATCAGGACA</td><td>>hg38_MT_+_1621_3279_RNR2_16S_50ntFlanks</td><td>1348</td><td>1385</td><td>38</td><td></td><td>rRF 12S and 16S</td><td>NO</td><td>0.00</td><td>0.00</td><td>33.89</td><td>0.00</td></tr>
<tr><td>>hg38_MT_+_1621_3279_RNR2_16S_50ntFlanks@1367.1396.30|TCGATGTTGGATCAGGACATCCCGATGGTG</td><td>>hg38_MT_+_1621_3279_RNR2_16S_50ntFlanks</td><td>1367</td><td>1396</td><td>30</td><td></td><td>rRF 12S and 16S</td><td>NO</td><td>0.00</td><td>0.00</td><td>61.39</td><td>0.00</td></tr>
<tr><td>>hg38_MT_+_1621_3279_RNR2_16S_50ntFlanks@1367.1401.35|TCGATGTTGGATCAGGACATCCCGATGGTGCAGCC</td><td>>hg38_MT_+_1621_3279_RNR2_16S_50ntFlanks</td><td>1367</td><td>1401</td><td>35</td><td></td><td>rRF 12S and 16S</td><td>NO</td><td>0.00</td><td>0.00</td><td>11.20</td><td>0.00</td></tr>
<tr><td>>hg38_MT_+_1621_3279_RNR2_16S_50ntFlanks@1368.1396.29|CGATGTTGGATCAGGACATCCCGATGGTG</td><td>>hg38_MT_+_1621_3279_RNR2_16S_50ntFlanks</td><td>1368</td><td>1396</td><td>29</td><td></td><td>rRF 12S and 16S</td><td>NO</td><td>0.00</td><td>0.00</td><td>12.38</td><td>0.00</td></tr>
<tr><td>>hg38_MT_+_1621_3279_RNR2_16S_50ntFlanks@1369.1396.28|GATGTTGGATCAGGACATCCCGATGGTG</td><td>>hg38_MT_+_1621_3279_RNR2_16S_50ntFlanks</td><td>1369</td><td>1396</td><td>28</td><td></td><td>rRF 12S and 16S</td><td>NO</td><td>0.00</td><td>0.00</td><td>34.62</td><td>0.00</td></tr>
<tr><td>>hg38_MT_+_1621_3279_RNR2_16S_50ntFlanks@1369.1401.33|GATGTTGGATCAGGACATCCCGATGGTGCAGCC</td><td>>hg38_MT_+_1621_3279_RNR2_16S_50ntFlanks</td><td>1369</td><td>1401</td><td>33</td><td></td><td>rRF 12S and 16S</td><td>NO</td><td>0.00</td><td>0.00</td><td>12.74</td><td>0.00</td></tr>
<tr><td>>hg38_MT_+_1621_3279_RNR2_16S_50ntFlanks@1370.1393.24|ATGTTGGATCAGGACATCCCGATG</td><td>>hg38_MT_+_1621_3279_RNR2_16S_50ntFlanks</td><td>1370</td><td>1393</td><td>24</td><td></td><td>rRF 12S and 16S</td><td>NO</td><td>0.00</td><td>0.00</td><td>54.55</td><td>0.00</td></tr>
<tr><td>>hg38_MT_+_1621_3279_RNR2_16S_50ntFlanks@1370.1395.26|ATGTTGGATCAGGACATCCCGATGGT</td><td>>hg38_MT_+_1621_3279_RNR2_16S_50ntFlanks</td><td>1370</td><td>1395</td><td>26</td><td></td><td>rRF 12S and 16S</td><td>NO</td><td>0.00</td><td>0.00</td><td>17.92</td><td>0.00</td></tr>
<tr><td>>hg38_MT_+_1621_3279_RNR2_16S_50ntFlanks@1370.1396.27|ATGTTGGATCAGGACATCCCGATGGTG</td><td>>hg38_MT_+_1621_3279_RNR2_16S_50ntFlanks</td><td>1370</td><td>1396</td><td>27</td><td></td><td>rRF 12S and 16S</td><td>NO</td><td>0.00</td><td>0.00</td><td>113.53</td><td>0.00</td></tr>
<tr><td>>hg38_MT_+_1621_3279_RNR2_16S_50ntFlanks@1370.1401.32|ATGTTGGATCAGGACATCCCGATGGTGCAGCC</td><td>>hg38_MT_+_1621_3279_RNR2_16S_50ntFlanks</td><td>1370</td><td>1401</td><td>32</td><td></td><td>rRF 12S and 16S</td><td>NO</td><td>0.00</td><td>0.00</td><td>12.36</td><td>0.00</td></tr>
<tr><td>>hg38_MT_+_1621_3279_RNR2_16S_50ntFlanks@1373.1396.24|TTGGATCAGGACATCCCGATGGTG</td><td>>hg38_MT_+_1621_3279_RNR2_16S_50ntFlanks</td><td>1373</td><td>1396</td><td>24</td><td></td><td>rRF 12S and 16S</td><td>NO</td><td>0.00</td><td>0.00</td><td>14.24</td><td>0.00</td></tr>
</table>

### MDA-MB-231

| | | | | | | | | | | |
|---|---|---|---|---|---|---|---|---|---|---|
| >hg38_MT_+_1621_3279_RNR2_16S_50ntFlanks@1380.1414.35\|AGGACATCCCGATGGTGCAGCCGCTATTAAAGGTT | >hg38_MT_+_1621_3279_RNR2_16S_50ntFlanks | 1380 | 1414 | 35 | rRF 12S and 16S | NO | 0.00 | 0.00 | 55.39 | 0.00 |
| >hg38_MT_+_1621_3279_RNR2_16S_50ntFlanks@1389.1414.26\|CGATGGTGCAGCCGCTATTAAAGGTT | >hg38_MT_+_1621_3279_RNR2_16S_50ntFlanks | 1389 | 1414 | 26 | rRF 12S and 16S | NO | 0.00 | 0.00 | 58.35 | 0.00 |
| >hg38_MT_+_1621_3279_RNR2_16S_50ntFlanks@1390.1414.25\|GATGGTGCAGCCGCTATTAAAGGTT | >hg38_MT_+_1621_3279_RNR2_16S_50ntFlanks | 1390 | 1414 | 25 | rRF 12S and 16S | NO | 0.00 | 0.00 | 12.86 | 0.00 |
| >multi-hg38_MT_+_1621_3279_RNR2_16S_50ntFlanks@51.70.20\|GCTAAACCTAGCCCCAAACC | >hg38_MT_+_1621_3279_RNR2_16S_50ntFlanks.multi | 51 | 70 | 20 | rRF 12S and 16S | NO | 0.00 | 0.00 | 290.39 | 22.17 |
| >multi-hg38_MT_+_1621_3279_RNR2_16S_50ntFlanks@51.71.21\|GCTAAACCTAGCCCCAAACCC | >hg38_MT_+_1621_3279_RNR2_16S_50ntFlanks.multi | 51 | 71 | 21 | rRF 12S and 16S | NO | 10.34 | 0.00 | 432.77 | 109.34 |
| >multi-hg38_MT_+_1621_3279_RNR2_16S_50ntFlanks@51.72.22\|GCTAAACCTAGCCCCAAACCCA | >hg38_MT_+_1621_3279_RNR2_16S_50ntFlanks.multi | 51 | 72 | 22 | rRF 12S and 16S | NO | 0.00 | 0.00 | 140.37 | 128.11 |
| >multi-hg38_MT_+_1621_3279_RNR2_16S_50ntFlanks@51.73.23\|GCTAAACCTAGCCCCAAACCCAC | >hg38_MT_+_1621_3279_RNR2_16S_50ntFlanks.multi | 51 | 73 | 23 | rRF 12S and 16S | NO | 0.00 | 0.00 | 96.17 | 72.92 |
| >multi-hg38_MT_+_1621_3279_RNR2_16S_50ntFlanks@51.74.24\|GCTAAACCTAGCCCCAAACCCACT | >hg38_MT_+_1621_3279_RNR2_16S_50ntFlanks.multi | 51 | 74 | 24 | rRF 12S and 16S | NO | 69.09 | 0.00 | 359.16 | 194.56 |
| >multi-hg38_MT_+_1621_3279_RNR2_16S_50ntFlanks@51.76.26\|GCTAAACCTAGCCCCAAACCCACTCC | >hg38_MT_+_1621_3279_RNR2_16S_50ntFlanks.multi | 51 | 76 | 26 | rRF 12S and 16S | NO | 0.00 | 0.00 | 62.39 | 15.12 |
| >multi-hg38_MT_+_1621_3279_RNR2_16S_50ntFlanks@51.79.29\|GCTAAACCTAGCCCCAAACCCACTCCACC | >hg38_MT_+_1621_3279_RNR2_16S_50ntFlanks.multi | 51 | 79 | 29 | rRF 12S and 16S | NO | 0.00 | 0.00 | 195.65 | 99.19 |
| >multi-hg38_MT_+_1621_3279_RNR2_16S_50ntFlanks@51.80.30\|GCTAAACCTAGCCCCAAACCCACTCCACCT | >hg38_MT_+_1621_3279_RNR2_16S_50ntFlanks.multi | 51 | 80 | 30 | rRF 12S and 16S | NO | 0.00 | 0.00 | 35.95 | 56.12 |
| >multi-hg38_MT_+_1621_3279_RNR2_16S_50ntFlanks@51.81.31\|GCTAAACCTAGCCCCAAACCCACTCCACCTT | >hg38_MT_+_1621_3279_RNR2_16S_50ntFlanks.multi | 51 | 81 | 31 | rRF 12S and 16S | NO | 0.00 | 0.00 | 34.44 | 17.78 |
| >multi-hg38_MT_+_1621_3279_RNR2_16S_50ntFlanks@51.82.32\|GCTAAACCTAGCCCCAAACCCACTCCACCTTA | >hg38_MT_+_1621_3279_RNR2_16S_50ntFlanks.multi | 51 | 82 | 32 | rRF 12S and 16S | NO | 0.00 | 0.00 | 99.14 | 18.58 |
| >multi-hg38_MT_+_1621_3279_RNR2_16S_50ntFlanks@51.84.34\|GCTAAACCTAGCCCCAAACCCACTCCACCTTACT | >hg38_MT_+_1621_3279_RNR2_16S_50ntFlanks.multi | 51 | 84 | 34 | rRF 12S and 16S | NO | 0.00 | 0.00 | 156.44 | 91.28 |
| >multi-hg38_MT_+_1621_3279_RNR2_16S_50ntFlanks@51.85.35\|GCTAAACCTAGCCCCAAACCCACTCCACCTTACTA | >hg38_MT_+_1621_3279_RNR2_16S_50ntFlanks.multi | 51 | 85 | 35 | rRF 12S and 16S | NO | 0.00 | 0.00 | 55.86 | 57.95 |
| >multi-hg38_MT_+_1621_3279_RNR2_16S_50ntFlanks@51.87.37\|GCTAAACCTAGCCCCAAACCCACTCCACCTTACTACC | >hg38_MT_+_1621_3279_RNR2_16S_50ntFlanks.multi | 51 | 87 | 37 | rRF 12S and 16S | NO | 51.71 | 0.00 | 353.68 | 116.26 |
| >multi-hg38_MT_+_1621_3279_RNR2_16S_50ntFlanks@51.88.38\|GCTAAACCTAGCCCCAAACCCACTCCACCTTACTACCA | >hg38_MT_+_1621_3279_RNR2_16S_50ntFlanks.multi | 51 | 88 | 38 | rRF 12S and 16S | NO | 63.67 | 0.00 | 522.40 | 152.55 |
| >multi-hg38_MT_+_1621_3279_RNR2_16S_50ntFlanks@194.214.21\|AAGCATAATATAGCAAGGACT | >hg38_MT_+_1621_3279_RNR2_16S_50ntFlanks.multi | 194 | 214 | 21 | rRF 12S and 16S | NO | 0.00 | 0.00 | 10.54 | 0.00 |
| >multi-hg38_MT_+_1621_3279_RNR2_16S_50ntFlanks@292.311.20\|AGACGAGCTACCTAAGAACA | >hg38_MT_+_1621_3279_RNR2_16S_50ntFlanks.multi | 292 | 311 | 20 | rRF 12S and 16S | NO | 0.00 | 0.00 | 12.81 | 0.00 |
| >multi-hg38_MT_+_1621_3279_RNR2_16S_50ntFlanks@459.492.34\|CTTGTAAATTTAACTGTTAGTCCAAAGAGGAACA | >hg38_MT_+_1621_3279_RNR2_16S_50ntFlanks.multi | 459 | 492 | 34 | rRF 12S and 16S | NO | 0.00 | 0.00 | 34.16 | 15.50 |
| >multi-hg38_MT_+_1621_3279_RNR2_16S_50ntFlanks@486.506.21\|AGGAACAGCTCTTTGGACACT | >hg38_MT_+_1621_3279_RNR2_16S_50ntFlanks.multi | 486 | 506 | 21 | rRF 12S and 16S | NO | 0.00 | 0.00 | 13.58 | 0.00 |
| >multi-hg38_MT_+_1621_3279_RNR2_16S_50ntFlanks@486.518.33\|AGGAACAGCTCTTTGGACACTAGGAAAAAACCT | >hg38_MT_+_1621_3279_RNR2_16S_50ntFlanks.multi | 486 | 518 | 33 | rRF 12S and 16S | NO | 0.00 | 0.00 | 13.58 | 0.00 |
| >multi-hg38_MT_+_1621_3279_RNR2_16S_50ntFlanks@493.517.25\|GCTCTTTGGACACTAGGAAAAAACC | >hg38_MT_+_1621_3279_RNR2_16S_50ntFlanks.multi | 493 | 517 | 25 | rRF 12S and 16S | NO | 0.00 | 0.00 | 11.96 | 19.87 |
| >multi-hg38_MT_+_1621_3279_RNR2_16S_50ntFlanks@493.518.26\|GCTCTTTGGACACTAGGAAAAAACCT | >hg38_MT_+_1621_3279_RNR2_16S_50ntFlanks.multi | 493 | 518 | 26 | rRF 12S and 16S | NO | 0.00 | 0.00 | 16.68 | 53.53 |
| >multi-hg38_MT_+_1621_3279_RNR2_16S_50ntFlanks@825.855.31\|TAAGGAAAGGTTAAAAAAAGTAAAAGGAACT | >hg38_MT_+_1621_3279_RNR2_16S_50ntFlanks.multi | 825 | 855 | 31 | rRF 12S and 16S | NO | 0.00 | 0.00 | 13.94 | 0.00 |
| >multi-hg38_MT_+_1621_3279_RNR2_16S_50ntFlanks@826.855.30\|AAGGAAAGGTTAAAAAAAGTAAAAGGAACT | >hg38_MT_+_1621_3279_RNR2_16S_50ntFlanks.multi | 826 | 855 | 30 | rRF 12S and 16S | NO | 0.00 | 0.00 | 35.05 | 0.00 |
| >multi-hg38_MT_+_1621_3279_RNR2_16S_50ntFlanks@1000.1017.18\|GGGACCTGTATGAATGGC | >hg38_MT_+_1621_3279_RNR2_16S_50ntFlanks.multi | 1000 | 1017 | 18 | rRF 12S and 16S | NO | 0.00 | 0.00 | 16.42 | 0.00 |
| >multi-hg38_MT_+_1621_3279_RNR2_16S_50ntFlanks@1095.1117.23\|AGACGAGAAGACCCTATGGAGCT | >hg38_MT_+_1621_3279_RNR2_16S_50ntFlanks.multi | 1095 | 1117 | 23 | rRF 12S and 16S | NO | 0.00 | 0.00 | 17.24 | 0.00 |
| >multi-hg38_MT_+_1621_3279_RNR2_16S_50ntFlanks@1095.1120.26\|AGACGAGAAGACCCTATGGAGCTTTA | >hg38_MT_+_1621_3279_RNR2_16S_50ntFlanks.multi | 1095 | 1120 | 26 | rRF 12S and 16S | NO | 0.00 | 0.00 | 55.43 | 0.00 |

### MDA-MB-231

| | | | | | | | | | | |
|---|---|---|---|---|---|---|---|---|---|---|
| >multi-hg38_MT_+_1621_3279_RNR2_16S_50ntFlanks@1349.1382.34\|ACAATAGGGTTTACGACCTCGATGTTGGATCAGG | >hg38_MT_+_1621_3279_RNR2_16S_50ntFlanks.multi | 1349 | 1382 | 34 | rRF 12S and 16S | NO | 0.00 | 0.00 | 14.95 | 0.00 |
| >multi-hg38_MT_+_1621_3279_RNR2_16S_50ntFlanks@1349.1385.37\|ACAATAGGGTTTACGACCTCGATGTTGGATCAGGACA | >hg38_MT_+_1621_3279_RNR2_16S_50ntFlanks.multi | 1349 | 1385 | 37 | rRF 12S and 16S | NO | 0.00 | 0.00 | 16.75 | 0.00 |
| >multi-hg38_MT_+_1621_3279_RNR2_16S_50ntFlanks@1349.1388.40\|ACAATAGGGTTTACGACCTCGATGTTGGATCAGGACATCC | >hg38_MT_+_1621_3279_RNR2_16S_50ntFlanks.multi | 1349 | 1388 | 40 | rRF 12S and 16S | NO | 0.00 | 0.00 | 73.17 | 15.91 |
| >multi-hg38_MT_+_1621_3279_RNR2_16S_50ntFlanks@1356.1388.33\|GGTTTACGACCTCGATGTTGGATCAGGACATCC | >hg38_MT_+_1621_3279_RNR2_16S_50ntFlanks.multi | 1356 | 1388 | 33 | rRF 12S and 16S | NO | 0.00 | 0.00 | 11.93 | 0.00 |
| >multi-hg38_MT_+_1621_3279_RNR2_16S_50ntFlanks@1357.1388.32\|GTTTACGACCTCGATGTTGGATCAGGACATCC | >hg38_MT_+_1621_3279_RNR2_16S_50ntFlanks.multi | 1357 | 1388 | 32 | rRF 12S and 16S | NO | 0.00 | 0.00 | 11.48 | 0.00 |
| >multi-hg38_MT_+_1621_3279_RNR2_16S_50ntFlanks@1363.1388.26\|GACCTCGATGTTGGATCAGGACATCC | >hg38_MT_+_1621_3279_RNR2_16S_50ntFlanks.multi | 1363 | 1388 | 26 | rRF 12S and 16S | NO | 0.00 | 0.00 | 62.41 | 0.00 |
| >multi-hg38_MT_+_1621_3279_RNR2_16S_50ntFlanks@1364.1388.25\|ACCTCGATGTTGGATCAGGACATCC | >hg38_MT_+_1621_3279_RNR2_16S_50ntFlanks.multi | 1364 | 1388 | 25 | rRF 12S and 16S | NO | 0.00 | 0.00 | 10.57 | 0.00 |
| >multi-hg38_MT_+_1621_3279_RNR2_16S_50ntFlanks@1367.1385.19\|TCGATGTTGGATCAGGACA | >hg38_MT_+_1621_3279_RNR2_16S_50ntFlanks.multi | 1367 | 1385 | 19 | rRF 12S and 16S | NO | 0.00 | 0.00 | 15.65 | 0.00 |
| >multi-hg38_MT_+_1621_3279_RNR2_16S_50ntFlanks@1370.1388.19\|ATGTTGGATCAGGACATCC | >hg38_MT_+_1621_3279_RNR2_16S_50ntFlanks.multi | 1370 | 1388 | 19 | rRF 12S and 16S | NO | 0.00 | 0.00 | 83.91 | 0.00 |
| >multi-hg38_MT_+_1621_3279_RNR2_16S_50ntFlanks@1370.1389.20\|ATGTTGGATCAGGACATCCC | >hg38_MT_+_1621_3279_RNR2_16S_50ntFlanks.multi | 1370 | 1389 | 20 | rRF 12S and 16S | NO | 0.00 | 0.00 | 11.58 | 0.00 |
| >multi-hg38_MT_+_1621_3279_RNR2_16S_50ntFlanks@1404.1421.18\|TATTAAAGGTTCGTTTGT | >hg38_MT_+_1621_3279_RNR2_16S_50ntFlanks.multi | 1404 | 1421 | 18 | rRF 12S and 16S | NO | 0.00 | 0.00 | 65.97 | 0.00 |
| >multi-hg38_MT_+_1621_3279_RNR2_16S_50ntFlanks@1416.1446.31\|GTTTGTTCAACGATTAAAGTCCTACGTGATC | >hg38_MT_+_1621_3279_RNR2_16S_50ntFlanks.multi | 1416 | 1446 | 31 | rRF 12S and 16S | NO | 0.00 | 0.00 | 13.34 | 0.00 |
| >multi-hg38_MT_+_1621_3279_RNR2_16S_50ntFlanks@1425.1451.27\|ACGATTAAAGTCCTACGTGATCTGAGT | >hg38_MT_+_1621_3279_RNR2_16S_50ntFlanks.multi | 1425 | 1451 | 27 | rRF 12S and 16S | NO | 0.00 | 0.00 | 11.06 | 0.00 |
| >multi-hg38_MT_+_1621_3279_RNR2_16S_50ntFlanks@1427.1448.22\|GATTAAAGTCCTACGTGATCTG | >hg38_MT_+_1621_3279_RNR2_16S_50ntFlanks.multi | 1427 | 1448 | 22 | rRF 12S and 16S | NO | 0.00 | 0.00 | 15.57 | 0.00 |
| >multi-hg38_MT_+_1621_3279_RNR2_16S_50ntFlanks@1427.1451.25\|GATTAAAGTCCTACGTGATCTGAGT | >hg38_MT_+_1621_3279_RNR2_16S_50ntFlanks.multi | 1427 | 1451 | 25 | rRF 12S and 16S | NO | 0.00 | 0.00 | 54.83 | 0.00 |
| >multi-hg38_MT_+_1621_3279_RNR2_16S_50ntFlanks@1427.1457.31\|GATTAAAGTCCTACGTGATCTGAGTTCAGAC | >hg38_MT_+_1621_3279_RNR2_16S_50ntFlanks.multi | 1427 | 1457 | 31 | rRF 12S and 16S | NO | 0.00 | 0.00 | 15.25 | 0.00 |
| >multi-hg38_MT_+_1621_3279_RNR2_16S_50ntFlanks@1427.1458.32\|GATTAAAGTCCTACGTGATCTGAGTTCAGACC | >hg38_MT_+_1621_3279_RNR2_16S_50ntFlanks.multi | 1427 | 1458 | 32 | rRF 12S and 16S | NO | 0.00 | 0.00 | 75.24 | 10.72 |
| >multi-hg38_MT_+_1621_3279_RNR2_16S_50ntFlanks@1428.1451.24\|ATTAAAGTCCTACGTGATCTGAGT | >hg38_MT_+_1621_3279_RNR2_16S_50ntFlanks.multi | 1428 | 1451 | 24 | rRF 12S and 16S | NO | 0.00 | 0.00 | 11.59 | 0.00 |
| >multi-hg38_MT_+_1621_3279_RNR2_16S_50ntFlanks@1428.1452.25\|ATTAAAGTCCTACGTGATCTGAGTT | >hg38_MT_+_1621_3279_RNR2_16S_50ntFlanks.multi | 1428 | 1452 | 25 | rRF 12S and 16S | NO | 0.00 | 0.00 | 13.99 | 0.00 |
| >multi-hg38_MT_+_1621_3279_RNR2_16S_50ntFlanks@1428.1453.26\|ATTAAAGTCCTACGTGATCTGAGTTC | >hg38_MT_+_1621_3279_RNR2_16S_50ntFlanks.multi | 1428 | 1453 | 26 | rRF 12S and 16S | NO | 0.00 | 0.00 | 55.97 | 0.00 |
| >multi-hg38_MT_+_1621_3279_RNR2_16S_50ntFlanks@1428.1457.30\|ATTAAAGTCCTACGTGATCTGAGTTCAGAC | >hg38_MT_+_1621_3279_RNR2_16S_50ntFlanks.multi | 1428 | 1457 | 30 | rRF 12S and 16S | NO | 0.00 | 0.00 | 11.11 | 0.00 |
| >multi-hg38_MT_+_1621_3279_RNR2_16S_50ntFlanks@1428.1458.31\|ATTAAAGTCCTACGTGATCTGAGTTCAGACC | >hg38_MT_+_1621_3279_RNR2_16S_50ntFlanks.multi | 1428 | 1458 | 31 | rRF 12S and 16S | NO | 0.00 | 0.00 | 61.29 | 11.86 |
| >multi-hg38_MT_+_1621_3279_RNR2_16S_50ntFlanks@1431.1451.21\|AAAGTCCTACGTGATCTGAGT | >hg38_MT_+_1621_3279_RNR2_16S_50ntFlanks.multi | 1431 | 1451 | 21 | rRF 12S and 16S | NO | 0.00 | 0.00 | 17.08 | 0.00 |
| >multi-hg38_MT_+_1621_3279_RNR2_16S_50ntFlanks@1431.1453.23\|AAAGTCCTACGTGATCTGAGTTC | >hg38_MT_+_1621_3279_RNR2_16S_50ntFlanks.multi | 1431 | 1453 | 23 | rRF 12S and 16S | NO | 0.00 | 0.00 | 35.24 | 0.00 |
| >multi-hg38_MT_+_1621_3279_RNR2_16S_50ntFlanks@1431.1458.28\|AAAGTCCTACGTGATCTGAGTTCAGACC | >hg38_MT_+_1621_3279_RNR2_16S_50ntFlanks.multi | 1431 | 1458 | 28 | rRF 12S and 16S | NO | 0.00 | 0.00 | 12.65 | 0.00 |
| >multi-hg38_MT_+_1621_3279_RNR2_16S_50ntFlanks@1432.1453.22\|AAGTCCTACGTGATCTGAGTTC | >hg38_MT_+_1621_3279_RNR2_16S_50ntFlanks.multi | 1432 | 1453 | 22 | rRF 12S and 16S | NO | 0.00 | 0.00 | 12.17 | 0.00 |
| >multi-hg38_MT_+_1621_3279_RNR2_16S_50ntFlanks@1432.1458.27\|AAGTCCTACGTGATCTGAGTTCAGACC | >hg38_MT_+_1621_3279_RNR2_16S_50ntFlanks.multi | 1432 | 1458 | 27 | rRF 12S and 16S | NO | 0.00 | 0.00 | 14.04 | 0.00 |

### MDA-MB-231

| | | | | | | | | | | |
|---|---|---|---|---|---|---|---|---|---|---|
| >multi-hg38_MT_+_1621_3279_RNR2_16S_50ntFlanks@1438.1457.20\|TACGTGATCTGAGTTCAGAC | >hg38_MT_+_1621_3279_RNR2_16S_50ntFlanks.multi | 1438 | 1457 | 20 | rRF 12S and 16S | NO | 0.00 | 0.00 | 58.47 | 0.00 |
| >multi-hg38_MT_+_1621_3279_RNR2_16S_50ntFlanks@1438.1458.21\|TACGTGATCTGAGTTCAGACC | >hg38_MT_+_1621_3279_RNR2_16S_50ntFlanks.multi | 1438 | 1458 | 21 | rRF 12S and 16S | NO | 0.00 | 0.00 | 13.92 | 0.00 |
| >multi-hg38_MT_+_1621_3279_RNR2_16S_50ntFlanks@1439.1457.19\|ACGTGATCTGAGTTCAGAC | >hg38_MT_+_1621_3279_RNR2_16S_50ntFlanks.multi | 1439 | 1457 | 19 | rRF 12S and 16S | NO | 0.00 | 0.00 | 16.75 | 0.00 |
| >multi-hg38_MT_+_598_1651_RNR1_12S_50ntFlanks@51.83.33\|AATAGGTTTGGTCCTAGCCTTTCTAT TAGCTCT | >hg38_MT_+_598_1651_RNR1_12S_50ntFlanks.multi | 51 | 83 | 33 | rRF 12S and 16S | NO | 0.00 | 0.00 | 17.62 | 0.00 |
| >multi-hg38_MT_+_598_1651_RNR1_12S_50ntFlanks@51.88.38\|AATAGGTTTGGTCCTAGCCTTTCTAT TAGCTCTTAGTA | >hg38_MT_+_598_1651_RNR1_12S_50ntFlanks.multi | 51 | 88 | 38 | rRF 12S and 16S | NO | 0.00 | 0.00 | 904.09 | 11.00 |
| >multi-hg38_MT_+_598_1651_RNR1_12S_50ntFlanks@52.77.26\|ATAGGTTTGGTCCTAGCCTTTCTATT | >hg38_MT_+_598_1651_RNR1_12S_50ntFlanks.multi | 52 | 77 | 26 | rRF 12S and 16S | NO | 0.00 | 0.00 | 573.55 | 84.02 |
| >multi-hg38_MT_+_598_1651_RNR1_12S_50ntFlanks@52.83.32\|ATAGGTTTGGTCCTAGCCTTTCTATT AGCTCT | >hg38_MT_+_598_1651_RNR1_12S_50ntFlanks.multi | 52 | 83 | 32 | rRF 12S and 16S | NO | 0.00 | 0.00 | 368.17 | 78.56 |
| >multi-hg38_MT_+_598_1651_RNR1_12S_50ntFlanks@52.84.33\|ATAGGTTTGGTCCTAGCCTTTCTATT AGCTCTT | >hg38_MT_+_598_1651_RNR1_12S_50ntFlanks.multi | 52 | 84 | 33 | rRF 12S and 16S | NO | 0.00 | 0.00 | 415.07 | 18.21 |
| >multi-hg38_MT_+_598_1651_RNR1_12S_50ntFlanks@52.87.36\|ATAGGTTTGGTCCTAGCCTTTCTATT AGCTCTTAGT | >hg38_MT_+_598_1651_RNR1_12S_50ntFlanks.multi | 52 | 87 | 36 | rRF 12S and 16S | NO | 0.00 | 0.00 | 289.60 | 63.86 |
| >multi-hg38_MT_+_598_1651_RNR1_12S_50ntFlanks@52.88.37\|ATAGGTTTGGTCCTAGCCTTTCTATT AGCTCTTAGTA | >hg38_MT_+_598_1651_RNR1_12S_50ntFlanks.multi | 52 | 88 | 37 | rRF 12S and 16S | YES | 53.25 | 0.00 | 5693.40 | 116.63 |
| >multi-hg38_MT_+_598_1651_RNR1_12S_50ntFlanks@52.90.39\|ATAGGTTTGGTCCTAGCCTTTCTATT AGCTCTTAGTAAG | >hg38_MT_+_598_1651_RNR1_12S_50ntFlanks.multi | 52 | 90 | 39 | rRF 12S and 16S | NO | 0.00 | 0.00 | 61.94 | 21.38 |
| >multi-hg38_MT_+_598_1651_RNR1_12S_50ntFlanks@52.93.42\|ATAGGTTTGGTCCTAGCCTTTCTATT AGCTCTTAGTAAGATT | >hg38_MT_+_598_1651_RNR1_12S_50ntFlanks.multi | 52 | 93 | 42 | rRF 12S and 16S | NO | 0.00 | 0.00 | 82.44 | 19.00 |
| >multi-hg38_MT_+_598_1651_RNR1_12S_50ntFlanks@368.404.37\|CAATAAAGCTAAAACTCACCTGAG TTGTAAAAAACTC | >hg38_MT_+_598_1651_RNR1_12S_50ntFlanks.multi | 368 | 404 | 37 | rRF 12S and 16S | NO | 0.00 | 0.00 | 13.45 | 17.96 |
| >multi-hg38_MT_+_598_1651_RNR1_12S_50ntFlanks@369.404.36\|AATAAAGCTAAAACTCACCTGAGT TGTAAAAAACTC | >hg38_MT_+_598_1651_RNR1_12S_50ntFlanks.multi | 369 | 404 | 36 | rRF 12S and 16S | NO | 0.00 | 0.00 | 32.16 | 18.54 |
| >multi-hg38_MT_+_598_1651_RNR1_12S_50ntFlanks@370.404.35\|ATAAAGCTAAAACTCACCTGAGTT GTAAAAAACTC | >hg38_MT_+_598_1651_RNR1_12S_50ntFlanks.multi | 370 | 404 | 35 | rRF 12S and 16S | NO | 0.00 | 0.00 | 16.02 | 19.91 |
| >multi-hg38_MT_+_598_1651_RNR1_12S_50ntFlanks@425.455.31\|ACGAAAGTGGCTTTAACATATCTG AACACAC | >hg38_MT_+_598_1651_RNR1_12S_50ntFlanks.multi | 425 | 455 | 31 | rRF 12S and 16S | NO | 0.00 | 0.00 | 10.37 | 0.00 |
| >multi-hg38_MT_+_598_1651_RNR1_12S_50ntFlanks@428.452.25\|AAAGTGGCTTTAACATATCTGAAC A | >hg38_MT_+_598_1651_RNR1_12S_50ntFlanks.multi | 428 | 452 | 25 | rRF 12S and 16S | NO | 0.00 | 0.00 | 15.26 | 0.00 |
| >multi-hg38_MT_+_598_1651_RNR1_12S_50ntFlanks@428.453.26\|AAAGTGGCTTTAACATATCTGAAC AC | >hg38_MT_+_598_1651_RNR1_12S_50ntFlanks.multi | 428 | 453 | 26 | rRF 12S and 16S | NO | 0.00 | 0.00 | 17.74 | 0.00 |
| >multi-hg38_MT_+_598_1651_RNR1_12S_50ntFlanks@428.454.27\|AAAGTGGCTTTAACATATCTGAAC ACA | >hg38_MT_+_598_1651_RNR1_12S_50ntFlanks.multi | 428 | 454 | 27 | rRF 12S and 16S | NO | 0.00 | 0.00 | 11.19 | 0.00 |
| >multi-hg38_MT_+_598_1651_RNR1_12S_50ntFlanks@428.455.28\|AAAGTGGCTTTAACATATCTGAAC ACAC | >hg38_MT_+_598_1651_RNR1_12S_50ntFlanks.multi | 428 | 455 | 28 | rRF 12S and 16S | NO | 0.00 | 0.00 | 75.67 | 10.95 |
| >multi-hg38_MT_+_598_1651_RNR1_12S_50ntFlanks@428.456.29\|AAAGTGGCTTTAACATATCTGAAC ACACA | >hg38_MT_+_598_1651_RNR1_12S_50ntFlanks.multi | 428 | 456 | 29 | rRF 12S and 16S | NO | 0.00 | 0.00 | 31.47 | 0.00 |
| >multi-hg38_MT_+_598_1651_RNR1_12S_50ntFlanks@428.462.35\|AAAGTGGCTTTAACATATCTGAAC ACACAATAGCT | >hg38_MT_+_598_1651_RNR1_12S_50ntFlanks.multi | 428 | 462 | 35 | rRF 12S and 16S | NO | 0.00 | 0.00 | 11.20 | 0.00 |
| >multi-hg38_MT_+_598_1651_RNR1_12S_50ntFlanks@428.465.38\|AAAGTGGCTTTAACATATCTGAAC ACACAATAGCTAAG | >hg38_MT_+_598_1651_RNR1_12S_50ntFlanks.multi | 428 | 465 | 38 | rRF 12S and 16S | NO | 0.00 | 0.00 | 11.57 | 0.00 |
| >multi-hg38_MT_+_598_1651_RNR1_12S_50ntFlanks@428.468.41\|AAAGTGGCTTTAACATATCTGAAC ACACAATAGCTAAGACC | >hg38_MT_+_598_1651_RNR1_12S_50ntFlanks.multi | 428 | 468 | 41 | rRF 12S and 16S | NO | 0.00 | 0.00 | 70.17 | 56.11 |
| >multi-hg38_MT_+_598_1651_RNR1_12S_50ntFlanks@428.469.42\|AAAGTGGCTTTAACATATCTGAAC ACACAATAGCTAAGACCC | >hg38_MT_+_598_1651_RNR1_12S_50ntFlanks.multi | 428 | 469 | 42 | rRF 12S and 16S | NO | 0.00 | 0.00 | 17.21 | 18.74 |
| >multi-hg38_MT_+_598_1651_RNR1_12S_50ntFlanks@439.469.31\|AACATATCTGAACACACAATAGCT AAGACCC | >hg38_MT_+_598_1651_RNR1_12S_50ntFlanks.multi | 439 | 469 | 31 | rRF 12S and 16S | NO | 0.00 | 0.00 | 10.51 | 0.00 |

### MDA-MB-231

| | | | | | | | | | | |
|---|---|---|---|---|---|---|---|---|---|---|
| >multi-hg38_MT_+_598_1651_RNR1_12S_50ntFlanks@442.480.39\|ATATCTGAACACACAATAGCTAAGACCCAAACTGGGATT | >hg38_MT_+_598_1651_RNR1_12S_50ntFlanks.multi | 442 | 480 | 39 | rRF 12S and 16S | NO | 0.00 | 0.00 | 15.87 | 12.35 |
| >multi-hg38_MT_+_598_1651_RNR1_12S_50ntFlanks@443.468.26\|TATCTGAACACACAATAGCTAAGACC | >hg38_MT_+_598_1651_RNR1_12S_50ntFlanks.multi | 443 | 468 | 26 | rRF 12S and 16S | NO | 0.00 | 0.00 | 13.04 | 0.00 |
| >multi-hg38_MT_+_598_1651_RNR1_12S_50ntFlanks@443.469.27\|TATCTGAACACACAATAGCTAAGACCC | >hg38_MT_+_598_1651_RNR1_12S_50ntFlanks.multi | 443 | 469 | 27 | rRF 12S and 16S | NO | 0.00 | 0.00 | 13.21 | 0.00 |
| >multi-hg38_MT_+_598_1651_RNR1_12S_50ntFlanks@443.480.38\|TATCTGAACACACAATAGCTAAGACCCAAACTGGGATT | >hg38_MT_+_598_1651_RNR1_12S_50ntFlanks.multi | 443 | 480 | 38 | rRF 12S and 16S | NO | 0.00 | 0.00 | 17.77 | 21.25 |
| >multi-hg38_MT_+_598_1651_RNR1_12S_50ntFlanks@608.633.26\|TGTTCTGTAATCGATAAACCCCGATC | >hg38_MT_+_598_1651_RNR1_12S_50ntFlanks.multi | 608 | 633 | 26 | rRF 12S and 16S | NO | 0.00 | 0.00 | 10.66 | 0.00 |
| >multi-hg38_MT_+_598_1651_RNR1_12S_50ntFlanks@634.665.32\|AACCTCACCACCTCTTGCTCAGCCTATATACC | >hg38_MT_+_598_1651_RNR1_12S_50ntFlanks.multi | 634 | 665 | 32 | rRF 12S and 16S | NO | 0.00 | 0.00 | 12.56 | 0.00 |
| >multi-hg38_MT_+_598_1651_RNR1_12S_50ntFlanks@718.744.27\|GTAAAGACGTTAGGTCAAGGTGTAGCC | >hg38_MT_+_598_1651_RNR1_12S_50ntFlanks.multi | 718 | 744 | 27 | rRF 12S and 16S | NO | 0.00 | 0.00 | 15.75 | 0.00 |
| >multi-hg38_MT_+_598_1651_RNR1_12S_50ntFlanks@719.744.26\|TAAAGACGTTAGGTCAAGGTGTAGCC | >hg38_MT_+_598_1651_RNR1_12S_50ntFlanks.multi | 719 | 744 | 26 | rRF 12S and 16S | NO | 0.00 | 0.00 | 35.79 | 10.74 |
| >multi-hg38_MT_+_598_1651_RNR1_12S_50ntFlanks@904.929.26\|AAGTATACTTCAAAGGACATTTAACT | >hg38_MT_+_598_1651_RNR1_12S_50ntFlanks.multi | 904 | 929 | 26 | rRF 12S and 16S | NO | 0.00 | 0.00 | 17.92 | 0.00 |
| >multi-hg38_MT_+_598_1651_RNR1_12S_50ntFlanks@905.929.25\|AGTATACTTCAAAGGACATTTAACT | >hg38_MT_+_598_1651_RNR1_12S_50ntFlanks.multi | 905 | 929 | 25 | rRF 12S and 16S | NO | 0.00 | 0.00 | 239.89 | 0.00 |
| >multi-hg38_MT_+_598_1651_RNR1_12S_50ntFlanks@905.935.31\|AGTATACTTCAAAGGACATTTAACTAAAACC | >hg38_MT_+_598_1651_RNR1_12S_50ntFlanks.multi | 905 | 935 | 31 | rRF 12S and 16S | NO | 0.00 | 0.00 | 13.81 | 0.00 |
| >multi-hg38_MT_+_598_1651_RNR1_12S_50ntFlanks@915.935.21\|AAAGGACATTTAACTAAAACC | >hg38_MT_+_598_1651_RNR1_12S_50ntFlanks.multi | 915 | 935 | 21 | rRF 12S and 16S | NO | 0.00 | 0.00 | 11.97 | 0.00 |
| >multi-hg38_MT_+_598_1651_RNR1_12S_50ntFlanks@916.935.20\|AAGGACATTTAACTAAAACC | >hg38_MT_+_598_1651_RNR1_12S_50ntFlanks.multi | 916 | 935 | 20 | rRF 12S and 16S | NO | 0.00 | 0.00 | 70.00 | 0.00 |
| >multi-hg38_MT_+_598_1651_RNR1_12S_50ntFlanks@917.935.19\|AGGACATTTAACTAAAACC | >hg38_MT_+_598_1651_RNR1_12S_50ntFlanks.multi | 917 | 935 | 19 | rRF 12S and 16S | NO | 0.00 | 0.00 | 58.83 | 0.00 |
| >multi-hg38_MT_+_598_1651_RNR1_12S_50ntFlanks@970.1004.35\|ATGGTAAGTGTACTGGAAAGTGCACTTGGACGAAC | >hg38_MT_+_598_1651_RNR1_12S_50ntFlanks.multi | 970 | 1004 | 35 | rRF 12S and 16S | NO | 0.00 | 0.00 | 10.95 | 10.84 |
| >multi-hg38_MT_+_598_1651_RNR1_12S_50ntFlanks@972.1004.33\|GGTAAGTGTACTGGAAAGTGCACTTGGACGAAC | >hg38_MT_+_598_1651_RNR1_12S_50ntFlanks.multi | 972 | 1004 | 33 | rRF 12S and 16S | NO | 0.00 | 0.00 | 17.38 | 13.72 |
| >hg38_21_+_8433172_8446622_RNA45SN1_45S_50ntFlanks@51.71.21\|GCTGACACGCTGTCCTCTGGC | >hg38_21_+_8433172_8446622_RNA45SN1_45S_50ntFlanks | 51 | 71 | 21 | rRF 45S | NO | 24.98 | 0.00 | 0.00 | 0.00 |
| >hg38_21_+_8433172_8446622_RNA45SN1_45S_50ntFlanks@51.74.24\|GCTGACACGCTGTCCTCTGGCGAC | >hg38_21_+_8433172_8446622_RNA45SN1_45S_50ntFlanks | 51 | 74 | 24 | rRF 45S | NO | 64.20 | 0.00 | 0.00 | 0.00 |
| >hg38_21_+_8433172_8446622_RNA45SN1_45S_50ntFlanks@474.496.23\|TCTTCGTGATCGATGTGGTGACG | >hg38_21_+_8433172_8446622_RNA45SN1_45S_50ntFlanks | 474 | 496 | 23 | rRF 45S | NO | 61.00 | 0.00 | 0.00 | 0.00 |
| >hg38_21_+_8433172_8446622_RNA45SN1_45S_50ntFlanks@474.497.24\|TCTTCGTGATCGATGTGGTGACGT | >hg38_21_+_8433172_8446622_RNA45SN1_45S_50ntFlanks | 474 | 497 | 24 | rRF 45S | NO | 204.37 | 0.00 | 11.95 | 11.83 |
| >hg38_21_+_8433172_8446622_RNA45SN1_45S_50ntFlanks@474.498.25\|TCTTCGTGATCGATGTGGTGACGTC | >hg38_21_+_8433172_8446622_RNA45SN1_45S_50ntFlanks | 474 | 498 | 25 | rRF 45S | NO | 59.70 | 0.00 | 0.00 | 14.50 |
| >hg38_21_+_8433172_8446622_RNA45SN1_45S_50ntFlanks@474.504.31\|TCTTCGTGATCGATGTGGTGACGTCGTGCTC | >hg38_21_+_8433172_8446622_RNA45SN1_45S_50ntFlanks | 474 | 504 | 31 | rRF 45S | NO | 78.81 | 0.00 | 0.00 | 0.00 |
| >hg38_21_+_8433172_8446622_RNA45SN1_45S_50ntFlanks@474.505.32\|TCTTCGTGATCGATGTGGTGACGTCGTGCTCT | >hg38_21_+_8433172_8446622_RNA45SN1_45S_50ntFlanks | 474 | 505 | 32 | rRF 45S | NO | 141.71 | 0.00 | 0.00 | 0.00 |
| >hg38_21_+_8433172_8446622_RNA45SN1_45S_50ntFlanks@474.506.33\|TCTTCGTGATCGATGTGGTGACGTCGTGCTCTC | >hg38_21_+_8433172_8446622_RNA45SN1_45S_50ntFlanks | 474 | 506 | 33 | rRF 45S | NO | 73.48 | 0.00 | 0.00 | 0.00 |
| >hg38_21_+_8433172_8446622_RNA45SN1_45S_50ntFlanks@474.516.43\|TCTTCGTGATCGATGTGGTGACGTCGTGCTCTCCCGGGCCGGG | >hg38_21_+_8433172_8446622_RNA45SN1_45S_50ntFlanks | 474 | 516 | 43 | rRF 45S | NO | 47.18 | 0.00 | 0.00 | 0.00 |
| >hg38_21_+_8433172_8446622_RNA45SN1_45S_50ntFlanks@475.492.18\|CTTCGTGATCGATGTGGT | >hg38_21_+_8433172_8446622_RNA45SN1_45S_50ntFlanks | 475 | 492 | 18 | rRF 45S | NO | 53.50 | 0.00 | 0.00 | 0.00 |
| >hg38_21_+_8433172_8446622_RNA45SN1_45S_50ntFlanks@475.494.20\|CTTCGTGATCGATGTGGTGA | >hg38_21_+_8433172_8446622_RNA45SN1_45S_50ntFlanks | 475 | 494 | 20 | rRF 45S | NO | 75.70 | 0.00 | 0.00 | 0.00 |
| >hg38_21_+_8433172_8446622_RNA45SN1_45S_50ntFlanks@475.495.21\|CTTCGTGATCGATGTGGTGAC | >hg38_21_+_8433172_8446622_RNA45SN1_45S_50ntFlanks | 475 | 495 | 21 | rRF 45S | NO | 369.84 | 0.00 | 12.95 | 16.98 |
| >hg38_21_+_8433172_8446622_RNA45SN1_45S_50ntFlanks@475.496.22\|CTTCGTGATCGATGTGGTGACG | >hg38_21_+_8433172_8446622_RNA45SN1_45S_50ntFlanks | 475 | 496 | 22 | rRF 45S | NO | 1100.88 | 13.21 | 136.40 | 109.15 |
| >hg38_21_+_8433172_8446622_RNA45SN1_45S_50ntFlanks@475.497.23\|CTTCGTGATCGATGTGGTGACGT | >hg38_21_+_8433172_8446622_RNA45SN1_45S_50ntFlanks | 475 | 497 | 23 | rRF 45S | NO | 2181.15 | 30.28 | 342.93 | 178.88 |
| >hg38_21_+_8433172_8446622_RNA45SN1_45S_50ntFlanks@475.498.24\|CTTCGTGATCGATGTGGTGACGTC | >hg38_21_+_8433172_8446622_RNA45SN1_45S_50ntFlanks | 475 | 498 | 24 | rRF 45S | NO | 1118.22 | 121.52 | 349.69 | 280.04 |
| >hg38_21_+_8433172_8446622_RNA45SN1_45S_50ntFlanks@475.499.25\|CTTCGTGATCGATGTGGTGACGTCG | >hg38_21_+_8433172_8446622_RNA45SN1_45S_50ntFlanks | 475 | 499 | 25 | rRF 45S | NO | 106.61 | 0.00 | 0.00 | 10.40 |

### MDA-MB-231

| | | | | | | | | | | |
|---|---|---|---|---|---|---|---|---|---|---|
| >hg38_21_+_8433172_8446622_RNA45SN1_45S_50ntFlanks@475.501.27\|CTTCGTGATCGATGTGGTGACGTCGTG | >hg38_21_+_8433172_8446622_RNA45SN1_45S_50ntFlanks | 475 | 501 | 27 | rRF 45S | NO | 86.36 | 0.00 | 11.87 | 16.31 |
| >hg38_21_+_8433172_8446622_RNA45SN1_45S_50ntFlanks@475.502.28\|CTTCGTGATCGATGTGGTGACGTCGTGC | >hg38_21_+_8433172_8446622_RNA45SN1_45S_50ntFlanks | 475 | 502 | 28 | rRF 45S | NO | 125.03 | 0.00 | 11.68 | 12.15 |
| >hg38_21_+_8433172_8446622_RNA45SN1_45S_50ntFlanks@475.503.29\|CTTCGTGATCGATGTGGTGACGTCGTGCT | >hg38_21_+_8433172_8446622_RNA45SN1_45S_50ntFlanks | 475 | 503 | 29 | rRF 45S | NO | 387.30 | 0.00 | 33.35 | 60.55 |
| >hg38_21_+_8433172_8446622_RNA45SN1_45S_50ntFlanks@475.504.30\|CTTCGTGATCGATGTGGTGACGTCGTGCTC | >hg38_21_+_8433172_8446622_RNA45SN1_45S_50ntFlanks | 475 | 504 | 30 | rRF 45S | NO | 1017.59 | 105.24 | 17.70 | 87.67 |
| >hg38_21_+_8433172_8446622_RNA45SN1_45S_50ntFlanks@475.505.31\|CTTCGTGATCGATGTGGTGACGTCGTGCTCT | >hg38_21_+_8433172_8446622_RNA45SN1_45S_50ntFlanks | 475 | 505 | 31 | rRF 45S | YES | 2023.25 | 250.54 | 67.18 | 136.18 |
| >hg38_21_+_8433172_8446622_RNA45SN1_45S_50ntFlanks@475.506.32\|CTTCGTGATCGATGTGGTGACGTCGTGCTCTC | >hg38_21_+_8433172_8446622_RNA45SN1_45S_50ntFlanks | 475 | 506 | 32 | rRF 45S | NO | 1062.90 | 204.77 | 0.00 | 91.85 |
| >hg38_21_+_8433172_8446622_RNA45SN1_45S_50ntFlanks@475.507.33\|CTTCGTGATCGATGTGGTGACGTCGTGCTCTCC | >hg38_21_+_8433172_8446622_RNA45SN1_45S_50ntFlanks | 475 | 507 | 33 | rRF 45S | NO | 250.24 | 109.08 | 0.00 | 65.93 |
| >hg38_21_+_8433172_8446622_RNA45SN1_45S_50ntFlanks@475.508.34\|CTTCGTGATCGATGTGGTGACGTCGTGCTCTCCC | >hg38_21_+_8433172_8446622_RNA45SN1_45S_50ntFlanks | 475 | 508 | 34 | rRF 45S | NO | 106.43 | 24.68 | 0.00 | 14.51 |
| >hg38_21_+_8433172_8446622_RNA45SN1_45S_50ntFlanks@475.509.35\|CTTCGTGATCGATGTGGTGACGTCGTGCTCTCCCG | >hg38_21_+_8433172_8446622_RNA45SN1_45S_50ntFlanks | 475 | 509 | 35 | rRF 45S | NO | 368.22 | 80.13 | 0.00 | 55.96 |
| >hg38_21_+_8433172_8446622_RNA45SN1_45S_50ntFlanks@475.510.36\|CTTCGTGATCGATGTGGTGACGTCGTGCTCTCCCGG | >hg38_21_+_8433172_8446622_RNA45SN1_45S_50ntFlanks | 475 | 510 | 36 | rRF 45S | NO | 458.96 | 100.48 | 0.00 | 65.77 |
| >hg38_21_+_8433172_8446622_RNA45SN1_45S_50ntFlanks@475.511.37\|CTTCGTGATCGATGTGGTGACGTCGTGCTCTCCCGGG | >hg38_21_+_8433172_8446622_RNA45SN1_45S_50ntFlanks | 475 | 511 | 37 | rRF 45S | NO | 282.26 | 58.24 | 0.00 | 17.38 |
| >hg38_21_+_8433172_8446622_RNA45SN1_45S_50ntFlanks@475.512.38\|CTTCGTGATCGATGTGGTGACGTCGTGCTCTCCCGGGC | >hg38_21_+_8433172_8446622_RNA45SN1_45S_50ntFlanks | 475 | 512 | 38 | rRF 45S | NO | 94.54 | 0.00 | 0.00 | 0.00 |
| >hg38_21_+_8433172_8446622_RNA45SN1_45S_50ntFlanks@475.513.39\|CTTCGTGATCGATGTGGTGACGTCGTGCTCTCCCGGGCC | >hg38_21_+_8433172_8446622_RNA45SN1_45S_50ntFlanks | 475 | 513 | 39 | rRF 45S | NO | 87.11 | 0.00 | 0.00 | 0.00 |
| >hg38_21_+_8433172_8446622_RNA45SN1_45S_50ntFlanks@475.514.40\|CTTCGTGATCGATGTGGTGACGTCGTGCTCTCCCGGGCCG | >hg38_21_+_8433172_8446622_RNA45SN1_45S_50ntFlanks | 475 | 514 | 40 | rRF 45S | NO | 280.08 | 13.30 | 0.00 | 0.00 |
| >hg38_21_+_8433172_8446622_RNA45SN1_45S_50ntFlanks@475.515.41\|CTTCGTGATCGATGTGGTGACGTCGTGCTCTCCCGGGCCGG | >hg38_21_+_8433172_8446622_RNA45SN1_45S_50ntFlanks | 475 | 515 | 41 | rRF 45S | NO | 240.20 | 0.00 | 0.00 | 0.00 |
| >hg38_21_+_8433172_8446622_RNA45SN1_45S_50ntFlanks@475.516.42\|CTTCGTGATCGATGTGGTGACGTCGTGCTCTCCCGGGCCGGG | >hg38_21_+_8433172_8446622_RNA45SN1_45S_50ntFlanks | 475 | 516 | 42 | rRF 45S | NO | 771.73 | 89.55 | 0.00 | 84.24 |
| >hg38_21_+_8433172_8446622_RNA45SN1_45S_50ntFlanks@475.517.43\|CTTCGTGATCGATGTGGTGACGTCGTGCTCTCCCGGGCCGGGT | >hg38_21_+_8433172_8446622_RNA45SN1_45S_50ntFlanks | 475 | 517 | 43 | rRF 45S | NO | 402.15 | 37.09 | 0.00 | 76.80 |
| >hg38_21_+_8433172_8446622_RNA45SN1_45S_50ntFlanks@475.518.44\|CTTCGTGATCGATGTGGTGACGTCGTGCTCTCCCGGGCCGGGTC | >hg38_21_+_8433172_8446622_RNA45SN1_45S_50ntFlanks | 475 | 518 | 44 | rRF 45S | NO | 245.67 | 29.99 | 0.00 | 57.19 |
| >hg38_21_+_8433172_8446622_RNA45SN1_45S_50ntFlanks@475.519.45\|CTTCGTGATCGATGTGGTGACGTCGTGCTCTCCCGGGCCGGGTCC | >hg38_21_+_8433172_8446622_RNA45SN1_45S_50ntFlanks | 475 | 519 | 45 | rRF 45S | NO | 85.72 | 0.00 | 0.00 | 0.00 |
| >hg38_21_+_8433172_8446622_RNA45SN1_45S_50ntFlanks@475.520.46\|CTTCGTGATCGATGTGGTGACGTCGTGCTCTCCCGGGCCGGGTCCG | >hg38_21_+_8433172_8446622_RNA45SN1_45S_50ntFlanks | 475 | 520 | 46 | rRF 45S | NO | 177.02 | 0.00 | 0.00 | 14.27 |
| >hg38_21_+_8433172_8446622_RNA45SN1_45S_50ntFlanks@475.521.47\|CTTCGTGATCGATGTGGTGACGTCGTGCTCTCCCGGGCCGGGTCCGA | >hg38_21_+_8433172_8446622_RNA45SN1_45S_50ntFlanks | 475 | 521 | 47 | rRF 45S | NO | 191.14 | 0.00 | 0.00 | 20.70 |
| >hg38_21_+_8433172_8446622_RNA45SN1_45S_50ntFlanks@475.522.48\|CTTCGTGATCGATGTGGTGACGTCGTGCTCTCCCGGGCCGGGTCCGAG | >hg38_21_+_8433172_8446622_RNA45SN1_45S_50ntFlanks | 475 | 522 | 48 | rRF 45S | NO | 136.42 | 0.00 | 0.00 | 0.00 |
| >hg38_21_+_8433172_8446622_RNA45SN1_45S_50ntFlanks@475.523.49\|CTTCGTGATCGATGTGGTGACGTCGTGCTCTCCCGGGCCGGGTCCGAGC | >hg38_21_+_8433172_8446622_RNA45SN1_45S_50ntFlanks | 475 | 523 | 49 | rRF 45S | NO | 148.19 | 0.00 | 0.00 | 0.00 |
| >hg38_21_+_8433172_8446622_RNA45SN1_45S_50ntFlanks@475.524.50\|CTTCGTGATCGATGTGGTGACGTCGTGCTCTCCCGGGCCGGGTCCGAGCC | >hg38_21_+_8433172_8446622_RNA45SN1_45S_50ntFlanks | 475 | 524 | 50 | rRF 45S | NO | 150.09 | 0.00 | 0.00 | 0.00 |
| >hg38_21_+_8433172_8446622_RNA45SN1_45S_50ntFlanks@476.494.19\|TTCGTGATCGATGTGGTGA | >hg38_21_+_8433172_8446622_RNA45SN1_45S_50ntFlanks | 476 | 494 | 19 | rRF 45S | NO | 74.52 | 0.00 | 0.00 | 0.00 |
| >hg38_21_+_8433172_8446622_RNA45SN1_45S_50ntFlanks@476.495.20\|TTCGTGATCGATGTGGTGAC | >hg38_21_+_8433172_8446622_RNA45SN1_45S_50ntFlanks | 476 | 495 | 20 | rRF 45S | NO | 381.17 | 0.00 | 0.00 | 12.96 |
| >hg38_21_+_8433172_8446622_RNA45SN1_45S_50ntFlanks@476.496.21\|TTCGTGATCGATGTGGTGACG | >hg38_21_+_8433172_8446622_RNA45SN1_45S_50ntFlanks | 476 | 496 | 21 | rRF 45S | NO | 296.43 | 0.00 | 0.00 | 16.67 |
| >hg38_21_+_8433172_8446622_RNA45SN1_45S_50ntFlanks@476.497.22\|TTCGTGATCGATGTGGTGACGT | >hg38_21_+_8433172_8446622_RNA45SN1_45S_50ntFlanks | 476 | 497 | 22 | rRF 45S | NO | 223.30 | 0.00 | 10.85 | 19.45 |
| >hg38_21_+_8433172_8446622_RNA45SN1_45S_50ntFlanks@476.498.23\|TTCGTGATCGATGTGGTGACGTC | >hg38_21_+_8433172_8446622_RNA45SN1_45S_50ntFlanks | 476 | 498 | 23 | rRF 45S | NO | 495.81 | 0.00 | 16.07 | 72.76 |
| >hg38_21_+_8433172_8446622_RNA45SN1_45S_50ntFlanks@476.499.24\|TTCGTGATCGATGTGGTGACGTCG | >hg38_21_+_8433172_8446622_RNA45SN1_45S_50ntFlanks | 476 | 499 | 24 | rRF 45S | NO | 386.05 | 27.73 | 33.31 | 88.79 |
| >hg38_21_+_8433172_8446622_RNA45SN1_45S_50ntFlanks@476.500.25\|TTCGTGATCGATGTGGTGACGTCGT | >hg38_21_+_8433172_8446622_RNA45SN1_45S_50ntFlanks | 476 | 500 | 25 | rRF 45S | NO | 24.67 | 0.00 | 0.00 | 0.00 |
| >hg38_21_+_8433172_8446622_RNA45SN1_45S_50ntFlanks@476.501.26\|TTCGTGATCGATGTGGTGACGTCGTG | >hg38_21_+_8433172_8446622_RNA45SN1_45S_50ntFlanks | 476 | 501 | 26 | rRF 45S | NO | 227.19 | 0.00 | 0.00 | 13.63 |
| >hg38_21_+_8433172_8446622_RNA45SN1_45S_50ntFlanks@476.502.27\|TTCGTGATCGATGTGGTGACGTCGTGC | >hg38_21_+_8433172_8446622_RNA45SN1_45S_50ntFlanks | 476 | 502 | 27 | rRF 45S | NO | 202.38 | 0.00 | 0.00 | 0.00 |
| >hg38_21_+_8433172_8446622_RNA45SN1_45S_50ntFlanks@476.503.28\|TTCGTGATCGATGTGGTGACGTCGTGCT | >hg38_21_+_8433172_8446622_RNA45SN1_45S_50ntFlanks | 476 | 503 | 28 | rRF 45S | NO | 406.57 | 0.00 | 0.00 | 14.16 |
| >hg38_21_+_8433172_8446622_RNA45SN1_45S_50ntFlanks@476.504.29\|TTCGTGATCGATGTGGTGACGTCGTGCTC | >hg38_21_+_8433172_8446622_RNA45SN1_45S_50ntFlanks | 476 | 504 | 29 | rRF 45S | NO | 544.20 | 0.00 | 0.00 | 19.50 |
| >hg38_21_+_8433172_8446622_RNA45SN1_45S_50ntFlanks@476.505.30\|TTCGTGATCGATGTGGTGACGTCGTGCTCT | >hg38_21_+_8433172_8446622_RNA45SN1_45S_50ntFlanks | 476 | 505 | 30 | rRF 45S | NO | 1057.43 | 63.28 | 0.00 | 77.21 |
| >hg38_21_+_8433172_8446622_RNA45SN1_45S_50ntFlanks@476.506.31\|TTCGTGATCGATGTGGTGACGTCGTGCTCTC | >hg38_21_+_8433172_8446622_RNA45SN1_45S_50ntFlanks | 476 | 506 | 31 | rRF 45S | NO | 609.64 | 62.64 | 0.00 | 22.07 |
| >hg38_21_+_8433172_8446622_RNA45SN1_45S_50ntFlanks@476.507.32\|TTCGTGATCGATGTGGTGACGTCGTGCTCTCC | >hg38_21_+_8433172_8446622_RNA45SN1_45S_50ntFlanks | 476 | 507 | 32 | rRF 45S | NO | 297.95 | 59.05 | 0.00 | 14.12 |

### MDA-MB-231

| | | | | | | | | | | |
|---|---|---|---|---|---|---|---|---|---|---|
| >hg38_21_+_8433172_8446622_RNA45SN1_45S_50ntFlanks@476.508.33\|TTCGTGATCGATGT<br>GGTGACGTCGTGCTCTCCC | >hg38_21_+_8433172_8446622_RNA45SN1_45S_50ntFlanks | 476 | 508 | 33 | rRF 45S | NO | 95.70 | 0.00 | 0.00 | 0.00 |
| >hg38_21_+_8433172_8446622_RNA45SN1_45S_50ntFlanks@476.509.34\|TTCGTGATCGATGT<br>GGTGACGTCGTGCTCTCCCG | >hg38_21_+_8433172_8446622_RNA45SN1_45S_50ntFlanks | 476 | 509 | 34 | rRF 45S | NO | 369.31 | 26.31 | 0.00 | 14.79 |
| >hg38_21_+_8433172_8446622_RNA45SN1_45S_50ntFlanks@476.510.35\|TTCGTGATCGATGT<br>GGTGACGTCGTGCTCTCCCGG | >hg38_21_+_8433172_8446622_RNA45SN1_45S_50ntFlanks | 476 | 510 | 35 | rRF 45S | NO | 296.40 | 24.71 | 0.00 | 14.42 |
| >hg38_21_+_8433172_8446622_RNA45SN1_45S_50ntFlanks@476.511.36\|TTCGTGATCGATGT<br>GGTGACGTCGTGCTCTCCCGGG | >hg38_21_+_8433172_8446622_RNA45SN1_45S_50ntFlanks | 476 | 511 | 36 | rRF 45S | NO | 196.01 | 13.32 | 0.00 | 0.00 |
| >hg38_21_+_8433172_8446622_RNA45SN1_45S_50ntFlanks@476.512.37\|TTCGTGATCGATGT<br>GGTGACGTCGTGCTCTCCCGGGC | >hg38_21_+_8433172_8446622_RNA45SN1_45S_50ntFlanks | 476 | 512 | 37 | rRF 45S | NO | 125.10 | 0.00 | 0.00 | 0.00 |
| >hg38_21_+_8433172_8446622_RNA45SN1_45S_50ntFlanks@476.513.38\|TTCGTGATCGATGT<br>GGTGACGTCGTGCTCTCCCGGGCC | >hg38_21_+_8433172_8446622_RNA45SN1_45S_50ntFlanks | 476 | 513 | 38 | rRF 45S | NO | 94.53 | 0.00 | 0.00 | 0.00 |
| >hg38_21_+_8433172_8446622_RNA45SN1_45S_50ntFlanks@476.514.39\|TTCGTGATCGATGT<br>GGTGACGTCGTGCTCTCCCGGGCCG | >hg38_21_+_8433172_8446622_RNA45SN1_45S_50ntFlanks | 476 | 514 | 39 | rRF 45S | NO | 65.19 | 0.00 | 0.00 | 0.00 |
| >hg38_21_+_8433172_8446622_RNA45SN1_45S_50ntFlanks@476.515.40\|TTCGTGATCGATGT<br>GGTGACGTCGTGCTCTCCCGGGCCGG | >hg38_21_+_8433172_8446622_RNA45SN1_45S_50ntFlanks | 476 | 515 | 40 | rRF 45S | NO | 74.95 | 0.00 | 0.00 | 0.00 |
| >hg38_21_+_8433172_8446622_RNA45SN1_45S_50ntFlanks@476.516.41\|TTCGTGATCGATGT<br>GGTGACGTCGTGCTCTCCCGGGCCGGG | >hg38_21_+_8433172_8446622_RNA45SN1_45S_50ntFlanks | 476 | 516 | 41 | rRF 45S | NO | 623.24 | 58.95 | 0.00 | 21.03 |
| >hg38_21_+_8433172_8446622_RNA45SN1_45S_50ntFlanks@476.517.42\|TTCGTGATCGATGT<br>GGTGACGTCGTGCTCTCCCGGGCCGGGT | >hg38_21_+_8433172_8446622_RNA45SN1_45S_50ntFlanks | 476 | 517 | 42 | rRF 45S | NO | 439.93 | 24.14 | 0.00 | 16.96 |
| >hg38_21_+_8433172_8446622_RNA45SN1_45S_50ntFlanks@476.518.43\|TTCGTGATCGATGT<br>GGTGACGTCGTGCTCTCCCGGGCCGGGTC | >hg38_21_+_8433172_8446622_RNA45SN1_45S_50ntFlanks | 476 | 518 | 43 | rRF 45S | NO | 276.61 | 0.00 | 0.00 | 0.00 |
| >hg38_21_+_8433172_8446622_RNA45SN1_45S_50ntFlanks@476.519.44\|TTCGTGATCGATGT<br>GGTGACGTCGTGCTCTCCCGGGCCGGGTCC | >hg38_21_+_8433172_8446622_RNA45SN1_45S_50ntFlanks | 476 | 519 | 44 | rRF 45S | NO | 116.04 | 0.00 | 0.00 | 0.00 |
| >hg38_21_+_8433172_8446622_RNA45SN1_45S_50ntFlanks@476.520.45\|TTCGTGATCGATGT<br>GGTGACGTCGTGCTCTCCCGGGCCGGGTCCG | >hg38_21_+_8433172_8446622_RNA45SN1_45S_50ntFlanks | 476 | 520 | 45 | rRF 45S | NO | 229.49 | 0.00 | 0.00 | 0.00 |
| >hg38_21_+_8433172_8446622_RNA45SN1_45S_50ntFlanks@476.521.46\|TTCGTGATCGATGT<br>GGTGACGTCGTGCTCTCCCGGGCCGGGTCCGA | >hg38_21_+_8433172_8446622_RNA45SN1_45S_50ntFlanks | 476 | 521 | 46 | rRF 45S | NO | 193.69 | 0.00 | 0.00 | 0.00 |
| >hg38_21_+_8433172_8446622_RNA45SN1_45S_50ntFlanks@476.522.47\|TTCGTGATCGATGT<br>GGTGACGTCGTGCTCTCCCGGGCCGGGTCCGAG | >hg38_21_+_8433172_8446622_RNA45SN1_45S_50ntFlanks | 476 | 522 | 47 | rRF 45S | NO | 139.66 | 0.00 | 0.00 | 0.00 |
| >hg38_21_+_8433172_8446622_RNA45SN1_45S_50ntFlanks@476.523.48\|TTCGTGATCGATGT<br>GGTGACGTCGTGCTCTCCCGGGCCGGGTCCGAGC | >hg38_21_+_8433172_8446622_RNA45SN1_45S_50ntFlanks | 476 | 523 | 48 | rRF 45S | NO | 307.18 | 0.00 | 0.00 | 0.00 |
| >hg38_21_+_8433172_8446622_RNA45SN1_45S_50ntFlanks@476.524.49\|TTCGTGATCGATGT<br>GGTGACGTCGTGCTCTCCCGGGCCGGGTCCGAGCC | >hg38_21_+_8433172_8446622_RNA45SN1_45S_50ntFlanks | 476 | 524 | 49 | rRF 45S | NO | 193.35 | 0.00 | 0.00 | 0.00 |
| >hg38_21_+_8433172_8446622_RNA45SN1_45S_50ntFlanks@476.525.50\|TTCGTGATCGATGT<br>GGTGACGTCGTGCTCTCCCGGGCCGGGTCCGAGCCG | >hg38_21_+_8433172_8446622_RNA45SN1_45S_50ntFlanks | 476 | 525 | 50 | rRF 45S | NO | 94.71 | 0.00 | 0.00 | 0.00 |
| >hg38_21_+_8433172_8446622_RNA45SN1_45S_50ntFlanks@480.497.18\|TGATCGATGTGGTG<br>ACGT | >hg38_21_+_8433172_8446622_RNA45SN1_45S_50ntFlanks | 480 | 497 | 18 | rRF 45S | NO | 51.32 | 0.00 | 0.00 | 0.00 |
| >hg38_21_+_8433172_8446622_RNA45SN1_45S_50ntFlanks@483.505.23\|TCGATGTGGTGAC<br>GTCGTGCTCT | >hg38_21_+_8433172_8446622_RNA45SN1_45S_50ntFlanks | 483 | 505 | 23 | rRF 45S | NO | 101.16 | 0.00 | 0.00 | 0.00 |
| >hg38_21_+_8433172_8446622_RNA45SN1_45S_50ntFlanks@483.506.24\|TCGATGTGGTGAC<br>GTCGTGCTCTC | >hg38_21_+_8433172_8446622_RNA45SN1_45S_50ntFlanks | 483 | 506 | 24 | rRF 45S | NO | 69.64 | 0.00 | 0.00 | 0.00 |
| >hg38_21_+_8433172_8446622_RNA45SN1_45S_50ntFlanks@483.507.25\|TCGATGTGGTGAC<br>GTCGTGCTCTCC | >hg38_21_+_8433172_8446622_RNA45SN1_45S_50ntFlanks | 483 | 507 | 25 | rRF 45S | NO | 53.76 | 0.00 | 0.00 | 0.00 |
| >hg38_21_+_8433172_8446622_RNA45SN1_45S_50ntFlanks@483.516.34\|TCGATGTGGTGAC<br>GTCGTGCTCTCCCGGGCCGGG | >hg38_21_+_8433172_8446622_RNA45SN1_45S_50ntFlanks | 483 | 516 | 34 | rRF 45S | NO | 59.78 | 0.00 | 0.00 | 0.00 |
| >hg38_21_+_8433172_8446622_RNA45SN1_45S_50ntFlanks@483.520.38\|TCGATGTGGTGAC<br>GTCGTGCTCTCCCGGGCCGGGTCCG | >hg38_21_+_8433172_8446622_RNA45SN1_45S_50ntFlanks | 483 | 520 | 38 | rRF 45S | NO | 50.28 | 0.00 | 0.00 | 0.00 |
| >hg38_21_+_8433172_8446622_RNA45SN1_45S_50ntFlanks@483.522.40\|TCGATGTGGTGAC<br>GTCGTGCTCTCCCGGGCCGGGTCCGAG | >hg38_21_+_8433172_8446622_RNA45SN1_45S_50ntFlanks | 483 | 522 | 40 | rRF 45S | NO | 72.25 | 0.00 | 0.00 | 0.00 |
| >hg38_21_+_8433172_8446622_RNA45SN1_45S_50ntFlanks@483.524.42\|TCGATGTGGTGAC<br>GTCGTGCTCTCCCGGGCCGGGTCCGAGCC | >hg38_21_+_8433172_8446622_RNA45SN1_45S_50ntFlanks | 483 | 524 | 42 | rRF 45S | NO | 108.40 | 0.00 | 0.00 | 0.00 |
| >hg38_21_+_8433172_8446622_RNA45SN1_45S_50ntFlanks@483.525.43\|TCGATGTGGTGAC<br>GTCGTGCTCTCCCGGGCCGGGTCCGAGCCG | >hg38_21_+_8433172_8446622_RNA45SN1_45S_50ntFlanks | 483 | 525 | 43 | rRF 45S | NO | 52.43 | 0.00 | 0.00 | 0.00 |
| >hg38_21_+_8433172_8446622_RNA45SN1_45S_50ntFlanks@483.529.47\|TCGATGTGGTGAC<br>GTCGTGCTCTCCCGGGCCGGGTCCGAGCCGCGAC | >hg38_21_+_8433172_8446622_RNA45SN1_45S_50ntFlanks | 483 | 529 | 47 | rRF 45S | NO | 75.27 | 0.00 | 0.00 | 0.00 |
| >hg38_21_+_8433172_8446622_RNA45SN1_45S_50ntFlanks@483.532.50\|TCGATGTGGTGAC<br>GTCGTGCTCTCCCGGGCCGGGTCCGAGCCGCGACGGG | >hg38_21_+_8433172_8446622_RNA45SN1_45S_50ntFlanks | 483 | 532 | 50 | rRF 45S | NO | 138.84 | 0.00 | 0.00 | 0.00 |
| >hg38_21_+_8433172_8446622_RNA45SN1_45S_50ntFlanks@485.504.20\|GATGTGGTGACGT<br>CGTGCTC | >hg38_21_+_8433172_8446622_RNA45SN1_45S_50ntFlanks | 485 | 504 | 20 | rRF 45S | NO | 61.61 | 0.00 | 0.00 | 0.00 |
| >hg38_21_+_8433172_8446622_RNA45SN1_45S_50ntFlanks@485.505.21\|GATGTGGTGACGT<br>CGTGCTCT | >hg38_21_+_8433172_8446622_RNA45SN1_45S_50ntFlanks | 485 | 505 | 21 | rRF 45S | NO | 262.31 | 0.00 | 35.58 | 0.00 |
| >hg38_21_+_8433172_8446622_RNA45SN1_45S_50ntFlanks@485.506.22\|GATGTGGTGACGT<br>CGTGCTCTC | >hg38_21_+_8433172_8446622_RNA45SN1_45S_50ntFlanks | 485 | 506 | 22 | rRF 45S | NO | 181.71 | 0.00 | 0.00 | 0.00 |
| >hg38_21_+_8433172_8446622_RNA45SN1_45S_50ntFlanks@485.507.23\|GATGTGGTGACGT<br>CGTGCTCTCC | >hg38_21_+_8433172_8446622_RNA45SN1_45S_50ntFlanks | 485 | 507 | 23 | rRF 45S | NO | 111.70 | 0.00 | 0.00 | 0.00 |
| >hg38_21_+_8433172_8446622_RNA45SN1_45S_50ntFlanks@485.508.24\|GATGTGGTGACGT<br>CGTGCTCTCCC | >hg38_21_+_8433172_8446622_RNA45SN1_45S_50ntFlanks | 485 | 508 | 24 | rRF 45S | NO | 55.67 | 0.00 | 0.00 | 0.00 |
| >hg38_21_+_8433172_8446622_RNA45SN1_45S_50ntFlanks@485.509.25\|GATGTGGTGACGT<br>CGTGCTCTCCCG | >hg38_21_+_8433172_8446622_RNA45SN1_45S_50ntFlanks | 485 | 509 | 25 | rRF 45S | NO | 63.34 | 0.00 | 0.00 | 0.00 |
| >hg38_21_+_8433172_8446622_RNA45SN1_45S_50ntFlanks@485.510.26\|GATGTGGTGACGT<br>CGTGCTCTCCCGG | >hg38_21_+_8433172_8446622_RNA45SN1_45S_50ntFlanks | 485 | 510 | 26 | rRF 45S | NO | 59.10 | 0.00 | 0.00 | 0.00 |
| >hg38_21_+_8433172_8446622_RNA45SN1_45S_50ntFlanks@485.511.27\|GATGTGGTGACGT<br>CGTGCTCTCCCGGG | >hg38_21_+_8433172_8446622_RNA45SN1_45S_50ntFlanks | 485 | 511 | 27 | rRF 45S | NO | 25.70 | 0.00 | 0.00 | 0.00 |
| >hg38_21_+_8433172_8446622_RNA45SN1_45S_50ntFlanks@485.514.30\|GATGTGGTGACGT<br>CGTGCTCTCCCGGGCCG | >hg38_21_+_8433172_8446622_RNA45SN1_45S_50ntFlanks | 485 | 514 | 30 | rRF 45S | NO | 82.50 | 0.00 | 0.00 | 0.00 |

### MDA-MB-231

| | | | | | | | | | | |
|---|---|---|---|---|---|---|---|---|---|---|
| >hg38_21_+_8433172_8446622_RNA45SN1_45S_50ntFlanks@485.515.31\|GATGTGGTGACGT CGTGCTCTCCCGGGCCGG | >hg38_21_+_8433172_8446622_RNA45SN1_45S_50ntFlanks | 485 | 515 | 31 | rRF 45S | NO | 73.47 | 0.00 | 0.00 | 0.00 |
| >hg38_21_+_8433172_8446622_RNA45SN1_45S_50ntFlanks@485.516.32\|GATGTGGTGACGT CGTGCTCTCCCGGGCCGGG | >hg38_21_+_8433172_8446622_RNA45SN1_45S_50ntFlanks | 485 | 516 | 32 | rRF 45S | NO | 89.66 | 0.00 | 0.00 | 0.00 |
| >hg38_21_+_8433172_8446622_RNA45SN1_45S_50ntFlanks@485.517.33\|GATGTGGTGACGT CGTGCTCTCCCGGGCCGGGT | >hg38_21_+_8433172_8446622_RNA45SN1_45S_50ntFlanks | 485 | 517 | 33 | rRF 45S | NO | 81.25 | 0.00 | 0.00 | 0.00 |
| >hg38_21_+_8433172_8446622_RNA45SN1_45S_50ntFlanks@485.518.34\|GATGTGGTGACGT CGTGCTCTCCCGGGCCGGGTC | >hg38_21_+_8433172_8446622_RNA45SN1_45S_50ntFlanks | 485 | 518 | 34 | rRF 45S | NO | 51.44 | 0.00 | 0.00 | 0.00 |
| >hg38_21_+_8433172_8446622_RNA45SN1_45S_50ntFlanks@485.519.35\|GATGTGGTGACGT CGTGCTCTCCCGGGCCGGGTCC | >hg38_21_+_8433172_8446622_RNA45SN1_45S_50ntFlanks | 485 | 519 | 35 | rRF 45S | NO | 61.39 | 0.00 | 0.00 | 0.00 |
| >hg38_21_+_8433172_8446622_RNA45SN1_45S_50ntFlanks@485.520.36\|GATGTGGTGACGT CGTGCTCTCCCGGGCCGGGTCCG | >hg38_21_+_8433172_8446622_RNA45SN1_45S_50ntFlanks | 485 | 520 | 36 | rRF 45S | NO | 116.66 | 0.00 | 0.00 | 0.00 |
| >hg38_21_+_8433172_8446622_RNA45SN1_45S_50ntFlanks@485.521.37\|GATGTGGTGACGT CGTGCTCTCCCGGGCCGGGTCCGA | >hg38_21_+_8433172_8446622_RNA45SN1_45S_50ntFlanks | 485 | 521 | 37 | rRF 45S | NO | 106.61 | 0.00 | 0.00 | 0.00 |
| >hg38_21_+_8433172_8446622_RNA45SN1_45S_50ntFlanks@485.522.38\|GATGTGGTGACGT CGTGCTCTCCCGGGCCGGGTCCGAG | >hg38_21_+_8433172_8446622_RNA45SN1_45S_50ntFlanks | 485 | 522 | 38 | rRF 45S | NO | 101.13 | 0.00 | 0.00 | 0.00 |
| >hg38_21_+_8433172_8446622_RNA45SN1_45S_50ntFlanks@485.524.40\|GATGTGGTGACGT CGTGCTCTCCCGGGCCGGGTCCGAGCC | >hg38_21_+_8433172_8446622_RNA45SN1_45S_50ntFlanks | 485 | 524 | 40 | rRF 45S | NO | 91.31 | 0.00 | 0.00 | 0.00 |
| >hg38_21_+_8433172_8446622_RNA45SN1_45S_50ntFlanks@485.525.41\|GATGTGGTGACGT CGTGCTCTCCCGGGCCGGGTCCGAGCCG | >hg38_21_+_8433172_8446622_RNA45SN1_45S_50ntFlanks | 485 | 525 | 41 | rRF 45S | NO | 100.61 | 0.00 | 0.00 | 0.00 |
| >hg38_21_+_8433172_8446622_RNA45SN1_45S_50ntFlanks@485.526.42\|GATGTGGTGACGT CGTGCTCTCCCGGGCCGGGTCCGAGCCGC | >hg38_21_+_8433172_8446622_RNA45SN1_45S_50ntFlanks | 485 | 526 | 42 | rRF 45S | NO | 55.59 | 0.00 | 0.00 | 0.00 |
| >hg38_21_+_8433172_8446622_RNA45SN1_45S_50ntFlanks@485.527.43\|GATGTGGTGACGT CGTGCTCTCCCGGGCCGGGTCCGAGCCGCG | >hg38_21_+_8433172_8446622_RNA45SN1_45S_50ntFlanks | 485 | 527 | 43 | rRF 45S | NO | 54.67 | 0.00 | 0.00 | 0.00 |
| >hg38_21_+_8433172_8446622_RNA45SN1_45S_50ntFlanks@485.529.45\|GATGTGGTGACGT CGTGCTCTCCCGGGCCGGGTCCGAGCCGCGAC | >hg38_21_+_8433172_8446622_RNA45SN1_45S_50ntFlanks | 485 | 529 | 45 | rRF 45S | NO | 26.08 | 0.00 | 0.00 | 0.00 |
| >hg38_21_+_8433172_8446622_RNA45SN1_45S_50ntFlanks@485.532.48\|GATGTGGTGACGT CGTGCTCTCCCGGGCCGGGTCCGAGCCGCGACGGG | >hg38_21_+_8433172_8446622_RNA45SN1_45S_50ntFlanks | 485 | 532 | 48 | rRF 45S | NO | 63.91 | 0.00 | 0.00 | 0.00 |
| >hg38_21_+_8433172_8446622_RNA45SN1_45S_50ntFlanks@3705.3722.18\|TACCTGGTTGAT CCTGCC | >hg38_21_+_8433172_8446622_RNA45SN1_45S_50ntFlanks | 3705 | 3722 | 18 | rRF 45S | NO | 144.17 | 1058.31 | 0.00 | 104.62 |
| >hg38_21_+_8433172_8446622_RNA45SN1_45S_50ntFlanks@3705.3723.19\|TACCTGGTTGAT CCTGCCA | >hg38_21_+_8433172_8446622_RNA45SN1_45S_50ntFlanks | 3705 | 3723 | 19 | rRF 45S | NO | 0.00 | 84.57 | 0.00 | 10.57 |
| >hg38_21_+_8433172_8446622_RNA45SN1_45S_50ntFlanks@3705.3725.21\|TACCTGGTTGAT CCTGCCAGT | >hg38_21_+_8433172_8446622_RNA45SN1_45S_50ntFlanks | 3705 | 3725 | 21 | rRF 45S | NO | 0.00 | 61.67 | 0.00 | 70.84 |
| >hg38_21_+_8433172_8446622_RNA45SN1_45S_50ntFlanks@3705.3728.24\|TACCTGGTTGAT CCTGCCAGTAGC | >hg38_21_+_8433172_8446622_RNA45SN1_45S_50ntFlanks | 3705 | 3728 | 24 | rRF 45S | NO | 0.00 | 92.83 | 0.00 | 0.00 |
| >hg38_21_+_8433172_8446622_RNA45SN1_45S_50ntFlanks@3705.3736.32\|TACCTGGTTGAT CCTGCCAGTAGCATATGCTT | >hg38_21_+_8433172_8446622_RNA45SN1_45S_50ntFlanks | 3705 | 3736 | 32 | rRF 45S | NO | 0.00 | 60.15 | 0.00 | 90.33 |
| >hg38_21_+_8433172_8446622_RNA45SN1_45S_50ntFlanks@3705.3739.35\|TACCTGGTTGAT CCTGCCAGTAGCATATGCTTGTC | >hg38_21_+_8433172_8446622_RNA45SN1_45S_50ntFlanks | 3705 | 3739 | 35 | rRF 45S | NO | 0.00 | 78.04 | 0.00 | 20.41 |
| >hg38_21_+_8433172_8446622_RNA45SN1_45S_50ntFlanks@3705.3741.37\|TACCTGGTTGAT CCTGCCAGTAGCATATGCTTGTCTC | >hg38_21_+_8433172_8446622_RNA45SN1_45S_50ntFlanks | 3705 | 3741 | 37 | rRF 45S | NO | 0.00 | 150.56 | 0.00 | 17.76 |
| >hg38_21_+_8433172_8446622_RNA45SN1_45S_50ntFlanks@3705.3742.38\|TACCTGGTTGAT CCTGCCAGTAGCATATGCTTGTCTCA | >hg38_21_+_8433172_8446622_RNA45SN1_45S_50ntFlanks | 3705 | 3742 | 38 | rRF 45S | NO | 0.00 | 110.37 | 0.00 | 15.95 |
| >hg38_21_+_8433172_8446622_RNA45SN1_45S_50ntFlanks@3705.3743.39\|TACCTGGTTGAT CCTGCCAGTAGCATATGCTTGTCTCAA | >hg38_21_+_8433172_8446622_RNA45SN1_45S_50ntFlanks | 3705 | 3743 | 39 | rRF 45S | NO | 0.00 | 215.55 | 0.00 | 14.45 |
| >hg38_21_+_8433172_8446622_RNA45SN1_45S_50ntFlanks@3705.3744.40\|TACCTGGTTGAT CCTGCCAGTAGCATATGCTTGTCTCAAA | >hg38_21_+_8433172_8446622_RNA45SN1_45S_50ntFlanks | 3705 | 3744 | 40 | rRF 45S | NO | 0.00 | 33.33 | 0.00 | 0.00 |
| >hg38_21_+_8433172_8446622_RNA45SN1_45S_50ntFlanks@3705.3745.41\|TACCTGGTTGAT CCTGCCAGTAGCATATGCTTGTCTCAAAG | >hg38_21_+_8433172_8446622_RNA45SN1_45S_50ntFlanks | 3705 | 3745 | 41 | rRF 45S | NO | 0.00 | 75.08 | 0.00 | 12.71 |
| >hg38_21_+_8433172_8446622_RNA45SN1_45S_50ntFlanks@3705.3746.42\|TACCTGGTTGAT CCTGCCAGTAGCATATGCTTGTCTCAAAGA | >hg38_21_+_8433172_8446622_RNA45SN1_45S_50ntFlanks | 3705 | 3746 | 42 | rRF 45S | NO | 0.00 | 110.32 | 0.00 | 81.55 |
| >hg38_21_+_8433172_8446622_RNA45SN1_45S_50ntFlanks@3705.3747.43\|TACCTGGTTGAT CCTGCCAGTAGCATATGCTTGTCTCAAAGAT | >hg38_21_+_8433172_8446622_RNA45SN1_45S_50ntFlanks | 3705 | 3747 | 43 | rRF 45S | NO | 0.00 | 36.34 | 0.00 | 0.00 |
| >hg38_21_+_8433172_8446622_RNA45SN1_45S_50ntFlanks@3705.3748.44\|TACCTGGTTGAT CCTGCCAGTAGCATATGCTTGTCTCAAAGATT | >hg38_21_+_8433172_8446622_RNA45SN1_45S_50ntFlanks | 3705 | 3748 | 44 | rRF 45S | NO | 0.00 | 136.13 | 0.00 | 14.16 |
| >hg38_21_+_8433172_8446622_RNA45SN1_45S_50ntFlanks@3840.3860.21\|CTCCTACTTGGA TAACTGTGG | >hg38_21_+_8433172_8446622_RNA45SN1_45S_50ntFlanks | 3840 | 3860 | 21 | rRF 45S | NO | 0.00 | 58.64 | 0.00 | 0.00 |
| >hg38_21_+_8433172_8446622_RNA45SN1_45S_50ntFlanks@3840.3868.29\|CTCCTACTTGGA TAACTGTGGTAATTCTA | >hg38_21_+_8433172_8446622_RNA45SN1_45S_50ntFlanks | 3840 | 3868 | 29 | rRF 45S | NO | 0.00 | 59.68 | 0.00 | 0.00 |
| >hg38_21_+_8433172_8446622_RNA45SN1_45S_50ntFlanks@3841.3868.28\|TCCTACTTGGAT AACTGTGGTAATTCTA | >hg38_21_+_8433172_8446622_RNA45SN1_45S_50ntFlanks | 3841 | 3868 | 28 | rRF 45S | NO | 0.00 | 13.02 | 0.00 | 0.00 |
| >hg38_21_+_8433172_8446622_RNA45SN1_45S_50ntFlanks@3842.3868.27\|CCTACTTGGATA ACTGTGGTAATTCTA | >hg38_21_+_8433172_8446622_RNA45SN1_45S_50ntFlanks | 3842 | 3868 | 27 | rRF 45S | NO | 0.00 | 26.56 | 0.00 | 0.00 |
| >hg38_21_+_8433172_8446622_RNA45SN1_45S_50ntFlanks@3844.3868.25\|TACTTGGATAAC TGTGGTAATTCTA | >hg38_21_+_8433172_8446622_RNA45SN1_45S_50ntFlanks | 3844 | 3868 | 25 | rRF 45S | NO | 0.00 | 24.81 | 0.00 | 0.00 |
| >hg38_21_+_8433172_8446622_RNA45SN1_45S_50ntFlanks@3845.3868.24\|ACTTGGATAACT GTGGTAATTCTA | >hg38_21_+_8433172_8446622_RNA45SN1_45S_50ntFlanks | 3845 | 3868 | 24 | rRF 45S | NO | 0.00 | 31.30 | 0.00 | 0.00 |
| >hg38_21_+_8433172_8446622_RNA45SN1_45S_50ntFlanks@3986.4023.38\|GGCTTTGGTGAC TCTAGATAACCTCGGGCCGATCGCAC | >hg38_21_+_8433172_8446622_RNA45SN1_45S_50ntFlanks | 3986 | 4023 | 38 | rRF 45S | NO | 0.00 | 35.76 | 0.00 | 0.00 |
| >hg38_21_+_8433172_8446622_RNA45SN1_45S_50ntFlanks@3988.4023.36\|CTTTGGTGACTC TAGATAACCTCGGGCCGATCGCAC | >hg38_21_+_8433172_8446622_RNA45SN1_45S_50ntFlanks | 3988 | 4023 | 36 | rRF 45S | NO | 0.00 | 79.14 | 0.00 | 0.00 |
| >hg38_21_+_8433172_8446622_RNA45SN1_45S_50ntFlanks@3990.4023.34\|TTGGTGACTCTA GATAACCTCGGGCCGATCGCAC | >hg38_21_+_8433172_8446622_RNA45SN1_45S_50ntFlanks | 3990 | 4023 | 34 | rRF 45S | NO | 0.00 | 63.18 | 0.00 | 0.00 |
| >hg38_21_+_8433172_8446622_RNA45SN1_45S_50ntFlanks@3991.4023.33\|TGGTGACTCTAG ATAACCTCGGGCCGATCGCAC | >hg38_21_+_8433172_8446622_RNA45SN1_45S_50ntFlanks | 3991 | 4023 | 33 | rRF 45S | NO | 0.00 | 13.01 | 0.00 | 0.00 |

### MDA-MB-231

| | | | | | | | | | | |
|---|---|---|---|---|---|---|---|---|---|---|
| >hg38_21_+_8433172_8446622_RNA45SN1_45S_50ntFlanks@3992.4023.32\|GGTGACTCTAGATAACCTCGGGCCGATCGCAC | >hg38_21_+_8433172_8446622_RNA45SN1_45S_50ntFlanks | 3992 | 4023 | 32 | rRF 45S | NO | 0.00 | 28.89 | 0.00 | 0.00 |
| >hg38_21_+_8433172_8446622_RNA45SN1_45S_50ntFlanks@3993.4023.31\|GTGACTCTAGATAACCTCGGGCCGATCGCAC | >hg38_21_+_8433172_8446622_RNA45SN1_45S_50ntFlanks | 3993 | 4023 | 31 | rRF 45S | NO | 24.35 | 73.49 | 0.00 | 0.00 |
| >hg38_21_+_8433172_8446622_RNA45SN1_45S_50ntFlanks@3993.4024.32\|GTGACTCTAGATAACCTCGGGCCGATCGCACG | >hg38_21_+_8433172_8446622_RNA45SN1_45S_50ntFlanks | 3993 | 4024 | 32 | rRF 45S | NO | 0.00 | 28.85 | 0.00 | 0.00 |
| >hg38_21_+_8433172_8446622_RNA45SN1_45S_50ntFlanks@4002.4023.22\|GATAACCTCGGGCCGATCGCAC | >hg38_21_+_8433172_8446622_RNA45SN1_45S_50ntFlanks | 4002 | 4023 | 22 | rRF 45S | NO | 0.00 | 30.06 | 0.00 | 0.00 |
| >hg38_21_+_8433172_8446622_RNA45SN1_45S_50ntFlanks@4005.4023.19\|AACCTCGGGCCGATCGCAC | >hg38_21_+_8433172_8446622_RNA45SN1_45S_50ntFlanks | 4005 | 4023 | 19 | rRF 45S | NO | 0.00 | 82.14 | 0.00 | 0.00 |
| >hg38_21_+_8433172_8446622_RNA45SN1_45S_50ntFlanks@4006.4023.18\|ACCTCGGGCCGATCGCAC | >hg38_21_+_8433172_8446622_RNA45SN1_45S_50ntFlanks | 4006 | 4023 | 18 | rRF 45S | NO | 0.00 | 57.72 | 0.00 | 0.00 |
| >hg38_21_+_8433172_8446622_RNA45SN1_45S_50ntFlanks@4045.4066.22\|CCATTCGAACGTCTGCCCTATC | >hg38_21_+_8433172_8446622_RNA45SN1_45S_50ntFlanks | 4045 | 4066 | 22 | rRF 45S | NO | 0.00 | 25.23 | 0.00 | 0.00 |
| >hg38_21_+_8433172_8446622_RNA45SN1_45S_50ntFlanks@4046.4066.21\|CATTCGAACGTCTGCCCTATC | >hg38_21_+_8433172_8446622_RNA45SN1_45S_50ntFlanks | 4046 | 4066 | 21 | rRF 45S | NO | 0.00 | 36.59 | 0.00 | 0.00 |
| >hg38_21_+_8433172_8446622_RNA45SN1_45S_50ntFlanks@4047.4066.20\|ATTCGAACGTCTGCCCTATC | >hg38_21_+_8433172_8446622_RNA45SN1_45S_50ntFlanks | 4047 | 4066 | 20 | rRF 45S | NO | 0.00 | 36.23 | 0.00 | 0.00 |
| >hg38_21_+_8433172_8446622_RNA45SN1_45S_50ntFlanks@4048.4066.19\|TTCGAACGTCTGCCCTATC | >hg38_21_+_8433172_8446622_RNA45SN1_45S_50ntFlanks | 4048 | 4066 | 19 | rRF 45S | NO | 0.00 | 32.99 | 0.00 | 0.00 |
| >hg38_21_+_8433172_8446622_RNA45SN1_45S_50ntFlanks@4049.4066.18\|TCGAACGTCTGCCCTATC | >hg38_21_+_8433172_8446622_RNA45SN1_45S_50ntFlanks | 4049 | 4066 | 18 | rRF 45S | NO | 0.00 | 35.45 | 0.00 | 0.00 |
| >hg38_21_+_8433172_8446622_RNA45SN1_45S_50ntFlanks@4453.4473.21\|TCGGCGCCCCCTCGATGCTCT | >hg38_21_+_8433172_8446622_RNA45SN1_45S_50ntFlanks | 4453 | 4473 | 21 | rRF 45S | NO | 0.00 | 55.89 | 0.00 | 0.00 |
| >hg38_21_+_8433172_8446622_RNA45SN1_45S_50ntFlanks@4585.4621.37\|GTTCTATTTTGTTGGTTTTCGGAACTGAGGCCATGAT | >hg38_21_+_8433172_8446622_RNA45SN1_45S_50ntFlanks | 4585 | 4621 | 37 | rRF 45S | NO | 0.00 | 28.84 | 0.00 | 0.00 |
| >hg38_21_+_8433172_8446622_RNA45SN1_45S_50ntFlanks@4585.4622.38\|GTTCTATTTTGTTGGTTTTCGGAACTGAGGCCATGATT | >hg38_21_+_8433172_8446622_RNA45SN1_45S_50ntFlanks | 4585 | 4622 | 38 | rRF 45S | NO | 0.00 | 26.19 | 0.00 | 0.00 |
| >hg38_21_+_8433172_8446622_RNA45SN1_45S_50ntFlanks@4678.4720.43\|CGGCGCAAGACGGACCAGAGCGAAAGCATTTGCCAAGAATGTT | >hg38_21_+_8433172_8446622_RNA45SN1_45S_50ntFlanks | 4678 | 4720 | 43 | rRF 45S | NO | 0.00 | 13.43 | 0.00 | 0.00 |
| >hg38_21_+_8433172_8446622_RNA45SN1_45S_50ntFlanks@4679.4720.42\|GGCGCAAGACGGACCAGAGCGAAAGCATTTGCCAAGAATGTT | >hg38_21_+_8433172_8446622_RNA45SN1_45S_50ntFlanks | 4679 | 4720 | 42 | rRF 45S | NO | 0.00 | 32.71 | 0.00 | 0.00 |
| >hg38_21_+_8433172_8446622_RNA45SN1_45S_50ntFlanks@4722.4759.38\|TCATTAATCAAGAACGAAAGTCGGAGGTTCGAAGACGA | >hg38_21_+_8433172_8446622_RNA45SN1_45S_50ntFlanks | 4722 | 4759 | 38 | rRF 45S | NO | 0.00 | 13.37 | 0.00 | 0.00 |
| >hg38_21_+_8433172_8446622_RNA45SN1_45S_50ntFlanks@4724.4758.35\|ATTAATCAAGAACGAAAGTCGGAGGTTCGAAGACG | >hg38_21_+_8433172_8446622_RNA45SN1_45S_50ntFlanks | 4724 | 4758 | 35 | rRF 45S | NO | 0.00 | 28.09 | 0.00 | 0.00 |
| >hg38_21_+_8433172_8446622_RNA45SN1_45S_50ntFlanks@4725.4758.34\|TTAATCAAGAACGAAAGTCGGAGGTTCGAAGACG | >hg38_21_+_8433172_8446622_RNA45SN1_45S_50ntFlanks | 4725 | 4758 | 34 | rRF 45S | NO | 0.00 | 13.63 | 0.00 | 0.00 |
| >hg38_21_+_8433172_8446622_RNA45SN1_45S_50ntFlanks@4725.4767.43\|TTAATCAAGAACGAAAGTCGGAGGTTCGAAGACGATCAGATAC | >hg38_21_+_8433172_8446622_RNA45SN1_45S_50ntFlanks | 4725 | 4767 | 43 | rRF 45S | NO | 0.00 | 71.10 | 0.00 | 0.00 |
| >hg38_21_+_8433172_8446622_RNA45SN1_45S_50ntFlanks@4726.4767.42\|TAATCAAGAACGAAAGTCGGAGGTTCGAAGACGATCAGATAC | >hg38_21_+_8433172_8446622_RNA45SN1_45S_50ntFlanks | 4726 | 4767 | 42 | rRF 45S | NO | 0.00 | 59.73 | 0.00 | 0.00 |
| >hg38_21_+_8433172_8446622_RNA45SN1_45S_50ntFlanks@4727.4758.32\|AATCAAGAACGAAAGTCGGAGGTTCGAAGACG | >hg38_21_+_8433172_8446622_RNA45SN1_45S_50ntFlanks | 4727 | 4758 | 32 | rRF 45S | NO | 0.00 | 37.73 | 35.55 | 0.00 |
| >hg38_21_+_8433172_8446622_RNA45SN1_45S_50ntFlanks@4727.4761.35\|AATCAAGAACGAAAGTCGGAGGTTCGAAGACGATC | >hg38_21_+_8433172_8446622_RNA45SN1_45S_50ntFlanks | 4727 | 4761 | 35 | rRF 45S | NO | 0.00 | 94.06 | 0.00 | 0.00 |
| >hg38_21_+_8433172_8446622_RNA45SN1_45S_50ntFlanks@4727.4765.39\|AATCAAGAACGAAAGTCGGAGGTTCGAAGACGATCAGAT | >hg38_21_+_8433172_8446622_RNA45SN1_45S_50ntFlanks | 4727 | 4765 | 39 | rRF 45S | NO | 0.00 | 175.25 | 0.00 | 0.00 |
| >hg38_21_+_8433172_8446622_RNA45SN1_45S_50ntFlanks@4727.4766.40\|AATCAAGAACGAAAGTCGGAGGTTCGAAGACGATCAGATA | >hg38_21_+_8433172_8446622_RNA45SN1_45S_50ntFlanks | 4727 | 4766 | 40 | rRF 45S | NO | 0.00 | 63.26 | 0.00 | 0.00 |
| >hg38_21_+_8433172_8446622_RNA45SN1_45S_50ntFlanks@4727.4767.41\|AATCAAGAACGAAAGTCGGAGGTTCGAAGACGATCAGATAC | >hg38_21_+_8433172_8446622_RNA45SN1_45S_50ntFlanks | 4727 | 4767 | 41 | rRF 45S | NO | 0.00 | 234.52 | 0.00 | 0.00 |
| >hg38_21_+_8433172_8446622_RNA45SN1_45S_50ntFlanks@4727.4768.42\|AATCAAGAACGAAAGTCGGAGGTTCGAAGACGATCAGATACC | >hg38_21_+_8433172_8446622_RNA45SN1_45S_50ntFlanks | 4727 | 4768 | 42 | rRF 45S | NO | 0.00 | 159.65 | 0.00 | 0.00 |
| >hg38_21_+_8433172_8446622_RNA45SN1_45S_50ntFlanks@4729.4767.39\|TCAAGAACGAAAGTCGGAGGTTCGAAGACGATCAGATAC | >hg38_21_+_8433172_8446622_RNA45SN1_45S_50ntFlanks | 4729 | 4767 | 39 | rRF 45S | NO | 0.00 | 32.01 | 0.00 | 0.00 |
| >hg38_21_+_8433172_8446622_RNA45SN1_45S_50ntFlanks@4730.4758.29\|CAAGAACGAAAGTCGGAGGTTCGAAGACG | >hg38_21_+_8433172_8446622_RNA45SN1_45S_50ntFlanks | 4730 | 4758 | 29 | rRF 45S | NO | 0.00 | 36.52 | 13.53 | 0.00 |
| >hg38_21_+_8433172_8446622_RNA45SN1_45S_50ntFlanks@4730.4761.32\|CAAGAACGAAAGTCGGAGGTTCGAAGACGATC | >hg38_21_+_8433172_8446622_RNA45SN1_45S_50ntFlanks | 4730 | 4761 | 32 | rRF 45S | NO | 0.00 | 30.94 | 0.00 | 0.00 |
| >hg38_21_+_8433172_8446622_RNA45SN1_45S_50ntFlanks@4730.4765.36\|CAAGAACGAAAGTCGGAGGTTCGAAGACGATCAGAT | >hg38_21_+_8433172_8446622_RNA45SN1_45S_50ntFlanks | 4730 | 4765 | 36 | rRF 45S | NO | 0.00 | 37.01 | 0.00 | 0.00 |
| >hg38_21_+_8433172_8446622_RNA45SN1_45S_50ntFlanks@4730.4766.37\|CAAGAACGAAAGTCGGAGGTTCGAAGACGATCAGATA | >hg38_21_+_8433172_8446622_RNA45SN1_45S_50ntFlanks | 4730 | 4766 | 37 | rRF 45S | NO | 0.00 | 25.89 | 0.00 | 0.00 |
| >hg38_21_+_8433172_8446622_RNA45SN1_45S_50ntFlanks@4730.4767.38\|CAAGAACGAAAGTCGGAGGTTCGAAGACGATCAGATAC | >hg38_21_+_8433172_8446622_RNA45SN1_45S_50ntFlanks | 4730 | 4767 | 38 | rRF 45S | NO | 0.00 | 106.70 | 0.00 | 0.00 |
| >hg38_21_+_8433172_8446622_RNA45SN1_45S_50ntFlanks@4730.4768.39\|CAAGAACGAAAGTCGGAGGTTCGAAGACGATCAGATACC | >hg38_21_+_8433172_8446622_RNA45SN1_45S_50ntFlanks | 4730 | 4768 | 39 | rRF 45S | NO | 0.00 | 61.22 | 0.00 | 0.00 |
| >hg38_21_+_8433172_8446622_RNA45SN1_45S_50ntFlanks@4731.4757.27\|AAGAACGAAAGTCGGAGGTTCGAAGAC | >hg38_21_+_8433172_8446622_RNA45SN1_45S_50ntFlanks | 4731 | 4757 | 27 | rRF 45S | NO | 0.00 | 13.18 | 0.00 | 0.00 |
| >hg38_21_+_8433172_8446622_RNA45SN1_45S_50ntFlanks@4731.4758.28\|AAGAACGAAAGTCGGAGGTTCGAAGACG | >hg38_21_+_8433172_8446622_RNA45SN1_45S_50ntFlanks | 4731 | 4758 | 28 | rRF 45S | NO | 0.00 | 161.22 | 31.29 | 0.00 |
| >hg38_21_+_8433172_8446622_RNA45SN1_45S_50ntFlanks@4731.4759.29\|AAGAACGAAAGTCGGAGGTTCGAAGACGA | >hg38_21_+_8433172_8446622_RNA45SN1_45S_50ntFlanks | 4731 | 4759 | 29 | rRF 45S | NO | 0.00 | 35.03 | 0.00 | 0.00 |
| >hg38_21_+_8433172_8446622_RNA45SN1_45S_50ntFlanks@4731.4761.31\|AAGAACGAAAGTCGGAGGTTCGAAGACGATC | >hg38_21_+_8433172_8446622_RNA45SN1_45S_50ntFlanks | 4731 | 4761 | 31 | rRF 45S | NO | 44.95 | 200.44 | 10.44 | 0.00 |

### MDA-MB-231

| | | | | | | | | | | |
|---|---|---|---|---|---|---|---|---|---|---|
| >hg38_21_+_8433172_8446622_RNA45SN1_45S_50ntFlanks@4731.4765.35\|AAGAACGAAAGTCGGAGGTTCGAAGACGATCAGAT | >hg38_21_+_8433172_8446622_RNA45SN1_45S_50ntFlanks | 4731 | 4765 | 35 | rRF 45S | NO | 0.00 | 144.68 | 10.35 | 0.00 |
| >hg38_21_+_8433172_8446622_RNA45SN1_45S_50ntFlanks@4731.4766.36\|AAGAACGAAAGTCGGAGGTTCGAAGACGATCAGATA | >hg38_21_+_8433172_8446622_RNA45SN1_45S_50ntFlanks | 4731 | 4766 | 36 | rRF 45S | NO | 0.00 | 91.38 | 0.00 | 0.00 |
| >hg38_21_+_8433172_8446622_RNA45SN1_45S_50ntFlanks@4731.4767.37\|AAGAACGAAAGTCGGAGGTTCGAAGACGATCAGATAC | >hg38_21_+_8433172_8446622_RNA45SN1_45S_50ntFlanks | 4731 | 4767 | 37 | rRF 45S | NO | 0.00 | 203.80 | 0.00 | 0.00 |
| >hg38_21_+_8433172_8446622_RNA45SN1_45S_50ntFlanks@4731.4768.38\|AAGAACGAAAGTCGGAGGTTCGAAGACGATCAGATACC | >hg38_21_+_8433172_8446622_RNA45SN1_45S_50ntFlanks | 4731 | 4768 | 38 | rRF 45S | NO | 0.00 | 145.78 | 0.00 | 0.00 |
| >hg38_21_+_8433172_8446622_RNA45SN1_45S_50ntFlanks@4732.4758.27\|AGAACGAAAGTCGGAGGTTCGAAGACG | >hg38_21_+_8433172_8446622_RNA45SN1_45S_50ntFlanks | 4732 | 4758 | 27 | rRF 45S | NO | 0.00 | 88.05 | 0.00 | 0.00 |
| >hg38_21_+_8433172_8446622_RNA45SN1_45S_50ntFlanks@4732.4761.30\|AGAACGAAAGTCGGAGGTTCGAAGACGATC | >hg38_21_+_8433172_8446622_RNA45SN1_45S_50ntFlanks | 4732 | 4761 | 30 | rRF 45S | NO | 0.00 | 106.99 | 0.00 | 0.00 |
| >hg38_21_+_8433172_8446622_RNA45SN1_45S_50ntFlanks@4732.4765.34\|AGAACGAAAGTCGGAGGTTCGAAGACGATCAGAT | >hg38_21_+_8433172_8446622_RNA45SN1_45S_50ntFlanks | 4732 | 4765 | 34 | rRF 45S | NO | 0.00 | 69.72 | 0.00 | 0.00 |
| >hg38_21_+_8433172_8446622_RNA45SN1_45S_50ntFlanks@4732.4766.35\|AGAACGAAAGTCGGAGGTTCGAAGACGATCAGATA | >hg38_21_+_8433172_8446622_RNA45SN1_45S_50ntFlanks | 4732 | 4766 | 35 | rRF 45S | NO | 0.00 | 59.84 | 0.00 | 0.00 |
| >hg38_21_+_8433172_8446622_RNA45SN1_45S_50ntFlanks@4732.4767.36\|AGAACGAAAGTCGGAGGTTCGAAGACGATCAGATAC | >hg38_21_+_8433172_8446622_RNA45SN1_45S_50ntFlanks | 4732 | 4767 | 36 | rRF 45S | NO | 0.00 | 114.07 | 0.00 | 0.00 |
| >hg38_21_+_8433172_8446622_RNA45SN1_45S_50ntFlanks@4732.4768.37\|AGAACGAAAGTCGGAGGTTCGAAGACGATCAGATACC | >hg38_21_+_8433172_8446622_RNA45SN1_45S_50ntFlanks | 4732 | 4768 | 37 | rRF 45S | NO | 0.00 | 75.22 | 0.00 | 0.00 |
| >hg38_21_+_8433172_8446622_RNA45SN1_45S_50ntFlanks@4733.4758.26\|GAACGAAAGTCGGAGGTTCGAAGACG | >hg38_21_+_8433172_8446622_RNA45SN1_45S_50ntFlanks | 4733 | 4758 | 26 | rRF 45S | NO | 0.00 | 71.07 | 0.00 | 0.00 |
| >hg38_21_+_8433172_8446622_RNA45SN1_45S_50ntFlanks@4733.4761.29\|GAACGAAAGTCGGAGGTTCGAAGACGATC | >hg38_21_+_8433172_8446622_RNA45SN1_45S_50ntFlanks | 4733 | 4761 | 29 | rRF 45S | NO | 0.00 | 96.77 | 0.00 | 0.00 |
| >hg38_21_+_8433172_8446622_RNA45SN1_45S_50ntFlanks@4733.4765.33\|GAACGAAAGTCGGAGGTTCGAAGACGATCAGAT | >hg38_21_+_8433172_8446622_RNA45SN1_45S_50ntFlanks | 4733 | 4765 | 33 | rRF 45S | NO | 0.00 | 68.04 | 0.00 | 0.00 |
| >hg38_21_+_8433172_8446622_RNA45SN1_45S_50ntFlanks@4733.4766.34\|GAACGAAAGTCGGAGGTTCGAAGACGATCAGATA | >hg38_21_+_8433172_8446622_RNA45SN1_45S_50ntFlanks | 4733 | 4766 | 34 | rRF 45S | NO | 0.00 | 34.21 | 0.00 | 0.00 |
| >hg38_21_+_8433172_8446622_RNA45SN1_45S_50ntFlanks@4733.4767.35\|GAACGAAAGTCGGAGGTTCGAAGACGATCAGATAC | >hg38_21_+_8433172_8446622_RNA45SN1_45S_50ntFlanks | 4733 | 4767 | 35 | rRF 45S | NO | 0.00 | 104.00 | 0.00 | 0.00 |
| >hg38_21_+_8433172_8446622_RNA45SN1_45S_50ntFlanks@4733.4768.36\|GAACGAAAGTCGGAGGTTCGAAGACGATCAGATACC | >hg38_21_+_8433172_8446622_RNA45SN1_45S_50ntFlanks | 4733 | 4768 | 36 | rRF 45S | NO | 0.00 | 32.50 | 0.00 | 0.00 |
| >hg38_21_+_8433172_8446622_RNA45SN1_45S_50ntFlanks@4734.4758.25\|AACGAAAGTCGGAGGTTCGAAGACG | >hg38_21_+_8433172_8446622_RNA45SN1_45S_50ntFlanks | 4734 | 4758 | 25 | rRF 45S | NO | 0.00 | 31.29 | 148.23 | 0.00 |
| >hg38_21_+_8433172_8446622_RNA45SN1_45S_50ntFlanks@4734.4761.28\|AACGAAAGTCGGAGGTTCGAAGACGATC | >hg38_21_+_8433172_8446622_RNA45SN1_45S_50ntFlanks | 4734 | 4761 | 28 | rRF 45S | NO | 0.00 | 35.59 | 0.00 | 0.00 |
| >hg38_21_+_8433172_8446622_RNA45SN1_45S_50ntFlanks@4734.4765.32\|AACGAAAGTCGGAGGTTCGAAGACGATCAGAT | >hg38_21_+_8433172_8446622_RNA45SN1_45S_50ntFlanks | 4734 | 4765 | 32 | rRF 45S | NO | 0.00 | 0.00 | 11.34 | 0.00 |
| >hg38_21_+_8433172_8446622_RNA45SN1_45S_50ntFlanks@4734.4767.34\|AACGAAAGTCGGAGGTTCGAAGACGATCAGATAC | >hg38_21_+_8433172_8446622_RNA45SN1_45S_50ntFlanks | 4734 | 4767 | 34 | rRF 45S | NO | 0.00 | 27.01 | 0.00 | 0.00 |
| >hg38_21_+_8433172_8446622_RNA45SN1_45S_50ntFlanks@4735.4758.24\|ACGAAAGTCGGAGGTTCGAAGACG | >hg38_21_+_8433172_8446622_RNA45SN1_45S_50ntFlanks | 4735 | 4758 | 24 | rRF 45S | NO | 0.00 | 82.09 | 0.00 | 0.00 |
| >hg38_21_+_8433172_8446622_RNA45SN1_45S_50ntFlanks@4735.4759.25\|ACGAAAGTCGGAGGTTCGAAGACGA | >hg38_21_+_8433172_8446622_RNA45SN1_45S_50ntFlanks | 4735 | 4759 | 25 | rRF 45S | NO | 0.00 | 26.09 | 0.00 | 0.00 |
| >hg38_21_+_8433172_8446622_RNA45SN1_45S_50ntFlanks@4735.4761.27\|ACGAAAGTCGGAGGTTCGAAGACGATC | >hg38_21_+_8433172_8446622_RNA45SN1_45S_50ntFlanks | 4735 | 4761 | 27 | rRF 45S | NO | 0.00 | 75.50 | 0.00 | 0.00 |
| >hg38_21_+_8433172_8446622_RNA45SN1_45S_50ntFlanks@4735.4767.33\|ACGAAAGTCGGAGGTTCGAAGACGATCAGATAC | >hg38_21_+_8433172_8446622_RNA45SN1_45S_50ntFlanks | 4735 | 4767 | 33 | rRF 45S | NO | 0.00 | 29.82 | 0.00 | 0.00 |
| >hg38_21_+_8433172_8446622_RNA45SN1_45S_50ntFlanks@4735.4768.34\|ACGAAAGTCGGAGGTTCGAAGACGATCAGATACC | >hg38_21_+_8433172_8446622_RNA45SN1_45S_50ntFlanks | 4735 | 4768 | 34 | rRF 45S | NO | 0.00 | 29.41 | 0.00 | 0.00 |
| >hg38_21_+_8433172_8446622_RNA45SN1_45S_50ntFlanks@4738.4757.20\|AAAGTCGGAGGTTCGAAGAC | >hg38_21_+_8433172_8446622_RNA45SN1_45S_50ntFlanks | 4738 | 4757 | 20 | rRF 45S | NO | 0.00 | 27.52 | 0.00 | 0.00 |
| >hg38_21_+_8433172_8446622_RNA45SN1_45S_50ntFlanks@4738.4758.21\|AAAGTCGGAGGTTCGAAGACG | >hg38_21_+_8433172_8446622_RNA45SN1_45S_50ntFlanks | 4738 | 4758 | 21 | rRF 45S | NO | 0.00 | 88.57 | 263.74 | 0.00 |
| >hg38_21_+_8433172_8446622_RNA45SN1_45S_50ntFlanks@4738.4761.24\|AAAGTCGGAGGTTCGAAGACGATC | >hg38_21_+_8433172_8446622_RNA45SN1_45S_50ntFlanks | 4738 | 4761 | 24 | rRF 45S | NO | 0.00 | 102.33 | 0.00 | 0.00 |
| >hg38_21_+_8433172_8446622_RNA45SN1_45S_50ntFlanks@4738.4765.28\|AAAGTCGGAGGTTCGAAGACGATCAGAT | >hg38_21_+_8433172_8446622_RNA45SN1_45S_50ntFlanks | 4738 | 4765 | 28 | rRF 45S | NO | 0.00 | 24.42 | 11.52 | 0.00 |
| >hg38_21_+_8433172_8446622_RNA45SN1_45S_50ntFlanks@4738.4766.29\|AAAGTCGGAGGTTCGAAGACGATCAGATA | >hg38_21_+_8433172_8446622_RNA45SN1_45S_50ntFlanks | 4738 | 4766 | 29 | rRF 45S | NO | 0.00 | 27.42 | 0.00 | 0.00 |
| >hg38_21_+_8433172_8446622_RNA45SN1_45S_50ntFlanks@4738.4767.30\|AAAGTCGGAGGTTCGAAGACGATCAGATAC | >hg38_21_+_8433172_8446622_RNA45SN1_45S_50ntFlanks | 4738 | 4767 | 30 | rRF 45S | NO | 0.00 | 62.92 | 0.00 | 0.00 |
| >hg38_21_+_8433172_8446622_RNA45SN1_45S_50ntFlanks@4738.4768.31\|AAAGTCGGAGGTTCGAAGACGATCAGATACC | >hg38_21_+_8433172_8446622_RNA45SN1_45S_50ntFlanks | 4738 | 4768 | 31 | rRF 45S | NO | 0.00 | 75.28 | 0.00 | 0.00 |
| >hg38_21_+_8433172_8446622_RNA45SN1_45S_50ntFlanks@4742.4759.18\|TCGGAGGTTCGAAGACGA | >hg38_21_+_8433172_8446622_RNA45SN1_45S_50ntFlanks | 4742 | 4759 | 18 | rRF 45S | NO | 24.57 | 0.00 | 0.00 | 0.00 |
| >hg38_21_+_8433172_8446622_RNA45SN1_45S_50ntFlanks@4823.4844.22\|ATGACCCGCCGGGCAGCTTCCG | >hg38_21_+_8433172_8446622_RNA45SN1_45S_50ntFlanks | 4823 | 4844 | 22 | rRF 45S | NO | 0.00 | 87.53 | 0.00 | 0.00 |
| >hg38_21_+_8433172_8446622_RNA45SN1_45S_50ntFlanks@4824.4844.21\|TGACCCGCCGGGCAGCTTCCG | >hg38_21_+_8433172_8446622_RNA45SN1_45S_50ntFlanks | 4824 | 4844 | 21 | rRF 45S | NO | 0.00 | 60.19 | 0.00 | 0.00 |
| >hg38_21_+_8433172_8446622_RNA45SN1_45S_50ntFlanks@5057.5081.25\|AGCGATTTGTCTGGTTAATTCCGAT | >hg38_21_+_8433172_8446622_RNA45SN1_45S_50ntFlanks | 5057 | 5081 | 25 | rRF 45S | NO | 0.00 | 0.00 | 16.79 | 0.00 |
| >hg38_21_+_8433172_8446622_RNA45SN1_45S_50ntFlanks@5078.5115.38\|CGATAACGAACGAGACTCTGGCATGCTAACTAGTTACG | >hg38_21_+_8433172_8446622_RNA45SN1_45S_50ntFlanks | 5078 | 5115 | 38 | rRF 45S | NO | 0.00 | 27.66 | 0.00 | 0.00 |
| >hg38_21_+_8433172_8446622_RNA45SN1_45S_50ntFlanks@5079.5112.34\|GATAACGAACGAGACTCTGGCATGCTAACTAGTT | >hg38_21_+_8433172_8446622_RNA45SN1_45S_50ntFlanks | 5079 | 5112 | 34 | rRF 45S | NO | 0.00 | 25.86 | 0.00 | 0.00 |

### MDA-MB-231

| | | | | | | | | | | |
|---|---|---|---|---|---|---|---|---|---|---|
| >hg38_21_+_8433172_8446622_RNA45SN1_45S_50ntFlanks@5082.5112.31\|AACGAACGAGA CTCTGGCATGCTAACTAGTT | >hg38_21_+_8433172_8446622_RNA45SN1_45S_50ntFlanks | 5082 | 5112 | 31 | rRF 45S | NO | 0.00 | 13.62 | 0.00 | 0.00 |
| >hg38_21_+_8433172_8446622_RNA45SN1_45S_50ntFlanks@5215.5233.19\|TCCGGGGCTGC ACGCGCGC | >hg38_21_+_8433172_8446622_RNA45SN1_45S_50ntFlanks | 5215 | 5233 | 19 | rRF 45S | NO | 0.00 | 28.06 | 0.00 | 0.00 |
| >hg38_21_+_8433172_8446622_RNA45SN1_45S_50ntFlanks@5216.5233.18\|CCGGGGCTGCA CGCGCGC | >hg38_21_+_8433172_8446622_RNA45SN1_45S_50ntFlanks | 5216 | 5233 | 18 | rRF 45S | NO | 0.00 | 79.85 | 0.00 | 0.00 |
| >hg38_21_+_8433172_8446622_RNA45SN1_45S_50ntFlanks@5422.5450.29\|GATTGGATGGTT TAGTGAGGCCCTCGGAT | >hg38_21_+_8433172_8446622_RNA45SN1_45S_50ntFlanks | 5422 | 5450 | 29 | rRF 45S | NO | 0.00 | 25.29 | 0.00 | 0.00 |
| >hg38_21_+_8433172_8446622_RNA45SN1_45S_50ntFlanks@5470.5516.47\|CCACGGCCCTG GCGGAGCGCTGAGAAGACGGTCGAACTTGACTATCT | >hg38_21_+_8433172_8446622_RNA45SN1_45S_50ntFlanks | 5470 | 5516 | 47 | rRF 45S | NO | 0.00 | 102.54 | 0.00 | 0.00 |
| >hg38_21_+_8433172_8446622_RNA45SN1_45S_50ntFlanks@5471.5516.46\|CACGGCCCTGG CGGAGCGCTGAGAAGACGGTCGAACTTGACTATCT | >hg38_21_+_8433172_8446622_RNA45SN1_45S_50ntFlanks | 5471 | 5516 | 46 | rRF 45S | NO | 0.00 | 485.93 | 0.00 | 0.00 |
| >hg38_21_+_8433172_8446622_RNA45SN1_45S_50ntFlanks@5472.5489.18\|ACGGCCCTGGC GGAGCGC | >hg38_21_+_8433172_8446622_RNA45SN1_45S_50ntFlanks | 5472 | 5489 | 18 | rRF 45S | NO | 23.82 | 150.56 | 11.47 | 10.97 |
| >hg38_21_+_8433172_8446622_RNA45SN1_45S_50ntFlanks@5472.5490.19\|ACGGCCCTGGC GGAGCGCT | >hg38_21_+_8433172_8446622_RNA45SN1_45S_50ntFlanks | 5472 | 5490 | 19 | rRF 45S | NO | 0.00 | 112.87 | 0.00 | 0.00 |
| >hg38_21_+_8433172_8446622_RNA45SN1_45S_50ntFlanks@5472.5491.20\|ACGGCCCTGGC GGAGCGCTG | >hg38_21_+_8433172_8446622_RNA45SN1_45S_50ntFlanks | 5472 | 5491 | 20 | rRF 45S | NO | 0.00 | 75.07 | 0.00 | 0.00 |
| >hg38_21_+_8433172_8446622_RNA45SN1_45S_50ntFlanks@5472.5494.23\|ACGGCCCTGGC GGAGCGCTGAGA | >hg38_21_+_8433172_8446622_RNA45SN1_45S_50ntFlanks | 5472 | 5494 | 23 | rRF 45S | NO | 0.00 | 162.46 | 0.00 | 0.00 |
| >hg38_21_+_8433172_8446622_RNA45SN1_45S_50ntFlanks@5472.5498.27\|ACGGCCCTGGC GGAGCGCTGAGAAGAC | >hg38_21_+_8433172_8446622_RNA45SN1_45S_50ntFlanks | 5472 | 5498 | 27 | rRF 45S | NO | 0.00 | 87.63 | 0.00 | 0.00 |
| >hg38_21_+_8433172_8446622_RNA45SN1_45S_50ntFlanks@5472.5499.28\|ACGGCCCTGGC GGAGCGCTGAGAAGACG | >hg38_21_+_8433172_8446622_RNA45SN1_45S_50ntFlanks | 5472 | 5499 | 28 | rRF 45S | NO | 24.45 | 347.46 | 0.00 | 0.00 |
| >hg38_21_+_8433172_8446622_RNA45SN1_45S_50ntFlanks@5472.5500.29\|ACGGCCCTGGC GGAGCGCTGAGAAGACGG | >hg38_21_+_8433172_8446622_RNA45SN1_45S_50ntFlanks | 5472 | 5500 | 29 | rRF 45S | NO | 0.00 | 186.95 | 0.00 | 0.00 |
| >hg38_21_+_8433172_8446622_RNA45SN1_45S_50ntFlanks@5472.5501.30\|ACGGCCCTGGC GGAGCGCTGAGAAGACGGT | >hg38_21_+_8433172_8446622_RNA45SN1_45S_50ntFlanks | 5472 | 5501 | 30 | rRF 45S | NO | 0.00 | 390.21 | 0.00 | 0.00 |
| >hg38_21_+_8433172_8446622_RNA45SN1_45S_50ntFlanks@5472.5502.31\|ACGGCCCTGGC GGAGCGCTGAGAAGACGGTC | >hg38_21_+_8433172_8446622_RNA45SN1_45S_50ntFlanks | 5472 | 5502 | 31 | rRF 45S | NO | 0.00 | 124.76 | 0.00 | 0.00 |
| >hg38_21_+_8433172_8446622_RNA45SN1_45S_50ntFlanks@5472.5506.35\|ACGGCCCTGGC GGAGCGCTGAGAAGACGGTCGAAC | >hg38_21_+_8433172_8446622_RNA45SN1_45S_50ntFlanks | 5472 | 5506 | 35 | rRF 45S | NO | 0.00 | 136.11 | 0.00 | 0.00 |
| >hg38_21_+_8433172_8446622_RNA45SN1_45S_50ntFlanks@5472.5507.36\|ACGGCCCTGGC GGAGCGCTGAGAAGACGGTCGAACT | >hg38_21_+_8433172_8446622_RNA45SN1_45S_50ntFlanks | 5472 | 5507 | 36 | rRF 45S | NO | 0.00 | 113.93 | 0.00 | 0.00 |
| >hg38_21_+_8433172_8446622_RNA45SN1_45S_50ntFlanks@5472.5509.38\|ACGGCCCTGGC GGAGCGCTGAGAAGACGGTCGAACTTG | >hg38_21_+_8433172_8446622_RNA45SN1_45S_50ntFlanks | 5472 | 5509 | 38 | rRF 45S | NO | 0.00 | 219.64 | 0.00 | 0.00 |
| >hg38_21_+_8433172_8446622_RNA45SN1_45S_50ntFlanks@5472.5510.39\|ACGGCCCTGGC GGAGCGCTGAGAAGACGGTCGAACTTGA | >hg38_21_+_8433172_8446622_RNA45SN1_45S_50ntFlanks | 5472 | 5510 | 39 | rRF 45S | NO | 0.00 | 76.14 | 0.00 | 0.00 |
| >hg38_21_+_8433172_8446622_RNA45SN1_45S_50ntFlanks@5472.5511.40\|ACGGCCCTGGC GGAGCGCTGAGAAGACGGTCGAACTTGAC | >hg38_21_+_8433172_8446622_RNA45SN1_45S_50ntFlanks | 5472 | 5511 | 40 | rRF 45S | NO | 0.00 | 99.67 | 0.00 | 0.00 |
| >hg38_21_+_8433172_8446622_RNA45SN1_45S_50ntFlanks@5472.5512.41\|ACGGCCCTGGC GGAGCGCTGAGAAGACGGTCGAACTTGACT | >hg38_21_+_8433172_8446622_RNA45SN1_45S_50ntFlanks | 5472 | 5512 | 41 | rRF 45S | NO | 108.64 | 538.36 | 0.00 | 0.00 |
| >hg38_21_+_8433172_8446622_RNA45SN1_45S_50ntFlanks@5472.5513.42\|ACGGCCCTGGC GGAGCGCTGAGAAGACGGTCGAACTTGACTA | >hg38_21_+_8433172_8446622_RNA45SN1_45S_50ntFlanks | 5472 | 5513 | 42 | rRF 45S | NO | 0.00 | 107.05 | 0.00 | 0.00 |
| >hg38_21_+_8433172_8446622_RNA45SN1_45S_50ntFlanks@5472.5514.43\|ACGGCCCTGGC GGAGCGCTGAGAAGACGGTCGAACTTGACTAT | >hg38_21_+_8433172_8446622_RNA45SN1_45S_50ntFlanks | 5472 | 5514 | 43 | rRF 45S | NO | 0.00 | 313.51 | 0.00 | 0.00 |
| >hg38_21_+_8433172_8446622_RNA45SN1_45S_50ntFlanks@5472.5515.44\|ACGGCCCTGGC GGAGCGCTGAGAAGACGGTCGAACTTGACTATC | >hg38_21_+_8433172_8446622_RNA45SN1_45S_50ntFlanks | 5472 | 5515 | 44 | rRF 45S | NO | 0.00 | 191.95 | 0.00 | 0.00 |
| >hg38_21_+_8433172_8446622_RNA45SN1_45S_50ntFlanks@5472.5516.45\|ACGGCCCTGGC GGAGCGCTGAGAAGACGGTCGAACTTGACTATCT | >hg38_21_+_8433172_8446622_RNA45SN1_45S_50ntFlanks | 5472 | 5516 | 45 | rRF 45S | YES | 77.27 | 3805.45 | 11.40 | 16.47 |
| >hg38_21_+_8433172_8446622_RNA45SN1_45S_50ntFlanks@5473.5490.18\|CGGCCCTGGCG GAGCGCT | >hg38_21_+_8433172_8446622_RNA45SN1_45S_50ntFlanks | 5473 | 5490 | 18 | rRF 45S | NO | 0.00 | 12.87 | 0.00 | 0.00 |
| >hg38_21_+_8433172_8446622_RNA45SN1_45S_50ntFlanks@5473.5499.27\|CGGCCCTGGCG GAGCGCTGAGAAGACG | >hg38_21_+_8433172_8446622_RNA45SN1_45S_50ntFlanks | 5473 | 5499 | 27 | rRF 45S | NO | 0.00 | 140.77 | 0.00 | 0.00 |
| >hg38_21_+_8433172_8446622_RNA45SN1_45S_50ntFlanks@5473.5512.40\|CGGCCCTGGCG GAGCGCTGAGAAGACGGTCGAACTTGACT | >hg38_21_+_8433172_8446622_RNA45SN1_45S_50ntFlanks | 5473 | 5512 | 40 | rRF 45S | NO | 0.00 | 120.00 | 0.00 | 0.00 |
| >hg38_21_+_8433172_8446622_RNA45SN1_45S_50ntFlanks@5473.5514.42\|CGGCCCTGGCG GAGCGCTGAGAAGACGGTCGAACTTGACTAT | >hg38_21_+_8433172_8446622_RNA45SN1_45S_50ntFlanks | 5473 | 5514 | 42 | rRF 45S | NO | 0.00 | 71.27 | 0.00 | 0.00 |
| >hg38_21_+_8433172_8446622_RNA45SN1_45S_50ntFlanks@5473.5516.44\|CGGCCCTGGCG GAGCGCTGAGAAGACGGTCGAACTTGACTATCT | >hg38_21_+_8433172_8446622_RNA45SN1_45S_50ntFlanks | 5473 | 5516 | 44 | rRF 45S | NO | 0.00 | 1380.75 | 0.00 | 0.00 |
| >hg38_21_+_8433172_8446622_RNA45SN1_45S_50ntFlanks@5476.5516.41\|CCCTGGCGGAG CGCTGAGAAGACGGTCGAACTTGACTATCT | >hg38_21_+_8433172_8446622_RNA45SN1_45S_50ntFlanks | 5476 | 5516 | 41 | rRF 45S | NO | 0.00 | 227.13 | 0.00 | 0.00 |
| >hg38_21_+_8433172_8446622_RNA45SN1_45S_50ntFlanks@5477.5516.40\|CCTGGCGGAGC GCTGAGAAGACGGTCGAACTTGACTATCT | >hg38_21_+_8433172_8446622_RNA45SN1_45S_50ntFlanks | 5477 | 5516 | 40 | rRF 45S | NO | 0.00 | 167.15 | 0.00 | 0.00 |
| >hg38_21_+_8433172_8446622_RNA45SN1_45S_50ntFlanks@5485.5516.32\|AGCGCTGAGAA GACGGTCGAACTTGACTATCT | >hg38_21_+_8433172_8446622_RNA45SN1_45S_50ntFlanks | 5485 | 5516 | 32 | rRF 45S | NO | 0.00 | 141.49 | 0.00 | 0.00 |
| >hg38_21_+_8433172_8446622_RNA45SN1_45S_50ntFlanks@5486.5516.31\|GCGCTGAGAAG ACGGTCGAACTTGACTATCT | >hg38_21_+_8433172_8446622_RNA45SN1_45S_50ntFlanks | 5486 | 5516 | 31 | rRF 45S | NO | 0.00 | 56.10 | 0.00 | 0.00 |
| >hg38_21_+_8433172_8446622_RNA45SN1_45S_50ntFlanks@5487.5516.30\|CGCTGAGAAGA CGGTCGAACTTGACTATCT | >hg38_21_+_8433172_8446622_RNA45SN1_45S_50ntFlanks | 5487 | 5516 | 30 | rRF 45S | NO | 0.00 | 279.08 | 0.00 | 0.00 |
| >hg38_21_+_8433172_8446622_RNA45SN1_45S_50ntFlanks@5488.5516.29\|GCTGAGAAGAC GGTCGAACTTGACTATCT | >hg38_21_+_8433172_8446622_RNA45SN1_45S_50ntFlanks | 5488 | 5516 | 29 | rRF 45S | NO | 0.00 | 112.58 | 0.00 | 0.00 |
| >hg38_21_+_8433172_8446622_RNA45SN1_45S_50ntFlanks@5489.5512.24\|CTGAGAAGACG GTCGAACTTGACT | >hg38_21_+_8433172_8446622_RNA45SN1_45S_50ntFlanks | 5489 | 5512 | 24 | rRF 45S | NO | 0.00 | 13.48 | 0.00 | 0.00 |
| >hg38_21_+_8433172_8446622_RNA45SN1_45S_50ntFlanks@5489.5516.28\|CTGAGAAGACG GTCGAACTTGACTATCT | >hg38_21_+_8433172_8446622_RNA45SN1_45S_50ntFlanks | 5489 | 5516 | 28 | rRF 45S | NO | 0.00 | 760.65 | 0.00 | 0.00 |

### MDA-MB-231

| | | | | | | | | | | |
|---|---|---|---|---|---|---|---|---|---|---|
| >hg38_21_+_8433172_8446622_RNA45SN1_45S_50ntFlanks@5489.5532.44\|CTGAGAAGACGGTCGAACTTGACTATCTAGAGGAAGTAAAAGTC | >hg38_21_+_8433172_8446622_RNA45SN1_45S_50ntFlanks | 5489 | 5532 | 44 | rRF 45S | NO | 0.00 | 35.31 | 0.00 | 0.00 |
| >hg38_21_+_8433172_8446622_RNA45SN1_45S_50ntFlanks@5490.5512.23\|TGAGAAGACGGTCGAACTTGACT | >hg38_21_+_8433172_8446622_RNA45SN1_45S_50ntFlanks | 5490 | 5512 | 23 | rRF 45S | NO | 0.00 | 27.42 | 0.00 | 0.00 |
| >hg38_21_+_8433172_8446622_RNA45SN1_45S_50ntFlanks@5490.5516.27\|TGAGAAGACGGTCGAACTTGACTATCT | >hg38_21_+_8433172_8446622_RNA45SN1_45S_50ntFlanks | 5490 | 5516 | 27 | rRF 45S | NO | 0.00 | 372.47 | 0.00 | 0.00 |
| >hg38_21_+_8433172_8446622_RNA45SN1_45S_50ntFlanks@5490.5532.43\|TGAGAAGACGGTCGAACTTGACTATCTAGAGGAAGTAAAAGTC | >hg38_21_+_8433172_8446622_RNA45SN1_45S_50ntFlanks | 5490 | 5532 | 43 | rRF 45S | NO | 0.00 | 96.94 | 0.00 | 0.00 |
| >hg38_21_+_8433172_8446622_RNA45SN1_45S_50ntFlanks@5491.5516.26\|GAGAAGACGGTCGAACTTGACTATCT | >hg38_21_+_8433172_8446622_RNA45SN1_45S_50ntFlanks | 5491 | 5516 | 26 | rRF 45S | NO | 0.00 | 500.93 | 13.03 | 0.00 |
| >hg38_21_+_8433172_8446622_RNA45SN1_45S_50ntFlanks@5491.5532.42\|GAGAAGACGGTCGAACTTGACTATCTAGAGGAAGTAAAAGTC | >hg38_21_+_8433172_8446622_RNA45SN1_45S_50ntFlanks | 5491 | 5532 | 42 | rRF 45S | NO | 0.00 | 61.77 | 0.00 | 0.00 |
| >hg38_21_+_8433172_8446622_RNA45SN1_45S_50ntFlanks@5492.5516.25\|AGAAGACGGTCGAACTTGACTATCT | >hg38_21_+_8433172_8446622_RNA45SN1_45S_50ntFlanks | 5492 | 5516 | 25 | rRF 45S | NO | 0.00 | 585.53 | 10.98 | 0.00 |
| >hg38_21_+_8433172_8446622_RNA45SN1_45S_50ntFlanks@5493.5516.24\|GAAGACGGTCGAACTTGACTATCT | >hg38_21_+_8433172_8446622_RNA45SN1_45S_50ntFlanks | 5493 | 5516 | 24 | rRF 45S | NO | 0.00 | 169.22 | 0.00 | 0.00 |
| >hg38_21_+_8433172_8446622_RNA45SN1_45S_50ntFlanks@5494.5516.23\|AAGACGGTCGAACTTGACTATCT | >hg38_21_+_8433172_8446622_RNA45SN1_45S_50ntFlanks | 5494 | 5516 | 23 | rRF 45S | NO | 0.00 | 291.15 | 0.00 | 0.00 |
| >hg38_21_+_8433172_8446622_RNA45SN1_45S_50ntFlanks@5495.5516.22\|AGACGGTCGAACTTGACTATCT | >hg38_21_+_8433172_8446622_RNA45SN1_45S_50ntFlanks | 5495 | 5516 | 22 | rRF 45S | NO | 0.00 | 81.60 | 0.00 | 0.00 |
| >hg38_21_+_8433172_8446622_RNA45SN1_45S_50ntFlanks@5497.5532.36\|ACGGTCGAACTTGACTATCTAGAGGAAGTAAAAGTC | >hg38_21_+_8433172_8446622_RNA45SN1_45S_50ntFlanks | 5497 | 5532 | 36 | rRF 45S | NO | 0.00 | 30.99 | 0.00 | 0.00 |
| >hg38_21_+_8433172_8446622_RNA45SN1_45S_50ntFlanks@6641.6683.43\|CGACCTCGTACGACTCTTAGCGGTGGATCACTCGGCTCGTGCG | >hg38_21_+_8433172_8446622_RNA45SN1_45S_50ntFlanks | 6641 | 6683 | 43 | rRF 45S | NO | 48.06 | 0.00 | 0.00 | 0.00 |
| >hg38_21_+_8433172_8446622_RNA45SN1_45S_50ntFlanks@6643.6683.41\|ACCTCGTACGACTCTTAGCGGTGGATCACTCGGCTCGTGCG | >hg38_21_+_8433172_8446622_RNA45SN1_45S_50ntFlanks | 6643 | 6683 | 41 | rRF 45S | NO | 66.08 | 0.00 | 0.00 | 0.00 |
| >hg38_21_+_8433172_8446622_RNA45SN1_45S_50ntFlanks@6644.6665.22\|CCTCGTACGACTCTTAGCGGTG | >hg38_21_+_8433172_8446622_RNA45SN1_45S_50ntFlanks | 6644 | 6665 | 22 | rRF 45S | NO | 49.61 | 0.00 | 0.00 | 54.90 |
| >hg38_21_+_8433172_8446622_RNA45SN1_45S_50ntFlanks@6644.6681.38\|CCTCGTACGACTCTTAGCGGTGGATCACTCGGCTCGTG | >hg38_21_+_8433172_8446622_RNA45SN1_45S_50ntFlanks | 6644 | 6681 | 38 | rRF 45S | NO | 73.17 | 0.00 | 0.00 | 0.00 |
| >hg38_21_+_8433172_8446622_RNA45SN1_45S_50ntFlanks@6644.6683.40\|CCTCGTACGACTCTTAGCGGTGGATCACTCGGCTCGTGCG | >hg38_21_+_8433172_8446622_RNA45SN1_45S_50ntFlanks | 6644 | 6683 | 40 | rRF 45S | NO | 85.58 | 0.00 | 0.00 | 0.00 |
| >hg38_21_+_8433172_8446622_RNA45SN1_45S_50ntFlanks@6645.6662.18\|CTCGTACGACTCTTAGCG | >hg38_21_+_8433172_8446622_RNA45SN1_45S_50ntFlanks | 6645 | 6662 | 18 | rRF 45S | NO | 222.71 | 299.78 | 61.67 | 165.29 |
| >hg38_21_+_8433172_8446622_RNA45SN1_45S_50ntFlanks@6645.6663.19\|CTCGTACGACTCTTAGCGG | >hg38_21_+_8433172_8446622_RNA45SN1_45S_50ntFlanks | 6645 | 6663 | 19 | rRF 45S | NO | 169.68 | 230.39 | 15.98 | 142.68 |
| >hg38_21_+_8433172_8446622_RNA45SN1_45S_50ntFlanks@6645.6664.20\|CTCGTACGACTCTTAGCGGT | >hg38_21_+_8433172_8446622_RNA45SN1_45S_50ntFlanks | 6645 | 6664 | 20 | rRF 45S | NO | 109.78 | 152.65 | 81.89 | 127.67 |
| >hg38_21_+_8433172_8446622_RNA45SN1_45S_50ntFlanks@6645.6665.21\|CTCGTACGACTCTTAGCGGTG | >hg38_21_+_8433172_8446622_RNA45SN1_45S_50ntFlanks | 6645 | 6665 | 21 | rRF 45S | NO | 208.85 | 227.97 | 68.75 | 196.82 |
| >hg38_21_+_8433172_8446622_RNA45SN1_45S_50ntFlanks@6645.6666.22\|CTCGTACGACTCTTAGCGGTGG | >hg38_21_+_8433172_8446622_RNA45SN1_45S_50ntFlanks | 6645 | 6666 | 22 | rRF 45S | NO | 119.54 | 116.04 | 0.00 | 139.44 |
| >hg38_21_+_8433172_8446622_RNA45SN1_45S_50ntFlanks@6645.6667.23\|CTCGTACGACTCTTAGCGGTGGA | >hg38_21_+_8433172_8446622_RNA45SN1_45S_50ntFlanks | 6645 | 6667 | 23 | rRF 45S | NO | 65.81 | 80.44 | 0.00 | 130.41 |
| >hg38_21_+_8433172_8446622_RNA45SN1_45S_50ntFlanks@6645.6668.24\|CTCGTACGACTCTTAGCGGTGGAT | >hg38_21_+_8433172_8446622_RNA45SN1_45S_50ntFlanks | 6645 | 6668 | 24 | rRF 45S | NO | 57.51 | 58.00 | 14.08 | 79.80 |
| >hg38_21_+_8433172_8446622_RNA45SN1_45S_50ntFlanks@6645.6669.25\|CTCGTACGACTCTTAGCGGTGGATC | >hg38_21_+_8433172_8446622_RNA45SN1_45S_50ntFlanks | 6645 | 6669 | 25 | rRF 45S | NO | 66.56 | 91.26 | 0.00 | 127.08 |
| >hg38_21_+_8433172_8446622_RNA45SN1_45S_50ntFlanks@6645.6672.28\|CTCGTACGACTCTTAGCGGTGGATCACT | >hg38_21_+_8433172_8446622_RNA45SN1_45S_50ntFlanks | 6645 | 6672 | 28 | rRF 45S | NO | 47.87 | 58.14 | 0.00 | 77.86 |
| >hg38_21_+_8433172_8446622_RNA45SN1_45S_50ntFlanks@6645.6674.30\|CTCGTACGACTCTTAGCGGTGGATCACTCG | >hg38_21_+_8433172_8446622_RNA45SN1_45S_50ntFlanks | 6645 | 6674 | 30 | rRF 45S | NO | 84.62 | 61.22 | 0.00 | 114.53 |
| >hg38_21_+_8433172_8446622_RNA45SN1_45S_50ntFlanks@6645.6677.33\|CTCGTACGACTCTTAGCGGTGGATCACTCGGCT | >hg38_21_+_8433172_8446622_RNA45SN1_45S_50ntFlanks | 6645 | 6677 | 33 | rRF 45S | NO | 25.29 | 0.00 | 0.00 | 20.91 |
| >hg38_21_+_8433172_8446622_RNA45SN1_45S_50ntFlanks@6645.6679.35\|CTCGTACGACTCTTAGCGGTGGATCACTCGGCTCG | >hg38_21_+_8433172_8446622_RNA45SN1_45S_50ntFlanks | 6645 | 6679 | 35 | rRF 45S | NO | 60.32 | 25.36 | 0.00 | 82.02 |
| >hg38_21_+_8433172_8446622_RNA45SN1_45S_50ntFlanks@6645.6680.36\|CTCGTACGACTCTTAGCGGTGGATCACTCGGCTCGT | >hg38_21_+_8433172_8446622_RNA45SN1_45S_50ntFlanks | 6645 | 6680 | 36 | rRF 45S | NO | 49.46 | 25.27 | 0.00 | 61.54 |
| >hg38_21_+_8433172_8446622_RNA45SN1_45S_50ntFlanks@6645.6681.37\|CTCGTACGACTCTTAGCGGTGGATCACTCGGCTCGTG | >hg38_21_+_8433172_8446622_RNA45SN1_45S_50ntFlanks | 6645 | 6681 | 37 | rRF 45S | NO | 109.25 | 27.52 | 0.00 | 56.51 |
| >hg38_21_+_8433172_8446622_RNA45SN1_45S_50ntFlanks@6646.6663.18\|TCGTACGACTCTTAGCGG | >hg38_21_+_8433172_8446622_RNA45SN1_45S_50ntFlanks | 6646 | 6663 | 18 | rRF 45S | NO | 591.58 | 664.90 | 129.63 | 385.07 |
| >hg38_21_+_8433172_8446622_RNA45SN1_45S_50ntFlanks@6646.6664.19\|TCGTACGACTCTTAGCGGT | >hg38_21_+_8433172_8446622_RNA45SN1_45S_50ntFlanks | 6646 | 6664 | 19 | rRF 45S | NO | 531.90 | 606.62 | 305.87 | 431.84 |
| >hg38_21_+_8433172_8446622_RNA45SN1_45S_50ntFlanks@6646.6665.20\|TCGTACGACTCTTAGCGGTG | >hg38_21_+_8433172_8446622_RNA45SN1_45S_50ntFlanks | 6646 | 6665 | 20 | rRF 45S | NO | 470.55 | 450.37 | 127.62 | 347.37 |
| >hg38_21_+_8433172_8446622_RNA45SN1_45S_50ntFlanks@6646.6666.21\|TCGTACGACTCTTAGCGGTGG | >hg38_21_+_8433172_8446622_RNA45SN1_45S_50ntFlanks | 6646 | 6666 | 21 | rRF 45S | NO | 273.63 | 208.32 | 73.97 | 260.95 |
| >hg38_21_+_8433172_8446622_RNA45SN1_45S_50ntFlanks@6646.6667.22\|TCGTACGACTCTTAGCGGTGGA | >hg38_21_+_8433172_8446622_RNA45SN1_45S_50ntFlanks | 6646 | 6667 | 22 | rRF 45S | NO | 142.47 | 135.96 | 89.60 | 269.92 |
| >hg38_21_+_8433172_8446622_RNA45SN1_45S_50ntFlanks@6646.6668.23\|TCGTACGACTCTTAGCGGTGGAT | >hg38_21_+_8433172_8446622_RNA45SN1_45S_50ntFlanks | 6646 | 6668 | 23 | rRF 45S | NO | 167.81 | 127.30 | 126.94 | 172.13 |
| >hg38_21_+_8433172_8446622_RNA45SN1_45S_50ntFlanks@6646.6669.24\|TCGTACGACTCTTAGCGGTGGATC | >hg38_21_+_8433172_8446622_RNA45SN1_45S_50ntFlanks | 6646 | 6669 | 24 | rRF 45S | NO | 207.28 | 240.05 | 63.11 | 282.04 |
| >hg38_21_+_8433172_8446622_RNA45SN1_45S_50ntFlanks@6646.6670.25\|TCGTACGACTCTTAGCGGTGGATCA | >hg38_21_+_8433172_8446622_RNA45SN1_45S_50ntFlanks | 6646 | 6670 | 25 | rRF 45S | NO | 26.68 | 24.25 | 0.00 | 82.61 |

### MDA-MB-231

| | | | | | | | | | | |
|---|---|---|---|---|---|---|---|---|---|---|
| >hg38_21_+_8433172_8446622_RNA45SN1_45S_50ntFlanks@6646.6672.27\|TCGTACGACTCTTAGCGGTGGATCACT | >hg38_21_+_8433172_8446622_RNA45SN1_45S_50ntFlanks | 6646 | 6672 | 27 | rRF 45S | NO | 158.04 | 164.18 | 0.00 | 194.82 |
| >hg38_21_+_8433172_8446622_RNA45SN1_45S_50ntFlanks@6646.6673.28\|TCGTACGACTCTTAGCGGTGGATCACTC | >hg38_21_+_8433172_8446622_RNA45SN1_45S_50ntFlanks | 6646 | 6673 | 28 | rRF 45S | NO | 67.83 | 63.91 | 0.00 | 74.00 |
| >hg38_21_+_8433172_8446622_RNA45SN1_45S_50ntFlanks@6646.6674.29\|TCGTACGACTCTTAGCGGTGGATCACTCG | >hg38_21_+_8433172_8446622_RNA45SN1_45S_50ntFlanks | 6646 | 6674 | 29 | rRF 45S | NO | 127.28 | 74.93 | 0.00 | 142.81 |
| >hg38_21_+_8433172_8446622_RNA45SN1_45S_50ntFlanks@6646.6675.30\|TCGTACGACTCTTAGCGGTGGATCACTCGG | >hg38_21_+_8433172_8446622_RNA45SN1_45S_50ntFlanks | 6646 | 6675 | 30 | rRF 45S | NO | 225.46 | 113.20 | 0.00 | 117.60 |
| >hg38_21_+_8433172_8446622_RNA45SN1_45S_50ntFlanks@6646.6676.31\|TCGTACGACTCTTAGCGGTGGATCACTCGGC | >hg38_21_+_8433172_8446622_RNA45SN1_45S_50ntFlanks | 6646 | 6676 | 31 | rRF 45S | NO | 115.23 | 64.91 | 34.73 | 140.97 |
| >hg38_21_+_8433172_8446622_RNA45SN1_45S_50ntFlanks@6646.6677.32\|TCGTACGACTCTTAGCGGTGGATCACTCGGCT | >hg38_21_+_8433172_8446622_RNA45SN1_45S_50ntFlanks | 6646 | 6677 | 32 | rRF 45S | NO | 148.78 | 73.87 | 11.53 | 118.80 |
| >hg38_21_+_8433172_8446622_RNA45SN1_45S_50ntFlanks@6646.6678.33\|TCGTACGACTCTTAGCGGTGGATCACTCGGCTC | >hg38_21_+_8433172_8446622_RNA45SN1_45S_50ntFlanks | 6646 | 6678 | 33 | rRF 45S | NO | 101.26 | 31.12 | 0.00 | 95.90 |
| >hg38_21_+_8433172_8446622_RNA45SN1_45S_50ntFlanks@6646.6679.34\|TCGTACGACTCTTAGCGGTGGATCACTCGGCTCG | >hg38_21_+_8433172_8446622_RNA45SN1_45S_50ntFlanks | 6646 | 6679 | 34 | rRF 45S | NO | 181.08 | 84.85 | 0.00 | 158.00 |
| >hg38_21_+_8433172_8446622_RNA45SN1_45S_50ntFlanks@6646.6680.35\|TCGTACGACTCTTAGCGGTGGATCACTCGGCTCGT | >hg38_21_+_8433172_8446622_RNA45SN1_45S_50ntFlanks | 6646 | 6680 | 35 | rRF 45S | NO | 188.46 | 98.77 | 0.00 | 143.57 |
| >hg38_21_+_8433172_8446622_RNA45SN1_45S_50ntFlanks@6646.6681.36\|TCGTACGACTCTTAGCGGTGGATCACTCGGCTCGTG | >hg38_21_+_8433172_8446622_RNA45SN1_45S_50ntFlanks | 6646 | 6681 | 36 | rRF 45S | NO | 1043.05 | 132.61 | 15.54 | 148.87 |
| >hg38_21_+_8433172_8446622_RNA45SN1_45S_50ntFlanks@6646.6683.38\|TCGTACGACTCTTAGCGGTGGATCACTCGGCTCGTGCG | >hg38_21_+_8433172_8446622_RNA45SN1_45S_50ntFlanks | 6646 | 6683 | 38 | rRF 45S | NO | 123.11 | 0.00 | 10.86 | 0.00 |
| >hg38_21_+_8433172_8446622_RNA45SN1_45S_50ntFlanks@6646.6685.40\|TCGTACGACTCTTAGCGGTGGATCACTCGGCTCGTGCGTC | >hg38_21_+_8433172_8446622_RNA45SN1_45S_50ntFlanks | 6646 | 6685 | 40 | rRF 45S | NO | 25.98 | 0.00 | 84.94 | 78.30 |
| >hg38_21_+_8433172_8446622_RNA45SN1_45S_50ntFlanks@6646.6689.44\|TCGTACGACTCTTAGCGGTGGATCACTCGGCTCGTGCGTCGATG | >hg38_21_+_8433172_8446622_RNA45SN1_45S_50ntFlanks | 6646 | 6689 | 44 | rRF 45S | NO | 91.74 | 12.93 | 17.41 | 111.48 |
| >hg38_21_+_8433172_8446622_RNA45SN1_45S_50ntFlanks@6646.6695.50\|TCGTACGACTCTTAGCGGTGGATCACTCGGCTCGTGCGTCGATGAAGAAC | >hg38_21_+_8433172_8446622_RNA45SN1_45S_50ntFlanks | 6646 | 6695 | 50 | rRF 45S | NO | 0.00 | 13.61 | 0.00 | 74.42 |
| >hg38_21_+_8433172_8446622_RNA45SN1_45S_50ntFlanks@6647.6664.18\|CGTACGACTCTTAGCGGT | >hg38_21_+_8433172_8446622_RNA45SN1_45S_50ntFlanks | 6647 | 6664 | 18 | rRF 45S | NO | 113.86 | 100.07 | 120.09 | 141.73 |
| >hg38_21_+_8433172_8446622_RNA45SN1_45S_50ntFlanks@6647.6665.19\|CGTACGACTCTTAGCGGTG | >hg38_21_+_8433172_8446622_RNA45SN1_45S_50ntFlanks | 6647 | 6665 | 19 | rRF 45S | NO | 134.18 | 95.09 | 61.62 | 137.75 |
| >hg38_21_+_8433172_8446622_RNA45SN1_45S_50ntFlanks@6647.6666.20\|CGTACGACTCTTAGCGGTGG | >hg38_21_+_8433172_8446622_RNA45SN1_45S_50ntFlanks | 6647 | 6666 | 20 | rRF 45S | NO | 73.44 | 27.99 | 12.74 | 82.34 |
| >hg38_21_+_8433172_8446622_RNA45SN1_45S_50ntFlanks@6647.6667.21\|CGTACGACTCTTAGCGGTGGA | >hg38_21_+_8433172_8446622_RNA45SN1_45S_50ntFlanks | 6647 | 6667 | 21 | rRF 45S | NO | 47.07 | 0.00 | 14.80 | 93.70 |
| >hg38_21_+_8433172_8446622_RNA45SN1_45S_50ntFlanks@6647.6668.22\|CGTACGACTCTTAGCGGTGGAT | >hg38_21_+_8433172_8446622_RNA45SN1_45S_50ntFlanks | 6647 | 6668 | 22 | rRF 45S | NO | 53.53 | 13.15 | 113.01 | 63.37 |
| >hg38_21_+_8433172_8446622_RNA45SN1_45S_50ntFlanks@6647.6669.23\|CGTACGACTCTTAGCGGTGGATC | >hg38_21_+_8433172_8446622_RNA45SN1_45S_50ntFlanks | 6647 | 6669 | 23 | rRF 45S | NO | 56.02 | 32.33 | 11.03 | 104.78 |
| >hg38_21_+_8433172_8446622_RNA45SN1_45S_50ntFlanks@6647.6672.26\|CGTACGACTCTTAGCGGTGGATCACT | >hg38_21_+_8433172_8446622_RNA45SN1_45S_50ntFlanks | 6647 | 6672 | 26 | rRF 45S | NO | 47.85 | 12.95 | 0.00 | 72.40 |
| >hg38_21_+_8433172_8446622_RNA45SN1_45S_50ntFlanks@6647.6676.30\|CGTACGACTCTTAGCGGTGGATCACTCGGC | >hg38_21_+_8433172_8446622_RNA45SN1_45S_50ntFlanks | 6647 | 6676 | 30 | rRF 45S | NO | 24.40 | 0.00 | 0.00 | 59.00 |
| >hg38_21_+_8433172_8446622_RNA45SN1_45S_50ntFlanks@6647.6677.31\|CGTACGACTCTTAGCGGTGGATCACTCGGCT | >hg38_21_+_8433172_8446622_RNA45SN1_45S_50ntFlanks | 6647 | 6677 | 31 | rRF 45S | NO | 48.81 | 0.00 | 0.00 | 19.94 |
| >hg38_21_+_8433172_8446622_RNA45SN1_45S_50ntFlanks@6647.6679.33\|CGTACGACTCTTAGCGGTGGATCACTCGGCTCG | >hg38_21_+_8433172_8446622_RNA45SN1_45S_50ntFlanks | 6647 | 6679 | 33 | rRF 45S | NO | 25.46 | 0.00 | 0.00 | 19.82 |
| >hg38_21_+_8433172_8446622_RNA45SN1_45S_50ntFlanks@6647.6680.34\|CGTACGACTCTTAGCGGTGGATCACTCGGCTCGT | >hg38_21_+_8433172_8446622_RNA45SN1_45S_50ntFlanks | 6647 | 6680 | 34 | rRF 45S | NO | 59.83 | 0.00 | 0.00 | 20.44 |
| >hg38_21_+_8433172_8446622_RNA45SN1_45S_50ntFlanks@6647.6681.35\|CGTACGACTCTTAGCGGTGGATCACTCGGCTCGTG | >hg38_21_+_8433172_8446622_RNA45SN1_45S_50ntFlanks | 6647 | 6681 | 35 | rRF 45S | NO | 451.48 | 0.00 | 0.00 | 58.54 |
| >hg38_21_+_8433172_8446622_RNA45SN1_45S_50ntFlanks@6648.6668.21\|GTACGACTCTTAGCGGTGGAT | >hg38_21_+_8433172_8446622_RNA45SN1_45S_50ntFlanks | 6648 | 6668 | 21 | rRF 45S | NO | 0.00 | 0.00 | 14.50 | 63.53 |
| >hg38_21_+_8433172_8446622_RNA45SN1_45S_50ntFlanks@6648.6676.29\|GTACGACTCTTAGCGGTGGATCACTCGGC | >hg38_21_+_8433172_8446622_RNA45SN1_45S_50ntFlanks | 6648 | 6676 | 29 | rRF 45S | NO | 74.20 | 0.00 | 0.00 | 113.00 |
| >hg38_21_+_8433172_8446622_RNA45SN1_45S_50ntFlanks@6648.6677.30\|GTACGACTCTTAGCGGTGGATCACTCGGCT | >hg38_21_+_8433172_8446622_RNA45SN1_45S_50ntFlanks | 6648 | 6677 | 30 | rRF 45S | NO | 52.26 | 0.00 | 0.00 | 57.81 |
| >hg38_21_+_8433172_8446622_RNA45SN1_45S_50ntFlanks@6648.6678.31\|GTACGACTCTTAGCGGTGGATCACTCGGCTC | >hg38_21_+_8433172_8446622_RNA45SN1_45S_50ntFlanks | 6648 | 6678 | 31 | rRF 45S | NO | 61.77 | 0.00 | 0.00 | 70.00 |
| >hg38_21_+_8433172_8446622_RNA45SN1_45S_50ntFlanks@6648.6679.32\|GTACGACTCTTAGCGGTGGATCACTCGGCTCG | >hg38_21_+_8433172_8446622_RNA45SN1_45S_50ntFlanks | 6648 | 6679 | 32 | rRF 45S | NO | 84.56 | 0.00 | 0.00 | 93.30 |
| >hg38_21_+_8433172_8446622_RNA45SN1_45S_50ntFlanks@6648.6680.33\|GTACGACTCTTAGCGGTGGATCACTCGGCTCGT | >hg38_21_+_8433172_8446622_RNA45SN1_45S_50ntFlanks | 6648 | 6680 | 33 | rRF 45S | NO | 69.06 | 0.00 | 0.00 | 68.50 |
| >hg38_21_+_8433172_8446622_RNA45SN1_45S_50ntFlanks@6648.6681.34\|GTACGACTCTTAGCGGTGGATCACTCGGCTCGTG | >hg38_21_+_8433172_8446622_RNA45SN1_45S_50ntFlanks | 6648 | 6681 | 34 | rRF 45S | NO | 394.18 | 0.00 | 0.00 | 62.97 |
| >hg38_21_+_8433172_8446622_RNA45SN1_45S_50ntFlanks@6648.6689.42\|GTACGACTCTTAGCGGTGGATCACTCGGCTCGTGCGTCGATG | >hg38_21_+_8433172_8446622_RNA45SN1_45S_50ntFlanks | 6648 | 6689 | 42 | rRF 45S | NO | 80.62 | 0.00 | 0.00 | 55.14 |
| >hg38_21_+_8433172_8446622_RNA45SN1_45S_50ntFlanks@6648.6692.45\|GTACGACTCTTAGCGGTGGATCACTCGGCTCGTGCGTCGATGAAG | >hg38_21_+_8433172_8446622_RNA45SN1_45S_50ntFlanks | 6648 | 6692 | 45 | rRF 45S | NO | 47.71 | 0.00 | 0.00 | 15.86 |
| >hg38_21_+_8433172_8446622_RNA45SN1_45S_50ntFlanks@6648.6695.48\|GTACGACTCTTAGCGGTGGATCACTCGGCTCGTGCGTCGATGAAGAAC | >hg38_21_+_8433172_8446622_RNA45SN1_45S_50ntFlanks | 6648 | 6695 | 48 | rRF 45S | NO | 24.77 | 24.33 | 0.00 | 60.78 |
| >hg38_21_+_8433172_8446622_RNA45SN1_45S_50ntFlanks@6648.6696.49\|GTACGACTCTTAGCGGTGGATCACTCGGCTCGTGCGTCGATGAAGAACG | >hg38_21_+_8433172_8446622_RNA45SN1_45S_50ntFlanks | 6648 | 6696 | 49 | rRF 45S | NO | 87.63 | 31.71 | 0.00 | 0.00 |
| >hg38_21_+_8433172_8446622_RNA45SN1_45S_50ntFlanks@6649.6679.31\|TACGACTCTTAGCGGTGGATCACTCGGCTCG | >hg38_21_+_8433172_8446622_RNA45SN1_45S_50ntFlanks | 6649 | 6679 | 31 | rRF 45S | NO | 55.02 | 0.00 | 0.00 | 82.28 |

### MDA-MB-231

| | | | | | | | | | | |
|---|---|---|---|---|---|---|---|---|---|---|
| >hg38_21_+_8433172_8446622_RNA45SN1_45S_50ntFlanks@6649.6681.33\|TACGACTCTTAG<br>CGGTGGATCACTCGGCTCGTG | >hg38_21_+_8433172_8446622_RNA45SN1_45S_50ntFlanks | 6649 | 6681 | 33 | rRF 45S | NO | 51.91 | 0.00 | 0.00 | 0.00 |
| >hg38_21_+_8433172_8446622_RNA45SN1_45S_50ntFlanks@6649.6683.35\|TACGACTCTTAG<br>CGGTGGATCACTCGGCTCGTGCG | >hg38_21_+_8433172_8446622_RNA45SN1_45S_50ntFlanks | 6649 | 6683 | 35 | rRF 45S | NO | 339.51 | 0.00 | 0.00 | 10.81 |
| >hg38_21_+_8433172_8446622_RNA45SN1_45S_50ntFlanks@6649.6689.41\|TACGACTCTTAG<br>CGGTGGATCACTCGGCTCGTGCGTCGATG | >hg38_21_+_8433172_8446622_RNA45SN1_45S_50ntFlanks | 6649 | 6689 | 41 | rRF 45S | NO | 10.50 | 0.00 | 0.00 | 15.16 |
| >hg38_21_+_8433172_8446622_RNA45SN1_45S_50ntFlanks@6650.6668.19\|ACGACTCTTAGC<br>GGTGGAT | >hg38_21_+_8433172_8446622_RNA45SN1_45S_50ntFlanks | 6650 | 6668 | 19 | rRF 45S | NO | 0.00 | 0.00 | 14.35 | 60.31 |
| >hg38_21_+_8433172_8446622_RNA45SN1_45S_50ntFlanks@6650.6676.27\|ACGACTCTTAGC<br>GGTGGATCACTCGGC | >hg38_21_+_8433172_8446622_RNA45SN1_45S_50ntFlanks | 6650 | 6676 | 27 | rRF 45S | NO | 48.86 | 0.00 | 0.00 | 86.33 |
| >hg38_21_+_8433172_8446622_RNA45SN1_45S_50ntFlanks@6650.6677.28\|ACGACTCTTAGC<br>GGTGGATCACTCGGCT | >hg38_21_+_8433172_8446622_RNA45SN1_45S_50ntFlanks | 6650 | 6677 | 28 | rRF 45S | NO | 55.00 | 0.00 | 0.00 | 65.94 |
| >hg38_21_+_8433172_8446622_RNA45SN1_45S_50ntFlanks@6650.6678.29\|ACGACTCTTAGC<br>GGTGGATCACTCGGCTC | >hg38_21_+_8433172_8446622_RNA45SN1_45S_50ntFlanks | 6650 | 6678 | 29 | rRF 45S | NO | 57.88 | 0.00 | 0.00 | 69.25 |
| >hg38_21_+_8433172_8446622_RNA45SN1_45S_50ntFlanks@6650.6679.30\|ACGACTCTTAGC<br>GGTGGATCACTCGGCTCG | >hg38_21_+_8433172_8446622_RNA45SN1_45S_50ntFlanks | 6650 | 6679 | 30 | rRF 45S | NO | 83.00 | 12.96 | 0.00 | 112.62 |
| >hg38_21_+_8433172_8446622_RNA45SN1_45S_50ntFlanks@6650.6681.32\|ACGACTCTTAGC<br>GGTGGATCACTCGGCTCGTG | >hg38_21_+_8433172_8446622_RNA45SN1_45S_50ntFlanks | 6650 | 6681 | 32 | rRF 45S | NO | 203.21 | 0.00 | 0.00 | 59.21 |
| >hg38_21_+_8433172_8446622_RNA45SN1_45S_50ntFlanks@6650.6683.34\|ACGACTCTTAGC<br>GGTGGATCACTCGGCTCGTGCG | >hg38_21_+_8433172_8446622_RNA45SN1_45S_50ntFlanks | 6650 | 6683 | 34 | rRF 45S | NO | 404.83 | 0.00 | 0.00 | 13.36 |
| >hg38_21_+_8433172_8446622_RNA45SN1_45S_50ntFlanks@6650.6689.40\|ACGACTCTTAGC<br>GGTGGATCACTCGGCTCGTGCGTCGATG | >hg38_21_+_8433172_8446622_RNA45SN1_45S_50ntFlanks | 6650 | 6689 | 40 | rRF 45S | NO | 59.68 | 0.00 | 0.00 | 19.33 |
| >hg38_21_+_8433172_8446622_RNA45SN1_45S_50ntFlanks@6650.6692.43\|ACGACTCTTAGC<br>GGTGGATCACTCGGCTCGTGCGTCGATGAAG | >hg38_21_+_8433172_8446622_RNA45SN1_45S_50ntFlanks | 6650 | 6692 | 43 | rRF 45S | NO | 61.63 | 0.00 | 0.00 | 19.17 |
| >hg38_21_+_8433172_8446622_RNA45SN1_45S_50ntFlanks@6650.6695.46\|ACGACTCTTAGC<br>GGTGGATCACTCGGCTCGTGCGTCGATGAAGAAC | >hg38_21_+_8433172_8446622_RNA45SN1_45S_50ntFlanks | 6650 | 6695 | 46 | rRF 45S | NO | 25.64 | 24.47 | 0.00 | 58.58 |
| >hg38_21_+_8433172_8446622_RNA45SN1_45S_50ntFlanks@6650.6696.47\|ACGACTCTTAGC<br>GGTGGATCACTCGGCTCGTGCGTCGATGAAGAACG | >hg38_21_+_8433172_8446622_RNA45SN1_45S_50ntFlanks | 6650 | 6696 | 47 | rRF 45S | NO | 99.12 | 25.40 | 0.00 | 62.03 |
| >hg38_21_+_8433172_8446622_RNA45SN1_45S_50ntFlanks@6650.6698.49\|ACGACTCTTAGC<br>GGTGGATCACTCGGCTCGTGCGTCGATGAAGAACGCA | >hg38_21_+_8433172_8446622_RNA45SN1_45S_50ntFlanks | 6650 | 6698 | 49 | rRF 45S | NO | 63.30 | 27.44 | 0.00 | 0.00 |
| >hg38_21_+_8433172_8446622_RNA45SN1_45S_50ntFlanks@6650.6699.50\|ACGACTCTTAGC<br>GGTGGATCACTCGGCTCGTGCGTCGATGAAGAACGCAG | >hg38_21_+_8433172_8446622_RNA45SN1_45S_50ntFlanks | 6650 | 6699 | 50 | rRF 45S | NO | 26.32 | 0.00 | 0.00 | 0.00 |
| >hg38_21_+_8433172_8446622_RNA45SN1_45S_50ntFlanks@6651.6668.18\|CGACTCTTAGCG<br>GTGGAT | >hg38_21_+_8433172_8446622_RNA45SN1_45S_50ntFlanks | 6651 | 6668 | 18 | rRF 45S | NO | 180.52 | 145.21 | 73.34 | 304.38 |
| >hg38_21_+_8433172_8446622_RNA45SN1_45S_50ntFlanks@6651.6669.19\|CGACTCTTAGCG<br>GTGGATC | >hg38_21_+_8433172_8446622_RNA45SN1_45S_50ntFlanks | 6651 | 6669 | 19 | rRF 45S | NO | 113.11 | 116.65 | 16.70 | 231.89 |
| >hg38_21_+_8433172_8446622_RNA45SN1_45S_50ntFlanks@6651.6672.22\|CGACTCTTAGCG<br>GTGGATCACT | >hg38_21_+_8433172_8446622_RNA45SN1_45S_50ntFlanks | 6651 | 6672 | 22 | rRF 45S | NO | 178.47 | 156.25 | 0.00 | 289.90 |
| >hg38_21_+_8433172_8446622_RNA45SN1_45S_50ntFlanks@6651.6673.23\|CGACTCTTAGCG<br>GTGGATCACTC | >hg38_21_+_8433172_8446622_RNA45SN1_45S_50ntFlanks | 6651 | 6673 | 23 | rRF 45S | NO | 134.98 | 134.77 | 0.00 | 187.21 |
| >hg38_21_+_8433172_8446622_RNA45SN1_45S_50ntFlanks@6651.6674.24\|CGACTCTTAGCG<br>GTGGATCACTCG | >hg38_21_+_8433172_8446622_RNA45SN1_45S_50ntFlanks | 6651 | 6674 | 24 | rRF 45S | NO | 171.19 | 125.82 | 0.00 | 233.44 |
| >hg38_21_+_8433172_8446622_RNA45SN1_45S_50ntFlanks@6651.6675.25\|CGACTCTTAGCG<br>GTGGATCACTCGG | >hg38_21_+_8433172_8446622_RNA45SN1_45S_50ntFlanks | 6651 | 6675 | 25 | rRF 45S | NO | 279.07 | 168.85 | 0.00 | 194.20 |
| >hg38_21_+_8433172_8446622_RNA45SN1_45S_50ntFlanks@6651.6676.26\|CGACTCTTAGCG<br>GTGGATCACTCGGC | >hg38_21_+_8433172_8446622_RNA45SN1_45S_50ntFlanks | 6651 | 6676 | 26 | rRF 45S | NO | 290.20 | 226.16 | 72.47 | 431.96 |
| >hg38_21_+_8433172_8446622_RNA45SN1_45S_50ntFlanks@6651.6677.27\|CGACTCTTAGCG<br>GTGGATCACTCGGCT | >hg38_21_+_8433172_8446622_RNA45SN1_45S_50ntFlanks | 6651 | 6677 | 27 | rRF 45S | NO | 300.25 | 192.23 | 14.04 | 307.73 |
| >hg38_21_+_8433172_8446622_RNA45SN1_45S_50ntFlanks@6651.6678.28\|CGACTCTTAGCG<br>GTGGATCACTCGGCTC | >hg38_21_+_8433172_8446622_RNA45SN1_45S_50ntFlanks | 6651 | 6678 | 28 | rRF 45S | NO | 348.72 | 242.18 | 11.00 | 337.04 |
| >hg38_21_+_8433172_8446622_RNA45SN1_45S_50ntFlanks@6651.6679.29\|CGACTCTTAGCG<br>GTGGATCACTCGGCTCG | >hg38_21_+_8433172_8446622_RNA45SN1_45S_50ntFlanks | 6651 | 6679 | 29 | rRF 45S | NO | 275.39 | 145.85 | 15.87 | 360.03 |
| >hg38_21_+_8433172_8446622_RNA45SN1_45S_50ntFlanks@6651.6680.30\|CGACTCTTAGCG<br>GTGGATCACTCGGCTCGT | >hg38_21_+_8433172_8446622_RNA45SN1_45S_50ntFlanks | 6651 | 6680 | 30 | rRF 45S | NO | 202.59 | 89.31 | 0.00 | 173.55 |
| >hg38_21_+_8433172_8446622_RNA45SN1_45S_50ntFlanks@6651.6681.31\|CGACTCTTAGCG<br>GTGGATCACTCGGCTCGTG | >hg38_21_+_8433172_8446622_RNA45SN1_45S_50ntFlanks | 6651 | 6681 | 31 | rRF 45S | NO | 1372.26 | 260.65 | 34.90 | 296.78 |
| >hg38_21_+_8433172_8446622_RNA45SN1_45S_50ntFlanks@6651.6682.32\|CGACTCTTAGCG<br>GTGGATCACTCGGCTCGTGC | >hg38_21_+_8433172_8446622_RNA45SN1_45S_50ntFlanks | 6651 | 6682 | 32 | rRF 45S | NO | 253.56 | 120.72 | 0.00 | 167.14 |
| >hg38_21_+_8433172_8446622_RNA45SN1_45S_50ntFlanks@6651.6683.33\|CGACTCTTAGCG<br>GTGGATCACTCGGCTCGTGCG | >hg38_21_+_8433172_8446622_RNA45SN1_45S_50ntFlanks | 6651 | 6683 | 33 | rRF 45S | NO | 1224.73 | 24.34 | 69.44 | 80.29 |
| >hg38_21_+_8433172_8446622_RNA45SN1_45S_50ntFlanks@6651.6684.34\|CGACTCTTAGCG<br>GTGGATCACTCGGCTCGTGCGT | >hg38_21_+_8433172_8446622_RNA45SN1_45S_50ntFlanks | 6651 | 6684 | 34 | rRF 45S | NO | 453.09 | 90.55 | 16.07 | 122.12 |
| >hg38_21_+_8433172_8446622_RNA45SN1_45S_50ntFlanks@6651.6685.35\|CGACTCTTAGCG<br>GTGGATCACTCGGCTCGTGCGTC | >hg38_21_+_8433172_8446622_RNA45SN1_45S_50ntFlanks | 6651 | 6685 | 35 | rRF 45S | NO | 102.53 | 0.00 | 91.22 | 77.35 |
| >hg38_21_+_8433172_8446622_RNA45SN1_45S_50ntFlanks@6651.6686.36\|CGACTCTTAGCG<br>GTGGATCACTCGGCTCGTGCGTCG | >hg38_21_+_8433172_8446622_RNA45SN1_45S_50ntFlanks | 6651 | 6686 | 36 | rRF 45S | NO | 57.64 | 0.00 | 0.00 | 13.06 |
| >hg38_21_+_8433172_8446622_RNA45SN1_45S_50ntFlanks@6651.6687.37\|CGACTCTTAGCG<br>GTGGATCACTCGGCTCGTGCGTCGA | >hg38_21_+_8433172_8446622_RNA45SN1_45S_50ntFlanks | 6651 | 6687 | 37 | rRF 45S | NO | 137.63 | 27.84 | 11.83 | 104.76 |
| >hg38_21_+_8433172_8446622_RNA45SN1_45S_50ntFlanks@6651.6688.38\|CGACTCTTAGCG<br>GTGGATCACTCGGCTCGTGCGTCGAT | >hg38_21_+_8433172_8446622_RNA45SN1_45S_50ntFlanks | 6651 | 6688 | 38 | rRF 45S | NO | 137.68 | 0.00 | 15.56 | 100.44 |
| >hg38_21_+_8433172_8446622_RNA45SN1_45S_50ntFlanks@6651.6689.39\|CGACTCTTAGCG<br>GTGGATCACTCGGCTCGTGCGTCGATG | >hg38_21_+_8433172_8446622_RNA45SN1_45S_50ntFlanks | 6651 | 6689 | 39 | rRF 45S | NO | 410.84 | 99.36 | 98.84 | 248.24 |
| >hg38_21_+_8433172_8446622_RNA45SN1_45S_50ntFlanks@6651.6690.40\|CGACTCTTAGCG<br>GTGGATCACTCGGCTCGTGCGTCGATGA | >hg38_21_+_8433172_8446622_RNA45SN1_45S_50ntFlanks | 6651 | 6690 | 40 | rRF 45S | NO | 150.98 | 32.24 | 14.15 | 102.83 |
| >hg38_21_+_8433172_8446622_RNA45SN1_45S_50ntFlanks@6651.6691.41\|CGACTCTTAGCG<br>GTGGATCACTCGGCTCGTGCGTCGATGAA | >hg38_21_+_8433172_8446622_RNA45SN1_45S_50ntFlanks | 6651 | 6691 | 41 | rRF 45S | NO | 58.57 | 0.00 | 0.00 | 18.29 |

### MDA-MB-231

| | | | | | | | | | | |
|---|---|---|---|---|---|---|---|---|---|---|
| >hg38_21_+_8433172_8446622_RNA45SN1_45S_50ntFlanks@6651.6692.42\|CGACTCTTAGCGGTGGATCACTCGGCTCGTGCGTCGATGAAG | >hg38_21_+_8433172_8446622_RNA45SN1_45S_50ntFlanks | 6651 | 6692 | 42 | rRF 45S | NO | 339.72 | 121.69 | 34.65 | 201.73 |
| >hg38_21_+_8433172_8446622_RNA45SN1_45S_50ntFlanks@6651.6693.43\|CGACTCTTAGCGGTGGATCACTCGGCTCGTGCGTCGATGAAGA | >hg38_21_+_8433172_8446622_RNA45SN1_45S_50ntFlanks | 6651 | 6693 | 43 | rRF 45S | NO | 201.88 | 100.67 | 12.45 | 190.93 |
| >hg38_21_+_8433172_8446622_RNA45SN1_45S_50ntFlanks@6651.6694.44\|CGACTCTTAGCGGTGGATCACTCGGCTCGTGCGTCGATGAAGAA | >hg38_21_+_8433172_8446622_RNA45SN1_45S_50ntFlanks | 6651 | 6694 | 44 | rRF 45S | NO | 146.14 | 66.13 | 0.00 | 119.03 |
| >hg38_21_+_8433172_8446622_RNA45SN1_45S_50ntFlanks@6651.6695.45\|CGACTCTTAGCGGTGGATCACTCGGCTCGTGCGTCGATGAAGAAC | >hg38_21_+_8433172_8446622_RNA45SN1_45S_50ntFlanks | 6651 | 6695 | 45 | rRF 45S | NO | 235.68 | 259.54 | 31.01 | 247.03 |
| >hg38_21_+_8433172_8446622_RNA45SN1_45S_50ntFlanks@6651.6696.46\|CGACTCTTAGCGGTGGATCACTCGGCTCGTGCGTCGATGAAGAACG | >hg38_21_+_8433172_8446622_RNA45SN1_45S_50ntFlanks | 6651 | 6696 | 46 | rRF 45S | NO | 808.69 | 371.28 | 12.94 | 333.75 |
| >hg38_21_+_8433172_8446622_RNA45SN1_45S_50ntFlanks@6651.6697.47\|CGACTCTTAGCGGTGGATCACTCGGCTCGTGCGTCGATGAAGAACGC | >hg38_21_+_8433172_8446622_RNA45SN1_45S_50ntFlanks | 6651 | 6697 | 47 | rRF 45S | NO | 184.81 | 213.39 | 0.00 | 265.45 |
| >hg38_21_+_8433172_8446622_RNA45SN1_45S_50ntFlanks@6651.6698.48\|CGACTCTTAGCGGTGGATCACTCGGCTCGTGCGTCGATGAAGAACGCA | >hg38_21_+_8433172_8446622_RNA45SN1_45S_50ntFlanks | 6651 | 6698 | 48 | rRF 45S | NO | 256.46 | 179.22 | 0.00 | 275.39 |
| >hg38_21_+_8433172_8446622_RNA45SN1_45S_50ntFlanks@6651.6699.49\|CGACTCTTAGCGGTGGATCACTCGGCTCGTGCGTCGATGAAGAACGCAG | >hg38_21_+_8433172_8446622_RNA45SN1_45S_50ntFlanks | 6651 | 6699 | 49 | rRF 45S | NO | 362.10 | 196.18 | 0.00 | 225.84 |
| >hg38_21_+_8433172_8446622_RNA45SN1_45S_50ntFlanks@6651.6700.50\|CGACTCTTAGCGGTGGATCACTCGGCTCGTGCGTCGATGAAGAACGCAGC | >hg38_21_+_8433172_8446622_RNA45SN1_45S_50ntFlanks | 6651 | 6700 | 50 | rRF 45S | NO | 116.01 | 190.55 | 0.00 | 265.08 |
| >hg38_21_+_8433172_8446622_RNA45SN1_45S_50ntFlanks@6652.6676.25\|GACTCTTAGCGGTGGATCACTCGGC | >hg38_21_+_8433172_8446622_RNA45SN1_45S_50ntFlanks | 6652 | 6676 | 25 | rRF 45S | NO | 74.60 | 78.63 | 12.82 | 93.05 |
| >hg38_21_+_8433172_8446622_RNA45SN1_45S_50ntFlanks@6652.6677.26\|GACTCTTAGCGGTGGATCACTCGGCT | >hg38_21_+_8433172_8446622_RNA45SN1_45S_50ntFlanks | 6652 | 6677 | 26 | rRF 45S | NO | 68.65 | 35.94 | 10.55 | 62.03 |
| >hg38_21_+_8433172_8446622_RNA45SN1_45S_50ntFlanks@6652.6678.27\|GACTCTTAGCGGTGGATCACTCGGCTC | >hg38_21_+_8433172_8446622_RNA45SN1_45S_50ntFlanks | 6652 | 6678 | 27 | rRF 45S | NO | 109.88 | 87.67 | 0.00 | 108.32 |
| >hg38_21_+_8433172_8446622_RNA45SN1_45S_50ntFlanks@6652.6679.28\|GACTCTTAGCGGTGGATCACTCGGCTCG | >hg38_21_+_8433172_8446622_RNA45SN1_45S_50ntFlanks | 6652 | 6679 | 28 | rRF 45S | NO | 85.25 | 30.31 | 0.00 | 95.79 |
| >hg38_21_+_8433172_8446622_RNA45SN1_45S_50ntFlanks@6652.6680.29\|GACTCTTAGCGGTGGATCACTCGGCTCGT | >hg38_21_+_8433172_8446622_RNA45SN1_45S_50ntFlanks | 6652 | 6680 | 29 | rRF 45S | NO | 24.52 | 0.00 | 0.00 | 13.50 |
| >hg38_21_+_8433172_8446622_RNA45SN1_45S_50ntFlanks@6652.6681.30\|GACTCTTAGCGGTGGATCACTCGGCTCGTG | >hg38_21_+_8433172_8446622_RNA45SN1_45S_50ntFlanks | 6652 | 6681 | 30 | rRF 45S | NO | 429.67 | 58.73 | 0.00 | 70.44 |
| >hg38_21_+_8433172_8446622_RNA45SN1_45S_50ntFlanks@6652.6682.31\|GACTCTTAGCGGTGGATCACTCGGCTCGTGC | >hg38_21_+_8433172_8446622_RNA45SN1_45S_50ntFlanks | 6652 | 6682 | 31 | rRF 45S | NO | 93.20 | 24.92 | 0.00 | 62.53 |
| >hg38_21_+_8433172_8446622_RNA45SN1_45S_50ntFlanks@6652.6683.32\|GACTCTTAGCGGTGGATCACTCGGCTCGTGCG | >hg38_21_+_8433172_8446622_RNA45SN1_45S_50ntFlanks | 6652 | 6683 | 32 | rRF 45S | NO | 957.42 | 0.00 | 0.00 | 18.47 |
| >hg38_21_+_8433172_8446622_RNA45SN1_45S_50ntFlanks@6652.6684.33\|GACTCTTAGCGGTGGATCACTCGGCTCGTGCGT | >hg38_21_+_8433172_8446622_RNA45SN1_45S_50ntFlanks | 6652 | 6684 | 33 | rRF 45S | NO | 128.09 | 13.20 | 0.00 | 13.45 |
| >hg38_21_+_8433172_8446622_RNA45SN1_45S_50ntFlanks@6652.6686.35\|GACTCTTAGCGGTGGATCACTCGGCTCGTGCGTCG | >hg38_21_+_8433172_8446622_RNA45SN1_45S_50ntFlanks | 6652 | 6686 | 35 | rRF 45S | NO | 171.73 | 0.00 | 0.00 | 19.38 |
| >hg38_21_+_8433172_8446622_RNA45SN1_45S_50ntFlanks@6652.6687.36\|GACTCTTAGCGGTGGATCACTCGGCTCGTGCGTCGA | >hg38_21_+_8433172_8446622_RNA45SN1_45S_50ntFlanks | 6652 | 6687 | 36 | rRF 45S | NO | 189.58 | 0.00 | 0.00 | 53.98 |
| >hg38_21_+_8433172_8446622_RNA45SN1_45S_50ntFlanks@6652.6688.37\|GACTCTTAGCGGTGGATCACTCGGCTCGTGCGTCGAT | >hg38_21_+_8433172_8446622_RNA45SN1_45S_50ntFlanks | 6652 | 6688 | 37 | rRF 45S | NO | 149.20 | 0.00 | 0.00 | 62.61 |
| >hg38_21_+_8433172_8446622_RNA45SN1_45S_50ntFlanks@6652.6689.38\|GACTCTTAGCGGTGGATCACTCGGCTCGTGCGTCGATG | >hg38_21_+_8433172_8446622_RNA45SN1_45S_50ntFlanks | 6652 | 6689 | 38 | rRF 45S | NO | 228.32 | 58.35 | 13.64 | 80.16 |
| >hg38_21_+_8433172_8446622_RNA45SN1_45S_50ntFlanks@6652.6690.39\|GACTCTTAGCGGTGGATCACTCGGCTCGTGCGTCGATGA | >hg38_21_+_8433172_8446622_RNA45SN1_45S_50ntFlanks | 6652 | 6690 | 39 | rRF 45S | NO | 68.45 | 0.00 | 0.00 | 12.26 |
| >hg38_21_+_8433172_8446622_RNA45SN1_45S_50ntFlanks@6652.6692.41\|GACTCTTAGCGGTGGATCACTCGGCTCGTGCGTCGATGAAG | >hg38_21_+_8433172_8446622_RNA45SN1_45S_50ntFlanks | 6652 | 6692 | 41 | rRF 45S | NO | 134.90 | 36.08 | 0.00 | 55.72 |
| >hg38_21_+_8433172_8446622_RNA45SN1_45S_50ntFlanks@6652.6693.42\|GACTCTTAGCGGTGGATCACTCGGCTCGTGCGTCGATGAAGA | >hg38_21_+_8433172_8446622_RNA45SN1_45S_50ntFlanks | 6652 | 6693 | 42 | rRF 45S | NO | 85.51 | 13.42 | 0.00 | 20.78 |
| >hg38_21_+_8433172_8446622_RNA45SN1_45S_50ntFlanks@6652.6694.43\|GACTCTTAGCGGTGGATCACTCGGCTCGTGCGTCGATGAAGAA | >hg38_21_+_8433172_8446622_RNA45SN1_45S_50ntFlanks | 6652 | 6694 | 43 | rRF 45S | NO | 58.53 | 0.00 | 0.00 | 14.89 |
| >hg38_21_+_8433172_8446622_RNA45SN1_45S_50ntFlanks@6652.6695.44\|GACTCTTAGCGGTGGATCACTCGGCTCGTGCGTCGATGAAGAAC | >hg38_21_+_8433172_8446622_RNA45SN1_45S_50ntFlanks | 6652 | 6695 | 44 | rRF 45S | NO | 105.87 | 98.62 | 12.54 | 80.98 |
| >hg38_21_+_8433172_8446622_RNA45SN1_45S_50ntFlanks@6652.6696.45\|GACTCTTAGCGGTGGATCACTCGGCTCGTGCGTCGATGAAGAACG | >hg38_21_+_8433172_8446622_RNA45SN1_45S_50ntFlanks | 6652 | 6696 | 45 | rRF 45S | NO | 210.90 | 108.01 | 0.00 | 77.63 |
| >hg38_21_+_8433172_8446622_RNA45SN1_45S_50ntFlanks@6652.6697.46\|GACTCTTAGCGGTGGATCACTCGGCTCGTGCGTCGATGAAGAACGC | >hg38_21_+_8433172_8446622_RNA45SN1_45S_50ntFlanks | 6652 | 6697 | 46 | rRF 45S | NO | 74.70 | 37.70 | 0.00 | 96.54 |
| >hg38_21_+_8433172_8446622_RNA45SN1_45S_50ntFlanks@6652.6698.47\|GACTCTTAGCGGTGGATCACTCGGCTCGTGCGTCGATGAAGAACGCA | >hg38_21_+_8433172_8446622_RNA45SN1_45S_50ntFlanks | 6652 | 6698 | 47 | rRF 45S | NO | 180.82 | 133.23 | 0.00 | 145.45 |
| >hg38_21_+_8433172_8446622_RNA45SN1_45S_50ntFlanks@6652.6699.48\|GACTCTTAGCGGTGGATCACTCGGCTCGTGCGTCGATGAAGAACGCAG | >hg38_21_+_8433172_8446622_RNA45SN1_45S_50ntFlanks | 6652 | 6699 | 48 | rRF 45S | NO | 114.39 | 35.08 | 0.00 | 68.54 |
| >hg38_21_+_8433172_8446622_RNA45SN1_45S_50ntFlanks@6652.6700.49\|GACTCTTAGCGGTGGATCACTCGGCTCGTGCGTCGATGAAGAACGCAGC | >hg38_21_+_8433172_8446622_RNA45SN1_45S_50ntFlanks | 6652 | 6700 | 49 | rRF 45S | NO | 10.64 | 36.30 | 0.00 | 87.38 |
| >hg38_21_+_8433172_8446622_RNA45SN1_45S_50ntFlanks@6652.6701.50\|GACTCTTAGCGGTGGATCACTCGGCTCGTGCGTCGATGAAGAACGCAGCT | >hg38_21_+_8433172_8446622_RNA45SN1_45S_50ntFlanks | 6652 | 6701 | 50 | rRF 45S | NO | 0.00 | 0.00 | 10.46 | 19.57 |
| >hg38_21_+_8433172_8446622_RNA45SN1_45S_50ntFlanks@6653.6676.24\|ACTCTTAGCGGTGGATCACTCGGC | >hg38_21_+_8433172_8446622_RNA45SN1_45S_50ntFlanks | 6653 | 6676 | 24 | rRF 45S | NO | 0.00 | 25.10 | 0.00 | 0.00 |
| >hg38_21_+_8433172_8446622_RNA45SN1_45S_50ntFlanks@6653.6681.29\|ACTCTTAGCGGTGGATCACTCGGCTCGTG | >hg38_21_+_8433172_8446622_RNA45SN1_45S_50ntFlanks | 6653 | 6681 | 29 | rRF 45S | NO | 154.01 | 0.00 | 0.00 | 0.00 |
| >hg38_21_+_8433172_8446622_RNA45SN1_45S_50ntFlanks@6653.6683.31\|ACTCTTAGCGGTGGATCACTCGGCTCGTGCG | >hg38_21_+_8433172_8446622_RNA45SN1_45S_50ntFlanks | 6653 | 6683 | 31 | rRF 45S | NO | 239.99 | 0.00 | 0.00 | 0.00 |
| >hg38_21_+_8433172_8446622_RNA45SN1_45S_50ntFlanks@6653.6686.34\|ACTCTTAGCGGTGGATCACTCGGCTCGTGCGTCG | >hg38_21_+_8433172_8446622_RNA45SN1_45S_50ntFlanks | 6653 | 6686 | 34 | rRF 45S | NO | 51.46 | 0.00 | 0.00 | 0.00 |
| >hg38_21_+_8433172_8446622_RNA45SN1_45S_50ntFlanks@6653.6687.35\|ACTCTTAGCGGTGGATCACTCGGCTCGTGCGTCGA | >hg38_21_+_8433172_8446622_RNA45SN1_45S_50ntFlanks | 6653 | 6687 | 35 | rRF 45S | NO | 65.96 | 0.00 | 0.00 | 0.00 |

### MDA-MB-231

| | | | | | | | | | | |
|---|---|---|---|---|---|---|---|---|---|---|
| >hg38_21_+_8433172_8446622_RNA45SN1_45S_50ntFlanks@6653.6688.36\|ACTCTTAGCGGTGGATCACTCGGCTCGTGCGTCGAT | >hg38_21_+_8433172_8446622_RNA45SN1_45S_50ntFlanks | 6653 | 6688 | 36 | rRF 45S | NO | 66.74 | 0.00 | 0.00 | 0.00 |
| >hg38_21_+_8433172_8446622_RNA45SN1_45S_50ntFlanks@6653.6689.37\|ACTCTTAGCGGTGGATCACTCGGCTCGTGCGTCGATG | >hg38_21_+_8433172_8446622_RNA45SN1_45S_50ntFlanks | 6653 | 6689 | 37 | rRF 45S | NO | 26.25 | 0.00 | 0.00 | 0.00 |
| >hg38_21_+_8433172_8446622_RNA45SN1_45S_50ntFlanks@6654.6681.28\|CTCTTAGCGGTGGATCACTCGGCTCGTG | >hg38_21_+_8433172_8446622_RNA45SN1_45S_50ntFlanks | 6654 | 6681 | 28 | rRF 45S | NO | 26.64 | 0.00 | 0.00 | 0.00 |
| >hg38_21_+_8433172_8446622_RNA45SN1_45S_50ntFlanks@6654.6683.30\|CTCTTAGCGGTGGATCACTCGGCTCGTGCG | >hg38_21_+_8433172_8446622_RNA45SN1_45S_50ntFlanks | 6654 | 6683 | 30 | rRF 45S | NO | 111.25 | 0.00 | 0.00 | 0.00 |
| >hg38_21_+_8433172_8446622_RNA45SN1_45S_50ntFlanks@6655.6681.27\|TCTTAGCGGTGGATCACTCGGCTCGTG | >hg38_21_+_8433172_8446622_RNA45SN1_45S_50ntFlanks | 6655 | 6681 | 27 | rRF 45S | NO | 23.85 | 0.00 | 0.00 | 0.00 |
| >hg38_21_+_8433172_8446622_RNA45SN1_45S_50ntFlanks@6655.6683.29\|TCTTAGCGGTGGATCACTCGGCTCGTGCG | >hg38_21_+_8433172_8446622_RNA45SN1_45S_50ntFlanks | 6655 | 6683 | 29 | rRF 45S | NO | 25.38 | 0.00 | 0.00 | 0.00 |
| >hg38_21_+_8433172_8446622_RNA45SN1_45S_50ntFlanks@6660.6683.24\|GCGGTGGATCACTCGGCTCGTGCG | >hg38_21_+_8433172_8446622_RNA45SN1_45S_50ntFlanks | 6660 | 6683 | 24 | rRF 45S | NO | 90.86 | 0.00 | 0.00 | 0.00 |
| >hg38_21_+_8433172_8446622_RNA45SN1_45S_50ntFlanks@6661.6681.21\|CGGTGGATCACTCGGCTCGTG | >hg38_21_+_8433172_8446622_RNA45SN1_45S_50ntFlanks | 6661 | 6681 | 21 | rRF 45S | NO | 108.94 | 0.00 | 0.00 | 0.00 |
| >hg38_21_+_8433172_8446622_RNA45SN1_45S_50ntFlanks@6661.6683.23\|CGGTGGATCACTCGGCTCGTGCG | >hg38_21_+_8433172_8446622_RNA45SN1_45S_50ntFlanks | 6661 | 6683 | 23 | rRF 45S | NO | 449.48 | 0.00 | 0.00 | 0.00 |
| >hg38_21_+_8433172_8446622_RNA45SN1_45S_50ntFlanks@6661.6686.26\|CGGTGGATCACTCGGCTCGTGCGTCG | >hg38_21_+_8433172_8446622_RNA45SN1_45S_50ntFlanks | 6661 | 6686 | 26 | rRF 45S | NO | 79.98 | 0.00 | 0.00 | 0.00 |
| >hg38_21_+_8433172_8446622_RNA45SN1_45S_50ntFlanks@6661.6687.27\|CGGTGGATCACTCGGCTCGTGCGTCGA | >hg38_21_+_8433172_8446622_RNA45SN1_45S_50ntFlanks | 6661 | 6687 | 27 | rRF 45S | NO | 79.81 | 0.00 | 0.00 | 0.00 |
| >hg38_21_+_8433172_8446622_RNA45SN1_45S_50ntFlanks@6661.6688.28\|CGGTGGATCACTCGGCTCGTGCGTCGAT | >hg38_21_+_8433172_8446622_RNA45SN1_45S_50ntFlanks | 6661 | 6688 | 28 | rRF 45S | NO | 60.60 | 0.00 | 0.00 | 0.00 |
| >hg38_21_+_8433172_8446622_RNA45SN1_45S_50ntFlanks@6661.6689.29\|CGGTGGATCACTCGGCTCGTGCGTCGATG | >hg38_21_+_8433172_8446622_RNA45SN1_45S_50ntFlanks | 6661 | 6689 | 29 | rRF 45S | NO | 62.38 | 0.00 | 0.00 | 12.93 |
| >hg38_21_+_8433172_8446622_RNA45SN1_45S_50ntFlanks@6661.6695.35\|CGGTGGATCACTCGGCTCGTGCGTCGATGAAGAAC | >hg38_21_+_8433172_8446622_RNA45SN1_45S_50ntFlanks | 6661 | 6695 | 35 | rRF 45S | NO | 0.00 | 0.00 | 10.41 | 10.99 |
| >hg38_21_+_8433172_8446622_RNA45SN1_45S_50ntFlanks@6661.6696.36\|CGGTGGATCACTCGGCTCGTGCGTCGATGAAGAACG | >hg38_21_+_8433172_8446622_RNA45SN1_45S_50ntFlanks | 6661 | 6696 | 36 | rRF 45S | NO | 54.21 | 0.00 | 0.00 | 0.00 |
| >hg38_21_+_8433172_8446622_RNA45SN1_45S_50ntFlanks@6661.6699.39\|CGGTGGATCACTCGGCTCGTGCGTCGATGAAGAACGCAG | >hg38_21_+_8433172_8446622_RNA45SN1_45S_50ntFlanks | 6661 | 6699 | 39 | rRF 45S | NO | 61.89 | 0.00 | 0.00 | 0.00 |
| >hg38_21_+_8433172_8446622_RNA45SN1_45S_50ntFlanks@6661.6701.41\|CGGTGGATCACTCGGCTCGTGCGTCGATGAAGAACGCAGCT | >hg38_21_+_8433172_8446622_RNA45SN1_45S_50ntFlanks | 6661 | 6701 | 41 | rRF 45S | NO | 10.71 | 0.00 | 13.35 | 16.36 |
| >hg38_21_+_8433172_8446622_RNA45SN1_45S_50ntFlanks@6697.6720.24\|CAGCTAGCTGCGAGAATTAATGTG | >hg38_21_+_8433172_8446622_RNA45SN1_45S_50ntFlanks | 6697 | 6720 | 24 | rRF 45S | NO | 0.00 | 83.98 | 0.00 | 0.00 |
| >hg38_21_+_8433172_8446622_RNA45SN1_45S_50ntFlanks@6698.6720.23\|AGCTAGCTGCGAGAATTAATGTG | >hg38_21_+_8433172_8446622_RNA45SN1_45S_50ntFlanks | 6698 | 6720 | 23 | rRF 45S | NO | 0.00 | 113.70 | 10.33 | 0.00 |
| >hg38_21_+_8433172_8446622_RNA45SN1_45S_50ntFlanks@6699.6720.22\|GCTAGCTGCGAGAATTAATGTG | >hg38_21_+_8433172_8446622_RNA45SN1_45S_50ntFlanks | 6699 | 6720 | 22 | rRF 45S | NO | 25.89 | 113.27 | 11.24 | 0.00 |
| >hg38_21_+_8433172_8446622_RNA45SN1_45S_50ntFlanks@6700.6720.21\|CTAGCTGCGAGAATTAATGTG | >hg38_21_+_8433172_8446622_RNA45SN1_45S_50ntFlanks | 6700 | 6720 | 21 | rRF 45S | NO | 0.00 | 67.46 | 0.00 | 0.00 |
| >hg38_21_+_8433172_8446622_RNA45SN1_45S_50ntFlanks@6732.6773.42\|ACATTGATCATCGACACTTCGAACGCACTTGCGGCCCCGGGT | >hg38_21_+_8433172_8446622_RNA45SN1_45S_50ntFlanks | 6732 | 6773 | 42 | rRF 45S | NO | 25.92 | 34.27 | 0.00 | 64.03 |
| >hg38_21_+_8433172_8446622_RNA45SN1_45S_50ntFlanks@6732.6774.43\|ACATTGATCATCGACACTTCGAACGCACTTGCGGCCCCGGGTT | >hg38_21_+_8433172_8446622_RNA45SN1_45S_50ntFlanks | 6732 | 6774 | 43 | rRF 45S | NO | 0.00 | 25.56 | 0.00 | 21.36 |
| >hg38_21_+_8433172_8446622_RNA45SN1_45S_50ntFlanks@6734.6773.40\|ATTGATCATCGACACTTCGAACGCACTTGCGGCCCCGGGT | >hg38_21_+_8433172_8446622_RNA45SN1_45S_50ntFlanks | 6734 | 6773 | 40 | rRF 45S | NO | 59.02 | 26.16 | 0.00 | 79.98 |
| >hg38_21_+_8433172_8446622_RNA45SN1_45S_50ntFlanks@6734.6774.41\|ATTGATCATCGACACTTCGAACGCACTTGCGGCCCCGGGTT | >hg38_21_+_8433172_8446622_RNA45SN1_45S_50ntFlanks | 6734 | 6774 | 41 | rRF 45S | NO | 10.64 | 24.61 | 0.00 | 60.97 |
| >hg38_21_+_8433172_8446622_RNA45SN1_45S_50ntFlanks@6759.6806.48\|CTTGCGGCCCCGGGTTCCTCCCGGGGCTACGCCTGTCTGAGCGTCGCT | >hg38_21_+_8433172_8446622_RNA45SN1_45S_50ntFlanks | 6759 | 6806 | 48 | rRF 45S | NO | 0.00 | 27.13 | 0.00 | 0.00 |
| >hg38_21_+_8433172_8446622_RNA45SN1_45S_50ntFlanks@6760.6806.47\|TTGCGGCCCCGGGTTCCTCCCGGGGCTACGCCTGTCTGAGCGTCGCT | >hg38_21_+_8433172_8446622_RNA45SN1_45S_50ntFlanks | 6760 | 6806 | 47 | rRF 45S | NO | 0.00 | 25.37 | 0.00 | 0.00 |
| >hg38_21_+_8433172_8446622_RNA45SN1_45S_50ntFlanks@6761.6806.46\|TGCGGCCCCGGGTTCCTCCCGGGGCTACGCCTGTCTGAGCGTCGCT | >hg38_21_+_8433172_8446622_RNA45SN1_45S_50ntFlanks | 6761 | 6806 | 46 | rRF 45S | NO | 0.00 | 13.47 | 0.00 | 14.52 |
| >hg38_21_+_8433172_8446622_RNA45SN1_45S_50ntFlanks@6772.6806.35\|GTTCCTCCCGGGGCTACGCCTGTCTGAGCGTCGCT | >hg38_21_+_8433172_8446622_RNA45SN1_45S_50ntFlanks | 6772 | 6806 | 35 | rRF 45S | NO | 0.00 | 29.31 | 0.00 | 12.86 |
| >hg38_21_+_8433172_8446622_RNA45SN1_45S_50ntFlanks@6773.6806.34\|TTCCTCCCGGGGCTACGCCTGTCTGAGCGTCGCT | >hg38_21_+_8433172_8446622_RNA45SN1_45S_50ntFlanks | 6773 | 6806 | 34 | rRF 45S | NO | 0.00 | 26.53 | 0.00 | 21.31 |
| >hg38_21_+_8433172_8446622_RNA45SN1_45S_50ntFlanks@6774.6806.33\|TCCTCCCGGGGCTACGCCTGTCTGAGCGTCGCT | >hg38_21_+_8433172_8446622_RNA45SN1_45S_50ntFlanks | 6774 | 6806 | 33 | rRF 45S | NO | 10.43 | 60.64 | 0.00 | 96.99 |
| >hg38_21_+_8433172_8446622_RNA45SN1_45S_50ntFlanks@6776.6802.27\|CTCCCGGGGCTACGCCTGTCTGAGCGT | >hg38_21_+_8433172_8446622_RNA45SN1_45S_50ntFlanks | 6776 | 6802 | 27 | rRF 45S | NO | 26.88 | 28.49 | 0.00 | 0.00 |
| >hg38_21_+_8433172_8446622_RNA45SN1_45S_50ntFlanks@6776.6806.31\|CTCCCGGGGCTACGCCTGTCTGAGCGTCGCT | >hg38_21_+_8433172_8446622_RNA45SN1_45S_50ntFlanks | 6776 | 6806 | 31 | rRF 45S | NO | 86.57 | 100.20 | 0.00 | 104.28 |
| >hg38_21_+_8433172_8446622_RNA45SN1_45S_50ntFlanks@6777.6806.30\|TCCCGGGGCTACGCCTGTCTGAGCGTCGCT | >hg38_21_+_8433172_8446622_RNA45SN1_45S_50ntFlanks | 6777 | 6806 | 30 | rRF 45S | NO | 0.00 | 28.57 | 0.00 | 18.93 |
| >hg38_21_+_8433172_8446622_RNA45SN1_45S_50ntFlanks@6778.6802.25\|CCCGGGGCTACGCCTGTCTGAGCGT | >hg38_21_+_8433172_8446622_RNA45SN1_45S_50ntFlanks | 6778 | 6802 | 25 | rRF 45S | NO | 73.12 | 159.27 | 0.00 | 15.28 |
| >hg38_21_+_8433172_8446622_RNA45SN1_45S_50ntFlanks@6778.6806.29\|CCCGGGGCTACGCCTGTCTGAGCGTCGCT | >hg38_21_+_8433172_8446622_RNA45SN1_45S_50ntFlanks | 6778 | 6806 | 29 | rRF 45S | NO | 64.30 | 83.32 | 16.91 | 85.84 |
| >hg38_21_+_8433172_8446622_RNA45SN1_45S_50ntFlanks@6779.6802.24\|CCGGGGCTACGCCTGTCTGAGCGT | >hg38_21_+_8433172_8446622_RNA45SN1_45S_50ntFlanks | 6779 | 6802 | 24 | rRF 45S | NO | 0.00 | 28.81 | 0.00 | 0.00 |
| >hg38_21_+_8433172_8446622_RNA45SN1_45S_50ntFlanks@6779.6806.28\|CCGGGGCTACGCCTGTCTGAGCGTCGCT | >hg38_21_+_8433172_8446622_RNA45SN1_45S_50ntFlanks | 6779 | 6806 | 28 | rRF 45S | NO | 0.00 | 31.76 | 33.69 | 19.77 |

### MDA-MB-231

| | | | | | | | | | | |
|---|---|---|---|---|---|---|---|---|---|---|
| >hg38_21_+_8433172_8446622_RNA45SN1_45S_50ntFlanks@6780.6802.23\|CGGGGCTACGC CTGTCTGAGCGT | >hg38_21_+_8433172_8446622_RNA45SN1_45S_50ntFlanks | 6780 | 6802 | 23 | rRF 45S | NO | 0.00 | 57.15 | 0.00 | 0.00 |
| >hg38_21_+_8433172_8446622_RNA45SN1_45S_50ntFlanks@6780.6806.27\|CGGGGCTACGC CTGTCTGAGCGTCGCT | >hg38_21_+_8433172_8446622_RNA45SN1_45S_50ntFlanks | 6780 | 6806 | 27 | rRF 45S | NO | 0.00 | 0.00 | 16.97 | 11.69 |
| >hg38_21_+_8433172_8446622_RNA45SN1_45S_50ntFlanks@6785.6806.22\|CTACGCCTGTCT GAGCGTCGCT | >hg38_21_+_8433172_8446622_RNA45SN1_45S_50ntFlanks | 6785 | 6806 | 22 | rRF 45S | NO | 52.03 | 95.56 | 63.65 | 101.87 |
| >hg38_21_+_8433172_8446622_RNA45SN1_45S_50ntFlanks@6786.6806.21\|TACGCCTGTCTG AGCGTCGCT | >hg38_21_+_8433172_8446622_RNA45SN1_45S_50ntFlanks | 6786 | 6806 | 21 | rRF 45S | NO | 71.98 | 128.85 | 34.14 | 127.01 |
| >hg38_21_+_8433172_8446622_RNA45SN1_45S_50ntFlanks@6787.6806.20\|ACGCCTGTCTGA GCGTCGCT | >hg38_21_+_8433172_8446622_RNA45SN1_45S_50ntFlanks | 6787 | 6806 | 20 | rRF 45S | NO | 84.27 | 131.08 | 65.03 | 98.64 |
| >hg38_21_+_8433172_8446622_RNA45SN1_45S_50ntFlanks@6788.6806.19\|CGCCTGTCTGA GCGTCGCT | >hg38_21_+_8433172_8446622_RNA45SN1_45S_50ntFlanks | 6788 | 6806 | 19 | rRF 45S | NO | 189.61 | 463.66 | 34.62 | 164.20 |
| >hg38_21_+_8433172_8446622_RNA45SN1_45S_50ntFlanks@6788.6807.20\|CGCCTGTCTGA GCGTCGCTT | >hg38_21_+_8433172_8446622_RNA45SN1_45S_50ntFlanks | 6788 | 6807 | 20 | rRF 45S | NO | 75.90 | 108.99 | 0.00 | 20.15 |
| >hg38_21_+_8433172_8446622_RNA45SN1_45S_50ntFlanks@6789.6806.18\|GCCTGTCTGAG CGTCGCT | >hg38_21_+_8433172_8446622_RNA45SN1_45S_50ntFlanks | 6789 | 6806 | 18 | rRF 45S | NO | 98.17 | 213.82 | 0.00 | 142.82 |
| >hg38_21_+_8433172_8446622_RNA45SN1_45S_50ntFlanks@6789.6807.19\|GCCTGTCTGAG CGTCGCTT | >hg38_21_+_8433172_8446622_RNA45SN1_45S_50ntFlanks | 6789 | 6807 | 19 | rRF 45S | NO | 0.00 | 33.72 | 0.00 | 17.65 |
| >hg38_21_+_8433172_8446622_RNA45SN1_45S_50ntFlanks@6790.6807.18\|CCTGTCTGAGC GTCGCTT | >hg38_21_+_8433172_8446622_RNA45SN1_45S_50ntFlanks | 6790 | 6807 | 18 | rRF 45S | NO | 79.07 | 111.73 | 0.00 | 70.63 |
| >hg38_21_+_8433172_8446622_RNA45SN1_45S_50ntFlanks@7968.7988.21\|TCCGAGACGCG ACCTCAGATC | >hg38_21_+_8433172_8446622_RNA45SN1_45S_50ntFlanks | 7968 | 7988 | 21 | rRF 45S | NO | 47.67 | 0.00 | 0.00 | 0.00 |
| >hg38_21_+_8433172_8446622_RNA45SN1_45S_50ntFlanks@7968.7994.27\|TCCGAGACGCG ACCTCAGATCAGACGT | >hg38_21_+_8433172_8446622_RNA45SN1_45S_50ntFlanks | 7968 | 7994 | 27 | rRF 45S | NO | 74.74 | 0.00 | 0.00 | 0.00 |
| >hg38_21_+_8433172_8446622_RNA45SN1_45S_50ntFlanks@7973.7991.19\|GACGCGACCTC AGATCAGA | >hg38_21_+_8433172_8446622_RNA45SN1_45S_50ntFlanks | 7973 | 7991 | 19 | rRF 45S | NO | 47.72 | 56.77 | 0.00 | 13.57 |
| >hg38_21_+_8433172_8446622_RNA45SN1_45S_50ntFlanks@7973.7992.20\|GACGCGACCTC AGATCAGAC | >hg38_21_+_8433172_8446622_RNA45SN1_45S_50ntFlanks | 7973 | 7992 | 20 | rRF 45S | NO | 62.99 | 60.95 | 0.00 | 17.51 |
| >hg38_21_+_8433172_8446622_RNA45SN1_45S_50ntFlanks@7973.7993.21\|GACGCGACCTC AGATCAGACG | >hg38_21_+_8433172_8446622_RNA45SN1_45S_50ntFlanks | 7973 | 7993 | 21 | rRF 45S | NO | 59.24 | 13.47 | 0.00 | 14.96 |
| >hg38_21_+_8433172_8446622_RNA45SN1_45S_50ntFlanks@7973.8008.36\|GACGCGACCTC AGATCAGACGTGGCGACCCGCTGAA | >hg38_21_+_8433172_8446622_RNA45SN1_45S_50ntFlanks | 7973 | 8008 | 36 | rRF 45S | NO | 100.92 | 65.92 | 0.00 | 96.82 |
| >hg38_21_+_8433172_8446622_RNA45SN1_45S_50ntFlanks@7973.8009.37\|GACGCGACCTC AGATCAGACGTGGCGACCCGCTGAAT | >hg38_21_+_8433172_8446622_RNA45SN1_45S_50ntFlanks | 7973 | 8009 | 37 | rRF 45S | NO | 98.10 | 146.47 | 113.29 | 96.49 |
| >hg38_21_+_8433172_8446622_RNA45SN1_45S_50ntFlanks@7973.8010.38\|GACGCGACCTC AGATCAGACGTGGCGACCCGCTGAATT | >hg38_21_+_8433172_8446622_RNA45SN1_45S_50ntFlanks | 7973 | 8010 | 38 | rRF 45S | NO | 109.38 | 116.63 | 0.00 | 113.87 |
| >hg38_21_+_8433172_8446622_RNA45SN1_45S_50ntFlanks@7973.8011.39\|GACGCGACCTC AGATCAGACGTGGCGACCCGCTGAATTT | >hg38_21_+_8433172_8446622_RNA45SN1_45S_50ntFlanks | 7973 | 8011 | 39 | rRF 45S | NO | 156.50 | 154.92 | 0.00 | 103.62 |
| >hg38_21_+_8433172_8446622_RNA45SN1_45S_50ntFlanks@7973.8012.40\|GACGCGACCTC AGATCAGACGTGGCGACCCGCTGAATTTA | >hg38_21_+_8433172_8446622_RNA45SN1_45S_50ntFlanks | 7973 | 8012 | 40 | rRF 45S | NO | 72.23 | 24.29 | 0.00 | 14.27 |
| >hg38_21_+_8433172_8446622_RNA45SN1_45S_50ntFlanks@7974.7991.18\|ACGCGACCTCA GATCAGA | >hg38_21_+_8433172_8446622_RNA45SN1_45S_50ntFlanks | 7974 | 7991 | 18 | rRF 45S | NO | 94.79 | 199.00 | 10.42 | 74.28 |
| >hg38_21_+_8433172_8446622_RNA45SN1_45S_50ntFlanks@7974.7992.19\|ACGCGACCTCA GATCAGAC | >hg38_21_+_8433172_8446622_RNA45SN1_45S_50ntFlanks | 7974 | 7992 | 19 | rRF 45S | NO | 80.24 | 109.75 | 0.00 | 81.59 |
| >hg38_21_+_8433172_8446622_RNA45SN1_45S_50ntFlanks@7974.7993.20\|ACGCGACCTCA GATCAGACG | >hg38_21_+_8433172_8446622_RNA45SN1_45S_50ntFlanks | 7974 | 7993 | 20 | rRF 45S | NO | 118.85 | 120.77 | 106.99 | 95.21 |
| >hg38_21_+_8433172_8446622_RNA45SN1_45S_50ntFlanks@7974.7994.21\|ACGCGACCTCA GATCAGACGT | >hg38_21_+_8433172_8446622_RNA45SN1_45S_50ntFlanks | 7974 | 7994 | 21 | rRF 45S | NO | 46.16 | 30.74 | 72.62 | 68.33 |
| >hg38_21_+_8433172_8446622_RNA45SN1_45S_50ntFlanks@7974.7995.22\|ACGCGACCTCA GATCAGACGTG | >hg38_21_+_8433172_8446622_RNA45SN1_45S_50ntFlanks | 7974 | 7995 | 22 | rRF 45S | NO | 91.62 | 125.17 | 16.40 | 99.57 |
| >hg38_21_+_8433172_8446622_RNA45SN1_45S_50ntFlanks@7974.7997.24\|ACGCGACCTCA GATCAGACGTGGC | >hg38_21_+_8433172_8446622_RNA45SN1_45S_50ntFlanks | 7974 | 7997 | 24 | rRF 45S | NO | 85.18 | 104.44 | 95.21 | 94.21 |
| >hg38_21_+_8433172_8446622_RNA45SN1_45S_50ntFlanks@7974.8004.31\|ACGCGACCTCA GATCAGACGTGGCGACCCGC | >hg38_21_+_8433172_8446622_RNA45SN1_45S_50ntFlanks | 7974 | 8004 | 31 | rRF 45S | NO | 23.94 | 31.64 | 0.00 | 59.15 |
| >hg38_21_+_8433172_8446622_RNA45SN1_45S_50ntFlanks@7974.8005.32\|ACGCGACCTCA GATCAGACGTGGCGACCCGCT | >hg38_21_+_8433172_8446622_RNA45SN1_45S_50ntFlanks | 7974 | 8005 | 32 | rRF 45S | NO | 83.89 | 83.94 | 16.64 | 173.36 |
| >hg38_21_+_8433172_8446622_RNA45SN1_45S_50ntFlanks@7974.8006.33\|ACGCGACCTCA GATCAGACGTGGCGACCCGCTG | >hg38_21_+_8433172_8446622_RNA45SN1_45S_50ntFlanks | 7974 | 8006 | 33 | rRF 45S | NO | 48.68 | 35.17 | 0.00 | 91.32 |
| >hg38_21_+_8433172_8446622_RNA45SN1_45S_50ntFlanks@7974.8007.34\|ACGCGACCTCA GATCAGACGTGGCGACCCGCTGA | >hg38_21_+_8433172_8446622_RNA45SN1_45S_50ntFlanks | 7974 | 8007 | 34 | rRF 45S | NO | 60.21 | 72.49 | 0.00 | 71.62 |
| >hg38_21_+_8433172_8446622_RNA45SN1_45S_50ntFlanks@7974.8008.35\|ACGCGACCTCA GATCAGACGTGGCGACCCGCTGAA | >hg38_21_+_8433172_8446622_RNA45SN1_45S_50ntFlanks | 7974 | 8008 | 35 | rRF 45S | NO | 259.99 | 414.13 | 90.98 | 349.22 |
| >hg38_21_+_8433172_8446622_RNA45SN1_45S_50ntFlanks@7974.8009.36\|ACGCGACCTCA GATCAGACGTGGCGACCCGCTGAAT | >hg38_21_+_8433172_8446622_RNA45SN1_45S_50ntFlanks | 7974 | 8009 | 36 | rRF 45S | NO | 276.53 | 828.93 | 441.66 | 337.25 |
| >hg38_21_+_8433172_8446622_RNA45SN1_45S_50ntFlanks@7974.8010.37\|ACGCGACCTCA GATCAGACGTGGCGACCCGCTGAATT | >hg38_21_+_8433172_8446622_RNA45SN1_45S_50ntFlanks | 7974 | 8010 | 37 | rRF 45S | NO | 347.96 | 652.88 | 93.38 | 391.24 |
| >hg38_21_+_8433172_8446622_RNA45SN1_45S_50ntFlanks@7974.8011.38\|ACGCGACCTCA GATCAGACGTGGCGACCCGCTGAATTT | >hg38_21_+_8433172_8446622_RNA45SN1_45S_50ntFlanks | 7974 | 8011 | 38 | rRF 45S | NO | 540.84 | 857.44 | 78.00 | 338.85 |
| >hg38_21_+_8433172_8446622_RNA45SN1_45S_50ntFlanks@7974.8012.39\|ACGCGACCTCA GATCAGACGTGGCGACCCGCTGAATTTA | >hg38_21_+_8433172_8446622_RNA45SN1_45S_50ntFlanks | 7974 | 8012 | 39 | rRF 45S | NO | 193.72 | 169.25 | 0.00 | 108.38 |
| >hg38_21_+_8433172_8446622_RNA45SN1_45S_50ntFlanks@7974.8013.40\|ACGCGACCTCA GATCAGACGTGGCGACCCGCTGAATTTAA | >hg38_21_+_8433172_8446622_RNA45SN1_45S_50ntFlanks | 7974 | 8013 | 40 | rRF 45S | NO | 103.21 | 113.93 | 0.00 | 77.46 |
| >hg38_21_+_8433172_8446622_RNA45SN1_45S_50ntFlanks@7974.8014.41\|ACGCGACCTCA GATCAGACGTGGCGACCCGCTGAATTTAAG | >hg38_21_+_8433172_8446622_RNA45SN1_45S_50ntFlanks | 7974 | 8014 | 41 | rRF 45S | NO | 57.54 | 72.34 | 0.00 | 21.92 |
| >hg38_21_+_8433172_8446622_RNA45SN1_45S_50ntFlanks@7974.8015.42\|ACGCGACCTCA GATCAGACGTGGCGACCCGCTGAATTTAAGC | >hg38_21_+_8433172_8446622_RNA45SN1_45S_50ntFlanks | 7974 | 8015 | 42 | rRF 45S | NO | 57.76 | 85.67 | 0.00 | 135.94 |

### MDA-MB-231

| | | | | | | | | | | |
|---|---|---|---|---|---|---|---|---|---|---|
| >hg38_21_+_8433172_8446622_RNA45SN1_45S_50ntFlanks@7974.8016.43\|ACGCGACCTCAGATCAGACGTGGCGACCCGCTGAATTTAAGCA | >hg38_21_+_8433172_8446622_RNA45SN1_45S_50ntFlanks | 7974 | 8016 | 43 | rRF 45S | NO | 62.02 | 117.19 | 0.00 | 216.10 |
| >hg38_21_+_8433172_8446622_RNA45SN1_45S_50ntFlanks@7975.7992.18\|CGCGACCTCAGATCAGAC | >hg38_21_+_8433172_8446622_RNA45SN1_45S_50ntFlanks | 7975 | 7992 | 18 | rRF 45S | NO | 279.03 | 558.81 | 76.72 | 194.52 |
| >hg38_21_+_8433172_8446622_RNA45SN1_45S_50ntFlanks@7975.7993.19\|CGCGACCTCAGATCAGACG | >hg38_21_+_8433172_8446622_RNA45SN1_45S_50ntFlanks | 7975 | 7993 | 19 | rRF 45S | NO | 346.46 | 457.55 | 268.86 | 237.98 |
| >hg38_21_+_8433172_8446622_RNA45SN1_45S_50ntFlanks@7975.7994.20\|CGCGACCTCAGATCAGACGT | >hg38_21_+_8433172_8446622_RNA45SN1_45S_50ntFlanks | 7975 | 7994 | 20 | rRF 45S | YES | 1032.89 | 2135.96 | 803.97 | 576.61 |
| >hg38_21_+_8433172_8446622_RNA45SN1_45S_50ntFlanks@7975.7995.21\|CGCGACCTCAGATCAGACGTG | >hg38_21_+_8433172_8446622_RNA45SN1_45S_50ntFlanks | 7975 | 7995 | 21 | rRF 45S | NO | 180.79 | 375.06 | 70.01 | 177.22 |
| >hg38_21_+_8433172_8446622_RNA45SN1_45S_50ntFlanks@7975.7996.22\|CGCGACCTCAGATCAGACGTGG | >hg38_21_+_8433172_8446622_RNA45SN1_45S_50ntFlanks | 7975 | 7996 | 22 | rRF 45S | NO | 166.14 | 251.61 | 58.42 | 131.22 |
| >hg38_21_+_8433172_8446622_RNA45SN1_45S_50ntFlanks@7975.7997.23\|CGCGACCTCAGATCAGACGTGGC | >hg38_21_+_8433172_8446622_RNA45SN1_45S_50ntFlanks | 7975 | 7997 | 23 | rRF 45S | NO | 311.57 | 548.64 | 260.27 | 308.00 |
| >hg38_21_+_8433172_8446622_RNA45SN1_45S_50ntFlanks@7975.8001.27\|CGCGACCTCAGATCAGACGTGGCGACC | >hg38_21_+_8433172_8446622_RNA45SN1_45S_50ntFlanks | 7975 | 8001 | 27 | rRF 45S | NO | 0.00 | 0.00 | 14.01 | 20.53 |
| >hg38_21_+_8433172_8446622_RNA45SN1_45S_50ntFlanks@7975.8002.28\|CGCGACCTCAGATCAGACGTGGCGACCC | >hg38_21_+_8433172_8446622_RNA45SN1_45S_50ntFlanks | 7975 | 8002 | 28 | rRF 45S | NO | 48.75 | 61.82 | 72.13 | 81.78 |
| >hg38_21_+_8433172_8446622_RNA45SN1_45S_50ntFlanks@7975.8003.29\|CGCGACCTCAGATCAGACGTGGCGACCCG | >hg38_21_+_8433172_8446622_RNA45SN1_45S_50ntFlanks | 7975 | 8003 | 29 | rRF 45S | NO | 54.90 | 98.00 | 0.00 | 85.12 |
| >hg38_21_+_8433172_8446622_RNA45SN1_45S_50ntFlanks@7975.8004.30\|CGCGACCTCAGATCAGACGTGGCGACCCGC | >hg38_21_+_8433172_8446622_RNA45SN1_45S_50ntFlanks | 7975 | 8004 | 30 | rRF 45S | NO | 171.86 | 252.31 | 59.76 | 176.50 |
| >hg38_21_+_8433172_8446622_RNA45SN1_45S_50ntFlanks@7975.8005.31\|CGCGACCTCAGATCAGACGTGGCGACCCGCT | >hg38_21_+_8433172_8446622_RNA45SN1_45S_50ntFlanks | 7975 | 8005 | 31 | rRF 45S | NO | 443.03 | 616.10 | 122.20 | 640.20 |
| >hg38_21_+_8433172_8446622_RNA45SN1_45S_50ntFlanks@7975.8006.32\|CGCGACCTCAGATCAGACGTGGCGACCCGCTG | >hg38_21_+_8433172_8446622_RNA45SN1_45S_50ntFlanks | 7975 | 8006 | 32 | rRF 45S | NO | 196.49 | 297.58 | 34.28 | 331.00 |
| >hg38_21_+_8433172_8446622_RNA45SN1_45S_50ntFlanks@7975.8007.33\|CGCGACCTCAGATCAGACGTGGCGACCCGCTGA | >hg38_21_+_8433172_8446622_RNA45SN1_45S_50ntFlanks | 7975 | 8007 | 33 | rRF 45S | NO | 238.36 | 411.00 | 61.50 | 241.92 |
| >hg38_21_+_8433172_8446622_RNA45SN1_45S_50ntFlanks@7975.8008.34\|CGCGACCTCAGATCAGACGTGGCGACCCGCTGAA | >hg38_21_+_8433172_8446622_RNA45SN1_45S_50ntFlanks | 7975 | 8008 | 34 | rRF 45S | YES | 962.94 | 2021.28 | 334.46 | 1062.75 |
| >hg38_21_+_8433172_8446622_RNA45SN1_45S_50ntFlanks@7975.8009.35\|CGCGACCTCAGATCAGACGTGGCGACCCGCTGAAT | >hg38_21_+_8433172_8446622_RNA45SN1_45S_50ntFlanks | 7975 | 8009 | 35 | rRF 45S | YES | 1069.53 | 4081.33 | 1071.39 | 1169.36 |
| >hg38_21_+_8433172_8446622_RNA45SN1_45S_50ntFlanks@7975.8010.36\|CGCGACCTCAGATCAGACGTGGCGACCCGCTGAATT | >hg38_21_+_8433172_8446622_RNA45SN1_45S_50ntFlanks | 7975 | 8010 | 36 | rRF 45S | YES | 1480.07 | 3630.41 | 373.11 | 1310.77 |
| >hg38_21_+_8433172_8446622_RNA45SN1_45S_50ntFlanks@7975.8011.37\|CGCGACCTCAGATCAGACGTGGCGACCCGCTGAATTT | >hg38_21_+_8433172_8446622_RNA45SN1_45S_50ntFlanks | 7975 | 8011 | 37 | rRF 45S | YES | 2462.17 | 4778.59 | 346.15 | 1150.53 |
| >hg38_21_+_8433172_8446622_RNA45SN1_45S_50ntFlanks@7975.8012.38\|CGCGACCTCAGATCAGACGTGGCGACCCGCTGAATTTA | >hg38_21_+_8433172_8446622_RNA45SN1_45S_50ntFlanks | 7975 | 8012 | 38 | rRF 45S | YES | 735.30 | 866.88 | 64.39 | 373.29 |
| >hg38_21_+_8433172_8446622_RNA45SN1_45S_50ntFlanks@7975.8013.39\|CGCGACCTCAGATCAGACGTGGCGACCCGCTGAATTTAA | >hg38_21_+_8433172_8446622_RNA45SN1_45S_50ntFlanks | 7975 | 8013 | 39 | rRF 45S | NO | 343.95 | 570.72 | 12.87 | 230.64 |
| >hg38_21_+_8433172_8446622_RNA45SN1_45S_50ntFlanks@7975.8014.40\|CGCGACCTCAGATCAGACGTGGCGACCCGCTGAATTTAAG | >hg38_21_+_8433172_8446622_RNA45SN1_45S_50ntFlanks | 7975 | 8014 | 40 | rRF 45S | NO | 271.44 | 440.69 | 0.00 | 217.22 |
| >hg38_21_+_8433172_8446622_RNA45SN1_45S_50ntFlanks@7975.8015.41\|CGCGACCTCAGATCAGACGTGGCGACCCGCTGAATTTAAGC | >hg38_21_+_8433172_8446622_RNA45SN1_45S_50ntFlanks | 7975 | 8015 | 41 | rRF 45S | NO | 256.23 | 411.87 | 69.56 | 429.20 |
| >hg38_21_+_8433172_8446622_RNA45SN1_45S_50ntFlanks@7975.8016.42\|CGCGACCTCAGATCAGACGTGGCGACCCGCTGAATTTAAGCA | >hg38_21_+_8433172_8446622_RNA45SN1_45S_50ntFlanks | 7975 | 8016 | 42 | rRF 45S | NO | 320.04 | 654.09 | 74.40 | 801.85 |
| >hg38_21_+_8433172_8446622_RNA45SN1_45S_50ntFlanks@7975.8017.43\|CGCGACCTCAGATCAGACGTGGCGACCCGCTGAATTTAAGCAT | >hg38_21_+_8433172_8446622_RNA45SN1_45S_50ntFlanks | 7975 | 8017 | 43 | rRF 45S | NO | 82.71 | 138.59 | 0.00 | 289.35 |
| >hg38_21_+_8433172_8446622_RNA45SN1_45S_50ntFlanks@7975.8018.44\|CGCGACCTCAGATCAGACGTGGCGACCCGCTGAATTTAAGCATA | >hg38_21_+_8433172_8446622_RNA45SN1_45S_50ntFlanks | 7975 | 8018 | 44 | rRF 45S | NO | 45.53 | 90.97 | 0.00 | 435.87 |
| >hg38_21_+_8433172_8446622_RNA45SN1_45S_50ntFlanks@7976.8008.33\|GCGACCTCAGATCAGACGTGGCGACCCGCTGAA | >hg38_21_+_8433172_8446622_RNA45SN1_45S_50ntFlanks | 7976 | 8008 | 33 | rRF 45S | NO | 0.00 | 58.56 | 0.00 | 0.00 |
| >hg38_21_+_8433172_8446622_RNA45SN1_45S_50ntFlanks@7976.8009.34\|GCGACCTCAGATCAGACGTGGCGACCCGCTGAAT | >hg38_21_+_8433172_8446622_RNA45SN1_45S_50ntFlanks | 7976 | 8009 | 34 | rRF 45S | NO | 0.00 | 90.46 | 10.45 | 0.00 |
| >hg38_21_+_8433172_8446622_RNA45SN1_45S_50ntFlanks@7976.8010.35\|GCGACCTCAGATCAGACGTGGCGACCCGCTGAATT | >hg38_21_+_8433172_8446622_RNA45SN1_45S_50ntFlanks | 7976 | 8010 | 35 | rRF 45S | NO | 0.00 | 75.70 | 0.00 | 0.00 |
| >hg38_21_+_8433172_8446622_RNA45SN1_45S_50ntFlanks@7976.8011.36\|GCGACCTCAGATCAGACGTGGCGACCCGCTGAATTT | >hg38_21_+_8433172_8446622_RNA45SN1_45S_50ntFlanks | 7976 | 8011 | 36 | rRF 45S | NO | 60.94 | 120.56 | 0.00 | 0.00 |
| >hg38_21_+_8433172_8446622_RNA45SN1_45S_50ntFlanks@7977.8009.33\|CGACCTCAGATCAGACGTGGCGACCCGCTGAAT | >hg38_21_+_8433172_8446622_RNA45SN1_45S_50ntFlanks | 7977 | 8009 | 33 | rRF 45S | NO | 0.00 | 37.24 | 0.00 | 0.00 |
| >hg38_21_+_8433172_8446622_RNA45SN1_45S_50ntFlanks@7977.8010.34\|CGACCTCAGATCAGACGTGGCGACCCGCTGAATT | >hg38_21_+_8433172_8446622_RNA45SN1_45S_50ntFlanks | 7977 | 8010 | 34 | rRF 45S | NO | 0.00 | 76.67 | 0.00 | 12.85 |
| >hg38_21_+_8433172_8446622_RNA45SN1_45S_50ntFlanks@7977.8011.35\|CGACCTCAGATCAGACGTGGCGACCCGCTGAATTT | >hg38_21_+_8433172_8446622_RNA45SN1_45S_50ntFlanks | 7977 | 8011 | 35 | rRF 45S | NO | 61.37 | 163.61 | 0.00 | 19.77 |
| >hg38_21_+_8433172_8446622_RNA45SN1_45S_50ntFlanks@7977.8012.36\|CGACCTCAGATCAGACGTGGCGACCCGCTGAATTTA | >hg38_21_+_8433172_8446622_RNA45SN1_45S_50ntFlanks | 7977 | 8012 | 36 | rRF 45S | NO | 0.00 | 28.68 | 0.00 | 0.00 |
| >hg38_21_+_8433172_8446622_RNA45SN1_45S_50ntFlanks@7977.8016.40\|CGACCTCAGATCAGACGTGGCGACCCGCTGAATTTAAGCA | >hg38_21_+_8433172_8446622_RNA45SN1_45S_50ntFlanks | 7977 | 8016 | 40 | rRF 45S | NO | 0.00 | 104.75 | 0.00 | 21.92 |
| >hg38_21_+_8433172_8446622_RNA45SN1_45S_50ntFlanks@7977.8017.41\|CGACCTCAGATCAGACGTGGCGACCCGCTGAATTTAAGCAT | >hg38_21_+_8433172_8446622_RNA45SN1_45S_50ntFlanks | 7977 | 8017 | 41 | rRF 45S | NO | 0.00 | 34.42 | 0.00 | 0.00 |
| >hg38_21_+_8433172_8446622_RNA45SN1_45S_50ntFlanks@7977.8019.43\|CGACCTCAGATCAGACGTGGCGACCCGCTGAATTTAAGCATAT | >hg38_21_+_8433172_8446622_RNA45SN1_45S_50ntFlanks | 7977 | 8019 | 43 | rRF 45S | NO | 0.00 | 29.73 | 0.00 | 0.00 |
| >hg38_21_+_8433172_8446622_RNA45SN1_45S_50ntFlanks@7978.8009.32\|GACCTCAGATCAGACGTGGCGACCCGCTGAAT | >hg38_21_+_8433172_8446622_RNA45SN1_45S_50ntFlanks | 7978 | 8009 | 32 | rRF 45S | NO | 0.00 | 27.75 | 0.00 | 0.00 |
| >hg38_21_+_8433172_8446622_RNA45SN1_45S_50ntFlanks@7978.8010.33\|GACCTCAGATCAGACGTGGCGACCCGCTGAATT | >hg38_21_+_8433172_8446622_RNA45SN1_45S_50ntFlanks | 7978 | 8010 | 33 | rRF 45S | NO | 0.00 | 35.21 | 0.00 | 0.00 |

### MDA-MB-231

| | | | | | | | | | | |
|---|---|---|---|---|---|---|---|---|---|---|
| >hg38_21_+_8433172_8446622_RNA45SN1_45S_50ntFlanks@7978.8011.34\|GACCTCAGATCA GACGTGGCGACCCGCTGAATTT | >hg38_21_+_8433172_8446622_RNA45SN1_45S_50ntFlanks | 7978 | 8011 | 34 | rRF 45S | NO | 0.00 | 109.72 | 0.00 | 14.94 |
| >hg38_21_+_8433172_8446622_RNA45SN1_45S_50ntFlanks@7978.8016.39\|GACCTCAGATCA GACGTGGCGACCCGCTGAATTTAAGCA | >hg38_21_+_8433172_8446622_RNA45SN1_45S_50ntFlanks | 7978 | 8016 | 39 | rRF 45S | NO | 0.00 | 30.12 | 0.00 | 0.00 |
| >hg38_21_+_8433172_8446622_RNA45SN1_45S_50ntFlanks@7988.8008.21\|CAGACGTGGCG ACCCGCTGAA | >hg38_21_+_8433172_8446622_RNA45SN1_45S_50ntFlanks | 7988 | 8008 | 21 | rRF 45S | NO | 0.00 | 67.08 | 0.00 | 0.00 |
| >hg38_21_+_8433172_8446622_RNA45SN1_45S_50ntFlanks@7988.8016.29\|CAGACGTGGCG ACCCGCTGAATTTAAGCA | >hg38_21_+_8433172_8446622_RNA45SN1_45S_50ntFlanks | 7988 | 8016 | 29 | rRF 45S | NO | 0.00 | 24.33 | 0.00 | 0.00 |
| >hg38_21_+_8433172_8446622_RNA45SN1_45S_50ntFlanks@7989.8008.20\|AGACGTGGCGA CCCGCTGAA | >hg38_21_+_8433172_8446622_RNA45SN1_45S_50ntFlanks | 7989 | 8008 | 20 | rRF 45S | NO | 24.91 | 83.80 | 0.00 | 0.00 |
| >hg38_21_+_8433172_8446622_RNA45SN1_45S_50ntFlanks@7989.8009.21\|AGACGTGGCGA CCCGCTGAAT | >hg38_21_+_8433172_8446622_RNA45SN1_45S_50ntFlanks | 7989 | 8009 | 21 | rRF 45S | NO | 83.73 | 141.26 | 95.92 | 17.85 |
| >hg38_21_+_8433172_8446622_RNA45SN1_45S_50ntFlanks@7989.8010.22\|AGACGTGGCGA CCCGCTGAATT | >hg38_21_+_8433172_8446622_RNA45SN1_45S_50ntFlanks | 7989 | 8010 | 22 | rRF 45S | NO | 89.07 | 128.17 | 0.00 | 22.05 |
| >hg38_21_+_8433172_8446622_RNA45SN1_45S_50ntFlanks@7989.8011.23\|AGACGTGGCGA CCCGCTGAATTT | >hg38_21_+_8433172_8446622_RNA45SN1_45S_50ntFlanks | 7989 | 8011 | 23 | rRF 45S | NO | 149.00 | 229.55 | 16.21 | 75.33 |
| >hg38_21_+_8433172_8446622_RNA45SN1_45S_50ntFlanks@7989.8012.24\|AGACGTGGCGA CCCGCTGAATTTA | >hg38_21_+_8433172_8446622_RNA45SN1_45S_50ntFlanks | 7989 | 8012 | 24 | rRF 45S | NO | 0.00 | 25.09 | 0.00 | 0.00 |
| >hg38_21_+_8433172_8446622_RNA45SN1_45S_50ntFlanks@7989.8016.28\|AGACGTGGCGA CCCGCTGAATTTAAGCA | >hg38_21_+_8433172_8446622_RNA45SN1_45S_50ntFlanks | 7989 | 8016 | 28 | rRF 45S | NO | 0.00 | 68.59 | 0.00 | 0.00 |
| >hg38_21_+_8433172_8446622_RNA45SN1_45S_50ntFlanks@7989.8017.29\|AGACGTGGCGA CCCGCTGAATTTAAGCAT | >hg38_21_+_8433172_8446622_RNA45SN1_45S_50ntFlanks | 7989 | 8017 | 29 | rRF 45S | NO | 0.00 | 35.74 | 0.00 | 0.00 |
| >hg38_21_+_8433172_8446622_RNA45SN1_45S_50ntFlanks@7989.8019.31\|AGACGTGGCGA CCCGCTGAATTTAAGCATAT | >hg38_21_+_8433172_8446622_RNA45SN1_45S_50ntFlanks | 7989 | 8019 | 31 | rRF 45S | NO | 0.00 | 57.85 | 0.00 | 0.00 |
| >hg38_21_+_8433172_8446622_RNA45SN1_45S_50ntFlanks@7990.8009.20\|GACGTGGCGAC CCGCTGAAT | >hg38_21_+_8433172_8446622_RNA45SN1_45S_50ntFlanks | 7990 | 8009 | 20 | rRF 45S | NO | 0.00 | 26.58 | 0.00 | 0.00 |
| >hg38_21_+_8433172_8446622_RNA45SN1_45S_50ntFlanks@7990.8010.21\|GACGTGGCGAC CCGCTGAATT | >hg38_21_+_8433172_8446622_RNA45SN1_45S_50ntFlanks | 7990 | 8010 | 21 | rRF 45S | NO | 0.00 | 24.13 | 0.00 | 0.00 |
| >hg38_21_+_8433172_8446622_RNA45SN1_45S_50ntFlanks@7990.8011.22\|GACGTGGCGAC CCGCTGAATTT | >hg38_21_+_8433172_8446622_RNA45SN1_45S_50ntFlanks | 7990 | 8011 | 22 | rRF 45S | NO | 54.78 | 82.34 | 0.00 | 15.02 |
| >hg38_21_+_8433172_8446622_RNA45SN1_45S_50ntFlanks@7990.8016.27\|GACGTGGCGAC CCGCTGAATTTAAGCA | >hg38_21_+_8433172_8446622_RNA45SN1_45S_50ntFlanks | 7990 | 8016 | 27 | rRF 45S | NO | 0.00 | 30.30 | 0.00 | 0.00 |
| >hg38_21_+_8433172_8446622_RNA45SN1_45S_50ntFlanks@7990.8019.30\|GACGTGGCGAC CCGCTGAATTTAAGCATAT | >hg38_21_+_8433172_8446622_RNA45SN1_45S_50ntFlanks | 7990 | 8019 | 30 | rRF 45S | NO | 0.00 | 31.80 | 0.00 | 0.00 |
| >hg38_21_+_8433172_8446622_RNA45SN1_45S_50ntFlanks@7991.8011.21\|ACGTGGCGACC CGCTGAATTT | >hg38_21_+_8433172_8446622_RNA45SN1_45S_50ntFlanks | 7991 | 8011 | 21 | rRF 45S | NO | 54.60 | 32.67 | 0.00 | 19.33 |
| >hg38_21_+_8433172_8446622_RNA45SN1_45S_50ntFlanks@7991.8016.26\|ACGTGGCGACC CGCTGAATTTAAGCA | >hg38_21_+_8433172_8446622_RNA45SN1_45S_50ntFlanks | 7991 | 8016 | 26 | rRF 45S | NO | 0.00 | 24.34 | 0.00 | 0.00 |
| >hg38_21_+_8433172_8446622_RNA45SN1_45S_50ntFlanks@7992.8011.20\|CGTGGCGACCC GCTGAATTT | >hg38_21_+_8433172_8446622_RNA45SN1_45S_50ntFlanks | 7992 | 8011 | 20 | rRF 45S | NO | 47.54 | 0.00 | 0.00 | 17.70 |
| >hg38_21_+_8433172_8446622_RNA45SN1_45S_50ntFlanks@7992.8016.25\|CGTGGCGACCC GCTGAATTTAAGCA | >hg38_21_+_8433172_8446622_RNA45SN1_45S_50ntFlanks | 7992 | 8016 | 25 | rRF 45S | NO | 0.00 | 33.09 | 0.00 | 0.00 |
| >hg38_21_+_8433172_8446622_RNA45SN1_45S_50ntFlanks@8020.8046.27\|TAGTCAGCGGA GGAGAAGAAACTAACC | >hg38_21_+_8433172_8446622_RNA45SN1_45S_50ntFlanks | 8020 | 8046 | 27 | rRF 45S | NO | 0.00 | 28.81 | 0.00 | 0.00 |
| >hg38_21_+_8433172_8446622_RNA45SN1_45S_50ntFlanks@8175.8202.28\|CCTTCTGATCGA GGCCCAGCCCGTGGAC | >hg38_21_+_8433172_8446622_RNA45SN1_45S_50ntFlanks | 8175 | 8202 | 28 | rRF 45S | NO | 0.00 | 24.06 | 0.00 | 0.00 |
| >hg38_21_+_8433172_8446622_RNA45SN1_45S_50ntFlanks@8175.8216.42\|CCTTCTGATCGA GGCCCAGCCCGTGGACGGTGTGAGGCCGGT | >hg38_21_+_8433172_8446622_RNA45SN1_45S_50ntFlanks | 8175 | 8216 | 42 | rRF 45S | NO | 0.00 | 0.00 | 12.03 | 0.00 |
| >hg38_21_+_8433172_8446622_RNA45SN1_45S_50ntFlanks@8176.8194.19\|CTTCTGATCGAG GCCCAGC | >hg38_21_+_8433172_8446622_RNA45SN1_45S_50ntFlanks | 8176 | 8194 | 19 | rRF 45S | NO | 0.00 | 32.30 | 0.00 | 0.00 |
| >hg38_21_+_8433172_8446622_RNA45SN1_45S_50ntFlanks@8176.8195.20\|CTTCTGATCGAG GCCCAGCC | >hg38_21_+_8433172_8446622_RNA45SN1_45S_50ntFlanks | 8176 | 8195 | 20 | rRF 45S | NO | 46.33 | 248.67 | 0.00 | 0.00 |
| >hg38_21_+_8433172_8446622_RNA45SN1_45S_50ntFlanks@8176.8196.21\|CTTCTGATCGAG GCCCAGCCC | >hg38_21_+_8433172_8446622_RNA45SN1_45S_50ntFlanks | 8176 | 8196 | 21 | rRF 45S | NO | 0.00 | 36.43 | 0.00 | 0.00 |
| >hg38_21_+_8433172_8446622_RNA45SN1_45S_50ntFlanks@8176.8202.27\|CTTCTGATCGAG GCCCAGCCCGTGGAC | >hg38_21_+_8433172_8446622_RNA45SN1_45S_50ntFlanks | 8176 | 8202 | 27 | rRF 45S | NO | 0.00 | 129.11 | 0.00 | 0.00 |
| >hg38_21_+_8433172_8446622_RNA45SN1_45S_50ntFlanks@8177.8195.19\|TTCTGATCGAGG CCCAGCC | >hg38_21_+_8433172_8446622_RNA45SN1_45S_50ntFlanks | 8177 | 8195 | 19 | rRF 45S | NO | 0.00 | 59.14 | 0.00 | 0.00 |
| >hg38_21_+_8433172_8446622_RNA45SN1_45S_50ntFlanks@8177.8203.27\|TTCTGATCGAGG CCCAGCCCGTGGACG | >hg38_21_+_8433172_8446622_RNA45SN1_45S_50ntFlanks | 8177 | 8203 | 27 | rRF 45S | NO | 0.00 | 26.83 | 0.00 | 0.00 |
| >hg38_21_+_8433172_8446622_RNA45SN1_45S_50ntFlanks@8195.8213.19\|CCGTGGACGGT GTGAGGCC | >hg38_21_+_8433172_8446622_RNA45SN1_45S_50ntFlanks | 8195 | 8213 | 19 | rRF 45S | NO | 0.00 | 143.96 | 0.00 | 0.00 |
| >hg38_21_+_8433172_8446622_RNA45SN1_45S_50ntFlanks@8196.8213.18\|CGTGGACGGTG TGAGGCC | >hg38_21_+_8433172_8446622_RNA45SN1_45S_50ntFlanks | 8196 | 8213 | 18 | rRF 45S | NO | 0.00 | 272.04 | 11.01 | 0.00 |
| >hg38_21_+_8433172_8446622_RNA45SN1_45S_50ntFlanks@8229.8251.23\|CGCGCCGGGCC CGGGTCTTCCCG | >hg38_21_+_8433172_8446622_RNA45SN1_45S_50ntFlanks | 8229 | 8251 | 23 | rRF 45S | NO | 0.00 | 56.64 | 0.00 | 0.00 |
| >hg38_21_+_8433172_8446622_RNA45SN1_45S_50ntFlanks@8342.8364.23\|CGTAAGGGAAA GTTGAAAAGAAC | >hg38_21_+_8433172_8446622_RNA45SN1_45S_50ntFlanks | 8342 | 8364 | 23 | rRF 45S | NO | 0.00 | 55.91 | 0.00 | 0.00 |
| >hg38_21_+_8433172_8446622_RNA45SN1_45S_50ntFlanks@8342.8365.24\|CGTAAGGGAAA GTTGAAAAGAACT | >hg38_21_+_8433172_8446622_RNA45SN1_45S_50ntFlanks | 8342 | 8365 | 24 | rRF 45S | NO | 0.00 | 183.46 | 17.67 | 0.00 |
| >hg38_21_+_8433172_8446622_RNA45SN1_45S_50ntFlanks@8342.8376.35\|CGTAAGGGAAA GTTGAAAAGAACTTTGAAGAGAGA | >hg38_21_+_8433172_8446622_RNA45SN1_45S_50ntFlanks | 8342 | 8376 | 35 | rRF 45S | NO | 0.00 | 26.23 | 0.00 | 0.00 |
| >hg38_21_+_8433172_8446622_RNA45SN1_45S_50ntFlanks@8342.8377.36\|CGTAAGGGAAA GTTGAAAAGAACTTTGAAGAGAGAG | >hg38_21_+_8433172_8446622_RNA45SN1_45S_50ntFlanks | 8342 | 8377 | 36 | rRF 45S | NO | 0.00 | 27.66 | 0.00 | 0.00 |
| >hg38_21_+_8433172_8446622_RNA45SN1_45S_50ntFlanks@8342.8378.37\|CGTAAGGGAAA GTTGAAAAGAACTTTGAAGAGAGAGT | >hg38_21_+_8433172_8446622_RNA45SN1_45S_50ntFlanks | 8342 | 8378 | 37 | rRF 45S | NO | 0.00 | 140.75 | 10.41 | 0.00 |

### MDA-MB-231

| | | | | | | | | | | |
|---|---|---|---|---|---|---|---|---|---|---|
| >hg38_21_+_8433172_8446622_RNA45SN1_45S_50ntFlanks@8342.8379.38\|CGTAAGGGAAAGTTGAAAAGAACTTTGAAGAGAGAGTT | >hg38_21_+_8433172_8446622_RNA45SN1_45S_50ntFlanks | 8342 | 8379 | 38 | rRF 45S | NO | 0.00 | 117.31 | 0.00 | 0.00 |
| >hg38_21_+_8433172_8446622_RNA45SN1_45S_50ntFlanks@8381.8406.26\|AAGAGGGCGTGAAACCGTTAAGAGGT | >hg38_21_+_8433172_8446622_RNA45SN1_45S_50ntFlanks | 8381 | 8406 | 26 | rRF 45S | NO | 0.00 | 26.81 | 13.09 | 0.00 |
| >hg38_21_+_8433172_8446622_RNA45SN1_45S_50ntFlanks@8686.8706.21\|CCGGCTCCGGGACGGCTGGGA | >hg38_21_+_8433172_8446622_RNA45SN1_45S_50ntFlanks | 8686 | 8706 | 21 | rRF 45S | NO | 0.00 | 87.77 | 0.00 | 0.00 |
| >hg38_21_+_8433172_8446622_RNA45SN1_45S_50ntFlanks@8687.8706.20\|CGGCTCCGGGACGGCTGGGA | >hg38_21_+_8433172_8446622_RNA45SN1_45S_50ntFlanks | 8687 | 8706 | 20 | rRF 45S | NO | 0.00 | 114.01 | 0.00 | 0.00 |
| >hg38_21_+_8433172_8446622_RNA45SN1_45S_50ntFlanks@8912.8936.25\|CGCCGAATCCCGGGGCCGAGGGAGC | >hg38_21_+_8433172_8446622_RNA45SN1_45S_50ntFlanks | 8912 | 8936 | 25 | rRF 45S | NO | 0.00 | 26.48 | 0.00 | 0.00 |
| >hg38_21_+_8433172_8446622_RNA45SN1_45S_50ntFlanks@9477.9497.21\|AGCGCACGTGTTAGGACCCGA | >hg38_21_+_8433172_8446622_RNA45SN1_45S_50ntFlanks | 9477 | 9497 | 21 | rRF 45S | NO | 0.00 | 116.85 | 0.00 | 0.00 |
| >hg38_21_+_8433172_8446622_RNA45SN1_45S_50ntFlanks@9478.9497.20\|GCGCACGTGTTAGGACCCGA | >hg38_21_+_8433172_8446622_RNA45SN1_45S_50ntFlanks | 9478 | 9497 | 20 | rRF 45S | NO | 0.00 | 77.52 | 0.00 | 0.00 |
| >hg38_21_+_8433172_8446622_RNA45SN1_45S_50ntFlanks@9526.9570.45\|GAAGCCAGAGGAAACTCTGGTGGAGGTCCGTAGCGGTCCTGACGT | >hg38_21_+_8433172_8446622_RNA45SN1_45S_50ntFlanks | 9526 | 9570 | 45 | rRF 45S | NO | 0.00 | 66.86 | 0.00 | 0.00 |
| >hg38_21_+_8433172_8446622_RNA45SN1_45S_50ntFlanks@9526.9571.46\|GAAGCCAGAGGAAACTCTGGTGGAGGTCCGTAGCGGTCCTGACGTG | >hg38_21_+_8433172_8446622_RNA45SN1_45S_50ntFlanks | 9526 | 9571 | 46 | rRF 45S | NO | 0.00 | 97.33 | 0.00 | 0.00 |
| >hg38_21_+_8433172_8446622_RNA45SN1_45S_50ntFlanks@9532.9571.40\|AGAGGAAACTCTGGTGGAGGTCCGTAGCGGTCCTGACGTG | >hg38_21_+_8433172_8446622_RNA45SN1_45S_50ntFlanks | 9532 | 9571 | 40 | rRF 45S | NO | 0.00 | 101.81 | 0.00 | 0.00 |
| >hg38_21_+_8433172_8446622_RNA45SN1_45S_50ntFlanks@9533.9553.21\|GAGGAAACTCTGGTGGAGGTC | >hg38_21_+_8433172_8446622_RNA45SN1_45S_50ntFlanks | 9533 | 9553 | 21 | rRF 45S | NO | 0.00 | 75.66 | 0.00 | 0.00 |
| >hg38_21_+_8433172_8446622_RNA45SN1_45S_50ntFlanks@9533.9570.38\|GAGGAAACTCTGGTGGAGGTCCGTAGCGGTCCTGACGT | >hg38_21_+_8433172_8446622_RNA45SN1_45S_50ntFlanks | 9533 | 9570 | 38 | rRF 45S | NO | 0.00 | 30.86 | 0.00 | 0.00 |
| >hg38_21_+_8433172_8446622_RNA45SN1_45S_50ntFlanks@9533.9571.39\|GAGGAAACTCTGGTGGAGGTCCGTAGCGGTCCTGACGTG | >hg38_21_+_8433172_8446622_RNA45SN1_45S_50ntFlanks | 9533 | 9571 | 39 | rRF 45S | NO | 0.00 | 110.91 | 0.00 | 0.00 |
| >hg38_21_+_8433172_8446622_RNA45SN1_45S_50ntFlanks@9534.9571.38\|AGGAAACTCTGGTGGAGGTCCGTAGCGGTCCTGACGTG | >hg38_21_+_8433172_8446622_RNA45SN1_45S_50ntFlanks | 9534 | 9571 | 38 | rRF 45S | NO | 0.00 | 92.35 | 0.00 | 0.00 |
| >hg38_21_+_8433172_8446622_RNA45SN1_45S_50ntFlanks@9535.9570.36\|GGAAACTCTGGTGGAGGTCCGTAGCGGTCCTGACGT | >hg38_21_+_8433172_8446622_RNA45SN1_45S_50ntFlanks | 9535 | 9570 | 36 | rRF 45S | NO | 0.00 | 28.86 | 0.00 | 0.00 |
| >hg38_21_+_8433172_8446622_RNA45SN1_45S_50ntFlanks@9535.9571.37\|GGAAACTCTGGTGGAGGTCCGTAGCGGTCCTGACGTG | >hg38_21_+_8433172_8446622_RNA45SN1_45S_50ntFlanks | 9535 | 9571 | 37 | rRF 45S | NO | 0.00 | 64.29 | 0.00 | 0.00 |
| >hg38_21_+_8433172_8446622_RNA45SN1_45S_50ntFlanks@9544.9571.28\|GGTGGAGGTCCGTAGCGGTCCTGACGTG | >hg38_21_+_8433172_8446622_RNA45SN1_45S_50ntFlanks | 9544 | 9571 | 28 | rRF 45S | NO | 0.00 | 93.76 | 0.00 | 0.00 |
| >hg38_21_+_8433172_8446622_RNA45SN1_45S_50ntFlanks@9544.9573.30\|GGTGGAGGTCCGTAGCGGTCCTGACGTGCA | >hg38_21_+_8433172_8446622_RNA45SN1_45S_50ntFlanks | 9544 | 9573 | 30 | rRF 45S | NO | 0.00 | 25.66 | 0.00 | 0.00 |
| >hg38_21_+_8433172_8446622_RNA45SN1_45S_50ntFlanks@9545.9571.27\|GTGGAGGTCCGTAGCGGTCCTGACGTG | >hg38_21_+_8433172_8446622_RNA45SN1_45S_50ntFlanks | 9545 | 9571 | 27 | rRF 45S | NO | 0.00 | 103.88 | 0.00 | 0.00 |
| >hg38_21_+_8433172_8446622_RNA45SN1_45S_50ntFlanks@9545.9572.28\|GTGGAGGTCCGTAGCGGTCCTGACGTGC | >hg38_21_+_8433172_8446622_RNA45SN1_45S_50ntFlanks | 9545 | 9572 | 28 | rRF 45S | NO | 0.00 | 24.74 | 0.00 | 0.00 |
| >hg38_21_+_8433172_8446622_RNA45SN1_45S_50ntFlanks@9546.9571.26\|TGGAGGTCCGTAGCGGTCCTGACGTG | >hg38_21_+_8433172_8446622_RNA45SN1_45S_50ntFlanks | 9546 | 9571 | 26 | rRF 45S | NO | 0.00 | 107.91 | 0.00 | 0.00 |
| >hg38_21_+_8433172_8446622_RNA45SN1_45S_50ntFlanks@9546.9572.27\|TGGAGGTCCGTAGCGGTCCTGACGTGC | >hg38_21_+_8433172_8446622_RNA45SN1_45S_50ntFlanks | 9546 | 9572 | 27 | rRF 45S | NO | 0.00 | 30.32 | 0.00 | 0.00 |
| >hg38_21_+_8433172_8446622_RNA45SN1_45S_50ntFlanks@9546.9573.28\|TGGAGGTCCGTAGCGGTCCTGACGTGCA | >hg38_21_+_8433172_8446622_RNA45SN1_45S_50ntFlanks | 9546 | 9573 | 28 | rRF 45S | NO | 0.00 | 29.53 | 0.00 | 0.00 |
| >hg38_21_+_8433172_8446622_RNA45SN1_45S_50ntFlanks@9548.9571.24\|GAGGTCCGTAGCGGTCCTGACGTG | >hg38_21_+_8433172_8446622_RNA45SN1_45S_50ntFlanks | 9548 | 9571 | 24 | rRF 45S | NO | 0.00 | 32.34 | 0.00 | 0.00 |
| >hg38_21_+_8433172_8446622_RNA45SN1_45S_50ntFlanks@9548.9573.26\|GAGGTCCGTAGCGGTCCTGACGTGCA | >hg38_21_+_8433172_8446622_RNA45SN1_45S_50ntFlanks | 9548 | 9573 | 26 | rRF 45S | NO | 0.00 | 25.99 | 0.00 | 0.00 |
| >hg38_21_+_8433172_8446622_RNA45SN1_45S_50ntFlanks@9551.9569.19\|GTCCGTAGCGGTCCTGACG | >hg38_21_+_8433172_8446622_RNA45SN1_45S_50ntFlanks | 9551 | 9569 | 19 | rRF 45S | NO | 0.00 | 37.41 | 0.00 | 0.00 |
| >hg38_21_+_8433172_8446622_RNA45SN1_45S_50ntFlanks@9551.9571.21\|GTCCGTAGCGGTCCTGACGTG | >hg38_21_+_8433172_8446622_RNA45SN1_45S_50ntFlanks | 9551 | 9571 | 21 | rRF 45S | NO | 0.00 | 30.71 | 0.00 | 0.00 |
| >hg38_21_+_8433172_8446622_RNA45SN1_45S_50ntFlanks@9552.9569.18\|TCCGTAGCGGTCCTGACG | >hg38_21_+_8433172_8446622_RNA45SN1_45S_50ntFlanks | 9552 | 9569 | 18 | rRF 45S | NO | 0.00 | 26.16 | 0.00 | 0.00 |
| >hg38_21_+_8433172_8446622_RNA45SN1_45S_50ntFlanks@9553.9570.18\|CCGTAGCGGTCCTGACGT | >hg38_21_+_8433172_8446622_RNA45SN1_45S_50ntFlanks | 9553 | 9570 | 18 | rRF 45S | NO | 0.00 | 25.98 | 0.00 | 0.00 |
| >hg38_21_+_8433172_8446622_RNA45SN1_45S_50ntFlanks@9554.9571.18\|CGTAGCGGTCCTGACGTG | >hg38_21_+_8433172_8446622_RNA45SN1_45S_50ntFlanks | 9554 | 9571 | 18 | rRF 45S | NO | 69.11 | 443.64 | 0.00 | 0.00 |
| >hg38_21_+_8433172_8446622_RNA45SN1_45S_50ntFlanks@9554.9572.19\|CGTAGCGGTCCTGACGTGC | >hg38_21_+_8433172_8446622_RNA45SN1_45S_50ntFlanks | 9554 | 9572 | 19 | rRF 45S | NO | 0.00 | 139.78 | 0.00 | 0.00 |
| >hg38_21_+_8433172_8446622_RNA45SN1_45S_50ntFlanks@9554.9573.20\|CGTAGCGGTCCTGACGTGCA | >hg38_21_+_8433172_8446622_RNA45SN1_45S_50ntFlanks | 9554 | 9573 | 20 | rRF 45S | NO | 25.59 | 225.52 | 0.00 | 0.00 |
| >hg38_21_+_8433172_8446622_RNA45SN1_45S_50ntFlanks@9554.9574.21\|CGTAGCGGTCCTGACGTGCAA | >hg38_21_+_8433172_8446622_RNA45SN1_45S_50ntFlanks | 9554 | 9574 | 21 | rRF 45S | NO | 0.00 | 57.41 | 0.00 | 0.00 |
| >hg38_21_+_8433172_8446622_RNA45SN1_45S_50ntFlanks@9554.9575.22\|CGTAGCGGTCCTGACGTGCAAA | >hg38_21_+_8433172_8446622_RNA45SN1_45S_50ntFlanks | 9554 | 9575 | 22 | rRF 45S | NO | 0.00 | 12.92 | 0.00 | 0.00 |
| >hg38_21_+_8433172_8446622_RNA45SN1_45S_50ntFlanks@9554.9576.23\|CGTAGCGGTCCTGACGTGCAAAT | >hg38_21_+_8433172_8446622_RNA45SN1_45S_50ntFlanks | 9554 | 9576 | 23 | rRF 45S | NO | 0.00 | 149.54 | 0.00 | 0.00 |
| >hg38_21_+_8433172_8446622_RNA45SN1_45S_50ntFlanks@9554.9577.24\|CGTAGCGGTCCTGACGTGCAAATC | >hg38_21_+_8433172_8446622_RNA45SN1_45S_50ntFlanks | 9554 | 9577 | 24 | rRF 45S | NO | 47.93 | 165.14 | 0.00 | 0.00 |
| >hg38_21_+_8433172_8446622_RNA45SN1_45S_50ntFlanks@9554.9581.28\|CGTAGCGGTCCTGACGTGCAAATCGGTC | >hg38_21_+_8433172_8446622_RNA45SN1_45S_50ntFlanks | 9554 | 9581 | 28 | rRF 45S | NO | 0.00 | 116.99 | 0.00 | 0.00 |
| >hg38_21_+_8433172_8446622_RNA45SN1_45S_50ntFlanks@9574.9603.30\|AATCGGTCGTCCGACCTGGGTATAGGGGCG | >hg38_21_+_8433172_8446622_RNA45SN1_45S_50ntFlanks | 9574 | 9603 | 30 | rRF 45S | NO | 50.04 | 37.44 | 17.13 | 15.05 |

### MDA-MB-231

| | | | | | | | | | | |
|---|---|---|---|---|---|---|---|---|---|---|
| >hg38_21_+_8433172_8446622_RNA45SN1_45S_50ntFlanks@10344.10373.30\|ATTCAAACGAGAACTTTGAAGGCCGAAGTG | >hg38_21_+_8433172_8446622_RNA45SN1_45S_50ntFlanks | 10344 | 10373 | 30 | rRF 45S | NO | 0.00 | 26.66 | 0.00 | 0.00 |
| >hg38_21_+_8433172_8446622_RNA45SN1_45S_50ntFlanks@10345.10373.29\|TTCAAACGAGAACTTTGAAGGCCGAAGTG | >hg38_21_+_8433172_8446622_RNA45SN1_45S_50ntFlanks | 10345 | 10373 | 29 | rRF 45S | NO | 0.00 | 35.12 | 0.00 | 0.00 |
| >hg38_21_+_8433172_8446622_RNA45SN1_45S_50ntFlanks@10345.10382.38\|TTCAAACGAGAACTTTGAAGGCCGAAGTGGAGAAGGGT | >hg38_21_+_8433172_8446622_RNA45SN1_45S_50ntFlanks | 10345 | 10382 | 38 | rRF 45S | NO | 0.00 | 28.40 | 0.00 | 0.00 |
| >hg38_21_+_8433172_8446622_RNA45SN1_45S_50ntFlanks@10346.10373.28\|TCAAACGAGAACTTTGAAGGCCGAAGTG | >hg38_21_+_8433172_8446622_RNA45SN1_45S_50ntFlanks | 10346 | 10373 | 28 | rRF 45S | NO | 0.00 | 24.53 | 0.00 | 0.00 |
| >hg38_21_+_8433172_8446622_RNA45SN1_45S_50ntFlanks@10346.10382.37\|TCAAACGAGAACTTTGAAGGCCGAAGTGGAGAAGGGT | >hg38_21_+_8433172_8446622_RNA45SN1_45S_50ntFlanks | 10346 | 10382 | 37 | rRF 45S | NO | 0.00 | 24.43 | 0.00 | 0.00 |
| >hg38_21_+_8433172_8446622_RNA45SN1_45S_50ntFlanks@10348.10367.20\|AAACGAGAACTTTGAAGGCC | >hg38_21_+_8433172_8446622_RNA45SN1_45S_50ntFlanks | 10348 | 10367 | 20 | rRF 45S | NO | 0.00 | 29.12 | 0.00 | 0.00 |
| >hg38_21_+_8433172_8446622_RNA45SN1_45S_50ntFlanks@10348.10369.22\|AAACGAGAACTTTGAAGGCCGA | >hg38_21_+_8433172_8446622_RNA45SN1_45S_50ntFlanks | 10348 | 10369 | 22 | rRF 45S | NO | 0.00 | 26.94 | 0.00 | 0.00 |
| >hg38_21_+_8433172_8446622_RNA45SN1_45S_50ntFlanks@10348.10371.24\|AAACGAGAACTTTGAAGGCCGAAG | >hg38_21_+_8433172_8446622_RNA45SN1_45S_50ntFlanks | 10348 | 10371 | 24 | rRF 45S | NO | 0.00 | 25.87 | 11.39 | 0.00 |
| >hg38_21_+_8433172_8446622_RNA45SN1_45S_50ntFlanks@10348.10372.25\|AAACGAGAACTTTGAAGGCCGAAGT | >hg38_21_+_8433172_8446622_RNA45SN1_45S_50ntFlanks | 10348 | 10372 | 25 | rRF 45S | NO | 49.83 | 32.65 | 15.40 | 0.00 |
| >hg38_21_+_8433172_8446622_RNA45SN1_45S_50ntFlanks@10348.10382.35\|AAACGAGAACTTTGAAGGCCGAAGTGGAGAAGGGT | >hg38_21_+_8433172_8446622_RNA45SN1_45S_50ntFlanks | 10348 | 10382 | 35 | rRF 45S | NO | 0.00 | 74.27 | 0.00 | 0.00 |
| >hg38_21_+_8433172_8446622_RNA45SN1_45S_50ntFlanks@10348.10383.36\|AAACGAGAACTTTGAAGGCCGAAGTGGAGAAGGGTT | >hg38_21_+_8433172_8446622_RNA45SN1_45S_50ntFlanks | 10348 | 10383 | 36 | rRF 45S | NO | 0.00 | 30.02 | 0.00 | 0.00 |
| >hg38_21_+_8433172_8446622_RNA45SN1_45S_50ntFlanks@10349.10367.19\|AACGAGAACTTTGAAGGCC | >hg38_21_+_8433172_8446622_RNA45SN1_45S_50ntFlanks | 10349 | 10367 | 19 | rRF 45S | NO | 0.00 | 32.28 | 0.00 | 0.00 |
| >hg38_21_+_8433172_8446622_RNA45SN1_45S_50ntFlanks@10349.10369.21\|AACGAGAACTTTGAAGGCCGA | >hg38_21_+_8433172_8446622_RNA45SN1_45S_50ntFlanks | 10349 | 10369 | 21 | rRF 45S | NO | 24.55 | 32.29 | 15.30 | 0.00 |
| >hg38_21_+_8433172_8446622_RNA45SN1_45S_50ntFlanks@10349.10371.23\|AACGAGAACTTTGAAGGCCGAAG | >hg38_21_+_8433172_8446622_RNA45SN1_45S_50ntFlanks | 10349 | 10371 | 23 | rRF 45S | NO | 10.58 | 26.79 | 15.67 | 0.00 |
| >hg38_21_+_8433172_8446622_RNA45SN1_45S_50ntFlanks@10349.10372.24\|AACGAGAACTTTGAAGGCCGAAGT | >hg38_21_+_8433172_8446622_RNA45SN1_45S_50ntFlanks | 10349 | 10372 | 24 | rRF 45S | NO | 59.94 | 35.71 | 56.18 | 12.20 |
| >hg38_21_+_8433172_8446622_RNA45SN1_45S_50ntFlanks@10349.10382.34\|AACGAGAACTTTGAAGGCCGAAGTGGAGAAGGGT | >hg38_21_+_8433172_8446622_RNA45SN1_45S_50ntFlanks | 10349 | 10382 | 34 | rRF 45S | NO | 0.00 | 106.53 | 11.03 | 0.00 |
| >hg38_21_+_8433172_8446622_RNA45SN1_45S_50ntFlanks@10349.10383.35\|AACGAGAACTTTGAAGGCCGAAGTGGAGAAGGGTT | >hg38_21_+_8433172_8446622_RNA45SN1_45S_50ntFlanks | 10349 | 10383 | 35 | rRF 45S | NO | 0.00 | 64.23 | 0.00 | 0.00 |
| >hg38_21_+_8433172_8446622_RNA45SN1_45S_50ntFlanks@10350.10382.33\|ACGAGAACTTTGAAGGCCGAAGTGGAGAAGGGT | >hg38_21_+_8433172_8446622_RNA45SN1_45S_50ntFlanks | 10350 | 10382 | 33 | rRF 45S | NO | 0.00 | 27.15 | 0.00 | 0.00 |
| >hg38_21_+_8433172_8446622_RNA45SN1_45S_50ntFlanks@10352.10369.18\|GAGAACTTTGAAGGCCGA | >hg38_21_+_8433172_8446622_RNA45SN1_45S_50ntFlanks | 10352 | 10369 | 18 | rRF 45S | NO | 10.49 | 0.00 | 12.85 | 0.00 |
| >hg38_21_+_8433172_8446622_RNA45SN1_45S_50ntFlanks@10380.10424.45\|GGTTCCATGTGAACAGCAGTTGAACATGGGTCAGTCGGTCCTGAG | >hg38_21_+_8433172_8446622_RNA45SN1_45S_50ntFlanks | 10380 | 10424 | 45 | rRF 45S | NO | 0.00 | 58.00 | 0.00 | 0.00 |
| >hg38_21_+_8433172_8446622_RNA45SN1_45S_50ntFlanks@10381.10424.44\|GTTCCATGTGAACAGCAGTTGAACATGGGTCAGTCGGTCCTGAG | >hg38_21_+_8433172_8446622_RNA45SN1_45S_50ntFlanks | 10381 | 10424 | 44 | rRF 45S | NO | 51.14 | 91.70 | 0.00 | 0.00 |
| >hg38_21_+_8433172_8446622_RNA45SN1_45S_50ntFlanks@10382.10424.43\|TTCCATGTGAACAGCAGTTGAACATGGGTCAGTCGGTCCTGAG | >hg38_21_+_8433172_8446622_RNA45SN1_45S_50ntFlanks | 10382 | 10424 | 43 | rRF 45S | NO | 62.44 | 133.36 | 0.00 | 0.00 |
| >hg38_21_+_8433172_8446622_RNA45SN1_45S_50ntFlanks@10383.10424.42\|TCCATGTGAACAGCAGTTGAACATGGGTCAGTCGGTCCTGAG | >hg38_21_+_8433172_8446622_RNA45SN1_45S_50ntFlanks | 10383 | 10424 | 42 | rRF 45S | NO | 100.66 | 233.08 | 0.00 | 19.65 |
| >hg38_21_+_8433172_8446622_RNA45SN1_45S_50ntFlanks@10384.10424.41\|CCATGTGAACAGCAGTTGAACATGGGTCAGTCGGTCCTGAG | >hg38_21_+_8433172_8446622_RNA45SN1_45S_50ntFlanks | 10384 | 10424 | 41 | rRF 45S | NO | 23.99 | 80.68 | 0.00 | 0.00 |
| >hg38_21_+_8433172_8446622_RNA45SN1_45S_50ntFlanks@10386.10424.39\|ATGTGAACAGCAGTTGAACATGGGTCAGTCGGTCCTGAG | >hg38_21_+_8433172_8446622_RNA45SN1_45S_50ntFlanks | 10386 | 10424 | 39 | rRF 45S | NO | 88.21 | 139.11 | 11.02 | 10.89 |
| >hg38_21_+_8433172_8446622_RNA45SN1_45S_50ntFlanks@10387.10424.38\|TGTGAACAGCAGTTGAACATGGGTCAGTCGGTCCTGAG | >hg38_21_+_8433172_8446622_RNA45SN1_45S_50ntFlanks | 10387 | 10424 | 38 | rRF 45S | NO | 79.77 | 120.48 | 0.00 | 11.72 |
| >hg38_21_+_8433172_8446622_RNA45SN1_45S_50ntFlanks@10388.10424.37\|GTGAACAGCAGTTGAACATGGGTCAGTCGGTCCTGAG | >hg38_21_+_8433172_8446622_RNA45SN1_45S_50ntFlanks | 10388 | 10424 | 37 | rRF 45S | NO | 70.03 | 122.90 | 16.38 | 0.00 |
| >hg38_21_+_8433172_8446622_RNA45SN1_45S_50ntFlanks@10389.10424.36\|TGAACAGCAGTTGAACATGGGTCAGTCGGTCCTGAG | >hg38_21_+_8433172_8446622_RNA45SN1_45S_50ntFlanks | 10389 | 10424 | 36 | rRF 45S | NO | 91.40 | 159.02 | 32.38 | 10.69 |
| >hg38_21_+_8433172_8446622_RNA45SN1_45S_50ntFlanks@10390.10424.35\|GAACAGCAGTTGAACATGGGTCAGTCGGTCCTGAG | >hg38_21_+_8433172_8446622_RNA45SN1_45S_50ntFlanks | 10390 | 10424 | 35 | rRF 45S | NO | 24.54 | 68.06 | 0.00 | 0.00 |
| >hg38_21_+_8433172_8446622_RNA45SN1_45S_50ntFlanks@10391.10424.34\|AACAGCAGTTGAACATGGGTCAGTCGGTCCTGAG | >hg38_21_+_8433172_8446622_RNA45SN1_45S_50ntFlanks | 10391 | 10424 | 34 | rRF 45S | NO | 65.99 | 98.57 | 15.53 | 10.83 |
| >hg38_21_+_8433172_8446622_RNA45SN1_45S_50ntFlanks@10392.10424.33\|ACAGCAGTTGAACATGGGTCAGTCGGTCCTGAG | >hg38_21_+_8433172_8446622_RNA45SN1_45S_50ntFlanks | 10392 | 10424 | 33 | rRF 45S | NO | 49.97 | 81.94 | 12.58 | 0.00 |
| >hg38_21_+_8433172_8446622_RNA45SN1_45S_50ntFlanks@10393.10424.32\|CAGCAGTTGAACATGGGTCAGTCGGTCCTGAG | >hg38_21_+_8433172_8446622_RNA45SN1_45S_50ntFlanks | 10393 | 10424 | 32 | rRF 45S | NO | 65.56 | 95.44 | 14.44 | 11.33 |
| >hg38_21_+_8433172_8446622_RNA45SN1_45S_50ntFlanks@10394.10424.31\|AGCAGTTGAACATGGGTCAGTCGGTCCTGAG | >hg38_21_+_8433172_8446622_RNA45SN1_45S_50ntFlanks | 10394 | 10424 | 31 | rRF 45S | NO | 55.00 | 72.33 | 55.93 | 0.00 |
| >hg38_21_+_8433172_8446622_RNA45SN1_45S_50ntFlanks@10395.10424.30\|GCAGTTGAACATGGGTCAGTCGGTCCTGAG | >hg38_21_+_8433172_8446622_RNA45SN1_45S_50ntFlanks | 10395 | 10424 | 30 | rRF 45S | NO | 68.15 | 93.31 | 32.57 | 0.00 |
| >hg38_21_+_8433172_8446622_RNA45SN1_45S_50ntFlanks@10396.10424.29\|CAGTTGAACATGGGTCAGTCGGTCCTGAG | >hg38_21_+_8433172_8446622_RNA45SN1_45S_50ntFlanks | 10396 | 10424 | 29 | rRF 45S | NO | 167.41 | 204.00 | 78.39 | 15.75 |
| >hg38_21_+_8433172_8446622_RNA45SN1_45S_50ntFlanks@10398.10424.27\|GTTGAACATGGGTCAGTCGGTCCTGAG | >hg38_21_+_8433172_8446622_RNA45SN1_45S_50ntFlanks | 10398 | 10424 | 27 | rRF 45S | NO | 96.22 | 99.76 | 32.83 | 0.00 |
| >hg38_21_+_8433172_8446622_RNA45SN1_45S_50ntFlanks@10400.10424.25\|TGAACATGGGTCAGTCGGTCCTGAG | >hg38_21_+_8433172_8446622_RNA45SN1_45S_50ntFlanks | 10400 | 10424 | 25 | rRF 45S | NO | 87.03 | 82.54 | 15.66 | 0.00 |
| >hg38_21_+_8433172_8446622_RNA45SN1_45S_50ntFlanks@10401.10424.24\|GAACATGGGTCAGTCGGTCCTGAG | >hg38_21_+_8433172_8446622_RNA45SN1_45S_50ntFlanks | 10401 | 10424 | 24 | rRF 45S | NO | 133.34 | 163.32 | 204.03 | 14.86 |

### MDA-MB-231

| | | | | | | | | | | |
|---|---|---|---|---|---|---|---|---|---|---|
| >hg38_21_+_8433172_8446622_RNA45SN1_45S_50ntFlanks@10402.10424.23\|AACATGGGTCAGTCGGTCCTGAG | >hg38_21_+_8433172_8446622_RNA45SN1_45S_50ntFlanks | 10402 | 10424 | 23 | rRF 45S | NO | 100.42 | 112.29 | 86.54 | 11.96 |
| >hg38_21_+_8433172_8446622_RNA45SN1_45S_50ntFlanks@10403.10424.22\|ACATGGGTCAGTCGGTCCTGAG | >hg38_21_+_8433172_8446622_RNA45SN1_45S_50ntFlanks | 10403 | 10424 | 22 | rRF 45S | NO | 72.83 | 75.18 | 64.13 | 0.00 |
| >hg38_21_+_8433172_8446622_RNA45SN1_45S_50ntFlanks@10404.10424.21\|CATGGGTCAGTCGGTCCTGAG | >hg38_21_+_8433172_8446622_RNA45SN1_45S_50ntFlanks | 10404 | 10424 | 21 | rRF 45S | NO | 76.61 | 87.77 | 269.43 | 0.00 |
| >hg38_21_+_8433172_8446622_RNA45SN1_45S_50ntFlanks@10405.10424.20\|ATGGGTCAGTCGGTCCTGAG | >hg38_21_+_8433172_8446622_RNA45SN1_45S_50ntFlanks | 10405 | 10424 | 20 | rRF 45S | NO | 85.18 | 82.20 | 865.87 | 0.00 |
| >hg38_21_+_8433172_8446622_RNA45SN1_45S_50ntFlanks@10406.10424.19\|TGGGTCAGTCGGTCCTGAG | >hg38_21_+_8433172_8446622_RNA45SN1_45S_50ntFlanks | 10406 | 10424 | 19 | rRF 45S | NO | 0.00 | 0.00 | 15.84 | 0.00 |
| >hg38_21_+_8433172_8446622_RNA45SN1_45S_50ntFlanks@10423.10456.34\|AGAGATGGGCGAGCGCCGTTCCGAAGGGACGGGC | >hg38_21_+_8433172_8446622_RNA45SN1_45S_50ntFlanks | 10423 | 10456 | 34 | rRF 45S | NO | 0.00 | 77.18 | 0.00 | 0.00 |
| >hg38_21_+_8433172_8446622_RNA45SN1_45S_50ntFlanks@10738.10756.19\|AAATCTCGCGCCGGGCCGT | >hg38_21_+_8433172_8446622_RNA45SN1_45S_50ntFlanks | 10738 | 10756 | 19 | rRF 45S | NO | 0.00 | 35.38 | 0.00 | 0.00 |
| >hg38_21_+_8433172_8446622_RNA45SN1_45S_50ntFlanks@10809.10830.22\|AATGTAGGTAAGGGAAGTCGGC | >hg38_21_+_8433172_8446622_RNA45SN1_45S_50ntFlanks | 10809 | 10830 | 22 | rRF 45S | NO | 0.00 | 27.25 | 0.00 | 0.00 |
| >hg38_21_+_8433172_8446622_RNA45SN1_45S_50ntFlanks@10834.10867.34\|CCGGATCCGTAACTTCGGGATAAGGATTGGCTCT | >hg38_21_+_8433172_8446622_RNA45SN1_45S_50ntFlanks | 10834 | 10867 | 34 | rRF 45S | NO | 0.00 | 72.99 | 0.00 | 0.00 |
| >hg38_21_+_8433172_8446622_RNA45SN1_45S_50ntFlanks@10835.10867.33\|CGGATCCGTAACTTCGGGATAAGGATTGGCTCT | >hg38_21_+_8433172_8446622_RNA45SN1_45S_50ntFlanks | 10835 | 10867 | 33 | rRF 45S | NO | 0.00 | 28.35 | 0.00 | 0.00 |
| >hg38_21_+_8433172_8446622_RNA45SN1_45S_50ntFlanks@10841.10867.27\|CGTAACTTCGGGATAAGGATTGGCTCT | >hg38_21_+_8433172_8446622_RNA45SN1_45S_50ntFlanks | 10841 | 10867 | 27 | rRF 45S | NO | 0.00 | 154.65 | 0.00 | 0.00 |
| >hg38_21_+_8433172_8446622_RNA45SN1_45S_50ntFlanks@10841.10868.28\|CGTAACTTCGGGATAAGGATTGGCTCTA | >hg38_21_+_8433172_8446622_RNA45SN1_45S_50ntFlanks | 10841 | 10868 | 28 | rRF 45S | NO | 0.00 | 27.66 | 0.00 | 0.00 |
| >hg38_21_+_8433172_8446622_RNA45SN1_45S_50ntFlanks@11326.11345.20\|CTCGTCGCGCGCGCGTCCGC | >hg38_21_+_8433172_8446622_RNA45SN1_45S_50ntFlanks | 11326 | 11345 | 20 | rRF 45S | NO | 61.69 | 33.91 | 0.00 | 0.00 |
| >hg38_21_+_8433172_8446622_RNA45SN1_45S_50ntFlanks@11538.11584.47\|CGCGCGGCGCCTCGCCTCGGCCGGCGCCTAGCAGCCGACTTAGAACT | >hg38_21_+_8433172_8446622_RNA45SN1_45S_50ntFlanks | 11538 | 11584 | 47 | rRF 45S | NO | 0.00 | 24.98 | 0.00 | 0.00 |
| >hg38_21_+_8433172_8446622_RNA45SN1_45S_50ntFlanks@11539.11583.45\|GCGCGGCGCCTCGCCTCGGCCGGCGCCTAGCAGCCGACTTAGAAC | >hg38_21_+_8433172_8446622_RNA45SN1_45S_50ntFlanks | 11539 | 11583 | 45 | rRF 45S | NO | 0.00 | 26.89 | 0.00 | 0.00 |
| >hg38_21_+_8433172_8446622_RNA45SN1_45S_50ntFlanks@11539.11584.46\|GCGCGGCGCCTCGCCTCGGCCGGCGCCTAGCAGCCGACTTAGAACT | >hg38_21_+_8433172_8446622_RNA45SN1_45S_50ntFlanks | 11539 | 11584 | 46 | rRF 45S | NO | 0.00 | 60.28 | 0.00 | 0.00 |
| >hg38_21_+_8433172_8446622_RNA45SN1_45S_50ntFlanks@11540.11582.43\|CGCGGCGCCTCGCCTCGGCCGGCGCCTAGCAGCCGACTTAGAA | >hg38_21_+_8433172_8446622_RNA45SN1_45S_50ntFlanks | 11540 | 11582 | 43 | rRF 45S | NO | 26.18 | 60.81 | 0.00 | 0.00 |
| >hg38_21_+_8433172_8446622_RNA45SN1_45S_50ntFlanks@11540.11583.44\|CGCGGCGCCTCGCCTCGGCCGGCGCCTAGCAGCCGACTTAGAAC | >hg38_21_+_8433172_8446622_RNA45SN1_45S_50ntFlanks | 11540 | 11583 | 44 | rRF 45S | NO | 48.32 | 124.01 | 0.00 | 10.50 |
| >hg38_21_+_8433172_8446622_RNA45SN1_45S_50ntFlanks@11540.11584.45\|CGCGGCGCCTCGCCTCGGCCGGCGCCTAGCAGCCGACTTAGAACT | >hg38_21_+_8433172_8446622_RNA45SN1_45S_50ntFlanks | 11540 | 11584 | 45 | rRF 45S | NO | 82.78 | 251.78 | 0.00 | 14.29 |
| >hg38_21_+_8433172_8446622_RNA45SN1_45S_50ntFlanks@11540.11586.47\|CGCGGCGCCTCGCCTCGGCCGGCGCCTAGCAGCCGACTTAGAACTGG | >hg38_21_+_8433172_8446622_RNA45SN1_45S_50ntFlanks | 11540 | 11586 | 47 | rRF 45S | NO | 0.00 | 27.36 | 0.00 | 0.00 |
| >hg38_21_+_8433172_8446622_RNA45SN1_45S_50ntFlanks@11541.11584.44\|GCGGCGCCTCGCCTCGGCCGGCGCCTAGCAGCCGACTTAGAACT | >hg38_21_+_8433172_8446622_RNA45SN1_45S_50ntFlanks | 11541 | 11584 | 44 | rRF 45S | NO | 0.00 | 24.91 | 0.00 | 0.00 |
| >hg38_21_+_8433172_8446622_RNA45SN1_45S_50ntFlanks@11542.11584.43\|CGGCGCCTCGCCTCGGCCGGCGCCTAGCAGCCGACTTAGAACT | >hg38_21_+_8433172_8446622_RNA45SN1_45S_50ntFlanks | 11542 | 11584 | 43 | rRF 45S | NO | 0.00 | 13.14 | 0.00 | 0.00 |
| >hg38_21_+_8433172_8446622_RNA45SN1_45S_50ntFlanks@11566.11583.18\|TAGCAGCCGACTTAGAAC | >hg38_21_+_8433172_8446622_RNA45SN1_45S_50ntFlanks | 11566 | 11583 | 18 | rRF 45S | NO | 0.00 | 57.92 | 0.00 | 0.00 |
| >hg38_21_+_8433172_8446622_RNA45SN1_45S_50ntFlanks@11566.11584.19\|TAGCAGCCGACTTAGAACT | >hg38_21_+_8433172_8446622_RNA45SN1_45S_50ntFlanks | 11566 | 11584 | 19 | rRF 45S | NO | 50.10 | 145.01 | 0.00 | 0.00 |
| >hg38_21_+_8433172_8446622_RNA45SN1_45S_50ntFlanks@11567.11584.18\|AGCAGCCGACTTAGAACT | >hg38_21_+_8433172_8446622_RNA45SN1_45S_50ntFlanks | 11567 | 11584 | 18 | rRF 45S | NO | 159.98 | 254.61 | 12.90 | 21.48 |
| >hg38_21_+_8433172_8446622_RNA45SN1_45S_50ntFlanks@11567.11585.19\|AGCAGCCGACTTAGAACTG | >hg38_21_+_8433172_8446622_RNA45SN1_45S_50ntFlanks | 11567 | 11585 | 19 | rRF 45S | NO | 69.23 | 59.18 | 0.00 | 12.18 |
| >hg38_21_+_8433172_8446622_RNA45SN1_45S_50ntFlanks@11567.11586.20\|AGCAGCCGACTTAGAACTGG | >hg38_21_+_8433172_8446622_RNA45SN1_45S_50ntFlanks | 11567 | 11586 | 20 | rRF 45S | NO | 66.14 | 36.80 | 12.73 | 12.32 |
| >hg38_21_+_8433172_8446622_RNA45SN1_45S_50ntFlanks@11568.11585.18\|GCAGCCGACTTAGAACTG | >hg38_21_+_8433172_8446622_RNA45SN1_45S_50ntFlanks | 11568 | 11585 | 18 | rRF 45S | NO | 24.38 | 25.55 | 0.00 | 0.00 |
| >hg38_21_+_8433172_8446622_RNA45SN1_45S_50ntFlanks@11568.11586.19\|GCAGCCGACTTAGAACTGG | >hg38_21_+_8433172_8446622_RNA45SN1_45S_50ntFlanks | 11568 | 11586 | 19 | rRF 45S | NO | 25.36 | 13.64 | 0.00 | 0.00 |
| >hg38_21_+_8433172_8446622_RNA45SN1_45S_50ntFlanks@11568.11611.44\|GCAGCCGACTTAGAACTGGTGCGGACCAGGGGAATCCGACTGTT | >hg38_21_+_8433172_8446622_RNA45SN1_45S_50ntFlanks | 11568 | 11611 | 44 | rRF 45S | NO | 0.00 | 27.08 | 0.00 | 16.97 |
| >hg38_21_+_8433172_8446622_RNA45SN1_45S_50ntFlanks@11573.11613.41\|CGACTTAGAACTGGTGCGGACCAGGGGAATCCGACTGTTTA | >hg38_21_+_8433172_8446622_RNA45SN1_45S_50ntFlanks | 11573 | 11613 | 41 | rRF 45S | NO | 0.00 | 32.16 | 0.00 | 0.00 |
| >hg38_21_+_8433172_8446622_RNA45SN1_45S_50ntFlanks@11580.11613.34\|GAACTGGTGCGGACCAGGGGAATCCGACTGTTTA | >hg38_21_+_8433172_8446622_RNA45SN1_45S_50ntFlanks | 11580 | 11613 | 34 | rRF 45S | NO | 0.00 | 37.42 | 0.00 | 0.00 |
| >hg38_21_+_8433172_8446622_RNA45SN1_45S_50ntFlanks@11582.11611.30\|ACTGGTGCGGACCAGGGGAATCCGACTGTT | >hg38_21_+_8433172_8446622_RNA45SN1_45S_50ntFlanks | 11582 | 11611 | 30 | rRF 45S | NO | 0.00 | 13.18 | 0.00 | 0.00 |
| >hg38_21_+_8433172_8446622_RNA45SN1_45S_50ntFlanks@11583.11611.29\|CTGGTGCGGACCAGGGGAATCCGACTGTT | >hg38_21_+_8433172_8446622_RNA45SN1_45S_50ntFlanks | 11583 | 11611 | 29 | rRF 45S | NO | 0.00 | 27.59 | 0.00 | 0.00 |
| >hg38_21_+_8433172_8446622_RNA45SN1_45S_50ntFlanks@11588.11611.24\|GCGGACCAGGGGAATCCGACTGTT | >hg38_21_+_8433172_8446622_RNA45SN1_45S_50ntFlanks | 11588 | 11611 | 24 | rRF 45S | NO | 0.00 | 25.63 | 0.00 | 0.00 |
| >hg38_21_+_8433172_8446622_RNA45SN1_45S_50ntFlanks@11595.11612.18\|AGGGGAATCCGACTGTTT | >hg38_21_+_8433172_8446622_RNA45SN1_45S_50ntFlanks | 11595 | 11612 | 18 | rRF 45S | NO | 0.00 | 24.84 | 0.00 | 0.00 |
| >hg38_21_+_8433172_8446622_RNA45SN1_45S_50ntFlanks@11665.11687.23\|TTCTGCCCAGTGCTCTGAATGTC | >hg38_21_+_8433172_8446622_RNA45SN1_45S_50ntFlanks | 11665 | 11687 | 23 | rRF 45S | NO | 0.00 | 60.61 | 0.00 | 0.00 |
| >hg38_21_+_8433172_8446622_RNA45SN1_45S_50ntFlanks@11665.11702.38\|TTCTGCCCAGTGCTCTGAATGTCAAAGTGAAGAAATTC | >hg38_21_+_8433172_8446622_RNA45SN1_45S_50ntFlanks | 11665 | 11702 | 38 | rRF 45S | NO | 0.00 | 98.38 | 0.00 | 0.00 |

### MDA-MB-231

| | | | | | | | | | | |
|---|---|---|---|---|---|---|---|---|---|---|
| >hg38_21_+_8433172_8446622_RNA45SN1_45S_50ntFlanks@11666.11687.22\|TCTGCCCAGTGCTCTGAATGTC | >hg38_21_+_8433172_8446622_RNA45SN1_45S_50ntFlanks | 11666 | 11687 | 22 | rRF 45S | NO | 0.00 | 125.23 | 0.00 | 0.00 |
| >hg38_21_+_8433172_8446622_RNA45SN1_45S_50ntFlanks@11666.11702.37\|TCTGCCCAGTGCTCTGAATGTCAAAGTGAAGAAATTC | >hg38_21_+_8433172_8446622_RNA45SN1_45S_50ntFlanks | 11666 | 11702 | 37 | rRF 45S | NO | 0.00 | 134.50 | 0.00 | 0.00 |
| >hg38_21_+_8433172_8446622_RNA45SN1_45S_50ntFlanks@11666.11703.38\|TCTGCCCAGTGCTCTGAATGTCAAAGTGAAGAAATTCA | >hg38_21_+_8433172_8446622_RNA45SN1_45S_50ntFlanks | 11666 | 11703 | 38 | rRF 45S | NO | 0.00 | 32.71 | 0.00 | 0.00 |
| >hg38_21_+_8433172_8446622_RNA45SN1_45S_50ntFlanks@11666.11706.41\|TCTGCCCAGTGCTCTGAATGTCAAAGTGAAGAAATTCAATG | >hg38_21_+_8433172_8446622_RNA45SN1_45S_50ntFlanks | 11666 | 11706 | 41 | rRF 45S | NO | 0.00 | 90.07 | 0.00 | 0.00 |
| >hg38_21_+_8433172_8446622_RNA45SN1_45S_50ntFlanks@11666.11707.42\|TCTGCCCAGTGCTCTGAATGTCAAAGTGAAGAAATTCAATGA | >hg38_21_+_8433172_8446622_RNA45SN1_45S_50ntFlanks | 11666 | 11707 | 42 | rRF 45S | NO | 0.00 | 33.79 | 0.00 | 0.00 |
| >hg38_21_+_8433172_8446622_RNA45SN1_45S_50ntFlanks@11667.11687.21\|CTGCCCAGTGCTCTGAATGTC | >hg38_21_+_8433172_8446622_RNA45SN1_45S_50ntFlanks | 11667 | 11687 | 21 | rRF 45S | NO | 0.00 | 87.18 | 0.00 | 0.00 |
| >hg38_21_+_8433172_8446622_RNA45SN1_45S_50ntFlanks@11667.11691.25\|CTGCCCAGTGCTCTGAATGTCAAAG | >hg38_21_+_8433172_8446622_RNA45SN1_45S_50ntFlanks | 11667 | 11691 | 25 | rRF 45S | NO | 0.00 | 26.93 | 0.00 | 0.00 |
| >hg38_21_+_8433172_8446622_RNA45SN1_45S_50ntFlanks@11667.11692.26\|CTGCCCAGTGCTCTGAATGTCAAAGT | >hg38_21_+_8433172_8446622_RNA45SN1_45S_50ntFlanks | 11667 | 11692 | 26 | rRF 45S | NO | 0.00 | 13.10 | 0.00 | 0.00 |
| >hg38_21_+_8433172_8446622_RNA45SN1_45S_50ntFlanks@11667.11702.36\|CTGCCCAGTGCTCTGAATGTCAAAGTGAAGAAATTC | >hg38_21_+_8433172_8446622_RNA45SN1_45S_50ntFlanks | 11667 | 11702 | 36 | rRF 45S | NO | 0.00 | 115.47 | 0.00 | 0.00 |
| >hg38_21_+_8433172_8446622_RNA45SN1_45S_50ntFlanks@11667.11703.37\|CTGCCCAGTGCTCTGAATGTCAAAGTGAAGAAATTCA | >hg38_21_+_8433172_8446622_RNA45SN1_45S_50ntFlanks | 11667 | 11703 | 37 | rRF 45S | NO | 0.00 | 28.61 | 0.00 | 0.00 |
| >hg38_21_+_8433172_8446622_RNA45SN1_45S_50ntFlanks@11667.11706.40\|CTGCCCAGTGCTCTGAATGTCAAAGTGAAGAAATTCAATG | >hg38_21_+_8433172_8446622_RNA45SN1_45S_50ntFlanks | 11667 | 11706 | 40 | rRF 45S | NO | 0.00 | 78.47 | 0.00 | 0.00 |
| >hg38_21_+_8433172_8446622_RNA45SN1_45S_50ntFlanks@11667.11707.41\|CTGCCCAGTGCTCTGAATGTCAAAGTGAAGAAATTCAATGA | >hg38_21_+_8433172_8446622_RNA45SN1_45S_50ntFlanks | 11667 | 11707 | 41 | rRF 45S | NO | 0.00 | 28.51 | 0.00 | 0.00 |
| >hg38_21_+_8433172_8446622_RNA45SN1_45S_50ntFlanks@11668.11685.18\|TGCCCAGTGCTCTGAATG | >hg38_21_+_8433172_8446622_RNA45SN1_45S_50ntFlanks | 11668 | 11685 | 18 | rRF 45S | NO | 0.00 | 13.12 | 0.00 | 0.00 |
| >hg38_21_+_8433172_8446622_RNA45SN1_45S_50ntFlanks@11668.11687.20\|TGCCCAGTGCTCTGAATGTC | >hg38_21_+_8433172_8446622_RNA45SN1_45S_50ntFlanks | 11668 | 11687 | 20 | rRF 45S | NO | 51.66 | 159.84 | 0.00 | 0.00 |
| >hg38_21_+_8433172_8446622_RNA45SN1_45S_50ntFlanks@11668.11688.21\|TGCCCAGTGCTCTGAATGTCA | >hg38_21_+_8433172_8446622_RNA45SN1_45S_50ntFlanks | 11668 | 11688 | 21 | rRF 45S | NO | 0.00 | 58.45 | 0.00 | 0.00 |
| >hg38_21_+_8433172_8446622_RNA45SN1_45S_50ntFlanks@11668.11691.24\|TGCCCAGTGCTCTGAATGTCAAAG | >hg38_21_+_8433172_8446622_RNA45SN1_45S_50ntFlanks | 11668 | 11691 | 24 | rRF 45S | NO | 0.00 | 31.19 | 0.00 | 0.00 |
| >hg38_21_+_8433172_8446622_RNA45SN1_45S_50ntFlanks@11668.11692.25\|TGCCCAGTGCTCTGAATGTCAAAGT | >hg38_21_+_8433172_8446622_RNA45SN1_45S_50ntFlanks | 11668 | 11692 | 25 | rRF 45S | NO | 0.00 | 26.05 | 0.00 | 0.00 |
| >hg38_21_+_8433172_8446622_RNA45SN1_45S_50ntFlanks@11668.11693.26\|TGCCCAGTGCTCTGAATGTCAAAGTG | >hg38_21_+_8433172_8446622_RNA45SN1_45S_50ntFlanks | 11668 | 11693 | 26 | rRF 45S | NO | 0.00 | 35.91 | 0.00 | 0.00 |
| >hg38_21_+_8433172_8446622_RNA45SN1_45S_50ntFlanks@11668.11701.34\|TGCCCAGTGCTCTGAATGTCAAAGTGAAGAAATT | >hg38_21_+_8433172_8446622_RNA45SN1_45S_50ntFlanks | 11668 | 11701 | 34 | rRF 45S | NO | 0.00 | 25.05 | 0.00 | 0.00 |
| >hg38_21_+_8433172_8446622_RNA45SN1_45S_50ntFlanks@11668.11702.35\|TGCCCAGTGCTCTGAATGTCAAAGTGAAGAAATTC | >hg38_21_+_8433172_8446622_RNA45SN1_45S_50ntFlanks | 11668 | 11702 | 35 | rRF 45S | NO | 54.81 | 314.16 | 0.00 | 0.00 |
| >hg38_21_+_8433172_8446622_RNA45SN1_45S_50ntFlanks@11668.11703.36\|TGCCCAGTGCTCTGAATGTCAAAGTGAAGAAATTCA | >hg38_21_+_8433172_8446622_RNA45SN1_45S_50ntFlanks | 11668 | 11703 | 36 | rRF 45S | NO | 0.00 | 135.43 | 0.00 | 0.00 |
| >hg38_21_+_8433172_8446622_RNA45SN1_45S_50ntFlanks@11668.11704.37\|TGCCCAGTGCTCTGAATGTCAAAGTGAAGAAATTCAA | >hg38_21_+_8433172_8446622_RNA45SN1_45S_50ntFlanks | 11668 | 11704 | 37 | rRF 45S | NO | 0.00 | 34.25 | 0.00 | 0.00 |
| >hg38_21_+_8433172_8446622_RNA45SN1_45S_50ntFlanks@11668.11705.38\|TGCCCAGTGCTCTGAATGTCAAAGTGAAGAAATTCAAT | >hg38_21_+_8433172_8446622_RNA45SN1_45S_50ntFlanks | 11668 | 11705 | 38 | rRF 45S | NO | 0.00 | 29.15 | 0.00 | 0.00 |
| >hg38_21_+_8433172_8446622_RNA45SN1_45S_50ntFlanks@11668.11706.39\|TGCCCAGTGCTCTGAATGTCAAAGTGAAGAAATTCAATG | >hg38_21_+_8433172_8446622_RNA45SN1_45S_50ntFlanks | 11668 | 11706 | 39 | rRF 45S | NO | 0.00 | 201.41 | 0.00 | 0.00 |
| >hg38_21_+_8433172_8446622_RNA45SN1_45S_50ntFlanks@11668.11707.40\|TGCCCAGTGCTCTGAATGTCAAAGTGAAGAAATTCAATGA | >hg38_21_+_8433172_8446622_RNA45SN1_45S_50ntFlanks | 11668 | 11707 | 40 | rRF 45S | NO | 0.00 | 139.07 | 0.00 | 0.00 |
| >hg38_21_+_8433172_8446622_RNA45SN1_45S_50ntFlanks@11668.11709.42\|TGCCCAGTGCTCTGAATGTCAAAGTGAAGAAATTCAATGAAG | >hg38_21_+_8433172_8446622_RNA45SN1_45S_50ntFlanks | 11668 | 11709 | 42 | rRF 45S | NO | 0.00 | 68.16 | 0.00 | 0.00 |
| >hg38_21_+_8433172_8446622_RNA45SN1_45S_50ntFlanks@11669.11687.19\|GCCCAGTGCTCTGAATGTC | >hg38_21_+_8433172_8446622_RNA45SN1_45S_50ntFlanks | 11669 | 11687 | 19 | rRF 45S | NO | 51.03 | 179.50 | 0.00 | 0.00 |
| >hg38_21_+_8433172_8446622_RNA45SN1_45S_50ntFlanks@11669.11688.20\|GCCCAGTGCTCTGAATGTCA | >hg38_21_+_8433172_8446622_RNA45SN1_45S_50ntFlanks | 11669 | 11688 | 20 | rRF 45S | NO | 0.00 | 35.68 | 0.00 | 0.00 |
| >hg38_21_+_8433172_8446622_RNA45SN1_45S_50ntFlanks@11669.11691.23\|GCCCAGTGCTCTGAATGTCAAAG | >hg38_21_+_8433172_8446622_RNA45SN1_45S_50ntFlanks | 11669 | 11691 | 23 | rRF 45S | NO | 0.00 | 26.94 | 0.00 | 0.00 |
| >hg38_21_+_8433172_8446622_RNA45SN1_45S_50ntFlanks@11669.11692.24\|GCCCAGTGCTCTGAATGTCAAAGT | >hg38_21_+_8433172_8446622_RNA45SN1_45S_50ntFlanks | 11669 | 11692 | 24 | rRF 45S | NO | 0.00 | 13.46 | 0.00 | 0.00 |
| >hg38_21_+_8433172_8446622_RNA45SN1_45S_50ntFlanks@11669.11693.25\|GCCCAGTGCTCTGAATGTCAAAGTG | >hg38_21_+_8433172_8446622_RNA45SN1_45S_50ntFlanks | 11669 | 11693 | 25 | rRF 45S | NO | 0.00 | 30.98 | 0.00 | 0.00 |
| >hg38_21_+_8433172_8446622_RNA45SN1_45S_50ntFlanks@11669.11702.34\|GCCCAGTGCTCTGAATGTCAAAGTGAAGAAATTC | >hg38_21_+_8433172_8446622_RNA45SN1_45S_50ntFlanks | 11669 | 11702 | 34 | rRF 45S | NO | 52.86 | 255.81 | 0.00 | 0.00 |
| >hg38_21_+_8433172_8446622_RNA45SN1_45S_50ntFlanks@11669.11703.35\|GCCCAGTGCTCTGAATGTCAAAGTGAAGAAATTCA | >hg38_21_+_8433172_8446622_RNA45SN1_45S_50ntFlanks | 11669 | 11703 | 35 | rRF 45S | NO | 0.00 | 99.00 | 0.00 | 0.00 |
| >hg38_21_+_8433172_8446622_RNA45SN1_45S_50ntFlanks@11669.11704.36\|GCCCAGTGCTCTGAATGTCAAAGTGAAGAAATTCAA | >hg38_21_+_8433172_8446622_RNA45SN1_45S_50ntFlanks | 11669 | 11704 | 36 | rRF 45S | NO | 0.00 | 27.48 | 0.00 | 0.00 |
| >hg38_21_+_8433172_8446622_RNA45SN1_45S_50ntFlanks@11669.11705.37\|GCCCAGTGCTCTGAATGTCAAAGTGAAGAAATTCAAT | >hg38_21_+_8433172_8446622_RNA45SN1_45S_50ntFlanks | 11669 | 11705 | 37 | rRF 45S | NO | 0.00 | 13.08 | 0.00 | 0.00 |
| >hg38_21_+_8433172_8446622_RNA45SN1_45S_50ntFlanks@11669.11706.38\|GCCCAGTGCTCTGAATGTCAAAGTGAAGAAATTCAATG | >hg38_21_+_8433172_8446622_RNA45SN1_45S_50ntFlanks | 11669 | 11706 | 38 | rRF 45S | NO | 0.00 | 189.44 | 0.00 | 0.00 |
| >hg38_21_+_8433172_8446622_RNA45SN1_45S_50ntFlanks@11669.11707.39\|GCCCAGTGCTCTGAATGTCAAAGTGAAGAAATTCAATGA | >hg38_21_+_8433172_8446622_RNA45SN1_45S_50ntFlanks | 11669 | 11707 | 39 | rRF 45S | NO | 0.00 | 122.22 | 0.00 | 0.00 |
| >hg38_21_+_8433172_8446622_RNA45SN1_45S_50ntFlanks@11669.11709.41\|GCCCAGTGCTCTGAATGTCAAAGTGAAGAAATTCAATGAAG | >hg38_21_+_8433172_8446622_RNA45SN1_45S_50ntFlanks | 11669 | 11709 | 41 | rRF 45S | NO | 0.00 | 74.05 | 0.00 | 0.00 |

### MDA-MB-231

| | | | | | | | | | | |
|---|---|---|---|---|---|---|---|---|---|---|
| >hg38_21_+_8433172_8446622_RNA45SN1_45S_50ntFlanks@11669.11710.42\|GCCCAGTGCT CTGAATGTCAAAGTGAAGAAATTCAATGAAGC | >hg38_21_+_8433172_8446622_RNA45SN1_45S_50ntFlanks | 11669 | 11710 | 42 | rRF 45S | NO | 0.00 | 57.02 | 0.00 | 0.00 |
| >hg38_21_+_8433172_8446622_RNA45SN1_45S_50ntFlanks@11672.11702.31\|CAGTGCTCTG AATGTCAAAGTGAAGAAATTC | >hg38_21_+_8433172_8446622_RNA45SN1_45S_50ntFlanks | 11672 | 11702 | 31 | rRF 45S | NO | 26.32 | 25.53 | 0.00 | 0.00 |
| >hg38_21_+_8433172_8446622_RNA45SN1_45S_50ntFlanks@11672.11706.35\|CAGTGCTCTG AATGTCAAAGTGAAGAAATTCAATG | >hg38_21_+_8433172_8446622_RNA45SN1_45S_50ntFlanks | 11672 | 11706 | 35 | rRF 45S | NO | 0.00 | 30.30 | 0.00 | 0.00 |
| >hg38_21_+_8433172_8446622_RNA45SN1_45S_50ntFlanks@11672.11707.36\|CAGTGCTCTG AATGTCAAAGTGAAGAAATTCAATGA | >hg38_21_+_8433172_8446622_RNA45SN1_45S_50ntFlanks | 11672 | 11707 | 36 | rRF 45S | NO | 0.00 | 33.51 | 0.00 | 0.00 |
| >hg38_21_+_8433172_8446622_RNA45SN1_45S_50ntFlanks@11672.11709.38\|CAGTGCTCTG AATGTCAAAGTGAAGAAATTCAATGAAG | >hg38_21_+_8433172_8446622_RNA45SN1_45S_50ntFlanks | 11672 | 11709 | 38 | rRF 45S | NO | 0.00 | 32.57 | 0.00 | 0.00 |
| >hg38_21_+_8433172_8446622_RNA45SN1_45S_50ntFlanks@11675.11702.28\|TGCTCTGAAT GTCAAAGTGAAGAAATTC | >hg38_21_+_8433172_8446622_RNA45SN1_45S_50ntFlanks | 11675 | 11702 | 28 | rRF 45S | NO | 59.21 | 75.42 | 0.00 | 0.00 |
| >hg38_21_+_8433172_8446622_RNA45SN1_45S_50ntFlanks@11675.11706.32\|TGCTCTGAAT GTCAAAGTGAAGAAATTCAATG | >hg38_21_+_8433172_8446622_RNA45SN1_45S_50ntFlanks | 11675 | 11706 | 32 | rRF 45S | NO | 0.00 | 56.22 | 0.00 | 0.00 |
| >hg38_21_+_8433172_8446622_RNA45SN1_45S_50ntFlanks@11675.11707.33\|TGCTCTGAAT GTCAAAGTGAAGAAATTCAATGA | >hg38_21_+_8433172_8446622_RNA45SN1_45S_50ntFlanks | 11675 | 11707 | 33 | rRF 45S | NO | 0.00 | 30.63 | 0.00 | 0.00 |
| >hg38_21_+_8433172_8446622_RNA45SN1_45S_50ntFlanks@11675.11709.35\|TGCTCTGAAT GTCAAAGTGAAGAAATTCAATGAAG | >hg38_21_+_8433172_8446622_RNA45SN1_45S_50ntFlanks | 11675 | 11709 | 35 | rRF 45S | NO | 0.00 | 26.42 | 0.00 | 0.00 |
| >hg38_21_+_8433172_8446622_RNA45SN1_45S_50ntFlanks@11703.11736.34\|AATGAAGCGC GGGTAAACGGCGGGAGTAACTATG | >hg38_21_+_8433172_8446622_RNA45SN1_45S_50ntFlanks | 11703 | 11736 | 34 | rRF 45S | NO | 0.00 | 25.32 | 0.00 | 0.00 |
| >hg38_21_+_8433172_8446622_RNA45SN1_45S_50ntFlanks@11710.11736.27\|CGCGGGTAAA CGGCGGGAGTAACTATG | >hg38_21_+_8433172_8446622_RNA45SN1_45S_50ntFlanks | 11710 | 11736 | 27 | rRF 45S | NO | 0.00 | 13.58 | 0.00 | 0.00 |
| >hg38_21_+_8433172_8446622_RNA45SN1_45S_50ntFlanks@11727.11744.18\|AGTAACTATG ACTCTCTT | >hg38_21_+_8433172_8446622_RNA45SN1_45S_50ntFlanks | 11727 | 11744 | 18 | rRF 45S | NO | 0.00 | 30.23 | 0.00 | 0.00 |
| >hg38_21_+_8433172_8446622_RNA45SN1_45S_50ntFlanks@11727.11745.19\|AGTAACTATG ACTCTCTTA | >hg38_21_+_8433172_8446622_RNA45SN1_45S_50ntFlanks | 11727 | 11745 | 19 | rRF 45S | NO | 0.00 | 13.01 | 0.00 | 0.00 |
| >hg38_21_+_8433172_8446622_RNA45SN1_45S_50ntFlanks@11727.11748.22\|AGTAACTATG ACTCTCTTAAGG | >hg38_21_+_8433172_8446622_RNA45SN1_45S_50ntFlanks | 11727 | 11748 | 22 | rRF 45S | NO | 0.00 | 13.08 | 0.00 | 0.00 |
| >hg38_21_+_8433172_8446622_RNA45SN1_45S_50ntFlanks@11729.11747.19\|TAACTATGAC TCTCTTAAG | >hg38_21_+_8433172_8446622_RNA45SN1_45S_50ntFlanks | 11729 | 11747 | 19 | rRF 45S | NO | 0.00 | 13.32 | 0.00 | 0.00 |
| >hg38_21_+_8433172_8446622_RNA45SN1_45S_50ntFlanks@11729.11748.20\|TAACTATGAC TCTCTTAAGG | >hg38_21_+_8433172_8446622_RNA45SN1_45S_50ntFlanks | 11729 | 11748 | 20 | rRF 45S | NO | 0.00 | 31.59 | 0.00 | 0.00 |
| >hg38_21_+_8433172_8446622_RNA45SN1_45S_50ntFlanks@11769.11794.26\|TAATTAGTGA CGCGCATGAATGGATG | >hg38_21_+_8433172_8446622_RNA45SN1_45S_50ntFlanks | 11769 | 11794 | 26 | rRF 45S | NO | 0.00 | 31.30 | 0.00 | 0.00 |
| >hg38_21_+_8433172_8446622_RNA45SN1_45S_50ntFlanks@11770.11794.25\|AATTAGTGAC GCGCATGAATGGATG | >hg38_21_+_8433172_8446622_RNA45SN1_45S_50ntFlanks | 11770 | 11794 | 25 | rRF 45S | NO | 0.00 | 99.50 | 0.00 | 0.00 |
| >hg38_21_+_8433172_8446622_RNA45SN1_45S_50ntFlanks@11771.11794.24\|ATTAGTGACG CGCATGAATGGATG | >hg38_21_+_8433172_8446622_RNA45SN1_45S_50ntFlanks | 11771 | 11794 | 24 | rRF 45S | NO | 0.00 | 56.05 | 0.00 | 0.00 |
| >hg38_21_+_8433172_8446622_RNA45SN1_45S_50ntFlanks@11772.11794.23\|TTAGTGACGC GCATGAATGGATG | >hg38_21_+_8433172_8446622_RNA45SN1_45S_50ntFlanks | 11772 | 11794 | 23 | rRF 45S | NO | 0.00 | 26.87 | 0.00 | 0.00 |
| >hg38_21_+_8433172_8446622_RNA45SN1_45S_50ntFlanks@11774.11794.21\|AGTGACGCGC ATGAATGGATG | >hg38_21_+_8433172_8446622_RNA45SN1_45S_50ntFlanks | 11774 | 11794 | 21 | rRF 45S | NO | 0.00 | 56.88 | 0.00 | 0.00 |
| >hg38_21_+_8433172_8446622_RNA45SN1_45S_50ntFlanks@11922.11950.29\|GAGAGGTGTA GAATAAGTGGGAGGCCCCC | >hg38_21_+_8433172_8446622_RNA45SN1_45S_50ntFlanks | 11922 | 11950 | 29 | rRF 45S | NO | 0.00 | 126.35 | 0.00 | 0.00 |
| >hg38_21_+_8433172_8446622_RNA45SN1_45S_50ntFlanks@12003.12029.27\|GCGGGCCGC CGGTGAAATACCACTACT | >hg38_21_+_8433172_8446622_RNA45SN1_45S_50ntFlanks | 12003 | 12029 | 27 | rRF 45S | NO | 0.00 | 82.09 | 0.00 | 0.00 |
| >hg38_21_+_8433172_8446622_RNA45SN1_45S_50ntFlanks@12003.12032.30\|GCGGGCCGC CGGTGAAATACCACTACTCTG | >hg38_21_+_8433172_8446622_RNA45SN1_45S_50ntFlanks | 12003 | 12032 | 30 | rRF 45S | NO | 0.00 | 13.45 | 0.00 | 0.00 |
| >hg38_21_+_8433172_8446622_RNA45SN1_45S_50ntFlanks@12003.12033.31\|GCGGGCCGC CGGTGAAATACCACTACTCTGA | >hg38_21_+_8433172_8446622_RNA45SN1_45S_50ntFlanks | 12003 | 12033 | 31 | rRF 45S | NO | 0.00 | 64.59 | 0.00 | 0.00 |
| >hg38_21_+_8433172_8446622_RNA45SN1_45S_50ntFlanks@12003.12034.32\|GCGGGCCGC CGGTGAAATACCACTACTCTGAT | >hg38_21_+_8433172_8446622_RNA45SN1_45S_50ntFlanks | 12003 | 12034 | 32 | rRF 45S | NO | 10.43 | 89.47 | 0.00 | 0.00 |
| >hg38_21_+_8433172_8446622_RNA45SN1_45S_50ntFlanks@12004.12029.26\|CGGGCCGCC GGTGAAATACCACTACT | >hg38_21_+_8433172_8446622_RNA45SN1_45S_50ntFlanks | 12004 | 12029 | 26 | rRF 45S | NO | 0.00 | 37.65 | 0.00 | 0.00 |
| >hg38_21_+_8433172_8446622_RNA45SN1_45S_50ntFlanks@12076.12095.20\|AGGGGCTCTC GCTTCTGGCG | >hg38_21_+_8433172_8446622_RNA45SN1_45S_50ntFlanks | 12076 | 12095 | 20 | rRF 45S | NO | 0.00 | 13.02 | 0.00 | 0.00 |
| >hg38_21_+_8433172_8446622_RNA45SN1_45S_50ntFlanks@12251.12271.21\|AAAGCTCGCT TGATCTTGATT | >hg38_21_+_8433172_8446622_RNA45SN1_45S_50ntFlanks | 12251 | 12271 | 21 | rRF 45S | NO | 0.00 | 13.13 | 0.00 | 0.00 |
| >hg38_21_+_8433172_8446622_RNA45SN1_45S_50ntFlanks@12251.12289.39\|AAAGCTCGCT TGATCTTGATTTTCAGTACGAATACAGAC | >hg38_21_+_8433172_8446622_RNA45SN1_45S_50ntFlanks | 12251 | 12289 | 39 | rRF 45S | NO | 0.00 | 24.85 | 0.00 | 0.00 |
| >hg38_21_+_8433172_8446622_RNA45SN1_45S_50ntFlanks@12388.12433.46\|CGTTCATAGC GACGTCGCTTTTTGATCCTTCGATGTCGGCTCTTCC | >hg38_21_+_8433172_8446622_RNA45SN1_45S_50ntFlanks | 12388 | 12433 | 46 | rRF 45S | NO | 0.00 | 141.14 | 0.00 | 0.00 |
| >hg38_21_+_8433172_8446622_RNA45SN1_45S_50ntFlanks@12388.12434.47\|CGTTCATAGC GACGTCGCTTTTTGATCCTTCGATGTCGGCTCTTCCT | >hg38_21_+_8433172_8446622_RNA45SN1_45S_50ntFlanks | 12388 | 12434 | 47 | rRF 45S | NO | 0.00 | 61.94 | 0.00 | 0.00 |
| >hg38_21_+_8433172_8446622_RNA45SN1_45S_50ntFlanks@12394.12414.21\|TAGCGACGTC GCTTTTTGATC | >hg38_21_+_8433172_8446622_RNA45SN1_45S_50ntFlanks | 12394 | 12414 | 21 | rRF 45S | NO | 0.00 | 62.69 | 0.00 | 0.00 |
| >hg38_21_+_8433172_8446622_RNA45SN1_45S_50ntFlanks@12395.12414.20\|AGCGACGTCG CTTTTTGATC | >hg38_21_+_8433172_8446622_RNA45SN1_45S_50ntFlanks | 12395 | 12414 | 20 | rRF 45S | NO | 49.50 | 80.45 | 0.00 | 0.00 |
| >hg38_21_+_8433172_8446622_RNA45SN1_45S_50ntFlanks@12396.12414.19\|GCGACGTCG CTTTTTGATC | >hg38_21_+_8433172_8446622_RNA45SN1_45S_50ntFlanks | 12396 | 12414 | 19 | rRF 45S | NO | 0.00 | 124.13 | 0.00 | 0.00 |
| >hg38_21_+_8433172_8446622_RNA45SN1_45S_50ntFlanks@12396.12418.23\|GCGACGTCG CTTTTTGATCCTTC | >hg38_21_+_8433172_8446622_RNA45SN1_45S_50ntFlanks | 12396 | 12418 | 23 | rRF 45S | NO | 0.00 | 26.33 | 0.00 | 0.00 |
| >hg38_21_+_8433172_8446622_RNA45SN1_45S_50ntFlanks@12397.12414.18\|CGACGTCGCT TTTTGATC | >hg38_21_+_8433172_8446622_RNA45SN1_45S_50ntFlanks | 12397 | 12414 | 18 | rRF 45S | NO | 0.00 | 57.00 | 0.00 | 0.00 |
| >hg38_21_+_8433172_8446622_RNA45SN1_45S_50ntFlanks@12397.12418.22\|CGACGTCGCT TTTTGATCCTTC | >hg38_21_+_8433172_8446622_RNA45SN1_45S_50ntFlanks | 12397 | 12418 | 22 | rRF 45S | NO | 0.00 | 138.25 | 0.00 | 0.00 |

### MDA-MB-231

| | | | | | | | | | | |
|---|---|---|---|---|---|---|---|---|---|---|
| >hg38_21_+_8433172_8446622_RNA45SN1_45S_50ntFlanks@12407.12433.27\|TTTTGATCCTTCGATGTCGGCTCTTCC | >hg38_21_+_8433172_8446622_RNA45SN1_45S_50ntFlanks | 12407 | 12433 | 27 | rRF 45S | NO | 0.00 | 76.95 | 0.00 | 0.00 |
| >hg38_21_+_8433172_8446622_RNA45SN1_45S_50ntFlanks@12407.12434.28\|TTTTGATCCTTCGATGTCGGCTCTTCCT | >hg38_21_+_8433172_8446622_RNA45SN1_45S_50ntFlanks | 12407 | 12434 | 28 | rRF 45S | NO | 0.00 | 31.49 | 0.00 | 0.00 |
| >hg38_21_+_8433172_8446622_RNA45SN1_45S_50ntFlanks@12407.12437.31\|TTTTGATCCTTCGATGTCGGCTCTTCCTATC | >hg38_21_+_8433172_8446622_RNA45SN1_45S_50ntFlanks | 12407 | 12437 | 31 | rRF 45S | NO | 0.00 | 12.88 | 0.00 | 0.00 |
| >hg38_21_+_8433172_8446622_RNA45SN1_45S_50ntFlanks@12407.12448.42\|TTTTGATCCTTCGATGTCGGCTCTTCCTATCATTGTGAAGCA | >hg38_21_+_8433172_8446622_RNA45SN1_45S_50ntFlanks | 12407 | 12448 | 42 | rRF 45S | NO | 0.00 | 13.19 | 0.00 | 0.00 |
| >hg38_21_+_8433172_8446622_RNA45SN1_45S_50ntFlanks@12408.12433.26\|TTTGATCCTTCGATGTCGGCTCTTCC | >hg38_21_+_8433172_8446622_RNA45SN1_45S_50ntFlanks | 12408 | 12433 | 26 | rRF 45S | NO | 0.00 | 34.62 | 0.00 | 0.00 |
| >hg38_21_+_8433172_8446622_RNA45SN1_45S_50ntFlanks@12409.12433.25\|TTGATCCTTCGATGTCGGCTCTTCC | >hg38_21_+_8433172_8446622_RNA45SN1_45S_50ntFlanks | 12409 | 12433 | 25 | rRF 45S | NO | 0.00 | 32.79 | 0.00 | 0.00 |
| >hg38_21_+_8433172_8446622_RNA45SN1_45S_50ntFlanks@12410.12433.24\|TGATCCTTCGATGTCGGCTCTTCC | >hg38_21_+_8433172_8446622_RNA45SN1_45S_50ntFlanks | 12410 | 12433 | 24 | rRF 45S | NO | 0.00 | 25.43 | 0.00 | 0.00 |
| >hg38_21_+_8433172_8446622_RNA45SN1_45S_50ntFlanks@12414.12433.20\|CCTTCGATGTCGGCTCTTCC | >hg38_21_+_8433172_8446622_RNA45SN1_45S_50ntFlanks | 12414 | 12433 | 20 | rRF 45S | NO | 0.00 | 33.45 | 0.00 | 0.00 |
| >hg38_21_+_8433172_8446622_RNA45SN1_45S_50ntFlanks@12414.12443.30\|CCTTCGATGTCGGCTCTTCCTATCATTGTG | >hg38_21_+_8433172_8446622_RNA45SN1_45S_50ntFlanks | 12414 | 12443 | 30 | rRF 45S | NO | 0.00 | 27.15 | 0.00 | 0.00 |
| >hg38_21_+_8433172_8446622_RNA45SN1_45S_50ntFlanks@12414.12444.31\|CCTTCGATGTCGGCTCTTCCTATCATTGTGA | >hg38_21_+_8433172_8446622_RNA45SN1_45S_50ntFlanks | 12414 | 12444 | 31 | rRF 45S | NO | 0.00 | 12.89 | 0.00 | 0.00 |
| >hg38_21_+_8433172_8446622_RNA45SN1_45S_50ntFlanks@12414.12447.34\|CCTTCGATGTCGGCTCTTCCTATCATTGTGAAGC | >hg38_21_+_8433172_8446622_RNA45SN1_45S_50ntFlanks | 12414 | 12447 | 34 | rRF 45S | NO | 0.00 | 13.38 | 0.00 | 0.00 |
| >hg38_21_+_8433172_8446622_RNA45SN1_45S_50ntFlanks@12414.12448.35\|CCTTCGATGTCGGCTCTTCCTATCATTGTGAAGCA | >hg38_21_+_8433172_8446622_RNA45SN1_45S_50ntFlanks | 12414 | 12448 | 35 | rRF 45S | NO | 0.00 | 64.00 | 0.00 | 0.00 |
| >hg38_21_+_8433172_8446622_RNA45SN1_45S_50ntFlanks@12415.12432.18\|CTTCGATGTCGGCTCTTC | >hg38_21_+_8433172_8446622_RNA45SN1_45S_50ntFlanks | 12415 | 12432 | 18 | rRF 45S | NO | 0.00 | 115.30 | 0.00 | 0.00 |
| >hg38_21_+_8433172_8446622_RNA45SN1_45S_50ntFlanks@12415.12433.19\|CTTCGATGTCGGCTCTTCC | >hg38_21_+_8433172_8446622_RNA45SN1_45S_50ntFlanks | 12415 | 12433 | 19 | rRF 45S | NO | 10.49 | 177.37 | 0.00 | 0.00 |
| >hg38_21_+_8433172_8446622_RNA45SN1_45S_50ntFlanks@12415.12434.20\|CTTCGATGTCGGCTCTTCCT | >hg38_21_+_8433172_8446622_RNA45SN1_45S_50ntFlanks | 12415 | 12434 | 20 | rRF 45S | NO | 54.34 | 150.76 | 0.00 | 0.00 |
| >hg38_21_+_8433172_8446622_RNA45SN1_45S_50ntFlanks@12415.12437.23\|CTTCGATGTCGGCTCTTCCTATC | >hg38_21_+_8433172_8446622_RNA45SN1_45S_50ntFlanks | 12415 | 12437 | 23 | rRF 45S | NO | 0.00 | 107.45 | 0.00 | 0.00 |
| >hg38_21_+_8433172_8446622_RNA45SN1_45S_50ntFlanks@12415.12438.24\|CTTCGATGTCGGCTCTTCCTATCA | >hg38_21_+_8433172_8446622_RNA45SN1_45S_50ntFlanks | 12415 | 12438 | 24 | rRF 45S | NO | 0.00 | 58.33 | 0.00 | 0.00 |
| >hg38_21_+_8433172_8446622_RNA45SN1_45S_50ntFlanks@12415.12439.25\|CTTCGATGTCGGCTCTTCCTATCAT | >hg38_21_+_8433172_8446622_RNA45SN1_45S_50ntFlanks | 12415 | 12439 | 25 | rRF 45S | NO | 0.00 | 33.81 | 0.00 | 0.00 |
| >hg38_21_+_8433172_8446622_RNA45SN1_45S_50ntFlanks@12415.12440.26\|CTTCGATGTCGGCTCTTCCTATCATT | >hg38_21_+_8433172_8446622_RNA45SN1_45S_50ntFlanks | 12415 | 12440 | 26 | rRF 45S | NO | 0.00 | 64.94 | 0.00 | 0.00 |
| >hg38_21_+_8433172_8446622_RNA45SN1_45S_50ntFlanks@12415.12441.27\|CTTCGATGTCGGCTCTTCCTATCATTG | >hg38_21_+_8433172_8446622_RNA45SN1_45S_50ntFlanks | 12415 | 12441 | 27 | rRF 45S | NO | 0.00 | 57.46 | 0.00 | 0.00 |
| >hg38_21_+_8433172_8446622_RNA45SN1_45S_50ntFlanks@12415.12442.28\|CTTCGATGTCGGCTCTTCCTATCATTGT | >hg38_21_+_8433172_8446622_RNA45SN1_45S_50ntFlanks | 12415 | 12442 | 28 | rRF 45S | NO | 0.00 | 78.30 | 0.00 | 0.00 |
| >hg38_21_+_8433172_8446622_RNA45SN1_45S_50ntFlanks@12415.12443.29\|CTTCGATGTCGGCTCTTCCTATCATTGTG | >hg38_21_+_8433172_8446622_RNA45SN1_45S_50ntFlanks | 12415 | 12443 | 29 | rRF 45S | NO | 0.00 | 81.87 | 0.00 | 0.00 |
| >hg38_21_+_8433172_8446622_RNA45SN1_45S_50ntFlanks@12415.12444.30\|CTTCGATGTCGGCTCTTCCTATCATTGTGA | >hg38_21_+_8433172_8446622_RNA45SN1_45S_50ntFlanks | 12415 | 12444 | 30 | rRF 45S | NO | 0.00 | 83.24 | 0.00 | 0.00 |
| >hg38_21_+_8433172_8446622_RNA45SN1_45S_50ntFlanks@12415.12446.32\|CTTCGATGTCGGCTCTTCCTATCATTGTGAAG | >hg38_21_+_8433172_8446622_RNA45SN1_45S_50ntFlanks | 12415 | 12446 | 32 | rRF 45S | NO | 0.00 | 34.75 | 0.00 | 0.00 |
| >hg38_21_+_8433172_8446622_RNA45SN1_45S_50ntFlanks@12415.12447.33\|CTTCGATGTCGGCTCTTCCTATCATTGTGAAGC | >hg38_21_+_8433172_8446622_RNA45SN1_45S_50ntFlanks | 12415 | 12447 | 33 | rRF 45S | NO | 0.00 | 76.47 | 0.00 | 0.00 |
| >hg38_21_+_8433172_8446622_RNA45SN1_45S_50ntFlanks@12415.12448.34\|CTTCGATGTCGGCTCTTCCTATCATTGTGAAGCA | >hg38_21_+_8433172_8446622_RNA45SN1_45S_50ntFlanks | 12415 | 12448 | 34 | rRF 45S | NO | 0.00 | 153.32 | 0.00 | 14.36 |
| >hg38_21_+_8433172_8446622_RNA45SN1_45S_50ntFlanks@12415.12449.35\|CTTCGATGTCGGCTCTTCCTATCATTGTGAAGCAG | >hg38_21_+_8433172_8446622_RNA45SN1_45S_50ntFlanks | 12415 | 12449 | 35 | rRF 45S | NO | 0.00 | 66.02 | 0.00 | 0.00 |
| >hg38_21_+_8433172_8446622_RNA45SN1_45S_50ntFlanks@12416.12433.18\|TTCGATGTCGGCTCTTCC | >hg38_21_+_8433172_8446622_RNA45SN1_45S_50ntFlanks | 12416 | 12433 | 18 | rRF 45S | NO | 45.49 | 468.99 | 0.00 | 0.00 |
| >hg38_21_+_8433172_8446622_RNA45SN1_45S_50ntFlanks@12416.12434.19\|TTCGATGTCGGCTCTTCCT | >hg38_21_+_8433172_8446622_RNA45SN1_45S_50ntFlanks | 12416 | 12434 | 19 | rRF 45S | NO | 47.18 | 332.71 | 0.00 | 0.00 |
| >hg38_21_+_8433172_8446622_RNA45SN1_45S_50ntFlanks@12416.12435.20\|TTCGATGTCGGCTCTTCCTA | >hg38_21_+_8433172_8446622_RNA45SN1_45S_50ntFlanks | 12416 | 12435 | 20 | rRF 45S | NO | 0.00 | 27.26 | 0.00 | 0.00 |
| >hg38_21_+_8433172_8446622_RNA45SN1_45S_50ntFlanks@12416.12436.21\|TTCGATGTCGGCTCTTCCTAT | >hg38_21_+_8433172_8446622_RNA45SN1_45S_50ntFlanks | 12416 | 12436 | 21 | rRF 45S | NO | 0.00 | 67.03 | 0.00 | 0.00 |
| >hg38_21_+_8433172_8446622_RNA45SN1_45S_50ntFlanks@12416.12437.22\|TTCGATGTCGGCTCTTCCTATC | >hg38_21_+_8433172_8446622_RNA45SN1_45S_50ntFlanks | 12416 | 12437 | 22 | rRF 45S | NO | 68.93 | 360.55 | 0.00 | 12.48 |
| >hg38_21_+_8433172_8446622_RNA45SN1_45S_50ntFlanks@12416.12438.23\|TTCGATGTCGGCTCTTCCTATCA | >hg38_21_+_8433172_8446622_RNA45SN1_45S_50ntFlanks | 12416 | 12438 | 23 | rRF 45S | NO | 57.40 | 215.04 | 0.00 | 0.00 |
| >hg38_21_+_8433172_8446622_RNA45SN1_45S_50ntFlanks@12416.12439.24\|TTCGATGTCGGCTCTTCCTATCAT | >hg38_21_+_8433172_8446622_RNA45SN1_45S_50ntFlanks | 12416 | 12439 | 24 | rRF 45S | NO | 0.00 | 169.65 | 0.00 | 0.00 |
| >hg38_21_+_8433172_8446622_RNA45SN1_45S_50ntFlanks@12416.12440.25\|TTCGATGTCGGCTCTTCCTATCATT | >hg38_21_+_8433172_8446622_RNA45SN1_45S_50ntFlanks | 12416 | 12440 | 25 | rRF 45S | NO | 24.28 | 216.05 | 0.00 | 0.00 |
| >hg38_21_+_8433172_8446622_RNA45SN1_45S_50ntFlanks@12416.12441.26\|TTCGATGTCGGCTCTTCCTATCATTG | >hg38_21_+_8433172_8446622_RNA45SN1_45S_50ntFlanks | 12416 | 12441 | 26 | rRF 45S | NO | 0.00 | 131.27 | 0.00 | 0.00 |
| >hg38_21_+_8433172_8446622_RNA45SN1_45S_50ntFlanks@12416.12442.27\|TTCGATGTCGGCTCTTCCTATCATTGT | >hg38_21_+_8433172_8446622_RNA45SN1_45S_50ntFlanks | 12416 | 12442 | 27 | rRF 45S | NO | 0.00 | 166.15 | 0.00 | 0.00 |
| >hg38_21_+_8433172_8446622_RNA45SN1_45S_50ntFlanks@12416.12443.28\|TTCGATGTCGGCTCTTCCTATCATTGTG | >hg38_21_+_8433172_8446622_RNA45SN1_45S_50ntFlanks | 12416 | 12443 | 28 | rRF 45S | NO | 0.00 | 145.03 | 0.00 | 0.00 |

### MDA-MB-231

| | | | | | | | | | | |
|---|---|---|---|---|---|---|---|---|---|---|
| >hg38_21_+_8433172_8446622_RNA45SN1_45S_50ntFlanks@12416.12444.29\|TTCGATGTCGGCTCTTCCTATCATTGTGA | >hg38_21_+_8433172_8446622_RNA45SN1_45S_50ntFlanks | 12416 | 12444 | 29 | rRF 45S | NO | 10.62 | 199.04 | 0.00 | 0.00 |
| >hg38_21_+_8433172_8446622_RNA45SN1_45S_50ntFlanks@12416.12445.30\|TTCGATGTCGGCTCTTCCTATCATTGTGAA | >hg38_21_+_8433172_8446622_RNA45SN1_45S_50ntFlanks | 12416 | 12445 | 30 | rRF 45S | NO | 0.00 | 81.47 | 0.00 | 0.00 |
| >hg38_21_+_8433172_8446622_RNA45SN1_45S_50ntFlanks@12416.12446.31\|TTCGATGTCGGCTCTTCCTATCATTGTGAAG | >hg38_21_+_8433172_8446622_RNA45SN1_45S_50ntFlanks | 12416 | 12446 | 31 | rRF 45S | NO | 0.00 | 175.90 | 0.00 | 0.00 |
| >hg38_21_+_8433172_8446622_RNA45SN1_45S_50ntFlanks@12416.12447.32\|TTCGATGTCGGCTCTTCCTATCATTGTGAAGC | >hg38_21_+_8433172_8446622_RNA45SN1_45S_50ntFlanks | 12416 | 12447 | 32 | rRF 45S | NO | 0.00 | 154.34 | 0.00 | 0.00 |
| >hg38_21_+_8433172_8446622_RNA45SN1_45S_50ntFlanks@12416.12448.33\|TTCGATGTCGGCTCTTCCTATCATTGTGAAGCA | >hg38_21_+_8433172_8446622_RNA45SN1_45S_50ntFlanks | 12416 | 12448 | 33 | rRF 45S | NO | 55.07 | 317.71 | 0.00 | 17.41 |
| >hg38_21_+_8433172_8446622_RNA45SN1_45S_50ntFlanks@12416.12449.34\|TTCGATGTCGGCTCTTCCTATCATTGTGAAGCAG | >hg38_21_+_8433172_8446622_RNA45SN1_45S_50ntFlanks | 12416 | 12449 | 34 | rRF 45S | NO | 0.00 | 80.82 | 0.00 | 12.35 |
| >hg38_21_+_8433172_8446622_RNA45SN1_45S_50ntFlanks@12416.12452.37\|TTCGATGTCGGCTCTTCCTATCATTGTGAAGCAGAAT | >hg38_21_+_8433172_8446622_RNA45SN1_45S_50ntFlanks | 12416 | 12452 | 37 | rRF 45S | NO | 0.00 | 28.35 | 0.00 | 0.00 |
| >hg38_21_+_8433172_8446622_RNA45SN1_45S_50ntFlanks@12416.12453.38\|TTCGATGTCGGCTCTTCCTATCATTGTGAAGCAGAATT | >hg38_21_+_8433172_8446622_RNA45SN1_45S_50ntFlanks | 12416 | 12453 | 38 | rRF 45S | NO | 0.00 | 27.01 | 0.00 | 0.00 |
| >hg38_21_+_8433172_8446622_RNA45SN1_45S_50ntFlanks@12416.12454.39\|TTCGATGTCGGCTCTTCCTATCATTGTGAAGCAGAATTC | >hg38_21_+_8433172_8446622_RNA45SN1_45S_50ntFlanks | 12416 | 12454 | 39 | rRF 45S | NO | 0.00 | 27.01 | 0.00 | 0.00 |
| >hg38_21_+_8433172_8446622_RNA45SN1_45S_50ntFlanks@12416.12457.42\|TTCGATGTCGGCTCTTCCTATCATTGTGAAGCAGAATTCACC | >hg38_21_+_8433172_8446622_RNA45SN1_45S_50ntFlanks | 12416 | 12457 | 42 | rRF 45S | NO | 0.00 | 159.41 | 0.00 | 0.00 |
| >hg38_21_+_8433172_8446622_RNA45SN1_45S_50ntFlanks@12416.12458.43\|TTCGATGTCGGCTCTTCCTATCATTGTGAAGCAGAATTCACCA | >hg38_21_+_8433172_8446622_RNA45SN1_45S_50ntFlanks | 12416 | 12458 | 43 | rRF 45S | NO | 0.00 | 60.55 | 0.00 | 0.00 |
| >hg38_21_+_8433172_8446622_RNA45SN1_45S_50ntFlanks@12416.12462.47\|TTCGATGTCGGCTCTTCCTATCATTGTGAAGCAGAATTCACCAAGCG | >hg38_21_+_8433172_8446622_RNA45SN1_45S_50ntFlanks | 12416 | 12462 | 47 | rRF 45S | NO | 0.00 | 29.11 | 0.00 | 0.00 |
| >hg38_21_+_8433172_8446622_RNA45SN1_45S_50ntFlanks@12417.12434.18\|TCGATGTCGGCTCTTCCT | >hg38_21_+_8433172_8446622_RNA45SN1_45S_50ntFlanks | 12417 | 12434 | 18 | rRF 45S | NO | 0.00 | 150.91 | 0.00 | 0.00 |
| >hg38_21_+_8433172_8446622_RNA45SN1_45S_50ntFlanks@12417.12437.21\|TCGATGTCGGCTCTTCCTATC | >hg38_21_+_8433172_8446622_RNA45SN1_45S_50ntFlanks | 12417 | 12437 | 21 | rRF 45S | NO | 0.00 | 151.48 | 0.00 | 0.00 |
| >hg38_21_+_8433172_8446622_RNA45SN1_45S_50ntFlanks@12417.12438.22\|TCGATGTCGGCTCTTCCTATCA | >hg38_21_+_8433172_8446622_RNA45SN1_45S_50ntFlanks | 12417 | 12438 | 22 | rRF 45S | NO | 0.00 | 96.96 | 0.00 | 0.00 |
| >hg38_21_+_8433172_8446622_RNA45SN1_45S_50ntFlanks@12417.12439.23\|TCGATGTCGGCTCTTCCTATCAT | >hg38_21_+_8433172_8446622_RNA45SN1_45S_50ntFlanks | 12417 | 12439 | 23 | rRF 45S | NO | 0.00 | 68.17 | 0.00 | 0.00 |
| >hg38_21_+_8433172_8446622_RNA45SN1_45S_50ntFlanks@12417.12440.24\|TCGATGTCGGCTCTTCCTATCATT | >hg38_21_+_8433172_8446622_RNA45SN1_45S_50ntFlanks | 12417 | 12440 | 24 | rRF 45S | NO | 0.00 | 81.74 | 0.00 | 0.00 |
| >hg38_21_+_8433172_8446622_RNA45SN1_45S_50ntFlanks@12417.12442.26\|TCGATGTCGGCTCTTCCTATCATTGT | >hg38_21_+_8433172_8446622_RNA45SN1_45S_50ntFlanks | 12417 | 12442 | 26 | rRF 45S | NO | 0.00 | 60.31 | 0.00 | 0.00 |
| >hg38_21_+_8433172_8446622_RNA45SN1_45S_50ntFlanks@12417.12443.27\|TCGATGTCGGCTCTTCCTATCATTGTG | >hg38_21_+_8433172_8446622_RNA45SN1_45S_50ntFlanks | 12417 | 12443 | 27 | rRF 45S | NO | 0.00 | 79.18 | 0.00 | 0.00 |
| >hg38_21_+_8433172_8446622_RNA45SN1_45S_50ntFlanks@12417.12444.28\|TCGATGTCGGCTCTTCCTATCATTGTGA | >hg38_21_+_8433172_8446622_RNA45SN1_45S_50ntFlanks | 12417 | 12444 | 28 | rRF 45S | NO | 0.00 | 84.31 | 0.00 | 0.00 |
| >hg38_21_+_8433172_8446622_RNA45SN1_45S_50ntFlanks@12417.12445.29\|TCGATGTCGGCTCTTCCTATCATTGTGAA | >hg38_21_+_8433172_8446622_RNA45SN1_45S_50ntFlanks | 12417 | 12445 | 29 | rRF 45S | NO | 0.00 | 26.58 | 0.00 | 0.00 |
| >hg38_21_+_8433172_8446622_RNA45SN1_45S_50ntFlanks@12417.12446.30\|TCGATGTCGGCTCTTCCTATCATTGTGAAG | >hg38_21_+_8433172_8446622_RNA45SN1_45S_50ntFlanks | 12417 | 12446 | 30 | rRF 45S | NO | 0.00 | 64.96 | 0.00 | 0.00 |
| >hg38_21_+_8433172_8446622_RNA45SN1_45S_50ntFlanks@12417.12447.31\|TCGATGTCGGCTCTTCCTATCATTGTGAAGC | >hg38_21_+_8433172_8446622_RNA45SN1_45S_50ntFlanks | 12417 | 12447 | 31 | rRF 45S | NO | 0.00 | 72.99 | 0.00 | 0.00 |
| >hg38_21_+_8433172_8446622_RNA45SN1_45S_50ntFlanks@12417.12448.32\|TCGATGTCGGCTCTTCCTATCATTGTGAAGCA | >hg38_21_+_8433172_8446622_RNA45SN1_45S_50ntFlanks | 12417 | 12448 | 32 | rRF 45S | NO | 0.00 | 162.60 | 0.00 | 0.00 |
| >hg38_21_+_8433172_8446622_RNA45SN1_45S_50ntFlanks@12417.12457.41\|TCGATGTCGGCTCTTCCTATCATTGTGAAGCAGAATTCACC | >hg38_21_+_8433172_8446622_RNA45SN1_45S_50ntFlanks | 12417 | 12457 | 41 | rRF 45S | NO | 0.00 | 35.75 | 0.00 | 0.00 |
| >hg38_21_+_8433172_8446622_RNA45SN1_45S_50ntFlanks@12418.12437.20\|CGATGTCGGCTCTTCCTATC | >hg38_21_+_8433172_8446622_RNA45SN1_45S_50ntFlanks | 12418 | 12437 | 20 | rRF 45S | NO | 0.00 | 110.64 | 0.00 | 0.00 |
| >hg38_21_+_8433172_8446622_RNA45SN1_45S_50ntFlanks@12418.12438.21\|CGATGTCGGCTCTTCCTATCA | >hg38_21_+_8433172_8446622_RNA45SN1_45S_50ntFlanks | 12418 | 12438 | 21 | rRF 45S | NO | 0.00 | 34.94 | 0.00 | 0.00 |
| >hg38_21_+_8433172_8446622_RNA45SN1_45S_50ntFlanks@12418.12439.22\|CGATGTCGGCTCTTCCTATCAT | >hg38_21_+_8433172_8446622_RNA45SN1_45S_50ntFlanks | 12418 | 12439 | 22 | rRF 45S | NO | 0.00 | 32.02 | 0.00 | 0.00 |
| >hg38_21_+_8433172_8446622_RNA45SN1_45S_50ntFlanks@12418.12440.23\|CGATGTCGGCTCTTCCTATCATT | >hg38_21_+_8433172_8446622_RNA45SN1_45S_50ntFlanks | 12418 | 12440 | 23 | rRF 45S | NO | 0.00 | 31.57 | 0.00 | 0.00 |
| >hg38_21_+_8433172_8446622_RNA45SN1_45S_50ntFlanks@12418.12442.25\|CGATGTCGGCTCTTCCTATCATTGT | >hg38_21_+_8433172_8446622_RNA45SN1_45S_50ntFlanks | 12418 | 12442 | 25 | rRF 45S | NO | 0.00 | 34.48 | 0.00 | 0.00 |
| >hg38_21_+_8433172_8446622_RNA45SN1_45S_50ntFlanks@12418.12443.26\|CGATGTCGGCTCTTCCTATCATTGTG | >hg38_21_+_8433172_8446622_RNA45SN1_45S_50ntFlanks | 12418 | 12443 | 26 | rRF 45S | NO | 0.00 | 65.35 | 0.00 | 0.00 |
| >hg38_21_+_8433172_8446622_RNA45SN1_45S_50ntFlanks@12418.12444.27\|CGATGTCGGCTCTTCCTATCATTGTGA | >hg38_21_+_8433172_8446622_RNA45SN1_45S_50ntFlanks | 12418 | 12444 | 27 | rRF 45S | NO | 0.00 | 56.61 | 0.00 | 0.00 |
| >hg38_21_+_8433172_8446622_RNA45SN1_45S_50ntFlanks@12418.12446.29\|CGATGTCGGCTCTTCCTATCATTGTGAAG | >hg38_21_+_8433172_8446622_RNA45SN1_45S_50ntFlanks | 12418 | 12446 | 29 | rRF 45S | NO | 0.00 | 30.54 | 0.00 | 0.00 |
| >hg38_21_+_8433172_8446622_RNA45SN1_45S_50ntFlanks@12418.12447.30\|CGATGTCGGCTCTTCCTATCATTGTGAAGC | >hg38_21_+_8433172_8446622_RNA45SN1_45S_50ntFlanks | 12418 | 12447 | 30 | rRF 45S | NO | 0.00 | 30.19 | 0.00 | 0.00 |
| >hg38_21_+_8433172_8446622_RNA45SN1_45S_50ntFlanks@12418.12448.31\|CGATGTCGGCTCTTCCTATCATTGTGAAGCA | >hg38_21_+_8433172_8446622_RNA45SN1_45S_50ntFlanks | 12418 | 12448 | 31 | rRF 45S | NO | 0.00 | 113.78 | 0.00 | 0.00 |
| >hg38_21_+_8433172_8446622_RNA45SN1_45S_50ntFlanks@12418.12449.32\|CGATGTCGGCTCTTCCTATCATTGTGAAGCAG | >hg38_21_+_8433172_8446622_RNA45SN1_45S_50ntFlanks | 12418 | 12449 | 32 | rRF 45S | NO | 0.00 | 29.28 | 0.00 | 0.00 |
| >hg38_21_+_8433172_8446622_RNA45SN1_45S_50ntFlanks@12418.12457.40\|CGATGTCGGCTCTTCCTATCATTGTGAAGCAGAATTCACC | >hg38_21_+_8433172_8446622_RNA45SN1_45S_50ntFlanks | 12418 | 12457 | 40 | rRF 45S | NO | 0.00 | 73.22 | 0.00 | 0.00 |
| >hg38_21_+_8433172_8446622_RNA45SN1_45S_50ntFlanks@12418.12458.41\|CGATGTCGGCTCTTCCTATCATTGTGAAGCAGAATTCACCA | >hg38_21_+_8433172_8446622_RNA45SN1_45S_50ntFlanks | 12418 | 12458 | 41 | rRF 45S | NO | 0.00 | 26.24 | 0.00 | 0.00 |

### MDA-MB-231

| | | | | | | | | | | |
|---|---|---|---|---|---|---|---|---|---|---|
| >hg38_21_+_8433172_8446622_RNA45SN1_45S_50ntFlanks@12419.12436.18\|GATGTCGGCTCTTCCTAT | >hg38_21_+_8433172_8446622_RNA45SN1_45S_50ntFlanks | 12419 | 12436 | 18 | rRF 45S | NO | 0.00 | 59.07 | 0.00 | 0.00 |
| >hg38_21_+_8433172_8446622_RNA45SN1_45S_50ntFlanks@12419.12437.19\|GATGTCGGCTCTTCCTATC | >hg38_21_+_8433172_8446622_RNA45SN1_45S_50ntFlanks | 12419 | 12437 | 19 | rRF 45S | NO | 120.02 | 468.04 | 0.00 | 15.90 |
| >hg38_21_+_8433172_8446622_RNA45SN1_45S_50ntFlanks@12419.12438.20\|GATGTCGGCTCTTCCTATCA | >hg38_21_+_8433172_8446622_RNA45SN1_45S_50ntFlanks | 12419 | 12438 | 20 | rRF 45S | NO | 85.59 | 224.38 | 0.00 | 0.00 |
| >hg38_21_+_8433172_8446622_RNA45SN1_45S_50ntFlanks@12419.12439.21\|GATGTCGGCTCTTCCTATCAT | >hg38_21_+_8433172_8446622_RNA45SN1_45S_50ntFlanks | 12419 | 12439 | 21 | rRF 45S | NO | 52.03 | 209.69 | 0.00 | 0.00 |
| >hg38_21_+_8433172_8446622_RNA45SN1_45S_50ntFlanks@12419.12440.22\|GATGTCGGCTCTTCCTATCATT | >hg38_21_+_8433172_8446622_RNA45SN1_45S_50ntFlanks | 12419 | 12440 | 22 | rRF 45S | NO | 0.00 | 110.43 | 0.00 | 0.00 |
| >hg38_21_+_8433172_8446622_RNA45SN1_45S_50ntFlanks@12419.12441.23\|GATGTCGGCTCTTCCTATCATTG | >hg38_21_+_8433172_8446622_RNA45SN1_45S_50ntFlanks | 12419 | 12441 | 23 | rRF 45S | NO | 10.69 | 159.14 | 0.00 | 0.00 |
| >hg38_21_+_8433172_8446622_RNA45SN1_45S_50ntFlanks@12419.12442.24\|GATGTCGGCTCTTCCTATCATTGT | >hg38_21_+_8433172_8446622_RNA45SN1_45S_50ntFlanks | 12419 | 12442 | 24 | rRF 45S | NO | 24.55 | 248.81 | 0.00 | 0.00 |
| >hg38_21_+_8433172_8446622_RNA45SN1_45S_50ntFlanks@12419.12443.25\|GATGTCGGCTCTTCCTATCATTGTG | >hg38_21_+_8433172_8446622_RNA45SN1_45S_50ntFlanks | 12419 | 12443 | 25 | rRF 45S | NO | 10.47 | 186.13 | 0.00 | 0.00 |
| >hg38_21_+_8433172_8446622_RNA45SN1_45S_50ntFlanks@12419.12444.26\|GATGTCGGCTCTTCCTATCATTGTGA | >hg38_21_+_8433172_8446622_RNA45SN1_45S_50ntFlanks | 12419 | 12444 | 26 | rRF 45S | NO | 0.00 | 163.62 | 0.00 | 0.00 |
| >hg38_21_+_8433172_8446622_RNA45SN1_45S_50ntFlanks@12419.12445.27\|GATGTCGGCTCTTCCTATCATTGTGAA | >hg38_21_+_8433172_8446622_RNA45SN1_45S_50ntFlanks | 12419 | 12445 | 27 | rRF 45S | NO | 0.00 | 64.15 | 0.00 | 0.00 |
| >hg38_21_+_8433172_8446622_RNA45SN1_45S_50ntFlanks@12419.12446.28\|GATGTCGGCTCTTCCTATCATTGTGAAG | >hg38_21_+_8433172_8446622_RNA45SN1_45S_50ntFlanks | 12419 | 12446 | 28 | rRF 45S | NO | 0.00 | 133.73 | 0.00 | 0.00 |
| >hg38_21_+_8433172_8446622_RNA45SN1_45S_50ntFlanks@12419.12447.29\|GATGTCGGCTCTTCCTATCATTGTGAAGC | >hg38_21_+_8433172_8446622_RNA45SN1_45S_50ntFlanks | 12419 | 12447 | 29 | rRF 45S | NO | 0.00 | 226.71 | 0.00 | 0.00 |
| >hg38_21_+_8433172_8446622_RNA45SN1_45S_50ntFlanks@12419.12448.30\|GATGTCGGCTCTTCCTATCATTGTGAAGCA | >hg38_21_+_8433172_8446622_RNA45SN1_45S_50ntFlanks | 12419 | 12448 | 30 | rRF 45S | NO | 24.87 | 400.63 | 0.00 | 11.51 |
| >hg38_21_+_8433172_8446622_RNA45SN1_45S_50ntFlanks@12419.12449.31\|GATGTCGGCTCTTCCTATCATTGTGAAGCAG | >hg38_21_+_8433172_8446622_RNA45SN1_45S_50ntFlanks | 12419 | 12449 | 31 | rRF 45S | NO | 0.00 | 176.73 | 0.00 | 0.00 |
| >hg38_21_+_8433172_8446622_RNA45SN1_45S_50ntFlanks@12419.12450.32\|GATGTCGGCTCTTCCTATCATTGTGAAGCAGA | >hg38_21_+_8433172_8446622_RNA45SN1_45S_50ntFlanks | 12419 | 12450 | 32 | rRF 45S | NO | 0.00 | 58.06 | 0.00 | 0.00 |
| >hg38_21_+_8433172_8446622_RNA45SN1_45S_50ntFlanks@12419.12452.34\|GATGTCGGCTCTTCCTATCATTGTGAAGCAGAAT | >hg38_21_+_8433172_8446622_RNA45SN1_45S_50ntFlanks | 12419 | 12452 | 34 | rRF 45S | NO | 0.00 | 97.19 | 0.00 | 0.00 |
| >hg38_21_+_8433172_8446622_RNA45SN1_45S_50ntFlanks@12419.12453.35\|GATGTCGGCTCTTCCTATCATTGTGAAGCAGAATT | >hg38_21_+_8433172_8446622_RNA45SN1_45S_50ntFlanks | 12419 | 12453 | 35 | rRF 45S | NO | 0.00 | 64.09 | 0.00 | 0.00 |
| >hg38_21_+_8433172_8446622_RNA45SN1_45S_50ntFlanks@12419.12454.36\|GATGTCGGCTCTTCCTATCATTGTGAAGCAGAATTC | >hg38_21_+_8433172_8446622_RNA45SN1_45S_50ntFlanks | 12419 | 12454 | 36 | rRF 45S | NO | 0.00 | 68.17 | 0.00 | 0.00 |
| >hg38_21_+_8433172_8446622_RNA45SN1_45S_50ntFlanks@12419.12455.37\|GATGTCGGCTCTTCCTATCATTGTGAAGCAGAATTCA | >hg38_21_+_8433172_8446622_RNA45SN1_45S_50ntFlanks | 12419 | 12455 | 37 | rRF 45S | NO | 0.00 | 13.56 | 0.00 | 0.00 |
| >hg38_21_+_8433172_8446622_RNA45SN1_45S_50ntFlanks@12419.12456.38\|GATGTCGGCTCTTCCTATCATTGTGAAGCAGAATTCAC | >hg38_21_+_8433172_8446622_RNA45SN1_45S_50ntFlanks | 12419 | 12456 | 38 | rRF 45S | NO | 0.00 | 82.49 | 0.00 | 0.00 |
| >hg38_21_+_8433172_8446622_RNA45SN1_45S_50ntFlanks@12419.12457.39\|GATGTCGGCTCTTCCTATCATTGTGAAGCAGAATTCACC | >hg38_21_+_8433172_8446622_RNA45SN1_45S_50ntFlanks | 12419 | 12457 | 39 | rRF 45S | NO | 24.01 | 176.35 | 0.00 | 0.00 |
| >hg38_21_+_8433172_8446622_RNA45SN1_45S_50ntFlanks@12419.12458.40\|GATGTCGGCTCTTCCTATCATTGTGAAGCAGAATTCACCA | >hg38_21_+_8433172_8446622_RNA45SN1_45S_50ntFlanks | 12419 | 12458 | 40 | rRF 45S | NO | 0.00 | 101.40 | 0.00 | 0.00 |
| >hg38_21_+_8433172_8446622_RNA45SN1_45S_50ntFlanks@12419.12459.41\|GATGTCGGCTCTTCCTATCATTGTGAAGCAGAATTCACCAA | >hg38_21_+_8433172_8446622_RNA45SN1_45S_50ntFlanks | 12419 | 12459 | 41 | rRF 45S | NO | 0.00 | 13.48 | 0.00 | 0.00 |
| >hg38_21_+_8433172_8446622_RNA45SN1_45S_50ntFlanks@12419.12460.42\|GATGTCGGCTCTTCCTATCATTGTGAAGCAGAATTCACCAAG | >hg38_21_+_8433172_8446622_RNA45SN1_45S_50ntFlanks | 12419 | 12460 | 42 | rRF 45S | NO | 0.00 | 25.00 | 0.00 | 0.00 |
| >hg38_21_+_8433172_8446622_RNA45SN1_45S_50ntFlanks@12419.12462.44\|GATGTCGGCTCTTCCTATCATTGTGAAGCAGAATTCACCAAGCG | >hg38_21_+_8433172_8446622_RNA45SN1_45S_50ntFlanks | 12419 | 12462 | 44 | rRF 45S | NO | 0.00 | 101.38 | 0.00 | 0.00 |
| >hg38_21_+_8433172_8446622_RNA45SN1_45S_50ntFlanks@12420.12437.18\|ATGTCGGCTCTTCCTATC | >hg38_21_+_8433172_8446622_RNA45SN1_45S_50ntFlanks | 12420 | 12437 | 18 | rRF 45S | NO | 51.00 | 136.11 | 0.00 | 0.00 |
| >hg38_21_+_8433172_8446622_RNA45SN1_45S_50ntFlanks@12420.12438.19\|ATGTCGGCTCTTCCTATCA | >hg38_21_+_8433172_8446622_RNA45SN1_45S_50ntFlanks | 12420 | 12438 | 19 | rRF 45S | NO | 23.85 | 65.19 | 0.00 | 0.00 |
| >hg38_21_+_8433172_8446622_RNA45SN1_45S_50ntFlanks@12420.12439.20\|ATGTCGGCTCTTCCTATCAT | >hg38_21_+_8433172_8446622_RNA45SN1_45S_50ntFlanks | 12420 | 12439 | 20 | rRF 45S | NO | 0.00 | 85.05 | 0.00 | 0.00 |
| >hg38_21_+_8433172_8446622_RNA45SN1_45S_50ntFlanks@12420.12440.21\|ATGTCGGCTCTTCCTATCATT | >hg38_21_+_8433172_8446622_RNA45SN1_45S_50ntFlanks | 12420 | 12440 | 21 | rRF 45S | NO | 0.00 | 85.56 | 0.00 | 0.00 |
| >hg38_21_+_8433172_8446622_RNA45SN1_45S_50ntFlanks@12420.12441.22\|ATGTCGGCTCTTCCTATCATTG | >hg38_21_+_8433172_8446622_RNA45SN1_45S_50ntFlanks | 12420 | 12441 | 22 | rRF 45S | NO | 0.00 | 34.45 | 0.00 | 0.00 |
| >hg38_21_+_8433172_8446622_RNA45SN1_45S_50ntFlanks@12420.12442.23\|ATGTCGGCTCTTCCTATCATTGT | >hg38_21_+_8433172_8446622_RNA45SN1_45S_50ntFlanks | 12420 | 12442 | 23 | rRF 45S | NO | 0.00 | 70.94 | 0.00 | 0.00 |
| >hg38_21_+_8433172_8446622_RNA45SN1_45S_50ntFlanks@12420.12443.24\|ATGTCGGCTCTTCCTATCATTGTG | >hg38_21_+_8433172_8446622_RNA45SN1_45S_50ntFlanks | 12420 | 12443 | 24 | rRF 45S | NO | 0.00 | 63.60 | 0.00 | 0.00 |
| >hg38_21_+_8433172_8446622_RNA45SN1_45S_50ntFlanks@12420.12444.25\|ATGTCGGCTCTTCCTATCATTGTGA | >hg38_21_+_8433172_8446622_RNA45SN1_45S_50ntFlanks | 12420 | 12444 | 25 | rRF 45S | NO | 0.00 | 66.91 | 0.00 | 0.00 |
| >hg38_21_+_8433172_8446622_RNA45SN1_45S_50ntFlanks@12420.12445.26\|ATGTCGGCTCTTCCTATCATTGTGAA | >hg38_21_+_8433172_8446622_RNA45SN1_45S_50ntFlanks | 12420 | 12445 | 26 | rRF 45S | NO | 0.00 | 13.09 | 0.00 | 0.00 |
| >hg38_21_+_8433172_8446622_RNA45SN1_45S_50ntFlanks@12420.12446.27\|ATGTCGGCTCTTCCTATCATTGTGAAG | >hg38_21_+_8433172_8446622_RNA45SN1_45S_50ntFlanks | 12420 | 12446 | 27 | rRF 45S | NO | 0.00 | 79.18 | 0.00 | 0.00 |
| >hg38_21_+_8433172_8446622_RNA45SN1_45S_50ntFlanks@12420.12447.28\|ATGTCGGCTCTTCCTATCATTGTGAAGC | >hg38_21_+_8433172_8446622_RNA45SN1_45S_50ntFlanks | 12420 | 12447 | 28 | rRF 45S | NO | 0.00 | 80.47 | 0.00 | 0.00 |
| >hg38_21_+_8433172_8446622_RNA45SN1_45S_50ntFlanks@12420.12448.29\|ATGTCGGCTCTTCCTATCATTGTGAAGCA | >hg38_21_+_8433172_8446622_RNA45SN1_45S_50ntFlanks | 12420 | 12448 | 29 | rRF 45S | NO | 0.00 | 153.72 | 0.00 | 73.81 |
| >hg38_21_+_8433172_8446622_RNA45SN1_45S_50ntFlanks@12420.12449.30\|ATGTCGGCTCTTCCTATCATTGTGAAGCAG | >hg38_21_+_8433172_8446622_RNA45SN1_45S_50ntFlanks | 12420 | 12449 | 30 | rRF 45S | NO | 0.00 | 56.41 | 0.00 | 19.36 |

### MDA-MB-231

| | | | | | | | | | | |
|---|---|---|---|---|---|---|---|---|---|---|
| >hg38_21_+_8433172_8446622_RNA45SN1_45S_50ntFlanks@12420.12457.38\|ATGTCGGCTCTTCCTATCATTGTGAAGCAGAATTCACC | >hg38_21_+_8433172_8446622_RNA45SN1_45S_50ntFlanks | 12420 | 12457 | 38 | rRF 45S | NO | 10.48 | 179.88 | 0.00 | 10.98 |
| >hg38_21_+_8433172_8446622_RNA45SN1_45S_50ntFlanks@12420.12458.39\|ATGTCGGCTCTTCCTATCATTGTGAAGCAGAATTCACCA | >hg38_21_+_8433172_8446622_RNA45SN1_45S_50ntFlanks | 12420 | 12458 | 39 | rRF 45S | NO | 0.00 | 87.18 | 0.00 | 12.94 |
| >hg38_21_+_8433172_8446622_RNA45SN1_45S_50ntFlanks@12420.12462.43\|ATGTCGGCTCTTCCTATCATTGTGAAGCAGAATTCACCAAGCG | >hg38_21_+_8433172_8446622_RNA45SN1_45S_50ntFlanks | 12420 | 12462 | 43 | rRF 45S | NO | 0.00 | 81.68 | 0.00 | 0.00 |
| >hg38_21_+_8433172_8446622_RNA45SN1_45S_50ntFlanks@12423.12457.35\|TCGGCTCTTCCTATCATTGTGAAGCAGAATTCACC | >hg38_21_+_8433172_8446622_RNA45SN1_45S_50ntFlanks | 12423 | 12457 | 35 | rRF 45S | NO | 0.00 | 57.51 | 0.00 | 0.00 |
| >hg38_21_+_8433172_8446622_RNA45SN1_45S_50ntFlanks@12438.12473.36\|ATTGTGAAGCAGAATTCACCAAGCGTTGGATTGTTC | >hg38_21_+_8433172_8446622_RNA45SN1_45S_50ntFlanks | 12438 | 12473 | 36 | rRF 45S | NO | 0.00 | 27.00 | 0.00 | 0.00 |
| >hg38_21_+_8433172_8446622_RNA45SN1_45S_50ntFlanks@12439.12473.35\|TTGTGAAGCAGAATTCACCAAGCGTTGGATTGTTC | >hg38_21_+_8433172_8446622_RNA45SN1_45S_50ntFlanks | 12439 | 12473 | 35 | rRF 45S | NO | 0.00 | 24.77 | 0.00 | 0.00 |
| >hg38_21_+_8433172_8446622_RNA45SN1_45S_50ntFlanks@12444.12480.37\|AAGCAGAATTCACCAAGCGTTGGATTGTTCACCCACT | >hg38_21_+_8433172_8446622_RNA45SN1_45S_50ntFlanks | 12444 | 12480 | 37 | rRF 45S | NO | 0.00 | 13.40 | 0.00 | 0.00 |
| >hg38_21_+_8433172_8446622_RNA45SN1_45S_50ntFlanks@12446.12480.35\|GCAGAATTCACCAAGCGTTGGATTGTTCACCCACT | >hg38_21_+_8433172_8446622_RNA45SN1_45S_50ntFlanks | 12446 | 12480 | 35 | rRF 45S | NO | 0.00 | 25.85 | 0.00 | 0.00 |
| >hg38_21_+_8433172_8446622_RNA45SN1_45S_50ntFlanks@12448.12480.33\|AGAATTCACCAAGCGTTGGATTGTTCACCCACT | >hg38_21_+_8433172_8446622_RNA45SN1_45S_50ntFlanks | 12448 | 12480 | 33 | rRF 45S | NO | 0.00 | 30.11 | 0.00 | 0.00 |
| >hg38_21_+_8433172_8446622_RNA45SN1_45S_50ntFlanks@12449.12473.25\|GAATTCACCAAGCGTTGGATTGTTC | >hg38_21_+_8433172_8446622_RNA45SN1_45S_50ntFlanks | 12449 | 12473 | 25 | rRF 45S | NO | 0.00 | 0.00 | 16.10 | 0.00 |
| >hg38_21_+_8433172_8446622_RNA45SN1_45S_50ntFlanks@12449.12480.32\|GAATTCACCAAGCGTTGGATTGTTCACCCACT | >hg38_21_+_8433172_8446622_RNA45SN1_45S_50ntFlanks | 12449 | 12480 | 32 | rRF 45S | NO | 0.00 | 66.21 | 0.00 | 0.00 |
| >hg38_21_+_8433172_8446622_RNA45SN1_45S_50ntFlanks@12450.12480.31\|AATTCACCAAGCGTTGGATTGTTCACCCACT | >hg38_21_+_8433172_8446622_RNA45SN1_45S_50ntFlanks | 12450 | 12480 | 31 | rRF 45S | NO | 0.00 | 36.27 | 0.00 | 0.00 |
| >hg38_21_+_8433172_8446622_RNA45SN1_45S_50ntFlanks@12453.12476.24\|TCACCAAGCGTTGGATTGTTCACC | >hg38_21_+_8433172_8446622_RNA45SN1_45S_50ntFlanks | 12453 | 12476 | 24 | rRF 45S | NO | 0.00 | 34.25 | 0.00 | 0.00 |
| >hg38_21_+_8433172_8446622_RNA45SN1_45S_50ntFlanks@12453.12477.25\|TCACCAAGCGTTGGATTGTTCACCC | >hg38_21_+_8433172_8446622_RNA45SN1_45S_50ntFlanks | 12453 | 12477 | 25 | rRF 45S | NO | 0.00 | 24.39 | 0.00 | 0.00 |
| >hg38_21_+_8433172_8446622_RNA45SN1_45S_50ntFlanks@12454.12476.23\|CACCAAGCGTTGGATTGTTCACC | >hg38_21_+_8433172_8446622_RNA45SN1_45S_50ntFlanks | 12454 | 12476 | 23 | rRF 45S | NO | 0.00 | 34.56 | 0.00 | 0.00 |
| >hg38_21_+_8433172_8446622_RNA45SN1_45S_50ntFlanks@12455.12476.22\|ACCAAGCGTTGGATTGTTCACC | >hg38_21_+_8433172_8446622_RNA45SN1_45S_50ntFlanks | 12455 | 12476 | 22 | rRF 45S | NO | 0.00 | 26.75 | 0.00 | 0.00 |
| >hg38_21_+_8433172_8446622_RNA45SN1_45S_50ntFlanks@12456.12476.21\|CCAAGCGTTGGATTGTTCACC | >hg38_21_+_8433172_8446622_RNA45SN1_45S_50ntFlanks | 12456 | 12476 | 21 | rRF 45S | NO | 0.00 | 25.94 | 0.00 | 0.00 |
| >hg38_21_+_8433172_8446622_RNA45SN1_45S_50ntFlanks@12457.12476.20\|CAAGCGTTGGATTGTTCACC | >hg38_21_+_8433172_8446622_RNA45SN1_45S_50ntFlanks | 12457 | 12476 | 20 | rRF 45S | NO | 0.00 | 58.17 | 0.00 | 0.00 |
| >hg38_21_+_8433172_8446622_RNA45SN1_45S_50ntFlanks@12458.12476.19\|AAGCGTTGGATTGTTCACC | >hg38_21_+_8433172_8446622_RNA45SN1_45S_50ntFlanks | 12458 | 12476 | 19 | rRF 45S | NO | 49.30 | 116.01 | 0.00 | 0.00 |
| >hg38_21_+_8433172_8446622_RNA45SN1_45S_50ntFlanks@12458.12477.20\|AAGCGTTGGATTGTTCACCC | >hg38_21_+_8433172_8446622_RNA45SN1_45S_50ntFlanks | 12458 | 12477 | 20 | rRF 45S | NO | 0.00 | 58.27 | 0.00 | 0.00 |
| >hg38_21_+_8433172_8446622_RNA45SN1_45S_50ntFlanks@12458.12496.39\|AAGCGTTGGATTGTTCACCCACTAATAGGGAACGTGAGC | >hg38_21_+_8433172_8446622_RNA45SN1_45S_50ntFlanks | 12458 | 12496 | 39 | rRF 45S | NO | 0.00 | 32.39 | 0.00 | 0.00 |
| >hg38_21_+_8433172_8446622_RNA45SN1_45S_50ntFlanks@12463.12496.34\|TTGGATTGTTCACCCACTAATAGGGAACGTGAGC | >hg38_21_+_8433172_8446622_RNA45SN1_45S_50ntFlanks | 12463 | 12496 | 34 | rRF 45S | NO | 0.00 | 33.53 | 0.00 | 0.00 |
| >hg38_21_+_8433172_8446622_RNA45SN1_45S_50ntFlanks@12473.12496.24\|CACCCACTAATAGGGAACGTGAGC | >hg38_21_+_8433172_8446622_RNA45SN1_45S_50ntFlanks | 12473 | 12496 | 24 | rRF 45S | NO | 0.00 | 24.95 | 0.00 | 0.00 |
| >hg38_21_+_8433172_8446622_RNA45SN1_45S_50ntFlanks@12474.12496.23\|ACCCACTAATAGGGAACGTGAGC | >hg38_21_+_8433172_8446622_RNA45SN1_45S_50ntFlanks | 12474 | 12496 | 23 | rRF 45S | NO | 0.00 | 85.77 | 0.00 | 0.00 |
| >hg38_21_+_8433172_8446622_RNA45SN1_45S_50ntFlanks@12474.12497.24\|ACCCACTAATAGGGAACGTGAGCT | >hg38_21_+_8433172_8446622_RNA45SN1_45S_50ntFlanks | 12474 | 12497 | 24 | rRF 45S | NO | 0.00 | 13.58 | 0.00 | 0.00 |
| >hg38_21_+_8433172_8446622_RNA45SN1_45S_50ntFlanks@12475.12496.22\|CCCACTAATAGGGAACGTGAGC | >hg38_21_+_8433172_8446622_RNA45SN1_45S_50ntFlanks | 12475 | 12496 | 22 | rRF 45S | NO | 0.00 | 58.64 | 0.00 | 0.00 |
| >hg38_21_+_8433172_8446622_RNA45SN1_45S_50ntFlanks@12475.12497.23\|CCCACTAATAGGGAACGTGAGCT | >hg38_21_+_8433172_8446622_RNA45SN1_45S_50ntFlanks | 12475 | 12497 | 23 | rRF 45S | NO | 0.00 | 26.41 | 0.00 | 0.00 |
| >hg38_21_+_8433172_8446622_RNA45SN1_45S_50ntFlanks@12477.12496.20\|CACTAATAGGGAACGTGAGC | >hg38_21_+_8433172_8446622_RNA45SN1_45S_50ntFlanks | 12477 | 12496 | 20 | rRF 45S | NO | 0.00 | 166.15 | 0.00 | 0.00 |
| >hg38_21_+_8433172_8446622_RNA45SN1_45S_50ntFlanks@12477.12497.21\|CACTAATAGGGAACGTGAGCT | >hg38_21_+_8433172_8446622_RNA45SN1_45S_50ntFlanks | 12477 | 12497 | 21 | rRF 45S | NO | 0.00 | 170.56 | 0.00 | 0.00 |
| >hg38_21_+_8433172_8446622_RNA45SN1_45S_50ntFlanks@12478.12496.19\|ACTAATAGGGAACGTGAGC | >hg38_21_+_8433172_8446622_RNA45SN1_45S_50ntFlanks | 12478 | 12496 | 19 | rRF 45S | NO | 0.00 | 90.71 | 0.00 | 0.00 |
| >hg38_21_+_8433172_8446622_RNA45SN1_45S_50ntFlanks@12478.12497.20\|ACTAATAGGGAACGTGAGCT | >hg38_21_+_8433172_8446622_RNA45SN1_45S_50ntFlanks | 12478 | 12497 | 20 | rRF 45S | NO | 0.00 | 142.43 | 0.00 | 0.00 |
| >hg38_21_+_8433172_8446622_RNA45SN1_45S_50ntFlanks@12479.12496.18\|CTAATAGGGAACGTGAGC | >hg38_21_+_8433172_8446622_RNA45SN1_45S_50ntFlanks | 12479 | 12496 | 18 | rRF 45S | NO | 0.00 | 101.60 | 0.00 | 0.00 |
| >hg38_21_+_8433172_8446622_RNA45SN1_45S_50ntFlanks@12479.12497.19\|CTAATAGGGAACGTGAGCT | >hg38_21_+_8433172_8446622_RNA45SN1_45S_50ntFlanks | 12479 | 12497 | 19 | rRF 45S | NO | 0.00 | 174.14 | 0.00 | 0.00 |
| >hg38_21_+_8433172_8446622_RNA45SN1_45S_50ntFlanks@12480.12497.18\|TAATAGGGAACGTGAGCT | >hg38_21_+_8433172_8446622_RNA45SN1_45S_50ntFlanks | 12480 | 12497 | 18 | rRF 45S | NO | 0.00 | 125.57 | 0.00 | 0.00 |
| >hg38_21_+_8433172_8446622_RNA45SN1_45S_50ntFlanks@12605.12624.20\|TATGTGCTTGGCTGAGGAGC | >hg38_21_+_8433172_8446622_RNA45SN1_45S_50ntFlanks | 12605 | 12624 | 20 | rRF 45S | NO | 0.00 | 28.35 | 0.00 | 0.00 |
| >hg38_21_+_8433172_8446622_RNA45SN1_45S_50ntFlanks@12607.12624.18\|TGTGCTTGGCTGAGGAGC | >hg38_21_+_8433172_8446622_RNA45SN1_45S_50ntFlanks | 12607 | 12624 | 18 | rRF 45S | NO | 0.00 | 31.86 | 0.00 | 0.00 |
| >hg38_21_+_8433172_8446622_RNA45SN1_45S_50ntFlanks@12625.12667.43\|CAATGGGGCGAAGCTACCATCTGTGGGATTATGACTGAACGCC | >hg38_21_+_8433172_8446622_RNA45SN1_45S_50ntFlanks | 12625 | 12667 | 43 | rRF 45S | NO | 0.00 | 28.64 | 0.00 | 0.00 |
| >hg38_21_+_8433172_8446622_RNA45SN1_45S_50ntFlanks@12626.12667.42\|AATGGGGCGAAGCTACCATCTGTGGGATTATGACTGAACGCC | >hg38_21_+_8433172_8446622_RNA45SN1_45S_50ntFlanks | 12626 | 12667 | 42 | rRF 45S | NO | 0.00 | 84.15 | 0.00 | 12.11 |

### MDA-MB-231

| | | | | | | | | | | |
|---|---|---|---|---|---|---|---|---|---|---|
| >hg38_21_+_8433172_8446622_RNA45SN1_45S_50ntFlanks@12635.12667.33\|AAGCTACCATCTGTGGGATTATGACTGAACGCC | >hg38_21_+_8433172_8446622_RNA45SN1_45S_50ntFlanks | 12635 | 12667 | 33 | rRF 45S | NO | 0.00 | 25.45 | 0.00 | 0.00 |
| >hg38_21_+_8433172_8446622_RNA45SN1_45S_50ntFlanks@12640.12667.28\|ACCATCTGTGGGATTATGACTGAACGCC | >hg38_21_+_8433172_8446622_RNA45SN1_45S_50ntFlanks | 12640 | 12667 | 28 | rRF 45S | NO | 0.00 | 26.74 | 0.00 | 0.00 |
| >hg38_21_+_8433172_8446622_RNA45SN1_45S_50ntFlanks@12640.12669.30\|ACCATCTGTGGGATTATGACTGAACGCCTC | >hg38_21_+_8433172_8446622_RNA45SN1_45S_50ntFlanks | 12640 | 12669 | 30 | rRF 45S | NO | 0.00 | 61.64 | 0.00 | 0.00 |
| >hg38_21_+_8433172_8446622_RNA45SN1_45S_50ntFlanks@12640.12670.31\|ACCATCTGTGGGATTATGACTGAACGCCTCT | >hg38_21_+_8433172_8446622_RNA45SN1_45S_50ntFlanks | 12640 | 12670 | 31 | rRF 45S | NO | 0.00 | 28.97 | 0.00 | 0.00 |
| >hg38_21_+_8433172_8446622_RNA45SN1_45S_50ntFlanks@12937.12957.21\|AACCATTCGTAGACGACCTGC | >hg38_21_+_8433172_8446622_RNA45SN1_45S_50ntFlanks | 12937 | 12957 | 21 | rRF 45S | NO | 0.00 | 24.92 | 0.00 | 0.00 |
| >hg38_21_+_8433172_8446622_RNA45SN1_45S_50ntFlanks@12941.12958.18\|ATTCGTAGACGACCTGCT | >hg38_21_+_8433172_8446622_RNA45SN1_45S_50ntFlanks | 12941 | 12958 | 18 | rRF 45S | NO | 0.00 | 25.83 | 0.00 | 0.00 |
| >hg38_21_+_8433172_8446622_RNA45SN1_45S_50ntFlanks@12941.12959.19\|ATTCGTAGACGACCTGCTT | >hg38_21_+_8433172_8446622_RNA45SN1_45S_50ntFlanks | 12941 | 12959 | 19 | rRF 45S | NO | 0.00 | 13.41 | 0.00 | 0.00 |
| >hg38_21_+_8433172_8446622_RNA45SN1_45S_50ntFlanks@12941.12960.20\|ATTCGTAGACGACCTGCTTC | >hg38_21_+_8433172_8446622_RNA45SN1_45S_50ntFlanks | 12941 | 12960 | 20 | rRF 45S | NO | 0.00 | 35.90 | 0.00 | 0.00 |
| >hg38_21_+_8433172_8446622_RNA45SN1_45S_50ntFlanks@12941.12961.21\|ATTCGTAGACGACCTGCTTCT | >hg38_21_+_8433172_8446622_RNA45SN1_45S_50ntFlanks | 12941 | 12961 | 21 | rRF 45S | NO | 0.00 | 13.53 | 0.00 | 0.00 |
| >hg38_21_+_8433172_8446622_RNA45SN1_45S_50ntFlanks@12945.12988.44\|GTAGACGACCTGCTTCTGGGTCGGGGTTTCGTACGTAGCAGAGC | >hg38_21_+_8433172_8446622_RNA45SN1_45S_50ntFlanks | 12945 | 12988 | 44 | rRF 45S | NO | 0.00 | 67.25 | 0.00 | 0.00 |
| >hg38_21_+_8433172_8446622_RNA45SN1_45S_50ntFlanks@12946.12988.43\|TAGACGACCTGCTTCTGGGTCGGGGTTTCGTACGTAGCAGAGC | >hg38_21_+_8433172_8446622_RNA45SN1_45S_50ntFlanks | 12946 | 12988 | 43 | rRF 45S | NO | 0.00 | 65.26 | 0.00 | 0.00 |
| >hg38_21_+_8433172_8446622_RNA45SN1_45S_50ntFlanks@12948.12988.41\|GACGACCTGCTTCTGGGTCGGGGTTTCGTACGTAGCAGAGC | >hg38_21_+_8433172_8446622_RNA45SN1_45S_50ntFlanks | 12948 | 12988 | 41 | rRF 45S | NO | 0.00 | 34.14 | 0.00 | 0.00 |
| >hg38_21_+_8433172_8446622_RNA45SN1_45S_50ntFlanks@12959.12993.35\|TCTGGGTCGGGGTTTCGTACGTAGCAGAGCAGCTC | >hg38_21_+_8433172_8446622_RNA45SN1_45S_50ntFlanks | 12959 | 12993 | 35 | rRF 45S | NO | 0.00 | 108.85 | 0.00 | 0.00 |
| >hg38_21_+_8433172_8446622_RNA45SN1_45S_50ntFlanks@12988.13013.26\|CAGCTCCCTCGCTGCGATCTATTGAA | >hg38_21_+_8433172_8446622_RNA45SN1_45S_50ntFlanks | 12988 | 13013 | 26 | rRF 45S | NO | 0.00 | 70.64 | 0.00 | 0.00 |
| >hg38_21_+_8433172_8446622_RNA45SN1_45S_50ntFlanks@12989.13013.25\|AGCTCCCTCGCTGCGATCTATTGAA | >hg38_21_+_8433172_8446622_RNA45SN1_45S_50ntFlanks | 12989 | 13013 | 25 | rRF 45S | NO | 0.00 | 25.95 | 0.00 | 0.00 |
| >hg38_21_+_8433172_8446622_RNA45SN1_45S_50ntFlanks@12990.13013.24\|GCTCCCTCGCTGCGATCTATTGAA | >hg38_21_+_8433172_8446622_RNA45SN1_45S_50ntFlanks | 12990 | 13013 | 24 | rRF 45S | NO | 0.00 | 67.56 | 0.00 | 0.00 |
| >hg38_21_+_8433172_8446622_RNA45SN1_45S_50ntFlanks@12990.13014.25\|GCTCCCTCGCTGCGATCTATTGAAA | >hg38_21_+_8433172_8446622_RNA45SN1_45S_50ntFlanks | 12990 | 13014 | 25 | rRF 45S | NO | 0.00 | 25.36 | 0.00 | 0.00 |
| >hg38_21_+_8433172_8446622_RNA45SN1_45S_50ntFlanks@12990.13017.28\|GCTCCCTCGCTGCGATCTATTGAAAGTC | >hg38_21_+_8433172_8446622_RNA45SN1_45S_50ntFlanks | 12990 | 13017 | 28 | rRF 45S | NO | 0.00 | 28.64 | 0.00 | 0.00 |
| >hg38_21_+_8433172_8446622_RNA45SN1_45S_50ntFlanks@12991.13039.49\|CTCCCTCGCTGCGATCTATTGAAAGTCAGCCCTCGACACAAGGGTTTGT | >hg38_21_+_8433172_8446622_RNA45SN1_45S_50ntFlanks | 12991 | 13039 | 49 | rRF 45S | NO | 0.00 | 24.41 | 0.00 | 0.00 |
| >hg38_21_+_8433172_8446622_RNA45SN1_45S_50ntFlanks@12992.13017.26\|TCCCTCGCTGCGATCTATTGAAAGTC | >hg38_21_+_8433172_8446622_RNA45SN1_45S_50ntFlanks | 12992 | 13017 | 26 | rRF 45S | NO | 0.00 | 13.56 | 0.00 | 0.00 |
| >hg38_21_+_8433172_8446622_RNA45SN1_45S_50ntFlanks@12992.13039.48\|TCCCTCGCTGCGATCTATTGAAAGTCAGCCCTCGACACAAGGGTTTGT | >hg38_21_+_8433172_8446622_RNA45SN1_45S_50ntFlanks | 12992 | 13039 | 48 | rRF 45S | NO | 0.00 | 64.36 | 0.00 | 0.00 |
| >hg38_21_+_8433172_8446622_RNA45SN1_45S_50ntFlanks@12993.13039.47\|CCCTCGCTGCGATCTATTGAAAGTCAGCCCTCGACACAAGGGTTTGT | >hg38_21_+_8433172_8446622_RNA45SN1_45S_50ntFlanks | 12993 | 13039 | 47 | rRF 45S | NO | 55.29 | 297.75 | 0.00 | 0.00 |
| >hg38_21_+_8433172_8446622_RNA45SN1_45S_50ntFlanks@12994.13036.43\|CCTCGCTGCGATCTATTGAAAGTCAGCCCTCGACACAAGGGTT | >hg38_21_+_8433172_8446622_RNA45SN1_45S_50ntFlanks | 12994 | 13036 | 43 | rRF 45S | NO | 0.00 | 25.73 | 0.00 | 0.00 |
| >hg38_21_+_8433172_8446622_RNA45SN1_45S_50ntFlanks@12994.13038.45\|CCTCGCTGCGATCTATTGAAAGTCAGCCCTCGACACAAGGGTTTG | >hg38_21_+_8433172_8446622_RNA45SN1_45S_50ntFlanks | 12994 | 13038 | 45 | rRF 45S | NO | 0.00 | 30.84 | 0.00 | 0.00 |
| >hg38_21_+_8433172_8446622_RNA45SN1_45S_50ntFlanks@12994.13039.46\|CCTCGCTGCGATCTATTGAAAGTCAGCCCTCGACACAAGGGTTTGT | >hg38_21_+_8433172_8446622_RNA45SN1_45S_50ntFlanks | 12994 | 13039 | 46 | rRF 45S | YES | 296.13 | 1795.13 | 0.00 | 318.17 |
| >hg38_21_+_8433172_8446622_RNA45SN1_45S_50ntFlanks@12995.13012.18\|CTCGCTGCGATCTATTGA | >hg38_21_+_8433172_8446622_RNA45SN1_45S_50ntFlanks | 12995 | 13012 | 18 | rRF 45S | NO | 0.00 | 69.04 | 0.00 | 0.00 |
| >hg38_21_+_8433172_8446622_RNA45SN1_45S_50ntFlanks@12995.13013.19\|CTCGCTGCGATCTATTGAA | >hg38_21_+_8433172_8446622_RNA45SN1_45S_50ntFlanks | 12995 | 13013 | 19 | rRF 45S | NO | 10.44 | 83.35 | 0.00 | 0.00 |
| >hg38_21_+_8433172_8446622_RNA45SN1_45S_50ntFlanks@12995.13014.20\|CTCGCTGCGATCTATTGAAA | >hg38_21_+_8433172_8446622_RNA45SN1_45S_50ntFlanks | 12995 | 13014 | 20 | rRF 45S | NO | 0.00 | 61.48 | 0.00 | 0.00 |
| >hg38_21_+_8433172_8446622_RNA45SN1_45S_50ntFlanks@12995.13015.21\|CTCGCTGCGATCTATTGAAAG | >hg38_21_+_8433172_8446622_RNA45SN1_45S_50ntFlanks | 12995 | 13015 | 21 | rRF 45S | NO | 10.52 | 72.07 | 0.00 | 0.00 |
| >hg38_21_+_8433172_8446622_RNA45SN1_45S_50ntFlanks@12995.13016.22\|CTCGCTGCGATCTATTGAAAGT | >hg38_21_+_8433172_8446622_RNA45SN1_45S_50ntFlanks | 12995 | 13016 | 22 | rRF 45S | NO | 50.20 | 88.30 | 0.00 | 0.00 |
| >hg38_21_+_8433172_8446622_RNA45SN1_45S_50ntFlanks@12995.13017.23\|CTCGCTGCGATCTATTGAAAGTC | >hg38_21_+_8433172_8446622_RNA45SN1_45S_50ntFlanks | 12995 | 13017 | 23 | rRF 45S | NO | 10.38 | 110.08 | 0.00 | 0.00 |
| >hg38_21_+_8433172_8446622_RNA45SN1_45S_50ntFlanks@12995.13035.41\|CTCGCTGCGATCTATTGAAAGTCAGCCCTCGACACAAGGGT | >hg38_21_+_8433172_8446622_RNA45SN1_45S_50ntFlanks | 12995 | 13035 | 41 | rRF 45S | NO | 0.00 | 59.05 | 0.00 | 0.00 |
| >hg38_21_+_8433172_8446622_RNA45SN1_45S_50ntFlanks@12995.13036.42\|CTCGCTGCGATCTATTGAAAGTCAGCCCTCGACACAAGGGTT | >hg38_21_+_8433172_8446622_RNA45SN1_45S_50ntFlanks | 12995 | 13036 | 42 | rRF 45S | NO | 59.33 | 398.34 | 0.00 | 60.28 |
| >hg38_21_+_8433172_8446622_RNA45SN1_45S_50ntFlanks@12995.13037.43\|CTCGCTGCGATCTATTGAAAGTCAGCCCTCGACACAAGGGTTT | >hg38_21_+_8433172_8446622_RNA45SN1_45S_50ntFlanks | 12995 | 13037 | 43 | rRF 45S | NO | 0.00 | 213.47 | 0.00 | 13.31 |
| >hg38_21_+_8433172_8446622_RNA45SN1_45S_50ntFlanks@12995.13038.44\|CTCGCTGCGATCTATTGAAAGTCAGCCCTCGACACAAGGGTTTG | >hg38_21_+_8433172_8446622_RNA45SN1_45S_50ntFlanks | 12995 | 13038 | 44 | rRF 45S | NO | 52.52 | 437.25 | 0.00 | 75.10 |
| >hg38_21_+_8433172_8446622_RNA45SN1_45S_50ntFlanks@12995.13039.45\|CTCGCTGCGATCTATTGAAAGTCAGCCCTCGACACAAGGGTTTGT | >hg38_21_+_8433172_8446622_RNA45SN1_45S_50ntFlanks | 12995 | 13039 | 45 | rRF 45S | YES | 2737.37 | 30716.47 | 116.31 | 1825.37 |
| >hg38_21_+_8433172_8446622_RNA45SN1_45S_50ntFlanks@12995.13040.46\|CTCGCTGCGATCTATTGAAAGTCAGCCCTCGACACAAGGGTTTGTC | >hg38_21_+_8433172_8446622_RNA45SN1_45S_50ntFlanks | 12995 | 13040 | 46 | rRF 45S | NO | 25.52 | 26.56 | 0.00 | 11.07 |
| >hg38_21_+_8433172_8446622_RNA45SN1_45S_50ntFlanks@12996.13013.18\|TCGCTGCGATCTATTGAA | >hg38_21_+_8433172_8446622_RNA45SN1_45S_50ntFlanks | 12996 | 13013 | 18 | rRF 45S | NO | 124.57 | 299.91 | 0.00 | 11.79 |

### MDA-MB-231

| | | | | | | | | | | |
|---|---|---|---|---|---|---|---|---|---|---|
| >hg38_21_+_8433172_8446622_RNA45SN1_45S_50ntFlanks@12996.13014.19\|TCGCTGCGATCTATTGAAA | >hg38_21_+_8433172_8446622_RNA45SN1_45S_50ntFlanks | 12996 | 13014 | 19 | rRF 45S | NO | 81.86 | 182.62 | 0.00 | 0.00 |
| >hg38_21_+_8433172_8446622_RNA45SN1_45S_50ntFlanks@12996.13015.20\|TCGCTGCGATCTATTGAAAG | >hg38_21_+_8433172_8446622_RNA45SN1_45S_50ntFlanks | 12996 | 13015 | 20 | rRF 45S | NO | 24.37 | 65.99 | 0.00 | 0.00 |
| >hg38_21_+_8433172_8446622_RNA45SN1_45S_50ntFlanks@12996.13016.21\|TCGCTGCGATCTATTGAAAGT | >hg38_21_+_8433172_8446622_RNA45SN1_45S_50ntFlanks | 12996 | 13016 | 21 | rRF 45S | NO | 0.00 | 30.35 | 0.00 | 0.00 |
| >hg38_21_+_8433172_8446622_RNA45SN1_45S_50ntFlanks@12996.13017.22\|TCGCTGCGATCTATTGAAAGTC | >hg38_21_+_8433172_8446622_RNA45SN1_45S_50ntFlanks | 12996 | 13017 | 22 | rRF 45S | NO | 61.53 | 161.98 | 0.00 | 0.00 |
| >hg38_21_+_8433172_8446622_RNA45SN1_45S_50ntFlanks@12996.13035.40\|TCGCTGCGATCTATTGAAAGTCAGCCCTCGACACAAGGGT | >hg38_21_+_8433172_8446622_RNA45SN1_45S_50ntFlanks | 12996 | 13035 | 40 | rRF 45S | NO | 0.00 | 24.02 | 0.00 | 0.00 |
| >hg38_21_+_8433172_8446622_RNA45SN1_45S_50ntFlanks@12996.13036.41\|TCGCTGCGATCTATTGAAAGTCAGCCCTCGACACAAGGGTT | >hg38_21_+_8433172_8446622_RNA45SN1_45S_50ntFlanks | 12996 | 13036 | 41 | rRF 45S | NO | 47.33 | 250.82 | 0.00 | 19.85 |
| >hg38_21_+_8433172_8446622_RNA45SN1_45S_50ntFlanks@12996.13037.42\|TCGCTGCGATCTATTGAAAGTCAGCCCTCGACACAAGGGTTT | >hg38_21_+_8433172_8446622_RNA45SN1_45S_50ntFlanks | 12996 | 13037 | 42 | rRF 45S | NO | 0.00 | 82.31 | 0.00 | 0.00 |
| >hg38_21_+_8433172_8446622_RNA45SN1_45S_50ntFlanks@12996.13038.43\|TCGCTGCGATCTATTGAAAGTCAGCCCTCGACACAAGGGTTTG | >hg38_21_+_8433172_8446622_RNA45SN1_45S_50ntFlanks | 12996 | 13038 | 43 | rRF 45S | NO | 10.50 | 235.44 | 0.00 | 21.77 |
| >hg38_21_+_8433172_8446622_RNA45SN1_45S_50ntFlanks@12996.13039.44\|TCGCTGCGATCTATTGAAAGTCAGCCCTCGACACAAGGGTTTGT | >hg38_21_+_8433172_8446622_RNA45SN1_45S_50ntFlanks | 12996 | 13039 | 44 | rRF 45S | YES | 1786.42 | 11638.96 | 166.66 | 1153.65 |
| >hg38_21_+_8433172_8446622_RNA45SN1_45S_50ntFlanks@12997.13014.18\|CGCTGCGATCTATTGAAA | >hg38_21_+_8433172_8446622_RNA45SN1_45S_50ntFlanks | 12997 | 13014 | 18 | rRF 45S | NO | 153.17 | 325.40 | 0.00 | 15.85 |
| >hg38_21_+_8433172_8446622_RNA45SN1_45S_50ntFlanks@12997.13015.19\|CGCTGCGATCTATTGAAAG | >hg38_21_+_8433172_8446622_RNA45SN1_45S_50ntFlanks | 12997 | 13015 | 19 | rRF 45S | NO | 53.79 | 71.81 | 0.00 | 0.00 |
| >hg38_21_+_8433172_8446622_RNA45SN1_45S_50ntFlanks@12997.13016.20\|CGCTGCGATCTATTGAAAGT | >hg38_21_+_8433172_8446622_RNA45SN1_45S_50ntFlanks | 12997 | 13016 | 20 | rRF 45S | NO | 10.38 | 25.16 | 0.00 | 0.00 |
| >hg38_21_+_8433172_8446622_RNA45SN1_45S_50ntFlanks@12997.13017.21\|CGCTGCGATCTATTGAAAGTC | >hg38_21_+_8433172_8446622_RNA45SN1_45S_50ntFlanks | 12997 | 13017 | 21 | rRF 45S | NO | 86.27 | 179.08 | 0.00 | 0.00 |
| >hg38_21_+_8433172_8446622_RNA45SN1_45S_50ntFlanks@12997.13035.39\|CGCTGCGATCTATTGAAAGTCAGCCCTCGACACAAGGGT | >hg38_21_+_8433172_8446622_RNA45SN1_45S_50ntFlanks | 12997 | 13035 | 39 | rRF 45S | NO | 0.00 | 35.00 | 0.00 | 0.00 |
| >hg38_21_+_8433172_8446622_RNA45SN1_45S_50ntFlanks@12997.13036.40\|CGCTGCGATCTATTGAAAGTCAGCCCTCGACACAAGGGTT | >hg38_21_+_8433172_8446622_RNA45SN1_45S_50ntFlanks | 12997 | 13036 | 40 | rRF 45S | NO | 63.32 | 238.90 | 0.00 | 60.77 |
| >hg38_21_+_8433172_8446622_RNA45SN1_45S_50ntFlanks@12997.13037.41\|CGCTGCGATCTATTGAAAGTCAGCCCTCGACACAAGGGTTT | >hg38_21_+_8433172_8446622_RNA45SN1_45S_50ntFlanks | 12997 | 13037 | 41 | rRF 45S | NO | 0.00 | 89.17 | 0.00 | 11.01 |
| >hg38_21_+_8433172_8446622_RNA45SN1_45S_50ntFlanks@12997.13038.42\|CGCTGCGATCTATTGAAAGTCAGCCCTCGACACAAGGGTTTG | >hg38_21_+_8433172_8446622_RNA45SN1_45S_50ntFlanks | 12997 | 13038 | 42 | rRF 45S | NO | 62.44 | 256.83 | 0.00 | 77.79 |
| >hg38_21_+_8433172_8446622_RNA45SN1_45S_50ntFlanks@12997.13039.43\|CGCTGCGATCTATTGAAAGTCAGCCCTCGACACAAGGGTTTGT | >hg38_21_+_8433172_8446622_RNA45SN1_45S_50ntFlanks | 12997 | 13039 | 43 | rRF 45S | YES | 2668.37 | 12381.66 | 389.96 | 1519.85 |
| >hg38_21_+_8433172_8446622_RNA45SN1_45S_50ntFlanks@12998.13039.42\|GCTGCGATCTATTGAAAGTCAGCCCTCGACACAAGGGTTTGT | >hg38_21_+_8433172_8446622_RNA45SN1_45S_50ntFlanks | 12998 | 13039 | 42 | rRF 45S | NO | 214.72 | 347.87 | 76.20 | 148.45 |
| >hg38_21_+_8433172_8446622_RNA45SN1_45S_50ntFlanks@12999.13027.29\|CTGCGATCTATTGAAAGTCAGCCCTCGAC | >hg38_21_+_8433172_8446622_RNA45SN1_45S_50ntFlanks | 12999 | 13027 | 29 | rRF 45S | NO | 0.00 | 28.45 | 0.00 | 0.00 |
| >hg38_21_+_8433172_8446622_RNA45SN1_45S_50ntFlanks@12999.13039.41\|CTGCGATCTATTGAAAGTCAGCCCTCGACACAAGGGTTTGT | >hg38_21_+_8433172_8446622_RNA45SN1_45S_50ntFlanks | 12999 | 13039 | 41 | rRF 45S | NO | 252.66 | 612.17 | 16.94 | 151.89 |
| >hg38_21_+_8433172_8446622_RNA45SN1_45S_50ntFlanks@13000.13039.40\|TGCGATCTATTGAAAGTCAGCCCTCGACACAAGGGTTTGT | >hg38_21_+_8433172_8446622_RNA45SN1_45S_50ntFlanks | 13000 | 13039 | 40 | rRF 45S | NO | 47.68 | 28.14 | 0.00 | 91.18 |
| >hg38_21_+_8433172_8446622_RNA45SN1_45S_50ntFlanks@13006.13039.34\|CTATTGAAAGTCAGCCCTCGACACAAGGGTTTGT | >hg38_21_+_8433172_8446622_RNA45SN1_45S_50ntFlanks | 13006 | 13039 | 34 | rRF 45S | NO | 10.64 | 13.56 | 0.00 | 13.99 |
| >hg38_21_+_8433172_8446622_RNA45SN1_45S_50ntFlanks@13007.13039.33\|TATTGAAAGTCAGCCCTCGACACAAGGGTTTGT | >hg38_21_+_8433172_8446622_RNA45SN1_45S_50ntFlanks | 13007 | 13039 | 33 | rRF 45S | NO | 69.33 | 87.40 | 0.00 | 35.47 |
| >hg38_21_+_8433172_8446622_RNA45SN1_45S_50ntFlanks@13008.13039.32\|ATTGAAAGTCAGCCCTCGACACAAGGGTTTGT | >hg38_21_+_8433172_8446622_RNA45SN1_45S_50ntFlanks | 13008 | 13039 | 32 | rRF 45S | NO | 64.12 | 36.23 | 11.66 | 15.57 |
| >hg38_21_+_8433172_8446622_RNA45SN1_45S_50ntFlanks@13022.13039.18\|CTCGACACAAGGGTTTGT | >hg38_21_+_8433172_8446622_RNA45SN1_45S_50ntFlanks | 13022 | 13039 | 18 | rRF 45S | NO | 45.01 | 63.25 | 10.89 | 20.42 |
| >hg38_1_-_228634819_228635039_RNA5S12_5S_50ntFlanks@51.75.25\|GTCTACGGCCATACCACCCTGAACG | >hg38_1_-_228634819_228635039_RNA5S12_5S_50ntFlanks | 51 | 75 | 25 | rRF 5S | NO | 0.00 | 86.35 | 0.00 | 0.00 |
| >hg38_1_-_228634819_228635039_RNA5S12_5S_50ntFlanks@51.79.29\|GTCTACGGCCATACCACCCTGAACGCGCC | >hg38_1_-_228634819_228635039_RNA5S12_5S_50ntFlanks | 51 | 79 | 29 | rRF 5S | NO | 73.28 | 145.06 | 0.00 | 10.34 |
| >hg38_1_-_228634819_228635039_RNA5S12_5S_50ntFlanks@51.80.30\|GTCTACGGCCATACCACCCTGAACGCGCCC | >hg38_1_-_228634819_228635039_RNA5S12_5S_50ntFlanks | 51 | 80 | 30 | rRF 5S | NO | 59.34 | 112.94 | 0.00 | 0.00 |
| >hg38_1_-_228634819_228635039_RNA5S12_5S_50ntFlanks@51.81.31\|GTCTACGGCCATACCACCCTGAACGCGCCCG | >hg38_1_-_228634819_228635039_RNA5S12_5S_50ntFlanks | 51 | 81 | 31 | rRF 5S | NO | 77.74 | 160.15 | 0.00 | 11.87 |
| >hg38_1_-_228634819_228635039_RNA5S12_5S_50ntFlanks@51.82.32\|GTCTACGGCCATACCACCCTGAACGCGCCCGA | >hg38_1_-_228634819_228635039_RNA5S12_5S_50ntFlanks | 51 | 82 | 32 | rRF 5S | NO | 77.26 | 147.56 | 0.00 | 13.93 |
| >hg38_1_-_228634819_228635039_RNA5S12_5S_50ntFlanks@51.83.33\|GTCTACGGCCATACCACCCTGAACGCGCCCGAT | >hg38_1_-_228634819_228635039_RNA5S12_5S_50ntFlanks | 51 | 83 | 33 | rRF 5S | NO | 94.08 | 189.62 | 0.00 | 12.26 |
| >hg38_1_-_228634819_228635039_RNA5S12_5S_50ntFlanks@51.84.34\|GTCTACGGCCATACCACCCTGAACGCGCCCGATC | >hg38_1_-_228634819_228635039_RNA5S12_5S_50ntFlanks | 51 | 84 | 34 | rRF 5S | NO | 208.50 | 957.22 | 62.91 | 72.26 |
| >hg38_1_-_228634819_228635039_RNA5S12_5S_50ntFlanks@51.85.35\|GTCTACGGCCATACCACCCTGAACGCGCCCGATCT | >hg38_1_-_228634819_228635039_RNA5S12_5S_50ntFlanks | 51 | 85 | 35 | rRF 5S | NO | 62.10 | 130.65 | 0.00 | 0.00 |

### MDA-MB-231

<table>
<tr><th></th><th></th><th></th><th></th><th></th><th></th><th></th><th></th><th></th><th></th><th></th></tr>
<tr><td>>hg38_1_-<br>_228634819_228635039_RNA5S12_5S_50ntFlanks@51.86.36|GTCTACGGCCATACCACCCTGA<br>ACGCGCCCGATCTC</td><td>>hg38_1_-_228634819_228635039_RNA5S12_5S_50ntFlanks</td><td>51</td><td>86</td><td>36</td><td>rRF 5S</td><td>NO</td><td>109.72</td><td>202.49</td><td>10.83</td><td>13.75</td></tr>
<tr><td>>hg38_1_-<br>_228634819_228635039_RNA5S12_5S_50ntFlanks@51.87.37|GTCTACGGCCATACCACCCTGA<br>ACGCGCCCGATCTCG</td><td>>hg38_1_-_228634819_228635039_RNA5S12_5S_50ntFlanks</td><td>51</td><td>87</td><td>37</td><td>rRF 5S</td><td>NO</td><td>63.91</td><td>135.58</td><td>0.00</td><td>13.84</td></tr>
<tr><td>>hg38_1_-<br>_228634819_228635039_RNA5S12_5S_50ntFlanks@51.88.38|GTCTACGGCCATACCACCCTGA<br>ACGCGCCCGATCTCGT</td><td>>hg38_1_-_228634819_228635039_RNA5S12_5S_50ntFlanks</td><td>51</td><td>88</td><td>38</td><td>rRF 5S</td><td>NO</td><td>122.28</td><td>331.85</td><td>32.11</td><td>35.45</td></tr>
<tr><td>>hg38_1_-<br>_228634819_228635039_RNA5S12_5S_50ntFlanks@51.89.39|GTCTACGGCCATACCACCCTGA<br>ACGCGCCCGATCTCGTC</td><td>>hg38_1_-_228634819_228635039_RNA5S12_5S_50ntFlanks</td><td>51</td><td>89</td><td>39</td><td>rRF 5S</td><td>YES</td><td>458.84</td><td>2410.09</td><td>232.67</td><td>247.62</td></tr>
<tr><td>>hg38_1_-<br>_228634819_228635039_RNA5S12_5S_50ntFlanks@51.90.40|GTCTACGGCCATACCACCCTGA<br>ACGCGCCCGATCTCGTCT</td><td>>hg38_1_-_228634819_228635039_RNA5S12_5S_50ntFlanks</td><td>51</td><td>90</td><td>40</td><td>rRF 5S</td><td>NO</td><td>0.00</td><td>80.15</td><td>0.00</td><td>0.00</td></tr>
<tr><td>>hg38_1_-<br>_228634819_228635039_RNA5S12_5S_50ntFlanks@51.91.41|GTCTACGGCCATACCACCCTGA<br>ACGCGCCCGATCTCGTCTG</td><td>>hg38_1_-_228634819_228635039_RNA5S12_5S_50ntFlanks</td><td>51</td><td>91</td><td>41</td><td>rRF 5S</td><td>NO</td><td>10.49</td><td>114.82</td><td>13.50</td><td>0.00</td></tr>
<tr><td>>hg38_1_-<br>_228634819_228635039_RNA5S12_5S_50ntFlanks@51.92.42|GTCTACGGCCATACCACCCTGA<br>ACGCGCCCGATCTCGTCTGA</td><td>>hg38_1_-_228634819_228635039_RNA5S12_5S_50ntFlanks</td><td>51</td><td>92</td><td>42</td><td>rRF 5S</td><td>NO</td><td>0.00</td><td>86.31</td><td>0.00</td><td>14.09</td></tr>
<tr><td>>hg38_1_-<br>_228634819_228635039_RNA5S12_5S_50ntFlanks@61.89.29|ATACCACCCTGAACGCGCCCG<br>ATCTCGTC</td><td>>hg38_1_-_228634819_228635039_RNA5S12_5S_50ntFlanks</td><td>61</td><td>89</td><td>29</td><td>rRF 5S</td><td>NO</td><td>0.00</td><td>82.25</td><td>0.00</td><td>0.00</td></tr>
<tr><td>>hg38_1_-<br>_228634819_228635039_RNA5S12_5S_50ntFlanks@89.123.35|CTGATCTCGGAAGCTAAGCAG<br>GGTCGGGCCTGGTT</td><td>>hg38_1_-_228634819_228635039_RNA5S12_5S_50ntFlanks</td><td>89</td><td>123</td><td>35</td><td>rRF 5S</td><td>NO</td><td>0.00</td><td>36.52</td><td>0.00</td><td>0.00</td></tr>
<tr><td>>hg38_1_-<br>_228634819_228635039_RNA5S12_5S_50ntFlanks@90.123.34|TGATCTCGGAAGCTAAGCAGG<br>GTCGGGCCTGGTT</td><td>>hg38_1_-_228634819_228635039_RNA5S12_5S_50ntFlanks</td><td>90</td><td>123</td><td>34</td><td>rRF 5S</td><td>NO</td><td>0.00</td><td>36.15</td><td>0.00</td><td>0.00</td></tr>
<tr><td>>hg38_1_-<br>_228634819_228635039_RNA5S12_5S_50ntFlanks@121.170.50|GTTAGTACTTGGATGGGAGAC<br>CGCCTGGGAATACCGGGTGCTGTAGGCTT</td><td>>hg38_1_-_228634819_228635039_RNA5S12_5S_50ntFlanks</td><td>121</td><td>170</td><td>50</td><td>rRF 5S</td><td>NO</td><td>0.00</td><td>31.71</td><td>0.00</td><td>0.00</td></tr>
<tr><td>>hg38_1_-<br>_228634819_228635039_RNA5S12_5S_50ntFlanks@122.170.49|TTAGTACTTGGATGGGAGACC<br>GCCTGGGAATACCGGGTGCTGTAGGCTT</td><td>>hg38_1_-_228634819_228635039_RNA5S12_5S_50ntFlanks</td><td>122</td><td>170</td><td>49</td><td>rRF 5S</td><td>NO</td><td>0.00</td><td>58.43</td><td>0.00</td><td>0.00</td></tr>
<tr><td>>hg38_1_-<br>_228634819_228635039_RNA5S12_5S_50ntFlanks@123.170.48|TAGTACTTGGATGGGAGACC<br>GCCTGGGAATACCGGGTGCTGTAGGCTT</td><td>>hg38_1_-_228634819_228635039_RNA5S12_5S_50ntFlanks</td><td>123</td><td>170</td><td>48</td><td>rRF 5S</td><td>NO</td><td>0.00</td><td>63.25</td><td>0.00</td><td>0.00</td></tr>
<tr><td>>hg38_1_-<br>_228634819_228635039_RNA5S12_5S_50ntFlanks@124.170.47|AGTACTTGGATGGGAGACCG<br>CCTGGGAATACCGGGTGCTGTAGGCTT</td><td>>hg38_1_-_228634819_228635039_RNA5S12_5S_50ntFlanks</td><td>124</td><td>170</td><td>47</td><td>rRF 5S</td><td>NO</td><td>0.00</td><td>77.37</td><td>0.00</td><td>0.00</td></tr>
<tr><td>>hg38_1_-<br>_228634819_228635039_RNA5S12_5S_50ntFlanks@124.171.48|AGTACTTGGATGGGAGACCG<br>CCTGGGAATACCGGGTGCTGTAGGCTTT</td><td>>hg38_1_-_228634819_228635039_RNA5S12_5S_50ntFlanks</td><td>124</td><td>171</td><td>48</td><td>rRF 5S</td><td>NO</td><td>0.00</td><td>29.96</td><td>0.00</td><td>0.00</td></tr>
<tr><td>>hg38_1_-<br>_228634819_228635039_RNA5S12_5S_50ntFlanks@125.170.46|GTACTTGGATGGGAGACCGC<br>CTGGGAATACCGGGTGCTGTAGGCTT</td><td>>hg38_1_-_228634819_228635039_RNA5S12_5S_50ntFlanks</td><td>125</td><td>170</td><td>46</td><td>rRF 5S</td><td>NO</td><td>0.00</td><td>70.01</td><td>0.00</td><td>0.00</td></tr>
<tr><td>>hg38_1_-<br>_228634819_228635039_RNA5S12_5S_50ntFlanks@125.171.47|GTACTTGGATGGGAGACCGC<br>CTGGGAATACCGGGTGCTGTAGGCTTT</td><td>>hg38_1_-_228634819_228635039_RNA5S12_5S_50ntFlanks</td><td>125</td><td>171</td><td>47</td><td>rRF 5S</td><td>NO</td><td>0.00</td><td>29.65</td><td>0.00</td><td>0.00</td></tr>
<tr><td>>hg38_1_-<br>_228634819_228635039_RNA5S12_5S_50ntFlanks@126.169.44|TACTTGGATGGGAGACCGCC<br>TGGGAATACCGGGTGCTGTAGGCT</td><td>>hg38_1_-_228634819_228635039_RNA5S12_5S_50ntFlanks</td><td>126</td><td>169</td><td>44</td><td>rRF 5S</td><td>NO</td><td>0.00</td><td>78.80</td><td>0.00</td><td>0.00</td></tr>
<tr><td>>hg38_1_-<br>_228634819_228635039_RNA5S12_5S_50ntFlanks@126.170.45|TACTTGGATGGGAGACCGCC<br>TGGGAATACCGGGTGCTGTAGGCTT</td><td>>hg38_1_-_228634819_228635039_RNA5S12_5S_50ntFlanks</td><td>126</td><td>170</td><td>45</td><td>rRF 5S</td><td>NO</td><td>0.00</td><td>285.17</td><td>0.00</td><td>0.00</td></tr>
<tr><td>>hg38_1_-<br>_228634819_228635039_RNA5S12_5S_50ntFlanks@126.171.46|TACTTGGATGGGAGACCGCC<br>TGGGAATACCGGGTGCTGTAGGCTTT</td><td>>hg38_1_-_228634819_228635039_RNA5S12_5S_50ntFlanks</td><td>126</td><td>171</td><td>46</td><td>rRF 5S</td><td>NO</td><td>0.00</td><td>200.40</td><td>0.00</td><td>0.00</td></tr>
<tr><td>>hg38_1_-<br>_228634819_228635039_RNA5S12_5S_50ntFlanks@127.169.43|ACTTGGATGGGAGACCGCCT<br>GGGAATACCGGGTGCTGTAGGCT</td><td>>hg38_1_-_228634819_228635039_RNA5S12_5S_50ntFlanks</td><td>127</td><td>169</td><td>43</td><td>rRF 5S</td><td>NO</td><td>0.00</td><td>68.13</td><td>0.00</td><td>0.00</td></tr>
<tr><td>>hg38_1_-<br>_228634819_228635039_RNA5S12_5S_50ntFlanks@127.170.44|ACTTGGATGGGAGACCGCCT<br>GGGAATACCGGGTGCTGTAGGCTT</td><td>>hg38_1_-_228634819_228635039_RNA5S12_5S_50ntFlanks</td><td>127</td><td>170</td><td>44</td><td>rRF 5S</td><td>NO</td><td>0.00</td><td>238.43</td><td>0.00</td><td>0.00</td></tr>
<tr><td>>hg38_1_-<br>_228634819_228635039_RNA5S12_5S_50ntFlanks@127.171.45|ACTTGGATGGGAGACCGCCT<br>GGGAATACCGGGTGCTGTAGGCTTT</td><td>>hg38_1_-_228634819_228635039_RNA5S12_5S_50ntFlanks</td><td>127</td><td>171</td><td>45</td><td>rRF 5S</td><td>NO</td><td>0.00</td><td>199.52</td><td>0.00</td><td>0.00</td></tr>
<tr><td>>hg38_1_-<br>_228634819_228635039_RNA5S12_5S_50ntFlanks@128.169.42|CTTGGATGGGAGACCGCCTG<br>GGAATACCGGGTGCTGTAGGCT</td><td>>hg38_1_-_228634819_228635039_RNA5S12_5S_50ntFlanks</td><td>128</td><td>169</td><td>42</td><td>rRF 5S</td><td>NO</td><td>0.00</td><td>37.76</td><td>0.00</td><td>0.00</td></tr>
<tr><td>>hg38_1_-<br>_228634819_228635039_RNA5S12_5S_50ntFlanks@128.170.43|CTTGGATGGGAGACCGCCTG<br>GGAATACCGGGTGCTGTAGGCTT</td><td>>hg38_1_-_228634819_228635039_RNA5S12_5S_50ntFlanks</td><td>128</td><td>170</td><td>43</td><td>rRF 5S</td><td>NO</td><td>0.00</td><td>201.49</td><td>0.00</td><td>0.00</td></tr>
<tr><td>>hg38_1_-<br>_228634819_228635039_RNA5S12_5S_50ntFlanks@128.171.44|CTTGGATGGGAGACCGCCTG<br>GGAATACCGGGTGCTGTAGGCTTT</td><td>>hg38_1_-_228634819_228635039_RNA5S12_5S_50ntFlanks</td><td>128</td><td>171</td><td>44</td><td>rRF 5S</td><td>NO</td><td>0.00</td><td>187.69</td><td>0.00</td><td>0.00</td></tr>
</table>

### MDA-MB-231

| | | | | | | | | | | |
|---|---|---|---|---|---|---|---|---|---|---|
| >hg38_1_-_228634819_228635039_RNA5S12_5S_50ntFlanks@129.169.41\|TTGGATGGGAGACCGCCTGGGAATACCGGGTGCTGTAGGCT | >hg38_1_-_228634819_228635039_RNA5S12_5S_50ntFlanks | 129 | 169 | 41 | rRF 5S | NO | 0.00 | 92.22 | 0.00 | 0.00 |
| >hg38_1_-_228634819_228635039_RNA5S12_5S_50ntFlanks@129.170.42\|TTGGATGGGAGACCGCCTGGGAATACCGGGTGCTGTAGGCTT | >hg38_1_-_228634819_228635039_RNA5S12_5S_50ntFlanks | 129 | 170 | 42 | rRF 5S | NO | 46.47 | 382.00 | 15.15 | 0.00 |
| >hg38_1_-_228634819_228635039_RNA5S12_5S_50ntFlanks@129.171.43\|TTGGATGGGAGACCGCCTGGGAATACCGGGTGCTGTAGGCTTT | >hg38_1_-_228634819_228635039_RNA5S12_5S_50ntFlanks | 129 | 171 | 43 | rRF 5S | NO | 0.00 | 304.35 | 0.00 | 0.00 |
| >hg38_1_-_228634819_228635039_RNA5S12_5S_50ntFlanks@130.169.40\|TGGATGGGAGACCGCCTGGGAATACCGGGTGCTGTAGGCT | >hg38_1_-_228634819_228635039_RNA5S12_5S_50ntFlanks | 130 | 169 | 40 | rRF 5S | NO | 0.00 | 95.11 | 0.00 | 0.00 |
| >hg38_1_-_228634819_228635039_RNA5S12_5S_50ntFlanks@130.170.41\|TGGATGGGAGACCGCCTGGGAATACCGGGTGCTGTAGGCTT | >hg38_1_-_228634819_228635039_RNA5S12_5S_50ntFlanks | 130 | 170 | 41 | rRF 5S | NO | 46.59 | 416.27 | 56.55 | 0.00 |
| >hg38_1_-_228634819_228635039_RNA5S12_5S_50ntFlanks@130.171.42\|TGGATGGGAGACCGCCTGGGAATACCGGGTGCTGTAGGCTTT | >hg38_1_-_228634819_228635039_RNA5S12_5S_50ntFlanks | 130 | 171 | 42 | rRF 5S | NO | 0.00 | 325.88 | 0.00 | 0.00 |
| >hg38_1_-_228634819_228635039_RNA5S12_5S_50ntFlanks@131.169.39\|GGATGGGAGACCGCCTGGGAATACCGGGTGCTGTAGGCT | >hg38_1_-_228634819_228635039_RNA5S12_5S_50ntFlanks | 131 | 169 | 39 | rRF 5S | NO | 0.00 | 64.54 | 0.00 | 0.00 |
| >hg38_1_-_228634819_228635039_RNA5S12_5S_50ntFlanks@131.170.40\|GGATGGGAGACCGCCTGGGAATACCGGGTGCTGTAGGCTT | >hg38_1_-_228634819_228635039_RNA5S12_5S_50ntFlanks | 131 | 170 | 40 | rRF 5S | NO | 0.00 | 254.68 | 15.67 | 0.00 |
| >hg38_1_-_228634819_228635039_RNA5S12_5S_50ntFlanks@131.171.41\|GGATGGGAGACCGCCTGGGAATACCGGGTGCTGTAGGCTTT | >hg38_1_-_228634819_228635039_RNA5S12_5S_50ntFlanks | 131 | 171 | 41 | rRF 5S | NO | 0.00 | 220.78 | 0.00 | 0.00 |
| >hg38_1_-_228634819_228635039_RNA5S12_5S_50ntFlanks@132.169.38\|GATGGGAGACCGCCTGGGAATACCGGGTGCTGTAGGCT | >hg38_1_-_228634819_228635039_RNA5S12_5S_50ntFlanks | 132 | 169 | 38 | rRF 5S | NO | 0.00 | 289.63 | 14.89 | 0.00 |
| >hg38_1_-_228634819_228635039_RNA5S12_5S_50ntFlanks@132.170.39\|GATGGGAGACCGCCTGGGAATACCGGGTGCTGTAGGCTT | >hg38_1_-_228634819_228635039_RNA5S12_5S_50ntFlanks | 132 | 170 | 39 | rRF 5S | NO | 345.13 | 2003.31 | 339.16 | 58.87 |
| >hg38_1_-_228634819_228635039_RNA5S12_5S_50ntFlanks@132.171.40\|GATGGGAGACCGCCTGGGAATACCGGGTGCTGTAGGCTTT | >hg38_1_-_228634819_228635039_RNA5S12_5S_50ntFlanks | 132 | 171 | 40 | rRF 5S | NO | 169.29 | 926.67 | 124.78 | 19.38 |
| >hg38_1_-_228634819_228635039_RNA5S12_5S_50ntFlanks@133.169.37\|ATGGGAGACCGCCTGGGAATACCGGGTGCTGTAGGCT | >hg38_1_-_228634819_228635039_RNA5S12_5S_50ntFlanks | 133 | 169 | 37 | rRF 5S | NO | 0.00 | 171.08 | 0.00 | 0.00 |
| >hg38_1_-_228634819_228635039_RNA5S12_5S_50ntFlanks@133.170.38\|ATGGGAGACCGCCTGGGAATACCGGGTGCTGTAGGCTT | >hg38_1_-_228634819_228635039_RNA5S12_5S_50ntFlanks | 133 | 170 | 38 | rRF 5S | NO | 259.86 | 679.15 | 143.68 | 17.99 |
| >hg38_1_-_228634819_228635039_RNA5S12_5S_50ntFlanks@133.171.39\|ATGGGAGACCGCCTGGGAATACCGGGTGCTGTAGGCTTT | >hg38_1_-_228634819_228635039_RNA5S12_5S_50ntFlanks | 133 | 171 | 39 | rRF 5S | NO | 147.77 | 502.70 | 57.74 | 15.86 |
| >hg38_1_-_228634819_228635039_RNA5S12_5S_50ntFlanks@134.168.35\|TGGGAGACCGCCTGGGAATACCGGGTGCTGTAGGC | >hg38_1_-_228634819_228635039_RNA5S12_5S_50ntFlanks | 134 | 168 | 35 | rRF 5S | NO | 0.00 | 35.84 | 0.00 | 0.00 |
| >hg38_1_-_228634819_228635039_RNA5S12_5S_50ntFlanks@134.169.36\|TGGGAGACCGCCTGGGAATACCGGGTGCTGTAGGCT | >hg38_1_-_228634819_228635039_RNA5S12_5S_50ntFlanks | 134 | 169 | 36 | rRF 5S | NO | 156.85 | 860.24 | 93.56 | 19.68 |
| >hg38_1_-_228634819_228635039_RNA5S12_5S_50ntFlanks@134.170.37\|TGGGAGACCGCCTGGGAATACCGGGTGCTGTAGGCTT | >hg38_1_-_228634819_228635039_RNA5S12_5S_50ntFlanks | 134 | 170 | 37 | rRF 5S | YES | 1553.51 | 4305.02 | 1048.99 | 260.88 |
| >hg38_1_-_228634819_228635039_RNA5S12_5S_50ntFlanks@134.171.38\|TGGGAGACCGCCTGGGAATACCGGGTGCTGTAGGCTTT | >hg38_1_-_228634819_228635039_RNA5S12_5S_50ntFlanks | 134 | 171 | 38 | rRF 5S | YES | 765.64 | 2133.17 | 458.12 | 210.01 |
| >hg38_1_-_228634819_228635039_RNA5S12_5S_50ntFlanks@134.172.39\|TGGGAGACCGCCTGGGAATACCGGGTGCTGTAGGCTTTT | >hg38_1_-_228634819_228635039_RNA5S12_5S_50ntFlanks | 134 | 172 | 39 | rRF 5S | NO | 0.00 | 93.22 | 0.00 | 0.00 |
| >hg38_1_-_228634819_228635039_RNA5S12_5S_50ntFlanks@135.169.35\|GGGAGACCGCCTGGGAATACCGGGTGCTGTAGGCT | >hg38_1_-_228634819_228635039_RNA5S12_5S_50ntFlanks | 135 | 169 | 35 | rRF 5S | NO | 10.55 | 131.95 | 15.41 | 0.00 |
| >hg38_1_-_228634819_228635039_RNA5S12_5S_50ntFlanks@135.170.36\|GGGAGACCGCCTGGGAATACCGGGTGCTGTAGGCTT | >hg38_1_-_228634819_228635039_RNA5S12_5S_50ntFlanks | 135 | 170 | 36 | rRF 5S | YES | 584.81 | 1243.11 | 594.86 | 95.19 |
| >hg38_1_-_228634819_228635039_RNA5S12_5S_50ntFlanks@135.171.37\|GGGAGACCGCCTGGGAATACCGGGTGCTGTAGGCTTT | >hg38_1_-_228634819_228635039_RNA5S12_5S_50ntFlanks | 135 | 171 | 37 | rRF 5S | YES | 335.78 | 833.13 | 238.93 | 87.09 |
| >hg38_1_-_228634819_228635039_RNA5S12_5S_50ntFlanks@135.172.38\|GGGAGACCGCCTGGGAATACCGGGTGCTGTAGGCTTTT | >hg38_1_-_228634819_228635039_RNA5S12_5S_50ntFlanks | 135 | 172 | 38 | rRF 5S | NO | 0.00 | 27.05 | 0.00 | 0.00 |
| >hg38_1_-_228634819_228635039_RNA5S12_5S_50ntFlanks@136.170.35\|GGAGACCGCCTGGGAATACCGGGTGCTGTAGGCTT | >hg38_1_-_228634819_228635039_RNA5S12_5S_50ntFlanks | 136 | 170 | 35 | rRF 5S | NO | 84.18 | 113.24 | 55.43 | 0.00 |
| >hg38_1_-_228634819_228635039_RNA5S12_5S_50ntFlanks@136.171.36\|GGAGACCGCCTGGGAATACCGGGTGCTGTAGGCTTT | >hg38_1_-_228634819_228635039_RNA5S12_5S_50ntFlanks | 136 | 171 | 36 | rRF 5S | NO | 131.87 | 186.27 | 55.69 | 16.99 |

### MDA-MB-231

| | | | | | | | | | | |
|---|---|---|---|---|---|---|---|---|---|---|
| >hg38_1_-_228634819_228635039_RNA5S12_5S_50ntFlanks@137.170.34\|GAGACCGCCTGGGAATACCGGGTGCTGTAGGCTT | >hg38_1_-_228634819_228635039_RNA5S12_5S_50ntFlanks | 137 | 170 | 34 | rRF 5S | NO | 0.00 | 64.76 | 10.79 | 0.00 |
| >hg38_1_-_228634819_228635039_RNA5S12_5S_50ntFlanks@137.171.35\|GAGACCGCCTGGGAATACCGGGTGCTGTAGGCTTT | >hg38_1_-_228634819_228635039_RNA5S12_5S_50ntFlanks | 137 | 171 | 35 | rRF 5S | NO | 82.26 | 169.90 | 33.17 | 10.46 |
| >hg38_1_-_228634819_228635039_RNA5S12_5S_50ntFlanks@138.169.32\|AGACCGCCTGGGAATACCGGGTGCTGTAGGCT | >hg38_1_-_228634819_228635039_RNA5S12_5S_50ntFlanks | 138 | 169 | 32 | rRF 5S | NO | 0.00 | 28.86 | 15.55 | 0.00 |
| >hg38_1_-_228634819_228635039_RNA5S12_5S_50ntFlanks@138.170.33\|AGACCGCCTGGGAATACCGGGTGCTGTAGGCTT | >hg38_1_-_228634819_228635039_RNA5S12_5S_50ntFlanks | 138 | 170 | 33 | rRF 5S | NO | 61.09 | 112.73 | 104.10 | 11.51 |
| >hg38_1_-_228634819_228635039_RNA5S12_5S_50ntFlanks@138.171.34\|AGACCGCCTGGGAATACCGGGTGCTGTAGGCTTT | >hg38_1_-_228634819_228635039_RNA5S12_5S_50ntFlanks | 138 | 171 | 34 | rRF 5S | NO | 60.75 | 145.99 | 83.40 | 16.49 |
| >hg38_1_-_228634819_228635039_RNA5S12_5S_50ntFlanks@139.169.31\|GACCGCCTGGGAATACCGGGTGCTGTAGGCT | >hg38_1_-_228634819_228635039_RNA5S12_5S_50ntFlanks | 139 | 169 | 31 | rRF 5S | NO | 0.00 | 63.68 | 0.00 | 0.00 |
| >hg38_1_-_228634819_228635039_RNA5S12_5S_50ntFlanks@139.170.32\|GACCGCCTGGGAATACCGGGTGCTGTAGGCTT | >hg38_1_-_228634819_228635039_RNA5S12_5S_50ntFlanks | 139 | 170 | 32 | rRF 5S | NO | 59.70 | 113.18 | 32.31 | 0.00 |
| >hg38_1_-_228634819_228635039_RNA5S12_5S_50ntFlanks@139.171.33\|GACCGCCTGGGAATACCGGGTGCTGTAGGCTTT | >hg38_1_-_228634819_228635039_RNA5S12_5S_50ntFlanks | 139 | 171 | 33 | rRF 5S | NO | 46.87 | 128.89 | 17.87 | 0.00 |
| >hg38_1_-_228634819_228635039_RNA5S12_5S_50ntFlanks@140.169.30\|ACCGCCTGGGAATACCGGGTGCTGTAGGCT | >hg38_1_-_228634819_228635039_RNA5S12_5S_50ntFlanks | 140 | 169 | 30 | rRF 5S | NO | 97.99 | 257.81 | 55.97 | 17.06 |
| >hg38_1_-_228634819_228635039_RNA5S12_5S_50ntFlanks@140.170.31\|ACCGCCTGGGAATACCGGGTGCTGTAGGCTT | >hg38_1_-_228634819_228635039_RNA5S12_5S_50ntFlanks | 140 | 170 | 31 | rRF 5S | NO | 416.74 | 788.09 | 175.23 | 90.40 |
| >hg38_1_-_228634819_228635039_RNA5S12_5S_50ntFlanks@140.171.32\|ACCGCCTGGGAATACCGGGTGCTGTAGGCTTT | >hg38_1_-_228634819_228635039_RNA5S12_5S_50ntFlanks | 140 | 171 | 32 | rRF 5S | NO | 181.66 | 425.25 | 91.97 | 21.43 |
| >hg38_1_-_228634819_228635039_RNA5S12_5S_50ntFlanks@141.169.29\|CCGCCTGGGAATACCGGGTGCTGTAGGCT | >hg38_1_-_228634819_228635039_RNA5S12_5S_50ntFlanks | 141 | 169 | 29 | rRF 5S | NO | 52.60 | 105.88 | 0.00 | 0.00 |
| >hg38_1_-_228634819_228635039_RNA5S12_5S_50ntFlanks@141.170.30\|CCGCCTGGGAATACCGGGTGCTGTAGGCTT | >hg38_1_-_228634819_228635039_RNA5S12_5S_50ntFlanks | 141 | 170 | 30 | rRF 5S | NO | 91.92 | 234.42 | 63.89 | 12.59 |
| >hg38_1_-_228634819_228635039_RNA5S12_5S_50ntFlanks@141.171.31\|CCGCCTGGGAATACCGGGTGCTGTAGGCTTT | >hg38_1_-_228634819_228635039_RNA5S12_5S_50ntFlanks | 141 | 171 | 31 | rRF 5S | NO | 91.12 | 314.05 | 73.44 | 15.38 |
| >hg38_1_-_228634819_228635039_RNA5S12_5S_50ntFlanks@142.170.29\|CGCCTGGGAATACCGGGTGCTGTAGGCTT | >hg38_1_-_228634819_228635039_RNA5S12_5S_50ntFlanks | 142 | 170 | 29 | rRF 5S | NO | 26.45 | 127.22 | 11.86 | 0.00 |
| >hg38_1_-_228634819_228635039_RNA5S12_5S_50ntFlanks@142.171.30\|CGCCTGGGAATACCGGGTGCTGTAGGCTTT | >hg38_1_-_228634819_228635039_RNA5S12_5S_50ntFlanks | 142 | 171 | 30 | rRF 5S | NO | 67.48 | 216.92 | 14.08 | 12.87 |
| >hg38_1_-_228634819_228635039_RNA5S12_5S_50ntFlanks@143.171.29\|GCCTGGGAATACCGGGTGCTGTAGGCTTT | >hg38_1_-_228634819_228635039_RNA5S12_5S_50ntFlanks | 143 | 171 | 29 | rRF 5S | NO | 0.00 | 13.35 | 0.00 | 0.00 |
| >hg38_1_-_228634819_228635039_RNA5S12_5S_50ntFlanks@144.169.26\|CCTGGGAATACCGGGTGCTGTAGGCT | >hg38_1_-_228634819_228635039_RNA5S12_5S_50ntFlanks | 144 | 169 | 26 | rRF 5S | NO | 0.00 | 28.59 | 0.00 | 0.00 |
| >hg38_1_-_228634819_228635039_RNA5S12_5S_50ntFlanks@144.170.27\|CCTGGGAATACCGGGTGCTGTAGGCTT | >hg38_1_-_228634819_228635039_RNA5S12_5S_50ntFlanks | 144 | 170 | 27 | rRF 5S | NO | 140.02 | 269.17 | 71.90 | 16.34 |
| >hg38_1_-_228634819_228635039_RNA5S12_5S_50ntFlanks@144.171.28\|CCTGGGAATACCGGGTGCTGTAGGCTTT | >hg38_1_-_228634819_228635039_RNA5S12_5S_50ntFlanks | 144 | 171 | 28 | rRF 5S | NO | 195.17 | 412.11 | 62.32 | 63.91 |
| >hg38_1_-_228634819_228635039_RNA5S12_5S_50ntFlanks@145.169.25\|CTGGGAATACCGGGTGCTGTAGGCT | >hg38_1_-_228634819_228635039_RNA5S12_5S_50ntFlanks | 145 | 169 | 25 | rRF 5S | NO | 0.00 | 75.64 | 0.00 | 0.00 |
| >hg38_1_-_228634819_228635039_RNA5S12_5S_50ntFlanks@145.170.26\|CTGGGAATACCGGGTGCTGTAGGCTT | >hg38_1_-_228634819_228635039_RNA5S12_5S_50ntFlanks | 145 | 170 | 26 | rRF 5S | NO | 25.12 | 261.47 | 12.30 | 0.00 |
| >hg38_1_-_228634819_228635039_RNA5S12_5S_50ntFlanks@145.171.27\|CTGGGAATACCGGGTGCTGTAGGCTTT | >hg38_1_-_228634819_228635039_RNA5S12_5S_50ntFlanks | 145 | 171 | 27 | rRF 5S | NO | 45.81 | 314.79 | 0.00 | 0.00 |
| >hg38_1_-_228634819_228635039_RNA5S12_5S_50ntFlanks@146.169.24\|TGGGAATACCGGGTGCTGTAGGCT | >hg38_1_-_228634819_228635039_RNA5S12_5S_50ntFlanks | 146 | 169 | 24 | rRF 5S | NO | 0.00 | 97.07 | 0.00 | 0.00 |
| >hg38_1_-_228634819_228635039_RNA5S12_5S_50ntFlanks@146.170.25\|TGGGAATACCGGGTGCTGTAGGCTT | >hg38_1_-_228634819_228635039_RNA5S12_5S_50ntFlanks | 146 | 170 | 25 | rRF 5S | NO | 59.10 | 386.46 | 18.04 | 0.00 |
| >hg38_1_-_228634819_228635039_RNA5S12_5S_50ntFlanks@146.171.26\|TGGGAATACCGGGTGCTGTAGGCTTT | >hg38_1_-_228634819_228635039_RNA5S12_5S_50ntFlanks | 146 | 171 | 26 | rRF 5S | NO | 48.09 | 327.34 | 11.08 | 0.00 |

### MDA-MB-231

<table>
<tr><td>>hg38_1_-_228634819_228635039_RNA5S12_5S_50ntFlanks@147.169.23|GGGAATACCGGGTGCTGTAGGCT</td><td>>hg38_1_-_228634819_228635039_RNA5S12_5S_50ntFlanks</td><td>147</td><td>169</td><td>23</td><td>rRF 5S</td><td>NO</td><td>0.00</td><td>124.67</td><td>10.53</td><td>0.00</td></tr>
<tr><td>>hg38_1_-_228634819_228635039_RNA5S12_5S_50ntFlanks@147.170.24|GGGAATACCGGGTGCTGTAGGCTT</td><td>>hg38_1_-_228634819_228635039_RNA5S12_5S_50ntFlanks</td><td>147</td><td>170</td><td>24</td><td>rRF 5S</td><td>NO</td><td>67.70</td><td>473.16</td><td>126.98</td><td>0.00</td></tr>
<tr><td>>hg38_1_-_228634819_228635039_RNA5S12_5S_50ntFlanks@147.171.25|GGGAATACCGGGTGCTGTAGGCTTT</td><td>>hg38_1_-_228634819_228635039_RNA5S12_5S_50ntFlanks</td><td>147</td><td>171</td><td>25</td><td>rRF 5S</td><td>NO</td><td>57.50</td><td>402.59</td><td>54.73</td><td>0.00</td></tr>
<tr><td>>hg38_1_-_228634819_228635039_RNA5S12_5S_50ntFlanks@148.169.22|GGAATACCGGGTGCTGTAGGCT</td><td>>hg38_1_-_228634819_228635039_RNA5S12_5S_50ntFlanks</td><td>148</td><td>169</td><td>22</td><td>rRF 5S</td><td>NO</td><td>0.00</td><td>93.58</td><td>0.00</td><td>0.00</td></tr>
<tr><td>>hg38_1_-_228634819_228635039_RNA5S12_5S_50ntFlanks@148.170.23|GGAATACCGGGTGCTGTAGGCTT</td><td>>hg38_1_-_228634819_228635039_RNA5S12_5S_50ntFlanks</td><td>148</td><td>170</td><td>23</td><td>rRF 5S</td><td>NO</td><td>50.30</td><td>299.31</td><td>32.29</td><td>0.00</td></tr>
<tr><td>>hg38_1_-_228634819_228635039_RNA5S12_5S_50ntFlanks@148.171.24|GGAATACCGGGTGCTGTAGGCTTT</td><td>>hg38_1_-_228634819_228635039_RNA5S12_5S_50ntFlanks</td><td>148</td><td>171</td><td>24</td><td>rRF 5S</td><td>NO</td><td>60.12</td><td>391.90</td><td>14.85</td><td>0.00</td></tr>
<tr><td>>hg38_1_-_228634819_228635039_RNA5S12_5S_50ntFlanks@149.169.21|GAATACCGGGTGCTGTAGGCT</td><td>>hg38_1_-_228634819_228635039_RNA5S12_5S_50ntFlanks</td><td>149</td><td>169</td><td>21</td><td>rRF 5S</td><td>NO</td><td>0.00</td><td>79.49</td><td>0.00</td><td>0.00</td></tr>
<tr><td>>hg38_1_-_228634819_228635039_RNA5S12_5S_50ntFlanks@149.170.22|GAATACCGGGTGCTGTAGGCTT</td><td>>hg38_1_-_228634819_228635039_RNA5S12_5S_50ntFlanks</td><td>149</td><td>170</td><td>22</td><td>rRF 5S</td><td>NO</td><td>55.45</td><td>325.92</td><td>17.90</td><td>0.00</td></tr>
<tr><td>>hg38_1_-_228634819_228635039_RNA5S12_5S_50ntFlanks@149.171.23|GAATACCGGGTGCTGTAGGCTTT</td><td>>hg38_1_-_228634819_228635039_RNA5S12_5S_50ntFlanks</td><td>149</td><td>171</td><td>23</td><td>rRF 5S</td><td>NO</td><td>67.76</td><td>401.98</td><td>13.20</td><td>0.00</td></tr>
<tr><td>>hg38_1_-_228634819_228635039_RNA5S12_5S_50ntFlanks@150.169.20|AATACCGGGTGCTGTAGGCT</td><td>>hg38_1_-_228634819_228635039_RNA5S12_5S_50ntFlanks</td><td>150</td><td>169</td><td>20</td><td>rRF 5S</td><td>NO</td><td>0.00</td><td>71.77</td><td>0.00</td><td>0.00</td></tr>
<tr><td>>hg38_1_-_228634819_228635039_RNA5S12_5S_50ntFlanks@150.170.21|AATACCGGGTGCTGTAGGCTT</td><td>>hg38_1_-_228634819_228635039_RNA5S12_5S_50ntFlanks</td><td>150</td><td>170</td><td>21</td><td>rRF 5S</td><td>NO</td><td>55.93</td><td>312.61</td><td>75.09</td><td>0.00</td></tr>
<tr><td>>hg38_1_-_228634819_228635039_RNA5S12_5S_50ntFlanks@150.171.22|AATACCGGGTGCTGTAGGCTTT</td><td>>hg38_1_-_228634819_228635039_RNA5S12_5S_50ntFlanks</td><td>150</td><td>171</td><td>22</td><td>rRF 5S</td><td>NO</td><td>75.36</td><td>391.25</td><td>36.05</td><td>0.00</td></tr>
<tr><td>>hg38_1_-_228634819_228635039_RNA5S12_5S_50ntFlanks@151.169.19|ATACCGGGTGCTGTAGGCT</td><td>>hg38_1_-_228634819_228635039_RNA5S12_5S_50ntFlanks</td><td>151</td><td>169</td><td>19</td><td>rRF 5S</td><td>NO</td><td>0.00</td><td>68.77</td><td>0.00</td><td>0.00</td></tr>
<tr><td>>hg38_1_-_228634819_228635039_RNA5S12_5S_50ntFlanks@151.170.20|ATACCGGGTGCTGTAGGCTT</td><td>>hg38_1_-_228634819_228635039_RNA5S12_5S_50ntFlanks</td><td>151</td><td>170</td><td>20</td><td>rRF 5S</td><td>NO</td><td>58.68</td><td>273.28</td><td>95.25</td><td>0.00</td></tr>
<tr><td>>hg38_1_-_228634819_228635039_RNA5S12_5S_50ntFlanks@151.171.21|ATACCGGGTGCTGTAGGCTTT</td><td>>hg38_1_-_228634819_228635039_RNA5S12_5S_50ntFlanks</td><td>151</td><td>171</td><td>21</td><td>rRF 5S</td><td>NO</td><td>67.15</td><td>320.66</td><td>32.67</td><td>0.00</td></tr>
<tr><td>>hg38_1_-_228634819_228635039_RNA5S12_5S_50ntFlanks@152.169.18|TACCGGGTGCTGTAGGCT</td><td>>hg38_1_-_228634819_228635039_RNA5S12_5S_50ntFlanks</td><td>152</td><td>169</td><td>18</td><td>rRF 5S</td><td>NO</td><td>0.00</td><td>95.33</td><td>0.00</td><td>0.00</td></tr>
<tr><td>>hg38_1_-_228634819_228635039_RNA5S12_5S_50ntFlanks@152.170.19|TACCGGGTGCTGTAGGCTT</td><td>>hg38_1_-_228634819_228635039_RNA5S12_5S_50ntFlanks</td><td>152</td><td>170</td><td>19</td><td>rRF 5S</td><td>NO</td><td>61.87</td><td>401.50</td><td>72.68</td><td>0.00</td></tr>
<tr><td>>hg38_1_-_228634819_228635039_RNA5S12_5S_50ntFlanks@152.171.20|TACCGGGTGCTGTAGGCTTT</td><td>>hg38_1_-_228634819_228635039_RNA5S12_5S_50ntFlanks</td><td>152</td><td>171</td><td>20</td><td>rRF 5S</td><td>NO</td><td>66.99</td><td>448.40</td><td>33.75</td><td>0.00</td></tr>
<tr><td>>hg38_1_-_228634819_228635039_RNA5S12_5S_50ntFlanks@153.170.18|ACCGGGTGCTGTAGGCTT</td><td>>hg38_1_-_228634819_228635039_RNA5S12_5S_50ntFlanks</td><td>153</td><td>170</td><td>18</td><td>rRF 5S</td><td>NO</td><td>249.12</td><td>1452.62</td><td>677.96</td><td>59.59</td></tr>
<tr><td>>hg38_1_-_228634819_228635039_RNA5S12_5S_50ntFlanks@153.171.19|ACCGGGTGCTGTAGGCTTT</td><td>>hg38_1_-_228634819_228635039_RNA5S12_5S_50ntFlanks</td><td>153</td><td>171</td><td>19</td><td>rRF 5S</td><td>NO</td><td>239.91</td><td>1515.24</td><td>442.32</td><td>68.07</td></tr>
<tr><td>>hg38_1_-_228634819_228635039_RNA5S12_5S_50ntFlanks@153.172.20|ACCGGGTGCTGTAGGCTTTT</td><td>>hg38_1_-_228634819_228635039_RNA5S12_5S_50ntFlanks</td><td>153</td><td>172</td><td>20</td><td>rRF 5S</td><td>NO</td><td>0.00</td><td>29.55</td><td>0.00</td><td>0.00</td></tr>
<tr><td>>hg38_1_-_228634819_228635039_RNA5S12_5S_50ntFlanks@154.171.18|CCGGGTGCTGTAGGCTTT</td><td>>hg38_1_-_228634819_228635039_RNA5S12_5S_50ntFlanks</td><td>154</td><td>171</td><td>18</td><td>rRF 5S</td><td>NO</td><td>62.85</td><td>465.44</td><td>62.20</td><td>0.00</td></tr>
<tr><td>>multi-trnaMT_GlnTTG_MT_-_4329_4400@1.26.26,trnalookalike2_GlnTTG_1_-_564879_564950@1.26.26|TAGGATGGGGTGTGATAGGTGGCACG</td><td>>GlnTTG.trnaMT_MT_-_4329_4400.multi</td><td>1</td><td>26</td><td>26</td><td>tRF lookalike</td><td>NO</td><td>0.00</td><td>0.00</td><td>14.11</td><td>0.00</td></tr>
<tr><td>>multi-trnaMT_GlnTTG_MT_-_4329_4400@1.30.30,trnalookalike2_GlnTTG_1_-_564879_564950@1.30.30|TAGGATGGGGTGTGATAGGTGGCACGGAGA</td><td>>GlnTTG.trnaMT_MT_-_4329_4400.multi</td><td>1</td><td>30</td><td>30</td><td>tRF lookalike</td><td>NO</td><td>0.00</td><td>0.00</td><td>33.38</td><td>0.00</td></tr>
<tr><td>>trnaMT_GluTTC_MT_-_14674_14742@31.72.42,trnalookalike8_GluTTC_5_-_93905172_93905240@31.72.42|TTCATATCATTGGTCGTGGTTGTAGTCCGTGCGAGAATACCA</td><td>>GluTTC.trnaMT_MT_-_14674_14742</td><td>31</td><td>72</td><td>42</td><td>tRF lookalike</td><td>NO</td><td>44.97</td><td>120.44</td><td>34.28</td><td>22.10</td></tr>
<tr><td>>trnaMT_GluTTC_MT_-_14674_14742@32.72.41,trnalookalike8_GluTTC_5_-_93905172_93905240@32.72.41|TCATATCATTGGTCGTGGTTGTAGTCCGTGCGAGAATACCA</td><td>>GluTTC.trnaMT_MT_-_14674_14742</td><td>32</td><td>72</td><td>41</td><td>tRF lookalike</td><td>NO</td><td>0.00</td><td>37.35</td><td>55.47</td><td>13.61</td></tr>
<tr><td>>trnaMT_GluTTC_MT_-_14674_14742@33.72.40,trnalookalike8_GluTTC_5_-_93905172_93905240@33.72.40|CATATCATTGGTCGTGGTTGTAGTCCGTGCGAGAATACCA</td><td>>GluTTC.trnaMT_MT_-_14674_14742</td><td>33</td><td>72</td><td>40</td><td>tRF lookalike</td><td>NO</td><td>0.00</td><td>32.90</td><td>34.72</td><td>13.70</td></tr>
<tr><td>>trnaMT_GluTTC_MT_-_14674_14742@34.71.38,trnalookalike8_GluTTC_5_-_93905172_93905240@34.71.38|ATATCATTGGTCGTGGTTGTAGTCCGTGCGAGAATACC</td><td>>GluTTC.trnaMT_MT_-_14674_14742</td><td>34</td><td>71</td><td>38</td><td>tRF lookalike</td><td>NO</td><td>0.00</td><td>25.90</td><td>82.14</td><td>14.00</td></tr>
<tr><td>>trnaMT_GluTTC_MT_-_14674_14742@34.72.39,trnalookalike8_GluTTC_5_-_93905172_93905240@34.72.39|ATATCATTGGTCGTGGTTGTAGTCCGTGCGAGAATACCA</td><td>>GluTTC.trnaMT_MT_-_14674_14742</td><td>34</td><td>72</td><td>39</td><td>tRF lookalike</td><td>NO</td><td>131.03</td><td>128.96</td><td>321.86</td><td>114.10</td></tr>
</table>

### MDA-MB-231

| | | | | | | | | | | |
|---|---|---|---|---|---|---|---|---|---|---|
| >trnaMT_GluTTC_MT_-_14674_14742@35.71.37,trnalookalike8_GluTTC_5_-_93905172_93905240@35.71.37\|TATCATTGGTCGTGGTTGTAGTCCGTGCGAGAATACC | >GluTTC.trnaMT_MT_-_14674_14742 | 35 | 71 | 37 | tRF lookalike | NO | 0.00 | 98.62 | 16.99 | 0.00 |
| >trnaMT_GluTTC_MT_-_14674_14742@35.72.38,trnalookalike8_GluTTC_5_-_93905172_93905240@35.72.38\|TATCATTGGTCGTGGTTGTAGTCCGTGCGAGAATACCA | >GluTTC.trnaMT_MT_-_14674_14742 | 35 | 72 | 38 | tRF lookalike | NO | 207.50 | 438.67 | 232.69 | 87.55 |
| >trnaMT_GluTTC_MT_-_14674_14742@37.71.35,trnalookalike8_GluTTC_5_-_93905172_93905240@37.71.35\|TCATTGGTCGTGGTTGTAGTCCGTGCGAGAATACC | >GluTTC.trnaMT_MT_-_14674_14742 | 37 | 71 | 35 | tRF lookalike | NO | 24.01 | 67.37 | 0.00 | 0.00 |
| >trnaMT_GluTTC_MT_-_14674_14742@37.72.36,trnalookalike8_GluTTC_5_-_93905172_93905240@37.72.36\|TCATTGGTCGTGGTTGTAGTCCGTGCGAGAATACCA | >GluTTC.trnaMT_MT_-_14674_14742 | 37 | 72 | 36 | tRF lookalike | NO | 278.40 | 292.08 | 97.83 | 20.07 |
| >trnaMT_GluTTC_MT_-_14674_14742@38.71.34,trnalookalike8_GluTTC_5_-_93905172_93905240@38.71.34\|CATTGGTCGTGGTTGTAGTCCGTGCGAGAATACC | >GluTTC.trnaMT_MT_-_14674_14742 | 38 | 71 | 34 | tRF lookalike | NO | 0.00 | 0.00 | 31.49 | 0.00 |
| >trnaMT_GluTTC_MT_-_14674_14742@38.72.35,trnalookalike8_GluTTC_5_-_93905172_93905240@38.72.35\|CATTGGTCGTGGTTGTAGTCCGTGCGAGAATACCA | >GluTTC.trnaMT_MT_-_14674_14742 | 38 | 72 | 35 | tRF lookalike | NO | 224.01 | 169.76 | 373.76 | 109.63 |
| >trnaMT_GluTTC_MT_-_14674_14742@39.71.33,trnalookalike8_GluTTC_5_-_93905172_93905240@39.71.33\|ATTGGTCGTGGTTGTAGTCCGTGCGAGAATACC | >GluTTC.trnaMT_MT_-_14674_14742 | 39 | 71 | 33 | tRF lookalike | NO | 104.78 | 92.08 | 180.57 | 21.50 |
| >trnaMT_GluTTC_MT_-_14674_14742@39.72.34,trnalookalike8_GluTTC_5_-_93905172_93905240@39.72.34\|ATTGGTCGTGGTTGTAGTCCGTGCGAGAATACCA | >GluTTC.trnaMT_MT_-_14674_14742 | 39 | 72 | 34 | tRF lookalike | NO | 719.23 | 432.89 | 1216.26 | 208.47 |
| >trnaMT_GluTTC_MT_-_14674_14742@40.71.32,trnalookalike8_GluTTC_5_-_93905172_93905240@40.71.32\|TTGGTCGTGGTTGTAGTCCGTGCGAGAATACC | >GluTTC.trnaMT_MT_-_14674_14742 | 40 | 71 | 32 | tRF lookalike | NO | 81.94 | 94.95 | 135.89 | 17.47 |
| >trnaMT_GluTTC_MT_-_14674_14742@40.72.33,trnalookalike8_GluTTC_5_-_93905172_93905240@40.72.33\|TTGGTCGTGGTTGTAGTCCGTGCGAGAATACCA | >GluTTC.trnaMT_MT_-_14674_14742 | 40 | 72 | 33 | tRF lookalike | YES | 652.30 | 633.56 | 1152.20 | 190.33 |
| >trnaMT_GluTTC_MT_-_14674_14742@41.71.31,trnalookalike8_GluTTC_5_-_93905172_93905240@41.71.31\|TGGTCGTGGTTGTAGTCCGTGCGAGAATACC | >GluTTC.trnaMT_MT_-_14674_14742 | 41 | 71 | 31 | tRF lookalike | NO | 0.00 | 30.36 | 14.56 | 0.00 |
| >trnaMT_GluTTC_MT_-_14674_14742@41.72.32,trnalookalike8_GluTTC_5_-_93905172_93905240@41.72.32\|TGGTCGTGGTTGTAGTCCGTGCGAGAATACCA | >GluTTC.trnaMT_MT_-_14674_14742 | 41 | 72 | 32 | tRF lookalike | NO | 292.35 | 344.53 | 332.79 | 78.08 |
| >trnaMT_GluTTC_MT_-_14674_14742@42.72.31,trnalookalike8_GluTTC_5_-_93905172_93905240@42.72.31\|GGTCGTGGTTGTAGTCCGTGCGAGAATACCA | >GluTTC.trnaMT_MT_-_14674_14742 | 42 | 72 | 31 | tRF lookalike | NO | 0.00 | 0.00 | 13.28 | 0.00 |
| >trnaMT_GluTTC_MT_-_14674_14742@44.71.28,trnalookalike8_GluTTC_5_-_93905172_93905240@44.71.28\|TCGTGGTTGTAGTCCGTGCGAGAATACC | >GluTTC.trnaMT_MT_-_14674_14742 | 44 | 71 | 28 | tRF lookalike | NO | 0.00 | 0.00 | 11.15 | 0.00 |
| >trnaMT_GluTTC_MT_-_14674_14742@44.72.29,trnalookalike8_GluTTC_5_-_93905172_93905240@44.72.29\|TCGTGGTTGTAGTCCGTGCGAGAATACCA | >GluTTC.trnaMT_MT_-_14674_14742 | 44 | 72 | 29 | tRF lookalike | NO | 50.78 | 59.93 | 150.37 | 21.31 |
| >trnaMT_GluTTC_MT_-_14674_14742@45.72.28,trnalookalike8_GluTTC_5_-_93905172_93905240@45.72.28\|CGTGGTTGTAGTCCGTGCGAGAATACCA | >GluTTC.trnaMT_MT_-_14674_14742 | 45 | 72 | 28 | tRF lookalike | NO | 46.57 | 32.23 | 33.98 | 12.16 |
| >trnaMT_GluTTC_MT_-_14674_14742@46.71.26,trnalookalike8_GluTTC_5_-_93905172_93905240@46.71.26\|GTGGTTGTAGTCCGTGCGAGAATACC | >GluTTC.trnaMT_MT_-_14674_14742 | 46 | 71 | 26 | tRF lookalike | NO | 0.00 | 0.00 | 16.14 | 0.00 |
| >trnaMT_GluTTC_MT_-_14674_14742@46.72.27,trnalookalike8_GluTTC_5_-_93905172_93905240@46.72.27\|GTGGTTGTAGTCCGTGCGAGAATACCA | >GluTTC.trnaMT_MT_-_14674_14742 | 46 | 72 | 27 | tRF lookalike | NO | 26.00 | 24.96 | 109.78 | 14.71 |
| >multi-trnaMT_GluTTC_MT_-_14674_14742@26.61.36,trnalookalike8_GluTTC_5_-_93905172_93905240@26.61.36\|GGTTTTTCATATCATTGGTCGTGGTTGTAGTCCGTG | >GluTTC.trnaMT_MT_-_14674_14742.multi | 26 | 61 | 36 | tRF lookalike | NO | 58.85 | 0.00 | 0.00 | 0.00 |
| >multi-trnaMT_GluTTC_MT_-_14674_14742@34.57.24,trnalookalike8_GluTTC_5_-_93905172_93905240@34.57.24\|ATATCATTGGTCGTGGTTGTAGTC | >GluTTC.trnaMT_MT_-_14674_14742.multi | 34 | 57 | 24 | tRF lookalike | NO | 0.00 | 0.00 | 11.06 | 0.00 |
| >multi-trnaMT_GluTTC_MT_-_14674_14742@39.57.19,trnalookalike8_GluTTC_5_-_93905172_93905240@39.57.19\|ATTGGTCGTGGTTGTAGTC | >GluTTC.trnaMT_MT_-_14674_14742.multi | 39 | 57 | 19 | tRF lookalike | NO | 0.00 | 0.00 | 33.04 | 0.00 |
| >multi-trnaMT_GluTTC_MT_-_14674_14742@39.58.20,trnalookalike8_GluTTC_5_-_93905172_93905240@39.58.20\|ATTGGTCGTGGTTGTAGTCC | >GluTTC.trnaMT_MT_-_14674_14742.multi | 39 | 58 | 20 | tRF lookalike | NO | 0.00 | 27.25 | 35.63 | 0.00 |
| >multi-trnaMT_GluTTC_MT_-_14674_14742@39.61.23,trnalookalike8_GluTTC_5_-_93905172_93905240@39.61.23\|ATTGGTCGTGGTTGTAGTCCGTG | >GluTTC.trnaMT_MT_-_14674_14742.multi | 39 | 61 | 23 | tRF lookalike | NO | 0.00 | 0.00 | 15.50 | 0.00 |
| >multi-trnaMT_GluTTC_MT_-_14674_14742@39.63.25,trnalookalike8_GluTTC_5_-_93905172_93905240@39.63.25\|ATTGGTCGTGGTTGTAGTCCGTGCG | >GluTTC.trnaMT_MT_-_14674_14742.multi | 39 | 63 | 25 | tRF lookalike | NO | 0.00 | 0.00 | 14.01 | 0.00 |
| >multi-trnaMT_GluTTC_MT_-_14674_14742@39.65.27,trnalookalike8_GluTTC_5_-_93905172_93905240@39.65.27\|ATTGGTCGTGGTTGTAGTCCGTGCGAG | >GluTTC.trnaMT_MT_-_14674_14742.multi | 39 | 65 | 27 | tRF lookalike | NO | 0.00 | 0.00 | 11.26 | 0.00 |
| >multi-trnaMT_GluTTC_MT_-_14674_14742@39.66.28,trnalookalike8_GluTTC_5_-_93905172_93905240@39.66.28\|ATTGGTCGTGGTTGTAGTCCGTGCGAGA | >GluTTC.trnaMT_MT_-_14674_14742.multi | 39 | 66 | 28 | tRF lookalike | NO | 0.00 | 0.00 | 12.59 | 0.00 |
| >multi-trnaMT_GluTTC_MT_-_14674_14742@40.57.18,trnalookalike8_GluTTC_5_-_93905172_93905240@40.57.18\|TTGGTCGTGGTTGTAGTC | >GluTTC.trnaMT_MT_-_14674_14742.multi | 40 | 57 | 18 | tRF lookalike | NO | 0.00 | 0.00 | 14.43 | 0.00 |
| >multi-trnaMT_GluTTC_MT_-_14674_14742@40.58.19,trnalookalike8_GluTTC_5_-_93905172_93905240@40.58.19\|TTGGTCGTGGTTGTAGTCC | >GluTTC.trnaMT_MT_-_14674_14742.multi | 40 | 58 | 19 | tRF lookalike | NO | 0.00 | 36.45 | 17.77 | 0.00 |
| >multi-trnaMT_GluTTC_MT_-_14674_14742@40.61.22,trnalookalike8_GluTTC_5_-_93905172_93905240@40.61.22\|TTGGTCGTGGTTGTAGTCCGTG | >GluTTC.trnaMT_MT_-_14674_14742.multi | 40 | 61 | 22 | tRF lookalike | NO | 0.00 | 0.00 | 13.14 | 0.00 |
| >multi-trnaMT_GluTTC_MT_-_14674_14742@40.63.24,trnalookalike8_GluTTC_5_-_93905172_93905240@40.63.24\|TTGGTCGTGGTTGTAGTCCGTGCG | >GluTTC.trnaMT_MT_-_14674_14742.multi | 40 | 63 | 24 | tRF lookalike | NO | 0.00 | 0.00 | 13.16 | 0.00 |
| >multi-trnaMT_GluTTC_MT_-_14674_14742@40.66.27,trnalookalike8_GluTTC_5_-_93905172_93905240@40.66.27\|TTGGTCGTGGTTGTAGTCCGTGCGAGA | >GluTTC.trnaMT_MT_-_14674_14742.multi | 40 | 66 | 27 | tRF lookalike | NO | 0.00 | 0.00 | 14.37 | 0.00 |
| >trnaMT_LysTTT_MT_+_8295_8364@37.73.37,trnalookalike7_LysTTT_1_+_568844_568913@37.73.37\|AAAGATTAAGAGAACCAACACCTCTTTACAGTGACCA | >LysTTT.trnaMT_MT_+_8295_8364 | 37 | 73 | 37 | tRF lookalike | NO | 0.00 | 0.00 | 58.97 | 0.00 |
| >trnaMT_LysTTT_MT_+_8295_8364@38.73.36,trnalookalike7_LysTTT_1_+_568844_568913@38.73.36\|AAGATTAAGAGAACCAACACCTCTTTACAGTGACCA | >LysTTT.trnaMT_MT_+_8295_8364 | 38 | 73 | 36 | tRF lookalike | NO | 0.00 | 0.00 | 68.63 | 0.00 |
| >multi-trnaMT_LysTTT_MT_+_8295_8364@1.20.20,trnalookalike7_LysTTT_1_+_568844_568913@1.20.20\|CACTGTAAAGCTAACTTAGC | >LysTTT.trnaMT_MT_+_8295_8364.multi | 1 | 20 | 20 | tRF lookalike | NO | 0.00 | 0.00 | 12.75 | 10.48 |
| >multi-trnaMT_LysTTT_MT_+_8295_8364@1.23.23,trnalookalike7_LysTTT_1_+_568844_568913@1.23.23\|CACTGTAAAGCTAACTTAGCATT | >LysTTT.trnaMT_MT_+_8295_8364.multi | 1 | 23 | 23 | tRF lookalike | NO | 0.00 | 0.00 | 15.81 | 0.00 |
| >multi-trnaMT_LysTTT_MT_+_8295_8364@1.26.26,trnalookalike7_LysTTT_1_+_568844_568913@1.26.26\|CACTGTAAAGCTAACTTAGCATTAAC | >LysTTT.trnaMT_MT_+_8295_8364.multi | 1 | 26 | 26 | tRF lookalike | NO | 0.00 | 0.00 | 106.68 | 13.17 |

### MDA-MB-231

| | | | | | | | | | | |
|---|---|---|---|---|---|---|---|---|---|---|
| >multi-trnaMT_LysTTT_MT_+_8295_8364@1.27.27,trnalookalike7_LysTTT_1_+_568844_568913@1.27.27\|CACTGTAAAGCTAACTTAGCATTAACC | >LysTTT.trnaMT_MT_+_8295_8364.multi | 1 | 27 | 27 | tRF lookalike | NO | 0.00 | 0.00 | 31.60 | 0.00 |
| >multi-trnaMT_LysTTT_MT_+_8295_8364@1.31.31,trnalookalike7_LysTTT_1_+_568844_568913@1.31.31\|CACTGTAAAGCTAACTTAGCATTAACCTTTT | >LysTTT.trnaMT_MT_+_8295_8364.multi | 1 | 31 | 31 | tRF lookalike | NO | 0.00 | 0.00 | 13.96 | 0.00 |
| >multi-trnaMT_LysTTT_MT_+_8295_8364@1.36.36,trnalookalike7_LysTTT_1_+_568844_568913@1.36.36\|CACTGTAAAGCTAACTTAGCATTAACCTTTTAAGTT | >LysTTT.trnaMT_MT_+_8295_8364.multi | 1 | 36 | 36 | tRF lookalike | NO | 0.00 | 0.00 | 62.30 | 13.57 |
| >multi-trnaMT_LysTTT_MT_+_8295_8364@1.38.38,trnalookalike7_LysTTT_1_+_568844_568913@1.38.38\|CACTGTAAAGCTAACTTAGCATTAACCTTTTAAGTTAA | >LysTTT.trnaMT_MT_+_8295_8364.multi | 1 | 38 | 38 | tRF lookalike | NO | 0.00 | 0.00 | 16.20 | 0.00 |
| >multi-am_trnaMT_LysTTT_MT_+_8295_8364@27.52.26,trnalookalike7_LysTTT_1_+_568844_568913@27.52.26\|CTTTTAAGTTAAAGATTAAGAGAACC | >LysTTT.trnaMT_MT_+_8295_8364.multi.am | 27 | 52 | 26 | tRF lookalike | NO | 0.00 | 0.00 | 10.73 | 0.00 |
| >multi-am_trnaMT_LysTTT_MT_+_8295_8364@34.52.19,trnalookalike7_LysTTT_1_+_568844_568913@34.52.19\|GTTAAAGATTAAGAGAACC | >LysTTT.trnaMT_MT_+_8295_8364.multi.am | 34 | 52 | 19 | tRF lookalike | NO | 0.00 | 0.00 | 10.77 | 0.00 |
| >multi-trnaMT_TrpTCA_MT_+_5512_5579@13.34.22,trnalookalike3_TrpTCA_1_+_566062_566129@13.34.22\|TAAATACAGACCAAGAGCCTTC | >TrpTCA.trnaMT_MT_+_5512_5579.multi | 13 | 34 | 22 | tRF lookalike | NO | 0.00 | 0.00 | 32.30 | 0.00 |
| >multi-trnaMT_TrpTCA_MT_+_5512_5579@18.35.18,trnalookalike3_TrpTCA_1_+_566062_566129@18.35.18\|ACAGACCAAGAGCCTTCA | >TrpTCA.trnaMT_MT_+_5512_5579.multi | 18 | 35 | 18 | tRF lookalike | NO | 0.00 | 0.00 | 15.04 | 0.00 |
| >trnaMT_HisGTG_MT_+_12138_12206@26.63.38\|AGATTGTGAATCTGACAACAGAGGCTTACGACCCCTTA | >HisGTG.trnaMT_MT_+_12138_12206 | 26 | 63 | 38 | tRF MT | NO | 0.00 | 0.00 | 11.03 | 0.00 |
| >trnaMT_HisGTG_MT_+_12138_12206@32.63.32\|TGAATCTGACAACAGAGGCTTACGACCCCTTA | >HisGTG.trnaMT_MT_+_12138_12206 | 32 | 63 | 32 | tRF MT | NO | 0.00 | 0.00 | 86.16 | 19.41 |
| >trnaMT_HisGTG_MT_+_12138_12206@32.69.38\|TGAATCTGACAACAGAGGCTTACGACCCCTTATTTACC | >HisGTG.trnaMT_MT_+_12138_12206 | 32 | 69 | 38 | tRF MT | NO | 0.00 | 0.00 | 65.30 | 103.84 |
| >trnaMT_HisGTG_MT_+_12138_12206@32.70.39\|TGAATCTGACAACAGAGGCTTACGACCCCTTATTTACCC | >HisGTG.trnaMT_MT_+_12138_12206 | 32 | 70 | 39 | tRF MT | NO | 73.77 | 24.37 | 217.74 | 167.54 |
| >trnaMT_HisGTG_MT_+_12138_12206@32.71.40\|TGAATCTGACAACAGAGGCTTACGACCCCTTATTTACCCC | >HisGTG.trnaMT_MT_+_12138_12206 | 32 | 71 | 40 | tRF MT | NO | 89.91 | 57.36 | 433.99 | 251.49 |
| >trnaMT_HisGTG_MT_+_12138_12206@32.72.41\|TGAATCTGACAACAGAGGCTTACGACCCCTTATTTACCCCA | >HisGTG.trnaMT_MT_+_12138_12206 | 32 | 72 | 41 | tRF MT | NO | 0.00 | 13.51 | 107.82 | 121.65 |
| >multi-am_trnaMT_HisGTG_MT_+_12138_12206@18.51.34\|AAAACATCAGATTGTGAATCTGACAACAGAGGCT | >HisGTG.trnaMT_MT_+_12138_12206.multi.am | 18 | 51 | 34 | tRF MT | NO | 0.00 | 0.00 | 16.19 | 0.00 |
| >multi-am_trnaMT_HisGTG_MT_+_12138_12206@19.51.33\|AAACATCAGATTGTGAATCTGACAACAGAGGCT | >HisGTG.trnaMT_MT_+_12138_12206.multi.am | 19 | 51 | 33 | tRF MT | NO | 0.00 | 0.00 | 16.25 | 0.00 |
| >multi-am_trnaMT_HisGTG_MT_+_12138_12206@26.44.19\|AGATTGTGAATCTGACAAC | >HisGTG.trnaMT_MT_+_12138_12206.multi.am | 26 | 44 | 19 | tRF MT | NO | 0.00 | 0.00 | 12.45 | 16.68 |
| >trnaMT_LeuTAA_MT_+_3230_3304@1.30.30\|GTTAAGATGGCAGAGCCCGGTAATCGCATA | >LeuTAA.trnaMT_MT_+_3230_3304 | 1 | 30 | 30 | tRF MT | NO | 0.00 | 0.00 | 66.65 | 15.50 |
| >trnaMT_MetCAT_MT_+_4402_4469@34.71.38\|ACCCCGAAAATGTTGGTTATACCCTTCCCGTACTACCA | >MetCAT.trnaMT_MT_+_4402_4469 | 34 | 71 | 38 | tRF MT | NO | 0.00 | 25.32 | 0.00 | 0.00 |
| >trnaMT_MetCAT_MT_+_4402_4469@36.71.36\|CCCGAAAATGTTGGTTATACCCTTCCCGTACTACCA | >MetCAT.trnaMT_MT_+_4402_4469 | 36 | 71 | 36 | tRF MT | NO | 10.48 | 24.76 | 0.00 | 0.00 |
| >trnaMT_MetCAT_MT_+_4402_4469@37.71.35\|CCGAAAATGTTGGTTATACCCTTCCCGTACTACCA | >MetCAT.trnaMT_MT_+_4402_4469 | 37 | 71 | 35 | tRF MT | NO | 0.00 | 32.94 | 12.31 | 0.00 |
| >trnaMT_MetCAT_MT_+_4402_4469@38.71.34\|CGAAAATGTTGGTTATACCCTTCCCGTACTACCA | >MetCAT.trnaMT_MT_+_4402_4469 | 38 | 71 | 34 | tRF MT | NO | 0.00 | 0.00 | 17.24 | 10.48 |
| >trnaMT_MetCAT_MT_+_4402_4469@39.71.33\|GAAAATGTTGGTTATACCCTTCCCGTACTACCA | >MetCAT.trnaMT_MT_+_4402_4469 | 39 | 71 | 33 | tRF MT | NO | 0.00 | 0.00 | 10.99 | 0.00 |
| >trnaMT_MetCAT_MT_+_4402_4469@40.71.32\|AAAATGTTGGTTATACCCTTCCCGTACTACCA | >MetCAT.trnaMT_MT_+_4402_4469 | 40 | 71 | 32 | tRF MT | NO | 0.00 | 0.00 | 10.61 | 0.00 |
| >trnaMT_MetCAT_MT_+_4402_4469@42.71.30\|AATGTTGGTTATACCCTTCCCGTACTACCA | >MetCAT.trnaMT_MT_+_4402_4469 | 42 | 71 | 30 | tRF MT | NO | 0.00 | 0.00 | 72.06 | 0.00 |
| >trnaMT_MetCAT_MT_+_4402_4469@43.71.29\|ATGTTGGTTATACCCTTCCCGTACTACCA | >MetCAT.trnaMT_MT_+_4402_4469 | 43 | 71 | 29 | tRF MT | NO | 0.00 | 0.00 | 12.91 | 0.00 |
| >trnaMT_MetCAT_MT_+_4402_4469@50.71.22\|TTATACCCTTCCCGTACTACCA | >MetCAT.trnaMT_MT_+_4402_4469 | 50 | 71 | 22 | tRF MT | NO | 0.00 | 26.47 | 0.00 | 0.00 |
| >trnaMT_MetCAT_MT_+_4402_4469@51.71.21\|TATACCCTTCCCGTACTACCA | >MetCAT.trnaMT_MT_+_4402_4469 | 51 | 71 | 21 | tRF MT | NO | 0.00 | 13.05 | 0.00 | 0.00 |
| >trnaMT_MetCAT_MT_+_4402_4469@52.70.19\|ATACCCTTCCCGTACTACC | >MetCAT.trnaMT_MT_+_4402_4469 | 52 | 70 | 19 | tRF MT | NO | 0.00 | 36.12 | 0.00 | 0.00 |
| >trnaMT_MetCAT_MT_+_4402_4469@52.71.20\|ATACCCTTCCCGTACTACCA | >MetCAT.trnaMT_MT_+_4402_4469 | 52 | 71 | 20 | tRF MT | NO | 49.29 | 96.66 | 0.00 | 0.00 |
| >trnaMT_MetCAT_MT_+_4402_4469@53.71.19\|TACCCTTCCCGTACTACCA | >MetCAT.trnaMT_MT_+_4402_4469 | 53 | 71 | 19 | tRF MT | NO | 0.00 | 25.80 | 0.00 | 0.00 |
| >multi-am_trnaMT_MetCAT_MT_+_4402_4469@1.18.18\|AGTAAGGTCAGCTAAATA | >MetCAT.trnaMT_MT_+_4402_4469.multi.am | 1 | 18 | 18 | tRF MT | NO | 0.00 | 0.00 | 35.12 | 0.00 |
| >multi-am_trnaMT_MetCAT_MT_+_4402_4469@1.21.21\|AGTAAGGTCAGCTAAATAAGC | >MetCAT.trnaMT_MT_+_4402_4469.multi.am | 1 | 21 | 21 | tRF MT | NO | 54.09 | 0.00 | 414.92 | 90.94 |
| >multi-am_trnaMT_MetCAT_MT_+_4402_4469@1.22.22\|AGTAAGGTCAGCTAAATAAGCT | >MetCAT.trnaMT_MT_+_4402_4469.multi.am | 1 | 22 | 22 | tRF MT | NO | 0.00 | 0.00 | 137.52 | 15.14 |
| >trnaMT_PheGAA_MT_+_577_647@1.18.18\|GTTTATGTAGCTTACCTC | >PheGAA.trnaMT_MT_+_577_647 | 1 | 18 | 18 | tRF MT | NO | 0.00 | 0.00 | 93.46 | 14.97 |
| >trnaMT_PheGAA_MT_+_577_647@1.19.19\|GTTTATGTAGCTTACCTCC | >PheGAA.trnaMT_MT_+_577_647 | 1 | 19 | 19 | tRF MT | NO | 46.32 | 0.00 | 60.84 | 53.54 |
| >trnaMT_PheGAA_MT_+_577_647@1.20.20\|GTTTATGTAGCTTACCTCCT | >PheGAA.trnaMT_MT_+_577_647 | 1 | 20 | 20 | tRF MT | NO | 0.00 | 0.00 | 12.98 | 12.16 |
| >trnaMT_PheGAA_MT_+_577_647@1.21.21\|GTTTATGTAGCTTACCTCCTC | >PheGAA.trnaMT_MT_+_577_647 | 1 | 21 | 21 | tRF MT | NO | 0.00 | 0.00 | 89.57 | 0.00 |
| >trnaMT_PheGAA_MT_+_577_647@1.34.34\|GTTTATGTAGCTTACCTCCTCAAAGCAATACACT | >PheGAA.trnaMT_MT_+_577_647 | 1 | 34 | 34 | tRF MT | NO | 0.00 | 0.00 | 84.88 | 55.72 |
| >trnaMT_PheGAA_MT_+_577_647@1.35.35\|GTTTATGTAGCTTACCTCCTCAAAGCAATACACTG | >PheGAA.trnaMT_MT_+_577_647 | 1 | 35 | 35 | tRF MT | NO | 0.00 | 0.00 | 31.56 | 14.15 |
| >trnaMT_PheGAA_MT_+_577_647@35.53.19\|GAAAATGTTTAGACGGGCT | >PheGAA.trnaMT_MT_+_577_647 | 35 | 53 | 19 | tRF MT | NO | 0.00 | 0.00 | 17.17 | 0.00 |
| >trnaMT_PheGAA_MT_+_577_647@35.54.20\|GAAAATGTTTAGACGGGCTC | >PheGAA.trnaMT_MT_+_577_647 | 35 | 54 | 20 | tRF MT | NO | 0.00 | 0.00 | 62.79 | 14.27 |
| >trnaMT_PheGAA_MT_+_577_647@35.55.21\|GAAAATGTTTAGACGGGCTCA | >PheGAA.trnaMT_MT_+_577_647 | 35 | 55 | 21 | tRF MT | NO | 0.00 | 0.00 | 64.06 | 0.00 |
| >trnaMT_PheGAA_MT_+_577_647@35.56.22\|GAAAATGTTTAGACGGGCTCAC | >PheGAA.trnaMT_MT_+_577_647 | 35 | 56 | 22 | tRF MT | NO | 0.00 | 0.00 | 33.72 | 12.54 |

### MDA-MB-231

| | | | | | | | | | | |
|---|---|---|---|---|---|---|---|---|---|---|
| >trnaMT_PheGAA_MT_+_577_647@35.57.23\|GAAAATGTTTAGACGGGCTCACA | >PheGAA.trnaMT_MT_+_577_647 | 35 | 57 | 23 | tRF MT | NO | 0.00 | 0.00 | 91.30 | 21.38 |
| >trnaMT_PheGAA_MT_+_577_647@35.59.25\|GAAAATGTTTAGACGGGCTCACATC | >PheGAA.trnaMT_MT_+_577_647 | 35 | 59 | 25 | tRF MT | NO | 0.00 | 0.00 | 11.45 | 11.05 |
| >trnaMT_PheGAA_MT_+_577_647@35.62.28\|GAAAATGTTTAGACGGGCTCACATCACC | >PheGAA.trnaMT_MT_+_577_647 | 35 | 62 | 28 | tRF MT | NO | 0.00 | 0.00 | 16.46 | 20.70 |
| >trnaMT_PheGAA_MT_+_577_647@36.55.20\|AAAATGTTTAGACGGGCTCA | >PheGAA.trnaMT_MT_+_577_647 | 36 | 55 | 20 | tRF MT | NO | 0.00 | 0.00 | 10.98 | 0.00 |
| >trnaMT_PheGAA_MT_+_577_647@36.57.22\|AAAATGTTTAGACGGGCTCACA | >PheGAA.trnaMT_MT_+_577_647 | 36 | 57 | 22 | tRF MT | NO | 0.00 | 0.00 | 14.79 | 11.77 |
| >trnaMT_PheGAA_MT_+_577_647@36.62.27\|AAAATGTTTAGACGGGCTCACATCACC | >PheGAA.trnaMT_MT_+_577_647 | 36 | 62 | 27 | tRF MT | NO | 0.00 | 0.00 | 11.15 | 18.61 |
| >trnaMT_PheGAA_MT_+_577_647@39.56.18\|ATGTTTAGACGGGCTCAC | >PheGAA.trnaMT_MT_+_577_647 | 39 | 56 | 18 | tRF MT | NO | 115.93 | 70.12 | 131.73 | 146.36 |
| >trnaMT_PheGAA_MT_+_577_647@39.57.19\|ATGTTTAGACGGGCTCACA | >PheGAA.trnaMT_MT_+_577_647 | 39 | 57 | 19 | tRF MT | NO | 0.00 | 0.00 | 17.54 | 20.29 |
| >trnaMT_PheGAA_MT_+_577_647@39.62.24\|ATGTTTAGACGGGCTCACATCACC | >PheGAA.trnaMT_MT_+_577_647 | 39 | 62 | 24 | tRF MT | NO | 0.00 | 0.00 | 10.56 | 20.64 |
| >trnaMT_PheGAA_MT_+_577_647@42.74.33\|TTTAGACGGGCTCACATCACCCCATAAACACCA | >PheGAA.trnaMT_MT_+_577_647 | 42 | 74 | 33 | tRF MT | NO | 0.00 | 0.00 | 62.44 | 12.02 |
| >trnaMT_PheGAA_MT_+_577_647@43.74.32\|TTAGACGGGCTCACATCACCCCATAAACACCA | >PheGAA.trnaMT_MT_+_577_647 | 43 | 74 | 32 | tRF MT | NO | 0.00 | 0.00 | 31.31 | 0.00 |
| >trnaMT_PheGAA_MT_+_577_647@44.74.31\|TAGACGGGCTCACATCACCCCATAAACACCA | >PheGAA.trnaMT_MT_+_577_647 | 44 | 74 | 31 | tRF MT | NO | 26.26 | 0.00 | 13.45 | 0.00 |
| >trnaMT_PheGAA_MT_+_577_647@53.74.22\|TCACATCACCCCATAAACACCA | >PheGAA.trnaMT_MT_+_577_647 | 53 | 74 | 22 | tRF MT | NO | 0.00 | 13.04 | 0.00 | 0.00 |
| >multi-am_trnaMT_PheGAA_MT_+_577_647@41.58.18\|GTTTAGACGGGCTCACAT | >PheGAA.trnaMT_MT_+_577_647.multi.am | 41 | 58 | 18 | tRF MT | NO | 24.11 | 0.00 | 17.41 | 71.02 |
| >multi-am_trnaMT_PheGAA_MT_+_577_647@41.62.22\|GTTTAGACGGGCTCACATCACC | >PheGAA.trnaMT_MT_+_577_647.multi.am | 41 | 62 | 22 | tRF MT | NO | 0.00 | 27.55 | 11.14 | 19.93 |
| >trnaMT_ProTGG_MT_-_15956_16023@1.30.30\|CAGAGAATAGTTTAAATTAGAATCTTAGCT | >ProTGG.trnaMT_MT_-_15956_16023 | 1 | 30 | 30 | tRF MT | NO | 0.00 | 0.00 | 31.89 | 0.00 |
| >trnaMT_ProTGG_MT_-_15956_16023@1.32.32\|CAGAGAATAGTTTAAATTAGAATCTTAGCTTT | >ProTGG.trnaMT_MT_-_15956_16023 | 1 | 32 | 32 | tRF MT | NO | 0.00 | 0.00 | 32.54 | 0.00 |
| >trnaMT_ProTGG_MT_-_15956_16023@39.58.20\|TAATGGTGGAGTTAAAGACT | >ProTGG.trnaMT_MT_-_15956_16023 | 39 | 58 | 20 | tRF MT | NO | 0.00 | 0.00 | 12.28 | 0.00 |
| >trnaMT_ProTGG_MT_-_15956_16023@40.58.19\|AATGGTGGAGTTAAAGACT | >ProTGG.trnaMT_MT_-_15956_16023 | 40 | 58 | 19 | tRF MT | NO | 0.00 | 0.00 | 105.81 | 0.00 |
| >trnaMT_ProTGG_MT_-_15956_16023@41.58.18\|ATGGTGGAGTTAAAGACT | >ProTGG.trnaMT_MT_-_15956_16023 | 41 | 58 | 18 | tRF MT | NO | 0.00 | 0.00 | 14.19 | 0.00 |
| >trnaMT_ProTGG_MT_-_15956_16023@44.70.27\|GTGGAGTTAAAGACTTTTTCTCTGACC | >ProTGG.trnaMT_MT_-_15956_16023 | 44 | 70 | 27 | tRF MT | NO | 0.00 | 0.00 | 76.38 | 0.00 |
| >trnaMT_ProTGG_MT_-_15956_16023@47.70.24\|GAGTTAAAGACTTTTTCTCTGACC | >ProTGG.trnaMT_MT_-_15956_16023 | 47 | 70 | 24 | tRF MT | NO | 0.00 | 0.00 | 13.41 | 0.00 |
| >trnaMT_ProTGG_MT_-_15956_16023@50.70.21\|TTAAAGACTTTTTCTCTGACC | >ProTGG.trnaMT_MT_-_15956_16023 | 50 | 70 | 21 | tRF MT | NO | 0.00 | 0.00 | 10.49 | 0.00 |
| >trnaMT_ProTGG_MT_-_15956_16023@50.71.22\|TTAAAGACTTTTTCTCTGACCA | >ProTGG.trnaMT_MT_-_15956_16023 | 50 | 71 | 22 | tRF MT | NO | 58.97 | 0.00 | 12.54 | 10.63 |
| >trnaMT_ProTGG_MT_-_15956_16023@51.71.21\|TAAAGACTTTTTCTCTGACCA | >ProTGG.trnaMT_MT_-_15956_16023 | 51 | 71 | 21 | tRF MT | NO | 46.56 | 0.00 | 11.18 | 0.00 |
| >multi-am_trnaMT_ProTGG_MT_-_15956_16023@52.70.19\|AAAGACTTTTTCTCTGACC | >ProTGG.trnaMT_MT_-_15956_16023.multi.am | 52 | 70 | 19 | tRF MT | NO | 45.46 | 0.00 | 160.42 | 17.64 |
| >multi-am_trnaMT_ProTGG_MT_-_15956_16023@52.71.20\|AAAGACTTTTTCTCTGACCA | >ProTGG.trnaMT_MT_-_15956_16023.multi.am | 52 | 71 | 20 | tRF MT | NO | 103.95 | 34.27 | 212.17 | 86.29 |
| >multi-am_trnaMT_ProTGG_MT_-_15956_16023@53.70.18\|AAGACTTTTTCTCTGACC | >ProTGG.trnaMT_MT_-_15956_16023.multi.am | 53 | 70 | 18 | tRF MT | NO | 0.00 | 0.00 | 130.94 | 17.37 |
| >multi-am_trnaMT_ProTGG_MT_-_15956_16023@53.71.19\|AAGACTTTTTCTCTGACCA | >ProTGG.trnaMT_MT_-_15956_16023.multi.am | 53 | 71 | 19 | tRF MT | NO | 91.98 | 75.02 | 178.39 | 95.54 |
| >multi-am_trnaMT_ProTGG_MT_-_15956_16023@54.71.18\|AGACTTTTTCTCTGACCA | >ProTGG.trnaMT_MT_-_15956_16023.multi.am | 54 | 71 | 18 | tRF MT | NO | 92.06 | 97.74 | 122.37 | 72.13 |
| >trnaMT_SerGCT_MT_+_12207_12265@1.18.18\|GAGAAAGCTCACAAGAAC | >SerGCT.trnaMT_MT_+_12207_12265 | 1 | 18 | 18 | tRF MT | YES | 313.37 | 0.00 | 7129.05 | 549.44 |
| >trnaMT_SerGCT_MT_+_12207_12265@1.19.19\|GAGAAAGCTCACAAGAACT | >SerGCT.trnaMT_MT_+_12207_12265 | 1 | 19 | 19 | tRF MT | NO | 90.20 | 0.00 | 1254.89 | 110.97 |
| >trnaMT_SerGCT_MT_+_12207_12265@1.20.20\|GAGAAAGCTCACAAGAACTG | >SerGCT.trnaMT_MT_+_12207_12265 | 1 | 20 | 20 | tRF MT | NO | 0.00 | 0.00 | 312.03 | 19.86 |
| >trnaMT_SerGCT_MT_+_12207_12265@1.21.21\|GAGAAAGCTCACAAGAACTGC | >SerGCT.trnaMT_MT_+_12207_12265 | 1 | 21 | 21 | tRF MT | NO | 0.00 | 0.00 | 1329.10 | 160.98 |
| >trnaMT_SerGCT_MT_+_12207_12265@1.22.22\|GAGAAAGCTCACAAGAACTGCT | >SerGCT.trnaMT_MT_+_12207_12265 | 1 | 22 | 22 | tRF MT | NO | 53.01 | 0.00 | 1434.28 | 132.58 |
| >trnaMT_SerGCT_MT_+_12207_12265@1.23.23\|GAGAAAGCTCACAAGAACTGCTA | >SerGCT.trnaMT_MT_+_12207_12265 | 1 | 23 | 23 | tRF MT | NO | 0.00 | 0.00 | 217.63 | 13.99 |
| >trnaMT_SerGCT_MT_+_12207_12265@1.24.24\|GAGAAAGCTCACAAGAACTGCTAA | >SerGCT.trnaMT_MT_+_12207_12265 | 1 | 24 | 24 | tRF MT | NO | 0.00 | 0.00 | 134.69 | 13.70 |
| >trnaMT_SerGCT_MT_+_12207_12265@1.25.25\|GAGAAAGCTCACAAGAACTGCTAAC | >SerGCT.trnaMT_MT_+_12207_12265 | 1 | 25 | 25 | tRF MT | YES | 162.70 | 31.03 | 3048.30 | 577.96 |
| >trnaMT_SerGCT_MT_+_12207_12265@1.26.26\|GAGAAAGCTCACAAGAACTGCTAACT | >SerGCT.trnaMT_MT_+_12207_12265 | 1 | 26 | 26 | tRF MT | NO | 113.50 | 0.00 | 1893.48 | 163.30 |
| >trnaMT_SerGCT_MT_+_12207_12265@14.40.27\|AGAACTGCTAACTCATGCCCCCATGTC | >SerGCT.trnaMT_MT_+_12207_12265 | 14 | 40 | 27 | tRF MT | NO | 0.00 | 0.00 | 11.18 | 0.00 |
| >trnaMT_SerGCT_MT_+_12207_12265@14.41.28\|AGAACTGCTAACTCATGCCCCCATGTCT | >SerGCT.trnaMT_MT_+_12207_12265 | 14 | 41 | 28 | tRF MT | NO | 0.00 | 0.00 | 11.87 | 0.00 |
| >trnaMT_SerGCT_MT_+_12207_12265@14.48.35\|AGAACTGCTAACTCATGCCCCCATGTCTAACAACA | >SerGCT.trnaMT_MT_+_12207_12265 | 14 | 48 | 35 | tRF MT | NO | 0.00 | 0.00 | 15.95 | 10.96 |
| >multi-am_trnaMT_SerGCT_MT_+_12207_12265@13.31.19\|AAGAACTGCTAACTCATGC | >SerGCT.trnaMT_MT_+_12207_12265.multi.am | 13 | 31 | 19 | tRF MT | NO | 0.00 | 0.00 | 11.05 | 0.00 |
| >multi-am_trnaMT_SerGCT_MT_+_12207_12265@14.31.18\|AGAACTGCTAACTCATGC | >SerGCT.trnaMT_MT_+_12207_12265.multi.am | 14 | 31 | 18 | tRF MT | NO | 0.00 | 0.00 | 14.23 | 0.00 |
| >trnaMT_ThrTGT_MT_+_15888_15953@31.69.39\|TTGTAAACCGGAGATGAAAACCTTTTTCCAAGGACACCA | >ThrTGT.trnaMT_MT_+_15888_15953 | 31 | 69 | 39 | tRF MT | NO | 0.00 | 0.00 | 31.72 | 0.00 |
| >trnaMT_ThrTGT_MT_+_15888_15953@34.69.36\|TAAACCGGAGATGAAAACCTTTTTCCAAGGACACCA | >ThrTGT.trnaMT_MT_+_15888_15953 | 34 | 69 | 36 | tRF MT | NO | 0.00 | 0.00 | 11.84 | 0.00 |
| >trnaMT_ThrTGT_MT_+_15888_15953@36.69.34\|AACCGGAGATGAAAACCTTTTTCCAAGGACACCA | >ThrTGT.trnaMT_MT_+_15888_15953 | 36 | 69 | 34 | tRF MT | NO | 0.00 | 0.00 | 16.56 | 0.00 |
| >trnaMT_ThrTGT_MT_+_15888_15953@37.69.33\|ACCGGAGATGAAAACCTTTTTCCAAGGACACCA | >ThrTGT.trnaMT_MT_+_15888_15953 | 37 | 69 | 33 | tRF MT | NO | 0.00 | 0.00 | 86.64 | 0.00 |
| >trnaMT_ThrTGT_MT_+_15888_15953@38.69.32\|CCGGAGATGAAAACCTTTTTCCAAGGACACCA | >ThrTGT.trnaMT_MT_+_15888_15953 | 38 | 69 | 32 | tRF MT | NO | 0.00 | 0.00 | 13.70 | 0.00 |
| >multi-am_trnaMT_TyrGTA_MT_-_5826_5891@1.31.31\|GGTAAAATGGCTGAGTGAAGCATTGGACTGT | >TyrGTA.trnaMT_MT_-_5826_5891.multi.am | 1 | 31 | 31 | tRF MT | NO | 0.00 | 0.00 | 13.57 | 10.99 |
| >multi-am_trnaMT_TyrGTA_MT_-_5826_5891@1.32.32\|GGTAAAATGGCTGAGTGAAGCATTGGACTGTA | >TyrGTA.trnaMT_MT_-_5826_5891.multi.am | 1 | 32 | 32 | tRF MT | NO | 0.00 | 0.00 | 17.86 | 54.66 |
| >multi-am_trnaMT_TyrGTA_MT_-_5826_5891@13.32.20\|GAGTGAAGCATTGGACTGTA | >TyrGTA.trnaMT_MT_-_5826_5891.multi.am | 13 | 32 | 20 | tRF MT | NO | 0.00 | 0.00 | 13.28 | 0.00 |
| >trnaMT_ValTAC_MT_+_1602_1670@29.72.44\|ACTTACACTTAGGAGATTTCAACTTAACTTGACCGCTCTGACCA | >ValTAC.trnaMT_MT_+_1602_1670 | 29 | 72 | 44 | tRF MT | NO | 0.00 | 0.00 | 11.93 | 19.04 |
| >trnaMT_ValTAC_MT_+_1602_1670@41.71.31\|GAGATTTCAACTTAACTTGACCGCTCTGACC | >ValTAC.trnaMT_MT_+_1602_1670 | 41 | 71 | 31 | tRF MT | NO | 0.00 | 0.00 | 61.75 | 16.48 |
| >trnaMT_ValTAC_MT_+_1602_1670@41.72.32\|GAGATTTCAACTTAACTTGACCGCTCTGACCA | >ValTAC.trnaMT_MT_+_1602_1670 | 41 | 72 | 32 | tRF MT | NO | 0.00 | 0.00 | 16.70 | 17.95 |
| >trnaMT_ValTAC_MT_+_1602_1670@42.71.30\|AGATTTCAACTTAACTTGACCGCTCTGACC | >ValTAC.trnaMT_MT_+_1602_1670 | 42 | 71 | 30 | tRF MT | NO | 103.19 | 0.00 | 395.45 | 250.18 |
| >trnaMT_ValTAC_MT_+_1602_1670@42.72.31\|AGATTTCAACTTAACTTGACCGCTCTGACCA | >ValTAC.trnaMT_MT_+_1602_1670 | 42 | 72 | 31 | tRF MT | NO | 300.53 | 0.00 | 260.96 | 390.97 |
| >trnaMT_ValTAC_MT_+_1602_1670@43.71.29\|GATTTCAACTTAACTTGACCGCTCTGACC | >ValTAC.trnaMT_MT_+_1602_1670 | 43 | 71 | 29 | tRF MT | NO | 0.00 | 0.00 | 56.52 | 57.54 |
| >trnaMT_ValTAC_MT_+_1602_1670@43.72.30\|GATTTCAACTTAACTTGACCGCTCTGACCA | >ValTAC.trnaMT_MT_+_1602_1670 | 43 | 72 | 30 | tRF MT | NO | 26.07 | 0.00 | 17.47 | 96.12 |
| >trnaMT_ValTAC_MT_+_1602_1670@44.71.28\|ATTTCAACTTAACTTGACCGCTCTGACC | >ValTAC.trnaMT_MT_+_1602_1670 | 44 | 71 | 28 | tRF MT | NO | 0.00 | 0.00 | 108.38 | 105.18 |
| >trnaMT_ValTAC_MT_+_1602_1670@44.72.29\|ATTTCAACTTAACTTGACCGCTCTGACCA | >ValTAC.trnaMT_MT_+_1602_1670 | 44 | 72 | 29 | tRF MT | NO | 63.05 | 0.00 | 60.40 | 134.67 |
| >trnaMT_ValTAC_MT_+_1602_1670@45.72.28\|TTTCAACTTAACTTGACCGCTCTGACCA | >ValTAC.trnaMT_MT_+_1602_1670 | 45 | 72 | 28 | tRF MT | NO | 49.54 | 0.00 | 0.00 | 19.46 |
| >trnaMT_ValTAC_MT_+_1602_1670@46.72.27\|TTCAACTTAACTTGACCGCTCTGACCA | >ValTAC.trnaMT_MT_+_1602_1670 | 46 | 72 | 27 | tRF MT | NO | 57.84 | 0.00 | 0.00 | 20.19 |
| >trnaMT_ValTAC_MT_+_1602_1670@47.72.26\|TCAACTTAACTTGACCGCTCTGACCA | >ValTAC.trnaMT_MT_+_1602_1670 | 47 | 72 | 26 | tRF MT | NO | 70.98 | 0.00 | 0.00 | 20.05 |
| >trnaMT_ValTAC_MT_+_1602_1670@48.71.24\|CAACTTAACTTGACCGCTCTGACC | >ValTAC.trnaMT_MT_+_1602_1670 | 48 | 71 | 24 | tRF MT | NO | 0.00 | 56.20 | 13.74 | 15.59 |

### MDA-MB-231

| | | | | | | | | | | |
|---|---|---|---|---|---|---|---|---|---|---|
| >trnaMT_ValTAC_MT_+_1602_1670@48.72.25\|CAACTTAACTTGACCGCTCTGACCA | >ValTAC.trnaMT_MT_+_1602_1670 | 48 | 72 | 25 | tRF MT | NO | 101.35 | 33.21 | 0.00 | 65.55 |
| >trnaMT_ValTAC_MT_+_1602_1670@49.72.24\|AACTTAACTTGACCGCTCTGACCA | >ValTAC.trnaMT_MT_+_1602_1670 | 49 | 72 | 24 | tRF MT | NO | 62.15 | 0.00 | 0.00 | 55.14 |
| >trnaMT_ValTAC_MT_+_1602_1670@50.71.22\|ACTTAACTTGACCGCTCTGACC | >ValTAC.trnaMT_MT_+_1602_1670 | 50 | 71 | 22 | tRF MT | NO | 0.00 | 0.00 | 12.15 | 14.84 |
| >trnaMT_ValTAC_MT_+_1602_1670@50.72.23\|ACTTAACTTGACCGCTCTGACCA | >ValTAC.trnaMT_MT_+_1602_1670 | 50 | 72 | 23 | tRF MT | NO | 102.50 | 13.03 | 14.13 | 74.69 |
| >trnaMT_ValTAC_MT_+_1602_1670@52.72.21\|TTAACTTGACCGCTCTGACCA | >ValTAC.trnaMT_MT_+_1602_1670 | 52 | 72 | 21 | tRF MT | NO | 70.85 | 0.00 | 0.00 | 20.28 |
| >trnaMT_ValTAC_MT_+_1602_1670@53.72.20\|TAACTTGACCGCTCTGACCA | >ValTAC.trnaMT_MT_+_1602_1670 | 53 | 72 | 20 | tRF MT | NO | 135.91 | 0.00 | 0.00 | 65.20 |
| >trnaMT_ValTAC_MT_+_1602_1670@54.71.18\|AACTTGACCGCTCTGACC | >ValTAC.trnaMT_MT_+_1602_1670 | 54 | 71 | 18 | tRF MT | NO | 55.89 | 0.00 | 0.00 | 12.10 |
| >trnaMT_ValTAC_MT_+_1602_1670@54.72.19\|AACTTGACCGCTCTGACCA | >ValTAC.trnaMT_MT_+_1602_1670 | 54 | 72 | 19 | tRF MT | NO | 181.56 | 0.00 | 10.83 | 64.21 |
| >trnaMT_ValTAC_MT_+_1602_1670@55.72.18\|ACTTGACCGCTCTGACCA | >ValTAC.trnaMT_MT_+_1602_1670 | 55 | 72 | 18 | tRF MT | NO | 181.61 | 0.00 | 10.90 | 97.99 |
| >multi-am_trnaMT_ValTAC_MT_+_1602_1670@28.57.30\|AACTTACACTTAGGAGATTTCAACTTAACT | >ValTAC.trnaMT_MT_+_1602_1670.multi.am | 28 | 57 | 30 | tRF MT | NO | 0.00 | 0.00 | 12.39 | 0.00 |
| >multi-am_trnaMT_ValTAC_MT_+_1602_1670@48.67.20\|CAACTTAACTTGACCGCTCT | >ValTAC.trnaMT_MT_+_1602_1670.multi.am | 48 | 67 | 20 | tRF MT | NO | 64.72 | 169.83 | 0.00 | 20.00 |
| >multi-am_trnaMT_ValTAC_MT_+_1602_1670@49.67.19\|AACTTAACTTGACCGCTCT | >ValTAC.trnaMT_MT_+_1602_1670.multi.am | 49 | 67 | 19 | tRF MT | NO | 0.00 | 13.61 | 0.00 | 0.00 |
| >multi-am_trnaMT_ValTAC_MT_+_1602_1670@50.67.18\|ACTTAACTTGACCGCTCT | >ValTAC.trnaMT_MT_+_1602_1670.multi.am | 50 | 67 | 18 | tRF MT | NO | 25.97 | 36.72 | 0.00 | 0.00 |
| >trna101_AlaAGC_6_-_28831462_28831533@58.75.18,trna102_AlaAGC_6_-_28806221_28806292@58.75.18,trna110_AlaTGC_6_-_28757547_28757618@58.75.18,trna113_AlaTGC_6_-_28726141_28726212@58.75.18,trna65_AlaAGC_6_+_28574933_28575004@58.75.18\|TCCCCGGCACCTCCACCA | >AlaAGC.trna101_6_-_28831462_28831533 | 58 | 75 | 18 | tRF nuclear | NO | 26.30 | 656.59 | 0.00 | 53.65 |
| >trna108_AlaAGC_6_-_28763741_28763812@58.75.18\|TCCCCAGTACCTCCACCA | >AlaAGC.trna108_6_-_28763741_28763812 | 58 | 75 | 18 | tRF nuclear | NO | 0.00 | 138.14 | 0.00 | 0.00 |
| >trna10_AlaCGC_6_+_26553731_26553802@1.18.18,trna119_AlaCGC_6_-_28641613_28641684@1.18.18,trna120_AlaAGC_6_-_28626014_28626085@1.18.18,trna13_AlaCGC_2_+_157257281_157257352@1.18.18,trna13_AlaTGC_12_-_125406301_125406372@1.18.18,trna66_AlaTGC_6_+_28611222_28611293@1.18.18,trna8_AlaTGC_12_+_125424512_125424583@1.18.18,trna8_AlaTGC_5_+_180633868_180633939@1.18.18\|GGGGATGTAGCTCAGTGG | >AlaCGC.trna10_6_+_26553731_26553802 | 1 | 18 | 18 | tRF nuclear | NO | 0.00 | 28.99 | 33.97 | 0.00 |
| >trna10_AlaCGC_6_+_26553731_26553802@1.19.19,trna119_AlaCGC_6_-_28641613_28641684@1.19.19,trna120_AlaAGC_6_-_28626014_28626085@1.19.19,trna13_AlaCGC_2_+_157257281_157257352@1.19.19,trna13_AlaTGC_12_-_125406301_125406372@1.19.19,trna66_AlaTGC_6_+_28611222_28611293@1.19.19,trna8_AlaTGC_12_+_125424512_125424583@1.19.19,trna8_AlaTGC_5_+_180633868_180633939@1.19.19\|GGGGATGTAGCTCAGTGGT | >AlaCGC.trna10_6_+_26553731_26553802 | 1 | 19 | 19 | tRF nuclear | NO | 0.00 | 58.67 | 17.62 | 0.00 |
| >trna10_AlaCGC_6_+_26553731_26553802@58.75.18,trna119_AlaCGC_6_-_28641613_28641684@58.75.18,trna13_AlaCGC_2_+_157257281_157257352@58.75.18,trna13_AlaTGC_12_-_125406301_125406372@58.75.18,trna66_AlaTGC_6_+_28611222_28611293@58.75.18,trna8_AlaTGC_12_+_125424512_125424583@58.75.18,trna8_AlaTGC_5_+_180633868_180633939@58.75.18\|TCCCCGGCATCTCCACCA | >AlaCGC.trna10_6_+_26553731_26553802 | 58 | 75 | 18 | tRF nuclear | NO | 0.00 | 508.85 | 0.00 | 15.93 |
| >trna12_ArgCCT_16_+_3243918_3243990@35.54.20,trna18_ArgCCT_17_+_73030001_73030073@35.54.20,trna21_ArgCCT_17_-_73030526_73030598@35.54.20,trna2_ArgCCT_16_+_3202901_3202973@35.54.20,trna3_ArgCCT_7_+_139025446_139025518@35.54.20\|CTAAGCCAGGGATTGTGGGT | >ArgCCT.trna12_16_+_3243918_3243990 | 35 | 54 | 20 | tRF nuclear | NO | 0.00 | 66.05 | 10.59 | 0.00 |
| >trna10_AspGTC_12_-_125424193_125424264@1.34.34,trna12_AspGTC_12_-_125411891_125411962@1.34.34,trna38_AspGTC_17_-_8125556_8125627@1.34.34,trna45_AspGTC_6_+_27447453_27447524@1.34.34,trna48_AspGTC_6_+_27471523_27471594@1.34.34,trna4_AspGTC_12_+_96429799_96429870@1.34.34,trna69_AspGTC_1_-_161440205_161440276@1.34.34,trna72_AspGTC_1_-_161432824_161432895@1.34.34,trna75_AspGTC_1_-_161425414_161425485@1.34.34,trna78_AspGTC_1_-_161418033_161418104@1.34.34,trna81_AspGTC_1_-_161410615_161410686@1.34.34\|TCCTCGTTAGTATAGTGGTGAGTATCCCCGCCTG | >AspGTC.trna10_12_-_125424193_125424264 | 1 | 34 | 34 | tRF nuclear | NO | 0.00 | 68.85 | 0.00 | 0.00 |
| >trna10_AspGTC_12_-_125424193_125424264@1.36.36,trna12_AspGTC_12_-_125411891_125411962@1.36.36,trna38_AspGTC_17_-_8125556_8125627@1.36.36,trna45_AspGTC_6_+_27447453_27447524@1.36.36,trna48_AspGTC_6_+_27471523_27471594@1.36.36,trna4_AspGTC_12_+_96429799_96429870@1.36.36,trna69_AspGTC_1_-_161440205_161440276@1.36.36,trna72_AspGTC_1_-_161432824_161432895@1.36.36,trna75_AspGTC_1_-_161425414_161425485@1.36.36,trna78_AspGTC_1_-_161418033_161418104@1.36.36,trna81_AspGTC_1_-_161410615_161410686@1.36.36\|TCCTCGTTAGTATAGTGGTGAGTATCCCCGCCTGTC | >AspGTC.trna10_12_-_125424193_125424264 | 1 | 36 | 36 | tRF nuclear | NO | 0.00 | 29.74 | 0.00 | 0.00 |
| >trna10_AspGTC_12_-_125424193_125424264@35.73.39,trna12_AspGTC_12_-_125411891_125411962@35.73.39,trna144_AspGTC_6_-_27551236_27551307@35.73.39,trna38_AspGTC_17_-_8125556_8125627@35.73.39,trna45_AspGTC_6_+_27447453_27447524@35.73.39,trna48_AspGTC_6_+_27471523_27471594@35.73.39,trna4_AspGTC_12_+_96429799_96429870@35.73.39,trna69_AspGTC_1_-_161440205_161440276@35.73.39,trna72_AspGTC_1_-_161432824_161432895@35.73.39,trna75_AspGTC_1_-_161425414_161425485@35.73.39,trna78_AspGTC_1_-_161418033_161418104@35.73.39,trna81_AspGTC_1_-_161410615_161410686@35.73.39\|TCACGCGGGAGACCGGGGTTCGATTCCCCGACGGGGAGC | >AspGTC.trna10_12_-_125424193_125424264 | 35 | 73 | 39 | tRF nuclear | NO | 0.00 | 190.83 | 0.00 | 0.00 |

### MDA-MB-231

| | | | | | | | | | | | |
|---|---|---|---|---|---|---|---|---|---|---|---|
| >trna10_AspGTC_12_-_125424193_125424264@35.74.40,trna12_AspGTC_12_-_125411891_125411962@35.74.40,trna144_AspGTC_6_-_27551236_27551307@35.74.40,trna38_AspGTC_17_-_8125556_8125627@35.74.40,trna45_AspGTC_6_+_27447453_27447524@35.74.40,trna48_AspGTC_6_+_27471523_27471594@35.74.40,trna4_AspGTC_12_+_96429799_96429870@35.74.40,trna69_AspGTC_1_-_161440205_161440276@35.74.40,trna72_AspGTC_1_-_161432824_161432895@35.74.40,trna75_AspGTC_1_-_161425414_161425485@35.74.40,trna78_AspGTC_1_-_161418033_161418104@35.74.40,trna81_AspGTC_1_-_161410615_161410686@35.74.40\|TCACGCGGGAGACCGGGGTTCGATTCCCCGACGGGGAGCC | >AspGTC.trna10_12_-_125424193_125424264 | 35 | 74 | 40 | | tRF nuclear | NO | 0.00 | 417.67 | 0.00 | 0.00 |
| >trna10_AspGTC_12_-_125424193_125424264@35.75.41,trna12_AspGTC_12_-_125411891_125411962@35.75.41,trna144_AspGTC_6_-_27551236_27551307@35.75.41,trna38_AspGTC_17_-_8125556_8125627@35.75.41,trna45_AspGTC_6_+_27447453_27447524@35.75.41,trna48_AspGTC_6_+_27471523_27471594@35.75.41,trna4_AspGTC_12_+_96429799_96429870@35.75.41,trna69_AspGTC_1_-_161440205_161440276@35.75.41,trna72_AspGTC_1_-_161432824_161432895@35.75.41,trna75_AspGTC_1_-_161425414_161425485@35.75.41,trna78_AspGTC_1_-_161418033_161418104@35.75.41,trna81_AspGTC_1_-_161410615_161410686@35.75.41\|TCACGCGGGAGACCGGGGTTCGATTCCCCGACGGGGAGCCA | >AspGTC.trna10_12_-_125424193_125424264 | 35 | 75 | 41 | | tRF nuclear | NO | 0.00 | 805.60 | 0.00 | 0.00 |
| >trna10_AspGTC_12_-_125424193_125424264@36.73.38,trna12_AspGTC_12_-_125411891_125411962@36.73.38,trna144_AspGTC_6_-_27551236_27551307@36.73.38,trna38_AspGTC_17_-_8125556_8125627@36.73.38,trna45_AspGTC_6_+_27447453_27447524@36.73.38,trna48_AspGTC_6_+_27471523_27471594@36.73.38,trna4_AspGTC_12_+_96429799_96429870@36.73.38,trna69_AspGTC_1_-_161440205_161440276@36.73.38,trna72_AspGTC_1_-_161432824_161432895@36.73.38,trna75_AspGTC_1_-_161425414_161425485@36.73.38,trna78_AspGTC_1_-_161418033_161418104@36.73.38,trna81_AspGTC_1_-_161410615_161410686@36.73.38\|CACGCGGGAGACCGGGGTTCGATTCCCCGACGGGGAGC | >AspGTC.trna10_12_-_125424193_125424264 | 36 | 73 | 38 | | tRF nuclear | NO | 0.00 | 256.97 | 0.00 | 0.00 |
| >trna10_AspGTC_12_-_125424193_125424264@36.74.39,trna12_AspGTC_12_-_125411891_125411962@36.74.39,trna144_AspGTC_6_-_27551236_27551307@36.74.39,trna38_AspGTC_17_-_8125556_8125627@36.74.39,trna45_AspGTC_6_+_27447453_27447524@36.74.39,trna48_AspGTC_6_+_27471523_27471594@36.74.39,trna4_AspGTC_12_+_96429799_96429870@36.74.39,trna69_AspGTC_1_-_161440205_161440276@36.74.39,trna72_AspGTC_1_-_161432824_161432895@36.74.39,trna75_AspGTC_1_-_161425414_161425485@36.74.39,trna78_AspGTC_1_-_161418033_161418104@36.74.39,trna81_AspGTC_1_-_161410615_161410686@36.74.39\|CACGCGGGAGACCGGGGTTCGATTCCCCGACGGGGAGCC | >AspGTC.trna10_12_-_125424193_125424264 | 36 | 74 | 39 | | tRF nuclear | NO | 0.00 | 408.17 | 0.00 | 0.00 |
| >trna10_AspGTC_12_-_125424193_125424264@36.75.40,trna12_AspGTC_12_-_125411891_125411962@36.75.40,trna144_AspGTC_6_-_27551236_27551307@36.75.40,trna38_AspGTC_17_-_8125556_8125627@36.75.40,trna45_AspGTC_6_+_27447453_27447524@36.75.40,trna48_AspGTC_6_+_27471523_27471594@36.75.40,trna4_AspGTC_12_+_96429799_96429870@36.75.40,trna69_AspGTC_1_-_161440205_161440276@36.75.40,trna72_AspGTC_1_-_161432824_161432895@36.75.40,trna75_AspGTC_1_-_161425414_161425485@36.75.40,trna78_AspGTC_1_-_161418033_161418104@36.75.40,trna81_AspGTC_1_-_161410615_161410686@36.75.40\|CACGCGGGAGACCGGGGTTCGATTCCCCGACGGGGAGCCA | >AspGTC.trna10_12_-_125424193_125424264 | 36 | 75 | 40 | | tRF nuclear | NO | 0.00 | 578.65 | 0.00 | 0.00 |
| >trna10_AspGTC_12_-_125424193_125424264@37.72.36,trna12_AspGTC_12_-_125411891_125411962@37.72.36,trna144_AspGTC_6_-_27551236_27551307@37.72.36,trna38_AspGTC_17_-_8125556_8125627@37.72.36,trna45_AspGTC_6_+_27447453_27447524@37.72.36,trna48_AspGTC_6_+_27471523_27471594@37.72.36,trna4_AspGTC_12_+_96429799_96429870@37.72.36,trna69_AspGTC_1_-_161440205_161440276@37.72.36,trna72_AspGTC_1_-_161432824_161432895@37.72.36,trna75_AspGTC_1_-_161425414_161425485@37.72.36,trna78_AspGTC_1_-_161418033_161418104@37.72.36,trna81_AspGTC_1_-_161410615_161410686@37.72.36\|ACGCGGGAGACCGGGGTTCGATTCCCCGACGGGGAG | >AspGTC.trna10_12_-_125424193_125424264 | 37 | 72 | 36 | | tRF nuclear | NO | 0.00 | 92.03 | 0.00 | 0.00 |
| >trna10_AspGTC_12_-_125424193_125424264@37.73.37,trna12_AspGTC_12_-_125411891_125411962@37.73.37,trna144_AspGTC_6_-_27551236_27551307@37.73.37,trna38_AspGTC_17_-_8125556_8125627@37.73.37,trna45_AspGTC_6_+_27447453_27447524@37.73.37,trna48_AspGTC_6_+_27471523_27471594@37.73.37,trna4_AspGTC_12_+_96429799_96429870@37.73.37,trna69_AspGTC_1_-_161440205_161440276@37.73.37,trna72_AspGTC_1_-_161432824_161432895@37.73.37,trna75_AspGTC_1_-_161425414_161425485@37.73.37,trna78_AspGTC_1_-_161418033_161418104@37.73.37,trna81_AspGTC_1_-_161410615_161410686@37.73.37\|ACGCGGGAGACCGGGGTTCGATTCCCCGACGGGGAGC | >AspGTC.trna10_12_-_125424193_125424264 | 37 | 73 | 37 | | tRF nuclear | NO | 0.00 | 736.95 | 10.92 | 0.00 |

### MDA-MB-231

| | | | | | | | | | | |
|---|---|---|---|---|---|---|---|---|---|---|
| >trna10_AspGTC_12_-_125424193_125424264@37.74.38,trna12_AspGTC_12_-_125411891_125411962@37.74.38,trna144_AspGTC_6_-_27551236_27551307@37.74.38,trna38_AspGTC_17_-_8125556_8125627@37.74.38,trna45_AspGTC_6_+_27447453_27447524@37.74.38,trna48_AspGTC_6_+_27471523_27471594@37.74.38,trna4_AspGTC_12_+_96429799_96429870@37.74.38,trna69_AspGTC_1_-_161440205_161440276@37.74.38,trna72_AspGTC_1_-_161432824_161432895@37.74.38,trna75_AspGTC_1_-_161425414_161425485@37.74.38,trna78_AspGTC_1_-_161418033_161418104@37.74.38,trna81_AspGTC_1_-_161410615_161410686@37.74.38\|ACGCGGGAGACCGGGGTTCGATTCCCCGACGGGGAGCC | >AspGTC.trna10_12_-_125424193_125424264 | 37 | 74 | 38 | tRF nuclear | NO | 0.00 | 504.19 | 0.00 | 0.00 |
| >trna10_AspGTC_12_-_125424193_125424264@37.75.39,trna12_AspGTC_12_-_125411891_125411962@37.75.39,trna144_AspGTC_6_-_27551236_27551307@37.75.39,trna38_AspGTC_17_-_8125556_8125627@37.75.39,trna45_AspGTC_6_+_27447453_27447524@37.75.39,trna48_AspGTC_6_+_27471523_27471594@37.75.39,trna4_AspGTC_12_+_96429799_96429870@37.75.39,trna69_AspGTC_1_-_161440205_161440276@37.75.39,trna72_AspGTC_1_-_161432824_161432895@37.75.39,trna75_AspGTC_1_-_161425414_161425485@37.75.39,trna78_AspGTC_1_-_161418033_161418104@37.75.39,trna81_AspGTC_1_-_161410615_161410686@37.75.39\|ACGCGGGAGACCGGGGTTCGATTCCCCGACGGGGAGCCA | >AspGTC.trna10_12_-_125424193_125424264 | 37 | 75 | 39 | tRF nuclear | NO | 0.00 | 383.13 | 0.00 | 0.00 |
| >trna10_AspGTC_12_-_125424193_125424264@38.73.36,trna12_AspGTC_12_-_125411891_125411962@38.73.36,trna144_AspGTC_6_-_27551236_27551307@38.73.36,trna38_AspGTC_17_-_8125556_8125627@38.73.36,trna45_AspGTC_6_+_27447453_27447524@38.73.36,trna48_AspGTC_6_+_27471523_27471594@38.73.36,trna4_AspGTC_12_+_96429799_96429870@38.73.36,trna69_AspGTC_1_-_161440205_161440276@38.73.36,trna72_AspGTC_1_-_161432824_161432895@38.73.36,trna75_AspGTC_1_-_161425414_161425485@38.73.36,trna78_AspGTC_1_-_161418033_161418104@38.73.36,trna81_AspGTC_1_-_161410615_161410686@38.73.36\|CGCGGGAGACCGGGGTTCGATTCCCCGACGGGGAGC | >AspGTC.trna10_12_-_125424193_125424264 | 38 | 73 | 36 | tRF nuclear | NO | 0.00 | 249.81 | 0.00 | 0.00 |
| >trna10_AspGTC_12_-_125424193_125424264@39.73.35,trna12_AspGTC_12_-_125411891_125411962@39.73.35,trna144_AspGTC_6_-_27551236_27551307@39.73.35,trna38_AspGTC_17_-_8125556_8125627@39.73.35,trna45_AspGTC_6_+_27447453_27447524@39.73.35,trna48_AspGTC_6_+_27471523_27471594@39.73.35,trna4_AspGTC_12_+_96429799_96429870@39.73.35,trna69_AspGTC_1_-_161440205_161440276@39.73.35,trna72_AspGTC_1_-_161432824_161432895@39.73.35,trna75_AspGTC_1_-_161425414_161425485@39.73.35,trna78_AspGTC_1_-_161418033_161418104@39.73.35,trna81_AspGTC_1_-_161410615_161410686@39.73.35\|GCGGGAGACCGGGGTTCGATTCCCCGACGGGGAGC | >AspGTC.trna10_12_-_125424193_125424264 | 39 | 73 | 35 | tRF nuclear | NO | 0.00 | 165.60 | 0.00 | 0.00 |
| >trna10_AspGTC_12_-_125424193_125424264@39.74.36,trna12_AspGTC_12_-_125411891_125411962@39.74.36,trna144_AspGTC_6_-_27551236_27551307@39.74.36,trna38_AspGTC_17_-_8125556_8125627@39.74.36,trna45_AspGTC_6_+_27447453_27447524@39.74.36,trna48_AspGTC_6_+_27471523_27471594@39.74.36,trna4_AspGTC_12_+_96429799_96429870@39.74.36,trna69_AspGTC_1_-_161440205_161440276@39.74.36,trna72_AspGTC_1_-_161432824_161432895@39.74.36,trna75_AspGTC_1_-_161425414_161425485@39.74.36,trna78_AspGTC_1_-_161418033_161418104@39.74.36,trna81_AspGTC_1_-_161410615_161410686@39.74.36\|GCGGGAGACCGGGGTTCGATTCCCCGACGGGGAGCC | >AspGTC.trna10_12_-_125424193_125424264 | 39 | 74 | 36 | tRF nuclear | NO | 0.00 | 155.10 | 0.00 | 0.00 |
| >trna10_AspGTC_12_-_125424193_125424264@39.75.37,trna12_AspGTC_12_-_125411891_125411962@39.75.37,trna144_AspGTC_6_-_27551236_27551307@39.75.37,trna38_AspGTC_17_-_8125556_8125627@39.75.37,trna45_AspGTC_6_+_27447453_27447524@39.75.37,trna48_AspGTC_6_+_27471523_27471594@39.75.37,trna4_AspGTC_12_+_96429799_96429870@39.75.37,trna69_AspGTC_1_-_161440205_161440276@39.75.37,trna72_AspGTC_1_-_161432824_161432895@39.75.37,trna75_AspGTC_1_-_161425414_161425485@39.75.37,trna78_AspGTC_1_-_161418033_161418104@39.75.37,trna81_AspGTC_1_-_161410615_161410686@39.75.37\|GCGGGAGACCGGGGTTCGATTCCCCGACGGGGAGCCA | >AspGTC.trna10_12_-_125424193_125424264 | 39 | 75 | 37 | tRF nuclear | NO | 0.00 | 195.64 | 0.00 | 0.00 |
| >trna5_AspGTC_12_+_98897281_98897352@35.74.40\|TCACGCGGGAGACCGGGGTTCAATTCCCCGACGGGGAGCC | >AspGTC.trna5_12_+_98897281_98897352 | 35 | 74 | 40 | tRF nuclear | NO | 0.00 | 122.95 | 0.00 | 0.00 |
| >trna5_AspGTC_12_+_98897281_98897352@35.75.41\|TCACGCGGGAGACCGGGGTTCAATTCCCCGACGGGGAGCCA | >AspGTC.trna5_12_+_98897281_98897352 | 35 | 75 | 41 | tRF nuclear | NO | 0.00 | 215.59 | 0.00 | 0.00 |
| >trna5_AspGTC_12_+_98897281_98897352@37.73.37\|ACGCGGGAGACCGGGGTTCAATTCCCCGACGGGGAGC | >AspGTC.trna5_12_+_98897281_98897352 | 37 | 73 | 37 | tRF nuclear | NO | 0.00 | 133.04 | 0.00 | 0.00 |
| >trna5_AspGTC_12_+_98897281_98897352@37.74.38\|ACGCGGGAGACCGGGGTTCAATTCCCCGACGGGGAGCC | >AspGTC.trna5_12_+_98897281_98897352 | 37 | 74 | 38 | tRF nuclear | NO | 0.00 | 110.50 | 0.00 | 0.00 |
| >trna10_GluCTC_3_-_103879524_103879595@16.42.27,trna116_GluCTC_1_-_145399233_145399304@16.42.27,trna59_GluCTC_1_+_249168447_249168518@16.42.27,trna71_GluCTC_1_-_161439189_161439260@16.42.27,trna74_GluCTC_1_-_161431809_161431880@16.42.27,trna77_GluCTC_1_-_161424398_161424469@16.42.27,trna77_GluCTC_6_+_28949976_28950047@16.42.27,trna80_GluCTC_1_-_161417018_161417089@16.42.27,trna87_GluCTC_6_-_126101393_126101464@16.42.27\|TGGTTAGGATTCGGCGCTCTCACCGCC | >GluCTC.trna10_3_-_103879524_103879595 | 16 | 42 | 27 | tRF nuclear | NO | 0.00 | 34.97 | 0.00 | 0.00 |

### MDA-MB-231

| | | | | | | | | | | |
|---|---|---|---|---|---|---|---|---|---|---|
| >trna116_GluCTC_1_-_145399233_145399304@1.31.31,trna59_GluCTC_1_+_249168447_249168518@1.31.31,trna71_GluCTC_1_-_161439189_161439260@1.31.31,trna74_GluCTC_1_-_161431809_161431880@1.31.31,trna77_GluCTC_1_-_161424398_161424469@1.31.31,trna77_GluCTC_6_+_28949976_28950047@1.31.31,trna80_GluCTC_1_-_161417018_161417089@1.31.31,trna87_GluCTC_6_-_126101393_126101464@1.31.31\|TCCCTGGTGGTCTAGTGGTTAGGATTCGGCG | >GluCTC.trna116_1_-_145399233_145399304 | 1 | 31 | 31 | tRF nuclear | NO | 77.05 | 704.84 | 119.38 | 63.77 |
| >trna116_GluCTC_1_-_145399233_145399304@1.32.32,trna59_GluCTC_1_+_249168447_249168518@1.32.32,trna71_GluCTC_1_-_161439189_161439260@1.32.32,trna74_GluCTC_1_-_161431809_161431880@1.32.32,trna77_GluCTC_1_-_161424398_161424469@1.32.32,trna77_GluCTC_6_+_28949976_28950047@1.32.32,trna80_GluCTC_1_-_161417018_161417089@1.32.32,trna87_GluCTC_6_-_126101393_126101464@1.32.32\|TCCCTGGTGGTCTAGTGGTTAGGATTCGGCGC | >GluCTC.trna116_1_-_145399233_145399304 | 1 | 32 | 32 | tRF nuclear | NO | 52.32 | 449.82 | 91.86 | 20.69 |
| >trna116_GluCTC_1_-_145399233_145399304@1.33.33,trna59_GluCTC_1_+_249168447_249168518@1.33.33,trna71_GluCTC_1_-_161439189_161439260@1.33.33,trna74_GluCTC_1_-_161431809_161431880@1.33.33,trna77_GluCTC_1_-_161424398_161424469@1.33.33,trna77_GluCTC_6_+_28949976_28950047@1.33.33,trna80_GluCTC_1_-_161417018_161417089@1.33.33,trna87_GluCTC_6_-_126101393_126101464@1.33.33\|TCCCTGGTGGTCTAGTGGTTAGGATTCGGCGCT | >GluCTC.trna116_1_-_145399233_145399304 | 1 | 33 | 33 | tRF nuclear | YES | 609.92 | 5954.90 | 221.44 | 150.42 |
| >trna116_GluCTC_1_-_145399233_145399304@1.34.34,trna59_GluCTC_1_+_249168447_249168518@1.34.34,trna71_GluCTC_1_-_161439189_161439260@1.34.34,trna74_GluCTC_1_-_161431809_161431880@1.34.34,trna77_GluCTC_1_-_161424398_161424469@1.34.34,trna77_GluCTC_6_+_28949976_28950047@1.34.34,trna80_GluCTC_1_-_161417018_161417089@1.34.34,trna87_GluCTC_6_-_126101393_126101464@1.34.34\|TCCCTGGTGGTCTAGTGGTTAGGATTCGGCGCTC | >GluCTC.trna116_1_-_145399233_145399304 | 1 | 34 | 34 | tRF nuclear | NO | 72.12 | 766.50 | 179.01 | 82.80 |
| >trna116_GluCTC_1_-_145399233_145399304@1.35.35,trna59_GluCTC_1_+_249168447_249168518@1.35.35,trna71_GluCTC_1_-_161439189_161439260@1.35.35,trna74_GluCTC_1_-_161431809_161431880@1.35.35,trna77_GluCTC_1_-_161424398_161424469@1.35.35,trna77_GluCTC_6_+_28949976_28950047@1.35.35,trna80_GluCTC_1_-_161417018_161417089@1.35.35,trna87_GluCTC_6_-_126101393_126101464@1.35.35\|TCCCTGGTGGTCTAGTGGTTAGGATTCGGCGCTCT | >GluCTC.trna116_1_-_145399233_145399304 | 1 | 35 | 35 | tRF nuclear | NO | 76.61 | 789.35 | 73.85 | 56.22 |
| >trna116_GluCTC_1_-_145399233_145399304@1.36.36,trna59_GluCTC_1_+_249168447_249168518@1.36.36,trna71_GluCTC_1_-_161439189_161439260@1.36.36,trna74_GluCTC_1_-_161431809_161431880@1.36.36,trna77_GluCTC_1_-_161424398_161424469@1.36.36,trna77_GluCTC_6_+_28949976_28950047@1.36.36,trna80_GluCTC_1_-_161417018_161417089@1.36.36,trna87_GluCTC_6_-_126101393_126101464@1.36.36\|TCCCTGGTGGTCTAGTGGTTAGGATTCGGCGCTCTC | >GluCTC.trna116_1_-_145399233_145399304 | 1 | 36 | 36 | tRF nuclear | NO | 78.96 | 953.45 | 63.53 | 65.47 |
| >trna116_GluCTC_1_-_145399233_145399304@1.39.39,trna59_GluCTC_1_+_249168447_249168518@1.39.39,trna71_GluCTC_1_-_161439189_161439260@1.39.39,trna74_GluCTC_1_-_161431809_161431880@1.39.39,trna77_GluCTC_1_-_161424398_161424469@1.39.39,trna77_GluCTC_6_+_28949976_28950047@1.39.39,trna80_GluCTC_1_-_161417018_161417089@1.39.39,trna87_GluCTC_6_-_126101393_126101464@1.39.39\|TCCCTGGTGGTCTAGTGGTTAGGATTCGGCGCTCTCACC | >GluCTC.trna116_1_-_145399233_145399304 | 1 | 39 | 39 | tRF nuclear | NO | 77.78 | 1539.99 | 0.00 | 14.01 |
| >trna116_GluCTC_1_-_145399233_145399304@1.42.42,trna59_GluCTC_1_+_249168447_249168518@1.42.42,trna71_GluCTC_1_-_161439189_161439260@1.42.42,trna74_GluCTC_1_-_161431809_161431880@1.42.42,trna77_GluCTC_1_-_161424398_161424469@1.42.42,trna77_GluCTC_6_+_28949976_28950047@1.42.42,trna80_GluCTC_1_-_161417018_161417089@1.42.42,trna87_GluCTC_6_-_126101393_126101464@1.42.42\|TCCCTGGTGGTCTAGTGGTTAGGATTCGGCGCTCTCACCGCC | >GluCTC.trna116_1_-_145399233_145399304 | 1 | 42 | 42 | tRF nuclear | NO | 0.00 | 112.23 | 0.00 | 0.00 |
| >trna116_GluCTC_1_-_145399233_145399304@3.33.31,trna59_GluCTC_1_+_249168447_249168518@3.33.31,trna71_GluCTC_1_-_161439189_161439260@3.33.31,trna74_GluCTC_1_-_161431809_161431880@3.33.31,trna77_GluCTC_1_-_161424398_161424469@3.33.31,trna77_GluCTC_6_+_28949976_28950047@3.33.31,trna80_GluCTC_1_-_161417018_161417089@3.33.31,trna87_GluCTC_6_-_126101393_126101464@3.33.31\|CCTGGTGGTCTAGTGGTTAGGATTCGGCGCT | >GluCTC.trna116_1_-_145399233_145399304 | 3 | 33 | 31 | tRF nuclear | NO | 0.00 | 33.69 | 0.00 | 14.47 |
| >trna116_GluCTC_1_-_145399233_145399304@35.74.40,trna134_GluTTC_1_-_16861774_16861845@35.74.40,trna71_GluCTC_1_-_161439189_161439260@35.74.40,trna74_GluCTC_1_-_161431809_161431880@35.74.40,trna77_GluCTC_1_-_161424398_161424469@35.74.40,trna77_GluCTC_6_+_28949976_28950047@35.74.40,trna80_GluCTC_1_-_161417018_161417089@35.74.40,trna84_GluTTC_1_-_161391883_161391954@35.74.40,trna87_GluCTC_6_-_126101393_126101464@35.74.40\|TCACCGCCGCGGCCCGGGTTCGATTCCCGGTCAGGGAACC | >GluCTC.trna116_1_-_145399233_145399304 | 35 | 74 | 40 | tRF nuclear | NO | 0.00 | 126.77 | 0.00 | 0.00 |

### MDA-MB-231

| | | | | | | | | | | |
|---|---|---|---|---|---|---|---|---|---|---|
| >trna116_GluCTC_1_-_145399233_145399304@35.75.41,trna134_GluTTC_1_-_16861774_16861845@35.75.41,trna71_GluCTC_1_-_161439189_161439260@35.75.41,trna74_GluCTC_1_-_161431809_161431880@35.75.41,trna77_GluCTC_1_-_161424398_161424469@35.75.41,trna77_GluCTC_6_+_28949976_28950047@35.75.41,trna80_GluCTC_1_-_161417018_161417089@35.75.41,trna84_GluTTC_1_-_161391883_161391954@35.75.41,trna87_GluCTC_6_-_126101393_126101464@35.75.41\|TCACCGCCGCGGCCCGGGTTCGATTCCCGGTCAGGGAACCA | >GluCTC.trna116_1_-_145399233_145399304 | 35 | 75 | 41 | tRF nuclear | NO | 0.00 | 189.78 | 0.00 | 0.00 |
| >trna116_GluCTC_1_-_145399233_145399304@36.74.39,trna134_GluTTC_1_-_16861774_16861845@36.74.39,trna71_GluCTC_1_-_161439189_161439260@36.74.39,trna74_GluCTC_1_-_161431809_161431880@36.74.39,trna77_GluCTC_1_-_161424398_161424469@36.74.39,trna77_GluCTC_6_+_28949976_28950047@36.74.39,trna80_GluCTC_1_-_161417018_161417089@36.74.39,trna84_GluTTC_1_-_161391883_161391954@36.74.39,trna87_GluCTC_6_-_126101393_126101464@36.74.39\|CACCGCCGCGGCCCGGGTTCGATTCCCGGTCAGGGAACC | >GluCTC.trna116_1_-_145399233_145399304 | 36 | 74 | 39 | tRF nuclear | NO | 0.00 | 142.79 | 0.00 | 0.00 |
| >trna116_GluCTC_1_-_145399233_145399304@36.75.40,trna134_GluTTC_1_-_16861774_16861845@36.75.40,trna71_GluCTC_1_-_161439189_161439260@36.75.40,trna74_GluCTC_1_-_161431809_161431880@36.75.40,trna77_GluCTC_1_-_161424398_161424469@36.75.40,trna77_GluCTC_6_+_28949976_28950047@36.75.40,trna80_GluCTC_1_-_161417018_161417089@36.75.40,trna84_GluTTC_1_-_161391883_161391954@36.75.40,trna87_GluCTC_6_-_126101393_126101464@36.75.40\|CACCGCCGCGGCCCGGGTTCGATTCCCGGTCAGGGAACCA | >GluCTC.trna116_1_-_145399233_145399304 | 36 | 75 | 40 | tRF nuclear | NO | 0.00 | 179.34 | 0.00 | 0.00 |
| >trna116_GluCTC_1_-_145399233_145399304@37.73.37,trna134_GluTTC_1_-_16861774_16861845@37.73.37,trna71_GluCTC_1_-_161439189_161439260@37.73.37,trna74_GluCTC_1_-_161431809_161431880@37.73.37,trna77_GluCTC_1_-_161424398_161424469@37.73.37,trna77_GluCTC_6_+_28949976_28950047@37.73.37,trna80_GluCTC_1_-_161417018_161417089@37.73.37,trna84_GluTTC_1_-_161391883_161391954@37.73.37,trna87_GluCTC_6_-_126101393_126101464@37.73.37\|ACCGCCGCGGCCCGGGTTCGATTCCCGGTCAGGGAAC | >GluCTC.trna116_1_-_145399233_145399304 | 37 | 73 | 37 | tRF nuclear | NO | 0.00 | 74.07 | 0.00 | 0.00 |
| >trna116_GluCTC_1_-_145399233_145399304@37.74.38,trna134_GluTTC_1_-_16861774_16861845@37.74.38,trna71_GluCTC_1_-_161439189_161439260@37.74.38,trna74_GluCTC_1_-_161431809_161431880@37.74.38,trna77_GluCTC_1_-_161424398_161424469@37.74.38,trna77_GluCTC_6_+_28949976_28950047@37.74.38,trna80_GluCTC_1_-_161417018_161417089@37.74.38,trna84_GluTTC_1_-_161391883_161391954@37.74.38,trna87_GluCTC_6_-_126101393_126101464@37.74.38\|ACCGCCGCGGCCCGGGTTCGATTCCCGGTCAGGGAACC | >GluCTC.trna116_1_-_145399233_145399304 | 37 | 74 | 38 | tRF nuclear | NO | 68.28 | 334.27 | 0.00 | 0.00 |
| >trna116_GluCTC_1_-_145399233_145399304@37.75.39,trna134_GluTTC_1_-_16861774_16861845@37.75.39,trna71_GluCTC_1_-_161439189_161439260@37.75.39,trna74_GluCTC_1_-_161431809_161431880@37.75.39,trna77_GluCTC_1_-_161424398_161424469@37.75.39,trna77_GluCTC_6_+_28949976_28950047@37.75.39,trna80_GluCTC_1_-_161417018_161417089@37.75.39,trna84_GluTTC_1_-_161391883_161391954@37.75.39,trna87_GluCTC_6_-_126101393_126101464@37.75.39\|ACCGCCGCGGCCCGGGTTCGATTCCCGGTCAGGGAACCA | >GluCTC.trna116_1_-_145399233_145399304 | 37 | 75 | 39 | tRF nuclear | NO | 85.60 | 319.83 | 0.00 | 0.00 |
| >trna116_GluCTC_1_-_145399233_145399304@38.74.37,trna134_GluTTC_1_-_16861774_16861845@38.74.37,trna71_GluCTC_1_-_161439189_161439260@38.74.37,trna74_GluCTC_1_-_161431809_161431880@38.74.37,trna77_GluCTC_1_-_161424398_161424469@38.74.37,trna77_GluCTC_6_+_28949976_28950047@38.74.37,trna80_GluCTC_1_-_161417018_161417089@38.74.37,trna84_GluTTC_1_-_161391883_161391954@38.74.37,trna87_GluCTC_6_-_126101393_126101464@38.74.37\|CCGCCGCGGCCCGGGTTCGATTCCCGGTCAGGGAACC | >GluCTC.trna116_1_-_145399233_145399304 | 38 | 74 | 37 | tRF nuclear | NO | 0.00 | 25.99 | 0.00 | 0.00 |
| >trna116_GluCTC_1_-_145399233_145399304@39.74.36,trna134_GluTTC_1_-_16861774_16861845@39.74.36,trna71_GluCTC_1_-_161439189_161439260@39.74.36,trna74_GluCTC_1_-_161431809_161431880@39.74.36,trna77_GluCTC_1_-_161424398_161424469@39.74.36,trna77_GluCTC_6_+_28949976_28950047@39.74.36,trna80_GluCTC_1_-_161417018_161417089@39.74.36,trna84_GluTTC_1_-_161391883_161391954@39.74.36,trna87_GluCTC_6_-_126101393_126101464@39.74.36\|CGCCGCGGCCCGGGTTCGATTCCCGGTCAGGGAACC | >GluCTC.trna116_1_-_145399233_145399304 | 39 | 74 | 36 | tRF nuclear | NO | 0.00 | 33.52 | 0.00 | 0.00 |
| >trna116_GluCTC_1_-_145399233_145399304@39.75.37,trna134_GluTTC_1_-_16861774_16861845@39.75.37,trna71_GluCTC_1_-_161439189_161439260@39.75.37,trna74_GluCTC_1_-_161431809_161431880@39.75.37,trna77_GluCTC_1_-_161424398_161424469@39.75.37,trna77_GluCTC_6_+_28949976_28950047@39.75.37,trna80_GluCTC_1_-_161417018_161417089@39.75.37,trna84_GluTTC_1_-_161391883_161391954@39.75.37,trna87_GluCTC_6_-_126101393_126101464@39.75.37\|CGCCGCGGCCCGGGTTCGATTCCCGGTCAGGGAACCA | >GluCTC.trna116_1_-_145399233_145399304 | 39 | 75 | 37 | tRF nuclear | NO | 0.00 | 29.78 | 0.00 | 0.00 |

### MDA-MB-231

<table>
<tr><td>>am_trna116_GluCTC_1_-_145399233_145399304@1.26.26,trna59_GluCTC_1_+_249168447_249168518@1.26.26,trna71_GluCTC_1_-_161439189_161439260@1.26.26,trna74_GluCTC_1_-_161431809_161431880@1.26.26,trna77_GluCTC_1_-_161424398_161424469@1.26.26,trna77_GluCTC_6_+_28949976_28950047@1.26.26,trna80_GluCTC_1_-_161417018_161417089@1.26.26,trna87_GluCTC_6_-_126101393_126101464@1.26.26|TCCCTGGTGGTCTAGTGGTTAGGATT</td><td>>GluCTC.trna116_1_-_145399233_145399304.am</td><td>1</td><td>26</td><td>26</td><td>tRF nuclear</td><td>NO</td><td>0.00</td><td>131.24</td><td>0.00</td><td>0.00</td></tr>
<tr><td>>am_trna116_GluCTC_1_-_145399233_145399304@1.27.27,trna59_GluCTC_1_+_249168447_249168518@1.27.27,trna71_GluCTC_1_-_161439189_161439260@1.27.27,trna74_GluCTC_1_-_161431809_161431880@1.27.27,trna77_GluCTC_1_-_161424398_161424469@1.27.27,trna77_GluCTC_6_+_28949976_28950047@1.27.27,trna80_GluCTC_1_-_161417018_161417089@1.27.27,trna87_GluCTC_6_-_126101393_126101464@1.27.27|TCCCTGGTGGTCTAGTGGTTAGGATTC</td><td>>GluCTC.trna116_1_-_145399233_145399304.am</td><td>1</td><td>27</td><td>27</td><td>tRF nuclear</td><td>NO</td><td>0.00</td><td>117.76</td><td>10.69</td><td>0.00</td></tr>
<tr><td>>am_trna116_GluCTC_1_-_145399233_145399304@1.28.28,trna59_GluCTC_1_+_249168447_249168518@1.28.28,trna71_GluCTC_1_-_161439189_161439260@1.28.28,trna74_GluCTC_1_-_161431809_161431880@1.28.28,trna77_GluCTC_1_-_161424398_161424469@1.28.28,trna77_GluCTC_6_+_28949976_28950047@1.28.28,trna80_GluCTC_1_-_161417018_161417089@1.28.28,trna87_GluCTC_6_-_126101393_126101464@1.28.28|TCCCTGGTGGTCTAGTGGTTAGGATTCG</td><td>>GluCTC.trna116_1_-_145399233_145399304.am</td><td>1</td><td>28</td><td>28</td><td>tRF nuclear</td><td>NO</td><td>0.00</td><td>112.12</td><td>0.00</td><td>0.00</td></tr>
<tr><td>>am_trna116_GluCTC_1_-_145399233_145399304@1.29.29,trna59_GluCTC_1_+_249168447_249168518@1.29.29,trna71_GluCTC_1_-_161439189_161439260@1.29.29,trna74_GluCTC_1_-_161431809_161431880@1.29.29,trna77_GluCTC_1_-_161424398_161424469@1.29.29,trna77_GluCTC_6_+_28949976_28950047@1.29.29,trna80_GluCTC_1_-_161417018_161417089@1.29.29,trna87_GluCTC_6_-_126101393_126101464@1.29.29|TCCCTGGTGGTCTAGTGGTTAGGATTCGG</td><td>>GluCTC.trna116_1_-_145399233_145399304.am</td><td>1</td><td>29</td><td>29</td><td>tRF nuclear</td><td>NO</td><td>0.00</td><td>265.22</td><td>10.73</td><td>0.00</td></tr>
<tr><td>>am_trna116_GluCTC_1_-_145399233_145399304@1.30.30,trna59_GluCTC_1_+_249168447_249168518@1.30.30,trna71_GluCTC_1_-_161439189_161439260@1.30.30,trna74_GluCTC_1_-_161431809_161431880@1.30.30,trna77_GluCTC_1_-_161424398_161424469@1.30.30,trna77_GluCTC_6_+_28949976_28950047@1.30.30,trna80_GluCTC_1_-_161417018_161417089@1.30.30,trna87_GluCTC_6_-_126101393_126101464@1.30.30|TCCCTGGTGGTCTAGTGGTTAGGATTCGGC</td><td>>GluCTC.trna116_1_-_145399233_145399304.am</td><td>1</td><td>30</td><td>30</td><td>tRF nuclear</td><td>NO</td><td>25.99</td><td>631.06</td><td>99.60</td><td>18.77</td></tr>
<tr><td>>trna11_GluTTC_15_-_26327381_26327452@1.29.29,trna3_GluTTC_13_-_45492062_45492133@1.29.29|TCCCACATGGTCTAGCGGTTAGGATTCCT</td><td>>GluTTC.trna11_15_-_26327381_26327452</td><td>1</td><td>29</td><td>29</td><td>tRF nuclear</td><td>NO</td><td>0.00</td><td>30.61</td><td>0.00</td><td>0.00</td></tr>
<tr><td>>trna11_GluTTC_15_-_26327381_26327452@1.31.31,trna3_GluTTC_13_-_45492062_45492133@1.31.31|TCCCACATGGTCTAGCGGTTAGGATTCCTGG</td><td>>GluTTC.trna11_15_-_26327381_26327452</td><td>1</td><td>31</td><td>31</td><td>tRF nuclear</td><td>NO</td><td>0.00</td><td>32.14</td><td>13.02</td><td>0.00</td></tr>
<tr><td>>trna11_GluTTC_15_-_26327381_26327452@1.32.32,trna3_GluTTC_13_-_45492062_45492133@1.32.32|TCCCACATGGTCTAGCGGTTAGGATTCCTGGT</td><td>>GluTTC.trna11_15_-_26327381_26327452</td><td>1</td><td>32</td><td>32</td><td>tRF nuclear</td><td>NO</td><td>0.00</td><td>60.43</td><td>15.31</td><td>11.47</td></tr>
<tr><td>>trna11_GluTTC_15_-_26327381_26327452@1.33.33,trna3_GluTTC_13_-_45492062_45492133@1.33.33|TCCCACATGGTCTAGCGGTTAGGATTCCTGGTT</td><td>>GluTTC.trna11_15_-_26327381_26327452</td><td>1</td><td>33</td><td>33</td><td>tRF nuclear</td><td>NO</td><td>73.17</td><td>468.95</td><td>145.79</td><td>113.01</td></tr>
<tr><td>>trna11_GluTTC_15_-_26327381_26327452@1.36.36,trna3_GluTTC_13_-_45492062_45492133@1.36.36|TCCCACATGGTCTAGCGGTTAGGATTCCTGGTTTTC</td><td>>GluTTC.trna11_15_-_26327381_26327452</td><td>1</td><td>36</td><td>36</td><td>tRF nuclear</td><td>NO</td><td>0.00</td><td>79.05</td><td>0.00</td><td>13.92</td></tr>
<tr><td>>trna134_GluTTC_1_-_16861774_16861845@1.33.33,trna5_GluTTC_1_+_17199078_17199149@1.33.33,trna84_GluTTC_1_-_161391883_161391954@1.33.33,trna94_GluTTC_1_-_149664355_149664427@1.33.33|TCCCTGGTGGTCTAGTGGCTAGGATTCGGCGCT</td><td>>GluTTC.trna134_1_-_16861774_16861845</td><td>1</td><td>33</td><td>33</td><td>tRF nuclear</td><td>NO</td><td>0.00</td><td>73.43</td><td>0.00</td><td>0.00</td></tr>
<tr><td>>trna20_GluTTC_2_-_131094701_131094772@1.31.31,trna5_GluTTC_13_-_41634874_41634945@1.31.31|TCCCATATGGTCTAGCGGTTAGGATTCCTGG</td><td>>GluTTC.trna20_2_-_131094701_131094772</td><td>1</td><td>31</td><td>31</td><td>tRF nuclear</td><td>NO</td><td>0.00</td><td>0.00</td><td>31.03</td><td>0.00</td></tr>
<tr><td>>trna20_GluTTC_2_-_131094701_131094772@1.32.32,trna5_GluTTC_13_-_41634874_41634945@1.32.32|TCCCATATGGTCTAGCGGTTAGGATTCCTGGT</td><td>>GluTTC.trna20_2_-_131094701_131094772</td><td>1</td><td>32</td><td>32</td><td>tRF nuclear</td><td>NO</td><td>0.00</td><td>0.00</td><td>17.47</td><td>0.00</td></tr>
<tr><td>>trna20_GluTTC_2_-_131094701_131094772@1.33.33,trna5_GluTTC_13_-_41634874_41634945@1.33.33|TCCCATATGGTCTAGCGGTTAGGATTCCTGGTT</td><td>>GluTTC.trna20_2_-_131094701_131094772</td><td>1</td><td>33</td><td>33</td><td>tRF nuclear</td><td>NO</td><td>0.00</td><td>206.14</td><td>183.88</td><td>54.17</td></tr>
<tr><td>>trna20_GluTTC_2_-_131094701_131094772@1.36.36,trna5_GluTTC_13_-_41634874_41634945@1.36.36|TCCCATATGGTCTAGCGGTTAGGATTCCTGGTTTTC</td><td>>GluTTC.trna20_2_-_131094701_131094772</td><td>1</td><td>36</td><td>36</td><td>tRF nuclear</td><td>NO</td><td>0.00</td><td>32.72</td><td>0.00</td><td>0.00</td></tr>
<tr><td>>am_trna128_GlyGCC_6_-_27870686_27870756@1.25.25,trna133_GlyCCC_1_-_16872434_16872504@1.25.25,trna18_GlyGCC_16_+_70822597_70822667@1.25.25,trna19_GlyGCC_16_+_70823410_70823480@1.25.25,trna19_GlyGCC_2_-_157257659_157257729@1.25.25,trna24_GlyGCC_16_-_70812942_70813012@1.25.25,trna25_GlyGCC_16_-_70812114_70812184@1.25.25,trna4_GlyCCC_1_+_17188416_17188486@1.25.25,trna5_GlyGCC_17_+_8029064_8029134@1.25.25,trna68_GlyGCC_1_-_161493637_161493707@1.25.25|GCATTGGTGGTTCAGTGGTAGAATT</td><td>>GlyGCC.trna128_6_-_27870686_27870756.am</td><td>1</td><td>25</td><td>25</td><td>tRF nuclear</td><td>NO</td><td>0.00</td><td>138.81</td><td>0.00</td><td>0.00</td></tr>
<tr><td>>am_trna128_GlyGCC_6_-_27870686_27870756@1.27.27,trna133_GlyCCC_1_-_16872434_16872504@1.27.27,trna18_GlyGCC_16_+_70822597_70822667@1.27.27,trna19_GlyGCC_16_+_70823410_70823480@1.27.27,trna19_GlyGCC_2_-_157257659_157257729@1.27.27,trna24_GlyGCC_16_-_70812942_70813012@1.27.27,trna25_GlyGCC_16_-_70812114_70812184@1.27.27,trna4_GlyCCC_1_+_17188416_17188486@1.27.27,trna5_GlyGCC_17_+_8029064_8029134@1.27.27,trna68_GlyGCC_1_-_161493637_161493707@1.27.27|GCATTGGTGGTTCAGTGGTAGAATTCT</td><td>>GlyGCC.trna128_6_-_27870686_27870756.am</td><td>1</td><td>27</td><td>27</td><td>tRF nuclear</td><td>NO</td><td>0.00</td><td>116.90</td><td>35.12</td><td>0.00</td></tr>
</table>

### MDA-MB-231

| | | | | | | | | | | |
|---|---|---|---|---|---|---|---|---|---|---|
| >am_trna128_GlyGCC_6_-_27870686_27870756@1.28.28,trna133_GlyCCC_1_-_16872434_16872504@1.28.28,trna18_GlyGCC_16_+_70822597_70822667@1.28.28,trna19_GlyGCC_16_+_70823410_70823480@1.28.28,trna19_GlyGCC_2_-_157257659_157257729@1.28.28,trna24_GlyGCC_16_-_70812942_70813012@1.28.28,trna25_GlyGCC_16_-_70812114_70812184@1.28.28,trna4_GlyCCC_1_+_17188416_17188486@1.28.28,trna5_GlyGCC_17_+_8029064_8029134@1.28.28,trna68_GlyGCC_1_-_161493637_161493707@1.28.28\|GCATTGGTGGTTCAGTGGTAGAATTCTC | >GlyGCC.trna128_6_-_27870686_27870756.am | 1 | 28 | 28 | tRF nuclear | NO | 0.00 | 288.13 | 82.24 | 0.00 |
| >am_trna128_GlyGCC_6_-_27870686_27870756@1.29.29,trna133_GlyCCC_1_-_16872434_16872504@1.29.29,trna18_GlyGCC_16_+_70822597_70822667@1.29.29,trna19_GlyGCC_16_+_70823410_70823480@1.29.29,trna19_GlyGCC_2_-_157257659_157257729@1.29.29,trna24_GlyGCC_16_-_70812942_70813012@1.29.29,trna25_GlyGCC_16_-_70812114_70812184@1.29.29,trna4_GlyCCC_1_+_17188416_17188486@1.29.29,trna5_GlyGCC_17_+_8029064_8029134@1.29.29,trna68_GlyGCC_1_-_161493637_161493707@1.29.29\|GCATTGGTGGTTCAGTGGTAGAATTCTCG | >GlyGCC.trna128_6_-_27870686_27870756.am | 1 | 29 | 29 | tRF nuclear | NO | 0.00 | 202.17 | 16.84 | 0.00 |
| >am_trna128_GlyGCC_6_-_27870686_27870756@1.30.30,trna133_GlyCCC_1_-_16872434_16872504@1.30.30,trna18_GlyGCC_16_+_70822597_70822667@1.30.30,trna19_GlyGCC_16_+_70823410_70823480@1.30.30,trna19_GlyGCC_2_-_157257659_157257729@1.30.30,trna24_GlyGCC_16_-_70812942_70813012@1.30.30,trna25_GlyGCC_16_-_70812114_70812184@1.30.30,trna4_GlyCCC_1_+_17188416_17188486@1.30.30,trna5_GlyGCC_17_+_8029064_8029134@1.30.30,trna68_GlyGCC_1_-_161493637_161493707@1.30.30\|GCATTGGTGGTTCAGTGGTAGAATTCTCGC | >GlyGCC.trna128_6_-_27870686_27870756.am | 1 | 30 | 30 | tRF nuclear | NO | 68.64 | 707.04 | 1339.97 | 75.94 |
| >am_trna128_GlyGCC_6_-_27870686_27870756@1.31.31,trna133_GlyCCC_1_-_16872434_16872504@1.31.31,trna18_GlyGCC_16_+_70822597_70822667@1.31.31,trna19_GlyGCC_16_+_70823410_70823480@1.31.31,trna19_GlyGCC_2_-_157257659_157257729@1.31.31,trna24_GlyGCC_16_-_70812942_70813012@1.31.31,trna25_GlyGCC_16_-_70812114_70812184@1.31.31,trna4_GlyCCC_1_+_17188416_17188486@1.31.31,trna5_GlyGCC_17_+_8029064_8029134@1.31.31,trna68_GlyGCC_1_-_161493637_161493707@1.31.31\|GCATTGGTGGTTCAGTGGTAGAATTCTCGCC | >GlyGCC.trna128_6_-_27870686_27870756.am | 1 | 31 | 31 | tRF nuclear | YES | 293.33 | 3111.49 | 1658.84 | 266.38 |
| >am_trna128_GlyGCC_6_-_27870686_27870756@1.32.32,trna133_GlyCCC_1_-_16872434_16872504@1.32.32,trna18_GlyGCC_16_+_70822597_70822667@1.32.32,trna19_GlyGCC_16_+_70823410_70823480@1.32.32,trna19_GlyGCC_2_-_157257659_157257729@1.32.32,trna24_GlyGCC_16_-_70812942_70813012@1.32.32,trna25_GlyGCC_16_-_70812114_70812184@1.32.32,trna4_GlyCCC_1_+_17188416_17188486@1.32.32,trna5_GlyGCC_17_+_8029064_8029134@1.32.32,trna68_GlyGCC_1_-_161493637_161493707@1.32.32\|GCATTGGTGGTTCAGTGGTAGAATTCTCGCCT | >GlyGCC.trna128_6_-_27870686_27870756.am | 1 | 32 | 32 | tRF nuclear | YES | 167.26 | 1191.57 | 921.10 | 176.84 |
| >am_trna128_GlyGCC_6_-_27870686_27870756@1.33.33,trna18_GlyGCC_16_+_70822597_70822667@1.33.33,trna19_GlyGCC_16_+_70823410_70823480@1.33.33,trna19_GlyGCC_2_-_157257659_157257729@1.33.33,trna24_GlyGCC_16_-_70812942_70813012@1.33.33,trna25_GlyGCC_16_-_70812114_70812184@1.33.33,trna5_GlyGCC_17_+_8029064_8029134@1.33.33,trna68_GlyGCC_1_-_161493637_161493707@1.33.33\|GCATTGGTGGTTCAGTGGTAGAATTCTCGCCTG | >GlyGCC.trna128_6_-_27870686_27870756.am | 1 | 33 | 33 | tRF nuclear | NO | 55.79 | 286.39 | 325.23 | 74.26 |
| >am_trna128_GlyGCC_6_-_27870686_27870756@1.34.34,trna18_GlyGCC_16_+_70822597_70822667@1.34.34,trna19_GlyGCC_16_+_70823410_70823480@1.34.34,trna19_GlyGCC_2_-_157257659_157257729@1.34.34,trna24_GlyGCC_16_-_70812942_70813012@1.34.34,trna25_GlyGCC_16_-_70812114_70812184@1.34.34,trna5_GlyGCC_17_+_8029064_8029134@1.34.34,trna68_GlyGCC_1_-_161493637_161493707@1.34.34\|GCATTGGTGGTTCAGTGGTAGAATTCTCGCCTGC | >GlyGCC.trna128_6_-_27870686_27870756.am | 1 | 34 | 34 | tRF nuclear | NO | 25.74 | 189.60 | 147.82 | 54.66 |
| >am_trna128_GlyGCC_6_-_27870686_27870756@1.35.35,trna18_GlyGCC_16_+_70822597_70822667@1.35.35,trna19_GlyGCC_16_+_70823410_70823480@1.35.35,trna19_GlyGCC_2_-_157257659_157257729@1.35.35,trna24_GlyGCC_16_-_70812942_70813012@1.35.35,trna25_GlyGCC_16_-_70812114_70812184@1.35.35,trna5_GlyGCC_17_+_8029064_8029134@1.35.35,trna68_GlyGCC_1_-_161493637_161493707@1.35.35\|GCATTGGTGGTTCAGTGGTAGAATTCTCGCCTGCC | >GlyGCC.trna128_6_-_27870686_27870756.am | 1 | 35 | 35 | tRF nuclear | NO | 98.51 | 581.42 | 303.94 | 87.28 |
| >am_trna128_GlyGCC_6_-_27870686_27870756@1.36.36,trna18_GlyGCC_16_+_70822597_70822667@1.36.36,trna19_GlyGCC_16_+_70823410_70823480@1.36.36,trna19_GlyGCC_2_-_157257659_157257729@1.36.36,trna24_GlyGCC_16_-_70812942_70813012@1.36.36,trna25_GlyGCC_16_-_70812114_70812184@1.36.36,trna5_GlyGCC_17_+_8029064_8029134@1.36.36,trna68_GlyGCC_1_-_161493637_161493707@1.36.36\|GCATTGGTGGTTCAGTGGTAGAATTCTCGCCTGCCA | >GlyGCC.trna128_6_-_27870686_27870756.am | 1 | 36 | 36 | tRF nuclear | NO | 0.00 | 111.05 | 34.02 | 0.00 |

### MDA-MB-231

| | | | | | | | | | | |
|---|---|---|---|---|---|---|---|---|---|---|
| >am_trna128_GlyGCC_6_-_27870686_27870756@3.35.33,trna18_GlyGCC_16_+_70822597_70822667@3.35.33,trna19_GlyGCC_16_+_70823410_70823480@3.35.33,trna19_GlyGCC_2_-_157257659_157257729@3.35.33,trna24_GlyGCC_16_-_70812942_70813012@3.35.33,trna25_GlyGCC_16_-_70812114_70812184@3.35.33,trna5_GlyGCC_17_+_8029064_8029134@3.35.33,trna68_GlyGCC_1_-_161493637_161493707@3.35.33\|ATTGGTGGTTCAGTGGTAGAATTCTCGCCTGCC | >GlyGCC.trna128_6_-_27870686_27870756.am | 3 | 35 | 33 | tRF nuclear | NO | 10.65 | 0.00 | 15.16 | 66.14 |
| >trna2_GlyGCC_21_-_18827107_18827177@1.31.31,trna35_GlyGCC_1_+_161413094_161413164@1.31.31,trna37_GlyGCC_1_+_161420467_161420537@1.31.31,trna39_GlyGCC_1_+_161427898_161427968@1.31.31,trna41_GlyGCC_1_+_161435258_161435328@1.31.31\|GCATGGGTGGTTCAGTGGTAGAATTCTCGCC | >GlyGCC.trna2_21_-_18827107_18827177 | 1 | 31 | 31 | tRF nuclear | NO | 0.00 | 84.08 | 63.15 | 0.00 |
| >trna2_GlyGCC_21_-_18827107_18827177@1.32.32,trna35_GlyGCC_1_+_161413094_161413164@1.32.32,trna37_GlyGCC_1_+_161420467_161420537@1.32.32,trna39_GlyGCC_1_+_161427898_161427968@1.32.32,trna41_GlyGCC_1_+_161435258_161435328@1.32.32\|GCATGGGTGGTTCAGTGGTAGAATTCTCGCCT | >GlyGCC.trna2_21_-_18827107_18827177 | 1 | 32 | 32 | tRF nuclear | NO | 0.00 | 12.97 | 18.10 | 0.00 |
| >trna111_HisGTG_1_-_147774845_147774916@1.30.30,trna118_HisGTG_1_-_145396881_145396952@1.30.30,trna16_HisGTG_1_+_146544773_146544844@1.30.30,trna1_HisGTG_15_+_45493349_45493420@1.30.30,trna21_HisGTG_1_+_147753471_147753542@1.30.30,trna33_HisGTG_6_+_27125906_27125977@1.30.30,trna7_HisGTG_9_-_14433938_14434009@1.30.30,trna8_HisGTG_15_-_45492611_45492682@1.30.30,trna9_HisGTG_15_-_45490804_45490875@1.30.30\|GCCGTGATCGTATAGTGGTTAGTACTCTGC | >HisGTG.trna111_1_-_147774845_147774916 | 1 | 30 | 30 | tRF nuclear | NO | 0.00 | 192.54 | 94.49 | 10.47 |
| >trna111_HisGTG_1_-_147774845_147774916@1.31.31,trna118_HisGTG_1_-_145396881_145396952@1.31.31,trna16_HisGTG_1_+_146544773_146544844@1.31.31,trna1_HisGTG_15_+_45493349_45493420@1.31.31,trna21_HisGTG_1_+_147753471_147753542@1.31.31,trna33_HisGTG_6_+_27125906_27125977@1.31.31,trna7_HisGTG_9_-_14433938_14434009@1.31.31,trna8_HisGTG_15_-_45492611_45492682@1.31.31,trna9_HisGTG_15_-_45490804_45490875@1.31.31\|GCCGTGATCGTATAGTGGTTAGTACTCTGCG | >HisGTG.trna111_1_-_147774845_147774916 | 1 | 31 | 31 | tRF nuclear | NO | 0.00 | 431.12 | 170.22 | 15.72 |
| >trna111_HisGTG_1_-_147774845_147774916@1.32.32,trna118_HisGTG_1_-_145396881_145396952@1.32.32,trna16_HisGTG_1_+_146544773_146544844@1.32.32,trna1_HisGTG_15_+_45493349_45493420@1.32.32,trna21_HisGTG_1_+_147753471_147753542@1.32.32,trna33_HisGTG_6_+_27125906_27125977@1.32.32,trna7_HisGTG_9_-_14433938_14434009@1.32.32,trna8_HisGTG_15_-_45492611_45492682@1.32.32,trna9_HisGTG_15_-_45490804_45490875@1.32.32\|GCCGTGATCGTATAGTGGTTAGTACTCTGCGT | >HisGTG.trna111_1_-_147774845_147774916 | 1 | 32 | 32 | tRF nuclear | NO | 0.00 | 85.60 | 15.31 | 0.00 |
| >trna111_HisGTG_1_-_147774845_147774916@1.33.33,trna118_HisGTG_1_-_145396881_145396952@1.33.33,trna16_HisGTG_1_+_146544773_146544844@1.33.33,trna1_HisGTG_15_+_45493349_45493420@1.33.33,trna21_HisGTG_1_+_147753471_147753542@1.33.33,trna33_HisGTG_6_+_27125906_27125977@1.33.33,trna7_HisGTG_9_-_14433938_14434009@1.33.33,trna8_HisGTG_15_-_45492611_45492682@1.33.33,trna9_HisGTG_15_-_45490804_45490875@1.33.33\|GCCGTGATCGTATAGTGGTTAGTACTCTGCGTT | >HisGTG.trna111_1_-_147774845_147774916 | 1 | 33 | 33 | tRF nuclear | NO | 47.27 | 354.18 | 82.27 | 0.00 |
| >trna111_HisGTG_1_-_147774845_147774916@1.34.34,trna118_HisGTG_1_-_145396881_145396952@1.34.34,trna16_HisGTG_1_+_146544773_146544844@1.34.34,trna1_HisGTG_15_+_45493349_45493420@1.34.34,trna21_HisGTG_1_+_147753471_147753542@1.34.34,trna33_HisGTG_6_+_27125906_27125977@1.34.34,trna7_HisGTG_9_-_14433938_14434009@1.34.34,trna8_HisGTG_15_-_45492611_45492682@1.34.34,trna9_HisGTG_15_-_45490804_45490875@1.34.34\|GCCGTGATCGTATAGTGGTTAGTACTCTGCGTTG | >HisGTG.trna111_1_-_147774845_147774916 | 1 | 34 | 34 | tRF nuclear | NO | 45.19 | 455.38 | 60.34 | 14.49 |
| >trna111_HisGTG_1_-_147774845_147774916@1.35.35,trna118_HisGTG_1_-_145396881_145396952@1.35.35,trna16_HisGTG_1_+_146544773_146544844@1.35.35,trna1_HisGTG_15_+_45493349_45493420@1.35.35,trna21_HisGTG_1_+_147753471_147753542@1.35.35,trna33_HisGTG_6_+_27125906_27125977@1.35.35,trna7_HisGTG_9_-_14433938_14434009@1.35.35,trna8_HisGTG_15_-_45492611_45492682@1.35.35,trna9_HisGTG_15_-_45490804_45490875@1.35.35\|GCCGTGATCGTATAGTGGTTAGTACTCTGCGTTGT | >HisGTG.trna111_1_-_147774845_147774916 | 1 | 35 | 35 | tRF nuclear | NO | 0.00 | 12.85 | 0.00 | 0.00 |
| >trna111_HisGTG_1_-_147774845_147774916@2.30.29,trna118_HisGTG_1_-_145396881_145396952@2.30.29,trna16_HisGTG_1_+_146544773_146544844@2.30.29,trna1_HisGTG_15_+_45493349_45493420@2.30.29,trna21_HisGTG_1_+_147753471_147753542@2.30.29,trna33_HisGTG_6_+_27125906_27125977@2.30.29,trna7_HisGTG_9_-_14433938_14434009@2.30.29,trna8_HisGTG_15_-_45492611_45492682@2.30.29,trna9_HisGTG_15_-_45490804_45490875@2.30.29\|CCGTGATCGTATAGTGGTTAGTACTCTGC | >HisGTG.trna111_1_-_147774845_147774916 | 2 | 30 | 29 | tRF nuclear | NO | 0.00 | 65.64 | 0.00 | 0.00 |
| >trna111_HisGTG_1_-_147774845_147774916@-1A.23.24,trna118_HisGTG_1_-_145396881_145396952@-1A.23.24,trna16_HisGTG_1_+_146544773_146544844@-1A.23.24,trna1_HisGTG_15_+_45493349_45493420@-1A.23.24,trna21_HisGTG_1_+_147753471_147753542@-1A.23.24,trna33_HisGTG_6_+_27125906_27125977@-1A.23.24,trna7_HisGTG_9_-_14433938_14434009@-1A.23.24,trna8_HisGTG_15_-_45492611_45492682@-1A.23.24,trna9_HisGTG_15_-_45490804_45490875@-1A.23.24\|AGCCGTGATCGTATAGTGGTTAGT | >HisGTG.trna111_1_-_147774845_147774916 | -1A | 23 | 24 | tRF nuclear | NO | 49.17 | 0.00 | 0.00 | 13.26 |

### MDA-MB-231

| | | | | | | | | | | |
|---|---|---|---|---|---|---|---|---|---|---|
| >trna111_HisGTG_1_-_147774845_147774916@-1A.26.27,trna118_HisGTG_1_-_145396881_145396952@-1A.26.27,trna16_HisGTG_1_+_146544773_146544844@-1A.26.27,trna1_HisGTG_15_+_45493349_45493420@-1A.26.27,trna21_HisGTG_1_+_147753471_147753542@-1A.26.27,trna33_HisGTG_6_+_27125906_27125977@-1A.26.27,trna7_HisGTG_9_-_14433938_14434009@-1A.26.27,trna8_HisGTG_15_-_45492611_45492682@-1A.26.27,trna9_HisGTG_15_-_45490804_45490875@-1A.26.27\|AGCCGTGATCGTATAGTGGTTAGTACT | >HisGTG.trna111_1_-_147774845_147774916 | -1A | 26 | 27 | tRF nuclear | NO | 0.00 | 28.68 | 0.00 | 10.66 |
| >trna111_HisGTG_1_-_147774845_147774916@-1A.33.34,trna118_HisGTG_1_-_145396881_145396952@-1A.33.34,trna16_HisGTG_1_+_146544773_146544844@-1A.33.34,trna1_HisGTG_15_+_45493349_45493420@-1A.33.34,trna21_HisGTG_1_+_147753471_147753542@-1A.33.34,trna33_HisGTG_6_+_27125906_27125977@-1A.33.34,trna7_HisGTG_9_-_14433938_14434009@-1A.33.34,trna8_HisGTG_15_-_45492611_45492682@-1A.33.34,trna9_HisGTG_15_-_45490804_45490875@-1A.33.34\|AGCCGTGATCGTATAGTGGTTAGTACTCTGCGTT | >HisGTG.trna111_1_-_147774845_147774916 | -1A | 33 | 34 | tRF nuclear | NO | 0.00 | 57.32 | 0.00 | 0.00 |
| >trna111_HisGTG_1_-_147774845_147774916@-1A.34.35,trna118_HisGTG_1_-_145396881_145396952@-1A.34.35,trna16_HisGTG_1_+_146544773_146544844@-1A.34.35,trna1_HisGTG_15_+_45493349_45493420@-1A.34.35,trna21_HisGTG_1_+_147753471_147753542@-1A.34.35,trna33_HisGTG_6_+_27125906_27125977@-1A.34.35,trna7_HisGTG_9_-_14433938_14434009@-1A.34.35,trna8_HisGTG_15_-_45492611_45492682@-1A.34.35,trna9_HisGTG_15_-_45490804_45490875@-1A.34.35\|AGCCGTGATCGTATAGTGGTTAGTACTCTGCGTTG | >HisGTG.trna111_1_-_147774845_147774916 | -1A | 34 | 35 | tRF nuclear | NO | 0.00 | 69.80 | 0.00 | 0.00 |
| >trna111_HisGTG_1_-_147774845_147774916@-1G.30.31,trna118_HisGTG_1_-_145396881_145396952@-1G.30.31,trna16_HisGTG_1_+_146544773_146544844@-1G.30.31,trna1_HisGTG_15_+_45493349_45493420@-1G.30.31,trna21_HisGTG_1_+_147753471_147753542@-1G.30.31,trna33_HisGTG_6_+_27125906_27125977@-1G.30.31,trna7_HisGTG_9_-_14433938_14434009@-1G.30.31,trna8_HisGTG_15_-_45492611_45492682@-1G.30.31,trna9_HisGTG_15_-_45490804_45490875@-1G.30.31\|GGCCGTGATCGTATAGTGGTTAGTACTCTGC | >HisGTG.trna111_1_-_147774845_147774916 | -1G | 30 | 31 | tRF nuclear | NO | 64.56 | 446.54 | 183.08 | 15.09 |
| >trna111_HisGTG_1_-_147774845_147774916@-1G.31.32,trna118_HisGTG_1_-_145396881_145396952@-1G.31.32,trna16_HisGTG_1_+_146544773_146544844@-1G.31.32,trna1_HisGTG_15_+_45493349_45493420@-1G.31.32,trna21_HisGTG_1_+_147753471_147753542@-1G.31.32,trna33_HisGTG_6_+_27125906_27125977@-1G.31.32,trna7_HisGTG_9_-_14433938_14434009@-1G.31.32,trna8_HisGTG_15_-_45492611_45492682@-1G.31.32,trna9_HisGTG_15_-_45490804_45490875@-1G.31.32\|GGCCGTGATCGTATAGTGGTTAGTACTCTGCG | >HisGTG.trna111_1_-_147774845_147774916 | -1G | 31 | 32 | tRF nuclear | NO | 106.49 | 818.73 | 567.18 | 78.03 |
| >trna111_HisGTG_1_-_147774845_147774916@-1G.32.33,trna118_HisGTG_1_-_145396881_145396952@-1G.32.33,trna16_HisGTG_1_+_146544773_146544844@-1G.32.33,trna1_HisGTG_15_+_45493349_45493420@-1G.32.33,trna21_HisGTG_1_+_147753471_147753542@-1G.32.33,trna33_HisGTG_6_+_27125906_27125977@-1G.32.33,trna7_HisGTG_9_-_14433938_14434009@-1G.32.33,trna8_HisGTG_15_-_45492611_45492682@-1G.32.33,trna9_HisGTG_15_-_45490804_45490875@-1G.32.33\|GGCCGTGATCGTATAGTGGTTAGTACTCTGCGT | >HisGTG.trna111_1_-_147774845_147774916 | -1G | 32 | 33 | tRF nuclear | NO | 74.79 | 339.06 | 260.55 | 20.84 |
| >trna111_HisGTG_1_-_147774845_147774916@-1G.33.34,trna118_HisGTG_1_-_145396881_145396952@-1G.33.34,trna16_HisGTG_1_+_146544773_146544844@-1G.33.34,trna1_HisGTG_15_+_45493349_45493420@-1G.33.34,trna21_HisGTG_1_+_147753471_147753542@-1G.33.34,trna33_HisGTG_6_+_27125906_27125977@-1G.33.34,trna7_HisGTG_9_-_14433938_14434009@-1G.33.34,trna8_HisGTG_15_-_45492611_45492682@-1G.33.34,trna9_HisGTG_15_-_45490804_45490875@-1G.33.34\|GGCCGTGATCGTATAGTGGTTAGTACTCTGCGTT | >HisGTG.trna111_1_-_147774845_147774916 | -1G | 33 | 34 | tRF nuclear | YES | 492.12 | 2851.44 | 907.81 | 170.55 |
| >trna111_HisGTG_1_-_147774845_147774916@-1G.34.35,trna118_HisGTG_1_-_145396881_145396952@-1G.34.35,trna16_HisGTG_1_+_146544773_146544844@-1G.34.35,trna1_HisGTG_15_+_45493349_45493420@-1G.34.35,trna21_HisGTG_1_+_147753471_147753542@-1G.34.35,trna33_HisGTG_6_+_27125906_27125977@-1G.34.35,trna7_HisGTG_9_-_14433938_14434009@-1G.34.35,trna8_HisGTG_15_-_45492611_45492682@-1G.34.35,trna9_HisGTG_15_-_45490804_45490875@-1G.34.35\|GGCCGTGATCGTATAGTGGTTAGTACTCTGCGTTG | >HisGTG.trna111_1_-_147774845_147774916 | -1G | 34 | 35 | tRF nuclear | YES | 194.32 | 1821.46 | 305.87 | 120.40 |
| >trna111_HisGTG_1_-_147774845_147774916@-1G.35.36,trna118_HisGTG_1_-_145396881_145396952@-1G.35.36,trna16_HisGTG_1_+_146544773_146544844@-1G.35.36,trna1_HisGTG_15_+_45493349_45493420@-1G.35.36,trna21_HisGTG_1_+_147753471_147753542@-1G.35.36,trna33_HisGTG_6_+_27125906_27125977@-1G.35.36,trna7_HisGTG_9_-_14433938_14434009@-1G.35.36,trna8_HisGTG_15_-_45492611_45492682@-1G.35.36,trna9_HisGTG_15_-_45490804_45490875@-1G.35.36\|GGCCGTGATCGTATAGTGGTTAGTACTCTGCGTTGT | >HisGTG.trna111_1_-_147774845_147774916 | -1G | 35 | 36 | tRF nuclear | NO | 51.49 | 226.07 | 31.14 | 15.35 |

### MDA-MB-231

| | | | | | | | | | | |
|---|---|---|---|---|---|---|---|---|---|---|
| >trna111_HisGTG_1_-_147774845_147774916@-1G.36.37,trna118_HisGTG_1_-_145396881_145396952@-1G.36.37,trna16_HisGTG_1_+_146544773_146544844@-1G.36.37,trna1_HisGTG_15_+_45493349_45493420@-1G.36.37,trna21_HisGTG_1_+_147753471_147753542@-1G.36.37,trna33_HisGTG_6_+_27125906_27125977@-1G.36.37,trna7_HisGTG_9_-_14433938_14434009@-1G.36.37,trna8_HisGTG_15_-_45492611_45492682@-1G.36.37,trna9_HisGTG_15_-_45490804_45490875@-1G.36.37\|GGCCGTGATCGTATAGTGGTTAGTACTCTGCGTTGTG | >HisGTG.trna111_1_-_147774845_147774916 | -1G | 36 | 37 | tRF nuclear | NO | 0.00 | 30.32 | 0.00 | 12.02 |
| >trna111_HisGTG_1_-_147774845_147774916@-1T.22.23,trna118_HisGTG_1_-_145396881_145396952@-1T.22.23,trna16_HisGTG_1_+_146544773_146544844@-1T.22.23,trna1_HisGTG_15_+_45493349_45493420@-1T.22.23,trna21_HisGTG_1_+_147753471_147753542@-1T.22.23,trna33_HisGTG_6_+_27125906_27125977@-1T.22.23,trna7_HisGTG_9_-_14433938_14434009@-1T.22.23,trna8_HisGTG_15_-_45492611_45492682@-1T.22.23,trna9_HisGTG_15_-_45490804_45490875@-1T.22.23\|TGCCGTGATCGTATAGTGGTTAG | >HisGTG.trna111_1_-_147774845_147774916 | -1T | 22 | 23 | tRF nuclear | NO | 0.00 | 0.00 | 15.44 | 0.00 |
| >trna111_HisGTG_1_-_147774845_147774916@-1T.23.24,trna118_HisGTG_1_-_145396881_145396952@-1T.23.24,trna16_HisGTG_1_+_146544773_146544844@-1T.23.24,trna1_HisGTG_15_+_45493349_45493420@-1T.23.24,trna21_HisGTG_1_+_147753471_147753542@-1T.23.24,trna33_HisGTG_6_+_27125906_27125977@-1T.23.24,trna7_HisGTG_9_-_14433938_14434009@-1T.23.24,trna8_HisGTG_15_-_45492611_45492682@-1T.23.24,trna9_HisGTG_15_-_45490804_45490875@-1T.23.24\|TGCCGTGATCGTATAGTGGTTAGT | >HisGTG.trna111_1_-_147774845_147774916 | -1T | 23 | 24 | tRF nuclear | NO | 275.32 | 116.31 | 89.57 | 117.59 |
| >trna111_HisGTG_1_-_147774845_147774916@-1T.25.26,trna118_HisGTG_1_-_145396881_145396952@-1T.25.26,trna16_HisGTG_1_+_146544773_146544844@-1T.25.26,trna1_HisGTG_15_+_45493349_45493420@-1T.25.26,trna21_HisGTG_1_+_147753471_147753542@-1T.25.26,trna33_HisGTG_6_+_27125906_27125977@-1T.25.26,trna7_HisGTG_9_-_14433938_14434009@-1T.25.26,trna8_HisGTG_15_-_45492611_45492682@-1T.25.26,trna9_HisGTG_15_-_45490804_45490875@-1T.25.26\|TGCCGTGATCGTATAGTGGTTAGTAC | >HisGTG.trna111_1_-_147774845_147774916 | -1T | 25 | 26 | tRF nuclear | NO | 23.95 | 0.00 | 0.00 | 10.92 |
| >trna111_HisGTG_1_-_147774845_147774916@-1T.26.27,trna118_HisGTG_1_-_145396881_145396952@-1T.26.27,trna16_HisGTG_1_+_146544773_146544844@-1T.26.27,trna1_HisGTG_15_+_45493349_45493420@-1T.26.27,trna21_HisGTG_1_+_147753471_147753542@-1T.26.27,trna33_HisGTG_6_+_27125906_27125977@-1T.26.27,trna7_HisGTG_9_-_14433938_14434009@-1T.26.27,trna8_HisGTG_15_-_45492611_45492682@-1T.26.27,trna9_HisGTG_15_-_45490804_45490875@-1T.26.27\|TGCCGTGATCGTATAGTGGTTAGTACT | >HisGTG.trna111_1_-_147774845_147774916 | -1T | 26 | 27 | tRF nuclear | NO | 183.71 | 201.39 | 12.33 | 101.53 |
| >trna111_HisGTG_1_-_147774845_147774916@-1T.28.29,trna118_HisGTG_1_-_145396881_145396952@-1T.28.29,trna16_HisGTG_1_+_146544773_146544844@-1T.28.29,trna1_HisGTG_15_+_45493349_45493420@-1T.28.29,trna21_HisGTG_1_+_147753471_147753542@-1T.28.29,trna33_HisGTG_6_+_27125906_27125977@-1T.28.29,trna7_HisGTG_9_-_14433938_14434009@-1T.28.29,trna8_HisGTG_15_-_45492611_45492682@-1T.28.29,trna9_HisGTG_15_-_45490804_45490875@-1T.28.29\|TGCCGTGATCGTATAGTGGTTAGTACTCT | >HisGTG.trna111_1_-_147774845_147774916 | -1T | 28 | 29 | tRF nuclear | NO | 26.17 | 60.07 | 0.00 | 12.78 |
| >trna111_HisGTG_1_-_147774845_147774916@-1T.30.31,trna118_HisGTG_1_-_145396881_145396952@-1T.30.31,trna16_HisGTG_1_+_146544773_146544844@-1T.30.31,trna1_HisGTG_15_+_45493349_45493420@-1T.30.31,trna21_HisGTG_1_+_147753471_147753542@-1T.30.31,trna33_HisGTG_6_+_27125906_27125977@-1T.30.31,trna7_HisGTG_9_-_14433938_14434009@-1T.30.31,trna8_HisGTG_15_-_45492611_45492682@-1T.30.31,trna9_HisGTG_15_-_45490804_45490875@-1T.30.31\|TGCCGTGATCGTATAGTGGTTAGTACTCTGC | >HisGTG.trna111_1_-_147774845_147774916 | -1T | 30 | 31 | tRF nuclear | NO | 0.00 | 64.89 | 0.00 | 0.00 |
| >trna111_HisGTG_1_-_147774845_147774916@-1T.31.32,trna118_HisGTG_1_-_145396881_145396952@-1T.31.32,trna16_HisGTG_1_+_146544773_146544844@-1T.31.32,trna1_HisGTG_15_+_45493349_45493420@-1T.31.32,trna21_HisGTG_1_+_147753471_147753542@-1T.31.32,trna33_HisGTG_6_+_27125906_27125977@-1T.31.32,trna7_HisGTG_9_-_14433938_14434009@-1T.31.32,trna8_HisGTG_15_-_45492611_45492682@-1T.31.32,trna9_HisGTG_15_-_45490804_45490875@-1T.31.32\|TGCCGTGATCGTATAGTGGTTAGTACTCTGCG | >HisGTG.trna111_1_-_147774845_147774916 | -1T | 31 | 32 | tRF nuclear | NO | 0.00 | 154.82 | 17.26 | 0.00 |
| >trna111_HisGTG_1_-_147774845_147774916@-1T.32.33,trna118_HisGTG_1_-_145396881_145396952@-1T.32.33,trna16_HisGTG_1_+_146544773_146544844@-1T.32.33,trna1_HisGTG_15_+_45493349_45493420@-1T.32.33,trna21_HisGTG_1_+_147753471_147753542@-1T.32.33,trna33_HisGTG_6_+_27125906_27125977@-1T.32.33,trna7_HisGTG_9_-_14433938_14434009@-1T.32.33,trna8_HisGTG_15_-_45492611_45492682@-1T.32.33,trna9_HisGTG_15_-_45490804_45490875@-1T.32.33\|TGCCGTGATCGTATAGTGGTTAGTACTCTGCGT | >HisGTG.trna111_1_-_147774845_147774916 | -1T | 32 | 33 | tRF nuclear | NO | 0.00 | 37.07 | 10.40 | 0.00 |

### MDA-MB-231

| | | | | | | | | | | |
|---|---|---|---|---|---|---|---|---|---|---|
| >trna111_HisGTG_1_-_147774845_147774916@-1T.33.34,trna118_HisGTG_1_-_145396881_145396952@-1T.33.34,trna16_HisGTG_1_+_146544773_146544844@-1T.33.34,trna1_HisGTG_15_+_45493349_45493420@-1T.33.34,trna21_HisGTG_1_+_147753471_147753542@-1T.33.34,trna33_HisGTG_6_+_27125906_27125977@-1T.33.34,trna7_HisGTG_9_-_14433938_14434009@-1T.33.34,trna8_HisGTG_15_-_45492611_45492682@-1T.33.34,trna9_HisGTG_15_-_45490804_45490875@-1T.33.34\|TGCCGTGATCGTATAGTGGTTAGTACTCTGCGTT | >HisGTG.trna111_1_-_147774845_147774916 | -1T | 33 | 34 | tRF nuclear | NO | 54.10 | 234.20 | 65.91 | 11.69 |
| >trna111_HisGTG_1_-_147774845_147774916@-1T.34.35,trna118_HisGTG_1_-_145396881_145396952@-1T.34.35,trna16_HisGTG_1_+_146544773_146544844@-1T.34.35,trna1_HisGTG_15_+_45493349_45493420@-1T.34.35,trna21_HisGTG_1_+_147753471_147753542@-1T.34.35,trna33_HisGTG_6_+_27125906_27125977@-1T.34.35,trna7_HisGTG_9_-_14433938_14434009@-1T.34.35,trna8_HisGTG_15_-_45492611_45492682@-1T.34.35,trna9_HisGTG_15_-_45490804_45490875@-1T.34.35\|TGCCGTGATCGTATAGTGGTTAGTACTCTGCGTTG | >HisGTG.trna111_1_-_147774845_147774916 | -1T | 34 | 35 | tRF nuclear | NO | 0.00 | 281.64 | 12.66 | 11.80 |
| >trna111_HisGTG_1_-_147774845_147774916@-1T.35.36,trna118_HisGTG_1_-_145396881_145396952@-1T.35.36,trna16_HisGTG_1_+_146544773_146544844@-1T.35.36,trna1_HisGTG_15_+_45493349_45493420@-1T.35.36,trna21_HisGTG_1_+_147753471_147753542@-1T.35.36,trna33_HisGTG_6_+_27125906_27125977@-1T.35.36,trna7_HisGTG_9_-_14433938_14434009@-1T.35.36,trna8_HisGTG_15_-_45492611_45492682@-1T.35.36,trna9_HisGTG_15_-_45490804_45490875@-1T.35.36\|TGCCGTGATCGTATAGTGGTTAGTACTCTGCGTTGT | >HisGTG.trna111_1_-_147774845_147774916 | -1T | 35 | 36 | tRF nuclear | NO | 0.00 | 24.93 | 0.00 | 0.00 |
| >am_trna111_HisGTG_1_-_147774845_147774916@1.26.26,trna118_HisGTG_1_-_145396881_145396952@1.26.26,trna16_HisGTG_1_+_146544773_146544844@1.26.26,trna1_HisGTG_15_+_45493349_45493420@1.26.26,trna21_HisGTG_1_+_147753471_147753542@1.26.26,trna33_HisGTG_6_+_27125906_27125977@1.26.26,trna7_HisGTG_9_-_14433938_14434009@1.26.26,trna8_HisGTG_15_-_45492611_45492682@1.26.26,trna9_HisGTG_15_-_45490804_45490875@1.26.26\|GCCGTGATCGTATAGTGGTTAGTACT | >HisGTG.trna111_1_-_147774845_147774916.am | 1 | 26 | 26 | tRF nuclear | NO | 0.00 | 28.42 | 0.00 | 0.00 |
| >am_trna111_HisGTG_1_-_147774845_147774916@1.29.29,trna118_HisGTG_1_-_145396881_145396952@1.29.29,trna16_HisGTG_1_+_146544773_146544844@1.29.29,trna1_HisGTG_15_+_45493349_45493420@1.29.29,trna21_HisGTG_1_+_147753471_147753542@1.29.29,trna33_HisGTG_6_+_27125906_27125977@1.29.29,trna7_HisGTG_9_-_14433938_14434009@1.29.29,trna8_HisGTG_15_-_45492611_45492682@1.29.29,trna9_HisGTG_15_-_45490804_45490875@1.29.29\|GCCGTGATCGTATAGTGGTTAGTACTCTG | >HisGTG.trna111_1_-_147774845_147774916.am | 1 | 29 | 29 | tRF nuclear | NO | 0.00 | 24.48 | 0.00 | 0.00 |
| >am_trna111_HisGTG_1_-_147774845_147774916@-1G.23.24,trna118_HisGTG_1_-_145396881_145396952@-1G.23.24,trna16_HisGTG_1_+_146544773_146544844@-1G.23.24,trna1_HisGTG_15_+_45493349_45493420@-1G.23.24,trna21_HisGTG_1_+_147753471_147753542@-1G.23.24,trna33_HisGTG_6_+_27125906_27125977@-1G.23.24,trna7_HisGTG_9_-_14433938_14434009@-1G.23.24,trna8_HisGTG_15_-_45492611_45492682@-1G.23.24,trna9_HisGTG_15_-_45490804_45490875@-1G.23.24\|GGCCGTGATCGTATAGTGGTTAGT | >HisGTG.trna111_1_-_147774845_147774916.am | -1G | 23 | 24 | tRF nuclear | NO | 100.56 | 64.67 | 33.69 | 17.23 |
| >am_trna111_HisGTG_1_-_147774845_147774916@-1G.26.27,trna118_HisGTG_1_-_145396881_145396952@-1G.26.27,trna16_HisGTG_1_+_146544773_146544844@-1G.26.27,trna1_HisGTG_15_+_45493349_45493420@-1G.26.27,trna21_HisGTG_1_+_147753471_147753542@-1G.26.27,trna33_HisGTG_6_+_27125906_27125977@-1G.26.27,trna7_HisGTG_9_-_14433938_14434009@-1G.26.27,trna8_HisGTG_15_-_45492611_45492682@-1G.26.27,trna9_HisGTG_15_-_45490804_45490875@-1G.26.27\|GGCCGTGATCGTATAGTGGTTAGTACT | >HisGTG.trna111_1_-_147774845_147774916.am | -1G | 26 | 27 | tRF nuclear | NO | 87.54 | 252.42 | 55.77 | 15.69 |
| >am_trna111_HisGTG_1_-_147774845_147774916@-1G.28.29,trna118_HisGTG_1_-_145396881_145396952@-1G.28.29,trna16_HisGTG_1_+_146544773_146544844@-1G.28.29,trna1_HisGTG_15_+_45493349_45493420@-1G.28.29,trna21_HisGTG_1_+_147753471_147753542@-1G.28.29,trna33_HisGTG_6_+_27125906_27125977@-1G.28.29,trna7_HisGTG_9_-_14433938_14434009@-1G.28.29,trna8_HisGTG_15_-_45492611_45492682@-1G.28.29,trna9_HisGTG_15_-_45490804_45490875@-1G.28.29\|GGCCGTGATCGTATAGTGGTTAGTACTCT | >HisGTG.trna111_1_-_147774845_147774916.am | -1G | 28 | 29 | tRF nuclear | NO | 10.52 | 149.16 | 17.77 | 0.00 |
| >am_trna111_HisGTG_1_-_147774845_147774916@-1G.29.30,trna118_HisGTG_1_-_145396881_145396952@-1G.29.30,trna16_HisGTG_1_+_146544773_146544844@-1G.29.30,trna1_HisGTG_15_+_45493349_45493420@-1G.29.30,trna21_HisGTG_1_+_147753471_147753542@-1G.29.30,trna33_HisGTG_6_+_27125906_27125977@-1G.29.30,trna7_HisGTG_9_-_14433938_14434009@-1G.29.30,trna8_HisGTG_15_-_45492611_45492682@-1G.29.30,trna9_HisGTG_15_-_45490804_45490875@-1G.29.30\|GGCCGTGATCGTATAGTGGTTAGTACTCTG | >HisGTG.trna111_1_-_147774845_147774916.am | -1G | 29 | 30 | tRF nuclear | NO | 0.00 | 65.01 | 12.97 | 0.00 |
| >am_trna17_LeuCAG_16_+_57333863_57333945@1.25.25,trna26_LeuCAG_16_-_57334392_57334474@1.25.25,trna34_LeuCAG_1_+_161411323_161411405@1.25.25,trna36_LeuCAG_1_+_161418741_161418823@1.25.25,trna38_LeuCAG_1_+_161426122_161426204@1.25.25,trna40_LeuCAG_1_+_161433532_161433614@1.25.25,trna42_LeuCAG_1_+_161440913_161440995@1.25.25,trna67_LeuCAG_1_-_161500132_161500214@1.25.25,trna7_LeuCAG_6_+_26521436_26521518@1.25.25\|GTCAGGATGGCCGAGCGGTCTAAGG | >LeuCAG.trna17_16_+_57333863_57333945.am | 1 | 25 | 25 | tRF nuclear | NO | 0.00 | 33.75 | 0.00 | 0.00 |

### MDA-MB-231

| | | | | | | | | | | |
|---|---|---|---|---|---|---|---|---|---|---|
| >trna10_LysCTT_16_+_3241501_3241573@1.33.33,trna13_LysCTT_14_-_58706613_58706685@1.33.33,trna2_LysCTT_15_+_79152904_79152976@1.33.33\|GCCCGGCTAGCTCAGTCGGTAGAGCATGGGACT | >LysCTT.trna10_16_+_3241501_3241573 | 1 | 33 | 33 | tRF nuclear | NO | 0.00 | 26.55 | 16.80 | 0.00 |
| >trna10_LysCTT_16_+_3241501_3241573@1.34.34,trna13_LysCTT_14_-_58706613_58706685@1.34.34,trna2_LysCTT_15_+_79152904_79152976@1.34.34\|GCCCGGCTAGCTCAGTCGGTAGAGCATGGGACTC | >LysCTT.trna10_16_+_3241501_3241573 | 1 | 34 | 34 | tRF nuclear | NO | 0.00 | 98.56 | 702.67 | 21.53 |
| >trna10_LysCTT_16_+_3241501_3241573@1.35.35,trna13_LysCTT_14_-_58706613_58706685@1.35.35,trna2_LysCTT_15_+_79152904_79152976@1.35.35\|GCCCGGCTAGCTCAGTCGGTAGAGCATGGGACTCT | >LysCTT.trna10_16_+_3241501_3241573 | 1 | 35 | 35 | tRF nuclear | NO | 24.90 | 541.53 | 201.51 | 99.39 |
| >trna10_LysCTT_16_+_3241501_3241573@1.36.36,trna13_LysCTT_14_-_58706613_58706685@1.36.36,trna2_LysCTT_15_+_79152904_79152976@1.36.36\|GCCCGGCTAGCTCAGTCGGTAGAGCATGGGACTCTT | >LysCTT.trna10_16_+_3241501_3241573 | 1 | 36 | 36 | tRF nuclear | NO | 0.00 | 55.80 | 316.31 | 76.06 |
| >trna10_LysCTT_16_+_3241501_3241573@3.33.31,trna13_LysCTT_14_-_58706613_58706685@3.33.31,trna2_LysCTT_15_+_79152904_79152976@3.33.31\|CCGGCTAGCTCAGTCGGTAGAGCATGGGACT | >LysCTT.trna10_16_+_3241501_3241573 | 3 | 33 | 31 | tRF nuclear | NO | 23.56 | 28.29 | 18.07 | 68.61 |
| >trna10_LysCTT_16_+_3241501_3241573@3.35.33,trna13_LysCTT_14_-_58706613_58706685@3.35.33,trna2_LysCTT_15_+_79152904_79152976@3.35.33\|CCGGCTAGCTCAGTCGGTAGAGCATGGGACTCT | >LysCTT.trna10_16_+_3241501_3241573 | 3 | 35 | 33 | tRF nuclear | NO | 0.00 | 28.09 | 64.23 | 71.35 |
| >trna10_LysCTT_16_+_3241501_3241573@3.36.34,trna13_LysCTT_14_-_58706613_58706685@3.36.34,trna2_LysCTT_15_+_79152904_79152976@3.36.34\|CCGGCTAGCTCAGTCGGTAGAGCATGGGACTCTT | >LysCTT.trna10_16_+_3241501_3241573 | 3 | 36 | 34 | tRF nuclear | NO | 0.00 | 0.00 | 31.10 | 11.22 |
| >trna10_LysCTT_16_+_3241501_3241573@4.33.30,trna13_LysCTT_14_-_58706613_58706685@4.33.30,trna2_LysCTT_15_+_79152904_79152976@4.33.30\|CGGCTAGCTCAGTCGGTAGAGCATGGGACT | >LysCTT.trna10_16_+_3241501_3241573 | 4 | 33 | 30 | tRF nuclear | NO | 57.80 | 0.00 | 62.07 | 68.26 |
| >trna10_LysCTT_16_+_3241501_3241573@4.35.32,trna13_LysCTT_14_-_58706613_58706685@4.35.32,trna2_LysCTT_15_+_79152904_79152976@4.35.32\|CGGCTAGCTCAGTCGGTAGAGCATGGGACTCT | >LysCTT.trna10_16_+_3241501_3241573 | 4 | 35 | 32 | tRF nuclear | NO | 0.00 | 0.00 | 14.58 | 73.32 |
| >trna10_LysCTT_16_+_3241501_3241573@4.36.33,trna13_LysCTT_14_-_58706613_58706685@4.36.33,trna2_LysCTT_15_+_79152904_79152976@4.36.33\|CGGCTAGCTCAGTCGGTAGAGCATGGGACTCTT | >LysCTT.trna10_16_+_3241501_3241573 | 4 | 36 | 33 | tRF nuclear | NO | 0.00 | 0.00 | 12.31 | 17.64 |
| >trna119_LysCTT_1_-_145395522_145395594@1.31.31,trna11_LysCTT_5_-_180648979_180649051@1.31.31,trna13_LysCTT_6_+_26556774_26556846@1.31.31,trna32_LysCTT_16_-_3207406_3207478@1.31.31,trna7_LysCTT_16_+_3225692_3225764@1.31.31,trna9_LysCTT_5_+_180634755_180634827@1.31.31\|GCCCGGCTAGCTCAGTCGGTAGAGCATGAGA | >LysCTT.trna119_1_-_145395522_145395594 | 1 | 31 | 31 | tRF nuclear | NO | 0.00 | 26.50 | 0.00 | 11.16 |
| >trna119_LysCTT_1_-_145395522_145395594@1.32.32,trna11_LysCTT_5_-_180648979_180649051@1.32.32,trna13_LysCTT_6_+_26556774_26556846@1.32.32,trna32_LysCTT_16_-_3207406_3207478@1.32.32,trna7_LysCTT_16_+_3225692_3225764@1.32.32,trna9_LysCTT_5_+_180634755_180634827@1.32.32\|GCCCGGCTAGCTCAGTCGGTAGAGCATGAGAC | >LysCTT.trna119_1_-_145395522_145395594 | 1 | 32 | 32 | tRF nuclear | NO | 25.97 | 204.22 | 75.49 | 69.04 |
| >trna119_LysCTT_1_-_145395522_145395594@1.33.33,trna11_LysCTT_5_-_180648979_180649051@1.33.33,trna13_LysCTT_6_+_26556774_26556846@1.33.33,trna7_LysCTT_16_+_3225692_3225764@1.33.33,trna9_LysCTT_5_+_180634755_180634827@1.33.33\|GCCCGGCTAGCTCAGTCGGTAGAGCATGAGACT | >LysCTT.trna119_1_-_145395522_145395594 | 1 | 33 | 33 | tRF nuclear | NO | 196.47 | 1231.12 | 178.85 | 125.74 |
| >trna119_LysCTT_1_-_145395522_145395594@1.34.34,trna11_LysCTT_5_-_180648979_180649051@1.34.34,trna13_LysCTT_6_+_26556774_26556846@1.34.34,trna7_LysCTT_16_+_3225692_3225764@1.34.34,trna9_LysCTT_5_+_180634755_180634827@1.34.34\|GCCCGGCTAGCTCAGTCGGTAGAGCATGAGACTC | >LysCTT.trna119_1_-_145395522_145395594 | 1 | 34 | 34 | tRF nuclear | NO | 117.12 | 415.93 | 126.40 | 94.45 |
| >trna119_LysCTT_1_-_145395522_145395594@1.35.35,trna11_LysCTT_5_-_180648979_180649051@1.35.35,trna13_LysCTT_6_+_26556774_26556846@1.35.35,trna7_LysCTT_16_+_3225692_3225764@1.35.35,trna9_LysCTT_5_+_180634755_180634827@1.35.35\|GCCCGGCTAGCTCAGTCGGTAGAGCATGAGACTCT | >LysCTT.trna119_1_-_145395522_145395594 | 1 | 35 | 35 | tRF nuclear | NO | 51.26 | 305.33 | 142.49 | 83.62 |
| >trna119_LysCTT_1_-_145395522_145395594@1.36.36,trna11_LysCTT_5_-_180648979_180649051@1.36.36,trna13_LysCTT_6_+_26556774_26556846@1.36.36,trna7_LysCTT_16_+_3225692_3225764@1.36.36,trna9_LysCTT_5_+_180634755_180634827@1.36.36\|GCCCGGCTAGCTCAGTCGGTAGAGCATGAGACTCTT | >LysCTT.trna119_1_-_145395522_145395594 | 1 | 36 | 36 | tRF nuclear | NO | 0.00 | 58.48 | 59.99 | 20.86 |
| >trna119_LysCTT_1_-_145395522_145395594@3.33.31,trna11_LysCTT_5_-_180648979_180649051@3.33.31,trna13_LysCTT_6_+_26556774_26556846@3.33.31,trna7_LysCTT_16_+_3225692_3225764@3.33.31,trna9_LysCTT_5_+_180634755_180634827@3.33.31\|CCGGCTAGCTCAGTCGGTAGAGCATGAGACT | >LysCTT.trna119_1_-_145395522_145395594 | 3 | 33 | 31 | tRF nuclear | NO | 60.79 | 35.83 | 12.97 | 21.19 |
| >trna119_LysCTT_1_-_145395522_145395594@3.35.33,trna11_LysCTT_5_-_180648979_180649051@3.35.33,trna13_LysCTT_6_+_26556774_26556846@3.35.33,trna7_LysCTT_16_+_3225692_3225764@3.35.33,trna9_LysCTT_5_+_180634755_180634827@3.35.33\|CCGGCTAGCTCAGTCGGTAGAGCATGAGACTCT | >LysCTT.trna119_1_-_145395522_145395594 | 3 | 35 | 33 | tRF nuclear | NO | 0.00 | 0.00 | 12.02 | 16.12 |
| >trna119_LysCTT_1_-_145395522_145395594@4.33.30,trna11_LysCTT_5_-_180648979_180649051@4.33.30,trna13_LysCTT_6_+_26556774_26556846@4.33.30,trna7_LysCTT_16_+_3225692_3225764@4.33.30,trna9_LysCTT_5_+_180634755_180634827@4.33.30\|CGGCTAGCTCAGTCGGTAGAGCATGAGACT | >LysCTT.trna119_1_-_145395522_145395594 | 4 | 33 | 30 | tRF nuclear | NO | 72.68 | 0.00 | 61.38 | 77.43 |
| >trna119_LysCTT_1_-_145395522_145395594@4.35.32,trna11_LysCTT_5_-_180648979_180649051@4.35.32,trna13_LysCTT_6_+_26556774_26556846@4.35.32,trna7_LysCTT_16_+_3225692_3225764@4.35.32,trna9_LysCTT_5_+_180634755_180634827@4.35.32\|CGGCTAGCTCAGTCGGTAGAGCATGAGACTCT | >LysCTT.trna119_1_-_145395522_145395594 | 4 | 35 | 32 | tRF nuclear | NO | 80.20 | 0.00 | 85.84 | 162.58 |
| >trna119_LysCTT_1_-_145395522_145395594@4.36.33,trna11_LysCTT_5_-_180648979_180649051@4.36.33,trna13_LysCTT_6_+_26556774_26556846@4.36.33,trna7_LysCTT_16_+_3225692_3225764@4.36.33,trna9_LysCTT_5_+_180634755_180634827@4.36.33\|CGGCTAGCTCAGTCGGTAGAGCATGAGACTCTT | >LysCTT.trna119_1_-_145395522_145395594 | 4 | 36 | 33 | tRF nuclear | NO | 23.60 | 0.00 | 76.84 | 59.33 |

### MDA-MB-231

<table>
<tr><th></th><th></th><th></th><th></th><th></th><th></th><th></th><th></th><th></th><th></th><th></th></tr>
<tr><td>>trna119_LysCTT_1_-_145395522_145395594@5.35.31,trna11_LysCTT_5_-_180648979_180649051@5.35.31,trna13_LysCTT_6_+_26556774_26556846@5.35.31,trna7_LysCTT_16_+_3225692_3225764@5.35.31,trna9_LysCTT_5_+_180634755_180634827@5.35.31|GGCTAGCTCAGTCGGTAGAGCATGAGACTCT</td><td>>LysCTT.trna119_1_-_145395522_145395594</td><td>5</td><td>35</td><td>31</td><td>tRF nuclear</td><td>NO</td><td>0.00</td><td>0.00</td><td>12.51</td><td>85.47</td></tr>
<tr><td>>trna119_LysCTT_1_-_145395522_145395594@5.36.32,trna11_LysCTT_5_-_180648979_180649051@5.36.32,trna13_LysCTT_6_+_26556774_26556846@5.36.32,trna7_LysCTT_16_+_3225692_3225764@5.36.32,trna9_LysCTT_5_+_180634755_180634827@5.36.32|GGCTAGCTCAGTCGGTAGAGCATGAGACTCTT</td><td>>LysCTT.trna119_1_-_145395522_145395594</td><td>5</td><td>36</td><td>32</td><td>tRF nuclear</td><td>NO</td><td>0.00</td><td>0.00</td><td>12.02</td><td>21.50</td></tr>
<tr><td>>am_trna119_LysCTT_1_-_145395522_145395594@13.33.21,trna11_LysCTT_5_-_180648979_180649051@13.33.21,trna128_LysCTT_1_-_55423542_55423614@13.33.21,trna13_LysCTT_6_+_26556774_26556846@13.33.21,trna6_LysCTT_19_-_52425396_52425468@13.33.21,trna7_LysCTT_16_+_3225692_3225764@13.33.21,trna9_LysCTT_5_+_180634755_180634827@13.33.21|CAGTCGGTAGAGCATGAGACT</td><td>>LysCTT.trna119_1_-_145395522_145395594.am</td><td>13</td><td>33</td><td>21</td><td>tRF nuclear</td><td>NO</td><td>0.00</td><td>0.00</td><td>12.45</td><td>14.93</td></tr>
<tr><td>>trna32_LysCTT_16_-_3207406_3207478@1.33.33|GCCCGGCTAGCTCAGTCGGTAGAGCATGAGACC</td><td>>LysCTT.trna32_16_-_3207406_3207478</td><td>1</td><td>33</td><td>33</td><td>tRF nuclear</td><td>NO</td><td>0.00</td><td>210.53</td><td>16.47</td><td>15.41</td></tr>
<tr><td>>trna14_LysTTT_11_-_59327808_59327880@1.19.19,trna2_LysTTT_17_+_8022473_8022545@1.19.19,trna54_LysTTT_1_+_204475655_204475727@1.19.19,trna5_LysTTT_11_+_59323902_59323974@1.19.19,trna62_LysTTT_1_-_204476158_204476230@1.19.19,trna76_LysTTT_6_+_28918806_28918878@1.19.19|GCCCGGATAGCTCAGTCGG</td><td>>LysTTT.trna14_11_-_59327808_59327880</td><td>1</td><td>19</td><td>19</td><td>tRF nuclear</td><td>NO</td><td>0.00</td><td>0.00</td><td>31.19</td><td>0.00</td></tr>
<tr><td>>trna14_LysTTT_11_-_59327808_59327880@1.32.32,trna2_LysTTT_17_+_8022473_8022545@1.32.32,trna54_LysTTT_1_+_204475655_204475727@1.32.32,trna5_LysTTT_11_+_59323902_59323974@1.32.32,trna62_LysTTT_1_-_204476158_204476230@1.32.32,trna76_LysTTT_6_+_28918806_28918878@1.32.32|GCCCGGATAGCTCAGTCGGTAGAGCATCAGAC</td><td>>LysTTT.trna14_11_-_59327808_59327880</td><td>1</td><td>32</td><td>32</td><td>tRF nuclear</td><td>NO</td><td>0.00</td><td>0.00</td><td>67.33</td><td>0.00</td></tr>
<tr><td>>trna14_LysTTT_11_-_59327808_59327880@1.33.33,trna2_LysTTT_17_+_8022473_8022545@1.33.33,trna54_LysTTT_1_+_204475655_204475727@1.33.33,trna5_LysTTT_11_+_59323902_59323974@1.33.33,trna62_LysTTT_1_-_204476158_204476230@1.33.33,trna76_LysTTT_6_+_28918806_28918878@1.33.33|GCCCGGATAGCTCAGTCGGTAGAGCATCAGACT</td><td>>LysTTT.trna14_11_-_59327808_59327880</td><td>1</td><td>33</td><td>33</td><td>tRF nuclear</td><td>NO</td><td>96.64</td><td>220.36</td><td>929.93</td><td>121.50</td></tr>
<tr><td>>trna143_LysTTT_6_-_27559593_27559665@14.31.18,trna149_LysTTT_6_-_27302769_27302841@14.31.18,trna14_LysTTT_11_-_59327808_59327880@14.31.18,trna2_LysTTT_17_+_8022473_8022545@14.31.18,trna54_LysTTT_1_+_204475655_204475727@14.31.18,trna5_LysTTT_11_+_59323902_59323974@14.31.18,trna62_LysTTT_1_-_204476158_204476230@14.31.18,trna76_LysTTT_6_+_28918806_28918878@14.31.18|AGTCGGTAGAGCATCAGA</td><td>>LysTTT.trna143_6_-_27559593_27559665</td><td>14</td><td>31</td><td>18</td><td>tRF nuclear</td><td>NO</td><td>0.00</td><td>24.15</td><td>14.59</td><td>0.00</td></tr>
<tr><td>>am_trna162_MetCAT_6_-_26758550_26758622@34.54.21,trna164_MetCAT_6_-_26735574_26735646@34.54.21,trna20_MetCAT_16_+_71460396_71460468@34.54.21,trna21_MetCAT_6_+_26701712_26701784@34.54.21,trna27_MetCAT_6_+_26766444_26766516@34.54.21,trna75_MetCAT_6_+_28912352_28912424@34.54.21,trna97_MetCAT_6_-_28921042_28921114@34.54.21|CATAATCTGAAGGTCCTGAGT</td><td>>MetCAT.trna162_6_-_26758550_26758622.am</td><td>34</td><td>54</td><td>21</td><td>tRF nuclear</td><td>NO</td><td>0.00</td><td>0.00</td><td>10.74</td><td>0.00</td></tr>
<tr><td>>trna16_TyrGTA_14_-_21131351_21131444@59.76.18,trna17_TyrGTA_14_-_21128117_21128210@59.76.18,trna19_TyrGTA_14_-_21121258_21121351@59.76.18,trna2_TyrGTA_2_+_27273650_27273738@59.76.18,trna4_TyrGTA_8_+_67025602_67025694@59.76.18,trna5_TyrGTA_14_+_21151432_21151520@59.76.18,trna5_TyrGTA_8_+_67026223_67026311@59.76.18|TTCCGGCTCGAAGGACCA</td><td>>TyrGTA.trna16_14_-_21131351_21131444</td><td>59</td><td>76</td><td>18</td><td>tRF nuclear</td><td>NO</td><td>0.00</td><td>63.13</td><td>31.43</td><td>0.00</td></tr>
<tr><td>>trna15_ValAAC_5_-_180615416_180615488@1.32.32|GTTTCCGTAGTGTAGTGGTCATCACGTTCGCC</td><td>>ValAAC.trna15_5_-_180615416_180615488</td><td>1</td><td>32</td><td>32</td><td>tRF nuclear</td><td>NO</td><td>0.00</td><td>65.78</td><td>14.03</td><td>11.14</td></tr>
<tr><td>>am_trna10_ValCAC_5_-_180649395_180649467@1.19.19,trna12_ValAAC_5_-_180645270_180645342@1.19.19,trna132_ValAAC_6_-_27721179_27721251@1.19.19,trna133_ValCAC_6_-_27696327_27696399@1.19.19,trna136_ValAAC_6_-_27648885_27648957@1.19.19,trna139_ValAAC_6_-_27618707_27618779@1.19.19,trna15_ValAAC_5_-_180615416_180615488@1.19.19,trna18_ValCAC_5_-_180529253_180529325@1.19.19,trna2_ValAAC_3_+_169490018_169490090@1.19.19,trna2_ValCAC_5_+_180524070_180524142@1.19.19,trna37_ValAAC_6_+_27203288_27203360@1.19.19,trna4_ValAAC_5_+_180591154_180591226@1.19.19,trna5_ValAAC_5_+_180596610_180596682@1.19.19,trna6_ValCAC_5_+_180600650_180600722@1.19.19,trna85_ValCAC_1_-_161369490_161369562@1.19.19,trna90_ValCAC_1_-_149684088_149684161@1.19.19,trna98_ValCAC_1_-_149298555_149298627@1.19.19,trna9_ValCAC_6_+_26538282_26538354@1.19.19|GTTTCCGTAGTGTAGTGGT</td><td>>ValCAC.trna10_5_-_180649395_180649467.am</td><td>1</td><td>19</td><td>19</td><td>tRF nuclear</td><td>NO</td><td>0.00</td><td>0.00</td><td>14.80</td><td>0.00</td></tr>
</table>

### MDA-MB-231

| | | | | | | | | | | |
|---|---|---|---|---|---|---|---|---|---|---|
| >am_trna10_ValCAC_5_-_180649395_180649467@1.23.23,trna12_ValAAC_5_-_180645270_180645342@1.23.23,trna132_ValAAC_6_-_27721179_27721251@1.23.23,trna136_ValAAC_6_-_27648885_27648957@1.23.23,trna139_ValAAC_6_-_27618707_27618779@1.23.23,trna18_ValCAC_5_-_180529253_180529325@1.23.23,trna2_ValAAC_3_+_169490018_169490090@1.23.23,trna2_ValCAC_5_+_180524070_180524142@1.23.23,trna37_ValAAC_6_+_27203288_27203360@1.23.23,trna4_ValAAC_5_+_180591154_180591226@1.23.23,trna5_ValAAC_5_+_180596610_180596682@1.23.23,trna6_ValCAC_5_+_180600650_180600722@1.23.23,trna85_ValCAC_1_-_161369490_161369562@1.23.23,trna90_ValCAC_1_-_149684088_149684161@1.23.23,trna98_ValCAC_1_-_149298555_149298627@1.23.23,trna9_ValCAC_6_+_26538282_26538354@1.23.23\|GTTTCCGTAGTGTAGTGGTTATC | >ValCAC.trna10_5_-_180649395_180649467.am | 1 | 23 | 23 | tRF nuclear | NO | 10.61 | 91.09 | 10.48 | 19.28 |
| >am_trna10_ValCAC_5_-_180649395_180649467@1.29.29,trna12_ValAAC_5_-_180645270_180645342@1.29.29,trna132_ValAAC_6_-_27721179_27721251@1.29.29,trna136_ValAAC_6_-_27648885_27648957@1.29.29,trna139_ValAAC_6_-_27618707_27618779@1.29.29,trna18_ValCAC_5_-_180529253_180529325@1.29.29,trna2_ValAAC_3_+_169490018_169490090@1.29.29,trna2_ValCAC_5_+_180524070_180524142@1.29.29,trna4_ValAAC_5_+_180591154_180591226@1.29.29,trna5_ValAAC_5_+_180596610_180596682@1.29.29,trna6_ValCAC_5_+_180600650_180600722@1.29.29,trna85_ValCAC_1_-_161369490_161369562@1.29.29,trna90_ValCAC_1_-_149684088_149684161@1.29.29,trna98_ValCAC_1_-_149298555_149298627@1.29.29,trna9_ValCAC_6_+_26538282_26538354@1.29.29\|GTTTCCGTAGTGTAGTGGTTATCACGTTC | >ValCAC.trna10_5_-_180649395_180649467.am | 1 | 29 | 29 | tRF nuclear | NO | 0.00 | 0.00 | 57.75 | 0.00 |
| >am_trna10_ValCAC_5_-_180649395_180649467@1.31.31,trna12_ValAAC_5_-_180645270_180645342@1.31.31,trna132_ValAAC_6_-_27721179_27721251@1.31.31,trna136_ValAAC_6_-_27648885_27648957@1.31.31,trna139_ValAAC_6_-_27618707_27618779@1.31.31,trna18_ValCAC_5_-_180529253_180529325@1.31.31,trna2_ValAAC_3_+_169490018_169490090@1.31.31,trna2_ValCAC_5_+_180524070_180524142@1.31.31,trna4_ValAAC_5_+_180591154_180591226@1.31.31,trna5_ValAAC_5_+_180596610_180596682@1.31.31,trna6_ValCAC_5_+_180600650_180600722@1.31.31,trna85_ValCAC_1_-_161369490_161369562@1.31.31,trna90_ValCAC_1_-_149684088_149684161@1.31.31,trna98_ValCAC_1_-_149298555_149298627@1.31.31,trna9_ValCAC_6_+_26538282_26538354@1.31.31\|GTTTCCGTAGTGTAGTGGTTATCACGTTCGC | >ValCAC.trna10_5_-_180649395_180649467.am | 1 | 31 | 31 | tRF nuclear | NO | 0.00 | 0.00 | 14.42 | 0.00 |
| >am_trna10_ValCAC_5_-_180649395_180649467@1.33.33,trna12_ValAAC_5_-_180645270_180645342@1.33.33,trna132_ValAAC_6_-_27721179_27721251@1.33.33,trna136_ValAAC_6_-_27648885_27648957@1.33.33,trna139_ValAAC_6_-_27618707_27618779@1.33.33,trna18_ValCAC_5_-_180529253_180529325@1.33.33,trna2_ValAAC_3_+_169490018_169490090@1.33.33,trna2_ValCAC_5_+_180524070_180524142@1.33.33,trna4_ValAAC_5_+_180591154_180591226@1.33.33,trna5_ValAAC_5_+_180596610_180596682@1.33.33,trna6_ValCAC_5_+_180600650_180600722@1.33.33,trna85_ValCAC_1_-_161369490_161369562@1.33.33,trna90_ValCAC_1_-_149684088_149684161@1.33.33,trna98_ValCAC_1_-_149298555_149298627@1.33.33,trna9_ValCAC_6_+_26538282_26538354@1.33.33\|GTTTCCGTAGTGTAGTGGTTATCACGTTCGCCT | >ValCAC.trna10_5_-_180649395_180649467.am | 1 | 33 | 33 | tRF nuclear | NO | 45.23 | 191.07 | 64.93 | 20.84 |
| >am_trna10_ValCAC_5_-_180649395_180649467@1.34.34,trna18_ValCAC_5_-_180529253_180529325@1.34.34,trna2_ValCAC_5_+_180524070_180524142@1.34.34,trna6_ValCAC_5_+_180600650_180600722@1.34.34,trna85_ValCAC_1_-_161369490_161369562@1.34.34,trna90_ValCAC_1_-_149684088_149684161@1.34.34,trna98_ValCAC_1_-_149298555_149298627@1.34.34,trna9_ValCAC_6_+_26538282_26538354@1.34.34\|GTTTCCGTAGTGTAGTGGTTATCACGTTCGCCTC | >ValCAC.trna10_5_-_180649395_180649467.am | 1 | 34 | 34 | tRF nuclear | NO | 0.00 | 131.85 | 15.24 | 0.00 |
| >am_trna10_ValCAC_5_-_180649395_180649467@1.36.36,trna18_ValCAC_5_-_180529253_180529325@1.36.36,trna2_ValCAC_5_+_180524070_180524142@1.36.36,trna6_ValCAC_5_+_180600650_180600722@1.36.36,trna85_ValCAC_1_-_161369490_161369562@1.36.36,trna90_ValCAC_1_-_149684088_149684161@1.36.36,trna98_ValCAC_1_-_149298555_149298627@1.36.36,trna9_ValCAC_6_+_26538282_26538354@1.36.36\|GTTTCCGTAGTGTAGTGGTTATCACGTTCGCCTCAC | >ValCAC.trna10_5_-_180649395_180649467.am | 1 | 36 | 36 | tRF nuclear | NO | 0.00 | 36.36 | 0.00 | 12.14 |
| >trna13_ValCAC_19_-_4724647_4724719@1.32.32\|GTTTCCGTAGTGTAGCGGTTATCACATTCGCC | >ValCAC.trna13_19_-_4724647_4724719 | 1 | 32 | 32 | tRF nuclear | NO | 0.00 | 0.00 | 15.07 | 0.00 |
| >trna13_ValCAC_19_-_4724647_4724719@1.34.34\|GTTTCCGTAGTGTAGCGGTTATCACATTCGCCTC | >ValCAC.trna13_19_-_4724647_4724719 | 1 | 34 | 34 | tRF nuclear | NO | 0.00 | 62.09 | 0.00 | 0.00 |
| >trna152_ValCAC_6_-_27248049_27248121@1.23.23\|GCTTCTGTAGTGTAGTGGTTATC | >ValCAC.trna152_6_-_27248049_27248121 | 1 | 23 | 23 | tRF nuclear | NO | 0.00 | 32.64 | 0.00 | 0.00 |
| >trna152_ValCAC_6_-_27248049_27248121@1.32.32\|GCTTCTGTAGTGTAGTGGTTATCACGTTCGCC | >ValCAC.trna152_6_-_27248049_27248121 | 1 | 32 | 32 | tRF nuclear | NO | 46.63 | 269.78 | 149.46 | 19.24 |
| >trna152_ValCAC_6_-_27248049_27248121@1.33.33\|GCTTCTGTAGTGTAGTGGTTATCACGTTCGCCT | >ValCAC.trna152_6_-_27248049_27248121 | 1 | 33 | 33 | tRF nuclear | NO | 0.00 | 149.65 | 76.02 | 10.66 |
| >trna152_ValCAC_6_-_27248049_27248121@1.34.34\|GCTTCTGTAGTGTAGTGGTTATCACGTTCGCCTC | >ValCAC.trna152_6_-_27248049_27248121 | 1 | 34 | 34 | tRF nuclear | NO | 82.88 | 424.58 | 84.52 | 13.62 |

### MDA-MB-231

<table>
<tr><td>>trna152_ValCAC_6_-_27248049_27248121@1.36.36|GCTTCTGTAGTGTAGTGGTTATCACGTTCGCCTCAC</td><td>>ValCAC.trna152_6_-_27248049_27248121</td><td>1</td><td>36</td><td>36</td><td>tRF nuclear</td><td>NO</td><td>0.00</td><td>119.22</td><td>0.00</td><td>14.04</td></tr>
<tr><td>>trna152_ValCAC_6_-_27248049_27248121@1.42.42|GCTTCTGTAGTGTAGTGGTTATCACGTTCGCCTCACACGCGA</td><td>>ValCAC.trna152_6_-_27248049_27248121</td><td>1</td><td>42</td><td>42</td><td>tRF nuclear</td><td>NO</td><td>0.00</td><td>30.11</td><td>0.00</td><td>0.00</td></tr>
<tr><td>>am_trna10_ValCAC_5_-_180649395_180649467@1.32.32,trna12_ValAAC_5_-_180645270_180645342@1.32.32,trna132_ValAAC_6_-_27721179_27721251@1.32.32,trna136_ValAAC_6_-_27648885_27648957@1.32.32,trna139_ValAAC_6_-_27618707_27618779@1.32.32,trna18_ValCAC_5_-_180529253_180529325@1.32.32,trna2_ValAAC_3_+_169490018_169490090@1.32.32,trna2_ValCAC_5_+_180524070_180524142@1.32.32,trna4_ValAAC_5_+_180591154_180591226@1.32.32,trna5_ValAAC_5_+_180596610_180596682@1.32.32,trna6_ValCAC_5_+_180600650_180600722@1.32.32,trna85_ValCAC_1_-_161369490_161369562@1.32.32,trna90_ValCAC_1_-_149684088_149684161@1.32.32,trna98_ValCAC_1_-_149298555_149298627@1.32.32,trna9_ValCAC_6_+_26538282_26538354@1.32.32|GTTTCCGTAGTGTAGTGGTTATCACGTTCGCC</td><td>>ValCAC.ttrna10_5_-_180649395_180649467.am</td><td>1</td><td>32</td><td>32</td><td>tRF nuclear</td><td>NO</td><td>102.64</td><td>628.12</td><td>112.41</td><td>110.49</td></tr>
<tr><td>>trna16_ValTAC_11_-_59318460_59318532@1.34.34|GGTTCCATAGTGTAGCGGTTATCACGTCTGCTTT</td><td>>ValTAC.trna16_11_-_59318460_59318532</td><td>1</td><td>34</td><td>34</td><td>tRF nuclear</td><td>NO</td><td>0.00</td><td>0.00</td><td>10.80</td><td>0.00</td></tr>
</table>
