## Supplementary material for "Single-nucleotide Differences and Cell Type Decide the Subcellular Localization of miRNA Isoforms (isomiRs), tRNA-derived Fragments (tRFs) and rRNA-derived Fragments (rRFs)": Supp. Table S9 (4 of 4)

### MDA-MB-468

<table>
<tr><th>Full Name</th><th>Short Name</th><th>Start</th><th>End</th><th>Length</th><th>Additional Nucleotides</th><th>group</th><th>Outlier</th><th>AVE NUC</th><th>AVE CYTO</th><th>AVE MITO</th><th>AVE MP</th></tr>
<tr><td>>MI0000061|hsa-let-7a-2&WithFlank&11|-|122146516|122146599@11.31.21(+1C);MI0000062|hsa-let-7a-3&WithFlank&22|+|46112743|46112828@10.30.21(+1C);MI0000060|hsa-let-7a-1&WithFlank&9|+|94175951|94176042@12.32.21(+1C)&[MIMAT0000062&hsa-let-7a-5p&offsets|0|-1(+1C);m-6&11|-|122146568|122146589&offsets|0|-1(+1C)];[MIMAT0000062_1&hsa-let-7a-5p&offsets|0|-1(+1C);m-5&22|+|46112752|46112773&offsets|0|-1(+1C)];[MIMAT0000062_2&hsa-let-7a-5p&offsets|0|-1(+1C);m-4&9|+|94175962|94175983&offsets|0|-1(+1C)]|TGAGGTAGTAGGTTGTATAGTC</td><td>>MI0000061|hsa-let-7a-2&WithFlank&11|-|122146516|122146599</td><td>11</td><td>31</td><td>21</td><td>(+1C)</td><td>isomiR</td><td>NO</td><td>30.98</td><td>0.00</td><td>0.00</td><td>0.00</td></tr>
<tr><td>>MI0000061|hsa-let-7a-2&WithFlank&11|-|122146516|122146599@11.31.21;MI0000062|hsa-let-7a-3&WithFlank&22|+|46112743|46112828@10.30.21;MI0000060|hsa-let-7a-1&WithFlank&9|+|94175951|94176042@12.32.21&[MIMAT0000062&hsa-let-7a-5p&offsets|0|-1;m-6&11|-|122146568|122146589&offsets|0|-1];[MIMAT0000062_1&hsa-let-7a-5p&offsets|0|-1;m-5&22|+|46112752|46112773&offsets|0|-1];[MIMAT0000062_2&hsa-let-7a-5p&offsets|0|-1;m-4&9|+|94175962|94175983&offsets|0|-1]|TGAGGTAGTAGGTTGTATAGT</td><td>>MI0000061|hsa-let-7a-2&WithFlank&11|-|122146516|122146599</td><td>11</td><td>31</td><td>21</td><td></td><td>isomiR</td><td>NO</td><td>13.10</td><td>0.00</td><td>0.00</td><td>0.00</td></tr>
<tr><td>>MI0000061|hsa-let-7a-2&WithFlank&11|-|122146516|122146599@11.32.22;MI0000062|hsa-let-7a-3&WithFlank&22|+|46112743|46112828@10.31.22;MI0000060|hsa-let-7a-1&WithFlank&9|+|94175951|94176042@12.33.22&[MIMAT0000062&hsa-let-7a-5p&offsets|0|0;m-6&11|-|122146568|122146589&offsets|0|0];[MIMAT0000062_1&hsa-let-7a-5p&offsets|0|0;m-5&22|+|46112752|46112773&offsets|0|0];[MIMAT0000062_2&hsa-let-7a-5p&offsets|0|0;m-4&9|+|94175962|94175983&offsets|0|0]|TGAGGTAGTAGGTTGTATAGTT</td><td>>MI0000061|hsa-let-7a-2&WithFlank&11|-|122146516|122146599</td><td>11</td><td>32</td><td>22</td><td></td><td>isomiR</td><td>NO</td><td>276.29</td><td>131.45</td><td>87.56</td><td>34.14</td></tr>
<tr><td>>MI0000063|hsa-let-7b&WithFlank&22|+|46113680|46113774@12.33.22&[MIMAT0000063&hsa-let-7b-5p&offsets|0|0;m-14&22|+|46113691|46113712&offsets|0|0]|TGAGGTAGTAGGTTGTGTGGTT</td><td>>MI0000063|hsa-let-7b&WithFlank&22|+|46113680|46113774</td><td>12</td><td>33</td><td>22</td><td></td><td>isomiR</td><td>NO</td><td>91.19</td><td>37.34</td><td>96.75</td><td>0.00</td></tr>
<tr><td>>MI0000064|hsa-let-7c&WithFlank&21|+|16539822|16539917@17.38.22&[MIMAT0000064&hsa-let-7c-5p&offsets|0|0;m-37&21|+|16539838|16539859&offsets|0|0]|TGAGGTAGTAGGTTGTATGGTT</td><td>>MI0000064|hsa-let-7c&WithFlank&21|+|16539822|16539917</td><td>17</td><td>38</td><td>22</td><td></td><td>isomiR</td><td>NO</td><td>47.05</td><td>28.27</td><td>37.19</td><td>0.00</td></tr>
<tr><td>>MI0000065|hsa-let-7d&WithFlank&9|+|94178828|94178926@14.35.22&[MIMAT0000065&hsa-let-7d-5p&offsets|0|0;m-90&9|+|94178841|94178862&offsets|0|0]|AGAGGTAGTAGGTTGCATAGTT</td><td>>MI0000065|hsa-let-7d&WithFlank&9|+|94178828|94178926</td><td>14</td><td>35</td><td>22</td><td></td><td>isomiR</td><td>NO</td><td>30.33</td><td>0.00</td><td>0.00</td><td>0.00</td></tr>
<tr><td>>MI0000066|hsa-let-7e&WithFlank&19|+|51692780|51692870@14.35.22&[MIMAT0000066&hsa-let-7e-5p&offsets|0|0;m-50&19|+|51692793|51692814&offsets|0|0]|TGAGGTAGGAGGTTGTATAGTT</td><td>>MI0000066|hsa-let-7e&WithFlank&19|+|51692780|51692870</td><td>14</td><td>35</td><td>22</td><td></td><td>isomiR</td><td>NO</td><td>52.26</td><td>0.00</td><td>0.00</td><td>0.00</td></tr>
<tr><td>>MI0000067|hsa-let-7f-1&WithFlank&9|+|94176341|94176439@13.33.21(+1C);MI0000068|hsa-let-7f-2&WithFlank&X|-|53557186|53557280@14.34.21(+1C)&[MIMAT0000067&hsa-let-7f-5p&offsets|0|-1(+1C);m-20&9|+|94176353|94176374&offsets|0|-1(+1C)];[MIMAT0000067_1&hsa-let-7f-5p&offsets|0|-1(+1C);m-19&X|-|53557246|53557267&offsets|0|-1(+1C)]|TGAGGTAGTAGATTGTATAGTC</td><td>>MI0000067|hsa-let-7f-1&WithFlank&9|+|94176341|94176439</td><td>13</td><td>33</td><td>21</td><td>(+1C)</td><td>isomiR</td><td>NO</td><td>58.25</td><td>33.48</td><td>6.93</td><td>0.00</td></tr>
<tr><td>>MI0000067|hsa-let-7f-1&WithFlank&9|+|94176341|94176439@13.33.21;MI0000068|hsa-let-7f-2&WithFlank&X|-|53557186|53557280@14.34.21&[MIMAT0000067&hsa-let-7f-5p&offsets|0|-1;m-20&9|+|94176353|94176374&offsets|0|-1];[MIMAT0000067_1&hsa-let-7f-5p&offsets|0|-1;m-19&X|-|53557246|53557267&offsets|0|-1]|TGAGGTAGTAGATTGTATAGT</td><td>>MI0000067|hsa-let-7f-1&WithFlank&9|+|94176341|94176439</td><td>13</td><td>33</td><td>21</td><td></td><td>isomiR</td><td>NO</td><td>40.09</td><td>0.00</td><td>0.00</td><td>0.00</td></tr>
<tr><td>>MI0000067|hsa-let-7f-1&WithFlank&9|+|94176341|94176439@13.34.22;MI0000068|hsa-let-7f-2&WithFlank&X|-|53557186|53557280@14.35.22&[MIMAT0000067&hsa-let-7f-5p&offsets|0|0;m-20&9|+|94176353|94176374&offsets|0|0];[MIMAT0000067_1&hsa-let-7f-5p&offsets|0|0;m-19&X|-|53557246|53557267&offsets|0|0]|TGAGGTAGTAGATTGTATAGTT</td><td>>MI0000067|hsa-let-7f-1&WithFlank&9|+|94176341|94176439</td><td>13</td><td>34</td><td>22</td><td></td><td>isomiR</td><td>NO</td><td>526.63</td><td>279.25</td><td>130.36</td><td>48.35</td></tr>
<tr><td>>MI0000070|hsa-mir-16-1&WithFlank&13|-|50048967|50049067@20.41.22;MI0000115|hsa-mir-16-2&WithFlank&3|+|160404739|160404831@16.37.22&[MIMAT0000069&hsa-miR-16-5p&offsets|0|0;m-62&13|-|50049027|50049048&offsets|0|0];[MIMAT0000069_1&hsa-miR-16-5p&offsets|0|0;m-61&3|+|160404754|160404775&offsets|0|0]|TAGCAGCACGTAAATATTGGCG</td><td>>MI0000070|hsa-mir-16-1&WithFlank&13|-|50048967|50049067</td><td>20</td><td>41</td><td>22</td><td></td><td>isomiR</td><td>NO</td><td>0.00</td><td>0.00</td><td>91.36</td><td>0.00</td></tr>
<tr><td>>MI0000070|hsa-mir-16-1&WithFlank&13|-|50048967|50049067@20.42.23;MI0000115|hsa-mir-16-2&WithFlank&3|+|160404739|160404831@16.38.23&[MIMAT0000069&hsa-miR-16-5p&offsets|0|+1;m-62&13|-|50049027|50049048&offsets|0|+1];[MIMAT0000069_1&hsa-miR-16-5p&offsets|0|+1;m-61&3|+|160404754|160404775&offsets|0|+1]|TAGCAGCACGTAAATATTGGCGT</td><td>>MI0000070|hsa-mir-16-1&WithFlank&13|-|50048967|50049067</td><td>20</td><td>42</td><td>23</td><td></td><td>isomiR</td><td>NO</td><td>0.00</td><td>0.00</td><td>31.57</td><td>0.00</td></tr>
<tr><td>>MI0000071|hsa-mir-17&WithFlank&13|+|91350599|91350694@20.42.23&[MIMAT0000070&hsa-miR-17-5p&offsets|0|0;m-88&13|+|91350618|91350640&offsets|0|0]|CAAAGTGCTTACAGTGCAGGTAG</td><td>>MI0000071|hsa-mir-17&WithFlank&13|+|91350599|91350694</td><td>20</td><td>42</td><td>23</td><td></td><td>isomiR</td><td>NO</td><td>21.34</td><td>12.29</td><td>89.22</td><td>0.00</td></tr>
<tr><td>>MI0000076|hsa-mir-20a&WithFlank&13|+|91351059|91351141@14.36.23&[MIMAT0000075&hsa-miR-20a-5p&offsets|0|0;m-96&13|+|91351072|91351094&offsets|0|0]|TAAAGTGCTTATAGTGCAGGTAG</td><td>>MI0000076|hsa-mir-20a&WithFlank&13|+|91351059|91351141</td><td>14</td><td>36</td><td>23</td><td></td><td>isomiR</td><td>NO</td><td>28.77</td><td>0.00</td><td>125.51</td><td>6.05</td></tr>
<tr><td>>MI0000077|hsa-mir-21&WithFlank&17|+|59841260|59841343@14.34.21&[MIMAT0000076&hsa-miR-21-5p&offsets|0|-1;m-1&17|+|59841273|59841294&offsets|0|-1]|TAGCTTATCAGACTGATGTTG</td><td>>MI0000077|hsa-mir-21&WithFlank&17|+|59841260|59841343</td><td>14</td><td>34</td><td>21</td><td></td><td>isomiR</td><td>NO</td><td>0.00</td><td>0.00</td><td>132.30</td><td>0.00</td></tr>
<tr><td>>MI0000077|hsa-mir-21&WithFlank&17|+|59841260|59841343@14.34.21(+1G)&[MIMAT0000076&hsa-miR-21-5p&offsets|0|-1(+1G);m-1&17|+|59841273|59841294&offsets|0|-1(+1G)]|TAGCTTATCAGACTGATGTTGG</td><td>>MI0000077|hsa-mir-21&WithFlank&17|+|59841260|59841343</td><td>14</td><td>34</td><td>21</td><td>(+1G)</td><td>isomiR</td><td>NO</td><td>0.00</td><td>0.00</td><td>67.49</td><td>0.00</td></tr>
</table>

### MDA-MB-468

<table>
<tr><th></th><th></th><th></th><th></th><th></th><th></th><th></th><th></th><th></th><th></th><th></th><th></th></tr>
<tr><td>>MI0000077|hsa-mir-21&WithFlank&17|+|59841260|59841343@14.35.22&[MIMAT0000076&hsa-miR-21-5p&offsets|0|0;m-1&17|+|59841273|59841294&offsets|0|0]|TAGCTTATCAGACTGATGTTGA</td><td>>MI0000077|hsa-mir-21&WithFlank&17|+|59841260|59841343</td><td>14</td><td>35</td><td>22</td><td></td><td>isomiR</td><td>NO</td><td>62.09</td><td>91.35</td><td>1530.61</td><td>39.61</td></tr>
<tr><td>>MI0000077|hsa-mir-21&WithFlank&17|+|59841260|59841343@14.35.22(+1A)&[MIMAT0000076&hsa-miR-21-5p&offsets|0|0(+1A);m-1&17|+|59841273|59841294&offsets|0|0(+1A)]|TAGCTTATCAGACTGATGTTGAA</td><td>>MI0000077|hsa-mir-21&WithFlank&17|+|59841260|59841343</td><td>14</td><td>35</td><td>22</td><td>(+1A)</td><td>isomiR</td><td>NO</td><td>0.00</td><td>0.00</td><td>44.75</td><td>0.00</td></tr>
<tr><td>>MI0000077|hsa-mir-21&WithFlank&17|+|59841260|59841343@14.35.22(+1U)&[MIMAT0000076&hsa-miR-21-5p&offsets|0|0(+1U);m-1&17|+|59841273|59841294&offsets|0|0(+1U)]|TAGCTTATCAGACTGATGTTGAT</td><td>>MI0000077|hsa-mir-21&WithFlank&17|+|59841260|59841343</td><td>14</td><td>35</td><td>22</td><td>(+1U)</td><td>isomiR</td><td>NO</td><td>44.10</td><td>62.39</td><td>124.47</td><td>25.71</td></tr>
<tr><td>>MI0000077|hsa-mir-21&WithFlank&17|+|59841260|59841343@14.36.23&[MIMAT0000076&hsa-miR-21-5p&offsets|0|+1;m-1&17|+|59841273|59841294&offsets|0|+1]|TAGCTTATCAGACTGATGTTGAC</td><td>>MI0000077|hsa-mir-21&WithFlank&17|+|59841260|59841343</td><td>14</td><td>36</td><td>23</td><td></td><td>isomiR</td><td>YES</td><td>568.48</td><td>800.25</td><td>791.24</td><td>219.12</td></tr>
<tr><td>>MI0000077|hsa-mir-21&WithFlank&17|+|59841260|59841343@14.36.23(+1A)&[MIMAT0000076&hsa-miR-21-5p&offsets|0|+1(+1A);m-1&17|+|59841273|59841294&offsets|0|+1(+1A)]|TAGCTTATCAGACTGATGTTGACA</td><td>>MI0000077|hsa-mir-21&WithFlank&17|+|59841260|59841343</td><td>14</td><td>36</td><td>23</td><td>(+1A)</td><td>isomiR</td><td>NO</td><td>150.19</td><td>206.40</td><td>74.91</td><td>58.95</td></tr>
<tr><td>>MI0000077|hsa-mir-21&WithFlank&17|+|59841260|59841343@14.36.23(+1C)&[MIMAT0000076&hsa-miR-21-5p&offsets|0|+1(+1C);m-1&17|+|59841273|59841294&offsets|0|+1(+1C)]|TAGCTTATCAGACTGATGTTGACC</td><td>>MI0000077|hsa-mir-21&WithFlank&17|+|59841260|59841343</td><td>14</td><td>36</td><td>23</td><td>(+1C)</td><td>isomiR</td><td>NO</td><td>20.10</td><td>29.37</td><td>0.00</td><td>0.00</td></tr>
<tr><td>>MI0000077|hsa-mir-21&WithFlank&17|+|59841260|59841343@14.36.23(+1G)&[MIMAT0000076&hsa-miR-21-5p&offsets|0|+1(+1G);m-1&17|+|59841273|59841294&offsets|0|+1(+1G)]|TAGCTTATCAGACTGATGTTGACG</td><td>>MI0000077|hsa-mir-21&WithFlank&17|+|59841260|59841343</td><td>14</td><td>36</td><td>23</td><td>(+1G)</td><td>isomiR</td><td>NO</td><td>0.00</td><td>27.26</td><td>0.00</td><td>0.00</td></tr>
<tr><td>>MI0000077|hsa-mir-21&WithFlank&17|+|59841260|59841343@14.37.24&[MIMAT0000076&hsa-miR-21-5p&offsets|0|+2;m-1&17|+|59841273|59841294&offsets|0|+2]|TAGCTTATCAGACTGATGTTGACT</td><td>>MI0000077|hsa-mir-21&WithFlank&17|+|59841260|59841343</td><td>14</td><td>37</td><td>24</td><td></td><td>isomiR</td><td>NO</td><td>138.28</td><td>151.57</td><td>77.29</td><td>51.72</td></tr>
<tr><td>>MI0000078|hsa-mir-22&WithFlank&17|-|1713897|1713993@59.80.22&[MIMAT0000077&hsa-miR-22-3p&offsets|0|0;m-2&17|-|1713914|1713935&offsets|0|0]|AAGCTGCCAGTTGAAGAACTGT</td><td>>MI0000078|hsa-mir-22&WithFlank&17|-|1713897|1713993</td><td>59</td><td>80</td><td>22</td><td></td><td>isomiR</td><td>NO</td><td>0.00</td><td>0.00</td><td>52.62</td><td>0.00</td></tr>
<tr><td>>MI0000081|hsa-mir-24-2&WithFlank&19|-|13836281|13836365@56.75.20;MI0000080|hsa-mir-24-1&WithFlank&9|+|95086015|95086094@50.69.20&[MIMAT0000080&hsa-miR-24-3p&offsets|0|-2;m-36&19|-|13836288|13836310&offsets|0|-3];[MIMAT0000080_1&hsa-miR-24-3p&offsets|0|-2;m-35&9|+|95086064|95086086&offsets|0|-3]|TGGCTCAGTTCAGCAGGAAC</td><td>>MI0000081|hsa-mir-24-2&WithFlank&19|-|13836281|13836365</td><td>56</td><td>75</td><td>20</td><td></td><td>isomiR</td><td>NO</td><td>37.42</td><td>27.75</td><td>113.36</td><td>0.00</td></tr>
<tr><td>>MI0000081|hsa-mir-24-2&WithFlank&19|-|13836281|13836365@56.76.21;MI0000080|hsa-mir-24-1&WithFlank&9|+|95086015|95086094@50.70.21&[MIMAT0000080&hsa-miR-24-3p&offsets|0|-1;m-36&19|-|13836288|13836310&offsets|0|-2];[MIMAT0000080_1&hsa-miR-24-3p&offsets|0|-1;m-35&9|+|95086064|95086086&offsets|0|-2]|TGGCTCAGTTCAGCAGGAACA</td><td>>MI0000081|hsa-mir-24-2&WithFlank&19|-|13836281|13836365</td><td>56</td><td>76</td><td>21</td><td></td><td>isomiR</td><td>NO</td><td>0.00</td><td>0.00</td><td>40.88</td><td>0.00</td></tr>
<tr><td>>MI0000081|hsa-mir-24-2&WithFlank&19|-|13836281|13836365@56.77.22(+1U);MI0000080|hsa-mir-24-1&WithFlank&9|+|95086015|95086094@50.71.22(+1U)&[MIMAT0000080&hsa-miR-24-3p&offsets|0|0(+1U);m-36&19|-|13836288|13836310&offsets|0|-1(+1U)];[MIMAT0000080_1&hsa-miR-24-3p&offsets|0|0(+1U);m-35&9|+|95086064|95086086&offsets|0|-1(+1U)]|TGGCTCAGTTCAGCAGGAACAGT</td><td>>MI0000081|hsa-mir-24-2&WithFlank&19|-|13836281|13836365</td><td>56</td><td>77</td><td>22</td><td>(+1U)</td><td>isomiR</td><td>NO</td><td>0.00</td><td>0.00</td><td>52.43</td><td>0.00</td></tr>
<tr><td>>MI0000081|hsa-mir-24-2&WithFlank&19|-|13836281|13836365@56.77.22;MI0000080|hsa-mir-24-1&WithFlank&9|+|95086015|95086094@50.71.22&[MIMAT0000080&hsa-miR-24-3p&offsets|0|0;m-36&19|-|13836288|13836310&offsets|0|-1];[MIMAT0000080_1&hsa-miR-24-3p&offsets|0|0;m-35&9|+|95086064|95086086&offsets|0|-1]|TGGCTCAGTTCAGCAGGAACAG</td><td>>MI0000081|hsa-mir-24-2&WithFlank&19|-|13836281|13836365</td><td>56</td><td>77</td><td>22</td><td></td><td>isomiR</td><td>NO</td><td>51.24</td><td>38.52</td><td>131.86</td><td>6.63</td></tr>
<tr><td>>MI0000082|hsa-mir-25&WithFlank&7|-|100093554|100093649@58.78.21&[MIMAT0000081&hsa-miR-25-3p&offsets|0|-1;m-23&7|-|100093571|100093592&offsets|0|-1]|CATTGCACTTGTCTCGGTCTG</td><td>>MI0000082|hsa-mir-25&WithFlank&7|-|100093554|100093649</td><td>58</td><td>78</td><td>21</td><td></td><td>isomiR</td><td>NO</td><td>21.49</td><td>0.00</td><td>0.00</td><td>0.00</td></tr>
<tr><td>>MI0000082|hsa-mir-25&WithFlank&7|-|100093554|100093649@58.79.22&[MIMAT0000081&hsa-miR-25-3p&offsets|0|0;m-23&7|-|100093571|100093592&offsets|0|0]|CATTGCACTTGTCTCGGTCTGA</td><td>>MI0000082|hsa-mir-25&WithFlank&7|-|100093554|100093649</td><td>58</td><td>79</td><td>22</td><td></td><td>isomiR</td><td>NO</td><td>193.74</td><td>55.96</td><td>73.32</td><td>6.41</td></tr>
<tr><td>>MI0000084|hsa-mir-26b&WithFlank&2|+|218402640|218402728@18.39.22&[MIMAT0000083&hsa-miR-26b-5p&offsets|0|+1;m-53&2|+|218402657|218402678&offsets|0|0]|TTCAAGTAATTCAGGATAGGTT</td><td>>MI0000084|hsa-mir-26b&WithFlank&2|+|218402640|218402728</td><td>18</td><td>39</td><td>22</td><td></td><td>isomiR</td><td>NO</td><td>0.00</td><td>0.00</td><td>27.72</td><td>0.00</td></tr>
<tr><td>>MI0000085|hsa-mir-27a&WithFlank&19|-|13836434|13836523@57.76.20&[MIMAT0000084&hsa-miR-27a-3p&offsets|0|-1;m-43&19|-|13836447|13836467&offsets|0|-1]|TTCACAGTGGCTAAGTTCCG</td><td>>MI0000085|hsa-mir-27a&WithFlank&19|-|13836434|13836523</td><td>57</td><td>76</td><td>20</td><td></td><td>isomiR</td><td>NO</td><td>6.10</td><td>0.00</td><td>82.31</td><td>14.12</td></tr>
<tr><td>>MI0000085|hsa-mir-27a&WithFlank&19|-|13836434|13836523@57.76.20(+1A)&[MIMAT0000084&hsa-miR-27a-3p&offsets|0|-1(+1A);m-43&19|-|13836447|13836467&offsets|0|-1(+1A)]|TTCACAGTGGCTAAGTTCCGA</td><td>>MI0000085|hsa-mir-27a&WithFlank&19|-|13836434|13836523</td><td>57</td><td>76</td><td>20</td><td>(+1A)</td><td>isomiR</td><td>NO</td><td>0.00</td><td>0.00</td><td>37.41</td><td>0.00</td></tr>
<tr><td>>MI0000085|hsa-mir-27a&WithFlank&19|-|13836434|13836523@57.76.20(+1U)&[MIMAT0000084&hsa-miR-27a-3p&offsets|0|-1(+1U);m-43&19|-|13836447|13836467&offsets|0|-1(+1U)]|TTCACAGTGGCTAAGTTCCGT</td><td>>MI0000085|hsa-mir-27a&WithFlank&19|-|13836434|13836523</td><td>57</td><td>76</td><td>20</td><td>(+1U)</td><td>isomiR</td><td>NO</td><td>0.00</td><td>0.00</td><td>74.61</td><td>0.00</td></tr>
</table>

### MDA-MB-468

<table>
<tr><th></th><th></th><th></th><th></th><th></th><th></th><th></th><th></th><th></th><th></th><th></th><th></th></tr>
<tr><td>>MI0000085|hsa-mir-27a&WithFlank&19|-|13836434|13836523@57.76.20(+2U)&[MIMAT0000084&hsa-miR-27a-3p&offsets|0|-1(+2U);m-43&19|-|13836447|13836467&offsets|0|-1(+2U)]|TTCACAGTGGCTAAGTTCCGTT</td><td>>MI0000085|hsa-mir-27a&WithFlank&19|-|13836434|13836523</td><td>57</td><td>76</td><td>20</td><td>(+2U)</td><td>isomiR</td><td>NO</td><td>0.00</td><td>0.00</td><td>41.00</td><td>0.00</td></tr>
<tr><td>>MI0000085|hsa-mir-27a&WithFlank&19|-|13836434|13836523@57.77.21&[MIMAT0000084&hsa-miR-27a-3p&offsets|0|0;m-43&19|-|13836447|13836467&offsets|0|0]|TTCACAGTGGCTAAGTTCCGC</td><td>>MI0000085|hsa-mir-27a&WithFlank&19|-|13836434|13836523</td><td>57</td><td>77</td><td>21</td><td></td><td>isomiR</td><td>NO</td><td>0.00</td><td>0.00</td><td>66.46</td><td>0.00</td></tr>
<tr><td>>MI0000088|hsa-mir-30a&WithFlank&6|-|71403545|71403627@12.33.22&[MIMAT0000087&hsa-miR-30a-5p&offsets|0|0;m-15&6|-|71403595|71403616&offsets|0|0]|TGTAAACATCCTCGACTGGAAG</td><td>>MI0000088|hsa-mir-30a&WithFlank&6|-|71403545|71403627</td><td>12</td><td>33</td><td>22</td><td></td><td>isomiR</td><td>NO</td><td>0.00</td><td>0.00</td><td>74.57</td><td>0.00</td></tr>
<tr><td>>MI0000088|hsa-mir-30a&WithFlank&6|-|71403545|71403627@12.33.22(+1U)&[MIMAT0000087&hsa-miR-30a-5p&offsets|0|0(+1U);m-15&6|-|71403595|71403616&offsets|0|0(+1U)]|TGTAAACATCCTCGACTGGAAGT</td><td>>MI0000088|hsa-mir-30a&WithFlank&6|-|71403545|71403627</td><td>12</td><td>33</td><td>22</td><td>(+1U)</td><td>isomiR</td><td>NO</td><td>0.00</td><td>0.00</td><td>46.30</td><td>0.00</td></tr>
<tr><td>>MI0000088|hsa-mir-30a&WithFlank&6|-|71403545|71403627@12.34.23&[MIMAT0000087&hsa-miR-30a-5p&offsets|0|+1;m-15&6|-|71403595|71403616&offsets|0|+1]|TGTAAACATCCTCGACTGGAAGC</td><td>>MI0000088|hsa-mir-30a&WithFlank&6|-|71403545|71403627</td><td>12</td><td>34</td><td>23</td><td></td><td>isomiR</td><td>NO</td><td>0.00</td><td>0.00</td><td>165.11</td><td>0.00</td></tr>
<tr><td>>MI0000088|hsa-mir-30a&WithFlank&6|-|71403545|71403627@12.34.23(+1A)&[MIMAT0000087&hsa-miR-30a-5p&offsets|0|+1(+1A);m-15&6|-|71403595|71403616&offsets|0|+1(+1A)]|TGTAAACATCCTCGACTGGAAGCA</td><td>>MI0000088|hsa-mir-30a&WithFlank&6|-|71403545|71403627</td><td>12</td><td>34</td><td>23</td><td>(+1A)</td><td>isomiR</td><td>NO</td><td>0.00</td><td>0.00</td><td>67.10</td><td>0.00</td></tr>
<tr><td>>MI0000088|hsa-mir-30a&WithFlank&6|-|71403545|71403627@12.34.23(+1C)&[MIMAT0000087&hsa-miR-30a-5p&offsets|0|+1(+1C);m-15&6|-|71403595|71403616&offsets|0|+1(+1C)]|TGTAAACATCCTCGACTGGAAGCC</td><td>>MI0000088|hsa-mir-30a&WithFlank&6|-|71403545|71403627</td><td>12</td><td>34</td><td>23</td><td>(+1C)</td><td>isomiR</td><td>NO</td><td>0.00</td><td>0.00</td><td>89.14</td><td>0.00</td></tr>
<tr><td>>MI0000088|hsa-mir-30a&WithFlank&6|-|71403545|71403627@12.34.23(+1G)&[MIMAT0000087&hsa-miR-30a-5p&offsets|0|+1(+1G);m-15&6|-|71403595|71403616&offsets|0|+1(+1G)]|TGTAAACATCCTCGACTGGAAGCG</td><td>>MI0000088|hsa-mir-30a&WithFlank&6|-|71403545|71403627</td><td>12</td><td>34</td><td>23</td><td>(+1G)</td><td>isomiR</td><td>NO</td><td>0.00</td><td>0.00</td><td>27.81</td><td>0.00</td></tr>
<tr><td>>MI0000088|hsa-mir-30a&WithFlank&6|-|71403545|71403627@12.35.24&[MIMAT0000087&hsa-miR-30a-5p&offsets|0|+2;m-15&6|-|71403595|71403616&offsets|0|+2]|TGTAAACATCCTCGACTGGAAGCT</td><td>>MI0000088|hsa-mir-30a&WithFlank&6|-|71403545|71403627</td><td>12</td><td>35</td><td>24</td><td></td><td>isomiR</td><td>NO</td><td>40.92</td><td>35.71</td><td>429.03</td><td>24.77</td></tr>
<tr><td>>MI0000088|hsa-mir-30a&WithFlank&6|-|71403545|71403627@12.35.24(+1U)&[MIMAT0000087&hsa-miR-30a-5p&offsets|0|+2(+1U);m-15&6|-|71403595|71403616&offsets|0|+2(+1U)]|TGTAAACATCCTCGACTGGAAGCTT</td><td>>MI0000088|hsa-mir-30a&WithFlank&6|-|71403545|71403627</td><td>12</td><td>35</td><td>24</td><td>(+1U)</td><td>isomiR</td><td>NO</td><td>0.00</td><td>0.00</td><td>27.76</td><td>0.00</td></tr>
<tr><td>>MI0000088|hsa-mir-30a&WithFlank&6|-|71403545|71403627@53.74.22&[MIMAT0000088&hsa-miR-30a-3p&offsets|0|0;m-38&6|-|71403554|71403575&offsets|0|0]|CTTTCAGTCGGATGTTTGCAGC</td><td>>MI0000088|hsa-mir-30a&WithFlank&6|-|71403545|71403627</td><td>53</td><td>74</td><td>22</td><td></td><td>isomiR</td><td>NO</td><td>38.21</td><td>0.00</td><td>27.34</td><td>0.00</td></tr>
<tr><td>>MI0000093|hsa-mir-92a-1&WithFlank&13|+|91351308|91351397@54.74.21(+1C);MI0000094|hsa-mir-92a-2&WithFlank&X|-|134169532|134169618@54.74.21(+1C)&[MIMAT0000092&hsa-miR-92a-3p&offsets|0|-1(+1C);m-9&13|+|91351361|91351382&offsets|0|-1(+1C)];[MIMAT0000092_1&hsa-miR-92a-3p&offsets|0|-1(+1C);m-11&X|-|134169544|134169565&offsets|0|-1(+1C)]|TATTGCACTTGTCCCGGCCTGC</td><td>>MI0000093|hsa-mir-92a-1&WithFlank&13|+|91351308|91351397</td><td>54</td><td>74</td><td>21</td><td>(+1C)</td><td>isomiR</td><td>NO</td><td>52.04</td><td>0.00</td><td>32.18</td><td>0.00</td></tr>
<tr><td>>MI0000093|hsa-mir-92a-1&WithFlank&13|+|91351308|91351397@54.75.22(+1A);MI0000094|hsa-mir-92a-2&WithFlank&X|-|134169532|134169618@54.75.22(+1A)&[MIMAT0000092&hsa-miR-92a-3p&offsets|0|0(+1A);m-9&13|+|91351361|91351382&offsets|0|0(+1A)];[MIMAT0000092_1&hsa-miR-92a-3p&offsets|0|0(+1A);m-11&X|-|134169544|134169565&offsets|0|0(+1A)]|TATTGCACTTGTCCCGGCCTGTA</td><td>>MI0000093|hsa-mir-92a-1&WithFlank&13|+|91351308|91351397</td><td>54</td><td>75</td><td>22</td><td>(+1A)</td><td>isomiR</td><td>NO</td><td>106.58</td><td>5.97</td><td>20.44</td><td>0.00</td></tr>
<tr><td>>MI0000093|hsa-mir-92a-1&WithFlank&13|+|91351308|91351397@54.75.22;MI0000094|hsa-mir-92a-2&WithFlank&X|-|134169532|134169618@54.75.22&[MIMAT0000092&hsa-miR-92a-3p&offsets|0|0;m-9&13|+|91351361|91351382&offsets|0|0];[MIMAT0000092_1&hsa-miR-92a-3p&offsets|0|0;m-11&X|-|134169544|134169565&offsets|0|0]|TATTGCACTTGTCCCGGCCTGT</td><td>>MI0000093|hsa-mir-92a-1&WithFlank&13|+|91351308|91351397</td><td>54</td><td>75</td><td>22</td><td></td><td>isomiR</td><td>NO</td><td>380.31</td><td>86.37</td><td>158.11</td><td>52.11</td></tr>
<tr><td>>MI0000093|hsa-mir-92a-1&WithFlank&13|+|91351308|91351397@54.76.23&[MIMAT0000092&hsa-miR-92a-3p&offsets|0|+1;m-9&13|+|91351361|91351382&offsets|0|+1]|TATTGCACTTGTCCCGGCCTGTT</td><td>>MI0000093|hsa-mir-92a-1&WithFlank&13|+|91351308|91351397</td><td>54</td><td>76</td><td>23</td><td></td><td>isomiR</td><td>NO</td><td>31.31</td><td>0.00</td><td>0.00</td><td>0.00</td></tr>
<tr><td>>MI0000095|hsa-mir-93&WithFlank&7|-|100093762|100093853@17.39.23&[MIMAT0000093&hsa-miR-93-5p&offsets|0|0;m-29&7|-|100093815|100093837&offsets|0|0]|CAAAGTGCTGTTCGTGCAGGTAG</td><td>>MI0000095|hsa-mir-93&WithFlank&7|-|100093762|100093853</td><td>17</td><td>39</td><td>23</td><td></td><td>isomiR</td><td>NO</td><td>30.67</td><td>27.45</td><td>107.11</td><td>0.00</td></tr>
<tr><td>>MI0000098|hsa-mir-96&WithFlank&7|-|129774686|129774775@15.37.23&[MIMAT0000095&hsa-miR-96-5p&offsets|0|0;m-248&7|-|129774739|129774761&offsets|0|0]|TTTGGCACTAGCACATTTTTGCT</td><td>>MI0000098|hsa-mir-96&WithFlank&7|-|129774686|129774775</td><td>15</td><td>37</td><td>23</td><td></td><td>isomiR</td><td>NO</td><td>0.00</td><td>0.00</td><td>28.65</td><td>0.00</td></tr>
<tr><td>>MI0000100|hsa-mir-98&WithFlank&X|-|53556217|53556347@28.49.22&[MIMAT0000096&hsa-miR-98-5p&offsets|0|0;m-143&X|-|53556299|53556320&offsets|0|0]|TGAGGTAGTAAGTTGTATTGTT</td><td>>MI0000100|hsa-mir-98&WithFlank&X|-|53556217|53556347</td><td>28</td><td>49</td><td>22</td><td></td><td>isomiR</td><td>NO</td><td>41.02</td><td>37.48</td><td>46.32</td><td>0.00</td></tr>
<tr><td>>MI0000101|hsa-mir-99a&WithFlank&21|+|16539083|16539175@19.39.21&[MIMAT0000097&hsa-miR-99a-5p&offsets|0|-1;m-64&21|+|16539101|16539121&offsets|0|0]|AACCCGTAGATCCGATCTTGT</td><td>>MI0000101|hsa-mir-99a&WithFlank&21|+|16539083|16539175</td><td>19</td><td>39</td><td>21</td><td></td><td>isomiR</td><td>NO</td><td>134.42</td><td>92.77</td><td>118.42</td><td>35.67</td></tr>
<tr><td>>MI0000101|hsa-mir-99a&WithFlank&21|+|16539083|16539175@19.40.22&[MIMAT0000097&hsa-miR-99a-5p&offsets|0|0;m-64&21|+|16539101|16539121&offsets|0|+1]|AACCCGTAGATCCGATCTTGTG</td><td>>MI0000101|hsa-mir-99a&WithFlank&21|+|16539083|16539175</td><td>19</td><td>40</td><td>22</td><td></td><td>isomiR</td><td>NO</td><td>64.66</td><td>65.43</td><td>63.45</td><td>14.34</td></tr>
</table>

### MDA-MB-468

<table>
<tr><th></th><th></th><th></th><th></th><th></th><th></th><th></th><th></th><th></th><th></th><th></th><th></th></tr>
<tr><td>>MI0000101|hsa-mir-99a&WithFlank&21|+|16539083|16539175@19.40.22(+1A)&[MIMAT0000097&hsa-miR-99a-5p&offsets|0|0(+1A);m-64&21|+|16539101|16539121&offsets|0|+1(+1A)]|AACCCGTAGATCCGATCTTGTGA</td><td>>MI0000101|hsa-mir-99a&WithFlank&21|+|16539083|16539175</td><td>19</td><td>40</td><td>22</td><td>(+1A)</td><td>isomiR</td><td>NO</td><td>71.40</td><td>61.28</td><td>42.33</td><td>0.00</td></tr>
<tr><td>>MI0000103|hsa-mir-101-1&WithFlank&1|-|65058428|65058514@52.73.22;MI0000739|hsa-mir-101-2&WithFlank&9|+|4850291|4850381@54.75.22&[MIMAT0000099&hsa-miR-101-3p&offsets|-1|0;m-17&1|-|65058443|65058462&offsets|-1|+1];[MIMAT0000099_1&hsa-miR-101-3p&offsets|-1|0;m-16&9|+|4850345|4850364&offsets|-1|+1]|GTACAGTACTGTGATAACTGAA</td><td>>MI0000103|hsa-mir-101-1&WithFlank&1|-|65058428|65058514</td><td>52</td><td>73</td><td>22</td><td></td><td>isomiR</td><td>NO</td><td>0.00</td><td>0.00</td><td>37.09</td><td>0.00</td></tr>
<tr><td>>MI0000103|hsa-mir-101-1&WithFlank&1|-|65058428|65058514@53.74.22;MI0000739|hsa-mir-101-2&WithFlank&9|+|4850291|4850381@55.76.22&[MIMAT0000099&hsa-miR-101-3p&offsets|0|+1;m-17&1|-|65058443|65058462&offsets|0|+2];[MIMAT0000099_1&hsa-miR-101-3p&offsets|0|+1;m-16&9|+|4850345|4850364&offsets|0|+2]|TACAGTACTGTGATAACTGAAG</td><td>>MI0000103|hsa-mir-101-1&WithFlank&1|-|65058428|65058514</td><td>53</td><td>74</td><td>22</td><td></td><td>isomiR</td><td>NO</td><td>0.00</td><td>0.00</td><td>20.70</td><td>0.00</td></tr>
<tr><td>>MI0000108|hsa-mir-103a-2&WithFlank&20|+|3917488|3917577@54.75.22;MI0000109|hsa-mir-103a-1&WithFlank&5|-|168560890|168560979@54.75.22&[MIMAT0000101&hsa-miR-103a-3p&offsets|0|-1;m-12&20|+|3917541|3917563&offsets|0|-1];[MIMAT0000101_1&hsa-miR-103a-3p&offsets|0|-1;m-13&5|-|168560904|168560926&offsets|0|-1;m-44220&5|-|168560908|168560926&offsets|0|+3]|AGCAGCATTGTACAGGGCTATG</td><td>>MI0000108|hsa-mir-103a-2&WithFlank&20|+|3917488|3917577</td><td>54</td><td>75</td><td>22</td><td></td><td>isomiR</td><td>NO</td><td>0.00</td><td>0.00</td><td>41.21</td><td>0.00</td></tr>
<tr><td>>MI0000108|hsa-mir-103a-2&WithFlank&20|+|3917488|3917577@54.76.23;MI0000109|hsa-mir-103a-1&WithFlank&5|-|168560890|168560979@54.76.23&[MIMAT0000101&hsa-miR-103a-3p&offsets|0|0;m-12&20|+|3917541|3917563&offsets|0|0];[MIMAT0000101_1&hsa-miR-103a-3p&offsets|0|0;m-13&5|-|168560904|168560926&offsets|0|0;m-44220&5|-|168560908|168560926&offsets|0|+4]|AGCAGCATTGTACAGGGCTATGA</td><td>>MI0000108|hsa-mir-103a-2&WithFlank&20|+|3917488|3917577</td><td>54</td><td>76</td><td>23</td><td></td><td>isomiR</td><td>NO</td><td>56.81</td><td>29.51</td><td>174.29</td><td>0.00</td></tr>
<tr><td>>MI0000234|hsa-mir-192&WithFlank&11|-|64891131|64891252@30.50.21&[MIMAT0000222&hsa-miR-192-5p&offsets|0|0;m-59&11|-|64891203|64891223&offsets|0|0]|CTGACCTATGAATTGACAGCC</td><td>>MI0000234|hsa-mir-192&WithFlank&11|-|64891131|64891252</td><td>30</td><td>50</td><td>21</td><td></td><td>isomiR</td><td>NO</td><td>0.00</td><td>0.00</td><td>31.27</td><td>0.00</td></tr>
<tr><td>>MI0000253|hsa-mir-148a&WithFlank&7|-|25949913|25949992@50.70.21(+1A)&[MIMAT0000243&hsa-miR-148a-3p&offsets|0|-1(+1A);m-7&7|-|25949922|25949943&offsets|0|-1(+1A)]|TCAGTGCACTACAGAACTTTGA</td><td>>MI0000253|hsa-mir-148a&WithFlank&7|-|25949913|25949992</td><td>50</td><td>70</td><td>21</td><td>(+1A)</td><td>isomiR</td><td>NO</td><td>0.00</td><td>0.00</td><td>41.89</td><td>0.00</td></tr>
<tr><td>>MI0000253|hsa-mir-148a&WithFlank&7|-|25949913|25949992@50.70.21(+1C)&[MIMAT0000243&hsa-miR-148a-3p&offsets|0|-1(+1C);m-7&7|-|25949922|25949943&offsets|0|-1(+1C)]|TCAGTGCACTACAGAACTTTGC</td><td>>MI0000253|hsa-mir-148a&WithFlank&7|-|25949913|25949992</td><td>50</td><td>70</td><td>21</td><td>(+1C)</td><td>isomiR</td><td>NO</td><td>34.70</td><td>0.00</td><td>165.30</td><td>0.00</td></tr>
<tr><td>>MI0000253|hsa-mir-148a&WithFlank&7|-|25949913|25949992@50.70.21(+1G)&[MIMAT0000243&hsa-miR-148a-3p&offsets|0|-1(+1G);m-7&7|-|25949922|25949943&offsets|0|-1(+1G)]|TCAGTGCACTACAGAACTTTGG</td><td>>MI0000253|hsa-mir-148a&WithFlank&7|-|25949913|25949992</td><td>50</td><td>70</td><td>21</td><td>(+1G)</td><td>isomiR</td><td>NO</td><td>0.00</td><td>0.00</td><td>39.79</td><td>0.00</td></tr>
<tr><td>>MI0000253|hsa-mir-148a&WithFlank&7|-|25949913|25949992@50.71.22&[MIMAT0000243&hsa-miR-148a-3p&offsets|0|0;m-7&7|-|25949922|25949943&offsets|0|0]|TCAGTGCACTACAGAACTTTGT</td><td>>MI0000253|hsa-mir-148a&WithFlank&7|-|25949913|25949992</td><td>50</td><td>71</td><td>22</td><td></td><td>isomiR</td><td>NO</td><td>293.61</td><td>185.03</td><td>888.79</td><td>58.54</td></tr>
<tr><td>>MI0000255|hsa-mir-30d&WithFlank&8|-|134804870|134804951@12.34.23&[MIMAT0000245&hsa-miR-30d-5p&offsets|0|+1;m-21&8|-|134804919|134804940&offsets|0|+1]|TGTAAACATCCCCGACTGGAAGC</td><td>>MI0000255|hsa-mir-30d&WithFlank&8|-|134804870|134804951</td><td>12</td><td>34</td><td>23</td><td></td><td>isomiR</td><td>NO</td><td>47.79</td><td>0.00</td><td>7.06</td><td>0.00</td></tr>
<tr><td>>MI0000255|hsa-mir-30d&WithFlank&8|-|134804870|134804951@12.35.24&[MIMAT0000245&hsa-miR-30d-5p&offsets|0|+2;m-21&8|-|134804919|134804940&offsets|0|+2]|TGTAAACATCCCCGACTGGAAGCT</td><td>>MI0000255|hsa-mir-30d&WithFlank&8|-|134804870|134804951</td><td>12</td><td>35</td><td>24</td><td></td><td>isomiR</td><td>NO</td><td>64.59</td><td>27.20</td><td>54.74</td><td>0.00</td></tr>
<tr><td>>MI0000264|hsa-mir-7-2&WithFlank&15|+|88611819|88611940@38.60.23;MI0000265|hsa-mir-7-3&WithFlank&19|+|4770664|4770785@37.59.23;MI0000263|hsa-mir-7-1&WithFlank&9|-|83969742|83969863@30.52.23&[MIMAT0000252&hsa-miR-7-5p&offsets|0|-1;m-400&15|+|88611856|88611879&offsets|0|-1];[MIMAT0000252_1&hsa-miR-7-5p&offsets|0|-1;m-402&19|+|4770700|4770723&offsets|0|-1];[MIMAT0000252_2&hsa-miR-7-5p&offsets|0|-1;m-401&9|-|83969811|83969834&offsets|0|-1]|TGGAAGACTAGTGATTTTGTTGT</td><td>>MI0000264|hsa-mir-7-2&WithFlank&15|+|88611819|88611940</td><td>38</td><td>60</td><td>23</td><td></td><td>isomiR</td><td>NO</td><td>0.00</td><td>0.00</td><td>36.42</td><td>0.00</td></tr>
<tr><td>>MI0000264|hsa-mir-7-2&WithFlank&15|+|88611819|88611940@38.61.24;MI0000265|hsa-mir-7-3&WithFlank&19|+|4770664|4770785@37.60.24;MI0000263|hsa-mir-7-1&WithFlank&9|-|83969742|83969863@30.53.24&[MIMAT0000252&hsa-miR-7-5p&offsets|0|0;m-400&15|+|88611856|88611879&offsets|0|0];[MIMAT0000252_1&hsa-miR-7-5p&offsets|0|0;m-402&19|+|4770700|4770723&offsets|0|0];[MIMAT0000252_2&hsa-miR-7-5p&offsets|0|0;m-401&9|-|83969811|83969834&offsets|0|0]|TGGAAGACTAGTGATTTTGTTGTT</td><td>>MI0000264|hsa-mir-7-2&WithFlank&15|+|88611819|88611940</td><td>38</td><td>61</td><td>24</td><td></td><td>isomiR</td><td>NO</td><td>96.58</td><td>127.70</td><td>100.58</td><td>23.14</td></tr>
<tr><td>>MI0000272|hsa-mir-182&WithFlank&7|-|129770377|129770498@29.52.24&[MIMAT0000259&hsa-miR-182-5p&offsets|0|0;m-22&7|-|129770449|129770470&offsets|0|+2]|TTTGGCAATGGTAGAACTCACACT</td><td>>MI0000272|hsa-mir-182&WithFlank&7|-|129770377|129770498</td><td>29</td><td>52</td><td>24</td><td></td><td>isomiR</td><td>NO</td><td>85.21</td><td>73.44</td><td>29.27</td><td>0.00</td></tr>
<tr><td>>MI0000273|hsa-mir-183&WithFlank&7|-|129774899|129775020@33.54.22&[MIMAT0000261&hsa-miR-183-5p&offsets|0|0;m-41&7|-|129774966|129774987&offsets|-1|-1]|TATGGCACTGGTAGAATTCACT</td><td>>MI0000273|hsa-mir-183&WithFlank&7|-|129774899|129775020</td><td>33</td><td>54</td><td>22</td><td></td><td>isomiR</td><td>NO</td><td>59.11</td><td>0.00</td><td>28.50</td><td>0.00</td></tr>
<tr><td>>MI0000285|hsa-mir-205&WithFlank&1|+|209432127|209432248@40.60.21&[MIMAT0000266&hsa-miR-205-5p&offsets|0|-1;m-351&1|+|209432166|209432187&offsets|0|-1]|TCCTTCATTCCACCGGAGTCT</td><td>>MI0000285|hsa-mir-205&WithFlank&1|+|209432127|209432248</td><td>40</td><td>60</td><td>21</td><td></td><td>isomiR</td><td>NO</td><td>0.00</td><td>0.00</td><td>49.60</td><td>0.00</td></tr>
<tr><td>>MI0000285|hsa-mir-205&WithFlank&1|+|209432127|209432248@40.60.21(+1A)&[MIMAT0000266&hsa-miR-205-5p&offsets|0|-1(+1A);m-351&1|+|209432166|209432187&offsets|0|-1(+1A)]|TCCTTCATTCCACCGGAGTCTA</td><td>>MI0000285|hsa-mir-205&WithFlank&1|+|209432127|209432248</td><td>40</td><td>60</td><td>21</td><td>(+1A)</td><td>isomiR</td><td>NO</td><td>0.00</td><td>0.00</td><td>39.07</td><td>0.00</td></tr>
</table>

### MDA-MB-468

<table>
<tr><td>>MI0000285|hsa-mir-205&WithFlank&1|+|209432127|209432248@40.61.22&[MIMAT0000266&hsa-miR-205-5p&offsets|0|0;m-351&1|+|209432166|209432187&offsets|0|0]|TCCTTCATTCCACCGGAGTCTG</td><td>>MI0000285|hsa-mir-205&WithFlank&1|+|209432127|209432248</td><td>40</td><td>61</td><td>22</td><td></td><td>isomiR</td><td>NO</td><td>64.66</td><td>70.12</td><td>211.98</td><td>36.39</td></tr>
<tr><td>>MI0000285|hsa-mir-205&WithFlank&1|+|209432127|209432248@40.61.22(+1A)&[MIMAT0000266&hsa-miR-205-5p&offsets|0|0(+1A);m-351&1|+|209432166|209432187&offsets|0|0(+1A)]|TCCTTCATTCCACCGGAGTCTGA</td><td>>MI0000285|hsa-mir-205&WithFlank&1|+|209432127|209432248</td><td>40</td><td>61</td><td>22</td><td>(+1A)</td><td>isomiR</td><td>NO</td><td>155.07</td><td>124.25</td><td>91.01</td><td>69.97</td></tr>
<tr><td>>MI0000285|hsa-mir-205&WithFlank&1|+|209432127|209432248@40.62.23&[MIMAT0000266&hsa-miR-205-5p&offsets|0|+1;m-351&1|+|209432166|209432187&offsets|0|+1]|TCCTTCATTCCACCGGAGTCTGT</td><td>>MI0000285|hsa-mir-205&WithFlank&1|+|209432127|209432248</td><td>40</td><td>62</td><td>23</td><td></td><td>isomiR</td><td>NO</td><td>77.62</td><td>87.27</td><td>80.92</td><td>26.44</td></tr>
<tr><td>>MI0000289|hsa-mir-181a-1&WithFlank&1|-|198859038|198859159@30.52.23;MI0000269|hsa-mir-181a-2&WithFlank&9|+|124692436|124692557@45.67.23&[MIMAT0000256&hsa-miR-181a-5p&offsets|0|0;m-32&1|-|198859108|198859130&offsets|0|0];[MIMAT0000256_1&hsa-miR-181a-5p&offsets|0|0;m-32&9|+|124692480|124692502&offsets|0|0]|AACATTCAACGCTGTCGGTGAGT</td><td>>MI0000289|hsa-mir-181a-1&WithFlank&1|-|198859038|19885915</td><td>30</td><td>52</td><td>23</td><td></td><td>isomiR</td><td>NO</td><td>0.00</td><td>0.00</td><td>50.61</td><td>0.00</td></tr>
<tr><td>>MI0000298|hsa-mir-221&WithFlank&X|-|45746151|45746272@71.92.22&[MIMAT0000278&hsa-miR-221-3p&offsets|0|-1;m-91&X|-|45746181|45746202&offsets|0|0]|AGCTACATTGTCTGCTGGGTTT</td><td>>MI0000298|hsa-mir-221&WithFlank&X|-|45746151|45746272</td><td>71</td><td>92</td><td>22</td><td></td><td>isomiR</td><td>NO</td><td>0.00</td><td>0.00</td><td>66.01</td><td>0.00</td></tr>
<tr><td>>MI0000298|hsa-mir-221&WithFlank&X|-|45746151|45746272@71.93.23&[MIMAT0000278&hsa-miR-221-3p&offsets|0|0;m-91&X|-|45746181|45746202&offsets|0|+1]|AGCTACATTGTCTGCTGGGTTTC</td><td>>MI0000298|hsa-mir-221&WithFlank&X|-|45746151|45746272</td><td>71</td><td>93</td><td>23</td><td></td><td>isomiR</td><td>NO</td><td>54.24</td><td>89.29</td><td>120.89</td><td>6.63</td></tr>
<tr><td>>MI0000299|hsa-mir-222&WithFlank&X|-|45747009|45747130@75.98.24&[MIMAT0000279&hsa-miR-222-3p&offsets|0|+3;m-140&X|-|45747033|45747056&offsets|0|0]|AGCTACATCTGGCTACTGGGTCTC</td><td>>MI0000299|hsa-mir-222&WithFlank&X|-|45747009|45747130</td><td>75</td><td>98</td><td>24</td><td></td><td>isomiR</td><td>NO</td><td>32.01</td><td>0.00</td><td>0.00</td><td>0.00</td></tr>
<tr><td>>MI0000299|hsa-mir-222&WithFlank&X|-|45747009|45747130@75.99.25&[MIMAT0000279&hsa-miR-222-3p&offsets|0|+4;m-140&X|-|45747033|45747056&offsets|0|+1]|AGCTACATCTGGCTACTGGGTCTCT</td><td>>MI0000299|hsa-mir-222&WithFlank&X|-|45747009|45747130</td><td>75</td><td>99</td><td>25</td><td></td><td>isomiR</td><td>NO</td><td>42.56</td><td>0.00</td><td>0.00</td><td>0.00</td></tr>
<tr><td>>MI0000342|hsa-mir-200b&WithFlank&1|+|1167098|1167204@63.83.21&[MIMAT0000318&hsa-miR-200b-3p&offsets|0|-1;m-93&1|+|1167160|1167182&offsets|0|-2]|TAATACTGCCTGGTAATGATG</td><td>>MI0000342|hsa-mir-200b&WithFlank&1|+|1167098|1167204</td><td>63</td><td>83</td><td>21</td><td></td><td>isomiR</td><td>NO</td><td>0.00</td><td>0.00</td><td>55.99</td><td>0.00</td></tr>
<tr><td>>MI0000342|hsa-mir-200b&WithFlank&1|+|1167098|1167204@63.84.22&[MIMAT0000318&hsa-miR-200b-3p&offsets|0|0;m-93&1|+|1167160|1167182&offsets|0|-1]|TAATACTGCCTGGTAATGATGA</td><td>>MI0000342|hsa-mir-200b&WithFlank&1|+|1167098|1167204</td><td>63</td><td>84</td><td>22</td><td></td><td>isomiR</td><td>NO</td><td>62.12</td><td>43.04</td><td>120.98</td><td>16.17</td></tr>
<tr><td>>MI0000342|hsa-mir-200b&WithFlank&1|+|1167098|1167204@63.84.22(+1U)&[MIMAT0000318&hsa-miR-200b-3p&offsets|0|0(+1U);m-93&1|+|1167160|1167182&offsets|0|-1(+1U)]|TAATACTGCCTGGTAATGATGAT</td><td>>MI0000342|hsa-mir-200b&WithFlank&1|+|1167098|1167204</td><td>63</td><td>84</td><td>22</td><td>(+1U)</td><td>isomiR</td><td>NO</td><td>12.81</td><td>0.00</td><td>0.00</td><td>0.00</td></tr>
<tr><td>>MI0000342|hsa-mir-200b&WithFlank&1|+|1167098|1167204@63.85.23&[MIMAT0000318&hsa-miR-200b-3p&offsets|0|+1;m-93&1|+|1167160|1167182&offsets|0|0]|TAATACTGCCTGGTAATGATGAC</td><td>>MI0000342|hsa-mir-200b&WithFlank&1|+|1167098|1167204</td><td>63</td><td>85</td><td>23</td><td></td><td>isomiR</td><td>NO</td><td>278.29</td><td>157.24</td><td>82.27</td><td>53.08</td></tr>
<tr><td>>MI0000342|hsa-mir-200b&WithFlank&1|+|1167098|1167204@63.85.23(+1A)&[MIMAT0000318&hsa-miR-200b-3p&offsets|0|+1(+1A);m-93&1|+|1167160|1167182&offsets|0|0(+1A)]|TAATACTGCCTGGTAATGATGACA</td><td>>MI0000342|hsa-mir-200b&WithFlank&1|+|1167098|1167204</td><td>63</td><td>85</td><td>23</td><td>(+1A)</td><td>isomiR</td><td>NO</td><td>43.89</td><td>44.36</td><td>13.66</td><td>6.05</td></tr>
<tr><td>>MI0000342|hsa-mir-200b&WithFlank&1|+|1167098|1167204@63.85.23(+1U)&[MIMAT0000318&hsa-miR-200b-3p&offsets|0|+1(+1U);m-93&1|+|1167160|1167182&offsets|0|0(+1U)]|TAATACTGCCTGGTAATGATGACT</td><td>>MI0000342|hsa-mir-200b&WithFlank&1|+|1167098|1167204</td><td>63</td><td>85</td><td>23</td><td>(+1U)</td><td>isomiR</td><td>NO</td><td>111.62</td><td>78.07</td><td>29.65</td><td>36.14</td></tr>
<tr><td>>MI0000342|hsa-mir-200b&WithFlank&1|+|1167098|1167204@64.85.22(+1U)&[MIMAT0000318&hsa-miR-200b-3p&offsets|+1|+1(+1U);m-93&1|+|1167160|1167182&offsets|+1|0(+1U)]|AATACTGCCTGGTAATGATGACT</td><td>>MI0000342|hsa-mir-200b&WithFlank&1|+|1167098|1167204</td><td>64</td><td>85</td><td>22</td><td>(+1U)</td><td>isomiR</td><td>NO</td><td>28.83</td><td>0.00</td><td>0.00</td><td>0.00</td></tr>
<tr><td>>MI0000433|hsa-let-7g&WithFlank&3|-|52268272|52268367@11.31.21(+1C)&[MIMAT0000414&hsa-let-7g-5p&offsets|0|-1(+1C);m-58&3|-|52268336|52268357&offsets|0|-1(+1C)]|TGAGGTAGTAGTTTGTACAGTC</td><td>>MI0000433|hsa-let-7g&WithFlank&3|-|52268272|52268367</td><td>11</td><td>31</td><td>21</td><td>(+1C)</td><td>isomiR</td><td>NO</td><td>35.12</td><td>0.00</td><td>0.00</td><td>0.00</td></tr>
<tr><td>>MI0000433|hsa-let-7g&WithFlank&3|-|52268272|52268367@11.32.22&[MIMAT0000414&hsa-let-7g-5p&offsets|0|0;m-58&3|-|52268336|52268357&offsets|0|0]|TGAGGTAGTAGTTTGTACAGTT</td><td>>MI0000433|hsa-let-7g&WithFlank&3|-|52268272|52268367</td><td>11</td><td>32</td><td>22</td><td></td><td>isomiR</td><td>NO</td><td>291.60</td><td>125.02</td><td>69.23</td><td>35.70</td></tr>
<tr><td>>MI0000434|hsa-let-7i&WithFlank&12|+|62603680|62603775@12.32.21(+1C)&[MIMAT0000415&hsa-let-7i-5p&offsets|0|-1(+1C);m-49&12|+|62603691|62603712&offsets|0|-1(+1C)]|TGAGGTAGTAGTTTGTGCTGTC</td><td>>MI0000434|hsa-let-7i&WithFlank&12|+|62603680|62603775</td><td>12</td><td>32</td><td>21</td><td>(+1C)</td><td>isomiR</td><td>NO</td><td>31.42</td><td>0.00</td><td>35.51</td><td>0.00</td></tr>
<tr><td>>MI0000434|hsa-let-7i&WithFlank&12|+|62603680|62603775@12.33.22&[MIMAT0000415&hsa-let-7i-5p&offsets|0|0;m-49&12|+|62603691|62603712&offsets|0|0]|TGAGGTAGTAGTTTGTGCTGTT</td><td>>MI0000434|hsa-let-7i&WithFlank&12|+|62603680|62603775</td><td>12</td><td>33</td><td>22</td><td></td><td>isomiR</td><td>NO</td><td>284.24</td><td>185.65</td><td>199.84</td><td>25.87</td></tr>
<tr><td>>MI0000434|hsa-let-7i&WithFlank&12|+|62603680|62603775@12.33.22(+1A)&[MIMAT0000415&hsa-let-7i-5p&offsets|0|0(+1A);m-49&12|+|62603691|62603712&offsets|0|0(+1A)]|TGAGGTAGTAGTTTGTGCTGTTA</td><td>>MI0000434|hsa-let-7i&WithFlank&12|+|62603680|62603775</td><td>12</td><td>33</td><td>22</td><td>(+1A)</td><td>isomiR</td><td>NO</td><td>0.00</td><td>0.00</td><td>36.71</td><td>0.00</td></tr>
</table>

### MDA-MB-468

<table>
<tr><td>>MI0000440|hsa-mir-27b&WithFlank&9|+|95085439|95085547@67.86.20&[MIMAT0000419&hsa-miR-27b-3p&offsets|0|-1;m-39&9|+|95085505|95085525&offsets|0|-1]|TTCACAGTGGCTAAGTTCTG</td><td>>MI0000440|hsa-mir-27b&WithFlank&9|+|95085439|95085547</td><td>67</td><td>86</td><td>20</td><td></td><td>isomiR</td><td>NO</td><td>0.00</td><td>0.00</td><td>13.56</td><td>0.00</td></tr>
<tr><td>>MI0000440|hsa-mir-27b&WithFlank&9|+|95085439|95085547@67.87.21&[MIMAT0000419&hsa-miR-27b-3p&offsets|0|0;m-39&9|+|95085505|95085525&offsets|0|0]|TTCACAGTGGCTAAGTTCTGC</td><td>>MI0000440|hsa-mir-27b&WithFlank&9|+|95085439|95085547</td><td>67</td><td>87</td><td>21</td><td></td><td>isomiR</td><td>NO</td><td>40.98</td><td>51.64</td><td>93.67</td><td>13.78</td></tr>
<tr><td>>MI0000447|hsa-mir-128-1&WithFlank&2|+|135665391|135665484@56.76.21;MI0000727|hsa-mir-128-2&WithFlank&3|+|35744470|35744565@58.78.21&[MIMAT0000424&hsa-miR-128-3p&offsets|0|0;m-108&2|+|135665446|135665466&offsets|0|0];[MIMAT0000424_1&hsa-miR-128-3p&offsets|0|0;m-117&3|+|35744527|35744547&offsets|0|0]|TCACAGTGAACCGGTCTCTTT</td><td>>MI0000447|hsa-mir-128-1&WithFlank&2|+|135665391|135665484</td><td>56</td><td>76</td><td>21</td><td></td><td>isomiR</td><td>NO</td><td>74.72</td><td>28.85</td><td>44.08</td><td>16.10</td></tr>
<tr><td>>MI0000447|hsa-mir-128-1&WithFlank&2|+|135665391|135665484@56.77.22&[MIMAT0000424&hsa-miR-128-3p&offsets|0|+1;m-108&2|+|135665446|135665466&offsets|0|+1]|TCACAGTGAACCGGTCTCTTTT</td><td>>MI0000447|hsa-mir-128-1&WithFlank&2|+|135665391|135665484</td><td>56</td><td>77</td><td>22</td><td></td><td>isomiR</td><td>NO</td><td>20.30</td><td>0.00</td><td>0.00</td><td>0.00</td></tr>
<tr><td>>MI0000459|hsa-mir-143&WithFlank&5|+|149428912|149429029@67.87.21&[MIMAT0000435&hsa-miR-143-3p&offsets|0|0;m-3&5|+|149428978|149428997&offsets|0|+1]|TGAGATGAAGCACTGTAGCTC</td><td>>MI0000459|hsa-mir-143&WithFlank&5|+|149428912|149429029</td><td>67</td><td>87</td><td>21</td><td></td><td>isomiR</td><td>NO</td><td>0.00</td><td>0.00</td><td>31.31</td><td>0.00</td></tr>
<tr><td>>MI0000465|hsa-mir-191&WithFlank&3|-|49020612|49020715@22.43.22&[MIMAT0000440&hsa-miR-191-5p&offsets|0|-1;m-48&3|-|49020672|49020694&offsets|0|-1]|CAACGGAATCCCAAAAGCAGCT</td><td>>MI0000465|hsa-mir-191&WithFlank&3|-|49020612|49020715</td><td>22</td><td>43</td><td>22</td><td></td><td>isomiR</td><td>NO</td><td>32.69</td><td>0.00</td><td>35.19</td><td>0.00</td></tr>
<tr><td>>MI0000465|hsa-mir-191&WithFlank&3|-|49020612|49020715@22.44.23&[MIMAT0000440&hsa-miR-191-5p&offsets|0|0;m-48&3|-|49020672|49020694&offsets|0|0]|CAACGGAATCCCAAAAGCAGCTG</td><td>>MI0000465|hsa-mir-191&WithFlank&3|-|49020612|49020715</td><td>22</td><td>44</td><td>23</td><td></td><td>isomiR</td><td>NO</td><td>47.10</td><td>5.99</td><td>59.35</td><td>0.00</td></tr>
<tr><td>>MI0000469|hsa-mir-125a&WithFlank&19|+|51693248|51693345@21.42.22&[MIMAT0000443&hsa-miR-125a-5p&offsets|0|-2;m-63&19|+|51693268|51693289&offsets|0|0]|TCCCTGAGACCCTTTAACCTGT</td><td>>MI0000469|hsa-mir-125a&WithFlank&19|+|51693248|51693345</td><td>21</td><td>42</td><td>22</td><td></td><td>isomiR</td><td>NO</td><td>19.73</td><td>0.00</td><td>0.00</td><td>0.00</td></tr>
<tr><td>>MI0000469|hsa-mir-125a&WithFlank&19|+|51693248|51693345@21.43.23&[MIMAT0000443&hsa-miR-125a-5p&offsets|0|-1;m-63&19|+|51693268|51693289&offsets|0|+1]|TCCCTGAGACCCTTTAACCTGTG</td><td>>MI0000469|hsa-mir-125a&WithFlank&19|+|51693248|51693345</td><td>21</td><td>43</td><td>23</td><td></td><td>isomiR</td><td>NO</td><td>80.66</td><td>28.89</td><td>0.00</td><td>6.59</td></tr>
<tr><td>>MI0000650|hsa-mir-200c&WithFlank&12|+|6963693|6963772@50.71.22&[MIMAT0000617&hsa-miR-200c-3p&offsets|0|-1;m-54&12|+|6963742|6963764&offsets|0|-1]|TAATACTGCCGGGTAATGATGG</td><td>>MI0000650|hsa-mir-200c&WithFlank&12|+|6963693|6963772</td><td>50</td><td>71</td><td>22</td><td></td><td>isomiR</td><td>NO</td><td>6.00</td><td>0.00</td><td>31.47</td><td>0.00</td></tr>
<tr><td>>MI0000650|hsa-mir-200c&WithFlank&12|+|6963693|6963772@50.72.23&[MIMAT0000617&hsa-miR-200c-3p&offsets|0|0;m-54&12|+|6963742|6963764&offsets|0|0]|TAATACTGCCGGGTAATGATGGA</td><td>>MI0000650|hsa-mir-200c&WithFlank&12|+|6963693|6963772</td><td>50</td><td>72</td><td>23</td><td></td><td>isomiR</td><td>NO</td><td>162.60</td><td>78.71</td><td>92.60</td><td>36.33</td></tr>
<tr><td>>MI0000734|hsa-mir-106b&WithFlank&7|-|100093987|100094080@58.79.22&[MIMAT0004672&hsa-miR-106b-3p&offsets|0|0;m-87&7|-|100094003|100094023&offsets|0|+1]|CCGCACTGTGGGTACTTGCTGC</td><td>>MI0000734|hsa-mir-106b&WithFlank&7|-|100093987|100094080</td><td>58</td><td>79</td><td>22</td><td></td><td>isomiR</td><td>NO</td><td>30.85</td><td>0.00</td><td>0.00</td><td>0.00</td></tr>
<tr><td>>MI0000736|hsa-mir-30c-1&WithFlank&1|+|40757278|40757378@23.45.23;MI0000254|hsa-mir-30c-2&WithFlank&6|-|71376954|71377037@13.35.23&[MIMAT0000244&hsa-miR-30c-5p&offsets|0|0;m-57&1|+|40757300|40757323&offsets|0|-1];[MIMAT0000244_1&hsa-miR-30c-5p&offsets|0|0;m-56&6|-|71377002|71377025&offsets|0|-1]|TGTAAACATCCTACACTCTCAGC</td><td>>MI0000736|hsa-mir-30c-1&WithFlank&1|+|40757278|40757378</td><td>23</td><td>45</td><td>23</td><td></td><td>isomiR</td><td>NO</td><td>113.68</td><td>30.69</td><td>0.00</td><td>15.31</td></tr>
<tr><td>>MI0000736|hsa-mir-30c-1&WithFlank&1|+|40757278|40757378@23.46.24;MI0000254|hsa-mir-30c-2&WithFlank&6|-|71376954|71377037@13.36.24&[MIMAT0000244&hsa-miR-30c-5p&offsets|0|+1;m-57&1|+|40757300|40757323&offsets|0|0];[MIMAT0000244_1&hsa-miR-30c-5p&offsets|0|+1;m-56&6|-|71377002|71377025&offsets|0|0]|TGTAAACATCCTACACTCTCAGCT</td><td>>MI0000736|hsa-mir-30c-1&WithFlank&1|+|40757278|40757378</td><td>23</td><td>46</td><td>24</td><td></td><td>isomiR</td><td>NO</td><td>145.67</td><td>45.06</td><td>6.75</td><td>39.83</td></tr>
<tr><td>>MI0000737|hsa-mir-200a&WithFlank&1|+|1167857|1167958@22.43.22&[MIMAT0001620&hsa-miR-200a-5p&offsets|0|0;m-154&1|+|1167878|1167899&offsets|0|0]|CATCTTACCGGACAGTGCTGGA</td><td>>MI0000737|hsa-mir-200a&WithFlank&1|+|1167857|1167958</td><td>22</td><td>43</td><td>22</td><td></td><td>isomiR</td><td>NO</td><td>0.00</td><td>0.00</td><td>47.07</td><td>0.00</td></tr>
<tr><td>>MI0000737|hsa-mir-200a&WithFlank&1|+|1167857|1167958@60.81.22&[MIMAT0000682&hsa-miR-200a-3p&offsets|0|0;m-142&1|+|1167916|1167938&offsets|0|-1]|TAACACTGTCTGGTAACGATGT</td><td>>MI0000737|hsa-mir-200a&WithFlank&1|+|1167857|1167958</td><td>60</td><td>81</td><td>22</td><td></td><td>isomiR</td><td>NO</td><td>0.00</td><td>0.00</td><td>72.55</td><td>0.00</td></tr>
<tr><td>>MI0000737|hsa-mir-200a&WithFlank&1|+|1167857|1167958@60.81.22(+1C)&[MIMAT0000682&hsa-miR-200a-3p&offsets|0|0(+1C);m-142&1|+|1167916|1167938&offsets|0|-1(+1C)]|TAACACTGTCTGGTAACGATGTC</td><td>>MI0000737|hsa-mir-200a&WithFlank&1|+|1167857|1167958</td><td>60</td><td>81</td><td>22</td><td>(+1C)</td><td>isomiR</td><td>NO</td><td>0.00</td><td>0.00</td><td>49.05</td><td>0.00</td></tr>
<tr><td>>MI0000737|hsa-mir-200a&WithFlank&1|+|1167857|1167958@60.82.23&[MIMAT0000682&hsa-miR-200a-3p&offsets|0|+1;m-142&1|+|1167916|1167938&offsets|0|0]|TAACACTGTCTGGTAACGATGTT</td><td>>MI0000737|hsa-mir-200a&WithFlank&1|+|1167857|1167958</td><td>60</td><td>82</td><td>23</td><td></td><td>isomiR</td><td>NO</td><td>0.00</td><td>12.77</td><td>259.73</td><td>14.65</td></tr>
<tr><td>>MI0000737|hsa-mir-200a&WithFlank&1|+|1167857|1167958@60.82.23(+1U)&[MIMAT0000682&hsa-miR-200a-3p&offsets|0|+1(+1U);m-142&1|+|1167916|1167938&offsets|0|0(+1U)]|TAACACTGTCTGGTAACGATGTTT</td><td>>MI0000737|hsa-mir-200a&WithFlank&1|+|1167857|1167958</td><td>60</td><td>82</td><td>23</td><td>(+1U)</td><td>isomiR</td><td>NO</td><td>0.00</td><td>0.00</td><td>58.06</td><td>0.00</td></tr>
<tr><td>>MI0000746|hsa-mir-99b&WithFlank&19|+|51692606|51692687@13.33.21&[MIMAT0000689&hsa-miR-99b-5p&offsets|0|-1;m-10&19|+|51692618|51692639&offsets|0|-1]|CACCCGTAGAACCGACCTTGC</td><td>>MI0000746|hsa-mir-99b&WithFlank&19|+|51692606|51692687</td><td>13</td><td>33</td><td>21</td><td></td><td>isomiR</td><td>NO</td><td>56.63</td><td>0.00</td><td>6.54</td><td>0.00</td></tr>
</table>

### MDA-MB-468

<table>
<tr><td>>MI0000746|hsa-mir-99b&WithFlank&19|+|51692606|51692687@13.34.22&[MIMAT0000689&hsa-miR-99b-5p&offsets|0|0;m-10&19|+|51692618|51692639&offsets|0|0]|CACCCGTAGAACCGACCTTGCG</td><td>>MI0000746|hsa-mir-99b&WithFlank&19|+|51692606|51692687</td><td>13</td><td>34</td><td>22</td><td></td><td>isomiR</td><td>NO</td><td>121.58</td><td>85.66</td><td>59.12</td><td>6.06</td></tr>
<tr><td>>MI0000749|hsa-mir-30e&WithFlank&1|+|40754349|40754452@23.45.23(+1A)&[MIMAT0000692&hsa-miR-30e-5p&offsets|0|+1(+1A);m-24&1|+|40754371|40754394&offsets|0|-1(+1A)]|TGTAAACATCCTTGACTGGAAGCA</td><td>>MI0000749|hsa-mir-30e&WithFlank&1|+|40754349|40754452</td><td>23</td><td>45</td><td>23</td><td>(+1A)</td><td>isomiR</td><td>YES</td><td>0.00</td><td>0.00</td><td>31.19</td><td>0.00</td></tr>
<tr><td>>MI0000749|hsa-mir-30e&WithFlank&1|+|40754349|40754452@23.45.23(+1C)&[MIMAT0000692&hsa-miR-30e-5p&offsets|0|+1(+1C);m-24&1|+|40754371|40754394&offsets|0|-1(+1C)]|TGTAAACATCCTTGACTGGAAGCC</td><td>>MI0000749|hsa-mir-30e&WithFlank&1|+|40754349|40754452</td><td>23</td><td>45</td><td>23</td><td>(+1C)</td><td>isomiR</td><td>NO</td><td>0.00</td><td>0.00</td><td>49.67</td><td>0.00</td></tr>
<tr><td>>MI0000749|hsa-mir-30e&WithFlank&1|+|40754349|40754452@23.46.24&[MIMAT0000692&hsa-miR-30e-5p&offsets|0|+2;m-24&1|+|40754371|40754394&offsets|0|0]|TGTAAACATCCTTGACTGGAAGCT</td><td>>MI0000749|hsa-mir-30e&WithFlank&1|+|40754349|40754452</td><td>23</td><td>46</td><td>24</td><td></td><td>isomiR</td><td>NO</td><td>0.00</td><td>0.00</td><td>246.80</td><td>0.00</td></tr>
<tr><td>>MI0000749|hsa-mir-30e&WithFlank&1|+|40754349|40754452@65.85.21(+1U)&[MIMAT0000693&hsa-miR-30e-3p&offsets|0|-1(+1U);m-25&1|+|40754413|40754434&offsets|0|-1(+1U)]|CTTTCAGTCGGATGTTTACAGT</td><td>>MI0000749|hsa-mir-30e&WithFlank&1|+|40754349|40754452</td><td>65</td><td>85</td><td>21</td><td>(+1U)</td><td>isomiR</td><td>NO</td><td>22.01</td><td>0.00</td><td>30.90</td><td>0.00</td></tr>
<tr><td>>MI0000749|hsa-mir-30e&WithFlank&1|+|40754349|40754452@65.86.22&[MIMAT0000693&hsa-miR-30e-3p&offsets|0|0;m-25&1|+|40754413|40754434&offsets|0|0]|CTTTCAGTCGGATGTTTACAGC</td><td>>MI0000749|hsa-mir-30e&WithFlank&1|+|40754349|40754452</td><td>65</td><td>86</td><td>22</td><td></td><td>isomiR</td><td>NO</td><td>35.51</td><td>0.00</td><td>32.28</td><td>0.00</td></tr>
<tr><td>>MI0000750|hsa-mir-26a-2&WithFlank&12|-|57824603|57824698@20.40.21(+1C);MI0000083|hsa-mir-26a-1&WithFlank&3|+|37969398|37969486@16.36.21(+1C)&[MIMAT0000082&hsa-miR-26a-5p&offsets|0|-1(+1C);m-33&12|-|57824658|57824679&offsets|0|-1(+1C)];[MIMAT0000082_1&hsa-miR-26a-5p&offsets|0|-1(+1C);m-34&3|+|37969413|37969434&offsets|0|-1(+1C)]|TTCAAGTAATCCAGGATAGGCC</td><td>>MI0000750|hsa-mir-26a-2&WithFlank&12|-|57824603|57824698</td><td>20</td><td>40</td><td>21</td><td>(+1C)</td><td>isomiR</td><td>NO</td><td>30.26</td><td>12.58</td><td>30.14</td><td>0.00</td></tr>
<tr><td>>MI0000750|hsa-mir-26a-2&WithFlank&12|-|57824603|57824698@20.41.22;MI0000083|hsa-mir-26a-1&WithFlank&3|+|37969398|37969486@16.37.22&[MIMAT0000082&hsa-miR-26a-5p&offsets|0|0;m-33&12|-|57824658|57824679&offsets|0|0];[MIMAT0000082_1&hsa-miR-26a-5p&offsets|0|0;m-34&3|+|37969413|37969434&offsets|0|0]|TTCAAGTAATCCAGGATAGGCT</td><td>>MI0000750|hsa-mir-26a-2&WithFlank&12|-|57824603|57824698</td><td>20</td><td>41</td><td>22</td><td></td><td>isomiR</td><td>NO</td><td>222.45</td><td>173.88</td><td>147.88</td><td>90.77</td></tr>
<tr><td>>MI0000764|hsa-mir-363&WithFlank&X|-|134169372|134169458@56.77.22&[MIMAT0000707&hsa-miR-363-3p&offsets|0|0;m-178&X|-|134169381|134169403&offsets|0|-1]|AATTGCACGGTATCCATCTGTA</td><td>>MI0000764|hsa-mir-363&WithFlank&X|-|134169372|134169458</td><td>56</td><td>77</td><td>22</td><td></td><td>isomiR</td><td>NO</td><td>0.00</td><td>0.00</td><td>20.99</td><td>0.00</td></tr>
<tr><td>>am_MI0000786|hsa-mir-378a&WithFlank&5|+|149732819|149732896@49.69.21&[MIMAT0000732&hsa-miR-378a-3p&offsets|0|-1;m-44&5|+|149732867|149732888&offsets|0|-1]|ACTGGACTTGGAGTCAGAAGG</td><td>>MI0000786|hsa-mir-378a&WithFlank&5|+|149732819|149732896.a</td><td>49</td><td>69</td><td>21</td><td></td><td>isomiR</td><td>NO</td><td>0.00</td><td>0.00</td><td>102.88</td><td>0.00</td></tr>
<tr><td>>am_MI0000786|hsa-mir-378a&WithFlank&5|+|149732819|149732896@49.69.21(+1A)&[MIMAT0000732&hsa-miR-378a-3p&offsets|0|-1(+1A);m-44&5|+|149732867|149732888&offsets|0|-1(+1A)]|ACTGGACTTGGAGTCAGAAGGA</td><td>>MI0000786|hsa-mir-378a&WithFlank&5|+|149732819|149732896.a</td><td>49</td><td>69</td><td>21</td><td>(+1A)</td><td>isomiR</td><td>NO</td><td>0.00</td><td>0.00</td><td>50.28</td><td>0.00</td></tr>
<tr><td>>am_MI0000786|hsa-mir-378a&WithFlank&5|+|149732819|149732896@49.69.21(+1U)&[MIMAT0000732&hsa-miR-378a-3p&offsets|0|-1(+1U);m-44&5|+|149732867|149732888&offsets|0|-1(+1U)]|ACTGGACTTGGAGTCAGAAGGT</td><td>>MI0000786|hsa-mir-378a&WithFlank&5|+|149732819|149732896.a</td><td>49</td><td>69</td><td>21</td><td>(+1U)</td><td>isomiR</td><td>NO</td><td>0.00</td><td>0.00</td><td>118.33</td><td>0.00</td></tr>
<tr><td>>am_MI0000786|hsa-mir-378a&WithFlank&5|+|149732819|149732896@49.70.22&[MIMAT0000732&hsa-miR-378a-3p&offsets|0|0;m-44&5|+|149732867|149732888&offsets|0|0]|ACTGGACTTGGAGTCAGAAGGC</td><td>>MI0000786|hsa-mir-378a&WithFlank&5|+|149732819|149732896.a</td><td>49</td><td>70</td><td>22</td><td></td><td>isomiR</td><td>NO</td><td>120.64</td><td>86.52</td><td>396.01</td><td>6.34</td></tr>
<tr><td>>am_MI0000786|hsa-mir-378a&WithFlank&5|+|149732819|149732896@49.70.22(+1A)&[MIMAT0000732&hsa-miR-378a-3p&offsets|0|0(+1A);m-44&5|+|149732867|149732888&offsets|0|0(+1A)]|ACTGGACTTGGAGTCAGAAGGCA</td><td>>MI0000786|hsa-mir-378a&WithFlank&5|+|149732819|149732896.a</td><td>49</td><td>70</td><td>22</td><td>(+1A)</td><td>isomiR</td><td>NO</td><td>0.00</td><td>0.00</td><td>69.66</td><td>0.00</td></tr>
<tr><td>>am_MI0000786|hsa-mir-378a&WithFlank&5|+|149732819|149732896@49.70.22(+1U)&[MIMAT0000732&hsa-miR-378a-3p&offsets|0|0(+1U);m-44&5|+|149732867|149732888&offsets|0|0(+1U)]|ACTGGACTTGGAGTCAGAAGGCT</td><td>>MI0000786|hsa-mir-378a&WithFlank&5|+|149732819|149732896.a</td><td>49</td><td>70</td><td>22</td><td>(+1U)</td><td>isomiR</td><td>NO</td><td>0.00</td><td>0.00</td><td>65.42</td><td>0.00</td></tr>
<tr><td>>am_MI0000786|hsa-mir-378a&WithFlank&5|+|149732819|149732896@50.70.21&[MIMAT0000732&hsa-miR-378a-3p&offsets|+1|0;m-44&5|+|149732867|149732888&offsets|+1|0]|CTGGACTTGGAGTCAGAAGGC</td><td>>MI0000786|hsa-mir-378a&WithFlank&5|+|149732819|149732896.a</td><td>50</td><td>70</td><td>21</td><td></td><td>isomiR</td><td>NO</td><td>0.00</td><td>0.00</td><td>69.77</td><td>0.00</td></tr>
<tr><td>>MI0000802|hsa-mir-340&WithFlank&5|-|180015297|180015403@22.43.22&[MIMAT0004692&hsa-miR-340-5p&offsets|0|0;m-220&5|-|180015361|180015382&offsets|0|0]|TTATAAAGCAATGAGACTGATT</td><td>>MI0000802|hsa-mir-340&WithFlank&5|-|180015297|180015403</td><td>22</td><td>43</td><td>22</td><td></td><td>isomiR</td><td>NO</td><td>0.00</td><td>0.00</td><td>43.56</td><td>0.00</td></tr>
<tr><td>>MI0000809|hsa-mir-151a&WithFlank&8|-|140732558|140732659@53.74.22&[MIMAT0000757&hsa-miR-151a-3p&offsets|0|+1;m-30&8|-|140732587|140732607&offsets|0|+1]|CTAGACTGAAGCTCCTTGAGGA</td><td>>MI0000809|hsa-mir-151a&WithFlank&8|-|140732558|140732659</td><td>53</td><td>74</td><td>22</td><td></td><td>isomiR</td><td>NO</td><td>21.46</td><td>0.00</td><td>13.61</td><td>0.00</td></tr>
<tr><td>>am_MI0000809|hsa-mir-151a&WithFlank&8|-|140732558|140732659@53.73.21&[MIMAT0000757&hsa-miR-151a-3p&offsets|0|0;m-30&8|-|140732587|140732607&offsets|0|0]|CTAGACTGAAGCTCCTTGAGG</td><td>>MI0000809|hsa-mir-151a&WithFlank&8|-|140732558|140732659.a</td><td>53</td><td>73</td><td>21</td><td></td><td>isomiR</td><td>NO</td><td>93.24</td><td>38.37</td><td>93.43</td><td>0.00</td></tr>
</table>

### MDA-MB-468

| | | | | | | | | | | |
|---|---|---|---|---|---|---|---|---|---|---|
| >MI0000810\|hsa-mir-135b&WithFlank&1\|-\|205448296\|205448404@22.44.23&[MIMAT0000758&hsa-miR-135b-5p&offsets\|0\|0;m-321&1\|-\|205448361\|205448383&offsets\|0\|0]\|TATGGCTTTTCATTCCTATGTGA | >MI0000810\|hsa-mir-135b&WithFlank&1\|-\|205448296\|205448404 | 22 | 44 | 23 | isomiR | NO | 0.00 | 0.00 | 20.98 | 0.00 |
| >MI0000811\|hsa-mir-148b&WithFlank&12\|+\|54337210\|54337320@69.90.22&[MIMAT0000759&hsa-miR-148b-3p&offsets\|0\|0;m-77&12\|+\|54337278\|54337299&offsets\|0\|0]\|TCAGTGCATCACAGAACTTTGT | >MI0000811\|hsa-mir-148b&WithFlank&12\|+\|54337210\|54337320 | 69 | 90 | 22 | isomiR | NO | 0.00 | 0.00 | 76.81 | 0.00 |
| >MI0001445\|hsa-mir-423&WithFlank&17\|+\|30117073\|30117178@23.45.23&[MIMAT0004748&hsa-miR-423-5p&offsets\|0\|0;m-141&17\|+\|30117095\|30117117&offsets\|0\|0]\|TGAGGGGCAGAGAGCGAGACTTT | >MI0001445\|hsa-mir-423&WithFlank&17\|+\|30117073\|30117178 | 23 | 45 | 23 | isomiR | NO | 0.00 | 0.00 | 29.83 | 0.00 |
| >MI0001445\|hsa-mir-423&WithFlank&17\|+\|30117073\|30117178@59.81.23&[MIMAT0001340&hsa-miR-423-3p&offsets\|0\|0;m-100&17\|+\|30117131\|30117153&offsets\|0\|0]\|AGCTCGGTCTGAGGCCCCTCAGT | >MI0001445\|hsa-mir-423&WithFlank&17\|+\|30117073\|30117178 | 59 | 81 | 23 | isomiR | NO | 20.29 | 0.00 | 0.00 | 0.00 |
| >MI0001641\|hsa-mir-429&WithFlank&1\|+\|1168999\|1169093@57.78.22&[MIMAT0001536&hsa-miR-429&offsets\|0\|0;m-207&1\|+\|1169055\|1169076&offsets\|0\|0]\|TAATACTGTCTGGTAAAACCGT | >MI0001641\|hsa-mir-429&WithFlank&1\|+\|1168999\|1169093 | 57 | 78 | 22 | isomiR | NO | 0.00 | 26.51 | 95.86 | 0.00 |
| >MI0003205\|hsa-mir-532&WithFlank&X\|+\|50003142\|50003244@26.47.22&[MIMAT0002888&hsa-miR-532-5p&offsets\|0\|0;m-65&X\|+\|50003167\|50003188&offsets\|0\|0]\|CATGCCTTGAGTGTAGGACCGT | >MI0003205\|hsa-mir-532&WithFlank&X\|+\|50003142\|50003244 | 26 | 47 | 22 | isomiR | NO | 0.00 | 0.00 | 38.56 | 0.00 |
| >am_MI0005763\|hsa-mir-941-1&WithFlank&20\|+\|63919443\|63919526@53.75.23;MI0005764\|hsa-mir-941-2&WithFlank&20\|+\|63919499\|63919582@53.75.23;MI0005765\|hsa-mir-941-3&WithFlank&20\|+\|63919555\|63919638@53.75.23;MI0005766\|hsa-mir-941-4&WithFlank&20\|+\|63919750\|63919833@53.75.23;MI0031520\|hsa-mir-941-5&WithFlank&20\|+\|63919862\|63919945@53.75.23&[MIMAT0004984&hsa-miR-941&offsets\|0\|0;m-263&20\|+\|63919495\|63919517&offsets\|0\|0];[MIMAT0004984_1&hsa-miR-941&offsets\|0\|0;m-263&20\|+\|63919551\|63919573&offsets\|0\|0];[MIMAT0004984_2&hsa-miR-941&offsets\|0\|0;m-263&20\|+\|63919607\|63919629&offsets\|0\|0];[MIMAT0004984_3&hsa-miR-941&offsets\|0\|0;m-263&20\|+\|63919802\|63919824&offsets\|0\|0];[MIMAT0004984_4&hsa-miR-941&offsets\|0\|0;m-263&20\|+\|63919914\|63919936&offsets\|0\|0]\|CACCCGGCTGTGTGCACATGTGC | >MI0005763\|hsa-mir-941-1&WithFlank&20\|+\|63919443\|63919526.a | 53 | 75 | 23 | isomiR | NO | 33.51 | 0.00 | 0.00 | 0.00 |
| >MI0025750\|hsa-mir-7974&WithFlank&19\|-\|11495538\|11495628@62.85.24&[MIMAT0031177&hsa-miR-7974&offsets\|0\|0;m-867&19\|-\|11495544\|11495567&offsets\|0\|0]\|AGGCTGTGATGCTCTCCTGAGCCC | >MI0025750\|hsa-mir-7974&WithFlank&19\|-\|11495538\|11495628 | 62 | 85 | 24 | isomiR | NO | 31.82 | 0.00 | 0.00 | 0.00 |
| >Antisense_hg38_21_-_8433172_8446622_RNA45SN1_45S_50ntFlanks@7674.7692.19\|CTCTCTCTCTCTCTCCCCC | >Antisense_hg38_21_-_8433172_8446622_RNA45SN1_45S_50n | 7674 | 7692 | 19 | rRF 45S | NO | 29.42 | 27.07 | 0.00 | 0.00 |
| >Antisense_hg38_21_-_8433172_8446622_RNA45SN1_45S_50ntFlanks@7674.7693.20\|CTCTCTCTCTCTCTCCCCCG | >Antisense_hg38_21_-_8433172_8446622_RNA45SN1_45S_50n | 7674 | 7693 | 20 | rRF 45S | NO | 66.55 | 77.56 | 0.00 | 0.00 |
| >hg38_21_+_8433172_8446622_RNA45SN1_45S_50ntFlanks@51.71.21\|GCTGACACGCTGTCCTCTGGC | >hg38_21_+_8433172_8446622_RNA45SN1_45S_50ntFlanks | 51 | 71 | 21 | rRF 45S | NO | 34.86 | 0.00 | 0.00 | 0.00 |
| >hg38_21_+_8433172_8446622_RNA45SN1_45S_50ntFlanks@51.72.22\|GCTGACACGCTGTCCTCTGGCG | >hg38_21_+_8433172_8446622_RNA45SN1_45S_50ntFlanks | 51 | 72 | 22 | rRF 45S | NO | 65.94 | 0.00 | 0.00 | 0.00 |
| >hg38_21_+_8433172_8446622_RNA45SN1_45S_50ntFlanks@475.496.22\|CTTCGTGATCGATGTGGTGACG | >hg38_21_+_8433172_8446622_RNA45SN1_45S_50ntFlanks | 475 | 496 | 22 | rRF 45S | NO | 48.97 | 0.00 | 0.00 | 0.00 |
| >hg38_21_+_8433172_8446622_RNA45SN1_45S_50ntFlanks@475.497.23\|CTTCGTGATCGATGTGGTGACGT | >hg38_21_+_8433172_8446622_RNA45SN1_45S_50ntFlanks | 475 | 497 | 23 | rRF 45S | NO | 54.35 | 0.00 | 0.00 | 0.00 |
| >hg38_21_+_8433172_8446622_RNA45SN1_45S_50ntFlanks@475.498.24\|CTTCGTGATCGATGTGGTGACGTC | >hg38_21_+_8433172_8446622_RNA45SN1_45S_50ntFlanks | 475 | 498 | 24 | rRF 45S | NO | 81.56 | 0.00 | 0.00 | 0.00 |
| >hg38_21_+_8433172_8446622_RNA45SN1_45S_50ntFlanks@475.503.29\|CTTCGTGATCGATGTGGTGACGTCGTGCT | >hg38_21_+_8433172_8446622_RNA45SN1_45S_50ntFlanks | 475 | 503 | 29 | rRF 45S | NO | 51.38 | 0.00 | 0.00 | 0.00 |
| >hg38_21_+_8433172_8446622_RNA45SN1_45S_50ntFlanks@475.504.30\|CTTCGTGATCGATGTGGTGACGTCGTGCTC | >hg38_21_+_8433172_8446622_RNA45SN1_45S_50ntFlanks | 475 | 504 | 30 | rRF 45S | NO | 127.18 | 0.00 | 0.00 | 0.00 |
| >hg38_21_+_8433172_8446622_RNA45SN1_45S_50ntFlanks@475.505.31\|CTTCGTGATCGATGTGGTGACGTCGTGCTCT | >hg38_21_+_8433172_8446622_RNA45SN1_45S_50ntFlanks | 475 | 505 | 31 | rRF 45S | NO | 319.03 | 0.00 | 0.00 | 0.00 |
| >hg38_21_+_8433172_8446622_RNA45SN1_45S_50ntFlanks@475.506.32\|CTTCGTGATCGATGTGGTGACGTCGTGCTCTC | >hg38_21_+_8433172_8446622_RNA45SN1_45S_50ntFlanks | 475 | 506 | 32 | rRF 45S | NO | 273.25 | 0.00 | 0.00 | 0.00 |
| >hg38_21_+_8433172_8446622_RNA45SN1_45S_50ntFlanks@475.507.33\|CTTCGTGATCGATGTGGTGACGTCGTGCTCTCC | >hg38_21_+_8433172_8446622_RNA45SN1_45S_50ntFlanks | 475 | 507 | 33 | rRF 45S | NO | 92.02 | 0.00 | 0.00 | 0.00 |
| >hg38_21_+_8433172_8446622_RNA45SN1_45S_50ntFlanks@475.509.35\|CTTCGTGATCGATGTGGTGACGTCGTGCTCTCCCG | >hg38_21_+_8433172_8446622_RNA45SN1_45S_50ntFlanks | 475 | 509 | 35 | rRF 45S | NO | 85.00 | 0.00 | 0.00 | 0.00 |
| >hg38_21_+_8433172_8446622_RNA45SN1_45S_50ntFlanks@475.510.36\|CTTCGTGATCGATGTGGTGACGTCGTGCTCTCCCGG | >hg38_21_+_8433172_8446622_RNA45SN1_45S_50ntFlanks | 475 | 510 | 36 | rRF 45S | NO | 149.49 | 0.00 | 0.00 | 0.00 |
| >hg38_21_+_8433172_8446622_RNA45SN1_45S_50ntFlanks@475.511.37\|CTTCGTGATCGATGTGGTGACGTCGTGCTCTCCCGGG | >hg38_21_+_8433172_8446622_RNA45SN1_45S_50ntFlanks | 475 | 511 | 37 | rRF 45S | NO | 62.33 | 0.00 | 0.00 | 0.00 |
| >hg38_21_+_8433172_8446622_RNA45SN1_45S_50ntFlanks@476.503.28\|TTCGTGATCGATGTGGTGACGTCGTGCT | >hg38_21_+_8433172_8446622_RNA45SN1_45S_50ntFlanks | 476 | 503 | 28 | rRF 45S | NO | 30.88 | 0.00 | 0.00 | 0.00 |

### MDA-MB-468

| | | | | | | | | | | |
|---|---|---|---|---|---|---|---|---|---|---|
| >hg38_21_+_8433172_8446622_RNA45SN1_45S_50ntFlanks@476.504.29\|TTCGTGATCGATGT GGTGACGTCGTGCTC | >hg38_21_+_8433172_8446622_RNA45SN1_45S_50ntFlanks | 476 | 504 | 29 | rRF 45S | NO | 30.67 | 0.00 | 0.00 | 0.00 |
| >hg38_21_+_8433172_8446622_RNA45SN1_45S_50ntFlanks@476.505.30\|TTCGTGATCGATGT GGTGACGTCGTGCTCT | >hg38_21_+_8433172_8446622_RNA45SN1_45S_50ntFlanks | 476 | 505 | 30 | rRF 45S | NO | 78.84 | 0.00 | 0.00 | 0.00 |
| >hg38_21_+_8433172_8446622_RNA45SN1_45S_50ntFlanks@476.506.31\|TTCGTGATCGATGT GGTGACGTCGTGCTCTC | >hg38_21_+_8433172_8446622_RNA45SN1_45S_50ntFlanks | 476 | 506 | 31 | rRF 45S | NO | 70.81 | 0.00 | 0.00 | 0.00 |
| >hg38_21_+_8433172_8446622_RNA45SN1_45S_50ntFlanks@482.505.24\|ATCGATGTGGTGA CGTCGTGCTCT | >hg38_21_+_8433172_8446622_RNA45SN1_45S_50ntFlanks | 482 | 505 | 24 | rRF 45S | NO | 55.86 | 0.00 | 0.00 | 0.00 |
| >hg38_21_+_8433172_8446622_RNA45SN1_45S_50ntFlanks@482.506.25\|ATCGATGTGGTGA CGTCGTGCTCTC | >hg38_21_+_8433172_8446622_RNA45SN1_45S_50ntFlanks | 482 | 506 | 25 | rRF 45S | NO | 56.35 | 0.00 | 0.00 | 0.00 |
| >hg38_21_+_8433172_8446622_RNA45SN1_45S_50ntFlanks@482.507.26\|ATCGATGTGGTGA CGTCGTGCTCTCC | >hg38_21_+_8433172_8446622_RNA45SN1_45S_50ntFlanks | 482 | 507 | 26 | rRF 45S | NO | 69.42 | 0.00 | 0.00 | 0.00 |
| >hg38_21_+_8433172_8446622_RNA45SN1_45S_50ntFlanks@974.992.19\|CGCGTGGGTCCTG AGGGAG | >hg38_21_+_8433172_8446622_RNA45SN1_45S_50ntFlanks | 974 | 992 | 19 | rRF 45S | NO | 12.95 | 0.00 | 0.00 | 0.00 |
| >hg38_21_+_8433172_8446622_RNA45SN1_45S_50ntFlanks@2959.2979.21\|CGACGTGTGGC GTGGGTCGAC | >hg38_21_+_8433172_8446622_RNA45SN1_45S_50ntFlanks | 2959 | 2979 | 21 | rRF 45S | NO | 30.70 | 0.00 | 0.00 | 0.00 |
| >hg38_21_+_8433172_8446622_RNA45SN1_45S_50ntFlanks@3593.3610.18\|CCCGTCCGTCC GTCCGTC | >hg38_21_+_8433172_8446622_RNA45SN1_45S_50ntFlanks | 3593 | 3610 | 18 | rRF 45S | NO | 0.00 | 39.47 | 0.00 | 0.00 |
| >hg38_21_+_8433172_8446622_RNA45SN1_45S_50ntFlanks@3705.3722.18\|TACCTGGTTGAT CCTGCC | >hg38_21_+_8433172_8446622_RNA45SN1_45S_50ntFlanks | 3705 | 3722 | 18 | rRF 45S | NO | 43.54 | 95.46 | 28.20 | 71.31 |
| >hg38_21_+_8433172_8446622_RNA45SN1_45S_50ntFlanks@3711.3737.27\|GTTGATCCTGCC AGTAGCATATGCTTG | >hg38_21_+_8433172_8446622_RNA45SN1_45S_50ntFlanks | 3711 | 3737 | 27 | rRF 45S | NO | 91.06 | 53.54 | 0.00 | 34.58 |
| >hg38_21_+_8433172_8446622_RNA45SN1_45S_50ntFlanks@3738.3756.19\|TCTCAAAGATTA AGCCATG | >hg38_21_+_8433172_8446622_RNA45SN1_45S_50ntFlanks | 3738 | 3756 | 19 | rRF 45S | NO | 0.00 | 0.00 | 58.27 | 0.00 |
| >hg38_21_+_8433172_8446622_RNA45SN1_45S_50ntFlanks@3769.3792.24\|CGCACGGCCGG TACAGTGAAACTG | >hg38_21_+_8433172_8446622_RNA45SN1_45S_50ntFlanks | 3769 | 3792 | 24 | rRF 45S | NO | 57.28 | 0.00 | 0.00 | 14.44 |
| >hg38_21_+_8433172_8446622_RNA45SN1_45S_50ntFlanks@3769.3794.26\|CGCACGGCCGG TACAGTGAAACTGCG | >hg38_21_+_8433172_8446622_RNA45SN1_45S_50ntFlanks | 3769 | 3794 | 26 | rRF 45S | NO | 19.68 | 0.00 | 0.00 | 0.00 |
| >hg38_21_+_8433172_8446622_RNA45SN1_45S_50ntFlanks@3771.3791.21\|CACGGCCGGTA CAGTGAAACT | >hg38_21_+_8433172_8446622_RNA45SN1_45S_50ntFlanks | 3771 | 3791 | 21 | rRF 45S | NO | 20.52 | 0.00 | 0.00 | 0.00 |
| >hg38_21_+_8433172_8446622_RNA45SN1_45S_50ntFlanks@3771.3792.22\|CACGGCCGGTA CAGTGAAACTG | >hg38_21_+_8433172_8446622_RNA45SN1_45S_50ntFlanks | 3771 | 3792 | 22 | rRF 45S | NO | 74.29 | 36.22 | 0.00 | 37.34 |
| >hg38_21_+_8433172_8446622_RNA45SN1_45S_50ntFlanks@3771.3794.24\|CACGGCCGGTA CAGTGAAACTGCG | >hg38_21_+_8433172_8446622_RNA45SN1_45S_50ntFlanks | 3771 | 3794 | 24 | rRF 45S | NO | 33.42 | 0.00 | 0.00 | 0.00 |
| >hg38_21_+_8433172_8446622_RNA45SN1_45S_50ntFlanks@3771.3799.29\|CACGGCCGGTA CAGTGAAACTGCGAATGG | >hg38_21_+_8433172_8446622_RNA45SN1_45S_50ntFlanks | 3771 | 3799 | 29 | rRF 45S | NO | 21.52 | 0.00 | 0.00 | 0.00 |
| >hg38_21_+_8433172_8446622_RNA45SN1_45S_50ntFlanks@3773.3792.20\|CGGCCGGTACA GTGAAACTG | >hg38_21_+_8433172_8446622_RNA45SN1_45S_50ntFlanks | 3773 | 3792 | 20 | rRF 45S | NO | 33.96 | 0.00 | 0.00 | 26.66 |
| >hg38_21_+_8433172_8446622_RNA45SN1_45S_50ntFlanks@3775.3792.18\|GCCGGTACAGT GAAACTG | >hg38_21_+_8433172_8446622_RNA45SN1_45S_50ntFlanks | 3775 | 3792 | 18 | rRF 45S | NO | 168.04 | 211.76 | 68.37 | 84.82 |
| >hg38_21_+_8433172_8446622_RNA45SN1_45S_50ntFlanks@3775.3794.20\|GCCGGTACAGT GAAACTGCG | >hg38_21_+_8433172_8446622_RNA45SN1_45S_50ntFlanks | 3775 | 3794 | 20 | rRF 45S | NO | 0.00 | 12.34 | 0.00 | 0.00 |
| >hg38_21_+_8433172_8446622_RNA45SN1_45S_50ntFlanks@3775.3798.24\|GCCGGTACAGT GAAACTGCGAATG | >hg38_21_+_8433172_8446622_RNA45SN1_45S_50ntFlanks | 3775 | 3798 | 24 | rRF 45S | NO | 0.00 | 19.58 | 0.00 | 0.00 |
| >hg38_21_+_8433172_8446622_RNA45SN1_45S_50ntFlanks@3776.3801.26\|CCGGTACAGTG AAACTGCGAATGGCT | >hg38_21_+_8433172_8446622_RNA45SN1_45S_50ntFlanks | 3776 | 3801 | 26 | rRF 45S | NO | 28.77 | 51.66 | 37.50 | 14.64 |
| >hg38_21_+_8433172_8446622_RNA45SN1_45S_50ntFlanks@3776.3805.30\|CCGGTACAGTG AAACTGCGAATGGCTCATT | >hg38_21_+_8433172_8446622_RNA45SN1_45S_50ntFlanks | 3776 | 3805 | 30 | rRF 45S | NO | 46.88 | 19.71 | 0.00 | 0.00 |
| >hg38_21_+_8433172_8446622_RNA45SN1_45S_50ntFlanks@3777.3801.25\|CGGTACAGTGAA ACTGCGAATGGCT | >hg38_21_+_8433172_8446622_RNA45SN1_45S_50ntFlanks | 3777 | 3801 | 25 | rRF 45S | NO | 71.02 | 54.67 | 32.69 | 39.92 |
| >hg38_21_+_8433172_8446622_RNA45SN1_45S_50ntFlanks@3777.3805.29\|CGGTACAGTGAA ACTGCGAATGGCTCATT | >hg38_21_+_8433172_8446622_RNA45SN1_45S_50ntFlanks | 3777 | 3805 | 29 | rRF 45S | NO | 20.03 | 0.00 | 0.00 | 0.00 |
| >hg38_21_+_8433172_8446622_RNA45SN1_45S_50ntFlanks@3782.3800.19\|CAGTGAAACTGC GAATGGC | >hg38_21_+_8433172_8446622_RNA45SN1_45S_50ntFlanks | 3782 | 3800 | 19 | rRF 45S | NO | 44.09 | 41.78 | 20.58 | 25.03 |
| >hg38_21_+_8433172_8446622_RNA45SN1_45S_50ntFlanks@3782.3801.20\|CAGTGAAACTGC GAATGGCT | >hg38_21_+_8433172_8446622_RNA45SN1_45S_50ntFlanks | 3782 | 3801 | 20 | rRF 45S | NO | 111.50 | 103.67 | 67.21 | 100.26 |
| >hg38_21_+_8433172_8446622_RNA45SN1_45S_50ntFlanks@3782.3802.21\|CAGTGAAACTGC GAATGGCTC | >hg38_21_+_8433172_8446622_RNA45SN1_45S_50ntFlanks | 3782 | 3802 | 21 | rRF 45S | NO | 20.31 | 0.00 | 0.00 | 0.00 |
| >hg38_21_+_8433172_8446622_RNA45SN1_45S_50ntFlanks@3782.3805.24\|CAGTGAAACTGC GAATGGCTCATT | >hg38_21_+_8433172_8446622_RNA45SN1_45S_50ntFlanks | 3782 | 3805 | 24 | rRF 45S | NO | 21.83 | 28.31 | 0.00 | 23.77 |
| >hg38_21_+_8433172_8446622_RNA45SN1_45S_50ntFlanks@3782.3810.29\|CAGTGAAACTGC GAATGGCTCATTAAATC | >hg38_21_+_8433172_8446622_RNA45SN1_45S_50ntFlanks | 3782 | 3810 | 29 | rRF 45S | NO | 29.75 | 5.93 | 0.00 | 0.00 |
| >hg38_21_+_8433172_8446622_RNA45SN1_45S_50ntFlanks@3782.3812.31\|CAGTGAAACTGC GAATGGCTCATTAAATCAG | >hg38_21_+_8433172_8446622_RNA45SN1_45S_50ntFlanks | 3782 | 3812 | 31 | rRF 45S | NO | 31.09 | 0.00 | 0.00 | 13.81 |
| >hg38_21_+_8433172_8446622_RNA45SN1_45S_50ntFlanks@3782.3814.33\|CAGTGAAACTGC GAATGGCTCATTAAATCAGTT | >hg38_21_+_8433172_8446622_RNA45SN1_45S_50ntFlanks | 3782 | 3814 | 33 | rRF 45S | NO | 30.50 | 0.00 | 0.00 | 0.00 |
| >hg38_21_+_8433172_8446622_RNA45SN1_45S_50ntFlanks@3783.3801.19\|AGTGAAACTGCG AATGGCT | >hg38_21_+_8433172_8446622_RNA45SN1_45S_50ntFlanks | 3783 | 3801 | 19 | rRF 45S | NO | 12.91 | 0.00 | 6.78 | 23.17 |

### MDA-MB-468

| | | | | | | | | | | |
|---|---|---|---|---|---|---|---|---|---|---|
| >hg38_21_+_8433172_8446622_RNA45SN1_45S_50ntFlanks@3785.3810.26\|TGAAACTGCGAATGGCTCATTAAATC | >hg38_21_+_8433172_8446622_RNA45SN1_45S_50ntFlanks | 3785 | 3810 | 26 | rRF 45S | NO | 39.00 | 0.00 | 0.00 | 0.00 |
| >hg38_21_+_8433172_8446622_RNA45SN1_45S_50ntFlanks@3785.3812.28\|TGAAACTGCGAATGGCTCATTAAATCAG | >hg38_21_+_8433172_8446622_RNA45SN1_45S_50ntFlanks | 3785 | 3812 | 28 | rRF 45S | NO | 45.58 | 0.00 | 0.00 | 6.39 |
| >hg38_21_+_8433172_8446622_RNA45SN1_45S_50ntFlanks@3787.3810.24\|AAACTGCGAATGGCTCATTAAATC | >hg38_21_+_8433172_8446622_RNA45SN1_45S_50ntFlanks | 3787 | 3810 | 24 | rRF 45S | NO | 81.73 | 46.06 | 0.00 | 33.32 |
| >hg38_21_+_8433172_8446622_RNA45SN1_45S_50ntFlanks@3787.3812.26\|AAACTGCGAATGGCTCATTAAATCAG | >hg38_21_+_8433172_8446622_RNA45SN1_45S_50ntFlanks | 3787 | 3812 | 26 | rRF 45S | NO | 34.38 | 0.00 | 0.00 | 16.36 |
| >hg38_21_+_8433172_8446622_RNA45SN1_45S_50ntFlanks@3788.3810.23\|AACTGCGAATGGCTCATTAAATC | >hg38_21_+_8433172_8446622_RNA45SN1_45S_50ntFlanks | 3788 | 3810 | 23 | rRF 45S | NO | 21.10 | 0.00 | 0.00 | 0.00 |
| >hg38_21_+_8433172_8446622_RNA45SN1_45S_50ntFlanks@3789.3810.22\|ACTGCGAATGGCTCATTAAATC | >hg38_21_+_8433172_8446622_RNA45SN1_45S_50ntFlanks | 3789 | 3810 | 22 | rRF 45S | NO | 40.52 | 29.38 | 0.00 | 14.06 |
| >hg38_21_+_8433172_8446622_RNA45SN1_45S_50ntFlanks@3802.3829.28\|CATTAAATCAGTTATGGTTCCTTTGGTC | >hg38_21_+_8433172_8446622_RNA45SN1_45S_50ntFlanks | 3802 | 3829 | 28 | rRF 45S | NO | 0.00 | 19.32 | 0.00 | 0.00 |
| >hg38_21_+_8433172_8446622_RNA45SN1_45S_50ntFlanks@3802.3834.33\|CATTAAATCAGTTATGGTTCCTTTGGTCGCTCG | >hg38_21_+_8433172_8446622_RNA45SN1_45S_50ntFlanks | 3802 | 3834 | 33 | rRF 45S | NO | 83.76 | 56.34 | 0.00 | 14.39 |
| >hg38_21_+_8433172_8446622_RNA45SN1_45S_50ntFlanks@3802.3836.35\|CATTAAATCAGTTATGGTTCCTTTGGTCGCTCGCT | >hg38_21_+_8433172_8446622_RNA45SN1_45S_50ntFlanks | 3802 | 3836 | 35 | rRF 45S | NO | 31.43 | 0.00 | 0.00 | 0.00 |
| >hg38_21_+_8433172_8446622_RNA45SN1_45S_50ntFlanks@3802.3837.36\|CATTAAATCAGTTATGGTTCCTTTGGTCGCTCGCTC | >hg38_21_+_8433172_8446622_RNA45SN1_45S_50ntFlanks | 3802 | 3837 | 36 | rRF 45S | NO | 41.30 | 0.00 | 0.00 | 0.00 |
| >hg38_21_+_8433172_8446622_RNA45SN1_45S_50ntFlanks@3811.3834.24\|AGTTATGGTTCCTTTGGTCGCTCG | >hg38_21_+_8433172_8446622_RNA45SN1_45S_50ntFlanks | 3811 | 3834 | 24 | rRF 45S | NO | 34.00 | 0.00 | 0.00 | 0.00 |
| >hg38_21_+_8433172_8446622_RNA45SN1_45S_50ntFlanks@3813.3834.22\|TTATGGTTCCTTTGGTCGCTCG | >hg38_21_+_8433172_8446622_RNA45SN1_45S_50ntFlanks | 3813 | 3834 | 22 | rRF 45S | NO | 34.90 | 32.90 | 0.00 | 0.00 |
| >hg38_21_+_8433172_8446622_RNA45SN1_45S_50ntFlanks@3813.3836.24\|TTATGGTTCCTTTGGTCGCTCGCT | >hg38_21_+_8433172_8446622_RNA45SN1_45S_50ntFlanks | 3813 | 3836 | 24 | rRF 45S | NO | 30.67 | 0.00 | 0.00 | 0.00 |
| >hg38_21_+_8433172_8446622_RNA45SN1_45S_50ntFlanks@3813.3837.25\|TTATGGTTCCTTTGGTCGCTCGCTC | >hg38_21_+_8433172_8446622_RNA45SN1_45S_50ntFlanks | 3813 | 3837 | 25 | rRF 45S | NO | 46.06 | 0.00 | 0.00 | 0.00 |
| >hg38_21_+_8433172_8446622_RNA45SN1_45S_50ntFlanks@3813.3842.30\|TTATGGTTCCTTTGGTCGCTCGCTCCTCTC | >hg38_21_+_8433172_8446622_RNA45SN1_45S_50ntFlanks | 3813 | 3842 | 30 | rRF 45S | NO | 13.02 | 0.00 | 0.00 | 6.38 |
| >hg38_21_+_8433172_8446622_RNA45SN1_45S_50ntFlanks@3815.3837.23\|ATGGTTCCTTTGGTCGCTCGCTC | >hg38_21_+_8433172_8446622_RNA45SN1_45S_50ntFlanks | 3815 | 3837 | 23 | rRF 45S | NO | 30.38 | 0.00 | 0.00 | 0.00 |
| >hg38_21_+_8433172_8446622_RNA45SN1_45S_50ntFlanks@3822.3852.31\|CTTTGGTCGCTCGCTCCTCTCCTACTTGGAT | >hg38_21_+_8433172_8446622_RNA45SN1_45S_50ntFlanks | 3822 | 3852 | 31 | rRF 45S | NO | 29.72 | 0.00 | 0.00 | 0.00 |
| >hg38_21_+_8433172_8446622_RNA45SN1_45S_50ntFlanks@3823.3852.30\|TTTGGTCGCTCGCTCCTCTCCTACTTGGAT | >hg38_21_+_8433172_8446622_RNA45SN1_45S_50ntFlanks | 3823 | 3852 | 30 | rRF 45S | NO | 35.02 | 0.00 | 0.00 | 0.00 |
| >hg38_21_+_8433172_8446622_RNA45SN1_45S_50ntFlanks@3830.3860.31\|GCTCGCTCCTCTCCTACTTGGATAACTGTGG | >hg38_21_+_8433172_8446622_RNA45SN1_45S_50ntFlanks | 3830 | 3860 | 31 | rRF 45S | NO | 56.12 | 0.00 | 0.00 | 0.00 |
| >hg38_21_+_8433172_8446622_RNA45SN1_45S_50ntFlanks@3831.3860.30\|CTCGCTCCTCTCCTACTTGGATAACTGTGG | >hg38_21_+_8433172_8446622_RNA45SN1_45S_50ntFlanks | 3831 | 3860 | 30 | rRF 45S | NO | 131.91 | 41.38 | 0.00 | 13.41 |
| >hg38_21_+_8433172_8446622_RNA45SN1_45S_50ntFlanks@3833.3860.28\|CGCTCCTCTCCTACTTGGATAACTGTGG | >hg38_21_+_8433172_8446622_RNA45SN1_45S_50ntFlanks | 3833 | 3860 | 28 | rRF 45S | NO | 34.47 | 0.00 | 0.00 | 0.00 |
| >hg38_21_+_8433172_8446622_RNA45SN1_45S_50ntFlanks@3835.3860.26\|CTCCTCTCCTACTTGGATAACTGTGG | >hg38_21_+_8433172_8446622_RNA45SN1_45S_50ntFlanks | 3835 | 3860 | 26 | rRF 45S | NO | 238.40 | 180.13 | 0.00 | 102.01 |
| >hg38_21_+_8433172_8446622_RNA45SN1_45S_50ntFlanks@3835.3861.27\|CTCCTCTCCTACTTGGATAACTGTGGT | >hg38_21_+_8433172_8446622_RNA45SN1_45S_50ntFlanks | 3835 | 3861 | 27 | rRF 45S | NO | 39.85 | 28.88 | 0.00 | 23.97 |
| >hg38_21_+_8433172_8446622_RNA45SN1_45S_50ntFlanks@3835.3868.34\|CTCCTCTCCTACTTGGATAACTGTGGTAATTCTA | >hg38_21_+_8433172_8446622_RNA45SN1_45S_50ntFlanks | 3835 | 3868 | 34 | rRF 45S | NO | 39.35 | 31.19 | 0.00 | 0.00 |
| >hg38_21_+_8433172_8446622_RNA45SN1_45S_50ntFlanks@3835.3871.37\|CTCCTCTCCTACTTGGATAACTGTGGTAATTCTAGAG | >hg38_21_+_8433172_8446622_RNA45SN1_45S_50ntFlanks | 3835 | 3871 | 37 | rRF 45S | NO | 61.44 | 49.13 | 0.00 | 0.00 |
| >hg38_21_+_8433172_8446622_RNA45SN1_45S_50ntFlanks@3835.3872.38\|CTCCTCTCCTACTTGGATAACTGTGGTAATTCTAGAGC | >hg38_21_+_8433172_8446622_RNA45SN1_45S_50ntFlanks | 3835 | 3872 | 38 | rRF 45S | NO | 30.95 | 0.00 | 0.00 | 0.00 |
| >hg38_21_+_8433172_8446622_RNA45SN1_45S_50ntFlanks@3837.3860.24\|CCTCTCCTACTTGGATAACTGTGG | >hg38_21_+_8433172_8446622_RNA45SN1_45S_50ntFlanks | 3837 | 3860 | 24 | rRF 45S | NO | 20.11 | 0.00 | 0.00 | 0.00 |
| >hg38_21_+_8433172_8446622_RNA45SN1_45S_50ntFlanks@3837.3861.25\|CCTCTCCTACTTGGATAACTGTGGT | >hg38_21_+_8433172_8446622_RNA45SN1_45S_50ntFlanks | 3837 | 3861 | 25 | rRF 45S | NO | 21.31 | 0.00 | 0.00 | 0.00 |
| >hg38_21_+_8433172_8446622_RNA45SN1_45S_50ntFlanks@3837.3868.32\|CCTCTCCTACTTGGATAACTGTGGTAATTCTA | >hg38_21_+_8433172_8446622_RNA45SN1_45S_50ntFlanks | 3837 | 3868 | 32 | rRF 45S | NO | 34.97 | 0.00 | 0.00 | 0.00 |
| >hg38_21_+_8433172_8446622_RNA45SN1_45S_50ntFlanks@3837.3871.35\|CCTCTCCTACTTGGATAACTGTGGTAATTCTAGAG | >hg38_21_+_8433172_8446622_RNA45SN1_45S_50ntFlanks | 3837 | 3871 | 35 | rRF 45S | NO | 33.76 | 12.41 | 0.00 | 0.00 |
| >hg38_21_+_8433172_8446622_RNA45SN1_45S_50ntFlanks@3838.3860.23\|CTCTCCTACTTGGATAACTGTGG | >hg38_21_+_8433172_8446622_RNA45SN1_45S_50ntFlanks | 3838 | 3860 | 23 | rRF 45S | NO | 113.63 | 107.13 | 0.00 | 23.64 |
| >hg38_21_+_8433172_8446622_RNA45SN1_45S_50ntFlanks@3838.3868.31\|CTCTCCTACTTGGATAACTGTGGTAATTCTA | >hg38_21_+_8433172_8446622_RNA45SN1_45S_50ntFlanks | 3838 | 3868 | 31 | rRF 45S | NO | 38.07 | 32.89 | 0.00 | 0.00 |
| >hg38_21_+_8433172_8446622_RNA45SN1_45S_50ntFlanks@3838.3871.34\|CTCTCCTACTTGGATAACTGTGGTAATTCTAGAG | >hg38_21_+_8433172_8446622_RNA45SN1_45S_50ntFlanks | 3838 | 3871 | 34 | rRF 45S | NO | 37.73 | 36.44 | 0.00 | 0.00 |
| >hg38_21_+_8433172_8446622_RNA45SN1_45S_50ntFlanks@3838.3872.35\|CTCTCCTACTTGGATAACTGTGGTAATTCTAGAGC | >hg38_21_+_8433172_8446622_RNA45SN1_45S_50ntFlanks | 3838 | 3872 | 35 | rRF 45S | NO | 12.51 | 0.00 | 0.00 | 0.00 |

### MDA-MB-468

| | | | | | | | | | | |
|---|---|---|---|---|---|---|---|---|---|---|
| >hg38_21_+_8433172_8446622_RNA45SN1_45S_50ntFlanks@3838.3873.36\|CTCTCCTACTTGGATAACTGTGGTAATTCTAGAGCT | >hg38_21_+_8433172_8446622_RNA45SN1_45S_50ntFlanks | 3838 | 3873 | 36 | rRF 45S | NO | 30.33 | 0.00 | 0.00 | 0.00 |
| >hg38_21_+_8433172_8446622_RNA45SN1_45S_50ntFlanks@3840.3860.21\|CTCCTACTTGGATAACTGTGG | >hg38_21_+_8433172_8446622_RNA45SN1_45S_50ntFlanks | 3840 | 3860 | 21 | rRF 45S | NO | 5.93 | 45.25 | 0.00 | 0.00 |
| >hg38_21_+_8433172_8446622_RNA45SN1_45S_50ntFlanks@3842.3859.18\|CCTACTTGGATAACTGTG | >hg38_21_+_8433172_8446622_RNA45SN1_45S_50ntFlanks | 3842 | 3859 | 18 | rRF 45S | NO | 0.00 | 40.63 | 0.00 | 0.00 |
| >hg38_21_+_8433172_8446622_RNA45SN1_45S_50ntFlanks@3842.3868.27\|CCTACTTGGATAACTGTGGTAATTCTA | >hg38_21_+_8433172_8446622_RNA45SN1_45S_50ntFlanks | 3842 | 3868 | 27 | rRF 45S | NO | 0.00 | 42.59 | 0.00 | 0.00 |
| >hg38_21_+_8433172_8446622_RNA45SN1_45S_50ntFlanks@3843.3860.18\|CTACTTGGATAACTGTGG | >hg38_21_+_8433172_8446622_RNA45SN1_45S_50ntFlanks | 3843 | 3860 | 18 | rRF 45S | NO | 0.00 | 31.66 | 0.00 | 0.00 |
| >hg38_21_+_8433172_8446622_RNA45SN1_45S_50ntFlanks@3843.3861.19\|CTACTTGGATAACTGTGGT | >hg38_21_+_8433172_8446622_RNA45SN1_45S_50ntFlanks | 3843 | 3861 | 19 | rRF 45S | NO | 37.58 | 104.21 | 0.00 | 15.42 |
| >hg38_21_+_8433172_8446622_RNA45SN1_45S_50ntFlanks@3843.3868.26\|CTACTTGGATAACTGTGGTAATTCTA | >hg38_21_+_8433172_8446622_RNA45SN1_45S_50ntFlanks | 3843 | 3868 | 26 | rRF 45S | NO | 0.00 | 64.11 | 0.00 | 25.25 |
| >hg38_21_+_8433172_8446622_RNA45SN1_45S_50ntFlanks@3843.3871.29\|CTACTTGGATAACTGTGGTAATTCTAGAG | >hg38_21_+_8433172_8446622_RNA45SN1_45S_50ntFlanks | 3843 | 3871 | 29 | rRF 45S | NO | 0.00 | 41.14 | 0.00 | 6.23 |
| >hg38_21_+_8433172_8446622_RNA45SN1_45S_50ntFlanks@3844.3868.25\|TACTTGGATAACTGTGGTAATTCTA | >hg38_21_+_8433172_8446622_RNA45SN1_45S_50ntFlanks | 3844 | 3868 | 25 | rRF 45S | NO | 0.00 | 29.10 | 0.00 | 0.00 |
| >hg38_21_+_8433172_8446622_RNA45SN1_45S_50ntFlanks@3845.3868.24\|ACTTGGATAACTGTGGTAATTCTA | >hg38_21_+_8433172_8446622_RNA45SN1_45S_50ntFlanks | 3845 | 3868 | 24 | rRF 45S | NO | 0.00 | 33.21 | 0.00 | 0.00 |
| >hg38_21_+_8433172_8446622_RNA45SN1_45S_50ntFlanks@3846.3868.23\|CTTGGATAACTGTGGTAATTCTA | >hg38_21_+_8433172_8446622_RNA45SN1_45S_50ntFlanks | 3846 | 3868 | 23 | rRF 45S | NO | 31.11 | 86.85 | 6.59 | 14.78 |
| >hg38_21_+_8433172_8446622_RNA45SN1_45S_50ntFlanks@3846.3869.24\|CTTGGATAACTGTGGTAATTCTAG | >hg38_21_+_8433172_8446622_RNA45SN1_45S_50ntFlanks | 3846 | 3869 | 24 | rRF 45S | NO | 13.07 | 63.17 | 7.19 | 13.70 |
| >hg38_21_+_8433172_8446622_RNA45SN1_45S_50ntFlanks@3850.3873.24\|GATAACTGTGGTAATTCTAGAGCT | >hg38_21_+_8433172_8446622_RNA45SN1_45S_50ntFlanks | 3850 | 3873 | 24 | rRF 45S | NO | 0.00 | 12.37 | 0.00 | 0.00 |
| >hg38_21_+_8433172_8446622_RNA45SN1_45S_50ntFlanks@3853.3873.21\|AACTGTGGTAATTCTAGAGCT | >hg38_21_+_8433172_8446622_RNA45SN1_45S_50ntFlanks | 3853 | 3873 | 21 | rRF 45S | NO | 0.00 | 0.00 | 32.96 | 0.00 |
| >hg38_21_+_8433172_8446622_RNA45SN1_45S_50ntFlanks@3853.3881.29\|AACTGTGGTAATTCTAGAGCTAATACATG | >hg38_21_+_8433172_8446622_RNA45SN1_45S_50ntFlanks | 3853 | 3881 | 29 | rRF 45S | NO | 34.10 | 42.04 | 20.98 | 33.61 |
| >hg38_21_+_8433172_8446622_RNA45SN1_45S_50ntFlanks@3854.3871.18\|ACTGTGGTAATTCTAGAG | >hg38_21_+_8433172_8446622_RNA45SN1_45S_50ntFlanks | 3854 | 3871 | 18 | rRF 45S | NO | 0.00 | 0.00 | 31.36 | 0.00 |
| >hg38_21_+_8433172_8446622_RNA45SN1_45S_50ntFlanks@3854.3872.19\|ACTGTGGTAATTCTAGAGC | >hg38_21_+_8433172_8446622_RNA45SN1_45S_50ntFlanks | 3854 | 3872 | 19 | rRF 45S | NO | 0.00 | 46.85 | 39.96 | 0.00 |
| >hg38_21_+_8433172_8446622_RNA45SN1_45S_50ntFlanks@3854.3873.20\|ACTGTGGTAATTCTAGAGCT | >hg38_21_+_8433172_8446622_RNA45SN1_45S_50ntFlanks | 3854 | 3873 | 20 | rRF 45S | NO | 0.00 | 49.48 | 56.37 | 0.00 |
| >hg38_21_+_8433172_8446622_RNA45SN1_45S_50ntFlanks@3854.3874.21\|ACTGTGGTAATTCTAGAGCTA | >hg38_21_+_8433172_8446622_RNA45SN1_45S_50ntFlanks | 3854 | 3874 | 21 | rRF 45S | NO | 0.00 | 27.76 | 28.00 | 0.00 |
| >hg38_21_+_8433172_8446622_RNA45SN1_45S_50ntFlanks@3854.3881.28\|ACTGTGGTAATTCTAGAGCTAATACATG | >hg38_21_+_8433172_8446622_RNA45SN1_45S_50ntFlanks | 3854 | 3881 | 28 | rRF 45S | NO | 99.71 | 113.41 | 39.72 | 45.22 |
| >hg38_21_+_8433172_8446622_RNA45SN1_45S_50ntFlanks@3854.3889.36\|ACTGTGGTAATTCTAGAGCTAATACATGCCGACGGG | >hg38_21_+_8433172_8446622_RNA45SN1_45S_50ntFlanks | 3854 | 3889 | 36 | rRF 45S | NO | 42.53 | 0.00 | 0.00 | 0.00 |
| >hg38_21_+_8433172_8446622_RNA45SN1_45S_50ntFlanks@3854.3898.45\|ACTGTGGTAATTCTAGAGCTAATACATGCCGACGGGCGCTGACCC | >hg38_21_+_8433172_8446622_RNA45SN1_45S_50ntFlanks | 3854 | 3898 | 45 | rRF 45S | YES | 65.77 | 42.24 | 0.00 | 25.13 |
| >hg38_21_+_8433172_8446622_RNA45SN1_45S_50ntFlanks@3861.3898.38\|TAATTCTAGAGCTAATACATGCCGACGGGCGCTGACCC | >hg38_21_+_8433172_8446622_RNA45SN1_45S_50ntFlanks | 3861 | 3898 | 38 | rRF 45S | NO | 0.00 | 30.95 | 0.00 | 0.00 |
| >hg38_21_+_8433172_8446622_RNA45SN1_45S_50ntFlanks@3866.3898.33\|CTAGAGCTAATACATGCCGACGGGCGCTGACCC | >hg38_21_+_8433172_8446622_RNA45SN1_45S_50ntFlanks | 3866 | 3898 | 33 | rRF 45S | NO | 0.00 | 31.43 | 0.00 | 39.38 |
| >hg38_21_+_8433172_8446622_RNA45SN1_45S_50ntFlanks@3870.3898.29\|AGCTAATACATGCCGACGGGCGCTGACCC | >hg38_21_+_8433172_8446622_RNA45SN1_45S_50ntFlanks | 3870 | 3898 | 29 | rRF 45S | NO | 0.00 | 32.52 | 0.00 | 39.22 |
| >hg38_21_+_8433172_8446622_RNA45SN1_45S_50ntFlanks@3872.3898.27\|CTAATACATGCCGACGGGCGCTGACCC | >hg38_21_+_8433172_8446622_RNA45SN1_45S_50ntFlanks | 3872 | 3898 | 27 | rRF 45S | NO | 76.91 | 129.21 | 0.00 | 101.93 |
| >hg38_21_+_8433172_8446622_RNA45SN1_45S_50ntFlanks@3872.3899.28\|CTAATACATGCCGACGGGCGCTGACCCC | >hg38_21_+_8433172_8446622_RNA45SN1_45S_50ntFlanks | 3872 | 3899 | 28 | rRF 45S | NO | 0.00 | 29.10 | 0.00 | 6.16 |
| >hg38_21_+_8433172_8446622_RNA45SN1_45S_50ntFlanks@3873.3898.26\|TAATACATGCCGACGGGCGCTGACCC | >hg38_21_+_8433172_8446622_RNA45SN1_45S_50ntFlanks | 3873 | 3898 | 26 | rRF 45S | NO | 114.37 | 213.55 | 0.00 | 120.87 |
| >hg38_21_+_8433172_8446622_RNA45SN1_45S_50ntFlanks@3873.3899.27\|TAATACATGCCGACGGGCGCTGACCCC | >hg38_21_+_8433172_8446622_RNA45SN1_45S_50ntFlanks | 3873 | 3899 | 27 | rRF 45S | NO | 12.80 | 44.02 | 0.00 | 14.36 |
| >hg38_21_+_8433172_8446622_RNA45SN1_45S_50ntFlanks@3874.3898.25\|AATACATGCCGACGGGCGCTGACCC | >hg38_21_+_8433172_8446622_RNA45SN1_45S_50ntFlanks | 3874 | 3898 | 25 | rRF 45S | NO | 186.19 | 302.90 | 49.53 | 148.98 |
| >hg38_21_+_8433172_8446622_RNA45SN1_45S_50ntFlanks@3874.3899.26\|AATACATGCCGACGGGCGCTGACCCC | >hg38_21_+_8433172_8446622_RNA45SN1_45S_50ntFlanks | 3874 | 3899 | 26 | rRF 45S | NO | 39.51 | 60.30 | 0.00 | 34.50 |
| >hg38_21_+_8433172_8446622_RNA45SN1_45S_50ntFlanks@3875.3898.24\|ATACATGCCGACGGGCGCTGACCC | >hg38_21_+_8433172_8446622_RNA45SN1_45S_50ntFlanks | 3875 | 3898 | 24 | rRF 45S | NO | 207.19 | 345.25 | 71.04 | 286.80 |
| >hg38_21_+_8433172_8446622_RNA45SN1_45S_50ntFlanks@3875.3899.25\|ATACATGCCGACGGGCGCTGACCCC | >hg38_21_+_8433172_8446622_RNA45SN1_45S_50ntFlanks | 3875 | 3899 | 25 | rRF 45S | NO | 43.93 | 79.52 | 0.00 | 51.13 |
| >hg38_21_+_8433172_8446622_RNA45SN1_45S_50ntFlanks@3876.3898.23\|TACATGCCGACGGGCGCTGACCC | >hg38_21_+_8433172_8446622_RNA45SN1_45S_50ntFlanks | 3876 | 3898 | 23 | rRF 45S | NO | 59.42 | 126.52 | 0.00 | 60.24 |

### MDA-MB-468

| | | | | | | | | | | |
|---|---|---|---|---|---|---|---|---|---|---|
| >hg38_21_+_8433172_8446622_RNA45SN1_45S_50ntFlanks@3876.3899.24\|TACATGCCGACGGGCGCTGACCCCC | >hg38_21_+_8433172_8446622_RNA45SN1_45S_50ntFlanks | 3876 | 3899 | 24 | rRF 45S | NO | 0.00 | 27.17 | 0.00 | 0.00 |
| >hg38_21_+_8433172_8446622_RNA45SN1_45S_50ntFlanks@3878.3898.21\|CATGCCGACGGGCGCTGACCC | >hg38_21_+_8433172_8446622_RNA45SN1_45S_50ntFlanks | 3878 | 3898 | 21 | rRF 45S | NO | 5.96 | 37.58 | 0.00 | 0.00 |
| >hg38_21_+_8433172_8446622_RNA45SN1_45S_50ntFlanks@3881.3898.18\|GCCGACGGGCGCTGACCC | >hg38_21_+_8433172_8446622_RNA45SN1_45S_50ntFlanks | 3881 | 3898 | 18 | rRF 45S | NO | 0.00 | 29.69 | 0.00 | 6.27 |
| >hg38_21_+_8433172_8446622_RNA45SN1_45S_50ntFlanks@3882.3899.18\|CCGACGGGCGCTGACCCC | >hg38_21_+_8433172_8446622_RNA45SN1_45S_50ntFlanks | 3882 | 3899 | 18 | rRF 45S | NO | 21.04 | 168.29 | 139.13 | 102.00 |
| >hg38_21_+_8433172_8446622_RNA45SN1_45S_50ntFlanks@3882.3915.34\|CCGACGGGCGCTGACCCCTTCGCGGGGGGGATG | >hg38_21_+_8433172_8446622_RNA45SN1_45S_50ntFlanks | 3882 | 3915 | 34 | rRF 45S | NO | 0.00 | 30.47 | 0.00 | 0.00 |
| >hg38_21_+_8433172_8446622_RNA45SN1_45S_50ntFlanks@3885.3915.31\|ACGGGCGCTGACCCCCTTCGCGGGGGGGATG | >hg38_21_+_8433172_8446622_RNA45SN1_45S_50ntFlanks | 3885 | 3915 | 31 | rRF 45S | NO | 6.04 | 50.43 | 0.00 | 0.00 |
| >hg38_21_+_8433172_8446622_RNA45SN1_45S_50ntFlanks@3899.3919.21\|CCTTCGCGGGGGGGATGCGTG | >hg38_21_+_8433172_8446622_RNA45SN1_45S_50ntFlanks | 3899 | 3919 | 21 | rRF 45S | NO | 110.60 | 133.73 | 151.60 | 154.99 |
| >hg38_21_+_8433172_8446622_RNA45SN1_45S_50ntFlanks@3899.3920.22\|CCTTCGCGGGGGGGATGCGTGC | >hg38_21_+_8433172_8446622_RNA45SN1_45S_50ntFlanks | 3899 | 3920 | 22 | rRF 45S | NO | 44.26 | 72.17 | 39.50 | 26.44 |
| >hg38_21_+_8433172_8446622_RNA45SN1_45S_50ntFlanks@3899.3927.29\|CCTTCGCGGGGGGGATGCGTGCATTTATC | >hg38_21_+_8433172_8446622_RNA45SN1_45S_50ntFlanks | 3899 | 3927 | 29 | rRF 45S | NO | 0.00 | 35.74 | 0.00 | 43.45 |
| >hg38_21_+_8433172_8446622_RNA45SN1_45S_50ntFlanks@3899.3937.39\|CCTTCGCGGGGGGGATGCGTGCATTTATCAGATCAAAAC | >hg38_21_+_8433172_8446622_RNA45SN1_45S_50ntFlanks | 3899 | 3937 | 39 | rRF 45S | YES | 12.92 | 29.07 | 0.00 | 37.65 |
| >hg38_21_+_8433172_8446622_RNA45SN1_45S_50ntFlanks@3899.3938.40\|CCTTCGCGGGGGGGATGCGTGCATTTATCAGATCAAAACC | >hg38_21_+_8433172_8446622_RNA45SN1_45S_50ntFlanks | 3899 | 3938 | 40 | rRF 45S | YES | 0.00 | 34.93 | 0.00 | 13.33 |
| >hg38_21_+_8433172_8446622_RNA45SN1_45S_50ntFlanks@3900.3919.20\|CTTCGCGGGGGGATGCGTG | >hg38_21_+_8433172_8446622_RNA45SN1_45S_50ntFlanks | 3900 | 3919 | 20 | rRF 45S | NO | 34.76 | 29.84 | 83.76 | 23.14 |
| >hg38_21_+_8433172_8446622_RNA45SN1_45S_50ntFlanks@3930.3959.30\|ATCAAAACCAACCCGGTCAGCCCCTCTCCG | >hg38_21_+_8433172_8446622_RNA45SN1_45S_50ntFlanks | 3930 | 3959 | 30 | rRF 45S | NO | 56.39 | 69.71 | 0.00 | 37.72 |
| >hg38_21_+_8433172_8446622_RNA45SN1_45S_50ntFlanks@3931.3959.29\|TCAAAACCAACCCGGTCAGCCCCTCTCCG | >hg38_21_+_8433172_8446622_RNA45SN1_45S_50ntFlanks | 3931 | 3959 | 29 | rRF 45S | NO | 37.56 | 46.76 | 0.00 | 25.83 |
| >hg38_21_+_8433172_8446622_RNA45SN1_45S_50ntFlanks@3939.3959.21\|AACCCGGTCAGCCCCTCTCCG | >hg38_21_+_8433172_8446622_RNA45SN1_45S_50ntFlanks | 3939 | 3959 | 21 | rRF 45S | NO | 47.63 | 81.70 | 0.00 | 100.69 |
| >hg38_21_+_8433172_8446622_RNA45SN1_45S_50ntFlanks@3940.3959.20\|ACCCGGTCAGCCCCTCTCCG | >hg38_21_+_8433172_8446622_RNA45SN1_45S_50ntFlanks | 3940 | 3959 | 20 | rRF 45S | NO | 0.00 | 30.38 | 0.00 | 6.07 |
| >hg38_21_+_8433172_8446622_RNA45SN1_45S_50ntFlanks@3984.4001.18\|GCGGCTTTGGTGACTCTA | >hg38_21_+_8433172_8446622_RNA45SN1_45S_50ntFlanks | 3984 | 4001 | 18 | rRF 45S | NO | 273.50 | 73.69 | 48.60 | 74.36 |
| >hg38_21_+_8433172_8446622_RNA45SN1_45S_50ntFlanks@3984.4002.19\|GCGGCTTTGGTGACTCTAG | >hg38_21_+_8433172_8446622_RNA45SN1_45S_50ntFlanks | 3984 | 4002 | 19 | rRF 45S | NO | 244.74 | 50.50 | 55.85 | 86.03 |
| >hg38_21_+_8433172_8446622_RNA45SN1_45S_50ntFlanks@3984.4004.21\|GCGGCTTTGGTGACTCTAGAT | >hg38_21_+_8433172_8446622_RNA45SN1_45S_50ntFlanks | 3984 | 4004 | 21 | rRF 45S | NO | 19.63 | 0.00 | 0.00 | 0.00 |
| >hg38_21_+_8433172_8446622_RNA45SN1_45S_50ntFlanks@3984.4005.22\|GCGGCTTTGGTGACTCTAGATA | >hg38_21_+_8433172_8446622_RNA45SN1_45S_50ntFlanks | 3984 | 4005 | 22 | rRF 45S | NO | 47.78 | 0.00 | 0.00 | 0.00 |
| >hg38_21_+_8433172_8446622_RNA45SN1_45S_50ntFlanks@3984.4010.27\|GCGGCTTTGGTGACTCTAGATAACCTC | >hg38_21_+_8433172_8446622_RNA45SN1_45S_50ntFlanks | 3984 | 4010 | 27 | rRF 45S | NO | 12.75 | 0.00 | 0.00 | 0.00 |
| >hg38_21_+_8433172_8446622_RNA45SN1_45S_50ntFlanks@3984.4024.41\|GCGGCTTTGGTGACTCTAGATAACCTCGGGCCGATCGCACG | >hg38_21_+_8433172_8446622_RNA45SN1_45S_50ntFlanks | 3984 | 4024 | 41 | rRF 45S | NO | 0.00 | 43.43 | 105.60 | 67.84 |
| >hg38_21_+_8433172_8446622_RNA45SN1_45S_50ntFlanks@3984.4026.43\|GCGGCTTTGGTGACTCTAGATAACCTCGGGCCGATCGCACGCC | >hg38_21_+_8433172_8446622_RNA45SN1_45S_50ntFlanks | 3984 | 4026 | 43 | rRF 45S | NO | 0.00 | 38.75 | 37.96 | 57.31 |
| >hg38_21_+_8433172_8446622_RNA45SN1_45S_50ntFlanks@3985.4002.18\|CGGCTTTGGTGACTCTAG | >hg38_21_+_8433172_8446622_RNA45SN1_45S_50ntFlanks | 3985 | 4002 | 18 | rRF 45S | NO | 402.59 | 217.61 | 135.55 | 155.91 |
| >hg38_21_+_8433172_8446622_RNA45SN1_45S_50ntFlanks@3985.4004.20\|CGGCTTTGGTGACTCTAGAT | >hg38_21_+_8433172_8446622_RNA45SN1_45S_50ntFlanks | 3985 | 4004 | 20 | rRF 45S | NO | 30.21 | 0.00 | 0.00 | 0.00 |
| >hg38_21_+_8433172_8446622_RNA45SN1_45S_50ntFlanks@3985.4005.21\|CGGCTTTGGTGACTCTAGATA | >hg38_21_+_8433172_8446622_RNA45SN1_45S_50ntFlanks | 3985 | 4005 | 21 | rRF 45S | NO | 87.47 | 6.03 | 37.66 | 26.52 |
| >hg38_21_+_8433172_8446622_RNA45SN1_45S_50ntFlanks@3985.4010.26\|CGGCTTTGGTGACTCTAGATAACCTC | >hg38_21_+_8433172_8446622_RNA45SN1_45S_50ntFlanks | 3985 | 4010 | 26 | rRF 45S | NO | 38.37 | 0.00 | 0.00 | 0.00 |
| >hg38_21_+_8433172_8446622_RNA45SN1_45S_50ntFlanks@3985.4013.29\|CGGCTTTGGTGACTCTAGATAACCTCGGG | >hg38_21_+_8433172_8446622_RNA45SN1_45S_50ntFlanks | 3985 | 4013 | 29 | rRF 45S | NO | 42.53 | 0.00 | 0.00 | 0.00 |
| >hg38_21_+_8433172_8446622_RNA45SN1_45S_50ntFlanks@3985.4014.30\|CGGCTTTGGTGACTCTAGATAACCTCGGGC | >hg38_21_+_8433172_8446622_RNA45SN1_45S_50ntFlanks | 3985 | 4014 | 30 | rRF 45S | NO | 30.47 | 0.00 | 0.00 | 0.00 |
| >hg38_21_+_8433172_8446622_RNA45SN1_45S_50ntFlanks@3985.4024.40\|CGGCTTTGGTGACTCTAGATAACCTCGGGCCGATCGCACG | >hg38_21_+_8433172_8446622_RNA45SN1_45S_50ntFlanks | 3985 | 4024 | 40 | rRF 45S | NO | 0.00 | 55.80 | 118.85 | 65.35 |
| >hg38_21_+_8433172_8446622_RNA45SN1_45S_50ntFlanks@3985.4026.42\|CGGCTTTGGTGACTCTAGATAACCTCGGGCCGATCGCACGCC | >hg38_21_+_8433172_8446622_RNA45SN1_45S_50ntFlanks | 3985 | 4026 | 42 | rRF 45S | NO | 57.59 | 92.98 | 47.42 | 104.31 |
| >hg38_21_+_8433172_8446622_RNA45SN1_45S_50ntFlanks@3988.4024.37\|CTTTGGTGACTCTAGATAACCTCGGGCCGATCGCACG | >hg38_21_+_8433172_8446622_RNA45SN1_45S_50ntFlanks | 3988 | 4024 | 37 | rRF 45S | NO | 46.35 | 0.00 | 21.12 | 16.30 |
| >hg38_21_+_8433172_8446622_RNA45SN1_45S_50ntFlanks@3990.4024.35\|TTGGTGACTCTAGATAACCTCGGGCCGATCGCACG | >hg38_21_+_8433172_8446622_RNA45SN1_45S_50ntFlanks | 3990 | 4024 | 35 | rRF 45S | NO | 41.35 | 0.00 | 0.00 | 0.00 |
| >hg38_21_+_8433172_8446622_RNA45SN1_45S_50ntFlanks@3991.4013.23\|TGGTGACTCTAGATAACCTCGGG | >hg38_21_+_8433172_8446622_RNA45SN1_45S_50ntFlanks | 3991 | 4013 | 23 | rRF 45S | NO | 13.00 | 0.00 | 0.00 | 0.00 |

### MDA-MB-468

| | | | | | | | | | | |
|---|---|---|---|---|---|---|---|---|---|---|
| >hg38_21_+_8433172_8446622_RNA45SN1_45S_50ntFlanks@3991.4024.34\|TGGTGACTCTAG ATAACCTCGGGCCGATCGCACG | >hg38_21_+_8433172_8446622_RNA45SN1_45S_50ntFlanks | 3991 | 4024 | 34 | rRF 45S | NO | 44.23 | 0.00 | 0.00 | 15.25 |
| >hg38_21_+_8433172_8446622_RNA45SN1_45S_50ntFlanks@3991.4026.36\|TGGTGACTCTAG ATAACCTCGGGCCGATCGCACGCC | >hg38_21_+_8433172_8446622_RNA45SN1_45S_50ntFlanks | 3991 | 4026 | 36 | rRF 45S | NO | 43.47 | 0.00 | 0.00 | 14.43 |
| >hg38_21_+_8433172_8446622_RNA45SN1_45S_50ntFlanks@3993.4013.21\|GTGACTCTAGAT AACCTCGGG | >hg38_21_+_8433172_8446622_RNA45SN1_45S_50ntFlanks | 3993 | 4013 | 21 | rRF 45S | NO | 75.35 | 57.15 | 6.62 | 13.66 |
| >hg38_21_+_8433172_8446622_RNA45SN1_45S_50ntFlanks@3993.4024.32\|GTGACTCTAGAT AACCTCGGGCCGATCGCACG | >hg38_21_+_8433172_8446622_RNA45SN1_45S_50ntFlanks | 3993 | 4024 | 32 | rRF 45S | NO | 57.59 | 0.00 | 13.89 | 69.99 |
| >hg38_21_+_8433172_8446622_RNA45SN1_45S_50ntFlanks@3993.4026.34\|GTGACTCTAGAT AACCTCGGGCCGATCGCACGCC | >hg38_21_+_8433172_8446622_RNA45SN1_45S_50ntFlanks | 3993 | 4026 | 34 | rRF 45S | NO | 41.31 | 0.00 | 0.00 | 41.66 |
| >hg38_21_+_8433172_8446622_RNA45SN1_45S_50ntFlanks@3996.4022.27\|ACTCTAGATAAC CTCGGGCCGATCGCA | >hg38_21_+_8433172_8446622_RNA45SN1_45S_50ntFlanks | 3996 | 4022 | 27 | rRF 45S | NO | 21.13 | 0.00 | 0.00 | 0.00 |
| >hg38_21_+_8433172_8446622_RNA45SN1_45S_50ntFlanks@3996.4023.28\|ACTCTAGATAAC CTCGGGCCGATCGCAC | >hg38_21_+_8433172_8446622_RNA45SN1_45S_50ntFlanks | 3996 | 4023 | 28 | rRF 45S | NO | 19.89 | 0.00 | 0.00 | 0.00 |
| >hg38_21_+_8433172_8446622_RNA45SN1_45S_50ntFlanks@3996.4024.29\|ACTCTAGATAAC CTCGGGCCGATCGCACG | >hg38_21_+_8433172_8446622_RNA45SN1_45S_50ntFlanks | 3996 | 4024 | 29 | rRF 45S | NO | 64.54 | 0.00 | 0.00 | 13.42 |
| >hg38_21_+_8433172_8446622_RNA45SN1_45S_50ntFlanks@3997.4024.28\|CTCTAGATAACC TCGGGCCGATCGCACG | >hg38_21_+_8433172_8446622_RNA45SN1_45S_50ntFlanks | 3997 | 4024 | 28 | rRF 45S | NO | 20.24 | 0.00 | 0.00 | 0.00 |
| >hg38_21_+_8433172_8446622_RNA45SN1_45S_50ntFlanks@3999.4024.26\|CTAGATAACCTC GGGCCGATCGCACG | >hg38_21_+_8433172_8446622_RNA45SN1_45S_50ntFlanks | 3999 | 4024 | 26 | rRF 45S | NO | 19.70 | 0.00 | 0.00 | 0.00 |
| >hg38_21_+_8433172_8446622_RNA45SN1_45S_50ntFlanks@4000.4023.24\|TAGATAACCTCG GGCCGATCGCAC | >hg38_21_+_8433172_8446622_RNA45SN1_45S_50ntFlanks | 4000 | 4023 | 24 | rRF 45S | NO | 20.57 | 0.00 | 0.00 | 0.00 |
| >hg38_21_+_8433172_8446622_RNA45SN1_45S_50ntFlanks@4000.4024.25\|TAGATAACCTCG GGCCGATCGCACG | >hg38_21_+_8433172_8446622_RNA45SN1_45S_50ntFlanks | 4000 | 4024 | 25 | rRF 45S | NO | 49.49 | 56.25 | 0.00 | 13.33 |
| >hg38_21_+_8433172_8446622_RNA45SN1_45S_50ntFlanks@4000.4042.43\|TAGATAACCTCG GGCCGATCGCACGCCCCCCGTGGCGGCGACG | >hg38_21_+_8433172_8446622_RNA45SN1_45S_50ntFlanks | 4000 | 4042 | 43 | rRF 45S | NO | 20.53 | 47.88 | 0.00 | 0.00 |
| >hg38_21_+_8433172_8446622_RNA45SN1_45S_50ntFlanks@4002.4020.19\|GATAACCTCGG GCCGATCG | >hg38_21_+_8433172_8446622_RNA45SN1_45S_50ntFlanks | 4002 | 4020 | 19 | rRF 45S | NO | 93.20 | 37.69 | 0.00 | 13.40 |
| >hg38_21_+_8433172_8446622_RNA45SN1_45S_50ntFlanks@4002.4022.21\|GATAACCTCGG GCCGATCGCA | >hg38_21_+_8433172_8446622_RNA45SN1_45S_50ntFlanks | 4002 | 4022 | 21 | rRF 45S | NO | 74.03 | 44.25 | 0.00 | 6.66 |
| >hg38_21_+_8433172_8446622_RNA45SN1_45S_50ntFlanks@4002.4023.22\|GATAACCTCGG GCCGATCGCAC | >hg38_21_+_8433172_8446622_RNA45SN1_45S_50ntFlanks | 4002 | 4023 | 22 | rRF 45S | NO | 109.83 | 46.59 | 0.00 | 6.29 |
| >hg38_21_+_8433172_8446622_RNA45SN1_45S_50ntFlanks@4002.4024.23\|GATAACCTCGG GCCGATCGCACG | >hg38_21_+_8433172_8446622_RNA45SN1_45S_50ntFlanks | 4002 | 4024 | 23 | rRF 45S | NO | 160.33 | 250.03 | 28.24 | 61.61 |
| >hg38_21_+_8433172_8446622_RNA45SN1_45S_50ntFlanks@4002.4025.24\|GATAACCTCGG GCCGATCGCACGC | >hg38_21_+_8433172_8446622_RNA45SN1_45S_50ntFlanks | 4002 | 4025 | 24 | rRF 45S | NO | 65.89 | 42.88 | 0.00 | 13.89 |
| >hg38_21_+_8433172_8446622_RNA45SN1_45S_50ntFlanks@4002.4026.25\|GATAACCTCGG GCCGATCGCACGCC | >hg38_21_+_8433172_8446622_RNA45SN1_45S_50ntFlanks | 4002 | 4026 | 25 | rRF 45S | NO | 13.14 | 34.88 | 0.00 | 0.00 |
| >hg38_21_+_8433172_8446622_RNA45SN1_45S_50ntFlanks@4002.4042.41\|GATAACCTCGG GCCGATCGCACGCCCCCCGTGGCGGCGACG | >hg38_21_+_8433172_8446622_RNA45SN1_45S_50ntFlanks | 4002 | 4042 | 41 | rRF 45S | NO | 45.96 | 97.49 | 0.00 | 0.00 |
| >hg38_21_+_8433172_8446622_RNA45SN1_45S_50ntFlanks@4003.4020.18\|ATAACCTCGGG CCGATCG | >hg38_21_+_8433172_8446622_RNA45SN1_45S_50ntFlanks | 4003 | 4020 | 18 | rRF 45S | NO | 185.68 | 61.19 | 0.00 | 47.94 |
| >hg38_21_+_8433172_8446622_RNA45SN1_45S_50ntFlanks@4003.4022.20\|ATAACCTCGGG CCGATCGCA | >hg38_21_+_8433172_8446622_RNA45SN1_45S_50ntFlanks | 4003 | 4022 | 20 | rRF 45S | NO | 124.84 | 59.04 | 0.00 | 39.53 |
| >hg38_21_+_8433172_8446622_RNA45SN1_45S_50ntFlanks@4003.4023.21\|ATAACCTCGGG CCGATCGCAC | >hg38_21_+_8433172_8446622_RNA45SN1_45S_50ntFlanks | 4003 | 4023 | 21 | rRF 45S | NO | 147.11 | 47.91 | 0.00 | 25.68 |
| >hg38_21_+_8433172_8446622_RNA45SN1_45S_50ntFlanks@4003.4024.22\|ATAACCTCGGG CCGATCGCACG | >hg38_21_+_8433172_8446622_RNA45SN1_45S_50ntFlanks | 4003 | 4024 | 22 | rRF 45S | NO | 200.77 | 357.54 | 45.69 | 109.88 |
| >hg38_21_+_8433172_8446622_RNA45SN1_45S_50ntFlanks@4003.4025.23\|ATAACCTCGGG CCGATCGCACGC | >hg38_21_+_8433172_8446622_RNA45SN1_45S_50ntFlanks | 4003 | 4025 | 23 | rRF 45S | NO | 101.84 | 82.42 | 0.00 | 52.30 |
| >hg38_21_+_8433172_8446622_RNA45SN1_45S_50ntFlanks@4003.4026.24\|ATAACCTCGGG CCGATCGCACGCC | >hg38_21_+_8433172_8446622_RNA45SN1_45S_50ntFlanks | 4003 | 4026 | 24 | rRF 45S | NO | 61.63 | 77.86 | 0.00 | 52.01 |
| >hg38_21_+_8433172_8446622_RNA45SN1_45S_50ntFlanks@4003.4039.37\|ATAACCTCGGG CCGATCGCACGCCCCCCGTGGCGGCG | >hg38_21_+_8433172_8446622_RNA45SN1_45S_50ntFlanks | 4003 | 4039 | 37 | rRF 45S | NO | 29.45 | 0.00 | 0.00 | 0.00 |
| >hg38_21_+_8433172_8446622_RNA45SN1_45S_50ntFlanks@4003.4042.40\|ATAACCTCGGG CCGATCGCACGCCCCCCGTGGCGGCGACG | >hg38_21_+_8433172_8446622_RNA45SN1_45S_50ntFlanks | 4003 | 4042 | 40 | rRF 45S | NO | 143.18 | 204.36 | 0.00 | 54.44 |
| >hg38_21_+_8433172_8446622_RNA45SN1_45S_50ntFlanks@4005.4022.18\|AACCTCGGGCC GATCGCA | >hg38_21_+_8433172_8446622_RNA45SN1_45S_50ntFlanks | 4005 | 4022 | 18 | rRF 45S | NO | 51.81 | 0.00 | 0.00 | 0.00 |
| >hg38_21_+_8433172_8446622_RNA45SN1_45S_50ntFlanks@4005.4023.19\|AACCTCGGGCC GATCGCAC | >hg38_21_+_8433172_8446622_RNA45SN1_45S_50ntFlanks | 4005 | 4023 | 19 | rRF 45S | NO | 49.32 | 0.00 | 0.00 | 0.00 |
| >hg38_21_+_8433172_8446622_RNA45SN1_45S_50ntFlanks@4005.4024.20\|AACCTCGGGCC GATCGCACG | >hg38_21_+_8433172_8446622_RNA45SN1_45S_50ntFlanks | 4005 | 4024 | 20 | rRF 45S | NO | 61.49 | 113.97 | 0.00 | 26.22 |
| >hg38_21_+_8433172_8446622_RNA45SN1_45S_50ntFlanks@4005.4025.21\|AACCTCGGGCC GATCGCACGC | >hg38_21_+_8433172_8446622_RNA45SN1_45S_50ntFlanks | 4005 | 4025 | 21 | rRF 45S | NO | 37.49 | 28.64 | 0.00 | 6.17 |
| >hg38_21_+_8433172_8446622_RNA45SN1_45S_50ntFlanks@4005.4026.22\|AACCTCGGGCC GATCGCACGCC | >hg38_21_+_8433172_8446622_RNA45SN1_45S_50ntFlanks | 4005 | 4026 | 22 | rRF 45S | NO | 6.10 | 28.97 | 0.00 | 6.12 |
| >hg38_21_+_8433172_8446622_RNA45SN1_45S_50ntFlanks@4005.4042.38\|AACCTCGGGCC GATCGCACGCCCCCCGTGGCGGCGACG | >hg38_21_+_8433172_8446622_RNA45SN1_45S_50ntFlanks | 4005 | 4042 | 38 | rRF 45S | NO | 49.98 | 57.19 | 0.00 | 6.23 |

### MDA-MB-468

| | | | | | | | | | | |
|---|---|---|---|---|---|---|---|---|---|---|
| >hg38_21_+_8433172_8446622_RNA45SN1_45S_50ntFlanks@4006.4023.18\|ACCTCGGGCCGATCGCAC | >hg38_21_+_8433172_8446622_RNA45SN1_45S_50ntFlanks | 4006 | 4023 | 18 | rRF 45S | NO | 71.52 | 28.01 | 0.00 | 0.00 |
| >hg38_21_+_8433172_8446622_RNA45SN1_45S_50ntFlanks@4006.4024.19\|ACCTCGGGCCGATCGCACG | >hg38_21_+_8433172_8446622_RNA45SN1_45S_50ntFlanks | 4006 | 4024 | 19 | rRF 45S | NO | 54.90 | 126.74 | 6.74 | 25.69 |
| >hg38_21_+_8433172_8446622_RNA45SN1_45S_50ntFlanks@4006.4025.20\|ACCTCGGGCCGATCGCACGC | >hg38_21_+_8433172_8446622_RNA45SN1_45S_50ntFlanks | 4006 | 4025 | 20 | rRF 45S | NO | 39.88 | 42.24 | 0.00 | 14.70 |
| >hg38_21_+_8433172_8446622_RNA45SN1_45S_50ntFlanks@4006.4026.21\|ACCTCGGGCCGATCGCACGCC | >hg38_21_+_8433172_8446622_RNA45SN1_45S_50ntFlanks | 4006 | 4026 | 21 | rRF 45S | NO | 0.00 | 32.04 | 0.00 | 0.00 |
| >hg38_21_+_8433172_8446622_RNA45SN1_45S_50ntFlanks@4006.4042.37\|ACCTCGGGCCGATCGCACGCCCCCCGTGGCGGCGACG | >hg38_21_+_8433172_8446622_RNA45SN1_45S_50ntFlanks | 4006 | 4042 | 37 | rRF 45S | NO | 48.54 | 74.06 | 0.00 | 0.00 |
| >hg38_21_+_8433172_8446622_RNA45SN1_45S_50ntFlanks@4007.4024.18\|CCTCGGGCCGATCGCACG | >hg38_21_+_8433172_8446622_RNA45SN1_45S_50ntFlanks | 4007 | 4024 | 18 | rRF 45S | NO | 6.04 | 76.31 | 30.91 | 23.10 |
| >hg38_21_+_8433172_8446622_RNA45SN1_45S_50ntFlanks@4007.4042.36\|CCTCGGGCCGATCGCACGCCCCCCGTGGCGGCGACG | >hg38_21_+_8433172_8446622_RNA45SN1_45S_50ntFlanks | 4007 | 4042 | 36 | rRF 45S | NO | 0.00 | 36.48 | 0.00 | 0.00 |
| >hg38_21_+_8433172_8446622_RNA45SN1_45S_50ntFlanks@4008.4025.18\|CTCGGGCCGATCGCACGC | >hg38_21_+_8433172_8446622_RNA45SN1_45S_50ntFlanks | 4008 | 4025 | 18 | rRF 45S | NO | 0.00 | 26.34 | 0.00 | 0.00 |
| >hg38_21_+_8433172_8446622_RNA45SN1_45S_50ntFlanks@4008.4042.35\|CTCGGGCCGATCGCACGCCCCCCGTGGCGGCGACG | >hg38_21_+_8433172_8446622_RNA45SN1_45S_50ntFlanks | 4008 | 4042 | 35 | rRF 45S | NO | 0.00 | 34.76 | 0.00 | 0.00 |
| >hg38_21_+_8433172_8446622_RNA45SN1_45S_50ntFlanks@4043.4063.21\|ACCCATTCGAACGTCTGCCCT | >hg38_21_+_8433172_8446622_RNA45SN1_45S_50ntFlanks | 4043 | 4063 | 21 | rRF 45S | NO | 34.96 | 0.00 | 0.00 | 0.00 |
| >hg38_21_+_8433172_8446622_RNA45SN1_45S_50ntFlanks@4045.4062.18\|CCATTCGAACGTCTGCCC | >hg38_21_+_8433172_8446622_RNA45SN1_45S_50ntFlanks | 4045 | 4062 | 18 | rRF 45S | NO | 42.45 | 67.47 | 0.00 | 0.00 |
| >hg38_21_+_8433172_8446622_RNA45SN1_45S_50ntFlanks@4045.4063.19\|CCATTCGAACGTCTGCCCT | >hg38_21_+_8433172_8446622_RNA45SN1_45S_50ntFlanks | 4045 | 4063 | 19 | rRF 45S | NO | 134.67 | 212.28 | 0.00 | 61.77 |
| >hg38_21_+_8433172_8446622_RNA45SN1_45S_50ntFlanks@4045.4066.22\|CCATTCGAACGTCTGCCCTATC | >hg38_21_+_8433172_8446622_RNA45SN1_45S_50ntFlanks | 4045 | 4066 | 22 | rRF 45S | NO | 88.48 | 43.81 | 0.00 | 49.58 |
| >hg38_21_+_8433172_8446622_RNA45SN1_45S_50ntFlanks@4046.4063.18\|CATTCGAACGTCTGCCCT | >hg38_21_+_8433172_8446622_RNA45SN1_45S_50ntFlanks | 4046 | 4063 | 18 | rRF 45S | NO | 183.52 | 346.37 | 41.57 | 84.36 |
| >hg38_21_+_8433172_8446622_RNA45SN1_45S_50ntFlanks@4046.4066.21\|CATTCGAACGTCTGCCCTATC | >hg38_21_+_8433172_8446622_RNA45SN1_45S_50ntFlanks | 4046 | 4066 | 21 | rRF 45S | NO | 144.58 | 86.36 | 0.00 | 78.79 |
| >hg38_21_+_8433172_8446622_RNA45SN1_45S_50ntFlanks@4046.4067.22\|CATTCGAACGTCTGCCCTATCA | >hg38_21_+_8433172_8446622_RNA45SN1_45S_50ntFlanks | 4046 | 4067 | 22 | rRF 45S | NO | 30.98 | 0.00 | 0.00 | 0.00 |
| >hg38_21_+_8433172_8446622_RNA45SN1_45S_50ntFlanks@4047.4064.18\|ATTCGAACGTCTGCCCTA | >hg38_21_+_8433172_8446622_RNA45SN1_45S_50ntFlanks | 4047 | 4064 | 18 | rRF 45S | NO | 0.00 | 37.27 | 0.00 | 13.79 |
| >hg38_21_+_8433172_8446622_RNA45SN1_45S_50ntFlanks@4047.4065.19\|ATTCGAACGTCTGCCCTAT | >hg38_21_+_8433172_8446622_RNA45SN1_45S_50ntFlanks | 4047 | 4065 | 19 | rRF 45S | NO | 32.57 | 45.79 | 0.00 | 0.00 |
| >hg38_21_+_8433172_8446622_RNA45SN1_45S_50ntFlanks@4047.4066.20\|ATTCGAACGTCTGCCCTATC | >hg38_21_+_8433172_8446622_RNA45SN1_45S_50ntFlanks | 4047 | 4066 | 20 | rRF 45S | NO | 235.48 | 155.98 | 40.55 | 140.79 |
| >hg38_21_+_8433172_8446622_RNA45SN1_45S_50ntFlanks@4047.4067.21\|ATTCGAACGTCTGCCCTATCA | >hg38_21_+_8433172_8446622_RNA45SN1_45S_50ntFlanks | 4047 | 4067 | 21 | rRF 45S | NO | 50.78 | 32.21 | 0.00 | 16.52 |
| >hg38_21_+_8433172_8446622_RNA45SN1_45S_50ntFlanks@4047.4073.27\|ATTCGAACGTCTGCCCTATCAACTTTC | >hg38_21_+_8433172_8446622_RNA45SN1_45S_50ntFlanks | 4047 | 4073 | 27 | rRF 45S | NO | 0.00 | 19.39 | 0.00 | 0.00 |
| >hg38_21_+_8433172_8446622_RNA45SN1_45S_50ntFlanks@4047.4081.35\|ATTCGAACGTCTGCCCTATCAACTTTCGATGGTAG | >hg38_21_+_8433172_8446622_RNA45SN1_45S_50ntFlanks | 4047 | 4081 | 35 | rRF 45S | NO | 0.00 | 26.45 | 0.00 | 39.93 |
| >hg38_21_+_8433172_8446622_RNA45SN1_45S_50ntFlanks@4048.4066.19\|TTCGAACGTCTGCCCTATC | >hg38_21_+_8433172_8446622_RNA45SN1_45S_50ntFlanks | 4048 | 4066 | 19 | rRF 45S | NO | 33.93 | 33.32 | 0.00 | 13.93 |
| >hg38_21_+_8433172_8446622_RNA45SN1_45S_50ntFlanks@4064.4081.18\|ATCAACTTTCGATGGTAG | >hg38_21_+_8433172_8446622_RNA45SN1_45S_50ntFlanks | 4064 | 4081 | 18 | rRF 45S | NO | 0.00 | 41.93 | 28.83 | 0.00 |
| >hg38_21_+_8433172_8446622_RNA45SN1_45S_50ntFlanks@4064.4098.35\|ATCAACTTTCGATGGTAGTCGCCGTGCCTACCATG | >hg38_21_+_8433172_8446622_RNA45SN1_45S_50ntFlanks | 4064 | 4098 | 35 | rRF 45S | NO | 0.00 | 26.01 | 0.00 | 0.00 |
| >hg38_21_+_8433172_8446622_RNA45SN1_45S_50ntFlanks@4064.4101.38\|ATCAACTTTCGATGGTAGTCGCCGTGCCTACCATGGTG | >hg38_21_+_8433172_8446622_RNA45SN1_45S_50ntFlanks | 4064 | 4101 | 38 | rRF 45S | NO | 0.00 | 56.64 | 0.00 | 0.00 |
| >hg38_21_+_8433172_8446622_RNA45SN1_45S_50ntFlanks@4069.4089.21\|CTTTCGATGGTAGTCGCCGTG | >hg38_21_+_8433172_8446622_RNA45SN1_45S_50ntFlanks | 4069 | 4089 | 21 | rRF 45S | NO | 0.00 | 12.66 | 7.28 | 37.62 |
| >hg38_21_+_8433172_8446622_RNA45SN1_45S_50ntFlanks@4069.4095.27\|CTTTCGATGGTAGTCGCCGTGCCTACC | >hg38_21_+_8433172_8446622_RNA45SN1_45S_50ntFlanks | 4069 | 4095 | 27 | rRF 45S | NO | 0.00 | 26.02 | 34.34 | 0.00 |
| >hg38_21_+_8433172_8446622_RNA45SN1_45S_50ntFlanks@4069.4098.30\|CTTTCGATGGTAGTCGCCGTGCCTACCATG | >hg38_21_+_8433172_8446622_RNA45SN1_45S_50ntFlanks | 4069 | 4098 | 30 | rRF 45S | NO | 0.00 | 38.92 | 0.00 | 0.00 |
| >hg38_21_+_8433172_8446622_RNA45SN1_45S_50ntFlanks@4069.4101.33\|CTTTCGATGGTAGTCGCCGTGCCTACCATGGTG | >hg38_21_+_8433172_8446622_RNA45SN1_45S_50ntFlanks | 4069 | 4101 | 33 | rRF 45S | NO | 0.00 | 55.99 | 0.00 | 0.00 |
| >hg38_21_+_8433172_8446622_RNA45SN1_45S_50ntFlanks@4099.4121.23\|GTGACCACGGGTGACGGGGAATC | >hg38_21_+_8433172_8446622_RNA45SN1_45S_50ntFlanks | 4099 | 4121 | 23 | rRF 45S | NO | 20.12 | 0.00 | 0.00 | 0.00 |
| >hg38_21_+_8433172_8446622_RNA45SN1_45S_50ntFlanks@4102.4120.19\|ACCACGGGTGACGGGGAAT | >hg38_21_+_8433172_8446622_RNA45SN1_45S_50ntFlanks | 4102 | 4120 | 19 | rRF 45S | NO | 0.00 | 27.32 | 0.00 | 0.00 |
| >hg38_21_+_8433172_8446622_RNA45SN1_45S_50ntFlanks@4102.4121.20\|ACCACGGGTGACGGGGAATC | >hg38_21_+_8433172_8446622_RNA45SN1_45S_50ntFlanks | 4102 | 4121 | 20 | rRF 45S | NO | 95.57 | 125.42 | 65.17 | 13.79 |
| >hg38_21_+_8433172_8446622_RNA45SN1_45S_50ntFlanks@4102.4127.26\|ACCACGGGTGACGGGGAATCAGGGTT | >hg38_21_+_8433172_8446622_RNA45SN1_45S_50ntFlanks | 4102 | 4127 | 26 | rRF 45S | NO | 47.36 | 0.00 | 0.00 | 0.00 |

### MDA-MB-468

| | | | | | | | | | | |
|---|---|---|---|---|---|---|---|---|---|---|
| >hg38_21_+_8433172_8446622_RNA45SN1_45S_50ntFlanks@4103.4121.19\|CCACGGGTGAC GGGGAATC | >hg38_21_+_8433172_8446622_RNA45SN1_45S_50ntFlanks | 4103 | 4121 | 19 | rRF 45S | NO | 51.96 | 64.66 | 39.82 | 0.00 |
| >hg38_21_+_8433172_8446622_RNA45SN1_45S_50ntFlanks@4122.4145.24\|AGGGTTCGATTC CGGAGAGGGAGC | >hg38_21_+_8433172_8446622_RNA45SN1_45S_50ntFlanks | 4122 | 4145 | 24 | rRF 45S | NO | 12.72 | 0.00 | 20.98 | 0.00 |
| >hg38_21_+_8433172_8446622_RNA45SN1_45S_50ntFlanks@4128.4162.35\|CGATTCCGGAG AGGGAGCCTGAGAAACGGCTACCA | >hg38_21_+_8433172_8446622_RNA45SN1_45S_50ntFlanks | 4128 | 4162 | 35 | rRF 45S | NO | 41.69 | 47.49 | 0.00 | 14.83 |
| >hg38_21_+_8433172_8446622_RNA45SN1_45S_50ntFlanks@4129.4162.34\|GATTCCGGAGA GGGAGCCTGAGAAACGGCTACCA | >hg38_21_+_8433172_8446622_RNA45SN1_45S_50ntFlanks | 4129 | 4162 | 34 | rRF 45S | NO | 32.86 | 46.58 | 0.00 | 0.00 |
| >hg38_21_+_8433172_8446622_RNA45SN1_45S_50ntFlanks@4130.4162.33\|ATTCCGGAGAG GGAGCCTGAGAAACGGCTACCA | >hg38_21_+_8433172_8446622_RNA45SN1_45S_50ntFlanks | 4130 | 4162 | 33 | rRF 45S | NO | 53.45 | 67.11 | 0.00 | 23.70 |
| >hg38_21_+_8433172_8446622_RNA45SN1_45S_50ntFlanks@4160.4182.23\|CCACATCCAAG GAAGGCAGCAGG | >hg38_21_+_8433172_8446622_RNA45SN1_45S_50ntFlanks | 4160 | 4182 | 23 | rRF 45S | NO | 29.69 | 0.00 | 0.00 | 0.00 |
| >hg38_21_+_8433172_8446622_RNA45SN1_45S_50ntFlanks@4160.4184.25\|CCACATCCAAG GAAGGCAGCAGGCG | >hg38_21_+_8433172_8446622_RNA45SN1_45S_50ntFlanks | 4160 | 4184 | 25 | rRF 45S | NO | 21.81 | 0.00 | 0.00 | 0.00 |
| >hg38_21_+_8433172_8446622_RNA45SN1_45S_50ntFlanks@4163.4182.20\|CATCCAAGGAA GGCAGCAGG | >hg38_21_+_8433172_8446622_RNA45SN1_45S_50ntFlanks | 4163 | 4182 | 20 | rRF 45S | NO | 103.70 | 54.72 | 27.53 | 0.00 |
| >hg38_21_+_8433172_8446622_RNA45SN1_45S_50ntFlanks@4163.4184.22\|CATCCAAGGAA GGCAGCAGGCG | >hg38_21_+_8433172_8446622_RNA45SN1_45S_50ntFlanks | 4163 | 4184 | 22 | rRF 45S | NO | 103.67 | 55.76 | 35.15 | 0.00 |
| >hg38_21_+_8433172_8446622_RNA45SN1_45S_50ntFlanks@4163.4186.24\|CATCCAAGGAA GGCAGCAGGCGCG | >hg38_21_+_8433172_8446622_RNA45SN1_45S_50ntFlanks | 4163 | 4186 | 24 | rRF 45S | NO | 52.94 | 36.83 | 0.00 | 0.00 |
| >hg38_21_+_8433172_8446622_RNA45SN1_45S_50ntFlanks@4201.4219.19\|CCGACCCGGGG AGGTAGTG | >hg38_21_+_8433172_8446622_RNA45SN1_45S_50ntFlanks | 4201 | 4219 | 19 | rRF 45S | NO | 0.00 | 0.00 | 34.02 | 0.00 |
| >hg38_21_+_8433172_8446622_RNA45SN1_45S_50ntFlanks@4202.4219.18\|CGACCCGGGGA GGTAGTG | >hg38_21_+_8433172_8446622_RNA45SN1_45S_50ntFlanks | 4202 | 4219 | 18 | rRF 45S | NO | 0.00 | 0.00 | 34.77 | 0.00 |
| >hg38_21_+_8433172_8446622_RNA45SN1_45S_50ntFlanks@4220.4242.23\|ACGAAAAATAAC AATACAGGACT | >hg38_21_+_8433172_8446622_RNA45SN1_45S_50ntFlanks | 4220 | 4242 | 23 | rRF 45S | NO | 21.04 | 75.40 | 68.72 | 47.14 |
| >hg38_21_+_8433172_8446622_RNA45SN1_45S_50ntFlanks@4223.4242.20\|AAAAATAACAAT ACAGGACT | >hg38_21_+_8433172_8446622_RNA45SN1_45S_50ntFlanks | 4223 | 4242 | 20 | rRF 45S | NO | 0.00 | 0.00 | 27.51 | 0.00 |
| >hg38_21_+_8433172_8446622_RNA45SN1_45S_50ntFlanks@4224.4242.19\|AAAATAACAATA CAGGACT | >hg38_21_+_8433172_8446622_RNA45SN1_45S_50ntFlanks | 4224 | 4242 | 19 | rRF 45S | NO | 12.77 | 56.38 | 62.61 | 24.01 |
| >hg38_21_+_8433172_8446622_RNA45SN1_45S_50ntFlanks@4225.4242.18\|AAATAACAATAC AGGACT | >hg38_21_+_8433172_8446622_RNA45SN1_45S_50ntFlanks | 4225 | 4242 | 18 | rRF 45S | NO | 0.00 | 34.89 | 55.13 | 13.79 |
| >hg38_21_+_8433172_8446622_RNA45SN1_45S_50ntFlanks@4240.4258.19\|ACTCTTTCGAGG CCCTGTA | >hg38_21_+_8433172_8446622_RNA45SN1_45S_50ntFlanks | 4240 | 4258 | 19 | rRF 45S | NO | 0.00 | 53.11 | 0.00 | 6.56 |
| >hg38_21_+_8433172_8446622_RNA45SN1_45S_50ntFlanks@4241.4258.18\|CTCTTTCGAGGC CCTGTA | >hg38_21_+_8433172_8446622_RNA45SN1_45S_50ntFlanks | 4241 | 4258 | 18 | rRF 45S | NO | 19.70 | 132.93 | 0.00 | 6.05 |
| >hg38_21_+_8433172_8446622_RNA45SN1_45S_50ntFlanks@4243.4263.21\|CTTTCGAGGCC CTGTAATTGG | >hg38_21_+_8433172_8446622_RNA45SN1_45S_50ntFlanks | 4243 | 4263 | 21 | rRF 45S | NO | 33.65 | 31.70 | 0.00 | 6.42 |
| >hg38_21_+_8433172_8446622_RNA45SN1_45S_50ntFlanks@4259.4282.24\|ATTGGAATGAGT CCACTTTAAATC | >hg38_21_+_8433172_8446622_RNA45SN1_45S_50ntFlanks | 4259 | 4282 | 24 | rRF 45S | NO | 0.00 | 0.00 | 39.28 | 0.00 |
| >hg38_21_+_8433172_8446622_RNA45SN1_45S_50ntFlanks@4279.4314.36\|AATCCTTTAACG AGGATCCATTGGAGGGCAAGTCTG | >hg38_21_+_8433172_8446622_RNA45SN1_45S_50ntFlanks | 4279 | 4314 | 36 | rRF 45S | NO | 30.11 | 0.00 | 0.00 | 0.00 |
| >hg38_21_+_8433172_8446622_RNA45SN1_45S_50ntFlanks@4283.4307.25\|CTTTAACGAGGA TCCATTGGAGGGC | >hg38_21_+_8433172_8446622_RNA45SN1_45S_50ntFlanks | 4283 | 4307 | 25 | rRF 45S | NO | 34.55 | 0.00 | 6.63 | 0.00 |
| >hg38_21_+_8433172_8446622_RNA45SN1_45S_50ntFlanks@4283.4314.32\|CTTTAACGAGGA TCCATTGGAGGGCAAGTCTG | >hg38_21_+_8433172_8446622_RNA45SN1_45S_50ntFlanks | 4283 | 4314 | 32 | rRF 45S | NO | 57.31 | 28.70 | 0.00 | 0.00 |
| >hg38_21_+_8433172_8446622_RNA45SN1_45S_50ntFlanks@4283.4317.35\|CTTTAACGAGGA TCCATTGGAGGGCAAGTCTGGTG | >hg38_21_+_8433172_8446622_RNA45SN1_45S_50ntFlanks | 4283 | 4317 | 35 | rRF 45S | NO | 337.23 | 135.57 | 74.58 | 45.03 |
| >hg38_21_+_8433172_8446622_RNA45SN1_45S_50ntFlanks@4283.4321.39\|CTTTAACGAGGA TCCATTGGAGGGCAAGTCTGGTGCCAG | >hg38_21_+_8433172_8446622_RNA45SN1_45S_50ntFlanks | 4283 | 4321 | 39 | rRF 45S | NO | 36.92 | 0.00 | 0.00 | 0.00 |
| >hg38_21_+_8433172_8446622_RNA45SN1_45S_50ntFlanks@4297.4314.18\|CATTGGAGGGC AAGTCTG | >hg38_21_+_8433172_8446622_RNA45SN1_45S_50ntFlanks | 4297 | 4314 | 18 | rRF 45S | NO | 36.33 | 33.16 | 0.00 | 0.00 |
| >hg38_21_+_8433172_8446622_RNA45SN1_45S_50ntFlanks@4297.4317.21\|CATTGGAGGGC AAGTCTGGTG | >hg38_21_+_8433172_8446622_RNA45SN1_45S_50ntFlanks | 4297 | 4317 | 21 | rRF 45S | NO | 28.70 | 55.81 | 55.59 | 13.32 |
| >hg38_21_+_8433172_8446622_RNA45SN1_45S_50ntFlanks@4297.4319.23\|CATTGGAGGGC AAGTCTGGTGCC | >hg38_21_+_8433172_8446622_RNA45SN1_45S_50ntFlanks | 4297 | 4319 | 23 | rRF 45S | NO | 0.00 | 19.50 | 0.00 | 0.00 |
| >hg38_21_+_8433172_8446622_RNA45SN1_45S_50ntFlanks@4297.4321.25\|CATTGGAGGGC AAGTCTGGTGCCAG | >hg38_21_+_8433172_8446622_RNA45SN1_45S_50ntFlanks | 4297 | 4321 | 25 | rRF 45S | NO | 19.50 | 40.87 | 0.00 | 0.00 |
| >hg38_21_+_8433172_8446622_RNA45SN1_45S_50ntFlanks@4353.4371.19\|TATTAAAGTTGC TGCAGTT | >hg38_21_+_8433172_8446622_RNA45SN1_45S_50ntFlanks | 4353 | 4371 | 19 | rRF 45S | NO | 0.00 | 30.15 | 0.00 | 0.00 |
| >hg38_21_+_8433172_8446622_RNA45SN1_45S_50ntFlanks@4367.4389.23\|CAGTTAAAAAGC TCGTAGTTGGA | >hg38_21_+_8433172_8446622_RNA45SN1_45S_50ntFlanks | 4367 | 4389 | 23 | rRF 45S | NO | 45.53 | 133.13 | 37.45 | 14.40 |
| >hg38_21_+_8433172_8446622_RNA45SN1_45S_50ntFlanks@4367.4391.25\|CAGTTAAAAAGC TCGTAGTTGGATC | >hg38_21_+_8433172_8446622_RNA45SN1_45S_50ntFlanks | 4367 | 4391 | 25 | rRF 45S | NO | 63.02 | 145.38 | 38.44 | 14.83 |
| >hg38_21_+_8433172_8446622_RNA45SN1_45S_50ntFlanks@4367.4396.30\|CAGTTAAAAAGC TCGTAGTTGGATCTTGGG | >hg38_21_+_8433172_8446622_RNA45SN1_45S_50ntFlanks | 4367 | 4396 | 30 | rRF 45S | NO | 32.47 | 0.00 | 0.00 | 0.00 |
| >hg38_21_+_8433172_8446622_RNA45SN1_45S_50ntFlanks@4367.4398.32\|CAGTTAAAAAGC TCGTAGTTGGATCTTGGGAG | >hg38_21_+_8433172_8446622_RNA45SN1_45S_50ntFlanks | 4367 | 4398 | 32 | rRF 45S | NO | 84.45 | 35.27 | 0.00 | 14.80 |

### MDA-MB-468

| | | | | | | | | | | |
|---|---|---|---|---|---|---|---|---|---|---|
| >hg38_21_+_8433172_8446622_RNA45SN1_45S_50ntFlanks@4370.4389.20\|TTAAAAAGCTCGTAGTTGGA | >hg38_21_+_8433172_8446622_RNA45SN1_45S_50ntFlanks | 4370 | 4389 | 20 | rRF 45S | NO | 0.00 | 19.50 | 0.00 | 0.00 |
| >hg38_21_+_8433172_8446622_RNA45SN1_45S_50ntFlanks@4370.4398.29\|TTAAAAAGCTCGTAGTTGGATCTTGGGAG | >hg38_21_+_8433172_8446622_RNA45SN1_45S_50ntFlanks | 4370 | 4398 | 29 | rRF 45S | NO | 20.61 | 0.00 | 0.00 | 0.00 |
| >hg38_21_+_8433172_8446622_RNA45SN1_45S_50ntFlanks@4380.4408.29\|CGTAGTTGGATCTTGGGAGCGGGCGGGCG | >hg38_21_+_8433172_8446622_RNA45SN1_45S_50ntFlanks | 4380 | 4408 | 29 | rRF 45S | NO | 12.62 | 0.00 | 0.00 | 0.00 |
| >hg38_21_+_8433172_8446622_RNA45SN1_45S_50ntFlanks@4399.4423.25\|CGGGCGGGCGGTCCGCCGCGAGGCG | >hg38_21_+_8433172_8446622_RNA45SN1_45S_50ntFlanks | 4399 | 4423 | 25 | rRF 45S | NO | 0.00 | 33.13 | 0.00 | 0.00 |
| >hg38_21_+_8433172_8446622_RNA45SN1_45S_50ntFlanks@4443.4470.28\|CCCTTGCCTCTCGGCGCCCCCTCGATGC | >hg38_21_+_8433172_8446622_RNA45SN1_45S_50ntFlanks | 4443 | 4470 | 28 | rRF 45S | NO | 12.92 | 0.00 | 0.00 | 14.34 |
| >hg38_21_+_8433172_8446622_RNA45SN1_45S_50ntFlanks@4443.4471.29\|CCCTTGCCTCTCGGCGCCCCCTCGATGCT | >hg38_21_+_8433172_8446622_RNA45SN1_45S_50ntFlanks | 4443 | 4471 | 29 | rRF 45S | NO | 91.18 | 122.17 | 0.00 | 103.80 |
| >hg38_21_+_8433172_8446622_RNA45SN1_45S_50ntFlanks@4443.4472.30\|CCCTTGCCTCTCGGCGCCCCCTCGATGCTC | >hg38_21_+_8433172_8446622_RNA45SN1_45S_50ntFlanks | 4443 | 4472 | 30 | rRF 45S | NO | 36.44 | 56.72 | 0.00 | 38.75 |
| >hg38_21_+_8433172_8446622_RNA45SN1_45S_50ntFlanks@4444.4471.28\|CCTTGCCTCTCGGCGCCCCCTCGATGCT | >hg38_21_+_8433172_8446622_RNA45SN1_45S_50ntFlanks | 4444 | 4471 | 28 | rRF 45S | NO | 12.83 | 30.73 | 0.00 | 16.30 |
| >hg38_21_+_8433172_8446622_RNA45SN1_45S_50ntFlanks@4445.4471.27\|CTTGCCTCTCGGCGCCCCCTCGATGCT | >hg38_21_+_8433172_8446622_RNA45SN1_45S_50ntFlanks | 4445 | 4471 | 27 | rRF 45S | NO | 46.16 | 65.94 | 0.00 | 35.68 |
| >hg38_21_+_8433172_8446622_RNA45SN1_45S_50ntFlanks@4445.4472.28\|CTTGCCTCTCGGCGCCCCCTCGATGCTC | >hg38_21_+_8433172_8446622_RNA45SN1_45S_50ntFlanks | 4445 | 4472 | 28 | rRF 45S | NO | 32.19 | 30.93 | 0.00 | 0.00 |
| >hg38_21_+_8433172_8446622_RNA45SN1_45S_50ntFlanks@4449.4471.23\|CCTCTCGGCGCCCCCTCGATGCT | >hg38_21_+_8433172_8446622_RNA45SN1_45S_50ntFlanks | 4449 | 4471 | 23 | rRF 45S | NO | 80.01 | 81.00 | 0.00 | 97.35 |
| >hg38_21_+_8433172_8446622_RNA45SN1_45S_50ntFlanks@4449.4472.24\|CCTCTCGGCGCCCCCTCGATGCTC | >hg38_21_+_8433172_8446622_RNA45SN1_45S_50ntFlanks | 4449 | 4472 | 24 | rRF 45S | NO | 75.18 | 59.20 | 0.00 | 61.64 |
| >hg38_21_+_8433172_8446622_RNA45SN1_45S_50ntFlanks@4472.4492.21\|CTTAGCTGAGTGTCCCGCGGG | >hg38_21_+_8433172_8446622_RNA45SN1_45S_50ntFlanks | 4472 | 4492 | 21 | rRF 45S | NO | 0.00 | 28.24 | 0.00 | 0.00 |
| >hg38_21_+_8433172_8446622_RNA45SN1_45S_50ntFlanks@4472.4493.22\|CTTAGCTGAGTGTCCCGCGGGG | >hg38_21_+_8433172_8446622_RNA45SN1_45S_50ntFlanks | 4472 | 4493 | 22 | rRF 45S | NO | 0.00 | 41.30 | 0.00 | 0.00 |
| >hg38_21_+_8433172_8446622_RNA45SN1_45S_50ntFlanks@4472.4494.23\|CTTAGCTGAGTGTCCCGCGGGGC | >hg38_21_+_8433172_8446622_RNA45SN1_45S_50ntFlanks | 4472 | 4494 | 23 | rRF 45S | NO | 38.53 | 101.44 | 29.97 | 36.95 |
| >hg38_21_+_8433172_8446622_RNA45SN1_45S_50ntFlanks@4472.4496.25\|CTTAGCTGAGTGTCCCGCGGGGCCC | >hg38_21_+_8433172_8446622_RNA45SN1_45S_50ntFlanks | 4472 | 4496 | 25 | rRF 45S | NO | 0.00 | 33.98 | 0.00 | 5.98 |
| >hg38_21_+_8433172_8446622_RNA45SN1_45S_50ntFlanks@4472.4497.26\|CTTAGCTGAGTGTCCCGCGGGGCCCG | >hg38_21_+_8433172_8446622_RNA45SN1_45S_50ntFlanks | 4472 | 4497 | 26 | rRF 45S | NO | 0.00 | 38.63 | 6.83 | 14.53 |
| >hg38_21_+_8433172_8446622_RNA45SN1_45S_50ntFlanks@4486.4511.26\|CCGCGGGGCCCGAAGCGTTTACTTTG | >hg38_21_+_8433172_8446622_RNA45SN1_45S_50ntFlanks | 4486 | 4511 | 26 | rRF 45S | NO | 20.36 | 0.00 | 0.00 | 0.00 |
| >hg38_21_+_8433172_8446622_RNA45SN1_45S_50ntFlanks@4547.4565.19\|CTGGATACCGCAGCTAGGA | >hg38_21_+_8433172_8446622_RNA45SN1_45S_50ntFlanks | 4547 | 4565 | 19 | rRF 45S | NO | 0.00 | 0.00 | 27.58 | 0.00 |
| >hg38_21_+_8433172_8446622_RNA45SN1_45S_50ntFlanks@4555.4572.18\|CGCAGCTAGGAATAATGG | >hg38_21_+_8433172_8446622_RNA45SN1_45S_50ntFlanks | 4555 | 4572 | 18 | rRF 45S | NO | 67.27 | 37.16 | 27.86 | 0.00 |
| >hg38_21_+_8433172_8446622_RNA45SN1_45S_50ntFlanks@4555.4573.19\|CGCAGCTAGGAATAATGGA | >hg38_21_+_8433172_8446622_RNA45SN1_45S_50ntFlanks | 4555 | 4573 | 19 | rRF 45S | NO | 36.61 | 0.00 | 6.77 | 0.00 |
| >hg38_21_+_8433172_8446622_RNA45SN1_45S_50ntFlanks@4555.4587.33\|CGCAGCTAGGAATAATGGAATAGGACCGCGGTT | >hg38_21_+_8433172_8446622_RNA45SN1_45S_50ntFlanks | 4555 | 4587 | 33 | rRF 45S | NO | 20.00 | 0.00 | 0.00 | 0.00 |
| >hg38_21_+_8433172_8446622_RNA45SN1_45S_50ntFlanks@4557.4587.31\|CAGCTAGGAATAATGGAATAGGACCGCGGTT | >hg38_21_+_8433172_8446622_RNA45SN1_45S_50ntFlanks | 4557 | 4587 | 31 | rRF 45S | NO | 39.52 | 31.94 | 0.00 | 0.00 |
| >hg38_21_+_8433172_8446622_RNA45SN1_45S_50ntFlanks@4560.4579.20\|CTAGGAATAATGGAATAGGA | >hg38_21_+_8433172_8446622_RNA45SN1_45S_50ntFlanks | 4560 | 4579 | 20 | rRF 45S | NO | 0.00 | 39.99 | 0.00 | 0.00 |
| >hg38_21_+_8433172_8446622_RNA45SN1_45S_50ntFlanks@4560.4587.28\|CTAGGAATAATGGAATAGGACCGCGGTT | >hg38_21_+_8433172_8446622_RNA45SN1_45S_50ntFlanks | 4560 | 4587 | 28 | rRF 45S | NO | 51.34 | 61.90 | 6.50 | 0.00 |
| >hg38_21_+_8433172_8446622_RNA45SN1_45S_50ntFlanks@4564.4587.24\|GAATAATGGAATAGGACCGCGGTT | >hg38_21_+_8433172_8446622_RNA45SN1_45S_50ntFlanks | 4564 | 4587 | 24 | rRF 45S | NO | 20.55 | 0.00 | 0.00 | 0.00 |
| >hg38_21_+_8433172_8446622_RNA45SN1_45S_50ntFlanks@4565.4587.23\|AATAATGGAATAGGACCGCGGTT | >hg38_21_+_8433172_8446622_RNA45SN1_45S_50ntFlanks | 4565 | 4587 | 23 | rRF 45S | NO | 74.11 | 49.89 | 0.00 | 0.00 |
| >hg38_21_+_8433172_8446622_RNA45SN1_45S_50ntFlanks@4566.4587.22\|ATAATGGAATAGGACCGCGGTT | >hg38_21_+_8433172_8446622_RNA45SN1_45S_50ntFlanks | 4566 | 4587 | 22 | rRF 45S | NO | 38.22 | 31.45 | 0.00 | 0.00 |
| >hg38_21_+_8433172_8446622_RNA45SN1_45S_50ntFlanks@4567.4587.21\|TAATGGAATAGGACCGCGGTT | >hg38_21_+_8433172_8446622_RNA45SN1_45S_50ntFlanks | 4567 | 4587 | 21 | rRF 45S | NO | 48.09 | 54.65 | 0.00 | 0.00 |
| >hg38_21_+_8433172_8446622_RNA45SN1_45S_50ntFlanks@4567.4589.23\|TAATGGAATAGGACCGCGGTTCT | >hg38_21_+_8433172_8446622_RNA45SN1_45S_50ntFlanks | 4567 | 4589 | 23 | rRF 45S | NO | 0.00 | 28.87 | 0.00 | 0.00 |
| >hg38_21_+_8433172_8446622_RNA45SN1_45S_50ntFlanks@4569.4587.19\|ATGGAATAGGACCGCGGTT | >hg38_21_+_8433172_8446622_RNA45SN1_45S_50ntFlanks | 4569 | 4587 | 19 | rRF 45S | NO | 38.45 | 26.42 | 0.00 | 0.00 |
| >hg38_21_+_8433172_8446622_RNA45SN1_45S_50ntFlanks@4570.4587.18\|TGGAATAGGACCGCGGTT | >hg38_21_+_8433172_8446622_RNA45SN1_45S_50ntFlanks | 4570 | 4587 | 18 | rRF 45S | NO | 48.65 | 34.42 | 0.00 | 0.00 |
| >hg38_21_+_8433172_8446622_RNA45SN1_45S_50ntFlanks@4588.4608.21\|CTATTTTGTTGGTTTTCGGAA | >hg38_21_+_8433172_8446622_RNA45SN1_45S_50ntFlanks | 4588 | 4608 | 21 | rRF 45S | NO | 0.00 | 36.09 | 0.00 | 0.00 |
| >hg38_21_+_8433172_8446622_RNA45SN1_45S_50ntFlanks@4588.4613.26\|CTATTTTGTTGGTTTTCGGAACTGAG | >hg38_21_+_8433172_8446622_RNA45SN1_45S_50ntFlanks | 4588 | 4613 | 26 | rRF 45S | NO | 0.00 | 0.00 | 36.75 | 0.00 |

### MDA-MB-468

| | | | | | | | | | | |
|---|---|---|---|---|---|---|---|---|---|---|
| >hg38_21_+_8433172_8446622_RNA45SN1_45S_50ntFlanks@4588.4614.27\|CTATTTTGTTGGTTTTCGGAACTGAGG | >hg38_21_+_8433172_8446622_RNA45SN1_45S_50ntFlanks | 4588 | 4614 | 27 | rRF 45S | NO | 44.88 | 55.65 | 31.08 | 0.00 |
| >hg38_21_+_8433172_8446622_RNA45SN1_45S_50ntFlanks@4588.4615.28\|CTATTTTGTTGGTTTTCGGAACTGAGGC | >hg38_21_+_8433172_8446622_RNA45SN1_45S_50ntFlanks | 4588 | 4615 | 28 | rRF 45S | NO | 12.76 | 12.65 | 0.00 | 0.00 |
| >hg38_21_+_8433172_8446622_RNA45SN1_45S_50ntFlanks@4588.4616.29\|CTATTTTGTTGGTTTCGGAACTGAGGCC | >hg38_21_+_8433172_8446622_RNA45SN1_45S_50ntFlanks | 4588 | 4616 | 29 | rRF 45S | NO | 132.70 | 207.60 | 55.45 | 44.53 |
| >hg38_21_+_8433172_8446622_RNA45SN1_45S_50ntFlanks@4588.4619.32\|CTATTTTGTTGGTTTCGGAACTGAGGCCATG | >hg38_21_+_8433172_8446622_RNA45SN1_45S_50ntFlanks | 4588 | 4619 | 32 | rRF 45S | NO | 0.00 | 28.11 | 0.00 | 0.00 |
| >hg38_21_+_8433172_8446622_RNA45SN1_45S_50ntFlanks@4590.4616.27\|ATTTTGTTGGTTTTCGGAACTGAGGCC | >hg38_21_+_8433172_8446622_RNA45SN1_45S_50ntFlanks | 4590 | 4616 | 27 | rRF 45S | NO | 30.84 | 38.24 | 28.71 | 0.00 |
| >hg38_21_+_8433172_8446622_RNA45SN1_45S_50ntFlanks@4657.4682.26\|CTAGAGGTGAAATTCTTGGACCGGCG | >hg38_21_+_8433172_8446622_RNA45SN1_45S_50ntFlanks | 4657 | 4682 | 26 | rRF 45S | NO | 103.30 | 51.68 | 0.00 | 38.07 |
| >hg38_21_+_8433172_8446622_RNA45SN1_45S_50ntFlanks@4657.4688.32\|CTAGAGGTGAAATTCTTGGACCGGCGCAAGAC | >hg38_21_+_8433172_8446622_RNA45SN1_45S_50ntFlanks | 4657 | 4688 | 32 | rRF 45S | NO | 32.98 | 26.23 | 0.00 | 61.85 |
| >hg38_21_+_8433172_8446622_RNA45SN1_45S_50ntFlanks@4657.4689.33\|CTAGAGGTGAAATTCTTGGACCGGCGCAAGACG | >hg38_21_+_8433172_8446622_RNA45SN1_45S_50ntFlanks | 4657 | 4689 | 33 | rRF 45S | NO | 248.81 | 290.95 | 112.60 | 437.24 |
| >hg38_21_+_8433172_8446622_RNA45SN1_45S_50ntFlanks@4683.4707.25\|CAAGACGGACCAGAGCGAAAGCATT | >hg38_21_+_8433172_8446622_RNA45SN1_45S_50ntFlanks | 4683 | 4707 | 25 | rRF 45S | NO | 30.57 | 29.57 | 21.20 | 37.93 |
| >hg38_21_+_8433172_8446622_RNA45SN1_45S_50ntFlanks@4685.4707.23\|AGACGGACCAGAGCGAAAGCATT | >hg38_21_+_8433172_8446622_RNA45SN1_45S_50ntFlanks | 4685 | 4707 | 23 | rRF 45S | NO | 13.04 | 0.00 | 0.00 | 6.22 |
| >hg38_21_+_8433172_8446622_RNA45SN1_45S_50ntFlanks@4722.4748.27\|TCATTAATCAAGAACGAAAGTCGGAGG | >hg38_21_+_8433172_8446622_RNA45SN1_45S_50ntFlanks | 4722 | 4748 | 27 | rRF 45S | NO | 21.91 | 44.77 | 0.00 | 24.31 |
| >hg38_21_+_8433172_8446622_RNA45SN1_45S_50ntFlanks@4722.4751.30\|TCATTAATCAAGAACGAAAGTCGGAGGTTC | >hg38_21_+_8433172_8446622_RNA45SN1_45S_50ntFlanks | 4722 | 4751 | 30 | rRF 45S | NO | 44.99 | 46.09 | 35.48 | 15.88 |
| >hg38_21_+_8433172_8446622_RNA45SN1_45S_50ntFlanks@4722.4752.31\|TCATTAATCAAGAACGAAAGTCGGAGGTTCG | >hg38_21_+_8433172_8446622_RNA45SN1_45S_50ntFlanks | 4722 | 4752 | 31 | rRF 45S | NO | 51.81 | 77.05 | 21.12 | 58.71 |
| >hg38_21_+_8433172_8446622_RNA45SN1_45S_50ntFlanks@4722.4758.37\|TCATTAATCAAGAACGAAAGTCGGAGGTTCGAAGACG | >hg38_21_+_8433172_8446622_RNA45SN1_45S_50ntFlanks | 4722 | 4758 | 37 | rRF 45S | NO | 714.77 | 387.54 | 122.22 | 321.95 |
| >hg38_21_+_8433172_8446622_RNA45SN1_45S_50ntFlanks@4722.4759.38\|TCATTAATCAAGAACGAAAGTCGGAGGTTCGAAGACGA | >hg38_21_+_8433172_8446622_RNA45SN1_45S_50ntFlanks | 4722 | 4759 | 38 | rRF 45S | NO | 280.45 | 206.59 | 46.09 | 136.22 |
| >hg38_21_+_8433172_8446622_RNA45SN1_45S_50ntFlanks@4723.4748.26\|CATTAATCAAGAACGAAAGTCGGAGG | >hg38_21_+_8433172_8446622_RNA45SN1_45S_50ntFlanks | 4723 | 4748 | 26 | rRF 45S | NO | 0.00 | 42.70 | 7.07 | 16.19 |
| >hg38_21_+_8433172_8446622_RNA45SN1_45S_50ntFlanks@4723.4751.29\|CATTAATCAAGAACGAAAGTCGGAGGTTC | >hg38_21_+_8433172_8446622_RNA45SN1_45S_50ntFlanks | 4723 | 4751 | 29 | rRF 45S | NO | 38.40 | 44.93 | 43.75 | 15.23 |
| >hg38_21_+_8433172_8446622_RNA45SN1_45S_50ntFlanks@4723.4752.30\|CATTAATCAAGAACGAAAGTCGGAGGTTCG | >hg38_21_+_8433172_8446622_RNA45SN1_45S_50ntFlanks | 4723 | 4752 | 30 | rRF 45S | NO | 44.15 | 70.73 | 38.74 | 60.73 |
| >hg38_21_+_8433172_8446622_RNA45SN1_45S_50ntFlanks@4723.4758.36\|CATTAATCAAGAACGAAAGTCGGAGGTTCGAAGACG | >hg38_21_+_8433172_8446622_RNA45SN1_45S_50ntFlanks | 4723 | 4758 | 36 | rRF 45S | NO | 555.52 | 274.80 | 155.46 | 394.90 |
| >hg38_21_+_8433172_8446622_RNA45SN1_45S_50ntFlanks@4723.4759.37\|CATTAATCAAGAACGAAAGTCGGAGGTTCGAAGACGA | >hg38_21_+_8433172_8446622_RNA45SN1_45S_50ntFlanks | 4723 | 4759 | 37 | rRF 45S | NO | 144.10 | 102.60 | 39.88 | 123.76 |
| >hg38_21_+_8433172_8446622_RNA45SN1_45S_50ntFlanks@4724.4748.25\|ATTAATCAAGAACGAAAGTCGGAGG | >hg38_21_+_8433172_8446622_RNA45SN1_45S_50ntFlanks | 4724 | 4748 | 25 | rRF 45S | NO | 21.46 | 40.40 | 29.99 | 16.15 |
| >hg38_21_+_8433172_8446622_RNA45SN1_45S_50ntFlanks@4724.4751.28\|ATTAATCAAGAACGAAAGTCGGAGGTTC | >hg38_21_+_8433172_8446622_RNA45SN1_45S_50ntFlanks | 4724 | 4751 | 28 | rRF 45S | NO | 48.66 | 50.57 | 59.85 | 15.62 |
| >hg38_21_+_8433172_8446622_RNA45SN1_45S_50ntFlanks@4724.4752.29\|ATTAATCAAGAACGAAAGTCGGAGGTTCG | >hg38_21_+_8433172_8446622_RNA45SN1_45S_50ntFlanks | 4724 | 4752 | 29 | rRF 45S | NO | 50.92 | 69.85 | 42.14 | 53.15 |
| >hg38_21_+_8433172_8446622_RNA45SN1_45S_50ntFlanks@4724.4758.35\|ATTAATCAAGAACGAAAGTCGGAGGTTCGAAGACG | >hg38_21_+_8433172_8446622_RNA45SN1_45S_50ntFlanks | 4724 | 4758 | 35 | rRF 45S | NO | 941.22 | 425.82 | 244.23 | 560.62 |
| >hg38_21_+_8433172_8446622_RNA45SN1_45S_50ntFlanks@4724.4759.36\|ATTAATCAAGAACGAAAGTCGGAGGTTCGAAGACGA | >hg38_21_+_8433172_8446622_RNA45SN1_45S_50ntFlanks | 4724 | 4759 | 36 | rRF 45S | NO | 303.83 | 166.50 | 69.42 | 197.60 |
| >hg38_21_+_8433172_8446622_RNA45SN1_45S_50ntFlanks@4725.4758.34\|TTAATCAAGAACGAAAGTCGGAGGTTCGAAGACG | >hg38_21_+_8433172_8446622_RNA45SN1_45S_50ntFlanks | 4725 | 4758 | 34 | rRF 45S | NO | 129.07 | 60.55 | 40.38 | 53.37 |
| >hg38_21_+_8433172_8446622_RNA45SN1_45S_50ntFlanks@4725.4759.35\|TTAATCAAGAACGAAAGTCGGAGGTTCGAAGACGA | >hg38_21_+_8433172_8446622_RNA45SN1_45S_50ntFlanks | 4725 | 4759 | 35 | rRF 45S | NO | 41.10 | 0.00 | 0.00 | 6.42 |
| >hg38_21_+_8433172_8446622_RNA45SN1_45S_50ntFlanks@4726.4752.27\|TAATCAAGAACGAAAGTCGGAGGTTCG | >hg38_21_+_8433172_8446622_RNA45SN1_45S_50ntFlanks | 4726 | 4752 | 27 | rRF 45S | NO | 6.08 | 32.03 | 0.00 | 0.00 |
| >hg38_21_+_8433172_8446622_RNA45SN1_45S_50ntFlanks@4726.4758.33\|TAATCAAGAACGAAAGTCGGAGGTTCGAAGACG | >hg38_21_+_8433172_8446622_RNA45SN1_45S_50ntFlanks | 4726 | 4758 | 33 | rRF 45S | NO | 320.60 | 174.17 | 86.50 | 120.94 |
| >hg38_21_+_8433172_8446622_RNA45SN1_45S_50ntFlanks@4726.4759.34\|TAATCAAGAACGAAAGTCGGAGGTTCGAAGACGA | >hg38_21_+_8433172_8446622_RNA45SN1_45S_50ntFlanks | 4726 | 4759 | 34 | rRF 45S | NO | 130.10 | 84.31 | 34.99 | 56.41 |
| >hg38_21_+_8433172_8446622_RNA45SN1_45S_50ntFlanks@4727.4758.32\|AATCAAGAACGAAAGTCGGAGGTTCGAAGACG | >hg38_21_+_8433172_8446622_RNA45SN1_45S_50ntFlanks | 4727 | 4758 | 32 | rRF 45S | NO | 44.29 | 29.20 | 41.23 | 26.90 |
| >hg38_21_+_8433172_8446622_RNA45SN1_45S_50ntFlanks@4728.4751.24\|ATCAAGAACGAAAGTCGGAGGTTC | >hg38_21_+_8433172_8446622_RNA45SN1_45S_50ntFlanks | 4728 | 4751 | 24 | rRF 45S | NO | 47.64 | 52.72 | 56.70 | 0.00 |
| >hg38_21_+_8433172_8446622_RNA45SN1_45S_50ntFlanks@4728.4752.25\|ATCAAGAACGAAAGTCGGAGGTTCG | >hg38_21_+_8433172_8446622_RNA45SN1_45S_50ntFlanks | 4728 | 4752 | 25 | rRF 45S | NO | 32.38 | 43.62 | 30.67 | 0.00 |
| >hg38_21_+_8433172_8446622_RNA45SN1_45S_50ntFlanks@4728.4758.31\|ATCAAGAACGAAAGTCGGAGGTTCGAAGACG | >hg38_21_+_8433172_8446622_RNA45SN1_45S_50ntFlanks | 4728 | 4758 | 31 | rRF 45S | NO | 245.35 | 129.51 | 161.33 | 71.20 |

### MDA-MB-468

| | | | | | | | | | | |
|---|---|---|---|---|---|---|---|---|---|---|
| >hg38_21_+_8433172_8446622_RNA45SN1_45S_50ntFlanks@4728.4759.32\|ATCAAGAACGAAAGTCGGAGGTTCGAAGACGA | >hg38_21_+_8433172_8446622_RNA45SN1_45S_50ntFlanks | 4728 | 4759 | 32 | rRF 45S | NO | 32.77 | 33.72 | 37.66 | 0.00 |
| >hg38_21_+_8433172_8446622_RNA45SN1_45S_50ntFlanks@4729.4751.23\|TCAAGAACGAAAGTCGGAGGTTC | >hg38_21_+_8433172_8446622_RNA45SN1_45S_50ntFlanks | 4729 | 4751 | 23 | rRF 45S | NO | 20.56 | 19.66 | 32.04 | 0.00 |
| >hg38_21_+_8433172_8446622_RNA45SN1_45S_50ntFlanks@4729.4758.30\|TCAAGAACGAAAGTCGGAGGTTCGAAGACG | >hg38_21_+_8433172_8446622_RNA45SN1_45S_50ntFlanks | 4729 | 4758 | 30 | rRF 45S | NO | 148.98 | 73.63 | 110.99 | 67.39 |
| >hg38_21_+_8433172_8446622_RNA45SN1_45S_50ntFlanks@4729.4759.31\|TCAAGAACGAAAGTCGGAGGTTCGAAGACGA | >hg38_21_+_8433172_8446622_RNA45SN1_45S_50ntFlanks | 4729 | 4759 | 31 | rRF 45S | NO | 41.97 | 33.64 | 36.23 | 13.43 |
| >hg38_21_+_8433172_8446622_RNA45SN1_45S_50ntFlanks@4730.4751.22\|CAAGAACGAAAGTCGGAGGTTC | >hg38_21_+_8433172_8446622_RNA45SN1_45S_50ntFlanks | 4730 | 4751 | 22 | rRF 45S | NO | 32.70 | 34.31 | 42.96 | 0.00 |
| >hg38_21_+_8433172_8446622_RNA45SN1_45S_50ntFlanks@4730.4752.23\|CAAGAACGAAAGTCGGAGGTTCG | >hg38_21_+_8433172_8446622_RNA45SN1_45S_50ntFlanks | 4730 | 4752 | 23 | rRF 45S | NO | 0.00 | 26.58 | 0.00 | 0.00 |
| >hg38_21_+_8433172_8446622_RNA45SN1_45S_50ntFlanks@4730.4758.29\|CAAGAACGAAAGTCGGAGGTTCGAAGACG | >hg38_21_+_8433172_8446622_RNA45SN1_45S_50ntFlanks | 4730 | 4758 | 29 | rRF 45S | NO | 197.38 | 81.36 | 153.15 | 102.18 |
| >hg38_21_+_8433172_8446622_RNA45SN1_45S_50ntFlanks@4730.4759.30\|CAAGAACGAAAGTCGGAGGTTCGAAGACGA | >hg38_21_+_8433172_8446622_RNA45SN1_45S_50ntFlanks | 4730 | 4759 | 30 | rRF 45S | NO | 48.22 | 26.71 | 53.19 | 25.84 |
| >hg38_21_+_8433172_8446622_RNA45SN1_45S_50ntFlanks@4731.4748.18\|AAGAACGAAAGTCGGAGG | >hg38_21_+_8433172_8446622_RNA45SN1_45S_50ntFlanks | 4731 | 4748 | 18 | rRF 45S | NO | 0.00 | 28.61 | 33.80 | 0.00 |
| >hg38_21_+_8433172_8446622_RNA45SN1_45S_50ntFlanks@4731.4751.21\|AAGAACGAAAGTCGGAGGTTC | >hg38_21_+_8433172_8446622_RNA45SN1_45S_50ntFlanks | 4731 | 4751 | 21 | rRF 45S | NO | 78.15 | 68.50 | 92.28 | 0.00 |
| >hg38_21_+_8433172_8446622_RNA45SN1_45S_50ntFlanks@4731.4758.28\|AAGAACGAAAGTCGGAGGTTCGAAGACG | >hg38_21_+_8433172_8446622_RNA45SN1_45S_50ntFlanks | 4731 | 4758 | 28 | rRF 45S | NO | 492.24 | 244.12 | 367.63 | 153.94 |
| >hg38_21_+_8433172_8446622_RNA45SN1_45S_50ntFlanks@4731.4759.29\|AAGAACGAAAGTCGGAGGTTCGAAGACGA | >hg38_21_+_8433172_8446622_RNA45SN1_45S_50ntFlanks | 4731 | 4759 | 29 | rRF 45S | NO | 152.61 | 92.30 | 152.85 | 55.04 |
| >hg38_21_+_8433172_8446622_RNA45SN1_45S_50ntFlanks@4732.4751.20\|AGAACGAAAGTCGGAGGTTC | >hg38_21_+_8433172_8446622_RNA45SN1_45S_50ntFlanks | 4732 | 4751 | 20 | rRF 45S | NO | 5.99 | 0.00 | 30.87 | 0.00 |
| >hg38_21_+_8433172_8446622_RNA45SN1_45S_50ntFlanks@4732.4758.27\|AGAACGAAAGTCGGAGGTTCGAAGACG | >hg38_21_+_8433172_8446622_RNA45SN1_45S_50ntFlanks | 4732 | 4758 | 27 | rRF 45S | NO | 123.97 | 62.51 | 115.21 | 44.22 |
| >hg38_21_+_8433172_8446622_RNA45SN1_45S_50ntFlanks@4732.4759.28\|AGAACGAAAGTCGGAGGTTCGAAGACGA | >hg38_21_+_8433172_8446622_RNA45SN1_45S_50ntFlanks | 4732 | 4759 | 28 | rRF 45S | NO | 39.72 | 0.00 | 51.02 | 0.00 |
| >hg38_21_+_8433172_8446622_RNA45SN1_45S_50ntFlanks@4733.4758.26\|GAACGAAAGTCGGAGGTTCGAAGACG | >hg38_21_+_8433172_8446622_RNA45SN1_45S_50ntFlanks | 4733 | 4758 | 26 | rRF 45S | NO | 66.47 | 37.46 | 66.96 | 16.60 |
| >hg38_21_+_8433172_8446622_RNA45SN1_45S_50ntFlanks@4733.4759.27\|GAACGAAAGTCGGAGGTTCGAAGACGA | >hg38_21_+_8433172_8446622_RNA45SN1_45S_50ntFlanks | 4733 | 4759 | 27 | rRF 45S | NO | 5.92 | 0.00 | 31.16 | 0.00 |
| >hg38_21_+_8433172_8446622_RNA45SN1_45S_50ntFlanks@4734.4751.18\|AACGAAAGTCGGAGGTTC | >hg38_21_+_8433172_8446622_RNA45SN1_45S_50ntFlanks | 4734 | 4751 | 18 | rRF 45S | NO | 56.90 | 39.04 | 39.38 | 0.00 |
| >hg38_21_+_8433172_8446622_RNA45SN1_45S_50ntFlanks@4734.4752.19\|AACGAAAGTCGGAGGTTCG | >hg38_21_+_8433172_8446622_RNA45SN1_45S_50ntFlanks | 4734 | 4752 | 19 | rRF 45S | NO | 0.00 | 36.20 | 20.26 | 0.00 |
| >hg38_21_+_8433172_8446622_RNA45SN1_45S_50ntFlanks@4734.4758.25\|AACGAAAGTCGGAGGTTCGAAGACG | >hg38_21_+_8433172_8446622_RNA45SN1_45S_50ntFlanks | 4734 | 4758 | 25 | rRF 45S | NO | 125.33 | 50.01 | 107.41 | 45.51 |
| >hg38_21_+_8433172_8446622_RNA45SN1_45S_50ntFlanks@4734.4759.26\|AACGAAAGTCGGAGGTTCGAAGACGA | >hg38_21_+_8433172_8446622_RNA45SN1_45S_50ntFlanks | 4734 | 4759 | 26 | rRF 45S | NO | 65.48 | 29.76 | 61.97 | 6.11 |
| >hg38_21_+_8433172_8446622_RNA45SN1_45S_50ntFlanks@4735.4758.24\|ACGAAAGTCGGAGGTTCGAAGACG | >hg38_21_+_8433172_8446622_RNA45SN1_45S_50ntFlanks | 4735 | 4758 | 24 | rRF 45S | NO | 91.21 | 19.52 | 39.87 | 0.00 |
| >hg38_21_+_8433172_8446622_RNA45SN1_45S_50ntFlanks@4735.4759.25\|ACGAAAGTCGGAGGTTCGAAGACGA | >hg38_21_+_8433172_8446622_RNA45SN1_45S_50ntFlanks | 4735 | 4759 | 25 | rRF 45S | NO | 35.33 | 0.00 | 0.00 | 0.00 |
| >hg38_21_+_8433172_8446622_RNA45SN1_45S_50ntFlanks@4736.4758.23\|CGAAAGTCGGAGGTTCGAAGACG | >hg38_21_+_8433172_8446622_RNA45SN1_45S_50ntFlanks | 4736 | 4758 | 23 | rRF 45S | NO | 49.06 | 0.00 | 29.82 | 0.00 |
| >hg38_21_+_8433172_8446622_RNA45SN1_45S_50ntFlanks@4738.4758.21\|AAAGTCGGAGGTTCGAAGACG | >hg38_21_+_8433172_8446622_RNA45SN1_45S_50ntFlanks | 4738 | 4758 | 21 | rRF 45S | NO | 211.24 | 142.62 | 280.42 | 82.33 |
| >hg38_21_+_8433172_8446622_RNA45SN1_45S_50ntFlanks@4738.4759.22\|AAAGTCGGAGGTTCGAAGACGA | >hg38_21_+_8433172_8446622_RNA45SN1_45S_50ntFlanks | 4738 | 4759 | 22 | rRF 45S | NO | 0.00 | 32.23 | 84.76 | 0.00 |
| >hg38_21_+_8433172_8446622_RNA45SN1_45S_50ntFlanks@4739.4758.20\|AAGTCGGAGGTTCGAAGACG | >hg38_21_+_8433172_8446622_RNA45SN1_45S_50ntFlanks | 4739 | 4758 | 20 | rRF 45S | NO | 0.00 | 0.00 | 42.18 | 0.00 |
| >hg38_21_+_8433172_8446622_RNA45SN1_45S_50ntFlanks@4742.4759.18\|TCGGAGGTTCGAAGACGA | >hg38_21_+_8433172_8446622_RNA45SN1_45S_50ntFlanks | 4742 | 4759 | 18 | rRF 45S | NO | 28.76 | 0.00 | 0.00 | 0.00 |
| >hg38_21_+_8433172_8446622_RNA45SN1_45S_50ntFlanks@4753.4782.30\|AAGACGATCAGATACCGTCGTAGTTCCGAC | >hg38_21_+_8433172_8446622_RNA45SN1_45S_50ntFlanks | 4753 | 4782 | 30 | rRF 45S | NO | 32.55 | 43.93 | 0.00 | 33.32 |
| >hg38_21_+_8433172_8446622_RNA45SN1_45S_50ntFlanks@4753.4783.31\|AAGACGATCAGATACCGTCGTAGTTCCGACC | >hg38_21_+_8433172_8446622_RNA45SN1_45S_50ntFlanks | 4753 | 4783 | 31 | rRF 45S | NO | 42.12 | 31.12 | 0.00 | 35.46 |
| >hg38_21_+_8433172_8446622_RNA45SN1_45S_50ntFlanks@4759.4778.20\|ATCAGATACCGTCGTAGTTC | >hg38_21_+_8433172_8446622_RNA45SN1_45S_50ntFlanks | 4759 | 4778 | 20 | rRF 45S | NO | 97.22 | 123.39 | 46.92 | 41.73 |
| >hg38_21_+_8433172_8446622_RNA45SN1_45S_50ntFlanks@4759.4779.21\|ATCAGATACCGTCGTAGTTCC | >hg38_21_+_8433172_8446622_RNA45SN1_45S_50ntFlanks | 4759 | 4779 | 21 | rRF 45S | NO | 71.66 | 74.19 | 0.00 | 14.80 |
| >hg38_21_+_8433172_8446622_RNA45SN1_45S_50ntFlanks@4759.4780.22\|ATCAGATACCGTCGTAGTTCCG | >hg38_21_+_8433172_8446622_RNA45SN1_45S_50ntFlanks | 4759 | 4780 | 22 | rRF 45S | YES | 550.07 | 802.88 | 120.07 | 212.01 |
| >hg38_21_+_8433172_8446622_RNA45SN1_45S_50ntFlanks@4759.4781.23\|ATCAGATACCGTCGTAGTTCCGA | >hg38_21_+_8433172_8446622_RNA45SN1_45S_50ntFlanks | 4759 | 4781 | 23 | rRF 45S | NO | 0.00 | 41.89 | 0.00 | 0.00 |

### MDA-MB-468

| | | | | | | | | | | |
|---|---|---|---|---|---|---|---|---|---|---|
| >hg38_21_+_8433172_8446622_RNA45SN1_45S_50ntFlanks@4759.4782.24\|ATCAGATACCGTCGTAGTTCCGAC | >hg38_21_+_8433172_8446622_RNA45SN1_45S_50ntFlanks | 4759 | 4782 | 24 | rRF 45S | NO | 36.95 | 164.31 | 140.79 | 84.22 |
| >hg38_21_+_8433172_8446622_RNA45SN1_45S_50ntFlanks@4759.4783.25\|ATCAGATACCGTCGTAGTTCCGACC | >hg38_21_+_8433172_8446622_RNA45SN1_45S_50ntFlanks | 4759 | 4783 | 25 | rRF 45S | NO | 50.70 | 110.31 | 56.43 | 67.00 |
| >hg38_21_+_8433172_8446622_RNA45SN1_45S_50ntFlanks@4759.4786.28\|ATCAGATACCGTCGTAGTTCCGACCATA | >hg38_21_+_8433172_8446622_RNA45SN1_45S_50ntFlanks | 4759 | 4786 | 28 | rRF 45S | NO | 0.00 | 50.96 | 44.75 | 49.31 |
| >hg38_21_+_8433172_8446622_RNA45SN1_45S_50ntFlanks@4759.4793.35\|ATCAGATACCGTCGTAGTTCCGACCATAAACGATG | >hg38_21_+_8433172_8446622_RNA45SN1_45S_50ntFlanks | 4759 | 4793 | 35 | rRF 45S | NO | 41.11 | 81.87 | 0.00 | 16.33 |
| >hg38_21_+_8433172_8446622_RNA45SN1_45S_50ntFlanks@4760.4778.19\|TCAGATACCGTCGTAGTTC | >hg38_21_+_8433172_8446622_RNA45SN1_45S_50ntFlanks | 4760 | 4778 | 19 | rRF 45S | NO | 39.26 | 59.99 | 6.51 | 6.30 |
| >hg38_21_+_8433172_8446622_RNA45SN1_45S_50ntFlanks@4760.4779.20\|TCAGATACCGTCGTAGTTCC | >hg38_21_+_8433172_8446622_RNA45SN1_45S_50ntFlanks | 4760 | 4779 | 20 | rRF 45S | NO | 32.57 | 43.72 | 0.00 | 0.00 |
| >hg38_21_+_8433172_8446622_RNA45SN1_45S_50ntFlanks@4760.4780.21\|TCAGATACCGTCGTAGTTCCG | >hg38_21_+_8433172_8446622_RNA45SN1_45S_50ntFlanks | 4760 | 4780 | 21 | rRF 45S | NO | 243.72 | 425.92 | 56.82 | 103.62 |
| >hg38_21_+_8433172_8446622_RNA45SN1_45S_50ntFlanks@4760.4782.23\|TCAGATACCGTCGTAGTTCCGAC | >hg38_21_+_8433172_8446622_RNA45SN1_45S_50ntFlanks | 4760 | 4782 | 23 | rRF 45S | NO | 31.24 | 115.69 | 53.23 | 45.83 |
| >hg38_21_+_8433172_8446622_RNA45SN1_45S_50ntFlanks@4760.4783.24\|TCAGATACCGTCGTAGTTCCGACC | >hg38_21_+_8433172_8446622_RNA45SN1_45S_50ntFlanks | 4760 | 4783 | 24 | rRF 45S | NO | 20.16 | 60.25 | 29.07 | 33.31 |
| >hg38_21_+_8433172_8446622_RNA45SN1_45S_50ntFlanks@4760.4786.27\|TCAGATACCGTCGTAGTTCCGACCATA | >hg38_21_+_8433172_8446622_RNA45SN1_45S_50ntFlanks | 4760 | 4786 | 27 | rRF 45S | NO | 0.00 | 29.14 | 28.28 | 15.73 |
| >hg38_21_+_8433172_8446622_RNA45SN1_45S_50ntFlanks@4760.4793.34\|TCAGATACCGTCGTAGTTCCGACCATAAACGATG | >hg38_21_+_8433172_8446622_RNA45SN1_45S_50ntFlanks | 4760 | 4793 | 34 | rRF 45S | NO | 29.65 | 42.45 | 0.00 | 0.00 |
| >hg38_21_+_8433172_8446622_RNA45SN1_45S_50ntFlanks@4761.4780.20\|CAGATACCGTCGTAGTTCCG | >hg38_21_+_8433172_8446622_RNA45SN1_45S_50ntFlanks | 4761 | 4780 | 20 | rRF 45S | NO | 0.00 | 12.48 | 0.00 | 0.00 |
| >hg38_21_+_8433172_8446622_RNA45SN1_45S_50ntFlanks@4762.4780.19\|AGATACCGTCGTAGTTCCG | >hg38_21_+_8433172_8446622_RNA45SN1_45S_50ntFlanks | 4762 | 4780 | 19 | rRF 45S | NO | 29.24 | 63.49 | 0.00 | 13.84 |
| >hg38_21_+_8433172_8446622_RNA45SN1_45S_50ntFlanks@4763.4780.18\|GATACCGTCGTAGTTCCG | >hg38_21_+_8433172_8446622_RNA45SN1_45S_50ntFlanks | 4763 | 4780 | 18 | rRF 45S | NO | 32.27 | 108.38 | 0.00 | 16.07 |
| >hg38_21_+_8433172_8446622_RNA45SN1_45S_50ntFlanks@4770.4793.24\|TCGTAGTTCCGACCATAAACGATG | >hg38_21_+_8433172_8446622_RNA45SN1_45S_50ntFlanks | 4770 | 4793 | 24 | rRF 45S | NO | 0.00 | 47.94 | 0.00 | 0.00 |
| >hg38_21_+_8433172_8446622_RNA45SN1_45S_50ntFlanks@4772.4793.22\|GTAGTTCCGACCATAAACGATG | >hg38_21_+_8433172_8446622_RNA45SN1_45S_50ntFlanks | 4772 | 4793 | 22 | rRF 45S | NO | 12.72 | 36.41 | 0.00 | 0.00 |
| >hg38_21_+_8433172_8446622_RNA45SN1_45S_50ntFlanks@4773.4793.21\|TAGTTCCGACCATAAACGATG | >hg38_21_+_8433172_8446622_RNA45SN1_45S_50ntFlanks | 4773 | 4793 | 21 | rRF 45S | NO | 121.57 | 180.79 | 30.28 | 36.61 |
| >hg38_21_+_8433172_8446622_RNA45SN1_45S_50ntFlanks@4853.4883.31\|AAGTCTTTGGGTTCCGGGGGGAGTATGGTTG | >hg38_21_+_8433172_8446622_RNA45SN1_45S_50ntFlanks | 4853 | 4883 | 31 | rRF 45S | NO | 46.96 | 0.00 | 0.00 | 0.00 |
| >hg38_21_+_8433172_8446622_RNA45SN1_45S_50ntFlanks@4971.4999.29\|CCGGCCCGGACACGGACAGGATTGACAGA | >hg38_21_+_8433172_8446622_RNA45SN1_45S_50ntFlanks | 4971 | 4999 | 29 | rRF 45S | NO | 0.00 | 0.00 | 43.69 | 23.66 |
| >hg38_21_+_8433172_8446622_RNA45SN1_45S_50ntFlanks@4975.4994.20\|CCCGGACACGGACAGGATTG | >hg38_21_+_8433172_8446622_RNA45SN1_45S_50ntFlanks | 4975 | 4994 | 20 | rRF 45S | NO | 0.00 | 0.00 | 40.97 | 0.00 |
| >hg38_21_+_8433172_8446622_RNA45SN1_45S_50ntFlanks@4983.5002.20\|CGGACAGGATTGACAGATTG | >hg38_21_+_8433172_8446622_RNA45SN1_45S_50ntFlanks | 4983 | 5002 | 20 | rRF 45S | NO | 0.00 | 0.00 | 33.55 | 16.55 |
| >hg38_21_+_8433172_8446622_RNA45SN1_45S_50ntFlanks@4995.5012.18\|ACAGATTGATAGCTCTTT | >hg38_21_+_8433172_8446622_RNA45SN1_45S_50ntFlanks | 4995 | 5012 | 18 | rRF 45S | NO | 0.00 | 0.00 | 27.93 | 0.00 |
| >hg38_21_+_8433172_8446622_RNA45SN1_45S_50ntFlanks@4995.5013.19\|ACAGATTGATAGCTCTTTC | >hg38_21_+_8433172_8446622_RNA45SN1_45S_50ntFlanks | 4995 | 5013 | 19 | rRF 45S | NO | 0.00 | 0.00 | 61.45 | 13.56 |
| >hg38_21_+_8433172_8446622_RNA45SN1_45S_50ntFlanks@4995.5022.28\|ACAGATTGATAGCTCTTTCTCGATTCCG | >hg38_21_+_8433172_8446622_RNA45SN1_45S_50ntFlanks | 4995 | 5022 | 28 | rRF 45S | NO | 37.13 | 0.00 | 0.00 | 23.10 |
| >hg38_21_+_8433172_8446622_RNA45SN1_45S_50ntFlanks@4995.5024.30\|ACAGATTGATAGCTCTTTCTCGATTCCGTG | >hg38_21_+_8433172_8446622_RNA45SN1_45S_50ntFlanks | 4995 | 5024 | 30 | rRF 45S | NO | 105.26 | 0.00 | 31.39 | 37.67 |
| >hg38_21_+_8433172_8446622_RNA45SN1_45S_50ntFlanks@5071.5097.27\|TTAATTCCGATAACGAACGAGACTCTG | >hg38_21_+_8433172_8446622_RNA45SN1_45S_50ntFlanks | 5071 | 5097 | 27 | rRF 45S | NO | 20.32 | 0.00 | 0.00 | 0.00 |
| >hg38_21_+_8433172_8446622_RNA45SN1_45S_50ntFlanks@5071.5098.28\|TTAATTCCGATAACGAACGAGACTCTGG | >hg38_21_+_8433172_8446622_RNA45SN1_45S_50ntFlanks | 5071 | 5098 | 28 | rRF 45S | NO | 65.85 | 0.00 | 6.53 | 14.41 |
| >hg38_21_+_8433172_8446622_RNA45SN1_45S_50ntFlanks@5073.5097.25\|AATTCCGATAACGAACGAGACTCTG | >hg38_21_+_8433172_8446622_RNA45SN1_45S_50ntFlanks | 5073 | 5097 | 25 | rRF 45S | NO | 42.62 | 0.00 | 0.00 | 13.31 |
| >hg38_21_+_8433172_8446622_RNA45SN1_45S_50ntFlanks@5073.5098.26\|AATTCCGATAACGAACGAGACTCTGG | >hg38_21_+_8433172_8446622_RNA45SN1_45S_50ntFlanks | 5073 | 5098 | 26 | rRF 45S | NO | 108.47 | 26.80 | 42.28 | 55.09 |
| >hg38_21_+_8433172_8446622_RNA45SN1_45S_50ntFlanks@5073.5102.30\|AATTCCGATAACGAACGAGACTCTGGCATG | >hg38_21_+_8433172_8446622_RNA45SN1_45S_50ntFlanks | 5073 | 5102 | 30 | rRF 45S | NO | 34.88 | 0.00 | 0.00 | 0.00 |
| >hg38_21_+_8433172_8446622_RNA45SN1_45S_50ntFlanks@5073.5110.38\|AATTCCGATAACGAACGAGACTCTGGCATGCTAACTAG | >hg38_21_+_8433172_8446622_RNA45SN1_45S_50ntFlanks | 5073 | 5110 | 38 | rRF 45S | NO | 37.65 | 0.00 | 0.00 | 42.41 |
| >hg38_21_+_8433172_8446622_RNA45SN1_45S_50ntFlanks@5074.5098.25\|ATTCCGATAACGAACGAGACTCTGG | >hg38_21_+_8433172_8446622_RNA45SN1_45S_50ntFlanks | 5074 | 5098 | 25 | rRF 45S | NO | 13.08 | 0.00 | 0.00 | 0.00 |
| >hg38_21_+_8433172_8446622_RNA45SN1_45S_50ntFlanks@5078.5097.20\|CGATAACGAACGAGACTCTG | >hg38_21_+_8433172_8446622_RNA45SN1_45S_50ntFlanks | 5078 | 5097 | 20 | rRF 45S | NO | 0.00 | 0.00 | 35.60 | 0.00 |
| >hg38_21_+_8433172_8446622_RNA45SN1_45S_50ntFlanks@5078.5098.21\|CGATAACGAACGAGACTCTGG | >hg38_21_+_8433172_8446622_RNA45SN1_45S_50ntFlanks | 5078 | 5098 | 21 | rRF 45S | NO | 0.00 | 0.00 | 35.27 | 0.00 |

### MDA-MB-468

| | | | | | | | | | | |
|---|---|---|---|---|---|---|---|---|---|---|
| >hg38_21_+_8433172_8446622_RNA45SN1_45S_50ntFlanks@5078.5102.25\|CGATAACGAACGAGACTCTGGCATG | >hg38_21_+_8433172_8446622_RNA45SN1_45S_50ntFlanks | 5078 | 5102 | 25 | rRF 45S | NO | 36.58 | 0.00 | 31.98 | 0.00 |
| >hg38_21_+_8433172_8446622_RNA45SN1_45S_50ntFlanks@5084.5102.19\|CGAACGAGACTCTGGCATG | >hg38_21_+_8433172_8446622_RNA45SN1_45S_50ntFlanks | 5084 | 5102 | 19 | rRF 45S | NO | 36.67 | 0.00 | 46.94 | 24.35 |
| >hg38_21_+_8433172_8446622_RNA45SN1_45S_50ntFlanks@5084.5105.22\|CGAACGAGACTCTGGCATGCTA | >hg38_21_+_8433172_8446622_RNA45SN1_45S_50ntFlanks | 5084 | 5105 | 22 | rRF 45S | NO | 31.29 | 0.00 | 13.57 | 6.53 |
| >hg38_21_+_8433172_8446622_RNA45SN1_45S_50ntFlanks@5084.5110.27\|CGAACGAGACTCTGGCATGCTAACTAG | >hg38_21_+_8433172_8446622_RNA45SN1_45S_50ntFlanks | 5084 | 5110 | 27 | rRF 45S | NO | 70.80 | 0.00 | 34.24 | 75.74 |
| >hg38_21_+_8433172_8446622_RNA45SN1_45S_50ntFlanks@5186.5205.20\|CAATAACAGGTCTGTGATGC | >hg38_21_+_8433172_8446622_RNA45SN1_45S_50ntFlanks | 5186 | 5205 | 20 | rRF 45S | NO | 12.85 | 0.00 | 0.00 | 0.00 |
| >hg38_21_+_8433172_8446622_RNA45SN1_45S_50ntFlanks@5237.5254.18\|ACTGACTGGCTCAGCGTG | >hg38_21_+_8433172_8446622_RNA45SN1_45S_50ntFlanks | 5237 | 5254 | 18 | rRF 45S | YES | 0.00 | 0.00 | 42.48 | 0.00 |
| >hg38_21_+_8433172_8446622_RNA45SN1_45S_50ntFlanks@5417.5445.29\|CTACCGATTGGATGGTTTAGTGAGGCCCT | >hg38_21_+_8433172_8446622_RNA45SN1_45S_50ntFlanks | 5417 | 5445 | 29 | rRF 45S | NO | 37.55 | 51.52 | 0.00 | 0.00 |
| >hg38_21_+_8433172_8446622_RNA45SN1_45S_50ntFlanks@5420.5443.24\|CCGATTGGATGGTTTAGTGAGGCC | >hg38_21_+_8433172_8446622_RNA45SN1_45S_50ntFlanks | 5420 | 5443 | 24 | rRF 45S | NO | 101.70 | 374.07 | 55.17 | 38.59 |
| >hg38_21_+_8433172_8446622_RNA45SN1_45S_50ntFlanks@5420.5444.25\|CCGATTGGATGGTTTAGTGAGGCCC | >hg38_21_+_8433172_8446622_RNA45SN1_45S_50ntFlanks | 5420 | 5444 | 25 | rRF 45S | NO | 20.61 | 77.84 | 0.00 | 0.00 |
| >hg38_21_+_8433172_8446622_RNA45SN1_45S_50ntFlanks@5420.5445.26\|CCGATTGGATGGTTTAGTGAGGCCCT | >hg38_21_+_8433172_8446622_RNA45SN1_45S_50ntFlanks | 5420 | 5445 | 26 | rRF 45S | NO | 68.13 | 193.69 | 7.09 | 34.21 |
| >hg38_21_+_8433172_8446622_RNA45SN1_45S_50ntFlanks@5421.5443.23\|CGATTGGATGGTTTAGTGAGGCC | >hg38_21_+_8433172_8446622_RNA45SN1_45S_50ntFlanks | 5421 | 5443 | 23 | rRF 45S | NO | 51.38 | 118.09 | 45.07 | 13.33 |
| >hg38_21_+_8433172_8446622_RNA45SN1_45S_50ntFlanks@5421.5445.25\|CGATTGGATGGTTTAGTGAGGCCCT | >hg38_21_+_8433172_8446622_RNA45SN1_45S_50ntFlanks | 5421 | 5445 | 25 | rRF 45S | NO | 33.42 | 62.38 | 6.49 | 15.95 |
| >hg38_21_+_8433172_8446622_RNA45SN1_45S_50ntFlanks@5423.5466.44\|ATTGGATGGTTTAGTGAGGCCCTCGGATCGGCCCCGCCGGGGTC | >hg38_21_+_8433172_8446622_RNA45SN1_45S_50ntFlanks | 5423 | 5466 | 44 | rRF 45S | YES | 73.66 | 70.68 | 0.00 | 0.00 |
| >hg38_21_+_8433172_8446622_RNA45SN1_45S_50ntFlanks@5436.5466.31\|GTGAGGCCCTCGGATCGGCCCCGCCGGGGTC | >hg38_21_+_8433172_8446622_RNA45SN1_45S_50ntFlanks | 5436 | 5466 | 31 | rRF 45S | NO | 34.33 | 31.09 | 0.00 | 0.00 |
| >hg38_21_+_8433172_8446622_RNA45SN1_45S_50ntFlanks@5442.5466.25\|CCCTCGGATCGGCCCCGCCGGGGTC | >hg38_21_+_8433172_8446622_RNA45SN1_45S_50ntFlanks | 5442 | 5466 | 25 | rRF 45S | NO | 36.49 | 35.58 | 0.00 | 15.50 |
| >hg38_21_+_8433172_8446622_RNA45SN1_45S_50ntFlanks@5443.5466.24\|CCTCGGATCGGCCCCGCCGGGGTC | >hg38_21_+_8433172_8446622_RNA45SN1_45S_50ntFlanks | 5443 | 5466 | 24 | rRF 45S | NO | 73.65 | 78.14 | 0.00 | 52.62 |
| >hg38_21_+_8433172_8446622_RNA45SN1_45S_50ntFlanks@5444.5465.22\|CTCGGATCGGCCCCGCCGGGGT | >hg38_21_+_8433172_8446622_RNA45SN1_45S_50ntFlanks | 5444 | 5465 | 22 | rRF 45S | NO | 21.01 | 0.00 | 0.00 | 0.00 |
| >hg38_21_+_8433172_8446622_RNA45SN1_45S_50ntFlanks@5444.5466.23\|CTCGGATCGGCCCCGCCGGGGTC | >hg38_21_+_8433172_8446622_RNA45SN1_45S_50ntFlanks | 5444 | 5466 | 23 | rRF 45S | YES | 1011.23 | 1693.70 | 685.90 | 520.51 |
| >hg38_21_+_8433172_8446622_RNA45SN1_45S_50ntFlanks@5444.5467.24\|CTCGGATCGGCCCCGCCGGGGTCG | >hg38_21_+_8433172_8446622_RNA45SN1_45S_50ntFlanks | 5444 | 5467 | 24 | rRF 45S | NO | 49.99 | 96.44 | 130.33 | 39.83 |
| >hg38_21_+_8433172_8446622_RNA45SN1_45S_50ntFlanks@5445.5466.22\|TCGGATCGGCCCCGCCGGGGTC | >hg38_21_+_8433172_8446622_RNA45SN1_45S_50ntFlanks | 5445 | 5466 | 22 | rRF 45S | NO | 236.86 | 488.32 | 368.97 | 152.36 |
| >hg38_21_+_8433172_8446622_RNA45SN1_45S_50ntFlanks@5445.5467.23\|TCGGATCGGCCCCGCCGGGGTCG | >hg38_21_+_8433172_8446622_RNA45SN1_45S_50ntFlanks | 5445 | 5467 | 23 | rRF 45S | NO | 0.00 | 26.40 | 37.59 | 0.00 |
| >hg38_21_+_8433172_8446622_RNA45SN1_45S_50ntFlanks@5446.5466.21\|CGGATCGGCCCCGCCGGGGTC | >hg38_21_+_8433172_8446622_RNA45SN1_45S_50ntFlanks | 5446 | 5466 | 21 | rRF 45S | NO | 354.77 | 719.44 | 869.57 | 403.80 |
| >hg38_21_+_8433172_8446622_RNA45SN1_45S_50ntFlanks@5446.5467.22\|CGGATCGGCCCCGCCGGGGTCG | >hg38_21_+_8433172_8446622_RNA45SN1_45S_50ntFlanks | 5446 | 5467 | 22 | rRF 45S | NO | 0.00 | 32.76 | 57.81 | 13.49 |
| >hg38_21_+_8433172_8446622_RNA45SN1_45S_50ntFlanks@5449.5466.18\|ATCGGCCCCGCCGGGGTC | >hg38_21_+_8433172_8446622_RNA45SN1_45S_50ntFlanks | 5449 | 5466 | 18 | rRF 45S | NO | 0.00 | 0.00 | 122.67 | 0.00 |
| >hg38_21_+_8433172_8446622_RNA45SN1_45S_50ntFlanks@5467.5498.32\|GGCCCACGGCCCTGGCGGAGCGCTGAGAAGAC | >hg38_21_+_8433172_8446622_RNA45SN1_45S_50ntFlanks | 5467 | 5498 | 32 | rRF 45S | NO | 21.04 | 0.00 | 0.00 | 0.00 |
| >hg38_21_+_8433172_8446622_RNA45SN1_45S_50ntFlanks@5467.5499.33\|GGCCCACGGCCCTGGCGGAGCGCTGAGAAGACG | >hg38_21_+_8433172_8446622_RNA45SN1_45S_50ntFlanks | 5467 | 5499 | 33 | rRF 45S | NO | 418.61 | 157.67 | 38.13 | 74.60 |
| >hg38_21_+_8433172_8446622_RNA45SN1_45S_50ntFlanks@5467.5514.48\|GGCCCACGGCCCTGGCGGAGCGCTGAGAAGACGGTCGAACTTGACTAT | >hg38_21_+_8433172_8446622_RNA45SN1_45S_50ntFlanks | 5467 | 5514 | 48 | rRF 45S | YES | 86.79 | 53.85 | 0.00 | 0.00 |
| >hg38_21_+_8433172_8446622_RNA45SN1_45S_50ntFlanks@5468.5499.32\|GCCCACGGCCCTGGCGGAGCGCTGAGAAGACG | >hg38_21_+_8433172_8446622_RNA45SN1_45S_50ntFlanks | 5468 | 5499 | 32 | rRF 45S | NO | 44.14 | 0.00 | 0.00 | 0.00 |
| >hg38_21_+_8433172_8446622_RNA45SN1_45S_50ntFlanks@5469.5499.31\|CCCACGGCCCTGGCGGAGCGCTGAGAAGACG | >hg38_21_+_8433172_8446622_RNA45SN1_45S_50ntFlanks | 5469 | 5499 | 31 | rRF 45S | NO | 49.22 | 35.82 | 0.00 | 0.00 |
| >hg38_21_+_8433172_8446622_RNA45SN1_45S_50ntFlanks@5470.5499.30\|CCACGGCCCTGGCGGAGCGCTGAGAAGACG | >hg38_21_+_8433172_8446622_RNA45SN1_45S_50ntFlanks | 5470 | 5499 | 30 | rRF 45S | NO | 65.80 | 77.55 | 30.23 | 14.35 |
| >hg38_21_+_8433172_8446622_RNA45SN1_45S_50ntFlanks@5471.5499.29\|CACGGCCCTGGCGGAGCGCTGAGAAGACG | >hg38_21_+_8433172_8446622_RNA45SN1_45S_50ntFlanks | 5471 | 5499 | 29 | rRF 45S | NO | 166.58 | 250.04 | 84.87 | 47.31 |
| >hg38_21_+_8433172_8446622_RNA45SN1_45S_50ntFlanks@5471.5514.44\|CACGGCCCTGGCGGAGCGCTGAGAAGACGGTCGAACTTGACTAT | >hg38_21_+_8433172_8446622_RNA45SN1_45S_50ntFlanks | 5471 | 5514 | 44 | rRF 45S | NO | 78.49 | 167.48 | 0.00 | 34.15 |
| >hg38_21_+_8433172_8446622_RNA45SN1_45S_50ntFlanks@5472.5499.28\|ACGGCCCTGGCGGAGCGCTGAGAAGACG | >hg38_21_+_8433172_8446622_RNA45SN1_45S_50ntFlanks | 5472 | 5499 | 28 | rRF 45S | NO | 108.98 | 138.93 | 43.82 | 41.74 |
| >hg38_21_+_8433172_8446622_RNA45SN1_45S_50ntFlanks@5472.5514.43\|ACGGCCCTGGCGGAGCGCTGAGAAGACGGTCGAACTTGACTAT | >hg38_21_+_8433172_8446622_RNA45SN1_45S_50ntFlanks | 5472 | 5514 | 43 | rRF 45S | NO | 57.30 | 100.77 | 0.00 | 14.31 |

### MDA-MB-468

| | | | | | | | | | | |
|---|---|---|---|---|---|---|---|---|---|---|
| >hg38_21_+_8433172_8446622_RNA45SN1_45S_50ntFlanks@5473.5499.27\|CGGCCCTGGCGGAGCGCTGAGAAGACG | >hg38_21_+_8433172_8446622_RNA45SN1_45S_50ntFlanks | 5473 | 5499 | 27 | rRF 45S | NO | 55.36 | 87.80 | 85.95 | 52.13 |
| >hg38_21_+_8433172_8446622_RNA45SN1_45S_50ntFlanks@5473.5514.42\|CGGCCCTGGCGGAGCGCTGAGAAGACGGTCGAACTTGACTAT | >hg38_21_+_8433172_8446622_RNA45SN1_45S_50ntFlanks | 5473 | 5514 | 42 | rRF 45S | NO | 35.83 | 61.93 | 0.00 | 25.49 |
| >hg38_21_+_8433172_8446622_RNA45SN1_45S_50ntFlanks@5482.5499.18\|CGGAGCGCTGAGAAGACG | >hg38_21_+_8433172_8446622_RNA45SN1_45S_50ntFlanks | 5482 | 5499 | 18 | rRF 45S | NO | 0.00 | 0.00 | 102.66 | 0.00 |
| >hg38_21_+_8433172_8446622_RNA45SN1_45S_50ntFlanks@5482.5514.33\|CGGAGCGCTGAGAAGACGGTCGAACTTGACTAT | >hg38_21_+_8433172_8446622_RNA45SN1_45S_50ntFlanks | 5482 | 5514 | 33 | rRF 45S | NO | 13.10 | 28.16 | 31.29 | 0.00 |
| >hg38_21_+_8433172_8446622_RNA45SN1_45S_50ntFlanks@5484.5509.26\|GAGCGCTGAGAAGACGGTCGAACTTG | >hg38_21_+_8433172_8446622_RNA45SN1_45S_50ntFlanks | 5484 | 5509 | 26 | rRF 45S | NO | 0.00 | 0.00 | 55.08 | 0.00 |
| >hg38_21_+_8433172_8446622_RNA45SN1_45S_50ntFlanks@5484.5514.31\|GAGCGCTGAGAAGACGGTCGAACTTGACTAT | >hg38_21_+_8433172_8446622_RNA45SN1_45S_50ntFlanks | 5484 | 5514 | 31 | rRF 45S | NO | 249.97 | 391.11 | 245.60 | 169.31 |
| >hg38_21_+_8433172_8446622_RNA45SN1_45S_50ntFlanks@5484.5518.35\|GAGCGCTGAGAAGACGGTCGAACTTGACTATCTAG | >hg38_21_+_8433172_8446622_RNA45SN1_45S_50ntFlanks | 5484 | 5518 | 35 | rRF 45S | NO | 0.00 | 31.95 | 0.00 | 0.00 |
| >hg38_21_+_8433172_8446622_RNA45SN1_45S_50ntFlanks@5485.5506.22\|AGCGCTGAGAAGACGGTCGAAC | >hg38_21_+_8433172_8446622_RNA45SN1_45S_50ntFlanks | 5485 | 5506 | 22 | rRF 45S | NO | 0.00 | 0.00 | 75.33 | 0.00 |
| >hg38_21_+_8433172_8446622_RNA45SN1_45S_50ntFlanks@5485.5509.25\|AGCGCTGAGAAGACGGTCGAACTTG | >hg38_21_+_8433172_8446622_RNA45SN1_45S_50ntFlanks | 5485 | 5509 | 25 | rRF 45S | NO | 0.00 | 28.71 | 127.99 | 15.73 |
| >hg38_21_+_8433172_8446622_RNA45SN1_45S_50ntFlanks@5485.5510.26\|AGCGCTGAGAAGACGGTCGAACTTGA | >hg38_21_+_8433172_8446622_RNA45SN1_45S_50ntFlanks | 5485 | 5510 | 26 | rRF 45S | NO | 0.00 | 0.00 | 75.09 | 0.00 |
| >hg38_21_+_8433172_8446622_RNA45SN1_45S_50ntFlanks@5485.5512.28\|AGCGCTGAGAAGACGGTCGAACTTGACT | >hg38_21_+_8433172_8446622_RNA45SN1_45S_50ntFlanks | 5485 | 5512 | 28 | rRF 45S | NO | 0.00 | 0.00 | 36.94 | 0.00 |
| >hg38_21_+_8433172_8446622_RNA45SN1_45S_50ntFlanks@5485.5513.29\|AGCGCTGAGAAGACGGTCGAACTTGACTA | >hg38_21_+_8433172_8446622_RNA45SN1_45S_50ntFlanks | 5485 | 5513 | 29 | rRF 45S | NO | 0.00 | 0.00 | 65.13 | 14.14 |
| >hg38_21_+_8433172_8446622_RNA45SN1_45S_50ntFlanks@5485.5514.30\|AGCGCTGAGAAGACGGTCGAACTTGACTAT | >hg38_21_+_8433172_8446622_RNA45SN1_45S_50ntFlanks | 5485 | 5514 | 30 | rRF 45S | NO | 158.18 | 441.98 | 396.04 | 203.51 |
| >hg38_21_+_8433172_8446622_RNA45SN1_45S_50ntFlanks@5485.5515.31\|AGCGCTGAGAAGACGGTCGAACTTGACTATC | >hg38_21_+_8433172_8446622_RNA45SN1_45S_50ntFlanks | 5485 | 5515 | 31 | rRF 45S | NO | 5.97 | 32.54 | 48.75 | 16.57 |
| >hg38_21_+_8433172_8446622_RNA45SN1_45S_50ntFlanks@5485.5516.32\|AGCGCTGAGAAGACGGTCGAACTTGACTATCT | >hg38_21_+_8433172_8446622_RNA45SN1_45S_50ntFlanks | 5485 | 5516 | 32 | rRF 45S | NO | 33.87 | 80.04 | 36.85 | 40.08 |
| >hg38_21_+_8433172_8446622_RNA45SN1_45S_50ntFlanks@5485.5518.34\|AGCGCTGAGAAGACGGTCGAACTTGACTATCTAG | >hg38_21_+_8433172_8446622_RNA45SN1_45S_50ntFlanks | 5485 | 5518 | 34 | rRF 45S | NO | 36.95 | 98.09 | 58.62 | 42.80 |
| >hg38_21_+_8433172_8446622_RNA45SN1_45S_50ntFlanks@5485.5526.42\|AGCGCTGAGAAGACGGTCGAACTTGACTATCTAGAGGAAGTA | >hg38_21_+_8433172_8446622_RNA45SN1_45S_50ntFlanks | 5485 | 5526 | 42 | rRF 45S | NO | 36.33 | 32.14 | 0.00 | 82.70 |
| >hg38_21_+_8433172_8446622_RNA45SN1_45S_50ntFlanks@5486.5514.29\|GCGCTGAGAAGACGGTCGAACTTGACTAT | >hg38_21_+_8433172_8446622_RNA45SN1_45S_50ntFlanks | 5486 | 5514 | 29 | rRF 45S | NO | 34.31 | 88.72 | 130.72 | 53.13 |
| >hg38_21_+_8433172_8446622_RNA45SN1_45S_50ntFlanks@5487.5509.23\|CGCTGAGAAGACGGTCGAACTTG | >hg38_21_+_8433172_8446622_RNA45SN1_45S_50ntFlanks | 5487 | 5509 | 23 | rRF 45S | NO | 0.00 | 0.00 | 71.83 | 0.00 |
| >hg38_21_+_8433172_8446622_RNA45SN1_45S_50ntFlanks@5487.5510.24\|CGCTGAGAAGACGGTCGAACTTGA | >hg38_21_+_8433172_8446622_RNA45SN1_45S_50ntFlanks | 5487 | 5510 | 24 | rRF 45S | NO | 0.00 | 0.00 | 45.57 | 0.00 |
| >hg38_21_+_8433172_8446622_RNA45SN1_45S_50ntFlanks@5487.5513.27\|CGCTGAGAAGACGGTCGAACTTGACTA | >hg38_21_+_8433172_8446622_RNA45SN1_45S_50ntFlanks | 5487 | 5513 | 27 | rRF 45S | NO | 0.00 | 0.00 | 42.04 | 0.00 |
| >hg38_21_+_8433172_8446622_RNA45SN1_45S_50ntFlanks@5487.5514.28\|CGCTGAGAAGACGGTCGAACTTGACTAT | >hg38_21_+_8433172_8446622_RNA45SN1_45S_50ntFlanks | 5487 | 5514 | 28 | rRF 45S | NO | 68.07 | 347.70 | 349.45 | 179.44 |
| >hg38_21_+_8433172_8446622_RNA45SN1_45S_50ntFlanks@5487.5515.29\|CGCTGAGAAGACGGTCGAACTTGACTATC | >hg38_21_+_8433172_8446622_RNA45SN1_45S_50ntFlanks | 5487 | 5515 | 29 | rRF 45S | NO | 0.00 | 0.00 | 59.84 | 16.60 |
| >hg38_21_+_8433172_8446622_RNA45SN1_45S_50ntFlanks@5487.5516.30\|CGCTGAGAAGACGGTCGAACTTGACTATCT | >hg38_21_+_8433172_8446622_RNA45SN1_45S_50ntFlanks | 5487 | 5516 | 30 | rRF 45S | NO | 6.12 | 73.78 | 34.29 | 24.30 |
| >hg38_21_+_8433172_8446622_RNA45SN1_45S_50ntFlanks@5487.5526.40\|CGCTGAGAAGACGGTCGAACTTGACTATCTAGAGGAAGTA | >hg38_21_+_8433172_8446622_RNA45SN1_45S_50ntFlanks | 5487 | 5526 | 40 | rRF 45S | NO | 31.85 | 48.76 | 13.65 | 84.00 |
| >hg38_21_+_8433172_8446622_RNA45SN1_45S_50ntFlanks@5489.5509.21\|CTGAGAAGACGGTCGAACTTG | >hg38_21_+_8433172_8446622_RNA45SN1_45S_50ntFlanks | 5489 | 5509 | 21 | rRF 45S | NO | 0.00 | 0.00 | 62.00 | 0.00 |
| >hg38_21_+_8433172_8446622_RNA45SN1_45S_50ntFlanks@5489.5510.22\|CTGAGAAGACGGTCGAACTTGA | >hg38_21_+_8433172_8446622_RNA45SN1_45S_50ntFlanks | 5489 | 5510 | 22 | rRF 45S | NO | 0.00 | 0.00 | 48.35 | 0.00 |
| >hg38_21_+_8433172_8446622_RNA45SN1_45S_50ntFlanks@5489.5514.26\|CTGAGAAGACGGTCGAACTTGACTAT | >hg38_21_+_8433172_8446622_RNA45SN1_45S_50ntFlanks | 5489 | 5514 | 26 | rRF 45S | NO | 0.00 | 70.67 | 250.97 | 47.99 |
| >hg38_21_+_8433172_8446622_RNA45SN1_45S_50ntFlanks@5489.5516.28\|CTGAGAAGACGGTCGAACTTGACTATCT | >hg38_21_+_8433172_8446622_RNA45SN1_45S_50ntFlanks | 5489 | 5516 | 28 | rRF 45S | NO | 0.00 | 45.52 | 27.83 | 15.22 |
| >hg38_21_+_8433172_8446622_RNA45SN1_45S_50ntFlanks@5492.5509.18\|AGAAGACGGTCGAACTTG | >hg38_21_+_8433172_8446622_RNA45SN1_45S_50ntFlanks | 5492 | 5509 | 18 | rRF 45S | NO | 0.00 | 35.27 | 51.75 | 0.00 |
| >hg38_21_+_8433172_8446622_RNA45SN1_45S_50ntFlanks@5492.5510.19\|AGAAGACGGTCGAACTTGA | >hg38_21_+_8433172_8446622_RNA45SN1_45S_50ntFlanks | 5492 | 5510 | 19 | rRF 45S | NO | 0.00 | 0.00 | 30.18 | 0.00 |
| >hg38_21_+_8433172_8446622_RNA45SN1_45S_50ntFlanks@5492.5513.22\|AGAAGACGGTCGAACTTGACTA | >hg38_21_+_8433172_8446622_RNA45SN1_45S_50ntFlanks | 5492 | 5513 | 22 | rRF 45S | NO | 0.00 | 0.00 | 51.73 | 0.00 |
| >hg38_21_+_8433172_8446622_RNA45SN1_45S_50ntFlanks@5492.5514.23\|AGAAGACGGTCGAACTTGACTAT | >hg38_21_+_8433172_8446622_RNA45SN1_45S_50ntFlanks | 5492 | 5514 | 23 | rRF 45S | NO | 0.00 | 103.10 | 293.01 | 57.35 |
| >hg38_21_+_8433172_8446622_RNA45SN1_45S_50ntFlanks@5492.5516.25\|AGAAGACGGTCGAACTTGACTATCT | >hg38_21_+_8433172_8446622_RNA45SN1_45S_50ntFlanks | 5492 | 5516 | 25 | rRF 45S | NO | 0.00 | 29.47 | 0.00 | 0.00 |

### MDA-MB-468

| | | | | | | | | | | |
|---|---|---|---|---|---|---|---|---|---|---|
| >hg38_21_+_8433172_8446622_RNA45SN1_45S_50ntFlanks@5492.5526.35\|AGAAGACGGTCGAACTTGACTATCTAGAGGAAGTA | >hg38_21_+_8433172_8446622_RNA45SN1_45S_50ntFlanks | 5492 | 5526 | 35 | rRF 45S | NO | 21.21 | 50.65 | 46.61 | 85.76 |
| >hg38_21_+_8433172_8446622_RNA45SN1_45S_50ntFlanks@5494.5514.21\|AAGACGGTCGAACTTGACTAT | >hg38_21_+_8433172_8446622_RNA45SN1_45S_50ntFlanks | 5494 | 5514 | 21 | rRF 45S | NO | 0.00 | 0.00 | 113.81 | 0.00 |
| >hg38_21_+_8433172_8446622_RNA45SN1_45S_50ntFlanks@5494.5526.33\|AAGACGGTCGAACTTGACTATCTAGAGGAAGTA | >hg38_21_+_8433172_8446622_RNA45SN1_45S_50ntFlanks | 5494 | 5526 | 33 | rRF 45S | NO | 0.00 | 0.00 | 34.99 | 36.46 |
| >hg38_21_+_8433172_8446622_RNA45SN1_45S_50ntFlanks@5515.5535.21\|CTAGAGGAAGTAAAAGTCGTA | >hg38_21_+_8433172_8446622_RNA45SN1_45S_50ntFlanks | 5515 | 5535 | 21 | rRF 45S | NO | 0.00 | 40.00 | 35.28 | 0.00 |
| >hg38_21_+_8433172_8446622_RNA45SN1_45S_50ntFlanks@5523.5550.28\|AGTAAAAGTCGTAACAAGGTTTCCGTAG | >hg38_21_+_8433172_8446622_RNA45SN1_45S_50ntFlanks | 5523 | 5550 | 28 | rRF 45S | NO | 71.48 | 0.00 | 0.00 | 0.00 |
| >hg38_21_+_8433172_8446622_RNA45SN1_45S_50ntFlanks@5524.5550.27\|GTAAAAGTCGTAACAAGGTTTCCGTAG | >hg38_21_+_8433172_8446622_RNA45SN1_45S_50ntFlanks | 5524 | 5550 | 27 | rRF 45S | NO | 41.83 | 27.01 | 0.00 | 0.00 |
| >hg38_21_+_8433172_8446622_RNA45SN1_45S_50ntFlanks@5525.5550.26\|TAAAAGTCGTAACAAGGTTTCCGTAG | >hg38_21_+_8433172_8446622_RNA45SN1_45S_50ntFlanks | 5525 | 5550 | 26 | rRF 45S | NO | 46.86 | 26.60 | 0.00 | 0.00 |
| >hg38_21_+_8433172_8446622_RNA45SN1_45S_50ntFlanks@5526.5550.25\|AAAAGTCGTAACAAGGTTTCCGTAG | >hg38_21_+_8433172_8446622_RNA45SN1_45S_50ntFlanks | 5526 | 5550 | 25 | rRF 45S | NO | 35.32 | 0.00 | 0.00 | 0.00 |
| >hg38_21_+_8433172_8446622_RNA45SN1_45S_50ntFlanks@5527.5549.23\|AAAGTCGTAACAAGGTTTCCGTA | >hg38_21_+_8433172_8446622_RNA45SN1_45S_50ntFlanks | 5527 | 5549 | 23 | rRF 45S | NO | 0.00 | 0.00 | 28.49 | 0.00 |
| >hg38_21_+_8433172_8446622_RNA45SN1_45S_50ntFlanks@5527.5550.24\|AAAGTCGTAACAAGGTTTCCGTAG | >hg38_21_+_8433172_8446622_RNA45SN1_45S_50ntFlanks | 5527 | 5550 | 24 | rRF 45S | NO | 125.37 | 66.41 | 44.58 | 5.99 |
| >hg38_21_+_8433172_8446622_RNA45SN1_45S_50ntFlanks@6645.6662.18\|CTCGTACGACTCTTAGCG | >hg38_21_+_8433172_8446622_RNA45SN1_45S_50ntFlanks | 6645 | 6662 | 18 | rRF 45S | NO | 39.43 | 78.16 | 43.67 | 25.69 |
| >hg38_21_+_8433172_8446622_RNA45SN1_45S_50ntFlanks@6645.6663.19\|CTCGTACGACTCTTAGCGG | >hg38_21_+_8433172_8446622_RNA45SN1_45S_50ntFlanks | 6645 | 6663 | 19 | rRF 45S | NO | 33.69 | 39.20 | 38.68 | 46.87 |
| >hg38_21_+_8433172_8446622_RNA45SN1_45S_50ntFlanks@6645.6665.21\|CTCGTACGACTCTTAGCGGTG | >hg38_21_+_8433172_8446622_RNA45SN1_45S_50ntFlanks | 6645 | 6665 | 21 | rRF 45S | NO | 34.96 | 37.84 | 31.83 | 42.43 |
| >hg38_21_+_8433172_8446622_RNA45SN1_45S_50ntFlanks@6645.6666.22\|CTCGTACGACTCTTAGCGGTGG | >hg38_21_+_8433172_8446622_RNA45SN1_45S_50ntFlanks | 6645 | 6666 | 22 | rRF 45S | NO | 43.39 | 48.44 | 0.00 | 56.68 |
| >hg38_21_+_8433172_8446622_RNA45SN1_45S_50ntFlanks@6645.6669.25\|CTCGTACGACTCTTAGCGGTGGATC | >hg38_21_+_8433172_8446622_RNA45SN1_45S_50ntFlanks | 6645 | 6669 | 25 | rRF 45S | NO | 53.96 | 35.03 | 40.16 | 145.39 |
| >hg38_21_+_8433172_8446622_RNA45SN1_45S_50ntFlanks@6645.6670.26\|CTCGTACGACTCTTAGCGGTGGATCA | >hg38_21_+_8433172_8446622_RNA45SN1_45S_50ntFlanks | 6645 | 6670 | 26 | rRF 45S | NO | 30.82 | 12.50 | 27.47 | 141.08 |
| >hg38_21_+_8433172_8446622_RNA45SN1_45S_50ntFlanks@6645.6679.35\|CTCGTACGACTCTTAGCGGTGGATCACTCGGCTCG | >hg38_21_+_8433172_8446622_RNA45SN1_45S_50ntFlanks | 6645 | 6679 | 35 | rRF 45S | NO | 69.41 | 0.00 | 0.00 | 157.50 |
| >hg38_21_+_8433172_8446622_RNA45SN1_45S_50ntFlanks@6645.6680.36\|CTCGTACGACTCTTAGCGGTGGATCACTCGGCTCGT | >hg38_21_+_8433172_8446622_RNA45SN1_45S_50ntFlanks | 6645 | 6680 | 36 | rRF 45S | NO | 21.88 | 0.00 | 0.00 | 66.41 |
| >hg38_21_+_8433172_8446622_RNA45SN1_45S_50ntFlanks@6645.6681.37\|CTCGTACGACTCTTAGCGGTGGATCACTCGGCTCGTG | >hg38_21_+_8433172_8446622_RNA45SN1_45S_50ntFlanks | 6645 | 6681 | 37 | rRF 45S | NO | 166.82 | 50.14 | 96.30 | 350.77 |
| >hg38_21_+_8433172_8446622_RNA45SN1_45S_50ntFlanks@6646.6663.18\|TCGTACGACTCTTAGCGG | >hg38_21_+_8433172_8446622_RNA45SN1_45S_50ntFlanks | 6646 | 6663 | 18 | rRF 45S | NO | 117.92 | 215.68 | 188.08 | 180.84 |
| >hg38_21_+_8433172_8446622_RNA45SN1_45S_50ntFlanks@6646.6664.19\|TCGTACGACTCTTAGCGGT | >hg38_21_+_8433172_8446622_RNA45SN1_45S_50ntFlanks | 6646 | 6664 | 19 | rRF 45S | NO | 137.75 | 72.36 | 69.25 | 178.19 |
| >hg38_21_+_8433172_8446622_RNA45SN1_45S_50ntFlanks@6646.6665.20\|TCGTACGACTCTTAGCGGTG | >hg38_21_+_8433172_8446622_RNA45SN1_45S_50ntFlanks | 6646 | 6665 | 20 | rRF 45S | NO | 94.60 | 110.88 | 80.29 | 112.47 |
| >hg38_21_+_8433172_8446622_RNA45SN1_45S_50ntFlanks@6646.6666.21\|TCGTACGACTCTTAGCGGTGG | >hg38_21_+_8433172_8446622_RNA45SN1_45S_50ntFlanks | 6646 | 6666 | 21 | rRF 45S | NO | 122.58 | 201.65 | 82.45 | 182.57 |
| >hg38_21_+_8433172_8446622_RNA45SN1_45S_50ntFlanks@6646.6667.22\|TCGTACGACTCTTAGCGGTGGA | >hg38_21_+_8433172_8446622_RNA45SN1_45S_50ntFlanks | 6646 | 6667 | 22 | rRF 45S | NO | 0.00 | 12.28 | 13.54 | 51.97 |
| >hg38_21_+_8433172_8446622_RNA45SN1_45S_50ntFlanks@6646.6668.23\|TCGTACGACTCTTAGCGGTGGAT | >hg38_21_+_8433172_8446622_RNA45SN1_45S_50ntFlanks | 6646 | 6668 | 23 | rRF 45S | NO | 33.09 | 38.41 | 48.22 | 58.30 |
| >hg38_21_+_8433172_8446622_RNA45SN1_45S_50ntFlanks@6646.6669.24\|TCGTACGACTCTTAGCGGTGGATC | >hg38_21_+_8433172_8446622_RNA45SN1_45S_50ntFlanks | 6646 | 6669 | 24 | rRF 45S | NO | 179.55 | 177.65 | 195.47 | 468.50 |
| >hg38_21_+_8433172_8446622_RNA45SN1_45S_50ntFlanks@6646.6670.25\|TCGTACGACTCTTAGCGGTGGATCA | >hg38_21_+_8433172_8446622_RNA45SN1_45S_50ntFlanks | 6646 | 6670 | 25 | rRF 45S | NO | 101.13 | 123.11 | 136.72 | 443.84 |
| >hg38_21_+_8433172_8446622_RNA45SN1_45S_50ntFlanks@6646.6675.30\|TCGTACGACTCTTAGCGGTGGATCACTCGG | >hg38_21_+_8433172_8446622_RNA45SN1_45S_50ntFlanks | 6646 | 6675 | 30 | rRF 45S | NO | 79.22 | 47.85 | 7.11 | 122.12 |
| >hg38_21_+_8433172_8446622_RNA45SN1_45S_50ntFlanks@6646.6677.32\|TCGTACGACTCTTAGCGGTGGATCACTCGGCT | >hg38_21_+_8433172_8446622_RNA45SN1_45S_50ntFlanks | 6646 | 6677 | 32 | rRF 45S | NO | 37.40 | 0.00 | 6.49 | 62.36 |
| >hg38_21_+_8433172_8446622_RNA45SN1_45S_50ntFlanks@6646.6678.33\|TCGTACGACTCTTAGCGGTGGATCACTCGGCTC | >hg38_21_+_8433172_8446622_RNA45SN1_45S_50ntFlanks | 6646 | 6678 | 33 | rRF 45S | NO | 28.66 | 0.00 | 0.00 | 51.49 |
| >hg38_21_+_8433172_8446622_RNA45SN1_45S_50ntFlanks@6646.6679.34\|TCGTACGACTCTTAGCGGTGGATCACTCGGCTCG | >hg38_21_+_8433172_8446622_RNA45SN1_45S_50ntFlanks | 6646 | 6679 | 34 | rRF 45S | NO | 215.52 | 76.33 | 57.07 | 431.08 |
| >hg38_21_+_8433172_8446622_RNA45SN1_45S_50ntFlanks@6646.6680.35\|TCGTACGACTCTTAGCGGTGGATCACTCGGCTCGT | >hg38_21_+_8433172_8446622_RNA45SN1_45S_50ntFlanks | 6646 | 6680 | 35 | rRF 45S | NO | 92.37 | 48.54 | 69.00 | 221.29 |
| >hg38_21_+_8433172_8446622_RNA45SN1_45S_50ntFlanks@6646.6681.36\|TCGTACGACTCTTAGCGGTGGATCACTCGGCTCGTG | >hg38_21_+_8433172_8446622_RNA45SN1_45S_50ntFlanks | 6646 | 6681 | 36 | rRF 45S | YES | 918.93 | 359.86 | 481.24 | 1650.08 |
| >hg38_21_+_8433172_8446622_RNA45SN1_45S_50ntFlanks@6646.6685.40\|TCGTACGACTCTTAGCGGTGGATCACTCGGCTCGTGCGTC | >hg38_21_+_8433172_8446622_RNA45SN1_45S_50ntFlanks | 6646 | 6685 | 40 | rRF 45S | NO | 0.00 | 0.00 | 27.72 | 36.33 |

### MDA-MB-468

| | | | | | | | | | | |
|---|---|---|---|---|---|---|---|---|---|---|
| >hg38_21_+_8433172_8446622_RNA45SN1_45S_50ntFlanks@6646.6689.44\|TCGTACGACTCTTAGCGGTGGATCACTCGGCTCGTGCGTCGATG | >hg38_21_+_8433172_8446622_RNA45SN1_45S_50ntFlanks | 6646 | 6689 | 44 | rRF 45S | NO | 54.96 | 45.91 | 153.55 | 979.13 |
| >hg38_21_+_8433172_8446622_RNA45SN1_45S_50ntFlanks@6647.6665.19\|CGTACGACTCTTAGCGGTG | >hg38_21_+_8433172_8446622_RNA45SN1_45S_50ntFlanks | 6647 | 6665 | 19 | rRF 45S | NO | 0.00 | 0.00 | 30.36 | 25.87 |
| >hg38_21_+_8433172_8446622_RNA45SN1_45S_50ntFlanks@6647.6666.20\|CGTACGACTCTTAGCGGTGG | >hg38_21_+_8433172_8446622_RNA45SN1_45S_50ntFlanks | 6647 | 6666 | 20 | rRF 45S | NO | 21.62 | 37.35 | 13.79 | 39.39 |
| >hg38_21_+_8433172_8446622_RNA45SN1_45S_50ntFlanks@6647.6669.23\|CGTACGACTCTTAGCGGTGGATC | >hg38_21_+_8433172_8446622_RNA45SN1_45S_50ntFlanks | 6647 | 6669 | 23 | rRF 45S | NO | 34.00 | 33.04 | 53.30 | 117.42 |
| >hg38_21_+_8433172_8446622_RNA45SN1_45S_50ntFlanks@6647.6670.24\|CGTACGACTCTTAGCGGTGGATCA | >hg38_21_+_8433172_8446622_RNA45SN1_45S_50ntFlanks | 6647 | 6670 | 24 | rRF 45S | NO | 0.00 | 0.00 | 30.53 | 106.55 |
| >hg38_21_+_8433172_8446622_RNA45SN1_45S_50ntFlanks@6647.6679.33\|CGTACGACTCTTAGCGGTGGATCACTCGGCTCG | >hg38_21_+_8433172_8446622_RNA45SN1_45S_50ntFlanks | 6647 | 6679 | 33 | rRF 45S | NO | 30.39 | 0.00 | 0.00 | 84.75 |
| >hg38_21_+_8433172_8446622_RNA45SN1_45S_50ntFlanks@6647.6681.35\|CGTACGACTCTTAGCGGTGGATCACTCGGCTCGTG | >hg38_21_+_8433172_8446622_RNA45SN1_45S_50ntFlanks | 6647 | 6681 | 35 | rRF 45S | NO | 188.23 | 76.47 | 147.07 | 436.42 |
| >hg38_21_+_8433172_8446622_RNA45SN1_45S_50ntFlanks@6647.6689.43\|CGTACGACTCTTAGCGGTGGATCACTCGGCTCGTGCGTCGATG | >hg38_21_+_8433172_8446622_RNA45SN1_45S_50ntFlanks | 6647 | 6689 | 43 | rRF 45S | NO | 0.00 | 0.00 | 29.67 | 142.91 |
| >hg38_21_+_8433172_8446622_RNA45SN1_45S_50ntFlanks@6648.6679.32\|GTACGACTCTTAGCGGTGGATCACTCGGCTCG | >hg38_21_+_8433172_8446622_RNA45SN1_45S_50ntFlanks | 6648 | 6679 | 32 | rRF 45S | NO | 65.80 | 0.00 | 0.00 | 157.50 |
| >hg38_21_+_8433172_8446622_RNA45SN1_45S_50ntFlanks@6648.6681.34\|GTACGACTCTTAGCGGTGGATCACTCGGCTCGTG | >hg38_21_+_8433172_8446622_RNA45SN1_45S_50ntFlanks | 6648 | 6681 | 34 | rRF 45S | NO | 138.05 | 74.34 | 111.87 | 470.65 |
| >hg38_21_+_8433172_8446622_RNA45SN1_45S_50ntFlanks@6648.6689.42\|GTACGACTCTTAGCGGTGGATCACTCGGCTCGTGCGTCGATG | >hg38_21_+_8433172_8446622_RNA45SN1_45S_50ntFlanks | 6648 | 6689 | 42 | rRF 45S | NO | 73.78 | 19.52 | 50.60 | 496.95 |
| >hg38_21_+_8433172_8446622_RNA45SN1_45S_50ntFlanks@6649.6679.31\|TACGACTCTTAGCGGTGGATCACTCGGCTCG | >hg38_21_+_8433172_8446622_RNA45SN1_45S_50ntFlanks | 6649 | 6679 | 31 | rRF 45S | NO | 50.67 | 12.64 | 0.00 | 121.27 |
| >hg38_21_+_8433172_8446622_RNA45SN1_45S_50ntFlanks@6649.6681.33\|TACGACTCTTAGCGGTGGATCACTCGGCTCGTG | >hg38_21_+_8433172_8446622_RNA45SN1_45S_50ntFlanks | 6649 | 6681 | 33 | rRF 45S | NO | 12.52 | 19.54 | 84.49 | 117.86 |
| >hg38_21_+_8433172_8446622_RNA45SN1_45S_50ntFlanks@6649.6689.41\|TACGACTCTTAGCGGTGGATCACTCGGCTCGTGCGTCGATG | >hg38_21_+_8433172_8446622_RNA45SN1_45S_50ntFlanks | 6649 | 6689 | 41 | rRF 45S | NO | 37.05 | 0.00 | 36.82 | 286.82 |
| >hg38_21_+_8433172_8446622_RNA45SN1_45S_50ntFlanks@6650.6679.30\|ACGACTCTTAGCGGTGGATCACTCGGCTCG | >hg38_21_+_8433172_8446622_RNA45SN1_45S_50ntFlanks | 6650 | 6679 | 30 | rRF 45S | NO | 60.89 | 33.25 | 0.00 | 184.09 |
| >hg38_21_+_8433172_8446622_RNA45SN1_45S_50ntFlanks@6650.6681.32\|ACGACTCTTAGCGGTGGATCACTCGGCTCGTG | >hg38_21_+_8433172_8446622_RNA45SN1_45S_50ntFlanks | 6650 | 6681 | 32 | rRF 45S | NO | 131.32 | 95.89 | 136.22 | 430.54 |
| >hg38_21_+_8433172_8446622_RNA45SN1_45S_50ntFlanks@6650.6689.40\|ACGACTCTTAGCGGTGGATCACTCGGCTCGTGCGTCGATG | >hg38_21_+_8433172_8446622_RNA45SN1_45S_50ntFlanks | 6650 | 6689 | 40 | rRF 45S | NO | 46.59 | 0.00 | 58.83 | 379.40 |
| >hg38_21_+_8433172_8446622_RNA45SN1_45S_50ntFlanks@6650.6698.49\|ACGACTCTTAGCGGTGGATCACTCGGCTCGTGCGTCGATGAAGAACGCA | >hg38_21_+_8433172_8446622_RNA45SN1_45S_50ntFlanks | 6650 | 6698 | 49 | rRF 45S | NO | 80.71 | 0.00 | 0.00 | 152.88 |
| >hg38_21_+_8433172_8446622_RNA45SN1_45S_50ntFlanks@6650.6699.50\|ACGACTCTTAGCGGTGGATCACTCGGCTCGTGCGTCGATGAAGAACGCAG | >hg38_21_+_8433172_8446622_RNA45SN1_45S_50ntFlanks | 6650 | 6699 | 50 | rRF 45S | NO | 47.26 | 0.00 | 0.00 | 98.61 |
| >hg38_21_+_8433172_8446622_RNA45SN1_45S_50ntFlanks@6651.6668.18\|CGACTCTTAGCGGTGGAT | >hg38_21_+_8433172_8446622_RNA45SN1_45S_50ntFlanks | 6651 | 6668 | 18 | rRF 45S | NO | 13.15 | 76.59 | 38.01 | 88.96 |
| >hg38_21_+_8433172_8446622_RNA45SN1_45S_50ntFlanks@6651.6669.19\|CGACTCTTAGCGGTGGATC | >hg38_21_+_8433172_8446622_RNA45SN1_45S_50ntFlanks | 6651 | 6669 | 19 | rRF 45S | NO | 37.13 | 116.84 | 128.36 | 314.47 |
| >hg38_21_+_8433172_8446622_RNA45SN1_45S_50ntFlanks@6651.6670.20\|CGACTCTTAGCGGTGGATCA | >hg38_21_+_8433172_8446622_RNA45SN1_45S_50ntFlanks | 6651 | 6670 | 20 | rRF 45S | NO | 0.00 | 43.93 | 68.26 | 268.58 |
| >hg38_21_+_8433172_8446622_RNA45SN1_45S_50ntFlanks@6651.6675.25\|CGACTCTTAGCGGTGGATCACTCGG | >hg38_21_+_8433172_8446622_RNA45SN1_45S_50ntFlanks | 6651 | 6675 | 25 | rRF 45S | NO | 102.33 | 100.29 | 31.66 | 195.15 |
| >hg38_21_+_8433172_8446622_RNA45SN1_45S_50ntFlanks@6651.6676.26\|CGACTCTTAGCGGTGGATCACTCGGC | >hg38_21_+_8433172_8446622_RNA45SN1_45S_50ntFlanks | 6651 | 6676 | 26 | rRF 45S | NO | 30.90 | 0.00 | 13.59 | 104.61 |
| >hg38_21_+_8433172_8446622_RNA45SN1_45S_50ntFlanks@6651.6677.27\|CGACTCTTAGCGGTGGATCACTCGGCT | >hg38_21_+_8433172_8446622_RNA45SN1_45S_50ntFlanks | 6651 | 6677 | 27 | rRF 45S | NO | 58.11 | 45.73 | 52.02 | 218.83 |
| >hg38_21_+_8433172_8446622_RNA45SN1_45S_50ntFlanks@6651.6678.28\|CGACTCTTAGCGGTGGATCACTCGGCTC | >hg38_21_+_8433172_8446622_RNA45SN1_45S_50ntFlanks | 6651 | 6678 | 28 | rRF 45S | NO | 87.84 | 47.87 | 31.66 | 242.29 |
| >hg38_21_+_8433172_8446622_RNA45SN1_45S_50ntFlanks@6651.6679.29\|CGACTCTTAGCGGTGGATCACTCGGCTCG | >hg38_21_+_8433172_8446622_RNA45SN1_45S_50ntFlanks | 6651 | 6679 | 29 | rRF 45S | NO | 202.63 | 206.72 | 160.16 | 1259.78 |
| >hg38_21_+_8433172_8446622_RNA45SN1_45S_50ntFlanks@6651.6680.30\|CGACTCTTAGCGGTGGATCACTCGGCTCGT | >hg38_21_+_8433172_8446622_RNA45SN1_45S_50ntFlanks | 6651 | 6680 | 30 | rRF 45S | NO | 121.53 | 80.60 | 115.01 | 426.63 |
| >hg38_21_+_8433172_8446622_RNA45SN1_45S_50ntFlanks@6651.6681.31\|CGACTCTTAGCGGTGGATCACTCGGCTCGTG | >hg38_21_+_8433172_8446622_RNA45SN1_45S_50ntFlanks | 6651 | 6681 | 31 | rRF 45S | YES | 1613.25 | 1092.18 | 1026.43 | 3967.73 |
| >hg38_21_+_8433172_8446622_RNA45SN1_45S_50ntFlanks@6651.6683.33\|CGACTCTTAGCGGTGGATCACTCGGCTCGTGCG | >hg38_21_+_8433172_8446622_RNA45SN1_45S_50ntFlanks | 6651 | 6683 | 33 | rRF 45S | NO | 49.22 | 0.00 | 58.99 | 163.53 |
| >hg38_21_+_8433172_8446622_RNA45SN1_45S_50ntFlanks@6651.6684.34\|CGACTCTTAGCGGTGGATCACTCGGCTCGTGCGT | >hg38_21_+_8433172_8446622_RNA45SN1_45S_50ntFlanks | 6651 | 6684 | 34 | rRF 45S | NO | 21.81 | 0.00 | 0.00 | 87.69 |
| >hg38_21_+_8433172_8446622_RNA45SN1_45S_50ntFlanks@6651.6685.35\|CGACTCTTAGCGGTGGATCACTCGGCTCGTGCGTC | >hg38_21_+_8433172_8446622_RNA45SN1_45S_50ntFlanks | 6651 | 6685 | 35 | rRF 45S | NO | 0.00 | 0.00 | 65.46 | 89.33 |
| >hg38_21_+_8433172_8446622_RNA45SN1_45S_50ntFlanks@6651.6686.36\|CGACTCTTAGCGGTGGATCACTCGGCTCGTGCGTCG | >hg38_21_+_8433172_8446622_RNA45SN1_45S_50ntFlanks | 6651 | 6686 | 36 | rRF 45S | NO | 0.00 | 0.00 | 68.16 | 93.04 |
| >hg38_21_+_8433172_8446622_RNA45SN1_45S_50ntFlanks@6651.6688.38\|CGACTCTTAGCGGTGGATCACTCGGCTCGTGCGTCGAT | >hg38_21_+_8433172_8446622_RNA45SN1_45S_50ntFlanks | 6651 | 6688 | 38 | rRF 45S | NO | 20.66 | 0.00 | 66.77 | 242.24 |

### MDA-MB-468

| | | | | | | | | | | |
|---|---|---|---|---|---|---|---|---|---|---|
| >hg38_21_+_8433172_8446622_RNA45SN1_45S_50ntFlanks@6651.6689.39\|CGACTCTTAGCG GTGGATCACTCGGCTCGTGCGTCGATG | >hg38_21_+_8433172_8446622_RNA45SN1_45S_50ntFlanks | 6651 | 6689 | 39 | rRF 45S | YES | 698.95 | 291.42 | 636.88 | 4434.62 |
| >hg38_21_+_8433172_8446622_RNA45SN1_45S_50ntFlanks@6651.6690.40\|CGACTCTTAGCG GTGGATCACTCGGCTCGTGCGTCGATGA | >hg38_21_+_8433172_8446622_RNA45SN1_45S_50ntFlanks | 6651 | 6690 | 40 | rRF 45S | NO | 28.81 | 0.00 | 0.00 | 115.19 |
| >hg38_21_+_8433172_8446622_RNA45SN1_45S_50ntFlanks@6651.6692.42\|CGACTCTTAGCG GTGGATCACTCGGCTCGTGCGTCGATGAAG | >hg38_21_+_8433172_8446622_RNA45SN1_45S_50ntFlanks | 6651 | 6692 | 42 | rRF 45S | NO | 89.49 | 28.76 | 52.62 | 417.50 |
| >hg38_21_+_8433172_8446622_RNA45SN1_45S_50ntFlanks@6651.6696.46\|CGACTCTTAGCG GTGGATCACTCGGCTCGTGCGTCGATGAAGAACG | >hg38_21_+_8433172_8446622_RNA45SN1_45S_50ntFlanks | 6651 | 6696 | 46 | rRF 45S | NO | 117.83 | 0.00 | 20.97 | 298.58 |
| >hg38_21_+_8433172_8446622_RNA45SN1_45S_50ntFlanks@6651.6697.47\|CGACTCTTAGCG GTGGATCACTCGGCTCGTGCGTCGATGAAGAACGC | >hg38_21_+_8433172_8446622_RNA45SN1_45S_50ntFlanks | 6651 | 6697 | 47 | rRF 45S | NO | 97.05 | 33.26 | 0.00 | 300.40 |
| >hg38_21_+_8433172_8446622_RNA45SN1_45S_50ntFlanks@6651.6698.48\|CGACTCTTAGCG GTGGATCACTCGGCTCGTGCGTCGATGAAGAACGCA | >hg38_21_+_8433172_8446622_RNA45SN1_45S_50ntFlanks | 6651 | 6698 | 48 | rRF 45S | NO | 257.02 | 71.07 | 28.09 | 901.53 |
| >hg38_21_+_8433172_8446622_RNA45SN1_45S_50ntFlanks@6651.6699.49\|CGACTCTTAGCG GTGGATCACTCGGCTCGTGCGTCGATGAAGAACGCAG | >hg38_21_+_8433172_8446622_RNA45SN1_45S_50ntFlanks | 6651 | 6699 | 49 | rRF 45S | NO | 611.22 | 165.92 | 43.40 | 1257.46 |
| >hg38_21_+_8433172_8446622_RNA45SN1_45S_50ntFlanks@6652.6678.27\|GACTCTTAGCGG TGGATCACTCGGCTC | >hg38_21_+_8433172_8446622_RNA45SN1_45S_50ntFlanks | 6652 | 6678 | 27 | rRF 45S | NO | 30.92 | 0.00 | 0.00 | 60.55 |
| >hg38_21_+_8433172_8446622_RNA45SN1_45S_50ntFlanks@6652.6679.28\|GACTCTTAGCGG TGGATCACTCGGCTCG | >hg38_21_+_8433172_8446622_RNA45SN1_45S_50ntFlanks | 6652 | 6679 | 28 | rRF 45S | NO | 79.70 | 41.95 | 0.00 | 230.07 |
| >hg38_21_+_8433172_8446622_RNA45SN1_45S_50ntFlanks@6652.6681.30\|GACTCTTAGCGG TGGATCACTCGGCTCGTG | >hg38_21_+_8433172_8446622_RNA45SN1_45S_50ntFlanks | 6652 | 6681 | 30 | rRF 45S | NO | 317.53 | 190.89 | 153.06 | 716.10 |
| >hg38_21_+_8433172_8446622_RNA45SN1_45S_50ntFlanks@6652.6683.32\|GACTCTTAGCGG TGGATCACTCGGCTCGTGCG | >hg38_21_+_8433172_8446622_RNA45SN1_45S_50ntFlanks | 6652 | 6683 | 32 | rRF 45S | NO | 45.87 | 0.00 | 0.00 | 64.20 |
| >hg38_21_+_8433172_8446622_RNA45SN1_45S_50ntFlanks@6652.6686.35\|GACTCTTAGCGG TGGATCACTCGGCTCGTGCGTCG | >hg38_21_+_8433172_8446622_RNA45SN1_45S_50ntFlanks | 6652 | 6686 | 35 | rRF 45S | NO | 48.39 | 0.00 | 21.03 | 221.30 |
| >hg38_21_+_8433172_8446622_RNA45SN1_45S_50ntFlanks@6652.6688.37\|GACTCTTAGCGG TGGATCACTCGGCTCGTGCGTCGAT | >hg38_21_+_8433172_8446622_RNA45SN1_45S_50ntFlanks | 6652 | 6688 | 37 | rRF 45S | NO | 37.37 | 0.00 | 0.00 | 151.84 |
| >hg38_21_+_8433172_8446622_RNA45SN1_45S_50ntFlanks@6652.6689.38\|GACTCTTAGCGG TGGATCACTCGGCTCGTGCGTCGATG | >hg38_21_+_8433172_8446622_RNA45SN1_45S_50ntFlanks | 6652 | 6689 | 38 | rRF 45S | NO | 571.66 | 120.30 | 110.84 | 1440.63 |
| >hg38_21_+_8433172_8446622_RNA45SN1_45S_50ntFlanks@6652.6692.41\|GACTCTTAGCGG TGGATCACTCGGCTCGTGCGTCGATGAAG | >hg38_21_+_8433172_8446622_RNA45SN1_45S_50ntFlanks | 6652 | 6692 | 41 | rRF 45S | NO | 41.16 | 0.00 | 0.00 | 108.87 |
| >hg38_21_+_8433172_8446622_RNA45SN1_45S_50ntFlanks@6652.6696.45\|GACTCTTAGCGG TGGATCACTCGGCTCGTGCGTCGATGAAGAACG | >hg38_21_+_8433172_8446622_RNA45SN1_45S_50ntFlanks | 6652 | 6696 | 45 | rRF 45S | NO | 21.45 | 0.00 | 0.00 | 56.97 |
| >hg38_21_+_8433172_8446622_RNA45SN1_45S_50ntFlanks@6652.6697.46\|GACTCTTAGCGG TGGATCACTCGGCTCGTGCGTCGATGAAGAACGC | >hg38_21_+_8433172_8446622_RNA45SN1_45S_50ntFlanks | 6652 | 6697 | 46 | rRF 45S | NO | 41.22 | 0.00 | 0.00 | 36.24 |
| >hg38_21_+_8433172_8446622_RNA45SN1_45S_50ntFlanks@6652.6698.47\|GACTCTTAGCGG TGGATCACTCGGCTCGTGCGTCGATGAAGAACGCA | >hg38_21_+_8433172_8446622_RNA45SN1_45S_50ntFlanks | 6652 | 6698 | 47 | rRF 45S | NO | 260.77 | 49.09 | 0.00 | 541.09 |
| >hg38_21_+_8433172_8446622_RNA45SN1_45S_50ntFlanks@6652.6699.48\|GACTCTTAGCGG TGGATCACTCGGCTCGTGCGTCGATGAAGAACGCAG | >hg38_21_+_8433172_8446622_RNA45SN1_45S_50ntFlanks | 6652 | 6699 | 48 | rRF 45S | NO | 120.82 | 31.84 | 0.00 | 320.97 |
| >hg38_21_+_8433172_8446622_RNA45SN1_45S_50ntFlanks@6653.6681.29\|ACTCTTAGCGGT GGATCACTCGGCTCGTG | >hg38_21_+_8433172_8446622_RNA45SN1_45S_50ntFlanks | 6653 | 6681 | 29 | rRF 45S | NO | 121.56 | 149.67 | 58.67 | 150.26 |
| >hg38_21_+_8433172_8446622_RNA45SN1_45S_50ntFlanks@6653.6686.34\|ACTCTTAGCGGT GGATCACTCGGCTCGTGCGTCG | >hg38_21_+_8433172_8446622_RNA45SN1_45S_50ntFlanks | 6653 | 6686 | 34 | rRF 45S | NO | 12.69 | 0.00 | 0.00 | 36.45 |
| >hg38_21_+_8433172_8446622_RNA45SN1_45S_50ntFlanks@6653.6688.36\|ACTCTTAGCGGT GGATCACTCGGCTCGTGCGTCGAT | >hg38_21_+_8433172_8446622_RNA45SN1_45S_50ntFlanks | 6653 | 6688 | 36 | rRF 45S | NO | 20.99 | 0.00 | 0.00 | 37.33 |
| >hg38_21_+_8433172_8446622_RNA45SN1_45S_50ntFlanks@6653.6689.37\|ACTCTTAGCGGT GGATCACTCGGCTCGTGCGTCGATG | >hg38_21_+_8433172_8446622_RNA45SN1_45S_50ntFlanks | 6653 | 6689 | 37 | rRF 45S | NO | 99.77 | 37.65 | 0.00 | 189.42 |
| >hg38_21_+_8433172_8446622_RNA45SN1_45S_50ntFlanks@6654.6681.28\|CTCTTAGCGGTG GATCACTCGGCTCGTG | >hg38_21_+_8433172_8446622_RNA45SN1_45S_50ntFlanks | 6654 | 6681 | 28 | rRF 45S | NO | 49.63 | 91.06 | 60.45 | 80.59 |
| >hg38_21_+_8433172_8446622_RNA45SN1_45S_50ntFlanks@6654.6689.36\|CTCTTAGCGGTG GATCACTCGGCTCGTGCGTCGATG | >hg38_21_+_8433172_8446622_RNA45SN1_45S_50ntFlanks | 6654 | 6689 | 36 | rRF 45S | NO | 54.76 | 0.00 | 0.00 | 78.54 |
| >hg38_21_+_8433172_8446622_RNA45SN1_45S_50ntFlanks@6655.6681.27\|TCTTAGCGGTGG ATCACTCGGCTCGTG | >hg38_21_+_8433172_8446622_RNA45SN1_45S_50ntFlanks | 6655 | 6681 | 27 | rRF 45S | NO | 104.95 | 142.19 | 51.46 | 115.13 |
| >hg38_21_+_8433172_8446622_RNA45SN1_45S_50ntFlanks@6655.6689.35\|TCTTAGCGGTGG ATCACTCGGCTCGTGCGTCGATG | >hg38_21_+_8433172_8446622_RNA45SN1_45S_50ntFlanks | 6655 | 6689 | 35 | rRF 45S | NO | 66.98 | 0.00 | 0.00 | 89.43 |
| >hg38_21_+_8433172_8446622_RNA45SN1_45S_50ntFlanks@6656.6681.26\|CTTAGCGGTGGA TCACTCGGCTCGTG | >hg38_21_+_8433172_8446622_RNA45SN1_45S_50ntFlanks | 6656 | 6681 | 26 | rRF 45S | NO | 109.74 | 179.89 | 124.14 | 146.74 |
| >hg38_21_+_8433172_8446622_RNA45SN1_45S_50ntFlanks@6656.6689.34\|CTTAGCGGTGGA TCACTCGGCTCGTGCGTCGATG | >hg38_21_+_8433172_8446622_RNA45SN1_45S_50ntFlanks | 6656 | 6689 | 34 | rRF 45S | NO | 46.60 | 0.00 | 27.86 | 53.15 |
| >hg38_21_+_8433172_8446622_RNA45SN1_45S_50ntFlanks@6657.6681.25\|TTAGCGGTGGAT CACTCGGCTCGTG | >hg38_21_+_8433172_8446622_RNA45SN1_45S_50ntFlanks | 6657 | 6681 | 25 | rRF 45S | NO | 30.96 | 45.70 | 51.98 | 44.03 |
| >hg38_21_+_8433172_8446622_RNA45SN1_45S_50ntFlanks@6659.6681.23\|AGCGGTGGATC ACTCGGCTCGTG | >hg38_21_+_8433172_8446622_RNA45SN1_45S_50ntFlanks | 6659 | 6681 | 23 | rRF 45S | NO | 31.20 | 43.62 | 81.58 | 79.19 |
| >hg38_21_+_8433172_8446622_RNA45SN1_45S_50ntFlanks@6659.6689.31\|AGCGGTGGATC ACTCGGCTCGTGCGTCGATG | >hg38_21_+_8433172_8446622_RNA45SN1_45S_50ntFlanks | 6659 | 6689 | 31 | rRF 45S | NO | 0.00 | 0.00 | 29.40 | 26.68 |
| >hg38_21_+_8433172_8446622_RNA45SN1_45S_50ntFlanks@6660.6681.22\|GCGGTGGATCA CTCGGCTCGTG | >hg38_21_+_8433172_8446622_RNA45SN1_45S_50ntFlanks | 6660 | 6681 | 22 | rRF 45S | NO | 36.71 | 27.32 | 51.04 | 52.76 |
| >hg38_21_+_8433172_8446622_RNA45SN1_45S_50ntFlanks@6660.6689.30\|GCGGTGGATCA CTCGGCTCGTGCGTCGATG | >hg38_21_+_8433172_8446622_RNA45SN1_45S_50ntFlanks | 6660 | 6689 | 30 | rRF 45S | NO | 36.20 | 0.00 | 46.57 | 60.99 |

### MDA-MB-468

| | | | | | | | | | | |
|---|---|---|---|---|---|---|---|---|---|---|
| >hg38_21_+_8433172_8446622_RNA45SN1_45S_50ntFlanks@6661.6681.21\|CGGTGGATCACTCGGCTCGTG | >hg38_21_+_8433172_8446622_RNA45SN1_45S_50ntFlanks | 6661 | 6681 | 21 | rRF 45S | NO | 121.25 | 76.71 | 108.53 | 111.36 |
| >hg38_21_+_8433172_8446622_RNA45SN1_45S_50ntFlanks@6661.6686.26\|CGGTGGATCACTCGGCTCGTGCGTCG | >hg38_21_+_8433172_8446622_RNA45SN1_45S_50ntFlanks | 6661 | 6686 | 26 | rRF 45S | NO | 39.85 | 0.00 | 31.69 | 95.46 |
| >hg38_21_+_8433172_8446622_RNA45SN1_45S_50ntFlanks@6661.6688.28\|CGGTGGATCACTCGGCTCGTGCGTCGAT | >hg38_21_+_8433172_8446622_RNA45SN1_45S_50ntFlanks | 6661 | 6688 | 28 | rRF 45S | NO | 42.65 | 0.00 | 34.70 | 65.07 |
| >hg38_21_+_8433172_8446622_RNA45SN1_45S_50ntFlanks@6661.6689.29\|CGGTGGATCACTCGGCTCGTGCGTCGATG | >hg38_21_+_8433172_8446622_RNA45SN1_45S_50ntFlanks | 6661 | 6689 | 29 | rRF 45S | NO | 261.72 | 78.45 | 255.60 | 462.28 |
| >hg38_21_+_8433172_8446622_RNA45SN1_45S_50ntFlanks@6661.6696.36\|CGGTGGATCACTCGGCTCGTGCGTCGATGAAGAACG | >hg38_21_+_8433172_8446622_RNA45SN1_45S_50ntFlanks | 6661 | 6696 | 36 | rRF 45S | NO | 0.00 | 0.00 | 32.05 | 36.37 |
| >hg38_21_+_8433172_8446622_RNA45SN1_45S_50ntFlanks@6661.6697.37\|CGGTGGATCACTCGGCTCGTGCGTCGATGAAGAACGC | >hg38_21_+_8433172_8446622_RNA45SN1_45S_50ntFlanks | 6661 | 6697 | 37 | rRF 45S | NO | 21.76 | 0.00 | 7.29 | 66.52 |
| >hg38_21_+_8433172_8446622_RNA45SN1_45S_50ntFlanks@6661.6699.39\|CGGTGGATCACTCGGCTCGTGCGTCGATGAAGAACGCAG | >hg38_21_+_8433172_8446622_RNA45SN1_45S_50ntFlanks | 6661 | 6699 | 39 | rRF 45S | NO | 98.76 | 56.21 | 56.63 | 206.12 |
| >hg38_21_+_8433172_8446622_RNA45SN1_45S_50ntFlanks@6663.6681.19\|GTGGATCACTCGGCTCGTG | >hg38_21_+_8433172_8446622_RNA45SN1_45S_50ntFlanks | 6663 | 6681 | 19 | rRF 45S | NO | 118.10 | 79.47 | 172.72 | 87.46 |
| >hg38_21_+_8433172_8446622_RNA45SN1_45S_50ntFlanks@6663.6686.24\|GTGGATCACTCGGCTCGTGCGTCG | >hg38_21_+_8433172_8446622_RNA45SN1_45S_50ntFlanks | 6663 | 6686 | 24 | rRF 45S | NO | 40.36 | 0.00 | 43.39 | 60.38 |
| >hg38_21_+_8433172_8446622_RNA45SN1_45S_50ntFlanks@6663.6688.26\|GTGGATCACTCGGCTCGTGCGTCGAT | >hg38_21_+_8433172_8446622_RNA45SN1_45S_50ntFlanks | 6663 | 6688 | 26 | rRF 45S | NO | 0.00 | 0.00 | 20.94 | 35.08 |
| >hg38_21_+_8433172_8446622_RNA45SN1_45S_50ntFlanks@6663.6689.27\|GTGGATCACTCGGCTCGTGCGTCGATG | >hg38_21_+_8433172_8446622_RNA45SN1_45S_50ntFlanks | 6663 | 6689 | 27 | rRF 45S | NO | 71.85 | 70.07 | 220.65 | 182.78 |
| >hg38_21_+_8433172_8446622_RNA45SN1_45S_50ntFlanks@6664.6681.18\|TGGATCACTCGGCTCGTG | >hg38_21_+_8433172_8446622_RNA45SN1_45S_50ntFlanks | 6664 | 6681 | 18 | rRF 45S | NO | 88.71 | 79.97 | 182.77 | 85.69 |
| >hg38_21_+_8433172_8446622_RNA45SN1_45S_50ntFlanks@6664.6686.23\|TGGATCACTCGGCTCGTGCGTCG | >hg38_21_+_8433172_8446622_RNA45SN1_45S_50ntFlanks | 6664 | 6686 | 23 | rRF 45S | NO | 0.00 | 0.00 | 13.86 | 39.01 |
| >hg38_21_+_8433172_8446622_RNA45SN1_45S_50ntFlanks@6664.6689.26\|TGGATCACTCGGCTCGTGCGTCGATG | >hg38_21_+_8433172_8446622_RNA45SN1_45S_50ntFlanks | 6664 | 6689 | 26 | rRF 45S | NO | 0.00 | 0.00 | 84.50 | 35.34 |
| >hg38_21_+_8433172_8446622_RNA45SN1_45S_50ntFlanks@6666.6689.24\|GATCACTCGGCTCGTGCGTCGATG | >hg38_21_+_8433172_8446622_RNA45SN1_45S_50ntFlanks | 6666 | 6689 | 24 | rRF 45S | NO | 0.00 | 0.00 | 53.39 | 6.06 |
| >hg38_21_+_8433172_8446622_RNA45SN1_45S_50ntFlanks@6667.6689.23\|ATCACTCGGCTCGTGCGTCGATG | >hg38_21_+_8433172_8446622_RNA45SN1_45S_50ntFlanks | 6667 | 6689 | 23 | rRF 45S | NO | 0.00 | 0.00 | 35.48 | 0.00 |
| >hg38_21_+_8433172_8446622_RNA45SN1_45S_50ntFlanks@6669.6686.18\|CACTCGGCTCGTGCGTCG | >hg38_21_+_8433172_8446622_RNA45SN1_45S_50ntFlanks | 6669 | 6686 | 18 | rRF 45S | NO | 0.00 | 0.00 | 27.94 | 15.34 |
| >hg38_21_+_8433172_8446622_RNA45SN1_45S_50ntFlanks@6670.6696.27\|ACTCGGCTCGTGCGTCGATGAAGAACG | >hg38_21_+_8433172_8446622_RNA45SN1_45S_50ntFlanks | 6670 | 6696 | 27 | rRF 45S | NO | 0.00 | 0.00 | 37.11 | 33.59 |
| >hg38_21_+_8433172_8446622_RNA45SN1_45S_50ntFlanks@6670.6699.30\|ACTCGGCTCGTGCGTCGATGAAGAACGCAG | >hg38_21_+_8433172_8446622_RNA45SN1_45S_50ntFlanks | 6670 | 6699 | 30 | rRF 45S | NO | 0.00 | 26.26 | 39.08 | 97.22 |
| >hg38_21_+_8433172_8446622_RNA45SN1_45S_50ntFlanks@6671.6696.26\|CTCGGCTCGTGCGTCGATGAAGAACG | >hg38_21_+_8433172_8446622_RNA45SN1_45S_50ntFlanks | 6671 | 6696 | 26 | rRF 45S | NO | 40.70 | 49.60 | 109.64 | 159.40 |
| >hg38_21_+_8433172_8446622_RNA45SN1_45S_50ntFlanks@6671.6697.27\|CTCGGCTCGTGCGTCGATGAAGAACGC | >hg38_21_+_8433172_8446622_RNA45SN1_45S_50ntFlanks | 6671 | 6697 | 27 | rRF 45S | NO | 31.59 | 40.60 | 39.58 | 127.47 |
| >hg38_21_+_8433172_8446622_RNA45SN1_45S_50ntFlanks@6671.6698.28\|CTCGGCTCGTGCGTCGATGAAGAACGCA | >hg38_21_+_8433172_8446622_RNA45SN1_45S_50ntFlanks | 6671 | 6698 | 28 | rRF 45S | NO | 44.28 | 74.82 | 41.24 | 243.63 |
| >hg38_21_+_8433172_8446622_RNA45SN1_45S_50ntFlanks@6671.6699.29\|CTCGGCTCGTGCGTCGATGAAGAACGCAG | >hg38_21_+_8433172_8446622_RNA45SN1_45S_50ntFlanks | 6671 | 6699 | 29 | rRF 45S | NO | 100.33 | 197.03 | 181.78 | 534.99 |
| >hg38_21_+_8433172_8446622_RNA45SN1_45S_50ntFlanks@6687.6720.34\|ATGAAGAACGCAGCTAGCTGCGAGAATTAATGTG | >hg38_21_+_8433172_8446622_RNA45SN1_45S_50ntFlanks | 6687 | 6720 | 34 | rRF 45S | NO | 0.00 | 28.63 | 0.00 | 0.00 |
| >hg38_21_+_8433172_8446622_RNA45SN1_45S_50ntFlanks@6689.6720.32\|GAAGAACGCAGCTAGCTGCGAGAATTAATGTG | >hg38_21_+_8433172_8446622_RNA45SN1_45S_50ntFlanks | 6689 | 6720 | 32 | rRF 45S | NO | 0.00 | 34.33 | 0.00 | 0.00 |
| >hg38_21_+_8433172_8446622_RNA45SN1_45S_50ntFlanks@6690.6720.31\|AAGAACGCAGCTAGCTGCGAGAATTAATGTG | >hg38_21_+_8433172_8446622_RNA45SN1_45S_50ntFlanks | 6690 | 6720 | 31 | rRF 45S | NO | 61.42 | 239.80 | 32.70 | 102.18 |
| >hg38_21_+_8433172_8446622_RNA45SN1_45S_50ntFlanks@6697.6718.22\|CAGCTAGCTGCGAGAATTAATG | >hg38_21_+_8433172_8446622_RNA45SN1_45S_50ntFlanks | 6697 | 6718 | 22 | rRF 45S | NO | 40.51 | 40.03 | 56.82 | 135.98 |
| >hg38_21_+_8433172_8446622_RNA45SN1_45S_50ntFlanks@6697.6720.24\|CAGCTAGCTGCGAGAATTAATGTG | >hg38_21_+_8433172_8446622_RNA45SN1_45S_50ntFlanks | 6697 | 6720 | 24 | rRF 45S | NO | 87.49 | 90.13 | 44.66 | 37.00 |
| >hg38_21_+_8433172_8446622_RNA45SN1_45S_50ntFlanks@6698.6717.20\|AGCTAGCTGCGAGAATTAAT | >hg38_21_+_8433172_8446622_RNA45SN1_45S_50ntFlanks | 6698 | 6717 | 20 | rRF 45S | NO | 32.22 | 0.00 | 0.00 | 0.00 |
| >hg38_21_+_8433172_8446622_RNA45SN1_45S_50ntFlanks@6698.6718.21\|AGCTAGCTGCGAGAATTAATG | >hg38_21_+_8433172_8446622_RNA45SN1_45S_50ntFlanks | 6698 | 6718 | 21 | rRF 45S | NO | 749.62 | 524.03 | 305.92 | 158.91 |
| >hg38_21_+_8433172_8446622_RNA45SN1_45S_50ntFlanks@6698.6719.22\|AGCTAGCTGCGAGAATTAATGT | >hg38_21_+_8433172_8446622_RNA45SN1_45S_50ntFlanks | 6698 | 6719 | 22 | rRF 45S | NO | 119.47 | 72.64 | 35.00 | 0.00 |
| >hg38_21_+_8433172_8446622_RNA45SN1_45S_50ntFlanks@6698.6720.23\|AGCTAGCTGCGAGAATTAATGTG | >hg38_21_+_8433172_8446622_RNA45SN1_45S_50ntFlanks | 6698 | 6720 | 23 | rRF 45S | YES | 1774.55 | 1342.09 | 482.38 | 229.23 |
| >hg38_21_+_8433172_8446622_RNA45SN1_45S_50ntFlanks@6698.6721.24\|AGCTAGCTGCGAGAATTAATGTGA | >hg38_21_+_8433172_8446622_RNA45SN1_45S_50ntFlanks | 6698 | 6721 | 24 | rRF 45S | NO | 19.94 | 0.00 | 0.00 | 0.00 |
| >hg38_21_+_8433172_8446622_RNA45SN1_45S_50ntFlanks@6699.6717.19\|GCTAGCTGCGAGAATTAAT | >hg38_21_+_8433172_8446622_RNA45SN1_45S_50ntFlanks | 6699 | 6717 | 19 | rRF 45S | NO | 68.88 | 44.76 | 33.65 | 14.44 |

### MDA-MB-468

| | | | | | | | | | | |
|---|---|---|---|---|---|---|---|---|---|---|
| >hg38_21_+_8433172_8446622_RNA45SN1_45S_50ntFlanks@6699.6718.20\|GCTAGCTGCGAGAATTAATG | >hg38_21_+_8433172_8446622_RNA45SN1_45S_50ntFlanks | 6699 | 6718 | 20 | rRF 45S | YES | 1444.87 | 1228.43 | 587.60 | 332.23 |
| >hg38_21_+_8433172_8446622_RNA45SN1_45S_50ntFlanks@6699.6719.21\|GCTAGCTGCGAGAATTAATGT | >hg38_21_+_8433172_8446622_RNA45SN1_45S_50ntFlanks | 6699 | 6719 | 21 | rRF 45S | NO | 199.95 | 154.61 | 67.02 | 39.48 |
| >hg38_21_+_8433172_8446622_RNA45SN1_45S_50ntFlanks@6699.6720.22\|GCTAGCTGCGAGAATTAATGTG | >hg38_21_+_8433172_8446622_RNA45SN1_45S_50ntFlanks | 6699 | 6720 | 22 | rRF 45S | YES | 2936.80 | 2798.41 | 902.73 | 444.06 |
| >hg38_21_+_8433172_8446622_RNA45SN1_45S_50ntFlanks@6699.6721.23\|GCTAGCTGCGAGAATTAATGTGA | >hg38_21_+_8433172_8446622_RNA45SN1_45S_50ntFlanks | 6699 | 6721 | 23 | rRF 45S | NO | 42.29 | 0.00 | 0.00 | 0.00 |
| >hg38_21_+_8433172_8446622_RNA45SN1_45S_50ntFlanks@6700.6717.18\|CTAGCTGCGAGAATTAAT | >hg38_21_+_8433172_8446622_RNA45SN1_45S_50ntFlanks | 6700 | 6717 | 18 | rRF 45S | NO | 22.06 | 12.71 | 0.00 | 0.00 |
| >hg38_21_+_8433172_8446622_RNA45SN1_45S_50ntFlanks@6700.6718.19\|CTAGCTGCGAGAATTAATG | >hg38_21_+_8433172_8446622_RNA45SN1_45S_50ntFlanks | 6700 | 6718 | 19 | rRF 45S | NO | 622.58 | 852.29 | 172.15 | 227.86 |
| >hg38_21_+_8433172_8446622_RNA45SN1_45S_50ntFlanks@6700.6719.20\|CTAGCTGCGAGAATTAATGT | >hg38_21_+_8433172_8446622_RNA45SN1_45S_50ntFlanks | 6700 | 6719 | 20 | rRF 45S | NO | 29.07 | 56.80 | 20.81 | 0.00 |
| >hg38_21_+_8433172_8446622_RNA45SN1_45S_50ntFlanks@6700.6720.21\|CTAGCTGCGAGAATTAATGTG | >hg38_21_+_8433172_8446622_RNA45SN1_45S_50ntFlanks | 6700 | 6720 | 21 | rRF 45S | YES | 498.53 | 1426.21 | 415.45 | 189.91 |
| >hg38_21_+_8433172_8446622_RNA45SN1_45S_50ntFlanks@6702.6720.19\|AGCTGCGAGAATTAATGTG | >hg38_21_+_8433172_8446622_RNA45SN1_45S_50ntFlanks | 6702 | 6720 | 19 | rRF 45S | NO | 0.00 | 31.81 | 57.72 | 0.00 |
| >hg38_21_+_8433172_8446622_RNA45SN1_45S_50ntFlanks@6703.6720.18\|GCTGCGAGAATTAATGTG | >hg38_21_+_8433172_8446622_RNA45SN1_45S_50ntFlanks | 6703 | 6720 | 18 | rRF 45S | NO | 0.00 | 0.00 | 28.72 | 0.00 |
| >hg38_21_+_8433172_8446622_RNA45SN1_45S_50ntFlanks@6719.6736.18\|TGAATTGCAGGACACATT | >hg38_21_+_8433172_8446622_RNA45SN1_45S_50ntFlanks | 6719 | 6736 | 18 | rRF 45S | NO | 279.43 | 157.44 | 42.72 | 39.75 |
| >hg38_21_+_8433172_8446622_RNA45SN1_45S_50ntFlanks@6719.6737.19\|TGAATTGCAGGACACATTG | >hg38_21_+_8433172_8446622_RNA45SN1_45S_50ntFlanks | 6719 | 6737 | 19 | rRF 45S | NO | 220.46 | 181.64 | 55.86 | 35.92 |
| >hg38_21_+_8433172_8446622_RNA45SN1_45S_50ntFlanks@6719.6739.21\|TGAATTGCAGGACACATTGAT | >hg38_21_+_8433172_8446622_RNA45SN1_45S_50ntFlanks | 6719 | 6739 | 21 | rRF 45S | NO | 86.04 | 63.69 | 0.00 | 0.00 |
| >hg38_21_+_8433172_8446622_RNA45SN1_45S_50ntFlanks@6719.6740.22\|TGAATTGCAGGACACATTGATC | >hg38_21_+_8433172_8446622_RNA45SN1_45S_50ntFlanks | 6719 | 6740 | 22 | rRF 45S | NO | 404.62 | 267.56 | 79.37 | 52.00 |
| >hg38_21_+_8433172_8446622_RNA45SN1_45S_50ntFlanks@6719.6741.23\|TGAATTGCAGGACACATTGATCA | >hg38_21_+_8433172_8446622_RNA45SN1_45S_50ntFlanks | 6719 | 6741 | 23 | rRF 45S | NO | 36.44 | 27.76 | 0.00 | 0.00 |
| >hg38_21_+_8433172_8446622_RNA45SN1_45S_50ntFlanks@6719.6742.24\|TGAATTGCAGGACACATTGATCAT | >hg38_21_+_8433172_8446622_RNA45SN1_45S_50ntFlanks | 6719 | 6742 | 24 | rRF 45S | NO | 53.33 | 33.81 | 0.00 | 0.00 |
| >hg38_21_+_8433172_8446622_RNA45SN1_45S_50ntFlanks@6719.6743.25\|TGAATTGCAGGACACATTGATCATC | >hg38_21_+_8433172_8446622_RNA45SN1_45S_50ntFlanks | 6719 | 6743 | 25 | rRF 45S | YES | 640.39 | 410.26 | 117.36 | 83.31 |
| >hg38_21_+_8433172_8446622_RNA45SN1_45S_50ntFlanks@6719.6744.26\|TGAATTGCAGGACACATTGATCATCG | >hg38_21_+_8433172_8446622_RNA45SN1_45S_50ntFlanks | 6719 | 6744 | 26 | rRF 45S | YES | 424.35 | 1499.56 | 93.80 | 105.72 |
| >hg38_21_+_8433172_8446622_RNA45SN1_45S_50ntFlanks@6719.6745.27\|TGAATTGCAGGACACATTGATCATCGA | >hg38_21_+_8433172_8446622_RNA45SN1_45S_50ntFlanks | 6719 | 6745 | 27 | rRF 45S | NO | 40.62 | 97.43 | 0.00 | 0.00 |
| >hg38_21_+_8433172_8446622_RNA45SN1_45S_50ntFlanks@6719.6746.28\|TGAATTGCAGGACACATTGATCATCGAC | >hg38_21_+_8433172_8446622_RNA45SN1_45S_50ntFlanks | 6719 | 6746 | 28 | rRF 45S | NO | 54.46 | 91.24 | 44.17 | 41.44 |
| >hg38_21_+_8433172_8446622_RNA45SN1_45S_50ntFlanks@6719.6747.29\|TGAATTGCAGGACACATTGATCATCGACA | >hg38_21_+_8433172_8446622_RNA45SN1_45S_50ntFlanks | 6719 | 6747 | 29 | rRF 45S | NO | 58.30 | 98.36 | 13.85 | 68.15 |
| >hg38_21_+_8433172_8446622_RNA45SN1_45S_50ntFlanks@6719.6749.31\|TGAATTGCAGGACACATTGATCATCGACACT | >hg38_21_+_8433172_8446622_RNA45SN1_45S_50ntFlanks | 6719 | 6749 | 31 | rRF 45S | NO | 48.71 | 27.25 | 0.00 | 6.37 |
| >hg38_21_+_8433172_8446622_RNA45SN1_45S_50ntFlanks@6719.6751.33\|TGAATTGCAGGACACATTGATCATCGACACTTC | >hg38_21_+_8433172_8446622_RNA45SN1_45S_50ntFlanks | 6719 | 6751 | 33 | rRF 45S | NO | 36.02 | 0.00 | 0.00 | 6.65 |
| >hg38_21_+_8433172_8446622_RNA45SN1_45S_50ntFlanks@6721.6739.19\|AATTGCAGGACACATTGAT | >hg38_21_+_8433172_8446622_RNA45SN1_45S_50ntFlanks | 6721 | 6739 | 19 | rRF 45S | NO | 100.26 | 126.62 | 28.94 | 14.92 |
| >hg38_21_+_8433172_8446622_RNA45SN1_45S_50ntFlanks@6721.6740.20\|AATTGCAGGACACATTGATC | >hg38_21_+_8433172_8446622_RNA45SN1_45S_50ntFlanks | 6721 | 6740 | 20 | rRF 45S | YES | 520.50 | 556.84 | 152.34 | 69.36 |
| >hg38_21_+_8433172_8446622_RNA45SN1_45S_50ntFlanks@6721.6741.21\|AATTGCAGGACACATTGATCA | >hg38_21_+_8433172_8446622_RNA45SN1_45S_50ntFlanks | 6721 | 6741 | 21 | rRF 45S | NO | 44.95 | 41.37 | 0.00 | 0.00 |
| >hg38_21_+_8433172_8446622_RNA45SN1_45S_50ntFlanks@6721.6742.22\|AATTGCAGGACACATTGATCAT | >hg38_21_+_8433172_8446622_RNA45SN1_45S_50ntFlanks | 6721 | 6742 | 22 | rRF 45S | NO | 49.27 | 43.83 | 0.00 | 0.00 |
| >hg38_21_+_8433172_8446622_RNA45SN1_45S_50ntFlanks@6721.6743.23\|AATTGCAGGACACATTGATCATC | >hg38_21_+_8433172_8446622_RNA45SN1_45S_50ntFlanks | 6721 | 6743 | 23 | rRF 45S | NO | 537.88 | 447.51 | 99.79 | 77.64 |
| >hg38_21_+_8433172_8446622_RNA45SN1_45S_50ntFlanks@6721.6744.24\|AATTGCAGGACACATTGATCATCG | >hg38_21_+_8433172_8446622_RNA45SN1_45S_50ntFlanks | 6721 | 6744 | 24 | rRF 45S | YES | 738.62 | 2304.14 | 120.33 | 137.40 |
| >hg38_21_+_8433172_8446622_RNA45SN1_45S_50ntFlanks@6721.6745.25\|AATTGCAGGACACATTGATCATCGA | >hg38_21_+_8433172_8446622_RNA45SN1_45S_50ntFlanks | 6721 | 6745 | 25 | rRF 45S | NO | 97.83 | 206.39 | 0.00 | 0.00 |
| >hg38_21_+_8433172_8446622_RNA45SN1_45S_50ntFlanks@6721.6746.26\|AATTGCAGGACACATTGATCATCGAC | >hg38_21_+_8433172_8446622_RNA45SN1_45S_50ntFlanks | 6721 | 6746 | 26 | rRF 45S | NO | 117.17 | 177.99 | 89.50 | 63.57 |
| >hg38_21_+_8433172_8446622_RNA45SN1_45S_50ntFlanks@6721.6747.27\|AATTGCAGGACACATTGATCATCGACA | >hg38_21_+_8433172_8446622_RNA45SN1_45S_50ntFlanks | 6721 | 6747 | 27 | rRF 45S | NO | 110.79 | 166.03 | 45.77 | 90.45 |
| >hg38_21_+_8433172_8446622_RNA45SN1_45S_50ntFlanks@6721.6749.29\|AATTGCAGGACACATTGATCATCGACACT | >hg38_21_+_8433172_8446622_RNA45SN1_45S_50ntFlanks | 6721 | 6749 | 29 | rRF 45S | NO | 20.65 | 0.00 | 0.00 | 0.00 |
| >hg38_21_+_8433172_8446622_RNA45SN1_45S_50ntFlanks@6721.6750.30\|AATTGCAGGACACATTGATCATCGACACTT | >hg38_21_+_8433172_8446622_RNA45SN1_45S_50ntFlanks | 6721 | 6750 | 30 | rRF 45S | NO | 35.95 | 0.00 | 0.00 | 13.18 |

### MDA-MB-468

| | | | | | | | | | | |
|---|---|---|---|---|---|---|---|---|---|---|
| >hg38_21_+_8433172_8446622_RNA45SN1_45S_50ntFlanks@6721.6752.32\|AATTGCAGGACACATTGATCATCGACACTTCG | >hg38_21_+_8433172_8446622_RNA45SN1_45S_50ntFlanks | 6721 | 6752 | 32 | rRF 45S | NO | 45.45 | 30.35 | 0.00 | 24.68 |
| >hg38_21_+_8433172_8446622_RNA45SN1_45S_50ntFlanks@6721.6756.36\|AATTGCAGGACACATTGATCATCGACACTTCGAACG | >hg38_21_+_8433172_8446622_RNA45SN1_45S_50ntFlanks | 6721 | 6756 | 36 | rRF 45S | NO | 50.60 | 46.41 | 27.46 | 39.84 |
| >hg38_21_+_8433172_8446622_RNA45SN1_45S_50ntFlanks@6721.6757.37\|AATTGCAGGACACATTGATCATCGACACTTCGAACGC | >hg38_21_+_8433172_8446622_RNA45SN1_45S_50ntFlanks | 6721 | 6757 | 37 | rRF 45S | NO | 20.99 | 12.57 | 0.00 | 0.00 |
| >hg38_21_+_8433172_8446622_RNA45SN1_45S_50ntFlanks@6721.6758.38\|AATTGCAGGACACATTGATCATCGACACTTCGAACGCA | >hg38_21_+_8433172_8446622_RNA45SN1_45S_50ntFlanks | 6721 | 6758 | 38 | rRF 45S | NO | 214.96 | 168.41 | 21.10 | 83.62 |
| >hg38_21_+_8433172_8446622_RNA45SN1_45S_50ntFlanks@6721.6760.40\|AATTGCAGGACACATTGATCATCGACACTTCGAACGCACT | >hg38_21_+_8433172_8446622_RNA45SN1_45S_50ntFlanks | 6721 | 6760 | 40 | rRF 45S | NO | 64.84 | 35.63 | 0.00 | 6.29 |
| >hg38_21_+_8433172_8446622_RNA45SN1_45S_50ntFlanks@6721.6770.50\|AATTGCAGGACACATTGATCATCGACACTTCGAACGCACTTGCGGCCCCG | >hg38_21_+_8433172_8446622_RNA45SN1_45S_50ntFlanks | 6721 | 6770 | 50 | rRF 45S | NO | 34.74 | 12.80 | 0.00 | 59.13 |
| >hg38_21_+_8433172_8446622_RNA45SN1_45S_50ntFlanks@6722.6740.19\|ATTGCAGGACACATTGATC | >hg38_21_+_8433172_8446622_RNA45SN1_45S_50ntFlanks | 6722 | 6740 | 19 | rRF 45S | NO | 70.73 | 62.74 | 6.96 | 0.00 |
| >hg38_21_+_8433172_8446622_RNA45SN1_45S_50ntFlanks@6722.6743.22\|ATTGCAGGACACATTGATCATC | >hg38_21_+_8433172_8446622_RNA45SN1_45S_50ntFlanks | 6722 | 6743 | 22 | rRF 45S | NO | 84.83 | 58.33 | 0.00 | 0.00 |
| >hg38_21_+_8433172_8446622_RNA45SN1_45S_50ntFlanks@6722.6744.23\|ATTGCAGGACACATTGATCATCG | >hg38_21_+_8433172_8446622_RNA45SN1_45S_50ntFlanks | 6722 | 6744 | 23 | rRF 45S | NO | 99.27 | 260.91 | 0.00 | 0.00 |
| >hg38_21_+_8433172_8446622_RNA45SN1_45S_50ntFlanks@6722.6745.24\|ATTGCAGGACACATTGATCATCGA | >hg38_21_+_8433172_8446622_RNA45SN1_45S_50ntFlanks | 6722 | 6745 | 24 | rRF 45S | NO | 0.00 | 12.55 | 0.00 | 0.00 |
| >hg38_21_+_8433172_8446622_RNA45SN1_45S_50ntFlanks@6723.6740.18\|TTGCAGGACACATTGATC | >hg38_21_+_8433172_8446622_RNA45SN1_45S_50ntFlanks | 6723 | 6740 | 18 | rRF 45S | NO | 21.55 | 19.32 | 0.00 | 0.00 |
| >hg38_21_+_8433172_8446622_RNA45SN1_45S_50ntFlanks@6723.6743.21\|TTGCAGGACACATTGATCATC | >hg38_21_+_8433172_8446622_RNA45SN1_45S_50ntFlanks | 6723 | 6743 | 21 | rRF 45S | NO | 38.87 | 27.61 | 0.00 | 0.00 |
| >hg38_21_+_8433172_8446622_RNA45SN1_45S_50ntFlanks@6723.6744.22\|TTGCAGGACACATTGATCATCG | >hg38_21_+_8433172_8446622_RNA45SN1_45S_50ntFlanks | 6723 | 6744 | 22 | rRF 45S | NO | 44.34 | 115.60 | 0.00 | 0.00 |
| >hg38_21_+_8433172_8446622_RNA45SN1_45S_50ntFlanks@6725.6743.19\|GCAGGACACATTGATCATC | >hg38_21_+_8433172_8446622_RNA45SN1_45S_50ntFlanks | 6725 | 6743 | 19 | rRF 45S | NO | 33.76 | 0.00 | 0.00 | 0.00 |
| >hg38_21_+_8433172_8446622_RNA45SN1_45S_50ntFlanks@6725.6744.20\|GCAGGACACATTGATCATCG | >hg38_21_+_8433172_8446622_RNA45SN1_45S_50ntFlanks | 6725 | 6744 | 20 | rRF 45S | NO | 21.16 | 12.68 | 0.00 | 0.00 |
| >hg38_21_+_8433172_8446622_RNA45SN1_45S_50ntFlanks@6726.6744.19\|CAGGACACATTGATCATCG | >hg38_21_+_8433172_8446622_RNA45SN1_45S_50ntFlanks | 6726 | 6744 | 19 | rRF 45S | NO | 0.00 | 29.85 | 0.00 | 0.00 |
| >hg38_21_+_8433172_8446622_RNA45SN1_45S_50ntFlanks@6729.6770.42\|GACACATTGATCATCGACACTTCGAACGCACTTGCGGCCCCG | >hg38_21_+_8433172_8446622_RNA45SN1_45S_50ntFlanks | 6729 | 6770 | 42 | rRF 45S | NO | 28.75 | 0.00 | 0.00 | 0.00 |
| >hg38_21_+_8433172_8446622_RNA45SN1_45S_50ntFlanks@6729.6771.43\|GACACATTGATCATCGACACTTCGAACGCACTTGCGGCCCCGG | >hg38_21_+_8433172_8446622_RNA45SN1_45S_50ntFlanks | 6729 | 6771 | 43 | rRF 45S | NO | 19.63 | 0.00 | 0.00 | 0.00 |
| >hg38_21_+_8433172_8446622_RNA45SN1_45S_50ntFlanks@6730.6770.41\|ACACATTGATCATCGACACTTCGAACGCACTTGCGGCCCCG | >hg38_21_+_8433172_8446622_RNA45SN1_45S_50ntFlanks | 6730 | 6770 | 41 | rRF 45S | NO | 35.72 | 29.24 | 0.00 | 42.39 |
| >hg38_21_+_8433172_8446622_RNA45SN1_45S_50ntFlanks@6730.6771.42\|ACACATTGATCATCGACACTTCGAACGCACTTGCGGCCCCGG | >hg38_21_+_8433172_8446622_RNA45SN1_45S_50ntFlanks | 6730 | 6771 | 42 | rRF 45S | NO | 32.83 | 0.00 | 0.00 | 51.46 |
| >hg38_21_+_8433172_8446622_RNA45SN1_45S_50ntFlanks@6731.6752.22\|CACATTGATCATCGACACTTCG | >hg38_21_+_8433172_8446622_RNA45SN1_45S_50ntFlanks | 6731 | 6752 | 22 | rRF 45S | NO | 21.93 | 61.27 | 0.00 | 14.47 |
| >hg38_21_+_8433172_8446622_RNA45SN1_45S_50ntFlanks@6731.6753.23\|CACATTGATCATCGACACTTCGA | >hg38_21_+_8433172_8446622_RNA45SN1_45S_50ntFlanks | 6731 | 6753 | 23 | rRF 45S | NO | 0.00 | 33.38 | 0.00 | 0.00 |
| >hg38_21_+_8433172_8446622_RNA45SN1_45S_50ntFlanks@6731.6756.26\|CACATTGATCATCGACACTTCGAACG | >hg38_21_+_8433172_8446622_RNA45SN1_45S_50ntFlanks | 6731 | 6756 | 26 | rRF 45S | NO | 46.74 | 67.06 | 20.48 | 26.48 |
| >hg38_21_+_8433172_8446622_RNA45SN1_45S_50ntFlanks@6731.6758.28\|CACATTGATCATCGACACTTCGAACGCA | >hg38_21_+_8433172_8446622_RNA45SN1_45S_50ntFlanks | 6731 | 6758 | 28 | rRF 45S | NO | 0.00 | 12.76 | 0.00 | 0.00 |
| >hg38_21_+_8433172_8446622_RNA45SN1_45S_50ntFlanks@6731.6770.40\|CACATTGATCATCGACACTTCGAACGCACTTGCGGCCCCG | >hg38_21_+_8433172_8446622_RNA45SN1_45S_50ntFlanks | 6731 | 6770 | 40 | rRF 45S | NO | 113.37 | 84.28 | 0.00 | 117.36 |
| >hg38_21_+_8433172_8446622_RNA45SN1_45S_50ntFlanks@6731.6771.41\|CACATTGATCATCGACACTTCGAACGCACTTGCGGCCCCGG | >hg38_21_+_8433172_8446622_RNA45SN1_45S_50ntFlanks | 6731 | 6771 | 41 | rRF 45S | NO | 87.57 | 31.05 | 0.00 | 126.79 |
| >hg38_21_+_8433172_8446622_RNA45SN1_45S_50ntFlanks@6731.6772.42\|CACATTGATCATCGACACTTCGAACGCACTTGCGGCCCCGGG | >hg38_21_+_8433172_8446622_RNA45SN1_45S_50ntFlanks | 6731 | 6772 | 42 | rRF 45S | NO | 21.22 | 0.00 | 0.00 | 0.00 |
| >hg38_21_+_8433172_8446622_RNA45SN1_45S_50ntFlanks@6732.6752.21\|ACATTGATCATCGACACTTCG | >hg38_21_+_8433172_8446622_RNA45SN1_45S_50ntFlanks | 6732 | 6752 | 21 | rRF 45S | NO | 0.00 | 38.92 | 0.00 | 0.00 |
| >hg38_21_+_8433172_8446622_RNA45SN1_45S_50ntFlanks@6732.6756.25\|ACATTGATCATCGACACTTCGAACG | >hg38_21_+_8433172_8446622_RNA45SN1_45S_50ntFlanks | 6732 | 6756 | 25 | rRF 45S | NO | 30.98 | 31.08 | 27.29 | 6.69 |
| >hg38_21_+_8433172_8446622_RNA45SN1_45S_50ntFlanks@6732.6770.39\|ACATTGATCATCGACACTTCGAACGCACTTGCGGCCCCG | >hg38_21_+_8433172_8446622_RNA45SN1_45S_50ntFlanks | 6732 | 6770 | 39 | rRF 45S | NO | 92.82 | 58.42 | 27.88 | 96.66 |
| >hg38_21_+_8433172_8446622_RNA45SN1_45S_50ntFlanks@6732.6771.40\|ACATTGATCATCGACACTTCGAACGCACTTGCGGCCCCGG | >hg38_21_+_8433172_8446622_RNA45SN1_45S_50ntFlanks | 6732 | 6771 | 40 | rRF 45S | NO | 90.88 | 29.34 | 0.00 | 128.58 |
| >hg38_21_+_8433172_8446622_RNA45SN1_45S_50ntFlanks@6733.6752.20\|CATTGATCATCGACACTTCG | >hg38_21_+_8433172_8446622_RNA45SN1_45S_50ntFlanks | 6733 | 6752 | 20 | rRF 45S | NO | 0.00 | 71.60 | 44.99 | 16.13 |
| >hg38_21_+_8433172_8446622_RNA45SN1_45S_50ntFlanks@6733.6753.21\|CATTGATCATCGACACTTCGA | >hg38_21_+_8433172_8446622_RNA45SN1_45S_50ntFlanks | 6733 | 6753 | 21 | rRF 45S | NO | 0.00 | 30.56 | 32.30 | 0.00 |
| >hg38_21_+_8433172_8446622_RNA45SN1_45S_50ntFlanks@6733.6756.24\|CATTGATCATCGACACTTCGAACG | >hg38_21_+_8433172_8446622_RNA45SN1_45S_50ntFlanks | 6733 | 6756 | 24 | rRF 45S | NO | 37.44 | 61.43 | 63.35 | 24.64 |

### MDA-MB-468

| | | | | | | | | | | |
|---|---|---|---|---|---|---|---|---|---|---|
| >hg38_21_+_8433172_8446622_RNA45SN1_45S_50ntFlanks@6733.6758.26\|CATTGATCATCGACACTTCGAACGCA | >hg38_21_+_8433172_8446622_RNA45SN1_45S_50ntFlanks | 6733 | 6758 | 26 | rRF 45S | NO | 0.00 | 19.33 | 13.61 | 14.54 |
| >hg38_21_+_8433172_8446622_RNA45SN1_45S_50ntFlanks@6733.6770.38\|CATTGATCATCGACACTTCGAACGCACTTGCGGCCCCG | >hg38_21_+_8433172_8446622_RNA45SN1_45S_50ntFlanks | 6733 | 6770 | 38 | rRF 45S | NO | 170.58 | 94.51 | 38.53 | 154.44 |
| >hg38_21_+_8433172_8446622_RNA45SN1_45S_50ntFlanks@6733.6771.39\|CATTGATCATCGACACTTCGAACGCACTTGCGGCCCCGG | >hg38_21_+_8433172_8446622_RNA45SN1_45S_50ntFlanks | 6733 | 6771 | 39 | rRF 45S | NO | 160.77 | 48.01 | 0.00 | 213.25 |
| >hg38_21_+_8433172_8446622_RNA45SN1_45S_50ntFlanks@6733.6772.40\|CATTGATCATCGACACTTCGAACGCACTTGCGGCCCCGGG | >hg38_21_+_8433172_8446622_RNA45SN1_45S_50ntFlanks | 6733 | 6772 | 40 | rRF 45S | NO | 22.04 | 0.00 | 0.00 | 51.20 |
| >hg38_21_+_8433172_8446622_RNA45SN1_45S_50ntFlanks@6734.6770.37\|ATTGATCATCGACACTTCGAACGCACTTGCGGCCCCG | >hg38_21_+_8433172_8446622_RNA45SN1_45S_50ntFlanks | 6734 | 6770 | 37 | rRF 45S | NO | 76.32 | 37.62 | 13.60 | 74.54 |
| >hg38_21_+_8433172_8446622_RNA45SN1_45S_50ntFlanks@6734.6771.38\|ATTGATCATCGACACTTCGAACGCACTTGCGGCCCCGG | >hg38_21_+_8433172_8446622_RNA45SN1_45S_50ntFlanks | 6734 | 6771 | 38 | rRF 45S | NO | 57.76 | 0.00 | 0.00 | 81.57 |
| >hg38_21_+_8433172_8446622_RNA45SN1_45S_50ntFlanks@6735.6770.36\|TTGATCATCGACACTTCGAACGCACTTGCGGCCCCG | >hg38_21_+_8433172_8446622_RNA45SN1_45S_50ntFlanks | 6735 | 6770 | 36 | rRF 45S | NO | 31.27 | 0.00 | 0.00 | 33.28 |
| >hg38_21_+_8433172_8446622_RNA45SN1_45S_50ntFlanks@6735.6771.37\|TTGATCATCGACACTTCGAACGCACTTGCGGCCCCGG | >hg38_21_+_8433172_8446622_RNA45SN1_45S_50ntFlanks | 6735 | 6771 | 37 | rRF 45S | NO | 31.68 | 0.00 | 0.00 | 45.25 |
| >hg38_21_+_8433172_8446622_RNA45SN1_45S_50ntFlanks@6737.6770.34\|GATCATCGACACTTCGAACGCACTTGCGGCCCCG | >hg38_21_+_8433172_8446622_RNA45SN1_45S_50ntFlanks | 6737 | 6770 | 34 | rRF 45S | NO | 44.28 | 30.32 | 0.00 | 54.72 |
| >hg38_21_+_8433172_8446622_RNA45SN1_45S_50ntFlanks@6737.6771.35\|GATCATCGACACTTCGAACGCACTTGCGGCCCCGG | >hg38_21_+_8433172_8446622_RNA45SN1_45S_50ntFlanks | 6737 | 6771 | 35 | rRF 45S | NO | 38.24 | 0.00 | 0.00 | 63.33 |
| >hg38_21_+_8433172_8446622_RNA45SN1_45S_50ntFlanks@6738.6756.19\|ATCATCGACACTTCGAACG | >hg38_21_+_8433172_8446622_RNA45SN1_45S_50ntFlanks | 6738 | 6756 | 19 | rRF 45S | NO | 0.00 | 49.87 | 32.92 | 0.00 |
| >hg38_21_+_8433172_8446622_RNA45SN1_45S_50ntFlanks@6738.6770.33\|ATCATCGACACTTCGAACGCACTTGCGGCCCCG | >hg38_21_+_8433172_8446622_RNA45SN1_45S_50ntFlanks | 6738 | 6770 | 33 | rRF 45S | NO | 173.30 | 179.67 | 127.92 | 253.85 |
| >hg38_21_+_8433172_8446622_RNA45SN1_45S_50ntFlanks@6738.6771.34\|ATCATCGACACTTCGAACGCACTTGCGGCCCCGG | >hg38_21_+_8433172_8446622_RNA45SN1_45S_50ntFlanks | 6738 | 6771 | 34 | rRF 45S | NO | 203.67 | 111.02 | 79.50 | 373.21 |
| >hg38_21_+_8433172_8446622_RNA45SN1_45S_50ntFlanks@6738.6772.35\|ATCATCGACACTTCGAACGCACTTGCGGCCCCGGG | >hg38_21_+_8433172_8446622_RNA45SN1_45S_50ntFlanks | 6738 | 6772 | 35 | rRF 45S | NO | 58.23 | 0.00 | 0.00 | 131.37 |
| >hg38_21_+_8433172_8446622_RNA45SN1_45S_50ntFlanks@6739.6770.32\|TCATCGACACTTCGAACGCACTTGCGGCCCCG | >hg38_21_+_8433172_8446622_RNA45SN1_45S_50ntFlanks | 6739 | 6770 | 32 | rRF 45S | NO | 6.04 | 19.31 | 6.49 | 47.71 |
| >hg38_21_+_8433172_8446622_RNA45SN1_45S_50ntFlanks@6739.6771.33\|TCATCGACACTTCGAACGCACTTGCGGCCCCGG | >hg38_21_+_8433172_8446622_RNA45SN1_45S_50ntFlanks | 6739 | 6771 | 33 | rRF 45S | NO | 20.08 | 0.00 | 0.00 | 63.78 |
| >hg38_21_+_8433172_8446622_RNA45SN1_45S_50ntFlanks@6740.6770.31\|CATCGACACTTCGAACGCACTTGCGGCCCCG | >hg38_21_+_8433172_8446622_RNA45SN1_45S_50ntFlanks | 6740 | 6770 | 31 | rRF 45S | NO | 80.49 | 91.40 | 79.52 | 128.76 |
| >hg38_21_+_8433172_8446622_RNA45SN1_45S_50ntFlanks@6740.6771.32\|CATCGACACTTCGAACGCACTTGCGGCCCCGG | >hg38_21_+_8433172_8446622_RNA45SN1_45S_50ntFlanks | 6740 | 6771 | 32 | rRF 45S | NO | 94.09 | 53.25 | 54.19 | 203.56 |
| >hg38_21_+_8433172_8446622_RNA45SN1_45S_50ntFlanks@6740.6772.33\|CATCGACACTTCGAACGCACTTGCGGCCCCGGG | >hg38_21_+_8433172_8446622_RNA45SN1_45S_50ntFlanks | 6740 | 6772 | 33 | rRF 45S | NO | 35.74 | 0.00 | 0.00 | 87.72 |
| >hg38_21_+_8433172_8446622_RNA45SN1_45S_50ntFlanks@6741.6770.30\|ATCGACACTTCGAACGCACTTGCGGCCCCG | >hg38_21_+_8433172_8446622_RNA45SN1_45S_50ntFlanks | 6741 | 6770 | 30 | rRF 45S | NO | 63.49 | 142.11 | 83.15 | 92.38 |
| >hg38_21_+_8433172_8446622_RNA45SN1_45S_50ntFlanks@6741.6771.31\|ATCGACACTTCGAACGCACTTGCGGCCCCGG | >hg38_21_+_8433172_8446622_RNA45SN1_45S_50ntFlanks | 6741 | 6771 | 31 | rRF 45S | NO | 67.19 | 57.50 | 56.78 | 129.85 |
| >hg38_21_+_8433172_8446622_RNA45SN1_45S_50ntFlanks@6741.6772.32\|ATCGACACTTCGAACGCACTTGCGGCCCCGGG | >hg38_21_+_8433172_8446622_RNA45SN1_45S_50ntFlanks | 6741 | 6772 | 32 | rRF 45S | NO | 21.42 | 0.00 | 0.00 | 67.64 |
| >hg38_21_+_8433172_8446622_RNA45SN1_45S_50ntFlanks@6742.6770.29\|TCGACACTTCGAACGCACTTGCGGCCCCG | >hg38_21_+_8433172_8446622_RNA45SN1_45S_50ntFlanks | 6742 | 6770 | 29 | rRF 45S | NO | 0.00 | 46.33 | 28.79 | 6.68 |
| >hg38_21_+_8433172_8446622_RNA45SN1_45S_50ntFlanks@6743.6770.28\|CGACACTTCGAACGCACTTGCGGCCCCG | >hg38_21_+_8433172_8446622_RNA45SN1_45S_50ntFlanks | 6743 | 6770 | 28 | rRF 45S | NO | 60.54 | 177.74 | 89.56 | 79.96 |
| >hg38_21_+_8433172_8446622_RNA45SN1_45S_50ntFlanks@6743.6771.29\|CGACACTTCGAACGCACTTGCGGCCCCGG | >hg38_21_+_8433172_8446622_RNA45SN1_45S_50ntFlanks | 6743 | 6771 | 29 | rRF 45S | NO | 61.16 | 83.99 | 65.15 | 110.71 |
| >hg38_21_+_8433172_8446622_RNA45SN1_45S_50ntFlanks@6744.6770.27\|GACACTTCGAACGCACTTGCGGCCCCG | >hg38_21_+_8433172_8446622_RNA45SN1_45S_50ntFlanks | 6744 | 6770 | 27 | rRF 45S | NO | 66.18 | 205.44 | 62.06 | 71.75 |
| >hg38_21_+_8433172_8446622_RNA45SN1_45S_50ntFlanks@6744.6771.28\|GACACTTCGAACGCACTTGCGGCCCCGG | >hg38_21_+_8433172_8446622_RNA45SN1_45S_50ntFlanks | 6744 | 6771 | 28 | rRF 45S | NO | 35.06 | 70.12 | 50.62 | 61.99 |
| >hg38_21_+_8433172_8446622_RNA45SN1_45S_50ntFlanks@6745.6769.25\|ACACTTCGAACGCACTTGCGGCCCC | >hg38_21_+_8433172_8446622_RNA45SN1_45S_50ntFlanks | 6745 | 6769 | 25 | rRF 45S | NO | 0.00 | 35.04 | 0.00 | 6.13 |
| >hg38_21_+_8433172_8446622_RNA45SN1_45S_50ntFlanks@6745.6770.26\|ACACTTCGAACGCACTTGCGGCCCCG | >hg38_21_+_8433172_8446622_RNA45SN1_45S_50ntFlanks | 6745 | 6770 | 26 | rRF 45S | NO | 149.35 | 502.62 | 261.91 | 261.41 |
| >hg38_21_+_8433172_8446622_RNA45SN1_45S_50ntFlanks@6745.6771.27\|ACACTTCGAACGCACTTGCGGCCCCGG | >hg38_21_+_8433172_8446622_RNA45SN1_45S_50ntFlanks | 6745 | 6771 | 27 | rRF 45S | NO | 103.36 | 224.66 | 170.59 | 240.48 |
| >hg38_21_+_8433172_8446622_RNA45SN1_45S_50ntFlanks@6745.6772.28\|ACACTTCGAACGCACTTGCGGCCCCGGG | >hg38_21_+_8433172_8446622_RNA45SN1_45S_50ntFlanks | 6745 | 6772 | 28 | rRF 45S | NO | 0.00 | 29.47 | 42.20 | 56.65 |
| >hg38_21_+_8433172_8446622_RNA45SN1_45S_50ntFlanks@6746.6770.25\|CACTTCGAACGCACTTGCGGCCCCG | >hg38_21_+_8433172_8446622_RNA45SN1_45S_50ntFlanks | 6746 | 6770 | 25 | rRF 45S | NO | 44.49 | 148.14 | 88.00 | 81.70 |
| >hg38_21_+_8433172_8446622_RNA45SN1_45S_50ntFlanks@6746.6771.26\|CACTTCGAACGCACTTGCGGCCCCGG | >hg38_21_+_8433172_8446622_RNA45SN1_45S_50ntFlanks | 6746 | 6771 | 26 | rRF 45S | NO | 20.54 | 61.35 | 60.67 | 68.50 |
| >hg38_21_+_8433172_8446622_RNA45SN1_45S_50ntFlanks@6747.6770.24\|ACTTCGAACGCACTTGCGGCCCCG | >hg38_21_+_8433172_8446622_RNA45SN1_45S_50ntFlanks | 6747 | 6770 | 24 | rRF 45S | NO | 0.00 | 71.10 | 56.55 | 40.49 |

### MDA-MB-468

| | | | | | | | | | | |
|---|---|---|---|---|---|---|---|---|---|---|
| >hg38_21_+_8433172_8446622_RNA45SN1_45S_50ntFlanks@6747.6771.25\|ACTTCGAACGCA<br>CTTGCGGCCCCGG | >hg38_21_+_8433172_8446622_RNA45SN1_45S_50ntFlanks | 6747 | 6771 | 25 | rRF 45S | NO | 0.00 | 29.60 | 44.81 | 23.72 |
| >hg38_21_+_8433172_8446622_RNA45SN1_45S_50ntFlanks@6748.6770.23\|CTTCGAACGCA<br>CTTGCGGCCCCG | >hg38_21_+_8433172_8446622_RNA45SN1_45S_50ntFlanks | 6748 | 6770 | 23 | rRF 45S | NO | 0.00 | 36.00 | 50.38 | 25.80 |
| >hg38_21_+_8433172_8446622_RNA45SN1_45S_50ntFlanks@6748.6771.24\|CTTCGAACGCA<br>CTTGCGGCCCCGG | >hg38_21_+_8433172_8446622_RNA45SN1_45S_50ntFlanks | 6748 | 6771 | 24 | rRF 45S | NO | 0.00 | 0.00 | 38.38 | 0.00 |
| >hg38_21_+_8433172_8446622_RNA45SN1_45S_50ntFlanks@6751.6770.20\|CGAACGCACTT<br>GCGGCCCCG | >hg38_21_+_8433172_8446622_RNA45SN1_45S_50ntFlanks | 6751 | 6770 | 20 | rRF 45S | NO | 0.00 | 0.00 | 36.79 | 0.00 |
| >hg38_21_+_8433172_8446622_RNA45SN1_45S_50ntFlanks@6771.6799.29\|GGTTCCTCCCG<br>GGGCTACGCCTGTCTGAG | >hg38_21_+_8433172_8446622_RNA45SN1_45S_50ntFlanks | 6771 | 6799 | 29 | rRF 45S | NO | 0.00 | 0.00 | 20.53 | 26.72 |
| >hg38_21_+_8433172_8446622_RNA45SN1_45S_50ntFlanks@6771.6806.36\|GGTTCCTCCCG<br>GGGCTACGCCTGTCTGAGCGTCGCT | >hg38_21_+_8433172_8446622_RNA45SN1_45S_50ntFlanks | 6771 | 6806 | 36 | rRF 45S | NO | 0.00 | 0.00 | 61.57 | 90.72 |
| >hg38_21_+_8433172_8446622_RNA45SN1_45S_50ntFlanks@6772.6799.28\|GTTCCTCCCGG<br>GGCTACGCCTGTCTGAG | >hg38_21_+_8433172_8446622_RNA45SN1_45S_50ntFlanks | 6772 | 6799 | 28 | rRF 45S | NO | 55.71 | 64.82 | 13.70 | 122.46 |
| >hg38_21_+_8433172_8446622_RNA45SN1_45S_50ntFlanks@6772.6803.32\|GTTCCTCCCGG<br>GGCTACGCCTGTCTGAGCGTC | >hg38_21_+_8433172_8446622_RNA45SN1_45S_50ntFlanks | 6772 | 6803 | 32 | rRF 45S | NO | 0.00 | 0.00 | 54.74 | 70.90 |
| >hg38_21_+_8433172_8446622_RNA45SN1_45S_50ntFlanks@6772.6806.35\|GTTCCTCCCGG<br>GGCTACGCCTGTCTGAGCGTCGCT | >hg38_21_+_8433172_8446622_RNA45SN1_45S_50ntFlanks | 6772 | 6806 | 35 | rRF 45S | NO | 46.56 | 29.53 | 71.83 | 202.89 |
| >hg38_21_+_8433172_8446622_RNA45SN1_45S_50ntFlanks@6772.6807.36\|GTTCCTCCCGG<br>GGCTACGCCTGTCTGAGCGTCGCTT | >hg38_21_+_8433172_8446622_RNA45SN1_45S_50ntFlanks | 6772 | 6807 | 36 | rRF 45S | NO | 39.28 | 0.00 | 0.00 | 141.73 |
| >hg38_21_+_8433172_8446622_RNA45SN1_45S_50ntFlanks@6773.6793.21\|TTCCTCCCGGG<br>GCTACGCCTG | >hg38_21_+_8433172_8446622_RNA45SN1_45S_50ntFlanks | 6773 | 6793 | 21 | rRF 45S | NO | 40.26 | 35.39 | 0.00 | 34.71 |
| >hg38_21_+_8433172_8446622_RNA45SN1_45S_50ntFlanks@6773.6799.27\|TTCCTCCCGGG<br>GCTACGCCTGTCTGAG | >hg38_21_+_8433172_8446622_RNA45SN1_45S_50ntFlanks | 6773 | 6799 | 27 | rRF 45S | NO | 0.00 | 33.02 | 0.00 | 51.36 |
| >hg38_21_+_8433172_8446622_RNA45SN1_45S_50ntFlanks@6773.6803.31\|TTCCTCCCGGG<br>GCTACGCCTGTCTGAGCGTC | >hg38_21_+_8433172_8446622_RNA45SN1_45S_50ntFlanks | 6773 | 6803 | 31 | rRF 45S | NO | 57.74 | 71.53 | 41.13 | 181.71 |
| >hg38_21_+_8433172_8446622_RNA45SN1_45S_50ntFlanks@6773.6806.34\|TTCCTCCCGGG<br>GCTACGCCTGTCTGAGCGTCGCT | >hg38_21_+_8433172_8446622_RNA45SN1_45S_50ntFlanks | 6773 | 6806 | 34 | rRF 45S | NO | 0.00 | 0.00 | 42.00 | 110.31 |
| >hg38_21_+_8433172_8446622_RNA45SN1_45S_50ntFlanks@6776.6803.28\|CTCCCGGGGCT<br>ACGCCTGTCTGAGCGTC | >hg38_21_+_8433172_8446622_RNA45SN1_45S_50ntFlanks | 6776 | 6803 | 28 | rRF 45S | NO | 44.60 | 51.12 | 0.00 | 117.61 |
| >hg38_21_+_8433172_8446622_RNA45SN1_45S_50ntFlanks@6779.6803.25\|CCGGGGCTACG<br>CCTGTCTGAGCGTC | >hg38_21_+_8433172_8446622_RNA45SN1_45S_50ntFlanks | 6779 | 6803 | 25 | rRF 45S | NO | 0.00 | 28.17 | 37.42 | 55.88 |
| >hg38_21_+_8433172_8446622_RNA45SN1_45S_50ntFlanks@6784.6803.20\|GCTACGCCTGT<br>CTGAGCGTC | >hg38_21_+_8433172_8446622_RNA45SN1_45S_50ntFlanks | 6784 | 6803 | 20 | rRF 45S | NO | 0.00 | 0.00 | 128.47 | 0.00 |
| >hg38_21_+_8433172_8446622_RNA45SN1_45S_50ntFlanks@6785.6806.22\|CTACGCCTGTCT<br>GAGCGTCGCT | >hg38_21_+_8433172_8446622_RNA45SN1_45S_50ntFlanks | 6785 | 6806 | 22 | rRF 45S | NO | 0.00 | 0.00 | 70.46 | 39.37 |
| >hg38_21_+_8433172_8446622_RNA45SN1_45S_50ntFlanks@7973.8006.34\|GACGCGACCTC<br>AGATCAGACGTGGCGACCCGCTG | >hg38_21_+_8433172_8446622_RNA45SN1_45S_50ntFlanks | 7973 | 8006 | 34 | rRF 45S | NO | 32.08 | 0.00 | 0.00 | 68.27 |
| >hg38_21_+_8433172_8446622_RNA45SN1_45S_50ntFlanks@7973.8008.36\|GACGCGACCTC<br>AGATCAGACGTGGCGACCCGCTGAA | >hg38_21_+_8433172_8446622_RNA45SN1_45S_50ntFlanks | 7973 | 8008 | 36 | rRF 45S | NO | 37.63 | 0.00 | 0.00 | 55.58 |
| >hg38_21_+_8433172_8446622_RNA45SN1_45S_50ntFlanks@7974.7991.18\|ACGCGACCTCA<br>GATCAGA | >hg38_21_+_8433172_8446622_RNA45SN1_45S_50ntFlanks | 7974 | 7991 | 18 | rRF 45S | NO | 34.06 | 0.00 | 0.00 | 6.09 |
| >hg38_21_+_8433172_8446622_RNA45SN1_45S_50ntFlanks@7974.7992.19\|ACGCGACCTCA<br>GATCAGAC | >hg38_21_+_8433172_8446622_RNA45SN1_45S_50ntFlanks | 7974 | 7992 | 19 | rRF 45S | NO | 44.51 | 0.00 | 0.00 | 25.86 |
| >hg38_21_+_8433172_8446622_RNA45SN1_45S_50ntFlanks@7974.7993.20\|ACGCGACCTCA<br>GATCAGACG | >hg38_21_+_8433172_8446622_RNA45SN1_45S_50ntFlanks | 7974 | 7993 | 20 | rRF 45S | NO | 55.21 | 32.87 | 36.97 | 68.62 |
| >hg38_21_+_8433172_8446622_RNA45SN1_45S_50ntFlanks@7974.7994.21\|ACGCGACCTCA<br>GATCAGACGT | >hg38_21_+_8433172_8446622_RNA45SN1_45S_50ntFlanks | 7974 | 7994 | 21 | rRF 45S | NO | 0.00 | 28.61 | 38.16 | 15.74 |
| >hg38_21_+_8433172_8446622_RNA45SN1_45S_50ntFlanks@7974.7995.22\|ACGCGACCTCA<br>GATCAGACGTG | >hg38_21_+_8433172_8446622_RNA45SN1_45S_50ntFlanks | 7974 | 7995 | 22 | rRF 45S | NO | 42.80 | 99.47 | 54.34 | 58.75 |
| >hg38_21_+_8433172_8446622_RNA45SN1_45S_50ntFlanks@7974.8006.33\|ACGCGACCTCA<br>GATCAGACGTGGCGACCCGCTG | >hg38_21_+_8433172_8446622_RNA45SN1_45S_50ntFlanks | 7974 | 8006 | 33 | rRF 45S | NO | 299.09 | 134.99 | 93.16 | 501.69 |
| >hg38_21_+_8433172_8446622_RNA45SN1_45S_50ntFlanks@7974.8007.34\|ACGCGACCTCA<br>GATCAGACGTGGCGACCCGCTGA | >hg38_21_+_8433172_8446622_RNA45SN1_45S_50ntFlanks | 7974 | 8007 | 34 | rRF 45S | NO | 44.21 | 0.00 | 0.00 | 73.22 |
| >hg38_21_+_8433172_8446622_RNA45SN1_45S_50ntFlanks@7974.8008.35\|ACGCGACCTCA<br>GATCAGACGTGGCGACCCGCTGAA | >hg38_21_+_8433172_8446622_RNA45SN1_45S_50ntFlanks | 7974 | 8008 | 35 | rRF 45S | NO | 243.35 | 47.26 | 61.36 | 332.17 |
| >hg38_21_+_8433172_8446622_RNA45SN1_45S_50ntFlanks@7974.8009.36\|ACGCGACCTCA<br>GATCAGACGTGGCGACCCGCTGAAT | >hg38_21_+_8433172_8446622_RNA45SN1_45S_50ntFlanks | 7974 | 8009 | 36 | rRF 45S | NO | 74.75 | 0.00 | 35.83 | 116.56 |
| >hg38_21_+_8433172_8446622_RNA45SN1_45S_50ntFlanks@7974.8010.37\|ACGCGACCTCA<br>GATCAGACGTGGCGACCCGCTGAATT | >hg38_21_+_8433172_8446622_RNA45SN1_45S_50ntFlanks | 7974 | 8010 | 37 | rRF 45S | NO | 44.66 | 0.00 | 0.00 | 73.03 |
| >hg38_21_+_8433172_8446622_RNA45SN1_45S_50ntFlanks@7974.8011.38\|ACGCGACCTCA<br>GATCAGACGTGGCGACCCGCTGAATTT | >hg38_21_+_8433172_8446622_RNA45SN1_45S_50ntFlanks | 7974 | 8011 | 38 | rRF 45S | NO | 76.52 | 0.00 | 0.00 | 98.26 |
| >hg38_21_+_8433172_8446622_RNA45SN1_45S_50ntFlanks@7974.8012.39\|ACGCGACCTCA<br>GATCAGACGTGGCGACCCGCTGAATTTA | >hg38_21_+_8433172_8446622_RNA45SN1_45S_50ntFlanks | 7974 | 8012 | 39 | rRF 45S | NO | 105.89 | 0.00 | 0.00 | 176.44 |
| >hg38_21_+_8433172_8446622_RNA45SN1_45S_50ntFlanks@7975.7992.18\|CGCGACCTCAG<br>ATCAGAC | >hg38_21_+_8433172_8446622_RNA45SN1_45S_50ntFlanks | 7975 | 7992 | 18 | rRF 45S | NO | 190.34 | 83.64 | 39.83 | 156.55 |
| >hg38_21_+_8433172_8446622_RNA45SN1_45S_50ntFlanks@7975.7993.19\|CGCGACCTCAG<br>ATCAGACG | >hg38_21_+_8433172_8446622_RNA45SN1_45S_50ntFlanks | 7975 | 7993 | 19 | rRF 45S | NO | 186.20 | 158.66 | 133.25 | 251.61 |

### MDA-MB-468

| | | | | | | | | | | |
|---|---|---|---|---|---|---|---|---|---|---|
| >hg38_21_+_8433172_8446622_RNA45SN1_45S_50ntFlanks@7975.7994.20\|CGCGACCTCAGATCAGACGT | >hg38_21_+_8433172_8446622_RNA45SN1_45S_50ntFlanks | 7975 | 7994 | 20 | rRF 45S | YES | 1017.33 | 1121.56 | 511.11 | 627.29 |
| >hg38_21_+_8433172_8446622_RNA45SN1_45S_50ntFlanks@7975.7995.21\|CGCGACCTCAGATCAGACGTG | >hg38_21_+_8433172_8446622_RNA45SN1_45S_50ntFlanks | 7975 | 7995 | 21 | rRF 45S | NO | 106.07 | 368.34 | 180.53 | 141.23 |
| >hg38_21_+_8433172_8446622_RNA45SN1_45S_50ntFlanks@7975.7996.22\|CGCGACCTCAGATCAGACGTGG | >hg38_21_+_8433172_8446622_RNA45SN1_45S_50ntFlanks | 7975 | 7996 | 22 | rRF 45S | NO | 80.63 | 176.44 | 108.82 | 132.24 |
| >hg38_21_+_8433172_8446622_RNA45SN1_45S_50ntFlanks@7975.7997.23\|CGCGACCTCAGATCAGACGTGGC | >hg38_21_+_8433172_8446622_RNA45SN1_45S_50ntFlanks | 7975 | 7997 | 23 | rRF 45S | NO | 66.64 | 39.40 | 61.17 | 89.07 |
| >hg38_21_+_8433172_8446622_RNA45SN1_45S_50ntFlanks@7975.8002.28\|CGCGACCTCAGATCAGACGTGGCGACCC | >hg38_21_+_8433172_8446622_RNA45SN1_45S_50ntFlanks | 7975 | 8002 | 28 | rRF 45S | NO | 0.00 | 0.00 | 49.13 | 0.00 |
| >hg38_21_+_8433172_8446622_RNA45SN1_45S_50ntFlanks@7975.8003.29\|CGCGACCTCAGATCAGACGTGGCGACCCG | >hg38_21_+_8433172_8446622_RNA45SN1_45S_50ntFlanks | 7975 | 8003 | 29 | rRF 45S | NO | 50.75 | 107.39 | 60.12 | 92.18 |
| >hg38_21_+_8433172_8446622_RNA45SN1_45S_50ntFlanks@7975.8004.30\|CGCGACCTCAGATCAGACGTGGCGACCCGC | >hg38_21_+_8433172_8446622_RNA45SN1_45S_50ntFlanks | 7975 | 8004 | 30 | rRF 45S | NO | 45.97 | 0.00 | 27.65 | 97.91 |
| >hg38_21_+_8433172_8446622_RNA45SN1_45S_50ntFlanks@7975.8005.31\|CGCGACCTCAGATCAGACGTGGCGACCCGCT | >hg38_21_+_8433172_8446622_RNA45SN1_45S_50ntFlanks | 7975 | 8005 | 31 | rRF 45S | NO | 183.00 | 124.06 | 68.22 | 327.74 |
| >hg38_21_+_8433172_8446622_RNA45SN1_45S_50ntFlanks@7975.8006.32\|CGCGACCTCAGATCAGACGTGGCGACCCGCTG | >hg38_21_+_8433172_8446622_RNA45SN1_45S_50ntFlanks | 7975 | 8006 | 32 | rRF 45S | YES | 1948.88 | 948.12 | 398.90 | 2983.36 |
| >hg38_21_+_8433172_8446622_RNA45SN1_45S_50ntFlanks@7975.8007.33\|CGCGACCTCAGATCAGACGTGGCGACCCGCTGA | >hg38_21_+_8433172_8446622_RNA45SN1_45S_50ntFlanks | 7975 | 8007 | 33 | rRF 45S | NO | 272.27 | 60.32 | 72.68 | 402.93 |
| >hg38_21_+_8433172_8446622_RNA45SN1_45S_50ntFlanks@7975.8008.34\|CGCGACCTCAGATCAGACGTGGCGACCCGCTGAA | >hg38_21_+_8433172_8446622_RNA45SN1_45S_50ntFlanks | 7975 | 8008 | 34 | rRF 45S | NO | 1299.19 | 224.33 | 235.08 | 1683.42 |
| >hg38_21_+_8433172_8446622_RNA45SN1_45S_50ntFlanks@7975.8009.35\|CGCGACCTCAGATCAGACGTGGCGACCCGCTGAAT | >hg38_21_+_8433172_8446622_RNA45SN1_45S_50ntFlanks | 7975 | 8009 | 35 | rRF 45S | NO | 444.66 | 75.12 | 123.21 | 730.78 |
| >hg38_21_+_8433172_8446622_RNA45SN1_45S_50ntFlanks@7975.8010.36\|CGCGACCTCAGATCAGACGTGGCGACCCGCTGAATT | >hg38_21_+_8433172_8446622_RNA45SN1_45S_50ntFlanks | 7975 | 8010 | 36 | rRF 45S | NO | 286.41 | 46.35 | 40.73 | 470.49 |
| >hg38_21_+_8433172_8446622_RNA45SN1_45S_50ntFlanks@7975.8011.37\|CGCGACCTCAGATCAGACGTGGCGACCCGCTGAATTT | >hg38_21_+_8433172_8446622_RNA45SN1_45S_50ntFlanks | 7975 | 8011 | 37 | rRF 45S | NO | 542.60 | 73.72 | 47.37 | 614.98 |
| >hg38_21_+_8433172_8446622_RNA45SN1_45S_50ntFlanks@7975.8012.38\|CGCGACCTCAGATCAGACGTGGCGACCCGCTGAATTTA | >hg38_21_+_8433172_8446622_RNA45SN1_45S_50ntFlanks | 7975 | 8012 | 38 | rRF 45S | NO | 724.72 | 92.03 | 65.04 | 1101.25 |
| >hg38_21_+_8433172_8446622_RNA45SN1_45S_50ntFlanks@7975.8013.39\|CGCGACCTCAGATCAGACGTGGCGACCCGCTGAATTTAA | >hg38_21_+_8433172_8446622_RNA45SN1_45S_50ntFlanks | 7975 | 8013 | 39 | rRF 45S | NO | 46.75 | 0.00 | 0.00 | 70.14 |
| >hg38_21_+_8433172_8446622_RNA45SN1_45S_50ntFlanks@7975.8014.40\|CGCGACCTCAGATCAGACGTGGCGACCCGCTGAATTTAAG | >hg38_21_+_8433172_8446622_RNA45SN1_45S_50ntFlanks | 7975 | 8014 | 40 | rRF 45S | NO | 78.68 | 0.00 | 0.00 | 137.55 |
| >hg38_21_+_8433172_8446622_RNA45SN1_45S_50ntFlanks@7975.8016.42\|CGCGACCTCAGATCAGACGTGGCGACCCGCTGAATTTAAGCA | >hg38_21_+_8433172_8446622_RNA45SN1_45S_50ntFlanks | 7975 | 8016 | 42 | rRF 45S | NO | 29.97 | 0.00 | 0.00 | 230.81 |
| >hg38_21_+_8433172_8446622_RNA45SN1_45S_50ntFlanks@7977.7994.18\|CGACCTCAGATCAGACGT | >hg38_21_+_8433172_8446622_RNA45SN1_45S_50ntFlanks | 7977 | 7994 | 18 | rRF 45S | NO | 0.00 | 19.61 | 0.00 | 0.00 |
| >hg38_21_+_8433172_8446622_RNA45SN1_45S_50ntFlanks@7977.7995.19\|CGACCTCAGATCAGACGTG | >hg38_21_+_8433172_8446622_RNA45SN1_45S_50ntFlanks | 7977 | 7995 | 19 | rRF 45S | NO | 0.00 | 27.22 | 0.00 | 0.00 |
| >hg38_21_+_8433172_8446622_RNA45SN1_45S_50ntFlanks@7977.8006.30\|CGACCTCAGATCAGACGTGGCGACCCGCTG | >hg38_21_+_8433172_8446622_RNA45SN1_45S_50ntFlanks | 7977 | 8006 | 30 | rRF 45S | NO | 135.86 | 41.68 | 51.34 | 123.23 |
| >hg38_21_+_8433172_8446622_RNA45SN1_45S_50ntFlanks@7977.8008.32\|CGACCTCAGATCAGACGTGGCGACCCGCTGAA | >hg38_21_+_8433172_8446622_RNA45SN1_45S_50ntFlanks | 7977 | 8008 | 32 | rRF 45S | NO | 41.60 | 0.00 | 7.13 | 43.04 |
| >hg38_21_+_8433172_8446622_RNA45SN1_45S_50ntFlanks@7978.8006.29\|GACCTCAGATCAGACGTGGCGACCCGCTG | >hg38_21_+_8433172_8446622_RNA45SN1_45S_50ntFlanks | 7978 | 8006 | 29 | rRF 45S | NO | 32.14 | 0.00 | 0.00 | 14.90 |
| >hg38_21_+_8433172_8446622_RNA45SN1_45S_50ntFlanks@7979.8003.25\|ACCTCAGATCAGACGTGGCGACCCG | >hg38_21_+_8433172_8446622_RNA45SN1_45S_50ntFlanks | 7979 | 8003 | 25 | rRF 45S | NO | 89.90 | 46.03 | 0.00 | 14.34 |
| >hg38_21_+_8433172_8446622_RNA45SN1_45S_50ntFlanks@7979.8005.27\|ACCTCAGATCAGACGTGGCGACCCGCT | >hg38_21_+_8433172_8446622_RNA45SN1_45S_50ntFlanks | 7979 | 8005 | 27 | rRF 45S | NO | 12.86 | 0.00 | 0.00 | 14.80 |
| >hg38_21_+_8433172_8446622_RNA45SN1_45S_50ntFlanks@7979.8006.28\|ACCTCAGATCAGACGTGGCGACCCGCTG | >hg38_21_+_8433172_8446622_RNA45SN1_45S_50ntFlanks | 7979 | 8006 | 28 | rRF 45S | NO | 512.03 | 196.58 | 167.00 | 424.35 |
| >hg38_21_+_8433172_8446622_RNA45SN1_45S_50ntFlanks@7979.8008.30\|ACCTCAGATCAGACGTGGCGACCCGCTGAA | >hg38_21_+_8433172_8446622_RNA45SN1_45S_50ntFlanks | 7979 | 8008 | 30 | rRF 45S | NO | 103.72 | 26.01 | 76.66 | 124.68 |
| >hg38_21_+_8433172_8446622_RNA45SN1_45S_50ntFlanks@7979.8009.31\|ACCTCAGATCAGACGTGGCGACCCGCTGAAT | >hg38_21_+_8433172_8446622_RNA45SN1_45S_50ntFlanks | 7979 | 8009 | 31 | rRF 45S | NO | 12.91 | 0.00 | 21.05 | 25.63 |
| >hg38_21_+_8433172_8446622_RNA45SN1_45S_50ntFlanks@7979.8012.34\|ACCTCAGATCAGACGTGGCGACCCGCTGAATTTA | >hg38_21_+_8433172_8446622_RNA45SN1_45S_50ntFlanks | 7979 | 8012 | 34 | rRF 45S | NO | 30.02 | 0.00 | 0.00 | 66.64 |
| >hg38_21_+_8433172_8446622_RNA45SN1_45S_50ntFlanks@7980.8003.24\|CCTCAGATCAGACGTGGCGACCCG | >hg38_21_+_8433172_8446622_RNA45SN1_45S_50ntFlanks | 7980 | 8003 | 24 | rRF 45S | NO | 51.93 | 27.98 | 0.00 | 0.00 |
| >hg38_21_+_8433172_8446622_RNA45SN1_45S_50ntFlanks@7980.8005.26\|CCTCAGATCAGACGTGGCGACCCGCT | >hg38_21_+_8433172_8446622_RNA45SN1_45S_50ntFlanks | 7980 | 8005 | 26 | rRF 45S | NO | 28.85 | 0.00 | 0.00 | 25.15 |
| >hg38_21_+_8433172_8446622_RNA45SN1_45S_50ntFlanks@7980.8006.27\|CCTCAGATCAGACGTGGCGACCCGCTG | >hg38_21_+_8433172_8446622_RNA45SN1_45S_50ntFlanks | 7980 | 8006 | 27 | rRF 45S | NO | 410.41 | 159.73 | 208.23 | 397.55 |
| >hg38_21_+_8433172_8446622_RNA45SN1_45S_50ntFlanks@7980.8008.29\|CCTCAGATCAGACGTGGCGACCCGCTGAA | >hg38_21_+_8433172_8446622_RNA45SN1_45S_50ntFlanks | 7980 | 8008 | 29 | rRF 45S | NO | 70.71 | 0.00 | 75.52 | 112.40 |
| >hg38_21_+_8433172_8446622_RNA45SN1_45S_50ntFlanks@7981.8003.23\|CTCAGATCAGACGTGGCGACCCG | >hg38_21_+_8433172_8446622_RNA45SN1_45S_50ntFlanks | 7981 | 8003 | 23 | rRF 45S | NO | 13.21 | 0.00 | 0.00 | 0.00 |

### MDA-MB-468

| | | | | | | | | | | |
|---|---|---|---|---|---|---|---|---|---|---|
| >hg38_21_+_8433172_8446622_RNA45SN1_45S_50ntFlanks@7981.8005.25\|CTCAGATCAGACGTGGCGACCCGCT | >hg38_21_+_8433172_8446622_RNA45SN1_45S_50ntFlanks | 7981 | 8005 | 25 | rRF 45S | NO | 19.53 | 0.00 | 0.00 | 15.84 |
| >hg38_21_+_8433172_8446622_RNA45SN1_45S_50ntFlanks@7981.8006.26\|CTCAGATCAGACGTGGCGACCCGCTG | >hg38_21_+_8433172_8446622_RNA45SN1_45S_50ntFlanks | 7981 | 8006 | 26 | rRF 45S | NO | 296.72 | 149.76 | 195.12 | 320.89 |
| >hg38_21_+_8433172_8446622_RNA45SN1_45S_50ntFlanks@7981.8008.28\|CTCAGATCAGACGTGGCGACCCGCTGAA | >hg38_21_+_8433172_8446622_RNA45SN1_45S_50ntFlanks | 7981 | 8008 | 28 | rRF 45S | NO | 39.40 | 0.00 | 64.29 | 90.53 |
| >hg38_21_+_8433172_8446622_RNA45SN1_45S_50ntFlanks@7982.8006.25\|TCAGATCAGACGTGGCGACCCGCTG | >hg38_21_+_8433172_8446622_RNA45SN1_45S_50ntFlanks | 7982 | 8006 | 25 | rRF 45S | NO | 131.48 | 72.56 | 141.58 | 244.70 |
| >hg38_21_+_8433172_8446622_RNA45SN1_45S_50ntFlanks@7982.8008.27\|TCAGATCAGACGTGGCGACCCGCTGAA | >hg38_21_+_8433172_8446622_RNA45SN1_45S_50ntFlanks | 7982 | 8008 | 27 | rRF 45S | NO | 12.80 | 0.00 | 58.08 | 78.19 |
| >hg38_21_+_8433172_8446622_RNA45SN1_45S_50ntFlanks@7983.8003.21\|CAGATCAGACGTGGCGACCCG | >hg38_21_+_8433172_8446622_RNA45SN1_45S_50ntFlanks | 7983 | 8003 | 21 | rRF 45S | NO | 52.07 | 35.71 | 37.33 | 14.86 |
| >hg38_21_+_8433172_8446622_RNA45SN1_45S_50ntFlanks@7983.8004.22\|CAGATCAGACGTGGCGACCCGC | >hg38_21_+_8433172_8446622_RNA45SN1_45S_50ntFlanks | 7983 | 8004 | 22 | rRF 45S | NO | 61.92 | 0.00 | 49.00 | 111.05 |
| >hg38_21_+_8433172_8446622_RNA45SN1_45S_50ntFlanks@7983.8005.23\|CAGATCAGACGTGGCGACCCGCT | >hg38_21_+_8433172_8446622_RNA45SN1_45S_50ntFlanks | 7983 | 8005 | 23 | rRF 45S | NO | 51.29 | 12.50 | 51.19 | 73.18 |
| >hg38_21_+_8433172_8446622_RNA45SN1_45S_50ntFlanks@7983.8006.24\|CAGATCAGACGTGGCGACCCGCTG | >hg38_21_+_8433172_8446622_RNA45SN1_45S_50ntFlanks | 7983 | 8006 | 24 | rRF 45S | YES | 368.75 | 236.77 | 437.88 | 665.38 |
| >hg38_21_+_8433172_8446622_RNA45SN1_45S_50ntFlanks@7983.8007.25\|CAGATCAGACGTGGCGACCCGCTGA | >hg38_21_+_8433172_8446622_RNA45SN1_45S_50ntFlanks | 7983 | 8007 | 25 | rRF 45S | NO | 0.00 | 0.00 | 41.58 | 62.06 |
| >hg38_21_+_8433172_8446622_RNA45SN1_45S_50ntFlanks@7983.8008.26\|CAGATCAGACGTGGCGACCCGCTGAA | >hg38_21_+_8433172_8446622_RNA45SN1_45S_50ntFlanks | 7983 | 8008 | 26 | rRF 45S | NO | 72.49 | 41.25 | 174.86 | 243.60 |
| >hg38_21_+_8433172_8446622_RNA45SN1_45S_50ntFlanks@7983.8009.27\|CAGATCAGACGTGGCGACCCGCTGAAT | >hg38_21_+_8433172_8446622_RNA45SN1_45S_50ntFlanks | 7983 | 8009 | 27 | rRF 45S | NO | 0.00 | 0.00 | 43.97 | 46.77 |
| >hg38_21_+_8433172_8446622_RNA45SN1_45S_50ntFlanks@7983.8012.30\|CAGATCAGACGTGGCGACCCGCTGAATTTA | >hg38_21_+_8433172_8446622_RNA45SN1_45S_50ntFlanks | 7983 | 8012 | 30 | rRF 45S | NO | 0.00 | 0.00 | 30.27 | 51.99 |
| >hg38_21_+_8433172_8446622_RNA45SN1_45S_50ntFlanks@7984.8003.20\|AGATCAGACGTGGCGACCCG | >hg38_21_+_8433172_8446622_RNA45SN1_45S_50ntFlanks | 7984 | 8003 | 20 | rRF 45S | NO | 35.57 | 5.98 | 31.03 | 0.00 |
| >hg38_21_+_8433172_8446622_RNA45SN1_45S_50ntFlanks@7984.8004.21\|AGATCAGACGTGGCGACCCGC | >hg38_21_+_8433172_8446622_RNA45SN1_45S_50ntFlanks | 7984 | 8004 | 21 | rRF 45S | NO | 12.78 | 0.00 | 39.94 | 48.46 |
| >hg38_21_+_8433172_8446622_RNA45SN1_45S_50ntFlanks@7984.8005.22\|AGATCAGACGTGGCGACCCGCT | >hg38_21_+_8433172_8446622_RNA45SN1_45S_50ntFlanks | 7984 | 8005 | 22 | rRF 45S | NO | 0.00 | 0.00 | 39.40 | 34.87 |
| >hg38_21_+_8433172_8446622_RNA45SN1_45S_50ntFlanks@7984.8006.23\|AGATCAGACGTGGCGACCCGCTG | >hg38_21_+_8433172_8446622_RNA45SN1_45S_50ntFlanks | 7984 | 8006 | 23 | rRF 45S | NO | 112.74 | 113.08 | 321.50 | 252.01 |
| >hg38_21_+_8433172_8446622_RNA45SN1_45S_50ntFlanks@7984.8007.24\|AGATCAGACGTGGCGACCCGCTGA | >hg38_21_+_8433172_8446622_RNA45SN1_45S_50ntFlanks | 7984 | 8007 | 24 | rRF 45S | NO | 0.00 | 0.00 | 39.72 | 34.40 |
| >hg38_21_+_8433172_8446622_RNA45SN1_45S_50ntFlanks@7984.8008.25\|AGATCAGACGTGGCGACCCGCTGAA | >hg38_21_+_8433172_8446622_RNA45SN1_45S_50ntFlanks | 7984 | 8008 | 25 | rRF 45S | NO | 37.54 | 0.00 | 183.70 | 149.81 |
| >hg38_21_+_8433172_8446622_RNA45SN1_45S_50ntFlanks@7984.8009.26\|AGATCAGACGTGGCGACCCGCTGAAT | >hg38_21_+_8433172_8446622_RNA45SN1_45S_50ntFlanks | 7984 | 8009 | 26 | rRF 45S | NO | 0.00 | 0.00 | 52.62 | 23.43 |
| >hg38_21_+_8433172_8446622_RNA45SN1_45S_50ntFlanks@7985.8004.20\|GATCAGACGTGGCGACCCGC | >hg38_21_+_8433172_8446622_RNA45SN1_45S_50ntFlanks | 7985 | 8004 | 20 | rRF 45S | NO | 0.00 | 0.00 | 31.40 | 25.21 |
| >hg38_21_+_8433172_8446622_RNA45SN1_45S_50ntFlanks@7985.8005.21\|GATCAGACGTGGCGACCCGCT | >hg38_21_+_8433172_8446622_RNA45SN1_45S_50ntFlanks | 7985 | 8005 | 21 | rRF 45S | NO | 0.00 | 0.00 | 31.96 | 15.22 |
| >hg38_21_+_8433172_8446622_RNA45SN1_45S_50ntFlanks@7985.8006.22\|GATCAGACGTGGCGACCCGCTG | >hg38_21_+_8433172_8446622_RNA45SN1_45S_50ntFlanks | 7985 | 8006 | 22 | rRF 45S | NO | 75.21 | 87.92 | 260.94 | 187.17 |
| >hg38_21_+_8433172_8446622_RNA45SN1_45S_50ntFlanks@7985.8007.23\|GATCAGACGTGGCGACCCGCTGA | >hg38_21_+_8433172_8446622_RNA45SN1_45S_50ntFlanks | 7985 | 8007 | 23 | rRF 45S | NO | 0.00 | 0.00 | 20.92 | 14.45 |
| >hg38_21_+_8433172_8446622_RNA45SN1_45S_50ntFlanks@7985.8008.24\|GATCAGACGTGGCGACCCGCTGAA | >hg38_21_+_8433172_8446622_RNA45SN1_45S_50ntFlanks | 7985 | 8008 | 24 | rRF 45S | NO | 0.00 | 0.00 | 90.00 | 62.82 |
| >hg38_21_+_8433172_8446622_RNA45SN1_45S_50ntFlanks@7985.8009.25\|GATCAGACGTGGCGACCCGCTGAAT | >hg38_21_+_8433172_8446622_RNA45SN1_45S_50ntFlanks | 7985 | 8009 | 25 | rRF 45S | NO | 0.00 | 0.00 | 32.14 | 0.00 |
| >hg38_21_+_8433172_8446622_RNA45SN1_45S_50ntFlanks@7985.8012.28\|GATCAGACGTGGCGACCCGCTGAATTTA | >hg38_21_+_8433172_8446622_RNA45SN1_45S_50ntFlanks | 7985 | 8012 | 28 | rRF 45S | NO | 0.00 | 0.00 | 29.89 | 38.60 |
| >hg38_21_+_8433172_8446622_RNA45SN1_45S_50ntFlanks@7986.8003.18\|ATCAGACGTGGCGACCCG | >hg38_21_+_8433172_8446622_RNA45SN1_45S_50ntFlanks | 7986 | 8003 | 18 | rRF 45S | NO | 146.37 | 164.14 | 92.85 | 81.88 |
| >hg38_21_+_8433172_8446622_RNA45SN1_45S_50ntFlanks@7986.8004.19\|ATCAGACGTGGCGACCCGC | >hg38_21_+_8433172_8446622_RNA45SN1_45S_50ntFlanks | 7986 | 8004 | 19 | rRF 45S | NO | 6.18 | 0.00 | 44.25 | 42.29 |
| >hg38_21_+_8433172_8446622_RNA45SN1_45S_50ntFlanks@7986.8005.20\|ATCAGACGTGGCGACCCGCT | >hg38_21_+_8433172_8446622_RNA45SN1_45S_50ntFlanks | 7986 | 8005 | 20 | rRF 45S | NO | 0.00 | 36.25 | 104.50 | 74.31 |
| >hg38_21_+_8433172_8446622_RNA45SN1_45S_50ntFlanks@7986.8006.21\|ATCAGACGTGGCGACCCGCTG | >hg38_21_+_8433172_8446622_RNA45SN1_45S_50ntFlanks | 7986 | 8006 | 21 | rRF 45S | NO | 69.93 | 137.42 | 523.05 | 299.24 |
| >hg38_21_+_8433172_8446622_RNA45SN1_45S_50ntFlanks@7986.8007.22\|ATCAGACGTGGCGACCCGCTGA | >hg38_21_+_8433172_8446622_RNA45SN1_45S_50ntFlanks | 7986 | 8007 | 22 | rRF 45S | NO | 0.00 | 0.00 | 37.19 | 13.81 |
| >hg38_21_+_8433172_8446622_RNA45SN1_45S_50ntFlanks@7986.8008.23\|ATCAGACGTGGCGACCCGCTGAA | >hg38_21_+_8433172_8446622_RNA45SN1_45S_50ntFlanks | 7986 | 8008 | 23 | rRF 45S | NO | 0.00 | 0.00 | 151.74 | 77.76 |
| >hg38_21_+_8433172_8446622_RNA45SN1_45S_50ntFlanks@7986.8009.24\|ATCAGACGTGGCGACCCGCTGAAT | >hg38_21_+_8433172_8446622_RNA45SN1_45S_50ntFlanks | 7986 | 8009 | 24 | rRF 45S | NO | 0.00 | 0.00 | 53.71 | 0.00 |

### MDA-MB-468

| | | | | | | | | | | |
|---|---|---|---|---|---|---|---|---|---|---|
| >hg38_21_+_8433172_8446622_RNA45SN1_45S_50ntFlanks@7986.8011.26\|ATCAGACGTGGCGACCCGCTGAATTT | >hg38_21_+_8433172_8446622_RNA45SN1_45S_50ntFlanks | 7986 | 8011 | 26 | rRF 45S | NO | 0.00 | 0.00 | 40.99 | 37.72 |
| >hg38_21_+_8433172_8446622_RNA45SN1_45S_50ntFlanks@7986.8012.27\|ATCAGACGTGGCGACCCGCTGAATTTA | >hg38_21_+_8433172_8446622_RNA45SN1_45S_50ntFlanks | 7986 | 8012 | 27 | rRF 45S | NO | 0.00 | 0.00 | 91.76 | 105.10 |
| >hg38_21_+_8433172_8446622_RNA45SN1_45S_50ntFlanks@7986.8014.29\|ATCAGACGTGGCGACCCGCTGAATTTAAG | >hg38_21_+_8433172_8446622_RNA45SN1_45S_50ntFlanks | 7986 | 8014 | 29 | rRF 45S | NO | 0.00 | 0.00 | 33.30 | 23.69 |
| >hg38_21_+_8433172_8446622_RNA45SN1_45S_50ntFlanks@7986.8016.31\|ATCAGACGTGGCGACCCGCTGAATTTAAGCA | >hg38_21_+_8433172_8446622_RNA45SN1_45S_50ntFlanks | 7986 | 8016 | 31 | rRF 45S | NO | 0.00 | 0.00 | 41.57 | 50.33 |
| >hg38_21_+_8433172_8446622_RNA45SN1_45S_50ntFlanks@7986.8017.32\|ATCAGACGTGGCGACCCGCTGAATTTAAGCAT | >hg38_21_+_8433172_8446622_RNA45SN1_45S_50ntFlanks | 7986 | 8017 | 32 | rRF 45S | NO | 0.00 | 0.00 | 39.93 | 58.05 |
| >hg38_21_+_8433172_8446622_RNA45SN1_45S_50ntFlanks@7987.8006.20\|TCAGACGTGGCGACCCGCTG | >hg38_21_+_8433172_8446622_RNA45SN1_45S_50ntFlanks | 7987 | 8006 | 20 | rRF 45S | NO | 0.00 | 0.00 | 43.55 | 6.66 |
| >hg38_21_+_8433172_8446622_RNA45SN1_45S_50ntFlanks@7987.8008.22\|TCAGACGTGGCGACCCGCTGAA | >hg38_21_+_8433172_8446622_RNA45SN1_45S_50ntFlanks | 7987 | 8008 | 22 | rRF 45S | NO | 0.00 | 0.00 | 27.75 | 5.96 |
| >hg38_21_+_8433172_8446622_RNA45SN1_45S_50ntFlanks@7988.8006.19\|CAGACGTGGCGACCCGCTG | >hg38_21_+_8433172_8446622_RNA45SN1_45S_50ntFlanks | 7988 | 8006 | 19 | rRF 45S | NO | 0.00 | 0.00 | 77.57 | 16.55 |
| >hg38_21_+_8433172_8446622_RNA45SN1_45S_50ntFlanks@7988.8007.20\|CAGACGTGGCGACCCGCTGA | >hg38_21_+_8433172_8446622_RNA45SN1_45S_50ntFlanks | 7988 | 8007 | 20 | rRF 45S | NO | 0.00 | 0.00 | 101.63 | 100.92 |
| >hg38_21_+_8433172_8446622_RNA45SN1_45S_50ntFlanks@7988.8008.21\|CAGACGTGGCGACCCGCTGAA | >hg38_21_+_8433172_8446622_RNA45SN1_45S_50ntFlanks | 7988 | 8008 | 21 | rRF 45S | NO | 74.61 | 49.25 | 441.37 | 380.10 |
| >hg38_21_+_8433172_8446622_RNA45SN1_45S_50ntFlanks@7988.8009.22\|CAGACGTGGCGACCCGCTGAAT | >hg38_21_+_8433172_8446622_RNA45SN1_45S_50ntFlanks | 7988 | 8009 | 22 | rRF 45S | NO | 0.00 | 0.00 | 141.65 | 97.36 |
| >hg38_21_+_8433172_8446622_RNA45SN1_45S_50ntFlanks@7988.8010.23\|CAGACGTGGCGACCCGCTGAATT | >hg38_21_+_8433172_8446622_RNA45SN1_45S_50ntFlanks | 7988 | 8010 | 23 | rRF 45S | NO | 0.00 | 0.00 | 45.41 | 43.10 |
| >hg38_21_+_8433172_8446622_RNA45SN1_45S_50ntFlanks@7988.8011.24\|CAGACGTGGCGACCCGCTGAATTT | >hg38_21_+_8433172_8446622_RNA45SN1_45S_50ntFlanks | 7988 | 8011 | 24 | rRF 45S | NO | 0.00 | 0.00 | 74.24 | 81.44 |
| >hg38_21_+_8433172_8446622_RNA45SN1_45S_50ntFlanks@7988.8012.25\|CAGACGTGGCGACCCGCTGAATTTA | >hg38_21_+_8433172_8446622_RNA45SN1_45S_50ntFlanks | 7988 | 8012 | 25 | rRF 45S | NO | 57.17 | 29.08 | 210.21 | 293.15 |
| >hg38_21_+_8433172_8446622_RNA45SN1_45S_50ntFlanks@7988.8014.27\|CAGACGTGGCGACCCGCTGAATTTAAG | >hg38_21_+_8433172_8446622_RNA45SN1_45S_50ntFlanks | 7988 | 8014 | 27 | rRF 45S | NO | 0.00 | 0.00 | 67.78 | 67.79 |
| >hg38_21_+_8433172_8446622_RNA45SN1_45S_50ntFlanks@7988.8016.29\|CAGACGTGGCGACCCGCTGAATTTAAGCA | >hg38_21_+_8433172_8446622_RNA45SN1_45S_50ntFlanks | 7988 | 8016 | 29 | rRF 45S | NO | 0.00 | 0.00 | 46.18 | 47.96 |
| >hg38_21_+_8433172_8446622_RNA45SN1_45S_50ntFlanks@7988.8017.30\|CAGACGTGGCGACCCGCTGAATTTAAGCAT | >hg38_21_+_8433172_8446622_RNA45SN1_45S_50ntFlanks | 7988 | 8017 | 30 | rRF 45S | NO | 0.00 | 0.00 | 40.00 | 52.45 |
| >hg38_21_+_8433172_8446622_RNA45SN1_45S_50ntFlanks@7988.8018.31\|CAGACGTGGCGACCCGCTGAATTTAAGCATA | >hg38_21_+_8433172_8446622_RNA45SN1_45S_50ntFlanks | 7988 | 8018 | 31 | rRF 45S | NO | 0.00 | 0.00 | 13.70 | 36.92 |
| >hg38_21_+_8433172_8446622_RNA45SN1_45S_50ntFlanks@7989.8006.18\|AGACGTGGCGACCCGCTG | >hg38_21_+_8433172_8446622_RNA45SN1_45S_50ntFlanks | 7989 | 8006 | 18 | rRF 45S | NO | 40.39 | 31.93 | 697.92 | 226.71 |
| >hg38_21_+_8433172_8446622_RNA45SN1_45S_50ntFlanks@7989.8007.19\|AGACGTGGCGACCCGCTGA | >hg38_21_+_8433172_8446622_RNA45SN1_45S_50ntFlanks | 7989 | 8007 | 19 | rRF 45S | NO | 0.00 | 0.00 | 269.51 | 105.48 |
| >hg38_21_+_8433172_8446622_RNA45SN1_45S_50ntFlanks@7989.8008.20\|AGACGTGGCGACCCGCTGAA | >hg38_21_+_8433172_8446622_RNA45SN1_45S_50ntFlanks | 7989 | 8008 | 20 | rRF 45S | NO | 72.44 | 30.79 | 916.71 | 283.99 |
| >hg38_21_+_8433172_8446622_RNA45SN1_45S_50ntFlanks@7989.8009.21\|AGACGTGGCGACCCGCTGAAT | >hg38_21_+_8433172_8446622_RNA45SN1_45S_50ntFlanks | 7989 | 8009 | 21 | rRF 45S | NO | 0.00 | 0.00 | 205.77 | 51.62 |
| >hg38_21_+_8433172_8446622_RNA45SN1_45S_50ntFlanks@7989.8010.22\|AGACGTGGCGACCCGCTGAATT | >hg38_21_+_8433172_8446622_RNA45SN1_45S_50ntFlanks | 7989 | 8010 | 22 | rRF 45S | NO | 0.00 | 0.00 | 47.83 | 24.77 |
| >hg38_21_+_8433172_8446622_RNA45SN1_45S_50ntFlanks@7989.8011.23\|AGACGTGGCGACCCGCTGAATTT | >hg38_21_+_8433172_8446622_RNA45SN1_45S_50ntFlanks | 7989 | 8011 | 23 | rRF 45S | NO | 0.00 | 0.00 | 78.09 | 61.75 |
| >hg38_21_+_8433172_8446622_RNA45SN1_45S_50ntFlanks@7989.8012.24\|AGACGTGGCGACCCGCTGAATTTA | >hg38_21_+_8433172_8446622_RNA45SN1_45S_50ntFlanks | 7989 | 8012 | 24 | rRF 45S | NO | 56.70 | 0.00 | 218.73 | 209.13 |
| >hg38_21_+_8433172_8446622_RNA45SN1_45S_50ntFlanks@7989.8014.26\|AGACGTGGCGACCCGCTGAATTTAAG | >hg38_21_+_8433172_8446622_RNA45SN1_45S_50ntFlanks | 7989 | 8014 | 26 | rRF 45S | NO | 0.00 | 0.00 | 59.63 | 42.17 |
| >hg38_21_+_8433172_8446622_RNA45SN1_45S_50ntFlanks@7989.8016.28\|AGACGTGGCGACCCGCTGAATTTAAGCA | >hg38_21_+_8433172_8446622_RNA45SN1_45S_50ntFlanks | 7989 | 8016 | 28 | rRF 45S | NO | 0.00 | 0.00 | 20.33 | 0.00 |
| >hg38_21_+_8433172_8446622_RNA45SN1_45S_50ntFlanks@7990.8007.18\|GACGTGGCGACCCGCTGA | >hg38_21_+_8433172_8446622_RNA45SN1_45S_50ntFlanks | 7990 | 8007 | 18 | rRF 45S | NO | 0.00 | 0.00 | 38.21 | 0.00 |
| >hg38_21_+_8433172_8446622_RNA45SN1_45S_50ntFlanks@7990.8008.19\|GACGTGGCGACCCGCTGAA | >hg38_21_+_8433172_8446622_RNA45SN1_45S_50ntFlanks | 7990 | 8008 | 19 | rRF 45S | NO | 0.00 | 0.00 | 207.89 | 93.64 |
| >hg38_21_+_8433172_8446622_RNA45SN1_45S_50ntFlanks@7990.8009.20\|GACGTGGCGACCCGCTGAAT | >hg38_21_+_8433172_8446622_RNA45SN1_45S_50ntFlanks | 7990 | 8009 | 20 | rRF 45S | NO | 0.00 | 0.00 | 58.41 | 6.21 |
| >hg38_21_+_8433172_8446622_RNA45SN1_45S_50ntFlanks@7990.8012.23\|GACGTGGCGACCCGCTGAATTTA | >hg38_21_+_8433172_8446622_RNA45SN1_45S_50ntFlanks | 7990 | 8012 | 23 | rRF 45S | NO | 0.00 | 0.00 | 47.60 | 58.22 |
| >hg38_21_+_8433172_8446622_RNA45SN1_45S_50ntFlanks@7991.8008.18\|ACGTGGCGACCCGCTGAA | >hg38_21_+_8433172_8446622_RNA45SN1_45S_50ntFlanks | 7991 | 8008 | 18 | rRF 45S | YES | 123.26 | 104.08 | 1100.04 | 888.03 |
| >hg38_21_+_8433172_8446622_RNA45SN1_45S_50ntFlanks@7991.8009.19\|ACGTGGCGACCCGCTGAAT | >hg38_21_+_8433172_8446622_RNA45SN1_45S_50ntFlanks | 7991 | 8009 | 19 | rRF 45S | NO | 38.21 | 12.45 | 392.84 | 212.76 |
| >hg38_21_+_8433172_8446622_RNA45SN1_45S_50ntFlanks@7991.8010.20\|ACGTGGCGACCCGCTGAATT | >hg38_21_+_8433172_8446622_RNA45SN1_45S_50ntFlanks | 7991 | 8010 | 20 | rRF 45S | NO | 0.00 | 0.00 | 57.10 | 97.10 |

### MDA-MB-468

| | | | | | | | | | | |
|---|---|---|---|---|---|---|---|---|---|---|
| >hg38_21_+_8433172_8446622_RNA45SN1_45S_50ntFlanks@7991.8011.21\|ACGTGGCGACCCGCTGAATTT | >hg38_21_+_8433172_8446622_RNA45SN1_45S_50ntFlanks | 7991 | 8011 | 21 | rRF 45S | NO | 30.47 | 0.00 | 90.86 | 217.71 |
| >hg38_21_+_8433172_8446622_RNA45SN1_45S_50ntFlanks@7991.8012.22\|ACGTGGCGACCCGCTGAATTTA | >hg38_21_+_8433172_8446622_RNA45SN1_45S_50ntFlanks | 7991 | 8012 | 22 | rRF 45S | NO | 83.42 | 43.87 | 181.72 | 578.24 |
| >hg38_21_+_8433172_8446622_RNA45SN1_45S_50ntFlanks@7991.8013.23\|ACGTGGCGACCCGCTGAATTTAA | >hg38_21_+_8433172_8446622_RNA45SN1_45S_50ntFlanks | 7991 | 8013 | 23 | rRF 45S | NO | 0.00 | 0.00 | 27.41 | 50.81 |
| >hg38_21_+_8433172_8446622_RNA45SN1_45S_50ntFlanks@7991.8014.24\|ACGTGGCGACCCGCTGAATTTAAG | >hg38_21_+_8433172_8446622_RNA45SN1_45S_50ntFlanks | 7991 | 8014 | 24 | rRF 45S | NO | 20.30 | 0.00 | 85.90 | 114.82 |
| >hg38_21_+_8433172_8446622_RNA45SN1_45S_50ntFlanks@7991.8016.26\|ACGTGGCGACCCGCTGAATTTAAGCA | >hg38_21_+_8433172_8446622_RNA45SN1_45S_50ntFlanks | 7991 | 8016 | 26 | rRF 45S | NO | 0.00 | 0.00 | 32.23 | 67.02 |
| >hg38_21_+_8433172_8446622_RNA45SN1_45S_50ntFlanks@7992.8009.18\|CGTGGCGACCCGCTGAAT | >hg38_21_+_8433172_8446622_RNA45SN1_45S_50ntFlanks | 7992 | 8009 | 18 | rRF 45S | NO | 0.00 | 0.00 | 57.71 | 55.00 |
| >hg38_21_+_8433172_8446622_RNA45SN1_45S_50ntFlanks@7992.8012.21\|CGTGGCGACCCGCTGAATTTA | >hg38_21_+_8433172_8446622_RNA45SN1_45S_50ntFlanks | 7992 | 8012 | 21 | rRF 45S | NO | 0.00 | 0.00 | 21.26 | 102.66 |
| >hg38_21_+_8433172_8446622_RNA45SN1_45S_50ntFlanks@8079.8103.25\|AAGAGCCCAGCGCCGAATCCCCGCC | >hg38_21_+_8433172_8446622_RNA45SN1_45S_50ntFlanks | 8079 | 8103 | 25 | rRF 45S | NO | 0.00 | 0.00 | 76.63 | 46.10 |
| >hg38_21_+_8433172_8446622_RNA45SN1_45S_50ntFlanks@8079.8104.26\|AAGAGCCCAGCGCCGAATCCCCGCCC | >hg38_21_+_8433172_8446622_RNA45SN1_45S_50ntFlanks | 8079 | 8104 | 26 | rRF 45S | NO | 0.00 | 0.00 | 119.03 | 59.06 |
| >hg38_21_+_8433172_8446622_RNA45SN1_45S_50ntFlanks@8104.8136.33\|CCGCGGCGGGGCGCGGGACATGTGGCGTACGGA | >hg38_21_+_8433172_8446622_RNA45SN1_45S_50ntFlanks | 8104 | 8136 | 33 | rRF 45S | YES | 0.00 | 0.00 | 44.85 | 0.00 |
| >hg38_21_+_8433172_8446622_RNA45SN1_45S_50ntFlanks@8104.8138.35\|CCGCGGCGGGGCGCGGGACATGTGGCGTACGGAAG | >hg38_21_+_8433172_8446622_RNA45SN1_45S_50ntFlanks | 8104 | 8138 | 35 | rRF 45S | NO | 13.02 | 0.00 | 129.90 | 47.57 |
| >hg38_21_+_8433172_8446622_RNA45SN1_45S_50ntFlanks@8104.8139.36\|CCGCGGCGGGGCGCGGGACATGTGGCGTACGGAAGA | >hg38_21_+_8433172_8446622_RNA45SN1_45S_50ntFlanks | 8104 | 8139 | 36 | rRF 45S | NO | 0.00 | 0.00 | 46.45 | 0.00 |
| >hg38_21_+_8433172_8446622_RNA45SN1_45S_50ntFlanks@8104.8140.37\|CCGCGGCGGGGCGCGGGACATGTGGCGTACGGAAGAC | >hg38_21_+_8433172_8446622_RNA45SN1_45S_50ntFlanks | 8104 | 8140 | 37 | rRF 45S | NO | 50.92 | 0.00 | 238.64 | 93.11 |
| >hg38_21_+_8433172_8446622_RNA45SN1_45S_50ntFlanks@8104.8141.38\|CCGCGGCGGGGCGCGGGACATGTGGCGTACGGAAGACC | >hg38_21_+_8433172_8446622_RNA45SN1_45S_50ntFlanks | 8104 | 8141 | 38 | rRF 45S | NO | 20.68 | 0.00 | 69.97 | 50.24 |
| >hg38_21_+_8433172_8446622_RNA45SN1_45S_50ntFlanks@8104.8143.40\|CCGCGGCGGGGCGCGGGACATGTGGCGTACGGAAGACCCG | >hg38_21_+_8433172_8446622_RNA45SN1_45S_50ntFlanks | 8104 | 8143 | 40 | rRF 45S | NO | 362.05 | 94.81 | 349.48 | 641.55 |
| >hg38_21_+_8433172_8446622_RNA45SN1_45S_50ntFlanks@8105.8136.32\|CGCGGCGGGGCGCGGGACATGTGGCGTACGGA | >hg38_21_+_8433172_8446622_RNA45SN1_45S_50ntFlanks | 8105 | 8136 | 32 | rRF 45S | YES | 0.00 | 0.00 | 92.23 | 6.04 |
| >hg38_21_+_8433172_8446622_RNA45SN1_45S_50ntFlanks@8105.8138.34\|CGCGGCGGGGCGCGGGACATGTGGCGTACGGAAG | >hg38_21_+_8433172_8446622_RNA45SN1_45S_50ntFlanks | 8105 | 8138 | 34 | rRF 45S | NO | 41.61 | 0.00 | 335.24 | 100.66 |
| >hg38_21_+_8433172_8446622_RNA45SN1_45S_50ntFlanks@8105.8139.35\|CGCGGCGGGGCGCGGGACATGTGGCGTACGGAAGA | >hg38_21_+_8433172_8446622_RNA45SN1_45S_50ntFlanks | 8105 | 8139 | 35 | rRF 45S | NO | 0.00 | 0.00 | 111.44 | 37.11 |
| >hg38_21_+_8433172_8446622_RNA45SN1_45S_50ntFlanks@8105.8140.36\|CGCGGCGGGGCGCGGGACATGTGGCGTACGGAAGAC | >hg38_21_+_8433172_8446622_RNA45SN1_45S_50ntFlanks | 8105 | 8140 | 36 | rRF 45S | NO | 55.89 | 0.00 | 379.58 | 131.30 |
| >hg38_21_+_8433172_8446622_RNA45SN1_45S_50ntFlanks@8105.8141.37\|CGCGGCGGGGCGCGGGACATGTGGCGTACGGAAGACC | >hg38_21_+_8433172_8446622_RNA45SN1_45S_50ntFlanks | 8105 | 8141 | 37 | rRF 45S | NO | 0.00 | 0.00 | 151.77 | 54.00 |
| >hg38_21_+_8433172_8446622_RNA45SN1_45S_50ntFlanks@8105.8142.38\|CGCGGCGGGGCGCGGGACATGTGGCGTACGGAAGACCC | >hg38_21_+_8433172_8446622_RNA45SN1_45S_50ntFlanks | 8105 | 8142 | 38 | rRF 45S | NO | 0.00 | 0.00 | 64.88 | 41.79 |
| >hg38_21_+_8433172_8446622_RNA45SN1_45S_50ntFlanks@8105.8143.39\|CGCGGCGGGGCGCGGGACATGTGGCGTACGGAAGACCCG | >hg38_21_+_8433172_8446622_RNA45SN1_45S_50ntFlanks | 8105 | 8143 | 39 | rRF 45S | NO | 590.26 | 134.81 | 801.39 | 602.75 |
| >hg38_21_+_8433172_8446622_RNA45SN1_45S_50ntFlanks@8106.8138.33\|GCGGCGGGGCGCGGGACATGTGGCGTACGGAAG | >hg38_21_+_8433172_8446622_RNA45SN1_45S_50ntFlanks | 8106 | 8138 | 33 | rRF 45S | YES | 0.00 | 0.00 | 44.29 | 0.00 |
| >hg38_21_+_8433172_8446622_RNA45SN1_45S_50ntFlanks@8106.8140.35\|GCGGCGGGGCGCGGGACATGTGGCGTACGGAAGAC | >hg38_21_+_8433172_8446622_RNA45SN1_45S_50ntFlanks | 8106 | 8140 | 35 | rRF 45S | NO | 0.00 | 0.00 | 53.25 | 0.00 |
| >hg38_21_+_8433172_8446622_RNA45SN1_45S_50ntFlanks@8106.8143.38\|GCGGCGGGGCGCGGGACATGTGGCGTACGGAAGACCCG | >hg38_21_+_8433172_8446622_RNA45SN1_45S_50ntFlanks | 8106 | 8143 | 38 | rRF 45S | NO | 0.00 | 0.00 | 68.67 | 40.85 |
| >hg38_21_+_8433172_8446622_RNA45SN1_45S_50ntFlanks@8107.8138.32\|CGGCGGGGCGCGGGACATGTGGCGTACGGAAG | >hg38_21_+_8433172_8446622_RNA45SN1_45S_50ntFlanks | 8107 | 8138 | 32 | rRF 45S | NO | 0.00 | 0.00 | 61.75 | 0.00 |
| >hg38_21_+_8433172_8446622_RNA45SN1_45S_50ntFlanks@8107.8140.34\|CGGCGGGGCGCGGGACATGTGGCGTACGGAAGAC | >hg38_21_+_8433172_8446622_RNA45SN1_45S_50ntFlanks | 8107 | 8140 | 34 | rRF 45S | YES | 0.00 | 0.00 | 56.74 | 0.00 |
| >hg38_21_+_8433172_8446622_RNA45SN1_45S_50ntFlanks@8107.8141.35\|CGGCGGGGCGCGGGACATGTGGCGTACGGAAGACC | >hg38_21_+_8433172_8446622_RNA45SN1_45S_50ntFlanks | 8107 | 8141 | 35 | rRF 45S | YES | 0.00 | 0.00 | 32.09 | 0.00 |
| >hg38_21_+_8433172_8446622_RNA45SN1_45S_50ntFlanks@8107.8143.37\|CGGCGGGGCGCGGGACATGTGGCGTACGGAAGACCCG | >hg38_21_+_8433172_8446622_RNA45SN1_45S_50ntFlanks | 8107 | 8143 | 37 | rRF 45S | NO | 0.00 | 0.00 | 82.49 | 34.60 |
| >hg38_21_+_8433172_8446622_RNA45SN1_45S_50ntFlanks@8110.8138.29\|CGGGGCGCGGGACATGTGGCGTACGGAAG | >hg38_21_+_8433172_8446622_RNA45SN1_45S_50ntFlanks | 8110 | 8138 | 29 | rRF 45S | YES | 0.00 | 0.00 | 35.08 | 0.00 |
| >hg38_21_+_8433172_8446622_RNA45SN1_45S_50ntFlanks@8110.8140.31\|CGGGGCGCGGGACATGTGGCGTACGGAAGAC | >hg38_21_+_8433172_8446622_RNA45SN1_45S_50ntFlanks | 8110 | 8140 | 31 | rRF 45S | YES | 0.00 | 0.00 | 32.61 | 0.00 |
| >hg38_21_+_8433172_8446622_RNA45SN1_45S_50ntFlanks@8113.8138.26\|GGCGCGGGACATGTGGCGTACGGAAG | >hg38_21_+_8433172_8446622_RNA45SN1_45S_50ntFlanks | 8113 | 8138 | 26 | rRF 45S | NO | 0.00 | 0.00 | 312.13 | 23.34 |
| >hg38_21_+_8433172_8446622_RNA45SN1_45S_50ntFlanks@8113.8139.27\|GGCGCGGGACATGTGGCGTACGGAAGA | >hg38_21_+_8433172_8446622_RNA45SN1_45S_50ntFlanks | 8113 | 8139 | 27 | rRF 45S | YES | 0.00 | 0.00 | 76.82 | 0.00 |
| >hg38_21_+_8433172_8446622_RNA45SN1_45S_50ntFlanks@8113.8140.28\|GGCGCGGGACATGTGGCGTACGGAAGAC | >hg38_21_+_8433172_8446622_RNA45SN1_45S_50ntFlanks | 8113 | 8140 | 28 | rRF 45S | NO | 0.00 | 0.00 | 216.70 | 14.96 |

### MDA-MB-468

| | | | | | | | | | | |
|---|---|---|---|---|---|---|---|---|---|---|
| >hg38_21_+_8433172_8446622_RNA45SN1_45S_50ntFlanks@8113.8141.29\|GGCGCGGGACATGTGGCGTACGGAAGACC | >hg38_21_+_8433172_8446622_RNA45SN1_45S_50ntFlanks | 8113 | 8141 | 29 | rRF 45S | NO | 0.00 | 0.00 | 88.13 | 0.00 |
| >hg38_21_+_8433172_8446622_RNA45SN1_45S_50ntFlanks@8113.8143.31\|GGCGCGGGACATGTGGCGTACGGAAGACCCG | >hg38_21_+_8433172_8446622_RNA45SN1_45S_50ntFlanks | 8113 | 8143 | 31 | rRF 45S | NO | 0.00 | 0.00 | 194.60 | 25.85 |
| >hg38_21_+_8433172_8446622_RNA45SN1_45S_50ntFlanks@8114.8136.23\|GCGCGGGACATGTGGCGTACGGA | >hg38_21_+_8433172_8446622_RNA45SN1_45S_50ntFlanks | 8114 | 8136 | 23 | rRF 45S | YES | 0.00 | 0.00 | 135.29 | 0.00 |
| >hg38_21_+_8433172_8446622_RNA45SN1_45S_50ntFlanks@8114.8138.25\|GCGCGGGACATGTGGCGTACGGAAG | >hg38_21_+_8433172_8446622_RNA45SN1_45S_50ntFlanks | 8114 | 8138 | 25 | rRF 45S | NO | 34.19 | 0.00 | 651.01 | 85.73 |
| >hg38_21_+_8433172_8446622_RNA45SN1_45S_50ntFlanks@8114.8139.26\|GCGCGGGACATGTGGCGTACGGAAGA | >hg38_21_+_8433172_8446622_RNA45SN1_45S_50ntFlanks | 8114 | 8139 | 26 | rRF 45S | NO | 0.00 | 0.00 | 162.59 | 14.29 |
| >hg38_21_+_8433172_8446622_RNA45SN1_45S_50ntFlanks@8114.8140.27\|GCGCGGGACATGTGGCGTACGGAAGAC | >hg38_21_+_8433172_8446622_RNA45SN1_45S_50ntFlanks | 8114 | 8140 | 27 | rRF 45S | NO | 31.27 | 0.00 | 471.31 | 90.50 |
| >hg38_21_+_8433172_8446622_RNA45SN1_45S_50ntFlanks@8114.8141.28\|GCGCGGGACATGTGGCGTACGGAAGACC | >hg38_21_+_8433172_8446622_RNA45SN1_45S_50ntFlanks | 8114 | 8141 | 28 | rRF 45S | NO | 0.00 | 0.00 | 246.84 | 54.59 |
| >hg38_21_+_8433172_8446622_RNA45SN1_45S_50ntFlanks@8114.8142.29\|GCGCGGGACATGTGGCGTACGGAAGACCC | >hg38_21_+_8433172_8446622_RNA45SN1_45S_50ntFlanks | 8114 | 8142 | 29 | rRF 45S | NO | 0.00 | 0.00 | 60.23 | 13.92 |
| >hg38_21_+_8433172_8446622_RNA45SN1_45S_50ntFlanks@8114.8143.30\|GCGCGGGACATGTGGCGTACGGAAGACCCG | >hg38_21_+_8433172_8446622_RNA45SN1_45S_50ntFlanks | 8114 | 8143 | 30 | rRF 45S | NO | 36.62 | 0.00 | 653.39 | 175.30 |
| >hg38_21_+_8433172_8446622_RNA45SN1_45S_50ntFlanks@8115.8138.24\|CGCGGGACATGTGGCGTACGGAAG | >hg38_21_+_8433172_8446622_RNA45SN1_45S_50ntFlanks | 8115 | 8138 | 24 | rRF 45S | NO | 0.00 | 0.00 | 303.23 | 50.10 |
| >hg38_21_+_8433172_8446622_RNA45SN1_45S_50ntFlanks@8115.8139.25\|CGCGGGACATGTGGCGTACGGAAGA | >hg38_21_+_8433172_8446622_RNA45SN1_45S_50ntFlanks | 8115 | 8139 | 25 | rRF 45S | YES | 0.00 | 0.00 | 74.82 | 0.00 |
| >hg38_21_+_8433172_8446622_RNA45SN1_45S_50ntFlanks@8115.8140.26\|CGCGGGACATGTGGCGTACGGAAGAC | >hg38_21_+_8433172_8446622_RNA45SN1_45S_50ntFlanks | 8115 | 8140 | 26 | rRF 45S | NO | 0.00 | 0.00 | 238.01 | 50.62 |
| >hg38_21_+_8433172_8446622_RNA45SN1_45S_50ntFlanks@8115.8141.27\|CGCGGGACATGTGGCGTACGGAAGACC | >hg38_21_+_8433172_8446622_RNA45SN1_45S_50ntFlanks | 8115 | 8141 | 27 | rRF 45S | NO | 0.00 | 0.00 | 157.72 | 40.39 |
| >hg38_21_+_8433172_8446622_RNA45SN1_45S_50ntFlanks@8115.8142.28\|CGCGGGACATGTGGCGTACGGAAGACCC | >hg38_21_+_8433172_8446622_RNA45SN1_45S_50ntFlanks | 8115 | 8142 | 28 | rRF 45S | NO | 0.00 | 0.00 | 66.79 | 23.06 |
| >hg38_21_+_8433172_8446622_RNA45SN1_45S_50ntFlanks@8115.8143.29\|CGCGGGACATGTGGCGTACGGAAGACCCG | >hg38_21_+_8433172_8446622_RNA45SN1_45S_50ntFlanks | 8115 | 8143 | 29 | rRF 45S | NO | 45.05 | 0.00 | 573.67 | 311.15 |
| >hg38_21_+_8433172_8446622_RNA45SN1_45S_50ntFlanks@8153.8181.29\|GCCGCTCGTGGGGGGCCCAAGTCCTTCTG | >hg38_21_+_8433172_8446622_RNA45SN1_45S_50ntFlanks | 8153 | 8181 | 29 | rRF 45S | NO | 21.10 | 0.00 | 0.00 | 68.31 |
| >hg38_21_+_8433172_8446622_RNA45SN1_45S_50ntFlanks@8154.8181.28\|CCGCTCGTGGGGGGCCCAAGTCCTTCTG | >hg38_21_+_8433172_8446622_RNA45SN1_45S_50ntFlanks | 8154 | 8181 | 28 | rRF 45S | NO | 120.78 | 12.45 | 40.06 | 332.15 |
| >hg38_21_+_8433172_8446622_RNA45SN1_45S_50ntFlanks@8155.8181.27\|CGCTCGTGGGGGGCCCAAGTCCTTCTG | >hg38_21_+_8433172_8446622_RNA45SN1_45S_50ntFlanks | 8155 | 8181 | 27 | rRF 45S | NO | 36.83 | 0.00 | 6.52 | 173.52 |
| >hg38_21_+_8433172_8446622_RNA45SN1_45S_50ntFlanks@8157.8178.22\|CTCGTGGGGGGCCCAAGTCCTT | >hg38_21_+_8433172_8446622_RNA45SN1_45S_50ntFlanks | 8157 | 8178 | 22 | rRF 45S | NO | 0.00 | 0.00 | 38.84 | 53.27 |
| >hg38_21_+_8433172_8446622_RNA45SN1_45S_50ntFlanks@8157.8181.25\|CTCGTGGGGGGCCCAAGTCCTTCTG | >hg38_21_+_8433172_8446622_RNA45SN1_45S_50ntFlanks | 8157 | 8181 | 25 | rRF 45S | NO | 0.00 | 0.00 | 74.96 | 102.65 |
| >hg38_21_+_8433172_8446622_RNA45SN1_45S_50ntFlanks@8168.8185.18\|CCCAAGTCCTTCTGATCG | >hg38_21_+_8433172_8446622_RNA45SN1_45S_50ntFlanks | 8168 | 8185 | 18 | rRF 45S | NO | 40.42 | 35.44 | 20.24 | 0.00 |
| >hg38_21_+_8433172_8446622_RNA45SN1_45S_50ntFlanks@8168.8186.19\|CCCAAGTCCTTCTGATCGA | >hg38_21_+_8433172_8446622_RNA45SN1_45S_50ntFlanks | 8168 | 8186 | 19 | rRF 45S | NO | 94.33 | 71.40 | 6.70 | 25.61 |
| >hg38_21_+_8433172_8446622_RNA45SN1_45S_50ntFlanks@8168.8187.20\|CCCAAGTCCTTCTGATCGAG | >hg38_21_+_8433172_8446622_RNA45SN1_45S_50ntFlanks | 8168 | 8187 | 20 | rRF 45S | NO | 70.92 | 29.02 | 0.00 | 0.00 |
| >hg38_21_+_8433172_8446622_RNA45SN1_45S_50ntFlanks@8168.8188.21\|CCCAAGTCCTTCTGATCGAGG | >hg38_21_+_8433172_8446622_RNA45SN1_45S_50ntFlanks | 8168 | 8188 | 21 | rRF 45S | NO | 0.00 | 26.99 | 6.87 | 0.00 |
| >hg38_21_+_8433172_8446622_RNA45SN1_45S_50ntFlanks@8168.8197.30\|CCCAAGTCCTTCTGATCGAGGCCCAGCCCG | >hg38_21_+_8433172_8446622_RNA45SN1_45S_50ntFlanks | 8168 | 8197 | 30 | rRF 45S | NO | 70.20 | 0.00 | 0.00 | 26.81 |
| >hg38_21_+_8433172_8446622_RNA45SN1_45S_50ntFlanks@8168.8203.36\|CCCAAGTCCTTCTGATCGAGGCCCAGCCCGTGGACG | >hg38_21_+_8433172_8446622_RNA45SN1_45S_50ntFlanks | 8168 | 8203 | 36 | rRF 45S | NO | 0.00 | 0.00 | 20.46 | 0.00 |
| >hg38_21_+_8433172_8446622_RNA45SN1_45S_50ntFlanks@8168.8212.45\|CCCAAGTCCTTCTGATCGAGGCCCAGCCCGTGGACGGTGTGAGGC | >hg38_21_+_8433172_8446622_RNA45SN1_45S_50ntFlanks | 8168 | 8212 | 45 | rRF 45S | NO | 65.02 | 0.00 | 0.00 | 0.00 |
| >hg38_21_+_8433172_8446622_RNA45SN1_45S_50ntFlanks@8169.8186.18\|CCAAGTCCTTCTGATCGA | >hg38_21_+_8433172_8446622_RNA45SN1_45S_50ntFlanks | 8169 | 8186 | 18 | rRF 45S | NO | 6.00 | 27.17 | 0.00 | 0.00 |
| >hg38_21_+_8433172_8446622_RNA45SN1_45S_50ntFlanks@8169.8197.29\|CCAAGTCCTTCTGATCGAGGCCCAGCCCG | >hg38_21_+_8433172_8446622_RNA45SN1_45S_50ntFlanks | 8169 | 8197 | 29 | rRF 45S | NO | 33.52 | 0.00 | 0.00 | 14.52 |
| >hg38_21_+_8433172_8446622_RNA45SN1_45S_50ntFlanks@8169.8212.44\|CCAAGTCCTTCTGATCGAGGCCCAGCCCGTGGACGGTGTGAGGC | >hg38_21_+_8433172_8446622_RNA45SN1_45S_50ntFlanks | 8169 | 8212 | 44 | rRF 45S | NO | 47.12 | 0.00 | 0.00 | 13.39 |
| >hg38_21_+_8433172_8446622_RNA45SN1_45S_50ntFlanks@8169.8214.46\|CCAAGTCCTTCTGATCGAGGCCCAGCCCGTGGACGGTGTGAGGCCG | >hg38_21_+_8433172_8446622_RNA45SN1_45S_50ntFlanks | 8169 | 8214 | 46 | rRF 45S | NO | 30.25 | 0.00 | 0.00 | 0.00 |
| >hg38_21_+_8433172_8446622_RNA45SN1_45S_50ntFlanks@8170.8187.18\|CAAGTCCTTCTGATCGAG | >hg38_21_+_8433172_8446622_RNA45SN1_45S_50ntFlanks | 8170 | 8187 | 18 | rRF 45S | NO | 43.99 | 52.76 | 6.55 | 6.49 |
| >hg38_21_+_8433172_8446622_RNA45SN1_45S_50ntFlanks@8170.8188.19\|CAAGTCCTTCTGATCGAGG | >hg38_21_+_8433172_8446622_RNA45SN1_45S_50ntFlanks | 8170 | 8188 | 19 | rRF 45S | NO | 0.00 | 40.99 | 35.97 | 0.00 |
| >hg38_21_+_8433172_8446622_RNA45SN1_45S_50ntFlanks@8170.8195.26\|CAAGTCCTTCTGATCGAGGCCCAGCC | >hg38_21_+_8433172_8446622_RNA45SN1_45S_50ntFlanks | 8170 | 8195 | 26 | rRF 45S | NO | 43.44 | 94.21 | 74.30 | 51.81 |

### MDA-MB-468

| | | | | | | | | | | |
|---|---|---|---|---|---|---|---|---|---|---|
| >hg38_21_+_8433172_8446622_RNA45SN1_45S_50ntFlanks@8170.8197.28\|CAAGTCCTTCTGATCGAGGCCCAGCCCG | >hg38_21_+_8433172_8446622_RNA45SN1_45S_50ntFlanks | 8170 | 8197 | 28 | rRF 45S | NO | 204.70 | 84.17 | 0.00 | 175.04 |
| >hg38_21_+_8433172_8446622_RNA45SN1_45S_50ntFlanks@8170.8201.32\|CAAGTCCTTCTGATCGAGGCCCAGCCCGTGGA | >hg38_21_+_8433172_8446622_RNA45SN1_45S_50ntFlanks | 8170 | 8201 | 32 | rRF 45S | NO | 21.77 | 0.00 | 0.00 | 0.00 |
| >hg38_21_+_8433172_8446622_RNA45SN1_45S_50ntFlanks@8170.8212.43\|CAAGTCCTTCTGATCGAGGCCCAGCCCGTGGACGGTGTGAGGC | >hg38_21_+_8433172_8446622_RNA45SN1_45S_50ntFlanks | 8170 | 8212 | 43 | rRF 45S | NO | 376.30 | 76.00 | 31.56 | 222.10 |
| >hg38_21_+_8433172_8446622_RNA45SN1_45S_50ntFlanks@8170.8213.44\|CAAGTCCTTCTGATCGAGGCCCAGCCCGTGGACGGTGTGAGGCC | >hg38_21_+_8433172_8446622_RNA45SN1_45S_50ntFlanks | 8170 | 8213 | 44 | rRF 45S | NO | 52.28 | 32.89 | 0.00 | 48.18 |
| >hg38_21_+_8433172_8446622_RNA45SN1_45S_50ntFlanks@8170.8214.45\|CAAGTCCTTCTGATCGAGGCCCAGCCCGTGGACGGTGTGAGGCCG | >hg38_21_+_8433172_8446622_RNA45SN1_45S_50ntFlanks | 8170 | 8214 | 45 | rRF 45S | NO | 176.44 | 129.60 | 0.00 | 144.97 |
| >hg38_21_+_8433172_8446622_RNA45SN1_45S_50ntFlanks@8170.8215.46\|CAAGTCCTTCTGATCGAGGCCCAGCCCGTGGACGGTGTGAGGCCGG | >hg38_21_+_8433172_8446622_RNA45SN1_45S_50ntFlanks | 8170 | 8215 | 46 | rRF 45S | NO | 114.25 | 70.47 | 0.00 | 90.55 |
| >hg38_21_+_8433172_8446622_RNA45SN1_45S_50ntFlanks@8171.8197.27\|AAGTCCTTCTGATCGAGGCCCAGCCCG | >hg38_21_+_8433172_8446622_RNA45SN1_45S_50ntFlanks | 8171 | 8197 | 27 | rRF 45S | NO | 55.20 | 5.94 | 0.00 | 34.57 |
| >hg38_21_+_8433172_8446622_RNA45SN1_45S_50ntFlanks@8171.8212.42\|AAGTCCTTCTGATCGAGGCCCAGCCCGTGGACGGTGTGAGGC | >hg38_21_+_8433172_8446622_RNA45SN1_45S_50ntFlanks | 8171 | 8212 | 42 | rRF 45S | NO | 159.39 | 28.33 | 0.00 | 35.71 |
| >hg38_21_+_8433172_8446622_RNA45SN1_45S_50ntFlanks@8171.8213.43\|AAGTCCTTCTGATCGAGGCCCAGCCCGTGGACGGTGTGAGGCC | >hg38_21_+_8433172_8446622_RNA45SN1_45S_50ntFlanks | 8171 | 8213 | 43 | rRF 45S | NO | 29.16 | 0.00 | 0.00 | 0.00 |
| >hg38_21_+_8433172_8446622_RNA45SN1_45S_50ntFlanks@8171.8214.44\|AAGTCCTTCTGATCGAGGCCCAGCCCGTGGACGGTGTGAGGCCG | >hg38_21_+_8433172_8446622_RNA45SN1_45S_50ntFlanks | 8171 | 8214 | 44 | rRF 45S | NO | 72.62 | 68.07 | 0.00 | 24.67 |
| >hg38_21_+_8433172_8446622_RNA45SN1_45S_50ntFlanks@8172.8212.41\|AGTCCTTCTGATCGAGGCCCAGCCCGTGGACGGTGTGAGGC | >hg38_21_+_8433172_8446622_RNA45SN1_45S_50ntFlanks | 8172 | 8212 | 41 | rRF 45S | NO | 70.23 | 0.00 | 0.00 | 0.00 |
| >hg38_21_+_8433172_8446622_RNA45SN1_45S_50ntFlanks@8176.8193.18\|CTTCTGATCGAGGCCCAG | >hg38_21_+_8433172_8446622_RNA45SN1_45S_50ntFlanks | 8176 | 8193 | 18 | rRF 45S | NO | 0.00 | 32.35 | 20.31 | 0.00 |
| >hg38_21_+_8433172_8446622_RNA45SN1_45S_50ntFlanks@8176.8195.20\|CTTCTGATCGAGGCCCAGCC | >hg38_21_+_8433172_8446622_RNA45SN1_45S_50ntFlanks | 8176 | 8195 | 20 | rRF 45S | NO | 164.02 | 105.75 | 34.39 | 47.50 |
| >hg38_21_+_8433172_8446622_RNA45SN1_45S_50ntFlanks@8176.8196.21\|CTTCTGATCGAGGCCCAGCCC | >hg38_21_+_8433172_8446622_RNA45SN1_45S_50ntFlanks | 8176 | 8196 | 21 | rRF 45S | NO | 40.63 | 5.95 | 0.00 | 0.00 |
| >hg38_21_+_8433172_8446622_RNA45SN1_45S_50ntFlanks@8176.8197.22\|CTTCTGATCGAGGCCCAGCCCG | >hg38_21_+_8433172_8446622_RNA45SN1_45S_50ntFlanks | 8176 | 8197 | 22 | rRF 45S | NO | 54.26 | 80.65 | 31.22 | 35.08 |
| >hg38_21_+_8433172_8446622_RNA45SN1_45S_50ntFlanks@8176.8201.26\|CTTCTGATCGAGGCCCAGCCCGTGGA | >hg38_21_+_8433172_8446622_RNA45SN1_45S_50ntFlanks | 8176 | 8201 | 26 | rRF 45S | NO | 33.74 | 44.18 | 35.79 | 0.00 |
| >hg38_21_+_8433172_8446622_RNA45SN1_45S_50ntFlanks@8176.8202.27\|CTTCTGATCGAGGCCCAGCCCGTGGAC | >hg38_21_+_8433172_8446622_RNA45SN1_45S_50ntFlanks | 8176 | 8202 | 27 | rRF 45S | NO | 50.92 | 44.35 | 0.00 | 0.00 |
| >hg38_21_+_8433172_8446622_RNA45SN1_45S_50ntFlanks@8176.8203.28\|CTTCTGATCGAGGCCCAGCCCGTGGACG | >hg38_21_+_8433172_8446622_RNA45SN1_45S_50ntFlanks | 8176 | 8203 | 28 | rRF 45S | NO | 248.39 | 242.89 | 214.03 | 80.80 |
| >hg38_21_+_8433172_8446622_RNA45SN1_45S_50ntFlanks@8176.8204.29\|CTTCTGATCGAGGCCCAGCCCGTGGACGG | >hg38_21_+_8433172_8446622_RNA45SN1_45S_50ntFlanks | 8176 | 8204 | 29 | rRF 45S | NO | 110.57 | 92.17 | 88.22 | 42.57 |
| >hg38_21_+_8433172_8446622_RNA45SN1_45S_50ntFlanks@8176.8205.30\|CTTCTGATCGAGGCCCAGCCCGTGGACGGT | >hg38_21_+_8433172_8446622_RNA45SN1_45S_50ntFlanks | 8176 | 8205 | 30 | rRF 45S | NO | 21.21 | 0.00 | 0.00 | 0.00 |
| >hg38_21_+_8433172_8446622_RNA45SN1_45S_50ntFlanks@8176.8206.31\|CTTCTGATCGAGGCCCAGCCCGTGGACGGTG | >hg38_21_+_8433172_8446622_RNA45SN1_45S_50ntFlanks | 8176 | 8206 | 31 | rRF 45S | NO | 92.43 | 73.52 | 39.32 | 6.64 |
| >hg38_21_+_8433172_8446622_RNA45SN1_45S_50ntFlanks@8176.8208.33\|CTTCTGATCGAGGCCCAGCCCGTGGACGGTGTG | >hg38_21_+_8433172_8446622_RNA45SN1_45S_50ntFlanks | 8176 | 8208 | 33 | rRF 45S | NO | 75.48 | 28.47 | 0.00 | 0.00 |
| >hg38_21_+_8433172_8446622_RNA45SN1_45S_50ntFlanks@8176.8212.37\|CTTCTGATCGAGGCCCAGCCCGTGGACGGTGTGAGGC | >hg38_21_+_8433172_8446622_RNA45SN1_45S_50ntFlanks | 8176 | 8212 | 37 | rRF 45S | NO | 0.00 | 0.00 | 37.47 | 0.00 |
| >hg38_21_+_8433172_8446622_RNA45SN1_45S_50ntFlanks@8176.8213.38\|CTTCTGATCGAGGCCCAGCCCGTGGACGGTGTGAGGCC | >hg38_21_+_8433172_8446622_RNA45SN1_45S_50ntFlanks | 8176 | 8213 | 38 | rRF 45S | NO | 0.00 | 0.00 | 27.74 | 0.00 |
| >hg38_21_+_8433172_8446622_RNA45SN1_45S_50ntFlanks@8176.8214.39\|CTTCTGATCGAGGCCCAGCCCGTGGACGGTGTGAGGCCG | >hg38_21_+_8433172_8446622_RNA45SN1_45S_50ntFlanks | 8176 | 8214 | 39 | rRF 45S | NO | 0.00 | 52.09 | 57.00 | 13.93 |
| >hg38_21_+_8433172_8446622_RNA45SN1_45S_50ntFlanks@8176.8215.40\|CTTCTGATCGAGGCCCAGCCCGTGGACGGTGTGAGGCCGG | >hg38_21_+_8433172_8446622_RNA45SN1_45S_50ntFlanks | 8176 | 8215 | 40 | rRF 45S | NO | 0.00 | 31.29 | 29.17 | 6.18 |
| >hg38_21_+_8433172_8446622_RNA45SN1_45S_50ntFlanks@8176.8220.45\|CTTCTGATCGAGGCCCAGCCCGTGGACGGTGTGAGGCCGGTAGCG | >hg38_21_+_8433172_8446622_RNA45SN1_45S_50ntFlanks | 8176 | 8220 | 45 | rRF 45S | NO | 19.93 | 30.94 | 0.00 | 13.18 |
| >hg38_21_+_8433172_8446622_RNA45SN1_45S_50ntFlanks@8176.8223.48\|CTTCTGATCGAGGCCCAGCCCGTGGACGGTGTGAGGCCGGTAGCGGCC | >hg38_21_+_8433172_8446622_RNA45SN1_45S_50ntFlanks | 8176 | 8223 | 48 | rRF 45S | NO | 54.09 | 0.00 | 0.00 | 0.00 |
| >hg38_21_+_8433172_8446622_RNA45SN1_45S_50ntFlanks@8177.8203.27\|TTCTGATCGAGGCCCAGCCCGTGGACG | >hg38_21_+_8433172_8446622_RNA45SN1_45S_50ntFlanks | 8177 | 8203 | 27 | rRF 45S | NO | 41.73 | 34.35 | 0.00 | 0.00 |
| >hg38_21_+_8433172_8446622_RNA45SN1_45S_50ntFlanks@8177.8204.28\|TTCTGATCGAGGCCCAGCCCGTGGACGG | >hg38_21_+_8433172_8446622_RNA45SN1_45S_50ntFlanks | 8177 | 8204 | 28 | rRF 45S | NO | 32.98 | 0.00 | 0.00 | 0.00 |
| >hg38_21_+_8433172_8446622_RNA45SN1_45S_50ntFlanks@8177.8206.30\|TTCTGATCGAGGCCCAGCCCGTGGACGGTG | >hg38_21_+_8433172_8446622_RNA45SN1_45S_50ntFlanks | 8177 | 8206 | 30 | rRF 45S | NO | 29.12 | 0.00 | 0.00 | 0.00 |
| >hg38_21_+_8433172_8446622_RNA45SN1_45S_50ntFlanks@8178.8203.26\|TCTGATCGAGGCCCAGCCCGTGGACG | >hg38_21_+_8433172_8446622_RNA45SN1_45S_50ntFlanks | 8178 | 8203 | 26 | rRF 45S | NO | 21.90 | 29.05 | 0.00 | 0.00 |
| >hg38_21_+_8433172_8446622_RNA45SN1_45S_50ntFlanks@8179.8208.30\|CTGATCGAGGCCCAGCCCGTGGACGGTGTG | >hg38_21_+_8433172_8446622_RNA45SN1_45S_50ntFlanks | 8179 | 8208 | 30 | rRF 45S | NO | 44.03 | 0.00 | 0.00 | 0.00 |
| >hg38_21_+_8433172_8446622_RNA45SN1_45S_50ntFlanks@8179.8211.33\|CTGATCGAGGCCCAGCCCGTGGACGGTGTGAGG | >hg38_21_+_8433172_8446622_RNA45SN1_45S_50ntFlanks | 8179 | 8211 | 33 | rRF 45S | NO | 0.00 | 0.00 | 38.78 | 0.00 |

### MDA-MB-468

| | | | | | | | | | | |
|---|---|---|---|---|---|---|---|---|---|---|
| >hg38_21_+_8433172_8446622_RNA45SN1_45S_50ntFlanks@8179.8212.34\|CTGATCGAGGCCCAGCCCGTGGACGGTGTGAGGC | >hg38_21_+_8433172_8446622_RNA45SN1_45S_50ntFlanks | 8179 | 8212 | 34 | rRF 45S | NO | 0.00 | 0.00 | 34.88 | 0.00 |
| >hg38_21_+_8433172_8446622_RNA45SN1_45S_50ntFlanks@8182.8203.22\|ATCGAGGCCCAGCCCGTGGACG | >hg38_21_+_8433172_8446622_RNA45SN1_45S_50ntFlanks | 8182 | 8203 | 22 | rRF 45S | NO | 6.14 | 36.82 | 7.13 | 16.55 |
| >hg38_21_+_8433172_8446622_RNA45SN1_45S_50ntFlanks@8182.8204.23\|ATCGAGGCCCAGCCCGTGGACGG | >hg38_21_+_8433172_8446622_RNA45SN1_45S_50ntFlanks | 8182 | 8204 | 23 | rRF 45S | NO | 29.44 | 0.00 | 0.00 | 0.00 |
| >hg38_21_+_8433172_8446622_RNA45SN1_45S_50ntFlanks@8182.8206.25\|ATCGAGGCCCAGCCCGTGGACGGTG | >hg38_21_+_8433172_8446622_RNA45SN1_45S_50ntFlanks | 8182 | 8206 | 25 | rRF 45S | NO | 37.55 | 0.00 | 20.40 | 0.00 |
| >hg38_21_+_8433172_8446622_RNA45SN1_45S_50ntFlanks@8182.8208.27\|ATCGAGGCCCAGCCCGTGGACGGTGTG | >hg38_21_+_8433172_8446622_RNA45SN1_45S_50ntFlanks | 8182 | 8208 | 27 | rRF 45S | NO | 46.91 | 0.00 | 0.00 | 0.00 |
| >hg38_21_+_8433172_8446622_RNA45SN1_45S_50ntFlanks@8182.8209.28\|ATCGAGGCCCAGCCCGTGGACGGTGTGA | >hg38_21_+_8433172_8446622_RNA45SN1_45S_50ntFlanks | 8182 | 8209 | 28 | rRF 45S | NO | 19.61 | 0.00 | 0.00 | 0.00 |
| >hg38_21_+_8433172_8446622_RNA45SN1_45S_50ntFlanks@8182.8210.29\|ATCGAGGCCCAGCCCGTGGACGGTGTGAG | >hg38_21_+_8433172_8446622_RNA45SN1_45S_50ntFlanks | 8182 | 8210 | 29 | rRF 45S | NO | 62.98 | 0.00 | 0.00 | 0.00 |
| >hg38_21_+_8433172_8446622_RNA45SN1_45S_50ntFlanks@8182.8211.30\|ATCGAGGCCCAGCCCGTGGACGGTGTGAGG | >hg38_21_+_8433172_8446622_RNA45SN1_45S_50ntFlanks | 8182 | 8211 | 30 | rRF 45S | NO | 36.71 | 0.00 | 0.00 | 0.00 |
| >hg38_21_+_8433172_8446622_RNA45SN1_45S_50ntFlanks@8182.8223.42\|ATCGAGGCCCAGCCCGTGGACGGTGTGAGGCCGGTAGCGGCC | >hg38_21_+_8433172_8446622_RNA45SN1_45S_50ntFlanks | 8182 | 8223 | 42 | rRF 45S | NO | 51.29 | 31.14 | 0.00 | 0.00 |
| >hg38_21_+_8433172_8446622_RNA45SN1_45S_50ntFlanks@8183.8223.41\|TCGAGGCCCAGCCCGTGGACGGTGTGAGGCCGGTAGCGGCC | >hg38_21_+_8433172_8446622_RNA45SN1_45S_50ntFlanks | 8183 | 8223 | 41 | rRF 45S | NO | 51.30 | 28.71 | 0.00 | 0.00 |
| >hg38_21_+_8433172_8446622_RNA45SN1_45S_50ntFlanks@8184.8210.27\|CGAGGCCCAGCCCGTGGACGGTGTGAG | >hg38_21_+_8433172_8446622_RNA45SN1_45S_50ntFlanks | 8184 | 8210 | 27 | rRF 45S | NO | 29.91 | 0.00 | 0.00 | 0.00 |
| >hg38_21_+_8433172_8446622_RNA45SN1_45S_50ntFlanks@8184.8223.40\|CGAGGCCCAGCCCGTGGACGGTGTGAGGCCGGTAGCGGCC | >hg38_21_+_8433172_8446622_RNA45SN1_45S_50ntFlanks | 8184 | 8223 | 40 | rRF 45S | NO | 21.40 | 0.00 | 0.00 | 0.00 |
| >hg38_21_+_8433172_8446622_RNA45SN1_45S_50ntFlanks@8189.8212.24\|CCCAGCCCGTGGACGGTGTGAGGC | >hg38_21_+_8433172_8446622_RNA45SN1_45S_50ntFlanks | 8189 | 8212 | 24 | rRF 45S | NO | 0.00 | 12.50 | 42.50 | 0.00 |
| >hg38_21_+_8433172_8446622_RNA45SN1_45S_50ntFlanks@8190.8212.23\|CCAGCCCGTGGACGGTGTGAGGC | >hg38_21_+_8433172_8446622_RNA45SN1_45S_50ntFlanks | 8190 | 8212 | 23 | rRF 45S | NO | 37.23 | 34.25 | 61.40 | 13.20 |
| >hg38_21_+_8433172_8446622_RNA45SN1_45S_50ntFlanks@8190.8213.24\|CCAGCCCGTGGACGGTGTGAGGCC | >hg38_21_+_8433172_8446622_RNA45SN1_45S_50ntFlanks | 8190 | 8213 | 24 | rRF 45S | NO | 21.08 | 0.00 | 0.00 | 0.00 |
| >hg38_21_+_8433172_8446622_RNA45SN1_45S_50ntFlanks@8191.8211.21\|CAGCCCGTGGACGGTGTGAGG | >hg38_21_+_8433172_8446622_RNA45SN1_45S_50ntFlanks | 8191 | 8211 | 21 | rRF 45S | NO | 0.00 | 0.00 | 39.56 | 0.00 |
| >hg38_21_+_8433172_8446622_RNA45SN1_45S_50ntFlanks@8191.8212.22\|CAGCCCGTGGACGGTGTGAGGC | >hg38_21_+_8433172_8446622_RNA45SN1_45S_50ntFlanks | 8191 | 8212 | 22 | rRF 45S | NO | 47.22 | 50.16 | 69.63 | 15.22 |
| >hg38_21_+_8433172_8446622_RNA45SN1_45S_50ntFlanks@8191.8213.23\|CAGCCCGTGGACGGTGTGAGGCC | >hg38_21_+_8433172_8446622_RNA45SN1_45S_50ntFlanks | 8191 | 8213 | 23 | rRF 45S | NO | 76.00 | 37.63 | 36.57 | 24.26 |
| >hg38_21_+_8433172_8446622_RNA45SN1_45S_50ntFlanks@8191.8214.24\|CAGCCCGTGGACGGTGTGAGGCCG | >hg38_21_+_8433172_8446622_RNA45SN1_45S_50ntFlanks | 8191 | 8214 | 24 | rRF 45S | NO | 102.81 | 44.25 | 27.93 | 26.75 |
| >hg38_21_+_8433172_8446622_RNA45SN1_45S_50ntFlanks@8191.8215.25\|CAGCCCGTGGACGGTGTGAGGCCGG | >hg38_21_+_8433172_8446622_RNA45SN1_45S_50ntFlanks | 8191 | 8215 | 25 | rRF 45S | NO | 144.00 | 102.29 | 39.28 | 44.39 |
| >hg38_21_+_8433172_8446622_RNA45SN1_45S_50ntFlanks@8194.8211.18\|CCCGTGGACGGTGTGAGG | >hg38_21_+_8433172_8446622_RNA45SN1_45S_50ntFlanks | 8194 | 8211 | 18 | rRF 45S | NO | 98.10 | 64.38 | 121.28 | 6.54 |
| >hg38_21_+_8433172_8446622_RNA45SN1_45S_50ntFlanks@8194.8212.19\|CCCGTGGACGGTGTGAGGC | >hg38_21_+_8433172_8446622_RNA45SN1_45S_50ntFlanks | 8194 | 8212 | 19 | rRF 45S | NO | 111.99 | 51.13 | 83.34 | 6.19 |
| >hg38_21_+_8433172_8446622_RNA45SN1_45S_50ntFlanks@8194.8213.20\|CCCGTGGACGGTGTGAGGCC | >hg38_21_+_8433172_8446622_RNA45SN1_45S_50ntFlanks | 8194 | 8213 | 20 | rRF 45S | NO | 157.38 | 76.10 | 52.12 | 35.53 |
| >hg38_21_+_8433172_8446622_RNA45SN1_45S_50ntFlanks@8194.8214.21\|CCCGTGGACGGTGTGAGGCCG | >hg38_21_+_8433172_8446622_RNA45SN1_45S_50ntFlanks | 8194 | 8214 | 21 | rRF 45S | NO | 109.10 | 95.72 | 35.13 | 15.95 |
| >hg38_21_+_8433172_8446622_RNA45SN1_45S_50ntFlanks@8194.8215.22\|CCCGTGGACGGTGTGAGGCCGG | >hg38_21_+_8433172_8446622_RNA45SN1_45S_50ntFlanks | 8194 | 8215 | 22 | rRF 45S | NO | 172.57 | 200.97 | 56.94 | 40.03 |
| >hg38_21_+_8433172_8446622_RNA45SN1_45S_50ntFlanks@8194.8225.32\|CCCGTGGACGGTGTGAGGCCGGTAGCGGCCCC | >hg38_21_+_8433172_8446622_RNA45SN1_45S_50ntFlanks | 8194 | 8225 | 32 | rRF 45S | NO | 0.00 | 42.61 | 0.00 | 0.00 |
| >hg38_21_+_8433172_8446622_RNA45SN1_45S_50ntFlanks@8196.8213.18\|CGTGGACGGTGTGAGGCC | >hg38_21_+_8433172_8446622_RNA45SN1_45S_50ntFlanks | 8196 | 8213 | 18 | rRF 45S | NO | 123.01 | 96.41 | 79.79 | 15.02 |
| >hg38_21_+_8433172_8446622_RNA45SN1_45S_50ntFlanks@8196.8214.19\|CGTGGACGGTGTGAGGCCG | >hg38_21_+_8433172_8446622_RNA45SN1_45S_50ntFlanks | 8196 | 8214 | 19 | rRF 45S | NO | 66.15 | 220.99 | 106.71 | 13.21 |
| >hg38_21_+_8433172_8446622_RNA45SN1_45S_50ntFlanks@8196.8215.20\|CGTGGACGGTGTGAGGCCGG | >hg38_21_+_8433172_8446622_RNA45SN1_45S_50ntFlanks | 8196 | 8215 | 20 | rRF 45S | NO | 79.20 | 343.87 | 115.89 | 13.38 |
| >hg38_21_+_8433172_8446622_RNA45SN1_45S_50ntFlanks@8196.8216.21\|CGTGGACGGTGTGAGGCCGGT | >hg38_21_+_8433172_8446622_RNA45SN1_45S_50ntFlanks | 8196 | 8216 | 21 | rRF 45S | NO | 0.00 | 43.17 | 33.27 | 0.00 |
| >hg38_21_+_8433172_8446622_RNA45SN1_45S_50ntFlanks@8196.8222.27\|CGTGGACGGTGTGAGGCCGGTAGCGGC | >hg38_21_+_8433172_8446622_RNA45SN1_45S_50ntFlanks | 8196 | 8222 | 27 | rRF 45S | NO | 0.00 | 35.77 | 6.49 | 0.00 |
| >hg38_21_+_8433172_8446622_RNA45SN1_45S_50ntFlanks@8196.8223.28\|CGTGGACGGTGTGAGGCCGGTAGCGGCC | >hg38_21_+_8433172_8446622_RNA45SN1_45S_50ntFlanks | 8196 | 8223 | 28 | rRF 45S | NO | 41.23 | 78.82 | 29.34 | 0.00 |
| >hg38_21_+_8433172_8446622_RNA45SN1_45S_50ntFlanks@8196.8224.29\|CGTGGACGGTGTGAGGCCGGTAGCGGCCC | >hg38_21_+_8433172_8446622_RNA45SN1_45S_50ntFlanks | 8196 | 8224 | 29 | rRF 45S | NO | 88.77 | 125.48 | 6.70 | 0.00 |
| >hg38_21_+_8433172_8446622_RNA45SN1_45S_50ntFlanks@8196.8225.30\|CGTGGACGGTGTGAGGCCGGTAGCGGCCCC | >hg38_21_+_8433172_8446622_RNA45SN1_45S_50ntFlanks | 8196 | 8225 | 30 | rRF 45S | NO | 66.31 | 94.94 | 0.00 | 0.00 |

### MDA-MB-468

| | | | | | | | | | | |
|---|---|---|---|---|---|---|---|---|---|---|
| >hg38_21_+_8433172_8446622_RNA45SN1_45S_50ntFlanks@8196.8226.31\|CGTGGACGGTGTGAGGCCGGTAGCGGCCCCC | >hg38_21_+_8433172_8446622_RNA45SN1_45S_50ntFlanks | 8196 | 8226 | 31 | rRF 45S | NO | 12.70 | 60.60 | 0.00 | 0.00 |
| >hg38_21_+_8433172_8446622_RNA45SN1_45S_50ntFlanks@8197.8214.18\|GTGGACGGTGTGAGGCCG | >hg38_21_+_8433172_8446622_RNA45SN1_45S_50ntFlanks | 8197 | 8214 | 18 | rRF 45S | NO | 0.00 | 12.80 | 0.00 | 0.00 |
| >hg38_21_+_8433172_8446622_RNA45SN1_45S_50ntFlanks@8197.8215.19\|GTGGACGGTGTGAGGCCGG | >hg38_21_+_8433172_8446622_RNA45SN1_45S_50ntFlanks | 8197 | 8215 | 19 | rRF 45S | NO | 0.00 | 34.85 | 0.00 | 0.00 |
| >hg38_21_+_8433172_8446622_RNA45SN1_45S_50ntFlanks@8198.8215.18\|TGGACGGTGTGAGGCCGG | >hg38_21_+_8433172_8446622_RNA45SN1_45S_50ntFlanks | 8198 | 8215 | 18 | rRF 45S | NO | 38.97 | 228.77 | 98.19 | 0.00 |
| >hg38_21_+_8433172_8446622_RNA45SN1_45S_50ntFlanks@8198.8223.26\|TGGACGGTGTGAGGCCGGTAGCGGCC | >hg38_21_+_8433172_8446622_RNA45SN1_45S_50ntFlanks | 8198 | 8223 | 26 | rRF 45S | NO | 56.27 | 58.48 | 6.59 | 0.00 |
| >hg38_21_+_8433172_8446622_RNA45SN1_45S_50ntFlanks@8198.8224.27\|TGGACGGTGTGAGGCCGGTAGCGGCCC | >hg38_21_+_8433172_8446622_RNA45SN1_45S_50ntFlanks | 8198 | 8224 | 27 | rRF 45S | NO | 29.41 | 48.60 | 0.00 | 0.00 |
| >hg38_21_+_8433172_8446622_RNA45SN1_45S_50ntFlanks@8198.8225.28\|TGGACGGTGTGAGGCCGGTAGCGGCCCC | >hg38_21_+_8433172_8446622_RNA45SN1_45S_50ntFlanks | 8198 | 8225 | 28 | rRF 45S | NO | 20.89 | 71.24 | 0.00 | 0.00 |
| >hg38_21_+_8433172_8446622_RNA45SN1_45S_50ntFlanks@8317.8356.40\|CACGAGACCGATAGTCAACAAGTACCGTAAGGGAAAGTTG | >hg38_21_+_8433172_8446622_RNA45SN1_45S_50ntFlanks | 8317 | 8356 | 40 | rRF 45S | NO | 45.58 | 37.35 | 0.00 | 41.58 |
| >hg38_21_+_8433172_8446622_RNA45SN1_45S_50ntFlanks@8319.8356.38\|CGAGACCGATAGTCAACAAGTACCGTAAGGGAAAGTTG | >hg38_21_+_8433172_8446622_RNA45SN1_45S_50ntFlanks | 8319 | 8356 | 38 | rRF 45S | NO | 0.00 | 42.95 | 0.00 | 6.41 |
| >hg38_21_+_8433172_8446622_RNA45SN1_45S_50ntFlanks@8324.8356.33\|CCGATAGTCAACAAGTACCGTAAGGGAAAGTTG | >hg38_21_+_8433172_8446622_RNA45SN1_45S_50ntFlanks | 8324 | 8356 | 33 | rRF 45S | NO | 0.00 | 73.08 | 0.00 | 0.00 |
| >hg38_21_+_8433172_8446622_RNA45SN1_45S_50ntFlanks@8356.8373.18\|GAAAAGAACTTTGAAGAG | >hg38_21_+_8433172_8446622_RNA45SN1_45S_50ntFlanks | 8356 | 8373 | 18 | rRF 45S | NO | 0.00 | 0.00 | 34.52 | 0.00 |
| >hg38_21_+_8433172_8446622_RNA45SN1_45S_50ntFlanks@8356.8375.20\|GAAAAGAACTTTGAAGAGAG | >hg38_21_+_8433172_8446622_RNA45SN1_45S_50ntFlanks | 8356 | 8375 | 20 | rRF 45S | NO | 6.00 | 0.00 | 86.75 | 0.00 |
| >hg38_21_+_8433172_8446622_RNA45SN1_45S_50ntFlanks@8356.8377.22\|GAAAAGAACTTTGAAGAGAGAG | >hg38_21_+_8433172_8446622_RNA45SN1_45S_50ntFlanks | 8356 | 8377 | 22 | rRF 45S | NO | 40.03 | 0.00 | 64.87 | 0.00 |
| >hg38_21_+_8433172_8446622_RNA45SN1_45S_50ntFlanks@8356.8378.23\|GAAAAGAACTTTGAAGAGAGAGT | >hg38_21_+_8433172_8446622_RNA45SN1_45S_50ntFlanks | 8356 | 8378 | 23 | rRF 45S | NO | 144.87 | 63.44 | 208.24 | 6.16 |
| >hg38_21_+_8433172_8446622_RNA45SN1_45S_50ntFlanks@8356.8379.24\|GAAAAGAACTTTGAAGAGAGAGTT | >hg38_21_+_8433172_8446622_RNA45SN1_45S_50ntFlanks | 8356 | 8379 | 24 | rRF 45S | YES | 674.24 | 419.60 | 366.48 | 125.26 |
| >hg38_21_+_8433172_8446622_RNA45SN1_45S_50ntFlanks@8357.8375.19\|AAAAGAACTTTGAAGAGAG | >hg38_21_+_8433172_8446622_RNA45SN1_45S_50ntFlanks | 8357 | 8375 | 19 | rRF 45S | YES | 149.32 | 137.37 | 425.54 | 74.27 |
| >hg38_21_+_8433172_8446622_RNA45SN1_45S_50ntFlanks@8357.8376.20\|AAAAGAACTTTGAAGAGAGA | >hg38_21_+_8433172_8446622_RNA45SN1_45S_50ntFlanks | 8357 | 8376 | 20 | rRF 45S | NO | 30.22 | 0.00 | 72.44 | 0.00 |
| >hg38_21_+_8433172_8446622_RNA45SN1_45S_50ntFlanks@8357.8377.21\|AAAAGAACTTTGAAGAGAGAG | >hg38_21_+_8433172_8446622_RNA45SN1_45S_50ntFlanks | 8357 | 8377 | 21 | rRF 45S | NO | 262.98 | 159.24 | 224.90 | 85.73 |
| >hg38_21_+_8433172_8446622_RNA45SN1_45S_50ntFlanks@8357.8378.22\|AAAAGAACTTTGAAGAGAGAGT | >hg38_21_+_8433172_8446622_RNA45SN1_45S_50ntFlanks | 8357 | 8378 | 22 | rRF 45S | YES | 696.75 | 378.06 | 657.77 | 97.44 |
| >hg38_21_+_8433172_8446622_RNA45SN1_45S_50ntFlanks@8357.8379.23\|AAAAGAACTTTGAAGAGAGAGTT | >hg38_21_+_8433172_8446622_RNA45SN1_45S_50ntFlanks | 8357 | 8379 | 23 | rRF 45S | YES | 1985.93 | 1626.03 | 1376.70 | 616.52 |
| >hg38_21_+_8433172_8446622_RNA45SN1_45S_50ntFlanks@8357.8387.31\|AAAAGAACTTTGAAGAGAGAGTTCAAGAGGG | >hg38_21_+_8433172_8446622_RNA45SN1_45S_50ntFlanks | 8357 | 8387 | 31 | rRF 45S | NO | 6.13 | 31.79 | 27.76 | 0.00 |
| >hg38_21_+_8433172_8446622_RNA45SN1_45S_50ntFlanks@8357.8391.35\|AAAAGAACTTTGAAGAGAGAGTTCAAGAGGGCGTG | >hg38_21_+_8433172_8446622_RNA45SN1_45S_50ntFlanks | 8357 | 8391 | 35 | rRF 45S | NO | 88.25 | 46.12 | 29.10 | 25.82 |
| >hg38_21_+_8433172_8446622_RNA45SN1_45S_50ntFlanks@8358.8378.21\|AAAGAACTTTGAAGAGAGAGT | >hg38_21_+_8433172_8446622_RNA45SN1_45S_50ntFlanks | 8358 | 8378 | 21 | rRF 45S | NO | 0.00 | 0.00 | 27.19 | 0.00 |
| >hg38_21_+_8433172_8446622_RNA45SN1_45S_50ntFlanks@8358.8379.22\|AAAGAACTTTGAAGAGAGAGTT | >hg38_21_+_8433172_8446622_RNA45SN1_45S_50ntFlanks | 8358 | 8379 | 22 | rRF 45S | NO | 49.03 | 35.45 | 40.38 | 5.99 |
| >hg38_21_+_8433172_8446622_RNA45SN1_45S_50ntFlanks@8359.8378.20\|AAGAACTTTGAAGAGAGAGT | >hg38_21_+_8433172_8446622_RNA45SN1_45S_50ntFlanks | 8359 | 8378 | 20 | rRF 45S | NO | 0.00 | 0.00 | 13.62 | 0.00 |
| >hg38_21_+_8433172_8446622_RNA45SN1_45S_50ntFlanks@8359.8379.21\|AAGAACTTTGAAGAGAGAGTT | >hg38_21_+_8433172_8446622_RNA45SN1_45S_50ntFlanks | 8359 | 8379 | 21 | rRF 45S | NO | 28.73 | 35.67 | 32.11 | 0.00 |
| >hg38_21_+_8433172_8446622_RNA45SN1_45S_50ntFlanks@8378.8402.25\|TTCAAGAGGGCGTGAAACCGTTAAG | >hg38_21_+_8433172_8446622_RNA45SN1_45S_50ntFlanks | 8378 | 8402 | 25 | rRF 45S | NO | 0.00 | 0.00 | 20.84 | 0.00 |
| >hg38_21_+_8433172_8446622_RNA45SN1_45S_50ntFlanks@8379.8402.24\|TCAAGAGGGCGTGAAACCGTTAAG | >hg38_21_+_8433172_8446622_RNA45SN1_45S_50ntFlanks | 8379 | 8402 | 24 | rRF 45S | NO | 0.00 | 0.00 | 36.23 | 0.00 |
| >hg38_21_+_8433172_8446622_RNA45SN1_45S_50ntFlanks@8380.8397.18\|CAAGAGGGCGTGAAACCG | >hg38_21_+_8433172_8446622_RNA45SN1_45S_50ntFlanks | 8380 | 8397 | 18 | rRF 45S | NO | 37.08 | 88.94 | 69.07 | 6.56 |
| >hg38_21_+_8433172_8446622_RNA45SN1_45S_50ntFlanks@8380.8399.20\|CAAGAGGGCGTGAAACCGTT | >hg38_21_+_8433172_8446622_RNA45SN1_45S_50ntFlanks | 8380 | 8399 | 20 | rRF 45S | NO | 0.00 | 54.65 | 62.32 | 15.49 |
| >hg38_21_+_8433172_8446622_RNA45SN1_45S_50ntFlanks@8380.8400.21\|CAAGAGGGCGTGAAACCGTTA | >hg38_21_+_8433172_8446622_RNA45SN1_45S_50ntFlanks | 8380 | 8400 | 21 | rRF 45S | NO | 37.00 | 220.95 | 145.51 | 91.68 |
| >hg38_21_+_8433172_8446622_RNA45SN1_45S_50ntFlanks@8380.8401.22\|CAAGAGGGCGTGAAACCGTTAA | >hg38_21_+_8433172_8446622_RNA45SN1_45S_50ntFlanks | 8380 | 8401 | 22 | rRF 45S | NO | 0.00 | 49.71 | 28.09 | 14.85 |
| >hg38_21_+_8433172_8446622_RNA45SN1_45S_50ntFlanks@8380.8402.23\|CAAGAGGGCGTGAAACCGTTAAG | >hg38_21_+_8433172_8446622_RNA45SN1_45S_50ntFlanks | 8380 | 8402 | 23 | rRF 45S | NO | 42.57 | 123.41 | 271.64 | 146.61 |
| >hg38_21_+_8433172_8446622_RNA45SN1_45S_50ntFlanks@8380.8403.24\|CAAGAGGGCGTGAAACCGTTAAGA | >hg38_21_+_8433172_8446622_RNA45SN1_45S_50ntFlanks | 8380 | 8403 | 24 | rRF 45S | NO | 6.04 | 58.85 | 75.31 | 72.38 |

### MDA-MB-468

| | | | | | | | | | | |
|---|---|---|---|---|---|---|---|---|---|---|
| >hg38_21_+_8433172_8446622_RNA45SN1_45S_50ntFlanks@8380.8404.25\|CAAGAGGGCGT GAAACCGTTAAGAG | >hg38_21_+_8433172_8446622_RNA45SN1_45S_50ntFlanks | 8380 | 8404 | 25 | rRF 45S | NO | 0.00 | 52.63 | 33.27 | 40.70 |
| >hg38_21_+_8433172_8446622_RNA45SN1_45S_50ntFlanks@8380.8405.26\|CAAGAGGGCGT GAAACCGTTAAGAGG | >hg38_21_+_8433172_8446622_RNA45SN1_45S_50ntFlanks | 8380 | 8405 | 26 | rRF 45S | NO | 75.90 | 145.09 | 151.52 | 96.74 |
| >hg38_21_+_8433172_8446622_RNA45SN1_45S_50ntFlanks@8380.8406.27\|CAAGAGGGCGT GAAACCGTTAAGAGGT | >hg38_21_+_8433172_8446622_RNA45SN1_45S_50ntFlanks | 8380 | 8406 | 27 | rRF 45S | NO | 41.53 | 45.85 | 84.24 | 62.12 |
| >hg38_21_+_8433172_8446622_RNA45SN1_45S_50ntFlanks@8381.8399.19\|AAGAGGGCGTG AAACCGTT | >hg38_21_+_8433172_8446622_RNA45SN1_45S_50ntFlanks | 8381 | 8399 | 19 | rRF 45S | NO | 0.00 | 0.00 | 40.28 | 0.00 |
| >hg38_21_+_8433172_8446622_RNA45SN1_45S_50ntFlanks@8381.8400.20\|AAGAGGGCGTG AAACCGTTA | >hg38_21_+_8433172_8446622_RNA45SN1_45S_50ntFlanks | 8381 | 8400 | 20 | rRF 45S | NO | 0.00 | 76.71 | 82.39 | 23.06 |
| >hg38_21_+_8433172_8446622_RNA45SN1_45S_50ntFlanks@8381.8402.22\|AAGAGGGCGTG AAACCGTTAAG | >hg38_21_+_8433172_8446622_RNA45SN1_45S_50ntFlanks | 8381 | 8402 | 22 | rRF 45S | NO | 39.53 | 93.29 | 232.17 | 72.03 |
| >hg38_21_+_8433172_8446622_RNA45SN1_45S_50ntFlanks@8381.8403.23\|AAGAGGGCGTG AAACCGTTAAGA | >hg38_21_+_8433172_8446622_RNA45SN1_45S_50ntFlanks | 8381 | 8403 | 23 | rRF 45S | NO | 0.00 | 12.49 | 49.19 | 13.52 |
| >hg38_21_+_8433172_8446622_RNA45SN1_45S_50ntFlanks@8381.8405.25\|AAGAGGGCGTG AAACCGTTAAGAGG | >hg38_21_+_8433172_8446622_RNA45SN1_45S_50ntFlanks | 8381 | 8405 | 25 | rRF 45S | NO | 29.74 | 55.00 | 85.12 | 14.41 |
| >hg38_21_+_8433172_8446622_RNA45SN1_45S_50ntFlanks@8381.8406.26\|AAGAGGGCGTG AAACCGTTAAGAGGT | >hg38_21_+_8433172_8446622_RNA45SN1_45S_50ntFlanks | 8381 | 8406 | 26 | rRF 45S | NO | 0.00 | 0.00 | 51.67 | 0.00 |
| >hg38_21_+_8433172_8446622_RNA45SN1_45S_50ntFlanks@8382.8402.21\|AGAGGGCGTGA AACCGTTAAG | >hg38_21_+_8433172_8446622_RNA45SN1_45S_50ntFlanks | 8382 | 8402 | 21 | rRF 45S | NO | 0.00 | 0.00 | 29.93 | 0.00 |
| >hg38_21_+_8433172_8446622_RNA45SN1_45S_50ntFlanks@8384.8402.19\|AGGGCGTGAAA CCGTTAAG | >hg38_21_+_8433172_8446622_RNA45SN1_45S_50ntFlanks | 8384 | 8402 | 19 | rRF 45S | NO | 0.00 | 0.00 | 13.70 | 0.00 |
| >hg38_21_+_8433172_8446622_RNA45SN1_45S_50ntFlanks@8388.8405.18\|CGTGAAACCGTT AAGAGG | >hg38_21_+_8433172_8446622_RNA45SN1_45S_50ntFlanks | 8388 | 8405 | 18 | rRF 45S | NO | 0.00 | 37.46 | 0.00 | 0.00 |
| >hg38_21_+_8433172_8446622_RNA45SN1_45S_50ntFlanks@8423.8442.20\|CGCAGTCCGCC CGGAGGATT | >hg38_21_+_8433172_8446622_RNA45SN1_45S_50ntFlanks | 8423 | 8442 | 20 | rRF 45S | NO | 0.00 | 0.00 | 98.93 | 218.66 |
| >hg38_21_+_8433172_8446622_RNA45SN1_45S_50ntFlanks@8472.8490.19\|CGGCCCGGCGG ATCTTTCC | >hg38_21_+_8433172_8446622_RNA45SN1_45S_50ntFlanks | 8472 | 8490 | 19 | rRF 45S | NO | 35.70 | 0.00 | 0.00 | 0.00 |
| >hg38_21_+_8433172_8446622_RNA45SN1_45S_50ntFlanks@8472.8491.20\|CGGCCCGGCGG ATCTTTCCC | >hg38_21_+_8433172_8446622_RNA45SN1_45S_50ntFlanks | 8472 | 8491 | 20 | rRF 45S | NO | 48.33 | 29.53 | 0.00 | 43.17 |
| >hg38_21_+_8433172_8446622_RNA45SN1_45S_50ntFlanks@8472.8492.21\|CGGCCCGGCGG ATCTTTCCCG | >hg38_21_+_8433172_8446622_RNA45SN1_45S_50ntFlanks | 8472 | 8492 | 21 | rRF 45S | NO | 35.83 | 37.18 | 0.00 | 88.06 |
| >hg38_21_+_8433172_8446622_RNA45SN1_45S_50ntFlanks@8528.8551.24\|CTTCCCCCGCC GCCCCTCCTCCTC | >hg38_21_+_8433172_8446622_RNA45SN1_45S_50ntFlanks | 8528 | 8551 | 24 | rRF 45S | NO | 0.00 | 19.69 | 0.00 | 0.00 |
| >hg38_21_+_8433172_8446622_RNA45SN1_45S_50ntFlanks@8552.8569.18\|CTCCCCGGAGG GGGCGGG | >hg38_21_+_8433172_8446622_RNA45SN1_45S_50ntFlanks | 8552 | 8569 | 18 | rRF 45S | NO | 0.00 | 68.48 | 70.39 | 35.91 |
| >hg38_21_+_8433172_8446622_RNA45SN1_45S_50ntFlanks@8552.8577.26\|CTCCCCGGAGG GGGCGGGCTCCGGCG | >hg38_21_+_8433172_8446622_RNA45SN1_45S_50ntFlanks | 8552 | 8577 | 26 | rRF 45S | NO | 29.66 | 51.77 | 0.00 | 15.53 |
| >hg38_21_+_8433172_8446622_RNA45SN1_45S_50ntFlanks@8570.8593.24\|CTCCGGCGGGT GCGGGGGTGGGCG | >hg38_21_+_8433172_8446622_RNA45SN1_45S_50ntFlanks | 8570 | 8593 | 24 | rRF 45S | NO | 19.75 | 0.00 | 0.00 | 0.00 |
| >hg38_21_+_8433172_8446622_RNA45SN1_45S_50ntFlanks@8737.8754.18\|CCCGTCCGTCC GTCCGTC | >hg38_21_+_8433172_8446622_RNA45SN1_45S_50ntFlanks | 8737 | 8754 | 18 | rRF 45S | NO | 0.00 | 39.47 | 0.00 | 0.00 |
| >hg38_21_+_8433172_8446622_RNA45SN1_45S_50ntFlanks@8737.8758.22\|CCCGTCCGTCC GTCCGTCCGTC | >hg38_21_+_8433172_8446622_RNA45SN1_45S_50ntFlanks | 8737 | 8758 | 22 | rRF 45S | NO | 0.00 | 31.24 | 0.00 | 0.00 |
| >hg38_21_+_8433172_8446622_RNA45SN1_45S_50ntFlanks@8799.8823.25\|CCTCGGGAGGG CGCGCGGGTCGGGG | >hg38_21_+_8433172_8446622_RNA45SN1_45S_50ntFlanks | 8799 | 8823 | 25 | rRF 45S | NO | 20.27 | 0.00 | 0.00 | 0.00 |
| >hg38_21_+_8433172_8446622_RNA45SN1_45S_50ntFlanks@9004.9026.23\|TCTCCCCCGCG GGGGCGCGCCGG | >hg38_21_+_8433172_8446622_RNA45SN1_45S_50ntFlanks | 9004 | 9026 | 23 | rRF 45S | NO | 77.14 | 0.00 | 0.00 | 185.44 |
| >hg38_21_+_8433172_8446622_RNA45SN1_45S_50ntFlanks@9032.9059.28\|CCTCGTGGGGG GGCCGGGCCACCCCTCC | >hg38_21_+_8433172_8446622_RNA45SN1_45S_50ntFlanks | 9032 | 9059 | 28 | rRF 45S | NO | 0.00 | 0.00 | 35.98 | 37.25 |
| >hg38_21_+_8433172_8446622_RNA45SN1_45S_50ntFlanks@9033.9056.24\|CTCGTGGGGGG GCCGGGCCACCCC | >hg38_21_+_8433172_8446622_RNA45SN1_45S_50ntFlanks | 9033 | 9056 | 24 | rRF 45S | YES | 0.00 | 0.00 | 60.28 | 14.04 |
| >hg38_21_+_8433172_8446622_RNA45SN1_45S_50ntFlanks@9033.9058.26\|CTCGTGGGGGG GCCGGGCCACCCCTC | >hg38_21_+_8433172_8446622_RNA45SN1_45S_50ntFlanks | 9033 | 9058 | 26 | rRF 45S | NO | 0.00 | 0.00 | 134.12 | 45.91 |
| >hg38_21_+_8433172_8446622_RNA45SN1_45S_50ntFlanks@9033.9059.27\|CTCGTGGGGGG GCCGGGCCACCCCTCC | >hg38_21_+_8433172_8446622_RNA45SN1_45S_50ntFlanks | 9033 | 9059 | 27 | rRF 45S | NO | 0.00 | 0.00 | 102.66 | 64.85 |
| >hg38_21_+_8433172_8446622_RNA45SN1_45S_50ntFlanks@9035.9059.25\|CGTGGGGGGGC CGGGCCACCCCTCC | >hg38_21_+_8433172_8446622_RNA45SN1_45S_50ntFlanks | 9035 | 9059 | 25 | rRF 45S | YES | 0.00 | 0.00 | 13.56 | 14.35 |
| >hg38_21_+_8433172_8446622_RNA45SN1_45S_50ntFlanks@9129.9152.24\|CGGGTCGGGGG GCGGGGCGGACTG | >hg38_21_+_8433172_8446622_RNA45SN1_45S_50ntFlanks | 9129 | 9152 | 24 | rRF 45S | NO | 0.00 | 0.00 | 72.64 | 93.62 |
| >hg38_21_+_8433172_8446622_RNA45SN1_45S_50ntFlanks@9166.9197.32\|CCGGGCGGGTC GCGCCGTCGGGCCCGGGGGAG | >hg38_21_+_8433172_8446622_RNA45SN1_45S_50ntFlanks | 9166 | 9197 | 32 | rRF 45S | NO | 33.54 | 0.00 | 59.71 | 91.96 |
| >hg38_21_+_8433172_8446622_RNA45SN1_45S_50ntFlanks@9167.9197.31\|CGGGCGGGTCG CGCCGTCGGGCCCGGGGGAG | >hg38_21_+_8433172_8446622_RNA45SN1_45S_50ntFlanks | 9167 | 9197 | 31 | rRF 45S | NO | 0.00 | 0.00 | 44.80 | 38.28 |
| >hg38_21_+_8433172_8446622_RNA45SN1_45S_50ntFlanks@9171.9197.27\|CGGGTCGCGCC GTCGGGCCCGGGGGAG | >hg38_21_+_8433172_8446622_RNA45SN1_45S_50ntFlanks | 9171 | 9197 | 27 | rRF 45S | NO | 0.00 | 0.00 | 43.02 | 38.56 |
| >hg38_21_+_8433172_8446622_RNA45SN1_45S_50ntFlanks@9178.9197.20\|CGCCGTCGGGC CCGGGGGAG | >hg38_21_+_8433172_8446622_RNA45SN1_45S_50ntFlanks | 9178 | 9197 | 20 | rRF 45S | NO | 0.00 | 0.00 | 13.77 | 50.01 |

### MDA-MB-468

| | | | | | | | | | | |
|---|---|---|---|---|---|---|---|---|---|---|
| >hg38_21_+_8433172_8446622_RNA45SN1_45S_50ntFlanks@9180.9197.18\|CCGTCGGGCCCGGGGGAG | >hg38_21_+_8433172_8446622_RNA45SN1_45S_50ntFlanks | 9180 | 9197 | 18 | rRF 45S | NO | 21.85 | 0.00 | 56.38 | 108.78 |
| >hg38_21_+_8433172_8446622_RNA45SN1_45S_50ntFlanks@9198.9218.21\|GTTCTCTCGGGGCCACGCGCG | >hg38_21_+_8433172_8446622_RNA45SN1_45S_50ntFlanks | 9198 | 9218 | 21 | rRF 45S | NO | 0.00 | 0.00 | 33.65 | 53.18 |
| >hg38_21_+_8433172_8446622_RNA45SN1_45S_50ntFlanks@9198.9222.25\|GTTCTCTCGGGGCCACGCGCGCGTC | >hg38_21_+_8433172_8446622_RNA45SN1_45S_50ntFlanks | 9198 | 9222 | 25 | rRF 45S | NO | 39.13 | 0.00 | 66.48 | 126.56 |
| >hg38_21_+_8433172_8446622_RNA45SN1_45S_50ntFlanks@9198.9223.26\|GTTCTCTCGGGGCCACGCGCGCGTCC | >hg38_21_+_8433172_8446622_RNA45SN1_45S_50ntFlanks | 9198 | 9223 | 26 | rRF 45S | NO | 0.00 | 0.00 | 32.74 | 24.14 |
| >hg38_21_+_8433172_8446622_RNA45SN1_45S_50ntFlanks@9201.9218.18\|CTCTCGGGGCCACGCGCG | >hg38_21_+_8433172_8446622_RNA45SN1_45S_50ntFlanks | 9201 | 9218 | 18 | rRF 45S | NO | 0.00 | 0.00 | 45.23 | 36.41 |
| >hg38_21_+_8433172_8446622_RNA45SN1_45S_50ntFlanks@9201.9220.20\|CTCTCGGGGCCACGCGCGCG | >hg38_21_+_8433172_8446622_RNA45SN1_45S_50ntFlanks | 9201 | 9220 | 20 | rRF 45S | YES | 0.00 | 0.00 | 28.43 | 26.02 |
| >hg38_21_+_8433172_8446622_RNA45SN1_45S_50ntFlanks@9201.9222.22\|CTCTCGGGGCCACGCGCGCGTC | >hg38_21_+_8433172_8446622_RNA45SN1_45S_50ntFlanks | 9201 | 9222 | 22 | rRF 45S | NO | 0.00 | 0.00 | 45.64 | 26.51 |
| >hg38_21_+_8433172_8446622_RNA45SN1_45S_50ntFlanks@9202.9220.19\|TCTCGGGGCCACGCGCGCG | >hg38_21_+_8433172_8446622_RNA45SN1_45S_50ntFlanks | 9202 | 9220 | 19 | rRF 45S | YES | 0.00 | 0.00 | 50.41 | 16.25 |
| >hg38_21_+_8433172_8446622_RNA45SN1_45S_50ntFlanks@9202.9222.21\|TCTCGGGGCCACGCGCGCGTC | >hg38_21_+_8433172_8446622_RNA45SN1_45S_50ntFlanks | 9202 | 9222 | 21 | rRF 45S | YES | 0.00 | 0.00 | 57.17 | 6.56 |
| >hg38_21_+_8433172_8446622_RNA45SN1_45S_50ntFlanks@9203.9225.23\|CTCGGGGCCACGCGCGCGTCCCC | >hg38_21_+_8433172_8446622_RNA45SN1_45S_50ntFlanks | 9203 | 9225 | 23 | rRF 45S | YES | 0.00 | 0.00 | 44.94 | 6.50 |
| >hg38_21_+_8433172_8446622_RNA45SN1_45S_50ntFlanks@9304.9336.33\|ACCAAGGAGTCTAACACGTGCGCGAGTCGGGGG | >hg38_21_+_8433172_8446622_RNA45SN1_45S_50ntFlanks | 9304 | 9336 | 33 | rRF 45S | YES | 0.00 | 0.00 | 33.77 | 0.00 |
| >hg38_21_+_8433172_8446622_RNA45SN1_45S_50ntFlanks@9304.9338.35\|ACCAAGGAGTCTAACACGTGCGCGAGTCGGGGGCT | >hg38_21_+_8433172_8446622_RNA45SN1_45S_50ntFlanks | 9304 | 9338 | 35 | rRF 45S | NO | 0.00 | 0.00 | 58.22 | 46.95 |
| >hg38_21_+_8433172_8446622_RNA45SN1_45S_50ntFlanks@9305.9334.30\|CCAAGGAGTCTAACACGTGCGCGAGTCGGG | >hg38_21_+_8433172_8446622_RNA45SN1_45S_50ntFlanks | 9305 | 9334 | 30 | rRF 45S | NO | 0.00 | 0.00 | 13.65 | 0.00 |
| >hg38_21_+_8433172_8446622_RNA45SN1_45S_50ntFlanks@9305.9335.31\|CCAAGGAGTCTAACACGTGCGCGAGTCGGGG | >hg38_21_+_8433172_8446622_RNA45SN1_45S_50ntFlanks | 9305 | 9335 | 31 | rRF 45S | NO | 0.00 | 0.00 | 43.06 | 13.52 |
| >hg38_21_+_8433172_8446622_RNA45SN1_45S_50ntFlanks@9305.9336.32\|CCAAGGAGTCTAACACGTGCGCGAGTCGGGGG | >hg38_21_+_8433172_8446622_RNA45SN1_45S_50ntFlanks | 9305 | 9336 | 32 | rRF 45S | NO | 0.00 | 0.00 | 83.43 | 24.36 |
| >hg38_21_+_8433172_8446622_RNA45SN1_45S_50ntFlanks@9305.9337.33\|CCAAGGAGTCTAACACGTGCGCGAGTCGGGGGC | >hg38_21_+_8433172_8446622_RNA45SN1_45S_50ntFlanks | 9305 | 9337 | 33 | rRF 45S | NO | 0.00 | 0.00 | 36.88 | 13.37 |
| >hg38_21_+_8433172_8446622_RNA45SN1_45S_50ntFlanks@9305.9338.34\|CCAAGGAGTCTAACACGTGCGCGAGTCGGGGGCT | >hg38_21_+_8433172_8446622_RNA45SN1_45S_50ntFlanks | 9305 | 9338 | 34 | rRF 45S | NO | 21.06 | 0.00 | 123.97 | 98.54 |
| >hg38_21_+_8433172_8446622_RNA45SN1_45S_50ntFlanks@9306.9323.18\|CAAGGAGTCTAACACGTG | >hg38_21_+_8433172_8446622_RNA45SN1_45S_50ntFlanks | 9306 | 9323 | 18 | rRF 45S | NO | 0.00 | 0.00 | 38.00 | 6.44 |
| >hg38_21_+_8433172_8446622_RNA45SN1_45S_50ntFlanks@9306.9335.30\|CAAGGAGTCTAACACGTGCGCGAGTCGGGG | >hg38_21_+_8433172_8446622_RNA45SN1_45S_50ntFlanks | 9306 | 9335 | 30 | rRF 45S | NO | 0.00 | 0.00 | 49.90 | 14.33 |
| >hg38_21_+_8433172_8446622_RNA45SN1_45S_50ntFlanks@9306.9336.31\|CAAGGAGTCTAACACGTGCGCGAGTCGGGGG | >hg38_21_+_8433172_8446622_RNA45SN1_45S_50ntFlanks | 9306 | 9336 | 31 | rRF 45S | NO | 0.00 | 0.00 | 60.53 | 0.00 |
| >hg38_21_+_8433172_8446622_RNA45SN1_45S_50ntFlanks@9306.9338.33\|CAAGGAGTCTAACACGTGCGCGAGTCGGGGGCT | >hg38_21_+_8433172_8446622_RNA45SN1_45S_50ntFlanks | 9306 | 9338 | 33 | rRF 45S | NO | 0.00 | 0.00 | 88.41 | 53.39 |
| >hg38_21_+_8433172_8446622_RNA45SN1_45S_50ntFlanks@9307.9338.32\|AAGGAGTCTAACACGTGCGCGAGTCGGGGGCT | >hg38_21_+_8433172_8446622_RNA45SN1_45S_50ntFlanks | 9307 | 9338 | 32 | rRF 45S | NO | 0.00 | 0.00 | 110.03 | 6.25 |
| >hg38_21_+_8433172_8446622_RNA45SN1_45S_50ntFlanks@9308.9336.29\|AGGAGTCTAACACGTGCGCGAGTCGGGGG | >hg38_21_+_8433172_8446622_RNA45SN1_45S_50ntFlanks | 9308 | 9336 | 29 | rRF 45S | NO | 0.00 | 0.00 | 120.31 | 0.00 |
| >hg38_21_+_8433172_8446622_RNA45SN1_45S_50ntFlanks@9308.9338.31\|AGGAGTCTAACACGTGCGCGAGTCGGGGGCT | >hg38_21_+_8433172_8446622_RNA45SN1_45S_50ntFlanks | 9308 | 9338 | 31 | rRF 45S | NO | 0.00 | 0.00 | 183.53 | 38.83 |
| >hg38_21_+_8433172_8446622_RNA45SN1_45S_50ntFlanks@9313.9338.26\|TCTAACACGTGCGCGAGTCGGGGGCT | >hg38_21_+_8433172_8446622_RNA45SN1_45S_50ntFlanks | 9313 | 9338 | 26 | rRF 45S | NO | 0.00 | 0.00 | 32.59 | 23.02 |
| >hg38_21_+_8433172_8446622_RNA45SN1_45S_50ntFlanks@9339.9367.29\|CGCACGAAAGCCGCCGTGGCGCAATGAAG | >hg38_21_+_8433172_8446622_RNA45SN1_45S_50ntFlanks | 9339 | 9367 | 29 | rRF 45S | NO | 0.00 | 0.00 | 30.16 | 0.00 |
| >hg38_21_+_8433172_8446622_RNA45SN1_45S_50ntFlanks@9339.9370.32\|CGCACGAAAGCCGCCGTGGCGCAATGAAGGTG | >hg38_21_+_8433172_8446622_RNA45SN1_45S_50ntFlanks | 9339 | 9370 | 32 | rRF 45S | NO | 0.00 | 0.00 | 316.71 | 25.10 |
| >hg38_21_+_8433172_8446622_RNA45SN1_45S_50ntFlanks@9343.9370.28\|CGAAAGCCGCCGTGGCGCAATGAAGGTG | >hg38_21_+_8433172_8446622_RNA45SN1_45S_50ntFlanks | 9343 | 9370 | 28 | rRF 45S | NO | 0.00 | 0.00 | 93.32 | 44.65 |
| >hg38_21_+_8433172_8446622_RNA45SN1_45S_50ntFlanks@9345.9370.26\|AAAGCCGCCGTGGCGCAATGAAGGTG | >hg38_21_+_8433172_8446622_RNA45SN1_45S_50ntFlanks | 9345 | 9370 | 26 | rRF 45S | NO | 0.00 | 0.00 | 45.66 | 14.62 |
| >hg38_21_+_8433172_8446622_RNA45SN1_45S_50ntFlanks@9352.9370.19\|CCGTGGCGCAATGAAGGTG | >hg38_21_+_8433172_8446622_RNA45SN1_45S_50ntFlanks | 9352 | 9370 | 19 | rRF 45S | NO | 43.95 | 0.00 | 172.87 | 127.54 |
| >hg38_21_+_8433172_8446622_RNA45SN1_45S_50ntFlanks@9353.9370.18\|CGTGGCGCAATGAAGGTG | >hg38_21_+_8433172_8446622_RNA45SN1_45S_50ntFlanks | 9353 | 9370 | 18 | rRF 45S | NO | 0.00 | 0.00 | 53.35 | 0.00 |
| >hg38_21_+_8433172_8446622_RNA45SN1_45S_50ntFlanks@9381.9409.29\|CGCTCGCCGGCCGAGGTGGGATCCCGAGG | >hg38_21_+_8433172_8446622_RNA45SN1_45S_50ntFlanks | 9381 | 9409 | 29 | rRF 45S | NO | 39.50 | 0.00 | 0.00 | 85.43 |
| >hg38_21_+_8433172_8446622_RNA45SN1_45S_50ntFlanks@9454.9488.35\|CCGCGCCGGGGAGGTGGAGCACGAGCGCACGTGTT | >hg38_21_+_8433172_8446622_RNA45SN1_45S_50ntFlanks | 9454 | 9488 | 35 | rRF 45S | NO | 0.00 | 0.00 | 42.21 | 25.41 |
| >hg38_21_+_8433172_8446622_RNA45SN1_45S_50ntFlanks@9454.9503.50\|CCGCGCCGGGGAGGTGGAGCACGAGCGCACGTGTTAGGACCCGAAAGATG | >hg38_21_+_8433172_8446622_RNA45SN1_45S_50ntFlanks | 9454 | 9503 | 50 | rRF 45S | NO | 148.90 | 0.00 | 7.28 | 188.91 |

### MDA-MB-468

| | | | | | | | | | | |
|---|---|---|---|---|---|---|---|---|---|---|
| >hg38_21_+_8433172_8446622_RNA45SN1_45S_50ntFlanks@9470.9503.34\|GAGCACGAGCGCACGTGTTAGGACCCGAAAGATG | >hg38_21_+_8433172_8446622_RNA45SN1_45S_50ntFlanks | 9470 | 9503 | 34 | rRF 45S | NO | 0.00 | 0.00 | 56.57 | 44.88 |
| >hg38_21_+_8433172_8446622_RNA45SN1_45S_50ntFlanks@9471.9503.33\|AGCACGAGCGCACGTGTTAGGACCCGAAAGATG | >hg38_21_+_8433172_8446622_RNA45SN1_45S_50ntFlanks | 9471 | 9503 | 33 | rRF 45S | NO | 0.00 | 0.00 | 61.02 | 58.72 |
| >hg38_21_+_8433172_8446622_RNA45SN1_45S_50ntFlanks@9503.9526.24\|GGTGAACTATGCCTGGGCAGGGCG | >hg38_21_+_8433172_8446622_RNA45SN1_45S_50ntFlanks | 9503 | 9526 | 24 | rRF 45S | NO | 0.00 | 0.00 | 27.81 | 0.00 |
| >hg38_21_+_8433172_8446622_RNA45SN1_45S_50ntFlanks@9504.9521.18\|GTGAACTATGCCTGGGCA | >hg38_21_+_8433172_8446622_RNA45SN1_45S_50ntFlanks | 9504 | 9521 | 18 | rRF 45S | NO | 105.53 | 222.15 | 41.16 | 33.53 |
| >hg38_21_+_8433172_8446622_RNA45SN1_45S_50ntFlanks@9504.9522.19\|GTGAACTATGCCTGGGCAG | >hg38_21_+_8433172_8446622_RNA45SN1_45S_50ntFlanks | 9504 | 9522 | 19 | rRF 45S | NO | 66.55 | 104.17 | 33.61 | 15.58 |
| >hg38_21_+_8433172_8446622_RNA45SN1_45S_50ntFlanks@9504.9523.20\|GTGAACTATGCCTGGGCAGG | >hg38_21_+_8433172_8446622_RNA45SN1_45S_50ntFlanks | 9504 | 9523 | 20 | rRF 45S | NO | 167.04 | 226.77 | 91.33 | 44.64 |
| >hg38_21_+_8433172_8446622_RNA45SN1_45S_50ntFlanks@9504.9524.21\|GTGAACTATGCCTGGGCAGGG | >hg38_21_+_8433172_8446622_RNA45SN1_45S_50ntFlanks | 9504 | 9524 | 21 | rRF 45S | NO | 37.87 | 58.01 | 34.20 | 0.00 |
| >hg38_21_+_8433172_8446622_RNA45SN1_45S_50ntFlanks@9504.9525.22\|GTGAACTATGCCTGGGCAGGGC | >hg38_21_+_8433172_8446622_RNA45SN1_45S_50ntFlanks | 9504 | 9525 | 22 | rRF 45S | NO | 34.08 | 105.33 | 93.69 | 36.49 |
| >hg38_21_+_8433172_8446622_RNA45SN1_45S_50ntFlanks@9504.9526.23\|GTGAACTATGCCTGGGCAGGGCG | >hg38_21_+_8433172_8446622_RNA45SN1_45S_50ntFlanks | 9504 | 9526 | 23 | rRF 45S | NO | 127.58 | 433.67 | 535.52 | 157.46 |
| >hg38_21_+_8433172_8446622_RNA45SN1_45S_50ntFlanks@9504.9527.24\|GTGAACTATGCCTGGGCAGGGCGA | >hg38_21_+_8433172_8446622_RNA45SN1_45S_50ntFlanks | 9504 | 9527 | 24 | rRF 45S | NO | 0.00 | 39.59 | 61.83 | 0.00 |
| >hg38_21_+_8433172_8446622_RNA45SN1_45S_50ntFlanks@9504.9529.26\|GTGAACTATGCCTGGGCAGGGCGAAG | >hg38_21_+_8433172_8446622_RNA45SN1_45S_50ntFlanks | 9504 | 9529 | 26 | rRF 45S | NO | 0.00 | 29.16 | 37.51 | 0.00 |
| >hg38_21_+_8433172_8446622_RNA45SN1_45S_50ntFlanks@9504.9533.30\|GTGAACTATGCCTGGGCAGGGCGAAGCCAG | >hg38_21_+_8433172_8446622_RNA45SN1_45S_50ntFlanks | 9504 | 9533 | 30 | rRF 45S | NO | 54.69 | 93.10 | 51.33 | 65.11 |
| >hg38_21_+_8433172_8446622_RNA45SN1_45S_50ntFlanks@9505.9526.22\|TGAACTATGCCTGGGCAGGGCG | >hg38_21_+_8433172_8446622_RNA45SN1_45S_50ntFlanks | 9505 | 9526 | 22 | rRF 45S | NO | 0.00 | 27.43 | 43.60 | 0.00 |
| >hg38_21_+_8433172_8446622_RNA45SN1_45S_50ntFlanks@9507.9526.20\|AACTATGCCTGGGCAGGGCG | >hg38_21_+_8433172_8446622_RNA45SN1_45S_50ntFlanks | 9507 | 9526 | 20 | rRF 45S | NO | 0.00 | 0.00 | 37.22 | 0.00 |
| >hg38_21_+_8433172_8446622_RNA45SN1_45S_50ntFlanks@9533.9550.18\|GAGGAAACTCTGGTGGAG | >hg38_21_+_8433172_8446622_RNA45SN1_45S_50ntFlanks | 9533 | 9550 | 18 | rRF 45S | NO | 12.81 | 26.32 | 0.00 | 0.00 |
| >hg38_21_+_8433172_8446622_RNA45SN1_45S_50ntFlanks@9533.9551.19\|GAGGAAACTCTGGTGGAGG | >hg38_21_+_8433172_8446622_RNA45SN1_45S_50ntFlanks | 9533 | 9551 | 19 | rRF 45S | NO | 22.09 | 54.06 | 0.00 | 0.00 |
| >hg38_21_+_8433172_8446622_RNA45SN1_45S_50ntFlanks@9533.9552.20\|GAGGAAACTCTGGTGGAGGT | >hg38_21_+_8433172_8446622_RNA45SN1_45S_50ntFlanks | 9533 | 9552 | 20 | rRF 45S | NO | 20.75 | 31.41 | 0.00 | 0.00 |
| >hg38_21_+_8433172_8446622_RNA45SN1_45S_50ntFlanks@9533.9553.21\|GAGGAAACTCTGGTGGAGGTC | >hg38_21_+_8433172_8446622_RNA45SN1_45S_50ntFlanks | 9533 | 9553 | 21 | rRF 45S | YES | 714.32 | 1275.03 | 98.13 | 37.37 |
| >hg38_21_+_8433172_8446622_RNA45SN1_45S_50ntFlanks@9533.9554.22\|GAGGAAACTCTGGTGGAGGTCC | >hg38_21_+_8433172_8446622_RNA45SN1_45S_50ntFlanks | 9533 | 9554 | 22 | rRF 45S | NO | 0.00 | 32.59 | 0.00 | 0.00 |
| >hg38_21_+_8433172_8446622_RNA45SN1_45S_50ntFlanks@9533.9561.29\|GAGGAAACTCTGGTGGAGGTCCGTAGCGG | >hg38_21_+_8433172_8446622_RNA45SN1_45S_50ntFlanks | 9533 | 9561 | 29 | rRF 45S | NO | 52.69 | 36.09 | 0.00 | 0.00 |
| >hg38_21_+_8433172_8446622_RNA45SN1_45S_50ntFlanks@9533.9562.30\|GAGGAAACTCTGGTGGAGGTCCGTAGCGGT | >hg38_21_+_8433172_8446622_RNA45SN1_45S_50ntFlanks | 9533 | 9562 | 30 | rRF 45S | NO | 39.75 | 28.70 | 0.00 | 0.00 |
| >hg38_21_+_8433172_8446622_RNA45SN1_45S_50ntFlanks@9533.9563.31\|GAGGAAACTCTGGTGGAGGTCCGTAGCGGTC | >hg38_21_+_8433172_8446622_RNA45SN1_45S_50ntFlanks | 9533 | 9563 | 31 | rRF 45S | NO | 259.89 | 229.56 | 44.46 | 0.00 |
| >hg38_21_+_8433172_8446622_RNA45SN1_45S_50ntFlanks@9533.9564.32\|GAGGAAACTCTGGTGGAGGTCCGTAGCGGTCC | >hg38_21_+_8433172_8446622_RNA45SN1_45S_50ntFlanks | 9533 | 9564 | 32 | rRF 45S | NO | 53.43 | 0.00 | 0.00 | 0.00 |
| >hg38_21_+_8433172_8446622_RNA45SN1_45S_50ntFlanks@9533.9566.34\|GAGGAAACTCTGGTGGAGGTCCGTAGCGGTCCTG | >hg38_21_+_8433172_8446622_RNA45SN1_45S_50ntFlanks | 9533 | 9566 | 34 | rRF 45S | NO | 39.18 | 26.50 | 0.00 | 0.00 |
| >hg38_21_+_8433172_8446622_RNA45SN1_45S_50ntFlanks@9533.9569.37\|GAGGAAACTCTGGTGGAGGTCCGTAGCGGTCCTGACG | >hg38_21_+_8433172_8446622_RNA45SN1_45S_50ntFlanks | 9533 | 9569 | 37 | rRF 45S | NO | 51.34 | 39.45 | 0.00 | 0.00 |
| >hg38_21_+_8433172_8446622_RNA45SN1_45S_50ntFlanks@9533.9570.38\|GAGGAAACTCTGGTGGAGGTCCGTAGCGGTCCTGACGT | >hg38_21_+_8433172_8446622_RNA45SN1_45S_50ntFlanks | 9533 | 9570 | 38 | rRF 45S | NO | 50.79 | 12.29 | 0.00 | 0.00 |
| >hg38_21_+_8433172_8446622_RNA45SN1_45S_50ntFlanks@9533.9571.39\|GAGGAAACTCTGGTGGAGGTCCGTAGCGGTCCTGACGTG | >hg38_21_+_8433172_8446622_RNA45SN1_45S_50ntFlanks | 9533 | 9571 | 39 | rRF 45S | NO | 624.51 | 187.73 | 34.92 | 39.98 |
| >hg38_21_+_8433172_8446622_RNA45SN1_45S_50ntFlanks@9533.9572.40\|GAGGAAACTCTGGTGGAGGTCCGTAGCGGTCCTGACGTGC | >hg38_21_+_8433172_8446622_RNA45SN1_45S_50ntFlanks | 9533 | 9572 | 40 | rRF 45S | NO | 85.21 | 0.00 | 0.00 | 0.00 |
| >hg38_21_+_8433172_8446622_RNA45SN1_45S_50ntFlanks@9534.9551.18\|AGGAAACTCTGGTGGAGG | >hg38_21_+_8433172_8446622_RNA45SN1_45S_50ntFlanks | 9534 | 9551 | 18 | rRF 45S | NO | 276.19 | 524.06 | 107.44 | 16.59 |
| >hg38_21_+_8433172_8446622_RNA45SN1_45S_50ntFlanks@9534.9552.19\|AGGAAACTCTGGTGGAGGT | >hg38_21_+_8433172_8446622_RNA45SN1_45S_50ntFlanks | 9534 | 9552 | 19 | rRF 45S | NO | 242.48 | 280.70 | 64.81 | 0.00 |
| >hg38_21_+_8433172_8446622_RNA45SN1_45S_50ntFlanks@9534.9553.20\|AGGAAACTCTGGTGGAGGTC | >hg38_21_+_8433172_8446622_RNA45SN1_45S_50ntFlanks | 9534 | 9553 | 20 | rRF 45S | YES | 4861.67 | 7658.77 | 703.85 | 239.08 |
| >hg38_21_+_8433172_8446622_RNA45SN1_45S_50ntFlanks@9534.9554.21\|AGGAAACTCTGGTGGAGGTCC | >hg38_21_+_8433172_8446622_RNA45SN1_45S_50ntFlanks | 9534 | 9554 | 21 | rRF 45S | NO | 0.00 | 36.08 | 0.00 | 0.00 |
| >hg38_21_+_8433172_8446622_RNA45SN1_45S_50ntFlanks@9534.9558.25\|AGGAAACTCTGGTGGAGGTCCGTAG | >hg38_21_+_8433172_8446622_RNA45SN1_45S_50ntFlanks | 9534 | 9558 | 25 | rRF 45S | NO | 0.00 | 44.94 | 34.64 | 0.00 |
| >hg38_21_+_8433172_8446622_RNA45SN1_45S_50ntFlanks@9534.9559.26\|AGGAAACTCTGGTGGAGGTCCGTAGC | >hg38_21_+_8433172_8446622_RNA45SN1_45S_50ntFlanks | 9534 | 9559 | 26 | rRF 45S | NO | 0.00 | 0.00 | 52.18 | 0.00 |

### MDA-MB-468

| | | | | | | | | | | |
|---|---|---|---|---|---|---|---|---|---|---|
| >hg38_21_+_8433172_8446622_RNA45SN1_45S_50ntFlanks@9534.9560.27\|AGGAAACTCTGGTGGAGGTCCGTAGCG | >hg38_21_+_8433172_8446622_RNA45SN1_45S_50ntFlanks | 9534 | 9560 | 27 | rRF 45S | NO | 43.67 | 63.18 | 46.74 | 0.00 |
| >hg38_21_+_8433172_8446622_RNA45SN1_45S_50ntFlanks@9534.9561.28\|AGGAAACTCTGGTGGAGGTCCGTAGCGG | >hg38_21_+_8433172_8446622_RNA45SN1_45S_50ntFlanks | 9534 | 9561 | 28 | rRF 45S | NO | 307.94 | 226.06 | 82.11 | 42.03 |
| >hg38_21_+_8433172_8446622_RNA45SN1_45S_50ntFlanks@9534.9562.29\|AGGAAACTCTGGTGGAGGTCCGTAGCGGT | >hg38_21_+_8433172_8446622_RNA45SN1_45S_50ntFlanks | 9534 | 9562 | 29 | rRF 45S | NO | 215.57 | 177.50 | 78.98 | 36.07 |
| >hg38_21_+_8433172_8446622_RNA45SN1_45S_50ntFlanks@9534.9563.30\|AGGAAACTCTGGTGGAGGTCCGTAGCGGTC | >hg38_21_+_8433172_8446622_RNA45SN1_45S_50ntFlanks | 9534 | 9563 | 30 | rRF 45S | YES | 1607.09 | 1494.83 | 304.59 | 146.44 |
| >hg38_21_+_8433172_8446622_RNA45SN1_45S_50ntFlanks@9534.9564.31\|AGGAAACTCTGGTGGAGGTCCGTAGCGGTCC | >hg38_21_+_8433172_8446622_RNA45SN1_45S_50ntFlanks | 9534 | 9564 | 31 | rRF 45S | NO | 328.81 | 145.40 | 37.28 | 14.18 |
| >hg38_21_+_8433172_8446622_RNA45SN1_45S_50ntFlanks@9534.9565.32\|AGGAAACTCTGGTGGAGGTCCGTAGCGGTCCT | >hg38_21_+_8433172_8446622_RNA45SN1_45S_50ntFlanks | 9534 | 9565 | 32 | rRF 45S | NO | 184.42 | 111.73 | 0.00 | 14.87 |
| >hg38_21_+_8433172_8446622_RNA45SN1_45S_50ntFlanks@9534.9566.33\|AGGAAACTCTGGTGGAGGTCCGTAGCGGTCCTG | >hg38_21_+_8433172_8446622_RNA45SN1_45S_50ntFlanks | 9534 | 9566 | 33 | rRF 45S | NO | 277.49 | 221.85 | 49.91 | 42.13 |
| >hg38_21_+_8433172_8446622_RNA45SN1_45S_50ntFlanks@9534.9567.34\|AGGAAACTCTGGTGGAGGTCCGTAGCGGTCCTGA | >hg38_21_+_8433172_8446622_RNA45SN1_45S_50ntFlanks | 9534 | 9567 | 34 | rRF 45S | NO | 131.53 | 89.93 | 0.00 | 0.00 |
| >hg38_21_+_8433172_8446622_RNA45SN1_45S_50ntFlanks@9534.9568.35\|AGGAAACTCTGGTGGAGGTCCGTAGCGGTCCTGAC | >hg38_21_+_8433172_8446622_RNA45SN1_45S_50ntFlanks | 9534 | 9568 | 35 | rRF 45S | NO | 47.41 | 36.46 | 0.00 | 0.00 |
| >hg38_21_+_8433172_8446622_RNA45SN1_45S_50ntFlanks@9534.9569.36\|AGGAAACTCTGGTGGAGGTCCGTAGCGGTCCTGACG | >hg38_21_+_8433172_8446622_RNA45SN1_45S_50ntFlanks | 9534 | 9569 | 36 | rRF 45S | NO | 438.91 | 339.30 | 46.47 | 56.02 |
| >hg38_21_+_8433172_8446622_RNA45SN1_45S_50ntFlanks@9534.9570.37\|AGGAAACTCTGGTGGAGGTCCGTAGCGGTCCTGACGT | >hg38_21_+_8433172_8446622_RNA45SN1_45S_50ntFlanks | 9534 | 9570 | 37 | rRF 45S | NO | 350.65 | 203.08 | 20.80 | 33.14 |
| >hg38_21_+_8433172_8446622_RNA45SN1_45S_50ntFlanks@9534.9571.38\|AGGAAACTCTGGTGGAGGTCCGTAGCGGTCCTGACGTG | >hg38_21_+_8433172_8446622_RNA45SN1_45S_50ntFlanks | 9534 | 9571 | 38 | rRF 45S | YES | 3886.51 | 2257.44 | 206.74 | 250.03 |
| >hg38_21_+_8433172_8446622_RNA45SN1_45S_50ntFlanks@9534.9572.39\|AGGAAACTCTGGTGGAGGTCCGTAGCGGTCCTGACGTGC | >hg38_21_+_8433172_8446622_RNA45SN1_45S_50ntFlanks | 9534 | 9572 | 39 | rRF 45S | NO | 531.46 | 284.56 | 28.86 | 51.17 |
| >hg38_21_+_8433172_8446622_RNA45SN1_45S_50ntFlanks@9534.9573.40\|AGGAAACTCTGGTGGAGGTCCGTAGCGGTCCTGACGTGCA | >hg38_21_+_8433172_8446622_RNA45SN1_45S_50ntFlanks | 9534 | 9573 | 40 | rRF 45S | NO | 203.14 | 90.17 | 0.00 | 34.61 |
| >hg38_21_+_8433172_8446622_RNA45SN1_45S_50ntFlanks@9534.9574.41\|AGGAAACTCTGGTGGAGGTCCGTAGCGGTCCTGACGTGCAA | >hg38_21_+_8433172_8446622_RNA45SN1_45S_50ntFlanks | 9534 | 9574 | 41 | rRF 45S | NO | 123.00 | 42.49 | 0.00 | 0.00 |
| >hg38_21_+_8433172_8446622_RNA45SN1_45S_50ntFlanks@9535.9553.19\|GGAAACTCTGGTGGAGGTC | >hg38_21_+_8433172_8446622_RNA45SN1_45S_50ntFlanks | 9535 | 9553 | 19 | rRF 45S | NO | 0.00 | 34.99 | 0.00 | 0.00 |
| >hg38_21_+_8433172_8446622_RNA45SN1_45S_50ntFlanks@9535.9571.37\|GGAAACTCTGGTGGAGGTCCGTAGCGGTCCTGACGTG | >hg38_21_+_8433172_8446622_RNA45SN1_45S_50ntFlanks | 9535 | 9571 | 37 | rRF 45S | NO | 97.17 | 39.32 | 0.00 | 0.00 |
| >hg38_21_+_8433172_8446622_RNA45SN1_45S_50ntFlanks@9536.9571.36\|GAAACTCTGGTGGAGGTCCGTAGCGGTCCTGACGTG | >hg38_21_+_8433172_8446622_RNA45SN1_45S_50ntFlanks | 9536 | 9571 | 36 | rRF 45S | NO | 89.75 | 37.99 | 0.00 | 0.00 |
| >hg38_21_+_8433172_8446622_RNA45SN1_45S_50ntFlanks@9537.9571.35\|AAACTCTGGTGGAGGTCCGTAGCGGTCCTGACGTG | >hg38_21_+_8433172_8446622_RNA45SN1_45S_50ntFlanks | 9537 | 9571 | 35 | rRF 45S | NO | 76.84 | 29.65 | 0.00 | 0.00 |
| >hg38_21_+_8433172_8446622_RNA45SN1_45S_50ntFlanks@9538.9571.34\|AACTCTGGTGGAGGTCCGTAGCGGTCCTGACGTG | >hg38_21_+_8433172_8446622_RNA45SN1_45S_50ntFlanks | 9538 | 9571 | 34 | rRF 45S | NO | 129.07 | 47.52 | 0.00 | 0.00 |
| >hg38_21_+_8433172_8446622_RNA45SN1_45S_50ntFlanks@9539.9563.25\|ACTCTGGTGGAGGTCCGTAGCGGTC | >hg38_21_+_8433172_8446622_RNA45SN1_45S_50ntFlanks | 9539 | 9563 | 25 | rRF 45S | NO | 32.43 | 0.00 | 0.00 | 0.00 |
| >hg38_21_+_8433172_8446622_RNA45SN1_45S_50ntFlanks@9539.9569.31\|ACTCTGGTGGAGGTCCGTAGCGGTCCTGACG | >hg38_21_+_8433172_8446622_RNA45SN1_45S_50ntFlanks | 9539 | 9569 | 31 | rRF 45S | NO | 36.13 | 0.00 | 0.00 | 0.00 |
| >hg38_21_+_8433172_8446622_RNA45SN1_45S_50ntFlanks@9539.9571.33\|ACTCTGGTGGAGGTCCGTAGCGGTCCTGACGTG | >hg38_21_+_8433172_8446622_RNA45SN1_45S_50ntFlanks | 9539 | 9571 | 33 | rRF 45S | NO | 240.81 | 94.67 | 0.00 | 5.94 |
| >hg38_21_+_8433172_8446622_RNA45SN1_45S_50ntFlanks@9540.9571.32\|CTCTGGTGGAGGTCCGTAGCGGTCCTGACGTG | >hg38_21_+_8433172_8446622_RNA45SN1_45S_50ntFlanks | 9540 | 9571 | 32 | rRF 45S | NO | 126.14 | 66.39 | 0.00 | 0.00 |
| >hg38_21_+_8433172_8446622_RNA45SN1_45S_50ntFlanks@9543.9571.29\|TGGTGGAGGTCCGTAGCGGTCCTGACGTG | >hg38_21_+_8433172_8446622_RNA45SN1_45S_50ntFlanks | 9543 | 9571 | 29 | rRF 45S | NO | 36.90 | 5.96 | 0.00 | 0.00 |
| >hg38_21_+_8433172_8446622_RNA45SN1_45S_50ntFlanks@9544.9563.20\|GGTGGAGGTCCGTAGCGGTC | >hg38_21_+_8433172_8446622_RNA45SN1_45S_50ntFlanks | 9544 | 9563 | 20 | rRF 45S | NO | 34.65 | 31.83 | 0.00 | 0.00 |
| >hg38_21_+_8433172_8446622_RNA45SN1_45S_50ntFlanks@9544.9569.26\|GGTGGAGGTCCGTAGCGGTCCTGACG | >hg38_21_+_8433172_8446622_RNA45SN1_45S_50ntFlanks | 9544 | 9569 | 26 | rRF 45S | NO | 0.00 | 12.72 | 0.00 | 0.00 |
| >hg38_21_+_8433172_8446622_RNA45SN1_45S_50ntFlanks@9544.9570.27\|GGTGGAGGTCCGTAGCGGTCCTGACGT | >hg38_21_+_8433172_8446622_RNA45SN1_45S_50ntFlanks | 9544 | 9570 | 27 | rRF 45S | NO | 32.67 | 31.98 | 0.00 | 0.00 |
| >hg38_21_+_8433172_8446622_RNA45SN1_45S_50ntFlanks@9544.9571.28\|GGTGGAGGTCCGTAGCGGTCCTGACGTG | >hg38_21_+_8433172_8446622_RNA45SN1_45S_50ntFlanks | 9544 | 9571 | 28 | rRF 45S | NO | 378.41 | 180.29 | 30.38 | 23.81 |
| >hg38_21_+_8433172_8446622_RNA45SN1_45S_50ntFlanks@9544.9572.29\|GGTGGAGGTCCGTAGCGGTCCTGACGTGC | >hg38_21_+_8433172_8446622_RNA45SN1_45S_50ntFlanks | 9544 | 9572 | 29 | rRF 45S | NO | 61.68 | 12.64 | 0.00 | 0.00 |
| >hg38_21_+_8433172_8446622_RNA45SN1_45S_50ntFlanks@9544.9573.30\|GGTGGAGGTCCGTAGCGGTCCTGACGTGCA | >hg38_21_+_8433172_8446622_RNA45SN1_45S_50ntFlanks | 9544 | 9573 | 30 | rRF 45S | NO | 38.26 | 0.00 | 0.00 | 0.00 |
| >hg38_21_+_8433172_8446622_RNA45SN1_45S_50ntFlanks@9544.9574.31\|GGTGGAGGTCCGTAGCGGTCCTGACGTGCAA | >hg38_21_+_8433172_8446622_RNA45SN1_45S_50ntFlanks | 9544 | 9574 | 31 | rRF 45S | NO | 12.59 | 0.00 | 0.00 | 0.00 |
| >hg38_21_+_8433172_8446622_RNA45SN1_45S_50ntFlanks@9545.9562.18\|GTGGAGGTCCGTAGCGGT | >hg38_21_+_8433172_8446622_RNA45SN1_45S_50ntFlanks | 9545 | 9562 | 18 | rRF 45S | NO | 37.41 | 29.98 | 0.00 | 0.00 |
| >hg38_21_+_8433172_8446622_RNA45SN1_45S_50ntFlanks@9545.9563.19\|GTGGAGGTCCGTAGCGGTC | >hg38_21_+_8433172_8446622_RNA45SN1_45S_50ntFlanks | 9545 | 9563 | 19 | rRF 45S | NO | 523.00 | 414.69 | 47.29 | 26.72 |

### MDA-MB-468

| | | | | | | | | | | |
|---|---|---|---|---|---|---|---|---|---|---|
| >hg38_21_+_8433172_8446622_RNA45SN1_45S_50ntFlanks@9545.9564.20\|GTGGAGGTCCGTAGCGGTCC | >hg38_21_+_8433172_8446622_RNA45SN1_45S_50ntFlanks | 9545 | 9564 | 20 | rRF 45S | NO | 68.67 | 68.85 | 0.00 | 0.00 |
| >hg38_21_+_8433172_8446622_RNA45SN1_45S_50ntFlanks@9545.9566.22\|GTGGAGGTCCGTAGCGGTCCTG | >hg38_21_+_8433172_8446622_RNA45SN1_45S_50ntFlanks | 9545 | 9566 | 22 | rRF 45S | NO | 65.66 | 233.14 | 35.30 | 6.10 |
| >hg38_21_+_8433172_8446622_RNA45SN1_45S_50ntFlanks@9545.9567.23\|GTGGAGGTCCGTAGCGGTCCTGA | >hg38_21_+_8433172_8446622_RNA45SN1_45S_50ntFlanks | 9545 | 9567 | 23 | rRF 45S | NO | 0.00 | 43.34 | 0.00 | 0.00 |
| >hg38_21_+_8433172_8446622_RNA45SN1_45S_50ntFlanks@9545.9569.25\|GTGGAGGTCCGTAGCGGTCCTGACG | >hg38_21_+_8433172_8446622_RNA45SN1_45S_50ntFlanks | 9545 | 9569 | 25 | rRF 45S | NO | 0.00 | 89.32 | 20.56 | 6.34 |
| >hg38_21_+_8433172_8446622_RNA45SN1_45S_50ntFlanks@9545.9570.26\|GTGGAGGTCCGTAGCGGTCCTGACGT | >hg38_21_+_8433172_8446622_RNA45SN1_45S_50ntFlanks | 9545 | 9570 | 26 | rRF 45S | NO | 47.71 | 197.93 | 13.55 | 13.28 |
| >hg38_21_+_8433172_8446622_RNA45SN1_45S_50ntFlanks@9545.9571.27\|GTGGAGGTCCGTAGCGGTCCTGACGTG | >hg38_21_+_8433172_8446622_RNA45SN1_45S_50ntFlanks | 9545 | 9571 | 27 | rRF 45S | YES | 789.06 | 2069.68 | 169.92 | 172.25 |
| >hg38_21_+_8433172_8446622_RNA45SN1_45S_50ntFlanks@9545.9572.28\|GTGGAGGTCCGTAGCGGTCCTGACGTGC | >hg38_21_+_8433172_8446622_RNA45SN1_45S_50ntFlanks | 9545 | 9572 | 28 | rRF 45S | NO | 141.99 | 244.64 | 0.00 | 34.69 |
| >hg38_21_+_8433172_8446622_RNA45SN1_45S_50ntFlanks@9545.9573.29\|GTGGAGGTCCGTAGCGGTCCTGACGTGCA | >hg38_21_+_8433172_8446622_RNA45SN1_45S_50ntFlanks | 9545 | 9573 | 29 | rRF 45S | NO | 132.27 | 147.37 | 0.00 | 45.28 |
| >hg38_21_+_8433172_8446622_RNA45SN1_45S_50ntFlanks@9545.9574.30\|GTGGAGGTCCGTAGCGGTCCTGACGTGCAA | >hg38_21_+_8433172_8446622_RNA45SN1_45S_50ntFlanks | 9545 | 9574 | 30 | rRF 45S | NO | 152.13 | 82.62 | 0.00 | 23.62 |
| >hg38_21_+_8433172_8446622_RNA45SN1_45S_50ntFlanks@9545.9575.31\|GTGGAGGTCCGTAGCGGTCCTGACGTGCAAA | >hg38_21_+_8433172_8446622_RNA45SN1_45S_50ntFlanks | 9545 | 9575 | 31 | rRF 45S | NO | 31.41 | 0.00 | 0.00 | 0.00 |
| >hg38_21_+_8433172_8446622_RNA45SN1_45S_50ntFlanks@9546.9571.26\|TGGAGGTCCGTAGCGGTCCTGACGTG | >hg38_21_+_8433172_8446622_RNA45SN1_45S_50ntFlanks | 9546 | 9571 | 26 | rRF 45S | NO | 45.08 | 62.86 | 0.00 | 0.00 |
| >hg38_21_+_8433172_8446622_RNA45SN1_45S_50ntFlanks@9548.9571.24\|GAGGTCCGTAGCGGTCCTGACGTG | >hg38_21_+_8433172_8446622_RNA45SN1_45S_50ntFlanks | 9548 | 9571 | 24 | rRF 45S | NO | 35.91 | 99.26 | 0.00 | 15.08 |
| >hg38_21_+_8433172_8446622_RNA45SN1_45S_50ntFlanks@9548.9572.25\|GAGGTCCGTAGCGGTCCTGACGTGC | >hg38_21_+_8433172_8446622_RNA45SN1_45S_50ntFlanks | 9548 | 9572 | 25 | rRF 45S | NO | 0.00 | 12.54 | 0.00 | 0.00 |
| >hg38_21_+_8433172_8446622_RNA45SN1_45S_50ntFlanks@9548.9573.26\|GAGGTCCGTAGCGGTCCTGACGTGCA | >hg38_21_+_8433172_8446622_RNA45SN1_45S_50ntFlanks | 9548 | 9573 | 26 | rRF 45S | NO | 12.64 | 0.00 | 0.00 | 0.00 |
| >hg38_21_+_8433172_8446622_RNA45SN1_45S_50ntFlanks@9549.9571.23\|AGGTCCGTAGCGGTCCTGACGTG | >hg38_21_+_8433172_8446622_RNA45SN1_45S_50ntFlanks | 9549 | 9571 | 23 | rRF 45S | NO | 0.00 | 46.94 | 0.00 | 0.00 |
| >hg38_21_+_8433172_8446622_RNA45SN1_45S_50ntFlanks@9551.9571.21\|GTCCGTAGCGGTCCTGACGTG | >hg38_21_+_8433172_8446622_RNA45SN1_45S_50ntFlanks | 9551 | 9571 | 21 | rRF 45S | NO | 0.00 | 12.51 | 0.00 | 0.00 |
| >hg38_21_+_8433172_8446622_RNA45SN1_45S_50ntFlanks@9552.9569.18\|TCCGTAGCGGTCCTGACG | >hg38_21_+_8433172_8446622_RNA45SN1_45S_50ntFlanks | 9552 | 9569 | 18 | rRF 45S | NO | 31.62 | 30.60 | 0.00 | 0.00 |
| >hg38_21_+_8433172_8446622_RNA45SN1_45S_50ntFlanks@9552.9571.20\|TCCGTAGCGGTCCTGACGTG | >hg38_21_+_8433172_8446622_RNA45SN1_45S_50ntFlanks | 9552 | 9571 | 20 | rRF 45S | NO | 39.61 | 80.07 | 0.00 | 0.00 |
| >hg38_21_+_8433172_8446622_RNA45SN1_45S_50ntFlanks@9552.9572.21\|TCCGTAGCGGTCCTGACGTGC | >hg38_21_+_8433172_8446622_RNA45SN1_45S_50ntFlanks | 9552 | 9572 | 21 | rRF 45S | NO | 0.00 | 12.43 | 0.00 | 0.00 |
| >hg38_21_+_8433172_8446622_RNA45SN1_45S_50ntFlanks@9554.9571.18\|CGTAGCGGTCCTGACGTG | >hg38_21_+_8433172_8446622_RNA45SN1_45S_50ntFlanks | 9554 | 9571 | 18 | rRF 45S | YES | 7186.36 | 14325.55 | 890.04 | 1201.74 |
| >hg38_21_+_8433172_8446622_RNA45SN1_45S_50ntFlanks@9554.9572.19\|CGTAGCGGTCCTGACGTGC | >hg38_21_+_8433172_8446622_RNA45SN1_45S_50ntFlanks | 9554 | 9572 | 19 | rRF 45S | YES | 798.98 | 1499.62 | 129.17 | 163.01 |
| >hg38_21_+_8433172_8446622_RNA45SN1_45S_50ntFlanks@9554.9573.20\|CGTAGCGGTCCTGACGTGCA | >hg38_21_+_8433172_8446622_RNA45SN1_45S_50ntFlanks | 9554 | 9573 | 20 | rRF 45S | YES | 704.93 | 989.98 | 110.46 | 180.63 |
| >hg38_21_+_8433172_8446622_RNA45SN1_45S_50ntFlanks@9554.9574.21\|CGTAGCGGTCCTGACGTGCAA | >hg38_21_+_8433172_8446622_RNA45SN1_45S_50ntFlanks | 9554 | 9574 | 21 | rRF 45S | NO | 156.15 | 252.80 | 63.14 | 50.03 |
| >hg38_21_+_8433172_8446622_RNA45SN1_45S_50ntFlanks@9554.9575.22\|CGTAGCGGTCCTGACGTGCAAA | >hg38_21_+_8433172_8446622_RNA45SN1_45S_50ntFlanks | 9554 | 9575 | 22 | rRF 45S | NO | 40.42 | 62.10 | 31.92 | 38.26 |
| >hg38_21_+_8433172_8446622_RNA45SN1_45S_50ntFlanks@9554.9576.23\|CGTAGCGGTCCTGACGTGCAAAT | >hg38_21_+_8433172_8446622_RNA45SN1_45S_50ntFlanks | 9554 | 9576 | 23 | rRF 45S | NO | 0.00 | 27.71 | 0.00 | 0.00 |
| >hg38_21_+_8433172_8446622_RNA45SN1_45S_50ntFlanks@9554.9577.24\|CGTAGCGGTCCTGACGTGCAAATC | >hg38_21_+_8433172_8446622_RNA45SN1_45S_50ntFlanks | 9554 | 9577 | 24 | rRF 45S | NO | 66.84 | 44.79 | 0.00 | 0.00 |
| >hg38_21_+_8433172_8446622_RNA45SN1_45S_50ntFlanks@9554.9579.26\|CGTAGCGGTCCTGACGTGCAAATCGG | >hg38_21_+_8433172_8446622_RNA45SN1_45S_50ntFlanks | 9554 | 9579 | 26 | rRF 45S | NO | 21.72 | 28.83 | 0.00 | 0.00 |
| >hg38_21_+_8433172_8446622_RNA45SN1_45S_50ntFlanks@9554.9580.27\|CGTAGCGGTCCTGACGTGCAAATCGGT | >hg38_21_+_8433172_8446622_RNA45SN1_45S_50ntFlanks | 9554 | 9580 | 27 | rRF 45S | NO | 29.24 | 33.23 | 0.00 | 0.00 |
| >hg38_21_+_8433172_8446622_RNA45SN1_45S_50ntFlanks@9554.9581.28\|CGTAGCGGTCCTGACGTGCAAATCGGTC | >hg38_21_+_8433172_8446622_RNA45SN1_45S_50ntFlanks | 9554 | 9581 | 28 | rRF 45S | NO | 48.61 | 32.33 | 0.00 | 0.00 |
| >hg38_21_+_8433172_8446622_RNA45SN1_45S_50ntFlanks@9554.9582.29\|CGTAGCGGTCCTGACGTGCAAATCGGTCG | >hg38_21_+_8433172_8446622_RNA45SN1_45S_50ntFlanks | 9554 | 9582 | 29 | rRF 45S | NO | 70.81 | 43.12 | 0.00 | 0.00 |
| >hg38_21_+_8433172_8446622_RNA45SN1_45S_50ntFlanks@9554.9584.31\|CGTAGCGGTCCTGACGTGCAAATCGGTCGTC | >hg38_21_+_8433172_8446622_RNA45SN1_45S_50ntFlanks | 9554 | 9584 | 31 | rRF 45S | NO | 261.11 | 234.29 | 0.00 | 51.40 |
| >hg38_21_+_8433172_8446622_RNA45SN1_45S_50ntFlanks@9554.9595.42\|CGTAGCGGTCCTGACGTGCAAATCGGTCGTCCGACCTGGGTA | >hg38_21_+_8433172_8446622_RNA45SN1_45S_50ntFlanks | 9554 | 9595 | 42 | rRF 45S | NO | 6.18 | 19.36 | 0.00 | 38.97 |
| >hg38_21_+_8433172_8446622_RNA45SN1_45S_50ntFlanks@9554.9597.44\|CGTAGCGGTCCTGACGTGCAAATCGGTCGTCCGACCTGGGTATA | >hg38_21_+_8433172_8446622_RNA45SN1_45S_50ntFlanks | 9554 | 9597 | 44 | rRF 45S | NO | 87.15 | 37.51 | 0.00 | 146.65 |
| >hg38_21_+_8433172_8446622_RNA45SN1_45S_50ntFlanks@9556.9573.18\|TAGCGGTCCTGACGTGCA | >hg38_21_+_8433172_8446622_RNA45SN1_45S_50ntFlanks | 9556 | 9573 | 18 | rRF 45S | NO | 0.00 | 36.03 | 0.00 | 0.00 |

### MDA-MB-468

<table>
<tr><td>>hg38_21_+_8433172_8446622_RNA45SN1_45S_50ntFlanks@9556.9582.27|TAGCGGTCCTGACGTGCAAATCGGTCG</td><td>>hg38_21_+_8433172_8446622_RNA45SN1_45S_50ntFlanks</td><td>9556</td><td>9582</td><td>27</td><td>rRF 45S</td><td>NO</td><td>53.80</td><td>44.40</td><td>0.00</td><td>0.00</td></tr>
<tr><td>>hg38_21_+_8433172_8446622_RNA45SN1_45S_50ntFlanks@9562.9595.34|TCCTGACGTGCAAATCGGTCGTCCGACCTGGGTA</td><td>>hg38_21_+_8433172_8446622_RNA45SN1_45S_50ntFlanks</td><td>9562</td><td>9595</td><td>34</td><td>rRF 45S</td><td>NO</td><td>0.00</td><td>27.46</td><td>0.00</td><td>33.77</td></tr>
<tr><td>>hg38_21_+_8433172_8446622_RNA45SN1_45S_50ntFlanks@9562.9596.35|TCCTGACGTGCAAATCGGTCGTCCGACCTGGGTAT</td><td>>hg38_21_+_8433172_8446622_RNA45SN1_45S_50ntFlanks</td><td>9562</td><td>9596</td><td>35</td><td>rRF 45S</td><td>NO</td><td>97.93</td><td>86.30</td><td>0.00</td><td>168.43</td></tr>
<tr><td>>hg38_21_+_8433172_8446622_RNA45SN1_45S_50ntFlanks@9562.9597.36|TCCTGACGTGCAAATCGGTCGTCCGACCTGGGTATA</td><td>>hg38_21_+_8433172_8446622_RNA45SN1_45S_50ntFlanks</td><td>9562</td><td>9597</td><td>36</td><td>rRF 45S</td><td>NO</td><td>50.56</td><td>51.03</td><td>0.00</td><td>124.98</td></tr>
<tr><td>>hg38_21_+_8433172_8446622_RNA45SN1_45S_50ntFlanks@9563.9596.34|CCTGACGTGCAAATCGGTCGTCCGACCTGGGTAT</td><td>>hg38_21_+_8433172_8446622_RNA45SN1_45S_50ntFlanks</td><td>9563</td><td>9596</td><td>34</td><td>rRF 45S</td><td>NO</td><td>120.64</td><td>98.97</td><td>0.00</td><td>210.82</td></tr>
<tr><td>>hg38_21_+_8433172_8446622_RNA45SN1_45S_50ntFlanks@9563.9597.35|CCTGACGTGCAAATCGGTCGTCCGACCTGGGTATA</td><td>>hg38_21_+_8433172_8446622_RNA45SN1_45S_50ntFlanks</td><td>9563</td><td>9597</td><td>35</td><td>rRF 45S</td><td>NO</td><td>44.75</td><td>42.38</td><td>0.00</td><td>114.84</td></tr>
<tr><td>>hg38_21_+_8433172_8446622_RNA45SN1_45S_50ntFlanks@9564.9592.29|CTGACGTGCAAATCGGTCGTCCGACCTGG</td><td>>hg38_21_+_8433172_8446622_RNA45SN1_45S_50ntFlanks</td><td>9564</td><td>9592</td><td>29</td><td>rRF 45S</td><td>NO</td><td>90.48</td><td>242.46</td><td>33.51</td><td>40.87</td></tr>
<tr><td>>hg38_21_+_8433172_8446622_RNA45SN1_45S_50ntFlanks@9564.9594.31|CTGACGTGCAAATCGGTCGTCCGACCTGGGT</td><td>>hg38_21_+_8433172_8446622_RNA45SN1_45S_50ntFlanks</td><td>9564</td><td>9594</td><td>31</td><td>rRF 45S</td><td>NO</td><td>69.94</td><td>50.26</td><td>0.00</td><td>57.22</td></tr>
<tr><td>>hg38_21_+_8433172_8446622_RNA45SN1_45S_50ntFlanks@9564.9595.32|CTGACGTGCAAATCGGTCGTCCGACCTGGGTA</td><td>>hg38_21_+_8433172_8446622_RNA45SN1_45S_50ntFlanks</td><td>9564</td><td>9595</td><td>32</td><td>rRF 45S</td><td>NO</td><td>195.19</td><td>175.02</td><td>27.98</td><td>159.90</td></tr>
<tr><td>>hg38_21_+_8433172_8446622_RNA45SN1_45S_50ntFlanks@9564.9596.33|CTGACGTGCAAATCGGTCGTCCGACCTGGGTAT</td><td>>hg38_21_+_8433172_8446622_RNA45SN1_45S_50ntFlanks</td><td>9564</td><td>9596</td><td>33</td><td>rRF 45S</td><td>YES</td><td>1872.98</td><td>1403.21</td><td>117.29</td><td>1515.46</td></tr>
<tr><td>>hg38_21_+_8433172_8446622_RNA45SN1_45S_50ntFlanks@9564.9597.34|CTGACGTGCAAATCGGTCGTCCGACCTGGGTATA</td><td>>hg38_21_+_8433172_8446622_RNA45SN1_45S_50ntFlanks</td><td>9564</td><td>9597</td><td>34</td><td>rRF 45S</td><td>NO</td><td>746.96</td><td>594.92</td><td>61.64</td><td>840.54</td></tr>
<tr><td>>hg38_21_+_8433172_8446622_RNA45SN1_45S_50ntFlanks@9565.9592.28|TGACGTGCAAATCGGTCGTCCGACCTGG</td><td>>hg38_21_+_8433172_8446622_RNA45SN1_45S_50ntFlanks</td><td>9565</td><td>9592</td><td>28</td><td>rRF 45S</td><td>NO</td><td>33.45</td><td>120.67</td><td>0.00</td><td>0.00</td></tr>
<tr><td>>hg38_21_+_8433172_8446622_RNA45SN1_45S_50ntFlanks@9565.9594.30|TGACGTGCAAATCGGTCGTCCGACCTGGGT</td><td>>hg38_21_+_8433172_8446622_RNA45SN1_45S_50ntFlanks</td><td>9565</td><td>9594</td><td>30</td><td>rRF 45S</td><td>NO</td><td>19.46</td><td>0.00</td><td>0.00</td><td>0.00</td></tr>
<tr><td>>hg38_21_+_8433172_8446622_RNA45SN1_45S_50ntFlanks@9565.9595.31|TGACGTGCAAATCGGTCGTCCGACCTGGGTA</td><td>>hg38_21_+_8433172_8446622_RNA45SN1_45S_50ntFlanks</td><td>9565</td><td>9595</td><td>31</td><td>rRF 45S</td><td>NO</td><td>83.82</td><td>85.94</td><td>0.00</td><td>44.53</td></tr>
<tr><td>>hg38_21_+_8433172_8446622_RNA45SN1_45S_50ntFlanks@9565.9596.32|TGACGTGCAAATCGGTCGTCCGACCTGGGTAT</td><td>>hg38_21_+_8433172_8446622_RNA45SN1_45S_50ntFlanks</td><td>9565</td><td>9596</td><td>32</td><td>rRF 45S</td><td>YES</td><td>612.62</td><td>570.25</td><td>65.19</td><td>385.00</td></tr>
<tr><td>>hg38_21_+_8433172_8446622_RNA45SN1_45S_50ntFlanks@9565.9597.33|TGACGTGCAAATCGGTCGTCCGACCTGGGTATA</td><td>>hg38_21_+_8433172_8446622_RNA45SN1_45S_50ntFlanks</td><td>9565</td><td>9597</td><td>33</td><td>rRF 45S</td><td>NO</td><td>243.71</td><td>231.05</td><td>33.96</td><td>216.42</td></tr>
<tr><td>>hg38_21_+_8433172_8446622_RNA45SN1_45S_50ntFlanks@9566.9592.27|GACGTGCAAATCGGTCGTCCGACCTGG</td><td>>hg38_21_+_8433172_8446622_RNA45SN1_45S_50ntFlanks</td><td>9566</td><td>9592</td><td>27</td><td>rRF 45S</td><td>NO</td><td>0.00</td><td>39.63</td><td>0.00</td><td>0.00</td></tr>
<tr><td>>hg38_21_+_8433172_8446622_RNA45SN1_45S_50ntFlanks@9566.9595.30|GACGTGCAAATCGGTCGTCCGACCTGGGTA</td><td>>hg38_21_+_8433172_8446622_RNA45SN1_45S_50ntFlanks</td><td>9566</td><td>9595</td><td>30</td><td>rRF 45S</td><td>NO</td><td>29.73</td><td>34.92</td><td>0.00</td><td>13.88</td></tr>
<tr><td>>hg38_21_+_8433172_8446622_RNA45SN1_45S_50ntFlanks@9566.9596.31|GACGTGCAAATCGGTCGTCCGACCTGGGTAT</td><td>>hg38_21_+_8433172_8446622_RNA45SN1_45S_50ntFlanks</td><td>9566</td><td>9596</td><td>31</td><td>rRF 45S</td><td>NO</td><td>216.61</td><td>235.40</td><td>37.10</td><td>202.19</td></tr>
<tr><td>>hg38_21_+_8433172_8446622_RNA45SN1_45S_50ntFlanks@9566.9597.32|GACGTGCAAATCGGTCGTCCGACCTGGGTATA</td><td>>hg38_21_+_8433172_8446622_RNA45SN1_45S_50ntFlanks</td><td>9566</td><td>9597</td><td>32</td><td>rRF 45S</td><td>NO</td><td>90.71</td><td>101.30</td><td>0.00</td><td>116.16</td></tr>
<tr><td>>hg38_21_+_8433172_8446622_RNA45SN1_45S_50ntFlanks@9567.9584.18|ACGTGCAAATCGGTCGTC</td><td>>hg38_21_+_8433172_8446622_RNA45SN1_45S_50ntFlanks</td><td>9567</td><td>9584</td><td>18</td><td>rRF 45S</td><td>NO</td><td>0.00</td><td>49.18</td><td>27.38</td><td>0.00</td></tr>
<tr><td>>hg38_21_+_8433172_8446622_RNA45SN1_45S_50ntFlanks@9567.9585.19|ACGTGCAAATCGGTCGTCC</td><td>>hg38_21_+_8433172_8446622_RNA45SN1_45S_50ntFlanks</td><td>9567</td><td>9585</td><td>19</td><td>rRF 45S</td><td>NO</td><td>0.00</td><td>26.96</td><td>0.00</td><td>0.00</td></tr>
<tr><td>>hg38_21_+_8433172_8446622_RNA45SN1_45S_50ntFlanks@9567.9586.20|ACGTGCAAATCGGTCGTCCG</td><td>>hg38_21_+_8433172_8446622_RNA45SN1_45S_50ntFlanks</td><td>9567</td><td>9586</td><td>20</td><td>rRF 45S</td><td>NO</td><td>0.00</td><td>54.61</td><td>0.00</td><td>0.00</td></tr>
<tr><td>>hg38_21_+_8433172_8446622_RNA45SN1_45S_50ntFlanks@9567.9592.26|ACGTGCAAATCGGTCGTCCGACCTGG</td><td>>hg38_21_+_8433172_8446622_RNA45SN1_45S_50ntFlanks</td><td>9567</td><td>9592</td><td>26</td><td>rRF 45S</td><td>NO</td><td>0.00</td><td>144.79</td><td>33.32</td><td>6.51</td></tr>
<tr><td>>hg38_21_+_8433172_8446622_RNA45SN1_45S_50ntFlanks@9567.9594.28|ACGTGCAAATCGGTCGTCCGACCTGGGT</td><td>>hg38_21_+_8433172_8446622_RNA45SN1_45S_50ntFlanks</td><td>9567</td><td>9594</td><td>28</td><td>rRF 45S</td><td>NO</td><td>12.67</td><td>58.57</td><td>20.75</td><td>37.25</td></tr>
<tr><td>>hg38_21_+_8433172_8446622_RNA45SN1_45S_50ntFlanks@9567.9595.29|ACGTGCAAATCGGTCGTCCGACCTGGGTA</td><td>>hg38_21_+_8433172_8446622_RNA45SN1_45S_50ntFlanks</td><td>9567</td><td>9595</td><td>29</td><td>rRF 45S</td><td>NO</td><td>104.06</td><td>311.09</td><td>56.23</td><td>121.95</td></tr>
<tr><td>>hg38_21_+_8433172_8446622_RNA45SN1_45S_50ntFlanks@9567.9596.30|ACGTGCAAATCGGTCGTCCGACCTGGGTAT</td><td>>hg38_21_+_8433172_8446622_RNA45SN1_45S_50ntFlanks</td><td>9567</td><td>9596</td><td>30</td><td>rRF 45S</td><td>NO</td><td>233.87</td><td>495.21</td><td>133.85</td><td>370.42</td></tr>
<tr><td>>hg38_21_+_8433172_8446622_RNA45SN1_45S_50ntFlanks@9567.9597.31|ACGTGCAAATCGGTCGTCCGACCTGGGTATA</td><td>>hg38_21_+_8433172_8446622_RNA45SN1_45S_50ntFlanks</td><td>9567</td><td>9597</td><td>31</td><td>rRF 45S</td><td>NO</td><td>170.51</td><td>366.44</td><td>104.25</td><td>363.80</td></tr>
<tr><td>>hg38_21_+_8433172_8446622_RNA45SN1_45S_50ntFlanks@9567.9603.37|ACGTGCAAATCGGTCGTCCGACCTGGGTATAGGGGCG</td><td>>hg38_21_+_8433172_8446622_RNA45SN1_45S_50ntFlanks</td><td>9567</td><td>9603</td><td>37</td><td>rRF 45S</td><td>NO</td><td>22.16</td><td>26.96</td><td>0.00</td><td>58.66</td></tr>
<tr><td>>hg38_21_+_8433172_8446622_RNA45SN1_45S_50ntFlanks@9568.9586.19|CGTGCAAATCGGTCGTCCG</td><td>>hg38_21_+_8433172_8446622_RNA45SN1_45S_50ntFlanks</td><td>9568</td><td>9586</td><td>19</td><td>rRF 45S</td><td>NO</td><td>0.00</td><td>35.63</td><td>0.00</td><td>0.00</td></tr>
<tr><td>>hg38_21_+_8433172_8446622_RNA45SN1_45S_50ntFlanks@9568.9592.25|CGTGCAAATCGGTCGTCCGACCTGG</td><td>>hg38_21_+_8433172_8446622_RNA45SN1_45S_50ntFlanks</td><td>9568</td><td>9592</td><td>25</td><td>rRF 45S</td><td>NO</td><td>0.00</td><td>103.72</td><td>35.32</td><td>6.46</td></tr>
<tr><td>>hg38_21_+_8433172_8446622_RNA45SN1_45S_50ntFlanks@9568.9594.27|CGTGCAAATCGGTCGTCCGACCTGGGT</td><td>>hg38_21_+_8433172_8446622_RNA45SN1_45S_50ntFlanks</td><td>9568</td><td>9594</td><td>27</td><td>rRF 45S</td><td>NO</td><td>12.76</td><td>48.53</td><td>30.70</td><td>37.88</td></tr>
<tr><td>>hg38_21_+_8433172_8446622_RNA45SN1_45S_50ntFlanks@9568.9595.28|CGTGCAAATCGGTCGTCCGACCTGGGTA</td><td>>hg38_21_+_8433172_8446622_RNA45SN1_45S_50ntFlanks</td><td>9568</td><td>9595</td><td>28</td><td>rRF 45S</td><td>NO</td><td>91.36</td><td>282.22</td><td>58.45</td><td>114.07</td></tr>
<tr><td>>hg38_21_+_8433172_8446622_RNA45SN1_45S_50ntFlanks@9568.9596.29|CGTGCAAATCGGTCGTCCGACCTGGGTAT</td><td>>hg38_21_+_8433172_8446622_RNA45SN1_45S_50ntFlanks</td><td>9568</td><td>9596</td><td>29</td><td>rRF 45S</td><td>NO</td><td>182.79</td><td>379.87</td><td>132.32</td><td>335.75</td></tr>
</table>

### MDA-MB-468

| | | | | | | | | | | |
|---|---|---|---|---|---|---|---|---|---|---|
| >hg38_21_+_8433172_8446622_RNA45SN1_45S_50ntFlanks@9568.9597.30\|CGTGCAAATCG GTCGTCCGACCTGGGTATA | >hg38_21_+_8433172_8446622_RNA45SN1_45S_50ntFlanks | 9568 | 9597 | 30 | rRF 45S | NO | 166.56 | 305.84 | 111.85 | 404.71 |
| >hg38_21_+_8433172_8446622_RNA45SN1_45S_50ntFlanks@9568.9603.36\|CGTGCAAATCG GTCGTCCGACCTGGGTATAGGGGCG | >hg38_21_+_8433172_8446622_RNA45SN1_45S_50ntFlanks | 9568 | 9603 | 36 | rRF 45S | NO | 46.49 | 32.28 | 0.00 | 79.03 |
| >hg38_21_+_8433172_8446622_RNA45SN1_45S_50ntFlanks@9569.9592.24\|GTGCAAATCGGT CGTCCGACCTGG | >hg38_21_+_8433172_8446622_RNA45SN1_45S_50ntFlanks | 9569 | 9592 | 24 | rRF 45S | NO | 0.00 | 29.40 | 0.00 | 0.00 |
| >hg38_21_+_8433172_8446622_RNA45SN1_45S_50ntFlanks@9569.9595.27\|GTGCAAATCGGT CGTCCGACCTGGGTA | >hg38_21_+_8433172_8446622_RNA45SN1_45S_50ntFlanks | 9569 | 9595 | 27 | rRF 45S | NO | 0.00 | 35.66 | 0.00 | 0.00 |
| >hg38_21_+_8433172_8446622_RNA45SN1_45S_50ntFlanks@9569.9596.28\|GTGCAAATCGGT CGTCCGACCTGGGTAT | >hg38_21_+_8433172_8446622_RNA45SN1_45S_50ntFlanks | 9569 | 9596 | 28 | rRF 45S | NO | 0.00 | 31.53 | 21.27 | 36.98 |
| >hg38_21_+_8433172_8446622_RNA45SN1_45S_50ntFlanks@9569.9597.29\|GTGCAAATCGGT CGTCCGACCTGGGTATA | >hg38_21_+_8433172_8446622_RNA45SN1_45S_50ntFlanks | 9569 | 9597 | 29 | rRF 45S | NO | 31.68 | 49.99 | 39.41 | 74.86 |
| >hg38_21_+_8433172_8446622_RNA45SN1_45S_50ntFlanks@9570.9594.25\|TGCAAATCGGTC GTCCGACCTGGGT | >hg38_21_+_8433172_8446622_RNA45SN1_45S_50ntFlanks | 9570 | 9594 | 25 | rRF 45S | NO | 0.00 | 37.15 | 38.59 | 13.45 |
| >hg38_21_+_8433172_8446622_RNA45SN1_45S_50ntFlanks@9570.9595.26\|TGCAAATCGGTC GTCCGACCTGGGTA | >hg38_21_+_8433172_8446622_RNA45SN1_45S_50ntFlanks | 9570 | 9595 | 26 | rRF 45S | NO | 55.45 | 248.11 | 75.09 | 99.87 |
| >hg38_21_+_8433172_8446622_RNA45SN1_45S_50ntFlanks@9570.9596.27\|TGCAAATCGGTC GTCCGACCTGGGTAT | >hg38_21_+_8433172_8446622_RNA45SN1_45S_50ntFlanks | 9570 | 9596 | 27 | rRF 45S | NO | 53.79 | 196.37 | 118.86 | 160.91 |
| >hg38_21_+_8433172_8446622_RNA45SN1_45S_50ntFlanks@9570.9597.28\|TGCAAATCGGTC GTCCGACCTGGGTATA | >hg38_21_+_8433172_8446622_RNA45SN1_45S_50ntFlanks | 9570 | 9597 | 28 | rRF 45S | NO | 42.81 | 144.23 | 92.26 | 182.04 |
| >hg38_21_+_8433172_8446622_RNA45SN1_45S_50ntFlanks@9570.9603.34\|TGCAAATCGGTC GTCCGACCTGGGTATAGGGGCG | >hg38_21_+_8433172_8446622_RNA45SN1_45S_50ntFlanks | 9570 | 9603 | 34 | rRF 45S | NO | 19.82 | 12.41 | 0.00 | 50.42 |
| >hg38_21_+_8433172_8446622_RNA45SN1_45S_50ntFlanks@9572.9592.21\|CAAATCGGTCGT CCGACCTGG | >hg38_21_+_8433172_8446622_RNA45SN1_45S_50ntFlanks | 9572 | 9592 | 21 | rRF 45S | NO | 0.00 | 29.04 | 50.82 | 0.00 |
| >hg38_21_+_8433172_8446622_RNA45SN1_45S_50ntFlanks@9572.9594.23\|CAAATCGGTCGT CCGACCTGGGT | >hg38_21_+_8433172_8446622_RNA45SN1_45S_50ntFlanks | 9572 | 9594 | 23 | rRF 45S | NO | 0.00 | 67.18 | 158.72 | 46.91 |
| >hg38_21_+_8433172_8446622_RNA45SN1_45S_50ntFlanks@9572.9595.24\|CAAATCGGTCGT CCGACCTGGGTA | >hg38_21_+_8433172_8446622_RNA45SN1_45S_50ntFlanks | 9572 | 9595 | 24 | rRF 45S | NO | 30.70 | 427.35 | 221.26 | 132.38 |
| >hg38_21_+_8433172_8446622_RNA45SN1_45S_50ntFlanks@9572.9596.25\|CAAATCGGTCGT CCGACCTGGGTAT | >hg38_21_+_8433172_8446622_RNA45SN1_45S_50ntFlanks | 9572 | 9596 | 25 | rRF 45S | YES | 76.68 | 454.24 | 523.40 | 404.56 |
| >hg38_21_+_8433172_8446622_RNA45SN1_45S_50ntFlanks@9572.9597.26\|CAAATCGGTCGT CCGACCTGGGTATA | >hg38_21_+_8433172_8446622_RNA45SN1_45S_50ntFlanks | 9572 | 9597 | 26 | rRF 45S | YES | 72.80 | 388.84 | 378.05 | 490.35 |
| >hg38_21_+_8433172_8446622_RNA45SN1_45S_50ntFlanks@9572.9601.30\|CAAATCGGTCGT CCGACCTGGGTATAGGGG | >hg38_21_+_8433172_8446622_RNA45SN1_45S_50ntFlanks | 9572 | 9601 | 30 | rRF 45S | NO | 0.00 | 0.00 | 37.82 | 43.91 |
| >hg38_21_+_8433172_8446622_RNA45SN1_45S_50ntFlanks@9572.9603.32\|CAAATCGGTCGT CCGACCTGGGTATAGGGGCG | >hg38_21_+_8433172_8446622_RNA45SN1_45S_50ntFlanks | 9572 | 9603 | 32 | rRF 45S | NO | 121.53 | 115.86 | 132.16 | 212.33 |
| >hg38_21_+_8433172_8446622_RNA45SN1_45S_50ntFlanks@9572.9605.34\|CAAATCGGTCGT CCGACCTGGGTATAGGGGCGAA | >hg38_21_+_8433172_8446622_RNA45SN1_45S_50ntFlanks | 9572 | 9605 | 34 | rRF 45S | NO | 12.94 | 0.00 | 0.00 | 54.83 |
| >hg38_21_+_8433172_8446622_RNA45SN1_45S_50ntFlanks@9573.9592.20\|AAATCGGTCGTC CGACCTGG | >hg38_21_+_8433172_8446622_RNA45SN1_45S_50ntFlanks | 9573 | 9592 | 20 | rRF 45S | NO | 0.00 | 0.00 | 31.91 | 0.00 |
| >hg38_21_+_8433172_8446622_RNA45SN1_45S_50ntFlanks@9573.9594.22\|AAATCGGTCGTC CGACCTGGGT | >hg38_21_+_8433172_8446622_RNA45SN1_45S_50ntFlanks | 9573 | 9594 | 22 | rRF 45S | NO | 0.00 | 12.26 | 85.27 | 0.00 |
| >hg38_21_+_8433172_8446622_RNA45SN1_45S_50ntFlanks@9573.9595.23\|AAATCGGTCGTC CGACCTGGGTA | >hg38_21_+_8433172_8446622_RNA45SN1_45S_50ntFlanks | 9573 | 9595 | 23 | rRF 45S | NO | 0.00 | 158.61 | 110.17 | 55.55 |
| >hg38_21_+_8433172_8446622_RNA45SN1_45S_50ntFlanks@9573.9596.24\|AAATCGGTCGTC CGACCTGGGTAT | >hg38_21_+_8433172_8446622_RNA45SN1_45S_50ntFlanks | 9573 | 9596 | 24 | rRF 45S | NO | 33.23 | 159.97 | 249.41 | 163.55 |
| >hg38_21_+_8433172_8446622_RNA45SN1_45S_50ntFlanks@9573.9597.25\|AAATCGGTCGTC CGACCTGGGTATA | >hg38_21_+_8433172_8446622_RNA45SN1_45S_50ntFlanks | 9573 | 9597 | 25 | rRF 45S | NO | 34.14 | 146.72 | 181.58 | 209.75 |
| >hg38_21_+_8433172_8446622_RNA45SN1_45S_50ntFlanks@9573.9603.31\|AAATCGGTCGTC CGACCTGGGTATAGGGGCG | >hg38_21_+_8433172_8446622_RNA45SN1_45S_50ntFlanks | 9573 | 9603 | 31 | rRF 45S | NO | 56.05 | 42.21 | 62.55 | 88.37 |
| >hg38_21_+_8433172_8446622_RNA45SN1_45S_50ntFlanks@9574.9592.19\|AATCGGTCGTCC GACCTGG | >hg38_21_+_8433172_8446622_RNA45SN1_45S_50ntFlanks | 9574 | 9592 | 19 | rRF 45S | NO | 0.00 | 0.00 | 44.73 | 0.00 |
| >hg38_21_+_8433172_8446622_RNA45SN1_45S_50ntFlanks@9574.9594.21\|AATCGGTCGTCC GACCTGGGT | >hg38_21_+_8433172_8446622_RNA45SN1_45S_50ntFlanks | 9574 | 9594 | 21 | rRF 45S | NO | 0.00 | 40.15 | 154.50 | 16.01 |
| >hg38_21_+_8433172_8446622_RNA45SN1_45S_50ntFlanks@9574.9595.22\|AATCGGTCGTCC GACCTGGGTA | >hg38_21_+_8433172_8446622_RNA45SN1_45S_50ntFlanks | 9574 | 9595 | 22 | rRF 45S | NO | 0.00 | 279.41 | 207.96 | 82.10 |
| >hg38_21_+_8433172_8446622_RNA45SN1_45S_50ntFlanks@9574.9596.23\|AATCGGTCGTCC GACCTGGGTAT | >hg38_21_+_8433172_8446622_RNA45SN1_45S_50ntFlanks | 9574 | 9596 | 23 | rRF 45S | NO | 56.19 | 306.25 | 479.12 | 254.57 |
| >hg38_21_+_8433172_8446622_RNA45SN1_45S_50ntFlanks@9574.9597.24\|AATCGGTCGTCC GACCTGGGTATA | >hg38_21_+_8433172_8446622_RNA45SN1_45S_50ntFlanks | 9574 | 9597 | 24 | rRF 45S | NO | 55.20 | 281.40 | 338.85 | 339.02 |
| >hg38_21_+_8433172_8446622_RNA45SN1_45S_50ntFlanks@9574.9601.28\|AATCGGTCGTCC GACCTGGGTATAGGGG | >hg38_21_+_8433172_8446622_RNA45SN1_45S_50ntFlanks | 9574 | 9601 | 28 | rRF 45S | NO | 0.00 | 0.00 | 41.49 | 33.26 |
| >hg38_21_+_8433172_8446622_RNA45SN1_45S_50ntFlanks@9574.9603.30\|AATCGGTCGTCC GACCTGGGTATAGGGGCG | >hg38_21_+_8433172_8446622_RNA45SN1_45S_50ntFlanks | 9574 | 9603 | 30 | rRF 45S | NO | 90.10 | 74.68 | 141.59 | 141.40 |
| >hg38_21_+_8433172_8446622_RNA45SN1_45S_50ntFlanks@9574.9605.32\|AATCGGTCGTCC GACCTGGGTATAGGGGCGAA | >hg38_21_+_8433172_8446622_RNA45SN1_45S_50ntFlanks | 9574 | 9605 | 32 | rRF 45S | NO | 0.00 | 0.00 | 36.63 | 38.33 |
| >hg38_21_+_8433172_8446622_RNA45SN1_45S_50ntFlanks@9575.9594.20\|ATCGGTCGTCC GACCTGGGT | >hg38_21_+_8433172_8446622_RNA45SN1_45S_50ntFlanks | 9575 | 9594 | 20 | rRF 45S | NO | 0.00 | 0.00 | 75.17 | 0.00 |

### MDA-MB-468

| | | | | | | | | | | |
|---|---|---|---|---|---|---|---|---|---|---|
| >hg38_21_+_8433172_8446622_RNA45SN1_45S_50ntFlanks@9575.9595.21\|ATCGGTCGTCCGACCTGGGTA | >hg38_21_+_8433172_8446622_RNA45SN1_45S_50ntFlanks | 9575 | 9595 | 21 | rRF 45S | NO | 0.00 | 97.38 | 106.52 | 34.61 |
| >hg38_21_+_8433172_8446622_RNA45SN1_45S_50ntFlanks@9575.9596.22\|ATCGGTCGTCCGACCTGGGTAT | >hg38_21_+_8433172_8446622_RNA45SN1_45S_50ntFlanks | 9575 | 9596 | 22 | rRF 45S | NO | 0.00 | 122.37 | 261.28 | 118.51 |
| >hg38_21_+_8433172_8446622_RNA45SN1_45S_50ntFlanks@9575.9597.23\|ATCGGTCGTCCGACCTGGGTATA | >hg38_21_+_8433172_8446622_RNA45SN1_45S_50ntFlanks | 9575 | 9597 | 23 | rRF 45S | NO | 0.00 | 119.32 | 170.39 | 157.47 |
| >hg38_21_+_8433172_8446622_RNA45SN1_45S_50ntFlanks@9575.9603.29\|ATCGGTCGTCCGACCTGGGTATAGGGGCG | >hg38_21_+_8433172_8446622_RNA45SN1_45S_50ntFlanks | 9575 | 9603 | 29 | rRF 45S | NO | 37.19 | 34.01 | 79.87 | 68.87 |
| >hg38_21_+_8433172_8446622_RNA45SN1_45S_50ntFlanks@9576.9596.21\|TCGGTCGTCCGACCTGGGTAT | >hg38_21_+_8433172_8446622_RNA45SN1_45S_50ntFlanks | 9576 | 9596 | 21 | rRF 45S | NO | 0.00 | 0.00 | 33.01 | 0.00 |
| >hg38_21_+_8433172_8446622_RNA45SN1_45S_50ntFlanks@9602.9622.21\|CGAAAGACTAATCGAACCATC | >hg38_21_+_8433172_8446622_RNA45SN1_45S_50ntFlanks | 9602 | 9622 | 21 | rRF 45S | NO | 0.00 | 52.51 | 29.37 | 0.00 |
| >hg38_21_+_8433172_8446622_RNA45SN1_45S_50ntFlanks@9604.9621.18\|AAAGACTAATCGAACCAT | >hg38_21_+_8433172_8446622_RNA45SN1_45S_50ntFlanks | 9604 | 9621 | 18 | rRF 45S | NO | 0.00 | 28.41 | 6.70 | 0.00 |
| >hg38_21_+_8433172_8446622_RNA45SN1_45S_50ntFlanks@9604.9622.19\|AAAGACTAATCGAACCATC | >hg38_21_+_8433172_8446622_RNA45SN1_45S_50ntFlanks | 9604 | 9622 | 19 | rRF 45S | NO | 73.14 | 118.34 | 83.64 | 24.67 |
| >hg38_21_+_8433172_8446622_RNA45SN1_45S_50ntFlanks@9605.9622.18\|AAGACTAATCGAACCATC | >hg38_21_+_8433172_8446622_RNA45SN1_45S_50ntFlanks | 9605 | 9622 | 18 | rRF 45S | NO | 36.97 | 61.65 | 42.87 | 0.00 |
| >hg38_21_+_8433172_8446622_RNA45SN1_45S_50ntFlanks@9714.9737.24\|CGAATGATTAGAGGTCTTGGGGCC | >hg38_21_+_8433172_8446622_RNA45SN1_45S_50ntFlanks | 9714 | 9737 | 24 | rRF 45S | NO | 60.32 | 66.56 | 37.53 | 138.93 |
| >hg38_21_+_8433172_8446622_RNA45SN1_45S_50ntFlanks@9716.9737.22\|AATGATTAGAGGTCTTGGGGCC | >hg38_21_+_8433172_8446622_RNA45SN1_45S_50ntFlanks | 9716 | 9737 | 22 | rRF 45S | NO | 6.09 | 31.35 | 20.82 | 34.56 |
| >hg38_21_+_8433172_8446622_RNA45SN1_45S_50ntFlanks@9720.9737.18\|ATTAGAGGTCTTGGGGCC | >hg38_21_+_8433172_8446622_RNA45SN1_45S_50ntFlanks | 9720 | 9737 | 18 | rRF 45S | NO | 0.00 | 0.00 | 50.00 | 23.41 |
| >hg38_21_+_8433172_8446622_RNA45SN1_45S_50ntFlanks@9736.9753.18\|CCGAAACGATCTCAACCT | >hg38_21_+_8433172_8446622_RNA45SN1_45S_50ntFlanks | 9736 | 9753 | 18 | rRF 45S | NO | 0.00 | 12.58 | 0.00 | 0.00 |
| >hg38_21_+_8433172_8446622_RNA45SN1_45S_50ntFlanks@9736.9754.19\|CCGAAACGATCTCAACCTA | >hg38_21_+_8433172_8446622_RNA45SN1_45S_50ntFlanks | 9736 | 9754 | 19 | rRF 45S | NO | 31.16 | 73.41 | 0.00 | 25.94 |
| >hg38_21_+_8433172_8446622_RNA45SN1_45S_50ntFlanks@9736.9755.20\|CCGAAACGATCTCAACCTAT | >hg38_21_+_8433172_8446622_RNA45SN1_45S_50ntFlanks | 9736 | 9755 | 20 | rRF 45S | NO | 13.09 | 32.95 | 0.00 | 0.00 |
| >hg38_21_+_8433172_8446622_RNA45SN1_45S_50ntFlanks@9736.9765.30\|CCGAAACGATCTCAACCTATTCTCAAACTT | >hg38_21_+_8433172_8446622_RNA45SN1_45S_50ntFlanks | 9736 | 9765 | 30 | rRF 45S | NO | 94.69 | 0.00 | 0.00 | 133.39 |
| >hg38_21_+_8433172_8446622_RNA45SN1_45S_50ntFlanks@9736.9767.32\|CCGAAACGATCTCAACCTATTCTCAAACTTTA | >hg38_21_+_8433172_8446622_RNA45SN1_45S_50ntFlanks | 9736 | 9767 | 32 | rRF 45S | NO | 20.76 | 0.00 | 0.00 | 41.20 |
| >hg38_21_+_8433172_8446622_RNA45SN1_45S_50ntFlanks@9738.9755.18\|GAAACGATCTCAACCTAT | >hg38_21_+_8433172_8446622_RNA45SN1_45S_50ntFlanks | 9738 | 9755 | 18 | rRF 45S | NO | 40.38 | 61.91 | 31.73 | 16.22 |
| >hg38_21_+_8433172_8446622_RNA45SN1_45S_50ntFlanks@9738.9757.20\|GAAACGATCTCAACCTATTC | >hg38_21_+_8433172_8446622_RNA45SN1_45S_50ntFlanks | 9738 | 9757 | 20 | rRF 45S | NO | 0.00 | 0.00 | 28.32 | 0.00 |
| >hg38_21_+_8433172_8446622_RNA45SN1_45S_50ntFlanks@9738.9758.21\|GAAACGATCTCAACCTATTCT | >hg38_21_+_8433172_8446622_RNA45SN1_45S_50ntFlanks | 9738 | 9758 | 21 | rRF 45S | NO | 117.28 | 286.07 | 70.21 | 185.68 |
| >hg38_21_+_8433172_8446622_RNA45SN1_45S_50ntFlanks@9738.9759.22\|GAAACGATCTCAACCTATTCTC | >hg38_21_+_8433172_8446622_RNA45SN1_45S_50ntFlanks | 9738 | 9759 | 22 | rRF 45S | NO | 53.06 | 75.70 | 20.43 | 133.65 |
| >hg38_21_+_8433172_8446622_RNA45SN1_45S_50ntFlanks@9738.9763.26\|GAAACGATCTCAACCTATTCTCAAAC | >hg38_21_+_8433172_8446622_RNA45SN1_45S_50ntFlanks | 9738 | 9763 | 26 | rRF 45S | NO | 39.75 | 0.00 | 0.00 | 0.00 |
| >hg38_21_+_8433172_8446622_RNA45SN1_45S_50ntFlanks@9738.9764.27\|GAAACGATCTCAACCTATTCTCAAACT | >hg38_21_+_8433172_8446622_RNA45SN1_45S_50ntFlanks | 9738 | 9764 | 27 | rRF 45S | NO | 31.63 | 0.00 | 0.00 | 23.00 |
| >hg38_21_+_8433172_8446622_RNA45SN1_45S_50ntFlanks@9738.9765.28\|GAAACGATCTCAACCTATTCTCAAACTT | >hg38_21_+_8433172_8446622_RNA45SN1_45S_50ntFlanks | 9738 | 9765 | 28 | rRF 45S | NO | 220.92 | 60.35 | 40.53 | 888.79 |
| >hg38_21_+_8433172_8446622_RNA45SN1_45S_50ntFlanks@9738.9767.30\|GAAACGATCTCAACCTATTCTCAAACTTTA | >hg38_21_+_8433172_8446622_RNA45SN1_45S_50ntFlanks | 9738 | 9767 | 30 | rRF 45S | NO | 82.91 | 0.00 | 30.42 | 284.71 |
| >hg38_21_+_8433172_8446622_RNA45SN1_45S_50ntFlanks@9739.9757.19\|AAACGATCTCAACCTATTC | >hg38_21_+_8433172_8446622_RNA45SN1_45S_50ntFlanks | 9739 | 9757 | 19 | rRF 45S | NO | 39.44 | 65.56 | 44.32 | 122.73 |
| >hg38_21_+_8433172_8446622_RNA45SN1_45S_50ntFlanks@9739.9758.20\|AAACGATCTCAACCTATTCT | >hg38_21_+_8433172_8446622_RNA45SN1_45S_50ntFlanks | 9739 | 9758 | 20 | rRF 45S | NO | 135.29 | 204.72 | 54.22 | 219.29 |
| >hg38_21_+_8433172_8446622_RNA45SN1_45S_50ntFlanks@9739.9759.21\|AAACGATCTCAACCTATTCTC | >hg38_21_+_8433172_8446622_RNA45SN1_45S_50ntFlanks | 9739 | 9759 | 21 | rRF 45S | NO | 64.78 | 47.12 | 7.20 | 139.54 |
| >hg38_21_+_8433172_8446622_RNA45SN1_45S_50ntFlanks@9739.9765.27\|AAACGATCTCAACCTATTCTCAAACTT | >hg38_21_+_8433172_8446622_RNA45SN1_45S_50ntFlanks | 9739 | 9765 | 27 | rRF 45S | NO | 163.74 | 30.07 | 40.30 | 625.79 |
| >hg38_21_+_8433172_8446622_RNA45SN1_45S_50ntFlanks@9739.9767.29\|AAACGATCTCAACCTATTCTCAAACTTTA | >hg38_21_+_8433172_8446622_RNA45SN1_45S_50ntFlanks | 9739 | 9767 | 29 | rRF 45S | NO | 54.03 | 0.00 | 0.00 | 158.11 |
| >hg38_21_+_8433172_8446622_RNA45SN1_45S_50ntFlanks@9740.9758.19\|AACGATCTCAACCTATTCT | >hg38_21_+_8433172_8446622_RNA45SN1_45S_50ntFlanks | 9740 | 9758 | 19 | rRF 45S | NO | 0.00 | 31.22 | 0.00 | 24.05 |
| >hg38_21_+_8433172_8446622_RNA45SN1_45S_50ntFlanks@9740.9765.26\|AACGATCTCAACCTATTCTCAAACTT | >hg38_21_+_8433172_8446622_RNA45SN1_45S_50ntFlanks | 9740 | 9765 | 26 | rRF 45S | NO | 12.59 | 0.00 | 0.00 | 98.23 |
| >hg38_21_+_8433172_8446622_RNA45SN1_45S_50ntFlanks@9806.9830.25\|CGTGGAATGCGAGTGCCTAGTGGGC | >hg38_21_+_8433172_8446622_RNA45SN1_45S_50ntFlanks | 9806 | 9830 | 25 | rRF 45S | NO | 0.00 | 0.00 | 38.60 | 0.00 |
| >hg38_21_+_8433172_8446622_RNA45SN1_45S_50ntFlanks@9806.9838.33\|CGTGGAATGCGAGTGCCTAGTGGGCCACTTTTG | >hg38_21_+_8433172_8446622_RNA45SN1_45S_50ntFlanks | 9806 | 9838 | 33 | rRF 45S | NO | 0.00 | 0.00 | 36.72 | 41.06 |

### MDA-MB-468

| | | | | | | | | | | |
|---|---|---|---|---|---|---|---|---|---|---|
| >hg38_21_+_8433172_8446622_RNA45SN1_45S_50ntFlanks@9821.9850.30\|CCTAGTGGGCC ACTTTTGGTAAGCAGAACT | >hg38_21_+_8433172_8446622_RNA45SN1_45S_50ntFlanks | 9821 | 9850 | 30 | rRF 45S | NO | 41.64 | 0.00 | 0.00 | 82.12 |
| >hg38_21_+_8433172_8446622_RNA45SN1_45S_50ntFlanks@9822.9850.29\|CTAGTGGGCCA CTTTTGGTAAGCAGAACT | >hg38_21_+_8433172_8446622_RNA45SN1_45S_50ntFlanks | 9822 | 9850 | 29 | rRF 45S | NO | 36.54 | 0.00 | 0.00 | 14.43 |
| >hg38_21_+_8433172_8446622_RNA45SN1_45S_50ntFlanks@9862.9893.32\|ATGAACCGAAC GCCGGGTTAAGGCGCCCGATG | >hg38_21_+_8433172_8446622_RNA45SN1_45S_50ntFlanks | 9862 | 9893 | 32 | rRF 45S | YES | 0.00 | 0.00 | 35.39 | 0.00 |
| >hg38_21_+_8433172_8446622_RNA45SN1_45S_50ntFlanks@10099.10117.19\|CGCGCGCGC GTGTGGTGTG | >hg38_21_+_8433172_8446622_RNA45SN1_45S_50ntFlanks | 10099 | 10117 | 19 | rRF 45S | NO | 0.00 | 37.20 | 133.37 | 53.51 |
| >hg38_21_+_8433172_8446622_RNA45SN1_45S_50ntFlanks@10101.10125.25\|CGCGCGCGT GTGGTGTGCGTCGGAG | >hg38_21_+_8433172_8446622_RNA45SN1_45S_50ntFlanks | 10101 | 10125 | 25 | rRF 45S | NO | 0.00 | 0.00 | 34.39 | 40.53 |
| >hg38_21_+_8433172_8446622_RNA45SN1_45S_50ntFlanks@10101.10126.26\|CGCGCGCGT GTGGTGTGCGTCGGAGG | >hg38_21_+_8433172_8446622_RNA45SN1_45S_50ntFlanks | 10101 | 10126 | 26 | rRF 45S | NO | 0.00 | 29.89 | 6.50 | 35.95 |
| >hg38_21_+_8433172_8446622_RNA45SN1_45S_50ntFlanks@10101.10127.27\|CGCGCGCGT GTGGTGTGCGTCGGAGGG | >hg38_21_+_8433172_8446622_RNA45SN1_45S_50ntFlanks | 10101 | 10127 | 27 | rRF 45S | NO | 41.23 | 60.99 | 29.06 | 60.62 |
| >hg38_21_+_8433172_8446622_RNA45SN1_45S_50ntFlanks@10107.10125.19\|CGTGTGGTGT GCGTCGGAG | >hg38_21_+_8433172_8446622_RNA45SN1_45S_50ntFlanks | 10107 | 10125 | 19 | rRF 45S | NO | 29.40 | 47.38 | 32.88 | 0.00 |
| >hg38_21_+_8433172_8446622_RNA45SN1_45S_50ntFlanks@10107.10127.21\|CGTGTGGTGT GCGTCGGAGGG | >hg38_21_+_8433172_8446622_RNA45SN1_45S_50ntFlanks | 10107 | 10127 | 21 | rRF 45S | NO | 33.70 | 43.30 | 33.32 | 6.52 |
| >hg38_21_+_8433172_8446622_RNA45SN1_45S_50ntFlanks@10116.10135.20\|TGCGTCGGAG GGCGGCGGCG | >hg38_21_+_8433172_8446622_RNA45SN1_45S_50ntFlanks | 10116 | 10135 | 20 | rRF 45S | NO | 0.00 | 38.16 | 0.00 | 13.49 |
| >hg38_21_+_8433172_8446622_RNA45SN1_45S_50ntFlanks@10116.10136.21\|TGCGTCGGAG GGCGGCGGCGG | >hg38_21_+_8433172_8446622_RNA45SN1_45S_50ntFlanks | 10116 | 10136 | 21 | rRF 45S | NO | 12.61 | 42.94 | 0.00 | 34.60 |
| >hg38_21_+_8433172_8446622_RNA45SN1_45S_50ntFlanks@10116.10139.24\|TGCGTCGGAG GGCGGCGGCGGCGG | >hg38_21_+_8433172_8446622_RNA45SN1_45S_50ntFlanks | 10116 | 10139 | 24 | rRF 45S | NO | 0.00 | 12.54 | 0.00 | 0.00 |
| >hg38_21_+_8433172_8446622_RNA45SN1_45S_50ntFlanks@10186.10207.22\|TCCTCCCCTC CTCCCGCCCACG | >hg38_21_+_8433172_8446622_RNA45SN1_45S_50ntFlanks | 10186 | 10207 | 22 | rRF 45S | NO | 67.06 | 38.09 | 0.00 | 0.00 |
| >hg38_21_+_8433172_8446622_RNA45SN1_45S_50ntFlanks@10186.10208.23\|TCCTCCCCTC CTCCCGCCCACGC | >hg38_21_+_8433172_8446622_RNA45SN1_45S_50ntFlanks | 10186 | 10208 | 23 | rRF 45S | NO | 33.38 | 0.00 | 0.00 | 0.00 |
| >hg38_21_+_8433172_8446622_RNA45SN1_45S_50ntFlanks@10186.10209.24\|TCCTCCCCTC CTCCCGCCCACGCC | >hg38_21_+_8433172_8446622_RNA45SN1_45S_50ntFlanks | 10186 | 10209 | 24 | rRF 45S | NO | 20.95 | 0.00 | 0.00 | 0.00 |
| >hg38_21_+_8433172_8446622_RNA45SN1_45S_50ntFlanks@10342.10361.20\|ATATTCAAAC GAGAACTTTG | >hg38_21_+_8433172_8446622_RNA45SN1_45S_50ntFlanks | 10342 | 10361 | 20 | rRF 45S | NO | 0.00 | 59.03 | 62.20 | 16.15 |
| >hg38_21_+_8433172_8446622_RNA45SN1_45S_50ntFlanks@10344.10361.18\|ATTCAAACGA GAACTTTG | >hg38_21_+_8433172_8446622_RNA45SN1_45S_50ntFlanks | 10344 | 10361 | 18 | rRF 45S | NO | 0.00 | 0.00 | 13.73 | 0.00 |
| >hg38_21_+_8433172_8446622_RNA45SN1_45S_50ntFlanks@10346.10368.23\|TCAAACGAGA ACTTTGAAGGCCG | >hg38_21_+_8433172_8446622_RNA45SN1_45S_50ntFlanks | 10346 | 10368 | 23 | rRF 45S | NO | 160.15 | 252.92 | 103.02 | 139.25 |
| >hg38_21_+_8433172_8446622_RNA45SN1_45S_50ntFlanks@10346.10369.24\|TCAAACGAGA ACTTTGAAGGCCGA | >hg38_21_+_8433172_8446622_RNA45SN1_45S_50ntFlanks | 10346 | 10369 | 24 | rRF 45S | NO | 12.82 | 32.49 | 57.72 | 36.10 |
| >hg38_21_+_8433172_8446622_RNA45SN1_45S_50ntFlanks@10346.10372.27\|TCAAACGAGA ACTTTGAAGGCCGAAGT | >hg38_21_+_8433172_8446622_RNA45SN1_45S_50ntFlanks | 10346 | 10372 | 27 | rRF 45S | NO | 0.00 | 0.00 | 20.54 | 0.00 |
| >hg38_21_+_8433172_8446622_RNA45SN1_45S_50ntFlanks@10346.10373.28\|TCAAACGAGA ACTTTGAAGGCCGAAGTG | >hg38_21_+_8433172_8446622_RNA45SN1_45S_50ntFlanks | 10346 | 10373 | 28 | rRF 45S | NO | 19.67 | 0.00 | 0.00 | 13.97 |
| >hg38_21_+_8433172_8446622_RNA45SN1_45S_50ntFlanks@10346.10374.29\|TCAAACGAGA ACTTTGAAGGCCGAAGTGG | >hg38_21_+_8433172_8446622_RNA45SN1_45S_50ntFlanks | 10346 | 10374 | 29 | rRF 45S | NO | 79.76 | 67.64 | 90.14 | 78.93 |
| >hg38_21_+_8433172_8446622_RNA45SN1_45S_50ntFlanks@10346.10383.38\|TCAAACGAGA ACTTTGAAGGCCGAAGTGGAGAAGGGTT | >hg38_21_+_8433172_8446622_RNA45SN1_45S_50ntFlanks | 10346 | 10383 | 38 | rRF 45S | NO | 34.82 | 0.00 | 0.00 | 33.26 |
| >hg38_21_+_8433172_8446622_RNA45SN1_45S_50ntFlanks@10346.10388.43\|TCAAACGAGA ACTTTGAAGGCCGAAGTGGAGAAGGGTTCCATG | >hg38_21_+_8433172_8446622_RNA45SN1_45S_50ntFlanks | 10346 | 10388 | 43 | rRF 45S | NO | 34.60 | 0.00 | 0.00 | 36.96 |
| >hg38_21_+_8433172_8446622_RNA45SN1_45S_50ntFlanks@10347.10368.22\|CAAACGAGAA CTTTGAAGGCCG | >hg38_21_+_8433172_8446622_RNA45SN1_45S_50ntFlanks | 10347 | 10368 | 22 | rRF 45S | NO | 22.10 | 54.00 | 0.00 | 0.00 |
| >hg38_21_+_8433172_8446622_RNA45SN1_45S_50ntFlanks@10347.10374.28\|CAAACGAGAA CTTTGAAGGCCGAAGTGG | >hg38_21_+_8433172_8446622_RNA45SN1_45S_50ntFlanks | 10347 | 10374 | 28 | rRF 45S | NO | 0.00 | 0.00 | 31.17 | 0.00 |
| >hg38_21_+_8433172_8446622_RNA45SN1_45S_50ntFlanks@10348.10366.19\|AAACGAGAAC TTTGAAGGC | >hg38_21_+_8433172_8446622_RNA45SN1_45S_50ntFlanks | 10348 | 10366 | 19 | rRF 45S | NO | 39.30 | 51.91 | 34.54 | 0.00 |
| >hg38_21_+_8433172_8446622_RNA45SN1_45S_50ntFlanks@10348.10367.20\|AAACGAGAAC TTTGAAGGCC | >hg38_21_+_8433172_8446622_RNA45SN1_45S_50ntFlanks | 10348 | 10367 | 20 | rRF 45S | NO | 0.00 | 26.53 | 0.00 | 0.00 |
| >hg38_21_+_8433172_8446622_RNA45SN1_45S_50ntFlanks@10348.10368.21\|AAACGAGAAC TTTGAAGGCCG | >hg38_21_+_8433172_8446622_RNA45SN1_45S_50ntFlanks | 10348 | 10368 | 21 | rRF 45S | NO | 79.33 | 114.68 | 69.77 | 24.41 |
| >hg38_21_+_8433172_8446622_RNA45SN1_45S_50ntFlanks@10348.10371.24\|AAACGAGAAC TTTGAAGGCCGAAG | >hg38_21_+_8433172_8446622_RNA45SN1_45S_50ntFlanks | 10348 | 10371 | 24 | rRF 45S | NO | 20.65 | 0.00 | 13.83 | 0.00 |
| >hg38_21_+_8433172_8446622_RNA45SN1_45S_50ntFlanks@10348.10374.27\|AAACGAGAAC TTTGAAGGCCGAAGTGG | >hg38_21_+_8433172_8446622_RNA45SN1_45S_50ntFlanks | 10348 | 10374 | 27 | rRF 45S | NO | 71.65 | 61.74 | 55.27 | 23.79 |
| >hg38_21_+_8433172_8446622_RNA45SN1_45S_50ntFlanks@10348.10377.30\|AAACGAGAAC TTTGAAGGCCGAAGTGGAGA | >hg38_21_+_8433172_8446622_RNA45SN1_45S_50ntFlanks | 10348 | 10377 | 30 | rRF 45S | NO | 19.69 | 0.00 | 0.00 | 0.00 |
| >hg38_21_+_8433172_8446622_RNA45SN1_45S_50ntFlanks@10348.10383.36\|AAACGAGAAC TTTGAAGGCCGAAGTGGAGAAGGGTT | >hg38_21_+_8433172_8446622_RNA45SN1_45S_50ntFlanks | 10348 | 10383 | 36 | rRF 45S | NO | 104.85 | 56.87 | 31.96 | 33.80 |
| >hg38_21_+_8433172_8446622_RNA45SN1_45S_50ntFlanks@10349.10366.18\|AACGAGAACT TTGAAGGC | >hg38_21_+_8433172_8446622_RNA45SN1_45S_50ntFlanks | 10349 | 10366 | 18 | rRF 45S | NO | 57.31 | 91.92 | 44.17 | 0.00 |

### MDA-MB-468

<table>
<tr><td>>hg38_21_+_8433172_8446622_RNA45SN1_45S_50ntFlanks@10349.10367.19|AACGAGAACTTTGAAGGCC</td><td>>hg38_21_+_8433172_8446622_RNA45SN1_45S_50ntFlanks</td><td>10349</td><td>10367</td><td>19</td><td>rRF 45S</td><td>NO</td><td>21.80</td><td>48.30</td><td>0.00</td><td>0.00</td></tr>
<tr><td>>hg38_21_+_8433172_8446622_RNA45SN1_45S_50ntFlanks@10349.10368.20|AACGAGAACTTTGAAGGCCG</td><td>>hg38_21_+_8433172_8446622_RNA45SN1_45S_50ntFlanks</td><td>10349</td><td>10368</td><td>20</td><td>rRF 45S</td><td>NO</td><td>165.06</td><td>197.78</td><td>101.28</td><td>45.19</td></tr>
<tr><td>>hg38_21_+_8433172_8446622_RNA45SN1_45S_50ntFlanks@10349.10371.23|AACGAGAACTTTGAAGGCCGAAG</td><td>>hg38_21_+_8433172_8446622_RNA45SN1_45S_50ntFlanks</td><td>10349</td><td>10371</td><td>23</td><td>rRF 45S</td><td>NO</td><td>59.29</td><td>34.54</td><td>48.75</td><td>6.50</td></tr>
<tr><td>>hg38_21_+_8433172_8446622_RNA45SN1_45S_50ntFlanks@10349.10374.26|AACGAGAACTTTGAAGGCCGAAGTGG</td><td>>hg38_21_+_8433172_8446622_RNA45SN1_45S_50ntFlanks</td><td>10349</td><td>10374</td><td>26</td><td>rRF 45S</td><td>NO</td><td>80.94</td><td>73.29</td><td>65.08</td><td>33.75</td></tr>
<tr><td>>hg38_21_+_8433172_8446622_RNA45SN1_45S_50ntFlanks@10349.10376.28|AACGAGAACTTTGAAGGCCGAAGTGGAG</td><td>>hg38_21_+_8433172_8446622_RNA45SN1_45S_50ntFlanks</td><td>10349</td><td>10376</td><td>28</td><td>rRF 45S</td><td>NO</td><td>79.33</td><td>56.43</td><td>35.64</td><td>0.00</td></tr>
<tr><td>>hg38_21_+_8433172_8446622_RNA45SN1_45S_50ntFlanks@10349.10377.29|AACGAGAACTTTGAAGGCCGAAGTGGAGA</td><td>>hg38_21_+_8433172_8446622_RNA45SN1_45S_50ntFlanks</td><td>10349</td><td>10377</td><td>29</td><td>rRF 45S</td><td>NO</td><td>128.42</td><td>69.40</td><td>73.69</td><td>0.00</td></tr>
<tr><td>>hg38_21_+_8433172_8446622_RNA45SN1_45S_50ntFlanks@10349.10378.30|AACGAGAACTTTGAAGGCCGAAGTGGAGAA</td><td>>hg38_21_+_8433172_8446622_RNA45SN1_45S_50ntFlanks</td><td>10349</td><td>10378</td><td>30</td><td>rRF 45S</td><td>NO</td><td>41.17</td><td>0.00</td><td>0.00</td><td>0.00</td></tr>
<tr><td>>hg38_21_+_8433172_8446622_RNA45SN1_45S_50ntFlanks@10349.10382.34|AACGAGAACTTTGAAGGCCGAAGTGGAGAAGGGT</td><td>>hg38_21_+_8433172_8446622_RNA45SN1_45S_50ntFlanks</td><td>10349</td><td>10382</td><td>34</td><td>rRF 45S</td><td>NO</td><td>37.97</td><td>27.44</td><td>6.65</td><td>0.00</td></tr>
<tr><td>>hg38_21_+_8433172_8446622_RNA45SN1_45S_50ntFlanks@10349.10383.35|AACGAGAACTTTGAAGGCCGAAGTGGAGAAGGGTT</td><td>>hg38_21_+_8433172_8446622_RNA45SN1_45S_50ntFlanks</td><td>10349</td><td>10383</td><td>35</td><td>rRF 45S</td><td>NO</td><td>596.67</td><td>361.42</td><td>127.68</td><td>89.72</td></tr>
<tr><td>>hg38_21_+_8433172_8446622_RNA45SN1_45S_50ntFlanks@10349.10384.36|AACGAGAACTTTGAAGGCCGAAGTGGAGAAGGGTTC</td><td>>hg38_21_+_8433172_8446622_RNA45SN1_45S_50ntFlanks</td><td>10349</td><td>10384</td><td>36</td><td>rRF 45S</td><td>NO</td><td>120.68</td><td>65.02</td><td>31.49</td><td>0.00</td></tr>
<tr><td>>hg38_21_+_8433172_8446622_RNA45SN1_45S_50ntFlanks@10349.10388.40|AACGAGAACTTTGAAGGCCGAAGTGGAGAAGGGTTCCATG</td><td>>hg38_21_+_8433172_8446622_RNA45SN1_45S_50ntFlanks</td><td>10349</td><td>10388</td><td>40</td><td>rRF 45S</td><td>NO</td><td>73.31</td><td>88.29</td><td>7.21</td><td>42.98</td></tr>
<tr><td>>hg38_21_+_8433172_8446622_RNA45SN1_45S_50ntFlanks@10349.10390.42|AACGAGAACTTTGAAGGCCGAAGTGGAGAAGGGTTCCATGTG</td><td>>hg38_21_+_8433172_8446622_RNA45SN1_45S_50ntFlanks</td><td>10349</td><td>10390</td><td>42</td><td>rRF 45S</td><td>NO</td><td>82.21</td><td>94.13</td><td>0.00</td><td>46.63</td></tr>
<tr><td>>hg38_21_+_8433172_8446622_RNA45SN1_45S_50ntFlanks@10350.10367.18|ACGAGAACTTTGAAGGCC</td><td>>hg38_21_+_8433172_8446622_RNA45SN1_45S_50ntFlanks</td><td>10350</td><td>10367</td><td>18</td><td>rRF 45S</td><td>NO</td><td>54.15</td><td>109.99</td><td>45.86</td><td>0.00</td></tr>
<tr><td>>hg38_21_+_8433172_8446622_RNA45SN1_45S_50ntFlanks@10350.10368.19|ACGAGAACTTTGAAGGCCG</td><td>>hg38_21_+_8433172_8446622_RNA45SN1_45S_50ntFlanks</td><td>10350</td><td>10368</td><td>19</td><td>rRF 45S</td><td>YES</td><td>341.51</td><td>446.99</td><td>242.79</td><td>113.52</td></tr>
<tr><td>>hg38_21_+_8433172_8446622_RNA45SN1_45S_50ntFlanks@10350.10369.20|ACGAGAACTTTGAAGGCCGA</td><td>>hg38_21_+_8433172_8446622_RNA45SN1_45S_50ntFlanks</td><td>10350</td><td>10369</td><td>20</td><td>rRF 45S</td><td>NO</td><td>32.40</td><td>33.50</td><td>37.80</td><td>0.00</td></tr>
<tr><td>>hg38_21_+_8433172_8446622_RNA45SN1_45S_50ntFlanks@10350.10370.21|ACGAGAACTTTGAAGGCCGAA</td><td>>hg38_21_+_8433172_8446622_RNA45SN1_45S_50ntFlanks</td><td>10350</td><td>10370</td><td>21</td><td>rRF 45S</td><td>NO</td><td>0.00</td><td>0.00</td><td>28.28</td><td>0.00</td></tr>
<tr><td>>hg38_21_+_8433172_8446622_RNA45SN1_45S_50ntFlanks@10350.10371.22|ACGAGAACTTTGAAGGCCGAAG</td><td>>hg38_21_+_8433172_8446622_RNA45SN1_45S_50ntFlanks</td><td>10350</td><td>10371</td><td>22</td><td>rRF 45S</td><td>NO</td><td>104.98</td><td>66.17</td><td>87.27</td><td>43.06</td></tr>
<tr><td>>hg38_21_+_8433172_8446622_RNA45SN1_45S_50ntFlanks@10350.10372.23|ACGAGAACTTTGAAGGCCGAAGT</td><td>>hg38_21_+_8433172_8446622_RNA45SN1_45S_50ntFlanks</td><td>10350</td><td>10372</td><td>23</td><td>rRF 45S</td><td>NO</td><td>31.16</td><td>29.34</td><td>35.85</td><td>0.00</td></tr>
<tr><td>>hg38_21_+_8433172_8446622_RNA45SN1_45S_50ntFlanks@10350.10373.24|ACGAGAACTTTGAAGGCCGAAGTG</td><td>>hg38_21_+_8433172_8446622_RNA45SN1_45S_50ntFlanks</td><td>10350</td><td>10373</td><td>24</td><td>rRF 45S</td><td>NO</td><td>34.10</td><td>47.58</td><td>45.06</td><td>0.00</td></tr>
<tr><td>>hg38_21_+_8433172_8446622_RNA45SN1_45S_50ntFlanks@10350.10374.25|ACGAGAACTTTGAAGGCCGAAGTGG</td><td>>hg38_21_+_8433172_8446622_RNA45SN1_45S_50ntFlanks</td><td>10350</td><td>10374</td><td>25</td><td>rRF 45S</td><td>NO</td><td>169.40</td><td>167.06</td><td>140.54</td><td>85.34</td></tr>
<tr><td>>hg38_21_+_8433172_8446622_RNA45SN1_45S_50ntFlanks@10350.10376.27|ACGAGAACTTTGAAGGCCGAAGTGGAG</td><td>>hg38_21_+_8433172_8446622_RNA45SN1_45S_50ntFlanks</td><td>10350</td><td>10376</td><td>27</td><td>rRF 45S</td><td>NO</td><td>154.81</td><td>108.98</td><td>77.68</td><td>26.78</td></tr>
<tr><td>>hg38_21_+_8433172_8446622_RNA45SN1_45S_50ntFlanks@10350.10377.28|ACGAGAACTTTGAAGGCCGAAGTGGAGA</td><td>>hg38_21_+_8433172_8446622_RNA45SN1_45S_50ntFlanks</td><td>10350</td><td>10377</td><td>28</td><td>rRF 45S</td><td>NO</td><td>272.11</td><td>127.26</td><td>160.25</td><td>39.05</td></tr>
<tr><td>>hg38_21_+_8433172_8446622_RNA45SN1_45S_50ntFlanks@10350.10378.29|ACGAGAACTTTGAAGGCCGAAGTGGAGAA</td><td>>hg38_21_+_8433172_8446622_RNA45SN1_45S_50ntFlanks</td><td>10350</td><td>10378</td><td>29</td><td>rRF 45S</td><td>NO</td><td>91.11</td><td>0.00</td><td>29.16</td><td>0.00</td></tr>
<tr><td>>hg38_21_+_8433172_8446622_RNA45SN1_45S_50ntFlanks@10350.10379.30|ACGAGAACTTTGAAGGCCGAAGTGGAGAAG</td><td>>hg38_21_+_8433172_8446622_RNA45SN1_45S_50ntFlanks</td><td>10350</td><td>10379</td><td>30</td><td>rRF 45S</td><td>NO</td><td>51.74</td><td>28.01</td><td>35.40</td><td>0.00</td></tr>
<tr><td>>hg38_21_+_8433172_8446622_RNA45SN1_45S_50ntFlanks@10350.10380.31|ACGAGAACTTTGAAGGCCGAAGTGGAGAAGG</td><td>>hg38_21_+_8433172_8446622_RNA45SN1_45S_50ntFlanks</td><td>10350</td><td>10380</td><td>31</td><td>rRF 45S</td><td>NO</td><td>20.06</td><td>0.00</td><td>0.00</td><td>0.00</td></tr>
<tr><td>>hg38_21_+_8433172_8446622_RNA45SN1_45S_50ntFlanks@10350.10381.32|ACGAGAACTTTGAAGGCCGAAGTGGAGAAGGG</td><td>>hg38_21_+_8433172_8446622_RNA45SN1_45S_50ntFlanks</td><td>10350</td><td>10381</td><td>32</td><td>rRF 45S</td><td>NO</td><td>51.77</td><td>26.95</td><td>27.22</td><td>0.00</td></tr>
<tr><td>>hg38_21_+_8433172_8446622_RNA45SN1_45S_50ntFlanks@10350.10382.33|ACGAGAACTTTGAAGGCCGAAGTGGAGAAGGGT</td><td>>hg38_21_+_8433172_8446622_RNA45SN1_45S_50ntFlanks</td><td>10350</td><td>10382</td><td>33</td><td>rRF 45S</td><td>NO</td><td>88.46</td><td>54.58</td><td>53.54</td><td>14.04</td></tr>
<tr><td>>hg38_21_+_8433172_8446622_RNA45SN1_45S_50ntFlanks@10350.10383.34|ACGAGAACTTTGAAGGCCGAAGTGGAGAAGGGTT</td><td>>hg38_21_+_8433172_8446622_RNA45SN1_45S_50ntFlanks</td><td>10350</td><td>10383</td><td>34</td><td>rRF 45S</td><td>YES</td><td>1273.12</td><td>596.22</td><td>276.50</td><td>186.84</td></tr>
<tr><td>>hg38_21_+_8433172_8446622_RNA45SN1_45S_50ntFlanks@10350.10384.35|ACGAGAACTTTGAAGGCCGAAGTGGAGAAGGGTTC</td><td>>hg38_21_+_8433172_8446622_RNA45SN1_45S_50ntFlanks</td><td>10350</td><td>10384</td><td>35</td><td>rRF 45S</td><td>NO</td><td>278.73</td><td>128.41</td><td>77.44</td><td>44.87</td></tr>
<tr><td>>hg38_21_+_8433172_8446622_RNA45SN1_45S_50ntFlanks@10350.10388.39|ACGAGAACTTTGAAGGCCGAAGTGGAGAAGGGTTCCATG</td><td>>hg38_21_+_8433172_8446622_RNA45SN1_45S_50ntFlanks</td><td>10350</td><td>10388</td><td>39</td><td>rRF 45S</td><td>NO</td><td>165.20</td><td>194.48</td><td>62.21</td><td>108.64</td></tr>
<tr><td>>hg38_21_+_8433172_8446622_RNA45SN1_45S_50ntFlanks@10350.10390.41|ACGAGAACTTTGAAGGCCGAAGTGGAGAAGGGTTCCATGTG</td><td>>hg38_21_+_8433172_8446622_RNA45SN1_45S_50ntFlanks</td><td>10350</td><td>10390</td><td>41</td><td>rRF 45S</td><td>NO</td><td>223.00</td><td>231.23</td><td>50.69</td><td>119.36</td></tr>
<tr><td>>hg38_21_+_8433172_8446622_RNA45SN1_45S_50ntFlanks@10350.10391.42|ACGAGAACTTTGAAGGCCGAAGTGGAGAAGGGTTCCATGTGA</td><td>>hg38_21_+_8433172_8446622_RNA45SN1_45S_50ntFlanks</td><td>10350</td><td>10391</td><td>42</td><td>rRF 45S</td><td>NO</td><td>30.31</td><td>0.00</td><td>0.00</td><td>0.00</td></tr>
<tr><td>>hg38_21_+_8433172_8446622_RNA45SN1_45S_50ntFlanks@10351.10368.18|CGAGAACTTTGAAGGCCG</td><td>>hg38_21_+_8433172_8446622_RNA45SN1_45S_50ntFlanks</td><td>10351</td><td>10368</td><td>18</td><td>rRF 45S</td><td>YES</td><td>869.99</td><td>1283.07</td><td>519.39</td><td>300.65</td></tr>
<tr><td>>hg38_21_+_8433172_8446622_RNA45SN1_45S_50ntFlanks@10351.10369.19|CGAGAACTTTGAAGGCCGA</td><td>>hg38_21_+_8433172_8446622_RNA45SN1_45S_50ntFlanks</td><td>10351</td><td>10369</td><td>19</td><td>rRF 45S</td><td>NO</td><td>38.02</td><td>56.05</td><td>56.59</td><td>14.44</td></tr>
</table>

### MDA-MB-468

| | | | | | | | | | | |
|---|---|---|---|---|---|---|---|---|---|---|
| >hg38_21_+_8433172_8446622_RNA45SN1_45S_50ntFlanks@10351.10370.20\|CGAGAACTTTGAAGGCCGAA | >hg38_21_+_8433172_8446622_RNA45SN1_45S_50ntFlanks | 10351 | 10370 | 20 | rRF 45S | NO | 0.00 | 26.94 | 44.02 | 0.00 |
| >hg38_21_+_8433172_8446622_RNA45SN1_45S_50ntFlanks@10351.10371.21\|CGAGAACTTTGAAGGCCGAAG | >hg38_21_+_8433172_8446622_RNA45SN1_45S_50ntFlanks | 10351 | 10371 | 21 | rRF 45S | NO | 63.78 | 62.69 | 73.60 | 49.10 |
| >hg38_21_+_8433172_8446622_RNA45SN1_45S_50ntFlanks@10351.10372.22\|CGAGAACTTTGAAGGCCGAAGT | >hg38_21_+_8433172_8446622_RNA45SN1_45S_50ntFlanks | 10351 | 10372 | 22 | rRF 45S | NO | 21.26 | 34.54 | 41.19 | 0.00 |
| >hg38_21_+_8433172_8446622_RNA45SN1_45S_50ntFlanks@10351.10373.23\|CGAGAACTTTGAAGGCCGAAGTG | >hg38_21_+_8433172_8446622_RNA45SN1_45S_50ntFlanks | 10351 | 10373 | 23 | rRF 45S | NO | 39.41 | 79.81 | 74.19 | 6.48 |
| >hg38_21_+_8433172_8446622_RNA45SN1_45S_50ntFlanks@10351.10374.24\|CGAGAACTTTGAAGGCCGAAGTGG | >hg38_21_+_8433172_8446622_RNA45SN1_45S_50ntFlanks | 10351 | 10374 | 24 | rRF 45S | NO | 183.77 | 207.93 | 192.52 | 111.66 |
| >hg38_21_+_8433172_8446622_RNA45SN1_45S_50ntFlanks@10351.10376.26\|CGAGAACTTTGAAGGCCGAAGTGGAG | >hg38_21_+_8433172_8446622_RNA45SN1_45S_50ntFlanks | 10351 | 10376 | 26 | rRF 45S | NO | 127.24 | 125.70 | 127.97 | 40.89 |
| >hg38_21_+_8433172_8446622_RNA45SN1_45S_50ntFlanks@10351.10377.27\|CGAGAACTTTGAAGGCCGAAGTGGAGA | >hg38_21_+_8433172_8446622_RNA45SN1_45S_50ntFlanks | 10351 | 10377 | 27 | rRF 45S | NO | 548.20 | 385.51 | 465.50 | 111.34 |
| >hg38_21_+_8433172_8446622_RNA45SN1_45S_50ntFlanks@10351.10378.28\|CGAGAACTTTGAAGGCCGAAGTGGAGAA | >hg38_21_+_8433172_8446622_RNA45SN1_45S_50ntFlanks | 10351 | 10378 | 28 | rRF 45S | NO | 166.67 | 57.39 | 70.65 | 14.84 |
| >hg38_21_+_8433172_8446622_RNA45SN1_45S_50ntFlanks@10351.10379.29\|CGAGAACTTTGAAGGCCGAAGTGGAGAAG | >hg38_21_+_8433172_8446622_RNA45SN1_45S_50ntFlanks | 10351 | 10379 | 29 | rRF 45S | NO | 84.95 | 55.81 | 70.06 | 34.85 |
| >hg38_21_+_8433172_8446622_RNA45SN1_45S_50ntFlanks@10351.10380.30\|CGAGAACTTTGAAGGCCGAAGTGGAGAAGG | >hg38_21_+_8433172_8446622_RNA45SN1_45S_50ntFlanks | 10351 | 10380 | 30 | rRF 45S | NO | 56.70 | 45.52 | 49.47 | 0.00 |
| >hg38_21_+_8433172_8446622_RNA45SN1_45S_50ntFlanks@10351.10381.31\|CGAGAACTTTGAAGGCCGAAGTGGAGAAGGG | >hg38_21_+_8433172_8446622_RNA45SN1_45S_50ntFlanks | 10351 | 10381 | 31 | rRF 45S | NO | 99.67 | 59.43 | 58.54 | 15.95 |
| >hg38_21_+_8433172_8446622_RNA45SN1_45S_50ntFlanks@10351.10382.32\|CGAGAACTTTGAAGGCCGAAGTGGAGAAGGGT | >hg38_21_+_8433172_8446622_RNA45SN1_45S_50ntFlanks | 10351 | 10382 | 32 | rRF 45S | NO | 159.66 | 107.07 | 103.48 | 47.41 |
| >hg38_21_+_8433172_8446622_RNA45SN1_45S_50ntFlanks@10351.10383.33\|CGAGAACTTTGAAGGCCGAAGTGGAGAAGGGTT | >hg38_21_+_8433172_8446622_RNA45SN1_45S_50ntFlanks | 10351 | 10383 | 33 | rRF 45S | YES | 2631.91 | 1470.32 | 655.13 | 408.37 |
| >hg38_21_+_8433172_8446622_RNA45SN1_45S_50ntFlanks@10351.10384.34\|CGAGAACTTTGAAGGCCGAAGTGGAGAAGGGTTC | >hg38_21_+_8433172_8446622_RNA45SN1_45S_50ntFlanks | 10351 | 10384 | 34 | rRF 45S | NO | 414.27 | 244.40 | 152.47 | 85.50 |
| >hg38_21_+_8433172_8446622_RNA45SN1_45S_50ntFlanks@10351.10385.35\|CGAGAACTTTGAAGGCCGAAGTGGAGAAGGGTTCC | >hg38_21_+_8433172_8446622_RNA45SN1_45S_50ntFlanks | 10351 | 10385 | 35 | rRF 45S | NO | 12.88 | 0.00 | 0.00 | 0.00 |
| >hg38_21_+_8433172_8446622_RNA45SN1_45S_50ntFlanks@10351.10388.38\|CGAGAACTTTGAAGGCCGAAGTGGAGAAGGGTTCCATG | >hg38_21_+_8433172_8446622_RNA45SN1_45S_50ntFlanks | 10351 | 10388 | 38 | rRF 45S | YES | 390.11 | 459.29 | 156.29 | 247.07 |
| >hg38_21_+_8433172_8446622_RNA45SN1_45S_50ntFlanks@10351.10390.40\|CGAGAACTTTGAAGGCCGAAGTGGAGAAGGGTTCCATGTG | >hg38_21_+_8433172_8446622_RNA45SN1_45S_50ntFlanks | 10351 | 10390 | 40 | rRF 45S | NO | 754.25 | 736.76 | 159.88 | 360.50 |
| >hg38_21_+_8433172_8446622_RNA45SN1_45S_50ntFlanks@10351.10391.41\|CGAGAACTTTGAAGGCCGAAGTGGAGAAGGGTTCCATGTGA | >hg38_21_+_8433172_8446622_RNA45SN1_45S_50ntFlanks | 10351 | 10391 | 41 | rRF 45S | NO | 73.66 | 30.75 | 0.00 | 45.58 |
| >hg38_21_+_8433172_8446622_RNA45SN1_45S_50ntFlanks@10353.10383.31\|AGAACTTTGAAGGCCGAAGTGGAGAAGGGTT | >hg38_21_+_8433172_8446622_RNA45SN1_45S_50ntFlanks | 10353 | 10383 | 31 | rRF 45S | NO | 42.36 | 0.00 | 0.00 | 0.00 |
| >hg38_21_+_8433172_8446622_RNA45SN1_45S_50ntFlanks@10353.10388.36\|AGAACTTTGAAGGCCGAAGTGGAGAAGGGTTCCATG | >hg38_21_+_8433172_8446622_RNA45SN1_45S_50ntFlanks | 10353 | 10388 | 36 | rRF 45S | NO | 6.14 | 19.34 | 0.00 | 0.00 |
| >hg38_21_+_8433172_8446622_RNA45SN1_45S_50ntFlanks@10353.10390.38\|AGAACTTTGAAGGCCGAAGTGGAGAAGGGTTCCATGTG | >hg38_21_+_8433172_8446622_RNA45SN1_45S_50ntFlanks | 10353 | 10390 | 38 | rRF 45S | NO | 29.01 | 29.25 | 0.00 | 0.00 |
| >hg38_21_+_8433172_8446622_RNA45SN1_45S_50ntFlanks@10355.10383.29\|AACTTTGAAGGCCGAAGTGGAGAAGGGTT | >hg38_21_+_8433172_8446622_RNA45SN1_45S_50ntFlanks | 10355 | 10383 | 29 | rRF 45S | NO | 41.10 | 0.00 | 0.00 | 0.00 |
| >hg38_21_+_8433172_8446622_RNA45SN1_45S_50ntFlanks@10355.10388.34\|AACTTTGAAGGCCGAAGTGGAGAAGGGTTCCATG | >hg38_21_+_8433172_8446622_RNA45SN1_45S_50ntFlanks | 10355 | 10388 | 34 | rRF 45S | NO | 50.84 | 43.06 | 0.00 | 0.00 |
| >hg38_21_+_8433172_8446622_RNA45SN1_45S_50ntFlanks@10355.10390.36\|AACTTTGAAGGCCGAAGTGGAGAAGGGTTCCATGTG | >hg38_21_+_8433172_8446622_RNA45SN1_45S_50ntFlanks | 10355 | 10390 | 36 | rRF 45S | NO | 31.32 | 29.85 | 0.00 | 0.00 |
| >hg38_21_+_8433172_8446622_RNA45SN1_45S_50ntFlanks@10357.10383.27\|CTTTGAAGGCCGAAGTGGAGAAGGGTT | >hg38_21_+_8433172_8446622_RNA45SN1_45S_50ntFlanks | 10357 | 10383 | 27 | rRF 45S | NO | 74.22 | 29.74 | 7.12 | 0.00 |
| >hg38_21_+_8433172_8446622_RNA45SN1_45S_50ntFlanks@10357.10388.32\|CTTTGAAGGCCGAAGTGGAGAAGGGTTCCATG | >hg38_21_+_8433172_8446622_RNA45SN1_45S_50ntFlanks | 10357 | 10388 | 32 | rRF 45S | NO | 50.95 | 41.66 | 0.00 | 0.00 |
| >hg38_21_+_8433172_8446622_RNA45SN1_45S_50ntFlanks@10357.10390.34\|CTTTGAAGGCCGAAGTGGAGAAGGGTTCCATGTG | >hg38_21_+_8433172_8446622_RNA45SN1_45S_50ntFlanks | 10357 | 10390 | 34 | rRF 45S | NO | 38.73 | 37.12 | 0.00 | 0.00 |
| >hg38_21_+_8433172_8446622_RNA45SN1_45S_50ntFlanks@10361.10383.23\|GAAGGCCGAAGTGGAGAAGGGTT | >hg38_21_+_8433172_8446622_RNA45SN1_45S_50ntFlanks | 10361 | 10383 | 23 | rRF 45S | NO | 62.07 | 36.48 | 28.63 | 0.00 |
| >hg38_21_+_8433172_8446622_RNA45SN1_45S_50ntFlanks@10361.10388.28\|GAAGGCCGAAGTGGAGAAGGGTTCCATG | >hg38_21_+_8433172_8446622_RNA45SN1_45S_50ntFlanks | 10361 | 10388 | 28 | rRF 45S | NO | 34.72 | 49.13 | 0.00 | 0.00 |
| >hg38_21_+_8433172_8446622_RNA45SN1_45S_50ntFlanks@10361.10390.30\|GAAGGCCGAAGTGGAGAAGGGTTCCATGTG | >hg38_21_+_8433172_8446622_RNA45SN1_45S_50ntFlanks | 10361 | 10390 | 30 | rRF 45S | NO | 13.14 | 38.13 | 0.00 | 0.00 |
| >hg38_21_+_8433172_8446622_RNA45SN1_45S_50ntFlanks@10362.10382.21\|AAGGCCGAAGTGGAGAAGGGT | >hg38_21_+_8433172_8446622_RNA45SN1_45S_50ntFlanks | 10362 | 10382 | 21 | rRF 45S | NO | 19.85 | 0.00 | 0.00 | 0.00 |
| >hg38_21_+_8433172_8446622_RNA45SN1_45S_50ntFlanks@10362.10383.22\|AAGGCCGAAGTGGAGAAGGGTT | >hg38_21_+_8433172_8446622_RNA45SN1_45S_50ntFlanks | 10362 | 10383 | 22 | rRF 45S | NO | 283.05 | 201.48 | 91.79 | 57.77 |
| >hg38_21_+_8433172_8446622_RNA45SN1_45S_50ntFlanks@10362.10384.23\|AAGGCCGAAGTGGAGAAGGGTTC | >hg38_21_+_8433172_8446622_RNA45SN1_45S_50ntFlanks | 10362 | 10384 | 23 | rRF 45S | NO | 72.32 | 58.72 | 42.01 | 0.00 |
| >hg38_21_+_8433172_8446622_RNA45SN1_45S_50ntFlanks@10362.10388.27\|AAGGCCGAAGTGGAGAAGGGTTCCATG | >hg38_21_+_8433172_8446622_RNA45SN1_45S_50ntFlanks | 10362 | 10388 | 27 | rRF 45S | NO | 193.16 | 222.20 | 79.88 | 43.22 |

### MDA-MB-468

| | | | | | | | | | | |
|---|---|---|---|---|---|---|---|---|---|---|
| >hg38_21_+_8433172_8446622_RNA45SN1_45S_50ntFlanks@10362.10390.29\|AAGGCCGAAGTGGAGAAGGGTTCCATGTG | >hg38_21_+_8433172_8446622_RNA45SN1_45S_50ntFlanks | 10362 | 10390 | 29 | rRF 45S | NO | 145.71 | 191.59 | 68.59 | 47.70 |
| >hg38_21_+_8433172_8446622_RNA45SN1_45S_50ntFlanks@10369.10407.39\|AAGTGGAGAAGGGTTCCATGTGAACAGCAGTTGAACATG | >hg38_21_+_8433172_8446622_RNA45SN1_45S_50ntFlanks | 10369 | 10407 | 39 | rRF 45S | NO | 61.72 | 93.82 | 0.00 | 0.00 |
| >hg38_21_+_8433172_8446622_RNA45SN1_45S_50ntFlanks@10375.10406.32\|AGAAGGGTTCCATGTGAACAGCAGTTGAACAT | >hg38_21_+_8433172_8446622_RNA45SN1_45S_50ntFlanks | 10375 | 10406 | 32 | rRF 45S | NO | 38.08 | 12.77 | 0.00 | 0.00 |
| >hg38_21_+_8433172_8446622_RNA45SN1_45S_50ntFlanks@10375.10407.33\|AGAAGGGTTCCATGTGAACAGCAGTTGAACATG | >hg38_21_+_8433172_8446622_RNA45SN1_45S_50ntFlanks | 10375 | 10407 | 33 | rRF 45S | NO | 30.82 | 50.07 | 0.00 | 0.00 |
| >hg38_21_+_8433172_8446622_RNA45SN1_45S_50ntFlanks@10384.10405.22\|CCATGTGAACAGCAGTTGAACA | >hg38_21_+_8433172_8446622_RNA45SN1_45S_50ntFlanks | 10384 | 10405 | 22 | rRF 45S | NO | 28.83 | 0.00 | 6.60 | 0.00 |
| >hg38_21_+_8433172_8446622_RNA45SN1_45S_50ntFlanks@10384.10424.41\|CCATGTGAACAGCAGTTGAACATGGGTCAGTCGGTCCTGAG | >hg38_21_+_8433172_8446622_RNA45SN1_45S_50ntFlanks | 10384 | 10424 | 41 | rRF 45S | NO | 68.63 | 0.00 | 0.00 | 0.00 |
| >hg38_21_+_8433172_8446622_RNA45SN1_45S_50ntFlanks@10387.10410.24\|TGTGAACAGCAGTTGAACATGGGT | >hg38_21_+_8433172_8446622_RNA45SN1_45S_50ntFlanks | 10387 | 10410 | 24 | rRF 45S | NO | 39.73 | 0.00 | 0.00 | 0.00 |
| >hg38_21_+_8433172_8446622_RNA45SN1_45S_50ntFlanks@10388.10410.23\|GTGAACAGCAGTTGAACATGGGT | >hg38_21_+_8433172_8446622_RNA45SN1_45S_50ntFlanks | 10388 | 10410 | 23 | rRF 45S | NO | 61.80 | 44.60 | 0.00 | 0.00 |
| >hg38_21_+_8433172_8446622_RNA45SN1_45S_50ntFlanks@10389.10407.19\|TGAACAGCAGTTGAACATG | >hg38_21_+_8433172_8446622_RNA45SN1_45S_50ntFlanks | 10389 | 10407 | 19 | rRF 45S | NO | 126.04 | 165.29 | 53.87 | 6.28 |
| >hg38_21_+_8433172_8446622_RNA45SN1_45S_50ntFlanks@10389.10409.21\|TGAACAGCAGTTGAACATGGG | >hg38_21_+_8433172_8446622_RNA45SN1_45S_50ntFlanks | 10389 | 10409 | 21 | rRF 45S | NO | 82.72 | 84.36 | 0.00 | 0.00 |
| >hg38_21_+_8433172_8446622_RNA45SN1_45S_50ntFlanks@10389.10410.22\|TGAACAGCAGTTGAACATGGGT | >hg38_21_+_8433172_8446622_RNA45SN1_45S_50ntFlanks | 10389 | 10410 | 22 | rRF 45S | NO | 245.32 | 180.48 | 39.15 | 0.00 |
| >hg38_21_+_8433172_8446622_RNA45SN1_45S_50ntFlanks@10389.10424.36\|TGAACAGCAGTTGAACATGGGTCAGTCGGTCCTGAG | >hg38_21_+_8433172_8446622_RNA45SN1_45S_50ntFlanks | 10389 | 10424 | 36 | rRF 45S | NO | 128.06 | 76.65 | 0.00 | 50.90 |
| >hg38_21_+_8433172_8446622_RNA45SN1_45S_50ntFlanks@10389.10429.41\|TGAACAGCAGTTGAACATGGGTCAGTCGGTCCTGAGAGATG | >hg38_21_+_8433172_8446622_RNA45SN1_45S_50ntFlanks | 10389 | 10429 | 41 | rRF 45S | NO | 54.45 | 49.41 | 0.00 | 0.00 |
| >hg38_21_+_8433172_8446622_RNA45SN1_45S_50ntFlanks@10389.10430.42\|TGAACAGCAGTTGAACATGGGTCAGTCGGTCCTGAGAGATGG | >hg38_21_+_8433172_8446622_RNA45SN1_45S_50ntFlanks | 10389 | 10430 | 42 | rRF 45S | NO | 19.89 | 0.00 | 0.00 | 0.00 |
| >hg38_21_+_8433172_8446622_RNA45SN1_45S_50ntFlanks@10391.10408.18\|AACAGCAGTTGAACATGG | >hg38_21_+_8433172_8446622_RNA45SN1_45S_50ntFlanks | 10391 | 10408 | 18 | rRF 45S | NO | 30.99 | 47.13 | 0.00 | 0.00 |
| >hg38_21_+_8433172_8446622_RNA45SN1_45S_50ntFlanks@10391.10409.19\|AACAGCAGTTGAACATGGG | >hg38_21_+_8433172_8446622_RNA45SN1_45S_50ntFlanks | 10391 | 10409 | 19 | rRF 45S | NO | 144.66 | 284.19 | 27.36 | 0.00 |
| >hg38_21_+_8433172_8446622_RNA45SN1_45S_50ntFlanks@10391.10410.20\|AACAGCAGTTGAACATGGGT | >hg38_21_+_8433172_8446622_RNA45SN1_45S_50ntFlanks | 10391 | 10410 | 20 | rRF 45S | NO | 355.05 | 394.24 | 58.59 | 37.24 |
| >hg38_21_+_8433172_8446622_RNA45SN1_45S_50ntFlanks@10391.10419.29\|AACAGCAGTTGAACATGGGTCAGTCGGTC | >hg38_21_+_8433172_8446622_RNA45SN1_45S_50ntFlanks | 10391 | 10419 | 29 | rRF 45S | NO | 53.34 | 48.26 | 0.00 | 0.00 |
| >hg38_21_+_8433172_8446622_RNA45SN1_45S_50ntFlanks@10391.10420.30\|AACAGCAGTTGAACATGGGTCAGTCGGTCC | >hg38_21_+_8433172_8446622_RNA45SN1_45S_50ntFlanks | 10391 | 10420 | 30 | rRF 45S | NO | 57.13 | 55.31 | 0.00 | 15.25 |
| >hg38_21_+_8433172_8446622_RNA45SN1_45S_50ntFlanks@10391.10422.32\|AACAGCAGTTGAACATGGGTCAGTCGGTCCTG | >hg38_21_+_8433172_8446622_RNA45SN1_45S_50ntFlanks | 10391 | 10422 | 32 | rRF 45S | NO | 35.13 | 40.68 | 0.00 | 0.00 |
| >hg38_21_+_8433172_8446622_RNA45SN1_45S_50ntFlanks@10391.10424.34\|AACAGCAGTTGAACATGGGTCAGTCGGTCCTGAG | >hg38_21_+_8433172_8446622_RNA45SN1_45S_50ntFlanks | 10391 | 10424 | 34 | rRF 45S | NO | 171.51 | 104.92 | 0.00 | 84.79 |
| >hg38_21_+_8433172_8446622_RNA45SN1_45S_50ntFlanks@10391.10426.36\|AACAGCAGTTGAACATGGGTCAGTCGGTCCTGAGAG | >hg38_21_+_8433172_8446622_RNA45SN1_45S_50ntFlanks | 10391 | 10426 | 36 | rRF 45S | NO | 34.90 | 27.18 | 0.00 | 0.00 |
| >hg38_21_+_8433172_8446622_RNA45SN1_45S_50ntFlanks@10391.10429.39\|AACAGCAGTTGAACATGGGTCAGTCGGTCCTGAGAGATG | >hg38_21_+_8433172_8446622_RNA45SN1_45S_50ntFlanks | 10391 | 10429 | 39 | rRF 45S | NO | 134.34 | 141.98 | 0.00 | 48.40 |
| >hg38_21_+_8433172_8446622_RNA45SN1_45S_50ntFlanks@10391.10430.40\|AACAGCAGTTGAACATGGGTCAGTCGGTCCTGAGAGATGG | >hg38_21_+_8433172_8446622_RNA45SN1_45S_50ntFlanks | 10391 | 10430 | 40 | rRF 45S | NO | 63.70 | 59.97 | 0.00 | 26.07 |
| >hg38_21_+_8433172_8446622_RNA45SN1_45S_50ntFlanks@10391.10433.43\|AACAGCAGTTGAACATGGGTCAGTCGGTCCTGAGAGATGGGCG | >hg38_21_+_8433172_8446622_RNA45SN1_45S_50ntFlanks | 10391 | 10433 | 43 | rRF 45S | NO | 21.59 | 35.39 | 0.00 | 0.00 |
| >hg38_21_+_8433172_8446622_RNA45SN1_45S_50ntFlanks@10391.10435.45\|AACAGCAGTTGAACATGGGTCAGTCGGTCCTGAGAGATGGGCGAG | >hg38_21_+_8433172_8446622_RNA45SN1_45S_50ntFlanks | 10391 | 10435 | 45 | rRF 45S | NO | 32.75 | 40.69 | 0.00 | 16.28 |
| >hg38_21_+_8433172_8446622_RNA45SN1_45S_50ntFlanks@10392.10409.18\|ACAGCAGTTGAACATGGG | >hg38_21_+_8433172_8446622_RNA45SN1_45S_50ntFlanks | 10392 | 10409 | 18 | rRF 45S | NO | 142.81 | 241.33 | 7.13 | 0.00 |
| >hg38_21_+_8433172_8446622_RNA45SN1_45S_50ntFlanks@10392.10410.19\|ACAGCAGTTGAACATGGGT | >hg38_21_+_8433172_8446622_RNA45SN1_45S_50ntFlanks | 10392 | 10410 | 19 | rRF 45S | YES | 368.10 | 328.33 | 92.07 | 40.14 |
| >hg38_21_+_8433172_8446622_RNA45SN1_45S_50ntFlanks@10392.10411.20\|ACAGCAGTTGAACATGGGTC | >hg38_21_+_8433172_8446622_RNA45SN1_45S_50ntFlanks | 10392 | 10411 | 20 | rRF 45S | NO | 33.82 | 27.04 | 0.00 | 0.00 |
| >hg38_21_+_8433172_8446622_RNA45SN1_45S_50ntFlanks@10392.10418.27\|ACAGCAGTTGAACATGGGTCAGTCGGT | >hg38_21_+_8433172_8446622_RNA45SN1_45S_50ntFlanks | 10392 | 10418 | 27 | rRF 45S | NO | 21.23 | 0.00 | 0.00 | 0.00 |
| >hg38_21_+_8433172_8446622_RNA45SN1_45S_50ntFlanks@10392.10419.28\|ACAGCAGTTGAACATGGGTCAGTCGGTC | >hg38_21_+_8433172_8446622_RNA45SN1_45S_50ntFlanks | 10392 | 10419 | 28 | rRF 45S | NO | 87.58 | 48.15 | 0.00 | 0.00 |
| >hg38_21_+_8433172_8446622_RNA45SN1_45S_50ntFlanks@10392.10420.29\|ACAGCAGTTGAACATGGGTCAGTCGGTCC | >hg38_21_+_8433172_8446622_RNA45SN1_45S_50ntFlanks | 10392 | 10420 | 29 | rRF 45S | NO | 59.49 | 38.91 | 0.00 | 0.00 |
| >hg38_21_+_8433172_8446622_RNA45SN1_45S_50ntFlanks@10392.10422.31\|ACAGCAGTTGAACATGGGTCAGTCGGTCCTG | >hg38_21_+_8433172_8446622_RNA45SN1_45S_50ntFlanks | 10392 | 10422 | 31 | rRF 45S | NO | 77.32 | 51.04 | 0.00 | 0.00 |
| >hg38_21_+_8433172_8446622_RNA45SN1_45S_50ntFlanks@10392.10424.33\|ACAGCAGTTGAACATGGGTCAGTCGGTCCTGAG | >hg38_21_+_8433172_8446622_RNA45SN1_45S_50ntFlanks | 10392 | 10424 | 33 | rRF 45S | NO | 195.04 | 103.82 | 29.05 | 83.63 |

### MDA-MB-468

| | | | | | | | | | | |
|---|---|---|---|---|---|---|---|---|---|---|
| >hg38_21_+_8433172_8446622_RNA45SN1_45S_50ntFlanks@10392.10426.35\|ACAGCAGTTGAACATGGTCAGTCGGTCCTGAGAG | >hg38_21_+_8433172_8446622_RNA45SN1_45S_50ntFlanks | 10392 | 10426 | 35 | rRF 45S | NO | 62.31 | 35.23 | 0.00 | 0.00 |
| >hg38_21_+_8433172_8446622_RNA45SN1_45S_50ntFlanks@10392.10428.37\|ACAGCAGTTGAACATGGTCAGTCGGTCCTGAGAGAT | >hg38_21_+_8433172_8446622_RNA45SN1_45S_50ntFlanks | 10392 | 10428 | 37 | rRF 45S | NO | 20.14 | 0.00 | 0.00 | 0.00 |
| >hg38_21_+_8433172_8446622_RNA45SN1_45S_50ntFlanks@10392.10429.38\|ACAGCAGTTGAACATGGTCAGTCGGTCCTGAGAGATG | >hg38_21_+_8433172_8446622_RNA45SN1_45S_50ntFlanks | 10392 | 10429 | 38 | rRF 45S | NO | 219.49 | 168.16 | 37.99 | 60.12 |
| >hg38_21_+_8433172_8446622_RNA45SN1_45S_50ntFlanks@10392.10430.39\|ACAGCAGTTGAACATGGTCAGTCGGTCCTGAGAGATGG | >hg38_21_+_8433172_8446622_RNA45SN1_45S_50ntFlanks | 10392 | 10430 | 39 | rRF 45S | NO | 100.32 | 64.97 | 0.00 | 33.53 |
| >hg38_21_+_8433172_8446622_RNA45SN1_45S_50ntFlanks@10392.10431.40\|ACAGCAGTTGAACATGGTCAGTCGGTCCTGAGAGATGGG | >hg38_21_+_8433172_8446622_RNA45SN1_45S_50ntFlanks | 10392 | 10431 | 40 | rRF 45S | NO | 31.99 | 6.02 | 0.00 | 0.00 |
| >hg38_21_+_8433172_8446622_RNA45SN1_45S_50ntFlanks@10392.10432.41\|ACAGCAGTTGAACATGGTCAGTCGGTCCTGAGAGATGGGC | >hg38_21_+_8433172_8446622_RNA45SN1_45S_50ntFlanks | 10392 | 10432 | 41 | rRF 45S | NO | 19.60 | 0.00 | 0.00 | 0.00 |
| >hg38_21_+_8433172_8446622_RNA45SN1_45S_50ntFlanks@10392.10433.42\|ACAGCAGTTGAACATGGTCAGTCGGTCCTGAGAGATGGGCG | >hg38_21_+_8433172_8446622_RNA45SN1_45S_50ntFlanks | 10392 | 10433 | 42 | rRF 45S | NO | 51.37 | 45.81 | 0.00 | 14.29 |
| >hg38_21_+_8433172_8446622_RNA45SN1_45S_50ntFlanks@10392.10435.44\|ACAGCAGTTGAACATGGTCAGTCGGTCCTGAGAGATGGGCGAG | >hg38_21_+_8433172_8446622_RNA45SN1_45S_50ntFlanks | 10392 | 10435 | 44 | rRF 45S | NO | 62.00 | 54.89 | 0.00 | 38.71 |
| >hg38_21_+_8433172_8446622_RNA45SN1_45S_50ntFlanks@10393.10410.18\|CAGCAGTTGAACATGGGT | >hg38_21_+_8433172_8446622_RNA45SN1_45S_50ntFlanks | 10393 | 10410 | 18 | rRF 45S | NO | 200.75 | 226.21 | 52.22 | 0.00 |
| >hg38_21_+_8433172_8446622_RNA45SN1_45S_50ntFlanks@10393.10419.27\|CAGCAGTTGAACATGGGTCAGTCGGTC | >hg38_21_+_8433172_8446622_RNA45SN1_45S_50ntFlanks | 10393 | 10419 | 27 | rRF 45S | NO | 47.55 | 28.46 | 0.00 | 0.00 |
| >hg38_21_+_8433172_8446622_RNA45SN1_45S_50ntFlanks@10393.10420.28\|CAGCAGTTGAACATGGGTCAGTCGGTCC | >hg38_21_+_8433172_8446622_RNA45SN1_45S_50ntFlanks | 10393 | 10420 | 28 | rRF 45S | NO | 31.21 | 0.00 | 0.00 | 0.00 |
| >hg38_21_+_8433172_8446622_RNA45SN1_45S_50ntFlanks@10393.10422.30\|CAGCAGTTGAACATGGGTCAGTCGGTCCTG | >hg38_21_+_8433172_8446622_RNA45SN1_45S_50ntFlanks | 10393 | 10422 | 30 | rRF 45S | NO | 44.43 | 33.48 | 0.00 | 0.00 |
| >hg38_21_+_8433172_8446622_RNA45SN1_45S_50ntFlanks@10393.10424.32\|CAGCAGTTGAACATGGGTCAGTCGGTCCTGAG | >hg38_21_+_8433172_8446622_RNA45SN1_45S_50ntFlanks | 10393 | 10424 | 32 | rRF 45S | NO | 127.01 | 60.98 | 0.00 | 62.52 |
| >hg38_21_+_8433172_8446622_RNA45SN1_45S_50ntFlanks@10393.10426.34\|CAGCAGTTGAACATGGGTCAGTCGGTCCTGAGAG | >hg38_21_+_8433172_8446622_RNA45SN1_45S_50ntFlanks | 10393 | 10426 | 34 | rRF 45S | NO | 36.91 | 0.00 | 0.00 | 0.00 |
| >hg38_21_+_8433172_8446622_RNA45SN1_45S_50ntFlanks@10393.10429.37\|CAGCAGTTGAACATGGGTCAGTCGGTCCTGAGAGATG | >hg38_21_+_8433172_8446622_RNA45SN1_45S_50ntFlanks | 10393 | 10429 | 37 | rRF 45S | NO | 141.94 | 102.21 | 30.23 | 44.25 |
| >hg38_21_+_8433172_8446622_RNA45SN1_45S_50ntFlanks@10393.10430.38\|CAGCAGTTGAACATGGGTCAGTCGGTCCTGAGAGATGG | >hg38_21_+_8433172_8446622_RNA45SN1_45S_50ntFlanks | 10393 | 10430 | 38 | rRF 45S | NO | 63.70 | 44.41 | 0.00 | 15.94 |
| >hg38_21_+_8433172_8446622_RNA45SN1_45S_50ntFlanks@10393.10433.41\|CAGCAGTTGAACATGGGTCAGTCGGTCCTGAGAGATGGGCG | >hg38_21_+_8433172_8446622_RNA45SN1_45S_50ntFlanks | 10393 | 10433 | 41 | rRF 45S | NO | 21.28 | 12.64 | 0.00 | 0.00 |
| >hg38_21_+_8433172_8446622_RNA45SN1_45S_50ntFlanks@10393.10435.43\|CAGCAGTTGAACATGGGTCAGTCGGTCCTGAGAGATGGGCGAG | >hg38_21_+_8433172_8446622_RNA45SN1_45S_50ntFlanks | 10393 | 10435 | 43 | rRF 45S | NO | 33.36 | 30.46 | 0.00 | 23.26 |
| >hg38_21_+_8433172_8446622_RNA45SN1_45S_50ntFlanks@10394.10424.31\|AGCAGTTGAACATGGGTCAGTCGGTCCTGAG | >hg38_21_+_8433172_8446622_RNA45SN1_45S_50ntFlanks | 10394 | 10424 | 31 | rRF 45S | NO | 19.55 | 0.00 | 0.00 | 0.00 |
| >hg38_21_+_8433172_8446622_RNA45SN1_45S_50ntFlanks@10394.10429.36\|AGCAGTTGAACATGGGTCAGTCGGTCCTGAGAGATG | >hg38_21_+_8433172_8446622_RNA45SN1_45S_50ntFlanks | 10394 | 10429 | 36 | rRF 45S | NO | 19.91 | 0.00 | 0.00 | 0.00 |
| >hg38_21_+_8433172_8446622_RNA45SN1_45S_50ntFlanks@10396.10424.29\|CAGTTGAACATGGGTCAGTCGGTCCTGAG | >hg38_21_+_8433172_8446622_RNA45SN1_45S_50ntFlanks | 10396 | 10424 | 29 | rRF 45S | NO | 43.91 | 0.00 | 0.00 | 0.00 |
| >hg38_21_+_8433172_8446622_RNA45SN1_45S_50ntFlanks@10401.10429.29\|GAACATGGGTCAGTCGGTCCTGAGAGATG | >hg38_21_+_8433172_8446622_RNA45SN1_45S_50ntFlanks | 10401 | 10429 | 29 | rRF 45S | NO | 55.12 | 37.84 | 33.83 | 0.00 |
| >hg38_21_+_8433172_8446622_RNA45SN1_45S_50ntFlanks@10401.10430.30\|GAACATGGGTCAGTCGGTCCTGAGAGATGG | >hg38_21_+_8433172_8446622_RNA45SN1_45S_50ntFlanks | 10401 | 10430 | 30 | rRF 45S | NO | 19.43 | 0.00 | 0.00 | 0.00 |
| >hg38_21_+_8433172_8446622_RNA45SN1_45S_50ntFlanks@10402.10419.18\|AACATGGGTCAGTCGGTC | >hg38_21_+_8433172_8446622_RNA45SN1_45S_50ntFlanks | 10402 | 10419 | 18 | rRF 45S | NO | 99.59 | 102.82 | 32.65 | 0.00 |
| >hg38_21_+_8433172_8446622_RNA45SN1_45S_50ntFlanks@10402.10420.19\|AACATGGGTCAGTCGGTCC | >hg38_21_+_8433172_8446622_RNA45SN1_45S_50ntFlanks | 10402 | 10420 | 19 | rRF 45S | NO | 57.69 | 164.71 | 70.97 | 66.65 |
| >hg38_21_+_8433172_8446622_RNA45SN1_45S_50ntFlanks@10402.10422.21\|AACATGGGTCAGTCGGTCCTG | >hg38_21_+_8433172_8446622_RNA45SN1_45S_50ntFlanks | 10402 | 10422 | 21 | rRF 45S | NO | 0.00 | 29.78 | 48.39 | 0.00 |
| >hg38_21_+_8433172_8446622_RNA45SN1_45S_50ntFlanks@10402.10424.23\|AACATGGGTCAGTCGGTCCTGAG | >hg38_21_+_8433172_8446622_RNA45SN1_45S_50ntFlanks | 10402 | 10424 | 23 | rRF 45S | NO | 178.27 | 107.35 | 59.51 | 94.09 |
| >hg38_21_+_8433172_8446622_RNA45SN1_45S_50ntFlanks@10402.10426.25\|AACATGGGTCAGTCGGTCCTGAGAG | >hg38_21_+_8433172_8446622_RNA45SN1_45S_50ntFlanks | 10402 | 10426 | 25 | rRF 45S | NO | 21.70 | 40.51 | 32.31 | 6.01 |
| >hg38_21_+_8433172_8446622_RNA45SN1_45S_50ntFlanks@10402.10428.27\|AACATGGGTCAGTCGGTCCTGAGAGAT | >hg38_21_+_8433172_8446622_RNA45SN1_45S_50ntFlanks | 10402 | 10428 | 27 | rRF 45S | NO | 12.79 | 0.00 | 0.00 | 0.00 |
| >hg38_21_+_8433172_8446622_RNA45SN1_45S_50ntFlanks@10402.10429.28\|AACATGGGTCAGTCGGTCCTGAGAGATG | >hg38_21_+_8433172_8446622_RNA45SN1_45S_50ntFlanks | 10402 | 10429 | 28 | rRF 45S | NO | 396.21 | 316.71 | 120.07 | 96.93 |
| >hg38_21_+_8433172_8446622_RNA45SN1_45S_50ntFlanks@10402.10430.29\|AACATGGGTCAGTCGGTCCTGAGAGATGG | >hg38_21_+_8433172_8446622_RNA45SN1_45S_50ntFlanks | 10402 | 10430 | 29 | rRF 45S | NO | 229.39 | 178.48 | 56.89 | 59.52 |
| >hg38_21_+_8433172_8446622_RNA45SN1_45S_50ntFlanks@10402.10431.30\|AACATGGGTCAGTCGGTCCTGAGAGATGGG | >hg38_21_+_8433172_8446622_RNA45SN1_45S_50ntFlanks | 10402 | 10431 | 30 | rRF 45S | NO | 158.86 | 120.34 | 21.17 | 14.58 |
| >hg38_21_+_8433172_8446622_RNA45SN1_45S_50ntFlanks@10402.10432.31\|AACATGGGTCAGTCGGTCCTGAGAGATGGGC | >hg38_21_+_8433172_8446622_RNA45SN1_45S_50ntFlanks | 10402 | 10432 | 31 | rRF 45S | NO | 40.21 | 0.00 | 0.00 | 0.00 |
| >hg38_21_+_8433172_8446622_RNA45SN1_45S_50ntFlanks@10402.10433.32\|AACATGGGTCAGTCGGTCCTGAGAGATGGGCG | >hg38_21_+_8433172_8446622_RNA45SN1_45S_50ntFlanks | 10402 | 10433 | 32 | rRF 45S | NO | 70.67 | 55.13 | 6.92 | 0.00 |

### MDA-MB-468

| | | | | | | | | | | |
|---|---|---|---|---|---|---|---|---|---|---|
| >hg38_21_+_8433172_8446622_RNA45SN1_45S_50ntFlanks@10402.10435.34\|AACATGGGTC AGTCGGTCCTGAGAGATGGGCGAG | >hg38_21_+_8433172_8446622_RNA45SN1_45S_50ntFlanks | 10402 | 10435 | 34 | rRF 45S | NO | 116.98 | 82.20 | 31.29 | 37.89 |
| >hg38_21_+_8433172_8446622_RNA45SN1_45S_50ntFlanks@10403.10429.27\|ACATGGGTCA GTCGGTCCTGAGAGATG | >hg38_21_+_8433172_8446622_RNA45SN1_45S_50ntFlanks | 10403 | 10429 | 27 | rRF 45S | NO | 62.86 | 42.57 | 0.00 | 0.00 |
| >hg38_21_+_8433172_8446622_RNA45SN1_45S_50ntFlanks@10403.10430.28\|ACATGGGTCA GTCGGTCCTGAGAGATGG | >hg38_21_+_8433172_8446622_RNA45SN1_45S_50ntFlanks | 10403 | 10430 | 28 | rRF 45S | NO | 32.31 | 0.00 | 0.00 | 0.00 |
| >hg38_21_+_8433172_8446622_RNA45SN1_45S_50ntFlanks@10404.10422.19\|CATGGGTCAG TCGGTCCTG | >hg38_21_+_8433172_8446622_RNA45SN1_45S_50ntFlanks | 10404 | 10422 | 19 | rRF 45S | NO | 0.00 | 27.74 | 20.92 | 0.00 |
| >hg38_21_+_8433172_8446622_RNA45SN1_45S_50ntFlanks@10404.10424.21\|CATGGGTCAG TCGGTCCTGAG | >hg38_21_+_8433172_8446622_RNA45SN1_45S_50ntFlanks | 10404 | 10424 | 21 | rRF 45S | NO | 50.99 | 27.05 | 0.00 | 15.41 |
| >hg38_21_+_8433172_8446622_RNA45SN1_45S_50ntFlanks@10404.10429.26\|CATGGGTCAG TCGGTCCTGAGAGATG | >hg38_21_+_8433172_8446622_RNA45SN1_45S_50ntFlanks | 10404 | 10429 | 26 | rRF 45S | NO | 146.62 | 118.50 | 49.42 | 43.48 |
| >hg38_21_+_8433172_8446622_RNA45SN1_45S_50ntFlanks@10404.10430.27\|CATGGGTCAG TCGGTCCTGAGAGATGG | >hg38_21_+_8433172_8446622_RNA45SN1_45S_50ntFlanks | 10404 | 10430 | 27 | rRF 45S | NO | 72.22 | 40.27 | 0.00 | 0.00 |
| >hg38_21_+_8433172_8446622_RNA45SN1_45S_50ntFlanks@10404.10431.28\|CATGGGTCAG TCGGTCCTGAGAGATGGG | >hg38_21_+_8433172_8446622_RNA45SN1_45S_50ntFlanks | 10404 | 10431 | 28 | rRF 45S | NO | 21.78 | 0.00 | 0.00 | 0.00 |
| >hg38_21_+_8433172_8446622_RNA45SN1_45S_50ntFlanks@10404.10435.32\|CATGGGTCAG TCGGTCCTGAGAGATGGGCGAG | >hg38_21_+_8433172_8446622_RNA45SN1_45S_50ntFlanks | 10404 | 10435 | 32 | rRF 45S | NO | 29.38 | 0.00 | 0.00 | 0.00 |
| >hg38_21_+_8433172_8446622_RNA45SN1_45S_50ntFlanks@10407.10429.23\|GGGTCAGTCG GTCCTGAGAGATG | >hg38_21_+_8433172_8446622_RNA45SN1_45S_50ntFlanks | 10407 | 10429 | 23 | rRF 45S | NO | 32.13 | 0.00 | 0.00 | 0.00 |
| >hg38_21_+_8433172_8446622_RNA45SN1_45S_50ntFlanks@10408.10425.18\|GGTCAGTCGG TCCTGAGA | >hg38_21_+_8433172_8446622_RNA45SN1_45S_50ntFlanks | 10408 | 10425 | 18 | rRF 45S | NO | 12.98 | 34.01 | 0.00 | 0.00 |
| >hg38_21_+_8433172_8446622_RNA45SN1_45S_50ntFlanks@10408.10426.19\|GGTCAGTCGG TCCTGAGAG | >hg38_21_+_8433172_8446622_RNA45SN1_45S_50ntFlanks | 10408 | 10426 | 19 | rRF 45S | NO | 82.61 | 63.81 | 0.00 | 0.00 |
| >hg38_21_+_8433172_8446622_RNA45SN1_45S_50ntFlanks@10408.10428.21\|GGTCAGTCGG TCCTGAGAGAT | >hg38_21_+_8433172_8446622_RNA45SN1_45S_50ntFlanks | 10408 | 10428 | 21 | rRF 45S | NO | 52.41 | 27.55 | 0.00 | 0.00 |
| >hg38_21_+_8433172_8446622_RNA45SN1_45S_50ntFlanks@10408.10429.22\|GGTCAGTCGG TCCTGAGAGATG | >hg38_21_+_8433172_8446622_RNA45SN1_45S_50ntFlanks | 10408 | 10429 | 22 | rRF 45S | NO | 380.90 | 271.51 | 38.76 | 23.42 |
| >hg38_21_+_8433172_8446622_RNA45SN1_45S_50ntFlanks@10408.10430.23\|GGTCAGTCGG TCCTGAGAGATGG | >hg38_21_+_8433172_8446622_RNA45SN1_45S_50ntFlanks | 10408 | 10430 | 23 | rRF 45S | NO | 209.12 | 172.78 | 6.82 | 0.00 |
| >hg38_21_+_8433172_8446622_RNA45SN1_45S_50ntFlanks@10408.10431.24\|GGTCAGTCGG TCCTGAGAGATGGG | >hg38_21_+_8433172_8446622_RNA45SN1_45S_50ntFlanks | 10408 | 10431 | 24 | rRF 45S | NO | 102.52 | 85.89 | 0.00 | 0.00 |
| >hg38_21_+_8433172_8446622_RNA45SN1_45S_50ntFlanks@10408.10432.25\|GGTCAGTCGG TCCTGAGAGATGGGC | >hg38_21_+_8433172_8446622_RNA45SN1_45S_50ntFlanks | 10408 | 10432 | 25 | rRF 45S | NO | 81.91 | 65.09 | 0.00 | 0.00 |
| >hg38_21_+_8433172_8446622_RNA45SN1_45S_50ntFlanks@10408.10433.26\|GGTCAGTCGG TCCTGAGAGATGGGCG | >hg38_21_+_8433172_8446622_RNA45SN1_45S_50ntFlanks | 10408 | 10433 | 26 | rRF 45S | NO | 117.34 | 189.74 | 27.64 | 0.00 |
| >hg38_21_+_8433172_8446622_RNA45SN1_45S_50ntFlanks@10408.10434.27\|GGTCAGTCGG TCCTGAGAGATGGGCGA | >hg38_21_+_8433172_8446622_RNA45SN1_45S_50ntFlanks | 10408 | 10434 | 27 | rRF 45S | NO | 0.00 | 12.41 | 0.00 | 0.00 |
| >hg38_21_+_8433172_8446622_RNA45SN1_45S_50ntFlanks@10408.10435.28\|GGTCAGTCGG TCCTGAGAGATGGGCGAG | >hg38_21_+_8433172_8446622_RNA45SN1_45S_50ntFlanks | 10408 | 10435 | 28 | rRF 45S | NO | 133.05 | 191.08 | 48.67 | 39.10 |
| >hg38_21_+_8433172_8446622_RNA45SN1_45S_50ntFlanks@10409.10429.21\|GTCAGTCGGT CCTGAGAGATG | >hg38_21_+_8433172_8446622_RNA45SN1_45S_50ntFlanks | 10409 | 10429 | 21 | rRF 45S | NO | 135.15 | 107.91 | 0.00 | 0.00 |
| >hg38_21_+_8433172_8446622_RNA45SN1_45S_50ntFlanks@10409.10430.22\|GTCAGTCGGT CCTGAGAGATGG | >hg38_21_+_8433172_8446622_RNA45SN1_45S_50ntFlanks | 10409 | 10430 | 22 | rRF 45S | NO | 83.63 | 73.17 | 0.00 | 0.00 |
| >hg38_21_+_8433172_8446622_RNA45SN1_45S_50ntFlanks@10409.10431.23\|GTCAGTCGGT CCTGAGAGATGGG | >hg38_21_+_8433172_8446622_RNA45SN1_45S_50ntFlanks | 10409 | 10431 | 23 | rRF 45S | NO | 41.95 | 32.49 | 0.00 | 0.00 |
| >hg38_21_+_8433172_8446622_RNA45SN1_45S_50ntFlanks@10409.10433.25\|GTCAGTCGGT CCTGAGAGATGGGCG | >hg38_21_+_8433172_8446622_RNA45SN1_45S_50ntFlanks | 10409 | 10433 | 25 | rRF 45S | NO | 0.00 | 29.06 | 0.00 | 0.00 |
| >hg38_21_+_8433172_8446622_RNA45SN1_45S_50ntFlanks@10410.10428.19\|TCAGTCGGTC CTGAGAGAT | >hg38_21_+_8433172_8446622_RNA45SN1_45S_50ntFlanks | 10410 | 10428 | 19 | rRF 45S | NO | 96.26 | 74.79 | 0.00 | 0.00 |
| >hg38_21_+_8433172_8446622_RNA45SN1_45S_50ntFlanks@10410.10429.20\|TCAGTCGGTC CTGAGAGATG | >hg38_21_+_8433172_8446622_RNA45SN1_45S_50ntFlanks | 10410 | 10429 | 20 | rRF 45S | YES | 628.34 | 633.71 | 85.78 | 86.54 |
| >hg38_21_+_8433172_8446622_RNA45SN1_45S_50ntFlanks@10410.10430.21\|TCAGTCGGTC CTGAGAGATGG | >hg38_21_+_8433172_8446622_RNA45SN1_45S_50ntFlanks | 10410 | 10430 | 21 | rRF 45S | NO | 501.47 | 558.71 | 56.57 | 65.38 |
| >hg38_21_+_8433172_8446622_RNA45SN1_45S_50ntFlanks@10410.10431.22\|TCAGTCGGTC CTGAGAGATGGG | >hg38_21_+_8433172_8446622_RNA45SN1_45S_50ntFlanks | 10410 | 10431 | 22 | rRF 45S | NO | 260.32 | 242.76 | 27.83 | 23.51 |
| >hg38_21_+_8433172_8446622_RNA45SN1_45S_50ntFlanks@10410.10432.23\|TCAGTCGGTC CTGAGAGATGGGC | >hg38_21_+_8433172_8446622_RNA45SN1_45S_50ntFlanks | 10410 | 10432 | 23 | rRF 45S | NO | 49.59 | 93.08 | 31.98 | 6.58 |
| >hg38_21_+_8433172_8446622_RNA45SN1_45S_50ntFlanks@10410.10433.24\|TCAGTCGGTC CTGAGAGATGGGCG | >hg38_21_+_8433172_8446622_RNA45SN1_45S_50ntFlanks | 10410 | 10433 | 24 | rRF 45S | NO | 20.46 | 106.10 | 64.53 | 25.47 |
| >hg38_21_+_8433172_8446622_RNA45SN1_45S_50ntFlanks@10410.10435.26\|TCAGTCGGTC CTGAGAGATGGGCGAG | >hg38_21_+_8433172_8446622_RNA45SN1_45S_50ntFlanks | 10410 | 10435 | 26 | rRF 45S | NO | 0.00 | 6.02 | 63.89 | 0.00 |
| >hg38_21_+_8433172_8446622_RNA45SN1_45S_50ntFlanks@10410.10442.33\|TCAGTCGGTC CTGAGAGATGGGCGAGCGCCGTT | >hg38_21_+_8433172_8446622_RNA45SN1_45S_50ntFlanks | 10410 | 10442 | 33 | rRF 45S | NO | 5.95 | 38.48 | 0.00 | 0.00 |
| >hg38_21_+_8433172_8446622_RNA45SN1_45S_50ntFlanks@10410.10443.34\|TCAGTCGGTC CTGAGAGATGGGCGAGCGCCGTTC | >hg38_21_+_8433172_8446622_RNA45SN1_45S_50ntFlanks | 10410 | 10443 | 34 | rRF 45S | NO | 21.53 | 53.01 | 35.43 | 0.00 |
| >hg38_21_+_8433172_8446622_RNA45SN1_45S_50ntFlanks@10410.10453.44\|TCAGTCGGTC CTGAGAGATGGGCGAGCGCCGTTCCGAAGGGACG | >hg38_21_+_8433172_8446622_RNA45SN1_45S_50ntFlanks | 10410 | 10453 | 44 | rRF 45S | NO | 36.40 | 51.04 | 0.00 | 16.60 |

### MDA-MB-468

| | | | | | | | | | | |
|---|---|---|---|---|---|---|---|---|---|---|
| >hg38_21_+_8433172_8446622_RNA45SN1_45S_50ntFlanks@10411.10428.18\|CAGTCGGTCCTGAGAGAT | >hg38_21_+_8433172_8446622_RNA45SN1_45S_50ntFlanks | 10411 | 10428 | 18 | rRF 45S | NO | 360.10 | 308.72 | 34.83 | 77.95 |
| >hg38_21_+_8433172_8446622_RNA45SN1_45S_50ntFlanks@10411.10429.19\|CAGTCGGTCCTGAGAGATG | >hg38_21_+_8433172_8446622_RNA45SN1_45S_50ntFlanks | 10411 | 10429 | 19 | rRF 45S | YES | 1055.83 | 1093.93 | 158.26 | 202.84 |
| >hg38_21_+_8433172_8446622_RNA45SN1_45S_50ntFlanks@10411.10430.20\|CAGTCGGTCCTGAGAGATGG | >hg38_21_+_8433172_8446622_RNA45SN1_45S_50ntFlanks | 10411 | 10430 | 20 | rRF 45S | YES | 570.57 | 974.86 | 132.04 | 132.19 |
| >hg38_21_+_8433172_8446622_RNA45SN1_45S_50ntFlanks@10411.10431.21\|CAGTCGGTCCTGAGAGATGGG | >hg38_21_+_8433172_8446622_RNA45SN1_45S_50ntFlanks | 10411 | 10431 | 21 | rRF 45S | NO | 353.26 | 495.89 | 73.70 | 62.17 |
| >hg38_21_+_8433172_8446622_RNA45SN1_45S_50ntFlanks@10411.10432.22\|CAGTCGGTCCTGAGAGATGGGC | >hg38_21_+_8433172_8446622_RNA45SN1_45S_50ntFlanks | 10411 | 10432 | 22 | rRF 45S | NO | 34.93 | 160.63 | 68.12 | 26.89 |
| >hg38_21_+_8433172_8446622_RNA45SN1_45S_50ntFlanks@10411.10433.23\|CAGTCGGTCCTGAGAGATGGGCG | >hg38_21_+_8433172_8446622_RNA45SN1_45S_50ntFlanks | 10411 | 10433 | 23 | rRF 45S | NO | 0.00 | 110.93 | 100.19 | 45.87 |
| >hg38_21_+_8433172_8446622_RNA45SN1_45S_50ntFlanks@10411.10435.25\|CAGTCGGTCCTGAGAGATGGGCGAG | >hg38_21_+_8433172_8446622_RNA45SN1_45S_50ntFlanks | 10411 | 10435 | 25 | rRF 45S | NO | 0.00 | 35.40 | 84.87 | 39.66 |
| >hg38_21_+_8433172_8446622_RNA45SN1_45S_50ntFlanks@10411.10440.30\|CAGTCGGTCCTGAGAGATGGGCGAGCGCCG | >hg38_21_+_8433172_8446622_RNA45SN1_45S_50ntFlanks | 10411 | 10440 | 30 | rRF 45S | NO | 0.00 | 26.81 | 0.00 | 0.00 |
| >hg38_21_+_8433172_8446622_RNA45SN1_45S_50ntFlanks@10411.10442.32\|CAGTCGGTCCTGAGAGATGGGCGAGCGCCGTT | >hg38_21_+_8433172_8446622_RNA45SN1_45S_50ntFlanks | 10411 | 10442 | 32 | rRF 45S | NO | 29.25 | 66.53 | 41.99 | 26.66 |
| >hg38_21_+_8433172_8446622_RNA45SN1_45S_50ntFlanks@10411.10443.33\|CAGTCGGTCCTGAGAGATGGGCGAGCGCCGTTC | >hg38_21_+_8433172_8446622_RNA45SN1_45S_50ntFlanks | 10411 | 10443 | 33 | rRF 45S | NO | 56.99 | 117.90 | 130.00 | 57.36 |
| >hg38_21_+_8433172_8446622_RNA45SN1_45S_50ntFlanks@10411.10453.43\|CAGTCGGTCCTGAGAGATGGGCGAGCGCCGTTCCGAAGGGACG | >hg38_21_+_8433172_8446622_RNA45SN1_45S_50ntFlanks | 10411 | 10453 | 43 | rRF 45S | NO | 61.19 | 99.81 | 0.00 | 80.26 |
| >hg38_21_+_8433172_8446622_RNA45SN1_45S_50ntFlanks@10411.10456.46\|CAGTCGGTCCTGAGAGATGGGCGAGCGCCGTTCCGAAGGGACGGGC | >hg38_21_+_8433172_8446622_RNA45SN1_45S_50ntFlanks | 10411 | 10456 | 46 | rRF 45S | NO | 0.00 | 33.46 | 0.00 | 50.41 |
| >hg38_21_+_8433172_8446622_RNA45SN1_45S_50ntFlanks@10412.10430.19\|AGTCGGTCCTGAGAGATGG | >hg38_21_+_8433172_8446622_RNA45SN1_45S_50ntFlanks | 10412 | 10430 | 19 | rRF 45S | NO | 0.00 | 31.74 | 0.00 | 0.00 |
| >hg38_21_+_8433172_8446622_RNA45SN1_45S_50ntFlanks@10414.10431.18\|TCGGTCCTGAGAGATGGG | >hg38_21_+_8433172_8446622_RNA45SN1_45S_50ntFlanks | 10414 | 10431 | 18 | rRF 45S | NO | 0.00 | 78.96 | 0.00 | 0.00 |
| >hg38_21_+_8433172_8446622_RNA45SN1_45S_50ntFlanks@10420.10443.24\|CTGAGAGATGGGCGAGCGCCGTTC | >hg38_21_+_8433172_8446622_RNA45SN1_45S_50ntFlanks | 10420 | 10443 | 24 | rRF 45S | NO | 37.08 | 67.35 | 0.00 | 0.00 |
| >hg38_21_+_8433172_8446622_RNA45SN1_45S_50ntFlanks@10420.10453.34\|CTGAGAGATGGGCGAGCGCCGTTCCGAAGGGACG | >hg38_21_+_8433172_8446622_RNA45SN1_45S_50ntFlanks | 10420 | 10453 | 34 | rRF 45S | NO | 116.55 | 132.86 | 0.00 | 0.00 |
| >hg38_21_+_8433172_8446622_RNA45SN1_45S_50ntFlanks@10420.10455.36\|CTGAGAGATGGGCGAGCGCCGTTCCGAAGGGACGGG | >hg38_21_+_8433172_8446622_RNA45SN1_45S_50ntFlanks | 10420 | 10455 | 36 | rRF 45S | NO | 20.69 | 36.65 | 0.00 | 0.00 |
| >hg38_21_+_8433172_8446622_RNA45SN1_45S_50ntFlanks@10420.10456.37\|CTGAGAGATGGGCGAGCGCCGTTCCGAAGGGACGGGC | >hg38_21_+_8433172_8446622_RNA45SN1_45S_50ntFlanks | 10420 | 10456 | 37 | rRF 45S | NO | 232.22 | 297.20 | 0.00 | 6.10 |
| >hg38_21_+_8433172_8446622_RNA45SN1_45S_50ntFlanks@10420.10457.38\|CTGAGAGATGGGCGAGCGCCGTTCCGAAGGGACGGGCG | >hg38_21_+_8433172_8446622_RNA45SN1_45S_50ntFlanks | 10420 | 10457 | 38 | rRF 45S | NO | 45.85 | 97.35 | 0.00 | 0.00 |
| >hg38_21_+_8433172_8446622_RNA45SN1_45S_50ntFlanks@10421.10453.33\|TGAGAGATGGGCGAGCGCCGTTCCGAAGGGACG | >hg38_21_+_8433172_8446622_RNA45SN1_45S_50ntFlanks | 10421 | 10453 | 33 | rRF 45S | NO | 56.52 | 64.33 | 0.00 | 0.00 |
| >hg38_21_+_8433172_8446622_RNA45SN1_45S_50ntFlanks@10421.10455.35\|TGAGAGATGGGCGAGCGCCGTTCCGAAGGGACGGG | >hg38_21_+_8433172_8446622_RNA45SN1_45S_50ntFlanks | 10421 | 10455 | 35 | rRF 45S | NO | 0.00 | 12.45 | 0.00 | 0.00 |
| >hg38_21_+_8433172_8446622_RNA45SN1_45S_50ntFlanks@10421.10456.36\|TGAGAGATGGGCGAGCGCCGTTCCGAAGGGACGGGC | >hg38_21_+_8433172_8446622_RNA45SN1_45S_50ntFlanks | 10421 | 10456 | 36 | rRF 45S | NO | 116.73 | 181.48 | 0.00 | 0.00 |
| >hg38_21_+_8433172_8446622_RNA45SN1_45S_50ntFlanks@10421.10457.37\|TGAGAGATGGGCGAGCGCCGTTCCGAAGGGACGGGCG | >hg38_21_+_8433172_8446622_RNA45SN1_45S_50ntFlanks | 10421 | 10457 | 37 | rRF 45S | NO | 33.77 | 72.74 | 0.00 | 0.00 |
| >hg38_21_+_8433172_8446622_RNA45SN1_45S_50ntFlanks@10423.10453.31\|AGAGATGGGCGAGCGCCGTTCCGAAGGGACG | >hg38_21_+_8433172_8446622_RNA45SN1_45S_50ntFlanks | 10423 | 10453 | 31 | rRF 45S | NO | 49.56 | 48.78 | 0.00 | 0.00 |
| >hg38_21_+_8433172_8446622_RNA45SN1_45S_50ntFlanks@10423.10456.34\|AGAGATGGGCGAGCGCCGTTCCGAAGGGACGGGC | >hg38_21_+_8433172_8446622_RNA45SN1_45S_50ntFlanks | 10423 | 10456 | 34 | rRF 45S | NO | 70.16 | 84.91 | 0.00 | 0.00 |
| >hg38_21_+_8433172_8446622_RNA45SN1_45S_50ntFlanks@10423.10457.35\|AGAGATGGGCGAGCGCCGTTCCGAAGGGACGGGCG | >hg38_21_+_8433172_8446622_RNA45SN1_45S_50ntFlanks | 10423 | 10457 | 35 | rRF 45S | NO | 42.00 | 67.98 | 0.00 | 0.00 |
| >hg38_21_+_8433172_8446622_RNA45SN1_45S_50ntFlanks@10425.10453.29\|AGATGGGCGAGCGCCGTTCCGAAGGGACG | >hg38_21_+_8433172_8446622_RNA45SN1_45S_50ntFlanks | 10425 | 10453 | 29 | rRF 45S | NO | 36.74 | 55.66 | 0.00 | 0.00 |
| >hg38_21_+_8433172_8446622_RNA45SN1_45S_50ntFlanks@10425.10456.32\|AGATGGGCGAGCGCCGTTCCGAAGGGACGGGC | >hg38_21_+_8433172_8446622_RNA45SN1_45S_50ntFlanks | 10425 | 10456 | 32 | rRF 45S | NO | 41.41 | 79.85 | 0.00 | 0.00 |
| >hg38_21_+_8433172_8446622_RNA45SN1_45S_50ntFlanks@10425.10457.33\|AGATGGGCGAGCGCCGTTCCGAAGGGACGGGCG | >hg38_21_+_8433172_8446622_RNA45SN1_45S_50ntFlanks | 10425 | 10457 | 33 | rRF 45S | NO | 0.00 | 52.11 | 0.00 | 0.00 |
| >hg38_21_+_8433172_8446622_RNA45SN1_45S_50ntFlanks@10427.10453.27\|ATGGGCGAGCGCCGTTCCGAAGGGACG | >hg38_21_+_8433172_8446622_RNA45SN1_45S_50ntFlanks | 10427 | 10453 | 27 | rRF 45S | NO | 59.94 | 94.78 | 0.00 | 0.00 |
| >hg38_21_+_8433172_8446622_RNA45SN1_45S_50ntFlanks@10427.10456.30\|ATGGGCGAGCGCCGTTCCGAAGGGACGGGC | >hg38_21_+_8433172_8446622_RNA45SN1_45S_50ntFlanks | 10427 | 10456 | 30 | rRF 45S | NO | 58.39 | 143.61 | 6.82 | 0.00 |
| >hg38_21_+_8433172_8446622_RNA45SN1_45S_50ntFlanks@10427.10457.31\|ATGGGCGAGCGCCGTTCCGAAGGGACGGGCG | >hg38_21_+_8433172_8446622_RNA45SN1_45S_50ntFlanks | 10427 | 10457 | 31 | rRF 45S | NO | 0.00 | 40.50 | 0.00 | 0.00 |
| >hg38_21_+_8433172_8446622_RNA45SN1_45S_50ntFlanks@10436.10453.18\|CGCCGTTCCGAAGGGACG | >hg38_21_+_8433172_8446622_RNA45SN1_45S_50ntFlanks | 10436 | 10453 | 18 | rRF 45S | NO | 0.00 | 48.53 | 7.14 | 0.00 |
| >hg38_21_+_8433172_8446622_RNA45SN1_45S_50ntFlanks@10441.10460.20\|TTCCGAAGGGACGGGCGATG | >hg38_21_+_8433172_8446622_RNA45SN1_45S_50ntFlanks | 10441 | 10460 | 20 | rRF 45S | NO | 34.41 | 0.00 | 0.00 | 0.00 |

### MDA-MB-468

| | | | | | | | | | | | |
|---|---|---|---|---|---|---|---|---|---|---|---|
| >hg38_21_+_8433172_8446622_RNA45SN1_45S_50ntFlanks@10444.10462.19\|CGAAGGGACGGGCGATGGC | >hg38_21_+_8433172_8446622_RNA45SN1_45S_50ntFlanks | 10444 | 10462 | 19 | | rRF 45S | NO | 37.19 | 29.99 | 0.00 | 0.00 |
| >hg38_21_+_8433172_8446622_RNA45SN1_45S_50ntFlanks@10458.10477.20\|ATGGCCTCCGTTGCCCTCGG | >hg38_21_+_8433172_8446622_RNA45SN1_45S_50ntFlanks | 10458 | 10477 | 20 | | rRF 45S | NO | 95.85 | 361.86 | 63.07 | 56.95 |
| >hg38_21_+_8433172_8446622_RNA45SN1_45S_50ntFlanks@10458.10478.21\|ATGGCCTCCGTTGCCCTCGGC | >hg38_21_+_8433172_8446622_RNA45SN1_45S_50ntFlanks | 10458 | 10478 | 21 | | rRF 45S | NO | 257.60 | 252.70 | 0.00 | 77.25 |
| >hg38_21_+_8433172_8446622_RNA45SN1_45S_50ntFlanks@10458.10484.27\|ATGGCCTCCGTTGCCCTCGGCCGATCG | >hg38_21_+_8433172_8446622_RNA45SN1_45S_50ntFlanks | 10458 | 10484 | 27 | | rRF 45S | NO | 12.51 | 51.79 | 0.00 | 0.00 |
| >hg38_21_+_8433172_8446622_RNA45SN1_45S_50ntFlanks@10458.10487.30\|ATGGCCTCCGTTGCCCTCGGCCGATCGAAA | >hg38_21_+_8433172_8446622_RNA45SN1_45S_50ntFlanks | 10458 | 10487 | 30 | | rRF 45S | NO | 43.36 | 34.03 | 0.00 | 15.82 |
| >hg38_21_+_8433172_8446622_RNA45SN1_45S_50ntFlanks@10458.10488.31\|ATGGCCTCCGTTGCCCTCGGCCGATCGAAAG | >hg38_21_+_8433172_8446622_RNA45SN1_45S_50ntFlanks | 10458 | 10488 | 31 | | rRF 45S | NO | 79.55 | 41.62 | 0.00 | 45.44 |
| >hg38_21_+_8433172_8446622_RNA45SN1_45S_50ntFlanks@10458.10497.40\|ATGGCCTCCGTTGCCCTCGGCCGATCGAAAGGGAGTCGGG | >hg38_21_+_8433172_8446622_RNA45SN1_45S_50ntFlanks | 10458 | 10497 | 40 | | rRF 45S | NO | 19.90 | 43.61 | 33.69 | 53.65 |
| >hg38_21_+_8433172_8446622_RNA45SN1_45S_50ntFlanks@10463.10497.35\|CTCCGTTGCCCTCGGCCGATCGAAAGGGAGTCGGG | >hg38_21_+_8433172_8446622_RNA45SN1_45S_50ntFlanks | 10463 | 10497 | 35 | | rRF 45S | NO | 0.00 | 27.55 | 6.52 | 0.00 |
| >hg38_21_+_8433172_8446622_RNA45SN1_45S_50ntFlanks@10466.10497.32\|CGTTGCCCTCGGCCGATCGAAAGGGAGTCGGG | >hg38_21_+_8433172_8446622_RNA45SN1_45S_50ntFlanks | 10466 | 10497 | 32 | | rRF 45S | NO | 64.85 | 83.68 | 27.59 | 51.21 |
| >hg38_21_+_8433172_8446622_RNA45SN1_45S_50ntFlanks@10468.10487.20\|TTGCCCTCGGCCGATCGAAA | >hg38_21_+_8433172_8446622_RNA45SN1_45S_50ntFlanks | 10468 | 10487 | 20 | | rRF 45S | NO | 30.08 | 0.00 | 0.00 | 0.00 |
| >hg38_21_+_8433172_8446622_RNA45SN1_45S_50ntFlanks@10468.10488.21\|TTGCCCTCGGCCGATCGAAAG | >hg38_21_+_8433172_8446622_RNA45SN1_45S_50ntFlanks | 10468 | 10488 | 21 | | rRF 45S | NO | 43.77 | 0.00 | 0.00 | 0.00 |
| >hg38_21_+_8433172_8446622_RNA45SN1_45S_50ntFlanks@10468.10497.30\|TTGCCCTCGGCCGATCGAAAGGGAGTCGGG | >hg38_21_+_8433172_8446622_RNA45SN1_45S_50ntFlanks | 10468 | 10497 | 30 | | rRF 45S | NO | 0.00 | 46.95 | 0.00 | 0.00 |
| >hg38_21_+_8433172_8446622_RNA45SN1_45S_50ntFlanks@10471.10488.18\|CCCTCGGCCGATCGAAAG | >hg38_21_+_8433172_8446622_RNA45SN1_45S_50ntFlanks | 10471 | 10488 | 18 | | rRF 45S | NO | 100.77 | 50.78 | 0.00 | 38.45 |
| >hg38_21_+_8433172_8446622_RNA45SN1_45S_50ntFlanks@10478.10512.35\|CCGATCGAAAGGGAGTCGGGTTCAGATCCCCGAAT | >hg38_21_+_8433172_8446622_RNA45SN1_45S_50ntFlanks | 10478 | 10512 | 35 | | rRF 45S | NO | 0.00 | 28.66 | 0.00 | 0.00 |
| >hg38_21_+_8433172_8446622_RNA45SN1_45S_50ntFlanks@10479.10497.19\|CGATCGAAAGGGAGTCGGG | >hg38_21_+_8433172_8446622_RNA45SN1_45S_50ntFlanks | 10479 | 10497 | 19 | | rRF 45S | NO | 0.00 | 76.43 | 58.15 | 0.00 |
| >hg38_21_+_8433172_8446622_RNA45SN1_45S_50ntFlanks@10479.10512.34\|CGATCGAAAGGGAGTCGGGTTCAGATCCCCGAAT | >hg38_21_+_8433172_8446622_RNA45SN1_45S_50ntFlanks | 10479 | 10512 | 34 | | rRF 45S | NO | 0.00 | 75.66 | 0.00 | 0.00 |
| >hg38_21_+_8433172_8446622_RNA45SN1_45S_50ntFlanks@10481.10498.18\|ATCGAAAGGGAGTCGGGT | >hg38_21_+_8433172_8446622_RNA45SN1_45S_50ntFlanks | 10481 | 10498 | 18 | | rRF 45S | NO | 0.00 | 31.05 | 0.00 | 0.00 |
| >hg38_21_+_8433172_8446622_RNA45SN1_45S_50ntFlanks@10481.10512.32\|ATCGAAAGGGAGTCGGGTTCAGATCCCCGAAT | >hg38_21_+_8433172_8446622_RNA45SN1_45S_50ntFlanks | 10481 | 10512 | 32 | | rRF 45S | NO | 0.00 | 28.06 | 0.00 | 0.00 |
| >hg38_21_+_8433172_8446622_RNA45SN1_45S_50ntFlanks@10482.10511.30\|TCGAAAGGGAGTCGGGTTCAGATCCCCGAA | >hg38_21_+_8433172_8446622_RNA45SN1_45S_50ntFlanks | 10482 | 10511 | 30 | | rRF 45S | NO | 0.00 | 37.39 | 0.00 | 0.00 |
| >hg38_21_+_8433172_8446622_RNA45SN1_45S_50ntFlanks@10482.10512.31\|TCGAAAGGGAGTCGGGTTCAGATCCCCGAAT | >hg38_21_+_8433172_8446622_RNA45SN1_45S_50ntFlanks | 10482 | 10512 | 31 | | rRF 45S | NO | 0.00 | 62.69 | 0.00 | 0.00 |
| >hg38_21_+_8433172_8446622_RNA45SN1_45S_50ntFlanks@10498.10516.19\|TTCAGATCCCCGAATCCGG | >hg38_21_+_8433172_8446622_RNA45SN1_45S_50ntFlanks | 10498 | 10516 | 19 | | rRF 45S | NO | 0.00 | 43.26 | 0.00 | 0.00 |
| >hg38_21_+_8433172_8446622_RNA45SN1_45S_50ntFlanks@10508.10533.26\|CGAATCCGGAGTGGCGGAGATGGGCG | >hg38_21_+_8433172_8446622_RNA45SN1_45S_50ntFlanks | 10508 | 10533 | 26 | | rRF 45S | NO | 0.00 | 0.00 | 56.47 | 0.00 |
| >hg38_21_+_8433172_8446622_RNA45SN1_45S_50ntFlanks@10509.10533.25\|GAATCCGGAGTGGCGGAGATGGGCG | >hg38_21_+_8433172_8446622_RNA45SN1_45S_50ntFlanks | 10509 | 10533 | 25 | | rRF 45S | NO | 0.00 | 0.00 | 47.86 | 0.00 |
| >hg38_21_+_8433172_8446622_RNA45SN1_45S_50ntFlanks@10510.10532.23\|AATCCGGAGTGGCGGAGATGGGC | >hg38_21_+_8433172_8446622_RNA45SN1_45S_50ntFlanks | 10510 | 10532 | 23 | | rRF 45S | NO | 0.00 | 0.00 | 29.59 | 0.00 |
| >hg38_21_+_8433172_8446622_RNA45SN1_45S_50ntFlanks@10510.10533.24\|AATCCGGAGTGGCGGAGATGGGCG | >hg38_21_+_8433172_8446622_RNA45SN1_45S_50ntFlanks | 10510 | 10533 | 24 | | rRF 45S | NO | 105.07 | 66.48 | 269.44 | 67.18 |
| >hg38_21_+_8433172_8446622_RNA45SN1_45S_50ntFlanks@10510.10534.25\|AATCCGGAGTGGCGGAGATGGGCGC | >hg38_21_+_8433172_8446622_RNA45SN1_45S_50ntFlanks | 10510 | 10534 | 25 | | rRF 45S | NO | 0.00 | 0.00 | 29.66 | 0.00 |
| >hg38_21_+_8433172_8446622_RNA45SN1_45S_50ntFlanks@10511.10533.23\|ATCCGGAGTGGCGGAGATGGGCG | >hg38_21_+_8433172_8446622_RNA45SN1_45S_50ntFlanks | 10511 | 10533 | 23 | | rRF 45S | NO | 5.96 | 0.00 | 75.67 | 0.00 |
| >hg38_21_+_8433172_8446622_RNA45SN1_45S_50ntFlanks@10512.10532.21\|TCCGGAGTGGCGGAGATGGGC | >hg38_21_+_8433172_8446622_RNA45SN1_45S_50ntFlanks | 10512 | 10532 | 21 | | rRF 45S | NO | 0.00 | 0.00 | 42.68 | 0.00 |
| >hg38_21_+_8433172_8446622_RNA45SN1_45S_50ntFlanks@10512.10533.22\|TCCGGAGTGGCGGAGATGGGCG | >hg38_21_+_8433172_8446622_RNA45SN1_45S_50ntFlanks | 10512 | 10533 | 22 | | rRF 45S | NO | 135.03 | 84.61 | 362.91 | 68.69 |
| >hg38_21_+_8433172_8446622_RNA45SN1_45S_50ntFlanks@10512.10534.23\|TCCGGAGTGGCGGAGATGGGCGC | >hg38_21_+_8433172_8446622_RNA45SN1_45S_50ntFlanks | 10512 | 10534 | 23 | | rRF 45S | NO | 0.00 | 0.00 | 75.15 | 0.00 |
| >hg38_21_+_8433172_8446622_RNA45SN1_45S_50ntFlanks@10513.10533.21\|CCGGAGTGGCGGAGATGGGCG | >hg38_21_+_8433172_8446622_RNA45SN1_45S_50ntFlanks | 10513 | 10533 | 21 | | rRF 45S | NO | 173.19 | 79.57 | 403.07 | 100.36 |
| >hg38_21_+_8433172_8446622_RNA45SN1_45S_50ntFlanks@10513.10534.22\|CCGGAGTGGCGGAGATGGGCGC | >hg38_21_+_8433172_8446622_RNA45SN1_45S_50ntFlanks | 10513 | 10534 | 22 | | rRF 45S | NO | 12.66 | 0.00 | 100.54 | 6.19 |
| >hg38_21_+_8433172_8446622_RNA45SN1_45S_50ntFlanks@10514.10533.20\|CGGAGTGGCGGAGATGGGCG | >hg38_21_+_8433172_8446622_RNA45SN1_45S_50ntFlanks | 10514 | 10533 | 20 | | rRF 45S | NO | 29.43 | 0.00 | 98.32 | 15.55 |
| >hg38_21_+_8433172_8446622_RNA45SN1_45S_50ntFlanks@10516.10533.18\|GAGTGGCGGAGATGGGCG | >hg38_21_+_8433172_8446622_RNA45SN1_45S_50ntFlanks | 10516 | 10533 | 18 | | rRF 45S | NO | 31.50 | 26.14 | 148.70 | 35.14 |

### MDA-MB-468

| | | | | | | | | | | |
|---|---|---|---|---|---|---|---|---|---|---|
| >hg38_21_+_8433172_8446622_RNA45SN1_45S_50ntFlanks@10516.10534.19\|GAGTGGCGGA GATGGGCGC | >hg38_21_+_8433172_8446622_RNA45SN1_45S_50ntFlanks | 10516 | 10534 | 19 | rRF 45S | YES | 0.00 | 0.00 | 46.54 | 0.00 |
| >hg38_21_+_8433172_8446622_RNA45SN1_45S_50ntFlanks@10517.10534.18\|AGTGGCGGAG ATGGGCGC | >hg38_21_+_8433172_8446622_RNA45SN1_45S_50ntFlanks | 10517 | 10534 | 18 | rRF 45S | NO | 0.00 | 0.00 | 71.04 | 0.00 |
| >hg38_21_+_8433172_8446622_RNA45SN1_45S_50ntFlanks@10534.10567.34\|CCGCGAGGC GTCCAGTGCGGTAACGCGACCGATC | >hg38_21_+_8433172_8446622_RNA45SN1_45S_50ntFlanks | 10534 | 10567 | 34 | rRF 45S | NO | 0.00 | 0.00 | 47.82 | 13.29 |
| >hg38_21_+_8433172_8446622_RNA45SN1_45S_50ntFlanks@10534.10570.37\|CCGCGAGGC GTCCAGTGCGGTAACGCGACCGATCCCG | >hg38_21_+_8433172_8446622_RNA45SN1_45S_50ntFlanks | 10534 | 10570 | 37 | rRF 45S | NO | 0.00 | 0.00 | 32.71 | 52.18 |
| >hg38_21_+_8433172_8446622_RNA45SN1_45S_50ntFlanks@10571.10608.38\|GAGAAGCCG GCGGGAGCCCCGGGGAGAGTTCTCTTTTC | >hg38_21_+_8433172_8446622_RNA45SN1_45S_50ntFlanks | 10571 | 10608 | 38 | rRF 45S | NO | 12.90 | 35.07 | 0.00 | 0.00 |
| >hg38_21_+_8433172_8446622_RNA45SN1_45S_50ntFlanks@10572.10607.36\|AGAAGCCGG CGGGAGCCCCGGGGAGAGTTCTCTTTT | >hg38_21_+_8433172_8446622_RNA45SN1_45S_50ntFlanks | 10572 | 10607 | 36 | rRF 45S | NO | 50.65 | 42.78 | 0.00 | 0.00 |
| >hg38_21_+_8433172_8446622_RNA45SN1_45S_50ntFlanks@10572.10608.37\|AGAAGCCGG CGGGAGCCCCGGGGAGAGTTCTCTTTTC | >hg38_21_+_8433172_8446622_RNA45SN1_45S_50ntFlanks | 10572 | 10608 | 37 | rRF 45S | NO | 83.11 | 118.20 | 0.00 | 6.25 |
| >hg38_21_+_8433172_8446622_RNA45SN1_45S_50ntFlanks@10574.10607.34\|AAGCCGGCG GGAGCCCCGGGGAGAGTTCTCTTTT | >hg38_21_+_8433172_8446622_RNA45SN1_45S_50ntFlanks | 10574 | 10607 | 34 | rRF 45S | NO | 12.97 | 36.07 | 0.00 | 0.00 |
| >hg38_21_+_8433172_8446622_RNA45SN1_45S_50ntFlanks@10574.10608.35\|AAGCCGGCG GGAGCCCCGGGGAGAGTTCTCTTTTC | >hg38_21_+_8433172_8446622_RNA45SN1_45S_50ntFlanks | 10574 | 10608 | 35 | rRF 45S | NO | 79.15 | 140.12 | 0.00 | 13.61 |
| >hg38_21_+_8433172_8446622_RNA45SN1_45S_50ntFlanks@10587.10607.21\|CCCCGGGGA GAGTTCTCTTTT | >hg38_21_+_8433172_8446622_RNA45SN1_45S_50ntFlanks | 10587 | 10607 | 21 | rRF 45S | NO | 0.00 | 26.51 | 0.00 | 0.00 |
| >hg38_21_+_8433172_8446622_RNA45SN1_45S_50ntFlanks@10587.10608.22\|CCCCGGGGA GAGTTCTCTTTTC | >hg38_21_+_8433172_8446622_RNA45SN1_45S_50ntFlanks | 10587 | 10608 | 22 | rRF 45S | NO | 0.00 | 12.61 | 0.00 | 0.00 |
| >hg38_21_+_8433172_8446622_RNA45SN1_45S_50ntFlanks@10663.10706.44\|CCTTGGAAAG CGTCGCGGTTCCGGCGGCGTCCGGTGAGCTCTCG | >hg38_21_+_8433172_8446622_RNA45SN1_45S_50ntFlanks | 10663 | 10706 | 44 | rRF 45S | NO | 41.40 | 0.00 | 0.00 | 0.00 |
| >hg38_21_+_8433172_8446622_RNA45SN1_45S_50ntFlanks@10664.10706.43\|CTTGGAAAGC GTCGCGGTTCCGGCGGCGTCCGGTGAGCTCTCG | >hg38_21_+_8433172_8446622_RNA45SN1_45S_50ntFlanks | 10664 | 10706 | 43 | rRF 45S | NO | 89.81 | 46.43 | 0.00 | 41.33 |
| >hg38_21_+_8433172_8446622_RNA45SN1_45S_50ntFlanks@10668.10706.39\|GAAAGCGTCG CGGTTCCGGCGGCGTCCGGTGAGCTCTCG | >hg38_21_+_8433172_8446622_RNA45SN1_45S_50ntFlanks | 10668 | 10706 | 39 | rRF 45S | NO | 67.24 | 37.18 | 0.00 | 0.00 |
| >hg38_21_+_8433172_8446622_RNA45SN1_45S_50ntFlanks@10669.10706.38\|AAAGCGTCGC GGTTCCGGCGGCGTCCGGTGAGCTCTCG | >hg38_21_+_8433172_8446622_RNA45SN1_45S_50ntFlanks | 10669 | 10706 | 38 | rRF 45S | NO | 57.38 | 34.23 | 0.00 | 0.00 |
| >hg38_21_+_8433172_8446622_RNA45SN1_45S_50ntFlanks@10677.10706.30\|GCGGTTCCGG CGGCGTCCGGTGAGCTCTCG | >hg38_21_+_8433172_8446622_RNA45SN1_45S_50ntFlanks | 10677 | 10706 | 30 | rRF 45S | NO | 0.00 | 19.39 | 0.00 | 0.00 |
| >hg38_21_+_8433172_8446622_RNA45SN1_45S_50ntFlanks@10677.10708.32\|GCGGTTCCGG CGGCGTCCGGTGAGCTCTCGCT | >hg38_21_+_8433172_8446622_RNA45SN1_45S_50ntFlanks | 10677 | 10708 | 32 | rRF 45S | NO | 33.13 | 0.00 | 0.00 | 0.00 |
| >hg38_21_+_8433172_8446622_RNA45SN1_45S_50ntFlanks@10677.10709.33\|GCGGTTCCGG CGGCGTCCGGTGAGCTCTCGCTG | >hg38_21_+_8433172_8446622_RNA45SN1_45S_50ntFlanks | 10677 | 10709 | 33 | rRF 45S | NO | 49.13 | 29.96 | 0.00 | 14.20 |
| >hg38_21_+_8433172_8446622_RNA45SN1_45S_50ntFlanks@10677.10710.34\|GCGGTTCCGG CGGCGTCCGGTGAGCTCTCGCTGG | >hg38_21_+_8433172_8446622_RNA45SN1_45S_50ntFlanks | 10677 | 10710 | 34 | rRF 45S | NO | 307.53 | 149.94 | 0.00 | 96.02 |
| >hg38_21_+_8433172_8446622_RNA45SN1_45S_50ntFlanks@10677.10711.35\|GCGGTTCCGG CGGCGTCCGGTGAGCTCTCGCTGGC | >hg38_21_+_8433172_8446622_RNA45SN1_45S_50ntFlanks | 10677 | 10711 | 35 | rRF 45S | NO | 81.71 | 43.41 | 0.00 | 44.91 |
| >hg38_21_+_8433172_8446622_RNA45SN1_45S_50ntFlanks@10677.10712.36\|GCGGTTCCGG CGGCGTCCGGTGAGCTCTCGCTGGCC | >hg38_21_+_8433172_8446622_RNA45SN1_45S_50ntFlanks | 10677 | 10712 | 36 | rRF 45S | NO | 57.52 | 39.75 | 0.00 | 35.71 |
| >hg38_21_+_8433172_8446622_RNA45SN1_45S_50ntFlanks@10678.10705.28\|CGGTTCCGGC GGCGTCCGGTGAGCTCTC | >hg38_21_+_8433172_8446622_RNA45SN1_45S_50ntFlanks | 10678 | 10705 | 28 | rRF 45S | NO | 45.97 | 97.67 | 45.58 | 25.49 |
| >hg38_21_+_8433172_8446622_RNA45SN1_45S_50ntFlanks@10678.10706.29\|CGGTTCCGGC GGCGTCCGGTGAGCTCTCG | >hg38_21_+_8433172_8446622_RNA45SN1_45S_50ntFlanks | 10678 | 10706 | 29 | rRF 45S | YES | 957.14 | 1623.82 | 691.53 | 672.22 |
| >hg38_21_+_8433172_8446622_RNA45SN1_45S_50ntFlanks@10678.10707.30\|CGGTTCCGGC GGCGTCCGGTGAGCTCTCGC | >hg38_21_+_8433172_8446622_RNA45SN1_45S_50ntFlanks | 10678 | 10707 | 30 | rRF 45S | NO | 51.32 | 33.47 | 0.00 | 50.94 |
| >hg38_21_+_8433172_8446622_RNA45SN1_45S_50ntFlanks@10678.10708.31\|CGGTTCCGGC GGCGTCCGGTGAGCTCTCGCT | >hg38_21_+_8433172_8446622_RNA45SN1_45S_50ntFlanks | 10678 | 10708 | 31 | rRF 45S | YES | 1953.18 | 896.41 | 163.42 | 926.54 |
| >hg38_21_+_8433172_8446622_RNA45SN1_45S_50ntFlanks@10678.10709.32\|CGGTTCCGGC GGCGTCCGGTGAGCTCTCGCTG | >hg38_21_+_8433172_8446622_RNA45SN1_45S_50ntFlanks | 10678 | 10709 | 32 | rRF 45S | YES | 2767.75 | 1507.22 | 311.54 | 1582.40 |
| >hg38_21_+_8433172_8446622_RNA45SN1_45S_50ntFlanks@10678.10710.33\|CGGTTCCGGC GGCGTCCGGTGAGCTCTCGCTGG | >hg38_21_+_8433172_8446622_RNA45SN1_45S_50ntFlanks | 10678 | 10710 | 33 | rRF 45S | YES | 17484.17 | 12789.80 | 1832.61 | 8541.82 |
| >hg38_21_+_8433172_8446622_RNA45SN1_45S_50ntFlanks@10678.10711.34\|CGGTTCCGGC GGCGTCCGGTGAGCTCTCGCTGGC | >hg38_21_+_8433172_8446622_RNA45SN1_45S_50ntFlanks | 10678 | 10711 | 34 | rRF 45S | YES | 4318.71 | 2904.79 | 314.64 | 2883.23 |
| >hg38_21_+_8433172_8446622_RNA45SN1_45S_50ntFlanks@10678.10712.35\|CGGTTCCGGC GGCGTCCGGTGAGCTCTCGCTGGCC | >hg38_21_+_8433172_8446622_RNA45SN1_45S_50ntFlanks | 10678 | 10712 | 35 | rRF 45S | YES | 1007.40 | 728.50 | 385.40 | 836.42 |
| >hg38_21_+_8433172_8446622_RNA45SN1_45S_50ntFlanks@10679.10705.27\|GGTTCCGGCG GCGTCCGGTGAGCTCTC | >hg38_21_+_8433172_8446622_RNA45SN1_45S_50ntFlanks | 10679 | 10705 | 27 | rRF 45S | NO | 0.00 | 47.35 | 29.83 | 0.00 |
| >hg38_21_+_8433172_8446622_RNA45SN1_45S_50ntFlanks@10679.10706.28\|GGTTCCGGCG GCGTCCGGTGAGCTCTCG | >hg38_21_+_8433172_8446622_RNA45SN1_45S_50ntFlanks | 10679 | 10706 | 28 | rRF 45S | NO | 215.43 | 398.92 | 275.60 | 160.44 |
| >hg38_21_+_8433172_8446622_RNA45SN1_45S_50ntFlanks@10679.10708.30\|GGTTCCGGCG GCGTCCGGTGAGCTCTCGCT | >hg38_21_+_8433172_8446622_RNA45SN1_45S_50ntFlanks | 10679 | 10708 | 30 | rRF 45S | NO | 550.02 | 346.85 | 62.33 | 384.85 |
| >hg38_21_+_8433172_8446622_RNA45SN1_45S_50ntFlanks@10679.10709.31\|GGTTCCGGCG GCGTCCGGTGAGCTCTCGCTG | >hg38_21_+_8433172_8446622_RNA45SN1_45S_50ntFlanks | 10679 | 10709 | 31 | rRF 45S | YES | 566.09 | 428.46 | 119.28 | 529.14 |
| >hg38_21_+_8433172_8446622_RNA45SN1_45S_50ntFlanks@10679.10710.32\|GGTTCCGGCG GCGTCCGGTGAGCTCTCGCTGG | >hg38_21_+_8433172_8446622_RNA45SN1_45S_50ntFlanks | 10679 | 10710 | 32 | rRF 45S | YES | 2199.40 | 2272.74 | 466.58 | 1524.11 |

### MDA-MB-468

| | | | | | | | | | | |
|---|---|---|---|---|---|---|---|---|---|---|
| >hg38_21_+_8433172_8446622_RNA45SN1_45S_50ntFlanks@10679.10711.33\|GGTTCCGGCGGCGTCCGGTGAGCTCTCGCTGGC | >hg38_21_+_8433172_8446622_RNA45SN1_45S_50ntFlanks | 10679 | 10711 | 33 | rRF 45S | NO | 446.04 | 449.65 | 89.49 | 483.34 |
| >hg38_21_+_8433172_8446622_RNA45SN1_45S_50ntFlanks@10679.10712.34\|GGTTCCGGCGGCGTCCGGTGAGCTCTCGCTGGCC | >hg38_21_+_8433172_8446622_RNA45SN1_45S_50ntFlanks | 10679 | 10712 | 34 | rRF 45S | NO | 31.73 | 33.47 | 90.75 | 26.73 |
| >hg38_21_+_8433172_8446622_RNA45SN1_45S_50ntFlanks@10680.10705.26\|GTTCCGGCGGCGTCCGGTGAGCTCTC | >hg38_21_+_8433172_8446622_RNA45SN1_45S_50ntFlanks | 10680 | 10705 | 26 | rRF 45S | NO | 49.67 | 82.29 | 0.00 | 0.00 |
| >hg38_21_+_8433172_8446622_RNA45SN1_45S_50ntFlanks@10680.10706.27\|GTTCCGGCGGCGTCCGGTGAGCTCTCG | >hg38_21_+_8433172_8446622_RNA45SN1_45S_50ntFlanks | 10680 | 10706 | 27 | rRF 45S | YES | 841.78 | 933.67 | 193.33 | 225.73 |
| >hg38_21_+_8433172_8446622_RNA45SN1_45S_50ntFlanks@10680.10708.29\|GTTCCGGCGGCGTCCGGTGAGCTCTCGCT | >hg38_21_+_8433172_8446622_RNA45SN1_45S_50ntFlanks | 10680 | 10708 | 29 | rRF 45S | YES | 1232.02 | 1246.41 | 159.25 | 719.63 |
| >hg38_21_+_8433172_8446622_RNA45SN1_45S_50ntFlanks@10680.10709.30\|GTTCCGGCGGCGTCCGGTGAGCTCTCGCTG | >hg38_21_+_8433172_8446622_RNA45SN1_45S_50ntFlanks | 10680 | 10709 | 30 | rRF 45S | NO | 182.44 | 309.13 | 83.05 | 241.93 |
| >hg38_21_+_8433172_8446622_RNA45SN1_45S_50ntFlanks@10680.10710.31\|GTTCCGGCGGCGTCCGGTGAGCTCTCGCTGG | >hg38_21_+_8433172_8446622_RNA45SN1_45S_50ntFlanks | 10680 | 10710 | 31 | rRF 45S | YES | 867.64 | 1564.14 | 320.55 | 699.79 |
| >hg38_21_+_8433172_8446622_RNA45SN1_45S_50ntFlanks@10680.10711.32\|GTTCCGGCGGCGTCCGGTGAGCTCTCGCTGGC | >hg38_21_+_8433172_8446622_RNA45SN1_45S_50ntFlanks | 10680 | 10711 | 32 | rRF 45S | YES | 519.01 | 802.48 | 80.51 | 430.14 |
| >hg38_21_+_8433172_8446622_RNA45SN1_45S_50ntFlanks@10680.10712.33\|GTTCCGGCGGCGTCCGGTGAGCTCTCGCTGGCC | >hg38_21_+_8433172_8446622_RNA45SN1_45S_50ntFlanks | 10680 | 10712 | 33 | rRF 45S | NO | 395.50 | 425.79 | 70.76 | 227.87 |
| >hg38_21_+_8433172_8446622_RNA45SN1_45S_50ntFlanks@10681.10706.26\|TTCCGGCGGCGTCCGGTGAGCTCTCG | >hg38_21_+_8433172_8446622_RNA45SN1_45S_50ntFlanks | 10681 | 10706 | 26 | rRF 45S | NO | 241.59 | 219.16 | 35.93 | 50.34 |
| >hg38_21_+_8433172_8446622_RNA45SN1_45S_50ntFlanks@10681.10708.28\|TTCCGGCGGCGTCCGGTGAGCTCTCGCT | >hg38_21_+_8433172_8446622_RNA45SN1_45S_50ntFlanks | 10681 | 10708 | 28 | rRF 45S | NO | 0.00 | 30.59 | 0.00 | 14.99 |
| >hg38_21_+_8433172_8446622_RNA45SN1_45S_50ntFlanks@10681.10710.30\|TTCCGGCGGCGTCCGGTGAGCTCTCGCTGG | >hg38_21_+_8433172_8446622_RNA45SN1_45S_50ntFlanks | 10681 | 10710 | 30 | rRF 45S | NO | 74.55 | 153.31 | 50.98 | 89.69 |
| >hg38_21_+_8433172_8446622_RNA45SN1_45S_50ntFlanks@10681.10711.31\|TTCCGGCGGCGTCCGGTGAGCTCTCGCTGGC | >hg38_21_+_8433172_8446622_RNA45SN1_45S_50ntFlanks | 10681 | 10711 | 31 | rRF 45S | NO | 64.57 | 104.03 | 0.00 | 51.48 |
| >hg38_21_+_8433172_8446622_RNA45SN1_45S_50ntFlanks@10681.10712.32\|TTCCGGCGGCGTCCGGTGAGCTCTCGCTGGCC | >hg38_21_+_8433172_8446622_RNA45SN1_45S_50ntFlanks | 10681 | 10712 | 32 | rRF 45S | NO | 192.58 | 184.96 | 0.00 | 85.09 |
| >hg38_21_+_8433172_8446622_RNA45SN1_45S_50ntFlanks@10682.10706.25\|TCCGGCGGCGTCCGGTGAGCTCTCG | >hg38_21_+_8433172_8446622_RNA45SN1_45S_50ntFlanks | 10682 | 10706 | 25 | rRF 45S | NO | 172.36 | 152.37 | 29.30 | 59.83 |
| >hg38_21_+_8433172_8446622_RNA45SN1_45S_50ntFlanks@10682.10712.31\|TCCGGCGGCGTCCGGTGAGCTCTCGCTGGCC | >hg38_21_+_8433172_8446622_RNA45SN1_45S_50ntFlanks | 10682 | 10712 | 31 | rRF 45S | NO | 21.42 | 31.74 | 0.00 | 14.43 |
| >hg38_21_+_8433172_8446622_RNA45SN1_45S_50ntFlanks@10683.10705.23\|CCGGCGGCGTCCGGTGAGCTCTC | >hg38_21_+_8433172_8446622_RNA45SN1_45S_50ntFlanks | 10683 | 10705 | 23 | rRF 45S | NO | 72.86 | 67.61 | 0.00 | 36.27 |
| >hg38_21_+_8433172_8446622_RNA45SN1_45S_50ntFlanks@10683.10706.24\|CCGGCGGCGTCCGGTGAGCTCTCG | >hg38_21_+_8433172_8446622_RNA45SN1_45S_50ntFlanks | 10683 | 10706 | 24 | rRF 45S | NO | 803.57 | 633.00 | 166.64 | 334.83 |
| >hg38_21_+_8433172_8446622_RNA45SN1_45S_50ntFlanks@10683.10709.27\|CCGGCGGCGTCCGGTGAGCTCTCGCTG | >hg38_21_+_8433172_8446622_RNA45SN1_45S_50ntFlanks | 10683 | 10709 | 27 | rRF 45S | NO | 0.00 | 55.46 | 39.06 | 101.76 |
| >hg38_21_+_8433172_8446622_RNA45SN1_45S_50ntFlanks@10683.10710.28\|CCGGCGGCGTCCGGTGAGCTCTCGCTGG | >hg38_21_+_8433172_8446622_RNA45SN1_45S_50ntFlanks | 10683 | 10710 | 28 | rRF 45S | NO | 0.00 | 12.50 | 0.00 | 14.77 |
| >hg38_21_+_8433172_8446622_RNA45SN1_45S_50ntFlanks@10683.10711.29\|CCGGCGGCGTCCGGTGAGCTCTCGCTGGC | >hg38_21_+_8433172_8446622_RNA45SN1_45S_50ntFlanks | 10683 | 10711 | 29 | rRF 45S | NO | 65.68 | 132.22 | 41.59 | 93.48 |
| >hg38_21_+_8433172_8446622_RNA45SN1_45S_50ntFlanks@10683.10712.30\|CCGGCGGCGTCCGGTGAGCTCTCGCTGGCC | >hg38_21_+_8433172_8446622_RNA45SN1_45S_50ntFlanks | 10683 | 10712 | 30 | rRF 45S | NO | 137.42 | 187.66 | 34.31 | 109.20 |
| >hg38_21_+_8433172_8446622_RNA45SN1_45S_50ntFlanks@10684.10705.22\|CGGCGGCGTCCGGTGAGCTCTC | >hg38_21_+_8433172_8446622_RNA45SN1_45S_50ntFlanks | 10684 | 10705 | 22 | rRF 45S | NO | 37.41 | 12.27 | 0.00 | 0.00 |
| >hg38_21_+_8433172_8446622_RNA45SN1_45S_50ntFlanks@10684.10706.23\|CGGCGGCGTCCGGTGAGCTCTCG | >hg38_21_+_8433172_8446622_RNA45SN1_45S_50ntFlanks | 10684 | 10706 | 23 | rRF 45S | NO | 604.91 | 466.83 | 108.79 | 238.32 |
| >hg38_21_+_8433172_8446622_RNA45SN1_45S_50ntFlanks@10684.10708.25\|CGGCGGCGTCCGGTGAGCTCTCGCT | >hg38_21_+_8433172_8446622_RNA45SN1_45S_50ntFlanks | 10684 | 10708 | 25 | rRF 45S | NO | 66.70 | 59.17 | 0.00 | 54.21 |
| >hg38_21_+_8433172_8446622_RNA45SN1_45S_50ntFlanks@10684.10709.26\|CGGCGGCGTCCGGTGAGCTCTCGCTG | >hg38_21_+_8433172_8446622_RNA45SN1_45S_50ntFlanks | 10684 | 10709 | 26 | rRF 45S | NO | 76.36 | 150.63 | 52.57 | 134.45 |
| >hg38_21_+_8433172_8446622_RNA45SN1_45S_50ntFlanks@10684.10710.27\|CGGCGGCGTCCGGTGAGCTCTCGCTGG | >hg38_21_+_8433172_8446622_RNA45SN1_45S_50ntFlanks | 10684 | 10710 | 27 | rRF 45S | YES | 722.83 | 740.93 | 83.42 | 376.23 |
| >hg38_21_+_8433172_8446622_RNA45SN1_45S_50ntFlanks@10684.10711.28\|CGGCGGCGTCCGGTGAGCTCTCGCTGGC | >hg38_21_+_8433172_8446622_RNA45SN1_45S_50ntFlanks | 10684 | 10711 | 28 | rRF 45S | NO | 380.10 | 406.37 | 44.17 | 193.87 |
| >hg38_21_+_8433172_8446622_RNA45SN1_45S_50ntFlanks@10684.10712.29\|CGGCGGCGTCCGGTGAGCTCTCGCTGGCC | >hg38_21_+_8433172_8446622_RNA45SN1_45S_50ntFlanks | 10684 | 10712 | 29 | rRF 45S | NO | 37.96 | 53.81 | 0.00 | 23.00 |
| >hg38_21_+_8433172_8446622_RNA45SN1_45S_50ntFlanks@10685.10706.22\|GGCGGCGTCCGGTGAGCTCTCG | >hg38_21_+_8433172_8446622_RNA45SN1_45S_50ntFlanks | 10685 | 10706 | 22 | rRF 45S | NO | 32.49 | 0.00 | 0.00 | 0.00 |
| >hg38_21_+_8433172_8446622_RNA45SN1_45S_50ntFlanks@10685.10710.26\|GGCGGCGTCCGGTGAGCTCTCGCTGG | >hg38_21_+_8433172_8446622_RNA45SN1_45S_50ntFlanks | 10685 | 10710 | 26 | rRF 45S | NO | 57.53 | 38.41 | 0.00 | 24.04 |
| >hg38_21_+_8433172_8446622_RNA45SN1_45S_50ntFlanks@10686.10706.21\|GCGGCGTCCGGTGAGCTCTCG | >hg38_21_+_8433172_8446622_RNA45SN1_45S_50ntFlanks | 10686 | 10706 | 21 | rRF 45S | NO | 101.50 | 72.41 | 31.82 | 38.54 |
| >hg38_21_+_8433172_8446622_RNA45SN1_45S_50ntFlanks@10686.10708.23\|GCGGCGTCCGGTGAGCTCTCGCT | >hg38_21_+_8433172_8446622_RNA45SN1_45S_50ntFlanks | 10686 | 10708 | 23 | rRF 45S | NO | 71.44 | 46.57 | 0.00 | 24.73 |
| >hg38_21_+_8433172_8446622_RNA45SN1_45S_50ntFlanks@10686.10709.24\|GCGGCGTCCGGTGAGCTCTCGCTG | >hg38_21_+_8433172_8446622_RNA45SN1_45S_50ntFlanks | 10686 | 10709 | 24 | rRF 45S | NO | 63.62 | 74.42 | 0.00 | 50.52 |

### MDA-MB-468

| | | | | | | | | | | |
|---|---|---|---|---|---|---|---|---|---|---|
| >hg38_21_+_8433172_8446622_RNA45SN1_45S_50ntFlanks@10686.10710.25\|GCGGCGTCCGGTGAGCTCTCGCTGG | >hg38_21_+_8433172_8446622_RNA45SN1_45S_50ntFlanks | 10686 | 10710 | 25 | rRF 45S | NO | 220.89 | 225.84 | 32.08 | 121.94 |
| >hg38_21_+_8433172_8446622_RNA45SN1_45S_50ntFlanks@10686.10711.26\|GCGGCGTCCGGTGAGCTCTCGCTGGC | >hg38_21_+_8433172_8446622_RNA45SN1_45S_50ntFlanks | 10686 | 10711 | 26 | rRF 45S | NO | 43.03 | 59.74 | 0.00 | 34.30 |
| >hg38_21_+_8433172_8446622_RNA45SN1_45S_50ntFlanks@10686.10712.27\|GCGGCGTCCGGTGAGCTCTCGCTGGCC | >hg38_21_+_8433172_8446622_RNA45SN1_45S_50ntFlanks | 10686 | 10712 | 27 | rRF 45S | NO | 28.76 | 36.25 | 0.00 | 6.24 |
| >hg38_21_+_8433172_8446622_RNA45SN1_45S_50ntFlanks@10687.10705.19\|CGGCGTCCGGTGAGCTCTC | >hg38_21_+_8433172_8446622_RNA45SN1_45S_50ntFlanks | 10687 | 10705 | 19 | rRF 45S | NO | 30.82 | 19.66 | 0.00 | 0.00 |
| >hg38_21_+_8433172_8446622_RNA45SN1_45S_50ntFlanks@10687.10706.20\|CGGCGTCCGGTGAGCTCTCG | >hg38_21_+_8433172_8446622_RNA45SN1_45S_50ntFlanks | 10687 | 10706 | 20 | rRF 45S | NO | 338.46 | 400.24 | 92.77 | 138.89 |
| >hg38_21_+_8433172_8446622_RNA45SN1_45S_50ntFlanks@10687.10708.22\|CGGCGTCCGGTGAGCTCTCGCT | >hg38_21_+_8433172_8446622_RNA45SN1_45S_50ntFlanks | 10687 | 10708 | 22 | rRF 45S | NO | 162.50 | 199.61 | 7.16 | 75.86 |
| >hg38_21_+_8433172_8446622_RNA45SN1_45S_50ntFlanks@10687.10709.23\|CGGCGTCCGGTGAGCTCTCGCTG | >hg38_21_+_8433172_8446622_RNA45SN1_45S_50ntFlanks | 10687 | 10709 | 23 | rRF 45S | NO | 145.08 | 287.20 | 52.13 | 128.73 |
| >hg38_21_+_8433172_8446622_RNA45SN1_45S_50ntFlanks@10687.10710.24\|CGGCGTCCGGTGAGCTCTCGCTGG | >hg38_21_+_8433172_8446622_RNA45SN1_45S_50ntFlanks | 10687 | 10710 | 24 | rRF 45S | NO | 309.36 | 621.41 | 116.47 | 189.77 |
| >hg38_21_+_8433172_8446622_RNA45SN1_45S_50ntFlanks@10687.10711.25\|CGGCGTCCGGTGAGCTCTCGCTGGC | >hg38_21_+_8433172_8446622_RNA45SN1_45S_50ntFlanks | 10687 | 10711 | 25 | rRF 45S | NO | 310.07 | 515.90 | 65.23 | 154.32 |
| >hg38_21_+_8433172_8446622_RNA45SN1_45S_50ntFlanks@10687.10712.26\|CGGCGTCCGGTGAGCTCTCGCTGGCC | >hg38_21_+_8433172_8446622_RNA45SN1_45S_50ntFlanks | 10687 | 10712 | 26 | rRF 45S | NO | 60.39 | 167.85 | 30.05 | 39.83 |
| >hg38_21_+_8433172_8446622_RNA45SN1_45S_50ntFlanks@10689.10706.18\|GCGTCCGGTGAGCTCTCG | >hg38_21_+_8433172_8446622_RNA45SN1_45S_50ntFlanks | 10689 | 10706 | 18 | rRF 45S | NO | 146.38 | 142.49 | 51.53 | 50.67 |
| >hg38_21_+_8433172_8446622_RNA45SN1_45S_50ntFlanks@10689.10708.20\|GCGTCCGGTGAGCTCTCGCT | >hg38_21_+_8433172_8446622_RNA45SN1_45S_50ntFlanks | 10689 | 10708 | 20 | rRF 45S | NO | 91.27 | 124.35 | 0.00 | 37.16 |
| >hg38_21_+_8433172_8446622_RNA45SN1_45S_50ntFlanks@10689.10709.21\|GCGTCCGGTGAGCTCTCGCTG | >hg38_21_+_8433172_8446622_RNA45SN1_45S_50ntFlanks | 10689 | 10709 | 21 | rRF 45S | NO | 58.90 | 74.79 | 0.00 | 24.68 |
| >hg38_21_+_8433172_8446622_RNA45SN1_45S_50ntFlanks@10690.10708.19\|CGTCCGGTGAGCTCTCGCT | >hg38_21_+_8433172_8446622_RNA45SN1_45S_50ntFlanks | 10690 | 10708 | 19 | rRF 45S | NO | 373.91 | 821.86 | 158.93 | 171.07 |
| >hg38_21_+_8433172_8446622_RNA45SN1_45S_50ntFlanks@10690.10709.20\|CGTCCGGTGAGCTCTCGCTG | >hg38_21_+_8433172_8446622_RNA45SN1_45S_50ntFlanks | 10690 | 10709 | 20 | rRF 45S | NO | 113.57 | 214.71 | 33.41 | 66.96 |
| >hg38_21_+_8433172_8446622_RNA45SN1_45S_50ntFlanks@10690.10710.21\|CGTCCGGTGAGCTCTCGCTGG | >hg38_21_+_8433172_8446622_RNA45SN1_45S_50ntFlanks | 10690 | 10710 | 21 | rRF 45S | NO | 19.71 | 107.74 | 48.40 | 14.70 |
| >hg38_21_+_8433172_8446622_RNA45SN1_45S_50ntFlanks@10694.10711.18\|CGGTGAGCTCTCGCTGGC | >hg38_21_+_8433172_8446622_RNA45SN1_45S_50ntFlanks | 10694 | 10711 | 18 | rRF 45S | NO | 0.00 | 65.29 | 46.99 | 33.70 |
| >hg38_21_+_8433172_8446622_RNA45SN1_45S_50ntFlanks@10713.10736.24\|CTTGAAAATCCGGGGGAGAGGGTG | >hg38_21_+_8433172_8446622_RNA45SN1_45S_50ntFlanks | 10713 | 10736 | 24 | rRF 45S | NO | 0.00 | 31.70 | 28.55 | 0.00 |
| >hg38_21_+_8433172_8446622_RNA45SN1_45S_50ntFlanks@10727.10752.26\|GGAGAGGGTGTAAATCTCGCGCCGGG | >hg38_21_+_8433172_8446622_RNA45SN1_45S_50ntFlanks | 10727 | 10752 | 26 | rRF 45S | NO | 0.00 | 0.00 | 39.27 | 0.00 |
| >hg38_21_+_8433172_8446622_RNA45SN1_45S_50ntFlanks@10727.10755.29\|GGAGAGGGTGTAAATCTCGCGCCGGGCCG | >hg38_21_+_8433172_8446622_RNA45SN1_45S_50ntFlanks | 10727 | 10755 | 29 | rRF 45S | NO | 112.23 | 63.15 | 48.58 | 13.27 |
| >hg38_21_+_8433172_8446622_RNA45SN1_45S_50ntFlanks@10727.10757.31\|GGAGAGGGTGTAAATCTCGCGCCGGGCCGTA | >hg38_21_+_8433172_8446622_RNA45SN1_45S_50ntFlanks | 10727 | 10757 | 31 | rRF 45S | NO | 61.80 | 31.82 | 0.00 | 0.00 |
| >hg38_21_+_8433172_8446622_RNA45SN1_45S_50ntFlanks@10728.10755.28\|GAGAGGGTGTAAATCTCGCGCCGGGCCG | >hg38_21_+_8433172_8446622_RNA45SN1_45S_50ntFlanks | 10728 | 10755 | 28 | rRF 45S | NO | 45.45 | 0.00 | 0.00 | 0.00 |
| >hg38_21_+_8433172_8446622_RNA45SN1_45S_50ntFlanks@10729.10755.27\|AGAGGGTGTAAATCTCGCGCCGGGCCG | >hg38_21_+_8433172_8446622_RNA45SN1_45S_50ntFlanks | 10729 | 10755 | 27 | rRF 45S | NO | 56.47 | 30.37 | 0.00 | 0.00 |
| >hg38_21_+_8433172_8446622_RNA45SN1_45S_50ntFlanks@10731.10752.22\|AGGGTGTAAATCTCGCGCCGGG | >hg38_21_+_8433172_8446622_RNA45SN1_45S_50ntFlanks | 10731 | 10752 | 22 | rRF 45S | NO | 0.00 | 33.93 | 27.85 | 0.00 |
| >hg38_21_+_8433172_8446622_RNA45SN1_45S_50ntFlanks@10731.10755.25\|AGGGTGTAAATCTCGCGCCGGGCCG | >hg38_21_+_8433172_8446622_RNA45SN1_45S_50ntFlanks | 10731 | 10755 | 25 | rRF 45S | NO | 66.98 | 37.66 | 6.73 | 0.00 |
| >hg38_21_+_8433172_8446622_RNA45SN1_45S_50ntFlanks@10734.10755.22\|GTGTAAATCTCGCGCCGGGCCG | >hg38_21_+_8433172_8446622_RNA45SN1_45S_50ntFlanks | 10734 | 10755 | 22 | rRF 45S | NO | 36.00 | 0.00 | 0.00 | 0.00 |
| >hg38_21_+_8433172_8446622_RNA45SN1_45S_50ntFlanks@10735.10752.18\|TGTAAATCTCGCGCCGGG | >hg38_21_+_8433172_8446622_RNA45SN1_45S_50ntFlanks | 10735 | 10752 | 18 | rRF 45S | NO | 0.00 | 29.42 | 0.00 | 0.00 |
| >hg38_21_+_8433172_8446622_RNA45SN1_45S_50ntFlanks@10735.10755.21\|TGTAAATCTCGCGCCGGGCCG | >hg38_21_+_8433172_8446622_RNA45SN1_45S_50ntFlanks | 10735 | 10755 | 21 | rRF 45S | NO | 218.87 | 89.06 | 0.00 | 35.51 |
| >hg38_21_+_8433172_8446622_RNA45SN1_45S_50ntFlanks@10735.10757.23\|TGTAAATCTCGCGCCGGGCCGTA | >hg38_21_+_8433172_8446622_RNA45SN1_45S_50ntFlanks | 10735 | 10757 | 23 | rRF 45S | NO | 46.16 | 0.00 | 0.00 | 0.00 |
| >hg38_21_+_8433172_8446622_RNA45SN1_45S_50ntFlanks@10735.10767.33\|TGTAAATCTCGCGCCGGGCCGTACCCATATCCG | >hg38_21_+_8433172_8446622_RNA45SN1_45S_50ntFlanks | 10735 | 10767 | 33 | rRF 45S | NO | 29.82 | 0.00 | 0.00 | 0.00 |
| >hg38_21_+_8433172_8446622_RNA45SN1_45S_50ntFlanks@10736.10755.20\|GTAAATCTCGCGCCGGGCCG | >hg38_21_+_8433172_8446622_RNA45SN1_45S_50ntFlanks | 10736 | 10755 | 20 | rRF 45S | NO | 43.62 | 0.00 | 0.00 | 0.00 |
| >hg38_21_+_8433172_8446622_RNA45SN1_45S_50ntFlanks@10737.10755.19\|TAAATCTCGCGCCGGGCCG | >hg38_21_+_8433172_8446622_RNA45SN1_45S_50ntFlanks | 10737 | 10755 | 19 | rRF 45S | NO | 264.07 | 76.36 | 0.00 | 49.11 |
| >hg38_21_+_8433172_8446622_RNA45SN1_45S_50ntFlanks@10737.10757.21\|TAAATCTCGCGCCGGGCCGTA | >hg38_21_+_8433172_8446622_RNA45SN1_45S_50ntFlanks | 10737 | 10757 | 21 | rRF 45S | NO | 56.40 | 0.00 | 0.00 | 0.00 |
| >hg38_21_+_8433172_8446622_RNA45SN1_45S_50ntFlanks@10737.10767.31\|TAAATCTCGCGCCGGGCCGTACCCATATCCG | >hg38_21_+_8433172_8446622_RNA45SN1_45S_50ntFlanks | 10737 | 10767 | 31 | rRF 45S | NO | 38.64 | 0.00 | 0.00 | 0.00 |

### MDA-MB-468

| | | | | | | | | | | |
|---|---|---|---|---|---|---|---|---|---|---|
| >hg38_21_+_8433172_8446622_RNA45SN1_45S_50ntFlanks@10738.10755.18\|AAATCTCGCGCCGGGCCG | >hg38_21_+_8433172_8446622_RNA45SN1_45S_50ntFlanks | 10738 | 10755 | 18 | rRF 45S | NO | 159.54 | 31.42 | 0.00 | 6.32 |
| >hg38_21_+_8433172_8446622_RNA45SN1_45S_50ntFlanks@10738.10757.20\|AAATCTCGCGCCGGGCCGTA | >hg38_21_+_8433172_8446622_RNA45SN1_45S_50ntFlanks | 10738 | 10757 | 20 | rRF 45S | NO | 43.76 | 0.00 | 0.00 | 0.00 |
| >hg38_21_+_8433172_8446622_RNA45SN1_45S_50ntFlanks@10768.10785.18\|CAGCAGGTCTCCAAGGTG | >hg38_21_+_8433172_8446622_RNA45SN1_45S_50ntFlanks | 10768 | 10785 | 18 | rRF 45S | NO | 34.56 | 51.10 | 34.20 | 33.22 |
| >hg38_21_+_8433172_8446622_RNA45SN1_45S_50ntFlanks@10770.10796.27\|GCAGGTCTCCAAGGTGAACAGCCTCTG | >hg38_21_+_8433172_8446622_RNA45SN1_45S_50ntFlanks | 10770 | 10796 | 27 | rRF 45S | NO | 38.90 | 49.89 | 0.00 | 14.05 |
| >hg38_21_+_8433172_8446622_RNA45SN1_45S_50ntFlanks@10771.10791.21\|CAGGTCTCCAAGGTGAACAGC | >hg38_21_+_8433172_8446622_RNA45SN1_45S_50ntFlanks | 10771 | 10791 | 21 | rRF 45S | NO | 21.17 | 0.00 | 0.00 | 0.00 |
| >hg38_21_+_8433172_8446622_RNA45SN1_45S_50ntFlanks@10771.10796.26\|CAGGTCTCCAAGGTGAACAGCCTCTG | >hg38_21_+_8433172_8446622_RNA45SN1_45S_50ntFlanks | 10771 | 10796 | 26 | rRF 45S | NO | 36.81 | 47.72 | 27.35 | 6.03 |
| >hg38_21_+_8433172_8446622_RNA45SN1_45S_50ntFlanks@10771.10797.27\|CAGGTCTCCAAGGTGAACAGCCTCTGG | >hg38_21_+_8433172_8446622_RNA45SN1_45S_50ntFlanks | 10771 | 10797 | 27 | rRF 45S | NO | 0.00 | 60.18 | 32.95 | 0.00 |
| >hg38_21_+_8433172_8446622_RNA45SN1_45S_50ntFlanks@10771.10805.35\|CAGGTCTCCAAGGTGAACAGCCTCTGGCATGTTGG | >hg38_21_+_8433172_8446622_RNA45SN1_45S_50ntFlanks | 10771 | 10805 | 35 | rRF 45S | NO | 88.19 | 74.60 | 13.72 | 48.80 |
| >hg38_21_+_8433172_8446622_RNA45SN1_45S_50ntFlanks@10771.10806.36\|CAGGTCTCCAAGGTGAACAGCCTCTGGCATGTTGGA | >hg38_21_+_8433172_8446622_RNA45SN1_45S_50ntFlanks | 10771 | 10806 | 36 | rRF 45S | NO | 58.48 | 54.54 | 6.49 | 47.52 |
| >hg38_21_+_8433172_8446622_RNA45SN1_45S_50ntFlanks@10771.10816.46\|CAGGTCTCCAAGGTGAACAGCCTCTGGCATGTTGGAACAATGTAGG | >hg38_21_+_8433172_8446622_RNA45SN1_45S_50ntFlanks | 10771 | 10816 | 46 | rRF 45S | YES | 36.88 | 28.61 | 0.00 | 0.00 |
| >hg38_21_+_8433172_8446622_RNA45SN1_45S_50ntFlanks@10771.10818.48\|CAGGTCTCCAAGGTGAACAGCCTCTGGCATGTTGGAACAATGTAGGTA | >hg38_21_+_8433172_8446622_RNA45SN1_45S_50ntFlanks | 10771 | 10818 | 48 | rRF 45S | YES | 30.75 | 53.60 | 0.00 | 0.00 |
| >hg38_21_+_8433172_8446622_RNA45SN1_45S_50ntFlanks@10772.10796.25\|AGGTCTCCAAGGTGAACAGCCTCTG | >hg38_21_+_8433172_8446622_RNA45SN1_45S_50ntFlanks | 10772 | 10796 | 25 | rRF 45S | NO | 42.85 | 30.23 | 0.00 | 0.00 |
| >hg38_21_+_8433172_8446622_RNA45SN1_45S_50ntFlanks@10777.10818.42\|TCCAAGGTGAACAGCCTCTGGCATGTTGGAACAATGTAGGTA | >hg38_21_+_8433172_8446622_RNA45SN1_45S_50ntFlanks | 10777 | 10818 | 42 | rRF 45S | NO | 0.00 | 31.64 | 0.00 | 0.00 |
| >hg38_21_+_8433172_8446622_RNA45SN1_45S_50ntFlanks@10778.10818.41\|CCAAGGTGAACAGCCTCTGGCATGTTGGAACAATGTAGGTA | >hg38_21_+_8433172_8446622_RNA45SN1_45S_50ntFlanks | 10778 | 10818 | 41 | rRF 45S | NO | 0.00 | 31.94 | 0.00 | 0.00 |
| >hg38_21_+_8433172_8446622_RNA45SN1_45S_50ntFlanks@10781.10801.21\|AGGTGAACAGCCTCTGGCATG | >hg38_21_+_8433172_8446622_RNA45SN1_45S_50ntFlanks | 10781 | 10801 | 21 | rRF 45S | NO | 0.00 | 71.62 | 47.96 | 0.00 |
| >hg38_21_+_8433172_8446622_RNA45SN1_45S_50ntFlanks@10781.10802.22\|AGGTGAACAGCCTCTGGCATGT | >hg38_21_+_8433172_8446622_RNA45SN1_45S_50ntFlanks | 10781 | 10802 | 22 | rRF 45S | NO | 0.00 | 34.94 | 0.00 | 0.00 |
| >hg38_21_+_8433172_8446622_RNA45SN1_45S_50ntFlanks@10781.10803.23\|AGGTGAACAGCCTCTGGCATGTT | >hg38_21_+_8433172_8446622_RNA45SN1_45S_50ntFlanks | 10781 | 10803 | 23 | rRF 45S | NO | 0.00 | 37.54 | 0.00 | 0.00 |
| >hg38_21_+_8433172_8446622_RNA45SN1_45S_50ntFlanks@10781.10804.24\|AGGTGAACAGCCTCTGGCATGTTG | >hg38_21_+_8433172_8446622_RNA45SN1_45S_50ntFlanks | 10781 | 10804 | 24 | rRF 45S | NO | 0.00 | 30.56 | 0.00 | 0.00 |
| >hg38_21_+_8433172_8446622_RNA45SN1_45S_50ntFlanks@10781.10805.25\|AGGTGAACAGCCTCTGGCATGTTGG | >hg38_21_+_8433172_8446622_RNA45SN1_45S_50ntFlanks | 10781 | 10805 | 25 | rRF 45S | NO | 0.00 | 40.04 | 0.00 | 0.00 |
| >hg38_21_+_8433172_8446622_RNA45SN1_45S_50ntFlanks@10781.10806.26\|AGGTGAACAGCCTCTGGCATGTTGGA | >hg38_21_+_8433172_8446622_RNA45SN1_45S_50ntFlanks | 10781 | 10806 | 26 | rRF 45S | NO | 0.00 | 31.31 | 0.00 | 0.00 |
| >hg38_21_+_8433172_8446622_RNA45SN1_45S_50ntFlanks@10781.10807.27\|AGGTGAACAGCCTCTGGCATGTTGGAA | >hg38_21_+_8433172_8446622_RNA45SN1_45S_50ntFlanks | 10781 | 10807 | 27 | rRF 45S | NO | 29.40 | 32.28 | 0.00 | 0.00 |
| >hg38_21_+_8433172_8446622_RNA45SN1_45S_50ntFlanks@10781.10817.37\|AGGTGAACAGCCTCTGGCATGTTGGAACAATGTAGGT | >hg38_21_+_8433172_8446622_RNA45SN1_45S_50ntFlanks | 10781 | 10817 | 37 | rRF 45S | NO | 0.00 | 26.98 | 0.00 | 0.00 |
| >hg38_21_+_8433172_8446622_RNA45SN1_45S_50ntFlanks@10781.10818.38\|AGGTGAACAGCCTCTGGCATGTTGGAACAATGTAGGTA | >hg38_21_+_8433172_8446622_RNA45SN1_45S_50ntFlanks | 10781 | 10818 | 38 | rRF 45S | NO | 65.85 | 198.67 | 30.45 | 77.92 |
| >hg38_21_+_8433172_8446622_RNA45SN1_45S_50ntFlanks@10782.10801.20\|GGTGAACAGCCTCTGGCATG | >hg38_21_+_8433172_8446622_RNA45SN1_45S_50ntFlanks | 10782 | 10801 | 20 | rRF 45S | NO | 37.87 | 63.58 | 0.00 | 5.99 |
| >hg38_21_+_8433172_8446622_RNA45SN1_45S_50ntFlanks@10782.10818.37\|GGTGAACAGCCTCTGGCATGTTGGAACAATGTAGGTA | >hg38_21_+_8433172_8446622_RNA45SN1_45S_50ntFlanks | 10782 | 10818 | 37 | rRF 45S | NO | 0.00 | 36.58 | 0.00 | 13.23 |
| >hg38_21_+_8433172_8446622_RNA45SN1_45S_50ntFlanks@10783.10801.19\|GTGAACAGCCTCTGGCATG | >hg38_21_+_8433172_8446622_RNA45SN1_45S_50ntFlanks | 10783 | 10801 | 19 | rRF 45S | NO | 0.00 | 31.21 | 28.39 | 0.00 |
| >hg38_21_+_8433172_8446622_RNA45SN1_45S_50ntFlanks@10783.10816.34\|GTGAACAGCCTCTGGCATGTTGGAACAATGTAGG | >hg38_21_+_8433172_8446622_RNA45SN1_45S_50ntFlanks | 10783 | 10816 | 34 | rRF 45S | NO | 0.00 | 12.71 | 0.00 | 0.00 |
| >hg38_21_+_8433172_8446622_RNA45SN1_45S_50ntFlanks@10783.10817.35\|GTGAACAGCCTCTGGCATGTTGGAACAATGTAGGT | >hg38_21_+_8433172_8446622_RNA45SN1_45S_50ntFlanks | 10783 | 10817 | 35 | rRF 45S | NO | 0.00 | 34.18 | 0.00 | 0.00 |
| >hg38_21_+_8433172_8446622_RNA45SN1_45S_50ntFlanks@10783.10818.36\|GTGAACAGCCTCTGGCATGTTGGAACAATGTAGGTA | >hg38_21_+_8433172_8446622_RNA45SN1_45S_50ntFlanks | 10783 | 10818 | 36 | rRF 45S | NO | 49.06 | 153.05 | 34.09 | 44.79 |
| >hg38_21_+_8433172_8446622_RNA45SN1_45S_50ntFlanks@10784.10818.35\|TGAACAGCCTCTGGCATGTTGGAACAATGTAGGTA | >hg38_21_+_8433172_8446622_RNA45SN1_45S_50ntFlanks | 10784 | 10818 | 35 | rRF 45S | NO | 0.00 | 46.78 | 0.00 | 15.54 |
| >hg38_21_+_8433172_8446622_RNA45SN1_45S_50ntFlanks@10786.10812.27\|AACAGCCTCTGGCATGTTGGAACAATG | >hg38_21_+_8433172_8446622_RNA45SN1_45S_50ntFlanks | 10786 | 10812 | 27 | rRF 45S | NO | 0.00 | 28.81 | 0.00 | 0.00 |
| >hg38_21_+_8433172_8446622_RNA45SN1_45S_50ntFlanks@10786.10815.30\|AACAGCCTCTGGCATGTTGGAACAATGTAG | >hg38_21_+_8433172_8446622_RNA45SN1_45S_50ntFlanks | 10786 | 10815 | 30 | rRF 45S | NO | 20.12 | 40.14 | 0.00 | 0.00 |
| >hg38_21_+_8433172_8446622_RNA45SN1_45S_50ntFlanks@10786.10816.31\|AACAGCCTCTGGCATGTTGGAACAATGTAGG | >hg38_21_+_8433172_8446622_RNA45SN1_45S_50ntFlanks | 10786 | 10816 | 31 | rRF 45S | NO | 80.59 | 223.39 | 27.76 | 15.45 |
| >hg38_21_+_8433172_8446622_RNA45SN1_45S_50ntFlanks@10786.10817.32\|AACAGCCTCTGGCATGTTGGAACAATGTAGGT | >hg38_21_+_8433172_8446622_RNA45SN1_45S_50ntFlanks | 10786 | 10817 | 32 | rRF 45S | NO | 32.41 | 79.79 | 59.77 | 41.57 |

### MDA-MB-468

| | | | | | | | | | | |
|---|---|---|---|---|---|---|---|---|---|---|
| >hg38_21_+_8433172_8446622_RNA45SN1_45S_50ntFlanks@10786.10818.33\|AACAGCCTCTGGCATGTTGGAACAATGTAGGTA | >hg38_21_+_8433172_8446622_RNA45SN1_45S_50ntFlanks | 10786 | 10818 | 33 | rRF 45S | YES | 90.06 | 409.10 | 177.93 | 194.24 |
| >hg38_21_+_8433172_8446622_RNA45SN1_45S_50ntFlanks@10786.10820.35\|AACAGCCTCTGGCATGTTGGAACAATGTAGGTAAG | >hg38_21_+_8433172_8446622_RNA45SN1_45S_50ntFlanks | 10786 | 10820 | 35 | rRF 45S | NO | 0.00 | 29.99 | 0.00 | 35.35 |
| >hg38_21_+_8433172_8446622_RNA45SN1_45S_50ntFlanks@10786.10821.36\|AACAGCCTCTGGCATGTTGGAACAATGTAGGTAAGG | >hg38_21_+_8433172_8446622_RNA45SN1_45S_50ntFlanks | 10786 | 10821 | 36 | rRF 45S | NO | 0.00 | 29.22 | 0.00 | 42.99 |
| >hg38_21_+_8433172_8446622_RNA45SN1_45S_50ntFlanks@10786.10822.37\|AACAGCCTCTGGCATGTTGGAACAATGTAGGTAAGGG | >hg38_21_+_8433172_8446622_RNA45SN1_45S_50ntFlanks | 10786 | 10822 | 37 | rRF 45S | NO | 30.31 | 48.62 | 32.56 | 73.28 |
| >hg38_21_+_8433172_8446622_RNA45SN1_45S_50ntFlanks@10786.10823.38\|AACAGCCTCTGGCATGTTGGAACAATGTAGGTAAGGGA | >hg38_21_+_8433172_8446622_RNA45SN1_45S_50ntFlanks | 10786 | 10823 | 38 | rRF 45S | NO | 22.06 | 31.82 | 7.31 | 73.51 |
| >hg38_21_+_8433172_8446622_RNA45SN1_45S_50ntFlanks@10786.10825.40\|AACAGCCTCTGGCATGTTGGAACAATGTAGGTAAGGGAAG | >hg38_21_+_8433172_8446622_RNA45SN1_45S_50ntFlanks | 10786 | 10825 | 40 | rRF 45S | NO | 33.32 | 38.17 | 0.00 | 74.96 |
| >hg38_21_+_8433172_8446622_RNA45SN1_45S_50ntFlanks@10786.10826.41\|AACAGCCTCTGGCATGTTGGAACAATGTAGGTAAGGGAAGT | >hg38_21_+_8433172_8446622_RNA45SN1_45S_50ntFlanks | 10786 | 10826 | 41 | rRF 45S | NO | 48.64 | 45.04 | 0.00 | 104.97 |
| >hg38_21_+_8433172_8446622_RNA45SN1_45S_50ntFlanks@10787.10816.30\|ACAGCCTCTGGCATGTTGGAACAATGTAGG | >hg38_21_+_8433172_8446622_RNA45SN1_45S_50ntFlanks | 10787 | 10816 | 30 | rRF 45S | NO | 37.98 | 48.16 | 0.00 | 0.00 |
| >hg38_21_+_8433172_8446622_RNA45SN1_45S_50ntFlanks@10787.10818.32\|ACAGCCTCTGGCATGTTGGAACAATGTAGGTA | >hg38_21_+_8433172_8446622_RNA45SN1_45S_50ntFlanks | 10787 | 10818 | 32 | rRF 45S | NO | 13.21 | 95.49 | 67.82 | 66.58 |
| >hg38_21_+_8433172_8446622_RNA45SN1_45S_50ntFlanks@10788.10806.19\|CAGCCTCTGGCATGTTGGA | >hg38_21_+_8433172_8446622_RNA45SN1_45S_50ntFlanks | 10788 | 10806 | 19 | rRF 45S | NO | 0.00 | 12.78 | 0.00 | 0.00 |
| >hg38_21_+_8433172_8446622_RNA45SN1_45S_50ntFlanks@10788.10815.28\|CAGCCTCTGGCATGTTGGAACAATGTAG | >hg38_21_+_8433172_8446622_RNA45SN1_45S_50ntFlanks | 10788 | 10815 | 28 | rRF 45S | NO | 48.20 | 41.29 | 0.00 | 0.00 |
| >hg38_21_+_8433172_8446622_RNA45SN1_45S_50ntFlanks@10788.10816.29\|CAGCCTCTGGCATGTTGGAACAATGTAGG | >hg38_21_+_8433172_8446622_RNA45SN1_45S_50ntFlanks | 10788 | 10816 | 29 | rRF 45S | NO | 42.15 | 83.68 | 0.00 | 0.00 |
| >hg38_21_+_8433172_8446622_RNA45SN1_45S_50ntFlanks@10788.10818.31\|CAGCCTCTGGCATGTTGGAACAATGTAGGTA | >hg38_21_+_8433172_8446622_RNA45SN1_45S_50ntFlanks | 10788 | 10818 | 31 | rRF 45S | NO | 0.00 | 112.45 | 104.96 | 100.17 |
| >hg38_21_+_8433172_8446622_RNA45SN1_45S_50ntFlanks@10789.10818.30\|AGCCTCTGGCATGTTGGAACAATGTAGGTA | >hg38_21_+_8433172_8446622_RNA45SN1_45S_50ntFlanks | 10789 | 10818 | 30 | rRF 45S | NO | 0.00 | 0.00 | 31.75 | 13.96 |
| >hg38_21_+_8433172_8446622_RNA45SN1_45S_50ntFlanks@10794.10816.23\|CTGGCATGTTGGAACAATGTAGG | >hg38_21_+_8433172_8446622_RNA45SN1_45S_50ntFlanks | 10794 | 10816 | 23 | rRF 45S | NO | 45.21 | 61.95 | 71.28 | 15.78 |
| >hg38_21_+_8433172_8446622_RNA45SN1_45S_50ntFlanks@10794.10817.24\|CTGGCATGTTGGAACAATGTAGGT | >hg38_21_+_8433172_8446622_RNA45SN1_45S_50ntFlanks | 10794 | 10817 | 24 | rRF 45S | NO | 0.00 | 19.41 | 36.88 | 14.42 |
| >hg38_21_+_8433172_8446622_RNA45SN1_45S_50ntFlanks@10794.10818.25\|CTGGCATGTTGGAACAATGTAGGTA | >hg38_21_+_8433172_8446622_RNA45SN1_45S_50ntFlanks | 10794 | 10818 | 25 | rRF 45S | NO | 56.11 | 179.16 | 80.74 | 88.76 |
| >hg38_21_+_8433172_8446622_RNA45SN1_45S_50ntFlanks@10794.10822.29\|CTGGCATGTTGGAACAATGTAGGTAAGGG | >hg38_21_+_8433172_8446622_RNA45SN1_45S_50ntFlanks | 10794 | 10822 | 29 | rRF 45S | NO | 0.00 | 30.45 | 21.25 | 6.46 |
| >hg38_21_+_8433172_8446622_RNA45SN1_45S_50ntFlanks@10794.10825.32\|CTGGCATGTTGGAACAATGTAGGTAAGGGAAG | >hg38_21_+_8433172_8446622_RNA45SN1_45S_50ntFlanks | 10794 | 10825 | 32 | rRF 45S | NO | 20.71 | 12.37 | 6.57 | 16.30 |
| >hg38_21_+_8433172_8446622_RNA45SN1_45S_50ntFlanks@10794.10826.33\|CTGGCATGTTGGAACAATGTAGGTAAGGGAAGT | >hg38_21_+_8433172_8446622_RNA45SN1_45S_50ntFlanks | 10794 | 10826 | 33 | rRF 45S | NO | 0.00 | 0.00 | 28.12 | 16.17 |
| >hg38_21_+_8433172_8446622_RNA45SN1_45S_50ntFlanks@10795.10818.24\|TGGCATGTTGGAACAATGTAGGTA | >hg38_21_+_8433172_8446622_RNA45SN1_45S_50ntFlanks | 10795 | 10818 | 24 | rRF 45S | NO | 22.11 | 97.64 | 7.23 | 23.04 |
| >hg38_21_+_8433172_8446622_RNA45SN1_45S_50ntFlanks@10796.10818.23\|GGCATGTTGGAACAATGTAGGTA | >hg38_21_+_8433172_8446622_RNA45SN1_45S_50ntFlanks | 10796 | 10818 | 23 | rRF 45S | NO | 0.00 | 54.32 | 29.01 | 0.00 |
| >hg38_21_+_8433172_8446622_RNA45SN1_45S_50ntFlanks@10797.10817.21\|GCATGTTGGAACAATGTAGGT | >hg38_21_+_8433172_8446622_RNA45SN1_45S_50ntFlanks | 10797 | 10817 | 21 | rRF 45S | NO | 0.00 | 35.89 | 38.10 | 0.00 |
| >hg38_21_+_8433172_8446622_RNA45SN1_45S_50ntFlanks@10797.10818.22\|GCATGTTGGAACAATGTAGGTA | >hg38_21_+_8433172_8446622_RNA45SN1_45S_50ntFlanks | 10797 | 10818 | 22 | rRF 45S | NO | 38.01 | 252.22 | 108.72 | 51.65 |
| >hg38_21_+_8433172_8446622_RNA45SN1_45S_50ntFlanks@10797.10822.26\|GCATGTTGGAACAATGTAGGTAAGGG | >hg38_21_+_8433172_8446622_RNA45SN1_45S_50ntFlanks | 10797 | 10822 | 26 | rRF 45S | NO | 0.00 | 0.00 | 36.38 | 0.00 |
| >hg38_21_+_8433172_8446622_RNA45SN1_45S_50ntFlanks@10797.10823.27\|GCATGTTGGAACAATGTAGGTAAGGGA | >hg38_21_+_8433172_8446622_RNA45SN1_45S_50ntFlanks | 10797 | 10823 | 27 | rRF 45S | NO | 0.00 | 0.00 | 30.30 | 0.00 |
| >hg38_21_+_8433172_8446622_RNA45SN1_45S_50ntFlanks@10797.10826.30\|GCATGTTGGAACAATGTAGGTAAGGGAAGT | >hg38_21_+_8433172_8446622_RNA45SN1_45S_50ntFlanks | 10797 | 10826 | 30 | rRF 45S | NO | 0.00 | 0.00 | 38.92 | 0.00 |
| >hg38_21_+_8433172_8446622_RNA45SN1_45S_50ntFlanks@10798.10816.19\|CATGTTGGAACAATGTAGG | >hg38_21_+_8433172_8446622_RNA45SN1_45S_50ntFlanks | 10798 | 10816 | 19 | rRF 45S | NO | 0.00 | 43.68 | 0.00 | 0.00 |
| >hg38_21_+_8433172_8446622_RNA45SN1_45S_50ntFlanks@10798.10817.20\|CATGTTGGAACAATGTAGGT | >hg38_21_+_8433172_8446622_RNA45SN1_45S_50ntFlanks | 10798 | 10817 | 20 | rRF 45S | NO | 0.00 | 60.64 | 30.87 | 0.00 |
| >hg38_21_+_8433172_8446622_RNA45SN1_45S_50ntFlanks@10798.10818.21\|CATGTTGGAACAATGTAGGTA | >hg38_21_+_8433172_8446622_RNA45SN1_45S_50ntFlanks | 10798 | 10818 | 21 | rRF 45S | NO | 32.74 | 303.18 | 62.33 | 48.91 |
| >hg38_21_+_8433172_8446622_RNA45SN1_45S_50ntFlanks@10798.10822.25\|CATGTTGGAACAATGTAGGTAAGGG | >hg38_21_+_8433172_8446622_RNA45SN1_45S_50ntFlanks | 10798 | 10822 | 25 | rRF 45S | NO | 0.00 | 0.00 | 29.25 | 0.00 |
| >hg38_21_+_8433172_8446622_RNA45SN1_45S_50ntFlanks@10798.10826.29\|CATGTTGGAACAATGTAGGTAAGGGAAGT | >hg38_21_+_8433172_8446622_RNA45SN1_45S_50ntFlanks | 10798 | 10826 | 29 | rRF 45S | NO | 0.00 | 0.00 | 28.10 | 0.00 |
| >hg38_21_+_8433172_8446622_RNA45SN1_45S_50ntFlanks@10800.10818.19\|TGTTGGAACAATGTAGGTA | >hg38_21_+_8433172_8446622_RNA45SN1_45S_50ntFlanks | 10800 | 10818 | 19 | rRF 45S | NO | 0.00 | 28.92 | 0.00 | 0.00 |
| >hg38_21_+_8433172_8446622_RNA45SN1_45S_50ntFlanks@10801.10818.18\|GTTGGAACAATGTAGGTA | >hg38_21_+_8433172_8446622_RNA45SN1_45S_50ntFlanks | 10801 | 10818 | 18 | rRF 45S | NO | 0.00 | 50.18 | 0.00 | 0.00 |

### MDA-MB-468

| | | | | | | | | | | |
|---|---|---|---|---|---|---|---|---|---|---|
| >hg38_21_+_8433172_8446622_RNA45SN1_45S_50ntFlanks@10805.10822.18\|GAACAATGTAGGTAAGGG | >hg38_21_+_8433172_8446622_RNA45SN1_45S_50ntFlanks | 10805 | 10822 | 18 | rRF 45S | NO | 0.00 | 31.82 | 0.00 | 0.00 |
| >hg38_21_+_8433172_8446622_RNA45SN1_45S_50ntFlanks@10805.10823.19\|GAACAATGTAGGTAAGGGA | >hg38_21_+_8433172_8446622_RNA45SN1_45S_50ntFlanks | 10805 | 10823 | 19 | rRF 45S | NO | 0.00 | 26.79 | 0.00 | 0.00 |
| >hg38_21_+_8433172_8446622_RNA45SN1_45S_50ntFlanks@10806.10823.18\|AACAATGTAGGTAAGGGA | >hg38_21_+_8433172_8446622_RNA45SN1_45S_50ntFlanks | 10806 | 10823 | 18 | rRF 45S | NO | 36.30 | 49.89 | 0.00 | 35.75 |
| >hg38_21_+_8433172_8446622_RNA45SN1_45S_50ntFlanks@10806.10825.20\|AACAATGTAGGTAAGGGAAG | >hg38_21_+_8433172_8446622_RNA45SN1_45S_50ntFlanks | 10806 | 10825 | 20 | rRF 45S | NO | 35.21 | 36.03 | 0.00 | 24.08 |
| >hg38_21_+_8433172_8446622_RNA45SN1_45S_50ntFlanks@10806.10826.21\|AACAATGTAGGTAAGGGAAGT | >hg38_21_+_8433172_8446622_RNA45SN1_45S_50ntFlanks | 10806 | 10826 | 21 | rRF 45S | NO | 36.68 | 34.47 | 0.00 | 35.97 |
| >hg38_21_+_8433172_8446622_RNA45SN1_45S_50ntFlanks@10806.10828.23\|AACAATGTAGGTAAGGGAAGTCG | >hg38_21_+_8433172_8446622_RNA45SN1_45S_50ntFlanks | 10806 | 10828 | 23 | rRF 45S | NO | 21.26 | 32.87 | 0.00 | 23.31 |
| >hg38_21_+_8433172_8446622_RNA45SN1_45S_50ntFlanks@10806.10840.35\|AACAATGTAGGTAAGGGAAGTCGGCAAGCCGGATC | >hg38_21_+_8433172_8446622_RNA45SN1_45S_50ntFlanks | 10806 | 10840 | 35 | rRF 45S | NO | 37.57 | 43.76 | 0.00 | 50.73 |
| >hg38_21_+_8433172_8446622_RNA45SN1_45S_50ntFlanks@10807.10825.19\|ACAATGTAGGTAAGGGAAG | >hg38_21_+_8433172_8446622_RNA45SN1_45S_50ntFlanks | 10807 | 10825 | 19 | rRF 45S | NO | 0.00 | 29.64 | 0.00 | 13.61 |
| >hg38_21_+_8433172_8446622_RNA45SN1_45S_50ntFlanks@10807.10826.20\|ACAATGTAGGTAAGGGAAGT | >hg38_21_+_8433172_8446622_RNA45SN1_45S_50ntFlanks | 10807 | 10826 | 20 | rRF 45S | NO | 0.00 | 26.21 | 0.00 | 16.23 |
| >hg38_21_+_8433172_8446622_RNA45SN1_45S_50ntFlanks@10807.10840.34\|ACAATGTAGGTAAGGGAAGTCGGCAAGCCGGATC | >hg38_21_+_8433172_8446622_RNA45SN1_45S_50ntFlanks | 10807 | 10840 | 34 | rRF 45S | NO | 0.00 | 12.66 | 0.00 | 36.93 |
| >hg38_21_+_8433172_8446622_RNA45SN1_45S_50ntFlanks@10808.10825.18\|CAATGTAGGTAAGGGAAG | >hg38_21_+_8433172_8446622_RNA45SN1_45S_50ntFlanks | 10808 | 10825 | 18 | rRF 45S | NO | 73.56 | 128.53 | 0.00 | 80.01 |
| >hg38_21_+_8433172_8446622_RNA45SN1_45S_50ntFlanks@10808.10826.19\|CAATGTAGGTAAGGGAAGT | >hg38_21_+_8433172_8446622_RNA45SN1_45S_50ntFlanks | 10808 | 10826 | 19 | rRF 45S | NO | 72.48 | 118.74 | 6.85 | 86.48 |
| >hg38_21_+_8433172_8446622_RNA45SN1_45S_50ntFlanks@10808.10827.20\|CAATGTAGGTAAGGGAAGTC | >hg38_21_+_8433172_8446622_RNA45SN1_45S_50ntFlanks | 10808 | 10827 | 20 | rRF 45S | NO | 30.48 | 49.68 | 0.00 | 33.39 |
| >hg38_21_+_8433172_8446622_RNA45SN1_45S_50ntFlanks@10808.10828.21\|CAATGTAGGTAAGGGAAGTCG | >hg38_21_+_8433172_8446622_RNA45SN1_45S_50ntFlanks | 10808 | 10828 | 21 | rRF 45S | NO | 57.56 | 99.80 | 0.00 | 72.97 |
| >hg38_21_+_8433172_8446622_RNA45SN1_45S_50ntFlanks@10808.10829.22\|CAATGTAGGTAAGGGAAGTCGG | >hg38_21_+_8433172_8446622_RNA45SN1_45S_50ntFlanks | 10808 | 10829 | 22 | rRF 45S | NO | 35.76 | 45.34 | 0.00 | 26.25 |
| >hg38_21_+_8433172_8446622_RNA45SN1_45S_50ntFlanks@10808.10830.23\|CAATGTAGGTAAGGGAAGTCGGC | >hg38_21_+_8433172_8446622_RNA45SN1_45S_50ntFlanks | 10808 | 10830 | 23 | rRF 45S | NO | 28.77 | 48.32 | 0.00 | 15.68 |
| >hg38_21_+_8433172_8446622_RNA45SN1_45S_50ntFlanks@10808.10840.33\|CAATGTAGGTAAGGGAAGTCGGCAAGCCGGATC | >hg38_21_+_8433172_8446622_RNA45SN1_45S_50ntFlanks | 10808 | 10840 | 33 | rRF 45S | NO | 97.10 | 117.86 | 56.76 | 122.26 |
| >hg38_21_+_8433172_8446622_RNA45SN1_45S_50ntFlanks@10816.10840.25\|GTAAGGGAAGTCGGCAAGCCGGATC | >hg38_21_+_8433172_8446622_RNA45SN1_45S_50ntFlanks | 10816 | 10840 | 25 | rRF 45S | NO | 12.75 | 12.25 | 0.00 | 0.00 |
| >hg38_21_+_8433172_8446622_RNA45SN1_45S_50ntFlanks@10817.10840.24\|TAAGGGAAGTCGGCAAGCCGGATC | >hg38_21_+_8433172_8446622_RNA45SN1_45S_50ntFlanks | 10817 | 10840 | 24 | rRF 45S | NO | 22.13 | 28.36 | 0.00 | 0.00 |
| >hg38_21_+_8433172_8446622_RNA45SN1_45S_50ntFlanks@10819.10837.19\|AGGGAAGTCGGCAAGCCGG | >hg38_21_+_8433172_8446622_RNA45SN1_45S_50ntFlanks | 10819 | 10837 | 19 | rRF 45S | NO | 0.00 | 46.21 | 40.53 | 0.00 |
| >hg38_21_+_8433172_8446622_RNA45SN1_45S_50ntFlanks@10819.10838.20\|AGGGAAGTCGGCAAGCCGGA | >hg38_21_+_8433172_8446622_RNA45SN1_45S_50ntFlanks | 10819 | 10838 | 20 | rRF 45S | NO | 0.00 | 0.00 | 30.48 | 0.00 |
| >hg38_21_+_8433172_8446622_RNA45SN1_45S_50ntFlanks@10819.10839.21\|AGGGAAGTCGGCAAGCCGGAT | >hg38_21_+_8433172_8446622_RNA45SN1_45S_50ntFlanks | 10819 | 10839 | 21 | rRF 45S | NO | 0.00 | 0.00 | 35.54 | 0.00 |
| >hg38_21_+_8433172_8446622_RNA45SN1_45S_50ntFlanks@10819.10840.22\|AGGGAAGTCGGCAAGCCGGATC | >hg38_21_+_8433172_8446622_RNA45SN1_45S_50ntFlanks | 10819 | 10840 | 22 | rRF 45S | NO | 375.33 | 414.30 | 212.58 | 89.26 |
| >hg38_21_+_8433172_8446622_RNA45SN1_45S_50ntFlanks@10820.10840.21\|GGGAAGTCGGCAAGCCGGATC | >hg38_21_+_8433172_8446622_RNA45SN1_45S_50ntFlanks | 10820 | 10840 | 21 | rRF 45S | NO | 106.72 | 127.79 | 63.50 | 24.32 |
| >hg38_21_+_8433172_8446622_RNA45SN1_45S_50ntFlanks@10821.10838.18\|GGAAGTCGGCAAGCCGGA | >hg38_21_+_8433172_8446622_RNA45SN1_45S_50ntFlanks | 10821 | 10838 | 18 | rRF 45S | NO | 33.44 | 59.52 | 41.27 | 0.00 |
| >hg38_21_+_8433172_8446622_RNA45SN1_45S_50ntFlanks@10821.10840.20\|GGAAGTCGGCAAGCCGGATC | >hg38_21_+_8433172_8446622_RNA45SN1_45S_50ntFlanks | 10821 | 10840 | 20 | rRF 45S | NO | 112.49 | 264.00 | 114.36 | 44.96 |
| >hg38_21_+_8433172_8446622_RNA45SN1_45S_50ntFlanks@10821.10865.45\|GGAAGTCGGCAAGCCGGATCCGTAACTTCGGGATAAGGATTGGCT | >hg38_21_+_8433172_8446622_RNA45SN1_45S_50ntFlanks | 10821 | 10865 | 45 | rRF 45S | NO | 41.50 | 43.46 | 0.00 | 0.00 |
| >hg38_21_+_8433172_8446622_RNA45SN1_45S_50ntFlanks@10822.10840.19\|GAAGTCGGCAAGCCGGATC | >hg38_21_+_8433172_8446622_RNA45SN1_45S_50ntFlanks | 10822 | 10840 | 19 | rRF 45S | NO | 0.00 | 32.37 | 0.00 | 0.00 |
| >hg38_21_+_8433172_8446622_RNA45SN1_45S_50ntFlanks@10823.10865.43\|AAGTCGGCAAGCCGGATCCGTAACTTCGGGATAAGGATTGGCT | >hg38_21_+_8433172_8446622_RNA45SN1_45S_50ntFlanks | 10823 | 10865 | 43 | rRF 45S | NO | 44.10 | 52.18 | 0.00 | 0.00 |
| >hg38_21_+_8433172_8446622_RNA45SN1_45S_50ntFlanks@10826.10844.19\|TCGGCAAGCCGGATCCGTA | >hg38_21_+_8433172_8446622_RNA45SN1_45S_50ntFlanks | 10826 | 10844 | 19 | rRF 45S | NO | 0.00 | 40.80 | 0.00 | 0.00 |
| >hg38_21_+_8433172_8446622_RNA45SN1_45S_50ntFlanks@10826.10858.33\|TCGGCAAGCCGGATCCGTAACTTCGGGATAAGG | >hg38_21_+_8433172_8446622_RNA45SN1_45S_50ntFlanks | 10826 | 10858 | 33 | rRF 45S | NO | 0.00 | 31.40 | 0.00 | 0.00 |
| >hg38_21_+_8433172_8446622_RNA45SN1_45S_50ntFlanks@10826.10865.40\|TCGGCAAGCCGGATCCGTAACTTCGGGATAAGGATTGGCT | >hg38_21_+_8433172_8446622_RNA45SN1_45S_50ntFlanks | 10826 | 10865 | 40 | rRF 45S | NO | 0.00 | 65.59 | 0.00 | 0.00 |
| >hg38_21_+_8433172_8446622_RNA45SN1_45S_50ntFlanks@10827.10865.39\|CGGCAAGCCGGATCCGTAACTTCGGGATAAGGATTGGCT | >hg38_21_+_8433172_8446622_RNA45SN1_45S_50ntFlanks | 10827 | 10865 | 39 | rRF 45S | NO | 0.00 | 37.98 | 0.00 | 0.00 |
| >hg38_21_+_8433172_8446622_RNA45SN1_45S_50ntFlanks@10829.10852.24\|GCAAGCCGGATCCGTAACTTCGGG | >hg38_21_+_8433172_8446622_RNA45SN1_45S_50ntFlanks | 10829 | 10852 | 24 | rRF 45S | NO | 0.00 | 28.90 | 0.00 | 0.00 |

### MDA-MB-468

| | | | | | | | | | | |
|---|---|---|---|---|---|---|---|---|---|---|
| >hg38_21_+_8433172_8446622_RNA45SN1_45S_50ntFlanks@10829.10858.30\|GCAAGCCGG ATCCGTAACTTCGGGATAAGG | >hg38_21_+_8433172_8446622_RNA45SN1_45S_50ntFlanks | 10829 | 10858 | 30 | rRF 45S | NO | 0.00 | 37.36 | 0.00 | 0.00 |
| >hg38_21_+_8433172_8446622_RNA45SN1_45S_50ntFlanks@10830.10851.22\|CAAGCCGGAT CCGTAACTTCGG | >hg38_21_+_8433172_8446622_RNA45SN1_45S_50ntFlanks | 10830 | 10851 | 22 | rRF 45S | NO | 0.00 | 27.53 | 0.00 | 0.00 |
| >hg38_21_+_8433172_8446622_RNA45SN1_45S_50ntFlanks@10830.10852.23\|CAAGCCGGAT CCGTAACTTCGGG | >hg38_21_+_8433172_8446622_RNA45SN1_45S_50ntFlanks | 10830 | 10852 | 23 | rRF 45S | NO | 0.00 | 26.37 | 0.00 | 0.00 |
| >hg38_21_+_8433172_8446622_RNA45SN1_45S_50ntFlanks@10830.10858.29\|CAAGCCGGAT CCGTAACTTCGGGATAAGG | >hg38_21_+_8433172_8446622_RNA45SN1_45S_50ntFlanks | 10830 | 10858 | 29 | rRF 45S | NO | 0.00 | 43.21 | 32.92 | 25.20 |
| >hg38_21_+_8433172_8446622_RNA45SN1_45S_50ntFlanks@10832.10862.31\|AGCCGGATCC GTAACTTCGGGATAAGGATTG | >hg38_21_+_8433172_8446622_RNA45SN1_45S_50ntFlanks | 10832 | 10862 | 31 | rRF 45S | NO | 0.00 | 35.64 | 0.00 | 0.00 |
| >hg38_21_+_8433172_8446622_RNA45SN1_45S_50ntFlanks@10835.10858.24\|CGGATCCGTA ACTTCGGGATAAGG | >hg38_21_+_8433172_8446622_RNA45SN1_45S_50ntFlanks | 10835 | 10858 | 24 | rRF 45S | NO | 0.00 | 32.79 | 0.00 | 0.00 |
| >hg38_21_+_8433172_8446622_RNA45SN1_45S_50ntFlanks@10838.10865.28\|ATCCGTAACT TCGGGATAAGGATTGGCT | >hg38_21_+_8433172_8446622_RNA45SN1_45S_50ntFlanks | 10838 | 10865 | 28 | rRF 45S | NO | 0.00 | 46.68 | 0.00 | 0.00 |
| >hg38_21_+_8433172_8446622_RNA45SN1_45S_50ntFlanks@10838.10868.31\|ATCCGTAACT TCGGGATAAGGATTGGCTCTA | >hg38_21_+_8433172_8446622_RNA45SN1_45S_50ntFlanks | 10838 | 10868 | 31 | rRF 45S | NO | 0.00 | 44.58 | 0.00 | 0.00 |
| >hg38_21_+_8433172_8446622_RNA45SN1_45S_50ntFlanks@10839.10865.27\|TCCGTAACTT CGGGATAAGGATTGGCT | >hg38_21_+_8433172_8446622_RNA45SN1_45S_50ntFlanks | 10839 | 10865 | 27 | rRF 45S | NO | 30.09 | 117.13 | 0.00 | 0.00 |
| >hg38_21_+_8433172_8446622_RNA45SN1_45S_50ntFlanks@10839.10868.30\|TCCGTAACTT CGGGATAAGGATTGGCTCTA | >hg38_21_+_8433172_8446622_RNA45SN1_45S_50ntFlanks | 10839 | 10868 | 30 | rRF 45S | NO | 20.53 | 28.29 | 0.00 | 0.00 |
| >hg38_21_+_8433172_8446622_RNA45SN1_45S_50ntFlanks@10840.10865.26\|CCGTAACTTC GGGATAAGGATTGGCT | >hg38_21_+_8433172_8446622_RNA45SN1_45S_50ntFlanks | 10840 | 10865 | 26 | rRF 45S | NO | 144.45 | 331.36 | 66.38 | 50.85 |
| >hg38_21_+_8433172_8446622_RNA45SN1_45S_50ntFlanks@10840.10867.28\|CCGTAACTTC GGGATAAGGATTGGCTCT | >hg38_21_+_8433172_8446622_RNA45SN1_45S_50ntFlanks | 10840 | 10867 | 28 | rRF 45S | NO | 28.87 | 29.91 | 0.00 | 0.00 |
| >hg38_21_+_8433172_8446622_RNA45SN1_45S_50ntFlanks@10840.10868.29\|CCGTAACTTC GGGATAAGGATTGGCTCTA | >hg38_21_+_8433172_8446622_RNA45SN1_45S_50ntFlanks | 10840 | 10868 | 29 | rRF 45S | NO | 78.92 | 50.54 | 0.00 | 0.00 |
| >hg38_21_+_8433172_8446622_RNA45SN1_45S_50ntFlanks@10841.10858.18\|CGTAACTTCG GGATAAGG | >hg38_21_+_8433172_8446622_RNA45SN1_45S_50ntFlanks | 10841 | 10858 | 18 | rRF 45S | NO | 0.00 | 114.27 | 6.54 | 0.00 |
| >hg38_21_+_8433172_8446622_RNA45SN1_45S_50ntFlanks@10841.10859.19\|CGTAACTTCG GGATAAGGA | >hg38_21_+_8433172_8446622_RNA45SN1_45S_50ntFlanks | 10841 | 10859 | 19 | rRF 45S | NO | 0.00 | 74.24 | 0.00 | 0.00 |
| >hg38_21_+_8433172_8446622_RNA45SN1_45S_50ntFlanks@10841.10860.20\|CGTAACTTCG GGATAAGGAT | >hg38_21_+_8433172_8446622_RNA45SN1_45S_50ntFlanks | 10841 | 10860 | 20 | rRF 45S | NO | 0.00 | 41.08 | 0.00 | 0.00 |
| >hg38_21_+_8433172_8446622_RNA45SN1_45S_50ntFlanks@10841.10861.21\|CGTAACTTCG GGATAAGGATT | >hg38_21_+_8433172_8446622_RNA45SN1_45S_50ntFlanks | 10841 | 10861 | 21 | rRF 45S | NO | 53.92 | 269.70 | 51.11 | 6.67 |
| >hg38_21_+_8433172_8446622_RNA45SN1_45S_50ntFlanks@10841.10862.22\|CGTAACTTCG GGATAAGGATTG | >hg38_21_+_8433172_8446622_RNA45SN1_45S_50ntFlanks | 10841 | 10862 | 22 | rRF 45S | NO | 143.22 | 462.57 | 79.87 | 26.87 |
| >hg38_21_+_8433172_8446622_RNA45SN1_45S_50ntFlanks@10841.10863.23\|CGTAACTTCG GGATAAGGATTGG | >hg38_21_+_8433172_8446622_RNA45SN1_45S_50ntFlanks | 10841 | 10863 | 23 | rRF 45S | NO | 146.91 | 462.42 | 97.03 | 6.36 |
| >hg38_21_+_8433172_8446622_RNA45SN1_45S_50ntFlanks@10841.10864.24\|CGTAACTTCG GGATAAGGATTGGC | >hg38_21_+_8433172_8446622_RNA45SN1_45S_50ntFlanks | 10841 | 10864 | 24 | rRF 45S | NO | 298.71 | 550.81 | 71.99 | 24.02 |
| >hg38_21_+_8433172_8446622_RNA45SN1_45S_50ntFlanks@10841.10865.25\|CGTAACTTCG GGATAAGGATTGGCT | >hg38_21_+_8433172_8446622_RNA45SN1_45S_50ntFlanks | 10841 | 10865 | 25 | rRF 45S | YES | 2699.69 | 6513.12 | 464.00 | 259.04 |
| >hg38_21_+_8433172_8446622_RNA45SN1_45S_50ntFlanks@10841.10866.26\|CGTAACTTCG GGATAAGGATTGGCTC | >hg38_21_+_8433172_8446622_RNA45SN1_45S_50ntFlanks | 10841 | 10866 | 26 | rRF 45S | NO | 120.74 | 128.68 | 0.00 | 0.00 |
| >hg38_21_+_8433172_8446622_RNA45SN1_45S_50ntFlanks@10841.10867.27\|CGTAACTTCG GGATAAGGATTGGCTCT | >hg38_21_+_8433172_8446622_RNA45SN1_45S_50ntFlanks | 10841 | 10867 | 27 | rRF 45S | NO | 242.38 | 258.41 | 13.75 | 0.00 |
| >hg38_21_+_8433172_8446622_RNA45SN1_45S_50ntFlanks@10841.10868.28\|CGTAACTTCG GGATAAGGATTGGCTCTA | >hg38_21_+_8433172_8446622_RNA45SN1_45S_50ntFlanks | 10841 | 10868 | 28 | rRF 45S | NO | 744.82 | 562.80 | 87.05 | 50.91 |
| >hg38_21_+_8433172_8446622_RNA45SN1_45S_50ntFlanks@10841.10871.31\|CGTAACTTCG GGATAAGGATTGGCTCTAAGG | >hg38_21_+_8433172_8446622_RNA45SN1_45S_50ntFlanks | 10841 | 10871 | 31 | rRF 45S | NO | 42.06 | 29.20 | 0.00 | 0.00 |
| >hg38_21_+_8433172_8446622_RNA45SN1_45S_50ntFlanks@10841.10872.32\|CGTAACTTCG GGATAAGGATTGGCTCTAAGGG | >hg38_21_+_8433172_8446622_RNA45SN1_45S_50ntFlanks | 10841 | 10872 | 32 | rRF 45S | NO | 75.38 | 89.11 | 0.00 | 0.00 |
| >hg38_21_+_8433172_8446622_RNA45SN1_45S_50ntFlanks@10841.10873.33\|CGTAACTTCG GGATAAGGATTGGCTCTAAGGGC | >hg38_21_+_8433172_8446622_RNA45SN1_45S_50ntFlanks | 10841 | 10873 | 33 | rRF 45S | NO | 79.25 | 47.03 | 0.00 | 0.00 |
| >hg38_21_+_8433172_8446622_RNA45SN1_45S_50ntFlanks@10841.10874.34\|CGTAACTTCG GGATAAGGATTGGCTCTAAGGGCT | >hg38_21_+_8433172_8446622_RNA45SN1_45S_50ntFlanks | 10841 | 10874 | 34 | rRF 45S | NO | 88.83 | 70.11 | 0.00 | 0.00 |
| >hg38_21_+_8433172_8446622_RNA45SN1_45S_50ntFlanks@10841.10875.35\|CGTAACTTCG GGATAAGGATTGGCTCTAAGGGCTG | >hg38_21_+_8433172_8446622_RNA45SN1_45S_50ntFlanks | 10841 | 10875 | 35 | rRF 45S | NO | 64.99 | 67.61 | 0.00 | 0.00 |
| >hg38_21_+_8433172_8446622_RNA45SN1_45S_50ntFlanks@10842.10865.24\|GTAACTTCGG GATAAGGATTGGCT | >hg38_21_+_8433172_8446622_RNA45SN1_45S_50ntFlanks | 10842 | 10865 | 24 | rRF 45S | NO | 0.00 | 53.73 | 0.00 | 0.00 |
| >hg38_21_+_8433172_8446622_RNA45SN1_45S_50ntFlanks@10843.10861.19\|TAACTTCGGG ATAAGGATT | >hg38_21_+_8433172_8446622_RNA45SN1_45S_50ntFlanks | 10843 | 10861 | 19 | rRF 45S | NO | 0.00 | 34.48 | 0.00 | 0.00 |
| >hg38_21_+_8433172_8446622_RNA45SN1_45S_50ntFlanks@10843.10862.20\|TAACTTCGGG ATAAGGATTG | >hg38_21_+_8433172_8446622_RNA45SN1_45S_50ntFlanks | 10843 | 10862 | 20 | rRF 45S | NO | 0.00 | 83.06 | 0.00 | 0.00 |
| >hg38_21_+_8433172_8446622_RNA45SN1_45S_50ntFlanks@10843.10863.21\|TAACTTCGGG ATAAGGATTGG | >hg38_21_+_8433172_8446622_RNA45SN1_45S_50ntFlanks | 10843 | 10863 | 21 | rRF 45S | NO | 0.00 | 62.05 | 0.00 | 0.00 |
| >hg38_21_+_8433172_8446622_RNA45SN1_45S_50ntFlanks@10843.10864.22\|TAACTTCGGG ATAAGGATTGGC | >hg38_21_+_8433172_8446622_RNA45SN1_45S_50ntFlanks | 10843 | 10864 | 22 | rRF 45S | NO | 0.00 | 30.10 | 0.00 | 0.00 |

### MDA-MB-468

| | | | | | | | | | | |
|---|---|---|---|---|---|---|---|---|---|---|
| >hg38_21_+_8433172_8446622_RNA45SN1_45S_50ntFlanks@10843.10865.23\|TAACTTCGGGATAAGGATTGGCT | >hg38_21_+_8433172_8446622_RNA45SN1_45S_50ntFlanks | 10843 | 10865 | 23 | rRF 45S | NO | 65.99 | 185.31 | 0.00 | 0.00 |
| >hg38_21_+_8433172_8446622_RNA45SN1_45S_50ntFlanks@10843.10867.25\|TAACTTCGGGATAAGGATTGGCTCT | >hg38_21_+_8433172_8446622_RNA45SN1_45S_50ntFlanks | 10843 | 10867 | 25 | rRF 45S | NO | 13.21 | 33.61 | 0.00 | 0.00 |
| >hg38_21_+_8433172_8446622_RNA45SN1_45S_50ntFlanks@10843.10868.26\|TAACTTCGGGATAAGGATTGGCTCTA | >hg38_21_+_8433172_8446622_RNA45SN1_45S_50ntFlanks | 10843 | 10868 | 26 | rRF 45S | NO | 81.07 | 67.21 | 0.00 | 0.00 |
| >hg38_21_+_8433172_8446622_RNA45SN1_45S_50ntFlanks@10844.10865.22\|AACTTCGGGATAAGGATTGGCT | >hg38_21_+_8433172_8446622_RNA45SN1_45S_50ntFlanks | 10844 | 10865 | 22 | rRF 45S | NO | 31.40 | 58.55 | 0.00 | 0.00 |
| >hg38_21_+_8433172_8446622_RNA45SN1_45S_50ntFlanks@10845.10862.18\|ACTTCGGGATAAGGATTG | >hg38_21_+_8433172_8446622_RNA45SN1_45S_50ntFlanks | 10845 | 10862 | 18 | rRF 45S | NO | 34.43 | 107.19 | 0.00 | 0.00 |
| >hg38_21_+_8433172_8446622_RNA45SN1_45S_50ntFlanks@10845.10863.19\|ACTTCGGGATAAGGATTGG | >hg38_21_+_8433172_8446622_RNA45SN1_45S_50ntFlanks | 10845 | 10863 | 19 | rRF 45S | NO | 28.97 | 110.42 | 0.00 | 0.00 |
| >hg38_21_+_8433172_8446622_RNA45SN1_45S_50ntFlanks@10845.10864.20\|ACTTCGGGATAAGGATTGGC | >hg38_21_+_8433172_8446622_RNA45SN1_45S_50ntFlanks | 10845 | 10864 | 20 | rRF 45S | NO | 48.93 | 99.13 | 0.00 | 0.00 |
| >hg38_21_+_8433172_8446622_RNA45SN1_45S_50ntFlanks@10845.10865.21\|ACTTCGGGATAAGGATTGGCT | >hg38_21_+_8433172_8446622_RNA45SN1_45S_50ntFlanks | 10845 | 10865 | 21 | rRF 45S | NO | 406.61 | 851.27 | 69.54 | 51.51 |
| >hg38_21_+_8433172_8446622_RNA45SN1_45S_50ntFlanks@10845.10866.22\|ACTTCGGGATAAGGATTGGCTC | >hg38_21_+_8433172_8446622_RNA45SN1_45S_50ntFlanks | 10845 | 10866 | 22 | rRF 45S | NO | 30.35 | 32.88 | 0.00 | 0.00 |
| >hg38_21_+_8433172_8446622_RNA45SN1_45S_50ntFlanks@10845.10867.23\|ACTTCGGGATAAGGATTGGCTCT | >hg38_21_+_8433172_8446622_RNA45SN1_45S_50ntFlanks | 10845 | 10867 | 23 | rRF 45S | NO | 38.13 | 44.40 | 0.00 | 0.00 |
| >hg38_21_+_8433172_8446622_RNA45SN1_45S_50ntFlanks@10845.10868.24\|ACTTCGGGATAAGGATTGGCTCTA | >hg38_21_+_8433172_8446622_RNA45SN1_45S_50ntFlanks | 10845 | 10868 | 24 | rRF 45S | NO | 108.92 | 78.44 | 6.65 | 0.00 |
| >hg38_21_+_8433172_8446622_RNA45SN1_45S_50ntFlanks@10846.10863.18\|CTTCGGGATAAGGATTGG | >hg38_21_+_8433172_8446622_RNA45SN1_45S_50ntFlanks | 10846 | 10863 | 18 | rRF 45S | NO | 0.00 | 79.73 | 0.00 | 0.00 |
| >hg38_21_+_8433172_8446622_RNA45SN1_45S_50ntFlanks@10846.10864.19\|CTTCGGGATAAGGATTGGC | >hg38_21_+_8433172_8446622_RNA45SN1_45S_50ntFlanks | 10846 | 10864 | 19 | rRF 45S | NO | 33.42 | 76.60 | 0.00 | 0.00 |
| >hg38_21_+_8433172_8446622_RNA45SN1_45S_50ntFlanks@10846.10865.20\|CTTCGGGATAAGGATTGGCT | >hg38_21_+_8433172_8446622_RNA45SN1_45S_50ntFlanks | 10846 | 10865 | 20 | rRF 45S | NO | 193.64 | 489.49 | 59.13 | 24.63 |
| >hg38_21_+_8433172_8446622_RNA45SN1_45S_50ntFlanks@10846.10867.22\|CTTCGGGATAAGGATTGGCTCT | >hg38_21_+_8433172_8446622_RNA45SN1_45S_50ntFlanks | 10846 | 10867 | 22 | rRF 45S | NO | 20.39 | 33.89 | 0.00 | 0.00 |
| >hg38_21_+_8433172_8446622_RNA45SN1_45S_50ntFlanks@10846.10868.23\|CTTCGGGATAAGGATTGGCTCTA | >hg38_21_+_8433172_8446622_RNA45SN1_45S_50ntFlanks | 10846 | 10868 | 23 | rRF 45S | NO | 82.39 | 61.02 | 0.00 | 0.00 |
| >hg38_21_+_8433172_8446622_RNA45SN1_45S_50ntFlanks@10848.10865.18\|TCGGGATAAGGATTGGCT | >hg38_21_+_8433172_8446622_RNA45SN1_45S_50ntFlanks | 10848 | 10865 | 18 | rRF 45S | NO | 0.00 | 32.26 | 0.00 | 0.00 |
| >hg38_21_+_8433172_8446622_RNA45SN1_45S_50ntFlanks@10849.10867.19\|CGGGATAAGGATTGGCTCT | >hg38_21_+_8433172_8446622_RNA45SN1_45S_50ntFlanks | 10849 | 10867 | 19 | rRF 45S | NO | 0.00 | 28.04 | 0.00 | 0.00 |
| >hg38_21_+_8433172_8446622_RNA45SN1_45S_50ntFlanks@10849.10868.20\|CGGGATAAGGATTGGCTCTA | >hg38_21_+_8433172_8446622_RNA45SN1_45S_50ntFlanks | 10849 | 10868 | 20 | rRF 45S | NO | 48.55 | 38.64 | 0.00 | 0.00 |
| >hg38_21_+_8433172_8446622_RNA45SN1_45S_50ntFlanks@10853.10873.21\|ATAAGGATTGGCTCTAAGGGC | >hg38_21_+_8433172_8446622_RNA45SN1_45S_50ntFlanks | 10853 | 10873 | 21 | rRF 45S | NO | 0.00 | 30.04 | 0.00 | 0.00 |
| >hg38_21_+_8433172_8446622_RNA45SN1_45S_50ntFlanks@10866.10883.18\|CTAAGGGCTGGGTCGGTC | >hg38_21_+_8433172_8446622_RNA45SN1_45S_50ntFlanks | 10866 | 10883 | 18 | rRF 45S | NO | 0.00 | 27.82 | 0.00 | 0.00 |
| >hg38_21_+_8433172_8446622_RNA45SN1_45S_50ntFlanks@10866.10884.19\|CTAAGGGCTGGGTCGGTCG | >hg38_21_+_8433172_8446622_RNA45SN1_45S_50ntFlanks | 10866 | 10884 | 19 | rRF 45S | NO | 58.36 | 81.35 | 13.91 | 0.00 |
| >hg38_21_+_8433172_8446622_RNA45SN1_45S_50ntFlanks@10866.10886.21\|CTAAGGGCTGGGTCGGTCGGG | >hg38_21_+_8433172_8446622_RNA45SN1_45S_50ntFlanks | 10866 | 10886 | 21 | rRF 45S | NO | 33.83 | 50.44 | 0.00 | 0.00 |
| >hg38_21_+_8433172_8446622_RNA45SN1_45S_50ntFlanks@10866.10889.24\|CTAAGGGCTGGGTCGGTCGGGCTG | >hg38_21_+_8433172_8446622_RNA45SN1_45S_50ntFlanks | 10866 | 10889 | 24 | rRF 45S | NO | 67.15 | 47.68 | 0.00 | 6.19 |
| >hg38_21_+_8433172_8446622_RNA45SN1_45S_50ntFlanks@10866.10890.25\|CTAAGGGCTGGGTCGGTCGGGCTGG | >hg38_21_+_8433172_8446622_RNA45SN1_45S_50ntFlanks | 10866 | 10890 | 25 | rRF 45S | NO | 58.93 | 78.08 | 6.85 | 15.02 |
| >hg38_21_+_8433172_8446622_RNA45SN1_45S_50ntFlanks@10866.10891.26\|CTAAGGGCTGGGTCGGTCGGGCTGGG | >hg38_21_+_8433172_8446622_RNA45SN1_45S_50ntFlanks | 10866 | 10891 | 26 | rRF 45S | NO | 41.12 | 62.08 | 34.78 | 26.05 |
| >hg38_21_+_8433172_8446622_RNA45SN1_45S_50ntFlanks@10866.10892.27\|CTAAGGGCTGGGTCGGTCGGGCTGGGG | >hg38_21_+_8433172_8446622_RNA45SN1_45S_50ntFlanks | 10866 | 10892 | 27 | rRF 45S | NO | 40.46 | 51.97 | 45.42 | 33.94 |
| >hg38_21_+_8433172_8446622_RNA45SN1_45S_50ntFlanks@10866.10894.29\|CTAAGGGCTGGGTCGGTCGGGCTGGGGCG | >hg38_21_+_8433172_8446622_RNA45SN1_45S_50ntFlanks | 10866 | 10894 | 29 | rRF 45S | NO | 57.40 | 37.71 | 34.58 | 26.28 |
| >hg38_21_+_8433172_8446622_RNA45SN1_45S_50ntFlanks@10866.10901.36\|CTAAGGGCTGGGTCGGTCGGGCTGGGGCGCGAAGCG | >hg38_21_+_8433172_8446622_RNA45SN1_45S_50ntFlanks | 10866 | 10901 | 36 | rRF 45S | NO | 114.79 | 54.52 | 0.00 | 47.04 |
| >hg38_21_+_8433172_8446622_RNA45SN1_45S_50ntFlanks@10866.10902.37\|CTAAGGGCTGGGTCGGTCGGGCTGGGGCGCGAAGCGG | >hg38_21_+_8433172_8446622_RNA45SN1_45S_50ntFlanks | 10866 | 10902 | 37 | rRF 45S | NO | 191.66 | 84.99 | 34.67 | 73.96 |
| >hg38_21_+_8433172_8446622_RNA45SN1_45S_50ntFlanks@10866.10904.39\|CTAAGGGCTGGGTCGGTCGGGCTGGGGCGCGAAGCGGGG | >hg38_21_+_8433172_8446622_RNA45SN1_45S_50ntFlanks | 10866 | 10904 | 39 | rRF 45S | NO | 50.58 | 30.63 | 0.00 | 0.00 |
| >hg38_21_+_8433172_8446622_RNA45SN1_45S_50ntFlanks@10869.10886.18\|AGGGCTGGGTCGGTCGGG | >hg38_21_+_8433172_8446622_RNA45SN1_45S_50ntFlanks | 10869 | 10886 | 18 | rRF 45S | NO | 33.04 | 54.65 | 20.37 | 0.00 |
| >hg38_21_+_8433172_8446622_RNA45SN1_45S_50ntFlanks@10869.10889.21\|AGGGCTGGGTCGGTCGGGCTG | >hg38_21_+_8433172_8446622_RNA45SN1_45S_50ntFlanks | 10869 | 10889 | 21 | rRF 45S | NO | 78.16 | 64.04 | 6.61 | 0.00 |
| >hg38_21_+_8433172_8446622_RNA45SN1_45S_50ntFlanks@10869.10890.22\|AGGGCTGGGTCGGTCGGGCTGG | >hg38_21_+_8433172_8446622_RNA45SN1_45S_50ntFlanks | 10869 | 10890 | 22 | rRF 45S | NO | 62.44 | 81.28 | 30.90 | 0.00 |

### MDA-MB-468

| | | | | | | | | | | |
|---|---|---|---|---|---|---|---|---|---|---|
| >hg38_21_+_8433172_8446622_RNA45SN1_45S_50ntFlanks@10869.10891.23\|AGGGCTGGGTCGGTCGGGCTGGG | >hg38_21_+_8433172_8446622_RNA45SN1_45S_50ntFlanks | 10869 | 10891 | 23 | rRF 45S | NO | 40.90 | 67.84 | 44.94 | 0.00 |
| >hg38_21_+_8433172_8446622_RNA45SN1_45S_50ntFlanks@10869.10892.24\|AGGGCTGGGTCGGTCGGGCTGGGG | >hg38_21_+_8433172_8446622_RNA45SN1_45S_50ntFlanks | 10869 | 10892 | 24 | rRF 45S | NO | 36.30 | 42.54 | 64.76 | 0.00 |
| >hg38_21_+_8433172_8446622_RNA45SN1_45S_50ntFlanks@10869.10894.26\|AGGGCTGGGTCGGTCGGGCTGGGGCG | >hg38_21_+_8433172_8446622_RNA45SN1_45S_50ntFlanks | 10869 | 10894 | 26 | rRF 45S | NO | 45.42 | 19.57 | 47.23 | 0.00 |
| >hg38_21_+_8433172_8446622_RNA45SN1_45S_50ntFlanks@10869.10901.33\|AGGGCTGGGTCGGTCGGGCTGGGGCGCGAAGCG | >hg38_21_+_8433172_8446622_RNA45SN1_45S_50ntFlanks | 10869 | 10901 | 33 | rRF 45S | NO | 98.39 | 49.88 | 31.79 | 24.24 |
| >hg38_21_+_8433172_8446622_RNA45SN1_45S_50ntFlanks@10869.10902.34\|AGGGCTGGGTCGGTCGGGCTGGGGCGCGAAGCGG | >hg38_21_+_8433172_8446622_RNA45SN1_45S_50ntFlanks | 10869 | 10902 | 34 | rRF 45S | NO | 209.76 | 90.77 | 69.42 | 53.38 |
| >hg38_21_+_8433172_8446622_RNA45SN1_45S_50ntFlanks@10869.10904.36\|AGGGCTGGGTCGGTCGGGCTGGGGCGCGAAGCGGGG | >hg38_21_+_8433172_8446622_RNA45SN1_45S_50ntFlanks | 10869 | 10904 | 36 | rRF 45S | NO | 58.07 | 35.73 | 0.00 | 0.00 |
| >hg38_21_+_8433172_8446622_RNA45SN1_45S_50ntFlanks@10873.10890.18\|CTGGGTCGGTCGGGCTGG | >hg38_21_+_8433172_8446622_RNA45SN1_45S_50ntFlanks | 10873 | 10890 | 18 | rRF 45S | NO | 28.84 | 60.15 | 0.00 | 0.00 |
| >hg38_21_+_8433172_8446622_RNA45SN1_45S_50ntFlanks@10873.10891.19\|CTGGGTCGGTCGGGCTGGG | >hg38_21_+_8433172_8446622_RNA45SN1_45S_50ntFlanks | 10873 | 10891 | 19 | rRF 45S | NO | 42.77 | 72.37 | 21.02 | 0.00 |
| >hg38_21_+_8433172_8446622_RNA45SN1_45S_50ntFlanks@10873.10892.20\|CTGGGTCGGTCGGGCTGGGG | >hg38_21_+_8433172_8446622_RNA45SN1_45S_50ntFlanks | 10873 | 10892 | 20 | rRF 45S | NO | 58.20 | 80.77 | 45.57 | 0.00 |
| >hg38_21_+_8433172_8446622_RNA45SN1_45S_50ntFlanks@10873.10894.22\|CTGGGTCGGTCGGGCTGGGGCG | >hg38_21_+_8433172_8446622_RNA45SN1_45S_50ntFlanks | 10873 | 10894 | 22 | rRF 45S | NO | 44.44 | 38.23 | 34.41 | 0.00 |
| >hg38_21_+_8433172_8446622_RNA45SN1_45S_50ntFlanks@10873.10901.29\|CTGGGTCGGTCGGGCTGGGGCGCGAAGCG | >hg38_21_+_8433172_8446622_RNA45SN1_45S_50ntFlanks | 10873 | 10901 | 29 | rRF 45S | NO | 28.99 | 12.39 | 0.00 | 0.00 |
| >hg38_21_+_8433172_8446622_RNA45SN1_45S_50ntFlanks@10873.10902.30\|CTGGGTCGGTCGGGCTGGGGCGCGAAGCGG | >hg38_21_+_8433172_8446622_RNA45SN1_45S_50ntFlanks | 10873 | 10902 | 30 | rRF 45S | NO | 81.66 | 66.52 | 38.81 | 6.05 |
| >hg38_21_+_8433172_8446622_RNA45SN1_45S_50ntFlanks@10873.10904.32\|CTGGGTCGGTCGGGCTGGGGCGCGAAGCGGGG | >hg38_21_+_8433172_8446622_RNA45SN1_45S_50ntFlanks | 10873 | 10904 | 32 | rRF 45S | NO | 32.52 | 32.03 | 0.00 | 0.00 |
| >hg38_21_+_8433172_8446622_RNA45SN1_45S_50ntFlanks@10878.10901.24\|TCGGTCGGGCTGGGGCGCGAAGCG | >hg38_21_+_8433172_8446622_RNA45SN1_45S_50ntFlanks | 10878 | 10901 | 24 | rRF 45S | NO | 41.37 | 37.94 | 0.00 | 0.00 |
| >hg38_21_+_8433172_8446622_RNA45SN1_45S_50ntFlanks@10878.10902.25\|TCGGTCGGGCTGGGGCGCGAAGCGG | >hg38_21_+_8433172_8446622_RNA45SN1_45S_50ntFlanks | 10878 | 10902 | 25 | rRF 45S | NO | 40.50 | 42.59 | 0.00 | 0.00 |
| >hg38_21_+_8433172_8446622_RNA45SN1_45S_50ntFlanks@10882.10902.21\|TCGGGCTGGGGCGCGAAGCGG | >hg38_21_+_8433172_8446622_RNA45SN1_45S_50ntFlanks | 10882 | 10902 | 21 | rRF 45S | NO | 12.77 | 30.42 | 0.00 | 0.00 |
| >hg38_21_+_8433172_8446622_RNA45SN1_45S_50ntFlanks@10905.10925.21\|CTGGGCGCGCGCCGCGGCTGG | >hg38_21_+_8433172_8446622_RNA45SN1_45S_50ntFlanks | 10905 | 10925 | 21 | rRF 45S | NO | 0.00 | 28.66 | 0.00 | 0.00 |
| >hg38_21_+_8433172_8446622_RNA45SN1_45S_50ntFlanks@10905.10926.22\|CTGGGCGCGCGCCGCGGCTGGA | >hg38_21_+_8433172_8446622_RNA45SN1_45S_50ntFlanks | 10905 | 10926 | 22 | rRF 45S | NO | 0.00 | 27.16 | 0.00 | 0.00 |
| >hg38_21_+_8433172_8446622_RNA45SN1_45S_50ntFlanks@10905.10932.28\|CTGGGCGCGCGCCGCGGCTGGACGAGGC | >hg38_21_+_8433172_8446622_RNA45SN1_45S_50ntFlanks | 10905 | 10932 | 28 | rRF 45S | NO | 21.13 | 0.00 | 0.00 | 0.00 |
| >hg38_21_+_8433172_8446622_RNA45SN1_45S_50ntFlanks@10905.10933.29\|CTGGGCGCGCGCCGCGGCTGGACGAGGCG | >hg38_21_+_8433172_8446622_RNA45SN1_45S_50ntFlanks | 10905 | 10933 | 29 | rRF 45S | NO | 40.15 | 12.79 | 0.00 | 0.00 |
| >hg38_21_+_8433172_8446622_RNA45SN1_45S_50ntFlanks@10905.10936.32\|CTGGGCGCGCGCCGCGGCTGGACGAGGCGCCG | >hg38_21_+_8433172_8446622_RNA45SN1_45S_50ntFlanks | 10905 | 10936 | 32 | rRF 45S | NO | 0.00 | 32.01 | 0.00 | 0.00 |
| >hg38_21_+_8433172_8446622_RNA45SN1_45S_50ntFlanks@10905.10941.37\|CTGGGCGCGCGCCGCGGCTGGACGAGGCGCCGCCGCC | >hg38_21_+_8433172_8446622_RNA45SN1_45S_50ntFlanks | 10905 | 10941 | 37 | rRF 45S | NO | 0.00 | 29.61 | 0.00 | 0.00 |
| >hg38_21_+_8433172_8446622_RNA45SN1_45S_50ntFlanks@10912.10931.20\|CGCGCCGCGGCTGGACGAGG | >hg38_21_+_8433172_8446622_RNA45SN1_45S_50ntFlanks | 10912 | 10931 | 20 | rRF 45S | NO | 32.22 | 0.00 | 0.00 | 0.00 |
| >hg38_21_+_8433172_8446622_RNA45SN1_45S_50ntFlanks@10912.10932.21\|CGCGCCGCGGCTGGACGAGGC | >hg38_21_+_8433172_8446622_RNA45SN1_45S_50ntFlanks | 10912 | 10932 | 21 | rRF 45S | NO | 0.00 | 34.43 | 30.13 | 0.00 |
| >hg38_21_+_8433172_8446622_RNA45SN1_45S_50ntFlanks@10914.10931.18\|CGCCGCGGCTGGACGAGG | >hg38_21_+_8433172_8446622_RNA45SN1_45S_50ntFlanks | 10914 | 10931 | 18 | rRF 45S | NO | 55.38 | 53.35 | 50.86 | 14.93 |
| >hg38_21_+_8433172_8446622_RNA45SN1_45S_50ntFlanks@10914.10932.19\|CGCCGCGGCTGGACGAGGC | >hg38_21_+_8433172_8446622_RNA45SN1_45S_50ntFlanks | 10914 | 10932 | 19 | rRF 45S | NO | 0.00 | 43.49 | 46.39 | 0.00 |
| >hg38_21_+_8433172_8446622_RNA45SN1_45S_50ntFlanks@10914.10936.23\|CGCCGCGGCTGGACGAGGCGCCG | >hg38_21_+_8433172_8446622_RNA45SN1_45S_50ntFlanks | 10914 | 10936 | 23 | rRF 45S | NO | 0.00 | 51.02 | 44.39 | 48.25 |
| >hg38_21_+_8433172_8446622_RNA45SN1_45S_50ntFlanks@10914.10946.33\|CGCCGCGGCTGGACGAGGCGCCGCCGCCCCCCC | >hg38_21_+_8433172_8446622_RNA45SN1_45S_50ntFlanks | 10914 | 10946 | 33 | rRF 45S | NO | 6.12 | 34.25 | 0.00 | 16.53 |
| >hg38_21_+_8433172_8446622_RNA45SN1_45S_50ntFlanks@10916.10933.18\|CCGCGGCTGGACGAGGCG | >hg38_21_+_8433172_8446622_RNA45SN1_45S_50ntFlanks | 10916 | 10933 | 18 | rRF 45S | NO | 36.36 | 48.59 | 29.70 | 0.00 |
| >hg38_21_+_8433172_8446622_RNA45SN1_45S_50ntFlanks@10916.10936.21\|CCGCGGCTGGACGAGGCGCCG | >hg38_21_+_8433172_8446622_RNA45SN1_45S_50ntFlanks | 10916 | 10936 | 21 | rRF 45S | NO | 0.00 | 36.53 | 0.00 | 0.00 |
| >hg38_21_+_8433172_8446622_RNA45SN1_45S_50ntFlanks@10921.10946.26\|GCTGGACGAGGCGCCGCCGCCCCCCC | >hg38_21_+_8433172_8446622_RNA45SN1_45S_50ntFlanks | 10921 | 10946 | 26 | rRF 45S | NO | 0.00 | 32.84 | 0.00 | 0.00 |
| >hg38_21_+_8433172_8446622_RNA45SN1_45S_50ntFlanks@10922.10939.18\|CTGGACGAGGCGCCGCCG | >hg38_21_+_8433172_8446622_RNA45SN1_45S_50ntFlanks | 10922 | 10939 | 18 | rRF 45S | NO | 90.36 | 92.29 | 56.77 | 67.95 |
| >hg38_21_+_8433172_8446622_RNA45SN1_45S_50ntFlanks@10922.10941.20\|CTGGACGAGGCGCCGCCGCC | >hg38_21_+_8433172_8446622_RNA45SN1_45S_50ntFlanks | 10922 | 10941 | 20 | rRF 45S | NO | 53.24 | 57.32 | 0.00 | 15.02 |
| >hg38_21_+_8433172_8446622_RNA45SN1_45S_50ntFlanks@10922.10945.24\|CTGGACGAGGCGCCGCCGCCCCCC | >hg38_21_+_8433172_8446622_RNA45SN1_45S_50ntFlanks | 10922 | 10945 | 24 | rRF 45S | NO | 44.06 | 101.34 | 39.21 | 13.84 |

### MDA-MB-468

| | | | | | | | | | | |
|---|---|---|---|---|---|---|---|---|---|---|
| >hg38_21_+_8433172_8446622_RNA45SN1_45S_50ntFlanks@10922.10946.25\|CTGGACGAGGCGCCGCCGCCCCCCC | >hg38_21_+_8433172_8446622_RNA45SN1_45S_50ntFlanks | 10922 | 10946 | 25 | rRF 45S | NO | 64.94 | 161.54 | 48.57 | 36.33 |
| >hg38_21_+_8433172_8446622_RNA45SN1_45S_50ntFlanks@10922.10947.26\|CTGGACGAGGCGCCGCCGCCCCCCCC | >hg38_21_+_8433172_8446622_RNA45SN1_45S_50ntFlanks | 10922 | 10947 | 26 | rRF 45S | NO | 0.00 | 27.78 | 0.00 | 0.00 |
| >hg38_21_+_8433172_8446622_RNA45SN1_45S_50ntFlanks@11027.11045.19\|CTCTCTCTCTCTCTCCCCC | >hg38_21_+_8433172_8446622_RNA45SN1_45S_50ntFlanks | 11027 | 11045 | 19 | rRF 45S | NO | 29.42 | 27.07 | 0.00 | 0.00 |
| >hg38_21_+_8433172_8446622_RNA45SN1_45S_50ntFlanks@11027.11046.20\|CTCTCTCTCTCTCTCCCCCG | >hg38_21_+_8433172_8446622_RNA45SN1_45S_50ntFlanks | 11027 | 11046 | 20 | rRF 45S | NO | 66.55 | 77.56 | 0.00 | 0.00 |
| >hg38_21_+_8433172_8446622_RNA45SN1_45S_50ntFlanks@11027.11048.22\|CTCTCTCTCTCTCTCCCCCGCT | >hg38_21_+_8433172_8446622_RNA45SN1_45S_50ntFlanks | 11027 | 11048 | 22 | rRF 45S | NO | 13.16 | 0.00 | 0.00 | 0.00 |
| >hg38_21_+_8433172_8446622_RNA45SN1_45S_50ntFlanks@11027.11049.23\|CTCTCTCTCTCTCTCCCCCGCTC | >hg38_21_+_8433172_8446622_RNA45SN1_45S_50ntFlanks | 11027 | 11049 | 23 | rRF 45S | NO | 69.18 | 81.24 | 0.00 | 14.20 |
| >hg38_21_+_8433172_8446622_RNA45SN1_45S_50ntFlanks@11027.11050.24\|CTCTCTCTCTCTCTCCCCCGCTCC | >hg38_21_+_8433172_8446622_RNA45SN1_45S_50ntFlanks | 11027 | 11050 | 24 | rRF 45S | NO | 101.98 | 133.32 | 0.00 | 36.11 |
| >hg38_21_+_8433172_8446622_RNA45SN1_45S_50ntFlanks@11027.11051.25\|CTCTCTCTCTCTCTCCCCCGCTCCC | >hg38_21_+_8433172_8446622_RNA45SN1_45S_50ntFlanks | 11027 | 11051 | 25 | rRF 45S | NO | 91.40 | 162.03 | 0.00 | 41.98 |
| >hg38_21_+_8433172_8446622_RNA45SN1_45S_50ntFlanks@11027.11053.27\|CTCTCTCTCTCTCTCCCCCGCTCCCCG | >hg38_21_+_8433172_8446622_RNA45SN1_45S_50ntFlanks | 11027 | 11053 | 27 | rRF 45S | NO | 93.67 | 112.94 | 0.00 | 0.00 |
| >hg38_21_+_8433172_8446622_RNA45SN1_45S_50ntFlanks@11027.11055.29\|CTCTCTCTCTCTCTCCCCCGCTCCCCGTC | >hg38_21_+_8433172_8446622_RNA45SN1_45S_50ntFlanks | 11027 | 11055 | 29 | rRF 45S | NO | 0.00 | 28.21 | 0.00 | 0.00 |
| >hg38_21_+_8433172_8446622_RNA45SN1_45S_50ntFlanks@11027.11059.33\|CTCTCTCTCTCTCTCCCCCGCTCCCCGTCCTCC | >hg38_21_+_8433172_8446622_RNA45SN1_45S_50ntFlanks | 11027 | 11059 | 33 | rRF 45S | NO | 20.57 | 0.00 | 0.00 | 0.00 |
| >hg38_21_+_8433172_8446622_RNA45SN1_45S_50ntFlanks@11027.11060.34\|CTCTCTCTCTCTCTCCCCCGCTCCCCGTCCTCCC | >hg38_21_+_8433172_8446622_RNA45SN1_45S_50ntFlanks | 11027 | 11060 | 34 | rRF 45S | NO | 150.25 | 100.08 | 0.00 | 89.06 |
| >hg38_21_+_8433172_8446622_RNA45SN1_45S_50ntFlanks@11027.11061.35\|CTCTCTCTCTCTCTCCCCCGCTCCCCGTCCTCCCC | >hg38_21_+_8433172_8446622_RNA45SN1_45S_50ntFlanks | 11027 | 11061 | 35 | rRF 45S | NO | 42.51 | 31.91 | 0.00 | 0.00 |
| >hg38_21_+_8433172_8446622_RNA45SN1_45S_50ntFlanks@11027.11075.49\|CTCTCTCTCTCTCTCCCCCGCTCCCCGTCCTCCCCCCTCCCCGGGGGAG | >hg38_21_+_8433172_8446622_RNA45SN1_45S_50ntFlanks | 11027 | 11075 | 49 | rRF 45S | YES | 42.79 | 26.56 | 0.00 | 0.00 |
| >hg38_21_+_8433172_8446622_RNA45SN1_45S_50ntFlanks@11029.11060.32\|CTCTCTCTCTCTCCCCCGCTCCCCGTCCTCCC | >hg38_21_+_8433172_8446622_RNA45SN1_45S_50ntFlanks | 11029 | 11060 | 32 | rRF 45S | NO | 12.67 | 0.00 | 0.00 | 0.00 |
| >hg38_21_+_8433172_8446622_RNA45SN1_45S_50ntFlanks@11035.11060.26\|CTCTCTCCCCCGCTCCCCGTCCTCCC | >hg38_21_+_8433172_8446622_RNA45SN1_45S_50ntFlanks | 11035 | 11060 | 26 | rRF 45S | YES | 20.01 | 0.00 | 0.00 | 0.00 |
| >hg38_21_+_8433172_8446622_RNA45SN1_45S_50ntFlanks@11037.11060.24\|CTCTCCCCCGCTCCCCGTCCTCCC | >hg38_21_+_8433172_8446622_RNA45SN1_45S_50ntFlanks | 11037 | 11060 | 24 | rRF 45S | NO | 90.75 | 0.00 | 0.00 | 24.66 |
| >hg38_21_+_8433172_8446622_RNA45SN1_45S_50ntFlanks@11037.11075.39\|CTCTCCCCCGCTCCCCGTCCTCCCCCCTCCCCGGGGGAG | >hg38_21_+_8433172_8446622_RNA45SN1_45S_50ntFlanks | 11037 | 11075 | 39 | rRF 45S | NO | 21.49 | 0.00 | 0.00 | 0.00 |
| >hg38_21_+_8433172_8446622_RNA45SN1_45S_50ntFlanks@11038.11060.23\|TCTCCCCCGCTCCCCGTCCTCCC | >hg38_21_+_8433172_8446622_RNA45SN1_45S_50ntFlanks | 11038 | 11060 | 23 | rRF 45S | NO | 61.09 | 26.87 | 0.00 | 0.00 |
| >hg38_21_+_8433172_8446622_RNA45SN1_45S_50ntFlanks@11051.11075.25\|CCGTCCTCCCCCCTCCCCGGGGGAG | >hg38_21_+_8433172_8446622_RNA45SN1_45S_50ntFlanks | 11051 | 11075 | 25 | rRF 45S | NO | 0.00 | 45.62 | 0.00 | 0.00 |
| >hg38_21_+_8433172_8446622_RNA45SN1_45S_50ntFlanks@11052.11075.24\|CGTCCTCCCCCCTCCCCGGGGGAG | >hg38_21_+_8433172_8446622_RNA45SN1_45S_50ntFlanks | 11052 | 11075 | 24 | rRF 45S | NO | 0.00 | 12.45 | 0.00 | 0.00 |
| >hg38_21_+_8433172_8446622_RNA45SN1_45S_50ntFlanks@11076.11103.28\|CGCCGCGTGGGGGCGGCGGCGGGGGGAG | >hg38_21_+_8433172_8446622_RNA45SN1_45S_50ntFlanks | 11076 | 11103 | 28 | rRF 45S | NO | 62.87 | 0.00 | 0.00 | 0.00 |
| >hg38_21_+_8433172_8446622_RNA45SN1_45S_50ntFlanks@11104.11121.18\|AAGGGTCGGGGCGGCAGG | >hg38_21_+_8433172_8446622_RNA45SN1_45S_50ntFlanks | 11104 | 11121 | 18 | rRF 45S | NO | 20.54 | 0.00 | 0.00 | 0.00 |
| >hg38_21_+_8433172_8446622_RNA45SN1_45S_50ntFlanks@11104.11123.20\|AAGGGTCGGGGCGGCAGGGG | >hg38_21_+_8433172_8446622_RNA45SN1_45S_50ntFlanks | 11104 | 11123 | 20 | rRF 45S | NO | 74.24 | 46.28 | 28.08 | 0.00 |
| >hg38_21_+_8433172_8446622_RNA45SN1_45S_50ntFlanks@11104.11133.30\|AAGGGTCGGGGCGGCAGGGGCCGGCGGCGG | >hg38_21_+_8433172_8446622_RNA45SN1_45S_50ntFlanks | 11104 | 11133 | 30 | rRF 45S | NO | 51.30 | 27.11 | 0.00 | 0.00 |
| >hg38_21_+_8433172_8446622_RNA45SN1_45S_50ntFlanks@11161.11178.18\|CACGGTCCCCCGCGAGGG | >hg38_21_+_8433172_8446622_RNA45SN1_45S_50ntFlanks | 11161 | 11178 | 18 | rRF 45S | NO | 110.62 | 219.31 | 48.30 | 58.71 |
| >hg38_21_+_8433172_8446622_RNA45SN1_45S_50ntFlanks@11161.11179.19\|CACGGTCCCCCGCGAGGGG | >hg38_21_+_8433172_8446622_RNA45SN1_45S_50ntFlanks | 11161 | 11179 | 19 | rRF 45S | NO | 63.00 | 135.44 | 35.56 | 23.73 |
| >hg38_21_+_8433172_8446622_RNA45SN1_45S_50ntFlanks@11161.11188.28\|CACGGTCCCCCGCGAGGGGGGCCCGGGC | >hg38_21_+_8433172_8446622_RNA45SN1_45S_50ntFlanks | 11161 | 11188 | 28 | rRF 45S | NO | 52.83 | 48.50 | 6.51 | 39.74 |
| >hg38_21_+_8433172_8446622_RNA45SN1_45S_50ntFlanks@11188.11225.38\|CACCCGGGGGGCCGGCGGCGGCGGCGACTCTGGACGCG | >hg38_21_+_8433172_8446622_RNA45SN1_45S_50ntFlanks | 11188 | 11225 | 38 | rRF 45S | NO | 13.22 | 29.44 | 0.00 | 0.00 |
| >hg38_21_+_8433172_8446622_RNA45SN1_45S_50ntFlanks@11189.11225.37\|ACCCGGGGGGCCGGCGGCGGCGGCGACTCTGGACGCG | >hg38_21_+_8433172_8446622_RNA45SN1_45S_50ntFlanks | 11189 | 11225 | 37 | rRF 45S | NO | 35.67 | 32.92 | 0.00 | 0.00 |
| >hg38_21_+_8433172_8446622_RNA45SN1_45S_50ntFlanks@11190.11225.36\|CCCGGGGGGCCGGCGGCGGCGGCGACTCTGGACGCG | >hg38_21_+_8433172_8446622_RNA45SN1_45S_50ntFlanks | 11190 | 11225 | 36 | rRF 45S | NO | 61.24 | 41.66 | 0.00 | 0.00 |
| >hg38_21_+_8433172_8446622_RNA45SN1_45S_50ntFlanks@11199.11220.22\|CCGGCGGCGGCGGCGACTCTGG | >hg38_21_+_8433172_8446622_RNA45SN1_45S_50ntFlanks | 11199 | 11220 | 22 | rRF 45S | NO | 46.05 | 50.46 | 0.00 | 0.00 |
| >hg38_21_+_8433172_8446622_RNA45SN1_45S_50ntFlanks@11199.11223.25\|CCGGCGGCGGCGGCGACTCTGGACG | >hg38_21_+_8433172_8446622_RNA45SN1_45S_50ntFlanks | 11199 | 11223 | 25 | rRF 45S | NO | 45.79 | 57.69 | 0.00 | 0.00 |
| >hg38_21_+_8433172_8446622_RNA45SN1_45S_50ntFlanks@11199.11224.26\|CCGGCGGCGGCGGCGACTCTGGACGC | >hg38_21_+_8433172_8446622_RNA45SN1_45S_50ntFlanks | 11199 | 11224 | 26 | rRF 45S | NO | 0.00 | 12.71 | 0.00 | 0.00 |

### MDA-MB-468

| | | | | | | | | | | |
|---|---|---|---|---|---|---|---|---|---|---|
| >hg38_21_+_8433172_8446622_RNA45SN1_45S_50ntFlanks@11199.11225.27\|CCGGCGGCGGCGGCGACTCTGGACGCG | >hg38_21_+_8433172_8446622_RNA45SN1_45S_50ntFlanks | 11199 | 11225 | 27 | rRF 45S | NO | 173.67 | 300.36 | 34.62 | 63.50 |
| >hg38_21_+_8433172_8446622_RNA45SN1_45S_50ntFlanks@11199.11227.29\|CCGGCGGCGGCGGCGACTCTGGACGCGAG | >hg38_21_+_8433172_8446622_RNA45SN1_45S_50ntFlanks | 11199 | 11227 | 29 | rRF 45S | NO | 59.55 | 97.30 | 0.00 | 6.47 |
| >hg38_21_+_8433172_8446622_RNA45SN1_45S_50ntFlanks@11199.11229.31\|CCGGCGGCGGCGGCGACTCTGGACGCGAGCC | >hg38_21_+_8433172_8446622_RNA45SN1_45S_50ntFlanks | 11199 | 11229 | 31 | rRF 45S | NO | 0.00 | 12.63 | 0.00 | 0.00 |
| >hg38_21_+_8433172_8446622_RNA45SN1_45S_50ntFlanks@11199.11233.35\|CCGGCGGCGGCGGCGACTCTGGACGCGAGCCGGGC | >hg38_21_+_8433172_8446622_RNA45SN1_45S_50ntFlanks | 11199 | 11233 | 35 | rRF 45S | NO | 0.00 | 30.25 | 0.00 | 0.00 |
| >hg38_21_+_8433172_8446622_RNA45SN1_45S_50ntFlanks@11200.11223.24\|CGGCGGCGGCGGCGACTCTGGACG | >hg38_21_+_8433172_8446622_RNA45SN1_45S_50ntFlanks | 11200 | 11223 | 24 | rRF 45S | NO | 32.13 | 31.66 | 0.00 | 0.00 |
| >hg38_21_+_8433172_8446622_RNA45SN1_45S_50ntFlanks@11200.11225.26\|CGGCGGCGGCGGCGACTCTGGACGCG | >hg38_21_+_8433172_8446622_RNA45SN1_45S_50ntFlanks | 11200 | 11225 | 26 | rRF 45S | NO | 52.74 | 137.42 | 36.56 | 26.22 |
| >hg38_21_+_8433172_8446622_RNA45SN1_45S_50ntFlanks@11200.11227.28\|CGGCGGCGGCGGCGACTCTGGACGCGAG | >hg38_21_+_8433172_8446622_RNA45SN1_45S_50ntFlanks | 11200 | 11227 | 28 | rRF 45S | NO | 0.00 | 34.08 | 0.00 | 0.00 |
| >hg38_21_+_8433172_8446622_RNA45SN1_45S_50ntFlanks@11200.11232.33\|CGGCGGCGGCGGCGACTCTGGACGCGAGCCGGG | >hg38_21_+_8433172_8446622_RNA45SN1_45S_50ntFlanks | 11200 | 11232 | 33 | rRF 45S | NO | 138.59 | 405.05 | 54.00 | 38.15 |
| >hg38_21_+_8433172_8446622_RNA45SN1_45S_50ntFlanks@11200.11233.34\|CGGCGGCGGCGGCGACTCTGGACGCGAGCCGGGC | >hg38_21_+_8433172_8446622_RNA45SN1_45S_50ntFlanks | 11200 | 11233 | 34 | rRF 45S | NO | 0.00 | 33.68 | 0.00 | 0.00 |
| >hg38_21_+_8433172_8446622_RNA45SN1_45S_50ntFlanks@11200.11234.35\|CGGCGGCGGCGGCGACTCTGGACGCGAGCCGGGCC | >hg38_21_+_8433172_8446622_RNA45SN1_45S_50ntFlanks | 11200 | 11234 | 35 | rRF 45S | NO | 0.00 | 39.37 | 0.00 | 0.00 |
| >hg38_21_+_8433172_8446622_RNA45SN1_45S_50ntFlanks@11203.11225.23\|CGGCGGCGGCGACTCTGGACGCG | >hg38_21_+_8433172_8446622_RNA45SN1_45S_50ntFlanks | 11203 | 11225 | 23 | rRF 45S | NO | 0.00 | 29.57 | 0.00 | 0.00 |
| >hg38_21_+_8433172_8446622_RNA45SN1_45S_50ntFlanks@11203.11232.30\|CGGCGGCGGCGACTCTGGACGCGAGCCGGG | >hg38_21_+_8433172_8446622_RNA45SN1_45S_50ntFlanks | 11203 | 11232 | 30 | rRF 45S | NO | 0.00 | 45.30 | 0.00 | 0.00 |
| >hg38_21_+_8433172_8446622_RNA45SN1_45S_50ntFlanks@11205.11225.21\|GCGGCGGCGACTCTGGACGCG | >hg38_21_+_8433172_8446622_RNA45SN1_45S_50ntFlanks | 11205 | 11225 | 21 | rRF 45S | NO | 0.00 | 51.91 | 31.16 | 0.00 |
| >hg38_21_+_8433172_8446622_RNA45SN1_45S_50ntFlanks@11205.11232.28\|GCGGCGGCGACTCTGGACGCGAGCCGGG | >hg38_21_+_8433172_8446622_RNA45SN1_45S_50ntFlanks | 11205 | 11232 | 28 | rRF 45S | NO | 32.91 | 85.80 | 20.68 | 0.00 |
| >hg38_21_+_8433172_8446622_RNA45SN1_45S_50ntFlanks@11206.11225.20\|CGGCGGCGACTCTGGACGCG | >hg38_21_+_8433172_8446622_RNA45SN1_45S_50ntFlanks | 11206 | 11225 | 20 | rRF 45S | NO | 0.00 | 26.44 | 0.00 | 0.00 |
| >hg38_21_+_8433172_8446622_RNA45SN1_45S_50ntFlanks@11206.11232.27\|CGGCGGCGACTCTGGACGCGAGCCGGG | >hg38_21_+_8433172_8446622_RNA45SN1_45S_50ntFlanks | 11206 | 11232 | 27 | rRF 45S | NO | 0.00 | 77.75 | 29.72 | 0.00 |
| >hg38_21_+_8433172_8446622_RNA45SN1_45S_50ntFlanks@11207.11225.19\|GGCGGCGACTCTGGACGCG | >hg38_21_+_8433172_8446622_RNA45SN1_45S_50ntFlanks | 11207 | 11225 | 19 | rRF 45S | NO | 0.00 | 60.73 | 44.74 | 0.00 |
| >hg38_21_+_8433172_8446622_RNA45SN1_45S_50ntFlanks@11207.11232.26\|GGCGGCGACTCTGGACGCGAGCCGGG | >hg38_21_+_8433172_8446622_RNA45SN1_45S_50ntFlanks | 11207 | 11232 | 26 | rRF 45S | NO | 41.49 | 80.25 | 0.00 | 0.00 |
| >hg38_21_+_8433172_8446622_RNA45SN1_45S_50ntFlanks@11207.11234.28\|GGCGGCGACTCTGGACGCGAGCCGGGCC | >hg38_21_+_8433172_8446622_RNA45SN1_45S_50ntFlanks | 11207 | 11234 | 28 | rRF 45S | NO | 0.00 | 12.26 | 0.00 | 0.00 |
| >hg38_21_+_8433172_8446622_RNA45SN1_45S_50ntFlanks@11208.11225.18\|GCGGCGACTCTGGACGCG | >hg38_21_+_8433172_8446622_RNA45SN1_45S_50ntFlanks | 11208 | 11225 | 18 | rRF 45S | NO | 52.17 | 575.65 | 411.15 | 64.57 |
| >hg38_21_+_8433172_8446622_RNA45SN1_45S_50ntFlanks@11208.11227.20\|GCGGCGACTCTGGACGCGAG | >hg38_21_+_8433172_8446622_RNA45SN1_45S_50ntFlanks | 11208 | 11227 | 20 | rRF 45S | NO | 47.72 | 232.47 | 100.84 | 33.95 |
| >hg38_21_+_8433172_8446622_RNA45SN1_45S_50ntFlanks@11208.11228.21\|GCGGCGACTCTGGACGCGAGC | >hg38_21_+_8433172_8446622_RNA45SN1_45S_50ntFlanks | 11208 | 11228 | 21 | rRF 45S | NO | 12.76 | 66.39 | 35.15 | 0.00 |
| >hg38_21_+_8433172_8446622_RNA45SN1_45S_50ntFlanks@11208.11229.22\|GCGGCGACTCTGGACGCGAGCC | >hg38_21_+_8433172_8446622_RNA45SN1_45S_50ntFlanks | 11208 | 11229 | 22 | rRF 45S | NO | 52.83 | 92.68 | 6.61 | 0.00 |
| >hg38_21_+_8433172_8446622_RNA45SN1_45S_50ntFlanks@11208.11230.23\|GCGGCGACTCTGGACGCGAGCCG | >hg38_21_+_8433172_8446622_RNA45SN1_45S_50ntFlanks | 11208 | 11230 | 23 | rRF 45S | NO | 94.92 | 127.91 | 40.42 | 15.92 |
| >hg38_21_+_8433172_8446622_RNA45SN1_45S_50ntFlanks@11208.11231.24\|GCGGCGACTCTGGACGCGAGCCGG | >hg38_21_+_8433172_8446622_RNA45SN1_45S_50ntFlanks | 11208 | 11231 | 24 | rRF 45S | NO | 134.20 | 188.32 | 30.07 | 24.07 |
| >hg38_21_+_8433172_8446622_RNA45SN1_45S_50ntFlanks@11208.11232.25\|GCGGCGACTCTGGACGCGAGCCGGG | >hg38_21_+_8433172_8446622_RNA45SN1_45S_50ntFlanks | 11208 | 11232 | 25 | rRF 45S | YES | 935.88 | 1272.39 | 230.06 | 161.56 |
| >hg38_21_+_8433172_8446622_RNA45SN1_45S_50ntFlanks@11208.11233.26\|GCGGCGACTCTGGACGCGAGCCGGGC | >hg38_21_+_8433172_8446622_RNA45SN1_45S_50ntFlanks | 11208 | 11233 | 26 | rRF 45S | NO | 274.08 | 536.87 | 68.18 | 66.89 |
| >hg38_21_+_8433172_8446622_RNA45SN1_45S_50ntFlanks@11208.11234.27\|GCGGCGACTCTGGACGCGAGCCGGGCC | >hg38_21_+_8433172_8446622_RNA45SN1_45S_50ntFlanks | 11208 | 11234 | 27 | rRF 45S | NO | 163.89 | 567.29 | 60.72 | 65.14 |
| >hg38_21_+_8433172_8446622_RNA45SN1_45S_50ntFlanks@11208.11235.28\|GCGGCGACTCTGGACGCGAGCCGGGCCC | >hg38_21_+_8433172_8446622_RNA45SN1_45S_50ntFlanks | 11208 | 11235 | 28 | rRF 45S | NO | 0.00 | 12.62 | 0.00 | 0.00 |
| >hg38_21_+_8433172_8446622_RNA45SN1_45S_50ntFlanks@11208.11238.31\|GCGGCGACTCTGGACGCGAGCCGGGCCCTTC | >hg38_21_+_8433172_8446622_RNA45SN1_45S_50ntFlanks | 11208 | 11238 | 31 | rRF 45S | NO | 0.00 | 39.44 | 0.00 | 0.00 |
| >hg38_21_+_8433172_8446622_RNA45SN1_45S_50ntFlanks@11208.11239.32\|GCGGCGACTCTGGACGCGAGCCGGGCCCTTCC | >hg38_21_+_8433172_8446622_RNA45SN1_45S_50ntFlanks | 11208 | 11239 | 32 | rRF 45S | NO | 0.00 | 58.99 | 0.00 | 0.00 |
| >hg38_21_+_8433172_8446622_RNA45SN1_45S_50ntFlanks@11208.11250.43\|GCGGCGACTCTGGACGCGAGCCGGGCCCTTCCCGTGGATCGCC | >hg38_21_+_8433172_8446622_RNA45SN1_45S_50ntFlanks | 11208 | 11250 | 43 | rRF 45S | NO | 48.14 | 44.39 | 0.00 | 0.00 |
| >hg38_21_+_8433172_8446622_RNA45SN1_45S_50ntFlanks@11208.11252.45\|GCGGCGACTCTGGACGCGAGCCGGGCCCTTCCCGTGGATCGCCCC | >hg38_21_+_8433172_8446622_RNA45SN1_45S_50ntFlanks | 11208 | 11252 | 45 | rRF 45S | NO | 108.78 | 68.71 | 0.00 | 23.88 |
| >hg38_21_+_8433172_8446622_RNA45SN1_45S_50ntFlanks@11208.11254.47\|GCGGCGACTCTGGACGCGAGCCGGGCCCTTCCCGTGGATCGCCCCAG | >hg38_21_+_8433172_8446622_RNA45SN1_45S_50ntFlanks | 11208 | 11254 | 47 | rRF 45S | NO | 1213.74 | 678.82 | 0.00 | 227.24 |

### MDA-MB-468

| | | | | | | | | | | |
|---|---|---|---|---|---|---|---|---|---|---|
| >hg38_21_+_8433172_8446622_RNA45SN1_45S_50ntFlanks@11209.11227.19\|CGGCGACTCTGGACGCGAG | >hg38_21_+_8433172_8446622_RNA45SN1_45S_50ntFlanks | 11209 | 11227 | 19 | rRF 45S | NO | 29.90 | 160.58 | 79.39 | 15.00 |
| >hg38_21_+_8433172_8446622_RNA45SN1_45S_50ntFlanks@11209.11229.21\|CGGCGACTCTGGACGCGAGCC | >hg38_21_+_8433172_8446622_RNA45SN1_45S_50ntFlanks | 11209 | 11229 | 21 | rRF 45S | NO | 35.23 | 62.30 | 7.23 | 0.00 |
| >hg38_21_+_8433172_8446622_RNA45SN1_45S_50ntFlanks@11209.11230.22\|CGGCGACTCTGGACGCGAGCCG | >hg38_21_+_8433172_8446622_RNA45SN1_45S_50ntFlanks | 11209 | 11230 | 22 | rRF 45S | NO | 46.81 | 104.65 | 47.53 | 23.49 |
| >hg38_21_+_8433172_8446622_RNA45SN1_45S_50ntFlanks@11209.11231.23\|CGGCGACTCTGGACGCGAGCCGG | >hg38_21_+_8433172_8446622_RNA45SN1_45S_50ntFlanks | 11209 | 11231 | 23 | rRF 45S | NO | 78.23 | 152.30 | 31.58 | 26.43 |
| >hg38_21_+_8433172_8446622_RNA45SN1_45S_50ntFlanks@11209.11232.24\|CGGCGACTCTGGACGCGAGCCGGG | >hg38_21_+_8433172_8446622_RNA45SN1_45S_50ntFlanks | 11209 | 11232 | 24 | rRF 45S | YES | 942.73 | 1733.99 | 361.24 | 282.38 |
| >hg38_21_+_8433172_8446622_RNA45SN1_45S_50ntFlanks@11209.11233.25\|CGGCGACTCTGGACGCGAGCCGGGC | >hg38_21_+_8433172_8446622_RNA45SN1_45S_50ntFlanks | 11209 | 11233 | 25 | rRF 45S | NO | 89.83 | 228.98 | 40.40 | 36.31 |
| >hg38_21_+_8433172_8446622_RNA45SN1_45S_50ntFlanks@11209.11234.26\|CGGCGACTCTGGACGCGAGCCGGGCC | >hg38_21_+_8433172_8446622_RNA45SN1_45S_50ntFlanks | 11209 | 11234 | 26 | rRF 45S | NO | 66.43 | 314.00 | 61.75 | 50.62 |
| >hg38_21_+_8433172_8446622_RNA45SN1_45S_50ntFlanks@11209.11252.44\|CGGCGACTCTGGACGCGAGCCGGGCCCTTCCCGTGGATCGCCCC | >hg38_21_+_8433172_8446622_RNA45SN1_45S_50ntFlanks | 11209 | 11252 | 44 | rRF 45S | NO | 95.45 | 47.93 | 0.00 | 16.37 |
| >hg38_21_+_8433172_8446622_RNA45SN1_45S_50ntFlanks@11209.11254.46\|CGGCGACTCTGGACGCGAGCCGGGCCCTTCCCGTGGATCGCCCCAG | >hg38_21_+_8433172_8446622_RNA45SN1_45S_50ntFlanks | 11209 | 11254 | 46 | rRF 45S | YES | 1539.07 | 785.47 | 0.00 | 326.94 |
| >hg38_21_+_8433172_8446622_RNA45SN1_45S_50ntFlanks@11210.11227.18\|GGCGACTCTGGACGCGAG | >hg38_21_+_8433172_8446622_RNA45SN1_45S_50ntFlanks | 11210 | 11227 | 18 | rRF 45S | NO | 0.00 | 12.78 | 0.00 | 0.00 |
| >hg38_21_+_8433172_8446622_RNA45SN1_45S_50ntFlanks@11210.11232.23\|GGCGACTCTGGACGCGAGCCGGG | >hg38_21_+_8433172_8446622_RNA45SN1_45S_50ntFlanks | 11210 | 11232 | 23 | rRF 45S | NO | 84.45 | 88.35 | 57.89 | 16.65 |
| >hg38_21_+_8433172_8446622_RNA45SN1_45S_50ntFlanks@11210.11233.24\|GGCGACTCTGGACGCGAGCCGGGC | >hg38_21_+_8433172_8446622_RNA45SN1_45S_50ntFlanks | 11210 | 11233 | 24 | rRF 45S | NO | 6.16 | 40.92 | 0.00 | 0.00 |
| >hg38_21_+_8433172_8446622_RNA45SN1_45S_50ntFlanks@11210.11254.45\|GGCGACTCTGGACGCGAGCCGGGCCCTTCCCGTGGATCGCCCCAG | >hg38_21_+_8433172_8446622_RNA45SN1_45S_50ntFlanks | 11210 | 11254 | 45 | rRF 45S | NO | 105.93 | 54.88 | 0.00 | 39.66 |
| >hg38_21_+_8433172_8446622_RNA45SN1_45S_50ntFlanks@11211.11232.22\|GCGACTCTGGACGCGAGCCGGG | >hg38_21_+_8433172_8446622_RNA45SN1_45S_50ntFlanks | 11211 | 11232 | 22 | rRF 45S | NO | 34.36 | 108.50 | 45.41 | 0.00 |
| >hg38_21_+_8433172_8446622_RNA45SN1_45S_50ntFlanks@11211.11254.44\|GCGACTCTGGACGCGAGCCGGGCCCTTCCCGTGGATCGCCCCAG | >hg38_21_+_8433172_8446622_RNA45SN1_45S_50ntFlanks | 11211 | 11254 | 44 | rRF 45S | NO | 33.75 | 29.33 | 0.00 | 6.13 |
| >hg38_21_+_8433172_8446622_RNA45SN1_45S_50ntFlanks@11212.11229.18\|CGACTCTGGACGCGAGCC | >hg38_21_+_8433172_8446622_RNA45SN1_45S_50ntFlanks | 11212 | 11229 | 18 | rRF 45S | NO | 0.00 | 43.07 | 0.00 | 0.00 |
| >hg38_21_+_8433172_8446622_RNA45SN1_45S_50ntFlanks@11212.11230.19\|CGACTCTGGACGCGAGCCG | >hg38_21_+_8433172_8446622_RNA45SN1_45S_50ntFlanks | 11212 | 11230 | 19 | rRF 45S | NO | 0.00 | 12.57 | 0.00 | 0.00 |
| >hg38_21_+_8433172_8446622_RNA45SN1_45S_50ntFlanks@11212.11231.20\|CGACTCTGGACGCGAGCCGG | >hg38_21_+_8433172_8446622_RNA45SN1_45S_50ntFlanks | 11212 | 11231 | 20 | rRF 45S | NO | 0.00 | 39.88 | 0.00 | 0.00 |
| >hg38_21_+_8433172_8446622_RNA45SN1_45S_50ntFlanks@11212.11232.21\|CGACTCTGGACGCGAGCCGGG | >hg38_21_+_8433172_8446622_RNA45SN1_45S_50ntFlanks | 11212 | 11232 | 21 | rRF 45S | NO | 30.40 | 310.77 | 74.06 | 16.34 |
| >hg38_21_+_8433172_8446622_RNA45SN1_45S_50ntFlanks@11212.11250.39\|CGACTCTGGACGCGAGCCGGGCCCTTCCCGTGGATCGCC | >hg38_21_+_8433172_8446622_RNA45SN1_45S_50ntFlanks | 11212 | 11250 | 39 | rRF 45S | NO | 31.61 | 37.71 | 0.00 | 0.00 |
| >hg38_21_+_8433172_8446622_RNA45SN1_45S_50ntFlanks@11212.11251.40\|CGACTCTGGACGCGAGCCGGGCCCTTCCCGTGGATCGCCC | >hg38_21_+_8433172_8446622_RNA45SN1_45S_50ntFlanks | 11212 | 11251 | 40 | rRF 45S | NO | 12.58 | 28.20 | 0.00 | 0.00 |
| >hg38_21_+_8433172_8446622_RNA45SN1_45S_50ntFlanks@11212.11252.41\|CGACTCTGGACGCGAGCCGGGCCCTTCCCGTGGATCGCCCC | >hg38_21_+_8433172_8446622_RNA45SN1_45S_50ntFlanks | 11212 | 11252 | 41 | rRF 45S | NO | 65.54 | 56.31 | 0.00 | 16.38 |
| >hg38_21_+_8433172_8446622_RNA45SN1_45S_50ntFlanks@11212.11253.42\|CGACTCTGGACGCGAGCCGGGCCCTTCCCGTGGATCGCCCCA | >hg38_21_+_8433172_8446622_RNA45SN1_45S_50ntFlanks | 11212 | 11253 | 42 | rRF 45S | NO | 33.83 | 29.16 | 0.00 | 0.00 |
| >hg38_21_+_8433172_8446622_RNA45SN1_45S_50ntFlanks@11212.11254.43\|CGACTCTGGACGCGAGCCGGGCCCTTCCCGTGGATCGCCCCAG | >hg38_21_+_8433172_8446622_RNA45SN1_45S_50ntFlanks | 11212 | 11254 | 43 | rRF 45S | NO | 31.26 | 38.01 | 0.00 | 25.98 |
| >hg38_21_+_8433172_8446622_RNA45SN1_45S_50ntFlanks@11215.11234.20\|CTCTGGACGCGAGCCGGGCC | >hg38_21_+_8433172_8446622_RNA45SN1_45S_50ntFlanks | 11215 | 11234 | 20 | rRF 45S | NO | 0.00 | 42.67 | 0.00 | 0.00 |
| >hg38_21_+_8433172_8446622_RNA45SN1_45S_50ntFlanks@11217.11234.18\|CTGGACGCGAGCCGGGCC | >hg38_21_+_8433172_8446622_RNA45SN1_45S_50ntFlanks | 11217 | 11234 | 18 | rRF 45S | NO | 36.17 | 214.05 | 45.04 | 34.02 |
| >hg38_21_+_8433172_8446622_RNA45SN1_45S_50ntFlanks@11217.11248.32\|CTGGACGCGAGCCGGGCCCTTCCCGTGGATCG | >hg38_21_+_8433172_8446622_RNA45SN1_45S_50ntFlanks | 11217 | 11248 | 32 | rRF 45S | NO | 19.41 | 0.00 | 0.00 | 0.00 |
| >hg38_21_+_8433172_8446622_RNA45SN1_45S_50ntFlanks@11217.11250.34\|CTGGACGCGAGCCGGGCCCTTCCCGTGGATCGCC | >hg38_21_+_8433172_8446622_RNA45SN1_45S_50ntFlanks | 11217 | 11250 | 34 | rRF 45S | NO | 0.00 | 26.58 | 0.00 | 0.00 |
| >hg38_21_+_8433172_8446622_RNA45SN1_45S_50ntFlanks@11233.11250.18\|CCCTTCCCGTGGATCGCC | >hg38_21_+_8433172_8446622_RNA45SN1_45S_50ntFlanks | 11233 | 11250 | 18 | rRF 45S | NO | 34.40 | 94.37 | 0.00 | 15.70 |
| >hg38_21_+_8433172_8446622_RNA45SN1_45S_50ntFlanks@11233.11252.20\|CCCTTCCCGTGGATCGCCCC | >hg38_21_+_8433172_8446622_RNA45SN1_45S_50ntFlanks | 11233 | 11252 | 20 | rRF 45S | NO | 44.48 | 41.41 | 0.00 | 24.08 |
| >hg38_21_+_8433172_8446622_RNA45SN1_45S_50ntFlanks@11233.11253.21\|CCCTTCCCGTGGATCGCCCCA | >hg38_21_+_8433172_8446622_RNA45SN1_45S_50ntFlanks | 11233 | 11253 | 21 | rRF 45S | NO | 20.08 | 32.53 | 0.00 | 16.49 |
| >hg38_21_+_8433172_8446622_RNA45SN1_45S_50ntFlanks@11233.11254.22\|CCCTTCCCGTGGATCGCCCCAG | >hg38_21_+_8433172_8446622_RNA45SN1_45S_50ntFlanks | 11233 | 11254 | 22 | rRF 45S | NO | 110.77 | 123.66 | 6.90 | 89.80 |
| >hg38_21_+_8433172_8446622_RNA45SN1_45S_50ntFlanks@11234.11254.21\|CCTTCCCGTGGATCGCCCCAG | >hg38_21_+_8433172_8446622_RNA45SN1_45S_50ntFlanks | 11234 | 11254 | 21 | rRF 45S | NO | 0.00 | 35.03 | 0.00 | 14.67 |
| >hg38_21_+_8433172_8446622_RNA45SN1_45S_50ntFlanks@11235.11252.18\|CTTCCCGTGGATCGCCCC | >hg38_21_+_8433172_8446622_RNA45SN1_45S_50ntFlanks | 11235 | 11252 | 18 | rRF 45S | NO | 130.85 | 181.78 | 6.71 | 64.96 |

### MDA-MB-468

<table>
<tr><td>>hg38_21_+_8433172_8446622_RNA45SN1_45S_50ntFlanks@11235.11253.19|CTTCCCGTGGATCGCCCCA</td><td>>hg38_21_+_8433172_8446622_RNA45SN1_45S_50ntFlanks</td><td>11235</td><td>11253</td><td>19</td><td>rRF 45S</td><td>NO</td><td>60.91</td><td>125.03</td><td>0.00</td><td>59.70</td></tr>
<tr><td>>hg38_21_+_8433172_8446622_RNA45SN1_45S_50ntFlanks@11235.11254.20|CTTCCCGTGGATCGCCCCAG</td><td>>hg38_21_+_8433172_8446622_RNA45SN1_45S_50ntFlanks</td><td>11235</td><td>11254</td><td>20</td><td>rRF 45S</td><td>NO</td><td>112.67</td><td>267.36</td><td>54.53</td><td>109.99</td></tr>
<tr><td>>hg38_21_+_8433172_8446622_RNA45SN1_45S_50ntFlanks@11235.11255.21|CTTCCCGTGGATCGCCCCAGC</td><td>>hg38_21_+_8433172_8446622_RNA45SN1_45S_50ntFlanks</td><td>11235</td><td>11255</td><td>21</td><td>rRF 45S</td><td>NO</td><td>28.80</td><td>32.54</td><td>0.00</td><td>0.00</td></tr>
<tr><td>>hg38_21_+_8433172_8446622_RNA45SN1_45S_50ntFlanks@11235.11268.34|CTTCCCGTGGATCGCCCCAGCTGCGGCGGGCGTC</td><td>>hg38_21_+_8433172_8446622_RNA45SN1_45S_50ntFlanks</td><td>11235</td><td>11268</td><td>34</td><td>rRF 45S</td><td>NO</td><td>28.66</td><td>0.00</td><td>0.00</td><td>13.58</td></tr>
<tr><td>>hg38_21_+_8433172_8446622_RNA45SN1_45S_50ntFlanks@11235.11269.35|CTTCCCGTGGATCGCCCCAGCTGCGGCGGGCGTCG</td><td>>hg38_21_+_8433172_8446622_RNA45SN1_45S_50ntFlanks</td><td>11235</td><td>11269</td><td>35</td><td>rRF 45S</td><td>NO</td><td>75.59</td><td>50.74</td><td>0.00</td><td>73.89</td></tr>
<tr><td>>hg38_21_+_8433172_8446622_RNA45SN1_45S_50ntFlanks@11312.11344.33|CCCCCACCCCACGTCTCGTCGCGCGCGCGTCCG</td><td>>hg38_21_+_8433172_8446622_RNA45SN1_45S_50ntFlanks</td><td>11312</td><td>11344</td><td>33</td><td>rRF 45S</td><td>NO</td><td>54.94</td><td>0.00</td><td>0.00</td><td>6.45</td></tr>
<tr><td>>hg38_21_+_8433172_8446622_RNA45SN1_45S_50ntFlanks@11313.11344.32|CCCCACCCCACGTCTCGTCGCGCGCGCGTCCG</td><td>>hg38_21_+_8433172_8446622_RNA45SN1_45S_50ntFlanks</td><td>11313</td><td>11344</td><td>32</td><td>rRF 45S</td><td>NO</td><td>42.64</td><td>0.00</td><td>0.00</td><td>0.00</td></tr>
<tr><td>>hg38_21_+_8433172_8446622_RNA45SN1_45S_50ntFlanks@11314.11342.29|CCCACCCCACGTCTCGTCGCGCGCGCGTC</td><td>>hg38_21_+_8433172_8446622_RNA45SN1_45S_50ntFlanks</td><td>11314</td><td>11342</td><td>29</td><td>rRF 45S</td><td>NO</td><td>31.28</td><td>0.00</td><td>0.00</td><td>0.00</td></tr>
<tr><td>>hg38_21_+_8433172_8446622_RNA45SN1_45S_50ntFlanks@11314.11344.31|CCCACCCCACGTCTCGTCGCGCGCGCGTCCG</td><td>>hg38_21_+_8433172_8446622_RNA45SN1_45S_50ntFlanks</td><td>11314</td><td>11344</td><td>31</td><td>rRF 45S</td><td>NO</td><td>162.31</td><td>60.08</td><td>0.00</td><td>54.50</td></tr>
<tr><td>>hg38_21_+_8433172_8446622_RNA45SN1_45S_50ntFlanks@11315.11332.18|CCACCCCACGTCTCGTCG</td><td>>hg38_21_+_8433172_8446622_RNA45SN1_45S_50ntFlanks</td><td>11315</td><td>11332</td><td>18</td><td>rRF 45S</td><td>NO</td><td>47.80</td><td>38.62</td><td>0.00</td><td>6.22</td></tr>
<tr><td>>hg38_21_+_8433172_8446622_RNA45SN1_45S_50ntFlanks@11315.11334.20|CCACCCCACGTCTCGTCGCG</td><td>>hg38_21_+_8433172_8446622_RNA45SN1_45S_50ntFlanks</td><td>11315</td><td>11334</td><td>20</td><td>rRF 45S</td><td>NO</td><td>12.67</td><td>33.41</td><td>0.00</td><td>0.00</td></tr>
<tr><td>>hg38_21_+_8433172_8446622_RNA45SN1_45S_50ntFlanks@11315.11336.22|CCACCCCACGTCTCGTCGCGCG</td><td>>hg38_21_+_8433172_8446622_RNA45SN1_45S_50ntFlanks</td><td>11315</td><td>11336</td><td>22</td><td>rRF 45S</td><td>NO</td><td>59.73</td><td>47.21</td><td>0.00</td><td>35.23</td></tr>
<tr><td>>hg38_21_+_8433172_8446622_RNA45SN1_45S_50ntFlanks@11315.11338.24|CCACCCCACGTCTCGTCGCGCGCG</td><td>>hg38_21_+_8433172_8446622_RNA45SN1_45S_50ntFlanks</td><td>11315</td><td>11338</td><td>24</td><td>rRF 45S</td><td>NO</td><td>78.55</td><td>54.13</td><td>0.00</td><td>39.37</td></tr>
<tr><td>>hg38_21_+_8433172_8446622_RNA45SN1_45S_50ntFlanks@11315.11342.28|CCACCCCACGTCTCGTCGCGCGCGCGTC</td><td>>hg38_21_+_8433172_8446622_RNA45SN1_45S_50ntFlanks</td><td>11315</td><td>11342</td><td>28</td><td>rRF 45S</td><td>NO</td><td>105.29</td><td>57.73</td><td>0.00</td><td>40.66</td></tr>
<tr><td>>hg38_21_+_8433172_8446622_RNA45SN1_45S_50ntFlanks@11315.11343.29|CCACCCCACGTCTCGTCGCGCGCGCGTCC</td><td>>hg38_21_+_8433172_8446622_RNA45SN1_45S_50ntFlanks</td><td>11315</td><td>11343</td><td>29</td><td>rRF 45S</td><td>NO</td><td>57.83</td><td>0.00</td><td>0.00</td><td>0.00</td></tr>
<tr><td>>hg38_21_+_8433172_8446622_RNA45SN1_45S_50ntFlanks@11315.11344.30|CCACCCCACGTCTCGTCGCGCGCGCGTCCG</td><td>>hg38_21_+_8433172_8446622_RNA45SN1_45S_50ntFlanks</td><td>11315</td><td>11344</td><td>30</td><td>rRF 45S</td><td>NO</td><td>552.64</td><td>214.99</td><td>0.00</td><td>185.67</td></tr>
<tr><td>>hg38_21_+_8433172_8446622_RNA45SN1_45S_50ntFlanks@11316.11334.19|CACCCCACGTCTCGTCGCG</td><td>>hg38_21_+_8433172_8446622_RNA45SN1_45S_50ntFlanks</td><td>11316</td><td>11334</td><td>19</td><td>rRF 45S</td><td>NO</td><td>49.33</td><td>128.73</td><td>0.00</td><td>53.21</td></tr>
<tr><td>>hg38_21_+_8433172_8446622_RNA45SN1_45S_50ntFlanks@11316.11336.21|CACCCCACGTCTCGTCGCGCG</td><td>>hg38_21_+_8433172_8446622_RNA45SN1_45S_50ntFlanks</td><td>11316</td><td>11336</td><td>21</td><td>rRF 45S</td><td>NO</td><td>169.28</td><td>198.44</td><td>0.00</td><td>136.05</td></tr>
<tr><td>>hg38_21_+_8433172_8446622_RNA45SN1_45S_50ntFlanks@11316.11338.23|CACCCCACGTCTCGTCGCGCGCG</td><td>>hg38_21_+_8433172_8446622_RNA45SN1_45S_50ntFlanks</td><td>11316</td><td>11338</td><td>23</td><td>rRF 45S</td><td>NO</td><td>280.64</td><td>254.75</td><td>0.00</td><td>171.41</td></tr>
<tr><td>>hg38_21_+_8433172_8446622_RNA45SN1_45S_50ntFlanks@11316.11340.25|CACCCCACGTCTCGTCGCGCGCGCG</td><td>>hg38_21_+_8433172_8446622_RNA45SN1_45S_50ntFlanks</td><td>11316</td><td>11340</td><td>25</td><td>rRF 45S</td><td>NO</td><td>76.35</td><td>43.38</td><td>0.00</td><td>81.77</td></tr>
<tr><td>>hg38_21_+_8433172_8446622_RNA45SN1_45S_50ntFlanks@11316.11342.27|CACCCCACGTCTCGTCGCGCGCGCGTC</td><td>>hg38_21_+_8433172_8446622_RNA45SN1_45S_50ntFlanks</td><td>11316</td><td>11342</td><td>27</td><td>rRF 45S</td><td>NO</td><td>511.32</td><td>319.47</td><td>6.71</td><td>252.09</td></tr>
<tr><td>>hg38_21_+_8433172_8446622_RNA45SN1_45S_50ntFlanks@11316.11343.28|CACCCCACGTCTCGTCGCGCGCGCGTCC</td><td>>hg38_21_+_8433172_8446622_RNA45SN1_45S_50ntFlanks</td><td>11316</td><td>11343</td><td>28</td><td>rRF 45S</td><td>NO</td><td>208.89</td><td>64.23</td><td>0.00</td><td>75.70</td></tr>
<tr><td>>hg38_21_+_8433172_8446622_RNA45SN1_45S_50ntFlanks@11316.11344.29|CACCCCACGTCTCGTCGCGCGCGCGTCCG</td><td>>hg38_21_+_8433172_8446622_RNA45SN1_45S_50ntFlanks</td><td>11316</td><td>11344</td><td>29</td><td>rRF 45S</td><td>YES</td><td>2177.55</td><td>931.77</td><td>31.17</td><td>862.20</td></tr>
<tr><td>>hg38_21_+_8433172_8446622_RNA45SN1_45S_50ntFlanks@11317.11334.18|ACCCCACGTCTCGTCGCG</td><td>>hg38_21_+_8433172_8446622_RNA45SN1_45S_50ntFlanks</td><td>11317</td><td>11334</td><td>18</td><td>rRF 45S</td><td>NO</td><td>0.00</td><td>47.25</td><td>0.00</td><td>15.59</td></tr>
<tr><td>>hg38_21_+_8433172_8446622_RNA45SN1_45S_50ntFlanks@11317.11336.20|ACCCCACGTCTCGTCGCGCG</td><td>>hg38_21_+_8433172_8446622_RNA45SN1_45S_50ntFlanks</td><td>11317</td><td>11336</td><td>20</td><td>rRF 45S</td><td>NO</td><td>80.08</td><td>83.52</td><td>0.00</td><td>70.84</td></tr>
<tr><td>>hg38_21_+_8433172_8446622_RNA45SN1_45S_50ntFlanks@11317.11338.22|ACCCCACGTCTCGTCGCGCGCG</td><td>>hg38_21_+_8433172_8446622_RNA45SN1_45S_50ntFlanks</td><td>11317</td><td>11338</td><td>22</td><td>rRF 45S</td><td>NO</td><td>146.62</td><td>108.90</td><td>0.00</td><td>94.38</td></tr>
<tr><td>>hg38_21_+_8433172_8446622_RNA45SN1_45S_50ntFlanks@11317.11340.24|ACCCCACGTCTCGTCGCGCGCGCG</td><td>>hg38_21_+_8433172_8446622_RNA45SN1_45S_50ntFlanks</td><td>11317</td><td>11340</td><td>24</td><td>rRF 45S</td><td>NO</td><td>41.29</td><td>0.00</td><td>0.00</td><td>40.80</td></tr>
<tr><td>>hg38_21_+_8433172_8446622_RNA45SN1_45S_50ntFlanks@11317.11342.26|ACCCCACGTCTCGTCGCGCGCGCGTC</td><td>>hg38_21_+_8433172_8446622_RNA45SN1_45S_50ntFlanks</td><td>11317</td><td>11342</td><td>26</td><td>rRF 45S</td><td>NO</td><td>246.17</td><td>148.78</td><td>0.00</td><td>116.31</td></tr>
<tr><td>>hg38_21_+_8433172_8446622_RNA45SN1_45S_50ntFlanks@11317.11343.27|ACCCCACGTCTCGTCGCGCGCGCGTCC</td><td>>hg38_21_+_8433172_8446622_RNA45SN1_45S_50ntFlanks</td><td>11317</td><td>11343</td><td>27</td><td>rRF 45S</td><td>NO</td><td>143.30</td><td>47.52</td><td>0.00</td><td>49.94</td></tr>
<tr><td>>hg38_21_+_8433172_8446622_RNA45SN1_45S_50ntFlanks@11317.11344.28|ACCCCACGTCTCGTCGCGCGCGCGTCCG</td><td>>hg38_21_+_8433172_8446622_RNA45SN1_45S_50ntFlanks</td><td>11317</td><td>11344</td><td>28</td><td>rRF 45S</td><td>NO</td><td>1397.90</td><td>552.26</td><td>0.00</td><td>497.16</td></tr>
<tr><td>>hg38_21_+_8433172_8446622_RNA45SN1_45S_50ntFlanks@11318.11336.19|CCCCACGTCTCGTCGCGCG</td><td>>hg38_21_+_8433172_8446622_RNA45SN1_45S_50ntFlanks</td><td>11318</td><td>11336</td><td>19</td><td>rRF 45S</td><td>NO</td><td>37.87</td><td>64.65</td><td>0.00</td><td>39.59</td></tr>
<tr><td>>hg38_21_+_8433172_8446622_RNA45SN1_45S_50ntFlanks@11318.11338.21|CCCCACGTCTCGTCGCGCGCG</td><td>>hg38_21_+_8433172_8446622_RNA45SN1_45S_50ntFlanks</td><td>11318</td><td>11338</td><td>21</td><td>rRF 45S</td><td>NO</td><td>56.47</td><td>74.27</td><td>0.00</td><td>48.39</td></tr>
<tr><td>>hg38_21_+_8433172_8446622_RNA45SN1_45S_50ntFlanks@11318.11342.25|CCCCACGTCTCGTCGCGCGCGCGTC</td><td>>hg38_21_+_8433172_8446622_RNA45SN1_45S_50ntFlanks</td><td>11318</td><td>11342</td><td>25</td><td>rRF 45S</td><td>NO</td><td>186.66</td><td>126.95</td><td>0.00</td><td>88.34</td></tr>
<tr><td>>hg38_21_+_8433172_8446622_RNA45SN1_45S_50ntFlanks@11318.11343.26|CCCCACGTCTCGTCGCGCGCGCGTCC</td><td>>hg38_21_+_8433172_8446622_RNA45SN1_45S_50ntFlanks</td><td>11318</td><td>11343</td><td>26</td><td>rRF 45S</td><td>NO</td><td>133.72</td><td>47.75</td><td>0.00</td><td>39.09</td></tr>
</table>

### MDA-MB-468

| | | | | | | | | | | |
|---|---|---|---|---|---|---|---|---|---|---|
| >hg38_21_+_8433172_8446622_RNA45SN1_45S_50ntFlanks@11318.11344.27\|CCCCACGTCTCGTCGCGCGCGCGTCCG | >hg38_21_+_8433172_8446622_RNA45SN1_45S_50ntFlanks | 11318 | 11344 | 27 | rRF 45S | NO | 982.82 | 506.86 | 0.00 | 398.99 |
| >hg38_21_+_8433172_8446622_RNA45SN1_45S_50ntFlanks@11319.11338.20\|CCCACGTCTCGTCGCGCGCG | >hg38_21_+_8433172_8446622_RNA45SN1_45S_50ntFlanks | 11319 | 11338 | 20 | rRF 45S | NO | 0.00 | 12.38 | 0.00 | 0.00 |
| >hg38_21_+_8433172_8446622_RNA45SN1_45S_50ntFlanks@11319.11342.24\|CCCACGTCTCGTCGCGCGCGCGTC | >hg38_21_+_8433172_8446622_RNA45SN1_45S_50ntFlanks | 11319 | 11342 | 24 | rRF 45S | NO | 67.45 | 80.23 | 0.00 | 24.81 |
| >hg38_21_+_8433172_8446622_RNA45SN1_45S_50ntFlanks@11319.11343.25\|CCCACGTCTCGTCGCGCGCGCGTCC | >hg38_21_+_8433172_8446622_RNA45SN1_45S_50ntFlanks | 11319 | 11343 | 25 | rRF 45S | NO | 43.36 | 0.00 | 0.00 | 0.00 |
| >hg38_21_+_8433172_8446622_RNA45SN1_45S_50ntFlanks@11319.11344.26\|CCCACGTCTCGTCGCGCGCGCGTCCG | >hg38_21_+_8433172_8446622_RNA45SN1_45S_50ntFlanks | 11319 | 11344 | 26 | rRF 45S | NO | 330.72 | 251.87 | 0.00 | 136.11 |
| >hg38_21_+_8433172_8446622_RNA45SN1_45S_50ntFlanks@11320.11338.19\|CCACGTCTCGTCGCGCGCG | >hg38_21_+_8433172_8446622_RNA45SN1_45S_50ntFlanks | 11320 | 11338 | 19 | rRF 45S | NO | 0.00 | 19.35 | 0.00 | 0.00 |
| >hg38_21_+_8433172_8446622_RNA45SN1_45S_50ntFlanks@11320.11342.23\|CCACGTCTCGTCGCGCGCGCGTC | >hg38_21_+_8433172_8446622_RNA45SN1_45S_50ntFlanks | 11320 | 11342 | 23 | rRF 45S | NO | 111.31 | 146.05 | 0.00 | 64.09 |
| >hg38_21_+_8433172_8446622_RNA45SN1_45S_50ntFlanks@11320.11343.24\|CCACGTCTCGTCGCGCGCGCGTCC | >hg38_21_+_8433172_8446622_RNA45SN1_45S_50ntFlanks | 11320 | 11343 | 24 | rRF 45S | NO | 62.19 | 35.47 | 0.00 | 6.31 |
| >hg38_21_+_8433172_8446622_RNA45SN1_45S_50ntFlanks@11320.11344.25\|CCACGTCTCGTCGCGCGCGCGTCCG | >hg38_21_+_8433172_8446622_RNA45SN1_45S_50ntFlanks | 11320 | 11344 | 25 | rRF 45S | NO | 447.42 | 404.24 | 0.00 | 236.05 |
| >hg38_21_+_8433172_8446622_RNA45SN1_45S_50ntFlanks@11321.11338.18\|CACGTCTCGTCGCGCGCG | >hg38_21_+_8433172_8446622_RNA45SN1_45S_50ntFlanks | 11321 | 11338 | 18 | rRF 45S | NO | 45.54 | 108.63 | 0.00 | 25.68 |
| >hg38_21_+_8433172_8446622_RNA45SN1_45S_50ntFlanks@11321.11340.20\|CACGTCTCGTCGCGCGCGCG | >hg38_21_+_8433172_8446622_RNA45SN1_45S_50ntFlanks | 11321 | 11340 | 20 | rRF 45S | NO | 29.04 | 33.04 | 0.00 | 14.10 |
| >hg38_21_+_8433172_8446622_RNA45SN1_45S_50ntFlanks@11321.11342.22\|CACGTCTCGTCGCGCGCGCGTC | >hg38_21_+_8433172_8446622_RNA45SN1_45S_50ntFlanks | 11321 | 11342 | 22 | rRF 45S | NO | 144.10 | 200.62 | 0.00 | 74.32 |
| >hg38_21_+_8433172_8446622_RNA45SN1_45S_50ntFlanks@11321.11343.23\|CACGTCTCGTCGCGCGCGCGTCC | >hg38_21_+_8433172_8446622_RNA45SN1_45S_50ntFlanks | 11321 | 11343 | 23 | rRF 45S | NO | 133.75 | 68.24 | 0.00 | 44.81 |
| >hg38_21_+_8433172_8446622_RNA45SN1_45S_50ntFlanks@11321.11344.24\|CACGTCTCGTCGCGCGCGCGTCCG | >hg38_21_+_8433172_8446622_RNA45SN1_45S_50ntFlanks | 11321 | 11344 | 24 | rRF 45S | NO | 693.80 | 619.00 | 37.25 | 333.29 |
| >hg38_21_+_8433172_8446622_RNA45SN1_45S_50ntFlanks@11322.11340.19\|ACGTCTCGTCGCGCGCGCG | >hg38_21_+_8433172_8446622_RNA45SN1_45S_50ntFlanks | 11322 | 11340 | 19 | rRF 45S | NO | 35.82 | 39.94 | 0.00 | 13.49 |
| >hg38_21_+_8433172_8446622_RNA45SN1_45S_50ntFlanks@11322.11342.21\|ACGTCTCGTCGCGCGCGCGTC | >hg38_21_+_8433172_8446622_RNA45SN1_45S_50ntFlanks | 11322 | 11342 | 21 | rRF 45S | NO | 166.95 | 200.02 | 20.42 | 63.30 |
| >hg38_21_+_8433172_8446622_RNA45SN1_45S_50ntFlanks@11322.11343.22\|ACGTCTCGTCGCGCGCGCGTCC | >hg38_21_+_8433172_8446622_RNA45SN1_45S_50ntFlanks | 11322 | 11343 | 22 | rRF 45S | NO | 212.21 | 109.73 | 0.00 | 55.51 |
| >hg38_21_+_8433172_8446622_RNA45SN1_45S_50ntFlanks@11322.11344.23\|ACGTCTCGTCGCGCGCGCGTCCG | >hg38_21_+_8433172_8446622_RNA45SN1_45S_50ntFlanks | 11322 | 11344 | 23 | rRF 45S | NO | 823.59 | 740.29 | 65.40 | 331.26 |
| >hg38_21_+_8433172_8446622_RNA45SN1_45S_50ntFlanks@11323.11340.18\|CGTCTCGTCGCGCGCGCG | >hg38_21_+_8433172_8446622_RNA45SN1_45S_50ntFlanks | 11323 | 11340 | 18 | rRF 45S | NO | 68.37 | 87.73 | 7.15 | 56.08 |
| >hg38_21_+_8433172_8446622_RNA45SN1_45S_50ntFlanks@11323.11342.20\|CGTCTCGTCGCGCGCGCGTC | >hg38_21_+_8433172_8446622_RNA45SN1_45S_50ntFlanks | 11323 | 11342 | 20 | rRF 45S | NO | 524.39 | 871.59 | 106.53 | 231.71 |
| >hg38_21_+_8433172_8446622_RNA45SN1_45S_50ntFlanks@11323.11343.21\|CGTCTCGTCGCGCGCGCGTCC | >hg38_21_+_8433172_8446622_RNA45SN1_45S_50ntFlanks | 11323 | 11343 | 21 | rRF 45S | NO | 840.19 | 550.18 | 13.71 | 268.05 |
| >hg38_21_+_8433172_8446622_RNA45SN1_45S_50ntFlanks@11323.11344.22\|CGTCTCGTCGCGCGCGCGTCCG | >hg38_21_+_8433172_8446622_RNA45SN1_45S_50ntFlanks | 11323 | 11344 | 22 | rRF 45S | YES | 1089.34 | 1403.31 | 210.15 | 578.20 |
| >hg38_21_+_8433172_8446622_RNA45SN1_45S_50ntFlanks@11324.11344.21\|GTCTCGTCGCGCGCGCGTCCG | >hg38_21_+_8433172_8446622_RNA45SN1_45S_50ntFlanks | 11324 | 11344 | 21 | rRF 45S | NO | 21.12 | 30.93 | 0.00 | 0.00 |
| >hg38_21_+_8433172_8446622_RNA45SN1_45S_50ntFlanks@11325.11344.20\|TCTCGTCGCGCGCGCGTCCG | >hg38_21_+_8433172_8446622_RNA45SN1_45S_50ntFlanks | 11325 | 11344 | 20 | rRF 45S | NO | 56.16 | 73.49 | 0.00 | 26.25 |
| >hg38_21_+_8433172_8446622_RNA45SN1_45S_50ntFlanks@11452.11471.20\|CTCCTCGGCGCGCGGCGGCG | >hg38_21_+_8433172_8446622_RNA45SN1_45S_50ntFlanks | 11452 | 11471 | 20 | rRF 45S | NO | 34.02 | 30.62 | 0.00 | 0.00 |
| >hg38_21_+_8433172_8446622_RNA45SN1_45S_50ntFlanks@11452.11472.21\|CTCCTCGGCGCGCGGCGGCGG | >hg38_21_+_8433172_8446622_RNA45SN1_45S_50ntFlanks | 11452 | 11472 | 21 | rRF 45S | NO | 55.91 | 60.96 | 0.00 | 0.00 |
| >hg38_21_+_8433172_8446622_RNA45SN1_45S_50ntFlanks@11452.11478.27\|CTCCTCGGCGCGCGGCGGCGGCGGCGG | >hg38_21_+_8433172_8446622_RNA45SN1_45S_50ntFlanks | 11452 | 11478 | 27 | rRF 45S | NO | 61.56 | 91.23 | 32.93 | 0.00 |
| >hg38_21_+_8433172_8446622_RNA45SN1_45S_50ntFlanks@11453.11477.25\|TCCTCGGCGCGCGGCGGCGGCGGCG | >hg38_21_+_8433172_8446622_RNA45SN1_45S_50ntFlanks | 11453 | 11477 | 25 | rRF 45S | NO | 0.00 | 26.04 | 0.00 | 0.00 |
| >hg38_21_+_8433172_8446622_RNA45SN1_45S_50ntFlanks@11515.11539.25\|GTCCGCCCCCGGGGCCGCGGTTCCG | >hg38_21_+_8433172_8446622_RNA45SN1_45S_50ntFlanks | 11515 | 11539 | 25 | rRF 45S | NO | 30.40 | 0.00 | 0.00 | 0.00 |
| >hg38_21_+_8433172_8446622_RNA45SN1_45S_50ntFlanks@11517.11539.23\|CCGCCCCCGGGGCCGCGGTTCCG | >hg38_21_+_8433172_8446622_RNA45SN1_45S_50ntFlanks | 11517 | 11539 | 23 | rRF 45S | NO | 39.20 | 26.71 | 0.00 | 0.00 |
| >hg38_21_+_8433172_8446622_RNA45SN1_45S_50ntFlanks@11534.11551.18\|GTTCCGCGCGGCGCCTCG | >hg38_21_+_8433172_8446622_RNA45SN1_45S_50ntFlanks | 11534 | 11551 | 18 | rRF 45S | NO | 230.00 | 121.89 | 0.00 | 68.45 |
| >hg38_21_+_8433172_8446622_RNA45SN1_45S_50ntFlanks@11534.11552.19\|GTTCCGCGCGGCGCCTCGC | >hg38_21_+_8433172_8446622_RNA45SN1_45S_50ntFlanks | 11534 | 11552 | 19 | rRF 45S | NO | 97.45 | 60.38 | 0.00 | 41.66 |
| >hg38_21_+_8433172_8446622_RNA45SN1_45S_50ntFlanks@11534.11553.20\|GTTCCGCGCGGCGCCTCGCC | >hg38_21_+_8433172_8446622_RNA45SN1_45S_50ntFlanks | 11534 | 11553 | 20 | rRF 45S | NO | 54.34 | 104.50 | 0.00 | 15.80 |
| >hg38_21_+_8433172_8446622_RNA45SN1_45S_50ntFlanks@11534.11555.22\|GTTCCGCGCGGCGCCTCGCCTC | >hg38_21_+_8433172_8446622_RNA45SN1_45S_50ntFlanks | 11534 | 11555 | 22 | rRF 45S | NO | 5.97 | 38.54 | 0.00 | 0.00 |

### MDA-MB-468

| | | | | | | | | | | |
|---|---|---|---|---|---|---|---|---|---|---|
| >hg38_21_+_8433172_8446622_RNA45SN1_45S_50ntFlanks@11534.11557.24\|GTTCCGCGCGGCGCCTCGCCTCGG | >hg38_21_+_8433172_8446622_RNA45SN1_45S_50ntFlanks | 11534 | 11557 | 24 | rRF 45S | NO | 0.00 | 26.37 | 0.00 | 0.00 |
| >hg38_21_+_8433172_8446622_RNA45SN1_45S_50ntFlanks@11534.11560.27\|GTTCCGCGCGGCGCCTCGCCTCGGCCG | >hg38_21_+_8433172_8446622_RNA45SN1_45S_50ntFlanks | 11534 | 11560 | 27 | rRF 45S | NO | 36.44 | 58.05 | 0.00 | 39.90 |
| >hg38_21_+_8433172_8446622_RNA45SN1_45S_50ntFlanks@11535.11552.18\|TTCCGCGCGGCGCCTCGC | >hg38_21_+_8433172_8446622_RNA45SN1_45S_50ntFlanks | 11535 | 11552 | 18 | rRF 45S | NO | 35.13 | 0.00 | 0.00 | 0.00 |
| >hg38_21_+_8433172_8446622_RNA45SN1_45S_50ntFlanks@11537.11560.24\|CCGCGCGGCGCCTCGCCTCGGCCG | >hg38_21_+_8433172_8446622_RNA45SN1_45S_50ntFlanks | 11537 | 11560 | 24 | rRF 45S | NO | 0.00 | 12.38 | 0.00 | 0.00 |
| >hg38_21_+_8433172_8446622_RNA45SN1_45S_50ntFlanks@11537.11580.44\|CCGCGCGGCGCCTCGCCTCGGCCGGCGCCTAGCAGCCGACTTAG | >hg38_21_+_8433172_8446622_RNA45SN1_45S_50ntFlanks | 11537 | 11580 | 44 | rRF 45S | NO | 33.97 | 0.00 | 0.00 | 0.00 |
| >hg38_21_+_8433172_8446622_RNA45SN1_45S_50ntFlanks@11540.11580.41\|CGCGGCGCCTCGCCTCGGCCGGCGCCTAGCAGCCGACTTAG | >hg38_21_+_8433172_8446622_RNA45SN1_45S_50ntFlanks | 11540 | 11580 | 41 | rRF 45S | NO | 57.48 | 0.00 | 0.00 | 0.00 |
| >hg38_21_+_8433172_8446622_RNA45SN1_45S_50ntFlanks@11552.11569.18\|CCTCGGCCGGCGCCTAGC | >hg38_21_+_8433172_8446622_RNA45SN1_45S_50ntFlanks | 11552 | 11569 | 18 | rRF 45S | NO | 46.22 | 31.76 | 0.00 | 41.00 |
| >hg38_21_+_8433172_8446622_RNA45SN1_45S_50ntFlanks@11552.11591.40\|CCTCGGCCGGCGCCTAGCAGCCGACTTAGAACTGGTGCGG | >hg38_21_+_8433172_8446622_RNA45SN1_45S_50ntFlanks | 11552 | 11591 | 40 | rRF 45S | NO | 46.59 | 36.10 | 0.00 | 0.00 |
| >hg38_21_+_8433172_8446622_RNA45SN1_45S_50ntFlanks@11552.11592.41\|CCTCGGCCGGCGCCTAGCAGCCGACTTAGAACTGGTGCGGA | >hg38_21_+_8433172_8446622_RNA45SN1_45S_50ntFlanks | 11552 | 11592 | 41 | rRF 45S | NO | 28.81 | 26.81 | 0.00 | 0.00 |
| >hg38_21_+_8433172_8446622_RNA45SN1_45S_50ntFlanks@11561.11585.25\|GCGCCTAGCAGCCGACTTAGAACTG | >hg38_21_+_8433172_8446622_RNA45SN1_45S_50ntFlanks | 11561 | 11585 | 25 | rRF 45S | NO | 111.61 | 112.95 | 66.02 | 113.57 |
| >hg38_21_+_8433172_8446622_RNA45SN1_45S_50ntFlanks@11561.11586.26\|GCGCCTAGCAGCCGACTTAGAACTGG | >hg38_21_+_8433172_8446622_RNA45SN1_45S_50ntFlanks | 11561 | 11586 | 26 | rRF 45S | NO | 51.35 | 31.94 | 49.01 | 67.50 |
| >hg38_21_+_8433172_8446622_RNA45SN1_45S_50ntFlanks@11561.11588.28\|GCGCCTAGCAGCCGACTTAGAACTGGTG | >hg38_21_+_8433172_8446622_RNA45SN1_45S_50ntFlanks | 11561 | 11588 | 28 | rRF 45S | NO | 20.89 | 99.96 | 41.46 | 26.72 |
| >hg38_21_+_8433172_8446622_RNA45SN1_45S_50ntFlanks@11561.11590.30\|GCGCCTAGCAGCCGACTTAGAACTGGTGCG | >hg38_21_+_8433172_8446622_RNA45SN1_45S_50ntFlanks | 11561 | 11590 | 30 | rRF 45S | NO | 0.00 | 19.37 | 0.00 | 0.00 |
| >hg38_21_+_8433172_8446622_RNA45SN1_45S_50ntFlanks@11561.11591.31\|GCGCCTAGCAGCCGACTTAGAACTGGTGCGG | >hg38_21_+_8433172_8446622_RNA45SN1_45S_50ntFlanks | 11561 | 11591 | 31 | rRF 45S | NO | 0.00 | 82.19 | 44.92 | 64.77 |
| >hg38_21_+_8433172_8446622_RNA45SN1_45S_50ntFlanks@11561.11592.32\|GCGCCTAGCAGCCGACTTAGAACTGGTGCGGA | >hg38_21_+_8433172_8446622_RNA45SN1_45S_50ntFlanks | 11561 | 11592 | 32 | rRF 45S | NO | 31.19 | 116.09 | 66.03 | 82.30 |
| >hg38_21_+_8433172_8446622_RNA45SN1_45S_50ntFlanks@11561.11593.33\|GCGCCTAGCAGCCGACTTAGAACTGGTGCGGAC | >hg38_21_+_8433172_8446622_RNA45SN1_45S_50ntFlanks | 11561 | 11593 | 33 | rRF 45S | NO | 0.00 | 43.30 | 0.00 | 25.20 |
| >hg38_21_+_8433172_8446622_RNA45SN1_45S_50ntFlanks@11561.11594.34\|GCGCCTAGCAGCCGACTTAGAACTGGTGCGGACC | >hg38_21_+_8433172_8446622_RNA45SN1_45S_50ntFlanks | 11561 | 11594 | 34 | rRF 45S | NO | 0.00 | 41.03 | 0.00 | 15.89 |
| >hg38_21_+_8433172_8446622_RNA45SN1_45S_50ntFlanks@11561.11600.40\|GCGCCTAGCAGCCGACTTAGAACTGGTGCGGACCAGGGGA | >hg38_21_+_8433172_8446622_RNA45SN1_45S_50ntFlanks | 11561 | 11600 | 40 | rRF 45S | NO | 112.85 | 32.78 | 0.00 | 45.44 |
| >hg38_21_+_8433172_8446622_RNA45SN1_45S_50ntFlanks@11561.11601.41\|GCGCCTAGCAGCCGACTTAGAACTGGTGCGGACCAGGGGAA | >hg38_21_+_8433172_8446622_RNA45SN1_45S_50ntFlanks | 11561 | 11601 | 41 | rRF 45S | NO | 212.95 | 61.58 | 7.03 | 136.39 |
| >hg38_21_+_8433172_8446622_RNA45SN1_45S_50ntFlanks@11561.11602.42\|GCGCCTAGCAGCCGACTTAGAACTGGTGCGGACCAGGGGAAT | >hg38_21_+_8433172_8446622_RNA45SN1_45S_50ntFlanks | 11561 | 11602 | 42 | rRF 45S | NO | 136.07 | 39.52 | 0.00 | 100.20 |
| >hg38_21_+_8433172_8446622_RNA45SN1_45S_50ntFlanks@11561.11603.43\|GCGCCTAGCAGCCGACTTAGAACTGGTGCGGACCAGGGGAATC | >hg38_21_+_8433172_8446622_RNA45SN1_45S_50ntFlanks | 11561 | 11603 | 43 | rRF 45S | NO | 201.11 | 43.59 | 0.00 | 225.85 |
| >hg38_21_+_8433172_8446622_RNA45SN1_45S_50ntFlanks@11562.11588.27\|CGCCTAGCAGCCGACTTAGAACTGGTG | >hg38_21_+_8433172_8446622_RNA45SN1_45S_50ntFlanks | 11562 | 11588 | 27 | rRF 45S | NO | 29.45 | 34.46 | 28.14 | 0.00 |
| >hg38_21_+_8433172_8446622_RNA45SN1_45S_50ntFlanks@11562.11592.31\|CGCCTAGCAGCCGACTTAGAACTGGTGCGGA | >hg38_21_+_8433172_8446622_RNA45SN1_45S_50ntFlanks | 11562 | 11592 | 31 | rRF 45S | NO | 0.00 | 78.30 | 94.62 | 41.46 |
| >hg38_21_+_8433172_8446622_RNA45SN1_45S_50ntFlanks@11564.11581.18\|CCTAGCAGCCGACTTAGA | >hg38_21_+_8433172_8446622_RNA45SN1_45S_50ntFlanks | 11564 | 11581 | 18 | rRF 45S | NO | 83.38 | 81.12 | 0.00 | 77.36 |
| >hg38_21_+_8433172_8446622_RNA45SN1_45S_50ntFlanks@11564.11582.19\|CCTAGCAGCCGACTTAGAA | >hg38_21_+_8433172_8446622_RNA45SN1_45S_50ntFlanks | 11564 | 11582 | 19 | rRF 45S | NO | 183.47 | 90.36 | 0.00 | 99.82 |
| >hg38_21_+_8433172_8446622_RNA45SN1_45S_50ntFlanks@11564.11585.22\|CCTAGCAGCCGACTTAGAACTG | >hg38_21_+_8433172_8446622_RNA45SN1_45S_50ntFlanks | 11564 | 11585 | 22 | rRF 45S | NO | 174.95 | 178.45 | 66.60 | 87.18 |
| >hg38_21_+_8433172_8446622_RNA45SN1_45S_50ntFlanks@11564.11586.23\|CCTAGCAGCCGACTTAGAACTGG | >hg38_21_+_8433172_8446622_RNA45SN1_45S_50ntFlanks | 11564 | 11586 | 23 | rRF 45S | NO | 46.57 | 46.38 | 48.12 | 43.19 |
| >hg38_21_+_8433172_8446622_RNA45SN1_45S_50ntFlanks@11564.11588.25\|CCTAGCAGCCGACTTAGAACTGGTG | >hg38_21_+_8433172_8446622_RNA45SN1_45S_50ntFlanks | 11564 | 11588 | 25 | rRF 45S | NO | 60.72 | 98.33 | 62.46 | 42.22 |
| >hg38_21_+_8433172_8446622_RNA45SN1_45S_50ntFlanks@11564.11590.27\|CCTAGCAGCCGACTTAGAACTGGTGCG | >hg38_21_+_8433172_8446622_RNA45SN1_45S_50ntFlanks | 11564 | 11590 | 27 | rRF 45S | NO | 0.00 | 0.00 | 31.55 | 0.00 |
| >hg38_21_+_8433172_8446622_RNA45SN1_45S_50ntFlanks@11564.11591.28\|CCTAGCAGCCGACTTAGAACTGGTGCGG | >hg38_21_+_8433172_8446622_RNA45SN1_45S_50ntFlanks | 11564 | 11591 | 28 | rRF 45S | NO | 52.68 | 86.04 | 89.17 | 88.92 |
| >hg38_21_+_8433172_8446622_RNA45SN1_45S_50ntFlanks@11564.11592.29\|CCTAGCAGCCGACTTAGAACTGGTGCGGA | >hg38_21_+_8433172_8446622_RNA45SN1_45S_50ntFlanks | 11564 | 11592 | 29 | rRF 45S | NO | 0.00 | 37.83 | 57.03 | 6.68 |
| >hg38_21_+_8433172_8446622_RNA45SN1_45S_50ntFlanks@11564.11600.37\|CCTAGCAGCCGACTTAGAACTGGTGCGGACCAGGGGA | >hg38_21_+_8433172_8446622_RNA45SN1_45S_50ntFlanks | 11564 | 11600 | 37 | rRF 45S | NO | 100.30 | 58.59 | 44.30 | 57.07 |
| >hg38_21_+_8433172_8446622_RNA45SN1_45S_50ntFlanks@11564.11601.38\|CCTAGCAGCCGACTTAGAACTGGTGCGGACCAGGGGAA | >hg38_21_+_8433172_8446622_RNA45SN1_45S_50ntFlanks | 11564 | 11601 | 38 | rRF 45S | NO | 275.30 | 136.49 | 77.95 | 221.07 |
| >hg38_21_+_8433172_8446622_RNA45SN1_45S_50ntFlanks@11564.11602.39\|CCTAGCAGCCGACTTAGAACTGGTGCGGACCAGGGGAAT | >hg38_21_+_8433172_8446622_RNA45SN1_45S_50ntFlanks | 11564 | 11602 | 39 | rRF 45S | NO | 159.98 | 80.42 | 40.71 | 161.31 |

### MDA-MB-468

| | | | | | | | | | | |
|---|---|---|---|---|---|---|---|---|---|---|
| >hg38_21_+_8433172_8446622_RNA45SN1_45S_50ntFlanks@11564.11603.40\|CCTAGCAGCC GACTTAGAACTGGTGCGGACCAGGGGAATC | >hg38_21_+_8433172_8446622_RNA45SN1_45S_50ntFlanks | 11564 | 11603 | 40 | rRF 45S | NO | 189.43 | 72.45 | 44.56 | 335.80 |
| >hg38_21_+_8433172_8446622_RNA45SN1_45S_50ntFlanks@11564.11605.42\|CCTAGCAGCC GACTTAGAACTGGTGCGGACCAGGGGAATCCG | >hg38_21_+_8433172_8446622_RNA45SN1_45S_50ntFlanks | 11564 | 11605 | 42 | rRF 45S | NO | 33.90 | 0.00 | 0.00 | 55.74 |
| >hg38_21_+_8433172_8446622_RNA45SN1_45S_50ntFlanks@11564.11611.48\|CCTAGCAGCC GACTTAGAACTGGTGCGGACCAGGGGAATCCGACTGTT | >hg38_21_+_8433172_8446622_RNA45SN1_45S_50ntFlanks | 11564 | 11611 | 48 | rRF 45S | NO | 53.88 | 40.33 | 0.00 | 137.38 |
| >hg38_21_+_8433172_8446622_RNA45SN1_45S_50ntFlanks@11565.11601.37\|CTAGCAGCCG ACTTAGAACTGGTGCGGACCAGGGGAA | >hg38_21_+_8433172_8446622_RNA45SN1_45S_50ntFlanks | 11565 | 11601 | 37 | rRF 45S | NO | 33.64 | 0.00 | 0.00 | 0.00 |
| >hg38_21_+_8433172_8446622_RNA45SN1_45S_50ntFlanks@11567.11585.19\|AGCAGCCGA CTTAGAACTG | >hg38_21_+_8433172_8446622_RNA45SN1_45S_50ntFlanks | 11567 | 11585 | 19 | rRF 45S | NO | 52.91 | 56.20 | 38.49 | 14.67 |
| >hg38_21_+_8433172_8446622_RNA45SN1_45S_50ntFlanks@11567.11586.20\|AGCAGCCGA CTTAGAACTGG | >hg38_21_+_8433172_8446622_RNA45SN1_45S_50ntFlanks | 11567 | 11586 | 20 | rRF 45S | NO | 0.00 | 0.00 | 28.32 | 0.00 |
| >hg38_21_+_8433172_8446622_RNA45SN1_45S_50ntFlanks@11567.11588.22\|AGCAGCCGA CTTAGAACTGGTG | >hg38_21_+_8433172_8446622_RNA45SN1_45S_50ntFlanks | 11567 | 11588 | 22 | rRF 45S | NO | 0.00 | 32.02 | 31.65 | 0.00 |
| >hg38_21_+_8433172_8446622_RNA45SN1_45S_50ntFlanks@11567.11592.26\|AGCAGCCGA CTTAGAACTGGTGCGGA | >hg38_21_+_8433172_8446622_RNA45SN1_45S_50ntFlanks | 11567 | 11592 | 26 | rRF 45S | NO | 0.00 | 0.00 | 27.37 | 0.00 |
| >hg38_21_+_8433172_8446622_RNA45SN1_45S_50ntFlanks@11567.11600.34\|AGCAGCCGA CTTAGAACTGGTGCGGACCAGGGGA | >hg38_21_+_8433172_8446622_RNA45SN1_45S_50ntFlanks | 11567 | 11600 | 34 | rRF 45S | NO | 50.23 | 0.00 | 0.00 | 0.00 |
| >hg38_21_+_8433172_8446622_RNA45SN1_45S_50ntFlanks@11567.11601.35\|AGCAGCCGA CTTAGAACTGGTGCGGACCAGGGGAA | >hg38_21_+_8433172_8446622_RNA45SN1_45S_50ntFlanks | 11567 | 11601 | 35 | rRF 45S | NO | 94.64 | 0.00 | 0.00 | 6.46 |
| >hg38_21_+_8433172_8446622_RNA45SN1_45S_50ntFlanks@11567.11602.36\|AGCAGCCGA CTTAGAACTGGTGCGGACCAGGGGAAT | >hg38_21_+_8433172_8446622_RNA45SN1_45S_50ntFlanks | 11567 | 11602 | 36 | rRF 45S | NO | 43.32 | 0.00 | 0.00 | 0.00 |
| >hg38_21_+_8433172_8446622_RNA45SN1_45S_50ntFlanks@11567.11603.37\|AGCAGCCGA CTTAGAACTGGTGCGGACCAGGGGAATC | >hg38_21_+_8433172_8446622_RNA45SN1_45S_50ntFlanks | 11567 | 11603 | 37 | rRF 45S | NO | 64.60 | 0.00 | 0.00 | 49.30 |
| >hg38_21_+_8433172_8446622_RNA45SN1_45S_50ntFlanks@11568.11585.18\|GCAGCCGACT TAGAACTG | >hg38_21_+_8433172_8446622_RNA45SN1_45S_50ntFlanks | 11568 | 11585 | 18 | rRF 45S | NO | 90.75 | 111.28 | 68.01 | 39.57 |
| >hg38_21_+_8433172_8446622_RNA45SN1_45S_50ntFlanks@11568.11586.19\|GCAGCCGACT TAGAACTGG | >hg38_21_+_8433172_8446622_RNA45SN1_45S_50ntFlanks | 11568 | 11586 | 19 | rRF 45S | NO | 21.03 | 33.88 | 46.22 | 6.06 |
| >hg38_21_+_8433172_8446622_RNA45SN1_45S_50ntFlanks@11568.11588.21\|GCAGCCGACT TAGAACTGGTG | >hg38_21_+_8433172_8446622_RNA45SN1_45S_50ntFlanks | 11568 | 11588 | 21 | rRF 45S | NO | 0.00 | 46.43 | 44.26 | 0.00 |
| >hg38_21_+_8433172_8446622_RNA45SN1_45S_50ntFlanks@11568.11592.25\|GCAGCCGACT TAGAACTGGTGCGGA | >hg38_21_+_8433172_8446622_RNA45SN1_45S_50ntFlanks | 11568 | 11592 | 25 | rRF 45S | NO | 0.00 | 31.31 | 47.65 | 0.00 |
| >hg38_21_+_8433172_8446622_RNA45SN1_45S_50ntFlanks@11568.11600.33\|GCAGCCGACT TAGAACTGGTGCGGACCAGGGGA | >hg38_21_+_8433172_8446622_RNA45SN1_45S_50ntFlanks | 11568 | 11600 | 33 | rRF 45S | NO | 69.05 | 0.00 | 6.87 | 0.00 |
| >hg38_21_+_8433172_8446622_RNA45SN1_45S_50ntFlanks@11568.11601.34\|GCAGCCGACT TAGAACTGGTGCGGACCAGGGGAA | >hg38_21_+_8433172_8446622_RNA45SN1_45S_50ntFlanks | 11568 | 11601 | 34 | rRF 45S | NO | 138.91 | 35.76 | 41.66 | 24.51 |
| >hg38_21_+_8433172_8446622_RNA45SN1_45S_50ntFlanks@11568.11602.35\|GCAGCCGACT TAGAACTGGTGCGGACCAGGGGAAT | >hg38_21_+_8433172_8446622_RNA45SN1_45S_50ntFlanks | 11568 | 11602 | 35 | rRF 45S | NO | 69.93 | 0.00 | 0.00 | 14.29 |
| >hg38_21_+_8433172_8446622_RNA45SN1_45S_50ntFlanks@11568.11603.36\|GCAGCCGACT TAGAACTGGTGCGGACCAGGGGAATC | >hg38_21_+_8433172_8446622_RNA45SN1_45S_50ntFlanks | 11568 | 11603 | 36 | rRF 45S | NO | 116.54 | 35.27 | 35.03 | 78.70 |
| >hg38_21_+_8433172_8446622_RNA45SN1_45S_50ntFlanks@11568.11605.38\|GCAGCCGACT TAGAACTGGTGCGGACCAGGGGAATCCG | >hg38_21_+_8433172_8446622_RNA45SN1_45S_50ntFlanks | 11568 | 11605 | 38 | rRF 45S | NO | 33.13 | 0.00 | 0.00 | 15.71 |
| >hg38_21_+_8433172_8446622_RNA45SN1_45S_50ntFlanks@11569.11586.18\|CAGCCGACTT AGAACTGG | >hg38_21_+_8433172_8446622_RNA45SN1_45S_50ntFlanks | 11569 | 11586 | 18 | rRF 45S | NO | 13.07 | 97.55 | 72.96 | 33.42 |
| >hg38_21_+_8433172_8446622_RNA45SN1_45S_50ntFlanks@11569.11590.22\|CAGCCGACTT AGAACTGGTGCG | >hg38_21_+_8433172_8446622_RNA45SN1_45S_50ntFlanks | 11569 | 11590 | 22 | rRF 45S | NO | 0.00 | 0.00 | 30.63 | 0.00 |
| >hg38_21_+_8433172_8446622_RNA45SN1_45S_50ntFlanks@11569.11591.23\|CAGCCGACTT AGAACTGGTGCGG | >hg38_21_+_8433172_8446622_RNA45SN1_45S_50ntFlanks | 11569 | 11591 | 23 | rRF 45S | NO | 0.00 | 0.00 | 33.26 | 0.00 |
| >hg38_21_+_8433172_8446622_RNA45SN1_45S_50ntFlanks@11569.11592.24\|CAGCCGACTT AGAACTGGTGCGGA | >hg38_21_+_8433172_8446622_RNA45SN1_45S_50ntFlanks | 11569 | 11592 | 24 | rRF 45S | NO | 0.00 | 33.57 | 94.18 | 0.00 |
| >hg38_21_+_8433172_8446622_RNA45SN1_45S_50ntFlanks@11569.11593.25\|CAGCCGACTT AGAACTGGTGCGGAC | >hg38_21_+_8433172_8446622_RNA45SN1_45S_50ntFlanks | 11569 | 11593 | 25 | rRF 45S | NO | 0.00 | 0.00 | 28.44 | 0.00 |
| >hg38_21_+_8433172_8446622_RNA45SN1_45S_50ntFlanks@11569.11594.26\|CAGCCGACTT AGAACTGGTGCGGACC | >hg38_21_+_8433172_8446622_RNA45SN1_45S_50ntFlanks | 11569 | 11594 | 26 | rRF 45S | NO | 0.00 | 37.09 | 32.33 | 0.00 |
| >hg38_21_+_8433172_8446622_RNA45SN1_45S_50ntFlanks@11569.11597.29\|CAGCCGACTT AGAACTGGTGCGGACCAGG | >hg38_21_+_8433172_8446622_RNA45SN1_45S_50ntFlanks | 11569 | 11597 | 29 | rRF 45S | NO | 0.00 | 27.97 | 0.00 | 0.00 |
| >hg38_21_+_8433172_8446622_RNA45SN1_45S_50ntFlanks@11569.11599.31\|CAGCCGACTT AGAACTGGTGCGGACCAGGGG | >hg38_21_+_8433172_8446622_RNA45SN1_45S_50ntFlanks | 11569 | 11599 | 31 | rRF 45S | NO | 46.12 | 40.26 | 0.00 | 0.00 |
| >hg38_21_+_8433172_8446622_RNA45SN1_45S_50ntFlanks@11569.11600.32\|CAGCCGACTT AGAACTGGTGCGGACCAGGGGA | >hg38_21_+_8433172_8446622_RNA45SN1_45S_50ntFlanks | 11569 | 11600 | 32 | rRF 45S | NO | 429.72 | 143.06 | 96.62 | 50.81 |
| >hg38_21_+_8433172_8446622_RNA45SN1_45S_50ntFlanks@11569.11601.33\|CAGCCGACTT AGAACTGGTGCGGACCAGGGGAA | >hg38_21_+_8433172_8446622_RNA45SN1_45S_50ntFlanks | 11569 | 11601 | 33 | rRF 45S | NO | 944.27 | 266.48 | 182.14 | 148.53 |
| >hg38_21_+_8433172_8446622_RNA45SN1_45S_50ntFlanks@11569.11602.34\|CAGCCGACTT AGAACTGGTGCGGACCAGGGGAAT | >hg38_21_+_8433172_8446622_RNA45SN1_45S_50ntFlanks | 11569 | 11602 | 34 | rRF 45S | NO | 512.42 | 155.54 | 98.32 | 129.43 |
| >hg38_21_+_8433172_8446622_RNA45SN1_45S_50ntFlanks@11569.11603.35\|CAGCCGACTT AGAACTGGTGCGGACCAGGGGAATC | >hg38_21_+_8433172_8446622_RNA45SN1_45S_50ntFlanks | 11569 | 11603 | 35 | rRF 45S | NO | 588.81 | 190.88 | 128.44 | 344.91 |
| >hg38_21_+_8433172_8446622_RNA45SN1_45S_50ntFlanks@11569.11605.37\|CAGCCGACTT AGAACTGGTGCGGACCAGGGGAATCCG | >hg38_21_+_8433172_8446622_RNA45SN1_45S_50ntFlanks | 11569 | 11605 | 37 | rRF 45S | NO | 97.48 | 38.12 | 28.56 | 84.63 |

### MDA-MB-468

| | | | | | | | | | | |
|---|---|---|---|---|---|---|---|---|---|---|
| >hg38_21_+_8433172_8446622_RNA45SN1_45S_50ntFlanks@11569.11609.41\|CAGCCGACTTAGAACTGGTGCGGACCAGGGGAATCCGACTG | >hg38_21_+_8433172_8446622_RNA45SN1_45S_50ntFlanks | 11569 | 11609 | 41 | rRF 45S | NO | 0.00 | 19.48 | 0.00 | 53.56 |
| >hg38_21_+_8433172_8446622_RNA45SN1_45S_50ntFlanks@11569.11611.43\|CAGCCGACTTAGAACTGGTGCGGACCAGGGGAATCCGACTGTT | >hg38_21_+_8433172_8446622_RNA45SN1_45S_50ntFlanks | 11569 | 11611 | 43 | rRF 45S | NO | 6.06 | 26.73 | 0.00 | 84.23 |
| >hg38_21_+_8433172_8446622_RNA45SN1_45S_50ntFlanks@11569.11612.44\|CAGCCGACTTAGAACTGGTGCGGACCAGGGGAATCCGACTGTTT | >hg38_21_+_8433172_8446622_RNA45SN1_45S_50ntFlanks | 11569 | 11612 | 44 | rRF 45S | NO | 84.07 | 49.72 | 0.00 | 142.85 |
| >hg38_21_+_8433172_8446622_RNA45SN1_45S_50ntFlanks@11570.11591.22\|AGCCGACTTAGAACTGGTGCGG | >hg38_21_+_8433172_8446622_RNA45SN1_45S_50ntFlanks | 11570 | 11591 | 22 | rRF 45S | NO | 0.00 | 33.75 | 43.30 | 0.00 |
| >hg38_21_+_8433172_8446622_RNA45SN1_45S_50ntFlanks@11570.11592.23\|AGCCGACTTAGAACTGGTGCGGA | >hg38_21_+_8433172_8446622_RNA45SN1_45S_50ntFlanks | 11570 | 11592 | 23 | rRF 45S | NO | 0.00 | 0.00 | 46.23 | 0.00 |
| >hg38_21_+_8433172_8446622_RNA45SN1_45S_50ntFlanks@11570.11600.31\|AGCCGACTTAGAACTGGTGCGGACCAGGGGA | >hg38_21_+_8433172_8446622_RNA45SN1_45S_50ntFlanks | 11570 | 11600 | 31 | rRF 45S | NO | 82.62 | 19.70 | 6.95 | 0.00 |
| >hg38_21_+_8433172_8446622_RNA45SN1_45S_50ntFlanks@11570.11601.32\|AGCCGACTTAGAACTGGTGCGGACCAGGGGAA | >hg38_21_+_8433172_8446622_RNA45SN1_45S_50ntFlanks | 11570 | 11601 | 32 | rRF 45S | NO | 210.80 | 58.72 | 56.82 | 44.63 |
| >hg38_21_+_8433172_8446622_RNA45SN1_45S_50ntFlanks@11570.11602.33\|AGCCGACTTAGAACTGGTGCGGACCAGGGGAAT | >hg38_21_+_8433172_8446622_RNA45SN1_45S_50ntFlanks | 11570 | 11602 | 33 | rRF 45S | NO | 107.29 | 32.61 | 31.41 | 36.12 |
| >hg38_21_+_8433172_8446622_RNA45SN1_45S_50ntFlanks@11570.11603.34\|AGCCGACTTAGAACTGGTGCGGACCAGGGGAATC | >hg38_21_+_8433172_8446622_RNA45SN1_45S_50ntFlanks | 11570 | 11603 | 34 | rRF 45S | NO | 140.42 | 38.53 | 46.63 | 104.83 |
| >hg38_21_+_8433172_8446622_RNA45SN1_45S_50ntFlanks@11570.11605.36\|AGCCGACTTAGAACTGGTGCGGACCAGGGGAATCCG | >hg38_21_+_8433172_8446622_RNA45SN1_45S_50ntFlanks | 11570 | 11605 | 36 | rRF 45S | NO | 19.66 | 0.00 | 0.00 | 13.96 |
| >hg38_21_+_8433172_8446622_RNA45SN1_45S_50ntFlanks@11571.11592.22\|GCCGACTTAGAACTGGTGCGGA | >hg38_21_+_8433172_8446622_RNA45SN1_45S_50ntFlanks | 11571 | 11592 | 22 | rRF 45S | NO | 0.00 | 0.00 | 37.16 | 0.00 |
| >hg38_21_+_8433172_8446622_RNA45SN1_45S_50ntFlanks@11571.11600.30\|GCCGACTTAGAACTGGTGCGGACCAGGGGA | >hg38_21_+_8433172_8446622_RNA45SN1_45S_50ntFlanks | 11571 | 11600 | 30 | rRF 45S | NO | 46.42 | 0.00 | 0.00 | 0.00 |
| >hg38_21_+_8433172_8446622_RNA45SN1_45S_50ntFlanks@11571.11601.31\|GCCGACTTAGAACTGGTGCGGACCAGGGGAA | >hg38_21_+_8433172_8446622_RNA45SN1_45S_50ntFlanks | 11571 | 11601 | 31 | rRF 45S | NO | 86.25 | 6.04 | 29.49 | 0.00 |
| >hg38_21_+_8433172_8446622_RNA45SN1_45S_50ntFlanks@11571.11602.32\|GCCGACTTAGAACTGGTGCGGACCAGGGGAAT | >hg38_21_+_8433172_8446622_RNA45SN1_45S_50ntFlanks | 11571 | 11602 | 32 | rRF 45S | NO | 43.06 | 0.00 | 0.00 | 0.00 |
| >hg38_21_+_8433172_8446622_RNA45SN1_45S_50ntFlanks@11571.11603.33\|GCCGACTTAGAACTGGTGCGGACCAGGGGAATC | >hg38_21_+_8433172_8446622_RNA45SN1_45S_50ntFlanks | 11571 | 11603 | 33 | rRF 45S | NO | 51.78 | 0.00 | 6.67 | 37.96 |
| >hg38_21_+_8433172_8446622_RNA45SN1_45S_50ntFlanks@11572.11589.18\|CCGACTTAGAACTGGTGC | >hg38_21_+_8433172_8446622_RNA45SN1_45S_50ntFlanks | 11572 | 11589 | 18 | rRF 45S | NO | 0.00 | 29.41 | 13.64 | 0.00 |
| >hg38_21_+_8433172_8446622_RNA45SN1_45S_50ntFlanks@11572.11590.19\|CCGACTTAGAACTGGTGCG | >hg38_21_+_8433172_8446622_RNA45SN1_45S_50ntFlanks | 11572 | 11590 | 19 | rRF 45S | NO | 177.40 | 215.16 | 119.24 | 16.57 |
| >hg38_21_+_8433172_8446622_RNA45SN1_45S_50ntFlanks@11572.11591.20\|CCGACTTAGAACTGGTGCGG | >hg38_21_+_8433172_8446622_RNA45SN1_45S_50ntFlanks | 11572 | 11591 | 20 | rRF 45S | YES | 115.54 | 382.76 | 334.64 | 90.79 |
| >hg38_21_+_8433172_8446622_RNA45SN1_45S_50ntFlanks@11572.11592.21\|CCGACTTAGAACTGGTGCGGA | >hg38_21_+_8433172_8446622_RNA45SN1_45S_50ntFlanks | 11572 | 11592 | 21 | rRF 45S | NO | 0.00 | 71.22 | 149.22 | 0.00 |
| >hg38_21_+_8433172_8446622_RNA45SN1_45S_50ntFlanks@11572.11593.22\|CCGACTTAGAACTGGTGCGGAC | >hg38_21_+_8433172_8446622_RNA45SN1_45S_50ntFlanks | 11572 | 11593 | 22 | rRF 45S | NO | 0.00 | 57.43 | 54.86 | 14.33 |
| >hg38_21_+_8433172_8446622_RNA45SN1_45S_50ntFlanks@11572.11594.23\|CCGACTTAGAACTGGTGCGGACC | >hg38_21_+_8433172_8446622_RNA45SN1_45S_50ntFlanks | 11572 | 11594 | 23 | rRF 45S | NO | 0.00 | 44.95 | 6.65 | 0.00 |
| >hg38_21_+_8433172_8446622_RNA45SN1_45S_50ntFlanks@11572.11597.26\|CCGACTTAGAACTGGTGCGGACCAGG | >hg38_21_+_8433172_8446622_RNA45SN1_45S_50ntFlanks | 11572 | 11597 | 26 | rRF 45S | NO | 0.00 | 48.61 | 0.00 | 0.00 |
| >hg38_21_+_8433172_8446622_RNA45SN1_45S_50ntFlanks@11572.11599.28\|CCGACTTAGAACTGGTGCGGACCAGGGG | >hg38_21_+_8433172_8446622_RNA45SN1_45S_50ntFlanks | 11572 | 11599 | 28 | rRF 45S | NO | 21.19 | 69.44 | 28.14 | 0.00 |
| >hg38_21_+_8433172_8446622_RNA45SN1_45S_50ntFlanks@11572.11600.29\|CCGACTTAGAACTGGTGCGGACCAGGGGA | >hg38_21_+_8433172_8446622_RNA45SN1_45S_50ntFlanks | 11572 | 11600 | 29 | rRF 45S | NO | 401.09 | 366.67 | 204.67 | 60.81 |
| >hg38_21_+_8433172_8446622_RNA45SN1_45S_50ntFlanks@11572.11601.30\|CCGACTTAGAACTGGTGCGGACCAGGGGAA | >hg38_21_+_8433172_8446622_RNA45SN1_45S_50ntFlanks | 11572 | 11601 | 30 | rRF 45S | YES | 1103.26 | 914.15 | 403.96 | 176.33 |
| >hg38_21_+_8433172_8446622_RNA45SN1_45S_50ntFlanks@11572.11602.31\|CCGACTTAGAACTGGTGCGGACCAGGGGAAT | >hg38_21_+_8433172_8446622_RNA45SN1_45S_50ntFlanks | 11572 | 11602 | 31 | rRF 45S | NO | 546.65 | 471.03 | 210.05 | 134.62 |
| >hg38_21_+_8433172_8446622_RNA45SN1_45S_50ntFlanks@11572.11603.32\|CCGACTTAGAACTGGTGCGGACCAGGGGAATC | >hg38_21_+_8433172_8446622_RNA45SN1_45S_50ntFlanks | 11572 | 11603 | 32 | rRF 45S | NO | 670.27 | 485.06 | 266.25 | 370.23 |
| >hg38_21_+_8433172_8446622_RNA45SN1_45S_50ntFlanks@11572.11604.33\|CCGACTTAGAACTGGTGCGGACCAGGGGAATCC | >hg38_21_+_8433172_8446622_RNA45SN1_45S_50ntFlanks | 11572 | 11604 | 33 | rRF 45S | NO | 21.33 | 0.00 | 0.00 | 0.00 |
| >hg38_21_+_8433172_8446622_RNA45SN1_45S_50ntFlanks@11572.11605.34\|CCGACTTAGAACTGGTGCGGACCAGGGGAATCCG | >hg38_21_+_8433172_8446622_RNA45SN1_45S_50ntFlanks | 11572 | 11605 | 34 | rRF 45S | NO | 343.99 | 239.24 | 72.68 | 201.85 |
| >hg38_21_+_8433172_8446622_RNA45SN1_45S_50ntFlanks@11572.11606.35\|CCGACTTAGAACTGGTGCGGACCAGGGGAATCCGA | >hg38_21_+_8433172_8446622_RNA45SN1_45S_50ntFlanks | 11572 | 11606 | 35 | rRF 45S | NO | 31.40 | 26.94 | 0.00 | 15.82 |
| >hg38_21_+_8433172_8446622_RNA45SN1_45S_50ntFlanks@11572.11609.38\|CCGACTTAGAACTGGTGCGGACCAGGGGAATCCGACTG | >hg38_21_+_8433172_8446622_RNA45SN1_45S_50ntFlanks | 11572 | 11609 | 38 | rRF 45S | NO | 34.23 | 28.32 | 0.00 | 47.68 |
| >hg38_21_+_8433172_8446622_RNA45SN1_45S_50ntFlanks@11572.11611.40\|CCGACTTAGAACTGGTGCGGACCAGGGGAATCCGACTGTT | >hg38_21_+_8433172_8446622_RNA45SN1_45S_50ntFlanks | 11572 | 11611 | 40 | rRF 45S | NO | 75.74 | 46.53 | 13.70 | 144.20 |
| >hg38_21_+_8433172_8446622_RNA45SN1_45S_50ntFlanks@11572.11612.41\|CCGACTTAGAACTGGTGCGGACCAGGGGAATCCGACTGTTT | >hg38_21_+_8433172_8446622_RNA45SN1_45S_50ntFlanks | 11572 | 11612 | 41 | rRF 45S | NO | 170.50 | 84.64 | 27.25 | 194.45 |
| >hg38_21_+_8433172_8446622_RNA45SN1_45S_50ntFlanks@11573.11591.19\|CGACTTAGAACTGGTGCGG | >hg38_21_+_8433172_8446622_RNA45SN1_45S_50ntFlanks | 11573 | 11591 | 19 | rRF 45S | NO | 0.00 | 31.34 | 28.39 | 0.00 |

### MDA-MB-468

| | | | | | | | | | | |
|---|---|---|---|---|---|---|---|---|---|---|
| >hg38_21_+_8433172_8446622_RNA45SN1_45S_50ntFlanks@11573.11592.20\|CGACTTAGAACTGGTGCGGA | >hg38_21_+_8433172_8446622_RNA45SN1_45S_50ntFlanks | 11573 | 11592 | 20 | rRF 45S | NO | 28.97 | 39.48 | 43.25 | 0.00 |
| >hg38_21_+_8433172_8446622_RNA45SN1_45S_50ntFlanks@11573.11593.21\|CGACTTAGAACTGGTGCGGAC | >hg38_21_+_8433172_8446622_RNA45SN1_45S_50ntFlanks | 11573 | 11593 | 21 | rRF 45S | NO | 21.91 | 28.11 | 0.00 | 0.00 |
| >hg38_21_+_8433172_8446622_RNA45SN1_45S_50ntFlanks@11573.11600.28\|CGACTTAGAACTGGTGCGGACCAGGGGA | >hg38_21_+_8433172_8446622_RNA45SN1_45S_50ntFlanks | 11573 | 11600 | 28 | rRF 45S | NO | 52.59 | 37.58 | 34.38 | 0.00 |
| >hg38_21_+_8433172_8446622_RNA45SN1_45S_50ntFlanks@11573.11601.29\|CGACTTAGAACTGGTGCGGACCAGGGGAA | >hg38_21_+_8433172_8446622_RNA45SN1_45S_50ntFlanks | 11573 | 11601 | 29 | rRF 45S | NO | 140.32 | 82.40 | 58.81 | 0.00 |
| >hg38_21_+_8433172_8446622_RNA45SN1_45S_50ntFlanks@11573.11602.30\|CGACTTAGAACTGGTGCGGACCAGGGGAAT | >hg38_21_+_8433172_8446622_RNA45SN1_45S_50ntFlanks | 11573 | 11602 | 30 | rRF 45S | NO | 79.28 | 49.78 | 30.36 | 0.00 |
| >hg38_21_+_8433172_8446622_RNA45SN1_45S_50ntFlanks@11573.11603.31\|CGACTTAGAACTGGTGCGGACCAGGGGAATC | >hg38_21_+_8433172_8446622_RNA45SN1_45S_50ntFlanks | 11573 | 11603 | 31 | rRF 45S | NO | 80.03 | 61.44 | 33.39 | 15.26 |
| >hg38_21_+_8433172_8446622_RNA45SN1_45S_50ntFlanks@11573.11605.33\|CGACTTAGAACTGGTGCGGACCAGGGGAATCCG | >hg38_21_+_8433172_8446622_RNA45SN1_45S_50ntFlanks | 11573 | 11605 | 33 | rRF 45S | NO | 46.19 | 43.43 | 0.00 | 0.00 |
| >hg38_21_+_8433172_8446622_RNA45SN1_45S_50ntFlanks@11575.11592.18\|ACTTAGAACTGGTGCGGA | >hg38_21_+_8433172_8446622_RNA45SN1_45S_50ntFlanks | 11575 | 11592 | 18 | rRF 45S | NO | 167.43 | 339.20 | 105.00 | 45.92 |
| >hg38_21_+_8433172_8446622_RNA45SN1_45S_50ntFlanks@11575.11593.19\|ACTTAGAACTGGTGCGGAC | >hg38_21_+_8433172_8446622_RNA45SN1_45S_50ntFlanks | 11575 | 11593 | 19 | rRF 45S | NO | 106.86 | 215.38 | 36.84 | 15.35 |
| >hg38_21_+_8433172_8446622_RNA45SN1_45S_50ntFlanks@11575.11594.20\|ACTTAGAACTGGTGCGGACC | >hg38_21_+_8433172_8446622_RNA45SN1_45S_50ntFlanks | 11575 | 11594 | 20 | rRF 45S | NO | 177.87 | 144.77 | 27.74 | 0.00 |
| >hg38_21_+_8433172_8446622_RNA45SN1_45S_50ntFlanks@11575.11599.25\|ACTTAGAACTGGTGCGGACCAGGGG | >hg38_21_+_8433172_8446622_RNA45SN1_45S_50ntFlanks | 11575 | 11599 | 25 | rRF 45S | NO | 0.00 | 30.51 | 0.00 | 0.00 |
| >hg38_21_+_8433172_8446622_RNA45SN1_45S_50ntFlanks@11575.11600.26\|ACTTAGAACTGGTGCGGACCAGGGGA | >hg38_21_+_8433172_8446622_RNA45SN1_45S_50ntFlanks | 11575 | 11600 | 26 | rRF 45S | NO | 50.61 | 77.39 | 7.32 | 0.00 |
| >hg38_21_+_8433172_8446622_RNA45SN1_45S_50ntFlanks@11575.11601.27\|ACTTAGAACTGGTGCGGACCAGGGGAA | >hg38_21_+_8433172_8446622_RNA45SN1_45S_50ntFlanks | 11575 | 11601 | 27 | rRF 45S | NO | 143.32 | 124.35 | 37.07 | 6.26 |
| >hg38_21_+_8433172_8446622_RNA45SN1_45S_50ntFlanks@11575.11602.28\|ACTTAGAACTGGTGCGGACCAGGGGAAT | >hg38_21_+_8433172_8446622_RNA45SN1_45S_50ntFlanks | 11575 | 11602 | 28 | rRF 45S | NO | 81.31 | 152.33 | 7.00 | 6.53 |
| >hg38_21_+_8433172_8446622_RNA45SN1_45S_50ntFlanks@11575.11603.29\|ACTTAGAACTGGTGCGGACCAGGGGAATC | >hg38_21_+_8433172_8446622_RNA45SN1_45S_50ntFlanks | 11575 | 11603 | 29 | rRF 45S | NO | 367.73 | 724.83 | 110.83 | 90.03 |
| >hg38_21_+_8433172_8446622_RNA45SN1_45S_50ntFlanks@11575.11605.31\|ACTTAGAACTGGTGCGGACCAGGGGAATCCG | >hg38_21_+_8433172_8446622_RNA45SN1_45S_50ntFlanks | 11575 | 11605 | 31 | rRF 45S | NO | 127.78 | 230.96 | 36.15 | 45.09 |
| >hg38_21_+_8433172_8446622_RNA45SN1_45S_50ntFlanks@11575.11606.32\|ACTTAGAACTGGTGCGGACCAGGGGAATCCGA | >hg38_21_+_8433172_8446622_RNA45SN1_45S_50ntFlanks | 11575 | 11606 | 32 | rRF 45S | NO | 0.00 | 32.83 | 0.00 | 0.00 |
| >hg38_21_+_8433172_8446622_RNA45SN1_45S_50ntFlanks@11575.11607.33\|ACTTAGAACTGGTGCGGACCAGGGGAATCCGAC | >hg38_21_+_8433172_8446622_RNA45SN1_45S_50ntFlanks | 11575 | 11607 | 33 | rRF 45S | NO | 22.00 | 57.80 | 0.00 | 0.00 |
| >hg38_21_+_8433172_8446622_RNA45SN1_45S_50ntFlanks@11575.11609.35\|ACTTAGAACTGGTGCGGACCAGGGGAATCCGACTG | >hg38_21_+_8433172_8446622_RNA45SN1_45S_50ntFlanks | 11575 | 11609 | 35 | rRF 45S | NO | 20.42 | 43.14 | 20.65 | 23.41 |
| >hg38_21_+_8433172_8446622_RNA45SN1_45S_50ntFlanks@11575.11611.37\|ACTTAGAACTGGTGCGGACCAGGGGAATCCGACTGTT | >hg38_21_+_8433172_8446622_RNA45SN1_45S_50ntFlanks | 11575 | 11611 | 37 | rRF 45S | NO | 32.16 | 27.36 | 29.15 | 69.12 |
| >hg38_21_+_8433172_8446622_RNA45SN1_45S_50ntFlanks@11575.11613.39\|ACTTAGAACTGGTGCGGACCAGGGGAATCCGACTGTTTA | >hg38_21_+_8433172_8446622_RNA45SN1_45S_50ntFlanks | 11575 | 11613 | 39 | rRF 45S | NO | 21.28 | 12.48 | 0.00 | 41.99 |
| >hg38_21_+_8433172_8446622_RNA45SN1_45S_50ntFlanks@11576.11593.18\|CTTAGAACTGGTGCGGAC | >hg38_21_+_8433172_8446622_RNA45SN1_45S_50ntFlanks | 11576 | 11593 | 18 | rRF 45S | NO | 76.33 | 160.77 | 6.56 | 0.00 |
| >hg38_21_+_8433172_8446622_RNA45SN1_45S_50ntFlanks@11576.11594.19\|CTTAGAACTGGTGCGGACC | >hg38_21_+_8433172_8446622_RNA45SN1_45S_50ntFlanks | 11576 | 11594 | 19 | rRF 45S | NO | 116.52 | 120.58 | 0.00 | 0.00 |
| >hg38_21_+_8433172_8446622_RNA45SN1_45S_50ntFlanks@11576.11600.25\|CTTAGAACTGGTGCGGACCAGGGGA | >hg38_21_+_8433172_8446622_RNA45SN1_45S_50ntFlanks | 11576 | 11600 | 25 | rRF 45S | NO | 34.88 | 65.83 | 0.00 | 0.00 |
| >hg38_21_+_8433172_8446622_RNA45SN1_45S_50ntFlanks@11576.11601.26\|CTTAGAACTGGTGCGGACCAGGGGAA | >hg38_21_+_8433172_8446622_RNA45SN1_45S_50ntFlanks | 11576 | 11601 | 26 | rRF 45S | NO | 122.00 | 117.41 | 28.30 | 0.00 |
| >hg38_21_+_8433172_8446622_RNA45SN1_45S_50ntFlanks@11576.11602.27\|CTTAGAACTGGTGCGGACCAGGGGAAT | >hg38_21_+_8433172_8446622_RNA45SN1_45S_50ntFlanks | 11576 | 11602 | 27 | rRF 45S | NO | 62.59 | 121.31 | 0.00 | 0.00 |
| >hg38_21_+_8433172_8446622_RNA45SN1_45S_50ntFlanks@11576.11603.28\|CTTAGAACTGGTGCGGACCAGGGGAATC | >hg38_21_+_8433172_8446622_RNA45SN1_45S_50ntFlanks | 11576 | 11603 | 28 | rRF 45S | NO | 264.67 | 483.97 | 82.41 | 57.30 |
| >hg38_21_+_8433172_8446622_RNA45SN1_45S_50ntFlanks@11576.11605.30\|CTTAGAACTGGTGCGGACCAGGGGAATCCG | >hg38_21_+_8433172_8446622_RNA45SN1_45S_50ntFlanks | 11576 | 11605 | 30 | rRF 45S | NO | 102.79 | 151.69 | 20.33 | 24.86 |
| >hg38_21_+_8433172_8446622_RNA45SN1_45S_50ntFlanks@11576.11607.32\|CTTAGAACTGGTGCGGACCAGGGGAATCCGAC | >hg38_21_+_8433172_8446622_RNA45SN1_45S_50ntFlanks | 11576 | 11607 | 32 | rRF 45S | NO | 19.55 | 42.92 | 0.00 | 0.00 |
| >hg38_21_+_8433172_8446622_RNA45SN1_45S_50ntFlanks@11576.11609.34\|CTTAGAACTGGTGCGGACCAGGGGAATCCGACTG | >hg38_21_+_8433172_8446622_RNA45SN1_45S_50ntFlanks | 11576 | 11609 | 34 | rRF 45S | NO | 36.83 | 60.30 | 27.70 | 16.10 |
| >hg38_21_+_8433172_8446622_RNA45SN1_45S_50ntFlanks@11576.11611.36\|CTTAGAACTGGTGCGGACCAGGGGAATCCGACTGTT | >hg38_21_+_8433172_8446622_RNA45SN1_45S_50ntFlanks | 11576 | 11611 | 36 | rRF 45S | NO | 6.13 | 32.70 | 0.00 | 47.98 |
| >hg38_21_+_8433172_8446622_RNA45SN1_45S_50ntFlanks@11576.11613.38\|CTTAGAACTGGTGCGGACCAGGGGAATCCGACTGTTTA | >hg38_21_+_8433172_8446622_RNA45SN1_45S_50ntFlanks | 11576 | 11613 | 38 | rRF 45S | NO | 0.00 | 26.17 | 0.00 | 15.80 |
| >hg38_21_+_8433172_8446622_RNA45SN1_45S_50ntFlanks@11577.11594.18\|TTAGAACTGGTGCGGACC | >hg38_21_+_8433172_8446622_RNA45SN1_45S_50ntFlanks | 11577 | 11594 | 18 | rRF 45S | NO | 12.89 | 36.61 | 0.00 | 0.00 |
| >hg38_21_+_8433172_8446622_RNA45SN1_45S_50ntFlanks@11577.11601.25\|TTAGAACTGGTGCGGACCAGGGGAA | >hg38_21_+_8433172_8446622_RNA45SN1_45S_50ntFlanks | 11577 | 11601 | 25 | rRF 45S | NO | 21.20 | 34.42 | 0.00 | 0.00 |

### MDA-MB-468

| | | | | | | | | | | |
|---|---|---|---|---|---|---|---|---|---|---|
| >hg38_21_+_8433172_8446622_RNA45SN1_45S_50ntFlanks@11577.11602.26\|TTAGAACTGG TGCGGACCAGGGGAAT | >hg38_21_+_8433172_8446622_RNA45SN1_45S_50ntFlanks | 11577 | 11602 | 26 | rRF 45S | NO | 0.00 | 28.84 | 0.00 | 0.00 |
| >hg38_21_+_8433172_8446622_RNA45SN1_45S_50ntFlanks@11577.11603.27\|TTAGAACTGG TGCGGACCAGGGGAATC | >hg38_21_+_8433172_8446622_RNA45SN1_45S_50ntFlanks | 11577 | 11603 | 27 | rRF 45S | NO | 72.44 | 184.08 | 34.39 | 14.58 |
| >hg38_21_+_8433172_8446622_RNA45SN1_45S_50ntFlanks@11577.11605.29\|TTAGAACTGG TGCGGACCAGGGGAATCCG | >hg38_21_+_8433172_8446622_RNA45SN1_45S_50ntFlanks | 11577 | 11605 | 29 | rRF 45S | NO | 46.01 | 74.50 | 0.00 | 0.00 |
| >hg38_21_+_8433172_8446622_RNA45SN1_45S_50ntFlanks@11578.11601.24\|TAGAACTGGT GCGGACCAGGGGAA | >hg38_21_+_8433172_8446622_RNA45SN1_45S_50ntFlanks | 11578 | 11601 | 24 | rRF 45S | NO | 6.15 | 12.58 | 0.00 | 0.00 |
| >hg38_21_+_8433172_8446622_RNA45SN1_45S_50ntFlanks@11578.11603.26\|TAGAACTGGT GCGGACCAGGGGAATC | >hg38_21_+_8433172_8446622_RNA45SN1_45S_50ntFlanks | 11578 | 11603 | 26 | rRF 45S | NO | 49.42 | 132.88 | 28.69 | 6.54 |
| >hg38_21_+_8433172_8446622_RNA45SN1_45S_50ntFlanks@11578.11605.28\|TAGAACTGGT GCGGACCAGGGGAATCCG | >hg38_21_+_8433172_8446622_RNA45SN1_45S_50ntFlanks | 11578 | 11605 | 28 | rRF 45S | NO | 40.02 | 60.18 | 0.00 | 0.00 |
| >hg38_21_+_8433172_8446622_RNA45SN1_45S_50ntFlanks@11579.11601.23\|AGAACTGGTG CGGACCAGGGGAA | >hg38_21_+_8433172_8446622_RNA45SN1_45S_50ntFlanks | 11579 | 11601 | 23 | rRF 45S | NO | 40.33 | 80.64 | 13.90 | 0.00 |
| >hg38_21_+_8433172_8446622_RNA45SN1_45S_50ntFlanks@11579.11602.24\|AGAACTGGTG CGGACCAGGGGAAT | >hg38_21_+_8433172_8446622_RNA45SN1_45S_50ntFlanks | 11579 | 11602 | 24 | rRF 45S | NO | 0.00 | 65.80 | 20.43 | 0.00 |
| >hg38_21_+_8433172_8446622_RNA45SN1_45S_50ntFlanks@11579.11603.25\|AGAACTGGTG CGGACCAGGGGAATC | >hg38_21_+_8433172_8446622_RNA45SN1_45S_50ntFlanks | 11579 | 11603 | 25 | rRF 45S | NO | 146.35 | 468.21 | 219.38 | 91.53 |
| >hg38_21_+_8433172_8446622_RNA45SN1_45S_50ntFlanks@11579.11605.27\|AGAACTGGTG CGGACCAGGGGAATCCG | >hg38_21_+_8433172_8446622_RNA45SN1_45S_50ntFlanks | 11579 | 11605 | 27 | rRF 45S | NO | 111.19 | 171.97 | 57.50 | 60.47 |
| >hg38_21_+_8433172_8446622_RNA45SN1_45S_50ntFlanks@11579.11607.29\|AGAACTGGTG CGGACCAGGGGAATCCGAC | >hg38_21_+_8433172_8446622_RNA45SN1_45S_50ntFlanks | 11579 | 11607 | 29 | rRF 45S | NO | 0.00 | 37.52 | 6.65 | 15.38 |
| >hg38_21_+_8433172_8446622_RNA45SN1_45S_50ntFlanks@11579.11609.31\|AGAACTGGTG CGGACCAGGGGAATCCGACTG | >hg38_21_+_8433172_8446622_RNA45SN1_45S_50ntFlanks | 11579 | 11609 | 31 | rRF 45S | NO | 0.00 | 27.85 | 45.98 | 23.96 |
| >hg38_21_+_8433172_8446622_RNA45SN1_45S_50ntFlanks@11579.11611.33\|AGAACTGGTG CGGACCAGGGGAATCCGACTGTT | >hg38_21_+_8433172_8446622_RNA45SN1_45S_50ntFlanks | 11579 | 11611 | 33 | rRF 45S | NO | 21.19 | 48.67 | 47.79 | 84.27 |
| >hg38_21_+_8433172_8446622_RNA45SN1_45S_50ntFlanks@11580.11600.21\|GAACTGGTGC GGACCAGGGGA | >hg38_21_+_8433172_8446622_RNA45SN1_45S_50ntFlanks | 11580 | 11600 | 21 | rRF 45S | NO | 0.00 | 54.93 | 0.00 | 0.00 |
| >hg38_21_+_8433172_8446622_RNA45SN1_45S_50ntFlanks@11580.11601.22\|GAACTGGTGC GGACCAGGGGAA | >hg38_21_+_8433172_8446622_RNA45SN1_45S_50ntFlanks | 11580 | 11601 | 22 | rRF 45S | NO | 46.45 | 115.55 | 38.88 | 5.92 |
| >hg38_21_+_8433172_8446622_RNA45SN1_45S_50ntFlanks@11580.11602.23\|GAACTGGTGC GGACCAGGGGAAT | >hg38_21_+_8433172_8446622_RNA45SN1_45S_50ntFlanks | 11580 | 11602 | 23 | rRF 45S | NO | 20.09 | 96.31 | 31.23 | 0.00 |
| >hg38_21_+_8433172_8446622_RNA45SN1_45S_50ntFlanks@11580.11603.24\|GAACTGGTGC GGACCAGGGGAATC | >hg38_21_+_8433172_8446622_RNA45SN1_45S_50ntFlanks | 11580 | 11603 | 24 | rRF 45S | NO | 152.18 | 637.07 | 208.45 | 84.04 |
| >hg38_21_+_8433172_8446622_RNA45SN1_45S_50ntFlanks@11580.11605.26\|GAACTGGTGC GGACCAGGGGAATCCG | >hg38_21_+_8433172_8446622_RNA45SN1_45S_50ntFlanks | 11580 | 11605 | 26 | rRF 45S | NO | 91.54 | 179.97 | 54.98 | 56.93 |
| >hg38_21_+_8433172_8446622_RNA45SN1_45S_50ntFlanks@11580.11607.28\|GAACTGGTGC GGACCAGGGGAATCCGAC | >hg38_21_+_8433172_8446622_RNA45SN1_45S_50ntFlanks | 11580 | 11607 | 28 | rRF 45S | NO | 0.00 | 31.56 | 0.00 | 6.33 |
| >hg38_21_+_8433172_8446622_RNA45SN1_45S_50ntFlanks@11580.11609.30\|GAACTGGTGC GGACCAGGGGAATCCGACTG | >hg38_21_+_8433172_8446622_RNA45SN1_45S_50ntFlanks | 11580 | 11609 | 30 | rRF 45S | NO | 34.19 | 59.86 | 63.28 | 40.28 |
| >hg38_21_+_8433172_8446622_RNA45SN1_45S_50ntFlanks@11580.11611.32\|GAACTGGTGC GGACCAGGGGAATCCGACTGTT | >hg38_21_+_8433172_8446622_RNA45SN1_45S_50ntFlanks | 11580 | 11611 | 32 | rRF 45S | NO | 54.80 | 107.33 | 85.97 | 138.11 |
| >hg38_21_+_8433172_8446622_RNA45SN1_45S_50ntFlanks@11580.11613.34\|GAACTGGTGC GGACCAGGGGAATCCGACTGTTTA | >hg38_21_+_8433172_8446622_RNA45SN1_45S_50ntFlanks | 11580 | 11613 | 34 | rRF 45S | NO | 19.52 | 53.33 | 0.00 | 61.86 |
| >hg38_21_+_8433172_8446622_RNA45SN1_45S_50ntFlanks@11581.11600.20\|AACTGGTGCG GACCAGGGGA | >hg38_21_+_8433172_8446622_RNA45SN1_45S_50ntFlanks | 11581 | 11600 | 20 | rRF 45S | NO | 0.00 | 75.21 | 7.05 | 0.00 |
| >hg38_21_+_8433172_8446622_RNA45SN1_45S_50ntFlanks@11581.11601.21\|AACTGGTGCG GACCAGGGGAA | >hg38_21_+_8433172_8446622_RNA45SN1_45S_50ntFlanks | 11581 | 11601 | 21 | rRF 45S | NO | 46.54 | 144.70 | 40.80 | 14.51 |
| >hg38_21_+_8433172_8446622_RNA45SN1_45S_50ntFlanks@11581.11602.22\|AACTGGTGCG GACCAGGGGAAT | >hg38_21_+_8433172_8446622_RNA45SN1_45S_50ntFlanks | 11581 | 11602 | 22 | rRF 45S | NO | 28.76 | 127.15 | 37.65 | 16.15 |
| >hg38_21_+_8433172_8446622_RNA45SN1_45S_50ntFlanks@11581.11603.23\|AACTGGTGCG GACCAGGGGAATC | >hg38_21_+_8433172_8446622_RNA45SN1_45S_50ntFlanks | 11581 | 11603 | 23 | rRF 45S | NO | 158.76 | 715.49 | 269.95 | 100.43 |
| >hg38_21_+_8433172_8446622_RNA45SN1_45S_50ntFlanks@11581.11605.25\|AACTGGTGCG GACCAGGGGAATCCG | >hg38_21_+_8433172_8446622_RNA45SN1_45S_50ntFlanks | 11581 | 11605 | 25 | rRF 45S | NO | 108.91 | 249.85 | 83.28 | 80.64 |
| >hg38_21_+_8433172_8446622_RNA45SN1_45S_50ntFlanks@11581.11606.26\|AACTGGTGCG GACCAGGGGAATCCGA | >hg38_21_+_8433172_8446622_RNA45SN1_45S_50ntFlanks | 11581 | 11606 | 26 | rRF 45S | NO | 0.00 | 19.57 | 0.00 | 0.00 |
| >hg38_21_+_8433172_8446622_RNA45SN1_45S_50ntFlanks@11581.11607.27\|AACTGGTGCG GACCAGGGGAATCCGAC | >hg38_21_+_8433172_8446622_RNA45SN1_45S_50ntFlanks | 11581 | 11607 | 27 | rRF 45S | NO | 0.00 | 33.50 | 0.00 | 6.42 |
| >hg38_21_+_8433172_8446622_RNA45SN1_45S_50ntFlanks@11581.11609.29\|AACTGGTGCG GACCAGGGGAATCCGACTG | >hg38_21_+_8433172_8446622_RNA45SN1_45S_50ntFlanks | 11581 | 11609 | 29 | rRF 45S | NO | 70.41 | 147.99 | 92.56 | 68.36 |
| >hg38_21_+_8433172_8446622_RNA45SN1_45S_50ntFlanks@11581.11611.31\|AACTGGTGCG GACCAGGGGAATCCGACTGTT | >hg38_21_+_8433172_8446622_RNA45SN1_45S_50ntFlanks | 11581 | 11611 | 31 | rRF 45S | NO | 40.31 | 93.72 | 74.52 | 112.77 |
| >hg38_21_+_8433172_8446622_RNA45SN1_45S_50ntFlanks@11581.11612.32\|AACTGGTGCG GACCAGGGGAATCCGACTGTTT | >hg38_21_+_8433172_8446622_RNA45SN1_45S_50ntFlanks | 11581 | 11612 | 32 | rRF 45S | NO | 60.87 | 142.89 | 66.12 | 134.99 |
| >hg38_21_+_8433172_8446622_RNA45SN1_45S_50ntFlanks@11582.11600.19\|ACTGGTGCGG ACCAGGGGA | >hg38_21_+_8433172_8446622_RNA45SN1_45S_50ntFlanks | 11582 | 11600 | 19 | rRF 45S | NO | 0.00 | 19.39 | 0.00 | 0.00 |
| >hg38_21_+_8433172_8446622_RNA45SN1_45S_50ntFlanks@11582.11601.20\|ACTGGTGCGG ACCAGGGGAA | >hg38_21_+_8433172_8446622_RNA45SN1_45S_50ntFlanks | 11582 | 11601 | 20 | rRF 45S | NO | 0.00 | 37.77 | 0.00 | 0.00 |

### MDA-MB-468

| | | | | | | | | | | |
|---|---|---|---|---|---|---|---|---|---|---|
| >hg38_21_+_8433172_8446622_RNA45SN1_45S_50ntFlanks@11582.11602.21\|ACTGGTGCGG ACCAGGGGAAT | >hg38_21_+_8433172_8446622_RNA45SN1_45S_50ntFlanks | 11582 | 11602 | 21 | rRF 45S | NO | 0.00 | 38.38 | 0.00 | 0.00 |
| >hg38_21_+_8433172_8446622_RNA45SN1_45S_50ntFlanks@11582.11603.22\|ACTGGTGCGG ACCAGGGGAATC | >hg38_21_+_8433172_8446622_RNA45SN1_45S_50ntFlanks | 11582 | 11603 | 22 | rRF 45S | NO | 45.13 | 117.38 | 74.13 | 6.37 |
| >hg38_21_+_8433172_8446622_RNA45SN1_45S_50ntFlanks@11582.11605.24\|ACTGGTGCGG ACCAGGGGAATCCG | >hg38_21_+_8433172_8446622_RNA45SN1_45S_50ntFlanks | 11582 | 11605 | 24 | rRF 45S | NO | 0.00 | 34.41 | 0.00 | 0.00 |
| >hg38_21_+_8433172_8446622_RNA45SN1_45S_50ntFlanks@11582.11611.30\|ACTGGTGCGG ACCAGGGGAATCCGACTGTT | >hg38_21_+_8433172_8446622_RNA45SN1_45S_50ntFlanks | 11582 | 11611 | 30 | rRF 45S | NO | 33.26 | 59.90 | 48.84 | 63.48 |
| >hg38_21_+_8433172_8446622_RNA45SN1_45S_50ntFlanks@11582.11612.31\|ACTGGTGCGG ACCAGGGGAATCCGACTGTTT | >hg38_21_+_8433172_8446622_RNA45SN1_45S_50ntFlanks | 11582 | 11612 | 31 | rRF 45S | NO | 0.00 | 12.66 | 0.00 | 15.83 |
| >hg38_21_+_8433172_8446622_RNA45SN1_45S_50ntFlanks@11583.11600.18\|CTGGTGCGGA CCAGGGGA | >hg38_21_+_8433172_8446622_RNA45SN1_45S_50ntFlanks | 11583 | 11600 | 18 | rRF 45S | NO | 5.97 | 71.13 | 0.00 | 0.00 |
| >hg38_21_+_8433172_8446622_RNA45SN1_45S_50ntFlanks@11583.11601.19\|CTGGTGCGGA CCAGGGGAA | >hg38_21_+_8433172_8446622_RNA45SN1_45S_50ntFlanks | 11583 | 11601 | 19 | rRF 45S | NO | 84.98 | 111.54 | 32.40 | 6.47 |
| >hg38_21_+_8433172_8446622_RNA45SN1_45S_50ntFlanks@11583.11602.20\|CTGGTGCGGA CCAGGGGAAT | >hg38_21_+_8433172_8446622_RNA45SN1_45S_50ntFlanks | 11583 | 11602 | 20 | rRF 45S | NO | 52.10 | 79.61 | 42.00 | 14.84 |
| >hg38_21_+_8433172_8446622_RNA45SN1_45S_50ntFlanks@11583.11603.21\|CTGGTGCGGA CCAGGGGAATC | >hg38_21_+_8433172_8446622_RNA45SN1_45S_50ntFlanks | 11583 | 11603 | 21 | rRF 45S | NO | 125.50 | 211.11 | 160.82 | 57.76 |
| >hg38_21_+_8433172_8446622_RNA45SN1_45S_50ntFlanks@11583.11605.23\|CTGGTGCGGA CCAGGGGAATCCG | >hg38_21_+_8433172_8446622_RNA45SN1_45S_50ntFlanks | 11583 | 11605 | 23 | rRF 45S | NO | 49.69 | 89.82 | 38.99 | 51.72 |
| >hg38_21_+_8433172_8446622_RNA45SN1_45S_50ntFlanks@11583.11609.27\|CTGGTGCGGA CCAGGGGAATCCGACTG | >hg38_21_+_8433172_8446622_RNA45SN1_45S_50ntFlanks | 11583 | 11609 | 27 | rRF 45S | NO | 29.44 | 55.66 | 37.42 | 40.47 |
| >hg38_21_+_8433172_8446622_RNA45SN1_45S_50ntFlanks@11583.11611.29\|CTGGTGCGGA CCAGGGGAATCCGACTGTT | >hg38_21_+_8433172_8446622_RNA45SN1_45S_50ntFlanks | 11583 | 11611 | 29 | rRF 45S | NO | 123.05 | 169.59 | 113.91 | 166.50 |
| >hg38_21_+_8433172_8446622_RNA45SN1_45S_50ntFlanks@11583.11612.30\|CTGGTGCGGA CCAGGGGAATCCGACTGTTT | >hg38_21_+_8433172_8446622_RNA45SN1_45S_50ntFlanks | 11583 | 11612 | 30 | rRF 45S | NO | 63.20 | 77.41 | 47.38 | 80.18 |
| >hg38_21_+_8433172_8446622_RNA45SN1_45S_50ntFlanks@11586.11603.18\|GTGCGGACCA GGGGAATC | >hg38_21_+_8433172_8446622_RNA45SN1_45S_50ntFlanks | 11586 | 11603 | 18 | rRF 45S | NO | 207.93 | 147.13 | 69.03 | 26.39 |
| >hg38_21_+_8433172_8446622_RNA45SN1_45S_50ntFlanks@11586.11605.20\|GTGCGGACCA GGGGAATCCG | >hg38_21_+_8433172_8446622_RNA45SN1_45S_50ntFlanks | 11586 | 11605 | 20 | rRF 45S | NO | 0.00 | 28.97 | 6.91 | 0.00 |
| >hg38_21_+_8433172_8446622_RNA45SN1_45S_50ntFlanks@11586.11609.24\|GTGCGGACCA GGGGAATCCGACTG | >hg38_21_+_8433172_8446622_RNA45SN1_45S_50ntFlanks | 11586 | 11609 | 24 | rRF 45S | NO | 0.00 | 51.44 | 29.60 | 0.00 |
| >hg38_21_+_8433172_8446622_RNA45SN1_45S_50ntFlanks@11586.11611.26\|GTGCGGACCA GGGGAATCCGACTGTT | >hg38_21_+_8433172_8446622_RNA45SN1_45S_50ntFlanks | 11586 | 11611 | 26 | rRF 45S | NO | 126.20 | 206.72 | 82.09 | 85.82 |
| >hg38_21_+_8433172_8446622_RNA45SN1_45S_50ntFlanks@11586.11612.27\|GTGCGGACCA GGGGAATCCGACTGTTT | >hg38_21_+_8433172_8446622_RNA45SN1_45S_50ntFlanks | 11586 | 11612 | 27 | rRF 45S | NO | 68.90 | 92.94 | 51.99 | 53.83 |
| >hg38_21_+_8433172_8446622_RNA45SN1_45S_50ntFlanks@11587.11611.25\|TGCGGACCAG GGGAATCCGACTGTT | >hg38_21_+_8433172_8446622_RNA45SN1_45S_50ntFlanks | 11587 | 11611 | 25 | rRF 45S | NO | 21.03 | 45.51 | 27.85 | 24.22 |
| >hg38_21_+_8433172_8446622_RNA45SN1_45S_50ntFlanks@11589.11609.21\|CGGACCAGG GGAATCCGACTG | >hg38_21_+_8433172_8446622_RNA45SN1_45S_50ntFlanks | 11589 | 11609 | 21 | rRF 45S | NO | 0.00 | 28.73 | 0.00 | 0.00 |
| >hg38_21_+_8433172_8446622_RNA45SN1_45S_50ntFlanks@11589.11611.23\|CGGACCAGG GGAATCCGACTGTT | >hg38_21_+_8433172_8446622_RNA45SN1_45S_50ntFlanks | 11589 | 11611 | 23 | rRF 45S | NO | 38.17 | 53.71 | 36.04 | 23.72 |
| >hg38_21_+_8433172_8446622_RNA45SN1_45S_50ntFlanks@11589.11612.24\|CGGACCAGG GGAATCCGACTGTTT | >hg38_21_+_8433172_8446622_RNA45SN1_45S_50ntFlanks | 11589 | 11612 | 24 | rRF 45S | NO | 0.00 | 19.60 | 0.00 | 0.00 |
| >hg38_21_+_8433172_8446622_RNA45SN1_45S_50ntFlanks@11592.11611.20\|ACCAGGGGAA TCCGACTGTT | >hg38_21_+_8433172_8446622_RNA45SN1_45S_50ntFlanks | 11592 | 11611 | 20 | rRF 45S | NO | 0.00 | 37.55 | 38.79 | 23.02 |
| >hg38_21_+_8433172_8446622_RNA45SN1_45S_50ntFlanks@11592.11612.21\|ACCAGGGGAA TCCGACTGTTT | >hg38_21_+_8433172_8446622_RNA45SN1_45S_50ntFlanks | 11592 | 11612 | 21 | rRF 45S | NO | 0.00 | 31.23 | 20.29 | 16.07 |
| >hg38_21_+_8433172_8446622_RNA45SN1_45S_50ntFlanks@11593.11611.19\|CCAGGGGAAT CCGACTGTT | >hg38_21_+_8433172_8446622_RNA45SN1_45S_50ntFlanks | 11593 | 11611 | 19 | rRF 45S | NO | 155.36 | 271.81 | 87.30 | 103.95 |
| >hg38_21_+_8433172_8446622_RNA45SN1_45S_50ntFlanks@11593.11612.20\|CCAGGGGAAT CCGACTGTTT | >hg38_21_+_8433172_8446622_RNA45SN1_45S_50ntFlanks | 11593 | 11612 | 20 | rRF 45S | NO | 67.55 | 105.37 | 44.51 | 53.86 |
| >hg38_21_+_8433172_8446622_RNA45SN1_45S_50ntFlanks@11593.11613.21\|CCAGGGGAAT CCGACTGTTTA | >hg38_21_+_8433172_8446622_RNA45SN1_45S_50ntFlanks | 11593 | 11613 | 21 | rRF 45S | NO | 34.95 | 34.22 | 0.00 | 36.55 |
| >hg38_21_+_8433172_8446622_RNA45SN1_45S_50ntFlanks@11594.11611.18\|CAGGGGAATC CGACTGTT | >hg38_21_+_8433172_8446622_RNA45SN1_45S_50ntFlanks | 11594 | 11611 | 18 | rRF 45S | NO | 118.75 | 185.51 | 54.77 | 94.02 |
| >hg38_21_+_8433172_8446622_RNA45SN1_45S_50ntFlanks@11594.11612.19\|CAGGGGAATC CGACTGTTT | >hg38_21_+_8433172_8446622_RNA45SN1_45S_50ntFlanks | 11594 | 11612 | 19 | rRF 45S | NO | 53.74 | 74.02 | 20.75 | 49.23 |
| >hg38_21_+_8433172_8446622_RNA45SN1_45S_50ntFlanks@11594.11613.20\|CAGGGGAATC CGACTGTTTA | >hg38_21_+_8433172_8446622_RNA45SN1_45S_50ntFlanks | 11594 | 11613 | 20 | rRF 45S | NO | 19.82 | 6.03 | 0.00 | 23.87 |
| >hg38_21_+_8433172_8446622_RNA45SN1_45S_50ntFlanks@11602.11619.18\|TCCGACTGTT TAATTAAA | >hg38_21_+_8433172_8446622_RNA45SN1_45S_50ntFlanks | 11602 | 11619 | 18 | rRF 45S | NO | 79.47 | 108.88 | 0.00 | 16.24 |
| >hg38_21_+_8433172_8446622_RNA45SN1_45S_50ntFlanks@11602.11620.19\|TCCGACTGTT TAATTAAAA | >hg38_21_+_8433172_8446622_RNA45SN1_45S_50ntFlanks | 11602 | 11620 | 19 | rRF 45S | NO | 88.40 | 82.88 | 0.00 | 13.68 |
| >hg38_21_+_8433172_8446622_RNA45SN1_45S_50ntFlanks@11602.11621.20\|TCCGACTGTT TAATTAAAAC | >hg38_21_+_8433172_8446622_RNA45SN1_45S_50ntFlanks | 11602 | 11621 | 20 | rRF 45S | NO | 0.00 | 36.38 | 0.00 | 0.00 |
| >hg38_21_+_8433172_8446622_RNA45SN1_45S_50ntFlanks@11603.11620.18\|CCGACTGTTT AATTAAAA | >hg38_21_+_8433172_8446622_RNA45SN1_45S_50ntFlanks | 11603 | 11620 | 18 | rRF 45S | NO | 46.61 | 48.67 | 0.00 | 0.00 |

### MDA-MB-468

| | | | | | | | | | | |
|---|---|---|---|---|---|---|---|---|---|---|
| >hg38_21_+_8433172_8446622_RNA45SN1_45S_50ntFlanks@11604.11621.18\|CGACTGTTTAATTAAAAC | >hg38_21_+_8433172_8446622_RNA45SN1_45S_50ntFlanks | 11604 | 11621 | 18 | rRF 45S | NO | 13.03 | 70.14 | 21.00 | 6.33 |
| >hg38_21_+_8433172_8446622_RNA45SN1_45S_50ntFlanks@11612.11640.29\|TAATTAAAACAAAGCATCGCGAAGGCCCG | >hg38_21_+_8433172_8446622_RNA45SN1_45S_50ntFlanks | 11612 | 11640 | 29 | rRF 45S | NO | 19.57 | 0.00 | 49.71 | 89.18 |
| >hg38_21_+_8433172_8446622_RNA45SN1_45S_50ntFlanks@11613.11640.28\|AATTAAAACAAAGCATCGCGAAGGCCCG | >hg38_21_+_8433172_8446622_RNA45SN1_45S_50ntFlanks | 11613 | 11640 | 28 | rRF 45S | NO | 29.79 | 0.00 | 124.29 | 114.14 |
| >hg38_21_+_8433172_8446622_RNA45SN1_45S_50ntFlanks@11614.11640.27\|ATTAAAACAAAGCATCGCGAAGGCCCG | >hg38_21_+_8433172_8446622_RNA45SN1_45S_50ntFlanks | 11614 | 11640 | 27 | rRF 45S | NO | 0.00 | 0.00 | 49.09 | 43.41 |
| >hg38_21_+_8433172_8446622_RNA45SN1_45S_50ntFlanks@11617.11640.24\|AAAACAAAGCATCGCGAAGGCCCG | >hg38_21_+_8433172_8446622_RNA45SN1_45S_50ntFlanks | 11617 | 11640 | 24 | rRF 45S | NO | 0.00 | 0.00 | 55.08 | 24.90 |
| >hg38_21_+_8433172_8446622_RNA45SN1_45S_50ntFlanks@11618.11640.23\|AAACAAAGCATCGCGAAGGCCCG | >hg38_21_+_8433172_8446622_RNA45SN1_45S_50ntFlanks | 11618 | 11640 | 23 | rRF 45S | NO | 0.00 | 0.00 | 88.60 | 47.93 |
| >hg38_21_+_8433172_8446622_RNA45SN1_45S_50ntFlanks@11619.11640.22\|AACAAAGCATCGCGAAGGCCCG | >hg38_21_+_8433172_8446622_RNA45SN1_45S_50ntFlanks | 11619 | 11640 | 22 | rRF 45S | NO | 0.00 | 0.00 | 106.46 | 58.18 |
| >hg38_21_+_8433172_8446622_RNA45SN1_45S_50ntFlanks@11620.11640.21\|ACAAAGCATCGCGAAGGCCCG | >hg38_21_+_8433172_8446622_RNA45SN1_45S_50ntFlanks | 11620 | 11640 | 21 | rRF 45S | NO | 0.00 | 0.00 | 156.73 | 96.10 |
| >hg38_21_+_8433172_8446622_RNA45SN1_45S_50ntFlanks@11621.11640.20\|CAAAGCATCGCGAAGGCCCG | >hg38_21_+_8433172_8446622_RNA45SN1_45S_50ntFlanks | 11621 | 11640 | 20 | rRF 45S | NO | 0.00 | 0.00 | 130.94 | 104.25 |
| >hg38_21_+_8433172_8446622_RNA45SN1_45S_50ntFlanks@11622.11640.19\|AAAGCATCGCGAAGGCCCG | >hg38_21_+_8433172_8446622_RNA45SN1_45S_50ntFlanks | 11622 | 11640 | 19 | rRF 45S | NO | 0.00 | 0.00 | 68.08 | 0.00 |
| >hg38_21_+_8433172_8446622_RNA45SN1_45S_50ntFlanks@11641.11658.18\|CGGCGGGTGTTGACGCGA | >hg38_21_+_8433172_8446622_RNA45SN1_45S_50ntFlanks | 11641 | 11658 | 18 | rRF 45S | YES | 0.00 | 0.00 | 58.59 | 6.51 |
| >hg38_21_+_8433172_8446622_RNA45SN1_45S_50ntFlanks@11641.11659.19\|CGGCGGGTGTTGACGCGAT | >hg38_21_+_8433172_8446622_RNA45SN1_45S_50ntFlanks | 11641 | 11659 | 19 | rRF 45S | NO | 0.00 | 0.00 | 143.57 | 60.90 |
| >hg38_21_+_8433172_8446622_RNA45SN1_45S_50ntFlanks@11641.11660.20\|CGGCGGGTGTTGACGCGATG | >hg38_21_+_8433172_8446622_RNA45SN1_45S_50ntFlanks | 11641 | 11660 | 20 | rRF 45S | YES | 162.59 | 147.00 | 4358.12 | 1117.47 |
| >hg38_21_+_8433172_8446622_RNA45SN1_45S_50ntFlanks@11641.11661.21\|CGGCGGGTGTTGACGCGATGT | >hg38_21_+_8433172_8446622_RNA45SN1_45S_50ntFlanks | 11641 | 11661 | 21 | rRF 45S | NO | 0.00 | 0.00 | 44.33 | 26.48 |
| >hg38_21_+_8433172_8446622_RNA45SN1_45S_50ntFlanks@11641.11662.22\|CGGCGGGTGTTGACGCGATGTG | >hg38_21_+_8433172_8446622_RNA45SN1_45S_50ntFlanks | 11641 | 11662 | 22 | rRF 45S | NO | 87.26 | 45.46 | 352.07 | 193.70 |
| >hg38_21_+_8433172_8446622_RNA45SN1_45S_50ntFlanks@11641.11663.23\|CGGCGGGTGTTGACGCGATGTGA | >hg38_21_+_8433172_8446622_RNA45SN1_45S_50ntFlanks | 11641 | 11663 | 23 | rRF 45S | NO | 21.94 | 0.00 | 73.45 | 44.72 |
| >hg38_21_+_8433172_8446622_RNA45SN1_45S_50ntFlanks@11641.11664.24\|CGGCGGGTGTTGACGCGATGTGAT | >hg38_21_+_8433172_8446622_RNA45SN1_45S_50ntFlanks | 11641 | 11664 | 24 | rRF 45S | NO | 61.45 | 27.39 | 102.56 | 72.20 |
| >hg38_21_+_8433172_8446622_RNA45SN1_45S_50ntFlanks@11641.11665.25\|CGGCGGGTGTTGACGCGATGTGATT | >hg38_21_+_8433172_8446622_RNA45SN1_45S_50ntFlanks | 11641 | 11665 | 25 | rRF 45S | NO | 210.26 | 102.62 | 320.82 | 308.65 |
| >hg38_21_+_8433172_8446622_RNA45SN1_45S_50ntFlanks@11641.11667.27\|CGGCGGGTGTTGACGCGATGTGATTTC | >hg38_21_+_8433172_8446622_RNA45SN1_45S_50ntFlanks | 11641 | 11667 | 27 | rRF 45S | NO | 41.89 | 0.00 | 62.82 | 47.00 |
| >hg38_21_+_8433172_8446622_RNA45SN1_45S_50ntFlanks@11641.11668.28\|CGGCGGGTGTTGACGCGATGTGATTTCT | >hg38_21_+_8433172_8446622_RNA45SN1_45S_50ntFlanks | 11641 | 11668 | 28 | rRF 45S | NO | 37.43 | 0.00 | 55.13 | 42.64 |
| >hg38_21_+_8433172_8446622_RNA45SN1_45S_50ntFlanks@11641.11669.29\|CGGCGGGTGTTGACGCGATGTGATTTCTG | >hg38_21_+_8433172_8446622_RNA45SN1_45S_50ntFlanks | 11641 | 11669 | 29 | rRF 45S | NO | 139.40 | 78.15 | 208.59 | 144.78 |
| >hg38_21_+_8433172_8446622_RNA45SN1_45S_50ntFlanks@11641.11670.30\|CGGCGGGTGTTGACGCGATGTGATTTCTGC | >hg38_21_+_8433172_8446622_RNA45SN1_45S_50ntFlanks | 11641 | 11670 | 30 | rRF 45S | NO | 228.65 | 77.62 | 153.75 | 233.58 |
| >hg38_21_+_8433172_8446622_RNA45SN1_45S_50ntFlanks@11641.11685.45\|CGGCGGGTGTTGACGCGATGTGATTTCTGCCCAGTGCTCTGAATG | >hg38_21_+_8433172_8446622_RNA45SN1_45S_50ntFlanks | 11641 | 11685 | 45 | rRF 45S | NO | 85.24 | 36.81 | 0.00 | 223.02 |
| >hg38_21_+_8433172_8446622_RNA45SN1_45S_50ntFlanks@11642.11660.19\|GGCGGGTGTTGACGCGATG | >hg38_21_+_8433172_8446622_RNA45SN1_45S_50ntFlanks | 11642 | 11660 | 19 | rRF 45S | NO | 0.00 | 0.00 | 59.08 | 0.00 |
| >hg38_21_+_8433172_8446622_RNA45SN1_45S_50ntFlanks@11643.11660.18\|GCGGGTGTTGACGCGATG | >hg38_21_+_8433172_8446622_RNA45SN1_45S_50ntFlanks | 11643 | 11660 | 18 | rRF 45S | NO | 0.00 | 0.00 | 189.24 | 34.01 |
| >hg38_21_+_8433172_8446622_RNA45SN1_45S_50ntFlanks@11660.11685.26\|GTGATTTCTGCCCAGTGCTCTGAATG | >hg38_21_+_8433172_8446622_RNA45SN1_45S_50ntFlanks | 11660 | 11685 | 26 | rRF 45S | NO | 59.03 | 38.53 | 28.11 | 105.44 |
| >hg38_21_+_8433172_8446622_RNA45SN1_45S_50ntFlanks@11661.11678.18\|TGATTTCTGCCCAGTGCT | >hg38_21_+_8433172_8446622_RNA45SN1_45S_50ntFlanks | 11661 | 11678 | 18 | rRF 45S | NO | 0.00 | 32.03 | 0.00 | 0.00 |
| >hg38_21_+_8433172_8446622_RNA45SN1_45S_50ntFlanks@11661.11679.19\|TGATTTCTGCCCAGTGCTC | >hg38_21_+_8433172_8446622_RNA45SN1_45S_50ntFlanks | 11661 | 11679 | 19 | rRF 45S | NO | 129.45 | 200.64 | 0.00 | 25.06 |
| >hg38_21_+_8433172_8446622_RNA45SN1_45S_50ntFlanks@11661.11680.20\|TGATTTCTGCCCAGTGCTCT | >hg38_21_+_8433172_8446622_RNA45SN1_45S_50ntFlanks | 11661 | 11680 | 20 | rRF 45S | NO | 12.87 | 41.42 | 0.00 | 0.00 |
| >hg38_21_+_8433172_8446622_RNA45SN1_45S_50ntFlanks@11661.11681.21\|TGATTTCTGCCCAGTGCTCTG | >hg38_21_+_8433172_8446622_RNA45SN1_45S_50ntFlanks | 11661 | 11681 | 21 | rRF 45S | NO | 430.49 | 447.14 | 28.17 | 83.76 |
| >hg38_21_+_8433172_8446622_RNA45SN1_45S_50ntFlanks@11661.11682.22\|TGATTTCTGCCCAGTGCTCTGA | >hg38_21_+_8433172_8446622_RNA45SN1_45S_50ntFlanks | 11661 | 11682 | 22 | rRF 45S | NO | 59.34 | 66.15 | 0.00 | 0.00 |
| >hg38_21_+_8433172_8446622_RNA45SN1_45S_50ntFlanks@11661.11685.25\|TGATTTCTGCCCAGTGCTCTGAATG | >hg38_21_+_8433172_8446622_RNA45SN1_45S_50ntFlanks | 11661 | 11685 | 25 | rRF 45S | NO | 430.74 | 577.08 | 137.93 | 181.20 |
| >hg38_21_+_8433172_8446622_RNA45SN1_45S_50ntFlanks@11663.11685.23\|ATTTCTGCCCAGTGCTCTGAATG | >hg38_21_+_8433172_8446622_RNA45SN1_45S_50ntFlanks | 11663 | 11685 | 23 | rRF 45S | NO | 6.11 | 61.64 | 32.57 | 0.00 |
| >hg38_21_+_8433172_8446622_RNA45SN1_45S_50ntFlanks@11664.11685.22\|TTTCTGCCCAGTGCTCTGAATG | >hg38_21_+_8433172_8446622_RNA45SN1_45S_50ntFlanks | 11664 | 11685 | 22 | rRF 45S | NO | 0.00 | 29.01 | 0.00 | 0.00 |

### MDA-MB-468

| | | | | | | | | | | |
|---|---|---|---|---|---|---|---|---|---|---|
| >hg38_21_+_8433172_8446622_RNA45SN1_45S_50ntFlanks@11665.11685.21\|TTCTGCCCAG TGCTCTGAATG | >hg38_21_+_8433172_8446622_RNA45SN1_45S_50ntFlanks | 11665 | 11685 | 21 | rRF 45S | NO | 0.00 | 42.97 | 0.00 | 0.00 |
| >hg38_21_+_8433172_8446622_RNA45SN1_45S_50ntFlanks@11666.11685.20\|TCTGCCCAGT GCTCTGAATG | >hg38_21_+_8433172_8446622_RNA45SN1_45S_50ntFlanks | 11666 | 11685 | 20 | rRF 45S | NO | 94.02 | 244.96 | 86.83 | 90.46 |
| >hg38_21_+_8433172_8446622_RNA45SN1_45S_50ntFlanks@11666.11702.37\|TCTGCCCAGT GCTCTGAATGTCAAAGTGAAGAAATTC | >hg38_21_+_8433172_8446622_RNA45SN1_45S_50ntFlanks | 11666 | 11702 | 37 | rRF 45S | NO | 30.00 | 0.00 | 0.00 | 0.00 |
| >hg38_21_+_8433172_8446622_RNA45SN1_45S_50ntFlanks@11667.11685.19\|CTGCCCAGTG CTCTGAATG | >hg38_21_+_8433172_8446622_RNA45SN1_45S_50ntFlanks | 11667 | 11685 | 19 | rRF 45S | NO | 35.07 | 50.24 | 0.00 | 15.45 |
| >hg38_21_+_8433172_8446622_RNA45SN1_45S_50ntFlanks@11668.11685.18\|TGCCCAGTGC TCTGAATG | >hg38_21_+_8433172_8446622_RNA45SN1_45S_50ntFlanks | 11668 | 11685 | 18 | rRF 45S | NO | 33.84 | 43.49 | 0.00 | 0.00 |
| >hg38_21_+_8433172_8446622_RNA45SN1_45S_50ntFlanks@11668.11701.34\|TGCCCAGTGC TCTGAATGTCAAAGTGAAGAAATT | >hg38_21_+_8433172_8446622_RNA45SN1_45S_50ntFlanks | 11668 | 11701 | 34 | rRF 45S | NO | 60.87 | 0.00 | 0.00 | 0.00 |
| >hg38_21_+_8433172_8446622_RNA45SN1_45S_50ntFlanks@11668.11702.35\|TGCCCAGTGC TCTGAATGTCAAAGTGAAGAAATTC | >hg38_21_+_8433172_8446622_RNA45SN1_45S_50ntFlanks | 11668 | 11702 | 35 | rRF 45S | NO | 176.19 | 44.78 | 0.00 | 41.35 |
| >hg38_21_+_8433172_8446622_RNA45SN1_45S_50ntFlanks@11669.11702.34\|GCCCAGTGCT CTGAATGTCAAAGTGAAGAAATTC | >hg38_21_+_8433172_8446622_RNA45SN1_45S_50ntFlanks | 11669 | 11702 | 34 | rRF 45S | NO | 20.79 | 0.00 | 0.00 | 0.00 |
| >hg38_21_+_8433172_8446622_RNA45SN1_45S_50ntFlanks@11670.11687.18\|CCCAGTGCTC TGAATGTC | >hg38_21_+_8433172_8446622_RNA45SN1_45S_50ntFlanks | 11670 | 11687 | 18 | rRF 45S | NO | 69.76 | 95.38 | 6.80 | 0.00 |
| >hg38_21_+_8433172_8446622_RNA45SN1_45S_50ntFlanks@11670.11688.19\|CCCAGTGCTC TGAATGTCA | >hg38_21_+_8433172_8446622_RNA45SN1_45S_50ntFlanks | 11670 | 11688 | 19 | rRF 45S | NO | 58.41 | 42.12 | 0.00 | 0.00 |
| >hg38_21_+_8433172_8446622_RNA45SN1_45S_50ntFlanks@11670.11691.22\|CCCAGTGCTC TGAATGTCAAAG | >hg38_21_+_8433172_8446622_RNA45SN1_45S_50ntFlanks | 11670 | 11691 | 22 | rRF 45S | NO | 84.66 | 36.48 | 0.00 | 16.09 |
| >hg38_21_+_8433172_8446622_RNA45SN1_45S_50ntFlanks@11670.11692.23\|CCCAGTGCTC TGAATGTCAAAGT | >hg38_21_+_8433172_8446622_RNA45SN1_45S_50ntFlanks | 11670 | 11692 | 23 | rRF 45S | NO | 40.21 | 0.00 | 0.00 | 0.00 |
| >hg38_21_+_8433172_8446622_RNA45SN1_45S_50ntFlanks@11670.11693.24\|CCCAGTGCTC TGAATGTCAAAGTG | >hg38_21_+_8433172_8446622_RNA45SN1_45S_50ntFlanks | 11670 | 11693 | 24 | rRF 45S | NO | 70.83 | 51.98 | 0.00 | 0.00 |
| >hg38_21_+_8433172_8446622_RNA45SN1_45S_50ntFlanks@11670.11698.29\|CCCAGTGCTC TGAATGTCAAAGTGAAGAA | >hg38_21_+_8433172_8446622_RNA45SN1_45S_50ntFlanks | 11670 | 11698 | 29 | rRF 45S | NO | 47.69 | 0.00 | 0.00 | 0.00 |
| >hg38_21_+_8433172_8446622_RNA45SN1_45S_50ntFlanks@11670.11699.30\|CCCAGTGCTC TGAATGTCAAAGTGAAGAAA | >hg38_21_+_8433172_8446622_RNA45SN1_45S_50ntFlanks | 11670 | 11699 | 30 | rRF 45S | NO | 50.29 | 0.00 | 0.00 | 0.00 |
| >hg38_21_+_8433172_8446622_RNA45SN1_45S_50ntFlanks@11670.11700.31\|CCCAGTGCTC TGAATGTCAAAGTGAAGAAAT | >hg38_21_+_8433172_8446622_RNA45SN1_45S_50ntFlanks | 11670 | 11700 | 31 | rRF 45S | NO | 72.22 | 0.00 | 0.00 | 0.00 |
| >hg38_21_+_8433172_8446622_RNA45SN1_45S_50ntFlanks@11670.11701.32\|CCCAGTGCTC TGAATGTCAAAGTGAAGAAATT | >hg38_21_+_8433172_8446622_RNA45SN1_45S_50ntFlanks | 11670 | 11701 | 32 | rRF 45S | NO | 195.60 | 43.15 | 0.00 | 0.00 |
| >hg38_21_+_8433172_8446622_RNA45SN1_45S_50ntFlanks@11670.11702.33\|CCCAGTGCTC TGAATGTCAAAGTGAAGAAATTC | >hg38_21_+_8433172_8446622_RNA45SN1_45S_50ntFlanks | 11670 | 11702 | 33 | rRF 45S | NO | 422.73 | 159.75 | 29.24 | 72.18 |
| >hg38_21_+_8433172_8446622_RNA45SN1_45S_50ntFlanks@11670.11703.34\|CCCAGTGCTC TGAATGTCAAAGTGAAGAAATTCA | >hg38_21_+_8433172_8446622_RNA45SN1_45S_50ntFlanks | 11670 | 11703 | 34 | rRF 45S | NO | 109.09 | 50.47 | 0.00 | 24.49 |
| >hg38_21_+_8433172_8446622_RNA45SN1_45S_50ntFlanks@11670.11704.35\|CCCAGTGCTC TGAATGTCAAAGTGAAGAAATTCAA | >hg38_21_+_8433172_8446622_RNA45SN1_45S_50ntFlanks | 11670 | 11704 | 35 | rRF 45S | NO | 53.83 | 0.00 | 0.00 | 0.00 |
| >hg38_21_+_8433172_8446622_RNA45SN1_45S_50ntFlanks@11670.11706.37\|CCCAGTGCTC TGAATGTCAAAGTGAAGAAATTCAATG | >hg38_21_+_8433172_8446622_RNA45SN1_45S_50ntFlanks | 11670 | 11706 | 37 | rRF 45S | NO | 37.92 | 0.00 | 0.00 | 0.00 |
| >hg38_21_+_8433172_8446622_RNA45SN1_45S_50ntFlanks@11671.11688.18\|CCAGTGCTCT GAATGTCA | >hg38_21_+_8433172_8446622_RNA45SN1_45S_50ntFlanks | 11671 | 11688 | 18 | rRF 45S | NO | 20.17 | 29.12 | 0.00 | 0.00 |
| >hg38_21_+_8433172_8446622_RNA45SN1_45S_50ntFlanks@11671.11691.21\|CCAGTGCTCT GAATGTCAAAG | >hg38_21_+_8433172_8446622_RNA45SN1_45S_50ntFlanks | 11671 | 11691 | 21 | rRF 45S | NO | 44.22 | 0.00 | 0.00 | 0.00 |
| >hg38_21_+_8433172_8446622_RNA45SN1_45S_50ntFlanks@11671.11692.22\|CCAGTGCTCT GAATGTCAAAGT | >hg38_21_+_8433172_8446622_RNA45SN1_45S_50ntFlanks | 11671 | 11692 | 22 | rRF 45S | NO | 20.14 | 0.00 | 0.00 | 0.00 |
| >hg38_21_+_8433172_8446622_RNA45SN1_45S_50ntFlanks@11671.11693.23\|CCAGTGCTCT GAATGTCAAAGTG | >hg38_21_+_8433172_8446622_RNA45SN1_45S_50ntFlanks | 11671 | 11693 | 23 | rRF 45S | NO | 54.10 | 50.54 | 0.00 | 0.00 |
| >hg38_21_+_8433172_8446622_RNA45SN1_45S_50ntFlanks@11671.11699.29\|CCAGTGCTCT GAATGTCAAAGTGAAGAAA | >hg38_21_+_8433172_8446622_RNA45SN1_45S_50ntFlanks | 11671 | 11699 | 29 | rRF 45S | NO | 20.04 | 0.00 | 0.00 | 0.00 |
| >hg38_21_+_8433172_8446622_RNA45SN1_45S_50ntFlanks@11671.11700.30\|CCAGTGCTCT GAATGTCAAAGTGAAGAAAT | >hg38_21_+_8433172_8446622_RNA45SN1_45S_50ntFlanks | 11671 | 11700 | 30 | rRF 45S | NO | 39.27 | 0.00 | 0.00 | 0.00 |
| >hg38_21_+_8433172_8446622_RNA45SN1_45S_50ntFlanks@11671.11701.31\|CCAGTGCTCT GAATGTCAAAGTGAAGAAATT | >hg38_21_+_8433172_8446622_RNA45SN1_45S_50ntFlanks | 11671 | 11701 | 31 | rRF 45S | NO | 83.85 | 32.82 | 0.00 | 0.00 |
| >hg38_21_+_8433172_8446622_RNA45SN1_45S_50ntFlanks@11671.11702.32\|CCAGTGCTCT GAATGTCAAAGTGAAGAAATTC | >hg38_21_+_8433172_8446622_RNA45SN1_45S_50ntFlanks | 11671 | 11702 | 32 | rRF 45S | NO | 224.94 | 104.79 | 7.32 | 24.38 |
| >hg38_21_+_8433172_8446622_RNA45SN1_45S_50ntFlanks@11671.11703.33\|CCAGTGCTCT GAATGTCAAAGTGAAGAAATTCA | >hg38_21_+_8433172_8446622_RNA45SN1_45S_50ntFlanks | 11671 | 11703 | 33 | rRF 45S | NO | 90.97 | 49.69 | 0.00 | 0.00 |
| >hg38_21_+_8433172_8446622_RNA45SN1_45S_50ntFlanks@11671.11704.34\|CCAGTGCTCT GAATGTCAAAGTGAAGAAATTCAA | >hg38_21_+_8433172_8446622_RNA45SN1_45S_50ntFlanks | 11671 | 11704 | 34 | rRF 45S | NO | 56.35 | 12.62 | 0.00 | 0.00 |
| >hg38_21_+_8433172_8446622_RNA45SN1_45S_50ntFlanks@11671.11706.36\|CCAGTGCTCT GAATGTCAAAGTGAAGAAATTCAATG | >hg38_21_+_8433172_8446622_RNA45SN1_45S_50ntFlanks | 11671 | 11706 | 36 | rRF 45S | NO | 68.27 | 28.85 | 0.00 | 0.00 |
| >hg38_21_+_8433172_8446622_RNA45SN1_45S_50ntFlanks@11672.11701.30\|CAGTGCTCTG AATGTCAAAGTGAAGAAATT | >hg38_21_+_8433172_8446622_RNA45SN1_45S_50ntFlanks | 11672 | 11701 | 30 | rRF 45S | NO | 41.04 | 0.00 | 0.00 | 0.00 |
| >hg38_21_+_8433172_8446622_RNA45SN1_45S_50ntFlanks@11672.11702.31\|CAGTGCTCTG AATGTCAAAGTGAAGAAATTC | >hg38_21_+_8433172_8446622_RNA45SN1_45S_50ntFlanks | 11672 | 11702 | 31 | rRF 45S | NO | 113.38 | 49.75 | 0.00 | 0.00 |

### MDA-MB-468

| | | | | | | | | | | |
|---|---|---|---|---|---|---|---|---|---|---|
| >hg38_21_+_8433172_8446622_RNA45SN1_45S_50ntFlanks@11672.11703.32\|CAGTGCTCTG AATGTCAAAGTGAAGAAATTCA | >hg38_21_+_8433172_8446622_RNA45SN1_45S_50ntFlanks | 11672 | 11703 | 32 | rRF 45S | NO | 38.26 | 0.00 | 0.00 | 0.00 |
| >hg38_21_+_8433172_8446622_RNA45SN1_45S_50ntFlanks@11672.11704.33\|CAGTGCTCTG AATGTCAAAGTGAAGAAATTCAA | >hg38_21_+_8433172_8446622_RNA45SN1_45S_50ntFlanks | 11672 | 11704 | 33 | rRF 45S | NO | 34.54 | 0.00 | 0.00 | 0.00 |
| >hg38_21_+_8433172_8446622_RNA45SN1_45S_50ntFlanks@11672.11706.35\|CAGTGCTCTG AATGTCAAAGTGAAGAAATTCAATG | >hg38_21_+_8433172_8446622_RNA45SN1_45S_50ntFlanks | 11672 | 11706 | 35 | rRF 45S | NO | 50.26 | 12.65 | 0.00 | 0.00 |
| >hg38_21_+_8433172_8446622_RNA45SN1_45S_50ntFlanks@11677.11702.26\|CTCTGAATGT CAAAGTGAAGAAATTC | >hg38_21_+_8433172_8446622_RNA45SN1_45S_50ntFlanks | 11677 | 11702 | 26 | rRF 45S | NO | 12.91 | 45.46 | 28.27 | 0.00 |
| >hg38_21_+_8433172_8446622_RNA45SN1_45S_50ntFlanks@11682.11702.21\|AATGTCAAAG TGAAGAAATTC | >hg38_21_+_8433172_8446622_RNA45SN1_45S_50ntFlanks | 11682 | 11702 | 21 | rRF 45S | NO | 0.00 | 0.00 | 34.57 | 0.00 |
| >hg38_21_+_8433172_8446622_RNA45SN1_45S_50ntFlanks@11683.11702.20\|ATGTCAAAGT GAAGAAATTC | >hg38_21_+_8433172_8446622_RNA45SN1_45S_50ntFlanks | 11683 | 11702 | 20 | rRF 45S | NO | 0.00 | 0.00 | 30.32 | 0.00 |
| >hg38_21_+_8433172_8446622_RNA45SN1_45S_50ntFlanks@11694.11717.24\|AAGAAATTCA ATGAAGCGCGGGTA | >hg38_21_+_8433172_8446622_RNA45SN1_45S_50ntFlanks | 11694 | 11717 | 24 | rRF 45S | NO | 0.00 | 12.80 | 0.00 | 0.00 |
| >hg38_21_+_8433172_8446622_RNA45SN1_45S_50ntFlanks@11694.11728.35\|AAGAAATTCA ATGAAGCGCGGGTAAACGGCGGGAG | >hg38_21_+_8433172_8446622_RNA45SN1_45S_50ntFlanks | 11694 | 11728 | 35 | rRF 45S | NO | 19.86 | 0.00 | 0.00 | 0.00 |
| >hg38_21_+_8433172_8446622_RNA45SN1_45S_50ntFlanks@11703.11726.24\|AATGAAGCGC GGGTAAACGGCGGG | >hg38_21_+_8433172_8446622_RNA45SN1_45S_50ntFlanks | 11703 | 11726 | 24 | rRF 45S | NO | 29.70 | 0.00 | 0.00 | 0.00 |
| >hg38_21_+_8433172_8446622_RNA45SN1_45S_50ntFlanks@11703.11728.26\|AATGAAGCGC GGGTAAACGGCGGGAG | >hg38_21_+_8433172_8446622_RNA45SN1_45S_50ntFlanks | 11703 | 11728 | 26 | rRF 45S | NO | 250.08 | 159.12 | 83.74 | 93.87 |
| >hg38_21_+_8433172_8446622_RNA45SN1_45S_50ntFlanks@11703.11736.34\|AATGAAGCGC GGGTAAACGGCGGGAGTAACTATG | >hg38_21_+_8433172_8446622_RNA45SN1_45S_50ntFlanks | 11703 | 11736 | 34 | rRF 45S | NO | 12.88 | 0.00 | 0.00 | 0.00 |
| >hg38_21_+_8433172_8446622_RNA45SN1_45S_50ntFlanks@11703.11740.38\|AATGAAGCGC GGGTAAACGGCGGGAGTAACTATGACTC | >hg38_21_+_8433172_8446622_RNA45SN1_45S_50ntFlanks | 11703 | 11740 | 38 | rRF 45S | NO | 33.07 | 0.00 | 0.00 | 0.00 |
| >hg38_21_+_8433172_8446622_RNA45SN1_45S_50ntFlanks@11704.11728.25\|ATGAAGCGCG GGTAAACGGCGGGAG | >hg38_21_+_8433172_8446622_RNA45SN1_45S_50ntFlanks | 11704 | 11728 | 25 | rRF 45S | NO | 134.02 | 67.61 | 45.51 | 54.67 |
| >hg38_21_+_8433172_8446622_RNA45SN1_45S_50ntFlanks@11705.11728.24\|TGAAGCGCGG GTAAACGGCGGGAG | >hg38_21_+_8433172_8446622_RNA45SN1_45S_50ntFlanks | 11705 | 11728 | 24 | rRF 45S | NO | 101.91 | 56.83 | 28.27 | 25.43 |
| >hg38_21_+_8433172_8446622_RNA45SN1_45S_50ntFlanks@11707.11728.22\|AAGCGCGGGT AAACGGCGGGAG | >hg38_21_+_8433172_8446622_RNA45SN1_45S_50ntFlanks | 11707 | 11728 | 22 | rRF 45S | NO | 42.06 | 28.49 | 0.00 | 0.00 |
| >hg38_21_+_8433172_8446622_RNA45SN1_45S_50ntFlanks@11707.11740.34\|AAGCGCGGGT AAACGGCGGGAGTAACTATGACTC | >hg38_21_+_8433172_8446622_RNA45SN1_45S_50ntFlanks | 11707 | 11740 | 34 | rRF 45S | NO | 12.66 | 0.00 | 0.00 | 0.00 |
| >hg38_21_+_8433172_8446622_RNA45SN1_45S_50ntFlanks@11710.11728.19\|CGCGGGTAAA CGGCGGGAG | >hg38_21_+_8433172_8446622_RNA45SN1_45S_50ntFlanks | 11710 | 11728 | 19 | rRF 45S | NO | 46.61 | 52.14 | 29.68 | 0.00 |
| >hg38_21_+_8433172_8446622_RNA45SN1_45S_50ntFlanks@11710.11740.31\|CGCGGGTAAA CGGCGGGAGTAACTATGACTC | >hg38_21_+_8433172_8446622_RNA45SN1_45S_50ntFlanks | 11710 | 11740 | 31 | rRF 45S | NO | 19.94 | 0.00 | 0.00 | 0.00 |
| >hg38_21_+_8433172_8446622_RNA45SN1_45S_50ntFlanks@11718.11736.19\|AACGGCGGG AGTAACTATG | >hg38_21_+_8433172_8446622_RNA45SN1_45S_50ntFlanks | 11718 | 11736 | 19 | rRF 45S | NO | 101.22 | 50.93 | 0.00 | 0.00 |
| >hg38_21_+_8433172_8446622_RNA45SN1_45S_50ntFlanks@11718.11737.20\|AACGGCGGG AGTAACTATGA | >hg38_21_+_8433172_8446622_RNA45SN1_45S_50ntFlanks | 11718 | 11737 | 20 | rRF 45S | NO | 19.65 | 0.00 | 0.00 | 0.00 |
| >hg38_21_+_8433172_8446622_RNA45SN1_45S_50ntFlanks@11718.11738.21\|AACGGCGGG AGTAACTATGAC | >hg38_21_+_8433172_8446622_RNA45SN1_45S_50ntFlanks | 11718 | 11738 | 21 | rRF 45S | NO | 88.05 | 44.95 | 0.00 | 0.00 |
| >hg38_21_+_8433172_8446622_RNA45SN1_45S_50ntFlanks@11718.11739.22\|AACGGCGGG AGTAACTATGACT | >hg38_21_+_8433172_8446622_RNA45SN1_45S_50ntFlanks | 11718 | 11739 | 22 | rRF 45S | NO | 35.92 | 0.00 | 0.00 | 0.00 |
| >hg38_21_+_8433172_8446622_RNA45SN1_45S_50ntFlanks@11718.11740.23\|AACGGCGGG AGTAACTATGACTC | >hg38_21_+_8433172_8446622_RNA45SN1_45S_50ntFlanks | 11718 | 11740 | 23 | rRF 45S | NO | 366.65 | 120.31 | 0.00 | 24.09 |
| >hg38_21_+_8433172_8446622_RNA45SN1_45S_50ntFlanks@11718.11742.25\|AACGGCGGG AGTAACTATGACTCTC | >hg38_21_+_8433172_8446622_RNA45SN1_45S_50ntFlanks | 11718 | 11742 | 25 | rRF 45S | NO | 47.70 | 288.92 | 39.85 | 6.15 |
| >hg38_21_+_8433172_8446622_RNA45SN1_45S_50ntFlanks@11718.11743.26\|AACGGCGGG AGTAACTATGACTCTCT | >hg38_21_+_8433172_8446622_RNA45SN1_45S_50ntFlanks | 11718 | 11743 | 26 | rRF 45S | NO | 0.00 | 44.40 | 0.00 | 0.00 |
| >hg38_21_+_8433172_8446622_RNA45SN1_45S_50ntFlanks@11718.11744.27\|AACGGCGGG AGTAACTATGACTCTCTT | >hg38_21_+_8433172_8446622_RNA45SN1_45S_50ntFlanks | 11718 | 11744 | 27 | rRF 45S | NO | 0.00 | 39.36 | 34.68 | 0.00 |
| >hg38_21_+_8433172_8446622_RNA45SN1_45S_50ntFlanks@11718.11751.34\|AACGGCGGG AGTAACTATGACTCTCTTAAGGTAG | >hg38_21_+_8433172_8446622_RNA45SN1_45S_50ntFlanks | 11718 | 11751 | 34 | rRF 45S | NO | 37.13 | 58.05 | 0.00 | 0.00 |
| >hg38_21_+_8433172_8446622_RNA45SN1_45S_50ntFlanks@11719.11740.22\|ACGGCGGGA GTAACTATGACTC | >hg38_21_+_8433172_8446622_RNA45SN1_45S_50ntFlanks | 11719 | 11740 | 22 | rRF 45S | NO | 21.18 | 0.00 | 0.00 | 0.00 |
| >hg38_21_+_8433172_8446622_RNA45SN1_45S_50ntFlanks@11727.11744.18\|AGTAACTATG ACTCTCTT | >hg38_21_+_8433172_8446622_RNA45SN1_45S_50ntFlanks | 11727 | 11744 | 18 | rRF 45S | NO | 50.65 | 26.92 | 0.00 | 0.00 |
| >hg38_21_+_8433172_8446622_RNA45SN1_45S_50ntFlanks@11727.11751.25\|AGTAACTATG ACTCTCTTAAGGTAG | >hg38_21_+_8433172_8446622_RNA45SN1_45S_50ntFlanks | 11727 | 11751 | 25 | rRF 45S | NO | 91.68 | 0.00 | 0.00 | 0.00 |
| >hg38_21_+_8433172_8446622_RNA45SN1_45S_50ntFlanks@11728.11745.18\|GTAACTATGA CTCTCTTA | >hg38_21_+_8433172_8446622_RNA45SN1_45S_50ntFlanks | 11728 | 11745 | 18 | rRF 45S | NO | 21.85 | 0.00 | 0.00 | 0.00 |
| >hg38_21_+_8433172_8446622_RNA45SN1_45S_50ntFlanks@11728.11747.20\|GTAACTATGA CTCTCTTAAG | >hg38_21_+_8433172_8446622_RNA45SN1_45S_50ntFlanks | 11728 | 11747 | 20 | rRF 45S | NO | 31.33 | 0.00 | 0.00 | 0.00 |
| >hg38_21_+_8433172_8446622_RNA45SN1_45S_50ntFlanks@11728.11748.21\|GTAACTATGA CTCTCTTAAGG | >hg38_21_+_8433172_8446622_RNA45SN1_45S_50ntFlanks | 11728 | 11748 | 21 | rRF 45S | NO | 12.57 | 0.00 | 0.00 | 0.00 |
| >hg38_21_+_8433172_8446622_RNA45SN1_45S_50ntFlanks@11728.11751.24\|GTAACTATGA CTCTCTTAAGGTAG | >hg38_21_+_8433172_8446622_RNA45SN1_45S_50ntFlanks | 11728 | 11751 | 24 | rRF 45S | NO | 148.50 | 0.00 | 0.00 | 0.00 |

### MDA-MB-468

| | | | | | | | | | | |
|---|---|---|---|---|---|---|---|---|---|---|
| >hg38_21_+_8433172_8446622_RNA45SN1_45S_50ntFlanks@11729.11746.18\|TAACTATGACTCTCTTAA | >hg38_21_+_8433172_8446622_RNA45SN1_45S_50ntFlanks | 11729 | 11746 | 18 | rRF 45S | NO | 0.00 | 12.67 | 0.00 | 0.00 |
| >hg38_21_+_8433172_8446622_RNA45SN1_45S_50ntFlanks@11729.11747.19\|TAACTATGACTCTCTTAAG | >hg38_21_+_8433172_8446622_RNA45SN1_45S_50ntFlanks | 11729 | 11747 | 19 | rRF 45S | NO | 433.42 | 471.44 | 0.00 | 73.19 |
| >hg38_21_+_8433172_8446622_RNA45SN1_45S_50ntFlanks@11729.11748.20\|TAACTATGACTCTCTTAAGG | >hg38_21_+_8433172_8446622_RNA45SN1_45S_50ntFlanks | 11729 | 11748 | 20 | rRF 45S | NO | 426.68 | 371.04 | 0.00 | 75.85 |
| >hg38_21_+_8433172_8446622_RNA45SN1_45S_50ntFlanks@11729.11749.21\|TAACTATGACTCTCTTAAGGT | >hg38_21_+_8433172_8446622_RNA45SN1_45S_50ntFlanks | 11729 | 11749 | 21 | rRF 45S | NO | 446.47 | 492.13 | 34.78 | 62.60 |
| >hg38_21_+_8433172_8446622_RNA45SN1_45S_50ntFlanks@11729.11750.22\|TAACTATGACTCTCTTAAGGTA | >hg38_21_+_8433172_8446622_RNA45SN1_45S_50ntFlanks | 11729 | 11750 | 22 | rRF 45S | NO | 100.35 | 165.79 | 0.00 | 0.00 |
| >hg38_21_+_8433172_8446622_RNA45SN1_45S_50ntFlanks@11729.11751.23\|TAACTATGACTCTCTTAAGGTAG | >hg38_21_+_8433172_8446622_RNA45SN1_45S_50ntFlanks | 11729 | 11751 | 23 | rRF 45S | YES | 4381.34 | 3462.43 | 119.24 | 379.80 |
| >hg38_21_+_8433172_8446622_RNA45SN1_45S_50ntFlanks@11729.11752.24\|TAACTATGACTCTCTTAAGGTAGC | >hg38_21_+_8433172_8446622_RNA45SN1_45S_50ntFlanks | 11729 | 11752 | 24 | rRF 45S | NO | 38.03 | 86.45 | 0.00 | 0.00 |
| >hg38_21_+_8433172_8446622_RNA45SN1_45S_50ntFlanks@11729.11753.25\|TAACTATGACTCTCTTAAGGTAGCC | >hg38_21_+_8433172_8446622_RNA45SN1_45S_50ntFlanks | 11729 | 11753 | 25 | rRF 45S | NO | 0.00 | 174.03 | 60.90 | 0.00 |
| >hg38_21_+_8433172_8446622_RNA45SN1_45S_50ntFlanks@11729.11761.33\|TAACTATGACTCTCTTAAGGTAGCCAAATGCCT | >hg38_21_+_8433172_8446622_RNA45SN1_45S_50ntFlanks | 11729 | 11761 | 33 | rRF 45S | NO | 0.00 | 42.18 | 0.00 | 0.00 |
| >hg38_21_+_8433172_8446622_RNA45SN1_45S_50ntFlanks@11729.11764.36\|TAACTATGACTCTCTTAAGGTAGCCAAATGCCTCGT | >hg38_21_+_8433172_8446622_RNA45SN1_45S_50ntFlanks | 11729 | 11764 | 36 | rRF 45S | NO | 29.23 | 127.70 | 0.00 | 0.00 |
| >hg38_21_+_8433172_8446622_RNA45SN1_45S_50ntFlanks@11729.11765.37\|TAACTATGACTCTCTTAAGGTAGCCAAATGCCTCGTC | >hg38_21_+_8433172_8446622_RNA45SN1_45S_50ntFlanks | 11729 | 11765 | 37 | rRF 45S | NO | 421.04 | 711.98 | 0.00 | 75.81 |
| >hg38_21_+_8433172_8446622_RNA45SN1_45S_50ntFlanks@11729.11766.38\|TAACTATGACTCTCTTAAGGTAGCCAAATGCCTCGTCA | >hg38_21_+_8433172_8446622_RNA45SN1_45S_50ntFlanks | 11729 | 11766 | 38 | rRF 45S | NO | 0.00 | 48.94 | 0.00 | 0.00 |
| >hg38_21_+_8433172_8446622_RNA45SN1_45S_50ntFlanks@11729.11767.39\|TAACTATGACTCTCTTAAGGTAGCCAAATGCCTCGTCAT | >hg38_21_+_8433172_8446622_RNA45SN1_45S_50ntFlanks | 11729 | 11767 | 39 | rRF 45S | NO | 101.72 | 326.24 | 35.08 | 49.62 |
| >hg38_21_+_8433172_8446622_RNA45SN1_45S_50ntFlanks@11731.11751.21\|ACTATGACTCTCTTAAGGTAG | >hg38_21_+_8433172_8446622_RNA45SN1_45S_50ntFlanks | 11731 | 11751 | 21 | rRF 45S | NO | 32.55 | 0.00 | 0.00 | 0.00 |
| >hg38_21_+_8433172_8446622_RNA45SN1_45S_50ntFlanks@11733.11751.19\|TATGACTCTCTTAAGGTAG | >hg38_21_+_8433172_8446622_RNA45SN1_45S_50ntFlanks | 11733 | 11751 | 19 | rRF 45S | NO | 12.97 | 0.00 | 0.00 | 0.00 |
| >hg38_21_+_8433172_8446622_RNA45SN1_45S_50ntFlanks@11739.11758.20\|TCTCTTAAGGTAGCCAAATG | >hg38_21_+_8433172_8446622_RNA45SN1_45S_50ntFlanks | 11739 | 11758 | 20 | rRF 45S | NO | 0.00 | 34.06 | 0.00 | 0.00 |
| >hg38_21_+_8433172_8446622_RNA45SN1_45S_50ntFlanks@11741.11758.18\|TCTTAAGGTAGCCAAATG | >hg38_21_+_8433172_8446622_RNA45SN1_45S_50ntFlanks | 11741 | 11758 | 18 | rRF 45S | NO | 0.00 | 26.94 | 0.00 | 0.00 |
| >hg38_21_+_8433172_8446622_RNA45SN1_45S_50ntFlanks@11752.11780.29\|CCAAATGCCTCGTCATCTAATTAGTGACG | >hg38_21_+_8433172_8446622_RNA45SN1_45S_50ntFlanks | 11752 | 11780 | 29 | rRF 45S | NO | 0.00 | 41.11 | 0.00 | 0.00 |
| >hg38_21_+_8433172_8446622_RNA45SN1_45S_50ntFlanks@11759.11777.19\|CCTCGTCATCTAATTAGTG | >hg38_21_+_8433172_8446622_RNA45SN1_45S_50ntFlanks | 11759 | 11777 | 19 | rRF 45S | NO | 0.00 | 38.63 | 0.00 | 0.00 |
| >hg38_21_+_8433172_8446622_RNA45SN1_45S_50ntFlanks@11766.11790.25\|ATCTAATTAGTGACGCGCATGAATG | >hg38_21_+_8433172_8446622_RNA45SN1_45S_50ntFlanks | 11766 | 11790 | 25 | rRF 45S | NO | 33.86 | 38.48 | 0.00 | 0.00 |
| >hg38_21_+_8433172_8446622_RNA45SN1_45S_50ntFlanks@11766.11791.26\|ATCTAATTAGTGACGCGCATGAATGG | >hg38_21_+_8433172_8446622_RNA45SN1_45S_50ntFlanks | 11766 | 11791 | 26 | rRF 45S | NO | 41.28 | 26.90 | 0.00 | 0.00 |
| >hg38_21_+_8433172_8446622_RNA45SN1_45S_50ntFlanks@11766.11803.38\|ATCTAATTAGTGACGCGCATGAATGGATGAACGAGATT | >hg38_21_+_8433172_8446622_RNA45SN1_45S_50ntFlanks | 11766 | 11803 | 38 | rRF 45S | NO | 33.34 | 12.58 | 0.00 | 0.00 |
| >hg38_21_+_8433172_8446622_RNA45SN1_45S_50ntFlanks@11767.11790.24\|TCTAATTAGTGACGCGCATGAATG | >hg38_21_+_8433172_8446622_RNA45SN1_45S_50ntFlanks | 11767 | 11790 | 24 | rRF 45S | NO | 13.18 | 31.20 | 0.00 | 0.00 |
| >hg38_21_+_8433172_8446622_RNA45SN1_45S_50ntFlanks@11767.11791.25\|TCTAATTAGTGACGCGCATGAATGG | >hg38_21_+_8433172_8446622_RNA45SN1_45S_50ntFlanks | 11767 | 11791 | 25 | rRF 45S | NO | 21.32 | 0.00 | 0.00 | 0.00 |
| >hg38_21_+_8433172_8446622_RNA45SN1_45S_50ntFlanks@11767.11794.28\|TCTAATTAGTGACGCGCATGAATGGATG | >hg38_21_+_8433172_8446622_RNA45SN1_45S_50ntFlanks | 11767 | 11794 | 28 | rRF 45S | NO | 185.88 | 104.55 | 6.63 | 0.00 |
| >hg38_21_+_8433172_8446622_RNA45SN1_45S_50ntFlanks@11767.11803.37\|TCTAATTAGTGACGCGCATGAATGGATGAACGAGATT | >hg38_21_+_8433172_8446622_RNA45SN1_45S_50ntFlanks | 11767 | 11803 | 37 | rRF 45S | NO | 52.94 | 39.04 | 0.00 | 0.00 |
| >hg38_21_+_8433172_8446622_RNA45SN1_45S_50ntFlanks@11768.11786.19\|CTAATTAGTGACGCGCATG | >hg38_21_+_8433172_8446622_RNA45SN1_45S_50ntFlanks | 11768 | 11786 | 19 | rRF 45S | NO | 50.61 | 97.23 | 0.00 | 6.63 |
| >hg38_21_+_8433172_8446622_RNA45SN1_45S_50ntFlanks@11768.11790.23\|CTAATTAGTGACGCGCATGAATG | >hg38_21_+_8433172_8446622_RNA45SN1_45S_50ntFlanks | 11768 | 11790 | 23 | rRF 45S | NO | 129.53 | 190.46 | 0.00 | 0.00 |
| >hg38_21_+_8433172_8446622_RNA45SN1_45S_50ntFlanks@11768.11791.24\|CTAATTAGTGACGCGCATGAATGG | >hg38_21_+_8433172_8446622_RNA45SN1_45S_50ntFlanks | 11768 | 11791 | 24 | rRF 45S | NO | 79.01 | 63.74 | 0.00 | 0.00 |
| >hg38_21_+_8433172_8446622_RNA45SN1_45S_50ntFlanks@11768.11792.25\|CTAATTAGTGACGCGCATGAATGGA | >hg38_21_+_8433172_8446622_RNA45SN1_45S_50ntFlanks | 11768 | 11792 | 25 | rRF 45S | NO | 234.32 | 203.61 | 31.42 | 14.84 |
| >hg38_21_+_8433172_8446622_RNA45SN1_45S_50ntFlanks@11768.11794.27\|CTAATTAGTGACGCGCATGAATGGATG | >hg38_21_+_8433172_8446622_RNA45SN1_45S_50ntFlanks | 11768 | 11794 | 27 | rRF 45S | NO | 439.05 | 245.96 | 71.10 | 35.97 |
| >hg38_21_+_8433172_8446622_RNA45SN1_45S_50ntFlanks@11768.11803.36\|CTAATTAGTGACGCGCATGAATGGATGAACGAGATT | >hg38_21_+_8433172_8446622_RNA45SN1_45S_50ntFlanks | 11768 | 11803 | 36 | rRF 45S | NO | 145.74 | 119.41 | 30.99 | 16.41 |
| >hg38_21_+_8433172_8446622_RNA45SN1_45S_50ntFlanks@11768.11804.37\|CTAATTAGTGACGCGCATGAATGGATGAACGAGATTC | >hg38_21_+_8433172_8446622_RNA45SN1_45S_50ntFlanks | 11768 | 11804 | 37 | rRF 45S | NO | 59.80 | 42.50 | 0.00 | 6.03 |
| >hg38_21_+_8433172_8446622_RNA45SN1_45S_50ntFlanks@11768.11806.39\|CTAATTAGTGACGCGCATGAATGGATGAACGAGATTCCC | >hg38_21_+_8433172_8446622_RNA45SN1_45S_50ntFlanks | 11768 | 11806 | 39 | rRF 45S | NO | 20.82 | 0.00 | 0.00 | 0.00 |

### MDA-MB-468

| | | | | | | | | | | |
|---|---|---|---|---|---|---|---|---|---|---|
| >hg38_21_+_8433172_8446622_RNA45SN1_45S_50ntFlanks@11769.11786.18\|TAATTAGTGACGCGCATG | >hg38_21_+_8433172_8446622_RNA45SN1_45S_50ntFlanks | 11769 | 11786 | 18 | rRF 45S | NO | 0.00 | 46.62 | 0.00 | 0.00 |
| >hg38_21_+_8433172_8446622_RNA45SN1_45S_50ntFlanks@11769.11790.22\|TAATTAGTGACGCGCATGAATG | >hg38_21_+_8433172_8446622_RNA45SN1_45S_50ntFlanks | 11769 | 11790 | 22 | rRF 45S | NO | 48.80 | 84.34 | 0.00 | 0.00 |
| >hg38_21_+_8433172_8446622_RNA45SN1_45S_50ntFlanks@11769.11791.23\|TAATTAGTGACGCGCATGAATGG | >hg38_21_+_8433172_8446622_RNA45SN1_45S_50ntFlanks | 11769 | 11791 | 23 | rRF 45S | NO | 129.14 | 95.35 | 0.00 | 0.00 |
| >hg38_21_+_8433172_8446622_RNA45SN1_45S_50ntFlanks@11769.11792.24\|TAATTAGTGACGCGCATGAATGGA | >hg38_21_+_8433172_8446622_RNA45SN1_45S_50ntFlanks | 11769 | 11792 | 24 | rRF 45S | NO | 102.46 | 69.19 | 0.00 | 0.00 |
| >hg38_21_+_8433172_8446622_RNA45SN1_45S_50ntFlanks@11769.11794.26\|TAATTAGTGACGCGCATGAATGGATG | >hg38_21_+_8433172_8446622_RNA45SN1_45S_50ntFlanks | 11769 | 11794 | 26 | rRF 45S | NO | 180.74 | 102.90 | 0.00 | 0.00 |
| >hg38_21_+_8433172_8446622_RNA45SN1_45S_50ntFlanks@11769.11803.35\|TAATTAGTGACGCGCATGAATGGATGAACGAGATT | >hg38_21_+_8433172_8446622_RNA45SN1_45S_50ntFlanks | 11769 | 11803 | 35 | rRF 45S | NO | 62.83 | 66.57 | 0.00 | 0.00 |
| >hg38_21_+_8433172_8446622_RNA45SN1_45S_50ntFlanks@11769.11804.36\|TAATTAGTGACGCGCATGAATGGATGAACGAGATTC | >hg38_21_+_8433172_8446622_RNA45SN1_45S_50ntFlanks | 11769 | 11804 | 36 | rRF 45S | NO | 69.68 | 61.77 | 0.00 | 0.00 |
| >hg38_21_+_8433172_8446622_RNA45SN1_45S_50ntFlanks@11770.11794.25\|AATTAGTGACGCGCATGAATGGATG | >hg38_21_+_8433172_8446622_RNA45SN1_45S_50ntFlanks | 11770 | 11794 | 25 | rRF 45S | NO | 36.66 | 0.00 | 0.00 | 0.00 |
| >hg38_21_+_8433172_8446622_RNA45SN1_45S_50ntFlanks@11771.11791.21\|ATTAGTGACGCGCATGAATGG | >hg38_21_+_8433172_8446622_RNA45SN1_45S_50ntFlanks | 11771 | 11791 | 21 | rRF 45S | NO | 20.39 | 0.00 | 0.00 | 0.00 |
| >hg38_21_+_8433172_8446622_RNA45SN1_45S_50ntFlanks@11771.11794.24\|ATTAGTGACGCGCATGAATGGATG | >hg38_21_+_8433172_8446622_RNA45SN1_45S_50ntFlanks | 11771 | 11794 | 24 | rRF 45S | NO | 67.61 | 0.00 | 0.00 | 0.00 |
| >hg38_21_+_8433172_8446622_RNA45SN1_45S_50ntFlanks@11772.11794.23\|TTAGTGACGCGCATGAATGGATG | >hg38_21_+_8433172_8446622_RNA45SN1_45S_50ntFlanks | 11772 | 11794 | 23 | rRF 45S | NO | 52.19 | 0.00 | 0.00 | 0.00 |
| >hg38_21_+_8433172_8446622_RNA45SN1_45S_50ntFlanks@11778.11803.26\|ACGCGCATGAATGGATGAACGAGATT | >hg38_21_+_8433172_8446622_RNA45SN1_45S_50ntFlanks | 11778 | 11803 | 26 | rRF 45S | NO | 0.00 | 30.16 | 40.78 | 0.00 |
| >hg38_21_+_8433172_8446622_RNA45SN1_45S_50ntFlanks@11778.11804.27\|ACGCGCATGAATGGATGAACGAGATTC | >hg38_21_+_8433172_8446622_RNA45SN1_45S_50ntFlanks | 11778 | 11804 | 27 | rRF 45S | NO | 0.00 | 0.00 | 48.09 | 0.00 |
| >hg38_21_+_8433172_8446622_RNA45SN1_45S_50ntFlanks@11781.11798.18\|CGCATGAATGGATGAACG | >hg38_21_+_8433172_8446622_RNA45SN1_45S_50ntFlanks | 11781 | 11798 | 18 | rRF 45S | NO | 54.74 | 68.20 | 29.69 | 0.00 |
| >hg38_21_+_8433172_8446622_RNA45SN1_45S_50ntFlanks@11781.11803.23\|CGCATGAATGGATGAACGAGATT | >hg38_21_+_8433172_8446622_RNA45SN1_45S_50ntFlanks | 11781 | 11803 | 23 | rRF 45S | NO | 203.55 | 74.35 | 66.46 | 0.00 |
| >hg38_21_+_8433172_8446622_RNA45SN1_45S_50ntFlanks@11781.11804.24\|CGCATGAATGGATGAACGAGATTC | >hg38_21_+_8433172_8446622_RNA45SN1_45S_50ntFlanks | 11781 | 11804 | 24 | rRF 45S | NO | 428.03 | 163.59 | 146.02 | 53.71 |
| >hg38_21_+_8433172_8446622_RNA45SN1_45S_50ntFlanks@11781.11805.25\|CGCATGAATGGATGAACGAGATTCC | >hg38_21_+_8433172_8446622_RNA45SN1_45S_50ntFlanks | 11781 | 11805 | 25 | rRF 45S | NO | 61.28 | 41.24 | 0.00 | 0.00 |
| >hg38_21_+_8433172_8446622_RNA45SN1_45S_50ntFlanks@11781.11806.26\|CGCATGAATGGATGAACGAGATTCCC | >hg38_21_+_8433172_8446622_RNA45SN1_45S_50ntFlanks | 11781 | 11806 | 26 | rRF 45S | NO | 61.23 | 53.55 | 34.51 | 34.26 |
| >hg38_21_+_8433172_8446622_RNA45SN1_45S_50ntFlanks@11782.11804.23\|GCATGAATGGATGAACGAGATTC | >hg38_21_+_8433172_8446622_RNA45SN1_45S_50ntFlanks | 11782 | 11804 | 23 | rRF 45S | NO | 32.02 | 0.00 | 0.00 | 0.00 |
| >hg38_21_+_8433172_8446622_RNA45SN1_45S_50ntFlanks@11783.11800.18\|CATGAATGGATGAACGAG | >hg38_21_+_8433172_8446622_RNA45SN1_45S_50ntFlanks | 11783 | 11800 | 18 | rRF 45S | NO | 63.08 | 32.48 | 34.21 | 0.00 |
| >hg38_21_+_8433172_8446622_RNA45SN1_45S_50ntFlanks@11783.11802.20\|CATGAATGGATGAACGAGAT | >hg38_21_+_8433172_8446622_RNA45SN1_45S_50ntFlanks | 11783 | 11802 | 20 | rRF 45S | NO | 31.34 | 0.00 | 30.64 | 0.00 |
| >hg38_21_+_8433172_8446622_RNA45SN1_45S_50ntFlanks@11783.11803.21\|CATGAATGGATGAACGAGATT | >hg38_21_+_8433172_8446622_RNA45SN1_45S_50ntFlanks | 11783 | 11803 | 21 | rRF 45S | YES | 1006.52 | 260.40 | 184.58 | 49.32 |
| >hg38_21_+_8433172_8446622_RNA45SN1_45S_50ntFlanks@11783.11804.22\|CATGAATGGATGAACGAGATTC | >hg38_21_+_8433172_8446622_RNA45SN1_45S_50ntFlanks | 11783 | 11804 | 22 | rRF 45S | YES | 2104.51 | 1312.18 | 523.81 | 224.63 |
| >hg38_21_+_8433172_8446622_RNA45SN1_45S_50ntFlanks@11783.11805.23\|CATGAATGGATGAACGAGATTCC | >hg38_21_+_8433172_8446622_RNA45SN1_45S_50ntFlanks | 11783 | 11805 | 23 | rRF 45S | NO | 128.50 | 138.57 | 65.47 | 38.88 |
| >hg38_21_+_8433172_8446622_RNA45SN1_45S_50ntFlanks@11783.11806.24\|CATGAATGGATGAACGAGATTCCC | >hg38_21_+_8433172_8446622_RNA45SN1_45S_50ntFlanks | 11783 | 11806 | 24 | rRF 45S | NO | 103.04 | 133.14 | 115.16 | 88.67 |
| >hg38_21_+_8433172_8446622_RNA45SN1_45S_50ntFlanks@11784.11803.20\|ATGAATGGATGAACGAGATT | >hg38_21_+_8433172_8446622_RNA45SN1_45S_50ntFlanks | 11784 | 11803 | 20 | rRF 45S | NO | 307.05 | 86.09 | 65.10 | 0.00 |
| >hg38_21_+_8433172_8446622_RNA45SN1_45S_50ntFlanks@11784.11804.21\|ATGAATGGATGAACGAGATTC | >hg38_21_+_8433172_8446622_RNA45SN1_45S_50ntFlanks | 11784 | 11804 | 21 | rRF 45S | NO | 613.49 | 353.94 | 193.75 | 48.83 |
| >hg38_21_+_8433172_8446622_RNA45SN1_45S_50ntFlanks@11784.11805.22\|ATGAATGGATGAACGAGATTCC | >hg38_21_+_8433172_8446622_RNA45SN1_45S_50ntFlanks | 11784 | 11805 | 22 | rRF 45S | NO | 36.43 | 38.20 | 29.75 | 0.00 |
| >hg38_21_+_8433172_8446622_RNA45SN1_45S_50ntFlanks@11784.11806.23\|ATGAATGGATGAACGAGATTCCC | >hg38_21_+_8433172_8446622_RNA45SN1_45S_50ntFlanks | 11784 | 11806 | 23 | rRF 45S | NO | 29.39 | 39.23 | 55.31 | 13.64 |
| >hg38_21_+_8433172_8446622_RNA45SN1_45S_50ntFlanks@11785.11803.19\|TGAATGGATGAACGAGATT | >hg38_21_+_8433172_8446622_RNA45SN1_45S_50ntFlanks | 11785 | 11803 | 19 | rRF 45S | NO | 396.66 | 103.62 | 64.56 | 0.00 |
| >hg38_21_+_8433172_8446622_RNA45SN1_45S_50ntFlanks@11785.11804.20\|TGAATGGATGAACGAGATTC | >hg38_21_+_8433172_8446622_RNA45SN1_45S_50ntFlanks | 11785 | 11804 | 20 | rRF 45S | YES | 777.48 | 473.14 | 198.58 | 55.99 |
| >hg38_21_+_8433172_8446622_RNA45SN1_45S_50ntFlanks@11785.11805.21\|TGAATGGATGAACGAGATTCC | >hg38_21_+_8433172_8446622_RNA45SN1_45S_50ntFlanks | 11785 | 11805 | 21 | rRF 45S | NO | 34.75 | 49.61 | 31.20 | 0.00 |
| >hg38_21_+_8433172_8446622_RNA45SN1_45S_50ntFlanks@11785.11806.22\|TGAATGGATGAACGAGATTCCC | >hg38_21_+_8433172_8446622_RNA45SN1_45S_50ntFlanks | 11785 | 11806 | 22 | rRF 45S | NO | 0.00 | 36.26 | 44.99 | 0.00 |
| >hg38_21_+_8433172_8446622_RNA45SN1_45S_50ntFlanks@11786.11803.18\|GAATGGATGAACGAGATT | >hg38_21_+_8433172_8446622_RNA45SN1_45S_50ntFlanks | 11786 | 11803 | 18 | rRF 45S | NO | 120.21 | 31.94 | 41.67 | 0.00 |

### MDA-MB-468

| | | | | | | | | | | |
|---|---|---|---|---|---|---|---|---|---|---|
| >hg38_21_+_8433172_8446622_RNA45SN1_45S_50ntFlanks@11786.11804.19\|GAATGGATGA ACGAGATTC | >hg38_21_+_8433172_8446622_RNA45SN1_45S_50ntFlanks | 11786 | 11804 | 19 | rRF 45S | NO | 76.48 | 56.16 | 64.85 | 0.00 |
| >hg38_21_+_8433172_8446622_RNA45SN1_45S_50ntFlanks@11787.11804.18\|AATGGATGAA CGAGATTC | >hg38_21_+_8433172_8446622_RNA45SN1_45S_50ntFlanks | 11787 | 11804 | 18 | rRF 45S | YES | 456.70 | 252.72 | 302.14 | 56.03 |
| >hg38_21_+_8433172_8446622_RNA45SN1_45S_50ntFlanks@11804.11821.18\|CCCACTGTCC CTACCTAC | >hg38_21_+_8433172_8446622_RNA45SN1_45S_50ntFlanks | 11804 | 11821 | 18 | rRF 45S | NO | 32.46 | 0.00 | 0.00 | 0.00 |
| >hg38_21_+_8433172_8446622_RNA45SN1_45S_50ntFlanks@11821.11844.24\|CTATCCAGCG AAACCACAGCCAAG | >hg38_21_+_8433172_8446622_RNA45SN1_45S_50ntFlanks | 11821 | 11844 | 24 | rRF 45S | NO | 12.63 | 0.00 | 27.32 | 6.53 |
| >hg38_21_+_8433172_8446622_RNA45SN1_45S_50ntFlanks@11843.11860.18\|AGGGAACGG GCTTGGCGG | >hg38_21_+_8433172_8446622_RNA45SN1_45S_50ntFlanks | 11843 | 11860 | 18 | rRF 45S | NO | 0.00 | 0.00 | 34.01 | 0.00 |
| >hg38_21_+_8433172_8446622_RNA45SN1_45S_50ntFlanks@11843.11872.30\|AGGGAACGG GCTTGGCGGAATCAGCGGGGA | >hg38_21_+_8433172_8446622_RNA45SN1_45S_50ntFlanks | 11843 | 11872 | 30 | rRF 45S | NO | 0.00 | 0.00 | 56.39 | 0.00 |
| >hg38_21_+_8433172_8446622_RNA45SN1_45S_50ntFlanks@11844.11872.29\|GGGAACGGG CTTGGCGGAATCAGCGGGGA | >hg38_21_+_8433172_8446622_RNA45SN1_45S_50ntFlanks | 11844 | 11872 | 29 | rRF 45S | NO | 0.00 | 0.00 | 48.52 | 0.00 |
| >hg38_21_+_8433172_8446622_RNA45SN1_45S_50ntFlanks@11845.11871.27\|GGAACGGGCT TGGCGGAATCAGCGGGG | >hg38_21_+_8433172_8446622_RNA45SN1_45S_50ntFlanks | 11845 | 11871 | 27 | rRF 45S | NO | 0.00 | 0.00 | 33.44 | 0.00 |
| >hg38_21_+_8433172_8446622_RNA45SN1_45S_50ntFlanks@11845.11872.28\|GGAACGGGCT TGGCGGAATCAGCGGGGA | >hg38_21_+_8433172_8446622_RNA45SN1_45S_50ntFlanks | 11845 | 11872 | 28 | rRF 45S | NO | 33.92 | 0.00 | 114.04 | 26.15 |
| >hg38_21_+_8433172_8446622_RNA45SN1_45S_50ntFlanks@11845.11873.29\|GGAACGGGCT TGGCGGAATCAGCGGGGAA | >hg38_21_+_8433172_8446622_RNA45SN1_45S_50ntFlanks | 11845 | 11873 | 29 | rRF 45S | NO | 0.00 | 0.00 | 42.78 | 0.00 |
| >hg38_21_+_8433172_8446622_RNA45SN1_45S_50ntFlanks@11846.11872.27\|GAACGGGCTT GGCGGAATCAGCGGGGA | >hg38_21_+_8433172_8446622_RNA45SN1_45S_50ntFlanks | 11846 | 11872 | 27 | rRF 45S | NO | 0.00 | 0.00 | 20.40 | 0.00 |
| >hg38_21_+_8433172_8446622_RNA45SN1_45S_50ntFlanks@11847.11871.25\|AACGGGCTTG GCGGAATCAGCGGGG | >hg38_21_+_8433172_8446622_RNA45SN1_45S_50ntFlanks | 11847 | 11871 | 25 | rRF 45S | NO | 0.00 | 0.00 | 13.69 | 0.00 |
| >hg38_21_+_8433172_8446622_RNA45SN1_45S_50ntFlanks@11847.11872.26\|AACGGGCTTG GCGGAATCAGCGGGGA | >hg38_21_+_8433172_8446622_RNA45SN1_45S_50ntFlanks | 11847 | 11872 | 26 | rRF 45S | NO | 37.64 | 0.00 | 77.81 | 24.90 |
| >hg38_21_+_8433172_8446622_RNA45SN1_45S_50ntFlanks@11847.11873.27\|AACGGGCTTG GCGGAATCAGCGGGGAA | >hg38_21_+_8433172_8446622_RNA45SN1_45S_50ntFlanks | 11847 | 11873 | 27 | rRF 45S | NO | 0.00 | 0.00 | 13.72 | 0.00 |
| >hg38_21_+_8433172_8446622_RNA45SN1_45S_50ntFlanks@11848.11872.25\|ACGGGCTTGG CGGAATCAGCGGGGA | >hg38_21_+_8433172_8446622_RNA45SN1_45S_50ntFlanks | 11848 | 11872 | 25 | rRF 45S | NO | 6.15 | 0.00 | 50.14 | 0.00 |
| >hg38_21_+_8433172_8446622_RNA45SN1_45S_50ntFlanks@11849.11872.24\|CGGGCTTGGC GGAATCAGCGGGGA | >hg38_21_+_8433172_8446622_RNA45SN1_45S_50ntFlanks | 11849 | 11872 | 24 | rRF 45S | NO | 47.95 | 27.93 | 72.87 | 37.18 |
| >hg38_21_+_8433172_8446622_RNA45SN1_45S_50ntFlanks@11872.11893.22\|AAAGAAGACC CTGTTGAGCTTG | >hg38_21_+_8433172_8446622_RNA45SN1_45S_50ntFlanks | 11872 | 11893 | 22 | rRF 45S | NO | 0.00 | 12.81 | 0.00 | 0.00 |
| >hg38_21_+_8433172_8446622_RNA45SN1_45S_50ntFlanks@11873.11890.18\|AAGAAGACCC TGTTGAGC | >hg38_21_+_8433172_8446622_RNA45SN1_45S_50ntFlanks | 11873 | 11890 | 18 | rRF 45S | NO | 101.71 | 75.10 | 52.25 | 13.34 |
| >hg38_21_+_8433172_8446622_RNA45SN1_45S_50ntFlanks@11873.11892.20\|AAGAAGACCC TGTTGAGCTT | >hg38_21_+_8433172_8446622_RNA45SN1_45S_50ntFlanks | 11873 | 11892 | 20 | rRF 45S | NO | 49.41 | 50.44 | 35.34 | 6.25 |
| >hg38_21_+_8433172_8446622_RNA45SN1_45S_50ntFlanks@11873.11893.21\|AAGAAGACCC TGTTGAGCTTG | >hg38_21_+_8433172_8446622_RNA45SN1_45S_50ntFlanks | 11873 | 11893 | 21 | rRF 45S | NO | 223.22 | 393.49 | 198.02 | 70.51 |
| >hg38_21_+_8433172_8446622_RNA45SN1_45S_50ntFlanks@11873.11897.25\|AAGAAGACCC TGTTGAGCTTGACTC | >hg38_21_+_8433172_8446622_RNA45SN1_45S_50ntFlanks | 11873 | 11897 | 25 | rRF 45S | NO | 51.87 | 62.24 | 0.00 | 6.48 |
| >hg38_21_+_8433172_8446622_RNA45SN1_45S_50ntFlanks@11873.11898.26\|AAGAAGACCC TGTTGAGCTTGACTCT | >hg38_21_+_8433172_8446622_RNA45SN1_45S_50ntFlanks | 11873 | 11898 | 26 | rRF 45S | NO | 114.51 | 111.23 | 31.00 | 41.56 |
| >hg38_21_+_8433172_8446622_RNA45SN1_45S_50ntFlanks@11873.11899.27\|AAGAAGACCC TGTTGAGCTTGACTCTA | >hg38_21_+_8433172_8446622_RNA45SN1_45S_50ntFlanks | 11873 | 11899 | 27 | rRF 45S | NO | 36.21 | 34.11 | 0.00 | 0.00 |
| >hg38_21_+_8433172_8446622_RNA45SN1_45S_50ntFlanks@11873.11900.28\|AAGAAGACCC TGTTGAGCTTGACTCTAG | >hg38_21_+_8433172_8446622_RNA45SN1_45S_50ntFlanks | 11873 | 11900 | 28 | rRF 45S | NO | 80.65 | 85.68 | 28.17 | 23.61 |
| >hg38_21_+_8433172_8446622_RNA45SN1_45S_50ntFlanks@11873.11902.30\|AAGAAGACCC TGTTGAGCTTGACTCTAGTC | >hg38_21_+_8433172_8446622_RNA45SN1_45S_50ntFlanks | 11873 | 11902 | 30 | rRF 45S | NO | 0.00 | 30.42 | 0.00 | 0.00 |
| >hg38_21_+_8433172_8446622_RNA45SN1_45S_50ntFlanks@11873.11903.31\|AAGAAGACCC TGTTGAGCTTGACTCTAGTCT | >hg38_21_+_8433172_8446622_RNA45SN1_45S_50ntFlanks | 11873 | 11903 | 31 | rRF 45S | NO | 0.00 | 0.00 | 13.60 | 15.09 |
| >hg38_21_+_8433172_8446622_RNA45SN1_45S_50ntFlanks@11873.11904.32\|AAGAAGACCC TGTTGAGCTTGACTCTAGTCTG | >hg38_21_+_8433172_8446622_RNA45SN1_45S_50ntFlanks | 11873 | 11904 | 32 | rRF 45S | NO | 0.00 | 31.90 | 106.99 | 37.21 |
| >hg38_21_+_8433172_8446622_RNA45SN1_45S_50ntFlanks@11873.11905.33\|AAGAAGACCC TGTTGAGCTTGACTCTAGTCTGG | >hg38_21_+_8433172_8446622_RNA45SN1_45S_50ntFlanks | 11873 | 11905 | 33 | rRF 45S | NO | 0.00 | 29.48 | 54.90 | 50.08 |
| >hg38_21_+_8433172_8446622_RNA45SN1_45S_50ntFlanks@11873.11906.34\|AAGAAGACCC TGTTGAGCTTGACTCTAGTCTGGC | >hg38_21_+_8433172_8446622_RNA45SN1_45S_50ntFlanks | 11873 | 11906 | 34 | rRF 45S | NO | 44.84 | 104.95 | 42.30 | 90.59 |
| >hg38_21_+_8433172_8446622_RNA45SN1_45S_50ntFlanks@11873.11907.35\|AAGAAGACCC TGTTGAGCTTGACTCTAGTCTGGCA | >hg38_21_+_8433172_8446622_RNA45SN1_45S_50ntFlanks | 11873 | 11907 | 35 | rRF 45S | NO | 0.00 | 30.88 | 0.00 | 25.69 |
| >hg38_21_+_8433172_8446622_RNA45SN1_45S_50ntFlanks@11873.11908.36\|AAGAAGACCC TGTTGAGCTTGACTCTAGTCTGGCAC | >hg38_21_+_8433172_8446622_RNA45SN1_45S_50ntFlanks | 11873 | 11908 | 36 | rRF 45S | NO | 0.00 | 29.56 | 0.00 | 40.24 |
| >hg38_21_+_8433172_8446622_RNA45SN1_45S_50ntFlanks@11874.11892.19\|AGAAGACCCT GTTGAGCTT | >hg38_21_+_8433172_8446622_RNA45SN1_45S_50ntFlanks | 11874 | 11892 | 19 | rRF 45S | NO | 31.64 | 44.70 | 29.84 | 0.00 |
| >hg38_21_+_8433172_8446622_RNA45SN1_45S_50ntFlanks@11874.11893.20\|AGAAGACCCT GTTGAGCTTG | >hg38_21_+_8433172_8446622_RNA45SN1_45S_50ntFlanks | 11874 | 11893 | 20 | rRF 45S | NO | 243.30 | 477.06 | 180.09 | 85.16 |
| >hg38_21_+_8433172_8446622_RNA45SN1_45S_50ntFlanks@11874.11897.24\|AGAAGACCCT GTTGAGCTTGACTC | >hg38_21_+_8433172_8446622_RNA45SN1_45S_50ntFlanks | 11874 | 11897 | 24 | rRF 45S | NO | 64.91 | 80.60 | 0.00 | 16.29 |

### MDA-MB-468

| | | | | | | | | | | |
|---|---|---|---|---|---|---|---|---|---|---|
| >hg38_21_+_8433172_8446622_RNA45SN1_45S_50ntFlanks@11874.11898.25\|AGAAGACCCTGTTGAGCTTGACTCT | >hg38_21_+_8433172_8446622_RNA45SN1_45S_50ntFlanks | 11874 | 11898 | 25 | rRF 45S | NO | 159.77 | 188.85 | 43.10 | 70.58 |
| >hg38_21_+_8433172_8446622_RNA45SN1_45S_50ntFlanks@11874.11899.26\|AGAAGACCCTGTTGAGCTTGACTCTA | >hg38_21_+_8433172_8446622_RNA45SN1_45S_50ntFlanks | 11874 | 11899 | 26 | rRF 45S | NO | 41.16 | 50.37 | 0.00 | 16.18 |
| >hg38_21_+_8433172_8446622_RNA45SN1_45S_50ntFlanks@11874.11900.27\|AGAAGACCCTGTTGAGCTTGACTCTAG | >hg38_21_+_8433172_8446622_RNA45SN1_45S_50ntFlanks | 11874 | 11900 | 27 | rRF 45S | NO | 64.79 | 92.66 | 20.37 | 33.94 |
| >hg38_21_+_8433172_8446622_RNA45SN1_45S_50ntFlanks@11874.11902.29\|AGAAGACCCTGTTGAGCTTGACTCTAGTC | >hg38_21_+_8433172_8446622_RNA45SN1_45S_50ntFlanks | 11874 | 11902 | 29 | rRF 45S | NO | 20.36 | 35.10 | 0.00 | 0.00 |
| >hg38_21_+_8433172_8446622_RNA45SN1_45S_50ntFlanks@11874.11903.30\|AGAAGACCCTGTTGAGCTTGACTCTAGTCT | >hg38_21_+_8433172_8446622_RNA45SN1_45S_50ntFlanks | 11874 | 11903 | 30 | rRF 45S | NO | 0.00 | 0.00 | 13.82 | 16.60 |
| >hg38_21_+_8433172_8446622_RNA45SN1_45S_50ntFlanks@11874.11904.31\|AGAAGACCCTGTTGAGCTTGACTCTAGTCTG | >hg38_21_+_8433172_8446622_RNA45SN1_45S_50ntFlanks | 11874 | 11904 | 31 | rRF 45S | NO | 0.00 | 5.97 | 103.33 | 39.83 |
| >hg38_21_+_8433172_8446622_RNA45SN1_45S_50ntFlanks@11874.11905.32\|AGAAGACCCTGTTGAGCTTGACTCTAGTCTGG | >hg38_21_+_8433172_8446622_RNA45SN1_45S_50ntFlanks | 11874 | 11905 | 32 | rRF 45S | NO | 0.00 | 0.00 | 56.71 | 52.11 |
| >hg38_21_+_8433172_8446622_RNA45SN1_45S_50ntFlanks@11874.11906.33\|AGAAGACCCTGTTGAGCTTGACTCTAGTCTGGC | >hg38_21_+_8433172_8446622_RNA45SN1_45S_50ntFlanks | 11874 | 11906 | 33 | rRF 45S | NO | 46.91 | 116.77 | 44.61 | 125.47 |
| >hg38_21_+_8433172_8446622_RNA45SN1_45S_50ntFlanks@11874.11907.34\|AGAAGACCCTGTTGAGCTTGACTCTAGTCTGGCA | >hg38_21_+_8433172_8446622_RNA45SN1_45S_50ntFlanks | 11874 | 11907 | 34 | rRF 45S | NO | 0.00 | 31.35 | 0.00 | 44.14 |
| >hg38_21_+_8433172_8446622_RNA45SN1_45S_50ntFlanks@11874.11908.35\|AGAAGACCCTGTTGAGCTTGACTCTAGTCTGGCAC | >hg38_21_+_8433172_8446622_RNA45SN1_45S_50ntFlanks | 11874 | 11908 | 35 | rRF 45S | NO | 20.33 | 42.13 | 0.00 | 60.76 |
| >hg38_21_+_8433172_8446622_RNA45SN1_45S_50ntFlanks@11874.11909.36\|AGAAGACCCTGTTGAGCTTGACTCTAGTCTGGCACG | >hg38_21_+_8433172_8446622_RNA45SN1_45S_50ntFlanks | 11874 | 11909 | 36 | rRF 45S | NO | 0.00 | 27.61 | 0.00 | 44.26 |
| >hg38_21_+_8433172_8446622_RNA45SN1_45S_50ntFlanks@11875.11893.19\|GAAGACCCTGTTGAGCTTG | >hg38_21_+_8433172_8446622_RNA45SN1_45S_50ntFlanks | 11875 | 11893 | 19 | rRF 45S | NO | 46.15 | 94.91 | 36.30 | 0.00 |
| >hg38_21_+_8433172_8446622_RNA45SN1_45S_50ntFlanks@11875.11898.24\|GAAGACCCTGTTGAGCTTGACTCT | >hg38_21_+_8433172_8446622_RNA45SN1_45S_50ntFlanks | 11875 | 11898 | 24 | rRF 45S | NO | 12.93 | 27.13 | 0.00 | 0.00 |
| >hg38_21_+_8433172_8446622_RNA45SN1_45S_50ntFlanks@11876.11893.18\|AAGACCCTGTTGAGCTTG | >hg38_21_+_8433172_8446622_RNA45SN1_45S_50ntFlanks | 11876 | 11893 | 18 | rRF 45S | NO | 90.04 | 347.23 | 112.42 | 42.55 |
| >hg38_21_+_8433172_8446622_RNA45SN1_45S_50ntFlanks@11876.11897.22\|AAGACCCTGTTGAGCTTGACTC | >hg38_21_+_8433172_8446622_RNA45SN1_45S_50ntFlanks | 11876 | 11897 | 22 | rRF 45S | NO | 38.63 | 63.25 | 0.00 | 0.00 |
| >hg38_21_+_8433172_8446622_RNA45SN1_45S_50ntFlanks@11876.11898.23\|AAGACCCTGTTGAGCTTGACTCT | >hg38_21_+_8433172_8446622_RNA45SN1_45S_50ntFlanks | 11876 | 11898 | 23 | rRF 45S | NO | 84.58 | 84.68 | 0.00 | 36.19 |
| >hg38_21_+_8433172_8446622_RNA45SN1_45S_50ntFlanks@11876.11899.24\|AAGACCCTGTTGAGCTTGACTCTA | >hg38_21_+_8433172_8446622_RNA45SN1_45S_50ntFlanks | 11876 | 11899 | 24 | rRF 45S | NO | 36.53 | 36.11 | 0.00 | 6.12 |
| >hg38_21_+_8433172_8446622_RNA45SN1_45S_50ntFlanks@11876.11900.25\|AAGACCCTGTTGAGCTTGACTCTAG | >hg38_21_+_8433172_8446622_RNA45SN1_45S_50ntFlanks | 11876 | 11900 | 25 | rRF 45S | NO | 35.20 | 40.00 | 0.00 | 0.00 |
| >hg38_21_+_8433172_8446622_RNA45SN1_45S_50ntFlanks@11876.11904.29\|AAGACCCTGTTGAGCTTGACTCTAGTCTG | >hg38_21_+_8433172_8446622_RNA45SN1_45S_50ntFlanks | 11876 | 11904 | 29 | rRF 45S | NO | 0.00 | 0.00 | 38.67 | 0.00 |
| >hg38_21_+_8433172_8446622_RNA45SN1_45S_50ntFlanks@11876.11905.30\|AAGACCCTGTTGAGCTTGACTCTAGTCTGG | >hg38_21_+_8433172_8446622_RNA45SN1_45S_50ntFlanks | 11876 | 11905 | 30 | rRF 45S | NO | 0.00 | 0.00 | 33.14 | 24.48 |
| >hg38_21_+_8433172_8446622_RNA45SN1_45S_50ntFlanks@11876.11906.31\|AAGACCCTGTTGAGCTTGACTCTAGTCTGGC | >hg38_21_+_8433172_8446622_RNA45SN1_45S_50ntFlanks | 11876 | 11906 | 31 | rRF 45S | NO | 0.00 | 30.61 | 0.00 | 44.30 |
| >hg38_21_+_8433172_8446622_RNA45SN1_45S_50ntFlanks@11881.11903.23\|CCTGTTGAGCTTGACTCTAGTCT | >hg38_21_+_8433172_8446622_RNA45SN1_45S_50ntFlanks | 11881 | 11903 | 23 | rRF 45S | NO | 0.00 | 42.53 | 0.00 | 0.00 |
| >hg38_21_+_8433172_8446622_RNA45SN1_45S_50ntFlanks@11881.11904.24\|CCTGTTGAGCTTGACTCTAGTCTG | >hg38_21_+_8433172_8446622_RNA45SN1_45S_50ntFlanks | 11881 | 11904 | 24 | rRF 45S | NO | 71.21 | 140.07 | 74.77 | 46.67 |
| >hg38_21_+_8433172_8446622_RNA45SN1_45S_50ntFlanks@11881.11906.26\|CCTGTTGAGCTTGACTCTAGTCTGGC | >hg38_21_+_8433172_8446622_RNA45SN1_45S_50ntFlanks | 11881 | 11906 | 26 | rRF 45S | NO | 0.00 | 44.96 | 0.00 | 0.00 |
| >hg38_21_+_8433172_8446622_RNA45SN1_45S_50ntFlanks@11882.11904.23\|CTGTTGAGCTTGACTCTAGTCTG | >hg38_21_+_8433172_8446622_RNA45SN1_45S_50ntFlanks | 11882 | 11904 | 23 | rRF 45S | NO | 35.20 | 90.30 | 46.19 | 34.77 |
| >hg38_21_+_8433172_8446622_RNA45SN1_45S_50ntFlanks@11885.11903.19\|TTGAGCTTGACTCTAGTCT | >hg38_21_+_8433172_8446622_RNA45SN1_45S_50ntFlanks | 11885 | 11903 | 19 | rRF 45S | NO | 0.00 | 52.50 | 0.00 | 0.00 |
| >hg38_21_+_8433172_8446622_RNA45SN1_45S_50ntFlanks@11885.11904.20\|TTGAGCTTGACTCTAGTCTG | >hg38_21_+_8433172_8446622_RNA45SN1_45S_50ntFlanks | 11885 | 11904 | 20 | rRF 45S | NO | 21.35 | 194.37 | 79.21 | 46.81 |
| >hg38_21_+_8433172_8446622_RNA45SN1_45S_50ntFlanks@11885.11905.21\|TTGAGCTTGACTCTAGTCTGG | >hg38_21_+_8433172_8446622_RNA45SN1_45S_50ntFlanks | 11885 | 11905 | 21 | rRF 45S | NO | 32.40 | 88.72 | 28.27 | 0.00 |
| >hg38_21_+_8433172_8446622_RNA45SN1_45S_50ntFlanks@11885.11912.28\|TTGAGCTTGACTCTAGTCTGGCACGGTG | >hg38_21_+_8433172_8446622_RNA45SN1_45S_50ntFlanks | 11885 | 11912 | 28 | rRF 45S | NO | 0.00 | 0.00 | 20.30 | 0.00 |
| >hg38_21_+_8433172_8446622_RNA45SN1_45S_50ntFlanks@11886.11903.18\|TGAGCTTGACTCTAGTCT | >hg38_21_+_8433172_8446622_RNA45SN1_45S_50ntFlanks | 11886 | 11903 | 18 | rRF 45S | NO | 0.00 | 26.49 | 0.00 | 0.00 |
| >hg38_21_+_8433172_8446622_RNA45SN1_45S_50ntFlanks@11886.11904.19\|TGAGCTTGACTCTAGTCTG | >hg38_21_+_8433172_8446622_RNA45SN1_45S_50ntFlanks | 11886 | 11904 | 19 | rRF 45S | NO | 0.00 | 108.33 | 46.41 | 23.87 |
| >hg38_21_+_8433172_8446622_RNA45SN1_45S_50ntFlanks@11886.11905.20\|TGAGCTTGACTCTAGTCTGG | >hg38_21_+_8433172_8446622_RNA45SN1_45S_50ntFlanks | 11886 | 11905 | 20 | rRF 45S | NO | 0.00 | 46.25 | 0.00 | 0.00 |
| >hg38_21_+_8433172_8446622_RNA45SN1_45S_50ntFlanks@11890.11907.18\|CTTGACTCTAGTCTGGCA | >hg38_21_+_8433172_8446622_RNA45SN1_45S_50ntFlanks | 11890 | 11907 | 18 | rRF 45S | NO | 0.00 | 108.20 | 58.45 | 15.82 |
| >hg38_21_+_8433172_8446622_RNA45SN1_45S_50ntFlanks@11890.11912.23\|CTTGACTCTAGTCTGGCACGGTG | >hg38_21_+_8433172_8446622_RNA45SN1_45S_50ntFlanks | 11890 | 11912 | 23 | rRF 45S | NO | 12.83 | 147.84 | 45.30 | 25.35 |

### MDA-MB-468

| | | | | | | | | | | |
|---|---|---|---|---|---|---|---|---|---|---|
| >hg38_21_+_8433172_8446622_RNA45SN1_45S_50ntFlanks@11891.11909.19\|TTGACTCTAG TCTGGCACG | >hg38_21_+_8433172_8446622_RNA45SN1_45S_50ntFlanks | 11891 | 11909 | 19 | rRF 45S | NO | 0.00 | 32.56 | 0.00 | 0.00 |
| >hg38_21_+_8433172_8446622_RNA45SN1_45S_50ntFlanks@11891.11910.20\|TTGACTCTAG TCTGGCACGG | >hg38_21_+_8433172_8446622_RNA45SN1_45S_50ntFlanks | 11891 | 11910 | 20 | rRF 45S | NO | 0.00 | 41.24 | 0.00 | 0.00 |
| >hg38_21_+_8433172_8446622_RNA45SN1_45S_50ntFlanks@11891.11912.22\|TTGACTCTAG TCTGGCACGGTG | >hg38_21_+_8433172_8446622_RNA45SN1_45S_50ntFlanks | 11891 | 11912 | 22 | rRF 45S | NO | 84.71 | 199.73 | 28.59 | 25.89 |
| >hg38_21_+_8433172_8446622_RNA45SN1_45S_50ntFlanks@11894.11912.19\|ACTCTAGTCT GGCACGGTG | >hg38_21_+_8433172_8446622_RNA45SN1_45S_50ntFlanks | 11894 | 11912 | 19 | rRF 45S | NO | 84.36 | 152.21 | 0.00 | 0.00 |
| >hg38_21_+_8433172_8446622_RNA45SN1_45S_50ntFlanks@11894.11916.23\|ACTCTAGTCT GGCACGGTGAAGA | >hg38_21_+_8433172_8446622_RNA45SN1_45S_50ntFlanks | 11894 | 11916 | 23 | rRF 45S | NO | 46.24 | 0.00 | 0.00 | 0.00 |
| >hg38_21_+_8433172_8446622_RNA45SN1_45S_50ntFlanks@11918.11948.31\|ACATGAGAGG TGTAGAATAAGTGGGAGGCCC | >hg38_21_+_8433172_8446622_RNA45SN1_45S_50ntFlanks | 11918 | 11948 | 31 | rRF 45S | NO | 0.00 | 63.15 | 58.16 | 0.00 |
| >hg38_21_+_8433172_8446622_RNA45SN1_45S_50ntFlanks@11918.11949.32\|ACATGAGAGG TGTAGAATAAGTGGGAGGCCCC | >hg38_21_+_8433172_8446622_RNA45SN1_45S_50ntFlanks | 11918 | 11949 | 32 | rRF 45S | NO | 0.00 | 45.89 | 0.00 | 0.00 |
| >hg38_21_+_8433172_8446622_RNA45SN1_45S_50ntFlanks@11918.11950.33\|ACATGAGAGG TGTAGAATAAGTGGGAGGCCCCC | >hg38_21_+_8433172_8446622_RNA45SN1_45S_50ntFlanks | 11918 | 11950 | 33 | rRF 45S | NO | 0.00 | 152.34 | 7.12 | 0.00 |
| >hg38_21_+_8433172_8446622_RNA45SN1_45S_50ntFlanks@11918.11951.34\|ACATGAGAGG TGTAGAATAAGTGGGAGGCCCCCG | >hg38_21_+_8433172_8446622_RNA45SN1_45S_50ntFlanks | 11918 | 11951 | 34 | rRF 45S | NO | 0.00 | 142.31 | 27.70 | 0.00 |
| >hg38_21_+_8433172_8446622_RNA45SN1_45S_50ntFlanks@11918.11952.35\|ACATGAGAGG TGTAGAATAAGTGGGAGGCCCCCGG | >hg38_21_+_8433172_8446622_RNA45SN1_45S_50ntFlanks | 11918 | 11952 | 35 | rRF 45S | NO | 0.00 | 38.05 | 0.00 | 0.00 |
| >hg38_21_+_8433172_8446622_RNA45SN1_45S_50ntFlanks@11918.11954.37\|ACATGAGAGG TGTAGAATAAGTGGGAGGCCCCCGGCG | >hg38_21_+_8433172_8446622_RNA45SN1_45S_50ntFlanks | 11918 | 11954 | 37 | rRF 45S | NO | 0.00 | 26.55 | 0.00 | 0.00 |
| >hg38_21_+_8433172_8446622_RNA45SN1_45S_50ntFlanks@11919.11950.32\|CATGAGAGGT GTAGAATAAGTGGGAGGCCCCC | >hg38_21_+_8433172_8446622_RNA45SN1_45S_50ntFlanks | 11919 | 11950 | 32 | rRF 45S | NO | 0.00 | 27.32 | 0.00 | 0.00 |
| >hg38_21_+_8433172_8446622_RNA45SN1_45S_50ntFlanks@11919.11951.33\|CATGAGAGGT GTAGAATAAGTGGGAGGCCCCCG | >hg38_21_+_8433172_8446622_RNA45SN1_45S_50ntFlanks | 11919 | 11951 | 33 | rRF 45S | NO | 0.00 | 30.08 | 0.00 | 0.00 |
| >hg38_21_+_8433172_8446622_RNA45SN1_45S_50ntFlanks@11920.11948.29\|ATGAGAGGTG TAGAATAAGTGGGAGGCCC | >hg38_21_+_8433172_8446622_RNA45SN1_45S_50ntFlanks | 11920 | 11948 | 29 | rRF 45S | NO | 0.00 | 26.38 | 61.67 | 0.00 |
| >hg38_21_+_8433172_8446622_RNA45SN1_45S_50ntFlanks@11920.11950.31\|ATGAGAGGTG TAGAATAAGTGGGAGGCCCCC | >hg38_21_+_8433172_8446622_RNA45SN1_45S_50ntFlanks | 11920 | 11950 | 31 | rRF 45S | NO | 0.00 | 42.22 | 7.33 | 0.00 |
| >hg38_21_+_8433172_8446622_RNA45SN1_45S_50ntFlanks@11920.11951.32\|ATGAGAGGTG TAGAATAAGTGGGAGGCCCCCG | >hg38_21_+_8433172_8446622_RNA45SN1_45S_50ntFlanks | 11920 | 11951 | 32 | rRF 45S | NO | 0.00 | 46.30 | 28.94 | 0.00 |
| >hg38_21_+_8433172_8446622_RNA45SN1_45S_50ntFlanks@11921.11948.28\|TGAGAGGTGT AGAATAAGTGGGAGGCCC | >hg38_21_+_8433172_8446622_RNA45SN1_45S_50ntFlanks | 11921 | 11948 | 28 | rRF 45S | NO | 0.00 | 0.00 | 50.53 | 0.00 |
| >hg38_21_+_8433172_8446622_RNA45SN1_45S_50ntFlanks@11921.11950.30\|TGAGAGGTGT AGAATAAGTGGGAGGCCCCC | >hg38_21_+_8433172_8446622_RNA45SN1_45S_50ntFlanks | 11921 | 11950 | 30 | rRF 45S | NO | 0.00 | 35.48 | 0.00 | 0.00 |
| >hg38_21_+_8433172_8446622_RNA45SN1_45S_50ntFlanks@11921.11951.31\|TGAGAGGTGT AGAATAAGTGGGAGGCCCCCG | >hg38_21_+_8433172_8446622_RNA45SN1_45S_50ntFlanks | 11921 | 11951 | 31 | rRF 45S | NO | 0.00 | 38.44 | 0.00 | 0.00 |
| >hg38_21_+_8433172_8446622_RNA45SN1_45S_50ntFlanks@11998.12032.35\|GCCCTGCGG GCCGCCGGTGAAATACCACTACTCTG | >hg38_21_+_8433172_8446622_RNA45SN1_45S_50ntFlanks | 11998 | 12032 | 35 | rRF 45S | NO | 31.35 | 0.00 | 0.00 | 6.30 |
| >hg38_21_+_8433172_8446622_RNA45SN1_45S_50ntFlanks@11999.12029.31\|CCCTGCGGG CCGCCGGTGAAATACCACTACT | >hg38_21_+_8433172_8446622_RNA45SN1_45S_50ntFlanks | 11999 | 12029 | 31 | rRF 45S | NO | 47.55 | 0.00 | 0.00 | 0.00 |
| >hg38_21_+_8433172_8446622_RNA45SN1_45S_50ntFlanks@11999.12030.32\|CCCTGCGGG CCGCCGGTGAAATACCACTACTC | >hg38_21_+_8433172_8446622_RNA45SN1_45S_50ntFlanks | 11999 | 12030 | 32 | rRF 45S | NO | 113.60 | 38.70 | 0.00 | 13.67 |
| >hg38_21_+_8433172_8446622_RNA45SN1_45S_50ntFlanks@11999.12031.33\|CCCTGCGGG CCGCCGGTGAAATACCACTACTCT | >hg38_21_+_8433172_8446622_RNA45SN1_45S_50ntFlanks | 11999 | 12031 | 33 | rRF 45S | NO | 224.98 | 127.94 | 0.00 | 78.78 |
| >hg38_21_+_8433172_8446622_RNA45SN1_45S_50ntFlanks@11999.12032.34\|CCCTGCGGG CCGCCGGTGAAATACCACTACTCTG | >hg38_21_+_8433172_8446622_RNA45SN1_45S_50ntFlanks | 11999 | 12032 | 34 | rRF 45S | YES | 1435.70 | 998.23 | 67.31 | 412.66 |
| >hg38_21_+_8433172_8446622_RNA45SN1_45S_50ntFlanks@11999.12033.35\|CCCTGCGGG CCGCCGGTGAAATACCACTACTCTGA | >hg38_21_+_8433172_8446622_RNA45SN1_45S_50ntFlanks | 11999 | 12033 | 35 | rRF 45S | NO | 55.41 | 35.34 | 0.00 | 38.70 |
| >hg38_21_+_8433172_8446622_RNA45SN1_45S_50ntFlanks@11999.12035.37\|CCCTGCGGG CCGCCGGTGAAATACCACTACTCTGATC | >hg38_21_+_8433172_8446622_RNA45SN1_45S_50ntFlanks | 11999 | 12035 | 37 | rRF 45S | NO | 30.96 | 0.00 | 0.00 | 36.87 |
| >hg38_21_+_8433172_8446622_RNA45SN1_45S_50ntFlanks@11999.12043.45\|CCCTGCGGG CCGCCGGTGAAATACCACTACTCTGATCGTTTTTTC | >hg38_21_+_8433172_8446622_RNA45SN1_45S_50ntFlanks | 11999 | 12043 | 45 | rRF 45S | NO | 96.18 | 34.11 | 0.00 | 103.28 |
| >hg38_21_+_8433172_8446622_RNA45SN1_45S_50ntFlanks@12000.12032.33\|CCTGCGGGC CGCCGGTGAAATACCACTACTCTG | >hg38_21_+_8433172_8446622_RNA45SN1_45S_50ntFlanks | 12000 | 12032 | 33 | rRF 45S | NO | 87.68 | 52.58 | 0.00 | 25.43 |
| >hg38_21_+_8433172_8446622_RNA45SN1_45S_50ntFlanks@12001.12030.30\|CTGCGGGCC GCCGGTGAAATACCACTACTC | >hg38_21_+_8433172_8446622_RNA45SN1_45S_50ntFlanks | 12001 | 12030 | 30 | rRF 45S | NO | 46.26 | 0.00 | 0.00 | 0.00 |
| >hg38_21_+_8433172_8446622_RNA45SN1_45S_50ntFlanks@12001.12031.31\|CTGCGGGCC GCCGGTGAAATACCACTACTCT | >hg38_21_+_8433172_8446622_RNA45SN1_45S_50ntFlanks | 12001 | 12031 | 31 | rRF 45S | NO | 48.25 | 47.79 | 0.00 | 15.14 |
| >hg38_21_+_8433172_8446622_RNA45SN1_45S_50ntFlanks@12001.12032.32\|CTGCGGGCC GCCGGTGAAATACCACTACTCTG | >hg38_21_+_8433172_8446622_RNA45SN1_45S_50ntFlanks | 12001 | 12032 | 32 | rRF 45S | NO | 272.80 | 315.09 | 63.65 | 131.46 |
| >hg38_21_+_8433172_8446622_RNA45SN1_45S_50ntFlanks@12002.12029.28\|TGCGGGCCG CCGGTGAAATACCACTACT | >hg38_21_+_8433172_8446622_RNA45SN1_45S_50ntFlanks | 12002 | 12029 | 28 | rRF 45S | NO | 19.66 | 0.00 | 0.00 | 0.00 |
| >hg38_21_+_8433172_8446622_RNA45SN1_45S_50ntFlanks@12002.12030.29\|TGCGGGCCG CCGGTGAAATACCACTACTC | >hg38_21_+_8433172_8446622_RNA45SN1_45S_50ntFlanks | 12002 | 12030 | 29 | rRF 45S | NO | 63.60 | 43.05 | 0.00 | 6.54 |
| >hg38_21_+_8433172_8446622_RNA45SN1_45S_50ntFlanks@12002.12031.30\|TGCGGGCCG CCGGTGAAATACCACTACTCT | >hg38_21_+_8433172_8446622_RNA45SN1_45S_50ntFlanks | 12002 | 12031 | 30 | rRF 45S | NO | 64.95 | 92.60 | 0.00 | 43.13 |

### MDA-MB-468

| | | | | | | | | | | |
|---|---|---|---|---|---|---|---|---|---|---|
| >hg38_21_+_8433172_8446622_RNA45SN1_45S_50ntFlanks@12002.12032.31\|TGCGGGCCGCCGGTGAAATACCACTACTCTG | >hg38_21_+_8433172_8446622_RNA45SN1_45S_50ntFlanks | 12002 | 12032 | 31 | rRF 45S | NO | 340.15 | 474.04 | 107.17 | 180.79 |
| >hg38_21_+_8433172_8446622_RNA45SN1_45S_50ntFlanks@12002.12043.42\|TGCGGGCCGCCGGTGAAATACCACTACTCTGATCGTTTTTTC | >hg38_21_+_8433172_8446622_RNA45SN1_45S_50ntFlanks | 12002 | 12043 | 42 | rRF 45S | NO | 101.48 | 34.49 | 0.00 | 74.48 |
| >hg38_21_+_8433172_8446622_RNA45SN1_45S_50ntFlanks@12003.12030.28\|GCGGGCCGCCGGTGAAATACCACTACTC | >hg38_21_+_8433172_8446622_RNA45SN1_45S_50ntFlanks | 12003 | 12030 | 28 | rRF 45S | NO | 41.07 | 0.00 | 0.00 | 13.63 |
| >hg38_21_+_8433172_8446622_RNA45SN1_45S_50ntFlanks@12003.12031.29\|GCGGGCCGCCGGTGAAATACCACTACTCT | >hg38_21_+_8433172_8446622_RNA45SN1_45S_50ntFlanks | 12003 | 12031 | 29 | rRF 45S | NO | 35.70 | 47.65 | 6.81 | 41.25 |
| >hg38_21_+_8433172_8446622_RNA45SN1_45S_50ntFlanks@12003.12032.30\|GCGGGCCGCCGGTGAAATACCACTACTCTG | >hg38_21_+_8433172_8446622_RNA45SN1_45S_50ntFlanks | 12003 | 12032 | 30 | rRF 45S | NO | 176.99 | 271.30 | 167.38 | 159.35 |
| >hg38_21_+_8433172_8446622_RNA45SN1_45S_50ntFlanks@12003.12043.41\|GCGGGCCGCCGGTGAAATACCACTACTCTGATCGTTTTTTC | >hg38_21_+_8433172_8446622_RNA45SN1_45S_50ntFlanks | 12003 | 12043 | 41 | rRF 45S | YES | 54.27 | 0.00 | 0.00 | 53.74 |
| >hg38_21_+_8433172_8446622_RNA45SN1_45S_50ntFlanks@12004.12032.29\|CGGGCCGCCGGTGAAATACCACTACTCTG | >hg38_21_+_8433172_8446622_RNA45SN1_45S_50ntFlanks | 12004 | 12032 | 29 | rRF 45S | NO | 0.00 | 26.32 | 60.63 | 16.19 |
| >hg38_21_+_8433172_8446622_RNA45SN1_45S_50ntFlanks@12006.12032.27\|GGCCGCCGGTGAAATACCACTACTCTG | >hg38_21_+_8433172_8446622_RNA45SN1_45S_50ntFlanks | 12006 | 12032 | 27 | rRF 45S | NO | 40.67 | 37.41 | 7.30 | 37.43 |
| >hg38_21_+_8433172_8446622_RNA45SN1_45S_50ntFlanks@12007.12029.23\|GCCGCCGGTGAAATACCACTACT | >hg38_21_+_8433172_8446622_RNA45SN1_45S_50ntFlanks | 12007 | 12029 | 23 | rRF 45S | NO | 0.00 | 0.00 | 32.46 | 115.35 |
| >hg38_21_+_8433172_8446622_RNA45SN1_45S_50ntFlanks@12007.12030.24\|GCCGCCGGTGAAATACCACTACTC | >hg38_21_+_8433172_8446622_RNA45SN1_45S_50ntFlanks | 12007 | 12030 | 24 | rRF 45S | NO | 31.05 | 0.00 | 6.82 | 119.62 |
| >hg38_21_+_8433172_8446622_RNA45SN1_45S_50ntFlanks@12007.12031.25\|GCCGCCGGTGAAATACCACTACTCT | >hg38_21_+_8433172_8446622_RNA45SN1_45S_50ntFlanks | 12007 | 12031 | 25 | rRF 45S | NO | 79.35 | 51.82 | 38.68 | 193.56 |
| >hg38_21_+_8433172_8446622_RNA45SN1_45S_50ntFlanks@12007.12032.26\|GCCGCCGGTGAAATACCACTACTCTG | >hg38_21_+_8433172_8446622_RNA45SN1_45S_50ntFlanks | 12007 | 12032 | 26 | rRF 45S | NO | 359.56 | 278.03 | 314.69 | 430.73 |
| >hg38_21_+_8433172_8446622_RNA45SN1_45S_50ntFlanks@12007.12043.37\|GCCGCCGGTGAAATACCACTACTCTGATCGTTTTTTC | >hg38_21_+_8433172_8446622_RNA45SN1_45S_50ntFlanks | 12007 | 12043 | 37 | rRF 45S | NO | 143.13 | 58.99 | 0.00 | 233.92 |
| >hg38_21_+_8433172_8446622_RNA45SN1_45S_50ntFlanks@12008.12029.22\|CCGCCGGTGAAATACCACTACT | >hg38_21_+_8433172_8446622_RNA45SN1_45S_50ntFlanks | 12008 | 12029 | 22 | rRF 45S | NO | 0.00 | 0.00 | 95.99 | 67.69 |
| >hg38_21_+_8433172_8446622_RNA45SN1_45S_50ntFlanks@12008.12030.23\|CCGCCGGTGAAATACCACTACTC | >hg38_21_+_8433172_8446622_RNA45SN1_45S_50ntFlanks | 12008 | 12030 | 23 | rRF 45S | NO | 37.64 | 89.10 | 190.42 | 203.31 |
| >hg38_21_+_8433172_8446622_RNA45SN1_45S_50ntFlanks@12008.12031.24\|CCGCCGGTGAAATACCACTACTCT | >hg38_21_+_8433172_8446622_RNA45SN1_45S_50ntFlanks | 12008 | 12031 | 24 | rRF 45S | NO | 73.77 | 208.10 | 242.36 | 248.86 |
| >hg38_21_+_8433172_8446622_RNA45SN1_45S_50ntFlanks@12008.12032.25\|CCGCCGGTGAAATACCACTACTCTG | >hg38_21_+_8433172_8446622_RNA45SN1_45S_50ntFlanks | 12008 | 12032 | 25 | rRF 45S | YES | 608.85 | 1296.90 | 1174.96 | 932.85 |
| >hg38_21_+_8433172_8446622_RNA45SN1_45S_50ntFlanks@12008.12033.26\|CCGCCGGTGAAATACCACTACTCTGA | >hg38_21_+_8433172_8446622_RNA45SN1_45S_50ntFlanks | 12008 | 12033 | 26 | rRF 45S | NO | 30.35 | 59.77 | 31.89 | 40.65 |
| >hg38_21_+_8433172_8446622_RNA45SN1_45S_50ntFlanks@12008.12034.27\|CCGCCGGTGAAATACCACTACTCTGAT | >hg38_21_+_8433172_8446622_RNA45SN1_45S_50ntFlanks | 12008 | 12034 | 27 | rRF 45S | NO | 0.00 | 55.23 | 0.00 | 13.70 |
| >hg38_21_+_8433172_8446622_RNA45SN1_45S_50ntFlanks@12008.12035.28\|CCGCCGGTGAAATACCACTACTCTGATC | >hg38_21_+_8433172_8446622_RNA45SN1_45S_50ntFlanks | 12008 | 12035 | 28 | rRF 45S | NO | 0.00 | 0.00 | 33.69 | 13.91 |
| >hg38_21_+_8433172_8446622_RNA45SN1_45S_50ntFlanks@12008.12037.30\|CCGCCGGTGAAATACCACTACTCTGATCGT | >hg38_21_+_8433172_8446622_RNA45SN1_45S_50ntFlanks | 12008 | 12037 | 30 | rRF 45S | NO | 12.77 | 0.00 | 0.00 | 13.46 |
| >hg38_21_+_8433172_8446622_RNA45SN1_45S_50ntFlanks@12008.12039.32\|CCGCCGGTGAAATACCACTACTCTGATCGTTT | >hg38_21_+_8433172_8446622_RNA45SN1_45S_50ntFlanks | 12008 | 12039 | 32 | rRF 45S | NO | 35.69 | 35.95 | 0.00 | 24.01 |
| >hg38_21_+_8433172_8446622_RNA45SN1_45S_50ntFlanks@12008.12041.34\|CCGCCGGTGAAATACCACTACTCTGATCGTTTTT | >hg38_21_+_8433172_8446622_RNA45SN1_45S_50ntFlanks | 12008 | 12041 | 34 | rRF 45S | NO | 37.30 | 12.74 | 0.00 | 34.22 |
| >hg38_21_+_8433172_8446622_RNA45SN1_45S_50ntFlanks@12008.12042.35\|CCGCCGGTGAAATACCACTACTCTGATCGTTTTTT | >hg38_21_+_8433172_8446622_RNA45SN1_45S_50ntFlanks | 12008 | 12042 | 35 | rRF 45S | NO | 51.81 | 43.04 | 0.00 | 61.65 |
| >hg38_21_+_8433172_8446622_RNA45SN1_45S_50ntFlanks@12008.12043.36\|CCGCCGGTGAAATACCACTACTCTGATCGTTTTTTC | >hg38_21_+_8433172_8446622_RNA45SN1_45S_50ntFlanks | 12008 | 12043 | 36 | rRF 45S | NO | 482.15 | 234.05 | 58.58 | 503.72 |
| >hg38_21_+_8433172_8446622_RNA45SN1_45S_50ntFlanks@12033.12053.21\|ATCGTTTTTTCACTGACCCGG | >hg38_21_+_8433172_8446622_RNA45SN1_45S_50ntFlanks | 12033 | 12053 | 21 | rRF 45S | NO | 0.00 | 37.26 | 13.82 | 60.76 |
| >hg38_21_+_8433172_8446622_RNA45SN1_45S_50ntFlanks@12044.12068.25\|ACTGACCCGGTGAGGCGGGGGGGCG | >hg38_21_+_8433172_8446622_RNA45SN1_45S_50ntFlanks | 12044 | 12068 | 25 | rRF 45S | NO | 50.82 | 6.03 | 38.79 | 69.43 |
| >hg38_21_+_8433172_8446622_RNA45SN1_45S_50ntFlanks@12044.12070.27\|ACTGACCCGGTGAGGCGGGGGGGCGAG | >hg38_21_+_8433172_8446622_RNA45SN1_45S_50ntFlanks | 12044 | 12070 | 27 | rRF 45S | NO | 96.94 | 29.39 | 40.78 | 88.23 |
| >hg38_21_+_8433172_8446622_RNA45SN1_45S_50ntFlanks@12044.12071.28\|ACTGACCCGGTGAGGCGGGGGGGCGAGC | >hg38_21_+_8433172_8446622_RNA45SN1_45S_50ntFlanks | 12044 | 12071 | 28 | rRF 45S | NO | 45.65 | 0.00 | 30.53 | 65.13 |
| >hg38_21_+_8433172_8446622_RNA45SN1_45S_50ntFlanks@12044.12072.29\|ACTGACCCGGTGAGGCGGGGGGGCGAGCC | >hg38_21_+_8433172_8446622_RNA45SN1_45S_50ntFlanks | 12044 | 12072 | 29 | rRF 45S | NO | 60.94 | 0.00 | 20.43 | 83.18 |
| >hg38_21_+_8433172_8446622_RNA45SN1_45S_50ntFlanks@12045.12070.26\|CTGACCCGGTGAGGCGGGGGGGCGAG | >hg38_21_+_8433172_8446622_RNA45SN1_45S_50ntFlanks | 12045 | 12070 | 26 | rRF 45S | NO | 34.39 | 0.00 | 0.00 | 16.02 |
| >hg38_21_+_8433172_8446622_RNA45SN1_45S_50ntFlanks@12046.12070.25\|TGACCCGGTGAGGCGGGGGGGCGAG | >hg38_21_+_8433172_8446622_RNA45SN1_45S_50ntFlanks | 12046 | 12070 | 25 | rRF 45S | NO | 12.96 | 0.00 | 0.00 | 0.00 |
| >hg38_21_+_8433172_8446622_RNA45SN1_45S_50ntFlanks@12053.12070.18\|GTGAGGCGGGGGGGCGAG | >hg38_21_+_8433172_8446622_RNA45SN1_45S_50ntFlanks | 12053 | 12070 | 18 | rRF 45S | NO | 13.21 | 0.00 | 0.00 | 0.00 |
| >hg38_21_+_8433172_8446622_RNA45SN1_45S_50ntFlanks@12054.12072.19\|TGAGGCGGGGGGGCGAGCC | >hg38_21_+_8433172_8446622_RNA45SN1_45S_50ntFlanks | 12054 | 12072 | 19 | rRF 45S | NO | 12.56 | 0.00 | 0.00 | 0.00 |

### MDA-MB-468

| | | | | | | | | | | |
|---|---|---|---|---|---|---|---|---|---|---|
| >hg38_21_+_8433172_8446622_RNA45SN1_45S_50ntFlanks@12071.12092.22\|CCCCGAGGGGCTCTCGCTTCTG | >hg38_21_+_8433172_8446622_RNA45SN1_45S_50ntFlanks | 12071 | 12092 | 22 | rRF 45S | NO | 0.00 | 33.19 | 0.00 | 0.00 |
| >hg38_21_+_8433172_8446622_RNA45SN1_45S_50ntFlanks@12071.12094.24\|CCCCGAGGGGCTCTCGCTTCTGGC | >hg38_21_+_8433172_8446622_RNA45SN1_45S_50ntFlanks | 12071 | 12094 | 24 | rRF 45S | NO | 0.00 | 51.33 | 0.00 | 0.00 |
| >hg38_21_+_8433172_8446622_RNA45SN1_45S_50ntFlanks@12071.12095.25\|CCCCGAGGGGCTCTCGCTTCTGGCG | >hg38_21_+_8433172_8446622_RNA45SN1_45S_50ntFlanks | 12071 | 12095 | 25 | rRF 45S | NO | 56.55 | 247.36 | 0.00 | 57.91 |
| >hg38_21_+_8433172_8446622_RNA45SN1_45S_50ntFlanks@12071.12096.26\|CCCCGAGGGGCTCTCGCTTCTGGCGC | >hg38_21_+_8433172_8446622_RNA45SN1_45S_50ntFlanks | 12071 | 12096 | 26 | rRF 45S | NO | 0.00 | 52.63 | 0.00 | 6.46 |
| >hg38_21_+_8433172_8446622_RNA45SN1_45S_50ntFlanks@12071.12097.27\|CCCCGAGGGGCTCTCGCTTCTGGCGCC | >hg38_21_+_8433172_8446622_RNA45SN1_45S_50ntFlanks | 12071 | 12097 | 27 | rRF 45S | NO | 6.11 | 33.91 | 0.00 | 0.00 |
| >hg38_21_+_8433172_8446622_RNA45SN1_45S_50ntFlanks@12071.12106.36\|CCCCGAGGGGCTCTCGCTTCTGGCGCCAAGCGCCCG | >hg38_21_+_8433172_8446622_RNA45SN1_45S_50ntFlanks | 12071 | 12106 | 36 | rRF 45S | NO | 21.99 | 28.45 | 0.00 | 0.00 |
| >hg38_21_+_8433172_8446622_RNA45SN1_45S_50ntFlanks@12072.12090.19\|CCCGAGGGGCTCTCGCTTC | >hg38_21_+_8433172_8446622_RNA45SN1_45S_50ntFlanks | 12072 | 12090 | 19 | rRF 45S | NO | 0.00 | 30.48 | 0.00 | 0.00 |
| >hg38_21_+_8433172_8446622_RNA45SN1_45S_50ntFlanks@12072.12092.21\|CCCGAGGGGCTCTCGCTTCTG | >hg38_21_+_8433172_8446622_RNA45SN1_45S_50ntFlanks | 12072 | 12092 | 21 | rRF 45S | NO | 0.00 | 57.41 | 0.00 | 0.00 |
| >hg38_21_+_8433172_8446622_RNA45SN1_45S_50ntFlanks@12072.12093.22\|CCCGAGGGGCTCTCGCTTCTGG | >hg38_21_+_8433172_8446622_RNA45SN1_45S_50ntFlanks | 12072 | 12093 | 22 | rRF 45S | NO | 0.00 | 26.16 | 0.00 | 0.00 |
| >hg38_21_+_8433172_8446622_RNA45SN1_45S_50ntFlanks@12072.12094.23\|CCCGAGGGGCTCTCGCTTCTGGC | >hg38_21_+_8433172_8446622_RNA45SN1_45S_50ntFlanks | 12072 | 12094 | 23 | rRF 45S | NO | 0.00 | 36.09 | 0.00 | 0.00 |
| >hg38_21_+_8433172_8446622_RNA45SN1_45S_50ntFlanks@12072.12095.24\|CCCGAGGGGCTCTCGCTTCTGGCG | >hg38_21_+_8433172_8446622_RNA45SN1_45S_50ntFlanks | 12072 | 12095 | 24 | rRF 45S | NO | 43.41 | 245.59 | 29.80 | 39.50 |
| >hg38_21_+_8433172_8446622_RNA45SN1_45S_50ntFlanks@12072.12096.25\|CCCGAGGGGCTCTCGCTTCTGGCGC | >hg38_21_+_8433172_8446622_RNA45SN1_45S_50ntFlanks | 12072 | 12096 | 25 | rRF 45S | NO | 0.00 | 59.47 | 0.00 | 0.00 |
| >hg38_21_+_8433172_8446622_RNA45SN1_45S_50ntFlanks@12072.12097.26\|CCCGAGGGGCTCTCGCTTCTGGCGCC | >hg38_21_+_8433172_8446622_RNA45SN1_45S_50ntFlanks | 12072 | 12097 | 26 | rRF 45S | NO | 41.01 | 61.66 | 0.00 | 25.48 |
| >hg38_21_+_8433172_8446622_RNA45SN1_45S_50ntFlanks@12073.12090.18\|CCGAGGGGCTCTCGCTTC | >hg38_21_+_8433172_8446622_RNA45SN1_45S_50ntFlanks | 12073 | 12090 | 18 | rRF 45S | NO | 0.00 | 57.89 | 7.17 | 6.39 |
| >hg38_21_+_8433172_8446622_RNA45SN1_45S_50ntFlanks@12073.12092.20\|CCGAGGGGCTCTCGCTTCTG | >hg38_21_+_8433172_8446622_RNA45SN1_45S_50ntFlanks | 12073 | 12092 | 20 | rRF 45S | NO | 0.00 | 79.29 | 0.00 | 0.00 |
| >hg38_21_+_8433172_8446622_RNA45SN1_45S_50ntFlanks@12073.12094.22\|CCGAGGGGCTCTCGCTTCTGGC | >hg38_21_+_8433172_8446622_RNA45SN1_45S_50ntFlanks | 12073 | 12094 | 22 | rRF 45S | NO | 0.00 | 68.55 | 0.00 | 0.00 |
| >hg38_21_+_8433172_8446622_RNA45SN1_45S_50ntFlanks@12073.12095.23\|CCGAGGGGCTCTCGCTTCTGGCG | >hg38_21_+_8433172_8446622_RNA45SN1_45S_50ntFlanks | 12073 | 12095 | 23 | rRF 45S | NO | 70.80 | 436.95 | 92.68 | 66.89 |
| >hg38_21_+_8433172_8446622_RNA45SN1_45S_50ntFlanks@12073.12096.24\|CCGAGGGGCTCTCGCTTCTGGCGC | >hg38_21_+_8433172_8446622_RNA45SN1_45S_50ntFlanks | 12073 | 12096 | 24 | rRF 45S | NO | 36.31 | 156.05 | 6.49 | 37.41 |
| >hg38_21_+_8433172_8446622_RNA45SN1_45S_50ntFlanks@12073.12097.25\|CCGAGGGGCTCTCGCTTCTGGCGCC | >hg38_21_+_8433172_8446622_RNA45SN1_45S_50ntFlanks | 12073 | 12097 | 25 | rRF 45S | NO | 131.23 | 199.03 | 0.00 | 100.23 |
| >hg38_21_+_8433172_8446622_RNA45SN1_45S_50ntFlanks@12073.12098.26\|CCGAGGGGCTCTCGCTTCTGGCGCCA | >hg38_21_+_8433172_8446622_RNA45SN1_45S_50ntFlanks | 12073 | 12098 | 26 | rRF 45S | NO | 0.00 | 31.48 | 0.00 | 0.00 |
| >hg38_21_+_8433172_8446622_RNA45SN1_45S_50ntFlanks@12074.12095.22\|CGAGGGGCTCTCGCTTCTGGCG | >hg38_21_+_8433172_8446622_RNA45SN1_45S_50ntFlanks | 12074 | 12095 | 22 | rRF 45S | NO | 39.97 | 105.78 | 27.53 | 44.25 |
| >hg38_21_+_8433172_8446622_RNA45SN1_45S_50ntFlanks@12075.12095.21\|GAGGGGCTCTCGCTTCTGGCG | >hg38_21_+_8433172_8446622_RNA45SN1_45S_50ntFlanks | 12075 | 12095 | 21 | rRF 45S | NO | 0.00 | 32.89 | 0.00 | 0.00 |
| >hg38_21_+_8433172_8446622_RNA45SN1_45S_50ntFlanks@12076.12095.20\|AGGGGCTCTCGCTTCTGGCG | >hg38_21_+_8433172_8446622_RNA45SN1_45S_50ntFlanks | 12076 | 12095 | 20 | rRF 45S | NO | 241.15 | 719.91 | 37.52 | 67.49 |
| >hg38_21_+_8433172_8446622_RNA45SN1_45S_50ntFlanks@12076.12096.21\|AGGGGCTCTCGCTTCTGGCGC | >hg38_21_+_8433172_8446622_RNA45SN1_45S_50ntFlanks | 12076 | 12096 | 21 | rRF 45S | NO | 21.17 | 46.13 | 0.00 | 0.00 |
| >hg38_21_+_8433172_8446622_RNA45SN1_45S_50ntFlanks@12076.12097.22\|AGGGGCTCTCGCTTCTGGCGCC | >hg38_21_+_8433172_8446622_RNA45SN1_45S_50ntFlanks | 12076 | 12097 | 22 | rRF 45S | NO | 0.00 | 36.41 | 0.00 | 0.00 |
| >hg38_21_+_8433172_8446622_RNA45SN1_45S_50ntFlanks@12077.12095.19\|GGGGCTCTCGCTTCTGGCG | >hg38_21_+_8433172_8446622_RNA45SN1_45S_50ntFlanks | 12077 | 12095 | 19 | rRF 45S | NO | 49.26 | 33.73 | 0.00 | 0.00 |
| >hg38_21_+_8433172_8446622_RNA45SN1_45S_50ntFlanks@12078.12095.18\|GGGCTCTCGCTTCTGGCG | >hg38_21_+_8433172_8446622_RNA45SN1_45S_50ntFlanks | 12078 | 12095 | 18 | rRF 45S | NO | 55.98 | 47.35 | 0.00 | 0.00 |
| >hg38_21_+_8433172_8446622_RNA45SN1_45S_50ntFlanks@12081.12106.26\|CTCTCGCTTCTGGCGCCAAGCGCCCG | >hg38_21_+_8433172_8446622_RNA45SN1_45S_50ntFlanks | 12081 | 12106 | 26 | rRF 45S | NO | 0.00 | 51.29 | 0.00 | 42.66 |
| >hg38_21_+_8433172_8446622_RNA45SN1_45S_50ntFlanks@12081.12107.27\|CTCTCGCTTCTGGCGCCAAGCGCCCGG | >hg38_21_+_8433172_8446622_RNA45SN1_45S_50ntFlanks | 12081 | 12107 | 27 | rRF 45S | NO | 0.00 | 54.11 | 0.00 | 39.76 |
| >hg38_21_+_8433172_8446622_RNA45SN1_45S_50ntFlanks@12081.12108.28\|CTCTCGCTTCTGGCGCCAAGCGCCCGGC | >hg38_21_+_8433172_8446622_RNA45SN1_45S_50ntFlanks | 12081 | 12108 | 28 | rRF 45S | NO | 0.00 | 51.65 | 0.00 | 44.85 |
| >hg38_21_+_8433172_8446622_RNA45SN1_45S_50ntFlanks@12081.12110.30\|CTCTCGCTTCTGGCGCCAAGCGCCCGGCCG | >hg38_21_+_8433172_8446622_RNA45SN1_45S_50ntFlanks | 12081 | 12110 | 30 | rRF 45S | NO | 20.45 | 87.23 | 27.19 | 74.62 |
| >hg38_21_+_8433172_8446622_RNA45SN1_45S_50ntFlanks@12126.12146.21\|CGACCCGCTCCGGGGACAGTG | >hg38_21_+_8433172_8446622_RNA45SN1_45S_50ntFlanks | 12126 | 12146 | 21 | rRF 45S | NO | 0.00 | 48.46 | 113.57 | 35.68 |
| >hg38_21_+_8433172_8446622_RNA45SN1_45S_50ntFlanks@12126.12150.25\|CGACCCGCTCCGGGGACAGTGCCAG | >hg38_21_+_8433172_8446622_RNA45SN1_45S_50ntFlanks | 12126 | 12150 | 25 | rRF 45S | NO | 35.26 | 57.46 | 7.26 | 84.38 |
| >hg38_21_+_8433172_8446622_RNA45SN1_45S_50ntFlanks@12133.12150.18\|CTCCGGGGACAGTGCCAG | >hg38_21_+_8433172_8446622_RNA45SN1_45S_50ntFlanks | 12133 | 12150 | 18 | rRF 45S | NO | 97.40 | 136.86 | 27.92 | 103.56 |

### MDA-MB-468

| | | | | | | | | | | |
|---|---|---|---|---|---|---|---|---|---|---|
| >hg38_21_+_8433172_8446622_RNA45SN1_45S_50ntFlanks@12151.12179.29\|GTGGGGAGTTTGACTGGGGCGGTACACCT | >hg38_21_+_8433172_8446622_RNA45SN1_45S_50ntFlanks | 12151 | 12179 | 29 | rRF 45S | NO | 0.00 | 0.00 | 28.57 | 0.00 |
| >hg38_21_+_8433172_8446622_RNA45SN1_45S_50ntFlanks@12151.12180.30\|GTGGGGAGTTTGACTGGGGCGGTACACCTG | >hg38_21_+_8433172_8446622_RNA45SN1_45S_50ntFlanks | 12151 | 12180 | 30 | rRF 45S | NO | 30.19 | 12.45 | 39.12 | 0.00 |
| >hg38_21_+_8433172_8446622_RNA45SN1_45S_50ntFlanks@12151.12182.32\|GTGGGGAGTTTGACTGGGGCGGTACACCTGTC | >hg38_21_+_8433172_8446622_RNA45SN1_45S_50ntFlanks | 12151 | 12182 | 32 | rRF 45S | NO | 20.15 | 0.00 | 38.80 | 6.16 |
| >hg38_21_+_8433172_8446622_RNA45SN1_45S_50ntFlanks@12154.12180.27\|GGGAGTTTGACTGGGGCGGTACACCTG | >hg38_21_+_8433172_8446622_RNA45SN1_45S_50ntFlanks | 12154 | 12180 | 27 | rRF 45S | NO | 0.00 | 0.00 | 37.35 | 0.00 |
| >hg38_21_+_8433172_8446622_RNA45SN1_45S_50ntFlanks@12154.12182.29\|GGGAGTTTGACTGGGGCGGTACACCTGTC | >hg38_21_+_8433172_8446622_RNA45SN1_45S_50ntFlanks | 12154 | 12182 | 29 | rRF 45S | NO | 0.00 | 0.00 | 39.57 | 0.00 |
| >hg38_21_+_8433172_8446622_RNA45SN1_45S_50ntFlanks@12157.12179.23\|AGTTTGACTGGGGCGGTACACCT | >hg38_21_+_8433172_8446622_RNA45SN1_45S_50ntFlanks | 12157 | 12179 | 23 | rRF 45S | NO | 0.00 | 29.65 | 91.49 | 37.17 |
| >hg38_21_+_8433172_8446622_RNA45SN1_45S_50ntFlanks@12157.12180.24\|AGTTTGACTGGGGCGGTACACCTG | >hg38_21_+_8433172_8446622_RNA45SN1_45S_50ntFlanks | 12157 | 12180 | 24 | rRF 45S | NO | 38.18 | 43.80 | 142.70 | 68.62 |
| >hg38_21_+_8433172_8446622_RNA45SN1_45S_50ntFlanks@12157.12181.25\|AGTTTGACTGGGGCGGTACACCTGT | >hg38_21_+_8433172_8446622_RNA45SN1_45S_50ntFlanks | 12157 | 12181 | 25 | rRF 45S | NO | 19.70 | 5.94 | 105.19 | 53.45 |
| >hg38_21_+_8433172_8446622_RNA45SN1_45S_50ntFlanks@12157.12182.26\|AGTTTGACTGGGGCGGTACACCTGTC | >hg38_21_+_8433172_8446622_RNA45SN1_45S_50ntFlanks | 12157 | 12182 | 26 | rRF 45S | NO | 0.00 | 0.00 | 145.59 | 52.84 |
| >hg38_21_+_8433172_8446622_RNA45SN1_45S_50ntFlanks@12157.12183.27\|AGTTTGACTGGGGCGGTACACCTGTCA | >hg38_21_+_8433172_8446622_RNA45SN1_45S_50ntFlanks | 12157 | 12183 | 27 | rRF 45S | NO | 0.00 | 0.00 | 36.10 | 0.00 |
| >hg38_21_+_8433172_8446622_RNA45SN1_45S_50ntFlanks@12158.12180.23\|GTTTGACTGGGGCGGTACACCTG | >hg38_21_+_8433172_8446622_RNA45SN1_45S_50ntFlanks | 12158 | 12180 | 23 | rRF 45S | NO | 0.00 | 0.00 | 21.06 | 0.00 |
| >hg38_21_+_8433172_8446622_RNA45SN1_45S_50ntFlanks@12159.12178.20\|TTTGACTGGGGCGGTACACC | >hg38_21_+_8433172_8446622_RNA45SN1_45S_50ntFlanks | 12159 | 12178 | 20 | rRF 45S | NO | 34.45 | 12.74 | 51.09 | 6.05 |
| >hg38_21_+_8433172_8446622_RNA45SN1_45S_50ntFlanks@12159.12179.21\|TTTGACTGGGGCGGTACACCT | >hg38_21_+_8433172_8446622_RNA45SN1_45S_50ntFlanks | 12159 | 12179 | 21 | rRF 45S | NO | 54.69 | 95.33 | 231.20 | 100.07 |
| >hg38_21_+_8433172_8446622_RNA45SN1_45S_50ntFlanks@12159.12180.22\|TTTGACTGGGGCGGTACACCTG | >hg38_21_+_8433172_8446622_RNA45SN1_45S_50ntFlanks | 12159 | 12180 | 22 | rRF 45S | NO | 44.77 | 64.95 | 252.01 | 89.71 |
| >hg38_21_+_8433172_8446622_RNA45SN1_45S_50ntFlanks@12159.12181.23\|TTTGACTGGGGCGGTACACCTGT | >hg38_21_+_8433172_8446622_RNA45SN1_45S_50ntFlanks | 12159 | 12181 | 23 | rRF 45S | NO | 28.97 | 32.84 | 183.56 | 74.59 |
| >hg38_21_+_8433172_8446622_RNA45SN1_45S_50ntFlanks@12159.12182.24\|TTTGACTGGGGCGGTACACCTGTC | >hg38_21_+_8433172_8446622_RNA45SN1_45S_50ntFlanks | 12159 | 12182 | 24 | rRF 45S | NO | 0.00 | 0.00 | 196.70 | 74.75 |
| >hg38_21_+_8433172_8446622_RNA45SN1_45S_50ntFlanks@12159.12183.25\|TTTGACTGGGGCGGTACACCTGTCA | >hg38_21_+_8433172_8446622_RNA45SN1_45S_50ntFlanks | 12159 | 12183 | 25 | rRF 45S | NO | 0.00 | 0.00 | 37.69 | 0.00 |
| >hg38_21_+_8433172_8446622_RNA45SN1_45S_50ntFlanks@12160.12179.20\|TTGACTGGGGCGGTACACCT | >hg38_21_+_8433172_8446622_RNA45SN1_45S_50ntFlanks | 12160 | 12179 | 20 | rRF 45S | NO | 0.00 | 0.00 | 20.68 | 0.00 |
| >hg38_21_+_8433172_8446622_RNA45SN1_45S_50ntFlanks@12160.12180.21\|TTGACTGGGGCGGTACACCTG | >hg38_21_+_8433172_8446622_RNA45SN1_45S_50ntFlanks | 12160 | 12180 | 21 | rRF 45S | NO | 0.00 | 0.00 | 27.40 | 0.00 |
| >hg38_21_+_8433172_8446622_RNA45SN1_45S_50ntFlanks@12161.12179.19\|TGACTGGGGCGGTACACCT | >hg38_21_+_8433172_8446622_RNA45SN1_45S_50ntFlanks | 12161 | 12179 | 19 | rRF 45S | NO | 0.00 | 0.00 | 61.79 | 15.60 |
| >hg38_21_+_8433172_8446622_RNA45SN1_45S_50ntFlanks@12161.12180.20\|TGACTGGGGCGGTACACCTG | >hg38_21_+_8433172_8446622_RNA45SN1_45S_50ntFlanks | 12161 | 12180 | 20 | rRF 45S | NO | 0.00 | 0.00 | 71.83 | 14.34 |
| >hg38_21_+_8433172_8446622_RNA45SN1_45S_50ntFlanks@12161.12181.21\|TGACTGGGGCGGTACACCTGT | >hg38_21_+_8433172_8446622_RNA45SN1_45S_50ntFlanks | 12161 | 12181 | 21 | rRF 45S | NO | 0.00 | 0.00 | 41.25 | 0.00 |
| >hg38_21_+_8433172_8446622_RNA45SN1_45S_50ntFlanks@12162.12179.18\|GACTGGGGCGGTACACCT | >hg38_21_+_8433172_8446622_RNA45SN1_45S_50ntFlanks | 12162 | 12179 | 18 | rRF 45S | NO | 0.00 | 0.00 | 33.61 | 0.00 |
| >hg38_21_+_8433172_8446622_RNA45SN1_45S_50ntFlanks@12162.12180.19\|GACTGGGGCGGTACACCTG | >hg38_21_+_8433172_8446622_RNA45SN1_45S_50ntFlanks | 12162 | 12180 | 19 | rRF 45S | NO | 0.00 | 0.00 | 85.95 | 16.39 |
| >hg38_21_+_8433172_8446622_RNA45SN1_45S_50ntFlanks@12162.12181.20\|GACTGGGGCGGTACACCTGT | >hg38_21_+_8433172_8446622_RNA45SN1_45S_50ntFlanks | 12162 | 12181 | 20 | rRF 45S | NO | 0.00 | 0.00 | 40.32 | 0.00 |
| >hg38_21_+_8433172_8446622_RNA45SN1_45S_50ntFlanks@12163.12180.18\|ACTGGGGCGGTACACCTG | >hg38_21_+_8433172_8446622_RNA45SN1_45S_50ntFlanks | 12163 | 12180 | 18 | rRF 45S | NO | 32.33 | 48.16 | 303.38 | 98.56 |
| >hg38_21_+_8433172_8446622_RNA45SN1_45S_50ntFlanks@12163.12181.19\|ACTGGGGCGGTACACCTGT | >hg38_21_+_8433172_8446622_RNA45SN1_45S_50ntFlanks | 12163 | 12181 | 19 | rRF 45S | NO | 0.00 | 12.80 | 204.50 | 89.99 |
| >hg38_21_+_8433172_8446622_RNA45SN1_45S_50ntFlanks@12163.12182.20\|ACTGGGGCGGTACACCTGTC | >hg38_21_+_8433172_8446622_RNA45SN1_45S_50ntFlanks | 12163 | 12182 | 20 | rRF 45S | NO | 46.59 | 47.80 | 345.88 | 203.88 |
| >hg38_21_+_8433172_8446622_RNA45SN1_45S_50ntFlanks@12163.12183.21\|ACTGGGGCGGTACACCTGTCA | >hg38_21_+_8433172_8446622_RNA45SN1_45S_50ntFlanks | 12163 | 12183 | 21 | rRF 45S | NO | 0.00 | 0.00 | 47.04 | 39.46 |
| >hg38_21_+_8433172_8446622_RNA45SN1_45S_50ntFlanks@12164.12182.19\|CTGGGGCGGTACACCTGTC | >hg38_21_+_8433172_8446622_RNA45SN1_45S_50ntFlanks | 12164 | 12182 | 19 | rRF 45S | NO | 0.00 | 0.00 | 20.54 | 0.00 |
| >hg38_21_+_8433172_8446622_RNA45SN1_45S_50ntFlanks@12202.12220.19\|CTAAGGCGAGCTCAGGGAG | >hg38_21_+_8433172_8446622_RNA45SN1_45S_50ntFlanks | 12202 | 12220 | 19 | rRF 45S | NO | 38.37 | 0.00 | 29.52 | 16.37 |
| >hg38_21_+_8433172_8446622_RNA45SN1_45S_50ntFlanks@12202.12221.20\|CTAAGGCGAGCTCAGGGAGG | >hg38_21_+_8433172_8446622_RNA45SN1_45S_50ntFlanks | 12202 | 12221 | 20 | rRF 45S | NO | 29.20 | 0.00 | 13.78 | 15.46 |
| >hg38_21_+_8433172_8446622_RNA45SN1_45S_50ntFlanks@12244.12268.25\|AAGGGCAAAAGCTCGCTTGATCTTG | >hg38_21_+_8433172_8446622_RNA45SN1_45S_50ntFlanks | 12244 | 12268 | 25 | rRF 45S | NO | 0.00 | 0.00 | 34.53 | 0.00 |
| >hg38_21_+_8433172_8446622_RNA45SN1_45S_50ntFlanks@12258.12284.27\|GCTTGATCTTGATTTTCAGTACGAATA | >hg38_21_+_8433172_8446622_RNA45SN1_45S_50ntFlanks | 12258 | 12284 | 27 | rRF 45S | NO | 51.60 | 0.00 | 0.00 | 14.85 |

### MDA-MB-468

| | | | | | | | | | | |
|---|---|---|---|---|---|---|---|---|---|---|
| >hg38_21_+_8433172_8446622_RNA45SN1_45S_50ntFlanks@12259.12282.24\|CTTGATCTTGATTTTCAGTACGAA | >hg38_21_+_8433172_8446622_RNA45SN1_45S_50ntFlanks | 12259 | 12282 | 24 | rRF 45S | NO | 0.00 | 0.00 | 31.87 | 15.93 |
| >hg38_21_+_8433172_8446622_RNA45SN1_45S_50ntFlanks@12259.12284.26\|CTTGATCTTGATTTTCAGTACGAATA | >hg38_21_+_8433172_8446622_RNA45SN1_45S_50ntFlanks | 12259 | 12284 | 26 | rRF 45S | NO | 51.52 | 0.00 | 62.07 | 58.54 |
| >hg38_21_+_8433172_8446622_RNA45SN1_45S_50ntFlanks@12274.12293.20\|CAGTACGAATACAGACCGTG | >hg38_21_+_8433172_8446622_RNA45SN1_45S_50ntFlanks | 12274 | 12293 | 20 | rRF 45S | NO | 0.00 | 0.00 | 40.86 | 0.00 |
| >hg38_21_+_8433172_8446622_RNA45SN1_45S_50ntFlanks@12324.12342.19\|TTGGGTTTTAAGCAGGAGG | >hg38_21_+_8433172_8446622_RNA45SN1_45S_50ntFlanks | 12324 | 12342 | 19 | rRF 45S | NO | 0.00 | 40.49 | 0.00 | 0.00 |
| >hg38_21_+_8433172_8446622_RNA45SN1_45S_50ntFlanks@12324.12343.20\|TTGGGTTTTAAGCAGGAGGT | >hg38_21_+_8433172_8446622_RNA45SN1_45S_50ntFlanks | 12324 | 12343 | 20 | rRF 45S | NO | 0.00 | 28.13 | 0.00 | 0.00 |
| >hg38_21_+_8433172_8446622_RNA45SN1_45S_50ntFlanks@12324.12344.21\|TTGGGTTTTAAGCAGGAGGTG | >hg38_21_+_8433172_8446622_RNA45SN1_45S_50ntFlanks | 12324 | 12344 | 21 | rRF 45S | NO | 96.43 | 295.11 | 65.17 | 23.81 |
| >hg38_21_+_8433172_8446622_RNA45SN1_45S_50ntFlanks@12324.12345.22\|TTGGGTTTTAAGCAGGAGGTGT | >hg38_21_+_8433172_8446622_RNA45SN1_45S_50ntFlanks | 12324 | 12345 | 22 | rRF 45S | NO | 0.00 | 66.35 | 0.00 | 0.00 |
| >hg38_21_+_8433172_8446622_RNA45SN1_45S_50ntFlanks@12324.12346.23\|TTGGGTTTTAAGCAGGAGGTGTC | >hg38_21_+_8433172_8446622_RNA45SN1_45S_50ntFlanks | 12324 | 12346 | 23 | rRF 45S | NO | 140.90 | 228.64 | 120.67 | 44.57 |
| >hg38_21_+_8433172_8446622_RNA45SN1_45S_50ntFlanks@12324.12348.25\|TTGGGTTTTAAGCAGGAGGTGTCAG | >hg38_21_+_8433172_8446622_RNA45SN1_45S_50ntFlanks | 12324 | 12348 | 25 | rRF 45S | NO | 43.52 | 84.58 | 37.45 | 0.00 |
| >hg38_21_+_8433172_8446622_RNA45SN1_45S_50ntFlanks@12329.12346.18\|TTTTAAGCAGGAGGTGTC | >hg38_21_+_8433172_8446622_RNA45SN1_45S_50ntFlanks | 12329 | 12346 | 18 | rRF 45S | NO | 32.81 | 0.00 | 0.00 | 0.00 |
| >hg38_21_+_8433172_8446622_RNA45SN1_45S_50ntFlanks@12347.12365.19\|AGAAAAGTTACCACAGGGA | >hg38_21_+_8433172_8446622_RNA45SN1_45S_50ntFlanks | 12347 | 12365 | 19 | rRF 45S | NO | 0.00 | 71.10 | 0.00 | 0.00 |
| >hg38_21_+_8433172_8446622_RNA45SN1_45S_50ntFlanks@12357.12375.19\|CCACAGGGATAACTGGCTT | >hg38_21_+_8433172_8446622_RNA45SN1_45S_50ntFlanks | 12357 | 12375 | 19 | rRF 45S | NO | 0.00 | 31.30 | 0.00 | 0.00 |
| >hg38_21_+_8433172_8446622_RNA45SN1_45S_50ntFlanks@12357.12376.20\|CCACAGGGATAACTGGCTTG | >hg38_21_+_8433172_8446622_RNA45SN1_45S_50ntFlanks | 12357 | 12376 | 20 | rRF 45S | NO | 19.49 | 53.05 | 46.03 | 15.94 |
| >hg38_21_+_8433172_8446622_RNA45SN1_45S_50ntFlanks@12360.12382.23\|CAGGGATAACTGGCTTGTGGCGG | >hg38_21_+_8433172_8446622_RNA45SN1_45S_50ntFlanks | 12360 | 12382 | 23 | rRF 45S | NO | 36.68 | 0.00 | 0.00 | 0.00 |
| >hg38_21_+_8433172_8446622_RNA45SN1_45S_50ntFlanks@12360.12384.25\|CAGGGATAACTGGCTTGTGGCGGCC | >hg38_21_+_8433172_8446622_RNA45SN1_45S_50ntFlanks | 12360 | 12384 | 25 | rRF 45S | NO | 0.00 | 28.65 | 0.00 | 0.00 |
| >hg38_21_+_8433172_8446622_RNA45SN1_45S_50ntFlanks@12363.12382.20\|GGATAACTGGCTTGTGGCGG | >hg38_21_+_8433172_8446622_RNA45SN1_45S_50ntFlanks | 12363 | 12382 | 20 | rRF 45S | NO | 34.04 | 36.60 | 0.00 | 0.00 |
| >hg38_21_+_8433172_8446622_RNA45SN1_45S_50ntFlanks@12365.12382.18\|ATAACTGGCTTGTGGCGG | >hg38_21_+_8433172_8446622_RNA45SN1_45S_50ntFlanks | 12365 | 12382 | 18 | rRF 45S | NO | 32.32 | 110.04 | 7.31 | 15.41 |
| >hg38_21_+_8433172_8446622_RNA45SN1_45S_50ntFlanks@12365.12383.19\|ATAACTGGCTTGTGGCGGC | >hg38_21_+_8433172_8446622_RNA45SN1_45S_50ntFlanks | 12365 | 12383 | 19 | rRF 45S | NO | 0.00 | 32.97 | 0.00 | 0.00 |
| >hg38_21_+_8433172_8446622_RNA45SN1_45S_50ntFlanks@12366.12383.18\|TAACTGGCTTGTGGCGGC | >hg38_21_+_8433172_8446622_RNA45SN1_45S_50ntFlanks | 12366 | 12383 | 18 | rRF 45S | NO | 31.81 | 185.09 | 32.54 | 34.97 |
| >hg38_21_+_8433172_8446622_RNA45SN1_45S_50ntFlanks@12366.12384.19\|TAACTGGCTTGTGGCGGCC | >hg38_21_+_8433172_8446622_RNA45SN1_45S_50ntFlanks | 12366 | 12384 | 19 | rRF 45S | NO | 0.00 | 73.03 | 31.42 | 16.50 |
| >hg38_21_+_8433172_8446622_RNA45SN1_45S_50ntFlanks@12367.12384.18\|AACTGGCTTGTGGCGGCC | >hg38_21_+_8433172_8446622_RNA45SN1_45S_50ntFlanks | 12367 | 12384 | 18 | rRF 45S | NO | 0.00 | 5.96 | 27.60 | 0.00 |
| >hg38_21_+_8433172_8446622_RNA45SN1_45S_50ntFlanks@12386.12414.29\|AGCGTTCATAGCGACGTCGCTTTTTGATC | >hg38_21_+_8433172_8446622_RNA45SN1_45S_50ntFlanks | 12386 | 12414 | 29 | rRF 45S | NO | 0.00 | 36.17 | 7.20 | 58.25 |
| >hg38_21_+_8433172_8446622_RNA45SN1_45S_50ntFlanks@12394.12414.21\|TAGCGACGTCGCTTTTTGATC | >hg38_21_+_8433172_8446622_RNA45SN1_45S_50ntFlanks | 12394 | 12414 | 21 | rRF 45S | NO | 75.80 | 115.58 | 0.00 | 75.95 |
| >hg38_21_+_8433172_8446622_RNA45SN1_45S_50ntFlanks@12394.12419.26\|TAGCGACGTCGCTTTTTGATCCTTCG | >hg38_21_+_8433172_8446622_RNA45SN1_45S_50ntFlanks | 12394 | 12419 | 26 | rRF 45S | NO | 54.51 | 57.92 | 0.00 | 68.29 |
| >hg38_21_+_8433172_8446622_RNA45SN1_45S_50ntFlanks@12396.12414.19\|GCGACGTCGCTTTTTGATC | >hg38_21_+_8433172_8446622_RNA45SN1_45S_50ntFlanks | 12396 | 12414 | 19 | rRF 45S | NO | 106.65 | 155.51 | 0.00 | 47.48 |
| >hg38_21_+_8433172_8446622_RNA45SN1_45S_50ntFlanks@12396.12416.21\|GCGACGTCGCTTTTTGATCCT | >hg38_21_+_8433172_8446622_RNA45SN1_45S_50ntFlanks | 12396 | 12416 | 21 | rRF 45S | NO | 35.48 | 0.00 | 0.00 | 6.02 |
| >hg38_21_+_8433172_8446622_RNA45SN1_45S_50ntFlanks@12396.12417.22\|GCGACGTCGCTTTTTGATCCTT | >hg38_21_+_8433172_8446622_RNA45SN1_45S_50ntFlanks | 12396 | 12417 | 22 | rRF 45S | NO | 98.15 | 33.78 | 0.00 | 49.70 |
| >hg38_21_+_8433172_8446622_RNA45SN1_45S_50ntFlanks@12397.12414.18\|CGACGTCGCTTTTTGATC | >hg38_21_+_8433172_8446622_RNA45SN1_45S_50ntFlanks | 12397 | 12414 | 18 | rRF 45S | NO | 51.86 | 181.48 | 38.45 | 73.28 |
| >hg38_21_+_8433172_8446622_RNA45SN1_45S_50ntFlanks@12397.12417.21\|CGACGTCGCTTTTTGATCCTT | >hg38_21_+_8433172_8446622_RNA45SN1_45S_50ntFlanks | 12397 | 12417 | 21 | rRF 45S | NO | 67.34 | 78.86 | 0.00 | 68.46 |
| >hg38_21_+_8433172_8446622_RNA45SN1_45S_50ntFlanks@12397.12418.22\|CGACGTCGCTTTTTGATCCTTC | >hg38_21_+_8433172_8446622_RNA45SN1_45S_50ntFlanks | 12397 | 12418 | 22 | rRF 45S | NO | 12.77 | 35.12 | 0.00 | 0.00 |
| >hg38_21_+_8433172_8446622_RNA45SN1_45S_50ntFlanks@12399.12419.21\|ACGTCGCTTTTTGATCCTTCG | >hg38_21_+_8433172_8446622_RNA45SN1_45S_50ntFlanks | 12399 | 12419 | 21 | rRF 45S | NO | 0.00 | 29.17 | 0.00 | 0.00 |
| >hg38_21_+_8433172_8446622_RNA45SN1_45S_50ntFlanks@12404.12431.28\|GCTTTTTGATCCTTCGATGTCGGCTCTT | >hg38_21_+_8433172_8446622_RNA45SN1_45S_50ntFlanks | 12404 | 12431 | 28 | rRF 45S | NO | 0.00 | 43.85 | 0.00 | 0.00 |
| >hg38_21_+_8433172_8446622_RNA45SN1_45S_50ntFlanks@12404.12432.29\|GCTTTTTGATCCTTCGATGTCGGCTCTTC | >hg38_21_+_8433172_8446622_RNA45SN1_45S_50ntFlanks | 12404 | 12432 | 29 | rRF 45S | NO | 0.00 | 36.13 | 0.00 | 0.00 |
| >hg38_21_+_8433172_8446622_RNA45SN1_45S_50ntFlanks@12404.12434.31\|GCTTTTTGATCCTTCGATGTCGGCTCTTCCT | >hg38_21_+_8433172_8446622_RNA45SN1_45S_50ntFlanks | 12404 | 12434 | 31 | rRF 45S | NO | 63.80 | 0.00 | 0.00 | 0.00 |

### MDA-MB-468

| | | | | | | | | | | |
|---|---|---|---|---|---|---|---|---|---|---|
| >hg38_21_+_8433172_8446622_RNA45SN1_45S_50ntFlanks@12404.12435.32\|GCTTTTTGATCCTTCGATGTCGGCTCTTCCTA | >hg38_21_+_8433172_8446622_RNA45SN1_45S_50ntFlanks | 12404 | 12435 | 32 | rRF 45S | NO | 34.54 | 0.00 | 0.00 | 0.00 |
| >hg38_21_+_8433172_8446622_RNA45SN1_45S_50ntFlanks@12404.12436.33\|GCTTTTTGATCCTTCGATGTCGGCTCTTCCTAT | >hg38_21_+_8433172_8446622_RNA45SN1_45S_50ntFlanks | 12404 | 12436 | 33 | rRF 45S | NO | 57.71 | 0.00 | 0.00 | 0.00 |
| >hg38_21_+_8433172_8446622_RNA45SN1_45S_50ntFlanks@12405.12422.18\|CTTTTTGATCCTTCGATG | >hg38_21_+_8433172_8446622_RNA45SN1_45S_50ntFlanks | 12405 | 12422 | 18 | rRF 45S | NO | 0.00 | 146.93 | 20.32 | 6.29 |
| >hg38_21_+_8433172_8446622_RNA45SN1_45S_50ntFlanks@12405.12423.19\|CTTTTTGATCCTTCGATGT | >hg38_21_+_8433172_8446622_RNA45SN1_45S_50ntFlanks | 12405 | 12423 | 19 | rRF 45S | NO | 0.00 | 46.99 | 0.00 | 0.00 |
| >hg38_21_+_8433172_8446622_RNA45SN1_45S_50ntFlanks@12405.12424.20\|CTTTTTGATCCTTCGATGTC | >hg38_21_+_8433172_8446622_RNA45SN1_45S_50ntFlanks | 12405 | 12424 | 20 | rRF 45S | NO | 41.70 | 201.40 | 42.55 | 14.40 |
| >hg38_21_+_8433172_8446622_RNA45SN1_45S_50ntFlanks@12405.12425.21\|CTTTTTGATCCTTCGATGTCG | >hg38_21_+_8433172_8446622_RNA45SN1_45S_50ntFlanks | 12405 | 12425 | 21 | rRF 45S | NO | 34.02 | 138.32 | 21.23 | 14.73 |
| >hg38_21_+_8433172_8446622_RNA45SN1_45S_50ntFlanks@12405.12426.22\|CTTTTTGATCCTTCGATGTCGG | >hg38_21_+_8433172_8446622_RNA45SN1_45S_50ntFlanks | 12405 | 12426 | 22 | rRF 45S | NO | 187.56 | 565.64 | 82.58 | 64.75 |
| >hg38_21_+_8433172_8446622_RNA45SN1_45S_50ntFlanks@12405.12427.23\|CTTTTTGATCCTTCGATGTCGGC | >hg38_21_+_8433172_8446622_RNA45SN1_45S_50ntFlanks | 12405 | 12427 | 23 | rRF 45S | NO | 0.00 | 19.53 | 0.00 | 0.00 |
| >hg38_21_+_8433172_8446622_RNA45SN1_45S_50ntFlanks@12405.12428.24\|CTTTTTGATCCTTCGATGTCGGCT | >hg38_21_+_8433172_8446622_RNA45SN1_45S_50ntFlanks | 12405 | 12428 | 24 | rRF 45S | NO | 0.00 | 28.94 | 0.00 | 0.00 |
| >hg38_21_+_8433172_8446622_RNA45SN1_45S_50ntFlanks@12405.12429.25\|CTTTTTGATCCTTCGATGTCGGCTC | >hg38_21_+_8433172_8446622_RNA45SN1_45S_50ntFlanks | 12405 | 12429 | 25 | rRF 45S | NO | 438.81 | 566.45 | 34.34 | 43.83 |
| >hg38_21_+_8433172_8446622_RNA45SN1_45S_50ntFlanks@12405.12430.26\|CTTTTTGATCCTTCGATGTCGGCTCT | >hg38_21_+_8433172_8446622_RNA45SN1_45S_50ntFlanks | 12405 | 12430 | 26 | rRF 45S | NO | 70.88 | 184.86 | 0.00 | 0.00 |
| >hg38_21_+_8433172_8446622_RNA45SN1_45S_50ntFlanks@12405.12431.27\|CTTTTTGATCCTTCGATGTCGGCTCTT | >hg38_21_+_8433172_8446622_RNA45SN1_45S_50ntFlanks | 12405 | 12431 | 27 | rRF 45S | YES | 691.59 | 1595.35 | 92.29 | 102.57 |
| >hg38_21_+_8433172_8446622_RNA45SN1_45S_50ntFlanks@12405.12432.28\|CTTTTTGATCCTTCGATGTCGGCTCTTC | >hg38_21_+_8433172_8446622_RNA45SN1_45S_50ntFlanks | 12405 | 12432 | 28 | rRF 45S | YES | 696.00 | 982.93 | 129.36 | 119.24 |
| >hg38_21_+_8433172_8446622_RNA45SN1_45S_50ntFlanks@12405.12433.29\|CTTTTTGATCCTTCGATGTCGGCTCTTCC | >hg38_21_+_8433172_8446622_RNA45SN1_45S_50ntFlanks | 12405 | 12433 | 29 | rRF 45S | NO | 785.17 | 340.39 | 30.00 | 86.57 |
| >hg38_21_+_8433172_8446622_RNA45SN1_45S_50ntFlanks@12405.12434.30\|CTTTTTGATCCTTCGATGTCGGCTCTTCCT | >hg38_21_+_8433172_8446622_RNA45SN1_45S_50ntFlanks | 12405 | 12434 | 30 | rRF 45S | YES | 2440.41 | 1142.70 | 50.08 | 323.98 |
| >hg38_21_+_8433172_8446622_RNA45SN1_45S_50ntFlanks@12405.12435.31\|CTTTTTGATCCTTCGATGTCGGCTCTTCCTA | >hg38_21_+_8433172_8446622_RNA45SN1_45S_50ntFlanks | 12405 | 12435 | 31 | rRF 45S | NO | 1065.86 | 372.46 | 40.70 | 191.74 |
| >hg38_21_+_8433172_8446622_RNA45SN1_45S_50ntFlanks@12405.12436.32\|CTTTTTGATCCTTCGATGTCGGCTCTTCCTAT | >hg38_21_+_8433172_8446622_RNA45SN1_45S_50ntFlanks | 12405 | 12436 | 32 | rRF 45S | YES | 2477.00 | 548.54 | 35.19 | 499.36 |
| >hg38_21_+_8433172_8446622_RNA45SN1_45S_50ntFlanks@12405.12437.33\|CTTTTTGATCCTTCGATGTCGGCTCTTCCTATC | >hg38_21_+_8433172_8446622_RNA45SN1_45S_50ntFlanks | 12405 | 12437 | 33 | rRF 45S | NO | 904.61 | 247.70 | 6.83 | 196.29 |
| >hg38_21_+_8433172_8446622_RNA45SN1_45S_50ntFlanks@12405.12438.34\|CTTTTTGATCCTTCGATGTCGGCTCTTCCTATCA | >hg38_21_+_8433172_8446622_RNA45SN1_45S_50ntFlanks | 12405 | 12438 | 34 | rRF 45S | NO | 96.31 | 0.00 | 0.00 | 55.38 |
| >hg38_21_+_8433172_8446622_RNA45SN1_45S_50ntFlanks@12405.12439.35\|CTTTTTGATCCTTCGATGTCGGCTCTTCCTATCAT | >hg38_21_+_8433172_8446622_RNA45SN1_45S_50ntFlanks | 12405 | 12439 | 35 | rRF 45S | NO | 77.46 | 0.00 | 0.00 | 0.00 |
| >hg38_21_+_8433172_8446622_RNA45SN1_45S_50ntFlanks@12405.12440.36\|CTTTTTGATCCTTCGATGTCGGCTCTTCCTATCATT | >hg38_21_+_8433172_8446622_RNA45SN1_45S_50ntFlanks | 12405 | 12440 | 36 | rRF 45S | NO | 129.49 | 41.20 | 0.00 | 6.34 |
| >hg38_21_+_8433172_8446622_RNA45SN1_45S_50ntFlanks@12405.12441.37\|CTTTTTGATCCTTCGATGTCGGCTCTTCCTATCATTG | >hg38_21_+_8433172_8446622_RNA45SN1_45S_50ntFlanks | 12405 | 12441 | 37 | rRF 45S | NO | 997.05 | 228.27 | 0.00 | 132.46 |
| >hg38_21_+_8433172_8446622_RNA45SN1_45S_50ntFlanks@12405.12442.38\|CTTTTTGATCCTTCGATGTCGGCTCTTCCTATCATTGT | >hg38_21_+_8433172_8446622_RNA45SN1_45S_50ntFlanks | 12405 | 12442 | 38 | rRF 45S | NO | 85.67 | 0.00 | 0.00 | 0.00 |
| >hg38_21_+_8433172_8446622_RNA45SN1_45S_50ntFlanks@12405.12443.39\|CTTTTTGATCCTTCGATGTCGGCTCTTCCTATCATTGTG | >hg38_21_+_8433172_8446622_RNA45SN1_45S_50ntFlanks | 12405 | 12443 | 39 | rRF 45S | YES | 2209.25 | 457.92 | 0.00 | 261.68 |
| >hg38_21_+_8433172_8446622_RNA45SN1_45S_50ntFlanks@12405.12444.40\|CTTTTTGATCCTTCGATGTCGGCTCTTCCTATCATTGTGA | >hg38_21_+_8433172_8446622_RNA45SN1_45S_50ntFlanks | 12405 | 12444 | 40 | rRF 45S | NO | 332.60 | 60.25 | 0.00 | 45.45 |
| >hg38_21_+_8433172_8446622_RNA45SN1_45S_50ntFlanks@12405.12445.41\|CTTTTTGATCCTTCGATGTCGGCTCTTCCTATCATTGTGAA | >hg38_21_+_8433172_8446622_RNA45SN1_45S_50ntFlanks | 12405 | 12445 | 41 | rRF 45S | NO | 47.20 | 0.00 | 0.00 | 0.00 |
| >hg38_21_+_8433172_8446622_RNA45SN1_45S_50ntFlanks@12405.12446.42\|CTTTTTGATCCTTCGATGTCGGCTCTTCCTATCATTGTGAAG | >hg38_21_+_8433172_8446622_RNA45SN1_45S_50ntFlanks | 12405 | 12446 | 42 | rRF 45S | NO | 343.36 | 121.17 | 0.00 | 74.64 |
| >hg38_21_+_8433172_8446622_RNA45SN1_45S_50ntFlanks@12406.12431.26\|TTTTTGATCCTTCGATGTCGGCTCTT | >hg38_21_+_8433172_8446622_RNA45SN1_45S_50ntFlanks | 12406 | 12431 | 26 | rRF 45S | NO | 6.00 | 63.47 | 0.00 | 0.00 |
| >hg38_21_+_8433172_8446622_RNA45SN1_45S_50ntFlanks@12406.12432.27\|TTTTTGATCCTTCGATGTCGGCTCTTC | >hg38_21_+_8433172_8446622_RNA45SN1_45S_50ntFlanks | 12406 | 12432 | 27 | rRF 45S | NO | 0.00 | 33.40 | 0.00 | 0.00 |
| >hg38_21_+_8433172_8446622_RNA45SN1_45S_50ntFlanks@12406.12433.28\|TTTTTGATCCTTCGATGTCGGCTCTTCC | >hg38_21_+_8433172_8446622_RNA45SN1_45S_50ntFlanks | 12406 | 12433 | 28 | rRF 45S | NO | 35.51 | 0.00 | 0.00 | 0.00 |
| >hg38_21_+_8433172_8446622_RNA45SN1_45S_50ntFlanks@12406.12434.29\|TTTTTGATCCTTCGATGTCGGCTCTTCCT | >hg38_21_+_8433172_8446622_RNA45SN1_45S_50ntFlanks | 12406 | 12434 | 29 | rRF 45S | NO | 137.51 | 54.30 | 0.00 | 0.00 |
| >hg38_21_+_8433172_8446622_RNA45SN1_45S_50ntFlanks@12406.12435.30\|TTTTTGATCCTTCGATGTCGGCTCTTCCTA | >hg38_21_+_8433172_8446622_RNA45SN1_45S_50ntFlanks | 12406 | 12435 | 30 | rRF 45S | NO | 49.05 | 0.00 | 0.00 | 0.00 |
| >hg38_21_+_8433172_8446622_RNA45SN1_45S_50ntFlanks@12406.12436.31\|TTTTTGATCCTTCGATGTCGGCTCTTCCTAT | >hg38_21_+_8433172_8446622_RNA45SN1_45S_50ntFlanks | 12406 | 12436 | 31 | rRF 45S | NO | 122.00 | 19.42 | 0.00 | 0.00 |
| >hg38_21_+_8433172_8446622_RNA45SN1_45S_50ntFlanks@12406.12437.32\|TTTTTGATCCTTCGATGTCGGCTCTTCCTATC | >hg38_21_+_8433172_8446622_RNA45SN1_45S_50ntFlanks | 12406 | 12437 | 32 | rRF 45S | NO | 49.61 | 0.00 | 0.00 | 0.00 |

### MDA-MB-468

| | | | | | | | | | | |
|---|---|---|---|---|---|---|---|---|---|---|
| >hg38_21_+_8433172_8446622_RNA45SN1_45S_50ntFlanks@12406.12441.36\|TTTTTGATCCT TCGATGTCGGCTCTTCCTATCATTG | >hg38_21_+_8433172_8446622_RNA45SN1_45S_50ntFlanks | 12406 | 12441 | 36 | rRF 45S | NO | 46.83 | 0.00 | 0.00 | 0.00 |
| >hg38_21_+_8433172_8446622_RNA45SN1_45S_50ntFlanks@12406.12443.38\|TTTTTGATCCT TCGATGTCGGCTCTTCCTATCATTGTG | >hg38_21_+_8433172_8446622_RNA45SN1_45S_50ntFlanks | 12406 | 12443 | 38 | rRF 45S | NO | 110.77 | 0.00 | 0.00 | 0.00 |
| >hg38_21_+_8433172_8446622_RNA45SN1_45S_50ntFlanks@12407.12426.20\|TTTTGATCCTT CGATGTCGG | >hg38_21_+_8433172_8446622_RNA45SN1_45S_50ntFlanks | 12407 | 12426 | 20 | rRF 45S | NO | 0.00 | 54.33 | 0.00 | 0.00 |
| >hg38_21_+_8433172_8446622_RNA45SN1_45S_50ntFlanks@12407.12429.23\|TTTTGATCCTT CGATGTCGGCTC | >hg38_21_+_8433172_8446622_RNA45SN1_45S_50ntFlanks | 12407 | 12429 | 23 | rRF 45S | NO | 0.00 | 41.55 | 0.00 | 0.00 |
| >hg38_21_+_8433172_8446622_RNA45SN1_45S_50ntFlanks@12407.12431.25\|TTTTGATCCTT CGATGTCGGCTCTT | >hg38_21_+_8433172_8446622_RNA45SN1_45S_50ntFlanks | 12407 | 12431 | 25 | rRF 45S | NO | 43.39 | 208.81 | 0.00 | 0.00 |
| >hg38_21_+_8433172_8446622_RNA45SN1_45S_50ntFlanks@12407.12432.26\|TTTTGATCCTT CGATGTCGGCTCTTC | >hg38_21_+_8433172_8446622_RNA45SN1_45S_50ntFlanks | 12407 | 12432 | 26 | rRF 45S | NO | 28.71 | 79.00 | 0.00 | 0.00 |
| >hg38_21_+_8433172_8446622_RNA45SN1_45S_50ntFlanks@12407.12433.27\|TTTTGATCCTT CGATGTCGGCTCTTCC | >hg38_21_+_8433172_8446622_RNA45SN1_45S_50ntFlanks | 12407 | 12433 | 27 | rRF 45S | NO | 88.94 | 52.05 | 0.00 | 0.00 |
| >hg38_21_+_8433172_8446622_RNA45SN1_45S_50ntFlanks@12407.12434.28\|TTTTGATCCTT CGATGTCGGCTCTTCCT | >hg38_21_+_8433172_8446622_RNA45SN1_45S_50ntFlanks | 12407 | 12434 | 28 | rRF 45S | NO | 254.41 | 131.53 | 0.00 | 59.05 |
| >hg38_21_+_8433172_8446622_RNA45SN1_45S_50ntFlanks@12407.12435.29\|TTTTGATCCTT CGATGTCGGCTCTTCCTA | >hg38_21_+_8433172_8446622_RNA45SN1_45S_50ntFlanks | 12407 | 12435 | 29 | rRF 45S | NO | 103.78 | 50.91 | 0.00 | 26.89 |
| >hg38_21_+_8433172_8446622_RNA45SN1_45S_50ntFlanks@12407.12436.30\|TTTTGATCCTT CGATGTCGGCTCTTCCTAT | >hg38_21_+_8433172_8446622_RNA45SN1_45S_50ntFlanks | 12407 | 12436 | 30 | rRF 45S | NO | 348.06 | 98.23 | 0.00 | 106.99 |
| >hg38_21_+_8433172_8446622_RNA45SN1_45S_50ntFlanks@12407.12437.31\|TTTTGATCCTT CGATGTCGGCTCTTCCTATC | >hg38_21_+_8433172_8446622_RNA45SN1_45S_50ntFlanks | 12407 | 12437 | 31 | rRF 45S | NO | 156.17 | 43.44 | 0.00 | 45.18 |
| >hg38_21_+_8433172_8446622_RNA45SN1_45S_50ntFlanks@12407.12441.35\|TTTTGATCCTT CGATGTCGGCTCTTCCTATCATTG | >hg38_21_+_8433172_8446622_RNA45SN1_45S_50ntFlanks | 12407 | 12441 | 35 | rRF 45S | NO | 140.61 | 26.89 | 0.00 | 16.26 |
| >hg38_21_+_8433172_8446622_RNA45SN1_45S_50ntFlanks@12407.12443.37\|TTTTGATCCTT CGATGTCGGCTCTTCCTATCATTGTG | >hg38_21_+_8433172_8446622_RNA45SN1_45S_50ntFlanks | 12407 | 12443 | 37 | rRF 45S | NO | 322.00 | 54.34 | 0.00 | 70.28 |
| >hg38_21_+_8433172_8446622_RNA45SN1_45S_50ntFlanks@12407.12444.38\|TTTTGATCCTT CGATGTCGGCTCTTCCTATCATTGTGA | >hg38_21_+_8433172_8446622_RNA45SN1_45S_50ntFlanks | 12407 | 12444 | 38 | rRF 45S | NO | 56.77 | 0.00 | 0.00 | 0.00 |
| >hg38_21_+_8433172_8446622_RNA45SN1_45S_50ntFlanks@12407.12446.40\|TTTTGATCCTT CGATGTCGGCTCTTCCTATCATTGTGAAG | >hg38_21_+_8433172_8446622_RNA45SN1_45S_50ntFlanks | 12407 | 12446 | 40 | rRF 45S | NO | 71.62 | 29.41 | 0.00 | 0.00 |
| >hg38_21_+_8433172_8446622_RNA45SN1_45S_50ntFlanks@12408.12431.24\|TTTGATCCTTC GATGTCGGCTCTT | >hg38_21_+_8433172_8446622_RNA45SN1_45S_50ntFlanks | 12408 | 12431 | 24 | rRF 45S | NO | 0.00 | 74.48 | 0.00 | 0.00 |
| >hg38_21_+_8433172_8446622_RNA45SN1_45S_50ntFlanks@12408.12432.25\|TTTGATCCTTC GATGTCGGCTCTTC | >hg38_21_+_8433172_8446622_RNA45SN1_45S_50ntFlanks | 12408 | 12432 | 25 | rRF 45S | NO | 0.00 | 27.73 | 0.00 | 0.00 |
| >hg38_21_+_8433172_8446622_RNA45SN1_45S_50ntFlanks@12408.12434.27\|TTTGATCCTTC GATGTCGGCTCTTCCT | >hg38_21_+_8433172_8446622_RNA45SN1_45S_50ntFlanks | 12408 | 12434 | 27 | rRF 45S | NO | 41.83 | 6.06 | 0.00 | 0.00 |
| >hg38_21_+_8433172_8446622_RNA45SN1_45S_50ntFlanks@12408.12436.29\|TTTGATCCTTC GATGTCGGCTCTTCCTAT | >hg38_21_+_8433172_8446622_RNA45SN1_45S_50ntFlanks | 12408 | 12436 | 29 | rRF 45S | NO | 41.63 | 0.00 | 0.00 | 0.00 |
| >hg38_21_+_8433172_8446622_RNA45SN1_45S_50ntFlanks@12408.12443.36\|TTTGATCCTTC GATGTCGGCTCTTCCTATCATTGTG | >hg38_21_+_8433172_8446622_RNA45SN1_45S_50ntFlanks | 12408 | 12443 | 36 | rRF 45S | NO | 54.65 | 0.00 | 0.00 | 0.00 |
| >hg38_21_+_8433172_8446622_RNA45SN1_45S_50ntFlanks@12409.12434.26\|TTGATCCTTC GATGTCGGCTCTTCCT | >hg38_21_+_8433172_8446622_RNA45SN1_45S_50ntFlanks | 12409 | 12434 | 26 | rRF 45S | NO | 12.50 | 0.00 | 0.00 | 0.00 |
| >hg38_21_+_8433172_8446622_RNA45SN1_45S_50ntFlanks@12411.12434.24\|GATCCTTCGA TGTCGGCTCTTCCT | >hg38_21_+_8433172_8446622_RNA45SN1_45S_50ntFlanks | 12411 | 12434 | 24 | rRF 45S | NO | 20.33 | 0.00 | 0.00 | 0.00 |
| >hg38_21_+_8433172_8446622_RNA45SN1_45S_50ntFlanks@12411.12436.26\|GATCCTTCGA TGTCGGCTCTTCCTAT | >hg38_21_+_8433172_8446622_RNA45SN1_45S_50ntFlanks | 12411 | 12436 | 26 | rRF 45S | NO | 21.53 | 0.00 | 0.00 | 0.00 |
| >hg38_21_+_8433172_8446622_RNA45SN1_45S_50ntFlanks@12411.12443.33\|GATCCTTCGA TGTCGGCTCTTCCTATCATTGTG | >hg38_21_+_8433172_8446622_RNA45SN1_45S_50ntFlanks | 12411 | 12443 | 33 | rRF 45S | NO | 30.51 | 0.00 | 0.00 | 0.00 |
| >hg38_21_+_8433172_8446622_RNA45SN1_45S_50ntFlanks@12412.12429.18\|ATCCTTCGAT GTCGGCTC | >hg38_21_+_8433172_8446622_RNA45SN1_45S_50ntFlanks | 12412 | 12429 | 18 | rRF 45S | NO | 0.00 | 27.29 | 0.00 | 0.00 |
| >hg38_21_+_8433172_8446622_RNA45SN1_45S_50ntFlanks@12412.12430.19\|ATCCTTCGAT GTCGGCTCT | >hg38_21_+_8433172_8446622_RNA45SN1_45S_50ntFlanks | 12412 | 12430 | 19 | rRF 45S | NO | 0.00 | 28.26 | 0.00 | 0.00 |
| >hg38_21_+_8433172_8446622_RNA45SN1_45S_50ntFlanks@12412.12431.20\|ATCCTTCGAT GTCGGCTCTT | >hg38_21_+_8433172_8446622_RNA45SN1_45S_50ntFlanks | 12412 | 12431 | 20 | rRF 45S | NO | 20.84 | 50.83 | 0.00 | 0.00 |
| >hg38_21_+_8433172_8446622_RNA45SN1_45S_50ntFlanks@12412.12432.21\|ATCCTTCGAT GTCGGCTCTTC | >hg38_21_+_8433172_8446622_RNA45SN1_45S_50ntFlanks | 12412 | 12432 | 21 | rRF 45S | NO | 115.52 | 112.69 | 0.00 | 15.42 |
| >hg38_21_+_8433172_8446622_RNA45SN1_45S_50ntFlanks@12412.12433.22\|ATCCTTCGAT GTCGGCTCTTCC | >hg38_21_+_8433172_8446622_RNA45SN1_45S_50ntFlanks | 12412 | 12433 | 22 | rRF 45S | NO | 38.84 | 0.00 | 0.00 | 0.00 |
| >hg38_21_+_8433172_8446622_RNA45SN1_45S_50ntFlanks@12412.12434.23\|ATCCTTCGAT GTCGGCTCTTCCT | >hg38_21_+_8433172_8446622_RNA45SN1_45S_50ntFlanks | 12412 | 12434 | 23 | rRF 45S | NO | 102.25 | 50.90 | 0.00 | 13.41 |
| >hg38_21_+_8433172_8446622_RNA45SN1_45S_50ntFlanks@12412.12435.24\|ATCCTTCGAT GTCGGCTCTTCCTA | >hg38_21_+_8433172_8446622_RNA45SN1_45S_50ntFlanks | 12412 | 12435 | 24 | rRF 45S | NO | 63.05 | 0.00 | 0.00 | 0.00 |
| >hg38_21_+_8433172_8446622_RNA45SN1_45S_50ntFlanks@12412.12436.25\|ATCCTTCGAT GTCGGCTCTTCCTAT | >hg38_21_+_8433172_8446622_RNA45SN1_45S_50ntFlanks | 12412 | 12436 | 25 | rRF 45S | NO | 129.23 | 35.99 | 0.00 | 38.54 |
| >hg38_21_+_8433172_8446622_RNA45SN1_45S_50ntFlanks@12412.12437.26\|ATCCTTCGAT GTCGGCTCTTCCTATC | >hg38_21_+_8433172_8446622_RNA45SN1_45S_50ntFlanks | 12412 | 12437 | 26 | rRF 45S | NO | 42.45 | 0.00 | 0.00 | 0.00 |
| >hg38_21_+_8433172_8446622_RNA45SN1_45S_50ntFlanks@12412.12441.30\|ATCCTTCGAT GTCGGCTCTTCCTATCATTG | >hg38_21_+_8433172_8446622_RNA45SN1_45S_50ntFlanks | 12412 | 12441 | 30 | rRF 45S | NO | 57.23 | 12.62 | 0.00 | 0.00 |

### MDA-MB-468

| | | | | | | | | | | |
|---|---|---|---|---|---|---|---|---|---|---|
| >hg38_21_+_8433172_8446622_RNA45SN1_45S_50ntFlanks@12412.12443.32\|ATCCTTCGATGTCGGCTCTTCCTATCATTGTG | >hg38_21_+_8433172_8446622_RNA45SN1_45S_50ntFlanks | 12412 | 12443 | 32 | rRF 45S | NO | 126.21 | 81.19 | 0.00 | 38.33 |
| >hg38_21_+_8433172_8446622_RNA45SN1_45S_50ntFlanks@12412.12444.33\|ATCCTTCGATGTCGGCTCTTCCTATCATTGTGA | >hg38_21_+_8433172_8446622_RNA45SN1_45S_50ntFlanks | 12412 | 12444 | 33 | rRF 45S | NO | 34.73 | 0.00 | 0.00 | 0.00 |
| >hg38_21_+_8433172_8446622_RNA45SN1_45S_50ntFlanks@12412.12446.35\|ATCCTTCGATGTCGGCTCTTCCTATCATTGTGAAG | >hg38_21_+_8433172_8446622_RNA45SN1_45S_50ntFlanks | 12412 | 12446 | 35 | rRF 45S | NO | 0.00 | 37.71 | 0.00 | 0.00 |
| >hg38_21_+_8433172_8446622_RNA45SN1_45S_50ntFlanks@12413.12431.19\|TCCTTCGATGTCGGCTCTT | >hg38_21_+_8433172_8446622_RNA45SN1_45S_50ntFlanks | 12413 | 12431 | 19 | rRF 45S | NO | 0.00 | 28.56 | 0.00 | 0.00 |
| >hg38_21_+_8433172_8446622_RNA45SN1_45S_50ntFlanks@12413.12432.20\|TCCTTCGATGTCGGCTCTTC | >hg38_21_+_8433172_8446622_RNA45SN1_45S_50ntFlanks | 12413 | 12432 | 20 | rRF 45S | NO | 67.15 | 64.12 | 0.00 | 0.00 |
| >hg38_21_+_8433172_8446622_RNA45SN1_45S_50ntFlanks@12413.12434.22\|TCCTTCGATGTCGGCTCTTCCT | >hg38_21_+_8433172_8446622_RNA45SN1_45S_50ntFlanks | 12413 | 12434 | 22 | rRF 45S | NO | 50.68 | 19.61 | 0.00 | 0.00 |
| >hg38_21_+_8433172_8446622_RNA45SN1_45S_50ntFlanks@12413.12435.23\|TCCTTCGATGTCGGCTCTTCCTA | >hg38_21_+_8433172_8446622_RNA45SN1_45S_50ntFlanks | 12413 | 12435 | 23 | rRF 45S | NO | 33.54 | 0.00 | 0.00 | 0.00 |
| >hg38_21_+_8433172_8446622_RNA45SN1_45S_50ntFlanks@12413.12436.24\|TCCTTCGATGTCGGCTCTTCCTAT | >hg38_21_+_8433172_8446622_RNA45SN1_45S_50ntFlanks | 12413 | 12436 | 24 | rRF 45S | NO | 62.73 | 0.00 | 0.00 | 0.00 |
| >hg38_21_+_8433172_8446622_RNA45SN1_45S_50ntFlanks@12413.12441.29\|TCCTTCGATGTCGGCTCTTCCTATCATTG | >hg38_21_+_8433172_8446622_RNA45SN1_45S_50ntFlanks | 12413 | 12441 | 29 | rRF 45S | NO | 29.45 | 0.00 | 0.00 | 0.00 |
| >hg38_21_+_8433172_8446622_RNA45SN1_45S_50ntFlanks@12413.12443.31\|TCCTTCGATGTCGGCTCTTCCTATCATTGTG | >hg38_21_+_8433172_8446622_RNA45SN1_45S_50ntFlanks | 12413 | 12443 | 31 | rRF 45S | NO | 54.43 | 40.47 | 0.00 | 0.00 |
| >hg38_21_+_8433172_8446622_RNA45SN1_45S_50ntFlanks@12415.12432.18\|CTTCGATGTCGGCTCTTC | >hg38_21_+_8433172_8446622_RNA45SN1_45S_50ntFlanks | 12415 | 12432 | 18 | rRF 45S | NO | 345.21 | 574.91 | 41.09 | 73.29 |
| >hg38_21_+_8433172_8446622_RNA45SN1_45S_50ntFlanks@12415.12433.19\|CTTCGATGTCGGCTCTTCC | >hg38_21_+_8433172_8446622_RNA45SN1_45S_50ntFlanks | 12415 | 12433 | 19 | rRF 45S | NO | 88.15 | 97.41 | 0.00 | 6.16 |
| >hg38_21_+_8433172_8446622_RNA45SN1_45S_50ntFlanks@12415.12434.20\|CTTCGATGTCGGCTCTTCCT | >hg38_21_+_8433172_8446622_RNA45SN1_45S_50ntFlanks | 12415 | 12434 | 20 | rRF 45S | NO | 330.32 | 323.16 | 6.89 | 95.81 |
| >hg38_21_+_8433172_8446622_RNA45SN1_45S_50ntFlanks@12415.12435.21\|CTTCGATGTCGGCTCTTCCTA | >hg38_21_+_8433172_8446622_RNA45SN1_45S_50ntFlanks | 12415 | 12435 | 21 | rRF 45S | NO | 327.95 | 209.15 | 0.00 | 111.81 |
| >hg38_21_+_8433172_8446622_RNA45SN1_45S_50ntFlanks@12415.12436.22\|CTTCGATGTCGGCTCTTCCTAT | >hg38_21_+_8433172_8446622_RNA45SN1_45S_50ntFlanks | 12415 | 12436 | 22 | rRF 45S | NO | 651.13 | 367.45 | 0.00 | 309.52 |
| >hg38_21_+_8433172_8446622_RNA45SN1_45S_50ntFlanks@12415.12437.23\|CTTCGATGTCGGCTCTTCCTATC | >hg38_21_+_8433172_8446622_RNA45SN1_45S_50ntFlanks | 12415 | 12437 | 23 | rRF 45S | NO | 208.05 | 165.49 | 0.00 | 136.92 |
| >hg38_21_+_8433172_8446622_RNA45SN1_45S_50ntFlanks@12415.12440.26\|CTTCGATGTCGGCTCTTCCTATCATT | >hg38_21_+_8433172_8446622_RNA45SN1_45S_50ntFlanks | 12415 | 12440 | 26 | rRF 45S | NO | 19.95 | 0.00 | 0.00 | 0.00 |
| >hg38_21_+_8433172_8446622_RNA45SN1_45S_50ntFlanks@12415.12441.27\|CTTCGATGTCGGCTCTTCCTATCATTG | >hg38_21_+_8433172_8446622_RNA45SN1_45S_50ntFlanks | 12415 | 12441 | 27 | rRF 45S | NO | 185.71 | 152.63 | 0.00 | 134.06 |
| >hg38_21_+_8433172_8446622_RNA45SN1_45S_50ntFlanks@12415.12443.29\|CTTCGATGTCGGCTCTTCCTATCATTGTG | >hg38_21_+_8433172_8446622_RNA45SN1_45S_50ntFlanks | 12415 | 12443 | 29 | rRF 45S | NO | 213.74 | 349.67 | 0.00 | 281.47 |
| >hg38_21_+_8433172_8446622_RNA45SN1_45S_50ntFlanks@12415.12444.30\|CTTCGATGTCGGCTCTTCCTATCATTGTGA | >hg38_21_+_8433172_8446622_RNA45SN1_45S_50ntFlanks | 12415 | 12444 | 30 | rRF 45S | NO | 71.82 | 88.93 | 0.00 | 65.35 |
| >hg38_21_+_8433172_8446622_RNA45SN1_45S_50ntFlanks@12415.12445.31\|CTTCGATGTCGGCTCTTCCTATCATTGTGAA | >hg38_21_+_8433172_8446622_RNA45SN1_45S_50ntFlanks | 12415 | 12445 | 31 | rRF 45S | NO | 29.32 | 37.97 | 0.00 | 33.77 |
| >hg38_21_+_8433172_8446622_RNA45SN1_45S_50ntFlanks@12415.12446.32\|CTTCGATGTCGGCTCTTCCTATCATTGTGAAG | >hg38_21_+_8433172_8446622_RNA45SN1_45S_50ntFlanks | 12415 | 12446 | 32 | rRF 45S | NO | 39.19 | 83.96 | 0.00 | 75.82 |
| >hg38_21_+_8433172_8446622_RNA45SN1_45S_50ntFlanks@12415.12455.41\|CTTCGATGTCGGCTCTTCCTATCATTGTGAAGCAGAATTCA | >hg38_21_+_8433172_8446622_RNA45SN1_45S_50ntFlanks | 12415 | 12455 | 41 | rRF 45S | NO | 0.00 | 28.77 | 0.00 | 101.90 |
| >hg38_21_+_8433172_8446622_RNA45SN1_45S_50ntFlanks@12416.12433.18\|TTCGATGTCGGCTCTTCC | >hg38_21_+_8433172_8446622_RNA45SN1_45S_50ntFlanks | 12416 | 12433 | 18 | rRF 45S | NO | 0.00 | 12.43 | 0.00 | 0.00 |
| >hg38_21_+_8433172_8446622_RNA45SN1_45S_50ntFlanks@12416.12434.19\|TTCGATGTCGGCTCTTCCT | >hg38_21_+_8433172_8446622_RNA45SN1_45S_50ntFlanks | 12416 | 12434 | 19 | rRF 45S | NO | 45.88 | 83.52 | 0.00 | 0.00 |
| >hg38_21_+_8433172_8446622_RNA45SN1_45S_50ntFlanks@12416.12435.20\|TTCGATGTCGGCTCTTCCTA | >hg38_21_+_8433172_8446622_RNA45SN1_45S_50ntFlanks | 12416 | 12435 | 20 | rRF 45S | NO | 70.28 | 92.71 | 0.00 | 0.00 |
| >hg38_21_+_8433172_8446622_RNA45SN1_45S_50ntFlanks@12416.12436.21\|TTCGATGTCGGCTCTTCCTAT | >hg38_21_+_8433172_8446622_RNA45SN1_45S_50ntFlanks | 12416 | 12436 | 21 | rRF 45S | NO | 109.28 | 108.95 | 0.00 | 43.15 |
| >hg38_21_+_8433172_8446622_RNA45SN1_45S_50ntFlanks@12416.12437.22\|TTCGATGTCGGCTCTTCCTATC | >hg38_21_+_8433172_8446622_RNA45SN1_45S_50ntFlanks | 12416 | 12437 | 22 | rRF 45S | NO | 37.46 | 44.81 | 0.00 | 0.00 |
| >hg38_21_+_8433172_8446622_RNA45SN1_45S_50ntFlanks@12416.12441.26\|TTCGATGTCGGCTCTTCCTATCATTG | >hg38_21_+_8433172_8446622_RNA45SN1_45S_50ntFlanks | 12416 | 12441 | 26 | rRF 45S | NO | 20.90 | 43.01 | 0.00 | 0.00 |
| >hg38_21_+_8433172_8446622_RNA45SN1_45S_50ntFlanks@12416.12443.28\|TTCGATGTCGGCTCTTCCTATCATTGTG | >hg38_21_+_8433172_8446622_RNA45SN1_45S_50ntFlanks | 12416 | 12443 | 28 | rRF 45S | NO | 30.50 | 84.89 | 0.00 | 26.20 |
| >hg38_21_+_8433172_8446622_RNA45SN1_45S_50ntFlanks@12416.12444.29\|TTCGATGTCGGCTCTTCCTATCATTGTGA | >hg38_21_+_8433172_8446622_RNA45SN1_45S_50ntFlanks | 12416 | 12444 | 29 | rRF 45S | NO | 0.00 | 28.01 | 0.00 | 0.00 |
| >hg38_21_+_8433172_8446622_RNA45SN1_45S_50ntFlanks@12416.12446.31\|TTCGATGTCGGCTCTTCCTATCATTGTGAAG | >hg38_21_+_8433172_8446622_RNA45SN1_45S_50ntFlanks | 12416 | 12446 | 31 | rRF 45S | NO | 0.00 | 32.68 | 0.00 | 0.00 |
| >hg38_21_+_8433172_8446622_RNA45SN1_45S_50ntFlanks@12417.12434.18\|TCGATGTCGGCTCTTCCT | >hg38_21_+_8433172_8446622_RNA45SN1_45S_50ntFlanks | 12417 | 12434 | 18 | rRF 45S | NO | 295.55 | 631.33 | 30.54 | 75.60 |
| >hg38_21_+_8433172_8446622_RNA45SN1_45S_50ntFlanks@12417.12435.19\|TCGATGTCGGCTCTTCCTA | >hg38_21_+_8433172_8446622_RNA45SN1_45S_50ntFlanks | 12417 | 12435 | 19 | rRF 45S | NO | 386.75 | 514.36 | 21.12 | 92.87 |

### MDA-MB-468

<table>
<tr><td>>hg38_21_+_8433172_8446622_RNA45SN1_45S_50ntFlanks@12417.12436.20|TCGATGTCGGCTCTTCCTAT</td><td>>hg38_21_+_8433172_8446622_RNA45SN1_45S_50ntFlanks</td><td>12417</td><td>12436</td><td>20</td><td>rRF 45S</td><td>NO</td><td>690.44</td><td>733.02</td><td>6.80</td><td>227.91</td></tr>
<tr><td>>hg38_21_+_8433172_8446622_RNA45SN1_45S_50ntFlanks@12417.12437.21|TCGATGTCGGCTCTTCCTATC</td><td>>hg38_21_+_8433172_8446622_RNA45SN1_45S_50ntFlanks</td><td>12417</td><td>12437</td><td>21</td><td>rRF 45S</td><td>NO</td><td>224.02</td><td>289.17</td><td>0.00</td><td>95.27</td></tr>
<tr><td>>hg38_21_+_8433172_8446622_RNA45SN1_45S_50ntFlanks@12417.12438.22|TCGATGTCGGCTCTTCCTATCA</td><td>>hg38_21_+_8433172_8446622_RNA45SN1_45S_50ntFlanks</td><td>12417</td><td>12438</td><td>22</td><td>rRF 45S</td><td>NO</td><td>33.27</td><td>6.05</td><td>0.00</td><td>49.72</td></tr>
<tr><td>>hg38_21_+_8433172_8446622_RNA45SN1_45S_50ntFlanks@12417.12439.23|TCGATGTCGGCTCTTCCTATCAT</td><td>>hg38_21_+_8433172_8446622_RNA45SN1_45S_50ntFlanks</td><td>12417</td><td>12439</td><td>23</td><td>rRF 45S</td><td>NO</td><td>6.17</td><td>27.74</td><td>0.00</td><td>0.00</td></tr>
<tr><td>>hg38_21_+_8433172_8446622_RNA45SN1_45S_50ntFlanks@12417.12440.24|TCGATGTCGGCTCTTCCTATCATT</td><td>>hg38_21_+_8433172_8446622_RNA45SN1_45S_50ntFlanks</td><td>12417</td><td>12440</td><td>24</td><td>rRF 45S</td><td>NO</td><td>21.42</td><td>36.32</td><td>0.00</td><td>0.00</td></tr>
<tr><td>>hg38_21_+_8433172_8446622_RNA45SN1_45S_50ntFlanks@12417.12441.25|TCGATGTCGGCTCTTCCTATCATTG</td><td>>hg38_21_+_8433172_8446622_RNA45SN1_45S_50ntFlanks</td><td>12417</td><td>12441</td><td>25</td><td>rRF 45S</td><td>NO</td><td>96.51</td><td>163.73</td><td>0.00</td><td>59.41</td></tr>
<tr><td>>hg38_21_+_8433172_8446622_RNA45SN1_45S_50ntFlanks@12417.12443.27|TCGATGTCGGCTCTTCCTATCATTGTG</td><td>>hg38_21_+_8433172_8446622_RNA45SN1_45S_50ntFlanks</td><td>12417</td><td>12443</td><td>27</td><td>rRF 45S</td><td>NO</td><td>199.84</td><td>667.30</td><td>0.00</td><td>168.24</td></tr>
<tr><td>>hg38_21_+_8433172_8446622_RNA45SN1_45S_50ntFlanks@12417.12444.28|TCGATGTCGGCTCTTCCTATCATTGTGA</td><td>>hg38_21_+_8433172_8446622_RNA45SN1_45S_50ntFlanks</td><td>12417</td><td>12444</td><td>28</td><td>rRF 45S</td><td>NO</td><td>70.26</td><td>182.20</td><td>0.00</td><td>43.98</td></tr>
<tr><td>>hg38_21_+_8433172_8446622_RNA45SN1_45S_50ntFlanks@12417.12445.29|TCGATGTCGGCTCTTCCTATCATTGTGAA</td><td>>hg38_21_+_8433172_8446622_RNA45SN1_45S_50ntFlanks</td><td>12417</td><td>12445</td><td>29</td><td>rRF 45S</td><td>NO</td><td>29.86</td><td>78.42</td><td>0.00</td><td>5.93</td></tr>
<tr><td>>hg38_21_+_8433172_8446622_RNA45SN1_45S_50ntFlanks@12417.12446.30|TCGATGTCGGCTCTTCCTATCATTGTGAAG</td><td>>hg38_21_+_8433172_8446622_RNA45SN1_45S_50ntFlanks</td><td>12417</td><td>12446</td><td>30</td><td>rRF 45S</td><td>NO</td><td>94.45</td><td>186.38</td><td>0.00</td><td>36.25</td></tr>
<tr><td>>hg38_21_+_8433172_8446622_RNA45SN1_45S_50ntFlanks@12417.12455.39|TCGATGTCGGCTCTTCCTATCATTGTGAAGCAGAATTCA</td><td>>hg38_21_+_8433172_8446622_RNA45SN1_45S_50ntFlanks</td><td>12417</td><td>12455</td><td>39</td><td>rRF 45S</td><td>NO</td><td>87.23</td><td>110.31</td><td>0.00</td><td>94.63</td></tr>
<tr><td>>hg38_21_+_8433172_8446622_RNA45SN1_45S_50ntFlanks@12418.12435.18|CGATGTCGGCTCTTCCTA</td><td>>hg38_21_+_8433172_8446622_RNA45SN1_45S_50ntFlanks</td><td>12418</td><td>12435</td><td>18</td><td>rRF 45S</td><td>YES</td><td>1093.19</td><td>1102.04</td><td>58.26</td><td>256.82</td></tr>
<tr><td>>hg38_21_+_8433172_8446622_RNA45SN1_45S_50ntFlanks@12418.12436.19|CGATGTCGGCTCTTCCTAT</td><td>>hg38_21_+_8433172_8446622_RNA45SN1_45S_50ntFlanks</td><td>12418</td><td>12436</td><td>19</td><td>rRF 45S</td><td>YES</td><td>1855.33</td><td>1558.28</td><td>52.61</td><td>563.99</td></tr>
<tr><td>>hg38_21_+_8433172_8446622_RNA45SN1_45S_50ntFlanks@12418.12437.20|CGATGTCGGCTCTTCCTATC</td><td>>hg38_21_+_8433172_8446622_RNA45SN1_45S_50ntFlanks</td><td>12418</td><td>12437</td><td>20</td><td>rRF 45S</td><td>NO</td><td>542.97</td><td>596.53</td><td>13.55</td><td>234.10</td></tr>
<tr><td>>hg38_21_+_8433172_8446622_RNA45SN1_45S_50ntFlanks@12418.12438.21|CGATGTCGGCTCTTCCTATCA</td><td>>hg38_21_+_8433172_8446622_RNA45SN1_45S_50ntFlanks</td><td>12418</td><td>12438</td><td>21</td><td>rRF 45S</td><td>NO</td><td>54.91</td><td>33.36</td><td>0.00</td><td>95.14</td></tr>
<tr><td>>hg38_21_+_8433172_8446622_RNA45SN1_45S_50ntFlanks@12418.12439.22|CGATGTCGGCTCTTCCTATCAT</td><td>>hg38_21_+_8433172_8446622_RNA45SN1_45S_50ntFlanks</td><td>12418</td><td>12439</td><td>22</td><td>rRF 45S</td><td>NO</td><td>48.72</td><td>48.94</td><td>0.00</td><td>15.93</td></tr>
<tr><td>>hg38_21_+_8433172_8446622_RNA45SN1_45S_50ntFlanks@12418.12440.23|CGATGTCGGCTCTTCCTATCATT</td><td>>hg38_21_+_8433172_8446622_RNA45SN1_45S_50ntFlanks</td><td>12418</td><td>12440</td><td>23</td><td>rRF 45S</td><td>NO</td><td>39.03</td><td>46.47</td><td>0.00</td><td>6.09</td></tr>
<tr><td>>hg38_21_+_8433172_8446622_RNA45SN1_45S_50ntFlanks@12418.12441.24|CGATGTCGGCTCTTCCTATCATTG</td><td>>hg38_21_+_8433172_8446622_RNA45SN1_45S_50ntFlanks</td><td>12418</td><td>12441</td><td>24</td><td>rRF 45S</td><td>NO</td><td>100.14</td><td>156.75</td><td>0.00</td><td>76.98</td></tr>
<tr><td>>hg38_21_+_8433172_8446622_RNA45SN1_45S_50ntFlanks@12418.12442.25|CGATGTCGGCTCTTCCTATCATTGT</td><td>>hg38_21_+_8433172_8446622_RNA45SN1_45S_50ntFlanks</td><td>12418</td><td>12442</td><td>25</td><td>rRF 45S</td><td>NO</td><td>20.81</td><td>31.58</td><td>0.00</td><td>0.00</td></tr>
<tr><td>>hg38_21_+_8433172_8446622_RNA45SN1_45S_50ntFlanks@12418.12443.26|CGATGTCGGCTCTTCCTATCATTGTG</td><td>>hg38_21_+_8433172_8446622_RNA45SN1_45S_50ntFlanks</td><td>12418</td><td>12443</td><td>26</td><td>rRF 45S</td><td>NO</td><td>882.18</td><td>808.48</td><td>38.22</td><td>298.93</td></tr>
<tr><td>>hg38_21_+_8433172_8446622_RNA45SN1_45S_50ntFlanks@12418.12444.27|CGATGTCGGCTCTTCCTATCATTGTGA</td><td>>hg38_21_+_8433172_8446622_RNA45SN1_45S_50ntFlanks</td><td>12418</td><td>12444</td><td>27</td><td>rRF 45S</td><td>NO</td><td>222.99</td><td>157.47</td><td>0.00</td><td>68.71</td></tr>
<tr><td>>hg38_21_+_8433172_8446622_RNA45SN1_45S_50ntFlanks@12418.12445.28|CGATGTCGGCTCTTCCTATCATTGTGAA</td><td>>hg38_21_+_8433172_8446622_RNA45SN1_45S_50ntFlanks</td><td>12418</td><td>12445</td><td>28</td><td>rRF 45S</td><td>NO</td><td>115.39</td><td>83.35</td><td>0.00</td><td>38.60</td></tr>
<tr><td>>hg38_21_+_8433172_8446622_RNA45SN1_45S_50ntFlanks@12418.12446.29|CGATGTCGGCTCTTCCTATCATTGTGAAG</td><td>>hg38_21_+_8433172_8446622_RNA45SN1_45S_50ntFlanks</td><td>12418</td><td>12446</td><td>29</td><td>rRF 45S</td><td>NO</td><td>230.70</td><td>288.41</td><td>29.91</td><td>83.89</td></tr>
<tr><td>>hg38_21_+_8433172_8446622_RNA45SN1_45S_50ntFlanks@12418.12449.32|CGATGTCGGCTCTTCCTATCATTGTGAAGCAG</td><td>>hg38_21_+_8433172_8446622_RNA45SN1_45S_50ntFlanks</td><td>12418</td><td>12449</td><td>32</td><td>rRF 45S</td><td>NO</td><td>31.85</td><td>47.07</td><td>0.00</td><td>6.44</td></tr>
<tr><td>>hg38_21_+_8433172_8446622_RNA45SN1_45S_50ntFlanks@12418.12454.37|CGATGTCGGCTCTTCCTATCATTGTGAAGCAGAATTC</td><td>>hg38_21_+_8433172_8446622_RNA45SN1_45S_50ntFlanks</td><td>12418</td><td>12454</td><td>37</td><td>rRF 45S</td><td>NO</td><td>20.25</td><td>26.27</td><td>0.00</td><td>6.26</td></tr>
<tr><td>>hg38_21_+_8433172_8446622_RNA45SN1_45S_50ntFlanks@12418.12455.38|CGATGTCGGCTCTTCCTATCATTGTGAAGCAGAATTCA</td><td>>hg38_21_+_8433172_8446622_RNA45SN1_45S_50ntFlanks</td><td>12418</td><td>12455</td><td>38</td><td>rRF 45S</td><td>NO</td><td>173.29</td><td>260.90</td><td>0.00</td><td>255.03</td></tr>
<tr><td>>hg38_21_+_8433172_8446622_RNA45SN1_45S_50ntFlanks@12418.12456.39|CGATGTCGGCTCTTCCTATCATTGTGAAGCAGAATTCAC</td><td>>hg38_21_+_8433172_8446622_RNA45SN1_45S_50ntFlanks</td><td>12418</td><td>12456</td><td>39</td><td>rRF 45S</td><td>NO</td><td>115.50</td><td>92.50</td><td>0.00</td><td>49.41</td></tr>
<tr><td>>hg38_21_+_8433172_8446622_RNA45SN1_45S_50ntFlanks@12419.12436.18|GATGTCGGCTCTTCCTAT</td><td>>hg38_21_+_8433172_8446622_RNA45SN1_45S_50ntFlanks</td><td>12419</td><td>12436</td><td>18</td><td>rRF 45S</td><td>NO</td><td>305.24</td><td>192.70</td><td>0.00</td><td>45.13</td></tr>
<tr><td>>hg38_21_+_8433172_8446622_RNA45SN1_45S_50ntFlanks@12419.12437.19|GATGTCGGCTCTTCCTATC</td><td>>hg38_21_+_8433172_8446622_RNA45SN1_45S_50ntFlanks</td><td>12419</td><td>12437</td><td>19</td><td>rRF 45S</td><td>NO</td><td>107.25</td><td>95.84</td><td>0.00</td><td>6.51</td></tr>
<tr><td>>hg38_21_+_8433172_8446622_RNA45SN1_45S_50ntFlanks@12419.12441.23|GATGTCGGCTCTTCCTATCATTG</td><td>>hg38_21_+_8433172_8446622_RNA45SN1_45S_50ntFlanks</td><td>12419</td><td>12441</td><td>23</td><td>rRF 45S</td><td>NO</td><td>55.59</td><td>45.63</td><td>0.00</td><td>0.00</td></tr>
<tr><td>>hg38_21_+_8433172_8446622_RNA45SN1_45S_50ntFlanks@12419.12443.25|GATGTCGGCTCTTCCTATCATTGTG</td><td>>hg38_21_+_8433172_8446622_RNA45SN1_45S_50ntFlanks</td><td>12419</td><td>12443</td><td>25</td><td>rRF 45S</td><td>NO</td><td>110.56</td><td>64.90</td><td>0.00</td><td>0.00</td></tr>
<tr><td>>hg38_21_+_8433172_8446622_RNA45SN1_45S_50ntFlanks@12419.12444.26|GATGTCGGCTCTTCCTATCATTGTGA</td><td>>hg38_21_+_8433172_8446622_RNA45SN1_45S_50ntFlanks</td><td>12419</td><td>12444</td><td>26</td><td>rRF 45S</td><td>NO</td><td>39.13</td><td>0.00</td><td>0.00</td><td>0.00</td></tr>
<tr><td>>hg38_21_+_8433172_8446622_RNA45SN1_45S_50ntFlanks@12419.12446.28|GATGTCGGCTCTTCCTATCATTGTGAAG</td><td>>hg38_21_+_8433172_8446622_RNA45SN1_45S_50ntFlanks</td><td>12419</td><td>12446</td><td>28</td><td>rRF 45S</td><td>NO</td><td>48.36</td><td>46.52</td><td>0.00</td><td>0.00</td></tr>
<tr><td>>hg38_21_+_8433172_8446622_RNA45SN1_45S_50ntFlanks@12419.12455.37|GATGTCGGCTCTTCCTATCATTGTGAAGCAGAATTCA</td><td>>hg38_21_+_8433172_8446622_RNA45SN1_45S_50ntFlanks</td><td>12419</td><td>12455</td><td>37</td><td>rRF 45S</td><td>NO</td><td>30.87</td><td>28.00</td><td>0.00</td><td>0.00</td></tr>
</table>

### MDA-MB-468

| | | | | | | | | | | |
|---|---|---|---|---|---|---|---|---|---|---|
| >hg38_21_+_8433172_8446622_RNA45SN1_45S_50ntFlanks@12420.12437.18\|ATGTCGGCTCTTCCTATC | >hg38_21_+_8433172_8446622_RNA45SN1_45S_50ntFlanks | 12420 | 12437 | 18 | rRF 45S | NO | 696.08 | 574.35 | 29.35 | 113.78 |
| >hg38_21_+_8433172_8446622_RNA45SN1_45S_50ntFlanks@12420.12438.19\|ATGTCGGCTCTTCCTATCA | >hg38_21_+_8433172_8446622_RNA45SN1_45S_50ntFlanks | 12420 | 12438 | 19 | rRF 45S | NO | 69.11 | 33.94 | 0.00 | 13.73 |
| >hg38_21_+_8433172_8446622_RNA45SN1_45S_50ntFlanks@12420.12439.20\|ATGTCGGCTCTTCCTATCAT | >hg38_21_+_8433172_8446622_RNA45SN1_45S_50ntFlanks | 12420 | 12439 | 20 | rRF 45S | NO | 80.65 | 99.92 | 0.00 | 24.04 |
| >hg38_21_+_8433172_8446622_RNA45SN1_45S_50ntFlanks@12420.12440.21\|ATGTCGGCTCTTCCTATCATT | >hg38_21_+_8433172_8446622_RNA45SN1_45S_50ntFlanks | 12420 | 12440 | 21 | rRF 45S | NO | 67.92 | 61.62 | 0.00 | 0.00 |
| >hg38_21_+_8433172_8446622_RNA45SN1_45S_50ntFlanks@12420.12441.22\|ATGTCGGCTCTTCCTATCATTG | >hg38_21_+_8433172_8446622_RNA45SN1_45S_50ntFlanks | 12420 | 12441 | 22 | rRF 45S | NO | 449.06 | 262.55 | 0.00 | 59.58 |
| >hg38_21_+_8433172_8446622_RNA45SN1_45S_50ntFlanks@12420.12442.23\|ATGTCGGCTCTTCCTATCATTGT | >hg38_21_+_8433172_8446622_RNA45SN1_45S_50ntFlanks | 12420 | 12442 | 23 | rRF 45S | NO | 38.36 | 28.15 | 0.00 | 0.00 |
| >hg38_21_+_8433172_8446622_RNA45SN1_45S_50ntFlanks@12420.12443.24\|ATGTCGGCTCTTCCTATCATTGTG | >hg38_21_+_8433172_8446622_RNA45SN1_45S_50ntFlanks | 12420 | 12443 | 24 | rRF 45S | NO | 691.88 | 435.23 | 34.77 | 92.29 |
| >hg38_21_+_8433172_8446622_RNA45SN1_45S_50ntFlanks@12420.12444.25\|ATGTCGGCTCTTCCTATCATTGTGA | >hg38_21_+_8433172_8446622_RNA45SN1_45S_50ntFlanks | 12420 | 12444 | 25 | rRF 45S | NO | 287.16 | 154.04 | 0.00 | 25.98 |
| >hg38_21_+_8433172_8446622_RNA45SN1_45S_50ntFlanks@12420.12445.26\|ATGTCGGCTCTTCCTATCATTGTGAA | >hg38_21_+_8433172_8446622_RNA45SN1_45S_50ntFlanks | 12420 | 12445 | 26 | rRF 45S | NO | 199.19 | 104.10 | 0.00 | 14.96 |
| >hg38_21_+_8433172_8446622_RNA45SN1_45S_50ntFlanks@12420.12446.27\|ATGTCGGCTCTTCCTATCATTGTGAAG | >hg38_21_+_8433172_8446622_RNA45SN1_45S_50ntFlanks | 12420 | 12446 | 27 | rRF 45S | NO | 649.80 | 797.65 | 56.08 | 80.36 |
| >hg38_21_+_8433172_8446622_RNA45SN1_45S_50ntFlanks@12420.12447.28\|ATGTCGGCTCTTCCTATCATTGTGAAGC | >hg38_21_+_8433172_8446622_RNA45SN1_45S_50ntFlanks | 12420 | 12447 | 28 | rRF 45S | NO | 12.64 | 0.00 | 0.00 | 0.00 |
| >hg38_21_+_8433172_8446622_RNA45SN1_45S_50ntFlanks@12420.12448.29\|ATGTCGGCTCTTCCTATCATTGTGAAGCA | >hg38_21_+_8433172_8446622_RNA45SN1_45S_50ntFlanks | 12420 | 12448 | 29 | rRF 45S | NO | 31.05 | 29.00 | 0.00 | 6.07 |
| >hg38_21_+_8433172_8446622_RNA45SN1_45S_50ntFlanks@12420.12449.30\|ATGTCGGCTCTTCCTATCATTGTGAAGCAG | >hg38_21_+_8433172_8446622_RNA45SN1_45S_50ntFlanks | 12420 | 12449 | 30 | rRF 45S | NO | 68.41 | 90.23 | 0.00 | 13.50 |
| >hg38_21_+_8433172_8446622_RNA45SN1_45S_50ntFlanks@12420.12450.31\|ATGTCGGCTCTTCCTATCATTGTGAAGCAGA | >hg38_21_+_8433172_8446622_RNA45SN1_45S_50ntFlanks | 12420 | 12450 | 31 | rRF 45S | NO | 0.00 | 30.98 | 0.00 | 0.00 |
| >hg38_21_+_8433172_8446622_RNA45SN1_45S_50ntFlanks@12420.12455.36\|ATGTCGGCTCTTCCTATCATTGTGAAGCAGAATTCA | >hg38_21_+_8433172_8446622_RNA45SN1_45S_50ntFlanks | 12420 | 12455 | 36 | rRF 45S | NO | 201.88 | 241.51 | 13.88 | 151.96 |
| >hg38_21_+_8433172_8446622_RNA45SN1_45S_50ntFlanks@12420.12456.37\|ATGTCGGCTCTTCCTATCATTGTGAAGCAGAATTCAC | >hg38_21_+_8433172_8446622_RNA45SN1_45S_50ntFlanks | 12420 | 12456 | 37 | rRF 45S | NO | 118.56 | 78.94 | 0.00 | 35.52 |
| >hg38_21_+_8433172_8446622_RNA45SN1_45S_50ntFlanks@12423.12441.19\|TCGGCTCTTCCTATCATTG | >hg38_21_+_8433172_8446622_RNA45SN1_45S_50ntFlanks | 12423 | 12441 | 19 | rRF 45S | NO | 21.88 | 0.00 | 0.00 | 0.00 |
| >hg38_21_+_8433172_8446622_RNA45SN1_45S_50ntFlanks@12423.12443.21\|TCGGCTCTTCCTATCATTGTG | >hg38_21_+_8433172_8446622_RNA45SN1_45S_50ntFlanks | 12423 | 12443 | 21 | rRF 45S | NO | 44.46 | 41.30 | 0.00 | 0.00 |
| >hg38_21_+_8433172_8446622_RNA45SN1_45S_50ntFlanks@12431.12455.25\|TCCTATCATTGTGAAGCAGAATTCA | >hg38_21_+_8433172_8446622_RNA45SN1_45S_50ntFlanks | 12431 | 12455 | 25 | rRF 45S | NO | 12.59 | 0.00 | 0.00 | 0.00 |
| >hg38_21_+_8433172_8446622_RNA45SN1_45S_50ntFlanks@12432.12455.24\|CCTATCATTGTGAAGCAGAATTCA | >hg38_21_+_8433172_8446622_RNA45SN1_45S_50ntFlanks | 12432 | 12455 | 24 | rRF 45S | NO | 32.26 | 0.00 | 0.00 | 0.00 |
| >hg38_21_+_8433172_8446622_RNA45SN1_45S_50ntFlanks@12433.12455.23\|CTATCATTGTGAAGCAGAATTCA | >hg38_21_+_8433172_8446622_RNA45SN1_45S_50ntFlanks | 12433 | 12455 | 23 | rRF 45S | NO | 0.00 | 0.00 | 13.67 | 0.00 |
| >hg38_21_+_8433172_8446622_RNA45SN1_45S_50ntFlanks@12434.12455.22\|TATCATTGTGAAGCAGAATTCA | >hg38_21_+_8433172_8446622_RNA45SN1_45S_50ntFlanks | 12434 | 12455 | 22 | rRF 45S | NO | 13.02 | 0.00 | 0.00 | 0.00 |
| >hg38_21_+_8433172_8446622_RNA45SN1_45S_50ntFlanks@12435.12454.20\|ATCATTGTGAAGCAGAATTC | >hg38_21_+_8433172_8446622_RNA45SN1_45S_50ntFlanks | 12435 | 12454 | 20 | rRF 45S | NO | 33.55 | 0.00 | 34.34 | 0.00 |
| >hg38_21_+_8433172_8446622_RNA45SN1_45S_50ntFlanks@12435.12455.21\|ATCATTGTGAAGCAGAATTCA | >hg38_21_+_8433172_8446622_RNA45SN1_45S_50ntFlanks | 12435 | 12455 | 21 | rRF 45S | NO | 76.88 | 94.99 | 79.94 | 26.08 |
| >hg38_21_+_8433172_8446622_RNA45SN1_45S_50ntFlanks@12436.12454.19\|TCATTGTGAAGCAGAATTC | >hg38_21_+_8433172_8446622_RNA45SN1_45S_50ntFlanks | 12436 | 12454 | 19 | rRF 45S | NO | 72.80 | 72.22 | 76.40 | 0.00 |
| >hg38_21_+_8433172_8446622_RNA45SN1_45S_50ntFlanks@12436.12455.20\|TCATTGTGAAGCAGAATTCA | >hg38_21_+_8433172_8446622_RNA45SN1_45S_50ntFlanks | 12436 | 12455 | 20 | rRF 45S | NO | 109.15 | 218.85 | 142.89 | 77.56 |
| >hg38_21_+_8433172_8446622_RNA45SN1_45S_50ntFlanks@12436.12473.38\|TCATTGTGAAGCAGAATTCACCAAGCGTTGGATTGTTC | >hg38_21_+_8433172_8446622_RNA45SN1_45S_50ntFlanks | 12436 | 12473 | 38 | rRF 45S | NO | 21.69 | 91.57 | 0.00 | 0.00 |
| >hg38_21_+_8433172_8446622_RNA45SN1_45S_50ntFlanks@12437.12454.18\|CATTGTGAAGCAGAATTC | >hg38_21_+_8433172_8446622_RNA45SN1_45S_50ntFlanks | 12437 | 12454 | 18 | rRF 45S | NO | 151.69 | 168.93 | 142.32 | 15.12 |
| >hg38_21_+_8433172_8446622_RNA45SN1_45S_50ntFlanks@12437.12455.19\|CATTGTGAAGCAGAATTCA | >hg38_21_+_8433172_8446622_RNA45SN1_45S_50ntFlanks | 12437 | 12455 | 19 | rRF 45S | YES | 397.72 | 1125.83 | 513.00 | 250.07 |
| >hg38_21_+_8433172_8446622_RNA45SN1_45S_50ntFlanks@12437.12472.36\|CATTGTGAAGCAGAATTCACCAAGCGTTGGATTGTT | >hg38_21_+_8433172_8446622_RNA45SN1_45S_50ntFlanks | 12437 | 12472 | 36 | rRF 45S | NO | 0.00 | 31.50 | 0.00 | 0.00 |
| >hg38_21_+_8433172_8446622_RNA45SN1_45S_50ntFlanks@12437.12473.37\|CATTGTGAAGCAGAATTCACCAAGCGTTGGATTGTTC | >hg38_21_+_8433172_8446622_RNA45SN1_45S_50ntFlanks | 12437 | 12473 | 37 | rRF 45S | NO | 113.78 | 490.91 | 52.24 | 16.20 |
| >hg38_21_+_8433172_8446622_RNA45SN1_45S_50ntFlanks@12437.12474.38\|CATTGTGAAGCAGAATTCACCAAGCGTTGGATTGTTCA | >hg38_21_+_8433172_8446622_RNA45SN1_45S_50ntFlanks | 12437 | 12474 | 38 | rRF 45S | NO | 0.00 | 46.05 | 0.00 | 0.00 |
| >hg38_21_+_8433172_8446622_RNA45SN1_45S_50ntFlanks@12438.12455.18\|ATTGTGAAGCAGAATTCA | >hg38_21_+_8433172_8446622_RNA45SN1_45S_50ntFlanks | 12438 | 12455 | 18 | rRF 45S | YES | 72.60 | 182.31 | 176.97 | 55.64 |
| >hg38_21_+_8433172_8446622_RNA45SN1_45S_50ntFlanks@12438.12473.36\|ATTGTGAAGCAGAATTCACCAAGCGTTGGATTGTTC | >hg38_21_+_8433172_8446622_RNA45SN1_45S_50ntFlanks | 12438 | 12473 | 36 | rRF 45S | NO | 20.59 | 95.25 | 0.00 | 0.00 |

### MDA-MB-468

| | | | | | | | | | | |
|---|---|---|---|---|---|---|---|---|---|---|
| >hg38_21_+_8433172_8446622_RNA45SN1_45S_50ntFlanks@12444.12476.33\|AAGCAGAATTCACCAAGCGTTGGATTGTTCACC | >hg38_21_+_8433172_8446622_RNA45SN1_45S_50ntFlanks | 12444 | 12476 | 33 | rRF 45S | NO | 0.00 | 26.02 | 28.11 | 0.00 |
| >hg38_21_+_8433172_8446622_RNA45SN1_45S_50ntFlanks@12444.12480.37\|AAGCAGAATTCACCAAGCGTTGGATTGTTCACCCACT | >hg38_21_+_8433172_8446622_RNA45SN1_45S_50ntFlanks | 12444 | 12480 | 37 | rRF 45S | NO | 12.87 | 0.00 | 0.00 | 0.00 |
| >hg38_21_+_8433172_8446622_RNA45SN1_45S_50ntFlanks@12444.12484.41\|AAGCAGAATTCACCAAGCGTTGGATTGTTCACCCACTAATA | >hg38_21_+_8433172_8446622_RNA45SN1_45S_50ntFlanks | 12444 | 12484 | 41 | rRF 45S | NO | 60.98 | 0.00 | 0.00 | 0.00 |
| >hg38_21_+_8433172_8446622_RNA45SN1_45S_50ntFlanks@12444.12485.42\|AAGCAGAATTCACCAAGCGTTGGATTGTTCACCCACTAATAG | >hg38_21_+_8433172_8446622_RNA45SN1_45S_50ntFlanks | 12444 | 12485 | 42 | rRF 45S | NO | 39.69 | 0.00 | 0.00 | 0.00 |
| >hg38_21_+_8433172_8446622_RNA45SN1_45S_50ntFlanks@12444.12486.43\|AAGCAGAATTCACCAAGCGTTGGATTGTTCACCCACTAATAGG | >hg38_21_+_8433172_8446622_RNA45SN1_45S_50ntFlanks | 12444 | 12486 | 43 | rRF 45S | NO | 40.50 | 0.00 | 0.00 | 0.00 |
| >hg38_21_+_8433172_8446622_RNA45SN1_45S_50ntFlanks@12455.12473.19\|ACCAAGCGTTGGATTGTTC | >hg38_21_+_8433172_8446622_RNA45SN1_45S_50ntFlanks | 12455 | 12473 | 19 | rRF 45S | NO | 0.00 | 0.00 | 46.52 | 0.00 |
| >hg38_21_+_8433172_8446622_RNA45SN1_45S_50ntFlanks@12455.12476.22\|ACCAAGCGTTGGATTGTTCACC | >hg38_21_+_8433172_8446622_RNA45SN1_45S_50ntFlanks | 12455 | 12476 | 22 | rRF 45S | NO | 0.00 | 47.36 | 20.42 | 0.00 |
| >hg38_21_+_8433172_8446622_RNA45SN1_45S_50ntFlanks@12456.12473.18\|CCAAGCGTTGGATTGTTC | >hg38_21_+_8433172_8446622_RNA45SN1_45S_50ntFlanks | 12456 | 12473 | 18 | rRF 45S | YES | 77.35 | 813.24 | 919.47 | 45.37 |
| >hg38_21_+_8433172_8446622_RNA45SN1_45S_50ntFlanks@12456.12474.19\|CCAAGCGTTGGATTGTTCA | >hg38_21_+_8433172_8446622_RNA45SN1_45S_50ntFlanks | 12456 | 12474 | 19 | rRF 45S | NO | 73.74 | 815.55 | 219.60 | 47.34 |
| >hg38_21_+_8433172_8446622_RNA45SN1_45S_50ntFlanks@12456.12475.20\|CCAAGCGTTGGATTGTTCAC | >hg38_21_+_8433172_8446622_RNA45SN1_45S_50ntFlanks | 12456 | 12475 | 20 | rRF 45S | YES | 376.94 | 1728.20 | 246.87 | 178.97 |
| >hg38_21_+_8433172_8446622_RNA45SN1_45S_50ntFlanks@12456.12476.21\|CCAAGCGTTGGATTGTTCACC | >hg38_21_+_8433172_8446622_RNA45SN1_45S_50ntFlanks | 12456 | 12476 | 21 | rRF 45S | YES | 847.47 | 2698.55 | 762.76 | 392.23 |
| >hg38_21_+_8433172_8446622_RNA45SN1_45S_50ntFlanks@12456.12477.22\|CCAAGCGTTGGATTGTTCACCC | >hg38_21_+_8433172_8446622_RNA45SN1_45S_50ntFlanks | 12456 | 12477 | 22 | rRF 45S | NO | 288.08 | 668.94 | 182.38 | 136.04 |
| >hg38_21_+_8433172_8446622_RNA45SN1_45S_50ntFlanks@12456.12478.23\|CCAAGCGTTGGATTGTTCACCCA | >hg38_21_+_8433172_8446622_RNA45SN1_45S_50ntFlanks | 12456 | 12478 | 23 | rRF 45S | YES | 638.99 | 909.91 | 170.63 | 236.16 |
| >hg38_21_+_8433172_8446622_RNA45SN1_45S_50ntFlanks@12456.12479.24\|CCAAGCGTTGGATTGTTCACCCAC | >hg38_21_+_8433172_8446622_RNA45SN1_45S_50ntFlanks | 12456 | 12479 | 24 | rRF 45S | NO | 28.69 | 46.31 | 52.26 | 16.40 |
| >hg38_21_+_8433172_8446622_RNA45SN1_45S_50ntFlanks@12456.12480.25\|CCAAGCGTTGGATTGTTCACCCACT | >hg38_21_+_8433172_8446622_RNA45SN1_45S_50ntFlanks | 12456 | 12480 | 25 | rRF 45S | NO | 58.67 | 204.79 | 124.88 | 46.41 |
| >hg38_21_+_8433172_8446622_RNA45SN1_45S_50ntFlanks@12456.12481.26\|CCAAGCGTTGGATTGTTCACCCACTA | >hg38_21_+_8433172_8446622_RNA45SN1_45S_50ntFlanks | 12456 | 12481 | 26 | rRF 45S | NO | 39.51 | 94.32 | 39.24 | 23.98 |
| >hg38_21_+_8433172_8446622_RNA45SN1_45S_50ntFlanks@12456.12483.28\|CCAAGCGTTGGATTGTTCACCCACTAAT | >hg38_21_+_8433172_8446622_RNA45SN1_45S_50ntFlanks | 12456 | 12483 | 28 | rRF 45S | NO | 159.60 | 115.96 | 37.26 | 54.08 |
| >hg38_21_+_8433172_8446622_RNA45SN1_45S_50ntFlanks@12456.12484.29\|CCAAGCGTTGGATTGTTCACCCACTAATA | >hg38_21_+_8433172_8446622_RNA45SN1_45S_50ntFlanks | 12456 | 12484 | 29 | rRF 45S | YES | 724.77 | 542.90 | 123.21 | 251.12 |
| >hg38_21_+_8433172_8446622_RNA45SN1_45S_50ntFlanks@12456.12485.30\|CCAAGCGTTGGATTGTTCACCCACTAATAG | >hg38_21_+_8433172_8446622_RNA45SN1_45S_50ntFlanks | 12456 | 12485 | 30 | rRF 45S | NO | 309.12 | 204.30 | 56.20 | 75.41 |
| >hg38_21_+_8433172_8446622_RNA45SN1_45S_50ntFlanks@12456.12486.31\|CCAAGCGTTGGATTGTTCACCCACTAATAGG | >hg38_21_+_8433172_8446622_RNA45SN1_45S_50ntFlanks | 12456 | 12486 | 31 | rRF 45S | YES | 761.47 | 504.38 | 131.37 | 161.02 |
| >hg38_21_+_8433172_8446622_RNA45SN1_45S_50ntFlanks@12456.12487.32\|CCAAGCGTTGGATTGTTCACCCACTAATAGGG | >hg38_21_+_8433172_8446622_RNA45SN1_45S_50ntFlanks | 12456 | 12487 | 32 | rRF 45S | YES | 3646.01 | 3258.44 | 376.85 | 393.97 |
| >hg38_21_+_8433172_8446622_RNA45SN1_45S_50ntFlanks@12456.12488.33\|CCAAGCGTTGGATTGTTCACCCACTAATAGGGA | >hg38_21_+_8433172_8446622_RNA45SN1_45S_50ntFlanks | 12456 | 12488 | 33 | rRF 45S | YES | 562.19 | 553.88 | 139.35 | 115.32 |
| >hg38_21_+_8433172_8446622_RNA45SN1_45S_50ntFlanks@12456.12489.34\|CCAAGCGTTGGATTGTTCACCCACTAATAGGGAA | >hg38_21_+_8433172_8446622_RNA45SN1_45S_50ntFlanks | 12456 | 12489 | 34 | rRF 45S | NO | 136.32 | 103.84 | 34.59 | 38.16 |
| >hg38_21_+_8433172_8446622_RNA45SN1_45S_50ntFlanks@12456.12490.35\|CCAAGCGTTGGATTGTTCACCCACTAATAGGGAAC | >hg38_21_+_8433172_8446622_RNA45SN1_45S_50ntFlanks | 12456 | 12490 | 35 | rRF 45S | NO | 109.04 | 95.35 | 38.83 | 46.19 |
| >hg38_21_+_8433172_8446622_RNA45SN1_45S_50ntFlanks@12456.12493.38\|CCAAGCGTTGGATTGTTCACCCACTAATAGGGAACGTG | >hg38_21_+_8433172_8446622_RNA45SN1_45S_50ntFlanks | 12456 | 12493 | 38 | rRF 45S | NO | 44.41 | 43.97 | 0.00 | 15.23 |
| >hg38_21_+_8433172_8446622_RNA45SN1_45S_50ntFlanks@12456.12496.41\|CCAAGCGTTGGATTGTTCACCCACTAATAGGGAACGTGAGC | >hg38_21_+_8433172_8446622_RNA45SN1_45S_50ntFlanks | 12456 | 12496 | 41 | rRF 45S | NO | 51.57 | 53.04 | 0.00 | 13.89 |
| >hg38_21_+_8433172_8446622_RNA45SN1_45S_50ntFlanks@12456.12499.44\|CCAAGCGTTGGATTGTTCACCCACTAATAGGGAACGTGAGCTGG | >hg38_21_+_8433172_8446622_RNA45SN1_45S_50ntFlanks | 12456 | 12499 | 44 | rRF 45S | NO | 0.00 | 49.19 | 0.00 | 0.00 |
| >hg38_21_+_8433172_8446622_RNA45SN1_45S_50ntFlanks@12457.12474.18\|CAAGCGTTGGATTGTTCA | >hg38_21_+_8433172_8446622_RNA45SN1_45S_50ntFlanks | 12457 | 12474 | 18 | rRF 45S | NO | 0.00 | 117.81 | 0.00 | 0.00 |
| >hg38_21_+_8433172_8446622_RNA45SN1_45S_50ntFlanks@12457.12475.19\|CAAGCGTTGGATTGTTCAC | >hg38_21_+_8433172_8446622_RNA45SN1_45S_50ntFlanks | 12457 | 12475 | 19 | rRF 45S | NO | 20.20 | 113.99 | 0.00 | 25.38 |
| >hg38_21_+_8433172_8446622_RNA45SN1_45S_50ntFlanks@12457.12476.20\|CAAGCGTTGGATTGTTCACC | >hg38_21_+_8433172_8446622_RNA45SN1_45S_50ntFlanks | 12457 | 12476 | 20 | rRF 45S | YES | 237.96 | 639.42 | 84.81 | 107.36 |
| >hg38_21_+_8433172_8446622_RNA45SN1_45S_50ntFlanks@12457.12477.21\|CAAGCGTTGGATTGTTCACCC | >hg38_21_+_8433172_8446622_RNA45SN1_45S_50ntFlanks | 12457 | 12477 | 21 | rRF 45S | NO | 188.32 | 328.82 | 31.99 | 53.95 |
| >hg38_21_+_8433172_8446622_RNA45SN1_45S_50ntFlanks@12457.12478.22\|CAAGCGTTGGATTGTTCACCCA | >hg38_21_+_8433172_8446622_RNA45SN1_45S_50ntFlanks | 12457 | 12478 | 22 | rRF 45S | NO | 203.32 | 301.65 | 33.64 | 80.75 |
| >hg38_21_+_8433172_8446622_RNA45SN1_45S_50ntFlanks@12457.12484.28\|CAAGCGTTGGATTGTTCACCCACTAATA | >hg38_21_+_8433172_8446622_RNA45SN1_45S_50ntFlanks | 12457 | 12484 | 28 | rRF 45S | NO | 33.60 | 34.91 | 0.00 | 35.15 |
| >hg38_21_+_8433172_8446622_RNA45SN1_45S_50ntFlanks@12457.12486.30\|CAAGCGTTGGATTGTTCACCCACTAATAGG | >hg38_21_+_8433172_8446622_RNA45SN1_45S_50ntFlanks | 12457 | 12486 | 30 | rRF 45S | NO | 34.86 | 0.00 | 0.00 | 0.00 |

### MDA-MB-468

| | | | | | | | | | | |
|---|---|---|---|---|---|---|---|---|---|---|
| >hg38_21_+_8433172_8446622_RNA45SN1_45S_50ntFlanks@12457.12487.31\|CAAGCGTTGGATTGTTCACCCACTAATAGGG | >hg38_21_+_8433172_8446622_RNA45SN1_45S_50ntFlanks | 12457 | 12487 | 31 | rRF 45S | NO | 191.49 | 148.72 | 0.00 | 40.12 |
| >hg38_21_+_8433172_8446622_RNA45SN1_45S_50ntFlanks@12457.12488.32\|CAAGCGTTGGATTGTTCACCCACTAATAGGGA | >hg38_21_+_8433172_8446622_RNA45SN1_45S_50ntFlanks | 12457 | 12488 | 32 | rRF 45S | NO | 70.87 | 47.30 | 0.00 | 0.00 |
| >hg38_21_+_8433172_8446622_RNA45SN1_45S_50ntFlanks@12458.12475.18\|AAGCGTTGGATTGTTCAC | >hg38_21_+_8433172_8446622_RNA45SN1_45S_50ntFlanks | 12458 | 12475 | 18 | rRF 45S | NO | 0.00 | 56.89 | 0.00 | 0.00 |
| >hg38_21_+_8433172_8446622_RNA45SN1_45S_50ntFlanks@12458.12476.19\|AAGCGTTGGATTGTTCACC | >hg38_21_+_8433172_8446622_RNA45SN1_45S_50ntFlanks | 12458 | 12476 | 19 | rRF 45S | NO | 114.72 | 307.17 | 46.75 | 36.96 |
| >hg38_21_+_8433172_8446622_RNA45SN1_45S_50ntFlanks@12458.12477.20\|AAGCGTTGGATTGTTCACCC | >hg38_21_+_8433172_8446622_RNA45SN1_45S_50ntFlanks | 12458 | 12477 | 20 | rRF 45S | NO | 56.88 | 106.80 | 0.00 | 0.00 |
| >hg38_21_+_8433172_8446622_RNA45SN1_45S_50ntFlanks@12458.12478.21\|AAGCGTTGGATTGTTCACCCA | >hg38_21_+_8433172_8446622_RNA45SN1_45S_50ntFlanks | 12458 | 12478 | 21 | rRF 45S | NO | 63.14 | 96.06 | 0.00 | 13.59 |
| >hg38_21_+_8433172_8446622_RNA45SN1_45S_50ntFlanks@12458.12487.30\|AAGCGTTGGATTGTTCACCCACTAATAGGG | >hg38_21_+_8433172_8446622_RNA45SN1_45S_50ntFlanks | 12458 | 12487 | 30 | rRF 45S | NO | 83.48 | 62.84 | 0.00 | 0.00 |
| >hg38_21_+_8433172_8446622_RNA45SN1_45S_50ntFlanks@12459.12476.18\|AGCGTTGGATTGTTCACC | >hg38_21_+_8433172_8446622_RNA45SN1_45S_50ntFlanks | 12459 | 12476 | 18 | rRF 45S | NO | 122.65 | 272.91 | 57.00 | 39.03 |
| >hg38_21_+_8433172_8446622_RNA45SN1_45S_50ntFlanks@12459.12477.19\|AGCGTTGGATTGTTCACCC | >hg38_21_+_8433172_8446622_RNA45SN1_45S_50ntFlanks | 12459 | 12477 | 19 | rRF 45S | NO | 39.63 | 75.53 | 0.00 | 0.00 |
| >hg38_21_+_8433172_8446622_RNA45SN1_45S_50ntFlanks@12459.12478.20\|AGCGTTGGATTGTTCACCCA | >hg38_21_+_8433172_8446622_RNA45SN1_45S_50ntFlanks | 12459 | 12478 | 20 | rRF 45S | NO | 33.34 | 65.64 | 0.00 | 0.00 |
| >hg38_21_+_8433172_8446622_RNA45SN1_45S_50ntFlanks@12459.12487.29\|AGCGTTGGATTGTTCACCCACTAATAGGG | >hg38_21_+_8433172_8446622_RNA45SN1_45S_50ntFlanks | 12459 | 12487 | 29 | rRF 45S | NO | 84.70 | 47.50 | 0.00 | 0.00 |
| >hg38_21_+_8433172_8446622_RNA45SN1_45S_50ntFlanks@12474.12496.23\|ACCCACTAATAGGGAACGTGAGC | >hg38_21_+_8433172_8446622_RNA45SN1_45S_50ntFlanks | 12474 | 12496 | 23 | rRF 45S | NO | 0.00 | 27.82 | 0.00 | 0.00 |
| >hg38_21_+_8433172_8446622_RNA45SN1_45S_50ntFlanks@12474.12498.25\|ACCCACTAATAGGGAACGTGAGCTG | >hg38_21_+_8433172_8446622_RNA45SN1_45S_50ntFlanks | 12474 | 12498 | 25 | rRF 45S | NO | 0.00 | 33.57 | 0.00 | 0.00 |
| >hg38_21_+_8433172_8446622_RNA45SN1_45S_50ntFlanks@12475.12496.22\|CCCACTAATAGGGAACGTGAGC | >hg38_21_+_8433172_8446622_RNA45SN1_45S_50ntFlanks | 12475 | 12496 | 22 | rRF 45S | NO | 0.00 | 31.23 | 0.00 | 0.00 |
| >hg38_21_+_8433172_8446622_RNA45SN1_45S_50ntFlanks@12475.12498.24\|CCCACTAATAGGGAACGTGAGCTG | >hg38_21_+_8433172_8446622_RNA45SN1_45S_50ntFlanks | 12475 | 12498 | 24 | rRF 45S | NO | 49.04 | 50.79 | 0.00 | 6.43 |
| >hg38_21_+_8433172_8446622_RNA45SN1_45S_50ntFlanks@12476.12493.18\|CCACTAATAGGGAACGTG | >hg38_21_+_8433172_8446622_RNA45SN1_45S_50ntFlanks | 12476 | 12493 | 18 | rRF 45S | NO | 0.00 | 42.27 | 0.00 | 0.00 |
| >hg38_21_+_8433172_8446622_RNA45SN1_45S_50ntFlanks@12476.12496.21\|CCACTAATAGGGAACGTGAGC | >hg38_21_+_8433172_8446622_RNA45SN1_45S_50ntFlanks | 12476 | 12496 | 21 | rRF 45S | NO | 137.86 | 158.29 | 72.86 | 58.06 |
| >hg38_21_+_8433172_8446622_RNA45SN1_45S_50ntFlanks@12476.12497.22\|CCACTAATAGGGAACGTGAGCT | >hg38_21_+_8433172_8446622_RNA45SN1_45S_50ntFlanks | 12476 | 12497 | 22 | rRF 45S | NO | 110.39 | 137.81 | 48.53 | 35.37 |
| >hg38_21_+_8433172_8446622_RNA45SN1_45S_50ntFlanks@12476.12498.23\|CCACTAATAGGGAACGTGAGCTG | >hg38_21_+_8433172_8446622_RNA45SN1_45S_50ntFlanks | 12476 | 12498 | 23 | rRF 45S | NO | 156.35 | 197.06 | 64.24 | 40.81 |
| >hg38_21_+_8433172_8446622_RNA45SN1_45S_50ntFlanks@12476.12499.24\|CCACTAATAGGGAACGTGAGCTGG | >hg38_21_+_8433172_8446622_RNA45SN1_45S_50ntFlanks | 12476 | 12499 | 24 | rRF 45S | NO | 21.32 | 39.36 | 0.00 | 0.00 |
| >hg38_21_+_8433172_8446622_RNA45SN1_45S_50ntFlanks@12476.12500.25\|CCACTAATAGGGAACGTGAGCTGGG | >hg38_21_+_8433172_8446622_RNA45SN1_45S_50ntFlanks | 12476 | 12500 | 25 | rRF 45S | NO | 71.92 | 66.75 | 0.00 | 0.00 |
| >hg38_21_+_8433172_8446622_RNA45SN1_45S_50ntFlanks@12477.12496.20\|CACTAATAGGGAACGTGAGC | >hg38_21_+_8433172_8446622_RNA45SN1_45S_50ntFlanks | 12477 | 12496 | 20 | rRF 45S | NO | 20.06 | 51.03 | 0.00 | 0.00 |
| >hg38_21_+_8433172_8446622_RNA45SN1_45S_50ntFlanks@12477.12497.21\|CACTAATAGGGAACGTGAGCT | >hg38_21_+_8433172_8446622_RNA45SN1_45S_50ntFlanks | 12477 | 12497 | 21 | rRF 45S | NO | 0.00 | 43.55 | 0.00 | 0.00 |
| >hg38_21_+_8433172_8446622_RNA45SN1_45S_50ntFlanks@12477.12498.22\|CACTAATAGGGAACGTGAGCTG | >hg38_21_+_8433172_8446622_RNA45SN1_45S_50ntFlanks | 12477 | 12498 | 22 | rRF 45S | NO | 85.39 | 144.59 | 0.00 | 0.00 |
| >hg38_21_+_8433172_8446622_RNA45SN1_45S_50ntFlanks@12477.12499.23\|CACTAATAGGGAACGTGAGCTGG | >hg38_21_+_8433172_8446622_RNA45SN1_45S_50ntFlanks | 12477 | 12499 | 23 | rRF 45S | NO | 0.00 | 41.37 | 0.00 | 0.00 |
| >hg38_21_+_8433172_8446622_RNA45SN1_45S_50ntFlanks@12478.12496.19\|ACTAATAGGGAACGTGAGC | >hg38_21_+_8433172_8446622_RNA45SN1_45S_50ntFlanks | 12478 | 12496 | 19 | rRF 45S | NO | 31.97 | 28.94 | 0.00 | 0.00 |
| >hg38_21_+_8433172_8446622_RNA45SN1_45S_50ntFlanks@12478.12497.20\|ACTAATAGGGAACGTGAGCT | >hg38_21_+_8433172_8446622_RNA45SN1_45S_50ntFlanks | 12478 | 12497 | 20 | rRF 45S | NO | 21.65 | 32.53 | 0.00 | 0.00 |
| >hg38_21_+_8433172_8446622_RNA45SN1_45S_50ntFlanks@12478.12498.21\|ACTAATAGGGAACGTGAGCTG | >hg38_21_+_8433172_8446622_RNA45SN1_45S_50ntFlanks | 12478 | 12498 | 21 | rRF 45S | NO | 59.15 | 60.23 | 0.00 | 0.00 |
| >hg38_21_+_8433172_8446622_RNA45SN1_45S_50ntFlanks@12478.12499.22\|ACTAATAGGGAACGTGAGCTGG | >hg38_21_+_8433172_8446622_RNA45SN1_45S_50ntFlanks | 12478 | 12499 | 22 | rRF 45S | NO | 30.11 | 28.70 | 0.00 | 0.00 |
| >hg38_21_+_8433172_8446622_RNA45SN1_45S_50ntFlanks@12479.12496.18\|CTAATAGGGAACGTGAGC | >hg38_21_+_8433172_8446622_RNA45SN1_45S_50ntFlanks | 12479 | 12496 | 18 | rRF 45S | NO | 48.19 | 67.57 | 0.00 | 0.00 |
| >hg38_21_+_8433172_8446622_RNA45SN1_45S_50ntFlanks@12479.12497.19\|CTAATAGGGAACGTGAGCT | >hg38_21_+_8433172_8446622_RNA45SN1_45S_50ntFlanks | 12479 | 12497 | 19 | rRF 45S | NO | 48.65 | 78.97 | 0.00 | 0.00 |
| >hg38_21_+_8433172_8446622_RNA45SN1_45S_50ntFlanks@12479.12498.20\|CTAATAGGGAACGTGAGCTG | >hg38_21_+_8433172_8446622_RNA45SN1_45S_50ntFlanks | 12479 | 12498 | 20 | rRF 45S | NO | 76.69 | 114.57 | 0.00 | 0.00 |
| >hg38_21_+_8433172_8446622_RNA45SN1_45S_50ntFlanks@12479.12499.21\|CTAATAGGGAACGTGAGCTGG | >hg38_21_+_8433172_8446622_RNA45SN1_45S_50ntFlanks | 12479 | 12499 | 21 | rRF 45S | NO | 38.98 | 47.34 | 0.00 | 0.00 |
| >hg38_21_+_8433172_8446622_RNA45SN1_45S_50ntFlanks@12479.12500.22\|CTAATAGGGAACGTGAGCTGGG | >hg38_21_+_8433172_8446622_RNA45SN1_45S_50ntFlanks | 12479 | 12500 | 22 | rRF 45S | NO | 38.22 | 42.28 | 0.00 | 0.00 |

### MDA-MB-468

| | | | | | | | | | | |
|---|---|---|---|---|---|---|---|---|---|---|
| >hg38_21_+_8433172_8446622_RNA45SN1_45S_50ntFlanks@12481.12498.18\|AATAGGGAACGTGAGCTG | >hg38_21_+_8433172_8446622_RNA45SN1_45S_50ntFlanks | 12481 | 12498 | 18 | rRF 45S | NO | 29.24 | 31.57 | 0.00 | 0.00 |
| >hg38_21_+_8433172_8446622_RNA45SN1_45S_50ntFlanks@12488.12505.18\|AACGTGAGCTGGGTTTAG | >hg38_21_+_8433172_8446622_RNA45SN1_45S_50ntFlanks | 12488 | 12505 | 18 | rRF 45S | NO | 12.66 | 0.00 | 0.00 | 0.00 |
| >hg38_21_+_8433172_8446622_RNA45SN1_45S_50ntFlanks@12488.12509.22\|AACGTGAGCTGGGTTTAGACCG | >hg38_21_+_8433172_8446622_RNA45SN1_45S_50ntFlanks | 12488 | 12509 | 22 | rRF 45S | NO | 341.19 | 336.69 | 64.97 | 16.52 |
| >hg38_21_+_8433172_8446622_RNA45SN1_45S_50ntFlanks@12488.12521.34\|AACGTGAGCTGGGTTTAGACCGTCGTGAGACAGG | >hg38_21_+_8433172_8446622_RNA45SN1_45S_50ntFlanks | 12488 | 12521 | 34 | rRF 45S | NO | 37.73 | 41.56 | 0.00 | 23.69 |
| >hg38_21_+_8433172_8446622_RNA45SN1_45S_50ntFlanks@12488.12522.35\|AACGTGAGCTGGGTTTAGACCGTCGTGAGACAGGT | >hg38_21_+_8433172_8446622_RNA45SN1_45S_50ntFlanks | 12488 | 12522 | 35 | rRF 45S | NO | 28.99 | 0.00 | 0.00 | 0.00 |
| >hg38_21_+_8433172_8446622_RNA45SN1_45S_50ntFlanks@12488.12524.37\|AACGTGAGCTGGGTTTAGACCGTCGTGAGACAGGTTA | >hg38_21_+_8433172_8446622_RNA45SN1_45S_50ntFlanks | 12488 | 12524 | 37 | rRF 45S | NO | 20.88 | 0.00 | 0.00 | 0.00 |
| >hg38_21_+_8433172_8446622_RNA45SN1_45S_50ntFlanks@12488.12525.38\|AACGTGAGCTGGGTTTAGACCGTCGTGAGACAGGTTAG | >hg38_21_+_8433172_8446622_RNA45SN1_45S_50ntFlanks | 12488 | 12525 | 38 | rRF 45S | NO | 21.26 | 0.00 | 0.00 | 0.00 |
| >hg38_21_+_8433172_8446622_RNA45SN1_45S_50ntFlanks@12489.12509.21\|ACGTGAGCTGGGTTTAGACCG | >hg38_21_+_8433172_8446622_RNA45SN1_45S_50ntFlanks | 12489 | 12509 | 21 | rRF 45S | NO | 44.66 | 65.34 | 0.00 | 0.00 |
| >hg38_21_+_8433172_8446622_RNA45SN1_45S_50ntFlanks@12510.12527.18\|TCGTGAGACAGGTTAGTT | >hg38_21_+_8433172_8446622_RNA45SN1_45S_50ntFlanks | 12510 | 12527 | 18 | rRF 45S | NO | 0.00 | 91.87 | 0.00 | 0.00 |
| >hg38_21_+_8433172_8446622_RNA45SN1_45S_50ntFlanks@12539.12557.19\|ATGATGTGTTGTTGCCATG | >hg38_21_+_8433172_8446622_RNA45SN1_45S_50ntFlanks | 12539 | 12557 | 19 | rRF 45S | NO | 0.00 | 0.00 | 21.13 | 13.67 |
| >hg38_21_+_8433172_8446622_RNA45SN1_45S_50ntFlanks@12539.12563.25\|ATGATGTGTTGTTGCCATGGTAATC | >hg38_21_+_8433172_8446622_RNA45SN1_45S_50ntFlanks | 12539 | 12563 | 25 | rRF 45S | NO | 52.08 | 41.27 | 28.46 | 13.28 |
| >hg38_21_+_8433172_8446622_RNA45SN1_45S_50ntFlanks@12539.12565.27\|ATGATGTGTTGTTGCCATGGTAATCCT | >hg38_21_+_8433172_8446622_RNA45SN1_45S_50ntFlanks | 12539 | 12565 | 27 | rRF 45S | NO | 29.09 | 0.00 | 0.00 | 0.00 |
| >hg38_21_+_8433172_8446622_RNA45SN1_45S_50ntFlanks@12539.12566.28\|ATGATGTGTTGTTGCCATGGTAATCCTG | >hg38_21_+_8433172_8446622_RNA45SN1_45S_50ntFlanks | 12539 | 12566 | 28 | rRF 45S | NO | 99.89 | 95.49 | 61.55 | 43.58 |
| >hg38_21_+_8433172_8446622_RNA45SN1_45S_50ntFlanks@12539.12567.29\|ATGATGTGTTGTTGCCATGGTAATCCTGC | >hg38_21_+_8433172_8446622_RNA45SN1_45S_50ntFlanks | 12539 | 12567 | 29 | rRF 45S | NO | 78.13 | 57.60 | 35.42 | 24.99 |
| >hg38_21_+_8433172_8446622_RNA45SN1_45S_50ntFlanks@12539.12568.30\|ATGATGTGTTGTTGCCATGGTAATCCTGCT | >hg38_21_+_8433172_8446622_RNA45SN1_45S_50ntFlanks | 12539 | 12568 | 30 | rRF 45S | NO | 0.00 | 0.00 | 32.67 | 0.00 |
| >hg38_21_+_8433172_8446622_RNA45SN1_45S_50ntFlanks@12539.12569.31\|ATGATGTGTTGTTGCCATGGTAATCCTGCTC | >hg38_21_+_8433172_8446622_RNA45SN1_45S_50ntFlanks | 12539 | 12569 | 31 | rRF 45S | NO | 244.37 | 174.88 | 58.73 | 93.32 |
| >hg38_21_+_8433172_8446622_RNA45SN1_45S_50ntFlanks@12539.12573.35\|ATGATGTGTTGTTGCCATGGTAATCCTGCTCAGTA | >hg38_21_+_8433172_8446622_RNA45SN1_45S_50ntFlanks | 12539 | 12573 | 35 | rRF 45S | NO | 6.18 | 0.00 | 31.22 | 34.03 |
| >hg38_21_+_8433172_8446622_RNA45SN1_45S_50ntFlanks@12539.12587.49\|ATGATGTGTTGTTGCCATGGTAATCCTGCTCAGTACGAGAGGAACCGCA | >hg38_21_+_8433172_8446622_RNA45SN1_45S_50ntFlanks | 12539 | 12587 | 49 | rRF 45S | YES | 21.38 | 0.00 | 0.00 | 0.00 |
| >hg38_21_+_8433172_8446622_RNA45SN1_45S_50ntFlanks@12567.12587.21\|CTCAGTACGAGAGGAACCGCA | >hg38_21_+_8433172_8446622_RNA45SN1_45S_50ntFlanks | 12567 | 12587 | 21 | rRF 45S | NO | 29.99 | 36.51 | 0.00 | 46.28 |
| >hg38_21_+_8433172_8446622_RNA45SN1_45S_50ntFlanks@12570.12587.18\|AGTACGAGAGGAACCGCA | >hg38_21_+_8433172_8446622_RNA45SN1_45S_50ntFlanks | 12570 | 12587 | 18 | rRF 45S | NO | 41.92 | 54.91 | 57.49 | 39.32 |
| >hg38_21_+_8433172_8446622_RNA45SN1_45S_50ntFlanks@12574.12604.31\|CGAGAGGAACCGCAGGTTCAGACATTTGGTG | >hg38_21_+_8433172_8446622_RNA45SN1_45S_50ntFlanks | 12574 | 12604 | 31 | rRF 45S | NO | 0.00 | 0.00 | 40.97 | 0.00 |
| >hg38_21_+_8433172_8446622_RNA45SN1_45S_50ntFlanks@12578.12604.27\|AGGAACCGCAGGTTCAGACATTTGGTG | >hg38_21_+_8433172_8446622_RNA45SN1_45S_50ntFlanks | 12578 | 12604 | 27 | rRF 45S | NO | 0.00 | 0.00 | 48.89 | 0.00 |
| >hg38_21_+_8433172_8446622_RNA45SN1_45S_50ntFlanks@12589.12614.26\|GTTCAGACATTTGGTGTATGTGCTTG | >hg38_21_+_8433172_8446622_RNA45SN1_45S_50ntFlanks | 12589 | 12614 | 26 | rRF 45S | NO | 0.00 | 0.00 | 35.79 | 0.00 |
| >hg38_21_+_8433172_8446622_RNA45SN1_45S_50ntFlanks@12602.12626.25\|GTGTATGTGCTTGGCTGAGGAGCCA | >hg38_21_+_8433172_8446622_RNA45SN1_45S_50ntFlanks | 12602 | 12626 | 25 | rRF 45S | NO | 0.00 | 26.01 | 0.00 | 0.00 |
| >hg38_21_+_8433172_8446622_RNA45SN1_45S_50ntFlanks@12605.12622.18\|TATGTGCTTGGCTGAGGA | >hg38_21_+_8433172_8446622_RNA45SN1_45S_50ntFlanks | 12605 | 12622 | 18 | rRF 45S | NO | 36.67 | 0.00 | 0.00 | 0.00 |
| >hg38_21_+_8433172_8446622_RNA45SN1_45S_50ntFlanks@12605.12623.19\|TATGTGCTTGGCTGAGGAG | >hg38_21_+_8433172_8446622_RNA45SN1_45S_50ntFlanks | 12605 | 12623 | 19 | rRF 45S | NO | 76.43 | 42.04 | 0.00 | 0.00 |
| >hg38_21_+_8433172_8446622_RNA45SN1_45S_50ntFlanks@12605.12624.20\|TATGTGCTTGGCTGAGGAGC | >hg38_21_+_8433172_8446622_RNA45SN1_45S_50ntFlanks | 12605 | 12624 | 20 | rRF 45S | NO | 393.48 | 111.68 | 0.00 | 5.97 |
| >hg38_21_+_8433172_8446622_RNA45SN1_45S_50ntFlanks@12605.12625.21\|TATGTGCTTGGCTGAGGAGCC | >hg38_21_+_8433172_8446622_RNA45SN1_45S_50ntFlanks | 12605 | 12625 | 21 | rRF 45S | YES | 564.81 | 590.36 | 80.70 | 61.19 |
| >hg38_21_+_8433172_8446622_RNA45SN1_45S_50ntFlanks@12605.12626.22\|TATGTGCTTGGCTGAGGAGCCA | >hg38_21_+_8433172_8446622_RNA45SN1_45S_50ntFlanks | 12605 | 12626 | 22 | rRF 45S | NO | 240.34 | 454.85 | 70.79 | 43.62 |
| >hg38_21_+_8433172_8446622_RNA45SN1_45S_50ntFlanks@12606.12624.19\|ATGTGCTTGGCTGAGGAGC | >hg38_21_+_8433172_8446622_RNA45SN1_45S_50ntFlanks | 12606 | 12624 | 19 | rRF 45S | NO | 50.06 | 0.00 | 0.00 | 0.00 |
| >hg38_21_+_8433172_8446622_RNA45SN1_45S_50ntFlanks@12606.12625.20\|ATGTGCTTGGCTGAGGAGCC | >hg38_21_+_8433172_8446622_RNA45SN1_45S_50ntFlanks | 12606 | 12625 | 20 | rRF 45S | NO | 54.83 | 50.03 | 0.00 | 0.00 |
| >hg38_21_+_8433172_8446622_RNA45SN1_45S_50ntFlanks@12606.12626.21\|ATGTGCTTGGCTGAGGAGCCA | >hg38_21_+_8433172_8446622_RNA45SN1_45S_50ntFlanks | 12606 | 12626 | 21 | rRF 45S | NO | 5.93 | 32.77 | 0.00 | 0.00 |
| >hg38_21_+_8433172_8446622_RNA45SN1_45S_50ntFlanks@12607.12625.19\|TGTGCTTGGCTGAGGAGCC | >hg38_21_+_8433172_8446622_RNA45SN1_45S_50ntFlanks | 12607 | 12625 | 19 | rRF 45S | NO | 20.11 | 27.57 | 0.00 | 0.00 |
| >hg38_21_+_8433172_8446622_RNA45SN1_45S_50ntFlanks@12615.12653.39\|GCTGAGGAGCCAATGGGGCGAAGCTACCATCTGTGGGAT | >hg38_21_+_8433172_8446622_RNA45SN1_45S_50ntFlanks | 12615 | 12653 | 39 | rRF 45S | NO | 108.27 | 91.54 | 36.22 | 123.91 |

### MDA-MB-468

| | | | | | | | | | | |
|---|---|---|---|---|---|---|---|---|---|---|
| >hg38_21_+_8433172_8446622_RNA45SN1_45S_50ntFlanks@12615.12654.40\|GCTGAGGAGCCAATGGGGCGAAGCTACCATCTGTGGGATT | >hg38_21_+_8433172_8446622_RNA45SN1_45S_50ntFlanks | 12615 | 12654 | 40 | rRF 45S | NO | 465.74 | 268.69 | 53.95 | 321.93 |
| >hg38_21_+_8433172_8446622_RNA45SN1_45S_50ntFlanks@12615.12661.47\|GCTGAGGAGCCAATGGGGCGAAGCTACCATCTGTGGGATTATGACTG | >hg38_21_+_8433172_8446622_RNA45SN1_45S_50ntFlanks | 12615 | 12661 | 47 | rRF 45S | NO | 41.90 | 0.00 | 0.00 | 62.14 |
| >hg38_21_+_8433172_8446622_RNA45SN1_45S_50ntFlanks@12616.12653.38\|CTGAGGAGCCAATGGGGCGAAGCTACCATCTGTGGGAT | >hg38_21_+_8433172_8446622_RNA45SN1_45S_50ntFlanks | 12616 | 12653 | 38 | rRF 45S | NO | 36.81 | 26.52 | 0.00 | 49.01 |
| >hg38_21_+_8433172_8446622_RNA45SN1_45S_50ntFlanks@12616.12654.39\|CTGAGGAGCCAATGGGGCGAAGCTACCATCTGTGGGATT | >hg38_21_+_8433172_8446622_RNA45SN1_45S_50ntFlanks | 12616 | 12654 | 39 | rRF 45S | NO | 96.67 | 57.55 | 6.98 | 111.66 |
| >hg38_21_+_8433172_8446622_RNA45SN1_45S_50ntFlanks@12617.12653.37\|TGAGGAGCCAATGGGGCGAAGCTACCATCTGTGGGAT | >hg38_21_+_8433172_8446622_RNA45SN1_45S_50ntFlanks | 12617 | 12653 | 37 | rRF 45S | NO | 41.53 | 34.67 | 0.00 | 35.90 |
| >hg38_21_+_8433172_8446622_RNA45SN1_45S_50ntFlanks@12617.12654.38\|TGAGGAGCCAATGGGGCGAAGCTACCATCTGTGGGATT | >hg38_21_+_8433172_8446622_RNA45SN1_45S_50ntFlanks | 12617 | 12654 | 38 | rRF 45S | NO | 101.35 | 73.79 | 13.64 | 82.44 |
| >hg38_21_+_8433172_8446622_RNA45SN1_45S_50ntFlanks@12619.12653.35\|AGGAGCCAATGGGGCGAAGCTACCATCTGTGGGAT | >hg38_21_+_8433172_8446622_RNA45SN1_45S_50ntFlanks | 12619 | 12653 | 35 | rRF 45S | NO | 56.21 | 52.36 | 33.24 | 43.75 |
| >hg38_21_+_8433172_8446622_RNA45SN1_45S_50ntFlanks@12619.12654.36\|AGGAGCCAATGGGGCGAAGCTACCATCTGTGGGATT | >hg38_21_+_8433172_8446622_RNA45SN1_45S_50ntFlanks | 12619 | 12654 | 36 | rRF 45S | NO | 132.40 | 109.46 | 41.00 | 105.76 |
| >hg38_21_+_8433172_8446622_RNA45SN1_45S_50ntFlanks@12624.12652.29\|CCAATGGGGCGAAGCTACCATCTGTGGGA | >hg38_21_+_8433172_8446622_RNA45SN1_45S_50ntFlanks | 12624 | 12652 | 29 | rRF 45S | NO | 0.00 | 0.00 | 27.24 | 0.00 |
| >hg38_21_+_8433172_8446622_RNA45SN1_45S_50ntFlanks@12624.12653.30\|CCAATGGGGCGAAGCTACCATCTGTGGGAT | >hg38_21_+_8433172_8446622_RNA45SN1_45S_50ntFlanks | 12624 | 12653 | 30 | rRF 45S | NO | 0.00 | 12.82 | 64.27 | 66.91 |
| >hg38_21_+_8433172_8446622_RNA45SN1_45S_50ntFlanks@12624.12654.31\|CCAATGGGGCGAAGCTACCATCTGTGGGATT | >hg38_21_+_8433172_8446622_RNA45SN1_45S_50ntFlanks | 12624 | 12654 | 31 | rRF 45S | NO | 98.87 | 173.10 | 247.58 | 450.91 |
| >hg38_21_+_8433172_8446622_RNA45SN1_45S_50ntFlanks@12624.12657.34\|CCAATGGGGCGAAGCTACCATCTGTGGGATTATG | >hg38_21_+_8433172_8446622_RNA45SN1_45S_50ntFlanks | 12624 | 12657 | 34 | rRF 45S | NO | 39.60 | 19.49 | 50.38 | 136.86 |
| >hg38_21_+_8433172_8446622_RNA45SN1_45S_50ntFlanks@12624.12661.38\|CCAATGGGGCGAAGCTACCATCTGTGGGATTATGACTG | >hg38_21_+_8433172_8446622_RNA45SN1_45S_50ntFlanks | 12624 | 12661 | 38 | rRF 45S | NO | 136.22 | 39.85 | 50.00 | 255.16 |
| >hg38_21_+_8433172_8446622_RNA45SN1_45S_50ntFlanks@12625.12653.29\|CAATGGGGCGAAGCTACCATCTGTGGGAT | >hg38_21_+_8433172_8446622_RNA45SN1_45S_50ntFlanks | 12625 | 12653 | 29 | rRF 45S | NO | 71.41 | 104.14 | 74.71 | 118.95 |
| >hg38_21_+_8433172_8446622_RNA45SN1_45S_50ntFlanks@12625.12654.30\|CAATGGGGCGAAGCTACCATCTGTGGGATT | >hg38_21_+_8433172_8446622_RNA45SN1_45S_50ntFlanks | 12625 | 12654 | 30 | rRF 45S | NO | 314.55 | 256.78 | 129.41 | 285.02 |
| >hg38_21_+_8433172_8446622_RNA45SN1_45S_50ntFlanks@12625.12661.37\|CAATGGGGCGAAGCTACCATCTGTGGGATTATGACTG | >hg38_21_+_8433172_8446622_RNA45SN1_45S_50ntFlanks | 12625 | 12661 | 37 | rRF 45S | NO | 13.17 | 0.00 | 6.86 | 63.44 |
| >hg38_21_+_8433172_8446622_RNA45SN1_45S_50ntFlanks@12626.12652.27\|AATGGGGCGAAGCTACCATCTGTGGGA | >hg38_21_+_8433172_8446622_RNA45SN1_45S_50ntFlanks | 12626 | 12652 | 27 | rRF 45S | NO | 0.00 | 19.38 | 47.16 | 6.51 |
| >hg38_21_+_8433172_8446622_RNA45SN1_45S_50ntFlanks@12626.12653.28\|AATGGGGCGAAGCTACCATCTGTGGGAT | >hg38_21_+_8433172_8446622_RNA45SN1_45S_50ntFlanks | 12626 | 12653 | 28 | rRF 45S | NO | 184.00 | 281.27 | 264.20 | 221.21 |
| >hg38_21_+_8433172_8446622_RNA45SN1_45S_50ntFlanks@12626.12654.29\|AATGGGGCGAAGCTACCATCTGTGGGATT | >hg38_21_+_8433172_8446622_RNA45SN1_45S_50ntFlanks | 12626 | 12654 | 29 | rRF 45S | YES | 567.59 | 525.83 | 313.42 | 425.89 |
| >hg38_21_+_8433172_8446622_RNA45SN1_45S_50ntFlanks@12626.12657.32\|AATGGGGCGAAGCTACCATCTGTGGGATTATG | >hg38_21_+_8433172_8446622_RNA45SN1_45S_50ntFlanks | 12626 | 12657 | 32 | rRF 45S | NO | 39.74 | 12.35 | 35.40 | 64.67 |
| >hg38_21_+_8433172_8446622_RNA45SN1_45S_50ntFlanks@12626.12659.34\|AATGGGGCGAAGCTACCATCTGTGGGATTATGAC | >hg38_21_+_8433172_8446622_RNA45SN1_45S_50ntFlanks | 12626 | 12659 | 34 | rRF 45S | NO | 73.26 | 31.33 | 40.63 | 101.21 |
| >hg38_21_+_8433172_8446622_RNA45SN1_45S_50ntFlanks@12626.12661.36\|AATGGGGCGAAGCTACCATCTGTGGGATTATGACTG | >hg38_21_+_8433172_8446622_RNA45SN1_45S_50ntFlanks | 12626 | 12661 | 36 | rRF 45S | NO | 94.18 | 46.81 | 95.33 | 180.63 |
| >hg38_21_+_8433172_8446622_RNA45SN1_45S_50ntFlanks@12627.12652.26\|ATGGGGCGAAGCTACCATCTGTGGGA | >hg38_21_+_8433172_8446622_RNA45SN1_45S_50ntFlanks | 12627 | 12652 | 26 | rRF 45S | NO | 0.00 | 32.88 | 41.82 | 16.51 |
| >hg38_21_+_8433172_8446622_RNA45SN1_45S_50ntFlanks@12627.12653.27\|ATGGGGCGAAGCTACCATCTGTGGGAT | >hg38_21_+_8433172_8446622_RNA45SN1_45S_50ntFlanks | 12627 | 12653 | 27 | rRF 45S | NO | 149.32 | 223.26 | 263.03 | 242.10 |
| >hg38_21_+_8433172_8446622_RNA45SN1_45S_50ntFlanks@12627.12654.28\|ATGGGGCGAAGCTACCATCTGTGGGATT | >hg38_21_+_8433172_8446622_RNA45SN1_45S_50ntFlanks | 12627 | 12654 | 28 | rRF 45S | NO | 467.55 | 414.06 | 304.28 | 469.14 |
| >hg38_21_+_8433172_8446622_RNA45SN1_45S_50ntFlanks@12627.12657.31\|ATGGGGCGAAGCTACCATCTGTGGGATTATG | >hg38_21_+_8433172_8446622_RNA45SN1_45S_50ntFlanks | 12627 | 12657 | 31 | rRF 45S | NO | 54.98 | 28.95 | 42.74 | 110.39 |
| >hg38_21_+_8433172_8446622_RNA45SN1_45S_50ntFlanks@12627.12659.33\|ATGGGGCGAAGCTACCATCTGTGGGATTATGAC | >hg38_21_+_8433172_8446622_RNA45SN1_45S_50ntFlanks | 12627 | 12659 | 33 | rRF 45S | NO | 131.24 | 48.52 | 65.66 | 203.97 |
| >hg38_21_+_8433172_8446622_RNA45SN1_45S_50ntFlanks@12627.12661.35\|ATGGGGCGAAGCTACCATCTGTGGGATTATGACTG | >hg38_21_+_8433172_8446622_RNA45SN1_45S_50ntFlanks | 12627 | 12661 | 35 | rRF 45S | NO | 59.58 | 35.48 | 101.28 | 187.91 |
| >hg38_21_+_8433172_8446622_RNA45SN1_45S_50ntFlanks@12628.12653.26\|TGGGGCGAAGCTACCATCTGTGGGAT | >hg38_21_+_8433172_8446622_RNA45SN1_45S_50ntFlanks | 12628 | 12653 | 26 | rRF 45S | NO | 53.42 | 86.83 | 72.01 | 86.35 |
| >hg38_21_+_8433172_8446622_RNA45SN1_45S_50ntFlanks@12628.12654.27\|TGGGGCGAAGCTACCATCTGTGGGATT | >hg38_21_+_8433172_8446622_RNA45SN1_45S_50ntFlanks | 12628 | 12654 | 27 | rRF 45S | NO | 223.61 | 169.93 | 92.81 | 193.27 |
| >hg38_21_+_8433172_8446622_RNA45SN1_45S_50ntFlanks@12628.12659.32\|TGGGGCGAAGCTACCATCTGTGGGATTATGAC | >hg38_21_+_8433172_8446622_RNA45SN1_45S_50ntFlanks | 12628 | 12659 | 32 | rRF 45S | NO | 37.37 | 0.00 | 0.00 | 55.76 |
| >hg38_21_+_8433172_8446622_RNA45SN1_45S_50ntFlanks@12628.12661.34\|TGGGGCGAAGCTACCATCTGTGGGATTATGACTG | >hg38_21_+_8433172_8446622_RNA45SN1_45S_50ntFlanks | 12628 | 12661 | 34 | rRF 45S | NO | 73.10 | 0.00 | 50.43 | 111.91 |
| >hg38_21_+_8433172_8446622_RNA45SN1_45S_50ntFlanks@12632.12653.22\|GCGAAGCTACCATCTGTGGGAT | >hg38_21_+_8433172_8446622_RNA45SN1_45S_50ntFlanks | 12632 | 12653 | 22 | rRF 45S | NO | 0.00 | 43.09 | 93.81 | 66.56 |
| >hg38_21_+_8433172_8446622_RNA45SN1_45S_50ntFlanks@12632.12654.23\|GCGAAGCTACCATCTGTGGGATT | >hg38_21_+_8433172_8446622_RNA45SN1_45S_50ntFlanks | 12632 | 12654 | 23 | rRF 45S | NO | 78.80 | 82.00 | 130.41 | 151.03 |

### MDA-MB-468

| | | | | | | | | | | |
|---|---|---|---|---|---|---|---|---|---|---|
| >hg38_21_+_8433172_8446622_RNA45SN1_45S_50ntFlanks@12632.12659.28\|GCGAAGCTACCATCTGTGGGATTATGAC | >hg38_21_+_8433172_8446622_RNA45SN1_45S_50ntFlanks | 12632 | 12659 | 28 | rRF 45S | NO | 21.14 | 0.00 | 47.77 | 72.02 |
| >hg38_21_+_8433172_8446622_RNA45SN1_45S_50ntFlanks@12632.12661.30\|GCGAAGCTACCATCTGTGGGATTATGACTG | >hg38_21_+_8433172_8446622_RNA45SN1_45S_50ntFlanks | 12632 | 12661 | 30 | rRF 45S | NO | 30.05 | 0.00 | 60.76 | 77.08 |
| >hg38_21_+_8433172_8446622_RNA45SN1_45S_50ntFlanks@12633.12651.19\|CGAAGCTACCATCTGTGGG | >hg38_21_+_8433172_8446622_RNA45SN1_45S_50ntFlanks | 12633 | 12651 | 19 | rRF 45S | NO | 20.04 | 52.43 | 71.20 | 24.81 |
| >hg38_21_+_8433172_8446622_RNA45SN1_45S_50ntFlanks@12633.12652.20\|CGAAGCTACCATCTGTGGGA | >hg38_21_+_8433172_8446622_RNA45SN1_45S_50ntFlanks | 12633 | 12652 | 20 | rRF 45S | NO | 0.00 | 40.37 | 93.30 | 39.31 |
| >hg38_21_+_8433172_8446622_RNA45SN1_45S_50ntFlanks@12633.12653.21\|CGAAGCTACCATCTGTGGGAT | >hg38_21_+_8433172_8446622_RNA45SN1_45S_50ntFlanks | 12633 | 12653 | 21 | rRF 45S | YES | 100.98 | 211.86 | 497.90 | 292.06 |
| >hg38_21_+_8433172_8446622_RNA45SN1_45S_50ntFlanks@12633.12654.22\|CGAAGCTACCATCTGTGGGATT | >hg38_21_+_8433172_8446622_RNA45SN1_45S_50ntFlanks | 12633 | 12654 | 22 | rRF 45S | YES | 175.99 | 278.87 | 607.36 | 525.78 |
| >hg38_21_+_8433172_8446622_RNA45SN1_45S_50ntFlanks@12633.12657.25\|CGAAGCTACCATCTGTGGGATTATG | >hg38_21_+_8433172_8446622_RNA45SN1_45S_50ntFlanks | 12633 | 12657 | 25 | rRF 45S | NO | 0.00 | 0.00 | 92.00 | 76.70 |
| >hg38_21_+_8433172_8446622_RNA45SN1_45S_50ntFlanks@12633.12659.27\|CGAAGCTACCATCTGTGGGATTATGAC | >hg38_21_+_8433172_8446622_RNA45SN1_45S_50ntFlanks | 12633 | 12659 | 27 | rRF 45S | NO | 59.11 | 31.57 | 99.26 | 189.41 |
| >hg38_21_+_8433172_8446622_RNA45SN1_45S_50ntFlanks@12633.12661.29\|CGAAGCTACCATCTGTGGGATTATGACTG | >hg38_21_+_8433172_8446622_RNA45SN1_45S_50ntFlanks | 12633 | 12661 | 29 | rRF 45S | NO | 95.85 | 49.82 | 223.95 | 284.01 |
| >hg38_21_+_8433172_8446622_RNA45SN1_45S_50ntFlanks@12634.12653.20\|GAAGCTACCATCTGTGGGAT | >hg38_21_+_8433172_8446622_RNA45SN1_45S_50ntFlanks | 12634 | 12653 | 20 | rRF 45S | NO | 0.00 | 0.00 | 33.62 | 0.00 |
| >hg38_21_+_8433172_8446622_RNA45SN1_45S_50ntFlanks@12634.12654.21\|GAAGCTACCATCTGTGGGATT | >hg38_21_+_8433172_8446622_RNA45SN1_45S_50ntFlanks | 12634 | 12654 | 21 | rRF 45S | NO | 0.00 | 0.00 | 53.26 | 14.05 |
| >hg38_21_+_8433172_8446622_RNA45SN1_45S_50ntFlanks@12634.12661.28\|GAAGCTACCATCTGTGGGATTATGACTG | >hg38_21_+_8433172_8446622_RNA45SN1_45S_50ntFlanks | 12634 | 12661 | 28 | rRF 45S | NO | 0.00 | 0.00 | 43.21 | 39.39 |
| >hg38_21_+_8433172_8446622_RNA45SN1_45S_50ntFlanks@12635.12652.18\|AAGCTACCATCTGTGGGA | >hg38_21_+_8433172_8446622_RNA45SN1_45S_50ntFlanks | 12635 | 12652 | 18 | rRF 45S | NO | 0.00 | 0.00 | 61.25 | 0.00 |
| >hg38_21_+_8433172_8446622_RNA45SN1_45S_50ntFlanks@12635.12653.19\|AAGCTACCATCTGTGGGAT | >hg38_21_+_8433172_8446622_RNA45SN1_45S_50ntFlanks | 12635 | 12653 | 19 | rRF 45S | NO | 0.00 | 66.79 | 325.25 | 89.92 |
| >hg38_21_+_8433172_8446622_RNA45SN1_45S_50ntFlanks@12635.12654.20\|AAGCTACCATCTGTGGGATT | >hg38_21_+_8433172_8446622_RNA45SN1_45S_50ntFlanks | 12635 | 12654 | 20 | rRF 45S | YES | 51.68 | 123.94 | 655.42 | 288.50 |
| >hg38_21_+_8433172_8446622_RNA45SN1_45S_50ntFlanks@12635.12657.23\|AAGCTACCATCTGTGGGATTATG | >hg38_21_+_8433172_8446622_RNA45SN1_45S_50ntFlanks | 12635 | 12657 | 23 | rRF 45S | NO | 0.00 | 0.00 | 108.30 | 68.28 |
| >hg38_21_+_8433172_8446622_RNA45SN1_45S_50ntFlanks@12635.12659.25\|AAGCTACCATCTGTGGGATTATGAC | >hg38_21_+_8433172_8446622_RNA45SN1_45S_50ntFlanks | 12635 | 12659 | 25 | rRF 45S | NO | 68.29 | 40.81 | 162.92 | 233.47 |
| >hg38_21_+_8433172_8446622_RNA45SN1_45S_50ntFlanks@12635.12660.26\|AAGCTACCATCTGTGGGATTATGACT | >hg38_21_+_8433172_8446622_RNA45SN1_45S_50ntFlanks | 12635 | 12660 | 26 | rRF 45S | NO | 0.00 | 0.00 | 32.85 | 25.25 |
| >hg38_21_+_8433172_8446622_RNA45SN1_45S_50ntFlanks@12635.12661.27\|AAGCTACCATCTGTGGGATTATGACTG | >hg38_21_+_8433172_8446622_RNA45SN1_45S_50ntFlanks | 12635 | 12661 | 27 | rRF 45S | NO | 137.73 | 77.03 | 446.95 | 432.37 |
| >hg38_21_+_8433172_8446622_RNA45SN1_45S_50ntFlanks@12636.12653.18\|AGCTACCATCTGTGGGAT | >hg38_21_+_8433172_8446622_RNA45SN1_45S_50ntFlanks | 12636 | 12653 | 18 | rRF 45S | NO | 0.00 | 0.00 | 97.69 | 35.80 |
| >hg38_21_+_8433172_8446622_RNA45SN1_45S_50ntFlanks@12636.12654.19\|AGCTACCATCTGTGGGATT | >hg38_21_+_8433172_8446622_RNA45SN1_45S_50ntFlanks | 12636 | 12654 | 19 | rRF 45S | NO | 0.00 | 49.08 | 211.67 | 113.22 |
| >hg38_21_+_8433172_8446622_RNA45SN1_45S_50ntFlanks@12636.12657.22\|AGCTACCATCTGTGGGATTATG | >hg38_21_+_8433172_8446622_RNA45SN1_45S_50ntFlanks | 12636 | 12657 | 22 | rRF 45S | NO | 0.00 | 0.00 | 46.57 | 25.58 |
| >hg38_21_+_8433172_8446622_RNA45SN1_45S_50ntFlanks@12636.12659.24\|AGCTACCATCTGTGGGATTATGAC | >hg38_21_+_8433172_8446622_RNA45SN1_45S_50ntFlanks | 12636 | 12659 | 24 | rRF 45S | NO | 31.38 | 0.00 | 79.05 | 126.02 |
| >hg38_21_+_8433172_8446622_RNA45SN1_45S_50ntFlanks@12636.12661.26\|AGCTACCATCTGTGGGATTATGACTG | >hg38_21_+_8433172_8446622_RNA45SN1_45S_50ntFlanks | 12636 | 12661 | 26 | rRF 45S | NO | 48.21 | 12.59 | 111.19 | 170.49 |
| >hg38_21_+_8433172_8446622_RNA45SN1_45S_50ntFlanks@12637.12654.18\|GCTACCATCTGTGGGATT | >hg38_21_+_8433172_8446622_RNA45SN1_45S_50ntFlanks | 12637 | 12654 | 18 | rRF 45S | NO | 0.00 | 0.00 | 150.96 | 51.47 |
| >hg38_21_+_8433172_8446622_RNA45SN1_45S_50ntFlanks@12637.12657.21\|GCTACCATCTGTGGGATTATG | >hg38_21_+_8433172_8446622_RNA45SN1_45S_50ntFlanks | 12637 | 12657 | 21 | rRF 45S | NO | 0.00 | 0.00 | 35.66 | 0.00 |
| >hg38_21_+_8433172_8446622_RNA45SN1_45S_50ntFlanks@12637.12659.23\|GCTACCATCTGTGGGATTATGAC | >hg38_21_+_8433172_8446622_RNA45SN1_45S_50ntFlanks | 12637 | 12659 | 23 | rRF 45S | NO | 0.00 | 0.00 | 69.72 | 71.69 |
| >hg38_21_+_8433172_8446622_RNA45SN1_45S_50ntFlanks@12637.12661.25\|GCTACCATCTGTGGGATTATGACTG | >hg38_21_+_8433172_8446622_RNA45SN1_45S_50ntFlanks | 12637 | 12661 | 25 | rRF 45S | NO | 6.18 | 0.00 | 86.85 | 93.33 |
| >hg38_21_+_8433172_8446622_RNA45SN1_45S_50ntFlanks@12638.12657.20\|CTACCATCTGTGGGATTATG | >hg38_21_+_8433172_8446622_RNA45SN1_45S_50ntFlanks | 12638 | 12657 | 20 | rRF 45S | NO | 0.00 | 0.00 | 67.17 | 23.16 |
| >hg38_21_+_8433172_8446622_RNA45SN1_45S_50ntFlanks@12638.12659.22\|CTACCATCTGTGGGATTATGAC | >hg38_21_+_8433172_8446622_RNA45SN1_45S_50ntFlanks | 12638 | 12659 | 22 | rRF 45S | NO | 0.00 | 0.00 | 86.27 | 131.78 |
| >hg38_21_+_8433172_8446622_RNA45SN1_45S_50ntFlanks@12638.12661.24\|CTACCATCTGTGGGATTATGACTG | >hg38_21_+_8433172_8446622_RNA45SN1_45S_50ntFlanks | 12638 | 12661 | 24 | rRF 45S | NO | 33.83 | 26.04 | 164.88 | 173.29 |
| >hg38_21_+_8433172_8446622_RNA45SN1_45S_50ntFlanks@12639.12661.23\|TACCATCTGTGGGATTATGACTG | >hg38_21_+_8433172_8446622_RNA45SN1_45S_50ntFlanks | 12639 | 12661 | 23 | rRF 45S | NO | 0.00 | 0.00 | 35.38 | 33.54 |
| >hg38_21_+_8433172_8446622_RNA45SN1_45S_50ntFlanks@12640.12659.20\|ACCATCTGTGGGATTATGAC | >hg38_21_+_8433172_8446622_RNA45SN1_45S_50ntFlanks | 12640 | 12659 | 20 | rRF 45S | NO | 0.00 | 0.00 | 49.64 | 59.29 |
| >hg38_21_+_8433172_8446622_RNA45SN1_45S_50ntFlanks@12640.12661.22\|ACCATCTGTGGGATTATGACTG | >hg38_21_+_8433172_8446622_RNA45SN1_45S_50ntFlanks | 12640 | 12661 | 22 | rRF 45S | NO | 0.00 | 0.00 | 109.88 | 57.00 |

### MDA-MB-468

| | | | | | | | | | | |
|---|---|---|---|---|---|---|---|---|---|---|
| >hg38_21_+_8433172_8446622_RNA45SN1_45S_50ntFlanks@12641.12659.19\|CCATCTGTGGGATTATGAC | >hg38_21_+_8433172_8446622_RNA45SN1_45S_50ntFlanks | 12641 | 12659 | 19 | rRF 45S | NO | 21.36 | 60.85 | 51.84 | 166.93 |
| >hg38_21_+_8433172_8446622_RNA45SN1_45S_50ntFlanks@12641.12661.21\|CCATCTGTGGGATTATGACTG | >hg38_21_+_8433172_8446622_RNA45SN1_45S_50ntFlanks | 12641 | 12661 | 21 | rRF 45S | NO | 20.34 | 28.08 | 83.13 | 114.67 |
| >hg38_21_+_8433172_8446622_RNA45SN1_45S_50ntFlanks@12642.12659.18\|CATCTGTGGGATTATGAC | >hg38_21_+_8433172_8446622_RNA45SN1_45S_50ntFlanks | 12642 | 12659 | 18 | rRF 45S | NO | 0.00 | 29.21 | 0.00 | 42.97 |
| >hg38_21_+_8433172_8446622_RNA45SN1_45S_50ntFlanks@12642.12661.20\|CATCTGTGGGATTATGACTG | >hg38_21_+_8433172_8446622_RNA45SN1_45S_50ntFlanks | 12642 | 12661 | 20 | rRF 45S | NO | 0.00 | 0.00 | 35.47 | 43.39 |
| >hg38_21_+_8433172_8446622_RNA45SN1_45S_50ntFlanks@12643.12661.19\|ATCTGTGGGATTATGACTG | >hg38_21_+_8433172_8446622_RNA45SN1_45S_50ntFlanks | 12643 | 12661 | 19 | rRF 45S | NO | 0.00 | 0.00 | 36.21 | 42.19 |
| >hg38_21_+_8433172_8446622_RNA45SN1_45S_50ntFlanks@12644.12661.18\|TCTGTGGGATTATGACTG | >hg38_21_+_8433172_8446622_RNA45SN1_45S_50ntFlanks | 12644 | 12661 | 18 | rRF 45S | NO | 0.00 | 30.25 | 76.84 | 70.27 |
| >hg38_21_+_8433172_8446622_RNA45SN1_45S_50ntFlanks@12654.12674.21\|TATGACTGAACGCCTCTAAGT | >hg38_21_+_8433172_8446622_RNA45SN1_45S_50ntFlanks | 12654 | 12674 | 21 | rRF 45S | NO | 154.54 | 79.21 | 0.00 | 0.00 |
| >hg38_21_+_8433172_8446622_RNA45SN1_45S_50ntFlanks@12654.12675.22\|TATGACTGAACGCCTCTAAGTC | >hg38_21_+_8433172_8446622_RNA45SN1_45S_50ntFlanks | 12654 | 12675 | 22 | rRF 45S | NO | 322.64 | 524.87 | 0.00 | 84.36 |
| >hg38_21_+_8433172_8446622_RNA45SN1_45S_50ntFlanks@12655.12673.19\|ATGACTGAACGCCTCTAAG | >hg38_21_+_8433172_8446622_RNA45SN1_45S_50ntFlanks | 12655 | 12673 | 19 | rRF 45S | NO | 81.33 | 19.38 | 0.00 | 0.00 |
| >hg38_21_+_8433172_8446622_RNA45SN1_45S_50ntFlanks@12655.12674.20\|ATGACTGAACGCCTCTAAGT | >hg38_21_+_8433172_8446622_RNA45SN1_45S_50ntFlanks | 12655 | 12674 | 20 | rRF 45S | NO | 576.99 | 269.35 | 27.98 | 56.14 |
| >hg38_21_+_8433172_8446622_RNA45SN1_45S_50ntFlanks@12655.12675.21\|ATGACTGAACGCCTCTAAGTC | >hg38_21_+_8433172_8446622_RNA45SN1_45S_50ntFlanks | 12655 | 12675 | 21 | rRF 45S | YES | 557.57 | 1332.44 | 34.28 | 182.00 |
| >hg38_21_+_8433172_8446622_RNA45SN1_45S_50ntFlanks@12656.12674.19\|TGACTGAACGCCTCTAAGT | >hg38_21_+_8433172_8446622_RNA45SN1_45S_50ntFlanks | 12656 | 12674 | 19 | rRF 45S | NO | 34.27 | 0.00 | 0.00 | 0.00 |
| >hg38_21_+_8433172_8446622_RNA45SN1_45S_50ntFlanks@12656.12675.20\|TGACTGAACGCCTCTAAGTC | >hg38_21_+_8433172_8446622_RNA45SN1_45S_50ntFlanks | 12656 | 12675 | 20 | rRF 45S | NO | 0.00 | 37.58 | 0.00 | 0.00 |
| >hg38_21_+_8433172_8446622_RNA45SN1_45S_50ntFlanks@12657.12674.18\|GACTGAACGCCTCTAAGT | >hg38_21_+_8433172_8446622_RNA45SN1_45S_50ntFlanks | 12657 | 12674 | 18 | rRF 45S | NO | 71.19 | 36.75 | 0.00 | 0.00 |
| >hg38_21_+_8433172_8446622_RNA45SN1_45S_50ntFlanks@12658.12675.18\|ACTGAACGCCTCTAAGTC | >hg38_21_+_8433172_8446622_RNA45SN1_45S_50ntFlanks | 12658 | 12675 | 18 | rRF 45S | NO | 98.39 | 67.20 | 0.00 | 37.24 |
| >hg38_21_+_8433172_8446622_RNA45SN1_45S_50ntFlanks@12676.12703.28\|AGAATCCCGCCCAGGCGGAACGATACGG | >hg38_21_+_8433172_8446622_RNA45SN1_45S_50ntFlanks | 12676 | 12703 | 28 | rRF 45S | NO | 0.00 | 36.23 | 112.32 | 14.62 |
| >hg38_21_+_8433172_8446622_RNA45SN1_45S_50ntFlanks@12676.12708.33\|AGAATCCCGCCCAGGCGGAACGATACGGCAGCG | >hg38_21_+_8433172_8446622_RNA45SN1_45S_50ntFlanks | 12676 | 12708 | 33 | rRF 45S | NO | 0.00 | 26.02 | 43.87 | 0.00 |
| >hg38_21_+_8433172_8446622_RNA45SN1_45S_50ntFlanks@12678.12703.26\|AATCCCGCCCAGGCGGAACGATACGG | >hg38_21_+_8433172_8446622_RNA45SN1_45S_50ntFlanks | 12678 | 12703 | 26 | rRF 45S | NO | 0.00 | 27.70 | 89.92 | 6.42 |
| >hg38_21_+_8433172_8446622_RNA45SN1_45S_50ntFlanks@12709.12742.34\|CCGCGGAGCCTCGGTTGGCCTCGGATAGCCGGTC | >hg38_21_+_8433172_8446622_RNA45SN1_45S_50ntFlanks | 12709 | 12742 | 34 | rRF 45S | NO | 0.00 | 0.00 | 36.33 | 47.93 |
| >hg38_21_+_8433172_8446622_RNA45SN1_45S_50ntFlanks@12710.12742.33\|CGCGGAGCCTCGGTTGGCCTCGGATAGCCGGTC | >hg38_21_+_8433172_8446622_RNA45SN1_45S_50ntFlanks | 12710 | 12742 | 33 | rRF 45S | NO | 0.00 | 0.00 | 37.02 | 0.00 |
| >hg38_21_+_8433172_8446622_RNA45SN1_45S_50ntFlanks@12712.12733.22\|CGGAGCCTCGGTTGGCCTCGGA | >hg38_21_+_8433172_8446622_RNA45SN1_45S_50ntFlanks | 12712 | 12733 | 22 | rRF 45S | NO | 0.00 | 0.00 | 32.84 | 0.00 |
| >hg38_21_+_8433172_8446622_RNA45SN1_45S_50ntFlanks@12712.12742.31\|CGGAGCCTCGGTTGGCCTCGGATAGCCGGTC | >hg38_21_+_8433172_8446622_RNA45SN1_45S_50ntFlanks | 12712 | 12742 | 31 | rRF 45S | NO | 0.00 | 0.00 | 188.95 | 14.14 |
| >hg38_21_+_8433172_8446622_RNA45SN1_45S_50ntFlanks@12712.12745.34\|CGGAGCCTCGGTTGGCCTCGGATAGCCGGTCCCC | >hg38_21_+_8433172_8446622_RNA45SN1_45S_50ntFlanks | 12712 | 12745 | 34 | rRF 45S | NO | 0.00 | 0.00 | 94.52 | 13.34 |
| >hg38_21_+_8433172_8446622_RNA45SN1_45S_50ntFlanks@12712.12747.36\|CGGAGCCTCGGTTGGCCTCGGATAGCCGGTCCCCCG | >hg38_21_+_8433172_8446622_RNA45SN1_45S_50ntFlanks | 12712 | 12747 | 36 | rRF 45S | NO | 0.00 | 0.00 | 75.63 | 26.60 |
| >hg38_21_+_8433172_8446622_RNA45SN1_45S_50ntFlanks@12712.12751.40\|CGGAGCCTCGGTTGGCCTCGGATAGCCGGTCCCCCGCCTG | >hg38_21_+_8433172_8446622_RNA45SN1_45S_50ntFlanks | 12712 | 12751 | 40 | rRF 45S | NO | 0.00 | 0.00 | 29.41 | 70.12 |
| >hg38_21_+_8433172_8446622_RNA45SN1_45S_50ntFlanks@12714.12747.34\|GAGCCTCGGTTGGCCTCGGATAGCCGGTCCCCCG | >hg38_21_+_8433172_8446622_RNA45SN1_45S_50ntFlanks | 12714 | 12747 | 34 | rRF 45S | NO | 0.00 | 0.00 | 39.52 | 13.33 |
| >hg38_21_+_8433172_8446622_RNA45SN1_45S_50ntFlanks@12717.12742.26\|CCTCGGTTGGCCTCGGATAGCCGGTC | >hg38_21_+_8433172_8446622_RNA45SN1_45S_50ntFlanks | 12717 | 12742 | 26 | rRF 45S | NO | 0.00 | 0.00 | 64.43 | 0.00 |
| >hg38_21_+_8433172_8446622_RNA45SN1_45S_50ntFlanks@12717.12747.31\|CCTCGGTTGGCCTCGGATAGCCGGTCCCCCG | >hg38_21_+_8433172_8446622_RNA45SN1_45S_50ntFlanks | 12717 | 12747 | 31 | rRF 45S | NO | 0.00 | 0.00 | 51.74 | 0.00 |
| >hg38_21_+_8433172_8446622_RNA45SN1_45S_50ntFlanks@12841.12862.22\|CGGAGTGCCCTTCGTCCTGGGA | >hg38_21_+_8433172_8446622_RNA45SN1_45S_50ntFlanks | 12841 | 12862 | 22 | rRF 45S | YES | 0.00 | 0.00 | 13.54 | 0.00 |
| >hg38_21_+_8433172_8446622_RNA45SN1_45S_50ntFlanks@12874.12891.18\|GCTGGAAAGGCGGCCGCC | >hg38_21_+_8433172_8446622_RNA45SN1_45S_50ntFlanks | 12874 | 12891 | 18 | rRF 45S | NO | 0.00 | 34.60 | 0.00 | 0.00 |
| >hg38_21_+_8433172_8446622_RNA45SN1_45S_50ntFlanks@12892.12932.41\|CCCTCGCCCGTCACGCACCGCACGTTCGTGGGGAACCTGGC | >hg38_21_+_8433172_8446622_RNA45SN1_45S_50ntFlanks | 12892 | 12932 | 41 | rRF 45S | NO | 44.56 | 0.00 | 0.00 | 252.38 |
| >hg38_21_+_8433172_8446622_RNA45SN1_45S_50ntFlanks@12893.12930.38\|CCTCGCCCGTCACGCACCGCACGTTCGTGGGGAACCTG | >hg38_21_+_8433172_8446622_RNA45SN1_45S_50ntFlanks | 12893 | 12930 | 38 | rRF 45S | NO | 0.00 | 0.00 | 40.95 | 121.00 |
| >hg38_21_+_8433172_8446622_RNA45SN1_45S_50ntFlanks@12893.12932.40\|CCTCGCCCGTCACGCACCGCACGTTCGTGGGGAACCTGGC | >hg38_21_+_8433172_8446622_RNA45SN1_45S_50ntFlanks | 12893 | 12932 | 40 | rRF 45S | NO | 34.67 | 0.00 | 28.07 | 183.37 |
| >hg38_21_+_8433172_8446622_RNA45SN1_45S_50ntFlanks@12893.12933.41\|CCTCGCCCGTCACGCACCGCACGTTCGTGGGGAACCTGGCG | >hg38_21_+_8433172_8446622_RNA45SN1_45S_50ntFlanks | 12893 | 12933 | 41 | rRF 45S | NO | 37.58 | 0.00 | 52.89 | 187.31 |

### MDA-MB-468

| | | | | | | | | | | |
|---|---|---|---|---|---|---|---|---|---|---|
| >hg38_21_+_8433172_8446622_RNA45SN1_45S_50ntFlanks@12907.12933.27\|CACCGCACGTTCGTGGGGAACCTGGCG | >hg38_21_+_8433172_8446622_RNA45SN1_45S_50ntFlanks | 12907 | 12933 | 27 | rRF 45S | NO | 54.91 | 30.79 | 32.70 | 82.19 |
| >hg38_21_+_8433172_8446622_RNA45SN1_45S_50ntFlanks@12912.12933.22\|CACGTTCGTGGGGAACCTGGCG | >hg38_21_+_8433172_8446622_RNA45SN1_45S_50ntFlanks | 12912 | 12933 | 22 | rRF 45S | NO | 13.04 | 41.48 | 38.36 | 38.29 |
| >hg38_21_+_8433172_8446622_RNA45SN1_45S_50ntFlanks@12914.12933.20\|CGTTCGTGGGGAACCTGGCG | >hg38_21_+_8433172_8446622_RNA45SN1_45S_50ntFlanks | 12914 | 12933 | 20 | rRF 45S | NO | 0.00 | 0.00 | 45.28 | 13.17 |
| >hg38_21_+_8433172_8446622_RNA45SN1_45S_50ntFlanks@12931.12956.26\|GCGCTAAACCATTCGTAGACGACCTG | >hg38_21_+_8433172_8446622_RNA45SN1_45S_50ntFlanks | 12931 | 12956 | 26 | rRF 45S | NO | 81.04 | 49.42 | 28.68 | 51.74 |
| >hg38_21_+_8433172_8446622_RNA45SN1_45S_50ntFlanks@12932.12956.25\|CGCTAAACCATTCGTAGACGACCTG | >hg38_21_+_8433172_8446622_RNA45SN1_45S_50ntFlanks | 12932 | 12956 | 25 | rRF 45S | NO | 76.62 | 58.39 | 31.18 | 40.68 |
| >hg38_21_+_8433172_8446622_RNA45SN1_45S_50ntFlanks@12932.12972.41\|CGCTAAACCATTCGTAGACGACCTGCTTCTGGGTCGGGGTT | >hg38_21_+_8433172_8446622_RNA45SN1_45S_50ntFlanks | 12932 | 12972 | 41 | rRF 45S | NO | 28.76 | 78.82 | 0.00 | 23.88 |
| >hg38_21_+_8433172_8446622_RNA45SN1_45S_50ntFlanks@12933.12956.24\|GCTAAACCATTCGTAGACGACCTG | >hg38_21_+_8433172_8446622_RNA45SN1_45S_50ntFlanks | 12933 | 12956 | 24 | rRF 45S | NO | 196.09 | 101.69 | 0.00 | 15.02 |
| >hg38_21_+_8433172_8446622_RNA45SN1_45S_50ntFlanks@12934.12951.18\|CTAAACCATTCGTAGACG | >hg38_21_+_8433172_8446622_RNA45SN1_45S_50ntFlanks | 12934 | 12951 | 18 | rRF 45S | NO | 32.73 | 133.41 | 37.16 | 16.50 |
| >hg38_21_+_8433172_8446622_RNA45SN1_45S_50ntFlanks@12934.12952.19\|CTAAACCATTCGTAGACGA | >hg38_21_+_8433172_8446622_RNA45SN1_45S_50ntFlanks | 12934 | 12952 | 19 | rRF 45S | NO | 0.00 | 80.05 | 6.88 | 0.00 |
| >hg38_21_+_8433172_8446622_RNA45SN1_45S_50ntFlanks@12934.12953.20\|CTAAACCATTCGTAGACGAC | >hg38_21_+_8433172_8446622_RNA45SN1_45S_50ntFlanks | 12934 | 12953 | 20 | rRF 45S | NO | 54.10 | 136.38 | 28.85 | 6.28 |
| >hg38_21_+_8433172_8446622_RNA45SN1_45S_50ntFlanks@12934.12954.21\|CTAAACCATTCGTAGACGACC | >hg38_21_+_8433172_8446622_RNA45SN1_45S_50ntFlanks | 12934 | 12954 | 21 | rRF 45S | NO | 13.17 | 44.88 | 0.00 | 0.00 |
| >hg38_21_+_8433172_8446622_RNA45SN1_45S_50ntFlanks@12934.12955.22\|CTAAACCATTCGTAGACGACCT | >hg38_21_+_8433172_8446622_RNA45SN1_45S_50ntFlanks | 12934 | 12955 | 22 | rRF 45S | NO | 65.24 | 61.33 | 0.00 | 0.00 |
| >hg38_21_+_8433172_8446622_RNA45SN1_45S_50ntFlanks@12934.12956.23\|CTAAACCATTCGTAGACGACCTG | >hg38_21_+_8433172_8446622_RNA45SN1_45S_50ntFlanks | 12934 | 12956 | 23 | rRF 45S | NO | 961.17 | 856.77 | 92.65 | 138.11 |
| >hg38_21_+_8433172_8446622_RNA45SN1_45S_50ntFlanks@12934.12957.24\|CTAAACCATTCGTAGACGACCTGC | >hg38_21_+_8433172_8446622_RNA45SN1_45S_50ntFlanks | 12934 | 12957 | 24 | rRF 45S | NO | 44.23 | 33.67 | 0.00 | 0.00 |
| >hg38_21_+_8433172_8446622_RNA45SN1_45S_50ntFlanks@12934.12958.25\|CTAAACCATTCGTAGACGACCTGCT | >hg38_21_+_8433172_8446622_RNA45SN1_45S_50ntFlanks | 12934 | 12958 | 25 | rRF 45S | NO | 51.68 | 12.78 | 0.00 | 0.00 |
| >hg38_21_+_8433172_8446622_RNA45SN1_45S_50ntFlanks@12934.12959.26\|CTAAACCATTCGTAGACGACCTGCTT | >hg38_21_+_8433172_8446622_RNA45SN1_45S_50ntFlanks | 12934 | 12959 | 26 | rRF 45S | NO | 133.61 | 48.05 | 0.00 | 16.07 |
| >hg38_21_+_8433172_8446622_RNA45SN1_45S_50ntFlanks@12934.12960.27\|CTAAACCATTCGTAGACGACCTGCTTC | >hg38_21_+_8433172_8446622_RNA45SN1_45S_50ntFlanks | 12934 | 12960 | 27 | rRF 45S | NO | 74.51 | 27.31 | 0.00 | 0.00 |
| >hg38_21_+_8433172_8446622_RNA45SN1_45S_50ntFlanks@12934.12961.28\|CTAAACCATTCGTAGACGACCTGCTTCT | >hg38_21_+_8433172_8446622_RNA45SN1_45S_50ntFlanks | 12934 | 12961 | 28 | rRF 45S | NO | 117.08 | 37.94 | 0.00 | 0.00 |
| >hg38_21_+_8433172_8446622_RNA45SN1_45S_50ntFlanks@12934.12962.29\|CTAAACCATTCGTAGACGACCTGCTTCTG | >hg38_21_+_8433172_8446622_RNA45SN1_45S_50ntFlanks | 12934 | 12962 | 29 | rRF 45S | NO | 64.36 | 31.33 | 0.00 | 0.00 |
| >hg38_21_+_8433172_8446622_RNA45SN1_45S_50ntFlanks@12934.12965.32\|CTAAACCATTCGTAGACGACCTGCTTCTGGGT | >hg38_21_+_8433172_8446622_RNA45SN1_45S_50ntFlanks | 12934 | 12965 | 32 | rRF 45S | NO | 43.40 | 86.14 | 0.00 | 0.00 |
| >hg38_21_+_8433172_8446622_RNA45SN1_45S_50ntFlanks@12934.12966.33\|CTAAACCATTCGTAGACGACCTGCTTCTGGGTC | >hg38_21_+_8433172_8446622_RNA45SN1_45S_50ntFlanks | 12934 | 12966 | 33 | rRF 45S | NO | 0.00 | 109.93 | 0.00 | 0.00 |
| >hg38_21_+_8433172_8446622_RNA45SN1_45S_50ntFlanks@12934.12969.36\|CTAAACCATTCGTAGACGACCTGCTTCTGGGTCGGG | >hg38_21_+_8433172_8446622_RNA45SN1_45S_50ntFlanks | 12934 | 12969 | 36 | rRF 45S | NO | 0.00 | 52.85 | 0.00 | 0.00 |
| >hg38_21_+_8433172_8446622_RNA45SN1_45S_50ntFlanks@12934.12970.37\|CTAAACCATTCGTAGACGACCTGCTTCTGGGTCGGGG | >hg38_21_+_8433172_8446622_RNA45SN1_45S_50ntFlanks | 12934 | 12970 | 37 | rRF 45S | NO | 0.00 | 56.53 | 0.00 | 0.00 |
| >hg38_21_+_8433172_8446622_RNA45SN1_45S_50ntFlanks@12934.12972.39\|CTAAACCATTCGTAGACGACCTGCTTCTGGGTCGGGGTT | >hg38_21_+_8433172_8446622_RNA45SN1_45S_50ntFlanks | 12934 | 12972 | 39 | rRF 45S | NO | 69.71 | 511.57 | 30.37 | 68.15 |
| >hg38_21_+_8433172_8446622_RNA45SN1_45S_50ntFlanks@12935.12953.19\|TAAACCATTCGTAGACGAC | >hg38_21_+_8433172_8446622_RNA45SN1_45S_50ntFlanks | 12935 | 12953 | 19 | rRF 45S | NO | 0.00 | 40.99 | 0.00 | 0.00 |
| >hg38_21_+_8433172_8446622_RNA45SN1_45S_50ntFlanks@12935.12956.22\|TAAACCATTCGTAGACGACCTG | >hg38_21_+_8433172_8446622_RNA45SN1_45S_50ntFlanks | 12935 | 12956 | 22 | rRF 45S | NO | 268.94 | 293.49 | 6.67 | 40.17 |
| >hg38_21_+_8433172_8446622_RNA45SN1_45S_50ntFlanks@12935.12959.25\|TAAACCATTCGTAGACGACCTGCTT | >hg38_21_+_8433172_8446622_RNA45SN1_45S_50ntFlanks | 12935 | 12959 | 25 | rRF 45S | NO | 21.10 | 0.00 | 0.00 | 0.00 |
| >hg38_21_+_8433172_8446622_RNA45SN1_45S_50ntFlanks@12935.12961.27\|TAAACCATTCGTAGACGACCTGCTTCT | >hg38_21_+_8433172_8446622_RNA45SN1_45S_50ntFlanks | 12935 | 12961 | 27 | rRF 45S | NO | 31.97 | 0.00 | 0.00 | 0.00 |
| >hg38_21_+_8433172_8446622_RNA45SN1_45S_50ntFlanks@12935.12966.32\|TAAACCATTCGTAGACGACCTGCTTCTGGGTC | >hg38_21_+_8433172_8446622_RNA45SN1_45S_50ntFlanks | 12935 | 12966 | 32 | rRF 45S | NO | 0.00 | 34.70 | 0.00 | 0.00 |
| >hg38_21_+_8433172_8446622_RNA45SN1_45S_50ntFlanks@12936.12953.18\|AAACCATTCGTAGACGAC | >hg38_21_+_8433172_8446622_RNA45SN1_45S_50ntFlanks | 12936 | 12953 | 18 | rRF 45S | NO | 39.70 | 65.40 | 0.00 | 0.00 |
| >hg38_21_+_8433172_8446622_RNA45SN1_45S_50ntFlanks@12936.12955.20\|AAACCATTCGTAGACGACCT | >hg38_21_+_8433172_8446622_RNA45SN1_45S_50ntFlanks | 12936 | 12955 | 20 | rRF 45S | NO | 53.28 | 35.22 | 0.00 | 0.00 |
| >hg38_21_+_8433172_8446622_RNA45SN1_45S_50ntFlanks@12936.12956.21\|AAACCATTCGTAGACGACCTG | >hg38_21_+_8433172_8446622_RNA45SN1_45S_50ntFlanks | 12936 | 12956 | 21 | rRF 45S | NO | 834.48 | 511.33 | 92.59 | 119.21 |
| >hg38_21_+_8433172_8446622_RNA45SN1_45S_50ntFlanks@12936.12957.22\|AAACCATTCGTAGACGACCTGC | >hg38_21_+_8433172_8446622_RNA45SN1_45S_50ntFlanks | 12936 | 12957 | 22 | rRF 45S | NO | 37.67 | 0.00 | 0.00 | 0.00 |
| >hg38_21_+_8433172_8446622_RNA45SN1_45S_50ntFlanks@12936.12958.23\|AAACCATTCGTAGACGACCTGCT | >hg38_21_+_8433172_8446622_RNA45SN1_45S_50ntFlanks | 12936 | 12958 | 23 | rRF 45S | NO | 39.53 | 0.00 | 0.00 | 0.00 |

### MDA-MB-468

| | | | | | | | | | | |
|---|---|---|---|---|---|---|---|---|---|---|
| >hg38_21_+_8433172_8446622_RNA45SN1_45S_50ntFlanks@12936.12959.24\|AAACCATTCGTAGACGACCTGCTT | >hg38_21_+_8433172_8446622_RNA45SN1_45S_50ntFlanks | 12936 | 12959 | 24 | rRF 45S | NO | 77.15 | 5.94 | 0.00 | 0.00 |
| >hg38_21_+_8433172_8446622_RNA45SN1_45S_50ntFlanks@12936.12960.25\|AAACCATTCGTAGACGACCTGCTTC | >hg38_21_+_8433172_8446622_RNA45SN1_45S_50ntFlanks | 12936 | 12960 | 25 | rRF 45S | NO | 62.33 | 0.00 | 0.00 | 0.00 |
| >hg38_21_+_8433172_8446622_RNA45SN1_45S_50ntFlanks@12936.12961.26\|AAACCATTCGTAGACGACCTGCTTCT | >hg38_21_+_8433172_8446622_RNA45SN1_45S_50ntFlanks | 12936 | 12961 | 26 | rRF 45S | NO | 80.85 | 0.00 | 0.00 | 0.00 |
| >hg38_21_+_8433172_8446622_RNA45SN1_45S_50ntFlanks@12936.12962.27\|AAACCATTCGTAGACGACCTGCTTCTG | >hg38_21_+_8433172_8446622_RNA45SN1_45S_50ntFlanks | 12936 | 12962 | 27 | rRF 45S | NO | 42.86 | 0.00 | 0.00 | 0.00 |
| >hg38_21_+_8433172_8446622_RNA45SN1_45S_50ntFlanks@12936.12966.31\|AAACCATTCGTAGACGACCTGCTTCTGGGTC | >hg38_21_+_8433172_8446622_RNA45SN1_45S_50ntFlanks | 12936 | 12966 | 31 | rRF 45S | NO | 0.00 | 50.87 | 0.00 | 0.00 |
| >hg38_21_+_8433172_8446622_RNA45SN1_45S_50ntFlanks@12937.12955.19\|AACCATTCGTAGACGACCT | >hg38_21_+_8433172_8446622_RNA45SN1_45S_50ntFlanks | 12937 | 12955 | 19 | rRF 45S | NO | 38.60 | 32.06 | 0.00 | 0.00 |
| >hg38_21_+_8433172_8446622_RNA45SN1_45S_50ntFlanks@12937.12956.20\|AACCATTCGTAGACGACCTG | >hg38_21_+_8433172_8446622_RNA45SN1_45S_50ntFlanks | 12937 | 12956 | 20 | rRF 45S | NO | 608.80 | 442.88 | 75.15 | 89.68 |
| >hg38_21_+_8433172_8446622_RNA45SN1_45S_50ntFlanks@12937.12957.21\|AACCATTCGTAGACGACCTGC | >hg38_21_+_8433172_8446622_RNA45SN1_45S_50ntFlanks | 12937 | 12957 | 21 | rRF 45S | NO | 22.11 | 0.00 | 0.00 | 0.00 |
| >hg38_21_+_8433172_8446622_RNA45SN1_45S_50ntFlanks@12937.12958.22\|AACCATTCGTAGACGACCTGCT | >hg38_21_+_8433172_8446622_RNA45SN1_45S_50ntFlanks | 12937 | 12958 | 22 | rRF 45S | NO | 21.44 | 0.00 | 0.00 | 0.00 |
| >hg38_21_+_8433172_8446622_RNA45SN1_45S_50ntFlanks@12937.12959.23\|AACCATTCGTAGACGACCTGCTT | >hg38_21_+_8433172_8446622_RNA45SN1_45S_50ntFlanks | 12937 | 12959 | 23 | rRF 45S | NO | 61.25 | 0.00 | 0.00 | 0.00 |
| >hg38_21_+_8433172_8446622_RNA45SN1_45S_50ntFlanks@12937.12960.24\|AACCATTCGTAGACGACCTGCTTC | >hg38_21_+_8433172_8446622_RNA45SN1_45S_50ntFlanks | 12937 | 12960 | 24 | rRF 45S | NO | 38.91 | 0.00 | 0.00 | 0.00 |
| >hg38_21_+_8433172_8446622_RNA45SN1_45S_50ntFlanks@12937.12961.25\|AACCATTCGTAGACGACCTGCTTCT | >hg38_21_+_8433172_8446622_RNA45SN1_45S_50ntFlanks | 12937 | 12961 | 25 | rRF 45S | NO | 63.54 | 0.00 | 0.00 | 0.00 |
| >hg38_21_+_8433172_8446622_RNA45SN1_45S_50ntFlanks@12937.12962.26\|AACCATTCGTAGACGACCTGCTTCTG | >hg38_21_+_8433172_8446622_RNA45SN1_45S_50ntFlanks | 12937 | 12962 | 26 | rRF 45S | NO | 31.43 | 0.00 | 0.00 | 0.00 |
| >hg38_21_+_8433172_8446622_RNA45SN1_45S_50ntFlanks@12937.12966.30\|AACCATTCGTAGACGACCTGCTTCTGGGTC | >hg38_21_+_8433172_8446622_RNA45SN1_45S_50ntFlanks | 12937 | 12966 | 30 | rRF 45S | NO | 0.00 | 49.75 | 0.00 | 0.00 |
| >hg38_21_+_8433172_8446622_RNA45SN1_45S_50ntFlanks@12938.12956.19\|ACCATTCGTAGACGACCTG | >hg38_21_+_8433172_8446622_RNA45SN1_45S_50ntFlanks | 12938 | 12956 | 19 | rRF 45S | NO | 194.12 | 205.80 | 34.45 | 38.10 |
| >hg38_21_+_8433172_8446622_RNA45SN1_45S_50ntFlanks@12938.12966.29\|ACCATTCGTAGACGACCTGCTTCTGGGTC | >hg38_21_+_8433172_8446622_RNA45SN1_45S_50ntFlanks | 12938 | 12966 | 29 | rRF 45S | NO | 0.00 | 12.58 | 0.00 | 0.00 |
| >hg38_21_+_8433172_8446622_RNA45SN1_45S_50ntFlanks@12939.12956.18\|CCATTCGTAGACGACCTG | >hg38_21_+_8433172_8446622_RNA45SN1_45S_50ntFlanks | 12939 | 12956 | 18 | rRF 45S | NO | 68.96 | 152.31 | 32.96 | 25.43 |
| >hg38_21_+_8433172_8446622_RNA45SN1_45S_50ntFlanks@12939.12966.28\|CCATTCGTAGACGACCTGCTTCTGGGTC | >hg38_21_+_8433172_8446622_RNA45SN1_45S_50ntFlanks | 12939 | 12966 | 28 | rRF 45S | NO | 0.00 | 26.55 | 0.00 | 0.00 |
| >hg38_21_+_8433172_8446622_RNA45SN1_45S_50ntFlanks@12940.12966.27\|CATTCGTAGACGACCTGCTTCTGGGTC | >hg38_21_+_8433172_8446622_RNA45SN1_45S_50ntFlanks | 12940 | 12966 | 27 | rRF 45S | NO | 0.00 | 125.29 | 0.00 | 0.00 |
| >hg38_21_+_8433172_8446622_RNA45SN1_45S_50ntFlanks@12940.12972.33\|CATTCGTAGACGACCTGCTTCTGGGTCGGGGTT | >hg38_21_+_8433172_8446622_RNA45SN1_45S_50ntFlanks | 12940 | 12972 | 33 | rRF 45S | NO | 0.00 | 53.12 | 0.00 | 0.00 |
| >hg38_21_+_8433172_8446622_RNA45SN1_45S_50ntFlanks@12941.12965.25\|ATTCGTAGACGACCTGCTTCTGGGT | >hg38_21_+_8433172_8446622_RNA45SN1_45S_50ntFlanks | 12941 | 12965 | 25 | rRF 45S | NO | 0.00 | 32.80 | 0.00 | 0.00 |
| >hg38_21_+_8433172_8446622_RNA45SN1_45S_50ntFlanks@12941.12966.26\|ATTCGTAGACGACCTGCTTCTGGGTC | >hg38_21_+_8433172_8446622_RNA45SN1_45S_50ntFlanks | 12941 | 12966 | 26 | rRF 45S | NO | 0.00 | 111.71 | 13.87 | 0.00 |
| >hg38_21_+_8433172_8446622_RNA45SN1_45S_50ntFlanks@12942.12959.18\|TTCGTAGACGACCTGCTT | >hg38_21_+_8433172_8446622_RNA45SN1_45S_50ntFlanks | 12942 | 12959 | 18 | rRF 45S | NO | 0.00 | 79.29 | 0.00 | 0.00 |
| >hg38_21_+_8433172_8446622_RNA45SN1_45S_50ntFlanks@12942.12960.19\|TTCGTAGACGACCTGCTTC | >hg38_21_+_8433172_8446622_RNA45SN1_45S_50ntFlanks | 12942 | 12960 | 19 | rRF 45S | NO | 0.00 | 38.20 | 0.00 | 0.00 |
| >hg38_21_+_8433172_8446622_RNA45SN1_45S_50ntFlanks@12942.12961.20\|TTCGTAGACGACCTGCTTCT | >hg38_21_+_8433172_8446622_RNA45SN1_45S_50ntFlanks | 12942 | 12961 | 20 | rRF 45S | NO | 6.10 | 31.34 | 0.00 | 0.00 |
| >hg38_21_+_8433172_8446622_RNA45SN1_45S_50ntFlanks@12942.12962.21\|TTCGTAGACGACCTGCTTCTG | >hg38_21_+_8433172_8446622_RNA45SN1_45S_50ntFlanks | 12942 | 12962 | 21 | rRF 45S | NO | 0.00 | 27.28 | 0.00 | 0.00 |
| >hg38_21_+_8433172_8446622_RNA45SN1_45S_50ntFlanks@12942.12965.24\|TTCGTAGACGACCTGCTTCTGGGT | >hg38_21_+_8433172_8446622_RNA45SN1_45S_50ntFlanks | 12942 | 12965 | 24 | rRF 45S | NO | 0.00 | 39.82 | 0.00 | 0.00 |
| >hg38_21_+_8433172_8446622_RNA45SN1_45S_50ntFlanks@12942.12966.25\|TTCGTAGACGACCTGCTTCTGGGTC | >hg38_21_+_8433172_8446622_RNA45SN1_45S_50ntFlanks | 12942 | 12966 | 25 | rRF 45S | NO | 0.00 | 90.04 | 0.00 | 0.00 |
| >hg38_21_+_8433172_8446622_RNA45SN1_45S_50ntFlanks@12942.12972.31\|TTCGTAGACGACCTGCTTCTGGGTCGGGGTT | >hg38_21_+_8433172_8446622_RNA45SN1_45S_50ntFlanks | 12942 | 12972 | 31 | rRF 45S | NO | 0.00 | 84.05 | 51.05 | 0.00 |
| >hg38_21_+_8433172_8446622_RNA45SN1_45S_50ntFlanks@12942.12973.32\|TTCGTAGACGACCTGCTTCTGGGTCGGGGTTT | >hg38_21_+_8433172_8446622_RNA45SN1_45S_50ntFlanks | 12942 | 12973 | 32 | rRF 45S | NO | 0.00 | 29.02 | 0.00 | 0.00 |
| >hg38_21_+_8433172_8446622_RNA45SN1_45S_50ntFlanks@12944.12964.21\|CGTAGACGACCTGCTTCTGGG | >hg38_21_+_8433172_8446622_RNA45SN1_45S_50ntFlanks | 12944 | 12964 | 21 | rRF 45S | NO | 36.12 | 0.00 | 0.00 | 0.00 |
| >hg38_21_+_8433172_8446622_RNA45SN1_45S_50ntFlanks@12944.12972.29\|CGTAGACGACCTGCTTCTGGGTCGGGGTT | >hg38_21_+_8433172_8446622_RNA45SN1_45S_50ntFlanks | 12944 | 12972 | 29 | rRF 45S | NO | 0.00 | 29.82 | 31.64 | 0.00 |
| >hg38_21_+_8433172_8446622_RNA45SN1_45S_50ntFlanks@12948.12982.35\|GACGACCTGCTTCTGGGTCGGGGTTTCGTACGTAG | >hg38_21_+_8433172_8446622_RNA45SN1_45S_50ntFlanks | 12948 | 12982 | 35 | rRF 45S | NO | 0.00 | 29.77 | 0.00 | 0.00 |
| >hg38_21_+_8433172_8446622_RNA45SN1_45S_50ntFlanks@12949.12982.34\|ACGACCTGCTTCTGGGTCGGGGTTTCGTACGTAG | >hg38_21_+_8433172_8446622_RNA45SN1_45S_50ntFlanks | 12949 | 12982 | 34 | rRF 45S | NO | 0.00 | 26.79 | 0.00 | 0.00 |

### MDA-MB-468

| | | | | | | | | | | |
|---|---|---|---|---|---|---|---|---|---|---|
| >hg38_21_+_8433172_8446622_RNA45SN1_45S_50ntFlanks@12949.12990.42\|ACGACCTGCTTCTGGGTCGGGGTTTCGTACGTAGCAGAGCAG | >hg38_21_+_8433172_8446622_RNA45SN1_45S_50ntFlanks | 12949 | 12990 | 42 | rRF 45S | NO | 45.75 | 0.00 | 0.00 | 0.00 |
| >hg38_21_+_8433172_8446622_RNA45SN1_45S_50ntFlanks@12954.12972.19\|CTGCTTCTGGGTCGGGGTT | >hg38_21_+_8433172_8446622_RNA45SN1_45S_50ntFlanks | 12954 | 12972 | 19 | rRF 45S | NO | 53.34 | 48.14 | 0.00 | 0.00 |
| >hg38_21_+_8433172_8446622_RNA45SN1_45S_50ntFlanks@12954.12982.29\|CTGCTTCTGGGTCGGGGTTTCGTACGTAG | >hg38_21_+_8433172_8446622_RNA45SN1_45S_50ntFlanks | 12954 | 12982 | 29 | rRF 45S | NO | 31.92 | 0.00 | 0.00 | 0.00 |
| >hg38_21_+_8433172_8446622_RNA45SN1_45S_50ntFlanks@12956.12982.27\|GCTTCTGGGTCGGGGTTTCGTACGTAG | >hg38_21_+_8433172_8446622_RNA45SN1_45S_50ntFlanks | 12956 | 12982 | 27 | rRF 45S | NO | 29.27 | 0.00 | 0.00 | 0.00 |
| >hg38_21_+_8433172_8446622_RNA45SN1_45S_50ntFlanks@12957.12975.19\|CTTCTGGGTCGGGGTTTCG | >hg38_21_+_8433172_8446622_RNA45SN1_45S_50ntFlanks | 12957 | 12975 | 19 | rRF 45S | NO | 13.06 | 0.00 | 0.00 | 0.00 |
| >hg38_21_+_8433172_8446622_RNA45SN1_45S_50ntFlanks@12957.12979.23\|CTTCTGGGTCGGGGTTTCGTACG | >hg38_21_+_8433172_8446622_RNA45SN1_45S_50ntFlanks | 12957 | 12979 | 23 | rRF 45S | NO | 156.54 | 64.47 | 0.00 | 34.42 |
| >hg38_21_+_8433172_8446622_RNA45SN1_45S_50ntFlanks@12957.12980.24\|CTTCTGGGTCGGGGTTTCGTACGT | >hg38_21_+_8433172_8446622_RNA45SN1_45S_50ntFlanks | 12957 | 12980 | 24 | rRF 45S | NO | 46.29 | 0.00 | 0.00 | 0.00 |
| >hg38_21_+_8433172_8446622_RNA45SN1_45S_50ntFlanks@12957.12981.25\|CTTCTGGGTCGGGGTTTCGTACGTA | >hg38_21_+_8433172_8446622_RNA45SN1_45S_50ntFlanks | 12957 | 12981 | 25 | rRF 45S | NO | 33.11 | 0.00 | 0.00 | 0.00 |
| >hg38_21_+_8433172_8446622_RNA45SN1_45S_50ntFlanks@12957.12982.26\|CTTCTGGGTCGGGGTTTCGTACGTAG | >hg38_21_+_8433172_8446622_RNA45SN1_45S_50ntFlanks | 12957 | 12982 | 26 | rRF 45S | NO | 371.55 | 162.34 | 31.87 | 79.32 |
| >hg38_21_+_8433172_8446622_RNA45SN1_45S_50ntFlanks@12957.12985.29\|CTTCTGGGTCGGGGTTTCGTACGTAGCAG | >hg38_21_+_8433172_8446622_RNA45SN1_45S_50ntFlanks | 12957 | 12985 | 29 | rRF 45S | NO | 21.52 | 0.00 | 0.00 | 0.00 |
| >hg38_21_+_8433172_8446622_RNA45SN1_45S_50ntFlanks@12957.12986.30\|CTTCTGGGTCGGGGTTTCGTACGTAGCAGA | >hg38_21_+_8433172_8446622_RNA45SN1_45S_50ntFlanks | 12957 | 12986 | 30 | rRF 45S | NO | 52.65 | 12.59 | 0.00 | 0.00 |
| >hg38_21_+_8433172_8446622_RNA45SN1_45S_50ntFlanks@12957.12987.31\|CTTCTGGGTCGGGGTTTCGTACGTAGCAGAG | >hg38_21_+_8433172_8446622_RNA45SN1_45S_50ntFlanks | 12957 | 12987 | 31 | rRF 45S | NO | 68.28 | 28.35 | 0.00 | 0.00 |
| >hg38_21_+_8433172_8446622_RNA45SN1_45S_50ntFlanks@12957.12988.32\|CTTCTGGGTCGGGGTTTCGTACGTAGCAGAGC | >hg38_21_+_8433172_8446622_RNA45SN1_45S_50ntFlanks | 12957 | 12988 | 32 | rRF 45S | NO | 33.92 | 0.00 | 0.00 | 0.00 |
| >hg38_21_+_8433172_8446622_RNA45SN1_45S_50ntFlanks@12957.12989.33\|CTTCTGGGTCGGGGTTTCGTACGTAGCAGAGCA | >hg38_21_+_8433172_8446622_RNA45SN1_45S_50ntFlanks | 12957 | 12989 | 33 | rRF 45S | NO | 45.90 | 0.00 | 0.00 | 0.00 |
| >hg38_21_+_8433172_8446622_RNA45SN1_45S_50ntFlanks@12957.12990.34\|CTTCTGGGTCGGGGTTTCGTACGTAGCAGAGCAG | >hg38_21_+_8433172_8446622_RNA45SN1_45S_50ntFlanks | 12957 | 12990 | 34 | rRF 45S | NO | 149.37 | 44.26 | 0.00 | 13.42 |
| >hg38_21_+_8433172_8446622_RNA45SN1_45S_50ntFlanks@12957.12991.35\|CTTCTGGGTCGGGGTTTCGTACGTAGCAGAGCAGC | >hg38_21_+_8433172_8446622_RNA45SN1_45S_50ntFlanks | 12957 | 12991 | 35 | rRF 45S | NO | 28.75 | 0.00 | 0.00 | 0.00 |
| >hg38_21_+_8433172_8446622_RNA45SN1_45S_50ntFlanks@12959.12982.24\|TCTGGGTCGGGTTTCGTACGTAG | >hg38_21_+_8433172_8446622_RNA45SN1_45S_50ntFlanks | 12959 | 12982 | 24 | rRF 45S | NO | 46.97 | 0.00 | 0.00 | 0.00 |
| >hg38_21_+_8433172_8446622_RNA45SN1_45S_50ntFlanks@12960.12982.23\|CTGGGTCGGGGTTTCGTACGTAG | >hg38_21_+_8433172_8446622_RNA45SN1_45S_50ntFlanks | 12960 | 12982 | 23 | rRF 45S | NO | 44.87 | 0.00 | 0.00 | 0.00 |
| >hg38_21_+_8433172_8446622_RNA45SN1_45S_50ntFlanks@12960.12990.31\|CTGGGTCGGGGTTTCGTACGTAGCAGAGCAG | >hg38_21_+_8433172_8446622_RNA45SN1_45S_50ntFlanks | 12960 | 12990 | 31 | rRF 45S | NO | 39.78 | 0.00 | 0.00 | 0.00 |
| >hg38_21_+_8433172_8446622_RNA45SN1_45S_50ntFlanks@12971.12988.18\|TTTCGTACGTAGCAGAGC | >hg38_21_+_8433172_8446622_RNA45SN1_45S_50ntFlanks | 12971 | 12988 | 18 | rRF 45S | NO | 21.77 | 0.00 | 0.00 | 0.00 |
| >hg38_21_+_8433172_8446622_RNA45SN1_45S_50ntFlanks@12971.12989.19\|TTTCGTACGTAGCAGAGCA | >hg38_21_+_8433172_8446622_RNA45SN1_45S_50ntFlanks | 12971 | 12989 | 19 | rRF 45S | NO | 38.41 | 33.47 | 0.00 | 0.00 |
| >hg38_21_+_8433172_8446622_RNA45SN1_45S_50ntFlanks@12971.12990.20\|TTTCGTACGTAGCAGAGCAG | >hg38_21_+_8433172_8446622_RNA45SN1_45S_50ntFlanks | 12971 | 12990 | 20 | rRF 45S | NO | 126.48 | 54.67 | 0.00 | 0.00 |
| >hg38_21_+_8433172_8446622_RNA45SN1_45S_50ntFlanks@12971.12991.21\|TTTCGTACGTAGCAGAGCAGC | >hg38_21_+_8433172_8446622_RNA45SN1_45S_50ntFlanks | 12971 | 12991 | 21 | rRF 45S | NO | 73.34 | 52.85 | 0.00 | 0.00 |
| >hg38_21_+_8433172_8446622_RNA45SN1_45S_50ntFlanks@12973.12990.18\|TCGTACGTAGCAGAGCAG | >hg38_21_+_8433172_8446622_RNA45SN1_45S_50ntFlanks | 12973 | 12990 | 18 | rRF 45S | YES | 1418.84 | 528.30 | 78.71 | 100.69 |
| >hg38_21_+_8433172_8446622_RNA45SN1_45S_50ntFlanks@12973.12991.19\|TCGTACGTAGCAGAGCAGC | >hg38_21_+_8433172_8446622_RNA45SN1_45S_50ntFlanks | 12973 | 12991 | 19 | rRF 45S | YES | 822.65 | 521.00 | 50.20 | 82.39 |
| >hg38_21_+_8433172_8446622_RNA45SN1_45S_50ntFlanks@12973.12992.20\|TCGTACGTAGCAGAGCAGCT | >hg38_21_+_8433172_8446622_RNA45SN1_45S_50ntFlanks | 12973 | 12992 | 20 | rRF 45S | NO | 134.50 | 118.77 | 0.00 | 13.81 |
| >hg38_21_+_8433172_8446622_RNA45SN1_45S_50ntFlanks@12973.13001.29\|TCGTACGTAGCAGAGCAGCTCCCTCGCTG | >hg38_21_+_8433172_8446622_RNA45SN1_45S_50ntFlanks | 12973 | 13001 | 29 | rRF 45S | NO | 48.15 | 38.42 | 0.00 | 0.00 |
| >hg38_21_+_8433172_8446622_RNA45SN1_45S_50ntFlanks@12974.12991.18\|CGTACGTAGCAGAGCAGC | >hg38_21_+_8433172_8446622_RNA45SN1_45S_50ntFlanks | 12974 | 12991 | 18 | rRF 45S | NO | 647.44 | 288.75 | 39.68 | 70.23 |
| >hg38_21_+_8433172_8446622_RNA45SN1_45S_50ntFlanks@12974.12992.19\|CGTACGTAGCAGAGCAGCT | >hg38_21_+_8433172_8446622_RNA45SN1_45S_50ntFlanks | 12974 | 12992 | 19 | rRF 45S | NO | 45.40 | 30.47 | 0.00 | 0.00 |
| >hg38_21_+_8433172_8446622_RNA45SN1_45S_50ntFlanks@12988.13013.26\|CAGCTCCCTCGCTGCGATCTATTGAA | >hg38_21_+_8433172_8446622_RNA45SN1_45S_50ntFlanks | 12988 | 13013 | 26 | rRF 45S | NO | 31.67 | 0.00 | 0.00 | 66.14 |
| >hg38_21_+_8433172_8446622_RNA45SN1_45S_50ntFlanks@12990.13012.23\|GCTCCCTCGCTGCGATCTATTGA | >hg38_21_+_8433172_8446622_RNA45SN1_45S_50ntFlanks | 12990 | 13012 | 23 | rRF 45S | NO | 81.15 | 0.00 | 0.00 | 0.00 |
| >hg38_21_+_8433172_8446622_RNA45SN1_45S_50ntFlanks@12990.13013.24\|GCTCCCTCGCTGCGATCTATTGAA | >hg38_21_+_8433172_8446622_RNA45SN1_45S_50ntFlanks | 12990 | 13013 | 24 | rRF 45S | NO | 393.67 | 85.74 | 0.00 | 69.11 |
| >hg38_21_+_8433172_8446622_RNA45SN1_45S_50ntFlanks@12990.13014.25\|GCTCCCTCGCTGCGATCTATTGAAA | >hg38_21_+_8433172_8446622_RNA45SN1_45S_50ntFlanks | 12990 | 13014 | 25 | rRF 45S | NO | 119.67 | 0.00 | 0.00 | 0.00 |
| >hg38_21_+_8433172_8446622_RNA45SN1_45S_50ntFlanks@12990.13015.26\|GCTCCCTCGCTGCGATCTATTGAAAG | >hg38_21_+_8433172_8446622_RNA45SN1_45S_50ntFlanks | 12990 | 13015 | 26 | rRF 45S | NO | 62.49 | 0.00 | 0.00 | 0.00 |

### MDA-MB-468

| | | | | | | | | | | |
|---|---|---|---|---|---|---|---|---|---|---|
| >hg38_21_+_8433172_8446622_RNA45SN1_45S_50ntFlanks@12991.13008.18\|CTCCCTCGCT GCGATCTA | >hg38_21_+_8433172_8446622_RNA45SN1_45S_50ntFlanks | 12991 | 13008 | 18 | rRF 45S | NO | 38.75 | 26.33 | 0.00 | 0.00 |
| >hg38_21_+_8433172_8446622_RNA45SN1_45S_50ntFlanks@12991.13009.19\|CTCCCTCGCT GCGATCTAT | >hg38_21_+_8433172_8446622_RNA45SN1_45S_50ntFlanks | 12991 | 13009 | 19 | rRF 45S | NO | 37.49 | 0.00 | 0.00 | 0.00 |
| >hg38_21_+_8433172_8446622_RNA45SN1_45S_50ntFlanks@12991.13010.20\|CTCCCTCGCT GCGATCTATT | >hg38_21_+_8433172_8446622_RNA45SN1_45S_50ntFlanks | 12991 | 13010 | 20 | rRF 45S | NO | 56.22 | 46.41 | 0.00 | 22.95 |
| >hg38_21_+_8433172_8446622_RNA45SN1_45S_50ntFlanks@12991.13011.21\|CTCCCTCGCT GCGATCTATTG | >hg38_21_+_8433172_8446622_RNA45SN1_45S_50ntFlanks | 12991 | 13011 | 21 | rRF 45S | NO | 294.83 | 126.91 | 0.00 | 64.21 |
| >hg38_21_+_8433172_8446622_RNA45SN1_45S_50ntFlanks@12991.13012.22\|CTCCCTCGCT GCGATCTATTGA | >hg38_21_+_8433172_8446622_RNA45SN1_45S_50ntFlanks | 12991 | 13012 | 22 | rRF 45S | NO | 653.02 | 198.70 | 0.00 | 69.20 |
| >hg38_21_+_8433172_8446622_RNA45SN1_45S_50ntFlanks@12991.13013.23\|CTCCCTCGCT GCGATCTATTGAA | >hg38_21_+_8433172_8446622_RNA45SN1_45S_50ntFlanks | 12991 | 13013 | 23 | rRF 45S | YES | 2270.37 | 600.84 | 34.43 | 251.52 |
| >hg38_21_+_8433172_8446622_RNA45SN1_45S_50ntFlanks@12991.13014.24\|CTCCCTCGCT GCGATCTATTGAAA | >hg38_21_+_8433172_8446622_RNA45SN1_45S_50ntFlanks | 12991 | 13014 | 24 | rRF 45S | NO | 828.93 | 152.97 | 0.00 | 84.81 |
| >hg38_21_+_8433172_8446622_RNA45SN1_45S_50ntFlanks@12991.13015.25\|CTCCCTCGCT GCGATCTATTGAAAG | >hg38_21_+_8433172_8446622_RNA45SN1_45S_50ntFlanks | 12991 | 13015 | 25 | rRF 45S | NO | 1022.61 | 205.28 | 0.00 | 193.02 |
| >hg38_21_+_8433172_8446622_RNA45SN1_45S_50ntFlanks@12991.13016.26\|CTCCCTCGCT GCGATCTATTGAAAGT | >hg38_21_+_8433172_8446622_RNA45SN1_45S_50ntFlanks | 12991 | 13016 | 26 | rRF 45S | NO | 168.84 | 43.40 | 0.00 | 6.03 |
| >hg38_21_+_8433172_8446622_RNA45SN1_45S_50ntFlanks@12991.13017.27\|CTCCCTCGCT GCGATCTATTGAAAGTC | >hg38_21_+_8433172_8446622_RNA45SN1_45S_50ntFlanks | 12991 | 13017 | 27 | rRF 45S | NO | 115.24 | 0.00 | 0.00 | 0.00 |
| >hg38_21_+_8433172_8446622_RNA45SN1_45S_50ntFlanks@12991.13018.28\|CTCCCTCGCT GCGATCTATTGAAAGTCA | >hg38_21_+_8433172_8446622_RNA45SN1_45S_50ntFlanks | 12991 | 13018 | 28 | rRF 45S | NO | 20.26 | 0.00 | 0.00 | 0.00 |
| >hg38_21_+_8433172_8446622_RNA45SN1_45S_50ntFlanks@12991.13019.29\|CTCCCTCGCT GCGATCTATTGAAAGTCAG | >hg38_21_+_8433172_8446622_RNA45SN1_45S_50ntFlanks | 12991 | 13019 | 29 | rRF 45S | NO | 133.27 | 35.53 | 0.00 | 6.36 |
| >hg38_21_+_8433172_8446622_RNA45SN1_45S_50ntFlanks@12991.13027.37\|CTCCCTCGCT GCGATCTATTGAAAGTCAGCCCTCGAC | >hg38_21_+_8433172_8446622_RNA45SN1_45S_50ntFlanks | 12991 | 13027 | 37 | rRF 45S | NO | 59.69 | 33.98 | 0.00 | 51.97 |
| >hg38_21_+_8433172_8446622_RNA45SN1_45S_50ntFlanks@12991.13028.38\|CTCCCTCGCT GCGATCTATTGAAAGTCAGCCCTCGACA | >hg38_21_+_8433172_8446622_RNA45SN1_45S_50ntFlanks | 12991 | 13028 | 38 | rRF 45S | NO | 32.49 | 27.43 | 0.00 | 36.37 |
| >hg38_21_+_8433172_8446622_RNA45SN1_45S_50ntFlanks@12991.13039.49\|CTCCCTCGCT GCGATCTATTGAAAGTCAGCCCTCGACACAAGGGTTTGT | >hg38_21_+_8433172_8446622_RNA45SN1_45S_50ntFlanks | 12991 | 13039 | 49 | rRF 45S | NO | 250.26 | 141.73 | 0.00 | 228.39 |
| >hg38_21_+_8433172_8446622_RNA45SN1_45S_50ntFlanks@12992.13011.20\|TCCCTCGCTG CGATCTATTG | >hg38_21_+_8433172_8446622_RNA45SN1_45S_50ntFlanks | 12992 | 13011 | 20 | rRF 45S | NO | 40.79 | 0.00 | 0.00 | 0.00 |
| >hg38_21_+_8433172_8446622_RNA45SN1_45S_50ntFlanks@12992.13012.21\|TCCCTCGCTG CGATCTATTGA | >hg38_21_+_8433172_8446622_RNA45SN1_45S_50ntFlanks | 12992 | 13012 | 21 | rRF 45S | NO | 80.80 | 34.77 | 0.00 | 0.00 |
| >hg38_21_+_8433172_8446622_RNA45SN1_45S_50ntFlanks@12992.13013.22\|TCCCTCGCTG CGATCTATTGAA | >hg38_21_+_8433172_8446622_RNA45SN1_45S_50ntFlanks | 12992 | 13013 | 22 | rRF 45S | NO | 271.01 | 85.76 | 0.00 | 35.72 |
| >hg38_21_+_8433172_8446622_RNA45SN1_45S_50ntFlanks@12992.13014.23\|TCCCTCGCTG CGATCTATTGAAA | >hg38_21_+_8433172_8446622_RNA45SN1_45S_50ntFlanks | 12992 | 13014 | 23 | rRF 45S | NO | 101.40 | 5.98 | 0.00 | 0.00 |
| >hg38_21_+_8433172_8446622_RNA45SN1_45S_50ntFlanks@12992.13015.24\|TCCCTCGCTG CGATCTATTGAAAG | >hg38_21_+_8433172_8446622_RNA45SN1_45S_50ntFlanks | 12992 | 13015 | 24 | rRF 45S | NO | 127.06 | 34.49 | 0.00 | 15.99 |
| >hg38_21_+_8433172_8446622_RNA45SN1_45S_50ntFlanks@12992.13039.48\|TCCCTCGCTG CGATCTATTGAAAGTCAGCCCTCGACACAAGGGTTTGT | >hg38_21_+_8433172_8446622_RNA45SN1_45S_50ntFlanks | 12992 | 13039 | 48 | rRF 45S | NO | 67.83 | 33.11 | 0.00 | 0.00 |
| >hg38_21_+_8433172_8446622_RNA45SN1_45S_50ntFlanks@12993.13012.20\|CCCTCGCTGC GATCTATTGA | >hg38_21_+_8433172_8446622_RNA45SN1_45S_50ntFlanks | 12993 | 13012 | 20 | rRF 45S | NO | 37.41 | 0.00 | 0.00 | 0.00 |
| >hg38_21_+_8433172_8446622_RNA45SN1_45S_50ntFlanks@12993.13013.21\|CCCTCGCTGC GATCTATTGAA | >hg38_21_+_8433172_8446622_RNA45SN1_45S_50ntFlanks | 12993 | 13013 | 21 | rRF 45S | NO | 118.01 | 34.28 | 0.00 | 0.00 |
| >hg38_21_+_8433172_8446622_RNA45SN1_45S_50ntFlanks@12993.13014.22\|CCCTCGCTGC GATCTATTGAAA | >hg38_21_+_8433172_8446622_RNA45SN1_45S_50ntFlanks | 12993 | 13014 | 22 | rRF 45S | NO | 41.31 | 0.00 | 0.00 | 0.00 |
| >hg38_21_+_8433172_8446622_RNA45SN1_45S_50ntFlanks@12993.13015.23\|CCCTCGCTGC GATCTATTGAAAG | >hg38_21_+_8433172_8446622_RNA45SN1_45S_50ntFlanks | 12993 | 13015 | 23 | rRF 45S | NO | 48.88 | 0.00 | 0.00 | 0.00 |
| >hg38_21_+_8433172_8446622_RNA45SN1_45S_50ntFlanks@12993.13039.47\|CCCTCGCTGC GATCTATTGAAAGTCAGCCCTCGACACAAGGGTTTGT | >hg38_21_+_8433172_8446622_RNA45SN1_45S_50ntFlanks | 12993 | 13039 | 47 | rRF 45S | NO | 60.84 | 0.00 | 0.00 | 0.00 |
| >hg38_21_+_8433172_8446622_RNA45SN1_45S_50ntFlanks@12994.13012.19\|CCTCGCTGCG ATCTATTGA | >hg38_21_+_8433172_8446622_RNA45SN1_45S_50ntFlanks | 12994 | 13012 | 19 | rRF 45S | NO | 46.10 | 31.55 | 0.00 | 0.00 |
| >hg38_21_+_8433172_8446622_RNA45SN1_45S_50ntFlanks@12994.13013.20\|CCTCGCTGCG ATCTATTGAA | >hg38_21_+_8433172_8446622_RNA45SN1_45S_50ntFlanks | 12994 | 13013 | 20 | rRF 45S | NO | 184.38 | 86.40 | 0.00 | 35.30 |
| >hg38_21_+_8433172_8446622_RNA45SN1_45S_50ntFlanks@12994.13014.21\|CCTCGCTGCG ATCTATTGAAA | >hg38_21_+_8433172_8446622_RNA45SN1_45S_50ntFlanks | 12994 | 13014 | 21 | rRF 45S | NO | 68.21 | 0.00 | 0.00 | 0.00 |
| >hg38_21_+_8433172_8446622_RNA45SN1_45S_50ntFlanks@12994.13015.22\|CCTCGCTGCG ATCTATTGAAAG | >hg38_21_+_8433172_8446622_RNA45SN1_45S_50ntFlanks | 12994 | 13015 | 22 | rRF 45S | NO | 76.32 | 28.75 | 0.00 | 0.00 |
| >hg38_21_+_8433172_8446622_RNA45SN1_45S_50ntFlanks@12994.13039.46\|CCTCGCTGCG ATCTATTGAAAGTCAGCCCTCGACACAAGGGTTTGT | >hg38_21_+_8433172_8446622_RNA45SN1_45S_50ntFlanks | 12994 | 13039 | 46 | rRF 45S | NO | 64.16 | 6.07 | 0.00 | 0.00 |
| >hg38_21_+_8433172_8446622_RNA45SN1_45S_50ntFlanks@12995.13013.19\|CTCGCTGCGA TCTATTGAA | >hg38_21_+_8433172_8446622_RNA45SN1_45S_50ntFlanks | 12995 | 13013 | 19 | rRF 45S | NO | 74.01 | 39.88 | 0.00 | 0.00 |
| >hg38_21_+_8433172_8446622_RNA45SN1_45S_50ntFlanks@12995.13014.20\|CTCGCTGCGA TCTATTGAAA | >hg38_21_+_8433172_8446622_RNA45SN1_45S_50ntFlanks | 12995 | 13014 | 20 | rRF 45S | NO | 29.62 | 0.00 | 0.00 | 0.00 |
| >hg38_21_+_8433172_8446622_RNA45SN1_45S_50ntFlanks@12995.13015.21\|CTCGCTGCGA TCTATTGAAAG | >hg38_21_+_8433172_8446622_RNA45SN1_45S_50ntFlanks | 12995 | 13015 | 21 | rRF 45S | NO | 29.54 | 0.00 | 0.00 | 0.00 |

### MDA-MB-468

| | | | | | | | | | | |
|---|---|---|---|---|---|---|---|---|---|---|
| >hg38_21_+_8433172_8446622_RNA45SN1_45S_50ntFlanks@12995.13039.45\|CTCGCTGCGATCTATTGAAAGTCAGCCCTCGACACAAGGGTTTGT | >hg38_21_+_8433172_8446622_RNA45SN1_45S_50ntFlanks | 12995 | 13039 | 45 | rRF 45S | NO | 216.71 | 125.38 | 0.00 | 81.80 |
| >hg38_21_+_8433172_8446622_RNA45SN1_45S_50ntFlanks@12996.13013.18\|TCGCTGCGATCTATTGAA | >hg38_21_+_8433172_8446622_RNA45SN1_45S_50ntFlanks | 12996 | 13013 | 18 | rRF 45S | NO | 48.52 | 41.00 | 0.00 | 0.00 |
| >hg38_21_+_8433172_8446622_RNA45SN1_45S_50ntFlanks@12996.13039.44\|TCGCTGCGATCTATTGAAAGTCAGCCCTCGACACAAGGGTTTGT | >hg38_21_+_8433172_8446622_RNA45SN1_45S_50ntFlanks | 12996 | 13039 | 44 | rRF 45S | NO | 163.78 | 114.04 | 0.00 | 81.31 |
| >hg38_21_+_8433172_8446622_RNA45SN1_45S_50ntFlanks@12997.13014.18\|CGCTGCGATCTATTGAAA | >hg38_21_+_8433172_8446622_RNA45SN1_45S_50ntFlanks | 12997 | 13014 | 18 | rRF 45S | NO | 46.98 | 27.88 | 0.00 | 0.00 |
| >hg38_21_+_8433172_8446622_RNA45SN1_45S_50ntFlanks@12997.13039.43\|CGCTGCGATCTATTGAAAGTCAGCCCTCGACACAAGGGTTTGT | >hg38_21_+_8433172_8446622_RNA45SN1_45S_50ntFlanks | 12997 | 13039 | 43 | rRF 45S | NO | 95.31 | 74.59 | 0.00 | 76.00 |
| >hg38_21_+_8433172_8446622_RNA45SN1_45S_50ntFlanks@12999.13027.29\|CTGCGATCTATTGAAAGTCAGCCCTCGAC | >hg38_21_+_8433172_8446622_RNA45SN1_45S_50ntFlanks | 12999 | 13027 | 29 | rRF 45S | NO | 89.82 | 52.70 | 0.00 | 23.39 |
| >hg38_21_+_8433172_8446622_RNA45SN1_45S_50ntFlanks@12999.13028.30\|CTGCGATCTATTGAAAGTCAGCCCTCGACA | >hg38_21_+_8433172_8446622_RNA45SN1_45S_50ntFlanks | 12999 | 13028 | 30 | rRF 45S | NO | 38.63 | 32.93 | 0.00 | 0.00 |
| >hg38_21_+_8433172_8446622_RNA45SN1_45S_50ntFlanks@12999.13039.41\|CTGCGATCTATTGAAAGTCAGCCCTCGACACAAGGGTTTGT | >hg38_21_+_8433172_8446622_RNA45SN1_45S_50ntFlanks | 12999 | 13039 | 41 | rRF 45S | NO | 0.00 | 12.56 | 0.00 | 23.92 |
| >hg38_21_+_8433172_8446622_RNA45SN1_45S_50ntFlanks@13000.13027.28\|TGCGATCTATTGAAAGTCAGCCCTCGAC | >hg38_21_+_8433172_8446622_RNA45SN1_45S_50ntFlanks | 13000 | 13027 | 28 | rRF 45S | NO | 57.40 | 40.00 | 0.00 | 0.00 |
| >hg38_21_+_8433172_8446622_RNA45SN1_45S_50ntFlanks@13001.13027.27\|GCGATCTATTGAAAGTCAGCCCTCGAC | >hg38_21_+_8433172_8446622_RNA45SN1_45S_50ntFlanks | 13001 | 13027 | 27 | rRF 45S | NO | 30.79 | 27.16 | 0.00 | 0.00 |
| >hg38_21_+_8433172_8446622_RNA45SN1_45S_50ntFlanks@13002.13020.19\|CGATCTATTGAAAGTCAGC | >hg38_21_+_8433172_8446622_RNA45SN1_45S_50ntFlanks | 13002 | 13020 | 19 | rRF 45S | NO | 0.00 | 34.36 | 0.00 | 0.00 |
| >hg38_21_+_8433172_8446622_RNA45SN1_45S_50ntFlanks@13002.13023.22\|CGATCTATTGAAAGTCAGCCCT | >hg38_21_+_8433172_8446622_RNA45SN1_45S_50ntFlanks | 13002 | 13023 | 22 | rRF 45S | NO | 54.67 | 72.93 | 32.13 | 0.00 |
| >hg38_21_+_8433172_8446622_RNA45SN1_45S_50ntFlanks@13002.13025.24\|CGATCTATTGAAAGTCAGCCCTCG | >hg38_21_+_8433172_8446622_RNA45SN1_45S_50ntFlanks | 13002 | 13025 | 24 | rRF 45S | NO | 0.00 | 59.52 | 0.00 | 0.00 |
| >hg38_21_+_8433172_8446622_RNA45SN1_45S_50ntFlanks@13002.13026.25\|CGATCTATTGAAAGTCAGCCCTCGA | >hg38_21_+_8433172_8446622_RNA45SN1_45S_50ntFlanks | 13002 | 13026 | 25 | rRF 45S | NO | 0.00 | 34.26 | 0.00 | 0.00 |
| >hg38_21_+_8433172_8446622_RNA45SN1_45S_50ntFlanks@13002.13027.26\|CGATCTATTGAAAGTCAGCCCTCGAC | >hg38_21_+_8433172_8446622_RNA45SN1_45S_50ntFlanks | 13002 | 13027 | 26 | rRF 45S | NO | 554.12 | 1049.29 | 44.76 | 182.25 |
| >hg38_21_+_8433172_8446622_RNA45SN1_45S_50ntFlanks@13002.13028.27\|CGATCTATTGAAAGTCAGCCCTCGACA | >hg38_21_+_8433172_8446622_RNA45SN1_45S_50ntFlanks | 13002 | 13028 | 27 | rRF 45S | NO | 266.91 | 495.94 | 36.03 | 117.21 |
| >hg38_21_+_8433172_8446622_RNA45SN1_45S_50ntFlanks@13002.13029.28\|CGATCTATTGAAAGTCAGCCCTCGACAC | >hg38_21_+_8433172_8446622_RNA45SN1_45S_50ntFlanks | 13002 | 13029 | 28 | rRF 45S | NO | 182.81 | 288.04 | 0.00 | 106.75 |
| >hg38_21_+_8433172_8446622_RNA45SN1_45S_50ntFlanks@13002.13039.38\|CGATCTATTGAAAGTCAGCCCTCGACACAAGGGTTTGT | >hg38_21_+_8433172_8446622_RNA45SN1_45S_50ntFlanks | 13002 | 13039 | 38 | rRF 45S | NO | 67.99 | 175.15 | 28.62 | 180.30 |
| >hg38_21_+_8433172_8446622_RNA45SN1_45S_50ntFlanks@13004.13023.20\|ATCTATTGAAAGTCAGCCCT | >hg38_21_+_8433172_8446622_RNA45SN1_45S_50ntFlanks | 13004 | 13023 | 20 | rRF 45S | NO | 0.00 | 28.46 | 0.00 | 0.00 |
| >hg38_21_+_8433172_8446622_RNA45SN1_45S_50ntFlanks@13004.13025.22\|ATCTATTGAAAGTCAGCCCTCG | >hg38_21_+_8433172_8446622_RNA45SN1_45S_50ntFlanks | 13004 | 13025 | 22 | rRF 45S | NO | 65.34 | 119.27 | 0.00 | 0.00 |
| >hg38_21_+_8433172_8446622_RNA45SN1_45S_50ntFlanks@13004.13027.24\|ATCTATTGAAAGTCAGCCCTCGAC | >hg38_21_+_8433172_8446622_RNA45SN1_45S_50ntFlanks | 13004 | 13027 | 24 | rRF 45S | NO | 138.10 | 223.79 | 0.00 | 42.27 |
| >hg38_21_+_8433172_8446622_RNA45SN1_45S_50ntFlanks@13004.13028.25\|ATCTATTGAAAGTCAGCCCTCGACA | >hg38_21_+_8433172_8446622_RNA45SN1_45S_50ntFlanks | 13004 | 13028 | 25 | rRF 45S | NO | 39.23 | 121.32 | 0.00 | 0.00 |
| >hg38_21_+_8433172_8446622_RNA45SN1_45S_50ntFlanks@13004.13029.26\|ATCTATTGAAAGTCAGCCCTCGACAC | >hg38_21_+_8433172_8446622_RNA45SN1_45S_50ntFlanks | 13004 | 13029 | 26 | rRF 45S | NO | 0.00 | 26.41 | 0.00 | 0.00 |
| >hg38_21_+_8433172_8446622_RNA45SN1_45S_50ntFlanks@13005.13025.21\|TCTATTGAAAGTCAGCCCTCG | >hg38_21_+_8433172_8446622_RNA45SN1_45S_50ntFlanks | 13005 | 13025 | 21 | rRF 45S | NO | 0.00 | 30.33 | 0.00 | 0.00 |
| >hg38_21_+_8433172_8446622_RNA45SN1_45S_50ntFlanks@13005.13027.23\|TCTATTGAAAGTCAGCCCTCGAC | >hg38_21_+_8433172_8446622_RNA45SN1_45S_50ntFlanks | 13005 | 13027 | 23 | rRF 45S | NO | 0.00 | 37.93 | 0.00 | 0.00 |
| >hg38_21_+_8433172_8446622_RNA45SN1_45S_50ntFlanks@13005.13028.24\|TCTATTGAAAGTCAGCCCTCGACA | >hg38_21_+_8433172_8446622_RNA45SN1_45S_50ntFlanks | 13005 | 13028 | 24 | rRF 45S | NO | 0.00 | 30.79 | 0.00 | 0.00 |
| >hg38_21_+_8433172_8446622_RNA45SN1_45S_50ntFlanks@13006.13025.20\|CTATTGAAAGTCAGCCCTCG | >hg38_21_+_8433172_8446622_RNA45SN1_45S_50ntFlanks | 13006 | 13025 | 20 | rRF 45S | NO | 53.07 | 87.16 | 0.00 | 0.00 |
| >hg38_21_+_8433172_8446622_RNA45SN1_45S_50ntFlanks@13006.13027.22\|CTATTGAAAGTCAGCCCTCGAC | >hg38_21_+_8433172_8446622_RNA45SN1_45S_50ntFlanks | 13006 | 13027 | 22 | rRF 45S | NO | 31.36 | 101.69 | 0.00 | 0.00 |
| >hg38_21_+_8433172_8446622_RNA45SN1_45S_50ntFlanks@13006.13028.23\|CTATTGAAAGTCAGCCCTCGACA | >hg38_21_+_8433172_8446622_RNA45SN1_45S_50ntFlanks | 13006 | 13028 | 23 | rRF 45S | NO | 0.00 | 93.89 | 0.00 | 0.00 |
| >hg38_21_+_8433172_8446622_RNA45SN1_45S_50ntFlanks@13007.13025.19\|TATTGAAAGTCAGCCCTCG | >hg38_21_+_8433172_8446622_RNA45SN1_45S_50ntFlanks | 13007 | 13025 | 19 | rRF 45S | NO | 52.43 | 93.60 | 0.00 | 0.00 |
| >hg38_21_+_8433172_8446622_RNA45SN1_45S_50ntFlanks@13007.13027.21\|TATTGAAAGTCAGCCCTCGAC | >hg38_21_+_8433172_8446622_RNA45SN1_45S_50ntFlanks | 13007 | 13027 | 21 | rRF 45S | NO | 0.00 | 57.66 | 0.00 | 0.00 |
| >hg38_21_+_8433172_8446622_RNA45SN1_45S_50ntFlanks@13007.13028.22\|TATTGAAAGTCAGCCCTCGACA | >hg38_21_+_8433172_8446622_RNA45SN1_45S_50ntFlanks | 13007 | 13028 | 22 | rRF 45S | NO | 0.00 | 73.26 | 0.00 | 0.00 |
| >hg38_21_+_8433172_8446622_RNA45SN1_45S_50ntFlanks@13008.13025.18\|ATTGAAAGTCAGCCCTCG | >hg38_21_+_8433172_8446622_RNA45SN1_45S_50ntFlanks | 13008 | 13025 | 18 | rRF 45S | NO | 69.50 | 113.90 | 0.00 | 0.00 |
| >hg38_21_+_8433172_8446622_RNA45SN1_45S_50ntFlanks@13008.13027.20\|ATTGAAAGTCAGCCCTCGAC | >hg38_21_+_8433172_8446622_RNA45SN1_45S_50ntFlanks | 13008 | 13027 | 20 | rRF 45S | NO | 0.00 | 56.69 | 0.00 | 0.00 |

### MDA-MB-468

| | | | | | | | | | | |
|---|---|---|---|---|---|---|---|---|---|---|
| >hg38_21_+_8433172_8446622_RNA45SN1_45S_50ntFlanks@13008.13028.21\|ATTGAAAGTCAGCCCTCGACA | >hg38_21_+_8433172_8446622_RNA45SN1_45S_50ntFlanks | 13008 | 13028 | 21 | rRF 45S | NO | 0.00 | 56.82 | 0.00 | 0.00 |
| >hg38_21_+_8433172_8446622_RNA45SN1_45S_50ntFlanks@13009.13027.19\|TTGAAAGTCAGCCCTCGAC | >hg38_21_+_8433172_8446622_RNA45SN1_45S_50ntFlanks | 13009 | 13027 | 19 | rRF 45S | NO | 0.00 | 36.75 | 0.00 | 0.00 |
| >hg38_1_-_228634819_228635039_RNA5S12_5S_50ntFlanks@51.71.21\|GTCTACGGCCATACCACCCTG | >hg38_1_-_228634819_228635039_RNA5S12_5S_50ntFlanks | 51 | 71 | 21 | rRF 5S | NO | 64.92 | 41.37 | 0.00 | 6.51 |
| >hg38_1_-_228634819_228635039_RNA5S12_5S_50ntFlanks@51.81.31\|GTCTACGGCCATACCACCCTGAACGCGCCCG | >hg38_1_-_228634819_228635039_RNA5S12_5S_50ntFlanks | 51 | 81 | 31 | rRF 5S | NO | 48.42 | 0.00 | 0.00 | 0.00 |
| >hg38_1_-_228634819_228635039_RNA5S12_5S_50ntFlanks@51.82.32\|GTCTACGGCCATACCACCCTGAACGCGCCCGA | >hg38_1_-_228634819_228635039_RNA5S12_5S_50ntFlanks | 51 | 82 | 32 | rRF 5S | NO | 32.78 | 0.00 | 0.00 | 0.00 |
| >hg38_1_-_228634819_228635039_RNA5S12_5S_50ntFlanks@51.83.33\|GTCTACGGCCATACCACCCTGAACGCGCCCGAT | >hg38_1_-_228634819_228635039_RNA5S12_5S_50ntFlanks | 51 | 83 | 33 | rRF 5S | NO | 734.01 | 112.58 | 0.00 | 72.73 |
| >hg38_1_-_228634819_228635039_RNA5S12_5S_50ntFlanks@51.84.34\|GTCTACGGCCATACCACCCTGAACGCGCCCGATC | >hg38_1_-_228634819_228635039_RNA5S12_5S_50ntFlanks | 51 | 84 | 34 | rRF 5S | NO | 120.54 | 27.41 | 0.00 | 0.00 |
| >hg38_1_-_228634819_228635039_RNA5S12_5S_50ntFlanks@84.101.18\|CTCGTCTGATCTCGGAAG | >hg38_1_-_228634819_228635039_RNA5S12_5S_50ntFlanks | 84 | 101 | 18 | rRF 5S | NO | 81.32 | 176.86 | 42.90 | 14.87 |
| >hg38_1_-_228634819_228635039_RNA5S12_5S_50ntFlanks@84.102.19\|CTCGTCTGATCTCGGAAGC | >hg38_1_-_228634819_228635039_RNA5S12_5S_50ntFlanks | 84 | 102 | 19 | rRF 5S | NO | 30.66 | 45.48 | 0.00 | 0.00 |
| >hg38_1_-_228634819_228635039_RNA5S12_5S_50ntFlanks@84.103.20\|CTCGTCTGATCTCGGAAGCT | >hg38_1_-_228634819_228635039_RNA5S12_5S_50ntFlanks | 84 | 103 | 20 | rRF 5S | NO | 75.41 | 100.45 | 0.00 | 0.00 |
| >hg38_1_-_228634819_228635039_RNA5S12_5S_50ntFlanks@84.112.29\|CTCGTCTGATCTCGGAAGCTAAGCAGGGT | >hg38_1_-_228634819_228635039_RNA5S12_5S_50ntFlanks | 84 | 112 | 29 | rRF 5S | NO | 66.52 | 27.15 | 0.00 | 0.00 |
| >hg38_1_-_228634819_228635039_RNA5S12_5S_50ntFlanks@124.147.24\|AGTACTTGGATGGGAGACCGCCTG | >hg38_1_-_228634819_228635039_RNA5S12_5S_50ntFlanks | 124 | 147 | 24 | rRF 5S | NO | 21.33 | 0.00 | 0.00 | 0.00 |
| >hg38_1_-_228634819_228635039_RNA5S12_5S_50ntFlanks@132.169.38\|GATGGGAGACCGCCTGGGAATACCGGGTGCTGTAGGCT | >hg38_1_-_228634819_228635039_RNA5S12_5S_50ntFlanks | 132 | 169 | 38 | rRF 5S | NO | 52.64 | 104.59 | 47.39 | 14.16 |
| >hg38_1_-_228634819_228635039_RNA5S12_5S_50ntFlanks@132.170.39\|GATGGGAGACCGCCTGGGAATACCGGGTGCTGTAGGCTT | >hg38_1_-_228634819_228635039_RNA5S12_5S_50ntFlanks | 132 | 170 | 39 | rRF 5S | NO | 428.59 | 719.13 | 256.82 | 169.87 |
| >hg38_1_-_228634819_228635039_RNA5S12_5S_50ntFlanks@132.171.40\|GATGGGAGACCGCCTGGGAATACCGGGTGCTGTAGGCTTT | >hg38_1_-_228634819_228635039_RNA5S12_5S_50ntFlanks | 132 | 171 | 40 | rRF 5S | NO | 93.65 | 168.91 | 71.83 | 58.16 |
| >hg38_1_-_228634819_228635039_RNA5S12_5S_50ntFlanks@133.169.37\|ATGGGAGACCGCCTGGGAATACCGGGTGCTGTAGGCT | >hg38_1_-_228634819_228635039_RNA5S12_5S_50ntFlanks | 133 | 169 | 37 | rRF 5S | NO | 0.00 | 30.06 | 6.90 | 0.00 |
| >hg38_1_-_228634819_228635039_RNA5S12_5S_50ntFlanks@133.170.38\|ATGGGAGACCGCCTGGGAATACCGGGTGCTGTAGGCTT | >hg38_1_-_228634819_228635039_RNA5S12_5S_50ntFlanks | 133 | 170 | 38 | rRF 5S | NO | 66.57 | 118.15 | 106.31 | 63.84 |
| >hg38_1_-_228634819_228635039_RNA5S12_5S_50ntFlanks@133.171.39\|ATGGGAGACCGCCTGGGAATACCGGGTGCTGTAGGCTTT | >hg38_1_-_228634819_228635039_RNA5S12_5S_50ntFlanks | 133 | 171 | 39 | rRF 5S | NO | 0.00 | 37.68 | 39.20 | 15.92 |
| >hg38_1_-_228634819_228635039_RNA5S12_5S_50ntFlanks@134.170.37\|TGGGAGACCGCCTGGGAATACCGGGTGCTGTAGGCTT | >hg38_1_-_228634819_228635039_RNA5S12_5S_50ntFlanks | 134 | 170 | 37 | rRF 5S | NO | 149.71 | 68.46 | 50.55 | 34.82 |
| >hg38_1_-_228634819_228635039_RNA5S12_5S_50ntFlanks@134.171.38\|TGGGAGACCGCCTGGGAATACCGGGTGCTGTAGGCTTT | >hg38_1_-_228634819_228635039_RNA5S12_5S_50ntFlanks | 134 | 171 | 38 | rRF 5S | NO | 34.53 | 0.00 | 0.00 | 0.00 |
| >hg38_1_-_228634819_228635039_RNA5S12_5S_50ntFlanks@135.170.36\|GGGAGACCGCCTGGGAATACCGGGTGCTGTAGGCTT | >hg38_1_-_228634819_228635039_RNA5S12_5S_50ntFlanks | 135 | 170 | 36 | rRF 5S | NO | 94.47 | 57.02 | 103.30 | 61.63 |
| >hg38_1_-_228634819_228635039_RNA5S12_5S_50ntFlanks@135.171.37\|GGGAGACCGCCTGGGAATACCGGGTGCTGTAGGCTTT | >hg38_1_-_228634819_228635039_RNA5S12_5S_50ntFlanks | 135 | 171 | 37 | rRF 5S | NO | 6.13 | 0.00 | 32.95 | 14.00 |
| >hg38_1_-_228634819_228635039_RNA5S12_5S_50ntFlanks@136.170.35\|GGAGACCGCCTGGGAATACCGGGTGCTGTAGGCTT | >hg38_1_-_228634819_228635039_RNA5S12_5S_50ntFlanks | 136 | 170 | 35 | rRF 5S | NO | 0.00 | 0.00 | 46.92 | 14.03 |
| >hg38_1_-_228634819_228635039_RNA5S12_5S_50ntFlanks@136.171.36\|GGAGACCGCCTGGGAATACCGGGTGCTGTAGGCTTT | >hg38_1_-_228634819_228635039_RNA5S12_5S_50ntFlanks | 136 | 171 | 36 | rRF 5S | NO | 0.00 | 0.00 | 40.07 | 15.82 |

### MDA-MB-468

| | | | | | | | | | | |
|---|---|---|---|---|---|---|---|---|---|---|
| >hg38_1_-_228634819_228635039_RNA5S12_5S_50ntFlanks@140.169.30\|ACCGCCTGGGAATACCGGGTGCTGTAGGCT | >hg38_1_-_228634819_228635039_RNA5S12_5S_50ntFlanks | 140 | 169 | 30 | rRF 5S | NO | 0.00 | 51.74 | 268.66 | 63.39 |
| >hg38_1_-_228634819_228635039_RNA5S12_5S_50ntFlanks@140.170.31\|ACCGCCTGGGAATACCGGGTGCTGTAGGCTT | >hg38_1_-_228634819_228635039_RNA5S12_5S_50ntFlanks | 140 | 170 | 31 | rRF 5S | NO | 47.27 | 140.55 | 540.75 | 190.17 |
| >hg38_1_-_228634819_228635039_RNA5S12_5S_50ntFlanks@140.171.32\|ACCGCCTGGGAATACCGGGTGCTGTAGGCTTT | >hg38_1_-_228634819_228635039_RNA5S12_5S_50ntFlanks | 140 | 171 | 32 | rRF 5S | NO | 0.00 | 33.08 | 225.12 | 48.13 |
| >hg38_1_-_228634819_228635039_RNA5S12_5S_50ntFlanks@141.169.29\|CCGCCTGGGAATACCGGGTGCTGTAGGCT | >hg38_1_-_228634819_228635039_RNA5S12_5S_50ntFlanks | 141 | 169 | 29 | rRF 5S | NO | 0.00 | 0.00 | 47.83 | 6.08 |
| >hg38_1_-_228634819_228635039_RNA5S12_5S_50ntFlanks@141.170.30\|CCGCCTGGGAATACCGGGTGCTGTAGGCTT | >hg38_1_-_228634819_228635039_RNA5S12_5S_50ntFlanks | 141 | 170 | 30 | rRF 5S | NO | 0.00 | 0.00 | 109.22 | 6.47 |
| >hg38_1_-_228634819_228635039_RNA5S12_5S_50ntFlanks@141.171.31\|CCGCCTGGGAATACCGGGTGCTGTAGGCTTT | >hg38_1_-_228634819_228635039_RNA5S12_5S_50ntFlanks | 141 | 171 | 31 | rRF 5S | NO | 0.00 | 0.00 | 104.22 | 0.00 |
| >hg38_1_-_228634819_228635039_RNA5S12_5S_50ntFlanks@142.170.29\|CGCCTGGGAATACCGGGTGCTGTAGGCTT | >hg38_1_-_228634819_228635039_RNA5S12_5S_50ntFlanks | 142 | 170 | 29 | rRF 5S | NO | 0.00 | 0.00 | 55.81 | 0.00 |
| >hg38_1_-_228634819_228635039_RNA5S12_5S_50ntFlanks@142.171.30\|CGCCTGGGAATACCGGGTGCTGTAGGCTTT | >hg38_1_-_228634819_228635039_RNA5S12_5S_50ntFlanks | 142 | 171 | 30 | rRF 5S | NO | 0.00 | 0.00 | 67.03 | 0.00 |
| >hg38_1_-_228634819_228635039_RNA5S12_5S_50ntFlanks@144.170.27\|CCTGGGAATACCGGGTGCTGTAGGCTT | >hg38_1_-_228634819_228635039_RNA5S12_5S_50ntFlanks | 144 | 170 | 27 | rRF 5S | NO | 0.00 | 0.00 | 140.33 | 25.14 |
| >hg38_1_-_228634819_228635039_RNA5S12_5S_50ntFlanks@144.171.28\|CCTGGGAATACCGGGTGCTGTAGGCTTT | >hg38_1_-_228634819_228635039_RNA5S12_5S_50ntFlanks | 144 | 171 | 28 | rRF 5S | NO | 0.00 | 0.00 | 162.28 | 41.88 |
| >hg38_1_-_228634819_228635039_RNA5S12_5S_50ntFlanks@147.170.24\|GGGAATACCGGGTGCTGTAGGCTT | >hg38_1_-_228634819_228635039_RNA5S12_5S_50ntFlanks | 147 | 170 | 24 | rRF 5S | NO | 13.19 | 0.00 | 102.18 | 14.18 |
| >hg38_1_-_228634819_228635039_RNA5S12_5S_50ntFlanks@147.171.25\|GGGAATACCGGGTGCTGTAGGCTTT | >hg38_1_-_228634819_228635039_RNA5S12_5S_50ntFlanks | 147 | 171 | 25 | rRF 5S | NO | 0.00 | 0.00 | 90.23 | 0.00 |
| >hg38_1_-_228634819_228635039_RNA5S12_5S_50ntFlanks@148.169.22\|GGAATACCGGGTGCTGTAGGCT | >hg38_1_-_228634819_228635039_RNA5S12_5S_50ntFlanks | 148 | 169 | 22 | rRF 5S | NO | 5.93 | 33.51 | 181.39 | 39.47 |
| >hg38_1_-_228634819_228635039_RNA5S12_5S_50ntFlanks@148.170.23\|GGAATACCGGGTGCTGTAGGCTT | >hg38_1_-_228634819_228635039_RNA5S12_5S_50ntFlanks | 148 | 170 | 23 | rRF 5S | NO | 48.95 | 63.90 | 444.12 | 97.97 |
| >hg38_1_-_228634819_228635039_RNA5S12_5S_50ntFlanks@148.171.24\|GGAATACCGGGTGCTGTAGGCTTT | >hg38_1_-_228634819_228635039_RNA5S12_5S_50ntFlanks | 148 | 171 | 24 | rRF 5S | NO | 43.52 | 74.04 | 502.88 | 87.48 |
| >hg38_1_-_228634819_228635039_RNA5S12_5S_50ntFlanks@149.170.22\|GAATACCGGGTGCTGTAGGCTT | >hg38_1_-_228634819_228635039_RNA5S12_5S_50ntFlanks | 149 | 170 | 22 | rRF 5S | NO | 0.00 | 0.00 | 139.31 | 22.97 |
| >hg38_1_-_228634819_228635039_RNA5S12_5S_50ntFlanks@149.171.23\|GAATACCGGGTGCTGTAGGCTTT | >hg38_1_-_228634819_228635039_RNA5S12_5S_50ntFlanks | 149 | 171 | 23 | rRF 5S | NO | 0.00 | 0.00 | 167.21 | 23.23 |
| >hg38_1_-_228634819_228635039_RNA5S12_5S_50ntFlanks@150.169.20\|AATACCGGGTGCTGTAGGCT | >hg38_1_-_228634819_228635039_RNA5S12_5S_50ntFlanks | 150 | 169 | 20 | rRF 5S | NO | 0.00 | 0.00 | 164.54 | 16.00 |
| >hg38_1_-_228634819_228635039_RNA5S12_5S_50ntFlanks@150.170.21\|AATACCGGGTGCTGTAGGCTT | >hg38_1_-_228634819_228635039_RNA5S12_5S_50ntFlanks | 150 | 170 | 21 | rRF 5S | NO | 0.00 | 26.63 | 446.99 | 95.69 |
| >hg38_1_-_228634819_228635039_RNA5S12_5S_50ntFlanks@150.171.22\|AATACCGGGTGCTGTAGGCTTT | >hg38_1_-_228634819_228635039_RNA5S12_5S_50ntFlanks | 150 | 171 | 22 | rRF 5S | NO | 0.00 | 31.81 | 639.38 | 103.92 |
| >hg38_1_-_228634819_228635039_RNA5S12_5S_50ntFlanks@151.170.20\|ATACCGGGTGCTGTAGGCTT | >hg38_1_-_228634819_228635039_RNA5S12_5S_50ntFlanks | 151 | 170 | 20 | rRF 5S | NO | 0.00 | 0.00 | 315.69 | 59.48 |
| >hg38_1_-_228634819_228635039_RNA5S12_5S_50ntFlanks@151.171.21\|ATACCGGGTGCTGTAGGCTTT | >hg38_1_-_228634819_228635039_RNA5S12_5S_50ntFlanks | 151 | 171 | 21 | rRF 5S | NO | 0.00 | 0.00 | 398.65 | 56.60 |
| >hg38_1_-_228634819_228635039_RNA5S12_5S_50ntFlanks@152.170.19\|TACCGGGTGCTGTAGGCTT | >hg38_1_-_228634819_228635039_RNA5S12_5S_50ntFlanks | 152 | 170 | 19 | rRF 5S | NO | 0.00 | 0.00 | 193.70 | 41.33 |

### MDA-MB-468

| | | | | | | | | | | |
|---|---|---|---|---|---|---|---|---|---|---|
| >hg38_1_-_228634819_228635039_RNA5S12_5S_50ntFlanks@152.171.20\|TACCGGGTGCTGTAGGCTTT | >hg38_1_-_228634819_228635039_RNA5S12_5S_50ntFlanks | 152 | 171 | 20 | rRF 5S | NO | 0.00 | 0.00 | 247.79 | 37.17 |
| >hg38_1_-_228634819_228635039_RNA5S12_5S_50ntFlanks@153.170.18\|ACCGGGTGCTGTAGGCTT | >hg38_1_-_228634819_228635039_RNA5S12_5S_50ntFlanks | 153 | 170 | 18 | rRF 5S | NO | 19.99 | 29.05 | 145.38 | 14.46 |
| >hg38_1_-_228634819_228635039_RNA5S12_5S_50ntFlanks@153.171.19\|ACCGGGTGCTGTAGGCTTT | >hg38_1_-_228634819_228635039_RNA5S12_5S_50ntFlanks | 153 | 171 | 19 | rRF 5S | NO | 0.00 | 5.93 | 138.71 | 13.55 |
| >hg38_1_-_228634819_228635039_RNA5S12_5S_50ntFlanks@154.171.18\|CCGGGTGCTGTAGGCTTT | >hg38_1_-_228634819_228635039_RNA5S12_5S_50ntFlanks | 154 | 171 | 18 | rRF 5S | NO | 0.00 | 31.14 | 563.11 | 79.82 |
| >trnaMT_GluTTC_MT_-_14674_14742@39.72.34,trnalookalike8_GluTTC_5_-_93905172_93905240@39.72.34\|ATTGGTCGTGGTTGTAGTCCGTGCGAGAATACCA | >GluTTC.trnaMT_MT_-_14674_14742 | 39 | 72 | 34 | tRF lookalike | NO | 44.21 | 0.00 | 69.28 | 16.15 |
| >trnaMT_GluTTC_MT_-_14674_14742@40.72.33,trnalookalike8_GluTTC_5_-_93905172_93905240@40.72.33\|TTGGTCGTGGTTGTAGTCCGTGCGAGAATACCA | >GluTTC.trnaMT_MT_-_14674_14742 | 40 | 72 | 33 | tRF lookalike | NO | 115.68 | 0.00 | 66.11 | 25.16 |
| >trnaMT_GluTTC_MT_-_14674_14742@41.72.32,trnalookalike8_GluTTC_5_-_93905172_93905240@41.72.32\|TGGTCGTGGTTGTAGTCCGTGCGAGAATACCA | >GluTTC.trnaMT_MT_-_14674_14742 | 41 | 72 | 32 | tRF lookalike | NO | 54.22 | 0.00 | 49.70 | 37.85 |
| >multi-trnaMT_GluTTC_MT_-_14674_14742@27.61.35,trnalookalike8_GluTTC_5_-_93905172_93905240@27.61.35\|GTTTTTCATATCATTGGTCGTGGTTGTAGTCCGTG | >GluTTC.trnaMT_MT_-_14674_14742.multi | 27 | 61 | 35 | tRF lookalike | YES | 36.34 | 0.00 | 0.00 | 0.00 |
| >trnaMT_TrpTCA_MT_+_5512_5579@53.71.19,trnalookalike3_TrpTCA_1_+_566062_566129@53.71.19\|CAATACTTAATTTCTGCCA | >TrpTCA.trnaMT_MT_+_5512_5579 | 53 | 71 | 19 | tRF lookalike | NO | 0.00 | 0.00 | 37.95 | 6.30 |
| >multi-am_trnaMT_ArgTCG_MT_+_10405_10469@28.48.21\|ATTTCGACTCATTAAATTATG | >ArgTCG.trnaMT_MT_+_10405_10469.multi.am | 28 | 48 | 21 | tRF MT | NO | 77.91 | 31.15 | 0.00 | 35.33 |
| >trnaMT_HisGTG_MT_+_12138_12206@32.71.40\|TGAATCTGACAACAGAGGCTTACGACCCCTTATTTACCCC | >HisGTG.trnaMT_MT_+_12138_12206 | 32 | 71 | 40 | tRF MT | NO | 0.00 | 0.00 | 43.39 | 160.33 |
| >trnaMT_LeuTAA_MT_+_3230_3304@16.44.29\|CCCGGTAATCGCATAAAACTTAAAACTTT | >LeuTAA.trnaMT_MT_+_3230_3304 | 16 | 44 | 29 | tRF MT | NO | 153.48 | 79.35 | 27.72 | 52.89 |
| >trnaMT_LeuTAA_MT_+_3230_3304@18.44.27\|CGGTAATCGCATAAAACTTAAAACTTT | >LeuTAA.trnaMT_MT_+_3230_3304 | 18 | 44 | 27 | tRF MT | NO | 22.09 | 5.97 | 7.24 | 15.63 |
| >multi-am_trnaMT_ProTGG_MT_-_15956_16023@53.71.19\|AAGACTTTTTCTCTGACCA | >ProTGG.trnaMT_MT_-_15956_16023.multi.am | 53 | 71 | 19 | tRF MT | NO | 0.00 | 0.00 | 35.28 | 76.55 |
| >trnaMT_SerGCT_MT_+_12207_12265@1.18.18\|GAGAAAGCTCACAAGAAC | >SerGCT.trnaMT_MT_+_12207_12265 | 1 | 18 | 18 | tRF MT | YES | 0.00 | 0.00 | 686.47 | 47.16 |
| >trnaMT_SerGCT_MT_+_12207_12265@1.19.19\|GAGAAAGCTCACAAGAACT | >SerGCT.trnaMT_MT_+_12207_12265 | 1 | 19 | 19 | tRF MT | NO | 0.00 | 0.00 | 246.41 | 35.33 |
| >trnaMT_SerGCT_MT_+_12207_12265@1.20.20\|GAGAAAGCTCACAAGAACTG | >SerGCT.trnaMT_MT_+_12207_12265 | 1 | 20 | 20 | tRF MT | NO | 0.00 | 0.00 | 83.85 | 26.61 |
| >trnaMT_SerGCT_MT_+_12207_12265@1.22.22\|GAGAAAGCTCACAAGAACTGCT | >SerGCT.trnaMT_MT_+_12207_12265 | 1 | 22 | 22 | tRF MT | NO | 0.00 | 0.00 | 200.86 | 41.20 |
| >trnaMT_SerGCT_MT_+_12207_12265@1.25.25\|GAGAAAGCTCACAAGAACTGCTAAC | >SerGCT.trnaMT_MT_+_12207_12265 | 1 | 25 | 25 | tRF MT | NO | 0.00 | 0.00 | 241.17 | 69.30 |
| >trnaMT_SerGCT_MT_+_12207_12265@1.26.26\|GAGAAAGCTCACAAGAACTGCTAACT | >SerGCT.trnaMT_MT_+_12207_12265 | 1 | 26 | 26 | tRF MT | NO | 0.00 | 0.00 | 166.68 | 42.94 |
| >trnaMT_SerGCT_MT_+_12207_12265@1.38.38\|GAGAAAGCTCACAAGAACTGCTAACTCATGCCCCCATG | >SerGCT.trnaMT_MT_+_12207_12265 | 1 | 38 | 38 | tRF MT | NO | 13.15 | 0.00 | 0.00 | 0.00 |
| >trnaMT_SerGCT_MT_+_12207_12265@7.38.32\|GCTCACAAGAACTGCTAACTCATGCCCCCATG | >SerGCT.trnaMT_MT_+_12207_12265 | 7 | 38 | 32 | tRF MT | YES | 33.09 | 0.00 | 0.00 | 0.00 |
| >trnaMT_SerGCT_MT_+_12207_12265@8.38.31\|CTCACAAGAACTGCTAACTCATGCCCCCATG | >SerGCT.trnaMT_MT_+_12207_12265 | 8 | 38 | 31 | tRF MT | NO | 105.09 | 0.00 | 0.00 | 39.21 |
| >trnaMT_SerGCT_MT_+_12207_12265@10.38.29\|CACAAGAACTGCTAACTCATGCCCCCATG | >SerGCT.trnaMT_MT_+_12207_12265 | 10 | 38 | 29 | tRF MT | NO | 81.17 | 0.00 | 0.00 | 26.35 |
| >multi-am_trnaMT_SerGCT_MT_+_12207_12265@11.38.28\|ACAAGAACTGCTAACTCATGCCCCCATG | >SerGCT.trnaMT_MT_+_12207_12265.multi.am | 11 | 38 | 28 | tRF MT | YES | 29.45 | 0.00 | 0.00 | 0.00 |
| >multi-am_trnaMT_SerGCT_MT_+_12207_12265@12.38.27\|CAAGAACTGCTAACTCATGCCCCCATG | >SerGCT.trnaMT_MT_+_12207_12265.multi.am | 12 | 38 | 27 | tRF MT | NO | 69.18 | 0.00 | 0.00 | 33.61 |
| >multi-am_trnaMT_SerGCT_MT_+_12207_12265@13.38.26\|AAGAACTGCTAACTCATGCCCCCATG | >SerGCT.trnaMT_MT_+_12207_12265.multi.am | 13 | 38 | 26 | tRF MT | YES | 57.23 | 0.00 | 0.00 | 0.00 |
| >multi-am_trnaMT_SerGCT_MT_+_12207_12265@16.38.23\|AACTGCTAACTCATGCCCCCATG | >SerGCT.trnaMT_MT_+_12207_12265.multi.am | 16 | 38 | 23 | tRF MT | NO | 46.30 | 0.00 | 0.00 | 0.00 |
| >trnaMT_ValTAC_MT_+_1602_1670@50.72.23\|ACTTAACTTGACCGCTCTGACCA | >ValTAC.trnaMT_MT_+_1602_1670 | 50 | 72 | 23 | tRF MT | NO | 136.50 | 0.00 | 0.00 | 37.20 |
| >trnaMT_ValTAC_MT_+_1602_1670@51.71.21\|CTTAACTTGACCGCTCTGACC | >ValTAC.trnaMT_MT_+_1602_1670 | 51 | 71 | 21 | tRF MT | NO | 30.18 | 0.00 | 20.76 | 41.70 |
| >trnaMT_ValTAC_MT_+_1602_1670@51.72.22\|CTTAACTTGACCGCTCTGACCA | >ValTAC.trnaMT_MT_+_1602_1670 | 51 | 72 | 22 | tRF MT | NO | 247.55 | 0.00 | 49.78 | 127.87 |
| >trnaMT_ValTAC_MT_+_1602_1670@55.72.18\|ACTTGACCGCTCTGACCA | >ValTAC.trnaMT_MT_+_1602_1670 | 55 | 72 | 18 | tRF MT | NO | 20.14 | 0.00 | 0.00 | 122.62 |
| >multi-am_trnaMT_ValTAC_MT_+_1602_1670@30.68.39\|CTTACACTTAGGAGATTTCAACTTAACTTGACCGCTCTG | >ValTAC.trnaMT_MT_+_1602_1670.multi.am | 30 | 68 | 39 | tRF MT | YES | 31.35 | 0.00 | 0.00 | 0.00 |
| >multi-am_trnaMT_ValTAC_MT_+_1602_1670@44.68.25\|ATTTCAACTTAACTTGACCGCTCTG | >ValTAC.trnaMT_MT_+_1602_1670.multi.am | 44 | 68 | 25 | tRF MT | NO | 35.68 | 0.00 | 0.00 | 0.00 |
| >multi-am_trnaMT_ValTAC_MT_+_1602_1670@48.68.21\|CAACTTAACTTGACCGCTCTG | >ValTAC.trnaMT_MT_+_1602_1670.multi.am | 48 | 68 | 21 | tRF MT | NO | 31.60 | 0.00 | 0.00 | 0.00 |
| >multi-am_trnaMT_ValTAC_MT_+_1602_1670@49.68.20\|AACTTAACTTGACCGCTCTG | >ValTAC.trnaMT_MT_+_1602_1670.multi.am | 49 | 68 | 20 | tRF MT | NO | 128.49 | 0.00 | 0.00 | 14.27 |
| >multi-am_trnaMT_ValTAC_MT_+_1602_1670@50.67.18\|ACTTAACTTGACCGCTCT | >ValTAC.trnaMT_MT_+_1602_1670.multi.am | 50 | 67 | 18 | tRF MT | NO | 22.00 | 0.00 | 0.00 | 0.00 |
| >multi-am_trnaMT_ValTAC_MT_+_1602_1670@50.68.19\|ACTTAACTTGACCGCTCTG | >ValTAC.trnaMT_MT_+_1602_1670.multi.am | 50 | 68 | 19 | tRF MT | NO | 848.72 | 173.72 | 85.48 | 91.93 |
| >multi-am_trnaMT_ValTAC_MT_+_1602_1670@51.68.18\|CTTAACTTGACCGCTCTG | >ValTAC.trnaMT_MT_+_1602_1670.multi.am | 51 | 68 | 18 | tRF MT | YES | 1459.38 | 899.94 | 213.96 | 350.59 |
| >trna12_ArgCCT_16_+_3243918_3243990@35.54.20,trna18_ArgCCT_17_+_73030001_73030073@35.54.20,trna21_ArgCCT_17_-_73030526_73030598@35.54.20,trna2_ArgCCT_16_+_3202901_3202973@35.54.20,trna3_ArgCCT_7_+_139025446_139025518@35.54.20\|CTAAGCCAGGGATTGTGGGT | >ArgCCT.trna12_16_+_3243918_3243990 | 35 | 54 | 20 | tRF nuclear | NO | 0.00 | 37.64 | 0.00 | 0.00 |

### MDA-MB-468

<table>
<tr><td>>trna12_ArgCCT_16_+_3243918_3243990@36.54.19,trna18_ArgCCT_17_+_73030001_73030073@36.54.19,trna21_ArgCCT_17_-_73030526_73030598@36.54.19,trna2_ArgCCT_16_+_3202901_3202973@36.54.19,trna3_ArgCCT_7_+_139025446_139025518@36.54.19|TAAGCCAGGGATTGTGGGT</td><td>>ArgCCT.trna12_16_+_3243918_3243990</td><td>36</td><td>54</td><td>19</td><td>tRF nuclear</td><td>NO</td><td>0.00</td><td>31.80</td><td>0.00</td><td>0.00</td></tr>
<tr><td>>am_trna2_ArgCCT_16_+_3202901_3202973@14.31.18,trna3_ArgCCT_7_+_139025446_139025518@14.31.18|AATGGATAAGGCATTGGC</td><td>>ArgCCT.trna2_16_+_3202901_3202973.am</td><td>14</td><td>31</td><td>18</td><td>tRF nuclear</td><td>NO</td><td>39.28</td><td>0.00</td><td>0.00</td><td>0.00</td></tr>
<tr><td>>trna9_ArgTCT_1_+_94313129_94313213@1.29.29|GGCTCCGTGGCGCAATGGATAGCGCATTG</td><td>>ArgTCT.trna9_1_+_94313129_94313213</td><td>1</td><td>29</td><td>29</td><td>tRF nuclear</td><td>NO</td><td>0.00</td><td>32.97</td><td>0.00</td><td>0.00</td></tr>
<tr><td>>trna10_AspGTC_12_-_125424193_125424264@1.29.29,trna12_AspGTC_12_-_125411891_125411962@1.29.29,trna38_AspGTC_17_-_8125556_8125627@1.29.29,trna45_AspGTC_6_+_27447453_27447524@1.29.29,trna48_AspGTC_6_+_27471523_27471594@1.29.29,trna4_AspGTC_12_+_96429799_96429870@1.29.29,trna69_AspGTC_1_-_161440205_161440276@1.29.29,trna72_AspGTC_1_-_161432824_161432895@1.29.29,trna75_AspGTC_1_-_161425414_161425485@1.29.29,trna78_AspGTC_1_-_161418033_161418104@1.29.29,trna81_AspGTC_1_-_161410615_161410686@1.29.29|TCCTCGTTAGTATAGTGGTGAGTATCCCC</td><td>>AspGTC.trna10_12_-_125424193_125424264</td><td>1</td><td>29</td><td>29</td><td>tRF nuclear</td><td>NO</td><td>0.00</td><td>53.68</td><td>0.00</td><td>0.00</td></tr>
<tr><td>>trna10_AspGTC_12_-_125424193_125424264@1.30.30,trna12_AspGTC_12_-_125411891_125411962@1.30.30,trna38_AspGTC_17_-_8125556_8125627@1.30.30,trna45_AspGTC_6_+_27447453_27447524@1.30.30,trna48_AspGTC_6_+_27471523_27471594@1.30.30,trna4_AspGTC_12_+_96429799_96429870@1.30.30,trna69_AspGTC_1_-_161440205_161440276@1.30.30,trna72_AspGTC_1_-_161432824_161432895@1.30.30,trna75_AspGTC_1_-_161425414_161425485@1.30.30,trna78_AspGTC_1_-_161418033_161418104@1.30.30,trna81_AspGTC_1_-_161410615_161410686@1.30.30|TCCTCGTTAGTATAGTGGTGAGTATCCCCG</td><td>>AspGTC.trna10_12_-_125424193_125424264</td><td>1</td><td>30</td><td>30</td><td>tRF nuclear</td><td>NO</td><td>42.28</td><td>90.82</td><td>0.00</td><td>0.00</td></tr>
<tr><td>>trna10_AspGTC_12_-_125424193_125424264@1.31.31,trna12_AspGTC_12_-_125411891_125411962@1.31.31,trna38_AspGTC_17_-_8125556_8125627@1.31.31,trna45_AspGTC_6_+_27447453_27447524@1.31.31,trna48_AspGTC_6_+_27471523_27471594@1.31.31,trna4_AspGTC_12_+_96429799_96429870@1.31.31,trna69_AspGTC_1_-_161440205_161440276@1.31.31,trna72_AspGTC_1_-_161432824_161432895@1.31.31,trna75_AspGTC_1_-_161425414_161425485@1.31.31,trna78_AspGTC_1_-_161418033_161418104@1.31.31,trna81_AspGTC_1_-_161410615_161410686@1.31.31|TCCTCGTTAGTATAGTGGTGAGTATCCCCGC</td><td>>AspGTC.trna10_12_-_125424193_125424264</td><td>1</td><td>31</td><td>31</td><td>tRF nuclear</td><td>NO</td><td>0.00</td><td>33.83</td><td>0.00</td><td>0.00</td></tr>
<tr><td>>trna10_AspGTC_12_-_125424193_125424264@31.72.42,trna12_AspGTC_12_-_125411891_125411962@31.72.42,trna38_AspGTC_17_-_8125556_8125627@31.72.42,trna45_AspGTC_6_+_27447453_27447524@31.72.42,trna48_AspGTC_6_+_27471523_27471594@31.72.42,trna4_AspGTC_12_+_96429799_96429870@31.72.42,trna69_AspGTC_1_-_161440205_161440276@31.72.42,trna72_AspGTC_1_-_161432824_161432895@31.72.42,trna75_AspGTC_1_-_161425414_161425485@31.72.42,trna78_AspGTC_1_-_161418033_161418104@31.72.42,trna81_AspGTC_1_-_161410615_161410686@31.72.42|CCTGTCACGCGGGAGACCGGGGTTCGATTCCCCGACGGGGAG</td><td>>AspGTC.trna10_12_-_125424193_125424264</td><td>31</td><td>72</td><td>42</td><td>tRF nuclear</td><td>NO</td><td>39.01</td><td>137.60</td><td>0.00</td><td>0.00</td></tr>
<tr><td>>trna10_AspGTC_12_-_125424193_125424264@32.70.39,trna12_AspGTC_12_-_125411891_125411962@32.70.39,trna144_AspGTC_6_-_27551236_27551307@32.70.39,trna38_AspGTC_17_-_8125556_8125627@32.70.39,trna45_AspGTC_6_+_27447453_27447524@32.70.39,trna48_AspGTC_6_+_27471523_27471594@32.70.39,trna4_AspGTC_12_+_96429799_96429870@32.70.39,trna69_AspGTC_1_-_161440205_161440276@32.70.39,trna72_AspGTC_1_-_161432824_161432895@32.70.39,trna75_AspGTC_1_-_161425414_161425485@32.70.39,trna78_AspGTC_1_-_161418033_161418104@32.70.39,trna81_AspGTC_1_-_161410615_161410686@32.70.39|CTGTCACGCGGGAGACCGGGGTTCGATTCCCCGACGGGG</td><td>>AspGTC.trna10_12_-_125424193_125424264</td><td>32</td><td>70</td><td>39</td><td>tRF nuclear</td><td>NO</td><td>33.03</td><td>53.05</td><td>0.00</td><td>0.00</td></tr>
<tr><td>>trna10_AspGTC_12_-_125424193_125424264@32.71.40,trna12_AspGTC_12_-_125411891_125411962@32.71.40,trna144_AspGTC_6_-_27551236_27551307@32.71.40,trna38_AspGTC_17_-_8125556_8125627@32.71.40,trna45_AspGTC_6_+_27447453_27447524@32.71.40,trna48_AspGTC_6_+_27471523_27471594@32.71.40,trna4_AspGTC_12_+_96429799_96429870@32.71.40,trna69_AspGTC_1_-_161440205_161440276@32.71.40,trna72_AspGTC_1_-_161432824_161432895@32.71.40,trna75_AspGTC_1_-_161425414_161425485@32.71.40,trna78_AspGTC_1_-_161418033_161418104@32.71.40,trna81_AspGTC_1_-_161410615_161410686@32.71.40|CTGTCACGCGGGAGACCGGGGTTCGATTCCCCGACGGGGA</td><td>>AspGTC.trna10_12_-_125424193_125424264</td><td>32</td><td>71</td><td>40</td><td>tRF nuclear</td><td>NO</td><td>0.00</td><td>12.36</td><td>0.00</td><td>0.00</td></tr>
</table>

### MDA-MB-468

| | | | | | | | | | | |
|---|---|---|---|---|---|---|---|---|---|---|
| >trna10_AspGTC_12_-_125424193_125424264@32.72.41,trna12_AspGTC_12_-_125411891_125411962@32.72.41,trna144_AspGTC_6_-_27551236_27551307@32.72.41,trna38_AspGTC_17_-_8125556_8125627@32.72.41,trna45_AspGTC_6_+_27447453_27447524@32.72.41,trna48_AspGTC_6_+_27471523_27471594@32.72.41,trna4_AspGTC_12_+_96429799_96429870@32.72.41,trna69_AspGTC_1_-_161440205_161440276@32.72.41,trna72_AspGTC_1_-_161432824_161432895@32.72.41,trna75_AspGTC_1_-_161425414_161425485@32.72.41,trna78_AspGTC_1_-_161418033_161418104@32.72.41,trna81_AspGTC_1_-_161410615_161410686@32.72.41\|CTGTCACGCGGGAGACCGGGGTTCGATTCCCCGACGGGGAG | >AspGTC.trna10_12_-_125424193_125424264 | 32 | 72 | 41 | tRF nuclear | NO | 79.61 | 170.40 | 0.00 | 0.00 |
| >trna10_AspGTC_12_-_125424193_125424264@32.73.42,trna12_AspGTC_12_-_125411891_125411962@32.73.42,trna144_AspGTC_6_-_27551236_27551307@32.73.42,trna38_AspGTC_17_-_8125556_8125627@32.73.42,trna45_AspGTC_6_+_27447453_27447524@32.73.42,trna48_AspGTC_6_+_27471523_27471594@32.73.42,trna4_AspGTC_12_+_96429799_96429870@32.73.42,trna69_AspGTC_1_-_161440205_161440276@32.73.42,trna72_AspGTC_1_-_161432824_161432895@32.73.42,trna75_AspGTC_1_-_161425414_161425485@32.73.42,trna78_AspGTC_1_-_161418033_161418104@32.73.42,trna81_AspGTC_1_-_161410615_161410686@32.73.42\|CTGTCACGCGGGAGACCGGGGTTCGATTCCCCGACGGGGAGC | >AspGTC.trna10_12_-_125424193_125424264 | 32 | 73 | 42 | tRF nuclear | NO | 6.03 | 36.66 | 0.00 | 0.00 |
| >trna10_AspGTC_12_-_125424193_125424264@35.72.38,trna12_AspGTC_12_-_125411891_125411962@35.72.38,trna144_AspGTC_6_-_27551236_27551307@35.72.38,trna38_AspGTC_17_-_8125556_8125627@35.72.38,trna45_AspGTC_6_+_27447453_27447524@35.72.38,trna48_AspGTC_6_+_27471523_27471594@35.72.38,trna4_AspGTC_12_+_96429799_96429870@35.72.38,trna69_AspGTC_1_-_161440205_161440276@35.72.38,trna72_AspGTC_1_-_161432824_161432895@35.72.38,trna75_AspGTC_1_-_161425414_161425485@35.72.38,trna78_AspGTC_1_-_161418033_161418104@35.72.38,trna81_AspGTC_1_-_161410615_161410686@35.72.38\|TCACGCGGGAGACCGGGGTTCGATTCCCCGACGGGGAG | >AspGTC.trna10_12_-_125424193_125424264 | 35 | 72 | 38 | tRF nuclear | NO | 0.00 | 0.00 | 28.76 | 0.00 |
| >trna10_AspGTC_12_-_125424193_125424264@37.70.34,trna12_AspGTC_12_-_125411891_125411962@37.70.34,trna144_AspGTC_6_-_27551236_27551307@37.70.34,trna38_AspGTC_17_-_8125556_8125627@37.70.34,trna45_AspGTC_6_+_27447453_27447524@37.70.34,trna48_AspGTC_6_+_27471523_27471594@37.70.34,trna4_AspGTC_12_+_96429799_96429870@37.70.34,trna69_AspGTC_1_-_161440205_161440276@37.70.34,trna72_AspGTC_1_-_161432824_161432895@37.70.34,trna75_AspGTC_1_-_161425414_161425485@37.70.34,trna78_AspGTC_1_-_161418033_161418104@37.70.34,trna81_AspGTC_1_-_161410615_161410686@37.70.34\|ACGCGGGAGACCGGGGTTCGATTCCCCGACGGGG | >AspGTC.trna10_12_-_125424193_125424264 | 37 | 70 | 34 | tRF nuclear | NO | 0.00 | 40.12 | 101.27 | 0.00 |
| >trna10_AspGTC_12_-_125424193_125424264@37.72.36,trna12_AspGTC_12_-_125411891_125411962@37.72.36,trna144_AspGTC_6_-_27551236_27551307@37.72.36,trna38_AspGTC_17_-_8125556_8125627@37.72.36,trna45_AspGTC_6_+_27447453_27447524@37.72.36,trna48_AspGTC_6_+_27471523_27471594@37.72.36,trna4_AspGTC_12_+_96429799_96429870@37.72.36,trna69_AspGTC_1_-_161440205_161440276@37.72.36,trna72_AspGTC_1_-_161432824_161432895@37.72.36,trna75_AspGTC_1_-_161425414_161425485@37.72.36,trna78_AspGTC_1_-_161418033_161418104@37.72.36,trna81_AspGTC_1_-_161410615_161410686@37.72.36\|ACGCGGGAGACCGGGGTTCGATTCCCCGACGGGGAG | >AspGTC.trna10_12_-_125424193_125424264 | 37 | 72 | 36 | tRF nuclear | NO | 0.00 | 67.46 | 6.49 | 0.00 |
| >trna10_AspGTC_12_-_125424193_125424264@38.72.35,trna12_AspGTC_12_-_125411891_125411962@38.72.35,trna144_AspGTC_6_-_27551236_27551307@38.72.35,trna38_AspGTC_17_-_8125556_8125627@38.72.35,trna45_AspGTC_6_+_27447453_27447524@38.72.35,trna48_AspGTC_6_+_27471523_27471594@38.72.35,trna4_AspGTC_12_+_96429799_96429870@38.72.35,trna69_AspGTC_1_-_161440205_161440276@38.72.35,trna72_AspGTC_1_-_161432824_161432895@38.72.35,trna75_AspGTC_1_-_161425414_161425485@38.72.35,trna78_AspGTC_1_-_161418033_161418104@38.72.35,trna81_AspGTC_1_-_161410615_161410686@38.72.35\|CGCGGGAGACCGGGGTTCGATTCCCCGACGGGGAG | >AspGTC.trna10_12_-_125424193_125424264 | 38 | 72 | 35 | tRF nuclear | NO | 0.00 | 51.88 | 6.64 | 0.00 |

### MDA-MB-468

| | | | | | | | | | | |
|---|---|---|---|---|---|---|---|---|---|---|
| >trna10_AspGTC_12_-_125424193_125424264@39.72.34,trna12_AspGTC_12_-_125411891_125411962@39.72.34,trna144_AspGTC_6_-_27551236_27551307@39.72.34,trna38_AspGTC_17_-_8125556_8125627@39.72.34,trna45_AspGTC_6_+_27447453_27447524@39.72.34,trna48_AspGTC_6_+_27471523_27471594@39.72.34,trna4_AspGTC_12_+_96429799_96429870@39.72.34,trna69_AspGTC_1_-_161440205_161440276@39.72.34,trna72_AspGTC_1_-_161432824_161432895@39.72.34,trna75_AspGTC_1_-_161425414_161425485@39.72.34,trna78_AspGTC_1_-_161418033_161418104@39.72.34,trna81_AspGTC_1_-_161410615_161410686@39.72.34\|GCGGGAGACCGGGGTTCGATTCCCCGACGGGGAG | >AspGTC.trna10_12_-_125424193_125424264 | 39 | 72 | 34 | tRF nuclear | YES | 0.00 | 46.03 | 0.00 | 0.00 |
| >trna112_GlnCTG_1_-_147737382_147737453@30.50.21,trna131_GlnCTG_6_-_27759135_27759206@30.50.21,trna146_GlnCTG_6_-_27515531_27515602@30.50.21,trna15_GlnCTG_1_+_145963304_145963375@30.50.21,trna19_GlnCTG_1_+_147505038_147505109@30.50.21,trna1_GlnCTG_6_+_18836402_18836473@30.50.21,trna28_GlnCTG_1_+_149186125_149186196@30.50.21,trna3_GlnCTG_17_+_8023070_8023141@30.50.21,trna49_GlnCTG_6_+_27487308_27487379@30.50.21,trna7_GlnCTG_15_-_66161400_66161471@30.50.21,trna99_GlnCTG_6_-_28909378_28909449@30.50.21\|GACTCTGAATCCAGCGATCCG | >GlnCTG.trna112_1_-_147737382_147737453 | 30 | 50 | 21 | tRF nuclear | NO | 19.97 | 26.07 | 0.00 | 0.00 |
| >trna146_GlnCTG_6_-_27515531_27515602@1.30.30,trna1_GlnCTG_6_+_18836402_18836473@1.30.30,trna3_GlnCTG_17_+_8023070_8023141@1.30.30,trna42_GlnCTG_6_+_27263212_27263283@1.30.30,trna49_GlnCTG_6_+_27487308_27487379@1.30.30,trna7_GlnCTG_15_-_66161400_66161471@1.30.30,trna99_GlnCTG_6_-_28909378_28909449@1.30.30\|GGTTCCATGGTGTAATGGTTAGCACTCTGG | >GlnCTG.trna146_6_-_27515531_27515602 | 1 | 30 | 30 | tRF nuclear | NO | 0.00 | 12.46 | 0.00 | 0.00 |
| >am_trna146_GlnCTG_6_-_27515531_27515602@1.29.29,trna1_GlnCTG_6_+_18836402_18836473@1.29.29,trna3_GlnCTG_17_+_8023070_8023141@1.29.29,trna42_GlnCTG_6_+_27263212_27263283@1.29.29,trna49_GlnCTG_6_+_27487308_27487379@1.29.29,trna7_GlnCTG_15_-_66161400_66161471@1.29.29,trna99_GlnCTG_6_-_28909378_28909449@1.29.29\|GGTTCCATGGTGTAATGGTTAGCACTCTG | >GlnCTG.trna146_6_-_27515531_27515602.am | 1 | 29 | 29 | tRF nuclear | YES | 183.29 | 1435.11 | 164.88 | 61.29 |
| >trna130_GlnTTG_6_-_27763640_27763711@1.29.29,trna173_GlnTTG_6_-_26311975_26312046@1.29.29,trna174_GlnTTG_6_-_26311424_26311495@1.29.29\|GGCCCCATGGTGTAATGGTTAGCACTCTG | >GlnTTG.trna130_6_-_27763640_27763711 | 1 | 29 | 29 | tRF nuclear | NO | 56.98 | 143.13 | 0.00 | 0.00 |
| >trna16_GlnTTG_17_+_47269890_47269961@1.29.29,trna64_GlnTTG_6_+_28557156_28557227@1.29.29,trna84_GlnTTG_6_+_145503859_145503930@1.29.29\|GGTCCCATGGTGTAATGGTTAGCACTCTG | >GlnTTG.trna16_17_+_47269890_47269961 | 1 | 29 | 29 | tRF nuclear | NO | 62.73 | 332.48 | 13.86 | 0.00 |
| >trna116_GluCTC_1_-_145399233_145399304@1.31.31,trna59_GluCTC_1_+_249168447_249168518@1.31.31,trna71_GluCTC_1_-_161439189_161439260@1.31.31,trna74_GluCTC_1_-_161431809_161431880@1.31.31,trna77_GluCTC_1_-_161424398_161424469@1.31.31,trna77_GluCTC_6_+_28949976_28950047@1.31.31,trna80_GluCTC_1_-_161417018_161417089@1.31.31,trna87_GluCTC_6_-_126101393_126101464@1.31.31\|TCCCTGGTGGTCTAGTGGTTAGGATTCGGCG | >GluCTC.trna116_1_-_145399233_145399304 | 1 | 31 | 31 | tRF nuclear | YES | 6127.60 | 19041.18 | 250.20 | 265.24 |
| >trna116_GluCTC_1_-_145399233_145399304@1.33.33,trna59_GluCTC_1_+_249168447_249168518@1.33.33,trna71_GluCTC_1_-_161439189_161439260@1.33.33,trna74_GluCTC_1_-_161431809_161431880@1.33.33,trna77_GluCTC_1_-_161424398_161424469@1.33.33,trna77_GluCTC_6_+_28949976_28950047@1.33.33,trna80_GluCTC_1_-_161417018_161417089@1.33.33,trna87_GluCTC_6_-_126101393_126101464@1.33.33\|TCCCTGGTGGTCTAGTGGTTAGGATTCGGCGCT | >GluCTC.trna116_1_-_145399233_145399304 | 1 | 33 | 33 | tRF nuclear | NO | 68.06 | 67.09 | 0.00 | 0.00 |
| >trna116_GluCTC_1_-_145399233_145399304@2.31.30,trna59_GluCTC_1_+_249168447_249168518@2.31.30,trna71_GluCTC_1_-_161439189_161439260@2.31.30,trna74_GluCTC_1_-_161431809_161431880@2.31.30,trna77_GluCTC_1_-_161424398_161424469@2.31.30,trna77_GluCTC_6_+_28949976_28950047@2.31.30,trna80_GluCTC_1_-_161417018_161417089@2.31.30,trna87_GluCTC_6_-_126101393_126101464@2.31.30\|CCCTGGTGGTCTAGTGGTTAGGATTCGGCG | >GluCTC.trna116_1_-_145399233_145399304 | 2 | 31 | 30 | tRF nuclear | NO | 19.56 | 43.68 | 0.00 | 0.00 |
| >trna116_GluCTC_1_-_145399233_145399304@13.31.19,trna25_GluCTC_2_-_71273488_71273560@13.31.19,trna59_GluCTC_1_+_249168447_249168518@13.31.19,trna71_GluCTC_1_-_161439189_161439260@13.31.19,trna74_GluCTC_1_-_161431809_161431880@13.31.19,trna77_GluCTC_1_-_161424398_161424469@13.31.19,trna77_GluCTC_6_+_28949976_28950047@13.31.19,trna80_GluCTC_1_-_161417018_161417089@13.31.19,trna87_GluCTC_6_-_126101393_126101464@13.31.19\|TAGTGGTTAGGATTCGGCG | >GluCTC.trna116_1_-_145399233_145399304 | 13 | 31 | 19 | tRF nuclear | NO | 0.00 | 12.38 | 0.00 | 0.00 |

### MDA-MB-468

<table>
<tr><th></th><th></th><th></th><th></th><th></th><th></th><th></th><th></th><th></th><th></th><th></th></tr>
<tr><td>>trna116_GluCTC_1_-_145399233_145399304@30.70.41,trna71_GluCTC_1_-_161439189_161439260@30.70.41,trna74_GluCTC_1_-_161431809_161431880@30.70.41,trna77_GluCTC_1_-_161424398_161424469@30.70.41,trna77_GluCTC_6_+_28949976_28950047@30.70.41,trna80_GluCTC_1_-_161417018_161417089@30.70.41,trna87_GluCTC_6_-_126101393_126101464@30.70.41|CGCTCTCACCGCCGCGGCCCGGGTTCGATTCCCGGTCAGGG</td><td>>GluCTC.trna116_1_-_145399233_145399304</td><td>30</td><td>70</td><td>41</td><td>tRF nuclear</td><td>YES</td><td>12.85</td><td>35.23</td><td>0.00</td><td>0.00</td></tr>
<tr><td>>trna116_GluCTC_1_-_145399233_145399304@31.70.40,trna71_GluCTC_1_-_161439189_161439260@31.70.40,trna74_GluCTC_1_-_161431809_161431880@31.70.40,trna77_GluCTC_1_-_161424398_161424469@31.70.40,trna77_GluCTC_6_+_28949976_28950047@31.70.40,trna80_GluCTC_1_-_161417018_161417089@31.70.40,trna87_GluCTC_6_-_126101393_126101464@31.70.40|GCTCTCACCGCCGCGGCCCGGGTTCGATTCCCGGTCAGGG</td><td>>GluCTC.trna116_1_-_145399233_145399304</td><td>31</td><td>70</td><td>40</td><td>tRF nuclear</td><td>NO</td><td>32.78</td><td>0.00</td><td>0.00</td><td>0.00</td></tr>
<tr><td>>trna116_GluCTC_1_-_145399233_145399304@32.69.38,trna59_GluCTC_1_+_249168447_249168518@32.69.38,trna71_GluCTC_1_-_161439189_161439260@32.69.38,trna74_GluCTC_1_-_161431809_161431880@32.69.38,trna77_GluCTC_1_-_161424398_161424469@32.69.38,trna77_GluCTC_6_+_28949976_28950047@32.69.38,trna80_GluCTC_1_-_161417018_161417089@32.69.38,trna87_GluCTC_6_-_126101393_126101464@32.69.38|CTCTCACCGCCGCGGCCCGGGTTCGATTCCCGGTCAGG</td><td>>GluCTC.trna116_1_-_145399233_145399304</td><td>32</td><td>69</td><td>38</td><td>tRF nuclear</td><td>NO</td><td>53.07</td><td>51.06</td><td>0.00</td><td>0.00</td></tr>
<tr><td>>trna116_GluCTC_1_-_145399233_145399304@32.70.39,trna71_GluCTC_1_-_161439189_161439260@32.70.39,trna74_GluCTC_1_-_161431809_161431880@32.70.39,trna77_GluCTC_1_-_161424398_161424469@32.70.39,trna77_GluCTC_6_+_28949976_28950047@32.70.39,trna80_GluCTC_1_-_161417018_161417089@32.70.39,trna87_GluCTC_6_-_126101393_126101464@32.70.39|CTCTCACCGCCGCGGCCCGGGTTCGATTCCCGGTCAGGG</td><td>>GluCTC.trna116_1_-_145399233_145399304</td><td>32</td><td>70</td><td>39</td><td>tRF nuclear</td><td>NO</td><td>312.21</td><td>551.46</td><td>0.00</td><td>46.81</td></tr>
<tr><td>>trna116_GluCTC_1_-_145399233_145399304@32.71.40,trna71_GluCTC_1_-_161439189_161439260@32.71.40,trna74_GluCTC_1_-_161431809_161431880@32.71.40,trna77_GluCTC_1_-_161424398_161424469@32.71.40,trna77_GluCTC_6_+_28949976_28950047@32.71.40,trna80_GluCTC_1_-_161417018_161417089@32.71.40,trna87_GluCTC_6_-_126101393_126101464@32.71.40|CTCTCACCGCCGCGGCCCGGGTTCGATTCCCGGTCAGGGA</td><td>>GluCTC.trna116_1_-_145399233_145399304</td><td>32</td><td>71</td><td>40</td><td>tRF nuclear</td><td>NO</td><td>55.83</td><td>170.47</td><td>0.00</td><td>0.00</td></tr>
<tr><td>>trna116_GluCTC_1_-_145399233_145399304@32.72.41,trna71_GluCTC_1_-_161439189_161439260@32.72.41,trna74_GluCTC_1_-_161431809_161431880@32.72.41,trna77_GluCTC_1_-_161424398_161424469@32.72.41,trna77_GluCTC_6_+_28949976_28950047@32.72.41,trna80_GluCTC_1_-_161417018_161417089@32.72.41,trna87_GluCTC_6_-_126101393_126101464@32.72.41|CTCTCACCGCCGCGGCCCGGGTTCGATTCCCGGTCAGGGAA</td><td>>GluCTC.trna116_1_-_145399233_145399304</td><td>32</td><td>72</td><td>41</td><td>tRF nuclear</td><td>NO</td><td>0.00</td><td>68.52</td><td>0.00</td><td>0.00</td></tr>
<tr><td>>trna116_GluCTC_1_-_145399233_145399304@32.73.42,trna71_GluCTC_1_-_161439189_161439260@32.73.42,trna74_GluCTC_1_-_161431809_161431880@32.73.42,trna77_GluCTC_1_-_161424398_161424469@32.73.42,trna77_GluCTC_6_+_28949976_28950047@32.73.42,trna80_GluCTC_1_-_161417018_161417089@32.73.42,trna87_GluCTC_6_-_126101393_126101464@32.73.42|CTCTCACCGCCGCGGCCCGGGTTCGATTCCCGGTCAGGGAAC</td><td>>GluCTC.trna116_1_-_145399233_145399304</td><td>32</td><td>73</td><td>42</td><td>tRF nuclear</td><td>NO</td><td>51.92</td><td>120.69</td><td>0.00</td><td>0.00</td></tr>
<tr><td>>trna116_GluCTC_1_-_145399233_145399304@32.75.44,trna71_GluCTC_1_-_161439189_161439260@32.75.44,trna74_GluCTC_1_-_161431809_161431880@32.75.44,trna77_GluCTC_1_-_161424398_161424469@32.75.44,trna77_GluCTC_6_+_28949976_28950047@32.75.44,trna80_GluCTC_1_-_161417018_161417089@32.75.44,trna87_GluCTC_6_-_126101393_126101464@32.75.44|CTCTCACCGCCGCGGCCCGGGTTCGATTCCCGGTCAGGGAACCA</td><td>>GluCTC.trna116_1_-_145399233_145399304</td><td>32</td><td>75</td><td>44</td><td>tRF nuclear</td><td>NO</td><td>67.26</td><td>93.02</td><td>0.00</td><td>0.00</td></tr>
<tr><td>>trna116_GluCTC_1_-_145399233_145399304@36.70.35,trna134_GluTTC_1_-_16861774_16861845@36.70.35,trna71_GluCTC_1_-_161439189_161439260@36.70.35,trna74_GluCTC_1_-_161431809_161431880@36.70.35,trna77_GluCTC_1_-_161424398_161424469@36.70.35,trna77_GluCTC_6_+_28949976_28950047@36.70.35,trna80_GluCTC_1_-_161417018_161417089@36.70.35,trna84_GluTTC_1_-_161391883_161391954@36.70.35,trna87_GluCTC_6_-_126101393_126101464@36.70.35|CACCGCCGCGGCCCGGGTTCGATTCCCGGTCAGGG</td><td>>GluCTC.trna116_1_-_145399233_145399304</td><td>36</td><td>70</td><td>35</td><td>tRF nuclear</td><td>NO</td><td>0.00</td><td>35.92</td><td>0.00</td><td>0.00</td></tr>
</table>

### MDA-MB-468

| | | | | | | | | | | |
|---|---|---|---|---|---|---|---|---|---|---|
| >am_trna116_GluCTC_1_-_145399233_145399304@1.26.26,trna59_GluCTC_1_+_249168447_249168518@1.26.26,trna71_GluCTC_1_-_161439189_161439260@1.26.26,trna74_GluCTC_1_-_161431809_161431880@1.26.26,trna77_GluCTC_1_-_161424398_161424469@1.26.26,trna77_GluCTC_6_+_28949976_28950047@1.26.26,trna80_GluCTC_1_-_161417018_161417089@1.26.26,trna87_GluCTC_6_-_126101393_126101464@1.26.26\|TCCCTGGTGGTCTAGTGGTTAGGATT | >GluCTC.trna116_1_-_145399233_145399304.am | 1 | 26 | 26 | tRF nuclear | NO | 38.33 | 71.39 | 0.00 | 0.00 |
| >am_trna116_GluCTC_1_-_145399233_145399304@1.27.27,trna59_GluCTC_1_+_249168447_249168518@1.27.27,trna71_GluCTC_1_-_161439189_161439260@1.27.27,trna74_GluCTC_1_-_161431809_161431880@1.27.27,trna77_GluCTC_1_-_161424398_161424469@1.27.27,trna77_GluCTC_6_+_28949976_28950047@1.27.27,trna80_GluCTC_1_-_161417018_161417089@1.27.27,trna87_GluCTC_6_-_126101393_126101464@1.27.27\|TCCCTGGTGGTCTAGTGGTTAGGATTC | >GluCTC.trna116_1_-_145399233_145399304.am | 1 | 27 | 27 | tRF nuclear | NO | 164.20 | 310.95 | 32.36 | 0.00 |
| >am_trna116_GluCTC_1_-_145399233_145399304@1.28.28,trna59_GluCTC_1_+_249168447_249168518@1.28.28,trna71_GluCTC_1_-_161439189_161439260@1.28.28,trna74_GluCTC_1_-_161431809_161431880@1.28.28,trna77_GluCTC_1_-_161424398_161424469@1.28.28,trna77_GluCTC_6_+_28949976_28950047@1.28.28,trna80_GluCTC_1_-_161417018_161417089@1.28.28,trna87_GluCTC_6_-_126101393_126101464@1.28.28\|TCCCTGGTGGTCTAGTGGTTAGGATTCG | >GluCTC.trna116_1_-_145399233_145399304.am | 1 | 28 | 28 | tRF nuclear | YES | 872.99 | 1735.59 | 105.32 | 91.77 |
| >am_trna116_GluCTC_1_-_145399233_145399304@1.29.29,trna59_GluCTC_1_+_249168447_249168518@1.29.29,trna71_GluCTC_1_-_161439189_161439260@1.29.29,trna74_GluCTC_1_-_161431809_161431880@1.29.29,trna77_GluCTC_1_-_161424398_161424469@1.29.29,trna77_GluCTC_6_+_28949976_28950047@1.29.29,trna80_GluCTC_1_-_161417018_161417089@1.29.29,trna87_GluCTC_6_-_126101393_126101464@1.29.29\|TCCCTGGTGGTCTAGTGGTTAGGATTCGG | >GluCTC.trna116_1_-_145399233_145399304.am | 1 | 29 | 29 | tRF nuclear | YES | 1437.52 | 5414.50 | 227.66 | 156.06 |
| >am_trna116_GluCTC_1_-_145399233_145399304@1.30.30,trna59_GluCTC_1_+_249168447_249168518@1.30.30,trna71_GluCTC_1_-_161439189_161439260@1.30.30,trna74_GluCTC_1_-_161431809_161431880@1.30.30,trna77_GluCTC_1_-_161424398_161424469@1.30.30,trna77_GluCTC_6_+_28949976_28950047@1.30.30,trna80_GluCTC_1_-_161417018_161417089@1.30.30,trna87_GluCTC_6_-_126101393_126101464@1.30.30\|TCCCTGGTGGTCTAGTGGTTAGGATTCGGC | >GluCTC.trna116_1_-_145399233_145399304.am | 1 | 30 | 30 | tRF nuclear | YES | 790.06 | 2322.35 | 91.63 | 63.91 |
| >trna59_GluCTC_1_+_249168447_249168518@32.70.39\|CTCTCACCGCCGCGGCCCGGGTTCGATTCCCGGTCAGGA | >GluCTC.trna59_1_+_249168447_249168518 | 32 | 70 | 39 | tRF nuclear | NO | 22.15 | 53.92 | 0.00 | 0.00 |
| >trna11_GluTTC_15_-_26327381_26327452@1.28.28,trna3_GluTTC_13_-_45492062_45492133@1.28.28\|TCCCACATGGTCTAGCGGTTAGGATTCC | >GluTTC.trna11_15_-_26327381_26327452 | 1 | 28 | 28 | tRF nuclear | NO | 0.00 | 26.24 | 0.00 | 0.00 |
| >trna11_GluTTC_15_-_26327381_26327452@1.29.29,trna3_GluTTC_13_-_45492062_45492133@1.29.29\|TCCCACATGGTCTAGCGGTTAGGATTCCT | >GluTTC.trna11_15_-_26327381_26327452 | 1 | 29 | 29 | tRF nuclear | NO | 83.84 | 375.05 | 7.16 | 68.85 |
| >trna11_GluTTC_15_-_26327381_26327452@1.30.30,trna3_GluTTC_13_-_45492062_45492133@1.30.30\|TCCCACATGGTCTAGCGGTTAGGATTCCTG | >GluTTC.trna11_15_-_26327381_26327452 | 1 | 30 | 30 | tRF nuclear | NO | 59.82 | 199.50 | 47.96 | 26.45 |
| >trna11_GluTTC_15_-_26327381_26327452@1.31.31,trna3_GluTTC_13_-_45492062_45492133@1.31.31\|TCCCACATGGTCTAGCGGTTAGGATTCCTGG | >GluTTC.trna11_15_-_26327381_26327452 | 1 | 31 | 31 | tRF nuclear | NO | 37.73 | 198.21 | 0.00 | 0.00 |
| >trna134_GluTTC_1_-_16861774_16861845@1.31.31,trna5_GluTTC_1_+_17199078_17199149@1.31.31,trna84_GluTTC_1_-_161391883_161391954@1.31.31,trna94_GluTTC_1_-_149664355_149664427@1.31.31\|TCCCTGGTGGTCTAGTGGCTAGGATTCGGCG | >GluTTC.trna134_1_-_16861774_16861845 | 1 | 31 | 31 | tRF nuclear | NO | 316.00 | 441.15 | 0.00 | 0.00 |
| >am_trna134_GluTTC_1_-_16861774_16861845@1.28.28,trna5_GluTTC_1_+_17199078_17199149@1.28.28,trna84_GluTTC_1_-_161391883_161391954@1.28.28,trna94_GluTTC_1_-_149664355_149664427@1.28.28\|TCCCTGGTGGTCTAGTGGCTAGGATTCG | >GluTTC.trna134_1_-_16861774_16861845.am | 1 | 28 | 28 | tRF nuclear | NO | 50.15 | 101.85 | 0.00 | 0.00 |
| >am_trna134_GluTTC_1_-_16861774_16861845@1.29.29,trna5_GluTTC_1_+_17199078_17199149@1.29.29,trna84_GluTTC_1_-_161391883_161391954@1.29.29,trna94_GluTTC_1_-_149664355_149664427@1.29.29\|TCCCTGGTGGTCTAGTGGCTAGGATTCGG | >GluTTC.trna134_1_-_16861774_16861845.am | 1 | 29 | 29 | tRF nuclear | NO | 151.66 | 364.69 | 0.00 | 0.00 |
| >am_trna134_GluTTC_1_-_16861774_16861845@1.30.30,trna5_GluTTC_1_+_17199078_17199149@1.30.30,trna84_GluTTC_1_-_161391883_161391954@1.30.30,trna94_GluTTC_1_-_149664355_149664427@1.30.30\|TCCCTGGTGGTCTAGTGGCTAGGATTCGGC | >GluTTC.trna134_1_-_16861774_16861845.am | 1 | 30 | 30 | tRF nuclear | NO | 76.55 | 103.96 | 0.00 | 0.00 |
| >trna20_GluTTC_2_-_131094701_131094772@1.29.29,trna5_GluTTC_13_-_41634874_41634945@1.29.29\|TCCCATATGGTCTAGCGGTTAGGATTCCT | >GluTTC.trna20_2_-_131094701_131094772 | 1 | 29 | 29 | tRF nuclear | NO | 0.00 | 31.74 | 0.00 | 0.00 |
| >trna20_GluTTC_2_-_131094701_131094772@1.30.30,trna5_GluTTC_13_-_41634874_41634945@1.30.30\|TCCCATATGGTCTAGCGGTTAGGATTCCTG | >GluTTC.trna20_2_-_131094701_131094772 | 1 | 30 | 30 | tRF nuclear | NO | 0.00 | 30.95 | 0.00 | 0.00 |
| >trna20_GluTTC_2_-_131094701_131094772@1.31.31,trna5_GluTTC_13_-_41634874_41634945@1.31.31\|TCCCATATGGTCTAGCGGTTAGGATTCCTGG | >GluTTC.trna20_2_-_131094701_131094772 | 1 | 31 | 31 | tRF nuclear | NO | 0.00 | 43.45 | 0.00 | 0.00 |

### MDA-MB-468

| | | | | | | | | | | |
|---|---|---|---|---|---|---|---|---|---|---|
| >am_trna128_GlyGCC_6_-_27870686_27870756@1.25.25,trna133_GlyCCC_1_-_16872434_16872504@1.25.25,trna18_GlyGCC_16_+_70822597_70822667@1.25.25,trna19_GlyGCC_16_+_70823410_70823480@1.25.25,trna19_GlyGCC_2_-_157257659_157257729@1.25.25,trna24_GlyGCC_16_-_70812942_70813012@1.25.25,trna25_GlyGCC_16_-_70812114_70812184@1.25.25,trna4_GlyCCC_1_+_17188416_17188486@1.25.25,trna5_GlyGCC_17_+_8029064_8029134@1.25.25,trna68_GlyGCC_1_-_161493637_161493707@1.25.25\|GCATTGGTGGTTCAGTGGTAGAATT | >GlyGCC.trna128_6_-_27870686_27870756.am | 1 | 25 | 25 | tRF nuclear | NO | 36.18 | 39.94 | 0.00 | 0.00 |
| >am_trna128_GlyGCC_6_-_27870686_27870756@1.26.26,trna133_GlyCCC_1_-_16872434_16872504@1.26.26,trna18_GlyGCC_16_+_70822597_70822667@1.26.26,trna19_GlyGCC_16_+_70823410_70823480@1.26.26,trna19_GlyGCC_2_-_157257659_157257729@1.26.26,trna24_GlyGCC_16_-_70812942_70813012@1.26.26,trna25_GlyGCC_16_-_70812114_70812184@1.26.26,trna4_GlyCCC_1_+_17188416_17188486@1.26.26,trna5_GlyGCC_17_+_8029064_8029134@1.26.26,trna68_GlyGCC_1_-_161493637_161493707@1.26.26\|GCATTGGTGGTTCAGTGGTAGAATTC | >GlyGCC.trna128_6_-_27870686_27870756.am | 1 | 26 | 26 | tRF nuclear | NO | 30.55 | 0.00 | 0.00 | 0.00 |
| >am_trna128_GlyGCC_6_-_27870686_27870756@1.27.27,trna133_GlyCCC_1_-_16872434_16872504@1.27.27,trna18_GlyGCC_16_+_70822597_70822667@1.27.27,trna19_GlyGCC_16_+_70823410_70823480@1.27.27,trna19_GlyGCC_2_-_157257659_157257729@1.27.27,trna24_GlyGCC_16_-_70812942_70813012@1.27.27,trna25_GlyGCC_16_-_70812114_70812184@1.27.27,trna4_GlyCCC_1_+_17188416_17188486@1.27.27,trna5_GlyGCC_17_+_8029064_8029134@1.27.27,trna68_GlyGCC_1_-_161493637_161493707@1.27.27\|GCATTGGTGGTTCAGTGGTAGAATTCT | >GlyGCC.trna128_6_-_27870686_27870756.am | 1 | 27 | 27 | tRF nuclear | NO | 130.02 | 131.31 | 0.00 | 0.00 |
| >am_trna128_GlyGCC_6_-_27870686_27870756@1.28.28,trna133_GlyCCC_1_-_16872434_16872504@1.28.28,trna18_GlyGCC_16_+_70822597_70822667@1.28.28,trna19_GlyGCC_16_+_70823410_70823480@1.28.28,trna19_GlyGCC_2_-_157257659_157257729@1.28.28,trna24_GlyGCC_16_-_70812942_70813012@1.28.28,trna25_GlyGCC_16_-_70812114_70812184@1.28.28,trna4_GlyCCC_1_+_17188416_17188486@1.28.28,trna5_GlyGCC_17_+_8029064_8029134@1.28.28,trna68_GlyGCC_1_-_161493637_161493707@1.28.28\|GCATTGGTGGTTCAGTGGTAGAATTCTC | >GlyGCC.trna128_6_-_27870686_27870756.am | 1 | 28 | 28 | tRF nuclear | YES | 2254.61 | 4827.38 | 398.28 | 184.83 |
| >am_trna128_GlyGCC_6_-_27870686_27870756@1.29.29,trna133_GlyCCC_1_-_16872434_16872504@1.29.29,trna18_GlyGCC_16_+_70822597_70822667@1.29.29,trna19_GlyGCC_16_+_70823410_70823480@1.29.29,trna19_GlyGCC_2_-_157257659_157257729@1.29.29,trna24_GlyGCC_16_-_70812942_70813012@1.29.29,trna25_GlyGCC_16_-_70812114_70812184@1.29.29,trna4_GlyCCC_1_+_17188416_17188486@1.29.29,trna5_GlyGCC_17_+_8029064_8029134@1.29.29,trna68_GlyGCC_1_-_161493637_161493707@1.29.29\|GCATTGGTGGTTCAGTGGTAGAATTCTCG | >GlyGCC.trna128_6_-_27870686_27870756.am | 1 | 29 | 29 | tRF nuclear | YES | 1036.20 | 3349.80 | 274.42 | 112.88 |
| >am_trna128_GlyGCC_6_-_27870686_27870756@1.30.30,trna133_GlyCCC_1_-_16872434_16872504@1.30.30,trna18_GlyGCC_16_+_70822597_70822667@1.30.30,trna19_GlyGCC_16_+_70823410_70823480@1.30.30,trna19_GlyGCC_2_-_157257659_157257729@1.30.30,trna24_GlyGCC_16_-_70812942_70813012@1.30.30,trna25_GlyGCC_16_-_70812114_70812184@1.30.30,trna4_GlyCCC_1_+_17188416_17188486@1.30.30,trna5_GlyGCC_17_+_8029064_8029134@1.30.30,trna68_GlyGCC_1_-_161493637_161493707@1.30.30\|GCATTGGTGGTTCAGTGGTAGAATTCTCGC | >GlyGCC.trna128_6_-_27870686_27870756.am | 1 | 30 | 30 | tRF nuclear | YES | 377.64 | 2273.25 | 190.84 | 44.24 |
| >am_trna128_GlyGCC_6_-_27870686_27870756@1.31.31,trna133_GlyCCC_1_-_16872434_16872504@1.31.31,trna18_GlyGCC_16_+_70822597_70822667@1.31.31,trna19_GlyGCC_16_+_70823410_70823480@1.31.31,trna19_GlyGCC_2_-_157257659_157257729@1.31.31,trna24_GlyGCC_16_-_70812942_70813012@1.31.31,trna25_GlyGCC_16_-_70812114_70812184@1.31.31,trna4_GlyCCC_1_+_17188416_17188486@1.31.31,trna5_GlyGCC_17_+_8029064_8029134@1.31.31,trna68_GlyGCC_1_-_161493637_161493707@1.31.31\|GCATTGGTGGTTCAGTGGTAGAATTCTCGCC | >GlyGCC.trna128_6_-_27870686_27870756.am | 1 | 31 | 31 | tRF nuclear | NO | 21.61 | 189.33 | 6.55 | 0.00 |
| >trna2_GlyGCC_21_-_18827107_18827177@1.28.28,trna35_GlyGCC_1_+_161413094_161413164@1.28.28,trna37_GlyGCC_1_+_161420467_161420537@1.28.28,trna39_GlyGCC_1_+_161427898_161427968@1.28.28,trna41_GlyGCC_1_+_161435258_161435328@1.28.28\|GCATGGGTGGTTCAGTGGTAGAATTCTC | >GlyGCC.trna2_21_-_18827107_18827177 | 1 | 28 | 28 | tRF nuclear | YES | 1077.32 | 3505.75 | 291.20 | 69.21 |
| >trna2_GlyGCC_21_-_18827107_18827177@1.29.29,trna35_GlyGCC_1_+_161413094_161413164@1.29.29,trna37_GlyGCC_1_+_161420467_161420537@1.29.29,trna39_GlyGCC_1_+_161427898_161427968@1.29.29,trna41_GlyGCC_1_+_161435258_161435328@1.29.29\|GCATGGGTGGTTCAGTGGTAGAATTCTCG | >GlyGCC.trna2_21_-_18827107_18827177 | 1 | 29 | 29 | tRF nuclear | YES | 845.75 | 3374.50 | 230.78 | 73.74 |

### MDA-MB-468

| | | | | | | | | | | |
|---|---|---|---|---|---|---|---|---|---|---|
| >trna2_GlyGCC_21_-_18827107_18827177@1.30.30,trna35_GlyGCC_1_+_161413094_161413164@1.30.30,trna37_GlyGCC_1_+_161420467_161420537@1.30.30,trna39_GlyGCC_1_+_161427898_161427968@1.30.30,trna41_GlyGCC_1_+_161435258_161435328@1.30.30\|GCATGGGTGGTTCAGTGGTAGAATTCTCGC | >GlyGCC.trna2_21_-_18827107_18827177 | 1 | 30 | 30 | tRF nuclear | NO | 237.77 | 1217.83 | 150.85 | 16.54 |
| >trna2_GlyGCC_21_-_18827107_18827177@1.31.31,trna35_GlyGCC_1_+_161413094_161413164@1.31.31,trna37_GlyGCC_1_+_161420467_161420537@1.31.31,trna39_GlyGCC_1_+_161427898_161427968@1.31.31,trna41_GlyGCC_1_+_161435258_161435328@1.31.31\|GCATGGGTGGTTCAGTGGTAGAATTCTCGCC | >GlyGCC.trna2_21_-_18827107_18827177 | 1 | 31 | 31 | tRF nuclear | NO | 0.00 | 92.50 | 0.00 | 0.00 |
| >am_trna2_GlyGCC_21_-_18827107_18827177@1.27.27,trna35_GlyGCC_1_+_161413094_161413164@1.27.27,trna37_GlyGCC_1_+_161420467_161420537@1.27.27,trna39_GlyGCC_1_+_161427898_161427968@1.27.27,trna41_GlyGCC_1_+_161435258_161435328@1.27.27\|GCATGGGTGGTTCAGTGGTAGAATTCT | >GlyGCC.trna2_21_-_18827107_18827177.am | 1 | 27 | 27 | tRF nuclear | NO | 71.72 | 81.03 | 0.00 | 0.00 |
| >trna10_GlyTCC_17_+_8124866_8124937@1.29.29\|GCGTTGGTGGTATAGTGGTAAGCATAGCT | >GlyTCC.trna10_17_+_8124866_8124937 | 1 | 29 | 29 | tRF nuclear | NO | 0.00 | 34.30 | 0.00 | 0.00 |
| >trna10_GlyTCC_17_+_8124866_8124937@1.30.30\|GCGTTGGTGGTATAGTGGTAAGCATAGCTG | >GlyTCC.trna10_17_+_8124866_8124937 | 1 | 30 | 30 | tRF nuclear | NO | 0.00 | 94.07 | 0.00 | 0.00 |
| >am_trna117_GlyTCC_1_-_145397864_145397935@1.29.29,trna45_GlyTCC_1_+_161500903_161500974@1.29.29,trna70_GlyTCC_1_-_161439547_161439618@1.29.29,trna73_GlyTCC_1_-_161432166_161432237@1.29.29,trna76_GlyTCC_1_-_161424756_161424827@1.29.29,trna79_GlyTCC_1_-_161417375_161417446@1.29.29\|GCGTTGGTGGTATAGTGGTGAGCATAGCT | >GlyTCC.trna117_1_-_145397864_145397935.am | 1 | 29 | 29 | tRF nuclear | NO | 102.87 | 311.71 | 39.52 | 0.00 |
| >am_trna117_GlyTCC_1_-_145397864_145397935@1.30.30,trna45_GlyTCC_1_+_161500903_161500974@1.30.30,trna70_GlyTCC_1_-_161439547_161439618@1.30.30,trna73_GlyTCC_1_-_161432166_161432237@1.30.30,trna76_GlyTCC_1_-_161424756_161424827@1.30.30,trna79_GlyTCC_1_-_161417375_161417446@1.30.30\|GCGTTGGTGGTATAGTGGTGAGCATAGCTG | >GlyTCC.trna117_1_-_145397864_145397935.am | 1 | 30 | 30 | tRF nuclear | NO | 105.79 | 516.35 | 68.74 | 26.11 |
| >am_trna2_GlyTCC_19_+_4724082_4724153@1.29.29\|GCGTTGGTGGTATAGTGGTTAGCATAGCT | >GlyTCC.trna2_19_+_4724082_4724153.am | 1 | 29 | 29 | tRF nuclear | NO | 0.00 | 42.33 | 0.00 | 0.00 |
| >am_trna2_GlyTCC_19_+_4724082_4724153@1.30.30\|GCGTTGGTGGTATAGTGGTTAGCATAGCTG | >GlyTCC.trna2_19_+_4724082_4724153.am | 1 | 30 | 30 | tRF nuclear | NO | 0.00 | 111.46 | 0.00 | 0.00 |
| >am_trna106_HisGTG_1_-_149155828_149155899@7.28.22,trna111_HisGTG_1_-_147774845_147774916@7.28.22,trna118_HisGTG_1_-_145396881_145396952@7.28.22,trna16_HisGTG_1_+_146544773_146544844@7.28.22,trna1_HisGTG_15_+_45493349_45493420@7.28.22,trna21_HisGTG_1_+_147753471_147753542@7.28.22,trna33_HisGTG_6_+_27125906_27125977@7.28.22,trna7_HisGTG_9_-_14433938_14434009@7.28.22,trna8_HisGTG_15_-_45492611_45492682@7.28.22,trna9_HisGTG_15_-_45490804_45490875@7.28.22\|ATCGTATAGTGGTTAGTACTCT | >HisGTG.trna106_1_-_149155828_149155899.am | 7 | 28 | 22 | tRF nuclear | NO | 0.00 | 59.61 | 0.00 | 0.00 |
| >am_trna106_HisGTG_1_-_149155828_149155899@7.29.23,trna111_HisGTG_1_-_147774845_147774916@7.29.23,trna118_HisGTG_1_-_145396881_145396952@7.29.23,trna16_HisGTG_1_+_146544773_146544844@7.29.23,trna1_HisGTG_15_+_45493349_45493420@7.29.23,trna21_HisGTG_1_+_147753471_147753542@7.29.23,trna33_HisGTG_6_+_27125906_27125977@7.29.23,trna7_HisGTG_9_-_14433938_14434009@7.29.23,trna8_HisGTG_15_-_45492611_45492682@7.29.23,trna9_HisGTG_15_-_45490804_45490875@7.29.23\|ATCGTATAGTGGTTAGTACTCTG | >HisGTG.trna106_1_-_149155828_149155899.am | 7 | 29 | 23 | tRF nuclear | NO | 0.00 | 46.11 | 27.25 | 0.00 |
| >trna111_HisGTG_1_-_147774845_147774916@1.30.30,trna118_HisGTG_1_-_145396881_145396952@1.30.30,trna16_HisGTG_1_+_146544773_146544844@1.30.30,trna1_HisGTG_15_+_45493349_45493420@1.30.30,trna21_HisGTG_1_+_147753471_147753542@1.30.30,trna33_HisGTG_6_+_27125906_27125977@1.30.30,trna7_HisGTG_9_-_14433938_14434009@1.30.30,trna8_HisGTG_15_-_45492611_45492682@1.30.30,trna9_HisGTG_15_-_45490804_45490875@1.30.30\|GCCGTGATCGTATAGTGGTTAGTACTCTGC | >HisGTG.trna111_1_-_147774845_147774916 | 1 | 30 | 30 | tRF nuclear | NO | 58.37 | 169.53 | 0.00 | 0.00 |
| >trna111_HisGTG_1_-_147774845_147774916@1.31.31,trna118_HisGTG_1_-_145396881_145396952@1.31.31,trna16_HisGTG_1_+_146544773_146544844@1.31.31,trna1_HisGTG_15_+_45493349_45493420@1.31.31,trna21_HisGTG_1_+_147753471_147753542@1.31.31,trna33_HisGTG_6_+_27125906_27125977@1.31.31,trna7_HisGTG_9_-_14433938_14434009@1.31.31,trna8_HisGTG_15_-_45492611_45492682@1.31.31,trna9_HisGTG_15_-_45490804_45490875@1.31.31\|GCCGTGATCGTATAGTGGTTAGTACTCTGCG | >HisGTG.trna111_1_-_147774845_147774916 | 1 | 31 | 31 | tRF nuclear | YES | 721.72 | 2379.47 | 160.39 | 50.60 |

### MDA-MB-468

| | | | | | | | | | | |
|---|---|---|---|---|---|---|---|---|---|---|
| >trna111_HisGTG_1_-_147774845_147774916@2.31.30,trna118_HisGTG_1_-_145396881_145396952@2.31.30,trna16_HisGTG_1_+_146544773_146544844@2.31.30,trna1_HisGTG_15_+_45493349_45493420@2.31.30,trna21_HisGTG_1_+_147753471_147753542@2.31.30,trna33_HisGTG_6_+_27125906_27125977@2.31.30,trna7_HisGTG_9_-_14433938_14434009@2.31.30,trna8_HisGTG_15_-_45492611_45492682@2.31.30,trna9_HisGTG_15_-_45490804_45490875@2.31.30\|CCGTGATCGTATAGTGGTTAGTACTCTGCG | >HisGTG.trna111_1_-_147774845_147774916 | 2 | 31 | 30 | tRF nuclear | NO | 0.00 | 19.65 | 0.00 | 0.00 |
| >trna111_HisGTG_1_-_147774845_147774916@3.31.29,trna118_HisGTG_1_-_145396881_145396952@3.31.29,trna16_HisGTG_1_+_146544773_146544844@3.31.29,trna1_HisGTG_15_+_45493349_45493420@3.31.29,trna21_HisGTG_1_+_147753471_147753542@3.31.29,trna33_HisGTG_6_+_27125906_27125977@3.31.29,trna7_HisGTG_9_-_14433938_14434009@3.31.29,trna8_HisGTG_15_-_45492611_45492682@3.31.29,trna9_HisGTG_15_-_45490804_45490875@3.31.29\|CGTGATCGTATAGTGGTTAGTACTCTGCG | >HisGTG.trna111_1_-_147774845_147774916 | 3 | 31 | 29 | tRF nuclear | NO | 0.00 | 28.69 | 0.00 | 0.00 |
| >trna111_HisGTG_1_-_147774845_147774916@-1A.29.30,trna118_HisGTG_1_-_145396881_145396952@-1A.29.30,trna16_HisGTG_1_+_146544773_146544844@-1A.29.30,trna1_HisGTG_15_+_45493349_45493420@-1A.29.30,trna21_HisGTG_1_+_147753471_147753542@-1A.29.30,trna33_HisGTG_6_+_27125906_27125977@-1A.29.30,trna7_HisGTG_9_-_14433938_14434009@-1A.29.30,trna8_HisGTG_15_-_45492611_45492682@-1A.29.30,trna9_HisGTG_15_-_45490804_45490875@-1A.29.30\|AGCCGTGATCGTATAGTGGTTAGTACTCTG | >HisGTG.trna111_1_-_147774845_147774916 | -1A | 29 | 30 | tRF nuclear | NO | 58.00 | 90.42 | 0.00 | 0.00 |
| >trna111_HisGTG_1_-_147774845_147774916@-1A.31.32,trna118_HisGTG_1_-_145396881_145396952@-1A.31.32,trna16_HisGTG_1_+_146544773_146544844@-1A.31.32,trna1_HisGTG_15_+_45493349_45493420@-1A.31.32,trna21_HisGTG_1_+_147753471_147753542@-1A.31.32,trna33_HisGTG_6_+_27125906_27125977@-1A.31.32,trna7_HisGTG_9_-_14433938_14434009@-1A.31.32,trna8_HisGTG_15_-_45492611_45492682@-1A.31.32,trna9_HisGTG_15_-_45490804_45490875@-1A.31.32\|AGCCGTGATCGTATAGTGGTTAGTACTCTGCG | >HisGTG.trna111_1_-_147774845_147774916 | -1A | 31 | 32 | tRF nuclear | NO | 113.83 | 207.84 | 32.57 | 0.00 |
| >trna111_HisGTG_1_-_147774845_147774916@-1G.30.31,trna118_HisGTG_1_-_145396881_145396952@-1G.30.31,trna16_HisGTG_1_+_146544773_146544844@-1G.30.31,trna1_HisGTG_15_+_45493349_45493420@-1G.30.31,trna21_HisGTG_1_+_147753471_147753542@-1G.30.31,trna33_HisGTG_6_+_27125906_27125977@-1G.30.31,trna7_HisGTG_9_-_14433938_14434009@-1G.30.31,trna8_HisGTG_15_-_45492611_45492682@-1G.30.31,trna9_HisGTG_15_-_45490804_45490875@-1G.30.31\|GGCCGTGATCGTATAGTGGTTAGTACTCTGC | >HisGTG.trna111_1_-_147774845_147774916 | -1G | 30 | 31 | tRF nuclear | NO | 124.01 | 152.43 | 0.00 | 0.00 |
| >trna111_HisGTG_1_-_147774845_147774916@-1G.31.32,trna118_HisGTG_1_-_145396881_145396952@-1G.31.32,trna16_HisGTG_1_+_146544773_146544844@-1G.31.32,trna1_HisGTG_15_+_45493349_45493420@-1G.31.32,trna21_HisGTG_1_+_147753471_147753542@-1G.31.32,trna33_HisGTG_6_+_27125906_27125977@-1G.31.32,trna7_HisGTG_9_-_14433938_14434009@-1G.31.32,trna8_HisGTG_15_-_45492611_45492682@-1G.31.32,trna9_HisGTG_15_-_45490804_45490875@-1G.31.32\|GGCCGTGATCGTATAGTGGTTAGTACTCTGCG | >HisGTG.trna111_1_-_147774845_147774916 | -1G | 31 | 32 | tRF nuclear | YES | 1463.63 | 1842.86 | 294.57 | 113.40 |
| >trna111_HisGTG_1_-_147774845_147774916@-1T.28.29,trna118_HisGTG_1_-_145396881_145396952@-1T.28.29,trna16_HisGTG_1_+_146544773_146544844@-1T.28.29,trna1_HisGTG_15_+_45493349_45493420@-1T.28.29,trna21_HisGTG_1_+_147753471_147753542@-1T.28.29,trna33_HisGTG_6_+_27125906_27125977@-1T.28.29,trna7_HisGTG_9_-_14433938_14434009@-1T.28.29,trna8_HisGTG_15_-_45492611_45492682@-1T.28.29,trna9_HisGTG_15_-_45490804_45490875@-1T.28.29\|TGCCGTGATCGTATAGTGGTTAGTACTCT | >HisGTG.trna111_1_-_147774845_147774916 | -1T | 28 | 29 | tRF nuclear | NO | 64.73 | 112.15 | 0.00 | 0.00 |
| >trna111_HisGTG_1_-_147774845_147774916@-1T.29.30,trna118_HisGTG_1_-_145396881_145396952@-1T.29.30,trna16_HisGTG_1_+_146544773_146544844@-1T.29.30,trna1_HisGTG_15_+_45493349_45493420@-1T.29.30,trna21_HisGTG_1_+_147753471_147753542@-1T.29.30,trna33_HisGTG_6_+_27125906_27125977@-1T.29.30,trna7_HisGTG_9_-_14433938_14434009@-1T.29.30,trna8_HisGTG_15_-_45492611_45492682@-1T.29.30,trna9_HisGTG_15_-_45490804_45490875@-1T.29.30\|TGCCGTGATCGTATAGTGGTTAGTACTCTG | >HisGTG.trna111_1_-_147774845_147774916 | -1T | 29 | 30 | tRF nuclear | NO | 364.36 | 848.75 | 63.78 | 53.90 |

### MDA-MB-468

| | | | | | | | | | | |
|---|---|---|---|---|---|---|---|---|---|---|
| >trna111_HisGTG_1_-_147774845_147774916@-1T.30.31,trna118_HisGTG_1_-_145396881_145396952@-1T.30.31,trna16_HisGTG_1_+_146544773_146544844@-1T.30.31,trna1_HisGTG_15_+_45493349_45493420@-1T.30.31,trna21_HisGTG_1_+_147753471_147753542@-1T.30.31,trna33_HisGTG_6_+_27125906_27125977@-1T.30.31,trna7_HisGTG_9_-_14433938_14434009@-1T.30.31,trna8_HisGTG_15_-_45492611_45492682@-1T.30.31,trna9_HisGTG_15_-_45490804_45490875@-1T.30.31\|TGCCGTGATCGTATAGTGGTTAGTACTCTGC | >HisGTG.trna111_1_-_147774845_147774916 | -1T | 30 | 31 | tRF nuclear | NO | 43.72 | 104.17 | 0.00 | 0.00 |
| >trna111_HisGTG_1_-_147774845_147774916@-1T.31.32,trna118_HisGTG_1_-_145396881_145396952@-1T.31.32,trna16_HisGTG_1_+_146544773_146544844@-1T.31.32,trna1_HisGTG_15_+_45493349_45493420@-1T.31.32,trna21_HisGTG_1_+_147753471_147753542@-1T.31.32,trna33_HisGTG_6_+_27125906_27125977@-1T.31.32,trna7_HisGTG_9_-_14433938_14434009@-1T.31.32,trna8_HisGTG_15_-_45492611_45492682@-1T.31.32,trna9_HisGTG_15_-_45490804_45490875@-1T.31.32\|TGCCGTGATCGTATAGTGGTTAGTACTCTGCG | >HisGTG.trna111_1_-_147774845_147774916 | -1T | 31 | 32 | tRF nuclear | YES | 530.80 | 1497.45 | 170.60 | 64.35 |
| >am_trna111_HisGTG_1_-_147774845_147774916@1.28.28,trna118_HisGTG_1_-_145396881_145396952@1.28.28,trna16_HisGTG_1_+_146544773_146544844@1.28.28,trna1_HisGTG_15_+_45493349_45493420@1.28.28,trna21_HisGTG_1_+_147753471_147753542@1.28.28,trna33_HisGTG_6_+_27125906_27125977@1.28.28,trna7_HisGTG_9_-_14433938_14434009@1.28.28,trna8_HisGTG_15_-_45492611_45492682@1.28.28,trna9_HisGTG_15_-_45490804_45490875@1.28.28\|GCCGTGATCGTATAGTGGTTAGTACTCT | >HisGTG.trna111_1_-_147774845_147774916.am | 1 | 28 | 28 | tRF nuclear | NO | 56.04 | 117.28 | 0.00 | 0.00 |
| >am_trna111_HisGTG_1_-_147774845_147774916@1.29.29,trna118_HisGTG_1_-_145396881_145396952@1.29.29,trna16_HisGTG_1_+_146544773_146544844@1.29.29,trna1_HisGTG_15_+_45493349_45493420@1.29.29,trna21_HisGTG_1_+_147753471_147753542@1.29.29,trna33_HisGTG_6_+_27125906_27125977@1.29.29,trna7_HisGTG_9_-_14433938_14434009@1.29.29,trna8_HisGTG_15_-_45492611_45492682@1.29.29,trna9_HisGTG_15_-_45490804_45490875@1.29.29\|GCCGTGATCGTATAGTGGTTAGTACTCTG | >HisGTG.trna111_1_-_147774845_147774916.am | 1 | 29 | 29 | tRF nuclear | YES | 534.14 | 1564.57 | 84.72 | 54.47 |
| >am_trna111_HisGTG_1_-_147774845_147774916@2.29.28,trna118_HisGTG_1_-_145396881_145396952@2.29.28,trna16_HisGTG_1_+_146544773_146544844@2.29.28,trna1_HisGTG_15_+_45493349_45493420@2.29.28,trna21_HisGTG_1_+_147753471_147753542@2.29.28,trna33_HisGTG_6_+_27125906_27125977@2.29.28,trna7_HisGTG_9_-_14433938_14434009@2.29.28,trna8_HisGTG_15_-_45492611_45492682@2.29.28,trna9_HisGTG_15_-_45490804_45490875@2.29.28\|CCGTGATCGTATAGTGGTTAGTACTCTG | >HisGTG.trna111_1_-_147774845_147774916.am | 2 | 29 | 28 | tRF nuclear | NO | 0.00 | 27.57 | 6.64 | 0.00 |
| >am_trna111_HisGTG_1_-_147774845_147774916@3.29.27,trna118_HisGTG_1_-_145396881_145396952@3.29.27,trna16_HisGTG_1_+_146544773_146544844@3.29.27,trna1_HisGTG_15_+_45493349_45493420@3.29.27,trna21_HisGTG_1_+_147753471_147753542@3.29.27,trna33_HisGTG_6_+_27125906_27125977@3.29.27,trna7_HisGTG_9_-_14433938_14434009@3.29.27,trna8_HisGTG_15_-_45492611_45492682@3.29.27,trna9_HisGTG_15_-_45490804_45490875@3.29.27\|CGTGATCGTATAGTGGTTAGTACTCTG | >HisGTG.trna111_1_-_147774845_147774916.am | 3 | 29 | 27 | tRF nuclear | NO | 0.00 | 19.65 | 28.45 | 0.00 |
| >am_trna111_HisGTG_1_-_147774845_147774916@-1G.28.29,trna118_HisGTG_1_-_145396881_145396952@-1G.28.29,trna16_HisGTG_1_+_146544773_146544844@-1G.28.29,trna1_HisGTG_15_+_45493349_45493420@-1G.28.29,trna21_HisGTG_1_+_147753471_147753542@-1G.28.29,trna33_HisGTG_6_+_27125906_27125977@-1G.28.29,trna7_HisGTG_9_-_14433938_14434009@-1G.28.29,trna8_HisGTG_15_-_45492611_45492682@-1G.28.29,trna9_HisGTG_15_-_45490804_45490875@-1G.28.29\|GGCCGTGATCGTATAGTGGTTAGTACTCT | >HisGTG.trna111_1_-_147774845_147774916.am | -1G | 28 | 29 | tRF nuclear | NO | 137.62 | 138.83 | 0.00 | 0.00 |
| >am_trna111_HisGTG_1_-_147774845_147774916@-1G.29.30,trna118_HisGTG_1_-_145396881_145396952@-1G.29.30,trna16_HisGTG_1_+_146544773_146544844@-1G.29.30,trna1_HisGTG_15_+_45493349_45493420@-1G.29.30,trna21_HisGTG_1_+_147753471_147753542@-1G.29.30,trna33_HisGTG_6_+_27125906_27125977@-1G.29.30,trna7_HisGTG_9_-_14433938_14434009@-1G.29.30,trna8_HisGTG_15_-_45492611_45492682@-1G.29.30,trna9_HisGTG_15_-_45490804_45490875@-1G.29.30\|GGCCGTGATCGTATAGTGGTTAGTACTCTG | >HisGTG.trna111_1_-_147774845_147774916.am | -1G | 29 | 30 | tRF nuclear | NO | 779.38 | 941.41 | 94.58 | 79.44 |
| >trna10_LysCTT_16_+_3241501_3241573@1.28.28,trna119_LysCTT_1_-_145395522_145395594@1.28.28,trna11_LysCTT_5_-_180648979_180649051@1.28.28,trna13_LysCTT_14_-_58706613_58706685@1.28.28,trna13_LysCTT_6_+_26556774_26556846@1.28.28,trna2_LysCTT_15_+_79152904_79152976@1.28.28,trna32_LysCTT_16_-_3207406_3207478@1.28.28,trna7_LysCTT_16_+_3225692_3225764@1.28.28,trna9_LysCTT_5_+_180634755_180634827@1.28.28\|GCCCGGCTAGCTCAGTCGGTAGAGCATG | >LysCTT.trna10_16_+_3241501_3241573 | 1 | 28 | 28 | tRF nuclear | NO | 108.84 | 449.13 | 30.24 | 87.52 |

### MDA-MB-468

| | | | | | | | | | | |
|---|---|---|---|---|---|---|---|---|---|---|
| >trna10_LysCTT_16_+_3241501_3241573@1.29.29,trna13_LysCTT_14_-_58706613_58706685@1.29.29,trna2_LysCTT_15_+_79152904_79152976@1.29.29\|GCCCGGCTAGCTCAGTCGGTAGAGCATGG | >LysCTT.trna10_16_+_3241501_3241573 | 1 | 29 | 29 | tRF nuclear | NO | 0.00 | 99.84 | 0.00 | 0.00 |
| >trna10_LysCTT_16_+_3241501_3241573@1.30.30,trna13_LysCTT_14_-_58706613_58706685@1.30.30,trna2_LysCTT_15_+_79152904_79152976@1.30.30\|GCCCGGCTAGCTCAGTCGGTAGAGCATGGG | >LysCTT.trna10_16_+_3241501_3241573 | 1 | 30 | 30 | tRF nuclear | NO | 0.00 | 52.14 | 0.00 | 0.00 |
| >trna10_LysCTT_16_+_3241501_3241573@14.31.18,trna13_LysCTT_14_-_58706613_58706685@14.31.18,trna2_LysCTT_15_+_79152904_79152976@14.31.18,trna4_LysCTT_18_-_43669271_43669343@14.31.18\|AGTCGGTAGAGCATGGGA | >LysCTT.trna10_16_+_3241501_3241573 | 14 | 31 | 18 | tRF nuclear | NO | 79.80 | 97.57 | 0.00 | 0.00 |
| >am_trna10_LysCTT_16_+_3241501_3241573@11.28.18,trna119_LysCTT_1_-_145395522_145395594@11.28.18,trna11_LysCTT_5_-_180648979_180649051@11.28.18,trna128_LysCTT_1_-_55423542_55423614@11.28.18,trna13_LysCTT_14_-_58706613_58706685@11.28.18,trna13_LysCTT_6_+_26556774_26556846@11.28.18,trna2_LysCTT_15_+_79152904_79152976@11.28.18,trna32_LysCTT_16_-_3207406_3207478@11.28.18,trna4_LysCTT_18_-_43669271_43669343@11.28.18,trna6_LysCTT_19_-_52425396_52425468@11.28.18,trna7_LysCTT_16_+_3225692_3225764@11.28.18,trna9_LysCTT_5_+_180634755_180634827@11.28.18\|CTCAGTCGGTAGAGCATG | >LysCTT.trna10_16_+_3241501_3241573.am | 11 | 28 | 18 | tRF nuclear | NO | 0.00 | 59.99 | 7.29 | 0.00 |
| >am_trna10_LysCTT_16_+_3241501_3241573@31.51.21,trna119_LysCTT_1_-_145395522_145395594@31.51.21,trna11_LysCTT_5_-_180648979_180649051@31.51.21,trna13_LysCTT_16_+_3246153_3246225@31.51.21,trna13_LysCTT_6_+_26556774_26556846@31.51.21,trna30_LysCTT_16_-_3230555_3230627@31.51.21,trna5_LysCTT_16_+_3214939_3215011@31.51.21,trna7_LysCTT_16_+_3225692_3225764@31.51.21,trna8_LysCTT_1_+_39970195_39970267@31.51.21,trna9_LysCTT_5_+_180634755_180634827@31.51.21\|ACTCTTAATCTCAGGGTCGTG | >LysCTT.trna10_16_+_3241501_3241573.am | 31 | 51 | 21 | tRF nuclear | NO | 0.00 | 80.73 | 0.00 | 0.00 |
| >trna119_LysCTT_1_-_145395522_145395594@1.29.29,trna11_LysCTT_5_-_180648979_180649051@1.29.29,trna13_LysCTT_6_+_26556774_26556846@1.29.29,trna32_LysCTT_16_-_3207406_3207478@1.29.29,trna7_LysCTT_16_+_3225692_3225764@1.29.29,trna9_LysCTT_5_+_180634755_180634827@1.29.29\|GCCCGGCTAGCTCAGTCGGTAGAGCATGA | >LysCTT.trna119_1_-_145395522_145395594 | 1 | 29 | 29 | tRF nuclear | NO | 63.35 | 224.80 | 0.00 | 23.46 |
| >trna119_LysCTT_1_-_145395522_145395594@1.30.30,trna11_LysCTT_5_-_180648979_180649051@1.30.30,trna13_LysCTT_6_+_26556774_26556846@1.30.30,trna32_LysCTT_16_-_3207406_3207478@1.30.30,trna7_LysCTT_16_+_3225692_3225764@1.30.30,trna9_LysCTT_5_+_180634755_180634827@1.30.30\|GCCCGGCTAGCTCAGTCGGTAGAGCATGAG | >LysCTT.trna119_1_-_145395522_145395594 | 1 | 30 | 30 | tRF nuclear | YES | 544.34 | 2681.82 | 78.55 | 215.80 |
| >trna119_LysCTT_1_-_145395522_145395594@1.31.31,trna11_LysCTT_5_-_180648979_180649051@1.31.31,trna13_LysCTT_6_+_26556774_26556846@1.31.31,trna32_LysCTT_16_-_3207406_3207478@1.31.31,trna7_LysCTT_16_+_3225692_3225764@1.31.31,trna9_LysCTT_5_+_180634755_180634827@1.31.31\|GCCCGGCTAGCTCAGTCGGTAGAGCATGAGA | >LysCTT.trna119_1_-_145395522_145395594 | 1 | 31 | 31 | tRF nuclear | NO | 123.58 | 1073.72 | 38.12 | 57.85 |
| >trna119_LysCTT_1_-_145395522_145395594@1.32.32,trna11_LysCTT_5_-_180648979_180649051@1.32.32,trna13_LysCTT_6_+_26556774_26556846@1.32.32,trna32_LysCTT_16_-_3207406_3207478@1.32.32,trna7_LysCTT_16_+_3225692_3225764@1.32.32,trna9_LysCTT_5_+_180634755_180634827@1.32.32\|GCCCGGCTAGCTCAGTCGGTAGAGCATGAGAC | >LysCTT.trna119_1_-_145395522_145395594 | 1 | 32 | 32 | tRF nuclear | NO | 0.00 | 113.24 | 0.00 | 0.00 |
| >am_trna119_LysCTT_1_-_145395522_145395594@14.31.18,trna11_LysCTT_5_-_180648979_180649051@14.31.18,trna128_LysCTT_1_-_55423542_55423614@14.31.18,trna13_LysCTT_6_+_26556774_26556846@14.31.18,trna32_LysCTT_16_-_3207406_3207478@14.31.18,trna6_LysCTT_19_-_52425396_52425468@14.31.18,trna7_LysCTT_16_+_3225692_3225764@14.31.18,trna9_LysCTT_5_+_180634755_180634827@14.31.18\|AGTCGGTAGAGCATGAGA | >LysCTT.trna119_1_-_145395522_145395594.am | 14 | 31 | 18 | tRF nuclear | NO | 40.28 | 0.00 | 0.00 | 0.00 |
| >am_trna13_LysCTT_14_-_58706613_58706685@31.51.21,trna2_LysCTT_15_+_79152904_79152976@31.51.21,trna4_LysCTT_18_-_43669271_43669343@31.51.21\|ACTCTTAATCCCAGGGTCGTG | >LysCTT.trna13_14_-_58706613_58706685.am | 31 | 51 | 21 | tRF nuclear | NO | 0.00 | 53.74 | 0.00 | 0.00 |
| >trna14_LysTTT_11_-_59327808_59327880@1.30.30,trna2_LysTTT_17_+_8022473_8022545@1.30.30,trna54_LysTTT_1_+_204475655_204475727@1.30.30,trna5_LysTTT_11_+_59323902_59323974@1.30.30,trna62_LysTTT_1_-_204476158_204476230@1.30.30,trna76_LysTTT_6_+_28918806_28918878@1.30.30\|GCCCGGATAGCTCAGTCGGTAGAGCATCAG | >LysTTT.trna14_11_-_59327808_59327880 | 1 | 30 | 30 | tRF nuclear | NO | 0.00 | 83.34 | 0.00 | 0.00 |
| >trna143_LysTTT_6_-_27559593_27559665@13.30.18,trna149_LysTTT_6_-_27302769_27302841@13.30.18,trna14_LysTTT_11_-_59327808_59327880@13.30.18,trna2_LysTTT_17_+_8022473_8022545@13.30.18,trna54_LysTTT_1_+_204475655_204475727@13.30.18,trna5_LysTTT_11_+_59323902_59323974@13.30.18,trna62_LysTTT_1_-_204476158_204476230@13.30.18,trna76_LysTTT_6_+_28918806_28918878@13.30.18\|CAGTCGGTAGAGCATCAG | >LysTTT.trna143_6_-_27559593_27559665 | 13 | 30 | 18 | tRF nuclear | NO | 0.00 | 77.22 | 0.00 | 0.00 |

### MDA-MB-468

| | | | | | | | | | | |
|---|---|---|---|---|---|---|---|---|---|---|
| >am_trna11_ProAGG_16_+_3241989_3242060@1.28.28,trna12_ProAGG_6_+_26555498_26555569@1.28.28,trna12_ProTGG_11_-_75946869_75946940@1.28.28,trna14_ProTGG_5_-_180615854_180615925@1.28.28,trna22_ProAGG_14_-_21081560_21081631@1.28.28,trna23_ProAGG_14_-_21077495_21077566@1.28.28,trna28_ProTGG_16_-_3234133_3234204@1.28.28,trna29_ProAGG_16_-_3232635_3232706@1.28.28,trna2_ProAGG_7_+_128423504_128423575@1.28.28,trna30_ProCGG_6_+_27059521_27059592@1.28.28,trna37_ProCGG_17_-_8126151_8126222@1.28.28,trna3_ProTGG_16_+_3208923_3208994@1.28.28,trna52_ProCGG_1_+_167683962_167684033@1.28.28,trna65_ProAGG_1_-_167684725_167684796@1.28.28,trna6_ProCGG_16_+_3222049_3222120@1.28.28,trna6_ProTGG_14_+_21152175_21152246@1.28.28,trna8_ProTGG_16_+_3238094_3238165@1.28.28,trna9_ProAGG_11_+_75946557_75946628@1.28.28,trna9_ProAGG_16_+_3239634_3239705@1.28.28\|GGCTCGTTGGTCTAGGGGTATGATTCTC | >ProAGG.trna11_16_+_3241989_3242060.am | 1 | 28 | 28 | tRF nuclear | NO | 125.88 | 244.90 | 0.00 | 0.00 |
| >am_trna11_ProAGG_16_+_3241989_3242060@1.29.29,trna12_ProAGG_6_+_26555498_26555569@1.29.29,trna12_ProTGG_11_-_75946869_75946940@1.29.29,trna14_ProTGG_5_-_180615854_180615925@1.29.29,trna22_ProAGG_14_-_21081560_21081631@1.29.29,trna23_ProAGG_14_-_21077495_21077566@1.29.29,trna28_ProTGG_16_-_3234133_3234204@1.29.29,trna29_ProAGG_16_-_3232635_3232706@1.29.29,trna2_ProAGG_7_+_128423504_128423575@1.29.29,trna30_ProCGG_6_+_27059521_27059592@1.29.29,trna37_ProCGG_17_-_8126151_8126222@1.29.29,trna3_ProTGG_16_+_3208923_3208994@1.29.29,trna52_ProCGG_1_+_167683962_167684033@1.29.29,trna65_ProAGG_1_-_167684725_167684796@1.29.29,trna6_ProCGG_16_+_3222049_3222120@1.29.29,trna6_ProTGG_14_+_21152175_21152246@1.29.29,trna8_ProTGG_16_+_3238094_3238165@1.29.29,trna9_ProAGG_11_+_75946557_75946628@1.29.29,trna9_ProAGG_16_+_3239634_3239705@1.29.29\|GGCTCGTTGGTCTAGGGGTATGATTCTCG | >ProAGG.trna11_16_+_3241989_3242060.am | 1 | 29 | 29 | tRF nuclear | NO | 171.53 | 594.22 | 0.00 | 6.42 |
| >trna12_TrpCCA_17_+_19411494_19411565@30.52.23,trna168_TrpCCA_6_-_26331672_26331743@30.52.23,trna170_TrpCCA_6_-_26319330_26319401@30.52.23,trna1_TrpCCA_7_+_99067307_99067378@30.52.23,trna39_TrpCCA_17_-_8124187_8124258@30.52.23,trna6_TrpCCA_17_+_8089676_8089747@30.52.23\|ACTCCAGATCAGAAGGTTGCGTG | >TrpCCA.trna12_17_+_19411494_19411565 | 30 | 52 | 23 | tRF nuclear | NO | 33.19 | 0.00 | 0.00 | 0.00 |
| >trna6_TrpCCA_12_+_98898030_98898101@30.52.23\|ACTCCAGATCAGAAGGCTGCGTG | >TrpCCA.trna6_12_+_98898030_98898101 | 30 | 52 | 23 | tRF nuclear | NO | 21.36 | 0.00 | 0.00 | 0.00 |
| >trna15_ValAAC_5_-_180615416_180615488@1.23.23\|GTTTCCGTAGTGTAGTGGTCATC | >ValAAC.trna15_5_-_180615416_180615488 | 1 | 23 | 23 | tRF nuclear | NO | 0.00 | 42.30 | 0.00 | 0.00 |
| >trna15_ValAAC_5_-_180615416_180615488@1.29.29\|GTTTCCGTAGTGTAGTGGTCATCACGTTC | >ValAAC.trna15_5_-_180615416_180615488 | 1 | 29 | 29 | tRF nuclear | NO | 35.25 | 178.60 | 133.86 | 6.14 |
| >trna37_ValAAC_6_+_27203288_27203360@1.29.29\|GTTTCCGTAGTGTAGTGGTTATCACGTTT | >ValAAC.trna37_6_+_27203288_27203360 | 1 | 29 | 29 | tRF nuclear | NO | 0.00 | 95.64 | 44.53 | 0.00 |
| >am_trna10_ValCAC_5_-_180649395_180649467@1.23.23,trna12_ValAAC_5_-_180645270_180645342@1.23.23,trna132_ValAAC_6_-_27721179_27721251@1.23.23,trna136_ValAAC_6_-_27648885_27648957@1.23.23,trna139_ValAAC_6_-_27618707_27618779@1.23.23,trna18_ValCAC_5_-_180529253_180529325@1.23.23,trna2_ValAAC_3_+_169490018_169490090@1.23.23,trna2_ValCAC_5_+_180524070_180524142@1.23.23,trna37_ValAAC_6_+_27203288_27203360@1.23.23,trna4_ValAAC_5_+_180591154_180591226@1.23.23,trna5_ValAAC_5_+_180596610_180596682@1.23.23,trna6_ValCAC_5_+_180600650_180600722@1.23.23,trna85_ValCAC_1_-_161369490_161369562@1.23.23,trna90_ValCAC_1_-_149684088_149684161@1.23.23,trna98_ValCAC_1_-_149298555_149298627@1.23.23,trna9_ValCAC_6_+_26538282_26538354@1.23.23\|GTTTCCGTAGTGTAGTGGTTATC | >ValCAC.trna10_5_-_180649395_180649467.am | 1 | 23 | 23 | tRF nuclear | NO | 43.62 | 249.00 | 0.00 | 6.04 |
| >am_trna10_ValCAC_5_-_180649395_180649467@1.29.29,trna12_ValAAC_5_-_180645270_180645342@1.29.29,trna132_ValAAC_6_-_27721179_27721251@1.29.29,trna136_ValAAC_6_-_27648885_27648957@1.29.29,trna139_ValAAC_6_-_27618707_27618779@1.29.29,trna18_ValCAC_5_-_180529253_180529325@1.29.29,trna2_ValAAC_3_+_169490018_169490090@1.29.29,trna2_ValCAC_5_+_180524070_180524142@1.29.29,trna4_ValAAC_5_+_180591154_180591226@1.29.29,trna5_ValAAC_5_+_180596610_180596682@1.29.29,trna6_ValCAC_5_+_180600650_180600722@1.29.29,trna85_ValCAC_1_-_161369490_161369562@1.29.29,trna90_ValCAC_1_-_149684088_149684161@1.29.29,trna98_ValCAC_1_-_149298555_149298627@1.29.29,trna9_ValCAC_6_+_26538282_26538354@1.29.29\|GTTTCCGTAGTGTAGTGGTTATCACGTTC | >ValCAC.trna10_5_-_180649395_180649467.am | 1 | 29 | 29 | tRF nuclear | YES | 320.57 | 1753.09 | 392.37 | 138.82 |

### MDA-MB-468

<table>
<tr><th></th><th></th><th></th><th></th><th></th><th></th><th></th><th></th><th></th><th></th><th></th></tr>
<tr><td>>am_trna10_ValCAC_5_-_180649395_180649467@1.31.31,trna12_ValAAC_5_-_180645270_180645342@1.31.31,trna132_ValAAC_6_-_27721179_27721251@1.31.31,trna136_ValAAC_6_-_27648885_27648957@1.31.31,trna139_ValAAC_6_-_27618707_27618779@1.31.31,trna18_ValCAC_5_-_180529253_180529325@1.31.31,trna2_ValAAC_3_+_169490018_169490090@1.31.31,trna2_ValCAC_5_+_180524070_180524142@1.31.31,trna4_ValAAC_5_+_180591154_180591226@1.31.31,trna5_ValAAC_5_+_180596610_180596682@1.31.31,trna6_ValCAC_5_+_180600650_180600722@1.31.31,trna85_ValCAC_1_-_161369490_161369562@1.31.31,trna90_ValCAC_1_-_149684088_149684161@1.31.31,trna98_ValCAC_1_-_149298555_149298627@1.31.31,trna9_ValCAC_6_+_26538282_26538354@1.31.31|GTTTCCGTAGTGTAGTGGTTATCACGTTCGC</td><td>>ValCAC.trna10_5_-_180649395_180649467.am</td><td>1</td><td>31</td><td>31</td><td>tRF nuclear</td><td>NO</td><td>20.62</td><td>62.07</td><td>0.00</td><td>0.00</td></tr>
<tr><td>>am_trna10_ValCAC_5_-_180649395_180649467@5.23.19,trna12_ValAAC_5_-_180645270_180645342@5.23.19,trna132_ValAAC_6_-_27721179_27721251@5.23.19,trna136_ValAAC_6_-_27648885_27648957@5.23.19,trna139_ValAAC_6_-_27618707_27618779@5.23.19,trna18_ValCAC_5_-_180529253_180529325@5.23.19,trna2_ValAAC_3_+_169490018_169490090@5.23.19,trna2_ValCAC_5_+_180524070_180524142@5.23.19,trna37_ValAAC_6_+_27203288_27203360@5.23.19,trna4_ValAAC_5_+_180591154_180591226@5.23.19,trna5_ValAAC_5_+_180596610_180596682@5.23.19,trna6_ValCAC_5_+_180600650_180600722@5.23.19,trna85_ValCAC_1_-_161369490_161369562@5.23.19,trna90_ValCAC_1_-_149684088_149684161@5.23.19,trna98_ValCAC_1_-_149298555_149298627@5.23.19,trna9_ValCAC_6_+_26538282_26538354@5.23.19|CCGTAGTGTAGTGGTTATC</td><td>>ValCAC.trna10_5_-_180649395_180649467.am</td><td>5</td><td>23</td><td>19</td><td>tRF nuclear</td><td>NO</td><td>0.00</td><td>12.52</td><td>0.00</td><td>0.00</td></tr>
<tr><td>>trna152_ValCAC_6_-_27248049_27248121@1.23.23|GCTTCTGTAGTGTAGTGGTTATC</td><td>>ValCAC.trna152_6_-_27248049_27248121</td><td>1</td><td>23</td><td>23</td><td>tRF nuclear</td><td>NO</td><td>0.00</td><td>44.67</td><td>0.00</td><td>0.00</td></tr>
<tr><td>>trna152_ValCAC_6_-_27248049_27248121@1.24.24|GCTTCTGTAGTGTAGTGGTTATCA</td><td>>ValCAC.trna152_6_-_27248049_27248121</td><td>1</td><td>24</td><td>24</td><td>tRF nuclear</td><td>NO</td><td>0.00</td><td>19.29</td><td>0.00</td><td>0.00</td></tr>
<tr><td>>trna152_ValCAC_6_-_27248049_27248121@1.28.28|GCTTCTGTAGTGTAGTGGTTATCACGTT</td><td>>ValCAC.trna152_6_-_27248049_27248121</td><td>1</td><td>28</td><td>28</td><td>tRF nuclear</td><td>NO</td><td>0.00</td><td>12.25</td><td>0.00</td><td>0.00</td></tr>
<tr><td>>trna152_ValCAC_6_-_27248049_27248121@1.29.29|GCTTCTGTAGTGTAGTGGTTATCACGTTC</td><td>>ValCAC.trna152_6_-_27248049_27248121</td><td>1</td><td>29</td><td>29</td><td>tRF nuclear</td><td>YES</td><td>337.06</td><td>2128.04</td><td>148.00</td><td>63.48</td></tr>
<tr><td>>trna152_ValCAC_6_-_27248049_27248121@1.30.30|GCTTCTGTAGTGTAGTGGTTATCACGTTCG</td><td>>ValCAC.trna152_6_-_27248049_27248121</td><td>1</td><td>30</td><td>30</td><td>tRF nuclear</td><td>NO</td><td>0.00</td><td>32.43</td><td>0.00</td><td>0.00</td></tr>
<tr><td>>trna16_ValTAC_11_-_59318460_59318532@1.30.30|GGTTCCATAGTGTAGCGGTTATCACGTCTG</td><td>>ValTAC.trna16_11_-_59318460_59318532</td><td>1</td><td>30</td><td>30</td><td>tRF nuclear</td><td>NO</td><td>0.00</td><td>30.13</td><td>0.00</td><td>0.00</td></tr>
<tr><td>>trna17_ValTAC_11_-_59318102_59318174@1.30.30,trna4_ValTAC_X_-_18693029_18693101@1.30.30|GGTTCCATAGTGTAGTGGTTATCACGTCTG</td><td>>ValTAC.trna17_11_-_59318102_59318174</td><td>1</td><td>30</td><td>30</td><td>tRF nuclear</td><td>NO</td><td>0.00</td><td>71.66</td><td>0.00</td><td>0.00</td></tr>
</table>
